# Identification of Cancer-Associated Fibroblasts in Glioblastoma and Defining Their Pro-tumoral Effects

**DOI:** 10.1101/2021.05.08.443250

**Authors:** Saket Jain, Jonathan W. Rick, Rushikesh Joshi, Angad Beniwal, Jordan Spatz, Alexander Chih-Chieh Chang, Alan T. Nguyen, Sweta Sudhir, Ankush Chandra, Alex Haddad, Harsh Wadhwa, Sumedh S. Shah, Serah Choi, Josie L. Hayes, Lin Wang, Garima Yagnik, Joseph F. Costello, Aaron Diaz, Manish K. Aghi

**Affiliations:** Department of Neurosurgery; University of California San Francisco (UCSF)

**Author notes:** Corresponding author: Manish K. Aghi, MD, PhD; Professor of Neurological Surgery; UCSF; Diller Cancer Research Building; 1450 Third Street Rm HD-465; San Francisco, CA 94158. Authors contributed equally to this work (Saket Jain, Jonathan W. Rick).

**Keywords:** fibroblasts, glioblastoma, stem cells, EDA fibronectin

## Abstract

Despite their identification in some cancers, pro-tumoral cancer-associated fibroblasts (CAFs) were presumed absent in glioblastoma given the lack of brain fibroblasts. Serial trypsinization of primary glioblastoma cultures yielded cells with CAF morphology, CAF transcriptomic profile, and mesenchymal lineage in single-cell RNA-seq. Glioblastoma CAFs were attracted to glioblastoma stem cells (GSCs) and CAFs enriched GSCs. We created a resource of inferred crosstalk by mapping expression of receptors to their cognate ligands, identifying PDGF-β and TGF-β as mediators of GSC effects on CAFs, and osteopontin and hepatocyte growth factor as mediators of CAF-induced GSC enrichment. Glioblastoma CAFs also induced M2 macrophage polarization by producing the EDA fibronectin variant. Glioblastoma CAFs were enriched in the subventricular zone which houses neural stem cells that produce GSCs. Including CAFs in GSC-derived xenografts induced *in vivo* growth. These findings are among the first to identify glioblastoma CAFs and their GSC interactions, making them an intriguing target.

## INTRODUCTION

Glioblastoma (GBM) is an aggressive primary brain cancer with a poor prognosis.^1^ Current therapies have failed in large part because they treat GBM cells in isolation and fail to account for the recent understanding that GBM is an organ with complex interplay between tumor cells and their microenvironment.^2^ In terms of the cellular makeup of the GBM microenvironment, while numerous studies have focused on endothelial cells^3^ and immune cells,^4^ little attention has been paid to whether cancer-associated fibroblasts (CAFs), a cell type described as the most important one in the stroma of carcinomas,^5^ exist in GBM. While many have presumed the lack of CAFs in GBM based on the lack of fibroblasts in the central nervous system,^6^ some studies have identified cells expressing markers associated with CAFs in GBM.^7–9^ However, these studies fail to comprehensively profile the effects of the identified cells on every component of GBM and its microenvironment. More importantly, the reliance of these studies on cell surface markers without comprehensive gene expression profiling raises the possibility that the identified cells could be other cells in the microenvironment such as pericytes, which share overlapping cell surface markers with fibroblasts.^10^

To directly address this knowledge gap, we used a serial trypsinization method described for isolating CAFs in other cancers^11^ and analyzed the resulting cells transcriptomically to verify that they were CAFs based on their gene expression profile. We used single-cell lineage trajectory analysis to define the origin of these cells. And, we comprehensively determined the effects of these cells on GBM cells and the GBM microenvironment *in vivo*.

## RESULTS

### Identifying CAFs in GBM by Serial Trypsinization

To determine whether a CAF-like population existed in GBM, we used an established serial trypsinization method^11^ in which dissociated GBM patient samples were cultured for five weeks in DMEM/F12 with 10% fetal bovine serum. Cells underwent media change every 4 days and serial trypsinization to remove less adherent tumor cells, resulting in retention of cells resistant to trypsinization that have been confirmed to be CAFs in other cancers.^11^ Within five weeks of culturing cells isolated from newly diagnosed patient GBM samples in this manner, a population of cells emerged that uniformly exhibited the large spindle-shaped morphology that has been described for CAFs and fibroblasts.^12^

We quantified the morphology of these cells by developing a modified version of visually-aided morpho-phenotyping recognition (VAMPIRE) analysis^13^ to classify and compare irregular cellular and nuclear shapes. By pairing nuclear and cytoplasm datasets by cell, we generated a 16 data-point profile for each cell. We then designed a machine learning logistic regression classifier utilizing breast cancer CAF data (**Supp. Fig. 1**) and GBM cell line data (**Supp. Fig. 2**) to achieve a nominal accuracy of 91% in distinguishing GBM cells from CAFs. Our classifier identified 77% of the cells isolated from serial trypsinization of patient GBMs as exhibiting CAF morphology. In contrast, when patient GBM samples were grown in culture without serial trypsinization, the classifier found GBM cells predominated at 82%, reducing the population of cells with CAF morphology to 18% (P<0.0001; **Figs. 1A-B**), supporting our hypothesis that these cells isolated by serial trypsinization were CAFs and underscoring the importance of serial trypsinization in isolating this CAF population from GBM.

**Figure 1.**
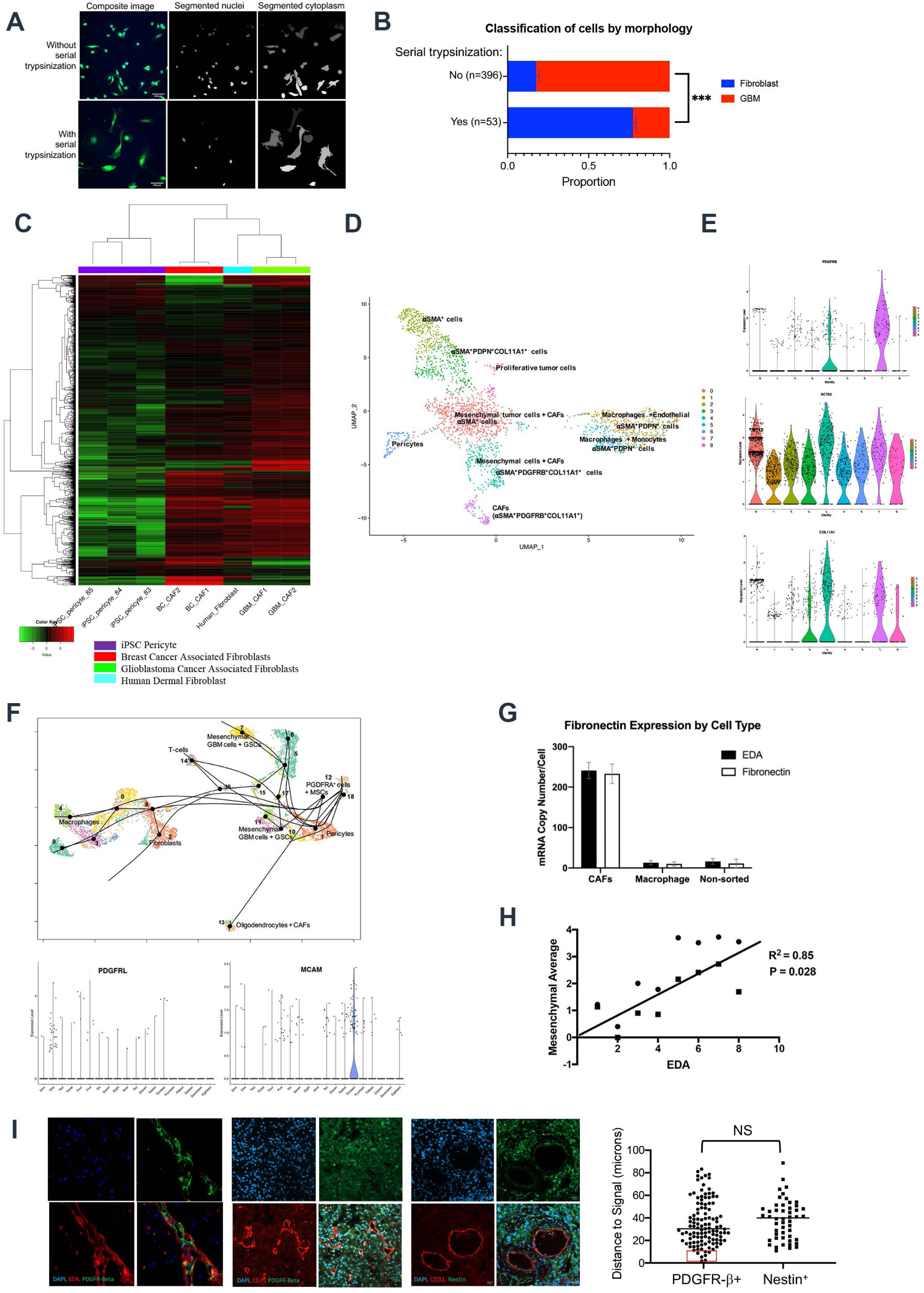
Identification of CAFs in GBM. (**A**) Representative segmented images of patient derived GBM cells with and without serial trypsinization. (**B**) Quantification of VAMPIRE analysis, revealing that nearly 77% of cells isolated from serial trypsinization of GBM exhibited fibroblast morphology, defined using 1997T and 2124T breast cancer-associated fibroblasts, compared to just 23% of these cells exhibiting GBM morphology, defined using GBM6, GBM43 and U251 cells (P<0.001). (**C**) Serially trypsinized cells from patient GBMs exhibited a transcriptomic profile similar to breast cancer CAFs and normal human dermal fibroblasts but distinct from brain pericytes as assessed by bulk RNA-seq. (**D**) Heterogeneous expression of markers expressed by CAFs from other cancers among 9 clusters identified by scRNA-seq of 7,276 serially trypsinized cells from patient GBM; and **(E)** Violin plots showing expression of three CAF markers in these clusters. **(F**) Neighbor clustering and a non-dimensional reduction technique UMAP were used to identify and visualize 18 robust cell clusters based on the 23386 most variable genes across scRNA-seq of 12074 cells from 8 patient GBMs. Shown are violin plots for expression of CAF marker MCAM and pericyte marker PDGFRL, along with results of a slingshot lineage trajectory analysis performed on the 18 clusters. (**G**) qPCR revealed elevated expression of total fibronectin and the EDA splice variant of fibronectin in CAF-like cells isolated by serial trypsinization of patient GBMs relative to (1) CD11b^+^ TAMs and (2) a population enriched for tumor cells obtained by flow sorting a freshly resected tumor to eliminate stromal cells expressing CD11b, CD31, and CD3 (n=3/group). (**H**) EDA fibronectin expression correlated with aggregate expression of five mesenchymal genes (**Supp. Table 6**) as assessed by qPCR of patient newly diagnosed GBM specimens (n=8; P=0.0012). (**I**) IF of patient GBMs revealed PDGFR-β staining in close proximity to EDA staining (left panel: red=EDA; green=PDGFR-β), with PDGFR-β^+^ cells also in comparable proximity to CD31^+^ vessels (middle panel: red=CD31; green=PDGFR-β) as Nestin^+^ GSCs (right panel: red=CD31; green=nestin), as confirmed by bar graph on the right (P=0.3=NS=not significant), with some PDGFR-β+ cells intimately associated with vessels (Red box), consistent with them being pericytes. 100x magnification; scale bar 20 μm.

We then performed bulk RNA-seq to analyze the gene expression profile of these CAF-like cells that we had identified in patient GBMs. Bulk RNA-seq revealed that these CAF-like cells in GBM exhibited a transcriptomic profile (**Supp. Table 1)** similar to the published profile of breast cancer CAFs^14^ but different from that of pericytes,^15^ a cell type with some overlap in morphology and cell surface marker expression with CAFs (**Fig. 1C**). We also compared these CAF-like cells to normal fibroblasts from 8 different tissues, revealing that these cells most closely resembled dermal fibroblasts^16^ (**Fig. 1C**) compared to normal fibroblasts from seven other tissues^17^ (**Supp. Fig. 3**). Together, these findings supported the hypothesis that these cells were GBM CAFs.

### Assessing GBM CAF heterogeneity and identifying CAFs in GBM using single cell-sequencing

We then carried out single-cell RNA sequencing (scRNA-seq) to assess the expression of CAF markers in 7,276 cells isolated from patient GBM by serial trypsinization. As has been shown with other cancers,^18^ CAF markers were expressed by large numbers of these cells isolated by serial trypsinization of patient GBMs but not in a uniform manner, including *ACTA2* (encodes *α*-SMA, 55% of cells), *TNC* (35% of cells), *PDGFRB* (12.5% of cells), *CO11A1* (24.6% of cells), and *PDPN* (27.8% of cells) (**Figs. 1D-E; Supp. Fig. 4)**, findings corroborated at the protein level by flow cytometry (**Supp. Fig. 5**). In contrast, markers expressed by cells sharing some lineage with CAFs but not by CAFs were absent from most of the cells isolated from GBM by serial trypsinization, including *EPCAM*, an epithelial cell marker only expressed by 0.07% of the cells; *SMTN*, a smooth muscle cell marker expressed by only 4% of the cells; and *PECAM1*, an endothelial cell marker expressed by only 10.7% of the cells.

Cluster analysis revealed 9 clusters within these cells arising through serial trypsinization of GBM (**Figs. 1D-E**). In terms of CAF markers from other cancers, *α*SMA was expressed in all 9 clusters, while PDGFRB and COL11A1 were expressed in Cluster 4 and 7. While most of these cells had a CAF-like profile, we observed a cluster representing pericytes and a separate cluster of macrophages and endothelial cells. We also observed a cluster containing mesenchymal cells and CAF-like cells, suggesting a mesenchymal lineage of CAFs.

We then sought to determine if population(s) of cells expressing these CAF markers could be identified in scRNA-seq of GBMs. To do so, we analyzed previously archived scRNA-seq results from 8 patient GBMs.^19^ Using the Seurat 10x genomic workflow to analyze the dataset, we identified the 23386 most variable genes across 12074 cells. We used shared nearest neighbor clustering and ran a non-dimensional reduction technique UMAP to identify and visualize 18 robust cell clusters (**Supp. Fig. 6**). Analysis of these clusters revealed dense expression of *MCAM*, whose expression in our CAF-like cells cultured by serial trypsinization was higher than in pericytes (log_2_(Fold Change)=7.7; P=4.2X10^-11^; **Supp. Table 1**), in cluster 13, and preferential expression of platelet-derived growth factor receptor-like (*PDGFRL*), a previously described pericyte marker^20^ whose expression in our CAF-like cells cultured by serial trypsinization was lower than in pericytes (log_2_(Fold Change)=-3.2; P=0.04; **Supp. Table 1**) in cluster 1 **(Fig. 1F).** These two clusters were part of a lineage trajectory when analyzed by Slingshot,^21^ a top-ranked trajectory inference tool^22^ (**Fig. 1F**). Slingshot lineage trajectory analysis also identified a link between these two clusters (1 and 13) and cluster 12, which contained cells expressing mesenchymal stem cell (MSC) markers THY1, BMPR2, and PDGFR-*α*^23^ (**Supp. Fig. 7**). These findings suggested possible shared origin of CAFs and pericytes from MSCs, with pericytes and CAFs in clusters 1 and 13 also expressing these MSC markers albeit not as frequently or strongly as the MSCs in cluster 12.

### CAF production of fibronectin in GBM

Because our RNA-seq analysis revealed that the ECM protein fibronectin (*FN1* gene) was differentially expressed in GBM (log_2_(Fold Change)=5.3; P=6.9X10^-22^) and breast cancer CAFs (log_2_(Fold Change)=2.6; P=9.9X10^-6^) relative to pericytes (**Supp. Table 1**), particularly the former, and because fibronectin has been shown to be the most abundant ECM protein in GBM,^24^ we then further analyzed CAF expression of *FN1*.

First, to verify that fibronectin was differentially expressed between GBM versus normal brain, using the GlioVis databank, we queried the expression of fibronectin (FN1) in GBM and non-tumor control brain tissue. We found that GBM had significantly higher expression of *FN1* than non-tumor brain samples (P<0.001, **Supp. Fig. 8A**). GBM also had much higher expression of *FN1* than low grade gliomas (P<0.001, **Supp. Fig. 8B**).

Because fibronectin lacking splice variants is not a component of cancer pathogenesis, we next analyzed expression of total fibronectin and its extra-domain A (EDA) splice variant in GBM CAFs, tumor-associated macrophages (TAMs), and tumor cells. qPCR revealed 32-fold elevation of total fibronectin and 16-fold elevation of the EDA splice variant in CAFs relative to TAMs (P=0.002-0.004; **Fig. 1G**) and tumor cells (P=0.002; **Fig. 1G**), suggesting that EDA is a more specific GBM CAF biomarker than the cell surface receptors described for other CAFs (**Fig. 1E**). Transcriptomic analysis also revealed a positive correlation between patient GBM expression of EDA and aggregate expression of the mesenchymal subtype genes *CHI3LI*, *TIMP1*, and *SPOCD1* that confer a worse prognosis (P=0.0012; **Fig. 1H**).^25^

In terms of microscopic intratumoral EDA distribution, immunofluorescence (IF) confirmed expression of the EDA splice variant of fibronectin in GBM patient specimens in close proximity to cells expressing PDGFR-β **(Fig. 1I; Supp. Fig. 9**), a CAF marker identified from other cancers that we also found to be expressed by a portion of the cells we isolated by serial trypsinization of patient GBMs (**Fig. 1E**). While the pattern of EDA staining resembled blood vessels structurally (**Fig. 1I**), co-staining for CD31 and EDA revealed distinct distribution of blood vessels from areas of EDA deposition (**Supp. Fig. 10**).

Interestingly, co-staining of PDGFR-β with CD31 revealed PDGFR-β^+^ cells to reside in the perivascular niche in the same degree of close proximity to blood vessels exhibited by tumor-initiating Nestin^+^ GBM stem-like cells (GSCs) that have been shown to give rise to GBM cells^26^ (P=0.3; **Fig. 1I**). Some PDGFR-β^+^ cells were intimately attached to vessels in a manner not seen with Nestin^+^ cells (**Fig. 1I**), consistent with some of these PDGFR-β^+^ cells being pericytes.

### CAFs induce pro-tumoral effects on GBM stem cells

In light of the location of these GBM CAFs in the perivascular niche alongside tumor-initiating GSCs, we then analyzed the effects of GBM CAFs on these GSCs. This was done by taking GSC-containing neurospheres derived from GBM6 GBM cells and culturing them in conditioned media (CM) from GBM CAFs for 72 hours. These cells were then transcriptomically assessed and compared to GBM6 neurospheres in control media using the NanoString nCounter platform and a 770 gene multiplex to analyze expression of genes from various step in the cancer progression process including angiogenesis, extracellular matrix (ECM) remodeling, epithelial-to-mesenchymal transition (EMT), and invasion. The analysis revealed that secreted factors from GBM CAFs upregulated the Ras, VEGF, MAPK, PI3K-Akt, and HIF-1 signaling pathways in GSCs (P<0.002; **Figs. 2A-C**).

**Figure 2.**
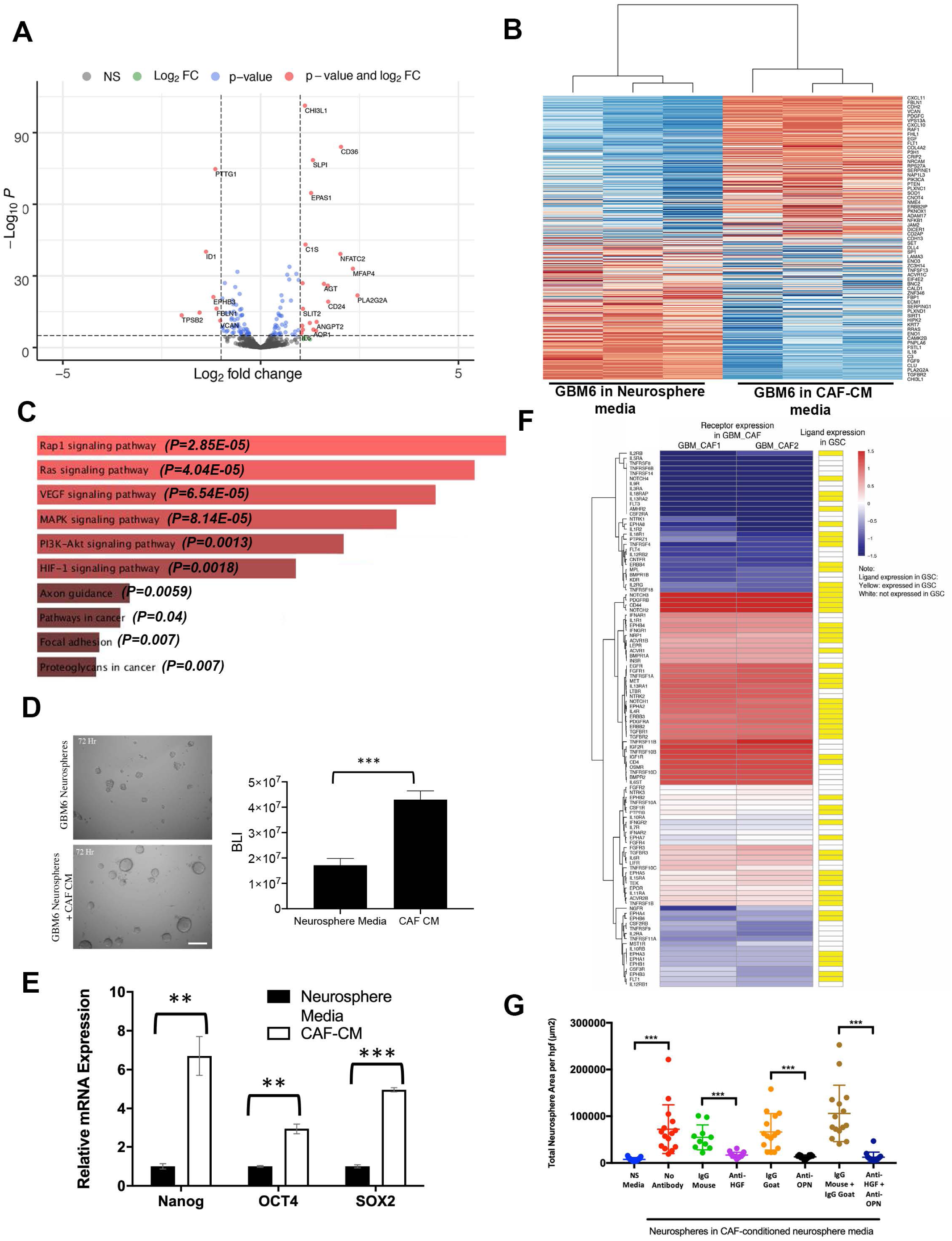
CAFs induce pro-tumoral effects of GBM stem cells. Multiplex transcriptomic analysis using the NanoString nCounter platform revealed genes in the cancer progression process upregulated by CAF CM in GBM6 stem cells, as seen by (**A**) Volcano plot to the left showing significantly (P<0.05) up- (to the right of rightmost vertical dashed line) and downregulated genes (to the left of leftmost vertical dashed line); (**B**) heatmap in the middle showing significantly (P<0.05) up- and downregulated genes; and (**C**) pathway analysis to the right showing that CAF CM upregulated Ras, VEGF, MAPK, PI3K-Akt, and HIF-1 signaling pathways in GBM stem cells (P<0.002). (**D**) Luciferase-expressing GBM43-derived neurospheres were grown in neurosphere media or CAF CM for 72 hours, after which bioluminescence (BLI) was measured, with CAF CM increasing the BLI significantly (P<0.001). (**E**) Compared to growth in neurosphere media, growth of neurospheres derived from DBTRG-05MG GBM cells in CAF CM for 24 hours elevated expression of stem cell genes Nanog 6.7-fold (P=0.009), Sox2 5.0-fold (P<0.001), and Oct4 3.0-fold (P=0.005) (n=3/group). (**F**) We mapped of the expression of receptors expressed by CAFs (**Supp. Table S2**) to that of their cognate ligands/agonists expressed by GBM6 neurospheres ^27^ based on a database of 491 known receptor-ligand interactions ^58^. Shown are cognate pairs that were co-expressed by GBM CAFs and GSCs for which the number of FPKM of the ligand is > 0.05 and read counts of the receptor is > 10, which represented 174 CAF ligands with receptors expressed by GBM stem cells. (**G**) 1000 GBM6 cells were seeded in a 12-well plate in triplicate with either neurosphere media or CAF CM with or without antibodies targeting osteopontin (OPN) and/or hepatocyte growth factor (HGF) for 72 hours. CAF CM induced neurosphere formation as measured by the total neurosphere area (P<0.001), effects that were mitigated by both anti-HGF (P<0.001) and anti-OPN (P<0.001), with the combination of anti-HGF and anti-OPN reducing the total neurosphere area more than either antibody alone (P<0.001) (n=15 hpf across 3 wells/group).

We then analyzed the functional consequences of these transcriptomic changes that CAF CM induced in the GSC-enriched GBM neurospheres. Culturing GSC- containing neurospheres derived from luciferase-expressing GBM43 GBM cells in CM from GBM CAFs for 72 hours led to increased bioluminescence (BLI) compared to growing these cells in neurosphere media (P<0.001; **Fig. 2D**). Consistent with these results, incubating GSC-containing neurospheres derived from DBTRG-05MG GBM cells in GBM CAF CM for 24 hours also increased the expression of GBM stem cell genes Nanog 6.7-fold (P=0.009), Sox2 5.0-fold (P<0.001), and Oct4 3.0-fold (P=0.005) (**Fig. 2E**).

To investigate potential mediators of CAF effects on GSCs, we created a resource of inferred crosstalk by mapping the expression of receptors expressed by GSCs to that of their cognate ligands/agonists expressed by CAF cells, using our RNA-seq results from GBM CAFs and published RNA-seq results from GBM6-derived neurospheres^27^ (**Fig. 2F; Supp. Table 2**). Based on these results, in investigating the mediators causing CAFs to promote GSC enrichment, we focused on osteopontin (OPN) and its receptor CD44 and hepatocyte growth factor (HGF) and its receptor c-Met which appeared in our GSC-CAF receptor-ligand analysis (**Supp. Table 2**). We therefore carried out a neurosphere formation assay in the presence of anti-OPN and/or anti-HGF antibodies. CAF CM increased the total neurosphere area, which accounts for both the total number and size of neurospheres, per high power field (P<0.001), effects mitigated by either anti-HGF (P<0.001) or anti-OPN (P<0.001), with the combination of anti-HGF and anti-OPN reducing the total neurosphere area more than either antibody alone (P<0.001) (**Fig. 2G**). These results suggest that the increased neurosphere formation in CAF CM is mediated through both the OPN-CD44 and HGF-cMET axes.

We then determined whether CAFs chemotactically attracted GSCs. We performed a chemotaxis assay comparing the migration of neurospheres derived from GBM6 cells towards control media or CAF CM and found no difference in the levels of chemotaxis between these conditions (P=0.1; **Supp. Fig. 11)**.

### GBM stem cells mediate CAF invasion and proliferation via PDGF and TGF-β pathways

Given that fibroblasts do not typically reside in the central nervous system, we sought to determine what factors attract CAFs to GBM. We hypothesized that GSCs may recruit CAFs to the perivascular niche of GBM. In order to ascertain if CAFs were attracted to GSCs, we assessed the trans-Matrigel chemotactic response of CAFs to GSC CM **(Fig. 3A)**. We found that GSC CM attracted CAFs 5-times as much as neurosphere control media (58.9 vs 12.8 cells per hpf; P<0.001; **Fig. 3B**).

**Figure 3.**
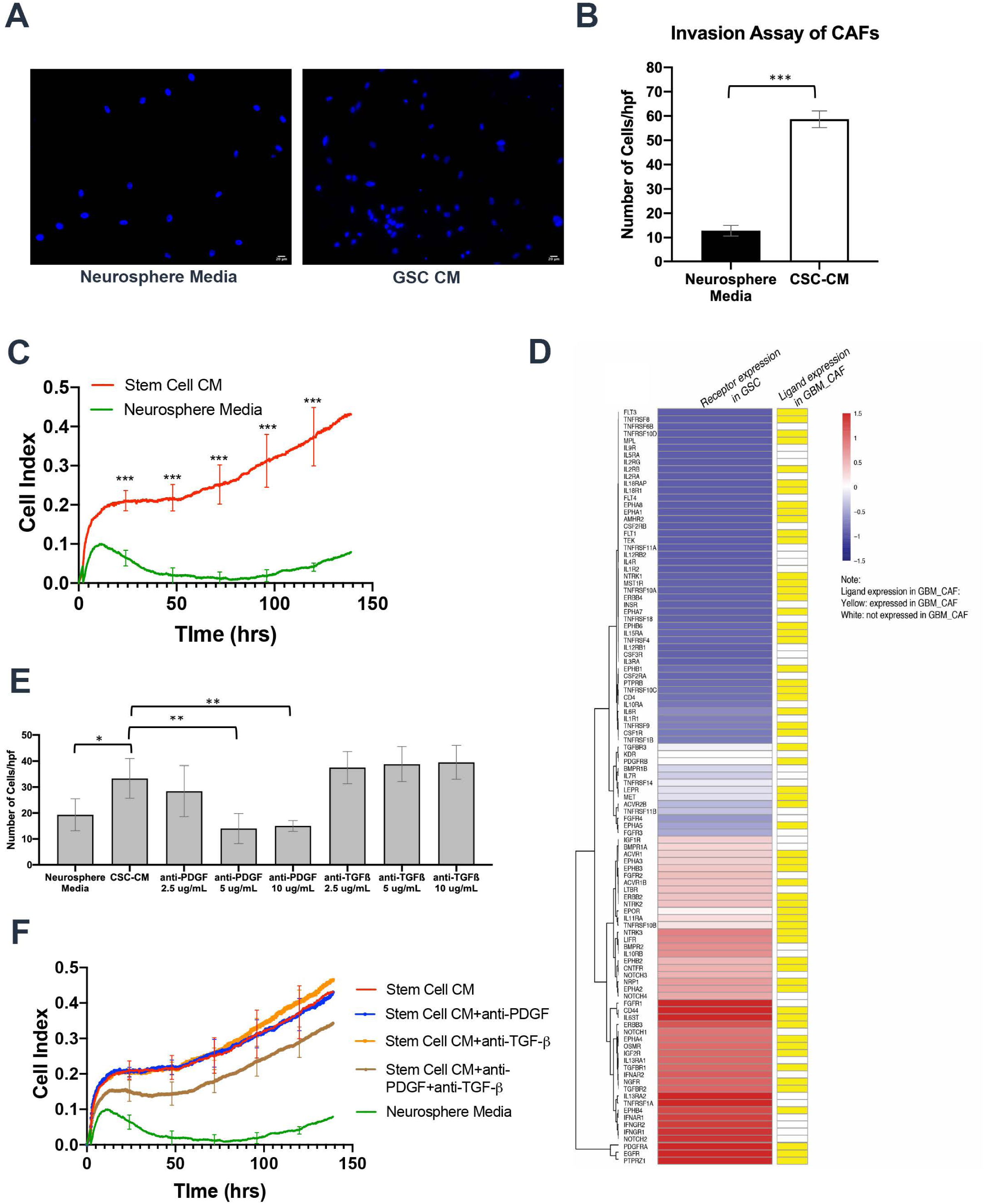
GBM stem cells mediate CAFs invasion and proliferation via PDGF and TGF-β pathways. Compared to neurosphere media, CM from GBM6 stem cell enriched neurospheres (**A-B**) attracted more cancer-associated fibroblasts (CAFs) in chemotaxis assays (P<0.001) and (**C**) stimulated CAF proliferation (P<0.001; n=5/group). (**D**) We mapped of the expression of receptors expressed by CAFs (**Supp. Table S2**) to that of their cognate ligands/agonists expressed by GBM6 neurospheres ^27^ based on a database of 491 known receptor-ligand interactions ^58^. Shown are cognate pairs that were co-expressed by GBM CAFs and GSCs for which the number of FPKM of the ligand is > 0.05 and read counts of the receptor is > 10, which represented 189 GBM stem cell ligands with receptors expressed by CAFs. (**E**) The chemotaxis of CAFs towards GBM6 neurosphere CM was abrogated by neutralizing antibodies against PDGF, but not TGF- β. TGF-β neutralizing antibodies did not abrogate invasion at 2.5-10 µg/mL (P=0.2-0.4). PDGF neutralizing antibodies reduced the number of invading cells at 5 and 10 µg/ml (P=0.002). (**F**) While neutralizing antibodies against PDGF (P=0.7-0.9) or TGF-β (P=0.5-0.9) did not affect GBM stem cell CM-induced CAF proliferation, the combination of neutralizing antibodies against PDGF and TGF-β reversed GBM stem cell-induced CAF proliferation (P=0.002-0.02) (n=5/group). * P<0.05; ** P<0.01; *** P<0.001.

We also wanted to determine if GSCs could contribute to the enrichment of CAFs via direct proliferation. We found that, at 120 hours, the number of CAFs grown in GSC CM increased over 4-fold, while CAFs grown in neurosphere control media did not grow during the same time interval (P<0.001; **Fig. 3C; Supp Fig. 12; Supp. Table 3**).

Then, to investigate potential mediators of these GSC effects on CAFs, we created the converse of our map between CAF ligands/agonists and GSC receptors (**Fig. 2D**) by mapping the expression of receptors expressed by CAFs to that of their cognate ligands/agonists expressed by GSCs, using the RNA-seq results described above (**Fig. 3D; Supp. Table 2**). Using this resource, to investigate the mediators enabling GSCs to chemotactically recruit CAFs and stimulate their proliferation, we focused on PDGF and TGF-β since both appeared in our GSC-CAF ligand-receptor analysis (**Fig. 3D**; **Supp. Table 2**) and both have receptors present on a majority of CAFs studied to date.^28^ Varying concentrations of neutralizing antibodies to TGF-β or PDGF were placed in the GSC CM before the Boyden chamber and CAFs were applied. The TGF-β neutralizing antibodies did not inhibit invasion at concentrations ranging from 2.5 to 10 µg/mL (P=0.2-0.4; **Fig. 3E**). PDGF neutralizing antibodies, however, significantly reduced the number of invading cells at 5 and 10 µg/ml (P=0.002; **Fig. 3E).** In terms of mediators of GSC-induced CAF proliferation, while 5 µg/mL neutralizing antibody against PDGF (P=0.7-0.9) or 1 µg/mL neutralizing antibody against TGF-β (P=0.5-0.9) did not affect GSC CM-induced CAF proliferation, combining neutralizing antibodies against PDGF and TGF-β at these concentrations reversed GSC-induced CAF proliferation (P=0.002-0.02; **Fig. 3F**).

### CAFs fail to induce pro-tumoral effects on non-stem GBM cells

We then analyzed whether GBM CAFs exerted protumoral effects on non-stem adherent GBM cells similar to the protumoral effects we found them to have on GBM stem cells. Addition of CM from GBM CAFs to non-stem adherent DBTRG-05MG cells caused no change in mesenchymal gene expression (P=0.8; **Supp. Fig. 13A**). There were also no phenotypic effects of GBM CAFs on non-stem GBM cells, as addition of CM from GBM CAFs to non-stem adherent GBM6 cells caused no change in tumor cell morphology as assessed by shape factor^29^ (P=0.06-0.8; **Supp. Fig. 13B),** invasion in Matrigel chambers (P=0.5; **Supp. Fig. 13C**), or proliferation (P=0.3-0.9; **Supp. Fig. 13D**) of the GBM cells. These results show that the pro-tumoral effects of GBM CAFs are specific to GSCs.

### Cultured GBM CAFs induce angiogenesis and M2 macrophage polarization via the EDA-TLR4 axis

We then investigated whether CAFs influence other cells in the tumor microenvironment. We started by investigating the effects of CAFs on endothelial cells because of our finding that CAF CM activated VEGF signaling in GSCs (**Fig. 2C**). We found that adding CAF CM to cultured HUVEC cells increased all three stages of angiogenesis: expansion of the network by tip cells, tubule formation, and fusion of the newly formed vessels.^30^ Specifically, CAF CM significantly increased expansion at 4 hours, extension at 8 hours, and mesh fusion at 16 hours (**Fig. 4A; Supp. Figs. 14-17**). Additionally, we found that a serial CM experiment in which HUVEC cells were grown in CM taken from GBM6 cells grown in CAF CM led to a significant increase in mesh formation and fusion compared to growing in HUVECs in CAF CM alone (**Fig. 4A; Supp. Figs. 14-17**), indicating an additional possible synergistic effect between CAFs and GBM cells in the process of angiogenesis.

**Figure 4.**
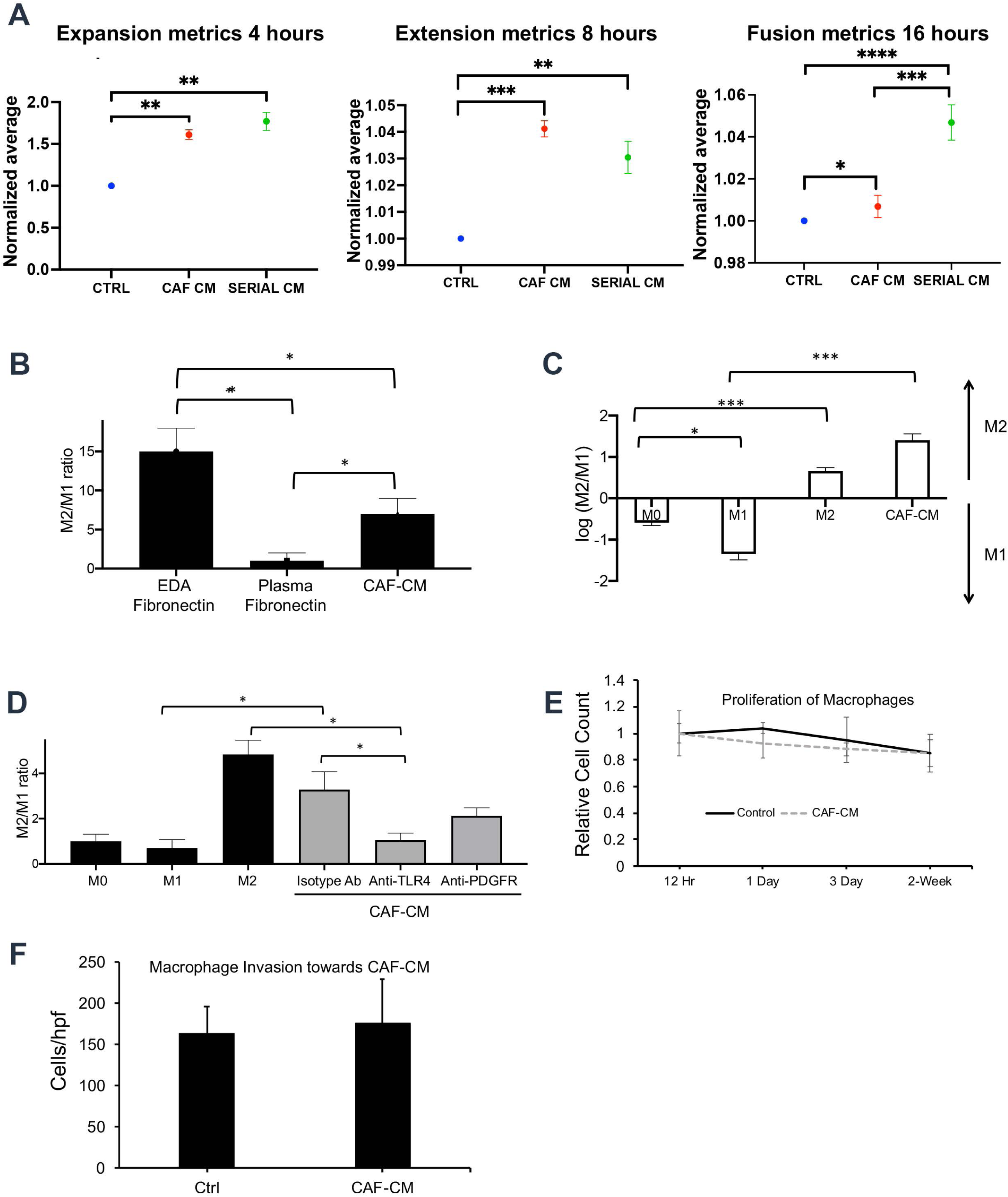
GBM CAFs induce angiogenesis and M2 macrophage polarization in culture. **(A)** Angiogenesis assays in cultured HUVEC cells revealed that CAF CM enhanced all stages of angiogenesis: expansion metrics at 4 hours (P=0.003); extension metrics at 8 hours (P<0.001), and fusion metrics at 16 hours (P=0.02). Serial CM taken from GBM cells grown in CAF CM enhanced fusion metrics at 16 hours more than CAF CM (P<0.001) (n=6/group). We then assessed the effect of CAF CM on macrophage polarization using ratio of gene expression assessed by qPCR of three M2 genes *(ARG1*, *TGFB1*, and *MMP9*) to three M1 genes (*NOS2*, *CXCL10*, and *IL1B*). **(B)** CAF CM and the EDA splice variant of fibronectin that they produce caused M2 polarization of cultured macrophages derived from human monocytes isolated from peripheral blood in a manner not seen with plasma fibronectin lacking the EDA splice variant (n=3/group; P=0.01). (**C**) When the THP-1 immortalized monocyte-like cell line was differentiated into macrophages followed by incubation in CAF CM, CAF CM drove more M2 polarization than achieved with a cytokine positive control known to drive M2 polarization (n=3/group; P=0.03). **(D)** The effects of CAF CM on M2 polarization of cultured macrophages derived from human monocytes isolated from peripheral blood were reversed by a blocking antibody against toll-like receptor 4 (TLR4), a known receptor for EDA fibronectin (n=3/group; P=0.01). CAFs did not induce (**E**) macrophage proliferation (n=3/group; P=0.3-0.9) or **(F)** chemotaxis (n=3/group; P=0.7). * P<0.05; ** P<0.01; *** P<0.001.

We then investigated the effects of CAFs on the macrophages that comprise up to 40% of the mass of a GBM.^31^ We found that CAF CM and the EDA splice variant of fibronectin that they produce cause M2 polarization of cultured macrophages derived from human monocytes isolated from peripheral blood in a manner not seen with plasma fibronectin lacking the EDA splice variant (P=0.01; **Fig. 4B**). Similarly, when the THP-1 immortalized monocyte-like cell line was differentiated into macrophages followed by incubation in CAF CM, CAF CM drove more M2 polarization than achieved with a cytokine positive control known to drive M2 polarization (P=0.03; **Fig. 4C**). The effects of CAF CM on M2 polarization of cultured macrophages derived from human monocytes isolated from peripheral blood were reversed by a blocking antibody against toll-like receptor 4 (TLR4), a known receptor for EDA fibronectin^32^ (P=0.01; **Fig. 4D**). While CAFs caused M2 macrophage polarization, CAFs did not induce macrophage proliferation (P=0.3-0.9; **Fig. 4E**) or chemotaxis (P=0.7; **Fig. 4F**).

### Regional variation in CAF localization in GBM

In order to evaluate variation in CAF localization between different tumor regions, we acquired site-directed biopsies from different regions of patient GBMs. We included regions of GBM described by our group^33^ and others:^34^ (1) tumor core; (2) leading edge of tumor enhancement; and (3) peritumoral brain zone (PBZ), the non-enhancing FLAIR bright regions surrounding the tumor **(Fig. 5A)**. Because of our finding that CAFs interact with tumor-initiating GSCs, we also sampled tissue from the subventricular zone (SVZ), the largest germinal zone in the brain found along the lateral walls of the lateral ventricles which houses the neural stem cells felt to give rise to GSCs,^35^ in cases where tumor involved this area. We then performed qPCR for fibronectin and its EDA and EDB splice variants, revealing that samples taken from SVZ GBM had 22-fold increased expression of CAF-specific EDA and 22-fold increased total FN expression, but just 5-fold increased EDB expression normalized relative to the tumor core (**Fig. 5B**). Consistent with these results, IF revealed SVZ GBM to be enriched for EDA fibronectin (**Fig. 5C**). These SVZ GBM areas were also enriched by flow cytometry for cells expressing *α*-SMA, a marker expressed by many of our cultured CAF cells (**Fig. 1E**), with 4.9% of cells from the tumor core expressing *α*-SMA compared to 13.4% of the cells from SVZ areas of GBM (P=0.02; **Fig. 5D**). IF for PDGFR-*α*, another marker we had found to be expressed by some of our cultured CAF cells (**Supp. Figs. 4-5**), also revealed more PDFR-*α*^+^ cells in SVZ areas of GBM relative to non-SVZ areas of GBM (**Fig. 5E**). In contrast, no staining for PDGFR-β or EDA (**Supp. Fig. 18**) and no detectable EDA mRNA above the threshold of accurate detection by qPCR (**Fig. 5G**) was observed in SVZ samples taken from non-tumor bearing patient specimens resected during epilepsy cases. Similarly, no staining for PDGFR-β or EDA was noted in SVZ samples taken from the autopsies of GBM patients whose tumors did not radiographically involve the SVZ (**Fig. 5F**).

**Figure 5.**
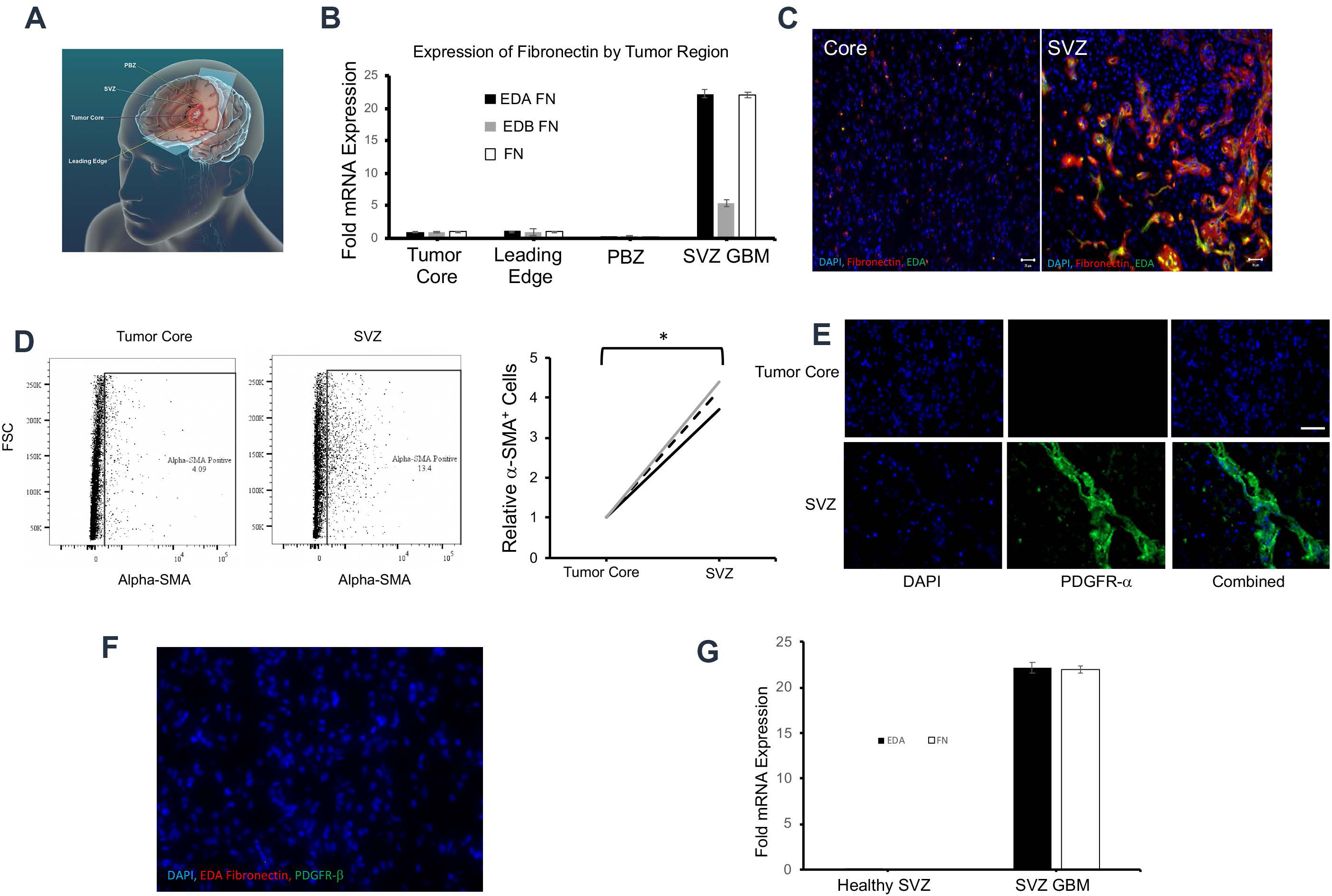
Regional variation of CAF localization in GBM. **(A)** Schema of where site-directed biopsies from patient GBMs were taken. **(B)** qPCR for total and EDA fibronectin revealed comparable elevation of both in SVZ GBM compared to the PBZ, leading edge, and tumor core (n=3/group). (**C**) IF confirmed elevated EDA (green) and total fibronectin (red) in SVZ GBM compared to the tumor core; **(D)** Flow cytometry for CAF marker *α*-SMA reveals elevation in the SVZ compared to the tumor core (n=3 paired specimens; P=0.02). (**E**) IF confirmed elevated staining of CAF marker PDGFR-*α* in the SVZ compared to the tumor core. (**F**) IF revealed no PDGFR-*α* or EDA staining in the SVZ of a GBM patient whose tumor did not involve the SVZ. 100x magnification; Scale bar 30 μM. * P<0.05; ** P<0.01; *** P<0.001. (**G**) Total and EDA fibronectin expression by qPCR was elevated in SVZ GBM but virtually undetectable in tumor-free SVZ from epilepsy surgery (n=3).

### Inclusion of CAFs with GBM stem cells induces tumor growth in vivo

To determine whether the pro-tumoral effects of CAFs on tumor-initiating GSCs we noted in cultured neurospheres also occurred *in vivo*, we intracranially implanted 40,000 GBM6 neurosphere cells, below the 100,000 neurosphere cell threshold reported to be needed to establish intracranial GBM6 tumors,^36^ into 10 athymic mice and 35,000 neurosphere cells mixed with 5,000 CAFs generated from a patient GBM by serial trypsinization into 10 athymic mice. The inclusion of CAFs with neurospheres enabled tumor growth to reach endpoint in the majority of mice, which did not occur in the absence of CAFs (P=0.03; **Fig. 6A**). In fact, addition of CAFs to GBM6 neurospheres caused mice with 35,000 GBM6 neurosphere cells to reach endpoint with the same time point as mice with 100,000 GBM6 neurosphere cells and no CAFs (P=0.4; **Supp. Fig. 19**), revealing that the tumor-promoting effects of CAFs on GSCs we noted in culture were also present *in vivo* in the intracranial tumor microenvironment. Analysis of these tumors at endpoint revealed that pro-tumoral effects of GBM CAFs noted in culture were also occurring *in vivo*. Consistent with our findings with cultured GSCs grown in CAF CM, transcriptomic profiling of tumors derived from GBM6 neurospheres grown alongside CAFs *in vivo* compared to GBM6 neurospheres grown without CAFs *in vivo* revealed increased expression of genes involved in HIF-1 signaling pathways, as well as central carbon metabolism, adherens junctions, and TGF-β signaling (P<0.003; **Figs. 6B-D**). IF revealed that CAFs caused GBM6 neurosphere-derived tumors to exhibit increased vessel diameter (P<0.001) (**Figs. 6E-F**) but with decreased vessel density (P<0.001) and decreased total vessel length (P=0.04) (**Supp. Fig. 20**), with the net effect of the former greater than the latter leading to increased total vessel surface area (P<0.001) (**Figs. 6E-F**). Moreover, flow cytometry analysis revealed that CAFs increased the percentage of macrophages that were CD206^+^ M2 pro-tumoral macrophages in GBM6 neurosphere-derived tumors (P=0.0096; **Fig. 6G**).

**Figure 6.**
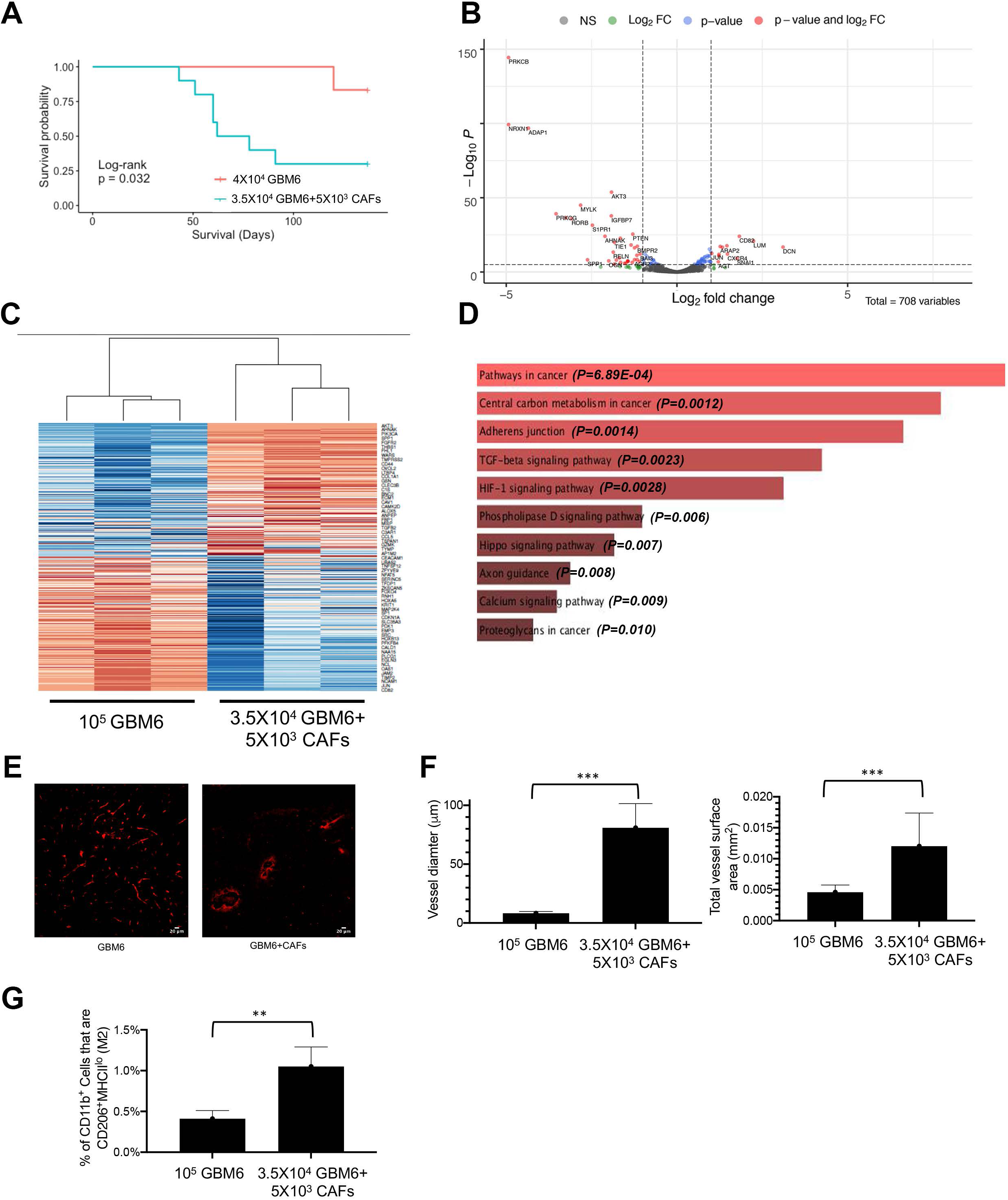
CAFs induce GBM tumor growth intracranially *in vivo*. (**A**) Kaplan-Meier curve showing intracranial implantation of 3.5X10^4^ GBM6 neurospheres with 5X10^3^ CAFs reduced survival compared to mice receiving 4.0X10^4^ GBM6 neurospheres, a threshold not associated with tumor formation in most mice (n=10/group; P=0.03). Compared to mice receiving 10^5^ GBM6 cells in neurospheres (higher number used to generate tumors), intracranial implantation of 3.5X10^4^ GBM6 neurospheres with 5X10^3^ CAFs led to transcriptional changes as determined by NanoString nCounter multiplex analysis, revealing genes in the cancer progression process upregulated by CAFs in GBM6 neurosphere-derived xenografts as seen by (**B**) Volcano plot to the left showing significantly (P<0.05) up- (to the right of rightmost vertical dashed line) and downregulated genes (to the left of leftmost vertical dashed line), (**C**) heatmap in the middle showing significantly (P<0.05) up- and downregulated genes, and (**D**) pathway analysis to the right showing that CAFs upregulated HIF-1, central carbon metabolism, adherens junctions, and TGF-β signaling pathways in GBM6 tumors (P<0.003); (**E-F**) led to increased vessel diameter (P<0.001) and increased total vessel surface area (P<0.001) (3 mice/group; 8 fields/mouse); and (**G**) increased the percentage of macrophages that were CD206^+^ M2 pro-tumoral macrophages in GBM6 neurosphere-derived tumors (P=0.0096). 100x magnification; Scale bar 20 μM. * P<0.05; ** P<0.01; *** P<0.001.

## DISCUSSION

GBMs derive much of their aggressive biology and treatment refractoriness from their regional tumor microenvironment.^2^ Compared to other solid tumors, it is currently unknown whether CAFs exist in GBM and the role that they play in GBM biology. The main argument made for a lack of CAFs in GBM is that, apart from a small amount in the blood vessels, there are no baseline fibroblasts in the brain.^6^ However, because of evidence suggesting that CAFs in other solid tumors arise from marrow-derived precursors rather than usurping of local normal fibroblasts in the organ where the tumor forms,^37–40^ it seems plausible that CAFs could exist in GBM. Indeed, recent studies have identified cells expressing markers associated with CAFs in GBM,^7–9^ but comprehensive gene expression profiling to prove these cells were CAFs and evidence for their role in GBM biology had been lacking, a knowledge gap that our current study addresses.

We began the process by determining if serial trypsinization of cells from a primary GBM specimen, a method that has been shown to generate CAFs in other cancers,^11^ could isolate CAF-like cells. Trypsin detaches cultured cells from the culture dish through proteolytic effects on cell surface integrins, and serial trypsinization takes advantage of the fact that primary GBM cells are less adherent and durable than CAFs. Most of these cells transcriptomically resembled CAFs in single-cell analysis and morphologic analysis revealed 77% of these cells to be CAFs, similar to the 79% found to be CAFs when the method was used in a murine lineage tracing study.^11^ Interestingly, the cells that emerged from GBM serial trypsinization did not uniformly express CAF markers, but instead our cluster analysis suggested possible subtypes of GBM CAFs based on patterns of marker expression. Such subtypes with distinct functionality have been described in CAFs from other cancers^41^ and further work will be needed to determine if that is the case with GBM.

Among the unique proteins we found to be expressed by GBM CAFs was the EDA splice variant of fibronectin. The EDA fibronectin splice variant arises at the 11^th^ Type III repeat (Extra-domain A; EDA). When fibronectin expresses the EDA domain, it is termed cellular fibronectin and has pivotal roles in wound healing, embryogenesis, and cancer – hence cellular fibronectin is sometimes referred to as oncofetal fibronectin.^42^ EDA containing fibronectin is thought to be principally produced by fibroblasts, and in malignancy, cancer associated fibroblasts (CAFs) are commonly the source.^42^ In contrast, fibronectin lacking splice variants is called plasma fibronectin and is produced by hepatocytes and is not a component of cancer pathogenesis.^42^

CAFs in systemic cancers have been shown to render the immune microenvironment more pro-tumoral by recruiting more monocytes and promoting their differentiation and polarization into M2 macrophages.^43^ We found similar effects of GBM CAFs, which drove M2 polarization of macrophages via toll-like receptor 4 (TLR4), a known receptor for the EDA fibronectin that we found to be produced by CAFs.

An additional profound effect we found that our novel CAF population exerted on the GBM microenvironment was on its microvasculature. The pathology of GBM is characterized by three findings: proliferation of atypical astrocytic neoplastic cells, tumor cell necrosis, and aberrant microvasculature composed of hypertrophied and glomeruloid blood vessels.^44^ Our finding that CAFs shifted the GBM vasculature to a larger, hypertrophied phenotype suggests that they play a role in establishing this defining feature of GBM. The unique architecture of GBM microvasculature has been postulated as one explanation for why GBMs are less responsive to anti-angiogenic therapies like bevacizumab,^45^ and it would be interesting to determine if CAFs play a role in this resistance by maintaining the unique vascular architecture of GBM.

Not only did we find an impact of GBM CAFs on the tumor vasculature, but we found that these cells were enriched in the perivascular niche in close proximity to blood vessels. Despite their reduced prevalence, our finding of CAF enrichment in the perivascular niche positions them to impact GBM biology. The GBM perivascular niche is defined as the area of the tumor that borders tumor vessels, which has garnered attention because it is a prime location for the GSCs whose recruitment of and nourishment by CAFs we demonstrated (**Figure 7**). Localization of CAFs to the perivascular niche empowers CAFs, despite their relatively low frequency in tumors, to maintain and nourish GSCs, another rare cell type that also resides in the perivascular niche.

**Figure 7.**
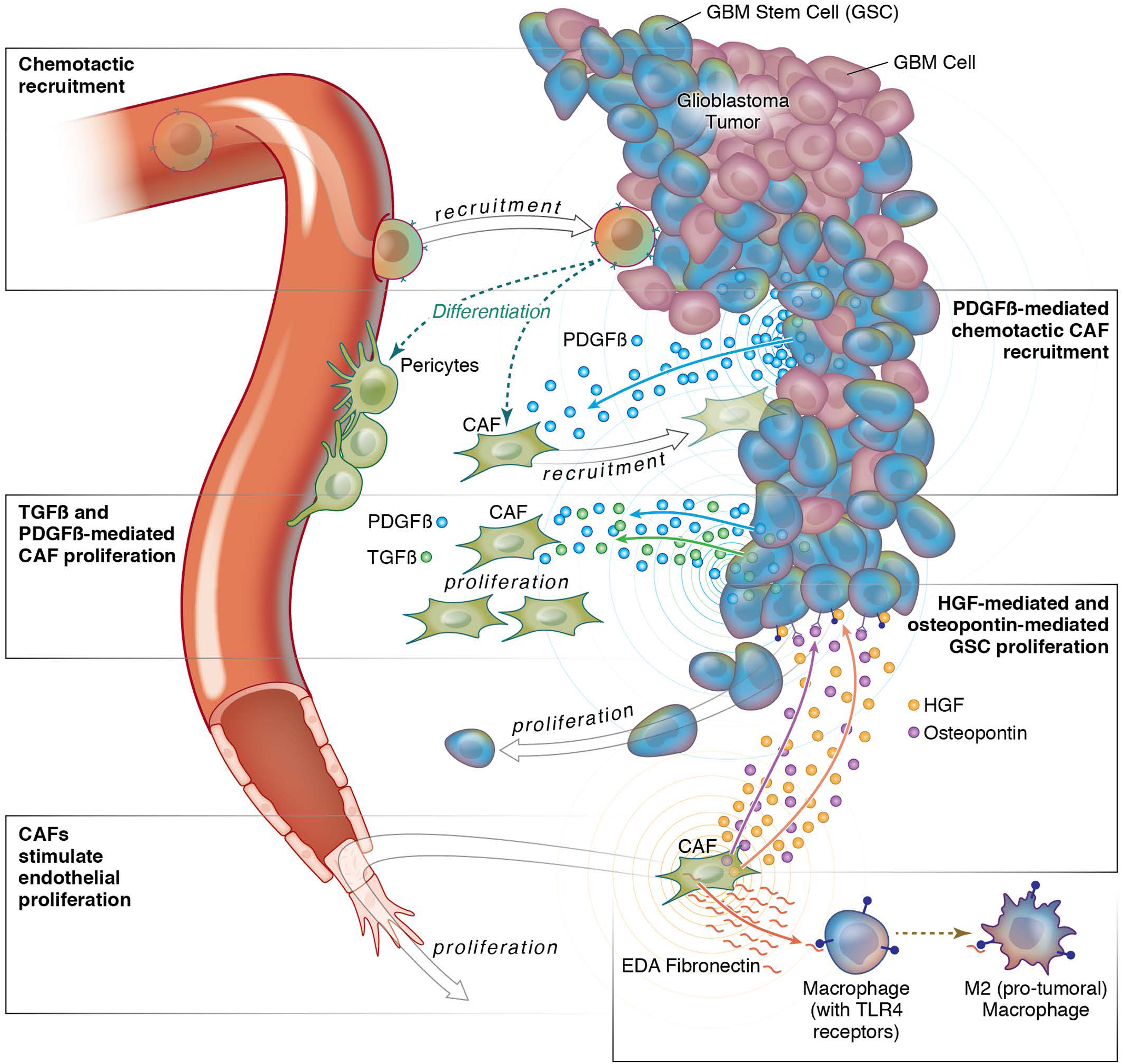
Summary of interactions between CAFs and GSCs in the perivascular niche of GBM. Shown are the interactions between CAFs and GSCs in the GBM perivascular niche that we demonstrated. GSCs recruit CAFs via PDGF-β secretion; CAFs promote GSC proliferation via HGF and osteopontin secretion; and GSCs promote CAF proliferation via TGFβ and PDGF-β secretion. CAFs also exert pro-tumoral effects on other cells in the GBM microenvironment by stimulating angiogenesis and M2 macrophage polarization.

We also found regional variation in these CAF-like cells in GBM, with cells expressing CAF markers being more prevalent in the GBM SVZ. This finding is of interest because GSCs are not housed uniformly throughout the GBM perivascular niche, but instead contact the vasculature at sites that lack astrocyte endfeet and pericyte coverage, a modification of the BBB unique to the SVZ. Interestingly, we found CAF enrichment in the SVZ of GBM patients but only when the SVZ contained tumor by imaging. Patients with GBMs that contact the SVZ have an overall survival less than those with tumors distantly located to SVZ.^46^ These differences in survival between SVZ-involved GBMs have been correlated with proteomic differences^47^ which suggest that tumor-initiating GSC enrichment in the SVZ of SVZ-involved GBMs are responsible for the poor prognosis of these patients. Our work expands upon these findings by defining the cellular makeup of the SVZ tumor microenvironment and how GSCs could recruit CAFs to the SVZ, as well as a potential role of CAF enrichment in GBMs in the SVZ in the worsened prognosis of GBMs involving the SVZ.

Another area of uncertainty that we attempted to address is the lineage of this novel CAF population we identified. While initial studies of CAFs in mouse models of other cancers have identified CAFs originating from local fibroblasts, endothelial cells, and vascular mural cells,^48–50^ other studies have implicated bone marrow-derived cells, most likely mesenchymal stem cells, as a source of CAFs.^37–40, 51^ Our single-cell lineage analysis suggested that CAFs and pericytes share a lineage and suggested that this shared lineage is from MSCs. Indeed, pericytes have been shown to derive from PDGFR^+^ myeloid progenitor cells, specifically mesenchymal stem cells (MSCs).^52^ MSCs are multipotent stem cells in bone marrow that make and repair skeletal tissues and co-exist with hematopoietic stem cells in the marrow that make blood cells. Breakdown of the blood-brain barrier (BBB) around GBM enables recruitment of endothelial and myeloid progenitor cells derived from hematopoietic stem cells in the marrow for neovascularization^53^ and establishment of tumor-associated macrophages,^4^ respectively. We hypothesize that BBB breakdown also allows the recruitment of MSCs to GBM which then differentiate into CAFs and pericytes.

There are, of course, some limitations to our work and potential areas of future study. Our hypotheses about GBM CAF lineage from single cell sequencing data are not a substitute for the traditional method of lineage tracing that involves genetic labeling of a cell followed by the tracking of its offspring. Unfortunately, studying GBM CAFs in mouse models proved challenging because we found that implanted murine GBM models do not produce CAF-like cells during serial trypsinization, suggesting that these cells were recruited to tumors that naturally formed like human GBM and would best be studied in transgenic mice that naturally form GBMs, a potential area of future study. The lack of ubiquitous CAF markers in GBM also made it impossible to quantify this population or reliably visualize it in tissue, a problem that also arises in other cancers.^54^ Further work will also be needed to develop a reliable protocol for that and to determine if those metrics can offer prognostic or therapeutic insights for GBM patients, as has been done for CAFs in other cancers.^55^ Overall, our findings provide compelling evidence that GBM CAFs play a significant role in creating a pro-tumoral GBM microenvironment, insight which we can now begin to exploit for therapeutic benefit.

## METHODS

### Cell Culture

DBTRG-05MG (ATCC), U251 (ATCC), GBM6 (Mayo Clinic), and GBM43 (Mayo Clinic) GBM cells; HUVEC cells (ATCC); and THP-1 human monocytes (ATCC) were verified using short tandem repeat (STR) profiling, passaged under 6 times, and confirmed to be Mycoplasma-free. Breast cancer CAFs were kindly provided by the Breast Cancer Now Tissue Bank (London, UK). GBM cells were cultured in DMEM/F-12 plus 10% FBS and 1% P/S at 37°C. HUVEC cells were grown in EGM-2 media (Lonza Cat # CC-3162). THP-1 cells were grown in complete RPMI with HEPES. To isolate and grow GBM CAFs in culture, the serial trypsinization method^11^ was used in which dissociated GBM patient samples were cultured in DMEM/F12 media with 10% fetal bovine serum and 1% Penicillin and streptomycin. Cells underwent media change every 4 days and serial trypsinization with 0.25% trypsin-EDTA. Because primary GBM cells are less adherent than tumor cells, detaching within 30-60 seconds of trypsinization compared to 10-15 minutes for CAFs (as assessed by microscopy), we would trypsinize for 30 seconds and discard the supernatant which had weakly adherent primary GBM cells, after which we would trypsinize for 15 minutes to detach the CAFs which were then transferred to a fresh plate. This serial trypsinization resulted in a cell population with consistent fibroblast morphology within five weeks. To generate GSC-containing neurospheres, GBM cells were grown in neurosphere media, consisting of DMEM/F12 (Gibco) supplemented with 20 ng/ml EGF (Peprotech), 20 ng/ml bFGF (Peprotech), and 2% GEM21/neuroplex (Gemini Bio-Products). When comparing the effects of CAF CM media vs Neurosphere media, CAF CM was generated by replacing the media of cultured CAF cells by neurosphere media for 72 hours, after which the media was collected, centrifuged at 300g for 5 mins followed by filtration through a 40 μm filter.

### Human GBM Tissue Acquisition

Site-directed biopsies were guided via BrainLab^TM^ interoperative MRI with IRB approval (approval #11-06160). Biopsy locations were confirmed by image capture at the time of biopsy. Samples were then transported while suspended in standard culture media on ice to the laboratory for processing. The regions were: (1) tumor core, the center of the tumor; tumor leading edge, the outer edge of the tumor enhancement as seen on MRI; and peri-tumoral brain zone (PBZ); the FLAIR bright region outside gadolinium enhancement on MRI.^34^ Samples were also obtained from the SVZ region when GBM tumors invaded this area. Additional SVZ samples were obtained as non-tumor controls from (1) patients undergoing surgical correction of epilepsy and (2) non-tumor autopsies.

### Sample Dissociation

In order to separate cells from the surrounding stroma, samples were placed on sterile culture plates and finely chopped with sterile scalpels. Tumor chunks were suspended in papain at 37°C for 30 minutes and vortexed to assure good mixture. After this incubation, the solution was applied to a 50 μm filter and rinsed with culture media. Cells were then centrifuged for 5 minutes at 500 g. Media was aspirated, and cells were treated with 1 ml of ACK RBC Lysis Buffer (Lonza) for 2 minutes. The RBC lysis reaction was halted by addition of 5 mL dPBS. Remaining cells were centrifuged for 5 minutes at 500 g, ACK lysis buffer/dPBS was aspirated, and cells were resuspended in fresh dPBS and counted.

### Morphology Analysis

15,000 cells/well were seeded in Permanox 2-chamber slides (Sigma #C6682). Cells were incubated overnight at 37°C, followed by staining with CytoTracker Green (ThermoFisher Scientific #C2925) supplemented media for 30 minutes, and then fixed using 4% paraformaldehyde solution in PBS (Thermo #J19943-K2). Cells were imaged at 20x on a Zeiss Cell Observer Spinning Disc Confocal microscope using ZEN Blue 2012 (Carl Zeiss) software. Images were segmented into their blue and green channels for DAPI and CellTrackerGreen staining using ImageJ. CellProfiler was used to identify nuclei as primary objects, and cytoplasm as secondary objects. Propagation and watershed methods were used, with thresholds manually adjusted and verified. Morphology analysis was done with the VAMPIRE analysis package^56^ with 100 coordinates and 100 shape modes as the settings and corresponding data was fed into our logistic regression. Data was compiled through four biological repeats, and statistical testing was done with an unpaired t-test by randomly splitting our total data set into three to check for consistent outcomes in each subpopulation. These splitting functions were also adjusted to test after normalizing for sample size and remained significant. These segmented nuclei and cytoplasm were then fed separately into the VAMPIRE pipeline to calculate their respective unique morphological features such as circularity. By pairing the nuclei and cytoplasm datasets by cell, we generated a 16 data-point profile for each cell. We then designed a machine learning logistic regression classifier utilizing breast cancer CAF data and GBM data from GBM6, GBM43 and U251 to achieve nominal accuracy of 91% using a 70%/30% train/test split of approximately 2704 cellular images.

### Neurosphere formation assay

To determine the effect of CAF CM on neurospheres, 10,000 GBM6 cells expressing luciferase were seeded in triplicate in a 12-well low attachment plate with either neurosphere media or CAF CM. We then assessed bioluminescence after 72 hours and imaged at 72 hours under 100x magnification using a Nikon D90 mounted on a Nikon Eclipse TS100 microscope, with 15 high power fields (hpf) from each condition analyzed using ImageJ. ROI manager function was used to measure the total area of neurospheres which accounted for the total number of spheres and sphere diameter.

### Nanostring Multiplex Transcriptomic Analysis

Using the RNeasy Mini kit (Qiagen), RNA was extracted from GBM6 neurospheres grown in neurosphere media or CAF CM and GBM6 xenografts grown with or without CAFs. A bioanalyzer was used to determine quantity and quality of the RNA sample. RNA (175 ng) from each sample was hybridized with the codeset for 18 hours. 30 μl of the reaction was loaded into the nCounter cartridge and run on the nCounter SPRINT Profiler. The raw data was then extracted followed by quality control and alignment using the Nanostring analysis software. Raw files were further processed and analyzed using the DESeq2 package in R to reveal differentially expressed genes.

### Cell proliferation assay

GBM CAFs were plated at 1000 cells per well in 96 well plates in neurosphere media or GSC CM. Proliferation was continuously assessed using the xCELLigence RTCA MP instrument (ACEA Biosciences) to measure impedance as a surrogate for cell count over 120 hours.^57^ First, 50 μL of media was added to each well of 96 well E-Plates (ACEA Biosciences) and the background impedance was measured and displayed as Cell Index. Dissociated adherent GBM CAF cells were seeded at 1000, 3000, 5000, or 7000 cells/well of the E-Plate in a volume of 100 μL and allowed to passively adhere on the electrode surface. Post seeding, the E-Plate was kept at ambient temperature inside a laminar flow hood for 30 minutes and then transferred to the RTCA MP instrument inside a cell culture incubator. Data recording was initiated immediately at 15-minute intervals for the entire duration of the experiment.

### RNA Extraction

RNA was extracted using the RNeasy^TM^ products supplied by Qiagen. This protocol was either applied to whole *ex vivo* samples or to dissociated cells as required by the experiment. Extracted RNA was stored at -80°C.

### Bulk RNA Sequencing

GBM CAF RNA libraries were prepared and Illumina HiSeq NGS preformed (UC Davis DNA Technologies Core, Davis, CA) per standard protocols. GBM CAF RNA-Seq datasets were aligned (BowTie2) and gene exons counted (FeatureCounts) with standard inputs using the Galaxy public server (https://usegalaxy.org/). iPSC-Pericyte (GSE117469 and GSM2790558),^15^ dermal fibroblast (GSM3124683),^16^ and breast cancer CAF (GSE106503)^14^ RNA-seq results were obtained from GEO and RNA-seq results from seven different types of normal human fibroblasts were kindly provided by Susan Thibeault (University of Wisconsin).^17^ Differential gene expression, heatmap, and sample cluster were performed by iDEP8.1 (http://bioinformatics.sdstate.edu/idep/). To infer the receptor-ligand interactions between GBM CAFs and GBM6 neurospheres, we compared the RNA-seq data we generated from a pair of GBM CAFs and published RNA-Seq from GBM6 neurospheres^27^ to a database of 491 known receptor-ligand interactions.^58^ Then we annotated cognate pairs that were co-expressed by GBM CAFs and GSCs for which the number of FPKM of the ligand is > 0.05 and read counts of the receptor is > 10. This produced 189 GSC ligands with receptors expressed by CAFs and 174 CAF ligands with receptors expressed by GSCs for further analysis.

### Single-cell RNA-Sequencing

Single-cell sequencing was carried out using the chromium Next GEM Single Cell 3’v3.1 protocol (10x genomics). Fresh tumor tissue post resection was collected in the Hiberbate^TM^ media and dissociated with an enzyme cocktail including 32 mg of Collagenase IV (Worthington #LS0042019); 10 mg of Deoxyribonuclease I (Worthington #LS002007) and 20 mg Soybean trypsin inhibitor (Worthington #LS003587) in 10 ml DPBS followed by RBC lysis to form a single cell suspension which was then serially trypsinized to generate CAFs. CAFs were used for library preparation for single-cell sequencing using the manufacturers protocol. Post library preparation cells were sequenced using the Illumina Novaseq. Raw data was preprocessed using Cell Ranger to obtain the matrix and count files. Data analysis was carried out using the Seurat algorithm on R. Single-cell RNA-Seq publicly available data from 8 GBM patients GBMs ^19^ was downloaded from the Broad institute database. The data was analyzed in R using scRNA-seq Seurat 10x genomics workflow. Low-quality/dying cells which often express high mitochondrial contamination were filtered by PercentageFeatureSet function. Cell doublets or cells expressing aberrantly high gene counts were filtered out. Data was normalized using LogNormalize, a global-scaling normalization method. We next calculated the highly variable genes followed by scaling the data using the ScaleData function. Next, we calculated the PCA to explore the heterogeneity within the dataset. We then clustered the cells using FindClusters function which applies modularity optimization techniques. We carried out non-linear dimensional reduction technique to generate UMAPs to visualize these datasets. FindMarkers function was used to identify makers of clustered cells. Lineage specificity analysis was done using the slingshot algorithm https://bustools.github.io/BUS_notebooks_R/slingshot.html.

### Quantitative Polymerase Chain Reactions

cDNA was created using qScript XLT cDNA Supermix (Quanta Bio), both following standard manufacturer’s protocol. cDNA was diluted to a constant concentration for all samples to ensure similar nucleic acid loading levels. Quantitative PCR was carried out using Power Syber Green Master Mix (Applied Biosystems) and primers described in **Supp. Table 4**. qPCR was performed on an Applied Biosystems StepOne Real-Time PCR cycler following recommended guidelines described by Applied Biosystems for Syber: 95° C for 10 min, followed by 40 cycles of 95° C for 15 sec and 60° C for 1 min. Ct values were calculated using the StepOne software accompanying the real-time cycler. Samples were prepared with three technical replicates for each primer pair.

### Immunofluorescence

Tissue from the operating room or from mouse tumors was promptly suspended in 4% paraformaldehyde in water for two hours. These samples were then transferred to a 30% sucrose solution for 20 hours. Samples were then submerged into Tissue-Plus Optimal Cooling Temperature (OCT) Compound TM (Fisher Scientific) and frozen at -80°C for 24 hours. The OCT tissue blocks were then sectioned into 10 μm thick slices using a Leica HM550 Cryostat. Slides were rinsed with acetone and phosphate buffered saline (PBS) solution. 5% blocking solution (TNB) was made by mixing blocking solution with tris buffered saline (TBS). Slides were coated in blocking solution for 2 hours followed by primary antibodies in TNB for 12 hours at 4°C. Slides were rinsed with PBS and then secondary antibody (in blocking solution) was applied for 2 hours. The solution was aspirated and the sample was allowed to dry before 4,6-diamidino-2-phenylindole (DAPI) was added and the coverslip was applied. Samples were kept in the dark before visualizing with a Zeiss M1 fluorescent microscope. Images were processed using Fiji’s ImageJ software. Antibodies used are in **Supp. Table 5**.

### Flow Cytometry and Fluorescence-Activated Cell Sorting (FACS)

Samples were prepared via manual mechanical separation and papain digestion. RBC lysis was performed. Samples were resuspended in DMEM, pelleted, and resuspended in FACS buffer with Fc-block (Human Seroblock, Bio-Rad). These samples were then re-pelleted and suspended in a cocktail of fluorophore-conjugated primary antibodies (**Supp. Table 5**). After incubation at 4 degrees, the samples were rinsed 3 times in FACS buffer and then suspended in FACS buffer for analysis and sorting with FACSARIA III (BD Biosciences). Living single cells were selected via FSC/SSC isolation.

### Invasion Assays

All invasion assays were completed using the Matrigel^TM^ (Corning, New York) matrix solution. Matrigel was placed on Boyden chamber membranes per the manufactures protocol. The test media was placed at the bottom of the Boyden chambers and invading cells were placed on the other surface. After 24 hours, non-invading cells were washed away, and invasion was quantified via DAPI staining. Invasion was reported as number of cells per high-power field (hpf; 40x magnification).

### Population-based Bioinformatics

Population-based bioinformatic data was obtained from GlioVis (gliovis.bioinfo.cnio.es), a composite database that collects genetic information from multiple repositories. We used the U133A array from the TCGA_GBM dataset. Statistics performed on these analyses were done using the Gliovis statistical tools.

### Angiogenesis Assay in Culture

Geltrex™ LDEV-Free Reduced Growth Factor Basement Membrane Matrix (ThermoFisher Cat # A1413202) was thawed overnight at 4°C. 120 μL of pure Geltrex was plated into 48-well tissue culture treated plates (Corning #353078), with plates tapped to spread the Geltrex, and incubated at 37℃ for 30 minutes. 40,000 HUVEC cells in EGM-2 with hydrocortisone, ascorbic acid, GA-1000, and heparin but without growth factors (bFGF-B, VEGF, R3-IGF-1, bEGF and bovine brain extract) were then added to each well. 100 μL of each condition of media was then added to each well. Plate was tilted in all directions to distribute cells. After 3 hours and 30 minutes, an additional 100 μL of EGM-2 without growth factor was added with 1.5 μL of 1 mg/mL calcein-am (ThermoFisher Cat # C1430). Cells were then imaged 30 minutes later at the 4-hour timepoint, and at 8, 16, and 24 hour timepoints. Cells were imaged at 2.5x on a Zeiss Cell Observer Spinning Disc Confocal microscope using ZEN Blue 2012 (Carl Zeiss) software, with full z-stacks and stitching used to capture the entire well in 3-dimensional space.

### Quantifying Angiogenesis in HUVEC culture assays and in vivo

Images were processed to produce a Max Intensity Z-projection that was then analyzed using the ImageJ Angiogenesis Analyzer package, available at http://image.bio.methods.free.fr/ImageJ/?Angiogenesis-Analyzer-for-ImageJ&lang=en#outil_sommaire_0. Default software settings were used with the exception of not suppressing isolated elements. For cell culture assays, we normalized and averaged expansion (Nb extrem. /Nb branches / Tot. branches length), extension (Tot. master segments length / Tot. length / Tot. branching length / Tot. segments length), and fusion (Nb Junctions / Nb master junction / Nb master segments / Nb meshes / Nb pieces / Nb segment) metrics. Composite figures were compiled based on all possible individual metrics that describe the biological phenomenon in order to avoid bias. Each datapoint was normalized to control and then compared using a paired t-test in the composite statistical test. For *in vivo* assessment, total vessel length was derived from the software and converted to microns from pixel values using scale bar calculations.

### Macrophage Studies

THP-1 cells and monocytes isolated from peripheral blood (AllCells) run through the MojoSort™ Human CD14 Selection Kit were treated with 50 ng/μL PMA (phorbol myristate acetate) for 4 days to allow cell adhesion to the plate and differentiation into resting M0 macrophages. Resulting M0 macrophages were then incubated with 20 ng/mL IFN-*γ* (for M1 polarization), 20 ng/mL IL-4 (for M2 polarization), or experimental conditions. Cells then underwent qPCR analysis of expression of three M1 (*NOS2*, *CXCL10*, and *IL1B*) and three M2 genes *(ARG1*, *TGFB1*, and *MMP9*), from which we derived an M2/M1 ratio of the expression of the three M2 markers divided by expression of the three M1 markers as we previously described.^33^ To assess macrophage proliferation, cells were plated into black 96-well clear bottom plates and analyzed using CyQuant cell proliferation assay kits (Thermo Scientific, c7026).

### Murine Intracranial Xenograft Tumors

Animal experiments were approved by the UCSF IACUC (approval #AN105170-02). Cells (either 40,000 or 100,000 GBM6 cells grown as neurospheres or 35,000 GBM6 cells grown as neurospheres mixed with 5,000 GBM CAFs) were implanted intracranially into the right frontal lobes of athymic mice (6-8 weeks, female) stereotactically.

### Statistics

Quantitative PCRs, invasion assays, cell proliferation assays, neurosphere formation assays were done with three technical and biological replicates. P-values were generated using the non-parametric two-tailed T-test to compare effects between two conditions. NanoString raw data was analyzed using the DESeq2 package in R. The DESeq2 package carries out an internal normalization where a geometric mean is calculated for each gene across replicates, the counts for a gene in each replicate is then divided by the mean. Count outliers were removed using Cook’s distance analysis. The Wald test is used for testing significance. Kaplan-Meier analysis was carried out for survival studies. Single-cell RNAseq analysis was carried out using the standard Seurat workflow.

### Data availability

The custom script used for our cell morphology analysis has been made available at https://github.com/alexanderchang1/GBM_CAF_open. Sequencing data that support the findings of this study have been deposited in the National Center for Biotechnology Information Gene Expression Omnibus (GEO) and are accessible through the GEO Series accession number <u>GSE132825</u>. All other relevant data are available from the corresponding author on request.

## ACKNOWLEDGEMENTS

M.K.A. was supported by the NIH (1R01CA227136 and 2R01NS079697) and the Uncle Kory Foundation. J.R., A.C., and S.S. were supported by Howard Hughes Medical Institute (HHMI) fellowships. A.C. was supported by Alpha Omega Alpha (AOA) Carolyn L. Kuckein Student Research Fellowship. J.R. was supported by UCSF School of Medicine (SOM) Pathways Explore Summer Grants. This study was supported in part by HDFCCC Laboratory for Cell Analysis Shared Resource Facility through grants from NIH (P30CA082103 and S10 OD021818-01).

