## Supplementary Table 1 for "Identification of Cancer-Associated Fibroblasts in Glioblastoma and Defining Their Pro-tumoral Effects"

**Supplementary Table 1: Results of CAF RNA-seq performed on serially trypinsized cells from two patient GBMs as compared to archived RNA-seq data from two breast cancer CAFs and iPSC-derived pericytes.**

| Row-names | Gene name | BC_CAF-iPSC_pericyte___log2FoldChange | BC_CAF-iPSC_pericyte___padj | GBM_CAF-iPSC_pericyte___log2FoldChange | GBM_CAF-iPSC_pericyte___padj | sum | iPSC_pericyte_83 | iPSC_pericyte_84 | iPSC_pericyte_85 | BC_CAF1 | BC_CAF2 | GBM_CAF1 | GBM_CAF2 |
|---|---|---|---|---|---|---|---|---|---|---|---|---|---|
| ENSG00000122641 | INHBA | 15.23475208 | 4.16E-30 | 13.26893946 | 2.87E-23 | 28.50369154 | 2 | 2 | 2 | 15.27039578 | 15.252111 | 13.63010036 | 12.86013896 |
| ENSG00000163453 | IGFBP7 | 13.19948976 | 2.76E-23 | 13.6552506 | 4.13E-25 | 26.85474036 | 3.327711234 | 2 | 2 | 14.17896925 | 14.1969236 | 14.95324776 | 14.24892143 |
| ENSG00000144810 | COL8A1 | 12.36155052 | 2.88E-21 | 13.87852371 | 7.98E-27 | 26.24007423 | 2 | 3.076665209 | 2 | 13.3331339 | 13.3672165 | 14.60522221 | 15.08830309 |
| ENSG00000164761 | TNFRSF11B | 12.76404613 | 3.65E-19 | 12.24862457 | 6.43E-18 | 25.0126707 | 2 | 2 | 2 | 12.78826861 | 12.7942119 | 11.47523074 | 12.78817145 |
| ENSG00000103888 | CEMIP | 11.259841 | 2.01E-18 | 13.72078662 | 3.68E-27 | 24.98062762 | 2 | 2 | 3.369317972 | 12.21177394 | 12.2856726 | 14.61641343 | 14.79632422 |
| ENSG00000133816 | MICAL2 | 11.95029075 | 1.85E-20 | 12.87016173 | 8.36E-24 | 24.82045248 | 2 | 2 | 2 | 11.96799313 | 11.9881291 | 12.78215604 | 13.00391932 |
| ENSG00000074416 | MGLL | 11.42509103 | 6.00E-18 | 13.36008498 | 2.09E-24 | 24.78517601 | 2 | 2 | 2 | 11.41586652 | 11.490225 | 13.08008705 | 13.63993691 |
| ENSG00000169429 | CXCL8 | 17.22511006 | 6.05E-23 | 6.983778924 | 0.000140535 | 24.20888898 | 2 | 2 | 2 | 17.27216181 | 17.2306876 | 4.806835018 | 7.893928381 |
| ENSG00000049540 | ELN | 10.31101304 | 1.57E-13 | 13.27734295 | 6.08E-22 | 23.58835599 | 2 | 2 | 2 | 10.34037819 | 10.3433038 | 13.74101055 | 12.67493222 |
| ENSG00000105887 | MTPN | 11.37783241 | 1.04E-18 | 11.85362949 | 2.00E-20 | 23.2314619 | 2 | 2 | 2 | 11.43320503 | 11.3789528 | 11.80202 | 11.95689865 |
| ENSG00000134531 | EMP1 | 10.75201886 | 1.21E-13 | 11.39984752 | 2.47E-15 | 22.15186638 | 2 | 2 | 2 | 10.78928396 | 10.7740351 | 10.57227185 | 11.96193261 |
| ENSG00000205413 | SAMD9 | 13.07258495 | 1.58E-24 | 8.876446682 | 6.18E-12 | 21.94903164 | 2 | 2 | 2 | 13.11647091 | 13.0825676 | 8.90036762 | 8.929158763 |
| ENSG00000118503 | TNFAIP3 | 11.56798782 | 4.62E-15 | 10.13474606 | 7.31E-12 | 21.70273387 | 3.327711234 | 2 | 2 | 12.52726555 | 12.5863798 | 10.03160495 | 11.74047552 |
| ENSG00000158747 | NBL1 | 10.41629231 | 6.94E-16 | 11.20882678 | 2.04E-18 | 21.62511909 | 2 | 2 | 2 | 10.40946091 | 10.4832045 | 11.2075668 | 11.26700563 |
| ENSG00000198542 | ITGBL1 | 11.28123233 | 1.79E-17 | 10.22603105 | 1.13E-14 | 21.50726338 | 2 | 2 | 2 | 11.29778296 | 11.3218808 | 9.952507193 | 10.5085588 |
| ENSG00000148516 | ZEB1 | 10.89154343 | 5.28E-17 | 10.54762294 | 3.56E-16 | 21.43916637 | 2 | 2 | 2 | 10.94606981 | 10.8952763 | 10.7065785 | 10.43629373 |
| ENSG00000108691 | CCL2 | 10.9971921 | 4.22E-12 | 10.43464146 | 4.84E-11 | 21.43183356 | 2 | 2 | 2 | 11.02737923 | 11.0253156 | 9.016300808 | 11.17325195 |
| ENSG00000163531 | NFASC | 9.219804288 | 7.50E-11 | 11.91083054 | 1.34E-17 | 21.13063483 | 2 | 2 | 2 | 9.250166145 | 9.2611121 | 11.2693135 | 12.39414002 |
| ENSG00000117228 | GBP1 | 11.38867332 | 6.80E-18 | 9.507845495 | 7.07E-13 | 20.89651881 | 2 | 2 | 2 | 11.4169563 | 11.4173641 | 9.261890418 | 9.77651925 |
| ENSG00000187134 | AKR1C1 | 13.16450662 | 1.69E-23 | 7.308886609 | 5.85E-08 | 20.47339323 | 2 | 2 | 2 | 13.1714883 | 13.2112425 | 7.071691703 | 7.618424749 |
| ENSG00000136235 | GPNMB | 11.06733702 | 1.54E-16 | 9.393327834 | 2.82E-12 | 20.46066485 | 2 | 2 | 3.369317972 | 12.0312718 | 12.0820105 | 10.70622743 | 9.973214279 |
| ENSG00000132274 | TRIM22 | 10.38544843 | 1.12E-19 | 9.978505426 | 1.94E-18 | 20.36395386 | 2 | 3.686303414 | 2 | 12.2506787 | 12.2240915 | 11.97684375 | 11.66847267 |
| ENSG00000133048 | CHI3L1 | 11.44508223 | 9.45E-09 | 8.829573988 | 1.33E-05 | 20.27465622 | 2 | 2 | 2 | 11.44969881 | 11.4969662 | 9.802049415 | 5.390901532 |
| ENSG00000119922 | IFIT2 | 14.56591798 | 2.56E-28 | 5.684368489 | 3.93E-05 | 20.25028647 | 2 | 3.076665209 | 2 | 15.55008331 | 15.5583383 | 6.363520437 | 7.019463152 |
| ENSG00000000971 | CFH | 10.39323397 | 1.80E-12 | 9.768149176 | 3.53E-11 | 20.16138315 | 3.327711234 | 2 | 2 | 11.35280306 | 11.4136982 | 9.714689414 | 11.35932363 |
| ENSG00000178860 | MSC | 10.98784246 | 1.27E-16 | 9.084770198 | 9.47E-12 | 20.07261266 | 2 | 2 | 2 | 11.00320274 | 11.0306754 | 8.825193025 | 9.367379581 |
| ENSG00000162783 | IER5 | 9.179123658 | 2.13E-12 | 10.65874127 | 1.60E-16 | 19.83786493 | 2 | 2 | 2 | 9.18717761 | 9.24275568 | 10.79309453 | 10.57593535 |
| ENSG00000173210 | ABLIM3 | 10.03820491 | 9.72E-15 | 9.759188532 | 4.18E-14 | 19.79739345 | 2 | 2 | 2 | 10.07805004 | 10.0618152 | 9.843290512 | 9.738960951 |
| ENSG00000151846 | PABPC3 | 9.049221252 | 6.75E-12 | 10.69040209 | 2.29E-16 | 19.73962335 | 2 | 2 | 2 | 9.093229054 | 9.07923325 | 10.53150331 | 10.88707283 |
| ENSG00000147883 | CDKN2B | 8.754122749 | 4.80E-11 | 10.97417692 | 6.17E-17 | 19.72829967 | 2 | 2 | 2 | 8.784246083 | 8.80280692 | 10.75048495 | 11.21848118 |
| ENSG00000073756 | PTGS2 | 14.20279878 | 1.26E-20 | 5.511656368 | 0.000648354 | 19.71445514 | 2 | 2 | 2 | 14.24911738 | 14.2096019 | 6.280431262 | 4.532869193 |
| ENSG00000154678 | PDE1C | 8.316084246 | 1.39E-09 | 11.39424826 | 2.60E-17 | 19.71033251 | 2 | 2 | 2 | 8.365459376 | 8.35477078 | 10.98841621 | 11.75610955 |
| ENSG00000180914 | OXTR | 5.67763549 | 3.00E-05 | 14.02151181 | 1.21E-27 | 19.6991473 | 2 | 2 | 2 | 5.720439317 | 5.89549381 | 14.17941331 | 13.903882 |
| ENSG00000133121 | STARD13 | 10.14445215 | 1.35E-14 | 9.542861265 | 3.97E-13 | 19.68731342 | 2 | 2 | 2 | 10.16727262 | 10.1843212 | 9.364829867 | 9.761669384 |
| ENSG00000119138 | KLF9 | 9.138482047 | 3.06E-12 | 10.51951401 | 4.83E-16 | 19.65799605 | 2 | 2 | 2 | 9.157231967 | 9.19229617 | 10.67475396 | 10.41281336 |
| ENSG00000140945 | CDH13 | 8.827190941 | 1.04E-11 | 10.76292001 | 4.37E-17 | 19.59011095 | 2 | 3.076665209 | 2 | 9.79991149 | 9.84325683 | 11.82235644 | 11.67962039 |
| ENSG00000113739 | STC2 | 8.818404267 | 6.69E-40 | 10.5213964 | 3.56E-57 | 19.33980067 | 4.467311984 | 4.11352228 | 3.369317972 | 12.46477506 | 12.5147103 | 14.02649715 | 14.34091864 |
| ENSG00000148677 | ANKRD1 | 8.209103465 | 1.88E-09 | 10.94031107 | 3.46E-16 | 19.14941454 | 2 | 2 | 2 | 8.260583448 | 8.248381 | 11.27737127 | 10.57769608 |
| ENSG00000111801 | BTN3A3 | 10.82075211 | 5.42E-17 | 8.310620088 | 1.74E-10 | 19.1313722 | 2 | 2 | 2 | 10.81199516 | 10.8875237 | 8.354863925 | 8.354591622 |
| ENSG00000091592 | NLRP1 | 8.254351061 | 1.43E-09 | 10.83694694 | 6.11E-16 | 19.091298 | 2 | 2 | 2 | 8.266399315 | 8.33118782 | 10.48750758 | 11.16624784 |
| ENSG00000156804 | FBXO32 | 9.548020166 | 1.92E-14 | 9.531478438 | 1.61E-14 | 19.0794986 | 2 | 3.076665209 | 3.369317972 | 11.41978589 | 11.4048184 | 11.86528459 | 10.69530541 |
| ENSG00000213923 | CSNK1E | 9.630241292 | 9.03E-14 | 9.437873222 | 2.32E-13 | 19.06811451 | 2 | 3.076665209 | 2 | 10.62961854 | 10.6144921 | 10.47258492 | 10.3865723 |
| ENSG00000182752 | PAPPA | 9.560164219 | 3.83E-23 | 9.447605902 | 7.54E-23 | 19.00777012 | 2 | 3.686303414 | 3.369317972 | 11.96188328 | 12.0424814 | 12.08435998 | 11.66599202 |
| ENSG00000140853 | NLRC5 | 10.21539356 | 5.49E-15 | 8.705728659 | 3.27E-11 | 18.92112222 | 2 | 2 | 2 | 10.19396441 | 10.2972338 | 8.612361808 | 8.867684177 |
| ENSG00000139278 | GLIPR1 | 9.253883196 | 1.95E-14 | 9.652343393 | 9.35E-16 | 18.90622659 | 2 | 3.686303414 | 2 | 11.09675311 | 11.0956404 | 11.06680613 | 11.82322502 |
| ENSG00000154175 | ABI3BP | 9.239273217 | 9.06E-13 | 9.666878124 | 5.46E-14 | 18.90615134 | 2 | 2 | 3.369317972 | 10.21165595 | 10.2525441 | 10.66635607 | 10.65083574 |
| ENSG00000135318 | NT5E | 9.858199049 | 1.63E-11 | 9.031998025 | 7.38E-10 | 18.89019707 | 2 | 5.946638857 | 2 | 14.18294647 | 14.1959195 | 13.23119452 | 13.48460951 |
| ENSG00000188219 | POTEE | 9.590275282 | 2.32E-12 | 9.245185667 | 1.29E-11 | 18.83546095 | 2 | 2 | 2 | 9.55186119 | 9.69266358 | 8.864319213 | 9.603790255 |
| ENSG00000146648 | EGFR | 9.105618931 | 5.89E-12 | 9.701555966 | 1.53E-13 | 18.8071749 | 3.327711234 | 2 | 2 | 10.04290398 | 10.1531049 | 10.44857879 | 10.90235372 |
| ENSG00000013588 | GPRC5A | 7.294522805 | 4.10E-08 | 11.50590051 | 5.93E-19 | 18.80042332 | 2 | 2 | 2 | 7.285019958 | 7.42465798 | 11.39429296 | 11.66177264 |
| ENSG00000112837 | TBX18 | 7.732695523 | 4.40E-09 | 11.04759459 | 1.05E-17 | 18.78029011 | 2 | 2 | 2 | 7.826439377 | 7.74329312 | 10.98195321 | 11.16550298 |
| ENSG00000133805 | AMPD3 | 10.6549223 | 1.56E-15 | 8.098957646 | 1.99E-09 | 18.75387995 | 2 | 2 | 2 | 10.66527283 | 10.7040796 | 7.825413909 | 8.408109518 |
| ENSG00000188375 | H3F3C | 10.43070934 | 1.45E-15 | 8.303199372 | 2.86E-10 | 18.73390871 | 2 | 2 | 2 | 10.46530827 | 10.4570431 | 8.195243072 | 8.485179747 |
| ENSG00000008517 | IL32 | 8.496873615 | 5.90E-11 | 10.23416791 | 1.44E-15 | 18.73104152 | 3.327711234 | 2 | 2 | 9.459264257 | 9.52609769 | 11.22425456 | 11.22523682 |
| ENSG00000253159 | PCDHGA12 | 10.14766264 | 2.97E-13 | 8.564899689 | 9.39E-10 | 18.71256233 | 2 | 2 | 2 | 10.14850007 | 10.2089163 | 8.061837451 | 9.0005841 |
| ENSG00000183023 | SLC8A1 | 7.900319979 | 6.87E-09 | 10.7762413 | 7.32E-16 | 18.67656128 | 2 | 2 | 2 | 7.987681034 | 7.91175216 | 10.45095555 | 11.09046346 |
| ENSG00000154027 | AK5 | 9.310228768 | 7.18E-12 | 9.324184924 | 5.62E-12 | 18.63441369 | 2 | 2 | 2 | 9.366644887 | 9.32403997 | 9.653364877 | 8.989564175 |
| ENSG00000163347 | CLDN1 | 9.419440255 | 6.84E-12 | 9.165794268 | 2.28E-11 | 18.58523452 | 2 | 2 | 2 | 9.521380925 | 9.38345569 | 8.772510235 | 9.532624709 |
| ENSG00000136869 | TLR4 | 8.317420613 | 4.15E-09 | 10.17636352 | 2.96E-13 | 18.49378413 | 2 | 3.076665209 | 2 | 9.338394921 | 9.29056951 | 10.49646885 | 11.62307742 |
| ENSG00000196611 | MMP1 | 13.04516608 | 3.79E-19 | 5.441822874 | 0.000400501 | 18.48698895 | 2 | 3.076665209 | 3.369317972 | 14.90812579 | 14.8999667 | 8.112172997 | 5.540219242 |
| ENSG00000136244 | IL6 | 13.9185218 | 1.08E-26 | 4.474859558 | 0.00134027 | 18.39338136 | 2 | 2 | 2 | 13.93497504 | 13.9554865 | 4.5389195 | 4.911532223 |
| ENSG00000163739 | CXCL1 | 13.71917198 | 8.26E-15 | 4.651016124 | 0.016089489 | 18.3701881 | 2 | 2 | 2 | 13.75359206 | 13.7383793 | 3.10571689 | 5.661099865 |
| ENSG00000167264 | DUS2 | 8.168931648 | 1.33E-09 | 10.14807202 | 2.04E-14 | 18.31700367 | 2 | 2 | 2 | 8.164192202 | 8.26379462 | 9.904757465 | 10.4101208 |
| ENSG00000135480 | KRT7 | 6.848888144 | 2.55E-07 | 11.45452737 | 5.71E-19 | 18.30341551 | 2 | 2 | 2 | 6.846474326 | 6.99676009 | 11.54710573 | 11.4157153 |
| ENSG00000090339 | ICAM1 | 10.09012651 | 1.82E-31 | 8.201913658 | 2.82E-21 | 18.29204017 | 2 | 4.710564154 | 4.058998049 | 13.69222093 | 13.741006 | 11.20657426 | 12.26354895 |
| ENSG00000163661 | PTX3 | 10.31366559 | 1.87E-15 | 7.965004277 | 1.18E-09 | 18.27866986 | 2 | 2 | 2 | 10.32606281 | 10.3626704 | 7.958717528 | 8.067092663 |
| ENSG00000144959 | NCEH1 | 9.499772933 | 1.01E-12 | 8.740478155 | 5.76E-11 | 18.24025109 | 2 | 3.076665209 | 2 | 10.49739553 | 10.4864929 | 9.422103197 | 9.992781936 |
| ENSG00000172716 | SLFN11 | 8.327530668 | 3.98E-10 | 9.893407033 | 5.27E-14 | 18.2209377 | 2 | 2 | 2 | 8.310225403 | 8.43013848 | 9.73525344 | 10.09400257 |
| ENSG00000177409 | SAMD9L | 11.0274062 | 2.31E-21 | 7.123511689 | 1.55E-09 | 18.15091789 | 2 | 3.686303414 | 2 | 12.88366224 | 12.8667447 | 8.722467118 | 9.202055802 |
| ENSG00000154127 | UBASH3B | 7.743307924 | 5.10E-09 | 10.30007287 | 2.33E-15 | 18.04338079 | 2 | 2 | 2 | 7.786523606 | 7.80517502 | 10.19848552 | 10.45226125 |
| ENSG00000137809 | ITGA11 | 8.695281512 | 3.89E-27 | 9.318137221 | 2.14E-31 | 18.01341873 | 3.327711234 | 4.442710129 | 2 | 11.82786535 | 11.8844426 | 12.66594881 | 12.26354895 |
| ENSG00000169991 | IFFO2 | 7.854233332 | 2.91E-09 | 10.14138067 | 6.21E-15 | 17.995614 | 2 | 2 | 2 | 7.885575888 | 7.92393105 | 10.28236597 | 10.05408645 |
| ENSG00000101347 | SAMHD1 | 10.30086729 | 1.27E-15 | 7.619911436 | 4.97E-09 | 17.92077873 | 3.327711234 | 2 | 2 | 11.28088322 | 11.3017217 | 8.621327277 | 8.624257514 |
| ENSG00000145337 | PYURF | 9.239548032 | 1.69E-12 | 8.671729884 | 3.54E-11 | 17.91127792 | 2 | 2 | 2 | 9.332862708 | 9.21527115 | 8.666300387 | 8.756196613 |
| ENSG00000134363 | FST | 9.734053505 | 1.83E-13 | 8.157688725 | 8.86E-10 | 17.89174223 | 3.327711234 | 2 | 2 | 10.71112597 | 10.739998 | 9.380756704 | 8.889499502 |
| ENSG00000135424 | ITGA7 | 4.548829003 | 0.001525395 | 13.33023503 | 6.97E-24 | 17.87906403 | 2 | 2 | 2 | 5.022023783 | 4.53541753 | 13.17471064 | 13.51908818 |
| ENSG00000141574 | SECTM1 | 10.84420053 | 1.12E-16 | 7.034504393 | 1.31E-07 | 17.87870492 | 2 | 2 | 2 | 10.83696223 | 10.9094332 | 6.936150978 | 7.253757517 |
| ENSG00000150938 | CRIM1 | 7.321373314 | 2.83E-10 | 10.55200025 | 1.91E-20 | 17.87337356 | 2 | 5.303523747 | 2 | 11.01044056 | 11.0396575 | 14.05584185 | 14.42690002 |
| ENSG00000054598 | FOXC1 | 5.99760648 | 7.61E-06 | 11.83697788 | 2.61E-20 | 17.83458436 | 2 | 2 | 2 | 6.078767801 | 6.14061114 | 11.89367234 | 11.83551901 |
| ENSG00000113578 | FGF1 | 7.157442139 | 7.49E-08 | 10.66029989 | 1.97E-16 | 17.81774203 | 2 | 2 | 2 | 7.292639438 | 7.14977896 | 10.58798261 | 10.78560504 |
| ENSG00000184584 | TMEM173 | 9.071734254 | 2.02E-13 | 8.682689487 | 1.90E-12 | 17.75442374 | 2 | 2 | 4.058998049 | 10.92446939 | 11.0102587 | 10.03141812 | 10.97629934 |
| ENSG00000178726 | THBD | 7.508283374 | 3.43E-08 | 10.24519032 | 1.45E-14 | 17.75347369 | 2 | 2 | 2 | 7.61176303 | 7.51759662 | 9.975607996 | 10.52491227 |
| ENSG00000135905 | DOCK10 | 9.556562366 | 3.81E-13 | 8.19635623 | 5.85E-10 | 17.7529186 | 2 | 2 | 2 | 9.632130776 | 9.54750898 | 8.37208139 | 8.098650814 |
| ENSG00000187688 | TRPV2 | 9.04455143 | 5.63E-12 | 8.689380777 | 3.48E-11 | 17.73393221 | 2 | 2 | 2 | 9.03990038 | 9.12214832 | 8.801285298 | 8.653693725 |
| ENSG00000091490 | SEL1L3 | 9.046877699 | 3.05E-15 | 8.653119756 | 3.81E-14 | 17.69999746 | 4.00722531 | 2 | 2 | 10.90316293 | 10.9466189 | 10.37200212 | 10.67645725 |
| ENSG00000146859 | TMEM140 | 10.46707858 | 2.40E-15 | 7.212115459 | 8.01E-08 | 17.67919404 | 2 | 2 | 2 | 10.49450782 | 10.5003855 | 7.475964489 | 7.043754524 |
| ENSG00000187678 | SPRY4 | 9.18346534 | 2.39E-11 | 8.44175099 | 8.95E-10 | 17.62521633 | 3.327711234 | 2 | 2 | 10.1289481 | 10.2227847 | 9.808823657 | 8.938158123 |
| ENSG00000005884 | ITGA3 | 7.996148344 | 7.04E-15 | 9.617186836 | 2.86E-21 | 17.61333518 | 5.330594134 | 2 | 3.369317972 | 11.71721007 | 11.7879618 | 12.94236789 | 13.70408423 |
| ENSG00000155816 | FMN2 | 8.740808399 | 7.13E-11 | 8.789098831 | 5.03E-11 | 17.52990723 | 2 | 2 | 2 | 8.809693772 | 8.75042728 | 8.585635246 | 9.035944835 |
| ENSG00000174348 | PODN | 7.820357985 | 1.21E-08 | 9.664394538 | 8.56E-13 | 17.48475252 | 2 | 2 | 2 | 7.932663244 | 7.80780625 | 9.346924687 | 9.979766365 |
| ENSG00000240771 | ARHGEF25 | 8.403320644 | 2.68E-10 | 9.044581588 | 7.64E-12 | 17.44790223 | 2 | 2 | 2 | 8.532433657 | 8.35477078 | 8.941912865 | 9.209095161 |
| ENSG00000118523 | CTGF | 6.60887738 | 9.99E-09 | 10.83488509 | 5.42E-22 | 17.44376247 | 4.00722531 | 6.899080544 | 3.369317972 | 12.13173051 | 12.1248442 | 16.41885044 | 16.28415028 |
| ENSG00000276547 | PCDHGB5 | 10.4948288 | 5.50E-16 | 6.925566537 | 1.55E-07 | 17.42039534 | 2 | 2 | 2 | 10.49945465 | 10.5503412 | 6.973973146 | 7.021346309 |
| ENSG00000197442 | MAP3K5 | 9.68726639 | 2.01E-13 | 7.724670244 | 6.45E-09 | 17.41193663 | 2 | 2 | 2 | 9.711725296 | 9.72921087 | 7.927733561 | 7.609689469 |
| ENSG00000102802 | MEDAG | 9.62706863 | 1.24E-13 | 7.777687297 | 2.91E-09 | 17.40475593 | 3.327711234 | 2 | 2 | 10.66527283 | 10.5710111 | 8.685899845 | 8.866637078 |
| ENSG00000118785 | SPP1 | 7.391376779 | 6.92E-05 | 10.00144922 | 3.00E-08 | 17.392826 | 2 | 2 | 2 | 7.412876874 | 7.48837134 | 10.88560796 | 7.672153316 |
| ENSG00000176046 | NUPR1 | 8.186907362 | 1.68E-11 | 9.205607373 | 2.15E-14 | 17.39251474 | 2 | 3.686303414 | 2 | 10.03156504 | 10.0323066 | 11.37343913 | 10.62660779 |
| ENSG00000023445 | BIRC3 | 10.11986278 | 5.14E-15 | 7.234629655 | 3.48E-08 | 17.35449244 | 2 | 2 | 3.369317972 | 11.09484773 | 11.1262675 | 8.317350121 | 8.162331091 |
| ENSG00000162692 | VCAM1 | 8.021905583 | 1.20E-08 | 9.268545422 | 2.76E-11 | 17.290451 | 2 | 3.686303414 | 2 | 9.823772619 | 9.84965517 | 9.923201804 | 11.71335694 |
| ENSG00000254598 | CSNK2A3 | 8.817349981 | 1.84E-11 | 8.459454335 | 1.12E-10 | 17.27680432 | 2 | 2 | 2 | 8.842507967 | 8.86981338 | 8.483898848 | 8.519561679 |
| ENSG00000111331 | OAS3 | 11.55000833 | 8.47E-19 | 5.723742851 | 2.35E-05 | 17.27375118 | 2 | 2 | 4.058998049 | 13.42975069 | 13.482197 | 6.791847192 | 8.196067955 |
| ENSG00000176485 | PLA2G16 | 7.886750452 | 3.89E-09 | 9.345923342 | 1.57E-12 | 17.23267379 | 2 | 2 | 2 | 7.954442493 | 7.91907183 | 9.192066532 | 9.548056138 |
| ENSG00000171310 | CHST11 | 9.228783263 | 2.33E-11 | 7.935547634 | 1.10E-08 | 17.1643309 | 2 | 2 | 2 | 9.212503151 | 9.31482024 | 8.315509958 | 7.554837457 |

| | | | | | | | | | | | | | |
|---|---|---|---|---|---|---|---|---|---|---|---|---|---|
| ENSG00000172985 | SH3RF3 | 8.046563591 | 3.23E-09 | 9.100578517 | 1.35E-11 | 17.14714211 | 2 | 2 | 2 | 8.017874397 | 8.16669309 | 9.370453824 | 8.858759519 |
| ENSG00000253873 | PCDHGA11 | 10.18460662 | 5.33E-15 | 6.957514663 | 1.53E-07 | 17.14212128 | 2 | 2 | 2 | 10.1837567 | 10.2472229 | 6.952030889 | 7.101862022 |
| ENSG00000104881 | PPP1R13L | 6.767861479 | 2.87E-07 | 10.37292166 | 5.94E-16 | 17.14078314 | 2 | 3.076665209 | 2 | 7.770242853 | 7.79460185 | 11.34943455 | 11.37698864 |
| ENSG00000186340 | THBS2 | 7.669158951 | 4.79E-23 | 9.465755718 | 3.82E-35 | 17.13491467 | 4.00722531 | 3.076665209 | 4.058998049 | 10.86502241 | 10.878791 | 12.46227734 | 12.84506654 |
| ENSG00000138166 | DUSP5 | 8.344137776 | 4.26E-10 | 8.755473594 | 4.54E-11 | 17.09961137 | 2 | 2 | 3.369317972 | 9.332862708 | 9.34954609 | 9.977742731 | 9.480189131 |
| ENSG00000224831 | TMEM183B | 8.635649523 | 5.06E-11 | 8.422582887 | 1.42E-10 | 17.05823241 | 2 | 2 | 2 | 8.683521494 | 8.66883353 | 8.464101787 | 8.466640307 |
| ENSG00000137959 | IFI44L | 13.02778908 | 3.52E-24 | 3.968812775 | 0.004363792 | 16.99660186 | 2 | 2 | 2 | 13.05661732 | 13.053069 | 4.331190017 | 4.30448772 |
| ENSG00000133106 | EPSTI1 | 11.05812323 | 8.69E-17 | 5.923652631 | 1.66E-05 | 16.98177586 | 2 | 2 | 2 | 11.07233814 | 11.1018181 | 5.758846871 | 6.276965131 |
| ENSG00000067798 | NAV3 | 8.907953406 | 1.25E-26 | 8.053810183 | 3.93E-22 | 16.96176359 | 4.00722531 | 2 | 4.058998049 | 11.76814309 | 11.8176385 | 11.02115097 | 10.85456511 |
| ENSG00000196954 | CASP4 | 10.00116352 | 2.98E-14 | 6.960147065 | 2.03E-07 | 16.96131058 | 2 | 2 | 2 | 10.03042623 | 10.0356777 | 7.182183216 | 6.862343163 |
| ENSG00000152784 | PRDM8 | 10.24244539 | 1.73E-14 | 6.698264715 | 9.16E-07 | 16.9407101 | 2 | 2 | 2 | 10.24983013 | 10.2967656 | 6.497238296 | 7.013798886 |
| ENSG00000049130 | KITLG | 9.093647403 | 1.14E-10 | 7.773647619 | 4.40E-08 | 16.86729502 | 2 | 2 | 2 | 9.155143657 | 9.10513517 | 8.218405973 | 7.283877355 |
| ENSG00000121691 | CAT | 8.521376892 | 3.04E-10 | 8.336701666 | 6.96E-10 | 16.85807856 | 2 | 2 | 2 | 8.535647908 | 8.59001038 | 8.112172997 | 8.606804877 |
| ENSG00000006016 | CRLF1 | 10.61783112 | 6.31E-13 | 6.213114885 | 4.72E-05 | 16.83094601 | 2 | 2 | 2 | 10.59498781 | 10.6987733 | 6.850634117 | 5.448021784 |
| ENSG00000137842 | TMEM62 | 9.69260104 | 2.78E-13 | 7.120023036 | 1.24E-07 | 16.81262408 | 2 | 2 | 2 | 9.726568065 | 9.72504082 | 7.368839097 | 6.977399345 |
| ENSG00000122861 | PLAU | 9.556955631 | 4.48E-26 | 7.237554812 | 1.60E-15 | 16.79451044 | 2 | 4.11352228 | 3.369317972 | 12.36497354 | 12.4351513 | 10.37877362 | 9.716470666 |
| ENSG00000149633 | KIAA1755 | 9.075152744 | 5.27E-12 | 7.645921628 | 7.73E-09 | 16.72107437 | 2 | 2 | 2 | 9.086669782 | 9.13687201 | 7.633939843 | 7.763868281 |
| ENSG00000138685 | FGF2 | 9.978536548 | 4.43E-25 | 6.697690981 | 6.16E-12 | 16.67622753 | 3.327711234 | 5.457120953 | 2 | 14.07182595 | 14.0003157 | 11.00468611 | 10.46182695 |
| ENSG00000185551 | NR2F2 | 6.661820644 | 5.31E-14 | 10.00597837 | 1.07E-30 | 16.66779901 | 2 | 4.442710129 | 2 | 9.490242771 | 9.51325794 | 12.71516678 | 12.95259962 |
| ENSG00000100342 | APOL1 | 9.587776216 | 1.33E-33 | 7.021158745 | 1.31E-18 | 16.60893496 | 4.00722531 | 3.076665209 | 4.058998049 | 12.78346024 | 12.7982697 | 9.927625703 | 10.47700137 |
| ENSG00000177425 | PAWR | 6.513479058 | 1.03E-06 | 10.01942027 | 9.74E-15 | 16.53289933 | 2 | 3.076665209 | 2 | 7.454228844 | 7.60793141 | 11.10121445 | 10.91342392 |
| ENSG00000134470 | IL15RA | 9.921760852 | 1.40E-13 | 6.585826992 | 1.57E-06 | 16.50758784 | 2 | 2 | 2 | 9.938102197 | 9.96965331 | 6.380048704 | 6.911910672 |
| ENSG00000131015 | ULBP2 | 8.862840499 | 1.73E-11 | 7.582267024 | 1.06E-08 | 16.44510752 | 2 | 2 | 2 | 8.852852814 | 8.94822691 | 7.629009275 | 7.646759583 |
| ENSG00000077238 | IL4R | 7.864479742 | 8.27E-12 | 8.497050272 | 9.93E-14 | 16.36153001 | 2 | 3.686303414 | 2 | 9.669187952 | 9.77497559 | 10.42416644 | 10.27852722 |
| ENSG00000174032 | SLC25A30 | 8.748323978 | 3.41E-11 | 7.610829741 | 9.70E-09 | 16.35915372 | 2 | 2 | 2 | 8.826850269 | 8.74768919 | 7.705944963 | 7.62463201 |
| ENSG00000117036 | ETV3 | 7.509993321 | 1.82E-08 | 8.844400412 | 1.89E-11 | 16.35439373 | 2 | 2 | 2 | 7.632926745 | 7.49817903 | 8.915024678 | 8.850309106 |
| ENSG00000117226 | GBP3 | 9.620498097 | 1.54E-23 | 6.723568803 | 4.25E-12 | 16.3440669 | 2 | 4.11352228 | 2 | 12.03198325 | 12.0687713 | 9.049523349 | 9.266987143 |
| ENSG00000197646 | PDCD1LG2 | 8.362630273 | 7.05E-09 | 7.974738178 | 3.35E-08 | 16.33736845 | 2 | 2 | 2 | 8.386974157 | 8.42500614 | 7.408510937 | 8.45344973 |
| ENSG00000147889 | CDKN2A | 9.104782745 | 5.23E-12 | 7.149957604 | 8.41E-08 | 16.25474035 | 2 | 2 | 2 | 9.16555511 | 9.11685323 | 7.121690475 | 7.30412922 |
| ENSG00000136542 | GALNT5 | 9.295470675 | 5.41E-27 | 6.884886065 | 2.15E-15 | 16.18035674 | 2 | 4.442710129 | 2 | 12.1230758 | 12.1403237 | 9.781089302 | 9.670110091 |
| ENSG00000155158 | TTC39B | 7.812198373 | 1.03E-08 | 8.36600404 | 6.65E-10 | 16.17820241 | 2 | 2 | 2 | 7.852449262 | 7.8745862 | 8.628262104 | 8.151194533 |
| ENSG00000103647 | CORO2B | 7.978183834 | 3.63E-09 | 8.157712437 | 1.38E-09 | 16.13589627 | 2 | 2 | 2 | 8.006336225 | 8.04707549 | 8.381493453 | 8.000872619 |
| ENSG00000214944 | ARHGEF28 | 7.343638323 | 8.91E-08 | 8.778289183 | 8.81E-11 | 16.12192751 | 2 | 2 | 2 | 7.326437704 | 7.47849652 | 9.042123971 | 8.55118937 |
| ENSG00000108771 | DHX58 | 9.486177668 | 4.08E-13 | 6.627774725 | 6.63E-07 | 16.11395239 | 2 | 2 | 2 | 9.50179306 | 9.53882419 | 6.715756428 | 6.704867638 |
| ENSG00000111799 | COL12A1 | 7.999418766 | 1.28E-10 | 8.1068074 | 6.38E-11 | 16.10622617 | 2 | 5.457120953 | 2 | 11.80507162 | 11.8920568 | 11.82656184 | 12.07566644 |
| ENSG00000156253 | RWDD2B | 7.594690618 | 2.18E-08 | 8.486367699 | 2.67E-10 | 16.08105832 | 2 | 2 | 2 | 7.626911583 | 7.67287587 | 8.322856568 | 8.708487604 |
| ENSG00000105974 | CAV1 | 6.58135894 | 9.98E-12 | 9.493125548 | 1.57E-23 | 16.07448449 | 3.327711234 | 5.457120953 | 2 | 10.64271979 | 10.6460801 | 13.30271973 | 13.76627258 |
| ENSG00000180660 | MAB21L1 | 8.156002274 | 1.16E-09 | 7.912987405 | 3.42E-09 | 16.06898968 | 2 | 2 | 2 | 8.199071993 | 8.20512425 | 8.092249024 | 7.820122783 |
| ENSG00000184110 | EIF3C | 10.23844932 | 1.32E-14 | 5.826818252 | 2.21E-05 | 16.06526757 | 2 | 2 | 2 | 10.26201025 | 10.2769623 | 5.711121288 | 6.155328188 |
| ENSG00000106366 | SERPINE1 | 7.476436617 | 9.27E-08 | 8.541136413 | 6.87E-10 | 16.01757303 | 8.674867122 | 4.93638933 | 4.058998049 | 14.66453201 | 14.7005231 | 15.92548369 | 15.54374762 |
| ENSG00000113389 | NPR3 | 6.100704484 | 6.07E-06 | 9.915165446 | 2.58E-14 | 16.01586993 | 2 | 3.076665209 | 2 | 7.165617905 | 7.09482005 | 10.7412617 | 11.05447682 |
| ENSG00000106991 | ENG | 8.30353082 | 1.13E-37 | 7.696301012 | 9.61E-33 | 15.99983183 | 3.327711234 | 4.710564154 | 4.058998049 | 12.12894005 | 12.1784379 | 11.39182091 | 11.68772122 |
| ENSG00000262576 | PCDHGA4 | 10.21892957 | 4.80E-13 | 5.780373237 | 8.13E-05 | 15.9993028 | 2 | 2 | 2 | 10.21967182 | 10.2798081 | 5.318934974 | 6.323497254 |
| ENSG00000162496 | DHRS3 | 9.523209818 | 1.29E-12 | 6.470521626 | 2.41E-06 | 15.99373144 | 2 | 2 | 2 | 9.574696759 | 9.53961588 | 6.779440836 | 6.298867018 |
| ENSG00000126778 | SIX1 | 6.952096722 | 1.41E-06 | 9.02758735 | 1.58E-10 | 15.97968407 | 2 | 2 | 2 | 7.119236776 | 6.9210099 | 8.536980328 | 9.450446099 |
| ENSG00000188158 | NHS | 7.499838004 | 2.18E-08 | 8.418917018 | 2.19E-10 | 15.91875502 | 2 | 2 | 2 | 7.527093191 | 7.58663873 | 8.388512422 | 8.531503464 |
| ENSG00000182253 | SYNM | 7.880557912 | 1.17E-08 | 7.986964212 | 6.49E-09 | 15.86752212 | 2 | 2 | 2 | 7.942383523 | 7.91907183 | 8.297599653 | 7.714671806 |
| ENSG00000100302 | RASD2 | 7.899894061 | 7.44E-08 | 7.954413572 | 5.36E-08 | 15.85430763 | 2 | 2 | 2 | 7.913023977 | 7.98565528 | 8.453542774 | 7.34530375 |
| ENSG00000175745 | NR2F1 | 6.342129786 | 3.56E-08 | 9.507789595 | 2.33E-17 | 15.84991938 | 2 | 3.076665209 | 3.369317972 | 8.201097719 | 8.25031674 | 11.31891342 | 11.42774592 |
| ENSG00000181588 | MEX3D | 7.711632835 | 5.04E-09 | 8.119528388 | 6.04E-10 | 15.83116122 | 2 | 3.076665209 | 2 | 8.636393836 | 8.78689171 | 9.147938916 | 9.087636837 |
| ENSG00000150527 | CTAGE5 | 9.024923139 | 2.62E-11 | 6.771812774 | 8.44E-07 | 15.79673591 | 2 | 2 | 2 | 9.080080551 | 9.04395769 | 7.078941352 | 6.574984961 |
| ENSG00000176692 | FOXC2 | 4.45785955 | 0.00182751 | 11.33343421 | 9.82E-18 | 15.79129376 | 2 | 2 | 2 | 4.76150579 | 4.68029874 | 11.09165896 | 11.58959617 |
| ENSG00000185022 | MAFF | 8.575479738 | 1.41E-20 | 7.21379253 | 5.37E-15 | 15.78927227 | 2 | 4.11352228 | 3.369317972 | 11.40294453 | 11.4332138 | 9.649231086 | 10.37906502 |
| ENSG00000128340 | RAC2 | 10.54296157 | 4.58E-11 | 5.244770427 | 0.001958798 | 15.787732 | 2 | 2 | 2 | 10.58066597 | 10.5655798 | 6.072436333 | 4.161535617 |
| ENSG00000206190 | ATP10A | 7.55682658 | 1.84E-10 | 8.224195055 | 2.77E-12 | 15.78102163 | 2 | 3.686303414 | 2 | 9.384581425 | 9.43712594 | 10.29159999 | 9.820713969 |
| ENSG00000176641 | RNF152 | 6.140207192 | 6.33E-06 | 9.574344448 | 3.18E-13 | 15.71455164 | 2 | 2 | 2 | 6.164317909 | 6.32089174 | 9.70112864 | 9.508660255 |
| ENSG00000101421 | CHMP4B | 7.798175668 | 1.32E-14 | 7.89563063 | 4.41E-15 | 15.6938063 | 2 | 2 | 4.523675391 | 10.28320062 | 10.2812288 | 10.13729849 | 10.58662664 |
| ENSG00000028277 | POU2F2 | 9.427391611 | 1.45E-10 | 6.244992575 | 3.59E-05 | 15.67238419 | 2 | 2 | 2 | 9.439666599 | 9.48394374 | 5.586442682 | 6.839042789 |
| ENSG00000157168 | NRG1 | 11.23958285 | 2.07E-14 | 4.430834848 | 0.005079404 | 15.6704177 | 2 | 2 | 2 | 11.25015134 | 11.2862384 | 5.19095825 | 3.940519005 |
| ENSG00000003137 | CYP26B1 | 5.997893483 | 1.63E-05 | 9.666906369 | 5.28E-13 | 15.66479985 | 2 | 2 | 2 | 6.130704144 | 6.08962627 | 9.421533019 | 9.933764315 |
| ENSG00000196139 | AKR1C3 | 9.819584212 | 3.83E-14 | 5.835082837 | 1.29E-05 | 15.65466705 | 2 | 3.076665209 | 2 | 10.72386293 | 10.8931098 | 6.900587441 | 6.845434848 |
| ENSG00000253910 | PCDHGB2 | 10.17762342 | 6.73E-14 | 5.427924336 | 0.00012485 | 15.60554775 | 2 | 2 | 2 | 10.18273194 | 10.2345667 | 5.22272385 | 5.867805474 |
| ENSG00000120217 | CD274 | 10.04037135 | 2.30E-14 | 5.536606374 | 4.95E-05 | 15.57697772 | 2 | 2 | 2 | 10.07252815 | 10.0717013 | 5.815498119 | 5.529673353 |
| ENSG00000283154 | IQCJ-SCHIP1 | 7.576066127 | 1.92E-08 | 7.976990385 | 2.61E-09 | 15.55305651 | 2 | 2 | 2 | 7.523859802 | 7.73225478 | 7.989819469 | 8.060697341 |
| ENSG00000182218 | HHIPL1 | 7.338493294 | 2.78E-08 | 8.196059906 | 3.77E-10 | 15.5345532 | 2 | 3.076665209 | 2 | 8.31397419 | 8.37442884 | 9.192400798 | 9.19581595 |
| ENSG00000070371 | CLTCL1 | 6.771629255 | 5.81E-07 | 8.748062711 | 4.53E-11 | 15.51969197 | 2 | 2 | 2 | 6.856790872 | 6.84106088 | 8.89667997 | 8.669391068 |
| ENSG00000125538 | IL1B | 14.32195919 | 1.23E-26 | 1.18429571 | 0.481883257 | 15.5062549 | 2 | 2 | 2 | 14.34346783 | 14.3537526 | 2.492221014 | 2.817676617 |
| ENSG00000078098 | FAP | 9.26323256 | 7.20E-14 | 6.235629274 | 7.90E-07 | 15.49886183 | 2 | 3.686303414 | 2 | 11.08938989 | 11.1114346 | 8.471282103 | 7.574209638 |
| ENSG00000134107 | BHLHE40 | 8.425802229 | 2.13E-25 | 7.070199799 | 2.25E-18 | 15.49600203 | 2 | 3.076665209 | 5.156626401 | 11.90336716 | 11.9204473 | 10.59305515 | 10.52341813 |
| ENSG00000141052 | MYOCD | 4.197644785 | 0.003288418 | 11.2797101 | 7.23E-18 | 15.47735489 | 2 | 2 | 2 | 4.574143677 | 4.43005853 | 11.12828449 | 11.46843325 |
| ENSG00000251322 | SHANK3 | 7.604786314 | 2.30E-07 | 7.87075027 | 7.16E-08 | 15.47553658 | 2 | 2 | 2 | 7.565338955 | 7.74878077 | 8.352472895 | 7.302581437 |
| ENSG00000004799 | PDK4 | 9.905885481 | 1.21E-09 | 5.553542327 | 0.001134029 | 15.45942781 | 2 | 2 | 2 | 9.942952235 | 9.93336537 | 6.375345632 | 4.365060851 |
| ENSG00000135454 | B4GALNT1 | 8.12427994 | 2.70E-09 | 7.327564663 | 8.96E-08 | 15.4518446 | 2 | 2 | 2 | 8.017874397 | 8.30907138 | 7.552954291 | 7.203223239 |
| ENSG00000121310 | ECHDC2 | 7.214382946 | 1.26E-07 | 8.215996542 | 1.14E-09 | 15.43037949 | 2 | 2 | 2 | 7.173892946 | 7.37584259 | 8.108635266 | 8.399447278 |
| ENSG00000164823 | OSGIN2 | 8.281955824 | 8.81E-19 | 7.125328613 | 2.69E-14 | 15.40728444 | 3.327711234 | 3.076665209 | 3.369317972 | 10.75935206 | 10.7163854 | 9.608982234 | 9.561707646 |
| ENSG00000106153 | CHCHD2 | 7.613167438 | 2.82E-15 | 7.793812376 | 4.11E-16 | 15.40697981 | 2 | 4.11352228 | 2 | 10.07031346 | 10.0261059 | 10.25973689 | 10.19631519 |
| ENSG00000101825 | MXRA5 | 10.9224266 | 1.23E-16 | 4.462669624 | 0.001487806 | 15.38509622 | 2 | 2 | 2 | 10.94788047 | 10.9555451 | 4.90146639 | 4.522269241 |
| ENSG00000105376 | ICAM5 | 5.402727663 | 9.53E-05 | 9.971712526 | 3.74E-14 | 15.37444019 | 2 | 2 | 2 | 5.37770462 | 5.71663161 | 9.998153538 | 10.00924387 |
| ENSG00000120875 | DUSP4 | 7.484565651 | 2.89E-08 | 7.882116774 | 4.12E-09 | 15.36668243 | 3.327711234 | 2 | 2 | 8.553199415 | 8.4215744 | 9.084881607 | 8.64700078 |
| ENSG00000159216 | RUNX1 | 8.433136852 | 3.45E-18 | 6.904846874 | 1.29E-12 | 15.33798373 | 2 | 4.11352228 | 2 | 10.85186662 | 10.8759728 | 9.230659332 | 9.444493373 |
| ENSG00000189320 | FAM180A | 8.038850326 | 2.26E-09 | 7.26275343 | 7.33E-08 | 15.30160376 | 2 | 2 | 2 | 8.010962573 | 8.15847021 | 7.244115921 | 7.402777233 |
| ENSG00000108932 | SLC16A6 | 11.64797754 | 9.99E-16 | 3.628800204 | 0.02347325 | 15.27677774 | 2 | 2 | 2 | 11.67720334 | 11.6750476 | 3.429858444 | 4.489993931 |
| ENSG00000131979 | GCH1 | 9.216167288 | 1.28E-11 | 6.058051462 | 1.45E-05 | 15.27421875 | 2 | 2 | 2 | 9.293531584 | 9.20931286 | 5.85994354 | 6.420873184 |
| ENSG00000233954 | UQCRHL | 7.359022784 | 3.76E-08 | 7.904323544 | 2.62E-09 | 15.26334633 | 3.327711234 | 2 | 2 | 8.401141138 | 8.32752521 | 8.740866084 | 9.051693903 |
| ENSG00000188483 | IER5L | 7.64479529 | 4.33E-08 | 7.59398741 | 4.84E-08 | 15.2387827 | 2 | 2 | 2 | 7.647855834 | 7.74878077 | 7.91644678 | 7.321046127 |
| ENSG00000188522 | FAM83G | 6.884989229 | 2.36E-07 | 8.348320164 | 1.95E-10 | 15.23330939 | 2 | 3.076665209 | 2 | 7.898117066 | 7.89700047 | 9.258701797 | 9.4271968 |
| ENSG00000130513 | GDF15 | 8.427785472 | 1.90E-08 | 6.801399962 | 7.63E-06 | 15.22918543 | 2 | 5.303523747 | 2 | 12.05134158 | 12.1252398 | 11.18431183 | 8.965802436 |
| ENSG00000184979 | USP18 | 10.92916137 | 1.66E-16 | 4.274930422 | 0.002609328 | 15.20409179 | 2 | 2 | 2 | 10.95600041 | 10.9608743 | 4.340871764 | 4.766376438 |
| ENSG00000138134 | STAMBPL1 | 9.464317437 | 4.51E-12 | 5.737698263 | 4.86E-05 | 15.2020157 | 2 | 2 | 2 | 9.516508835 | 9.48064953 | 6.148830293 | 5.513708555 |
| ENSG00000105976 | MET | 7.224611017 | 4.53E-14 | 7.974427665 | 4.37E-17 | 15.19903868 | 2 | 2 | 4.874587708 | 10.13796435 | 10.1458442 | 10.74730333 | 11.01921956 |
| ENSG00000128284 | APOL3 | 7.684612857 | 2.41E-08 | 7.505557164 | 4.76E-08 | 15.19017002 | 2 | 2 | 3.369317972 | 8.664561167 | 8.70880006 | 8.121330705 | 8.815461059 |
| ENSG00000137266 | SLC22A23 | 6.400944072 | 5.70E-08 | 8.768567553 | 2.96E-14 | 15.16951163 | 3.327711234 | 3.076665209 | 2 | 8.332573284 | 8.22887947 | 10.40713106 | 10.83046661 |
| ENSG00000164161 | HHIP | 4.734297451 | 0.000676627 | 10.42810508 | 1.21E-15 | 15.16240253 | 2 | 2 | 2 | 5.040242897 | 4.87352701 | 10.51047766 | 10.40472054 |
| ENSG00000123096 | SSPN | 6.874300857 | 4.70E-07 | 8.284503889 | 6.91E-10 | 15.15880475 | 2 | 2 | 2 | 6.946463121 | 6.94988431 | 8.460220609 | 8.184349581 |
| ENSG00000261934 | PCDHGA9 | 10.13713138 | 1.54E-14 | 5.01972191 | 0.000282184 | 15.15685329 | 2 | 2 | 2 | 10.14060551 | 10.1959181 | 5.04460184 | 5.361470313 |
| ENSG00000093010 | COMT | 7.681166332 | 1.30E-09 | 7.465241816 | 3.60E-09 | 15.14640815 | 5.330594134 | 2 | 2 | 11.18537258 | 11.2610183 | 10.72748832 | 11.24299785 |
| ENSG00000164619 | BMPER | 7.74583155 | 2.01E-07 | 7.393011799 | 7.04E-07 | 15.13884335 | 2 | 3.076665209 | 2 | 8.747293529 | 8.74768919 | 7.511762185 | 8.943630224 |
| ENSG00000214193 | SH3D21 | 6.310659996 | 4.11E-06 | 8.827213561 | 3.64E-11 | 15.13787356 | 2 | 2 | 2 | 6.252910529 | 6.54560135 | 8.853791658 | 8.878113577 |
| ENSG00000176928 | GCNT4 | 6.076952483 | 3.19E-05 | 9.026969493 | 1.63E-10 | 15.10392198 | 2 | 2 | 2 | 6.096288482 | 6.26836889 | 9.445569009 | 8.544002356 |
| ENSG00000132334 | PTPRE | 8.638193896 | 1.58E-10 | 6.450274621 | 2.69E-06 | 15.08846852 | 2 | 2 | 2 | 8.618316353 | 8.73668456 | 6.713894915 | 6.341697848 |
| ENSG00000179862 | CITED4 | 5.969874025 | 2.29E-05 | 9.056174793 | 2.91E-11 | 15.02604882 | 2 | 2 | 2 | 6.024891728 | 6.14061114 | 8.7818697 | 9.349127298 |
| ENSG00000149596 | JPH2 | 4.19614938 | 0.00341118 | 10.80012163 | 2.30E-16 | 14.99627101 | 2 | 2 | 2 | 4.299492648 | 4.68029874 | 10.80854976 | 10.8504662 |
| ENSG00000137573 | SULF1 | 7.218065709 | 1.02E-06 | 7.739230696 | 1.27E-07 | 14.95729641 | 3.327711234 | 6.78762695 | 2 | 12.46372082 | 12.5019715 | 12.99447815 | 13.01308674 |
| ENSG00000115290 | GRB14 | 9.714462343 | 1.20E-13 | 5.242445231 | 0.000121751 | 14.95690757 | 2 | 2 | 2 | 9.789178534 | 9.70471389 | 5.4588802 | 5.361470313 |
| ENSG00000183486 | MX2 | 12.28646142 | 9.04E-17 | 2.658180048 | 0.115813971 | 14.94464147 | 2 | 2 | 2 | 12.30065678 | 12.3271867 | 3.782262616 | 2.817676617 |
| ENSG00000147251 | DOCK11 | 7.433108751 | 3.49E-07 | 7.501373983 | 2.42E-07 | 14.93448273 | 2 | 2 | 2 | 7.580980221 | 7.39696604 | 7.946086551 | 7.026981076 |
| ENSG00000166401 | SERPINB8 | 8.455540409 | 5.36E-09 | 6.471507249 | 1.14E-05 | 14.92704766 | 2 | 2 | 3.369317972 | 9.449924671 | 9.45402153 | 6.741568503 | 7.98359008 |

| | | | | | | | | | | | | | |
|---|---|---|---|---|---|---|---|---|---|---|---|---|---|
| ENSG00000008441 | NFIX | 6.256515292 | 3.13E-08 | 8.634499079 | 6.16E-15 | 14.89101437 | 2 | 5.131603625 | 2 | 9.794554993 | 9.78702715 | 11.79949881 | 12.45392616 |
| ENSG00000171812 | COL8A2 | 5.35569622 | 5.87E-07 | 9.522169315 | 3.85E-20 | 14.87786553 | 2 | 4.11352228 | 2 | 7.794595599 | 7.7919464 | 11.53621767 | 12.24704688 |
| ENSG00000149571 | KIRREL3 | 5.991284318 | 1.94E-05 | 8.880687345 | 6.20E-11 | 14.87197166 | 2 | 3.076665209 | 2 | 6.877205252 | 7.15805139 | 9.515898077 | 10.16237085 |
| ENSG00000241978 | AKAP2 | 11.96533226 | 5.82E-20 | 2.903961037 | 0.048142698 | 14.8692933 | 2 | 2 | 2 | 11.96710062 | 12.0186877 | 3.411599105 | 3.660199087 |
| ENSG00000148841 | ITPRIP | 8.845312785 | 5.86E-34 | 5.995410793 | 3.04E-16 | 14.84072358 | 4.815609471 | 3.686303414 | 2 | 12.27263705 | 12.3259253 | 9.346924687 | 9.558793178 |
| ENSG00000197971 | MBP | 6.515712628 | 1.06E-06 | 8.324315508 | 2.05E-10 | 14.84002814 | 2 | 3.076665209 | 2 | 7.565338955 | 7.50468064 | 9.311972588 | 9.33102078 |
| ENSG00000188211 | NCR3LG1 | 8.11500381 | 1.43E-08 | 6.719211803 | 3.44E-06 | 14.83421561 | 2 | 2 | 2 | 8.223194971 | 8.09740995 | 6.322573937 | 7.156006254 |
| ENSG00000078401 | EDN1 | 6.649513771 | 1.06E-06 | 8.182619018 | 9.81E-10 | 14.83213279 | 2 | 2 | 2 | 6.744602355 | 6.71776877 | 8.156706039 | 8.296681143 |
| ENSG00000253506 | NACA2 | 8.777887517 | 3.78E-11 | 6.04972769 | 8.54E-06 | 14.82761521 | 2 | 2 | 2 | 8.756965077 | 8.87484109 | 6.118193494 | 6.199243628 |
| ENSG00000111335 | OAS2 | 11.06584019 | 2.92E-19 | 3.696813311 | 0.005582023 | 14.7626535 | 4.00722531 | 2 | 2 | 12.94659359 | 12.9593671 | 5.158477451 | 6.092227206 |
| ENSG00000134899 | ERCC5 | 9.569735008 | 3.50E-13 | 5.190094993 | 0.000153911 | 14.75983 | 2 | 2 | 2 | 9.595637371 | 9.61140668 | 5.269098128 | 5.453611367 |
| ENSG00000112769 | LAMA4 | 7.594891898 | 1.58E-21 | 7.143787682 | 2.35E-19 | 14.73867958 | 4.467311984 | 4.11352228 | 6.073378139 | 12.50078528 | 12.5634025 | 12.5050204 | 11.47994833 |
| ENSG00000073910 | FRY | 5.397317071 | 0.000146384 | 9.341278642 | 5.70E-12 | 14.73859571 | 2 | 2 | 2 | 5.639097847 | 5.45984938 | 9.075851216 | 9.624999381 |
| ENSG00000006210 | CX3CL1 | 1.599854907 | 0.390522475 | 13.11117997 | 2.83E-19 | 14.71103488 | 2 | 2 | 2 | 3.162721474 | 2.38539787 | 13.00940021 | 13.25640611 |
| ENSG00000111110 | PPM1H | 4.868690713 | 0.001915398 | 9.826405338 | 2.30E-11 | 14.69509605 | 2 | 2 | 2 | 4.965943888 | 5.18011485 | 9.042123971 | 10.37741184 |
| ENSG00000076706 | MCAM | 7.020728799 | 2.08E-09 | 7.669660597 | 4.24E-11 | 14.6903894 | 4.00722531 | 3.076665209 | 2 | 9.464333207 | 9.50192864 | 10.70130335 | 9.167185134 |
| ENSG00000102287 | GABRE | 7.115720141 | 4.72E-07 | 7.535195713 | 7.60E-08 | 14.65091585 | 2 | 2 | 2 | 7.186217188 | 7.17852729 | 7.258234779 | 7.862758575 |
| ENSG00000196639 | HRH1 | 6.455199689 | 2.14E-08 | 8.158073287 | 6.04E-13 | 14.61327298 | 2 | 3.076665209 | 3.369317972 | 8.354580458 | 8.32017198 | 10.04499344 | 10.0106665 |
| ENSG00000169432 | SCN9A | 8.442527715 | 1.33E-10 | 6.16774534 | 4.16E-06 | 14.61027305 | 3.327711234 | 2 | 2 | 9.476928096 | 9.40011621 | 7.121690475 | 7.266516281 |
| ENSG00000089127 | OAS1 | 12.11453806 | 7.27E-18 | 2.489400316 | 0.123275594 | 14.60393837 | 2 | 2 | 2 | 12.14809842 | 12.1362731 | 3.584558147 | 2.885414003 |
| ENSG00000158301 | GPRASP2 | 6.846868502 | 5.55E-07 | 7.754509131 | 9.42E-09 | 14.60137763 | 2 | 2 | 2 | 6.936770382 | 6.90635305 | 7.718996793 | 7.889811111 |
| ENSG00000135828 | RNASEL | 7.179294241 | 1.33E-07 | 7.399000589 | 4.53E-08 | 14.57829483 | 2 | 2 | 2 | 7.246306984 | 7.24216308 | 7.415399995 | 7.500519913 |
| ENSG00000051382 | PIK3CB | 7.257595836 | 5.25E-08 | 7.320217753 | 3.56E-08 | 14.57781359 | 3.327711234 | 2 | 2 | 8.182762967 | 8.34212018 | 8.369718748 | 8.281832879 |
| ENSG00000147459 | DOCK5 | 7.146691462 | 2.90E-19 | 7.422439378 | 6.41E-21 | 14.56913084 | 3.327711234 | 3.686303414 | 4.058998049 | 10.33669748 | 10.3437572 | 10.34082931 | 10.84569152 |
| ENSG00000167785 | ZNF558 | 6.994663538 | 1.32E-06 | 7.573453347 | 1.24E-07 | 14.56811689 | 2 | 2 | 2 | 6.912249745 | 7.20272083 | 7.951626215 | 7.213143202 |
| ENSG00000057657 | PRDM1 | 9.921878738 | 2.55E-12 | 4.622334841 | 0.002135746 | 14.54421358 | 2 | 2 | 2 | 9.959800107 | 9.94834802 | 5.248671144 | 4.329022995 |
| ENSG00000184675 | AMER1 | 6.688641392 | 1.57E-06 | 7.852149147 | 1.04E-08 | 14.54079054 | 2 | 2 | 2 | 6.622948477 | 6.90143415 | 7.738355777 | 8.050589918 |
| ENSG00000156642 | NPTN | 7.251733481 | 3.24E-07 | 7.276247254 | 2.69E-07 | 14.52798073 | 2 | 2 | 5.59507254 | 11.07923811 | 11.0764016 | 10.94185947 | 11.24662458 |
| ENSG00000253485 | PCDHGA5 | 10.13839488 | 1.05E-14 | 4.388851436 | 0.001648343 | 14.52724632 | 2 | 2 | 2 | 10.13690654 | 10.2019318 | 4.563936166 | 4.757365452 |
| ENSG00000168404 | MLKL | 9.020994699 | 1.09E-11 | 5.497460614 | 6.18E-05 | 14.51845531 | 2 | 2 | 2 | 9.073461088 | 9.04284131 | 5.54096877 | 5.740981724 |
| ENSG00000101333 | PLCB4 | 6.886272462 | 3.76E-07 | 7.605804823 | 1.42E-08 | 14.49207728 | 2 | 2 | 3.369317972 | 7.90310317 | 7.89452713 | 8.787633372 | 8.40450657 |
| ENSG00000119681 | LTBP2 | 7.078309556 | 5.84E-07 | 7.381190925 | 1.59E-07 | 14.45950048 | 2 | 2 | 5.59507254 | 10.86054366 | 10.9469174 | 11.14176121 | 11.26879021 |
| ENSG00000113070 | HBEGF | 8.052448369 | 1.37E-08 | 6.390311855 | 9.00E-06 | 14.44276022 | 2 | 2 | 2 | 8.096156705 | 8.10384772 | 6.072436333 | 6.802276223 |
| ENSG00000006327 | TNFRSF12A | 7.229321172 | 1.21E-11 | 7.211576627 | 1.18E-11 | 14.4408978 | 3.327711234 | 5.72254323 | 2 | 11.4916791 | 11.5486124 | 11.46772494 | 11.53684257 |
| ENSG00000137801 | THBS1 | 6.713056322 | 2.00E-06 | 7.708393902 | 3.21E-08 | 14.42145022 | 10.15425007 | 8.469613413 | 4.058998049 | 15.65859347 | 15.6962604 | 16.69977347 | 16.64533661 |
| ENSG00000228672 | PROB1 | 6.725089719 | 2.25E-06 | 7.692871881 | 4.09E-08 | 14.4179616 | 2 | 2 | 2 | 6.710878132 | 6.89154576 | 8.007404089 | 7.427130611 |
| ENSG00000164465 | DCBLD1 | 7.359613594 | 3.17E-14 | 7.052884194 | 3.06E-13 | 14.41249779 | 2 | 4.11352228 | 2 | 9.809901631 | 9.77967427 | 9.45312092 | 9.525344379 |
| ENSG00000204956 | PCDHGA1 | 10.15456767 | 7.57E-14 | 4.251128495 | 0.003453531 | 14.40569616 | 2 | 2 | 2 | 10.15530726 | 10.2158672 | 4.230621763 | 4.810605986 |
| ENSG00000261609 | GAN | 5.859625691 | 0.00010005 | 8.535071444 | 4.36E-09 | 14.39469714 | 2 | 2 | 2 | 6.024891728 | 5.93454154 | 9.011762258 | 7.950369405 |
| ENSG00000179588 | ZFPM1 | 6.49009794 | 2.28E-06 | 7.897678118 | 4.76E-09 | 14.38777606 | 2 | 2 | 2 | 6.535745081 | 6.61935067 | 7.890311949 | 8.002780183 |
| ENSG00000006468 | ETV1 | 8.912993781 | 2.62E-11 | 5.474163754 | 7.50E-05 | 14.38715754 | 2 | 2 | 2 | 8.979812467 | 8.92175022 | 5.755231204 | 5.475755622 |
| ENSG00000107201 | DDX58 | 9.900908506 | 4.82E-26 | 4.465416267 | 4.44E-06 | 14.36632477 | 2 | 4.93638933 | 2 | 13.24904661 | 13.2653088 | 7.660277586 | 8.011333147 |
| ENSG00000262209 | PCDHGB3 | 10.17333512 | 6.59E-15 | 4.179832038 | 0.002761417 | 14.35316716 | 2 | 2 | 2 | 10.1755382 | 10.2330992 | 4.530483275 | 4.445805334 |
| ENSG00000099998 | GGT5 | 5.578841382 | 7.67E-05 | 8.770990819 | 9.84E-11 | 14.3498322 | 2 | 2 | 2 | 5.829220145 | 5.60003091 | 8.599312524 | 8.994365793 |
| ENSG00000188641 | DPYD | 8.602660628 | 8.09E-11 | 5.737696239 | 2.43E-05 | 14.34035687 | 2 | 2 | 3.369317972 | 9.536701552 | 9.65735634 | 6.621594001 | 6.922026307 |
| ENSG00000106546 | AHR | 7.481983079 | 5.78E-39 | 6.857273907 | 4.20E-33 | 14.33925699 | 4.815609471 | 5.303523747 | 5.39244166 | 12.5470107 | 12.4794558 | 11.43822139 | 12.23269239 |
| ENSG00000152977 | ZIC1 | 3.088849543 | 0.167558017 | 11.24250808 | 5.09E-09 | 14.33135762 | 2 | 2 | 2 | 4.358765249 | 2.2680111 | 11.30228498 | 11.23946309 |
| ENSG00000164171 | ITGA2 | 9.931527026 | 5.05E-40 | 4.387951093 | 1.39E-08 | 14.31947812 | 3.327711234 | 4.11352228 | 5.39244166 | 14.14727587 | 14.1454883 | 8.983840816 | 8.124285971 |
| ENSG00000078269 | SYNJ2 | 7.516431156 | 1.65E-14 | 6.802692513 | 3.86E-12 | 14.31912367 | 2 | 4.93638933 | 2 | 10.87647575 | 10.8592642 | 9.939423255 | 10.34450826 |
| ENSG00000138061 | CYP1B1 | 7.94619651 | 6.97E-07 | 6.352168295 | 9.47E-05 | 14.29836481 | 3.327711234 | 7.232742069 | 2 | 13.6491886 | 13.6246674 | 12.06368006 | 12.02371203 |
| ENSG00000157654 | PALM2-AKAF | 11.9985539 | 3.26E-20 | 2.298509911 | 0.126314768 | 14.29706381 | 2 | 2 | 2 | 12.00208679 | 12.050141 | 3.235948948 | 3.100202476 |
| ENSG00000164342 | TLR3 | 8.802754409 | 3.33E-11 | 5.479230293 | 6.44E-05 | 14.2819847 | 2 | 2 | 2 | 8.882187604 | 8.80016655 | 5.661762729 | 5.59182244 |
| ENSG00000105711 | SCN1B | 6.983819774 | 5.89E-07 | 7.240886195 | 1.88E-07 | 14.22470597 | 2 | 2 | 2 | 7.012547947 | 7.09482005 | 7.497765909 | 7.08034504 |
| ENSG00000109738 | GLRB | 7.174963059 | 1.81E-07 | 7.026374986 | 3.03E-07 | 14.20133804 | 2 | 2 | 2 | 7.186217188 | 7.29187696 | 6.998648055 | 7.186536869 |
| ENSG00000161281 | COX7A1 | 7.307782327 | 1.69E-07 | 6.86146273 | 9.31E-07 | 14.16924506 | 2 | 2 | 2 | 7.363085424 | 7.37584259 | 6.69514648 | 7.142226902 |
| ENSG00000157227 | MMP14 | 7.961989602 | 9.08E-25 | 6.200047337 | 1.43E-15 | 14.16203694 | 5.330594134 | 5.946638857 | 7.600047325 | 14.48432121 | 14.541477 | 12.96826794 | 12.49706716 |
| ENSG00000152689 | RASGRP3 | 7.98181187 | 8.50E-09 | 6.160312539 | 1.24E-05 | 14.14212441 | 2 | 2 | 2 | 8.040677544 | 8.02009991 | 6.48637478 | 5.999739876 |
| ENSG00000144596 | GRIP2 | 8.593095709 | 5.46E-10 | 5.548512162 | 0.000102519 | 14.14160787 | 2 | 2 | 2 | 8.595396228 | 8.67172529 | 5.954417431 | 5.367404784 |
| ENSG00000240764 | PCDHGC5 | 10.1543297 | 2.21E-14 | 3.98045929 | 0.00548702 | 14.13478899 | 2 | 2 | 2 | 10.15791688 | 10.2128923 | 4.133696865 | 4.500832793 |
| ENSG00000168874 | ATOH8 | 8.323676331 | 2.51E-09 | 5.802623994 | 5.02E-05 | 14.12630033 | 2 | 2 | 2 | 8.264463296 | 8.46389529 | 5.60262961 | 6.192573806 |
| ENSG00000122862 | SRGN | 5.992311569 | 1.14E-06 | 8.106876745 | 1.69E-11 | 14.09918831 | 5.866944937 | 2 | 3.369317972 | 10.28749693 | 10.3037729 | 12.30044439 | 12.50562677 |
| ENSG00000197961 | ZNF121 | 6.304632261 | 6.38E-06 | 7.788115495 | 1.26E-08 | 14.09274776 | 2 | 2 | 2 | 6.483440992 | 6.31350454 | 7.973574582 | 7.692387724 |
| ENSG00000169629 | RGPD8 | 10.43410229 | 1.48E-15 | 3.633023638 | 0.01085929 | 14.06712593 | 2 | 2 | 2 | 10.47329248 | 10.4557854 | 4.133696865 | 3.972018587 |
| ENSG00000148082 | SHC3 | 9.940464915 | 6.33E-13 | 4.124537439 | 0.005519013 | 14.06500235 | 2 | 2 | 2 | 9.891822146 | 10.0490837 | 4.742549839 | 4.062589736 |
| ENSG00000157680 | DGKI | 6.988831791 | 1.76E-07 | 7.067510973 | 1.13E-07 | 14.05634276 | 2 | 2 | 3.369317972 | 8.047448792 | 7.95036812 | 8.087224739 | 8.067092663 |
| ENSG00000105738 | SIPA1L3 | 6.97231752 | 3.44E-14 | 7.034573352 | 1.52E-14 | 14.00689087 | 2 | 4.710564154 | 2 | 10.09666793 | 10.098275 | 9.943797765 | 10.34713831 |
| ENSG00000115461 | IGFBP5 | 6.059605911 | 1.56E-25 | 7.936491466 | 6.61E-44 | 13.99609738 | 8.107517749 | 9.470501606 | 9.541233192 | 15.2092316 | 15.2364035 | 16.78321454 | 17.35901814 |
| ENSG00000080573 | COL5A3 | 7.456904998 | 2.84E-08 | 6.518084443 | 1.44E-06 | 13.97498944 | 2 | 3.076665209 | 2 | 8.397612413 | 8.5227358 | 7.667989909 | 7.39408282 |
| ENSG00000165891 | E2F7 | 7.707391411 | 8.42E-10 | 6.266366559 | 7.99E-07 | 13.97375797 | 2 | 3.686303414 | 2 | 9.548682898 | 9.54829592 | 8.491524932 | 7.619668339 |
| ENSG00000185532 | PRKG1 | 7.619601049 | 6.47E-08 | 6.353383137 | 8.31E-06 | 13.97298419 | 2 | 3.076665209 | 2 | 8.654247601 | 8.59001038 | 7.741100262 | 6.887339785 |
| ENSG00000035862 | TIMP2 | 6.67793799 | 1.73E-10 | 7.267303634 | 2.61E-12 | 13.94524162 | 7.218819781 | 8.053983481 | 4.058998049 | 13.78072469 | 13.8490983 | 14.27041633 | 14.52724307 |
| ENSG00000172123 | SLFN12 | 7.185178603 | 1.21E-07 | 6.754695965 | 6.73E-07 | 13.93987457 | 2 | 2 | 3.369317972 | 8.221200057 | 8.16464176 | 7.559188434 | 7.953331607 |
| ENSG00000119508 | NR4A3 | 7.218669792 | 2.76E-05 | 6.71820432 | 9.94E-05 | 13.93687411 | 2 | 2 | 2 | 7.250225364 | 7.31425651 | 7.479255779 | 5.459179378 |
| ENSG00000120738 | EGR1 | 7.881109764 | 1.06E-12 | 6.038847147 | 6.99E-08 | 13.91995691 | 2 | 4.710564154 | 4.058998049 | 11.4613657 | 11.5111625 | 10.33690382 | 8.28888532 |
| ENSG00000113657 | DPYSL3 | 5.789680994 | 4.55E-15 | 8.088992354 | 8.75E-29 | 13.87867335 | 4.467311984 | 6.228547981 | 4.058998049 | 10.98303584 | 10.9103517 | 13.19345143 | 13.29308713 |
| ENSG00000182379 | NXPH4 | 4.258478958 | 0.002380789 | 9.619888033 | 1.55E-13 | 13.87836699 | 3.327711234 | 2 | 2 | 5.289065324 | 5.48637102 | 10.53940814 | 10.68063396 |
| ENSG00000116194 | ANGPTL1 | 9.395871856 | 2.36E-11 | 4.477090073 | 0.002725828 | 13.87296193 | 2 | 2 | 2 | 9.42153757 | 9.43967292 | 4.291799915 | 5.080008298 |
| ENSG00000154237 | LRRK1 | 6.797632679 | 7.15E-07 | 7.045745794 | 2.35E-07 | 13.84337847 | 2 | 3.076665209 | 2 | 7.789219291 | 7.83385794 | 8.266822833 | 7.808173621 |
| ENSG00000123095 | BHLHE41 | 6.537982947 | 2.34E-06 | 7.303167547 | 9.27E-08 | 13.84115049 | 2 | 2 | 2 | 6.610808945 | 6.63721363 | 7.317051196 | 7.411419562 |
| ENSG00000254245 | PCDHGA3 | 10.27941858 | 2.21E-13 | 3.55310053 | 0.021103887 | 13.83251911 | 2 | 2 | 2 | 10.28797351 | 10.3323804 | 3.56816602 | 4.329022995 |
| ENSG00000251493 | FOXD1 | 7.118376566 | 1.08E-09 | 6.712093257 | 9.11E-09 | 13.83046982 | 2 | 3.686303414 | 2 | 8.926937992 | 9.03050377 | 8.639092784 | 8.512214961 |
| ENSG00000124875 | CXCL6 | 11.95237304 | 5.82E-20 | 1.876244137 | 0.224818109 | 13.82861718 | 2 | 2 | 2 | 12.01365886 | 11.9459029 | 3.011868911 | 2.885414003 |
| ENSG00000092969 | TGFB2 | 4.264136308 | 3.86E-06 | 9.552393745 | 2.31E-27 | 13.81653005 | 4.00722531 | 2 | 4.058998049 | 7.454228844 | 6.88657603 | 12.44277645 | 12.452313 |
| ENSG00000135114 | OASL | 13.25936858 | 4.21E-23 | 0.53672737 | 0.767692243 | 13.79609595 | 2 | 2 | 2 | 13.26565594 | 13.3067071 | 2.560105256 | 2.326754077 |
| ENSG00000162520 | SYNC | 6.933305946 | 1.79E-10 | 6.860120219 | 2.59E-10 | 13.79342616 | 2 | 4.710564154 | 2 | 10.04177408 | 10.0039039 | 9.366015681 | 10.36467412 |
| ENSG00000177606 | JUN | 7.08028268 | 9.95E-16 | 6.659606652 | 3.79E-14 | 13.73988933 | 2 | 5.595927442 | 4.058998049 | 11.39212402 | 11.4591793 | 11.07271919 | 10.93618637 |
| ENSG00000111341 | MGP | 2.565356494 | 0.048428417 | 11.16712765 | 7.27E-23 | 13.73248414 | 2 | 3.686303414 | 2 | 4.694064037 | 4.63360314 | 12.94685733 | 13.08746641 |
| ENSG00000180801 | ARSJ | 5.239001167 | 2.60E-05 | 8.486609031 | 1.55E-12 | 13.7256102 | 2 | 5.303523747 | 2 | 8.907220097 | 8.98007065 | 11.81456027 | 12.47327371 |
| ENSG00000156265 | MAP3K7CL | 4.215614527 | 0.000512022 | 9.50562439 | 6.46E-17 | 13.72123892 | 4.00722531 | 2 | 2 | 6.139181293 | 6.20590375 | 11.26407555 | 11.49256057 |
| ENSG00000167972 | ABCA3 | 3.47444478 | 0.036622122 | 10.23382622 | 4.44E-12 | 13.708271 | 2 | 2 | 2 | 4.387512679 | 3.28037825 | 10.38053488 | 10.1391448 |
| ENSG00000115107 | STEAP3 | 6.302534688 | 8.40E-11 | 7.400620234 | 1.22E-14 | 13.70315492 | 2 | 3.076665209 | 4.058998049 | 8.799033614 | 8.74219737 | 9.730197405 | 9.982668878 |
| ENSG00000144369 | FAM171B | 7.265449374 | 1.09E-06 | 6.429813421 | 1.84E-05 | 13.69526279 | 2 | 2 | 2 | 7.409376773 | 7.24216308 | 6.911998488 | 5.980509018 |
| ENSG00000188112 | C6orf132 | 4.845725432 | 0.000881385 | 8.844103539 | 1.42E-10 | 13.68982897 | 2 | 2 | 3.369317972 | 5.775854763 | 6.0722229 | 9.393896061 | 10.17807184 |
| ENSG00000166446 | CDYL2 | 8.096723313 | 1.06E-06 | 5.575808004 | 0.001131917 | 13.67253132 | 2 | 2 | 2 | 8.149581172 | 8.13770558 | 6.353989995 | 4.543391831 |
| ENSG00000137752 | CASP1 | 9.061125551 | 7.42E-12 | 4.603104221 | 0.000949913 | 13.66422977 | 2 | 2 | 2 | 9.083378928 | 9.11260313 | 4.820739789 | 4.870334174 |
| ENSG00000167702 | KIFC2 | 6.645885023 | 1.13E-06 | 7.002647166 | 2.39E-07 | 13.64853219 | 2 | 3.076665209 | 2 | 7.549526251 | 7.76782485 | 8.154648976 | 7.856442979 |
| ENSG00000116675 | DNAJC6 | 4.746168883 | 0.001122184 | 8.902014015 | 8.65E-11 | 13.6481829 | 2 | 2 | 2 | 5.194621444 | 4.70309173 | 8.798653052 | 9.069094714 |
| ENSG00000110852 | CLEC2B | 9.468026559 | 6.93E-13 | 4.158708717 | 0.003156428 | 13.62673528 | 2 | 2 | 2 | 9.521380925 | 9.48312089 | 4.5389195 | 4.400220394 |
| ENSG00000099822 | HCN2 | 5.002601144 | 0.000506245 | 8.59782554 | 3.13E-10 | 13.60042668 | 2 | 2 | 2 | 5.242616075 | 5.14724905 | 8.853368954 | 8.386355612 |
| ENSG00000167363 | FN3K | 5.797329359 | 3.82E-05 | 7.780849627 | 1.28E-08 | 13.57817899 | 2 | 2 | 2 | 5.808110275 | 6.02777321 | 7.765568476 | 7.897008641 |
| ENSG00000254221 | PCDHGB1 | 10.14510539 | 1.15E-13 | 3.411494743 | 0.023508946 | 13.55660013 | 2 | 2 | 2 | 10.14744996 | 10.2049293 | 3.584558147 | 4.147809227 |
| ENSG00000102575 | ACP5 | 6.46078331 | 0.002810253 | 7.094736666 | 0.000864919 | 13.55551998 | 2 | 2 | 2 | 6.529309835 | 6.57060569 | 8.059640467 | 4.292061816 |
| ENSG00000152953 | STK32B | 4.132508721 | 0.00623234 | 9.42122094 | 1.24E-11 | 13.55372966 | 2 | 2 | 2 | 4.470469939 | 4.43005853 | 9.003023259 | 9.800020464 |
| ENSG00000206538 | VGLL3 | 4.682725976 | 3.90E-12 | 8.854140034 | 1.31E-41 | 13.53686601 | 3.327711234 | 5.131603625 | 3.369317972 | 8.727753724 | 8.65574839 | 12.88907816 | 12.81060063 |
| ENSG00000153234 | NR4A2 | 8.696347292 | 8.82E-11 | 4.835972119 | 0.000564093 | 13.53231941 | 2 | 2 | 2 | 8.716468139 | 8.75588792 | 5.158477451 | 4.927687581 |
| ENSG00000160013 | PTGIR | 7.395100476 | 1.84E-07 | 6.122198649 | 1.98E-05 | 13.51729913 | 2 | 2 | 2 | 7.461007055 | 7.44845993 | 6.468821191 | 5.937276556 |
| ENSG00000198598 | MMP17 | 8.065669592 | 4.37E-09 | 5.4424983 | 0.000118706 | 13.50816789 | 2 | 2 | 2 | 8.132699624 | 8.09310209 | 5.766051132 | 5.396716407 |

| | | | | | | | | | | | | | |
|---|---|---|---|---|---|---|---|---|---|---|---|---|---|
| ENSG00000074855 | ANO8 | 5.98476352 | 1.09E-05 | 7.513693259 | 1.65E-08 | 13.49845678 | 2 | 2 | 3.369317972 | 6.970413838 | 7.06433609 | 8.573367846 | 8.459712934 |
| ENSG00000110107 | PRPF19 | 7.149202388 | 5.49E-10 | 6.308324793 | 5.06E-08 | 13.45752718 | 2 | 3.076665209 | 3.369317972 | 9.042162213 | 9.00665321 | 8.157391075 | 8.226587404 |
| ENSG00000136999 | NOV | 6.798032807 | 1.01E-06 | 6.628032515 | 1.80E-06 | 13.42606532 | 2 | 2 | 2 | 6.887305128 | 6.86146731 | 6.732402829 | 6.688368862 |
| ENSG00000087116 | ADAMTS2 | 5.168893171 | 2.10E-12 | 8.248290787 | 2.01E-30 | 13.41718396 | 2 | 5.303523747 | 4.523675391 | 9.436230969 | 9.45988854 | 12.68338458 | 12.33617257 |
| ENSG00000102007 | PLP2 | 6.581459187 | 5.11E-12 | 6.806513437 | 7.07E-13 | 13.38797262 | 3.327711234 | 5.457120953 | 2 | 10.6209439 | 10.6719453 | 10.77189113 | 10.96411291 |
| ENSG00000100292 | HMOX1 | 8.675230074 | 2.91E-09 | 4.707377669 | 0.002199945 | 13.38260774 | 2 | 3.686303414 | 2 | 10.48331504 | 10.4741226 | 7.237652294 | 5.282014932 |
| ENSG00000187479 | C11orf96 | 6.490459016 | 1.52E-11 | 6.889357202 | 5.69E-13 | 13.37981622 | 2 | 4.93638933 | 2 | 9.747510575 | 9.94596123 | 10.21173728 | 10.28245849 |
| ENSG00000170624 | SGCD | 5.289393798 | 0.000272372 | 8.089698854 | 6.62E-09 | 13.37909265 | 2 | 2 | 2 | 5.461211713 | 5.44640346 | 7.826275727 | 8.392188801 |
| ENSG00000178951 | ZBTB7A | 6.275960856 | 3.69E-06 | 7.098992757 | 1.15E-07 | 13.37495361 | 2 | 5.595927442 | 2 | 10.18478072 | 10.3364864 | 11.17122857 | 10.99030784 |
| ENSG00000164070 | HSPA4L | 5.064245722 | 0.000453925 | 8.307950406 | 1.58E-09 | 13.37219613 | 2 | 2 | 2 | 5.210797711 | 5.28958453 | 8.568740544 | 8.096865967 |
| ENSG00000235376 | RPEL1 | 6.675651473 | 2.28E-06 | 6.69220034 | 1.97E-06 | 13.36785181 | 2 | 2 | 2 | 6.820355514 | 6.68951263 | 6.615624322 | 6.913939481 |
| ENSG00000137033 | IL33 | 14.01558365 | 1.90E-21 | -0.653185163 | 0.748118922 | 13.36239849 | 2 | 2 | 2 | 14.06846635 | 14.0156896 | 2.34604626 | 2.059860178 |
| ENSG00000107984 | DKK1 | 9.247193409 | 6.19E-13 | 4.112153131 | 0.002650534 | 13.35934654 | 3.327711234 | 3.076665209 | 2 | 11.1032668 | 11.1106357 | 6.564892306 | 5.269393099 |
| ENSG00000157601 | MX1 | 10.77000438 | 6.15E-30 | 2.573677512 | 0.013923728 | 13.3436819 | 2 | 4.710564154 | 2 | 13.87406507 | 13.887268 | 5.4588802 | 6.06327132 |
| ENSG00000198482 | ZNF808 | 7.335447684 | 2.46E-07 | 6.007850076 | 3.01E-05 | 13.34329776 | 2 | 2 | 2 | 7.45762193 | 7.33264456 | 6.351597515 | 5.842465082 |
| ENSG00000134954 | ETS1 | 6.7648362 | 7.17E-10 | 6.57785291 | 2.00E-09 | 13.34268911 | 2 | 5.72254323 | 3.369317972 | 11.06734964 | 11.0471929 | 10.6402526 | 11.06934348 |
| ENSG00000105889 | STEAP1B | 8.856319832 | 6.51E-11 | 4.481731227 | 0.001775206 | 13.33805106 | 2 | 2 | 2 | 8.912174893 | 8.87734839 | 4.530483275 | 4.927687581 |
| ENSG00000010704 | HFE | 6.687427044 | 1.45E-06 | 6.645562109 | 1.57E-06 | 13.33298915 | 2 | 3.076665209 | 2 | 7.66556918 | 7.74054144 | 7.372383768 | 7.90417044 |
| ENSG00000178776 | C5orf46 | 5.011699957 | 0.000899661 | 8.313907304 | 6.78E-09 | 13.32560726 | 2 | 2 | 2 | 5.258265913 | 5.14724905 | 7.862014947 | 8.726491728 |
| ENSG00000134769 | DTNA | 6.297014054 | 1.82E-05 | 7.016520559 | 1.27E-06 | 13.31353461 | 2 | 2 | 2 | 6.436065776 | 6.35006802 | 6.717615541 | 7.379474672 |
| ENSG00000153904 | DDAH1 | 5.566723621 | 2.92E-06 | 7.731928235 | 2.81E-11 | 13.29865186 | 3.327711234 | 5.946638857 | 2 | 10.08135305 | 10.0345549 | 12.29212498 | 12.14224092 |
| ENSG00000115267 | IFIH1 | 9.535034437 | 2.06E-28 | 3.757544321 | 3.17E-05 | 13.29257876 | 2 | 4.442710129 | 2 | 12.37946895 | 12.363074 | 6.683779344 | 6.617932662 |
| ENSG00000109452 | INPP4B | 7.135815646 | 5.33E-06 | 6.155573789 | 0.000100443 | 13.29138943 | 2 | 2 | 2 | 7.075737784 | 7.31795292 | 6.743394671 | 5.524371359 |
| ENSG00000163565 | IFI16 | 8.491629413 | 8.19E-10 | 4.796542071 | 0.000904021 | 13.28817148 | 2 | 5.72254323 | 2 | 12.58957959 | 12.6104106 | 9.056150495 | 8.760712793 |
| ENSG00000183801 | OLFML1 | 4.927831053 | 0.005313564 | 8.325688201 | 5.97E-07 | 13.25351925 | 2 | 2 | 2 | 5.128039203 | 5.13053105 | 6.994053541 | 9.060878381 |
| ENSG00000115641 | FHL2 | 5.611328668 | 2.33E-07 | 7.625961766 | 6.87E-13 | 13.23729043 | 3.327711234 | 5.72254323 | 2 | 9.879225887 | 9.93276281 | 11.90309722 | 11.92945766 |
| ENSG00000172493 | AFF1 | 6.600194919 | 4.88E-14 | 6.637093156 | 2.66E-14 | 13.23728808 | 2 | 4.710564154 | 2 | 9.741955564 | 9.73406074 | 9.817517875 | 9.730640067 |
| ENSG00000112183 | RBM24 | 4.57175992 | 0.001312777 | 8.664026258 | 9.20E-11 | 13.23578618 | 2 | 2 | 2 | 4.76150579 | 4.87352701 | 8.723392661 | 8.684919451 |
| ENSG00000117586 | TNFSF4 | 9.414383255 | 2.77E-09 | 3.818737604 | 0.026949319 | 13.23312086 | 2 | 2 | 2 | 9.453327884 | 9.44475345 | 4.742549839 | 3.311948089 |
| ENSG00000185950 | IRS2 | 6.198387481 | 1.45E-10 | 7.018576531 | 2.29E-13 | 13.21696401 | 2 | 2 | 4.874587708 | 9.081180848 | 9.16073939 | 9.907406852 | 9.966387947 |
| ENSG00000124508 | BTN2A2 | 6.745945437 | 2.10E-13 | 6.462112274 | 1.87E-12 | 13.20805771 | 2 | 4.442710129 | 2 | 9.615519136 | 9.53803206 | 9.138949083 | 9.43604749 |
| ENSG00000153944 | MSI2 | 6.135070214 | 2.95E-07 | 7.067440856 | 2.23E-09 | 13.20251107 | 2 | 3.686303414 | 2 | 8.074220386 | 7.93118934 | 8.781425385 | 9.05767046 |
| ENSG00000151632 | AKR1C2 | 9.947846482 | 1.68E-23 | 3.230727436 | 0.002564517 | 13.17857392 | 2 | 2 | 4.523675391 | 12.41491918 | 12.4381264 | 5.598599872 | 6.003555476 |
| ENSG00000134321 | RSAD2 | 13.39828359 | 4.52E-17 | -0.223249482 | 0.920567538 | 13.17503411 | 2 | 3.076665209 | 3.369317972 | 15.26494408 | 15.2457575 | 3.296906311 | 2.117335275 |
| ENSG00000124785 | NRN1 | 8.412188813 | 1.68E-07 | 4.729914713 | 0.005277336 | 13.14210353 | 2 | 2 | 2 | 8.447952961 | 8.46222608 | 5.528313928 | 3.987514168 |
| ENSG00000128567 | PODXL | 5.418654136 | 4.50E-07 | 7.717671677 | 1.74E-13 | 13.13632581 | 6.138360429 | 3.686303414 | 3.369317972 | 10.18119344 | 10.1515521 | 11.6770541 | 12.96674738 |
| ENSG00000180530 | NRIP1 | 7.236459462 | 1.65E-11 | 5.889172181 | 5.61E-08 | 13.12563164 | 2 | 2 | 4.523675391 | 9.777686595 | 9.68838639 | 8.685899845 | 8.034828728 |
| ENSG00000253537 | PCDHGA7 | 9.220312596 | 2.16E-12 | 3.899408699 | 0.005658731 | 13.1197213 | 2 | 3.076665209 | 2 | 10.18170646 | 10.244312 | 5.185595313 | 4.943664032 |
| ENSG00000226742 | HSBP1L1 | 6.733812829 | 1.89E-06 | 6.372621001 | 6.69E-06 | 13.10643383 | 2 | 2 | 2 | 6.809774134 | 6.81514011 | 6.597565628 | 6.320441373 |
| ENSG00000081041 | CXCL2 | 12.14473503 | 6.54E-17 | 0.956052317 | 0.608101226 | 13.10078734 | 2 | 2 | 2 | 12.21566187 | 12.1277429 | 2.3070652 | 2.817676617 |
| ENSG00000145242 | EPHA5 | 6.502828051 | 3.42E-05 | 6.576689466 | 2.49E-05 | 13.07951752 | 2 | 2 | 2 | 6.522845756 | 6.65485813 | 5.982522014 | 7.12130792 |
| ENSG00000277203 | F8A1 | 8.415558423 | 2.68E-10 | 4.64638498 | 0.000872927 | 13.0619434 | 2 | 2 | 3.369317972 | 9.353044608 | 9.46906021 | 5.574182174 | 5.900914815 |
| ENSG00000121350 | PYROXD1 | 7.163240802 | 2.74E-07 | 5.850705358 | 3.41E-05 | 13.01394616 | 2 | 2 | 2 | 7.246306984 | 7.21069602 | 5.883320383 | 6.055940621 |
| ENSG00000163814 | CDCP1 | 8.228032737 | 1.24E-18 | 4.767534142 | 6.34E-07 | 12.99556688 | 4.00722531 | 2 | 4.874587708 | 11.73741575 | 11.7703643 | 8.72385521 | 7.727417522 |
| ENSG00000065534 | MYLK | 6.009407532 | 2.29E-05 | 6.973271693 | 5.94E-07 | 12.98267922 | 3.327711234 | 7.051611315 | 3.369317972 | 11.57546597 | 11.6152815 | 13.16171034 | 11.5021165 |
| ENSG00000198841 | KTI12 | 7.121659715 | 5.37E-07 | 5.857359901 | 4.71E-05 | 12.97901962 | 2 | 2 | 2 | 7.132034771 | 7.24216308 | 6.167993056 | 5.759151667 |
| ENSG00000185567 | AHNAK2 | 5.317952351 | 1.32E-05 | 7.623099693 | 1.27E-10 | 12.94105204 | 4.815609471 | 6.535001748 | 2 | 10.54203082 | 10.6435047 | 12.32812587 | 13.30214374 |
| ENSG00000177989 | ODF3B | 7.562354798 | 7.17E-08 | 5.369991598 | 0.000193253 | 12.9323464 | 2 | 2 | 2 | 7.523859802 | 7.70710533 | 5.703636713 | 5.32534152 |
| ENSG00000115415 | STAT1 | 7.404061525 | 2.97E-14 | 5.522747535 | 2.15E-08 | 12.92680906 | 7.159102577 | 8.276714465 | 4.874587708 | 14.71318337 | 14.7019162 | 13.18543446 | 12.34846028 |
| ENSG00000154556 | SORBS2 | 4.846844978 | 0.000497308 | 8.076640171 | 1.14E-09 | 12.92348515 | 3.327711234 | 2 | 2 | 5.978405189 | 5.88556448 | 9.168133494 | 8.97651941 |
| ENSG00000241399 | CD302 | 6.824274748 | 1.18E-06 | 6.090904354 | 1.64E-05 | 12.9151791 | 2 | 2 | 2 | 6.970413838 | 6.82556437 | 6.200262696 | 6.195912571 |
| ENSG00000142552 | RCN3 | 6.216191329 | 1.10E-07 | 6.686390069 | 8.52E-09 | 12.9025814 | 4.815609471 | 6.602424674 | 2 | 11.53166763 | 11.5617274 | 12.00470931 | 12.02805033 |
| ENSG00000171617 | ENC1 | 7.273854582 | 1.65E-13 | 5.623434517 | 1.72E-08 | 12.8972891 | 2 | 4.710564154 | 2 | 10.46993609 | 10.2892535 | 9.012519675 | 8.406669418 |
| ENSG00000167779 | IGFBP6 | 8.379586388 | 1.73E-12 | 4.508419928 | 0.0002746 | 12.88800632 | 2 | 3.686303414 | 6.203069917 | 13.045527 | 13.0670016 | 9.083080045 | 9.297612054 |
| ENSG00000072657 | TRHDE | 6.678920468 | 5.03E-06 | 6.20829868 | 2.30E-05 | 12.88721915 | 2 | 2 | 2 | 6.892328675 | 6.61334686 | 6.022174708 | 6.549111181 |
| ENSG00000172403 | SYNPO2 | 4.328192943 | 9.81E-07 | 8.537437011 | 1.15E-23 | 12.86562995 | 4.00722531 | 5.457120953 | 5.156626401 | 9.169698743 | 9.1566168 | 14.00882487 | 12.16264744 |
| ENSG00000139083 | ETV6 | 5.896375837 | 1.27E-11 | 6.962031175 | 5.74E-16 | 12.85840701 | 2 | 4.11352228 | 3.369317972 | 8.737556707 | 8.7802082 | 9.940418627 | 9.683551156 |
| ENSG00000174332 | GLIS1 | 5.289472578 | 0.000218113 | 7.562651669 | 4.19E-08 | 12.85212425 | 2 | 2 | 2 | 5.474671438 | 5.43283104 | 7.716209895 | 7.513957873 |
| ENSG00000105767 | CADM4 | 5.396728635 | 0.000195383 | 7.446667798 | 1.06E-07 | 12.84339643 | 2 | 2 | 2 | 5.540148523 | 5.56313034 | 7.686143116 | 7.297928101 |
| ENSG00000169083 | AR | 6.494130249 | 2.20E-06 | 6.342960969 | 3.67E-06 | 12.83709122 | 2 | 3.076665209 | 2 | 7.507583254 | 7.52080763 | 7.207540845 | 7.509939612 |
| ENSG00000180138 | CSNK1A1L | 6.758212703 | 1.78E-06 | 6.073331379 | 1.96E-05 | 12.83154408 | 2 | 2 | 2 | 6.836083475 | 6.83591385 | 6.290458506 | 6.06327132 |
| ENSG00000074527 | NTN4 | 3.062546602 | 0.014298818 | 9.765141439 | 5.61E-18 | 12.82768804 | 2 | 5.303523747 | 2 | 6.750147204 | 6.88158913 | 13.4572701 | 13.47939587 |
| ENSG00000163412 | EIF4E3 | 5.107556054 | 0.000400716 | 7.683412054 | 2.78E-08 | 12.79096811 | 2 | 2 | 2 | 5.334065616 | 5.24368216 | 7.872900025 | 7.586981404 |
| ENSG00000141013 | GAS8 | 6.47763442 | 7.72E-06 | 6.306200362 | 1.29E-05 | 12.78383478 | 2 | 2 | 2 | 6.61689148 | 6.51372389 | 6.173421703 | 6.600402875 |
| ENSG00000127528 | KLF2 | 5.243336524 | 7.80E-05 | 7.50991184 | 5.04E-09 | 12.75324836 | 2 | 3.686303414 | 2 | 7.194375246 | 7.02868983 | 9.76049665 | 8.765214879 |
| ENSG00000176438 | SYNE3 | 4.607304901 | 0.001569431 | 8.137916546 | 3.21E-09 | 12.74522145 | 2 | 2 | 2 | 4.670864177 | 5.00774437 | 8.124136806 | 8.242012195 |
| ENSG00000071575 | TRIB2 | 4.312302687 | 2.48E-05 | 8.423795092 | 6.10E-18 | 12.73609778 | 2 | 4.710564154 | 2 | 7.510853288 | 7.38644297 | 11.82779956 | 11.15439097 |
| ENSG00000110455 | ACCS | 6.480258129 | 3.06E-05 | 6.240682143 | 5.82E-05 | 12.72094027 | 2 | 2 | 2 | 6.509830055 | 6.6253296 | 5.755231204 | 6.755525668 |
| ENSG00000070882 | OSBPL3 | 6.926050283 | 2.32E-08 | 5.792467049 | 3.76E-06 | 12.71851733 | 4.00722531 | 2 | 2 | 8.88471049 | 8.75861051 | 8.008923105 | 7.317985043 |
| ENSG00000162892 | IL24 | 11.5577434 | 2.95E-09 | 1.154038378 | 0.638141963 | 12.71178178 | 2 | 2 | 2 | 11.58964598 | 11.5824281 | 3.059555873 | 2.059860178 |
| ENSG00000184470 | TXNRD2 | 5.879927846 | 2.19E-05 | 6.82466941 | 5.41E-07 | 12.70459726 | 2 | 2 | 3.369317972 | 6.917187326 | 6.91614081 | 7.680435163 | 7.980689421 |
| ENSG00000128342 | LIF | 7.356325667 | 1.25E-09 | 5.34639163 | 1.52E-05 | 12.7027173 | 6.138360429 | 3.686303414 | 2 | 11.97155766 | 12.0084303 | 9.477737996 | 10.35872838 |
| ENSG00000159208 | CIART | 6.400646238 | 7.03E-06 | 6.295992307 | 9.41E-06 | 12.69663854 | 2 | 2 | 2 | 6.5612028 | 6.42052719 | 6.34920106 | 6.435070131 |
| ENSG00000137878 | GCOM1 | 9.161417346 | 7.60E-11 | 3.518252897 | 0.022469646 | 12.67967024 | 2 | 2 | 2 | 9.252121432 | 9.14105132 | 3.600766115 | 4.26688425 |
| ENSG00000001617 | SEMA3F | 6.804804388 | 9.44E-10 | 5.864648191 | 1.61E-07 | 12.66945258 | 2 | 3.686303414 | 3.369317972 | 9.252121432 | 9.23593348 | 7.633939843 | 8.772500938 |
| ENSG00000198133 | TMEM229B | 5.626469571 | 0.000104489 | 7.031495788 | 6.63E-07 | 12.65796536 | 2 | 2 | 2 | 5.829220145 | 5.69405321 | 7.241533944 | 6.944034147 |
| ENSG00000185278 | ZBTB37 | 4.97439965 | 0.000555287 | 7.656863959 | 2.69E-08 | 12.63126361 | 2 | 2 | 2 | 5.093559143 | 5.24368216 | 7.760166845 | 7.66011679 |
| ENSG00000138378 | STAT4 | 5.542024556 | 0.00135052 | 7.076470944 | 2.44E-05 | 12.6184955 | 2 | 2 | 2 | 5.818703821 | 5.53799497 | 7.753839274 | 6.066922748 |
| ENSG00000041353 | RAB27B | 8.531848592 | 1.02E-13 | 4.080564461 | 0.000735092 | 12.61241305 | 2 | 3.686303414 | 2 | 10.40902283 | 10.3662449 | 5.99483933 | 6.052261256 |
| ENSG00000102524 | TNFSF13B | 10.05051427 | 2.38E-11 | 2.559424727 | 0.135257068 | 12.609939 | 2 | 2 | 2 | 10.12521908 | 10.0379207 | 3.709392871 | 2.782576693 |
| ENSG00000140403 | DNAJA4 | 5.540379592 | 7.57E-05 | 7.060859212 | 2.29E-07 | 12.6012388 | 2 | 3.076665209 | 2 | 6.394182228 | 6.76185814 | 8.022522757 | 8.117259579 |
| ENSG00000204941 | PSG5 | 5.245811773 | 0.000459654 | 7.336998916 | 3.69E-07 | 12.58281069 | 2 | 2 | 2 | 5.686138681 | 5.07918227 | 7.411959578 | 7.383872668 |
| ENSG00000276910 | | 8.264783965 | 1.10E-09 | 4.315107821 | 0.002615937 | 12.57989179 | 2 | 2 | 2 | 8.276040626 | 8.34212018 | 4.504874511 | 4.692661645 |
| ENSG00000134245 | WNT2B | 4.363223706 | 0.003871463 | 8.193886086 | 6.68E-09 | 12.55710979 | 2 | 2 | 2 | 4.846787657 | 4.40247105 | 7.984424805 | 8.456236734 |
| ENSG00000167767 | KRT80 | 2.012384905 | 0.221153215 | 10.54444285 | 5.04E-15 | 12.55682776 | 2 | 2 | 2 | 3.162721474 | 2.86123014 | 10.4114447 | 10.72117 |
| ENSG00000163735 | CXCL5 | 13.62562716 | 6.29E-21 | -1.112161687 | 0.571127612 | 12.51346547 | 2 | 2 | 2 | 13.67524369 | 13.6293234 | 2.048074405 | 2.277179783 |
| ENSG00000264364 | DYNLL2 | 6.379367104 | 1.92E-12 | 6.111805021 | 1.44E-11 | 12.49117213 | 2 | 4.442710129 | 2 | 9.231457548 | 9.20133 | 8.883491454 | 9.015305099 |
| ENSG00000144036 | EXOC6B | 5.844987291 | 1.96E-08 | 6.644637234 | 1.10E-10 | 12.48962453 | 2 | 4.710564154 | 2 | 8.95000826 | 8.95417718 | 9.326055204 | 10.07214007 |
| ENSG00000106688 | SLC1A1 | 6.799934133 | 1.30E-09 | 5.688618705 | 4.90E-07 | 12.48855284 | 2 | 2 | 4.874587708 | 9.771565364 | 9.72295127 | 8.963225048 | 8.233102165 |
| ENSG00000175899 | A2M | 4.435797002 | 0.052158739 | 8.048177417 | 0.000140835 | 12.48397442 | 3.327711234 | 2 | 2 | 6.456560224 | 2.59489879 | 9.420391988 | 8.542691771 |
| ENSG00000183337 | BCOR | 5.546887283 | 2.66E-08 | 6.92024309 | 1.73E-12 | 12.46713037 | 2 | 4.442710129 | 2 | 8.258639605 | 8.47883193 | 9.432328044 | 9.98604778 |
| ENSG00000013619 | MAMLD1 | 5.389289155 | 0.000113829 | 7.072675474 | 1.91E-07 | 12.46196463 | 2 | 2 | 3.369317972 | 6.422239259 | 6.46787796 | 7.92612656 | 8.223318958 |
| ENSG00000253731 | PCDHGA6 | 8.346014586 | 2.24E-11 | 4.095588627 | 0.001915611 | 12.44160321 | 2 | 2 | 4.058998049 | 10.2136641 | 10.2764875 | 5.574182174 | 6.451924088 |
| ENSG00000136830 | FAM129B | 6.25240834 | 5.87E-13 | 6.18732931 | 8.72E-13 | 12.43973765 | 5.532353208 | 7.73146271 | 5.39244166 | 12.85170056 | 12.8802457 | 12.69764229 | 12.89743356 |
| ENSG00000148926 | ADM | 6.319606769 | 8.02E-08 | 6.109821694 | 2.08E-07 | 12.42942846 | 4.467311984 | 6.464272742 | 2 | 11.44500551 | 11.4766471 | 11.10583548 | 11.38368346 |
| ENSG00000102452 | NALCN | 4.483303997 | 0.00209302 | 7.935433057 | 7.27E-09 | 12.41873705 | 2 | 2 | 2 | 4.647285135 | 4.83276841 | 7.918872821 | 8.047821014 |
| ENSG00000131981 | LGALS3 | 6.298892608 | 1.05E-21 | 6.114744884 | 9.93E-21 | 12.41363749 | 3.327711234 | 4.11352228 | 5.59507254 | 10.67003398 | 10.6372309 | 10.45318893 | 10.48674576 |
| ENSG00000128203 | ASPHD2 | 6.619593552 | 4.23E-06 | 5.775464118 | 6.88E-05 | 12.39505767 | 2 | 2 | 2 | 6.598566398 | 6.79936111 | 6.016144783 | 5.777095613 |
| ENSG00000213799 | ZNF845 | 6.38489564 | 8.90E-06 | 6.000074753 | 3.05E-05 | 12.38497039 | 3.327711234 | 2 | 2 | 7.546342806 | 7.2537885 | 7.336538937 | 6.64749689 |
| ENSG00000184922 | FMNL1 | 7.610863002 | 7.44E-07 | 4.772662136 | 0.002930366 | 12.38352514 | 2 | 2 | 2 | 7.593372273 | 7.73502229 | 5.427419529 | 4.365060851 |
| ENSG00000143127 | ITGA10 | 4.047210226 | 0.007642363 | 8.333236132 | 2.80E-09 | 12.38044636 | 2 | 2 | 2 | 4.415698455 | 4.3456612 | 8.033758684 | 8.654300638 |
| ENSG00000115339 | GALNT3 | 6.795342121 | 4.53E-06 | 5.582791148 | 0.000203796 | 12.37813327 | 2 | 2 | 2 | 6.872128674 | 6.8715633 | 6.007052374 | 5.367404784 |
| ENSG00000112139 | MDGA1 | 6.286129377 | 2.40E-09 | 6.09069067 | 7.07E-09 | 12.37682005 | 3.327711234 | 3.076665209 | 3.369317972 | 8.71221318 | 8.79752133 | 8.056705941 | 8.935663927 |
| ENSG00000106246 | PTCD1 | 7.349603154 | 1.31E-07 | 5.00289973 | 0.00049334 | 12.35250288 | 2 | 2 | 2 | 7.440576128 | 7.37938471 | 5.258920789 | 5.12973572 |
| ENSG00000261115 | TMEM178B | 2.374407678 | 0.164591992 | 9.973805487 | 6.45E-12 | 12.34821316 | 3.327711234 | 2 | 2 | 4.205751501 | 3.33976443 | 10.45249137 | 11.34219318 |
| ENSG00000171246 | NPTX1 | 6.626231649 | 0.001337504 | 5.709121645 | 0.006337615 | 12.33535329 | 2 | 2 | 2 | 6.739036112 | 6.67805337 | 6.658838928 | 3.753786374 |
| ENSG00000162889 | MAPKAPK2 | 6.077907173 | 4.81E-07 | 6.255484551 | 1.92E-07 | 12.33339172 | 2 | 6.311476754 | 4.058998049 | 10.96317997 | 11.0510847 | 11.07770369 | 11.28494885 |

| | | | | | | | | | | | | | |
|---|---|---|---|---|---|---|---|---|---|---|---|---|---|
| ENSG00000177283 | FZD8 | 7.117599289 | 2.90E-11 | 5.212261649 | 1.70E-06 | 12.32986094 | 2 | 4.11352228 | 2 | 9.491898484 | 9.57325401 | 7.946879235 | 7.272853598 |
| ENSG00000162944 | RFTN2 | 6.333454194 | 3.38E-05 | 5.994711648 | 8.72E-05 | 12.32816584 | 2 | 2 | 2 | 6.456560224 | 6.39974839 | 6.444330878 | 5.66592325 |
| ENSG00000153246 | PLA2R1 | 6.181537594 | 6.24E-09 | 6.136536616 | 7.32E-09 | 12.31807421 | 2 | 4.11352228 | 2 | 8.607665593 | 8.60672783 | 8.245719321 | 8.822498333 |
| ENSG00000104205 | SGK3 | 5.787915971 | 9.38E-05 | 6.528422075 | 7.59E-06 | 12.31633805 | 2 | 2 | 2 | 5.987823101 | 5.83486613 | 6.382394503 | 6.815359915 |
| ENSG00000081853 | PCDHGA2 | 8.349722268 | 4.08E-13 | 3.962721148 | 0.001120882 | 12.31244342 | 2 | 3.686303414 | 2 | 10.17347624 | 10.2370092 | 5.86330637 | 5.96101835 |
| ENSG00000151967 | SCHIP1 | 7.577610105 | 4.59E-08 | 4.732787331 | 0.001029882 | 12.31039744 | 2 | 2 | 2 | 7.530319349 | 7.72948195 | 5.002522427 | 4.911532223 |
| ENSG00000107957 | SH3PXD2A | 5.812156255 | 5.96E-05 | 6.494362698 | 5.46E-06 | 12.30651895 | 4.815609471 | 7.393647515 | 2 | 11.79538311 | 11.8174753 | 12.44330319 | 12.53145936 |
| ENSG00000161956 | SENP3 | 6.612782873 | 3.45E-11 | 5.677542088 | 1.56E-08 | 12.29032496 | 4.467311984 | 2 | 2 | 9.037634995 | 9.14417788 | 8.177792357 | 8.154630332 |
| ENSG00000115963 | RND3 | 7.591753554 | 1.39E-07 | 4.691555825 | 0.001786849 | 12.28330938 | 8.009475818 | 6.728495673 | 2 | 14.48660668 | 14.4773038 | 12.03186006 | 10.92905675 |
| ENSG00000168062 | BATF2 | 8.902939506 | 3.36E-11 | 3.37762757 | 0.021742881 | 12.28056708 | 2 | 2 | 2 | 8.904736303 | 8.97656697 | 3.968654369 | 3.753786374 |
| ENSG00000137449 | CPEB2 | 5.052551984 | 4.99E-06 | 7.227998301 | 1.91E-11 | 12.28055029 | 2 | 5.131603625 | 2 | 8.682071821 | 8.51792306 | 10.69754037 | 10.83167443 |
| ENSG00000149346 | SLX4IP | 4.912909845 | 0.000645538 | 7.365799354 | 9.59E-08 | 12.2787092 | 2 | 3.076665209 | 2 | 6.061031724 | 5.92487824 | 8.62380783 | 8.065268319 |
| ENSG00000162745 | OLFML2B | 3.658670053 | 0.016459729 | 8.615778848 | 6.37E-10 | 12.2744489 | 2 | 3.076665209 | 2 | 4.907595604 | 4.811949 | 9.986250224 | 9.103279088 |
| ENSG00000172572 | PDE3A | 3.993730553 | 0.010492446 | 8.277867241 | 8.63E-09 | 12.27159779 | 2 | 2 | 2 | 4.415698455 | 4.2560182 | 8.676850263 | 7.851158781 |
| ENSG00000066697 | MSANTD3 | 6.43182894 | 4.48E-10 | 5.828873616 | 1.76E-08 | 12.26070256 | 2 | 4.93638933 | 2 | 9.800579656 | 9.75398117 | 8.929534325 | 9.389783813 |
| ENSG00000258643 | BCL2L2-PABI | 10.19431583 | 1.44E-12 | 2.059801539 | 0.220786883 | 12.25411737 | 2 | 2 | 2 | 10.23853288 | 10.2123959 | 2.68698273 | 3.33635788 |
| ENSG00000170458 | CD14 | 5.274443218 | 0.000312209 | 6.978650187 | 8.54E-07 | 12.25309341 | 2 | 2 | 2 | 5.44762523 | 5.43283104 | 6.911998488 | 7.174740504 |
| ENSG00000166396 | SERPINB7 | 8.063666707 | 7.27E-09 | 4.176803589 | 0.004736945 | 12.2404703 | 2 | 2 | 2 | 8.160032699 | 8.06037647 | 4.261535101 | 4.683176719 |
| ENSG00000090376 | IRAK3 | 8.264979763 | 2.98E-09 | 3.971460605 | 0.007602279 | 12.23644037 | 2 | 2 | 3.369317972 | 9.28496215 | 9.23983583 | 5.4588802 | 4.711446447 |
| ENSG00000177932 | ZNF354C | 5.956340904 | 5.96E-05 | 6.278970348 | 1.93E-05 | 12.23531125 | 2 | 3.076665209 | 2 | 6.970413838 | 7.01053089 | 7.705008156 | 6.746446439 |
| ENSG00000174600 | CMKLR1 | 4.064095652 | 0.005817392 | 8.167508143 | 2.26E-09 | 12.2316038 | 2 | 2 | 2 | 4.329433326 | 4.45712836 | 8.109343506 | 8.310608339 |
| ENSG00000242419 | PCDHGC4 | 10.1386649 | 2.00E-13 | 2.065675297 | 0.198633428 | 12.2043402 | 2 | 2 | 2 | 10.14007766 | 10.1994291 | 2.803598391 | 3.261853426 |
| ENSG00000131378 | RFTN1 | 5.504963302 | 1.93E-09 | 6.696055619 | 1.30E-13 | 12.20101892 | 2 | 5.303523747 | 3.369317972 | 9.437090644 | 9.45318143 | 10.78635657 | 10.45818207 |
| ENSG00000255529 | POLR2M | 6.743426855 | 1.01E-11 | 5.453311745 | 4.92E-08 | 12.1967386 | 2 | 4.11352228 | 2 | 9.234427714 | 9.12003262 | 7.888662695 | 7.916365698 |
| ENSG00000156535 | CD109 | 7.348525958 | 4.30E-29 | 4.839693817 | 2.91E-13 | 12.18821977 | 3.327711234 | 4.11352228 | 4.523675391 | 11.03449708 | 11.0139665 | 8.544331623 | 8.512214961 |
| ENSG00000133069 | TMCC2 | 4.430879119 | 0.00254798 | 7.754264669 | 2.01E-08 | 12.18514379 | 2 | 2 | 2 | 4.694064037 | 4.70309173 | 7.924517766 | 7.678134128 |
| ENSG00000151414 | NEK7 | 5.433943034 | 2.71E-05 | 6.721390202 | 1.16E-07 | 12.15533324 | 2 | 6.728495673 | 4.523675391 | 10.83989045 | 10.7710044 | 11.9932919 | 12.18324251 |
| ENSG00000006459 | KDM7A | 6.343241413 | 2.48E-08 | 5.793326897 | 3.87E-07 | 12.13656831 | 2 | 4.11352228 | 2 | 8.717883675 | 8.78689171 | 8.594261972 | 7.681710751 |
| ENSG00000163734 | CXCL3 | 12.91730977 | 1.15E-21 | -0.781904958 | 0.675642045 | 12.13540481 | 2 | 2 | 2 | 12.96593307 | 12.9225886 | 2.267001569 | 2.117335275 |
| ENSG00000159231 | CBR3 | 6.068929369 | 3.17E-05 | 6.066279197 | 2.90E-05 | 12.13520857 | 2 | 2 | 2 | 6.197166296 | 6.1572134 | 6.057834398 | 6.283256815 |
| ENSG00000197808 | ZNF461 | 6.090594234 | 5.11E-05 | 6.016336933 | 5.89E-05 | 12.10693117 | 2 | 2 | 2 | 6.24507773 | 6.14893615 | 6.380048704 | 5.821002687 |
| ENSG00000140961 | OSGIN1 | 8.032480815 | 1.60E-08 | 4.057818861 | 0.007388815 | 12.09029968 | 2 | 2 | 2 | 8.056427969 | 8.10384772 | 4.644324387 | 4.077147165 |
| ENSG00000083857 | FAT1 | 5.414341658 | 6.94E-05 | 6.675394276 | 5.39E-07 | 12.08973593 | 6.366709671 | 7.60566235 | 2 | 11.91575529 | 11.9163384 | 13.25854844 | 13.08889818 |
| ENSG00000136514 | RTP4 | 8.966750681 | 2.45E-11 | 3.116318349 | 0.035697933 | 12.08306903 | 2 | 2 | 2 | 9.010169232 | 8.99861459 | 3.56816602 | 3.789585079 |
| ENSG00000197774 | EME2 | 4.722961074 | 0.001259444 | 7.353936805 | 1.33E-07 | 12.07689788 | 2 | 2 | 2 | 4.965943888 | 4.93258395 | 7.347386207 | 7.478755384 |
| ENSG00000155629 | PIK3AP1 | 5.642825884 | 0.00184965 | 6.40584129 | 0.000313297 | 12.04866717 | 2 | 2 | 2 | 5.639097847 | 5.90535527 | 7.143982415 | 5.294527297 |
| ENSG00000159733 | ZFYVE28 | 5.610629062 | 5.96E-05 | 6.416245672 | 3.07E-06 | 12.02687473 | 2 | 2 | 3.369317972 | 6.716553866 | 6.59518404 | 7.538302335 | 7.330190599 |
| ENSG00000221963 | APOL6 | 7.050143105 | 2.47E-29 | 4.973034014 | 3.23E-15 | 12.02317712 | 3.327711234 | 4.93638933 | 5.59507254 | 11.73479622 | 11.7190443 | 9.856639061 | 9.421147165 |
| ENSG00000007202 | KIAA0100 | 5.624140788 | 9.94E-10 | 6.383756907 | 2.44E-12 | 12.00789769 | 6.366709671 | 4.442710129 | 3.369317972 | 10.67878272 | 10.7365497 | 11.17993582 | 11.70488663 |
| ENSG00000180233 | ZNRF2 | 6.417248447 | 1.13E-05 | 5.574599335 | 0.000155741 | 11.99184778 | 2 | 2 | 2 | 6.490083648 | 6.52655952 | 5.866661379 | 5.545463418 |
| ENSG00000169213 | RAB3B | 4.31754499 | 5.36E-10 | 7.648338629 | 1.31E-29 | 11.96588362 | 5.330594134 | 4.93638933 | 6.882775709 | 10.15791688 | 10.1853333 | 13.48511211 | 13.51071303 |
| ENSG00000065320 | NTN1 | 6.561455193 | 1.47E-05 | 5.356494604 | 0.000488129 | 11.9179498 | 2 | 2 | 2 | 6.907295206 | 6.32824132 | 5.304163849 | 5.699242824 |
| ENSG00000031081 | ARHGAP31 | 5.639190288 | 7.67E-14 | 6.273123309 | 4.56E-17 | 11.9123136 | 2 | 4.442710129 | 4.058998049 | 9.021676912 | 9.14105132 | 9.577337433 | 9.836576475 |
| ENSG00000262406 | MMP12 | 5.869460779 | 0.046404119 | 5.991000344 | 0.039274911 | 11.86046112 | 2 | 2 | 2 | 6.01571348 | 5.96314905 | 7.05561307 | 2.172608095 |
| ENSG00000125657 | TNFSF9 | 6.467791727 | 3.55E-05 | 5.390463432 | 0.00067229 | 11.85825516 | 2 | 2 | 2 | 6.604700657 | 6.507263 | 5.038665118 | 5.921228818 |
| ENSG00000147119 | CHST7 | 5.32867275 | 1.15E-05 | 6.528027858 | 4.13E-08 | 11.85670061 | 2 | 3.686303414 | 2 | 7.157295126 | 7.26914476 | 8.306273771 | 8.472158355 |
| ENSG00000112096 | SOD2 | 8.394878984 | 1.42E-75 | 3.44999721 | 1.85E-13 | 11.84487619 | 7.904282243 | 6.844429755 | 7.934808822 | 15.99212855 | 15.9937102 | 10.91507154 | 11.17462754 |
| ENSG00000182890 | GLUD2 | 5.243031407 | 2.03E-07 | 6.601777187 | 2.78E-11 | 11.84480859 | 2 | 4.93638933 | 2 | 8.62888906 | 8.57927024 | 9.823356991 | 10.07372892 |
| ENSG00000140379 | BCL2A1 | 8.159361065 | 1.89E-05 | 3.682552932 | 0.075024979 | 11.841914 | 2 | 2 | 2 | 8.258639605 | 8.15020018 | 4.785723309 | 2.70970617 |
| ENSG00000188042 | ARL4C | 5.355503976 | 0.000225083 | 6.475041275 | 5.29E-06 | 11.83054525 | 2 | 5.72254323 | 2 | 9.477763976 | 9.46156046 | 10.55547888 | 10.61355952 |
| ENSG00000111424 | VDR | 6.151853212 | 2.67E-07 | 5.6693141 | 2.29E-06 | 11.82116731 | 3.327711234 | 2 | 3.369317972 | 8.083034979 | 8.02463101 | 7.749302529 | 7.388257299 |
| ENSG00000168060 | NAALADL1 | 4.974391091 | 0.000777117 | 6.83916125 | 1.61E-06 | 11.81355234 | 2 | 2 | 2 | 5.093559143 | 5.24368216 | 6.910373854 | 6.91798856 |
| ENSG00000185633 | NDUFA4L2 | 0.402374303 | 0.811969254 | 11.40814573 | 2.47E-21 | 11.81052003 | 2 | 3.686303414 | 2 | 3.289987342 | 2.94009902 | 12.97484098 | 13.47570932 |
| ENSG00000041982 | TNC | 6.969284358 | 7.46E-19 | 4.829619972 | 1.29E-09 | 11.79890433 | 2 | 4.710564154 | 3.369317972 | 10.50929771 | 10.2510948 | 8.181164698 | 8.336553984 |
| ENSG00000129355 | CDKN2D | 5.345610914 | 0.000145175 | 6.445519183 | 2.87E-06 | 11.7911301 | 2 | 3.076665209 | 2 | 6.328984825 | 6.47451726 | 7.559188434 | 7.36768071 |
| ENSG00000091039 | OSBPL8 | 6.610814149 | 2.20E-06 | 5.165665385 | 0.000283328 | 11.77647953 | 3.327711234 | 6.464272742 | 2 | 11.58189337 | 11.5601677 | 10.36193351 | 9.851472775 |
| ENSG00000181754 | AMIGO1 | 3.418024553 | 0.024824384 | 8.358316443 | 1.18E-09 | 11.776341 | 2 | 2 | 2 | 3.997744386 | 3.7864959 | 8.52588267 | 8.266036253 |
| ENSG00000136630 | HLX | 6.608617742 | 2.54E-06 | 5.155961167 | 0.000317146 | 11.76457891 | 2 | 3.076665209 | 2 | 7.602596956 | 7.64959655 | 6.453284622 | 5.949196291 |
| ENSG00000112619 | PRPH2 | 5.369075304 | 0.000416219 | 6.393057686 | 1.70E-05 | 11.76213299 | 2 | 2 | 2 | 5.552894743 | 5.49945122 | 6.712030998 | 6.215784511 |
| ENSG00000072121 | ZFYVE26 | 6.32981128 | 2.01E-07 | 5.415057565 | 1.06E-05 | 11.74486885 | 2 | 5.303523747 | 2 | 9.999333534 | 10.0596091 | 9.253905612 | 8.972144811 |
| ENSG00000168811 | IL12A | 8.541827338 | 2.32E-10 | 3.200113698 | 0.030786735 | 11.74194104 | 2 | 2 | 2 | 8.559529208 | 8.60672783 | 3.796405078 | 3.67940853 |
| ENSG00000172731 | LRRC20 | 5.038066911 | 0.000702835 | 6.697128987 | 3.18E-06 | 11.7351959 | 2 | 2 | 2 | 5.210797711 | 5.24368216 | 6.681876084 | 6.866539459 |
| ENSG00000128881 | TTBK2 | 5.318857644 | 7.78E-07 | 6.415534404 | 1.39E-09 | 11.73439205 | 2 | 4.11352228 | 2 | 7.691737974 | 7.81828345 | 9.08380094 | 8.543347212 |
| ENSG00000142798 | HSPG2 | 5.113012101 | 9.45E-11 | 6.613096789 | 1.81E-17 | 11.72610889 | 6.653098118 | 6.464272742 | 8.536706852 | 12.57305446 | 12.6628552 | 13.97758897 | 14.24620657 |
| ENSG00000149289 | ZC3H12C | 7.398496028 | 3.34E-10 | 4.318685772 | 0.000425805 | 11.7171818 | 2 | 5.303523747 | 2 | 11.13115783 | 11.0670971 | 7.971238879 | 8.103992141 |
| ENSG00000123643 | SLC36A1 | 5.027946002 | 2.54E-05 | 6.677937314 | 9.44E-09 | 11.70588332 | 2 | 4.93638933 | 2 | 8.291333947 | 8.42500614 | 9.41724949 | 10.4104801 |
| ENSG00000069667 | RORA | 6.519337782 | 4.17E-12 | 5.181331258 | 4.92E-08 | 11.70066904 | 4.467311984 | 3.076665209 | 2 | 9.437090644 | 9.36838404 | 8.362013331 | 7.72395257 |
| ENSG00000184378 | ACTRT3 | 5.607224019 | 0.000333419 | 6.077974753 | 8.07E-05 | 11.68519877 | 2 | 2 | 2 | 5.552894743 | 5.91514978 | 5.843010798 | 6.463051659 |
| ENSG00000154589 | LY96 | 7.494959764 | 1.28E-07 | 4.189544496 | 0.005165222 | 11.68450426 | 2 | 2 | 2 | 7.514115926 | 7.58969985 | 4.675266036 | 4.292061816 |
| ENSG00000133657 | ATP13A3 | 6.638815729 | 0.001939481 | 5.032820634 | 0.022450111 | 11.67163636 | 2 | 7.790474954 | 2 | 12.84507664 | 12.8140605 | 11.11026486 | 11.3317026 |
| ENSG00000184985 | SORCS2 | 7.000241637 | 2.39E-07 | 4.665566867 | 0.000873364 | 11.6658085 | 2 | 2 | 3.369317972 | 8.006336225 | 8.01555453 | 5.790986459 | 5.736403255 |
| ENSG00000249471 | ZNF324B | 5.787859202 | 0.000137041 | 5.874166648 | 9.77E-05 | 11.66202585 | 2 | 2 | 2 | 5.978405189 | 5.84514933 | 6.205571486 | 5.745545708 |
| ENSG00000135801 | TAF5L | 5.546802302 | 4.28E-08 | 6.106829549 | 1.14E-09 | 11.65363185 | 2 | 4.11352228 | 2 | 7.997038782 | 7.98797735 | 8.600320517 | 8.487906181 |
| ENSG00000185634 | SHC4 | 4.828600826 | 0.000552035 | 6.807705503 | 4.41E-07 | 11.63630633 | 3.327711234 | 3.076665209 | 2 | 6.699459234 | 6.77805036 | 9.231960665 | 7.764992659 |
| ENSG00000168453 | HR | 5.541331347 | 0.000273049 | 6.075500064 | 5.15E-05 | 11.61683141 | 2 | 2 | 2 | 5.709096143 | 5.65950892 | 5.932173402 | 6.397863414 |
| ENSG00000175600 | SUGCT | 3.935714364 | 0.008033366 | 7.670483561 | 2.78E-08 | 11.60619793 | 2 | 3.076665209 | 2 | 4.846787657 | 5.3193943 | 8.843612414 | 8.479026367 |
| ENSG00000163659 | TIPARP | 6.356305336 | 1.68E-10 | 5.249055358 | 1.74E-07 | 11.60536069 | 2 | 6.228547981 | 5.59507254 | 11.63493198 | 11.6660124 | 10.62572272 | 10.46026599 |
| ENSG00000134259 | NGF | 2.082487458 | 0.200921344 | 9.518357362 | 1.60E-12 | 11.60084482 | 2 | 2 | 2 | 3.162721474 | 2.94009902 | 9.51429464 | 9.589581388 |
| ENSG00000142871 | CYR61 | 4.198399171 | 0.021458981 | 7.400862084 | 1.48E-05 | 11.59926125 | 8.653385977 | 5.131603625 | 2 | 11.32078887 | 11.399377 | 14.67186756 | 14.44113699 |
| ENSG00000106819 | ASPN | 1.777301931 | 0.335944632 | 9.817608792 | 4.92E-11 | 11.59491072 | 2 | 2 | 2 | 2.947924294 | 2.86123014 | 9.01856474 | 10.37446816 |
| ENSG00000142583 | SLC2A5 | 6.068297715 | 0.000247921 | 5.52293584 | 0.000909649 | 11.59123356 | 2 | 2 | 2 | 6.087554739 | 6.26070706 | 6.129409568 | 4.982847849 |
| ENSG00000161217 | PCYT1A | 5.013681978 | 0.000661823 | 6.558661576 | 4.50E-06 | 11.57234355 | 2 | 5.72254323 | 2 | 9.144656522 | 9.11366683 | 10.48546314 | 10.828989 |
| ENSG00000115232 | ITGA4 | 6.989554019 | 1.44E-05 | 4.582246683 | 0.006305222 | 11.5718007 | 2 | 2 | 2 | 7.210554257 | 6.89154576 | 5.274160001 | 4.175132639 |
| ENSG00000173391 | OLR1 | 2.777319919 | 0.088103631 | 8.75991591 | 5.42E-10 | 11.53723583 | 2 | 2 | 2 | 3.515101962 | 3.39680241 | 8.403026507 | 9.109931338 |
| ENSG00000151835 | SACS | 4.978994169 | 0.000535791 | 6.556551024 | 2.65E-06 | 11.53554519 | 2 | 6.464272742 | 3.369317972 | 10.00688028 | 9.88372429 | 11.74214602 | 11.25794964 |
| ENSG00000072840 | EVC | 6.24153683 | 0.000353647 | 5.283515407 | 0.002856714 | 11.52505224 | 2 | 6.464272742 | 2 | 11.02652273 | 11.1590942 | 10.09769543 | 10.17870477 |
| ENSG00000197312 | DDI2 | 5.403246559 | 2.69E-07 | 6.116398647 | 3.84E-09 | 11.51964521 | 2 | 4.11352228 | 2 | 7.783822874 | 7.90193448 | 8.72431761 | 8.345600992 |
| ENSG00000258102 | MAP1LC3B2 | 6.227748798 | 2.33E-18 | 5.291710068 | 1.18E-13 | 11.51945887 | 4.00722531 | 3.686303414 | 4.058998049 | 9.708880707 | 9.67042697 | 8.660996377 | 8.852955181 |
| ENSG00000078018 | MAP2 | 3.57829697 | 0.005204952 | 7.902479106 | 2.95E-11 | 11.48077608 | 4.467311984 | 2 | 2 | 6.598566398 | 5.51241389 | 10.58976003 | 10.18249657 |
| ENSG00000115252 | PDE1A | 2.859811552 | 0.114510897 | 8.619998221 | 4.60E-08 | 11.47980977 | 2 | 2 | 2 | 3.566290607 | 3.45167086 | 7.625052652 | 9.257431793 |
| ENSG00000176749 | CDK5R1 | 5.418226877 | 0.001152927 | 6.049327564 | 0.000220783 | 11.46755444 | 2 | 2 | 2 | 5.709096143 | 5.41912973 | 6.597565628 | 5.524371359 |
| ENSG00000114270 | COL7A1 | 6.831084582 | 7.08E-14 | 4.618574946 | 6.95E-07 | 11.44965953 | 6.89194353 | 3.686303414 | 5.772716771 | 12.65338764 | 12.6844922 | 10.44170614 | 10.47751587 |
| ENSG00000130052 | STARD8 | 5.739298207 | 0.000121932 | 5.709484055 | 0.00012244 | 11.44878226 | 2 | 2 | 2 | 5.870535235 | 5.86549842 | 5.912830251 | 5.76365862 |
| ENSG00000106823 | ECM2 | 2.535488634 | 0.101148097 | 8.902406661 | 2.87E-11 | 11.4378953 | 2 | 2 | 3.369317972 | 4.034562184 | 3.86943969 | 9.691228581 | 10.07644859 |
| ENSG00000211448 | DIO2 | 4.804403686 | 0.005436694 | 6.629671065 | 6.66E-05 | 11.43407475 | 2 | 2 | 6.073378139 | 9.184108452 | 9.17507637 | 11.02124505 | 10.97374872 |
| ENSG00000164105 | SAP30 | 5.399408467 | 0.000203086 | 6.013155197 | 2.66E-05 | 11.41256366 | 2 | 3.076665209 | 2 | 6.336376195 | 6.56439512 | 6.756113529 | 7.287011591 |
| ENSG00000175592 | FOSL1 | 6.732406483 | 9.83E-16 | 4.677196923 | 3.92E-08 | 11.40960341 | 2 | 4.11352228 | 5.59507254 | 10.92600118 | 10.9892592 | 8.766685705 | 9.043840859 |
| ENSG00000138311 | ZNF365 | 5.39037471 | 0.000453925 | 6.015568296 | 6.84E-05 | 11.40594301 | 2 | 2 | 2 | 5.627094009 | 5.45984938 | 5.899789217 | 6.323497254 |
| ENSG00000134775 | FHOD3 | 7.296428699 | 3.96E-14 | 4.102524968 | 4.03E-05 | 11.39895367 | 3.327711234 | 3.686303414 | 2 | 9.701744597 | 9.78435771 | 6.516588833 | 6.685996416 |
| ENSG00000130751 | NPAS1 | 3.984923144 | 0.009315299 | 7.402720929 | 2.22E-07 | 11.38764407 | 2 | 2 | 3.369317972 | 5.128039203 | 5.16377554 | 7.8927823 | 8.786963584 |
| ENSG00000197044 | ZNF441 | 5.521491071 | 0.000129797 | 5.857900152 | 4.07E-05 | 11.37939122 | 2 | 3.076665209 | 2 | 6.470063213 | 6.66650236 | 7.111828383 | 6.640162436 |
| ENSG00000134668 | SPOCD1 | 5.893497214 | 4.80E-09 | 5.473435492 | 5.69E-08 | 11.36693271 | 2 | 4.93638933 | 2 | 9.201414645 | 9.29338856 | 8.883077364 | 8.777523662 |
| ENSG00000120899 | PTK2B | 5.451806831 | 0.000114314 | 5.890373966 | 2.44E-05 | 11.3421808 | 2 | 3.076665209 | 2 | 6.442929658 | 6.56439512 | 7.001702955 | 6.847559257 |
| ENSG00000137819 | PAQR5 | 4.987391995 | 6.56E-06 | 6.351352782 | 4.44E-09 | 11.33874478 | 3.327711234 | 3.686303414 | 2 | 7.461007055 | 7.43830714 | 8.262372054 | 9.180729263 |
| ENSG00000101160 | CTSZ | 6.21095136 | 4.41E-08 | 5.114649462 | 8.49E-06 | 11.32560082 | 2 | 5.946638857 | 6.718950348 | 11.91945108 | 11.9583902 | 10.99110861 | 10.68108075 |
| ENSG00000130038 | CRACR2A | 5.48339505 | 0.000418165 | 5.789477204 | 0.000164092 | 11.27287225 | 2 | 2 | 2 | 5.552894743 | 5.70538658 | 5.646229369 | 6.141542327 |
| ENSG00000205683 | DPF3 | 6.771568117 | 2.49E-06 | 4.499045386 | 0.002591993 | 11.2706135 | 2 | 2 | 2 | 6.846474326 | 6.85130017 | 4.799832052 | 4.711446447 |

| | | | | | | | | | | | | | |
|---|---|---|---|---|---|---|---|---|---|---|---|---|---|
| ENSG00000166920 | C15orf48 | 11.5690917 | 2.32E-08 | -0.305551318 | 0.915553996 | 11.26354038 | 2 | 2 | 2 | 11.58790527 | 11.6068075 | 2.492221014 | 2 |
| ENSG00000127824 | TUBA4A | 4.667757694 | 2.40E-05 | 6.588488736 | 8.15E-10 | 11.25624643 | 4.815609471 | 2 | 2 | 7.602596956 | 7.61995902 | 9.190394041 | 9.771772716 |
| ENSG00000065308 | TRAM2 | 4.468777569 | 0.000742187 | 6.780669714 | 9.65E-08 | 11.24944728 | 2 | 6.899080544 | 7.163363831 | 10.87457316 | 10.8896987 | 12.90512493 | 13.43049852 |
| ENSG00000196850 | PPTC7 | 5.430571601 | 7.44E-09 | 5.809834551 | 4.70E-10 | 11.24040615 | 2 | 4.442710129 | 2 | 8.291333947 | 8.25031674 | 8.559959383 | 8.727068782 |
| ENSG00000172935 | MRGPRF | 4.355506542 | 0.000620294 | 6.883355449 | 1.59E-08 | 11.23886199 | 2 | 5.131603625 | 2 | 7.883054517 | 7.90193448 | 9.852414792 | 10.79623028 |
| ENSG00000168283 | BMI1 | 6.447989679 | 1.32E-08 | 4.775339081 | 3.68E-05 | 11.22332876 | 2 | 4.11352228 | 2 | 8.869506572 | 8.84440862 | 7.578754018 | 6.716538017 |
| ENSG00000099860 | GADD45B | 4.599562783 | 1.24E-05 | 6.600671675 | 1.05E-10 | 11.20023446 | 4.00722531 | 5.303523747 | 2 | 8.660150093 | 8.71862106 | 11.2093021 | 9.836041609 |
| ENSG00000125843 | AP5S1 | 4.140193442 | 0.000887822 | 7.029744923 | 2.84E-09 | 11.16993837 | 2 | 3.686303414 | 2 | 5.978405189 | 6.16544343 | 8.912592153 | 8.864016001 |
| ENSG00000163874 | ZC3H12A | 7.229784796 | 3.12E-17 | 3.91417973 | 9.70E-06 | 11.14396453 | 4.467311984 | 4.710564154 | 2 | 11.04665822 | 11.0621467 | 7.139828829 | 8.19690135 |
| ENSG00000187583 | PLEKHN1 | 7.048572396 | 4.58E-06 | 4.0875162 | 0.01187557 | 11.1360886 | 2 | 2 | 2 | 7.007926688 | 7.21862736 | 4.749835737 | 3.972018587 |
| ENSG00000171951 | SCG2 | 6.890158699 | 4.27E-07 | 4.226221142 | 0.002992839 | 11.11637984 | 2 | 3.076665209 | 2 | 7.925329813 | 7.87959733 | 5.385935071 | 5.337485258 |
| ENSG00000077157 | PPP1R12B | 2.708821257 | 1.51E-06 | 8.386220862 | 3.19E-56 | 11.09504212 | 4.815609471 | 5.131603625 | 4.058998049 | 7.288834728 | 7.30683517 | 13.04393834 | 12.82549673 |
| ENSG00000171992 | SYNPO | 5.059931338 | 3.98E-07 | 6.024697421 | 9.02E-10 | 11.08462876 | 2 | 3.076665209 | 5.39244166 | 8.845101141 | 8.78422202 | 9.7566591 | 9.790113704 |
| ENSG00000166592 | RRAD | 5.935758823 | 3.92E-07 | 5.132692664 | 1.37E-05 | 11.06845149 | 2 | 4.11352228 | 2 | 8.369067528 | 8.32017198 | 7.953205078 | 7.0024032 |
| ENSG00000182621 | PLCB1 | 5.139523725 | 3.67E-06 | 5.925898384 | 6.08E-08 | 11.06542211 | 2 | 4.442710129 | 2 | 7.982979299 | 7.91663606 | 8.215119669 | 9.103723527 |
| ENSG00000188559 | RALGAPA2 | 4.914510747 | 5.43E-05 | 6.124139232 | 2.61E-07 | 11.03864998 | 3.327711234 | 3.076665209 | 2 | 6.766655021 | 6.88657603 | 8.100109146 | 7.910280955 |
| ENSG00000109743 | BST1 | 4.5979203 | 1.77E-09 | 6.416496775 | 9.68E-18 | 11.01441707 | 4.815609471 | 3.686303414 | 2 | 8.087422161 | 8.05816814 | 9.959986891 | 9.786796275 |
| ENSG00000152601 | MBNL1 | 5.279630877 | 0.01073585 | 5.734295167 | 0.004914121 | 11.01392604 | 2 | 7.315436877 | 2 | 11.04468547 | 10.9391381 | 11.53041218 | 11.35792713 |
| ENSG00000100918 | REC8 | 6.376095146 | 1.69E-05 | 4.635221512 | 0.002359129 | 11.01131666 | 2 | 2 | 2 | 6.496695859 | 6.44101097 | 4.764298067 | 4.975095763 |
| ENSG00000160201 | U2AF1 | 8.547838207 | 5.10E-09 | 2.447316431 | 0.139938782 | 10.99515464 | 2 | 2 | 2 | 8.663092307 | 8.51148098 | 2.911488689 | 3.539239711 |
| ENSG00000198945 | L3MBTL3 | 6.110562738 | 5.39E-07 | 4.877909298 | 8.21E-05 | 10.98847204 | 2 | 3.686303414 | 2 | 7.847284664 | 8.09525763 | 6.838725184 | 6.707209278 |
| ENSG00000138650 | PCDH10 | 5.944544487 | 2.04E-06 | 5.040740583 | 6.79E-05 | 10.98528507 | 3.327711234 | 3.076665209 | 2 | 7.753776273 | 7.90193448 | 7.240241221 | 6.580104496 |
| ENSG00000177576 | C18orf32 | 7.235385817 | 5.81E-10 | 3.745791518 | 0.002391435 | 10.98117733 | 2 | 3.076665209 | 3.369317972 | 9.162439546 | 9.05507424 | 5.825876956 | 5.611557759 |
| ENSG00000130590 | SAMD10 | 4.471117014 | 0.00360969 | 6.50686092 | 9.12E-06 | 10.97797793 | 2 | 2 | 2 | 4.783303513 | 4.68029874 | 6.654963382 | 6.530719415 |
| ENSG00000114529 | C3orf52 | 6.089069341 | 1.40E-05 | 4.881478659 | 0.000611597 | 10.970548 | 3.327711234 | 2 | 2 | 7.026323566 | 7.20671394 | 5.925754458 | 6.003555476 |
| ENSG00000132000 | PODNL1 | 7.389895539 | 6.13E-07 | 3.557958006 | 0.025915975 | 10.94785355 | 2 | 2 | 2 | 7.454228844 | 7.4450836 | 4.271694079 | 3.660199087 |
| ENSG00000119689 | DLST | 5.263637208 | 0.001152409 | 5.67851475 | 0.000383457 | 10.94215196 | 2 | 6.04685525 | 2 | 9.715981692 | 9.68338024 | 10.16515422 | 10.05867839 |
| ENSG00000163864 | NMNAT3 | 5.601874889 | 0.000253258 | 5.328465976 | 0.000499008 | 10.93034087 | 2 | 2 | 2 | 5.662809983 | 5.81407719 | 5.565950232 | 5.408276315 |
| ENSG00000182575 | NXPH3 | 4.678224501 | 0.013191227 | 6.25057209 | 0.000556851 | 10.92879659 | 2 | 2 | 2 | 4.927308402 | 4.89348248 | 6.983275763 | 5.191251321 |
| ENSG00000253626 | EIF5AL1 | 6.246091011 | 2.39E-19 | 4.6806931 | 2.27E-11 | 10.92678411 | 5.095959458 | 4.11352228 | 4.058998049 | 10.41383446 | 10.4881343 | 8.464101787 | 9.225115665 |
| ENSG00000155980 | KIF5A | 5.002609039 | 0.001272804 | 5.920780405 | 9.29E-05 | 10.92338944 | 2 | 2 | 2 | 5.242616075 | 5.14724905 | 6.162543904 | 5.900914815 |
| ENSG00000198794 | SCAMP5 | 5.298754233 | 0.000334643 | 5.621954092 | 0.000119053 | 10.92070832 | 2 | 3.076665209 | 2 | 6.5612028 | 6.12381559 | 6.423979439 | 6.876977108 |
| ENSG00000187815 | ZFP69 | 5.03835279 | 0.001598865 | 5.881915897 | 0.000161735 | 10.92026869 | 2 | 2 | 2 | 5.258265913 | 5.19627118 | 6.260165241 | 5.685057129 |
| ENSG00000067066 | SP100 | 6.66859587 | 1.72E-24 | 4.248512842 | 1.36E-10 | 10.91710871 | 4.467311984 | 4.93638933 | 6.43197674 | 12.04374227 | 12.0377259 | 9.546552631 | 9.702454181 |
| ENSG00000196878 | LAMB3 | 6.012780149 | 6.19E-09 | 4.862284359 | 3.48E-06 | 10.87506451 | 4.815609471 | 2 | 2 | 8.908460392 | 8.95655043 | 7.930140711 | 7.649197382 |
| ENSG00000276497 | | 5.407280655 | 0.00015131 | 5.466262816 | 0.000116623 | 10.87354347 | 2 | 3.076665209 | 2 | 6.490083648 | 6.43421531 | 6.412547789 | 6.617932662 |
| ENSG00000140451 | PIF1 | 5.157218495 | 0.000824293 | 5.707983961 | 0.000165601 | 10.86520246 | 2 | 2 | 2 | 5.363304456 | 5.30456641 | 5.805044075 | 5.871985934 |
| ENSG00000172613 | RAD9A | 5.410350889 | 0.000176723 | 5.451990966 | 0.000144706 | 10.86234186 | 2 | 3.076665209 | 2 | 6.415275989 | 6.51372389 | 6.678062013 | 6.308152703 |
| ENSG00000184226 | PCDH9 | 7.531614929 | 2.14E-07 | 3.326259784 | 0.035010376 | 10.85787471 | 2 | 2 | 2 | 7.559034656 | 7.61696151 | 4.053534413 | 3.560117069 |
| ENSG00000108370 | RGS9 | 2.642628143 | 0.113003183 | 8.205551545 | 1.24E-08 | 10.84817969 | 2 | 2 | 2 | 3.227757282 | 3.50452879 | 8.542759496 | 7.884647945 |
| ENSG00000137218 | FRS3 | 5.326381895 | 0.000623727 | 5.513297654 | 0.000346886 | 10.83967955 | 2 | 2 | 2 | 5.540148523 | 5.43283104 | 5.794513725 | 5.508347454 |
| ENSG00000185862 | EVI2B | 6.803323177 | 0.000344238 | 4.023798155 | 0.044848802 | 10.82712133 | 2 | 2 | 2 | 6.907295206 | 6.85130017 | 5.008609438 | 3.15612297 |
| ENSG00000151490 | PTPRO | 5.509235681 | 0.000456779 | 5.307647886 | 0.000715544 | 10.81688357 | 2 | 2 | 2 | 5.590471342 | 5.71663161 | 5.238348159 | 5.670730563 |
| ENSG00000133083 | DCLK1 | 3.626557482 | 0.098953314 | 7.182976432 | 0.000326774 | 10.80953391 | 2 | 2 | 2 | 4.739373666 | 2.68924543 | 7.009312024 | 7.452476174 |
| ENSG00000169604 | ANTXR1 | 5.662394594 | 1.09E-08 | 5.141394561 | 2.33E-07 | 10.80378915 | 4.00722531 | 7.669933291 | 6.882775709 | 12.43151429 | 12.4430008 | 11.82747678 | 12.00074925 |
| ENSG00000121898 | CPXM2 | 10.33316856 | 2.69E-14 | 0.465586146 | 0.800845428 | 10.79875471 | 2 | 2 | 2 | 10.34359113 | 10.3839861 | 2.526562374 | 2.326754077 |
| ENSG00000166073 | GPR176 | 5.138532416 | 9.51E-06 | 5.658593993 | 8.09E-07 | 10.79712641 | 4.467311984 | 7.051611315 | 3.369317972 | 10.80768086 | 10.8906298 | 11.31170051 | 11.42383583 |
| ENSG00000010818 | HIVEP2 | 5.744685245 | 1.93E-10 | 5.050587506 | 2.58E-08 | 10.79527275 | 3.327711234 | 5.303523747 | 6.629488387 | 11.23490719 | 11.2465449 | 10.3054184 | 10.75579885 |
| ENSG00000075223 | SEMA3C | 6.456987294 | 0.00021155 | 4.331481463 | 0.017146254 | 10.78846876 | 2 | 6.464272742 | 2 | 11.34457954 | 11.2732076 | 9.287461761 | 9.088535264 |
| ENSG00000138347 | MYPN | 7.844585263 | 1.40E-11 | 2.942135878 | 0.02039551 | 10.78672114 | 3.327711234 | 3.076665209 | 2 | 9.743346323 | 9.68338024 | 5.119635168 | 4.861951484 |
| ENSG00000104951 | IL4I1 | 9.046143397 | 0.000978424 | 1.736889595 | 0.591567059 | 10.78303299 | 2 | 2 | 2 | 9.046675269 | 9.1189736 | 3.429858444 | 2 |
| ENSG00000262045 | | 7.641282711 | 5.12E-08 | 3.139146295 | 0.04071117 | 10.78042901 | 2 | 2 | 2 | 7.791909949 | 7.59275449 | 3.600766115 | 3.789585079 |
| ENSG00000185652 | NTF3 | 4.678226271 | 0.00284647 | 6.101190341 | 5.47E-05 | 10.77941661 | 2 | 2 | 2 | 4.927308402 | 4.89348248 | 6.353989995 | 6.044874254 |
| ENSG00000058085 | LAMC2 | 4.544613792 | 3.15E-08 | 6.223498924 | 9.55E-15 | 10.76811272 | 4.00722531 | 3.076665209 | 4.523675391 | 8.045195239 | 8.04484669 | 10.0231736 | 9.306133598 |
| ENSG00000157625 | TAB3 | 6.014572741 | 4.08E-14 | 4.751931877 | 3.27E-09 | 10.76650462 | 3.327711234 | 4.93638933 | 2 | 9.621581886 | 9.65370449 | 8.474034275 | 8.290447859 |
| ENSG00000113361 | CDH6 | 4.106186497 | 4.45E-11 | 6.65904277 | 7.45E-28 | 10.76522927 | 4.00722531 | 4.442710129 | 4.874587708 | 8.334420183 | 8.29413645 | 11.04040399 | 10.63554893 |
| ENSG00000117016 | RIMS3 | 2.689912473 | 0.121631897 | 8.058535206 | 9.04E-08 | 10.74844768 | 2 | 2 | 2 | 3.462030052 | 3.33976443 | 8.550081477 | 7.460826701 |
| ENSG00000105339 | DENND3 | 4.314502192 | 7.88E-05 | 6.422020238 | 1.12E-09 | 10.73652243 | 2 | 4.11352228 | 2 | 6.604700657 | 6.93551934 | 8.659546439 | 9.011520488 |
| ENSG00000077092 | RARB | 4.408953842 | 3.98E-05 | 6.324182854 | 1.15E-09 | 10.7331367 | 2 | 4.11352228 | 2 | 6.79375511 | 6.94988431 | 8.915024678 | 8.569323645 |
| ENSG00000153162 | BMP6 | 3.600613829 | 0.005178798 | 7.114696114 | 3.18E-09 | 10.71530994 | 2 | 3.686303414 | 2 | 5.590471342 | 5.55061739 | 9.143105199 | 8.771941776 |
| ENSG00000115993 | TRAK2 | 5.715366676 | 0.000121908 | 4.993264582 | 0.000885779 | 10.70863126 | 3.327711234 | 6.602424674 | 2 | 10.75280717 | 10.8564076 | 9.622937045 | 10.43523469 |
| ENSG00000248546 | ANP32C | 5.236922051 | 0.000789893 | 5.467817922 | 0.000393634 | 10.70473997 | 2 | 2 | 2 | 5.501219927 | 5.30456641 | 5.665620077 | 5.566251551 |
| ENSG00000130775 | THEMIS2 | 7.521719479 | 5.21E-11 | 3.174431597 | 0.010215226 | 10.69615108 | 2 | 4.11352228 | 2 | 9.932626344 | 9.90722553 | 6.092633892 | 5.108633494 |
| ENSG00000204397 | CARD16 | 8.282130891 | 6.80E-09 | 2.412746179 | 0.13571439 | 10.69487707 | 2 | 2 | 2 | 8.369067528 | 8.28283292 | 3.44788957 | 2.981406154 |
| ENSG00000154319 | FAM167A | 10.45711719 | 1.87E-11 | 0.236654862 | 0.914220926 | 10.69377205 | 2 | 2 | 2 | 10.43808429 | 10.5353368 | 2.139668783 | 2.592996859 |
| ENSG00000162104 | ADCY9 | 4.089651108 | 0.002690684 | 6.584261618 | 3.86E-07 | 10.67391273 | 3.327711234 | 6.228547981 | 2 | 8.836004565 | 8.84952549 | 11.39000052 | 11.26114065 |
| ENSG00000163131 | CTSS | 7.028239332 | 1.76E-12 | 3.644790896 | 0.000478057 | 10.67303023 | 5.330594134 | 4.11352228 | 5.59507254 | 11.95904309 | 11.9815895 | 9.366015681 | 6.862343163 |
| ENSG00000121966 | CXCR4 | 6.977876896 | 0.000205988 | 3.692450517 | 0.065664664 | 10.67032741 | 4.00722531 | 2 | 2 | 8.880924505 | 8.96835847 | 6.601598251 | 3.287118176 |
| ENSG00000228049 | POLR2J2 | 9.434607059 | 7.76E-12 | 1.22892022 | 0.470226835 | 10.66352728 | 2 | 2 | 2 | 9.455875035 | 9.48229757 | 2.526562374 | 2.817676617 |
| ENSG00000240891 | PLCXD2 | 4.984350042 | 0.00141248 | 5.678353634 | 0.000203796 | 10.66270368 | 2 | 2 | 2 | 5.22679461 | 5.13053105 | 5.83618166 | 5.785984581 |
| ENSG00000198743 | SLC5A3 | 5.072280526 | 0.000195008 | 5.589684428 | 3.19E-05 | 10.66196495 | 3.327711234 | 6.844429755 | 3.369317972 | 10.47077592 | 10.4749506 | 11.51325565 | 10.15724173 |
| ENSG00000154188 | ANGPT1 | 6.192472058 | 1.19E-09 | 4.466411596 | 1.75E-05 | 10.65888365 | 2 | 4.11352228 | 2 | 8.700804887 | 8.54970888 | 6.892381078 | 6.975457902 |
| ENSG00000180357 | ZNF609 | 5.331380177 | 0.002435814 | 5.30724326 | 0.002454553 | 10.63862344 | 2 | 6.389896633 | 2 | 10.08355087 | 10.1385468 | 10.14836757 | 10.02340748 |
| ENSG00000215041 | NEURL4 | 5.415291939 | 8.41E-08 | 5.217980383 | 2.35E-07 | 10.63327232 | 2 | 3.076665209 | 4.058998049 | 7.877998516 | 7.91419618 | 7.838287442 | 7.552234734 |
| ENSG00000114251 | WNT5A | 7.284095001 | 1.17E-19 | 3.344094837 | 6.57E-05 | 10.62818984 | 4.815609471 | 7.393647515 | 6.203069917 | 13.75272914 | 13.7132496 | 10.06685284 | 9.470227873 |
| ENSG00000111961 | SASH1 | 5.128805429 | 3.22E-05 | 5.49766166 | 6.84E-06 | 10.62646709 | 3.327711234 | 7.431218269 | 4.874587708 | 11.19633074 | 11.23119 | 11.43094503 | 11.71902466 |
| ENSG00000065413 | ANKRD44 | 5.062708401 | 0.000553262 | 5.561732193 | 0.000116602 | 10.62444059 | 2 | 2 | 3.369317972 | 6.078767801 | 6.18985553 | 6.805896464 | 6.38329369 |
| ENSG00000162645 | GBP2 | 5.158161374 | 4.23E-05 | 5.465387913 | 1.20E-05 | 10.62354929 | 2 | 5.303523747 | 2 | 8.915879855 | 8.80675844 | 9.221844492 | 9.110373733 |
| ENSG00000124942 | AHNAK | 3.825010446 | 1.02E-08 | 6.782750644 | 1.47E-25 | 10.60776109 | 9.174465797 | 9.608085847 | 7.777116083 | 12.83116471 | 12.8825876 | 15.47033881 | 16.09161663 |
| ENSG00000143384 | MCL1 | 5.862753052 | 1.87E-06 | 4.738464244 | 0.000148669 | 10.6012173 | 3.327711234 | 7.761270542 | 8.206335842 | 13.28792539 | 13.2609349 | 12.25288089 | 12.04110352 |
| ENSG00000099260 | PALMD | 2.150635865 | 0.224142044 | 8.433407748 | 1.28E-08 | 10.58404361 | 2 | 2 | 2 | 3.289987342 | 2.86123014 | 8.822604079 | 8.018893446 |
| ENSG00000145777 | TSLP | 5.107930615 | 0.001199225 | 5.460767704 | 0.000443325 | 10.56869832 | 2 | 2 | 2 | 5.391962469 | 5.18011485 | 5.677130623 | 5.540219242 |
| ENSG00000103037 | SETD6 | 5.012885554 | 0.000565281 | 5.526829163 | 0.000112508 | 10.53971472 | 2 | 3.076665209 | 2 | 6.061031724 | 6.11534392 | 6.710164669 | 6.429408106 |
| ENSG00000121690 | DEPDC7 | 5.546975605 | 0.000748006 | 4.990322323 | 0.002581427 | 10.53729793 | 2 | 2 | 2 | 5.614989456 | 5.76075522 | 5.502666283 | 4.77533149 |
| ENSG00000115008 | IL1A | 9.72218958 | 4.79E-13 | 0.81360867 | 0.639891142 | 10.53579825 | 2 | 2 | 2 | 9.759931695 | 9.75056618 | 2.592885947 | 2.466008509 |
| ENSG00000159200 | RCAN1 | 4.563362809 | 0.03932955 | 5.971156161 | 0.005045064 | 10.53451897 | 2 | 7.669933291 | 2 | 10.61259734 | 10.6577951 | 11.96035313 | 12.11714577 |
| ENSG00000105516 | DBP | 3.975078927 | 0.016635932 | 6.555492183 | 2.44E-05 | 10.53057111 | 2 | 2 | 2 | 4.358765249 | 4.2865223 | 6.972416862 | 6.209190898 |
| ENSG00000144891 | AGTR1 | 7.204979756 | 3.73E-07 | 3.314137975 | 0.030925245 | 10.51911773 | 2 | 2 | 2 | 7.242377933 | 7.29563107 | 3.782262616 | 3.858622973 |
| ENSG00000268043 | NBPF12 | 8.502373814 | 4.99E-25 | 2.011430947 | 0.02816633 | 10.51380476 | 3.327711234 | 4.710564154 | 2 | 11.86826293 | 11.9424648 | 5.780352619 | 5.256659864 |
| ENSG00000129473 | BCL2L2 | 5.715984363 | 0.001807112 | 4.7965099 | 0.009956274 | 10.51249426 | 2 | 6.602424674 | 2 | 10.67550816 | 10.7410309 | 9.517766503 | 10.02270259 |
| ENSG00000101384 | JAG1 | 1.510630705 | 0.347244214 | 8.981894419 | 1.41E-11 | 10.49252512 | 2 | 5.595927442 | 2 | 5.775854763 | 5.43283104 | 13.1475794 | 12.75272692 |
| ENSG00000172780 | RAB43 | 7.793587208 | 9.95E-16 | 2.695294881 | 0.010790371 | 10.48888209 | 4.00722531 | 2 | 3.369317972 | 10.22067068 | 10.3093543 | 5.445480974 | 5.171036168 |
| ENSG00000147526 | TACC1 | 4.512495707 | 1.60E-05 | 5.972655136 | 4.82E-09 | 10.48515084 | 5.095959458 | 6.951736442 | 3.369317972 | 10.22615207 | 10.2859546 | 11.66698686 | 11.75858124 |
| ENSG00000164099 | PRSS12 | 5.149275579 | 0.000139918 | 5.331874574 | 7.13E-05 | 10.48115015 | 2 | 5.457120953 | 2 | 9.006698952 | 8.99630959 | 9.061649957 | 9.294500788 |
| ENSG00000170160 | CCDC144A | 4.699678514 | 0.003425349 | 5.760840738 | 0.000218364 | 10.46051925 | 2 | 2 | 2 | 4.76150579 | 5.07918227 | 5.979426183 | 5.79040861 |
| ENSG00000143643 | TTC13 | 5.108037382 | 4.02E-07 | 5.349614473 | 8.93E-08 | 10.45765185 | 2 | 3.076665209 | 4.058998049 | 7.552702688 | 7.63485389 | 7.745662833 | 7.912312056 |
| ENSG00000234444 | ZNF736 | 5.736159256 | 5.25E-05 | 4.711433848 | 0.001067759 | 10.4475931 | 3.327711234 | 2 | 2 | 6.79375511 | 6.76185814 | 5.83960027 | 5.772630506 |
| ENSG00000173227 | SYT12 | 3.643909142 | 0.020787524 | 6.792288365 | 3.02E-06 | 10.43619751 | 2 | 2 | 2 | 4.105493497 | 4.02226117 | 6.860764111 | 6.876977108 |
| ENSG00000272031 | ANKRD34A | 4.093117478 | 0.007427744 | 6.33901228 | 1.22E-05 | 10.43212976 | 3.327711234 | 2 | 2 | 5.093559143 | 5.37722731 | 7.018390248 | 7.640647 |
| ENSG00000248643 | RBM14-RBM | 8.582905017 | 3.95E-10 | 1.847069631 | 0.250975661 | 10.42997465 | 2 | 2 | 2 | 8.654247601 | 8.59306436 | 2.831336763 | 3.042026731 |
| ENSG00000145945 | FAM50B | 4.244436116 | 0.004301255 | 6.172427606 | 1.39E-05 | 10.41686372 | 2 | 3.076665209 | 2 | 5.552894743 | 5.18011485 | 7.125896536 | 7.271271876 |
| ENSG00000188167 | TMPPE | 5.081515641 | 0.001330054 | 5.334227366 | 0.000647675 | 10.41574301 | 2 | 2 | 2 | 5.22679461 | 5.30456641 | 5.413723492 | 5.571402115 |
| ENSG00000158296 | SLC13A3 | 6.114807307 | 0.000113586 | 4.268629392 | 0.009338878 | 10.3834367 | 2 | 2 | 2 | 6.061031724 | 6.36443784 | 4.806835018 | 4.26688425 |
| ENSG00000142627 | EPHA2 | 4.486411981 | 3.42E-09 | 5.893203321 | 2.62E-15 | 10.3796153 | 2 | 5.457120953 | 4.874587708 | 9.082280307 | 8.92660044 | 10.43748218 | 10.37446816 |
| ENSG00000110080 | ST3GAL4 | 5.402895735 | 1.92E-11 | 4.976310044 | 6.78E-10 | 10.37920578 | 3.327711234 | 4.93638933 | 2 | 8.99156386 | 9.06500642 | 8.660513226 | 8.549232824 |
| ENSG00000169245 | CXCL10 | 12.5294375 | 1.85E-20 | -2.158680165 | 0.253232241 | 10.37075733 | 2 | 2 | 2 | 12.57842157 | 12.5348018 | 2.048074405 | 2.117335275 |
| ENSG00000101307 | SIRPB1 | 7.444122588 | 1.20E-07 | 2.921102697 | 0.058832994 | 10.36522528 | 2 | 2 | 2 | 7.467753569 | 7.53675633 | 3.600766115 | 3.496556178 |

| | | | | | | | | | | | | | |
|---|---|---|---|---|---|---|---|---|---|---|---|---|---|
| ENSG00000110090 | CPT1A | 4.636314178 | 1.70E-14 | 5.72542173 | 8.87E-22 | 10.36173591 | 4.467311984 | 5.303523747 | 6.43197674 | 10.11933966 | 10.1196108 | 11.18909341 | 11.22237466 |
| ENSG00000039139 | DNAH5 | 4.430875509 | 0.005775587 | 5.923726453 | 0.000121103 | 10.35460196 | 2 | 2 | 2 | 4.694064037 | 4.70309173 | 5.883320383 | 6.182510897 |
| ENSG00000117594 | HSD11B1 | 9.863461927 | 3.60E-05 | 0.491027451 | 0.876760915 | 10.35448938 | 2 | 2 | 2 | 9.860122857 | 9.93095363 | 2.775316226 | 2 |
| ENSG00000169252 | ADRB2 | 5.516272079 | 0.000496545 | 4.83396541 | 0.002485754 | 10.35023749 | 2 | 2 | 2 | 5.709096143 | 5.6121243 | 5.196301326 | 4.878668439 |
| ENSG00000177311 | ZBTB38 | 5.379409309 | 0.002523465 | 4.960429568 | 0.00559508 | 10.33983888 | 3.327711234 | 7.60566235 | 2 | 11.42152442 | 11.445516 | 11.135169 | 10.88448866 |
| ENSG00000163249 | CCNYL1 | 4.826120981 | 1.56E-08 | 5.501395719 | 6.99E-11 | 10.3275167 | 4.467311984 | 3.686303414 | 2 | 7.990026167 | 8.07574085 | 8.450751194 | 8.913527044 |
| ENSG00000082482 | KCNK2 | 5.970952763 | 7.33E-07 | 4.356377122 | 0.000424437 | 10.32732989 | 3.327711234 | 3.076665209 | 2 | 7.872924734 | 7.84158259 | 6.157074092 | 6.418016952 |
| ENSG00000155367 | PPM1J | 4.469138829 | 0.00686426 | 5.857964304 | 0.000228036 | 10.32710313 | 2 | 2 | 2 | 4.443344104 | 4.97065356 | 6.063692919 | 5.888587685 |
| ENSG00000137502 | RAB30 | 4.834325478 | 0.003061431 | 5.47430682 | 0.000633362 | 10.3086323 | 2 | 5.946638857 | 2 | 9.20242622 | 9.13791797 | 10.13138331 | 9.387595483 |
| ENSG00000177675 | CD163L1 | 6.029376308 | 0.000128328 | 4.258573328 | 0.008973742 | 10.28794964 | 2 | 2 | 2 | 6.155987605 | 6.12381559 | 4.281782022 | 4.784231301 |
| ENSG00000141664 | ZCCHC2 | 5.86505253 | 2.65E-06 | 4.38838563 | 0.000597178 | 10.25343816 | 2 | 4.93638933 | 2 | 9.206465443 | 9.14937381 | 8.17644121 | 7.045606259 |
| ENSG00000114200 | BCHE | 6.490916412 | 1.26E-05 | 3.754267178 | 0.017060366 | 10.24518359 | 2 | 2 | 2 | 6.676346669 | 6.47451726 | 4.177580368 | 4.119958326 |
| ENSG00000145390 | USP53 | 4.461244088 | 0.008799219 | 5.780617737 | 0.000440009 | 10.24186183 | 2 | 6.140560021 | 2 | 8.957217624 | 9.03275482 | 10.14370816 | 10.45557294 |
| ENSG00000099999 | RNF215 | 4.901678349 | 3.79E-06 | 5.33999117 | 3.56E-07 | 10.24166952 | 2 | 4.11352228 | 2 | 7.234487582 | 7.46188679 | 7.786089373 | 7.77730333 |
| ENSG00000170276 | HSPB2 | 6.072701241 | 0.000102491 | 4.160038738 | 0.010418755 | 10.23273998 | 2 | 2 | 2 | 6.104969671 | 6.25300433 | 4.240999983 | 4.673629022 |
| ENSG00000149968 | MMP3 | 12.32614326 | 3.78E-16 | -2.105632411 | 0.304706002 | 10.22051085 | 2 | 2 | 2 | 12.34266693 | 12.3645268 | 2 | 2.172608095 |
| ENSG00000198113 | TOR4A | 5.980050313 | 1.43E-17 | 4.236441957 | 2.38E-09 | 10.21649227 | 3.327711234 | 3.686303414 | 4.874587708 | 9.675758322 | 9.66970392 | 8.05817395 | 7.823364542 |
| ENSG00000168386 | FILIP1L | 4.493304266 | 6.49E-08 | 5.715920155 | 2.61E-12 | 10.20922442 | 6.563806279 | 6.899080544 | 4.058998049 | 10.71821594 | 10.6891726 | 12.17379072 | 11.62268983 |
| ENSG00000162775 | RBM15 | 4.863005299 | 3.91E-06 | 5.335175921 | 3.01E-07 | 10.19818122 | 2 | 4.11352228 | 2 | 7.281195074 | 7.35080119 | 7.717139459 | 7.837328594 |
| ENSG00000143458 | GABPB2 | 4.399165724 | 0.000613773 | 5.798840646 | 3.16E-06 | 10.19800637 | 2 | 2 | 4.058998049 | 6.314087501 | 6.40670796 | 7.896069536 | 7.511280276 |
| ENSG00000212864 | RNF208 | 4.127521263 | 0.00581032 | 6.068101059 | 2.08E-05 | 10.19562232 | 2 | 3.076665209 | 2 | 5.093559143 | 5.43283104 | 7.007793418 | 7.184857568 |
| ENSG00000138411 | HECW2 | 3.956502369 | 0.000319037 | 6.238038698 | 3.29E-09 | 10.19454107 | 2 | 4.93638933 | 2 | 7.250225364 | 7.40045674 | 9.376938504 | 9.7580035 |
| ENSG00000198431 | TXNRD1 | 6.350192041 | 6.51E-17 | 3.836671686 | 8.19E-07 | 10.18686373 | 6.138360429 | 8.540087354 | 7.858115942 | 14.13292277 | 14.1571423 | 11.36011126 | 11.86249151 |
| ENSG00000261150 | EPPK1 | 4.433566182 | 0.001787884 | 5.75287232 | 2.83E-05 | 10.1864385 | 2 | 3.686303414 | 2 | 6.456560224 | 6.18176397 | 8.028526139 | 6.990916783 |
| ENSG00000155666 | KDM8 | 4.839337368 | 0.003057116 | 5.345974525 | 0.000860079 | 10.18531189 | 2 | 2 | 2 | 5.076005082 | 5.0259382 | 5.294231737 | 5.689801209 |
| ENSG00000166189 | HPS6 | 4.568852222 | 2.59E-05 | 5.614510767 | 1.29E-07 | 10.18336299 | 2 | 4.93638933 | 2 | 7.875463856 | 7.97632944 | 8.812201585 | 9.094361437 |
| ENSG00000134243 | SORT1 | 2.920942758 | 0.010240552 | 7.261386584 | 3.61E-12 | 10.18232934 | 3.327711234 | 6.228547981 | 3.369317972 | 7.872924734 | 7.74329312 | 11.97242248 | 12.26359867 |
| ENSG00000154553 | PDLIM3 | 3.9941632 | 0.058745872 | 6.186388449 | 0.001802707 | 10.18055165 | 2 | 2 | 2 | 4.470469939 | 4.1930041 | 7.019897747 | 4.729989799 |
| ENSG00000106785 | TRIM14 | 6.220604582 | 1.11E-19 | 3.955451039 | 1.38E-08 | 10.17605562 | 2 | 4.710564154 | 5.39244166 | 10.52636311 | 10.5702365 | 8.256629373 | 8.338065764 |
| ENSG00000095951 | HIVEP1 | 4.528056225 | 0.01185213 | 5.642947911 | 0.001217827 | 10.17100414 | 2 | 6.389896633 | 2 | 9.298270478 | 9.32495871 | 10.28440567 | 10.54697728 |
| ENSG00000160445 | ZER1 | 4.609522631 | 1.48E-08 | 5.56085422 | 4.18E-12 | 10.17037685 | 2 | 5.595927442 | 5.39244166 | 9.374743937 | 9.48723044 | 10.45806243 | 10.29613416 |
| ENSG00000120129 | DUSP1 | 4.502317542 | 0.001807112 | 5.657132369 | 5.53E-05 | 10.15944991 | 3.327711234 | 7.572420273 | 4.058998049 | 10.6367454 | 10.6065803 | 12.17725686 | 11.21385686 |
| ENSG00000105825 | TFPI2 | 8.70548735 | 1.00E-07 | 1.448242356 | 0.457022495 | 10.15372971 | 8.493334308 | 5.595927442 | 2 | 15.78127422 | 15.7487679 | 9.001879477 | 7.802709293 |
| ENSG00000177465 | ACOT4 | 4.273636829 | 0.009987576 | 5.873402564 | 0.000210517 | 10.14703939 | 2 | 2 | 2 | 4.443344104 | 4.68029874 | 5.744329515 | 6.205882756 |
| ENSG00000212710 | CTAGE1 | 6.318874479 | 2.84E-05 | 3.828102735 | 0.016148442 | 10.14697721 | 2 | 2 | 2 | 6.429169081 | 6.39974839 | 4.088436536 | 4.316807515 |
| ENSG00000165752 | STK32C | 4.243678226 | 0.003967676 | 5.897130032 | 3.02E-05 | 10.14080826 | 3.327711234 | 2 | 2 | 5.433909581 | 5.3193943 | 6.910373854 | 6.955898408 |
| ENSG00000172260 | NEGR1 | 3.978870286 | 0.025627053 | 6.160641307 | 0.000260431 | 10.13951159 | 2 | 2 | 6.073378139 | 8.411675819 | 8.31092742 | 10.46111737 | 10.59613413 |
| ENSG00000196535 | MYO18A | 5.412807314 | 4.78E-07 | 4.723238592 | 1.32E-05 | 10.13604591 | 2 | 5.595927442 | 3.369317972 | 9.573133444 | 9.62191107 | 8.7818697 | 9.032680928 |
| ENSG00000169184 | MN1 | 3.691651357 | 0.001101338 | 6.443969569 | 1.91E-09 | 10.13562093 | 2 | 4.93638933 | 2 | 7.106324235 | 7.02417148 | 9.484031995 | 10.02716108 |
| ENSG00000183615 | FAM167B | 4.483544779 | 0.006679598 | 5.649276178 | 0.000399175 | 10.13282096 | 2 | 2 | 2 | 4.694064037 | 4.79082474 | 5.536762811 | 5.995914157 |
| ENSG00000260916 | CCPG1 | 6.267787801 | 3.23E-35 | 3.864039551 | 4.64E-14 | 10.13182735 | 5.70932728 | 5.303523747 | 4.058998049 | 11.33723019 | 11.2569323 | 8.893805266 | 8.913527044 |
| ENSG00000114698 | PLSCR4 | 6.18499999 | 2.61E-18 | 3.946381472 | 4.53E-08 | 10.13138146 | 2 | 5.131603625 | 4.523675391 | 10.40023308 | 10.2683912 | 7.991357103 | 8.226587404 |
| ENSG00000146376 | ARHGAP18 | 5.863527082 | 4.43E-06 | 4.267557228 | 0.001141619 | 10.13108431 | 5.095959458 | 2 | 2 | 9.089953145 | 9.19028095 | 7.160478198 | 7.881541153 |
| ENSG00000138356 | AOX1 | 4.848578844 | 0.001149949 | 5.281727354 | 0.000329491 | 10.1303062 | 2 | 5.131603625 | 2 | 8.29513208 | 8.40602977 | 7.657374802 | 9.403927624 |
| ENSG00000105894 | PTN | 5.909938552 | 1.64E-14 | 4.215238492 | 6.86E-08 | 10.12517704 | 3.327711234 | 4.710564154 | 4.058998049 | 9.754424396 | 9.75193315 | 7.606109069 | 8.425990514 |
| ENSG00000255150 | EID3 | 5.956108663 | 0.000110351 | 4.162285004 | 0.00911236 | 10.11839367 | 2 | 2 | 2 | 6.052081244 | 6.08962627 | 4.555645384 | 4.388595494 |
| ENSG00000129667 | RHBDF2 | 5.892457838 | 1.60E-09 | 4.217831575 | 2.37E-05 | 10.11028941 | 2 | 4.710564154 | 2 | 9.012478123 | 9.01236778 | 7.506395059 | 7.20156326 |
| ENSG00000204147 | ASAH2B | 6.729995082 | 1.55E-06 | 3.377641548 | 0.024591077 | 10.10763663 | 2 | 3.076665209 | 2 | 7.662632018 | 7.82349365 | 4.785723309 | 4.434543419 |
| ENSG00000136689 | IL1RN | 6.145024745 | 0.018875043 | 3.961441321 | 0.144697175 | 10.10646607 | 2 | 2 | 2 | 6.29903474 | 6.19790195 | 5.130840083 | 2.225841198 |
| ENSG00000132359 | RAP1GAP2 | 4.11664322 | 0.014880133 | 5.987799408 | 0.000187366 | 10.10444263 | 2 | 2 | 2 | 4.574143677 | 4.2865223 | 6.370627179 | 5.768151538 |
| ENSG00000105963 | ADAP1 | 5.027068052 | 0.000567369 | 5.071366993 | 0.000461699 | 10.09843505 | 2 | 2 | 3.369317972 | 6.139181293 | 6.06344184 | 6.216130818 | 6.066922748 |
| ENSG00000116260 | QSOX1 | 4.938349612 | 0.004382954 | 5.159168772 | 0.002657411 | 10.09751838 | 9.397585735 | 9.28800521 | 2 | 13.65085696 | 13.7273466 | 13.71677233 | 14.08098316 |
| ENSG00000178015 | GPR150 | 4.608276584 | 0.008438864 | 5.48742961 | 0.001245769 | 10.09570619 | 2 | 2 | 2 | 4.846787657 | 4.85329164 | 6.019162896 | 5.108633494 |
| ENSG00000198824 | CHAMP1 | 4.687370942 | 8.04E-05 | 5.403412324 | 3.81E-06 | 10.09078327 | 2 | 5.131603625 | 2 | 8.192977693 | 8.2657098 | 8.83678613 | 9.033147652 |
| ENSG00000122863 | CHST3 | 3.700540162 | 0.053839803 | 6.389190655 | 0.000350509 | 10.08973082 | 2 | 6.602424674 | 2 | 8.693628619 | 8.71442024 | 11.38839668 | 11.37349133 |
| ENSG00000149380 | P4HA3 | 7.125850819 | 6.51E-09 | 2.960469145 | 0.026657325 | 10.08631996 | 3.327711234 | 2 | 3.369317972 | 9.052296807 | 8.99400091 | 5.333556395 | 4.654341953 |
| ENSG00000170054 | SERPINA9 | 2.213792519 | 0.186455215 | 7.87204008 | 2.63E-08 | 10.0858326 | 2 | 2 | 2 | 3.227757282 | 3.01487888 | 8.022522757 | 7.814703611 |
| ENSG00000164694 | FNDC1 | 7.455830825 | 1.80E-07 | 2.603167801 | 0.099916399 | 10.05898863 | 2 | 2 | 5.39244166 | 11.0319387 | 11.0712094 | 5.614651724 | 6.723495198 |
| ENSG00000158258 | CLSTN2 | -0.686502473 | 0.723460473 | 10.74284512 | 1.61E-15 | 10.05634265 | 3.327711234 | 2 | 2 | 2.498691189 | 2.2680111 | 11.56082781 | 11.88625902 |
| ENSG00000117115 | PADI2 | 0.693574828 | 0.751812007 | 9.337248726 | 5.79E-09 | 10.03082355 | 2 | 2 | 2 | 2.784612485 | 2.14022021 | 9.498161492 | 9.234480927 |
| ENSG00000118515 | SGK1 | 6.044624108 | 7.85E-14 | 3.982196768 | 1.46E-06 | 10.02682088 | 3.327711234 | 5.303523747 | 3.369317972 | 10.13478858 | 10.1843212 | 8.394916616 | 7.762743025 |
| ENSG00000123570 | RAB9B | 4.678597778 | 0.005365654 | 5.335871471 | 0.001156865 | 10.01446925 | 2 | 2 | 2 | 4.984880435 | 4.83276841 | 5.201624686 | 5.740981724 |
| ENSG00000184194 | GPR173 | 5.101320308 | 8.04E-05 | 4.91172111 | 0.000143774 | 10.01304142 | 2 | 3.686303414 | 2 | 7.066878356 | 6.89154576 | 6.556607032 | 7.006211772 |
| ENSG00000132718 | SYT11 | 4.383794096 | 1.21E-17 | 5.622752907 | 8.74E-29 | 10.006547 | 5.095959458 | 4.93638933 | 4.874587708 | 9.229474035 | 9.10192275 | 10.28832008 | 10.50418883 |
| ENSG00000198517 | MAFK | 5.167954634 | 0.001877118 | 4.83515801 | 0.003724409 | 10.00311264 | 6.96351253 | 3.076665209 | 2 | 10.5295909 | 10.5609081 | 10.1025056 | 10.31659934 |
| ENSG00000179630 | LACC1 | 6.239040206 | 1.34E-17 | 3.760620649 | 4.94E-07 | 9.999660855 | 4.467311984 | 4.11352228 | 2 | 9.699596862 | 9.67763759 | 7.249266052 | 7.234405279 |
| ENSG00000196526 | AFAP1 | 3.932163029 | 0.001734936 | 6.066460719 | 4.31E-07 | 9.998623747 | 6.73718411 | 8.077869079 | 3.369317972 | 10.88374584 | 10.9406373 | 12.74694674 | 13.29096652 |
| ENSG00000125965 | GDF5 | 4.838777421 | 0.005488003 | 5.154287069 | 0.002690113 | 9.993064489 | 2 | 2 | 2 | 4.984880435 | 5.11361705 | 4.854926465 | 5.689801209 |
| ENSG00000111674 | ENO2 | 4.695461659 | 0.033493947 | 5.297245343 | 0.014352343 | 9.992707002 | 2 | 2 | 7.398165132 | 10.43808429 | 10.499165 | 10.87325587 | 11.24188786 |
| ENSG00000158869 | FCER1G | 4.665867641 | 0.015592762 | 5.323874194 | 0.004720857 | 9.989741835 | 2 | 2 | 2 | 4.76150579 | 5.0259382 | 6.046045499 | 4.553838275 |
| ENSG00000119318 | RAD23B | 4.684979733 | 2.23E-12 | 5.296831376 | 1.11E-15 | 9.981811108 | 4.00722531 | 6.228547981 | 6.322058544 | 10.44877988 | 10.4140861 | 10.98378895 | 11.09784436 |
| ENSG00000170485 | NPAS2 | 3.908188356 | 6.42E-05 | 6.067865182 | 1.14E-10 | 9.976053538 | 2 | 4.710564154 | 2 | 7.012547947 | 7.1204461 | 9.167793558 | 9.218564658 |
| ENSG00000095383 | TBC1D2 | 3.925819185 | 0.002410798 | 6.046698472 | 9.83E-07 | 9.972517657 | 2 | 5.303523747 | 2 | 7.596453725 | 7.6957855 | 9.734335478 | 9.756591061 |
| ENSG00000150281 | CTF1 | 5.004475735 | 0.00103094 | 4.964904478 | 0.001062609 | 9.969380213 | 2 | 3.076665209 | 2 | 6.197166296 | 5.9536761 | 5.696113106 | 6.323497254 |
| ENSG00000114670 | NEK11 | 5.354853428 | 0.000792713 | 4.613133766 | 0.004247726 | 9.967987194 | 2 | 2 | 2 | 5.527288686 | 5.49945122 | 4.854926465 | 4.853519801 |
| ENSG00000163145 | C1QTNF7 | 1.496142873 | 0.397553222 | 8.458330535 | 1.64E-09 | 9.954473408 | 2 | 2 | 2 | 2.695456494 | 2.86123014 | 8.652274805 | 8.331250243 |
| ENSG00000107863 | ARHGAP21 | 4.899767137 | 6.83E-05 | 5.053510207 | 3.51E-05 | 9.953277344 | 7.031698319 | 6.464272742 | 2 | 11.04524939 | 11.0446855 | 11.16664251 | 11.22951945 |
| ENSG00000214226 | C17orf67 | 5.87864461 | 0.000176081 | 4.072488553 | 0.012349323 | 9.951133163 | 2 | 2 | 2 | 5.890757403 | 6.09824986 | 4.513461373 | 4.279527956 |
| ENSG00000186493 | C5orf38 | 6.27871189 | 0.000142503 | 3.67115678 | 0.03552107 | 9.949868669 | 2 | 2 | 2 | 6.470063213 | 6.27599024 | 4.434275199 | 3.620995423 |
| ENSG00000103174 | NAGPA | 4.987177624 | 5.23E-06 | 4.952299047 | 5.66E-06 | 9.93947667 | 2 | 4.11352228 | 2 | 7.461007055 | 7.39696604 | 7.17948788 | 7.583161732 |
| ENSG00000111276 | CDKN1B | 4.101900065 | 0.004901099 | 5.836555511 | 2.99E-05 | 9.938455576 | 2 | 5.595927442 | 2 | 8.134820652 | 8.06698125 | 9.811872597 | 9.830413512 |
| ENSG00000131634 | TMEM204 | 5.202928159 | 0.000376911 | 4.721050805 | 0.001326409 | 9.923978964 | 2 | 3.076665209 | 2 | 6.18902405 | 6.34282908 | 5.711121288 | 5.913137508 |
| ENSG00000139645 | ANKRD52 | 4.027507384 | 0.001119359 | 5.895463286 | 6.98E-07 | 9.92297067 | 7.940205782 | 6.228547981 | 3.369317972 | 10.76552522 | 10.7571169 | 12.48827223 | 12.75329358 |
| ENSG00000198835 | GJC2 | 4.099329107 | 0.019482624 | 5.818264329 | 0.000482998 | 9.917593436 | 2 | 2 | 2 | 4.497095146 | 4.3456612 | 5.449961224 | 6.308152703 |
| ENSG00000125740 | FOSB | 5.93148272 | 0.000251401 | 3.981274425 | 0.018528026 | 9.912757145 | 2 | 3.076665209 | 2 | 6.836083475 | 7.08617588 | 5.654016955 | 4.341135913 |
| ENSG00000253846 | PCDHGA10 | 6.700069293 | 1.62E-09 | 3.211261844 | 0.006717774 | 9.911331137 | 2 | 5.131603625 | 2 | 10.20056091 | 10.2688688 | 6.880813179 | 6.70252219 |
| ENSG00000145743 | FBXL17 | 4.689708732 | 2.20E-06 | 5.221054949 | 9.64E-08 | 9.910763681 | 2 | 4.710564154 | 2 | 7.756533788 | 7.88956768 | 8.342266573 | 8.353844539 |
| ENSG00000151651 | ADAM8 | 5.140717324 | 0.037143257 | 4.758850618 | 0.052981698 | 9.899567942 | 2 | 2 | 2 | 5.334065616 | 5.30456641 | 5.805044075 | 2.851942824 |
| ENSG00000091409 | ITGA6 | 5.222553211 | 7.95E-07 | 4.663904446 | 1.18E-05 | 9.886457657 | 4.00722531 | 4.11352228 | 2 | 8.360030171 | 8.42329129 | 6.952030889 | 8.387816122 |
| ENSG00000135540 | NHSL1 | 4.418726403 | 0.000535144 | 5.457772712 | 1.14E-05 | 9.876499116 | 4.00722531 | 2 | 2 | 6.449761039 | 6.29111322 | 7.284811203 | 7.460826701 |
| ENSG00000151474 | FRMD4A | 5.386792743 | 0.001902897 | 4.486201787 | 0.011008233 | 9.87299453 | 2 | 6.311476754 | 2 | 10.11558569 | 10.0585048 | 9.255186154 | 9.124459205 |
| ENSG00000169583 | CLIC3 | 4.796957066 | 0.011041916 | 5.073891144 | 0.006426257 | 9.870848211 | 2 | 2 | 2 | 4.846787657 | 5.16377554 | 4.478802948 | 5.76365862 |
| ENSG00000134070 | IRAK2 | 6.82924899 | 4.57E-21 | 3.039971767 | 6.08E-05 | 9.869220757 | 4.00722531 | 3.686303414 | 4.058998049 | 10.28367862 | 10.2953601 | 6.387074688 | 6.711881188 |
| ENSG00000164176 | EDIL3 | 3.75652613 | 2.04E-07 | 6.101353265 | 3.52E-18 | 9.857879395 | 5.532353208 | 7.701025996 | 6.718950348 | 10.67550816 | 10.5932967 | 13.17282734 | 12.75010323 |
| ENSG00000169957 | ZNF768 | 4.405285788 | 0.000517545 | 5.439178187 | 1.12E-05 | 9.844463975 | 2 | 5.131603625 | 2 | 7.975897842 | 7.90685167 | 9.260615815 | 8.589138522 |
| ENSG00000145040 | UCN2 | 6.086010481 | 4.52E-06 | 3.753993897 | 0.006933647 | 9.840004378 | 2 | 3.686303414 | 2 | 7.860161659 | 8.00642058 | 5.227950776 | 6.048572483 |
| ENSG00000157933 | SKI | 4.36584549 | 0.012278946 | 5.469725777 | 0.001201483 | 9.835871267 | 2 | 7.315436877 | 3.369317972 | 10.14797511 | 10.1385468 | 11.28913979 | 11.19823048 |
| ENSG00000198805 | PNP | 6.739020229 | 1.34E-10 | 3.096116242 | 0.005764691 | 9.835136471 | 2 | 4.93638933 | 2 | 10.03894548 | 10.1185514 | 6.710164669 | 6.245088435 |
| ENSG00000081277 | PKP1 | 0.856590506 | 0.674378575 | 8.9725828 | 2.94E-09 | 9.829173305 | 2 | 2 | 2 | 2.270786764 | 2.77779907 | 8.73537099 | 9.24096022 |
| ENSG00000143387 | CTSK | 6.19281176 | 1.99E-05 | 3.636095999 | 0.017626246 | 9.82890776 | 3.327711234 | 6.602424674 | 2 | 11.25722675 | 11.308705 | 8.589193677 | 8.872386739 |
| ENSG00000084112 | SSH1 | 4.612004484 | 3.62E-08 | 5.198123732 | 3.61E-10 | 9.810128216 | 4.815609471 | 7.538394143 | 7.398165132 | 11.54408973 | 11.5956213 | 11.99669313 | 12.29866876 |
| ENSG00000020633 | RUNX3 | 4.063842474 | 0.061048542 | 5.744906714 | 0.005243018 | 9.808749188 | 2 | 2 | 2 | 4.299492648 | 4.4836996 | 6.601598251 | 4.329022995 |
| ENSG00000184156 | KCNQ3 | 4.228891835 | 0.021105714 | 5.579141944 | 0.001502148 | 9.808033778 | 2 | 2 | 2 | 4.574143677 | 4.4836996 | 5.056402493 | 6.172377306 |
| ENSG00000205090 | TMEM240 | 3.83869444 | 0.021503596 | 5.968331317 | 0.00014242 | 9.807025757 | 2 | 2 | 2 | 4.139692846 | 4.2865223 | 6.249924465 | 5.892708443 |
| ENSG00000117069 | ST6GALNAC5 | 6.461278565 | 1.49E-05 | 3.338581844 | 0.036665023 | 9.799860409 | 2 | 2 | 2 | 6.610808945 | 6.48770488 | 3.851628042 | 3.824516949 |
| ENSG00000152223 | EPG5 | 4.973646975 | 8.87E-08 | 4.805057692 | 2.33E-07 | 9.778704666 | 3.327711234 | 6.728495673 | 5.59507254 | 10.589972 | 10.7560955 | 10.55911238 | 10.45348223 |
| ENSG00000139197 | PEX5 | 4.906136734 | 4.16E-05 | 4.872419878 | 4.38E-05 | 9.778556612 | 2 | 5.131603625 | 2 | 8.395844809 | 8.49034333 | 8.329558253 | 8.487225055 |

| | | | | | | | | | | | | | |
|---|---|---|---|---|---|---|---|---|---|---|---|---|---|
| ENSG00000253276 | CCDC71L | 6.289782742 | 1.60E-14 | 3.473714428 | 4.22E-05 | 9.76349717 | 4.815609471 | 5.838937808 | 7.285536013 | 12.53169913 | 12.5382719 | 10.07194845 | 9.266590264 |
| ENSG00000126785 | RHOJ | 1.918112081 | 0.08648314 | 7.835491978 | 4.47E-16 | 9.75360406 | 3.327711234 | 4.11352228 | 2 | 5.058234798 | 4.83276841 | 10.17618144 | 11.03852583 |
| ENSG00000168970 | JMJD7-PLA2( | 5.417017768 | 0.001788712 | 4.335546148 | 0.014425739 | 9.752563915 | 2 | 2 | 2 | 5.514313187 | 5.62411716 | 4.971696071 | 4.147809227 |
| ENSG00000128805 | ARHGAP22 | 5.741075765 | 2.95E-05 | 4.003550646 | 0.004852454 | 9.744626411 | 5.330594134 | 2 | 2 | 9.286870873 | 9.30367823 | 7.796681167 | 7.322574236 |
| ENSG00000134594 | RAB33A | 3.388773887 | 0.050844567 | 6.354825786 | 6.29E-05 | 9.743599673 | 2 | 2 | 2 | 3.709785322 | 4.02226117 | 6.734240628 | 6.092227206 |
| ENSG00000125730 | C3 | 6.570393024 | 4.36E-06 | 3.166082088 | 0.039545549 | 9.736475112 | 2 | 4.442710129 | 2 | 9.283050899 | 9.33957954 | 6.678062013 | 4.634793542 |
| ENSG00000230667 | SETSIP | 7.558452893 | 1.12E-05 | 2.173201486 | 0.267968334 | 9.73165438 | 2 | 2 | 2 | 7.650823203 | 7.57741621 | 2.526562374 | 3.518055793 |
| ENSG00000174740 | PABPC5 | 4.79754584 | 0.0040697 | 4.91807645 | 0.002942813 | 9.715622291 | 2 | 2 | 2 | 4.946755474 | 5.07918227 | 5.258920789 | 4.967301796 |
| ENSG00000117245 | KIF17 | 3.115581199 | 0.061794627 | 6.582233802 | 1.22E-05 | 9.697815002 | 2 | 2 | 2 | 3.663521627 | 3.69849104 | 6.823267818 | 6.490500786 |
| ENSG00000134201 | GSTM5 | 2.594998485 | 0.140051343 | 7.102579209 | 3.54E-06 | 9.697577694 | 2 | 2 | 2 | 3.162721474 | 3.50452879 | 6.802396951 | 7.462213768 |
| ENSG00000168502 | MTCL1 | 3.682707154 | 2.58E-05 | 6.014773788 | 8.75E-13 | 9.697480942 | 2 | 4.93638933 | 3.369317972 | 7.311513918 | 7.34356593 | 9.816433958 | 9.417219134 |
| ENSG00000140511 | HAPLN3 | 4.759620503 | 1.04E-05 | 4.933346669 | 4.21E-06 | 9.692967171 | 4.00722531 | 3.076665209 | 6.322058544 | 9.693134408 | 9.64563775 | 9.840947095 | 9.844043851 |
| ENSG00000148426 | PROSER2 | 2.370490653 | 0.092407051 | 7.315395622 | 2.95E-09 | 9.685886275 | 4.00722531 | 2 | 2 | 4.497095146 | 4.56059747 | 8.852946125 | 9.495172628 |
| ENSG00000198728 | LDB1 | 5.046869711 | 1.10E-07 | 4.63290054 | 1.17E-06 | 9.67977025 | 4.467311984 | 5.838937808 | 2 | 9.635130631 | 9.76417792 | 9.262208893 | 9.316140048 |
| ENSG00000185164 | NOMO2 | 7.571759675 | 5.14E-46 | 2.104461046 | 0.000215968 | 9.676220721 | 4.467311984 | 6.04685525 | 5.156626401 | 12.82920138 | 12.8778999 | 7.3389565 | 7.496463996 |
| ENSG00000152503 | TRIM36 | 4.484682994 | 0.034068642 | 5.191504983 | 0.011775054 | 9.676187976 | 2 | 2 | 2 | 4.867343068 | 4.60967607 | 6.034159474 | 4.077147165 |
| ENSG00000115828 | QPCT | 4.912551649 | 6.85E-11 | 4.751631685 | 2.67E-10 | 9.664183334 | 3.327711234 | 4.442710129 | 3.369317972 | 8.402902269 | 8.32017198 | 8.291371768 | 8.109313767 |
| ENSG00000010438 | PRSS3 | 3.213684752 | 0.065966526 | 6.450171519 | 4.72E-05 | 9.663856271 | 2 | 2 | 2 | 3.840289928 | 3.65239003 | 6.121005698 | 6.864442837 |
| ENSG00000069424 | KCNAB2 | 4.851761512 | 1.87E-05 | 4.809477188 | 2.06E-05 | 9.6612387 | 2 | 4.710564154 | 2 | 7.794595594 | 8.10170499 | 8.209845962 | 7.543088047 |
| ENSG00000105499 | PLA2G4C | 5.475634328 | 2.75E-08 | 4.181636009 | 3.06E-05 | 9.657270337 | 2 | 4.710564154 | 2 | 8.530823842 | 8.66303256 | 7.260787065 | 7.388257299 |
| ENSG00000103044 | HAS3 | 3.937408428 | 0.018276173 | 5.712081093 | 0.000300843 | 9.649489521 | 2 | 2 | 2 | 4.299492648 | 4.2865223 | 5.755231204 | 5.925257518 |
| ENSG00000197779 | ZNF81 | 4.799881637 | 2.84E-05 | 4.849493651 | 2.10E-05 | 9.649375288 | 2 | 4.11352228 | 2 | 7.178012736 | 7.29563107 | 7.550870243 | 6.96179415 |
| ENSG00000074755 | ZZEF1 | 4.404990897 | 0.001013885 | 5.24352046 | 6.32E-05 | 9.648511357 | 4.00722531 | 6.535001748 | 2 | 9.526236618 | 9.52849248 | 10.34278806 | 10.38181612 |
| ENSG00000170498 | KISS1 | 0.013475643 | 0.995777461 | 9.62855468 | 3.93E-11 | 9.642030322 | 2 | 2 | 2 | 2.498691189 | 2.14022021 | 9.852626299 | 9.442386533 |
| ENSG00000158023 | WDR66 | 4.572789645 | 0.007975336 | 5.068302221 | 0.002694218 | 9.641091866 | 2 | 2 | 2 | 4.927308402 | 4.70309173 | 4.977914261 | 5.486701579 |
| ENSG00000144681 | STAC | 2.083293554 | 0.242536361 | 7.552463654 | 4.71E-07 | 9.635757208 | 2 | 2 | 2 | 3.227757282 | 2.86123014 | 7.86621124 | 7.294817518 |
| ENSG00000135362 | PRR5L | 4.018233434 | 0.000905022 | 5.616297898 | 1.52E-06 | 9.634531332 | 2 | 5.131603625 | 2 | 7.59952861 | 7.54308677 | 9.0649396 | 9.228787577 |
| ENSG00000253368 | TRNP1 | 4.027268062 | 0.00224118 | 5.605953233 | 1.01E-05 | 9.633221295 | 2 | 5.303523747 | 2 | 7.772969103 | 7.7127321 | 9.170171434 | 9.426131059 |
| ENSG00000004468 | CD38 | 7.473499398 | 2.20E-06 | 2.156307426 | 0.231087488 | 9.629806825 | 2 | 2 | 2 | 7.43025113 | 7.62593544 | 3.393105702 | 2.70970617 |
| ENSG00000182095 | TNRC18 | 4.490270688 | 0.002137885 | 5.138495153 | 0.000341963 | 9.628765841 | 2 | 5.595927442 | 7.600047325 | 10.74657923 | 10.8020184 | 11.46440721 | 11.37744818 |
| ENSG00000184785 | SMIM10 | 5.031464951 | 0.000876133 | 4.592437016 | 0.002499057 | 9.623901966 | 2 | 3.076665209 | 2 | 6.113598934 | 6.09824986 | 5.413723492 | 5.933281326 |
| ENSG00000184260 | HIST2H2AC | 9.272260933 | 1.09E-11 | 0.345701207 | 0.854663923 | 9.61796214 | 2 | 2 | 2 | 9.262828394 | 9.35135081 | 2.34604626 | 2.466008509 |
| ENSG00000151376 | ME3 | 4.364944925 | 0.000634997 | 5.252941183 | 2.52E-05 | 9.617886108 | 4.00722531 | 2 | 2 | 6.365568715 | 6.27599024 | 7.134272047 | 7.216434756 |
| ENSG00000186564 | FOXD2 | 3.303747627 | 0.047844479 | 6.300060645 | 3.58E-05 | 9.603808272 | 2 | 2 | 2 | 3.754611435 | 3.86943969 | 6.490730008 | 6.29575846 |
| ENSG00000116183 | PAPPA2 | 5.132786454 | 0.001862035 | 4.46467139 | 0.007405859 | 9.597457844 | 2 | 2 | 2 | 5.37770462 | 5.24368216 | 4.813804156 | 4.634793542 |
| ENSG00000258659 | TRIM34 | 7.315552352 | 4.49E-07 | 2.263490551 | 0.167292121 | 9.579042903 | 2 | 2 | 2 | 7.447418636 | 7.30311013 | 3.27687201 | 3.012034821 |
| ENSG00000120690 | ELF1 | 5.688112584 | 1.81E-15 | 3.886175538 | 9.11E-08 | 9.574288123 | 2 | 5.131603625 | 5.39244166 | 10.26249532 | 10.2158672 | 8.540660658 | 8.351600966 |
| ENSG00000149503 | INCENP | 4.481391235 | 6.39E-05 | 5.087187817 | 3.97E-06 | 9.568579052 | 2 | 4.11352228 | 2 | 7.136275664 | 6.70088158 | 7.504242605 | 7.544398272 |
| ENSG00000175315 | CST6 | 2.389392098 | 0.191457657 | 7.17727049 | 5.11E-06 | 9.566662588 | 2 | 2 | 2 | 3.162721474 | 3.28037825 | 6.62753908 | 7.672153316 |
| ENSG00000133107 | TRPC4 | 2.56045088 | 0.121367017 | 6.991171485 | 1.52E-06 | 9.551622364 | 2 | 3.076665209 | 2 | 3.95996233 | 3.98554648 | 8.373261261 | 7.500519913 |
| ENSG00000115419 | GLS | 4.05166088 | 2.04E-07 | 5.49809559 | 5.77E-13 | 9.549756469 | 5.532353208 | 7.099070191 | 4.874587708 | 10.25715053 | 10.0479713 | 11.75011531 | 11.43146841 |
| ENSG00000131019 | ULBP3 | 5.578252554 | 0.000993283 | 3.967817821 | 0.023750038 | 9.546070375 | 2 | 2 | 2 | 5.731694 | 5.70538658 | 3.943456545 | 4.614976609 |
| ENSG00000171262 | FAM98B | 4.449388618 | 2.01E-07 | 5.092840625 | 1.66E-09 | 9.542229243 | 3.327711234 | 4.442710129 | 2 | 7.635924947 | 7.6318872 | 8.15052603 | 8.374617996 |
| ENSG00000138623 | SEMA7A | 5.395305635 | 0.003175086 | 4.141813945 | 0.027327995 | 9.53711958 | 2 | 2 | 6.322058544 | 9.978816937 | 10.0945086 | 8.797335126 | 8.786410001 |
| ENSG00000158560 | DYNC1I1 | 4.907956502 | 0.003293724 | 4.623909948 | 0.005658731 | 9.53186645 | 2 | 2 | 2 | 5.003571641 | 5.21224859 | 4.881704255 | 4.845038551 |
| ENSG00000197956 | S100A6 | 5.090320588 | 3.99E-11 | 4.437394857 | 1.02E-08 | 9.527715445 | 9.097018025 | 7.051611315 | 7.502634626 | 13.21868427 | 13.2461576 | 12.22525492 | 12.86434166 |
| ENSG00000050165 | DKK3 | 4.699973595 | 2.33E-08 | 4.809668242 | 9.37E-09 | 9.509641838 | 6.366709671 | 8.848535039 | 9.031722608 | 13.14167328 | 13.1776882 | 13.29176405 | 13.24675504 |
| ENSG00000164211 | STARD4 | 5.020873136 | 1.44E-06 | 4.488296362 | 1.86E-05 | 9.509169498 | 6.366709671 | 5.457120953 | 2 | 10.38783677 | 10.2788601 | 9.611484085 | 9.97345748 |
| ENSG00000130758 | MAP3K10 | 4.219945678 | 4.70E-06 | 5.284915715 | 4.90E-09 | 9.504861393 | 3.327711234 | 5.131603625 | 2 | 7.913023977 | 8.11238697 | 9.222826582 | 8.897724096 |
| ENSG00000188372 | ZP3 | 4.89211992 | 0.00149867 | 4.608902462 | 0.002796225 | 9.501022382 | 2 | 3.076665209 | 2 | 5.900763176 | 6.04571775 | 5.988693817 | 5.367404784 |
| ENSG00000096433 | ITPR3 | 6.663477447 | 1.01E-12 | 2.831125258 | 0.004649784 | 9.494602705 | 3.327711234 | 4.11352228 | 6.203069917 | 11.45458917 | 11.5791589 | 7.367655602 | 7.989373961 |
| ENSG00000067715 | SYT1 | 7.985600531 | 2.55E-06 | 1.507607642 | 0.454406107 | 9.493208173 | 3.327711234 | 2 | 2 | 8.985700128 | 8.98472903 | 2.457042231 | 3.789585079 |
| ENSG00000141526 | SLC16A3 | 5.209859184 | 1.35E-06 | 4.280251617 | 9.03E-05 | 9.490110801 | 8.87372952 | 6.04685525 | 5.930865648 | 12.77948335 | 12.8444124 | 11.6021717 | 12.11868584 |
| ENSG00000205609 | EIF3CL | 10.23675981 | 3.34E-14 | -0.7508763 | 0.685511159 | 9.485883514 | 2 | 2 | 2 | 10.26006833 | 10.2755373 | 2.183373728 | 2.225841198 |
| ENSG00000114473 | IQCG | 4.334295744 | 0.001101338 | 5.14231486 | 7.38E-05 | 9.476610603 | 2 | 3.686303414 | 2 | 6.197166296 | 6.29111322 | 6.770513376 | 7.232780814 |
| ENSG00000185745 | IFIT1 | 8.719931752 | 1.23E-11 | 0.756471555 | 0.639891142 | 9.476403307 | 7.579279349 | 4.442710129 | 3.369317972 | 14.85650684 | 14.8459655 | 6.529346487 | 7.252154709 |
| ENSG00000226784 | PGAM4 | 11.56603534 | 4.30E-14 | -2.103206978 | 0.3091698 | 9.462828359 | 2 | 2 | 2 | 11.57917009 | 11.6092606 | 2 | 2.172608095 |
| ENSG00000048052 | HDAC9 | 5.160710871 | 2.33E-07 | 4.298254194 | 2.06E-05 | 9.458965066 | 2 | 4.710564154 | 2 | 8.327018759 | 8.24255817 | 7.406207255 | 7.469129172 |
| ENSG00000171877 | FRMD5 | 5.609066422 | 0.000555458 | 3.847510372 | 0.022853959 | 9.456576795 | 2 | 2 | 2 | 5.870535235 | 5.6121243 | 4.340871764 | 4.09155917 |
| ENSG00000159082 | SYNJ1 | 4.36379392 | 5.18E-09 | 5.088899036 | 5.18E-12 | 9.452692956 | 4.00722531 | 4.710564154 | 2 | 7.985332082 | 8.06917614 | 8.68304821 | 8.802925779 |
| ENSG00000168014 | C2CD3 | 4.055916603 | 0.000672752 | 5.396209813 | 3.09E-06 | 9.452126416 | 4.00722531 | 6.140560021 | 2 | 8.828161587 | 8.85716692 | 10.12141194 | 10.22744137 |
| ENSG00000157111 | TMEM171 | 5.64379386 | 7.06E-08 | 3.784973952 | 0.000459733 | 9.428767811 | 2 | 4.11352228 | 2 | 8.132699624 | 8.02689124 | 6.320128742 | 6.232137896 |
| ENSG00000118473 | SGIP1 | 4.429422363 | 0.001919616 | 4.99817985 | 0.000364719 | 9.427602212 | 5.330594134 | 2 | 2 | 7.949630994 | 8.04484669 | 8.722004124 | 8.376090431 |
| ENSG00000100100 | PIK3IP1 | 5.372946357 | 9.16E-06 | 4.054522649 | 0.001071222 | 9.427469006 | 2 | 5.131603625 | 2 | 8.848982184 | 8.95417718 | 7.782541424 | 7.395535532 |
| ENSG00000157064 | NMNAT2 | 3.7728109 | 0.004958412 | 5.652264968 | 1.00E-05 | 9.425075868 | 3.327711234 | 2 | 3.369317972 | 5.674521485 | 5.86549842 | 7.16184437 | 7.893928381 |
| ENSG00000066923 | STAG3 | 4.305151708 | 0.016178162 | 5.11663929 | 0.003232449 | 9.421790998 | 2 | 2 | 2 | 4.739373666 | 4.43005853 | 5.586442682 | 4.935697922 |
| ENSG00000164023 | SGMS2 | 3.260896055 | 0.008082884 | 6.160299144 | 8.54E-08 | 9.421195198 | 2 | 5.131603625 | 2 | 6.917187326 | 6.75096147 | 9.56215648 | 9.810132015 |
| ENSG00000168056 | LTBP3 | 4.436701143 | 0.05663139 | 4.978095552 | 0.028692244 | 9.414796695 | 2 | 8.029695761 | 2 | 10.83304867 | 10.9079011 | 11.30120057 | 11.51370137 |
| ENSG00000138593 | SECISBP2L | 4.770018765 | 0.000875927 | 4.642948896 | 0.001171005 | 9.412967661 | 2 | 7.315436877 | 5.156626401 | 10.77813214 | 10.7299752 | 10.78923234 | 10.44526431 |
| ENSG00000171428 | NAT1 | 4.871980654 | 0.001047492 | 4.538212815 | 0.002318491 | 9.410193469 | 2 | 3.076665209 | 2 | 5.900763176 | 6.00960265 | 5.684753614 | 5.606950214 |
| ENSG00000184995 | IFNE | 7.169095511 | 2.93E-06 | 2.239483032 | 0.196332247 | 9.408578543 | 2 | 2 | 2 | 7.214570819 | 7.2537885 | 3.374372158 | 2.851942824 |
| ENSG00000182685 | BRICD5 | 3.548766083 | 0.042452833 | 5.853033332 | 0.000314836 | 9.401799415 | 2 | 2 | 2 | 4.17310023 | 3.7864959 | 6.140539117 | 5.785984581 |
| ENSG00000149201 | CCDC81 | 1.05059529 | 0.548089791 | 8.35101951 | 9.92E-10 | 9.4016148 | 2 | 3.076665209 | 2 | 3.162721474 | 2.86123014 | 9.214621994 | 9.470227873 |
| ENSG00000081386 | ZNF510 | 4.575667798 | 2.66E-06 | 4.805909766 | 6.68E-07 | 9.381577565 | 3.327711234 | 4.11352228 | 2 | 7.433701013 | 7.45518898 | 7.925322387 | 7.358771532 |
| ENSG00000057019 | DCBLD2 | 5.882240238 | 1.06E-44 | 3.489229222 | 2.37E-16 | 9.37146946 | 7.031698319 | 7.874700259 | 7.029880454 | 13.24291444 | 13.230921 | 10.95777532 | 10.72579699 |
| ENSG00000151388 | ADAMTS12 | 4.480102774 | 0.000767754 | 4.886709519 | 0.000199805 | 9.366812294 | 2 | 7.145017454 | 6.43197674 | 10.68747873 | 10.6916678 | 10.61491551 | 11.45508498 |
| ENSG00000171100 | MTM1 | 4.70887209 | 1.36E-05 | 4.656523614 | 1.56E-05 | 9.365395703 | 2 | 4.11352228 | 2 | 7.132034771 | 7.19067478 | 7.171371525 | 7.04745562 |
| ENSG00000163689 | C3orf67 | 5.559358378 | 0.001451898 | 3.804002805 | 0.035956933 | 9.363361183 | 2 | 2 | 2 | 5.627094009 | 5.77157844 | 4.530483275 | 3.735548302 |
| ENSG00000054965 | FAM168A | 3.993934831 | 0.055580779 | 5.369102421 | 0.007227479 | 9.363037252 | 2 | 7.099070191 | 2 | 9.469384409 | 9.51728265 | 10.79066808 | 10.93331374 |
| ENSG00000166033 | HTRA1 | 4.165111451 | 1.37E-08 | 5.18434938 | 7.06E-13 | 9.349460831 | 7.667647223 | 8.806266215 | 6.629488387 | 12.07983178 | 12.1570582 | 13.27104463 | 12.99040357 |
| ENSG00000142409 | ZNF787 | 4.507547727 | 0.00063259 | 4.840170155 | 0.000203725 | 9.347717881 | 2 | 5.303523747 | 2 | 8.139053376 | 8.284723 | 8.516835903 | 8.567391554 |
| ENSG00000157353 | FUK | 3.874385028 | 0.003295139 | 5.468880876 | 1.52E-05 | 9.343265904 | 2 | 3.686303414 | 2 | 5.959382894 | 5.65950892 | 7.367655602 | 7.314917451 |
| ENSG00000260428 | SCX | 5.564725446 | 0.00151176 | 3.769156546 | 0.038665093 | 9.333881992 | 2 | 2 | 2 | 5.48800675 | 5.89549381 | 4.487545924 | 3.735548302 |
| ENSG00000197106 | SLC6A17 | 3.414022188 | 0.090465939 | 5.918811937 | 0.001434587 | 9.332834126 | 2 | 2 | 2 | 3.349643765 | 4.2865223 | 4.668821191 | 5.414021719 |
| ENSG00000081923 | ATP8B1 | 3.000444915 | 2.48E-05 | 6.327044162 | 9.73E-21 | 9.327489077 | 6.257048221 | 4.11352228 | 5.156626401 | 8.29513208 | 8.24255817 | 11.45119913 | 11.69592421 |
| ENSG00000101596 | SMCHD1 | 5.33287477 | 2.00E-05 | 3.993650253 | 0.00183886 | 9.326525023 | 4.467311984 | 6.535001748 | 2 | 10.54681762 | 10.5189709 | 9.447810736 | 8.899772939 |
| ENSG00000205517 | RGL3 | 4.535724254 | 0.010654158 | 4.785207901 | 0.006279821 | 9.320932154 | 2 | 2 | 2 | 4.907595604 | 4.65713986 | 5.258920789 | 4.692661645 |
| ENSG00000129048 | ACKR4 | 6.964812535 | 1.05E-08 | 2.345660989 | 0.083161849 | 9.310473524 | 2 | 3.686303414 | 2 | 8.857997553 | 8.77886778 | 4.281782022 | 4.644600858 |
| ENSG00000121281 | ADCY7 | 5.688117803 | 2.33E-12 | 3.619583761 | 1.42E-05 | 9.307701564 | 2 | 4.93638933 | 3.369317972 | 9.308641573 | 9.3166689 | 7.231159579 | 7.313381205 |
| ENSG00000182492 | BGN | 3.832407867 | 0.00024625 | 5.474583188 | 6.09E-08 | 9.306991055 | 9.886101849 | 7.701025996 | 6.718950348 | 12.49501766 | 12.5683571 | 14.05464728 | 14.28359817 |
| ENSG00000130024 | PHF10 | 3.977742492 | 0.000425977 | 5.322786409 | 1.15E-06 | 9.3005289 | 3.327711234 | 5.595927442 | 2 | 8.195011988 | 8.14188248 | 9.699953617 | 9.271345616 |
| ENSG00000138172 | CALHM2 | 3.877292691 | 0.001742493 | 5.422613057 | 5.41E-06 | 9.299905748 | 2 | 5.131603625 | 2 | 7.388195945 | 7.47187556 | 9.075488822 | 8.820877391 |
| ENSG00000078053 | AMPH | 5.012451268 | 0.001326022 | 4.284376767 | 0.006766036 | 9.296828035 | 2 | 3.686303414 | 2 | 6.815074525 | 6.90143415 | 5.169385838 | 6.746446439 |
| ENSG00000163625 | WDFY3 | 4.448472221 | 0.010175586 | 4.845854042 | 0.004483519 | 9.294326263 | 2 | 7.274681896 | 3.369317972 | 10.1755382 | 10.1969221 | 10.52713712 | 10.6357026 |
| ENSG00000100393 | EP300 | 4.77325981 | 0.000140189 | 4.518825882 | 0.000318465 | 9.292085691 | 2 | 5.946638857 | 7.029880454 | 10.68204987 | 10.7454982 | 10.51355748 | 10.40508119 |
| ENSG00000105643 | ARRDC2 | 5.459323749 | 0.000179763 | 3.828108585 | 0.011275691 | 9.287432334 | 2 | 5.595927442 | 2 | 9.376537567 | 9.50436385 | 8.086505553 | 7.512619696 |
| ENSG00000221995 | TIAF1 | 6.444132427 | 6.79E-06 | 2.839835025 | 0.06836276 | 9.283967452 | 2 | 3.076665209 | 2 | 7.555872146 | 7.36873214 | 3.968654369 | 4.376876162 |
| ENSG00000118762 | PKD2 | 3.563125295 | 3.04E-07 | 5.711844276 | 2.68E-17 | 9.274969571 | 6.009025997 | 7.431218269 | 5.930865648 | 10.17141133 | 10.1639279 | 11.83979864 | 12.66796169 |
| ENSG00000179456 | ZBTB18 | 3.666653424 | 0.000413591 | 5.607111147 | 1.75E-08 | 9.273764571 | 2 | 4.442710129 | 2 | 6.658765962 | 6.40670796 | 8.522164399 | 8.335797499 |
| ENSG00000060140 | STYK1 | 4.288815185 | 0.030317415 | 4.980456948 | 0.009741936 | 9.269272132 | 2 | 2 | 2 | 4.497095146 | 4.65713986 | 4.360042493 | 5.694529741 |
| ENSG00000178773 | CPNE7 | 5.907971112 | 3.98E-05 | 3.358922264 | 0.027424614 | 9.266893376 | 2 | 5.303523747 | 2 | 9.561354269 | 9.60839132 | 6.231826305 | 7.599641096 |
| ENSG00000137203 | TFAP2A | 3.508384158 | 0.000968694 | 5.754190144 | 1.27E-08 | 9.262574303 | 2 | 4.442710129 | 2 | 6.496695859 | 6.26070706 | 8.3832514 | 8.741991736 |
| ENSG00000181619 | GPR135 | 4.14467377 | 0.007220064 | 5.105905741 | 0.000630205 | 9.250579511 | 2 | 3.076665209 | 2 | 5.144975053 | 5.41912973 | 6.031172603 | 6.308152703 |
| ENSG00000170373 | CST1 | 3.993161482 | 0.028289069 | 5.248902369 | 0.002615599 | 9.242063851 | 2 | 2 | 2 | 4.329433326 | 4.3456612 | 4.984105765 | 5.7500953 |
| ENSG00000198722 | UNC13B | 3.864817549 | 0.001319542 | 5.370158688 | 3.73E-06 | 9.234976237 | 3.327711234 | 5.72254323 | 2 | 8.153770873 | 8.17690623 | 9.342112202 | 9.916412852 |
| ENSG00000083454 | P2RX5 | 6.073705754 | 0.000934242 | 3.161106199 | 0.107213033 | 9.234811953 | 2 | 2 | 2 | 6.276156955 | 6.08095082 | 4.166733998 | 3.042026731 |
| ENSG00000185158 | LRRC37B | 5.21439679 | 0.000170463 | 4.013736191 | 0.004739329 | 9.228132981 | 4.00722531 | 2 | 2 | 7.161462517 | 7.14145882 | 6.387074688 | 5.475755622 |

| | | | | | | | | | | | | | |
|---|---|---|---|---|---|---|---|---|---|---|---|---|---|
| ENSG00000167978 | SRRM2 | 4.213683134 | 7.32E-14 | 5.007891421 | 2.42E-19 | 9.221574555 | 8.257427002 | 8.929529568 | 7.342949392 | 12.5192709 | 12.5228406 | 13.29033993 | 13.33894516 |
| ENSG00000111199 | TRPV4 | 4.826350842 | 0.000209355 | 4.391213526 | 0.000791158 | 9.217564368 | 2 | 3.686303414 | 2 | 6.542151749 | 6.8715633 | 6.344396176 | 6.254725615 |
| ENSG00000112425 | EPM2A | 4.249269982 | 0.001105783 | 4.968264008 | 9.61E-05 | 9.217533991 | 2 | 3.686303414 | 2 | 6.104969671 | 6.22964624 | 6.804147769 | 6.905807071 |
| ENSG00000188906 | LRRK2 | 5.421562237 | 1.59E-05 | 3.795889199 | 0.003429913 | 9.217451436 | 2 | 3.686303414 | 2 | 7.273514749 | 7.32163988 | 5.692336538 | 5.799216178 |
| ENSG00000137198 | GMPR | 6.414743772 | 1.91E-10 | 2.789903486 | 0.010112025 | 9.204647258 | 2 | 4.442710129 | 2 | 9.219515356 | 9.23788597 | 5.427419529 | 5.945233978 |
| ENSG00000178607 | ERN1 | 4.793814136 | 5.90E-08 | 4.40623318 | 6.65E-07 | 9.200047316 | 2 | 3.686303414 | 4.523675391 | 8.04293816 | 7.97398853 | 7.865372957 | 7.343799539 |
| ENSG00000196754 | S100A2 | 4.74410065 | 0.008829144 | 4.454237696 | 0.014029183 | 9.198338346 | 2 | 2 | 2 | 4.867343068 | 5.06165168 | 4.311629396 | 5.035976547 |
| ENSG00000187801 | ZFP69B | 4.897221216 | 0.001721482 | 4.296014639 | 0.006440115 | 9.193235856 | 2 | 3.076665209 | 2 | 5.920568914 | 6.03677338 | 5.725974948 | 5.04340913 |
| ENSG00000181773 | GPR3 | 4.69989795 | 0.013126122 | 4.492002871 | 0.017626246 | 9.191900821 | 2 | 2 | 2 | 4.804776789 | 5.04390545 | 4.155805465 | 5.171036168 |
| ENSG00000141753 | IGFBP4 | 4.356853012 | 3.02E-24 | 4.834898145 | 4.91E-30 | 9.191751157 | 9.579430841 | 9.566483142 | 8.699419242 | 13.66216008 | 13.7119771 | 14.35551441 | 13.94591334 |
| ENSG00000122254 | HS3ST2 | 2.688097229 | 0.167050868 | 6.498999846 | 0.000153586 | 9.187097075 | 2 | 2 | 2 | 3.227757282 | 3.55551834 | 7.087593115 | 5.812327524 |
| ENSG00000184374 | COLEC10 | 5.369307178 | 0.001440368 | 3.813946263 | 0.028788809 | 9.18325344 | 2 | 2 | 2 | 5.590471342 | 5.45984938 | 4.388329244 | 3.972018587 |
| ENSG00000108846 | ABCC3 | 6.969398152 | 3.18E-09 | 2.210397383 | 0.092310672 | 9.179795535 | 2 | 2 | 5.156626401 | 10.25178588 | 10.2779115 | 5.673303969 | 5.571402115 |
| ENSG00000189058 | APOD | 4.361743477 | 0.033609824 | 4.817221442 | 0.01663167 | 9.178964919 | 2 | 2 | 2 | 4.623314311 | 4.65713986 | 4.076896059 | 5.601925227 |
| ENSG00000198752 | CDC42BPB | 4.714503213 | 2.56E-11 | 4.459548247 | 2.86E-10 | 9.17405146 | 4.467311984 | 6.844429755 | 6.534109959 | 10.91709408 | 10.9155453 | 10.69965825 | 10.62320178 |
| ENSG00000159212 | CLIC6 | 3.080716141 | 0.110520142 | 6.092501167 | 0.000488026 | 9.173217308 | 2 | 2 | 2 | 3.515101962 | 3.7864959 | 5.472156123 | 6.681239787 |
| ENSG00000187957 | DNER | 8.059874221 | 1.31E-14 | 1.109816135 | 0.376931525 | 9.169690355 | 2 | 4.11352228 | 2 | 10.50274314 | 10.4477945 | 3.694366604 | 4.175132639 |
| ENSG00000166741 | NNMT | 5.713820895 | 0.000312208 | 3.454605337 | 0.038310303 | 9.168426232 | 5.330594134 | 2 | 2 | 9.268635235 | 9.28774494 | 7.657374802 | 5.995914157 |
| ENSG00000156802 | ATAD2 | 3.949460218 | 5.30E-05 | 5.216292322 | 4.03E-08 | 9.16575254 | 3.327711234 | 4.442710129 | 2 | 7.210554257 | 7.04216061 | 8.697250351 | 7.949380652 |
| ENSG00000106780 | MEGF9 | 4.871502098 | 5.10E-18 | 4.289926009 | 2.42E-14 | 9.161428106 | 4.467311984 | 5.131603625 | 4.874587708 | 9.512436154 | 9.56938265 | 9.156530478 | 8.740277714 |
| ENSG00000105612 | DNASE2 | 4.943154684 | 2.51E-11 | 4.19926985 | 1.78E-08 | 9.142424534 | 4.467311984 | 5.838937808 | 3.369317972 | 9.69385389 | 9.76350037 | 9.068585983 | 8.906411656 |
| ENSG00000064726 | BTBD1 | 4.399666174 | 2.30E-13 | 4.736437884 | 1.82E-15 | 9.136104058 | 4.467311984 | 6.228547981 | 5.156626401 | 9.805913872 | 9.78368958 | 10.19532025 | 10.06211278 |
| ENSG00000170027 | YWHAG | 4.25201471 | 1.92E-14 | 4.871323451 | 8.03E-19 | 9.123338161 | 6.89194353 | 7.051611315 | 8.206335842 | 11.72109746 | 11.7227089 | 12.1422582 | 12.51497152 |
| ENSG00000128739 | SNRPN | 5.649836859 | 4.38E-18 | 3.473106729 | 1.82E-07 | 9.122943587 | 3.327711234 | 4.442710129 | 4.523675391 | 9.57078529 | 9.46239569 | 7.313367756 | 7.421437279 |
| ENSG00000164266 | SPINK1 | 2.936187977 | 0.178578057 | 6.184080836 | 0.001643664 | 9.120268813 | 2 | 2 | 2 | 3.406930997 | 3.69849104 | 6.984820384 | 4.878668439 |
| ENSG00000137501 | SYTL2 | 3.590470427 | 0.021105714 | 5.527422273 | 0.000171403 | 9.1178927 | 2 | 3.686303414 | 2 | 5.552894743 | 5.49945122 | 7.900986387 | 6.512090157 |
| ENSG00000096696 | DSP | 2.476191041 | 0.003147211 | 6.630262886 | 1.18E-17 | 9.106453927 | 6.563806279 | 7.701025996 | 9.014156722 | 10.53039672 | 10.56791 | 14.72627191 | 14.67264228 |
| ENSG00000163053 | SLC16A14 | 3.918402359 | 0.036751137 | 5.17895821 | 0.003863831 | 9.097360569 | 2 | 2 | 2 | 4.299492648 | 4.2560182 | 5.73334482 | 4.836507146 |
| ENSG00000158406 | HIST1H4H | 5.97031586 | 0.001880748 | 3.12221573 | 0.127753231 | 9.09253159 | 2 | 2 | 2 | 6.104969671 | 6.06344184 | 2.911488689 | 4.188602707 |
| ENSG00000176842 | IRX5 | 6.515388372 | 1.51E-05 | 2.574987365 | 0.121083356 | 9.090375737 | 2 | 2 | 2 | 6.496695859 | 6.70088158 | 3.374372158 | 3.287118176 |
| ENSG00000138688 | KIAA1109 | 4.232180426 | 4.00E-05 | 4.835912491 | 1.89E-06 | 9.068092917 | 3.327711234 | 6.899080544 | 5.39244166 | 9.972308137 | 10.0272353 | 10.7919921 | 10.38273198 |
| ENSG00000167470 | MIDN | 4.26033537 | 0.000594915 | 4.804576212 | 8.21E-05 | 9.064911581 | 2 | 6.228547981 | 4.058998049 | 9.083378928 | 9.16587611 | 9.556194935 | 9.768412794 |
| ENSG00000025708 | TYMP | 6.41876226 | 7.62E-11 | 2.64612842 | 0.013130749 | 9.064890679 | 2 | 4.442710129 | 5.930865648 | 11.02223255 | 11.0469145 | 7.561260509 | 6.97156716 |
| ENSG00000165084 | C8orf34 | 3.560303972 | 0.031536406 | 5.502775705 | 0.000409205 | 9.063079676 | 2 | 2 | 3.369317972 | 4.739373666 | 4.811949 | 6.010089548 | 6.947999751 |
| ENSG00000113083 | LOX | 4.880424182 | 0.000353266 | 4.18001283 | 0.002518254 | 9.060437012 | 8.777722034 | 8.702538795 | 3.369317972 | 13.04475171 | 13.0269828 | 12.37356765 | 12.29721225 |
| ENSG00000206560 | ANKRD28 | 4.938183536 | 0.000613202 | 4.117673061 | 0.004914836 | 9.055856597 | 4.00722531 | 6.844429755 | 2 | 10.33485359 | 10.3319235 | 9.628876603 | 9.394150543 |
| ENSG00000164308 | ERAP2 | 3.969864118 | 0.000200919 | 5.079476025 | 1.05E-06 | 9.049340143 | 4.00722531 | 3.076665209 | 6.203069917 | 8.769305409 | 8.78955647 | 9.683639816 | 10.05638425 |
| ENSG00000131480 | AOC2 | 4.776691808 | 0.038740151 | 4.27099541 | 0.064751221 | 9.047687218 | 2 | 2 | 2 | 4.965943888 | 5.0259382 | 5.269098128 | 3.128433649 |
| ENSG00000056736 | IL17RB | 4.560451638 | 0.011297503 | 4.484289849 | 0.012268798 | 9.044741488 | 2 | 2 | 2 | 4.907595604 | 4.70309173 | 4.990270811 | 4.445805334 |
| ENSG00000153993 | SEMA3D | 4.691606872 | 0.003501197 | 4.352610195 | 0.006975079 | 9.044217066 | 2 | 2 | 4.058998049 | 6.804454198 | 6.54560135 | 5.289239922 | 6.963754056 |
| ENSG00000173846 | PLK3 | 5.358424429 | 0.000880586 | 3.684280805 | 0.027686483 | 9.042705234 | 2 | 5.946638857 | 2 | 9.677214349 | 9.70612498 | 8.252146986 | 7.773956279 |
| ENSG00000183111 | ARHGEF37 | 2.700789913 | 0.017734312 | 6.337430973 | 1.06E-09 | 9.038220886 | 4.467311984 | 2 | 2 | 5.258265913 | 5.39133044 | 8.791168836 | 8.856651551 |
| ENSG00000023171 | GRAMD1B | 4.4004533 | 3.31E-05 | 4.617024456 | 1.12E-05 | 9.017477756 | 2 | 4.710564154 | 2 | 7.504305792 | 7.55253065 | 7.857806412 | 7.613439639 |
| ENSG00000169047 | IRS1 | 4.272685452 | 0.000900252 | 4.741108817 | 0.000182815 | 9.013794269 | 2 | 5.946638857 | 3.369317972 | 8.733363582 | 8.80675844 | 9.448650487 | 8.984746522 |
| ENSG00000233822 | HIST1H2BN | 6.276751247 | 8.24E-06 | 2.735817246 | 0.074484213 | 9.012568493 | 3.327711234 | 2 | 2 | 7.352188525 | 7.24992376 | 4.076896059 | 4.133950982 |
| ENSG00000124788 | ATXN1 | 3.796772796 | 0.090096447 | 5.215768466 | 0.014623638 | 9.012541263 | 2 | 7.538394143 | 2 | 9.80191506 | 9.67331554 | 11.14478699 | 11.16273299 |
| ENSG00000171051 | FPR1 | 6.281310078 | 4.22E-05 | 2.726609731 | 0.103755371 | 9.007919809 | 2 | 2 | 2 | 6.276156955 | 6.47451726 | 3.374372158 | 3.474731315 |
| ENSG00000213160 | KLHL23 | 1.778869165 | 0.362317951 | 7.226800848 | 7.76E-06 | 9.005670012 | 2 | 2 | 2 | 3.09461492 | 2.68924543 | 7.702194078 | 6.711881188 |
| ENSG00000136237 | RAPGEF5 | 3.585400667 | 0.027691078 | 5.412342745 | 0.00042866 | 8.997743411 | 3.327711234 | 2 | 2 | 4.76150579 | 4.83276841 | 6.804147769 | 6.026239558 |
| ENSG00000132294 | EFR3A | 5.141287956 | 0.013783233 | 3.840496178 | 0.072675764 | 8.981784135 | 2 | 7.315436877 | 2 | 10.86054366 | 10.8487621 | 9.661838508 | 9.446947471 |
| ENSG00000189337 | KAZN | 3.105431582 | 0.042682089 | 5.868211478 | 3.37E-05 | 8.97364306 | 2 | 5.595927442 | 2 | 7.106324235 | 7.14145882 | 9.658938866 | 10.02058586 |
| ENSG00000056586 | RC3H2 | 3.85195384 | 0.000170463 | 5.118724625 | 2.77E-07 | 8.970678465 | 4.00722531 | 6.899080544 | 4.523675391 | 9.490242771 | 9.52849248 | 10.73013842 | 10.81195485 |
| ENSG00000163960 | UBXN7 | 3.693180416 | 0.000170033 | 5.276578681 | 2.79E-08 | 8.969759097 | 2 | 5.838937808 | 5.930865648 | 8.902248225 | 8.9350493 | 10.586203 | 10.39585643 |
| ENSG00000186470 | BTN3A2 | 5.514567362 | 1.10E-09 | 3.451641634 | 0.000226682 | 8.966208995 | 3.327711234 | 6.728495673 | 5.930865648 | 11.31238029 | 11.3353933 | 9.185700723 | 9.346125282 |
| ENSG00000006606 | CCL26 | 4.947124716 | 0.004419898 | 4.012780719 | 0.023554727 | 8.959905435 | 2 | 2 | 2 | 5.194621444 | 5.09650239 | 4.521997428 | 4.161535617 |
| ENSG00000249992 | TMEM158 | 7.718838195 | 1.84E-11 | 1.240106639 | 0.363255554 | 8.958944834 | 6.366709671 | 4.442710129 | 2 | 12.6386094 | 12.7873956 | 6.645228724 | 5.871985934 |
| ENSG00000188157 | AGRN | 4.286140565 | 8.16E-08 | 4.668578928 | 3.84E-09 | 8.954719493 | 9.2903348 | 8.414402626 | 6.80318718 | 12.74770046 | 12.7758512 | 12.87005887 | 13.37427865 |
| ENSG00000111912 | NCOA7 | 4.7472518 | 8.06E-22 | 4.202885665 | 1.61E-17 | 8.950137466 | 6.138360429 | 7.051611315 | 6.322058544 | 11.32427804 | 11.2159175 | 10.40063634 | 10.99438813 |
| ENSG00000113805 | CNTN3 | 7.231325156 | 1.35E-09 | 1.715914515 | 0.211573817 | 8.947239671 | 2 | 2 | 4.058998049 | 9.157231967 | 9.09439916 | 4.133696865 | 3.875378318 |
| ENSG00000152409 | JMY | 3.956101377 | 0.000155707 | 4.991100099 | 1.01E-06 | 8.947201475 | 4.00722531 | 5.595927442 | 2 | 8.300810597 | 8.29037844 | 9.661114144 | 8.875513287 |
| ENSG00000241697 | TMEFF1 | 6.635392849 | 1.09E-05 | 2.311546548 | 0.170499056 | 8.946939397 | 2 | 2 | 2 | 6.804454198 | 6.6253296 | 3.27687201 | 3.071407821 |
| ENSG00000119917 | IFIT3 | 8.16509734 | 2.82E-07 | 0.768823264 | 0.699688818 | 8.933920604 | 2 | 5.838937808 | 8.380957451 | 15.14069964 | 15.1709914 | 7.182183216 | 8.210170621 |
| ENSG00000109321 | AREG | 9.125028253 | 4.75E-11 | -0.218224377 | 0.910973565 | 8.906803876 | 2 | 3.076665209 | 2 | 10.11665925 | 10.1201401 | 2.34604626 | 2.671842496 |
| ENSG00000135914 | HTR2B | 1.690223964 | 0.361833497 | 7.216223311 | 2.00E-06 | 8.906447275 | 2 | 2 | 2 | 2.947924294 | 2.77779907 | 6.932953913 | 7.56002886 |
| ENSG00000111266 | DUSP16 | 4.857319668 | 5.62E-05 | 4.043167396 | 0.000945348 | 8.900487064 | 3.327711234 | 5.838937808 | 2 | 9.257971441 | 9.24663965 | 8.517369639 | 8.369452607 |
| ENSG00000187942 | LDLRAD2 | 3.899565296 | 0.034437161 | 5.000204583 | 0.004698955 | 8.899769879 | 2 | 2 | 2 | 4.358765249 | 4.16043394 | 4.868377487 | 5.459179378 |
| ENSG00000008294 | SPAG9 | 4.773769694 | 3.42E-11 | 4.114380198 | 1.36E-08 | 8.888149892 | 6.366709671 | 7.819099893 | 5.772716771 | 11.66459642 | 11.6718015 | 11.0292199 | 10.99018765 |
| ENSG00000198053 | SIRPA | 4.194106351 | 0.000939403 | 4.693283818 | 0.00016843 | 8.88739017 | 7.031698319 | 5.946638857 | 2 | 10.08684133 | 10.1984268 | 10.40294892 | 10.8459572 |
| ENSG00000152049 | KCNE4 | 3.20527256 | 0.00056585 | 5.677365834 | 1.34E-10 | 8.882638394 | 5.532353208 | 3.076665209 | 5.772716771 | 8.139053376 | 8.25224987 | 11.04993501 | 10.10247774 |
| ENSG00000142611 | PRDM16 | 3.75755215 | 0.052169147 | 5.123505645 | 0.005357536 | 8.881057795 | 2 | 2 | 2 | 4.415698455 | 3.82856379 | 5.626574484 | 4.886954834 |
| ENSG00000186174 | BCL9L | 4.333012226 | 0.014995326 | 4.54247202 | 0.009917482 | 8.875484247 | 2 | 4.710564154 | 8.236946975 | 11.03165415 | 11.100209 | 11.21128275 | 11.33681566 |
| ENSG00000159399 | HK2 | 5.060373974 | 0.000341058 | 3.80732126 | 0.008757632 | 8.867695234 | 7.096806373 | 7.790474954 | 2 | 11.91235914 | 11.9600175 | 10.50913655 | 10.84236645 |
| ENSG00000160460 | SPTBN4 | 2.69035537 | 0.127930027 | 6.176350878 | 8.09E-05 | 8.866706248 | 2 | 2 | 2 | 3.515101962 | 3.28037825 | 6.265258498 | 6.29264319 |
| ENSG00000168679 | SLC16A4 | 5.269751219 | 2.54E-23 | 3.595242251 | 1.85E-11 | 8.86499347 | 4.467311984 | 4.710564154 | 5.59507254 | 10.08738902 | 10.0311812 | 8.331987532 | 8.459018364 |
| ENSG00000008086 | CDKL5 | 3.657144508 | 4.62E-07 | 5.203413808 | 1.76E-13 | 8.860558316 | 2 | 4.442710129 | 5.156626401 | 7.734324667 | 7.70710533 | 9.323615907 | 9.170158797 |
| ENSG00000048471 | SNX29 | 4.357269646 | 5.91E-08 | 4.499147832 | 1.78E-08 | 8.856417477 | 2 | 4.442710129 | 5.156626401 | 8.432517176 | 8.38855922 | 8.245719321 | 8.802925779 |
| ENSG00000115902 | SLC1A4 | 1.876951314 | 0.363985262 | 6.974728426 | 8.35E-05 | 8.851679739 | 2 | 6.602424674 | 2 | 7.075737784 | 6.73446058 | 12.03768528 | 11.8900644 |
| ENSG00000180257 | ZNF816 | 4.403750004 | 0.001160148 | 4.43236426 | 0.000997654 | 8.836114263 | 2 | 3.686303414 | 2 | 6.321555392 | 6.29111322 | 6.560755617 | 6.06327132 |
| ENSG00000153714 | LURAP1L | 5.500080598 | 1.98E-11 | 3.328905846 | 8.70E-05 | 8.828986444 | 3.327711234 | 4.93638933 | 3.369317972 | 9.279220784 | 9.35944426 | 7.508544306 | 6.753261212 |
| ENSG00000164506 | STXBP5 | 4.943824931 | 0.008424242 | 3.882546272 | 0.042592362 | 8.826371204 | 2 | 6.602424674 | 2 | 9.960996031 | 9.91578828 | 9.023457772 | 8.72879856 |
| ENSG00000274290 | HIST1H2BE | 7.706299672 | 6.07E-08 | 1.118303045 | 0.523326453 | 8.824602717 | 2 | 2 | 2 | 7.770242853 | 7.74878077 | 2.560105256 | 2.70970617 |
| ENSG00000134516 | DOCK2 | 4.830339166 | 0.00023646 | 3.993386186 | 0.002779311 | 8.823725353 | 4.00722531 | 2 | 2 | 6.682159616 | 6.8563927 | 5.798032389 | 6.13459968 |
| ENSG00000175946 | KLHL38 | 2.012037713 | 0.265703867 | 6.809802605 | 8.10E-06 | 8.821840319 | 2 | 2 | 2 | 3.09461492 | 2.94009902 | 7.091899617 | 6.645056213 |
| ENSG00000150556 | LYPD6B | 4.743744471 | 0.007283609 | 4.067666523 | 0.022954732 | 8.811410994 | 2 | 2 | 2 | 4.804776789 | 5.11361705 | 4.311629396 | 4.479073019 |
| ENSG00000027075 | PRKCH | 1.588461057 | 0.143712254 | 7.221267023 | 5.01E-15 | 8.809728081 | 3.327711234 | 4.442710129 | 2 | 5.022023783 | 4.83276841 | 9.887932872 | 10.72261753 |
| ENSG00000239704 | CDRT4 | 7.772291328 | 6.03E-08 | 1.028217102 | 0.564110709 | 8.80050843 | 2 | 2 | 2 | 7.711768301 | 7.92877397 | 2.68698273 | 2.509591799 |
| ENSG00000161955 | TNFSF13 | 5.148183554 | 0.002492025 | 3.639273535 | 0.038718339 | 8.787457089 | 2 | 2 | 2 | 5.210797711 | 5.43283104 | 4.029788235 | 4.09155917 |
| ENSG00000179772 | FOXS1 | -1.684535475 | 0.428119634 | 10.46953201 | 2.11E-12 | 8.784996532 | 3.327711234 | 2 | 2 | 2.3892346 | 2 | 11.78416757 | 11.04050035 |
| ENSG00000100055 | CYTH4 | 4.132509496 | 0.054329477 | 4.652256268 | 0.026219867 | 8.784765764 | 2 | 2 | 2 | 4.470469939 | 4.43005853 | 5.47655442 | 3.875378318 |
| ENSG00000130589 | HELZ2 | 6.012493091 | 4.63E-10 | 2.769822063 | 0.007277007 | 8.782315155 | 5.095959458 | 7.701025996 | 5.156626401 | 12.47045468 | 12.5434121 | 9.307967678 | 9.237319199 |
| ENSG00000174004 | NRROS | 5.693927948 | 0.003801199 | 3.083717045 | 0.14026642 | 8.777644993 | 2 | 2 | 2 | 5.627094009 | 6.00043082 | 4.144793515 | 2.918126211 |
| ENSG00000132357 | CARD6 | 5.46942958 | 1.54E-12 | 3.306124352 | 3.44E-05 | 8.775553931 | 3.327711234 | 3.686303414 | 5.39244166 | 9.54788723 | 9.55457603 | 7.141214687 | 7.644317657 |
| ENSG00000141905 | NFIC | 4.228418833 | 1.18E-13 | 4.545449851 | 9.39E-16 | 8.773868684 | 7.096806373 | 6.04685525 | 7.646393253 | 11.21699976 | 11.260058 | 11.40695016 | 11.68972084 |
| ENSG00000138604 | GLCE | 4.82261625 | 0.000673951 | 3.944971549 | 0.006315896 | 8.7675878 | 2 | 5.457120953 | 2 | 8.709369553 | 8.63367258 | 7.610117916 | 7.978752403 |
| ENSG00000163635 | ATXN7 | 4.577930579 | 4.30E-08 | 4.183105897 | 6.07E-07 | 8.761036475 | 2 | 5.595927442 | 4.874587708 | 9.172798677 | 9.20032903 | 8.836358413 | 8.752233325 |
| ENSG00000198648 | STK39 | 3.964476128 | 0.003618428 | 4.793675344 | 0.000308673 | 8.758151471 | 3.327711234 | 6.140560021 | 2 | 8.556367783 | 8.71301725 | 9.391568904 | 9.524680706 |
| ENSG00000068654 | POLR1A | 3.974597393 | 0.009844814 | 4.782326074 | 0.001419617 | 8.756923467 | 3.327711234 | 6.602424674 | 2 | 9.00785664 | 9.1347778 | 9.751677657 | 9.990140122 |
| ENSG00000061337 | LZTS1 | 1.377678148 | 0.334988875 | 7.371895952 | 5.10E-10 | 8.7495741 | 2 | 2 | 5.156626401 | 4.965943888 | 4.79082474 | 10.49971375 | 10.81927984 |
| ENSG00000180855 | ZNF443 | 4.723324601 | 0.011100462 | 4.015939843 | 0.032745113 | 8.739264444 | 2 | 2 | 2 | 5.022023783 | 4.87352701 | 4.667592615 | 3.956354764 |
| ENSG00000163485 | ADORA1 | 5.63166593 | 0.000525107 | 3.104415889 | 0.072475386 | 8.736081819 | 2 | 2 | 2 | 5.720439317 | 5.81407719 | 3.648325314 | 3.698365557 |
| ENSG00000052795 | FNIP2 | 4.127242846 | 0.044781658 | 4.602149567 | 0.022444642 | 8.729392413 | 2 | 7.002537973 | 2 | 9.575477782 | 9.48064953 | 10.06009163 | 9.940497614 |
| ENSG00000151276 | MAGI1 | 3.7386577 | 0.000167805 | 4.982258644 | 2.42E-07 | 8.720916344 | 3.327711234 | 5.303523747 | 2 | 7.671425626 | 7.70428369 | 8.997677834 | 8.827888359 |
| ENSG00000177575 | CD163 | 1.935682413 | 0.307741842 | 6.781936302 | 2.05E-05 | 8.717618715 | 2 | 2 | 2 | 2.784612485 | 3.15372745 | 7.139828829 | 6.509409088 |
| ENSG00000107130 | NCS1 | 4.264739673 | 0.000798129 | 4.451372046 | 0.000410013 | 8.716111719 | 2 | 6.899080544 | 5.59507254 | 9.977635701 | 10.0722485 | 10.0536666 | 10.35406616 |
| ENSG00000139625 | MAP3K12 | 4.544606119 | 1.54E-07 | 4.161442703 | 1.66E-06 | 8.706048823 | 5.330594134 | 3.686303414 | 2 | 8.485003881 | 8.40776525 | 8.085786009 | 8.051511707 |
| ENSG00000146416 | AIG1 | 4.150213419 | 0.00590766 | 4.555734894 | 0.002143773 | 8.705948312 | 2 | 5.595927442 | 2 | 8.14116509 | 8.14812525 | 8.438403662 | 8.644559262 |
| ENSG00000182578 | CSF1R | 3.244230898 | 0.078987729 | 5.457595397 | 0.001364214 | 8.701826295 | 2 | 2 | 2 | 3.79808662 | 3.74316441 | 5.85994354 | 5.300743019 |

| | | | | | | | | | | | | | |
|---|---|---|---|---|---|---|---|---|---|---|---|---|---|
| ENSG00000166750 | SLFN5 | 5.645927178 | 0.001311774 | 3.051054841 | 0.102207077 | 8.696982019 | 8.609439662 | 5.131603625 | 2 | 12.76808443 | 12.7631125 | 10.36682853 | 9.953377568 |
| ENSG00000174943 | KCTD13 | 4.464865671 | 5.21E-08 | 4.230147505 | 2.48E-07 | 8.695013176 | 4.467311984 | 4.11352228 | 2 | 7.872924734 | 7.97866656 | 7.899349298 | 7.459438299 |
| ENSG00000162511 | LAPTM5 | 1.889106343 | 0.31619153 | 6.804277836 | 2.44E-05 | 8.693384178 | 3.327711234 | 3.076665209 | 2 | 3.881293667 | 4.31639476 | 9.481845891 | 6.605433158 |
| ENSG00000153029 | MR1 | 3.843200016 | 2.59E-16 | 4.844771988 | 1.16E-25 | 8.687972003 | 4.815609471 | 5.595927442 | 5.39244166 | 9.03081742 | 9.03162973 | 10.07231173 | 9.980734518 |
| ENSG00000166016 | ABTB2 | 5.030963174 | 3.48E-10 | 3.656702346 | 7.78E-06 | 8.687665521 | 4.00722531 | 3.686303414 | 3.369317972 | 8.192977693 | 8.28094035 | 6.977080688 | 6.804465095 |
| ENSG00000110218 | PANX1 | 4.924683509 | 3.39E-05 | 3.762447747 | 0.00197542 | 8.687131256 | 6.653098118 | 6.602424674 | 2 | 10.92262909 | 10.9213279 | 9.542625156 | 9.954856344 |
| ENSG00000134013 | LOXL2 | 3.910757536 | 4.55E-07 | 4.776289054 | 3.92E-10 | 8.68704659 | 9.895281535 | 9.726146444 | 7.600047325 | 13.24586113 | 13.3136663 | 14.05496899 | 14.23054703 |
| ENSG00000186765 | FSCN2 | 2.735469455 | 0.175994512 | 5.942462043 | 0.001005841 | 8.677931498 | 2 | 2 | 2 | 3.615725078 | 3.2184422 | 6.544089217 | 5.319231103 |
| ENSG00000126882 | FAM78A | 3.548820592 | 0.110727349 | 5.127683283 | 0.01461872 | 8.676503875 | 2 | 2 | 2 | 4.17310023 | 3.7864959 | 5.941748584 | 4.147809227 |
| ENSG00000242221 | PSG2 | 2.391008583 | 0.206839295 | 6.281485465 | 0.000142458 | 8.672494048 | 2 | 2 | 2 | 3.349643765 | 3.08597292 | 5.925754458 | 6.723495198 |
| ENSG00000169499 | PLEKHA2 | 3.918321376 | 1.02E-13 | 4.751461885 | 6.52E-20 | 8.669783261 | 5.70932728 | 4.442710129 | 5.59507254 | 9.085573664 | 9.06500642 | 9.821413244 | 9.981702022 |
| ENSG00000119900 | OGFRL1 | 5.082680168 | 1.35E-23 | 3.586693001 | 2.39E-12 | 8.669373168 | 4.467311984 | 5.72254323 | 5.39244166 | 10.27264434 | 10.2355442 | 8.646918883 | 8.877593894 |
| ENSG00000011454 | RABGAP1 | 3.863894114 | 0.009737709 | 4.802950072 | 0.000925327 | 8.666844186 | 4.467311984 | 7.099070191 | 2 | 9.50916971 | 9.59626647 | 10.67247936 | 10.27813351 |
| ENSG00000136040 | PLXNC1 | 3.115211289 | 0.118797449 | 5.549694002 | 0.002463699 | 8.664905291 | 2 | 2 | 2 | 3.615725078 | 3.74316441 | 6.165271053 | 4.982847849 |
| ENSG00000182134 | TDRKH | 3.282596501 | 0.042407869 | 5.38155109 | 0.000354475 | 8.664147591 | 2 | 3.076665209 | 2 | 4.598938466 | 4.4836996 | 6.162543904 | 6.669279147 |
| ENSG00000134463 | ECHDC3 | 3.213368236 | 0.074785353 | 5.441540968 | 0.00106239 | 8.654909204 | 2 | 2 | 2 | 3.79808662 | 3.69849104 | 5.65789504 | 5.524371359 |
| ENSG00000164309 | CMYA5 | 6.550094872 | 2.17E-05 | 2.104505109 | 0.227259568 | 8.654599981 | 2 | 2 | 2 | 6.5612028 | 6.70653264 | 2.911488689 | 3.209956562 |
| ENSG00000121289 | CEP89 | 3.573552177 | 0.002199247 | 5.079409191 | 5.90E-06 | 8.652961367 | 2 | 4.93638933 | 2 | 7.030886362 | 6.8563927 | 8.416251718 | 8.426701165 |
| ENSG00000135766 | EGLN1 | 4.21150811 | 0.000734564 | 4.438947428 | 0.000325136 | 8.650455538 | 4.00722531 | 7.35507212 | 4.058998049 | 10.163122 | 10.2094139 | 10.31312388 | 10.50671159 |
| ENSG00000170634 | ACYP2 | 4.209434957 | 8.24E-05 | 4.436160618 | 2.75E-05 | 8.645595574 | 3.327711234 | 3.686303414 | 2 | 6.682159616 | 6.71776877 | 6.768721236 | 7.05482945 |
| ENSG00000135363 | LMO2 | 3.776712715 | 0.046133395 | 4.865065777 | 0.007316377 | 8.641778492 | 2 | 2 | 2 | 4.139692846 | 4.1930041 | 4.705557986 | 5.367404784 |
| ENSG00000177352 | CCDC71 | 3.793507094 | 0.001214951 | 4.848091067 | 2.04E-05 | 8.641598162 | 2 | 4.93638933 | 2 | 7.127781374 | 7.18663697 | 8.092249024 | 8.28340307 |
| ENSG00000123700 | KCNJ2 | 4.198925526 | 0.031426015 | 4.435206346 | 0.020987656 | 8.634131871 | 2 | 2 | 2 | 4.739373666 | 4.22485516 | 5.020706951 | 4.292061816 |
| ENSG00000196943 | NOP9 | 4.514739579 | 2.27E-09 | 4.118787746 | 5.33E-08 | 8.633527325 | 5.095959458 | 5.457120953 | 2 | 9.20242622 | 9.07159009 | 8.763089655 | 8.728222198 |
| ENSG00000151952 | TMEM132D | 1.39154689 | 0.556481113 | 7.241566745 | 0.000163335 | 8.633113635 | 2 | 2 | 2 | 2.868578001 | 2.59489879 | 5.582367412 | 8.068003971 |
| ENSG00000173517 | PEAK1 | 3.762709178 | 0.001267977 | 4.866623243 | 1.75E-05 | 8.629332421 | 5.866944937 | 7.954278496 | 4.058998049 | 10.43808429 | 10.5201748 | 11.54344182 | 11.61849721 |
| ENSG00000125686 | MED1 | 4.189417566 | 0.047220426 | 4.436335062 | 0.032697033 | 8.625752627 | 2 | 7.189546432 | 2 | 9.759244431 | 9.79831747 | 10.00178175 | 10.04740207 |
| ENSG00000104756 | KCTD9 | 4.611056131 | 0.002344951 | 4.012587793 | 0.00887817 | 8.623643924 | 3.327711234 | 6.602424674 | 2 | 9.797905134 | 9.60612566 | 8.975319061 | 9.233668966 |
| ENSG00000047365 | ARAP2 | 4.081212359 | 0.101317769 | 4.542129765 | 0.060381002 | 8.623342124 | 2 | 2 | 2 | 4.329433326 | 4.4836996 | 5.545162502 | 3.128433649 |
| ENSG00000095203 | EPB41L4B | 2.859406534 | 0.112536896 | 5.752258743 | 0.000393732 | 8.611665277 | 2 | 2 | 2 | 3.515101962 | 3.50452879 | 5.758846871 | 5.992078266 |
| ENSG00000177508 | IRX3 | 5.031313049 | 0.001032336 | 3.57540954 | 0.024293372 | 8.606722589 | 2 | 3.076665209 | 2 | 6.087554739 | 6.12381559 | 4.990270811 | 4.553838275 |
| ENSG00000112343 | TRIM38 | 5.592867965 | 8.78E-09 | 3.011134606 | 0.003301386 | 8.604002571 | 4.467311984 | 6.844429755 | 4.058998049 | 11.20493678 | 11.195718 | 8.69913345 | 8.55898913 |
| ENSG00000127399 | LRRC61 | 6.749806174 | 2.13E-05 | 1.847242261 | 0.310818746 | 8.597048435 | 2 | 2 | 2 | 6.772115916 | 6.88158913 | 2.68698273 | 3.15612297 |
| ENSG00000137628 | DDX60 | 6.51145857 | 2.68E-22 | 2.084283292 | 0.00396904 | 8.595741861 | 4.815609471 | 5.72254323 | 3.369317972 | 11.30956651 | 11.3218808 | 7.116063252 | 6.725806822 |
| ENSG00000171055 | FEZ2 | 4.453093635 | 4.93E-11 | 4.142576536 | 1.03E-09 | 8.595670171 | 4.00722531 | 6.04685525 | 5.156626401 | 9.677941812 | 9.62862374 | 9.577849253 | 9.066361132 |
| ENSG00000100558 | PLEK2 | 5.390969793 | 2.68E-05 | 3.204445527 | 0.018019059 | 8.59541532 | 2 | 3.686303414 | 2 | 7.261917051 | 7.26532099 | 5.04460184 | 5.349527629 |
| ENSG00000176155 | CCDC57 | 4.795786217 | 1.21E-07 | 3.798705265 | 3.64E-05 | 8.594491482 | 2 | 5.131603625 | 3.369317972 | 8.593855195 | 8.58388295 | 7.590975897 | 7.622152309 |
| ENSG00000123689 | G0S2 | 7.332003286 | 8.87E-21 | 1.26006099 | 0.165797053 | 8.592064276 | 2 | 3.686303414 | 4.874587708 | 10.79398911 | 10.8013584 | 5.096960836 | 4.729989799 |
| ENSG00000162616 | DNAJB4 | 3.951509478 | 3.57E-06 | 4.636696202 | 3.34E-08 | 8.588205679 | 4.00722531 | 6.78762695 | 5.59507254 | 9.845303961 | 9.6580856 | 10.63251333 | 10.211653 |
| ENSG00000138395 | CDK15 | 3.586725007 | 0.025019077 | 4.9999964 | 0.001038388 | 8.586721406 | 3.327711234 | 2 | 2 | 4.965943888 | 4.60967607 | 6.189586148 | 5.953147751 |
| ENSG00000127603 | MACF1 | 3.357913025 | 3.09E-09 | 5.221786007 | 3.88E-21 | 8.579699032 | 8.16936288 | 9.687856435 | 8.865353883 | 12.36993409 | 12.4157698 | 14.27801527 | 14.23390067 |
| ENSG00000229937 | PRPS1L1 | 3.973668015 | 0.040174325 | 4.600007476 | 0.014521931 | 8.573675491 | 2 | 2 | 2 | 4.139692846 | 4.4836996 | 4.45225155 | 5.150533729 |
| ENSG00000169683 | LRRC45 | 3.979876955 | 0.002187892 | 4.590077009 | 0.000303676 | 8.569953964 | 3.327711234 | 3.076665209 | 2 | 5.900763176 | 5.9725602 | 6.619606851 | 6.400759785 |
| ENSG00000114268 | PFKFB4 | 4.99585869 | 0.004382954 | 3.565287954 | 0.048832636 | 8.561146644 | 2 | 6.228547981 | 2 | 9.576258382 | 9.64931004 | 7.864534187 | 8.463873348 |
| ENSG00000152154 | TMEM178A | 4.800546312 | 0.011612142 | 3.749098355 | 0.053196051 | 8.549644666 | 2 | 2 | 2 | 5.37770462 | 4.53541753 | 4.311629396 | 3.956354764 |
| ENSG00000105971 | CAV2 | 3.231382538 | 0.00129249 | 5.309331137 | 2.87E-08 | 8.540713675 | 4.00722531 | 5.457120953 | 7.225742779 | 9.263797825 | 9.27353851 | 11.44714298 | 11.22309073 |
| ENSG00000172137 | CALB2 | 8.605979182 | 5.56E-10 | -0.06745967 | 0.97269175 | 8.538519512 | 2 | 2 | 2 | 8.633396609 | 8.6601233 | 2.34604626 | 2.277179783 |
| ENSG00000126464 | PRR12 | 4.126970846 | 0.00015212 | 4.399831248 | 4.51E-05 | 8.526802095 | 5.095959458 | 6.311476754 | 2 | 9.270565666 | 9.36213202 | 9.600443341 | 9.57619232 |
| ENSG00000174529 | TMEM81 | 3.547727235 | 0.058360787 | 4.97872866 | 0.004973982 | 8.526455895 | 2 | 2 | 2 | 4.034562184 | 3.94787299 | 5.390603804 | 4.919632516 |
| ENSG00000183087 | GAS6 | 5.494411386 | 2.52E-06 | 3.030213822 | 0.014425801 | 8.524625208 | 7.384433967 | 7.847167909 | 3.369317972 | 12.51021088 | 12.5811024 | 10.3504468 | 9.762514036 |
| ENSG00000125898 | FAM110A | 4.385896319 | 0.004693545 | 4.138081795 | 0.007684751 | 8.523978114 | 2 | 3.076665209 | 2 | 5.514313187 | 5.49945122 | 5.385935071 | 5.171036168 |
| ENSG00000134318 | ROCK2 | 3.788277668 | 0.001047374 | 4.735616215 | 2.57E-05 | 8.523893883 | 3.327711234 | 7.274681896 | 5.39244166 | 9.787831283 | 9.89180734 | 10.76763523 | 10.8022669 |
| ENSG00000184923 | NUTM2A | 6.695347911 | 4.30E-05 | 1.826909171 | 0.331750585 | 8.522257081 | 2 | 2 | 2 | 6.825617242 | 6.72335419 | 2.62493832 | 3.183290849 |
| ENSG00000128422 | KRT17 | 2.821542772 | 0.15015685 | 5.697386356 | 0.001258335 | 8.518929128 | 2 | 2 | 2 | 3.79808662 | 3.08597292 | 5.744329515 | 5.909074772 |
| ENSG00000102554 | KLF5 | 3.016885928 | 0.011204778 | 5.495435499 | 7.70E-07 | 8.512321427 | 2 | 4.93638933 | 2 | 6.5612028 | 6.24526024 | 8.86850882 | 8.797990868 |
| ENSG00000172432 | GTPBP2 | 4.367510099 | 0.001011359 | 4.143447163 | 0.00183886 | 8.510957262 | 7.159102577 | 5.595927442 | 2 | 10.26055405 | 10.3455691 | 10.27985161 | 9.848823977 |
| ENSG00000099364 | FBXL19 | 4.244683475 | 0.013556653 | 4.265152558 | 0.012599871 | 8.509836033 | 5.866944937 | 2 | 2 | 8.442826028 | 8.38855922 | 8.50665722 | 8.362041249 |
| ENSG00000033867 | SLC4A7 | 3.229295081 | 9.45E-13 | 5.279245243 | 6.93E-33 | 8.508540324 | 5.866944937 | 6.464272742 | 7.029880454 | 9.6581704 | 9.73198422 | 11.86481281 | 11.6038049 |
| ENSG00000082641 | NFE2L1 | 4.212162052 | 8.01E-19 | 4.293358353 | 1.01E-19 | 8.505520406 | 7.667647223 | 8.916342327 | 8.486635645 | 12.62008676 | 12.6749758 | 12.7696937 | 12.68699458 |
| ENSG00000204634 | TBC1D8 | 3.65919245 | 0.007262014 | 4.844398467 | 0.000218044 | 8.503590916 | 2 | 3.686303414 | 2 | 5.639097847 | 5.58783529 | 6.577231679 | 6.870723585 |
| ENSG00000110844 | PRPF40B | 4.292586615 | 0.001220428 | 4.206280166 | 0.001463478 | 8.49886678 | 2 | 3.686303414 | 2 | 6.237202172 | 6.17362678 | 6.078235975 | 6.168983563 |
| ENSG00000178104 | PDE4DIP | 5.008717999 | 0.000112427 | 3.487912106 | 0.009445149 | 8.496630105 | 4.00722531 | 8.376385542 | 5.59507254 | 12.0064372 | 12.0433189 | 10.56428727 | 10.44684159 |
| ENSG00000225614 | ZNF469 | 3.275091567 | 0.146749658 | 5.205189508 | 0.01401918 | 8.480281074 | 2 | 7.431218269 | 2 | 9.111653326 | 9.11366683 | 10.69589097 | 11.30943053 |
| ENSG00000273173 | SNURF | 9.480468985 | 1.65E-10 | -1.002012766 | 0.614177 | 8.478456219 | 2 | 2 | 2 | 9.565291356 | 9.46239569 | 2.267001569 | 2.059860178 |
| ENSG00000111052 | LIN7A | 2.535726363 | 0.076658392 | 5.936887369 | 3.98E-06 | 8.472613731 | 2 | 4.11352228 | 2 | 5.178261742 | 4.98931816 | 8.838922815 | 7.520630205 |
| ENSG00000130402 | ACTN4 | 3.032743133 | 0.001405029 | 5.432451405 | 1.58E-09 | 8.465194538 | 9.751047882 | 7.874700259 | 7.029880454 | 11.67556519 | 11.7141436 | 13.95211632 | 14.2218472 |
| ENSG00000197506 | SLC28A3 | 3.497936115 | 0.063878162 | 4.953053537 | 0.005506115 | 8.450989652 | 2 | 2 | 2 | 4.105493497 | 3.7864959 | 4.977914261 | 5.306932076 |
| ENSG00000135604 | STX11 | 3.52627378 | 0.000350662 | 4.922719529 | 2.31E-07 | 8.448993309 | 2 | 3.076665209 | 4.523675391 | 6.554880361 | 6.39275508 | 7.958717528 | 7.692387724 |
| ENSG00000170442 | KRT86 | 1.689483666 | 0.4198367 | 6.757311064 | 9.43E-05 | 8.44679473 | 2 | 2 | 2 | 2.868578001 | 2.86123014 | 7.395794919 | 5.905000563 |
| ENSG00000112715 | VEGFA | 5.149069036 | 5.54E-07 | 3.295228685 | 0.002061106 | 8.444297721 | 6.138360429 | 8.191702463 | 4.874587708 | 12.14652252 | 12.1562841 | 10.39091191 | 10.2048333 |
| ENSG00000198855 | FICD | 3.820079318 | 0.000637206 | 4.621494851 | 2.29E-05 | 8.441574169 | 2 | 4.710564154 | 2 | 7.066878356 | 6.83591385 | 7.619097297 | 7.846917442 |
| ENSG00000147437 | GNRH1 | 4.756040247 | 0.007796496 | 3.681366835 | 0.043775504 | 8.437407081 | 2 | 2 | 2 | 5.058234798 | 4.89348248 | 4.177580368 | 4.002845082 |
| ENSG00000196159 | FAT4 | 3.707925812 | 0.054329477 | 4.727966788 | 0.010905332 | 8.4358926 | 2 | 6.535001748 | 2 | 8.593855195 | 8.69038176 | 9.718175864 | 9.590215873 |
| ENSG00000174945 | AMZ1 | 1.7756677 | 0.443101353 | 6.645197601 | 0.000596608 | 8.4208653 | 2 | 2 | 2 | 2.784612485 | 3.01487888 | 7.44824109 | 5.217771644 |
| ENSG00000124813 | RUNX2 | 6.053255828 | 1.14E-06 | 2.367114942 | 0.08323504 | 8.42037077 | 2 | 2 | 4.874587708 | 9.020530266 | 9.02259731 | 5.815498119 | 4.990558504 |
| ENSG00000087903 | RFX2 | 3.498830889 | 0.017592591 | 4.914373278 | 0.000468869 | 8.413204167 | 2 | 3.686303414 | 2 | 5.44762523 | 5.45984938 | 6.292954465 | 7.150854407 |
| ENSG00000185591 | SP1 | 3.866497077 | 0.001939227 | 4.529560624 | 0.000207819 | 8.396057701 | 6.468621203 | 6.899080544 | 2 | 9.946579093 | 9.94237362 | 10.71148464 | 10.49082905 |
| ENSG00000162852 | CNST | 4.088282043 | 0.007712736 | 4.307741101 | 0.004514133 | 8.396023143 | 2 | 6.535001748 | 3.369317972 | 9.135152476 | 9.11472974 | 9.623185015 | 8.994845077 |
| ENSG00000171940 | ZNF217 | 5.077010019 | 0.001072632 | 3.315270843 | 0.040591855 | 8.392280862 | 2 | 7.189546432 | 4.058998049 | 10.78591371 | 10.797723 | 9.003023259 | 9.071823126 |
| ENSG00000121879 | PIK3CA | 4.767898268 | 0.000473948 | 3.624074383 | 0.009668661 | 8.391972651 | 3.327711234 | 6.140560021 | 2 | 9.384581425 | 9.47817393 | 8.58359791 | 7.942440336 |
| ENSG00000067840 | PDZD4 | 0.571869157 | 0.776233807 | 7.815927214 | 2.65E-07 | 8.387796371 | 2 | 3.686303414 | 2 | 3.615725078 | 2.59489879 | 8.885973502 | 10.09757713 |
| ENSG00000181019 | NQO1 | 5.77326988 | 3.25E-28 | 2.607611387 | 1.54E-06 | 8.380881267 | 6.366709671 | 6.311476754 | 6.629488387 | 12.14822967 | 12.1383651 | 8.402448739 | 9.403205664 |
| ENSG00000133794 | ARNTL | 5.263476455 | 2.25E-09 | 3.116434945 | 0.000679074 | 8.3799114 | 3.327711234 | 4.442710129 | 2 | 8.418656444 | 8.44034872 | 6.514451546 | 6.13807518 |
| ENSG00000177453 | NIM1K | 4.974563799 | 0.013104298 | 3.400061511 | 0.100917556 | 8.374625309 | 2 | 2 | 2 | 5.128039203 | 5.21224859 | 3.235948948 | 4.329022995 |
| ENSG00000087095 | NLK | 4.368440605 | 1.67E-06 | 4.001111538 | 1.25E-05 | 8.369552142 | 4.00722531 | 5.303523747 | 2 | 8.486665614 | 8.4705529 | 8.367944224 | 7.81035357 |
| ENSG00000114302 | PRKAR2A | 3.352450204 | 0.052698164 | 5.015726417 | 0.002179419 | 8.368176621 | 2 | 7.051611315 | 3.369317972 | 8.948803189 | 8.81332035 | 10.45569736 | 10.61496321 |
| ENSG00000257365 | FNTB | 4.798983141 | 0.001854643 | 3.567715117 | 0.024667973 | 8.366698258 | 5.532353208 | 2 | 2 | 8.522747711 | 8.66157866 | 7.336538937 | 7.427130611 |
| ENSG00000143515 | ATP8B2 | 4.027789445 | 1.92E-09 | 4.331039304 | 7.96E-11 | 8.358828749 | 7.384433967 | 5.946638857 | 5.930865648 | 10.53200702 | 10.5760362 | 10.43041464 | 11.18537541 |
| ENSG00000138641 | HERC3 | 4.582030096 | 4.78E-06 | 3.774959631 | 0.000201497 | 8.356989728 | 3.327711234 | 5.303523747 | 2 | 8.535647908 | 8.4936155 | 7.564363052 | 7.859604233 |
| ENSG00000122877 | EGR2 | 5.09847937 | 0.016034926 | 3.25498839 | 0.139084171 | 8.353467761 | 2 | 2 | 3.369317972 | 6.197166296 | 6.14061114 | 5.284230775 | 2.782576693 |
| ENSG00000125618 | PAX8 | 4.153025464 | 0.014151141 | 4.19773519 | 0.012393734 | 8.350760654 | 2 | 2 | 3.369317972 | 5.144975053 | 5.43283104 | 4.799832052 | 5.727202496 |
| ENSG00000186088 | GSAP | 6.576324517 | 8.91E-06 | 1.768990809 | 0.295440666 | 8.345315326 | 2 | 2 | 4.058998049 | 8.567402595 | 8.46222608 | 4.588526421 | 3.287118176 |
| ENSG00000151690 | MFSD6 | 3.37350894 | 1.85E-05 | 4.96301614 | 7.26E-11 | 8.33652508 | 2 | 4.93638933 | 4.058998049 | 7.273514749 | 7.19470132 | 8.871434321 | 8.72128783 |
| ENSG00000213949 | ITGA1 | 4.647910424 | 0.000421918 | 3.687424558 | 0.006183393 | 8.335334982 | 2 | 7.002537973 | 5.930865648 | 10.60458486 | 10.5632459 | 10.03197853 | 9.062251023 |
| ENSG00000185262 | UBALD2 | 4.813755985 | 5.20E-08 | 3.519099315 | 9.94E-05 | 8.3328553 | 4.467311984 | 5.595927442 | 2 | 9.333786218 | 9.24178305 | 8.153962636 | 7.843728236 |
| ENSG00000130653 | PNPLA7 | 3.859404043 | 0.039042892 | 4.466767144 | 0.013950213 | 8.326171187 | 2 | 2 | 2 | 4.237680128 | 4.22485516 | 4.799832052 | 4.654341953 |
| ENSG00000112378 | PERP | 3.547917517 | 6.03E-16 | 4.777411636 | 1.88E-28 | 8.325329153 | 6.816637977 | 6.844429755 | 6.534109959 | 10.24934078 | 10.2345667 | 11.09309185 | 11.76653247 |
| ENSG00000043143 | JADE2 | 5.22490407 | 1.36E-15 | 3.095100706 | 4.17E-06 | 8.320004776 | 3.327711234 | 5.72254323 | 5.59507254 | 10.27841188 | 10.3373973 | 8.319187939 | 8.056111835 |
| ENSG00000121577 | POPDC2 | 2.548700716 | 0.176945672 | 5.768803074 | 0.00056294 | 8.31750379 | 2 | 2 | 2 | 3.462030052 | 3.15372745 | 6.060766632 | 5.708623051 |
| ENSG00000163395 | IGFN1 | 2.49683765 | 0.281095622 | 5.817045798 | 0.004719809 | 8.313883449 | 2 | 2 | 2 | 3.289987342 | 3.28037825 | 4.487545924 | 6.649933445 |
| ENSG00000164463 | CREBRF | 5.131996233 | 0.000203981 | 3.179888796 | 0.028543748 | 8.311885029 | 2 | 6.228547981 | 3.369317972 | 9.918537335 | 9.83876104 | 8.224302521 | 7.597118029 |
| ENSG00000162630 | B3GALT2 | 1.500815324 | 0.50971472 | 6.811047416 | 0.000247567 | 8.31186274 | 2 | 2 | 2 | 3.09461492 | 2.38539787 | 7.375919751 | 6.141542327 |
| ENSG00000244405 | ETV5 | 4.054207174 | 5.65E-16 | 4.253764943 | 1.07E-17 | 8.307972117 | 4.467311984 | 5.303523747 | 5.772716771 | 9.142549916 | 9.21427981 | 9.354710897 | 9.399228411 |
| ENSG00000166924 | NYAP1 | 3.692786932 | 0.023196476 | 4.610800796 | 0.00331552 | 8.303587727 | 2 | 2 | 3.369317972 | 4.783303513 | 4.98931816 | 5.9289675 | 5.459179378 |
| ENSG00000183715 | OPCML | 5.267094572 | 0.037231944 | 3.028416182 | 0.253721513 | 8.295510753 | 2 | 2 | 2 | 5.474671438 | 5.39133044 | 4.291799915 | 2.326754077 |
| ENSG00000130489 | SCO2 | 5.317096923 | 1.93E-14 | 2.975742509 | 3.49E-05 | 8.292839432 | 5.095959458 | 4.442710129 | 2 | 9.363030664 | 9.34230452 | 7.11465301 | 6.975457902 |
| ENSG00000000938 | FGR | 3.275430812 | 0.20396079 | 5.015586062 | 0.037934228 | 8.291016874 | 2 | 2 | 2 | 3.921164147 | 3.65239003 | 5.99483933 | 3.33635788 |

| | | | | | | | | | | | | | |
|---|---|---|---|---|---|---|---|---|---|---|---|---|---|
| ENSG00000106100 | NOD1 | 3.625809444 | 0.007885856 | 4.660603955 | 0.000397713 | 8.286413399 | 3.327711234 | 3.076665209 | 2 | 5.614989456 | 5.60003091 | 6.845542321 | 6.254725615 |
| ENSG00000227124 | ZNF717 | 4.149801872 | 0.00360402 | 4.135500475 | 0.003541393 | 8.285302347 | 2 | 3.686303414 | 2 | 6.104969671 | 6.00960265 | 6.346800618 | 5.661099865 |
| ENSG00000101608 | MYL12A | 3.660120009 | 7.29E-48 | 4.625061695 | 6.35E-77 | 8.285181703 | 8.981092835 | 8.607278467 | 8.742726175 | 12.44525263 | 12.4115594 | 13.35833349 | 13.42672912 |
| ENSG00000152242 | C18orf25 | 3.828697786 | 0.007024937 | 4.4548156 | 0.001326915 | 8.283513385 | 3.327711234 | 6.228547981 | 2 | 8.651287275 | 8.50824913 | 9.37340502 | 9.00820079 |
| ENSG00000179820 | MYADM | 3.843302968 | 0.005370243 | 4.439856422 | 0.001021668 | 8.283159389 | 3.327711234 | 7.954278496 | 8.721235202 | 11.61113422 | 11.675768 | 11.99089185 | 12.45192034 |
| ENSG00000101236 | RNF24 | 4.830482721 | 0.001200331 | 3.450935345 | 0.025360922 | 8.281418066 | 3.327711234 | 7.467835385 | 3.369317972 | 10.81365108 | 10.797723 | 9.318112363 | 9.536251143 |
| ENSG00000221994 | ZNF630 | 3.835074198 | 0.020143473 | 4.437392512 | 0.005795548 | 8.27246671 | 3.327711234 | 2 | 2 | 4.825935136 | 5.18011485 | 5.80853718 | 5.243813244 |
| ENSG00000187556 | NANOS3 | 3.732211093 | 0.056266195 | 4.539508237 | 0.016135868 | 8.27171933 | 2 | 2 | 2 | 3.881293667 | 4.3456612 | 4.580376151 | 4.975095763 |
| ENSG00000141682 | PMAIP1 | 6.663398997 | 6.13E-05 | 1.604927199 | 0.402280751 | 8.268326196 | 2 | 3.076665209 | 6.958202216 | 12.05596597 | 11.9962262 | 7.470462265 | 6.33565646 |
| ENSG00000169826 | CSGALNACT2 | 4.673093807 | 0.0019028 | 3.594939886 | 0.020035793 | 8.268033693 | 2 | 7.538394143 | 5.156626401 | 10.8531554 | 10.8394695 | 9.7848625 | 9.758285822 |
| ENSG00000048828 | FAM120A | 3.75647436 | 2.99E-06 | 4.509541666 | 1.19E-08 | 8.266016026 | 5.330594134 | 7.819099893 | 6.629488387 | 10.68132448 | 10.6309298 | 11.25865892 | 11.54324631 |
| ENSG00000176697 | BDNF | 3.916589394 | 4.54E-05 | 4.334788548 | 4.83E-06 | 8.251377941 | 2 | 5.595927442 | 5.930865648 | 9.066811112 | 8.96128525 | 9.547075488 | 9.303427654 |
| ENSG00000179715 | PCED1B | 3.424951881 | 0.013666189 | 4.818276585 | 0.000275564 | 8.243228466 | 2 | 3.686303414 | 2 | 5.289065324 | 5.49945122 | 6.835304498 | 6.559516442 |
| ENSG00000166825 | ANPEP | 5.966012516 | 0.000116017 | 2.275150353 | 0.181490912 | 8.241162869 | 8.468948322 | 7.315436877 | 2 | 13.32873996 | 13.3982649 | 9.073675485 | 10.10492182 |
| ENSG00000185129 | PURA | 3.516793028 | 0.021758694 | 4.721893817 | 0.001338547 | 8.238686845 | 4.00722531 | 6.899080544 | 2 | 8.980991923 | 8.95536429 | 10.36549517 | 9.936261795 |
| ENSG00000114127 | XRN1 | 5.015682045 | 4.18E-05 | 3.216914373 | 0.011903882 | 8.232596418 | 4.00722531 | 7.35507212 | 4.058998049 | 11.00087894 | 10.9761604 | 9.251341112 | 9.1401427 |
| ENSG00000243716 | NPIPB5 | 7.290568365 | 1.62E-31 | 0.936030687 | 0.203235525 | 8.226599052 | 5.330594134 | 3.686303414 | 4.058998049 | 11.5128316 | 11.4889959 | 5.227950776 | 5.367404784 |
| ENSG00000047188 | YTHDC2 | 4.046718313 | 9.32E-07 | 4.174603332 | 3.54E-07 | 8.221321646 | 2 | 5.131603625 | 4.058998049 | 8.020170999 | 8.06258142 | 8.275683409 | 8.050589918 |
| ENSG00000164091 | WDR82 | 3.779453233 | 2.45E-06 | 4.441540451 | 1.89E-08 | 8.220993684 | 5.095959458 | 7.315436877 | 5.59507254 | 10.06920484 | 10.0804323 | 10.74103322 | 10.72882541 |
| ENSG00000005469 | CROT | 4.630939842 | 1.27E-07 | 3.587671061 | 5.74E-05 | 8.218610904 | 2 | 4.442710129 | 3.369317972 | 7.77569021 | 7.8441483 | 6.875826972 | 6.707209278 |
| ENSG00000171533 | MAP6 | 2.824771867 | 0.060296866 | 5.391002286 | 9.86E-05 | 8.215774154 | 2 | 5.457120953 | 2 | 6.699459234 | 6.73446058 | 9.049523349 | 9.402844548 |
| ENSG00000114859 | CLCN2 | 5.090023711 | 0.003959538 | 3.124667276 | 0.09228972 | 8.214690988 | 2 | 2 | 2 | 5.22679461 | 5.3193943 | 3.838019203 | 3.518055793 |
| ENSG00000075643 | MOCOS | 4.727931341 | 0.007409183 | 3.4827898 | 0.054660707 | 8.210721141 | 2 | 2 | 3.369317972 | 5.870535235 | 5.77157844 | 5.227950776 | 3.8919413 |
| ENSG00000115808 | STRN | 4.09068888 | 0.027679774 | 4.11969899 | 0.025482471 | 8.21038787 | 2 | 6.389896633 | 2 | 8.935480545 | 8.81462916 | 9.005688562 | 8.796891925 |
| ENSG00000130997 | POLN | 4.927933462 | 0.005269294 | 3.280538323 | 0.074166356 | 8.208471784 | 2 | 2 | 2 | 5.144975053 | 5.11361705 | 3.838019203 | 3.753786374 |
| ENSG00000106003 | LFNG | 3.95677742 | 0.058415094 | 4.243613723 | 0.037791254 | 8.200391143 | 2 | 2 | 2 | 4.387512679 | 4.22485516 | 3.904814898 | 4.982847849 |
| ENSG00000187372 | PCDHB13 | 4.012586919 | 0.036823894 | 4.184454704 | 0.026879191 | 8.197041623 | 2 | 2 | 2 | 4.523237872 | 4.16043394 | 4.360042493 | 4.614976609 |
| ENSG00000188647 | PTAR1 | 3.800557613 | 0.030593626 | 4.394853803 | 0.01045232 | 8.195411415 | 2 | 7.189546432 | 3.369317972 | 9.517321994 | 9.39837151 | 10.28315082 | 9.773170377 |
| ENSG00000172061 | LRRC15 | 6.347925878 | 3.71E-05 | 1.846733131 | 0.293334688 | 8.19465901 | 2 | 2 | 2 | 6.490083648 | 6.39275508 | 2.987421008 | 2.885414003 |
| ENSG00000182584 | ACTL10 | 3.597145927 | 0.059410678 | 4.59431599 | 0.011923495 | 8.191461916 | 2 | 2 | 2 | 4.139692846 | 3.90918929 | 4.778616857 | 4.895193906 |
| ENSG00000116711 | PLA2G4A | 7.173589768 | 1.28E-11 | 1.016536039 | 0.427095076 | 8.190125807 | 2 | 4.442710129 | 2 | 9.926517587 | 10.0153322 | 4.470006665 | 3.789585079 |
| ENSG00000102466 | FGF14 | 2.151478169 | 0.27946754 | 6.03778076 | 0.000434519 | 8.189258929 | 2 | 2 | 2 | 3.349643765 | 2.77779907 | 6.010089548 | 6.27380897 |
| ENSG00000108599 | AKAP10 | 4.292120726 | 1.89E-10 | 3.890877478 | 8.48E-09 | 8.182998204 | 3.327711234 | 5.131603625 | 5.59507254 | 9.060130342 | 9.07923325 | 8.896269648 | 8.405948829 |
| ENSG00000185404 | SP140L | 4.267620293 | 5.90E-10 | 3.911205079 | 1.45E-08 | 8.178825372 | 5.095959458 | 4.11352228 | 3.369317972 | 8.293234263 | 8.36015869 | 7.812424466 | 8.122532579 |
| ENSG00000186150 | UBL4B | 1.390400159 | 0.494744255 | 6.786991611 | 3.36E-05 | 8.17739177 | 2 | 2 | 2 | 2.784612485 | 2.68924543 | 6.262714117 | 7.287011591 |
| ENSG00000136925 | TSTD2 | 4.523548691 | 2.57E-10 | 3.63744358 | 4.95E-07 | 8.160992271 | 4.467311984 | 4.710564154 | 2 | 8.369067528 | 8.3920703 | 7.525623979 | 7.492396644 |
| ENSG00000173442 | EHBP1L1 | 3.188683664 | 0.025590506 | 4.97181313 | 0.000232764 | 8.160496794 | 2 | 7.189546432 | 7.225742779 | 9.764733396 | 9.79897887 | 11.27540049 | 11.79831821 |
| ENSG00000275338 | | 7.379553318 | 6.79E-08 | 0.778200119 | 0.649722124 | 8.157753437 | 2 | 3.686303414 | 2 | 9.178978633 | 9.23104065 | 3.679182183 | 2.851942824 |
| ENSG00000142910 | TINAGL1 | -0.844394841 | 0.49749462 | 9.001312748 | 3.58E-22 | 8.156917907 | 3.327711234 | 3.076665209 | 4.058998049 | 2.868578001 | 3.15372745 | 11.47790684 | 12.20053827 |
| ENSG00000167522 | ANKRD11 | 3.917916742 | 0.006512679 | 4.228129007 | 0.002900244 | 8.146045749 | 3.327711234 | 8.639735838 | 6.322058544 | 11.22174696 | 11.260058 | 11.65071957 | 11.44336891 |
| ENSG00000138760 | SCARB2 | 4.403237424 | 2.65E-05 | 3.738882362 | 0.000420382 | 8.142119786 | 9.128497779 | 9.125613817 | 5.59507254 | 12.99606985 | 12.9927763 | 12.31341061 | 12.34733473 |
| ENSG00000225855 | RUSC1-AS1 | 2.899136394 | 0.127934297 | 5.242075933 | 0.002585025 | 8.141212327 | 2 | 2 | 2 | 3.566290607 | 3.50452879 | 5.217477917 | 5.581648407 |
| ENSG00000164124 | TMEM144 | 4.483245877 | 0.018992676 | 3.657147793 | 0.058715617 | 8.140393671 | 2 | 2 | 2 | 4.647285135 | 4.83276841 | 4.311629396 | 3.789585079 |
| ENSG00000139318 | DUSP6 | 6.381625656 | 3.07E-08 | 1.758349133 | 0.17846657 | 8.13997479 | 4.467311984 | 2 | 3.369317972 | 9.333786218 | 9.30274583 | 5.399896186 | 4.105828626 |
| ENSG00000125650 | PSPN | 1.595973537 | 0.397020618 | 6.541291901 | 2.24E-05 | 8.137265438 | 2 | 2 | 2 | 2.784612485 | 2.86123014 | 6.726875343 | 6.520103598 |
| ENSG00000090539 | CHRD | 3.328624741 | 0.044397422 | 4.807086522 | 0.002132231 | 8.135711262 | 2 | 3.076665209 | 2 | 4.739373666 | 4.40247105 | 6.075339068 | 5.689801209 |
| ENSG00000213145 | CRIP1 | 7.440982399 | 2.32E-07 | 0.693761795 | 0.708536423 | 8.134744195 | 2 | 2 | 2 | 7.501020868 | 7.49817903 | 2.526562374 | 2.466008509 |
| ENSG00000079387 | SENP1 | 4.127691222 | 6.33E-10 | 4.006752149 | 1.82E-09 | 8.13444337 | 4.00722531 | 4.11352228 | 4.523675391 | 8.087422161 | 7.94798467 | 7.962642137 | 7.835189056 |
| ENSG00000138738 | PRDM5 | 4.041209899 | 0.002888636 | 4.089965702 | 0.002381928 | 8.1311756 | 2 | 3.686303414 | 2 | 6.034011955 | 5.89549381 | 5.966975997 | 6.055940621 |
| ENSG00000152804 | HHEX | 5.217223641 | 9.33E-05 | 2.907929202 | 0.040183283 | 8.125152842 | 4.00722531 | 2 | 2 | 7.148924055 | 7.14977896 | 5.233158834 | 4.702084619 |
| ENSG00000171132 | PRKCE | 4.219219065 | 0.000106595 | 3.902050377 | 0.000349813 | 8.121269443 | 2 | 4.442710129 | 2 | 6.941624891 | 7.16627666 | 6.788313392 | 6.714211481 |
| ENSG00000271425 | NBPF10 | 8.416205419 | 6.39E-75 | -0.301634614 | 0.61592295 | 8.114570804 | 5.095959458 | 5.457120953 | 5.59507254 | 13.61227351 | 13.7312934 | 4.977914261 | 5.269393099 |
| ENSG00000168661 | ZNF30 | 3.866284949 | 0.022954373 | 4.24160811 | 0.011002269 | 8.107893059 | 2 | 3.076665209 | 2 | 4.887609717 | 5.18011485 | 5.725974948 | 4.911532223 |
| ENSG00000183840 | GPR39 | 6.005900179 | 0.000406094 | 2.098758643 | 0.269325152 | 8.104658821 | 2 | 2 | 2 | 5.949776857 | 6.26836889 | 2.803598391 | 3.287118176 |
| ENSG00000068724 | TTC7A | 3.761417506 | 0.051722704 | 4.34300901 | 0.021313338 | 8.104426516 | 2 | 6.535001748 | 2 | 8.705093576 | 8.68609779 | 9.163026016 | 9.374395329 |
| ENSG00000085433 | WDR47 | 4.238951812 | 0.002112811 | 3.860631288 | 0.005396684 | 8.099583101 | 2 | 6.140560021 | 3.369317972 | 8.939126212 | 8.88234995 | 8.739951687 | 8.299011699 |
| ENSG00000138792 | ENPEP | 3.710750922 | 0.078653566 | 4.378679339 | 0.031664471 | 8.089430261 | 2 | 2 | 2 | 3.95996233 | 4.2560182 | 5.079718116 | 4.047883918 |
| ENSG00000276966 | HIST1H4E | 5.490983653 | 0.004215891 | 2.588222596 | 0.21149879 | 8.079206249 | 2 | 2 | 2 | 5.742861563 | 5.52526112 | 2.831336763 | 3.717076715 |
| ENSG00000146950 | SHROOM2 | 1.109721726 | 0.509773777 | 6.967623362 | 2.41E-07 | 8.077345088 | 3.327711234 | 3.076665209 | 2 | 3.754611435 | 3.33976443 | 8.04860508 | 9.342363956 |
| ENSG00000162729 | IGSF8 | 4.529374396 | 0.003438359 | 3.544371864 | 0.025418276 | 8.073746261 | 4.00722531 | 7.099070191 | 2 | 10.10372352 | 10.2094139 | 9.047307518 | 9.296834866 |
| ENSG00000151176 | PLBD2 | 3.263224998 | 4.32E-10 | 4.809095612 | 5.92E-21 | 8.07232061 | 7.27616306 | 7.60566235 | 6.203069917 | 10.34771157 | 10.4188272 | 11.74812711 | 12.08565654 |
| ENSG00000154114 | TBCEL | 4.04947814 | 0.003951987 | 4.021205091 | 0.004075293 | 8.070683231 | 3.327711234 | 6.228547981 | 2 | 8.826850269 | 8.77349359 | 8.817412208 | 8.725914443 |
| ENSG00000161013 | MGAT4B | 4.204949726 | 0.003504466 | 3.856702823 | 0.007768297 | 8.061652549 | 6.563806279 | 7.761270542 | 2 | 10.83924025 | 10.9176784 | 10.54557014 | 10.51742604 |
| ENSG00000133321 | RARRES3 | 5.27517315 | 1.02E-11 | 2.785326799 | 0.000607542 | 8.060499949 | 4.00722531 | 4.710564154 | 2 | 8.920805049 | 8.92538942 | 6.310306306 | 6.645056213 |
| ENSG00000281429 | | 4.218223822 | 0.001966935 | 3.841560988 | 0.005092678 | 8.05978481 | 3.327711234 | 6.951736442 | 3.369317972 | 9.706030497 | 9.73613428 | 9.355905054 | 9.337081571 |
| ENSG00000135272 | MDFIC | 5.199143103 | 4.70E-09 | 2.858504515 | 0.002185545 | 8.057647618 | 3.327711234 | 6.228547981 | 4.523675391 | 10.30549718 | 10.2232775 | 7.905886539 | 7.979721237 |
| ENSG00000135090 | TAOK3 | 4.33986567 | 0.029985966 | 3.713869079 | 0.065273313 | 8.053734749 | 2 | 6.844429755 | 2 | 9.590237441 | 9.57170671 | 9.011762258 | 8.904372226 |
| ENSG00000102543 | CDADC1 | 4.675091177 | 3.61E-05 | 3.375267487 | 0.003775209 | 8.050358664 | 2 | 4.11352228 | 2 | 7.127781374 | 7.11195456 | 5.982522014 | 5.768151538 |
| ENSG00000130559 | CAMSAP1 | 3.162350605 | 0.124169408 | 4.88786415 | 0.011612184 | 8.050214755 | 2 | 6.78762695 | 2 | 8.468280417 | 8.23475766 | 10.12998797 | 10.00615667 |
| ENSG00000177076 | ACER2 | 3.362057223 | 0.097955385 | 4.686111842 | 0.014701036 | 8.048169064 | 2 | 2 | 2 | 3.881293667 | 3.82856379 | 5.258920789 | 4.468068808 |
| ENSG00000100307 | CBX7 | 3.360684086 | 1.71E-05 | 4.680448885 | 6.81E-10 | 8.041132971 | 2 | 4.710564154 | 4.523675391 | 7.148924055 | 7.32531744 | 8.692059118 | 8.362041249 |
| ENSG00000136158 | SPRY2 | 5.805158017 | 0.001165874 | 2.235293719 | 0.256352368 | 8.040451735 | 2 | 6.464272742 | 2 | 10.64569773 | 10.6701389 | 7.344982741 | 6.868633039 |
| ENSG00000171916 | LGALS9C | 5.410762981 | 0.002162092 | 2.624232049 | 0.16716103 | 8.034995031 | 2 | 2 | 2 | 5.602782483 | 5.52526112 | 3.10571689 | 3.580696614 |
| ENSG00000140323 | DISP2 | 2.775635056 | 0.150440253 | 5.258632734 | 0.002666452 | 8.03426779 | 2 | 2 | 2 | 3.289987342 | 3.60476711 | 5.47655442 | 5.373314945 |
| ENSG00000275395 | FCGBP | 4.654042263 | 0.01717456 | 3.379777154 | 0.091924093 | 8.033819418 | 2 | 2 | 2 | 4.694064037 | 5.06165168 | 3.500668998 | 4.161535617 |
| ENSG00000166439 | RNF169 | 4.269148573 | 0.017100337 | 3.762838571 | 0.036779004 | 8.031987144 | 2 | 6.228547981 | 2 | 8.935480545 | 8.84568954 | 8.459109781 | 8.317521841 |
| ENSG00000143226 | FCGR2A | 2.33424278 | 0.299070637 | 5.690178799 | 0.004075293 | 8.02442158 | 2 | 2 | 2 | 3.289987342 | 3.08597292 | 6.442083733 | 4.711446447 |
| ENSG00000173166 | RAPH1 | 4.522589275 | 2.40E-07 | 3.497510596 | 8.76E-05 | 8.020099871 | 2 | 4.93638933 | 5.772716771 | 9.263797825 | 9.15867957 | 8.096541671 | 8.291228494 |
| ENSG00000184860 | SDR42E1 | 4.521947976 | 0.014871185 | 3.498028198 | 0.064654885 | 8.019976174 | 2 | 2 | 2 | 4.71689672 | 4.83276841 | 4.029788235 | 3.875378318 |
| ENSG00000043355 | ZIC2 | -1.313532109 | 0.596370107 | 9.332501553 | 1.74E-08 | 8.018969444 | 2 | 2 | 2 | 2 | 2.2680111 | 8.702892291 | 9.820984284 |
| ENSG00000106617 | PRKAG2 | 3.242943566 | 3.01E-05 | 4.771340448 | 2.08E-10 | 8.014284014 | 4.467311984 | 4.442710129 | 2 | 7.066878356 | 6.80989959 | 8.462439703 | 8.420292672 |
| ENSG00000185187 | SIGIRR | 3.370776409 | 8.93E-05 | 4.642127463 | 2.46E-08 | 8.012903872 | 4.00722531 | 4.11352228 | 2 | 6.598566398 | 6.60731796 | 7.857806412 | 7.820122783 |
| ENSG00000107159 | CA9 | 4.620082934 | 0.011889226 | 3.39184837 | 0.072085279 | 8.011931304 | 2 | 2 | 2 | 4.846787657 | 4.87352701 | 3.810410249 | 3.940519005 |
| ENSG00000120708 | TGFBI | 4.981548779 | 5.41E-06 | 3.027809542 | 0.008414663 | 8.009358322 | 11.88152073 | 9.680074956 | 9.735974718 | 15.79016427 | 15.8181284 | 14.62132509 | 12.08486888 |
| ENSG00000184307 | ZDHHC23 | 5.273937827 | 0.002252871 | 2.735021446 | 0.138306822 | 8.008959273 | 2 | 2 | 2 | 5.363304456 | 5.51241389 | 3.517841605 | 3.33635788 |
| ENSG00000110092 | CCND1 | 4.379159459 | 4.82E-26 | 3.621990402 | 2.62E-18 | 8.001149861 | 9.90440318 | 10.22575081 | 9.990713521 | 14.40261977 | 14.4407672 | 13.14527188 | 14.04602869 |
| ENSG00000243710 | CFAP57 | 3.860722499 | 0.072220677 | 4.139909037 | 0.048259223 | 8.000631536 | 2 | 2 | 2 | 4.415698455 | 4.02226117 | 3.824280769 | 4.895193906 |
| ENSG00000164283 | ESM1 | 7.18619485 | 1.55E-05 | 0.812086443 | 0.692464461 | 7.998281293 | 5.866944937 | 2 | 2 | 11.39655036 | 11.2888305 | 5.642319755 | 4.365060851 |
| ENSG00000150991 | UBC | 5.107471661 | 2.21E-21 | 2.886053696 | 1.59E-07 | 7.993525357 | 11.80808675 | 10.74900803 | 10.49365885 | 16.26186415 | 16.2178222 | 13.99870688 | 14.03874037 |
| ENSG00000146757 | ZNF92 | 5.500236622 | 1.26E-06 | 2.492826421 | 0.042401898 | 7.993063043 | 2 | 4.442710129 | 2 | 8.406418094 | 8.185025 | 5.72966459 | 5.013446246 |
| ENSG00000241839 | PLEKHO2 | 3.495947156 | 0.044742745 | 4.494682434 | 0.007374689 | 7.990629591 | 2 | 7.099070191 | 3.369317972 | 9.029678018 | 9.10940731 | 9.98740646 | 10.13567188 |
| ENSG00000138675 | FGF5 | 4.630532146 | 1.08E-10 | 3.355808581 | 4.48E-06 | 7.986340727 | 5.866944937 | 4.11352228 | 6.203069917 | 10.1390214 | 10.0787993 | 8.512558892 | 9.109931338 |
| ENSG00000085831 | TTC39A | 3.418422052 | 0.081057036 | 4.567380847 | 0.014257344 | 7.985802899 | 2 | 2 | 2 | 4.034562184 | 3.74316441 | 4.894909044 | 4.729989799 |
| ENSG00000183671 | GPR1 | 2.7904203 | 0.054717846 | 5.189418883 | 0.00010205 | 7.979839182 | 2 | 2 | 4.058998049 | 4.694064037 | 5.07918227 | 6.928144991 | 7.311843322 |
| ENSG00000259305 | ZHX1-C8orf7 | 6.815631771 | 1.62E-06 | 1.163404679 | 0.495779443 | 7.97903645 | 2 | 3.076665209 | 2 | 7.683067677 | 7.96222667 | 2.937246078 | 3.209956562 |
| ENSG00000101210 | EEF1A2 | 2.645389014 | 0.214129738 | 5.33334526 | 0.005583444 | 7.978734273 | 2 | 2 | 2 | 3.566290607 | 3.15372745 | 5.951260634 | 4.819291477 |
| ENSG00000181788 | SIAH2 | 3.288270443 | 5.54E-05 | 4.688505409 | 2.95E-09 | 7.976775852 | 4.815609471 | 5.946638857 | 3.369317972 | 8.219202381 | 8.28094035 | 9.903328854 | 9.314605104 |
| ENSG00000074370 | ATP2A3 | 3.21337828 | 0.11657228 | 4.762383412 | 0.013281894 | 7.975761692 | 2 | 2 | 2 | 3.79808662 | 3.69849104 | 5.338397457 | 4.511590831 |
| ENSG00000162825 | NBPF20 | 9.128144727 | 1.30E-70 | -1.168058261 | 0.051202725 | 7.960086465 | 5.095959458 | 4.710564154 | 6.073378139 | 14.28879814 | 14.4042595 | 4.406883583 | 4.316807515 |
| ENSG00000157193 | LRP8 | 4.317119515 | 0.004529009 | 3.626925772 | 0.019007831 | 7.944045287 | 3.327711234 | 2 | 6.203069917 | 8.936696791 | 8.96364685 | 7.781653072 | 8.629822748 |
| ENSG00000172986 | GXYLT2 | 2.961273953 | 0.044744796 | 4.982083986 | 0.000281972 | 7.94335794 | 2 | 3.686303414 | 2 | 5.110902178 | 4.83276841 | 7.121690475 | 6.525421271 |
| ENSG00000196914 | ARHGEF12 | 3.936133398 | 2.52E-13 | 4.007139101 | 7.09E-14 | 7.943272499 | 6.563806279 | 8.029695761 | 7.502634626 | 11.40360438 | 11.401556 | 11.30754056 | 11.62230214 |
| ENSG00000184363 | PKP3 | 3.147521088 | 0.109887072 | 4.794448059 | 0.009113066 | 7.941969147 | 2 | 2 | 2 | 3.515101962 | 3.86943969 | 5.002522427 | 5.020995508 |
| ENSG00000126945 | HNRNPH2 | 3.85506626 | 3.52E-08 | 4.083886781 | 4.07E-09 | 7.938953041 | 6.563806279 | 4.93638933 | 4.874587708 | 9.363935069 | 9.46239569 | 9.40316733 | 9.846170307 |
| ENSG00000125968 | ID1 | 6.103521504 | 5.74E-06 | 1.834589651 | 0.227161376 | 7.938111155 | 5.330594134 | 2 | 2 | 9.660380652 | 9.64857633 | 5.390603804 | 5.621921006 |
| ENSG00000163291 | PAQR3 | 3.990789124 | 0.004333061 | 3.945505717 | 0.004644445 | 7.936294841 | 2 | 5.131603625 | 2 | 7.590284224 | 7.44845993 | 7.815899769 | 7.032593922 |
| ENSG00000082438 | COBLL1 | 2.153886206 | 0.315977132 | 5.771841878 | 0.002483378 | 7.925728084 | 2 | 6.844429755 | 2 | 7.337530098 | 7.5014335 | 11.20060466 | 10.786851 |

| | | | | | | | | | | | | | |
|---|---|---|---|---|---|---|---|---|---|---|---|---|---|
| ENSG00000187118 | CMC1 | 4.502164297 | 0.001019301 | 3.421371872 | 0.015113682 | 7.923536169 | 2 | 3.686303414 | 2 | 6.483440992 | 6.30607931 | 5.565950232 | 5.191251321 |
| ENSG00000189195 | BTBD8 | 2.445189841 | 0.231434543 | 5.470671262 | 0.002657411 | 7.915861103 | 2 | 2 | 2 | 3.349643765 | 3.15372745 | 6.007052374 | 5.08721802 |
| ENSG00000136379 | ABHD17C | 4.850537942 | 3.29E-10 | 3.055482082 | 0.000127227 | 7.906020024 | 4.00722531 | 5.303523747 | 3.369317972 | 9.137269904 | 9.10727282 | 6.983275763 | 7.646759583 |
| ENSG00000153815 | CMIP | 4.293605271 | 0.005775587 | 3.607879551 | 0.022516836 | 7.901484822 | 6.468621203 | 3.076665209 | 2 | 9.175891964 | 9.18219179 | 8.365574792 | 8.621777169 |
| ENSG00000079819 | EPB41L2 | 3.229695597 | 0.031289994 | 4.669673712 | 0.001065455 | 7.899369309 | 2 | 7.431218269 | 7.285536013 | 9.965769838 | 9.99180603 | 10.83683561 | 11.82693096 |
| ENSG00000129566 | TEP1 | 3.877560457 | 1.62E-13 | 4.015548432 | 1.49E-14 | 7.893108889 | 5.095959458 | 5.131603625 | 6.203069917 | 9.277301907 | 9.32403997 | 9.288400199 | 9.573627872 |
| ENSG00000273820 | USP27X | 3.673514098 | 0.009982485 | 4.219154605 | 0.002441241 | 7.892668703 | 2 | 3.686303414 | 2 | 5.363304456 | 5.83486613 | 6.18690462 | 6.06327132 |
| ENSG00000152133 | GPATCH11 | 3.134102391 | 0.004533018 | 4.756763845 | 5.97E-06 | 7.890866236 | 3.327711234 | 4.11352228 | 2 | 6.01571348 | 6.08095082 | 7.975129619 | 7.123062801 |
| ENSG00000140199 | SLC12A6 | 3.60423027 | 6.87E-07 | 4.285237533 | 2.08E-09 | 7.889467803 | 4.467311984 | 5.838937808 | 6.718950348 | 9.476928206 | 9.42432312 | 10.39846491 | 9.796725763 |
| ENSG00000150636 | CCDC102B | 4.678975799 | 0.013552546 | 3.210163922 | 0.101122291 | 7.889139721 | 2 | 2 | 2 | 5.040242897 | 4.76938658 | 3.943456545 | 3.518055793 |
| ENSG00000170417 | TMEM182 | 3.150314916 | 0.125476823 | 4.737348422 | 0.013968105 | 7.887663339 | 2 | 2 | 2 | 3.881293667 | 3.50452879 | 5.253805068 | 4.594883674 |
| ENSG00000090554 | FLT3LG | 6.628415357 | 1.35E-05 | 1.257592209 | 0.490286193 | 7.886007566 | 2 | 2 | 2 | 6.604700657 | 6.80989959 | 2.62493832 | 2.746601469 |
| ENSG00000163823 | CCR1 | 2.444086818 | 0.295066592 | 5.441695623 | 0.009007577 | 7.88578244 | 2 | 2 | 2 | 3.227757282 | 3.28037825 | 6.252491482 | 4.341135913 |
| ENSG00000100403 | ZC3H7B | 3.419653484 | 0.098091497 | 4.451999595 | 0.024694885 | 7.871653079 | 8.892187472 | 2 | 4.874587708 | 10.7662095 | 10.8158091 | 11.45091976 | 12.11670545 |
| ENSG00000185955 | C7orf61 | 3.596105637 | 0.08781545 | 4.273768727 | 0.03515611 | 7.869874364 | 2 | 2 | 2 | 3.997744386 | 4.05806467 | 4.04171018 | 4.951586397 |
| ENSG00000102032 | RENBP | 4.093671598 | 0.013079091 | 3.769493009 | 0.022726528 | 7.863164607 | 2 | 3.076665209 | 2 | 5.194621444 | 5.28958453 | 5.212212839 | 4.644600858 |
| ENSG00000113580 | NR3C1 | 4.844288223 | 8.66E-05 | 3.017184654 | 0.019926216 | 7.861472876 | 8.16936288 | 7.73146271 | 3.369317972 | 12.22651752 | 12.2242146 | 10.49511466 | 10.30098708 |
| ENSG00000115414 | FN1 | 2.550145312 | 9.94E-06 | 5.308509995 | 6.92E-22 | 7.858655307 | 13.80200664 | 15.22492386 | 14.22230326 | 17.07667487 | 17.1124152 | 19.85784669 | 19.84810262 |
| ENSG00000225830 | ERCC6 | 1.731686895 | 0.184583015 | 6.117307888 | 6.43E-08 | 7.848994783 | 6.816637977 | 7.73146271 | 3.369317972 | 8.50645854 | 8.51792306 | 12.47218794 | 13.20189837 |
| ENSG00000165118 | C9orf64 | 4.296723709 | 9.87E-06 | 3.547282791 | 0.000316727 | 7.8440065 | 2 | 3.076665209 | 4.874587708 | 7.474468681 | 7.62295033 | 6.952030889 | 6.678855578 |
| ENSG00000125746 | EML2 | 3.650926775 | 6.01E-05 | 4.192280808 | 2.81E-06 | 7.843207583 | 5.70932728 | 4.442710129 | 2 | 8.14116509 | 8.1187584 | 8.59173005 | 8.734549554 |
| ENSG00000140545 | MFGE8 | 4.315491043 | 4.84E-06 | 3.524109664 | 0.000234908 | 7.839600707 | 10.75779627 | 9.079034257 | 7.777116083 | 13.98613001 | 14.0263193 | 13.23375446 | 13.19646542 |
| ENSG00000101194 | SLC17A9 | 4.434096283 | 0.016051741 | 3.40499464 | 0.070302637 | 7.839090923 | 2 | 6.389896633 | 2 | 9.246247604 | 9.18725282 | 8.248936733 | 8.145162126 |
| ENSG00000115540 | MOB4 | 4.174568437 | 1.02E-13 | 3.651810213 | 8.66E-11 | 7.82637865 | 4.467311984 | 4.710564154 | 5.59507254 | 8.983347946 | 8.96128525 | 8.586652837 | 8.309067496 |
| ENSG00000122515 | ZMIZ2 | 3.701710124 | 0.010778206 | 4.124631016 | 0.00381062 | 7.82634114 | 2 | 7.315436877 | 5.39244166 | 9.733582734 | 9.73889436 | 10.05219399 | 10.2555122 |
| ENSG00000225932 | CTAGE4 | 6.255592151 | 8.98E-05 | 1.569969387 | 0.397192742 | 7.825561538 | 2 | 2 | 2 | 6.372775472 | 6.33555364 | 2.937246078 | 2.671842496 |
| ENSG00000181625 | SLX1B | 8.95786565 | 3.72E-10 | -1.14221716 | 0.548817763 | 7.81564849 | 2 | 2 | 2 | 8.977450658 | 9.01350799 | 2.094598369 | 2.225841198 |
| ENSG00000126803 | HSPA2 | 1.643847027 | 0.309143132 | 6.170818812 | 7.73E-06 | 7.814665839 | 2 | 2 | 5.39244166 | 5.406080789 | 5.34860058 | 9.803800654 | 9.725739902 |
| ENSG00000064763 | FAR2 | 3.4828275 | 0.000167216 | 4.330632229 | 1.60E-06 | 7.813459729 | 4.815609471 | 3.686303414 | 2 | 7.110641269 | 6.84618961 | 7.513903457 | 8.058864887 |
| ENSG00000178222 | RNF212 | 3.815033538 | 0.018412016 | 3.994352657 | 0.012343956 | 7.809386194 | 2 | 3.076665209 | 2 | 4.984880435 | 5.00774437 | 5.073924441 | 5.230851202 |
| ENSG00000263155 | MYZAP | 5.631459281 | 0.002669901 | 2.175042336 | 0.293035077 | 7.806501617 | 2 | 2 | 2 | 5.686138881 | 5.84514933 | 2.68698273 | 3.430065404 |
| ENSG00000160877 | NACC1 | 3.547474472 | 0.000344238 | 4.258343239 | 1.15E-05 | 7.805817711 | 5.095959458 | 8.053983481 | 6.322058544 | 10.49161431 | 10.5189709 | 10.79210238 | 11.54111489 |
| ENSG00000100345 | MYH9 | 2.629286489 | 0.008881252 | 5.174472469 | 3.36E-08 | 7.803758958 | 8.285626337 | 10.87022386 | 11.4884794 | 13.34084865 | 13.3502258 | 15.81354649 | 15.96311156 |
| ENSG00000128512 | DOCK4 | 5.416093234 | 2.66E-08 | 2.386423175 | 0.023658021 | 7.802516409 | 2 | 4.11352228 | 3.369317972 | 8.192977693 | 8.34574597 | 5.614651724 | 5.072762365 |
| ENSG00000134686 | PHC2 | 3.946770212 | 0.109430874 | 3.852540258 | 0.112945603 | 7.79931047 | 2 | 8.395519306 | 2 | 10.69648168 | 10.8013584 | 10.56221954 | 10.74315987 |
| ENSG00000107816 | LZTS2 | 3.514772876 | 0.00938604 | 4.281818359 | 0.001131917 | 7.796591235 | 2 | 6.464272742 | 7.225742779 | 9.774969254 | 9.78702715 | 10.56854263 | 10.52175615 |
| ENSG00000196932 | TMEM26 | 4.571578677 | 0.014868925 | 3.224631572 | 0.095146591 | 7.796210249 | 2 | 2 | 2 | 4.739373666 | 4.89348248 | 3.648325314 | 3.858622973 |
| ENSG00000160218 | TRAPPC10 | 4.299315599 | 0.000245132 | 3.495703571 | 0.003420748 | 7.79501917 | 2 | 5.131603625 | 6.534109959 | 9.640365415 | 9.58633944 | 8.736288294 | 8.890530144 |
| ENSG00000033327 | GAB2 | 2.904284387 | 0.127194913 | 4.890558998 | 0.005830103 | 7.794843384 | 2 | 6.311476754 | 2 | 7.626911583 | 7.62891439 | 9.653364877 | 9.528327138 |
| ENSG00000151778 | SERP2 | 3.467767398 | 0.043810684 | 4.318274834 | 0.00911236 | 7.786042232 | 2 | 3.076665209 | 2 | 4.470469939 | 4.89348248 | 5.102662916 | 5.722580018 |
| ENSG00000058063 | ATP11B | 4.020211732 | 0.0142187 | 3.764431999 | 0.022137903 | 7.784643731 | 4.467311984 | 7.572420273 | 2 | 10.12521908 | 10.1422001 | 9.896164504 | 9.861494087 |
| ENSG00000088387 | DOCK9 | 3.443413842 | 1.71E-05 | 4.340351749 | 2.84E-08 | 7.783765591 | 4.00722531 | 4.93638933 | 2 | 7.333842102 | 7.25764291 | 8.111466145 | 8.237969002 |
| ENSG00000038219 | BOD1L1 | 3.847867957 | 0.015565291 | 3.929123728 | 0.012788451 | 7.776991685 | 3.327711234 | 8.077869079 | 4.058998049 | 10.41993524 | 10.4507437 | 10.61778047 | 10.4074232 |
| ENSG00000048405 | ZNF800 | 3.876761441 | 0.024179818 | 3.891569965 | 0.022869703 | 7.768331405 | 2 | 5.946638857 | 2 | 8.221200057 | 8.21902896 | 8.445712697 | 7.987448576 |
| ENSG00000164574 | GALNT10 | 2.575052524 | 0.000171546 | 5.17721411 | 1.70E-15 | 7.752266634 | 6.96351253 | 8.234834549 | 6.43197674 | 9.95740528 | 10.0010327 | 12.5158451 | 12.63521027 |
| ENSG00000158373 | HIST1H2BD | 5.751358837 | 1.25E-10 | 1.994733228 | 0.04303466 | 7.746092064 | 3.327711234 | 4.710564154 | 2 | 9.155143657 | 9.14937381 | 5.201624686 | 5.777095613 |
| ENSG00000107551 | RASSF4 | 2.075188279 | 0.093856973 | 5.668794227 | 2.26E-07 | 7.743982506 | 2 | 4.710564154 | 2 | 5.48800675 | 5.11361705 | 8.984998983 | 8.509534124 |
| ENSG00000221823 | PPP3R1 | 4.133460082 | 8.75E-09 | 3.609888444 | 5.96E-07 | 7.743348526 | 4.00722531 | 6.464272742 | 6.203069917 | 10.00514223 | 9.89490419 | 9.4014335 | 9.458113344 |
| ENSG00000169933 | FRMPD4 | 4.886213837 | 0.002602476 | 2.854294738 | 0.095264998 | 7.740508575 | 2 | 4.442710129 | 2 | 7.501020868 | 7.74878077 | 4.220168345 | 6.397863414 |
| ENSG00000198774 | RASSF9 | 2.150847631 | 0.28400482 | 5.588922417 | 0.001382281 | 7.739770048 | 2 | 2 | 2 | 3.289987342 | 2.86123014 | 5.766051132 | 5.689801209 |
| ENSG00000139209 | SLC38A4 | 3.418722624 | 0.094962877 | 4.320772398 | 0.027228112 | 7.739495022 | 2 | 2 | 2 | 4.070463721 | 3.69849104 | 4.785723309 | 4.400220394 |
| ENSG00000160584 | SIK3 | 4.086010875 | 0.028911985 | 3.651528112 | 0.0515772 | 7.737538987 | 2 | 6.389896633 | 2 | 8.822909147 | 8.91810182 | 8.28637003 | 8.580862167 |
| ENSG00000162390 | ACOT11 | 4.098022861 | 0.049030905 | 3.638747112 | 0.080103167 | 7.736769972 | 2 | 2 | 2 | 4.299492648 | 4.53541753 | 3.616794016 | 4.400220394 |
| ENSG00000075391 | RASAL2 | 3.75217178 | 0.000927691 | 3.981319419 | 0.00038106 | 7.7334912 | 7.435667355 | 4.93638933 | 4.058998049 | 9.879858319 | 9.85666073 | 10.1379928 | 10.05385647 |
| ENSG00000150551 | LYPD1 | 6.405009125 | 0.004752131 | 1.324548647 | 0.618981618 | 7.729557773 | 2 | 2 | 2 | 6.710878132 | 6.24526024 | 3.12825509 | 2.117335275 |
| ENSG00000196189 | SEMA4A | 2.975077761 | 0.133011196 | 4.752426353 | 0.009778296 | 7.727504115 | 2 | 2 | 2 | 3.615725078 | 3.55551834 | 5.079718116 | 4.861951484 |
| ENSG00000165475 | CRYL1 | 3.34802026 | 0.013546981 | 4.378989221 | 0.00080388 | 7.727009481 | 2 | 5.131603625 | 2 | 6.872128674 | 6.93551934 | 8.044167181 | 7.762743025 |
| ENSG00000138448 | ITGAV | 3.51610778 | 0.00358975 | 4.207034823 | 0.000357883 | 7.723142603 | 4.467311984 | 8.747909548 | 7.285536013 | 11.17355701 | 11.145897 | 12.08377451 | 11.57027009 |
| ENSG00000040199 | PHLPP2 | 3.79165093 | 1.71E-08 | 3.930278737 | 4.06E-09 | 7.721929666 | 4.467311984 | 5.457120953 | 3.369317972 | 8.372666678 | 8.27524772 | 8.391426944 | 8.528858263 |
| ENSG00000080200 | CRYBG3 | 4.1573085 | 1.48E-13 | 3.562705974 | 2.90E-10 | 7.720014474 | 5.70932728 | 5.303523747 | 4.058998049 | 9.198375657 | 9.18725282 | 8.749069679 | 8.440841157 |
| ENSG00000130363 | RSPH3 | 3.795298184 | 5.55E-07 | 3.923109672 | 1.91E-07 | 7.718407856 | 4.815609471 | 5.303523747 | 2 | 8.241026137 | 8.17690623 | 8.294489071 | 8.375354401 |
| ENSG00000185760 | KCNQ5 | 6.181903684 | 5.54E-06 | 1.527028089 | 0.331063165 | 7.708931774 | 2 | 2 | 4.058998049 | 8.178656739 | 8.03813956 | 3.44788957 | 4.228274364 |
| ENSG00000161682 | FAM171A2 | 2.502821509 | 0.063823695 | 5.206037609 | 2.22E-05 | 7.708859117 | 2 | 4.11352228 | 2 | 5.022023783 | 5.11361705 | 7.985196707 | 7.124815549 |
| ENSG00000072952 | MRVI1 | 1.35351924 | 0.527194576 | 6.354747267 | 0.000381431 | 7.708266507 | 2 | 3.076665209 | 7.552162999 | 7.352188525 | 7.34718809 | 12.37080139 | 12.25951588 |
| ENSG00000158457 | TSPAN33 | 2.494673508 | 0.255434807 | 5.209707605 | 0.00836966 | 7.704381114 | 2 | 2 | 2 | 3.023133209 | 3.50452879 | 5.832754931 | 4.720747915 |
| ENSG00000139178 | C1RL | 4.258534669 | 4.16E-05 | 3.445464597 | 0.00111041 | 7.703999265 | 2 | 6.04685525 | 5.59507254 | 9.424141421 | 9.45821469 | 8.328342078 | 8.886919671 |
| ENSG00000184887 | BTBD6 | 3.298285651 | 0.019232909 | 4.403488605 | 0.001143848 | 7.701774256 | 4.00722531 | 6.535001748 | 2 | 8.404661252 | 8.45217004 | 9.526277199 | 9.523352442 |
| ENSG00000071242 | RPS6KA2 | 1.674283714 | 0.128350341 | 6.024894564 | 3.25E-10 | 7.699178278 | 4.815609471 | 6.78762695 | 4.058998049 | 7.477814553 | 7.16216989 | 11.08420293 | 12.04729732 |
| ENSG00000174738 | NR1D2 | 3.859667609 | 0.062809849 | 3.833296534 | 0.061534934 | 7.692964143 | 2 | 6.951736442 | 2 | 9.225498809 | 9.19732195 | 9.302094275 | 9.058129171 |
| ENSG00000211455 | STK38L | 2.93561061 | 0.000690859 | 4.756575115 | 8.75E-09 | 7.692185725 | 6.816637977 | 7.847167909 | 5.156626401 | 9.954406146 | 9.82583819 | 11.92035242 | 11.45925874 |
| ENSG00000058866 | DGKG | 5.416941938 | 0.00154394 | 2.271353055 | 0.225799741 | 7.688294993 | 2 | 2 | 2 | 5.501219927 | 5.63601114 | 3.235948948 | 3.071407821 |
| ENSG00000130592 | LSP1 | 4.003122896 | 0.074838025 | 3.683860214 | 0.098657948 | 7.68698311 | 3.327711234 | 2 | 2 | 5.058234798 | 5.2591458 | 5.669467138 | 3.012034821 |
| ENSG00000167549 | CORO6 | 2.253529971 | 0.067543282 | 5.433180548 | 8.58E-07 | 7.68671052 | 2 | 4.11352228 | 2 | 4.76150579 | 4.97065356 | 7.730090858 | 7.98359008 |
| ENSG00000055332 | EIF2AK2 | 4.383236353 | 0.000174604 | 3.301433038 | 0.005942742 | 7.684669392 | 6.89194353 | 7.819099893 | 3.369317972 | 11.22995516 | 11.2370588 | 10.23317759 | 10.07100411 |
| ENSG00000135605 | TEC | 3.331780942 | 0.102509199 | 4.351550327 | 0.025215033 | 7.683331269 | 2 | 2 | 2 | 3.615725078 | 4.02226117 | 4.705557986 | 4.553838275 |
| ENSG00000197483 | ZNF628 | 3.267239111 | 0.001032776 | 4.412598143 | 4.37E-06 | 7.679837253 | 3.327711234 | 4.442710129 | 2 | 6.336376195 | 6.59518404 | 7.291067412 | 7.822284765 |
| ENSG00000158711 | ELK4 | 3.722573332 | 7.29E-17 | 3.948551853 | 4.49E-19 | 7.671125186 | 5.095959458 | 6.04685525 | 5.930865648 | 9.423273993 | 9.35675148 | 9.665695653 | 9.562677828 |
| ENSG00000004866 | ST7 | 3.175216663 | 0.070101138 | 4.493717652 | 0.007066912 | 7.668934315 | 3.327711234 | 7.051611315 | 2 | 8.695066731 | 8.71301725 | 10.19532025 | 9.807133274 |
| ENSG00000100139 | MICALL1 | 2.486778635 | 0.100010522 | 5.177808704 | 0.000167739 | 7.66458734 | 3.327711234 | 7.761270542 | 4.523675391 | 8.83470036 | 8.82115545 | 11.62791697 | 11.37763195 |
| ENSG00000151470 | C4orf33 | 4.497190771 | 0.000105898 | 3.16637398 | 0.008341427 | 7.663564751 | 2 | 4.11352228 | 2 | 6.912249745 | 6.97351258 | 5.801542492 | 5.5506886 |
| ENSG00000185304 | RGPD2 | 9.191187738 | 2.41E-07 | -1.531443747 | 0.505767117 | 7.659743991 | 2 | 2 | 2 | 9.217515344 | 9.23691006 | 2.225793514 | 2 |
| ENSG00000275793 | RIMBP3 | 5.655514789 | 0.000990822 | 1.987812727 | 0.299605222 | 7.643327517 | 2 | 2 | 2 | 5.839660368 | 5.73886209 | 3.172301926 | 2.817676617 |
| ENSG00000278087 | | 12.82964265 | 2.59E-20 | -5.191149786 | 0.022002471 | 7.638492869 | 2 | 2 | 2 | 12.834431 | 12.8788387 | 2 | 2 |
| ENSG00000196110 | ZNF699 | 3.180030469 | 0.006978751 | 4.455588863 | 7.83E-05 | 7.635619331 | 2 | 4.710564154 | 2 | 6.351046291 | 6.32089174 | 7.689935911 | 7.437040198 |
| ENSG00000166823 | MESP1 | 2.390486296 | 0.240656755 | 5.244923154 | 0.003905932 | 7.63540945 | 2 | 2 | 2 | 3.289987342 | 3.15372745 | 5.707383854 | 5.04340913 |
| ENSG00000163840 | DTX3L | 4.88205459 | 1.01E-12 | 2.749619062 | 0.000110145 | 7.631673652 | 5.095959458 | 6.728495673 | 7.342949392 | 11.44969881 | 11.5103551 | 9.451446131 | 9.251026218 |
| ENSG00000197256 | KANK2 | 3.119952073 | 0.003412706 | 4.511395231 | 1.03E-05 | 7.631347304 | 4.815609471 | 8.433041974 | 7.225742779 | 10.52515083 | 10.5667454 | 11.96143224 | 11.90836987 |
| ENSG00000145623 | OSMR | 3.747923683 | 0.006332639 | 3.88091809 | 0.004330689 | 7.628841772 | 4.467311984 | 6.04685525 | 8.978369997 | 11.33307966 | 11.3510373 | 11.46468397 | 11.48498941 |
| ENSG00000159871 | LYPD5 | 2.012858346 | 0.350878999 | 5.614784097 | 0.002663586 | 7.627642443 | 2 | 2 | 2 | 3.162721474 | 2.86123014 | 6.205571486 | 5.08721802 |
| ENSG00000198753 | PLXNB3 | 4.676561152 | 1.30E-05 | 2.950892864 | 0.008495313 | 7.627454016 | 2 | 4.442710129 | 2 | 7.580980221 | 7.42465798 | 5.812021848 | 5.888587685 |
| ENSG00000119314 | PTBP3 | 3.79211763 | 0.001734709 | 3.833328002 | 0.001437647 | 7.625445632 | 6.009025997 | 6.728495673 | 2 | 9.597176503 | 9.56082893 | 9.802049415 | 9.411845434 |
| ENSG00000179388 | EGR3 | 4.867856792 | 0.002189164 | 2.755531516 | 0.101324269 | 7.623388308 | 3.327711234 | 2 | 2 | 6.061031724 | 5.83486613 | 4.340871764 | 3.858622973 |
| ENSG00000112658 | SRF | 2.953023063 | 0.086709662 | 4.668825289 | 0.003843291 | 7.621848352 | 4.00722531 | 7.35507212 | 2 | 8.812346549 | 8.84055903 | 10.41158826 | 10.64580898 |
| ENSG00000029534 | ANK1 | 0.69075686 | 0.745023679 | 6.928631153 | 1.18E-05 | 7.619388012 | 2 | 2 | 2 | 2.600425781 | 2.38539787 | 7.229857522 | 6.728114749 |
| ENSG00000180263 | FGD6 | 3.926971903 | 1.97E-10 | 3.688394811 | 2.27E-09 | 7.615366714 | 4.467311984 | 4.710564154 | 4.058998049 | 8.207157886 | 8.04037874 | 7.8927823 | 7.889811111 |
| ENSG00000115561 | CHMP3 | 3.840555027 | 0.004238809 | 3.772916475 | 0.004854847 | 7.613471501 | 7.384433967 | 4.442710129 | 3.369317972 | 9.802582299 | 9.80162141 | 9.289338026 | 10.07463605 |
| ENSG00000126709 | IFI6 | 7.167089072 | 6.78E-15 | 0.445860761 | 0.705642182 | 7.612949833 | 5.532353208 | 8.124485672 | 8.699419242 | 15.06164194 | 15.122211 | 8.047866378 | 8.663373824 |
| ENSG00000137216 | TMEM63B | 4.292837833 | 7.99E-05 | 3.319124694 | 0.002864384 | 7.611962527 | 4.815609471 | 6.899080544 | 3.369317972 | 9.941741253 | 9.93757622 | 8.986541761 | 8.956482366 |
| ENSG00000203867 | RBM20 | -0.949754075 | 0.638134514 | 8.556281998 | 2.95E-09 | 7.606527923 | 2 | 3.076665209 | 2 | 2.3892346 | 2.2680111 | 9.060551741 | 9.918182687 |
| ENSG00000184992 | BRI3BP | 2.680571936 | 0.061396429 | 4.920407824 | 0.000187114 | 7.60097976 | 2 | 3.686303414 | 2 | 4.825935136 | 4.65713986 | 6.768721236 | 6.836905794 |
| ENSG00000173530 | TNFRSF10D | 2.209350163 | 0.090904877 | 5.387704956 | 4.34E-06 | 7.597055119 | 2 | 6.728495673 | 6.203069917 | 8.017874397 | 8.14604733 | 11.10494797 | 11.36841437 |
| ENSG00000117318 | ID3 | 3.120496692 | 6.24E-05 | 4.475352413 | 3.10E-09 | 7.595849105 | 8.19931886 | 6.311476754 | 6.43197674 | 10.29936444 | 10.3782441 | 11.8938265 | 11.45604253 |
| ENSG00000132669 | RIN2 | 3.667421312 | 1.35E-07 | 3.928120356 | 1.25E-08 | 7.595541668 | 5.866944937 | 5.303523747 | 7.163363831 | 9.877326927 | 9.92974626 | 9.919572134 | 10.372072 |
| ENSG00000156453 | PCDH1 | 0.29836381 | 0.870417637 | 7.292692023 | 3.29E-08 | 7.591055833 | 4.467311984 | 2 | 2 | 3.615725078 | 3.28037825 | 10.37023041 | 8.864016001 |
| ENSG00000248592 | TMEM110-M | 6.995547237 | 7.30E-06 | 0.593259835 | 0.768959881 | 7.588807072 | 2 | 2 | 2 | 7.075737784 | 7.05550676 | 2.592885947 | 2.326754077 |
| ENSG00000048392 | RRM2B | 3.312820157 | 0.022492512 | 4.27288224 | 0.002294707 | 7.585702397 | 2 | 7.315436877 | 6.882775709 | 9.858198568 | 9.76756088 | 11.18456388 | 10.18753685 |
| ENSG00000111802 | TDP2 | 4.553110608 | 0.003256601 | 3.028670632 | 0.05973596 | 7.58178124 | 3.327711234 | 6.666836837 | 2 | 9.698163261 | 9.71736428 | 8.33380681 | 8.044120856 |

| | | | | | | | | | | | | | |
|---|---|---|---|---|---|---|---|---|---|---|---|---|---|
| ENSG00000146833 | TRIM4 | 3.491473863 | 0.000690666 | 4.088678657 | 4.96E-05 | 7.580152519 | 3.327711234 | 3.686303414 | 6.073378139 | 8.130575473 | 8.20911072 | 8.904453984 | 8.601149963 |
| ENSG00000265817 | FSBP | 5.609066294 | 0.000800798 | 1.96731063 | 0.292598927 | 7.576376924 | 2 | 2 | 2 | 5.870535235 | 5.6121243 | 3.011868911 | 2.981406154 |
| ENSG00000213199 | ASIC3 | 3.180110954 | 0.160294059 | 4.393582479 | 0.039954567 | 7.573693433 | 2 | 2 | 2 | 3.566290607 | 3.86943969 | 5.147485955 | 3.940519005 |
| ENSG00000116539 | ASH1L | 3.765275161 | 9.68E-05 | 3.807176369 | 7.43E-05 | 7.57245153 | 4.00722531 | 6.899080544 | 7.398165132 | 10.4133977 | 10.3378525 | 10.46679345 | 10.36726769 |
| ENSG00000156381 | ANKRD9 | 2.801762768 | 0.001483636 | 4.769637588 | 1.15E-08 | 7.571400356 | 5.532353208 | 4.442710129 | 2 | 7.190301983 | 7.14562489 | 9.280561171 | 8.906921063 |
| ENSG00000175556 | LONRF3 | 4.484969351 | 0.044941351 | 3.086070495 | 0.178705011 | 7.571039846 | 2 | 2 | 2 | 4.907595604 | 4.56059747 | 4.133696865 | 2.950113106 |
| ENSG00000176148 | TCP11L1 | 3.386774524 | 0.002505663 | 4.183163794 | 0.000124829 | 7.569938318 | 4.00722531 | 5.838937808 | 2 | 7.927778434 | 7.93600799 | 8.780536345 | 8.653086557 |
| ENSG00000103404 | USP31 | 3.769281769 | 0.00128085 | 3.799889924 | 0.001089398 | 7.569171694 | 3.327711234 | 5.595927442 | 2 | 7.947219213 | 7.97164382 | 8.086505553 | 7.885682058 |
| ENSG00000115844 | DLX2 | 5.739410546 | 0.001424077 | 1.82831572 | 0.367174663 | 7.567726267 | 2 | 2 | 2 | 5.890757403 | 5.84514933 | 2.62493832 | 3.183290849 |
| ENSG00000085465 | OVGP1 | 3.973653387 | 0.061258008 | 3.590991928 | 0.089476325 | 7.564645314 | 2 | 2 | 2 | 4.139692846 | 4.4836996 | 4.350488971 | 3.600986722 |
| ENSG00000129315 | CCNT1 | 4.002785425 | 0.062310616 | 3.56128226 | 0.096925701 | 7.564067685 | 2 | 7.145017454 | 2 | 9.572351151 | 9.52289845 | 9.473343311 | 8.620535396 |
| ENSG00000162139 | NEU3 | 2.731587147 | 0.024129772 | 4.826598526 | 1.90E-05 | 7.558185673 | 2 | 4.710564154 | 2 | 5.720439317 | 6.08095082 | 8.035995406 | 7.819040576 |
| ENSG00000124201 | ZNFX1 | 4.552686663 | 1.50E-05 | 3.002286521 | 0.006037433 | 7.554973184 | 5.330594134 | 8.777382925 | 8.960138144 | 12.83793404 | 12.9288719 | 11.20035539 | 11.45847708 |
| ENSG00000173918 | C1QTNF1 | 4.710741917 | 0.003177064 | 2.838573521 | 0.090878546 | 7.549315438 | 4.00722531 | 7.232742069 | 2 | 10.44536591 | 10.4770185 | 8.247007147 | 8.88330014 |
| ENSG00000158714 | SLAMF8 | 4.828951605 | 0.055881142 | 2.717769255 | 0.303576997 | 7.546720859 | 2 | 2 | 2 | 5.040242897 | 5.04390545 | 4.017766949 | 2.374681325 |
| ENSG00000125148 | MT2A | 6.407060603 | 1.60E-13 | 1.139420581 | 0.26164227 | 7.546481184 | 8.285626337 | 7.572420273 | 5.39244166 | 13.8152717 | 13.9436096 | 8.944296493 | 8.220862755 |
| ENSG00000186417 | GLDN | 3.418726327 | 0.102212351 | 4.126967663 | 0.039942724 | 7.545693991 | 2 | 2 | 2 | 4.070463721 | 3.69849104 | 4.644324387 | 4.215171317 |
| ENSG00000185519 | FAM131C | 3.181160802 | 0.122984674 | 4.362248004 | 0.025584749 | 7.543408806 | 2 | 2 | 2 | 3.709785322 | 3.74316441 | 4.854926465 | 4.388595494 |
| ENSG00000158887 | MPZ | 3.180813036 | 0.137755718 | 4.361191917 | 0.031648409 | 7.542004954 | 2 | 2 | 2 | 3.663521627 | 3.7864959 | 4.990270811 | 4.175132639 |
| ENSG00000175564 | UCP3 | 4.163332285 | 0.039266418 | 3.373767598 | 0.097996916 | 7.537099882 | 2 | 2 | 2 | 4.17310023 | 4.72553021 | 3.917810786 | 3.807156736 |
| ENSG00000137478 | FCHSD2 | 2.711963179 | 0.228046677 | 4.824695066 | 0.02108503 | 7.536658245 | 2 | 7.274681896 | 2 | 8.350935846 | 8.44373613 | 10.56738332 | 10.42424552 |
| ENSG00000115648 | MLPH | 5.070932377 | 5.36E-09 | 2.465633025 | 0.007862246 | 7.536565402 | 2 | 4.11352228 | 4.523675391 | 8.532433657 | 8.52753255 | 6.295446113 | 5.631815919 |
| ENSG00000171873 | ADRA1D | 4.699497129 | 0.060057468 | 2.836764058 | 0.274683967 | 7.536261187 | 2 | 2 | 2 | 4.739373666 | 5.09650239 | 4.088436536 | 2.466008509 |
| ENSG00000178996 | SNX18 | 2.130164232 | 0.214848439 | 5.405356275 | 0.000410013 | 7.535520507 | 4.00722531 | 7.145017454 | 2 | 7.862723323 | 7.77859536 | 11.33504966 | 10.75353482 |
| ENSG00000135823 | STX6 | 3.016848168 | 0.062403761 | 4.513709512 | 0.003194404 | 7.53055768 | 4.00722531 | 7.051611315 | 2 | 8.610716699 | 8.61125371 | 9.935834217 | 10.2442687 |
| ENSG00000178075 | GRAMD1C | 4.16961602 | 0.019121386 | 3.355993757 | 0.063125672 | 7.525609777 | 2 | 3.076665209 | 2 | 5.144975053 | 5.45984938 | 5.032703865 | 3.987514168 |
| ENSG00000224531 | SMIM13 | 3.433254972 | 0.075515059 | 4.09219879 | 0.028865749 | 7.525453763 | 2 | 6.389896633 | 2 | 8.221200057 | 8.22887947 | 9.145869306 | 8.545966004 |
| ENSG00000172216 | CEBPB | 4.430886458 | 3.68E-09 | 3.082184506 | 6.32E-05 | 7.513070964 | 6.009025997 | 7.701025996 | 8.408111533 | 12.04993732 | 12.1001021 | 10.80593075 | 10.64687673 |
| ENSG00000020577 | SAMD4A | 2.804066637 | 0.009617767 | 4.705541026 | 4.12E-06 | 7.509607663 | 7.031698319 | 7.669933291 | 4.058998049 | 9.641111693 | 9.64637295 | 11.10796328 | 11.8719151 |
| ENSG00000137177 | KIF13A | 3.682379034 | 3.01E-18 | 3.825502873 | 7.88E-20 | 7.507881907 | 7.384433967 | 6.728495673 | 6.322058544 | 10.46193323 | 10.5616878 | 10.65560768 | 10.65523878 |
| ENSG00000145246 | ATP10D | 3.371111502 | 0.022108229 | 4.136767086 | 0.003803056 | 7.507878588 | 2 | 5.838937808 | 7.502634626 | 9.610955295 | 9.66245344 | 10.54570096 | 10.23699419 |
| ENSG00000179111 | HES7 | 4.064223735 | 0.062918791 | 3.440072984 | 0.116333358 | 7.504296719 | 2 | 2 | 2 | 4.358765249 | 4.43005853 | 4.321442857 | 3.33635788 |
| ENSG00000254852 | NPIPA2 | 11.23534759 | 4.39E-16 | -3.744722507 | 0.101374362 | 7.490625088 | 2 | 2 | 2 | 11.26329852 | 11.2648535 | 2 | 2.059860178 |
| ENSG00000257594 | GALNT4 | 6.27619247 | 1.35E-05 | 1.214222082 | 0.478299157 | 7.490414552 | 2 | 3.076665209 | 2 | 7.25413313 | 7.34718809 | 3.215043717 | 2.981406154 |
| ENSG00000077150 | NFKB2 | 4.011487073 | 0.003482895 | 3.477957869 | 0.012368406 | 7.489444942 | 4.00722531 | 7.701025996 | 4.058998049 | 10.27889147 | 10.3005071 | 9.463959918 | 10.00258629 |
| ENSG00000142623 | PADI1 | 1.4987618 | 0.487075206 | 5.98487083 | 0.000780302 | 7.48363263 | 2 | 2 | 2 | 2.947924294 | 2.59489879 | 5.598599872 | 6.468583424 |
| ENSG00000196460 | RFX8 | 8.209931355 | 1.73E-08 | -0.727281878 | 0.711605352 | 7.482649476 | 2 | 2 | 2 | 8.235106869 | 8.27524772 | 2.139668783 | 2.277179783 |
| ENSG00000184481 | FOXO4 | 2.989536499 | 0.000119872 | 4.491543627 | 1.85E-09 | 7.481080127 | 4.467311984 | 4.442710129 | 2 | 6.70517998 | 6.6952083 | 8.204552906 | 8.126037235 |
| ENSG00000176454 | LPCAT4 | 3.309433968 | 0.00052729 | 4.17110557 | 7.22E-06 | 7.480539538 | 2 | 5.72254323 | 5.156626401 | 8.001694993 | 8.22494733 | 9.148283563 | 8.766338209 |
| ENSG00000122691 | TWIST1 | 5.288927208 | 1.99E-09 | 2.1858045 | 0.022620058 | 7.474731708 | 4.00722531 | 4.710564154 | 2 | 8.874592369 | 8.96718201 | 6.239610476 | 5.470251342 |
| ENSG00000198853 | RUSC2 | 2.842806896 | 0.017939507 | 4.628576928 | 4.36E-05 | 7.471383824 | 7.485143553 | 8.337339476 | 4.058998049 | 10.19853443 | 10.3126002 | 12.08454008 | 11.99219313 |
| ENSG00000050327 | ARHGEF5 | 4.739401809 | 1.55E-08 | 2.726372975 | 0.001899142 | 7.465774784 | 4.467311984 | 2 | 4.874587708 | 8.469961514 | 8.5130942 | 6.459963649 | 6.590289384 |
| ENSG00000138246 | DNAJC13 | 4.203235238 | 0.011917779 | 3.261781339 | 0.055805422 | 7.465016578 | 4.467311984 | 7.701025996 | 2 | 10.4372252 | 10.4308254 | 9.580405632 | 9.407171982 |
| ENSG00000148842 | CNNM2 | 1.811933407 | 0.183141741 | 5.650823555 | 2.04E-06 | 7.462756962 | 4.467311984 | 6.311476754 | 2 | 6.965655398 | 6.77267312 | 11.02068045 | 10.19109642 |
| ENSG00000101935 | AMMECR1 | 3.716675033 | 0.000483867 | 3.743679224 | 0.000403865 | 7.460354258 | 3.327711234 | 5.303523747 | 2 | 7.725907304 | 7.5835711 | 7.51817649 | 7.830900438 |
| ENSG00000101335 | MYL9 | 2.041738462 | 4.17E-05 | 5.416981309 | 1.17E-30 | 7.458719771 | 10.44422623 | 10.22039562 | 9.272377397 | 12.10655496 | 12.0929736 | 15.33288762 | 15.60222779 |
| ENSG00000152332 | UHMK1 | 3.89149463 | 0.007379211 | 3.564639218 | 0.014671069 | 7.456133848 | 7.579279349 | 7.538394143 | 2 | 10.8903519 | 10.7920865 | 10.43748218 | 10.59043716 |
| ENSG00000164125 | FAM198B | 3.812485021 | 5.28E-05 | 3.641994132 | 0.000110991 | 7.454479153 | 3.327711234 | 4.710564154 | 2 | 7.307758725 | 7.16627666 | 7.258234779 | 6.860240429 |
| ENSG00000149115 | TNKS1BP1 | 3.277288373 | 0.000432249 | 4.17698842 | 4.12E-06 | 7.454276793 | 5.330594134 | 8.147241016 | 8.266922072 | 10.93607039 | 11.0094017 | 11.77710915 | 11.96021567 |
| ENSG00000182326 | C1S | 5.022304587 | 7.00E-06 | 2.429928172 | 0.043006389 | 7.452232759 | 5.532353208 | 9.514922632 | 9.099922433 | 13.78119023 | 13.8184383 | 10.77323249 | 11.54414712 |
| ENSG00000172927 | MYEOV | 2.391693452 | 0.324077768 | 5.055365338 | 0.021023289 | 7.44705879 | 2 | 2 | 2 | 3.406930997 | 3.01487888 | 4.088436536 | 5.876154317 |
| ENSG00000132613 | MTSS1L | 3.277662631 | 0.007863092 | 4.16859713 | 0.000478334 | 7.446259761 | 2 | 5.946638857 | 6.718950348 | 8.954818504 | 9.07377798 | 9.785748878 | 10.00829467 |
| ENSG00000278318 | ZNF229 | 4.078374474 | 3.96E-08 | 3.365118452 | 7.46E-06 | 7.443492926 | 5.330594134 | 4.93638933 | 2 | 8.517880242 | 8.5227358 | 7.907516231 | 7.720479276 |
| ENSG00000196834 | POTEI | 9.590271172 | 8.18E-12 | -2.15516918 | 0.263504222 | 7.435101992 | 2 | 2 | 2 | 9.55186119 | 9.69266358 | 2.048074405 | 2.117335275 |
| ENSG00000100629 | CEP128 | 3.0453365 | 0.148151369 | 4.383901602 | 0.026905364 | 7.429238103 | 2 | 2 | 2 | 3.406930997 | 3.82856379 | 4.521997428 | 4.784231301 |
| ENSG00000102221 | JADE3 | 3.300383865 | 0.034024524 | 4.12880432 | 0.006072155 | 7.429188185 | 2 | 5.303523747 | 2 | 6.984595707 | 7.04216061 | 8.227567998 | 7.256957802 |
| ENSG00000176273 | SLC35G1 | 3.012182896 | 0.010924161 | 4.414206648 | 8.78E-05 | 7.426389544 | 2 | 4.442710129 | 2 | 5.987823101 | 5.80356929 | 7.08038692 | 7.382408159 |
| ENSG00000116514 | RNF19B | 4.100192263 | 0.018991992 | 3.326140156 | 0.060701932 | 7.426332419 | 2 | 6.04685525 | 2 | 8.519504557 | 8.56222967 | 7.908330388 | 7.634508409 |
| ENSG00000108622 | ICAM2 | 3.148558051 | 0.137320752 | 4.277606407 | 0.03302563 | 7.426164457 | 2 | 2 | 2 | 3.663521627 | 3.74316441 | 4.230621763 | 4.845038551 |
| ENSG00000181751 | C5orf30 | 3.637647867 | 3.06E-07 | 3.78416694 | 8.27E-08 | 7.421814807 | 6.009025997 | 5.303523747 | 3.369317972 | 8.797695539 | 8.70739158 | 8.765787532 | 9.020021944 |
| ENSG00000180777 | ANKRD30B | 5.045281219 | 8.78E-05 | 2.369149985 | 0.088401773 | 7.414431204 | 2 | 4.11352228 | 2 | 7.464384256 | 7.44169935 | 4.416072028 | 5.331426166 |
| ENSG00000173110 | HSPA6 | 5.666112195 | 0.001534589 | 1.745133011 | 0.388182097 | 7.411245206 | 2 | 2 | 2 | 5.697663076 | 5.89549381 | 3.10571689 | 2.632958241 |
| ENSG00000173905 | GOLIM4 | 3.666427024 | 0.001296732 | 3.744125805 | 0.000940881 | 7.410552829 | 4.467311984 | 7.73146271 | 5.156626401 | 10.1651988 | 10.0755276 | 10.00159102 | 10.371703 |
| ENSG00000254788 | CKLF-CMTM1 | 5.883876011 | 0.000275437 | 1.525476148 | 0.416362316 | 7.409352159 | 2 | 2 | 2 | 5.949776857 | 6.05460702 | 2.803598391 | 2.782576693 |
| ENSG00000072110 | ACTN1 | 2.788584999 | 1.88E-08 | 4.620747411 | 1.06E-21 | 7.40933241 | 9.689059027 | 9.785372135 | 8.585098385 | 12.20913422 | 12.2509268 | 14.08406122 | 14.03853699 |
| ENSG00000152291 | TGOLN2 | 3.665718382 | 3.53E-06 | 3.741767008 | 1.97E-06 | 7.40748539 | 7.096806373 | 9.338312924 | 7.896971941 | 12.05512627 | 12.0847252 | 11.97178976 | 12.30138363 |
| ENSG00000065809 | FAM107B | 3.228864661 | 0.002123869 | 4.178433353 | 4.08E-05 | 7.407298013 | 4.815609471 | 6.666836837 | 3.369317972 | 8.729158235 | 8.67605209 | 9.444446837 | 9.822335095 |
| ENSG00000188266 | HYKK | 2.63457679 | 0.122761253 | 4.772305538 | 0.002281644 | 7.406882328 | 2 | 3.076665209 | 2 | 4.034562184 | 4.02226117 | 6.010089548 | 5.699242824 |
| ENSG00000157657 | ZNF618 | 4.448810538 | 0.000273699 | 2.954377145 | 0.020693942 | 7.403187683 | 2 | 6.602424674 | 5.156626401 | 9.844008139 | 9.88372429 | 8.559441176 | 8.176765766 |
| ENSG00000107077 | KDM4C | 4.03360758 | 0.021123194 | 3.361693325 | 0.05738556 | 7.395300904 | 2 | 6.04685525 | 2 | 8.474993083 | 8.47552601 | 7.755649985 | 7.868000534 |
| ENSG00000196369 | SRGAP2B | 6.831623359 | 9.16E-12 | 0.559242485 | 0.661268816 | 7.390865844 | 4.815609471 | 2 | 3.369317972 | 10.11504861 | 10.0501952 | 4.487545924 | 3.753786374 |
| ENSG00000119682 | AREL1 | 3.528922327 | 7.60E-05 | 3.861710382 | 1.17E-05 | 7.390632709 | 5.095959458 | 7.002537973 | 4.523675391 | 9.463489617 | 9.39312469 | 9.632330087 | 9.876916845 |
| ENSG00000170469 | SPATA24 | 2.977468897 | 0.181130407 | 4.412705086 | 0.034523766 | 7.390173983 | 2 | 2 | 2 | 3.881293667 | 3.2184422 | 4.959178705 | 4.341135913 |
| ENSG00000120322 | PCDHB8 | 4.522783711 | 0.017920625 | 2.865134761 | 0.149193138 | 7.387918472 | 2 | 2 | 2 | 4.846787657 | 4.70309173 | 3.465698119 | 3.560117069 |
| ENSG00000114019 | AMOTL2 | 2.290129305 | 0.076570922 | 5.097641836 | 1.38E-05 | 7.387771141 | 4.00722531 | 7.819099893 | 8.536706852 | 9.948991943 | 9.93637437 | 12.79607961 | 12.6923973 |
| ENSG00000149177 | PTPRJ | 2.713266121 | 1.86E-05 | 4.671236946 | 1.65E-14 | 7.384503067 | 5.532353208 | 7.274681896 | 6.80318718 | 9.358500114 | 9.41399842 | 11.05674387 | 11.57340217 |
| ENSG00000100580 | TMED8 | 3.37331142 | 0.000123283 | 4.008946988 | 3.28E-06 | 7.382258408 | 4.467311984 | 3.686303414 | 6.203069917 | 8.358215887 | 8.34212018 | 8.973764228 | 8.985229012 |
| ENSG00000072364 | AFF4 | 3.619344728 | 0.016568695 | 3.75613387 | 0.01202378 | 7.375478598 | 9.667789716 | 7.051611315 | 4.058998049 | 11.95289514 | 11.9020956 | 12.0083211 | 12.11830098 |
| ENSG00000130147 | SH3BP4 | 3.24018063 | 0.013191227 | 4.129721985 | 0.001089398 | 7.369902615 | 3.327711234 | 8.053983481 | 6.629488387 | 10.16052179 | 10.1858391 | 10.74991732 | 11.31587024 |
| ENSG00000138028 | CGREF1 | 1.444434559 | 0.362218166 | 5.925239184 | 8.16E-06 | 7.369673743 | 2 | 3.686303414 | 2 | 3.615725078 | 3.90918929 | 8.055971376 | 7.429968874 |
| ENSG00000172164 | SNTB1 | 3.307045293 | 0.000124409 | 4.058124793 | 1.45E-06 | 7.365170086 | 2 | 5.457120953 | 4.874587708 | 7.708923796 | 7.94798467 | 8.648868792 | 8.495377414 |
| ENSG00000010030 | ETV7 | 4.714492707 | 0.00020089 | 2.649701998 | 0.04875895 | 7.364194705 | 2 | 2 | 4.874587708 | 7.617841552 | 7.77591026 | 5.626574484 | 5.807970301 |
| ENSG00000070731 | ST6GALNAC2 | 4.327352479 | 0.00692683 | 3.032895919 | 0.066788326 | 7.360248398 | 2 | 3.076665209 | 2 | 5.420062284 | 5.48637102 | 4.406883583 | 4.26688425 |
| ENSG00000123500 | COL10A1 | 8.776980359 | 5.95E-10 | -1.417516949 | 0.453396071 | 7.359463409 | 2 | 2 | 2 | 8.832088406 | 8.80016655 | 2.094598369 | 2.172608095 |
| ENSG00000139304 | PTPRQ | 3.571713969 | 0.120948604 | 3.783434219 | 0.092291365 | 7.355148188 | 2 | 2 | 2 | 3.95996233 | 4.05806467 | 3.465698119 | 4.644600858 |
| ENSG00000130066 | SAT1 | 4.926976378 | 2.72E-14 | 2.423220296 | 0.000354082 | 7.350196674 | 8.39321512 | 7.099070191 | 7.285536013 | 12.62471618 | 12.5906835 | 10.54740056 | 9.474014382 |
| ENSG00000160299 | PCNT | 3.302237014 | 0.03878829 | 4.044692574 | 0.008995102 | 7.346929588 | 2 | 6.602424674 | 3.369317972 | 8.319579168 | 8.4936155 | 9.182675508 | 9.10594367 |
| ENSG00000174370 | C11orf45 | 4.079870386 | 0.101901089 | 3.266398909 | 0.192069539 | 7.346269295 | 2 | 2 | 2 | 4.105493497 | 4.65713986 | 4.360042493 | 2.817676617 |
| ENSG00000196498 | NCOR2 | 3.062873503 | 0.029091374 | 4.282502156 | 0.001407621 | 7.345375659 | 3.327711234 | 8.451443573 | 8.434763955 | 10.90471749 | 10.9382377 | 12.13331984 | 12.14564413 |
| ENSG00000140526 | ABHD2 | 3.520431063 | 0.00084619 | 3.822843421 | 0.000240226 | 7.343274484 | 3.327711234 | 6.389896633 | 7.163363831 | 9.755113959 | 9.75670736 | 9.960968181 | 10.14714405 |
| ENSG00000081052 | COL4A4 | -0.208684324 | 0.908905358 | 7.551796196 | 1.83E-09 | 7.343111872 | 2 | 3.686303414 | 2 | 3.023133209 | 2.59489879 | 9.385734536 | 9.401037612 |
| ENSG00000104635 | SLC39A14 | 4.79258068 | 2.97E-06 | 2.547651632 | 0.019771557 | 7.340232312 | 5.330594134 | 8.993716073 | 8.296287027 | 12.93963415 | 12.9555902 | 10.73381744 | 10.67645725 |
| ENSG00000179431 | FJX1 | 3.790146787 | 0.033889281 | 3.548761401 | 0.046421006 | 7.338908188 | 2 | 6.04685525 | 2 | 8.201097719 | 8.2657098 | 7.682340325 | 8.253273225 |
| ENSG00000138135 | CH25H | 5.834501132 | 0.000451072 | 1.502676835 | 0.435247515 | 7.337177968 | 2 | 2 | 2 | 6.024891728 | 5.88556448 | 2.885263063 | 2.671842496 |
| ENSG00000173276 | ZBTB21 | 4.230065652 | 0.046591905 | 3.10601288 | 0.151419628 | 7.336078532 | 2 | 7.189546432 | 2 | 9.79991149 | 9.83811764 | 8.832074253 | 8.560285 |
| ENSG00000134802 | SLC43A3 | 6.224579334 | 2.37E-10 | 1.111449232 | 0.339130027 | 7.336028566 | 2 | 4.710564154 | 2 | 9.311457162 | 9.36838404 | 4.620674722 | 4.365060851 |
| ENSG00000136048 | DRAM1 | 3.256129734 | 3.52E-07 | 4.076905733 | 8.68E-11 | 7.333035467 | 8.716893051 | 7.73146271 | 6.958202216 | 11.2069543 | 11.2228348 | 11.84522962 | 12.20204378 |
| ENSG00000103269 | RHBDL1 | 2.658055954 | 0.127502931 | 4.673947578 | 0.003551201 | 7.332003532 | 2 | 3.076665209 | 2 | 3.997744386 | 4.0930011 | 6.013120342 | 5.481238982 |
| ENSG00000164106 | SCRG1 | 1.372926523 | 0.484134885 | 5.954595254 | 0.000242523 | 7.327521776 | 2 | 3.076665209 | 2 | 3.349643765 | 3.01487888 | 6.242195894 | 7.478755384 |
| ENSG00000102024 | PLS3 | 2.760994088 | 0.000660428 | 4.561845628 | 3.85E-09 | 7.322839716 | 9.174465797 | 8.993716073 | 6.80318718 | 11.380775 | 11.4074231 | 13.20212973 | 13.18477655 |
| ENSG00000143776 | CDC42BPA | 3.08634527 | 1.18E-09 | 4.227799196 | 1.88E-17 | 7.314144466 | 5.866944937 | 7.232742069 | 6.534109959 | 9.685919789 | 9.73682479 | 10.83169479 | 10.86666148 |
| ENSG00000172650 | AGAP5 | 7.776846802 | 1.33E-07 | -0.463010793 | 0.818792684 | 7.313836009 | 2 | 2 | 2 | 7.821180652 | 7.83643742 | 2.3070652 | 2.172608095 |
| ENSG00000130707 | ASS1 | 2.755359231 | 0.034922085 | 4.551961909 | 0.000201678 | 7.30732114 | 5.532353208 | 6.666836837 | 2 | 8.349110081 | 8.33666436 | 9.60772968 | 10.50771945 |
| ENSG00000087510 | TFAP2C | 1.754986582 | 0.166572561 | 5.552206848 | 4.75E-07 | 7.30719343 | 2 | 4.93638933 | 2 | 5.22679461 | 5.2591458 | 8.834646275 | 8.946110698 |
| ENSG00000160685 | ZBTB7B | 3.137116684 | 0.169849097 | 4.169984963 | 0.055706281 | 7.307101647 | 2 | 7.467835385 | 2 | 9.004380786 | 9.02033034 | 10.04017303 | 10.03882852 |
| ENSG00000163888 | CAMK2N2 | 3.975283781 | 0.050350232 | 3.326786254 | 0.102751946 | 7.302070034 | 2 | 2 | 2 | 4.387512679 | 4.2560182 | 3.917810786 | 3.735548302 |
| ENSG00000154654 | NCAM2 | 6.329000542 | 0.000586428 | 0.969062781 | 0.666867266 | 7.298063323 | 2 | 2 | 2 | 6.306580752 | 6.53293476 | 2.831336763 | 2.277179783 |

| | | | | | | | | | | | | | |
|---|---|---|---|---|---|---|---|---|---|---|---|---|---|
| ENSG00000144395 | CCDC150 | 3.011023436 | 0.148866715 | 4.285191095 | 0.029111318 | 7.296214531 | 2 | 2 | 2 | 3.566290607 | 3.65239003 | 4.461156421 | 4.683176719 |
| ENSG00000198756 | COLGALT2 | 2.724014486 | 0.053129029 | 4.568827205 | 0.000492089 | 7.29284169 | 2 | 4.442710129 | 2 | 5.818703821 | 5.34860058 | 6.770513376 | 7.776188509 |
| ENSG00000112541 | PDE10A | 0.276588494 | 0.878583032 | 7.010675373 | 4.46E-08 | 7.287263867 | 2 | 3.686303414 | 2 | 2.784612485 | 3.28037825 | 8.923506363 | 8.778637453 |
| ENSG00000121743 | GJA3 | 2.819852287 | 0.177650086 | 4.466676831 | 0.021719442 | 7.286529118 | 2 | 2 | 2 | 3.615725078 | 3.33976443 | 4.727866721 | 4.729989799 |
| ENSG00000162734 | PEA15 | 2.89806883 | 2.65E-05 | 4.386257963 | 5.04E-11 | 7.284326793 | 8.075571721 | 9.328390744 | 10.02619908 | 12.21691381 | 12.258881 | 13.53349039 | 13.89487424 |
| ENSG00000157766 | ACAN | -0.893518308 | 0.552070286 | 8.176877342 | 2.60E-12 | 7.283359033 | 2 | 5.131603625 | 2 | 3.566290607 | 3.08597292 | 11.67280806 | 11.72040189 |
| ENSG00000143469 | SYT14 | 6.467457802 | 6.79E-05 | 0.810403595 | 0.689153776 | 7.277861397 | 2 | 2 | 2 | 6.548530093 | 6.56439512 | 2.384001718 | 2.671842496 |
| ENSG00000182957 | SPATA13 | 4.5774309 | 2.86E-09 | 2.699722514 | 0.000778051 | 7.277153414 | 5.095959458 | 4.11352228 | 2 | 8.508095759 | 8.40776525 | 6.693258166 | 6.55171954 |
| ENSG00000269313 | MAGIX | 3.333646724 | 0.113679396 | 3.938947329 | 0.052055276 | 7.272594053 | 2 | 2 | 2 | 3.881293667 | 3.7864959 | 4.111244182 | 4.456980017 |
| ENSG00000166963 | MAP1A | 3.208800099 | 1.67E-06 | 4.063503982 | 6.50E-10 | 7.272304081 | 8.493334308 | 9.695596167 | 8.007629717 | 12.09175474 | 12.1430612 | 12.68025986 | 13.21414468 |
| ENSG00000106327 | TFR2 | 3.669898089 | 0.100833309 | 3.600156373 | 0.101255863 | 7.270054462 | 2 | 2 | 2 | 4.470469939 | 3.55551834 | 3.904814898 | 4.147809227 |
| ENSG00000158186 | MRAS | 2.633843194 | 4.23E-05 | 4.627970767 | 5.85E-14 | 7.261813961 | 6.009025997 | 7.538394143 | 6.718950348 | 9.496029474 | 9.50355257 | 11.05729453 | 11.81917129 |
| ENSG00000172059 | KLF11 | 3.202369987 | 0.002137885 | 4.058777008 | 6.10E-05 | 7.261146995 | 3.327711234 | 5.838937808 | 3.369317972 | 7.799951945 | 7.72392024 | 8.839776604 | 8.328971229 |
| ENSG00000055208 | TAB2 | 3.638775437 | 0.059087638 | 3.619765716 | 0.057302601 | 7.258541154 | 4.815609471 | 8.671479029 | 2 | 10.81265775 | 10.797723 | 10.99149283 | 10.54681386 |
| ENSG00000156925 | ZIC3 | 1.019042799 | 0.778887556 | 6.237754085 | 0.036451497 | 7.256796883 | 2 | 2 | 2 | 3.023133209 | 2 | 6.092633892 | 6.546498097 |
| ENSG00000106683 | LIMK1 | 4.42576042 | 2.63E-29 | 2.830542578 | 1.20E-12 | 7.256302998 | 6.468621203 | 6.535001748 | 5.930865648 | 10.6671059 | 10.7478979 | 8.983840816 | 9.244188987 |
| ENSG00000113448 | PDE4D | 5.506463548 | 7.03E-06 | 1.749526342 | 0.204157205 | 7.25598989 | 2 | 4.11352228 | 2 | 7.870381136 | 7.95512324 | 4.764298067 | 3.987514168 |
| ENSG00000149639 | SOGA1 | 2.606993285 | 1.60E-05 | 4.647993458 | 1.07E-15 | 7.254986743 | 6.009025997 | 7.35507212 | 7.552162999 | 9.649295375 | 9.72990471 | 11.41699656 | 11.98014078 |
| ENSG00000106772 | PRUNE2 | -3.752916103 | 0.531143362 | 11.00549732 | 0.028438282 | 7.252581214 | 2 | 2 | 2 | 2 | 2 | 10.26164954 | 11.53536074 |
| ENSG00000174106 | LEMD3 | 3.564029911 | 2.40E-05 | 3.685059989 | 1.09E-05 | 7.2490899 | 2 | 5.131603625 | 5.59507254 | 8.199071993 | 8.284723 | 8.362013331 | 8.362041249 |
| ENSG00000109320 | NFKB1 | 4.19387434 | 0.059805354 | 3.052872323 | 0.17862881 | 7.246746663 | 2 | 7.503546079 | 2 | 10.06086293 | 10.136455 | 8.62331206 | 9.240151898 |
| ENSG00000119771 | KLHL29 | 2.513856288 | 0.010502522 | 4.730024455 | 2.45E-07 | 7.243880743 | 2 | 5.303523747 | 5.59507254 | 7.202487431 | 7.38644297 | 9.087759434 | 9.795350748 |
| ENSG00000167207 | NOD2 | 3.771481084 | 0.028728494 | 3.469198385 | 0.043887828 | 7.240679469 | 3.327711234 | 2 | 2 | 4.867343068 | 5.04390545 | 4.984105765 | 4.341135913 |
| ENSG00000173193 | PARP14 | 5.128672735 | 1.99E-05 | 2.11032117 | 0.108033027 | 7.238993905 | 4.467311984 | 9.00621767 | 7.552162999 | 13.01713496 | 13.0340228 | 10.20843574 | 9.78485758 |
| ENSG00000051341 | POLQ | 3.548274734 | 0.116163431 | 3.684300299 | 0.094939611 | 7.232575033 | 2 | 2 | 2 | 4.105493497 | 3.86943969 | 3.5515855 | 4.489993931 |
| ENSG00000197780 | TAF13 | 3.708342731 | 0.070288814 | 3.52006932 | 0.083321465 | 7.228412051 | 2 | 2 | 6.534109959 | 8.572106084 | 8.60521604 | 8.271259924 | 8.522888769 |
| ENSG00000121552 | CSTA | 5.437607043 | 0.026843757 | 1.787206801 | 0.513707623 | 7.224813844 | 2 | 2 | 2 | 5.590471342 | 5.58783529 | 3.393105702 | 2.172608095 |
| ENSG00000104154 | SLC30A4 | 2.468611917 | 0.001318023 | 4.753766017 | 4.55E-11 | 7.222377934 | 4.467311984 | 3.686303414 | 4.523675391 | 6.586219071 | 6.23747437 | 8.852100097 | 8.421719243 |
| ENSG00000117013 | KCNQ4 | 0.858796749 | 0.684667709 | 6.35974563 | 8.20E-05 | 7.218542379 | 2 | 2 | 2 | 2.498691189 | 2.59489879 | 6.625560108 | 6.257923755 |
| ENSG00000171790 | SLFNL1 | 0.272692498 | 0.911517345 | 6.944313062 | 4.65E-05 | 7.21700556 | 2 | 2 | 2 | 2.270786764 | 2.49394783 | 7.463831776 | 6.362645675 |
| ENSG00000137962 | ARHGAP29 | 1.801497033 | 0.20822859 | 5.414893977 | 1.61E-05 | 7.216391009 | 6.89194353 | 6.899080544 | 2 | 8.102673083 | 8.08227582 | 11.34184575 | 11.96425948 |
| ENSG00000115548 | KDM3A | 4.319142138 | 0.00281989 | 2.892201893 | 0.05435683 | 7.211344031 | 2 | 7.145017454 | 4.874587708 | 10.13319806 | 10.0858626 | 8.839349773 | 8.526871175 |
| ENSG00000254996 | ANKHD1-EIF4 | 7.021681409 | 4.58E-19 | 0.186687095 | 0.862702263 | 7.208368504 | 4.00722531 | 4.11352228 | 2 | 10.19039992 | 10.2123959 | 3.796405078 | 3.875378318 |
| ENSG00000076513 | ANKRD13A | 2.778331682 | 5.07E-07 | 4.429043181 | 1.48E-16 | 7.207374863 | 6.366709671 | 7.35507212 | 5.930865648 | 9.427605927 | 9.44644299 | 11.06762631 | 11.09639353 |
| ENSG00000171109 | MFN1 | 3.828013466 | 0.079373234 | 3.374537077 | 0.121142926 | 7.202550543 | 2 | 7.274681896 | 2 | 9.506715015 | 9.50274083 | 9.030205459 | 9.077717044 |
| ENSG00000132688 | NES | 1.927496978 | 0.188460431 | 5.273343721 | 4.74E-05 | 7.200840699 | 4.815609471 | 8.004992147 | 4.058998049 | 8.814994458 | 8.23280093 | 11.62445289 | 12.10559801 |
| ENSG00000282637 | | 2.120012726 | 0.108674252 | 5.076619508 | 1.73E-05 | 7.196632234 | 2 | 4.710564154 | 2 | 5.289065324 | 5.39133044 | 7.761068522 | 8.485179747 |
| ENSG00000163235 | TGFA | 6.168884709 | 9.95E-05 | 1.023739846 | 0.591655943 | 7.192624555 | 2 | 2 | 3.369317972 | 7.261917051 | 7.12888795 | 3.296906311 | 2.632958241 |
| ENSG00000154864 | PIEZO2 | 0.901266606 | 0.661500335 | 6.290655063 | 0.000109903 | 7.191921669 | 2 | 3.076665209 | 2 | 3.289987342 | 2.49394783 | 7.117472117 | 7.489678691 |
| ENSG00000153253 | SCN3A | 5.354282511 | 0.011041916 | 1.835252001 | 0.434458206 | 7.189534512 | 2 | 2 | 2 | 5.433909581 | 5.58783529 | 2.42098416 | 3.311948089 |
| ENSG00000173120 | KDM2A | 3.791443751 | 0.000619313 | 3.396428122 | 0.002339673 | 7.187871873 | 4.00722531 | 7.901717006 | 7.691296561 | 10.99183871 | 11.0615956 | 10.63841368 | 10.62753531 |
| ENSG00000009413 | REV3L | 3.773255152 | 0.00024102 | 3.413563906 | 0.000966346 | 7.186819058 | 4.00722531 | 7.145017454 | 5.156626401 | 9.793212756 | 9.66027117 | 9.488666496 | 9.24176809 |
| ENSG00000197142 | ACSL5 | 5.491250308 | 1.83E-05 | 1.689943535 | 0.243517965 | 7.181193843 | 2 | 2 | 4.058998049 | 7.497728447 | 7.33629421 | 3.917810786 | 4.077147165 |
| ENSG00000181652 | ATG9B | 4.150350771 | 0.042255933 | 3.030015804 | 0.14563462 | 7.180366575 | 2 | 2 | 2 | 4.670864177 | 4.22485516 | 3.584558147 | 3.660199087 |
| ENSG00000131725 | WDR44 | 3.57931774 | 7.67E-05 | 3.59774275 | 6.46E-05 | 7.177060491 | 2 | 5.595927442 | 5.39244166 | 8.30834735 | 8.49850989 | 8.544331623 | 8.294346816 |
| ENSG00000250254 | PTTG2 | 3.471550796 | 0.104383657 | 3.704968003 | 0.075016235 | 7.176518799 | 2 | 2 | 2 | 4.034562184 | 3.82856379 | 3.904814898 | 4.292061816 |
| ENSG00000140465 | CYP1A1 | 4.114055548 | 0.078971311 | 3.061049818 | 0.19694971 | 7.175105367 | 2 | 2 | 2 | 4.17310023 | 4.65713986 | 2.962551659 | 4.105828626 |
| ENSG00000181195 | PENK | 6.933700553 | 2.18E-06 | 0.236614424 | 0.904062298 | 7.170314977 | 2 | 5.595927442 | 2 | 10.9250823 | 10.8900091 | 4.894909044 | 3.924507493 |
| ENSG00000174282 | ZBTB4 | 2.752031432 | 0.063911628 | 4.416030162 | 0.001529662 | 7.168061594 | 3.327711234 | 8.981105197 | 7.934808822 | 10.68892298 | 10.7523441 | 12.47675767 | 12.28192772 |
| ENSG00000115904 | SOS1 | 3.53496833 | 0.011191826 | 3.631375125 | 0.008618924 | 7.166343456 | 6.138360429 | 7.315436877 | 2 | 9.771565364 | 9.75670736 | 10.11013328 | 9.557495966 |
| ENSG00000173757 | STAT5B | 2.839686165 | 0.040073481 | 4.32450111 | 0.000934116 | 7.164187274 | 2 | 7.099070191 | 5.930865648 | 8.882187604 | 8.8250571 | 10.23852871 | 10.41728984 |
| ENSG00000006607 | FARP2 | 1.822550436 | 0.211727163 | 5.340695751 | 3.07E-05 | 7.163246188 | 4.00722531 | 6.389896633 | 2 | 6.851641821 | 6.86652414 | 10.0366402 | 10.57561499 |
| ENSG00000154429 | CCSAP | 2.991703724 | 0.000698097 | 4.169997409 | 9.51E-07 | 7.161701134 | 4.815609471 | 5.131603625 | 2 | 7.423326516 | 7.15805139 | 8.746340327 | 8.087908463 |
| ENSG00000151914 | DST | 3.237516714 | 8.99E-05 | 3.92333883 | 1.27E-06 | 7.160855543 | 10.26462289 | 8.916342327 | 7.818184403 | 12.55227704 | 12.5779266 | 13.18885183 | 13.30984437 |
| ENSG00000167971 | CASKIN1 | 2.387612241 | 0.273627021 | 4.767484298 | 0.015667908 | 7.155096539 | 2 | 2 | 2 | 2.947924294 | 3.45167086 | 5.125248504 | 4.836507146 |
| ENSG00000181284 | TMEM102 | 3.083146923 | 0.01291029 | 4.06523846 | 0.000637466 | 7.148385382 | 2 | 4.11352228 | 2 | 5.565529336 | 5.63601114 | 6.475428841 | 6.569847194 |
| ENSG00000160741 | CRTC2 | 3.57687149 | 0.075654914 | 3.561297305 | 0.072951675 | 7.138168795 | 2 | 6.666836837 | 2 | 8.630393142 | 8.65866647 | 8.678282924 | 8.578306014 |
| ENSG00000173821 | RNF213 | 4.446528568 | 1.68E-17 | 2.68872402 | 4.84E-07 | 7.135252588 | 9.410448395 | 8.717821467 | 8.007629717 | 13.22864867 | 13.2860975 | 11.59105918 | 11.40547739 |
| ENSG00000182175 | RGMA | 2.555463744 | 0.551538856 | 4.579378793 | 0.251602132 | 7.134842537 | 2 | 2 | 2 | 3.997744386 | 2 | 4.778616857 | 4.870334174 |
| ENSG00000179954 | SSC5D | 2.770940849 | 0.033290369 | 4.363195214 | 0.00037493 | 7.134136063 | 5.532353208 | 6.666836837 | 2 | 8.378048624 | 8.33848526 | 9.457021219 | 10.29943592 |
| ENSG00000125817 | CENPB | 3.024221293 | 0.054799589 | 4.109421294 | 0.006373412 | 7.133642586 | 8.042902253 | 7.669933291 | 2 | 10.19904132 | 10.3751428 | 11.35802693 | 11.38688232 |
| ENSG00000114480 | GBE1 | 3.420396981 | 0.024746731 | 3.712341746 | 0.013403649 | 7.132738727 | 7.579279349 | 7.901717006 | 2 | 10.56816382 | 10.5644133 | 10.75818265 | 10.95046432 |
| ENSG00000272333 | KMT2B | 3.353699635 | 0.121326135 | 3.778020727 | 0.072863536 | 7.131720362 | 2 | 7.099070191 | 2 | 8.793673851 | 8.91810182 | 9.373110172 | 9.176932811 |
| ENSG00000167202 | TBC1D2B | 3.130476712 | 0.192379762 | 3.997448234 | 0.082542158 | 7.127924947 | 2 | 7.847167909 | 2 | 9.354865403 | 9.41658653 | 10.52342208 | 9.910074228 |
| ENSG00000198399 | ITSN2 | 3.276583718 | 0.103576285 | 3.850911914 | 0.048101168 | 7.127495632 | 2 | 6.602424674 | 2 | 8.270263574 | 8.2960118 | 8.916644086 | 8.783084035 |
| ENSG00000111252 | SH2B3 | 4.266270714 | 0.00023647 | 2.85899777 | 0.018167791 | 7.125268484 | 6.468621203 | 8.469613413 | 4.523675391 | 11.50034187 | 11.5421087 | 9.88710712 | 10.31583232 |
| ENSG00000114023 | FAM162A | 3.269492314 | 0.00621802 | 3.854939322 | 0.000948646 | 7.124431637 | 6.366709671 | 4.710564154 | 2 | 8.30834735 | 8.3920703 | 8.923506363 | 8.937160962 |
| ENSG00000112078 | KCTD20 | 1.649689172 | 0.102464016 | 5.474288926 | 6.80E-10 | 7.123978098 | 7.709880939 | 9.018611864 | 6.073378139 | 9.72797373 | 9.65735634 | 13.61012915 | 13.40498581 |
| ENSG00000116199 | FAM20B | 2.635524718 | 0.000170716 | 4.478100077 | 2.51E-11 | 7.113624794 | 6.468621203 | 6.728495673 | 4.523675391 | 8.801706047 | 8.74219737 | 10.47327283 | 10.72623001 |
| ENSG00000169981 | ZNF35 | 3.49348253 | 0.015288584 | 3.619880014 | 0.011100946 | 7.113362544 | 2 | 5.131603625 | 2 | 7.127781374 | 6.93551934 | 6.838725184 | 7.418582165 |
| ENSG00000077264 | PAK3 | 4.766228498 | 0.014403895 | 2.330410594 | 0.264448359 | 7.096639092 | 2 | 2 | 2 | 4.984880435 | 4.98931816 | 3.393105702 | 2.950113106 |
| ENSG00000139289 | PHLDA1 | 5.558719374 | 2.16E-44 | 1.532842195 | 0.000319758 | 7.091561569 | 7.53297911 | 7.467835385 | 8.236946975 | 13.31874609 | 13.3075788 | 9.462297671 | 9.107717328 |
| ENSG00000172058 | SERF1A | 8.278040659 | 7.83E-09 | -1.18744399 | 0.532750733 | 7.09059667 | 2 | 2 | 2 | 8.304583895 | 8.34030387 | 2.183373728 | 2.117335275 |
| ENSG00000168961 | LGALS9 | 5.207392534 | 3.85E-05 | 1.882503368 | 0.179033043 | 7.089895902 | 2 | 4.93638933 | 2 | 8.529212228 | 8.5227358 | 5.677130623 | 4.886954834 |
| ENSG00000141497 | ZMYND15 | 3.046463071 | 0.161240809 | 4.043029878 | 0.049620411 | 7.089492949 | 2 | 2 | 2 | 3.566290607 | 3.69849104 | 4.122514202 | 4.594883674 |
| ENSG00000109063 | MYH3 | 4.01120481 | 0.049833933 | 3.070546395 | 0.138210579 | 7.081751205 | 2 | 2 | 2 | 4.329433326 | 4.37434574 | 3.600766115 | 3.698365557 |
| ENSG00000172183 | ISG20 | 5.599378006 | 4.78E-05 | 1.480808025 | 0.347140092 | 7.080186032 | 2 | 6.311476754 | 3.369317972 | 10.40067384 | 10.4444166 | 6.231826305 | 6.498634707 |
| ENSG00000100097 | LGALS1 | 3.867594037 | 3.83E-06 | 3.201605637 | 0.000161199 | 7.069199674 | 10.55263183 | 8.57407404 | 8.585098385 | 13.39748587 | 13.4440161 | 12.33389445 | 13.08096347 |
| ENSG00000115009 | CCL20 | 7.578465761 | 5.71E-06 | -0.511052057 | 0.820068239 | 7.067413705 | 2 | 2 | 2 | 7.68596357 | 7.58049694 | 2.094598369 | 2.374681325 |
| ENSG00000249884 | RNF103-CHM | 9.779844158 | 4.18E-11 | -2.712987522 | 0.194894738 | 7.066856637 | 2 | 2 | 2 | 9.80923777 | 9.81606952 | 2 | 2.117335275 |
| ENSG00000152495 | CAMK4 | 4.180617221 | 0.046457783 | 2.885804541 | 0.179747748 | 7.066421762 | 2 | 2 | 2 | 4.387512679 | 4.58534546 | 3.767980144 | 3.236138336 |
| ENSG00000196233 | LCOR | 2.78006534 | 0.040340218 | 4.286163549 | 0.000804245 | 7.066228889 | 4.467311984 | 6.535001748 | 2 | 8.056427969 | 7.93841129 | 9.451725397 | 9.52733357 |
| ENSG00000168298 | HIST1H1E | 4.319232539 | 0.031747935 | 2.744561561 | 0.188337529 | 7.0637941 | 2 | 2 | 2 | 4.670864177 | 4.53541753 | 3.27687201 | 3.580696614 |
| ENSG00000053438 | NNAT | 4.304670729 | 0.042473193 | 2.757015011 | 0.209143226 | 7.06168574 | 2 | 2 | 2 | 4.670864177 | 4.50979031 | 3.738983888 | 3.071407821 |
| ENSG00000174136 | RGMB | 4.529982179 | 8.15E-11 | 2.52414992 | 0.000524836 | 7.054132099 | 6.009025997 | 7.928237109 | 6.534109959 | 11.58441754 | 11.553128 | 9.647281667 | 9.485655697 |
| ENSG00000111328 | CDK2AP1 | 3.967486498 | 3.85E-33 | 3.082175817 | 1.55E-20 | 7.049662315 | 7.096806373 | 6.844429755 | 6.958202216 | 10.91122844 | 10.862748 | 9.880069077 | 10.11906036 |
| ENSG00000149308 | NPAT | 3.619797894 | 0.051517521 | 3.428069921 | 0.063487602 | 7.047867815 | 2 | 6.228547981 | 2 | 8.264463296 | 8.23084153 | 8.07857076 | 8.038552764 |
| ENSG00000127329 | PTPRB | 4.398178356 | 3.84E-05 | 2.648500109 | 0.018649379 | 7.046678466 | 3.327711234 | 3.076665209 | 4.058998049 | 7.326437704 | 7.31795292 | 6.075339068 | 5.0581604 |
| ENSG00000063127 | SLC6A16 | 2.080764065 | 0.328511143 | 4.964901421 | 0.008306252 | 7.045665486 | 2 | 2 | 2 | 2.947924294 | 3.15372745 | 5.357600875 | 4.935697922 |
| ENSG00000131018 | SYNE1 | 3.108843219 | 0.003101637 | 3.934744306 | 0.000115824 | 7.043587525 | 5.330594134 | 8.414402626 | 8.825624954 | 11.18332463 | 11.2399842 | 12.20539422 | 11.84578422 |
| ENSG00000143450 | OAZ3 | 3.71269421 | 0.080939257 | 3.328678999 | 0.115085799 | 7.041373208 | 2 | 2 | 2 | 4.237680128 | 3.98554648 | 3.694366604 | 3.956354764 |
| ENSG00000115756 | HPCAL1 | 4.020607589 | 0.001424077 | 3.017034999 | 0.020100516 | 7.037642589 | 6.96351253 | 6.464272742 | 2 | 10.09449002 | 10.1587842 | 9.041010784 | 9.211159041 |
| ENSG00000102271 | KLHL4 | 3.879625073 | 0.062735523 | 3.153677427 | 0.132250448 | 7.0333025 | 2 | 2 | 2 | 4.299492648 | 4.1930041 | 3.600766115 | 3.807156736 |
| ENSG00000137403 | | 12.22300053 | 2.68E-18 | -5.191150116 | 0.022468449 | 7.031850415 | 2 | 2 | 2 | 12.16350377 | 12.3326784 | 2 | 2 |
| ENSG00000082781 | ITGB5 | 3.612813598 | 1.10E-06 | 3.417975451 | 4.18E-06 | 7.03078905 | 8.696033109 | 8.777382925 | 6.629488387 | 11.87367064 | 11.9476934 | 11.71192115 | 11.72112623 |
| ENSG00000170214 | ADRA1B | 0.313586387 | 0.885376787 | 6.716363218 | 1.49E-05 | 7.029949604 | 2 | 3.076665209 | 2 | 2.784612485 | 2.59489879 | 7.05119671 | 8.192729549 |
| ENSG00000110987 | BCL7A | 3.318966262 | 0.000595251 | 3.710809158 | 9.25E-05 | 7.02977542 | 3.327711234 | 4.442710129 | 2 | 6.542151749 | 6.507263 | 7.045287119 | 6.741885308 |
| ENSG00000083093 | PALB2 | 3.843314315 | 0.008808181 | 3.177735533 | 0.032897253 | 7.021049849 | 2 | 5.303523747 | 2 | 7.565338955 | 7.53039799 | 7.054142452 | 6.728114749 |
| ENSG00000120451 | SNX19 | 3.236269806 | 0.082560548 | 3.782791706 | 0.036346775 | 7.019061513 | 5.330594134 | 8.607278467 | 2 | 10.33439225 | 10.4338094 | 10.89385283 | 10.96545847 |
| ENSG00000031691 | CENPQ | 3.701626997 | 0.011105632 | 3.314119944 | 0.02391519 | 7.015746941 | 2 | 3.076665209 | 3.369317972 | 5.839660368 | 5.49945122 | 5.661762729 | 4.886954834 |
| ENSG00000204381 | LAYN | 3.093650003 | 0.021356951 | 3.920490942 | 0.002561312 | 7.014140945 | 6.653098118 | 7.099070191 | 2 | 9.319871109 | 9.42346554 | 10.21749687 | 10.1734218 |
| ENSG00000029153 | ARNTL2 | 3.709477397 | 0.000536632 | 3.299600178 | 0.002254154 | 7.009077574 | 5.866944937 | 3.076665209 | 6.629488387 | 9.305820479 | 9.43201844 | 9.232285816 | 8.63228931 |
| ENSG00000163430 | FSTL1 | 2.602805657 | 2.48E-07 | 4.406115134 | 2.14E-19 | 7.008920792 | 11.20805444 | 12.08448117 | 12.35768921 | 14.55571806 | 14.5688082 | 16.09506566 | 16.59301128 |
| ENSG00000130720 | FIBCD1 | 2.95495031 | 0.080067967 | 4.05342139 | 0.011575145 | 7.0083717 | 2 | 3.686303414 | 2 | 4.946755474 | 4.95174433 | 6.430795201 | 5.20457242 |
| ENSG00000184384 | MAML2 | 3.259035589 | 0.008961459 | 3.749098087 | 0.002132217 | 7.008133677 | 6.468621203 | 6.728495673 | 2 | 9.271529914 | 9.21030762 | 9.702302705 | 9.754328277 |
| ENSG00000050405 | LIMA1 | 3.096326461 | 2.74E-06 | 3.910200332 | 1.58E-09 | 7.006526794 | 7.159102577 | 8.356994601 | 9.031722608 | 11.43385118 | 11.4700176 | 12.44498745 | 12.0593766 |
| ENSG00000146535 | GNA12 | 2.712172898 | 0.000860731 | 4.293117396 | 3.45E-08 | 7.005290293 | 6.257048221 | 8.717821467 | 7.777116083 | 10.56816382 | 10.6316726 | 12.20311522 | 12.15392944 |

| | | | | | | | | | | | | | |
|---|---|---|---|---|---|---|---|---|---|---|---|---|---|
| ENSG00000118369 | USP35 | 2.88787372 | 0.1083703 | 4.117193529 | 0.016000913 | 7.005067249 | 2 | 5.946638857 | 2 | 7.173892946 | 7.32163988 | 8.336833852 | 8.568035872 |
| ENSG00000107249 | GLIS3 | 3.112243425 | 0.02449614 | 3.891652318 | 0.00367982 | 7.003895743 | 3.327711234 | 5.131603625 | 7.646393253 | 9.420668575 | 9.36392108 | 9.554115566 | 10.5978704 |
| ENSG00000180008 | SOCS4 | 3.525681562 | 0.004196673 | 3.477936183 | 0.004611432 | 7.003617745 | 3.327711234 | 6.78762695 | 4.058998049 | 9.029678018 | 8.88733423 | 9.089196199 | 8.711989853 |
| ENSG00000136828 | RALGPS1 | 2.100184869 | 0.167050868 | 4.903131145 | 0.000285449 | 7.003316014 | 2 | 3.686303414 | 2 | 4.237680128 | 4.2865223 | 6.959905775 | 6.577546999 |
| ENSG00000107263 | RAPGEF1 | 2.858499539 | 0.063038247 | 4.143687183 | 0.004524025 | 7.002186723 | 4.00722531 | 7.669933291 | 3.369317972 | 9.021676912 | 9.12214832 | 10.42459331 | 10.27478258 |
| ENSG00000180776 | ZDHHC20 | 3.268715852 | 0.00011428 | 3.731851735 | 7.67E-06 | 7.000567586 | 4.467311984 | 7.099070191 | 5.772716771 | 9.404936476 | 9.35405368 | 9.851780083 | 9.829070247 |
| ENSG00000104419 | NDRG1 | 4.174265519 | 0.006207698 | 2.822079567 | 0.074468007 | 6.996345086 | 8.138771685 | 7.051611315 | 2 | 11.24377823 | 11.2751103 | 9.646549955 | 10.13436737 |
| ENSG00000134326 | CMPK2 | 9.307518351 | 4.93E-21 | -2.315769499 | 0.048314828 | 6.991748852 | 2 | 4.442710129 | 2 | 12.08902254 | 12.1387571 | 2.526562374 | 2.326754077 |
| ENSG00000123575 | FAM199X | 2.950947649 | 0.112649338 | 4.040763335 | 0.02294093 | 6.991710984 | 3.327711234 | 7.315436877 | 2 | 8.801706047 | 8.66303256 | 9.987213818 | 9.623140479 |
| ENSG00000182742 | HOXB4 | 7.352935807 | 7.99E-06 | -0.361708631 | 0.872376417 | 6.991227176 | 2 | 2 | 2 | 7.405868159 | 7.42122542 | 2.384001718 | 2.117335275 |
| ENSG00000183668 | PSG9 | 1.497853592 | 0.527022694 | 5.492304838 | 0.00562232 | 6.990158431 | 2 | 2 | 2 | 2.868578001 | 2.68924543 | 4.749835737 | 6.185873005 |
| ENSG00000187098 | MITF | 2.518537148 | 0.003498455 | 4.469766418 | 3.49E-08 | 6.988303566 | 4.00722531 | 4.710564154 | 2 | 6.147608921 | 6.30607931 | 7.975906509 | 8.254874792 |
| ENSG00000063601 | MTMR1 | 2.927543491 | 0.151113687 | 4.059263922 | 0.036333082 | 6.986807413 | 2 | 6.602424674 | 2 | 7.942383523 | 7.93600799 | 8.951423836 | 9.157371148 |
| ENSG00000144827 | ABHD10 | 3.126198895 | 0.002731351 | 3.859808163 | 0.000142658 | 6.986007058 | 4.467311984 | 5.72254323 | 2 | 7.68596357 | 7.72392024 | 8.676850263 | 8.126037235 |
| ENSG00000039560 | RAI14 | 3.844135864 | 1.65E-06 | 3.141054069 | 0.00011376 | 6.985189933 | 7.031698319 | 8.540087354 | 9.541233192 | 12.52827438 | 12.5324175 | 11.80202 | 11.85327028 |
| ENSG00000134242 | PTPN22 | 5.01168299 | 0.055895663 | 1.969540872 | 0.488789509 | 6.981223863 | 2 | 2 | 2 | 5.258265913 | 5.14724905 | 3.517841605 | 2.172608095 |
| ENSG00000135862 | LAMC1 | 2.345118471 | 0.010398313 | 4.631907945 | 5.53E-08 | 6.977026416 | 10.18451821 | 11.29895829 | 8.561105505 | 12.72235977 | 12.7690318 | 14.9059816 | 15.14789346 |
| ENSG00000100647 | SUSD6 | 3.818082363 | 0.038583311 | 3.158834485 | 0.090151257 | 6.976916848 | 2 | 8.414402626 | 4.874587708 | 10.71715466 | 10.776052 | 10.14163246 | 10.03557113 |
| ENSG00000170525 | PFKFB3 | 3.7704302 | 0.000130097 | 3.200219432 | 0.001343175 | 6.970649631 | 6.563806279 | 6.228547981 | 8.884815469 | 11.45882823 | 11.5018493 | 10.7250644 | 11.07615369 |
| ENSG00000109762 | SNX25 | 3.247202842 | 0.051734578 | 3.718587895 | 0.022521058 | 6.965790737 | 3.327711234 | 6.78762695 | 2 | 8.504819462 | 8.54498542 | 9.01024623 | 8.973604485 |
| ENSG00000145949 | MYLK4 | 3.244565412 | 0.134476647 | 3.719726475 | 0.075349452 | 6.964291887 | 2 | 2 | 2 | 3.840289928 | 3.69849104 | 3.993419637 | 4.241259475 |
| ENSG00000151012 | SLC7A11 | 5.502337118 | 6.82E-08 | 1.461265756 | 0.207497621 | 6.963602874 | 5.095959458 | 7.145017454 | 8.677268317 | 13.08794963 | 13.066109 | 8.947864566 | 9.138408485 |
| ENSG00000161921 | CXCL16 | 2.702773711 | 3.17E-05 | 4.260471183 | 9.13E-12 | 6.963244894 | 3.327711234 | 5.303523747 | 5.156626401 | 7.307758725 | 7.46188679 | 8.831215892 | 9.00677572 |
| ENSG00000162545 | CAMK2N1 | 2.822195125 | 0.001659602 | 4.13421516 | 1.58E-06 | 6.956410284 | 5.330594134 | 6.464272742 | 7.934808822 | 9.694573013 | 9.77026155 | 10.700246 | 11.3162538 |
| ENSG00000204010 | IFIT1B | 9.663443238 | 8.84E-11 | -2.711962854 | 0.196453617 | 6.951480384 | 2 | 2 | 2 | 9.690973808 | 9.70259467 | 2 | 2.117335275 |
| ENSG00000186635 | ARAP1 | 3.771376435 | 0.023509926 | 3.179239271 | 0.058676271 | 6.950615706 | 2 | 7.901717006 | 5.156626401 | 10.21115348 | 10.3135262 | 9.771053529 | 9.568807213 |
| ENSG00000180881 | CAPS2 | 2.89955963 | 0.188248635 | 4.044993774 | 0.051489483 | 6.944553404 | 2 | 2 | 2 | 3.615725078 | 3.45167086 | 4.588526421 | 4.133950982 |
| ENSG00000139970 | RTN1 | 0.42852882 | 0.833848549 | 6.515220466 | 1.25E-05 | 6.943749286 | 2 | 3.076665209 | 2 | 2.784612485 | 2.68924543 | 7.1328795 | 7.84903967 |
| ENSG00000107796 | ACTA2 | 1.006423158 | 0.521983587 | 5.936594945 | 3.96E-06 | 6.943018103 | 8.009475818 | 9.318399851 | 4.058998049 | 9.278261665 | 9.22120493 | 14.24028087 | 14.09820361 |
| ENSG00000125484 | GTF3C4 | 3.22065677 | 0.059838231 | 3.722042383 | 0.025556146 | 6.942699153 | 4.00722531 | 7.35507212 | 2 | 9.122382231 | 9.06060056 | 9.568866087 | 9.61099888 |
| ENSG00000187642 | PERM1 | 3.667383715 | 0.112319087 | 3.273705736 | 0.152745431 | 6.941089451 | 2 | 2 | 2 | 4.17310023 | 3.98554648 | 4.177580368 | 3.261853426 |
| ENSG00000005238 | FAM214B | 3.590311237 | 0.047154938 | 3.349213213 | 0.063066737 | 6.93952445 | 4.467311984 | 8.029695761 | 2 | 10.10101391 | 10.1546561 | 9.740063182 | 10.02199736 |
| ENSG00000088256 | GNA11 | 2.829133407 | 0.299070637 | 4.108806512 | 0.109843769 | 6.937939918 | 2 | 9.079034257 | 2 | 10.29130517 | 10.3491863 | 11.6102001 | 11.58474907 |
| ENSG00000162433 | AK4 | 2.733280611 | 0.026147826 | 4.202571659 | 0.000304705 | 6.93585227 | 4.00722531 | 6.78762695 | 8.266922072 | 9.856271709 | 9.89118718 | 11.21284883 | 11.45534619 |
| ENSG00000149657 | LSM14B | 2.911407748 | 0.172910901 | 4.021174525 | 0.047681033 | 6.932582274 | 6.653098118 | 2 | 2 | 7.852449262 | 8.01099479 | 9.011762258 | 9.04984997 |
| ENSG00000213085 | CFAP45 | 3.839333468 | 0.069491099 | 3.09092108 | 0.146121333 | 6.930254548 | 2 | 2 | 2 | 4.237680128 | 4.1930041 | 3.796405078 | 3.518055793 |
| ENSG00000171150 | SOCS5 | 3.760373457 | 0.048272272 | 3.168314955 | 0.098094243 | 6.928688412 | 4.00722531 | 8.147241016 | 2 | 10.38472097 | 10.3715902 | 9.837317946 | 9.737529735 |
| ENSG00000107317 | PTGDS | 0.170255443 | 0.934841508 | 6.757919971 | 7.48E-06 | 6.928175414 | 2 | 3.686303414 | 2 | 3.227757282 | 2.68924543 | 7.613117257 | 9.139275853 |
| ENSG00000104332 | SFRP1 | 5.324816833 | 2.53E-09 | 1.598862978 | 0.110078372 | 6.923679811 | 5.70932728 | 4.710564154 | 5.59507254 | 10.57090794 | 10.5648023 | 7.581819208 | 5.497565112 |
| ENSG00000065600 | TMEM206 | 3.886433455 | 0.016316848 | 3.036710244 | 0.06554535 | 6.923143699 | 2 | 5.595927442 | 2 | 7.900612272 | 7.85436579 | 7.319501608 | 6.711881188 |
| ENSG00000123338 | NCKAP1L | 1.275646484 | 0.621394099 | 5.643850455 | 0.008475154 | 6.919496939 | 2 | 2 | 2 | 2.784612485 | 2.59489879 | 6.482006365 | 4.353147976 |
| ENSG00000112561 | TFEB | 3.146834581 | 0.013266834 | 3.770495855 | 0.002274706 | 6.917330436 | 2 | 4.710564154 | 2 | 6.221320894 | 6.35727082 | 7.043805932 | 6.714211481 |
| ENSG00000168016 | TRANK1 | 4.54810141 | 1.52E-13 | 2.368813938 | 0.000230354 | 6.916915347 | 5.70932728 | 7.503546079 | 6.80318718 | 11.34480861 | 11.3855729 | 9.369567287 | 8.993406747 |
| ENSG00000008394 | MGST1 | 4.290238632 | 2.22E-12 | 2.624208877 | 3.11E-05 | 6.914447509 | 7.384433967 | 5.72254323 | 6.43197674 | 10.93058681 | 10.8519526 | 9.033940582 | 9.406811858 |
| ENSG00000063244 | U2AF2 | 3.409572142 | 1.93E-08 | 3.50271027 | 6.62E-09 | 6.912282412 | 7.159102577 | 5.595927442 | 7.225742779 | 10.15739533 | 10.1807731 | 10.21979429 | 10.30292368 |
| ENSG00000173482 | PTPRM | 2.76955821 | 0.161589056 | 4.141818928 | 0.026558049 | 6.911377138 | 2 | 8.777382925 | 5.156626401 | 10.02185637 | 10.1009593 | 11.35780343 | 11.50042979 |
| ENSG00000135837 | CEP350 | 3.577227925 | 0.011803936 | 3.331651676 | 0.019496236 | 6.908879601 | 4.00722531 | 8.077869079 | 4.523675391 | 10.19752012 | 10.2583269 | 9.942804722 | 10.02317256 |
| ENSG00000165417 | GTF2A1 | 3.407418064 | 0.044940119 | 3.498058642 | 0.036484987 | 6.905476706 | 3.327711234 | 6.899080544 | 2 | 8.89725916 | 8.67172529 | 8.920281122 | 8.835935827 |
| ENSG00000108256 | NUFIP2 | 3.570829643 | 9.25E-05 | 3.333740444 | 0.000271524 | 6.904570087 | 7.53297911 | 8.376385542 | 5.39244166 | 11.11674327 | 11.0725776 | 11.05976992 | 10.62382165 |
| ENSG00000125637 | PSD4 | 2.240048484 | 0.011707809 | 4.663984205 | 1.21E-08 | 6.904032689 | 3.327711234 | 4.442710129 | 3.369317972 | 5.742861563 | 5.80356929 | 7.865372957 | 8.317521841 |
| ENSG00000203797 | DDO | 5.923206793 | 0.009469798 | 0.979532975 | 0.719706986 | 6.902739769 | 2 | 2 | 2 | 6.069927018 | 6.00960265 | 2.937246078 | 2.117335275 |
| ENSG00000133065 | SLC41A1 | 3.082573895 | 0.003820004 | 3.819591558 | 0.000229589 | 6.902165453 | 4.467311984 | 7.790474954 | 5.930865648 | 9.72797373 | 9.70330142 | 10.39324012 | 10.50452546 |
| ENSG00000164930 | FZD6 | 3.6778466 | 0.112536896 | 3.222182716 | 0.162850128 | 6.900029315 | 2 | 7.701025996 | 2 | 9.80923777 | 9.75806852 | 9.470865405 | 9.175242287 |
| ENSG00000167114 | SLC27A4 | 2.988478166 | 0.05824713 | 3.910722387 | 0.009837756 | 6.899200553 | 3.327711234 | 6.464272742 | 2 | 7.966401501 | 7.96929529 | 8.720150661 | 9.022374596 |
| ENSG00000068366 | ACSL4 | 4.753955195 | 7.60E-05 | 2.139511711 | 0.100111403 | 6.893466906 | 4.467311984 | 8.124485672 | 9.06622548 | 12.85250628 | 12.8630334 | 10.11402027 | 10.36930224 |
| ENSG00000148411 | NACC2 | 2.391589617 | 0.000157675 | 4.500802437 | 7.72E-14 | 6.892392054 | 5.330594134 | 6.844429755 | 7.225742779 | 9.044420506 | 8.98240172 | 11.05398739 | 11.17229884 |
| ENSG00000259330 | INAFM2 | 1.69392158 | 0.000341709 | 5.198085402 | 1.55E-32 | 6.892006982 | 5.866944937 | 5.72254323 | 5.156626401 | 7.190301983 | 7.28055563 | 10.78856921 | 10.61620978 |
| ENSG00000116903 | EXOC8 | 3.649301244 | 5.22E-07 | 3.239551458 | 9.57E-06 | 6.888852702 | 3.327711234 | 4.93638933 | 5.772716771 | 8.491639362 | 8.40429221 | 8.24055649 | 7.816873723 |
| ENSG00000143970 | ASXL2 | 3.066552838 | 1.82E-11 | 3.820824391 | 1.85E-17 | 6.887377229 | 6.816637977 | 5.946638857 | 5.772716771 | 9.215512556 | 9.26686067 | 9.971718548 | 10.01161415 |
| ENSG00000137693 | YAP1 | 3.486548563 | 0.000136335 | 3.400001924 | 0.000194399 | 6.886550487 | 7.53297911 | 8.414402626 | 5.39244166 | 11.048347 | 11.0193685 | 10.88447059 | 11.079855 |
| ENSG00000197102 | DYNC1H1 | 3.221003615 | 0.004115118 | 3.656886327 | 0.00090549 | 6.877889942 | 6.563806279 | 10.46189757 | 9.589475645 | 12.75840141 | 12.8065979 | 13.11073939 | 13.31850957 |
| ENSG00000161888 | SPC24 | 3.012579551 | 0.197226444 | 3.858294934 | 0.083094108 | 6.870874485 | 2 | 2 | 2 | 3.754611435 | 3.45167086 | 3.724264246 | 4.604965121 |
| ENSG00000181982 | CCDC149 | 3.126727382 | 0.000167279 | 3.741782678 | 4.17E-06 | 6.86851006 | 4.467311984 | 4.93638933 | 2 | 7.053486362 | 7.24992376 | 7.937338157 | 7.548321824 |
| ENSG00000100156 | SLC16A8 | 2.275064302 | 0.297706831 | 4.580549106 | 0.019950506 | 6.855613408 | 2 | 2 | 2 | 3.227757282 | 3.08597292 | 5.002522427 | 4.624919101 |
| ENSG00000110880 | CORO1C | 2.440879815 | 0.045753442 | 4.413224832 | 9.96E-05 | 6.854104647 | 5.70932728 | 9.828248506 | 8.845626171 | 11.3151886 | 11.3278511 | 12.95249272 | 13.56695364 |
| ENSG00000145723 | GIN1 | 3.720437925 | 0.02162448 | 3.132891384 | 0.055208449 | 6.853329308 | 2 | 3.686303414 | 2 | 5.590471342 | 5.6826301 | 5.536762811 | 4.489993931 |
| ENSG00000164849 | GPR146 | 4.727372448 | 0.002681847 | 2.125306579 | 0.213653476 | 6.852679027 | 2 | 3.076665209 | 2 | 5.775854763 | 5.86549842 | 3.584558147 | 3.735548302 |
| ENSG00000113319 | RASGRF2 | 4.673604958 | 0.000556 | 2.178289618 | 0.135501339 | 6.851894576 | 6.257048221 | 2 | 4.058998049 | 9.463489617 | 9.50030288 | 7.134272047 | 6.903766785 |
| ENSG00000180917 | CMTR2 | 3.864205747 | 0.034408534 | 2.98614017 | 0.10784488 | 6.850345917 | 2 | 6.140560021 | 2 | 8.439397922 | 8.36015869 | 7.772739333 | 7.253757517 |
| ENSG00000145365 | TIFA | 3.361428392 | 4.80E-05 | 3.486825349 | 2.07E-05 | 6.848253741 | 3.327711234 | 4.442710129 | 3.369317972 | 6.872128674 | 6.80463998 | 6.860764111 | 7.052989523 |
| ENSG00000203814 | HIST2H2BF | 6.958915563 | 8.69E-07 | -0.11248077 | 0.95448384 | 6.846434792 | 2 | 3.076665209 | 2 | 7.980622673 | 7.95986274 | 2.457042231 | 2.632958241 |
| ENSG00000157542 | KCNJ6 | -0.724548903 | 0.752249153 | 7.568088807 | 1.07E-06 | 6.843539905 | 2 | 2 | 2 | 2.270786764 | 2.14022021 | 7.846806418 | 7.360260221 |
| ENSG00000162542 | TMCO4 | 2.640238434 | 0.00087139 | 4.199848698 | 2.70E-08 | 6.840087132 | 2 | 4.93638933 | 4.523675391 | 6.670510205 | 6.72335419 | 8.261097876 | 8.176765766 |
| ENSG00000213906 | LTB4R2 | 4.371457102 | 0.013378187 | 2.465704348 | 0.185599465 | 6.83716145 | 2 | 3.076665209 | 2 | 5.48800675 | 5.49945122 | 4.281782022 | 3.383972333 |
| ENSG00000101639 | CEP192 | 3.540152727 | 0.001471851 | 3.296048447 | 0.003154055 | 6.836201174 | 2 | 5.946638857 | 4.523675391 | 8.297027403 | 8.28283292 | 8.218405973 | 7.8574975 |
| ENSG00000166971 | AKTIP | 2.844274831 | 0.003720972 | 3.988274141 | 2.23E-05 | 6.832548972 | 3.327711234 | 4.93638933 | 2 | 6.470063213 | 6.53293476 | 7.697491716 | 7.517964974 |
| ENSG00000173480 | ZNF417 | 4.439550672 | 0.000613741 | 2.38913565 | 0.083245787 | 6.828686322 | 2 | 5.838937808 | 3.369317972 | 8.852852814 | 8.81724322 | 7.131485609 | 6.432241896 |
| ENSG00000109458 | GAB1 | 2.041104352 | 0.237601833 | 4.786561489 | 0.002077008 | 6.827665841 | 3.327711234 | 6.728495673 | 2 | 7.292639438 | 7.27676203 | 9.796563052 | 10.1753259 |
| ENSG00000153233 | PTPRR | 4.457289799 | 0.028243263 | 2.370140214 | 0.270737833 | 6.827430013 | 2 | 2 | 2 | 4.670864177 | 4.76938658 | 2.987421008 | 3.407202941 |
| ENSG00000146067 | FAM193B | 2.969153569 | 0.000902883 | 3.856769236 | 8.89E-06 | 6.825922806 | 3.327711234 | 6.228547981 | 5.156626401 | 8.211183901 | 8.21506984 | 9.317805994 | 8.823577951 |
| ENSG00000176853 | FAM91A1 | 3.460337303 | 0.008386015 | 3.363562987 | 0.010235235 | 6.82390029 | 2 | 6.951736442 | 5.930865648 | 9.383689875 | 9.34954609 | 9.369567287 | 9.16420533 |
| ENSG00000087494 | PTHLH | 4.431349699 | 0.000240382 | 2.389591042 | 0.063104764 | 6.820940741 | 3.327711234 | 3.686303414 | 4.058998049 | 7.638916932 | 7.66709116 | 4.350488971 | 6.38329369 |
| ENSG00000132932 | ATP8A2 | 1.274445776 | 0.562605288 | 5.545208786 | 0.00218059 | 6.819654562 | 2 | 2 | 2 | 2.695456494 | 2.68924543 | 5.258920789 | 6.018717758 |
| ENSG00000021300 | PLEKHB1 | 1.012622956 | 0.641177228 | 5.805608199 | 0.000743299 | 6.818231155 | 2 | 2 | 2 | 2.695456494 | 2.49394783 | 5.805044075 | 6.044874254 |
| ENSG00000150594 | ADRA2A | 0.857221333 | 0.727489842 | 5.955775681 | 0.002130827 | 6.812997014 | 2 | 2 | 2 | 2.3892346 | 2.68924543 | 5.185595313 | 6.61544141 |
| ENSG00000136802 | LRRC8A | 3.523261794 | 0.002887069 | 3.287501004 | 0.005604071 | 6.810762798 | 7.031698319 | 7.73146271 | 3.369317972 | 10.29652513 | 10.4389106 | 10.22028611 | 10.04323752 |
| ENSG00000138286 | FAM149B1 | 3.233619719 | 0.001152009 | 3.577093105 | 0.000257739 | 6.810712824 | 4.00722531 | 5.457120953 | 2 | 7.514115926 | 7.4450836 | 7.926930284 | 7.697107799 |
| ENSG00000198315 | ZKSCAN8 | 3.561212774 | 0.010243646 | 3.247311935 | 0.020011794 | 6.808524709 | 4.467311984 | 7.538394143 | 3.369317972 | 9.697445926 | 9.6919516 | 9.558789939 | 9.181571564 |
| ENSG00000205339 | IPO7 | 3.458259319 | 1.07E-08 | 3.348657202 | 2.95E-08 | 6.806916522 | 6.009025997 | 7.572420273 | 7.691296561 | 10.69684063 | 10.689886 | 10.70388473 | 10.45330786 |
| ENSG00000114978 | MOB1A | 3.737792873 | 2.98E-14 | 3.067600753 | 5.88E-10 | 6.805393626 | 7.485143553 | 7.638155693 | 6.322058544 | 10.9941771 | 10.9493026 | 10.36356703 | 10.24063603 |
| ENSG00000090857 | PDPR | 3.498793147 | 0.005182914 | 3.305321883 | 0.008424324 | 6.80411503 | 4.467311984 | 7.503546079 | 4.058998049 | 9.58946437 | 9.70471389 | 9.482119335 | 9.428971287 |
| ENSG00000179104 | TMTC2 | 3.248234319 | 0.011460822 | 3.55440453 | 0.00485155 | 6.802638849 | 2 | 4.442710129 | 2 | 6.283823264 | 5.89549381 | 6.184218099 | 6.574984961 |
| ENSG00000282214 | | 3.525985164 | 0.024581743 | 3.275325487 | 0.036849285 | 6.801310652 | 2 | 6.951736442 | 4.058998049 | 8.960808841 | 9.09547637 | 8.906900259 | 8.643336952 |
| ENSG00000166578 | IQCD | 1.863568792 | 0.459530549 | 4.934330364 | 0.024930503 | 6.797899156 | 2 | 2 | 2 | 3.289987342 | 2.49394783 | 4.659878161 | 5.492143571 |
| ENSG00000162004 | CCDC78 | 2.390511696 | 0.325100102 | 4.407196375 | 0.047258258 | 6.797708071 | 2 | 2 | 2 | 3.289987342 | 3.15372745 | 5.19095825 | 3.875378318 |
| ENSG00000129675 | ARHGEF6 | 2.590408802 | 0.016876187 | 4.205187545 | 3.81E-05 | 6.795596347 | 2 | 5.838937808 | 4.523675391 | 7.300218887 | 7.25764291 | 8.834646275 | 8.901818877 |
| ENSG00000103257 | SLC7A5 | 2.712990547 | 6.19E-05 | 4.077956508 | 4.63E-10 | 6.790947054 | 8.138771685 | 7.431218269 | 6.629488387 | 10.18988999 | 10.2374972 | 12.01230258 | 10.94712477 |
| ENSG00000102218 | RP2 | 3.489169917 | 9.51E-08 | 3.30030614 | 4.50E-07 | 6.789476057 | 5.532353208 | 6.04685525 | 4.058998049 | 8.813671111 | 8.85589616 | 8.403026507 | 8.857178832 |
| ENSG00000143473 | KCNH1 | 4.54717839 | 0.020944825 | 2.241167181 | 0.286960302 | 6.788345571 | 2 | 2 | 2 | 4.76150579 | 4.83276841 | 3.256555576 | 3.012034821 |
| ENSG00000131781 | FMO5 | 2.645912701 | 0.252177378 | 4.141965527 | 0.053922041 | 6.787878228 | 2 | 2 | 2 | 3.615725078 | 3.08597292 | 4.675266036 | 4.201948172 |
| ENSG00000156113 | KCNMA1 | 3.418038313 | 0.012001607 | 3.365839001 | 0.013077994 | 6.783877314 | 3.327711234 | 3.686303414 | 6.958202216 | 8.870779703 | 8.93384535 | 8.769824912 | 8.927653397 |
| ENSG00000197632 | SERPINB2 | 11.97408823 | 1.09E-17 | -5.191150361 | 0.022295807 | 6.782937869 | 2 | 2 | 2 | 12.0000521 | 12.0037045 | 2 | 2 |
| ENSG00000163171 | CDC42EP3 | 2.435872328 | 0.123290993 | 4.345997728 | 0.002975696 | 6.781870057 | 8.836089686 | 5.303523747 | 4.058998049 | 9.836208668 | 9.81083239 | 11.74784286 | 11.70993099 |
| ENSG00000109381 | ELF2 | 3.247481578 | 0.004804831 | 3.534262262 | 0.001845662 | 6.78174384 | 3.327711234 | 6.140560021 | 3.369317972 | 8.013270196 | 8.09094333 | 8.513629334 | 8.132149972 |
| ENSG00000149781 | FERMT3 | 2.97391488 | 0.089940832 | 3.806427063 | 0.023485564 | 6.780341943 | 2 | 3.686303414 | 2 | 5.003571641 | 4.91316568 | 6.237020416 | 4.861951484 |
| ENSG00000196865 | NHLRC2 | 2.797403188 | 2.53E-05 | 3.976683057 | 6.49E-10 | 6.774086246 | 3.327711234 | 5.595927442 | 5.39244166 | 7.751013478 | 7.7377845 | 8.874770504 | 8.94164277 |
| ENSG00000076641 | PAG1 | 2.816975801 | 1.22E-05 | 3.953900508 | 2.46E-10 | 6.770876308 | 4.815609471 | 4.11352228 | 5.59507254 | 7.527093191 | 7.58663873 | 8.841908869 | 8.491306993 |

| | | | | | | | | | | | | | |
|---|---|---|---|---|---|---|---|---|---|---|---|---|---|
| ENSG00000196337 | CGB7 | 4.063065477 | 0.070101138 | 2.706345289 | 0.240957826 | 6.769410766 | 2 | 2 | 2 | 4.17310023 | 4.58534546 | 3.724264246 | 3.012034821 |
| ENSG00000120756 | PLS1 | 1.187115691 | 0.524942496 | 5.582172558 | 0.000240832 | 6.769288249 | 2 | 3.076665209 | 2 | 3.09461492 | 3.08597292 | 6.382394503 | 6.841176624 |
| ENSG00000141337 | ARSG | 3.847945381 | 5.10E-07 | 2.920136689 | 0.000187368 | 6.76808207 | 2 | 4.93638933 | 4.874587708 | 7.90310317 | 8.14604733 | 7.113241388 | 7.130061054 |
| ENSG00000162078 | ZG16B | 1.014024556 | 0.665819919 | 5.753120077 | 0.002140974 | 6.767144633 | 2 | 2 | 2 | 2.784612485 | 2.38539787 | 5.472156123 | 6.199243628 |
| ENSG00000110395 | CBL | 3.245470639 | 0.022825231 | 3.520694458 | 0.012183874 | 6.766165097 | 5.095959458 | 7.928237109 | 3.369317972 | 9.848538434 | 9.73406074 | 10.274494 | 9.823414835 |
| ENSG00000152952 | PLOD2 | 4.642233504 | 0.000222301 | 2.119058652 | 0.119536754 | 6.761292156 | 5.330594134 | 10.16572116 | 9.212024683 | 13.87521525 | 13.8218999 | 10.98378895 | 11.60482568 |
| ENSG00000170577 | SIX2 | -1.313350828 | 0.590651191 | 8.073728458 | 8.10E-07 | 6.760377631 | 2 | 2 | 2 | 2 | 2.2680111 | 8.249579355 | 7.979721237 |
| ENSG00000146676 | PURB | 3.250361919 | 4.36E-08 | 3.508925295 | 2.54E-09 | 6.759287215 | 5.866944937 | 7.315436877 | 6.073378139 | 9.80923777 | 9.78034427 | 10.02879997 | 10.0755426 |
| ENSG00000068781 | STON1-GTF2 | 6.617191483 | 0.000337602 | 0.141684102 | 0.954947148 | 6.758875585 | 2 | 2 | 2 | 6.699459234 | 6.70088158 | 2.139668783 | 2.551896972 |
| ENSG00000149582 | TMEM25 | 2.192398011 | 0.231366574 | 4.565986976 | 0.006248071 | 6.758384987 | 2 | 5.946638857 | 2 | 6.573765157 | 6.58294721 | 8.908528807 | 8.896185552 |
| ENSG00000135932 | CAB39 | 3.602908284 | 0.016039408 | 3.155417988 | 0.036346775 | 6.758326272 | 6.468621203 | 7.819099893 | 2 | 10.30314152 | 10.2807554 | 9.905980863 | 9.783748583 |
| ENSG00000185742 | C11orf87 | 2.494629492 | 0.303181423 | 4.25763593 | 0.055927555 | 6.752265422 | 2 | 2 | 2 | 3.023133209 | 3.50452879 | 3.956110469 | 4.975095763 |
| ENSG00000155962 | CLIC2 | 2.975080278 | 0.221769007 | 3.774079409 | 0.104457634 | 6.749159687 | 2 | 2 | 2 | 3.615725078 | 3.55551834 | 4.628601098 | 3.474731315 |
| ENSG00000127743 | IL17B | 2.774647091 | 0.248162973 | 3.973560306 | 0.078790818 | 6.748207396 | 2 | 2 | 2 | 3.162721474 | 3.69849104 | 4.690491514 | 3.824516949 |
| ENSG00000136153 | LMO7 | 2.532852309 | 0.039713176 | 4.213415107 | 0.000259014 | 6.746267417 | 6.138360429 | 9.357954881 | 10.22252928 | 11.85111104 | 11.8451168 | 12.95483431 | 13.93649465 |
| ENSG00000196700 | ZNF512B | 2.878532033 | 0.015001375 | 3.867711618 | 0.000670543 | 6.746243651 | 6.366709671 | 6.311476754 | 2 | 8.551612617 | 8.62623826 | 9.45200461 | 9.681170767 |
| ENSG00000115306 | SPTBN1 | 2.989734532 | 0.000905307 | 3.750551898 | 1.92E-05 | 6.74028643 | 7.750913381 | 10.48441188 | 9.516493218 | 12.58764526 | 12.6467792 | 13.03616105 | 13.65407799 |
| ENSG00000079156 | OSBPL6 | 4.465254108 | 0.003554578 | 2.271707022 | 0.165246475 | 6.736961129 | 2 | 3.076665209 | 4.058998049 | 6.979883891 | 6.99214046 | 5.634468599 | 3.580696614 |
| ENSG00000066739 | ATG2B | 3.466700034 | 0.057486219 | 3.270199934 | 0.071384845 | 6.736899969 | 2 | 7.315436877 | 3.369317972 | 9.264766605 | 9.23104065 | 9.02571051 | 9.079525723 |
| ENSG00000060237 | WNK1 | 3.097848028 | 3.12E-22 | 3.629892467 | 1.37E-30 | 6.727740495 | 8.257427002 | 8.557180776 | 8.143093325 | 11.39544505 | 11.4429793 | 12.11712511 | 11.76273823 |
| ENSG00000047457 | CP | 2.97629186 | 0.2361363 | 3.751091213 | 0.117977654 | 6.727383074 | 2 | 2 | 2 | 3.754611435 | 3.39680241 | 4.652122235 | 3.36036153 |
| ENSG00000171862 | PTEN | 3.199783026 | 0.000645674 | 3.526733398 | 0.000136151 | 6.726516424 | 6.653098118 | 7.847167909 | 4.874587708 | 10.12521908 | 10.0345549 | 10.65148032 | 10.11112449 |
| ENSG00000121904 | CSMD2 | 3.658402043 | 0.001197467 | 3.064962093 | 0.007494621 | 6.723364136 | 3.327711234 | 2 | 4.523675391 | 6.610808945 | 6.67805337 | 6.351597515 | 5.745545708 |
| ENSG00000110429 | FBXO3 | 2.902121863 | 0.052698164 | 3.820702689 | 0.007862246 | 6.722824552 | 2 | 6.228547981 | 3.369317972 | 7.708923796 | 7.6318872 | 8.599312524 | 8.552492262 |
| ENSG00000119227 | PIGZ | 3.042423922 | 0.060267353 | 3.680206608 | 0.019086058 | 6.72263053 | 2 | 5.457120953 | 2 | 6.872128674 | 6.95938211 | 7.245405178 | 7.779530392 |
| ENSG00000186994 | KANK3 | 1.690389006 | 0.472807027 | 5.031838986 | 0.012952329 | 6.722227992 | 2 | 2 | 2 | 2.947924294 | 2.77779907 | 5.680947153 | 4.543391831 |
| ENSG00000136141 | LRCH1 | 3.60863038 | 0.003020234 | 3.11309842 | 0.011549101 | 6.7217288 | 2 | 6.464272742 | 5.156626401 | 8.924487942 | 8.92538942 | 8.601327805 | 8.245238611 |
| ENSG00000147164 | SNX12 | 3.025142263 | 0.117406202 | 3.693535934 | 0.047554519 | 6.718678197 | 3.327711234 | 7.572420273 | 2 | 9.04103174 | 9.06939887 | 9.512957081 | 9.900385864 |
| ENSG00000270898 | GPR75-ASB3 | 7.882161789 | 6.95E-08 | -1.16446949 | 0.545860427 | 6.717692299 | 2 | 2 | 2 | 7.786523606 | 8.06478301 | 2.139668783 | 2.172608095 |
| ENSG00000176871 | WSB2 | 3.041989234 | 0.000769315 | 3.673207505 | 3.22E-05 | 6.715196739 | 5.532353208 | 8.276714465 | 7.029880454 | 10.37891658 | 10.3514424 | 10.69895264 | 11.24037286 |
| ENSG00000155287 | SLC25A28 | 3.913086838 | 0.002673268 | 2.794827598 | 0.037844164 | 6.707914436 | 2 | 4.93638933 | 6.718950348 | 9.29542901 | 9.35044873 | 8.389095798 | 8.017950573 |
| ENSG00000240184 | PCDHGC3 | 3.776647898 | 0.038521438 | 2.925324421 | 0.114540794 | 6.701972319 | 4.467311984 | 8.101365653 | 2 | 10.35637181 | 10.4049916 | 9.216266648 | 9.792596779 |
| ENSG00000171608 | PIK3CD | 2.177922735 | 0.000866952 | 4.522925131 | 1.74E-13 | 6.700847866 | 5.532353208 | 5.303523747 | 6.073378139 | 7.656739692 | 7.8208909 | 9.477737996 | 10.48025681 |
| ENSG00000139618 | BRCA2 | 5.045888383 | 6.66E-06 | 1.654950995 | 0.187386354 | 6.700839378 | 2 | 4.11352228 | 2 | 7.536750104 | 7.42465798 | 4.406883583 | 4.329022995 |
| ENSG00000187987 | ZSCAN23 | 3.561260315 | 0.037885253 | 3.138588207 | 0.067466122 | 6.699848522 | 2 | 3.076665209 | 2 | 4.887609717 | 4.65713986 | 4.425202323 | 4.4231929 |
| ENSG00000171827 | ZNF570 | 3.174923644 | 0.00305872 | 3.522953655 | 0.000823875 | 6.697877299 | 3.327711234 | 5.131603625 | 2 | 6.993973425 | 6.95938211 | 7.519242774 | 7.076727469 |
| ENSG00000082512 | TRAF5 | 2.531958139 | 0.238863041 | 4.165477204 | 0.038278239 | 6.697435343 | 2 | 6.844429755 | 2 | 7.694616532 | 7.88209639 | 9.479108595 | 9.331400327 |
| ENSG00000148541 | FAM13C | 3.283629631 | 0.150709752 | 3.413625362 | 0.125922464 | 6.697254993 | 2 | 3.076665209 | 2 | 4.739373666 | 4.31639476 | 5.381251181 | 3.100202476 |
| ENSG00000205177 | C11orf91 | 5.978747771 | 0.000264903 | 0.71698913 | 0.726496359 | 6.695736901 | 2 | 2 | 2 | 5.968925393 | 6.20590375 | 2.457042231 | 2.551896972 |
| ENSG00000165029 | ABCA1 | 4.717553116 | 8.87E-30 | 1.977023143 | 4.95E-06 | 6.694576259 | 7.624139768 | 6.899080544 | 6.629488387 | 11.76848464 | 11.8038653 | 8.98885284 | 9.118313725 |
| ENSG00000140464 | PML | 3.618641366 | 4.34E-07 | 3.074961611 | 2.12E-05 | 6.693602977 | 8.493334308 | 8.414402626 | 6.43197674 | 11.59966248 | 11.6791849 | 11.04836626 | 11.14427326 |
| ENSG00000010327 | STAB1 | 1.274445254 | 0.558024285 | 5.416869472 | 0.002481102 | 6.691314726 | 2 | 2 | 2 | 2.695456494 | 2.68924543 | 5.328699035 | 5.777095613 |
| ENSG00000160886 | LY6K | 3.993352998 | 0.074741742 | 2.696784584 | 0.240873009 | 6.690137582 | 2 | 2 | 2 | 4.358765249 | 4.31639476 | 3.011868911 | 3.717076715 |
| ENSG00000197496 | SLC2A10 | 2.940715106 | 0.006444915 | 3.748610898 | 0.000334778 | 6.689326004 | 4.00722531 | 7.60566235 | 6.203069917 | 9.452477833 | 9.49050967 | 10.2641958 | 10.28774876 |
| ENSG00000073712 | FERMT2 | 2.245825321 | 1.11E-14 | 4.441770381 | 6.36E-56 | 6.687595703 | 8.009475818 | 8.317412869 | 7.858115942 | 10.28272246 | 10.3455691 | 12.49041464 | 12.5231767 |
| ENSG00000171388 | APLN | 1.815555109 | 0.312172567 | 4.865662446 | 0.001918418 | 6.681217555 | 3.327711234 | 2 | 2 | 3.515101962 | 3.39680241 | 6.043083161 | 5.85096143 |
| ENSG00000155393 | HEATR3 | 3.398616747 | 0.038177023 | 3.282060146 | 0.043831981 | 6.680676893 | 2 | 5.595927442 | 2 | 7.501020868 | 7.29187696 | 7.329261848 | 7.242500282 |
| ENSG00000196605 | ZNF846 | 3.307271811 | 0.012174173 | 3.369215573 | 0.009915802 | 6.676487383 | 2 | 4.11352228 | 2 | 5.697663076 | 5.88556448 | 6.043083161 | 5.631815919 |
| ENSG00000123143 | PKN1 | 2.766440651 | 0.162667495 | 3.908272456 | 0.037527109 | 6.674713107 | 5.095959458 | 8.791896851 | 2 | 10.08190282 | 10.054079 | 11.24438305 | 11.16848011 |
| ENSG00000180758 | GPR157 | 2.026650048 | 0.174256431 | 4.645967717 | 0.000501618 | 6.672617765 | 2 | 2 | 4.874587708 | 5.128039203 | 5.19627118 | 7.87872738 | 7.354296226 |
| ENSG00000216490 | IFI30 | 9.031942134 | 2.43E-15 | -2.36353864 | 0.083660985 | 6.668403494 | 2 | 4.11352228 | 2 | 11.40799569 | 11.4431908 | 2.457042231 | 2.172608095 |
| ENSG00000165731 | RET | 2.467990535 | 0.104145312 | 4.19954176 | 0.002768615 | 6.667532295 | 2 | 2 | 4.058998049 | 4.694064037 | 4.56059747 | 6.231826305 | 6.141542327 |
| ENSG00000169057 | MECP2 | 3.067822962 | 0.051305942 | 3.598649426 | 0.018882709 | 6.666472389 | 8.107517749 | 7.051611315 | 2 | 10.09829921 | 10.1695649 | 10.61429194 | 10.7112881 |
| ENSG00000103260 | METRN | 2.737151656 | 0.034673548 | 3.928078106 | 0.001437419 | 6.665229762 | 4.467311984 | 6.311476754 | 2 | 7.694616532 | 7.84926609 | 9.163708059 | 8.699106494 |
| ENSG00000158683 | PKD1L1 | 2.441806182 | 0.307380329 | 4.220088375 | 0.054660707 | 6.661894557 | 2 | 2 | 2 | 2.947924294 | 3.50452879 | 4.82764224 | 4.133950982 |
| ENSG00000171206 | TRIM8 | 3.330764778 | 0.000953251 | 3.326387523 | 0.000915639 | 6.657152301 | 7.27616306 | 8.505280662 | 5.156626401 | 10.78523872 | 10.894967 | 10.69871735 | 10.9627661 |
| ENSG00000180543 | TSPYL5 | 2.824209316 | 0.177920442 | 3.832317682 | 0.054838286 | 6.656526999 | 2 | 6.728495673 | 2 | 7.959237999 | 7.96694292 | 8.958123037 | 8.960904642 |
| ENSG00000110427 | KIAA1549L | 3.602372588 | 0.022954373 | 3.050300876 | 0.056438674 | 6.652673465 | 2 | 3.686303414 | 6.80318718 | 8.872051712 | 8.87860041 | 8.319800025 | 8.338065764 |
| ENSG00000108021 | FAM208B | 3.211361392 | 0.017806575 | 3.44034869 | 0.010049511 | 6.651710083 | 6.366709671 | 8.213429689 | 3.369317972 | 10.19853443 | 10.2084185 | 10.63608101 | 10.19360379 |
| ENSG00000009335 | UBE3C | 2.904296539 | 0.025535393 | 3.742108727 | 0.002868555 | 6.646405266 | 3.327711234 | 8.053983481 | 7.502634626 | 10.14060551 | 10.1148377 | 10.87043852 | 11.0505632 |
| ENSG00000183060 | LYSMD4 | 2.414977476 | 0.076743239 | 4.231387166 | 0.00080315 | 6.646364642 | 2 | 4.93638933 | 2 | 5.775854763 | 5.87556634 | 7.613117257 | 7.521960976 |
| ENSG00000130487 | KLHDC7B | 2.897841727 | 0.359800435 | 3.747421414 | 0.214267271 | 6.645263141 | 2 | 2 | 2 | 3.406930997 | 3.65239003 | 4.914492341 | 2.374681325 |
| ENSG00000151532 | VTI1A | 3.560133845 | 3.15E-09 | 3.085093078 | 3.32E-07 | 6.645226923 | 4.815609471 | 5.595927442 | 4.058998049 | 8.365459376 | 8.44204342 | 7.829717863 | 8.034828728 |
| ENSG00000031003 | FAM13B | 4.564411499 | 7.51E-08 | 2.080333391 | 0.023172228 | 6.64474489 | 3.327711234 | 5.838937808 | 4.058998049 | 9.217515344 | 9.26207178 | 6.870823472 | 6.723495198 |
| ENSG00000244165 | P2RY11 | 3.772990929 | 0.002468582 | 2.870165375 | 0.024934862 | 6.643156304 | 2 | 4.710564154 | 2 | 7.021746293 | 6.74548212 | 6.028179536 | 6.029985803 |
| ENSG00000139514 | SLC7A1 | 2.205045471 | 0.277947155 | 4.436572097 | 0.017129277 | 6.641617568 | 4.815609471 | 8.655694736 | 2 | 9.360314042 | 9.36392108 | 11.43892588 | 11.7207641 |
| ENSG00000116032 | GRIN3B | 3.733624046 | 0.111579909 | 2.902094201 | 0.219716772 | 6.635718247 | 2 | 2 | 2 | 4.105493497 | 4.16043394 | 3.904814898 | 3.042026731 |
| ENSG00000149187 | CELF1 | 3.771086599 | 0.010057832 | 2.862911878 | 0.056113002 | 6.633998477 | 2 | 7.701025996 | 6.718950348 | 10.42167359 | 10.4832045 | 9.592360305 | 9.502606422 |
| ENSG00000137463 | MGARP | 2.287334733 | 0.095959724 | 4.34657288 | 0.000521666 | 6.633907613 | 2 | 4.11352228 | 2 | 4.76150579 | 4.98931816 | 6.505870593 | 7.013798886 |
| ENSG00000065613 | SLK | 3.378085042 | 3.31E-07 | 3.254596603 | 8.58E-07 | 6.632681644 | 5.532353208 | 7.467835385 | 7.098165388 | 10.25325094 | 10.2564019 | 10.31004663 | 9.928254593 |
| ENSG00000075426 | FOSL2 | 2.182747727 | 2.06E-11 | 4.449342492 | 3.22E-45 | 6.632090219 | 7.485143553 | 7.819099893 | 8.143093325 | 9.966365455 | 10.0345549 | 12.29329414 | 12.23203189 |
| ENSG00000145214 | DGKQ | 3.036216329 | 0.019509132 | 3.594834547 | 0.004534024 | 6.631050876 | 4.467311984 | 6.389896633 | 2 | 8.076429088 | 8.185025 | 8.801285298 | 8.555744353 |
| ENSG00000108582 | CPD | 2.930468851 | 0.000719561 | 3.699658679 | 1.17E-05 | 6.63012753 | 6.138360429 | 8.834582582 | 8.35328246 | 11.06484891 | 11.075856 | 11.86743185 | 11.80885057 |
| ENSG00000174405 | LIG4 | 3.194471367 | 0.099101291 | 3.435348976 | 0.06941259 | 6.629820344 | 2 | 6.311476754 | 2 | 7.885575888 | 7.93600799 | 8.260460366 | 8.026414333 |
| ENSG00000072786 | STK10 | 3.358093793 | 2.52E-05 | 3.270795573 | 3.91E-05 | 6.628889367 | 6.816637977 | 6.228547981 | 4.058998049 | 9.323594948 | 9.43031191 | 9.330617798 | 9.250624924 |
| ENSG00000185972 | CCIN | 3.691457431 | 0.103405461 | 2.932426618 | 0.197522413 | 6.623884049 | 2 | 2 | 2 | 4.358765249 | 3.7864959 | 3.355392153 | 3.735548302 |
| ENSG00000096968 | JAK2 | 3.983114804 | 0.009351711 | 2.637033617 | 0.097308454 | 6.620148421 | 2 | 6.464272742 | 3.369317972 | 8.965583267 | 8.94225201 | 7.851893762 | 7.357281305 |
| ENSG00000091157 | WDR7 | 2.931868387 | 0.148258746 | 3.68649736 | 0.058167426 | 6.618365747 | 2 | 6.535001748 | 2 | 7.810605345 | 7.93841129 | 8.61485781 | 8.626114978 |
| ENSG00000230413 | | 11.80880312 | 3.88E-17 | -5.191150432 | 0.02246952 | 6.617652689 | 2 | 2 | 2 | 11.78512278 | 11.8866222 | 2 | 2 |
| ENSG00000127663 | KDM4B | 2.573404422 | 0.045544105 | 4.04223205 | 0.000846027 | 6.615636472 | 3.327711234 | 7.761270542 | 6.203069917 | 9.186155283 | 9.18523054 | 10.69471167 | 10.60007717 |
| ENSG00000103056 | SMPD3 | 0.686380815 | 0.777377621 | 5.925474024 | 0.001442563 | 6.61185484 | 2 | 2 | 2 | 2.270786764 | 2.68924543 | 6.249924465 | 5.799216178 |
| ENSG00000139192 | TAPBPL | 4.203312448 | 0.000588482 | 2.406591689 | 0.062792893 | 6.609904137 | 3.327711234 | 5.72254323 | 2 | 8.48998335 | 8.50338773 | 6.651077397 | 6.821857532 |
| ENSG00000065923 | SLC9A7 | 2.824031389 | 4.92E-09 | 3.777637803 | 1.47E-15 | 6.601669192 | 6.816637977 | 7.669933291 | 6.534109959 | 9.924066823 | 9.8787275 | 10.91253766 | 10.78933969 |
| ENSG00000197386 | HTT | 2.941630093 | 0.027016433 | 3.657961923 | 0.004541723 | 6.599592016 | 3.327711234 | 8.053983481 | 6.43197674 | 9.768836458 | 9.88060333 | 10.39918908 | 10.66761654 |
| ENSG00000129911 | KLF16 | 2.906850529 | 0.001810688 | 3.687445914 | 4.57E-05 | 6.594296442 | 3.327711234 | 5.457120953 | 6.534109959 | 8.36365191 | 8.43865202 | 9.002642099 | 9.32912155 |
| ENSG00000282107 | | 4.790482846 | 0.000862526 | 1.80327906 | 0.256970059 | 6.593761907 | 2 | 3.686303414 | 2 | 6.628980152 | 6.6952083 | 3.679182183 | 4.30448772 |
| ENSG00000258484 | SPESP1 | 1.861070434 | 0.456251283 | 4.730173802 | 0.030900875 | 6.591244236 | 2 | 2 | 2 | 3.09461492 | 2.77779907 | 5.48093935 | 4.119958326 |
| ENSG00000198924 | DCLRE1A | 2.707015332 | 0.037504648 | 3.883870881 | 0.001646167 | 6.590886213 | 2 | 4.442710129 | 2 | 5.602782483 | 5.60003091 | 6.931352719 | 6.449128733 |
| ENSG00000259030 | FPGT-TNNI3 | 3.936767379 | 0.087461944 | 2.651622658 | 0.260827638 | 6.588390037 | 2 | 2 | 2 | 4.205751501 | 4.37434574 | 2.937246078 | 3.717076715 |
| ENSG00000118518 | RNF146 | 3.531845266 | 0.018380127 | 3.055784173 | 0.043002955 | 6.587629439 | 4.00722531 | 6.728495673 | 2 | 8.839910123 | 8.81854848 | 8.557366484 | 8.126912071 |
| ENSG00000188611 | ASAH2 | 4.984858058 | 0.002812393 | 1.599491673 | 0.393071188 | 6.584349732 | 2 | 2 | 3.369317972 | 6.034011955 | 6.08962627 | 3.616794016 | 2.950113106 |
| ENSG00000115839 | RAB3GAP1 | 3.101889655 | 0.01740322 | 3.481017815 | 0.006513019 | 6.58290747 | 6.89194353 | 8.169643011 | 3.369317972 | 10.18529247 | 10.2014316 | 10.52938802 | 10.61215446 |
| ENSG00000082684 | SEMA5B | 1.867348823 | 0.662327642 | 4.71441858 | 0.219692889 | 6.581767402 | 2 | 2 | 2 | 3.515101962 | 2 | 5.002522427 | 4.878668439 |
| ENSG00000081189 | MEF2C | 3.482524503 | 0.002014131 | 3.09794356 | 0.006424701 | 6.580468064 | 2 | 3.686303414 | 4.523675391 | 6.820355514 | 6.64311914 | 6.818078339 | 5.703940561 |
| ENSG00000154529 | CNTNAP3B | 3.047982349 | 0.182152175 | 3.531665905 | 0.108113626 | 6.579648254 | 2 | 2 | 2 | 3.754611435 | 3.50452879 | 4.133696865 | 3.807156736 |
| ENSG00000132109 | TRIM21 | 4.349012075 | 0.000227241 | 2.227065528 | 0.077869736 | 6.576077603 | 6.73718411 | 6.140560021 | 2 | 10.19039992 | 10.1670053 | 8.041200979 | 8.104880443 |
| ENSG00000047056 | WDR37 | 3.495872057 | 0.086812219 | 3.079249981 | 0.129694026 | 6.575122038 | 2 | 6.666836837 | 2 | 8.554784469 | 8.57309694 | 7.937338157 | 8.339575961 |
| ENSG00000137393 | RNF144B | 3.041618323 | 0.056953525 | 3.532976221 | 0.023237554 | 6.574594544 | 6.257048221 | 2 | 3.369317972 | 7.805288479 | 7.71553728 | 8.138086022 | 8.344097092 |
| ENSG00000221843 | C2orf16 | 1.310980885 | 0.492404429 | 5.263297584 | 0.000926448 | 6.574278469 | 2 | 2 | 3.369317972 | 2.947924294 | 3.33976443 | 6.533574091 | 6.081436739 |
| ENSG00000178209 | PLEC | 2.875426082 | 1.73E-18 | 3.693616396 | 2.94E-30 | 6.569042478 | 10.25038534 | 9.962611189 | 9.778213818 | 12.86261997 | 12.8987213 | 13.45045629 | 13.91034311 |
| ENSG00000150093 | ITGB1 | 3.735732169 | 8.91E-12 | 2.827256813 | 3.54E-07 | 6.562988982 | 10.68025843 | 12.130888 | 11.78368112 | 15.38856894 | 15.3839076 | 14.52891921 | 14.42496738 |
| ENSG00000114166 | KAT2B | 2.79762879 | 0.004686679 | 3.765167133 | 7.79E-05 | 6.562795923 | 2 | 5.595927442 | 4.523675391 | 7.300218887 | 7.30683517 | 8.152588976 | 8.347103327 |
| ENSG00000204103 | MAFB | 3.669345828 | 0.021398494 | 2.892571815 | 0.074588837 | 6.561917642 | 2 | 4.93638933 | 2 | 7.062428156 | 6.90635305 | 6.795372357 | 5.331426166 |
| ENSG00000068383 | INPP5A | 2.589486662 | 0.017388264 | 3.970246511 | 0.000118328 | 6.559733174 | 2 | 6.04685525 | 5.39244166 | 7.77569021 | 7.64076905 | 8.950238388 | 9.187034567 |
| ENSG00000151468 | CCDC3 | -1.970914457 | 0.114510897 | 8.528194679 | 1.32E-17 | 6.557280222 | 4.467311984 | 3.076665209 | 4.058998049 | 2.947924294 | 2.59489879 | 10.99953788 | 12.6187847 |
| ENSG00000222047 | C10orf55 | 3.974263736 | 0.062077767 | 2.582001897 | 0.240345235 | 6.556265633 | 2 | 2 | 2 | 4.237680128 | 4.40247105 | 3.235948948 | 3.430065404 |
| ENSG00000011422 | PLAUR | 4.572034112 | 3.24E-05 | 1.982268736 | 0.097369201 | 6.554302848 | 4.00722531 | 7.847167909 | 7.818184403 | 11.79956717 | 11.8745935 | 9.01781049 | 9.459850232 |

| | | | | | | | | | | | | | |
|---|---|---|---|---|---|---|---|---|---|---|---|---|---|
| ENSG00000179922 | ZNF784 | 2.674345306 | 0.035422866 | 3.874474966 | 0.001290688 | 6.548820271 | 2 | 4.442710129 | 2 | 5.420062284 | 5.71663161 | 6.693258166 | 6.718860807 |
| ENSG00000160813 | PPP1R35 | 3.447295308 | 0.000334546 | 3.098520648 | 0.00134027 | 6.545815956 | 2 | 4.442710129 | 3.369317972 | 6.604700657 | 6.6952083 | 6.305369896 | 6.329589667 |
| ENSG00000005249 | PRKAR2B | 5.198849104 | 4.16E-05 | 1.346771376 | 0.356359551 | 6.54562048 | 3.327711234 | 3.076665209 | 2 | 7.19843704 | 6.99676009 | 3.679182183 | 3.771796761 |
| ENSG00000109927 | TECTA | 4.595296216 | 0.020435724 | 1.949451494 | 0.364963349 | 6.54474771 | 2 | 2 | 2 | 4.670864177 | 4.98931816 | 3.082820997 | 2.885414003 |
| ENSG00000119397 | CNTRL | 4.109430629 | 0.00103094 | 2.434829009 | 0.064928899 | 6.544259638 | 2 | 4.11352228 | 2 | 6.592405944 | 6.52015598 | 5.185595313 | 4.810605986 |
| ENSG00000167925 | GHDC | 3.316691099 | 0.002846809 | 3.227383735 | 0.003624465 | 6.544074834 | 3.327711234 | 6.04685525 | 3.369317972 | 7.999368766 | 8.08661608 | 8.063300251 | 7.839464964 |
| ENSG00000099814 | CEP170B | 2.927409656 | 0.059854337 | 3.615517188 | 0.016349295 | 6.542926844 | 2 | 7.393647515 | 7.858115942 | 9.938102197 | 9.99006945 | 10.72275211 | 10.57385172 |
| ENSG00000144843 | ADPRH | 2.3555514 | 0.077127293 | 4.180936791 | 0.000676414 | 6.536488191 | 2 | 4.710564154 | 2 | 5.44762523 | 5.64780787 | 7.08038692 | 7.464983909 |
| ENSG00000168490 | PHYHIP | 3.213038073 | 0.161654871 | 3.317359814 | 0.137443915 | 6.530397887 | 2 | 2 | 2 | 3.754611435 | 3.74316441 | 4.029788235 | 3.580696614 |
| ENSG00000111886 | GABRR2 | 4.132322139 | 0.061450217 | 2.395664499 | 0.298391409 | 6.527986638 | 2 | 2 | 2 | 4.443344104 | 4.45712836 | 3.500668998 | 2.885414003 |
| ENSG00000126461 | SCAF1 | 2.692668961 | 0.045443019 | 3.827825096 | 0.002799381 | 6.520494057 | 7.53297911 | 8.297207175 | 3.369317972 | 10.10372352 | 10.0551867 | 11.1203133 | 11.29801859 |
| ENSG00000122547 | EEPD1 | 2.546703445 | 0.105084134 | 3.966504052 | 0.006924386 | 6.513207496 | 2 | 3.686303414 | 2 | 4.598938466 | 4.63360314 | 6.106889539 | 5.611957759 |
| ENSG00000139200 | PIANP | 1.864214691 | 0.164314918 | 4.645670045 | 9.38E-05 | 6.509884736 | 2 | 4.710564154 | 2 | 5.22679461 | 4.98931816 | 7.510690356 | 7.945418846 |
| ENSG00000196196 | HRCT1 | 1.935477082 | 0.397539867 | 4.571354566 | 0.023994345 | 6.506831648 | 2 | 2 | 2 | 2.784612485 | 3.15372745 | 4.834511823 | 4.80186789 |
| ENSG00000183793 | NPIPA5 | 10.24908125 | 2.86E-13 | -3.743754742 | 0.103805123 | 6.505326512 | 2 | 2 | 2 | 10.28749693 | 10.2726831 | 2 | 2.059860178 |
| ENSG00000156599 | ZDHHC5 | 3.246118577 | 0.070433128 | 3.258055917 | 0.065664664 | 6.504174494 | 8.737455672 | 6.140560021 | 2 | 10.56934051 | 10.6264653 | 10.5520967 | 10.66581178 |
| ENSG00000079308 | TNS1 | 2.936134843 | 0.012711443 | 3.567302274 | 0.001852113 | 6.503437116 | 9.579430841 | 9.574900212 | 5.772716771 | 11.94460653 | 11.996947 | 12.92076824 | 12.19131528 |
| ENSG00000231094 | | 5.418222878 | 0.007764718 | 1.084805896 | 0.651103372 | 6.503028775 | 2 | 2 | 2 | 5.709096143 | 5.41912973 | 2.3070652 | 2.885414003 |
| ENSG00000101752 | MIB1 | 2.859316204 | 0.241959192 | 3.637628268 | 0.120921014 | 6.496944472 | 2 | 7.901717006 | 2 | 9.239364442 | 9.09977716 | 10.22911045 | 9.591800868 |
| ENSG00000223802 | CERS1 | 4.311902383 | 0.008465185 | 2.184190768 | 0.211685601 | 6.496093151 | 2 | 3.076665209 | 2 | 5.319220934 | 5.55061739 | 3.724264246 | 3.67940853 |
| ENSG00000165059 | PRKACG | 3.244232234 | 0.150422971 | 3.250323815 | 0.140213693 | 6.494556049 | 2 | 2 | 2 | 3.79808662 | 3.74316441 | 3.724264246 | 3.824516949 |
| ENSG00000120549 | KIAA1217 | 2.95965392 | 0.156702886 | 3.527912322 | 0.080230773 | 6.487566241 | 2 | 3.686303414 | 8.236946975 | 9.652259787 | 9.62041512 | 9.510546344 | 10.66791712 |
| ENSG00000175395 | ZNF25 | 2.547219261 | 0.121156062 | 3.939168201 | 0.010890162 | 6.486387463 | 2 | 6.464272742 | 3.369317972 | 7.562190249 | 7.51115308 | 9.249414741 | 8.463873348 |
| ENSG00000174080 | CTSF | 3.1720076 | 0.044825118 | 3.311940909 | 0.033432537 | 6.483948508 | 7.624139768 | 5.303523747 | 2 | 9.439666599 | 9.42003014 | 9.581426916 | 9.556522292 |
| ENSG00000173040 | EVC2 | 3.530914144 | 0.000680216 | 2.94989298 | 0.005175579 | 6.480807124 | 4.00722531 | 2 | 4.874587708 | 7.136275664 | 7.07311172 | 6.807643042 | 6.222348126 |
| ENSG00000278211 | | 6.082413083 | 0.000191933 | 0.397961587 | 0.852566533 | 6.48037467 | 2 | 2 | 2 | 6.29903474 | 6.0722229 | 2.457042231 | 2.374681325 |
| ENSG00000123908 | AGO2 | 2.234311756 | 0.048417421 | 4.244543405 | 4.85E-05 | 6.478855161 | 7.27616306 | 6.602424674 | 3.369317972 | 8.627383409 | 8.64695856 | 10.75059845 | 10.51258091 |
| ENSG00000163083 | INHBB | 4.766464661 | 0.015772075 | 1.710501748 | 0.434458206 | 6.476966409 | 2 | 2 | 2 | 5.022023783 | 4.95174433 | 2.68698273 | 3.042026731 |
| ENSG00000188707 | ZBED6CL | 5.646742537 | 0.000406306 | 0.829967377 | 0.670620849 | 6.476709914 | 3.327711234 | 2 | 2 | 6.733448311 | 6.64900057 | 2.62493832 | 3.15612297 |
| ENSG00000225967 | | 11.66763464 | 8.91E-17 | -5.191150548 | 0.022456739 | 6.476484095 | 2 | 2 | 2 | 11.68717402 | 11.7042919 | 2 | 2 |
| ENSG00000141068 | KSR1 | 1.683164237 | 0.334772114 | 4.792832494 | 0.001787956 | 6.475996731 | 2 | 5.595927442 | 2 | 5.742861563 | 5.78232107 | 9.048785116 | 8.43731913 |
| ENSG00000112787 | FBRSL1 | 2.803029538 | 0.019331325 | 3.672512249 | 0.001455406 | 6.475541786 | 2 | 6.311476754 | 5.156626401 | 7.961629787 | 8.0713677 | 9.004546892 | 8.736843548 |
| ENSG00000205129 | C4orf47 | 2.937815303 | 0.198526732 | 3.533627547 | 0.106686868 | 6.47144285 | 2 | 2 | 2 | 3.615725078 | 3.50452879 | 4.065262523 | 3.8919413 |
| ENSG00000156313 | RPGR | 4.260248306 | 0.000823063 | 2.207702812 | 0.104869869 | 6.467951118 | 2 | 4.11352228 | 2 | 6.682159616 | 6.71216164 | 4.521997428 | 5.050803619 |
| ENSG00000152127 | MGAT5 | 2.816047145 | 0.248162973 | 3.651855693 | 0.117745175 | 6.467902838 | 2 | 7.901717006 | 2 | 9.081180848 | 9.17405701 | 9.964103831 | 9.955102659 |
| ENSG00000091527 | CDV3 | 2.817176336 | 0.105584609 | 3.648517957 | 0.029020898 | 6.465694293 | 7.53297911 | 8.820493869 | 2 | 10.51174802 | 10.5632459 | 11.18758515 | 11.52718335 |
| ENSG00000166340 | TPP1 | 3.507600157 | 0.00677234 | 2.954123461 | 0.024616215 | 6.461723617 | 8.075571721 | 8.94259736 | 4.058998049 | 11.5005475 | 11.5214183 | 10.82922498 | 11.0770593 |
| ENSG00000038945 | MSR1 | 3.976615816 | 0.007764718 | 2.484692149 | 0.111079115 | 6.461307965 | 5.532353208 | 2 | 5.930865648 | 8.993902695 | 9.0282492 | 8.291371768 | 5.876154317 |
| ENSG00000120647 | CCDC77 | 3.702815615 | 0.004961216 | 2.758187184 | 0.041666835 | 6.461002799 | 2 | 4.710564154 | 2 | 6.846474326 | 6.77805036 | 6.167993056 | 5.606950214 |
| ENSG00000182158 | CREB3L2 | 3.056512833 | 0.03505329 | 3.395395386 | 0.017054345 | 6.45190822 | 7.940205782 | 9.566483142 | 4.058998049 | 11.42737655 | 11.4826289 | 11.84623902 | 11.73918884 |
| ENSG00000260314 | MRC1 | 1.860225968 | 0.429847898 | 4.589651478 | 0.02674158 | 6.449877445 | 2 | 2 | 2 | 3.023133209 | 2.86123014 | 5.180212365 | 4.376876162 |
| ENSG00000175445 | LPL | 1.320572128 | 0.714968044 | 5.126937466 | 0.100920521 | 6.447509594 | 3.327711234 | 2 | 2 | 3.79808662 | 2 | 7.05119671 | 3.807156736 |
| ENSG00000138379 | MSTN | 0.855699976 | 0.734406956 | 5.591317167 | 0.005313728 | 6.447017143 | 2 | 2 | 2 | 2.270786764 | 2.77779907 | 5.376552034 | 6.014942101 |
| ENSG00000196116 | TDRD7 | 4.075609328 | 0.063652316 | 2.36973438 | 0.300233814 | 6.445343708 | 2 | 2 | 7.098165388 | 9.523810814 | 9.5600488 | 7.770054437 | 7.932467322 |
| ENSG00000205426 | KRT81 | 3.332378729 | 0.167819097 | 3.110864196 | 0.190983933 | 6.443242924 | 2 | 2 | 2 | 3.709785322 | 3.94787299 | 4.029788235 | 3.209956562 |
| ENSG00000112893 | MAN2A1 | 2.804021099 | 0.000760977 | 3.636995337 | 6.96E-06 | 6.441016436 | 6.366709671 | 7.874700259 | 5.59507254 | 9.703174645 | 9.72016048 | 10.28597271 | 10.758765 |
| ENSG00000119737 | GPR75 | 4.145582195 | 0.012427993 | 2.293179636 | 0.190459822 | 6.438761832 | 2 | 3.076665209 | 2 | 5.304221918 | 5.27444544 | 3.694366604 | 3.858622973 |
| ENSG00000158158 | CNNM4 | 4.172749248 | 0.001230164 | 2.264871327 | 0.098726147 | 6.437620575 | 2 | 6.228547981 | 4.058998049 | 8.959612762 | 9.10727282 | 7.364099281 | 6.915965441 |
| ENSG00000151229 | SLC2A13 | 3.559644917 | 0.009351472 | 2.877944868 | 0.038778125 | 6.437589784 | 4.00722531 | 6.389896633 | 2 | 8.621345041 | 8.49524881 | 7.980559086 | 7.787298144 |
| ENSG00000061938 | TNK2 | 2.487564732 | 1.56E-05 | 3.940316829 | 1.18E-12 | 6.42788156 | 5.70932728 | 5.838937808 | 4.874587708 | 7.975897842 | 7.90193448 | 9.654578447 | 9.033147652 |
| ENSG00000128298 | BAIAP2L2 | 3.273125484 | 0.15309083 | 3.154043831 | 0.160572676 | 6.427169315 | 2 | 2 | 2 | 3.615725078 | 3.94787299 | 3.753554863 | 3.660199087 |
| ENSG00000152558 | TMEM123 | 3.593452491 | 2.38E-17 | 2.832431255 | 3.20E-11 | 6.425883746 | 7.940205782 | 8.968383114 | 8.460932916 | 12.13040239 | 12.0764293 | 11.2843655 | 11.40042151 |
| ENSG00000276349 | | 8.269516572 | 4.81E-06 | -1.846443244 | 0.426694614 | 6.423073328 | 2 | 2 | 2 | 8.287525788 | 8.34030387 | 2.183373728 | 2 |
| ENSG00000101605 | MYOM1 | 1.251527823 | 0.506916165 | 5.171232162 | 0.000918547 | 6.422759985 | 2 | 2 | 3.369317972 | 3.162721474 | 3.08597292 | 6.430795201 | 6.014942101 |
| ENSG00000164983 | TMEM65 | 2.829303322 | 0.145913141 | 3.589495816 | 0.054407162 | 6.418799139 | 6.009025997 | 2 | 2 | 7.277360022 | 7.0774796 | 7.985968197 | 7.860656447 |
| ENSG00000277075 | HIST1H2AE | 4.964290496 | 0.019164101 | 1.453106966 | 0.543480551 | 6.417397462 | 2 | 2 | 2 | 4.946755474 | 5.34860058 | 2.42098416 | 3.042026731 |
| ENSG00000086015 | MAST2 | 3.011336201 | 0.020000883 | 3.405385838 | 0.007283244 | 6.416722039 | 3.327711234 | 7.819099893 | 6.073378139 | 9.569217728 | 9.69550802 | 10.14509029 | 9.896287008 |
| ENSG00000087250 | MT3 | 3.497393865 | 0.182333448 | 2.919241808 | 0.265028723 | 6.416635673 | 2 | 2 | 2 | 4.034562184 | 3.86943969 | 2.746468522 | 4.062589736 |
| ENSG00000196712 | NF1 | 3.362256073 | 4.80E-22 | 3.0514345 | 1.48E-18 | 6.413690573 | 7.940205782 | 7.431218269 | 7.225742779 | 10.90254063 | 10.8893881 | 10.53150331 | 10.63831254 |
| ENSG00000114554 | PLXNA1 | 3.092285977 | 9.76E-06 | 3.319102663 | 1.64E-06 | 6.41138864 | 7.218819781 | 8.88960062 | 7.342949392 | 11.1032668 | 11.1286372 | 11.45091976 | 11.2251347 |
| ENSG00000119280 | C1orf198 | 0.49410387 | 0.418545189 | 5.911300893 | 6.72E-34 | 6.405404763 | 8.653385977 | 9.090820894 | 7.777116083 | 9.09649754 | 9.10513517 | 14.41570354 | 14.5942172 |
| ENSG00000205078 | SYCE1L | 3.081048508 | 0.179212705 | 3.324116159 | 0.134438751 | 6.405164667 | 2 | 2 | 2 | 3.566290607 | 3.74316441 | 3.796405078 | 3.858622973 |
| ENSG00000102606 | ARHGEF7 | 2.730244862 | 0.047425404 | 3.669207368 | 0.005421366 | 6.39945223 | 3.327711234 | 8.053983481 | 6.073378139 | 9.500148669 | 9.57248057 | 10.37921413 | 10.55918138 |
| ENSG00000196604 | POTEF | 7.635431549 | 4.71E-08 | -1.243565637 | 0.480855236 | 6.391865912 | 3.327711234 | 2 | 2 | 8.529212228 | 8.7394436 | 2.225793514 | 2.326754077 |
| ENSG00000116717 | GADD45A | 2.52846033 | 0.008801191 | 3.861825749 | 2.56E-05 | 6.390286079 | 7.750913381 | 4.93638933 | 6.203069917 | 9.17383051 | 9.23886123 | 10.37465563 | 10.67780108 |
| ENSG00000115762 | PLEKHB2 | 3.08026489 | 0.030837796 | 3.30561267 | 0.0187503 | 6.38587756 | 8.75772933 | 8.433041974 | 3.369317972 | 11.10137001 | 11.1007456 | 11.26494987 | 11.38459814 |
| ENSG00000158825 | CDA | 4.16565343 | 0.045577007 | 2.219835317 | 0.312041119 | 6.385488746 | 2 | 2 | 2 | 4.54891529 | 4.40247105 | 3.12825509 | 3.128433649 |
| ENSG00000151240 | DIP2C | 2.918666772 | 0.02326726 | 3.466413091 | 0.005685328 | 6.385079863 | 5.330594134 | 7.538394143 | 3.369317972 | 9.147810675 | 9.20332986 | 9.800076727 | 9.636102773 |
| ENSG00000137076 | TLN1 | 2.83656522 | 0.089599878 | 3.546107998 | 0.027546367 | 6.382673218 | 4.00722531 | 10.893964 | 9.907528187 | 12.68945875 | 12.786312 | 13.27312014 | 13.60376928 |
| ENSG00000065526 | SPEN | 2.817701673 | 0.047964022 | 3.56018276 | 0.009627368 | 6.377884433 | 3.327711234 | 6.728495673 | 8.486635645 | 10.05583453 | 10.1211983 | 10.72587282 | 10.92566722 |
| ENSG00000197714 | ZNF460 | 3.249064996 | 0.024456892 | 3.123914594 | 0.03002915 | 6.372979591 | 2 | 4.710564154 | 2 | 6.336376195 | 6.39275508 | 6.631488899 | 5.717942682 |
| ENSG00000129250 | KIF1C | 2.361305455 | 0.055403903 | 4.00908084 | 0.000487614 | 6.370386295 | 5.866944937 | 8.124485672 | 9.868959282 | 11.06067137 | 11.0934854 | 12.48011561 | 12.93103613 |
| ENSG00000138660 | AP1AR | 3.817276996 | 7.43E-05 | 2.551919242 | 0.010895983 | 6.369196238 | 3.327711234 | 3.076665209 | 5.156626401 | 7.629922299 | 7.52720828 | 6.653021698 | 5.980509018 |
| ENSG00000171860 | C3AR1 | 3.535658969 | 0.035913463 | 2.831122633 | 0.096882542 | 6.366781602 | 4.467311984 | 2 | 2 | 6.052081244 | 6.22964624 | 6.05195199 | 4.553838275 |
| ENSG00000166173 | LARP6 | 2.692126572 | 0.00192982 | 3.668457535 | 1.21E-05 | 6.360584108 | 5.70932728 | 8.234834549 | 6.958202216 | 9.940529154 | 10.0373603 | 11.18489989 | 10.70133804 |
| ENSG00000196659 | TTC30B | 4.105938697 | 0.002373085 | 2.251950145 | 0.116476087 | 6.357888843 | 2 | 3.076665209 | 3.369317972 | 5.949776857 | 6.1572134 | 4.369533167 | 4.456980017 |
| ENSG00000139675 | HNRNPA1L2 | 5.976342497 | 9.56E-07 | 0.377529503 | 0.810974431 | 6.353872 | 3.327711234 | 5.946638857 | 2 | 10.47371148 | 10.4524262 | 5.169385838 | 4.911532223 |
| ENSG00000271079 | CTAGE15 | 5.91344464 | 0.000827265 | 0.439299218 | 0.847708847 | 6.352743857 | 2 | 2 | 2 | 6.034011955 | 6.02777321 | 2.560105256 | 2.277179783 |
| ENSG00000130779 | CLIP1 | 2.798165715 | 6.77E-24 | 3.549349925 | 1.32E-38 | 6.34751564 | 7.975256505 | 7.790474954 | 7.600047325 | 10.58299811 | 10.5640243 | 11.34161973 | 11.30450932 |
| ENSG00000100485 | SOS2 | 3.139707374 | 0.002436283 | 3.206225991 | 0.001829498 | 6.345933365 | 4.467311984 | 7.051611315 | 4.523675391 | 8.95841569 | 8.99284518 | 9.0649396 | 9.018137056 |
| ENSG00000198829 | SUCNR1 | 1.860271891 | 0.551575085 | 4.482046599 | 0.109237936 | 6.34231849 | 2 | 2 | 2 | 3.023133209 | 2.86123014 | 5.553513572 | 2.70970617 |
| ENSG00000163697 | APBB2 | 3.02584228 | 0.091688187 | 3.316369935 | 0.057771995 | 6.342212214 | 3.327711234 | 8.623598428 | 4.058998049 | 10.14692461 | 10.1153688 | 10.57085845 | 10.25331056 |
| ENSG00000118985 | ELL2 | 3.944304117 | 4.42E-11 | 2.397233267 | 0.00010896 | 6.341537384 | 7.159102577 | 8.623598428 | 8.763901704 | 12.28187806 | 12.2569567 | 10.87408959 | 10.55788452 |
| ENSG00000075461 | CACNG4 | 0.836004157 | 0.795137033 | 5.504430379 | 0.037980527 | 6.340434536 | 2 | 4.11352228 | 2 | 4.387512679 | 2 | 7.57465694 | 7.953331607 |
| ENSG00000099937 | SERPIND1 | 4.928672011 | 0.010846156 | 1.409965459 | 0.521530272 | 6.33863747 | 2 | 2 | 2 | 5.258265913 | 4.98931816 | 2.592885947 | 2.885414003 |
| ENSG00000135503 | ACVR1B | 3.164267245 | 0.001104484 | 3.17399027 | 0.000997654 | 6.338257515 | 5.70932728 | 5.72254323 | 2 | 8.166267464 | 8.29225867 | 8.117815385 | 8.353097069 |
| ENSG00000130517 | PGPEP1 | 3.120404645 | 2.36E-05 | 3.216496908 | 1.15E-05 | 6.336901553 | 5.532353208 | 5.946638857 | 7.502634626 | 9.623848865 | 9.66099896 | 9.739377057 | 9.736956851 |
| ENSG00000157106 | SMG1 | 3.691729125 | 0.004232874 | 2.644737687 | 0.047077556 | 6.336466812 | 5.866944937 | 8.968383114 | 5.156626401 | 11.29090415 | 11.3191169 | 10.40164855 | 10.10403354 |
| ENSG00000113595 | TRIM23 | 3.076948193 | 0.002281184 | 3.258711197 | 0.001075078 | 6.335659391 | 5.532353208 | 5.131603625 | 2 | 7.783822874 | 7.69294162 | 8.306273771 | 7.383872668 |
| ENSG00000120051 | CFAP58 | 4.100832317 | 0.059311619 | 2.23480162 | 0.328120774 | 6.335633937 | 2 | 2 | 2 | 4.694064037 | 4.0930011 | 3.059555873 | 3.209956562 |
| ENSG00000071967 | CYBRD1 | 2.737813032 | 2.71E-16 | 3.596826432 | 1.04E-27 | 6.334639464 | 8.855032352 | 8.590771778 | 8.325066185 | 11.36279078 | 11.3010215 | 11.98530831 | 12.36968045 |
| ENSG00000177169 | ULK1 | 2.51022061 | 0.019562832 | 3.822287906 | 0.000178123 | 6.332508516 | 4.467311984 | 7.901717006 | 6.534109959 | 9.378328971 | 9.35315328 | 10.85953478 | 10.45783446 |
| ENSG00000165322 | ARHGAP12 | 3.59871718 | 0.001801872 | 2.733366903 | 0.021176021 | 6.332084083 | 6.257048221 | 6.389896633 | 2 | 9.287824288 | 9.29432703 | 8.488261545 | 8.376090431 |
| ENSG00000183287 | CCBE1 | 3.018868067 | 0.001148305 | 3.312079144 | 0.000292817 | 6.330947211 | 4.00722531 | 4.442710129 | 6.43197674 | 8.203120604 | 8.2561084 | 8.260460366 | 8.7391339 |
| ENSG00000105146 | AURKC | 3.148191506 | 0.182411111 | 3.18112633 | 0.167543084 | 6.329317836 | 2 | 2 | 2 | 3.615725078 | 3.7864959 | 3.956110469 | 3.45257121 |
| ENSG00000140396 | NCOA2 | 3.542966138 | 0.002346365 | 2.778115599 | 0.019945307 | 6.321081738 | 2 | 6.389896633 | 6.073378139 | 9.109497937 | 9.17914662 | 8.457441933 | 8.312916523 |
| ENSG00000174564 | IL20RB | 2.702385035 | 0.159914848 | 3.61434833 | 0.047321841 | 6.316733365 | 2 | 3.076665209 | 2 | 3.663521627 | 4.40247105 | 4.612704558 | 5.005857272 |
| ENSG00000146242 | TPBG | 4.968673395 | 5.26E-13 | 1.345897957 | 0.081368703 | 6.314571352 | 8.285626337 | 6.311476754 | 8.110401118 | 12.72491396 | 12.7391892 | 9.259978091 | 8.962375727 |
| ENSG00000172399 | MYOZ2 | -0.310504651 | 0.917283156 | 6.623134479 | 0.002070948 | 6.312629829 | 2 | 2 | 2 | 2.270786764 | 2.2680111 | 4.644324387 | 7.521960976 |
| ENSG00000133275 | CSNK1G2 | 3.00341222 | 0.085411097 | 3.308853728 | 0.051766586 | 6.312265949 | 4.00722531 | 8.433041974 | 3.369317972 | 9.897454779 | 9.9554848 | 10.11895226 | 10.33507589 |
| ENSG00000125633 | CCDC93 | 3.700083876 | 6.55E-05 | 2.610404304 | 0.006426257 | 6.31048818 | 4.00722531 | 7.051611315 | 6.958202216 | 10.15426209 | 10.1592994 | 9.242652114 | 8.88019044 |
| ENSG00000186660 | ZFP91 | 3.587205888 | 4.23E-05 | 2.721834742 | 0.002429239 | 6.30904063 | 5.532353208 | 7.901717006 | 8.380957451 | 11.25088489 | 11.2494513 | 10.39018357 | 10.38328123 |
| ENSG00000213064 | SFT2D2 | 1.563793955 | 0.197587957 | 4.742488391 | 7.90E-06 | 6.306282346 | 4.467311984 | 5.838937808 | 2 | 6.394182228 | 6.14893615 | 8.955762164 | 9.719663687 |
| ENSG00000092607 | TBX15 | 3.398117033 | 0.1005893 | 2.907777199 | 0.159343355 | 6.305894232 | 2 | 6.535001748 | 2 | 8.287525788 | 8.37797443 | 8.345877055 | 7.089349481 |
| ENSG00000167105 | TMEM92 | 2.427248941 | 0.200555174 | 3.875728698 | 0.028223199 | 6.302977639 | 2 | 3.076665209 | 2 | 3.754611435 | 3.98554648 | 4.652122235 | 5.361470313 |
| ENSG00000154262 | ABCA6 | 6.746784176 | 1.37E-05 | -0.44482059 | 0.832938355 | 6.301963586 | 2 | 2 | 2 | 6.710878132 | 6.93069905 | 2.267001569 | 2.225841198 |
| ENSG00000167216 | KATNAL2 | 2.494072687 | 0.307672642 | 3.804905541 | 0.094717057 | 6.298978227 | 2 | 2 | 2 | 2.947924294 | 3.55551834 | 4.291799915 | 4.077147165 |

| | | | | | | | | | | | | | |
|---|---|---|---|---|---|---|---|---|---|---|---|---|---|
| ENSG00000139263 | LRIG3 | 3.21462642 | 0.015276125 | 3.082852392 | 0.019964283 | 6.297478812 | 4.467311984 | 6.464272742 | 2 | 8.349110081 | 8.38680047 | 8.328950294 | 8.140837768 |
| ENSG00000149591 | TAGLN | 2.395130635 | 0.021070779 | 3.898777465 | 6.75E-05 | 6.2939081 | 11.54780176 | 10.20962523 | 8.296287027 | 12.92821476 | 12.9521752 | 14.45351417 | 14.43323509 |
| ENSG00000126860 | EVI2A | 4.738217158 | 0.010240552 | 1.554036979 | 0.451083615 | 6.292254136 | 2 | 3.076665209 | 2 | 5.720439317 | 5.93454154 | 3.709392871 | 2.70970617 |
| ENSG00000101079 | NDRG3 | 2.880648239 | 0.032370417 | 3.411402999 | 0.009226918 | 6.292051237 | 4.467311984 | 7.274681896 | 3.369317972 | 8.767939462 | 8.81201036 | 9.12745764 | 9.484290996 |
| ENSG00000198198 | SZT2 | 3.323927874 | 2.11E-13 | 2.965039368 | 6.27E-11 | 6.288967242 | 5.095959458 | 6.04685525 | 6.073378139 | 9.029678018 | 9.06500642 | 8.707109313 | 8.675981178 |
| ENSG00000163349 | HIPK1 | 2.953087218 | 0.103167922 | 3.333313672 | 0.058285878 | 6.28640089 | 4.00722531 | 8.687092496 | 3.369317972 | 10.11504861 | 10.1217271 | 10.61191991 | 10.37299407 |
| ENSG00000170006 | TMEM154 | 5.519519833 | 3.37E-07 | 0.765031331 | 0.562234526 | 6.284551163 | 2 | 4.442710129 | 2 | 8.291333947 | 8.36552655 | 3.753554863 | 4.175132639 |
| ENSG00000122674 | CCZ1 | 6.901370272 | 4.48E-14 | -0.617541778 | 0.596177968 | 6.283828493 | 4.00722531 | 3.076665209 | 3.369317972 | 9.860122857 | 9.70471389 | 3.316666208 | 2.918126211 |
| ENSG00000259075 | POC1B-GALN | 7.235914361 | 1.45E-06 | -0.952773198 | 0.630693854 | 6.283141163 | 2 | 2 | 2 | 7.226553841 | 7.36873214 | 2.183373728 | 2.172608095 |
| ENSG00000183580 | FBXL7 | 2.98530754 | 0.174608987 | 3.292955993 | 0.123227604 | 6.278263532 | 2 | 7.051611315 | 2 | 8.537252353 | 8.34574597 | 8.756774919 | 8.741420621 |
| ENSG00000109854 | HTATIP2 | 3.648235386 | 0.006781203 | 2.628922124 | 0.058029621 | 6.27715751 | 2 | 4.93638933 | 6.80318718 | 9.044420506 | 9.21030762 | 8.200570331 | 8.036691948 |
| ENSG00000113742 | CPEB4 | 3.799205756 | 9.91E-07 | 2.475656906 | 0.002138016 | 6.274862662 | 6.563806279 | 5.131603625 | 7.502634626 | 10.47622292 | 10.3666911 | 9.243297541 | 8.952540041 |
| ENSG00000213658 | LAT | 4.700389565 | 0.01506708 | 1.57380015 | 0.466152464 | 6.274189716 | 2 | 2 | 2 | 4.887609717 | 4.97065356 | 2.775316226 | 2.851942824 |
| ENSG00000126746 | ZNF384 | 3.67424972 | 0.007764718 | 2.595755622 | 0.06838385 | 6.270005342 | 2 | 6.666836837 | 4.523675391 | 9.031955922 | 8.96718201 | 7.995960198 | 7.869046644 |
| ENSG00000163378 | EOGT | 3.518892963 | 0.008000873 | 2.744406657 | 0.042845232 | 6.263299619 | 2 | 6.951736442 | 5.930865648 | 9.483600848 | 9.36302682 | 8.409366736 | 8.866637078 |
| ENSG00000115525 | ST3GAL5 | 3.410493016 | 1.75E-06 | 2.851101467 | 7.81E-05 | 6.261594483 | 3.327711234 | 5.457120953 | 4.523675391 | 8.001694993 | 7.89204953 | 7.334117317 | 7.462213768 |
| ENSG00000026508 | CD44 | 3.767844057 | 5.48E-70 | 2.49322201 | 5.61E-31 | 6.261066066 | 10.35389212 | 10.18234215 | 10.26770388 | 14.02560113 | 14.0380274 | 12.6485434 | 12.85918508 |
| ENSG00000162723 | SLAMF9 | 3.993531291 | 0.232704499 | 2.266738357 | 0.515187862 | 6.260269649 | 2 | 2 | 2 | 4.387512679 | 4.2865223 | 3.767980144 | 2.059860178 |
| ENSG00000178096 | BOLA1 | 2.501991805 | 0.060355042 | 3.755839105 | 0.002766242 | 6.257830911 | 2 | 4.710564154 | 2 | 5.662809983 | 5.70538658 | 6.947285242 | 6.791281771 |
| ENSG00000148154 | UGCG | 3.234930177 | 7.84E-06 | 3.021510255 | 3.09E-05 | 6.256440432 | 7.384433967 | 7.002537973 | 5.156626401 | 9.991161874 | 9.96965331 | 9.945583522 | 9.565261796 |
| ENSG00000188234 | AGAP4 | 5.429660284 | 3.17E-09 | 0.822763757 | 0.457717047 | 6.252424041 | 3.327711234 | 4.11352228 | 2 | 8.272191829 | 8.30721295 | 4.099885429 | 4.018014792 |
| ENSG00000213347 | MXD3 | 2.883517588 | 0.066569578 | 3.367969244 | 0.027078632 | 6.251486833 | 2 | 3.686303414 | 2 | 4.846787657 | 4.95174433 | 5.524070859 | 5.101530266 |
| ENSG00000135631 | RAB11FIP5 | 2.384887431 | 0.000408059 | 3.866127923 | 2.15E-09 | 6.251015354 | 8.009475818 | 8.169643011 | 6.322058544 | 10.06309216 | 10.0864045 | 11.43271451 | 11.66328098 |
| ENSG00000198162 | MAN1A2 | 2.284076047 | 0.371250007 | 3.965421594 | 0.094841246 | 6.249497641 | 2 | 8.147241016 | 2 | 8.866956935 | 8.8250571 | 10.40957704 | 10.62040904 |
| ENSG00000155100 | OTUD6B | 2.995405596 | 0.009301554 | 3.253839023 | 0.004074912 | 6.249244618 | 3.327711234 | 5.303523747 | 2 | 7.066878356 | 6.81514011 | 7.052670333 | 7.32867054 |
| ENSG00000127081 | ZNF484 | 3.485599946 | 0.023045186 | 2.762629414 | 0.076431446 | 6.24822936 | 2 | 5.303523747 | 2 | 7.178012736 | 7.21069602 | 6.649130473 | 6.326546677 |
| ENSG00000150477 | KIAA1328 | 2.757969598 | 0.0324034 | 3.489996132 | 0.004927027 | 6.24796573 | 2 | 4.442710129 | 2 | 5.651002631 | 5.64780787 | 6.419417643 | 6.245088435 |
| ENSG00000170962 | PDGFD | 2.861159916 | 0.315977132 | 3.386792057 | 0.21749096 | 6.247951973 | 2 | 2 | 2 | 3.709785322 | 3.28037825 | 2.775316226 | 4.489993931 |
| ENSG00000187720 | THSD4 | 1.95656353 | 0.139117554 | 4.289495866 | 0.000334043 | 6.246059396 | 2 | 6.464272742 | 5.39244166 | 7.43025113 | 7.30311013 | 10.00006426 | 9.248616776 |
| ENSG00000143816 | WNT9A | 1.27091119 | 0.609720423 | 4.97244723 | 0.017198851 | 6.24335842 | 2 | 2 | 2 | 2.3892346 | 2.94009902 | 5.206928476 | 5.12973572 |
| ENSG00000188643 | S100A16 | 2.58201302 | 7.13E-06 | 3.660961726 | 6.11E-11 | 6.242974746 | 8.674867122 | 7.761270542 | 7.896971941 | 10.70292919 | 10.7598372 | 11.20607774 | 12.23269239 |
| ENSG00000170113 | NIPA1 | 2.915919346 | 0.018178424 | 3.326305576 | 0.005911789 | 6.242224922 | 2 | 6.389896633 | 6.322058544 | 8.584574266 | 8.69180693 | 9.242652114 | 8.818171767 |
| ENSG00000123610 | TNFAIP6 | 8.546681066 | 6.43E-06 | -2.305640583 | 0.287678243 | 6.241040483 | 2 | 6.311476754 | 2 | 13.21893422 | 13.2489436 | 3.724264246 | 2.374681325 |
| ENSG00000143847 | PPFIA4 | 2.994622115 | 0.00021565 | 3.246232843 | 4.88E-05 | 6.240854959 | 3.327711234 | 6.140560021 | 5.772716771 | 8.385193462 | 8.41641145 | 8.609360894 | 8.68907177 |
| ENSG00000118432 | CNR1 | 1.154970505 | 0.697209163 | 5.084849603 | 0.04093731 | 6.239820108 | 2 | 2 | 2 | 3.023133209 | 2.14022021 | 4.927401733 | 5.545463418 |
| ENSG00000164180 | TMEM161B | 3.494448889 | 0.019477382 | 2.744119814 | 0.071516817 | 6.238568703 | 2 | 5.131603625 | 2 | 7.088925745 | 6.96881778 | 6.577231679 | 5.980509018 |
| ENSG00000176624 | MEX3C | 3.026719552 | 0.040445365 | 3.207232611 | 0.02745771 | 6.233952163 | 6.257048221 | 7.538394143 | 2 | 9.432787137 | 9.47403846 | 9.797661995 | 9.447297716 |
| ENSG00000064205 | WISP2 | -1.305489008 | 0.672951696 | 7.53692807 | 0.000876588 | 6.231439061 | 2 | 2 | 2 | 2.270786764 | 2 | 8.334412727 | 5.98823215 |
| ENSG00000122912 | SLC25A16 | 2.712716093 | 0.111386568 | 3.518359019 | 0.031231601 | 6.231075112 | 2 | 5.595927442 | 2 | 6.733448311 | 6.73446058 | 7.479255779 | 7.552234734 |
| ENSG00000161914 | ZNF653 | 2.955754666 | 0.020812654 | 3.272608288 | 0.008995102 | 6.228362954 | 2 | 4.442710129 | 2 | 5.839660368 | 5.8245091 | 6.084012396 | 6.172377306 |
| ENSG00000115020 | PIKFYVE | 3.38581335 | 2.06E-07 | 2.841147308 | 1.62E-05 | 6.226960659 | 5.866944937 | 7.232742069 | 5.59507254 | 9.76610238 | 9.78836002 | 9.314431634 | 9.152224182 |
| ENSG00000123094 | RASSF8 | 4.207705001 | 0.000644627 | 2.017787724 | 0.128084739 | 6.225492725 | 8.042902253 | 7.431218269 | 3.369317972 | 11.39897907 | 11.383149 | 9.443604635 | 8.92815536 |
| ENSG00000167065 | DUSP18 | 3.395045366 | 0.000214195 | 2.830377466 | 0.002345349 | 6.225422832 | 4.00722531 | 4.11352228 | 2 | 6.652857793 | 6.5890786 | 6.040114727 | 6.127623461 |
| ENSG00000163431 | LMOD1 | 0.490577839 | 0.841484908 | 5.734656108 | 0.002880079 | 6.225233947 | 8.009475818 | 3.076665209 | 2 | 7.044488801 | 6.8715633 | 12.34020583 | 11.95141501 |
| ENSG00000137831 | UACA | 1.836287071 | 0.081673461 | 4.388624686 | 3.83E-06 | 6.224911758 | 5.532353208 | 8.762721501 | 7.734844356 | 9.650037049 | 9.70541961 | 12.37393608 | 12.05771452 |
| ENSG00000134030 | CTIF | 1.5673442 | 0.103494474 | 4.655824799 | 4.37E-08 | 6.223168999 | 6.009025997 | 7.638155693 | 8.742726175 | 9.467702639 | 9.32403997 | 12.21874707 | 12.69866275 |
| ENSG00000107185 | RGP1 | 1.792894998 | 0.349647791 | 4.425981132 | 0.009967186 | 6.21887613 | 2 | 7.189546432 | 3.369317972 | 7.366699513 | 7.57741621 | 10.03067056 | 10.13066486 |
| ENSG00000083168 | KAT6A | 2.898462613 | 0.00551247 | 3.319862158 | 0.001171726 | 6.218324772 | 6.009025997 | 7.467835385 | 4.058998049 | 9.274418797 | 9.30181283 | 9.787520004 | 9.622210129 |
| ENSG00000160223 | ICOSLG | 4.430703453 | 0.068175471 | 1.7867648 | 0.496500125 | 6.217468253 | 2 | 2 | 2 | 4.670864177 | 4.72553021 | 3.296906311 | 2.374681325 |
| ENSG00000050344 | NFE2L3 | 4.414978197 | 2.54E-08 | 1.800353449 | 0.036866149 | 6.215331646 | 6.366709671 | 5.946638857 | 4.058998049 | 10.05807153 | 10.0722485 | 7.898530056 | 6.881131115 |
| ENSG00000109610 | SOD3 | 1.71914596 | 0.352977549 | 4.492557899 | 0.006315896 | 6.211703859 | 2 | 5.72254323 | 2 | 5.775854763 | 6.03677338 | 8.963225048 | 8.117259579 |
| ENSG00000132563 | REEP2 | -3.753168213 | 0.122079915 | 9.961365229 | 9.38E-12 | 6.208197016 | 2 | 2 | 2 | 2 | 2 | 9.888345571 | 10.09131582 |
| ENSG00000105576 | TNPO2 | 2.933768252 | 3.48E-20 | 3.271652633 | 3.36E-25 | 6.205420885 | 7.218819781 | 6.899080544 | 7.163363831 | 9.973493741 | 9.99815569 | 10.3869745 | 10.25511215 |
| ENSG00000146094 | DOK3 | 3.627768937 | 0.004232874 | 2.576530499 | 0.049027138 | 6.204299436 | 2 | 4.442710129 | 2 | 6.476767609 | 6.44777477 | 5.65789504 | 5.275717819 |
| ENSG00000150630 | VEGFC | 5.834104801 | 6.99E-05 | 0.369516888 | 0.84468287 | 6.203621688 | 7.904282243 | 5.946638857 | 2 | 12.42470411 | 12.4639002 | 7.200910877 | 6.821857532 |
| ENSG00000162482 | AKR7A3 | 3.571442141 | 0.12645146 | 2.628913066 | 0.266884496 | 6.200355207 | 2 | 2 | 2 | 3.921164147 | 4.0930011 | 3.082820997 | 3.600986722 |
| ENSG00000074181 | NOTCH3 | 1.320545039 | 0.002273448 | 4.878301729 | 7.28E-35 | 6.198846768 | 8.444143041 | 9.256956408 | 8.608698767 | 10.10155624 | 10.1664929 | 13.83562452 | 13.52227091 |
| ENSG00000204219 | TCEA3 | 2.35911222 | 0.037885253 | 3.838491573 | 0.000306801 | 6.197603793 | 2 | 5.131603625 | 3.369317972 | 6.043074888 | 6.33555364 | 7.802823992 | 7.421437279 |
| ENSG00000158163 | DZIP1L | 2.898440262 | 0.00251516 | 3.298067002 | 0.000443025 | 6.196507264 | 4.00722531 | 4.442710129 | 2 | 6.5612028 | 6.17362678 | 6.546183077 | 6.951954485 |
| ENSG00000163491 | NEK10 | 3.61381902 | 0.009718181 | 2.57903915 | 0.072993837 | 6.19285817 | 3.327711234 | 3.076665209 | 2 | 5.651002631 | 5.53799497 | 4.580376151 | 4.77533149 |
| ENSG00000163820 | FYCO1 | 1.966165714 | 0.085906155 | 4.22350523 | 4.79E-05 | 6.189670944 | 4.00722531 | 6.78762695 | 7.777116083 | 8.707945635 | 8.83927354 | 11.02321941 | 11.02075007 |
| ENSG00000047644 | WWC3 | 2.568275238 | 0.017140458 | 3.619285098 | 0.000439159 | 6.187560336 | 3.327711234 | 6.311476754 | 7.029880454 | 8.664561167 | 8.7297638 | 9.925616522 | 9.529981565 |
| ENSG00000267697 | LUZP6 | 11.37782338 | 6.62E-16 | -5.191150709 | 0.022650204 | 6.186672676 | 2 | 2 | 2 | 11.43320503 | 11.3789528 | 2 | 2 |
| ENSG00000131730 | CKMT2 | 2.08072363 | 0.386456011 | 4.105733342 | 0.058167426 | 6.186456972 | 2 | 2 | 2 | 2.947924294 | 3.15372745 | 4.133696865 | 4.673629022 |
| ENSG00000172456 | FGGY | 3.597711774 | 0.042507581 | 2.587437628 | 0.152650368 | 6.185149402 | 2 | 3.076665209 | 2 | 4.76150579 | 4.85329164 | 3.767980144 | 4.188602707 |
| ENSG00000165807 | PPP1R36 | 3.27312122 | 0.163554675 | 2.910994154 | 0.210663463 | 6.184115373 | 2 | 2 | 2 | 3.615725078 | 3.94787299 | 3.648325314 | 3.430065404 |
| ENSG00000075275 | CELSR1 | 1.689166646 | 0.388531091 | 4.487943586 | 0.009759057 | 6.177110232 | 2 | 2 | 5.59507254 | 6.104969671 | 4.97065356 | 8.372671446 | 8.312916523 |
| ENSG00000168389 | MFSD2A | 4.838241404 | 2.85E-08 | 1.335781555 | 0.176958539 | 6.174022959 | 3.327711234 | 4.710564154 | 2 | 8.304583895 | 8.20112674 | 4.868377487 | 4.998228167 |
| ENSG00000086991 | NOX4 | 3.781542615 | 0.000132612 | 2.390650889 | 0.021303089 | 6.172193504 | 3.327711234 | 4.11352228 | 4.058998049 | 7.186217188 | 7.32531744 | 6.398709156 | 5.217771644 |
| ENSG00000175294 | CATSPER1 | 2.938233356 | 0.267665409 | 3.231918637 | 0.208181348 | 6.170151993 | 2 | 2 | 2 | 3.663521627 | 3.45167086 | 3.011868911 | 4.254128754 |
| ENSG00000166822 | TMEM170A | 3.755429525 | 4.83E-05 | 2.41452044 | 0.012405533 | 6.169949965 | 4.00722531 | 3.076665209 | 5.39244166 | 7.880528731 | 7.89700047 | 6.900587441 | 6.179140936 |
| ENSG00000136167 | LCP1 | 0.052330339 | 0.984043262 | 6.117453105 | 0.001915899 | 6.169783444 | 2 | 3.076665209 | 2 | 2.784612485 | 2.38539787 | 7.959503304 | 5.108633494 |
| ENSG00000132207 | SLX1A | 8.957862302 | 1.61E-09 | -2.79188483 | 0.184524743 | 6.165977471 | 2 | 2 | 2 | 8.977450658 | 9.01350799 | 2.094598369 | 2 |
| ENSG00000042286 | AIFM2 | 3.975571649 | 8.10E-14 | 2.189870025 | 7.42E-05 | 6.165441674 | 5.095959458 | 5.838937808 | 5.59507254 | 9.411949708 | 9.42260746 | 7.336538937 | 7.911296863 |
| ENSG00000160469 | BRSK1 | 3.409634008 | 0.004777904 | 2.755371609 | 0.025405862 | 6.165005617 | 2 | 4.442710129 | 6.203069917 | 8.235106869 | 8.30535212 | 7.631969638 | 7.622152309 |
| ENSG00000078804 | TP53INP2 | 2.689028545 | 3.20E-09 | 3.475563277 | 6.75E-15 | 6.164591822 | 7.579279349 | 7.874700259 | 7.098165388 | 10.1970127 | 10.2530269 | 10.65220953 | 11.2961726 |
| ENSG00000113719 | ERGIC1 | 2.75788183 | 0.020652733 | 3.40519288 | 0.003212389 | 6.16307471 | 8.285626337 | 5.838937808 | 9.798878489 | 11.42953799 | 11.4842747 | 11.85159294 | 12.31828471 |
| ENSG00000160703 | NLRX1 | 2.477452083 | 0.23731318 | 3.685607493 | 0.062813847 | 6.163059576 | 2 | 6.602424674 | 2 | 7.337530098 | 7.64076905 | 8.660996377 | 8.712572735 |
| ENSG00000022267 | FHL1 | -2.147887483 | 0.185646208 | 8.310923966 | 2.50E-09 | 6.163036482 | 8.009475818 | 7.315436877 | 2 | 5.144975053 | 5.09650239 | 15.12381879 | 15.6380693 |
| ENSG00000123684 | LPGAT1 | 2.572892658 | 0.002065909 | 3.587163082 | 8.27E-06 | 6.16005574 | 6.009025997 | 7.503546079 | 5.156626401 | 9.067921572 | 9.12952895 | 10.05127285 | 10.16258417 |
| ENSG00000146826 | C7orf43 | 3.189914467 | 0.033156281 | 2.969324128 | 0.046894072 | 6.159238595 | 3.327711234 | 6.228547981 | 2 | 7.93509946 | 7.96222667 | 7.612118169 | 7.842663598 |
| ENSG00000102781 | KATNAL1 | 2.379095017 | 1.47E-05 | 3.775130933 | 9.66E-13 | 6.15422595 | 5.095959458 | 6.535001748 | 6.629488387 | 8.578353644 | 8.54498542 | 9.996814524 | 9.898081691 |
| ENSG00000203812 | HIST2H2AA3 | 11.34428912 | 8.54E-16 | -5.19115072 | 0.022702853 | 6.153138398 | 2 | 2 | 2 | 11.3344645 | 11.4102394 | 2 | 2 |
| ENSG00000272196 | HIST2H2AA4 | 11.34428912 | 8.54E-16 | -5.19115072 | 0.022702853 | 6.153138398 | 2 | 2 | 2 | 11.3344645 | 11.4102394 | 2 | 2 |
| ENSG00000198146 | ZNF770 | 3.261943757 | 8.92E-05 | 2.880909522 | 0.000593789 | 6.142853279 | 4.467311984 | 6.602424674 | 5.39244166 | 8.990393018 | 8.90095288 | 8.96596483 | 8.018893446 |
| ENSG00000001561 | ENPP4 | 1.050262246 | 0.592855432 | 5.09001386 | 0.001529662 | 6.140276106 | 2 | 3.076665209 | 2 | 3.09461492 | 2.94009902 | 6.433059984 | 5.825320785 |
| ENSG00000187566 | NHLRC1 | 2.935248432 | 0.066092631 | 3.199598706 | 0.040098237 | 6.134847137 | 2 | 2 | 4.058998049 | 5.003571641 | 5.07918227 | 5.008609438 | 5.502966356 |
| ENSG00000100888 | CHD8 | 2.960590317 | 0.019088294 | 3.174218237 | 0.010836307 | 6.134808554 | 5.095959458 | 8.590771778 | 5.59507254 | 10.1568736 | 10.290665 | 10.4929453 | 10.38144961 |
| ENSG00000140575 | IQGAP1 | 3.135228821 | 5.26E-05 | 2.998668176 | 0.0001103 | 6.133896997 | 8.042902253 | 10.21502048 | 8.845626171 | 12.43270008 | 12.4615001 | 12.22272076 | 12.39364033 |
| ENSG00000171817 | ZNF540 | 2.973445569 | 0.29251725 | 3.157435879 | 0.249742787 | 6.130881448 | 2 | 2 | 2 | 3.406930997 | 3.74316441 | 4.281782022 | 2.746601469 |
| ENSG00000133030 | MPRIP | 2.355622558 | 0.066342136 | 3.770045937 | 0.00174607 | 6.125668494 | 4.815609471 | 9.079034257 | 9.06622548 | 10.84021544 | 10.8974396 | 12.26318033 | 12.29978441 |
| ENSG00000130477 | UNC13A | 1.358942678 | 0.196475406 | 4.765537198 | 1.49E-07 | 6.124479876 | 3.327711234 | 3.686303414 | 5.772716771 | 5.720439317 | 5.98191036 | 8.932738969 | 9.374763636 |
| ENSG00000155111 | CDK19 | 3.028440123 | 0.03281527 | 3.095183509 | 0.027475652 | 6.123623632 | 2 | 6.389896633 | 4.058998049 | 8.109160159 | 7.96929529 | 8.266822833 | 7.926450253 |
| ENSG00000104447 | TRPS1 | 3.373578742 | 5.46E-06 | 2.749140206 | 0.000261617 | 6.122718948 | 4.815609471 | 6.844429755 | 5.39244166 | 9.237391778 | 9.30833119 | 8.702422971 | 8.603665995 |
| ENSG00000169641 | LUZP1 | 3.123050695 | 0.017260781 | 2.999552232 | 0.02221024 | 6.122602928 | 4.00722531 | 8.806266215 | 7.552162999 | 10.84960847 | 10.884411 | 10.57483812 | 10.89518287 |
| ENSG00000146858 | ZC3HAV1L | 2.424295804 | 0.010714204 | 3.698112377 | 3.87E-05 | 6.122408181 | 4.467311984 | 3.076665209 | 4.058998049 | 6.213314212 | 5.74985018 | 7.463831776 | 6.926052784 |
| ENSG00000206455 | | 4.68720374 | 0.003695722 | 1.435203289 | 0.432125179 | 6.12240703 | 3.327711234 | 2 | 2 | 5.709096143 | 5.85535975 | 3.082820997 | 3.36036153 |
| ENSG00000159363 | ATP13A2 | 3.190185484 | 0.036927252 | 2.930968936 | 0.054822702 | 6.12115442 | 2 | 7.393647515 | 5.39244166 | 9.286870873 | 9.29432703 | 8.813940551 | 9.2234807 |
| ENSG00000165272 | AQP3 | 2.73647328 | 0.26922137 | 3.382773976 | 0.152201542 | 6.119247256 | 2 | 2 | 2 | 3.709785322 | 3.08597292 | 3.796405078 | 3.940519005 |
| ENSG00000166225 | FRS2 | 3.102861168 | 0.054064283 | 3.014365784 | 0.058825076 | 6.117226953 | 4.815609471 | 7.431218269 | 2 | 9.208480821 | 9.06390621 | 9.110931391 | 8.986675514 |
| ENSG00000115956 | PLEK | 0.010133594 | 0.997264364 | 6.101480583 | 0.007467316 | 6.111614177 | 2 | 2 | 2 | 2.270786764 | 2.38539787 | 7.027411703 | 4.119958326 |
| ENSG00000101224 | CDC25B | 3.153180195 | 0.178994785 | 2.958319296 | 0.201973614 | 6.111499491 | 2 | 7.572420273 | 2 | 9.136211578 | 9.12952895 | 8.557885437 | 9.24096022 |
| ENSG00000159176 | CSRP1 | 2.22481964 | 0.003921045 | 3.884673196 | 9.88E-08 | 6.109492836 | 9.567995545 | 7.874700259 | 7.934808822 | 10.90409587 | 10.8971308 | 12.85044134 | 12.19183793 |
| ENSG00000170265 | ZNF282 | 2.904538895 | 0.21164903 | 3.203191472 | 0.156449615 | 6.107730367 | 2 | 7.467835385 | 2 | 8.773395507 | 8.78955647 | 9.225442227 | 8.913019964 |
| ENSG00000139370 | SLC15A4 | 3.429601219 | 0.010028324 | 2.676907156 | 0.048890727 | 6.106508375 | 2 | 6.140560021 | 7.029880454 | 9.425874714 | 9.46406471 | 8.864738722 | 8.509534124 |
| ENSG00000156795 | WDYHV1 | 2.350909734 | 0.104695101 | 3.754209358 | 0.005570045 | 6.105119091 | 2 | 4.93638933 | 2 | 5.85002558 | 5.65950892 | 6.964610156 | 7.211494604 |
| ENSG00000038532 | CLEC16A | 2.448978171 | 0.185413931 | 3.655996781 | 0.036320638 | 6.104974952 | 3.327711234 | 7.099070191 | 2 | 7.97352961 | 8.08661608 | 8.927929329 | 9.472294474 |

| | | | | | | | | | | | | | |
|---|---|---|---|---|---|---|---|---|---|---|---|---|---|
| ENSG00000141858 | SAMD1 | 2.538846335 | 0.000553577 | 3.563988517 | 4.86E-07 | 6.102834852 | 4.00722531 | 4.710564154 | 5.930865648 | 7.395290829 | 7.53675633 | 8.688271909 | 8.22740336 |
| ENSG00000168826 | ZBTB49 | 3.244200067 | 0.01448403 | 2.855146191 | 0.032443941 | 6.099346258 | 2 | 4.11352228 | 2 | 5.797438367 | 5.67111583 | 5.454427604 | 5.300743019 |
| ENSG00000170571 | EMB | 4.948607569 | 0.000729237 | 1.147847756 | 0.499266159 | 6.096455325 | 4.00722531 | 2 | 2 | 6.989292186 | 6.80463998 | 3.917810786 | 3.236138336 |
| ENSG00000198455 | ZXDB | 2.845407184 | 0.123195888 | 3.249417557 | 0.06987157 | 6.094824741 | 2 | 5.946638857 | 2 | 7.127781374 | 7.28055563 | 7.624061799 | 7.576773052 |
| ENSG00000197776 | KLHDC1 | 3.313965611 | 0.0960601 | 2.779023133 | 0.161994098 | 6.092988743 | 2 | 3.076665209 | 2 | 4.76150579 | 4.3456612 | 4.580376151 | 3.496556178 |
| ENSG00000139926 | FRMD6 | 4.054876088 | 6.36E-07 | 2.036765167 | 0.019469114 | 6.091641256 | 6.138360429 | 8.451443573 | 8.561105505 | 12.08271877 | 12.1014432 | 9.813394654 | 10.30156833 |
| ENSG00000184227 | ACOT1 | 3.875630924 | 1.86E-05 | 2.215959653 | 0.020780792 | 6.091590577 | 4.00722531 | 3.686303414 | 5.156626401 | 8.001694993 | 7.95749494 | 5.677130623 | 6.834765628 |
| ENSG00000123411 | IKZF4 | 2.727556021 | 0.100196808 | 3.363494898 | 0.035315165 | 6.091050919 | 2 | 5.457120953 | 2 | 6.598566398 | 6.6253296 | 7.118879607 | 7.327148877 |
| ENSG00000100226 | GTPBP1 | 3.059099934 | 2.47E-05 | 3.031181975 | 2.74E-05 | 6.090281909 | 8.257427002 | 7.701025996 | 6.073378139 | 10.57833006 | 10.6877448 | 10.46845053 | 10.73199123 |
| ENSG00000048707 | VPS13D | 2.738825349 | 0.006353097 | 3.349209341 | 0.000597178 | 6.08803469 | 4.467311984 | 6.951736442 | 7.818184403 | 9.636628223 | 9.66172639 | 10.22503182 | 10.28853085 |
| ENSG00000103710 | RASL12 | 2.367679863 | 0.177357792 | 3.719784818 | 0.023412451 | 6.087464681 | 2 | 3.686303414 | 2 | 4.574143677 | 4.31639476 | 6.060766632 | 5.020995508 |
| ENSG00000082497 | SERTAD4 | -2.311083228 | 0.371950735 | 8.398403621 | 3.19E-08 | 6.087320393 | 2 | 2 | 2 | 2 | 2.14022021 | 8.592743352 | 8.272375674 |
| ENSG00000133704 | IPO8 | 2.968055741 | 0.000260005 | 3.118667058 | 0.000108468 | 6.086722799 | 4.815609471 | 5.946638857 | 7.285536013 | 9.239364442 | 9.26015177 | 9.379289361 | 9.419363022 |
| ENSG00000170759 | KIF5B | 2.849324163 | 0.001065505 | 3.233884891 | 0.000156598 | 6.083209055 | 7.975256505 | 9.71856931 | 7.342949392 | 11.58267051 | 11.5238208 | 11.90615729 | 11.96847501 |
| ENSG00000170385 | SLC30A1 | 3.231671785 | 0.162428131 | 2.851455366 | 0.214965079 | 6.083127151 | 2 | 7.467835385 | 2 | 9.03990038 | 9.16894939 | 9.014789544 | 8.37166861 |
| ENSG00000170142 | UBE2E1 | 3.460878138 | 0.000392827 | 2.620980549 | 0.008953574 | 6.081858688 | 4.00722531 | 7.189546432 | 7.225742779 | 10.12468558 | 10.1036386 | 9.295574678 | 9.26102247 |
| ENSG00000001461 | NIPAL3 | 1.869196158 | 0.283079066 | 4.208068764 | 0.007459944 | 6.077264922 | 4.815609471 | 8.968383114 | 4.058998049 | 9.322664889 | 9.39749836 | 11.49474119 | 11.86616355 |
| ENSG00000142192 | APP | 3.214465921 | 6.16E-08 | 2.861961899 | 1.63E-06 | 6.076427821 | 9.473095744 | 9.424669273 | 10.70548138 | 13.21373885 | 13.1973249 | 12.5744683 | 13.08673594 |
| ENSG00000171448 | ZBTB26 | 2.719056788 | 0.047700201 | 3.357105265 | 0.011354296 | 6.076162053 | 2 | 4.710564154 | 2 | 5.910700032 | 5.84514933 | 6.637393426 | 6.30506411 |
| ENSG00000001629 | ANKIB1 | 3.099463387 | 0.016178162 | 2.974706814 | 0.021001623 | 6.074170201 | 4.00722531 | 8.337339476 | 8.65477198 | 11.02280532 | 11.0390977 | 10.8358731 | 10.97374872 |
| ENSG00000243708 | PLA2G4B | 5.098579721 | 0.005893523 | 0.975515079 | 0.658188823 | 6.0740948 | 2 | 2 | 2 | 5.242616075 | 5.3193943 | 2.656294034 | 2.509591799 |
| ENSG00000184185 | KCNJ12 | 1.054158706 | 0.394151052 | 5.019611174 | 8.64E-07 | 6.073769881 | 2 | 3.076665209 | 4.523675391 | 4.070463721 | 4.45712836 | 7.742927022 | 8.100433454 |
| ENSG00000166349 | RAG1 | 2.494032362 | 0.358758119 | 3.578896849 | 0.160899275 | 6.072929211 | 2 | 2 | 2 | 2.947924294 | 3.55551834 | 4.496236235 | 3.287118176 |
| ENSG00000213020 | ZNF611 | 3.555595066 | 1.98E-07 | 2.512111855 | 0.000343479 | 6.067706922 | 4.467311984 | 3.686303414 | 5.39244166 | 7.88809286 | 7.99492122 | 6.939340974 | 6.907844475 |
| ENSG00000070404 | FSTL3 | 1.911812009 | 0.019331325 | 4.155259572 | 3.39E-08 | 6.067071582 | 8.19931886 | 7.503546079 | 9.624627483 | 10.5946026 | 10.6387096 | 12.72467338 | 12.97888139 |
| ENSG00000108406 | DHX40 | 3.328794141 | 7.15E-05 | 2.737727709 | 0.001294981 | 6.06652185 | 4.815609471 | 5.303523747 | 7.098165388 | 9.395237275 | 9.24566964 | 8.644966335 | 8.821417907 |
| ENSG00000084731 | KIF3C | 2.775062318 | 0.047622861 | 3.291083531 | 0.015805954 | 6.066145849 | 6.563806279 | 7.232742069 | 2 | 9.078979414 | 9.12320501 | 9.6467939 | 9.580669163 |
| ENSG00000090530 | P3H2 | 2.384890014 | 0.069351955 | 3.67971874 | 0.002915078 | 6.064608754 | 6.816637977 | 6.04685525 | 2 | 8.170409056 | 8.30721295 | 9.671701874 | 9.359213231 |
| ENSG00000183621 | ZNF438 | 2.751740555 | 0.095756389 | 3.309874715 | 0.037646364 | 6.06161527 | 2 | 2 | 5.156626401 | 6.260701031 | 6.04571775 | 6.658838928 | 6.718860807 |
| ENSG00000147854 | UHRF2 | 3.278585168 | 0.002216582 | 2.780186477 | 0.01057194 | 6.058771645 | 6.73718411 | 7.051611315 | 3.369317972 | 9.626112287 | 9.58863636 | 9.390112523 | 8.767460665 |
| ENSG00000136935 | GOLGA1 | 2.957605775 | 0.025431454 | 3.098130744 | 0.017970055 | 6.055736519 | 2 | 6.389896633 | 4.523675391 | 8.008651253 | 8.11451391 | 8.1339154 | 8.265241863 |
| ENSG00000186350 | RXRA | 2.677041884 | 0.066450453 | 3.37434884 | 0.016614757 | 6.051390724 | 7.435667355 | 6.311476754 | 2 | 8.966774408 | 9.10620439 | 9.671701874 | 9.788455943 |
| ENSG00000083799 | CYLD | 3.309087402 | 0.023540207 | 2.740283947 | 0.064089806 | 6.049371349 | 9.247966931 | 6.04685525 | 4.523675391 | 11.18818376 | 11.100209 | 10.53743599 | 10.61605402 |
| ENSG00000105939 | ZC3HAV1 | 4.14344369 | 0.000411397 | 1.905180222 | 0.132382182 | 6.048623912 | 4.467311984 | 8.762721501 | 8.325066185 | 12.14455022 | 12.1401932 | 9.998344725 | 9.813396275 |
| ENSG00000224712 | NPIPA3 | 11.23582915 | 1.65E-15 | -5.191150797 | 0.022726528 | 6.044678354 | 2 | 2 | 2 | 11.26402542 | 11.2650929 | 2 | 2 |
| ENSG00000115896 | PLCL1 | 2.33302308 | 0.392606579 | 3.711301304 | 0.142587272 | 6.044324384 | 2 | 2 | 2 | 3.162721474 | 3.2184422 | 4.667592615 | 3.209956562 |
| ENSG00000163297 | ANTXR2 | 4.875945914 | 8.44E-06 | 1.166995108 | 0.355406319 | 6.042941023 | 6.366709671 | 7.002537973 | 9.624627483 | 13.22230432 | 13.2509995 | 9.642885818 | 9.418291476 |
| ENSG00000198915 | RASGEF1A | 2.270572932 | 0.385331055 | 3.771849892 | 0.117860642 | 6.042422824 | 2 | 2 | 2 | 2.695456494 | 3.50452879 | 4.230621763 | 4.09155917 |
| ENSG00000164949 | GEM | 3.897143001 | 0.006549526 | 2.141862256 | 0.157487659 | 6.039005258 | 2 | 7.189546432 | 5.930865648 | 9.945371153 | 9.9912274 | 8.698192208 | 7.521960976 |
| ENSG00000116132 | PRRX1 | 3.367804535 | 2.47E-29 | 2.667260335 | 5.94E-19 | 6.03506487 | 8.107517749 | 8.433041974 | 7.896971941 | 11.51710644 | 11.530009 | 10.85806066 | 10.78961595 |
| ENSG00000176399 | DMRTA1 | 4.133423449 | 0.061854072 | 1.899365042 | 0.423484962 | 6.032788492 | 2 | 2 | 2 | 4.598938466 | 4.2865223 | 3.1504466 | 2.746601469 |
| ENSG00000176463 | SLCO3A1 | 4.604967402 | 0.007224622 | 1.4236499 | 0.458667613 | 6.028617301 | 2 | 5.595927442 | 2 | 8.581467308 | 8.57464274 | 6.081127077 | 4.56420962 |
| ENSG00000165474 | GJB2 | 4.778111699 | 0.019324738 | 1.247289795 | 0.593027095 | 6.025401495 | 2 | 2 | 2 | 5.178261742 | 4.79082474 | 2.492221014 | 2.851942824 |
| ENSG00000197696 | NMB | 4.750507803 | 1.63E-05 | 1.274306805 | 0.310228532 | 6.024814608 | 6.009025997 | 4.11352228 | 2 | 9.34848254 | 9.3808074 | 6.037140174 | 5.909074772 |
| ENSG00000103642 | LACTB | 3.433400421 | 7.82E-07 | 2.589118279 | 0.00026611 | 6.0225187 | 4.815609471 | 6.951736442 | 6.203069917 | 9.610193249 | 9.63159714 | 8.642032548 | 8.910481888 |
| ENSG00000172548 | NIPAL4 | 0.689486832 | 0.790784372 | 5.332734687 | 0.008595717 | 6.022221519 | 2 | 2 | 2 | 2.498691189 | 2.49394783 | 6.016144783 | 4.664017718 |
| ENSG00000116852 | KIF21B | 2.273800221 | 0.374366032 | 3.747416652 | 0.113061522 | 6.021216872 | 2 | 2 | 2 | 3.09461492 | 3.2184422 | 4.547306681 | 3.580696614 |
| ENSG00000137877 | SPTBN5 | 5.057189104 | 0.001872533 | 0.962833754 | 0.616400319 | 6.020022858 | 2 | 3.686303414 | 2 | 6.799114572 | 6.98285659 | 2.803598391 | 3.858622973 |
| ENSG00000100644 | HIF1A | 2.976305113 | 0.000580679 | 3.043013578 | 0.000395937 | 6.019318691 | 8.138771685 | 10.36815316 | 10.77492406 | 13.10522462 | 13.0935265 | 13.12849113 | 13.20267654 |
| ENSG00000185052 | SLC24A3 | -1.688040961 | 0.404229173 | 7.706417123 | 6.18E-08 | 6.018376162 | 2 | 3.076665209 | 2 | 2.141737246 | 2.2680111 | 8.545902039 | 8.854540499 |
| ENSG00000054967 | RELT | 3.332610944 | 0.00059294 | 2.685738321 | 0.006620286 | 6.018349265 | 5.866944937 | 4.93638933 | 2 | 8.080836374 | 8.15020018 | 7.442631903 | 7.517964974 |
| ENSG00000171757 | LRRC34 | 3.204030721 | 0.075282483 | 2.811435577 | 0.11706113 | 6.015466298 | 3.327711234 | 2 | 2 | 4.574143677 | 4.37434574 | 4.053534413 | 4.26688425 |
| ENSG00000185697 | MYBL1 | 2.387162618 | 0.037134263 | 3.628170294 | 0.000758226 | 6.015332912 | 3.327711234 | 2 | 4.523675391 | 5.391962469 | 5.49945122 | 6.625560108 | 6.592824411 |
| ENSG00000050748 | MAPK9 | 2.888458564 | 0.000529129 | 3.125426843 | 0.00014468 | 6.013885408 | 5.866944937 | 6.78762695 | 4.058998049 | 8.767939462 | 8.81069917 | 9.035431928 | 9.013414035 |
| ENSG00000241360 | PDXP | 4.015898441 | 0.009415065 | 1.997257932 | 0.226994232 | 6.013156373 | 2 | 5.457120953 | 2 | 7.870381136 | 7.86451142 | 6.028179536 | 5.807970301 |
| ENSG00000061656 | SPAG4 | 2.78699126 | 0.129712892 | 3.225707667 | 0.070507478 | 6.012698928 | 2 | 2 | 5.59507254 | 6.836083475 | 6.507263 | 7.241533944 | 6.953927794 |
| ENSG00000141298 | SSH2 | 2.609175172 | 0.260754859 | 3.401909252 | 0.125372994 | 6.011084424 | 2 | 7.35507212 | 2 | 8.406418094 | 8.34574597 | 8.977260248 | 9.325315571 |
| ENSG00000085117 | CD82 | 4.241239554 | 1.06E-09 | 1.768448095 | 0.018944745 | 6.009687649 | 7.096806373 | 7.503546079 | 5.39244166 | 11.08829583 | 11.1490131 | 8.408216037 | 8.870820921 |
| ENSG00000174485 | DENND4A | 3.343727804 | 2.90E-05 | 2.664916642 | 0.001064564 | 6.008644446 | 4.467311984 | 5.457120953 | 6.882775709 | 9.162439556 | 9.19229617 | 8.591223132 | 8.412421212 |
| ENSG00000239305 | RNF103 | 3.239543246 | 0.151721189 | 2.762815487 | 0.220786883 | 6.002358733 | 2 | 7.315436877 | 2 | 9.028537716 | 8.88981993 | 8.586652837 | 8.383430148 |
| ENSG00000108854 | SMURF2 | 2.797182886 | 0.001110684 | 3.205103911 | 0.000140544 | 6.002286797 | 6.138360429 | 8.376385542 | 6.629488387 | 10.17244415 | 10.1623867 | 10.35597781 | 10.76355445 |
| ENSG00000164935 | DCSTAMP | 2.014562453 | 0.566415954 | 3.987168608 | 0.215756786 | 6.001731061 | 2 | 2 | 2 | 3.289987342 | 2.68924543 | 5.147485955 | 2.277179783 |
| ENSG00000164220 | F2RL2 | 4.396514425 | 3.52E-10 | 1.605063669 | 0.037080714 | 6.001578093 | 3.327711234 | 4.11352228 | 4.874587708 | 8.272191829 | 8.30162324 | 5.528313928 | 5.670730563 |
| ENSG00000173917 | HOXB2 | 5.439397426 | 0.001373482 | 0.562143408 | 0.789642976 | 6.001540834 | 2 | 2 | 5.930865648 | 9.615519136 | 9.69550802 | 5.096960836 | 4.827924991 |
| ENSG00000160781 | PAQR6 | 2.729369702 | 0.14612275 | 3.269738387 | 0.069822896 | 5.99910809 | 3.327711234 | 2 | 2 | 4.329433326 | 3.82856379 | 4.612704558 | 4.445805334 |
| ENSG00000119844 | AFTPH | 3.412519634 | 0.007180368 | 2.58617839 | 0.046628793 | 5.998698025 | 4.00722531 | 6.844429755 | 3.369317972 | 8.912174893 | 8.87358581 | 8.299462788 | 7.811442311 |
| ENSG00000116525 | TRIM62 | 2.505587247 | 0.182039968 | 3.488904327 | 0.050274744 | 5.994491573 | 2 | 5.946638857 | 2 | 6.809774134 | 6.94032357 | 7.668951058 | 7.984555672 |
| ENSG00000131773 | KHDRBS3 | 2.486726867 | 0.163957612 | 3.507709908 | 0.038301117 | 5.994436775 | 2 | 5.72254323 | 2 | 6.70517998 | 6.57060569 | 7.6993745 | 7.558732759 |
| ENSG00000249931 | GOLGA8K | 7.469271978 | 2.14E-06 | -1.476031839 | 0.470865845 | 5.993240139 | 2 | 2 | 2 | 7.549526251 | 7.50468064 | 2.183373728 | 2.059860178 |
| ENSG00000139438 | FAM222A | 3.212350354 | 0.17595559 | 2.780805188 | 0.238575668 | 5.993155542 | 2 | 2 | 2 | 3.663521627 | 3.82856379 | 3.5515855 | 3.36036153 |
| ENSG00000197208 | SLC22A4 | 5.399654523 | 4.96E-08 | 0.592908605 | 0.630920961 | 5.992563127 | 2 | 3.076665209 | 4.874587708 | 8.589222193 | 8.69038176 | 3.904814898 | 4.411752372 |
| ENSG00000138435 | CHRNA1 | 3.740204137 | 0.199271942 | 2.251183441 | 0.454855955 | 5.991387579 | 2 | 2 | 3.369317972 | 5.128039203 | 4.70309173 | 4.504874511 | 2.059860178 |
| ENSG00000163359 | COL6A3 | 3.946248851 | 1.40E-09 | 2.035519861 | 0.003115315 | 5.981768712 | 11.18203661 | 11.26034006 | 9.541233192 | 14.76074113 | 14.8187889 | 13.05031454 | 12.6865867 |
| ENSG00000011478 | QPCTL | 2.838047984 | 3.72E-05 | 3.143095517 | 3.65E-06 | 5.981143501 | 5.866944937 | 4.11352228 | 5.930865648 | 8.192977693 | 8.17894021 | 8.313052751 | 8.642725409 |
| ENSG00000197969 | VPS13A | 2.869547716 | 0.001128625 | 3.108259432 | 0.000350141 | 5.977807148 | 7.709880939 | 5.131603625 | 6.322058544 | 9.669187952 | 9.45653888 | 9.739377057 | 9.866478759 |
| ENSG00000105855 | ITGB8 | 5.541755181 | 1.31E-08 | 0.432151156 | 0.728068232 | 5.973906336 | 2 | 4.93638933 | 4.058998049 | 9.369349625 | 9.32035914 | 3.943456545 | 4.911532223 |
| ENSG00000104321 | TRPA1 | 7.780651506 | 2.47E-07 | -1.811091662 | 0.300664025 | 5.969559844 | 2 | 5.946638857 | 2 | 12.11504048 | 12.1075961 | 3.374372158 | 3.183290849 |
| ENSG00000123977 | DAW1 | 3.011795051 | 0.302944922 | 2.953823836 | 0.301212939 | 5.965618888 | 2 | 2 | 2 | 3.663521627 | 3.55551834 | 2.560105256 | 4.161535617 |
| ENSG00000118276 | B4GALT6 | 3.42608518 | 0.00680255 | 2.539092811 | 0.050514234 | 5.96517799 | 2 | 4.442710129 | 2 | 6.276156955 | 6.26836889 | 5.463319095 | 5.431121909 |
| ENSG00000126012 | KDM5C | 2.849582857 | 0.131914752 | 3.109087171 | 0.092258249 | 5.958670028 | 2 | 9.396454291 | 7.285536013 | 10.92201514 | 10.9843247 | 11.26947193 | 11.15321065 |
| ENSG00000185722 | ANKFY1 | 3.196177305 | 6.34E-17 | 2.759603351 | 5.56E-13 | 5.955780656 | 7.667647223 | 7.954278496 | 7.225742779 | 10.78962055 | 10.8671698 | 10.20462959 | 10.56047708 |
| ENSG00000130940 | CASZ1 | 1.803556345 | 0.112165694 | 4.150545955 | 4.18E-05 | 5.9541023 | 3.327711234 | 4.11352228 | 2 | 4.804776789 | 4.89348248 | 7.170014349 | 6.866539459 |
| ENSG00000153207 | AHCTF1 | 3.573591631 | 0.002523465 | 2.379539116 | 0.053196051 | 5.953130748 | 5.095959458 | 7.145017454 | 3.369317972 | 9.479434065 | 9.45905186 | 8.551645649 | 7.959237827 |
| ENSG00000107771 | CCSER2 | 2.577159257 | 1.11E-13 | 3.373664261 | 5.42E-23 | 5.950823518 | 6.816637977 | 7.315436877 | 7.342949392 | 9.803915852 | 9.64563775 | 10.5938145 | 10.4433342 |
| ENSG00000197324 | LRP10 | 3.030775841 | 9.75E-09 | 2.917333826 | 3.32E-08 | 5.948109667 | 10.50545418 | 10.54135596 | 9.287079172 | 13.21837177 | 13.2696695 | 13.24378775 | 13.00831746 |
| ENSG00000119321 | FKBP15 | 3.070096685 | 0.016495651 | 2.875426673 | 0.024934862 | 5.945523358 | 4.467311984 | 7.761270542 | 4.523675391 | 9.441381351 | 9.51808625 | 9.203388342 | 9.365527647 |
| ENSG00000180155 | LYNX1 | 1.004099921 | 0.435671309 | 4.940850989 | 3.91E-06 | 5.94495091 | 2 | 6.311476754 | 5.772716771 | 6.401247806 | 6.56439512 | 10.27275639 | 10.44298299 |
| ENSG00000180787 | ZFP3 | 2.287993543 | 0.032097652 | 3.654196788 | 0.00025706 | 5.94219033 | 3.327711234 | 4.442710129 | 2 | 5.44762523 | 5.64780787 | 6.99251878 | 6.669279147 |
| ENSG00000176994 | SMCR8 | 3.301888984 | 1.52E-06 | 2.638366772 | 0.000157435 | 5.940255756 | 7.624139768 | 6.535001748 | 5.59507254 | 10.04459716 | 10.0955857 | 9.482666068 | 9.333296562 |
| ENSG00000118496 | FBXO30 | 3.476106324 | 0.135689274 | 2.459430412 | 0.299147391 | 5.935536736 | 2 | 7.638155693 | 2 | 9.541506037 | 9.49786081 | 8.711314045 | 8.278687361 |
| ENSG00000134287 | ARF3 | 2.899097204 | 0.003712692 | 3.035499387 | 0.002170653 | 5.93459659 | 6.96351253 | 9.386925327 | 6.958202216 | 11.10921196 | 11.1863961 | 10.99972888 | 11.5222047 |
| ENSG00000122126 | OCRL | 2.565755622 | 0.319889927 | 3.366743365 | 0.172175696 | 5.932498987 | 2 | 8.356994601 | 2 | 9.356683903 | 9.31389502 | 10.15747031 | 10.10714012 |
| ENSG00000142279 | WTIP | 2.074267255 | 0.051696866 | 3.857411459 | 8.82E-05 | 5.931678714 | 2 | 5.72254323 | 5.39244166 | 6.941624891 | 7.06433609 | 9.004546892 | 8.45344973 |
| ENSG00000127418 | FGFRL1 | 2.681969191 | 0.181657578 | 3.247072603 | 0.093525732 | 5.929041794 | 4.467311984 | 8.469613413 | 2 | 9.610955295 | 9.66535801 | 9.908424552 | 10.44438731 |
| ENSG00000119927 | GPAM | 3.190679513 | 0.119128865 | 2.736471904 | 0.18059707 | 5.927151417 | 2 | 6.464272742 | 2 | 8.027038952 | 8.08878133 | 8.015738956 | 7.049302614 |
| ENSG00000181381 | DDX60L | 4.818851628 | 0.031171927 | 1.104640878 | 0.666867266 | 5.923492506 | 2 | 7.73146271 | 2 | 10.95719949 | 10.9469174 | 7.131485609 | 7.401331798 |
| ENSG00000170891 | CYTL1 | -0.313248726 | 0.911524164 | 6.236730233 | 0.001732414 | 5.923481507 | 2 | 2 | 2 | 2.141737246 | 2.38539787 | 5.399896186 | 6.899677537 |
| ENSG00000126821 | SGPP1 | 2.344046073 | 0.302845914 | 3.578398312 | 0.094088158 | 5.922444385 | 2 | 7.099070191 | 2 | 7.862723323 | 7.85690889 | 9.200067611 | 8.948586914 |
| ENSG00000198121 | LPAR1 | 3.345414805 | 0.001006517 | 2.574550067 | 0.013658585 | 5.919964872 | 5.095959458 | 8.505280662 | 8.076950877 | 11.14516677 | 11.1026219 | 10.356127 | 10.35387936 |
| ENSG00000105429 | MEGF8 | 2.279541318 | 0.096844942 | 3.638089335 | 0.004641907 | 5.917630653 | 3.327711234 | 7.874700259 | 8.076950877 | 9.607141034 | 9.74371185 | 10.92536393 | 11.12994454 |
| ENSG00000067141 | NEO1 | 2.170277047 | 0.20811252 | 3.74604617 | 0.019679078 | 5.916323217 | 2 | 8.213429689 | 6.43197674 | 9.166592134 | 9.14209426 | 10.563383 | 10.86744686 |
| ENSG00000134046 | MBD2 | 3.183282449 | 0.065440948 | 2.731607821 | 0.114871963 | 5.91489027 | 2 | 4.93638933 | 7.858115942 | 9.575477782 | 9.60612566 | 8.994997676 | 9.275296471 |
| ENSG00000136021 | SCYL2 | 3.258479898 | 0.007752174 | 2.655267129 | 0.032913606 | 5.913747027 | 4.00722531 | 8.004992147 | 5.930865648 | 10.00572181 | 10.043513 | 9.567578199 | 9.265398973 |
| ENSG00000100311 | PDGFB | 1.276809956 | 0.610415582 | 4.636624124 | 0.028655643 | 5.913434079 | 2 | 2 | 2 | 2.868578001 | 2.49394783 | 5.108342547 | 4.594883674 |
| ENSG00000153179 | RASSF3 | 2.418292721 | 2.77E-07 | 3.494846124 | 2.44E-14 | 5.913138845 | 6.009025997 | 7.051611315 | 7.163363831 | 9.193296408 | 9.22514727 | 10.3869745 | 10.16599298 |
| ENSG00000073060 | SCARB1 | 1.86662175 | 0.352079152 | 4.046334955 | 0.026558185 | 5.912956704 | 3.327711234 | 7.467835385 | 2 | 7.694616532 | 7.91663606 | 9.751450819 | 10.15873961 |
| ENSG00000137941 | TTLL7 | 2.725817057 | 0.005157524 | 3.185969928 | 0.000801746 | 5.911786986 | 4.467311984 | 3.076665209 | 4.058998049 | 6.321555392 | 6.25300433 | 7.086154752 | 6.248307987 |

| | | | | | | | | | | | | | |
|---|---|---|---|---|---|---|---|---|---|---|---|---|---|
| ENSG00000104267 | CA2 | 1.494607862 | 0.683257266 | 4.412645119 | 0.170346935 | 5.907252981 | 2 | 2 | 2 | 2.600425781 | 2.94009902 | 5.545162502 | 2.225841198 |
| ENSG00000068305 | MEF2A | 2.349257773 | 0.152743092 | 3.55593196 | 0.021881678 | 5.905189732 | 7.384433967 | 8.169643011 | 2 | 9.590237441 | 9.57248057 | 10.81094636 | 10.7560816 |
| ENSG00000115594 | IL1R1 | 4.604721595 | 5.26E-07 | 1.299970236 | 0.211023502 | 5.90469183 | 5.330594134 | 7.60566235 | 8.486635645 | 12.19878022 | 12.2084944 | 8.888451287 | 8.93066256 |
| ENSG00000204217 | BMPR2 | 2.772322312 | 0.002123869 | 3.130273687 | 0.000412201 | 5.902595998 | 7.384433967 | 9.192779271 | 6.718950348 | 10.9554005 | 10.8890776 | 11.31638534 | 11.24249341 |
| ENSG00000162614 | NEXN | 2.541537893 | 0.002036933 | 3.354437516 | 2.58E-05 | 5.895975409 | 7.485143553 | 7.002537973 | 4.874587708 | 9.344822471 | 9.28397021 | 10.09608847 | 10.15080456 |
| ENSG00000125249 | RAP2A | 2.796585355 | 0.003671412 | 3.099131879 | 0.001066317 | 5.895717234 | 6.653098118 | 6.311476754 | 3.369317972 | 8.70652031 | 8.72559535 | 9.215280081 | 8.784747976 |
| ENSG00000127947 | PTPN12 | 4.001090442 | 2.04E-07 | 1.893493481 | 0.022344999 | 5.894583923 | 6.653098118 | 8.451443573 | 6.629488387 | 11.48981601 | 11.5153943 | 9.646549955 | 9.106387289 |
| ENSG00000243156 | MICAL3 | 2.300318278 | 0.109150595 | 3.592299439 | 0.007741661 | 5.892617718 | 2 | 7.099070191 | 6.629488387 | 8.567402595 | 8.57772939 | 9.649231086 | 10.03603692 |
| ENSG00000243649 | CFB | 11.07974309 | 5.21E-15 | -5.191150855 | 0.022950398 | 5.88859223 | 2 | 2 | 2 | 11.06401437 | 11.1521225 | 2 | 2 |
| ENSG00000164296 | TIGD6 | 3.07832547 | 0.04116967 | 2.809491032 | 0.061814261 | 5.887816502 | 2 | 5.131603625 | 2 | 6.622948477 | 6.63721363 | 6.353989995 | 6.392053169 |
| ENSG00000167700 | MFSD3 | 2.651890779 | 0.095299137 | 3.235355547 | 0.034772335 | 5.887246326 | 3.327711234 | 6.311476754 | 2 | 7.348537858 | 7.62891439 | 7.800194554 | 8.294346816 |
| ENSG00000173706 | HEG1 | 1.767576402 | 0.210659147 | 4.119549466 | 0.001171726 | 5.887125868 | 4.467311984 | 9.267380431 | 8.784770914 | 10.24443813 | 10.253992 | 12.29485156 | 12.8473242 |
| ENSG00000133818 | RRAS2 | 2.305716272 | 0.063703589 | 3.580783744 | 0.002227172 | 5.886500015 | 7.33131396 | 7.538394143 | 3.369317972 | 9.213506984 | 9.12531607 | 10.14664362 | 10.68242029 |
| ENSG00000166707 | ZCCHC18 | 2.595332244 | 0.294006549 | 3.287979782 | 0.16118623 | 5.883312026 | 2 | 2 | 2 | 3.227757282 | 3.45167086 | 3.930690651 | 3.660199087 |
| ENSG00000170185 | USP38 | 3.007978735 | 0.100023481 | 2.8740986 | 0.111705957 | 5.882077335 | 2 | 7.638155693 | 4.058998049 | 9.159317259 | 9.13791797 | 9.092065441 | 8.93516457 |
| ENSG00000165997 | ARL5B | 3.169004983 | 0.005967521 | 2.70901109 | 0.020479608 | 5.878016072 | 3.327711234 | 6.844429755 | 4.874587708 | 8.81895723 | 8.74905888 | 8.582068017 | 8.02265879 |
| ENSG00000147614 | ATP6V0D2 | 5.216132875 | 0.009388115 | 0.661446867 | 0.785803002 | 5.877579742 | 3.327711234 | 2 | 4.058998049 | 7.764774849 | 7.85944752 | 4.487545924 | 2.117335275 |
| ENSG00000073331 | ALPK1 | 2.913539041 | 3.37E-05 | 2.963151474 | 2.10E-05 | 5.876690516 | 4.815609471 | 4.11352228 | 4.058998049 | 6.965655398 | 7.03768431 | 7.172727426 | 6.915965441 |
| ENSG00000170832 | USP32 | 3.296694244 | 0.03895919 | 2.577986874 | 0.111493496 | 5.874681119 | 2 | 7.874700259 | 5.930865648 | 9.891822146 | 9.90230962 | 9.121501891 | 9.240556116 |
| ENSG00000154229 | PRKCA | 1.392685249 | 0.088331632 | 4.479636238 | 4.59E-10 | 5.872321486 | 5.095959458 | 7.145017454 | 6.629488387 | 7.860161659 | 7.91175216 | 10.48764377 | 11.30189717 |
| ENSG00000131845 | ZNF304 | 2.737293584 | 0.004826128 | 3.133450689 | 0.000982659 | 5.870744273 | 3.327711234 | 5.946638857 | 4.058998049 | 7.433701013 | 7.58663873 | 7.927733561 | 7.873223515 |
| ENSG00000137075 | RNF38 | 2.774525904 | 0.058330657 | 3.09394138 | 0.030778033 | 5.868467283 | 3.327711234 | 6.951736442 | 3.369317972 | 8.310225403 | 8.2657098 | 8.693004369 | 8.508863136 |
| ENSG00000165244 | ZNF367 | 2.081434466 | 0.421293524 | 3.774888307 | 0.109758105 | 5.856322773 | 2 | 2 | 2 | 3.023133209 | 3.08597292 | 3.632645806 | 4.553838275 |
| ENSG00000099812 | MISP | 1.392695656 | 0.580922984 | 4.461242112 | 0.039231993 | 5.853937768 | 2 | 2 | 2 | 2.947924294 | 2.49394783 | 4.727866721 | 4.720747915 |
| ENSG00000147059 | SPIN2A | 4.631269401 | 0.020403301 | 1.218325855 | 0.593116818 | 5.849595256 | 2 | 2 | 2 | 4.76150579 | 4.97065356 | 2.592885947 | 2.746601469 |
| ENSG00000197892 | KIF13B | 2.816345506 | 0.030340464 | 3.032717791 | 0.017911792 | 5.849063297 | 2 | 4.710564154 | 6.43197674 | 7.805288479 | 8.07355593 | 8.10721774 | 8.208518621 |
| ENSG00000104689 | TNFRSF10A | 4.025529351 | 4.10E-06 | 1.822898921 | 0.053995487 | 5.848428273 | 3.327711234 | 3.076665209 | 5.156626401 | 7.697489359 | 7.84670947 | 5.684753614 | 5.641643428 |
| ENSG00000065060 | UHRF1BP1 | 2.229328004 | 0.026657522 | 3.618867056 | 0.000131309 | 5.84819506 | 5.095959458 | 7.145017454 | 4.523675391 | 8.321442665 | 8.15847021 | 9.743945072 | 9.484290996 |
| ENSG00000077420 | APBB1IP | 4.247776626 | 0.00464393 | 1.595577343 | 0.33487913 | 5.843553969 | 4.00722531 | 3.076665209 | 2 | 6.722207358 | 6.79406287 | 4.971696071 | 3.287118176 |
| ENSG00000124882 | EREG | 6.349544106 | 1.76E-30 | -0.507425928 | 0.462478098 | 5.842118178 | 5.532353208 | 4.11352228 | 4.874587708 | 11.06039244 | 11.0379776 | 4.461156421 | 4.489993931 |
| ENSG00000162757 | C1orf74 | 2.824351946 | 0.040592042 | 3.015224656 | 0.025868544 | 5.839576602 | 2 | 4.11352228 | 2 | 5.304221918 | 5.39133044 | 5.606648123 | 5.431121909 |
| ENSG00000183530 | PRR14L | 2.659643464 | 0.019452444 | 3.17946183 | 0.004097253 | 5.839105294 | 4.00722531 | 7.761270542 | 6.322058544 | 9.265734736 | 9.39662469 | 9.937031556 | 9.756873659 |
| ENSG00000263464 | PPIAL4C | 7.901226884 | 5.51E-05 | -2.06401995 | 0.405294552 | 5.837206934 | 2 | 2 | 2 | 7.939959586 | 7.96222667 | 2 | 2.172608095 |
| ENSG00000130988 | RGN | -0.72409335 | 0.778992908 | 6.559111261 | 0.000322101 | 5.835017911 | 2 | 2 | 2 | 2.270786764 | 2.14022021 | 5.919306827 | 7.124815549 |
| ENSG00000078369 | GNB1 | 2.613953308 | 0.000485712 | 3.215577907 | 1.12E-05 | 5.829531214 | 8.285626337 | 10.39216648 | 10.11951029 | 12.45409166 | 12.4656716 | 12.99790049 | 13.12177191 |
| ENSG00000116819 | TFAP2E | 2.898715294 | 0.253395005 | 2.930395393 | 0.237344892 | 5.829110687 | 2 | 2 | 2 | 3.515101962 | 3.55551834 | 3.851628042 | 3.183290849 |
| ENSG00000140548 | ZNF710 | 2.937295545 | 0.044706135 | 2.890810076 | 0.046006971 | 5.828105621 | 3.327711234 | 6.951736442 | 3.369317972 | 8.491639362 | 8.40429221 | 8.607356811 | 8.164036804 |
| ENSG00000138642 | HERC6 | 6.454049373 | 5.44E-06 | -0.627129984 | 0.72430187 | 5.826919388 | 3.327711234 | 6.535001748 | 2 | 11.47879537 | 11.4830405 | 4.652122235 | 4.644600858 |
| ENSG00000197134 | ZNF257 | 2.214045646 | 0.392182361 | 3.612566179 | 0.130326951 | 5.826611824 | 2 | 2 | 2 | 3.227757282 | 3.01487888 | 4.388329244 | 3.580696614 |
| ENSG00000075702 | WDR62 | 2.61230771 | 0.199709404 | 3.214269274 | 0.099251185 | 5.826576983 | 3.327711234 | 2 | 2 | 4.268917409 | 3.69849104 | 3.930690651 | 4.886954834 |
| ENSG00000213999 | MEF2B | 5.503574515 | 0.001640812 | 0.321158918 | 0.888753558 | 5.824733434 | 2 | 2 | 2 | 5.709096143 | 5.58783529 | 2.42098416 | 2.374681325 |
| ENSG00000120328 | PCDHB12 | 2.939474669 | 0.248121056 | 2.884474021 | 0.247980941 | 5.82394869 | 2 | 2 | 2 | 3.79808662 | 3.28037825 | 3.256555576 | 3.753786374 |
| ENSG00000197081 | IGF2R | 2.99987186 | 8.12E-05 | 2.823244421 | 0.000213322 | 5.823116281 | 10.02639697 | 8.077869079 | 8.434763955 | 12.09937748 | 12.1001021 | 11.82446067 | 12.01576379 |
| ENSG00000178297 | TMPRSS9 | 3.243893455 | 0.183184126 | 2.579037228 | 0.290945022 | 5.822930683 | 2 | 2 | 2 | 3.754611435 | 3.7864959 | 3.534812207 | 3.100202476 |
| ENSG00000062582 | MRPS24 | 4.405278527 | 9.11E-08 | 1.416682774 | 0.123888817 | 5.8219613 | 4.815609471 | 4.710564154 | 4.058998049 | 8.75144642 | 8.71161289 | 5.06226682 | 6.33565646 |
| ENSG00000204172 | AGAP9 | 4.365074667 | 0.001280898 | 1.456502569 | 0.336305876 | 5.821577236 | 2 | 5.131603625 | 2 | 7.930222906 | 7.85181819 | 5.108342547 | 5.1574003 |
| ENSG00000172379 | ARNT2 | 0.450268134 | 0.691119414 | 5.368131248 | 1.00E-09 | 5.818399382 | 4.467311984 | 6.844429755 | 4.874587708 | 6.328984825 | 6.17362678 | 10.72933238 | 11.38843349 |
| ENSG00000146373 | RNF217 | 2.329821478 | 0.323812042 | 3.487988312 | 0.117523606 | 5.817809791 | 2 | 7.35507212 | 2 | 8.197043419 | 7.99722844 | 9.555675374 | 8.854012253 |
| ENSG00000169504 | CLIC4 | 2.434116611 | 0.002123425 | 3.379900085 | 9.67E-06 | 5.814016696 | 8.696033109 | 11.00402286 | 10.21486071 | 12.72860514 | 12.6363624 | 13.61559199 | 13.64144911 |
| ENSG00000168675 | LDLRAD4 | 0.212583897 | 0.925360182 | 5.60116025 | 0.00097184 | 5.813744147 | 2 | 4.11352228 | 2 | 3.840289928 | 2.38539787 | 8.189226297 | 7.64187159 |
| ENSG00000073008 | PVR | 2.436187983 | 2.42E-05 | 3.376555406 | 1.76E-09 | 5.812743389 | 7.159102577 | 8.540087354 | 8.35328246 | 10.55911051 | 10.5554386 | 11.203095 | 11.7390458 |
| ENSG00000128274 | A4GALT | 2.169811609 | 0.333515024 | 3.642594602 | 0.081866442 | 5.812406212 | 2 | 3.076665209 | 8.206335842 | 8.759716509 | 8.84055903 | 10.11154794 | 10.40399898 |
| ENSG00000135677 | GNS | 3.012561827 | 3.62E-06 | 2.796413439 | 1.82E-05 | 5.808975266 | 8.653385977 | 10.19331737 | 8.784770914 | 12.38047484 | 12.4206128 | 12.15680513 | 12.21207488 |
| ENSG00000126456 | IRF3 | 3.141237377 | 0.051321923 | 2.667682949 | 0.09933964 | 5.808920326 | 5.095959458 | 7.572420273 | 2 | 9.275380474 | 9.40272932 | 8.856747125 | 8.883817772 |
| ENSG00000158859 | ADAMTS4 | 4.709533829 | 5.90E-19 | 1.096846141 | 0.065488624 | 5.806379969 | 5.095959458 | 6.666836837 | 6.322058544 | 10.80535243 | 10.8420389 | 7.332904981 | 7.154291015 |
| ENSG00000173267 | SNCG | 3.073446104 | 0.059053709 | 2.728867184 | 0.093025295 | 5.802313287 | 2 | 3.686303414 | 2 | 4.739373666 | 5.3193943 | 4.659878161 | 4.853519801 |
| ENSG00000184292 | TACSTD2 | 2.938642954 | 0.232847072 | 2.85852276 | 0.237776037 | 5.797165714 | 2 | 2 | 2 | 3.709785322 | 3.39680241 | 3.411599105 | 3.600986722 |
| ENSG00000143322 | ABL2 | 3.834974644 | 2.54E-08 | 1.961458758 | 0.007316377 | 5.796433402 | 8.042902253 | 8.876041558 | 6.882775709 | 11.95139166 | 11.9952166 | 9.880484031 | 10.29710605 |
| ENSG00000154930 | ACSS1 | 2.367139927 | 0.046591905 | 3.428834062 | 0.002286691 | 5.795973989 | 2 | 4.11352228 | 3.369317972 | 5.334065616 | 5.34860058 | 6.615624322 | 5.964937606 |
| ENSG00000198554 | WDHD1 | 3.247707878 | 0.001493487 | 2.543171185 | 0.015108153 | 5.790879063 | 3.327711234 | 4.710564154 | 2 | 6.693715713 | 6.67228943 | 6.176128386 | 5.829625997 |
| ENSG00000100201 | DDX17 | 2.711253137 | 0.000139786 | 3.079551563 | 1.13E-05 | 5.7908047 | 7.940205782 | 9.949724236 | 9.083172337 | 11.92329082 | 11.9103827 | 12.41400745 | 12.14288977 |
| ENSG00000187498 | COL4A1 | 1.634787047 | 0.038349318 | 4.153876789 | 6.19E-09 | 5.788663837 | 11.09308737 | 12.91590856 | 11.51057165 | 13.69244595 | 13.7010937 | 16.49368867 | 15.87054846 |
| ENSG00000163935 | SFMBT1 | 3.118196222 | 0.029640097 | 2.670134479 | 0.064397771 | 5.788330702 | 2 | 4.93638933 | 2 | 6.542151749 | 6.40670796 | 6.043083161 | 6.059610627 |
| ENSG00000169258 | GPRIN1 | 3.063531247 | 0.078250731 | 2.721497317 | 0.116627441 | 5.785028564 | 2 | 5.595927442 | 2 | 6.902323594 | 7.21862736 | 6.975527754 | 6.454714037 |
| ENSG00000144218 | AFF3 | 1.810346532 | 0.094524523 | 3.971292992 | 4.86E-05 | 5.781639524 | 2 | 5.303523747 | 5.930865648 | 6.777556219 | 6.84106088 | 8.795135904 | 9.053535482 |
| ENSG00000159921 | GNE | 2.746171259 | 1.96E-11 | 3.035078946 | 6.55E-14 | 5.781250205 | 6.138360429 | 5.838937808 | 6.322058544 | 8.740345368 | 8.7935444 | 8.960872516 | 9.141009026 |
| ENSG00000166147 | FBN1 | 2.22421739 | 0.030773929 | 3.556347823 | 0.000245067 | 5.780565213 | 10.48735708 | 12.47802568 | 9.624627483 | 13.57375396 | 13.6090065 | 14.84878204 | 14.99421534 |
| ENSG00000110046 | ATG2A | 3.407096321 | 0.000107002 | 2.372733448 | 0.009191319 | 5.77982977 | 6.009025997 | 6.951736442 | 4.058998049 | 9.385472425 | 9.51245165 | 8.211166197 | 8.610562542 |
| ENSG00000124635 | HIST1H2BJ | 5.301595587 | 2.04E-06 | 0.477543959 | 0.735593063 | 5.779139547 | 2 | 4.11352228 | 2 | 7.756533788 | 7.70710533 | 3.429858444 | 3.580696614 |
| ENSG00000158201 | ABHD3 | 3.622423252 | 0.011003457 | 2.156041648 | 0.148400754 | 5.778464901 | 2 | 4.442710129 | 2 | 6.548530093 | 6.31350454 | 5.445480974 | 4.594883674 |
| ENSG00000133216 | EPHB2 | 5.211800756 | 2.39E-07 | 0.566545163 | 0.651103372 | 5.778345919 | 6.257048221 | 5.303523747 | 2 | 10.37802153 | 10.404557 | 5.951260634 | 5.73181021 |
| ENSG00000152583 | SPARCL1 | 2.861160538 | 0.327927671 | 2.914807801 | 0.30681403 | 5.77596834 | 2 | 2 | 2 | 3.709785322 | 3.28037825 | 4.099885429 | 2.632958241 |
| ENSG00000162636 | FAM102B | 2.915113487 | 0.127032028 | 2.860181378 | 0.128290491 | 5.775294866 | 2 | 6.04685525 | 2 | 7.494428494 | 7.23826702 | 7.542503814 | 7.052989523 |
| ENSG00000176165 | FOXG1 | 5.97491672 | 0.056914368 | -0.201051524 | 0.959149058 | 5.773865196 | 2 | 2 | 2 | 6.113598934 | 6.06344184 | 2.526562374 | 2 |
| ENSG00000182979 | MTA1 | 2.659250883 | 0.036049949 | 3.114568802 | 0.011726998 | 5.773819685 | 3.327711234 | 7.393647515 | 5.39244166 | 8.859280876 | 8.76132797 | 9.326359826 | 9.196649491 |
| ENSG00000114999 | TTL | 2.44045415 | 0.037564908 | 3.332080035 | 0.002992423 | 5.772534185 | 7.384433967 | 7.099070191 | 3.369317972 | 9.074566443 | 9.15041075 | 10.04035872 | 9.958055169 |
| ENSG00000198814 | GK | 6.866756812 | 1.61E-06 | -1.096499397 | 0.524822542 | 5.770257414 | 2 | 5.72254323 | 2 | 11.00610223 | 10.9424344 | 3.517841605 | 3.660199087 |
| ENSG00000171346 | KRT15 | 3.570888536 | 0.175625164 | 2.197497131 | 0.419359474 | 5.768385668 | 2 | 2 | 2 | 3.840289928 | 4.16043394 | 2.560105256 | 3.518055793 |
| ENSG00000165030 | NFIL3 | 3.196756798 | 0.045765337 | 2.570519391 | 0.111954789 | 5.767276189 | 4.00722531 | 6.951736442 | 2 | 8.697938662 | 8.69749358 | 8.164223608 | 7.988411589 |
| ENSG00000215472 | RPL17-C18or | 10.95339604 | 1.38E-14 | -5.191150878 | 0.023227301 | 5.762245166 | 2 | 2 | 2 | 11.05396204 | 10.9075945 | 2 | 2 |
| ENSG00000162923 | WDR26 | 3.03455677 | 0.001104484 | 2.724867992 | 0.003635849 | 5.759424762 | 6.138360429 | 8.88960062 | 8.978369997 | 11.44179676 | 11.5006301 | 11.13212295 | 11.19186952 |
| ENSG00000147454 | SLC25A37 | 3.80552222 | 6.67E-06 | 1.952306112 | 0.030734544 | 5.757828332 | 5.330594134 | 8.029695761 | 7.734844356 | 11.19759955 | 11.207224 | 9.395058233 | 9.314988993 |
| ENSG00000100364 | KIAA0930 | 2.735092031 | 0.006672564 | 3.022721244 | 0.002294707 | 5.757813275 | 7.667647223 | 7.73146271 | 4.523675391 | 9.910512694 | 9.89118718 | 10.19064296 | 10.18417862 |
| TMEM44 | | 2.973032158 | 0.036596823 | 2.783158572 | 0.049531896 | 5.756190729 | 2 | 4.93638933 | 6.882775709 | 8.559529208 | 8.4936155 | 8.303800769 | 8.373881215 |
| ENSG00000144355 | DLX1 | 5.522779477 | 0.005659011 | 0.230092795 | 0.930138959 | 5.752872272 | 2 | 2 | 2 | 5.742861563 | 5.58783529 | 2.183373728 | 2.551896972 |
| ENSG00000137094 | DNAJB5 | 2.567271209 | 0.022996195 | 3.181311626 | 0.00364531 | 5.748582835 | 4.467311984 | 4.11352228 | 6.882775709 | 8.063125866 | 8.25224987 | 8.354863925 | 9.088535264 |
| ENSG00000271147 | ARMCX5-GPI | 4.155408393 | 6.14E-05 | 1.590491059 | 0.16435224 | 5.745899452 | 2 | 3.686303414 | 4.058998049 | 7.048994596 | 7.06873058 | 4.933813368 | 4.400220394 |
| ENSG00000172046 | USP19 | 2.72501915 | 0.116680459 | 3.018608495 | 0.074520332 | 5.743627645 | 4.467311984 | 7.538394143 | 2 | 8.792330793 | 8.83669913 | 9.249735981 | 8.946110698 |
| ENSG00000107036 | RIC1 | 2.383495329 | 0.063345312 | 3.359203354 | 0.005993894 | 5.742698683 | 3.327711234 | 7.35507212 | 7.777116083 | 9.340234287 | 9.42346554 | 10.21848193 | 10.47768733 |
| ENSG00000003402 | CFLAR | 2.83145974 | 2.38E-05 | 2.910945783 | 1.24E-05 | 5.742405524 | 8.075571721 | 7.669933291 | 9.330306409 | 11.35621574 | 11.350135 | 11.20077081 | 11.63219444 |
| ENSG00000072062 | PRKACA | 2.312155079 | 0.004214141 | 3.429141488 | 9.16E-06 | 5.741296567 | 5.532353208 | 7.60566235 | 7.896971941 | 9.577818317 | 9.63233954 | 10.54922865 | 10.86966979 |
| ENSG00000106733 | NMRK1 | 3.46404664 | 0.02530452 | 2.275430503 | 0.155557022 | 5.739477143 | 2 | 5.303523747 | 2 | 7.19843704 | 7.14562489 | 6.049001768 | 6.018717758 |
| ENSG00000085982 | USP40 | 3.21416886 | 9.14E-05 | 2.522074019 | 0.002647154 | 5.736242879 | 5.095959458 | 6.666836837 | 4.523675391 | 8.859280876 | 8.90710092 | 7.897710349 | 8.44785952 |
| ENSG00000110756 | HPS5 | 2.084421352 | 0.093501828 | 3.649971246 | 0.001511468 | 5.734392598 | 5.70932728 | 7.73146271 | 4.058998049 | 8.595396228 | 8.6202631 | 9.783088106 | 10.46442481 |
| ENSG00000180938 | ZNF572 | 2.7336337 | 0.27254373 | 3.000726037 | 0.212779129 | 5.734359737 | 2 | 2 | 2 | 3.406930997 | 3.45167086 | 3.738983888 | 3.45257121 |
| ENSG00000137414 | FAM8A1 | 2.349379819 | 0.223030005 | 3.381278064 | 0.064416773 | 5.730657883 | 4.815609471 | 8.276714465 | 2 | 9.16866395 | 9.1347778 | 10.43083967 | 9.872489883 |
| ENSG00000133937 | GSC | 5.038445434 | 0.006880061 | 0.691754231 | 0.761301217 | 5.730199665 | 2 | 2 | 2 | 5.273747807 | 5.18011485 | 2.526562374 | 2.466008509 |
| ENSG00000184402 | SS18L1 | 2.94451219 | 0.000827513 | 2.782033699 | 0.001598036 | 5.726545889 | 4.467311984 | 6.228547981 | 4.058998049 | 8.128448191 | 8.08878133 | 8.206540078 | 7.634508409 |
| ENSG00000120306 | CYSTM1 | 2.603717031 | 0.09775828 | 3.12091107 | 0.039779899 | 5.724628101 | 5.330594134 | 7.431218269 | 2 | 8.722121967 | 8.73530306 | 9.228053138 | 9.255433104 |
| ENSG00000197694 | SPTAN1 | 2.807911015 | 0.004062414 | 2.915644207 | 0.00262317 | 5.723555223 | 6.89194353 | 10.00059544 | 9.385998894 | 12.0444476 | 12.036184 | 11.97417317 | 12.30302946 |
| ENSG00000164741 | DLC1 | 1.938587204 | 0.298439087 | 3.784266443 | 0.026791677 | 5.722853647 | 3.327711234 | 9.00621767 | 4.874587708 | 9.476928206 | 9.42174886 | 11.47316881 | 11.07852969 |
| ENSG00000158470 | B4GALT5 | 3.133790741 | 0.000211037 | 2.58862914 | 0.002611125 | 5.722419881 | 5.330594134 | 7.701025996 | 7.97167868 | 10.48289882 | 10.4848496 | 9.947565107 | 9.933514329 |
| ENSG00000176532 | PRR15 | 1.274449266 | 0.656481504 | 4.446228641 | 0.070664362 | 5.720677906 | 2 | 2 | 2 | 2.695456494 | 2.68924543 | 5.367107572 | 3.474731315 |
| ENSG00000175518 | UBQLNL | 3.181502112 | 0.188518397 | 2.533718569 | 0.295652928 | 5.715220681 | 2 | 2 | 2 | 3.754611435 | 3.69849104 | 3.374372158 | 3.236138336 |
| ENSG00000145901 | TNIP1 | 3.132466063 | 5.11E-12 | 2.576989944 | 1.75E-08 | 5.709456007 | 9.097018025 | 9.225224669 | 9.344432377 | 12.32361727 | 12.3748777 | 11.19677777 | 12.21593427 |
| ENSG00000120137 | PANK3 | 1.945919721 | 0.351874922 | 3.762469346 | 0.050135255 | 5.708389068 | 4.00722531 | 8.276714465 | 2 | 8.657201865 | 8.73806474 | 10.59987492 | 10.40291596 |
| ENSG00000007933 | FMO3 | 1.586482027 | 0.370269344 | 4.118769091 | 0.008770914 | 5.705251118 | 2 | 4.710564154 | 2 | 4.694064037 | 4.95174433 | 7.786089373 | 6.124122661 |

| | | | | | | | | | | | | | |
|---|---|---|---|---|---|---|---|---|---|---|---|---|---|
| ENSG00000129657 | SEC14L1 | 1.978139911 | 0.184717964 | 3.721841555 | 0.006750332 | 5.699981466 | 3.327711234 | 7.928237109 | 8.585098385 | 9.706743577 | 9.67187198 | 11.00991086 | 11.75207515 |
| ENSG00000111269 | CREBL2 | 2.728266545 | 0.058804128 | 2.970717133 | 0.035621047 | 5.698983678 | 7.33131396 | 6.140560021 | 2 | 8.970341944 | 8.95060997 | 9.030205459 | 9.353992291 |
| ENSG00000170396 | ZNF804A | 7.140845344 | 3.05E-06 | -1.442201161 | 0.468753938 | 5.698644183 | 2 | 2 | 2 | 7.242377933 | 7.17037177 | 2.139668783 | 2.117335275 |
| ENSG00000129538 | RNASE1 | -0.7313856 | 0.818399248 | 6.41697834 | 0.006572253 | 5.685592741 | 2 | 2 | 2 | 2 | 2.38539787 | 6.693258166 | 6.29575846 |
| ENSG00000124225 | PMEPA1 | 1.527130177 | 0.056262369 | 4.157930164 | 6.56E-09 | 5.685060341 | 5.095959458 | 6.951736442 | 7.398165132 | 8.370868225 | 8.13979554 | 10.91618505 | 10.83234501 |
| ENSG00000178919 | FOXE1 | 4.405994612 | 0.001807656 | 1.276663541 | 0.425335565 | 5.682658153 | 2 | 4.93638933 | 2 | 7.66556918 | 7.77322015 | 5.163941955 | 4.279527956 |
| ENSG00000124208 | TMEM189-U | 10.87265855 | 2.05E-14 | -5.191150959 | 0.023163678 | 5.681507594 | 2 | 2 | 2 | 10.85861994 | 10.9442293 | 2 | 2 |
| ENSG00000262418 | | 1.389352449 | 0.620673635 | 4.291808583 | 0.077889486 | 5.681161032 | 2 | 2 | 2 | 2.695456494 | 2.77779907 | 5.180212365 | 3.539239711 |
| ENSG00000085449 | WDFY1 | 2.913905438 | 0.053258105 | 2.763671404 | 0.064987892 | 5.677576841 | 3.327711234 | 8.777382925 | 6.629488387 | 10.41601627 | 10.395402 | 10.27922234 | 10.23232731 |
| ENSG00000172159 | FRMD3 | 4.119131334 | 0.043612511 | 1.557800054 | 0.483865109 | 5.676931388 | 2 | 3.076665209 | 2 | 5.128039203 | 5.39133044 | 3.767980144 | 2.592996859 |
| ENSG00000116406 | EDEM3 | 3.348143684 | 0.006716746 | 2.319752523 | 0.06941259 | 5.667896208 | 5.095959458 | 8.505280662 | 5.59507254 | 10.51500864 | 10.5655798 | 9.80401941 | 9.156086127 |
| ENSG00000117500 | TMED5 | 3.579668181 | 0.000178186 | 2.088063833 | 0.038440494 | 5.667732014 | 5.330594134 | 8.029695761 | 6.203069917 | 10.58416277 | 10.4528466 | 9.338190203 | 8.651871454 |
| ENSG00000095752 | IL11 | 4.09474139 | 0.000183325 | 1.566545523 | 0.193576 | 5.661286912 | 7.790810999 | 6.311476754 | 4.058998049 | 10.74623244 | 10.7874282 | 7.811554331 | 8.592942424 |
| ENSG00000049246 | PER3 | 2.255840863 | 0.007490024 | 3.402847196 | 2.10E-05 | 5.658688059 | 4.815609471 | 4.710564154 | 2 | 6.268449691 | 6.39974839 | 7.317051196 | 7.558732759 |
| ENSG00000213139 | CRYGS | 3.049919775 | 0.245300741 | 2.606857701 | 0.318173125 | 5.656777476 | 2 | 2 | 2 | 3.95996233 | 3.2184422 | 3.584558147 | 3.071407821 |
| ENSG00000225828 | FAM229A | 2.541294475 | 0.12116737 | 3.113294565 | 0.047946397 | 5.65458904 | 2 | 3.686303414 | 2 | 4.623314311 | 4.58534546 | 5.279204176 | 4.886954834 |
| ENSG00000170471 | RALGAPB | 2.406662217 | 0.062355471 | 3.247828276 | 0.008609548 | 5.654490493 | 4.815609471 | 8.557180776 | 5.930865648 | 9.685196338 | 9.64710779 | 10.65694047 | 10.33356098 |
| ENSG00000282230 | | 2.933899382 | 0.03884132 | 2.719490378 | 0.054838286 | 5.65338976 | 4.467311984 | 9.814097497 | 9.06622548 | 11.88757367 | 11.8120795 | 11.36330635 | 11.86550852 |
| ENSG00000047936 | ROS1 | 2.209989299 | 0.435112068 | 3.443201777 | 0.190011047 | 5.653191075 | 2 | 2 | 2 | 2.784612485 | 3.39680241 | 3.256555576 | 4.365060851 |
| ENSG00000174705 | SH3PXD2B | 3.214799262 | 0.02248445 | 2.433423726 | 0.091103426 | 5.648222989 | 6.009025997 | 9.357954881 | 5.156626401 | 11.18971483 | 11.1670574 | 10.18477494 | 10.58519509 |
| ENSG00000271605 | MILR1 | 4.544557479 | 0.00113016 | 1.103529743 | 0.493413792 | 5.648087222 | 2 | 2 | 4.058998049 | 6.529309835 | 6.50077304 | 3.679182183 | 3.496556178 |
| ENSG00000105509 | HAS1 | 5.844374058 | 0.000699054 | -0.200552797 | 0.932012683 | 5.643821262 | 2 | 2 | 2 | 5.987823101 | 5.94414054 | 2.34604626 | 2.225841198 |
| ENSG00000224470 | ATXN1L | 2.541358187 | 0.130186997 | 3.097609919 | 0.055730923 | 5.638968106 | 6.73718411 | 8.297207175 | 2 | 9.618553696 | 9.70330142 | 10.17584339 | 10.25411154 |
| ENSG00000155380 | SLC16A1 | 3.467886792 | 0.000211964 | 2.167459181 | 0.027439899 | 5.635345973 | 5.330594134 | 8.395519306 | 7.342949392 | 10.97595466 | 10.8983658 | 9.7324978 | 9.544459294 |
| ENSG00000198742 | SMURF1 | 2.963774581 | 6.81E-08 | 2.670364072 | 1.28E-06 | 5.634138653 | 6.257048221 | 7.538394143 | 6.43197674 | 9.778365132 | 9.83360578 | 9.511886139 | 9.516692718 |
| ENSG00000088899 | LZTS3 | 1.834573145 | 0.341039924 | 3.798008134 | 0.03100898 | 5.632581279 | 3.327711234 | 7.145017454 | 2 | 7.487805859 | 7.45518898 | 9.03282106 | 9.709771424 |
| ENSG00000197713 | RPE | 3.021683493 | 0.085482239 | 2.610315281 | 0.136341225 | 5.631998773 | 4.00722531 | 7.393647515 | 2 | 8.941551548 | 8.91810182 | 8.5805365 | 8.461794641 |
| ENSG00000260873 | | 2.19199165 | 0.005020885 | 3.439745829 | 3.70E-06 | 5.631737479 | 5.330594134 | 6.951736442 | 7.646393253 | 9.117027752 | 9.03500236 | 10.48982111 | 10.12323121 |
| ENSG00000168916 | ZNF608 | 1.733502694 | 0.489660234 | 3.897990894 | 0.083545957 | 5.631493587 | 2 | 3.076665209 | 2 | 3.95996233 | 2.49394783 | 5.565950232 | 4.30448772 |
| ENSG00000103335 | PIEZO1 | 2.717391927 | 0.058485499 | 2.9112882 | 0.038684826 | 5.628680127 | 9.435834703 | 8.834582582 | 4.058998049 | 11.28279734 | 11.3321984 | 11.37586927 | 11.61670776 |
| ENSG00000081377 | CDC14B | 2.434161315 | 0.117408639 | 3.193748953 | 0.032070986 | 5.627910268 | 3.327711234 | 7.145017454 | 3.369317972 | 8.087422161 | 8.16464176 | 8.701014095 | 9.035013043 |
| ENSG00000075073 | TACR2 | 2.331786228 | 0.367465205 | 3.29543827 | 0.17543064 | 5.627224498 | 2 | 2 | 2 | 3.023133209 | 3.33976443 | 4.017766949 | 3.560117069 |
| ENSG00000178623 | GPR35 | 2.973017037 | 0.356981191 | 2.649978549 | 0.40736792 | 5.622995585 | 2 | 2 | 2 | 3.349643765 | 3.7864959 | 3.993419637 | 2.277179783 |
| ENSG00000110367 | DDX6 | 2.893727756 | 5.66E-18 | 2.727712449 | 2.86E-16 | 5.621440206 | 8.042902253 | 7.503546079 | 8.143093325 | 10.75073417 | 10.8131925 | 10.63140433 | 10.6018087 |
| ENSG00000078081 | LAMP3 | 2.935635988 | 0.17616338 | 2.685448169 | 0.211125211 | 5.621084157 | 2 | 2 | 4.058998049 | 5.110902178 | 5.06165168 | 5.598599872 | 3.311948089 |
| ENSG00000104365 | IKBKB | 2.896582202 | 0.150100354 | 2.722632223 | 0.170797377 | 5.619214425 | 3.327711234 | 7.790474954 | 2 | 9.122382231 | 9.15558431 | 8.977260248 | 8.955497794 |
| ENSG00000175073 | VCPIP1 | 3.008453781 | 0.025684466 | 2.609605731 | 0.054660118 | 5.618059512 | 2 | 6.666836837 | 6.882775709 | 9.160358775 | 9.16485023 | 8.914619542 | 8.602408528 |
| ENSG00000184151 | ICE1 | 2.649276038 | 0.021794347 | 2.962617153 | 0.008934398 | 5.61189319 | 6.257048221 | 7.847167909 | 4.058998049 | 9.401417034 | 9.34230452 | 9.768589685 | 9.593700566 |
| ENSG00000101972 | STAG2 | 3.366707267 | 0.114452129 | 2.242889143 | 0.304089967 | 5.60959641 | 3.327711234 | 8.414402626 | 2 | 10.21617036 | 10.2188359 | 9.256785236 | 8.922624134 |
| ENSG00000165724 | ZMYND19 | 2.449623303 | 0.150422971 | 3.157641846 | 0.052769782 | 5.607265149 | 3.327711234 | 6.602424674 | 2 | 7.593372273 | 7.53675633 | 8.469076579 | 8.018893446 |
| ENSG00000166855 | CLPX | 2.91358492 | 0.221769007 | 2.691354644 | 0.255544364 | 5.604939564 | 2 | 7.669933291 | 2 | 9.006698952 | 8.97890371 | 8.865577373 | 8.672989404 |
| ENSG00000155313 | USP25 | 2.301322347 | 0.27254373 | 3.30165937 | 0.096358916 | 5.602981716 | 2 | 7.901717006 | 3.369317972 | 8.752828065 | 8.5575471 | 9.879446421 | 9.381010503 |
| ENSG00000120049 | KCNIP2 | 2.815147938 | 0.129045271 | 2.787000424 | 0.125279255 | 5.602148362 | 3.327711234 | 2 | 2 | 4.17310023 | 4.16043394 | 4.155805465 | 4.133950982 |
| ENSG00000107331 | ABCA2 | 2.05885584 | 0.16074342 | 3.541646658 | 0.009371668 | 5.600502498 | 3.327711234 | 6.844429755 | 8.460932916 | 9.286870873 | 9.39487575 | 10.89467473 | 10.73901752 |
| ENSG00000214753 | HNRNPUL2 | 3.353748635 | 0.001883357 | 2.246698126 | 0.045247931 | 5.600446761 | 7.53297911 | 7.761270542 | 4.058998049 | 10.45090952 | 10.4465287 | 9.237154294 | 9.44799795 |
| ENSG00000074054 | CLASP1 | 2.929962661 | 1.20E-11 | 2.669971733 | 6.91E-10 | 5.599934395 | 7.579279349 | 6.602424674 | 7.285536013 | 10.09449002 | 10.0842356 | 9.948159052 | 9.702160718 |
| ENSG00000155085 | AK9 | 2.27937321 | 0.036303182 | 3.319362898 | 0.001236986 | 5.598736109 | 4.00722531 | 3.686303414 | 2 | 5.210797711 | 5.36298494 | 6.234425698 | 6.267475858 |
| ENSG00000184524 | CEND1 | 2.691435164 | 0.301726535 | 2.907140995 | 0.249167678 | 5.598576159 | 2 | 2 | 2 | 3.615725078 | 3.15372745 | 3.316666208 | 3.735548302 |
| ENSG00000170802 | FOXN2 | 3.273723562 | 2.66E-06 | 2.323939382 | 0.001196943 | 5.597662944 | 3.327711234 | 5.303523747 | 5.772716771 | 8.270263574 | 8.20711886 | 7.358152527 | 7.253757517 |
| ENSG00000023516 | AKAP11 | 2.661252011 | 0.000258801 | 2.934779274 | 4.45E-05 | 5.596031285 | 8.257427002 | 6.228547981 | 7.342949392 | 10.12735113 | 10.1233123 | 10.58442118 | 10.18375829 |
| ENSG00000049656 | CLPTM1L | 3.448094924 | 0.001384172 | 2.146982464 | 0.057294878 | 5.595077388 | 6.257048221 | 8.213429689 | 4.874587708 | 10.49161431 | 10.4914114 | 8.956549551 | 9.401760658 |
| ENSG00000175137 | SH3BP5L | 2.413317257 | 0.190541209 | 3.179306781 | 0.071210753 | 5.592624037 | 2 | 5.72254323 | 8.434763955 | 9.413697701 | 9.47155148 | 10.05550524 | 10.34130817 |
| ENSG00000184206 | GOLGA6L4 | 6.62100781 | 4.08E-06 | -1.030000294 | 0.569078707 | 5.591007516 | 2 | 2 | 3.369317972 | 7.650823203 | 7.62593544 | 2.34604626 | 2.277179783 |
| ENSG00000133063 | CHIT1 | -3.753087267 | 0.350187134 | 9.344072956 | 0.003068505 | 5.590985689 | 2 | 2 | 2 | 2 | 2 | 10.35403689 | 4.522269241 |
| ENSG00000151327 | FAM177A1 | 2.729347674 | 0.000546124 | 2.860647245 | 0.000252904 | 5.589994918 | 5.330594134 | 7.701025996 | 6.718950348 | 9.590237441 | 9.58096568 | 9.641172698 | 9.787626348 |
| ENSG00000145675 | PIK3R1 | 2.976007056 | 0.0296999 | 2.613292597 | 0.057302601 | 5.589299653 | 2 | 6.602424674 | 4.874587708 | 8.360030171 | 8.29788471 | 8.207201861 | 7.690021882 |
| ENSG00000088836 | SLC4A11 | 2.560743749 | 0.103124682 | 3.02808097 | 0.045706083 | 5.588824719 | 2 | 2 | 4.523675391 | 5.022023783 | 5.45984938 | 5.586442682 | 5.772630506 |
| ENSG00000141084 | RANBP10 | 2.931205783 | 0.106775299 | 2.65750796 | 0.140749923 | 5.588713743 | 2 | 7.099070191 | 3.369317972 | 8.486665614 | 8.5323134 | 8.278834759 | 8.198566699 |
| ENSG00000006534 | ALDH3B1 | 3.414418268 | 0.000735288 | 2.17214124 | 0.040046791 | 5.586559508 | 2 | 5.72254323 | 4.874587708 | 8.017874397 | 8.20112674 | 7.039353222 | 6.741885308 |
| ENSG00000134198 | TSPAN2 | 0.184200165 | 0.938887874 | 5.401221162 | 0.002674594 | 5.585421326 | 3.327711234 | 2 | 2 | 2.3892346 | 2.86123014 | 6.989444349 | 5.601925227 |
| ENSG00000169118 | CSNK1G1 | 3.251719729 | 1.30E-12 | 2.332255394 | 5.63E-07 | 5.583975123 | 5.866944937 | 6.78762695 | 6.322058544 | 9.58946437 | 9.58250309 | 8.845739021 | 8.476969394 |
| ENSG00000264522 | OTUD7B | 2.910478405 | 0.036655526 | 2.672457496 | 0.054639343 | 5.582935901 | 2 | 6.844429755 | 7.029880454 | 9.167628414 | 9.29150981 | 8.947072423 | 9.040594787 |
| ENSG00000175471 | MCTP1 | 4.841905864 | 0.025688178 | 0.740958587 | 0.774520328 | 5.582864451 | 2 | 2 | 3.369317972 | 6.01571348 | 5.83486613 | 3.336159116 | 2.172608095 |
| ENSG00000138119 | MYOF | 1.892786721 | 3.82E-05 | 3.688880545 | 4.02E-17 | 5.581667265 | 9.423197387 | 9.415325457 | 10.17589429 | 11.58461153 | 11.6216511 | 13.08110281 | 13.65771468 |
| ENSG00000182472 | CAPN12 | 3.081791088 | 0.210222627 | 2.499783037 | 0.308943128 | 5.581574126 | 2 | 2 | 2 | 3.663521627 | 3.65239003 | 3.235948948 | 3.33635788 |
| ENSG00000010671 | BTK | 2.733626332 | 0.337258691 | 2.846344344 | 0.303980057 | 5.579970676 | 2 | 2 | 2 | 3.406930997 | 3.45167086 | 3.993419637 | 2.746601469 |
| ENSG00000144283 | PKP4 | 1.305403598 | 0.524942496 | 4.27378479 | 0.016877206 | 5.579188389 | 4.00722531 | 7.847167909 | 2 | 7.794595599 | 7.51115308 | 10.43423541 | 10.75424271 |
| ENSG00000101096 | NFATC2 | 3.172165154 | 0.101896165 | 2.405818139 | 0.219476281 | 5.577983293 | 2 | 3.076665209 | 2 | 4.670864177 | 4.1930041 | 4.133696865 | 3.518055793 |
| ENSG00000165312 | OTUD1 | 2.074448594 | 0.157322534 | 3.503336984 | 0.010210539 | 5.577785578 | 2 | 5.838937808 | 3.369317972 | 6.610808945 | 6.42052719 | 8.166267073 | 7.592058619 |
| ENSG00000096717 | SIRT1 | 2.815860417 | 0.053222658 | 2.758288345 | 0.055572955 | 5.574148762 | 3.327711234 | 6.844429755 | 3.369317972 | 8.227176542 | 8.23671175 | 8.492068113 | 7.768360547 |
| ENSG00000114346 | ECT2 | 3.291901158 | 1.42E-10 | 2.277617201 | 1.44E-05 | 5.569518359 | 5.095959458 | 5.838937808 | 5.772716771 | 8.909699622 | 8.68609779 | 7.985196707 | 7.589522245 |
| ENSG00000099331 | MYO9B | 2.879869086 | 0.001137138 | 2.685649506 | 0.002488036 | 5.565518592 | 6.009025997 | 8.433041974 | 8.721235202 | 10.94938761 | 11.0071139 | 10.8686618 | 10.69560027 |
| ENSG00000206559 | ZCWPW2 | 4.09951821 | 0.05993045 | 1.464982238 | 0.540166698 | 5.564500448 | 2 | 2 | 2 | 4.523237872 | 4.31639476 | 2.717032197 | 2.817676617 |
| ENSG00000023287 | RB1CC1 | 2.702032789 | 0.000856796 | 2.856626364 | 0.000366438 | 5.558659153 | 6.468621203 | 8.607278467 | 7.225742779 | 10.40551335 | 10.3936516 | 10.83982598 | 10.19652354 |
| ENSG00000171368 | TPPP | 0.495926608 | 0.854188222 | 5.061578326 | 0.013923728 | 5.557504933 | 2 | 2 | 2 | 2.3892346 | 2.49394783 | 5.677130623 | 4.634793542 |
| ENSG00000100221 | JOSD1 | 2.200952461 | 0.086717068 | 3.354612336 | 0.00554415 | 5.555564797 | 4.815609471 | 8.607278467 | 6.203069917 | 9.520570052 | 9.56082893 | 10.53914534 | 10.82697164 |
| ENSG00000159403 | C1R | 3.618032772 | 1.56E-06 | 1.934602302 | 0.015816392 | 5.552635074 | 7.940205782 | 7.874700259 | 9.658943166 | 12.32698367 | 12.3696559 | 10.38170786 | 10.90592582 |
| ENSG00000115165 | CYTIP | 2.897844812 | 0.330348729 | 2.653856994 | 0.366255977 | 5.551701805 | 2 | 2 | 2 | 3.406930997 | 3.65239003 | 3.917810786 | 2.509591799 |
| ENSG00000121060 | TRIM25 | 3.613433879 | 0.001384172 | 1.938050801 | 0.106833944 | 5.55148468 | 8.928409675 | 6.04685525 | 5.930865648 | 11.23367077 | 11.289066 | 9.532099314 | 9.648340472 |
| ENSG00000166170 | BAG5 | 2.496921937 | 0.121290859 | 3.052724042 | 0.049497365 | 5.549645979 | 4.00722531 | 7.790474954 | 3.369317972 | 8.81895723 | 8.82894823 | 9.276467981 | 9.468503442 |
| ENSG00000249751 | ECSCR | 1.500040599 | 0.586949095 | 4.048838889 | 0.094671932 | 5.548879488 | 2 | 2 | 2 | 3.023133209 | 2.49394783 | 3.891700877 | 4.74829783 |
| ENSG00000063660 | GPC1 | 1.653900322 | 0.238657623 | 3.892539811 | 0.00205263 | 5.546440133 | 5.095959458 | 9.532315021 | 9.809100877 | 10.7322922 | 10.805643 | 12.97729211 | 13.03308733 |
| ENSG00000013563 | DNASE1L1 | 3.425892114 | 0.011026701 | 2.118529155 | 0.131976284 | 5.544421268 | 2 | 6.389896633 | 7.029880454 | 9.497678564 | 9.57942664 | 7.95714469 | 8.481764443 |
| ENSG00000196367 | TRRAP | 2.727396513 | 4.03E-15 | 2.81614 | 3.01E-16 | 5.543536513 | 6.816637977 | 7.467835385 | 7.225742779 | 9.875425464 | 9.92793329 | 9.947565107 | 10.03183938 |
| ENSG00000173230 | GOLGB1 | 2.577932112 | NA | 2.964064584 | NA | 5.541996696 | 4.815609471 | 10.38739569 | 5.39244166 | 11.4505505 | 11.4421327 | 11.91327231 | 11.74582436 |
| ENSG00000170835 | CEL | 4.163121868 | 0.084156162 | 1.378753002 | 0.603818931 | 5.541874871 | 2 | 2 | 2 | 4.139692846 | 4.74762503 | 2.987421008 | 2.421067459 |
| ENSG00000146122 | DAAM2 | -1.12503576 | 0.44518671 | 6.662929251 | 2.74E-09 | 5.537893491 | 3.327711234 | 3.686303414 | 2 | 2.600425781 | 2.77779907 | 9.471416419 | 8.594208165 |
| ENSG00000166548 | TK2 | 2.945557797 | 0.03177678 | 2.59025197 | 0.059960937 | 5.535809767 | 6.816637977 | 7.051611315 | 2 | 9.244284335 | 9.29620212 | 8.917453109 | 8.918082759 |
| ENSG00000130558 | OLFM1 | 2.610979635 | 0.170726471 | 2.924434427 | 0.111958462 | 5.535414063 | 3.327711234 | 2 | 2 | 4.105493497 | 3.90918929 | 4.088436536 | 4.400220394 |
| ENSG00000062524 | LTK | 1.692295777 | 0.524785881 | 3.840343301 | 0.104270596 | 5.532639078 | 2 | 2 | 2 | 3.09461492 | 2.59489879 | 4.122514202 | 4.30448772 |
| ENSG00000174437 | ATP2A2 | 2.703154829 | 2.27E-10 | 2.828553095 | 2.59E-11 | 5.531707924 | 8.797441463 | 9.799806309 | 9.301632639 | 12.05638564 | 12.0580365 | 12.09427683 | 12.26563573 |
| ENSG00000198930 | CSAG1 | 0.68600976 | 0.812768397 | 4.84562937 | 0.035498586 | 5.531639129 | 2 | 2 | 2 | 2.270786764 | 2.68924543 | 5.553513572 | 4.292061816 |
| ENSG00000141503 | MINK1 | 2.234302798 | 0.0437622 | 3.296404495 | 0.001696648 | 5.530707292 | 5.532353208 | 8.762721501 | 6.80318718 | 9.782429661 | 9.90722553 | 10.84163854 | 10.96570298 |
| ENSG00000214595 | EML6 | 3.011795918 | 0.238959909 | 2.51754657 | 0.323735478 | 5.529342488 | 2 | 2 | 2 | 3.663521627 | 3.55551834 | 3.534812207 | 3.012034821 |
| ENSG00000108604 | SMARCD2 | 2.80091221 | 0.102386938 | 2.727564534 | 0.106905881 | 5.528476745 | 4.467311984 | 7.467835385 | 2 | 8.716468139 | 8.92781046 | 8.863899583 | 8.633521012 |
| ENSG00000115504 | EHBP1 | 2.997137721 | 0.00405304 | 2.530368381 | 0.017000808 | 5.527506102 | 4.00722531 | 6.951736442 | 7.342949392 | 9.622337941 | 9.58250309 | 9.507594406 | 8.641501544 |
| ENSG00000185499 | MUC1 | 1.947067754 | 0.20711259 | 3.57924467 | 0.011950999 | 5.526312424 | 2 | 5.838937808 | 3.369317972 | 6.314087501 | 6.46787796 | 7.310906894 | 8.432373803 |
| ENSG00000162595 | DIRAS3 | 6.27082008 | 0.000120485 | -0.745807928 | 0.725350511 | 5.525012152 | 2 | 2 | 2 | 6.401247806 | 6.33555364 | 2.183373728 | 2.225841198 |
| ENSG00000152413 | HOMER1 | 3.164063129 | 3.57E-05 | 2.36094193 | 0.002637744 | 5.525005059 | 3.327711234 | 5.838937808 | 5.156626401 | 8.113468729 | 8.19711812 | 7.24282551 | 7.482861322 |
| ENSG00000213445 | SIPA1 | 2.447889533 | 0.013097086 | 3.076793723 | 0.001269607 | 5.524683256 | 3.327711234 | 6.602424674 | 5.772716771 | 8.16211395 | 8.11451391 | 8.636638396 | 8.87395086 |
| ENSG00000173744 | AGFG1 | 2.726985891 | 0.024500134 | 2.797686515 | 0.020011794 | 5.524672406 | 3.327711234 | 7.315436877 | 7.502634626 | 9.548682898 | 9.56394525 | 9.633561473 | 9.619726259 |
| ENSG00000130449 | ZSWIM6 | 3.20407876 | 0.010286652 | 2.318534936 | 0.071123944 | 5.522613696 | 2 | 6.535001748 | 5.39244166 | 8.610716699 | 8.68036596 | 7.848504194 | 7.692387724 |
| ENSG00000173621 | LRFN4 | 2.2220966 | 0.194855335 | 3.298720075 | 0.041382233 | 5.520816675 | 2 | 6.602424674 | 3.369317972 | 7.437142666 | 7.24992376 | 8.4151065 | 8.393643421 |
| ENSG00000041515 | MYO16 | 0.68785149 | 0.790662721 | 4.831893387 | 0.018228014 | 5.519744877 | 2 | 2 | 2 | 2.3892346 | 2.59489879 | 4.778616857 | 5.269393099 |
| ENSG00000070950 | RAD18 | 1.977772244 | 0.156896095 | 3.541644188 | 0.006021251 | 5.519416433 | 3.327711234 | 5.72254323 | 2 | 6.252910529 | 6.39275508 | 7.945293431 | 7.726263462 |
| ENSG00000141384 | TAF4B | 2.372126109 | 0.103759715 | 3.145380685 | 0.023910744 | 5.517506794 | 2 | 4.710564154 | 2 | 5.37770462 | 5.70538658 | 6.344396176 | 6.195912571 |
| ENSG00000164236 | ANKRD33B | 1.226571436 | 0.499380069 | 4.288543146 | 0.006284419 | 5.515114581 | 3.327711234 | 6.602424674 | 2 | 6.358325776 | 6.40670796 | 8.964399877 | 9.706263779 |

| | | | | | | | | | | | | | |
|---|---|---|---|---|---|---|---|---|---|---|---|---|---|
| ENSG00000085063 | CD59 | 2.894523491 | 1.91E-30 | 2.619480507 | 2.46E-25 | 5.514003997 | 10.81695923 | 10.72273565 | 10.64969256 | 13.60617143 | 13.6397447 | 13.14923767 | 13.52287338 |
| ENSG00000137413 | TAF8 | 2.935440768 | 2.00E-06 | 2.578471146 | 3.35E-05 | 5.513911915 | 6.009025997 | 4.710564154 | 5.156626401 | 8.229163214 | 8.1187584 | 8.035995406 | 7.574209638 |
| ENSG00000213859 | KCTD11 | 2.233791027 | 0.012308962 | 3.275708455 | 0.000117075 | 5.509499482 | 4.815609471 | 7.315436877 | 6.073378139 | 8.556367783 | 8.66303256 | 9.308892875 | 9.917424452 |
| ENSG00000185010 | F8 | 2.769220555 | 0.091117194 | 2.735738088 | 0.090145733 | 5.504958643 | 2 | 5.303523747 | 2 | 6.503277904 | 6.49425375 | 6.558682816 | 6.368575325 |
| ENSG00000196873 | CBWD3 | 7.927594045 | 2.69E-08 | -2.425390044 | 0.176769831 | 5.502204001 | 2 | 3.076665209 | 2 | 8.869506572 | 8.98356585 | 2.183373728 | 2.059860178 |
| ENSG00000127481 | UBR4 | 2.986241918 | 0.032193991 | 2.514522851 | 0.073848527 | 5.500764768 | 4.815609471 | 9.988044813 | 9.658943166 | 12.21465954 | 12.2824838 | 11.92080638 | 11.61880819 |
| ENSG00000058404 | CAMK2B | 2.333025254 | 0.366470827 | 3.167661276 | 0.193719441 | 5.50068653 | 2 | 2 | 2 | 3.162721474 | 3.2184422 | 3.891700877 | 3.518055793 |
| ENSG00000163815 | CLEC3B | 0.858631528 | 0.757769467 | 4.641014625 | 0.042085537 | 5.499646152 | 2 | 2 | 2 | 2.498691189 | 2.59489879 | 3.956110469 | 5.436777225 |
| ENSG00000127526 | SLC35E1 | 2.263278319 | 0.027309548 | 3.234385761 | 0.000941358 | 5.49766408 | 6.366709671 | 8.834582582 | 6.43197674 | 10.01265873 | 9.89614107 | 10.80593075 | 11.03281937 |
| ENSG00000164620 | RELL2 | 3.217101944 | 0.02572938 | 2.278994294 | 0.124033936 | 5.496096238 | 2 | 4.93638933 | 2 | 6.529309835 | 6.60731796 | 5.642319755 | 5.727202496 |
| ENSG00000133789 | SWAP70 | 2.394907539 | 0.031675145 | 3.100361006 | 0.003875548 | 5.495268545 | 5.532353208 | 8.004992147 | 5.156626401 | 9.176921588 | 9.20432875 | 9.634299801 | 10.11112449 |
| ENSG00000150712 | MTMR12 | 2.220415018 | 1.69E-05 | 3.270168357 | 5.48E-11 | 5.490583375 | 5.095959458 | 5.72254323 | 6.43197674 | 7.966401501 | 7.99029569 | 9.012519675 | 9.020021944 |
| ENSG00000137101 | CD72 | 2.813760808 | 0.167895393 | 2.676598198 | 0.181647286 | 5.490359007 | 3.327711234 | 2 | 2 | 3.95996233 | 4.3456612 | 4.45225155 | 3.518055793 |
| ENSG00000140948 | ZCCHC14 | 2.427148232 | 0.111655472 | 3.062435017 | 0.036695546 | 5.489583249 | 4.00722531 | 7.467835385 | 3.369317972 | 8.432517176 | 8.49198035 | 9.214621994 | 8.957957965 |
| ENSG00000123384 | LRP1 | 2.474409639 | 1.43E-06 | 3.014481284 | 2.38E-09 | 5.488890924 | 11.86992646 | 11.30403016 | 10.60345421 | 13.78321995 | 13.8597565 | 14.21286338 | 14.49704798 |
| ENSG00000033178 | UBA6 | 3.66686899 | 0.015236142 | 1.821562934 | 0.258826994 | 5.488431925 | 5.095959458 | 8.317412869 | 3.369317972 | 10.54761388 | 10.5628565 | 8.812201585 | 8.621156416 |
| ENSG00000155304 | HSPA13 | 2.960225964 | 0.106055103 | 2.528113709 | 0.167157761 | 5.488339674 | 5.866944937 | 8.623598428 | 2 | 10.21266037 | 10.1551727 | 9.961556635 | 9.51100768 |
| ENSG00000110651 | CD81 | 2.570449212 | 2.84E-06 | 2.916949514 | 7.42E-08 | 5.487398726 | 9.233563228 | 10.51097332 | 10.40208303 | 12.67030265 | 12.7675965 | 12.83409244 | 13.26589377 |
| ENSG00000147050 | KDM6A | 3.382059104 | 0.007464915 | 2.102685481 | 0.111330307 | 5.484744585 | 4.815609471 | 7.274681896 | 3.369317972 | 9.372049302 | 9.2990102 | 8.1689872 | 7.962181894 |
| ENSG00000108187 | PBLD | 2.745242651 | 0.017140458 | 2.735991764 | 0.016815901 | 5.481234415 | 2 | 4.93638933 | 3.369317972 | 6.358325776 | 6.42052719 | 6.043083161 | 6.654794245 |
| ENSG00000184060 | ADAP2 | 0.312646556 | 0.886720338 | 5.16646026 | 0.001313772 | 5.479106815 | 2 | 2 | 2 | 2.695456494 | 2.68924543 | 6.505870593 | 5.896817464 |
| ENSG00000058056 | USP13 | 2.290625527 | 9.84E-05 | 3.188297763 | 2.16E-08 | 5.47892329 | 5.330594134 | 6.899080544 | 6.203069917 | 8.481674661 | 8.59611189 | 9.314124483 | 9.538225362 |
| ENSG00000177363 | LRRN4CL | 0.564937038 | 0.666023939 | 4.910062942 | 1.62E-06 | 5.47499998 | 4.467311984 | 3.076665209 | 3.369317972 | 4.070463721 | 4.22485516 | 7.422256313 | 8.62735196 |
| ENSG00000168769 | TET2 | 2.963479293 | 0.019862551 | 2.510613623 | 0.050724086 | 5.474092916 | 6.73718411 | 5.838937808 | 2 | 8.693628619 | 8.66883353 | 8.411091064 | 8.031095054 |
| ENSG00000164713 | BRI3 | 3.010812637 | 3.54E-11 | 2.462338596 | 8.03E-08 | 5.473151233 | 7.27616306 | 8.376385542 | 7.934808822 | 10.84993128 | 11.0071139 | 10.26530837 | 10.49286637 |
| ENSG00000051596 | THOC3 | 3.747204424 | 4.56E-08 | 1.719646309 | 0.020018787 | 5.466850733 | 4.467311984 | 4.93638933 | 2 | 7.908072101 | 7.95512324 | 6.078235975 | 5.863612864 |
| ENSG00000134109 | EDEM1 | 3.394792868 | 1.74E-24 | 2.070960513 | 8.66E-10 | 5.46575338 | 7.031698319 | 7.099070191 | 6.958202216 | 10.36045596 | 10.4179663 | 8.935537205 | 9.197066081 |
| ENSG00000162704 | ARPC5 | 2.522091804 | 3.19E-08 | 2.941562621 | 6.63E-11 | 5.463654425 | 8.468948322 | 9.114109157 | 8.076950877 | 11.18486086 | 11.0853755 | 11.40802991 | 11.68801764 |
| ENSG00000104833 | TUBB4A | 5.142499084 | 0.000173358 | 0.32091598 | 0.856270394 | 5.463415064 | 3.327711234 | 3.076665209 | 2 | 7.012547947 | 7.07311172 | 2.803598391 | 3.33635788 |
| ENSG00000133135 | RNF128 | 5.173308602 | 0.044742745 | 0.288439355 | 0.929171616 | 5.461747957 | 2 | 2 | 2 | 5.363304456 | 5.33407135 | 2.656294034 | 2.059860178 |
| ENSG00000143126 | CELSR2 | 0.590709385 | 0.713873535 | 4.870684933 | 0.000158892 | 5.461394317 | 2 | 4.442710129 | 6.43197674 | 6.139181293 | 5.19627118 | 9.769486116 | 10.05155464 |
| ENSG00000073417 | PDE8A | 3.320752865 | 0.008045241 | 2.140189903 | 0.100850653 | 5.460942768 | 3.327711234 | 7.538394143 | 5.59507254 | 9.604847628 | 9.63900396 | 8.550603056 | 8.343344553 |
| ENSG00000108239 | TBC1D12 | 2.158997807 | 0.135211488 | 3.301322654 | 0.015400933 | 5.46032046 | 2 | 6.535001748 | 4.523675391 | 7.454228844 | 7.33629421 | 8.604345457 | 8.430249176 |
| ENSG00000197046 | SIGLEC15 | 2.936704802 | 0.184884539 | 2.522271791 | 0.253508506 | 5.458976593 | 2 | 3.076665209 | 2 | 4.205751501 | 4.31639476 | 4.504874511 | 3.012034821 |
| ENSG00000105329 | TGFB1 | 2.47117129 | 0.004754286 | 2.983697162 | 0.000469751 | 5.454868452 | 7.27616306 | 9.018611864 | 9.926432745 | 11.55323749 | 11.5710504 | 11.991084 | 12.15264135 |
| ENSG00000269713 | NBPF9 | 5.787830879 | 1.43E-28 | -0.334827077 | 0.614217941 | 5.453003802 | 6.138360429 | 6.535001748 | 7.552162999 | 12.56776544 | 12.6222308 | 6.497238296 | 6.572418365 |
| ENSG00000101152 | DNAJC5 | 2.281672594 | 0.027042585 | 3.169358695 | 0.001326571 | 5.451031289 | 7.867441304 | 9.246456519 | 6.073378139 | 10.46235554 | 10.5655798 | 11.26526767 | 11.52461319 |
| ENSG00000106686 | SPATA6L | 3.495974669 | 0.143386081 | 1.954433183 | 0.432815399 | 5.450407852 | 2 | 2 | 2 | 3.840289928 | 4.05806467 | 2.937246078 | 3.042026731 |
| ENSG00000147324 | MFHAS1 | 2.649072062 | 0.227801845 | 2.800351543 | 0.191758699 | 5.449423604 | 2 | 6.535001748 | 2 | 7.641902724 | 7.5462416 | 8.348878903 | 6.678855578 |
| ENSG00000078747 | ITCH | 2.573812601 | 2.92E-06 | 2.874033382 | 1.25E-07 | 5.447845982 | 6.366709671 | 7.669933291 | 7.225742779 | 9.723047898 | 9.74919792 | 10.28753805 | 9.728912497 |
| ENSG00000174197 | MGA | 2.907059644 | 0.000344331 | 2.538695432 | 0.001976772 | 5.445755076 | 6.138360429 | 7.847167909 | 5.930865648 | 9.734282329 | 9.86046766 | 9.724430319 | 9.065448786 |
| ENSG00000169896 | ITGAM | 0.274454513 | 0.927650433 | 5.170231574 | 0.024765019 | 5.444686087 | 2 | 2 | 2 | 2.3892346 | 2.38539787 | 6.040114727 | 3.956354764 |
| ENSG00000099994 | SUSD2 | 0.604850471 | 0.611488116 | 4.839308517 | 2.35E-07 | 5.444158988 | 4.00722531 | 2 | 4.523675391 | 4.358765249 | 4.02226117 | 8.010440524 | 8.187707412 |
| ENSG00000142233 | NTN5 | 1.150943712 | 0.701695664 | 4.291782511 | 0.093960346 | 5.442726223 | 2 | 2 | 2 | 2.784612485 | 2.49394783 | 5.233158834 | 3.36036153 |
| ENSG00000105219 | CNTD2 | 2.441176317 | 0.367367297 | 3.00028854 | 0.246966535 | 5.441464857 | 2 | 2 | 2 | 2.868578001 | 3.55551834 | 3.500668998 | 3.698365557 |
| ENSG00000133138 | TBC1D8B | 2.659459529 | 0.152145189 | 2.781650241 | 0.125418384 | 5.44110977 | 3.327711234 | 7.145017454 | 2 | 8.427334995 | 8.12087597 | 8.44402928 | 8.360554398 |
| ENSG00000115935 | WIPF1 | 2.96104282 | 0.025780112 | 2.475491403 | 0.065354655 | 5.436534222 | 3.327711234 | 7.874700259 | 8.042706615 | 10.34771157 | 10.3278045 | 10.0527464 | 9.623450463 |
| ENSG00000221968 | FADS3 | 2.073721174 | 0.000262222 | 3.361482086 | 6.91E-10 | 5.43520326 | 7.867441304 | 6.951736442 | 8.380957451 | 9.874156428 | 9.90538043 | 10.99733953 | 11.33141801 |
| ENSG00000274180 | NATD1 | 1.506649228 | 0.101137093 | 3.928282252 | 1.61E-06 | 5.43493148 | 5.095959458 | 7.189546432 | 5.772716771 | 7.821180652 | 7.72670378 | 10.57470991 | 9.619415474 |
| ENSG00000162851 | TFB2M | 2.983436939 | 0.020848047 | 2.450372458 | 0.060842073 | 5.433809397 | 2 | 5.595927442 | 3.369317972 | 7.202487431 | 7.17037177 | 6.708295923 | 6.632790505 |
| ENSG00000176383 | B3GNT4 | 2.497388939 | 0.342054331 | 2.930208374 | 0.246132276 | 5.427597313 | 2 | 2 | 2 | 3.349643765 | 3.2184422 | 3.782262616 | 3.287118176 |
| ENSG00000151067 | CACNA1C | 1.67392066 | 0.004986804 | 3.751369507 | 9.98E-12 | 5.425290167 | 4.815609471 | 6.04685525 | 5.156626401 | 7.048994596 | 7.08183429 | 9.233585684 | 8.974090715 |
| ENSG00000075292 | ZNF638 | 2.882194767 | 0.001788082 | 2.5412719 | 0.006389948 | 5.423466667 | 5.330594134 | 8.234834549 | 7.451345288 | 10.30737895 | 10.2707772 | 10.15969431 | 9.703334212 |
| ENSG00000117868 | ESYT2 | 1.904272384 | 0.086760878 | 3.516755737 | 0.000593789 | 5.421028121 | 5.532353208 | 8.376385542 | 9.049077186 | 10.11773201 | 10.1463641 | 11.73470634 | 11.74753181 |
| ENSG00000071127 | WDR1 | 1.49150699 | 4.72E-05 | 3.928974218 | 7.00E-30 | 5.420481208 | 10.34051281 | 9.896993485 | 9.849281147 | 11.52359623 | 11.5434907 | 13.69695759 | 14.19857858 |
| ENSG00000170579 | DLGAP1 | 0.28191919 | 0.945086484 | 5.138339023 | 0.122752719 | 5.420258213 | 2 | 2 | 2 | 2.695456494 | 2 | 4.881704255 | 5.651404445 |
| ENSG00000100399 | CHADL | 2.011409107 | 0.437765112 | 3.405585524 | 0.151541851 | 5.41699463 | 2 | 2 | 2 | 3.023133209 | 3.01487888 | 3.796405078 | 3.972018587 |
| ENSG00000177051 | FBXO46 | 1.882593899 | 0.046185884 | 3.532565859 | 4.87E-05 | 5.415159757 | 4.815609471 | 5.457120953 | 2 | 6.456560224 | 6.42052719 | 8.038228667 | 8.046896864 |
| ENSG00000137462 | TLR2 | 1.597880247 | 0.562650016 | 3.815463293 | 0.12122066 | 5.41334354 | 2 | 2 | 2 | 2.947924294 | 2.68924543 | 4.636484163 | 3.560117069 |
| ENSG00000164715 | LMTK2 | 2.264274195 | 0.213542763 | 3.147651539 | 0.069071834 | 5.411925735 | 2 | 4.11352228 | 7.451345288 | 8.211183901 | 8.21705076 | 8.978035992 | 9.191222844 |
| ENSG00000181804 | SLC9A9 | 3.812646118 | 0.007212171 | 1.599206258 | 0.299320086 | 5.411852376 | 2 | 4.93638933 | 2 | 7.136275664 | 7.14977896 | 4.771475226 | 5.319231103 |
| ENSG00000143067 | ZNF697 | 4.18292092 | 0.008608975 | 1.228315036 | 0.492372773 | 5.411235956 | 3.327711234 | 6.389896633 | 2 | 9.081180848 | 9.06939887 | 6.730562686 | 5.319231103 |
| ENSG00000124151 | NCOA3 | 2.670558104 | 0.0056444 | 2.739494593 | 0.004207821 | 5.410052696 | 5.095959458 | 7.954278496 | 8.076950877 | 10.12788365 | 10.2123959 | 10.30124027 | 10.17490298 |
| ENSG00000101938 | CHRDL1 | 2.148910113 | 0.448379821 | 3.258585084 | 0.216645944 | 5.407495196 | 2 | 2 | 2 | 3.09461492 | 3.08597292 | 4.240999983 | 3.100202476 |
| ENSG00000175662 | TOM1L2 | 1.861094751 | 0.299070637 | 3.545303627 | 0.031372002 | 5.406398378 | 2 | 8.451443573 | 6.80318718 | 9.125585405 | 9.11047337 | 10.64294545 | 10.9326885 |
| ENSG00000100346 | CACNA1I | 4.130992283 | 0.192660633 | 1.273345313 | 0.715242599 | 5.404337596 | 2 | 2 | 2 | 4.237680128 | 4.63360314 | 3.12825509 | 2.059860178 |
| ENSG00000156103 | MMP16 | 2.669819916 | 0.004515748 | 2.73225631 | 0.003390575 | 5.402076226 | 5.866944937 | 5.72254323 | 3.369317972 | 7.883054517 | 7.92635454 | 8.43671169 | 7.263337153 |
| ENSG00000151553 | FAM160B1 | 2.444538204 | 0.335980269 | 2.953856332 | 0.227974555 | 5.398394536 | 2 | 8.101365653 | 2 | 8.971529165 | 8.94703391 | 9.543411506 | 9.382476431 |
| ENSG00000116127 | ALMS1 | 2.311372033 | 0.327520219 | 3.086440291 | 0.170499056 | 5.397812324 | 2 | 7.35507212 | 2 | 8.045195239 | 8.1187584 | 8.728011488 | 8.959923085 |
| ENSG00000176208 | ATAD5 | 3.452476183 | 0.016039408 | 1.941588122 | 0.198118401 | 5.394064305 | 3.327711234 | 3.076665209 | 2 | 5.540148523 | 5.34860058 | 4.122514202 | 4.201948172 |
| ENSG00000162419 | GMEB1 | 2.284703441 | 0.000196909 | 3.10778202 | 1.58E-07 | 5.392485462 | 4.467311984 | 5.131603625 | 4.523675391 | 6.825617242 | 6.94032357 | 7.656405908 | 7.714671806 |
| ENSG00000115183 | TANC1 | 2.133923372 | 0.075173015 | 3.253501015 | 0.003975743 | 5.387424387 | 4.815609471 | 5.303523747 | 7.934808822 | 8.817637515 | 8.74357228 | 9.907203225 | 9.879774129 |
| ENSG00000166145 | SPINT1 | 1.665308784 | 0.300950806 | 3.721006907 | 0.009972043 | 5.386315691 | 4.467311984 | 2 | 2 | 4.329433326 | 4.58534546 | 6.535683256 | 6.029985803 |
| ENSG00000166479 | TMX3 | 3.054108486 | 0.057349984 | 2.327156174 | 0.153080856 | 5.381264661 | 2 | 8.029695761 | 7.818184403 | 10.40814626 | 10.3509915 | 9.592613586 | 9.71618004 |
| ENSG00000133059 | DSTYK | 2.338574328 | 0.035431212 | 3.041203607 | 0.004478253 | 5.379777935 | 4.815609471 | 7.847167909 | 5.59507254 | 9.011324139 | 8.9648262 | 9.77953276 | 9.587040651 |
| ENSG00000008118 | CAMK1G | 4.390214635 | 0.049461281 | 0.989073682 | 0.699544446 | 5.379288317 | 2 | 2 | 2 | 4.71689672 | 4.60967607 | 2.775316226 | 2.374681325 |
| ENSG00000116641 | DOCK7 | 2.618764995 | 0.075503547 | 2.75983479 | 0.0559313 | 5.378599785 | 6.816637977 | 8.623598428 | 3.369317972 | 9.966365455 | 10.0880289 | 9.856639061 | 10.42566819 |
| ENSG00000175216 | CKAP5 | 2.619152529 | 0.01784745 | 2.759438829 | 0.011575145 | 5.378591358 | 5.330594134 | 8.903033433 | 7.451345288 | 10.48248248 | 10.4350864 | 10.7065785 | 10.48248 |
| ENSG00000138071 | ACTR2 | 2.241337435 | 0.057431985 | 3.135089759 | 0.005293456 | 5.376427194 | 5.70932728 | 9.318399851 | 9.491321578 | 11.13672482 | 11.0815752 | 11.92508678 | 12.07492997 |
| ENSG00000205903 | ZNF316 | 2.644182449 | 0.002887069 | 2.730157897 | 0.001920201 | 5.374340346 | 4.00722531 | 6.602424674 | 6.882775709 | 8.842507967 | 8.86350401 | 8.951818769 | 8.924637944 |
| ENSG00000138669 | PRKG2 | 6.852127766 | 4.25E-05 | -1.481166354 | 0.487891953 | 5.370961412 | 2 | 2 | 2 | 6.979883891 | 6.8715633 | 2.183373728 | 2.059860178 |
| ENSG00000063438 | AHRR | 2.716034074 | 0.087305384 | 2.654462099 | 0.090359864 | 5.370496174 | 6.009025997 | 7.761270542 | 2 | 9.268635235 | 9.22613117 | 9.167113444 | 9.20537272 |
| ENSG00000198948 | MFAP3L | 1.615623527 | 0.03679441 | 3.754290265 | 9.32E-08 | 5.369913792 | 6.89194353 | 5.303523747 | 5.39244166 | 7.728718552 | 7.46188679 | 9.98393497 | 9.389054739 |
| ENSG00000176678 | FOXL1 | 1.301760167 | 0.644603834 | 4.064883623 | 0.103371184 | 5.36664379 | 8.418903792 | 2 | 2 | 8.130575473 | 8.14396641 | 10.5006588 | 11.18464037 |
| ENSG00000127920 | GNG11 | 4.412826109 | 1.23E-11 | 0.951807122 | 0.204070923 | 5.364633231 | 7.82963488 | 6.844429755 | 8.721235202 | 12.40440824 | 12.3454117 | 8.810895984 | 9.030812517 |
| ENSG00000158113 | LRRC43 | 2.54282498 | 0.377368801 | 2.818692227 | 0.310621084 | 5.361517207 | 2 | 2 | 2 | 2.695456494 | 3.74316441 | 3.616794016 | 3.33635788 |
| ENSG00000081014 | AP4E1 | 2.547839818 | 6.96E-06 | 2.8113683 | 5.09E-07 | 5.359208118 | 5.330594134 | 6.535001748 | 6.322058544 | 8.710792067 | 8.57772939 | 8.581047186 | 9.172702781 |
| ENSG00000103222 | ABCC1 | 6.139052554 | 6.06E-09 | -0.780909427 | 0.544666992 | 5.358143127 | 2 | 5.946638857 | 6.322058544 | 11.59119151 | 11.6549123 | 4.675266036 | 5.108633494 |
| ENSG00000133858 | ZFC3H1 | 2.933528219 | 0.053285833 | 2.420506489 | 0.113283635 | 5.354034708 | 7.435667355 | 7.638155693 | 2 | 9.897454779 | 9.84004698 | 9.426088142 | 9.287866987 |
| ENSG00000105835 | NAMPT | 5.415931016 | 5.39E-67 | -0.062038928 | 0.888380576 | 5.353892087 | 7.904282243 | 8.147241016 | 7.691296561 | 13.3498354 | 13.3065908 | 8.017249226 | 7.715835172 |
| ENSG00000135709 | KIAA0513 | 1.864996942 | 0.001066135 | 3.488271378 | 8.61E-11 | 5.35326832 | 6.366709671 | 5.131603625 | 6.203069917 | 7.723090567 | 7.7919464 | 9.537108665 | 9.165057331 |
| ENSG00000243896 | OR2A7 | 3.859837642 | 0.089428115 | 1.491394362 | 0.544635534 | 5.351232004 | 2 | 2 | 2 | 4.299492648 | 4.16043394 | 2.775316226 | 2.782576693 |
| ENSG00000157551 | KCNJ15 | 6.758835774 | 2.90E-05 | -1.408714414 | 0.496385053 | 5.35012136 | 2 | 2 | 2 | 6.941624891 | 6.72335419 | 2.094598369 | 2.172608095 |
| ENSG00000089902 | RCOR1 | 2.126052346 | 0.192916453 | 3.222256071 | 0.036232528 | 5.348308417 | 2 | 7.669933291 | 5.930865648 | 8.600009476 | 8.54340749 | 9.457577544 | 9.837645613 |
| ENSG00000149506 | ZP1 | 3.211653393 | 0.203854133 | 2.134262012 | 0.409012973 | 5.345915405 | 2 | 2 | 2 | 3.566290607 | 3.90918929 | 2.911488689 | 3.236138336 |
| ENSG00000104228 | TRIM35 | 2.641820563 | 0.040404199 | 2.699941311 | 0.033924497 | 5.341761874 | 5.866944937 | 6.602424674 | 2 | 8.304583895 | 8.29975519 | 8.643500187 | 8.005636809 |
| ENSG00000065970 | FOXJ2 | 1.967305803 | 0.174753955 | 3.371913488 | 0.012370521 | 5.339219291 | 2 | 7.051611315 | 6.534109959 | 8.14116509 | 8.20911072 | 9.698306977 | 9.423997212 |
| ENSG00000147234 | FRMPD3 | 1.389352484 | 0.619039988 | 3.946579654 | 0.106137285 | 5.335932138 | 2 | 2 | 2 | 2.695456494 | 2.77779907 | 3.56816602 | 4.784231301 |
| ENSG00000108679 | LGALS3BP | 3.204307339 | 8.47E-05 | 2.131536817 | 0.012057282 | 5.335844156 | 11.29732276 | 8.820493869 | 10.39531398 | 13.66921984 | 13.7104399 | 12.55282788 | 12.67974691 |
| ENSG00000179295 | PTPN11 | 2.276154318 | 1.22E-10 | 3.057731529 | 1.37E-18 | 5.333885847 | 8.16936288 | 8.607278467 | 8.865353883 | 10.83891504 | 10.8388264 | 11.46710344 | 11.75667488 |
| ENSG00000106560 | GIMAP2 | 3.281427504 | 0.098785844 | 2.049186219 | 0.316737831 | 5.330613723 | 3.327711234 | 2 | 2 | 4.415698455 | 4.65713986 | 3.968654369 | 3.128433649 |
| ENSG00000147027 | TMEM47 | 1.931983571 | 0.310383334 | 3.398565477 | 0.054125008 | 5.330549048 | 2 | 7.145017454 | 9.06622548 | 9.779043349 | 9.69834686 | 11.22351878 | 11.17916827 |
| ENSG00000185989 | RASA3 | 2.88278534 | 0.024475005 | 2.44595384 | 0.058553075 | 5.32873918 | 3.327711234 | 7.701025996 | 7.734844356 | 9.989404766 | 10.0451865 | 9.363939866 | 9.772611475 |
| ENSG00000164164 | OTUD4 | 2.694278278 | 0.10892118 | 2.633423721 | 0.111962102 | 5.327702 | 7.096806373 | 8.414402626 | 2 | 10.0321341 | 9.96199598 | 9.996814524 | 9.874314395 |
| ENSG00000102921 | N4BP1 | 3.045738375 | 0.013831602 | 2.281378797 | 0.071976668 | 5.327117172 | 4.00722531 | 8.395519306 | 7.098165388 | 10.36678618 | 10.3800133 | 9.658938866 | 9.563647359 |
| ENSG00000121068 | TBX2 | 3.139403656 | 0.036469923 | 2.186115358 | 0.155557022 | 5.325519014 | 6.138360429 | 7.467835385 | 2 | 9.445659335 | 9.51245165 | 8.91583461 | 8.012280354 |
| ENSG00000164291 | ARSK | 1.532336117 | 0.003324438 | 3.790421694 | 3.01E-15 | 5.322757811 | 5.866944937 | 5.303523747 | 5.772716771 | 7.144720234 | 7.05550676 | 9.497620601 | 9.129269498 |

| | | | | | | | | | | | | | |
|---|---|---|---|---|---|---|---|---|---|---|---|---|---|
| ENSG00000106714 | CNTNAP3 | 2.598455667 | 0.329996554 | 2.723873406 | 0.293334688 | 5.322329073 | 2 | 2 | 2 | 3.566290607 | 3.08597292 | 3.56816602 | 3.261853426 |
| ENSG00000164142 | FAM160A1 | 2.936153342 | 0.256740427 | 2.384628069 | 0.357240472 | 5.32078141 | 2 | 2 | 2 | 3.406930997 | 3.69849104 | 3.035909414 | 3.383972333 |
| ENSG00000198208 | RPS6KL1 | 2.011728946 | 0.32587825 | 3.307893372 | 0.081233304 | 5.319622318 | 2 | 3.076665209 | 2 | 3.406930997 | 3.74316441 | 4.868377487 | 4.175132639 |
| ENSG00000167964 | RAB26 | 3.243214407 | 0.203115525 | 2.072546261 | 0.428771225 | 5.315760668 | 2 | 2 | 2 | 3.663521627 | 3.86943969 | 2.803598391 | 3.261853426 |
| ENSG00000142327 | RNPEPL1 | 1.822070188 | 0.397020048 | 3.492993963 | 0.07747944 | 5.31506415 | 4.00722531 | 8.433041974 | 2 | 8.630393142 | 8.81462916 | 10.33327078 | 10.4387618 |
| ENSG00000117472 | TSPAN1 | 2.442338589 | 0.362156704 | 2.871594719 | 0.264665124 | 5.313933308 | 2 | 2 | 2 | 3.023133209 | 3.45167086 | 3.336159116 | 3.67940853 |
| ENSG00000189420 | ZFP92 | -1.306003541 | 0.649972951 | 6.618768172 | 0.001615626 | 5.31276463 | 2 | 2 | 2 | 2.270786764 | 2 | 6.934553331 | 6.423723772 |
| ENSG00000166128 | RAB8B | 3.70415574 | 0.000375915 | 1.608549897 | 0.155092367 | 5.312705637 | 4.815609471 | 7.431218269 | 4.874587708 | 9.917305676 | 9.92490661 | 8.084345843 | 7.557435494 |
| ENSG00000123080 | CDKN2C | 1.937891283 | 0.211843193 | 3.371092447 | 0.019079173 | 5.308983729 | 2 | 4.442710129 | 2 | 4.927308402 | 4.85329164 | 5.737015686 | 6.543880272 |
| ENSG00000168137 | SETD5 | 2.531668593 | 0.003344383 | 2.776069862 | 0.001089398 | 5.307738455 | 6.366709671 | 8.356994601 | 8.99637432 | 10.7774535 | 10.7784015 | 11.11547411 | 10.92163967 |
| ENSG00000114439 | BBX | 2.50017414 | 0.029907596 | 2.806998161 | 0.01288167 | 5.307172301 | 5.70932728 | 8.337339476 | 9.553445883 | 11.04750286 | 11.0320825 | 11.30568783 | 11.38542086 |
| ENSG00000013364 | MVP | 2.392200974 | 0.225247916 | 2.91052693 | 0.12583058 | 5.302727904 | 9.98400217 | 8.607278467 | 2 | 11.23144255 | 11.2789083 | 11.47598604 | 12.01912506 |
| ENSG00000124249 | KCNK15 | -1.305316762 | 0.681389915 | 6.605989017 | 0.005488246 | 5.300672254 | 2 | 2 | 2 | 2.270786764 | 2 | 7.308441828 | 5.581648407 |
| ENSG00000154065 | ANKRD29 | 4.546029466 | 0.118824374 | 0.753533914 | 0.823417076 | 5.29956338 | 2 | 2 | 2 | 4.574143677 | 4.98931816 | 2.858551861 | 2.059860178 |
| ENSG00000075539 | FRYL | 2.442801492 | 0.071895607 | 2.855436104 | 0.030430098 | 5.298237596 | 3.327711234 | 8.077869079 | 6.718950348 | 9.453327884 | 9.38785872 | 9.703944794 | 9.948684752 |
| ENSG00000107779 | BMPR1A | 2.73529087 | 3.83E-11 | 2.562658539 | 5.74E-10 | 5.297949409 | 6.009025997 | 6.04685525 | 5.772716771 | 8.545247918 | 8.673169 | 8.48662708 | 8.391460941 |
| ENSG00000066248 | NGEF | -0.723936751 | 0.790629813 | 6.019714113 | 0.002211198 | 5.295777362 | 2 | 2 | 2 | 2.270786764 | 2.14022021 | 5.318934974 | 6.64749689 |
| ENSG00000203896 | LIME1 | 3.586325322 | 0.012675026 | 1.709285689 | 0.267993165 | 5.295611011 | 2 | 2 | 4.523675391 | 6.24507773 | 6.14893615 | 4.604690117 | 4.4231929 |
| ENSG00000100242 | SUN2 | 2.467886049 | 0.165072736 | 2.827435083 | 0.100639252 | 5.295321132 | 8.609439662 | 6.602424674 | 2 | 9.767470067 | 9.82128771 | 10.09733848 | 10.20607564 |
| ENSG00000265190 | ANXA8 | 1.690418739 | 0.521219774 | 3.604181959 | 0.125703081 | 5.294600698 | 2 | 2 | 2 | 2.947924294 | 2.77779907 | 3.943456545 | 4.119958326 |
| ENSG00000110422 | HIPK3 | 2.474830727 | 0.003242657 | 2.819178567 | 0.00062762 | 5.294009294 | 7.940205782 | 7.467835385 | 5.39244166 | 9.750280081 | 9.69550802 | 10.24675978 | 9.859390114 |
| ENSG00000128596 | CCDC136 | 2.636359197 | 0.159971389 | 2.655400156 | 0.148260019 | 5.291759353 | 2 | 5.838937808 | 2 | 6.956091174 | 6.83074839 | 7.078941352 | 6.725806822 |
| ENSG00000282077 | | 4.26058699 | 0.052219697 | 1.031018608 | 0.681460704 | 5.291605598 | 2 | 2 | 2 | 4.739373666 | 4.3456612 | 2.492221014 | 2.70970617 |
| ENSG00000183684 | ALYREF | 2.94661578 | 0.004929756 | 2.344007469 | 0.028590946 | 5.290623249 | 3.327711234 | 6.311476754 | 4.523675391 | 8.051945366 | 8.11663771 | 7.198250335 | 7.737762724 |
| ENSG00000093183 | SEC22C | 2.313219991 | 0.000636648 | 2.974613282 | 6.20E-06 | 5.287833273 | 5.330594134 | 7.232742069 | 6.80318718 | 8.948803189 | 8.92660044 | 9.395638968 | 9.769813713 |
| ENSG00000137656 | BUD13 | 2.482119852 | 0.195669275 | 2.805649075 | 0.131096155 | 5.287768926 | 2 | 5.946638857 | 2 | 6.965655398 | 6.72891807 | 7.190239142 | 7.140495192 |
| ENSG00000142949 | PTPRF | 2.102374739 | 0.101674945 | 3.183894193 | 0.008569098 | 5.286268932 | 4.467311984 | 8.820493869 | 8.236946975 | 10.12521908 | 10.0809763 | 11.24067069 | 11.12086139 |
| ENSG00000155561 | NUP205 | 2.801720956 | 0.094156907 | 2.482854504 | 0.135518423 | 5.28457546 | 5.532353208 | 7.874700259 | 2 | 9.304878886 | 9.35675148 | 9.055415647 | 8.972144811 |
| ENSG00000015676 | NUDCD3 | 2.367340976 | 0.088711396 | 2.911149186 | 0.030006203 | 5.278490162 | 8.228665467 | 8.101365653 | 3.369317972 | 9.906793906 | 9.99873154 | 10.49009305 | 10.50081829 |
| ENSG00000107968 | MAP3K8 | 2.747445154 | 0.076258148 | 2.530893966 | 0.100093385 | 5.27833912 | 3.327711234 | 6.04685525 | 2 | 7.29643414 | 7.39346687 | 6.579278019 | 7.535201607 |
| ENSG00000006118 | TMEM132A | 3.636291041 | 0.000253038 | 1.640583741 | 0.127435542 | 5.276874782 | 5.866944937 | 9.102512015 | 8.941672936 | 12.13013662 | 12.1533131 | 10.15815498 | 10.14152759 |
| ENSG00000095587 | TLL2 | 3.147453096 | 0.244748361 | 2.128167559 | 0.440774103 | 5.275620655 | 2 | 2 | 2 | 3.515101962 | 3.86943969 | 2.656294034 | 3.407202941 |
| ENSG00000140563 | MCTP2 | 2.625286019 | 2.31E-05 | 2.650180951 | 1.68E-05 | 5.27546697 | 4.00722531 | 5.131603625 | 5.772716771 | 7.580980221 | 7.64371756 | 7.684242973 | 7.588252384 |
| ENSG00000060642 | PIGV | 3.075754089 | 0.070252457 | 2.198748185 | 0.204625268 | 5.274502274 | 2 | 6.728495673 | 3.369317972 | 8.260583448 | 8.34212018 | 7.304736313 | 7.563910189 |
| ENSG00000122008 | POLK | 3.155792929 | 0.037404281 | 2.117230849 | 0.175071314 | 5.273023779 | 2 | 7.274681896 | 5.39244166 | 9.190240253 | 9.12847688 | 8.423103999 | 7.766116162 |
| ENSG00000015568 | RGPD5 | 10.46403953 | 2.82E-13 | -5.191151153 | 0.023574771 | 5.272888379 | 2 | 2 | 2 | 10.50109983 | 10.4877241 | 2 | 2 |
| ENSG00000183054 | RGPD6 | 10.46403953 | 2.82E-13 | -5.191151153 | 0.023574771 | 5.272888379 | 2 | 2 | 2 | 10.50109983 | 10.4877241 | 2 | 2 |
| ENSG00000135842 | FAM129A | 2.699332003 | 0.026904691 | 2.573336137 | 0.034537867 | 5.27266814 | 6.257048221 | 7.467835385 | 4.874587708 | 9.198375657 | 9.23886123 | 9.879654003 | 7.234405279 |
| ENSG00000125845 | BMP2 | 4.932519785 | 0.00562177 | 0.338655386 | 0.880889575 | 5.27117517 | 7.940205782 | 5.131603625 | 2 | 11.44179676 | 11.4338527 | 7.647656456 | 5.20457242 |
| ENSG00000183578 | TNFAIP8L3 | 3.477391059 | 0.002290805 | 1.792799132 | 0.14042775 | 5.270190191 | 2 | 4.442710129 | 5.59507254 | 7.942383523 | 7.76511961 | 6.664632795 | 5.59182244 |
| ENSG00000198087 | CD2AP | 2.781812188 | 0.015572998 | 2.488250465 | 0.031551126 | 5.270062654 | 6.468621203 | 7.35507212 | 3.369317972 | 9.277301907 | 9.10192275 | 9.081998027 | 8.693802688 |
| ENSG00000164916 | FOXK1 | 1.944395126 | 0.001370344 | 3.320414389 | 8.54E-09 | 5.264809515 | 6.468621203 | 7.60566235 | 8.007629717 | 9.351221511 | 9.47486651 | 11.0025909 | 10.52905457 |
| ENSG00000187626 | ZKSCAN4 | 2.458914819 | 0.129733292 | 2.80556442 | 0.074759029 | 5.264479239 | 2 | 5.131603625 | 2 | 5.978405189 | 6.08962627 | 6.533574091 | 6.168983563 |
| ENSG00000214022 | REPIN1 | 2.584577143 | 0.014636321 | 2.679600202 | 0.010581937 | 5.264177345 | 5.70932728 | 8.522788974 | 6.203069917 | 9.960398193 | 9.90046182 | 10.00540087 | 10.04508992 |
| ENSG00000008277 | ADAM22 | 1.599895999 | 0.567865457 | 3.664077464 | 0.143706589 | 5.263973464 | 2 | 2 | 2 | 3.09461492 | 2.49394783 | 3.930690651 | 4.215171317 |
| ENSG00000131504 | DIAPH1 | 2.43044703 | 0.239601493 | 2.828836859 | 0.157060567 | 5.259283889 | 4.815609471 | 8.834582582 | 2 | 9.693134408 | 9.81476202 | 9.995665807 | 10.2930197 |
| ENSG00000047346 | FAM214A | 3.162665874 | 0.035415609 | 2.09138067 | 0.177532791 | 5.254046544 | 4.467311984 | 6.899080544 | 2 | 8.668958796 | 8.69465306 | 7.523500045 | 7.72163797 |
| ENSG00000120262 | CCDC170 | 2.012515229 | 0.30391278 | 3.241113355 | 0.073543326 | 5.253628584 | 2 | 2 | 3.369317972 | 3.515101962 | 3.65239003 | 4.461156421 | 4.553838275 |
| ENSG00000163449 | TMEM169 | 2.773988867 | 0.199271942 | 2.476398348 | 0.247980941 | 5.250387215 | 2 | 2 | 3.369317972 | 4.387512679 | 3.82856379 | 4.378961816 | 3.209956562 |
| ENSG00000169908 | TM4SF1 | 3.159118368 | 0.002904929 | 2.090928249 | 0.058149869 | 5.250046617 | 8.313285031 | 5.303523747 | 8.742726175 | 11.15146597 | 11.1825997 | 9.815783218 | 10.34017703 |
| ENSG00000161082 | CELF5 | 2.861150031 | 0.277466813 | 2.385603381 | 0.363109023 | 5.246753412 | 2 | 2 | 2 | 3.709785322 | 3.28037825 | 3.355392153 | 3.071407821 |
| ENSG00000145555 | MYO10 | 2.941264334 | 7.04E-08 | 2.300750089 | 3.33E-05 | 5.242014423 | 8.285626337 | 9.338312924 | 8.076950877 | 11.57488023 | 11.655095 | 11.08348224 | 10.85996842 |
| ENSG00000128285 | MCHR1 | 2.213376551 | 0.403701418 | 3.027019801 | 0.225372001 | 5.240396353 | 2 | 2 | 2 | 3.162721474 | 3.08597292 | 3.517841605 | 3.717076715 |
| ENSG00000213889 | PPM1N | 2.68947476 | 0.30391278 | 2.54966853 | 0.320161353 | 5.23914329 | 2 | 2 | 2 | 3.406930997 | 3.39680241 | 3.193831101 | 3.430065404 |
| ENSG00000179542 | SLITRK4 | 7.992334969 | 7.44E-08 | -2.753271311 | 0.186976308 | 5.239063658 | 2 | 2 | 2 | 8.087422161 | 7.99261031 | 2.048074405 | 2.059860178 |
| ENSG00000147010 | SH3KBP1 | 3.740009227 | 0.021998462 | 1.497688814 | 0.39901599 | 5.237698041 | 3.327711234 | 6.535001748 | 9.439621147 | 11.77036178 | 11.7718811 | 9.249414741 | 9.773170377 |
| ENSG00000138030 | KHK | 1.152227925 | 0.681613489 | 4.080808531 | 0.088401773 | 5.233036457 | 2 | 2 | 2 | 2.868578001 | 2.38539787 | 4.628601098 | 4.147809227 |
| ENSG00000151503 | NCAPD3 | 2.301224637 | 0.185410261 | 2.931138801 | 0.078715146 | 5.232363438 | 3.327711234 | 6.666836837 | 2 | 7.477814553 | 7.48179565 | 7.964208996 | 8.220862755 |
| ENSG00000111647 | UHRF1BP1L | 3.28660371 | 2.48E-05 | 1.945353204 | 0.017959609 | 5.231956915 | 4.00722531 | 6.389896633 | 6.534109959 | 9.231457548 | 9.20332986 | 7.951626215 | 7.827675587 |
| ENSG00000108439 | PNPO | 2.945934744 | 0.002537365 | 2.285465144 | 0.02246952 | 5.231399888 | 4.00722531 | 4.11352228 | 6.322058544 | 8.008651253 | 8.01099479 | 7.117472117 | 7.571641661 |
| ENSG00000179195 | ZNF664 | 1.591870014 | 0.005233903 | 3.637735593 | 5.88E-12 | 5.229605607 | 7.82963488 | 8.004992147 | 6.629488387 | 9.263797825 | 9.10299435 | 11.07199274 | 11.36146232 |
| ENSG00000136867 | SLC31A2 | 6.351005583 | 2.95E-08 | -1.122524023 | 0.42197184 | 5.22848156 | 3.327711234 | 3.686303414 | 2 | 8.7747563 | 8.80016655 | 2.911488689 | 2.421067459 |
| ENSG00000158079 | PTPDC1 | 2.575605227 | 0.064574411 | 2.65094195 | 0.053081508 | 5.226547177 | 3.327711234 | 5.72254323 | 2 | 6.902323594 | 6.88657603 | 7.16729616 | 6.735016458 |
| ENSG00000133740 | E2F5 | 1.935510386 | 0.217054056 | 3.289494536 | 0.023768024 | 5.225004922 | 4.467311984 | 2 | 2 | 4.694064037 | 4.68029874 | 5.966975997 | 5.825320785 |
| ENSG00000270136 | MINOS1-NBI | 10.41609146 | 4.03E-13 | -5.191151159 | 0.023706691 | 5.224940302 | 2 | 2 | 2 | 10.4133977 | 10.4790834 | 2 | 2 |
| ENSG00000122367 | LDB3 | -1.183882387 | 0.410884798 | 6.406962954 | 6.13E-09 | 5.223080566 | 4.00722531 | 2 | 3.369317972 | 2.868578001 | 2.49394783 | 9.092781861 | 8.689663984 |
| ENSG00000141255 | SPATA22 | 2.819431653 | 0.386043706 | 2.400077308 | 0.457596529 | 5.219508961 | 2 | 2 | 2 | 3.566290607 | 3.39680241 | 3.796405078 | 2.277179783 |
| ENSG00000186130 | ZBTB6 | 2.580528467 | 0.215761172 | 2.638601874 | 0.196222912 | 5.219130341 | 2 | 6.464272742 | 2 | 7.447418636 | 7.47187556 | 7.637872201 | 7.383872668 |
| ENSG00000158008 | EXTL1 | 1.037626851 | 0.487332808 | 4.180908336 | 0.000909742 | 5.218535187 | 2 | 4.710564154 | 3.369317972 | 4.497095146 | 4.76938658 | 6.560755617 | 8.146888246 |
| ENSG00000179041 | RRS1 | 2.469462047 | 0.098765799 | 2.748807325 | 0.058606472 | 5.218269372 | 3.327711234 | 5.946638857 | 2 | 6.951285179 | 7.02417148 | 6.958334233 | 7.5085977 |
| ENSG00000148341 | SH3GLB2 | 1.900459201 | 0.211593503 | 3.315028329 | 0.019191632 | 5.21548753 | 7.33131396 | 7.145017454 | 2 | 8.52598359 | 8.55127993 | 9.885040669 | 9.999723616 |
| ENSG00000188199 | NUTM2B | 6.448043195 | 0.000360365 | -1.234657172 | 0.587973908 | 5.213386022 | 2 | 2 | 2 | 6.604700657 | 6.46787796 | 2.225793514 | 2.059860178 |
| ENSG00000163877 | SNIP1 | 2.910585576 | 0.000969079 | 2.299880973 | 0.010756531 | 5.210466549 | 4.815609471 | 6.728495673 | 4.523675391 | 8.639384849 | 8.49198035 | 7.844256003 | 8.076179938 |
| ENSG00000100906 | NFKBIA | 3.321857081 | 5.16E-07 | 1.886952162 | 0.00682286 | 5.208809243 | 8.836089686 | 7.467835385 | 8.380957451 | 11.61835275 | 11.6488716 | 9.491385735 | 10.67585959 |
| ENSG00000116604 | MEF2D | 2.026386615 | 2.89E-11 | 3.181373988 | 1.19E-26 | 5.207760603 | 7.975256505 | 7.701025996 | 8.175061096 | 9.938102197 | 9.98484711 | 11.04198142 | 11.18159118 |
| ENSG00000266074 | BAHCC1 | 2.293224566 | 0.120787522 | 2.914372325 | 0.040105266 | 5.207596892 | 6.468621203 | 7.35507212 | 2 | 8.579911316 | 8.74905888 | 9.194070966 | 9.367009385 |
| ENSG00000197555 | SIPA1L1 | 3.723017218 | 1.23E-05 | 1.480199426 | 0.112527071 | 5.203216645 | 6.468621203 | 8.777382925 | 6.958202216 | 11.4340665 | 11.5095472 | 9.097786864 | 9.364044387 |
| ENSG00000169679 | BUB1 | 2.598957481 | 0.076258148 | 2.604175901 | 0.070716745 | 5.203133383 | 2 | 4.11352228 | 2 | 5.194621444 | 5.07918227 | 5.026717879 | 5.250250853 |
| ENSG00000131389 | SLC6A6 | 2.922701605 | 2.80E-06 | 2.275708628 | 0.000345801 | 5.198410233 | 7.096806373 | 7.979858146 | 8.805342542 | 11.00726039 | 11.0432906 | 10.53427493 | 10.20711012 |
| ENSG00000075151 | EIF4G3 | 2.664360981 | 0.002388547 | 2.53317796 | 0.00391914 | 5.197538941 | 6.73718411 | 9.225224669 | 7.691296561 | 10.90813165 | 10.8986743 | 10.68832675 | 10.85205426 |
| ENSG00000169248 | CXCL11 | 10.38726814 | 4.94E-13 | -5.191151162 | 0.023768024 | 5.196116975 | 2 | 2 | 2 | 10.4577033 | 10.3769158 | 2 | 2 |
| ENSG00000059804 | SLC2A3 | 2.482273802 | 5.66E-09 | 2.713235937 | 1.35E-10 | 5.195509739 | 8.716893051 | 7.979858146 | 8.236946975 | 10.80568529 | 10.8076163 | 11.29350887 | 10.725075 |
| ENSG00000187634 | SAMD11 | 2.805951894 | 0.074779862 | 2.389417369 | 0.128777677 | 5.195369263 | 2 | 3.076665209 | 4.058998049 | 5.565529336 | 5.2591458 | 5.622611168 | 4.077147165 |
| ENSG00000058804 | NDC1 | 2.683610608 | 0.191106627 | 2.511430305 | 0.215861 | 5.195040914 | 2 | 6.389896633 | 2 | 7.507583254 | 7.46522407 | 7.358152527 | 7.277588383 |
| ENSG00000214940 | NPIPA8 | 10.38455242 | 4.80E-13 | -5.191151179 | 0.023719047 | 5.193401241 | 2 | 2 | 2 | 10.39802725 | 10.432105 | 2 | 2 |
| ENSG00000214967 | NPIPA7 | 10.38455242 | 4.80E-13 | -5.191151179 | 0.023719047 | 5.193401241 | 2 | 2 | 2 | 10.39802725 | 10.432105 | 2 | 2 |
| ENSG00000103966 | EHD4 | 2.363495566 | 0.039654564 | 2.829257908 | 0.011254194 | 5.192753475 | 4.467311984 | 7.467835385 | 7.934808822 | 9.554240322 | 9.52849248 | 9.270464658 | 10.48946923 |
| ENSG00000261796 | ISY1-RAB43 | 10.3779691 | 5.71E-13 | -5.191151138 | 0.023890695 | 5.18681796 | 2 | 2 | 2 | 10.34404954 | 10.4703907 | 2 | 2 |
| ENSG00000123892 | RAB38 | 4.200838112 | 0.003614241 | 0.982683879 | 0.556098014 | 5.183521991 | 3.327711234 | 3.076665209 | 2 | 6.155987605 | 6.12381559 | 3.738983888 | 3.15612297 |
| ENSG00000140743 | CDR2 | 2.431988598 | 0.341594707 | 2.750799257 | 0.267661148 | 5.182787856 | 2 | 8.101365653 | 2 | 8.965583267 | 8.92781046 | 9.56215648 | 8.885886446 |
| ENSG00000123685 | BATF3 | 3.194994601 | 0.062735523 | 1.985730096 | 0.264144129 | 5.180724697 | 2 | 3.686303414 | 2 | 5.304221918 | 4.98931816 | 4.461156421 | 3.735548302 |
| ENSG00000163602 | RYBP | 3.039524859 | 0.009118461 | 2.140669425 | 0.075047762 | 5.180194284 | 3.327711234 | 6.535001748 | 7.451345288 | 9.56371781 | 9.46323044 | 8.766685705 | 8.461794641 |
| ENSG00000167632 | TRAPPC9 | 1.672146707 | 0.26684385 | 3.507689262 | 0.010549918 | 5.179835969 | 3.327711234 | 6.78762695 | 3.369317972 | 6.86703417 | 7.23436042 | 8.769376872 | 8.957466266 |
| ENSG00000164188 | RANBP3L | 0.495933931 | 0.85245466 | 4.681122192 | 0.02246952 | 5.177056124 | 2 | 2 | 2 | 2.3892346 | 2.49394783 | 4.881704255 | 4.943664032 |
| ENSG00000143756 | FBXO28 | 2.760284548 | 0.059583449 | 2.414454496 | 0.099599264 | 5.174739044 | 6.653098118 | 7.431218269 | 2 | 9.228481255 | 9.26398924 | 9.050261203 | 8.7391339 |
| ENSG00000100836 | PABPN1 | 3.096396012 | 2.86E-07 | 2.076967362 | 0.000862769 | 5.173363374 | 6.366709671 | 6.311476754 | 7.691296561 | 10.02013627 | 9.93817677 | 8.83678613 | 9.084938196 |
| ENSG00000196154 | S100A4 | 2.40691025 | 0.159728332 | 2.763788392 | 0.095263386 | 5.170698642 | 2 | 6.602424674 | 7.691296561 | 9.000896538 | 9.07814384 | 8.147770839 | 10.05316631 |
| ENSG00000166454 | ATMIN | 2.633959272 | 0.080939257 | 2.533094622 | 0.08995084 | 5.167053893 | 5.330594134 | 8.607278467 | 4.058998049 | 9.865241782 | 9.79434273 | 9.773513173 | 9.684739879 |
| ENSG00000160007 | ARHGAP35 | 1.760206001 | 6.38E-08 | 3.406560063 | 2.36E-27 | 5.166766064 | 8.75772933 | 8.522788974 | 8.884815469 | 10.4577033 | 10.4926385 | 11.89156381 | 12.31521775 |
| ENSG00000144647 | POMGNT2 | 1.49314763 | 0.036228406 | 3.670744415 | 1.13E-08 | 5.163892045 | 5.532353208 | 5.457120953 | 3.369317972 | 6.463327516 | 6.50077304 | 8.642521927 | 8.57382182 |
| ENSG00000095787 | WAC | 2.63030871 | 0.000207528 | 2.533290081 | 0.000351347 | 5.163598791 | 7.53297911 | 9.00621767 | 7.342949392 | 10.781859 | 10.7963987 | 10.88653786 | 10.46788136 |
| ENSG00000160216 | AGPAT3 | 2.45353413 | 2.17E-20 | 2.70893288 | 5.68E-25 | 5.16246701 | 8.042902253 | 8.029695761 | 7.858115942 | 10.38070504 | 10.4528466 | 10.62473233 | 10.71768999 |
| ENSG00000166169 | POLL | 1.996774262 | 0.214567154 | 3.165341153 | 0.035964483 | 5.162115415 | 4.00722531 | 6.666836837 | 2 | 7.182120794 | 7.33993465 | 8.591223132 | 8.204380326 |
| ENSG00000144560 | VGLL4 | 2.549689169 | 6.56E-10 | 2.611286634 | 2.05E-10 | 5.160975803 | 7.096806373 | 6.844429755 | 7.285536013 | 9.619311339 | 9.56160864 | 9.899034515 | 9.353618642 |
| ENSG00000172738 | TMEM217 | 2.975968555 | 0.004961216 | 2.184940552 | 0.044818092 | 5.160909106 | 4.00722531 | 2 | 4.874587708 | 6.640968371 | 6.49425375 | 5.818966034 | 5.825320785 |
| ENSG00000171621 | SPSB1 | 2.502438763 | 0.12806343 | 2.658293193 | 0.098271797 | 5.160731956 | 2 | 6.899080544 | 8.110401118 | 9.511620236 | 9.53327015 | 9.618963711 | 9.733801916 |
| ENSG00000095139 | ARCN1 | 2.38912447 | 0.033038824 | 2.769554243 | 0.011351964 | 5.158678714 | 9.015175615 | 10.5327403 | 6.958202216 | 11.85384977 | 11.8503907 | 12.1907878 | 12.27242173 |

| | | | | | | | | | | | | | |
|---|---|---|---|---|---|---|---|---|---|---|---|---|---|
| ENSG00000028137 | TNFRSF1B | 2.974356081 | 2.34E-05 | 2.183437642 | 0.002486145 | 5.157793722 | 3.327711234 | 4.93638933 | 5.39244166 | 7.565338955 | 7.56813436 | 6.937746858 | 6.6426114 |
| ENSG00000196155 | PLEKHG4 | 1.484333622 | 0.390209318 | 3.666821895 | 0.017730665 | 5.151155517 | 6.468621203 | 3.686303414 | 2 | 6.449761039 | 6.54560135 | 7.843404862 | 9.136672183 |
| ENSG00000198435 | NRARP | 1.939424255 | 0.479723716 | 3.209680182 | 0.204683486 | 5.149104437 | 2 | 2 | 2 | 3.162721474 | 2.77779907 | 3.878466556 | 3.600986722 |
| ENSG00000121454 | LHX4 | 0.42763088 | 0.852752459 | 4.721431192 | 0.007480866 | 5.149062072 | 2 | 3.076665209 | 2 | 2.695456494 | 2.77779907 | 5.141958644 | 6.27380897 |
| ENSG00000172071 | EIF2AK3 | 2.833493078 | 0.099708601 | 2.315329593 | 0.179625742 | 5.148822671 | 3.327711234 | 6.728495673 | 2 | 8.100504225 | 8.01782901 | 7.66606569 | 7.425709383 |
| ENSG00000197063 | MAFG | 2.916041429 | 6.36E-17 | 2.229374755 | 2.31E-10 | 5.145416184 | 7.867441304 | 7.467835385 | 8.206335842 | 10.75591108 | 10.7635692 | 10.06648818 | 10.08367608 |
| ENSG00000102878 | HSF4 | 3.562951209 | 0.010828316 | 1.581271309 | 0.294849354 | 5.144222518 | 2 | 4.710564154 | 2 | 6.676346669 | 6.66069199 | 5.079718116 | 4.56420962 |
| ENSG00000213967 | ZNF726 | 4.029219648 | 0.076034873 | 1.114809732 | 0.663448601 | 5.144029379 | 2 | 2 | 2 | 4.358765249 | 4.37434574 | 2.746468522 | 2.509591799 |
| ENSG00000148358 | GPR107 | 2.289527739 | 0.015836831 | 2.853767986 | 0.001919305 | 5.143295725 | 6.257048221 | 9.159587375 | 8.65477198 | 10.71325662 | 10.7523441 | 11.35735632 | 11.23206261 |
| ENSG00000166289 | PLEKHF1 | 2.665886509 | 0.12734523 | 2.475398677 | 0.152273554 | 5.141285187 | 2 | 5.457120953 | 2 | 6.535745081 | 6.55815769 | 6.579278019 | 6.113568999 |
| ENSG00000139572 | GPR84 | 6.2663893 | 0.000320814 | -1.125903753 | 0.610519493 | 5.140485548 | 2 | 2 | 2 | 6.283823264 | 6.44101097 | 2.094598369 | 2.225841198 |
| ENSG00000248905 | FMN1 | 4.001876443 | 0.024397609 | 1.138216699 | 0.568885801 | 5.140093142 | 2 | 2 | 5.59507254 | 7.789219291 | 7.93600799 | 5.494014735 | 4.692661645 |
| ENSG00000133878 | DUSP26 | 1.014354529 | 0.736178 | 4.120167769 | 0.107186013 | 5.134522297 | 2 | 2 | 2 | 2.784612485 | 2.38539787 | 4.990270811 | 3.539239711 |
| ENSG00000143878 | RHOB | 1.7918509 | 0.014995326 | 3.339404933 | 1.17E-06 | 5.131255833 | 9.128497779 | 8.234834549 | 7.163363831 | 10.1568736 | 10.1761984 | 11.8995701 | 11.49357837 |
| ENSG00000101311 | FERMT1 | 5.01199917 | 0.010010073 | 0.11803254 | 0.963624158 | 5.130031709 | 2 | 2 | 2 | 5.304221918 | 5.09650239 | 2.42098416 | 2.277179783 |
| ENSG00000171488 | LRRC8C | 3.660041145 | 0.008716756 | 1.468770963 | 0.33487913 | 5.128812108 | 2 | 4.442710129 | 6.322058544 | 8.657201865 | 8.58694992 | 7.013858277 | 5.631815919 |
| ENSG00000163029 | SMC6 | 2.412115533 | 0.086021641 | 2.714400283 | 0.046914405 | 5.126515816 | 4.467311984 | 7.189546432 | 3.369317972 | 8.29513208 | 8.20911072 | 8.953397421 | 7.992257228 |
| ENSG00000186026 | ZNF284 | 3.144609612 | 0.030874483 | 1.980661426 | 0.189987418 | 5.125271038 | 2 | 4.710564154 | 2 | 6.34372989 | 6.18176397 | 5.413723492 | 4.927687581 |
| ENSG00000054793 | ATP9A | 1.68137001 | 0.325849593 | 3.443154401 | 0.02799101 | 5.124524411 | 7.975256505 | 8.029695761 | 2 | 9.111653326 | 9.06830201 | 10.77323249 | 10.91228265 |
| ENSG00000179598 | PLD6 | 3.400011944 | 0.006529818 | 1.723276994 | 0.19604595 | 5.123288939 | 4.00722531 | 2 | 3.369317972 | 5.860316854 | 6.1068222 | 4.301748724 | 4.673629022 |
| ENSG00000187079 | TEAD1 | 1.928728878 | 0.004384186 | 3.19350787 | 6.53E-07 | 5.122236747 | 8.138771685 | 9.523645035 | 9.945092785 | 11.35143572 | 11.2588566 | 12.48349989 | 12.65034854 |
| ENSG00000187474 | FPR3 | -0.30732972 | 0.924945325 | 5.429398606 | 0.026356479 | 5.122068886 | 2 | 2 | 2 | 2.3892346 | 2.14022021 | 6.332313483 | 3.924507493 |
| ENSG00000146433 | TMEM181 | 2.419535852 | 0.01404794 | 2.70072789 | 0.005236775 | 5.120263742 | 5.330594134 | 8.356994601 | 7.029880454 | 9.795895983 | 9.76282251 | 10.04832124 | 10.07054948 |
| ENSG00000146426 | TIAM2 | 2.741151165 | 0.113850795 | 2.377328202 | 0.169003976 | 5.118479367 | 2 | 2 | 5.156626401 | 6.164317909 | 6.14061114 | 5.916072173 | 5.699242824 |
| ENSG00000281670 | | 2.258728545 | 0.082560548 | 2.857655601 | 0.022614967 | 5.116384146 | 7.904282243 | 7.274681896 | 3.369317972 | 9.262828394 | 9.34230452 | 9.778642552 | 10.00995536 |
| ENSG00000171385 | KCND3 | 1.69329442 | 0.555426444 | 3.416951758 | 0.190459822 | 5.110246178 | 2 | 2 | 2 | 3.162721474 | 2.49394783 | 3.865109708 | 3.924507493 |
| ENSG00000137869 | CYP19A1 | 3.666870847 | 0.164311333 | 1.441851432 | 0.61179806 | 5.108722279 | 2 | 2 | 2 | 4.105493497 | 4.05806467 | 3.059555873 | 2.374681325 |
| ENSG00000123104 | ITPR2 | 2.700600503 | 0.085360963 | 2.408019194 | 0.123018585 | 5.108619697 | 2 | 7.274681896 | 5.156626401 | 8.654247601 | 8.66738547 | 8.220374168 | 8.508191835 |
| ENSG00000155254 | MARVELD1 | 2.023135504 | 0.035422866 | 3.084017395 | 0.000695136 | 5.107152899 | 8.444143041 | 9.246456519 | 6.322058544 | 10.45176049 | 10.4423014 | 11.44763312 | 11.56164184 |
| ENSG00000130723 | PRRC2B | 2.121520886 | 0.010117882 | 2.984515568 | 0.000159121 | 5.106036454 | 7.27616306 | 9.461450882 | 9.516493218 | 11.13831145 | 11.193959 | 12.10838349 | 11.94342963 |
| ENSG00000114450 | GNB4 | 2.626108812 | 1.29E-08 | 2.479767928 | 7.86E-08 | 5.10587674 | 7.096806373 | 7.979858146 | 7.098165388 | 10.10480594 | 10.0356777 | 10.06301929 | 9.772052356 |
| ENSG00000250151 | ARPC4-TTLL3 | 8.86548825 | 1.42E-09 | -3.761123294 | 0.108737429 | 5.104364956 | 2 | 2 | 2 | 8.88471049 | 8.9229643 | 2.048074405 | 2 |
| ENSG00000170848 | PSG6 | 2.27703585 | 0.415158868 | 2.824975009 | 0.287937041 | 5.102010859 | 2 | 2 | 2 | 3.406930997 | 2.86123014 | 3.316666208 | 3.640730416 |
| ENSG00000166250 | CLMP | 4.967995761 | 0.003618428 | 0.132796533 | 0.952165908 | 5.100792294 | 2 | 6.228547981 | 8.721235202 | 12.29568877 | 12.333934 | 7.834009015 | 7.09293558 |
| ENSG00000170903 | MSANTD4 | 2.45092095 | 0.301707894 | 2.648979628 | 0.251602132 | 5.099900578 | 2 | 7.393647515 | 2 | 8.274117511 | 8.24255817 | 8.580025633 | 8.315988368 |
| ENSG00000170667 | RASA4B | 7.88103133 | 5.74E-07 | -2.785644032 | 0.158705253 | 5.095387298 | 2 | 3.076665209 | 2 | 8.737556707 | 9.01236778 | 2.183373728 | 2 |
| ENSG00000171943 | SRGAP2C | 3.966180648 | 4.81E-17 | 1.128818009 | 0.031069705 | 5.094998657 | 6.138360429 | 5.595927442 | 6.718950348 | 10.11612257 | 10.0501952 | 7.379447089 | 7.169655242 |
| ENSG00000147124 | ZNF41 | 3.114832526 | 0.147934794 | 1.979899455 | 0.370178383 | 5.094731981 | 2 | 6.844429755 | 2 | 8.386974157 | 8.34212018 | 7.191577434 | 7.307219812 |
| ENSG00000167081 | PBX3 | 2.960817349 | 0.000972978 | 2.133701338 | 0.021537163 | 5.094518687 | 3.327711234 | 5.72254323 | 6.073378139 | 8.343618883 | 8.22494733 | 7.821097076 | 7.011905845 |
| ENSG00000143178 | TBX19 | 2.365876789 | 0.189011771 | 2.72439038 | 0.117356112 | 5.090267168 | 2 | 4.11352228 | 2 | 4.804776789 | 4.98931816 | 5.755231204 | 4.341135913 |
| ENSG00000270629 | NBPF14 | 6.304119002 | 1.23E-23 | -1.215780007 | 0.089915373 | 5.088338995 | 5.866944937 | 7.790474954 | 7.451345288 | 13.49276167 | 13.5472161 | 6.016144783 | 6.155328188 |
| ENSG00000004700 | RECQL | 3.188873668 | 0.008031314 | 1.899136651 | 0.131991984 | 5.088010319 | 3.327711234 | 7.638155693 | 6.958202216 | 9.950196857 | 9.94476635 | 8.785419299 | 8.536120927 |
| ENSG00000068971 | PPP2R5B | 2.06912673 | 0.046720281 | 3.018250869 | 0.002233581 | 5.087377599 | 4.467311984 | 7.761270542 | 7.163363831 | 9.05005082 | 9.02486072 | 9.88958296 | 10.06759085 |
| ENSG00000179119 | SPTY2D1 | 2.58159614 | 0.014627781 | 2.500983888 | 0.017860434 | 5.082580028 | 7.624139768 | 6.728495673 | 4.058998049 | 9.276341511 | 9.27353851 | 9.239097094 | 9.149213283 |
| ENSG00000109674 | NEIL3 | 3.215455832 | 0.259817515 | 1.863654172 | 0.528078406 | 5.079110004 | 2 | 2 | 2 | 4.034562184 | 3.39680241 | 2.457042231 | 3.311948089 |
| ENSG00000101849 | TBL1X | 2.059447669 | 0.431634218 | 3.018933693 | 0.221243146 | 5.078381363 | 2 | 8.191702463 | 2 | 8.619831492 | 8.71442024 | 9.603714182 | 9.63702421 |
| ENSG00000127585 | FBXL16 | 0.69117069 | 0.799285627 | 4.386376926 | 0.043660941 | 5.077547615 | 2 | 2 | 2 | 2.600425781 | 2.38539787 | 4.742549839 | 4.574506939 |
| ENSG00000109667 | SLC2A9 | 1.152241343 | 0.685765481 | 3.921752218 | 0.107708201 | 5.073993561 | 2 | 2 | 2 | 2.868578001 | 2.38539787 | 4.122514202 | 4.4231929 |
| ENSG00000123240 | OPTN | 2.497219703 | 1.54E-05 | 2.575958263 | 7.20E-06 | 5.073177966 | 9.579430841 | 8.124485672 | 9.344432377 | 11.61189576 | 11.6394858 | 11.56670583 | 11.83009033 |
| ENSG00000066336 | SPI1 | -0.727421237 | 0.812768397 | 5.797567426 | 0.011254194 | 5.07014619 | 2 | 2 | 2 | 2.141737246 | 2.2680111 | 6.670403488 | 4.279527956 |
| ENSG00000087253 | LPCAT2 | 3.102998467 | 6.97E-06 | 1.964246078 | 0.006425146 | 5.067244545 | 5.532353208 | 6.228547981 | 4.523675391 | 8.702235867 | 8.5339035 | 7.803699407 | 7.112501355 |
| ENSG00000144791 | LIMD1 | 2.428541327 | 0.022341921 | 2.638383722 | 0.011703917 | 5.06692505 | 7.709880939 | 5.457120953 | 4.874587708 | 8.894758143 | 8.96128525 | 9.150005562 | 9.123144518 |
| ENSG00000105137 | SYDE1 | 1.861628336 | 0.000309993 | 3.204886362 | 7.87E-11 | 5.066514699 | 7.624139768 | 7.790474954 | 8.677268317 | 9.896204982 | 9.99988257 | 11.04845858 | 11.49459545 |
| ENSG00000144837 | PLA1A | 4.289629241 | 0.044487261 | 0.776243094 | 0.756508981 | 5.065872335 | 2 | 2 | 2 | 4.623314311 | 4.53541753 | 2.492221014 | 2.551896972 |
| ENSG00000163686 | ABHD6 | 2.463072939 | 0.101317769 | 2.595985253 | 0.077558619 | 5.059058193 | 3.327711234 | 5.946638857 | 2 | 7.007926688 | 6.95464102 | 7.364099281 | 6.802276223 |
| ENSG00000237765 | FAM200B | 2.274589408 | 0.003554638 | 2.783764315 | 0.00024308 | 5.058353723 | 6.009025997 | 5.457120953 | 3.369317972 | 7.494428494 | 7.45518898 | 8.058907395 | 7.884647945 |
| ENSG00000110203 | FOLR3 | 4.534593248 | 0.080939257 | 0.523103403 | 0.866209511 | 5.057696651 | 2 | 2 | 2 | 4.739373666 | 4.83276841 | 2.717032197 | 2.117335275 |
| ENSG00000122035 | RASL11A | 0.992303719 | 0.437031139 | 4.063517604 | 0.000132575 | 5.055821322 | 4.00722531 | 3.686303414 | 2 | 3.95996233 | 4.40247105 | 7.164572841 | 6.725806822 |
| ENSG00000197461 | PDGFA | 1.083689289 | 0.05874363 | 3.970908093 | 1.88E-15 | 5.054597382 | 5.532353208 | 4.93638933 | 5.59507254 | 6.229283386 | 6.48112613 | 9.220861732 | 9.150074181 |
| ENSG00000130943 | PKDREJ | 0.272349571 | 0.922914669 | 4.771027746 | 0.024667973 | 5.043377317 | 2 | 2 | 2 | 2.270786764 | 2.49394783 | 5.212212839 | 4.729989799 |
| ENSG00000102879 | CORO1A | 1.497904385 | 0.584591586 | 3.543578477 | 0.145407903 | 5.041482862 | 2 | 2 | 2 | 2.868578001 | 2.68924543 | 3.738983888 | 4.201948172 |
| ENSG00000145391 | SETD7 | 1.559182741 | 0.38549543 | 3.479496095 | 0.032370656 | 5.038678835 | 3.327711234 | 9.532315021 | 6.80318718 | 9.682298904 | 9.75056618 | 11.53694171 | 11.72134345 |
| ENSG00000180353 | HCLS1 | 0.694990781 | 0.837724165 | 4.343404927 | 0.116087069 | 5.038395708 | 2 | 2 | 2 | 2.784612485 | 2.14022021 | 5.163941955 | 3.753786374 |
| ENSG00000117751 | PPP1R8 | 2.538368044 | 0.011477015 | 2.49891976 | 0.012503529 | 5.037287804 | 4.00722531 | 7.274681896 | 6.322058544 | 8.905978734 | 8.86223882 | 8.784532718 | 8.903351429 |
| ENSG00000204176 | SYT15 | 1.716690029 | 0.182549331 | 3.31875885 | 0.004768505 | 5.035448879 | 5.095959458 | 3.076665209 | 2 | 5.501219927 | 5.19627118 | 6.977080688 | 6.735016458 |
| ENSG00000118680 | MYL12B | 1.743302761 | 0.011618622 | 3.287800669 | 3.47E-07 | 5.03110343 | 11.08909278 | 9.549500232 | 9.945092785 | 12.10059882 | 12.0761565 | 13.25401198 | 13.93132404 |
| ENSG00000142330 | CAPN10 | 2.181425281 | 0.151541791 | 2.845802743 | 0.050274744 | 5.027228024 | 4.00722531 | 6.464272742 | 2 | 7.29643414 | 7.23826702 | 7.873733947 | 7.961201206 |
| ENSG00000157224 | CLDN12 | 3.083407972 | 0.000882392 | 1.942789982 | 0.04529991 | 5.026197954 | 4.467311984 | 6.951736442 | 7.342949392 | 9.703174645 | 9.73267673 | 8.909342392 | 8.16744219 |
| ENSG00000172350 | ABCG4 | 0.012663238 | 0.997041173 | 5.005894059 | 0.023327067 | 5.018557296 | 2 | 2 | 2 | 2.3892346 | 2.2680111 | 5.737015686 | 4.329022995 |
| ENSG00000091986 | CCDC80 | 0.934463112 | 0.060422825 | 4.083154975 | 7.50E-21 | 5.017618087 | 10.84091191 | 10.42046463 | 10.74328839 | 11.59119151 | 11.6307818 | 14.20024542 | 15.15950973 |
| ENSG00000104177 | MYEF2 | 1.532554256 | 0.013676247 | 3.483149161 | 9.42E-10 | 5.015703417 | 5.866944937 | 5.303523747 | 4.523675391 | 6.699459234 | 6.84106088 | 8.490981547 | 8.852426354 |
| ENSG00000109861 | CTSC | 3.278431218 | 0.000217049 | 1.737230637 | 0.066158892 | 5.015661855 | 5.866944937 | 7.189546432 | 8.536706852 | 10.81431291 | 10.8474839 | 9.003404318 | 9.538828836 |
| ENSG00000067900 | ROCK1 | 2.651962488 | 2.38E-05 | 2.363623081 | 0.000181868 | 5.015585569 | 7.624139768 | 8.414402626 | 6.718950348 | 10.4290383 | 10.3437572 | 10.24595488 | 9.936760772 |
| ENSG00000039600 | SOX30 | 2.549348297 | 0.349396138 | 2.465404749 | 0.355912095 | 5.014753046 | 2 | 2 | 2 | 3.515101962 | 3.08597292 | 3.193831101 | 3.33635788 |
| ENSG00000178078 | STAP2 | 2.180179278 | 0.024886643 | 2.832817996 | 0.002385567 | 5.012997274 | 2 | 4.11352228 | 4.523675391 | 5.808110275 | 5.65950892 | 6.275391383 | 6.418016952 |
| ENSG00000183426 | NPIPA1 | 10.20385195 | 1.59E-12 | -5.191151239 | 0.024005483 | 5.012700712 | 2 | 2 | 2 | 10.22863675 | 10.2413953 | 2 | 2 |
| ENSG00000165434 | PGM2L1 | 2.358822732 | 7.03E-05 | 2.653295876 | 5.60E-06 | 5.012118609 | 6.009025997 | 6.311476754 | 6.718950348 | 8.690748087 | 8.65574839 | 9.381049994 | 8.37829627 |
| ENSG00000121680 | PEX16 | 2.64662678 | 0.000553874 | 2.362964357 | 0.002167809 | 5.009591137 | 3.327711234 | 4.710564154 | 4.874587708 | 6.634986713 | 7.0774796 | 6.514451546 | 6.681239787 |
| ENSG00000118507 | AKAP7 | 2.94524553 | 0.010416238 | 2.06126126 | 0.082028804 | 5.00650679 | 2 | 3.686303414 | 4.058998049 | 5.959382894 | 5.84514933 | 4.778616857 | 5.361470313 |
| ENSG00000132825 | PPP1R3D | 2.23213361 | 0.282224272 | 2.77233745 | 0.163643498 | 5.004471061 | 2 | 6.311476754 | 2 | 6.922108067 | 7.0150921 | 7.45939446 | 7.529919928 |
| ENSG00000125812 | GZF1 | 2.224587234 | 0.346349702 | 2.775187292 | 0.221243146 | 4.999774526 | 2 | 7.232742069 | 2 | 7.855024645 | 7.89452713 | 8.714110407 | 8.044120856 |
| ENSG00000177034 | MTX3 | 2.626699417 | 0.000917782 | 2.372752351 | 0.002916752 | 4.999451768 | 6.96351253 | 6.04685525 | 4.523675391 | 8.70652031 | 8.72280968 | 8.647894167 | 8.251669878 |
| ENSG00000127124 | HIVEP3 | 3.070347033 | 0.000111745 | 1.921371333 | 0.021235761 | 4.991718366 | 4.00722531 | 4.93638933 | 4.058998049 | 7.363085426 | 7.15392111 | 6.464399218 | 5.768151538 |
| ENSG00000189134 | NKAPL | 3.638215144 | 0.021706093 | 1.352128413 | 0.43754457 | 4.990343556 | 2 | 2 | 4.058998049 | 5.786686927 | 5.56313034 | 3.56816602 | 3.956354764 |
| ENSG00000134508 | CABLES1 | 2.813326782 | 0.067002675 | 2.176481799 | 0.161393405 | 4.989808581 | 2 | 7.002537973 | 5.39244166 | 8.570539957 | 8.63663561 | 7.137053113 | 8.502136032 |
| ENSG00000179097 | HTR1F | 3.116992736 | 0.540598702 | 1.872578133 | 0.721716302 | 4.989570869 | 2 | 2 | 2 | 3.840289928 | 3.50452879 | 3.517841605 | 2 |
| ENSG00000103121 | CMC2 | 2.17404781 | 0.075361599 | 2.811823887 | 0.016667916 | 4.985871697 | 6.138360429 | 4.710564154 | 2 | 7.093295078 | 7.10341274 | 7.617106703 | 7.817957556 |
| ENSG00000151718 | WWC2 | 2.066066269 | 0.026395472 | 2.915073008 | 0.001037164 | 4.981139277 | 5.532353208 | 6.728495673 | 8.206335842 | 9.222510184 | 9.27923788 | 10.12701838 | 10.06416949 |
| ENSG00000154305 | MIA3 | 2.84822169 | 0.004559857 | 2.132911125 | 0.038609931 | 4.981132815 | 5.532353208 | 8.777382925 | 7.691296561 | 10.66930252 | 10.6955802 | 10.10090399 | 9.824763374 |
| ENSG00000005471 | ABCB4 | 1.428686499 | 0.497158772 | 3.552261433 | 0.055590322 | 4.980947932 | 2 | 3.076665209 | 2 | 3.289987342 | 3.15372745 | 5.06226682 | 4.400220394 |
| ENSG00000198363 | ASPH | 3.082061351 | 8.87E-17 | 1.891977357 | 5.99E-07 | 4.974038707 | 9.460782017 | 8.955547846 | 9.372276026 | 12.38482562 | 12.3131338 | 10.84695645 | 11.41839745 |
| ENSG00000152229 | PSTPIP2 | 1.938236235 | 0.024764604 | 3.035186799 | 0.000189522 | 4.973423033 | 3.327711234 | 5.457120953 | 4.874587708 | 6.592405944 | 6.64311914 | 8.001311979 | 7.274433587 |
| ENSG00000129028 | THAP10 | 2.630501498 | 0.098666313 | 2.342422965 | 0.13767291 | 4.972924463 | 3.327711234 | 3.076665209 | 2 | 4.71689672 | 4.72553021 | 4.749835737 | 4.147809227 |
| ENSG00000197093 | GAL3ST4 | 1.826781321 | 0.132570607 | 3.143202082 | 0.005270568 | 4.969983403 | 2 | 5.595927442 | 4.058998049 | 6.087554739 | 6.35727082 | 7.487996019 | 7.5085977 |
| ENSG00000080819 | CPOX | 2.580309381 | 0.236852832 | 2.389553393 | 0.268850661 | 4.969862774 | 2 | 6.666836837 | 2 | 7.614805492 | 7.70428369 | 7.832294085 | 6.994755759 |
| ENSG00000165806 | CASP7 | 3.203971871 | 0.065184591 | 1.764200198 | 0.332600633 | 4.96817207 | 5.095959458 | 7.979858146 | 2 | 9.781753035 | 9.73820484 | 8.171023934 | 8.476283084 |
| ENSG00000107404 | DVL1 | 2.105057516 | 0.217270166 | 2.85996514 | 0.078740418 | 4.965022656 | 8.16936288 | 8.029695761 | 2 | 9.585592789 | 9.63382319 | 10.58899854 | 10.09198797 |
| | | 10.15614525 | 2.22E-12 | -5.191151248 | 0.024112824 | 4.964993998 | 2 | 2 | 2 | 10.16675444 | 10.2079206 | 2 | 2 |
| ENSG00000125347 | IRF1 | 3.227885286 | 4.13E-05 | 1.73652762 | 0.038112383 | 4.964412906 | 5.70932728 | 7.051611315 | 7.552162999 | 10.08300173 | 10.1989281 | 7.867049036 | 9.16888512 |
| ENSG00000114315 | HES1 | 1.659454898 | 0.250900803 | 3.304087056 | 0.012430294 | 4.963541954 | 4.00722531 | 5.838937808 | 2 | 6.503277904 | 5.92487824 | 8.269993575 | 7.208191747 |
| ENSG00000186352 | ANKRD37 | 2.621065768 | 0.13602062 | 2.342208114 | 0.179503712 | 4.963273882 | 2 | 2 | 5.156626401 | 6.113598934 | 5.96314905 | 5.696113106 | 5.859408034 |
| ENSG00000151461 | UPF2 | 2.741992269 | 0.00086982 | 2.221012126 | 0.008154016 | 4.963004394 | 5.330594134 | 7.669933291 | 6.43197674 | 9.460955886 | 9.5348592 | 9.182338982 | 8.746552532 |
| ENSG00000102034 | ELF4 | 2.39061265 | 0.136308666 | 2.571675612 | 0.100639252 | 4.962288262 | 7.867441304 | 6.464272742 | 2 | 8.984524517 | 9.22514727 | 9.185029001 | 9.387230438 |
| ENSG00000172380 | GNG12 | 1.798193227 | 0.096174103 | 3.15917945 | 0.001597541 | 4.957372677 | 7.579279349 | 10.68823825 | 9.083172337 | 11.45182709 | 11.4065554 | 12.54306469 | 12.99917931 |
| ENSG00000143669 | LYST | 3.102674493 | 0.002868772 | 1.854345534 | 0.090367254 | 4.957020028 | 4.00722531 | 7.35507212 | 6.43197674 | 9.55977642 | 9.50030288 | 8.642521927 | 7.829826288 |
| ENSG00000108352 | RAPGEFL1 | 1.855113589 | 0.110962 | 3.100403753 | 0.004067454 | 4.955517342 | 3.327711234 | 4.93638933 | 2 | 5.674521485 | 5.43283104 | 6.750676314 | 6.711881188 |
| ENSG00000180370 | PAK2 | 2.114853833 | 0.005935174 | 2.836011969 | 0.000128792 | 4.950865802 | 6.138360429 | 8.337339476 | 7.896971941 | 9.889311677 | 9.74714308 | 10.54713921 | 10.53004695 |

| | | | | | | | | | | | | | |
|---|---|---|---|---|---|---|---|---|---|---|---|---|---|
| ENSG00000165240 | ATP7A | 2.791309886 | 0.001200331 | 2.159128842 | 0.014597277 | 4.950438728 | 4.00722531 | 6.78762695 | 6.073378139 | 8.756965077 | 8.74494589 | 8.035995406 | 8.210995912 |
| ENSG00000186496 | ZNF396 | 0.276561518 | 0.922182516 | 4.673717032 | 0.029347467 | 4.95027855 | 2 | 2 | 2 | 2.498691189 | 2.2680111 | 5.056402493 | 4.739172856 |
| ENSG00000178074 | C2orf69 | 2.295935229 | 0.153238484 | 2.65254086 | 0.088390629 | 4.94847609 | 3.327711234 | 6.228547981 | 2 | 7.101994243 | 7.04216061 | 7.605105115 | 7.206537478 |
| ENSG00000204186 | ZDBF2 | 2.505529969 | 4.45E-05 | 2.442769274 | 6.58E-05 | 4.948299243 | 5.532353208 | 6.728495673 | 5.59507254 | 8.543652346 | 8.51952909 | 8.708044758 | 8.183508901 |
| ENSG00000146587 | RBAK | 2.306407115 | 0.342054331 | 2.641250071 | 0.261724935 | 4.947657185 | 2 | 7.538394143 | 2 | 8.246921218 | 8.27334517 | 8.798653052 | 8.348604099 |
| ENSG00000124613 | ZNF391 | 1.887051431 | 0.131912603 | 3.05828406 | 0.008662669 | 4.945335491 | 3.327711234 | 4.11352228 | 2 | 4.927308402 | 4.89348248 | 6.231826305 | 5.651404445 |
| ENSG00000148634 | HERC4 | 3.664594483 | 6.89E-19 | 1.277623226 | 0.004009929 | 4.94221771 | 7.384433967 | 8.169643011 | 8.296287027 | 11.65834356 | 11.6394858 | 9.416391256 | 9.10594367 |
| ENSG00000004897 | CDC27 | 3.01657372 | 0.013491212 | 1.920333316 | 0.130402547 | 4.936907036 | 6.653098118 | 8.806266215 | 4.874587708 | 10.61867235 | 10.5573944 | 9.45116681 | 9.540196884 |
| ENSG00000152104 | PTPN14 | 2.12386618 | 0.017335408 | 2.812779827 | 0.001057274 | 4.936646007 | 9.219014269 | 6.602424674 | 8.699419242 | 10.60343658 | 10.6697774 | 11.31960213 | 11.3294243 |
| ENSG00000116701 | NCF2 | 4.667650948 | 0.006771419 | 0.26591026 | 0.90314443 | 4.933561209 | 2 | 3.076665209 | 6.322058544 | 9.383689875 | 9.41054037 | 5.843010798 | 3.807156736 |
| ENSG00000137960 | GIPC2 | 1.497911908 | 0.596370107 | 3.434482572 | 0.173666446 | 4.93239448 | 2 | 2 | 2 | 2.868578001 | 2.68924543 | 3.517841605 | 4.215171317 |
| ENSG00000134343 | ANO3 | 0.495892678 | 0.862168375 | 4.436180789 | 0.045408424 | 4.932073467 | 2 | 2 | 2 | 2.3892346 | 2.49394783 | 5.04460184 | 4.254128754 |
| ENSG00000178852 | EFCAB13 | 2.659319641 | 0.203115525 | 2.272059925 | 0.274284885 | 4.931379565 | 2 | 2 | 3.369317972 | 4.17310023 | 3.90918929 | 4.133696865 | 3.261853426 |
| ENSG00000114120 | SLC25A36 | 2.588699801 | 0.11131837 | 2.338675263 | 0.147069388 | 4.927375063 | 2 | 7.928237109 | 7.451345288 | 9.682298904 | 9.70964668 | 9.73525344 | 9.087187414 |
| ENSG00000138180 | CEP55 | 2.143912168 | 0.300919213 | 2.783078113 | 0.157231541 | 4.926990281 | 2 | 3.686303414 | 2 | 4.070463721 | 4.40247105 | 3.796405078 | 5.349527629 |
| ENSG00000169926 | KLF13 | 5.18169143 | 3.30E-10 | -0.256392758 | 0.813974819 | 4.925298672 | 4.467311984 | 5.457120953 | 2 | 9.55977642 | 9.55222419 | 4.487545924 | 4.329022995 |
| ENSG00000167548 | KMT2D | 2.544942105 | 4.73E-07 | 2.378645864 | 2.65E-06 | 4.923587968 | 8.564112515 | 7.73146271 | 8.978369997 | 10.97536299 | 11.0945633 | 10.99264488 | 10.73644041 |
| ENSG00000026950 | BTN3A1 | 3.761824852 | 7.34E-17 | 1.159497799 | 0.019298939 | 4.921322651 | 7.159102577 | 6.844429755 | 7.858115942 | 11.05983441 | 11.0701139 | 8.627767864 | 8.30675314 |
| ENSG00000103657 | HERC1 | 2.216971659 | 0.152654576 | 2.700848001 | 0.07056968 | 4.917819661 | 3.327711234 | 8.607278467 | 6.322058544 | 9.524619869 | 9.51245165 | 10.09483737 | 9.898850157 |
| ENSG00000101191 | DIDO1 | 2.253054856 | 0.183753738 | 2.664038309 | 0.103880499 | 4.917093166 | 7.384433967 | 8.356994601 | 2 | 9.548682898 | 9.66535801 | 10.00197246 | 10.03253982 |
| ENSG00000173011 | TADA2B | 2.064678435 | 0.007676211 | 2.851494784 | 0.000123257 | 4.916173219 | 5.095959458 | 6.951736442 | 5.39244166 | 8.074220386 | 8.11238697 | 8.848711014 | 8.89361766 |
| ENSG00000151364 | KCTD14 | 3.442289685 | 0.17253254 | 1.47356142 | 0.584265208 | 4.915851105 | 2 | 2 | 2 | 3.615725078 | 4.16043394 | 2.68698273 | 2.851942824 |
| ENSG00000101773 | RBBP8 | 2.921787413 | 0.176804511 | 1.991011699 | 0.366230344 | 4.912799113 | 2 | 6.78762695 | 2 | 8.104838684 | 8.12931531 | 7.34858644 | 7.045606259 |
| ENSG00000198771 | RCSD1 | 2.447761313 | 0.008715311 | 2.464839645 | 0.007665374 | 4.912600958 | 4.467311984 | 4.442710129 | 2 | 6.229283386 | 6.11534392 | 6.016144783 | 6.344709081 |
| ENSG00000163755 | HPS3 | 3.419150131 | 0.00186218 | 1.490910645 | 0.211435731 | 4.910060776 | 3.327711234 | 6.951736442 | 7.163363831 | 9.883647099 | 9.90599812 | 8.055971376 | 7.907228933 |
| ENSG00000184083 | FAM120C | 2.554921135 | 0.005891845 | 2.352944224 | 0.011512049 | 4.907865359 | 5.330594134 | 5.303523747 | 2 | 7.186217188 | 7.24216308 | 6.864125031 | 7.157719456 |
| ENSG00000144749 | LRIG1 | 1.95490387 | 0.15865445 | 2.95240892 | 0.023955844 | 4.907312789 | 3.327711234 | 6.602424674 | 8.042706615 | 8.913410938 | 8.81724322 | 9.801172998 | 9.911598039 |
| ENSG00000160710 | ADAR | 3.039091046 | 0.000242028 | 1.867362577 | 0.03200445 | 4.906453623 | 8.816894989 | 10.94030351 | 9.149036566 | 12.9569702 | 13.0309294 | 11.66451928 | 11.96682692 |
| ENSG00000161813 | LARP4 | 2.678620081 | 0.004298315 | 2.22729533 | 0.019679078 | 4.905915411 | 4.815609471 | 7.790474954 | 7.285536013 | 9.755803193 | 9.70330142 | 9.40489908 | 9.145332871 |
| ENSG00000163545 | NUAK2 | 1.206523553 | 0.065393662 | 3.698516579 | 1.49E-10 | 4.905040132 | 4.00722531 | 5.457120953 | 5.772716771 | 6.408278948 | 6.35727082 | 8.828207602 | 8.80566012 |
| ENSG00000111615 | KRR1 | 3.577367754 | 1.31E-08 | 1.325821959 | 0.054930338 | 4.903189712 | 6.816637977 | 6.602424674 | 4.874587708 | 9.822457304 | 9.85983387 | 7.705944963 | 7.513957873 |
| ENSG00000167968 | DNASE1L2 | 2.691939554 | 0.343022212 | 2.210320044 | 0.435298592 | 4.902259599 | 2 | 2 | 2 | 3.663521627 | 3.08597292 | 3.374372158 | 2.817676617 |
| ENSG00000154642 | C21orf91 | 3.344810831 | 0.095959724 | 1.556806012 | 0.465241848 | 4.901616843 | 2 | 6.311476754 | 2 | 8.093978021 | 8.01782901 | 6.693258166 | 5.816671626 |
| ENSG00000163623 | NKX6-1 | -3.75316742 | 0.137326507 | 8.65435048 | 5.09E-08 | 4.90118306 | 2 | 2 | 2 | 2 | 2 | 8.743149543 | 8.644559262 |
| ENSG00000175155 | YPEL2 | 2.640044764 | 0.045589294 | 2.261052308 | 0.088390629 | 4.901097072 | 6.653098118 | 5.946638857 | 2 | 8.360030171 | 8.33666436 | 8.324077374 | 7.512619696 |
| ENSG00000088808 | PPP1R13B | 1.992910475 | 0.074800869 | 2.907190259 | 0.005874607 | 4.900100733 | 5.095959458 | 3.686303414 | 2 | 5.818703821 | 5.76075522 | 6.865802559 | 6.406535141 |
| ENSG00000155760 | FZD7 | 1.415148616 | 0.373567499 | 3.482781308 | 0.014716406 | 4.897929923 | 3.327711234 | 7.145017454 | 8.784770914 | 9.034230235 | 9.00665321 | 10.90450119 | 11.23581828 |
| ENSG00000162813 | BPNT1 | 2.322878729 | 0.00381227 | 2.570687511 | 0.001121109 | 4.89356624 | 3.327711234 | 5.72254323 | 5.930865648 | 7.491120977 | 7.67575955 | 7.862014947 | 7.796124663 |
| ENSG00000184677 | ZBTB40 | 2.455910803 | 0.131645633 | 2.436085252 | 0.128442152 | 4.891996055 | 6.138360429 | 7.819099893 | 2 | 9.060130342 | 9.06830201 | 9.309201142 | 8.720128855 |
| ENSG00000131115 | ZNF227 | 2.768843127 | 0.05130801 | 2.1210329 | 0.140636706 | 4.889876027 | 3.327711234 | 5.72254323 | 2 | 7.062428156 | 7.09482005 | 6.757921389 | 6.070564958 |
| ENSG00000102981 | PARD6A | 0.728535674 | 0.742254487 | 4.158365625 | 0.022192774 | 4.886901299 | 3.327711234 | 2 | 2 | 2.947924294 | 2.77779907 | 5.684753614 | 4.77533149 |
| ENSG00000165457 | FOLR2 | 0.684028022 | 0.834689821 | 4.201771236 | 0.115977634 | 4.885799258 | 2 | 2 | 2 | 2.141737246 | 2.77779907 | 4.713032626 | 4.26688425 |
| ENSG00000155966 | AFF2 | 2.393556069 | 0.446900339 | 2.49052849 | 0.415168199 | 4.88408456 | 2 | 2 | 2 | 3.566290607 | 2.77779907 | 2.656294034 | 3.717076715 |
| ENSG00000091136 | LAMB1 | 2.388907341 | 0.002939392 | 2.495174123 | 0.001719657 | 4.884081463 | 8.517314938 | 10.74528399 | 9.670202709 | 12.28510445 | 12.3186792 | 12.12734459 | 12.64314327 |
| ENSG00000226650 | KIF4B | 1.59892001 | 0.594260626 | 3.28482377 | 0.227519745 | 4.88374378 | 2 | 2 | 2 | 3.023133209 | 2.59489879 | 3.215043717 | 4.215171317 |
| ENSG00000138386 | NAB1 | 2.805281096 | 0.192332182 | 2.076838505 | 0.340665827 | 4.882119601 | 3.327711234 | 8.276714465 | 2 | 9.561354269 | 9.48394374 | 8.957336509 | 8.623017875 |
| ENSG00000172020 | GAP43 | 2.504840654 | 0.631147757 | 2.375988829 | 0.64532651 | 4.880829483 | 2 | 2 | 2 | 3.95996233 | 2 | 3.296906311 | 3.128433649 |
| ENSG00000119899 | SLC17A5 | 2.531419026 | 6.33E-07 | 2.349272143 | 4.03E-06 | 4.880691169 | 7.750913381 | 7.928237109 | 6.629488387 | 10.05359404 | 10.059057 | 9.947565107 | 9.798374054 |
| ENSG00000157240 | FZD1 | 2.969110281 | 0.053978339 | 1.910946716 | 0.229961504 | 4.880056996 | 4.467311984 | 6.951736442 | 2 | 8.537252353 | 8.5339035 | 7.41654498 | 7.567781104 |
| ENSG00000157450 | RNF111 | 2.499654145 | 0.070386148 | 2.379031688 | 0.082535125 | 4.878685833 | 5.70932728 | 7.874700259 | 3.369317972 | 9.133031936 | 9.06170328 | 9.08380094 | 8.864016001 |
| ENSG00000112531 | QKI | 2.002418905 | 0.001234563 | 2.875565897 | 1.42E-06 | 4.877984802 | 7.82963488 | 9.055168103 | 7.646393253 | 10.37174052 | 10.2664796 | 11.16842771 | 11.21354805 |
| ENSG00000130881 | LRP3 | 2.659138349 | 0.280324246 | 2.218695452 | 0.366255977 | 4.877833801 | 2 | 7.819099893 | 2 | 8.888486565 | 8.89106117 | 8.398397867 | 8.506848296 |
| ENSG00000042445 | RETSAT | 2.596174964 | 0.027389043 | 2.280500972 | 0.053903774 | 4.876675935 | 4.467311984 | 6.78762695 | 8.35328246 | 9.751662842 | 9.88434767 | 9.444727462 | 9.563647359 |
| ENSG00000183496 | MEX3B | 1.228887207 | 0.200963739 | 3.647441595 | 1.41E-05 | 4.876328802 | 5.095959458 | 5.595927442 | 3.369317972 | 6.229283386 | 5.99120031 | 8.871434321 | 7.894955865 |
| ENSG00000154122 | ANKH | 1.050445753 | 0.187221604 | 3.825473836 | 2.80E-08 | 4.875919588 | 7.33131396 | 7.274681896 | 8.922968322 | 9.069031178 | 9.11791381 | 11.70970445 | 11.99548996 |
| ENSG00000169174 | PCSK9 | 1.026543418 | 0.649972951 | 3.846154296 | 0.049924592 | 4.872697714 | 2 | 5.595927442 | 2 | 5.022023783 | 5.2591458 | 6.05195199 | 8.588503563 |
| ENSG00000175595 | ERCC4 | 2.695805489 | 0.000213156 | 2.176098037 | 0.003334494 | 4.871903526 | 4.00722531 | 5.457120953 | 6.073378139 | 7.997038782 | 7.96222667 | 7.727325329 | 7.156006254 |
| ENSG00000058668 | ATP2B4 | 1.474238251 | 0.000674855 | 3.395336229 | 6.92E-17 | 4.86957448 | 9.081016759 | 9.726146444 | 9.849281147 | 11.05760014 | 11.06132 | 12.6962582 | 13.21458197 |
| ENSG00000115457 | IGFBP2 | -2.460715785 | 0.009775319 | 7.328206961 | 1.51E-17 | 4.867491176 | 7.579279349 | 7.232742069 | 7.777116083 | 5.949776857 | 3.65239003 | 14.54529467 | 15.08748176 |
| ENSG00000182585 | EPGN | 7.86979128 | 5.02E-11 | -3.004225267 | 0.043497213 | 4.865566013 | 2 | 3.076665209 | 3.369317972 | 9.750280081 | 9.72921087 | 2.139668783 | 2.172608095 |
| ENSG00000118922 | KLF12 | 2.392858002 | 0.018020475 | 2.472007239 | 0.013658003 | 4.864865242 | 4.467311984 | 6.228547981 | 7.600047325 | 9.027396512 | 8.84055903 | 9.148283563 | 8.869253401 |
| ENSG00000177485 | ZBTB33 | 2.718798168 | 0.060978041 | 2.145940165 | 0.143233501 | 4.864738333 | 2 | 7.099070191 | 5.930865648 | 8.747293529 | 8.71862106 | 8.466867678 | 7.787298144 |
| ENSG00000144445 | KANSL1L | 2.818425244 | 0.000151121 | 2.045929135 | 0.00760211 | 4.86435438 | 5.095959458 | 5.131603625 | 3.369317972 | 7.43025113 | 7.34718809 | 6.884127769 | 6.347714043 |
| ENSG00000180535 | BHLHA15 | 1.93618411 | 0.508390119 | 2.926007849 | 0.28326297 | 4.862191959 | 2 | 2 | 2 | 2.868578001 | 3.08597292 | 3.082820997 | 3.908316285 |
| ENSG00000143179 | UCK2 | 2.086702697 | 0.009976623 | 2.771360206 | 0.000378704 | 4.858062903 | 6.563806279 | 7.790474954 | 5.59507254 | 8.978632046 | 9.00894176 | 9.462851966 | 9.858600332 |
| ENSG00000154493 | C10orf90 | 2.148907564 | 0.474821199 | 2.708112679 | 0.343881844 | 4.857020243 | 2 | 2 | 2 | 3.09461492 | 3.08597292 | 2.803598391 | 3.841670744 |
| ENSG00000184845 | DRD1 | -0.719010549 | 0.846636453 | 5.573362126 | 0.052942339 | 4.854351577 | 2 | 2 | 2 | 2.3892346 | 2 | 6.18690462 | 5.005857272 |
| ENSG00000131323 | TRAF3 | 2.141148845 | 0.109443181 | 2.712012956 | 0.034924845 | 4.853161801 | 3.327711234 | 7.60566235 | 7.858115942 | 9.256998085 | 9.33776001 | 9.734335478 | 9.986288828 |
| ENSG00000119042 | SATB2 | 3.298972655 | 0.006549526 | 1.550433926 | 0.235050017 | 4.849406581 | 6.009025997 | 6.311476754 | 2 | 8.874592369 | 8.7882247 | 7.472665674 | 6.632790505 |
| ENSG00000011198 | ABHD5 | 2.759273634 | 0.148420579 | 2.090042425 | 0.278555341 | 4.849316059 | 2 | 7.315436877 | 3.369317972 | 8.559529208 | 8.5323134 | 8.035995406 | 7.718159092 |
| ENSG00000090905 | TNRC6A | 2.587158585 | 0.001576415 | 2.262059038 | 0.0062626 | 4.849217622 | 7.096806373 | 8.540087354 | 6.322058544 | 10.1651988 | 10.2193301 | 10.0366402 | 9.678488127 |
| | | 10.04014075 | 5.00E-12 | -5.191151262 | 0.024397067 | 4.848989483 | 2 | 2 | 2 | 10.03270295 | 10.1100487 | 2 | 2 |
| ENSG00000127511 | SIN3B | 2.279647042 | 0.010409823 | 2.56666106 | 0.0032699 | 4.846308102 | 5.330594134 | 7.874700259 | 7.896971941 | 9.669187952 | 9.68122943 | 10.00654186 | 9.913880743 |
| ENSG00000184254 | ALDH1A3 | 2.624610763 | 0.158647869 | 2.216774328 | 0.232107541 | 4.841385091 | 2 | 3.686303414 | 2 | 4.623314311 | 4.68029874 | 4.720468739 | 3.753786374 |
| ENSG00000107897 | ACBD5 | 2.66114483 | 0.063226377 | 2.179471978 | 0.130474522 | 4.840616809 | 3.327711234 | 7.790474954 | 5.156626401 | 9.133031936 | 9.06390621 | 8.640073371 | 8.602408528 |
| ENSG00000139793 | MBNL2 | 2.733650716 | 0.07209881 | 2.106804991 | 0.170631309 | 4.840455707 | 2 | 7.572420273 | 6.718950348 | 9.380118153 | 9.28585881 | 8.852523173 | 8.555744353 |
| ENSG00000067057 | PFKP | 2.624311401 | 2.93E-06 | 2.212709375 | 9.77E-05 | 4.837020776 | 9.2903348 | 8.762721501 | 7.858115942 | 11.33699991 | 11.392601 | 10.7229835 | 11.15342532 |
| ENSG00000122870 | BICC1 | 2.219292907 | 1.51E-05 | 2.610580646 | 2.21E-07 | 4.829873554 | 7.384433967 | 7.761270542 | 8.296287027 | 10.07363422 | 10.0451865 | 10.00939041 | 10.78629737 |
| ENSG00000147650 | LRP12 | 2.557362577 | 0.122925492 | 2.272207839 | 0.168397072 | 4.829570416 | 2 | 8.029695761 | 7.97167868 | 9.93140666 | 9.98484711 | 9.65506359 | 9.693624218 |
| ENSG00000010404 | IDS | 2.326556374 | 1.70E-06 | 2.502964084 | 2.06E-07 | 4.829520458 | 8.928409675 | 8.762721501 | 7.858115942 | 10.89724025 | 10.9121868 | 10.82675092 | 11.2967558 |
| ENSG00000143013 | LMO4 | 3.710566359 | 6.96E-06 | 1.11660835 | 0.230307325 | 4.827174709 | 7.27616306 | 7.73146271 | 5.59507254 | 10.7879368 | 10.80663 | 8.716436576 | 7.455265057 |
| ENSG00000103540 | CCP110 | 2.693317485 | 0.076989387 | 2.133106352 | 0.16480724 | 4.826423837 | 2 | 6.464272742 | 4.058998049 | 7.794595599 | 7.7515168 | 7.524562403 | 6.847559257 |
| ENSG00000186111 | PIP5K1C | 1.711234209 | 0.168832359 | 3.11383577 | 0.006699223 | 4.825069978 | 5.70932728 | 8.557180776 | 9.658943166 | 10.35318724 | 10.4235527 | 11.45231606 | 12.06132282 |
| ENSG00000205364 | MT1M | 3.571145236 | 0.231366574 | 1.253401423 | 0.699127293 | 4.82454666 | 2 | 2 | 2 | 3.881293667 | 4.12711147 | 3.059555873 | 2.172608095 |
| ENSG00000100425 | BRD1 | 2.261747741 | 0.244284615 | 2.560447902 | 0.174059362 | 4.822195644 | 2 | 7.790474954 | 4.058998049 | 8.62888906 | 8.4705529 | 8.940719573 | 8.74825912 |
| ENSG00000163132 | MSX1 | 1.706284893 | 0.016446525 | 3.110446584 | 2.78E-06 | 4.816731477 | 6.366709671 | 7.274681896 | 5.59507254 | 8.310225403 | 8.21109983 | 9.285896341 | 9.946951964 |
| ENSG00000060558 | GNA15 | 1.85766975 | 0.558508397 | 2.958562374 | 0.318708894 | 4.816232124 | 2 | 2 | 2 | 2.784612485 | 3.08597292 | 4.111244182 | 2.70970617 |
| ENSG00000187097 | ENTPD5 | 2.603366888 | 4.78E-07 | 2.212772867 | 2.19E-05 | 4.816139755 | 5.70932728 | 5.131603625 | 5.930865648 | 8.067573917 | 8.12931531 | 7.538302335 | 7.87218043 |
| ENSG00000070444 | MNT | 2.323334258 | 0.182906893 | 2.492555376 | 0.142685271 | 4.815889633 | 3.327711234 | 7.73146271 | 3.369317972 | 8.508095759 | 8.60672783 | 8.490981547 | 8.927653397 |
| ENSG00000055813 | CCDC85A | 3.283522776 | 0.062977541 | 1.531345992 | 0.416586577 | 4.814868768 | 2 | 2 | 4.058998049 | 5.319220934 | 5.41912973 | 3.429858444 | 4.279527956 |
| ENSG00000270800 | RPS10-NUDT | 10.00314705 | 6.10E-12 | -5.191151287 | 0.024411167 | 4.811995768 | 2 | 2 | 2 | 10.03383996 | 10.0362388 | 2 | 2 |
| ENSG00000171988 | JMJD1C | 2.951420754 | 0.000334613 | 1.85959874 | 0.031149835 | 4.811019494 | 5.70932728 | 8.234834549 | 7.818184403 | 10.52636311 | 10.52338 | 9.4692111 | 9.405009888 |
| ENSG00000134294 | SLC38A2 | 2.849246014 | 0.002273448 | 1.959875466 | 0.042922706 | 4.809121481 | 8.19931886 | 11.05905903 | 10.28977157 | 13.10421017 | 13.1090105 | 12.49360536 | 11.87647387 |
| ENSG00000145819 | ARHGAP26 | 3.593856344 | 3.83E-12 | 1.215261815 | 0.032897253 | 4.809118159 | 5.095959458 | 5.946638857 | 5.930865648 | 9.239364442 | 9.18219179 | 7.090465544 | 6.61544141 |
| ENSG00000135686 | KLHL36 | 1.292351984 | 0.273065021 | 3.513620738 | 0.000749648 | 4.805972722 | 7.435667355 | 8.169643011 | 4.523675391 | 8.60460802 | 8.62325378 | 10.86667347 | 10.77921995 |
| ENSG00000065054 | SLC9A3R2 | 1.479252429 | 0.247900443 | 3.324195243 | 0.004035407 | 4.803447672 | 4.815609471 | 8.557180776 | 6.629488387 | 8.874592369 | 8.84312657 | 10.39991289 | 10.93967577 |
| ENSG00000157368 | IL34 | -0.191363728 | 0.902307277 | 4.99414876 | 9.20E-06 | 4.802785032 | 2 | 4.442710129 | 3.369317972 | 3.79808662 | 3.2184422 | 7.552954291 | 8.54661996 |
| ENSG00000119714 | GPR68 | 6.522618445 | 9.77E-05 | -1.720115929 | 0.370546992 | 4.802502516 | 6.009025997 | 2 | 2 | 10.7993477 | 10.8703199 | 3.517841605 | 3.100202476 |
| ENSG00000093167 | LRRFIP2 | 2.278335641 | 5.79E-06 | 2.524073363 | 3.67E-07 | 4.802409004 | 7.579279349 | 7.701025996 | 6.43197674 | 9.624603733 | 9.57788596 | 9.834108158 | 9.857810118 |
| ENSG00000174640 | SLCO2A1 | 2.372155284 | 0.232847072 | 2.428018112 | 0.210772821 | 4.800173396 | 2 | 3.076665209 | 2 | 3.921164147 | 3.74316441 | 3.891700877 | 3.858622973 |
| ENSG00000157540 | DYRK1A | 2.394609959 | 0.292274211 | 2.403991123 | 0.280132344 | 4.798601083 | 3.327711234 | 8.623598428 | 2 | 9.405814997 | 9.50355257 | 9.454515095 | 9.475045344 |
| ENSG00000186432 | KPNA4 | 2.869912299 | 0.000957099 | 1.928030189 | 0.032925752 | 4.797942488 | 5.866944937 | 8.234834549 | 8.511888463 | 10.79533062 | 10.7375851 | 9.911676379 | 9.738960951 |
| ENSG00000166889 | PATL1 | 2.868687053 | 0.084412675 | 1.927363274 | 0.258077792 | 4.796050327 | 2 | 8.317412869 | 7.734844356 | 10.34588171 | 10.3033068 | 9.341509511 | 9.432159874 |
| ENSG00000138639 | ARHGAP24 | 3.601021371 | 0.001899922 | 1.19468277 | 0.355912095 | 4.795704141 | 7.940205782 | 6.04685525 | 4.058998049 | 10.31675113 | 10.3296366 | 8.09796972 | 7.753709246 |
| ENSG00000187792 | ZNF70 | 1.448363288 | 0.099215121 | 3.345815592 | 2.06E-05 | 4.794178879 | 5.330594134 | 3.686303414 | 4.058998049 | 5.731694 | 5.8245091 | 7.802823992 | 7.357281305 |
| ENSG00000185305 | ARL15 | 2.296679603 | 0.188069595 | 2.495207393 | 0.141446273 | 4.791886996 | 2 | 5.303523747 | 2 | 6.213314212 | 5.85535975 | 6.339575235 | 6.117095472 |
| ENSG00000196569 | LAMA2 | 1.4195239 | 0.286782071 | 3.367576905 | 0.004848864 | 4.787100805 | 4.467311984 | 8.671479029 | 7.734844356 | 9.12771691 | 9.17405701 | 10.64490073 | 11.43217636 |

| | | | | | | | | | | | | | |
|---|---|---|---|---|---|---|---|---|---|---|---|---|---|
| ENSG00000050820 | BCAR1 | 2.47057639 | 0.000211512 | 2.316410856 | 0.000528448 | 4.786987246 | 8.042902253 | 8.213429689 | 9.61300511 | 11.21449494 | 11.3100976 | 11.03808111 | 11.17695249 |
| ENSG00000174306 | ZHX3 | 1.531618879 | 4.80E-05 | 3.25203491 | 9.01E-20 | 4.783653789 | 7.384433967 | 7.431218269 | 7.818184403 | 9.026254405 | 9.09977716 | 10.55911238 | 10.96521391 |
| ENSG00000115380 | EFEMP1 | 2.998621371 | 0.029383982 | 1.784919299 | 0.214737844 | 4.78354067 | 10.24321353 | 5.303523747 | 8.35328246 | 12.02313643 | 12.0305632 | 10.75908557 | 10.86875488 |
| ENSG00000198576 | ARC | 3.469809768 | 0.181904381 | 1.311782873 | 0.640863647 | 4.781592641 | 2 | 2 | 2 | 3.79808662 | 4.05806467 | 2.911488689 | 2.466008509 |
| ENSG00000113269 | RNF130 | 2.865844886 | 5.98E-05 | 1.915138735 | 0.009905697 | 4.780983621 | 6.73718411 | 7.051611315 | 4.874587708 | 9.357592295 | 9.26877178 | 8.443467705 | 8.295903454 |
| ENSG00000153560 | UBP1 | 2.846276786 | 0.079381692 | 1.934370142 | 0.244978858 | 4.780646927 | 8.138771685 | 7.790474954 | 2 | 10.2360652 | 10.2044302 | 9.365126412 | 9.258230494 |
| ENSG00000257529 | RPL36A-HNR | 9.971675083 | 8.02E-12 | -5.191151269 | 0.024561444 | 4.780523814 | 2 | 2 | 2 | 9.95800436 | 10.0479713 | 2 | 2 |
| ENSG00000130158 | DOCK6 | 2.43712143 | 9.89E-10 | 2.342013856 | 3.85E-09 | 4.779135285 | 6.009025997 | 6.04685525 | 6.322058544 | 8.473317843 | 8.5130942 | 8.44402928 | 8.353097069 |
| ENSG00000112186 | CAP2 | 1.255452465 | 0.33027738 | 3.523659916 | 0.001950899 | 4.779112381 | 3.327711234 | 5.131603625 | 7.029880454 | 7.206526483 | 6.88657603 | 9.144833386 | 9.419720027 |
| ENSG00000118482 | PHF3 | 2.551283859 | 0.146327881 | 2.223725421 | 0.203846251 | 4.77500928 | 2 | 7.467835385 | 8.65477198 | 10.11987514 | 10.1411573 | 10.06192211 | 9.49042201 |
| ENSG00000138594 | TMOD3 | 2.090231227 | 0.010985106 | 2.682969455 | 0.000733749 | 4.773200682 | 6.366709671 | 8.623598428 | 8.380957451 | 10.18733762 | 10.1222557 | 10.48122876 | 10.96997524 |
| ENSG00000197016 | ZNF470 | 2.659936215 | 0.035079261 | 2.111457041 | 0.098726147 | 4.771393256 | 3.327711234 | 3.686303414 | 6.43197674 | 7.626911583 | 7.72113133 | 7.284811203 | 6.977399345 |
| ENSG00000104866 | PPP1R37 | 2.301297364 | 0.118963488 | 2.468563181 | 0.087222372 | 4.769860545 | 4.00722531 | 4.11352228 | 7.691296561 | 8.543652346 | 8.5606705 | 8.743605802 | 8.691439168 |
| ENSG00000137193 | PIM1 | 1.060097271 | 0.18784302 | 3.709734681 | 8.99E-08 | 4.769831952 | 4.815609471 | 5.72254323 | 5.39244166 | 6.394182228 | 6.32824132 | 9.40691683 | 8.280260977 |
| ENSG00000188993 | LRRC66 | 3.642355167 | 0.214894239 | 1.127263972 | 0.726496359 | 4.769619139 | 2 | 2 | 2 | 3.881293667 | 4.22485516 | 2.987421008 | 2.172608095 |
| ENSG00000144233 | AMMECR1L | 2.118831784 | 0.112817684 | 2.65038275 | 0.039226073 | 4.769214534 | 3.327711234 | 6.78762695 | 7.934808822 | 9.054539304 | 8.95773559 | 9.556714309 | 9.513016723 |
| ENSG00000196862 | RGPD4 | 9.957208482 | 8.41E-12 | -5.19115129 | 0.02453583 | 4.766057192 | 2 | 2 | 2 | 10.01092762 | 9.96730153 | 2 | 2 |
| ENSG00000115738 | ID2 | 3.856845812 | 0.002192022 | 0.908207025 | 0.531253192 | 4.765052837 | 6.89194353 | 6.389896633 | 2 | 9.860122857 | 9.9170074 | 6.91847879 | 7.040043908 |
| ENSG00000104885 | DOT1L | 1.636004215 | 0.056298013 | 3.128688962 | 7.45E-05 | 4.764693177 | 5.532353208 | 7.928237109 | 6.882775709 | 8.671883119 | 8.74494589 | 10.254464 | 10.12804559 |
| ENSG00000144747 | TMF1 | 2.869627795 | 0.000400793 | 1.892596788 | 0.025292481 | 4.762224583 | 7.790810999 | 8.234834549 | 5.772716771 | 10.43335299 | 10.4482162 | 9.596154868 | 9.326839168 |
| ENSG00000174099 | MSRB3 | 1.227681112 | 0.370269344 | 3.533171386 | 0.003415182 | 4.760852498 | 9.410448395 | 7.638155693 | 5.156626401 | 9.492725629 | 9.44559847 | 11.76676501 | 11.76989677 |
| ENSG00000114353 | GNAI2 | 1.944140792 | 0.000372061 | 2.813366071 | 9.35E-08 | 4.757506863 | 10.65885801 | 10.42046463 | 9.385998894 | 12.20067981 | 12.1847779 | 12.83730227 | 13.2553564 |
| ENSG00000008710 | PKD1 | 2.163144042 | 0.002577478 | 2.590610002 | 0.000218902 | 4.753754045 | 9.613202746 | 8.687092496 | 10.49998282 | 11.8899302 | 11.9862429 | 12.67177419 | 11.97741506 |
| ENSG00000170100 | ZNF778 | 1.50183249 | 0.482249983 | 3.251759843 | 0.093750012 | 4.753592333 | 2 | 6.228547981 | 2 | 6.104969671 | 6.26836889 | 7.600074836 | 8.118139752 |
| ENSG00000182359 | KBTBD3 | 1.788924073 | 0.337673274 | 2.96097947 | 0.08700755 | 4.749903543 | 2 | 3.686303414 | 2 | 4.105493497 | 3.86943969 | 5.274160001 | 4.522269241 |
| ENSG00000112877 | CEP72 | 1.938988358 | 0.360070875 | 2.810438802 | 0.156791365 | 4.74942716 | 2 | 3.076665209 | 2 | 3.754611435 | 3.28037825 | 4.378961816 | 3.908316285 |
| ENSG00000174839 | DENND6A | 2.461772544 | 6.62E-05 | 2.286907519 | 0.000211681 | 4.748680063 | 4.815609471 | 5.838937808 | 4.523675391 | 7.590284224 | 7.53039799 | 7.408510937 | 7.370638252 |
| ENSG00000148660 | CAMK2G | 1.525206429 | 0.115460825 | 3.223180199 | 0.000251101 | 4.748386628 | 4.815609471 | 7.60566235 | 7.398165132 | 8.501535708 | 8.5606705 | 10.15232487 | 10.28186948 |
| ENSG00000198863 | RUNDC1 | 1.757552263 | 0.017287716 | 2.989836208 | 1.59E-05 | 4.747388471 | 5.330594134 | 7.35507212 | 6.43197674 | 8.388752656 | 8.27714776 | 9.546029585 | 9.565907066 |
| ENSG00000176225 | RTTN | 3.987047516 | 3.31E-05 | 0.758465213 | 0.503061508 | 4.745512729 | 5.532353208 | 5.131603625 | 2 | 8.625876185 | 8.64842725 | 5.85994354 | 5.08721802 |
| ENSG00000170677 | SOCS6 | 2.666976517 | 0.082560548 | 2.077863144 | 0.179747748 | 4.744839662 | 6.009025997 | 7.538394143 | 2 | 9.087765067 | 8.9648262 | 8.553208128 | 8.325926932 |
| ENSG00000157510 | AFAP1L1 | 0.666686287 | 0.606448179 | 4.07734749 | 0.000113649 | 4.744033777 | 3.327711234 | 4.93638933 | 6.80318718 | 6.328984825 | 6.21386141 | 9.495184079 | 9.723717339 |
| ENSG00000172175 | MALT1 | 2.637780564 | 0.06531578 | 2.106162001 | 0.144649282 | 4.743942565 | 2 | 6.844429755 | 7.098165388 | 8.959612762 | 9.03387903 | 8.480070636 | 8.461101072 |
| ENSG00000151151 | IPMK | 2.658120579 | 0.145039687 | 2.085699804 | 0.25649001 | 4.743820382 | 2 | 5.595927442 | 2 | 6.693715713 | 6.65485813 | 6.344396176 | 5.876154317 |
| ENSG00000163683 | SMIM14 | 1.595527996 | 0.002937529 | 3.146073905 | 4.18E-10 | 4.741601901 | 8.777722034 | 7.979858146 | 7.502634626 | 9.788505066 | 9.74164918 | 11.21466007 | 11.40213888 |
| ENSG00000115365 | LANCL1 | 1.934307186 | 0.01984582 | 2.802401205 | 0.00039686 | 4.736708391 | 7.867441304 | 7.874700259 | 5.59507254 | 9.350309099 | 9.33867006 | 10.29596165 | 10.155387 |
| ENSG00000122335 | SERAC1 | 1.152697247 | 0.423308674 | 3.583316511 | 0.004287245 | 4.736013759 | 6.816637977 | 5.838937808 | 2 | 6.907295206 | 7.00595521 | 9.193069097 | 9.491101628 |
| ENSG00000186812 | ZNF397 | 1.591942856 | 0.493419043 | 3.143683357 | 0.139684521 | 4.735626213 | 6.563806279 | 2 | 2 | 6.580005553 | 6.53928196 | 8.347678913 | 7.727417522 |
| ENSG00000198846 | TOX | 1.019291148 | 0.839558946 | 3.716181037 | 0.398978105 | 4.735472185 | 2 | 2 | 2 | 3.023133209 | 2 | 3.429858444 | 4.584731282 |
| ENSG00000141564 | RPTOR | 2.426331014 | 0.113542889 | 2.304591782 | 0.128348113 | 4.730922796 | 7.384433967 | 7.467835385 | 2 | 9.258944141 | 9.23886123 | 9.027210383 | 9.222253259 |
| ENSG00000112406 | HECA | 2.082849161 | 0.000150081 | 2.646874148 | 7.60E-07 | 4.729723309 | 7.159102577 | 6.04685525 | 6.203069917 | 8.622857004 | 8.53549186 | 9.333045295 | 8.914540669 |
| ENSG00000118655 | DCLRE1B | 1.544614728 | 0.428119634 | 3.184465291 | 0.072608488 | 4.729080019 | 2 | 5.72254323 | 2 | 5.731694 | 5.76075522 | 7.121690475 | 7.478755384 |
| ENSG00000104695 | PPP2CB | 2.201526865 | 0.000290549 | 2.525133361 | 2.35E-05 | 4.726660226 | 8.285626337 | 8.213429689 | 6.718950348 | 10.0774988 | 10.0901921 | 10.39687046 | 10.41585888 |
| ENSG00000253953 | PCDHGB4 | 4.501297502 | 0.000655225 | 0.223670094 | 0.895844052 | 4.724967595 | 3.327711234 | 4.442710129 | 7.342949392 | 10.39138945 | 10.4372122 | 5.790986459 | 6.538630327 |
| ENSG00000124177 | CHD6 | 2.30492783 | 0.02131845 | 2.41986879 | 0.014538236 | 4.72479662 | 8.517314938 | 9.00621767 | 5.930865648 | 10.58338644 | 10.5940591 | 10.44717886 | 10.92088325 |
| ENSG00000141294 | LRRC46 | 2.389914625 | 0.391949897 | 2.333363493 | 0.391460007 | 4.723278118 | 2 | 2 | 2 | 3.227757282 | 3.2184422 | 3.059555873 | 3.311948089 |
| ENSG00000071794 | HLTF | 1.46257296 | 0.318121128 | 3.260647481 | 0.014179007 | 4.723220441 | 2 | 6.951736442 | 6.322058544 | 7.590284224 | 7.47518984 | 9.299924348 | 9.319205044 |
| ENSG00000269067 | ZNF728 | 4.444206617 | 0.037659821 | 0.276625081 | 0.918154036 | 4.720831698 | 2 | 2 | 2 | 4.694064037 | 4.72553021 | 2.3070652 | 2.466008509 |
| ENSG00000175182 | FAM131A | 1.989529539 | 0.101224495 | 2.730093153 | 0.018317011 | 4.719622692 | 2 | 5.946638857 | 5.930865648 | 7.341208691 | 7.24992376 | 8.258546141 | 7.738907626 |
| ENSG00000117758 | STX12 | 2.455303158 | 0.098944573 | 2.263676718 | 0.12483688 | 4.718979877 | 7.096806373 | 8.747909548 | 3.369317972 | 10.03724565 | 10.0124836 | 9.737316722 | 9.924737419 |
| ENSG00000015532 | XYLT2 | 2.304372354 | 0.022979492 | 2.413307627 | 0.016137174 | 4.717679981 | 4.815609471 | 7.538394143 | 8.042706615 | 9.579376566 | 9.60234163 | 9.753037939 | 9.643457842 |
| ENSG00000185896 | LAMP1 | 2.467794043 | 0.000177625 | 2.249353078 | 0.000679074 | 4.717147121 | 10.00110988 | 8.876041558 | 10.66102425 | 12.47045468 | 12.4902458 | 12.11174512 | 12.39826717 |
| ENSG00000169756 | LIMS1 | 2.463767611 | 1.22E-14 | 2.250135684 | 1.78E-12 | 4.713903294 | 8.674867122 | 8.297207175 | 8.143093325 | 10.8713966 | 10.8030078 | 10.7387464 | 10.50031203 |
| ENSG00000179869 | ABCA13 | 2.867963601 | 0.000218272 | 1.84259875 | 0.023328609 | 4.710562352 | 4.815609471 | 4.93638933 | 5.39244166 | 7.764774849 | 7.77859536 | 6.031172603 | 7.261744957 |
| ENSG00000278461 | | 3.014261933 | 8.24E-06 | 1.695242991 | 0.017946326 | 4.709504924 | 4.00722531 | 5.303523747 | 4.523675391 | 7.691737974 | 7.45182838 | 6.173421703 | 6.420873184 |
| ENSG00000132640 | BTBD3 | 1.451749851 | 0.588240473 | 3.256740151 | 0.181428238 | 4.708490001 | 2 | 8.147241016 | 2 | 8.010962573 | 8.0358969 | 9.657971023 | 9.952144095 |
| ENSG00000135241 | PNPLA8 | 3.092272547 | 0.014980237 | 1.615821967 | 0.231005457 | 4.708094515 | 6.96351253 | 8.004992147 | 3.369317972 | 10.09394503 | 10.0907323 | 8.645942939 | 8.604294318 |
| ENSG00000173372 | C1QA | -2.311499221 | 0.415158868 | 7.018722266 | 0.00011303 | 4.707223045 | 2 | 2 | 2 | 2 | 2.14022021 | 7.399274054 | 6.690737414 |
| ENSG00000124466 | LYPD3 | 4.168470658 | 0.001516222 | 0.53861101 | 0.736592879 | 4.707081668 | 2 | 4.11352228 | 2 | 6.687949235 | 6.51372389 | 3.336159116 | 3.698365557 |
| ENSG00000130311 | DDA1 | 1.93107328 | 0.011671303 | 2.775995215 | 0.000147509 | 4.707068495 | 7.940205782 | 8.213429689 | 6.322058544 | 9.599482126 | 9.63011121 | 9.978130526 | 10.815215 |
| ENSG00000182446 | NPLOC4 | 2.196413452 | 0.003108658 | 2.508251783 | 0.000574909 | 4.704665235 | 8.631580145 | 9.338312924 | 7.225742779 | 10.82486117 | 10.8288217 | 11.11450432 | 11.16123925 |
| ENSG00000265972 | TXNIP | 2.43016199 | 0.000809887 | 2.2727605 | 0.001788445 | 4.70292249 | 11.39439513 | 10.22039562 | 9.658943166 | 13.00776242 | 13.0681681 | 13.22911886 | 12.42109226 |
| ENSG00000187796 | CARD9 | 0.72685174 | 0.738608861 | 3.975287714 | 0.026249826 | 4.702139454 | 3.327711234 | 2 | 2 | 2.784612485 | 2.94009902 | 5.385935071 | 4.836507146 |
| ENSG00000143412 | ANXA9 | -1.31051372 | 0.609315815 | 6.011610918 | 0.000843969 | 4.701097198 | 2 | 2 | 2 | 2.141737246 | 2.14022021 | 5.951260634 | 6.276965131 |
| ENSG00000131037 | EPS8L1 | 2.667086297 | 0.087812072 | 2.032813627 | 0.196453617 | 4.699899924 | 3.327711234 | 2 | 3.369317972 | 4.76150579 | 4.83276841 | 4.434275199 | 4.09155917 |
| ENSG00000116685 | KIAA2013 | 2.364688365 | 0.172285908 | 2.331022268 | 0.170945512 | 4.695710633 | 8.564112515 | 8.029695761 | 2 | 10.10155624 | 10.0777096 | 10.07684508 | 10.03510519 |
| ENSG00000173262 | SLC2A14 | 7.87240424 | 1.40E-07 | -3.180102844 | 0.101051926 | 4.692337396 | 2 | 2 | 3.369317972 | 8.899755849 | 8.84568954 | 2.139668783 | 2 |
| ENSG00000235098 | ANKRD65 | 2.082628425 | 0.148587465 | 2.60903543 | 0.058149869 | 4.691663855 | 2 | 3.076665209 | 4.058998049 | 4.984880435 | 4.53541753 | 4.881704255 | 5.524371359 |
| ENSG00000206140 | TMEM191C | 3.546620212 | 0.168573196 | 1.137300979 | 0.687792014 | 4.683921191 | 2 | 2 | 2 | 3.881293667 | 4.0930011 | 2.831336763 | 2.421067459 |
| ENSG00000153165 | RGPD3 | 9.874894914 | 1.41E-11 | -5.191151308 | 0.024699035 | 4.683743607 | 2 | 2 | 2 | 9.903687576 | 9.91090151 | 2 | 2 |
| ENSG00000108509 | CAMTA2 | 2.266074758 | 0.006973831 | 2.416663798 | 0.003546828 | 4.682738555 | 5.532353208 | 7.701025996 | 8.007629717 | 9.580933135 | 9.72573666 | 9.866098684 | 9.741247946 |
| ENSG00000091831 | ESR1 | 1.858549446 | 0.519421794 | 2.822232137 | 0.292236517 | 4.680781583 | 2 | 2 | 2 | 2.868578001 | 3.01487888 | 3.679182183 | 3.261853426 |
| ENSG00000270181 | BIVM-ERCC5 | 9.867533259 | 1.49E-11 | -5.191151307 | 0.024725305 | 4.676381952 | 2 | 2 | 2 | 9.9123685 | 9.88746056 | 2 | 2 |
| ENSG00000158427 | TMSB15B | 3.819206528 | 0.106744126 | 0.857035831 | 0.74998849 | 4.676242359 | 2 | 2 | 2 | 4.268917409 | 4.12711147 | 2.457042231 | 2.632958241 |
| ENSG00000120662 | MTRF1 | 2.491149602 | 0.133069745 | 2.184276196 | 0.185851802 | 4.675425797 | 2 | 5.131603625 | 2 | 6.043074888 | 6.08095082 | 5.696113106 | 5.85096143 |
| ENSG00000125952 | MAX | 2.836824107 | 8.10E-07 | 1.837694776 | 0.002094646 | 4.674518883 | 8.107517749 | 6.78762695 | 6.882775709 | 10.17502299 | 10.2094139 | 9.163367078 | 9.23204367 |
| ENSG00000079432 | CIC | 1.967570814 | 0.238207033 | 2.703394439 | 0.089199351 | 4.670965253 | 3.327711234 | 8.169643011 | 9.61300511 | 10.4372252 | 10.4587184 | 10.98899358 | 11.35251454 |
| ENSG00000171606 | ZNF274 | 2.377915927 | 0.177300996 | 2.293026132 | 0.186484076 | 4.670942058 | 4.00722531 | 7.099070191 | 2 | 8.03159949 | 8.01099479 | 8.305655918 | 7.442672408 |
| ENSG00000196437 | ZNF569 | 2.473383903 | 0.185725588 | 2.195620297 | 0.237878752 | 4.6690042 | 2 | 5.595927442 | 2 | 6.522845756 | 6.46787796 | 6.550361701 | 5.821002687 |
| ENSG00000167641 | PPP1R14A | -1.96494491 | 0.05126002 | 6.633263113 | 4.09E-15 | 4.668318203 | 4.815609471 | 5.457120953 | 2 | 3.09461492 | 3.45167086 | 11.30777198 | 10.91012447 |
| ENSG00000090975 | PITPNM2 | 2.229003061 | 0.107251275 | 2.436874049 | 0.070958342 | 4.665877111 | 2 | 6.602424674 | 5.156626401 | 7.677258395 | 7.65252709 | 7.913205701 | 7.823364542 |
| ENSG00000073670 | ADAM11 | 0.858624886 | 0.761711449 | 3.806692124 | 0.108864366 | 4.66531701 | 2 | 2 | 2 | 2.498691189 | 2.59489879 | 4.378961816 | 3.972018587 |
| ENSG00000038002 | AGA | 2.711919859 | 0.10980137 | 1.953236653 | 0.257630995 | 4.665156512 | 3.327711234 | 6.464272742 | 2 | 7.683067677 | 7.70145652 | 6.437578914 | 7.331709059 |
| ENSG00000162066 | AMDHD2 | 3.648696788 | 0.003512781 | 1.015110784 | 0.474513246 | 4.663807572 | 3.327711234 | 5.595927442 | 2 | 7.697489359 | 7.95749494 | 5.31402805 | 5.337485258 |
| ENSG00000159263 | SIM2 | 0.279018914 | 0.930366193 | 4.381707389 | 0.079386828 | 4.660726303 | 2 | 2 | 2 | 2.600425781 | 2.14022021 | 4.620674722 | 4.692661645 |
| ENSG00000175895 | PLEKHF2 | 2.316448096 | 0.043740293 | 2.344018533 | 0.038597696 | 4.660466629 | 4.815609471 | 5.838937808 | 2 | 7.026323566 | 7.1744553 | 7.3389565 | 6.883203642 |
| ENSG00000168038 | ULK4 | 2.144167381 | 0.170194337 | 2.514449294 | 0.094858183 | 4.658616674 | 2 | 4.442710129 | 2 | 4.907595604 | 5.21224859 | 5.196301326 | 5.576534358 |
| ENSG00000130827 | PLXNA3 | 2.193679761 | 0.007925031 | 2.464694479 | 0.002373092 | 4.658374239 | 6.816637977 | 8.191702463 | 9.180874355 | 10.50725261 | 10.5675219 | 10.60176359 | 10.98850403 |
| ENSG00000105063 | PPP6R1 | 2.298179606 | 0.025564784 | 2.359971456 | 0.020815791 | 4.658151062 | 5.095959458 | 8.376385542 | 7.734844356 | 9.864602909 | 9.88372429 | 9.931435456 | 9.939999927 |
| ENSG00000139718 | SETD1B | 2.344107313 | 0.120721992 | 2.312917027 | 0.120253602 | 4.65702434 | 7.485143553 | 6.389896633 | 2 | 8.782894215 | 8.75316018 | 8.787633372 | 8.684919451 |
| ENSG00000121797 | CCRL2 | 0.858622531 | 0.763635326 | 3.798138503 | 0.112770502 | 4.656761034 | 2 | 2 | 2 | 2.498691189 | 2.59489879 | 4.425202323 | 3.8919413 |
| ENSG00000115760 | BIRC6 | 2.424258428 | 0.028967364 | 2.232022951 | 0.044399174 | 4.656281379 | 5.866944937 | 9.461450882 | 9.049077186 | 11.12929738 | 11.2037318 | 10.91081207 | 11.03666497 |
| ENSG00000104312 | RIPK2 | 3.379558245 | 3.80E-11 | 1.275480082 | 0.022326117 | 4.655038327 | 8.042902253 | 7.189546432 | 6.80318718 | 10.78995706 | 10.7847595 | 8.506119506 | 8.860338475 |
| ENSG00000129317 | PUS7L | 1.694608589 | 0.341002462 | 2.960354592 | 0.073190118 | 4.654963181 | 3.327711234 | 6.602424674 | 2 | 6.820355514 | 6.84106088 | 8.092249024 | 8.040411184 |
| ENSG00000178105 | DDX10 | 1.982316489 | 0.091493251 | 2.671733701 | 0.017257852 | 4.65405019 | 7.159102577 | 6.728495673 | 3.369317972 | 8.312101014 | 8.42842973 | 8.720150661 | 9.325315571 |
| ENSG00000144857 | BOC | 2.041465191 | 0.110115082 | 2.609986731 | 0.03302563 | 4.651451921 | 2 | 5.72254323 | 4.058998049 | 6.664650034 | 6.37866593 | 7.021403672 | 7.130061054 |
| ENSG00000107338 | SHB | 2.624976303 | 0.052851591 | 2.025738468 | 0.140205611 | 4.650714771 | 3.327711234 | 7.790474954 | 5.930865648 | 9.217515344 | 9.17405701 | 8.234726135 | 8.894131604 |
| ENSG00000174151 | CYB561D1 | 2.356590949 | 0.026235442 | 2.293302156 | 0.029927201 | 4.649893105 | 3.327711234 | 6.728495673 | 5.59507254 | 8.074220386 | 8.07355593 | 8.125537812 | 7.887748062 |
| ENSG00000110330 | BIRC2 | 3.077421387 | 0.009582037 | 1.57171565 | 0.21425456 | 4.649137036 | 4.815609471 | 8.981105197 | 7.896971941 | 11.08446011 | 11.0544122 | 9.703475817 | 9.420076944 |
| ENSG00000112367 | FIG4 | 1.735619492 | 0.452840011 | 2.91168602 | 0.174638772 | 4.647305512 | 2 | 6.844429755 | 2 | 6.846474326 | 7.16216989 | 8.196576732 | 8.130406117 |
| ENSG00000134780 | DAGLA | 1.705395527 | 0.112733934 | 2.941228176 | 0.003132042 | 4.646623703 | 2 | 4.710564154 | 5.59507254 | 6.147608921 | 6.33555364 | 7.297296608 | 7.562617572 |
| ENSG00000146411 | SLC2A12 | -0.269907868 | 0.845522872 | 4.914543319 | 2.05E-06 | 4.644635452 | 6.138360429 | 5.303523747 | 2 | 5.040242897 | 4.98931816 | 9.60873181 | 10.33090604 |
| ENSG00000189120 | SP6 | 2.333643931 | 0.440702227 | 2.310744596 | 0.433987431 | 4.644388527 | 2 | 2 | 2 | 3.227757282 | 3.15372745 | 3.5515855 | 2.671842496 |
| ENSG00000213516 | RBMXL1 | 2.725522521 | 0.019173548 | 1.915673247 | 0.109758105 | 4.641195768 | 4.00722531 | 7.572420273 | 5.772716771 | 9.16555511 | 9.09547637 | 7.977459036 | 8.611187869 |
| ENSG00000256087 | ZNF432 | 2.755480421 | 0.023372934 | 1.883162516 | 0.132614409 | 4.638642937 | 2 | 6.228547981 | 6.073378139 | 8.235106869 | 8.30721295 | 7.594015257 | 7.20984412 |
| ENSG00000181894 | ZNF329 | 1.992607967 | 0.015889499 | 2.645733562 | 0.00085525 | 4.638341529 | 3.327711234 | 5.595927442 | 5.930865648 | 7.234487582 | 7.17037177 | 7.980559086 | 7.687652155 |
| ENSG00000138399 | FASTKD1 | 1.676997459 | 0.075317795 | 2.960424016 | 0.000709386 | 4.637421475 | 7.53297911 | 10.24169813 | 9.528916236 | 11.18792842 | 11.1077025 | 12.27036227 | 12.57426012 |

| | | | | | | | | | | | | | |
|---|---|---|---|---|---|---|---|---|---|---|---|---|---|
| ENSG00000104783 | KCNN4 | 0.092420682 | 0.966177728 | 4.544785493 | 0.007602279 | 4.637206176 | 5.70932728 | 2 | 2 | 4.470469939 | 4.2865223 | 8.768928693 | 8.272375674 |
| ENSG00000101574 | METTL4 | 2.472055089 | 0.228871904 | 2.163034257 | 0.290390881 | 4.635089346 | 2 | 6.228547981 | 2 | 6.984595707 | 7.24216308 | 7.019897747 | 6.587749895 |
| ENSG00000165152 | TMEM246 | 2.238510944 | 0.340614989 | 2.395475552 | 0.295009548 | 4.633986496 | 2 | 7.189546432 | 2 | 7.826439377 | 7.86451142 | 7.854430723 | 8.132149972 |
| ENSG00000188277 | C15orf62 | 1.014336527 | 0.731092219 | 3.618129302 | 0.153081117 | 4.632465829 | 2 | 2 | 2 | 2.784612485 | 2.38539787 | 4.155805465 | 3.924507493 |
| ENSG00000100410 | PHF5A | 2.276763331 | 0.011761813 | 2.355551093 | 0.008462275 | 4.632314424 | 6.563806279 | 4.93638933 | 4.058998049 | 7.677258395 | 7.77859536 | 7.80632247 | 7.805990373 |
| ENSG00000010318 | PHF7 | 2.100297411 | 0.17960083 | 2.530480908 | 0.092183627 | 4.630778319 | 2 | 4.11352228 | 2 | 4.739373666 | 4.65713986 | 5.180212365 | 4.951586397 |
| ENSG00000164733 | CTSB | 2.832325181 | 4.02E-09 | 1.79831437 | 0.00030473 | 4.630639551 | 12.49796133 | 11.76214832 | 11.62203638 | 14.83208287 | 14.8581549 | 14.118582 | 13.42052145 |
| ENSG00000189190 | ZNF600 | 2.747083529 | 0.04265939 | 1.88075426 | 0.176369212 | 4.627837789 | 2 | 6.04685525 | 4.058998049 | 7.444001439 | 7.5014335 | 6.818078339 | 6.420873184 |
| ENSG00000163121 | NEURL3 | 4.433857648 | 0.008153101 | 0.193885068 | 0.929171616 | 4.627742716 | 2 | 2 | 3.369317972 | 5.348759107 | 5.72778967 | 2.62493832 | 2.671842496 |
| ENSG00000144655 | CSRNP1 | 2.295984228 | 0.019121386 | 2.328381143 | 0.016731819 | 4.624365371 | 5.330594134 | 7.819099893 | 5.772716771 | 8.980991923 | 9.04507321 | 8.914214292 | 9.165908829 |
| ENSG00000181904 | C5orf24 | 1.565580316 | 1.57E-07 | 3.057486267 | 2.01E-26 | 4.623066584 | 8.042902253 | 7.847167909 | 8.042706615 | 9.601017164 | 9.44559847 | 11.11176776 | 10.90528859 |
| ENSG00000125753 | VASP | 1.804799693 | 0.087057774 | 2.815104861 | 0.004461713 | 4.619904554 | 5.70932728 | 8.191702463 | 8.922968322 | 9.852410253 | 9.9489441 | 10.7202043 | 11.07547412 |
| ENSG00000157426 | AASDH | 2.13912798 | 0.050098285 | 2.479999996 | 0.019580896 | 4.619127977 | 2 | 5.457120953 | 4.523675391 | 6.529309835 | 6.5890786 | 7.07314455 | 6.674075308 |
| ENSG00000145780 | FEM1C | 2.410474806 | 0.202020098 | 2.204301086 | 0.24003034 | 4.614775893 | 4.467311984 | 7.928237109 | 2 | 8.877128559 | 8.84312657 | 8.800408416 | 8.495377414 |
| ENSG00000047621 | C12orf4 | 2.389596536 | 0.035285398 | 2.220908737 | 0.049717407 | 4.610505273 | 3.327711234 | 6.666836837 | 4.874587708 | 7.913023977 | 7.8441483 | 7.613117257 | 7.80708241 |
| ENSG00000163156 | SCNM1 | 2.82649748 | 1.45E-05 | 1.783254097 | 0.008792063 | 4.609751577 | 5.330594134 | 4.442710129 | 6.073378139 | 8.058664058 | 8.1187584 | 6.898949899 | 7.219718817 |
| ENSG00000175793 | SFN | 1.778306598 | 0.560280703 | 2.83134867 | 0.320348511 | 4.609655268 | 2 | 2 | 2 | 3.023133209 | 2.77779907 | 2.937246078 | 3.8919413 |
| ENSG00000134996 | OSTF1 | 2.744378563 | 0.078977124 | 1.863847028 | 0.244680076 | 4.608225591 | 4.00722531 | 6.666836837 | 2 | 8.063125866 | 7.92393105 | 7.08038692 | 7.184857568 |
| ENSG00000115841 | RMDN2 | 3.383164901 | 0.000657972 | 1.221282107 | 0.267803599 | 4.604447008 | 2 | 3.686303414 | 4.523675391 | 6.70517998 | 6.56439512 | 4.764298067 | 4.553838275 |
| ENSG00000144674 | GOLGA4 | 2.406675371 | 1.03E-06 | 2.196540665 | 9.02E-06 | 4.603216036 | 8.313285031 | 9.13702746 | 8.007629717 | 11.00233175 | 10.938838 | 10.85879791 | 10.65660252 |
| ENSG00000070770 | CSNK2A2 | 1.459765371 | 0.347773435 | 3.141527708 | 0.026364101 | 4.601293079 | 7.384433967 | 6.78762695 | 2 | 7.93753157 | 8.01327646 | 9.651421039 | 9.632103066 |
| ENSG00000119684 | MLH3 | 2.695488514 | 0.093060766 | 1.904905627 | 0.244428647 | 4.600394141 | 4.815609471 | 7.189546432 | 2 | 8.581467308 | 8.46556258 | 8.031518488 | 7.388257299 |
| ENSG00000156510 | HKDC1 | 2.151756149 | 0.485453797 | 2.447322834 | 0.410308224 | 4.599078983 | 2 | 2 | 2 | 3.349643765 | 2.77779907 | 2.803598391 | 3.600986722 |
| ENSG00000197860 | SGTB | 2.053696164 | 0.03605895 | 2.54084682 | 0.007353946 | 4.594542984 | 4.00722531 | 7.002537973 | 6.534109959 | 8.302698479 | 8.32752521 | 9.020825125 | 8.532164007 |
| ENSG00000105967 | TFEC | 0.689489289 | 0.823523332 | 3.903113734 | 0.122897491 | 4.592603024 | 2 | 2 | 2 | 2.498691189 | 2.49394783 | 4.764298067 | 3.496556178 |
| ENSG00000104219 | ZDHHC2 | 2.181343352 | 0.167549493 | 2.410331359 | 0.116655583 | 4.591674711 | 4.00722531 | 7.538394143 | 3.369317972 | 8.341783829 | 8.22100445 | 8.309975344 | 8.683730876 |
| ENSG00000124074 | ENKD1 | 2.163759701 | 0.257710703 | 2.427498348 | 0.190345876 | 4.591258049 | 2 | 5.72254323 | 2 | 6.463327516 | 6.17362678 | 6.637393426 | 6.514766253 |
| ENSG00000073849 | ST6GAL1 | 0.016502355 | 0.991315828 | 4.56823799 | 3.36E-05 | 4.584740345 | 4.00722531 | 5.838937808 | 2 | 4.739373666 | 4.79082474 | 9.194070966 | 8.995324202 |
| ENSG00000166801 | FAM111A | 2.70970505 | 0.01749407 | 1.873234064 | 0.111040183 | 4.582939113 | 3.327711234 | 7.189546432 | 6.073378139 | 8.883449599 | 8.87484109 | 8.038228667 | 8.06618078 |
| ENSG00000275214 | | 11.57769263 | 6.99E-21 | -6.99822544 | 0.000718107 | 4.579467186 | 2 | 3.686303414 | 2 | 13.4043705 | 13.4189574 | 2 | 2 |
| ENSG00000121671 | CRY2 | 1.749419715 | 0.219370406 | 2.828735061 | 0.033929555 | 4.578154776 | 5.70932728 | 6.844429755 | 2 | 7.477814553 | 7.58663873 | 8.748160468 | 8.425990514 |
| ENSG00000146232 | NFKBIE | 2.270999171 | 0.135425564 | 2.304760314 | 0.122835348 | 4.575759485 | 2 | 7.189546432 | 6.43197674 | 8.471640654 | 8.56844956 | 7.985968197 | 8.960904642 |
| ENSG00000177464 | GPR4 | 3.608170841 | 0.054422065 | 0.962783204 | 0.64966917 | 4.570954045 | 3.327711234 | 2 | 2 | 4.443344104 | 5.11361705 | 3.059555873 | 2.885414003 |
| ENSG00000110446 | SLC15A3 | -3.753164585 | 0.151557124 | 8.323647244 | 9.48E-07 | 4.570482659 | 2 | 2 | 2 | 2 | 2 | 8.551124447 | 8.157201821 |
| ENSG00000155090 | KLF10 | 2.023207184 | 0.000282479 | 2.547083008 | 2.79E-06 | 4.570290192 | 8.228665467 | 7.393647515 | 7.285536013 | 9.695291778 | 9.70330142 | 10.56712557 | 9.764764027 |
| ENSG00000139597 | N4BP2L1 | 1.461837607 | 0.436838812 | 3.102218533 | 0.068532826 | 4.56405614 | 2 | 5.303523747 | 2 | 5.363304456 | 5.18011485 | 7.184873526 | 6.301968891 |
| ENSG00000101974 | ATP11C | 1.742432707 | 0.160411864 | 2.820799844 | 0.015161691 | 4.56323255 | 4.00722531 | 7.819099893 | 6.322058544 | 8.534041678 | 8.39031583 | 9.259659123 | 9.763920691 |
| ENSG00000197323 | TRIM33 | 2.839267278 | 0.044631453 | 1.722595792 | 0.241877934 | 4.56186307 | 3.327711234 | 8.029695761 | 5.772716771 | 9.614759498 | 9.52369893 | 8.619836896 | 8.284971554 |
| ENSG00000182902 | SLC25A18 | 1.150926308 | 0.693061884 | 3.407996869 | 0.179503712 | 4.558923176 | 2 | 2 | 2 | 2.784612485 | 2.49394783 | 3.917810786 | 3.858622973 |
| ENSG00000156671 | SAMD8 | 2.466272479 | 0.000431391 | 2.091180704 | 0.003223268 | 4.557453183 | 7.709880939 | 7.901717006 | 5.930865648 | 9.85562885 | 9.85666073 | 9.547336845 | 9.415430124 |
| ENSG00000102804 | TSC22D1 | 2.829671901 | 5.33E-06 | 1.726467688 | 0.008092771 | 4.556139589 | 10.79750658 | 9.856140801 | 9.132850258 | 12.9227779 | 12.9054164 | 11.84225044 | 11.7806379 |
| ENSG00000187961 | KLHL17 | 1.874392119 | 0.178652578 | 2.678461431 | 0.04234099 | 4.55285355 | 4.467311984 | 6.228547981 | 2 | 6.861921612 | 6.86652414 | 7.891135869 | 7.352801367 |
| ENSG00000205571 | SMN2 | 9.739807391 | 4.22E-11 | -5.191151232 | 0.025435 | 4.548656159 | 2 | 2 | 2 | 9.685196338 | 9.85538952 | 2 | 2 |
| ENSG00000172062 | SMN1 | 9.739469131 | 4.22E-11 | -5.191151233 | 0.025435 | 4.548317898 | 2 | 2 | 2 | 9.685196338 | 9.8547535 | 2 | 2 |
| ENSG00000175175 | PPM1E | -0.303340129 | 0.941709668 | 4.848779903 | 0.145407903 | 4.545439774 | 2 | 2 | 2 | 2.498691189 | 2 | 5.333556395 | 4.720747915 |
| ENSG00000167553 | TUBA1C | 2.993677919 | 9.39E-08 | 1.551470345 | 0.009202275 | 4.545148264 | 8.816894989 | 8.469613413 | 8.206335842 | 11.52096327 | 11.4772671 | 9.300854716 | 10.5556122 |
| ENSG00000119915 | ELOVL3 | 3.332028241 | 0.280963509 | 1.212355966 | 0.716509282 | 4.544384207 | 2 | 2 | 2 | 3.663521627 | 3.98554648 | 3.035909414 | 2.172608095 |
| ENSG00000013288 | MAN2B2 | 2.233181757 | 0.034724134 | 2.310332443 | 0.027071029 | 4.5435142 | 5.866944937 | 9.170736405 | 8.763901704 | 10.7018566 | 10.6994819 | 10.6478287 | 10.89672248 |
| ENSG00000196975 | ANXA4 | 3.031015746 | 7.22E-07 | 1.512398611 | 0.021023289 | 4.543414357 | 7.27616306 | 5.595927442 | 6.534109959 | 9.516508835 | 9.64710779 | 8.035250217 | 8.120777053 |
| ENSG00000126561 | STAT5A | 2.603073336 | 0.053580304 | 1.940147357 | 0.156889836 | 4.543220693 | 2 | 4.93638933 | 6.534109959 | 7.865280446 | 7.89204953 | 7.346184975 | 7.103640702 |
| ENSG00000169715 | MT1E | 4.702696723 | 0.008406966 | -0.159534351 | 0.944101725 | 4.543162373 | 7.031698319 | 2 | 3.369317972 | 10.14218788 | 10.1969221 | 5.899789217 | 4.766376438 |
| ENSG00000169247 | SH3TC2 | 0.977614717 | 0.564719306 | 3.564660514 | 0.014458835 | 4.54227523 | 3.327711234 | 3.076665209 | 3.369317972 | 3.515101962 | 4.1930041 | 5.002522427 | 6.735016458 |
| ENSG00000165355 | FBXO33 | 2.292705991 | 3.39E-05 | 2.249221854 | 4.43E-05 | 4.541927845 | 5.330594134 | 6.666836837 | 5.930865648 | 8.321442665 | 8.35116763 | 8.392591106 | 8.186868687 |
| ENSG00000100994 | PYGB | 1.365729408 | 0.256781312 | 3.172249205 | 0.00344396 | 4.537978613 | 8.418903792 | 9.497318007 | 5.772716771 | 9.867156706 | 9.94057646 | 11.72642042 | 11.6863866 |
| ENSG00000259571 | BLID | 3.116287235 | 0.233747682 | 1.421198117 | 0.609265042 | 4.537485352 | 2 | 2 | 2 | 3.754611335 | 3.60476711 | 2.717032197 | 2.782576693 |
| ENSG00000184014 | DENND5A | 2.470894845 | 0.000138719 | 2.06326463 | 0.001719657 | 4.534159475 | 8.855032352 | 9.680074956 | 7.934808822 | 11.45076334 | 11.4679397 | 10.91243621 | 11.18001149 |
| ENSG00000116678 | LEPR | 1.381987874 | 0.355101172 | 3.151001143 | 0.019679078 | 4.532989017 | 4.467311984 | 6.464272742 | 2 | 6.529309835 | 6.6253296 | 8.281350892 | 8.325926932 |
| ENSG00000109846 | CRYAB | -0.276020581 | 0.728428075 | 4.806918306 | 2.13E-15 | 4.530897725 | 8.257427002 | 7.393647515 | 8.865353883 | 8.036145657 | 7.99029569 | 13.43597286 | 12.59103903 |
| ENSG00000197629 | MPEG1 | -1.30515308 | 0.688878059 | 5.835817726 | 0.019545711 | 4.530664646 | 2 | 2 | 2 | 2.270786764 | 2 | 6.275391383 | 5.5506886 |
| ENSG00000196371 | FUT4 | 2.62609483 | 0.007362713 | 1.902868529 | 0.058813982 | 4.528963359 | 3.327711234 | 6.389896633 | 5.156626401 | 7.905589775 | 8.05595642 | 7.283556699 | 7.264927592 |
| ENSG00000013374 | NUB1 | 2.514679366 | 1.39E-09 | 2.012650357 | 1.67E-06 | 4.527329724 | 7.940205782 | 8.147241016 | 7.163363831 | 10.28797351 | 10.3378525 | 9.795903284 | 9.829876356 |
| ENSG00000187608 | ISG15 | 5.400089827 | 9.28E-14 | -0.877162753 | 0.303668356 | 4.522927074 | 9.2903348 | 7.232742069 | 9.06622548 | 14.14029863 | 14.1713234 | 7.580798201 | 8.165740501 |
| ENSG00000198914 | POU3F3 | 2.866948486 | 0.453973535 | 1.653663415 | 0.675209517 | 4.520611901 | 2 | 2 | 2 | 4.205751501 | 2.14022021 | 3.1504466 | 2.466008509 |
| ENSG00000135966 | TGFBRAP1 | 1.86720493 | 0.018700929 | 2.652776259 | 0.000461516 | 4.519981189 | 5.330594134 | 7.503546079 | 6.43197674 | 8.488325436 | 8.56844956 | 9.184693023 | 9.421503729 |
| ENSG00000105664 | COMP | 3.64416849 | 0.229538248 | 0.8757536 | 0.796725981 | 4.51992209 | 2 | 2 | 2 | 4.139692846 | 3.98554648 | 2.885263063 | 2.117335275 |
| ENSG00000066777 | ARFGEF1 | 2.633091534 | 0.002023832 | 1.886203102 | 0.0323856 | 4.519294636 | 7.435667355 | 8.053983481 | 5.59507254 | 9.933844998 | 9.98018912 | 9.543411506 | 8.786963584 |
| ENSG00000163346 | PBXIP1 | 2.173395419 | 0.001407766 | 2.344629182 | 0.000481509 | 4.518024601 | 8.468948322 | 8.702538795 | 6.882775709 | 10.34496592 | 10.4049916 | 10.41345332 | 10.66746623 |
| ENSG00000166068 | SPRED1 | 1.723638793 | 0.000146231 | 2.793810278 | 1.36E-10 | 4.517449071 | 7.33131396 | 7.503546079 | 6.534109959 | 8.966774408 | 8.81069917 | 10.08208599 | 9.814754207 |
| ENSG00000039319 | ZFYVE16 | 2.826510321 | 3.88E-11 | 1.690692991 | 0.000135456 | 4.517203312 | 6.816637977 | 7.669933291 | 7.163363831 | 10.10696834 | 10.0306182 | 9.108453756 | 8.748827534 |
| ENSG00000125457 | MIF4GD | 0.848049931 | 0.695215075 | 3.668234398 | 0.047681033 | 4.516284329 | 2 | 6.04685525 | 2 | 5.161714397 | 5.62411716 | 8.359038717 | 7.809264007 |
| ENSG00000164066 | INTU | 0.824637671 | 0.553979465 | 3.690645994 | 0.001690432 | 4.515283665 | 2 | 5.595927442 | 4.058998049 | 5.210797711 | 5.37722731 | 8.408216037 | 7.515294811 |
| ENSG00000137496 | IL18BP | 2.978467708 | 0.057973134 | 1.536185302 | 0.354201102 | 4.51465301 | 4.467311984 | 6.951736442 | 2 | 8.619831492 | 8.46389529 | 7.439255893 | 6.730418989 |
| ENSG00000171823 | FBXL14 | 2.319335158 | 0.230806091 | 2.193888372 | 0.252105582 | 4.51322353 | 2 | 5.838937808 | 2 | 6.628980152 | 6.54560135 | 6.453284622 | 6.482320745 |
| ENSG00000115226 | FNDC4 | 1.559509382 | 0.449705292 | 2.952845556 | 0.118891631 | 4.512354938 | 3.327711234 | 7.431218269 | 2 | 7.562190249 | 7.37938471 | 8.764439224 | 8.924637944 |
| ENSG00000244617 | ASPRV1 | 1.858525064 | 0.597963567 | 2.653665371 | 0.426580825 | 4.512190435 | 2 | 2 | 2 | 2.868578001 | 3.01487888 | 3.981090143 | 2.326754077 |
| ENSG00000158290 | CUL4B | 3.04918256 | 0.00515506 | 1.462155866 | 0.211603388 | 4.511338426 | 5.095959458 | 8.791896851 | 7.734844356 | 10.90533885 | 10.8529084 | 9.380756704 | 9.211159041 |
| ENSG00000177294 | FBXO39 | 3.59715341 | 0.182411111 | 0.914153706 | 0.76219473 | 4.511307116 | 2 | 2 | 2 | 4.139692846 | 3.90918929 | 2.803598391 | 2.277179783 |
| ENSG00000166123 | GPT2 | 1.784816194 | 0.10022545 | 2.725283379 | 0.007667651 | 4.510099574 | 5.866944937 | 7.761270542 | 4.874587708 | 8.418656444 | 8.44204342 | 9.602960031 | 9.074546388 |
| ENSG00000110274 | CEP164 | 2.420802535 | 0.006743859 | 2.086925449 | 0.021068527 | 4.507727984 | 4.00722531 | 6.78762695 | 6.203069917 | 8.451360823 | 8.40776525 | 8.275052313 | 7.90110545 |
| ENSG00000175054 | ATR | 2.66203026 | 0.076244071 | 1.845480495 | 0.229495532 | 4.507510754 | 2 | 7.274681896 | 5.772716771 | 8.652768197 | 8.86097251 | 7.985196707 | 7.922424886 |
| ENSG00000262607 | | 0.346727922 | 0.883560872 | 4.157308547 | 0.024419064 | 4.504036468 | 2 | 2 | 4.058998049 | 3.09461492 | 3.2184422 | 4.90146639 | 6.862343163 |
| ENSG00000184898 | RBM43 | 3.330452516 | 2.07E-07 | 1.170349373 | 0.099416364 | 4.500801889 | 5.532353208 | 4.442710129 | 6.203069917 | 8.76657222 | 8.6932307 | 6.723178551 | 6.501335855 |
| ENSG00000277224 | HIST1H2BF | 3.333959738 | 0.211918038 | 1.165160935 | 0.688459235 | 4.499120674 | 2 | 2 | 2 | 3.921164147 | 3.74316441 | 2.42098416 | 2.851942824 |
| ENSG00000172915 | NBEA | 2.201109161 | 0.028717201 | 2.297687199 | 0.020800484 | 4.49879636 | 2 | 5.131603625 | 5.39244166 | 6.79375511 | 6.79406287 | 6.666558925 | 7.078537388 |
| ENSG00000100985 | MMP9 | -3.273856792 | 0.31862506 | 7.771405272 | 0.00125484 | 4.49754848 | 2 | 3.076665209 | 2 | 2 | 2.14022021 | 9.738690605 | 4.353147976 |
| ENSG00000120063 | GNA13 | 2.151488964 | 0.084559691 | 2.342441124 | 0.054227008 | 4.493930089 | 4.467311984 | 8.234834549 | 8.536706852 | 10.02128323 | 9.95489142 | 10.39730548 | 9.920959478 |
| ENSG00000198168 | SVIP | 2.623748729 | 0.193809048 | 1.867898473 | 0.362474125 | 4.491647201 | 5.866944937 | 2 | 2 | 6.841288255 | 6.82556437 | 6.280431262 | 5.921228818 |
| ENSG00000139044 | B4GALNT3 | 0.279088054 | 0.93222913 | 4.209362084 | 0.106037661 | 4.488450138 | 2 | 2 | 2 | 2.600425781 | 2.14022021 | 4.166733998 | 4.793076545 |
| ENSG00000145794 | MEGF10 | 0.863476532 | 0.78336925 | 3.621910536 | 0.172416263 | 4.485387068 | 2 | 2 | 2 | 2.784612485 | 2.2680111 | 4.017766949 | 4.077147165 |
| ENSG00000277377 | | 4.417690307 | 0.037736766 | 0.06575594 | 0.980518967 | 4.483446247 | 2 | 2 | 2 | 4.739373666 | 4.63360314 | 2.3070652 | 2.374681325 |
| ENSG00000123243 | ITIH5 | -4.157567537 | 2.12E-05 | 8.639601772 | 3.14E-27 | 4.482034234 | 5.095959458 | 4.11352228 | 6.203069917 | 2.784612485 | 2.38539787 | 13.48928411 | 14.11736845 |
| ENSG00000136114 | THSD1 | 3.058979725 | 0.031675145 | 1.420138274 | 0.352050398 | 4.479117999 | 2 | 4.442710129 | 2 | 5.900763176 | 5.91514978 | 4.547306681 | 4.365060851 |
| ENSG00000107566 | ERLIN1 | 2.369767068 | 0.007296304 | 2.108680534 | 0.01795557 | 4.478447602 | 7.096806373 | 7.874700259 | 5.156626401 | 9.460955886 | 9.40620612 | 9.091707098 | 9.25263028 |
| ENSG00000149634 | SPATA25 | 2.016102074 | 0.531143362 | 2.461699069 | 0.425433438 | 4.477801143 | 2 | 2 | 2 | 3.406930997 | 2.49394783 | 3.500668998 | 2.981406154 |
| ENSG00000165626 | BEND7 | 1.124323933 | 0.516556851 | 3.353143987 | 0.026967588 | 4.477467921 | 2 | 4.93638933 | 2 | 4.867343068 | 4.40247105 | 6.828438696 | 6.535998174 |
| ENSG00000170248 | PDCD6IP | 2.288026277 | 0.00012178 | 2.189409771 | 0.000235994 | 4.477436048 | 8.797441463 | 8.981105197 | 7.451345288 | 10.84021544 | 10.8239557 | 10.6406201 | 10.82117286 |
| ENSG00000125355 | TMEM255A | 0.282075773 | 0.950628899 | 4.189882878 | 0.279259581 | 4.471958651 | 2 | 2 | 2 | 2.695456494 | 2 | 4.461156421 | 4.532869193 |
| ENSG00000158467 | AHCYL2 | 1.411858859 | 0.14556217 | 3.059000066 | 0.000486858 | 4.470858925 | 6.73718411 | 6.844429755 | 4.058998049 | 7.723090567 | 7.66128317 | 9.120096958 | 9.496527079 |
| ENSG00000240344 | PPIL3 | 2.653800486 | 0.060426137 | 1.816849579 | 0.210056903 | 4.470650065 | 4.467311984 | 6.464272742 | 2 | 7.797276258 | 7.8286851 | 7.182183216 | 6.782425482 |
| ENSG00000112419 | PHACTR2 | 0.923372171 | 0.197969988 | 3.544743656 | 9.58E-09 | 4.468115826 | 8.313285031 | 9.842262059 | 9.25752426 | 10.15216947 | 10.2247551 | 12.36480811 | 13.14363006 |
| ENSG00000104888 | SLC17A7 | 0.495876252 | 0.866101534 | 3.97169921 | 0.087078455 | 4.467575462 | 2 | 2 | 2 | 2.3892346 | 2.49394783 | 4.360042493 | 4.279527956 |
| ENSG00000119929 | CUTC | 2.214721831 | 0.085392635 | 2.25116235 | 0.074947938 | 4.46588418 | 4.467311984 | 6.04685525 | 2 | 6.931899482 | 7.16627666 | 7.15911073 | 7.015689447 |
| ENSG00000085514 | PILRA | 1.697734472 | 0.351613795 | 2.767894354 | 0.102455592 | 4.465628825 | 2 | 3.076665209 | 4.058998049 | 4.358765249 | 4.56059747 | 6.075339068 | 4.033026654 |
| ENSG00000183576 | SETD3 | 1.967922546 | 0.000770044 | 2.49500032 | 1.17E-05 | 4.462922866 | 6.138360429 | 7.538394143 | 7.600047325 | 9.129845271 | 9.21824109 | 9.795243215 | 9.595281739 |
| ENSG00000181418 | DDN | 1.012879912 | 0.73397139 | 3.449992171 | 0.179708528 | 4.462872083 | 2 | 2 | 2 | 2.695456494 | 2.49394783 | 3.56816602 | 4.201948172 |
| ENSG00000128591 | FLNC | 1.443149457 | 0.003480256 | 3.019069041 | 6.35E-11 | 4.462218498 | 9.497412313 | 10.59613751 | 10.30429804 | 11.60349653 | 11.6842056 | 13.06706733 | 13.35669676 |
| ENSG00000187244 | BCAM | 0.954622079 | 0.569254933 | 3.507549863 | 0.016033481 | 4.462171942 | 7.624139768 | 6.602424674 | 2 | 7.330144653 | 7.76511961 | 10.18174778 | 9.994939827 |

| | | | | | | | | | | | | | |
|---|---|---|---|---|---|---|---|---|---|---|---|---|---|
| ENSG00000179627 | ZBTB42 | 2.175234945 | 0.045106267 | 2.285351212 | 0.032219121 | 4.460586158 | 2 | 4.442710129 | 5.59507254 | 6.598566398 | 6.57678965 | 6.691367378 | 6.695462882 |
| ENSG00000182240 | BACE2 | 1.747772875 | 0.025627053 | 2.710814932 | 0.000252233 | 4.458587808 | 8.468948322 | 6.464272742 | 7.225742779 | 9.314267267 | 9.37460909 | 10.15266847 | 10.43999426 |
| ENSG00000165684 | SNAPC4 | 2.402816313 | 0.12734523 | 2.055723441 | 0.191066494 | 4.458539754 | 3.327711234 | 3.076665209 | 6.882775709 | 7.697489359 | 7.86198169 | 7.401588825 | 7.481493974 |
| ENSG00000136854 | STXBP1 | 1.932722512 | 0.259806896 | 2.524286948 | 0.12381131 | 4.45700946 | 7.435667355 | 8.297207175 | 2 | 9.231457548 | 9.30367823 | 9.705819181 | 9.992781936 |
| ENSG00000182263 | FIGN | 2.440084145 | 0.010021807 | 2.013139632 | 0.036087858 | 4.453223777 | 4.467311984 | 5.131603625 | 2 | 6.5612028 | 6.67228943 | 6.148830293 | 6.270645889 |
| ENSG00000244486 | SCARF2 | 1.349517772 | 0.231258748 | 3.103345825 | 0.002260394 | 4.452863597 | 6.138360429 | 8.276714465 | 9.083172337 | 9.551067273 | 9.65224116 | 11.89104907 | 10.47837298 |
| ENSG00000081320 | STK17B | 2.244837065 | 0.000587551 | 2.206962456 | 0.000694887 | 4.451799521 | 5.532353208 | 6.951736442 | 7.398165132 | 9.057896526 | 8.98821297 | 9.07802367 | 8.887436006 |
| ENSG00000154545 | MAGED4 | 9.642540498 | 6.47E-11 | -5.191151323 | 0.025401936 | 4.451389175 | 2 | 2 | 2 | 9.678668908 | 9.67331554 | 2 | 2 |
| ENSG00000187243 | MAGED4B | 9.642540498 | 6.47E-11 | -5.191151323 | 0.025401936 | 4.451389175 | 2 | 2 | 2 | 9.678668908 | 9.67331554 | 2 | 2 |
| ENSG00000166311 | SMPD1 | 2.048837934 | 0.134855166 | 2.397783063 | 0.070852767 | 4.446620997 | 4.815609471 | 9.540933215 | 9.099922433 | 10.79633594 | 10.8506773 | 11.10459281 | 11.23642639 |
| ENSG00000165935 | SMCO2 | 2.820338664 | 0.339862734 | 1.623569323 | 0.594804601 | 4.443907988 | 2 | 2 | 2 | 3.663521627 | 3.28037825 | 3.12825509 | 2.466008509 |
| ENSG00000104856 | RELB | 2.264191814 | 0.202526124 | 2.179698307 | 0.213719558 | 4.443890121 | 8.418903792 | 6.311476754 | 2 | 9.359407363 | 9.39837151 | 9.31719306 | 9.272532009 |
| ENSG00000166816 | LDHD | -1.315777443 | 0.684355643 | 5.758544897 | 0.019926216 | 4.442767454 | 2 | 2 | 2 | 2 | 2.2680111 | 5.832754931 | 5.937276556 |
| ENSG00000151726 | ACSL1 | 1.97049231 | 0.0190625 | 2.465024571 | 0.002440403 | 4.435516881 | 5.532353208 | 7.315436877 | 5.39244166 | 8.258639605 | 8.36731141 | 8.892160003 | 8.708487604 |
| ENSG00000002834 | LASP1 | 1.226145656 | 0.030568134 | 3.208995352 | 4.31E-10 | 4.435141008 | 10.89224845 | 11.12072186 | 9.819251342 | 11.90383364 | 11.9697428 | 13.9522767 | 13.8847597 |
| ENSG00000168411 | RFWD3 | 2.098874821 | 0.14406389 | 2.334590084 | 0.094670586 | 4.433464905 | 3.327711234 | 7.503546079 | 4.874587708 | 8.166267464 | 8.34936267 | 8.50665722 | 8.480396055 |
| ENSG00000162434 | JAK1 | 2.440051864 | 0.129911918 | 1.99253965 | 0.21842725 | 4.432591514 | 4.00722531 | 10.16572116 | 8.380957451 | 11.37674805 | 11.4162868 | 10.96091792 | 10.93880421 |
| ENSG00000188290 | HES4 | 2.172889715 | 0.22173333 | 2.258002404 | 0.194278246 | 4.430892118 | 8.19931886 | 5.838937808 | 2 | 9.025111392 | 9.01464729 | 9.1990699 | 9.002968638 |
| ENSG00000170370 | EMX2 | -0.718700302 | 0.861382316 | 5.146520932 | 0.112361418 | 4.42782063 | 2 | 2 | 2 | 2.3892346 | 2 | 5.963846589 | 4.147809227 |
| ENSG00000159899 | NPR2 | 2.534080135 | 0.103857132 | 1.893483742 | 0.230468038 | 4.427563876 | 6.468621203 | 7.701025996 | 2 | 9.077877436 | 9.20532695 | 8.528003102 | 8.487906181 |
| ENSG00000180089 | TMEM86B | 2.5371604 | 0.142232651 | 1.890004159 | 0.279023454 | 4.427164558 | 2 | 3.686303414 | 2 | 4.574143677 | 4.60967607 | 4.155805465 | 3.987514168 |
| ENSG00000204442 | FAM155A | 5.005142756 | 0.002167981 | -0.57851108 | 0.773343718 | 4.426631675 | 2 | 2 | 5.59507254 | 8.873322599 | 8.82765235 | 3.981090143 | 3.496556178 |
| ENSG00000048649 | RSF1 | 2.065181641 | 0.041980392 | 2.361070593 | 0.017309633 | 4.426252234 | 8.674867122 | 8.414402626 | 5.59507254 | 10.10534684 | 10.1089823 | 10.45430433 | 10.34788887 |
| ENSG00000213996 | TM6SF2 | 2.834731596 | 0.156125244 | 1.58471474 | 0.445656681 | 4.419446335 | 2 | 3.076665209 | 2 | 3.95996233 | 4.37434574 | 3.411599105 | 3.209956562 |
| ENSG00000101255 | TRIB3 | 1.532807668 | 0.387432577 | 2.884498959 | 0.077151749 | 4.417306626 | 2 | 7.467835385 | 6.629488387 | 7.917958914 | 8.15847021 | 10.05770853 | 8.067092663 |
| ENSG00000135018 | UBQLN1 | 2.200887476 | 4.83E-07 | 2.21564729 | 3.66E-07 | 4.416534765 | 7.940205782 | 8.834582582 | 8.922968322 | 10.8110007 | 10.829793 | 10.90806718 | 10.75834164 |
| ENSG00000196547 | MAN2A2 | 1.641634485 | 0.177844856 | 2.774849816 | 0.014545829 | 4.416484301 | 7.709880939 | 7.701025996 | 4.058998049 | 8.799033614 | 8.79487126 | 10.20975726 | 9.564939053 |
| ENSG00000118960 | HS1BP3 | 1.820340569 | 0.456251283 | 2.595194584 | 0.260775363 | 4.415535153 | 2 | 7.315436877 | 2 | 7.559034656 | 7.56191311 | 8.413960373 | 8.224954107 |
| ENSG00000162601 | MYSM1 | 2.597769835 | 0.043022105 | 1.817650377 | 0.166932988 | 4.415420211 | 3.327711234 | 6.389896633 | 7.734844356 | 9.226493643 | 9.21824109 | 8.652274805 | 8.212645079 |
| ENSG00000137672 | TRPC6 | 0.515189383 | 0.793804216 | 3.899750903 | 0.01477161 | 4.414940287 | 2 | 5.303523747 | 2 | 4.54891529 | 4.37434574 | 7.067324363 | 7.97389845 |
| ENSG00000189001 | SBSN | 3.621642239 | 0.148210562 | 0.792855281 | 0.781073497 | 4.41449752 | 2 | 2 | 2 | 4.237680128 | 3.82856379 | 2.62493832 | 2.421067459 |
| ENSG00000162591 | MEGF6 | -1.924240815 | 0.438437558 | 6.336052598 | 0.00313113 | 4.411811783 | 2 | 7.669933291 | 2 | 4.670864177 | 4.16043394 | 11.93560775 | 12.7588774 |
| ENSG00000108510 | MED13 | 2.466552612 | 0.175086607 | 1.944287291 | 0.287569328 | 4.410839903 | 2 | 8.791896851 | 8.825624954 | 10.68241243 | 10.6813023 | 10.28424887 | 10.02645802 |
| ENSG00000120337 | TNFSF18 | 5.716547896 | 0.052953075 | -1.305878904 | 0.706382819 | 4.410668992 | 2 | 2 | 2 | 5.720439317 | 5.96314905 | 2.267001569 | 2 |
| ENSG00000130584 | ZBTB46 | -0.47572303 | 0.812947514 | 4.886200295 | 0.001927913 | 4.410477265 | 2 | 5.72254323 | 2 | 3.881293667 | 4.16043394 | 8.96165712 | 9.027536965 |
| ENSG00000181555 | SETD2 | 2.163160337 | 0.050378597 | 2.246129777 | 0.038950505 | 4.409290114 | 5.095959458 | 8.255926468 | 8.608698767 | 10.01150489 | 10.1317374 | 10.13242893 | 10.17828285 |
| ENSG00000143199 | ADCY10 | 2.899992654 | 0.289703301 | 1.508921226 | 0.597377354 | 4.40891388 | 2 | 2 | 2 | 3.663521627 | 3.39680241 | 2.858551861 | 2.70970617 |
| ENSG00000145244 | CORIN | 1.008937075 | 0.456962569 | 3.399404944 | 0.003252545 | 4.408342019 | 2 | 3.686303414 | 4.058998049 | 4.387512679 | 4.02226117 | 5.960710379 | 6.62538071 |
| ENSG00000107290 | SETX | 2.59142962 | 6.32E-13 | 1.816616787 | 7.39E-07 | 4.408046407 | 8.493334308 | 8.337339476 | 9.06622548 | 11.23069904 | 11.2496932 | 10.41445658 | 10.51792633 |
| ENSG00000156711 | MAPK13 | 3.312713252 | 4.13E-09 | 1.092978467 | 0.080745253 | 4.405691719 | 6.73718411 | 5.595927442 | 6.203069917 | 9.473580277 | 9.47569407 | 7.575682301 | 6.91798856 |
| ENSG00000170191 | NANP | 1.922472875 | 0.387605511 | 2.482485845 | 0.243517965 | 4.40495872 | 2 | 6.602424674 | 2 | 6.984595707 | 6.92586261 | 7.540404604 | 7.459438299 |
| ENSG00000011347 | SYT7 | 3.420206639 | 0.079639272 | 0.984038486 | 0.650307428 | 4.404245125 | 5.70932728 | 2 | 2 | 7.440576128 | 7.43830714 | 5.610655473 | 4.445805334 |
| ENSG00000131381 | RBSN | 2.163075179 | 2.92E-05 | 2.240070326 | 1.30E-05 | 4.403145506 | 7.750913381 | 7.60566235 | 6.43197674 | 9.474417988 | 9.55457603 | 9.609733245 | 9.573306996 |
| ENSG00000153201 | RANBP2 | 2.5505255 | 4.10E-17 | 1.85257965 | 1.51E-09 | 4.40310515 | 8.564112515 | 9.030900487 | 8.845626171 | 11.34067986 | 11.4004669 | 10.80385399 | 10.53186455 |
| ENSG00000155903 | RASA2 | 2.670129547 | 0.053560884 | 1.732223801 | 0.225078882 | 4.402353348 | 6.009025997 | 7.002537973 | 2 | 8.601543953 | 8.68895519 | 7.79843893 | 7.63942137 |
| ENSG00000119596 | YLPM1 | 1.801475253 | 0.070841918 | 2.600838521 | 0.005996167 | 4.402313775 | 8.313285031 | 8.147241016 | 5.39244166 | 9.531881013 | 9.53088329 | 10.27780546 | 10.37538871 |
| ENSG00000145715 | RASA1 | 2.998669115 | 0.066585047 | 1.402598248 | 0.42122258 | 4.401267363 | 4.467311984 | 8.255926468 | 3.369317972 | 9.764048416 | 9.79963996 | 8.247007147 | 8.14947356 |
| ENSG00000222038 | POTEJ | 9.590269497 | 1.05E-10 | -5.191151262 | 0.025794745 | 4.399118235 | 2 | 2 | 2 | 9.55265467 | 9.6919516 | 2 | 2 |
| ENSG00000102786 | INTS6 | 2.564919756 | 0.025631643 | 1.833622892 | 0.120387966 | 4.398542649 | 3.327711234 | 7.051611315 | 6.958202216 | 8.967964567 | 9.0169232 | 8.498029643 | 7.996092644 |
| ENSG00000069966 | GNB5 | 1.814598426 | 5.20E-05 | 2.582362478 | 2.53E-09 | 4.396960904 | 6.009025997 | 6.04685525 | 6.80318718 | 8.04293816 | 8.11238697 | 8.850830124 | 8.823038243 |
| ENSG00000243147 | MRPL33 | 1.908069132 | 0.216950087 | 2.488601613 | 0.092611393 | 4.396670745 | 7.667647223 | 3.076665209 | 5.156626401 | 8.343618883 | 8.06917614 | 8.59173005 | 8.95894086 |
| ENSG00000143442 | POGZ | 2.258890333 | 0.127775345 | 2.136271071 | 0.14497505 | 4.395161403 | 3.327711234 | 8.522788974 | 6.882775709 | 9.587142668 | 9.62638965 | 9.702067969 | 9.22919499 |
| ENSG00000169018 | FEM1B | 1.356566964 | 0.009715897 | 3.037703048 | 3.69E-10 | 4.394270012 | 7.435667355 | 8.655694736 | 8.35328246 | 9.60637697 | 9.56472327 | 11.36982317 | 11.14470525 |
| ENSG00000141446 | ESCO1 | 2.484318086 | 0.096071046 | 1.908865335 | 0.205129503 | 4.393183421 | 5.095959458 | 6.899080544 | 2 | 8.170409056 | 8.09094333 | 7.872065621 | 7.176431616 |
| ENSG00000168890 | TMEM150A | 1.652156203 | 0.480384146 | 2.738377795 | 0.209907098 | 4.390533998 | 2 | 6.899080544 | 2 | 6.912249745 | 7.05550676 | 7.938932725 | 8.148612304 |
| ENSG00000019144 | PHLDB1 | 2.423349925 | 0.015831771 | 1.965973334 | 0.053853036 | 4.389323259 | 6.73718411 | 8.655694736 | 9.868959282 | 11.2965993 | 11.3514882 | 10.78724204 | 10.9440257 |
| ENSG00000101187 | SLCO4A1 | 3.521307695 | 0.071965527 | 0.86679076 | 0.695304679 | 4.388098455 | 3.327711234 | 2 | 2 | 4.623314311 | 4.85329164 | 3.172301926 | 2.632958241 |
| ENSG00000198815 | FOXJ3 | 2.367358191 | 0.032621666 | 2.019992523 | 0.070331611 | 4.387350713 | 5.866944937 | 8.147241016 | 5.156626401 | 9.30110636 | 9.34230452 | 9.083440537 | 8.861915706 |
| ENSG00000185104 | FAF1 | 1.811294217 | 0.001702654 | 2.574440111 | 3.35E-06 | 4.385734328 | 5.532353208 | 5.595927442 | 6.718950348 | 7.68596357 | 7.8720741 | 8.476781207 | 8.589773203 |
| ENSG00000239474 | KLHL41 | 1.274446705 | 0.688362385 | 3.110738614 | 0.272520906 | 4.385185319 | 2 | 2 | 2 | 2.695456494 | 2.68924543 | 4.144793515 | 2.981406154 |
| ENSG00000173404 | INSM1 | 1.282428516 | 0.837050918 | 3.101871805 | 0.584589142 | 4.384300321 | 2 | 2 | 2 | 3.162721474 | 2 | 4.425202323 | 2 |
| ENSG00000063854 | HAGH | 2.266267375 | 0.157563342 | 2.116002352 | 0.181344746 | 4.382269727 | 7.790810999 | 6.389896633 | 2 | 8.918344554 | 8.89972012 | 8.699603841 | 8.818171767 |
| ENSG00000174840 | PDE12 | 1.873960932 | 0.113862445 | 2.507545811 | 0.026981937 | 4.381506743 | 4.00722531 | 7.638155693 | 6.073378139 | 8.461536372 | 8.34393422 | 9.018941717 | 9.043840859 |
| ENSG00000001084 | GCLC | 3.361800744 | 0.039063396 | 1.017548336 | 0.574400364 | 4.379349081 | 2 | 6.666836837 | 8.236946975 | 10.39094584 | 10.4196875 | 8.209185391 | 7.934467448 |
| ENSG00000196422 | PPP1R26 | 2.032134984 | 0.232496722 | 2.34348691 | 0.155092367 | 4.375621894 | 2 | 7.701025996 | 2 | 8.349110081 | 8.5130942 | 8.920281122 | 8.537437492 |
| ENSG00000153922 | CHD1 | 2.654369115 | 0.102115752 | 1.720119292 | 0.301852744 | 4.374488407 | 2 | 7.701025996 | 5.930865648 | 9.142549916 | 9.09655277 | 8.456328963 | 7.875307424 |
| ENSG00000123358 | NR4A1 | 1.96337904 | 0.03398501 | 2.407252194 | 0.00729012 | 4.370631235 | 4.815609471 | 7.051611315 | 5.39244166 | 7.883054517 | 8.12510181 | 8.709446788 | 8.126037235 |
| ENSG00000196391 | ZNF774 | 0.859970152 | 0.615680633 | 3.507727115 | 0.016000913 | 4.367697267 | 2 | 4.442710129 | 2 | 4.299492648 | 3.65239003 | 6.375345632 | 6.267475858 |
| ENSG00000196890 | HIST3H2BB | 3.62007234 | 0.148005951 | 0.746366936 | 0.794186359 | 4.366439275 | 2 | 2 | 2 | 4.034562184 | 4.05806467 | 2.384001718 | 2.632958241 |
| ENSG00000184916 | JAG2 | 0.621609719 | 0.644603834 | 3.742013931 | 0.000583068 | 4.363623651 | 3.327711234 | 4.11352228 | 2 | 4.034562184 | 3.74316441 | 6.726875343 | 6.514766253 |
| ENSG00000108819 | PPP1R9B | 1.678881628 | 0.088518928 | 2.683944692 | 0.00361429 | 4.36282632 | 9.435834703 | 8.433041974 | 6.534109959 | 10.2141657 | 10.2394476 | 11.1725846 | 11.28426283 |
| ENSG00000131016 | AKAP12 | 0.664237014 | 0.343511582 | 3.696623728 | 2.46E-10 | 4.360860742 | 9.940323746 | 8.655694736 | 8.631919287 | 9.84659862 | 9.89984536 | 12.73924589 | 13.04543027 |
| ENSG00000070476 | ZXDC | 1.801786951 | 0.223962299 | 2.558426795 | 0.068912293 | 4.360213746 | 3.327711234 | 7.538394143 | 4.874587708 | 7.961629787 | 8.0314011 | 8.995380861 | 8.440137439 |
| ENSG00000117408 | IPO13 | 2.023644768 | 0.341002462 | 2.335900207 | 0.256784608 | 4.359544976 | 4.00722531 | 8.376385542 | 2 | 8.937912013 | 8.80016655 | 9.221844492 | 9.137974605 |
| ENSG00000141568 | FOXK2 | 1.6479922 | 0.262397112 | 2.70842538 | 0.048844314 | 4.356417581 | 3.327711234 | 8.101365653 | 8.266922072 | 9.244284335 | 9.26686067 | 10.45137457 | 10.15574229 |
| ENSG00000010072 | SPRTN | 2.010406268 | 0.364510704 | 2.34560461 | 0.273901976 | 4.356010878 | 2 | 6.602424674 | 2 | 6.946463121 | 7.12888795 | 7.483632518 | 7.239267729 |
| ENSG00000089057 | SLC23A2 | 1.792501542 | 0.490463824 | 2.561648318 | 0.297700044 | 4.354149861 | 2 | 7.928237109 | 2 | 8.136938566 | 8.14604733 | 8.71690136 | 9.066361132 |
| ENSG00000058091 | CDK14 | 2.952484641 | 9.83E-05 | 1.401613647 | 0.086347448 | 4.354098288 | 5.095959458 | 7.35507212 | 7.029880454 | 9.641111693 | 9.74233706 | 7.98287976 | 8.309067496 |
| ENSG00000176903 | PNMA1 | 1.628799547 | 2.31E-06 | 2.724263008 | 2.47E-16 | 4.353062555 | 7.975256505 | 7.35507212 | 7.777116083 | 9.302993856 | 9.34230452 | 10.46111737 | 10.36411774 |
| ENSG00000222009 | BTBD19 | 1.873716398 | 0.241482575 | 2.479335739 | 0.104869869 | 4.353052137 | 2 | 4.93638933 | 7.029880454 | 7.584088227 | 7.55253065 | 7.855275386 | 8.417435289 |
| ENSG00000254122 | PCDHGB7 | 3.606487754 | 1.99E-06 | 0.746343465 | 0.399833462 | 4.352831218 | 4.815609471 | 7.051611315 | 7.163363831 | 10.19039992 | 10.2554384 | 7.38764416 | 7.398436576 |
| ENSG00000175573 | C11orf68 | 1.710714624 | 0.184030119 | 2.641406072 | 0.029385896 | 4.352120696 | 4.00722531 | 7.701025996 | 8.236946975 | 9.115954467 | 9.16690126 | 10.05752505 | 10.07735401 |
| ENSG00000099904 | ZDHHC8 | 1.843113001 | 0.090381529 | 2.508014996 | 0.015805954 | 4.351127997 | 7.904282243 | 7.572420273 | 4.523675391 | 9.010169232 | 9.10406516 | 9.71701465 | 9.720533286 |
| ENSG00000136193 | SCRN1 | 1.863457192 | 0.044940119 | 2.487633011 | 0.005240878 | 4.351090203 | 8.855032352 | 10.3138733 | 7.734844356 | 11.18537258 | 11.2225883 | 11.82580793 | 11.82901558 |
| ENSG00000196083 | IL1RAP | 2.715530384 | 2.09E-08 | 1.634636166 | 0.001191054 | 4.350166549 | 5.330594134 | 6.140560021 | 5.59507254 | 8.439397922 | 8.3220138 | 7.237652294 | 7.398436576 |
| ENSG00000185046 | ANKS1B | 1.153728358 | 0.725192089 | 3.196326471 | 0.271421671 | 4.350054828 | 2 | 2 | 2 | 2.947924294 | 2.2680111 | 3.584558147 | 3.875378318 |
| ENSG00000060491 | OGFR | 2.947678434 | 1.44E-12 | 1.402311585 | 0.001432681 | 4.349990019 | 8.257427002 | 7.790474954 | 7.225742779 | 10.70756787 | 10.7810819 | 9.174577117 | 9.236508834 |
| ENSG00000167447 | SMG8 | 2.297016138 | 0.177474856 | 2.051667378 | 0.225372001 | 4.348683516 | 2 | 7.790474954 | 5.59507254 | 8.788294101 | 8.75042728 | 8.571313099 | 8.480396055 |
| ENSG00000034152 | MAP2K3 | 1.76890476 | 0.004683603 | 2.579484739 | 1.65E-05 | 4.348389498 | 8.816894989 | 7.701025996 | 7.896971941 | 9.927741409 | 10.0227123 | 10.25109847 | 11.1725107 |
| ENSG00000197622 | CDC42SE1 | 2.364962281 | 1.27E-10 | 1.983314407 | 8.68E-08 | 4.348276688 | 8.009475818 | 7.669933291 | 8.460932916 | 10.40111447 | 10.4384862 | 10.04443806 | 10.0346391 |
| ENSG00000110888 | CAPRIN2 | 2.217955595 | 0.084982806 | 2.127203552 | 0.094541013 | 4.345159147 | 3.327711234 | 6.78762695 | 7.734844356 | 8.941551548 | 9.00091592 | 8.970259716 | 8.786963584 |
| ENSG00000143367 | TUFT1 | 0.983915534 | 0.248121056 | 3.361035634 | 4.77E-06 | 4.344951168 | 4.00722531 | 5.595927442 | 6.43197674 | 6.573765157 | 6.56439512 | 8.802599622 | 8.984263871 |
| ENSG00000165521 | EML5 | 4.340921089 | 0.013143309 | 0.00378445 | 0.998969048 | 4.344705539 | 2 | 3.076665209 | 2 | 5.210797711 | 5.6826301 | 2.68698273 | 2.466008509 |
| ENSG00000180423 | HARBI1 | 2.059256814 | 0.21101766 | 2.284441371 | 0.152011997 | 4.343698185 | 2 | 4.710564154 | 2 | 5.161714397 | 5.33407135 | 5.684753614 | 5.177806074 |
| ENSG00000104218 | CSPP1 | 2.18380629 | 0.319942561 | 2.159163129 | 0.317458054 | 4.342969419 | 2 | 6.602424674 | 2 | 7.277360022 | 7.13728069 | 7.223329552 | 7.145684098 |
| ENSG00000111981 | ULBP1 | 0.803523706 | 0.423328228 | 3.538783967 | 2.39E-05 | 4.342307673 | 4.467311984 | 5.457120953 | 4.058998049 | 5.363304456 | 5.70538658 | 8.618842453 | 7.51663051 |
| ENSG00000101986 | ABCD1 | 2.600146421 | 0.000203189 | 1.74097549 | 0.017043752 | 4.341121911 | 5.532353208 | 6.228547981 | 7.451345288 | 9.161399541 | 9.16690126 | 8.351275892 | 8.273956186 |
| ENSG00000065357 | DGKA | 1.390903188 | 0.492446389 | 2.947229176 | 0.110137106 | 4.338132364 | 4.467311984 | 8.029695761 | 2 | 7.805288479 | 8.07574085 | 9.45312092 | 9.521357751 |
| ENSG00000022567 | SLC45A4 | 1.601157292 | 0.120909818 | 2.736424961 | 0.004198848 | 4.337582253 | 2 | 4.710564154 | 5.156626401 | 5.818703821 | 5.93454177 | 7.180836177 | 6.695462882 |
| ENSG00000074590 | NUAK1 | 1.693170838 | 0.335781482 | 2.644368944 | 0.109372081 | 4.337539782 | 4.00722531 | 4.11352228 | 8.460932916 | 8.689305661 | 8.63367258 | 9.853683372 | 9.306905794 |
| ENSG00000128165 | ADM2 | 1.028291973 | 0.439229935 | 3.306846439 | 0.004137928 | 4.335138412 | 4.00722531 | 7.002537973 | 5.39244166 | 6.927012081 | 6.99214046 | 9.831964324 | 8.054273543 |
| ENSG00000163681 | SLMAP | 2.175966264 | 0.146700672 | 2.155441676 | 0.143504721 | 4.33140794 | 2 | 6.899080544 | 8.585098385 | 9.621581886 | 9.50760441 | 9.736400072 | 9.324934421 |
| ENSG00000162714 | ZNF496 | 1.522082097 | 0.232251697 | 2.809305831 | 0.017180311 | 4.331387928 | 3.327711234 | 7.431218269 | 6.43197674 | 7.915493556 | 8.04037874 | 9.278673449 | 9.22674878 |
| ENSG00000175352 | NRIP3 | 2.136313201 | 0.013800383 | 2.194484868 | 0.010690228 | 4.330798069 | 4.467311984 | 5.838937808 | 7.029880454 | 8.122047456 | 8.25418043 | 8.327733606 | 8.161477479 |
| ENSG00000186197 | EDARADD | 7.904218508 | 5.69E-07 | -3.573834001 | 0.05519677 | 4.330384507 | 2 | 4.11352228 | 2 | 10.23853288 | 10.2291785 | 2.267001569 | 2 |
| ENSG00000111670 | GNPTAB | 1.336559917 | 0.001086736 | 2.992904793 | 4.99E-15 | 4.32946471 | 7.904282243 | 7.274681896 | 8.110401118 | 9.101928616 | 9.12109086 | 10.82211411 | 10.6966318 |
| ENSG00000119729 | RHOQ | 2.525954475 | 0.002821664 | 1.803364532 | 0.038931255 | 4.329319007 | 5.70932728 | 8.191702463 | 8.007629717 | 10.17037776 | 10.1406356 | 9.371930177 | 9.49788026 |

| | | | | | | | | | | | | | |
|---|---|---|---|---|---|---|---|---|---|---|---|---|---|
| ENSG00000196776 | CD47 | 2.916737894 | 0.01213142 | 1.407027397 | 0.258826994 | 4.323765291 | 9.143983763 | 7.847167909 | 5.156626401 | 11.00000655 | 11.026728 | 9.623185015 | 9.384672535 |
| ENSG00000198205 | ZXDA | 2.548522148 | 0.167369545 | 1.775070364 | 0.345152391 | 4.323592511 | 2 | 5.595927442 | 2 | 6.483440992 | 6.64900057 | 5.899789217 | 5.772630506 |
| ENSG00000073737 | DHRS9 | 2.496810989 | 0.523459664 | 1.826619643 | 0.642308336 | 4.323430633 | 2 | 2 | 2 | 3.289987342 | 3.28037825 | 3.465698119 | 2.059860178 |
| ENSG00000197879 | MYO1C | 1.068694035 | 0.094901214 | 3.253641074 | 8.27E-09 | 4.322335109 | 9.015175615 | 10.32389548 | 10.44859742 | 11.09075629 | 11.1562579 | 13.19651665 | 13.4092619 |
| ENSG00000126775 | ATG14 | 2.558481994 | 0.005070133 | 1.763441856 | 0.062126304 | 4.32192385 | 5.330594134 | 7.315436877 | 5.156626401 | 8.887228971 | 8.74494589 | 8.243785424 | 7.783974227 |
| ENSG00000085491 | SLC25A24 | 2.449142445 | 1.72E-10 | 1.86956576 | 1.54E-06 | 4.318708204 | 7.031698319 | 6.666836837 | 6.322058544 | 9.166592134 | 9.04395769 | 8.569255422 | 8.494021883 |
| ENSG00000129646 | QRICH2 | 2.31980206 | 0.174904784 | 1.995742366 | 0.241998457 | 4.315544427 | 3.327711234 | 2 | 5.39244166 | 6.139181293 | 6.29861567 | 6.550361701 | 4.757365452 |
| ENSG00000099204 | ABLIM1 | 0.972236936 | 0.262982679 | 3.343160596 | 8.39E-06 | 4.315397532 | 4.467311984 | 6.844429755 | 6.43197674 | 7.210554257 | 7.12888795 | 9.390986528 | 9.617238105 |
| ENSG00000108389 | MTMR4 | 0.962298544 | 0.698178574 | 3.352699953 | 0.123013154 | 4.314998497 | 3.327711234 | 8.717821467 | 2 | 8.14116509 | 8.11663771 | 10.26324148 | 10.70778414 |
| ENSG00000157617 | C2CD2 | 1.212672222 | 0.006321713 | 3.101483837 | 2.74E-14 | 4.314156059 | 6.89194353 | 7.73146271 | 7.398165132 | 8.652768197 | 8.51631523 | 10.6126694 | 10.29807728 |
| ENSG00000103111 | MON1B | 1.675898949 | 0.530264818 | 2.635524314 | 0.292216612 | 4.311423262 | 2 | 8.147241016 | 2 | 8.180711314 | 8.30535212 | 9.048785116 | 9.327219816 |
| ENSG00000165886 | UBTD1 | 2.057135241 | 1.06E-07 | 2.253309153 | 3.81E-09 | 4.310444394 | 6.816637977 | 6.844429755 | 6.203069917 | 8.697938662 | 8.65720817 | 8.919473684 | 8.823038243 |
| ENSG00000131095 | GFAP | 1.111715244 | 0.685365166 | 3.197329005 | 0.19011257 | 4.309044248 | 2 | 2 | 6.073378139 | 6.476767609 | 3.15372745 | 7.793159202 | 7.392628644 |
| ENSG00000122299 | ZC3H7A | 1.989402518 | 0.063965155 | 2.31923775 | 0.026467216 | 4.308640268 | 5.532353208 | 8.213429689 | 5.930865648 | 9.044420506 | 9.04841459 | 9.259020976 | 9.480531399 |
| ENSG00000132646 | PCNA | 1.940153717 | 0.272669235 | 2.364929723 | 0.164528915 | 4.30508344 | 4.467311984 | 7.35507212 | 2 | 7.783822874 | 7.95986274 | 8.415679223 | 8.158913605 |
| ENSG00000115977 | AAK1 | 1.909514811 | 0.000106004 | 2.394452083 | 6.41E-07 | 4.303966894 | 8.716893051 | 7.761270542 | 7.691296561 | 10.02357442 | 10.0306182 | 10.58734728 | 10.42921874 |
| ENSG00000049883 | PTCD2 | 1.752834916 | 0.28829652 | 2.550535788 | 0.101181698 | 4.303370705 | 3.327711234 | 6.140560021 | 2 | 6.535745081 | 6.39974839 | 7.443755487 | 7.000495137 |
| ENSG00000071054 | MAP4K4 | 3.08411984 | 6.71E-10 | 1.216323733 | 0.025387429 | 4.300443572 | 9.473095744 | 10.64891478 | 9.757248847 | 13.10109474 | 13.1642759 | 11.21433092 | 11.31769121 |
| ENSG00000163220 | S100A9 | 0.497864743 | 0.881014624 | 3.80176741 | 0.152741385 | 4.299632153 | 2 | 2 | 2 | 2.498691189 | 2.38539787 | 4.73522696 | 3.287118176 |
| ENSG00000156860 | FBRS | 2.000298597 | 0.105778838 | 2.296844324 | 0.055557322 | 4.297142921 | 7.579279349 | 9.443177293 | 5.59507254 | 10.28080824 | 10.2731592 | 10.54962008 | 10.59534423 |
| ENSG00000167323 | STIM1 | 1.632227523 | 0.269395698 | 2.663305225 | 0.053894075 | 4.295532748 | 7.159102577 | 8.451443573 | 3.369317972 | 9.060130342 | 8.95060977 | 10.10960242 | 9.949426738 |
| ENSG00000178878 | APOLD1 | 0.036865314 | 0.977083699 | 4.258551747 | 6.69E-06 | 4.295417061 | 6.138360429 | 4.442710129 | 6.073378139 | 5.697663076 | 5.7929843 | 8.539610093 | 10.54059012 |
| ENSG00000138496 | PARP9 | 3.824040474 | 3.99E-05 | 0.46998313 | 0.681431908 | 4.294023605 | 8.19931886 | 5.946638857 | 8.978369997 | 11.97066735 | 11.9540913 | 8.466314923 | 8.761839632 |
| ENSG00000128641 | MYO1B | 2.080657803 | 3.30E-09 | 2.212195986 | 2.45E-10 | 4.292853789 | 10.09970294 | 9.770792087 | 9.61300511 | 11.92880215 | 11.9068589 | 12.28418861 | 11.76849593 |
| ENSG00000183246 | RIMBP3C | 5.695499803 | 0.00189134 | -1.403130722 | 0.536670773 | 4.292369082 | 2 | 2 | 2 | 5.900763176 | 5.74985018 | 2.094598369 | 2.172608095 |
| ENSG00000166529 | ZSCAN21 | 2.066959841 | 0.260797132 | 2.222389 | 0.21477409 | 4.289348842 | 2 | 5.303523747 | 2 | 5.602782483 | 6.01871653 | 6.231826305 | 5.651404445 |
| ENSG00000250312 | ZNF718 | 2.505865235 | 0.052932111 | 1.783159471 | 0.177027586 | 4.289024706 | 2 | 4.710564154 | 3.369317972 | 6.052081244 | 5.85535975 | 5.673303969 | 4.77533149 |
| ENSG00000101336 | HCK | 0.269719039 | 0.937710599 | 4.018566073 | 0.146799193 | 4.288285111 | 2 | 2 | 2 | 2.141737246 | 2.59489879 | 4.834511823 | 3.640730416 |
| ENSG00000281742 | | 2.154036419 | 0.163051497 | 2.133941819 | 0.159212693 | 4.287978237 | 3.327711234 | 7.60566235 | 4.523675391 | 8.378048624 | 8.34212018 | 7.924517766 | 8.662770717 |
| ENSG00000135317 | SNX14 | 2.81673845 | 0.028112563 | 1.467200721 | 0.280137585 | 4.283939171 | 5.095959458 | 8.376385542 | 5.156626401 | 9.897454779 | 9.82063646 | 8.621823729 | 8.409548182 |
| ENSG00000127951 | FGL2 | 3.486121733 | 0.001087557 | 0.793803328 | 0.522353469 | 4.279925061 | 2 | 3.686303414 | 4.058998049 | 6.470063213 | 6.35727082 | 3.917810786 | 4.175132639 |
| ENSG00000170684 | ZNF296 | 2.750171206 | 0.162619057 | 1.529601617 | 0.455138882 | 4.279772823 | 3.327711234 | 2 | 2 | 4.139692846 | 4.0930011 | 3.27687201 | 3.287118176 |
| ENSG00000188672 | RHCE | 1.937762106 | 0.385331055 | 2.339180744 | 0.272104511 | 4.27694285 | 2 | 3.076665209 | 2 | 3.615725078 | 3.45167086 | 4.166733998 | 3.33635788 |
| ENSG00000136848 | DAB2IP | 1.002022132 | 0.483401635 | 3.273169572 | 0.00856295 | 4.275191704 | 8.631580145 | 7.638155693 | 4.058998049 | 8.633396609 | 8.69038176 | 11.07062965 | 10.75622296 |
| ENSG00000151422 | FER | 1.725001812 | 6.79E-05 | 2.546927946 | 1.07E-09 | 4.271929758 | 6.89194353 | 7.145017454 | 6.534109959 | 8.550024073 | 8.61576544 | 9.159610958 | 9.603790255 |
| ENSG00000133687 | TMTC1 | 1.629651486 | 0.237913826 | 2.642168315 | 0.040936541 | 4.271819801 | 6.366709671 | 5.838937808 | 2 | 7.144720234 | 7.12888795 | 7.47706242 | 8.575745325 |
| ENSG00000233493 | TMEM238 | 2.594781454 | 0.366381671 | 1.676078547 | 0.567684646 | 4.270860002 | 2 | 2 | 2 | 3.162721474 | 3.50452879 | 2.962551659 | 2.746601469 |
| ENSG00000166266 | CUL5 | 2.306410389 | 0.076743239 | 1.964182416 | 0.131858917 | 4.270592805 | 7.53297911 | 7.790474954 | 3.369317972 | 9.434510081 | 9.36034073 | 9.232935896 | 8.858759519 |
| ENSG00000139364 | TMEM132B | 4.761199703 | 2.65E-12 | -0.492299789 | 0.564054374 | 4.268899914 | 5.095959458 | 4.11352228 | 3.369317972 | 8.792330793 | 8.83798691 | 3.943456545 | 4.002845082 |
| ENSG00000128872 | TMOD2 | 2.499068622 | 0.005710292 | 1.769440179 | 0.057699308 | 4.268508802 | 4.815609471 | 6.04685525 | 3.369317972 | 7.587189552 | 7.48837134 | 6.913621294 | 6.739599323 |
| ENSG00000197070 | ARRDC1 | 1.462176587 | 0.461825521 | 2.805644991 | 0.122968358 | 4.267821578 | 2 | 7.099070191 | 3.369317972 | 7.075737784 | 7.05992818 | 8.505043475 | 8.25407423 |
| ENSG00000181027 | FKRP | 2.043181932 | 0.016750741 | 2.223989605 | 0.008135238 | 4.267171537 | 5.532353208 | 7.467835385 | 5.59507254 | 8.47833774 | 8.56378716 | 8.751793877 | 8.647610514 |
| ENSG00000104826 | LHB | 2.005881962 | 0.548198478 | 2.260740213 | 0.484174515 | 4.266622174 | 2 | 2 | 2 | 2.3892346 | 3.45167086 | 3.27687201 | 3.012034821 |
| ENSG00000135312 | HTR1B | 1.499055968 | 0.671487898 | 2.767283975 | 0.394823109 | 4.266339943 | 2 | 2 | 2 | 2.947924294 | 2.59489879 | 4.029788235 | 2.466008509 |
| ENSG00000128564 | VGF | 2.973845441 | 0.309469647 | 1.292362137 | 0.67791597 | 4.266207578 | 2 | 2 | 2 | 3.462030052 | 3.69849104 | 2.962551659 | 2.374681325 |
| ENSG00000172667 | ZMAT3 | 0.682097526 | 0.212680082 | 3.583487469 | 8.65E-15 | 4.265584995 | 7.624139768 | 8.276714465 | 8.699419242 | 9.010169232 | 8.86476809 | 12.03680136 | 11.58959617 |
| ENSG00000100065 | CARD10 | -0.182483795 | 0.895265695 | 4.444174254 | 1.30E-05 | 4.261690459 | 2 | 5.595927442 | 5.39244166 | 4.670864177 | 4.98931816 | 8.729394257 | 9.643763491 |
| ENSG00000235034 | C19orf81 | 1.486820593 | 0.441376957 | 2.773712462 | 0.114946876 | 4.260533054 | 2 | 3.686303414 | 2 | 3.754611435 | 3.7864959 | 5.085488617 | 4.400220394 |
| ENSG00000184271 | POU6F1 | 1.705646082 | 0.050999384 | 2.553652235 | 0.001937731 | 4.259298317 | 4.815609471 | 5.838937808 | 4.058998049 | 6.628980152 | 6.80989959 | 7.909144085 | 7.05850228 |
| ENSG00000109466 | KLHL2 | 2.486742583 | 0.181399395 | 1.772080648 | 0.348797592 | 4.25882323 | 3.327711234 | 7.002537973 | 2 | 7.93753157 | 8.01327646 | 7.514972902 | 6.988993457 |
| ENSG00000166004 | CEP295 | 2.271068987 | 0.318138811 | 1.980521362 | 0.381442321 | 4.251590349 | 2 | 6.844429755 | 2 | 7.584088227 | 7.48179565 | 7.552954291 | 6.866539459 |
| ENSG00000132740 | IGHMBP2 | 1.794369412 | 0.215468947 | 2.455804753 | 0.075024979 | 4.250174165 | 2 | 5.72254323 | 6.882775709 | 7.510853288 | 7.64959655 | 8.368535975 | 8.081604954 |
| ENSG00000070214 | SLC44A1 | 3.623737086 | 0.001296825 | 0.624389161 | 0.640695034 | 4.248126246 | 5.866944937 | 6.844429755 | 9.242516611 | 11.62402619 | 11.6150937 | 8.82044307 | 8.417435289 |
| ENSG00000075711 | DLG1 | 1.480835121 | 0.001328581 | 2.7667544 | 2.08E-10 | 4.247589521 | 7.384433967 | 7.901717006 | 6.882775709 | 9.021676912 | 8.82245717 | 10.16787644 | 10.24002969 |
| ENSG00000130433 | CACNG6 | 5.236575891 | 0.007634516 | -0.989333288 | 0.683334854 | 4.247242603 | 2 | 2 | 2 | 5.44762523 | 5.36298494 | 2.225793514 | 2.117335275 |
| ENSG00000173702 | MUC13 | 4.690417399 | 0.213876745 | -0.443285963 | 0.921262078 | 4.247131435 | 2 | 2 | 2 | 5.076005082 | 4.74762503 | 2.457042231 | 2 |
| ENSG00000197363 | ZNF517 | 1.300309689 | 0.498453223 | 2.946110092 | 0.089458136 | 4.246419781 | 2 | 5.457120953 | 2 | 5.178261742 | 5.36298494 | 7.000176314 | 6.600402875 |
| ENSG00000169760 | NLGN1 | 2.596386719 | 0.08193596 | 1.649747011 | 0.282680363 | 4.24613373 | 2 | 3.686303414 | 3.369317972 | 5.22679461 | 5.13053105 | 3.782262616 | 4.793076545 |
| ENSG00000226763 | SRRM5 | 2.442900379 | 0.403894818 | 1.798474787 | 0.540942209 | 4.241375166 | 2 | 2 | 2 | 3.09461492 | 3.39680241 | 3.035909414 | 2.782576693 |
| ENSG00000164631 | ZNF12 | 1.793735085 | 0.277947155 | 2.447348759 | 0.119536754 | 4.241083844 | 6.563806279 | 7.790474954 | 2 | 8.534041678 | 8.46722793 | 9.404322061 | 8.838074258 |
| ENSG00000129493 | HEATR5A | 1.799658024 | 0.001434462 | 2.441052085 | 7.53E-06 | 4.240710108 | 6.366709671 | 7.669933291 | 6.718950348 | 8.822909147 | 8.81201036 | 9.504905576 | 9.401760658 |
| ENSG00000124839 | RAB17 | 1.727686389 | 0.415598099 | 2.512579706 | 0.205097324 | 4.240266095 | 2 | 3.076665209 | 2 | 3.406930997 | 3.39680241 | 3.981090143 | 3.8919413 |
| ENSG00000174010 | KLHL15 | 2.461268624 | 0.063911628 | 1.777335398 | 0.189290362 | 4.238604022 | 2 | 4.93638933 | 6.43197674 | 7.759286043 | 7.55253065 | 7.02890981 | 6.955898408 |
| ENSG00000144040 | SFXN5 | 2.914706941 | 0.007907751 | 1.321134151 | 0.264665124 | 4.235841092 | 6.009025997 | 4.93638933 | 2 | 7.818544084 | 7.7866208 | 6.275391383 | 6.245088435 |
| ENSG00000116690 | PRG4 | 1.012885833 | 0.737721421 | 3.222218571 | 0.221458956 | 4.235104404 | 2 | 2 | 2 | 2.695456494 | 2.49394783 | 3.584558147 | 3.908316285 |
| ENSG00000146410 | MTFR2 | 3.471550595 | 0.167941338 | 0.763141351 | 0.789642976 | 4.234691946 | 2 | 2 | 2 | 4.034562184 | 3.82856379 | 2.526562374 | 2.509591799 |
| ENSG00000018408 | WWTR1 | 2.061608369 | 0.004263331 | 2.172585968 | 0.002338722 | 4.234194336 | 8.042902253 | 7.701025996 | 9.452720984 | 10.69144695 | 10.6189937 | 10.73805965 | 10.79416656 |
| ENSG00000173542 | MOB1B | 2.319263892 | 0.01380515 | 1.914299681 | 0.045052759 | 4.233563573 | 4.815609471 | 7.274681896 | 7.734844356 | 9.408447353 | 9.24566964 | 9.032447693 | 8.813832186 |
| ENSG00000164543 | STK17A | 1.960271648 | 0.000131113 | 2.272614269 | 6.38E-06 | 4.232885917 | 7.031698319 | 7.002537973 | 8.076950877 | 9.384581425 | 9.38785872 | 9.719104163 | 9.674305191 |
| ENSG00000244414 | CFHR1 | 7.995311529 | 2.68E-07 | -3.763576298 | 0.118337719 | 4.231735231 | 2 | 2 | 2 | 8.013270196 | 8.07355593 | 2.048074405 | 2 |
| ENSG00000160746 | ANO10 | 2.233570972 | 0.042924135 | 1.997998317 | 0.070335607 | 4.231569289 | 7.031698319 | 8.57407404 | 5.156626401 | 9.702459798 | 9.75874862 | 9.416105065 | 9.572665028 |
| ENSG00000076924 | XAB2 | 1.999499984 | 0.038085116 | 2.231987533 | 0.018144183 | 4.231487517 | 6.138360429 | 7.60566235 | 4.874587708 | 8.58612524 | 8.6021877 | 8.934738262 | 8.704976832 |
| ENSG00000269955 | C7orf55-LUC | 9.422318555 | 2.88E-10 | -5.191151298 | 0.026199962 | 4.231167256 | 2 | 2 | 2 | 9.436230969 | 9.47734778 | 2 | 2 |
| ENSG00000165105 | RASEF | 4.485977378 | 0.067479075 | -0.256198508 | 0.933434132 | 4.22977887 | 2 | 2 | 2 | 5.040242897 | 4.37434574 | 2.42098416 | 2.117335275 |
| ENSG00000161791 | FMNL3 | 2.364711015 | 8.27E-06 | 1.864811103 | 0.000554619 | 4.229522118 | 7.27616306 | 7.572420273 | 8.486635645 | 10.1842688 | 10.2409086 | 9.883799376 | 9.523684623 |
| ENSG00000196821 | C6orf106 | 2.157694542 | 0.003538216 | 2.071418461 | 0.005132356 | 4.229113003 | 7.750913381 | 8.862353853 | 9.809100877 | 11.14621855 | 11.2257892 | 11.04956604 | 11.14955624 |
| ENSG00000064607 | SUGP2 | 1.900077013 | 0.103634704 | 2.326859029 | 0.038665093 | 4.226936042 | 5.095959458 | 8.57407404 | 8.699419242 | 9.968745466 | 10.0306182 | 10.4078509 | 10.44263169 |
| ENSG00000203485 | INF2 | 1.579431484 | 0.001761901 | 2.644721783 | 3.51E-08 | 4.224153267 | 9.423197387 | 8.234834549 | 8.960138144 | 10.44877988 | 10.5852675 | 11.52052886 | 11.64071835 |
| ENSG00000184178 | SCFD2 | 2.169803856 | 0.364510704 | 2.051259341 | 0.384577928 | 4.221063197 | 2 | 7.274681896 | 2 | 7.910550164 | 7.81305436 | 7.50961773 | 7.949380652 |
| ENSG00000006831 | ADIPOR2 | 1.907191553 | 0.29492176 | 2.312306727 | 0.186045154 | 4.219498281 | 2 | 8.623598428 | 8.408111533 | 9.802582299 | 9.86489641 | 10.14871212 | 10.32138402 |
| ENSG00000154040 | CABYR | 1.164517985 | 0.517016221 | 3.054556038 | 0.054985297 | 4.219074023 | 2 | 5.131603625 | 2 | 4.907595604 | 4.811949 | 6.67615119 | 6.514766253 |
| ENSG00000165804 | ZNF219 | 1.514826708 | 0.026715109 | 2.702664407 | 2.18E-05 | 4.217491115 | 6.138360429 | 6.951736442 | 5.156626401 | 7.802622679 | 7.70992146 | 9.002260838 | 8.853483814 |
| ENSG00000251192 | ZNF674 | 2.424508787 | 0.033910747 | 1.792861367 | 0.123992425 | 4.217370154 | 2 | 4.11352228 | 5.39244166 | 6.503277904 | 6.60731796 | 6.213498222 | 5.646532192 |
| ENSG00000132694 | ARHGEF11 | 2.428987444 | 0.095066141 | 1.787805708 | 0.225799741 | 4.216793152 | 6.009025997 | 4.11352228 | 8.65477198 | 9.723047898 | 9.73475225 | 8.991160222 | 9.185355845 |
| ENSG00000136874 | STX17 | 2.755266564 | 0.003631933 | 1.461400591 | 0.146527599 | 4.216667155 | 4.815609471 | 7.315436877 | 5.39244166 | 8.95000826 | 8.99976571 | 7.811554331 | 7.574209638 |
| ENSG00000162607 | USP1 | 1.723819894 | 0.488639525 | 2.492381404 | 0.289726676 | 4.216201298 | 2 | 7.393647515 | 2 | 7.568480804 | 7.51759662 | 8.60986148 | 7.902127837 |
| ENSG00000160551 | TAOK1 | 1.981734107 | 4.48E-07 | 2.2343267 | 8.49E-09 | 4.216060808 | 8.797441463 | 8.88960062 | 9.516493218 | 11.08856943 | 11.0615956 | 11.48303928 | 11.15267381 |
| ENSG00000198887 | SMC5 | 2.016401093 | 0.003842733 | 2.198327397 | 0.001374482 | 4.21472849 | 5.70932728 | 7.145017454 | 5.59507254 | 8.386974157 | 8.26953254 | 8.553208128 | 8.463873348 |
| ENSG00000204060 | FOXO6 | 0.86351266 | 0.790662721 | 3.350298923 | 0.228061414 | 4.213811583 | 2 | 2 | 2 | 2.784612485 | 2.2680111 | 3.851628042 | 3.841670744 |
| ENSG00000135063 | FAM189A2 | -0.732281651 | 0.854305977 | 4.938402635 | 0.117842585 | 4.206120984 | 2 | 2 | 2 | 2 | 2.38539787 | 5.422868615 | 4.784231301 |
| ENSG00000175048 | ZDHHC14 | 2.258896471 | 0.313691383 | 1.946621284 | 0.382824967 | 4.205517755 | 2 | 6.728495673 | 2 | 7.444001439 | 7.36873214 | 7.349785676 | 6.804465095 |
| ENSG00000198182 | ZNF607 | 1.843347093 | 0.20811252 | 2.359476359 | 0.092298578 | 4.202823452 | 2 | 5.457120953 | 3.369317972 | 5.959382894 | 5.9725602 | 6.807643042 | 5.984375752 |
| ENSG00000076984 | MAP2K7 | 2.078348622 | 0.099215121 | 2.123745318 | 0.086209421 | 4.202093939 | 4.467311984 | 8.077869079 | 5.772716771 | 8.88597028 | 8.93143442 | 9.058352796 | 8.841275974 |
| ENSG00000105722 | ERF | 1.921447173 | 0.005467646 | 2.280094957 | 0.000727777 | 4.20154213 | 7.624139768 | 6.04685525 | 7.646393253 | 9.131970496 | 9.17099461 | 9.203056613 | 9.759696604 |
| ENSG00000182325 | FBXL6 | 2.221894978 | 0.297505663 | 1.97543189 | 0.350123893 | 4.197326868 | 2 | 6.311476754 | 2 | 6.993973425 | 6.9210099 | 6.457740741 | 6.942047249 |
| ENSG00000183718 | TRIM52 | 2.450570516 | 0.046457783 | 1.74242721 | 0.165909543 | 4.192997726 | 4.467311984 | 5.946638857 | 2 | 7.230526165 | 7.18258783 | 6.57313027 | 6.471341372 |
| ENSG00000143079 | CTTNBP2NL | 1.659356313 | 0.00410097 | 2.530733215 | 4.48E-06 | 4.190089528 | 8.75772933 | 7.572420273 | 7.552162999 | 9.748895993 | 9.68767229 | 10.60465476 | 10.56887086 |
| ENSG00000077235 | GTF3C1 | 1.947630487 | 5.71E-05 | 2.239260134 | 2.56E-06 | 4.186890621 | 8.107517749 | 8.968383114 | 7.97167868 | 10.3445078 | 10.385308 | 10.51703111 | 10.78255483 |
| ENSG00000107438 | PDLIM1 | 1.396113426 | 0.004683802 | 2.790360728 | 1.50E-09 | 4.186474154 | 7.904282243 | 8.702538795 | 7.777116083 | 9.558196845 | 9.60839132 | 10.76179094 | 11.1573911 |
| ENSG00000197121 | PGAP1 | 2.428428662 | 0.013540843 | 1.755817595 | 0.082334289 | 4.184246258 | 4.815609471 | 6.228547981 | 3.369317972 | 7.751013478 | 7.41433569 | 6.924930115 | 6.949978473 |
| ENSG00000178764 | ZHX2 | 1.592487124 | 0.301084135 | 2.591227043 | 0.071431314 | 4.183714167 | 6.009025997 | 7.189546432 | 2 | 7.725907304 | 7.6986238 | 8.443467705 | 8.913019964 |
| ENSG00000104983 | CCDC61 | 1.203030837 | 0.481130174 | 2.979531236 | 0.050033331 | 4.182562073 | 2 | 4.710564154 | 2 | 4.268917409 | 4.72553021 | 6.022174708 | 6.175763085 |
| ENSG00000139977 | NAA30 | 2.191281588 | 0.097979988 | 1.991240376 | 0.12941073 | 4.182521963 | 4.00722531 | 7.572420273 | 4.874587708 | 8.483340231 | 8.41813449 | 8.395497408 | 8.095972716 |
| ENSG00000184575 | XPOT | 1.701279734 | 0.243498812 | 2.479192547 | 0.072675764 | 4.18047228 | 4.467311984 | 9.443177293 | 8.677268317 | 10.26055405 | 10.2330992 | 11.36137528 | 10.57961443 |
| ENSG00000160714 | UBE2Q1 | 2.314980905 | 0.031200591 | 1.865308766 | 0.08678054 | 4.180289671 | 7.485143553 | 8.57407404 | 5.156626401 | 9.930185944 | 9.93817677 | 9.26856367 | 9.676099363 |
| ENSG00000153064 | BANK1 | 0.661560295 | 0.669226282 | 3.518715193 | 0.005828245 | 4.180275488 | 4.467311984 | 3.076665209 | 2 | 3.663521627 | 4.22485516 | 6.013120342 | 6.81752904 |
| ENSG00000092929 | UNC13D | 1.819414369 | 0.375778872 | 2.360815804 | 0.227601689 | 4.180230173 | 3.327711234 | 2 | 3.369317972 | 4.105493497 | 4.16043394 | 5.227950776 | 3.311948089 |

| | | | | | | | | | | | | | |
|---|---|---|---|---|---|---|---|---|---|---|---|---|---|
| ENSG00000259112 | NDUFC2-KCT | 9.370376944 | 4.33E-10 | -5.191151263 | 0.026510255 | 4.179225681 | 2 | 2 | 2 | 9.354865403 | 9.45402153 | 2 | 2 |
| ENSG00000003393 | ALS2 | 1.660283156 | 0.131737146 | 2.507894616 | 0.015985371 | 4.168177772 | 4.815609471 | 8.169643011 | 7.552162999 | 9.014783325 | 9.05507424 | 9.670022651 | 10.05936592 |
| ENSG00000150637 | CD226 | 2.495120064 | 0.437785917 | 1.669623792 | 0.608860386 | 4.164743856 | 2 | 2 | 2 | 3.09461492 | 3.45167086 | 3.215043717 | 2.374681325 |
| ENSG00000067225 | PKM | 2.17050867 | 8.51E-07 | 1.993360956 | 6.72E-06 | 4.163869625 | 12.54538064 | 12.24604664 | 12.48269311 | 14.57461638 | 14.6246954 | 13.84967977 | 14.83181639 |
| ENSG00000127325 | BEST3 | 3.472129694 | 0.172072167 | 0.691075062 | 0.811558361 | 4.163204755 | 2 | 2 | 2 | 4.105493497 | 3.74316441 | 2.526562374 | 2.466008509 |
| ENSG00000116663 | FBXO6 | 3.338388334 | 1.47E-08 | 0.824667618 | 0.222125563 | 4.163055952 | 5.532353208 | 4.442710129 | 5.772716771 | 8.535647908 | 8.48213028 | 6.197600957 | 5.925257518 |
| ENSG00000109756 | RAPGEF2 | 2.217262731 | 0.000108426 | 1.945254781 | 0.000758118 | 4.162517512 | 7.031698319 | 6.464272742 | 7.896971941 | 9.435370781 | 9.41227044 | 9.072949511 | 9.230009473 |
| ENSG00000197837 | HIST4H4 | 1.144078058 | 0.731338839 | 3.015333907 | 0.309952764 | 4.159411965 | 2 | 2 | 2 | 2.270786764 | 2.94009902 | 3.584558147 | 3.640730416 |
| ENSG00000156787 | TBC1D31 | 3.480709218 | 0.001532124 | 0.67694343 | 0.59982426 | 4.157652648 | 4.815609471 | 5.946638857 | 2 | 8.300810597 | 8.34212018 | 5.812021848 | 5.402507939 |
| ENSG00000265763 | ZNF488 | -0.723819587 | 0.800057969 | 4.876795264 | 0.02251104 | 4.152975676 | 2 | 2 | 2 | 2.270786764 | 2.14022021 | 5.014670875 | 5.150533729 |
| ENSG00000099330 | OCEL1 | 1.711589419 | 0.137305271 | 2.440488883 | 0.025468435 | 4.152078302 | 3.327711234 | 6.389896633 | 4.523675391 | 6.936770382 | 6.93551934 | 7.585896024 | 7.704159082 |
| ENSG00000123609 | NMI | 3.49158149 | 0.020971449 | 0.656442448 | 0.709826973 | 4.148023938 | 2 | 4.710564154 | 7.098165388 | 9.178978633 | 9.18523054 | 6.329884754 | 6.479583725 |
| ENSG00000021645 | NRXN3 | -0.366149336 | 0.535148173 | 4.51400768 | 2.25E-23 | 4.147858345 | 5.70932728 | 5.303523747 | 5.930865648 | 5.334065616 | 5.28958453 | 9.948950598 | 10.13414984 |
| ENSG00000164112 | TMEM155 | 6.969837767 | 0.00017384 | -2.822347402 | 0.246641741 | 4.147490365 | 2 | 2 | 2 | 6.979883891 | 7.0991228 | 2.094598369 | 2 |
| ENSG00000141040 | ZNF287 | 1.666865963 | 0.253975538 | 2.479425666 | 0.071647452 | 4.146291629 | 3.327711234 | 5.595927442 | 2 | 5.910700032 | 5.92487824 | 6.510167449 | 6.847559257 |
| ENSG00000171714 | ANO5 | 1.151019437 | 0.756818484 | 2.995075656 | 0.368954964 | 4.146095092 | 2 | 2 | 2 | 2.784612485 | 2.49394783 | 4.251304078 | 2.374681325 |
| ENSG00000177459 | ERICH5 | 0.856943564 | 0.794276136 | 3.287313909 | 0.242584119 | 4.144257473 | 2 | 2 | 2 | 2.3892346 | 2.68924543 | 3.193831101 | 4.228274364 |
| ENSG00000145860 | RNF145 | 1.971264372 | 8.52E-08 | 2.171908112 | 2.50E-09 | 4.143172485 | 8.540903468 | 8.077869079 | 7.777116083 | 10.11558569 | 10.1285838 | 10.23220252 | 10.40670299 |
| ENSG00000173575 | CHD2 | 2.409241404 | 0.005150692 | 1.733037327 | 0.050915017 | 4.142278731 | 6.468621203 | 8.94259736 | 8.784770914 | 10.80867762 | 10.810899 | 10.23788114 | 10.0252855 |
| ENSG00000175229 | GAL3ST3 | -2.311823937 | 0.458287513 | 6.452201368 | 0.002554195 | 4.140377432 | 2 | 2 | 2 | 2 | 2.14022021 | 5.747972572 | 7.051147246 |
| ENSG00000214655 | ZSWIM8 | 2.349034657 | 0.01537677 | 1.790612794 | 0.070728218 | 4.139647452 | 5.532353208 | 8.557180776 | 7.934808822 | 10.14902484 | 10.1143063 | 9.589063591 | 9.562354507 |
| ENSG00000136147 | PHF11 | 3.027548457 | 0.014409744 | 1.111032089 | 0.414485452 | 4.138580546 | 4.00722531 | 4.93638933 | 7.552162999 | 9.203437087 | 9.26686067 | 7.402744819 | 7.285445324 |
| ENSG00000226174 | TEX22 | 1.008440995 | 0.747712189 | 3.129527967 | 0.255172892 | 4.137968961 | 2 | 2 | 2 | 2.3892346 | 2.77779907 | 3.738983888 | 3.640730416 |
| ENSG00000164080 | RAD54L2 | 1.896561271 | 0.383636346 | 2.239670481 | 0.285451923 | 4.136231752 | 2 | 4.442710129 | 8.742726175 | 9.074566443 | 9.13791797 | 9.287461761 | 9.591484008 |
| ENSG00000112697 | TMEM30A | 1.84002324 | 0.003801199 | 2.294696364 | 0.000205536 | 4.134719604 | 8.777722034 | 10.3486505 | 9.478569125 | 11.5358871 | 11.4945185 | 12.14662045 | 11.76751454 |
| ENSG00000196358 | NTNG2 | 2.806529512 | 0.011761813 | 1.32289909 | 0.268600882 | 4.129428602 | 4.815609471 | 5.838937808 | 2 | 7.549526251 | 7.61094772 | 6.320128742 | 5.957088417 |
| ENSG00000072163 | LIMS2 | -0.112649604 | 0.944373487 | 4.241975349 | 0.000620166 | 4.129325745 | 8.107517749 | 6.899080544 | 3.369317972 | 6.998639523 | 6.94032357 | 11.46703437 | 11.06489955 |
| ENSG00000170871 | KIAA0232 | 1.4816217 | 0.011067153 | 2.645215264 | 1.29E-06 | 4.126836964 | 7.384433967 | 6.602424674 | 8.042706615 | 8.89725916 | 8.91200057 | 10.04351195 | 10.07938913 |
| ENSG00000131386 | GALNT15 | 2.600697687 | 0.046904744 | 1.523933258 | 0.26431896 | 4.124630945 | 2 | 3.076665209 | 4.874587708 | 5.949776857 | 5.93454154 | 5.227950776 | 4.644600858 |
| ENSG00000010610 | CD4 | -4.00803025 | 0.013021922 | 8.130483429 | 8.22E-12 | 4.122453179 | 2 | 4.93638933 | 2 | 2.141737246 | 2.2680111 | 11.41191034 | 11.4680018 |
| ENSG00000153094 | BCL2L11 | 2.251452502 | 0.091310742 | 1.870710802 | 0.160681517 | 4.122163304 | 3.327711234 | 4.93638933 | 4.874587708 | 6.676346669 | 6.52015598 | 6.998648055 | 4.522269241 |
| ENSG00000162194 | LBHD1 | 4.046151037 | 0.009852067 | 0.073492717 | 0.970602478 | 4.119643754 | 2 | 5.457120953 | 2 | 7.952038749 | 7.83643742 | 4.240999983 | 4.241259475 |
| ENSG00000122728 | TAF1L | 1.979918942 | 0.000706847 | 2.133846003 | 0.000205791 | 4.113764945 | 5.330594134 | 4.710564154 | 5.39244166 | 6.998639523 | 6.99676009 | 7.254397864 | 7.030725399 |
| ENSG00000100024 | UPB1 | 0.012690705 | 0.997041173 | 4.100351053 | 0.078044162 | 4.113041758 | 2 | 2 | 2 | 2.3892346 | 2.2680111 | 4.378961816 | 4.468068808 |
| ENSG00000101473 | ACOT8 | 1.457307622 | 0.292205343 | 2.652357323 | 0.037378894 | 4.109664944 | 4.815609471 | 6.311476754 | 2 | 6.640968371 | 6.58294721 | 7.792277365 | 7.754841567 |
| ENSG00000196968 | FUT11 | 1.364406184 | 0.491651383 | 2.744359437 | 0.130332141 | 4.108765622 | 8.609439662 | 5.838937808 | 2 | 8.616799622 | 8.52912793 | 9.88234985 | 10.00353926 |
| ENSG00000142875 | PRKACB | 1.376934383 | 0.278205251 | 2.731203277 | 0.018970092 | 4.10813766 | 6.96351253 | 8.029695761 | 4.058998049 | 8.504819462 | 8.35116763 | 9.976966829 | 9.537896513 |
| ENSG00000165071 | TMEM71 | 2.691942469 | 0.359851119 | 1.413429579 | 0.644563698 | 4.105372048 | 2 | 2 | 2 | 3.663521627 | 3.08597292 | 2.885263063 | 2.592996859 |
| ENSG00000204899 | MZT1 | 1.915875965 | 0.279417377 | 2.1890011 | 0.201181835 | 4.104877064 | 3.327711234 | 6.535001748 | 2 | 7.053486362 | 6.90635305 | 7.41654498 | 7.05482945 |
| ENSG00000152818 | UTRN | 1.46745059 | 0.015653107 | 2.637381922 | 3.42E-06 | 4.104832512 | 7.485143553 | 8.993716073 | 8.460932916 | 9.889311677 | 9.91517834 | 10.9525553 | 11.17653005 |
| ENSG00000163126 | ANKRD23 | 3.611539498 | 0.077966393 | 0.491051841 | 0.839388001 | 4.102591339 | 2 | 3.076665209 | 2 | 5.003571641 | 4.60967607 | 3.035909414 | 2.421067459 |
| ENSG00000180626 | ZNF594 | 2.91648701 | 4.19E-05 | 1.186065888 | 0.127844617 | 4.102552898 | 4.815609471 | 4.11352228 | 4.523675391 | 7.169761359 | 7.14977896 | 5.699879814 | 5.313094696 |
| ENSG00000072134 | EPN2 | 1.744714861 | 0.354286428 | 2.354943866 | 0.189186264 | 4.099658727 | 5.866944937 | 8.414402626 | 2 | 8.852852814 | 8.73253609 | 9.465896789 | 9.330261388 |
| ENSG00000122786 | CALD1 | 0.499761288 | 0.390679629 | 3.599119735 | 3.67E-14 | 4.098881023 | 11.72557631 | 12.86962125 | 12.5344401 | 12.97335982 | 12.9289475 | 16.21963215 | 15.857818 |
| ENSG00000115159 | GPD2 | 2.774277959 | 0.018963331 | 1.323712586 | 0.294912398 | 4.097990546 | 3.327711234 | 7.393647515 | 6.80318718 | 9.369349625 | 9.28207914 | 7.713417604 | 8.049667541 |
| ENSG00000114771 | AADAC | 6.752797127 | 0.001424077 | -2.65802783 | 0.315949328 | 4.094769296 | 2 | 2 | 2 | 6.83085985 | 6.83074839 | 2 | 2.117335275 |
| ENSG00000116984 | MTR | 1.786425337 | 0.003540321 | 2.3074622 | 0.000100034 | 4.093887537 | 6.816637977 | 8.101365653 | 6.882775709 | 9.16555511 | 9.18523054 | 9.817084406 | 9.558144718 |
| ENSG00000228623 | ZNF883 | 0.329463551 | 0.867562397 | 3.764163669 | 0.014619621 | 4.09362722 | 4.467311984 | 2 | 2 | 3.566290607 | 3.39680241 | 5.886629224 | 6.704867638 |
| ENSG00000165632 | TAF3 | 2.039626962 | 0.132711491 | 2.053493331 | 0.123279471 | 4.093120292 | 2 | 6.602424674 | 5.930865648 | 7.737119576 | 7.72113133 | 7.872900025 | 7.599641096 |
| ENSG00000170776 | AKAP13 | 1.725405029 | 9.76E-11 | 2.36723813 | 1.48E-19 | 4.092643159 | 8.855032352 | 8.57407404 | 8.585098385 | 10.33991861 | 10.4389106 | 10.95669348 | 11.10085297 |
| ENSG00000173402 | DAG1 | 0.753824512 | 0.271511354 | 3.337280599 | 9.54E-09 | 4.091105111 | 8.517314938 | 9.936721131 | 9.491321578 | 10.16156243 | 10.19993 | 12.38000169 | 13.0607906 |
| ENSG00000109323 | MANBA | 2.879294249 | 4.98E-07 | 1.211006168 | 0.051887684 | 4.090300416 | 6.009025997 | 7.538394143 | 7.398165132 | 9.974678372 | 9.98193763 | 8.409366736 | 8.229848462 |
| ENSG00000122420 | PTGFR | 2.924914678 | 0.142927959 | 1.165018797 | 0.586886119 | 4.089933475 | 4.00722531 | 8.169643011 | 2 | 9.564504798 | 9.57015775 | 8.018757917 | 7.602159758 |
| ENSG00000105605 | CACNG7 | 4.936744913 | 0.036966061 | -0.848515493 | 0.766427996 | 4.08822942 | 2 | 2 | 2 | 5.040242897 | 5.22805098 | 2.3070652 | 2.059860178 |
| ENSG00000149311 | ATM | 2.113430366 | 0.01977227 | 1.974333269 | 0.029695601 | 4.087763635 | 6.366709671 | 8.557180776 | 9.06622548 | 10.50766186 | 10.4397591 | 10.38887163 | 10.27990439 |
| ENSG00000141429 | GALNT1 | 0.666664562 | 0.136159792 | 3.418854663 | 3.54E-19 | 4.085519226 | 8.39321512 | 9.203675681 | 8.677268317 | 9.489414201 | 9.44390794 | 12.08526024 | 12.32863534 |
| ENSG00000189143 | CLDN4 | 0.977507391 | 0.671105744 | 3.107853434 | 0.118374111 | 4.085360825 | 2 | 2 | 3.369317972 | 3.162721474 | 2.77779907 | 4.652122235 | 4.09155917 |
| ENSG00000107581 | EIF3A | 1.746526271 | 0.000161536 | 2.336443624 | 2.07E-07 | 4.082969895 | 8.946185348 | 9.923599762 | 9.935792933 | 11.39699225 | 11.4287336 | 11.95937142 | 12.04371163 |
| ENSG00000106692 | FKTN | 1.650807606 | 0.003349323 | 2.430259052 | 6.26E-06 | 4.081066658 | 5.70932728 | 7.145017454 | 6.629488387 | 8.252792309 | 8.22887947 | 9.006068919 | 9.018608509 |
| ENSG00000270757 | HSPE1-MOB4 | 9.269271308 | 8.12E-10 | -5.191151265 | 0.026848553 | 4.078120043 | 2 | 2 | 2 | 9.316137634 | 9.29338856 | 2 | 2 |
| ENSG00000172201 | ID4 | -0.072187182 | 0.973612043 | 4.147475756 | 0.016029651 | 4.075288574 | 2 | 3.686303414 | 7.285536013 | 6.043074888 | 5.43283104 | 9.989523821 | 9.774287532 |
| ENSG00000198380 | GFPT1 | 1.98936392 | 8.34E-05 | 2.080646279 | 3.32E-05 | 4.0700102 | 8.836089686 | 9.763446433 | 8.65477198 | 11.17123391 | 11.146157 | 11.27469035 | 11.22462398 |
| ENSG00000133800 | LYVE1 | -2.311756458 | 0.448255827 | 6.378816374 | 0.002048422 | 4.067059917 | 2 | 2 | 2 | 2 | 2.14022021 | 5.938563912 | 6.860240429 |
| ENSG00000151458 | ANKRD50 | 2.008734087 | 1.11E-05 | 2.057895365 | 5.79E-06 | 4.066629452 | 6.89194353 | 7.467835385 | 6.80318718 | 9.108419034 | 9.04730166 | 9.363346227 | 8.843938654 |
| ENSG00000121964 | GTDC1 | 1.222257799 | 0.415495349 | 2.842918484 | 0.035360505 | 4.065176283 | 4.00722531 | 7.002537973 | 3.369317972 | 6.856790872 | 6.89649843 | 8.505581591 | 8.421006134 |
| ENSG00000152661 | GJA1 | 1.156704498 | 0.049729255 | 2.907205982 | 4.64E-08 | 4.06391048 | 10.87842776 | 10.71894309 | 9.647595056 | 11.69886939 | 11.6279928 | 13.13511385 | 13.64610971 |
| ENSG00000167778 | SPRYD3 | 1.170201333 | 0.471597639 | 2.893633039 | 0.046605398 | 4.063834371 | 3.327711234 | 7.60566235 | 8.884815469 | 8.926937992 | 9.01919552 | 10.64892515 | 10.72839317 |
| ENSG00000110876 | SELPLG | 2.420599233 | 0.199759863 | 1.642716176 | 0.393178386 | 4.063315409 | 2 | 5.595927442 | 2 | 6.276156955 | 6.59518404 | 5.880003936 | 5.524371359 |
| ENSG00000123124 | WWP1 | 2.187132244 | 7.60E-05 | 1.875248301 | 0.000778051 | 4.062380545 | 5.70932728 | 6.602424674 | 5.39244166 | 8.109160159 | 8.19108423 | 7.913205701 | 7.770601445 |
| ENSG00000135205 | CCDC146 | 2.438721787 | 0.063840574 | 1.623350349 | 0.229804148 | 4.062072136 | 4.467311984 | 3.076665209 | 2 | 5.44762523 | 5.47317115 | 5.008609438 | 4.411752372 |
| ENSG00000166140 | ZFYVE19 | 1.942865952 | 0.238681925 | 2.118791892 | 0.186976308 | 4.061657844 | 4.00722531 | 6.602424674 | 2 | 6.965655398 | 7.31795292 | 7.614115653 | 6.959831579 |
| ENSG00000115295 | CLIP4 | 1.745813976 | 0.223356242 | 2.315758843 | 0.091236017 | 4.061578818 | 3.327711234 | 8.191702463 | 6.882775709 | 8.828161587 | 8.88234995 | 9.415532511 | 9.4271968 |
| ENSG00000121741 | ZMYM2 | 2.162701664 | 0.112962085 | 1.892457134 | 0.164128485 | 4.055158798 | 4.00722531 | 8.522788974 | 8.380957451 | 10.07529173 | 10.0187431 | 10.09215279 | 9.376235922 |
| ENSG00000100503 | NIN | 1.882194432 | 0.002012733 | 2.172430516 | 0.000273878 | 4.054624948 | 6.366709671 | 7.901717006 | 7.818184403 | 9.423273993 | 9.32862783 | 9.646061941 | 9.68444279 |
| ENSG00000136378 | ADAMTS7 | -0.752028641 | 0.463646059 | 4.806407171 | 8.21E-09 | 4.05437853 | 8.009475818 | 7.979858146 | 5.39244166 | 6.755670824 | 6.84106088 | 12.14260421 | 12.45240024 |
| ENSG00000117475 | BLZF1 | 2.86888614 | 1.20E-13 | 1.18375915 | 0.004265603 | 4.05264529 | 7.384433967 | 7.393647515 | 6.629488387 | 9.974086178 | 10.0717013 | 8.429356794 | 8.267623722 |
| ENSG00000227051 | C14orf132 | 1.753544367 | 0.218891141 | 2.298942736 | 0.092079873 | 4.052487104 | 3.327711234 | 8.191702463 | 7.098165388 | 8.894758143 | 8.95891978 | 9.553595256 | 9.378808813 |
| ENSG00000108825 | PTGES3L-AA | 6.326026666 | 0.005941499 | -2.274386791 | 0.414555096 | 4.051639876 | 2 | 2 | 2 | 6.422239259 | 6.42052719 | 2.139668783 | 2 |
| ENSG00000255072 | PIGY | 9.239537298 | 1.08E-09 | -5.191151214 | 0.02716304 | 4.048386084 | 2 | 2 | 2 | 9.332862708 | 9.21527115 | 2 | 2 |
| ENSG00000165868 | HSPA12A | 1.55729252 | 0.394622747 | 2.487103237 | 0.145662728 | 4.044395757 | 3.327711234 | 6.602424674 | 2 | 6.722207358 | 6.67228943 | 7.792277365 | 7.379474672 |
| ENSG00000076928 | ARHGEF1 | 2.186103139 | 0.000615007 | 1.85787644 | 0.0040438 | 4.043979579 | 7.485143553 | 6.535001748 | 5.772716771 | 8.863124011 | 8.95180002 | 8.48225943 | 8.677176152 |
| ENSG00000182903 | ZNF721 | 2.274647798 | 0.177613024 | 1.768386197 | 0.297432059 | 4.043033995 | 2 | 7.099070191 | 4.523675391 | 7.985332082 | 7.98332947 | 7.781653072 | 7.117791742 |
| ENSG00000179071 | CCDC89 | 2.675165863 | 0.167382231 | 1.36611685 | 0.500751055 | 4.041282713 | 2 | 3.686303414 | 2 | 4.739373666 | 4.63360314 | 3.1504466 | 4.062589736 |
| ENSG00000107165 | TYRP1 | 4.375095859 | 0.084532433 | -0.334805001 | 0.914469974 | 4.040290858 | 2 | 2 | 2 | 4.523237872 | 4.76938658 | 2.094598369 | 2.421067459 |
| ENSG00000106392 | C1GALT1 | 2.123533601 | 0.02714508 | 1.915053984 | 0.046875269 | 4.038587584 | 4.467311984 | 7.315436877 | 6.073378139 | 8.522747711 | 8.40429221 | 8.245074976 | 8.272375674 |
| ENSG00000115355 | CCDC88A | 2.080989142 | 0.061157352 | 1.956721723 | 0.076411855 | 4.037710866 | 6.89194353 | 8.806266215 | 5.59507254 | 9.768836458 | 9.72712735 | 9.647281667 | 9.60127444 |
| ENSG00000149231 | CCDC82 | 2.343185699 | 0.001927216 | 1.694319613 | 0.029895597 | 4.037505312 | 5.330594134 | 7.232742069 | 6.073378139 | 8.778830996 | 8.70032852 | 8.351874517 | 7.791718153 |
| ENSG00000097033 | SH3GLB1 | 1.934965142 | 0.088857833 | 2.102450294 | 0.057993418 | 4.037415436 | 9.602033055 | 9.523645035 | 6.203069917 | 10.99680327 | 10.9537642 | 11.23167307 | 11.04767876 |
| ENSG00000175984 | DENND2C | 1.931559669 | 0.161064485 | 2.100651691 | 0.115858711 | 4.032211361 | 4.467311984 | 2 | 3.369317972 | 4.825935136 | 5.24368216 | 5.032703865 | 5.349527629 |
| ENSG00000126705 | AHDC1 | 0.917210481 | 0.159815785 | 3.114892014 | 3.70E-08 | 4.032102495 | 7.53297911 | 7.274681896 | 6.073378139 | 7.90310317 | 8.0713677 | 9.915730882 | 10.38529333 |
| ENSG00000177192 | PUS1 | 1.960827681 | 0.361980007 | 2.070503377 | 0.324040096 | 4.031331058 | 2 | 6.311476754 | 2 | 6.750147204 | 6.66069199 | 6.865802559 | 6.755525668 |
| ENSG00000162383 | SLC1A7 | -2.311645111 | 0.43323806 | 6.342038716 | 0.001216799 | 4.030393605 | 2 | 2 | 2 | 2 | 2.14022021 | 6.446574529 | 6.426568739 |
| ENSG00000074935 | TUBE1 | 2.363379723 | 0.299422571 | 1.667010298 | 0.470583967 | 4.03039002 | 2 | 6.464272742 | 2 | 7.190301983 | 7.29563107 | 7.186216802 | 5.475755622 |
| ENSG00000197361 | FBXL22 | 0.015496265 | 0.997041173 | 4.014055787 | 0.129312271 | 4.029552052 | 2 | 2 | 2 | 2.498691189 | 2.14022021 | 4.675266036 | 3.940519005 |
| ENSG00000164040 | PGRMC2 | 2.00784454 | 0.000106536 | 2.020905742 | 8.82E-05 | 4.028750282 | 7.624139768 | 8.903033433 | 8.65477198 | 10.49078653 | 10.4918206 | 10.53493405 | 10.47305071 |
| ENSG00000173641 | HSPB7 | 0.243515554 | 0.898893943 | 3.78402272 | 0.01182329 | 4.027538273 | 7.975256505 | 7.189546432 | 2 | 7.218576229 | 7.38291816 | 10.71031802 | 10.90286456 |
| ENSG00000188827 | SLX4 | 1.811335489 | 0.002335798 | 2.215627473 | 0.000125997 | 4.026962962 | 5.532353208 | 4.93638933 | 6.203069917 | 7.388195945 | 7.28433928 | 7.841701071 | 7.614687532 |
| ENSG00000108448 | TRIM16L | 3.582916858 | 8.58E-09 | 0.443095681 | 0.562406198 | 4.026012539 | 4.00722531 | 4.710564154 | 5.156626401 | 7.966401501 | 8.14188248 | 5.102662916 | 5.050803619 |
| ENSG00000149054 | ZNF215 | 2.650648555 | 0.102405989 | 1.374349305 | 0.420838949 | 4.024997861 | 2 | 4.93638933 | 2 | 6.078767801 | 5.9536761 | 4.920961476 | 4.80186789 |
| ENSG00000150457 | LATS2 | 0.939491323 | 0.173748267 | 3.085400466 | 2.87E-07 | 4.024891789 | 7.384433967 | 8.862353853 | 7.691296561 | 9.018234236 | 9.11579188 | 11.26701432 | 11.1410292 |
| ENSG00000187627 | RGPD1 | 9.214372382 | 1.18E-09 | -5.191151245 | 0.027118898 | 4.023221137 | 2 | 2 | 2 | 9.236404433 | 9.26398924 | 2 | 2 |
| ENSG00000197608 | ZNF841 | 2.494274658 | 0.000952027 | 1.526878193 | 0.053856478 | 4.02115085 | 5.330594134 | 6.535001748 | 4.874587708 | 8.166267464 | 8.23475766 | 7.567458938 | 6.847559257 |
| ENSG00000073614 | KDM5A | 2.22178311 | 0.098102627 | 1.799228277 | 0.181719899 | 4.021011387 | 4.467311984 | 9.079034257 | 8.721235202 | 10.54920508 | 10.5883315 | 10.12946437 | 10.16556732 |
| ENSG00000256977 | LIMS3 | 9.211192851 | 1.22E-09 | -5.191151235 | 0.027169109 | 4.020041616 | 2 | 2 | 2 | 9.277301907 | 9.21626181 | 2 | 2 |
| ENSG00000130783 | CCDC62 | 2.860715546 | 0.407755293 | 1.159216542 | 0.751485356 | 4.019932088 | 2 | 2 | 2 | 3.663521627 | 3.33976443 | 3.035909414 | 2.117335275 |
| ENSG00000196449 | YRDC | 2.083505786 | 0.044893175 | 1.935558165 | 0.061325919 | 4.019063951 | 4.467311984 | 7.145017454 | 5.156626401 | 8.03159949 | 8.16669309 | 7.993660487 | 7.915353355 |
| ENSG00000142347 | MYO1F | 0.691793707 | 0.760287901 | 3.326687697 | 0.082542158 | 4.018481404 | 4.00722531 | 2 | 2 | 3.162721474 | 3.50452879 | 5.976323695 | 4.4231929 |

| | | | | | | | | | | | | | |
|---|---|---|---|---|---|---|---|---|---|---|---|---|---|
| ENSG00000051825 | MPHOSPH9 | 1.98491133 | 0.389955811 | 2.032910796 | 0.366696857 | 4.017822125 | 2 | 6.844429755 | 2 | 7.363085426 | 7.13728069 | 7.215456797 | 7.382408159 |
| ENSG00000176170 | SPHK1 | 2.155072207 | 0.245027882 | 1.861493913 | 0.313731738 | 4.01656612 | 8.816894989 | 6.899080544 | 2 | 9.693134408 | 9.72990471 | 9.735024004 | 9.015305099 |
| ENSG00000182759 | MAFA | -0.307357422 | 0.924039274 | 4.321943988 | 0.079932646 | 4.014586566 | 2 | 2 | 2 | 2.3892346 | 2.14022021 | 4.946551782 | 4.161535617 |
| ENSG00000174899 | PQLC2L | 4.362046499 | 0.054872798 | -0.347533605 | 0.902389555 | 4.014512893 | 2 | 2 | 2 | 4.670864177 | 4.60967607 | 2.34604626 | 2.172608095 |
| ENSG00000215717 | TMEM167B | 2.00532718 | 0.079057979 | 2.002770183 | 0.075111972 | 4.008097363 | 4.815609471 | 8.433041974 | 7.342949392 | 9.425008328 | 9.51406378 | 9.599435435 | 9.3226454 |
| ENSG00000122477 | LRRC39 | 2.689468836 | 0.348815215 | 1.318510434 | 0.662784961 | 4.00797927 | 2 | 2 | 2 | 3.406930997 | 3.39680241 | 2.592885947 | 2.817676617 |
| ENSG00000198633 | ZNF534 | 1.709404053 | 0.310604101 | 2.298555335 | 0.149193138 | 4.007959388 | 2 | 4.11352228 | 2 | 4.470469939 | 4.2560182 | 4.977914261 | 4.729989799 |
| ENSG00000184619 | KRBA2 | 2.747136273 | 0.100138144 | 1.259285049 | 0.478214219 | 4.006421322 | 3.327711234 | 3.076665209 | 2 | 4.783303513 | 4.85329164 | 3.968654369 | 3.261853426 |
| ENSG00000127863 | TNFRSF19 | 4.256767635 | 0.004847695 | -0.253111148 | 0.89539314 | 4.003656487 | 2 | 5.303523747 | 2 | 7.990026167 | 7.90439517 | 4.088436536 | 3.620995423 |
| ENSG00000088888 | MAVS | 1.430525863 | 0.095909493 | 2.572245664 | 0.001180736 | 4.002771527 | 7.159102577 | 9.514922632 | 8.742726175 | 10.18273194 | 10.1949133 | 11.29537729 | 11.35960277 |
| ENSG00000152359 | POC5 | 2.393091584 | 0.185239825 | 1.608786305 | 0.382493233 | 4.001877889 | 2 | 5.303523747 | 2 | 6.113598934 | 6.14893615 | 5.646229369 | 5.115701921 |
| ENSG00000121039 | RDH10 | 2.992569752 | 0.022108229 | 1.008761815 | 0.484862967 | 4.001331566 | 3.327711234 | 7.73146271 | 7.896971941 | 10.23408802 | 10.2153718 | 7.957931324 | 8.501461594 |
| ENSG00000121621 | KIF18A | 2.268716152 | 0.159857485 | 1.731764474 | 0.285691035 | 4.000480626 | 3.327711234 | 3.686303414 | 2 | 4.946755474 | 4.811949 | 4.927401733 | 3.660199087 |
| ENSG00000240038 | AMY2B | 4.536107985 | 0.013985031 | -0.538029497 | 0.813331018 | 3.998078488 | 2 | 3.076665209 | 2 | 5.818703821 | 5.44640346 | 2.592885947 | 2.225841198 |
| ENSG00000120616 | EPC1 | 2.592761511 | 9.45E-09 | 1.404026771 | 0.00315168 | 3.996788282 | 5.532353208 | 6.389896633 | 6.322058544 | 8.68062069 | 8.65428713 | 7.5848779 | 7.402777233 |
| ENSG00000248333 | CDK11B | 3.056332831 | 0.00492344 | 0.938907334 | 0.441681135 | 3.995240165 | 7.790810999 | 4.710564154 | 5.59507254 | 9.620825434 | 9.61591793 | 7.464938976 | 7.583161732 |
| ENSG00000081177 | EXD2 | 1.540845865 | 0.167659551 | 2.452291107 | 0.019238354 | 3.993136971 | 5.330594134 | 7.73146271 | 5.156626401 | 8.178656739 | 8.03813956 | 9.075851216 | 8.944622926 |
| ENSG00000167840 | ZNF232 | 1.879526622 | 0.097789727 | 2.113373991 | 0.05457653 | 3.992900614 | 4.00722531 | 5.131603625 | 2 | 5.978405189 | 5.85535975 | 6.310306306 | 5.941260754 |
| ENSG00000071051 | NCK2 | 1.642103102 | 0.00017253 | 2.350431392 | 2.50E-08 | 3.992534494 | 6.816637977 | 6.899080544 | 7.098165388 | 8.553199415 | 8.54656162 | 9.512153949 | 8.93466504 |
| ENSG00000172939 | OXSR1 | 1.609675243 | 4.52E-07 | 2.382680987 | 1.55E-14 | 3.992356231 | 8.564112515 | 8.191702463 | 8.677268317 | 10.11987514 | 10.0451865 | 10.73312834 | 10.96521391 |
| ENSG00000164440 | TXLNB | 4.196290716 | 0.066039055 | -0.204719335 | 0.94282113 | 3.991571381 | 2 | 2 | 2 | 4.358765249 | 4.63360314 | 2.34604626 | 2.225841198 |
| ENSG00000176919 | C8G | 2.214057564 | 0.464304361 | 1.777420576 | 0.553713666 | 3.99147814 | 2 | 2 | 2 | 3.227757282 | 3.01487888 | 2.987421008 | 2.817676617 |
| ENSG00000173960 | UBXN2A | 1.783208833 | 0.003013039 | 2.206816159 | 0.000157435 | 3.990024992 | 7.096806373 | 5.838937808 | 6.43197674 | 8.239055745 | 8.2960118 | 8.947864566 | 8.367233194 |
| ENSG00000146463 | ZMYM4 | 2.273414827 | 0.088904811 | 1.715221885 | 0.204625268 | 3.988636712 | 4.00722531 | 8.639735838 | 8.076950877 | 10.10588754 | 10.0771644 | 9.5209639 | 9.550666404 |
| ENSG00000064999 | ANKS1A | 1.317161766 | 0.54318775 | 2.670749155 | 0.179837111 | 3.987910922 | 4.00722531 | 8.213429689 | 2 | 8.001694993 | 8.0314011 | 9.523622995 | 9.167610318 |
| ENSG00000215193 | PEX26 | 1.203580496 | 0.254759301 | 2.784194833 | 0.003409966 | 3.987775329 | 7.867441304 | 4.93638933 | 7.398165132 | 8.315844936 | 8.36909406 | 9.729736885 | 10.07282122 |
| ENSG00000068831 | RASGRP2 | -1.632406642 | 0.315223626 | 5.619836101 | 1.43E-05 | 3.987429459 | 3.327711234 | 2 | 4.058998049 | 2.270786764 | 2.77779907 | 7.631969638 | 8.538752857 |
| ENSG00000163785 | RYK | 1.951523651 | 4.16E-08 | 2.035367283 | 8.06E-09 | 3.986890934 | 6.96351253 | 7.315436877 | 6.80318718 | 9.014783325 | 8.94225201 | 9.119393978 | 9.002492046 |
| ENSG00000197372 | ZNF675 | 2.40445282 | 0.049351523 | 1.582077108 | 0.209143226 | 3.986529928 | 2 | 4.93638933 | 3.369317972 | 6.01571348 | 6.1068222 | 5.47655442 | 5.108633494 |
| ENSG00000162927 | PUS10 | 1.9912333 | 0.065284486 | 1.995155413 | 0.06000113 | 3.986388714 | 4.00722531 | 3.076665209 | 4.058998049 | 5.348759107 | 5.36298494 | 5.60262961 | 5.065479856 |
| ENSG00000165097 | KDM1B | 1.546189058 | 0.30511035 | 2.436463072 | 0.084387372 | 3.98265213 | 2 | 5.131603625 | 6.718950348 | 7.140504127 | 6.94032357 | 8.164905085 | 7.609689469 |
| ENSG00000173013 | CCDC96 | 2.091222849 | 0.244964234 | 1.891370552 | 0.28796405 | 3.982593401 | 2 | 2 | 4.523675391 | 4.825935136 | 4.89348248 | 4.416072028 | 4.919632516 |
| ENSG00000198729 | PPP1R14C | 1.279569437 | 0.714145675 | 2.701921349 | 0.396311091 | 3.981490787 | 2 | 2 | 2 | 3.023133209 | 2.2680111 | 3.663836244 | 3.100202476 |
| ENSG00000167523 | SPATA33 | 1.612857579 | 0.237679353 | 2.368449245 | 0.066070408 | 3.981306823 | 3.327711234 | 5.303523747 | 2 | 5.627094009 | 5.62411716 | 6.202919533 | 6.451924088 |
| ENSG00000121361 | KCNJ8 | 2.565164756 | 0.117408064 | 1.415067735 | 0.40849289 | 3.980232491 | 3.327711234 | 6.140560021 | 2 | 7.337530098 | 7.15805139 | 6.550361701 | 5.581648407 |
| ENSG00000132356 | PRKAA1 | 2.020577336 | 0.052853599 | 1.95928951 | 0.058034497 | 3.979866846 | 8.313285031 | 8.147241016 | 5.156626401 | 9.76541805 | 9.70330142 | 9.643130384 | 9.703627437 |
| ENSG00000175787 | ZNF169 | 1.25756401 | 0.524970355 | 2.721888807 | 0.125049462 | 3.979452816 | 2 | 3.686303414 | 2 | 3.840289928 | 3.33976443 | 4.834511823 | 4.634793542 |
| ENSG00000137975 | CLCA2 | 0.497871438 | 0.882991782 | 3.480268575 | 0.205101914 | 3.978140013 | 2 | 2 | 2 | 2.498691189 | 2.38539787 | 3.215043717 | 4.4231929 |
| ENSG00000143248 | RGS5 | -3.412758367 | 0.016504635 | 7.390698486 | 1.47E-08 | 3.977940119 | 8.892187472 | 7.954278496 | 3.369317972 | 4.825935136 | 4.63360314 | 15.40760851 | 15.18675872 |
| ENSG00000159259 | CHAF1B | 1.207146053 | 0.349641507 | 2.770024042 | 0.016350264 | 3.977170094 | 4.00722531 | 3.686303414 | 2 | 4.54891529 | 4.16043394 | 5.751606453 | 5.713290392 |
| ENSG00000019995 | ZRANB1 | 2.229422518 | 0.120981575 | 1.744488457 | 0.229217512 | 3.973910975 | 6.468621203 | 8.255926468 | 3.369317972 | 9.262828394 | 9.2990102 | 8.99844268 | 8.569323645 |
| ENSG00000182329 | KIAA2012 | 1.59688831 | 0.649878555 | 2.376948523 | 0.474346524 | 3.973836834 | 2 | 2 | 2 | 2.868578001 | 2.77779907 | 3.724264246 | 2.421067459 |
| ENSG00000099381 | SETD1A | 1.777838787 | 0.433814242 | 2.195725069 | 0.312608952 | 3.973563856 | 3.327711234 | 8.276714465 | 2 | 8.442826028 | 8.56222967 | 8.77875662 | 9.044303988 |
| ENSG00000162402 | USP24 | 2.051045699 | 0.168292291 | 1.92134335 | 0.191124222 | 3.972389049 | 3.327711234 | 8.557180776 | 7.098165388 | 9.44052423 | 9.51808625 | 9.35261874 | 9.349127298 |
| ENSG00000117152 | RGS4 | 0.084829216 | 0.88968542 | 3.886555059 | 4.15E-19 | 3.971384275 | 9.143983763 | 9.148351514 | 9.839340503 | 9.496854254 | 9.49867529 | 12.93004339 | 13.58122643 |
| ENSG00000124762 | CDKN1A | 0.569826231 | 0.643838836 | 3.401369374 | 0.000826214 | 3.971195605 | 12.98008639 | 10.95004268 | 10.2528023 | 12.42805917 | 12.4626484 | 15.75086129 | 14.56374084 |
| ENSG00000186834 | HEXIM1 | 2.529237251 | 0.031442498 | 1.44153827 | 0.243017249 | 3.970775521 | 8.696033109 | 7.051611315 | 4.874587708 | 10.04290398 | 10.135408 | 8.937532627 | 9.075453001 |
| ENSG00000164400 | CSF2 | 9.161904005 | 1.72E-09 | -5.19115121 | 0.027439899 | 3.970752795 | 2 | 2 | 2 | 9.161399541 | 9.23397834 | 2 | 2 |
| ENSG00000089094 | KDM2B | 2.115554878 | 0.142046401 | 1.854205551 | 0.19591446 | 3.96976043 | 4.467311984 | 7.315436877 | 3.369317972 | 7.987681034 | 8.14396641 | 7.6993745 | 7.914340301 |
| ENSG00000142687 | KIAA0319L | 1.72671269 | 0.174638808 | 2.24259072 | 0.065343113 | 3.96930341 | 5.532353208 | 9.018611864 | 6.534109959 | 9.480268385 | 9.50517467 | 9.929431577 | 10.07916314 |
| ENSG00000140534 | TICRR | 1.910529589 | 0.145397751 | 2.058130233 | 0.106931842 | 3.968659821 | 2 | 5.131603625 | 3.369317972 | 5.651002631 | 5.86549842 | 5.751606453 | 6.033722345 |
| ENSG00000099810 | MTAP | 1.985796858 | 0.005771922 | 1.981722623 | 0.005607289 | 3.96751948 | 5.095959458 | 7.099070191 | 6.80318718 | 8.519504557 | 8.50663048 | 8.625789205 | 8.381965189 |
| ENSG00000157107 | FCHO2 | 2.173046876 | 0.123897637 | 1.79311895 | 0.205383605 | 3.966165825 | 7.485143553 | 4.93638933 | 3.369317972 | 8.354580458 | 8.24644267 | 8.15052603 | 7.662532142 |
| ENSG00000124789 | NUP153 | 1.95411257 | 0.06013597 | 2.009947571 | 0.049513463 | 3.964060142 | 6.653098118 | 8.671479029 | 5.930865648 | 9.452477833 | 9.5678312 | 9.604970226 | 9.527664835 |
| ENSG00000184743 | ATL3 | 1.423062936 | 0.21977808 | 2.540257985 | 0.018389876 | 3.963320921 | 5.866944937 | 8.487557258 | 9.413059136 | 9.893075744 | 9.96494592 | 10.91456512 | 11.16177291 |
| ENSG00000030304 | MUSK | 1.014404791 | 0.754153092 | 2.948282904 | 0.302076863 | 3.962687695 | 2 | 2 | 2 | 2.784612485 | 2.38539787 | 3.355392153 | 3.753786374 |
| ENSG00000165617 | DACT1 | 1.818696993 | 0.308260189 | 2.138166469 | 0.214486601 | 3.956863462 | 7.485143553 | 8.337339476 | 2 | 9.172798677 | 9.22317745 | 9.959790553 | 8.870820921 |
| ENSG00000172057 | ORMDL3 | 1.601385503 | 0.373813246 | 2.351542826 | 0.166636971 | 3.952928329 | 9.48530526 | 5.595927442 | 4.058998049 | 9.6094308 | 9.61140668 | 10.25702296 | 10.45156308 |
| ENSG00000181074 | OR52N4 | 1.275726885 | 0.714005062 | 2.676305764 | 0.399502541 | 3.95203265 | 2 | 2 | 2 | 2.784612485 | 2.59489879 | 3.904814898 | 2.592996859 |
| ENSG00000117411 | B4GALT2 | 1.321824368 | 0.254759301 | 2.626233451 | 0.013611554 | 3.948057819 | 6.138360429 | 9.13702746 | 6.958202216 | 9.315202754 | 9.27828955 | 10.56931498 | 10.62180609 |
| ENSG00000107186 | MPDZ | 1.76598435 | 0.140821478 | 2.180372103 | 0.058715617 | 3.946356453 | 5.095959458 | 8.451443573 | 6.322058544 | 8.99156386 | 9.05839257 | 9.401144325 | 9.471950246 |
| ENSG00000116830 | TTF2 | 1.408824307 | 0.539839197 | 2.534984585 | 0.236787259 | 3.943808893 | 2 | 6.602424674 | 2 | 6.260701031 | 6.63721363 | 7.504242605 | 7.597118029 |
| ENSG00000138835 | RGS3 | 2.277342002 | 0.111902276 | 1.664866853 | 0.25288003 | 3.942208855 | 9.159305284 | 6.228547981 | 4.874587708 | 10.08245238 | 10.059057 | 9.377526577 | 9.540196884 |
| ENSG00000115568 | ZNF142 | 1.566927434 | 0.335922911 | 2.374913579 | 0.121465318 | 3.941841013 | 2 | 7.431218269 | 6.073378139 | 7.910550164 | 7.84158259 | 8.898729828 | 8.407389648 |
| ENSG00000163166 | IWS1 | 1.758235374 | 0.343732241 | 2.183377004 | 0.221346372 | 3.941612378 | 6.138360429 | 8.433041974 | 2 | 8.772033429 | 8.95417718 | 9.183011957 | 9.385769334 |
| ENSG00000055609 | KMT2C | 1.912748759 | 0.016043704 | 2.026419798 | 0.009779302 | 3.939168556 | 8.228665467 | 8.777382925 | 6.534109959 | 9.978816937 | 10.0579523 | 10.22208805 | 10.03626976 |
| ENSG00000184486 | POU3F2 | 3.339399653 | 0.282118771 | 0.599627194 | 0.864701858 | 3.939026846 | 2 | 2 | 2 | 4.470469939 | 2.68924543 | 2.560105256 | 2.374681325 |
| ENSG00000178226 | PRSS36 | 1.594829058 | 0.61206249 | 2.343554694 | 0.428528777 | 3.938383752 | 2 | 2 | 2 | 2.695456494 | 2.94009902 | 3.336159116 | 3.042026731 |
| ENSG00000165630 | PRPF18 | 3.307194222 | 0.000161834 | 0.631082667 | 0.542075443 | 3.938276889 | 2 | 5.131603625 | 4.874587708 | 7.656739692 | 7.57432889 | 4.990270811 | 5.18454436 |
| ENSG00000005889 | ZFX | 1.973565339 | 5.31E-09 | 1.960256032 | 5.84E-09 | 3.933821371 | 7.940205782 | 7.572420273 | 7.285536013 | 9.571568433 | 9.58019637 | 9.578360891 | 9.546749232 |
| ENSG00000159110 | IFNAR2 | 3.76166134 | 1.44E-09 | 0.17190437 | 0.835180221 | 3.933565709 | 5.330594134 | 5.595927442 | 4.058998049 | 8.81895723 | 8.73392024 | 5.096960836 | 5.524371359 |
| ENSG00000186832 | KRT16 | -0.303158672 | 0.946210441 | 4.23523163 | 0.258826994 | 3.932072958 | 2 | 2 | 2 | 2.498691189 | 2 | 4.848153631 | 4.133950982 |
| ENSG00000115363 | EVA1A | 2.409501782 | 0.004330368 | 1.519411116 | 0.085343919 | 3.928912898 | 7.904282243 | 5.838937808 | 6.322058544 | 9.313331174 | 9.35675148 | 8.091532339 | 8.741420621 |
| ENSG00000179532 | DNHD1 | 2.462163631 | 0.051696866 | 1.464457394 | 0.266306871 | 3.926621025 | 2 | 6.228547981 | 5.930865648 | 7.942383523 | 7.88956768 | 7.21150425 | 6.610445962 |
| ENSG00000114573 | ATP6V1A | 2.291182427 | 0.042589119 | 1.635014077 | 0.156803554 | 3.926196504 | 5.330594134 | 8.820493869 | 8.825264954 | 10.5602946 | 10.5974847 | 9.882142656 | 9.966632301 |
| ENSG00000188015 | S100A3 | 0.554604805 | 0.744767011 | 3.367909491 | 0.016078695 | 3.922514296 | 3.327711234 | 3.686303414 | 2 | 3.615725078 | 3.50452879 | 5.201624686 | 6.338680316 |
| ENSG00000274166 | | 7.064429395 | 1.11E-06 | -3.143713684 | 0.094537496 | 3.920715711 | 2 | 2 | 3.369317972 | 8.124184191 | 8.02236724 | 2.094598369 | 2.059860178 |
| ENSG00000256235 | SMIM3 | 2.764872092 | 2.42E-05 | 1.154696701 | 0.106137285 | 3.919568793 | 8.367060731 | 6.666836837 | 7.225742779 | 10.3495391 | 10.3093543 | 8.800846924 | 8.652479133 |
| ENSG00000198513 | ATL1 | 2.285945541 | 0.270109437 | 1.633344623 | 0.437547935 | 3.919290164 | 2 | 6.140560021 | 2 | 6.777556219 | 6.9210099 | 6.098353065 | 6.344709081 |
| ENSG00000164241 | C5orf63 | 2.139886124 | 0.203854133 | 1.778732054 | 0.289422154 | 3.918618178 | 3.327711234 | 2 | 3.369317972 | 4.105493497 | 4.58534546 | 4.209638631 | 3.940519005 |
| ENSG00000111843 | TMEM14C | 1.985839664 | 0.000759047 | 1.931189404 | 0.001031466 | 3.917029068 | 7.82963488 | 8.356994601 | 6.80318718 | 9.741259681 | 9.80689202 | 9.77752902 | 9.660173108 |
| ENSG00000113615 | SEC24A | 2.23980086 | 0.219632896 | 1.676009717 | 0.3644148 | 3.915810577 | 2 | 8.94259736 | 8.380957451 | 10.33669748 | 10.335119 | 9.797442274 | 9.750643668 |
| ENSG00000168172 | HOOK3 | 1.603339496 | 5.11E-05 | 2.31240017 | 1.58E-09 | 3.915739666 | 8.444143041 | 8.639735838 | 7.777116083 | 9.927741409 | 9.93517152 | 10.6333753 | 10.64305971 |
| ENSG00000112659 | CUL9 | 1.862196999 | 0.223543123 | 2.052911929 | 0.167672785 | 3.915108929 | 7.384433967 | 6.535001748 | 2 | 8.262524677 | 8.28661061 | 8.51523351 | 8.408829029 |
| ENSG00000196396 | PTPN1 | 1.751800426 | 0.334567146 | 2.16033079 | 0.215861 | 3.912131216 | 8.138771685 | 8.655694736 | 2 | 9.56056556 | 9.60006644 | 9.769486116 | 10.17617136 |
| ENSG00000242220 | TCP10L | 1.690450301 | 0.613127293 | 2.220811585 | 0.484665142 | 3.911261886 | 2 | 2 | 2 | 2.947924294 | 2.77779907 | 3.517841605 | 2.592996859 |
| ENSG00000132185 | FCRLA | 3.010996438 | 0.281735266 | 0.899615761 | 0.769071028 | 3.910612199 | 2 | 2 | 2 | 3.566290607 | 3.65239003 | 2.68698273 | 2.421067459 |
| ENSG00000197948 | FCHSD1 | 1.908303732 | 0.015099487 | 1.999254338 | 0.009971111 | 3.907558069 | 5.095959458 | 6.140560021 | 7.285536013 | 8.279879181 | 8.29975519 | 8.350677018 | 8.408109518 |
| ENSG00000111737 | RAB35 | 1.872837425 | 0.067435031 | 2.034606382 | 0.042390554 | 3.907443807 | 8.517314938 | 5.457120953 | 7.646393253 | 9.424141421 | 9.59930725 | 9.660389416 | 9.690077045 |
| ENSG00000153827 | TRIP12 | 2.256792155 | 1.24E-07 | 1.650453288 | 0.000157435 | 3.907245443 | 9.2903348 | 9.695596167 | 10.28245311 | 12.05260425 | 12.0764293 | 11.46059623 | 11.45726032 |
| ENSG00000171509 | RXFP1 | 1.690445815 | 0.59518254 | 2.215299096 | 0.464147051 | 3.905744912 | 2 | 2 | 2 | 2.947924294 | 2.77779907 | 3.355392153 | 2.851942824 |
| ENSG00000186814 | ZSCAN30 | 1.50564585 | 0.479224001 | 2.398700354 | 0.228768931 | 3.904346205 | 3.327711234 | 7.60566235 | 2 | 7.626911583 | 7.5462416 | 8.522164399 | 8.408109518 |
| ENSG00000122565 | CBX3 | 2.255883912 | 0.00010208 | 1.646183374 | 0.005906677 | 3.902067285 | 7.031698319 | 8.433041974 | 7.451345288 | 10.04290398 | 9.97727021 | 9.491657377 | 9.309990451 |
| ENSG00000101665 | SMAD7 | 2.248879609 | 0.217503328 | 1.65309165 | 0.370178383 | 3.901971259 | 6.009025997 | 8.433041974 | 2 | 9.36754703 | 9.29526488 | 8.780980934 | 8.699106494 |
| ENSG00000065243 | PKN2 | 2.248992834 | 0.09785965 | 1.65234966 | 0.230540721 | 3.901342494 | 4.815609471 | 8.414402626 | 5.39244166 | 9.359407363 | 9.29432703 | 8.964399877 | 8.462487876 |
| ENSG00000135324 | MRAP2 | -2.308732211 | 0.442830958 | 6.204309841 | 0.00226213 | 3.89557763 | 2 | 2 | 2 | 2.141737246 | 2 | 6.433059984 | 6.165581817 |
| ENSG00000134504 | KCTD1 | 1.914459697 | 0.309774396 | 1.980472064 | 0.281992691 | 3.894931761 | 2 | 6.844429755 | 3.369317972 | 7.230526165 | 7.30683517 | 7.600074836 | 7.006211772 |
| ENSG00000237172 | B3GNT9 | 1.59403503 | 0.194136557 | 2.300528882 | 0.047681033 | 3.894563913 | 7.709880939 | 7.874700259 | 4.058998049 | 8.838609445 | 8.83669913 | 9.450887435 | 9.622520313 |
| ENSG00000115598 | IL1RL2 | 3.114061087 | 0.26497168 | 0.779109435 | 0.802048583 | 3.893170522 | 2 | 2 | 2 | 3.462030052 | 3.86943969 | 2.656294034 | 2.374681325 |
| ENSG00000167074 | TEF | 1.369573053 | 0.379217697 | 2.521855017 | 0.079263128 | 3.89142807 | 6.138360429 | 7.189546432 | 2 | 7.398825229 | 7.65837038 | 8.475133675 | 8.840742848 |
| ENSG00000183624 | HMCES | 1.683027878 | 0.277913797 | 2.208061904 | 0.135257068 | 3.891089782 | 2 | 7.145017454 | 7.029880454 | 8.157948442 | 8.17894021 | 8.536980328 | 8.823577951 |
| ENSG00000123473 | STIL | 1.865041603 | 0.049361415 | 2.025503674 | 0.02901671 | 3.890545277 | 4.815609471 | 4.93638933 | 2 | 6.113598934 | 6.0722229 | 6.221381643 | 6.267475858 |
| ENSG00000135269 | TES | 0.83789391 | 0.565892094 | 3.051741934 | 0.016163983 | 3.889635843 | 8.517314938 | 8.451443573 | 4.058998049 | 8.744518279 | 8.78154738 | 10.85690136 | 11.06843302 |
| ENSG00000176658 | MYO1D | 1.116777974 | 0.44238177 | 2.771699121 | 0.03297173 | 3.888477094 | 4.467311984 | 7.761270542 | 9.116480286 | 9.148860528 | 9.15455108 | 10.58365688 | 10.98754108 |
| ENSG00000103042 | SLC38A7 | 2.241072812 | 0.195613485 | 1.647038766 | 0.348235366 | 3.888111578 | 2 | 7.189546432 | 8.408111533 | 9.576258382 | 9.54829592 | 8.936734789 | 9.00677572 |
| ENSG00000151348 | EXT2 | 2.398865362 | 0.078490009 | 1.487782223 | 0.29074063 | 3.886647585 | 10.00958825 | 9.267380431 | 5.156626401 | 11.51507238 | 11.5302082 | 10.39396692 | 10.80363534 |
| ENSG00000278558 | TMEM191B | 2.609393652 | 0.222533677 | 1.271382925 | 0.5723508 | 3.880776577 | 2 | 2 | 3.369317972 | 3.881293667 | 4.12711147 | 2.803598391 | 3.407202941 |

| | | | | | | | | | | | | | |
|---|---|---|---|---|---|---|---|---|---|---|---|---|---|
| ENSG00000197302 | ZNF720 | 1.849724522 | 0.158007309 | 2.030067467 | 0.110621519 | 3.879791989 | 5.532353208 | 2 | 5.59507254 | 6.652857793 | 6.77805036 | 6.91847879 | 6.862343163 |
| ENSG00000214517 | PPME1 | 0.468905141 | 0.706963346 | 3.408396091 | 0.000758118 | 3.877301233 | 9.635285994 | 9.616264339 | 6.43197674 | 9.596407142 | 9.57402703 | 12.33048163 | 12.68428561 |
| ENSG00000162129 | CLPB | 1.322351636 | 0.601636131 | 2.553792643 | 0.274904665 | 3.87614428 | 2 | 7.393647515 | 2 | 7.210554257 | 7.09050444 | 8.417967842 | 8.299011699 |
| ENSG00000134256 | CD101 | 1.384547394 | 0.695516458 | 2.49112559 | 0.446711602 | 3.875672983 | 2 | 2 | 2 | 2.270786764 | 3.08597292 | 3.059555873 | 3.474731315 |
| ENSG00000135452 | TSPAN31 | 2.144614413 | 0.002796453 | 1.730705067 | 0.018121893 | 3.87531948 | 5.70932728 | 7.274681896 | 5.772716771 | 8.598473365 | 8.54024645 | 8.020265032 | 8.28888532 |
| ENSG00000112276 | BVES | 2.851114639 | 0.005610729 | 1.020504935 | 0.368897909 | 3.871619574 | 4.467311984 | 6.666836837 | 7.691296561 | 9.601017164 | 9.56860713 | 8.127636775 | 7.302581437 |
| ENSG00000113721 | PDGFRB | 0.546736728 | 0.430113744 | 3.324563343 | 5.04E-09 | 3.871300071 | 9.497412313 | 10.79296235 | 9.767769415 | 10.66893665 | 10.6923799 | 13.70188182 | 13.15825212 |
| ENSG00000064989 | CALCRL | 0.55361321 | 0.689610102 | 3.317509787 | 0.003344821 | 3.871122997 | 4.00722531 | 4.442710129 | 2 | 4.598938466 | 3.86943969 | 7.078941352 | 6.362645675 |
| ENSG00000188763 | FZD9 | 0.227802372 | 0.902061374 | 3.643219219 | 0.010541206 | 3.871021591 | 2 | 4.710564154 | 2 | 3.79808662 | 3.74316441 | 6.433059984 | 6.994755759 |
| ENSG00000197226 | TBC1D9B | 1.631750367 | 1.00E-07 | 2.235869475 | 8.25E-14 | 3.867619842 | 9.159305284 | 9.497318007 | 9.031722608 | 10.85926146 | 10.8881455 | 11.34463044 | 11.59798046 |
| ENSG00000130810 | PPAN | 2.972178563 | 0.010826309 | 0.893913026 | 0.49395309 | 3.866091588 | 2 | 5.303523747 | 6.203069917 | 8.157948442 | 8.08010078 | 5.988693817 | 6.215784511 |
| ENSG00000187605 | TET3 | 0.914114326 | 0.590107068 | 2.951480803 | 0.047464741 | 3.865595129 | 6.138360429 | 7.538394143 | 2 | 7.25803034 | 7.4074129 | 9.364236594 | 9.324934421 |
| ENSG00000075407 | ZNF37A | 1.934716663 | 0.026360838 | 1.930549605 | 0.025734513 | 3.865266268 | 5.866944937 | 7.874700259 | 6.073378139 | 8.832088406 | 8.82245717 | 9.004546892 | 8.615557586 |
| ENSG00000151748 | SAV1 | 2.224509528 | 0.190049203 | 1.640643202 | 0.340182675 | 3.86515273 | 2 | 7.572420273 | 8.236946975 | 9.56371781 | 9.56860713 | 9.009866973 | 8.961885532 |
| ENSG00000120526 | NUDCD1 | 2.512768606 | 0.000367264 | 1.351815671 | 0.07180717 | 3.864584277 | 5.330594134 | 5.946638857 | 7.163363831 | 8.737556707 | 8.84440862 | 7.81936672 | 7.451079708 |
| ENSG00000178691 | SUZ12 | 2.167011133 | 0.000449271 | 1.697099415 | 0.007198919 | 3.864110548 | 5.70932728 | 6.951736442 | 7.398165132 | 9.023967474 | 8.91810182 | 8.632209947 | 8.368713182 |
| ENSG00000109511 | ANXA10 | 4.416628852 | 0.044311542 | -0.557392851 | 0.836438942 | 3.859236001 | 2 | 2 | 2 | 4.574143677 | 4.79082474 | 2.183373728 | 2.277179783 |
| ENSG00000128016 | ZFP36 | 1.594506055 | 0.237182775 | 2.263433052 | 0.077151749 | 3.857939107 | 7.485143553 | 7.761270542 | 3.369317972 | 8.657201865 | 8.65136017 | 9.473343311 | 9.143604888 |
| ENSG00000168610 | STAT3 | 1.431155922 | 0.004873329 | 2.426557809 | 4.48E-07 | 3.857713731 | 8.716893051 | 9.532315021 | 9.897982195 | 10.8713966 | 10.9036026 | 11.76485583 | 11.98936987 |
| ENSG00000185838 | GNB1L | 1.913479453 | 0.298900033 | 1.942553013 | 0.279801848 | 3.856032466 | 2 | 4.93638933 | 2 | 5.258265913 | 5.39133044 | 5.740677235 | 4.819291477 |
| ENSG00000223953 | C1QTNF5 | 6.604761574 | 8.17E-05 | -2.75116766 | 0.219885804 | 3.853593915 | 2 | 2 | 2 | 6.592405944 | 6.77805036 | 2.048074405 | 2.059860178 |
| ENSG00000170558 | CDH2 | 3.01980793 | 5.21E-06 | 0.833420424 | 0.268600882 | 3.853228354 | 9.521325812 | 8.791896851 | 7.818184403 | 11.87684221 | 11.8819476 | 10.01997556 | 9.283950453 |
| ENSG00000166780 | C16orf45 | 3.951720011 | 0.002208346 | -0.099070654 | 0.953066893 | 3.852649357 | 6.96351253 | 6.464272742 | 2 | 10.0321341 | 10.0836929 | 6.315225883 | 5.821002687 |
| ENSG00000235162 | C12orf75 | 1.548254703 | 0.00601843 | 2.304381855 | 1.84E-05 | 3.852636557 | 7.579279349 | 8.702538795 | 8.922968322 | 10.0321341 | 10.0645682 | 10.92315021 | 10.67046947 |
| ENSG00000078043 | PIAS2 | 1.965224316 | 0.131051356 | 1.88722739 | 0.141662309 | 3.852451706 | 3.327711234 | 7.35507212 | 5.59507254 | 8.205140657 | 8.06478301 | 8.087943566 | 8.031095054 |
| ENSG00000113916 | BCL6 | 2.004211213 | 0.288045778 | 1.846222942 | 0.321893604 | 3.850434155 | 8.998234871 | 7.189546432 | 2 | 9.786482773 | 9.74988221 | 9.851568452 | 9.322263543 |
| ENSG00000156504 | FAM122B | 2.161312927 | 0.053389543 | 1.688260638 | 0.135544734 | 3.849573566 | 6.816637977 | 5.303523747 | 3.369317972 | 7.939959586 | 7.71553728 | 7.272216802 | 7.459438299 |
| ENSG00000113600 | C9 | 4.06595011 | 0.109507808 | -0.216859972 | 0.944157185 | 3.849090139 | 2 | 2 | 2 | 4.598938466 | 4.16043394 | 2.139668783 | 2.421067459 |
| ENSG00000167077 | MEI1 | 2.139668156 | 0.199554781 | 1.709381387 | 0.305685651 | 3.849049543 | 2 | 4.11352228 | 2 | 4.647285135 | 4.79082474 | 4.188345803 | 4.522269241 |
| ENSG00000183479 | TREX2 | 0.493769106 | 0.883547191 | 3.35521693 | 0.22200216 | 3.848986036 | 2 | 2 | 2 | 2.270786764 | 2.59489879 | 4.076896059 | 3.580696614 |
| | WTH3DI | 9.038786388 | 3.86E-09 | -5.191151161 | 0.028077653 | 3.847635226 | 2 | 2 | 2 | 9.091045942 | 9.06060056 | 2 | 2 |
| ENSG00000100014 | SPECC1L | 1.913376227 | 0.184360992 | 1.933352071 | 0.171451798 | 3.846728298 | 4.815609471 | 8.981105197 | 6.073378139 | 9.60255057 | 9.48723044 | 9.58830174 | 9.543149124 |
| ENSG00000130725 | UBE2M | 1.789832368 | 0.019984214 | 2.055790145 | 0.006279543 | 3.845622513 | 7.096806373 | 8.234834549 | 6.203069917 | 9.138327453 | 9.24954577 | 9.385734536 | 9.529320022 |
| ENSG00000082126 | MPP4 | 1.796421316 | 0.454376509 | 2.049010497 | 0.375401268 | 3.845431813 | 5.532353208 | 2 | 2 | 5.614989456 | 5.7929843 | 4.177580368 | 6.711881188 |
| ENSG00000135722 | FBXL8 | 2.433680645 | 0.184268122 | 1.411164752 | 0.457954601 | 3.844845397 | 2 | 5.457120953 | 2 | 6.34372989 | 6.29861567 | 5.381251181 | 5.361470313 |
| ENSG00000246922 | UBAP1L | 1.177500794 | 0.557663675 | 2.667102872 | 0.141298013 | 3.844603666 | 2 | 4.11352228 | 2 | 3.840289928 | 4.02226117 | 5.684753614 | 4.316807515 |
| ENSG00000037749 | MFAP3 | 2.051857837 | 0.386972978 | 1.792112626 | 0.446711602 | 3.843970463 | 2 | 8.862353853 | 3.369317972 | 9.398771806 | 9.30087923 | 9.121501891 | 9.063165393 |
| ENSG00000008283 | CYB561 | 2.031022707 | 0.265448614 | 1.812458965 | 0.31713805 | 3.843481672 | 8.609439662 | 6.78762695 | 2 | 9.345738359 | 9.45149977 | 8.79689555 | 9.486337564 |
| ENSG00000160392 | C19orf47 | 1.543142711 | 0.067215792 | 2.299811026 | 0.003805056 | 3.842953737 | 5.532353208 | 5.838937808 | 3.369317972 | 6.79375511 | 6.65485813 | 7.34858644 | 7.563910189 |
| ENSG00000104517 | UBR5 | 2.21635571 | 0.017319385 | 1.62403783 | 0.089230416 | 3.840393541 | 6.257048221 | 8.590771778 | 9.031722608 | 10.5602946 | 10.5648023 | 10.13399593 | 9.789837542 |
| ENSG00000004139 | SARM1 | 2.298685981 | 0.302610403 | 1.540862191 | 0.496738835 | 3.839548171 | 2 | 6.666836837 | 2 | 7.497728447 | 7.26148706 | 6.643273885 | 6.657218517 |
| ENSG00000066422 | ZBTB11 | 1.866358618 | 0.228635685 | 1.971880222 | 0.192814682 | 3.83823884 | 7.096806373 | 7.232742069 | 2 | 8.446246007 | 8.41468635 | 8.82044307 | 8.178454506 |
| ENSG00000106665 | CLIP2 | 2.111796345 | 0.155215303 | 1.725447896 | 0.247204345 | 3.837244241 | 7.867441304 | 9.377333006 | 4.058998049 | 10.350452 | 10.3581897 | 10.0231736 | 9.913373788 |
| ENSG00000188396 | TCTEX1D4 | 1.384548723 | 0.694728637 | 2.452649284 | 0.453575633 | 3.837198007 | 2 | 2 | 2 | 2.270786764 | 3.08597292 | 3.355392153 | 3.15612297 |
| ENSG00000228454 | | 9.025510646 | 4.27E-09 | -5.191151148 | 0.02816633 | 3.834359498 | 2 | 2 | 2 | 9.087765067 | 9.03724641 | 2 | 2 |
| ENSG00000233312 | | 9.025510646 | 4.27E-09 | -5.191151148 | 0.02816633 | 3.834359498 | 2 | 2 | 2 | 9.087765067 | 9.03724641 | 2 | 2 |
| ENSG00000198894 | CIPC | 1.707915088 | 0.363034555 | 2.126400785 | 0.239257685 | 3.834315873 | 2 | 7.315436877 | 4.058998049 | 7.632926745 | 7.4949172 | 8.047866378 | 7.90110545 |
| ENSG00000177119 | ANO6 | 1.547512202 | 7.89E-07 | 2.286208696 | 6.02E-14 | 3.833720898 | 8.946185348 | 8.395519306 | 8.536706852 | 10.19345574 | 10.1629006 | 10.91497027 | 10.9149442 |
| ENSG00000158286 | RNF207 | 1.881140986 | 0.000674334 | 1.952576054 | 0.000352745 | 3.83371704 | 5.532353208 | 5.838937808 | 4.874587708 | 7.281195074 | 7.28811304 | 7.229857522 | 7.469129172 |
| ENSG00000169180 | XPO6 | 1.434132286 | 0.220372027 | 2.399435599 | 0.028282877 | 3.833567885 | 7.384433967 | 9.988044813 | 7.225742779 | 10.22465921 | 10.2262311 | 11.06141783 | 11.30508916 |
| ENSG00000176490 | DIRAS1 | 1.125023859 | 0.4962262 | 2.707194619 | 0.068373122 | 3.832218478 | 3.327711234 | 6.844429755 | 3.369317972 | 6.628980152 | 6.53293476 | 8.51523351 | 7.583161732 |
| ENSG00000138459 | SLC35A5 | 2.198547622 | 0.013143309 | 1.631467984 | 0.072675764 | 3.830015606 | 4.815609471 | 7.393647515 | 7.285536013 | 9.048926513 | 9.03612482 | 8.579514585 | 8.374617996 |
| ENSG00000115137 | DNAJC27 | 1.905929406 | 0.131015998 | 1.923228319 | 0.120415896 | 3.829157725 | 4.00722531 | 5.457120953 | 2 | 6.328984825 | 6.01871653 | 6.419417643 | 5.937276556 |
| ENSG00000085382 | HACE1 | 1.620032112 | 0.264978658 | 2.208528355 | 0.110369822 | 3.828560466 | 4.00722531 | 6.899080544 | 3.369317972 | 7.136275664 | 7.21069602 | 8.019511671 | 7.415721389 |
| ENSG00000173511 | VEGFB | 1.56870199 | 0.414528109 | 2.259457346 | 0.213653476 | 3.828159335 | 8.963744667 | 7.099070191 | 2 | 9.244284335 | 9.3304589 | 9.937430448 | 10.01208774 |
| ENSG00000179941 | BBS10 | 1.994280328 | 0.400992689 | 1.833105681 | 0.435161964 | 3.827386009 | 2 | 7.002537973 | 2 | 7.300218887 | 7.53039799 | 7.590975897 | 6.836905794 |
| ENSG00000145014 | TMEM44 | 1.925354088 | 0.00020695 | 1.900092628 | 0.000238991 | 3.825446716 | 6.816637977 | 7.954278496 | 8.007629717 | 9.623093601 | 9.5678312 | 9.428360321 | 9.700104804 |
| ENSG00000165458 | INPPL1 | 1.702496812 | 3.75E-09 | 2.12080255 | 8.36E-14 | 3.823299362 | 8.910412251 | 9.415325457 | 9.227351203 | 10.86342445 | 10.9343299 | 11.35227943 | 11.28053306 |
| ENSG00000135596 | MICAL1 | 1.362374518 | 0.311059736 | 2.460917333 | 0.048008107 | 3.823291851 | 4.00722531 | 8.234834549 | 8.042706615 | 8.883449599 | 9.00321357 | 9.982196088 | 10.08952187 |
| ENSG00000012124 | CD22 | 0.68948932 | 0.841054398 | 3.132766088 | 0.280017687 | 3.822255408 | 2 | 2 | 2 | 2.498691189 | 2.49394783 | 4.166733998 | 2.981406154 |
| ENSG00000114648 | KLHL18 | 1.608646446 | 0.158698358 | 2.206079153 | 0.042028392 | 3.814725599 | 4.00722531 | 7.002537973 | 7.552162999 | 8.365459376 | 8.36195019 | 9.070769397 | 8.829501448 |
| ENSG00000047230 | CTPS2 | 0.94352578 | 0.610039424 | 2.870386237 | 0.078715146 | 3.813912017 | 3.327711234 | 7.538394143 | 3.369317972 | 7.106324235 | 6.94032357 | 8.739036711 | 9.079977539 |
| ENSG00000269897 | COMMD3-BM | 9.004531104 | 4.87E-09 | -5.191151143 | 0.028266186 | 3.813379961 | 2 | 2 | 2 | 9.035366048 | 9.04841459 | 2 | 2 |
| ENSG00000122591 | FAM126A | 1.938297099 | 0.006763681 | 1.873016965 | 0.008764582 | 3.811314064 | 8.285626337 | 8.57407404 | 6.629488387 | 9.992332092 | 9.95489142 | 9.90251187 | 9.91489412 |
| ENSG00000250565 | ATP6V1E2 | 0.927847624 | 0.635885449 | 2.881626026 | 0.091607485 | 3.80947365 | 2 | 3.686303414 | 2 | 3.515101962 | 3.28037825 | 4.875056259 | 4.886954834 |
| ENSG00000144134 | RABL2A | 2.588700803 | 0.000460886 | 1.218952708 | 0.125850973 | 3.807653511 | 5.095959458 | 4.710564154 | 3.369317972 | 6.815074525 | 7.07311172 | 5.65789504 | 5.646532192 |
| ENSG00000141519 | CCDC40 | 1.681361362 | 0.338195259 | 2.126066254 | 0.205129503 | 3.807427616 | 2 | 4.710564154 | 2 | 5.003571641 | 4.811949 | 5.565950232 | 4.982847849 |
| ENSG00000275074 | NUDT18 | 1.343575016 | 0.566735645 | 2.463155327 | 0.259170136 | 3.806730343 | 2 | 6.728495673 | 2 | 6.503277904 | 6.53928196 | 7.560224843 | 7.65163107 |
| ENSG00000143324 | XPR1 | 1.757619468 | 0.283681035 | 2.048160658 | 0.195441403 | 3.805780126 | 4.00722531 | 5.72254323 | 8.960138144 | 9.246247604 | 9.33957954 | 9.522559945 | 9.640091423 |
| ENSG00000118200 | CAMSAP2 | 1.997623074 | 2.24E-11 | 1.807731834 | 1.51E-09 | 3.805354908 | 8.228665467 | 8.356994601 | 8.175061096 | 10.2793709 | 10.2054283 | 10.20661666 | 9.882366758 |
| | FAM196B | 8.995063304 | 5.87E-09 | -5.191151069 | 0.028636604 | 3.803912236 | 2 | 2 | 2 | 8.957217624 | 9.10406516 | 2 | 2 |
| ENSG00000124181 | PLCG1 | 1.932837716 | 0.186609 | 1.87035574 | 0.194491628 | 3.803193457 | 4.815609471 | 6.951736442 | 9.465702942 | 10.05527473 | 10.0977376 | 10.02260976 | 10.00615667 |
| ENSG00000138050 | THUMPD2 | 1.90848151 | 0.362456823 | 1.894699132 | 0.357657197 | 3.803180642 | 2 | 6.04685525 | 2 | 6.321555392 | 6.46787796 | 6.544089217 | 6.202567011 |
| ENSG00000179454 | KLHL28 | 1.689393596 | 0.131720426 | 2.113029519 | 0.049732463 | 3.802423115 | 4.815609471 | 7.002537973 | 4.523675391 | 7.596453725 | 7.51437845 | 8.333200639 | 7.488317792 |
| ENSG00000159256 | MORC3 | 2.200611086 | 0.009001216 | 1.598880122 | 0.064916066 | 3.799491208 | 5.330594134 | 7.002537973 | 7.734844356 | 9.205456697 | 9.09870316 | 8.830786521 | 8.2175813 |
| ENSG00000143353 | LYPLAL1 | 2.128483569 | 0.090021545 | 1.669475095 | 0.186478723 | 3.797958665 | 5.866944937 | 4.442710129 | 2 | 6.960881212 | 6.53928196 | 6.495072127 | 6.13459968 |
| ENSG00000164535 | DAGLB | 1.01390443 | 0.592369314 | 2.781812144 | 0.099466732 | 3.795716574 | 2 | 5.595927442 | 7.97167868 | 7.571615825 | 7.70428369 | 9.246842239 | 9.514354532 |
| ENSG00000083312 | TNPO1 | 2.108539217 | 1.81E-12 | 1.686955404 | 2.29E-08 | 3.795494622 | 9.015175615 | 9.34816733 | 9.180874355 | 11.32822224 | 11.2545233 | 11.04540869 | 10.67256802 |
| ENSG00000147679 | UTP23 | 2.399687087 | 0.072707408 | 1.394831785 | 0.316040074 | 3.794518872 | 4.467311984 | 7.051611315 | 3.369317972 | 8.136938566 | 8.11451391 | 7.268416942 | 7.0024032 |
| ENSG00000164707 | SLC13A4 | -1.060697213 | 0.571859046 | 4.854083778 | 0.001582235 | 3.793386565 | 4.815609471 | 2 | 2 | 3.227757282 | 2.59489879 | 8.144319434 | 7.425709383 |
| ENSG00000219607 | PPP1R3G | -0.003595342 | 0.998368875 | 3.795712391 | 0.030515328 | 3.792117048 | 2 | 5.838937808 | 2 | 4.329433326 | 4.63360314 | 7.988280194 | 8.067092663 |
| ENSG00000159433 | STARD9 | 1.447375651 | 0.188774353 | 2.341563013 | 0.023193105 | 3.788938663 | 3.327711234 | 6.464272742 | 6.073378139 | 7.110641269 | 7.2537885 | 8.392009143 | 7.627107456 |
| ENSG00000138592 | USP8 | 2.20264868 | 0.000763767 | 1.585566246 | 0.019165507 | 3.788214926 | 6.96351253 | 8.234834549 | 6.718950348 | 9.715981692 | 9.60536965 | 9.160294617 | 8.927151259 |
| ENSG00000151006 | PRSS53 | 1.526883649 | 0.399455299 | 2.260807349 | 0.184116925 | 3.787690998 | 2 | 4.93638933 | 2 | 5.194621444 | 4.74762503 | 5.801542492 | 5.475755622 |
| ENSG00000100030 | MAPK1 | 1.440512384 | 0.112649338 | 2.345768729 | 0.005734903 | 3.786281113 | 7.33131396 | 9.756063186 | 9.589475645 | 10.66233506 | 10.6610729 | 11.50392713 | 11.62416212 |
| ENSG00000186951 | PPARA | 1.260541306 | 0.234303473 | 2.524516938 | 0.009322992 | 3.785058243 | 5.330594134 | 8.077869079 | 8.35328246 | 8.942762689 | 9.01122668 | 10.29518375 | 10.17151518 |
| ENSG00000049860 | HEXB | 2.761326217 | 2.71E-10 | 1.023674638 | 0.032436902 | 3.785000855 | 7.867441304 | 8.834582582 | 8.825624954 | 11.330306 | 11.3221109 | 9.665454884 | 9.518360533 |
| ENSG00000134852 | CLOCK | 1.295737468 | 0.397539867 | 2.488516697 | 0.077151749 | 3.784254165 | 7.27616306 | 8.540087354 | 3.369317972 | 8.740345368 | 8.79619691 | 9.989139077 | 9.917677242 |
| ENSG00000123200 | ZC3H13 | 1.932699263 | 0.060764772 | 1.851273064 | 0.070288397 | 3.783972326 | 5.70932728 | 8.451443573 | 8.825624954 | 10.02013627 | 10.1280575 | 9.993749244 | 9.994220888 |
| ENSG00000164647 | STEAP1 | 7.578431545 | 1.56E-05 | -3.794939964 | 0.060834366 | 3.783491582 | 4.467311984 | 2 | 2 | 10.13319806 | 10.1057785 | 2.267001569 | 2 |
| ENSG00000100418 | DESI1 | 1.858491574 | 2.47E-05 | 1.923720159 | 1.10E-05 | 3.782211733 | 7.867441304 | 8.876041558 | 8.65477198 | 10.37846912 | 10.3800133 | 10.50362495 | 10.38236571 |
| ENSG00000178917 | ZNF852 | 1.877470295 | 0.177037027 | 1.902587534 | 0.161125537 | 3.78005783 | 4.00722531 | 4.11352228 | 2 | 5.040242897 | 5.3193943 | 5.699879814 | 4.456980017 |
| ENSG00000167395 | ZNF646 | 1.76167294 | 0.301934025 | 2.01717627 | 0.221484241 | 3.77884921 | 8.107517749 | 6.951736442 | 2 | 8.78154108 | 8.8250571 | 9.135128763 | 8.974090715 |
| ENSG00000021826 | CPS1 | 1.726301402 | 0.003288934 | 2.048824268 | 0.00033279 | 3.77512567 | 5.866944937 | 4.93638933 | 5.156626401 | 6.89733479 | 7.05550676 | 7.215456797 | 7.364717093 |
| ENSG00000180447 | GAS1 | 2.245774907 | 0.172055101 | 1.526230519 | 0.363325139 | 3.772005426 | 7.624139768 | 5.72254323 | 2 | 8.562683721 | 8.62325378 | 7.558151279 | 8.148612304 |
| ENSG00000262180 | OCLM | 2.643918238 | 0.387577122 | 1.125949141 | 0.727622186 | 3.769867379 | 2 | 2 | 2 | 3.406930997 | 3.33976443 | 2.858551861 | 2.374681325 |
| ENSG00000166762 | CATSPER2 | 1.049615791 | 0.641058771 | 2.71928983 | 0.17152891 | 3.768905621 | 2 | 3.076665209 | 2 | 3.023133209 | 3.01487888 | 4.065262523 | 4.119958326 |
| ENSG00000040531 | CTNS | 2.002882436 | 0.083653941 | 1.765433365 | 0.125782712 | 3.768315801 | 4.815609471 | 7.274681896 | 4.523675391 | 7.954442493 | 8.17486938 | 8.025527571 | 7.615934347 |
| ENSG00000173546 | CSPG4 | 0.114594227 | 0.942177035 | 3.652464792 | 0.002886376 | 3.767059019 | 10.14658276 | 7.467835385 | 6.43197674 | 8.923261356 | 9.0259911 | 12.84765862 | 12.04897144 |
| ENSG00000179833 | SERTAD2 | 1.98942365 | 8.94E-06 | 1.777183396 | 7.98E-05 | 3.766607046 | 7.940205782 | 7.874700259 | 8.677268317 | 10.15530726 | 10.2044302 | 9.732267925 | 10.17172715 |
| ENSG00000196628 | TCF4 | 2.533805805 | 0.001130382 | 1.231976761 | 0.140213693 | 3.765782565 | 8.493334308 | 8.88960062 | 6.629488387 | 10.80967368 | 10.8020184 | 9.422388201 | 9.590215873 |
| ENSG00000177697 | CD151 | 1.935090659 | 4.27E-07 | 1.828991123 | 1.81E-06 | 3.764081782 | 11.22635546 | 10.43440824 | 10.63827116 | 12.7143154 | 12.7616713 | 12.60099151 | 12.66265504 |
| ENSG00000121057 | AKAP1 | 1.105579359 | 0.405447051 | 2.655996407 | 0.026005702 | 3.761575766 | 7.27616306 | 4.710564154 | 4.058998049 | 7.097651217 | 7.08617588 | 8.683523874 | 8.541379994 |
| ENSG00000174125 | TLR1 | 0.486172233 | 0.785737459 | 3.275291813 | 0.024188368 | 3.761464046 | 2 | 4.11352228 | 2 | 3.462030052 | 3.50452879 | 5.661762729 | 5.825320785 |
| ENSG00000114841 | DNAH1 | 2.300830975 | 0.000123202 | 1.46037762 | 0.019857693 | 3.761208595 | 5.330594134 | 4.93638933 | 4.523675391 | 7.053486362 | 7.1744553 | 6.287958221 | 6.326546677 |
| ENSG00000010292 | NCAPD2 | 1.704838617 | 0.304685989 | 2.056362404 | 0.197633952 | 3.761201021 | 4.467311984 | 7.928237109 | 3.369317972 | 8.201097719 | 8.19912382 | 8.486081846 | 8.606177646 |
| ENSG00000169862 | CTNND2 | 2.603263373 | 0.471298431 | 1.156518261 | 0.760366765 | 3.759781634 | 2 | 2 | 2 | 3.95996233 | 2.2680111 | 2.62493832 | 2.671842496 |

| | | | | | | | | | | | | | |
|---|---|---|---|---|---|---|---|---|---|---|---|---|---|
| ENSG00000167701 | GPT | -0.723710535 | 0.808916091 | 4.482845591 | 0.049413588 | 3.759135056 | 2 | 2 | 2 | 2.270786764 | 2.14022021 | 4.690491514 | 4.793076545 |
| ENSG00000137098 | SPAG8 | 0.497858951 | 0.879726254 | 3.260275465 | 0.222125563 | 3.758134415 | 2 | 2 | 2 | 2.498691189 | 2.38539787 | 3.584558147 | 3.956354764 |
| ENSG00000127946 | HIP1 | 1.539801765 | 0.112267143 | 2.216561539 | 0.016024153 | 3.756363304 | 7.867441304 | 8.029695761 | 5.39244166 | 8.980991923 | 9.03612482 | 9.946772799 | 9.355859082 |
| ENSG00000133961 | NUMB | 2.252234963 | 1.86E-18 | 1.504017304 | 7.86E-09 | 3.756252267 | 8.892187472 | 8.88960062 | 9.212024683 | 11.2267272 | 11.2638957 | 10.55027224 | 10.44543965 |
| ENSG00000169032 | MAP2K1 | 1.667982611 | 0.020106114 | 2.087839776 | 0.002656384 | 3.755822387 | 9.460782017 | 7.60566235 | 8.460932916 | 10.36090904 | 10.3572918 | 10.75739214 | 10.79801646 |
| PPP1R12C | | 1.879381024 | 0.225247916 | 1.873297113 | 0.21938176 | 3.752678137 | 3.327711234 | 8.623598428 | 8.784770914 | 9.967555951 | 10.0396007 | 10.0182796 | 9.977343134 |
| ENSG00000165476 | REEP3 | 1.880660042 | 8.28E-09 | 1.87150613 | 8.64E-09 | 3.752166172 | 8.468948322 | 9.067150531 | 9.014156722 | 10.80035023 | 10.6966453 | 10.71055142 | 10.76944897 |
| ENSG00000132467 | UTP3 | 1.93971207 | 0.288829199 | 1.811515413 | 0.314949963 | 3.751227482 | 8.39321512 | 6.311476754 | 2 | 9.028537716 | 9.04284131 | 8.505043475 | 9.2234807 |
| ENSG00000198909 | MAP3K3 | 1.434152385 | 0.430098143 | 2.316179479 | 0.171763347 | 3.750331864 | 2 | 8.124485672 | 8.296287027 | 8.998569022 | 9.10834046 | 9.994516174 | 9.865692849 |
| ENSG00000214827 | MTCP1 | 3.428925614 | 0.022553657 | 0.320890204 | 0.861870179 | 3.749815818 | 2 | 4.11352228 | 2 | 5.775854763 | 5.98191036 | 3.584558147 | 3.128433649 |
| ENSG00000117362 | APH1A | 2.113223079 | 0.010063091 | 1.635331615 | 0.05114311 | 3.748554694 | 8.418903792 | 6.389896633 | 8.631919287 | 10.17399201 | 10.2039308 | 9.5642243 | 9.848028389 |
| ENSG00000161681 | SHANK1 | 2.635063569 | 0.108024028 | 1.112900592 | 0.525881687 | 3.747964161 | 5.095959458 | 2 | 7.029880454 | 8.31771326 | 8.37087451 | 6.499401217 | 7.140495192 |
| ENSG00000139737 | SLAIN1 | 2.018537754 | 0.611381475 | 1.727864402 | NA | 3.746402156 | 2 | 2 | 2 | 3.566290607 | 2.14022021 | 3.1504466 | 2.551896972 |
| ENSG00000102699 | PARP4 | 3.177328631 | 1.87E-05 | 0.567799909 | 0.519247447 | 3.745128539 | 7.435667355 | 7.099070191 | 8.99637432 | 11.24574219 | 11.2496932 | 8.569770117 | 8.726491728 |
| ENSG00000205143 | ARID3C | 0.49371813 | 0.88968542 | 3.250862887 | 0.266977444 | 3.744581018 | 2 | 2 | 2 | 2.270786764 | 2.59489879 | 4.220168345 | 3.128433649 |
| ENSG00000183340 | JRKL | 2.019141983 | 0.140555028 | 1.725308409 | 0.207119046 | 3.744450392 | 6.468621203 | 6.535001748 | 2 | 7.947219213 | 7.86703672 | 7.840848421 | 7.357281305 |
| ENSG00000139985 | ADAM21 | 0.858593391 | 0.789422857 | 2.884980633 | 0.299323387 | 3.743574024 | 2 | 2 | 2 | 2.498691189 | 2.59489879 | 3.534812207 | 3.518055793 |
| ENSG00000101577 | LPIN2 | 1.658962035 | 0.085333416 | 2.084170059 | 0.024699035 | 3.743132094 | 7.33131396 | 6.78762695 | 9.180874355 | 9.78107609 | 9.79633147 | 10.0712216 | 10.34074271 |
| ENSG00000165376 | CLDN2 | 0.689488183 | 0.831318691 | 3.053522525 | 0.26116563 | 3.743010708 | 2 | 2 | 2 | 2.498691189 | 2.49394783 | 3.694366604 | 3.580696614 |
| ENSG00000177189 | RPS6KA3 | 2.466136234 | 0.000285776 | 1.276169153 | 0.079091424 | 3.742305387 | 6.257048221 | 8.053983481 | 8.042706615 | 10.11451133 | 10.0955857 | 8.961264871 | 8.881227752 |
| ENSG00000196458 | ZNF605 | 1.958779011 | 0.466801761 | 1.781676375 | 0.501916413 | 3.740455386 | 2 | 8.255926468 | 2 | 8.642369674 | 8.62175922 | 8.827777334 | 7.95628774 |
| ENSG00000127483 | HP1BP3 | 1.800178115 | 0.007382667 | 1.939494076 | 0.003409966 | 3.739672191 | 7.82963488 | 9.497318007 | 9.242516611 | 10.82879697 | 10.7980539 | 10.71020131 | 11.15995768 |
| ENSG00000175970 | UNC119B | 1.549561175 | 0.454376509 | 2.189638428 | 0.263585743 | 3.739199602 | 5.095959458 | 8.57407404 | 2 | 8.657201865 | 8.65574839 | 9.379582949 | 9.197482551 |
| ENSG00000187105 | HEATR4 | 1.014440637 | 0.768549381 | 2.724111321 | 0.376042598 | 3.738551958 | 2 | 2 | 2 | 2.784612485 | 2.38539787 | 3.796405078 | 2.918126211 |
| ENSG00000178188 | SH2B1 | 1.580223507 | 0.169907773 | 2.157615809 | 0.048821317 | 3.737839317 | 4.467311984 | 8.004992147 | 7.552162999 | 8.829471715 | 8.85589616 | 9.444166158 | 9.387230438 |
| ENSG00000172869 | DMXL1 | 1.512620296 | 0.374656805 | 2.223616413 | 0.166829592 | 3.736236709 | 2 | 7.819099893 | 7.552162999 | 8.575233246 | 8.64254348 | 9.439386236 | 9.177777331 |
| ENSG00000153933 | DGKE | 2.285123542 | 0.003922264 | 1.451094162 | 0.079481787 | 3.736217704 | 4.815609471 | 4.442710129 | 6.322058544 | 7.620871236 | 7.52080763 | 6.730562686 | 6.791281771 |
| ENSG00000157214 | STEAP2 | 4.7391478 | 2.42E-10 | -1.005787395 | 0.250703287 | 3.733360405 | 4.00722531 | 5.457120953 | 5.59507254 | 9.777686595 | 9.74782835 | 4.720468739 | 3.789585079 |
| ENSG00000100003 | SEC14L2 | 2.397497345 | 0.12454222 | 1.333805739 | 0.41298492 | 3.731303084 | 2 | 5.72254323 | 7.342949392 | 8.496596021 | 8.53707846 | 7.17948788 | 7.712342256 |
| ENSG00000241322 | CDRT1 | 3.974453865 | 0.103494474 | -0.243155574 | 0.935446753 | 3.73129829 | 2 | 2 | 2 | 4.268917409 | 4.37434574 | 2.183373728 | 2.374681325 |
| ENSG00000197951 | ZNF71 | 1.483526004 | 0.536732782 | 2.246924464 | 0.323301531 | 3.730450468 | 2 | 6.951736442 | 2 | 6.755670824 | 6.98285659 | 7.815031728 | 7.386797235 |
| ENSG00000131080 | EDA2R | 1.887929307 | 0.181069609 | 1.841768587 | 0.184486831 | 3.729697894 | 6.816637977 | 6.140560021 | 2 | 7.794595599 | 7.78127547 | 7.954782214 | 7.493753702 |
| ENSG00000156136 | DCK | 1.541469513 | 0.249182594 | 2.18811344 | 0.084474028 | 3.729582953 | 6.257048221 | 6.228547981 | 2 | 7.169761359 | 7.18258783 | 8.039715588 | 7.532563185 |
| ENSG00000165323 | FAT3 | 0.922394504 | 0.461270807 | 2.80672631 | 0.009915802 | 3.729120814 | 4.00722531 | 3.686303414 | 5.59507254 | 5.433909581 | 5.48637102 | 7.803699407 | 6.323497254 |
| ENSG00000163605 | PPP4R2 | 2.320180052 | 0.007747348 | 1.408820019 | 0.122622819 | 3.729000071 | 8.631580145 | 6.464272742 | 6.80318718 | 9.968150831 | 9.88434767 | 9.086680922 | 8.951552775 |
| ENSG00000197355 | UAP1L1 | 1.770925407 | 0.18204098 | 1.953856671 | 0.129621144 | 3.724782079 | 7.435667355 | 7.60566235 | 3.369317972 | 8.689305661 | 8.75588792 | 8.556847345 | 9.184095521 |
| ENSG00000147133 | TAF1 | 1.967161568 | 0.000112837 | 1.753806639 | 0.000619607 | 3.720968208 | 7.384433967 | 6.228547981 | 6.629488387 | 8.744518279 | 8.7394436 | 8.610361892 | 8.446458576 |
| ENSG00000112146 | FBXO9 | 2.091831983 | 0.003639034 | 1.628657631 | 0.027126817 | 3.720489613 | 8.009475818 | 6.844429755 | 6.43197674 | 9.283050899 | 9.34864288 | 8.380906995 | 9.213631806 |
| ENSG00000170921 | TANC2 | 1.917391453 | 0.00063387 | 1.800230824 | 0.001356792 | 3.717622277 | 7.624139768 | 7.051611315 | 8.408111533 | 9.675758322 | 9.70259467 | 9.691228581 | 9.443440338 |
| ENSG00000188783 | PRELP | -1.086671818 | 0.266515549 | 4.801576717 | 2.64E-09 | 3.714904899 | 4.00722531 | 4.11352228 | 5.930865648 | 3.997744386 | 4.0930011 | 9.199402547 | 9.81909102 |
| ENSG00000081818 | PCDHB4 | 2.373737522 | 0.304276217 | 1.340113903 | 0.574460139 | 3.713851425 | 2 | 3.076665209 | 2 | 4.105493497 | 3.50452879 | 3.500668998 | 2.746601469 |
| ENSG00000144357 | UBR3 | 1.748715457 | 0.326774611 | 1.965053587 | 0.257062588 | 3.713769043 | 2 | 8.451443573 | 7.398165132 | 9.142549916 | 9.20532695 | 9.443042894 | 9.333296562 |
| ENSG00000205423 | CNEP1R1 | 1.913173316 | 0.009852067 | 1.799979832 | 0.015286757 | 3.713153147 | 5.095959458 | 6.666836837 | 5.156626401 | 7.72026832 | 7.7127321 | 7.692773977 | 7.512619696 |
| ENSG00000064115 | TM7SF3 | 1.836578556 | 0.068005105 | 1.876483139 | 0.058167426 | 3.713061695 | 5.330594134 | 8.356994601 | 7.858115942 | 9.484432763 | 9.44644299 | 9.457299408 | 9.551644037 |
| ENSG00000137145 | DENND4C | 2.078286051 | 0.010174598 | 1.633162686 | 0.047681033 | 3.711448736 | 8.009475818 | 8.077869079 | 5.930865648 | 9.689531608 | 9.66898051 | 9.470865405 | 8.9569744 |
| ENSG00000164327 | RICTOR | 3.234725717 | 3.55E-05 | 0.475667395 | 0.614612732 | 3.710393112 | 6.89194353 | 9.04308532 | 7.777116083 | 11.41826296 | 11.375853 | 8.91056191 | 8.332767588 |
| ENSG00000102265 | TIMP1 | 3.888523253 | 2.36E-18 | -0.180553879 | 0.755865023 | 3.707969374 | 12.59833413 | 11.82700743 | 11.61331269 | 15.9367421 | 15.9921748 | 12.01740554 | 11.76576038 |
| ENSG00000143514 | TP53BP2 | 1.527643867 | 0.375347947 | 2.17934904 | 0.18139849 | 3.706992907 | 6.009025997 | 9.640525262 | 4.523675391 | 9.713145491 | 9.73682479 | 10.19415235 | 10.53433941 |
| ENSG00000198952 | SMG5 | 1.863739276 | 0.056662136 | 1.842773415 | 0.056553913 | 3.706512691 | 6.257048221 | 8.732943944 | 9.083172337 | 10.26733709 | 10.3236737 | 10.48587226 | 10.02762959 |
| ENSG00000136448 | NMT1 | 1.840016501 | 0.129733292 | 1.865588719 | 0.117956298 | 3.70560522 | 5.532353208 | 9.461450882 | 8.742726175 | 10.41121192 | 10.4901834 | 10.51623025 | 10.43611728 |
| ENSG00000267508 | ZNF285 | 1.735073995 | 0.361413515 | 1.968494242 | 0.284195618 | 3.703568238 | 2 | 4.93638933 | 2 | 5.304221918 | 5.00774437 | 5.798032389 | 4.766376438 |
| ENSG00000101298 | SNPH | 1.840318881 | 0.510757053 | 1.861506566 | 0.49597846 | 3.701825446 | 2 | 2 | 8.434763955 | 8.658676732 | 8.70880006 | 8.90526987 | 8.472158355 |
| ENSG00000177733 | HNRNPA0 | 1.361340924 | 0.299725694 | 2.339273983 | 0.054822741 | 3.700614906 | 9.423197387 | 7.819099893 | 5.39244166 | 9.645581273 | 9.68051177 | 10.39846491 | 10.84263274 |
| ENSG00000133641 | C12orf29 | 1.821788504 | 0.292677052 | 1.878365578 | 0.266744705 | 3.700154082 | 2 | 6.78762695 | 4.058998049 | 7.218576229 | 7.18663697 | 7.535143176 | 6.913939481 |
| ENSG00000149930 | TAOK2 | 1.616757055 | 0.013230416 | 2.08314964 | 0.000947011 | 3.699906695 | 9.048471759 | 7.819099893 | 7.552162999 | 9.905552176 | 9.88994604 | 10.33251275 | 10.39185883 |
| ENSG00000106829 | TLE4 | 2.155983083 | 0.061614336 | 1.541996537 | 0.190242338 | 3.69797962 | 5.70932728 | 9.386925327 | 8.721235202 | 10.69000523 | 10.7526855 | 9.791055745 | 10.37022608 |
| ENSG00000104967 | NOVA2 | 2.33872267 | 0.514344567 | 1.358883852 | NA | 3.697606522 | 2 | 2 | 2 | 3.663521627 | 2.49394783 | 3.035909414 | 2.326754077 |
| ENSG00000104643 | MTMR9 | 1.79615764 | 0.016930176 | 1.900539541 | 0.010456421 | 3.696697181 | 5.532353208 | 7.572420273 | 6.629488387 | 8.561107327 | 8.60672783 | 8.700544163 | 8.674186858 |
| ENSG00000112079 | STK38 | 1.15297535 | 0.016667868 | 2.540488138 | 9.78E-09 | 3.693463488 | 7.667647223 | 8.655694736 | 8.007629717 | 9.281137113 | 9.36213202 | 10.81616151 | 10.58264661 |
| ENSG00000186976 | EFCAB6 | 1.49903845 | 0.649972951 | 2.194409952 | NA | 3.693448402 | 2 | 2 | 2 | 2.947924294 | 2.59489879 | 3.336159116 | 2.851942824 |
| ENSG00000100241 | SBF1 | 1.156366766 | 0.42327046 | 2.537018043 | 0.052381201 | 3.693384809 | 5.095959458 | 9.549500232 | 9.61300511 | 10.18170646 | 10.1772163 | 11.69314688 | 11.4067386 |
| ENSG00000072803 | FBXW11 | 1.480642584 | 0.001734936 | 2.211223514 | 1.02E-06 | 3.691866098 | 8.340423424 | 7.538394143 | 8.561105505 | 9.667723798 | 9.66608324 | 10.51369123 | 10.26488154 |
| ENSG00000163001 | CFAP36 | 2.423002447 | 0.001762985 | 1.26879576 | 0.124086277 | 3.691798207 | 4.815609471 | 7.051611315 | 6.073378139 | 8.619831492 | 8.65136017 | 7.508544306 | 7.489678691 |
| ENSG00000108797 | CNTNAP1 | 1.214402292 | 0.077982082 | 2.473789388 | 8.37E-05 | 3.68819168 | 7.904282243 | 8.191702463 | 6.629488387 | 8.873322599 | 8.97305476 | 9.835606948 | 10.45208674 |
| ENSG00000066629 | EML1 | 2.729435543 | 0.173153636 | 0.956782814 | 0.661209052 | 3.686218357 | 4.00722531 | 8.124485672 | 2 | 9.306761457 | 9.35225233 | 7.718996793 | 7.422862719 |
| ENSG00000101190 | TCFL5 | 1.526260006 | 0.379713551 | 2.159789054 | 0.189077353 | 3.68604906 | 2 | 7.051611315 | 4.523675391 | 7.222570549 | 7.19871665 | 8.018757917 | 7.613439639 |
| ENSG00000166949 | SMAD3 | 1.735980908 | 0.002281839 | 1.947918984 | 0.000492889 | 3.683899891 | 8.39321512 | 9.424669273 | 8.236946975 | 10.48206602 | 10.5585666 | 10.53044605 | 10.90923487 |
| ENSG00000196365 | LONP1 | 1.619628388 | 0.100882434 | 2.062842234 | 0.029695601 | 3.682470622 | 5.866944937 | 8.747909548 | 8.175061096 | 9.59332559 | 9.65954302 | 9.999109217 | 10.13393228 |
| ENSG00000257446 | ZNF878 | 3.049542221 | 0.304185458 | 0.632569292 | 0.849613217 | 3.682111513 | 2 | 2 | 2 | 3.921164147 | 3.28037825 | 2.656294034 | 2.277179783 |
| ENSG00000145936 | KCNMB1 | -3.753147707 | 0.220358877 | 7.434961295 | 0.000837372 | 3.681813588 | 2 | 2 | 2 | 2 | 2 | 7.900986387 | 6.92404095 |
| ENSG00000180044 | C3orf80 | 7.447651245 | 6.02E-06 | -3.765866893 | 0.128150716 | 3.681784352 | 2 | 2 | 2 | 7.507583254 | 7.50468064 | 2.048074405 | 2 |
| ENSG00000113716 | HMGXB3 | 1.787325627 | 0.194794952 | 1.893984078 | 0.1588785 | 3.681309704 | 6.257048221 | 6.311476754 | 9.658943166 | 10.11128345 | 10.0874876 | 10.15284024 | 10.25671168 |
| ENSG00000158125 | XDH | 2.900428658 | 0.097131191 | 0.777498999 | 0.690278517 | 3.677927657 | 2 | 2 | 4.058998049 | 5.128039203 | 4.91316568 | 3.296906311 | 3.496556178 |
| ENSG00000089159 | PXN | 2.142418081 | 0.017649875 | 1.533800942 | 0.098356149 | 3.676219023 | 7.384433967 | 9.267380431 | 9.990713521 | 11.33238675 | 11.395009 | 10.43804609 | 11.01733363 |
| ENSG00000130787 | HIP1R | 2.155472997 | 0.140659286 | 1.517396707 | 0.307557173 | 3.672869704 | 5.532353208 | 6.951736442 | 2 | 7.855024645 | 8.05152279 | 7.328045423 | 7.334741191 |
| ENSG00000174780 | SRP72 | 1.771216099 | 0.129861227 | 1.900379863 | 0.096243075 | 3.671595962 | 8.653385977 | 8.687092496 | 5.156626401 | 9.919767943 | 9.89428535 | 10.12526871 | 9.940248792 |
| ENSG00000166233 | ARIH1 | 1.821153328 | 2.75E-07 | 1.845662927 | 1.64E-07 | 3.666816255 | 8.19931886 | 8.147241016 | 8.325066185 | 10.00803782 | 10.0529704 | 10.29829283 | 9.76223254 |
| ENSG00000113441 | LNPEP | 1.630284279 | 0.196836305 | 2.036162154 | 0.093560552 | 3.666446433 | 8.816894989 | 7.232742069 | 4.874587708 | 9.342071312 | 9.31111579 | 9.597165068 | 9.852266467 |
| ENSG00000115556 | PLCD4 | 1.496206818 | 0.448944973 | 2.168359625 | 0.244375349 | 3.664566443 | 2 | 5.457120953 | 2 | 5.304221918 | 5.5755357 | 6.221381643 | 5.896817464 |
| ENSG00000073711 | PPP2R3A | 2.060652454 | 0.182837442 | 1.602644941 | 0.303025886 | 3.663297394 | 2 | 7.393647515 | 6.80318718 | 8.615281294 | 8.55284928 | 8.05302937 | 8.206864727 |
| ENSG00000180537 | RNF182 | -0.028813158 | 0.98413673 | 3.691899043 | 0.001006265 | 3.663085885 | 3.327711234 | 5.131603625 | 2 | 4.205751501 | 3.98554648 | 7.272216802 | 7.644317657 |
| ENSG00000167460 | TPM4 | 1.206034897 | 0.004834581 | 2.456909105 | 8.26E-10 | 3.662944002 | 12.64261488 | 11.84277663 | 11.90062666 | 13.37147027 | 13.3921961 | 14.59460754 | 14.66928705 |
| ENSG00000215251 | FASTKD5 | 1.862225852 | 0.200007734 | 1.798302171 | 0.209322464 | 3.660528022 | 5.532353208 | 7.819099893 | 3.369317972 | 8.381625466 | 8.39906692 | 8.125537812 | 8.504157459 |
| ENSG00000172264 | MACROD2 | 0.689488151 | 0.832844524 | 2.970138428 | 0.281038046 | 3.659626579 | 2 | 2 | 2 | 2.498691189 | 2.49394783 | 3.584558147 | 3.580696614 |
| ENSG00000119820 | YIPF4 | 1.622515454 | 0.065635381 | 2.036946638 | 0.016735344 | 3.659462092 | 5.866944937 | 7.847167909 | 8.325066185 | 9.272493518 | 9.26686067 | 9.876121046 | 9.457070207 |
| ENSG00000171503 | ETFDH | 1.508135773 | 0.323937413 | 2.150785495 | 0.137340944 | 3.658921267 | 2 | 7.099070191 | 6.534109959 | 7.668500375 | 7.83127384 | 8.344674564 | 8.424568162 |
| ENSG00000177374 | HIC1 | 0.769789225 | 0.633071268 | 2.885857858 | 0.038931255 | 3.655647083 | 6.816637977 | 7.099070191 | 2 | 7.035434774 | 7.26148706 | 9.04915428 | 9.403927624 |
| ENSG00000196743 | GM2A | 0.837689681 | 0.467543151 | 2.816272135 | 0.004738564 | 3.653961815 | 5.532353208 | 8.57407404 | 7.029880454 | 8.343618883 | 8.40255255 | 10.51235311 | 10.14174401 |
| ENSG00000172469 | MANEA | 1.80175155 | 0.007359418 | 1.852070346 | 0.005357536 | 3.653821896 | 4.815609471 | 5.838937808 | 6.629488387 | 7.742693204 | 7.5835711 | 7.799317009 | 7.62463201 |
| ENSG00000198466 | ZNF587 | 2.665358069 | 3.56E-08 | 0.987008852 | 0.06346313 | 3.652366921 | 7.33131396 | 6.464272742 | 7.451345288 | 9.764733396 | 9.74919792 | 8.296356225 | 7.856442979 |
| ENSG00000114331 | ACAP2 | 2.139193 | 1.25E-07 | 1.512471335 | 0.000268289 | 3.651664335 | 7.218819781 | 7.761270542 | 7.858115942 | 9.78445764 | 9.73129138 | 9.347224935 | 8.885886446 |
| ENSG00000162413 | KLHL21 | 2.116078209 | 0.265702905 | 1.535566072 | 0.426645605 | 3.651644281 | 9.410448395 | 8.213429689 | 2 | 10.44065847 | 10.4716357 | 9.95467636 | 9.796450865 |
| ENSG00000139651 | ZNF740 | 1.850650111 | 0.012834002 | 1.798707399 | 0.015373036 | 3.64935751 | 5.330594134 | 7.315436877 | 7.225742779 | 8.598473365 | 8.76809931 | 8.704768045 | 8.560285 |
| ENSG00000158480 | SPATA2 | 1.690286678 | 0.221271682 | 1.957722473 | 0.143141086 | 3.648009151 | 6.138360429 | 6.666836837 | 2 | 7.530319349 | 7.4949172 | 7.774526492 | 7.775072826 |
| ENSG00000023892 | DEF6 | 1.957355793 | 0.303980957 | 1.690643961 | 0.369090445 | 3.647999754 | 2 | 3.686303414 | 2 | 3.95996233 | 4.2560182 | 4.029788235 | 3.789585079 |
| ENSG00000018510 | AGPS | 2.260856807 | 0.017769551 | 1.387112855 | 0.163224785 | 3.647969662 | 7.667647223 | 8.687092496 | 5.772716771 | 10.12041043 | 9.98077219 | 9.234884382 | 9.129269498 |
| ENSG00000080561 | MID2 | 2.654866322 | 0.073563931 | 0.989946442 | 0.538569152 | 3.644812764 | 6.653098118 | 7.431218269 | 2 | 9.18513223 | 9.09655277 | 7.528804029 | 7.467748741 |
| ENSG00000198585 | NUDT16 | 1.467258145 | 0.218891141 | 2.177468171 | 0.053220172 | 3.644726316 | 4.00722531 | 7.538394143 | 7.451345288 | 8.427334995 | 8.40429221 | 8.889688585 | 9.31767336 |
| ENSG00000120279 | MYCT1 | 3.219183929 | 0.096972333 | 0.425506341 | 0.852507508 | 3.64469027 | 3.327711234 | 2 | 2 | 4.443344104 | 4.53541753 | 2.831336763 | 2.632958241 |
| ENSG00000102572 | STK24 | 2.351724385 | 7.58E-05 | 1.291756847 | 0.040597375 | 3.643481232 | 7.27616306 | 8.747909548 | 7.734844356 | 10.37936391 | 10.4205473 | 9.330921459 | 9.358468538 |
| ENSG00000145730 | PAM | 1.82970223 | 1.73E-09 | 1.811736057 | 2.29E-09 | 3.641438287 | 10.43795805 | 10.64091981 | 10.5677762 | 12.38192656 | 12.3758755 | 12.07985022 | 12.59614149 |
| ENSG00000123106 | CCDC91 | 2.380127198 | 0.014133546 | 1.260842454 | 0.220251675 | 3.640969652 | 6.009025997 | 6.535001748 | 3.369317972 | 8.14748175 | 8.09955907 | 6.931352719 | 7.119550903 |
| ENSG00000149782 | PLCB3 | 1.549283729 | 0.108536572 | 2.090825845 | 0.023427954 | 3.640109574 | 7.218819781 | 7.701025996 | 4.874587708 | 8.568972129 | 8.51148098 | 9.033940582 | 9.119193255 |
| ENSG00000110906 | KCTD10 | 1.250499998 | 0.057132864 | 2.3895414 | 7.98E-05 | 3.640041398 | 9.473095744 | 8.469613413 | 8.007629717 | 10.05303338 | 9.99815569 | 10.9216389 | 11.36776693 |
| ENSG00000187123 | LYPD6 | 4.838985764 | 0.019426473 | -1.199808284 | 0.630693854 | 3.63917748 | 2 | 2 | 2 | 5.022023783 | 5.07918227 | 2.183373728 | 2.117335275 |
| ENSG00000117528 | ABCD3 | 1.715816763 | 0.233103112 | 1.922797554 | 0.169025837 | 3.638614317 | 4.00722531 | 7.819099893 | 4.874587708 | 8.27796118 | 8.09094333 | 8.454100442 | 8.326688609 |
| ENSG00000178409 | BEND3 | 1.525993675 | 0.174897897 | 2.111241589 | 0.047215475 | 3.637235264 | 2 | 4.11352228 | 5.156626401 | 5.602782483 | 5.47317115 | 6.025180246 | 6.13807518 |

| | | | | | | | | | | | | | |
|---|---|---|---|---|---|---|---|---|---|---|---|---|---|
| ENSG00000082701 | GSK3B | 1.263642183 | 0.484301277 | 2.372755616 | 0.153698217 | 3.636397798 | 9.710019321 | 7.431218269 | 3.369317972 | 9.6388717 | 9.68695783 | 10.69648025 | 10.83689669 |
| ENSG00000222014 | RAB6C | 8.82662083 | 1.71E-08 | -5.19115099 | 0.029583545 | 3.63546984 | 2 | 2 | 2 | 8.902248225 | 8.82765235 | 2 | 2 |
| ENSG00000137802 | MAPKBP1 | 1.818732797 | 0.105302052 | 1.813423473 | 0.100794035 | 3.632156271 | 7.940205782 | 5.946638857 | 4.874587708 | 8.514626115 | 8.65574839 | 8.714575941 | 8.435201779 |
| ENSG00000173457 | PPP1R14B | 1.90504113 | 0.000144567 | 1.725298382 | 0.000614839 | 3.630339512 | 7.031698319 | 8.101365653 | 7.451345288 | 9.44052423 | 9.5427783 | 9.099926561 | 9.500245276 |
| ENSG00000151466 | SCLT1 | 2.3318898 | 0.003882515 | 1.298012847 | 0.128417272 | 3.629902647 | 3.327711234 | 5.457120953 | 4.874587708 | 7.026323566 | 6.97351258 | 6.181526565 | 5.821002687 |
| ENSG00000124795 | DEK | 2.038818495 | 0.007885856 | 1.587730044 | 0.04279221 | 3.626548539 | 7.485143553 | 8.981105197 | 7.163363831 | 10.17913955 | 10.1014956 | 9.871536104 | 9.485996671 |
| ENSG00000147065 | MSN | 1.324220322 | 0.215468947 | 2.30221188 | 0.02080403 | 3.626432202 | 8.946185348 | 11.98850994 | 11.18913795 | 12.4744381 | 12.5033925 | 13.13119617 | 13.73835164 |
| ENSG00000008256 | CYTH3 | 0.324063107 | 0.850996577 | 3.301787113 | 0.016654612 | 3.62585022 | 9.33149386 | 7.954278496 | 4.058998049 | 8.570539957 | 8.55598286 | 11.55739329 | 11.49721959 |
| ENSG00000130475 | FCHO1 | -0.286492525 | 0.864318577 | 3.909727128 | 0.001909905 | 3.623234604 | 2 | 4.11352228 | 3.369317972 | 3.349643765 | 3.2184422 | 6.046045499 | 7.255358547 |
| ENSG00000089916 | GPATCH2L | 1.657301695 | 0.159990977 | 1.963673113 | 0.084642081 | 3.620974808 | 5.095959458 | 8.356994601 | 8.699419242 | 9.678668908 | 9.64342987 | 10.16089043 | 9.74096227 |
| ENSG00000198862 | LTN1 | 1.968779097 | 0.006540456 | 1.652042769 | 0.024545065 | 3.620821866 | 7.159102577 | 8.213429689 | 6.534109959 | 9.439666599 | 9.41917001 | 9.440512349 | 8.693212171 |
| ENSG00000198689 | SLC9A6 | 1.5218079 | 0.374672148 | 2.09884885 | 0.197276691 | 3.620656749 | 2 | 7.73146271 | 6.073378139 | 8.049698831 | 8.06258142 | 8.517369639 | 8.727068782 |
| ENSG00000131023 | LATS1 | 1.546168424 | 0.05186708 | 2.074281655 | 0.006426257 | 3.620450079 | 7.624139768 | 7.901717006 | 5.772716771 | 8.883449599 | 8.89230135 | 9.558530648 | 9.249420371 |
| ENSG00000144026 | ZNF514 | 1.974743127 | 0.210990211 | 1.645208262 | 0.296894493 | 3.619951389 | 2 | 6.666836837 | 4.523675391 | 7.391747748 | 7.24992376 | 7.238947338 | 6.718860807 |
| ENSG00000141034 | GID4 | 2.042083539 | 0.226514714 | 1.577806122 | 0.35456714 | 3.619889661 | 2 | 6.535001748 | 7.97167868 | 8.85414072 | 8.86223882 | 8.60534994 | 8.157201821 |
| ENSG00000003400 | CASP10 | 1.519550726 | 0.116925716 | 2.100067596 | 0.022260314 | 3.619618322 | 3.327711234 | 3.686303414 | 4.874587708 | 5.406080789 | 5.3193943 | 5.912830251 | 5.88031069 |
| ENSG00000145545 | SRD5A1 | 1.408415344 | 0.415078879 | 2.210817026 | 0.171451798 | 3.619232371 | 7.667647223 | 5.72254323 | 2 | 7.781117077 | 7.82349365 | 8.695364791 | 8.484497333 |
| ENSG00000179046 | TRIML2 | 3.148890295 | 0.298769516 | 0.469759838 | 0.89230957 | 3.618650134 | 2 | 2 | 2 | 3.709785322 | 3.69849104 | 2.656294034 | 2.172608095 |
| ENSG00000263141 | | 2.487016298 | 0.002227636 | 1.129720929 | 0.198274928 | 3.616737227 | 6.653098118 | 5.946638857 | 4.523675391 | 8.406418094 | 8.31463236 | 7.407359556 | 6.520103598 |
| ENSG00000154760 | SLFN13 | 1.737220298 | 0.044788115 | 1.879172045 | 0.026212112 | 3.616392344 | 4.00722531 | 3.686303414 | 4.874587708 | 5.720439317 | 5.76075522 | 5.805044075 | 5.937276556 |
| ENSG00000205238 | SPDYE2 | 3.937391889 | 0.098417871 | -0.321416484 | 0.912368677 | 3.615975405 | 2 | 2 | 2 | 4.299492648 | 4.2865223 | 2.3070652 | 2.225841198 |
| ENSG00000168792 | ABHD15 | 1.436859392 | 0.049492187 | 2.178432884 | 0.001567678 | 3.615292276 | 4.467311984 | 6.311476754 | 6.43197674 | 7.352188525 | 7.36158648 | 8.131128273 | 8.035760639 |
| ENSG00000135740 | SLC9A5 | 1.149423383 | 0.496953626 | 2.462986632 | 0.109152121 | 3.612410014 | 4.00722531 | 6.535001748 | 2 | 6.306580752 | 6.35727082 | 7.512833218 | 7.68646583 |
| ENSG00000162267 | ITIH3 | -2.308256067 | 0.472536264 | 5.920309442 | 0.009193782 | 3.612053375 | 2 | 2 | 2 | 2.141737246 | 2 | 6.437578914 | 5.481238982 |
| ENSG00000171793 | CTPS1 | 0.435262512 | 0.760337788 | 3.176026944 | 0.006625005 | 3.611289456 | 5.330594134 | 8.557180776 | 9.385998894 | 8.951212326 | 8.90955283 | 11.97242248 | 11.26382758 |
| ENSG00000158483 | FAM86C1 | 2.429177637 | 0.007711612 | 1.181554901 | 0.226109663 | 3.610732539 | 3.327711234 | 4.93638933 | 4.058998049 | 6.522845756 | 6.44101697 | 4.996409624 | 5.586744392 |
| ENSG00000144136 | SLC20A1 | 3.159733509 | 0.001783365 | 0.449217874 | 0.712785235 | 3.608951382 | 8.418903792 | 9.656475478 | 6.43197674 | 11.81320634 | 11.8618308 | 8.804350194 | 9.405370462 |
| ENSG00000065361 | ERBB3 | -1.930947148 | 0.033219225 | 5.538848946 | 4.09E-13 | 3.607901798 | 4.815609471 | 3.686303414 | 5.39244166 | 3.566290607 | 3.01487888 | 9.756885121 | 10.29516161 |
| ENSG00000198610 | AKR1C4 | 8.798497025 | 6.32E-08 | -5.191150239 | 0.032903971 | 3.607346786 | 2 | 2 | 2 | 8.567402595 | 9.06500642 | 2 | 2 |
| ENSG00000125954 | CHURC1-FNT | 8.797582656 | 2.20E-08 | -5.191150929 | 0.029960917 | 3.606431726 | 2 | 2 | 2 | 8.772033429 | 8.8984863 | 2 | 2 |
| ENSG00000164185 | ZNF474 | 2.334272268 | 0.460540532 | 1.271854422 | NA | 3.60612669 | 2 | 2 | 2 | 3.289987342 | 3.08597292 | 2.885263063 | 2.466008509 |
| ENSG00000198467 | TPM2 | 1.443154361 | 0.011803936 | 2.162852611 | 7.38E-05 | 3.606006973 | 11.78357814 | 10.40638494 | 11.12511995 | 12.63421185 | 12.6707361 | 13.44241619 | 13.29782891 |
| ENSG00000177303 | CASKIN2 | 0.693507541 | 0.697239244 | 2.9094224 | 0.057634332 | 3.602929941 | 6.468621203 | 7.761270542 | 2 | 7.281195074 | 7.45518898 | 9.698306977 | 9.405009888 |
| ENSG00000130695 | CEP85 | 0.87580611 | 0.35418186 | 2.725201425 | 0.00089394 | 3.601007535 | 4.815609471 | 6.228547981 | 4.058998049 | 6.268449691 | 6.08962627 | 8.054501123 | 7.885682058 |
| ENSG00000128617 | OPN1SW | -3.753148157 | 0.2198682 | 7.351342842 | 0.000951235 | 3.598194685 | 2 | 2 | 2 | 2 | 2 | 6.937746858 | 7.768360547 |
| ENSG00000181072 | CHRM2 | 2.557206839 | 0.058610905 | 1.040138536 | 0.476093865 | 3.597345375 | 2 | 5.303523747 | 6.629488387 | 7.944803394 | 8.08010078 | 6.508020621 | 6.567271433 |
| ENSG00000142207 | URB1 | 1.435864604 | 0.191858509 | 2.16035339 | 0.037170481 | 3.596217993 | 4.815609471 | 8.053983481 | 6.958202216 | 8.485003881 | 8.60067114 | 9.393314624 | 9.118313725 |
| ENSG00000101463 | SYNDIG1 | -1.27004269 | 0.581333492 | 4.86590097 | 0.006504795 | 3.59585828 | 2 | 3.076665209 | 2 | 2.3892346 | 2.14022021 | 5.630526941 | 6.212491471 |
| ENSG00000164050 | PLXNB1 | 0.797991071 | 0.552458672 | 2.796722062 | 0.017084929 | 3.594713133 | 8.418903792 | 6.228547981 | 4.874587708 | 8.022463952 | 7.96929529 | 9.729736885 | 10.18942242 |
| ENSG00000182405 | PGBD4 | 1.16067629 | 0.540424469 | 2.432237172 | 0.160349824 | 3.592913461 | 2 | 5.131603625 | 2 | 4.76150579 | 4.93258395 | 6.197600957 | 5.768151538 |
| ENSG00000171729 | TMEM51 | 2.988601398 | 0.027435278 | 0.603198919 | 0.700098898 | 3.591800317 | 6.563806279 | 6.951736442 | 2 | 9.104095335 | 9.18016239 | 6.931352719 | 6.654794245 |
| ENSG00000119862 | LGALSL | -0.060544662 | 0.976835952 | 3.652094703 | 0.025542054 | 3.591550041 | 2 | 5.457120953 | 2 | 4.070463721 | 4.1930041 | 7.414254101 | 7.589522245 |
| ENSG00000112137 | PHACTR1 | 2.719878773 | 0.149717516 | 0.871460752 | 0.674724562 | 3.591339525 | 2 | 2 | 4.523675391 | 5.474671438 | 5.37722731 | 4.301748724 | 3.311948089 |
| ENSG00000143768 | LEFTY2 | 2.784455792 | 0.479814084 | 0.806315915 | 0.850336885 | 3.590771707 | 2 | 2 | 2 | 4.139692846 | 2.14022021 | 2.746468522 | 2.277179783 |
| ENSG00000085832 | EPS15 | 1.661896915 | 3.46E-07 | 1.928557735 | 1.92E-09 | 3.59045465 | 8.107517749 | 7.790474954 | 7.552162999 | 9.545497589 | 9.40967456 | 9.728815405 | 9.759978595 |
| ENSG00000165671 | NSD1 | 1.418222696 | 0.349265407 | 2.172103282 | 0.127258741 | 3.590325978 | 9.699577239 | 9.34816733 | 4.523675391 | 10.38605714 | 10.3813388 | 11.12215672 | 11.14934098 |
| ENSG00000132405 | TBC1D14 | 1.89672453 | 0.172055101 | 1.692225082 | 0.22004974 | 3.588949612 | 5.330594134 | 7.538394143 | 3.369317972 | 8.160032699 | 8.16669309 | 7.966556098 | 7.957271773 |
| ENSG00000142303 | ADAMTS10 | 0.521443173 | 0.84416062 | 3.06744605 | 0.170183679 | 3.588889223 | 2 | 7.189546432 | 2 | 6.069927018 | 6.29111322 | 8.515767839 | 8.797441501 |
| ENSG00000182827 | ACBD3 | 1.713018645 | 0.232251697 | 1.875200015 | 0.179198641 | 3.58821866 | 4.467311984 | 9.357954881 | 7.934808822 | 9.985884116 | 9.95192082 | 10.17110242 | 10.08884856 |
| ENSG00000188092 | GPR89B | 3.60855708 | 0.002803895 | -0.020950021 | 0.989200675 | 3.587607059 | 2 | 5.838937808 | 4.058998049 | 8.126317766 | 8.17282964 | 4.894909044 | 4.553838275 |
| ENSG00000080493 | SLC4A4 | 3.30322085 | 0.013507859 | 0.283442403 | 0.86358453 | 3.586663253 | 3.327711234 | 5.946638857 | 3.369317972 | 7.97352961 | 7.88209639 | 5.654016955 | 4.062589736 |
| ENSG00000170340 | B3GNT2 | 2.165574599 | 0.002318605 | 1.420056838 | 0.055057455 | 3.585631437 | 5.70932728 | 7.572420273 | 7.552162999 | 9.350309099 | 9.25726695 | 8.692059118 | 8.425990514 |
| ENSG00000101882 | NKAP | 1.739945689 | 0.10022545 | 1.844828809 | 0.074545495 | 3.584774498 | 4.467311984 | 6.728495673 | 4.523675391 | 7.300218887 | 7.40393901 | 7.674704537 | 7.196571833 |
| ENSG00000065883 | CDK13 | 1.598528014 | 0.026712193 | 1.985956285 | 0.004489235 | 3.584484298 | 8.313285031 | 8.077869079 | 6.43197674 | 9.382797773 | 9.41486164 | 9.910864109 | 9.645596027 |
| ENSG00000116922 | C1orf109 | 1.423906438 | 0.251883859 | 2.15891793 | 0.064830738 | 3.582824368 | 3.327711234 | 6.78762695 | 5.156626401 | 7.144720234 | 7.02417148 | 8.038228667 | 7.523290521 |
| ENSG00000116698 | SMG7 | 2.035976112 | 0.280396747 | 1.546737187 | 0.416362316 | 3.582713299 | 2 | 8.505280662 | 9.180874355 | 10.31908469 | 10.3319235 | 9.770158073 | 9.902941748 |
| ENSG00000153487 | ING1 | 1.75759886 | 0.254248233 | 1.820529461 | 0.227580218 | 3.578128322 | 4.00722531 | 7.232742069 | 3.369317972 | 7.587189552 | 7.60187978 | 7.852739912 | 7.428550441 |
| ENSG00000135956 | TMEM127 | 1.609653759 | 0.101754679 | 1.968177296 | 0.037955813 | 3.577831055 | 9.590776211 | 8.968383114 | 6.718950348 | 10.41033668 | 10.4786707 | 10.7701007 | 10.83422097 |
| ENSG00000117000 | RLF | 1.677345608 | 0.355050514 | 1.899847329 | 0.279798466 | 3.577192937 | 2 | 6.666836837 | 8.35328246 | 8.772033429 | 8.86476809 | 9.24845059 | 8.795792144 |
| ENSG00000155592 | ZKSCAN2 | 1.649013296 | 0.391405869 | 1.926744449 | 0.298395371 | 3.575757744 | 3.327711234 | 6.899080544 | 2 | 7.119236776 | 6.99676009 | 7.379447089 | 7.277588383 |
| ENSG00000171044 | XKR6 | 2.247500462 | 0.181478081 | 1.328182096 | 0.444850169 | 3.575682558 | 2 | 4.442710129 | 2 | 5.161714397 | 5.13053105 | 4.604690117 | 4.077147165 |
| ENSG00000183773 | AIFM3 | 2.083656679 | 0.51121293 | 1.491414887 | NA | 3.575071566 | 2 | 2 | 2 | 3.227757282 | 2.86123014 | 2.775316226 | 2.782576693 |
| ENSG00000175832 | ETV4 | 1.81056548 | 0.195586526 | 1.763992303 | 0.199377759 | 3.574557783 | 6.138360429 | 3.076665209 | 3.369317972 | 6.535745081 | 6.53293476 | 6.508020621 | 6.471341372 |
| ENSG00000133985 | TTC9 | -3.112005397 | 0.0683337 | 6.68578516 | 3.78E-06 | 3.573779762 | 2 | 5.457120953 | 2 | 2.498691189 | 2.49394783 | 9.931034903 | 10.93343876 |
| ENSG00000087088 | BAX | 1.568217745 | 0.000812241 | 2.004849116 | 1.05E-05 | 3.573066861 | 8.285626337 | 7.274681896 | 8.175061096 | 9.50179306 | 9.5348592 | 10.00444935 | 9.899618215 |
| ENSG00000168300 | PCMTD1 | 1.956511459 | 7.94E-06 | 1.614124309 | 0.000274788 | 3.570635768 | 7.709880939 | 7.467835385 | 7.342949392 | 9.52946469 | 9.36034073 | 9.40720485 | 8.727645605 |
| ENSG00000111321 | LTBR | 1.864222396 | 0.00491763 | 1.706081977 | 0.010511932 | 3.570304372 | 8.653385977 | 8.147241016 | 9.778213818 | 10.8531554 | 10.9026799 | 10.67176032 | 10.76734655 |
| ENSG00000176022 | B3GALT6 | 1.152490167 | 0.599989824 | 2.416914567 | 0.23044932 | 3.569404733 | 2 | 8.487557258 | 4.523675391 | 8.130575473 | 8.15434112 | 9.184693023 | 9.578432478 |
| ENSG00000130227 | XPO7 | 1.716068347 | 0.348810364 | 1.852887971 | 0.298467142 | 3.568956319 | 8.777722034 | 7.928237109 | 2 | 9.510803856 | 9.5631668 | 9.74508482 | 9.597808016 |
| ENSG00000124313 | IQSEC2 | 1.742383027 | 0.09775828 | 1.826469724 | 0.076006128 | 3.568852751 | 4.467311984 | 6.78762695 | 7.691296561 | 8.476666381 | 8.57154947 | 8.609360894 | 8.606804877 |
| ENSG00000080845 | DLGAP4 | 0.917484935 | 0.020792238 | 2.650779505 | 1.53E-13 | 3.56826444 | 9.657036304 | 9.883504052 | 9.25752426 | 10.52434207 | 10.5542639 | 12.03167326 | 12.47443408 |
| ENSG00000172318 | B3GALT1 | 2.646995536 | 0.41861794 | 0.921035823 | 0.79387618 | 3.568031359 | 2 | 2 | 2 | 3.709785322 | 2.94009902 | 2.267001569 | 2.817676617 |
| ENSG00000161638 | ITGA5 | 3.137959036 | 5.38E-09 | 0.429444685 | 0.509227321 | 3.567403721 | 10.65885801 | 11.24986486 | 12.02204252 | 14.51866036 | 14.5886674 | 11.71646058 | 11.96652151 |
| ENSG00000102349 | KLF8 | 0.430925973 | 0.738128035 | 3.134702412 | 0.002597654 | 3.565628385 | 3.327711234 | 3.076665209 | 5.39244166 | 4.54891529 | 4.70309173 | 7.199581219 | 7.034460028 |
| ENSG00000163872 | YEATS2 | 1.848510128 | 0.000223501 | 1.716911308 | 0.000631256 | 3.565421435 | 8.564112515 | 8.101365653 | 8.511889463 | 10.23161272 | 10.2467381 | 10.51569609 | 9.537567589 |
| ENSG00000185658 | BRWD1 | 1.860261666 | 0.000587551 | 1.704396993 | 0.001715093 | 3.564658658 | 8.009475818 | 8.395519306 | 7.163363831 | 9.80724435 | 9.78235238 | 9.867355293 | 9.369229141 |
| ENSG00000158555 | GDPD5 | 0.100689587 | 0.968286211 | 3.462555513 | 0.094151017 | 3.5632451 | 2 | 6.728495673 | 2 | 5.273747807 | 5.44640346 | 8.65615757 | 8.524881346 |
| ENSG00000116584 | ARHGEF2 | 1.687504616 | 0.20047706 | 1.874934596 | 0.143348036 | 3.562439212 | 5.532353208 | 9.640525262 | 8.076950877 | 10.2141657 | 10.2213052 | 10.76696209 | 9.919445528 |
| ENSG00000126804 | ZBTB1 | 2.226823746 | 2.38E-07 | 1.335522894 | 0.003033425 | 3.56234664 | 7.218819781 | 8.124485672 | 7.691296561 | 9.929575199 | 9.95370391 | 9.213963607 | 8.877074023 |
| ENSG00000150760 | DOCK1 | 1.791687544 | 0.02516631 | 1.770604013 | 0.026185977 | 3.562291557 | 6.653098118 | 8.791896851 | 7.691296561 | 9.722342833 | 9.77900397 | 9.576313249 | 9.868833922 |
| ENSG00000047634 | SCML1 | 1.988743476 | 0.264475702 | 1.572879709 | 0.379145564 | 3.561623184 | 4.00722531 | 7.051611315 | 2 | 7.574744049 | 7.61696151 | 7.401588825 | 6.942047249 |
| ENSG00000270617 | URGCP-MRP | 8.750868743 | 2.79E-08 | -5.191150933 | 0.03010975 | 3.559717811 | 2 | 2 | 2 | 8.804373539 | 8.77618319 | 2 | 2 |
| ENSG00000169507 | SLC38A11 | -3.753134638 | 0.262397112 | 7.312837345 | 0.004028686 | 3.559702707 | 2 | 2 | 2 | 2 | 2 | 7.98365249 | 6.298867018 |
| ENSG00000152749 | GPR180 | 2.05487971 | 0.077163483 | 1.503129971 | 0.203570906 | 3.55800968 | 6.009025997 | 7.954278496 | 4.874587708 | 8.893506006 | 8.83412012 | 8.742236593 | 7.714671806 |
| ENSG00000137200 | CMTR1 | 2.551144078 | 4.99E-06 | 1.005355732 | 0.100850653 | 3.55649981 | 8.540903468 | 7.901717006 | 7.029880454 | 10.48414712 | 10.4827929 | 8.90036762 | 8.989083133 |
| ENSG00000121903 | ZSCAN20 | 1.962307352 | 0.362261257 | 1.591563843 | 0.460972389 | 3.553871195 | 2 | 6.140560021 | 2 | 6.687949235 | 6.37156942 | 6.482006365 | 5.812327524 |
| ENSG00000185245 | GP1BA | -1.310505255 | 0.645571902 | 4.863660437 | 0.023508946 | 3.553155182 | 2 | 2 | 2 | 2.141737246 | 2.14022021 | 4.990270811 | 5.150533729 |
| ENSG00000188486 | H2AFX | 1.507510665 | 0.159032929 | 2.045467834 | 0.044582925 | 3.552978499 | 4.815609471 | 6.140560021 | 7.777116083 | 8.221200057 | 8.16053033 | 8.656642182 | 8.786410001 |
| ENSG00000126001 | CEP250 | 1.894275223 | 0.253228122 | 1.658252604 | 0.314385216 | 3.552527827 | 7.867441304 | 7.572420273 | 2 | 9.018234236 | 9.0282492 | 8.684949926 | 8.886403151 |
| ENSG00000109189 | USP46 | 1.981382358 | 0.001547707 | 1.570032761 | 0.014055845 | 3.551415119 | 5.70932728 | 6.666836837 | 5.156626401 | 7.971157485 | 7.89452713 | 7.541454591 | 7.517964974 |
| ENSG00000183891 | TTC32 | 1.84417828 | 0.153618856 | 1.706702241 | 0.179539773 | 3.550880521 | 3.327711234 | 4.11352228 | 2 | 4.867343068 | 4.87352701 | 4.68289886 | 4.819291477 |
| ENSG00000069275 | NUCKS1 | 1.142590798 | 0.036736378 | 2.407080914 | 1.61E-06 | 3.549671711 | 9.143983763 | 10.15453315 | 9.116480286 | 10.71325662 | 10.6873877 | 11.84730078 | 12.06982099 |
| ENSG00000165661 | QSOX2 | 1.441691014 | 0.306881895 | 2.107831747 | 0.113226946 | 3.549522761 | 4.467311984 | 5.303523747 | 8.143093325 | 8.151677543 | 8.32935768 | 8.788075779 | 9.00820079 |
| ENSG00000120784 | ZFP30 | 1.768868496 | 0.38484122 | 1.780638023 | 0.370592016 | 3.549506519 | 3.327711234 | 2 | 6.958202216 | 7.330144653 | 7.03319408 | 7.470462265 | 6.868633039 |
| ENSG00000092148 | HECTD1 | 1.472197762 | 0.132505099 | 2.076664377 | 0.025734513 | 3.54886214 | 7.53297911 | 10.13758676 | 8.561105505 | 10.62321188 | 10.5902432 | 11.27988996 | 11.13658383 |
| ENSG00000110047 | EHD1 | 0.798758374 | 0.243302388 | 2.748880971 | 3.58E-06 | 3.547639345 | 8.444143041 | 8.88960062 | 9.839340503 | 9.896204982 | 10.0423963 | 11.59984264 | 12.17686777 |
| ENSG00000137273 | FOXF2 | 0.852945632 | 0.626071633 | 2.694123412 | 0.080856643 | 3.547069044 | 2 | 5.131603625 | 7.225742779 | 6.733448311 | 6.82036165 | 8.705705008 | 8.442950253 |
| ENSG00000173041 | ZNF680 | 2.424471092 | 0.054569925 | 1.119638228 | 0.405601197 | 3.54410932 | 2 | 4.11352228 | 5.59507254 | 6.777556219 | 6.65485813 | 5.801542492 | 5.094391891 |
| ENSG00000143952 | VPS54 | 1.87136267 | 0.064736855 | 1.670881175 | 0.098450928 | 3.542243845 | 4.467311984 | 7.503546079 | 6.43197674 | 8.415170354 | 8.46222608 | 8.278834759 | 8.20189164 |
| ENSG00000100121 | GGTLC2 | 2.386858132 | 0.457964922 | 1.155338379 | NA | 3.54219651 | 2 | 2 | 2 | 2.868578001 | 3.50452879 | 2.457042231 | 2.817676617 |
| ENSG00000126767 | ELK1 | 1.797143361 | 0.252106496 | 1.7426535 | 0.259868009 | 3.539796862 | 9.27634997 | 6.951736442 | 4.058998049 | 9.732182526 | 9.79831747 | 9.405764176 | 9.963452459 |
| ENSG00000132953 | XPO4 | 1.788249823 | 0.290539585 | 1.749466527 | 0.291783001 | 3.53771635 | 2 | 7.847167909 | 6.43197674 | 8.511364632 | 8.48377664 | 8.3832514 | 8.531503464 |
| ENSG00000173432 | SAA1 | 3.957393374 | 0.096765227 | -0.419764834 | 0.885335383 | 3.53762854 | 2 | 2 | 2 | 4.470469939 | 4.12711147 | 2.225793514 | 2.277179783 |
| ENSG00000034510 | TMSB10 | 1.92401128 | 0.000256078 | 1.610916517 | 0.002580504 | 3.534927797 | 12.32193393 | 11.1798568 | 11.6816684 | 13.70168635 | 13.7498842 | 13.10104672 | 13.66933964 |
| ENSG00000183092 | BEGAIN | 0.188919815 | 0.939180462 | 3.345237539 | 0.088680604 | 3.534157354 | 2 | 2 | 3.369317972 | 2.784612485 | 2.49394783 | 4.340871764 | 4.810605986 |
| ENSG00000146574 | CCZ1B | 5.944292744 | 1.40E-14 | -2.410233427 | 0.00588506 | 3.534059317 | 4.467311984 | 4.710564154 | 2 | 9.856914282 | 9.70400783 | 2.62493832 | 2.851942824 |
| ENSG00000105321 | CCDC9 | 1.483144511 | 0.046523239 | 2.049339885 | 0.003865737 | 3.532484396 | 4.467311984 | 6.228547981 | 6.534109959 | 7.43025113 | 7.39346687 | 8.040458475 | 7.889811111 |

| | | | | | | | | | | | | | |
|---|---|---|---|---|---|---|---|---|---|---|---|---|---|
| ENSG00000173369 | C1QB | -3.753145913 | 0.228046677 | 7.284362014 | 0.001446259 | 3.531216101 | 2 | 2 | 2 | 2 | 2 | 7.728247761 | 6.826173076 |
| ENSG00000182853 | VMO1 | 0.272040666 | 0.940940174 | 3.258483055 | 0.283648588 | 3.530523721 | 2 | 2 | 2 | 2.270786764 | 2.49394783 | 4.340871764 | 2.851942824 |
| ENSG00000009954 | BAZ1B | 1.851448643 | 0.000740256 | 1.677846031 | 0.002375091 | 3.529294674 | 9.544849385 | 8.395519306 | 8.486635645 | 10.75453239 | 10.7437816 | 10.70435357 | 10.43505811 |
| ENSG00000145284 | SCD5 | -0.566728648 | 0.598764326 | 4.095304118 | 2.40E-06 | 3.52857547 | 5.532353208 | 6.844429755 | 8.236946975 | 6.856790872 | 6.55815769 | 11.15202309 | 11.47669218 |
| ENSG00000126883 | NUP214 | 1.87655858 | 0.310197758 | 1.650665691 | 0.36789886 | 3.527224272 | 3.327711234 | 8.687092496 | 4.523675391 | 9.062360705 | 9.07705362 | 8.828637742 | 8.862441066 |
| ENSG00000160193 | WDR4 | 1.954481436 | 0.33914729 | 1.570994813 | 0.443400867 | 3.525476249 | 2 | 5.838937808 | 2 | 6.213314212 | 6.26070706 | 5.696113106 | 6.037449234 |
| ENSG00000181450 | ZNF678 | 1.917128513 | 0.065154219 | 1.60742141 | 0.122910117 | 3.524549923 | 4.815609471 | 4.93638933 | 2 | 6.213314212 | 6.05460702 | 6.072436333 | 5.576534358 |
| ENSG00000027001 | MIPEP | 1.876426931 | 0.101350025 | 1.647977797 | 0.147500239 | 3.524404729 | 2 | 5.457120953 | 4.523675391 | 6.358325776 | 6.25300433 | 6.063692919 | 6.117095472 |
| ENSG00000074657 | ZNF532 | 1.455802484 | 0.350662169 | 2.068261665 | 0.161518911 | 3.524064148 | 4.467311984 | 9.680074956 | 7.646393253 | 9.886796832 | 9.89366625 | 10.55872351 | 10.43999426 |
| ENSG00000003249 | DBNDD1 | -0.765903048 | 0.370378063 | 4.287718464 | 5.62E-10 | 3.521815415 | 4.00722531 | 4.93638933 | 5.930865648 | 4.574143677 | 4.40247105 | 8.993080225 | 9.504290592 |
| ENSG00000003436 | TFPI | 2.638889693 | 3.46E-23 | 0.882741418 | 0.001957141 | 3.521631111 | 8.39321512 | 8.777382925 | 8.536706852 | 11.23638949 | 11.1853847 | 9.478834579 | 9.442386533 |
| ENSG00000162458 | FBLIM1 | 0.663252114 | 0.053329277 | 2.857656607 | 2.16E-21 | 3.520908721 | 9.204317094 | 9.532315021 | 8.978369997 | 9.856271709 | 9.98484711 | 12.15954314 | 12.06229494 |
| ENSG00000072518 | MARK2 | 1.700620978 | 4.21E-05 | 1.815098758 | 9.76E-06 | 3.515719736 | 6.89194353 | 7.503546079 | 6.718950348 | 8.745906571 | 8.79619691 | 8.887212926 | 8.882264318 |
| ENSG00000169871 | TRIM56 | 1.28907695 | 0.323747566 | 2.22559581 | 0.067085237 | 3.51467276 | 9.473095744 | 8.147241016 | 5.39244166 | 9.685919789 | 9.73613428 | 10.81051091 | 10.45679112 |
| ENSG00000101745 | ANKRD12 | 2.220207915 | 2.33E-12 | 1.292393592 | 8.02E-05 | 3.512601507 | 8.138771685 | 8.414402626 | 8.042706615 | 10.43894286 | 10.4002048 | 9.644841182 | 9.32912155 |
| ENSG00000186462 | NAP1L2 | 2.818518668 | 0.352516309 | 0.693240494 | 0.836549691 | 3.511759162 | 2 | 2 | 2 | 3.462030052 | 3.50452879 | 2.68698273 | 2.277179783 |
| ENSG00000179304 | FAM156B | 8.700469613 | 4.03E-08 | -5.191150868 | 0.030595573 | 3.509318745 | 2 | 2 | 2 | 8.709369553 | 8.77079897 | 2 | 2 |
| ENSG00000109790 | KLHL5 | 1.689216353 | 0.002646347 | 1.818383686 | 0.001044308 | 3.507600039 | 8.586954094 | 9.488434476 | 9.935792933 | 11.13407655 | 11.1101028 | 11.05159418 | 11.42632529 |
| ENSG00000017797 | RALBP1 | 1.578897175 | 0.013238784 | 1.925380753 | 0.001882006 | 3.504277928 | 7.53297911 | 9.102512015 | 8.266922072 | 10.02471865 | 10.0084861 | 10.22241543 | 10.48912908 |
| ENSG00000255526 | NEDD8-MDP | 8.695047998 | 4.09E-08 | -5.191150874 | 0.030591756 | 3.503897124 | 2 | 2 | 2 | 8.736160353 | 8.73392024 | 2 | 2 |
| ENSG00000254470 | AP5B1 | 2.268667918 | 0.032248782 | 1.234298901 | 0.268896089 | 3.502966819 | 5.095959458 | 7.60566235 | 8.486635645 | 9.800579656 | 9.91273599 | 8.796015995 | 8.862966236 |
| ENSG00000130224 | LRCH2 | 2.379073767 | 0.163782083 | 1.123250904 | 0.533552261 | 3.502324671 | 2 | 6.389896633 | 3.369317972 | 7.292639438 | 7.31055061 | 6.332313483 | 5.812327524 |
| ENSG00000177694 | NAALADL2 | 1.034706808 | 0.529862889 | 2.465482259 | 0.095140371 | 3.500189067 | 6.009025997 | 3.686303414 | 2 | 5.709096143 | 5.62411716 | 6.537789343 | 7.388257299 |
| ENSG00000198604 | BAZ1A | 2.591126045 | 0.073335148 | 0.908970036 | 0.565009493 | 3.500096081 | 3.327711234 | 8.639735838 | 7.858115942 | 10.31160396 | 10.3107463 | 8.732155826 | 8.53941009 |
| ENSG00000186591 | UBE2H | 1.5043178 | 0.016404216 | 1.995527797 | 0.000939774 | 3.499845596 | 10.30651143 | 10.30378101 | 8.865353883 | 11.49993054 | 11.4321483 | 11.92739794 | 11.98611928 |
| ENSG00000108587 | GOSR1 | 1.809084169 | 0.005532065 | 1.689854471 | 0.00975996 | 3.49893864 | 6.816637977 | 7.002537973 | 8.296287027 | 9.347568393 | 9.27068036 | 9.236830239 | 9.144037078 |
| ENSG00000135913 | USP37 | 2.266103908 | 0.156314863 | 1.23279514 | 0.459659586 | 3.498899048 | 2 | 6.844429755 | 4.523675391 | 7.831679003 | 7.67863747 | 6.807643042 | 6.693102083 |
| ENSG00000164828 | SUN1 | 0.762307692 | 0.186180631 | 2.735165911 | 4.20E-08 | 3.497473603 | 8.836089686 | 9.030900487 | 7.777116083 | 9.428470756 | 9.3816907 | 11.34545729 | 11.39761508 |
| ENSG00000155850 | SLC26A2 | 1.111860517 | 0.530264818 | 2.382410079 | 0.138711164 | 3.494270596 | 4.467311984 | 8.469613413 | 4.058998049 | 8.153770873 | 8.11238697 | 9.07476376 | 9.651383746 |
| ENSG00000015153 | YAF2 | 2.165351173 | 0.00023626 | 1.328332053 | 0.031839501 | 3.493683226 | 6.009025997 | 6.666836837 | 5.156626401 | 8.254744042 | 8.15020018 | 7.44824109 | 7.311843322 |
| ENSG00000163558 | PRKCI | 1.348954037 | 0.131883676 | 2.144699751 | 0.010661152 | 3.493653788 | 8.16936288 | 7.189546432 | 5.772716771 | 8.695066731 | 8.65428713 | 9.705116572 | 9.175665104 |
| ENSG00000165055 | METTL2B | 1.342639847 | 0.00692683 | 2.150405926 | 4.55E-06 | 3.493045773 | 6.366709671 | 6.535001748 | 5.59507254 | 7.64488235 | 7.42122542 | 8.296978073 | 8.365751687 |
| ENSG00000169777 | TAS2R1 | -1.30428344 | 0.736220139 | 4.794405526 | 0.127753231 | 3.490122086 | 2 | 2 | 2 | 2.270786764 | 2 | 4.406883583 | 5.436777225 |
| ENSG00000152684 | PELO | 1.959067364 | 0.28945592 | 1.53063052 | 0.41000811 | 3.489697883 | 8.998234871 | 8.053983481 | 2 | 9.923453481 | 10.0102007 | 9.544459308 | 9.537567589 |
| ENSG00000065802 | ASB1 | 1.302293506 | 0.429843212 | 2.187143625 | 0.153259447 | 3.489437132 | 3.327711234 | 8.213429689 | 9.116480286 | 9.404936476 | 9.50598504 | 10.25526417 | 10.41514286 |
| ENSG00000136367 | ZFHX2 | 0.012452852 | 0.992812774 | 3.476568643 | 0.000328492 | 3.489021495 | 4.00722531 | 4.93638933 | 2 | 4.299492648 | 4.02226117 | 7.451596164 | 7.142226902 |
| ENSG00000136643 | RPS6KC1 | 2.275062609 | 0.001730605 | 1.213946302 | 0.115961049 | 3.489008911 | 8.009475818 | 8.213429689 | 6.203069917 | 9.982354854 | 9.95963168 | 8.977648172 | 8.851897333 |
| ENSG00000168899 | VAMP5 | 1.46734928 | 0.214681289 | 2.020969105 | 0.07263329 | 3.488318386 | 5.866944937 | 6.464272742 | 8.631919287 | 8.902248225 | 8.98007065 | 8.757226889 | 9.97588724 |
| ENSG00000185220 | PGBD2 | 1.837750374 | 0.150495614 | 1.650100078 | 0.191572761 | 3.487850452 | 2 | 4.710564154 | 3.369317972 | 5.348759107 | 5.34860058 | 5.348031117 | 4.990558504 |
| ENSG00000100815 | TRIP11 | 1.735123931 | 0.163554675 | 1.752032262 | 0.150827473 | 3.487156194 | 8.107517749 | 8.451443573 | 4.523675391 | 9.461800958 | 9.49378147 | 9.707457275 | 9.244995051 |
| ENSG00000183665 | TRMT12 | 1.500436373 | 0.317999975 | 1.986693652 | 0.165909543 | 3.487130025 | 4.467311984 | 6.389896633 | 2 | 6.682159616 | 6.57678965 | 7.138441638 | 7.060335194 |
| ENSG00000196663 | TECPR2 | 1.266336495 | 0.485504025 | 2.219609941 | 0.187718342 | 3.485946436 | 2 | 7.979858146 | 8.110401118 | 8.715051214 | 8.72698617 | 9.922396 | 9.358096047 |
| ENSG00000185437 | SH3BGR | 3.665856255 | 0.141361699 | -0.180655174 | 0.95264385 | 3.485201081 | 2 | 2 | 2 | 3.95996233 | 4.1930041 | 2.3070652 | 2.277179783 |
| ENSG00000089060 | SLC8B1 | 1.402862729 | 0.331987296 | 2.082206125 | 0.126883017 | 3.485068855 | 7.27616306 | 4.710564154 | 3.369317972 | 7.206526483 | 7.45182838 | 7.977459036 | 8.02265879 |
| ENSG00000164808 | SPIDR | 1.41017389 | 0.24668092 | 2.073110285 | 0.071494314 | 3.483284175 | 6.138360429 | 7.145017454 | 3.369317972 | 7.484483106 | 7.69294162 | 8.151901655 | 8.331250243 |
| ENSG00000080371 | RAB21 | 1.754901318 | 0.000799417 | 1.728370128 | 0.000915639 | 3.483271446 | 7.031698319 | 7.819099893 | 8.35328246 | 9.594867189 | 9.53882419 | 9.577849253 | 9.502606422 |
| ENSG00000157014 | TATDN2 | 2.244725709 | 0.139433008 | 1.238162522 | 0.434687247 | 3.482888231 | 3.327711234 | 6.464272742 | 8.511888463 | 9.482768454 | 9.48147379 | 8.232777435 | 8.697340724 |
| ENSG00000138002 | IFT172 | 1.280202627 | 0.570234127 | 2.201368533 | 0.296233541 | 3.481571161 | 2 | 6.311476754 | 2 | 6.061031724 | 6.05460702 | 6.842137779 | 7.026981076 |
| ENSG00000177427 | MIEF2 | 0.681480327 | 0.685840372 | 2.79992398 | 0.052695388 | 3.481404307 | 2 | 6.602424674 | 4.523675391 | 5.949776857 | 6.1068222 | 8.113585664 | 8.044120856 |
| ENSG00000198160 | MIER1 | 1.990803486 | 0.007541767 | 1.48994492 | 0.051047807 | 3.480748406 | 6.009025997 | 7.35507212 | 8.110401118 | 9.392580682 | 9.3185152 | 8.956549551 | 8.75393321 |
| ENSG00000170881 | RNF139 | 1.699171049 | 0.275154513 | 1.777484847 | 0.243286625 | 3.476655896 | 3.327711234 | 8.101365653 | 5.59507254 | 8.395844809 | 8.54024645 | 8.703830473 | 8.37166861 |
| ENSG00000119723 | COQ6 | 2.552390503 | 0.004523149 | 0.923473776 | 0.352922604 | 3.475864279 | 3.327711234 | 5.457120953 | 4.058998049 | 6.902323594 | 7.03319408 | 5.318934974 | 5.540219242 |
| ENSG00000159374 | M1AP | 0.015595214 | 0.997041173 | 3.459162295 | 0.226906648 | 3.474757509 | 2 | 2 | 2 | 2.498691189 | 2.14022021 | 4.065262523 | 3.771796761 |
| ENSG00000081842 | PCDHA6 | 2.394168084 | 0.464304361 | 1.079753107 | NA | 3.473921191 | 2 | 2 | 2 | 3.615725078 | 2.68924543 | 2.68698273 | 2.551896972 |
| ENSG00000115310 | RTN4 | 1.640375464 | 1.92E-08 | 1.832590331 | 2.34E-10 | 3.472965755 | 11.04852648 | 11.53726652 | 11.53541251 | 13.02801696 | 13.0336013 | 13.20295944 | 13.24264946 |
| ENSG00000185112 | FAM43A | 1.455054709 | 0.315574141 | 2.014302705 | 0.143374446 | 3.469357414 | 2 | 6.311476754 | 6.43197674 | 7.300218887 | 7.12888795 | 8.117111293 | 7.294817518 |
| ENSG00000107372 | ZFAND5 | 1.289201606 | 0.163887287 | 2.175750551 | 0.011666318 | 3.464952157 | 10.1311248 | 10.27307349 | 7.858115942 | 11.06429261 | 11.0173806 | 11.77722057 | 12.06143722 |
| ENSG00000076108 | BAZ2A | 1.844682252 | 1.05E-08 | 1.618250956 | 5.96E-07 | 3.462933207 | 9.159305284 | 9.018611864 | 8.561105505 | 10.73753559 | 10.8089302 | 10.62906029 | 10.46165359 |
| ENSG00000070778 | PTPN21 | 1.348254286 | 0.479038245 | 2.113999591 | 0.23691073 | 3.462253877 | 2 | 8.732943944 | 7.225742779 | 8.966774408 | 8.88608977 | 9.698306977 | 9.677294239 |
| ENSG00000129003 | VPS13C | 2.083325284 | 0.00104027 | 1.377808501 | 0.037254341 | 3.461133784 | 7.096806373 | 7.503546079 | 8.65477198 | 9.982354854 | 9.95429779 | 9.209346464 | 9.321499526 |
| ENSG00000165219 | GAPVD1 | 1.700001036 | 0.241525716 | 1.758304585 | 0.216349452 | 3.45830562 | 4.00722531 | 8.395519306 | 9.014156722 | 9.906793906 | 9.82388974 | 10.01280126 | 9.829339 |
| ENSG00000164187 | LMBRD2 | 1.506454626 | 0.162158714 | 1.950672829 | 0.058080925 | 3.457127455 | 7.031698319 | 4.11352228 | 6.43197674 | 7.781117077 | 7.67287587 | 8.475683062 | 7.768360547 |
| ENSG00000133874 | RNF122 | 2.623926435 | 0.000487609 | 0.833171918 | 0.323536602 | 3.457098354 | 5.330594134 | 5.72254323 | 3.369317972 | 7.605658789 | 7.65837038 | 5.893224226 | 5.929275 |
| ENSG00000065615 | CYB5R4 | 2.319709965 | 1.40E-05 | 1.137068461 | 0.047012018 | 3.456778426 | 5.095959458 | 5.303523747 | 5.59507254 | 7.54315232 | 7.53992502 | 6.280431262 | 6.506723027 |
| ENSG00000198836 | OPA1 | 1.799593132 | 0.211077851 | 1.656260976 | 0.245873893 | 3.455854107 | 8.285626337 | 7.954278496 | 3.369317972 | 9.412823969 | 9.28680218 | 9.164389779 | 9.251828472 |
| ENSG00000164855 | TMEM184A | 1.210637026 | 0.591930254 | 2.242861272 | 0.280327402 | 3.453498298 | 2 | 3.686303414 | 2 | 3.709785322 | 3.39680241 | 4.813804156 | 3.560117069 |
| ENSG00000073150 | PANX2 | 2.281327889 | 0.196009997 | 1.17126754 | 0.526432906 | 3.452595528 | 4.00722531 | 6.951736442 | 2 | 7.797276258 | 7.7866208 | 7.131485609 | 6.117095472 |
| ENSG00000068650 | ATP11A | 1.654219603 | 0.414405236 | 1.798140637 | 0.361385021 | 3.452360239 | 4.815609471 | 8.297207175 | 2 | 8.43768081 | 8.52433649 | 8.62380783 | 8.624257514 |
| ENSG00000163932 | PRKCD | 2.183912777 | 0.079842495 | 1.268186981 | 0.328187708 | 3.452099757 | 8.107517749 | 6.535001748 | 4.058998049 | 9.15827499 | 9.14521857 | 8.327733606 | 8.157201821 |
| ENSG00000080503 | SMARCA2 | 1.62856316 | 0.369452745 | 1.823493334 | 0.299990634 | 3.452056494 | 2 | 8.469613413 | 7.098165388 | 8.937912013 | 9.01578569 | 9.161319498 | 9.180307928 |
| ENSG00000179761 | PIPOX | 1.217794822 | 0.526407754 | 2.232529353 | 0.203778744 | 3.450324175 | 2 | 3.076665209 | 3.369317972 | 3.921164147 | 3.28037825 | 4.68289886 | 4.033026654 |
| ENSG00000188994 | ZNF292 | 1.921311156 | 0.258819968 | 1.527429592 | 0.370755996 | 3.448740748 | 6.138360429 | 9.018611864 | 3.369317972 | 9.617795654 | 9.47652116 | 9.305806573 | 8.991966981 |
| ENSG00000169967 | MAP3K2 | 1.945796078 | 0.023002202 | 1.501381332 | 0.085501287 | 3.44717741 | 7.790810999 | 8.848535039 | 6.43197674 | 9.960398193 | 9.91761657 | 9.625909881 | 9.355112655 |
| ENSG00000175105 | ZNF654 | 2.157756176 | 0.154170396 | 1.28892813 | 0.410543204 | 3.446684307 | 2 | 6.728495673 | 7.098165388 | 8.504819462 | 8.43013848 | 7.942911451 | 7.18317631 |
| ENSG00000109680 | TBC1D19 | 2.506424594 | 0.061530373 | 0.939812734 | 0.519728384 | 3.446237328 | 2 | 5.838937808 | 4.058998049 | 7.114945424 | 7.0150921 | 5.343222328 | 5.785984581 |
| ENSG00000170775 | GPR37 | 4.897175552 | 0.003484322 | -1.453817022 | 0.451026653 | 3.44335853 | 2 | 4.11352228 | 2 | 7.300218887 | 7.21466714 | 2.831336763 | 2.172608095 |
| ENSG00000185640 | KRT79 | 4.319709069 | 0.074800869 | -0.877454815 | 0.759454549 | 3.442254254 | 2 | 2 | 2 | 4.739373666 | 4.45712836 | 2.094598369 | 2.277179783 |
| ENSG00000135999 | EPC2 | 2.114988183 | 0.390019856 | 1.326465979 | 0.598500734 | 3.441454163 | 2 | 7.431218269 | 2 | 7.982979299 | 7.94559728 | 7.249266052 | 7.133547492 |
| ENSG00000116977 | LGALS8 | 2.726676957 | 4.73E-06 | 0.712947531 | 0.294558331 | 3.439624488 | 9.048471759 | 8.395519306 | 7.451345288 | 11.13936824 | 11.1652631 | 8.971039232 | 9.302653594 |
| ENSG00000138757 | G3BP2 | 1.530352294 | 2.53E-13 | 1.907424873 | 2.24E-20 | 3.437777167 | 9.460782017 | 9.514922632 | 9.541233192 | 11.07565424 | 10.9854873 | 11.37402863 | 11.44064487 |
| ENSG00000134108 | ARL8B | 2.670247802 | 0.137084233 | 0.766800676 | 0.699334721 | 3.437048478 | 8.138771685 | 9.04308532 | 2 | 10.77881047 | 10.6952249 | 9.009108162 | 8.656119846 |
| ENSG00000108861 | DUSP3 | 1.492272668 | 0.053811959 | 1.944443358 | 0.008947043 | 3.436716026 | 9.790941832 | 9.338312924 | 7.734844356 | 10.67841925 | 10.6657943 | 11.03528871 | 11.20632352 |
| ENSG00000107742 | SPOCK2 | 2.214745958 | 0.498491454 | 1.221343176 | NA | 3.436089135 | 2 | 2 | 2 | 3.289987342 | 2.94009902 | 2.885263063 | 2.421067459 |
| ENSG00000115425 | PECR | 0.968794954 | 0.477022398 | 2.466947033 | 0.040591855 | 3.435741987 | 2 | 4.710564154 | 3.369317972 | 4.76150579 | 4.40247105 | 5.83960027 | 6.011156536 |
| ENSG00000111254 | AKAP3 | 0.272098667 | 0.936411459 | 3.159502559 | 0.259652636 | 3.431601225 | 2 | 2 | 2 | 2.270786764 | 2.49394783 | 3.865109708 | 3.539239711 |
| ENSG00000131873 | CHSY1 | 1.248740485 | 0.219547821 | 2.181386843 | 0.021222059 | 3.430127329 | 6.468621203 | 9.28800521 | 8.042706615 | 9.580155061 | 9.60385643 | 10.4390324 | 10.59881657 |
| ENSG00000118564 | FBXL5 | 0.821819696 | 0.536398142 | 2.605670516 | 0.026039832 | 3.427490212 | 5.330594134 | 9.067150531 | 9.099922433 | 9.371149971 | 9.36392108 | 11.3748388 | 10.8718893 |
| ENSG00000163064 | EN1 | 3.04642094 | 0.344298341 | 0.380426941 | 0.918213588 | 3.42684788 | 2 | 2 | 2 | 3.566290607 | 3.69849104 | 2.656294034 | 2.117335275 |
| ENSG00000110911 | SLC11A2 | 2.610345833 | 0.012491705 | 0.816205276 | 0.483694039 | 3.42655111 | 5.532353208 | 8.777382925 | 7.285536013 | 10.32141448 | 10.3491863 | 8.442343897 | 8.654300638 |
| ENSG00000100034 | PPM1F | 1.17726834 | 0.472493119 | 2.248141127 | 0.134649149 | 3.425409467 | 4.00722531 | 9.125613817 | 6.629488387 | 8.967964567 | 8.99168852 | 10.05752505 | 10.03183938 |
| ENSG00000168917 | SLC35G2 | 3.251988164 | 0.053572957 | 0.172252297 | 0.93387742 | 3.42424046 | 2 | 4.442710129 | 7.398165132 | 9.16866395 | 9.18624204 | 5.99483933 | 6.326546677 |
| ENSG00000160213 | CSTB | 1.768537516 | 0.075478678 | 1.655188279 | 0.093761478 | 3.423725795 | 10.13887448 | 7.701025996 | 8.175061096 | 10.80734846 | 10.8718924 | 10.41445658 | 10.9835621 |
| ENSG00000198677 | TTC37 | 1.761246557 | 0.047659578 | 1.661805177 | 0.06022343 | 3.423051734 | 8.009475818 | 9.74864196 | 7.552162999 | 10.52434207 | 10.5221789 | 10.47739344 | 10.36874764 |
| ENSG00000139132 | FGD4 | 0.717587394 | 0.431518756 | 2.704430729 | 0.000501743 | 3.422018124 | 5.70932728 | 7.954278496 | 7.777116083 | 8.168339746 | 8.12510181 | 9.989716154 | 10.23658897 |
| ENSG00000108061 | SHOC2 | 1.468991129 | 0.263164637 | 1.952013247 | 0.119669999 | 3.421004376 | 4.467311984 | 8.655694736 | 8.236946975 | 9.397888985 | 9.35764963 | 9.945186878 | 9.766168491 |
| ENSG00000173451 | THAP2 | 1.650011816 | 0.220999164 | 1.770535357 | 0.177528888 | 3.420547173 | 2 | 5.303523747 | 4.874587708 | 6.113598934 | 6.1068222 | 6.750676314 | 5.385063124 |
| ENSG00000166793 | YPEL4 | 2.043335023 | 0.211460922 | 1.375758144 | 0.40972733 | 3.419093167 | 2 | 3.686303414 | 4.058998049 | 5.22679461 | 4.91316568 | 5.102662916 | 3.474731315 |
| ENSG00000198373 | WWP2 | 1.544888061 | 0.314797955 | 1.873627623 | 0.205709574 | 3.418515684 | 3.327711234 | 6.535001748 | 8.408111533 | 8.707945635 | 8.73115061 | 8.97958623 | 9.10860334 |
| ENSG00000104093 | DMXL2 | 1.891679514 | 0.272777026 | 1.526277904 | 0.376725096 | 3.417957418 | 2 | 7.638155693 | 8.076950877 | 9.174861605 | 9.15867957 | 9.090272833 | 8.447159218 |
| ENSG00000100890 | KIAA0391 | 2.778430346 | 0.002395404 | 0.638759899 | 0.545022271 | 3.417190245 | 4.00722531 | 6.728495673 | 5.39244166 | 8.543652346 | 8.44373613 | 6.387074688 | 6.429408106 |
| ENSG00000183495 | EP400 | 1.636761231 | 0.002559868 | 1.77878949 | 0.000883053 | 3.415550722 | 7.096806373 | 7.954278496 | 8.460932916 | 9.566077486 | 9.55457603 | 9.767692698 | 9.63241113 |
| ENSG00000121892 | PDS5A | 1.588313232 | 0.026020592 | 1.825962356 | 0.00889175 | 3.414275589 | 7.904282243 | 9.470501606 | 8.110401118 | 10.25909638 | 10.2602493 | 10.68167537 | 10.28422407 |
| ENSG00000168350 | DEGS2 | 0.012819211 | 0.997163678 | 3.399352251 | 0.239702507 | 3.412171462 | 2 | 2 | 2 | 2.3892346 | 2.2680111 | 4.397636241 | 3.071407821 |
| ENSG00000105136 | ZNF419 | 1.806635513 | 0.203704581 | 1.605535119 | 0.256352368 | 3.412170632 | 2 | 6.535001748 | 5.59507254 | 7.273514749 | 7.39346687 | 7.435871964 | 6.764548117 |
| ENSG00000274600 | RIMBP3B | 5.695468939 | 0.030320381 | -2.284001069 | 0.458652814 | 3.41146787 | 2 | 2 | 2 | 5.900763176 | 5.74985018 | 2.139668783 | 2 |
| ENSG00000259024 | TVP23C-CDR' | 8.601227482 | 8.43E-08 | -5.191150712 | 0.031629908 | 3.41007677 | 2 | 2 | 2 | 8.583021623 | 8.69891174 | 2 | 2 |
| ENSG00000102547 | CAB39L | 1.740821355 | 0.503519889 | 1.669096175 | 0.515079411 | 3.40991753 | 2 | 7.790474954 | 2 | 7.939959586 | 7.96694292 | 7.623070264 | 8.103103292 |

| | | | | | | | | | | | | | |
|---|---|---|---|---|---|---|---|---|---|---|---|---|---|
| ENSG00000154222 | CC2D1B | 2.068570494 | 0.07702728 | 1.340758554 | 0.265688503 | 3.409329048 | 6.138360429 | 4.710564154 | 8.042706615 | 8.872051712 | 8.98123666 | 7.995960198 | 8.392188801 |
| ENSG00000174243 | DDX23 | 1.742986566 | 0.037693949 | 1.665741646 | 0.045767852 | 3.408728211 | 8.367060731 | 8.191702463 | 6.073378139 | 9.555032495 | 9.59474367 | 9.404610599 | 9.586086719 |
| ENSG00000081087 | OSTM1 | 1.679995665 | 0.001116395 | 1.72807689 | 0.000721942 | 3.408072555 | 7.82963488 | 8.540087354 | 7.398165132 | 9.72797373 | 9.6241521 | 9.558789939 | 9.873793342 |
| ENSG00000131094 | C1QL1 | 0.919676794 | 0.649272397 | 2.485979592 | 0.166578095 | 3.405656386 | 2 | 4.11352228 | 2 | 4.034562184 | 3.39680241 | 5.352823931 | 4.532869193 |
| ENSG00000180596 | HIST1H2BC | 4.320484731 | 1.31E-06 | -0.915357652 | 0.384545195 | 3.40512708 | 4.00722531 | 3.686303414 | 5.156626401 | 8.408172799 | 8.42842973 | 3.215043717 | 4.062589736 |
| ENSG00000272391 | POM121C | 2.602506947 | 2.06E-14 | 0.802316478 | 0.03268044 | 3.404823424 | 8.228665467 | 7.669933291 | 8.175061096 | 10.60649663 | 10.6316726 | 8.704299335 | 8.94213989 |
| ENSG00000119938 | PPP1R3C | 0.729984381 | 0.49620218 | 2.674793663 | 0.003705475 | 3.404778044 | 9.304185368 | 7.572420273 | 6.80318718 | 8.945181925 | 9.05949698 | 11.21786533 | 10.59534423 |
| ENSG00000110931 | CAMKK2 | 1.24965622 | 3.70E-05 | 2.153040276 | 1.01E-13 | 3.402696496 | 7.940205782 | 7.638155693 | 7.818184403 | 9.015934546 | 9.04730166 | 9.99643172 | 9.860179463 |
| ENSG00000144306 | SCRN3 | 1.319082723 | 0.371698295 | 2.083496556 | 0.131743628 | 3.402579279 | 3.327711234 | 6.899080544 | 4.058998049 | 6.979883891 | 6.78340763 | 7.64960537 | 7.608437243 |
| ENSG00000088543 | C3orf18 | 1.398159903 | 0.453666628 | 1.999258611 | 0.256697952 | 3.397418514 | 2 | 7.393647515 | 4.523675391 | 7.333842102 | 7.4314987 | 8.000548654 | 7.944426693 |
| ENSG00000182308 | DCAF4L1 | 2.387473346 | 0.446075855 | 1.009085259 | NA | 3.396558605 | 2 | 2 | 2 | 2.947924294 | 3.45167086 | 2.560105256 | 2.632958241 |
| ENSG00000179085 | DPM3 | 1.546425323 | 0.112649338 | 1.845853264 | 0.049503707 | 3.392278587 | 5.095959458 | 3.686303414 | 6.203069917 | 6.64692533 | 6.82036165 | 6.995586672 | 7.05482945 |
| ENSG00000164877 | MICALL2 | 1.136889993 | 0.615473512 | 2.255043031 | 0.280004532 | 3.391933024 | 8.468948322 | 2 | 4.523675391 | 8.058664058 | 8.13770558 | 9.078385428 | 9.322263543 |
| ENSG00000035403 | VCL | 0.315607148 | 0.726269433 | 3.075088089 | 1.70E-05 | 3.390695237 | 9.720386367 | 11.66299451 | 10.68342405 | 11.19937403 | 11.2319249 | 13.64177426 | 14.24245265 |
| ENSG00000183579 | ZNRF3 | 1.738356523 | 0.02413324 | 1.651305209 | 0.031331843 | 3.389661732 | 4.00722531 | 4.442710129 | 5.39244166 | 6.229283386 | 6.33555364 | 6.332313483 | 6.055940621 |
| ENSG00000065325 | GLP2R | -0.317990197 | 0.948248384 | 3.707175828 | 0.38169184 | 3.389185631 | 2 | 2 | 2 | 2 | 2.49394783 | 4.644324387 | 3.261853426 |
| ENSG00000178966 | RMI1 | 1.595134135 | 0.452687885 | 1.791513121 | 0.38150221 | 3.386647256 | 2 | 5.72254323 | 2 | 5.930370745 | 5.62411716 | 6.346800618 | 5.453611367 |
| ENSG00000132436 | FIGNL1 | 1.786897853 | 0.383862151 | 1.599502375 | 0.431616031 | 3.386400228 | 2 | 5.72254323 | 2 | 5.930370745 | 6.00043082 | 6.016144783 | 5.524371359 |
| ENSG00000155975 | VPS37A | 1.953629597 | 0.281556878 | 1.432544683 | 0.435144797 | 3.38617428 | 8.631580145 | 7.099070191 | 2 | 9.40053583 | 9.42945789 | 9.133040681 | 8.616180753 |
| ENSG00000135974 | C2orf49 | 1.636786211 | 0.178578057 | 1.749312484 | 0.140060743 | 3.386098695 | 7.624139768 | 5.303523747 | 4.523675391 | 8.04293816 | 7.99722844 | 8.351275892 | 7.871136591 |
| ENSG00000184905 | TCEAL2 | 3.362358 | 0.20486278 | 0.023426224 | 0.994111049 | 3.385784224 | 2 | 2 | 2 | 3.921164147 | 3.7864959 | 2.384001718 | 2.277179783 |
| ENSG00000075240 | GRAMD4 | 1.529554442 | 0.524284348 | 1.853812847 | 0.422430971 | 3.383367289 | 2 | 6.844429755 | 2 | 6.851641821 | 6.77267312 | 7.331691625 | 6.887339785 |
| ENSG00000100852 | ARHGAP5 | 2.011583805 | 0.18460402 | 1.370065752 | 0.375326773 | 3.381649557 | 4.00722531 | 9.367676479 | 7.858115942 | 10.26782038 | 10.2188359 | 9.867145934 | 9.283558215 |
| ENSG00000087074 | PPP1R15A | 2.74943733 | 0.002429018 | 0.631007747 | 0.546187635 | 3.380445077 | 7.27616306 | 9.308339287 | 10.10298738 | 12.00455363 | 12.0735623 | 9.868819956 | 9.980250523 |
| ENSG00000159423 | ALDH4A1 | 0.438524799 | 0.787199307 | 2.936899527 | 0.028740291 | 3.375424326 | 3.327711234 | 6.951736442 | 4.058998049 | 6.024891728 | 6.14893615 | 8.333200639 | 8.676578788 |
| ENSG00000144468 | RHBDD1 | 1.443370898 | 0.17057763 | 1.929823483 | 0.054562067 | 3.373194381 | 4.815609471 | 7.847167909 | 7.342949392 | 8.567402595 | 8.54656162 | 9.264754164 | 8.771382398 |
| ENSG00000219481 | NBPF1 | 4.013015945 | 6.94E-07 | -0.641279242 | 0.507862588 | 3.371736703 | 6.89194353 | 8.337339476 | 6.073378139 | 11.34961059 | 11.4541495 | 7.06586564 | 6.471341372 |
| ENSG00000101493 | ZNF516 | 1.603732389 | 0.201491557 | 1.767995429 | 0.147348187 | 3.371727819 | 4.467311984 | 8.297207175 | 6.718950348 | 8.756965077 | 8.82375772 | 9.087759434 | 8.80566012 |
| ENSG00000169962 | TAS1R3 | 0.007224108 | 0.99798804 | 3.364485949 | 0.247980941 | 3.371710057 | 2 | 2 | 2 | 2.141737246 | 2.49394783 | 4.029788235 | 3.660199087 |
| ENSG00000142459 | EVI5L | 0.80142163 | 0.613329998 | 2.570040711 | 0.065337615 | 3.371462341 | 6.73718411 | 8.356994601 | 3.369317972 | 7.942383523 | 8.06698125 | 9.661355639 | 9.84988408 |
| ENSG00000100505 | TRIM9 | 2.649567841 | 0.440574326 | 0.721164812 | 0.847276751 | 3.370732653 | 2 | 2 | 2 | 3.921164147 | 2.49394783 | 2.62493832 | 2.374681325 |
| ENSG00000139211 | AMIGO2 | 1.576904625 | 0.073765025 | 1.793568233 | 0.036652015 | 3.370472858 | 8.653385977 | 8.522788974 | 6.534109959 | 9.701744597 | 9.76823652 | 10.34924812 | 9.399228411 |
| ENSG00000040933 | INPP4A | 1.336697441 | 0.317344658 | 2.032534831 | 0.106808076 | 3.369232272 | 3.327711234 | 7.315436877 | 7.502634626 | 8.093978021 | 8.2580338 | 8.855903324 | 8.877074023 |
| ENSG00000146063 | TRIM41 | 1.746944156 | 0.305856687 | 1.618792953 | 0.337037541 | 3.365737109 | 8.107517749 | 7.189546432 | 2 | 8.836004565 | 8.89972012 | 8.772063028 | 8.709655965 |
| ENSG00000162654 | GBP4 | 5.734635802 | 6.38E-09 | -2.370906063 | 0.028948196 | 3.363729739 | 5.532353208 | 3.686303414 | 6.43197674 | 11.2046844 | 11.1596093 | 4.065262523 | 3.012034821 |
| ENSG00000204149 | AGAP6 | 3.095094872 | 0.027688126 | 0.267479697 | 0.877195052 | 3.362574569 | 2 | 4.442710129 | 6.534109959 | 8.203120604 | 8.284723 | 5.594558848 | 5.459179378 |
| ENSG00000161800 | RACGAP1 | 1.386926014 | 0.311279445 | 1.974854189 | 0.127639854 | 3.361780204 | 5.866944937 | 6.464272742 | 2 | 6.946463121 | 7.02417148 | 7.663174541 | 7.445480288 |
| ENSG00000142599 | RERE | 1.276897833 | 0.155953136 | 2.084623302 | 0.013226967 | 3.361521134 | 7.031698319 | 9.470501606 | 8.511888463 | 9.89995113 | 9.9465583 | 10.75863418 | 10.69795699 |
| ENSG00000188419 | CHM | 2.038229774 | 0.007413593 | 1.323270383 | 0.094621254 | 3.361500157 | 7.27616306 | 7.538394143 | 5.39244166 | 9.036500968 | 8.98821297 | 8.380320298 | 8.224954107 |
| ENSG00000131503 | ANKHD1 | 4.816655689 | 1.04E-18 | -1.45787615 | 0.016837458 | 3.358779539 | 6.138360429 | 5.131603625 | 4.874587708 | 10.14850007 | 10.1741606 | 4.350488971 | 4.09155917 |
| ENSG00000107560 | RAB11FIP2 | 1.890352807 | 0.118659477 | 1.468217222 | 0.229961504 | 3.358570029 | 3.327711234 | 7.189546432 | 6.43197674 | 8.132699624 | 8.23866319 | 7.968118715 | 7.543088047 |
| ENSG00000107643 | MAPK8 | 2.126900491 | 0.178652578 | 1.231059546 | 0.453051282 | 3.357960038 | 7.27616306 | 7.60566235 | 2 | 8.952415387 | 8.99976571 | 8.094396944 | 8.086110269 |
| ENSG00000097007 | ABL1 | 1.495004305 | 0.12639307 | 1.862035464 | 0.047854089 | 3.357039769 | 7.750913381 | 9.994333773 | 10.54960077 | 11.28518243 | 11.3595809 | 11.64101736 | 11.73618212 |
| ENSG00000127870 | RNF6 | 1.509272035 | 0.440574326 | 1.847531774 | 0.324961434 | 3.356803809 | 2 | 7.315436877 | 9.06622548 | 9.331938606 | 9.3808074 | 9.723042782 | 9.66228668 |
| ENSG00000146425 | DYNLT1 | 1.865017444 | 0.004829424 | 1.490368984 | 0.027424614 | 3.355386428 | 9.143983763 | 7.572420273 | 7.97167868 | 10.24542 | 10.2326097 | 9.732038014 | 9.98821576 |
| ENSG00000140992 | PDPK1 | 1.922345888 | 3.66E-05 | 1.429048411 | 0.002775982 | 3.351394299 | 7.790810999 | 8.213429689 | 7.098165388 | 9.690252889 | 9.68338024 | 9.258063226 | 9.13188655 |
| ENSG00000099917 | MED15 | 1.446593211 | 0.097990086 | 1.902266969 | 0.023148373 | 3.34886018 | 6.563806279 | 8.968383114 | 8.35328246 | 9.690973808 | 9.71736428 | 10.06173917 | 10.2482943 |
| ENSG00000015285 | WAS | 0.27207001 | 0.938309757 | 3.076788287 | 0.291783001 | 3.348858297 | 2 | 2 | 2 | 2.270786764 | 2.49394783 | 3.993419637 | 3.209956562 |
| ENSG00000166670 | MMP10 | 8.538358019 | 1.30E-07 | -5.191150625 | 0.032232657 | 3.347207394 | 2 | 2 | 2 | 8.634896 | 8.5227358 | 2 | 2 |
| ENSG00000196967 | ZNF585A | 2.169133567 | 0.027857303 | 1.176685786 | 0.258406162 | 3.345819353 | 6.257048221 | 6.140560021 | 4.058998049 | 7.839502934 | 7.89700047 | 7.434742221 | 6.041166521 |
| ENSG00000119986 | AVPI1 | 1.556417066 | 0.545415898 | 1.788287041 | 0.475468809 | 3.344704107 | 2 | 7.638155693 | 2 | 7.523859802 | 7.7127321 | 7.847655556 | 7.850099615 |
| ENSG00000111962 | UST | 1.221101671 | 0.208574574 | 2.123494073 | 0.018761048 | 3.344595744 | 6.138360429 | 8.607278467 | 7.552162999 | 8.935480545 | 9.0020652 | 9.387195342 | 10.22519644 |
| ENSG00000171223 | JUNB | 1.930019485 | 0.012630673 | 1.414037493 | 0.075077708 | 3.344056978 | 9.740899611 | 7.73146271 | 9.049077186 | 10.96138842 | 10.9898386 | 10.76066432 | 10.08209816 |
| ENSG00000196653 | ZNF502 | 1.62826827 | 0.281547017 | 1.715562842 | 0.244635807 | 3.343831112 | 3.327711234 | 5.595927442 | 2 | 5.959382894 | 5.7929843 | 6.013120342 | 5.905000563 |
| ENSG00000075303 | SLC25A40 | 1.525309076 | 0.069582907 | 1.817000766 | 0.025221004 | 3.342309841 | 3.327711234 | 5.303523747 | 4.874587708 | 6.078767801 | 6.18985553 | 6.537789343 | 6.27380897 |
| ENSG00000101442 | ACTR5 | 1.195662785 | 0.324473681 | 2.146485146 | 0.05484371 | 3.342147931 | 2 | 5.131603625 | 4.058998049 | 5.210797711 | 5.37722731 | 6.132200015 | 6.215784511 |
| ENSG00000242498 | ARPIN | 1.091032559 | 0.017866632 | 2.250947262 | 1.06E-07 | 3.341979821 | 6.366709671 | 6.844429755 | 6.203069917 | 7.697489359 | 7.4450836 | 8.691586261 | 8.747690481 |
| ENSG00000104361 | NIPAL2 | 1.910704697 | 0.090432691 | 1.430547503 | 0.210610076 | 3.3412522 | 5.532353208 | 7.099070191 | 8.825624954 | 9.634381252 | 9.59930725 | 8.927126162 | 9.324934421 |
| ENSG00000197863 | ZNF790 | 1.945995729 | 0.129843791 | 1.395038697 | 0.284984329 | 3.341034426 | 5.70932728 | 4.11352228 | 2 | 6.422239259 | 6.31350454 | 6.013120342 | 5.66592325 |
| ENSG00000088247 | KHSRP | 1.644872607 | 0.020681631 | 1.694806541 | 0.016160038 | 3.339679149 | 7.159102577 | 8.876041558 | 8.884815469 | 10.11343618 | 10.1572376 | 10.03682636 | 10.31985465 |
| ENSG00000177873 | ZNF619 | 1.218947667 | 0.142769855 | 2.116864693 | 0.005694583 | 3.33581236 | 5.095959458 | 4.442710129 | 4.058998049 | 5.731694 | 5.60003091 | 6.726875343 | 6.257923755 |
| ENSG00000196705 | ZNF431 | 1.748406212 | 0.044940119 | 1.586001351 | 0.068074655 | 3.334407563 | 3.327711234 | 5.838937808 | 5.39244166 | 6.882264028 | 6.82556437 | 6.78654324 | 6.602920209 |
| ENSG00000141425 | RPRD1A | 1.560100195 | 0.296977641 | 1.768459259 | 0.221520727 | 3.328559455 | 3.327711234 | 8.234834549 | 8.206335842 | 9.196346103 | 9.20931286 | 9.515363796 | 9.29605726 |
| ENSG00000198783 | ZNF830 | 1.68770209 | 0.474361527 | 1.639634987 | 0.478795283 | 3.327337077 | 2 | 6.78762695 | 2 | 6.979883891 | 6.83591385 | 6.830158212 | 6.895576665 |
| ENSG00000256525 | POLG2 | 1.407353188 | 0.279318358 | 1.919872459 | 0.119249183 | 3.327225647 | 2 | 5.131603625 | 4.058998049 | 5.420062284 | 5.53799497 | 6.280431262 | 5.529673353 |
| ENSG00000153071 | DAB2 | 1.249928089 | 0.035146883 | 2.076419429 | 0.000188322 | 3.326347517 | 10.79260216 | 10.71514053 | 10.20715117 | 11.83668751 | 11.8476762 | 11.9157543 | 13.15993594 |
| ENSG00000012223 | LTF | 0.01278131 | 0.9971266 | 3.311979747 | 0.221353293 | 3.324761056 | 2 | 2 | 2 | 2.3892346 | 2.2680111 | 3.648325314 | 3.972018587 |
| ENSG00000188747 | NOXA1 | -0.310504433 | 0.925672996 | 3.633863867 | 0.164988406 | 3.323359434 | 2 | 2 | 2 | 2.270786764 | 2.2680111 | 4.378961816 | 3.640730416 |
| ENSG00000175066 | GK5 | 1.445366762 | 0.119221214 | 1.876266104 | 0.034894638 | 3.321632866 | 5.532353208 | 6.228547981 | 7.858115942 | 8.304583895 | 8.2618769 | 8.778311345 | 8.637209821 |
| ENSG00000178694 | NSUN3 | 1.916432387 | 0.096878945 | 1.40065029 | 0.231333139 | 3.317082677 | 2 | 5.303523747 | 5.772716771 | 6.744602355 | 6.88657603 | 6.414841377 | 6.22561877 |
| ENSG00000198429 | ZNF69 | 1.476023984 | 0.072447018 | 1.840349916 | 0.019563833 | 3.3163739 | 4.815609471 | 4.710564154 | 3.369317972 | 5.709096143 | 5.80356929 | 6.095496313 | 6.099376274 |
| ENSG00000185201 | IFITM2 | 2.199501432 | 1.06E-08 | 1.116615205 | 0.006238495 | 3.316116637 | 10.80726569 | 10.64492263 | 10.82615578 | 12.90548945 | 13.0088736 | 11.42170857 | 12.22106411 |
| ENSG00000108468 | CBX1 | 1.492144699 | 0.219632896 | 1.823292509 | 0.119536754 | 3.315437209 | 8.87372952 | 6.951736442 | 5.39244166 | 9.144656522 | 9.24954577 | 9.375172844 | 9.664698405 |
| ENSG00000121957 | GPSM2 | 1.394561466 | 0.53868734 | 1.920563646 | 0.374841596 | 3.315125112 | 2 | 6.311476754 | 2 | 6.147608921 | 6.18176397 | 6.934553331 | 6.33565646 |
| ENSG00000172943 | PHF8 | 1.34282615 | 0.123384912 | 1.97113327 | 0.016889842 | 3.31395942 | 4.815609471 | 6.602424674 | 7.225742779 | 7.852449262 | 7.78928606 | 8.612361808 | 8.245238611 |
| ENSG00000107798 | LIPA | 2.1735307 | 0.001190589 | 1.140072489 | 0.110628724 | 3.313603189 | 7.750913381 | 9.148351514 | 7.934808822 | 10.5732559 | 10.6024186 | 9.881106239 | 9.141874832 |
| ENSG00000111530 | CAND1 | 1.447522376 | 0.261766667 | 1.865428034 | 0.130735409 | 3.312950411 | 9.27634997 | 10.07368926 | 5.930865648 | 10.63412381 | 10.6435047 | 11.13029223 | 10.97726982 |
| ENSG00000147588 | PMP2 | 3.221029076 | 0.544857088 | 0.091472225 | 0.987740261 | 3.312501301 | 2 | 2 | 2 | 4.523237872 | 2 | 2.457042231 | 2.225841198 |
| ENSG00000251349 | MSANTD3-TI | 8.50262631 | 1.62E-07 | -5.191150591 | 0.032497074 | 3.31147572 | 2 | 2 | 2 | 8.584574266 | 8.50338773 | 2 | 2 |
| ENSG00000111679 | PTPN6 | 0.316332522 | 0.916384376 | 2.99509129 | 0.227412477 | 3.311423812 | 3.327711234 | 2 | 2 | 2.947924294 | 2.38539787 | 4.971696071 | 3.042026731 |
| ENSG00000169627 | BOLA2B | 8.50185586 | 1.65E-07 | -5.191150583 | 0.032532189 | 3.310705278 | 2 | 2 | 2 | 8.589222193 | 8.49688027 | 2 | 2 |
| ENSG00000130193 | THEM6 | 0.881715739 | 0.529996014 | 2.428253465 | 0.050406299 | 3.309969204 | 6.257048221 | 5.303523747 | 2 | 6.18902405 | 6.08962627 | 7.302260672 | 7.900082338 |
| ENSG00000130962 | PRRG1 | 1.642587211 | 0.146700672 | 1.666742037 | 0.13261638 | 3.309329248 | 3.327711234 | 6.728495673 | 6.629488387 | 7.807949361 | 7.70992146 | 7.69371876 | 7.868000534 |
| ENSG00000136152 | COG3 | 1.99785036 | 0.051682313 | 1.311291674 | 0.215092713 | 3.309142035 | 4.815609471 | 7.701025996 | 6.322058544 | 8.66162195 | 8.72420319 | 8.295111724 | 7.667350749 |
| ENSG00000119392 | GLE1 | 1.413696196 | 0.38028167 | 1.894149091 | 0.217050348 | 3.307845287 | 7.435667355 | 6.464272742 | 2 | 7.728718552 | 7.94559728 | 8.122734438 | 8.483814596 |
| ENSG00000161671 | EMC10 | 1.880939635 | 0.004752131 | 1.42682627 | 0.036771903 | 3.307765905 | 9.533135543 | 8.169643011 | 8.266922072 | 10.62811365 | 10.7079584 | 9.861061271 | 10.50031203 |
| ENSG00000178935 | ZNF552 | 1.792208506 | 0.171506539 | 1.514652682 | 0.246643226 | 3.306861188 | 5.095959458 | 2 | 4.058998049 | 5.764940652 | 5.58783529 | 5.343222328 | 5.502966356 |
| ENSG00000129116 | PALLD | -0.06708444 | 0.91483416 | 3.373825505 | 8.15E-14 | 3.306741066 | 10.62615013 | 11.15761406 | 10.12770132 | 10.68059872 | 10.583733 | 13.85056693 | 14.26144901 |
| ENSG00000136682 | CBWD2 | 3.780661339 | 0.000665039 | -0.474658576 | 0.726349422 | 3.306002763 | 6.257048221 | 5.457120953 | 2 | 8.967964567 | 9.08358266 | 4.720468739 | 5.164234344 |
| ENSG00000118160 | SLC8A2 | 2.690456475 | 0.36407141 | 0.614958872 | 0.852211698 | 3.305415347 | 2 | 2 | 2 | 3.515101962 | 3.28037825 | 2.526562374 | 2.421067459 |
| ENSG00000136205 | TNS3 | 1.959165154 | 0.006834427 | 1.344405988 | 0.072934705 | 3.303571142 | 8.009475818 | 9.608085847 | 9.757248847 | 11.25892943 | 11.2665282 | 10.95373885 | 10.26269416 |
| ENSG00000118308 | LRMP | 1.27444235 | 0.725609862 | 2.028257093 | NA | 3.302699443 | 2 | 2 | 2 | 2.695456494 | 2.68924543 | 3.429858444 | 2.466008509 |
| ENSG00000119720 | NRDE2 | 1.701589844 | 0.111942166 | 1.599987847 | 0.130241043 | 3.301577691 | 5.532353208 | 7.051611315 | 4.058998049 | 7.584088227 | 7.79989812 | 7.586913431 | 7.605929526 |
| ENSG00000130150 | MOSPD2 | 1.152585811 | 0.516281765 | 2.148327779 | 0.188579572 | 3.30091359 | 2 | 7.928237109 | 6.629488387 | 7.994705029 | 7.97398853 | 9.005308106 | 8.931163478 |
| ENSG00000132840 | BHMT2 | 0.685023454 | 0.492643658 | 2.615740129 | 0.002064011 | 3.300763582 | 4.815609471 | 4.442710129 | 6.073378139 | 6.043074888 | 5.70538658 | 7.187558829 | 8.139971339 |
| ENSG00000125144 | MT1G | 8.020928413 | 1.90E-08 | -4.721352482 | 0.036346499 | 3.299575931 | 2 | 2 | 3.369317972 | 8.989221226 | 9.05063789 | 2.048074405 | 2 |
| ENSG00000090615 | GOLGA3 | 1.231472863 | 9.96E-05 | 2.065335865 | 9.68E-12 | 3.296808728 | 9.174465797 | 9.558016677 | 9.767769415 | 10.70435807 | 10.7890936 | 11.57807074 | 11.58140223 |
| ENSG00000173991 | TCAP | 0.01279932 | 0.997163678 | 3.283196652 | 0.240873009 | 3.295995972 | 2 | 2 | 2 | 2.3892346 | 2.2680111 | 4.133696865 | 3.36036153 |
| ENSG00000197771 | MCMBP | 1.820516675 | 0.034515932 | 1.474902538 | 0.090709877 | 3.295419213 | 7.579279349 | 8.255926468 | 5.772716771 | 9.344822471 | 9.3166689 | 9.059452688 | 8.912005269 |
| ENSG00000136807 | CDK9 | 1.392309583 | 0.025474975 | 1.901649888 | 0.001438346 | 3.293959472 | 6.257048221 | 7.701025996 | 7.734844356 | 8.690748087 | 8.80675844 | 9.22151698 | 9.288258056 |
| ENSG00000135482 | ZC3H10 | 1.290546862 | 0.56072077 | 2.002546334 | 0.339728351 | 3.293093196 | 2 | 6.140560021 | 2 | 5.968925393 | 5.83486613 | 6.656902456 | 6.493217192 |
| ENSG00000026751 | SLAMF7 | -3.725402998 | 0.02361388 | 7.015993643 | 2.07E-07 | 3.290590645 | 3.327711234 | 3.076665209 | 5.156626401 | 2.498691189 | 2.14022021 | 8.840629888 | 11.58498783 |
| ENSG00000145107 | TM4SF19 | 3.399630251 | 0.119911452 | -0.109145643 | 0.967718357 | 3.290484608 | 2 | 2 | 3.369317972 | 4.574143677 | 4.70309173 | 2.803598391 | 2.225841198 |
| ENSG00000137814 | HAUS2 | 1.40112673 | 0.018985894 | 1.889046189 | 0.000946600 | 3.290172919 | 5.532353208 | 5.946638857 | 6.629488387 | 7.559034656 | 7.32163988 | 8.140859753 | 7.667350749 |
| ENSG00000279389 | | 2.014286207 | 0.035203087 | 1.274979328 | 0.198028433 | 3.289265534 | 5.532353208 | 7.847167909 | 5.930865648 | 8.833394974 | 8.78422202 | 8.264917037 | 7.863808491 |
| ENSG00000197296 | FITM2 | -0.146404889 | 0.936681124 | 3.43524688 | 0.018858407 | 3.288841991 | 2 | 4.93638933 | 7.029880454 | 5.731694 | 5.53799497 | 8.897090174 | 9.294500788 |
| ENSG00000121858 | TNFSF10 | 5.904868845 | 0.00059577 | -2.621812825 | 0.164236009 | 3.28305602 | 6.009025997 | 2 | 2 | 10.27601154 | 10.1613584 | 3.011868911 | 2.671842496 |

| | | | | | | | | | | | | | |
|---|---|---|---|---|---|---|---|---|---|---|---|---|---|
| ENSG00000271880 | AGAP11 | 5.613045543 | 0.001118566 | -2.330358267 | 0.255544364 | 3.282687275 | 2 | 2 | 3.369317972 | 6.610808945 | 6.70653264 | 2.048074405 | 2.225841198 |
| ENSG00000136275 | C7orf69 | 0.274456092 | 0.939900699 | 3.007014159 | 0.323451531 | 3.281470251 | 2 | 2 | 2 | 2.3892346 | 2.38539787 | 4.099885429 | 2.851942824 |
| ENSG00000188554 | NBR1 | 1.545204187 | 0.010826309 | 1.735876565 | 0.003524935 | 3.281080752 | 9.081016759 | 10.38739569 | 9.227351203 | 11.25015134 | 11.2213553 | 11.39072895 | 11.4608208 |
| ENSG00000244115 | DNAJC25-GN | 8.471144183 | 2.12E-07 | -5.191150503 | 0.032980585 | 3.27999368 | 2 | 2 | 2 | 8.576794289 | 8.44711562 | 2 | 2 |
| ENSG00000187555 | USP7 | 1.623281294 | 0.011059186 | 1.65453515 | 0.009052045 | 3.277816443 | 8.444143041 | 8.395519306 | 6.882775709 | 9.693134408 | 9.66390646 | 9.697600698 | 9.721691937 |
| ENSG00000023608 | SNAPC1 | 2.397177531 | 0.074912837 | 0.879842527 | 0.547042483 | 3.277020058 | 3.327711234 | 7.232742069 | 4.874587708 | 8.330724217 | 8.32935768 | 6.989444349 | 6.685996416 |
| ENSG00000047578 | KIAA0556 | 1.465289188 | 0.116476863 | 1.808849819 | 0.044161511 | 3.274139007 | 7.485143553 | 7.315436877 | 4.874587708 | 8.356399318 | 8.40429221 | 8.639092784 | 8.797441501 |
| ENSG00000124120 | TTPAL | 0.707503143 | 0.530030976 | 2.566385533 | 0.008125446 | 3.273888676 | 5.330594134 | 8.029695761 | 6.322058544 | 7.683067677 | 7.70992146 | 9.263163896 | 9.763920691 |
| ENSG00000154803 | FLCN | 2.406104917 | 1.05E-05 | 0.867166721 | 0.150582065 | 3.273271638 | 5.70932728 | 7.099070191 | 6.718950348 | 9.002058889 | 8.98705259 | 7.600074836 | 7.342293759 |
| ENSG00000110497 | AMBRA1 | 1.800556154 | 0.048401835 | 1.472568236 | 0.109372081 | 3.27312439 | 7.27616306 | 7.819099893 | 5.156626401 | 8.81895723 | 8.97656697 | 8.465761957 | 8.676578788 |
| ENSG00000166922 | SCG5 | 8.839096306 | 1.18E-12 | -5.566038269 | 0.007570639 | 3.273058037 | 2 | 3.686303414 | 2 | 10.67076507 | 10.682378 | 2.048074405 | 2 |
| ENSG00000240857 | RDH14 | 1.417162562 | 0.063833268 | 1.853646762 | 0.011498073 | 3.270809324 | 6.009025997 | 7.232742069 | 5.39244166 | 7.815902688 | 7.83127384 | 8.217092349 | 8.28888532 |
| ENSG00000177565 | TBL1XR1 | 1.015862686 | 0.062718919 | 2.253885329 | 5.51E-06 | 3.269748015 | 9.143983763 | 8.169643011 | 8.143093325 | 9.664056897 | 9.46989111 | 10.91992417 | 10.67765183 |
| ENSG00000136295 | TTYH3 | 1.546000906 | 6.03E-05 | 1.723375201 | 5.79E-06 | 3.269376108 | 9.975371636 | 9.256956408 | 9.849281147 | 11.23391814 | 11.2935316 | 11.27870987 | 11.58713493 |
| ENSG00000276180 | HIST1H4I | -2.311964322 | 0.478464462 | 5.580879938 | 0.017373749 | 3.268915616 | 2 | 2 | 2 | 2 | 2.14022021 | 5.515547099 | 5.900914815 |
| ENSG00000142178 | SIK1 | 8.45901032 | 2.20E-07 | -5.191150516 | 0.03297173 | 3.267859804 | 2 | 2 | 2 | 8.458152491 | 8.54340749 | 2 | 2 |
| ENSG00000070759 | TESK2 | 1.023891733 | 0.617138584 | 2.241945116 | 0.229720627 | 3.265836849 | 2 | 5.457120953 | 2 | 4.965943888 | 5.07918227 | 5.985611216 | 6.27380897 |
| ENSG00000170325 | PRDM10 | 1.478362602 | 0.203115525 | 1.786579432 | 0.110104573 | 3.264942034 | 6.653098118 | 5.131603625 | 3.369317972 | 7.127781374 | 6.86146731 | 7.470462265 | 7.107191496 |
| ENSG00000132478 | UNK | 1.682988223 | 0.335558745 | 1.581601188 | 0.359814853 | 3.264589411 | 2 | 7.819099893 | 5.930865648 | 8.16211395 | 8.33484115 | 7.973574582 | 8.309067496 |
| ENSG00000134864 | GGACT | 1.183492611 | 0.280090671 | 2.080935945 | 0.038900682 | 3.264428556 | 4.467311984 | 4.710564154 | 2 | 5.022023783 | 5.30456641 | 5.916072173 | 6.06327132 |
| ENSG00000106459 | NRF1 | 1.593644647 | 0.208574574 | 1.669332153 | 0.177265 | 3.262976799 | 2 | 5.946638857 | 5.930865648 | 6.877205252 | 6.94511186 | 7.009312024 | 6.959831579 |
| ENSG00000257315 | ZBED6 | 2.92017094 | 0.060865068 | 0.342020052 | 0.853341364 | 3.262190992 | 2 | 7.35507212 | 5.39244166 | 8.929383887 | 9.04952667 | 6.34920106 | 6.577546999 |
| ENSG00000158615 | PPP1R15B | 1.411843954 | 0.043881335 | 1.848115962 | 0.006017652 | 3.259959916 | 8.16936288 | 8.487557258 | 6.718950348 | 9.384581425 | 9.36570792 | 9.949148417 | 9.654117214 |
| ENSG00000186566 | GPATCH8 | 1.350765896 | 0.160378969 | 1.909095568 | 0.036346775 | 3.259861464 | 6.468621203 | 9.159587375 | 8.266922072 | 9.632881324 | 9.72504082 | 10.13173193 | 10.33280292 |
| ENSG00000189339 | SLC35E2B | 1.940510468 | 0.015305913 | 1.318905967 | 0.110602847 | 3.259416434 | 6.257048221 | 8.004992147 | 8.536706852 | 9.734981585 | 9.85856545 | 9.198071497 | 9.161646304 |
| ENSG00000147100 | SLC16A2 | 1.134360936 | 0.000879733 | 2.124903058 | 3.90E-11 | 3.259263994 | 7.579279349 | 7.467835385 | 7.858115942 | 8.813671111 | 8.6932307 | 9.637249338 | 9.833364301 |
| ENSG00000029725 | RABEP1 | 1.886770318 | 0.029070155 | 1.372434755 | 0.121056843 | 3.259205074 | 6.563806279 | 8.993716073 | 7.896971941 | 10.02242928 | 10.0033302 | 9.421247846 | 9.576512555 |
| ENSG00000134313 | KIDINS220 | 1.582193416 | 0.000135408 | 1.6764341 | 4.39E-05 | 3.258627516 | 8.042902253 | 8.955547846 | 8.511888463 | 10.13690654 | 10.1243681 | 10.25686316 | 10.19151462 |
| ENSG00000027697 | IFNGR1 | 1.725201628 | 0.321721272 | 1.532674362 | 0.375035076 | 3.25787599 | 8.075571721 | 8.053983481 | 2 | 9.205456697 | 9.18624204 | 9.143796722 | 8.850838709 |
| ENSG00000152377 | SPOCK1 | 2.440537816 | 8.99E-06 | 0.817111968 | 0.181719899 | 3.257649784 | 9.602033055 | 9.672251278 | 10.78531698 | 12.54062562 | 12.5849424 | 10.80166469 | 11.06900213 |
| ENSG00000100626 | GALNT16 | -1.086111438 | 0.450183579 | 4.343693455 | 0.000162517 | 3.257582017 | 3.327711234 | 3.076665209 | 4.058998049 | 3.227757282 | 2.49394783 | 7.076045862 | 7.448282715 |
| ENSG00000116039 | ATP6V1B1 | 3.388980517 | 0.206233223 | -0.131556533 | 0.967718357 | 3.257423984 | 2 | 2 | 2 | 3.754611435 | 3.98554648 | 2.225793514 | 2.374681325 |
| ENSG00000116731 | PRDM2 | 1.592527772 | 0.012350658 | 1.664297286 | 0.008211245 | 3.256825058 | 6.89194353 | 7.701025996 | 8.511888463 | 9.386362875 | 9.45570025 | 9.584486439 | 9.394877047 |
| ENSG00000197343 | ZNF655 | 1.76613515 | 0.088075356 | 1.487867869 | 0.150474848 | 3.254003018 | 5.70932728 | 8.820493869 | 8.175061096 | 9.856271709 | 9.74645749 | 9.55749302 | 9.492799274 |
| ENSG00000100401 | RANGAP1 | 1.381469255 | 0.011026701 | 1.872463393 | 0.000337252 | 3.253932648 | 8.963744667 | 8.762721501 | 9.907528187 | 10.62849002 | 10.7188341 | 11.06196671 | 11.25994485 |
| ENSG00000107104 | KANK1 | 1.088881824 | 0.446977367 | 2.164580502 | 0.098016923 | 3.253462326 | 3.327711234 | 6.951736442 | 7.858115942 | 7.97352961 | 8.01099479 | 9.219878303 | 8.871343049 |
| ENSG00000142166 | IFNAR1 | 1.972546925 | 0.02795781 | 1.273815432 | 0.170653805 | 3.246362358 | 6.138360429 | 8.717821467 | 8.408111533 | 10.12841597 | 10.0193108 | 9.423527655 | 9.334054358 |
| ENSG00000099968 | BCL2L13 | 1.65816204 | 0.048419679 | 1.588138066 | 0.056802649 | 3.246300106 | 9.460782017 | 9.277729677 | 7.225742779 | 10.56226593 | 10.6246011 | 10.38038819 | 10.6544806 |
| ENSG00000198018 | ENTPD7 | 2.90665826 | 1.01E-08 | 0.335544721 | 0.590927058 | 3.242202981 | 6.89194353 | 6.899080544 | 7.934808822 | 10.22067068 | 10.1659803 | 7.481445808 | 7.795024296 |
| ENSG00000151320 | AKAP6 | -0.609729714 | 0.786127952 | 3.850765817 | 0.037996695 | 3.241036102 | 2 | 5.72254323 | 2 | 4.299492648 | 3.39680241 | 8.408216037 | 7.299480881 |
| ENSG00000067704 | IARS2 | 1.552605022 | 0.04039121 | 1.679866054 | 0.024026933 | 3.232471076 | 8.674867122 | 9.04308532 | 7.029880454 | 10.02300196 | 10.0198783 | 10.14024701 | 10.15617086 |
| ENSG00000179241 | LDLRAD3 | 1.721638084 | 0.31167516 | 1.509845056 | 0.371461791 | 3.23148314 | 4.467311984 | 7.002537973 | 2 | 7.366699513 | 7.33629421 | 7.039353222 | 7.244113847 |
| ENSG00000152137 | HSPB8 | 0.922563799 | 0.1808519 | 2.307568417 | 0.000173235 | 3.230132215 | 8.946185348 | 7.315436877 | 8.511888463 | 9.246247604 | 9.37815424 | 10.68452972 | 10.70163167 |
| ENSG00000163328 | GPR155 | 0.796223173 | 0.140821478 | 2.431580462 | 2.67E-07 | 3.227803636 | 6.96351253 | 7.051611315 | 5.930865648 | 7.474468681 | 7.56502709 | 9.195072139 | 9.070914229 |
| ENSG00000110583 | NAA40 | 1.29275132 | 0.282799717 | 1.934886631 | 0.08797541 | 3.227637951 | 5.532353208 | 8.101365653 | 5.39244166 | 8.085230238 | 8.30535212 | 8.651788724 | 8.996760622 |
| ENSG00000134250 | NOTCH2 | 1.225266896 | 0.083598237 | 1.998535319 | 0.002527641 | 3.223802215 | 9.895281535 | 11.67086841 | 11.08782089 | 12.26720893 | 12.2949627 | 13.0065992 | 13.09977244 |
| ENSG00000180354 | MTURN | 1.231700612 | 0.194489031 | 1.991002584 | 0.024727678 | 3.222703195 | 4.467311984 | 6.535001748 | 4.874587708 | 6.907295206 | 6.66650236 | 7.421115854 | 7.630812678 |
| ENSG00000105464 | GRIN2D | 0.68764422 | 0.730082566 | 2.53503807 | 0.14438651 | 3.22268229 | 2 | 5.131603625 | 2 | 4.647285135 | 4.22485516 | 6.075339068 | 6.113568999 |
| ENSG00000147421 | HMBOX1 | 1.307913202 | 0.49634303 | 1.914697592 | 0.291256234 | 3.222610795 | 2 | 7.232742069 | 4.058998049 | 7.035434774 | 7.16216989 | 7.824551575 | 7.547015159 |
| ENSG00000116266 | STXBP3 | 2.165957404 | 0.051020395 | 1.056166229 | 0.369090445 | 3.222123632 | 7.579279349 | 6.311476754 | 4.058998049 | 8.776115811 | 8.61876544 | 7.667028121 | 7.54177663 |
| ENSG00000172331 | BPGM | 1.884353966 | 1.47E-08 | 1.337308578 | 8.53E-05 | 3.221662544 | 8.313285031 | 8.276714465 | 8.631919287 | 10.25812379 | 10.3093543 | 9.525746748 | 9.92498893 |
| ENSG00000121486 | TRMT1L | 2.129193784 | 0.146475623 | 1.092045767 | 0.477862788 | 3.221239551 | 4.00722531 | 7.393647515 | 4.058998049 | 8.207157886 | 8.09094333 | 7.541454591 | 6.564691065 |
| ENSG00000105327 | BBC3 | 1.116903749 | 0.283441752 | 2.103986076 | 0.02803302 | 3.220889825 | 6.96351253 | 5.72254323 | 4.523675391 | 7.053486362 | 7.20272083 | 8.526943275 | 7.480125329 |
| ENSG00000177494 | ZBED2 | 2.211310859 | 0.502051288 | 1.007790487 | NA | 3.219101345 | 2 | 2 | 2 | 2.947924294 | 3.28037825 | 2.384001718 | 2.782576693 |
| ENSG00000036257 | CUL3 | 1.615137922 | 0.170042974 | 1.603668356 | 0.165345082 | 3.218806278 | 5.866944937 | 9.079034257 | 6.958202216 | 9.523001306 | 9.5188894 | 9.548120632 | 9.469883152 |
| ENSG00000174233 | ADCY6 | 1.015279618 | 0.374685886 | 2.202779184 | 0.034100279 | 3.218058802 | 6.563806279 | 8.213429689 | 9.735974718 | 9.663322397 | 9.74027243 | 10.8739854 | 10.89774798 |
| ENSG00000205045 | SLFN12L | 3.665346315 | 0.147447249 | -0.448419963 | 0.883149421 | 3.216926352 | 2 | 2 | 2 | 3.881293667 | 4.2560182 | 2.267001569 | 2.225841198 |
| ENSG00000214274 | ANG | 1.852947647 | 0.14259015 | 1.361979449 | 0.287006757 | 3.214927096 | 2 | 4.710564154 | 5.930865648 | 6.503277904 | 6.72891807 | 6.167993056 | 6.13807518 |
| ENSG00000061676 | NCKAP1 | 1.840330157 | 0.00515506 | 1.37364196 | 0.042119905 | 3.213972118 | 8.836089686 | 10.33384851 | 9.212024683 | 11.47440531 | 11.4175794 | 11.21251927 | 10.70353881 |
| ENSG00000165899 | OTOGL | 0.497902331 | 0.89458451 | 2.715591547 | 0.390470087 | 3.213493878 | 2 | 2 | 2 | 2.498691189 | 2.38539787 | 2.746468522 | 3.875378318 |
| ENSG00000160211 | G6PD | 1.978325781 | 0.12078466 | 1.233484174 | 0.348797592 | 3.211809955 | 10.05124847 | 7.145017454 | 6.882775709 | 10.72351064 | 10.7740351 | 9.549426 | 10.35294498 |
| ENSG00000134278 | SPIRE1 | 1.525711641 | 0.348069747 | 1.681773262 | 0.287145127 | 3.207484903 | 3.327711234 | 8.191702463 | 9.149036566 | 9.657432896 | 9.72643218 | 9.825944587 | 9.869095369 |
| ENSG00000103126 | AXIN1 | 1.126977345 | 0.425073921 | 2.08029856 | 0.109843769 | 3.207275905 | 7.435667355 | 7.761270542 | 3.369317972 | 8.149581172 | 8.19108423 | 9.350523544 | 8.831112735 |
| ENSG00000168944 | CEP120 | 1.796618131 | 0.494607899 | 1.409852552 | 0.592571552 | 3.206470683 | 2 | 7.847167909 | 2 | 7.992367495 | 8.13351654 | 8.067679772 | 7.161139769 |
| ENSG00000136052 | SLC41A2 | 2.773917665 | 0.001021144 | 0.428207414 | 0.673751517 | 3.202125079 | 5.532353208 | 4.11352228 | 6.43197674 | 8.291333947 | 8.26379462 | 6.242195894 | 5.722580018 |
| ENSG00000115275 | MOGS | 1.005622917 | 0.598708746 | 2.195779027 | 0.208181348 | 3.201401944 | 3.327711234 | 6.844429755 | 9.69246155 | 9.257971441 | 9.35044873 | 10.68654814 | 10.2622961 |
| ENSG00000135926 | TMBIM1 | 1.729050359 | 4.18E-06 | 1.472289804 | 0.000105356 | 3.201340163 | 10.11549941 | 9.338312924 | 9.926432745 | 11.49848994 | 11.6009408 | 11.25057591 | 11.33662661 |
| ENSG00000156587 | UBE2L6 | 2.771235371 | 1.07E-10 | 0.429636171 | 0.400966753 | 3.200871542 | 9.448362284 | 8.451443573 | 8.978369997 | 11.7599213 | 11.785461 | 9.32239471 | 9.54478665 |
| ENSG00000235109 | ZSCAN31 | 2.712266924 | 0.156635009 | 0.487526146 | 0.82326635 | 3.19979307 | 2 | 5.72254323 | 2 | 6.877205252 | 6.82036165 | 4.959178705 | 4.614976609 |
| ENSG00000182010 | RTKN2 | 0.555327983 | 0.781636058 | 2.642223993 | 0.118459494 | 3.197551976 | 2 | 4.11352228 | 2 | 3.754611435 | 3.2184422 | 4.952879058 | 5.319231103 |
| ENSG00000154133 | ROBO4 | 0.828347031 | 0.71541993 | 2.366368427 | 0.241080855 | 3.194715458 | 4.00722531 | 2 | 2 | 3.349643765 | 3.50452879 | 3.878466556 | 5.028505473 |
| ENSG00000092964 | DPYSL2 | 0.348082994 | 0.611410824 | 2.844214067 | 2.13E-07 | 3.192297061 | 9.33149386 | 9.540933215 | 9.344432377 | 9.80923777 | 9.70047233 | 11.41205386 | 12.77179957 |
| ENSG00000058453 | CROCC | 1.689883544 | 0.223753194 | 1.501954196 | 0.276627835 | 3.191837741 | 2 | 6.311476754 | 6.43197674 | 7.467753569 | 7.42465798 | 7.41654498 | 7.091143645 |
| ENSG00000047849 | MAP4 | 0.85643116 | 0.075173015 | 2.335106717 | 5.48E-08 | 3.191537876 | 10.81212061 | 11.46438781 | 11.81191322 | 12.2506787 | 12.2992933 | 13.65806653 | 13.84229602 |
| ENSG00000135924 | DNAJB2 | 1.75862777 | 0.000301529 | 1.432123083 | 0.003816216 | 3.190750852 | 7.096806373 | 7.35507212 | 7.97167868 | 9.20040236 | 9.30367823 | 9.177956973 | 8.62735196 |
| ENSG00000181104 | F2R | 1.726928389 | 0.200056839 | 1.463397099 | 0.277052271 | 3.190325488 | 8.777722034 | 7.60566235 | 4.523675391 | 9.520570052 | 9.44390794 | 9.664973224 | 8.575745325 |
| ENSG00000184371 | CSF1 | 1.636827642 | 0.000370047 | 1.552157628 | 0.000743084 | 3.188985269 | 9.567995545 | 8.876041558 | 9.287079172 | 10.88532146 | 10.9118811 | 10.39788531 | 11.13690957 |
| ENSG00000118804 | STBD1 | 3.029130793 | 0.019173548 | 0.159527005 | 0.922180661 | 3.188657798 | 6.257048221 | 4.442710129 | 2 | 7.852449262 | 8.05152279 | 5.233158834 | 5.217771644 |
| ENSG00000278812 | | 4.239394658 | 0.000129021 | -1.051620405 | 0.419522977 | 3.187774253 | 3.327711234 | 2 | 4.523675391 | 7.106324235 | 7.29937545 | 2.831336763 | 3.042026731 |
| ENSG00000137760 | ALKBH8 | 1.915034605 | 0.106975585 | 1.270944327 | 0.295788824 | 3.185978932 | 2 | 5.72254323 | 5.772716771 | 7.017154451 | 7.03319408 | 6.464399218 | 6.344709081 |
| ENSG00000132003 | ZSWIM4 | 1.810474987 | 0.181669225 | 1.374746603 | 0.314561325 | 3.18522159 | 4.00722531 | 5.72254323 | 7.934808822 | 8.43768081 | 8.50176361 | 7.678527483 | 8.328971229 |
| ENSG00000169992 | NLGN2 | 1.064438208 | 0.039785638 | 2.120657946 | 8.65E-06 | 3.185096155 | 8.981092835 | 8.469613413 | 9.589475645 | 10.13584795 | 10.1432423 | 11.06990214 | 11.30663426 |
| ENSG00000176619 | LMNB2 | 1.29018044 | 0.079160897 | 1.893450308 | 0.006446703 | 3.183630749 | 7.218819781 | 8.590771778 | 7.098165388 | 9.005540335 | 9.17609501 | 9.444446837 | 9.9049832 |
| ENSG00000251369 | ZNF550 | 1.3949007 | 0.214356852 | 1.787794796 | 0.096172658 | 3.182695497 | 3.327711234 | 6.228547981 | 5.156626401 | 6.755670824 | 6.57678965 | 7.403899888 | 6.574984961 |
| ENSG00000262862 | | 8.373406524 | 3.93E-07 | -5.191150377 | 0.033995019 | 3.182256146 | 2 | 2 | 2 | 8.43768081 | 8.39557285 | 2 | 2 |
| ENSG00000155744 | FAM126B | 1.731634634 | 0.085637804 | 1.448483333 | 0.15050827 | 3.180117967 | 4.467311984 | 7.315436877 | 6.203069917 | 8.134820652 | 8.09094333 | 8.057440132 | 7.570355956 |
| ENSG00000140937 | CDH11 | 1.049783193 | 0.1047992 | 2.130254815 | 0.000309103 | 3.180038009 | 9.435834703 | 10.81799725 | 9.703463497 | 11.18460494 | 11.1490131 | 12.27301174 | 12.21860406 |
| ENSG00000124370 | MCEE | 0.98953587 | 0.561910399 | 2.189810476 | 0.158634895 | 3.179346346 | 3.327711234 | 5.946638857 | 2 | 5.697663076 | 5.45984938 | 6.828438696 | 6.590289384 |
| ENSG00000110851 | PRDM4 | 1.771598147 | 0.160124234 | 1.405663451 | 0.267319871 | 3.177261598 | 4.00722531 | 7.60566235 | 8.076950877 | 9.037634995 | 9.10406516 | 8.666781604 | 8.748827534 |
| ENSG00000084764 | MAPRE3 | 1.792407453 | 0.187213731 | 1.379075319 | 0.314029219 | 3.171482772 | 3.327711234 | 5.838937808 | 7.600047325 | 8.14748175 | 8.21506984 | 7.709686122 | 7.837328594 |
| ENSG00000131791 | PRKAB2 | 0.91606495 | 0.629529096 | 2.255249125 | 0.189186264 | 3.171314075 | 2 | 8.169643011 | 7.691296561 | 8.258639605 | 8.29975519 | 10.08838597 | 8.88019044 |
| ENSG00000139505 | MTMR6 | 1.765519368 | 0.316926582 | 1.405303912 | 0.428859931 | 3.17082328 | 2 | 7.979858146 | 8.236946975 | 9.35577494 | 9.20831741 | 9.061649957 | 8.776966445 |
| ENSG00000034693 | PEX3 | 2.454654629 | 0.126821984 | 0.715214433 | 0.687732077 | 3.169869062 | 4.00722531 | 6.602424674 | 2 | 7.748245381 | 7.54938955 | 5.722275846 | 6.195912571 |
| ENSG00000143337 | TOR1AIP1 | 1.747383287 | 0.063359512 | 1.422392221 | 0.133187948 | 3.169775508 | 6.138360429 | 8.623598428 | 8.763901704 | 9.969934001 | 9.95726349 | 9.719336144 | 9.556197588 |
| ENSG00000163513 | TGFBR2 | 2.296062866 | 0.009222762 | 0.872647727 | 0.366230344 | 3.168710593 | 7.485143553 | 9.424669273 | 10.09465441 | 11.63155624 | 11.649055 | 10.0549539 | 10.36800786 |
| ENSG00000213214 | ARHGEF35 | 2.795738851 | 0.170233898 | 0.372113393 | 0.87553967 | 3.167852245 | 2 | 2 | 5.772716771 | 6.887305128 | 6.88657603 | 4.360042493 | 4.911532223 |
| ENSG00000112874 | NUDT12 | 1.062058495 | 0.646239362 | 2.10472761 | 0.325173109 | 3.166786105 | 2 | 6.311476754 | 2 | 5.94010643 | 5.76075522 | 7.0047514 | 6.662054875 |
| ENSG00000196924 | FLNA | 0.226305148 | 0.787571227 | 2.940116882 | 5.89E-06 | 3.16642203 | 12.39170213 | 13.27674723 | 14.17615456 | 13.66999705 | 13.7025084 | 16.32484542 | 16.47101995 |
| ENSG00000196456 | ZNF775 | 1.29018586 | 0.553674554 | 1.876047422 | 0.364727545 | 3.166233282 | 2 | 5.946638857 | 2 | 5.602782483 | 5.8245091 | 6.442083733 | 6.066922748 |
| ENSG00000134698 | AGO4 | 1.684221933 | 0.292538621 | 1.481432204 | 0.351097101 | 3.165654137 | 6.653098118 | 7.503546079 | 2 | 8.23312837 | 8.22100445 | 8.169666431 | 7.869046644 |
| ENSG00000198663 | C6orf89 | 1.206528573 | 3.23E-07 | 1.95844029 | 1.02E-17 | 3.164968863 | 9.031919741 | 9.067150531 | 9.049077186 | 10.23062141 | 10.2698233 | 11.0815887 | 10.91418426 |
| ENSG00000162444 | RBP7 | 0.49580458 | 0.888389865 | 2.667841846 | NA | 3.163646426 | 2 | 2 | 2 | 2.3892346 | 2.49394783 | 3.296906311 | 3.474731315 |
| ENSG00000104765 | BNIP3L | 1.169517402 | 0.004862033 | 1.992151526 | 3.87E-07 | 3.161668929 | 9.602033055 | 9.936721131 | 10.23015729 | 11.22772118 | 10.9849062 | 11.64438086 | 12.17327288 |
| ENSG00000156471 | PTDSS1 | 1.587010714 | 0.10807059 | 1.573405455 | 0.105765803 | 3.160416169 | 8.367060731 | 8.94259736 | 6.073378139 | 9.825086737 | 9.75670736 | 9.688861377 | 9.861494087 |
| ENSG00000105245 | NUMBL | 0.531084102 | 0.550985513 | 2.628638951 | 0.000420382 | 3.159723052 | 8.285626337 | 6.535001748 | 6.629488387 | 7.815902688 | 7.99029569 | 9.913907765 | 10.05224558 |
| ENSG00000242252 | BGLAP | 3.18149027 | 0.274207949 | -0.021934301 | 0.994944393 | 3.159555968 | 2 | 2 | 2 | 3.754611435 | 3.69849104 | 2.457042231 | 2.172608095 |
| ENSG00000154380 | ENAH | 0.326926092 | 0.472437308 | 2.832107976 | 7.92E-15 | 3.159034067 | 9.949165863 | 10.3486505 | 9.553445883 | 10.38561189 | 10.2428544 | 12.7687974 | 12.86450558 |

| | | | | | | | | | | | | | |
|---|---|---|---|---|---|---|---|---|---|---|---|---|---|
| ENSG00000135049 | AGTPBP1 | 1.245402354 | 0.39960716 | 1.910616191 | 0.168562587 | 3.156018545 | 2 | 6.666836837 | 5.772716771 | 6.965655398 | 6.91125523 | 7.452712791 | 7.708840864 |
| ENSG00000187239 | FNBP1 | 0.313532654 | 0.823523332 | 2.842409233 | 0.01185928 | 3.155941887 | 5.330594134 | 8.903033433 | 7.029880454 | 8.063125866 | 8.08661608 | 10.52408619 | 10.64535113 |
| ENSG00000169710 | FASN | 0.709820827 | 0.217903746 | 2.445873053 | 9.73E-07 | 3.15569388 | 9.761125267 | 10.62479571 | 9.868959282 | 10.83761346 | 10.8598983 | 12.15744732 | 12.91067024 |
| ENSG00000162144 | CYB561A3 | 1.890177232 | 0.000620266 | 1.264945766 | 0.027818382 | 3.155122998 | 7.435667355 | 7.189546432 | 8.380957451 | 9.596407142 | 9.65370449 | 8.808281231 | 9.176510366 |
| ENSG00000188229 | TUBB4B | 0.736145645 | 0.610764042 | 2.41855295 | 0.056285331 | 3.154698595 | 10.14658276 | 9.930175364 | 6.073378139 | 10.20157308 | 10.2707772 | 11.18632706 | 12.39785952 |
| ENSG00000164989 | CCDC171 | 2.052677102 | 0.298875829 | 1.101544266 | 0.59080501 | 3.154221367 | 2 | 3.686303414 | 2 | 4.139692846 | 4.22485516 | 3.724264246 | 3.236138336 |
| ENSG00000168234 | TTC39C | 1.850742811 | 0.001743699 | 1.298847874 | 0.033210265 | 3.149590685 | 5.095959458 | 5.131603625 | 5.39244166 | 6.946463121 | 6.93069905 | 6.564892306 | 6.232137896 |
| ENSG00000104497 | SNX16 | 1.617924239 | 0.461620063 | 1.529456903 | 0.478337468 | 3.147381142 | 2 | 6.04685525 | 2 | 6.237202172 | 5.99120031 | 6.290458506 | 5.727202496 |
| ENSG00000131127 | ZNF141 | 1.376220595 | 0.27007879 | 1.769675727 | 0.136351095 | 3.145896321 | 2 | 4.11352228 | 5.156626401 | 5.348759107 | 5.45984938 | 6.037140174 | 5.431121909 |
| ENSG00000023909 | GCLM | 2.17811235 | 0.023406655 | 0.965439718 | 0.351483224 | 3.143552068 | 5.532353208 | 7.467835385 | 8.460932916 | 9.747510575 | 9.72504082 | 8.402448739 | 8.651871454 |
| ENSG00000138771 | SHROOM3 | -0.502078289 | 0.488943881 | 3.644632636 | 3.80E-10 | 3.142554347 | 7.33131396 | 6.602424674 | 5.59507254 | 6.237202172 | 6.1572134 | 10.25462407 | 10.28186948 |
| ENSG00000052749 | RRP12 | 1.529845544 | 0.080997278 | 1.611530832 | 0.060381002 | 3.141376377 | 4.815609471 | 6.602424674 | 7.285536013 | 8.029321023 | 8.04707549 | 8.28197924 | 7.934467448 |
| ENSG00000019582 | CD74 | 0.595590002 | 0.722734802 | 2.545398029 | 0.078757712 | 3.14098803 | 3.327711234 | 6.311476754 | 8.206335842 | 7.546342806 | 7.61094772 | 9.953296359 | 8.855068552 |
| ENSG00000071537 | SEL1L | 1.65779824 | 0.038776418 | 1.483013292 | 0.064830738 | 3.140811532 | 7.624139768 | 9.733683991 | 9.553445883 | 10.93576629 | 10.8259041 | 10.74171855 | 10.67241823 |
| ENSG00000178385 | PLEKHM3 | 1.070037607 | 0.329903713 | 2.070509522 | 0.040228096 | 3.140547129 | 4.815609471 | 6.464272742 | 7.858115942 | 7.852449262 | 7.9408106 | 8.931137537 | 8.838608371 |
| ENSG00000151883 | PARP8 | 2.245778966 | 0.139563643 | 0.894667073 | 0.583522797 | 3.140446039 | 7.27616306 | 7.145017454 | 2 | 8.868232317 | 8.83412012 | 7.366471135 | 7.657697387 |
| ENSG00000145934 | TENM2 | 0.602496184 | 0.762843325 | 2.537206672 | 0.140420203 | 3.139702856 | 2 | 4.93638933 | 2 | 4.299492648 | 4.12711147 | 6.205571486 | 5.513708555 |
| ENSG00000152061 | RABGAP1L | 1.205999673 | 0.030197122 | 1.929745814 | 0.000214172 | 3.135745487 | 5.330594134 | 6.228547981 | 6.629488387 | 7.359462263 | 7.27295842 | 8.019511671 | 8.032963099 |
| ENSG00000156026 | MCU | 0.817652958 | 0.514396922 | 2.31684913 | 0.036346775 | 3.134502087 | 6.653098118 | 5.131603625 | 8.434763955 | 8.219202381 | 8.01327646 | 9.645817872 | 9.564293351 |
| ENSG00000188171 | ZNF626 | 1.780263211 | 0.016876187 | 1.351096154 | 0.075115142 | 3.131359365 | 4.467311984 | 5.131603625 | 5.156626401 | 6.722207358 | 6.47451726 | 6.490730008 | 5.821002687 |
| ENSG00000126391 | FRMD8 | 1.335496608 | 0.39305959 | 1.794143135 | 0.228416028 | 3.129639742 | 3.327711234 | 8.124485672 | 8.408111533 | 9.037634995 | 9.02486072 | 8.958123037 | 9.873011408 |
| ENSG00000140262 | TCF12 | 1.865944446 | 0.020534986 | 1.262610452 | 0.128777677 | 3.128554898 | 8.367060731 | 9.479495904 | 7.285536013 | 10.51378677 | 10.4971286 | 9.975607996 | 9.829070247 |
| ENSG00000171603 | CLSTN1 | 1.493908468 | 0.000723989 | 1.632825692 | 0.000179948 | 3.126734161 | 9.435834703 | 10.24169813 | 10.34701993 | 11.54707907 | 11.5582157 | 11.55836616 | 11.81320514 |
| ENSG00000163884 | KLF15 | -2.312004462 | 0.48457279 | 5.437613979 | 0.023854618 | 3.125609517 | 2 | 2 | 2 | 2 | 2.14022021 | 5.545162502 | 5.631815919 |
| ENSG00000099365 | STX1B | 2.315991592 | 0.226093212 | 0.807337786 | 0.696969597 | 3.123329377 | 2 | 5.457120953 | 2 | 6.221320894 | 6.18985553 | 4.5389195 | 5.080008298 |
| ENSG00000261587 | TMEM249 | 3.936331056 | 0.119879274 | -0.813188228 | 0.784238084 | 3.123142827 | 2 | 2 | 2 | 4.139692846 | 4.43005853 | 2.267001569 | 2.117335275 |
| ENSG00000123983 | ACSL3 | 1.904024408 | 0.05610284 | 1.218726581 | 0.237056099 | 3.122750989 | 6.468621203 | 9.377333006 | 8.236946975 | 10.37712591 | 10.3332939 | 9.42067733 | 9.887279953 |
| ENSG00000131507 | NDFIP1 | 1.757273518 | 9.86E-08 | 1.364266033 | 4.68E-05 | 3.121539551 | 8.468948322 | 9.090820894 | 8.99637432 | 10.64904058 | 10.6095994 | 10.27670249 | 10.19735668 |
| ENSG00000179546 | HTR1D | 0.725878283 | 0.771079266 | 2.393851486 | 0.272841778 | 3.119729769 | 2 | 2 | 3.369317972 | 2.695456494 | 3.01487888 | 4.133696865 | 3.496556178 |
| ENSG00000172575 | RASGRP1 | -2.512714196 | 0.034445258 | 5.630001759 | 7.62E-09 | 3.117287563 | 5.095959458 | 3.686303414 | 2 | 2.695456494 | 2.59489879 | 9.650204809 | 8.91098986 |
| ENSG00000212719 | C17orf51 | 1.28933874 | 0.441837321 | 1.827746458 | 0.249115502 | 3.117085198 | 6.366709671 | 7.431218269 | 2 | 7.555872146 | 7.81828345 | 8.593249736 | 7.722795734 |
| ENSG00000167861 | HID1 | -2.272313756 | 0.318802002 | 5.388292613 | 0.001147244 | 3.115978857 | 2 | 3.076665209 | 2 | 2.141737246 | 2.14022021 | 6.670403488 | 6.175763085 |
| ENSG00000153317 | ASAP1 | 1.215145914 | 0.344412784 | 1.899543476 | 0.115663273 | 3.11468939 | 6.366709671 | 9.956182102 | 10.28977157 | 10.83272206 | 10.7887607 | 11.3875211 | 11.59331975 |
| ENSG00000087008 | ACOX3 | 0.694260029 | 0.52990438 | 2.419589281 | 0.01125466 | 3.11384931 | 4.00722531 | 6.844429755 | 6.958202216 | 7.097651217 | 7.07311172 | 8.757226889 | 8.804020138 |
| ENSG00000111371 | SLC38A1 | 1.215754325 | 0.042514667 | 1.897463897 | 0.00077242 | 3.113218222 | 7.904282243 | 8.433041974 | 9.301632639 | 9.847245515 | 9.88808233 | 10.54085267 | 10.55349898 |
| ENSG00000173064 | HECTD4 | 1.338988539 | 0.024718288 | 1.774177207 | 0.001976772 | 3.113165747 | 9.27634997 | 8.101365653 | 8.110401118 | 9.909274163 | 9.96140527 | 10.49050086 | 10.23679159 |
| ENSG00000110172 | CHORDC1 | 1.748694895 | 0.356811536 | 1.364257323 | 0.474542652 | 3.112952218 | 2 | 7.189546432 | 4.058998049 | 7.507583254 | 7.47187556 | 7.535143176 | 6.520103598 |
| ENSG00000255112 | CHMP1B | 2.075936778 | 0.003866084 | 1.035444671 | 0.177968622 | 3.111381449 | 9.112843761 | 7.274681896 | 7.896971941 | 10.36136198 | 10.3568427 | 9.261890418 | 9.382110089 |
| ENSG00000167258 | CDK12 | 1.703701439 | 0.00072502 | 1.407570492 | 0.006022874 | 3.111271931 | 8.075571721 | 7.874700259 | 8.960138144 | 10.03554381 | 10.1014956 | 9.842651775 | 9.701867195 |
| ENSG00000120334 | CENPL | 1.448339037 | 0.267906024 | 1.662020766 | 0.188368879 | 3.110359803 | 3.327711234 | 5.946638857 | 3.369317972 | 6.229283386 | 6.05460702 | 6.442083733 | 6.245088435 |
| ENSG00000033030 | ZCCHC8 | 1.610344423 | 0.105038947 | 1.495251461 | 0.128390822 | 3.105595883 | 4.467311984 | 6.140560021 | 7.225742779 | 7.772969103 | 7.99953197 | 8.042684843 | 7.452476174 |
| ENSG00000130522 | JUND | 0.982717236 | 0.067009569 | 2.122408156 | 1.40E-05 | 3.105125392 | 8.75772933 | 9.856140801 | 9.647595056 | 10.44194384 | 10.5056622 | 11.40348951 | 11.79350523 |
| ENSG00000221890 | NPTXR | 0.367646493 | 0.877879848 | 2.73649753 | 0.165909543 | 3.104144023 | 3.327711234 | 7.638155693 | 2 | 6.516352584 | 6.51372389 | 8.723392661 | 8.933165411 |
| ENSG00000189410 | SH2D5 | 1.397532436 | 0.319054857 | 1.705675355 | 0.206700396 | 3.103207791 | 3.327711234 | 4.11352228 | 6.718950348 | 6.699459234 | 6.81514011 | 6.907119087 | 7.191563076 |
| ENSG00000103034 | NDRG4 | 3.087594726 | 0.127598657 | 0.015520842 | 0.994944393 | 3.103115568 | 2 | 6.228547981 | 2 | 7.772969103 | 7.66419009 | 5.102662916 | 4.511590831 |
| ENSG00000175305 | CCNE2 | 1.77137633 | 0.458136023 | 1.33006145 | 0.576450932 | 3.101437779 | 2 | 3.076665209 | 2 | 3.349643765 | 3.50452879 | 3.44788957 | 2.817676617 |
| ENSG00000162222 | TTC9C | 1.742105539 | 0.33524083 | 1.358228695 | 0.455315905 | 3.100334234 | 4.815609471 | 7.538394143 | 2 | 7.947219213 | 7.8208909 | 7.667989909 | 7.330190599 |
| ENSG00000108557 | RAI1 | 1.65881768 | 0.410884798 | 1.441231161 | 0.47216327 | 3.100048842 | 2 | 7.954278496 | 9.589475645 | 10.02529042 | 10.0923519 | 9.658455026 | 10.00568113 |
| ENSG00000101350 | KIF3B | 1.321676713 | 0.129712892 | 1.776313661 | 0.03302563 | 3.097990375 | 8.009475818 | 9.726146444 | 7.896971941 | 10.07252815 | 10.1772163 | 10.22094162 | 10.86574466 |
| ENSG00000082898 | XPO1 | 2.132119181 | 0.003707011 | 0.964536394 | 0.224818109 | 3.096655575 | 7.82963488 | 9.849218119 | 9.358421373 | 11.40931048 | 11.3114888 | 10.32567245 | 10.0543164 |
| ENSG00000165733 | BMS1 | 1.814578119 | 0.003642609 | 1.28121448 | 0.047023433 | 3.095792599 | 6.653098118 | 8.124485672 | 8.110401118 | 9.565291356 | 9.58019637 | 8.860117405 | 9.204958522 |
| ENSG00000128607 | KLHDC10 | 1.281418267 | 0.370174297 | 1.813345371 | 0.180586512 | 3.094763638 | 4.00722531 | 8.687092496 | 7.734844356 | 9.080080551 | 8.93625225 | 9.55463569 | 9.522022954 |
| ENSG00000274611 | TBC1D3 | 8.283594897 | 7.42E-07 | -5.191150183 | 0.035315165 | 3.092444713 | 2 | 2 | 2 | 8.302698479 | 8.35297033 | 2 | 2 |
| ENSG00000068394 | GPKOW | 1.531667536 | 0.487909731 | 1.560652422 | 0.470583967 | 3.092319958 | 3.327711234 | 7.847167909 | 2 | 7.821180652 | 7.86703672 | 7.684242973 | 8.039482273 |
| ENSG00000169313 | P2RY12 | 4.083114711 | 0.094962877 | -0.994016814 | 0.726496359 | 3.089097897 | 2 | 2 | 2 | 4.598938466 | 4.1930041 | 2.225793514 | 2.117335275 |
| ENSG00000105559 | PLEKHA4 | 2.801995215 | 3.35E-05 | 0.286699249 | 0.732333373 | 3.088694465 | 8.816894989 | 7.35507212 | 9.099922433 | 11.37876293 | 11.3862332 | 8.56564741 | 9.133628617 |
| ENSG00000114013 | CD86 | -1.310497508 | 0.677345605 | 4.399110958 | 0.073198426 | 3.088613449 | 2 | 2 | 2 | 2.141737246 | 2.14022021 | 5.119635168 | 4.018014792 |
| ENSG00000205572 | SERF1B | 8.278026389 | 7.69E-07 | -5.191150175 | 0.035365736 | 3.086876214 | 2 | 2 | 2 | 8.304583895 | 8.34030387 | 2 | 2 |
| ENSG00000187630 | DHRS4L2 | 1.941512121 | 0.177844856 | 1.144727819 | 0.442622978 | 3.08623994 | 2 | 4.93638933 | 6.43197674 | 7.097651217 | 7.20671394 | 6.022174708 | 6.676467423 |
| ENSG00000277974 | | 8.275165251 | 7.89E-07 | -5.191150164 | 0.035434326 | 3.084015088 | 2 | 2 | 2 | 8.293234263 | 8.34574597 | 2 | 2 |
| ENSG00000278299 | TBC1D3C | 8.275165251 | 7.89E-07 | -5.191150164 | 0.035434326 | 3.084015088 | 2 | 2 | 2 | 8.293234263 | 8.34574597 | 2 | 2 |
| ENSG00000011405 | PIK3C2A | 1.695122148 | 0.045753442 | 1.388714652 | 0.104797108 | 3.0838368 | 6.366709671 | 8.687092496 | 8.325066185 | 9.783781964 | 9.75874862 | 9.621944737 | 9.292162928 |
| ENSG00000174059 | CD34 | 1.119640448 | 0.680321745 | 1.963435452 | 0.435161964 | 3.0830759 | 2 | 2 | 3.369317972 | 3.023133209 | 3.08597292 | 2.68698273 | 4.09155917 |
| ENSG00000115825 | PRKD3 | 2.017413346 | 0.002635445 | 1.064437907 | 0.135544734 | 3.081851253 | 8.981092835 | 7.393647515 | 7.691296561 | 10.15164584 | 10.2360327 | 9.242329292 | 9.249822 |
| ENSG00000070159 | PTPN3 | -2.424045793 | 0.099653945 | 5.504713225 | 7.53E-07 | 3.080667432 | 2 | 3.076665209 | 4.058998049 | 2.270786764 | 2.38539787 | 8.04490778 | 7.930464419 |
| ENSG00000156869 | FRRS1 | 2.07026914 | 0.12909603 | 1.008562002 | 0.483213396 | 3.078831142 | 2 | 3.076665209 | 4.874587708 | 5.461211713 | 5.44640346 | 4.596630906 | 4.456980017 |
| ENSG00000138380 | CARF | 1.922716366 | 0.005958463 | 1.155803344 | 0.114203801 | 3.078519709 | 5.095959458 | 4.710564154 | 4.874587708 | 6.687949235 | 6.63721363 | 6.181526565 | 5.636738041 |
| ENSG00000282087 | | 8.269521542 | 8.20E-07 | -5.191150151 | 0.035498586 | 3.078371391 | 2 | 2 | 2 | 8.287525788 | 8.34030387 | 2 | 2 |
| ENSG00000141424 | SLC39A6 | 1.817909711 | 0.119309305 | 1.260079379 | 0.290138932 | 3.07798909 | 6.138360429 | 9.842262059 | 8.922968322 | 10.77711405 | 10.7271979 | 10.16345022 | 10.22805302 |
| ENSG00000011028 | MRC2 | 1.913487605 | 6.77E-27 | 1.164091828 | 1.22E-10 | 3.077579433 | 11.26581873 | 11.40661699 | 11.38635888 | 13.23212001 | 13.3004149 | 12.53234403 | 12.50284941 |
| ENSG00000154727 | GABPA | 1.819864845 | 0.239927272 | 1.257690958 | 0.426580825 | 3.077555802 | 2 | 6.728495673 | 7.225742779 | 8.209172298 | 8.20711886 | 7.872065621 | 7.399884913 |
| ENSG00000275153 | | 8.268573456 | 8.26E-07 | -5.191150148 | 0.035498785 | 3.077423308 | 2 | 2 | 2 | 8.285617933 | 8.34030387 | 2 | 2 |
| | LOC1010603 | 8.268573456 | 8.26E-07 | -5.191150148 | 0.035498785 | 3.077423308 | 2 | 2 | 2 | 8.285617933 | 8.34030387 | 2 | 2 |
| ENSG00000143479 | DYRK3 | 2.504819184 | 0.165298109 | 0.57071095 | 0.778729259 | 3.075530134 | 8.16936288 | 5.457120953 | 2 | 9.258944141 | 9.27258642 | 7.426809156 | 7.285445324 |
| ENSG00000143162 | CREG1 | 2.201017869 | 5.60E-05 | 0.872724827 | 0.144905041 | 3.073742697 | 8.963744667 | 7.901717006 | 7.818184403 | 10.50438458 | 10.5189709 | 9.356203439 | 9.0005841 |
| ENSG00000069869 | NEDD4 | 2.132358276 | 0.200056839 | 0.940604686 | 0.594921579 | 3.072962962 | 7.867441304 | 6.535001748 | 2 | 8.846395982 | 8.90095288 | 8.032265607 | 7.261744957 |
| ENSG00000119537 | KDSR | 1.862210167 | 2.17E-12 | 1.209028442 | 8.59E-06 | 3.071238609 | 8.228665467 | 8.276714465 | 8.434763955 | 10.22615207 | 10.096662 | 9.561380283 | 9.461931741 |
| ENSG00000147874 | HAUS6 | 1.795062027 | 0.100196808 | 1.275743301 | 0.251602132 | 3.070805328 | 4.00722531 | 6.04685525 | 7.285536013 | 8.091796043 | 7.99953197 | 7.675661224 | 7.382408159 |
| ENSG00000168488 | ATXN2L | 1.511308239 | 1.87E-09 | 1.559127061 | 4.43E-10 | 3.070435301 | 8.716893051 | 8.671479029 | 8.884815469 | 10.23458257 | 10.2845385 | 10.37656901 | 10.23476408 |
| ENSG00000148484 | RSU1 | 1.035327784 | 0.000888639 | 2.032293662 | 4.35E-12 | 3.067621447 | 8.609439662 | 8.834582582 | 8.904018013 | 9.829021922 | 9.80557618 | 10.63066452 | 10.97155887 |
| ENSG00000090470 | PDCD7 | 1.359872287 | 0.16375923 | 1.705967992 | 0.068587835 | 3.065840279 | 4.815609471 | 7.002537973 | 7.502634626 | 8.145379268 | 8.14188248 | 8.751340201 | 8.158913605 |
| ENSG00000198793 | MTOR | 1.223706216 | 0.000107077 | 1.841882538 | 1.39E-09 | 3.065588754 | 8.042902253 | 8.57407404 | 8.325066185 | 9.562930393 | 9.54040713 | 10.14336241 | 10.19067811 |
| ENSG00000075399 | VPS9D1 | 1.698964941 | 0.292000423 | 1.360409629 | 0.398956117 | 3.05937457 | 7.579279349 | 6.535001748 | 2 | 8.254744042 | 8.23084153 | 7.88783736 | 7.930464419 |
| ENSG00000107758 | PPP3CB | 0.891548837 | 0.433930421 | 2.167519725 | 0.033633667 | 3.059068562 | 5.330594134 | 8.540087354 | 7.734844356 | 8.634896 | 8.54182783 | 9.764548842 | 9.942486645 |
| ENSG00000182197 | EXT1 | 1.40258299 | 0.012710848 | 1.656465928 | 0.002554195 | 3.059048918 | 10.59831544 | 10.23108619 | 9.287079172 | 11.52136866 | 11.5517552 | 11.61737522 | 11.94454653 |
| ENSG00000068976 | PYGM | -0.464372904 | 0.802617873 | 3.522924727 | 0.017580275 | 3.058551823 | 2 | 4.710564154 | 2 | 3.09461492 | 3.50452879 | 6.615624322 | 6.622902297 |
| ENSG00000258644 | SYNJ2BP-CO | 8.248820207 | 9.40E-07 | -5.191150112 | 0.035746075 | 3.057670095 | 2 | 2 | 2 | 8.29892024 | 8.28849576 | 2 | 2 |
| ENSG00000167182 | SP2 | 1.136415155 | 0.231366574 | 1.920982143 | 0.03002915 | 3.057397298 | 6.89194353 | 6.844429755 | 4.523675391 | 7.517371202 | 7.54308677 | 7.912394292 | 8.60805852 |
| ENSG00000103599 | IQCH | 1.435999456 | 0.481906026 | 1.621306168 | 0.409695273 | 3.057305624 | 2 | 3.686303414 | 2 | 3.79808662 | 3.65239003 | 4.017766949 | 3.67940853 |
| ENSG00000146021 | KLHL3 | 3.337599882 | 0.003692538 | -0.280483606 | 0.845736528 | 3.057116276 | 3.327711234 | 5.131603625 | 2 | 7.097651217 | 7.14977896 | 4.005644654 | 3.807156736 |
| ENSG00000215187 | FAM166B | -1.316847075 | 0.744846249 | 4.373650241 | 0.191114239 | 3.056803166 | 2 | 2 | 2 | 2 | 2.2680111 | 4.785723309 | 4.500832793 |
| ENSG00000247626 | MARS2 | 1.581667375 | 0.330982566 | 1.47437911 | 0.359439631 | 3.056046485 | 2 | 5.838937808 | 3.369317972 | 6.061031724 | 6.02777321 | 6.132200015 | 5.727202496 |
| ENSG00000159228 | CBR1 | 1.744931257 | 0.000373498 | 1.310982387 | 0.009285622 | 3.055913644 | 7.485143553 | 8.487557258 | 8.631919287 | 10.04685164 | 9.98775074 | 9.589571269 | 9.581626682 |
| ENSG00000126368 | NR1D1 | 2.499315824 | 0.085143446 | 0.555281488 | 0.734896874 | 3.054597311 | 9.081016759 | 5.303523747 | 5.39244166 | 10.13372843 | 10.2084185 | 8.247007147 | 8.236348546 |
| ENSG00000173852 | DPY19L1 | 1.822552495 | 0.140480229 | 1.230582365 | 0.330230765 | 3.05313486 | 5.095959458 | 8.505280662 | 6.203069917 | 9.1138055 | 9.09224234 | 8.432756021 | 8.595472797 |
| | NARR | 8.244231739 | 9.99E-07 | -5.191150075 | 0.035936624 | 3.053081664 | 2 | 2 | 2 | 8.328872678 | 8.248381 | 2 | 2 |
| ENSG00000135218 | CD36 | 0.151052059 | 0.954430302 | 2.90144188 | 0.185697679 | 3.052493939 | 3.327711234 | 2 | 3.369317972 | 3.162721474 | 2.94009902 | 5.870008604 | 2.918126211 |
| ENSG00000109079 | TNFAIP1 | 0.062077921 | 0.96300973 | 2.989986037 | 0.004649784 | 3.052063959 | 9.940323746 | 9.433952963 | 6.43197674 | 9.225498809 | 9.29244949 | 12.02479168 | 12.31842831 |
| ENSG00000162817 | C1orf115 | 0.342943731 | 0.842753717 | 2.708350396 | 0.054562067 | 3.051294127 | 2 | 5.595927442 | 4.058998049 | 4.907595604 | 4.811949 | 6.126613714 | 7.617180085 |
| ENSG00000162924 | REL | 1.871703693 | 0.017393053 | 1.178624655 | 0.149212315 | 3.050328348 | 4.467311984 | 6.464272742 | 5.39244166 | 7.605658789 | 7.37229174 | 6.867478139 | 6.771278116 |
| ENSG00000266094 | RASSF5 | 0.791701709 | 0.645184945 | 2.257872853 | 0.139643037 | 3.049574561 | 3.327711234 | 5.838937808 | 2 | 5.540148523 | 5.04390545 | 6.625560108 | 6.739599323 |
| ENSG00000163701 | IL17RE | 0.340532129 | 0.871671212 | 2.708252683 | 0.112770502 | 3.048784812 | 2 | 4.11352228 | 2 | 3.462030052 | 3.28037825 | 4.720468739 | 5.566251551 |
| ENSG00000116871 | MAP7D1 | 1.090396929 | 0.035977572 | 1.958376397 | 5.08E-05 | 3.048773325 | 9.358295023 | 9.792607274 | 10.49998282 | 11.00233175 | 11.0915972 | 11.90263765 | 11.92606907 |
| ENSG00000139218 | SCAF11 | 0.96351038 | 0.157701036 | 2.08394333 | 0.000732621 | 3.047453709 | 8.653385977 | 10.25223258 | 9.465702942 | 10.55158859 | 10.5760362 | 11.56683474 | 11.7890907 |
| ENSG00000174579 | MSL2 | 1.450908327 | 0.453416873 | 1.595575369 | 0.393866241 | 3.046483696 | 2 | 4.710564154 | 7.734844356 | 7.650823203 | 7.8208909 | 8.030023089 | 7.713507501 |
| ENSG00000184216 | IRAK1 | 1.668925781 | 0.001995992 | 1.375625725 | 0.012347695 | 3.044551506 | 9.2903348 | 9.067150531 | 10.23015729 | 11.23614254 | 11.3278511 | 10.84981999 | 11.11767423 |
| ENSG00000166012 | TAF1D | 1.33857129 | 0.094280197 | 1.701974389 | 0.026587498 | 3.04054568 | 7.624139768 | 6.04685525 | 5.930865648 | 8.060896687 | 8.04707549 | 8.547470747 | 8.26365177 |

| | | | | | | | | | | | | | |
|---|---|---|---|---|---|---|---|---|---|---|---|---|---|
| ENSG00000218336 | TENM3 | 2.056129334 | 0.311473389 | 0.98015118 | 0.646868487 | 3.036280513 | 5.095959458 | 8.623598428 | 2 | 9.225498809 | 9.18320541 | 7.861174222 | 8.372406522 |
| ENSG00000110031 | LPXN | 4.871090537 | 0.001952423 | -1.835219961 | 0.289910617 | 3.035870576 | 4.00722531 | 4.11352228 | 8.434763955 | 11.78326477 | 11.8212241 | 5.737015686 | 4.522269241 |
| ENSG00000104687 | GSR | 1.945657821 | 0.349594603 | 1.090114584 | 0.61339599 | 3.035772405 | 8.998234871 | 5.457120953 | 2 | 9.484432763 | 9.44221542 | 8.681143984 | 8.544657203 |
| ENSG00000005810 | MYCBP2 | 1.979897114 | 0.007220064 | 1.050608998 | 0.17880563 | 3.030506112 | 8.816894989 | 8.876041558 | 6.958202216 | 10.35864221 | 10.4716357 | 9.51268942 | 9.468503442 |
| ENSG00000186318 | BACE1 | 1.49181805 | 0.000460725 | 1.538438352 | 0.000269051 | 3.030256401 | 8.797441463 | 8.916342327 | 8.042706615 | 10.0993857 | 10.144804 | 10.28315082 | 10.04439554 |
| ENSG00000177679 | SRRM3 | 0.301026642 | 0.858274786 | 2.729177194 | 0.044174878 | 3.030203836 | 5.866944937 | 3.686303414 | 2 | 4.907595604 | 4.87352701 | 7.042323222 | 7.252154709 |
| ENSG00000124343 | XG | 8.220939248 | 1.15E-06 | -5.191150033 | 0.036232528 | 3.029789215 | 2 | 2 | 2 | 8.291333947 | 8.240612 | 2 | 2 |
| ENSG00000274791 | F8A2 | 8.220466611 | 1.21E-06 | -5.191149991 | 0.036373066 | 3.029316619 | 2 | 2 | 2 | 8.207157886 | 8.3220138 | 2 | 2 |
| ENSG00000277150 | F8A3 | 8.220466611 | 1.21E-06 | -5.191149991 | 0.036373066 | 3.029316619 | 2 | 2 | 2 | 8.207157886 | 8.3220138 | 2 | 2 |
| ENSG00000042832 | TG | 3.846210474 | 0.067133789 | -0.817590789 | 0.73782085 | 3.028619684 | 2 | 3.076665209 | 2 | 4.965943888 | 5.07918227 | 2.139668783 | 2.551896972 |
| ENSG00000186847 | KRT14 | -3.229474305 | 0.156783493 | 6.2577098 | 0.000264451 | 3.028235495 | 2 | 2 | 4.058998049 | 2.270786764 | 2 | 7.44824109 | 8.72879856 |
| ENSG00000134285 | FKBP11 | 2.670839676 | 0.000124594 | 0.357069745 | 0.674724562 | 3.027909421 | 7.667647223 | 7.099070191 | 5.59507254 | 9.640365415 | 9.66898051 | 7.319501608 | 7.417152486 |
| ENSG00000152767 | FARP1 | 0.657285673 | 0.667164929 | 2.370106338 | 0.075077708 | 3.02739201 | 6.816637977 | 9.863030423 | 6.073378139 | 9.216514298 | 9.1617682 | 10.97536401 | 10.81059428 |
| ENSG00000213390 | ARHGAP19 | 1.180166259 | 0.487140735 | 1.846314164 | 0.246641741 | 3.026480423 | 2 | 5.946638857 | 3.369317972 | 5.786686927 | 5.73886209 | 6.353989995 | 6.423723772 |
| ENSG00000078814 | MYH7B | 0.429740808 | 0.867562397 | 2.596671108 | 0.220732836 | 3.026411916 | 2 | 3.076665209 | 2 | 2.868578001 | 2.59489879 | 4.155805465 | 3.824516949 |
| ENSG00000153237 | CCDC148 | 0.015684726 | 0.997041173 | 3.009269496 | 0.331063165 | 3.024954222 | 2 | 2 | 2 | 2.498691189 | 2.14022021 | 3.44788957 | 3.753786374 |
| ENSG00000156172 | C8orf37 | 0.725760001 | 0.514678379 | 2.298906217 | 0.016733631 | 3.024666218 | 4.815609471 | 3.686303414 | 3.369317972 | 4.470469939 | 4.83276841 | 5.922534244 | 6.215784511 |
| ENSG00000131089 | ARHGEF9 | -0.178082364 | 0.911633612 | 3.199558121 | 0.010279412 | 3.021475757 | 3.327711234 | 7.232742069 | 7.858115942 | 6.882264028 | 6.83591385 | 10.05219399 | 10.31813217 |
| ENSG00000136059 | VILL | 1.733217968 | 0.214218894 | 1.288001911 | 0.360920918 | 3.02121988 | 3.327711234 | 4.93638933 | 2 | 5.194621444 | 5.62411716 | 5.248671144 | 4.766376438 |
| ENSG00000182667 | NTM | 2.372084227 | 0.205285997 | 0.648800587 | 0.75370958 | 3.020884815 | 6.257048221 | 2 | 7.934808822 | 9.074566443 | 9.08900105 | 8.008923105 | 6.251520369 |
| ENSG00000138463 | DIRC2 | 1.427352618 | 0.466982135 | 1.592226903 | 0.401725514 | 3.019579521 | 2 | 7.701025996 | 4.523675391 | 7.671425626 | 7.70145652 | 7.732851095 | 7.955303035 |
| ENSG00000166886 | NAB2 | 1.518505827 | 0.002152817 | 1.500463068 | 0.002335708 | 3.018968895 | 7.904282243 | 7.467835385 | 8.561105505 | 9.508351943 | 9.57634363 | 9.40316733 | 9.63702421 |
| ENSG00000126858 | RHOT1 | 1.598548807 | 0.398338237 | 1.420157104 | 0.449551169 | 3.018705912 | 6.138360429 | 8.376385542 | 2 | 8.710792067 | 8.60823803 | 8.818278819 | 8.047821014 |
| ENSG00000073146 | MOV10L1 | 1.4834135 | 0.52563759 | 1.534255736 | 0.496731642 | 3.017669237 | 2 | 3.076665209 | 2 | 3.349643765 | 3.15372745 | 3.355392153 | 3.209956562 |
| ENSG00000165494 | PCF11 | 2.041286071 | 0.000236772 | 0.975246255 | 0.103004472 | 3.016532327 | 7.709880939 | 7.538394143 | 6.80318718 | 9.381905119 | 9.45821469 | 8.731695932 | 7.868000534 |
| ENSG00000139899 | CBLN3 | 1.791792236 | 0.45900055 | 1.224674195 | 0.617024792 | 3.016466431 | 2 | 7.002537973 | 2 | 7.178012736 | 7.26532099 | 6.52722803 | 6.806650651 |
| ENSG00000213341 | CHUK | 1.969665388 | 6.05E-06 | 1.046706362 | 0.023911633 | 3.01637175 | 6.653098118 | 6.602424674 | 6.882775709 | 8.70652031 | 8.57000035 | 7.953993861 | 7.463599504 |
| ENSG00000169136 | ATF5 | 2.610689615 | 3.96E-06 | 0.405128474 | 0.550387484 | 3.01581809 | 8.797441463 | 7.431218269 | 7.777116083 | 10.71112597 | 10.7114757 | 8.443467705 | 8.589138522 |
| ENSG00000179242 | CDH4 | 4.73959828 | 9.76E-06 | -1.723822982 | 0.160349824 | 3.015775299 | 3.327711234 | 2 | 4.523675391 | 7.745471964 | 7.63485389 | 2.746468522 | 2.551896972 |
| ENSG00000243207 | PPAN-P2RY1 | 8.205535819 | 1.40E-06 | -5.19114992 | 0.036768558 | 3.014385899 | 2 | 2 | 2 | 8.323303759 | 8.17486938 | 2 | 2 |
| ENSG00000159459 | UBR1 | 1.718805332 | 0.352202552 | 1.292700591 | 0.487618515 | 3.011505923 | 2 | 7.73146271 | 8.763901704 | 9.455026484 | 9.46822882 | 9.130251861 | 8.940150381 |
| ENSG00000179562 | GCC1 | 1.410737864 | 0.620468344 | 1.600010763 | 0.564099325 | 3.010748626 | 2 | 8.687092496 | 2 | 8.48998335 | 8.54813611 | 8.829927393 | 8.571895748 |
| ENSG00000068097 | HEATR6 | 1.187280873 | 0.476737643 | 1.822429944 | 0.245051346 | 3.009710817 | 4.00722531 | 7.874700259 | 4.058998049 | 7.590284224 | 7.6986238 | 8.301943231 | 8.237969002 |
| ENSG00000054523 | KIF1B | 1.239287775 | 8.10E-07 | 1.770395413 | 4.21E-13 | 3.009683188 | 8.716893051 | 8.862353853 | 8.585098385 | 9.954406146 | 9.97141463 | 10.49403038 | 10.49048921 |
| ENSG00000105397 | TYK2 | 1.338482146 | 0.48903311 | 1.669877245 | 0.367889318 | 3.008359391 | 8.313285031 | 9.067150531 | 2 | 9.429335066 | 9.54672161 | 9.810131135 | 9.82772573 |
| ENSG00000134247 | PTGFRN | 0.515132849 | 0.698766691 | 2.490654101 | 0.027497225 | 3.00578695 | 5.330594134 | 8.834582582 | 8.904018013 | 8.81102077 | 8.90095288 | 10.58734728 | 11.02533189 |
| ENSG00000131398 | KCNC3 | -3.753139074 | 0.256437356 | 6.758319752 | 0.007323777 | 3.005180679 | 2 | 2 | 2 | 2 | 2 | 6.950450741 | 6.711881188 |
| ENSG00000165490 | DDIAS | 2.060358502 | 0.307879292 | 0.943492214 | 0.658783502 | 3.003850716 | 2 | 3.686303414 | 2 | 4.497095146 | 3.7864959 | 3.500668998 | 3.287118176 |
| ENSG00000198353 | HOXC4 | 4.844761182 | 2.60E-05 | -1.844702684 | 0.155606188 | 3.000058498 | 4.467311984 | 4.11352228 | 2 | 8.304583895 | 8.26762243 | 3.12825509 | 2.374681325 |
| ENSG00000133612 | AGAP3 | 1.325540086 | 0.486515971 | 1.674084493 | 0.359872083 | 2.999624579 | 8.797441463 | 8.469613413 | 2 | 9.319871109 | 9.43542545 | 9.8644215 | 9.571701541 |
| ENSG00000272325 | NUDT3 | 2.735165094 | 0.01977227 | 0.261345905 | 0.855356794 | 2.996510999 | 2 | 6.04685525 | 5.59507254 | 7.939959586 | 7.92150349 | 5.390603804 | 5.713290392 |
| ENSG00000196935 | SRGAP1 | 2.074609006 | 0.059194088 | 0.921819938 | 0.434090596 | 2.996428944 | 8.340423424 | 7.145017454 | 4.874587708 | 9.374743937 | 9.45234084 | 8.254069715 | 8.28888532 |
| ENSG00000106351 | AGFG2 | 0.688830115 | 0.651205609 | 2.307091323 | 0.084128005 | 2.995921438 | 2 | 6.04685525 | 4.523675391 | 5.540148523 | 5.6826301 | 6.865802559 | 7.386797235 |
| ENSG00000134020 | PEBP4 | -0.727649172 | 0.837764017 | 3.722083119 | 0.189633354 | 2.994433947 | 2 | 2 | 2 | 2.141737246 | 2.2680111 | 4.659878161 | 3.261853426 |
| ENSG00000126353 | CCR7 | -1.310500242 | 0.668579277 | 4.30478236 | 0.070233601 | 2.994282119 | 2 | 2 | 2 | 2.141737246 | 2.14022021 | 4.406883583 | 4.757365452 |
| ENSG00000119403 | PHF19 | 0.265866542 | 0.885649929 | 2.727173883 | 0.066744424 | 2.993040425 | 5.532353208 | 7.232742069 | 2 | 6.463327516 | 6.18985553 | 8.75224741 | 8.716646332 |
| ENSG00000120008 | WDR11 | 1.402012553 | 0.372007399 | 1.590932431 | 0.295637434 | 2.992944984 | 3.327711234 | 8.687092496 | 8.296287027 | 9.245266304 | 9.40880824 | 9.518299895 | 9.516025051 |
| ENSG00000271321 | CTAGE6 | 5.839248142 | 0.013180916 | -2.84661821 | 0.320401275 | 2.992629932 | 2 | 2 | 2 | 5.978405189 | 5.94414054 | 2.094598369 | 2 |
| ENSG00000174456 | C12orf76 | 1.039912515 | 0.517016221 | 1.950113329 | 0.185017104 | 2.990025844 | 2 | 5.595927442 | 3.369317972 | 5.44762523 | 5.22805098 | 6.21875862 | 6.148451724 |
| ENSG00000151693 | ASAP2 | 1.637379061 | 0.10807059 | 1.351887694 | 0.185412187 | 2.989266755 | 5.70932728 | 6.844429755 | 8.486635645 | 9.018234236 | 9.10620439 | 8.823035893 | 8.735123394 |
| ENSG00000082196 | C1QTNF3 | 1.987246121 | 0.259802912 | 1.001367237 | 0.587346695 | 2.988613358 | 2 | 3.076665209 | 3.369317972 | 4.299492648 | 4.0930011 | 3.709392871 | 3.236138336 |
| ENSG00000166228 | PCBD1 | 0.714424984 | 0.559383109 | 2.27396168 | 0.03472132 | 2.988386664 | 4.467311984 | 7.901717006 | 6.882775709 | 7.725907304 | 7.65252709 | 9.044347772 | 9.395603187 |
| ENSG00000102854 | MSLN | 0.279381352 | 0.943520433 | 2.708508551 | NA | 2.987889903 | 2 | 2 | 2 | 2.600425781 | 2.14022021 | 2.937246078 | 3.771796761 |
| ENSG00000277443 | MARCKS | 1.596454329 | 0.036506808 | 1.390921256 | 0.069687163 | 2.987375585 | 9.081016759 | 10.80015962 | 10.29705308 | 11.82638973 | 11.8063345 | 11.00639811 | 12.03593134 |
| ENSG00000282185 | | -0.323181186 | 0.863478741 | 3.310292978 | 0.024438946 | 2.987111793 | 2 | 4.710564154 | 2 | 3.406930997 | 3.39680241 | 6.351597515 | 6.482320745 |
| ENSG00000015171 | ZMYND11 | 1.254422611 | 0.459689613 | 1.732404009 | 0.280587322 | 2.98682662 | 3.327711234 | 8.147241016 | 9.301632639 | 9.53348965 | 9.48723044 | 10.08298767 | 9.882625765 |
| ENSG00000078304 | PPP2R5C | 1.104527194 | 0.27893875 | 1.8822516 | 0.047119088 | 2.986778794 | 7.624139768 | 8.94259736 | 6.203069917 | 9.125585405 | 9.0561812 | 9.951322636 | 9.772331943 |
| ENSG00000146909 | NOM1 | 1.150050318 | 0.509715065 | 1.836126306 | 0.260870186 | 2.986176624 | 3.327711234 | 5.131603625 | 8.296287027 | 8.056427969 | 7.98565528 | 8.785862385 | 8.606804877 |
| ENSG00000168676 | KCTD19 | 0.012816003 | 0.997163678 | 2.972891288 | 0.307389297 | 2.985707291 | 2 | 2 | 2 | 2.3892346 | 2.2680111 | 3.296906311 | 3.824516949 |
| ENSG00000111490 | TBC1D30 | -0.307135341 | 0.93146871 | 3.292818809 | 0.25649001 | 2.985683468 | 2 | 2 | 2 | 2.3892346 | 2.14022021 | 3.968654369 | 3.620995423 |
| ENSG00000085377 | PREP | 1.653046962 | 0.167653704 | 1.331125875 | 0.268158791 | 2.984172837 | 4.815609471 | 8.395519306 | 6.882775709 | 8.918344554 | 9.01805981 | 8.431623835 | 8.839676004 |
| ENSG00000091127 | PUS7 | 1.421968731 | 0.181069609 | 1.561771204 | 0.130056692 | 2.983739935 | 3.327711234 | 5.838937808 | 6.534109959 | 7.03996889 | 7.13309042 | 7.196918221 | 7.24894374 |
| ENSG00000006451 | RALA | 2.641968903 | 2.16E-09 | 0.341712395 | 0.524228701 | 2.983681298 | 8.696033109 | 8.101365653 | 8.805342542 | 11.23144255 | 11.1448568 | 9.189724501 | 8.532164007 |
| ENSG00000144909 | OSBPL11 | 1.507105566 | 0.179604546 | 1.475828495 | 0.181024624 | 2.98293406 | 5.70932728 | 8.029695761 | 5.39244166 | 8.361842177 | 8.40255255 | 8.471832957 | 8.220043091 |
| ENSG00000120868 | APAF1 | 2.056712236 | 0.11481363 | 0.925714857 | 0.504387978 | 2.982427092 | 7.384433967 | 5.946638857 | 3.369317972 | 8.449657898 | 8.16874151 | 7.603095107 | 6.657218517 |
| ENSG00000140836 | ZFHX3 | 0.638704827 | 0.686208373 | 2.343372984 | 0.088390629 | 2.982077812 | 5.095959458 | 9.549500232 | 7.098165388 | 8.880924505 | 8.91566441 | 10.46914042 | 10.71012106 |
| ENSG00000204160 | ZDHHC18 | 1.326354984 | 0.343731455 | 1.655598575 | 0.218712264 | 2.98195356 | 8.564112515 | 7.145017454 | 4.058998049 | 8.780186674 | 8.79884455 | 9.202060967 | 9.024723418 |
| ENSG00000080824 | HSP90AA1 | 1.396408003 | 9.99E-10 | 1.58481909 | 2.44E-12 | 2.981227093 | 11.48534568 | 11.72102722 | 11.38976845 | 12.93879979 | 12.9321918 | 13.10118024 | 13.14611335 |
| ENSG00000214013 | GANC | 1.781680448 | 0.291638623 | 1.197745587 | 0.485369749 | 2.979426035 | 7.82963488 | 6.311476754 | 2 | 8.458152491 | 8.41468635 | 7.667989909 | 8.031095054 |
| ENSG00000156639 | ZFAND3 | 1.097543782 | 0.053991261 | 1.879930899 | 0.000395857 | 2.977474682 | 9.384607364 | 9.328390744 | 8.143093325 | 10.1220151 | 10.1741606 | 10.91689319 | 10.94017357 |
| ENSG00000100941 | PNN | 1.892537972 | 0.001161823 | 1.083787728 | 0.078447389 | 2.976325701 | 8.257427002 | 8.791896851 | 7.285536013 | 10.15112202 | 10.0971999 | 9.352020423 | 9.286301649 |
| ENSG00000132155 | RAF1 | 1.256474995 | 0.008781104 | 1.718431523 | 0.000188952 | 2.974906518 | 8.836089686 | 8.213429689 | 9.25752426 | 10.10642804 | 10.0390409 | 10.41072665 | 10.64687673 |
| ENSG00000101109 | STK4 | 1.650908382 | 0.178186326 | 1.321483892 | 0.28290231 | 2.972392274 | 4.815609471 | 8.687092496 | 8.143093325 | 9.55265467 | 9.54986851 | 9.306733157 | 9.135803248 |
| ENSG00000100079 | LGALS2 | -2.307876711 | 0.495905043 | 5.279750861 | 0.034674478 | 2.97187415 | 2 | 2 | 2 | 2.141737246 | 2 | 5.390603804 | 5.497565112 |
| ENSG00000105778 | AVL9 | 1.682661917 | 0.00326405 | 1.288692248 | 0.028067889 | 2.971354165 | 8.313285031 | 7.503546079 | 6.958202216 | 9.383689875 | 9.34048844 | 9.12500825 | 8.798540026 |
| ENSG00000261221 | ZNF865 | 0.587797955 | 0.298074813 | 2.381944371 | 6.27E-07 | 2.969742326 | 7.579279349 | 6.78762695 | 6.629488387 | 7.539954763 | 7.7127321 | 9.222499293 | 9.562031113 |
| ENSG00000146678 | IGFBP1 | -2.072956259 | 0.307942408 | 5.042484217 | 0.001703637 | 2.969527958 | 2 | 3.686303414 | 2 | 2.270786764 | 2.2680111 | 6.134985075 | 7.370638252 |
| ENSG00000012232 | EXTL3 | 0.973658171 | 0.271011111 | 1.995697363 | 0.013555764 | 2.969355533 | 9.544849385 | 7.638155693 | 7.858115942 | 9.580155061 | 9.58173459 | 10.44409424 | 10.73916056 |
| ENSG00000155265 | GOLGA7B | 0.497949615 | 0.915839444 | 2.470592243 | NA | 2.968541858 | 2 | 2 | 2 | 2.498691189 | 2.38539787 | 3.917810786 | 2.059860178 |
| ENSG00000240694 | PNMA2 | -2.634480682 | 0.01733931 | 5.603018159 | 1.73E-08 | 2.968537477 | 4.815609471 | 5.595927442 | 6.718950348 | 4.237680128 | 2.86123014 | 10.67080105 | 11.91008607 |
| ENSG00000186212 | SOWAHB | -2.307857601 | 0.496953626 | 5.275409601 | 0.03552107 | 2.967552 | 2 | 2 | 2 | 2.141737246 | 2 | 5.304163849 | 5.566251551 |
| | ATP5J2-PTCD | 8.158179723 | 1.81E-06 | -5.191149865 | 0.037234619 | 2.967029858 | 2 | 2 | 2 | 8.229163214 | 8.17894021 | 2 | 2 |
| ENSG00000148331 | ASB6 | 1.365954596 | 0.225370868 | 1.601039251 | 0.141569199 | 2.966993846 | 4.815609471 | 7.467835385 | 8.206335842 | 8.657201865 | 8.79221631 | 8.943899495 | 8.974576781 |
| ENSG00000247077 | PGAM5 | 0.757789377 | 0.379217697 | 2.208466303 | 0.003474236 | 2.96625568 | 5.866944937 | 6.228547981 | 7.734844356 | 7.552702688 | 7.59884443 | 9.069678103 | 8.94213989 |
| ENSG00000175213 | ZNF408 | 1.407611178 | 0.460238382 | 1.557728687 | 0.397767254 | 2.965339864 | 4.00722531 | 7.189546432 | 2 | 7.127781374 | 7.17852729 | 7.385306888 | 7.208191747 |
| ENSG00000183439 | TRIM61 | 3.212285742 | 0.088950597 | -0.247130029 | 0.914220926 | 2.965155713 | 2 | 3.686303414 | 2 | 5.058234798 | 5.24368216 | 3.035909414 | 2.466008509 |
| ENSG00000128791 | TWSG1 | 1.450916905 | 0.011158168 | 1.51410836 | 0.00747349 | 2.965025265 | 8.367060731 | 9.574900212 | 9.61300511 | 10.76244194 | 10.7069016 | 10.6229975 | 10.95404265 |
| ENSG00000165801 | ARHGEF40 | 1.127698759 | 0.237333582 | 1.837286557 | 0.039733569 | 2.964985316 | 7.750913381 | 9.608085847 | 7.552162999 | 9.691694368 | 9.80557618 | 10.12246481 | 10.7279608 |
| ENSG00000182541 | LIMK2 | 2.117863655 | 0.000122541 | 0.846326309 | 0.161125537 | 2.964189963 | 8.797441463 | 7.572420273 | 7.734844356 | 10.17347624 | 10.3042388 | 8.976484087 | 8.974576781 |
| ENSG00000278326 | | 8.151191053 | 1.97E-06 | -5.191149816 | 0.037509667 | 2.960041237 | 2 | 2 | 2 | 8.248880905 | 8.14396641 | 2 | 2 |
| ENSG00000171155 | C1GALT1C1 | 2.113652168 | 0.005097116 | 0.846387752 | 0.304133236 | 2.96003992 | 7.096806373 | 6.535001748 | 8.434763955 | 9.680848002 | 9.64710779 | 8.265552582 | 8.535462193 |
| ENSG00000156049 | GNA14 | -1.137865377 | 0.431178842 | 4.096821087 | 0.000445233 | 2.95895571 | 3.327711234 | 3.686303414 | 3.369317972 | 2.695456494 | 3.01487888 | 6.287958221 | 7.467748741 |
| ENSG00000152492 | CCDC50 | 1.149239406 | 0.138334654 | 1.807897097 | 0.013077994 | 2.957136503 | 9.858207046 | 9.28800521 | 7.896971941 | 10.3905021 | 10.3428503 | 10.84344883 | 11.18432523 |
| ENSG00000214814 | FER1L6 | 4.046953687 | 0.318639797 | -1.092861105 | 0.809485785 | 2.954092582 | 2 | 2 | 2 | 4.387512679 | 4.37434574 | 2.3070652 | 2 |
| ENSG00000239836 | | 9.10590246 | 1.12E-10 | -6.152947053 | 0.005662518 | 2.952955407 | 3.327711234 | 2 | 2 | 10.04685164 | 10.1499977 | 2 | 2 |
| ENSG00000053254 | FOXN3 | 0.540246659 | 0.756618096 | 2.410972976 | 0.108572237 | 2.951219635 | 4.00722531 | 8.655694736 | 9.767769415 | 9.299216392 | 9.27068036 | 11.16621714 | 11.13157999 |
| ENSG00000164323 | CFAP97 | 0.834980095 | 0.483350437 | 2.113693147 | 0.047119088 | 2.948673242 | 7.031698319 | 5.72254323 | 8.805342542 | 8.621345041 | 8.43695333 | 9.831105898 | 9.771493021 |
| ENSG00000067167 | TRAM1 | 1.918718128 | 0.001679195 | 1.029874519 | 0.112494976 | 2.948592646 | 9.015175615 | 9.896993485 | 10.5916593 | 11.94324566 | 11.8270724 | 11.04910471 | 10.94662937 |
| ENSG00000163946 | FAM208A | 1.244455741 | 0.08383018 | 1.70255654 | 0.013108075 | 2.947012281 | 7.624139768 | 9.030900487 | 7.646393253 | 9.566077486 | 9.43967292 | 10.17093281 | 9.713270561 |
| ENSG00000117533 | VAMP4 | 1.261219659 | 0.312962462 | 1.685656317 | 0.156803554 | 2.946875975 | 4.00722531 | 5.72254323 | 7.398165132 | 7.510853288 | 7.52080763 | 8.281350892 | 7.470508284 |
| ENSG00000129968 | ABHD17A | 1.335718102 | 0.012810571 | 1.610386298 | 0.002019672 | 2.9461044 | 7.096806373 | 7.431218269 | 8.296287027 | 8.989221226 | 9.03387903 | 9.267612236 | 9.301104226 |
| ENSG00000083444 | PLOD1 | 0.993388892 | 0.040278645 | 1.950375076 | 1.27E-05 | 2.943763968 | 12.13890018 | 11.48015755 | 11.12102291 | 12.62961284 | 12.6409809 | 13.58778272 | 13.59591238 |
| ENSG00000138316 | ADAMTS14 | 0.128812861 | 0.949002608 | 2.812114867 | 0.093670473 | 2.940927728 | 2 | 6.389896633 | 4.058998049 | 5.37770462 | 5.14724905 | 6.446574529 | 8.508191835 |
| ENSG00000157303 | SUSD3 | 2.401179135 | 0.269864545 | 0.539672239 | 0.82433351 | 2.940851375 | 2 | 3.076665209 | 2 | 3.997744386 | 3.69849104 | 2.885263063 | 2.671842496 |
| ENSG00000160789 | LMNA | 1.047226017 | 0.393187917 | 1.890279003 | 0.095521719 | 2.93750502 | 11.17828117 | 10.94355721 | 7.691296561 | 11.55759209 | 11.6302244 | 12.13170852 | 12.68814343 |
| ENSG00000069849 | ATP1B3 | 1.794420511 | 2.14E-06 | 1.141652159 | 0.003792422 | 2.93607267 | 9.097018025 | 9.488434476 | 9.746650996 | 11.2472134 | 11.2658107 | 10.35493302 | 10.8183324 |
| ENSG00000156052 | GNAQ | 1.30485017 | 0.346271967 | 1.630049107 | 0.220537149 | 2.934899277 | 5.095959458 | 9.523645035 | 8.076950877 | 9.778365132 | 9.69337522 | 10.08982211 | 10.0302037 |

| | | | | | | | | | | | | | |
|---|---|---|---|---|---|---|---|---|---|---|---|---|---|
| ENSG00000085719 | CPNE3 | 1.276910497 | 0.004297518 | 1.657791213 | 0.000126297 | 2.93470171 | 9.112843761 | 8.505280662 | 8.143093325 | 9.926517587 | 9.89552277 | 10.2461159 | 10.33375043 |
| ENSG00000080031 | PTPRH | 0.72858269 | 0.764673299 | 2.20304881 | 0.30263573 | 2.931631499 | 3.327711234 | 2 | 2 | 2.947924294 | 2.77779907 | 3.767980144 | 3.660199087 |
| ENSG00000116117 | PARD3B | -0.6900269 | 0.749166874 | 3.621321002 | 0.047708245 | 2.931294102 | 2 | 5.131603625 | 8.486635645 | 6.387081877 | 6.37866593 | 10.55599851 | 10.6947155 |
| ENSG00000163125 | RPRD2 | 1.287897482 | 0.418287941 | 1.643181522 | 0.28010057 | 2.931079004 | 3.327711234 | 8.762721501 | 8.076950877 | 9.133031936 | 9.21030762 | 9.556714309 | 9.493138564 |
| ENSG00000139734 | DIAPH3 | -1.427692635 | 0.420476172 | 4.357308364 | 0.004497948 | 2.929615729 | 2 | 6.535001748 | 3.369317972 | 3.921164147 | 4.0930011 | 9.272995418 | 9.500920284 |
| ENSG00000092529 | CAPN3 | 4.378714258 | 0.001386931 | -1.453049573 | 0.354405417 | 2.925664685 | 4.467311984 | 2 | 2 | 6.946463121 | 6.91614081 | 2.68698273 | 2.509591799 |
| ENSG00000172757 | CFL1 | 1.084165028 | 0.20823704 | 1.841262599 | 0.022093349 | 2.925427626 | 12.21602152 | 10.74900803 | 10.31150679 | 12.39989575 | 12.4351513 | 12.79907327 | 13.47179816 |
| ENSG00000109171 | SLAIN2 | 1.265291806 | 0.359143464 | 1.658199357 | 0.208570841 | 2.923491162 | 8.946185348 | 8.505280662 | 4.523675391 | 9.449924671 | 9.45234084 | 9.875912956 | 9.807406144 |
| ENSG00000244682 | FCGR2C | 0.429780277 | 0.874101085 | 2.493622671 | 0.267777028 | 2.923402949 | 2 | 3.076665209 | 2 | 2.868578001 | 2.59489879 | 4.301748724 | 3.407202941 |
| ENSG00000188779 | SKOR1 | 0.26663352 | 0.962060833 | 2.656548799 | NA | 2.923182319 | 2 | 2 | 2 | 2 | 2.68924543 | 3.256555576 | 3.496556178 |
| ENSG00000005339 | CREBBP | 1.427992015 | 0.015763734 | 1.494309202 | 0.010593672 | 2.922301217 | 9.143983763 | 7.73146271 | 8.380957451 | 9.925905286 | 9.96612421 | 10.07485216 | 9.946704253 |
| ENSG00000111145 | ELK3 | 1.663884875 | 0.023888254 | 1.258291463 | 0.094431932 | 2.922176338 | 7.096806373 | 9.055168103 | 8.325066185 | 10.01150489 | 10.0356777 | 9.490026756 | 9.738960951 |
| ENSG00000118058 | KMT2A | 1.423154445 | 0.078125311 | 1.498058272 | 0.058395723 | 2.921212717 | 8.564112515 | 9.799806309 | 7.777116083 | 10.350452 | 10.4019472 | 10.58021248 | 10.3092961 |
| ENSG00000104812 | GYS1 | 1.224767423 | 0.347067822 | 1.694369015 | 0.170827212 | 2.919136438 | 10.32020829 | 8.255926468 | 6.882775709 | 10.35136432 | 10.3813388 | 10.10534852 | 11.3170206 |
| | | 8.108156379 | 2.59E-06 | -5.191149723 | 0.038130367 | 2.917006657 | 2 | 2 | 2 | 8.126317766 | 8.18299958 | 2 | 2 |
| ENSG00000134758 | RNF138 | 1.997292401 | 0.023633746 | 0.918334499 | 0.332076786 | 2.9156269 | 5.095959458 | 5.72254323 | 3.369317972 | 7.012547947 | 6.83591385 | 6.118193494 | 5.641643428 |
| ENSG00000171466 | ZNF562 | 1.678941526 | 8.29E-06 | 1.235913944 | 0.001354312 | 2.91485547 | 6.73718411 | 7.274681896 | 7.451345288 | 8.868232317 | 8.80807321 | 8.427654172 | 8.37166861 |
| ENSG00000282750 | | 8.105987317 | 2.65E-06 | -5.191149706 | 0.038217881 | 2.914837611 | 2 | 2 | 2 | 8.113468729 | 8.19108423 | 2 | 2 |
| ENSG00000183779 | ZNF703 | 1.299128413 | 0.452192995 | 1.614185269 | 0.329209308 | 2.913313682 | 7.867441304 | 7.002537973 | 2 | 8.176599235 | 8.22691474 | 8.72755027 | 8.261263339 |
| ENSG00000102908 | NFAT5 | 2.063860479 | 0.044161456 | 0.847361536 | 0.444451639 | 2.911222015 | 7.218819781 | 9.367676479 | 10.36791565 | 11.57429426 | 11.4832463 | 10.41087029 | 10.21350733 |
| ENSG00000160255 | ITGB2 | -0.944792845 | 0.688658118 | 3.855947898 | 0.054642188 | 2.911155053 | 5.330594134 | 2 | 2 | 3.162721474 | 3.50452879 | 8.219062336 | 5.727202496 |
| ENSG00000115084 | SLC35F5 | 2.061029893 | 5.36E-07 | 0.850101691 | 0.057679184 | 2.911131584 | 7.790810999 | 8.147241016 | 8.380957451 | 10.18529247 | 10.1567216 | 9.231310145 | 8.642113606 |
| ENSG00000176407 | KCMF1 | 1.169698718 | 0.019862551 | 1.741243301 | 0.000261128 | 2.910942019 | 7.709880939 | 8.791896851 | 8.325066185 | 9.505076228 | 9.51325794 | 10.22143306 | 9.918940524 |
| ENSG00000205835 | GMNC | 0.010004217 | 0.997481474 | 2.900577038 | 0.348689353 | 2.910581255 | 2 | 2 | 2 | 2.270786764 | 2.38539787 | 2.911488689 | 3.972018587 |
| ENSG00000163577 | EIF5A2 | 1.310483156 | 0.003591958 | 1.600043453 | 0.000246393 | 2.91052661 | 6.73718411 | 6.140560021 | 6.629488387 | 7.831679003 | 7.72392024 | 8.181838221 | 7.934467448 |
| ENSG00000089692 | LAG3 | 0.497905282 | 0.895675696 | 2.411798403 | NA | 2.909703685 | 2 | 2 | 2 | 2.498691189 | 2.38539787 | 2.746468522 | 3.600986722 |
| ENSG00000168264 | IRF2BP2 | 1.05826229 | 0.392721273 | 1.850750879 | 0.106982382 | 2.909013169 | 6.653098118 | 9.377333006 | 6.718950348 | 9.207473484 | 9.26494702 | 10.30850554 | 9.671609742 |
| ENSG00000180891 | CUEDC1 | 0.541030609 | 0.725318811 | 2.367871685 | 0.073990493 | 2.908902294 | 8.75772933 | 7.467835385 | 4.058998049 | 8.221200057 | 8.2561084 | 9.967428009 | 10.13153689 |
| ENSG00000106012 | IQCE | 1.415943709 | 0.036071417 | 1.492538044 | 0.024788756 | 2.908481753 | 7.485143553 | 6.04685525 | 6.203069917 | 8.072008297 | 8.13141746 | 8.151901655 | 8.202721679 |
| ENSG00000124496 | TRERF1 | 0.903041054 | 0.621621222 | 2.0052456 | 0.230071506 | 2.908286654 | 5.095959458 | 7.431218269 | 2 | 7.030886362 | 7.00136498 | 7.985968197 | 8.195234077 |
| ENSG00000104722 | NEFM | 1.912322275 | 0.385406506 | 0.994962842 | 0.665769105 | 2.907285118 | 4.467311984 | 2 | 8.408111533 | 8.778830996 | 8.8250571 | 8.372671446 | 7.179807904 |
| ENSG00000182179 | UBA7 | 1.840788536 | 0.333078915 | 1.065972507 | 0.58769378 | 2.906761043 | 3.327711234 | 5.838937808 | 9.601288346 | 9.903687576 | 10.0136237 | 8.996912583 | 9.358468538 |
| ENSG00000163161 | ERCC3 | 1.459658133 | 0.141653394 | 1.446911344 | 0.138280206 | 2.906569477 | 6.366709671 | 8.747909548 | 6.80318718 | 9.123450746 | 9.18320541 | 9.100995221 | 9.17988647 |
| ENSG00000187908 | DMBT1 | 3.594462417 | 0.233603426 | -0.690089515 | 0.842587044 | 2.904372902 | 2 | 2 | 2 | 3.754611435 | 4.2560182 | 2.34604626 | 2.059860178 |
| ENSG00000148773 | MKI67 | 1.13744563 | 0.610493708 | 1.76685207 | 0.401339976 | 2.9042977 | 3.327711234 | 6.311476754 | 2 | 6.34372989 | 5.6121243 | 4.434275199 | 7.453871289 |
| ENSG00000105656 | ELL | 1.829122755 | 0.253735228 | 1.074743324 | 0.517007889 | 2.903866079 | 7.624139768 | 6.844429755 | 2 | 8.451360823 | 8.56844956 | 7.733770002 | 7.800517754 |
| ENSG00000148798 | INA | -0.144572707 | 0.938309757 | 3.046931005 | 0.041995639 | 2.902358298 | 4.467311984 | 6.78762695 | 2 | 5.37770462 | 5.43283104 | 8.725241966 | 8.16062336 |
| ENSG00000143797 | MBOAT2 | 1.314593438 | 0.0316272 | 1.587496711 | 0.007499883 | 2.902090149 | 8.075571721 | 7.232742069 | 8.143093325 | 9.218515697 | 9.10727282 | 9.833465342 | 8.882782322 |
| ENSG00000144589 | STK11IP | 0.969913071 | 0.556828814 | 1.93169378 | 0.204191294 | 2.901606852 | 5.095959458 | 6.899080544 | 2 | 6.693715713 | 6.61334686 | 7.378272268 | 7.770601445 |
| ENSG00000196715 | VKORC1L1 | 0.973460856 | 0.43698019 | 1.926391067 | 0.092679263 | 2.899851923 | 5.866944937 | 7.232742069 | 9.227351203 | 9.025111392 | 9.04060595 | 10.03085749 | 9.928254593 |
| ENSG00000163354 | DCST2 | 0.49578662 | 0.895265695 | 2.404046802 | NA | 2.899833422 | 2 | 2 | 2 | 2.3892346 | 2.49394783 | 3.5515855 | 2.817676617 |
| ENSG00000166436 | TRIM66 | 2.287547961 | 0.00023626 | 0.61106281 | 0.389304796 | 2.898610771 | 5.095959458 | 4.710564154 | 4.874587708 | 6.912249745 | 7.10768997 | 5.47655442 | 5.385063124 |
| ENSG00000163069 | SGCB | 1.543930637 | 1.41E-08 | 1.353748536 | 7.51E-07 | 2.897679174 | 9.423197387 | 9.338312924 | 9.149036566 | 10.8774261 | 10.8161359 | 10.5076599 | 10.7930647 |
| ENSG00000172890 | NADSYN1 | 1.334933896 | 0.030870877 | 1.560044109 | 0.009596225 | 2.894978005 | 7.218819781 | 8.077869079 | 6.629488387 | 8.799033614 | 8.72698617 | 8.900776778 | 9.067728571 |
| ENSG00000134982 | APC | 1.216166105 | 0.124010482 | 1.678055851 | 0.02597163 | 2.894221955 | 6.96351253 | 7.874700259 | 8.960138144 | 9.363030664 | 9.35764963 | 9.742804424 | 9.892948091 |
| ENSG00000140416 | TPM1 | 0.637968274 | 0.234421046 | 2.255751845 | 1.19E-06 | 2.893720119 | 12.62340907 | 11.50239087 | 12.26789926 | 12.82090962 | 12.858361 | 14.47704462 | 14.43668992 |
| ENSG00000154813 | DPH3 | 1.294127454 | 0.215656888 | 1.598460937 | 0.111748606 | 2.892588391 | 6.73718411 | 8.469613413 | 5.772716771 | 8.738951712 | 8.68609779 | 9.089196199 | 8.935663927 |
| ENSG00000116786 | PLEKHM2 | 1.465741702 | 4.71E-07 | 1.426758341 | 8.90E-07 | 2.892500043 | 9.128497779 | 9.514922632 | 9.06622548 | 10.70542879 | 10.7271979 | 10.72194193 | 10.63154769 |
| ENSG00000054277 | OPN3 | 0.577340645 | 0.698378828 | 2.315075569 | 0.07118462 | 2.892416214 | 2 | 4.710564154 | 6.073378139 | 5.552894743 | 5.49945122 | 6.897310495 | 7.405663765 |
| ENSG00000072310 | SREBF1 | 1.006370113 | 0.357081274 | 1.882635898 | 0.061692379 | 2.889006011 | 8.075571721 | 8.124485672 | 5.59507254 | 8.660150093 | 8.60672783 | 8.733075176 | 10.0028246 |
| ENSG00000141837 | CACNA1A | 1.783746604 | 0.317344658 | 1.103300299 | 0.547274868 | 2.887046903 | 2 | 5.303523747 | 7.646393253 | 8.054188409 | 8.08661608 | 6.981729486 | 7.728570659 |
| ENSG00000189308 | LIN54 | 1.547403489 | 0.141109656 | 1.339231476 | 0.20113831 | 2.886634965 | 4.467311984 | 7.145017454 | 5.59507254 | 7.688853661 | 7.63485389 | 7.730090858 | 7.123062801 |
| ENSG00000130856 | ZNF236 | 1.256620858 | 0.634315819 | 1.629448933 | 0.522581231 | 2.886069791 | 2 | 7.638155693 | 2 | 7.409376773 | 7.24216308 | 7.905886539 | 7.442672408 |
| ENSG00000124831 | LRRFIP1 | 1.095593514 | 0.362915467 | 1.78686846 | 0.112712865 | 2.882461974 | 9.922475255 | 8.929529568 | 6.322058544 | 10.0950348 | 10.08857 | 10.7425177 | 10.81860316 |
| ENSG00000137275 | RIPK1 | 1.538776737 | 0.025736029 | 1.341485744 | 0.053383916 | 2.880262481 | 6.816637977 | 8.376385542 | 8.561105505 | 9.617037214 | 9.63530531 | 9.357992454 | 9.498556374 |
| ENSG00000182132 | KCNIP1 | -1.265414385 | 0.719366965 | 4.144088735 | 0.164866748 | 2.87867435 | 3.327711234 | 2 | 2 | 2.498691189 | 2 | 3.235948948 | 6.102937566 |
| ENSG00000059588 | TARBP1 | 1.472343416 | 0.325101958 | 1.404983578 | 0.341761606 | 2.877326994 | 2 | 6.535001748 | 6.80318718 | 7.517371202 | 7.56191311 | 7.622078048 | 7.308762629 |
| ENSG00000119139 | TJP2 | 3.626360193 | 1.10E-06 | -0.749284856 | 0.394959847 | 2.877075337 | 4.467311984 | 4.11352228 | 4.874587708 | 7.852449262 | 7.86451142 | 3.600766115 | 4.161535617 |
| ENSG00000134853 | PDGFRA | 2.429635281 | 0.017875395 | 0.444768554 | 0.710952496 | 2.874403835 | 7.904282243 | 10.96612994 | 9.330306409 | 12.32756329 | 12.3434862 | 10.2021419 | 10.49133865 |
| ENSG00000100614 | PPM1A | 1.343161685 | 0.321826745 | 1.530684477 | 0.245064763 | 2.873846162 | 4.467311984 | 8.747909548 | 8.296287027 | 9.329162745 | 9.32954365 | 9.743945072 | 9.244592076 |
| ENSG00000162836 | ACP6 | -3.753133088 | 0.276585553 | 6.626294153 | 0.013315484 | 2.873161065 | 2 | 2 | 2 | 2 | 2 | 6.972416862 | 6.38621943 |
| ENSG00000185477 | GPRIN3 | 0.303711777 | 0.911798376 | 2.568776732 | 0.257812226 | 2.872488509 | 2 | 3.686303414 | 2 | 3.09461492 | 2.94009902 | 5.243518884 | 3.261853426 |
| ENSG00000282110 | | 1.397893884 | 0.821277934 | 1.474322103 | NA | 2.872215987 | 2 | 2 | 2 | 3.227757282 | 2 | 2.831336763 | 2.70970617 |
| ENSG00000105866 | SP4 | 0.975100895 | 0.54007403 | 1.896468801 | 0.195422681 | 2.871569696 | 2 | 5.457120953 | 3.369317972 | 4.907595604 | 5.39133044 | 6.106889539 | 5.913137508 |
| ENSG00000078140 | UBE2K | 1.516872063 | 0.311071331 | 1.354604018 | 0.362520034 | 2.871476081 | 9.233563228 | 8.505280662 | 4.058998049 | 9.952603669 | 9.76282251 | 9.724430319 | 9.673706637 |
| ENSG00000101966 | XIAP | 1.512994155 | 0.07809295 | 1.358347504 | 0.112145055 | 2.871341659 | 9.204317094 | 7.538394143 | 7.225742779 | 9.851120801 | 9.67835667 | 9.767692698 | 9.44062848 |
| ENSG00000105298 | CACTIN | 1.561499709 | 0.219703759 | 1.307066029 | 0.303728756 | 2.868565739 | 3.327711234 | 7.35507212 | 6.882775709 | 8.111316053 | 8.15640714 | 7.880388025 | 7.88671543 |
| ENSG00000119950 | MXI1 | 1.196041064 | 0.011144863 | 1.672271152 | 0.000207352 | 2.868312215 | 7.27616306 | 7.538394143 | 8.175061096 | 8.857997553 | 8.91810182 | 9.527072508 | 9.17354978 |
| ENSG00000087448 | KLHL42 | 0.973783029 | 0.001343782 | 1.893085046 | 3.08E-11 | 2.866868074 | 7.790810999 | 7.954278496 | 7.646393253 | 8.75144642 | 8.79221631 | 9.675532774 | 9.694509651 |
| ENSG00000136231 | IGF2BP3 | 1.304893142 | 0.527512895 | 1.560767111 | 0.432736935 | 2.865660253 | 4.00722531 | 7.701025996 | 2 | 7.530319349 | 7.51437845 | 7.801071566 | 7.744618542 |
| ENSG00000118412 | CASP8AP2 | 1.485455148 | 0.357081274 | 1.379748605 | 0.385870817 | 2.865203753 | 7.33131396 | 6.844429755 | 2 | 8.083034979 | 7.89700047 | 8.21050623 | 7.474637727 |
| ENSG00000114867 | EIF4G1 | 1.331037302 | 2.18E-05 | 1.533997148 | 6.63E-07 | 2.86503445 | 10.70657014 | 11.05905903 | 11.23155829 | 12.29106025 | 12.3962337 | 12.417014 | 12.66702221 |
| ENSG00000146007 | ZMAT2 | 1.421701693 | 4.45E-05 | 1.442809771 | 2.97E-05 | 2.864511464 | 7.667647223 | 7.638155693 | 8.042706615 | 9.207473484 | 9.17914662 | 9.05761907 | 9.355859082 |
| ENSG00000274641 | HIST1H2BO | 6.619714578 | 2.54E-05 | -3.759275672 | 0.072505978 | 2.860438906 | 3.327711234 | 2 | 2 | 7.688853661 | 7.5835711 | 2.094598369 | 2 |
| ENSG00000099219 | ERMP1 | 0.849719255 | 0.575968182 | 2.010169922 | 0.143374446 | 2.859889177 | 2 | 6.311476754 | 6.534109959 | 6.542151749 | 6.81514011 | 8.033012338 | 7.55613706 |
| ENSG00000164136 | IL15 | 1.918708666 | 0.284626966 | 0.940727841 | 0.617051806 | 2.859436507 | 6.138360429 | 2 | 3.369317972 | 6.516352584 | 6.60126376 | 5.85994354 | 5.379200993 |
| ENSG00000131149 | GSE1 | 1.252808981 | 0.437181027 | 1.606125414 | 0.29637811 | 2.858934395 | 4.467311984 | 9.018611864 | 5.930865648 | 8.88471049 | 8.8984863 | 9.358290408 | 9.116112551 |
| ENSG00000164122 | ASB5 | -1.272311879 | 0.588479045 | 4.12969454 | 0.027228112 | 2.85738266 | 2 | 3.076665209 | 2 | 2.270786764 | 2.2680111 | 5.590506472 | 4.870334174 |
| ENSG00000180398 | MCFD2 | 1.746793893 | 6.74E-05 | 1.11049032 | 0.015545758 | 2.857284213 | 10.09970294 | 10.87363932 | 11.06240811 | 12.48060115 | 12.4782175 | 11.74926356 | 11.93227541 |
| ENSG00000118046 | STK11 | 0.945916076 | 0.217903135 | 1.911308249 | 0.006424889 | 2.857224324 | 8.444143041 | 8.124485672 | 6.534109959 | 8.795015659 | 8.89354047 | 9.757111106 | 9.850148984 |
| ENSG00000104320 | NBN | 2.310456695 | 0.001663588 | 0.546659127 | 0.518789265 | 2.857115822 | 8.468948322 | 8.747909548 | 6.718950348 | 10.508071 | 10.5173643 | 8.878929913 | 8.626733602 |
| ENSG00000126218 | F10 | -0.343823244 | 0.863551136 | 3.199959587 | 0.046571766 | 2.856136343 | 4.00722531 | 6.951736442 | 2 | 5.348759107 | 5.24368216 | 8.674459329 | 8.725336927 |
| ENSG00000206340 | | 8.047083063 | 3.97E-06 | -5.191149543 | 0.039270477 | 2.85593352 | 2 | 2 | 2 | 8.115618198 | 8.07355593 | 2 | 2 |
| ENSG00000166272 | WBP1L | 0.811022465 | 0.064313177 | 2.044444216 | 2.11E-07 | 2.85546668 | 8.517314938 | 9.170736405 | 8.380957451 | 9.498502402 | 9.59245646 | 10.7588599 | 10.79141033 |
| ENSG00000116580 | GON4L | 1.708051563 | 5.57E-08 | 1.146611631 | 0.000397696 | 2.854663193 | 8.313285031 | 7.790474954 | 8.007629717 | 9.72797373 | 9.74782835 | 9.152756493 | 9.20620076 |
| ENSG00000122512 | PMS2 | 2.191834163 | 0.105206276 | 0.662568289 | 0.65836129 | 2.854402452 | 6.653098118 | 6.04685525 | 2 | 7.956842238 | 7.93841129 | 6.615624322 | 6.289521179 |
| ENSG00000139223 | ANP32D | 2.389904445 | 0.458595738 | 0.463608915 | NA | 2.853513361 | 2 | 2 | 2 | 3.227757282 | 3.2184422 | 2.3070652 | 2.551896972 |
| ENSG00000159202 | UBE2Z | 1.390626654 | 0.033756605 | 1.461636056 | 0.023809617 | 2.85226271 | 9.317904228 | 9.632483534 | 10.79563557 | 11.4400825 | 11.4577141 | 11.45029098 | 11.58618106 |
| ENSG00000271303 | SRXN1 | 4.647453335 | 0.013410886 | -1.796957671 | 0.381806697 | 2.850495664 | 3.327711234 | 7.790474954 | 2 | 10.8767926 | 10.8890776 | 4.628601098 | 4.729989799 |
| ENSG00000124215 | CDH26 | 2.015355299 | 0.596379651 | 0.834379539 | NA | 2.849734838 | 2 | 2 | 2 | 3.349643765 | 2.59489879 | 2.831336763 | 2.172608095 |
| ENSG00000158109 | TPRG1L | 0.883794463 | 0.14612275 | 1.965736024 | 0.000339822 | 2.849530488 | 9.204317094 | 8.297207175 | 7.896971941 | 9.4022977 | 9.49378147 | 10.44409424 | 10.60416654 |
| ENSG00000143398 | PIP5K1A | 0.979355878 | 0.056923475 | 1.866622143 | 8.15E-05 | 2.845978021 | 8.998234871 | 8.777382925 | 8.042706615 | 9.62912467 | 9.64121862 | 10.22388773 | 10.76411688 |
| ENSG00000106266 | SNX8 | 2.152674458 | 0.055618402 | 0.69253238 | 0.576450932 | 2.845206838 | 4.467311984 | 7.002537973 | 8.076950877 | 9.222510184 | 9.27353851 | 7.761068522 | 7.846917442 |
| ENSG00000120694 | HSPH1 | 2.249233505 | 1.61E-05 | 0.595796776 | 0.31653454 | 2.845030281 | 9.233563228 | 8.762721501 | 8.076950877 | 10.99037529 | 11.0213536 | 9.624671944 | 9.031747025 |
| ENSG00000119669 | IRF2BPL | 1.75163113 | 0.101896165 | 1.093381555 | 0.322038974 | 2.845012685 | 7.867441304 | 7.393647515 | 10.08627303 | 10.67332098 | 10.7170855 | 10.10570349 | 9.969317474 |
| ENSG00000129933 | MAU2 | 1.466601571 | 0.02248445 | 1.377252536 | 0.032067896 | 2.843854107 | 7.33131396 | 8.655694736 | 7.451345288 | 9.372948073 | 9.44306193 | 9.47746372 | 9.142307541 |
| ENSG00000234284 | ZNF879 | 1.507954291 | 0.483180449 | 1.335810784 | 0.530234743 | 2.843765074 | 2 | 5.838937808 | 2 | 5.808110275 | 5.8245091 | 5.805044075 | 5.492143571 |
| ENSG00000151929 | BAG3 | 1.003702538 | 0.311446617 | 1.83982764 | 0.044684078 | 2.843530178 | 9.751047882 | 7.928237109 | 7.451345288 | 9.695291778 | 9.76891184 | 10.39889946 | 10.7152199 |
| ENSG00000187288 | CIDEC | 2.329169315 | 0.517280172 | 0.513942789 | NA | 2.843112104 | 2 | 2 | 2 | 2.695456494 | 3.55551834 | 2.183373728 | 2.671842496 |
| ENSG00000160678 | S100A1 | -0.727654445 | 0.838100556 | 3.570689841 | 0.212471855 | 2.843035396 | 2 | 2 | 2 | 2.141737246 | 2.2680111 | 4.487545924 | 3.287118176 |
| ENSG00000100813 | ACIN1 | 1.545296846 | 3.14E-09 | 1.297535621 | 7.98E-07 | 2.842832466 | 9.27634997 | 9.298208073 | 8.978369997 | 10.72913699 | 10.7382749 | 10.56480374 | 10.40381853 |
| ENSG00000282607 | | 0.472225567 | 0.801541753 | 2.370294611 | 0.133814471 | 2.842520177 | 3.327711234 | 2 | 4.874587708 | 4.139692846 | 4.16043394 | 4.888321757 | 6.338680316 |
| ENSG00000149428 | HYOU1 | 1.648959525 | 0.041269777 | 1.19313316 | 0.147829677 | 2.842092685 | 8.107517749 | 10.07368926 | 9.301632639 | 10.98685701 | 11.0432906 | 10.03235202 | 10.9463816 |
| ENSG00000101306 | MYLK2 | 0.015720062 | 0.997041173 | 2.825459542 | 0.376931525 | 2.841179603 | 2 | 2 | 2 | 2.498691189 | 2.14022021 | 3.663836244 | 3.287118176 |
| ENSG00000198961 | PJA2 | 1.031169668 | 0.519532942 | 1.807097244 | 0.223166355 | 2.838266911 | 5.095959458 | 10.50658023 | 9.926432745 | 10.72562309 | 10.6923799 | 11.54455506 | 11.41991513 |
| ENSG00000183281 | PLGLB1 | 5.470641317 | 0.048431027 | -2.633351554 | 0.413811187 | 2.837289762 | 2 | 2 | 2 | 5.602782483 | 5.63601114 | 2 | 2.117335275 |
| ENSG00000111880 | RNGTT | 1.240267006 | 0.441837321 | 1.596244278 | 0.298915693 | 2.836511284 | 7.031698319 | 7.232742069 | 2 | 7.872924734 | 7.68723697 | 8.121330705 | 8.145162126 |

| | | | | | | | | | | | | | |
|---|---|---|---|---|---|---|---|---|---|---|---|---|---|
| ENSG00000136824 | SMC2 | 1.55258824 | 0.127668185 | 1.28224159 | 0.209153239 | 2.834829829 | 4.00722531 | 6.951736442 | 6.629488387 | 7.810605345 | 7.85181819 | 7.710619897 | 7.405663765 |
| ENSG00000198720 | ANKRD13B | 0.308209421 | 0.82459995 | 2.525119688 | 0.024879166 | 2.833329109 | 2 | 5.72254323 | 5.772716771 | 5.433909581 | 5.5755357 | 7.628021136 | 7.612190665 |
| ENSG00000130508 | PXDN | 0.824567426 | 0.033290369 | 2.008323781 | 1.18E-08 | 2.832891207 | 11.43272766 | 12.10496808 | 11.63358652 | 12.56462203 | 12.5888679 | 13.62095662 | 13.8868593 |
| ENSG00000168575 | SLC20A2 | 0.075762959 | 0.947432314 | 2.756708697 | 0.002325611 | 2.832471656 | 7.867441304 | 8.191702463 | 10.10298738 | 9.208480821 | 9.09870316 | 11.37785453 | 12.16903209 |
| ENSG00000136531 | SCN2A | 2.325581883 | 0.252142759 | 0.506671096 | 0.823417076 | 2.832252979 | 4.00722531 | 2 | 2 | 4.54891529 | 4.50979031 | 3.534812207 | 2.851942824 |
| ENSG00000168781 | PPIP5K1 | 1.19921197 | 0.385295588 | 1.632808293 | 0.213016766 | 2.832020263 | 7.218819781 | 4.442710129 | 4.058998049 | 7.136275664 | 7.06433609 | 7.354572674 | 7.674548617 |
| | TNFAIP8L2-S | 8.02130996 | 4.78E-06 | -5.191149458 | 0.039834475 | 2.830160502 | 2 | 2 | 2 | 8.045195239 | 8.09310209 | 2 | 2 |
| ENSG00000107611 | CUBN | 2.655294295 | 0.113756797 | 0.173841927 | 0.932122597 | 2.829136221 | 3.327711234 | 2 | 3.369317972 | 4.647285135 | 4.93258395 | 2.831336763 | 3.236138336 |
| ENSG00000085662 | AKR1B1 | 3.839895367 | 7.45E-05 | -1.010764034 | 0.361796964 | 2.829131333 | 10.37373127 | 10.10308486 | 7.225742779 | 13.55263966 | 13.6081105 | 8.612861354 | 8.862441066 |
| ENSG00000188021 | UBQLN2 | 1.115475623 | 0.568958143 | 1.713503637 | 0.355608544 | 2.82897926 | 8.836089686 | 8.876041558 | 2 | 9.430198858 | 9.33684939 | 9.968013835 | 9.990620812 |
| ENSG00000175820 | CCDC168 | 2.079153501 | 0.534782548 | 0.749273083 | NA | 2.828426584 | 2 | 2 | 2 | 2.784612485 | 3.28037825 | 2.560105256 | 2.466008509 |
| ENSG00000159214 | CCDC24 | 1.044628963 | 0.612008031 | 1.781997866 | 0.354312588 | 2.826626829 | 2 | 5.303523747 | 2 | 4.694064037 | 5.07918227 | 5.489669427 | 5.621921006 |
| ENSG00000109686 | SH3D19 | 1.002502974 | 0.020115865 | 1.823078833 | 5.77E-06 | 2.825581807 | 7.940205782 | 8.505280662 | 8.784770914 | 9.459264257 | 9.43116542 | 10.40063634 | 10.10913368 |
| ENSG00000079435 | LIPE | 0.78447177 | 0.691448868 | 2.040495998 | 0.248858956 | 2.824967767 | 2 | 4.442710129 | 2 | 4.205751501 | 3.60476711 | 4.965450964 | 4.943664032 |
| ENSG00000173678 | SPDYE2B | 3.937386335 | 0.120832449 | -1.11303236 | 0.702841297 | 2.824353975 | 2 | 2 | 2 | 4.299492648 | 4.2865223 | 2.094598369 | 2.225841198 |
| ENSG00000169914 | OTUD3 | 1.066907905 | 0.342227985 | 1.754556322 | 0.094172652 | 2.821464227 | 4.815609471 | 7.099070191 | 4.874587708 | 7.101994243 | 7.03768431 | 7.439255893 | 7.991296779 |
| ENSG00000213853 | EMP2 | 1.472604686 | 0.104022412 | 1.34687985 | 0.133542516 | 2.819484537 | 6.257048221 | 8.255926468 | 8.721235202 | 9.562142546 | 9.48805095 | 9.154129994 | 9.61099888 |
| ENSG00000075568 | TMEM131 | 1.848850241 | 0.069445337 | 0.969989336 | 0.364791599 | 2.818839577 | 7.096806373 | 9.181799937 | 6.629488387 | 9.963384909 | 9.89675912 | 9.080915197 | 9.031747025 |
| ENSG00000140553 | UNC45A | 1.146996467 | 0.403335693 | 1.669361475 | 0.197276691 | 2.816357942 | 5.532353208 | 9.599860728 | 7.646393253 | 9.528658349 | 9.58173459 | 10.09430085 | 10.05638425 |
| ENSG00000117479 | SLC19A2 | 2.035774245 | 0.114685272 | 0.780272858 | 0.575511775 | 2.816047103 | 6.366709671 | 5.838937808 | 2 | 7.52061915 | 7.5462416 | 6.202919533 | 6.429408106 |
| ENSG00000081026 | MAGI3 | 0.701616206 | 0.674035246 | 2.113061856 | 0.152092183 | 2.814678061 | 2 | 4.93638933 | 6.718950348 | 6.229283386 | 6.14061114 | 7.733770002 | 7.331709059 |
| ENSG00000186666 | BCDIN3D | 1.555128814 | 0.365734008 | 1.255422738 | 0.466314777 | 2.810551552 | 2 | 5.946638857 | 3.369317972 | 6.096288482 | 6.12381559 | 6.192262702 | 5.343519008 |
| ENSG00000170734 | POLH | 0.392747294 | 0.708629645 | 2.416413322 | 0.005377605 | 2.809160617 | 5.866944937 | 7.232742069 | 8.434763955 | 7.842101511 | 7.97866656 | 9.975996365 | 9.856492132 |
| ENSG00000203747 | FCGR3A | -3.75310509 | 0.347620176 | 6.561632913 | 0.043929255 | 2.808527823 | 2 | 2 | 2 | 2 | 2 | 7.406207255 | 4.943664032 |
| ENSG00000172348 | RCAN2 | -1.714574544 | 0.197249313 | 4.522488304 | 6.66E-05 | 2.807913759 | 2 | 5.303523747 | 4.058998049 | 3.406930997 | 2.94009902 | 8.015738956 | 9.035013043 |
| ENSG00000177054 | ZDHHC13 | 2.009608423 | 0.203194975 | 0.798202706 | 0.638087371 | 2.807811129 | 4.00722531 | 6.389896633 | 2 | 6.970413838 | 7.09482005 | 5.951260634 | 5.803599879 |
| ENSG00000105613 | MAST1 | 0.183876679 | 0.947256262 | 2.621302847 | 0.264448359 | 2.805179526 | 2 | 2 | 3.369317972 | 2.3892346 | 2.86123014 | 4.478802948 | 3.33635788 |
| ENSG00000128829 | EIF2AK4 | 1.128386994 | 0.38727917 | 1.676757001 | 0.172175696 | 2.805143995 | 5.330594134 | 8.505280662 | 9.372276026 | 9.600249849 | 9.58480612 | 10.13729849 | 10.13979504 |
| ENSG00000158473 | CD1D | 3.045259312 | 0.301996555 | -0.241216246 | 0.944101725 | 2.804043067 | 2 | 2 | 2 | 3.406930997 | 3.82856379 | 2.183373728 | 2.374681325 |
| ENSG00000176783 | RUFY1 | 1.300577444 | 0.460626621 | 1.500587689 | 0.37615226 | 2.801165133 | 7.975256505 | 7.73146271 | 2 | 8.534041678 | 8.59915299 | 8.789844052 | 8.73970592 |
| ENSG00000100266 | PACSIN2 | 1.185240983 | 0.403737922 | 1.615912333 | 0.231087488 | 2.801153316 | 7.485143553 | 9.00621767 | 4.523675391 | 9.060130342 | 9.09439916 | 9.498161492 | 9.512347353 |
| ENSG00000119787 | ATL2 | 1.81447857 | 0.113324339 | 0.985008399 | 0.41126962 | 2.799486968 | 4.00722531 | 7.315436877 | 6.80318718 | 8.485003881 | 8.22494733 | 7.934143724 | 7.010010317 |
| ENSG00000115464 | USP34 | 1.790023304 | 0.119919599 | 1.009093757 | 0.399869187 | 2.799117061 | 6.366709671 | 9.828248506 | 9.725218844 | 11.01965231 | 11.0665479 | 10.30896804 | 10.21823539 |
| ENSG00000168159 | RNF187 | 0.934878691 | 0.133409766 | 1.864113711 | 0.001063556 | 2.798992403 | 8.696033109 | 9.405920729 | 10.11951029 | 10.42037002 | 10.4827929 | 11.48720025 | 11.26243498 |
| ENSG00000187193 | MT1X | 3.6615396 | 0.000101566 | -0.862683757 | 0.430112177 | 2.798855844 | 6.366709671 | 6.228547981 | 3.369317972 | 9.422406043 | 9.34683476 | 5.371837532 | 4.604965121 |
| ENSG00000060749 | QSER1 | 1.153712923 | 0.119776467 | 1.644186545 | 0.019679078 | 2.797899468 | 7.750913381 | 8.213429689 | 6.322058544 | 8.847689662 | 8.69465306 | 9.279302964 | 9.240151898 |
| ENSG00000139668 | WDFY2 | 2.001133626 | 0.287567214 | 0.79592993 | 0.691793928 | 2.797063555 | 3.327711234 | 8.077869079 | 3.369317972 | 8.519504557 | 8.61125371 | 7.444878196 | 7.313381205 |
| ENSG00000176095 | IP6K1 | 1.065961396 | 0.478780358 | 1.730712906 | 0.218585328 | 2.796674302 | 6.89194353 | 9.923599762 | 5.930865648 | 9.682298904 | 9.60914575 | 10.2683239 | 10.34713831 |
| ENSG00000124557 | BTN1A1 | 3.305309898 | 0.256719105 | -0.510113158 | 0.881429417 | 2.79519674 | 2 | 2 | 2 | 3.95996233 | 3.65239003 | 2.34604626 | 2.117335275 |
| ENSG00000130382 | MLLT1 | 1.192985989 | 0.117455955 | 1.600556673 | 0.028078154 | 2.793542663 | 8.39321512 | 9.386925327 | 7.502634626 | 9.763363111 | 9.87371334 | 10.1071225 | 10.33507589 |
| ENSG00000116497 | S100PBP | 1.129905457 | 0.47627622 | 1.663198898 | 0.2660773 | 2.793104355 | 6.653098118 | 8.395519306 | 3.369317972 | 8.310225403 | 8.36731141 | 9.138255231 | 8.532164007 |
| ENSG00000137494 | ANKRD42 | 1.473998306 | 0.395630265 | 1.31882697 | 0.443133621 | 2.792825277 | 2 | 6.04685525 | 7.734844356 | 8.033874364 | 7.94559728 | 7.949254677 | 7.7169976 |
| ENSG00000158411 | MITD1 | 1.750203409 | 0.007222848 | 1.041435024 | 0.126314768 | 2.791638433 | 5.532353208 | 4.710564154 | 4.874587708 | 6.622948477 | 6.75096147 | 6.126613714 | 5.892708443 |
| ENSG00000145362 | ANK2 | -0.848018358 | 0.499737617 | 3.63935418 | 0.000559099 | 2.791335821 | 3.327711234 | 6.228547981 | 6.882775709 | 5.614989456 | 4.89348248 | 9.397089785 | 9.845904672 |
| ENSG00000132623 | ANKEF1 | 0.719244279 | 0.762684238 | 2.069811962 | 0.329610478 | 2.789056242 | 2 | 3.686303414 | 2 | 3.289987342 | 3.2184422 | 4.652122235 | 3.474731315 |
| ENSG00000184661 | CDCA2 | 2.805923552 | 0.010849998 | -0.01978254 | 0.988654005 | 2.786141012 | 3.327711234 | 5.595927442 | 3.369317972 | 7.161462517 | 7.17852729 | 4.281782022 | 4.810605986 |
| ENSG00000175414 | ARL10 | -0.362929574 | 0.85249814 | 3.148349855 | 0.047119088 | 2.785420281 | 8.777722034 | 5.595927442 | 3.369317972 | 7.030886362 | 7.00595521 | 10.57214341 | 10.39984297 |
| ENSG00000157999 | ANKRD61 | -0.313903303 | 0.931247849 | 3.098989004 | 0.298214368 | 2.785085701 | 2 | 2 | 2 | 2.141737246 | 2.38539787 | 3.679182183 | 3.660199087 |
| ENSG00000176915 | ANKLE2 | 1.351211071 | 5.93E-05 | 1.433873987 | 1.66E-05 | 2.785085059 | 9.371511179 | 9.28800521 | 8.742726175 | 10.52838133 | 10.4873138 | 10.61914866 | 10.56112449 |
| ENSG00000119685 | TTLL5 | 1.215139655 | 0.174428066 | 1.564350031 | 0.067386662 | 2.779489685 | 4.467311984 | 6.78762695 | 6.718950348 | 7.491120977 | 7.4949172 | 7.683291963 | 7.972925696 |
| ENSG00000237190 | CDKN2AIPNL | 0.878605881 | 0.564421351 | 1.899346125 | 0.172231671 | 2.777952006 | 2 | 6.464272742 | 5.156626401 | 6.283823264 | 6.22964624 | 7.146744843 | 7.325625608 |
| ENSG00000132471 | WBP2 | 1.031872246 | 8.80E-05 | 1.745761235 | 4.21E-12 | 2.777633482 | 9.497412313 | 9.632483534 | 9.61300511 | 10.57755058 | 10.6438729 | 11.16817282 | 11.46385329 |
| ENSG00000135315 | CEP162 | 1.522224911 | 0.387998502 | 1.254646785 | 0.475883236 | 2.776871696 | 3.327711234 | 6.228547981 | 2 | 6.379946408 | 6.26836889 | 6.151583464 | 5.98823215 |
| ENSG00000112365 | ZBTB24 | 1.336626944 | 0.381296261 | 1.440091386 | 0.332115168 | 2.77671833 | 3.327711234 | 7.002537973 | 4.058998049 | 6.979883891 | 6.99676009 | 7.183528998 | 6.987067564 |
| ENSG00000185621 | LMLN | 1.932493928 | 0.003085869 | 0.842076037 | 0.23478853 | 2.774569966 | 5.095959458 | 6.535001748 | 5.930865648 | 8.027038952 | 7.66998641 | 6.583362013 | 6.983208048 |
| ENSG00000148700 | ADD3 | 1.747515187 | 0.014968435 | 1.025874811 | 0.172755691 | 2.773389999 | 9.371511179 | 9.856140801 | 7.934808822 | 11.02051291 | 10.9793733 | 10.23187735 | 10.32710477 |
| ENSG00000149948 | HMGA2 | 2.564335009 | 3.00E-05 | 0.208826321 | 0.788857412 | 2.77316133 | 6.816637977 | 8.255926468 | 8.143093325 | 10.44750059 | 10.3857483 | 8.354863925 | 7.736616914 |
| ENSG00000148356 | LRSAM1 | 1.104322747 | 0.57816817 | 1.667132923 | 0.374392974 | 2.77145567 | 4.815609471 | 7.874700259 | 2 | 7.461007055 | 7.64666005 | 8.070592084 | 8.147750532 |
| ENSG00000183647 | ZNF530 | 1.14522636 | 0.421750953 | 1.625243748 | 0.225861304 | 2.770470107 | 2 | 4.93638933 | 3.369317972 | 5.003571641 | 4.811949 | 5.427419529 | 5.23734678 |
| ENSG00000251201 | TMED7-TICA | 8.922970381 | 3.03E-10 | -6.15294715 | 0.005714661 | 2.770023231 | 2 | 2 | 3.369317972 | 9.911750163 | 9.92248069 | 2 | 2 |
| ENSG00000136026 | CKAP4 | 1.748271279 | 1.18E-05 | 1.021679778 | 0.015233955 | 2.769951057 | 11.67849893 | 12.43359305 | 12.12382753 | 13.8324849 | 13.885323 | 12.89278527 | 13.33854399 |
| ENSG00000174137 | FAM53A | 1.090947744 | 0.564770025 | 1.677954217 | 0.347715703 | 2.768901961 | 2 | 4.710564154 | 2 | 4.443344104 | 4.37434574 | 4.996409624 | 4.80186789 |
| ENSG00000062370 | ZNF112 | 2.011641434 | 0.104928256 | 0.757183 | 0.572430102 | 2.768824434 | 4.467311984 | 5.595927442 | 2 | 6.529309835 | 6.507263 | 5.056402493 | 5.59182244 |
| ENSG00000159714 | ZDHHC1 | 1.488202593 | 0.313991908 | 1.280096897 | 0.384020069 | 2.768299491 | 6.468621203 | 6.666836837 | 2 | 7.337530098 | 7.56813436 | 7.541454591 | 6.895576665 |
| ENSG00000198042 | MAK16 | 1.949981094 | 0.224835356 | 0.817845983 | 0.633808927 | 2.767827077 | 2 | 7.35507212 | 5.772716771 | 8.126317766 | 8.10812366 | 7.024410815 | 6.994755759 |
| ENSG00000152484 | USP12 | 1.671765098 | 0.356532724 | 1.095260672 | 0.553726324 | 2.76702577 | 2 | 7.901717006 | 8.511888463 | 9.339314897 | 9.29150981 | 8.832932104 | 8.648219991 |
| ENSG00000163873 | GRIK3 | 2.867387832 | 0.607619146 | -0.101338069 | 0.986936055 | 2.766049763 | 2 | 2 | 2 | 4.237680128 | 2 | 2.384001718 | 2.225841198 |
| ENSG00000140521 | POLG | 0.766696349 | 0.416762166 | 1.99880831 | 0.017463001 | 2.765504658 | 8.631580145 | 6.311476754 | 7.552162999 | 8.621345041 | 8.4504872 | 9.908424552 | 9.59812349 |
| ENSG00000169592 | INO80E | 1.462706723 | 0.000263663 | 1.301602073 | 0.001237837 | 2.764308796 | 7.218819781 | 6.78762695 | 6.534109959 | 8.325162455 | 8.28283292 | 8.083625221 | 8.204380326 |
| ENSG00000163807 | KIAA1143 | 1.257307223 | 0.002222495 | 1.50618855 | 0.000167667 | 2.763495773 | 6.816637977 | 7.274681896 | 7.646393253 | 8.509731121 | 8.52593541 | 8.824330559 | 8.701457496 |
| ENSG00000166432 | ZMAT1 | 1.296378455 | 0.351836711 | 1.464015248 | 0.275776462 | 2.760393703 | 4.00722531 | 4.442710129 | 2 | 4.965943888 | 4.811949 | 5.454427604 | 4.445805334 |
| ENSG00000166685 | COG1 | 1.28743921 | 0.38727917 | 1.471270643 | 0.306126091 | 2.758709854 | 8.257427002 | 7.847167909 | 3.369317972 | 8.702235867 | 8.85462427 | 9.073312544 | 8.842874172 |
| ENSG00000060069 | CTDP1 | 2.025637517 | 0.113077307 | 0.732236112 | 0.598176643 | 2.757873629 | 3.327711234 | 6.464272742 | 7.552162999 | 8.47833774 | 8.6202631 | 7.289818338 | 7.269688419 |
| ENSG00000153936 | HS2ST1 | 1.965688948 | 0.037719519 | 0.78796485 | 0.44254216 | 2.753653798 | 6.366709671 | 6.728495673 | 8.677268317 | 9.595637371 | 9.5561418 | 8.558404203 | 8.239587639 |
| ENSG00000184304 | PRKD1 | 1.133883377 | 0.539294232 | 1.619100855 | 0.357994057 | 2.752984232 | 8.468948322 | 7.638155693 | 2 | 8.666028533 | 8.64548836 | 9.01441148 | 9.249822 |
| ENSG00000127990 | SGCE | 1.433984388 | 0.010605581 | 1.318937742 | 0.019302872 | 2.752922129 | 8.39321512 | 8.124485672 | 7.029880454 | 9.357592295 | 9.40533771 | 9.202392925 | 9.329501596 |
| ENSG00000133773 | CCDC59 | 1.636040299 | 0.143639472 | 1.113411858 | 0.329610478 | 2.749452157 | 6.468621203 | 6.78762695 | 3.369317972 | 7.703217906 | 7.7377845 | 7.325609491 | 7.083953563 |
| ENSG00000166046 | TCP11L2 | 1.392181144 | 0.621742885 | 1.356024708 | 0.627083786 | 2.748205852 | 8.257427002 | 2 | 2 | 8.047448792 | 8.08227582 | 8.14016682 | 7.909264332 |
| ENSG00000147799 | ARHGAP39 | 1.869292484 | 0.157705645 | 0.878594434 | 0.529438052 | 2.747886918 | 2 | 6.04685525 | 4.874587708 | 6.755670824 | 6.89154576 | 5.883320383 | 5.884455123 |
| ENSG00000152284 | TCF7L1 | 0.788308943 | 0.455839475 | 1.959527919 | 0.037602746 | 2.747836862 | 5.330594134 | 7.73146271 | 5.930865648 | 7.440576128 | 7.5462416 | 8.660513226 | 8.634751663 |
| ENSG00000141582 | CBX4 | 1.155647655 | 0.325307017 | 1.591790899 | 0.153932236 | 2.747438554 | 8.285626337 | 7.274681896 | 4.874587708 | 8.583021623 | 8.42671895 | 9.200067611 | 8.619914108 |
| ENSG00000107140 | TESK1 | 1.098174494 | 0.437912435 | 1.648892318 | 0.215287817 | 2.747066812 | 4.467311984 | 7.901717006 | 8.960138144 | 9.043291801 | 9.09332116 | 9.714689414 | 9.509666762 |
| ENSG00000004660 | CAMKK1 | 1.630854221 | 0.292091049 | 1.115970017 | 0.477543645 | 2.746824238 | 3.327711234 | 5.595927442 | 2 | 5.978405189 | 5.77157844 | 5.436478516 | 5.373314945 |
| ENSG00000140400 | MAN2C1 | 0.947657014 | 0.521741285 | 1.798936732 | 0.185049898 | 2.746593746 | 4.00722531 | 8.395519306 | 6.534109959 | 8.115618198 | 8.18906731 | 8.967137431 | 9.022374596 |
| ENSG00000157978 | LDLRAP1 | -0.489452467 | 0.805509199 | 3.235407096 | 0.049334117 | 2.745954629 | 7.485143553 | 8.487557258 | 2 | 7.012547947 | 7.03768431 | 10.35791611 | 10.99030784 |
| ENSG00000066855 | MTFR1 | 1.32415491 | 0.20606912 | 1.420864021 | 0.163643498 | 2.74501893 | 4.00722531 | 6.464272742 | 7.029880454 | 7.565338955 | 7.57741621 | 7.846806418 | 7.459438299 |
| ENSG00000145414 | NAF1 | 1.040943586 | 0.251528741 | 1.70387763 | 0.0439318 | 2.744821216 | 4.815609471 | 4.93638933 | 6.629488387 | 6.529309835 | 6.80463998 | 7.232460462 | 7.39408282 |
| ENSG00000204946 | ZNF783 | 1.32849268 | 0.337673274 | 1.414459558 | 0.294688747 | 2.742952239 | 5.095959458 | 3.686303414 | 7.342949392 | 7.337530098 | 7.4314987 | 7.447120996 | 7.491038307 |
| ENSG00000186891 | TNFRSF18 | 2.079149751 | 0.540424469 | 0.661664827 | NA | 2.740814579 | 2 | 2 | 2 | 2.784612485 | 3.28037825 | 2.592885947 | 2.374681325 |
| ENSG00000179832 | MROH1 | 1.713337853 | 0.091677988 | 1.027447214 | 0.328380953 | 2.740785067 | 7.031698319 | 5.457120953 | 8.325066185 | 9.088859521 | 9.0020652 | 8.364981826 | 8.368713182 |
| ENSG00000197535 | MYO5A | 0.697839044 | 0.667312826 | 2.042031404 | 0.157060567 | 2.739870449 | 4.00722531 | 8.806266215 | 9.116480286 | 9.062360705 | 9.13582529 | 10.12456825 | 10.69427292 |
| ENSG00000165732 | DDX21 | 1.271314794 | 0.076622525 | 1.468463096 | 0.035337738 | 2.73977789 | 7.940205782 | 9.687856435 | 9.516493218 | 10.51174802 | 10.4856715 | 10.72136295 | 10.66896863 |
| ENSG00000046889 | PREX2 | -1.269416704 | 0.637611669 | 4.008638154 | 0.06941864 | 2.73922145 | 2 | 2 | 3.369317972 | 2.3892346 | 2.14022021 | 5.815498119 | 3.956354764 |
| ENSG00000047932 | GOPC | 1.653517548 | 0.137415848 | 1.085565799 | 0.342050085 | 2.739083347 | 8.340423424 | 8.777382925 | 5.39244166 | 9.746123826 | 9.67547818 | 9.163708059 | 9.129269498 |
| ENSG00000117298 | ECE1 | 0.611853258 | 0.114890958 | 2.124435107 | 3.85E-10 | 2.736288365 | 10.1311248 | 10.74900803 | 10.23774519 | 10.96675639 | 11.0555197 | 12.42250978 | 12.6161781 |
| ENSG00000152778 | IFIT5 | 2.360511616 | 0.00037188 | 0.374881416 | 0.637246531 | 2.735393032 | 6.816637977 | 8.297207175 | 8.631919287 | 10.43765481 | 10.4541068 | 8.385592001 | 8.554444396 |
| ENSG00000157077 | ZFYVE9 | 1.359350186 | 0.039642635 | 1.375324082 | 0.035171121 | 2.734674269 | 6.816637977 | 6.728495673 | 8.110401118 | 8.697938862 | 8.68752719 | 8.726627392 | 8.690255954 |
| ENSG00000105668 | UPK1A | -0.687348191 | 0.780246302 | 3.420751441 | 0.083002579 | 2.73340325 | 2 | 2 | 3.369317972 | 2.3892346 | 2.38539787 | 4.301748724 | 4.943664032 |
| ENSG00000184602 | SNN | -0.202395157 | 0.906452235 | 2.935454527 | 0.030273608 | 2.733059369 | 4.467311984 | 7.274681896 | 3.369317972 | 5.829220145 | 5.77157844 | 8.852523173 | 8.834865421 |
| ENSG00000232653 | GOLGA8N | 7.923222943 | 9.74E-06 | -5.191149092 | 0.042128548 | 2.732073851 | 2 | 2 | 2 | 8.006336225 | 7.93841129 | 2 | 2 |
| ENSG00000144339 | TMEFF2 | 2.11075326 | 0.313991908 | 0.620277279 | 0.787507734 | 2.73103054 | 2 | 3.686303414 | 2 | 4.623314311 | 3.65239003 | 3.082820997 | 3.311948089 |
| ENSG00000188786 | MTF1 | 1.813673055 | 0.191729965 | 0.91592059 | 0.529629876 | 2.729593645 | 4.815609471 | 6.602424674 | 8.99637432 | 9.505895854 | 9.5300868 | 8.763539652 | 8.476283084 |
| ENSG00000180182 | MED14 | 1.171890363 | 0.433017846 | 1.555658549 | 0.273747396 | 2.727548913 | 4.00722531 | 8.791896851 | 7.285536013 | 8.875861021 | 8.80807321 | 9.325140951 | 9.113466702 |
| ENSG00000128159 | TUBGCP6 | 0.964174018 | 0.63904641 | 1.762763648 | 0.358092441 | 2.726937665 | 2 | 6.602424674 | 8.922968322 | 8.562683721 | 8.55441692 | 9.332135462 | 9.368859419 |
| ENSG00000183723 | CMTM4 | -0.400865562 | 0.486993352 | 3.122661312 | 1.37E-11 | 2.72179575 | 8.042902253 | 6.951736442 | 7.398165132 | 7.161462517 | 7.10341274 | 10.72240494 | 10.50587116 |
| ENSG00000206127 | GOLGA8O | 7.912392298 | 1.05E-05 | -5.191149049 | 0.042390794 | 2.721243249 | 2 | 2 | 2 | 7.994705029 | 7.92877397 | 2 | 2 |
| ENSG00000065518 | NDUFB4 | 1.870897843 | 0.002715259 | 0.84999279 | 0.207460653 | 2.720890634 | 8.418903792 | 7.638155693 | 6.80318718 | 9.584041242 | 9.65077635 | 8.480070636 | 8.718968947 |

| | | | | | | | | | | | | | |
|---|---|---|---|---|---|---|---|---|---|---|---|---|---|
| ENSG00000278567 | | 7.911711469 | 1.14E-05 | -5.191148959 | 0.042837614 | 2.720562511 | 2 | 2 | 2 | 7.872924734 | 8.04484669 | 2 | 2 |
| ENSG00000150347 | ARID5B | 0.644441967 | 0.311485333 | 2.075203831 | 0.000179121 | 2.719645798 | 9.247966931 | 10.3778065 | 10.60345421 | 10.84151467 | 10.8161359 | 12.37308854 | 12.13214626 |
| ENSG00000196652 | ZKSCAN5 | 1.432712851 | 0.223670516 | 1.286883826 | 0.271141542 | 2.719596677 | 4.467311984 | 7.928237109 | 6.629488387 | 8.30646685 | 8.37442884 | 8.364981826 | 8.006587762 |
| ENSG00000006837 | CDKL3 | 2.534170854 | 0.125092709 | 0.184862267 | 0.926455594 | 2.719033121 | 2 | 4.11352228 | 2 | 5.110902178 | 5.00774437 | 3.336159116 | 3.236138336 |
| ENSG00000130340 | SNX9 | 2.277146905 | 4.55E-07 | 0.440341668 | 0.406781563 | 2.717488573 | 8.910412251 | 9.497318007 | 9.999667074 | 11.77852445 | 11.8337041 | 10.02035217 | 9.925491821 |
| ENSG00000155034 | FBXL18 | 1.17438553 | 0.010955982 | 1.543052184 | 0.000501994 | 2.717437715 | 7.435667355 | 6.78762695 | 6.80318718 | 8.215198712 | 8.15227214 | 8.348878903 | 8.723602991 |
| ENSG00000013810 | TACC3 | 1.442976149 | 0.298925954 | 1.273863148 | 0.354405417 | 2.716839297 | 4.467311984 | 5.838937808 | 2 | 6.147608921 | 6.1572134 | 5.669467138 | 6.257923755 |
| ENSG00000116473 | RAP1A | 1.617517759 | 6.67E-05 | 1.098823591 | 0.008939862 | 2.71634135 | 6.96351253 | 6.899080544 | 7.600047325 | 8.78154108 | 8.75316018 | 8.317350121 | 8.18854565 |
| ENSG00000154845 | PPP4R1 | 1.269502587 | 0.008153101 | 1.446571412 | 0.002075724 | 2.716073999 | 8.468948322 | 8.702538795 | 7.734844356 | 9.665524776 | 9.58096568 | 9.588555735 | 9.984600648 |
| ENSG00000131943 | C19orf12 | 1.521052903 | 0.021456873 | 1.194789446 | 0.075777971 | 2.715842349 | 7.33131396 | 8.732943944 | 7.451345288 | 9.481935579 | 9.52849248 | 9.071859866 | 9.282773417 |
| ENSG00000157637 | SLC38A10 | 1.275514678 | 0.000285776 | 1.440251073 | 3.13E-05 | 2.715765751 | 9.689059027 | 9.574900212 | 9.014156722 | 10.69791695 | 10.7571169 | 10.85119671 | 10.93206299 |
| ENSG00000162409 | PRKAA2 | -0.453130259 | 0.769854407 | 3.167111114 | 0.012869116 | 2.713980855 | 2 | 6.535001748 | 5.39244166 | 5.161714397 | 5.13053105 | 8.682572389 | 8.53018147 |
| ENSG00000131446 | MGAT1 | 0.648162013 | 0.234880504 | 2.065133075 | 1.44E-05 | 2.713295088 | 10.88765625 | 10.80015962 | 9.935792933 | 11.25625289 | 11.2397407 | 12.37463584 | 12.90408303 |
| ENSG00000148396 | SEC16A | 1.032292346 | 0.001737311 | 1.678848743 | 8.59E-08 | 2.711141089 | 9.635285994 | 10.01303782 | 9.452720984 | 10.73054016 | 10.7747077 | 11.47783828 | 11.31375888 |
| ENSG00000106034 | CPED1 | 0.37120571 | 0.808679443 | 2.339793206 | 0.067094792 | 2.710998916 | 4.815609471 | 8.834582582 | 6.718950348 | 7.966401501 | 8.01782901 | 9.681736371 | 10.16578017 |
| ENSG00000184007 | PTP4A2 | 1.230214989 | 0.013448518 | 1.48072325 | 0.002239989 | 2.710938239 | 9.317904228 | 9.785372135 | 10.36791565 | 11.10109884 | 11.1257404 | 11.16204184 | 11.54037635 |
| ENSG00000135299 | ANKRD6 | -0.923545272 | 0.558941971 | 3.634376952 | 0.007051903 | 2.710831679 | 3.327711234 | 6.602424674 | 3.369317972 | 4.623314311 | 4.43005853 | 8.337438499 | 9.190804563 |
| ENSG00000124459 | ZNF45 | 1.326929285 | 0.39383571 | 1.383212314 | 0.362782971 | 2.710141599 | 6.96351253 | 6.951736442 | 2 | 7.68016596 | 7.6957855 | 7.814163164 | 7.66855289 |
| ENSG00000136449 | MYCBPAP | 1.27173153 | 0.547559684 | 1.43593974 | 0.482681185 | 2.707671271 | 2 | 3.686303414 | 2 | 3.663521627 | 3.55551834 | 3.917810786 | 3.496556178 |
| ENSG00000141252 | VPS53 | 1.189571563 | 0.000120617 | 1.516241998 | 4.74E-07 | 2.705813562 | 8.340423424 | 8.101365653 | 8.408111533 | 9.436230969 | 9.48723044 | 9.651907244 | 9.910582344 |
| ENSG00000119396 | RAB14 | 1.472365635 | 0.145397751 | 1.232427883 | 0.223084039 | 2.704793518 | 6.653098118 | 9.479495904 | 9.272377397 | 10.42471067 | 10.3051703 | 10.16889595 | 10.08390135 |
| ENSG00000153066 | TXNDC11 | 1.248975624 | 0.44518671 | 1.455213756 | 0.356290523 | 2.70418938 | 8.963744667 | 7.638155693 | 3.369317972 | 9.138327453 | 9.09977716 | 9.085961465 | 9.528327138 |
| ENSG00000156298 | TSPAN7 | -0.31396534 | 0.93407822 | 3.017927837 | 0.337194476 | 2.703962497 | 2 | 2 | 2 | 2.141737246 | 2.38539787 | 3.082820997 | 4.002845082 |
| ENSG00000158715 | SLC45A3 | 1.505109494 | 0.486454906 | 1.198558861 | 0.579500907 | 2.703668355 | 2 | 5.72254323 | 2 | 5.552894743 | 5.83486613 | 5.737015686 | 5.013446246 |
| ENSG00000167996 | FTH1 | 3.645269536 | 1.35E-06 | -0.941740737 | 0.274614244 | 2.703528799 | 14.02224794 | 12.20053628 | 12.26418878 | 16.70361438 | 16.7763081 | 12.29446237 | 11.99973474 |
| ENSG00000198715 | GLMP | 2.003086876 | 0.032193991 | 0.699505671 | 0.49591259 | 2.702592547 | 6.009025997 | 8.57407404 | 8.65477198 | 10.14797511 | 10.1190812 | 8.810024926 | 8.864540598 |
| ENSG00000197937 | ZNF347 | 0.749686183 | 0.393232855 | 1.94918618 | 0.01243418 | 2.698872362 | 5.095959458 | 6.951736442 | 5.59507254 | 6.892328675 | 6.80989959 | 8.161494476 | 7.869046644 |
| ENSG00000117222 | RBBP5 | 1.359031525 | 0.040747488 | 1.339720137 | 0.041517795 | 2.698751662 | 7.384433967 | 6.844429755 | 5.930865648 | 8.239055745 | 8.09310209 | 8.438967212 | 7.78508305 |
| ENSG00000165168 | CYBB | -2.307535798 | 0.518188676 | 5.004968725 | 0.063842507 | 2.697432926 | 2 | 2 | 2 | 2.141737246 | 2 | 5.436478516 | 4.911532223 |
| ENSG00000160321 | ZNF208 | 3.825406485 | 0.09741056 | -1.128591071 | 0.666867266 | 2.696815414 | 2 | 2 | 3.369317972 | 4.965943888 | 5.04390545 | 2.492221014 | 2.059860178 |
| ENSG00000135636 | DYSF | 0.526742625 | 0.57541332 | 2.169932554 | 0.006709168 | 2.696675179 | 4.00722531 | 5.946638857 | 6.322058544 | 6.213314212 | 6.21386141 | 7.994427465 | 7.581886257 |
| ENSG00000198019 | FCGR1B | 2.391122996 | 0.505231655 | 0.303858541 | NA | 2.694981537 | 2 | 2 | 2 | 3.349643765 | 3.08597292 | 2.62493832 | 2.117335275 |
| ENSG00000127415 | IDUA | 1.448205846 | 0.082711278 | 1.243668653 | 0.135257068 | 2.691874499 | 7.218819781 | 5.303523747 | 7.398165132 | 8.262524677 | 8.32752521 | 7.953205078 | 8.221681954 |
| ENSG00000011258 | MBTD1 | 1.649366555 | 0.011067153 | 1.042455474 | 0.122835348 | 2.691822029 | 4.815609471 | 6.04685525 | 6.203069917 | 7.450827758 | 7.33264456 | 7.012344451 | 6.559516442 |
| ENSG00000117602 | RCAN3 | 0.078997334 | 0.972415697 | 2.612704728 | 0.169351211 | 2.691702062 | 3.327711234 | 7.274681896 | 2 | 5.775854763 | 6.02777321 | 8.480618146 | 8.226587404 |
| ENSG00000132275 | RRP8 | 0.658538731 | 0.624897619 | 2.032532519 | 0.087371458 | 2.69107125 | 4.00722531 | 7.274681896 | 5.156626401 | 6.739036112 | 6.77805036 | 8.247650629 | 7.934467448 |
| ENSG00000110237 | ARHGEF17 | 0.367453575 | 0.745038229 | 2.32335018 | 0.013586383 | 2.690803755 | 7.53297911 | 9.616264339 | 10.42884603 | 9.922226014 | 10.031744 | 11.84735385 | 12.00753405 |
| ENSG00000168310 | IRF2 | 1.735573809 | 0.160777328 | 0.953568785 | 0.45939756 | 2.689142594 | 8.313285031 | 6.728495673 | 4.523675391 | 8.896009193 | 8.95655043 | 8.318575593 | 7.966097995 |
| ENSG00000197782 | ZNF780A | 2.204626412 | 0.015668605 | 0.48249955 | 0.646493605 | 2.687125963 | 4.00722531 | 6.464272742 | 6.073378139 | 8.056427969 | 7.90193448 | 6.651077397 | 5.863612864 |
| ENSG00000235194 | PPP1R3E | 0.940454805 | 0.688364462 | 1.745953082 | 0.424370697 | 2.686407887 | 2 | 6.228547981 | 2 | 5.709096143 | 5.6121243 | 6.453284622 | 6.380362005 |
| ENSG00000117505 | DR1 | 0.902390521 | 0.03191533 | 1.783457641 | 4.24E-06 | 2.685848162 | 8.009475818 | 8.147241016 | 8.742726175 | 9.256998085 | 9.19229617 | 10.21585362 | 9.977585641 |
| ENSG00000128272 | ATF4 | 1.257935558 | 0.254759301 | 1.426745434 | 0.182472647 | 2.684680992 | 11.59645304 | 10.35843478 | 8.677268317 | 11.90523218 | 11.905324 | 12.52778173 | 11.4079987 |
| ENSG00000173226 | IQCB1 | 0.977973036 | 0.522431564 | 1.706661253 | 0.227974555 | 2.684634288 | 4.00722531 | 6.844429755 | 3.369317972 | 6.503277904 | 6.51372389 | 7.382965824 | 7.017577533 |
| ENSG00000133703 | KRAS | 0.935335497 | 0.040893562 | 1.748344497 | 3.45E-05 | 2.683679993 | 7.624139768 | 7.232742069 | 8.143093325 | 8.68062069 | 8.56534297 | 9.403744811 | 9.457418003 |
| ENSG00000182504 | CEP97 | 1.486511731 | 0.012898585 | 1.196968583 | 0.048314828 | 2.683480314 | 6.468621203 | 6.04685525 | 5.930865648 | 7.708923796 | 7.47849652 | 7.645704907 | 6.885273196 |
| ENSG00000085224 | ATRX | 1.372507745 | 0.001997826 | 1.310309429 | 0.003174989 | 2.682817175 | 8.367060731 | 9.28800521 | 8.99637432 | 10.29841863 | 10.3065663 | 10.38974639 | 10.07395575 |
| ENSG00000163818 | LZTFL1 | 1.542985139 | 0.299070637 | 1.139137403 | 0.447404663 | 2.682122543 | 6.468621203 | 6.899080544 | 2 | 7.650823203 | 7.63485389 | 7.274744493 | 7.221358049 |
| ENSG00000180113 | TDRD6 | -0.314150389 | 0.943613626 | 2.994915704 | 0.433184926 | 2.680765315 | 2 | 2 | 2 | 2.141737246 | 2.38539787 | 4.301748724 | 2.172608095 |
| ENSG00000049245 | VAMP3 | 1.060260805 | 0.004700034 | 1.620134281 | 5.85E-06 | 2.680395087 | 8.910412251 | 9.497318007 | 9.624627483 | 10.49986612 | 10.3622229 | 10.98995534 | 10.99246943 |
| ENSG00000163626 | COX18 | 0.892411519 | 0.197017026 | 1.787963263 | 0.004469801 | 2.680374782 | 5.866944937 | 5.131603625 | 6.43197674 | 6.86703417 | 6.55815769 | 7.75021102 | 7.411419562 |
| ENSG00000043514 | TRIT1 | 1.520871805 | 0.354199679 | 1.159215915 | 0.483045217 | 2.68008772 | 7.096806373 | 5.131603625 | 2 | 7.307758725 | 7.34356593 | 6.812870125 | 7.119550903 |
| ENSG00000162522 | KIAA1522 | 0.7244839 | 0.492298378 | 1.952589319 | 0.037315852 | 2.677073219 | 9.112843761 | 8.487557258 | 6.322058544 | 9.012478123 | 9.1617682 | 10.12596884 | 10.47511327 |
| ENSG00000120733 | KDM3B | 1.322209443 | 0.318322589 | 1.352936174 | 0.297469212 | 2.675145617 | 5.70932728 | 9.599860728 | 7.552162999 | 9.728676049 | 9.71245786 | 9.697365195 | 9.80303401 |
| ENSG00000143369 | ECM1 | 0.967472598 | 0.152981087 | 1.706595965 | 0.006499774 | 2.674068563 | 10.7426185 | 9.608085847 | 9.636156974 | 11.0364838 | 11.0883544 | 11.3315637 | 12.15414401 |
| ENSG00000169621 | APLF | 1.243007645 | 0.561101819 | 1.42930552 | 0.49125968 | 2.672313166 | 2 | 5.595927442 | 2 | 5.527288686 | 5.13053105 | 5.699879814 | 5.294527297 |
| ENSG00000123552 | USP45 | 1.606482441 | 0.226793445 | 1.065233203 | 0.434090596 | 2.671715644 | 6.257048221 | 5.595927442 | 2 | 6.984595707 | 6.90635305 | 6.651077397 | 6.158754168 |
| ENSG00000164190 | NIPBL | 1.686982353 | 0.279279457 | 0.983993473 | 0.541021035 | 2.670975826 | 4.00722531 | 9.488434476 | 8.206335842 | 10.11182193 | 10.0907323 | 9.656275732 | 9.092123385 |
| ENSG00000178295 | GEN1 | 1.766442103 | 0.169077527 | 0.904419629 | 0.501207541 | 2.670861732 | 3.327711234 | 6.140560021 | 4.058998049 | 6.733448311 | 6.67228943 | 6.287958221 | 5.32534152 |
| ENSG00000142634 | EFHD2 | 1.728479323 | 0.002054399 | 0.937311808 | 0.115419908 | 2.665791131 | 7.384433967 | 7.393647515 | 8.536706852 | 9.595637371 | 9.56938265 | 8.761288263 | 8.831649431 |
| ENSG00000067369 | TP53BP1 | 1.417411793 | 0.029261156 | 1.247866721 | 0.055892984 | 2.665278514 | 7.790810999 | 9.338312924 | 9.242516611 | 10.35546265 | 10.3514424 | 10.15866828 | 10.20979627 |
| ENSG00000095564 | BTAF1 | 1.612292098 | 0.263419443 | 1.052978316 | 0.476093865 | 2.665270415 | 4.00722531 | 8.687092496 | 8.65477198 | 9.730079663 | 9.69763767 | 9.357098224 | 8.926648946 |
| ENSG00000187650 | VMAC | 1.137179203 | 0.407386256 | 1.526832416 | 0.241370875 | 2.664011619 | 2 | 4.11352228 | 5.59507254 | 5.540148523 | 5.47317115 | 5.751606453 | 5.97274432 |
| ENSG00000135835 | KIAA1614 | 0.974697615 | 0.171596185 | 1.688530319 | 0.010410997 | 2.663227934 | 6.138360429 | 4.93638933 | 6.43197674 | 6.970413838 | 6.73998187 | 7.671830666 | 7.425709383 |
| ENSG00000111731 | C2CD5 | 1.689549335 | 0.26495819 | 0.973450133 | 0.535268829 | 2.662999467 | 2 | 6.951736442 | 6.073378139 | 7.677258395 | 7.64959655 | 7.172727426 | 6.723495198 |
| ENSG00000161202 | DVL3 | 0.742359942 | 0.041992389 | 1.920289348 | 5.40E-09 | 2.66264929 | 9.189468652 | 9.77810053 | 9.330306409 | 10.16934346 | 10.2281967 | 11.46440721 | 11.27807399 |
| ENSG00000140443 | IGF1R | 0.779484532 | 0.149100125 | 1.882257119 | 9.95E-05 | 2.661741651 | 8.963744667 | 10.02537381 | 9.452720984 | 10.33761854 | 10.3135262 | 11.62184936 | 11.19927059 |
| ENSG00000187398 | LUZP2 | -3.509010171 | 0.033794082 | 6.169764805 | 4.48E-07 | 2.660754634 | 2 | 4.11352228 | 3.369317972 | 2.141737246 | 2.2680111 | 8.056705941 | 9.551969768 |
| ENSG00000162909 | CAPN2 | 0.881955897 | 0.037940998 | 1.776511717 | 5.60E-06 | 2.658467614 | 10.35389212 | 10.65686569 | 11.11279376 | 11.60655646 | 11.6367136 | 12.31329535 | 12.69269276 |
| ENSG00000166833 | NAV2 | 2.006471123 | 0.286042726 | 0.650372099 | 0.749694888 | 2.656843223 | 2 | 8.791896851 | 6.882775709 | 9.465176303 | 9.62191107 | 8.402448739 | 7.968052066 |
| ENSG00000141367 | CLTC | 1.210729687 | 0.041005338 | 1.446075035 | 0.012003587 | 2.656804722 | 10.93293345 | 12.0991443 | 10.9788475 | 12.65956989 | 12.6507234 | 12.76913358 | 13.00219207 |
| ENSG00000133739 | LRRCC1 | 2.015965816 | 0.170143299 | 0.637452859 | 0.691626719 | 2.653418674 | 3.327711234 | 4.710564154 | 2 | 5.552894743 | 5.41912973 | 4.705557986 | 3.735548302 |
| ENSG00000043093 | DCUN1D1 | 1.446082695 | 0.000365863 | 1.206230188 | 0.003365603 | 2.652312884 | 7.096806373 | 7.847167909 | 7.734844356 | 9.065699797 | 8.99400091 | 8.903637637 | 8.671790956 |
| ENSG00000152404 | CWF19L2 | 1.783069235 | 0.175251415 | 0.868256417 | 0.530574359 | 2.651325652 | 3.327711234 | 7.315436877 | 5.772716771 | 8.067573917 | 7.86198169 | 7.296052918 | 6.811011855 |
| ENSG00000021762 | OSBPL5 | 1.27021009 | 0.315574141 | 1.380423196 | 0.263184603 | 2.650633287 | 4.467311984 | 8.297207175 | 7.734844356 | 8.796356221 | 8.75179438 | 9.148628127 | 8.558340758 |
| ENSG00000155097 | ATP6V1C1 | 2.102017892 | 2.87E-10 | 0.547963962 | 0.146477581 | 2.649981855 | 8.138771685 | 8.791896851 | 8.434763955 | 10.56777138 | 10.5879489 | 9.062381638 | 8.999151483 |
| ENSG00000117139 | KDM5B | 1.174790195 | 0.099215121 | 1.473908514 | 0.03159717 | 2.648698709 | 8.493334308 | 10.12043894 | 10.08627303 | 10.89536487 | 10.9234525 | 11.33027328 | 11.07445415 |
| ENSG00000166908 | PIP4K2C | 0.965156699 | 0.216076248 | 1.683539008 | 0.020364713 | 2.648695707 | 7.031698319 | 7.393647515 | 8.721235202 | 8.864402784 | 8.84824797 | 9.356800024 | 9.755460113 |
| ENSG00000213648 | SULT1A4 | 7.839314907 | 1.74E-05 | -5.19114876 | 0.044185492 | 2.648166147 | 2 | 2 | 2 | 7.917958914 | 7.86198169 | 2 | 2 |
| ENSG00000261052 | SULT1A3 | 7.839314907 | 1.74E-05 | -5.19114876 | 0.044185492 | 2.648166147 | 2 | 2 | 2 | 7.917958914 | 7.86198169 | 2 | 2 |
| ENSG00000101846 | STS | 0.385943899 | 0.849097902 | 2.261509726 | 0.186667231 | 2.647453625 | 2 | 7.979858146 | 8.296287027 | 7.999368766 | 7.90193448 | 9.427508673 | 10.11200841 |
| ENSG00000136381 | IREB2 | 1.328409677 | 0.062735523 | 1.315930785 | 0.062080537 | 2.644340461 | 7.485143553 | 9.067150531 | 7.777116083 | 9.590237441 | 9.62936766 | 9.4949131 | 9.693328953 |
| ENSG00000155961 | RAB39B | 0.274454648 | 0.941409766 | 2.367105464 | NA | 2.641560113 | 2 | 2 | 2 | 2.3892346 | 2.38539787 | 3.429858444 | 2.950113106 |
| ENSG00000092295 | TGM1 | -0.185862332 | 0.936649074 | 2.82709693 | 0.140416036 | 2.641234598 | 5.095959458 | 2 | 2 | 3.566290607 | 3.74316441 | 5.4091291 | 6.709547124 |
| ENSG00000143951 | WDPCP | 0.945955489 | 0.663228562 | 1.692671387 | 0.403003915 | 2.638626876 | 2 | 5.595927442 | 2 | 5.22679461 | 4.91316568 | 5.740677235 | 5.768151538 |
| ENSG00000123560 | PLP1 | 0.914715675 | 0.822767511 | 1.72269462 | 0.65155864 | 2.637410294 | 4.00722531 | 2 | 2 | 4.237680128 | 2 | 3.5515855 | 4.479073019 |
| ENSG00000125971 | DYNLRB1 | 0.948663349 | 0.090124275 | 1.688744277 | 0.001093475 | 2.637407626 | 9.397585735 | 8.276714465 | 8.699419242 | 9.75855684 | 9.85029345 | 10.31281645 | 10.74087591 |
| ENSG00000144834 | TAGLN3 | 5.382677 | 0.006730413 | -2.748740215 | 0.268972233 | 2.633936785 | 2 | 2 | 2 | 5.501219927 | 5.5755357 | 2.048074405 | 2.059860178 |
| ENSG00000108306 | FBXL20 | 1.2615036 | 0.042589119 | 1.372180394 | 0.024545065 | 2.633683994 | 8.16936288 | 6.951736442 | 8.266922072 | 9.135152476 | 9.15144695 | 9.463959918 | 9.006300384 |
| ENSG00000137267 | TUBB2A | 1.126149674 | 0.400687122 | 1.507270483 | 0.238341966 | 2.633420157 | 8.946185348 | 7.145017454 | 5.156626401 | 8.983347946 | 8.84312657 | 8.792933324 | 9.664095852 |
| ENSG00000186399 | GOLGA8R | 7.824182883 | 2.01E-05 | -5.191148647 | 0.044848802 | 2.633034235 | 2 | 2 | 2 | 7.93509946 | 7.81305436 | 2 | 2 |
| ENSG00000189367 | KIAA0408 | 7.824023956 | 1.94E-05 | -5.191148689 | 0.044648908 | 2.632875266 | 2 | 2 | 2 | 7.908072101 | 7.84158259 | 2 | 2 |
| ENSG00000130699 | TAF4 | 2.113665643 | 0.315369213 | 0.519153485 | 0.823280183 | 2.632819127 | 2 | 5.946638857 | 2 | 6.496695859 | 6.48770488 | 5.264018433 | 4.729989799 |
| ENSG00000111358 | GTF2H3 | 1.670847243 | 0.058804128 | 0.960793281 | 0.296967694 | 2.631640524 | 4.815609471 | 7.232742069 | 6.43197674 | 8.076429088 | 8.13979554 | 7.17543541 | 7.614687532 |
| ENSG00000260097 | SPDYE6 | 2.273109036 | 0.486515971 | 0.356263819 | NA | 2.629372855 | 2 | 2 | 2 | 3.023133209 | 3.28037825 | 2.34604626 | 2.466008509 |
| ENSG00000175265 | GOLGA8A | 2.995602965 | 8.23E-09 | -0.368619795 | 0.562234526 | 2.626983171 | 7.904282243 | 8.862353853 | 7.646393253 | 11.24304106 | 11.2104591 | 7.959503304 | 7.805990373 |
| ENSG00000147036 | LANCL3 | -1.317080176 | 0.761841469 | 3.942291544 | 0.282983706 | 2.625211368 | 2 | 2 | 2 | 2 | 2.2680111 | 4.301748724 | 4.292061816 |
| ENSG00000171116 | HSFX1 | 3.148163868 | 0.280922232 | -0.523007381 | 0.877195052 | 2.625156487 | 2 | 2 | 2 | 3.615725078 | 3.7864959 | 2.139668783 | 2.326754077 |
| ENSG00000074047 | GLI2 | 1.507969014 | 0.41134066 | 1.116865386 | 0.545217341 | 2.6248344 | 2 | 8.029695761 | 6.073378139 | 8.264463296 | 8.28661061 | 8.189226297 | 7.512619696 |
| ENSG00000058335 | RASGRF1 | 0.271993944 | 0.944373487 | 2.351960904 | NA | 2.623954847 | 2 | 2 | 2 | 2.270786764 | 2.49394783 | 2.68698273 | 3.580696614 |
| ENSG00000140873 | ADAMTS18 | -0.302885678 | 0.955377328 | 2.926666065 | 0.542075443 | 2.623780387 | 2 | 2 | 2 | 2.498691189 | 2 | 3.796405078 | 3.261853426 |
| ENSG00000186815 | TPCN1 | 0.415328255 | 0.817831695 | 2.208198329 | 0.147401264 | 2.623526583 | 4.467311984 | 7.874700259 | 9.990713521 | 9.09649754 | 9.18421833 | 10.76830805 | 11.06922971 |
| ENSG00000108511 | HOXB6 | 5.45365659 | 0.000530987 | -2.830269874 | 0.111750609 | 2.623386715 | 2 | 4.11352228 | 2 | 7.844695415 | 7.77859536 | 2.384001718 | 2.059860178 |
| ENSG00000169169 | CPT1C | 1.548161696 | 0.239494508 | 1.074691805 | 0.423751061 | 2.622853501 | 7.159102577 | 7.538394143 | 3.369317972 | 8.332573384 | 8.35477078 | 7.633939843 | 8.085210331 |
| ENSG00000276802 | | 2.731243082 | 0.021534651 | -0.108592474 | 0.942222441 | 2.622650608 | 6.257048221 | 9.067150531 | 5.930865648 | 10.50397439 | 10.5542639 | 7.608114885 | 7.811442311 |
| ENSG00000215695 | RSC1A1 | 7.813253426 | 2.13E-05 | -5.191148622 | 0.045052759 | 2.622104804 | 2 | 2 | 2 | 7.815902688 | 7.91175216 | 2 | 2 |
| ENSG00000143321 | HDGF | 1.154053059 | 0.144865788 | 1.46697271 | 0.053383916 | 2.621025769 | 8.674867122 | 10.33384851 | 10.66665689 | 11.24008859 | 11.2904776 | 11.49263913 | 11.65852437 |
| ENSG00000197566 | ZNF624 | 0.8275107 | 0.634493636 | 1.791986234 | 0.260698796 | 2.619496934 | 2 | 5.838937808 | 3.369317972 | 5.289065324 | 5.40529704 | 6.423979439 | 6.033722345 |

| | | | | | | | | | | | | | |
|---|---|---|---|---|---|---|---|---|---|---|---|---|---|
| ENSG00000101654 | RNMT | 1.43278937 | 0.041269777 | 1.18541715 | 0.094023077 | 2.61820652 | 7.031698319 | 8.848535039 | 8.296287027 | 9.648553319 | 9.68266366 | 9.402011675 | 9.437105934 |
| ENSG00000119471 | HSDL2 | 0.850381466 | 0.6915899 | 1.767075949 | 0.370584029 | 2.617457416 | 2 | 8.762721501 | 5.772716771 | 8.205140657 | 8.19711812 | 9.162002347 | 9.053075307 |
| ENSG00000125551 | PLGLB2 | 5.470650263 | 0.033330344 | -2.853750004 | 0.349664874 | 2.61690026 | 2 | 2 | 2 | 5.602782483 | 5.63601114 | 2.094598369 | 2 |
| ENSG00000135632 | SMYD5 | 1.243603718 | 0.585257988 | 1.371208582 | 0.536695429 | 2.6148123 | 2 | 7.954278496 | 3.369317972 | 7.653784481 | 7.67863747 | 7.765568476 | 7.816873723 |
| ENSG00000155545 | MIER3 | 1.666752732 | 0.011919917 | 0.947738912 | 0.173968415 | 2.614491644 | 6.366709671 | 7.572420273 | 6.43197674 | 8.557949361 | 8.55598286 | 8.142936557 | 7.476011586 |
| ENSG00000160293 | VAV2 | 1.457189068 | 0.412532734 | 1.155962042 | 0.516505072 | 2.61315111 | 8.009475818 | 6.311476754 | 2 | 8.139053376 | 8.35836495 | 8.082904238 | 7.816873723 |
| ENSG00000204262 | COL5A2 | 1.327112453 | 0.092649431 | 1.28480411 | 0.098958811 | 2.611916563 | 11.65437133 | 13.45660594 | 12.03516535 | 13.91902576 | 13.9406531 | 13.70141217 | 14.05244606 |
| ENSG00000146834 | MEPCE | 1.235805372 | 0.502698475 | 1.3756491 | 0.44235966 | 2.611454472 | 2 | 8.004992147 | 8.296287027 | 8.807036108 | 8.79752133 | 9.036921734 | 8.838074258 |
| ENSG00000256374 | PPIAL4D | 7.800514757 | 2.26E-05 | -5.1911486 | 0.045228764 | 2.609366158 | 2 | 2 | 2 | 7.847284664 | 7.85690889 | 2 | 2 |
| ENSG00000271567 | PPIAL4E | 7.800514757 | 2.26E-05 | -5.1911486 | 0.045228764 | 2.609366158 | 2 | 2 | 2 | 7.847284664 | 7.85690889 | 2 | 2 |
| ENSG00000275974 | | 7.800514757 | 2.26E-05 | -5.1911486 | 0.045228764 | 2.609366158 | 2 | 2 | 2 | 7.847284664 | 7.85690889 | 2 | 2 |
| ENSG00000133114 | GPALPP1 | 1.06184983 | 0.060429834 | 1.547300531 | 0.003794368 | 2.60915036 | 6.468621203 | 7.189546432 | 6.43197674 | 7.842101511 | 7.74329312 | 8.53381827 | 7.950369405 |
| ENSG00000167110 | GOLGA2 | 1.181272242 | 0.022811457 | 1.425481202 | 0.004716327 | 2.606753443 | 9.689059027 | 8.557180776 | 9.478569125 | 10.46530827 | 10.5149509 | 10.64428999 | 10.8183324 |
| ENSG00000132819 | RBM38 | 0.129642725 | 0.88983543 | 2.476584016 | 0.00040416 | 2.606226741 | 4.467311984 | 5.595927442 | 6.203069917 | 5.614989456 | 5.78232107 | 7.637872201 | 8.216759768 |
| ENSG00000006576 | PHTF2 | 0.723389123 | 0.685840372 | 1.882243648 | 0.242785116 | 2.605632771 | 4.467311984 | 8.955547846 | 5.156626401 | 8.180711314 | 8.31278108 | 9.52945581 | 9.250223518 |
| ENSG00000153790 | C7orf31 | 1.100908968 | 0.34242241 | 1.504155948 | 0.170620999 | 2.605064916 | 5.095959458 | 4.710564154 | 2 | 5.602782483 | 5.16377554 | 5.83618166 | 5.699242824 |
| ENSG00000127364 | TAS2R4 | 0.855108633 | NA | 1.749858381 | NA | 2.604967015 | 2 | 2 | 2 | 2.270786764 | 2.77779907 | 3.296906311 | 2.326754077 |
| ENSG00000106367 | AP1S1 | 1.033844826 | 0.299078862 | 1.569820974 | 0.093567925 | 2.6036658 | 9.174465797 | 7.874700259 | 6.534109959 | 9.250166145 | 9.27068036 | 9.75507596 | 9.830950469 |
| ENSG00000173273 | TNKS | 1.413252573 | 0.254264315 | 1.189321379 | 0.337150716 | 2.602573952 | 5.095959458 | 8.791896851 | 8.699419242 | 9.650037049 | 9.59092963 | 9.518566517 | 9.266590264 |
| ENSG00000263528 | IKBKE | 2.534468276 | 0.215468947 | 0.066297591 | 0.977728724 | 2.600765866 | 3.327711234 | 7.572420273 | 2 | 8.573670512 | 8.56222967 | 5.626574484 | 6.559516442 |
| ENSG00000124222 | STX16 | 1.498094994 | 0.217082182 | 1.102165807 | 0.369236619 | 2.600260801 | 8.816894989 | 9.574900212 | 5.772716771 | 10.19853443 | 10.2335885 | 9.861691911 | 9.780694439 |
| ENSG00000062716 | VMP1 | 1.731862494 | 0.006082629 | 0.864924034 | 0.19944136 | 2.596786528 | 10.57565445 | 9.45234302 | 10.76445573 | 12.08888579 | 12.1023812 | 11.58483423 | 10.75777696 |
| ENSG00000272414 | FAM47E-STB | 7.786890439 | 2.65E-05 | -5.191148445 | 0.046085237 | 2.595741994 | 2 | 2 | 2 | 7.739909082 | 7.93118934 | 2 | 2 |
| ENSG00000168481 | LGI3 | -0.310502485 | 0.931034311 | 2.904513777 | 0.323536602 | 2.594011292 | 2 | 2 | 2 | 2.270786764 | 2.2680111 | 3.679182183 | 3.383972333 |
| ENSG00000172888 | ZNF621 | 1.121226836 | 0.562222129 | 1.469413257 | 0.430383523 | 2.590640092 | 2 | 7.761270542 | 8.845626171 | 8.914645924 | 8.95180002 | 9.388946358 | 9.159937761 |
| ENSG00000127666 | TICAM1 | 0.916365612 | 0.628420441 | 1.673802696 | 0.343240742 | 2.590168308 | 9.064836028 | 5.72254323 | 8.699419242 | 8.592312513 | 8.48870446 | 9.059452688 | 9.493138564 |
| ENSG00000146083 | RNF44 | 0.905325365 | 0.375529496 | 1.68479109 | 0.073836453 | 2.590116454 | 6.468621203 | 8.505280662 | 6.322058544 | 8.372666678 | 8.37442884 | 9.180655169 | 9.109931338 |
| ENSG00000078142 | PIK3C3 | 1.684925839 | 0.00948606 | 0.904859811 | 0.188307274 | 2.58978565 | 7.485143553 | 8.414402626 | 6.882775709 | 9.41544358 | 9.40880824 | 8.408216037 | 8.836470732 |
| ENSG00000261862 | | 7.776944904 | 2.63E-05 | -5.1911485 | 0.045860622 | 2.585796404 | 2 | 2 | 2 | 7.842101511 | 7.81567127 | 2 | 2 |
| ENSG00000145012 | LPP | 0.145169276 | 0.722229354 | 2.440179756 | 4.77E-15 | 2.585349032 | 9.848787621 | 9.583268459 | 10.14394518 | 10.02414665 | 10.0119132 | 12.43082064 | 12.17507145 |
| ENSG00000067596 | DHX8 | 1.281368151 | 0.273690508 | 1.30359691 | 0.25635543 | 2.584965061 | 5.532353208 | 8.981105197 | 8.561105505 | 9.52946469 | 9.56705484 | 9.409506941 | 9.715307813 |
| ENSG00000214711 | CAPN14 | 0.010001629 | 0.997481474 | 2.574723422 | NA | 2.58472505 | 2 | 2 | 2 | 2.270786764 | 2.38539787 | 3.5515855 | 3.071407821 |
| ENSG00000185924 | RTN4RL1 | -0.723399098 | 0.841155419 | 3.307633494 | 0.259517738 | 2.584234396 | 2 | 2 | 2 | 2.270786764 | 2.14022021 | 4.188345803 | 3.311948089 |
| ENSG00000064313 | TAF2 | 1.371088271 | 0.207448076 | 1.212827346 | 0.262333756 | 2.583915617 | 5.095959458 | 7.274681896 | 8.296287027 | 8.777474042 | 8.69749358 | 8.541710459 | 8.619914108 |
| ENSG00000197375 | SLC22A5 | 0.909563536 | 0.705604924 | 1.673354985 | 0.456551187 | 2.582918521 | 2 | 6.389896633 | 2 | 5.890757403 | 5.67111583 | 6.647180917 | 6.347714043 |
| ENSG00000169155 | ZBTB43 | 1.681598136 | 0.000326631 | 0.901126222 | 0.070466789 | 2.582724358 | 7.667647223 | 7.467835385 | 6.718950348 | 9.005540335 | 8.99861459 | 8.018757917 | 8.413855585 |
| ENSG00000123091 | RNF11 | 0.815993495 | 0.371365629 | 1.765690331 | 0.033376067 | 2.581683826 | 9.940323746 | 9.763446433 | 7.691296561 | 10.25471453 | 10.2173523 | 11.08384263 | 11.27679362 |
| ENSG00000204446 | C9orf170 | 2.444644759 | 0.440624206 | 0.135666309 | NA | 2.580311068 | 2 | 2 | 2 | 3.289987342 | 3.2184422 | 2.384001718 | 2.326754077 |
| ENSG00000173221 | GLRX | 2.388424651 | 1.42E-06 | 0.190855488 | 0.761845241 | 2.579280139 | 8.228665467 | 7.002537973 | 7.691296561 | 10.04910261 | 10.1111143 | 7.902621621 | 7.90110545 |
| ENSG00000024422 | EHD2 | 0.290813673 | 0.721168354 | 2.288357336 | 0.000505952 | 2.579171009 | 12.05742222 | 10.67655334 | 10.7904855 | 11.58169902 | 11.6378232 | 13.19868131 | 13.92358827 |
| ENSG00000136169 | SETDB2 | 1.961157148 | 0.190847995 | 0.609854535 | 0.710603737 | 2.571011683 | 2 | 6.844429755 | 6.718950348 | 8.22518713 | 8.03365075 | 7.090465544 | 6.468583424 |
| ENSG00000151338 | MIPOL1 | 0.922626262 | 0.693189026 | 1.647809572 | 0.451377576 | 2.570435835 | 2 | 6.04685525 | 2 | 5.363304456 | 5.5755357 | 6.473229652 | 5.722580018 |
| ENSG00000188026 | RILPL1 | 1.0572236 | 0.253880504 | 1.512533031 | 0.084716522 | 2.569756631 | 7.867441304 | 6.844429755 | 5.39244166 | 7.992367495 | 8.12087597 | 8.434452636 | 8.577026237 |
| ENSG00000138658 | ZGRF1 | 0.993995515 | 0.483854018 | 1.575195068 | 0.233466011 | 2.569190583 | 3.327711234 | 4.710564154 | 2 | 4.670864177 | 4.53541753 | 5.19095825 | 5.013446246 |
| ENSG00000135744 | AGT | -2.30028285 | 0.211593503 | 4.868289281 | 0.002344194 | 2.568006431 | 2 | 2 | 5.39244166 | 2.695456494 | 2.68924543 | 8.119222541 | 8.777523662 |
| ENSG00000285437 | POLR2J3 | 4.601493261 | 2.69E-08 | -2.034036654 | 0.025714394 | 2.567456607 | 4.815609471 | 3.686303414 | 5.930865648 | 9.452477833 | 9.47652116 | 3.172301926 | 3.717076715 |
| ENSG00000137965 | IFI44 | -3.753125373 | 0.302544659 | 6.32026359 | 0.028809061 | 2.567138217 | 2 | 2 | 2 | 2 | 2 | 6.396389756 | 6.435070131 |
| ENSG00000153767 | GTF2E1 | 1.666923177 | 0.228980535 | 0.898902049 | 0.534155222 | 2.565825226 | 3.327711234 | 7.467835385 | 5.59507254 | 7.925329813 | 7.93118934 | 7.377096489 | 6.944034147 |
| ENSG00000127914 | AKAP9 | 1.745130342 | 0.010140946 | 0.820306821 | 0.260698796 | 2.565437164 | 7.384433967 | 9.170736405 | 8.65477198 | 10.34359113 | 10.290665 | 9.475542327 | 9.313068526 |
| ENSG00000186453 | FAM228A | -0.103112999 | 0.968293784 | 2.668294011 | 0.204564701 | 2.565181012 | 3.327711234 | 2 | 2 | 2.498691189 | 2.59489879 | 4.251304078 | 3.824516949 |
| ENSG00000189319 | FAM53B | 0.957511745 | 0.579274794 | 1.606441988 | 0.320401275 | 2.563953733 | 2 | 7.431218269 | 7.225742779 | 7.608714138 | 7.7866208 | 8.589193677 | 8.036691948 |
| ENSG00000130956 | HABP4 | 0.743630791 | 0.688362385 | 1.818964001 | 0.278961643 | 2.562594791 | 8.138771685 | 7.145017454 | 2 | 7.875463856 | 7.88956768 | 8.985384832 | 8.903351429 |
| ENSG00000181885 | CLDN7 | 2.291414727 | 0.261829517 | 0.270617475 | 0.909261712 | 2.562032202 | 4.00722531 | 2 | 2 | 4.139692846 | 4.79082474 | 3.316666208 | 2.851942824 |
| ENSG00000132824 | SERINC3 | 1.154034432 | 9.21E-06 | 1.405904066 | 3.59E-08 | 2.559938498 | 10.4253394 | 10.62884366 | 10.26027233 | 11.63211941 | 11.5677554 | 11.76575458 | 11.93315092 |
| ENSG00000196220 | SRGAP3 | 1.345042367 | 0.52794944 | 1.212943004 | 0.563495697 | 2.557985371 | 2 | 4.442710129 | 2 | 4.54891529 | 4.12711147 | 4.720468739 | 3.539239711 |
| ENSG00000123607 | TTC21B | 1.516897314 | 0.219156141 | 1.039882744 | 0.409012973 | 2.556780057 | 3.327711234 | 7.099070191 | 6.629488387 | 7.872924734 | 7.80517502 | 7.570548195 | 7.142226902 |
| ENSG00000110841 | PPFIBP1 | 1.538255894 | 0.062284456 | 1.018067363 | 0.2306006 | 2.556323257 | 7.218819781 | 9.338312924 | 9.272377397 | 10.42037002 | 10.4062943 | 9.913096751 | 9.876656812 |
| ENSG00000196843 | ARID5A | 1.298725242 | 0.419943377 | 1.257299995 | 0.42731388 | 2.556025238 | 8.777722034 | 7.145017454 | 3.369317972 | 8.902248225 | 8.90218458 | 8.696779192 | 9.0086755 |
| ENSG00000163170 | BOLA3 | 1.49909005 | 0.07045483 | 1.056459374 | 0.211464999 | 2.555549425 | 6.366709671 | 6.04685525 | 4.058998049 | 7.2696592 | 7.18663697 | 6.67615119 | 6.913939481 |
| ENSG00000179387 | ELMOD2 | 1.086624195 | 0.532648303 | 1.468721601 | 0.377510751 | 2.555346006 | 2 | 7.572420273 | 6.203069917 | 7.59952861 | 7.48508724 | 8.175765162 | 7.604674031 |
| ENSG00000084628 | NKAIN1 | 2.975834417 | 0.562661877 | -0.421393043 | 0.94131204 | 2.554441374 | 2 | 2 | 2 | 3.709785322 | 3.45167086 | 2 | 2.466008509 |
| ENSG00000139112 | GABARAPL1 | 1.325307733 | 6.09E-05 | 1.228839949 | 0.00020825 | 2.554147683 | 10.27168932 | 9.74864196 | 9.703463497 | 11.22970711 | 11.2743971 | 11.16459958 | 11.14718668 |
| ENSG00000213988 | ZNF90 | 1.624378619 | 0.392606579 | 0.92870637 | 0.635038067 | 2.553084989 | 3.327711234 | 3.076665209 | 2 | 4.139692846 | 3.65239003 | 3.694366604 | 3.128433649 |
| ENSG00000155307 | SAMSN1 | -0.313998298 | 0.935584164 | 2.865793269 | 0.376931525 | 2.551794971 | 2 | 2 | 2 | 2.141737246 | 2.38539787 | 3.917810786 | 2.950113106 |
| ENSG00000183153 | GJD3 | 3.301116068 | 0.006082261 | -0.750293682 | 0.593027095 | 2.550822387 | 3.327711234 | 4.442710129 | 2 | 6.463327516 | 6.49425375 | 2.911488689 | 3.430065404 |
| ENSG00000125844 | RRBP1 | 0.91323918 | 0.001774792 | 1.63620575 | 3.24E-09 | 2.54944493 | 10.81695923 | 10.56689847 | 11.06667478 | 11.72409419 | 11.7574918 | 12.45002848 | 12.47623726 |
| ENSG00000128245 | YWHAH | 0.424797486 | 0.439449514 | 2.124373735 | 3.29E-06 | 2.549171221 | 8.696033109 | 9.246456519 | 9.798878489 | 9.746817367 | 9.7285167 | 11.46031868 | 11.40430528 |
| ENSG00000124134 | KCNS1 | -1.303810926 | 0.768597193 | 3.852061139 | 0.305716361 | 2.548250213 | 2 | 2 | 2 | 2.270786764 | 2 | 4.04171018 | 4.388595494 |
| ENSG00000013392 | RWDD2A | 0.84586136 | 0.082516252 | 1.702059951 | 0.000118641 | 2.54792131 | 6.009025997 | 6.844429755 | 6.629488387 | 7.423326516 | 7.31425651 | 8.166947586 | 8.252471774 |
| ENSG00000172824 | CES4A | 1.701061056 | 0.384580714 | 0.846831014 | 0.679226319 | 2.547892069 | 2 | 4.93638933 | 2 | 5.289065324 | 4.95174433 | 4.04171018 | 4.692661645 |
| ENSG00000141279 | NPEPPS | 1.760894541 | 0.069176139 | 0.78634686 | 0.447739474 | 2.547241401 | 8.340423424 | 10.06773758 | 7.502634626 | 10.76003928 | 10.8333485 | 9.80554977 | 9.845638987 |
| ENSG00000064393 | HIPK2 | 1.114461615 | 0.23620818 | 1.432592354 | 0.112072333 | 2.547053969 | 7.485143553 | 9.835272298 | 9.601288346 | 10.3851665 | 10.4036876 | 10.32506285 | 11.01638973 |
| ENSG00000143319 | ISG20L2 | 0.737044267 | 0.068175471 | 1.808398 | 5.97E-07 | 2.545442266 | 7.096806373 | 7.701025996 | 7.225742779 | 8.115618198 | 8.09310209 | 9.164389779 | 9.165057331 |
| ENSG00000087470 | DNM1L | 1.537643742 | 0.349323187 | 1.005855687 | 0.548363823 | 2.543499429 | 5.866944937 | 9.883504052 | 5.59507254 | 9.95740528 | 10.0102007 | 9.472242545 | 9.437105934 |
| ENSG00000121644 | DESI2 | 0.860629953 | 0.171930246 | 1.681501387 | 0.003528509 | 2.54213134 | 8.777722034 | 8.607278467 | 7.502634626 | 9.273456479 | 9.23006009 | 10.31895264 | 9.765606869 |
| ENSG00000003987 | MTMR7 | -1.303813348 | 0.768464398 | 3.845444016 | 0.306126091 | 2.541630668 | 2 | 2 | 2 | 2.270786764 | 2 | 4.301748724 | 4.133950982 |
| ENSG00000164674 | SYTL3 | 2.58059919 | 0.00141337 | -0.040851049 | 0.969279885 | 2.539748141 | 4.467311984 | 5.303523747 | 3.369317972 | 6.936770382 | 7.0991228 | 4.504874511 | 4.711446447 |
| ENSG00000178202 | KDELC2 | 0.53954224 | 0.574668905 | 1.997651743 | 0.016599301 | 2.537193983 | 8.340423424 | 9.214490408 | 6.882775709 | 9.017084849 | 8.92660044 | 10.15764151 | 10.64763892 |
| ENSG00000101216 | GMEB2 | 0.806738536 | 0.56521797 | 1.728468153 | 0.177968622 | 2.535206689 | 7.096806373 | 7.638155693 | 3.369317972 | 7.656739692 | 7.63485389 | 8.464101787 | 8.638437331 |
| ENSG00000169442 | CD52 | -1.279450278 | 0.745616581 | 3.814446666 | 0.264665347 | 2.534996389 | 3.327711234 | 2 | 2 | 2 | 2.49394783 | 5.873348082 | 2.374681325 |
| ENSG00000107281 | NPDC1 | 1.493609048 | 0.320964241 | 1.040700652 | 0.496230402 | 2.5343097 | 4.815609471 | 8.169643011 | 9.897982195 | 10.19345574 | 10.2252472 | 9.792379418 | 9.72342817 |
| ENSG00000131848 | ZSCAN5A | 0.836427041 | 0.689206562 | 1.697005253 | 0.377073141 | 2.533432294 | 2 | 5.131603625 | 2 | 4.54891529 | 4.58534546 | 5.008609438 | 5.566251551 |
| ENSG00000177666 | PNPLA2 | 1.078955795 | 0.225518037 | 1.453762856 | 0.087242653 | 2.532718651 | 9.922475255 | 8.717821467 | 7.777116083 | 10.13690654 | 10.1458442 | 10.13138331 | 10.81779073 |
| ENSG00000143622 | RIT1 | 1.585418608 | 0.391687888 | 0.94708529 | 0.61896509 | 2.532503898 | 8.674867122 | 8.297207175 | 2 | 9.50916971 | 9.47486651 | 8.89667997 | 8.818713298 |
| ENSG00000162733 | DDR2 | 0.254337744 | 0.741606385 | 2.278049059 | 0.000224741 | 2.532386803 | 9.820154628 | 8.747909548 | 10.38167985 | 10.02357442 | 10.0700583 | 12.17700352 | 11.94702543 |
| ENSG00000084636 | COL16A1 | 1.179449462 | 0.095026336 | 1.352578163 | 0.048281896 | 2.532027625 | 9.174465797 | 8.522788974 | 10.09465441 | 10.56305371 | 10.5971045 | 10.41603173 | 11.025801 |
| ENSG00000110514 | MADD | 1.47390807 | 0.257133882 | 1.057977615 | 0.423741616 | 2.531885684 | 4.467311984 | 8.505280662 | 8.236946975 | 9.235416412 | 9.36213202 | 8.68637457 | 9.063622361 |
| ENSG00000183474 | GTF2H2C | 5.909137245 | 4.06E-07 | -3.378556965 | 0.015050726 | 2.53058028 | 3.327711234 | 2 | 4.058998049 | 8.429064458 | 8.41985547 | 2.094598369 | 2.277179783 |
| ENSG00000146963 | LUC7L2 | 1.983153886 | 3.36E-05 | 0.546578297 | 0.314407502 | 2.529732183 | 7.940205782 | 6.844429755 | 7.646393253 | 9.446513412 | 9.5372395 | 7.992893101 | 8.142569067 |
| ENSG00000108179 | PPIF | 2.19491778 | 4.77E-08 | 0.334323244 | 0.487891953 | 2.529241024 | 8.39321512 | 8.981105197 | 8.699419242 | 10.86534179 | 10.9331253 | 8.720150661 | 9.313452824 |
| ENSG00000169031 | COL4A3 | -2.665163286 | 0.252535231 | 5.193943043 | 0.005963489 | 2.528779757 | 2 | 2 | 4.058998049 | 2 | 2.38539787 | 7.218085825 | 7.171352322 |
| ENSG00000100811 | YY1 | 0.52500514 | 0.231434376 | 2.003157498 | 1.08E-07 | 2.528162638 | 9.233563278 | 9.540933215 | 8.763901704 | 9.696010185 | 9.78502553 | 11.33497397 | 11.08417113 |
| ENSG00000048991 | R3HDM1 | 1.145139374 | 0.428693153 | 1.380051718 | 0.320805825 | 2.525191093 | 4.00722531 | 8.433041974 | 6.882775709 | 8.485003881 | 8.42329129 | 8.374440168 | 8.943630224 |
| ENSG00000143190 | POU2F1 | 1.468575201 | 0.077266405 | 1.055013595 | 0.212493403 | 2.523588796 | 5.866944937 | 7.928237109 | 6.629488387 | 8.540455898 | 8.50013767 | 8.191234678 | 8.028288444 |
| ENSG00000167280 | ENGASE | 1.091304147 | 0.517792902 | 1.432142201 | 0.372318292 | 2.523446348 | 2 | 6.535001748 | 7.451345288 | 7.481152682 | 7.62891439 | 8.014983227 | 7.755972999 |
| ENSG00000147894 | C9orf72 | 2.012818281 | 0.271053897 | 0.509362906 | 0.801100986 | 2.522181187 | 4.00722531 | 7.099070191 | 2 | 7.638916932 | 7.68437616 | 6.520853931 | 5.812327524 |
| ENSG00000112357 | PEX7 | 1.571106967 | 0.190366567 | 0.950429059 | 0.442284492 | 2.521536026 | 2 | 4.710564154 | 5.59507254 | 6.139181293 | 6.09824986 | 5.50697268 | 5.576534358 |
| ENSG00000131469 | RPL27 | 0.703091552 | 0.531143362 | 1.817369016 | 0.071495019 | 2.520460568 | 10.81212061 | 8.655694736 | 8.266922072 | 10.40375541 | 10.4114935 | 11.55927357 | 11.47883521 |
| ENSG00000119013 | NDUFB3 | 1.21038275 | 0.001829326 | 1.30802218 | 0.000603619 | 2.51840493 | 6.89194353 | 6.728495673 | 7.029880454 | 8.102673083 | 8.02009991 | 8.038228667 | 8.269209446 |
| ENSG00000185585 | OLFML2A | -2.235460683 | 0.013585291 | 4.752668595 | 6.73E-09 | 2.517207912 | 5.70932728 | 7.274681896 | 5.156626401 | 4.670864177 | 3.98554648 | 10.83029933 | 11.23835038 |
| ENSG00000175455 | CCDC14 | 1.572189539 | 0.254887129 | 0.932910458 | 0.513742625 | 2.505099997 | 4.00722531 | 8.356994601 | 8.110401118 | 9.309580714 | 9.18421833 | 8.939525293 | 8.193564875 |
| ENSG00000278760 | | 7.695995495 | 4.72E-05 | -5.191148055 | 0.048550418 | 2.50484744 | 2 | 2 | 2 | 7.671425626 | 7.82349365 | 2 | 2 |
| ENSG00000185483 | ROR1 | -0.816466534 | 0.581462748 | 3.320506394 | 0.007450401 | 2.504039861 | 3.327711234 | 5.303523747 | 2 | 3.79808662 | 3.45167086 | 7.20223931 | 7.296373648 |
| ENSG00000023572 | GLRX2 | 1.28881391 | 0.249770351 | 1.215183407 | 0.271243655 | 2.503997318 | 6.653098118 | 5.595927442 | 3.369317972 | 7.017154451 | 6.89154576 | 6.958334233 | 6.806650651 |
| ENSG00000010310 | GIPR | 0.236667037 | 0.915905242 | 2.265820918 | 0.226351497 | 2.502487955 | 2 | 5.303523747 | 2 | 3.997744386 | 4.43005853 | 6.132200015 | 5.867805474 |
| ENSG00000196151 | WDSUB1 | 1.067123085 | 0.444480686 | 1.43523671 | 0.278276141 | 2.502359795 | 3.327711234 | 4.11352228 | 6.534109959 | 6.18083559 | 6.3857277 | 6.651077397 | 6.62538071 |
| ENSG00000125945 | ZNF436 | 0.74086527 | 0.717844348 | 1.761163083 | 0.347741804 | 2.502028354 | 8.444143041 | 5.838937808 | 2 | 7.877998516 | 7.74878077 | 8.737663157 | 8.901818877 |

| | | | | | | | | | | | | | |
|---|---|---|---|---|---|---|---|---|---|---|---|---|---|
| ENSG00000189060 | H1F0 | 1.431559757 | 0.048733067 | 1.069711277 | 0.147317101 | 2.501271034 | 10.74769552 | 9.514922632 | 11.37608156 | 12.13716306 | 12.186169 | 11.66216814 | 11.92650878 |
| ENSG00000156304 | SCAF4 | 1.293957564 | 0.482463711 | 1.206412065 | 0.506531437 | 2.500369628 | 7.579279349 | 8.487557258 | 2 | 8.785596684 | 8.83153649 | 8.676850263 | 8.765776654 |
| ENSG00000187790 | FANCM | 1.629638197 | 0.076551676 | 0.870217543 | 0.365505781 | 2.49985574 | 3.327711234 | 4.442710129 | 5.156626401 | 6.052081244 | 5.78232107 | 5.385935071 | 5.050803619 |
| ENSG00000153339 | TRAPPC8 | 1.495933512 | 0.207531949 | 1.003241164 | 0.408534991 | 2.499174677 | 8.042902253 | 7.874700259 | 4.523675391 | 8.908460392 | 8.93143442 | 8.63860224 | 8.191057445 |
| ENSG00000181192 | DHTKD1 | 1.0337798 | 0.25067706 | 1.464634602 | 0.085942697 | 2.498414401 | 5.70932728 | 6.535001748 | 7.777116083 | 7.917958914 | 7.93360068 | 7.943705881 | 8.667588528 |
| ENSG00000065457 | ADAT1 | 1.189132353 | 0.179882373 | 1.309187903 | 0.128338636 | 2.498320256 | 5.532353208 | 7.847167909 | 6.80318718 | 8.219202381 | 8.17078703 | 8.322856568 | 8.304435067 |
| ENSG00000104626 | ERI1 | 0.746256918 | 0.29276914 | 1.750095194 | 0.005691812 | 2.496352112 | 5.095959458 | 6.464272742 | 5.59507254 | 6.628980152 | 6.51372389 | 7.652523812 | 7.429968874 |
| ENSG00000130821 | SLC6A8 | 2.001015341 | 0.198333509 | 0.494824257 | 0.775687249 | 2.495839597 | 4.467311984 | 8.820493869 | 5.39244166 | 9.349396108 | 9.47072153 | 8.050081351 | 7.778417291 |
| ENSG00000116106 | EPHA4 | 3.684371466 | 0.020403301 | -1.188599506 | 0.503356521 | 2.49577196 | 2 | 5.303523747 | 2 | 7.481152682 | 7.27676203 | 3.27687201 | 3.236138336 |
| ENSG00000145632 | PLK2 | 1.872170586 | 0.413198702 | 0.623213664 | 0.799962733 | 2.495384249 | 10.11549941 | 6.464272742 | 2 | 10.50970639 | 10.5040407 | 9.810566697 | 8.37093032 |
| ENSG00000124664 | SPDEF | 2.215432512 | 0.511909846 | 0.278914973 | NA | 2.494347485 | 2 | 2 | 2 | 3.349643765 | 2.86123014 | 2.3070652 | 2.466008509 |
| ENSG00000166813 | KIF7 | 1.066525759 | 0.040530546 | 1.427119586 | 0.004135001 | 2.493645345 | 7.031698319 | 6.535001748 | 7.646393253 | 8.15586117 | 8.18299958 | 8.52906215 | 8.522223965 |
| ENSG00000173548 | SNX33 | 0.434060427 | 0.808247156 | 2.059537274 | 0.179215774 | 2.493597701 | 4.467311984 | 8.077869079 | 10.05225195 | 9.206465443 | 9.28774494 | 10.75705321 | 10.9686339 |
| ENSG00000135074 | ADAM19 | 1.304036262 | 0.046986259 | 1.189527545 | 0.069532318 | 2.493563808 | 9.667789716 | 9.540933215 | 10.86587029 | 11.44329506 | 11.4685634 | 11.5630268 | 11.07999859 |
| ENSG00000186480 | INSIG1 | 2.166083015 | 0.017108279 | 0.325758971 | 0.762594103 | 2.491841986 | 8.418903792 | 10.75272247 | 10.31867969 | 12.28593974 | 12.2856726 | 9.739148276 | 10.92189172 |
| ENSG00000108175 | ZMIZ1 | 1.200454181 | 0.244599958 | 1.290918513 | 0.199257862 | 2.491327695 | 7.904282243 | 10.62073637 | 10.44204362 | 11.24843827 | 11.2703488 | 10.99331648 | 11.63534878 |
| ENSG00000174373 | RALGAPA1 | 1.260683513 | 0.341930387 | 1.229694764 | 0.346574255 | 2.490378277 | 3.327711234 | 7.051611315 | 7.398165132 | 8.008651253 | 7.84158259 | 8.025527571 | 7.755972999 |
| ENSG00000008324 | SS18L2 | 1.613313993 | 0.022108229 | 0.875650652 | 0.238915711 | 2.488964645 | 5.70932728 | 5.457120953 | 4.058998049 | 6.761173376 | 6.76185814 | 6.170709933 | 5.937276556 |
| ENSG00000124766 | SOX4 | 3.098312539 | 2.36E-05 | -0.610264177 | 0.47862258 | 2.488048362 | 8.737455672 | 7.503546079 | 7.163363831 | 11.04016618 | 11.0485841 | 7.808940772 | 6.730418989 |
| ENSG00000185386 | MAPK11 | -3.753121909 | 0.311453673 | 6.240599322 | 0.035923719 | 2.487477412 | 2 | 2 | 2 | 2 | 2 | 6.492902701 | 6.168983563 |
| ENSG00000153046 | CDYL | 0.843555674 | 0.016138255 | 1.64369842 | 3.93E-07 | 2.487254094 | 7.485143553 | 7.60566235 | 7.600047325 | 8.48998335 | 8.2960118 | 9.080915197 | 9.289821273 |
| ENSG00000146872 | TLK2 | 1.679598658 | 0.163759565 | 0.806919066 | 0.525667607 | 2.486517724 | 7.709880939 | 8.337339476 | 4.523675391 | 9.18717761 | 9.20931286 | 8.324077374 | 8.343344553 |
| ENSG00000104904 | OAZ1 | 1.069526125 | 0.002717218 | 1.415969572 | 4.04E-05 | 2.485495697 | 11.69698977 | 11.05003091 | 11.39656351 | 12.46319341 | 12.4818262 | 12.68103423 | 12.94449676 |
| ENSG00000213096 | ZNF254 | 1.884778024 | 0.001311774 | 0.600601609 | 0.360717166 | 2.485379633 | 5.330594134 | 5.72254323 | 6.43197674 | 7.753776273 | 7.64666005 | 6.656902456 | 6.222348126 |
| ENSG00000179855 | GIPC3 | -0.904009055 | 0.633283628 | 3.389106825 | 0.03824337 | 2.48509777 | 4.00722531 | 2 | 6.718950348 | 4.329433326 | 4.79082474 | 8.770720575 | 8.456236734 |
| ENSG00000095015 | MAP3K1 | 0.372792669 | 0.832844524 | 2.109713268 | 0.154214134 | 2.482505937 | 2 | 7.051611315 | 5.930865648 | 6.604700657 | 6.18176397 | 8.141552353 | 8.051511707 |
| ENSG00000005700 | IBTK | 1.746478042 | 0.000552485 | 0.735830003 | 0.181128882 | 2.482308045 | 7.867441304 | 9.00621767 | 8.825624954 | 10.39846869 | 10.3733676 | 9.21856602 | 9.524680706 |
| ENSG00000099942 | CRKL | 0.910901593 | 0.551800049 | 1.569740805 | 0.273134268 | 2.480642399 | 4.815609471 | 9.433952963 | 9.735974718 | 9.856914282 | 10.0187431 | 10.61778047 | 10.57545478 |
| ENSG00000234616 | JRK | 0.584544044 | 0.620673635 | 1.894815599 | 0.067784655 | 2.479359643 | 7.031698319 | 7.189546432 | 4.058998049 | 7.106324235 | 7.26532099 | 8.620333861 | 8.312147538 |
| ENSG00000121895 | TMEM156 | 3.220678467 | 0.000172096 | -0.74200312 | 0.45939756 | 2.478675347 | 4.467311984 | 4.442710129 | 4.874587708 | 7.54315232 | 7.6318872 | 3.429858444 | 4.4231929 |
| ENSG00000134644 | PUM1 | 1.217741635 | 0.011872202 | 1.259477679 | 0.008608574 | 2.477219314 | 8.737455672 | 9.566483142 | 8.608698767 | 10.24099634 | 10.2683912 | 10.30973854 | 10.28285103 |
| ENSG00000091513 | TF | 0.009998476 | 0.997481474 | 2.465808835 | NA | 2.475807311 | 2 | 2 | 2 | 2.270786764 | 2.38539787 | 3.393105702 | 3.128433649 |
| ENSG00000136122 | BORA | 2.127473214 | 0.263300675 | 0.347848986 | 0.872577887 | 2.4753222 | 2 | 5.303523747 | 2 | 5.808110275 | 5.94414054 | 4.271694079 | 4.365060851 |
| ENSG00000127980 | PEX1 | 1.252104972 | 0.4623472 | 1.223057643 | 0.463451338 | 2.475162614 | 5.095959458 | 7.051611315 | 2 | 7.017154451 | 7.07311172 | 7.278527746 | 6.697819825 |
| ENSG00000168564 | CDKN2AIP | 1.240394697 | 0.285861007 | 1.232472249 | 0.279466521 | 2.472866946 | 4.00722531 | 7.051611315 | 7.285536013 | 7.797276258 | 7.95274764 | 8.122032742 | 7.562617572 |
| ENSG00000166747 | AP1G1 | 1.016028867 | 0.000801566 | 1.456445392 | 5.93E-07 | 2.472474259 | 9.189468652 | 9.591588447 | 9.636156974 | 10.48206602 | 10.5161581 | 10.8420647 | 11.02908049 |
| ENSG00000070366 | SMG6 | 1.507760942 | 0.209654079 | 0.964260603 | 0.434758717 | 2.472021545 | 4.815609471 | 8.451443573 | 8.236946975 | 9.271529914 | 9.36392108 | 8.753154049 | 8.804567006 |
| ENSG00000128965 | CHAC1 | 0.697611415 | 0.745616581 | 1.772721154 | 0.365400061 | 2.470332569 | 2 | 6.140560021 | 8.076950877 | 7.467753569 | 7.56813436 | 9.352020423 | 6.786860422 |
| ENSG00000167555 | ZNF528 | 1.148438995 | 0.313233304 | 1.321623977 | 0.230545768 | 2.470062972 | 7.53297911 | 5.131603625 | 5.156626401 | 7.517371202 | 7.51437845 | 7.836577593 | 7.51663051 |
| ENSG00000079263 | SP140 | 3.74034959 | 0.024807493 | -1.270348852 | 0.500318204 | 2.470000738 | 3.327711234 | 2 | 3.369317972 | 5.697663076 | 5.84514933 | 2.68698273 | 2.225841198 |
| ENSG00000197381 | ADARB1 | -1.001646519 | 0.424331051 | 3.470721552 | 0.001279761 | 2.469075033 | 7.867441304 | 8.255926468 | 4.523675391 | 6.61689148 | 6.55189318 | 10.90419513 | 11.08709598 |
| ENSG00000101000 | PROCR | 2.899772903 | 0.031612654 | -0.431391148 | 0.787507734 | 2.468381755 | 7.904282243 | 5.838937808 | 3.369317972 | 9.539105795 | 9.51083771 | 6.159811591 | 6.359671687 |
| ENSG00000076201 | PTPN23 | 0.941661613 | 0.102251384 | 1.52640918 | 0.004583447 | 2.468070793 | 8.107517749 | 9.067150531 | 7.97167868 | 9.403177828 | 9.42088976 | 9.836890387 | 10.13567188 |
| ENSG00000163848 | ZNF148 | 1.040250431 | 0.262504755 | 1.427427064 | 0.106512505 | 2.467677495 | 9.015175615 | 8.297207175 | 6.534109959 | 9.375641031 | 9.21922971 | 9.817951213 | 9.537567589 |
| ENSG00000132535 | DLG4 | 1.436259864 | 0.399184147 | 1.03132221 | 0.548805363 | 2.467582074 | 9.081016759 | 6.728495673 | 3.369317972 | 9.18103275 | 9.20931286 | 8.832503242 | 8.753366804 |
| ENSG00000116044 | NFE2L2 | 1.627523515 | 0.007030767 | 0.839950964 | 0.190829323 | 2.46747448 | 9.128497779 | 9.308339287 | 10.40208303 | 11.34824023 | 11.3526149 | 10.76314172 | 10.33393985 |
| ENSG00000198265 | HELZ | 0.884874463 | 0.140909508 | 1.580406765 | 0.004461713 | 2.465281228 | 8.586954094 | 9.214490408 | 7.858115942 | 9.532685556 | 9.55143939 | 10.31542752 | 10.14908313 |
| ENSG00000167123 | CERCAM | 1.477743315 | 0.041225744 | 0.986422399 | 0.185611962 | 2.464165714 | 11.26581873 | 10.45279253 | 9.439621147 | 12.035819 | 12.0534707 | 11.74539597 | 11.33302995 |
| ENSG00000109819 | PPARGC1A | 2.644938647 | 0.391746117 | -0.180950602 | NA | 2.463988045 | 2 | 2 | 2 | 3.515101962 | 3.2184422 | 2.3070652 | 2.277179783 |
| ENSG00000161011 | SQSTM1 | 3.412778277 | 1.13E-09 | -0.94917777 | 0.132407854 | 2.463600507 | 12.02014453 | 11.73423878 | 10.54349105 | 14.94887343 | 14.9920283 | 10.5812339 | 10.64244805 |
| ENSG00000102886 | GDPD3 | 0.173692358 | 0.942525992 | 2.289528528 | 0.259652636 | 2.463220886 | 2 | 4.11352228 | 2 | 3.663521627 | 2.68924543 | 5.096960836 | 4.456980017 |
| ENSG00000135763 | URB2 | 0.703467952 | 0.564402109 | 1.759447454 | 0.109209308 | 2.462915406 | 3.327711234 | 6.728495673 | 6.073378139 | 6.744602355 | 6.48112613 | 7.701254831 | 7.58443608 |
| ENSG00000138035 | PNPT1 | 2.902007495 | 8.64E-17 | -0.441070221 | 0.284420805 | 2.460937274 | 7.27616306 | 7.60566235 | 7.342949392 | 10.29082969 | 10.3023742 | 6.805896464 | 7.162846889 |
| ENSG00000093000 | NUP50 | 1.316692809 | 0.007453224 | 1.144175514 | 0.021448254 | 2.460868323 | 7.159102577 | 7.819099893 | 8.296287027 | 9.147810675 | 9.11260313 | 8.878514511 | 9.035479014 |
| ENSG00000114021 | NIT2 | 1.525891421 | 0.012730716 | 0.934756455 | 0.143321997 | 2.460647876 | 6.257048221 | 5.946638857 | 7.285536013 | 8.049698831 | 8.11238697 | 7.438128798 | 7.561323797 |
| ENSG00000124198 | ARFGEF2 | 1.242417649 | 0.35327495 | 1.217027963 | 0.35551459 | 2.459445612 | 5.330594134 | 9.558016677 | 8.978369997 | 10.02471865 | 9.96317668 | 10.0116652 | 9.92498893 |
| ENSG00000096401 | CDC5L | 1.52867965 | 0.341899875 | 0.930119219 | 0.573981168 | 2.458798869 | 9.081016759 | 7.874700259 | 3.369317972 | 9.571568433 | 9.52609769 | 8.98885284 | 8.91909319 |
| ENSG00000186866 | POFUT2 | 1.010762326 | 0.00854774 | 1.447639954 | 8.09E-05 | 2.458402279 | 8.816894989 | 9.443177293 | 8.904018013 | 10.08355087 | 10.1041739 | 10.65111557 | 10.39640069 |
| ENSG00000131558 | EXOC4 | 0.706063269 | 0.05155675 | 1.752315556 | 7.77E-08 | 2.458378824 | 8.138771685 | 8.276714465 | 8.266922072 | 8.939126212 | 8.91810182 | 9.737316722 | 10.16833188 |
| ENSG00000108784 | NAGLU | 1.314365332 | 0.428693153 | 1.143519785 | 0.487108958 | 2.457885117 | 3.327711234 | 8.993716073 | 8.825624954 | 9.645581273 | 9.64857633 | 9.301164706 | 9.633950463 |
| ENSG00000135334 | AKIRIN2 | 1.194524485 | 0.108276603 | 1.263145911 | 0.082516727 | 2.457670396 | 8.517314938 | 7.819099893 | 6.629488387 | 8.952415387 | 9.10299435 | 9.22151698 | 8.962375727 |
| ENSG00000158445 | KCNB1 | -2.312396552 | 0.561910399 | 4.769951586 | 0.133361322 | 2.457555034 | 2 | 2 | 2 | 2 | 2.14022021 | 3.679182183 | 5.66592325 |
| ENSG00000058272 | PPP1R12A | 0.898698645 | 0.131061576 | 1.557896051 | 0.004816017 | 2.456594696 | 8.696033109 | 9.648522413 | 8.561105505 | 10.0132353 | 9.89180734 | 10.82632022 | 10.35742448 |
| ENSG00000130164 | LDLR | 1.282150194 | 0.372282912 | 1.1719926 | 0.410543204 | 2.454142795 | 6.138360429 | 10.54135596 | 10.11951029 | 11.07012316 | 11.0867303 | 10.17888295 | 11.47591991 |
| ENSG00000135002 | RFK | 1.027422038 | 0.535148173 | 1.426650042 | 0.366590687 | 2.45407208 | 7.33131396 | 7.099070191 | 2 | 7.753776273 | 7.55253065 | 8.111466145 | 7.98359008 |
| ENSG00000188283 | ZNF383 | 0.926443987 | 0.713873535 | 1.526777102 | 0.52282666 | 2.453221088 | 2 | 6.728495673 | 2 | 6.197166296 | 6.04571775 | 7.081831041 | 6.165581817 |
| ENSG00000142892 | PIGK | 1.233587955 | 0.498655693 | 1.218003713 | 0.495887113 | 2.451591668 | 8.313285031 | 7.847167909 | 2 | 8.799033614 | 8.68180105 | 8.690166757 | 8.761276323 |
| ENSG00000171680 | PLEKHG5 | -0.128643718 | 0.943594867 | 2.578437275 | 0.084015928 | 2.449793556 | 8.19931886 | 5.457120953 | 3.369317972 | 6.772115916 | 6.68951263 | 9.273627415 | 9.500920284 |
| ENSG00000169855 | ROBO1 | 2.872921056 | 2.92E-05 | -0.424320531 | 0.609295543 | 2.448600525 | 7.904282243 | 9.648522413 | 9.757248847 | 12.16972358 | 12.1875587 | 8.839776604 | 8.943133617 |
| ENSG00000021776 | AQR | 1.448220104 | 0.301217705 | 0.999613183 | 0.482248188 | 2.447833288 | 7.159102577 | 8.57407404 | 4.058998049 | 8.951212326 | 8.90710092 | 8.398397867 | 8.566746948 |
| ENSG00000100368 | CSF2RB | 0.00363709 | 0.999290064 | 2.443872465 | NA | 2.447509555 | 2 | 2 | 2 | 2 | 2.59489879 | 3.648325314 | 2.70970617 |
| ENSG00000112033 | PPARD | 0.83804147 | 0.296072769 | 1.605577185 | 0.029202998 | 2.443618655 | 8.107517749 | 7.538394143 | 9.344432377 | 9.32080297 | 9.39662469 | 10.00521062 | 10.23212406 |
| ENSG00000141314 | RHBDL3 | 2.083662902 | 0.546147534 | 0.359327793 | NA | 2.442990695 | 2 | 2 | 2 | 3.227757282 | 2.86123014 | 2.526562374 | 2.277179783 |
| ENSG00000133422 | MORC2 | 1.358122524 | 0.208280231 | 1.084587963 | 0.316802663 | 2.442710488 | 5.866944937 | 6.602424674 | 8.561105505 | 8.754208387 | 8.85462427 | 8.704299335 | 8.341838298 |
| ENSG00000088726 | TMEM40 | -2.312259723 | 0.531143362 | 4.754065177 | 0.096882542 | 2.441805454 | 2 | 2 | 2 | 2 | 2.14022021 | 4.667592615 | 5.230851202 |
| ENSG00000111058 | ACSS3 | 0.26977722 | 0.8692818 | 2.170780244 | 0.10548229 | 2.440557463 | 6.138360429 | 4.442710129 | 2 | 5.289065324 | 5.18011485 | 6.940933331 | 7.094725293 |
| ENSG00000122642 | FKBP9 | 1.413531148 | 8.04E-09 | 1.023387983 | 4.33E-05 | 2.436919131 | 10.22148166 | 10.03760521 | 10.36791565 | 11.58111582 | 11.6638355 | 11.16204184 | 11.30160663 |
| ENSG00000079805 | DNM2 | 0.790971047 | 0.313530525 | 1.642070046 | 0.022211412 | 2.433041093 | 8.228665467 | 10.10889278 | 9.778213818 | 10.34221503 | 10.3795712 | 11.25306166 | 11.16592866 |
| ENSG00000177731 | FLII | 1.180609335 | 0.105550682 | 1.250859183 | 0.079732946 | 2.431468518 | 9.992581382 | 9.672251278 | 8.266922072 | 10.60726063 | 10.7061966 | 10.45346786 | 10.95798087 |
| ENSG00000024862 | CCDC28A | 1.858680979 | 0.171070032 | 0.570673943 | 0.701499184 | 2.429354922 | 3.327711234 | 4.710564154 | 7.029880454 | 7.477814553 | 7.64371756 | 6.439833081 | 6.185873005 |
| ENSG00000196576 | PLXNB2 | 0.712798147 | 0.101674945 | 1.715830792 | 1.15E-05 | 2.428628939 | 11.0444064 | 10.44362967 | 10.23774519 | 11.29920208 | 11.3564391 | 12.37511444 | 12.28344877 |
| ENSG00000100300 | TSPO | 0.99977257 | 0.143386081 | 1.427795461 | 0.027684956 | 2.42756803 | 10.36714848 | 8.762721501 | 9.491321578 | 10.66417187 | 10.6998361 | 11.12688615 | 11.09136013 |
| ENSG00000120800 | UTP20 | 1.427459356 | 0.019135943 | 0.998974479 | 0.110655463 | 2.426433834 | 6.257048221 | 7.274681896 | 6.073378139 | 8.132699624 | 7.97164382 | 7.490172831 | 7.763868281 |
| ENSG00000102871 | TRADD | 1.363113742 | 0.466535804 | 1.062343679 | 0.570544354 | 2.425457421 | 8.257427002 | 6.228547981 | 2 | 8.254744042 | 8.41641145 | 7.851893762 | 8.208518621 |
| ENSG00000069974 | RAB27A | 1.452270037 | 0.001624359 | 0.967903223 | 0.042824635 | 2.42017326 | 6.653098118 | 6.04685525 | 6.43197674 | 7.867833045 | 7.70710533 | 7.426809156 | 7.194904178 |
| ENSG00000128594 | LRRC4 | 1.25288781 | 0.610659792 | 1.166837603 | 0.629414332 | 2.419725413 | 3.327711234 | 2 | 2 | 3.289987342 | 2.94009902 | 2.937246078 | 3.209956562 |
| ENSG00000102904 | TSNAXIP1 | 1.638659485 | 0.366658864 | 0.780733676 | 0.682314785 | 2.419393161 | 2 | 3.076665209 | 3.369317972 | 3.921164147 | 3.94787299 | 3.172301926 | 3.518055793 |
| ENSG00000175274 | TP53I11 | -1.835184867 | 0.074921525 | 4.252430441 | 4.67E-06 | 2.417245574 | 8.910412251 | 8.862353853 | 5.930865648 | 6.567497652 | 6.63721363 | 12.67330171 | 12.58675327 |
| ENSG00000188215 | DCUN1D3 | 0.926770603 | 0.757909059 | 1.490307571 | 0.603099089 | 2.417078174 | 8.716893051 | 2 | 2 | 8.049698831 | 8.08010078 | 8.693949001 | 8.545311752 |
| ENSG00000164743 | C8orf48 | 1.380372518 | 0.489368203 | 1.036698521 | 0.605522215 | 2.417071039 | 2 | 4.93638933 | 2 | 4.825935136 | 4.87352701 | 4.82764224 | 4.228274364 |
| ENSG00000100473 | COCH | -0.306986923 | 0.937842924 | 2.72400912 | NA | 2.417022197 | 2 | 2 | 2 | 2.3892346 | 2.14022021 | 3.796405078 | 2.918126211 |
| ENSG00000066455 | GOLGA5 | 1.179926175 | 2.16E-05 | 1.235685971 | 6.95E-06 | 2.415612147 | 8.609439662 | 8.655694736 | 8.296287027 | 9.718105191 | 9.69337522 | 9.779087725 | 9.743246098 |
| ENSG00000136560 | TANK | 1.517132713 | 0.422347566 | 0.897363835 | 0.644306757 | 2.414496549 | 8.87372952 | 7.638155693 | 2 | 9.310519243 | 9.30647181 | 8.591223132 | 8.789175794 |
| ENSG00000106211 | HSPB1 | -0.026639446 | 0.970008162 | 2.439734766 | 4.16E-06 | 2.41309532 | 11.59645304 | 10.59199661 | 10.38167985 | 10.91524435 | 10.9496004 | 13.44313619 | 13.34825857 |
| ENSG00000144824 | PHLDB2 | 0.47825396 | 0.712692238 | 1.934420458 | 0.085387145 | 2.412674417 | 9.097018025 | 9.267380431 | 5.772716771 | 9.1138055 | 9.17099461 | 10.64977738 | 10.53252493 |
| ENSG00000118873 | RAB3GAP2 | 1.387714128 | 0.276359671 | 1.024727998 | 0.427095076 | 2.412442126 | 5.095959458 | 9.079034257 | 8.35328246 | 9.638124262 | 9.58403885 | 9.402878503 | 9.079977539 |
| ENSG00000143147 | GPR161 | 1.090941266 | 0.39383571 | 1.32134025 | 0.282899381 | 2.412281516 | 6.138360429 | 8.862353853 | 5.930865648 | 8.705093576 | 8.73806474 | 9.088477995 | 8.796891925 |
| ENSG00000140044 | JDP2 | 1.221298346 | 0.199255138 | 1.190278528 | 0.203829413 | 2.411576874 | 6.009025997 | 7.638155693 | 7.646393253 | 8.486665614 | 8.46556258 | 9.035431928 | 7.429968874 |
| ENSG00000178397 | FAM220A | 0.299683633 | 0.844391096 | 2.110543088 | 0.095340194 | 2.410226721 | 3.327711234 | 6.389896633 | 7.502634626 | 6.722207358 | 6.88158913 | 8.843186716 | 8.251669878 |
| ENSG00000111300 | NAA25 | 1.204810003 | 0.45539264 | 1.204942268 | 0.445080176 | 2.409752271 | 3.327711234 | 6.899080544 | 8.608698767 | 8.576794289 | 8.66157866 | 8.710847458 | 8.522888769 |
| ENSG00000044115 | CTNNA1 | 0.749868986 | 0.154731945 | 1.656530369 | 0.000499008 | 2.406399355 | 11.33491623 | 11.46212086 | 10.44204362 | 11.90243375 | 11.8880216 | 12.63449703 | 12.9506245 |
| ENSG00000109089 | CDR2L | 1.224139693 | 0.009891083 | 1.179284116 | 0.012796144 | 2.403423809 | 8.16936288 | 8.903033433 | 8.143093325 | 9.591782342 | 9.75056618 | 9.830891212 | 9.393423672 |
| ENSG00000164978 | NUDT2 | 0.945497291 | 0.38669733 | 1.45779412 | 0.153789199 | 2.403291411 | 6.257048221 | 7.099070191 | 4.058998049 | 7.226553841 | 7.15805139 | 7.605105115 | 7.776188509 |
| ENSG00000151292 | CSNK1G3 | 1.422694697 | 0.182333448 | 0.978956788 | 0.367118564 | 2.401651485 | 7.096806373 | 8.376385542 | 5.39244166 | 8.837307593 | 8.81332035 | 8.600824249 | 8.133891722 |
| ENSG00000107443 | CCNJ | 1.691681187 | 0.056014362 | 0.709933359 | 0.456432004 | 2.401614545 | 4.467311984 | 6.78762695 | 5.59507254 | 7.539954763 | 7.60187978 | 6.639356243 | 6.597881141 |
| ENSG00000131759 | RARA | 1.090017762 | 0.170346386 | 1.311535018 | 0.086940568 | 2.40155278 | 8.042902253 | 7.874700259 | 6.073378139 | 8.633396609 | 8.64401667 | 8.732155826 | 8.974090715 |
| ENSG00000123975 | CKS2 | 2.127330793 | 0.14155575 | 0.272946428 | 0.871954133 | 2.400277221 | 6.89194353 | 5.595927442 | 2 | 7.883054517 | 7.87959733 | 6.157074092 | 6.018717758 |

| | | | | | | | | | | | | | |
|---|---|---|---|---|---|---|---|---|---|---|---|---|---|
| ENSG00000152193 | RNF219 | 1.909097033 | 0.188525669 | 0.48942734 | 0.761935984 | 2.398524373 | 5.70932728 | 6.78762695 | 2 | 7.66556918 | 7.6318872 | 6.462183138 | 6.06327132 |
| ENSG00000164970 | FAM219A | 1.184712402 | 0.079842495 | 1.212456576 | 0.068184653 | 2.397168978 | 7.940205782 | 7.874700259 | 9.25752426 | 9.706030497 | 9.65077635 | 9.673139654 | 9.738674821 |
| ENSG00000182606 | TRAK1 | -0.465060508 | 0.782975197 | 2.86181977 | 0.042053226 | 2.396759262 | 9.371511179 | 8.469613413 | 4.058998049 | 7.975897842 | 7.96458672 | 10.9714739 | 11.52776307 |
| ENSG00000164975 | SNAPC3 | 0.728129768 | 0.060346208 | 1.668412708 | 1.96E-06 | 2.396542476 | 8.19931886 | 8.487557258 | 8.380957451 | 9.072354885 | 9.09762836 | 10.24756422 | 9.750075969 |
| ENSG00000164209 | SLC25A46 | 1.14513461 | 0.324734879 | 1.249755038 | 0.269883074 | 2.394889648 | 8.257427002 | 8.505280662 | 5.156626401 | 9.004380786 | 9.02146427 | 9.198737176 | 9.029877404 |
| ENSG00000180509 | KCNE1 | 2.153028853 | 0.385151567 | 0.24144371 | 0.932122597 | 2.394472564 | 2 | 3.076665209 | 2 | 3.566290607 | 3.7864959 | 2.937246078 | 2.326754077 |
| ENSG00000160326 | SLC2A6 | 1.640229233 | 0.232267301 | 0.752867475 | 0.605340888 | 2.393096708 | 5.70932728 | 8.732943944 | 5.39244166 | 9.002058889 | 9.12426093 | 7.742927022 | 8.524881346 |
| ENSG00000152475 | ZNF837 | 0.364376059 | 0.865682896 | 2.026770574 | 0.265638215 | 2.391146633 | 2 | 4.93638933 | 2 | 4.105493497 | 3.94787299 | 5.618636935 | 5.20457242 |
| ENSG00000233608 | TWIST2 | 3.274516134 | 2.43E-06 | -0.884377645 | 0.265797026 | 2.390138489 | 6.89194353 | 6.389896633 | 5.39244166 | 9.535899247 | 9.59016561 | 4.940196634 | 5.945233978 |
| ENSG00000185219 | ZNF445 | 0.591274265 | 0.440185202 | 1.797454004 | 0.007204583 | 2.388728269 | 6.653098118 | 8.147241016 | 7.098165388 | 8.006336225 | 8.06258142 | 9.484578004 | 8.916565785 |
| ENSG00000169946 | ZFPM2 | 1.28501208 | 0.55785681 | 1.10331627 | 0.614923831 | 2.38832835 | 2 | 8.191702463 | 4.058998049 | 7.980622673 | 7.95274764 | 7.761068522 | 7.814703611 |
| ENSG00000197122 | SRC | 1.041587689 | 0.497513447 | 1.346217386 | 0.359814853 | 2.387805075 | 4.00722531 | 9.04308532 | 8.042706615 | 9.070139931 | 9.14105132 | 9.470314181 | 9.344245845 |
| ENSG00000108443 | RPS6KB1 | 1.681557781 | 0.200555174 | 0.706227225 | 0.614975927 | 2.387785006 | 8.042902253 | 8.276714465 | 4.058998049 | 9.333786218 | 9.23788597 | 8.291995768 | 8.347853908 |
| ENSG00000114796 | KLHL24 | 1.318996074 | 0.219370406 | 1.068237413 | 0.321488956 | 2.387233487 | 6.009025997 | 7.819099893 | 8.884815469 | 9.293531584 | 9.29432703 | 9.45200461 | 8.475596448 |
| ENSG00000163564 | PYHIN1 | 4.583262978 | 0.043170473 | -2.196422407 | 0.407814557 | 2.386840572 | 2 | 2 | 2 | 4.670864177 | 4.97065356 | 2.094598369 | 2.059860178 |
| ENSG00000129128 | SPCS3 | 0.764207024 | 0.038349318 | 1.622216774 | 1.58E-06 | 2.386423798 | 9.544849385 | 9.923599762 | 9.31604076 | 10.44322807 | 10.3190701 | 11.34658404 | 11.12173937 |
| ENSG00000070495 | JMJD6 | 1.325205189 | 0.203871564 | 1.060195126 | 0.311050695 | 2.385400315 | 8.16936288 | 7.315436877 | 5.156626401 | 8.575233246 | 8.6932307 | 8.226262694 | 8.506848296 |
| ENSG00000171444 | MCC | 0.339923643 | 0.728428075 | 2.044136516 | 0.011903882 | 2.384060159 | 4.467311984 | 6.666836837 | 5.59507254 | 6.24507773 | 6.14061114 | 7.857806412 | 7.826599033 |
| ENSG00000065150 | IPO5 | 0.724179564 | 0.397020048 | 1.658542585 | 0.031841861 | 2.382722148 | 10.16187685 | 10.69597594 | 8.677268317 | 10.72878599 | 10.8468444 | 11.41585194 | 11.9741374 |
| ENSG00000154451 | GBP5 | 5.370405629 | 2.29E-05 | -2.987965947 | 0.030448685 | 2.382439682 | 5.532353208 | 3.076665209 | 4.058998049 | 9.678668908 | 9.62713473 | 3.059555873 | 2.172608095 |
| ENSG00000160957 | RECQL4 | 0.749063758 | 0.676328124 | 1.632437128 | 0.318708894 | 2.381500886 | 3.327711234 | 5.946638857 | 2 | 5.334065616 | 5.39133044 | 6.192262702 | 6.168983563 |
| ENSG00000051523 | CYBA | 1.986448028 | 0.003480353 | 0.394065173 | 0.620474069 | 2.380513202 | 9.358295023 | 7.538394143 | 8.763901704 | 10.66893665 | 10.739998 | 9.138949083 | 9.09971864 |
| ENSG00000124782 | RREB1 | 0.966831602 | 0.1808519 | 1.413589647 | 0.038255118 | 2.380421249 | 7.82963488 | 8.376385542 | 6.629488387 | 8.692189072 | 8.79221631 | 9.042123971 | 9.317290185 |
| ENSG00000005007 | UPF1 | 0.973475844 | 0.001970076 | 1.406735544 | 3.14E-06 | 2.380211388 | 9.081016759 | 8.848535039 | 9.196533591 | 9.971714969 | 10.0529704 | 10.59634277 | 10.27517721 |
| ENSG00000132481 | TRIM47 | 0.319277679 | 0.880215531 | 2.060753695 | 0.247087402 | 2.380031374 | 8.285626337 | 6.04685525 | 2 | 7.318995108 | 7.28433928 | 8.842760893 | 9.173126343 |
| ENSG00000075420 | FNDC3B | 1.451216499 | 0.011919917 | 0.9281119 | 0.122378239 | 2.379328399 | 9.533135543 | 10.93049815 | 10.27509734 | 11.82589752 | 11.7866286 | 11.23597623 | 11.33046897 |
| ENSG00000152076 | CCDC74B | 2.68559618 | 0.028202356 | -0.308581604 | 0.835379021 | 2.377014576 | 2 | 4.710564154 | 3.369317972 | 6.096288482 | 6.16544343 | 3.429858444 | 3.789585079 |
| ENSG00000183354 | KIAA2026 | 1.358817989 | 0.01600303 | 1.01655093 | 0.078156821 | 2.37536892 | 8.493334308 | 7.572420273 | 7.398165132 | 9.214510118 | 9.27734059 | 9.123256136 | 8.651263518 |
| ENSG00000198464 | ZNF480 | 1.692626053 | 0.347244214 | 0.680927171 | 0.722957397 | 2.373553224 | 3.327711234 | 7.60566235 | 3.369317972 | 7.821180652 | 7.81567127 | 6.984820384 | 6.662054875 |
| ENSG00000154447 | SH3RF1 | 0.206462846 | 0.820270883 | 2.166665198 | 0.002482931 | 2.373128044 | 7.709880939 | 8.297207175 | 9.491321578 | 8.869506572 | 8.92417737 | 10.55742653 | 11.09023921 |
| ENSG00000012174 | MBTPS2 | 1.16763028 | 0.539874611 | 1.205479271 | 0.520317876 | 2.37310955 | 2 | 8.639735838 | 8.266922072 | 9.064587625 | 9.02372946 | 8.952213594 | 9.200809988 |
| ENSG00000135899 | SP110 | 2.952432247 | 0.001150989 | -0.579678786 | 0.586432904 | 2.372753461 | 8.16936288 | 5.457120953 | 7.451345288 | 10.26540233 | 10.3604317 | 6.497238296 | 7.09829807 |
| ENSG00000096654 | ZNF184 | 0.868514619 | 0.728012667 | 1.502327736 | 0.523757284 | 2.370842355 | 2 | 6.535001748 | 2 | 5.997179931 | 5.76075522 | 6.887434761 | 5.917188835 |
| ENSG00000110944 | IL23A | 1.162087386 | 0.591410858 | 1.208501607 | 0.566009975 | 2.370588993 | 2 | 3.686303414 | 2 | 3.615725078 | 3.45167086 | 3.709392871 | 3.407202941 |
| ENSG00000055917 | PUM2 | 1.39880682 | 0.299070637 | 0.971354989 | 0.477355787 | 2.370161809 | 5.330594134 | 9.523645035 | 9.31604076 | 10.32374052 | 10.2198241 | 10.03887247 | 9.625308966 |
| ENSG00000132406 | TMEM128 | 1.147845584 | 0.170554882 | 1.222171474 | 0.133443484 | 2.370017059 | 7.031698319 | 5.457120953 | 5.39244166 | 7.341208691 | 7.18663697 | 7.514972902 | 7.137025524 |
| ENSG00000184787 | UBE2G2 | 0.818184408 | 0.525545049 | 1.550901796 | 0.190242338 | 2.369086205 | 5.866944937 | 9.599860728 | 8.865353883 | 9.56056556 | 9.58940119 | 10.06849268 | 10.50771945 |
| ENSG00000163637 | PRICKLE2 | 1.561318462 | 0.380355326 | 0.807746487 | 0.664743846 | 2.369064948 | 2 | 7.979858146 | 8.110401118 | 8.998569022 | 9.0282492 | 8.289498148 | 8.245238611 |
| ENSG00000282650 | | -1.310495096 | 0.691746353 | 3.679377362 | 0.166987227 | 2.368882266 | 2 | 2 | 2 | 2.141737246 | 2.14022021 | 4.240999983 | 3.924507493 |
| ENSG00000077782 | FGFR1 | 1.328875491 | 0.143140149 | 1.039811088 | 0.256088094 | 2.368686579 | 7.790810999 | 10.04973379 | 10.06936278 | 10.92783718 | 10.956731 | 10.43593029 | 10.84329825 |
| ENSG00000126878 | AIF1L | -2.311950223 | 0.069585809 | 4.680592777 | 3.31E-05 | 2.368642554 | 5.866944937 | 5.131603625 | 3.369317972 | 3.881293667 | 2.49394783 | 9.408931763 | 9.818008038 |
| ENSG00000164010 | ERMAP | 1.721581315 | 0.239652463 | 0.646961026 | 0.682571547 | 2.368542341 | 6.468621203 | 6.728495673 | 2 | 7.59952861 | 7.83127384 | 6.928144991 | 6.368575325 |
| ENSG00000167771 | RCOR2 | 0.66778214 | 0.61090653 | 1.700685814 | 0.144465782 | 2.368467955 | 2 | 4.442710129 | 4.058998049 | 4.415698455 | 4.3456612 | 5.376552034 | 5.14363432 |
| ENSG00000172086 | KRCC1 | 0.845979953 | 0.127962856 | 1.521592188 | 0.002976633 | 2.367572141 | 7.096806373 | 7.669933291 | 6.882775709 | 8.115618198 | 8.08227582 | 9.049892323 | 8.413855585 |
| ENSG00000170917 | NUDT6 | 0.558337223 | 0.785383233 | 1.804546931 | 0.316598154 | 2.362884154 | 2 | 4.11352228 | 2 | 3.566290607 | 3.45167086 | 4.478802948 | 4.376876162 |
| ENSG00000250741 | NT5C1B-RDH | 7.553581144 | 0.000114143 | -5.191147348 | 0.053140582 | 2.362433795 | 2 | 2 | 2 | 7.608714138 | 7.61094772 | 2 | 2 |
| ENSG00000133318 | RTN3 | 0.866872767 | 0.138486261 | 1.495318548 | 0.005942742 | 2.362191314 | 10.22148166 | 8.981105197 | 9.358421373 | 10.48373114 | 10.4724652 | 11.09067303 | 11.11943354 |
| ENSG00000198331 | HYLS1 | 0.860542047 | 0.588479045 | 1.498873837 | 0.311050695 | 2.359415883 | 2 | 5.946638857 | 4.058998049 | 5.540148523 | 5.63601114 | 6.126613714 | 6.238627697 |
| ENSG00000080823 | MOK | 0.131443194 | 0.929816715 | 2.22577788 | 0.063181453 | 2.357221074 | 3.327711234 | 6.666836837 | 7.285536013 | 6.449761039 | 6.74548212 | 8.850830124 | 8.424568162 |
| ENSG00000121988 | ZRANB3 | 0.742643237 | 0.422586005 | 1.614501665 | 0.051229661 | 2.357144902 | 4.00722531 | 4.11352228 | 4.874587708 | 4.965943888 | 5.00774437 | 5.801542492 | 5.745545708 |
| ENSG00000096070 | BRPF3 | 1.31009197 | 0.519874318 | 1.046628698 | 0.607213739 | 2.356720668 | 2 | 9.090820894 | 9.413059136 | 9.965173975 | 9.99988257 | 9.877784692 | 9.543476778 |
| ENSG00000180346 | TIGD2 | 1.12886893 | 0.625679022 | 1.227367511 | 0.587145381 | 2.356236441 | 2 | 6.04685525 | 2 | 5.764940652 | 5.55061739 | 6.049001768 | 5.379200993 |
| ENSG00000063180 | CA11 | -0.72731905 | 0.719117881 | 3.082355302 | 0.073412065 | 2.355036252 | 2 | 5.303523747 | 2 | 3.709785322 | 3.33976443 | 6.736076089 | 6.841176624 |
| ENSG00000103266 | STUB1 | 1.268047467 | 0.215393228 | 1.086263786 | 0.287317112 | 2.354311252 | 6.468621203 | 8.862353853 | 9.330306409 | 9.91915277 | 9.88247671 | 9.881313582 | 9.538225362 |
| ENSG00000165675 | ENOX2 | 1.042916535 | 0.695713377 | 1.309583814 | 0.61151596 | 2.352500349 | 2 | 7.503546079 | 2 | 6.927012081 | 7.04662306 | 7.220710071 | 7.272853598 |
| ENSG00000198646 | NCOA6 | 0.924106431 | 0.236885537 | 1.427654858 | 0.051981925 | 2.351761288 | 6.89194353 | 8.395519306 | 8.742726175 | 9.069031178 | 9.16587611 | 9.840733868 | 9.35623215 |
| ENSG00000146054 | TRIM7 | 0.698504224 | 0.495803512 | 1.652796091 | 0.070347683 | 2.351300316 | 4.467311984 | 3.686303414 | 4.874587708 | 5.003571641 | 4.93258395 | 6.089765781 | 5.513708555 |
| ENSG00000104774 | MAN2B1 | 1.605804676 | 0.127670293 | 0.745021843 | 0.504623054 | 2.350826519 | 6.257048221 | 9.267380431 | 7.858115942 | 9.844656195 | 9.88932507 | 8.899548956 | 9.115231141 |
| ENSG00000165813 | CCDC186 | 0.983612907 | 0.583878202 | 1.367042616 | 0.425523578 | 2.350655523 | 2 | 7.761270542 | 7.285536013 | 7.975897842 | 7.89946959 | 8.660513226 | 7.862758575 |
| ENSG00000124226 | RNF114 | 1.671921042 | 0.000106412 | 0.675914562 | 0.151862412 | 2.347835604 | 9.533135543 | 8.57407404 | 9.212024683 | 10.79901337 | 10.8388264 | 9.887726478 | 9.76195099 |
| ENSG00000103047 | TANGO6 | 0.622331497 | 0.760903016 | 1.724689923 | 0.351097101 | 2.34702142 | 3.327711234 | 3.076665209 | 7.646393253 | 6.766655021 | 6.77805036 | 7.657374802 | 8.013226939 |
| ENSG00000112981 | NME5 | -0.310501882 | 0.932714208 | 2.65656894 NA | | 2.346067059 | 2 | 2 | 2 | 2.270786764 | 2.2680111 | 3.355392153 | 3.407202941 |
| ENSG00000185753 | CXorf38 | 0.836915159 | 0.306087183 | 1.508940377 | 0.045614369 | 2.345855536 | 6.653098118 | 6.666836837 | 8.266922072 | 8.205140657 | 8.23866319 | 8.891336668 | 8.882264318 |
| ENSG00000205726 | ITSN1 | 0.761685713 | 0.374329883 | 1.583722188 | 0.042742908 | 2.345407901 | 6.653098118 | 8.276714465 | 8.585098385 | 8.855427477 | 8.74905888 | 9.204383073 | 9.941741079 |
| ENSG00000179134 | SAMD4B | 0.864157677 | 0.16153158 | 1.481122249 | 0.00978288 | 2.345279925 | 8.696033109 | 8.929529568 | 9.963514551 | 10.14218788 | 10.1868502 | 10.76381664 | 10.79581778 |
| ENSG00000020426 | MNAT1 | 1.425921215 | 0.154641078 | 0.919192852 | 0.370178383 | 2.345114067 | 7.435667355 | 6.602424674 | 4.523675391 | 8.083034979 | 7.91663606 | 7.489084836 | 7.520630205 |
| ENSG00000143772 | ITPKB | 0.198806608 | 0.905370978 | 2.146069809 | 0.112945603 | 2.344876416 | 6.468621203 | 6.602424674 | 2 | 6.205262847 | 6.1572134 | 8.167627778 | 7.962181894 |
| ENSG00000104164 | BLOC1S6 | 1.003150477 | 0.30391278 | 1.340056142 | 0.14848394 | 2.343206619 | 7.667647223 | 9.461450882 | 7.225742779 | 9.508351943 | 9.41830936 | 9.683639816 | 9.906257642 |
| ENSG00000112701 | SENP6 | 1.40914275 | 0.354199679 | 0.933802937 | 0.546828054 | 2.342945687 | 4.00722531 | 8.276714465 | 9.149036566 | 9.6094308 | 9.62191107 | 9.415818816 | 8.806752407 |
| ENSG00000084676 | NCOA1 | 1.003007619 | 0.466535804 | 1.339752029 | 0.30559948 | 2.342759649 | 6.73718411 | 8.762721501 | 4.874587708 | 8.512996295 | 8.6202631 | 8.897500263 | 8.904882354 |
| ENSG00000113722 | CDX1 | -0.307017014 | 0.936463188 | 2.6489926 NA | | 2.341975586 | 2 | 2 | 2 | 2.3892346 | 2.14022021 | 3.393105702 | 3.36036153 |
| ENSG00000197579 | TOPORS | 1.027465915 | 0.555006373 | 1.31321203 | 0.434435201 | 2.340677945 | 3.327711234 | 9.055168103 | 6.882775709 | 8.765203682 | 8.8354102 | 9.092781861 | 9.075453001 |
| ENSG00000204406 | MBD5 | 1.210827225 | 0.410137517 | 1.128172472 | 0.436457359 | 2.338999697 | 4.00722531 | 6.728495673 | 8.561105505 | 8.508095759 | 8.62474679 | 8.500731308 | 8.47078082 |
| ENSG00000165424 | ZCCHC24 | 0.652294456 | 0.197034134 | 1.686160209 | 0.000175366 | 2.338454665 | 8.564112515 | 9.574900212 | 9.358421373 | 9.870978946 | 9.88683852 | 10.8308362 | 10.98440704 |
| ENSG00000150403 | TMCO3 | 0.309634389 | 0.514390461 | 2.025065989 | 1.42E-07 | 2.334700378 | 9.219014269 | 8.540087354 | 8.536706852 | 9.09649754 | 9.11685323 | 10.69069483 | 10.92918213 |
| ENSG00000136144 | RCBTB1 | 1.321192923 | 0.531161556 | 1.01222991 | 0.635038067 | 2.333422832 | 4.467311984 | 8.213429689 | 2 | 8.063125866 | 8.03365075 | 7.79843893 | 7.688837505 |
| ENSG00000198682 | PAPSS2 | 0.612836879 | 0.040383064 | 1.720522629 | 1.37E-10 | 2.333359508 | 8.963744667 | 9.13702746 | 9.242516611 | 9.706030497 | 9.74988221 | 10.69600884 | 10.95662832 |
| ENSG00000078808 | SDF4 | 1.23024544 | 0.054876729 | 1.102572067 | 0.085204406 | 2.332817507 | 11.31453223 | 10.22039562 | 9.888372618 | 11.81700892 | 11.8573927 | 11.68592437 | 11.73331274 |
| ENSG00000137460 | FHDC1 | 0.718084867 | 0.649032625 | 1.614257551 | 0.261053473 | 2.332342419 | 4.00722531 | 2 | 5.39244166 | 5.040242897 | 4.79082474 | 5.711121288 | 5.740981724 |
| ENSG00000143195 | ILDR2 | 0.66509634 | 0.838427142 | 1.666535507 | 0.574861708 | 2.331631846 | 3.327711234 | 4.11352228 | 2 | 4.647285135 | 2 | 4.764298067 | 4.522269241 |
| ENSG00000143924 | EML4 | 0.656557467 | 0.596166767 | 1.67470297 | 0.131976284 | 2.331260437 | 6.73718411 | 9.488434476 | 7.029880454 | 9.00785664 | 8.9350493 | 9.932236229 | 10.034406 |
| ENSG00000108100 | CCNY | 0.709297824 | 0.279279457 | 1.621069622 | 0.006021251 | 2.330367446 | 8.042902253 | 7.761270542 | 9.06622548 | 9.104095335 | 9.09762836 | 9.843290512 | 10.15531359 |
| ENSG00000163568 | AIM2 | 7.520834048 | 0.000141304 | -5.19114714 | 0.054407162 | 2.329686908 | 2 | 2 | 2 | 7.59952861 | 7.55566492 | 2 | 2 |
| ENSG00000144815 | NXPE3 | 0.837345016 | 0.495345811 | 1.491241596 | 0.189415864 | 2.328586612 | 4.815609471 | 7.790474954 | 8.325066185 | 8.379838154 | 8.40949864 | 9.139989235 | 8.937659629 |
| ENSG00000155330 | C16orf87 | 1.26516708 | 0.000854905 | 1.063171067 | 0.005585099 | 2.328338147 | 7.218819781 | 6.899080544 | 6.60318718 | 8.252792309 | 8.18704757 | 8.10721774 | 7.929461924 |
| ENSG00000135476 | ESPL1 | 1.299826981 | 0.443997681 | 1.028485828 | 0.543853497 | 2.328312809 | 2 | 5.838937808 | 3.369317972 | 5.900763176 | 5.64780787 | 5.498346994 | 5.561082532 |
| ENSG00000069482 | GAL | 0.391192469 | 0.88199664 | 1.936481853 | 0.389444033 | 2.327674322 | 3.327711234 | 3.686303414 | 2 | 3.754611435 | 3.01487888 | 5.371837532 | 2.632958241 |
| ENSG00000162298 | SYVN1 | 1.275279045 | 0.199759863 | 1.052382543 | 0.290417779 | 2.327661588 | 6.138360429 | 8.702538795 | 8.805342542 | 9.508351943 | 9.58863636 | 9.284015592 | 9.369598769 |
| ENSG00000005100 | DHX33 | 0.836421048 | 0.201194682 | 1.490981899 | 0.013866293 | 2.327402947 | 6.89194353 | 6.389896633 | 7.777116083 | 7.980622673 | 7.89452713 | 8.701014095 | 8.456932644 |
| ENSG00000176018 | LYSMD3 | 0.911569847 | 0.548376437 | 1.41548108 | 0.323969847 | 2.327050927 | 3.327711234 | 7.901717006 | 7.818184403 | 8.289431124 | 8.11238697 | 8.997677834 | 8.328971229 |
| ENSG00000133392 | MYH11 | 4.247663782 | 0.003295139 | -1.921559035 | 0.243248502 | 2.326104747 | 2 | 3.076665209 | 3.369317972 | 6.197166296 | 6.18176397 | 2.384001718 | 2.225841198 |
| ENSG00000185271 | KLHL33 | -2.312308415 | 0.540667714 | 4.636420665 | 0.119124184 | 2.32411225 | 2 | 2 | 2 | 2 | 2.14022021 | 4.434275199 | 5.211187127 |
| ENSG00000116191 | RALGPS2 | 0.724352198 | 0.711973842 | 1.599143011 | 0.373931636 | 2.32349521 | 8.468948322 | 6.78762695 | 2 | 7.992367495 | 8.00183183 | 8.994231002 | 8.71548362 |
| ENSG00000213047 | DENND1B | 1.430116838 | 0.35327495 | 0.893289162 | 0.571694567 | 2.323405999 | 2 | 6.844429755 | 5.772716771 | 7.344877929 | 7.1204461 | 6.882471426 | 6.530719415 |
| ENSG00000132361 | CLUH | 1.020675648 | 0.576726175 | 1.300492786 | 0.460189911 | 2.321168434 | 3.327711234 | 6.666836837 | 9.301632639 | 8.94033939 | 8.97539718 | 9.271414214 | 9.199147229 |
| ENSG00000164305 | CASP3 | 0.207215705 | 0.806463377 | 2.113621463 | 0.001567325 | 2.320837168 | 9.33149386 | 8.356994601 | 7.552162999 | 8.756965077 | 8.84055903 | 10.73324321 | 10.65826756 |
| ENSG00000218823 | PAPOLB | 1.499555752 | 0.435748344 | 0.821187643 | 0.679613476 | 2.320743396 | 2 | 4.11352228 | 2 | 4.070463721 | 4.2865223 | 3.796405078 | 3.560117069 |
| ENSG00000155511 | GRIA1 | -1.06138023 | 0.703860951 | 3.381313373 | 0.167205719 | 2.319933143 | 5.70932728 | 2 | 2 | 4.205751501 | 2.2680111 | 6.433059984 | 7.986484919 |
| ENSG00000157992 | KRTCAP3 | 1.480585034 | 0.547023674 | 0.837157974 | 0.739591934 | 2.317743008 | 2 | 3.076665209 | 2 | 3.023133209 | 3.45167086 | 3.035909414 | 2.782576693 |
| ENSG00000070367 | EXOC5 | 1.143132015 | 0.462285707 | 1.173995587 | 0.438543047 | 2.317127602 | 4.00722531 | 9.079034257 | 8.825624954 | 9.592554172 | 9.46739696 | 9.645573761 | 9.473326665 |
| ENSG00000164038 | SLC9B2 | 2.086456631 | 0.104768499 | 0.22539473 | 0.883235347 | 2.311851361 | 8.564112515 | 6.140560021 | 4.874587708 | 9.373846284 | 9.37105521 | 7.64960537 | 7.408544534 |
| ENSG00000105176 | URI1 | 0.852687418 | 0.569331332 | 1.457866831 | 0.298467142 | 2.310554249 | 4.00722531 | 8.395519306 | 8.677268317 | 8.83470036 | 8.83024294 | 9.461743163 | 9.404288469 |
| ENSG00000080345 | RIF1 | 1.255650283 | 0.015155122 | 1.053108888 | 0.043968507 | 2.308759171 | 8.737455672 | 7.761270542 | 8.721235202 | 9.695291778 | 9.72712735 | 9.704647975 | 9.284342586 |
| ENSG00000134987 | WDR36 | 1.361799511 | 0.150716856 | 0.946100291 | 0.327012123 | 2.307899802 | 5.866944937 | 8.147241016 | 6.534109959 | 8.578353644 | 8.48213028 | 8.293866149 | 7.923432281 |
| ENSG00000187013 | C17orf82 | 1.252890733 | 0.61318036 | 1.05465232 | 0.668986346 | 2.307543053 | 2 | 2 | 3.369317972 | 3.289987342 | 2.94009902 | 3.172301926 | 2.851942824 |
| ENSG00000076716 | GPC4 | 0.391889328 | 0.844353006 | 1.915471533 | 0.264665124 | 2.307360861 | 8.19931886 | 6.666836837 | 2 | 7.559034656 | 7.30683517 | 8.740866084 | 9.115231141 |
| ENSG00000120889 | TNFRSF10B | 1.002288575 | 1.28E-07 | 1.303258547 | 2.39E-12 | 2.305547122 | 10.46903009 | 10.44821838 | 10.57378414 | 11.47335809 | 11.5206165 | 11.79510361 | 11.80044455 |

| | | | | | | | | | | | | | |
|---|---|---|---|---|---|---|---|---|---|---|---|---|---|
| ENSG00000108799 | EZH1 | 0.868406056 | 0.641177228 | 1.436744769 | 0.413358122 | 2.305150825 | 8.257427002 | 7.431218269 | 2 | 8.190940526 | 8.17486938 | 8.961264871 | 8.48994763 |
| ENSG00000169246 | NPIPB3 | 5.44931766 | 6.69E-30 | -3.145845775 | 7.01E-10 | 2.303471885 | 6.366709671 | 6.728495673 | 5.39244166 | 11.67428979 | 11.6634724 | 3.663836244 | 3.600986722 |
| ENSG00000089050 | RBBP9 | 1.417786575 | 0.13915257 | 0.884969515 | 0.369236619 | 2.30275609 | 7.384433967 | 7.503546079 | 4.874587708 | 8.392303091 | 8.35836495 | 7.787860081 | 7.910280955 |
| ENSG00000141756 | FKBP10 | 0.737654234 | 0.251050327 | 1.563476168 | 0.007494256 | 2.301130402 | 12.26464645 | 11.09754095 | 12.48269311 | 12.77192248 | 12.826278 | 13.73492812 | 13.50561406 |
| ENSG00000150394 | CDH8 | -0.737231445 | 0.688378388 | 3.03786246 | 0.053922041 | 2.300631014 | 2 | 4.710564154 | 7.098165388 | 5.022023783 | 5.21224859 | 9.079831553 | 8.274745794 |
| ENSG00000072501 | SMC1A | 0.627710597 | 0.223475046 | 1.669022462 | 0.000259929 | 2.296733059 | 9.031919741 | 9.00621767 | 8.143093325 | 9.345738359 | 9.46906021 | 10.24160068 | 10.62444126 |
| ENSG00000011566 | MAP4K3 | 2.563194483 | 3.24E-06 | -0.267335605 | 0.696157407 | 2.295858878 | 6.563806279 | 7.874700259 | 7.858115942 | 10.08190282 | 10.0993493 | 7.440382108 | 7.126566171 |
| ENSG00000171867 | PRNP | 2.066344773 | 0.016788447 | 0.228749321 | 0.827299212 | 2.295094094 | 11.50346787 | 9.981728319 | 9.242516611 | 12.63561678 | 12.6047477 | 10.99024375 | 10.5456694 |
| ENSG00000214413 | BBIP1 | 1.360718295 | 0.36589956 | 0.933754478 | 0.541713981 | 2.294472773 | 6.89194353 | 6.389896633 | 2 | 7.344877929 | 7.49164798 | 6.940933331 | 7.065820006 |
| ENSG00000111186 | WNT5B | 0.113639527 | 0.888172933 | 2.179731972 | 0.000343839 | 2.293371499 | 7.709880939 | 8.981105197 | 9.31604076 | 8.918344554 | 8.9398551 | 11.04947379 | 10.92000025 |
| ENSG00000155858 | LSM11 | 0.268124543 | 0.701852817 | 2.024798334 | 0.000313211 | 2.292922877 | 6.468621203 | 6.602424674 | 6.80318718 | 7.017154451 | 6.74548212 | 9.000353022 | 8.071643455 |
| ENSG00000103479 | RBL2 | 1.215311903 | 0.329061986 | 1.074178547 | 0.384709619 | 2.28949045 | 5.330594134 | 9.00621767 | 8.845626171 | 9.562142546 | 9.6544356 | 9.518299895 | 9.417219134 |
| ENSG00000115520 | COQ10B | 1.458023393 | 0.157714101 | 0.830792877 | 0.437993223 | 2.288816269 | 9.081016759 | 8.414402626 | 6.073378139 | 9.835556805 | 9.66898051 | 9.149317009 | 9.112583673 |
| ENSG00000162777 | DENND2D | 0.243561862 | 0.865682896 | 2.045228439 | 0.077151749 | 2.288790301 | 4.467311984 | 4.11352228 | 2 | 4.034562184 | 4.0930011 | 5.853194273 | 5.282014932 |
| ENSG00000130529 | TRPM4 | 0.228462299 | 0.849223293 | 2.059994301 | 0.034146284 | 2.288456601 | 5.70932728 | 7.60566235 | 8.677268317 | 7.982979299 | 8.0087095 | 9.795903284 | 9.826110653 |
| ENSG00000074695 | LMAN1 | 1.084201768 | 0.03803836 | 1.202088253 | 0.019063806 | 2.286290021 | 10.63713535 | 10.51097332 | 9.516493218 | 11.39677132 | 11.3707459 | 11.46655078 | 11.53569017 |
| ENSG00000141437 | SLC25A52 | 4.582529336 | 0.174292193 | -2.297129444 | 0.537958482 | 2.285399893 | 2 | 2 | 2 | 4.54891529 | 5.06165168 | 2.139668783 | 2 |
| ENSG00000141569 | TRIM65 | 0.540439672 | 0.55375014 | 1.741416429 | 0.029917935 | 2.281856101 | 7.790810999 | 6.899080544 | 5.59507254 | 7.539954763 | 7.55253065 | 8.929133243 | 8.498084662 |
| ENSG00000029364 | SLC39A9 | 0.875208041 | 0.035558155 | 1.406517692 | 0.000321467 | 2.281725733 | 8.892187472 | 9.405920729 | 8.608698767 | 9.912986572 | 9.85538952 | 10.3722972 | 10.45435372 |
| ENSG00000184967 | NOC4L | 1.343871037 | 0.478334777 | 0.935078014 | 0.626461244 | 2.278949051 | 4.00722531 | 7.051611315 | 2 | 6.931899482 | 7.00136498 | 6.346800618 | 6.769038269 |
| ENSG00000163945 | UVSSA | 0.535893442 | 0.720558899 | 1.742581466 | 0.188722009 | 2.278474908 | 4.467311984 | 4.93638933 | 7.818184403 | 7.062428156 | 7.01963893 | 8.264281212 | 8.180141273 |
| ENSG00000112159 | MDN1 | 0.688468532 | 0.628451704 | 1.588984615 | 0.220081795 | 2.277453147 | 4.467311984 | 7.874700259 | 8.763901704 | 8.550024073 | 8.50338773 | 9.481298847 | 9.35623215 |
| ENSG00000108688 | CCL7 | 6.051420788 | 0.003930403 | -3.774546937 | 0.180505068 | 2.276873851 | 2 | 2 | 2 | 6.172600389 | 6.14893615 | 2.048074405 | 2 |
| ENSG00000171492 | LRRC8D | 2.138935195 | 0.145483541 | 0.13778175 | 0.937449278 | 2.276716945 | 3.327711234 | 7.35507212 | 8.536706852 | 9.651519255 | 9.58403885 | 7.596037946 | 7.680519528 |
| ENSG00000245848 | CEBPA | -0.688466766 | 0.803473818 | 2.964221236 | 0.201514215 | 2.27575447 | 2 | 3.686303414 | 2 | 2.695456494 | 2.49394783 | 5.646229369 | 3.261853426 |
| ENSG00000169895 | SYAP1 | 0.822875375 | 0.00491763 | 1.45137291 | 1.32E-07 | 2.274248285 | 8.816894989 | 8.834582582 | 8.677268317 | 9.607904693 | 9.58633944 | 10.07267492 | 10.35928683 |
| ENSG00000152102 | FAM168B | 0.243885491 | 0.895675696 | 2.028763373 | 0.189944554 | 2.272648864 | 4.467311984 | 10.28850895 | 9.945092785 | 9.826399658 | 9.77228374 | 11.51966333 | 11.63765248 |
| ENSG00000164494 | PDSS2 | 0.841496862 | 0.321145767 | 1.430780489 | 0.069800729 | 2.272277351 | 5.330594134 | 7.274681896 | 6.073378139 | 7.307758725 | 7.26914476 | 7.871230734 | 7.859604233 |
| ENSG00000174132 | FAM174A | 1.189166966 | 0.488975574 | 1.082958818 | 0.523941689 | 2.272125784 | 2 | 6.899080544 | 7.691296561 | 7.944803394 | 7.93600799 | 7.890311949 | 7.779530392 |
| ENSG00000135776 | ABCB10 | 0.982360736 | 0.552722287 | 1.28871532 | 0.418932978 | 2.271076057 | 2 | 6.389896633 | 7.285536013 | 7.307758725 | 7.27295842 | 7.644728141 | 7.532563185 |
| ENSG00000185267 | CDNF | 1.206204884 | 0.588327109 | 1.064718773 | 0.629414332 | 2.270923657 | 2 | 3.686303414 | 2 | 3.515101962 | 3.60476711 | 3.724264246 | 3.15612297 |
| ENSG00000130822 | PNCK | -0.718098908 | 0.895265695 | 2.988492177 | 0.525662412 | 2.270393268 | 2 | 2 | 2 | 2.3892346 | 2 | 3.930690651 | 3.15612297 |
| ENSG00000103978 | TMEM87A | 2.02167045 | 0.06340191 | 0.247700786 | 0.84812366 | 2.269371236 | 8.313285031 | 9.018611864 | 5.59507254 | 10.21014801 | 10.2227847 | 8.513629334 | 8.392188801 |
| ENSG00000177106 | EPS8L2 | 0.499594327 | 0.85245466 | 1.768284915 | 0.458817894 | 2.267879242 | 9.730679447 | 4.442710129 | 2 | 8.621345041 | 8.7394436 | 9.566031214 | 10.24002969 |
| ENSG00000033100 | CHPF2 | 0.958147825 | 0.013556653 | 1.308993957 | 0.000439228 | 2.267141782 | 9.689059027 | 10.27307349 | 10.43545992 | 11.11136778 | 11.1336271 | 11.40522087 | 11.53741843 |
| ENSG00000108262 | GIT1 | 0.600867924 | 0.765831172 | 1.666009922 | 0.361988871 | 2.266877846 | 8.468948322 | 8.702538795 | 2 | 8.509731121 | 8.70032852 | 9.869029072 | 9.428616564 |
| ENSG00000278742 | | 7.455553301 | 0.000218272 | -5.191146668 | 0.057347766 | 2.264406633 | 2 | 2 | 2 | 7.447418636 | 7.57741621 | 2 | 2 |
| ENSG00000156414 | TDRD9 | -3.753101612 | 0.362134662 | 6.017239333 | 0.078163215 | 2.264137721 | 2 | 2 | 2 | 2 | 2 | 5.119635168 | 6.716538017 |
| ENSG00000120265 | PCMT1 | 1.087765772 | 0.43457188 | 1.175115594 | 0.383915641 | 2.262881366 | 8.75772933 | 8.607278467 | 4.523675391 | 9.231457548 | 9.20532695 | 9.449489749 | 9.144901069 |
| ENSG00000131263 | RLIM | 1.266869209 | 0.187284818 | 0.994256456 | 0.303223054 | 2.261125665 | 9.820154628 | 8.762721501 | 7.163363831 | 10.225157 | 10.1908874 | 9.950927567 | 9.921967897 |
| ENSG00000007047 | MARK4 | 1.013033425 | 0.368308689 | 1.246752992 | 0.250438486 | 2.259786418 | 6.009025997 | 8.607278467 | 9.212024683 | 9.437090544 | 9.46739696 | 9.796563052 | 9.563647359 |
| ENSG00000169398 | PTK2 | 0.783383229 | 0.434884889 | 1.474313461 | 0.109780104 | 2.257696691 | 7.384433967 | 9.616264339 | 9.935792933 | 10.12788365 | 10.0928914 | 10.60364979 | 10.97180235 |
| ENSG00000258588 | TRIM6-TRIM | 7.448061742 | 0.000231352 | -5.191146595 | 0.057776246 | 2.256915147 | 2 | 2 | 2 | 7.580980221 | 7.4280824 | 2 | 2 |
| ENSG00000090060 | PAPOLA | 1.515736122 | 0.048471712 | 0.740924767 | 0.363979257 | 2.256660889 | 8.138771685 | 10.17682307 | 9.503962296 | 11.03591644 | 10.9808314 | 10.32506285 | 10.14066157 |
| ENSG00000109118 | PHF12 | 0.945266424 | 0.312962462 | 1.311387657 | 0.140295302 | 2.256654081 | 8.009475818 | 8.395519306 | 5.930865648 | 8.713632895 | 8.70880006 | 9.026460641 | 9.120511546 |
| ENSG00000090238 | YPEL3 | 0.541251753 | 0.707528996 | 1.714817322 | 0.178997619 | 2.256069075 | 4.00722531 | 7.847167909 | 8.110401118 | 7.971157485 | 7.98332947 | 9.348425302 | 8.89156005 |
| ENSG00000166326 | TRIM44 | 0.697623481 | 0.092286577 | 1.557874507 | 3.15E-05 | 2.255497987 | 9.646202137 | 10.35843478 | 9.839340503 | 10.66893665 | 10.6891726 | 11.41370331 | 11.65200579 |
| ENSG00000112763 | BTN2A1 | 1.746408691 | 0.00247249 | 0.508651381 | 0.435186101 | 2.255060072 | 9.689059027 | 8.276714465 | 9.31604076 | 10.95689981 | 10.9276923 | 9.815783218 | 9.592751029 |
| ENSG00000163964 | PIGX | 0.654371904 | 0.515257886 | 1.600453023 | 0.075349452 | 2.254824926 | 7.27616306 | 7.393647515 | 4.874587708 | 7.454228844 | 7.58969985 | 8.448514034 | 8.46040717 |
| ENSG00000007384 | RHBDF1 | 1.300844946 | 0.002487197 | 0.953653233 | 0.031386333 | 2.254498179 | 8.107517749 | 8.747909548 | 8.408111533 | 9.727271068 | 9.7560263 | 9.092423696 | 9.647426237 |
| ENSG00000178789 | CD300LB | -1.303268636 | 0.814508396 | 3.556201101 | 0.462938093 | 2.252932465 | 2 | 2 | 2 | 2.270786764 | 2 | 4.778616857 | 2.172608095 |
| ENSG00000183735 | TBK1 | 1.777920034 | 0.001515995 | 0.472442827 | 0.459674606 | 2.250362861 | 7.750913381 | 6.899080544 | 8.236946975 | 9.474417988 | 9.47569407 | 8.325906648 | 8.020777347 |
| ENSG00000157483 | MYO1E | 0.652357625 | 0.293988413 | 1.597070096 | 0.004002525 | 2.249427721 | 9.189468652 | 8.297207175 | 9.439236147 | 9.690252889 | 9.70188757 | 10.92345228 | 10.28029762 |
| ENSG00000187535 | IFT140 | 0.970824938 | 0.634148538 | 1.278218088 | 0.515353465 | 2.249043027 | 2 | 7.954278496 | 4.874587708 | 7.514115926 | 7.49164798 | 7.786974998 | 7.821204179 |
| ENSG00000117118 | SDHB | 1.41034914 | 0.198081092 | 0.8376604 | 0.459893021 | 2.24800954 | 8.981092835 | 7.145017454 | 6.073378139 | 9.314267267 | 9.26207178 | 8.550081477 | 8.872386739 |
| ENSG00000198081 | ZBTB14 | 0.939101304 | 0.531266648 | 1.308818796 | 0.360920918 | 2.2479201 | 6.257048221 | 6.602424674 | 2 | 6.809774134 | 6.77805036 | 7.293562319 | 6.994755759 |
| ENSG00000128513 | POT1 | 1.34709315 | 0.112649338 | 0.900401855 | 0.299350397 | 2.247495005 | 6.96351253 | 6.140560021 | 5.156626401 | 7.52061915 | 7.61395775 | 7.538302335 | 6.564691065 |
| ENSG00000167676 | PLIN4 | -0.727708903 | 0.844549702 | 2.973506641 | 0.332960127 | 2.245797738 | 2 | 2 | 2 | 2.141737246 | 2.2680111 | 3.865109708 | 3.236138336 |
| ENSG00000214026 | MRPL23 | 2.589643211 | 0.014438702 | -0.344475398 | 0.787237277 | 2.245167813 | 4.00722531 | 4.11352228 | 6.43197674 | 7.68016596 | 7.81567127 | 4.888321757 | 5.072762365 |
| ENSG00000137936 | BCAR3 | 1.090387397 | 0.061614336 | 1.153316643 | 0.04392157 | 2.24370404 | 7.53297911 | 8.732943944 | 7.896971941 | 9.228481255 | 9.23983583 | 9.470589819 | 9.098827153 |
| ENSG00000102053 | ZC3H12B | -0.167771671 | 0.91483416 | 2.410374805 | 0.053383916 | 2.242603134 | 2 | 4.11352228 | 5.39244166 | 4.105493497 | 4.37434574 | 6.098353065 | 6.887339785 |
| ENSG00000104324 | CPQ | 0.468101597 | 0.643120989 | 1.773344691 | 0.042824635 | 2.241446288 | 9.204317094 | 7.761270542 | 6.882775709 | 8.758341449 | 8.70032852 | 9.830032147 | 10.19985319 |
| ENSG00000065989 | PDE4A | 0.43308049 | 0.756699054 | 1.80792994 | 0.135544734 | 2.24101043 | 8.87372952 | 7.928237109 | 4.874587708 | 8.395844809 | 8.36015869 | 9.658696966 | 9.821254547 |
| ENSG00000163104 | SMARCAD1 | 1.749134507 | 0.190391386 | 0.490115373 | 0.738292369 | 2.23924988 | 4.00722531 | 8.276714465 | 6.80318718 | 8.977450658 | 8.8597051 | 7.774526492 | 7.572926221 |
| ENSG00000092439 | TRPM7 | 1.643741281 | 0.067435031 | 0.593530551 | 0.544635534 | 2.237271832 | 7.579279349 | 7.638155693 | 9.589475645 | 10.24198055 | 10.2089163 | 9.257743834 | 9.098827153 |
| ENSG00000152926 | ZNF117 | 1.561713645 | 0.022695801 | 0.67543045 | 0.361214879 | 2.237144095 | 4.815609471 | 6.04685525 | 6.43197674 | 7.450827758 | 7.36873214 | 6.7543034 | 6.317379004 |
| ENSG00000065000 | AP3D1 | 0.997358352 | 0.019549332 | 1.239760841 | 0.002731113 | 2.237119192 | 9.423197387 | 10.27823701 | 9.757248847 | 10.83076085 | 10.8903195 | 11.03789512 | 11.16486422 |
| ENSG00000151743 | AMN1 | 1.538281276 | 0.485453797 | 0.696023767 | 0.763713936 | 2.234305043 | 2 | 5.946638857 | 2 | 5.94010643 | 5.9536761 | 5.085488617 | 5.263040529 |
| ENSG00000117394 | SLC2A1 | 1.154690535 | 0.100527553 | 1.078057185 | 0.121709811 | 2.23274772 | 10.88304938 | 10.30378101 | 9.25752426 | 11.44457809 | 11.4469936 | 11.03463637 | 11.64071835 |
| ENSG00000154124 | OTULIN | 0.874070998 | 0.345449366 | 1.358288077 | 0.118418414 | 2.232359075 | 6.468621203 | 7.274681896 | 8.677268317 | 8.625876185 | 8.61876544 | 9.267294952 | 8.916059773 |
| ENSG00000174236 | REP15 | 3.009791982 | 0.305279886 | -0.77786129 | 0.813732645 | 2.231930693 | 2 | 2 | 2 | 3.406930997 | 3.7864959 | 2.225793514 | 2.172608095 |
| ENSG00000104936 | DMPK | -0.19762642 | 0.710155984 | 2.429553625 | 3.63E-09 | 2.231927205 | 9.204317094 | 8.337339476 | 9.083172337 | 8.737556707 | 8.70315789 | 11.20690519 | 11.45473662 |
| ENSG00000187210 | GCNT1 | 1.008738042 | 0.609550266 | 1.219823761 | 0.52282666 | 2.228561803 | 2 | 7.431218269 | 4.523675391 | 7.101994243 | 6.96881778 | 6.85908071 | 7.544398272 |
| ENSG00000107651 | SEC23IP | 1.240125569 | 0.307710975 | 0.988210598 | 0.418322864 | 2.228336167 | 6.366709671 | 9.558016677 | 7.398165132 | 9.616278374 | 9.63159714 | 9.202392925 | 9.526339317 |
| ENSG00000171121 | KCNMB3 | 3.110063055 | 0.05126002 | -0.881877375 | 0.62358047 | 2.22818568 | 2 | 4.93638933 | 2 | 6.415275989 | 6.48770488 | 3.12825509 | 3.287118176 |
| ENSG00000184719 | RNLS | 1.261817373 | 0.574255931 | 0.962664209 | 0.672113452 | 2.224481582 | 2 | 5.946638857 | 2 | 5.627094009 | 5.74985018 | 5.323825264 | 5.502966356 |
| ENSG00000120254 | MTHFD1L | 0.534564434 | 0.64508255 | 1.689648955 | 0.099251185 | 2.224213389 | 9.435834703 | 7.790474954 | 6.534109959 | 8.965583267 | 8.86981338 | 10.1449176 | 9.984600648 |
| ENSG00000115170 | ACVR1 | 1.690804058 | 0.047187606 | 0.53155749 | 0.572841244 | 2.222361549 | 7.940205782 | 6.464272742 | 8.805342542 | 9.658907527 | 9.72504082 | 8.624798858 | 8.45344973 |
| ENSG00000174576 | NPAS4 | -0.723374178 | 0.844052269 | 2.945493218 | 0.331905931 | 2.22211904 | 2 | 2 | 2 | 2.270786764 | 2.14022021 | 3.632645806 | 3.496556178 |
| ENSG00000068878 | PSME4 | 1.375046689 | 4.40E-06 | 0.846783891 | 0.006873339 | 2.22183058 | 8.892187472 | 9.00621767 | 9.344432377 | 10.41296081 | 10.505257 | 9.839880645 | 10.02129178 |
| ENSG00000081019 | RSBN1 | 1.126538129 | 0.528648153 | 1.093250027 | 0.533161677 | 2.219788156 | 7.975256505 | 7.002537973 | 2 | 8.149581172 | 8.05374131 | 8.317962987 | 7.769481431 |
| ENSG00000172830 | SSH3 | 0.363983274 | 0.688378388 | 1.855305416 | 0.0150633 | 2.219288691 | 5.866944937 | 6.311476754 | 7.734844356 | 7.25803034 | 7.16216989 | 8.69913345 | 8.654300638 |
| ENSG00000184557 | SOCS3 | 1.17466905 | 0.361180507 | 1.044243214 | 0.4140361 | 2.218912264 | 6.89194353 | 8.255926468 | 4.523675391 | 8.383410567 | 8.38503958 | 8.575932171 | 7.842663598 |
| ENSG00000213401 | MAGEA12 | -2.306780167 | 0.562496388 | 4.525257085 | 0.156315531 | 2.218476917 | 2 | 2 | 2 | 2.141737246 | 2 | 5.323825264 | 3.8919413 |
| ENSG00000197586 | ENTPD6 | 0.882987688 | 0.638882507 | 1.33525392 | 0.455061132 | 2.218241607 | 7.790810999 | 8.376385542 | 2 | 8.458152491 | 8.36195019 | 8.912592153 | 8.802378289 |
| ENSG00000119326 | CTNNAL1 | -0.208747818 | 0.833633118 | 2.424964205 | 0.001792503 | 2.216216387 | 6.73718411 | 8.791896851 | 7.552162999 | 7.762033056 | 7.72948195 | 10.0867686 | 10.58423994 |
| ENSG00000231924 | PSG1 | 0.694332923 | 0.759550096 | 1.520729472 | 0.467385744 | 2.215062395 | 2 | 2 | 5.156626401 | 4.237680128 | 4.45712836 | 4.875056259 | 5.20457242 |
| ENSG00000110079 | MS4A4A | -2.312353283 | 0.551211363 | 4.527326948 | 0.142012462 | 2.214973664 | 2 | 2 | 2 | 2 | 2.14022021 | 5.185595313 | 4.215171317 |
| ENSG00000143373 | ZNF687 | 0.629963618 | 0.795427588 | 1.584287128 | 0.47448492 | 2.214250746 | 4.815609471 | 9.125613817 | 2 | 8.235106869 | 8.2561084 | 9.167453541 | 9.213631806 |
| ENSG00000163995 | ABLIM2 | -2.312355123 | 0.551575085 | 4.526286499 | 0.142685271 | 2.213931376 | 2 | 2 | 2 | 2 | 2.14022021 | 5.19095825 | 4.201948172 |
| ENSG00000176401 | EID2B | 1.01369802 | 0.623226161 | 1.197938258 | 0.547523809 | 2.211636278 | 2 | 4.710564154 | 2 | 4.17310023 | 4.4836996 | 4.111244182 | 4.784231301 |
| ENSG00000147166 | ITGB1BP2 | -1.310494048 | 0.697008225 | 3.520945257 | 0.197561623 | 2.21045121 | 2 | 2 | 2 | 2.141737246 | 2.14022021 | 3.891700877 | 4.047883918 |
| ENSG00000171931 | FBXW10 | 3.272401786 | 0.30635441 | -1.062825412 | 0.764659891 | 2.209576374 | 2 | 2 | 2 | 3.515101962 | 4.02226117 | 2.048074405 | 2.277179783 |
| ENSG00000102385 | DRP2 | 0.674714874 | 0.765893437 | 1.534509789 | 0.462478098 | 2.209224663 | 2 | 5.595927442 | 2 | 4.946755474 | 4.72553021 | 5.367107572 | 5.812327524 |
| ENSG00000070269 | TMEM260 | 1.610925986 | 0.391746117 | 0.598244476 | 0.765896408 | 2.209170461 | 4.467311984 | 7.467835385 | 2 | 7.565338955 | 7.73225478 | 6.631488899 | 6.70252219 |
| ENSG00000173868 | PHOSPHO1 | -1.269611831 | 0.617831455 | 3.477712143 | 0.096104184 | 2.208100312 | 3.327711234 | 2 | 2 | 2.3892346 | 2.14022021 | 4.965450964 | 4.388595494 |
| ENSG00000069493 | CLEC2D | 4.450966765 | 3.76E-05 | -2.243056047 | 0.05509797 | 2.207910718 | 2 | 5.946638857 | 5.59507254 | 9.595637371 | 9.54119795 | 3.753554863 | 3.183290849 |
| ENSG00000177463 | NR2C2 | 1.036848313 | 0.326820527 | 1.170590951 | 0.25494197 | 2.207439263 | 9.544849385 | 7.099070191 | 7.502634626 | 9.512436154 | 9.48312089 | 9.743260791 | 9.508995835 |
| ENSG00000104375 | STK3 | 0.80380305 | 0.397404116 | 1.403441869 | 0.110621519 | 2.207244919 | 6.009025997 | 7.73146271 | 5.772716771 | 7.523859802 | 7.65545169 | 8.313667445 | 8.033896215 |
| ENSG00000106397 | PLOD3 | 1.339447667 | 0.477431438 | 0.867412905 | 0.651899269 | 2.206860572 | 9.975371636 | 10.64091981 | 3.369317972 | 11.06734964 | 11.1380772 | 10.60452918 | 10.65917495 |
| ENSG00000118197 | DDX59 | 1.140580059 | 0.236944877 | 1.06507941 | 0.26431896 | 2.20565947 | 4.815609471 | 7.274681896 | 6.073378139 | 7.555872146 | 7.45854177 | 7.566427714 | 7.288576158 |
| ENSG00000111540 | RAB5B | 0.474040238 | 0.535332813 | 1.73137772 | 0.008995102 | 2.205417959 | 9.097018025 | 10.27307349 | 10.8211137 | 10.68386176 | 10.7139326 | 11.81499449 | 12.08131906 |
| ENSG00000076067 | RBMS2 | 0.881829005 | 0.012336386 | 1.323420019 | 7.80E-05 | 2.205249025 | 9.87686338 | 9.770792087 | 9.372276026 | 10.55119161 | 10.5871833 | 10.81692048 | 11.18011685 |
| ENSG00000216588 | IGSF23 | -2.312322002 | 0.544079031 | 4.516778239 | 0.133193465 | 2.204456237 | 2 | 2 | 2 | 2 | 2.14022021 | 4.555645384 | 4.959465495 |
| ENSG00000138768 | USO1 | 1.375035242 | 0.001224561 | 0.828559793 | 0.063908241 | 2.203595035 | 8.737455672 | 9.470501606 | 9.647595056 | 10.72491929 | 10.6866731 | 10.20346921 | 10.1173006 |
| ENSG00000204611 | ZNF616 | 1.100075151 | 0.479960202 | 1.102409508 | 0.470407499 | 2.202484659 | 3.327711234 | 4.93638933 | 7.398165132 | 7.242377933 | 7.09050444 | 7.534088584 | 6.683620062 |
| ENSG00000086289 | EPDR1 | 0.546703924 | 0.35926735 | 1.654063983 | 0.001488322 | 2.200767907 | 7.790810999 | 7.051611315 | 8.143093325 | 8.188900477 | 8.32017198 | 9.098500449 | 9.569772634 |

| | | | | | | | | | | | | | |
|---|---|---|---|---|---|---|---|---|---|---|---|---|---|
| ENSG00000198730 | CTR9 | 1.250822554 | 0.261480902 | 0.949770045 | 0.397984664 | 2.200592599 | 6.009025997 | 9.079034257 | 7.552162999 | 9.302993856 | 9.27923788 | 8.800846924 | 9.161219358 |
| ENSG00000198796 | ALPK2 | 0.414499648 | 0.85527092 | 1.785334928 | 0.366670893 | 2.199834576 | 2 | 5.457120953 | 8.35328246 | 7.409376773 | 7.31795292 | 7.778984728 | 9.276085345 |
| ENSG00000164162 | ANAPC10 | 1.268298292 | 0.346404872 | 0.931143315 | 0.493153856 | 2.199441607 | 5.330594134 | 6.04685525 | 2 | 6.205262847 | 6.58294721 | 6.176128386 | 5.992078266 |
| ENSG00000167566 | NCKAP5L | 0.580074427 | 0.675598697 | 1.619316503 | 0.190484764 | 2.19939093 | 5.095959458 | 8.687092496 | 9.083172337 | 8.904736303 | 8.97305476 | 9.987021151 | 9.957809357 |
| ENSG00000143164 | DCAF6 | 1.080334628 | 2.34E-05 | 1.117150934 | 1.01E-05 | 2.197485562 | 8.855032352 | 8.903033433 | 8.825624954 | 9.979407193 | 9.89304688 | 9.877992513 | 10.06325576 |
| ENSG00000104432 | IL7 | 3.470607319 | 0.449678944 | -1.274101357 | 0.797193705 | 2.196505962 | 2 | 2 | 2 | 3.921164147 | 3.94787299 | 2 | 2.277179783 |
| ENSG00000108515 | ENO3 | 0.756748919 | 0.60852038 | 1.438819195 | 0.287769738 | 2.195568115 | 4.00722531 | 4.442710129 | 2 | 4.415698455 | 4.45712836 | 5.4091291 | 4.468068808 |
| ENSG00000144152 | FBLN7 | 0.893354197 | 0.717276749 | 1.301977508 | 0.580976488 | 2.195331705 | 2 | 6.535001748 | 2 | 5.808110275 | 6.00043082 | 6.428526856 | 6.141542327 |
| ENSG00000085871 | MGST2 | 1.42691189 | 0.183576111 | 0.767809654 | 0.49246368 | 2.194721544 | 7.159102577 | 5.457120953 | 4.523675391 | 7.412876874 | 7.57432889 | 6.510167449 | 7.133547492 |
| ENSG00000102038 | SMARCA1 | 1.059633281 | 0.180631866 | 1.133552576 | 0.1415706 | 2.193185857 | 8.009475818 | 10.019219 | 9.426401271 | 10.45855028 | 10.3940894 | 10.46042363 | 10.53927643 |
| ENSG00000103342 | GSPT1 | 0.950964451 | 0.07257246 | 1.240510775 | 0.01474777 | 2.191475226 | 9.781071289 | 9.616264339 | 8.65477198 | 10.40199532 | 10.3532448 | 10.58937934 | 10.73987554 |
| ENSG00000116005 | PCYOX1 | -0.112585724 | 0.883684699 | 2.303712203 | 6.33E-05 | 2.191126478 | 8.836089686 | 10.12617756 | 8.960138144 | 9.361220151 | 9.28774494 | 11.7437434 | 11.72401992 |
| ENSG00000136810 | TXN | 2.21967697 | 8.92E-05 | -0.030095589 | 0.968502972 | 2.189581381 | 9.820154628 | 8.395519306 | 9.049077186 | 11.40931048 | 11.4175794 | 9.088477995 | 9.250223518 |
| ENSG00000165115 | KIF27 | 2.140021933 | 0.108187743 | 0.049334939 | 0.974984507 | 2.189356873 | 2 | 4.442710129 | 4.058998049 | 5.627094009 | 5.6826301 | 4.271694079 | 3.383972333 |
| ENSG00000140829 | DHX38 | 1.259424096 | 0.453919948 | 0.929855064 | 0.58225381 | 2.189279159 | 3.327711234 | 7.73146271 | 9.132850258 | 9.275380474 | 9.27448997 | 8.766685705 | 9.10860334 |
| ENSG00000111897 | SERINC1 | 1.225885033 | 0.184268122 | 0.962961978 | 0.299579278 | 2.188847011 | 9.262228249 | 11.59012602 | 11.60159843 | 12.39271275 | 12.3511729 | 12.12283589 | 12.09580137 |
| ENSG00000109501 | WFS1 | -0.013752453 | 0.990899695 | 2.200707821 | 0.019084503 | 2.186955368 | 9.689059027 | 8.806266215 | 6.882775709 | 8.804373539 | 8.88234995 | 10.78181003 | 11.27343957 |
| ENSG00000122484 | RPAP2 | 0.665199535 | 0.680957539 | 1.520458532 | 0.301260958 | 2.185658067 | 6.563806279 | 6.78762695 | 2 | 6.89733479 | 6.6253296 | 7.807195765 | 7.355789538 |
| ENSG00000147576 | ADHFE1 | 1.756592586 | 0.340147427 | 0.428327528 | 0.833553126 | 2.184920114 | 2 | 4.442710129 | 2 | 4.71689672 | 4.70309173 | 3.810410249 | 3.539239711 |
| ENSG00000128595 | CALU | 0.522318521 | 0.305223019 | 1.662540458 | 0.000174595 | 2.184858979 | 12.48433893 | 13.3410766 | 12.55584475 | 13.36578153 | 13.3766069 | 14.37571671 | 14.6349142 |
| ENSG00000176014 | TUBB6 | 0.203316404 | 0.803640574 | 1.980347806 | 0.002293883 | 2.183664211 | 10.98115017 | 10.08551947 | 9.658943166 | 10.52393752 | 10.5810438 | 11.76350665 | 12.73131499 |
| ENSG00000125885 | MCM8 | 0.783400726 | 0.463670664 | 1.400240071 | 0.153651094 | 2.183640797 | 5.330594134 | 6.04685525 | 3.369317972 | 6.252910529 | 5.7929843 | 6.611630769 | 6.6426114 |
| ENSG00000158458 | NRG2 | 1.533939055 | 0.536685094 | 0.64688277 | 0.804608418 | 2.180821826 | 2 | 2 | 3.369317972 | 3.162721474 | 3.39680241 | 3.059555873 | 2.551896972 |
| ENSG00000172785 | CBWD1 | 3.153188618 | 0.015021195 | -0.973593346 | 0.501722792 | 2.179595272 | 6.73718411 | 6.389896633 | 2 | 9.031955922 | 9.16279627 | 4.965450964 | 5.28828468 |
| ENSG00000139547 | RDH16 | 2.690951034 | 0.412898781 | -0.511450235 | NA | 2.179500799 | 2 | 2 | 2 | 3.566290607 | 3.2184422 | 2.34604626 | 2.117335275 |
| ENSG00000147224 | PRPS1 | 0.843802214 | 0.322921865 | 1.335117654 | 0.095474849 | 2.178919868 | 9.397585735 | 7.503546079 | 8.296287027 | 9.404936476 | 9.47652116 | 9.51429464 | 10.25250913 |
| ENSG00000181722 | ZBTB20 | 2.06918468 | 0.292000423 | 0.108286792 | 0.96272717 | 2.177471472 | 2 | 6.951736442 | 9.196533591 | 9.933844998 | 9.9554848 | 7.805448645 | 8.172535245 |
| ENSG00000137807 | KIF23 | 0.787010275 | 0.633108815 | 1.388702188 | 0.366387369 | 2.175712463 | 4.467311984 | 6.389896633 | 2 | 5.949776857 | 5.9536761 | 6.414841377 | 6.620419619 |
| ENSG00000100600 | LGMN | 1.372869502 | 0.027892519 | 0.802353879 | 0.219885804 | 2.175223381 | 10.07567912 | 8.57407404 | 9.681375057 | 10.90533885 | 10.965595 | 10.48532674 | 10.23739929 |
| ENSG00000105698 | USF2 | 1.123391612 | 0.44980498 | 1.051015419 | 0.472698621 | 2.174407032 | 4.815609471 | 8.077869079 | 9.589475645 | 9.632130776 | 9.54672161 | 9.576569363 | 9.457070207 |
| ENSG00000239713 | APOBEC3G | 1.881235425 | 0.124042603 | 0.292619756 | 0.836490254 | 2.173855181 | 3.327711234 | 6.844429755 | 7.225742779 | 8.394075037 | 8.31092742 | 6.7543034 | 6.843307307 |
| ENSG00000144488 | ESPNL | 1.325321978 | 0.583890106 | 0.845337276 | 0.731701381 | 2.170659254 | 2 | 2 | 6.203069917 | 5.968925393 | 5.90535527 | 5.688550058 | 5.269393099 |
| ENSG00000152402 | GUCY1A2 | -0.306792081 | 0.947190051 | 2.474906812 | NA | 2.168114731 | 2 | 2 | 2 | 2.3892346 | 2.14022021 | 3.904814898 | 2.117335275 |
| ENSG00000137857 | DUOX1 | -1.310492654 | 0.702149071 | 3.476628704 | 0.21425456 | 2.16613605 | 2 | 2 | 2 | 2.141737246 | 2.14022021 | 4.199031499 | 3.620995423 |
| ENSG00000100243 | CYB5R3 | 0.177926023 | 0.833090562 | 1.986578946 | 0.00275976 | 2.164504969 | 12.54683767 | 11.6107392 | 10.87563042 | 11.97939826 | 12.0529163 | 13.58455511 | 14.0297935 |
| ENSG00000101413 | RPRD1B | 0.779605691 | 0.618091142 | 1.384471553 | 0.343161384 | 2.164077244 | 7.940205782 | 8.276714465 | 3.369317972 | 8.327018759 | 8.33484115 | 8.974541854 | 8.88330014 |
| ENSG00000125107 | CNOT1 | 1.195998688 | 0.024264667 | 0.967802786 | 0.072427994 | 2.163801474 | 9.371511179 | 10.44821838 | 9.528916236 | 11.06873707 | 11.0541353 | 10.89498282 | 10.76972906 |
| ENSG00000135338 | LCA5 | 0.792671252 | 0.593368805 | 1.370352353 | 0.322128031 | 2.163023606 | 2 | 6.140560021 | 6.073378139 | 6.29903474 | 6.32824132 | 7.1328795 | 6.541257688 |
| ENSG00000116096 | SPR | 1.306511771 | 0.451325678 | 0.856155112 | 0.627647664 | 2.162666883 | 2 | 7.393647515 | 5.59507254 | 7.444001439 | 7.5014335 | 7.130090369 | 6.932071508 |
| ENSG00000131473 | ACLY | 1.014753101 | 0.453973535 | 1.147629836 | 0.379839399 | 2.162382937 | 7.27616306 | 11.20718648 | 11.17734896 | 11.64445391 | 11.690457 | 11.48501894 | 12.05897558 |
| ENSG00000205208 | C4orf46 | 1.187485266 | 0.561800205 | 0.974829048 | 0.635400451 | 2.162314314 | 3.327711234 | 7.002537973 | 2 | 6.693715713 | 6.72335419 | 6.442083733 | 6.564691065 |
| ENSG00000102384 | CENPI | 0.439859469 | 0.843094589 | 1.721323467 | 0.374613381 | 2.161182936 | 2 | 4.93638933 | 2 | 3.881293667 | 4.2560182 | 4.764298067 | 5.448021784 |
| ENSG00000143771 | CNIH4 | 1.123941149 | 0.05248835 | 1.036586233 | 0.072440685 | 2.160527382 | 8.674867122 | 7.572420273 | 7.734844356 | 9.175891964 | 9.19932737 | 9.302713655 | 8.866113243 |
| ENSG00000170561 | IRX2 | 3.002047866 | 0.175108303 | -0.841904842 | 0.730630046 | 2.160143023 | 5.866944937 | 2 | 2 | 7.355829978 | 7.03319408 | 4.350488971 | 2.918126211 |
| ENSG00000176171 | BNIP3 | 1.417001985 | 0.034893697 | 0.742974259 | 0.294688747 | 2.159976244 | 10.38028415 | 9.159587375 | 8.941672936 | 11.0584384 | 11.0477495 | 10.15935238 | 10.57256798 |
| ENSG00000125868 | DSTN | 0.213268889 | 0.652839859 | 1.946352273 | 2.41E-07 | 2.159621162 | 11.18578229 | 10.62884366 | 10.40882047 | 11.02280532 | 10.9599875 | 12.79040486 | 12.65194484 |
| ENSG00000182831 | C16orf72 | 0.957091233 | 0.118250471 | 1.200001186 | 0.041683697 | 2.157092419 | 9.204317094 | 8.356994601 | 7.858115942 | 9.530270581 | 9.53167935 | 9.896369694 | 9.637331225 |
| ENSG00000148175 | STOM | 0.721167 | 0.112132289 | 1.434681298 | 0.00053967 | 2.155848298 | 10.29268328 | 10.27307349 | 10.84614969 | 11.22124799 | 11.2074731 | 12.19425512 | 11.59853243 |
| ENSG00000171552 | BCL2L1 | 0.348999595 | 0.628892599 | 1.806260295 | 0.002801737 | 2.15525989 | 9.635285994 | 8.671479029 | 9.926432745 | 9.81982307 | 9.87308535 | 10.83148019 | 11.65306896 |
| ENSG00000100697 | DICER1 | 1.023339536 | 0.192776283 | 1.128320242 | 0.139340009 | 2.151659778 | 7.53297911 | 9.405920729 | 8.460932916 | 9.672111811 | 9.69692814 | 9.995474265 | 9.548056138 |
| ENSG00000229809 | ZNF688 | 1.175930997 | 0.114471804 | 0.975505162 | 0.189949684 | 2.151436159 | 6.468621203 | 4.93638933 | 5.39244166 | 6.750147204 | 6.9210099 | 6.693258166 | 6.595354992 |
| ENSG00000218891 | ZNF579 | 0.230873267 | 0.909988721 | 1.918362864 | 0.260698796 | 2.149236131 | 2 | 7.099070191 | 8.206335842 | 7.437142666 | 7.37938471 | 9.155502188 | 8.984263871 |
| ENSG00000174238 | PITPNA | 0.828709484 | 0.042415488 | 1.320466184 | 0.000568123 | 2.149175668 | 8.737455672 | 8.806266215 | 9.426401271 | 9.842062221 | 9.84709924 | 10.23949951 | 10.42371165 |
| ENSG00000049239 | H6PD | 1.155266461 | 0.02537741 | 0.993481693 | 0.056299816 | 2.148748154 | 8.586954094 | 9.424669273 | 9.714382178 | 10.44536591 | 10.4869034 | 10.14853986 | 10.44614079 |
| ENSG00000153048 | CARHSP1 | 0.525811014 | 0.762607571 | 1.622720581 | 0.297427124 | 2.148531595 | 8.87372952 | 7.315436877 | 3.369317972 | 8.197043419 | 8.30348889 | 9.114816239 | 9.531303742 |
| ENSG00000116350 | SRSF4 | 0.933273804 | 0.575357254 | 1.215154416 | 0.448059647 | 2.14842822 | 9.781071289 | 8.981105197 | 4.058998049 | 9.832946403 | 9.75806852 | 10.10942542 | 10.04323752 |
| ENSG00000197548 | ATG7 | 1.702141255 | 0.241411697 | 0.446278469 | 0.781965112 | 2.148419724 | 3.327711234 | 8.191702463 | 7.858115942 | 9.110576034 | 9.20732128 | 7.821097076 | 8.008487789 |
| ENSG00000100596 | SPTLC2 | 1.312285848 | 2.16E-05 | 0.835498776 | 0.009547143 | 2.147784624 | 8.928409675 | 8.732943944 | 9.272377397 | 10.30219818 | 10.2873694 | 9.798760102 | 9.840581661 |
| ENSG00000185811 | IKZF1 | -1.310489421 | 0.712294566 | 3.457239919 | 0.238534046 | 2.146750498 | 2 | 2 | 2 | 2.141737246 | 2.14022021 | 4.388329244 | 3.236138336 |
| ENSG00000169221 | TBC1D10B | 0.751418883 | 0.016164965 | 1.394133701 | 1.58E-06 | 2.145552584 | 8.367060731 | 8.505280662 | 8.825624954 | 9.290680758 | 9.35044873 | 9.952309835 | 9.968341625 |
| ENSG00000152642 | GPD1L | -0.617600419 | 0.714145675 | 2.761944318 | 0.05471533 | 2.144343898 | 3.327711234 | 6.78762695 | 3.369317972 | 4.984880435 | 4.87352701 | 8.088662035 | 8.165740501 |
| ENSG00000078070 | MCCC1 | 0.544060145 | 0.832844524 | 1.599817227 | 0.489336405 | 2.143877373 | 2 | 8.433041974 | 3.369317972 | 7.433701013 | 7.45182838 | 8.075674526 | 8.797441501 |
| ENSG00000158042 | MRPL17 | 0.940534771 | 0.013480111 | 1.203323984 | 0.001053111 | 2.143858755 | 8.418903792 | 8.639735838 | 8.960138144 | 9.617037214 | 9.62713473 | 9.664250432 | 10.07418255 |
| ENSG00000197415 | VEPH1 | 3.862299748 | 0.018782812 | -1.720028368 | 0.333565204 | 2.14227138 | 6.468621203 | 2 | 3.369317972 | 8.769305409 | 8.7882247 | 3.810410249 | 3.600986722 |
| ENSG00000055130 | CUL1 | 1.409680826 | 0.353082226 | 0.727374985 | 0.646493605 | 2.137055811 | 8.946185348 | 9.148351514 | 4.058998049 | 9.894328254 | 9.88559363 | 9.2270746 | 9.195398998 |
| ENSG00000171631 | P2RY6 | -1.825784293 | 0.309957701 | 3.962743357 | 0.008764582 | 2.136959064 | 2 | 2 | 4.523675391 | 2.498691189 | 2.49394783 | 6.453284622 | 6.671679221 |
| ENSG00000005175 | RPAP3 | 1.504681019 | 0.111579909 | 0.631212177 | 0.532750733 | 2.135893196 | 5.330594134 | 7.847167909 | 6.534109959 | 8.427334995 | 8.3920703 | 7.596037946 | 7.503217537 |
| ENSG00000196782 | MAML3 | 1.549166801 | 0.000756769 | 0.584800361 | 0.248420606 | 2.133967161 | 6.89194353 | 6.140560021 | 6.534109959 | 8.085230238 | 8.00412803 | 6.977080688 | 7.216434756 |
| ENSG00000205542 | TMSB4X | 1.545809455 | 0.230169878 | 0.587589585 | 0.673347742 | 2.13339904 | 13.38954194 | 10.2574711 | 10.37481416 | 13.64291503 | 13.6712421 | 12.50279851 | 12.87228671 |
| ENSG00000167034 | NKX3-1 | 2.554818388 | 0.310002526 | -0.421514527 | 0.883235347 | 2.133303861 | 7.579279349 | 2 | 2 | 8.534041678 | 8.52593541 | 5.598599872 | 5.713290392 |
| ENSG00000158006 | PAFAH2 | 1.09851974 | 0.164155144 | 1.034625514 | 0.183338933 | 2.133145254 | 4.815609471 | 6.602424674 | 6.80318718 | 7.288834728 | 7.4450836 | 7.335328635 | 7.277588383 |
| ENSG00000166965 | RCCD1 | 0.129305505 | 0.930223809 | 2.003586846 | 0.09467842 | 2.132892352 | 3.327711234 | 5.838937808 | 7.163363831 | 6.221320894 | 6.27599024 | 8.110758945 | 8.021718375 |
| ENSG00000185798 | WDR53 | 1.014669265 | 0.686884062 | 1.117790682 | 0.649707664 | 2.132459947 | 2 | 6.78762695 | 2 | 6.155987605 | 6.36443784 | 6.419417643 | 6.301968891 |
| ENSG00000094975 | SUCO | 2.178283292 | 4.83E-06 | -0.046604406 | 0.942222441 | 2.131678886 | 7.904282243 | 8.777382925 | 7.97167868 | 10.45897359 | 10.437637 | 8.450751194 | 7.989373961 |
| ENSG00000159658 | EFCAB14 | 0.885505418 | 0.598361649 | 1.243089391 | 0.438092003 | 2.128594809 | 4.815609471 | 9.497318007 | 10.60345421 | 10.4615108 | 10.4753644 | 10.72979303 | 10.91456428 |
| ENSG00000104611 | SH2D4A | 1.100412822 | 0.552383336 | 1.027402343 | 0.575949683 | 2.127815164 | 7.82963488 | 5.595927442 | 2 | 7.593372273 | 7.62295033 | 7.595026956 | 7.476011586 |
| ENSG00000139629 | GALNT6 | 0.825125251 | 0.535581235 | 1.301734213 | 0.299116044 | 2.126859464 | 4.815609471 | 6.951736442 | 5.39244166 | 6.766655021 | 6.8715633 | 8.000548654 | 5.785984581 |
| ENSG00000167491 | GATAD2A | 0.787996563 | 0.211852193 | 1.338231584 | 0.023059327 | 2.126228147 | 8.928409675 | 9.566483142 | 8.143093325 | 9.787157185 | 9.77564777 | 10.24063129 | 10.41335128 |
| ENSG00000164692 | COL1A2 | 1.235715598 | 0.003820004 | 0.889991914 | 0.043497213 | 2.125707512 | 15.2112892 | 16.03971608 | 15.34601607 | 16.7869716 | 16.843947 | 16.52762301 | 16.41003914 |
| ENSG00000105717 | PBX4 | 0.343496344 | 0.886013357 | 1.781386183 | 0.37934489 | 2.124882526 | 2 | 3.686303414 | 2 | 2.947924294 | 3.15372745 | 3.838019203 | 4.09155917 |
| ENSG00000169410 | PTPN9 | 0.996895769 | 0.079429462 | 1.126042535 | 0.041645457 | 2.122938305 | 8.39321512 | 9.04308532 | 7.858115942 | 9.44822005 | 9.56860713 | 9.42949507 | 9.820443605 |
| ENSG00000169718 | DUS1L | 0.661835494 | 0.162215294 | 1.460580477 | 0.000597178 | 2.122415971 | 7.159102577 | 7.847167909 | 7.098165388 | 8.027038952 | 8.12510181 | 8.897500263 | 8.834865421 |
| ENSG00000112144 | ICK | 0.576350975 | 0.577039334 | 1.544706479 | 0.094399166 | 2.121057453 | 7.82963488 | 6.951736442 | 5.156626401 | 7.481152682 | 7.66128317 | 8.431623835 | 8.617426278 |
| ENSG00000159579 | RSPRY1 | 1.255664223 | 0.128350341 | 0.865018298 | 0.303980057 | 2.120682522 | 6.009025997 | 7.847167909 | 8.175061096 | 8.80037045 | 8.88608977 | 8.47458408 | 8.439433377 |
| ENSG00000172794 | RAB37 | -1.303609501 | 0.785196727 | 3.423661538 | 0.407675561 | 2.120052037 | 2 | 2 | 2 | 2.270786764 | 2 | 3.753554863 | 4.033026654 |
| ENSG00000167962 | ZNF598 | 0.569490076 | 0.594985066 | 1.55055171 | 0.104869869 | 2.120041787 | 8.946185348 | 7.145017454 | 6.534109959 | 8.444537031 | 8.51631523 | 9.584486439 | 9.310375569 |
| ENSG00000173218 | VANGL1 | 0.826358247 | 0.659761643 | 1.293526872 | 0.466865178 | 2.119885118 | 8.257427002 | 7.874700259 | 2 | 8.266399315 | 8.36909406 | 8.768480374 | 8.793039018 |
| ENSG00000178922 | HYI | 7.310913843 | 0.000544227 | -5.191145552 | 0.064663947 | 2.119768291 | 2 | 2 | 2 | 7.277360022 | 7.46188679 | 2 | 2 |
| ENSG00000198707 | CEP290 | 1.538513348 | 0.010550428 | 0.581054632 | 0.377325083 | 2.11956798 | 6.563806279 | 6.389896633 | 7.398165132 | 8.413424144 | 8.27524772 | 7.70407074 | 7.026981076 |
| ENSG00000157827 | FMNL2 | 1.601873532 | 5.08E-05 | 0.516317938 | 0.244087001 | 2.11819147 | 8.075571721 | 8.862353853 | 8.825624954 | 10.2136641 | 10.2384727 | 9.194070966 | 9.096149382 |
| ENSG00000118193 | KIF14 | 1.721529087 | 0.383405777 | 0.394856696 | 0.855899236 | 2.116385783 | 2 | 4.442710129 | 3.369317972 | 5.058234798 | 4.91316568 | 2.656294034 | 4.574506939 |
| ENSG00000162396 | PARS2 | 0.622459673 | 0.80586146 | 1.491978265 | 0.519967781 | 2.114437938 | 2 | 6.464272742 | 2 | 5.709096143 | 5.45984938 | 6.583362013 | 6.199243628 |
| ENSG00000105227 | PRX | 0.050800721 | 0.970008162 | 2.063092197 | 0.059117014 | 2.113892918 | 5.095959458 | 3.686303414 | 6.322058544 | 5.44762523 | 5.39133044 | 7.774526492 | 6.674075308 |
| ENSG00000103932 | RPAP1 | 0.693426144 | 0.557986927 | 1.420235183 | 0.191114239 | 2.113661327 | 4.467311984 | 7.701025996 | 6.882775709 | 7.494428494 | 7.59275449 | 8.425380879 | 8.070734444 |
| ENSG00000248167 | TRIM39-RPP | 7.304108672 | 0.000536632 | -5.191145622 | 0.064342105 | 2.11296305 | 2 | 2 | 2 | 7.366699513 | 7.36516374 | 2 | 2 |
| ENSG00000111676 | ATN1 | 0.589393105 | 0.299726211 | 1.523514912 | 0.002547818 | 2.112908017 | 9.710019321 | 10.83211025 | 10.02619908 | 10.87266806 | 10.8474839 | 11.76749432 | 11.81761433 |
| ENSG00000175548 | ALG10B | 0.974855736 | 0.52971614 | 1.136945908 | 0.446952646 | 2.111801643 | 2 | 4.93638933 | 6.322058544 | 6.096288482 | 6.18985553 | 6.577231679 | 5.949196291 |
| ENSG00000183475 | ASB7 | 1.191054147 | 0.056624958 | 0.918483126 | 0.145949245 | 2.109537273 | 6.89194353 | 7.274681896 | 5.772716771 | 7.956842238 | 7.95274764 | 7.759264603 | 7.610940609 |
| ENSG00000163376 | KBTBD8 | 2.129030575 | 0.307499144 | -0.022709082 | 0.992767357 | 2.106321493 | 2 | 5.838937808 | 2 | 6.456560224 | 6.34282908 | 4.652122235 | 4.254128754 |
| ENSG00000126790 | L3HYPDH | 1.780330451 | 0.329205175 | 0.325694062 | 0.874266884 | 2.106024513 | 2 | 7.572420273 | 8.631919287 | 9.371149971 | 9.39049411 | 7.805448645 | 8.065268319 |
| ENSG00000143819 | EPHX1 | 1.362412843 | 0.137222257 | 0.742887558 | 0.437636581 | 2.105300401 | 7.031698319 | 9.479495904 | 8.561105505 | 10.00514223 | 10.051861 | 9.133040681 | 9.645901223 |
| ENSG00000156467 | UQCRB | 1.026367173 | 0.253648116 | 1.077103351 | 0.220279485 | 2.103470523 | 9.344956677 | 7.431218269 | 7.552162999 | 9.432787137 | 9.38785872 | 9.405475868 | 9.514354532 |
| ENSG00000119408 | NEK6 | 1.293232827 | 0.02572938 | 0.809185808 | 0.179357289 | 2.102418635 | 8.696033109 | 8.234834549 | 7.398165132 | 9.531881013 | 9.44136842 | 8.749978316 | 9.224298414 |
| ENSG00000167671 | UBXN6 | 1.06148861 | 0.100196808 | 1.040081423 | 0.102066442 | 2.101570033 | 10.06758145 | 9.377333006 | 8.536706852 | 10.47454911 | 10.5546556 | 10.3964353 | 10.58503594 |
| ENSG00000111261 | MANSC1 | 0.095188805 | 0.947256262 | 2.006367875 | 0.094088158 | 2.101556681 | 7.709880939 | 6.140560021 | 4.058998049 | 6.676346669 | 6.75642009 | 8.181164698 | 8.903861917 |
| ENSG00000261794 | GOLGA8H | 7.289289823 | 0.000584335 | -5.191145507 | 0.064982565 | 2.098144316 | 2 | 2 | 2 | 7.348537858 | 7.35440525 | 2 | 2 |
| ENSG00000184208 | C22orf46 | 1.367525311 | 0.478780358 | 0.730503765 | 0.715215591 | 2.098029076 | 2 | 6.666836837 | 8.561105505 | 8.570539957 | 8.77214691 | 7.702194078 | 8.322876198 |

| | | | | | | | | | | | | | |
|---|---|---|---|---|---|---|---|---|---|---|---|---|---|
| ENSG00000032444 | PNPLA6 | 0.742322965 | 0.499134136 | 1.355206025 | 0.181428238 | 2.09752899 | 9.613202746 | 8.732943944 | 6.629488387 | 9.466861018 | 9.5427783 | 10.02204569 | 10.20317517 |
| ENSG00000196814 | MVB12B | 0.472514452 | 0.786259522 | 1.624572995 | 0.291093097 | 2.097087448 | 6.009025997 | 7.189546432 | 2 | 6.542151749 | 6.70088158 | 7.677572696 | 7.805990373 |
| ENSG00000033170 | FUT8 | 2.111948023 | 0.224881984 | -0.015629172 | 0.993931149 | 2.096318851 | 7.624139768 | 8.451443573 | 2 | 9.633631483 | 9.59321927 | 7.475964489 | 7.544398272 |
| ENSG00000035687 | ADSS | 1.740770814 | 0.074138279 | 0.355001497 | 0.749691042 | 2.095772311 | 8.75772933 | 7.669933291 | 5.930865648 | 9.587916984 | 9.58096568 | 8.068408401 | 8.340330468 |
| ENSG00000148498 | PARD3 | 0.969397607 | 0.570297901 | 1.126108591 | 0.497145211 | 2.095506198 | 5.330594134 | 8.968383114 | 4.523675391 | 8.52598359 | 8.50824913 | 8.728472558 | 8.613686471 |
| ENSG00000144229 | THSD7B | -1.310484826 | 0.725904626 | 3.405260271 | 0.276194973 | 2.094775445 | 2 | 2 | 2 | 2.141737246 | 2.14022021 | 2.831336763 | 4.489993931 |
| ENSG00000119431 | HDHD3 | 0.529363649 | 0.799141442 | 1.565366821 | 0.400363179 | 2.09473047 | 3.327711234 | 6.602424674 | 2 | 5.720439317 | 5.72778967 | 6.908747388 | 6.463051659 |
| ENSG00000170242 | USP47 | 0.47034009 | 0.630948994 | 1.6242741 | 0.057634332 | 2.094614191 | 7.790810999 | 10.17682307 | 9.681375057 | 10.01035013 | 9.96612421 | 11.26733166 | 10.99857641 |
| ENSG00000188610 | FAM72B | 3.975605547 | 0.032131199 | -1.8835802 | 0.369040004 | 2.092025347 | 2 | 3.076665209 | 2 | 5.144975053 | 5.13053105 | 2.139668783 | 2.225841198 |
| ENSG00000198523 | PLN | -0.302794136 | 0.958996611 | 2.394498538 | NA | 2.091704402 | 2 | 2 | 2 | 2.498691189 | 2 | 3.584558147 | 2.746601469 |
| ENSG00000114861 | FOXP1 | 1.247134941 | 0.046544775 | 0.844206088 | 0.189633354 | 2.091341029 | 8.564112515 | 7.701025996 | 9.212024683 | 9.863963753 | 9.83811764 | 9.489210754 | 9.410049744 |
| ENSG00000117525 | F3 | 1.257342908 | 0.283345486 | 0.83378505 | 0.485029668 | 2.091127959 | 7.82963488 | 8.655694736 | 10.81097638 | 10.90440671 | 10.9195043 | 10.75048495 | 10.17045486 |
| ENSG00000144476 | ACKR3 | -1.705607266 | 0.312710697 | 3.796040524 | 0.007955752 | 2.090433258 | 2 | 4.442710129 | 2 | 2.498691189 | 2.68924543 | 6.685680098 | 6.506723027 |
| ENSG00000063587 | ZNF275 | 0.599152545 | 0.684522822 | 1.490321074 | 0.263504222 | 2.089473619 | 4.467311984 | 8.356994601 | 8.865353883 | 8.658676732 | 8.7017439 | 9.510546344 | 9.617549359 |
| ENSG00000163923 | RPL39L | 0.658298662 | 0.523956356 | 1.430538929 | 0.125265786 | 2.088837591 | 4.467311984 | 5.457120953 | 6.882775709 | 6.529309835 | 6.56439512 | 7.196918221 | 7.383872668 |
| ENSG00000102109 | PCSK1N | -1.478187369 | 0.479762551 | 3.566701523 | 0.040078938 | 2.088514154 | 2 | 3.686303414 | 2 | 2.498691189 | 2.2680111 | 5.376552034 | 5.581648407 |
| ENSG00000205356 | TECPR1 | 0.357912054 | 0.759044086 | 1.729908263 | 0.083021331 | 2.087820317 | 5.330594134 | 7.638155693 | 8.35328246 | 7.947219213 | 7.8745862 | 9.284329221 | 9.25263028 |
| ENSG00000176771 | NCKAP5 | 0.708357214 | 0.569536355 | 1.375695775 | 0.229859615 | 2.084052988 | 5.330594134 | 4.710564154 | 2 | 5.110902178 | 5.24368216 | 5.912830251 | 5.646532192 |
| ENSG00000116962 | NID1 | 1.273360454 | 0.052328744 | 0.809913956 | 0.232791888 | 2.08327441 | 9.992581382 | 11.43463583 | 11.43667516 | 12.34427175 | 12.3883446 | 11.73900314 | 12.05104696 |
| ENSG00000100578 | KIAA0586 | 1.598995805 | 0.110752836 | 0.483987996 | 0.66285442 | 2.082983801 | 7.218819781 | 5.72254323 | 4.523675391 | 7.794595599 | 7.74329312 | 6.759726988 | 6.605433158 |
| ENSG00000114349 | GNAT1 | 3.97418412 | 0.311367551 | -1.892859025 | 0.658045217 | 2.081325096 | 2 | 2 | 2 | 4.237680128 | 4.40247105 | 2.183373728 | 2 |
| ENSG00000142784 | WDTC1 | 0.204889618 | 0.864559929 | 1.876227703 | 0.053436739 | 2.081117321 | 6.73718411 | 8.806266215 | 9.670207709 | 8.980991923 | 9.09008229 | 10.59432051 | 10.79911455 |
| ENSG00000188177 | ZC3H6 | 0.285820131 | 0.852386951 | 1.793123873 | 0.16325414 | 2.078494004 | 4.467311984 | 4.93638933 | 7.646393253 | 6.610808945 | 6.70653264 | 8.393172835 | 7.805990373 |
| ENSG00000079950 | STX7 | 0.81817604 | 0.211593503 | 1.260594199 | 0.040591855 | 2.078770239 | 7.53297911 | 7.572420273 | 8.763901704 | 8.896009193 | 8.86223882 | 9.443323792 | 9.181571564 |
| ENSG00000159708 | LRRC36 | -0.723328282 | 0.850026272 | 2.801721455 | NA | 2.078393173 | 2 | 2 | 2 | 2.270786764 | 2.14022021 | 3.767980144 | 3.100202476 |
| ENSG00000153832 | FBXO36 | 0.219314878 | 0.892703066 | 1.858793286 | 0.165125488 | 2.078108164 | 4.815609471 | 6.04685525 | 2 | 5.242616075 | 5.28958453 | 7.07314455 | 6.463051659 |
| ENSG00000091656 | ZFHX4 | 0.761280191 | 0.673950159 | 1.315811245 | 0.43784645 | 2.077091436 | 3.327711234 | 7.669933291 | 9.358421373 | 8.956018563 | 8.90710092 | 9.17593001 | 9.735236833 |
| ENSG00000129925 | TMEM8A | 1.011536175 | 0.000651291 | 1.064534128 | 0.000282431 | 2.076070303 | 8.928409675 | 8.955547846 | 9.180874355 | 9.963981511 | 10.0928914 | 10.21815365 | 9.933014227 |
| ENSG00000132510 | KDM6B | 1.063331231 | 0.04914446 | 1.012512125 | 0.058936535 | 2.075843355 | 8.075571721 | 9.055168103 | 8.110401118 | 9.524619869 | 9.58096568 | 9.641417554 | 9.348752388 |
| ENSG00000147316 | MCPH1 | 0.99471337 | 0.03160337 | 1.081041314 | 0.017106546 | 2.075754684 | 7.031698319 | 6.899080544 | 6.203069917 | 7.805288479 | 7.65545169 | 7.832294085 | 7.80161394 |
| ENSG00000127720 | METTL25 | 1.685673245 | 0.412705524 | 0.388512835 | 0.86358453 | 2.074186081 | 2 | 5.457120953 | 2 | 5.674521485 | 5.55061739 | 4.563936166 | 4.388595494 |
| ENSG00000149577 | SIDT2 | 0.262045899 | 0.865152618 | 1.811856615 | 0.158126095 | 2.073902515 | 5.095959458 | 9.433952963 | 8.434763955 | 8.765203682 | 8.72141482 | 10.32932464 | 10.23800673 |
| ENSG00000215375 | MYL5 | 0.291595036 | 0.869037712 | 1.781711608 | 0.230812916 | 2.073306644 | 6.366709671 | 3.076665209 | 3.369317972 | 5.304221918 | 5.33407135 | 6.645228724 | 6.777976866 |
| ENSG00000132952 | USPL1 | 1.25192505 | 0.438223548 | 0.82094495 | 0.617362697 | 2.07287 | 5.330594134 | 7.979858146 | 3.369317972 | 7.875463856 | 7.90685167 | 7.610117916 | 7.311843322 |
| ENSG00000145349 | CAMK2D | 0.105868856 | 0.841743726 | 1.966529087 | 7.45E-07 | 2.072397944 | 9.27634997 | 8.639735838 | 8.460932916 | 8.95841569 | 8.92053511 | 10.86761565 | 10.71157971 |
| ENSG00000075785 | RAB7A | 1.191584289 | 3.95E-05 | 0.880697515 | 0.003096188 | 2.072281804 | 11.01943573 | 10.70750522 | 10.92346452 | 12.07293465 | 12.0829612 | 11.56367292 | 11.94591045 |
| ENSG00000088280 | ASAP3 | 0.137132493 | 0.942460221 | 1.935037258 | 0.22807891 | 2.072169751 | 4.00722531 | 8.029695761 | 4.058998049 | 6.722207358 | 6.79936111 | 8.503966641 | 8.537437492 |
| ENSG00000165879 | FRAT1 | 1.649309323 | 0.393187917 | 0.420733766 | 0.842253868 | 2.070043089 | 2 | 4.93638933 | 2 | 5.093559143 | 5.07918227 | 3.956110469 | 4.175132639 |
| ENSG00000116237 | ICMT | 0.217117669 | 0.564310859 | 1.852325354 | 8.02E-10 | 2.069443023 | 9.497412313 | 9.159587375 | 9.703463497 | 9.664791023 | 9.69834686 | 11.23953894 | 11.38121091 |
| ENSG00000030419 | IKZF2 | 1.652441248 | 0.144835476 | 0.416910017 | 0.740394508 | 2.069351265 | 7.031698319 | 4.11352228 | 7.285536013 | 8.270263574 | 8.25418043 | 7.264607048 | 6.802276223 |
| ENSG00000125388 | GRK4 | 0.078559941 | 0.963296195 | 1.990331221 | 0.163471043 | 2.068891161 | 5.70932728 | 3.686303414 | 2 | 4.523237872 | 4.63360314 | 6.512311087 | 6.055940621 |
| ENSG00000187775 | DNAH17 | 0.95353929 | 0.634148538 | 1.115076707 | 0.565593677 | 2.068615997 | 2 | 4.710564154 | 2 | 4.299492648 | 4.2865223 | 4.388329244 | 4.456980017 |
| ENSG00000140299 | BNIP2 | 1.072717989 | 0.017649875 | 0.995650734 | 0.027841943 | 2.068368723 | 9.262228249 | 8.671479029 | 8.325066185 | 9.899327447 | 9.84197374 | 9.865050672 | 9.719663687 |
| ENSG00000130045 | NXNL2 | 1.348833847 | 0.468732877 | 0.719504684 | 0.707381508 | 2.068338532 | 3.327711234 | 2 | 3.369317972 | 3.709785322 | 3.82856379 | 3.374372158 | 3.33635788 |
| ENSG00000142556 | ZNF614 | 0.864472727 | 0.208583912 | 1.20383812 | 0.064420602 | 2.068310847 | 5.70932728 | 6.78762695 | 5.59507254 | 6.936770382 | 7.05550676 | 7.464938976 | 7.174740504 |
| ENSG00000126458 | RRAS | 0.125842831 | 0.857281815 | 1.941775241 | 0.000274788 | 2.067618071 | 9.624286623 | 8.317412869 | 9.301632639 | 9.260887574 | 9.33776001 | 11.1676629 | 11.04825611 |
| ENSG00000105483 | CARD8 | 1.162410655 | 0.194213675 | 0.904980874 | 0.314431097 | 2.067391529 | 4.815609471 | 6.728495673 | 7.029880454 | 7.647855834 | 7.53675633 | 7.621085148 | 6.994755759 |
| ENSG00000111364 | DDX55 | 1.297176716 | 0.268205982 | 0.769863359 | 0.524378192 | 2.067040075 | 3.327711234 | 6.602424674 | 6.629488387 | 7.402350992 | 7.31055061 | 6.99251878 | 6.681239787 |
| ENSG00000111834 | RSPH4A | 2.687966558 | 0.392020561 | -0.621070025 | NA | 2.066896533 | 2 | 2 | 2 | 3.227757282 | 3.55551834 | 2.267001569 | 2.172608095 |
| ENSG00000167635 | ZNF146 | 1.018763598 | 0.065760624 | 1.045770613 | 0.05485499 | 2.064534211 | 8.340423424 | 9.443177293 | 8.677268317 | 9.941741253 | 9.88808233 | 10.12876594 | 9.727471273 |
| ENSG00000102181 | CD99L2 | 0.618665536 | 0.065340768 | 1.44445115 | 1.90E-06 | 2.063116686 | 9.031919741 | 9.34816733 | 8.922968322 | 9.720225568 | 9.74577157 | 10.42331231 | 10.67735328 |
| ENSG00000205209 | SCGB2B2 | 2.646475826 | 0.406678307 | -0.58751271 | NA | 2.058963116 | 2 | 2 | 2 | 3.663521627 | 3.01487888 | 2.225793514 | 2.225841198 |
| ENSG00000157593 | SLC35B2 | 1.160213427 | 0.174135326 | 0.898464736 | 0.295507516 | 2.058678163 | 10.44422623 | 8.862353853 | 8.486635645 | 10.67696444 | 10.6920239 | 10.41660409 | 10.4302822 |
| ENSG00000134574 | DDB2 | -0.46952849 | 0.751812007 | 2.527512477 | 0.044161511 | 2.057983987 | 4.815609471 | 6.602424674 | 8.784770914 | 7.097651217 | 7.0991228 | 9.939024914 | 10.16811941 |
| ENSG00000133639 | BTG1 | 2.057505175 | 4.32E-06 | -0.000586523 | 0.999263508 | 2.056918652 | 9.317904228 | 9.883504052 | 9.116480286 | 11.52844449 | 11.5359724 | 9.755754661 | 9.139275853 |
| ENSG00000125834 | STK35 | 0.826380415 | 0.259012717 | 1.228198194 | 0.075285273 | 2.054578609 | 8.493334308 | 7.928237109 | 6.718950348 | 8.631895657 | 8.77079897 | 9.027959735 | 9.17143135 |
| ENSG00000139508 | SLC46A3 | 0.234320866 | 0.892887789 | 1.819801372 | 0.209657689 | 2.054122238 | 6.96351253 | 6.311476754 | 2 | 6.321555392 | 6.36443784 | 8.037484631 | 7.704159082 |
| ENSG00000136068 | FLNB | 0.789900989 | 0.231292431 | 1.260700946 | 0.041745906 | 2.050601935 | 9.730679447 | 10.87704672 | 11.18128933 | 11.49126529 | 11.5214183 | 11.78261435 | 12.14769299 |
| ENSG00000139687 | RB1 | 0.782164367 | 0.610603213 | 1.268152464 | 0.37875047 | 2.050316831 | 4.00722531 | 8.639735838 | 8.65477198 | 8.829471715 | 8.90095288 | 9.435155466 | 9.255433104 |
| ENSG00000078124 | ACER3 | 0.937784627 | 0.084457614 | 1.109467848 | 0.034537867 | 2.047252475 | 6.89143753 | 7.790474954 | 6.80318718 | 8.213192703 | 8.13351654 | 8.362013331 | 8.324402372 |
| ENSG00000167967 | E4F1 | 0.82642216 | 0.716973839 | 1.220110514 | 0.575342928 | 2.046532674 | 4.00722531 | 8.191702463 | 2 | 7.487805859 | 7.53992502 | 8.032265607 | 7.754841567 |
| ENSG00000144677 | CTDSPL | 0.221245882 | 0.833848549 | 1.825136933 | 0.033221064 | 2.046382815 | 7.33131396 | 8.848535039 | 6.629488387 | 8.074220386 | 8.19711812 | 9.752357958 | 9.701279971 |
| ENSG00000165914 | TTC7B | 0.597345931 | 0.738885038 | 1.448602268 | 0.376394793 | 2.045948199 | 7.667647223 | 7.232742069 | 2 | 7.497728447 | 7.46522407 | 8.337438499 | 8.299011699 |
| ENSG00000182141 | ZNF708 | 1.375307317 | 0.061315658 | 0.670203979 | 0.387886116 | 2.045511296 | 4.467311984 | 5.457120953 | 5.59507254 | 6.592405944 | 6.48770488 | 6.123812431 | 5.571402115 |
| ENSG00000068024 | HDAC4 | 0.92948396 | 0.351836711 | 1.115045024 | 0.246860689 | 2.044528983 | 8.107517749 | 6.389896633 | 5.772716771 | 8.065351606 | 7.96222667 | 8.189896068 | 8.205208936 |
| ENSG00000204962 | PCDHA8 | 2.394791035 | 0.502627905 | -0.351846149 | NA | 2.042944886 | 2 | 2 | 2 | 3.663521627 | 2.59489879 | 2.34604626 | 2.172608095 |
| ENSG00000170606 | HSPA4 | 0.521783279 | 0.243498812 | 1.518407118 | 0.000112402 | 2.040190397 | 10.32020829 | 9.672251278 | 10.37481416 | 10.67405041 | 10.6723063 | 11.48009892 | 11.8343144 |
| ENSG00000117425 | PTCH2 | 1.007616749 | 0.61882206 | 1.032529331 | 0.602505282 | 2.04014608 | 2 | 4.442710129 | 2 | 3.840289928 | 4.31639476 | 4.155805465 | 4.077147165 |
| ENSG00000198040 | ZNF84 | 0.976125375 | 0.579599436 | 1.063817107 | 0.536100907 | 2.039942482 | 6.366709671 | 8.848535039 | 3.369317972 | 8.50645854 | 8.48542112 | 8.888863837 | 8.193564875 |
| ENSG00000118260 | CREB1 | 1.005470001 | 0.370206288 | 1.032536461 | 0.34763435 | 2.038006463 | 8.928409675 | 7.099070191 | 6.073378139 | 8.857997553 | 8.82115545 | 9.009866973 | 8.707318295 |
| ENSG00000178809 | TRIM73 | 3.146338608 | 0.298074813 | -1.109995635 | 0.738292369 | 2.036342974 | 2 | 2 | 2 | 3.349643765 | 3.98554648 | 2.094598369 | 2.225841198 |
| ENSG00000105287 | PRKD2 | 1.64553101 | 0.180388625 | 0.388006168 | 0.778107221 | 2.033537178 | 5.330594134 | 7.874700259 | 9.180874355 | 9.772927884 | 9.79765578 | 8.563064723 | 8.5102048 |
| ENSG00000158623 | COPG2 | 0.529217728 | 0.788534399 | 1.502396899 | 0.397192742 | 2.031614627 | 4.815609471 | 7.393647515 | 2 | 6.522845756 | 6.63721363 | 7.438128798 | 7.605929526 |
| ENSG00000113273 | ARSB | -0.04275236 | 0.96053419 | 2.07355431 | 0.002204294 | 2.03080195 | 8.981092835 | 8.213429689 | 9.926432745 | 9.161399541 | 9.16382362 | 11.01246883 | 11.49043785 |
| ENSG00000148344 | PTGES | 2.472976157 | 0.016358013 | -0.443783557 | 0.713381003 | 2.0291926 | 9.317904228 | 6.728495673 | 8.99637432 | 11.11566977 | 11.2164127 | 7.241533944 | 8.859812348 |
| ENSG00000100625 | SIX4 | 0.892093948 | 0.201147319 | 1.136893421 | 0.089227181 | 2.028987369 | 6.366709671 | 7.002537973 | 7.896971941 | 8.098332103 | 8.08878133 | 8.10153367 | 8.536120927 |
| ENSG00000187860 | CCDC157 | 1.541048918 | 0.465537342 | 0.485539685 | 0.83120048 | 2.026588603 | 2 | 5.457120953 | 2 | 5.406080789 | 5.55061739 | 4.301748724 | 4.77533149 |
| ENSG00000183044 | ABAT | -0.133894872 | 0.959837734 | 2.160401778 | 0.351928367 | 2.026506906 | 2 | 6.464272742 | 2 | 5.210797711 | 4.53541753 | 6.040114727 | 7.632045641 |
| ENSG00000116741 | RGS2 | 3.094439944 | 0.115966618 | -1.067995374 | 0.620242104 | 2.02644457 | 2 | 6.04685525 | 2 | 7.507583254 | 7.58049694 | 3.993419637 | 3.640730416 |
| ENSG00000129534 | MIS18BP1 | 1.424095042 | 0.406678307 | 0.602313496 | 0.738341738 | 2.026408538 | 7.53297911 | 5.946638857 | 2 | 7.805288479 | 7.72392024 | 7.277267764 | 6.562106074 |
| ENSG00000141232 | TOB1 | 0.692896367 | 0.278800879 | 1.33332274 | 0.02331086 | 2.026219107 | 8.696033109 | 7.35507212 | 7.896971941 | 8.782894215 | 8.75042728 | 9.262527297 | 9.529981565 |
| ENSG00000125414 | MYH2 | -0.727730262 | 0.847740162 | 2.752531592 | NA | 2.02480133 | 2 | 2 | 2 | 2.141737246 | 2.2680111 | 3.600766115 | 3.261853426 |
| ENSG00000156140 | ADAMTS3 | -0.542821492 | 0.79462063 | 2.567120897 | 0.143469849 | 2.024299404 | 2 | 4.93638933 | 2 | 3.615725078 | 3.15372745 | 5.856572853 | 5.992078266 |
| ENSG00000168710 | AHCYL1 | 0.650427441 | 0.65319082 | 1.373441676 | 0.302366413 | 2.023869118 | 6.138360429 | 10.35843478 | 10.32581712 | 10.4457931 | 10.4431479 | 11.20533263 | 11.12524595 |
| ENSG00000102531 | FNDC3A | 1.682117034 | 0.036619822 | 0.339069233 | 0.715181868 | 2.021186267 | 8.75772933 | 10.51971956 | 8.784770914 | 11.29209249 | 11.2860026 | 9.872996531 | 10.02223248 |
| ENSG00000171469 | ZNF561 | 0.829315169 | 0.710155984 | 1.188861007 | 0.576686766 | 2.018176176 | 5.532353208 | 8.929529568 | 2 | 8.29513208 | 8.3183278 | 8.902003555 | 8.374617996 |
| ENSG00000124212 | PTGIS | -2.967908405 | 1.71E-05 | 4.982329361 | 3.42E-14 | 2.014420956 | 8.998234871 | 7.538394143 | 9.132850258 | 5.639097847 | 6.00043082 | 13.22463176 | 14.02033476 |
| ENSG00000140320 | BAHD1 | -0.024035888 | 0.992479093 | 2.038109097 | 0.326076859 | 2.014073209 | 8.716893051 | 4.710564154 | 2 | 7.161462517 | 7.24992376 | 9.327881973 | 9.143172568 |
| ENSG00000056558 | TRAF1 | 0.714654931 | 0.737269626 | 1.298986754 | 0.51714034 | 2.013641686 | 9.174465797 | 6.899080544 | 2 | 8.622857004 | 8.52113334 | 9.050629989 | 9.247008243 |
| ENSG00000214402 | LCNL1 | -3.753100702 | 0.367367297 | 5.7656261 | 0.097288553 | 2.012525398 | 2 | 2 | 2 | 2 | 2 | 5.436478516 | 6.238627697 |
| ENSG00000167106 | FAM102A | 0.993174344 | 0.153699077 | 1.019262224 | 0.134226978 | 2.012436569 | 7.096806373 | 8.169643011 | 8.65477198 | 9.073461088 | 9.10727282 | 9.379876477 | 8.793590064 |
| ENSG00000120925 | RNF170 | 1.441182039 | 0.282237791 | 0.570850055 | 0.690155642 | 2.012032094 | 5.70932728 | 8.101365653 | 4.523675391 | 8.302698479 | 8.2790453 | 7.611118389 | 7.236027917 |
| ENSG00000134184 | GSTM1 | 1.879597568 | 0.044020715 | 0.130781314 | 0.909359225 | 2.010378882 | 3.327711234 | 5.303523747 | 5.930865648 | 6.917187326 | 7.00595521 | 5.079718116 | 5.524371359 |
| ENSG00000130762 | ARHGEF16 | -3.651647412 | 0.050971196 | 5.661862532 | 1.57E-05 | 2.01021512 | 2 | 4.11352228 | 2 | 2.141737246 | 2.14022021 | 8.342868948 | 7.685278529 |
| ENSG00000113658 | SMAD5 | 0.823612406 | 0.183172174 | 1.184655633 | 0.043076236 | 2.008268039 | 7.975256505 | 9.13702746 | 8.143093325 | 9.37922384 | 9.2990102 | 9.880691463 | 9.49042201 |
| ENSG00000203965 | EFCAB7 | 1.412224052 | 0.238722314 | 0.594761866 | 0.641715954 | 2.006985918 | 2 | 4.93638933 | 4.523675391 | 5.48800675 | 5.55061739 | 4.977914261 | 4.584731282 |
| ENSG00000125398 | SOX9 | 2.492994734 | 0.084338909 | -0.488039148 | 0.767692243 | 2.004955585 | 2 | 6.844429755 | 6.882775709 | 8.78964092 | 8.68752719 | 5.740677235 | 5.949196291 |
| ENSG00000145911 | N4BP3 | 0.932948161 | 0.596379651 | 1.070943515 | 0.527954208 | 2.003891675 | 4.00722531 | 3.076665209 | 2 | 3.997744386 | 3.69849104 | 4.240999983 | 3.600986722 |
| ENSG00000103569 | AQP9 | -2.306085432 | 0.608215887 | 4.308584281 | 0.248121319 | 2.00249885 | 2 | 2 | 2 | 2.141737246 | 2 | 5.362362054 | 2.851942824 |
| ENSG00000181274 | FRAT2 | 1.068447869 | 0.474064658 | 0.93276735 | 0.527954208 | 2.001215219 | 3.327711234 | 6.728495673 | 4.058998049 | 6.496695859 | 6.507263 | 6.439833081 | 6.301968891 |
| ENSG00000120278 | PLEKHG1 | 0.059358047 | 0.977133743 | 1.941358321 | 0.263352567 | 2.000716367 | 4.00722531 | 3.076665209 | 4.058998049 | 4.470469939 | 2.49394783 | 5.801542492 | 4.654341953 |
| ENSG00000157954 | WIPI2 | 1.138662932 | 0.005531762 | 0.861177185 | 0.040634846 | 1.999840117 | 8.674867122 | 9.357954881 | 9.491321578 | 10.29699874 | 10.4028177 | 10.0143146 | 10.13197271 |
| ENSG00000124145 | SDC4 | 1.389759114 | 0.045746811 | 0.609097571 | 0.415216121 | 1.998856685 | 10.46286899 | 9.994333773 | 10.69449487 | 11.77954155 | 11.8176385 | 10.02373723 | 11.60270478 |
| ENSG00000167604 | NFKBID | 1.059044179 | 0.41446225 | 0.939768417 | 0.46274242 | 1.998812596 | 4.00722531 | 4.710564154 | 2 | 4.804776789 | 4.93258395 | 4.588526421 | 4.927687581 |
| ENSG00000013297 | CLDN11 | 0.36503847 | 0.747163598 | 1.633195902 | 0.094043218 | 1.998234372 | 9.233563228 | 7.232742069 | 6.882775709 | 8.509731121 | 8.5668971 | 9.983548733 | 9.584176963 |
| ENSG00000171522 | PTGER4 | -0.17050714 | 0.8177838 | 2.168222171 | 0.000117699 | 1.997715031 | 6.257048221 | 5.303523747 | 6.322058544 | 5.731694 | 5.94414054 | 7.909957323 | 8.26365177 |
| ENSG00000123066 | MED13L | 0.975460253 | 0.325172387 | 1.020877253 | 0.292072257 | 1.996337506 | 7.218819781 | 9.835272298 | 9.491321578 | 10.1698607 | 10.2039308 | 10.34849843 | 10.10580955 |

| | | | | | | | | | | | | | |
|---|---|---|---|---|---|---|---|---|---|---|---|---|---|
| ENSG00000124444 | ZNF576 | 1.009877852 | 0.432702068 | 0.98626107 | 0.434435201 | 1.996138922 | 5.866944937 | 7.051611315 | 3.369317972 | 7.169761359 | 6.93551934 | 6.975527754 | 7.091143645 |
| ENSG00000198900 | TOP1 | 1.892943283 | 5.06E-05 | 0.10316678 | 0.864675088 | 1.996110063 | 8.042902253 | 9.125613817 | 8.608698767 | 10.51744927 | 10.5783495 | 8.854214238 | 8.675981178 |
| ENSG00000180573 | HIST1H2AC | 2.423975732 | 0.069296116 | -0.427940406 | 0.78199021 | 1.996035325 | 8.228665467 | 6.04685525 | 4.058998049 | 9.387252775 | 9.37105521 | 6.305369896 | 6.811011855 |
| ENSG00000100084 | HIRA | 2.159059129 | 0.217270166 | -0.163562477 | 0.936599654 | 1.995496652 | 2 | 6.535001748 | 8.236946975 | 9.243301698 | 9.10406516 | 6.831875681 | 6.949978473 |
| ENSG00000025800 | KPNA6 | 0.592841996 | 0.529125418 | 1.401132362 | 0.098395659 | 1.993974358 | 8.517314938 | 10.74155032 | 9.478569125 | 10.42557724 | 10.4844385 | 11.16101747 | 11.35606301 |
| ENSG00000151692 | RNF144A | 2.57479024 | 0.07368721 | -0.580918544 | 0.72430187 | 1.993871695 | 3.327711234 | 3.686303414 | 3.369317972 | 5.48800675 | 5.6121243 | 3.5515855 | 2.551896972 |
| ENSG00000140525 | FANCI | 1.357596551 | 0.495803512 | 0.635527999 | 0.760366765 | 1.99312455 | 3.327711234 | 6.844429755 | 2 | 6.777556219 | 6.67228943 | 6.069527745 | 6.007361012 |
| ENSG00000273542 | HIST1H4K | 4.113472272 | 0.110962 | -2.120377681 | 0.467466101 | 1.993094591 | 2 | 2 | 2 | 4.070463721 | 4.72553021 | 2.048074405 | 2.117335275 |
| ENSG00000186063 | AIDA | 2.402091618 | 2.47E-06 | -0.409142676 | 0.50032005 | 1.992948942 | 7.624139768 | 8.834582582 | 8.763901704 | 10.91771013 | 10.8737771 | 8.164223608 | 8.038552764 |
| ENSG00000179938 | GOLGA8J | 7.183212894 | 0.001052791 | -5.191144637 | 0.070759925 | 1.992068258 | 2 | 2 | 2 | 7.242377933 | 7.2537885 | 2 | 2 |
| ENSG00000106483 | SFRP4 | -0.317065049 | 0.908151255 | 2.308499981 | 0.3091698 | 1.991434932 | 4.00722531 | 2 | 2 | 3.09461492 | 2.49394783 | 5.056402493 | 3.67940853 |
| ENSG00000087995 | METTL2A | 0.955415107 | 0.085357884 | 1.035720315 | 0.054916044 | 1.991135422 | 6.653098118 | 5.946638857 | 6.958202216 | 7.620871236 | 7.32531744 | 7.514972902 | 7.602159758 |
| ENSG00000213903 | LTB4R | 1.60284347 | 0.244378928 | 0.386931947 | 0.800472699 | 1.989775417 | 4.467311984 | 6.78762695 | 3.369317972 | 7.071314871 | 7.18258783 | 6.265258498 | 5.586744392 |
| ENSG00000142961 | MOB3C | 1.16733765 | 0.531143362 | 0.822401268 | 0.664743846 | 1.989738918 | 2 | 6.728495673 | 8.266922072 | 8.168339746 | 8.35477078 | 7.982106617 | 7.864857644 |
| ENSG00000159461 | AMFR | 1.161429693 | 0.450144124 | 0.826479648 | 0.594338258 | 1.987909341 | 5.095959458 | 9.695596167 | 10.23774519 | 10.54602092 | 10.6286993 | 10.21239669 | 10.2943831 |
| ENSG00000167291 | TBC1D16 | 0.554275226 | 0.386935633 | 1.43284042 | 0.011978839 | 1.987115645 | 8.313285031 | 8.88960062 | 9.670202709 | 9.591782342 | 9.6360458 | 10.56931498 | 10.40454019 |
| ENSG00000182087 | TMEM259 | 1.113588749 | 0.167448969 | 0.872890905 | 0.282010983 | 1.986479654 | 8.468948322 | 9.77810053 | 10.53119344 | 10.90782161 | 10.9516838 | 10.80647677 | 10.56274175 |
| ENSG00000144597 | EAF1 | 1.678974475 | 7.14E-07 | 0.304608198 | 0.446952646 | 1.983582673 | 7.790810999 | 8.376385542 | 7.97167868 | 9.746817367 | 9.73820484 | 8.424811996 | 8.331250243 |
| ENSG00000033627 | ATP6V0A1 | 0.705848517 | 0.361961292 | 1.277567849 | 0.074535792 | 1.983416365 | 7.218819781 | 9.055168103 | 8.561105505 | 9.118100239 | 9.21527115 | 9.641907141 | 9.824224109 |
| ENSG00000126070 | AGO3 | 0.263460219 | 0.895265695 | 1.718757087 | 0.311175193 | 1.982217305 | 3.327711234 | 8.077869079 | 4.523675391 | 6.887305128 | 6.96410766 | 8.620830655 | 8.018893446 |
| ENSG00000159884 | CCDC107 | 1.903588051 | 0.000157844 | 0.078213856 | 0.906201712 | 1.981801906 | 6.468621203 | 5.595927442 | 5.930865648 | 7.865280446 | 7.85690889 | 6.004008792 | 6.185873005 |
| ENSG00000155849 | ELMO1 | -0.949250998 | 0.705707601 | 2.929538429 | 0.158640059 | 1.980287431 | 2 | 2 | 3.369317972 | 2.3892346 | 2.2680111 | 4.166733998 | 4.341135913 |
| ENSG00000118454 | ANKRD13C | 0.962395522 | 0.410440873 | 1.016959244 | 0.370886214 | 1.979354765 | 5.095959458 | 8.414402626 | 7.398165132 | 8.425603456 | 8.48870446 | 8.525352075 | 8.497408326 |
| ENSG00000236334 | PPIAL4G | 8.131637665 | 2.47E-08 | -6.152947346 | 0.006279821 | 1.978690319 | 2 | 2 | 3.369317972 | 9.118100239 | 9.14209426 | 2 | 2 |
| ENSG00000143106 | PSMA5 | 1.715082549 | 0.000164731 | 0.263419042 | 0.630693854 | 1.978501591 | 8.107517749 | 7.60566235 | 8.631919287 | 9.837511512 | 9.89490419 | 8.362607519 | 8.485861839 |
| ENSG00000160563 | MED27 | 0.966173185 | 0.452168923 | 1.011813425 | 0.418932978 | 1.97798661 | 3.327711234 | 6.899080544 | 6.882775709 | 7.318995108 | 7.28811304 | 7.192914486 | 7.488317792 |
| ENSG00000164134 | NAA15 | 1.087526297 | 0.384241406 | 0.888888282 | 0.476444675 | 1.976414579 | 8.285626337 | 8.88960062 | 5.156626401 | 9.248208205 | 9.10620439 | 9.065304654 | 8.893103533 |
| ENSG00000196550 | FAM72A | 4.228078017 | 0.116554196 | -2.251841313 | 0.44325556 | 1.976236704 | 2 | 3.076665209 | 2 | 5.461211713 | 5.2591458 | 2 | 2.277179783 |
| ENSG00000069943 | PIGB | 1.801684997 | 0.175359664 | 0.174251143 | 0.91151969 | 1.975936139 | 7.435667355 | 5.72254323 | 3.369317972 | 8.054188409 | 8.0358969 | 6.247352872 | 6.649933445 |
| ENSG00000197724 | PHF2 | 0.695739744 | 0.117053593 | 1.278262607 | 0.00175921 | 1.974002351 | 8.609439662 | 8.053983481 | 7.777116083 | 8.875861021 | 8.87609528 | 9.513224692 | 9.393060099 |
| ENSG00000059122 | FLYWCH1 | 0.621251494 | 0.408406753 | 1.352075789 | 0.047025411 | 1.973327283 | 9.435834703 | 8.101365653 | 8.110401118 | 9.331938606 | 9.28585881 | 10.27575641 | 9.7489399 |
| ENSG00000197217 | ENTPD4 | 0.590165445 | 0.181562967 | 1.382535764 | 0.000480809 | 1.972701209 | 9.590776211 | 9.377333006 | 8.742726175 | 9.889939704 | 9.84709924 | 10.66659669 | 10.65022737 |
| ENSG00000108176 | DNAJC12 | 1.527874574 | 0.42279064 | 0.444534421 | 0.830158879 | 1.972408995 | 2 | 4.442710129 | 2 | 4.443344104 | 4.58534546 | 3.584558147 | 3.789585079 |
| ENSG00000108091 | CCDC6 | 0.805953489 | 0.015213259 | 1.165326286 | 0.000221542 | 1.971279775 | 8.257427002 | 8.147241016 | 8.266922072 | 9.021676912 | 9.01805981 | 9.538686959 | 9.194147421 |
| ENSG00000183726 | TMEM50A | 1.543837906 | 0.020447291 | 0.424907773 | 0.571127612 | 1.968745679 | 10.72212986 | 9.664384941 | 9.196533591 | 11.5430919 | 11.5505775 | 10.17296681 | 10.64840071 |
| ENSG00000198642 | KLHL9 | 0.825875243 | 0.038004723 | 1.142049101 | 0.00262317 | 1.967924344 | 8.107517749 | 8.848535039 | 8.699419242 | 9.433648866 | 9.38609912 | 9.752584654 | 9.695984164 |
| ENSG00000119285 | HEATR1 | 1.220800339 | 0.486454906 | 0.74690313 | 0.678808079 | 1.967703469 | 6.366709671 | 8.993716073 | 3.369317972 | 8.883449599 | 8.83798691 | 8.408791501 | 8.374617996 |
| ENSG00000167325 | RRM1 | 0.909580293 | 0.629335817 | 1.057573967 | 0.564110709 | 1.967154261 | 8.340423424 | 8.053983481 | 2 | 8.532433657 | 8.52113334 | 8.597294425 | 8.745413684 |
| ENSG00000044446 | PHKA2 | 0.46178078 | 0.801844483 | 1.505103633 | 0.356564979 | 1.966884413 | 6.009025997 | 7.467835385 | 2 | 6.772115916 | 6.84618961 | 7.939729349 | 7.702986259 |
| ENSG00000276276 | | 7.157683347 | 0.00128827 | -5.191144223 | 0.073251024 | 1.966539123 | 2 | 2 | 2 | 7.337530098 | 7.0991228 | 2 | 2 |
| ENSG00000228284 | | -3.753094472 | 0.381747878 | 5.718950646 | 0.112356605 | 1.965856174 | 2 | 2 | 2 | 2 | 2 | 6.341987719 | 5.094391891 |
| ENSG00000163527 | STT3B | 1.58782863 | 0.226367389 | 0.378020398 | 0.796658198 | 1.965849028 | 6.563806279 | 10.49332014 | 10.71094342 | 11.66752959 | 11.6300386 | 10.64709726 | 10.20151514 |
| ENSG00000181852 | RNF41 | 0.117366349 | 0.840490657 | 1.847778853 | 2.89E-05 | 1.965145203 | 8.816894989 | 8.077869079 | 9.083172337 | 8.793673851 | 8.8597051 | 10.51903133 | 10.57753611 |
| ENSG00000111877 | MCM9 | 0.867795267 | 0.519874318 | 1.097029585 | 0.393412229 | 1.964824852 | 5.70932728 | 5.838937808 | 2 | 6.104969671 | 6.01871653 | 6.146071859 | 6.400759785 |
| ENSG00000131462 | TUBG1 | 0.82391365 | 0.542787593 | 1.140854813 | 0.376931525 | 1.964768463 | 8.340423424 | 8.395519306 | 4.523675391 | 8.639384849 | 8.65866647 | 8.728933481 | 9.164631393 |
| ENSG00000153012 | LGI2 | -1.303522209 | 0.792847991 | 3.268045143 | 0.44874768 | 1.964522934 | 2 | 2 | 2 | 2.270786764 | 2 | 4.088436536 | 3.407202941 |
| ENSG00000008952 | SEC62 | 1.263753667 | 0.000289255 | 0.699954298 | 0.058364332 | 1.963707965 | 9.957954117 | 9.599860728 | 10.27509734 | 11.22224576 | 11.2324146 | 10.73312834 | 10.59281363 |
| ENSG00000107854 | TNKS2 | 1.219327687 | 0.138060742 | 0.743923919 | 0.379855077 | 1.963251606 | 7.485143553 | 9.540933215 | 9.099922433 | 10.19294689 | 10.1158997 | 9.920984758 | 9.394150543 |
| ENSG00000061936 | SFSWAP | 0.810995383 | 0.588403528 | 1.150583449 | 0.420944803 | 1.961578832 | 4.00722531 | 8.607278467 | 7.225742779 | 8.341783829 | 8.33301564 | 8.748160468 | 8.594208165 |
| ENSG00000178401 | DNAJC22 | 1.256708848 | 0.241959192 | 0.704386343 | 0.527535357 | 1.961095191 | 4.815609471 | 3.686303414 | 6.203069917 | 6.328984825 | 6.43421531 | 5.787450547 | 5.933281326 |
| ENSG00000138709 | LARP1B | 0.301656791 | 0.804733172 | 1.6589647 | 0.106711001 | 1.960621491 | 7.53297911 | 6.228547981 | 4.523675391 | 6.931899482 | 6.76185814 | 8.228872122 | 8.120777053 |
| ENSG00000172315 | TP53RK | 1.159928615 | 0.200892735 | 0.799585857 | 0.38602169 | 1.959514472 | 6.468621203 | 8.124485672 | 6.073378139 | 8.319579168 | 8.34936267 | 7.952415862 | 8.005636809 |
| ENSG00000198959 | TGM2 | 2.20590557 | 0.075674649 | -0.246848989 | 0.866860327 | 1.95905658 | 11.46391134 | 9.328390744 | 8.007629717 | 12.46477506 | 12.4950495 | 10.54975054 | 9.215689211 |
| ENSG00000175782 | SLC35E3 | 0.916315489 | 0.53071634 | 1.042734412 | 0.461118553 | 1.959049901 | 4.00722531 | 6.464272742 | 8.076950877 | 7.852449262 | 7.85436579 | 7.301021256 | 8.436613692 |
| ENSG00000197056 | ZMYM1 | 1.099523608 | 0.355390531 | 0.858813637 | 0.472358061 | 1.958337245 | 3.327711234 | 6.311476754 | 6.203069917 | 6.892328675 | 6.73998187 | 6.975527754 | 6.052261256 |
| ENSG00000101888 | NXT2 | 1.328329736 | 0.499226274 | 0.626616888 | 0.760440198 | 1.954946624 | 3.327711234 | 6.728495673 | 2 | 6.640968371 | 6.53928196 | 5.966975997 | 5.884455123 |
| ENSG00000165282 | PIGO | 0.805863993 | 0.669214488 | 1.14790757 | 0.523553954 | 1.953771563 | 2 | 7.099070191 | 8.206335842 | 7.930222906 | 8.01327646 | 8.275683409 | 8.342591622 |
| ENSG00000103994 | ZNF106 | 0.394527257 | 0.439157025 | 1.55790027 | 0.000319142 | 1.952427527 | 9.90440318 | 9.664384941 | 9.287079172 | 10.04007758 | 10.0277996 | 10.7983745 | 11.50464287 |
| ENSG00000116688 | MFN2 | 0.688286267 | 0.56825652 | 1.263252969 | 0.260527422 | 1.951539236 | 7.159102577 | 9.277729677 | 10.51254821 | 10.19090967 | 10.2462532 | 10.7395472 | 10.84303208 |
| ENSG00000086062 | B4GALT1 | 1.289695298 | 0.054670527 | 0.661768212 | 0.350940523 | 1.95146351 | 9.556468883 | 10.76380876 | 11.11279376 | 11.88411046 | 11.9216624 | 10.98658636 | 11.5167917 |
| ENSG00000139233 | LLPH | 0.991728033 | 0.359332006 | 0.959003985 | 0.367118564 | 1.950732018 | 7.904282243 | 7.051611315 | 4.874587708 | 7.908072101 | 8.15434112 | 7.940525532 | 8.065268319 |
| ENSG00000182952 | HMGN4 | 0.957413978 | 0.000598253 | 0.992084101 | 0.000320664 | 1.949498079 | 8.75772933 | 8.487557258 | 8.536706852 | 9.569217728 | 9.52209753 | 9.508400079 | 9.648949641 |
| ENSG00000155918 | RAET1L | 7.139715022 | 0.001475449 | -5.191143944 | 0.075016235 | 1.948571078 | 2 | 2 | 2 | 7.044488801 | 7.35080119 | 2 | 2 |
| ENSG00000174827 | PDZK1 | 2.068138782 | 0.300919213 | -0.120175854 | 0.958888695 | 1.947962928 | 2 | 2 | 4.058998049 | 4.268917409 | 4.40247105 | 2.885263063 | 2.950113106 |
| ENSG00000171365 | CLCN5 | 1.36186197 | 0.39960716 | 0.58568699 | 0.731331093 | 1.94754896 | 4.00722531 | 7.189546432 | 3.369317972 | 7.066878356 | 7.26532099 | 6.717615541 | 6.048572483 |
| ENSG00000117620 | SLC35A3 | 0.997748243 | 0.065697131 | 0.949021507 | 0.077151749 | 1.94676975 | 6.563806279 | 7.145017454 | 7.734844356 | 8.279879181 | 8.11451391 | 8.23926289 | 8.057947787 |
| ENSG00000035681 | NSMAF | 1.484870452 | 0.022233888 | 0.461620545 | 0.523553954 | 1.946490997 | 8.16936288 | 7.928237109 | 6.534109959 | 9.18513223 | 9.16073939 | 8.103667822 | 8.214292363 |
| ENSG00000084733 | RAB10 | 0.818999057 | 0.156553121 | 1.126456341 | 0.040371077 | 1.945455399 | 8.777722034 | 9.506147172 | 10.06083273 | 10.37936391 | 10.3310092 | 10.73289857 | 10.58742133 |
| ENSG00000243667 | WDR92 | 1.203640099 | 0.032193991 | 0.741495279 | 0.202759883 | 1.945135378 | 5.330594134 | 6.140560021 | 5.59507254 | 6.917187326 | 6.88158913 | 6.497238296 | 6.412287469 |
| ENSG00000101782 | RIOK3 | 1.108915401 | 0.079057979 | 0.836136236 | 0.19081938 | 1.945051637 | 8.009475818 | 9.45234302 | 8.561105505 | 9.895579677 | 9.91822548 | 9.638721848 | 9.632103066 |
| ENSG00000188626 | GOLGA8M | 7.135864817 | 0.001421933 | -5.191144077 | 0.074313018 | 1.94472074 | 2 | 2 | 2 | 7.093295078 | 7.30311013 | 2 | 2 |
| ENSG00000189079 | ARID2 | 0.856583248 | 0.151636445 | 1.087275979 | 0.057300582 | 1.943859227 | 7.975256505 | 7.954278496 | 6.80318718 | 8.589222193 | 8.45217004 | 8.926322547 | 8.550537483 |
| ENSG00000166200 | COPS2 | 1.41352353 | 0.18647032 | 0.529580402 | 0.646525284 | 1.943103933 | 6.468621203 | 9.599860728 | 8.699419242 | 10.16623608 | 10.1248958 | 9.35261874 | 9.173973093 |
| ENSG00000108828 | VAT1 | 0.50873371 | 0.183786544 | 1.433190187 | 2.19E-05 | 1.941923896 | 10.92399098 | 11.23400805 | 10.66102425 | 11.45373986 | 11.4826289 | 12.20981764 | 12.5533032 |
| ENSG00000012822 | CALCOCO1 | 1.070995946 | 0.017885637 | 0.869858499 | 0.057854079 | 1.940854445 | 9.204317094 | 9.357954881 | 10.03493575 | 10.59575793 | 10.6909553 | 10.59077471 | 10.27951104 |
| ENSG00000164403 | SHROOM1 | 0.747042284 | 0.533299715 | 1.192917244 | 0.290333557 | 1.939959528 | 7.485143553 | 5.72254323 | 4.523675391 | 7.062428156 | 7.18258783 | 7.746573618 | 7.342293759 |
| ENSG00000198046 | ZNF667 | 0.634669213 | 0.704230942 | 1.303510691 | 0.397528734 | 1.938179904 | 6.138360429 | 7.051611315 | 2 | 6.716553866 | 6.73998187 | 7.53303322 | 7.20156326 |
| ENSG00000102081 | FMR1 | 1.464137511 | 0.009798993 | 0.471612016 | 0.455371809 | 1.935749526 | 7.031698319 | 8.101365653 | 8.408111533 | 9.433648866 | 9.37903917 | 8.352472895 | 8.489267468 |
| ENSG00000198252 | STYX | 0.616957886 | 0.362270839 | 1.318679664 | 0.032017989 | 1.935637551 | 6.257048221 | 7.503546079 | 6.43197674 | 7.510853288 | 7.41778467 | 8.235375117 | 8.068003971 |
| ENSG00000157741 | UBN2 | 1.592560436 | 0.019814274 | 0.342926909 | 0.664743846 | 1.935487345 | 7.82963488 | 7.73146271 | 6.629488387 | 9.098672425 | 9.03162973 | 8.256629373 | 7.22299542 |
| ENSG00000135547 | HEY2 | -4.140665176 | 0.065574814 | 6.075882984 | 4.15E-06 | 1.935217808 | 3.327711234 | 3.076665209 | 2 | 2.141737246 | 2 | 7.908330388 | 8.002780183 |
| ENSG00000162378 | ZYG11B | 0.289297311 | 0.56778367 | 1.644152988 | 8.27E-05 | 1.933450299 | 8.540903468 | 8.191702463 | 9.049077186 | 8.899755849 | 8.93384535 | 10.39469335 | 10.12125705 |
| ENSG00000131626 | PPFIA1 | 1.086203592 | 0.06221272 | 0.845445916 | 0.15099294 | 1.931649508 | 8.816894989 | 9.479495904 | 8.143093325 | 10.05135008 | 9.94775169 | 9.706989438 | 9.812580902 |
| ENSG00000161091 | MFSD12 | 1.125340386 | 0.407755293 | 0.803652279 | 0.558480691 | 1.928992665 | 9.521325812 | 5.838937808 | 9.926432745 | 10.26878646 | 10.3666911 | 10.29565054 | 9.622520313 |
| ENSG00000177700 | POLR2L | 0.684915144 | 0.477619809 | 1.243264183 | 0.162230124 | 1.928179327 | 9.98400217 | 8.376385542 | 8.042706615 | 9.676486519 | 9.81279854 | 9.812090132 | 10.66776684 |
| ENSG00000023902 | PLEKHO1 | 1.178515208 | 0.434235922 | 0.749169306 | 0.626611066 | 1.927684514 | 4.00722531 | 8.876041558 | 8.460932916 | 9.250166145 | 9.33957954 | 9.060185483 | 8.649438172 |
| ENSG00000120159 | CAAP1 | 0.625445844 | 0.528827603 | 1.302200348 | 0.148716503 | 1.927646192 | 5.70932728 | 7.393647515 | 5.39244166 | 7.144720234 | 7.01053089 | 7.57260403 | 7.889811111 |
| ENSG00000116095 | PLEKHA3 | 1.002506444 | 0.597757024 | 0.924203044 | 0.624146383 | 1.926709487 | 2 | 7.274681896 | 8.434763955 | 8.392303091 | 8.36552655 | 8.474034275 | 8.105768199 |
| ENSG00000163762 | TM4SF18 | -1.310490463 | 0.711621313 | 3.23494391 | 0.2712184 | 1.924453448 | 2 | 2 | 2 | 2.141737246 | 2.14022021 | 3.411599105 | 4.047883918 |
| ENSG00000126822 | PLEKHG3 | -1.29854033 | 0.47271232 | 3.222582404 | 0.045860622 | 1.924042074 | 4.467311984 | 8.276714465 | 3.369317972 | 5.651002631 | 5.58783529 | 9.835606948 | 10.19067811 |
| ENSG00000128346 | C22orf23 | -2.398062482 | 0.152438771 | 4.31846721 | 0.002694218 | 1.920404728 | 2 | 4.93638933 | 2 | 2.498691189 | 2.59489879 | 7.87872738 | 7.331709059 |
| ENSG00000144043 | TEX261 | 0.806489411 | 0.129712892 | 1.113895105 | 0.028077653 | 1.920384516 | 9.613202746 | 8.862353853 | 8.699419242 | 9.879225887 | 9.95251543 | 9.950137104 | 10.45173765 |
| ENSG00000168398 | BDKRB2 | 5.27218442 | 0.000312209 | -3.352415574 | 0.030925245 | 1.919768846 | 4.00722531 | 6.666836837 | 2 | 10.45727963 | 10.5479824 | 3.215043717 | 2.592996859 |
| ENSG00000167005 | NUDT21 | 0.988887806 | 0.237601833 | 0.928162853 | 0.262807198 | 1.917050659 | 8.87372952 | 8.671479029 | 6.80318718 | 9.393466757 | 9.29994502 | 9.391277746 | 9.174819346 |
| ENSG00000165138 | ANKS6 | 0.434458782 | 0.8692818 | 1.482258435 | 0.529629876 | 1.916717217 | 3.327711234 | 8.590771778 | 2 | 7.580980221 | 7.38644297 | 8.457998097 | 8.575745325 |
| ENSG00000128482 | RNF112 | 0.47893797 | 0.752212984 | 1.435786676 | 0.285392525 | 1.914724646 | 4.00722531 | 4.11352228 | 2 | 4.070463721 | 3.98554648 | 4.416072028 | 5.108633494 |
| ENSG00000171604 | CXXC5 | -0.827870442 | 0.104928256 | 2.741504743 | 7.92E-10 | 1.913634301 | 8.836089686 | 8.916342327 | 7.97167868 | 7.920420067 | 7.71553728 | 11.19227235 | 11.52361706 |
| ENSG00000187391 | MAGI2 | 0.455316747 | 0.776973345 | 1.457370186 | 0.308265999 | 1.912686933 | 3.327711234 | 5.595927442 | 7.600047325 | 6.877205252 | 6.76727577 | 7.446000032 | 8.083408769 |
| ENSG00000169762 | TAPT1 | 1.352838821 | 0.387432577 | 0.559669833 | 0.735204809 | 1.912508655 | 8.285626337 | 5.303523747 | 4.058998049 | 8.27796118 | 8.240612 | 7.714348968 | 7.199901369 |
| ENSG00000160867 | FGFR4 | -0.524557064 | 0.760985399 | 2.436999835 | 0.087028008 | 1.912442771 | 3.327711234 | 3.686303414 | 2 | 3.023133209 | 2.86123014 | 4.894909044 | 5.164234344 |
| ENSG00000148218 | ALAD | 0.630803541 | 0.548066193 | 1.280744959 | 0.184467558 | 1.9115485 | 7.384433967 | 6.602424674 | 9.031722608 | 8.674801527 | 8.63070345 | 9.096715824 | 9.474014382 |
| ENSG00000169439 | SDC2 | -1.265349062 | 0.163640595 | 3.176349494 | 8.57E-05 | 1.911000432 | 10.11549941 | 9.318399851 | 7.691296561 | 8.164192202 | 8.02236724 | 12.25879808 | 12.73497751 |
| ENSG00000005893 | LAMP2 | 1.349915776 | 9.47E-05 | 0.560298158 | 0.137293355 | 1.910213933 | 9.624286623 | 10.28338212 | 10.04361983 | 11.38590424 | 11.3294542 | 10.47794197 | 10.65645106 |
| ENSG00000108960 | MMD | 0.718666891 | 0.723460473 | 1.191526712 | 0.535217469 | 1.910193603 | 4.00722531 | 7.051611315 | 2 | 6.490083648 | 6.22964624 | 7.210184325 | 6.280114403 |
| ENSG00000121542 | SEC22A | 0.844751962 | 0.531143362 | 1.065371108 | 0.410154943 | 1.91012307 | 3.327711234 | 7.232742069 | 6.43197674 | 7.169761359 | 7.20671394 | 7.562295432 | 7.224630935 |
| ENSG00000164776 | PHKG1 | 0.923977962 | 0.687785088 | 0.985932074 | 0.660564083 | 1.909910036 | 2 | 2 | 4.058998049 | 3.462030052 | 3.55551834 | 3.753554863 | 3.311948089 |
| ENSG00000167232 | ZNF91 | 1.229232395 | 0.514736001 | 0.68003481 | 0.726284264 | 1.909267205 | 2 | 8.213429689 | 6.718950348 | 8.422134132 | 8.14812525 | 8.123435792 | 7.247335571 |

| | | | | | | | | | | | | | |
|---|---|---|---|---|---|---|---|---|---|---|---|---|---|
| ENSG00000259207 | ITGB3 | 2.603260998 | 0.348943914 | -0.695657657 | 0.81957996 | 1.907603341 | 8.042902253 | 2 | 2 | 9.04103174 | 9.05396643 | 6.619606851 | 4.033026654 |
| ENSG00000147364 | FBXO25 | 0.254962654 | 0.900708025 | 1.652209824 | 0.344818335 | 1.907172478 | 7.904282243 | 5.838937808 | 2 | 6.820355514 | 6.95938211 | 8.159444235 | 8.350852333 |
| ENSG00000133731 | IMPA1 | 1.376960153 | 0.274972023 | 0.528356153 | 0.695554779 | 1.905316306 | 7.218819781 | 6.844429755 | 3.369317972 | 7.971157485 | 7.73502229 | 7.156371901 | 6.889403418 |
| ENSG00000198382 | UVRAG | 0.775157547 | 0.388769665 | 1.129646397 | 0.18225392 | 1.904803945 | 6.257048221 | 8.255926468 | 6.80318718 | 8.093978021 | 8.18097132 | 8.437839892 | 8.53677936 |
| ENSG00000122574 | WIPF3 | -2.983882974 | 0.213019329 | 4.888341312 | 0.014755667 | 1.904458338 | 2 | 4.11352228 | 2 | 2.3892346 | 2 | 7.747483829 | 6.371531035 |
| ENSG00000149658 | YTHDF1 | 0.645889421 | 0.395163317 | 1.257406922 | 0.071096291 | 1.903296343 | 8.19931886 | 9.159587375 | 7.451345288 | 9.013631184 | 9.15558431 | 9.755980824 | 9.629636184 |
| ENSG00000074266 | EED | 1.39183833 | 0.112243988 | 0.510679182 | 0.590583035 | 1.902517512 | 6.009025997 | 6.728495673 | 4.523675391 | 7.288834728 | 7.45518898 | 6.763331418 | 6.232137896 |
| ENSG00000204116 | CHIC1 | 1.146306395 | 0.47537973 | 0.755853802 | 0.643465101 | 1.902160198 | 2 | 7.002537973 | 6.073378139 | 7.261917051 | 7.06433609 | 6.833591108 | 6.741885308 |
| ENSG00000174292 | TNK1 | 0.442652377 | 0.840490657 | 1.45890352 | 0.451396209 | 1.901555897 | 2 | 4.11352228 | 2 | 3.515101962 | 3.33976443 | 4.301748724 | 3.972018587 |
| ENSG00000139436 | GIT2 | 1.479890151 | 0.03929762 | 0.421583223 | 0.599804676 | 1.901473375 | 8.107517749 | 8.234834549 | 6.43197674 | 9.287824288 | 9.23006009 | 8.201234858 | 8.220862755 |
| ENSG00000106591 | MRPL32 | 0.934484227 | 0.115211747 | 0.965949127 | 0.09634677 | 1.900433354 | 6.816637977 | 8.004992147 | 7.163363831 | 8.347282003 | 8.35656898 | 8.514698983 | 8.237969002 |
| ENSG00000067208 | EVI5 | 0.541599575 | 0.105778838 | 1.358821482 | 5.09E-06 | 1.900421057 | 7.975256505 | 7.874700259 | 8.007629717 | 8.496596021 | 8.47221251 | 9.177956973 | 9.402844548 |
| ENSG00000089048 | ESF1 | 1.532364861 | 0.023843846 | 0.367426197 | 0.635038067 | 1.899791058 | 7.384433967 | 7.954278496 | 6.203069917 | 8.808365552 | 8.93143442 | 7.79139499 | 7.647978997 |
| ENSG00000181472 | ZBTB2 | 1.474471667 | 0.023668321 | 0.425033011 | 0.560994904 | 1.899504678 | 7.624139768 | 7.232742069 | 5.930865648 | 8.568972129 | 8.51631523 | 7.481445808 | 7.537835213 |
| ENSG00000117010 | ZNF684 | 0.301503568 | 0.896917828 | 1.597398189 | 0.428364332 | 1.898901757 | 2 | 5.131603625 | 2 | 4.205751501 | 4.05806467 | 5.54934408 | 4.793076545 |
| ENSG00000162772 | ATF3 | 1.146381957 | 0.182039968 | 0.751165571 | 0.392356999 | 1.897547528 | 8.653385977 | 7.189546432 | 6.80318718 | 8.912174893 | 8.90955283 | 8.815677424 | 8.146025444 |
| ENSG00000059728 | MXD1 | 1.168038975 | 0.012450706 | 0.729195801 | 0.133193465 | 1.897234775 | 6.366709671 | 7.145017454 | 7.225742779 | 8.136938566 | 8.08010078 | 7.783429229 | 7.562617572 |
| ENSG00000082458 | DLG3 | 0.951392095 | 0.431799068 | 0.943715856 | 0.426131057 | 1.895107951 | 4.815609471 | 7.431218269 | 5.156626401 | 7.292639438 | 7.18663697 | 7.411959578 | 7.028854452 |
| ENSG00000157764 | BRAF | -0.136285158 | 0.891774766 | 2.029000629 | 0.009269015 | 1.892715471 | 6.138360429 | 8.029695761 | 6.534109959 | 6.975156635 | 7.08183429 | 9.154129994 | 9.165908829 |
| ENSG00000153214 | TMEM87B | 1.669242437 | 0.395924587 | 0.221821638 | 0.920635926 | 1.891064075 | 9.33149386 | 7.73146271 | 2 | 9.797905134 | 9.84133177 | 8.541185654 | 8.206037069 |
| ENSG00000126759 | CFP | 2.013763328 | NA | -0.124892997 | NA | 1.888870332 | 2 | 2 | 2 | 3.227757282 | 2.77779907 | 2.42098416 | 2.172608095 |
| ENSG00000165934 | CPSF2 | 0.716330331 | 0.228106188 | 1.172438209 | 0.035030931 | 1.888768539 | 8.009475818 | 8.639735838 | 9.287079172 | 9.471064221 | 9.42346554 | 10.02767645 | 9.762795477 |
| ENSG00000163781 | TOPBP1 | 0.916676303 | 0.632871056 | 0.971521755 | 0.605992978 | 1.888198059 | 3.327711234 | 8.639735838 | 4.874587708 | 8.160032699 | 8.0314011 | 8.247650629 | 8.047821014 |
| ENSG00000164292 | RHOBTB3 | 0.768906019 | 0.378155417 | 1.117254418 | 0.175036042 | 1.886160437 | 9.247966931 | 9.079034257 | 7.398165132 | 9.583264843 | 9.50436385 | 10.27812044 | 9.36032955 |
| ENSG00000148908 | RGS10 | 1.281865026 | 0.505521211 | 0.601590356 | 0.764974927 | 1.883455382 | 7.750913381 | 4.93638933 | 2 | 7.694616532 | 7.53675633 | 6.877490957 | 7.028854452 |
| ENSG00000204352 | C9orf129 | 7.073258558 | 0.001861972 | -5.191143651 | 0.077151749 | 1.882114907 | 2 | 2 | 2 | 7.169761359 | 7.11195456 | 2 | 2 |
| ENSG00000161940 | BCL6B | 0.897611828 | 0.725904626 | 0.981416451 | 0.692863851 | 1.879028279 | 3.327711234 | 2 | 2 | 2.947924294 | 2.94009902 | 3.10571689 | 2.851942824 |
| ENSG00000163704 | PRRT3 | 0.450554277 | 0.790303792 | 1.428319054 | 0.343471869 | 1.878873332 | 3.327711234 | 7.002537973 | 4.058998049 | 6.096288482 | 6.18176397 | 6.913621294 | 7.22299542 |
| ENSG00000101150 | TPD52L2 | 0.336044484 | 0.691501477 | 1.540480633 | 0.032896134 | 1.876525118 | 10.7426185 | 9.849218119 | 8.845626171 | 10.31300956 | 10.3782441 | 11.51506069 | 11.58004535 |
| ENSG00000118900 | UBN1 | 0.500765632 | 0.731288333 | 1.37548542 | 0.295571175 | 1.876251051 | 5.532353208 | 8.791896851 | 9.849211147 | 9.36574218 | 9.38433737 | 10.34534549 | 10.13892798 |
| ENSG00000273802 | HIST1H2BG | 3.745268006 | 0.055746187 | -1.870021459 | 0.368897909 | 1.875246547 | 6.009025997 | 2 | 2 | 8.078634414 | 8.08010078 | 2.987421008 | 3.45257121 |
| ENSG00000026025 | VIM | 1.37811376 | 0.036049949 | 0.493564062 | 0.492815255 | 1.871677822 | 13.96181853 | 13.49758156 | 12.33487604 | 14.79555473 | 14.7831527 | 14.00582069 | 13.79656997 |
| ENSG00000106701 | FSD1L | 1.318257443 | 0.292677052 | 0.553037321 | 0.67823435 | 1.871294764 | 2 | 4.93638933 | 5.59507254 | 6.078767801 | 5.86549842 | 5.180212365 | 5.361470313 |
| ENSG00000100949 | RABGGTA | 1.127620498 | 0.5253082 | 0.743084337 | 0.681279661 | 1.870704835 | 7.624139768 | 7.701025996 | 2 | 8.126317766 | 8.27144011 | 7.47156439 | 8.103992141 |
| ENSG00000153006 | SREK1IP1 | 0.893284845 | 0.036051738 | 0.976842785 | 0.019287415 | 1.87012763 | 7.33131396 | 7.790474954 | 7.502634626 | 8.463225341 | 8.41985547 | 8.729854886 | 8.284187525 |
| ENSG00000145358 | DDIT4L | 3.938628018 | 0.082799505 | -2.06871339 | 0.388256033 | 1.869914628 | 4.815609471 | 4.11352228 | 2 | 7.632926745 | 7.60187978 | 3.316666208 | 2 |
| ENSG00000061987 | MON2 | 1.012320805 | 0.085477895 | 0.857590652 | 0.144697175 | 1.869911457 | 7.867441304 | 8.981105197 | 7.896971941 | 9.352133347 | 9.36927498 | 9.231960665 | 9.181571564 |
| ENSG00000275031 | | 4.513142663 | 1.55E-14 | -2.64647408 | 1.87E-05 | 1.866668583 | 5.70932728 | 6.844429755 | 6.203069917 | 10.77167208 | 10.8115546 | 3.648325314 | 4.341135913 |
| ENSG00000203772 | SPRN | 1.358325998 | 0.552657351 | 0.503713332 | 0.836549691 | 1.86203933 | 2 | 5.838937808 | 2 | 5.786686927 | 5.55061739 | 5.318934974 | 4.329022995 |
| ENSG00000171295 | ZNF440 | 1.234180521 | 0.304198642 | 0.627763435 | 0.616846858 | 1.861943956 | 4.00722531 | 6.951736442 | 5.156626401 | 7.119236776 | 7.03319408 | 6.736076089 | 6.199243628 |
| ENSG00000178093 | TSSK6 | 0.669735175 | 0.759294561 | 1.192175293 | 0.561301735 | 1.861910467 | 2 | 4.710564154 | 2 | 4.105493497 | 4.02226117 | 4.834511823 | 4.002845082 |
| ENSG00000166262 | FAM227B | 0.16797141 | 0.929751797 | 1.693022808 | 0.28326297 | 1.860994218 | 3.327711234 | 3.686303414 | 2 | 3.162721474 | 3.45167086 | 4.644324387 | 4.105828626 |
| ENSG00000041880 | PARP3 | 1.240763293 | 0.03191533 | 0.620188936 | 0.311844973 | 1.860952229 | 7.790810999 | 6.602424674 | 7.691296561 | 8.676258519 | 8.64254348 | 7.861174222 | 8.210995912 |
| ENSG00000124523 | SIRT5 | 0.911756411 | 0.467848857 | 0.948519486 | 0.43787928 | 1.860275897 | 3.327711234 | 6.844429755 | 5.930865648 | 6.872128674 | 6.80989959 | 6.99251878 | 6.750993196 |
| ENSG00000120705 | ETF1 | 0.816984245 | 0.434235922 | 1.041983247 | 0.296112508 | 1.858967492 | 8.009475818 | 9.338312924 | 10.71094342 | 10.58610178 | 10.53732 | 10.71985653 | 10.84993646 |
| ENSG00000135655 | USP15 | 1.391776388 | 0.002659741 | 0.467143382 | 0.364515211 | 1.85891977 | 8.631580145 | 8.414402626 | 7.600047325 | 9.696010185 | 9.62936766 | 8.812636523 | 8.673588256 |
| ENSG00000072415 | MPP5 | 0.981657857 | 0.167897149 | 0.876533899 | 0.214740154 | 1.858191756 | 7.27616306 | 8.317412869 | 6.718950348 | 8.637890117 | 8.50824913 | 8.416251718 | 8.522888769 |
| ENSG00000114742 | WDR48 | 1.063363453 | 0.292532122 | 0.794205202 | 0.435384719 | 1.857568654 | 8.468948322 | 9.102512015 | 6.322058544 | 9.42153757 | 9.41227044 | 9.176605981 | 9.121828633 |
| ENSG00000215126 | CBWD6 | 4.588257983 | 8.16E-09 | -2.731257175 | 0.001820976 | 1.857000808 | 2 | 4.710564154 | 5.156626401 | 8.729158235 | 8.82635531 | 2.717032197 | 2.782576693 |
| ENSG00000123374 | CDK2 | 0.676552374 | 0.717263579 | 1.179439315 | 0.501916413 | 1.855991689 | 2 | 7.431218269 | 8.007629717 | 7.885575888 | 7.78928606 | 8.243140215 | 8.418864688 |
| ENSG00000177963 | RIC8A | 0.420657521 | 0.465054573 | 1.431573445 | 0.003946525 | 1.852230966 | 9.790941832 | 10.07961649 | 8.99637432 | 10.08135305 | 10.1427213 | 10.92445873 | 11.29335049 |
| ENSG00000197601 | FAR1 | 0.368739111 | 0.800325517 | 1.483389554 | 0.242330228 | 1.852128666 | 8.16936288 | 8.590771778 | 4.523675391 | 8.297027403 | 8.15227214 | 9.364533261 | 9.29605726 |
| ENSG00000125755 | SYMPK | 1.201233202 | 0.153219079 | 0.649476341 | 0.459476825 | 1.850709543 | 7.750913381 | 7.979858146 | 9.61300511 | 9.854985705 | 9.93095363 | 9.341810888 | 9.34725177 |
| ENSG00000165288 | BRWD3 | 1.081867233 | 0.062696835 | 0.768449349 | 0.194623596 | 1.850316582 | 7.579279349 | 6.78762695 | 7.858115942 | 8.537252353 | 8.51631523 | 8.48499076 | 7.887748062 |
| ENSG00000136827 | TOR1A | 1.291833851 | 0.000202427 | 0.554693466 | 0.142226777 | 1.846527317 | 8.963744667 | 9.102512015 | 8.585098385 | 10.18835911 | 10.1903833 | 9.267612236 | 9.621899879 |
| ENSG00000196507 | TCEAL3 | 0.140279331 | 0.896732205 | 1.705732919 | 0.04836641 | 1.84601225 | 8.892187472 | 6.535001748 | 7.858115942 | 8.229163214 | 8.15227214 | 9.531042494 | 9.928756347 |
| ENSG00000142208 | AKT1 | 1.268821073 | 0.412971533 | 0.5765566 | 0.723575557 | 1.845377674 | 10.27168932 | 10.23108619 | 5.156626401 | 10.90533885 | 10.9990778 | 10.10357235 | 10.40562199 |
| ENSG00000120594 | PLXDC2 | 0.405505548 | 0.609032994 | 1.439793986 | 0.035847057 | 1.845299534 | 6.366709671 | 6.728495673 | 6.882775709 | 6.941624891 | 7.18258783 | 8.573367846 | 7.321046127 |
| ENSG00000196876 | SCN8A | 2.716539415 | 0.066281011 | -0.871937728 | 0.593116818 | 1.844601686 | 2 | 7.315436877 | 6.534109959 | 9.088859521 | 9.07487069 | 5.630526941 | 5.581648407 |
| ENSG00000070614 | NDST1 | 0.563622702 | 0.546572519 | 1.279032617 | 0.1322507 | 1.84265532 | 9.913467515 | 9.785372135 | 11.61913434 | 11.22573252 | 11.2975154 | 11.64651719 | 12.24498338 |
| ENSG00000167721 | TSR1 | 1.040336636 | 0.098946169 | 0.802305742 | 0.207181946 | 1.842642377 | 8.674867122 | 8.732943944 | 7.342949392 | 9.399654088 | 9.42518018 | 9.08740002 | 9.259427717 |
| ENSG00000167333 | TRIM68 | 0.611503726 | 0.748108458 | 1.23101484 | 0.487308062 | 1.842518566 | 2 | 7.393647515 | 5.39244166 | 6.622948477 | 6.87658492 | 7.711553069 | 6.883203642 |
| ENSG00000078674 | PCM1 | 1.020512758 | 0.147002974 | 0.821753161 | 0.245421661 | 1.842265919 | 7.750913381 | 9.396454291 | 9.031722608 | 9.886167436 | 9.90783885 | 9.876537138 | 9.496865493 |
| ENSG00000148158 | SNX30 | 0.856229675 | 0.434435575 | 0.982663381 | 0.352922604 | 1.838893056 | 8.042902253 | 7.928237109 | 5.156626401 | 8.381625466 | 8.31092742 | 8.377971121 | 8.559637211 |
| ENSG00000172466 | ZNF24 | 0.954516605 | 0.522605424 | 0.881998629 | 0.548117081 | 1.836515234 | 5.095959458 | 9.514922632 | 9.636156974 | 10.01553929 | 9.93035007 | 10.20180989 | 9.521357751 |
| ENSG00000155252 | PI4K2A | 1.313290143 | 0.24500653 | 0.522853577 | 0.667109583 | 1.83614372 | 6.73718411 | 8.777382925 | 10.01740919 | 10.33623673 | 10.3599836 | 9.342112202 | 9.750927434 |
| ENSG00000100068 | LRP5L | 1.65163554 | 0.437566006 | 0.184099464 | 0.93862683 | 1.835735004 | 5.095959458 | 2 | 2 | 5.391962469 | 4.79082474 | 3.981090143 | 3.858622973 |
| ENSG00000141458 | NPC1 | 1.585602776 | 0.117971256 | 0.249171742 | 0.832879767 | 1.834774518 | 7.904282243 | 8.147241016 | 10.30429804 | 10.77371523 | 10.8171157 | 9.534737983 | 9.389783813 |
| ENSG00000101955 | SRPX | 1.549342349 | 0.478179842 | 0.285076922 | 0.906201712 | 1.834419271 | 3.327711234 | 9.125613817 | 3.369317972 | 9.141495458 | 9.11260313 | 7.79139499 | 7.958255135 |
| ENSG00000184828 | ZBTB7C | -1.303410978 | 0.802441015 | 3.137816698 | 0.491297476 | 1.834405719 | 2 | 2 | 2 | 2.270786764 | 2 | 4.199031499 | 2.918126211 |
| ENSG00000138495 | COX17 | 1.142188102 | 0.051185896 | 0.69196198 | 0.254317355 | 1.834150083 | 6.009025997 | 6.140560021 | 5.156626401 | 6.951285179 | 6.92586261 | 6.375345632 | 6.62538071 |
| ENSG00000105982 | RNF32 | 0.223521848 | 0.933881218 | 1.609739051 | 0.493182797 | 1.833260899 | 2 | 3.686303414 | 2 | 2.600425781 | 3.28037825 | 4.188345803 | 3.33635788 |
| ENSG00000266412 | NCOA4 | 1.067192167 | 0.04337239 | 0.76570091 | 0.155705957 | 1.832893077 | 9.590776211 | 10.76012279 | 10.18377222 | 11.3151886 | 11.330827 | 10.93098358 | 11.10773798 |
| ENSG00000197841 | ZNF181 | 1.243468323 | 0.436019907 | 0.587680084 | 0.724656426 | 1.831148406 | 6.366709671 | 6.728495673 | 2 | 7.19843704 | 7.21466714 | 7.02591204 | 5.909074772 |
| ENSG00000077254 | USP33 | 1.375837802 | 1.30E-05 | 0.454481203 | 0.195994401 | 1.830319004 | 8.564112515 | 8.522788974 | 8.960138144 | 10.01783961 | 10.0977376 | 9.242329292 | 9.033614226 |
| ENSG00000097021 | ACOT7 | 1.051644237 | 0.525295603 | 0.776008956 | 0.641507692 | 1.827653193 | 4.815609471 | 6.78762695 | 9.624627483 | 9.275380474 | 9.34864288 | 8.621823729 | 9.361816632 |
| ENSG00000137745 | MMP13 | 7.017586499 | 0.002488209 | -5.191143654 | 0.081042066 | 1.826443445 | 2 | 2 | 2 | 7.003290579 | 7.16627666 | 2 | 2 |
| ENSG00000219545 | UMAD1 | 1.02232181 | 0.538701364 | 0.802901589 | 0.632223509 | 1.8252234 | 4.815609471 | 6.666836837 | 2 | 6.408278948 | 6.51372389 | 6.004008792 | 6.465820193 |
| ENSG00000061273 | HDAC7 | 0.393558994 | 0.438437558 | 1.431177274 | 0.000991504 | 1.824736268 | 9.544849385 | 9.514922632 | 8.65477198 | 9.696728234 | 9.67763759 | 10.66370661 | 10.77657437 |
| ENSG00000109920 | FNBP4 | 1.11092149 | 0.279628699 | 0.710802317 | 0.498871333 | 1.821723807 | 8.737455672 | 9.148351514 | 6.322058544 | 9.568433308 | 9.61215953 | 9.252303334 | 9.130578617 |
| ENSG00000054392 | HHAT | 0.826428048 | 0.686228546 | 0.995024488 | 0.615427844 | 1.821452536 | 3.327711234 | 6.666836837 | 2 | 6.078767801 | 6.03677338 | 5.979426183 | 6.420873184 |
| ENSG00000176248 | ANAPC2 | 0.728187609 | 0.089730458 | 1.091346087 | 0.006751128 | 1.819533696 | 7.709880939 | 7.002537973 | 7.451345288 | 8.054188409 | 8.17894021 | 8.455215133 | 8.497408326 |
| ENSG00000151491 | EPS8 | -0.517095951 | 0.53177344 | 2.336033001 | 0.000833175 | 1.81893705 | 8.075571721 | 9.079034257 | 9.999667074 | 8.70652031 | 8.77618319 | 11.48201426 | 11.67783059 |
| ENSG00000106624 | AEBP1 | -0.323160872 | 0.682951928 | 2.141384438 | 0.000909649 | 1.818223566 | 10.45046729 | 11.26034006 | 12.15421689 | 11.10489064 | 11.1547085 | 13.55226621 | 13.63154322 |
| ENSG00000124493 | GRM4 | -0.727756714 | 0.851528683 | 2.545746269 | NA | 1.817989554 | 2 | 2 | 2 | 2.141737246 | 2.2680111 | 3.215043717 | 3.407202941 |
| ENSG00000049618 | ARID1B | 0.941019489 | 0.495694274 | 0.87529216 | 0.521153279 | 1.81631165 | 5.532353208 | 9.018611864 | 9.798878489 | 9.836208668 | 9.8787275 | 9.846692337 | 9.735523645 |
| ENSG00000089597 | GANAB | 0.930074645 | 0.154687626 | 0.885354353 | 0.169698897 | 1.815428998 | 10.80239439 | 11.36354124 | 12.2511264 | 12.50345529 | 12.5485341 | 12.32267771 | 12.62452591 |
| ENSG00000165271 | NOL6 | 0.80968831 | 0.514865066 | 1.005326195 | 0.398667475 | 1.815014505 | 5.330594134 | 8.540087354 | 8.784770914 | 8.934263273 | 8.97422645 | 9.007209385 | 9.277267848 |
| ENSG00000118217 | ATF6 | 0.71037993 | 0.097618305 | 1.102962958 | 0.006105215 | 1.813342888 | 8.87372952 | 9.04308532 | 9.647595056 | 9.963384909 | 9.89861166 | 10.27796296 | 10.36578622 |
| ENSG00000259332 | ST20-MTHFS | 7.004362544 | 0.002590896 | -5.191143003 | 0.081460409 | 1.813219541 | 2 | 2 | 2 | 7.044488801 | 7.10341274 | 2 | 2 |
| ENSG00000138092 | CENPO | 1.218954499 | 0.403701418 | 0.59347817 | 0.696976163 | 1.812432669 | 6.563806279 | 5.946638857 | 2 | 6.912249745 | 6.89154576 | 6.173421703 | 6.420873184 |
| ENSG00000171786 | NHLH1 | 2.060110941 | 0.313991908 | -0.247936522 | 0.916470603 | 1.812174419 | 2 | 3.686303414 | 2 | 4.070463721 | 4.2865223 | 2.62493832 | 2.918126211 |
| ENSG00000164880 | INTS1 | 0.602137651 | 0.671105744 | 1.207522821 | 0.356353281 | 1.809660472 | 10.57565445 | 7.761270542 | 7.225742779 | 9.851765671 | 9.94536391 | 10.46278097 | 10.54141057 |
| ENSG00000170004 | CHD3 | 0.514153223 | 0.544293669 | 1.292569277 | 0.090878546 | 1.8067225 | 8.855032352 | 10.52407288 | 10.45512158 | 10.59498781 | 10.6715842 | 10.9230495 | 11.77450991 |
| ENSG00000144426 | NBEAL1 | 0.84954597 | 0.365703512 | 0.955819972 | 0.293820393 | 1.805365942 | 8.816894989 | 7.232742069 | 6.718950348 | 8.673343061 | 8.75724985 | 9.029457273 | 8.577666268 |
| ENSG00000196961 | AP2A1 | 0.60874537 | 0.292553183 | 1.195945937 | 0.024032158 | 1.804691306 | 10.1076228 | 10.03760521 | 8.978369997 | 10.38650225 | 10.4132224 | 10.8706474 | 11.09225624 |
| ENSG00000106086 | PLEKHA8 | 0.947618085 | 0.037226271 | 0.855901662 | 0.059084307 | 1.803519748 | 6.96351253 | 7.701025996 | 7.502634626 | 8.370868225 | 8.35297033 | 8.439530542 | 8.0807022 |
| ENSG00000114423 | CBLB | 1.85259731 | 0.095126919 | -0.05021032 | 0.97020102 | 1.802386991 | 8.631580145 | 6.78762695 | 5.59507254 | 9.389030931 | 9.3304589 | 7.743839536 | 7.159430626 |
| ENSG00000155729 | KCTD18 | 0.388618136 | 0.833540602 | 1.412914998 | 0.384828556 | 1.801533134 | 7.579279349 | 6.666836837 | 2 | 7.080147179 | 6.92586261 | 7.886185273 | 8.117259579 |
| ENSG00000103353 | UBFD1 | 0.632787121 | 0.632327358 | 1.168045701 | 0.343161384 | 1.800832823 | 6.653098118 | 10.47544827 | 9.636156974 | 10.21917213 | 10.2340777 | 10.63251333 | 10.87787797 |
| ENSG00000113790 | EHHADH | -0.12596852 | 0.934841508 | 1.925824172 | 0.12649658 | 1.799855652 | 2 | 5.946638857 | 5.156626401 | 4.846787657 | 5.13053105 | 6.940933331 | 6.864442837 |
| ENSG00000183421 | RIPK4 | -2.654230762 | 0.18763416 | 4.453225096 | 0.010530438 | 1.798994334 | 4.815609471 | 2 | 2 | 2.600425781 | 2.14022021 | 7.309674888 | 7.570355956 |
| ENSG00000111790 | FGFR1OP2 | 0.952285579 | 0.459782955 | 0.846439653 | 0.505528206 | 1.798725233 | 8.257427002 | 7.572420273 | 4.523675391 | 8.404661252 | 8.34212018 | 8.548515606 | 7.922424886 |
| ENSG00000164649 | CDCA7L | 0.663722755 | 0.66611095 | 1.133013237 | 0.431244286 | 1.796735992 | 4.00722531 | 5.595927442 | 2 | 5.258265913 | 5.00774437 | 5.622611168 | 5.497565112 |
| ENSG00000149451 | ADAM33 | 1.691772579 | 0.261480902 | 0.101184304 | 0.954416752 | 1.792956883 | 6.96351253 | 6.666836837 | 2 | 7.818544084 | 7.99261031 | 6.221381643 | 6.495928492 |
| ENSG00000088002 | SULT2B1 | -1.303475717 | 0.796759656 | 3.096171555 | 0.484010617 | 1.792695839 | 2 | 2 | 2 | 2.270786764 | 2 | 3.694366604 | 3.640730416 |
| ENSG00000163219 | ARHGAP25 | -2.312443603 | 0.573719152 | 4.104630145 | 0.221372771 | 1.792186542 | 2 | 2 | 2 | 2 | 2.14022021 | 4.727866721 | 4.047883918 |

| | | | | | | | | | | | | | |
|---|---|---|---|---|---|---|---|---|---|---|---|---|---|
| ENSG00000166734 | CASC4 | 1.117142762 | 0.404897934 | 0.674978336 | 0.624569223 | 1.792121098 | 6.138360429 | 10.13758676 | 10.08627303 | 10.70507197 | 10.6668817 | 10.31450651 | 10.17236288 |
| ENSG00000178467 | P4HTM | 0.835469199 | 0.586697428 | 0.955241852 | 0.522645705 | 1.790711051 | 7.435667355 | 8.876041558 | 4.058998049 | 8.649804831 | 8.57000035 | 8.663891891 | 8.791384617 |
| ENSG00000243056 | EIF4EBP3 | 3.423983486 | 0.03057436 | -1.633377028 | 0.345260656 | 1.790606458 | 2 | 2 | 4.523675391 | 6.12217689 | 5.99120031 | 2.560105256 | 2.551896972 |
| ENSG00000018610 | CXorf56 | 0.982000291 | 0.094108586 | 0.808457617 | 0.167430714 | 1.790457908 | 6.563806279 | 6.951736442 | 5.772716771 | 7.437142666 | 7.5240115 | 7.454943454 | 7.152573734 |
| ENSG00000171456 | ASXL1 | 0.09154634 | 0.899933851 | 1.697561314 | 0.002384213 | 1.789107654 | 8.444143041 | 9.470501606 | 9.819251342 | 9.392580682 | 9.49296422 | 10.95846335 | 11.12491757 |
| ENSG00000103381 | CPPED1 | 0.718096036 | 0.370967351 | 1.070861701 | 0.15655242 | 1.788957737 | 8.609439662 | 8.213429689 | 6.718950348 | 8.693628619 | 8.81985255 | 8.904045868 | 9.286693143 |
| ENSG00000011275 | RNF216 | 0.177734422 | 0.711059078 | 1.610582106 | 2.28E-05 | 1.788316527 | 8.963744667 | 9.632483534 | 8.960138144 | 9.437090644 | 9.37105521 | 10.78558135 | 10.87683825 |
| ENSG00000112282 | MED23 | 1.332867249 | 0.300898696 | 0.455089935 | 0.742539931 | 1.787957183 | 5.70932728 | 9.018611864 | 6.534109959 | 9.115954467 | 9.11366683 | 8.385007206 | 8.091498142 |
| ENSG00000141469 | SLC14A1 | 1.106584767 | 0.581814479 | 0.680112932 | 0.738292369 | 1.786697699 | 3.327711234 | 2 | 3.369317972 | 3.515101962 | 3.69849104 | 3.600766115 | 3.012034821 |
| ENSG00000138778 | CENPE | 0.910755563 | 0.40768846 | 0.875539234 | 0.417447017 | 1.786294797 | 4.467311984 | 6.04685525 | 4.058998049 | 6.034011955 | 5.98191036 | 6.305369896 | 5.545463418 |
| ENSG00000256229 | ZNF486 | 0.895284273 | 0.284862917 | 0.890650422 | 0.275628117 | 1.785934695 | 5.70932728 | 4.11352228 | 5.59507254 | 6.034011955 | 6.12381559 | 6.170709933 | 5.97274432 |
| ENSG00000103855 | CD276 | 1.373925565 | 0.236772262 | 0.411915236 | 0.745570382 | 1.785840802 | 11.55650736 | 10.88722087 | 8.076950877 | 12.10263209 | 12.1465732 | 10.92697178 | 11.36748936 |
| ENSG00000131899 | LLGL1 | 1.010608601 | 0.467936122 | 0.772715614 | 0.579833528 | 1.783324215 | 8.653385977 | 4.93638933 | 9.132850258 | 9.389919188 | 9.35225233 | 8.939525293 | 9.306905794 |
| ENSG00000178163 | ZNF518B | 0.690272654 | 0.743410913 | 1.092911278 | 0.586432904 | 1.783183932 | 2 | 6.844429755 | 8.960138144 | 8.321442665 | 8.40081078 | 8.990775914 | 8.485179747 |
| ENSG00000182324 | KCNJ14 | 0.763828567 | 0.734475554 | 1.019185924 | 0.638604205 | 1.783014492 | 2 | 4.710564154 | 2 | 3.840289928 | 4.37434574 | 4.73522696 | 3.771796761 |
| ENSG00000077984 | CST7 | -0.314243794 | NA | 2.096937505 | NA | 1.782693711 | 2 | 2 | 2 | 2.141737246 | 2.38539787 | 3.648325314 | 2.059860178 |
| ENSG00000141556 | TBCD | 1.187051235 | 0.415814926 | 0.592309949 | 0.696686193 | 1.779361183 | 4.00722531 | 8.255926468 | 6.203069917 | 8.174538791 | 8.2561084 | 7.440382108 | 7.799420735 |
| ENSG00000131100 | ATP6V1E1 | 1.204763605 | 0.23034597 | 0.573544063 | 0.58833395 | 1.778307668 | 10.59268348 | 9.102512015 | 7.934808822 | 10.76928475 | 10.8430013 | 10.05624005 | 10.28872631 |
| ENSG00000146776 | ATXN7L1 | 1.570127391 | 0.281556878 | 0.206575743 | 0.901842972 | 1.776703134 | 2 | 5.72254323 | 6.629488387 | 7.153115662 | 7.22651534 | 5.801542492 | 5.968846243 |
| ENSG00000161835 | GRASP | -0.436094991 | 0.85051447 | 2.210299884 | 0.260738589 | 1.774204893 | 2 | 5.303523747 | 2 | 3.515101962 | 3.90918929 | 6.320128742 | 5.448021784 |
| ENSG00000256294 | ZNF225 | 0.939272193 | 0.405878812 | 0.833655425 | 0.456352327 | 1.772927618 | 3.327711234 | 6.311476754 | 6.322058544 | 6.676346669 | 6.76185814 | 6.613628927 | 6.622902297 |
| ENSG00000139437 | TCHP | 0.675931733 | 0.695919199 | 1.095981466 | 0.501628498 | 1.771913199 | 6.653098118 | 7.393647515 | 2 | 7.214570819 | 7.1204461 | 7.555035333 | 7.600900977 |
| ENSG00000205189 | ZBTB10 | 0.307868617 | 0.896153568 | 1.463770994 | 0.480226672 | 1.771639611 | 4.467311984 | 8.213429689 | 2 | 7.114945424 | 6.99676009 | 8.397818242 | 7.944426693 |
| ENSG00000170946 | DNAJC24 | 1.190666075 | 0.196009997 | 0.580646402 | 0.548363823 | 1.771312477 | 7.709880939 | 5.946638857 | 5.772716771 | 8.006336225 | 7.81828345 | 7.493431902 | 7.117791742 |
| ENSG00000117335 | CD46 | 1.116896131 | 0.153513211 | 0.65199915 | 0.422109347 | 1.768895281 | 8.340423424 | 10.28338212 | 9.888372618 | 10.82814175 | 10.8281739 | 10.50833129 | 10.20462614 |
| ENSG00000122203 | KIAA1191 | 0.610587974 | 0.12465991 | 1.156764709 | 0.001542477 | 1.767352683 | 9.689059027 | 10.12617756 | 10.42220168 | 10.71786226 | 10.7188341 | 11.19786031 | 11.32619054 |
| ENSG00000125733 | TRIP10 | 0.516524689 | 0.205222597 | 1.249961086 | 0.000591862 | 1.766485775 | 8.493334308 | 8.053983481 | 8.511888463 | 8.839910123 | 8.90341524 | 9.776191646 | 9.400314204 |
| ENSG00000126870 | WDR60 | 1.093607353 | 0.538070994 | 0.671489612 | 0.713730905 | 1.765096965 | 2 | 7.503546079 | 7.858115942 | 8.190940526 | 8.19510963 | 7.839995266 | 7.712342256 |
| ENSG00000125827 | TMX4 | 1.38302151 | 0.085117333 | 0.382050982 | 0.670218628 | 1.765072492 | 9.384607364 | 9.018611864 | 7.342949392 | 10.16882602 | 10.2109057 | 8.94032159 | 9.409690338 |
| ENSG00000137992 | DBT | 0.620825829 | 0.525902595 | 1.144035889 | 0.204625268 | 1.764861718 | 6.138360429 | 8.124485672 | 8.175061096 | 8.409925372 | 8.28283292 | 9.272046904 | 8.297458414 |
| ENSG00000150764 | DIXDC1 | 0.264639182 | 0.863874307 | 1.499903342 | 0.252589462 | 1.764542524 | 4.467311984 | 8.124485672 | 8.763901704 | 8.197043419 | 8.20112674 | 9.526807454 | 9.311145499 |
| ENSG00000155428 | TRIM74 | 3.210583301 | 0.267665409 | -1.447786938 | 0.649857772 | 1.762796363 | 2 | 2 | 2 | 3.406930997 | 4.02226117 | 2.139668783 | 2.117335275 |
| ENSG00000013016 | EHD3 | -0.721653629 | 0.118548292 | 2.484356572 | 6.37E-10 | 1.762702943 | 8.564112515 | 8.806266215 | 7.896971941 | 7.739909082 | 7.79460185 | 10.87554743 | 11.02027932 |
| ENSG00000141580 | WDR45B | 0.508091864 | 0.137498142 | 1.254144501 | 4.02E-05 | 1.762236365 | 9.204317094 | 9.479495904 | 9.681375057 | 9.985296505 | 9.96140527 | 10.58403908 | 10.8390337 |
| ENSG00000125741 | OPA3 | 0.286778506 | 0.766720309 | 1.474914211 | 0.072952984 | 1.761692717 | 6.563806279 | 8.297207175 | 8.763901704 | 8.408172799 | 8.43013848 | 9.354113447 | 9.806314352 |
| ENSG00000104442 | ARMC1 | 0.752600093 | 0.69344525 | 1.008036289 | 0.582641607 | 1.760636383 | 2 | 8.053983481 | 8.076950877 | 8.285617933 | 8.17690623 | 8.655672796 | 8.290447859 |
| ENSG00000110315 | RNF141 | -0.88909083 | 0.271602878 | 2.645844078 | 0.000180075 | 1.756753248 | 6.73718411 | 8.623598428 | 8.585098385 | 7.370304568 | 7.29937545 | 10.72829538 | 10.93843053 |
| ENSG00000166847 | DCTN5 | 0.579669376 | 0.433038514 | 1.176988842 | 0.080856643 | 1.756658218 | 10.18451821 | 8.522788974 | 9.372276026 | 10.06197797 | 10.1153688 | 10.69341333 | 10.67541118 |
| ENSG00000158220 | ESYT3 | -1.317468484 | 0.795427588 | 3.07358351 | 0.489950838 | 1.756115026 | 2 | 2 | 2 | 2 | 2.2680111 | 3.810410249 | 3.474731315 |
| ENSG00000116954 | RRAGC | 1.767001526 | 0.000538028 | -0.011626713 | 0.986196197 | 1.755374813 | 8.418903792 | 7.274681896 | 8.236946975 | 9.875425464 | 9.71666438 | 8.120628326 | 7.944426693 |
| ENSG00000136720 | HS6ST1 | 0.468928341 | 0.821640487 | 1.285900093 | 0.493153856 | 1.754828434 | 2 | 6.728495673 | 8.511888463 | 7.756533788 | 7.77052502 | 8.373850835 | 8.740849281 |
| ENSG00000088826 | SMOX | 3.503687942 | 0.000676627 | -1.748885556 | 0.114259051 | 1.754802386 | 8.19931886 | 6.728495673 | 4.874587708 | 10.63936224 | 10.6126122 | 5.758846871 | 5.191251321 |
| ENSG00000138802 | SEC24B | 1.100646987 | 0.215813666 | 0.64853403 | 0.480834472 | 1.749181018 | 6.96351253 | 9.277729677 | 8.325066185 | 9.555032495 | 9.58633944 | 9.245876366 | 8.985229012 |
| ENSG00000168994 | PXDC1 | -0.084593405 | 0.942177035 | 1.833205535 | 0.051887684 | 1.74861213 | 10.13887448 | 8.540087354 | 7.734844356 | 9.031955922 | 9.11153864 | 11.34267419 | 10.5004808 |
| ENSG00000230797 | YY2 | 0.404071721 | 0.854219249 | 1.344103576 | 0.489342742 | 1.748175297 | 2 | 4.442710129 | 2 | 3.79808662 | 3.50452879 | 4.504874511 | 4.201948172 |
| ENSG00000117280 | RAB29 | 0.523644412 | 0.718375041 | 1.224443382 | 0.357283417 | 1.748087794 | 8.564112515 | 4.710564154 | 8.805342542 | 8.652768197 | 8.673169 | 9.284015592 | 9.428971287 |
| ENSG00000100280 | AP1B1 | 0.481718746 | 0.686208373 | 1.266252281 | 0.237913947 | 1.747971027 | 9.810482605 | 9.74864196 | 6.882775709 | 9.769519169 | 9.76891184 | 10.207113 | 10.82845131 |
| ENSG00000163536 | SERPINI1 | 2.337666297 | 0.22692986 | -0.590148978 | 0.784860167 | 1.747517318 | 2 | 5.457120953 | 2 | 6.113598934 | 6.32824132 | 3.838019203 | 3.620995423 |
| ENSG00000140265 | ZSCAN29 | 1.063806919 | 0.153169017 | 0.682172042 | 0.370890738 | 1.745978961 | 6.138360429 | 7.928237109 | 7.451345288 | 8.369067528 | 8.43865202 | 8.202562993 | 7.828751338 |
| ENSG00000213079 | SCAF8 | 1.04791842 | 0.018176036 | 0.695769467 | 0.128956336 | 1.743687887 | 7.53297911 | 8.147241016 | 8.486635645 | 9.139384228 | 9.14833612 | 8.811331315 | 8.777523662 |
| ENSG00000126890 | CTAG2 | -0.7277976 | 0.857804601 | 2.470474711 | NA | 1.74267711 | 2 | 2 | 2 | 2.141737246 | 2.2680111 | 3.632645806 | 2.782576693 |
| ENSG00000164087 | POC1A | 0.625574074 | 0.78979414 | 1.1166413 | 0.611923598 | 1.742215374 | 2 | 5.595927442 | 2 | 4.497095146 | 5.04390545 | 5.201624686 | 5.250250853 |
| ENSG00000275594 | | 2.898265153 | 0.334766555 | -1.156717004 | 0.725350511 | 1.741548149 | 2 | 2 | 2 | 3.462030052 | 3.60476711 | 2.139668783 | 2.172608095 |
| ENSG00000185189 | NRBP2 | 0.764531662 | 0.53122754 | 0.976961841 | 0.406566361 | 1.741493503 | 7.27616306 | 7.051611315 | 9.778213818 | 9.316137634 | 9.39662469 | 9.540526122 | 9.595281739 |
| ENSG00000118513 | MYB | 2.332284341 | 0.686502469 | -0.591060112 | NA | 1.741224229 | 2 | 2 | 2 | 3.09461492 | 3.28037825 | 2 | 2.421067459 |
| ENSG00000185619 | PCGF3 | 0.919994104 | 0.064504706 | 0.820482349 | 0.098450928 | 1.740476453 | 9.304185368 | 8.395519306 | 8.434763955 | 9.667723798 | 9.70471389 | 9.614979403 | 9.559117299 |
| ENSG00000100350 | FOXRED2 | 0.609720413 | 0.521312957 | 1.129613943 | 0.195650955 | 1.739334356 | 5.095959458 | 7.35507212 | 6.629488387 | 7.300218887 | 7.16627666 | 7.511762185 | 7.942440336 |
| ENSG00000172845 | SP3 | 0.789041043 | 0.399039732 | 0.949491019 | 0.291402385 | 1.738532062 | 7.27616306 | 9.246456519 | 9.61300511 | 9.818504147 | 9.76823652 | 10.12491852 | 9.758568089 |
| ENSG00000112902 | SEMA5A | -0.124722214 | 0.895675696 | 1.862852672 | 0.012605537 | 1.738130459 | 9.710019321 | 10.00683004 | 11.25418149 | 10.36090904 | 10.3649055 | 11.74494028 | 12.77082042 |
| ENSG00000198561 | CTNND1 | 0.47580392 | 0.621621222 | 1.261915956 | 0.144252411 | 1.737719877 | 8.836089686 | 10.56267266 | 11.19694409 | 10.96377665 | 10.9670671 | 11.63500472 | 11.85709766 |
| ENSG00000075089 | ACTR6 | 1.461293785 | 0.047051348 | 0.276378319 | 0.744221494 | 1.737672104 | 6.563806279 | 6.389896633 | 4.874587708 | 7.574744049 | 7.4949172 | 6.635427934 | 6.092227206 |
| ENSG00000153558 | FBXL2 | 0.230000872 | 0.923810945 | 1.506606659 | 0.475075841 | 1.736607531 | 2 | 7.503546079 | 3.369317972 | 6.18083559 | 6.33555364 | 7.417689057 | 7.562617572 |
| ENSG00000133812 | SBF2 | 0.607126667 | 0.700598482 | 1.129318381 | 0.443373306 | 1.736445049 | 4.00722531 | 8.732943944 | 8.65477198 | 8.786946023 | 8.68895519 | 9.376644377 | 9.126210259 |
| ENSG00000085185 | BCORL1 | 0.146895616 | 0.913792484 | 1.589222878 | 0.154026394 | 1.736118495 | 4.815609471 | 7.979858146 | 8.175061096 | 7.668500375 | 7.75969386 | 9.120448319 | 9.158655025 |
| ENSG00000196757 | ZNF700 | 1.130140196 | 0.478107313 | 0.603010284 | 0.715036782 | 1.733150479 | 2 | 6.899080544 | 6.322058544 | 7.161462517 | 7.19067478 | 6.874161066 | 6.423723772 |
| ENSG00000166086 | JAM3 | 0.597375461 | 0.577257973 | 1.133358851 | 0.25426536 | 1.730734312 | 6.96351253 | 9.170736405 | 9.624627483 | 9.522191343 | 9.57788596 | 9.692174376 | 10.39167686 |
| ENSG00000135679 | MDM2 | 1.52073751 | 5.48E-06 | 0.209918677 | 0.605467095 | 1.730656186 | 9.602033055 | 9.849218119 | 9.849281147 | 11.29399181 | 11.283169 | 10.22682784 | 9.684739879 |
| ENSG00000186017 | ZNF566 | 0.731332716 | 0.398162916 | 0.999300884 | 0.222928155 | 1.730633601 | 4.815609471 | 6.464272742 | 5.772716771 | 6.640968371 | 6.45450701 | 7.121690475 | 6.406535141 |
| ENSG00000164576 | SAP30L | 0.575839456 | 0.194620623 | 1.154539342 | 0.004283896 | 1.730378799 | 8.285626337 | 7.819099893 | 8.007629717 | 8.583021623 | 8.64401667 | 9.419535622 | 8.91098986 |
| ENSG00000123636 | BAZ2B | 1.475147749 | 0.088652437 | 0.250241491 | 0.802048583 | 1.725389239 | 6.138360429 | 8.414402626 | 8.007629717 | 9.219515356 | 9.30925999 | 8.271892681 | 7.79392309 |
| ENSG00000113812 | ACTR8 | 1.148191606 | 0.48962595 | 0.576879331 | 0.737950893 | 1.725070938 | 8.797441463 | 6.844429755 | 3.369317972 | 8.734762645 | 8.66593596 | 8.2308261 | 8.034828728 |
| ENSG00000185344 | ATP6V0A2 | 0.826886042 | 0.568597763 | 0.898042806 | 0.526566702 | 1.724928848 | 5.330594134 | 8.540087354 | 5.39244166 | 8.054188409 | 8.07574085 | 8.261097876 | 7.997049907 |
| ENSG00000122257 | RBBP6 | 1.017467377 | 0.095026872 | 0.70712894 | 0.255721676 | 1.724596316 | 8.517314938 | 8.522788974 | 9.670202709 | 9.970527901 | 10.0678647 | 9.799857373 | 9.61630394 |
| ENSG00000066084 | DIP2B | 1.331288503 | 0.509240304 | 0.391357872 | 0.856534247 | 1.722646375 | 2 | 9.298208073 | 9.06622548 | 9.939922873 | 9.92005068 | 9.009487618 | 8.981364568 |
| ENSG00000184635 | ZNF93 | 0.870236725 | 0.719117881 | 0.851331203 | 0.722362352 | 1.721567928 | 2 | 6.228547981 | 2 | 5.590471342 | 5.60003091 | 5.553513572 | 5.601925227 |
| ENSG00000141867 | BRD4 | 0.543076889 | 0.232704499 | 1.178047086 | 0.004306487 | 1.721123975 | 9.657036304 | 8.955547846 | 8.941672936 | 9.755113959 | 9.77093593 | 10.28440567 | 10.49896112 |
| ENSG00000149679 | CABLES2 | 0.480257466 | 0.831692971 | 1.239458972 | 0.544751469 | 1.719716438 | 3.327711234 | 7.099070191 | 2 | 6.18083559 | 6.06344184 | 6.752490997 | 6.938065226 |
| ENSG00000176788 | BASP1 | 1.052477479 | 0.212500963 | 0.665100054 | 0.443633913 | 1.717577533 | 11.74855118 | 9.664384941 | 10.83618737 | 12.05344541 | 12.0166989 | 11.34538214 | 11.89841387 |
| ENSG00000110013 | SIAE | 1.133515142 | 0.057080264 | 0.583443363 | 0.352922604 | 1.716958505 | 7.218819781 | 7.35507212 | 6.073378139 | 8.113468729 | 8.10170499 | 7.722704302 | 7.392628644 |
| ENSG00000253767 | PCDHGA8 | 4.577231877 | 0.003100969 | -2.861558149 | 0.079386828 | 1.715673729 | 2 | 4.11352228 | 7.098165388 | 10.14797511 | 10.2029317 | 3.483289517 | 3.33635788 |
| ENSG00000029363 | BCLAF1 | 1.550725747 | 0.175563893 | 0.164940305 | 0.902362275 | 1.715666052 | 6.653098118 | 9.969011754 | 8.460932916 | 10.47329248 | 10.4637324 | 8.813506006 | 9.320735104 |
| ENSG00000151917 | BEND6 | 0.262091201 | 0.923796154 | 1.453080171 | 0.550340609 | 1.715171372 | 2 | 6.899080544 | 2 | 5.461211713 | 5.84514933 | 6.756113529 | 6.826173076 |
| ENSG00000198231 | DDX42 | 0.821120167 | 0.185774297 | 0.893580179 | 0.139114355 | 1.714700347 | 8.631580145 | 9.687856435 | 10.0085654 | 10.33807884 | 10.4006406 | 10.31818704 | 10.55593703 |
| ENSG00000151116 | UEVLD | 0.708528612 | 0.60637008 | 1.006029182 | 0.441586799 | 1.714557795 | 4.467311984 | 6.535001748 | 8.266922072 | 7.890605449 | 7.75969386 | 8.192572047 | 8.042267212 |
| ENSG00000066135 | KDM4A | 1.677439278 | 0.000102491 | 0.03562567 | 0.950358432 | 1.713064948 | 7.709880939 | 7.669933291 | 6.80318718 | 9.103012382 | 9.12637045 | 7.510690356 | 7.478755384 |
| ENSG00000136710 | CCDC115 | 0.79133698 | 0.552070286 | 0.921405089 | 0.476444675 | 1.712742069 | 4.00722531 | 7.901717006 | 7.285536013 | 7.954442493 | 7.79989812 | 7.967337618 | 8.045972122 |
| ENSG00000184343 | SRPK3 | -1.303435209 | 0.800325517 | 3.015417766 | 0.505167085 | 1.711982558 | 2 | 2 | 2 | 2.270786764 | 2 | 3.810410249 | 3.383972333 |
| ENSG00000103319 | EEF2K | 0.175771088 | 0.887688028 | 1.535772249 | 0.131017641 | 1.711543337 | 5.095959458 | 7.051611315 | 8.110401118 | 7.398825229 | 7.35080119 | 8.723392661 | 8.703804674 |
| ENSG00000182963 | GJC1 | 0.286610445 | 0.792847991 | 1.423592759 | 0.12513586 | 1.710203203 | 7.435667355 | 7.315436877 | 9.413059136 | 8.709369553 | 8.64989445 | 9.937829229 | 9.668007958 |
| ENSG00000146856 | AGBL3 | 0.187771121 | 0.946524933 | 1.522326214 | 0.540120633 | 1.710097335 | 2 | 2 | 3.369317972 | 2.695456494 | 2.59489879 | 3.584558147 | 2.885414003 |
| ENSG00000116209 | TMEM59 | 1.526189282 | 0.000238144 | 0.182617313 | 0.720364135 | 1.708806595 | 11.08909278 | 10.20420974 | 10.51254821 | 12.19611659 | 12.1473525 | 10.80996641 | 10.85033379 |
| ENSG00000136383 | ALPK3 | -0.903931393 | 0.570747269 | 2.612147445 | 0.063922883 | 1.708216053 | 2 | 5.72254323 | 6.718950348 | 4.867343068 | 5.00774437 | 8.68637457 | 7.690021882 |
| ENSG00000139350 | NEDD1 | 1.154315438 | 0.200056839 | 0.553395215 | 0.560395372 | 1.707710654 | 7.159102577 | 6.464272742 | 8.631919287 | 8.852852814 | 8.82635531 | 8.208524517 | 8.281047142 |
| ENSG00000132466 | ANKRD17 | 0.867297121 | 0.067189131 | 0.83862424 | 0.073994474 | 1.705921361 | 9.460782017 | 10.42046463 | 10.2528023 | 10.95089319 | 10.9846155 | 10.98330608 | 10.89377011 |
| ENSG00000170145 | SIK2 | 0.900979197 | 0.007742105 | 0.804498108 | 0.018178256 | 1.705477306 | 8.493334308 | 8.791896851 | 8.884815469 | 9.643348214 | 9.612912 | 9.694536155 | 9.349127298 |
| ENSG00000263001 | GTF2I | 2.59837292 | 0.017562521 | -0.89350443 | 0.460053783 | 1.70486849 | 6.468621203 | 8.702538795 | 10.01740919 | 11.57566116 | 11.6103915 | 8.018757917 | 8.215115301 |
| ENSG00000128487 | SPECC1 | 1.821668538 | 0.119484573 | -0.11743758 | 0.93349971 | 1.704230958 | 5.330594134 | 7.60566235 | 8.805342542 | 9.578597652 | 9.68122943 | 8.089380147 | 7.199901369 |
| ENSG00000150401 | DCUN1D2 | 0.233216328 | 0.821674232 | 1.470857051 | 0.086540628 | 1.704073379 | 5.330594134 | 7.099070191 | 5.59507254 | 6.535745081 | 6.42738748 | 7.967337618 | 7.327148877 |
| ENSG00000021355 | SERPINB1 | 1.651166782 | 0.001135193 | 0.052097607 | 0.937350796 | 1.703264388 | 8.009475818 | 7.315436877 | 7.691296561 | 9.354865403 | 9.29432703 | 7.304736313 | 8.0807022 |
| ENSG00000131373 | HACL1 | 0.66277102 | 0.436914894 | 1.040214259 | 0.190601024 | 1.702985279 | 7.218819781 | 5.457120953 | 6.073378139 | 7.030886362 | 7.09050444 | 7.218085825 | 7.612190665 |
| ENSG00000091542 | ALKBH5 | 0.489876863 | 0.329903713 | 1.212363833 | 0.007109557 | 1.702240695 | 10.25038534 | 9.357954881 | 9.926432745 | 10.27745223 | 10.4678974 | 10.91162437 | 11.26054287 |
| ENSG00000087586 | AURKA | 1.186019722 | 0.128090043 | 0.514982637 | 0.534560654 | 1.701002359 | 4.00722531 | 5.303523747 | 5.772716771 | 6.24507773 | 6.34282908 | 5.618636935 | 5.713290392 |
| ENSG00000112242 | E2F3 | 1.274518199 | 0.419141353 | 0.425626669 | 0.801603223 | 1.700144868 | 3.327711234 | 7.73146271 | 8.65477198 | 8.967964567 | 8.95536429 | 8.131128273 | 8.112850643 |
| ENSG00000160072 | ATAD3B | 1.239739037 | 0.110962 | 0.459544204 | 0.584645012 | 1.699283241 | 7.624139768 | 7.232742069 | 5.59507254 | 8.197043419 | 8.32935768 | 7.474865722 | 7.520630205 |
| ENSG00000166478 | ZNF143 | 0.991173746 | 0.472579685 | 0.706764051 | 0.611923598 | 1.697937797 | 7.53297911 | 7.099070191 | 3.369317972 | 7.77569021 | 7.74878077 | 7.621085148 | 7.331709059 |
| ENSG00000135047 | CTSL | 2.930331982 | 0.003161093 | -1.232557583 | 0.255070887 | 1.697774399 | 11.72557631 | 9.34816733 | 9.372276026 | 13.5295502 | 13.5511473 | 9.730427609 | 8.931664222 |
| ENSG00000196453 | ZNF777 | 0.755849822 | 0.706288309 | 0.940490892 | 0.628768646 | 1.696340714 | 8.444143041 | 6.311476754 | 2 | 7.855024645 | 7.95749494 | 8.157391075 | 8.017007082 |

| | | | | | | | | | | | | | |
|---|---|---|---|---|---|---|---|---|---|---|---|---|---|
| ENSG00000134369 | NAV1 | 0.231700443 | 0.626623393 | 1.464370629 | 0.000174595 | 1.696071072 | 11.12074405 | 11.49575678 | 10.91875185 | 11.42521186 | 11.4368305 | 12.34622686 | 12.92177157 |
| ENSG00000159840 | ZYX | -0.065875158 | 0.927489437 | 1.758259788 | 0.001514467 | 1.69238463 | 11.75864596 | 10.49332014 | 11.02342937 | 11.08829583 | 11.1492725 | 12.69425333 | 13.15230292 |
| ENSG00000073921 | PICALM | 0.860836774 | 0.306865781 | 0.831491923 | 0.315344239 | 1.692328697 | 9.174465797 | 10.30378101 | 11.26536143 | 11.37226044 | 11.3168068 | 11.37902969 | 11.24903735 |
| ENSG00000153879 | CEBPG | 0.262254474 | 0.838100556 | 1.429815407 | 0.188337529 | 1.692069881 | 6.138360429 | 9.00621767 | 7.646393253 | 8.300810597 | 8.29225867 | 9.910457803 | 8.781973674 |
| ENSG00000175550 | DRAP1 | 1.159436679 | 0.030199232 | 0.53249983 | 0.353010925 | 1.69193651 | 8.737455672 | 8.929529568 | 9.819251342 | 10.39227626 | 10.3962764 | 9.639212351 | 9.889343672 |
| ENSG00000174516 | PELI3 | 0.258201495 | 0.91292494 | 1.433555793 | 0.488684721 | 1.691757288 | 2 | 8.169643011 | 4.523675391 | 7.075737784 | 6.86146731 | 8.362013331 | 7.834118096 |
| ENSG00000068885 | IFT80 | 1.782638111 | 0.417782282 | -0.094013459 | 0.96989602 | 1.688624652 | 3.327711234 | 7.847167909 | 2 | 8.033874364 | 8.14604733 | 6.028179536 | 6.476841503 |
| ENSG00000197045 | GMFB | 1.451048163 | 0.12992063 | 0.236428046 | 0.831469687 | 1.687476208 | 8.107517749 | 8.791896851 | 6.073378139 | 9.528658349 | 9.42518018 | 8.245074976 | 8.302113262 |
| ENSG00000164048 | ZNF589 | 0.839924885 | 0.712294566 | 0.847332407 | 0.705796484 | 1.687257292 | 2 | 7.467835385 | 3.369317972 | 6.841288255 | 6.77805036 | 7.194250299 | 6.30506411 |
| ENSG00000153044 | CENPH | 0.581457228 | 0.710155984 | 1.103808891 | 0.44445996 | 1.68526612 | 4.467311984 | 3.686303414 | 2 | 4.415698455 | 3.7864959 | 4.720468739 | 4.365060851 |
| ENSG00000012211 | PRICKLE3 | 0.960214344 | 0.431243113 | 0.72230079 | 0.554788482 | 1.682515134 | 4.467311984 | 7.35507212 | 5.39244166 | 7.101994243 | 7.28055563 | 6.763331418 | 7.138761401 |
| ENSG00000198865 | CCDC152 | 3.218783623 | 0.310450907 | -1.537717382 | 0.65106688 | 1.681066242 | 3.327711234 | 2 | 2 | 4.387512679 | 4.58534546 | 2.42098416 | 2 |
| ENSG00000196455 | PIK3R4 | 0.931343823 | 0.30261211 | 0.749277178 | 0.407381942 | 1.680621001 | 7.667647223 | 8.414402626 | 6.073378139 | 8.645348336 | 8.55284928 | 8.348279032 | 8.487906181 |
| ENSG00000179051 | RCC2 | 1.086860776 | 0.033252373 | 0.59260517 | 0.270018177 | 1.679465947 | 9.590776211 | 8.57407404 | 9.589475645 | 10.34633939 | 10.4541068 | 9.908628006 | 9.908803154 |
| ENSG00000124275 | MTRR | 1.288641417 | 0.400196775 | 0.390818115 | 0.81390919 | 1.679459532 | 3.327711234 | 8.414402626 | 7.451345288 | 8.744518279 | 8.71301725 | 7.854430723 | 7.830900438 |
| ENSG00000131374 | TBC1D5 | 0.52782196 | 0.596606509 | 1.151427229 | 0.205709574 | 1.67929419 | 8.493334308 | 9.030900487 | 6.534109959 | 8.80304041 | 8.94105405 | 9.446970496 | 9.537567589 |
| ENSG00000188070 | C11orf95 | 0.340530758 | 0.840490657 | 1.338438289 | 0.365630894 | 1.678969047 | 4.00722531 | 8.993716073 | 8.206335842 | 8.459845423 | 8.41122995 | 9.731578082 | 9.035944835 |
| ENSG00000130818 | ZNF426 | 0.908912266 | 0.537027555 | 0.769789892 | 0.601793533 | 1.678702158 | 5.70932728 | 8.717821467 | 5.156626401 | 8.341783829 | 8.27714776 | 8.313667445 | 8.016062975 |
| ENSG00000179632 | MAF1 | 0.536197354 | 0.33517337 | 1.142108837 | 0.024149698 | 1.678306192 | 8.367060731 | 9.34816733 | 9.465702942 | 9.683748348 | 9.65881449 | 10.29968974 | 10.24949979 |
| ENSG00000123064 | DDX54 | 0.823283121 | 0.384414708 | 0.853640492 | 0.35551459 | 1.676923613 | 6.138360429 | 8.505280662 | 8.143093325 | 8.720710586 | 8.71442024 | 8.530120421 | 8.936662123 |
| ENSG00000112977 | DAP | 0.192643861 | 0.857804601 | 1.484197708 | 0.093533047 | 1.676841569 | 11.30078111 | 10.71132792 | 8.845626171 | 10.7521165 | 10.8336713 | 11.87479202 | 12.26508948 |
| ENSG00000182919 | C11orf54 | 0.128922828 | 0.863599094 | 1.547421878 | 0.00811335 | 1.676344706 | 6.816637977 | 6.464272742 | 5.39244166 | 6.372775472 | 6.56439512 | 7.888662695 | 7.808173621 |
| ENSG00000140749 | IGSF6 | -3.753079805 | 0.413653291 | 5.42867747 | 0.166693658 | 1.675597666 | 2 | 2 | 2 | 2 | 2 | 6.126613714 | 4.692661645 |
| ENSG00000014123 | UFL1 | 1.03541828 | 0.593517792 | 0.638220399 | 0.747455247 | 1.673638679 | 2 | 7.874700259 | 8.677268317 | 8.839910123 | 8.71020716 | 8.614358955 | 8.108428191 |
| ENSG00000181938 | GINS3 | 1.928720422 | 0.176419446 | -0.256147299 | 0.877983632 | 1.672573123 | 4.00722531 | 5.838937808 | 2 | 6.522845756 | 6.47451726 | 4.504874511 | 4.543391831 |
| ENSG00000161509 | GRIN2C | -2.312458114 | 0.577663983 | 3.983867454 | 0.243272299 | 1.67140934 | 2 | 2 | 2 | 2 | 2.14022021 | 4.425202323 | 4.228274364 |
| ENSG00000271092 | TMEM56-RW | 4.076651028 | 0.025536658 | -2.405995245 | 0.247980941 | 1.670655783 | 2 | 2 | 3.369317972 | 5.273747807 | 5.18011485 | 2.139668783 | 2.117335275 |
| ENSG00000163939 | PBRM1 | 0.813091611 | 0.559250209 | 0.854973898 | 0.531253192 | 1.668065509 | 5.095959458 | 8.955547846 | 9.25752426 | 9.381011913 | 9.37992356 | 9.323921045 | 9.514020196 |
| ENSG00000087460 | GNAS | 0.66147735 | 0.177667207 | 1.006392488 | 0.029583545 | 1.667869839 | 12.19573173 | 12.50249943 | 11.55983287 | 12.79322888 | 12.8042115 | 12.97462236 | 13.29462079 |
| ENSG00000160305 | DIP2A | 0.42244107 | 0.620033622 | 1.244196597 | 0.099323089 | 1.666637668 | 8.418903792 | 7.761270542 | 6.43197674 | 8.213192703 | 8.1187584 | 9.078747095 | 8.873429675 |
| ENSG00000110697 | PITPNM1 | 0.106952757 | 0.910881185 | 1.559624581 | 0.038741655 | 1.666577338 | 7.624139768 | 8.029695761 | 9.399592459 | 8.636393836 | 8.69180693 | 9.911270301 | 10.28108376 |
| ENSG00000047410 | TPR | 1.279819395 | 0.089945888 | 0.385784417 | 0.643465101 | 1.665603812 | 9.556468883 | 10.43902632 | 8.536706852 | 10.96556523 | 11.0116859 | 10.20827046 | 9.976858 |
| ENSG00000162702 | ZNF281 | 1.316406222 | 0.002491297 | 0.347520258 | 0.483213396 | 1.663926481 | 9.233563228 | 8.639735838 | 9.014156722 | 10.27601154 | 10.3014409 | 9.613232802 | 8.962375727 |
| ENSG00000103264 | FBXO31 | 0.344236409 | 0.681389915 | 1.319514954 | 0.06883389 | 1.663751363 | 8.493334308 | 8.702538795 | 6.882775709 | 8.483340231 | 8.64548836 | 9.611983935 | 9.45254121 |
| ENSG00000106636 | YKT6 | 0.518346363 | 0.304613133 | 1.145253811 | 0.0121807 | 1.663600174 | 9.497412313 | 9.415325457 | 10.26770388 | 10.29747219 | 10.290665 | 10.79815488 | 11.0311848 |
| ENSG00000169554 | ZEB2 | 1.001810441 | 0.29623641 | 0.659890321 | 0.499253973 | 1.661700762 | 8.75772933 | 10.76748534 | 8.763901704 | 10.75073417 | 10.7824203 | 10.46416585 | 10.3860243 |
| ENSG00000105497 | ZNF175 | -4.714961636 | 0.0591539 | 6.376095851 | 0.000147017 | 1.661134214 | 2 | 3.076665209 | 2 | 2 | 2 | 7.358152527 | 7.438450313 |
| ENSG00000014914 | MTMR11 | 1.003446234 | 0.60852038 | 0.657066801 | 0.740394508 | 1.660513035 | 2 | 6.140560021 | 8.296287027 | 7.954442493 | 8.0358969 | 7.673747216 | 7.636966981 |
| ENSG00000176182 | MYPOP | 0.682971386 | 0.590107068 | 0.977329206 | 0.418666323 | 1.660300592 | 5.70932728 | 5.595927442 | 8.266922072 | 7.781117077 | 7.7127321 | 8.116406856 | 7.952344882 |
| ENSG00000177548 | RABEP2 | 1.252688487 | 0.386166537 | 0.407599832 | 0.794186359 | 1.660288319 | 5.70932728 | 8.077869079 | 4.058998049 | 8.03841338 | 8.07792246 | 7.27600668 | 7.18317631 |
| ENSG00000122042 | UBL3 | 1.072414376 | 0.539282524 | 0.587770778 | 0.74490284 | 1.660185153 | 3.327711234 | 8.414402626 | 9.516493218 | 9.5407064 | 9.57942664 | 9.132692373 | 9.022374596 |
| ENSG00000114544 | SLC41A3 | 0.258391167 | 0.60106289 | 1.401704186 | 0.000584048 | 1.660095354 | 7.904282243 | 7.761270542 | 7.285536013 | 7.898117066 | 7.96458672 | 8.841056341 | 9.253832157 |
| ENSG00000198837 | DENND4B | 0.599957924 | 0.652929267 | 1.058637257 | 0.396684362 | 1.65859518 | 4.815609471 | 8.623598428 | 7.934808822 | 8.365459376 | 8.41985547 | 8.960480054 | 8.723602991 |
| ENSG00000180332 | KCTD4 | 0.727687405 | 0.786927084 | 0.928640424 | 0.718605505 | 1.65632783 | 2 | 3.076665209 | 2 | 2.868578001 | 2.86123014 | 2.803598391 | 3.100202476 |
| ENSG00000153989 | NUS1 | 1.613860488 | 0.008858946 | 0.041541349 | 0.958081539 | 1.655401837 | 7.624139768 | 7.979858146 | 9.049077186 | 10.00919444 | 9.88434767 | 8.363201462 | 8.410985412 |
| ENSG00000113356 | POLR3G | 0.283033323 | 0.856469262 | 1.372315582 | 0.303728756 | 1.655348905 | 4.00722531 | 5.131603625 | 2 | 4.299492648 | 4.65713986 | 5.654016955 | 5.177806074 |
| ENSG00000163110 | PDLIM5 | -0.400975475 | 0.770883416 | 2.055769801 | 0.080413293 | 1.654794327 | 7.750913381 | 11.19631665 | 11.58682058 | 10.49161431 | 10.4603916 | 12.98609149 | 12.87062285 |
| ENSG00000170364 | SETMAR | -0.048210558 | 0.966538801 | 1.700898449 | 0.066536768 | 1.652687891 | 4.467311984 | 6.04685525 | 4.058998049 | 5.058234798 | 5.21224859 | 6.68757835 | 6.841176624 |
| ENSG00000197619 | ZNF615 | 0.930605078 | 0.521753772 | 0.72181086 | 0.619812492 | 1.652415937 | 4.00722531 | 7.35507212 | 4.874587708 | 7.157295126 | 6.90635305 | 7.182183216 | 6.37742435 |
| ENSG00000066279 | ASPM | 1.050047958 | 0.563707037 | 0.602323105 | 0.747588951 | 1.652371062 | 3.327711234 | 5.595927442 | 2 | 5.363304456 | 5.30456641 | 4.209638631 | 5.419744333 |
| ENSG00000145050 | MANF | 1.210561686 | 0.244396401 | 0.439208478 | 0.695304679 | 1.649770164 | 9.435834703 | 6.951736442 | 7.691296561 | 9.653739712 | 9.5678312 | 8.416823986 | 9.175242287 |
| ENSG00000082996 | RNF13 | 1.523204258 | 0.406678307 | 0.126397912 | 0.951751695 | 1.64960217 | 7.975256505 | 8.540087354 | 2 | 9.255049401 | 9.17914662 | 7.962642137 | 7.698285408 |
| ENSG00000128335 | APOL2 | 1.606955258 | 0.025425486 | 0.041102498 | 0.964424173 | 1.648057757 | 8.836089686 | 9.591588447 | 10.60345421 | 11.44136838 | 11.4780933 | 9.638721848 | 10.11774074 |
| ENSG00000198056 | PRIM1 | 1.874424026 | 0.40298478 | -0.226507331 | 0.929596218 | 1.647916696 | 2 | 3.686303414 | 2 | 3.95996233 | 4.0930011 | 3.035909414 | 2.466008509 |
| ENSG00000248672 | LY75-CD302 | 6.837015545 | 0.005498029 | -5.191141267 | 0.09315428 | 1.645874278 | 2 | 2 | 2 | 6.984595707 | 6.83591385 | 2 | 2 |
| ENSG00000155008 | APOOL | 0.086827905 | 0.942525992 | 1.558541114 | 0.117467726 | 1.645369019 | 5.532353208 | 7.051611315 | 8.35328246 | 7.53353831 | 7.42122542 | 9.092423696 | 8.745413684 |
| ENSG00000164073 | MFSD8 | 1.45080403 | 0.26540262 | 0.194542859 | 0.896638566 | 1.645346889 | 7.435667355 | 7.099070191 | 3.369317972 | 8.258639605 | 8.05595642 | 6.981729486 | 6.881131115 |
| ENSG00000056277 | ZNF280C | 1.279188332 | 0.123978923 | 0.366003025 | 0.690278517 | 1.645191358 | 4.00722531 | 5.131603625 | 5.39244166 | 6.29903474 | 5.93454154 | 5.47655442 | 5.065479856 |
| ENSG00000106244 | PDAP1 | 0.48647754 | 0.489850205 | 1.158685065 | 0.067503656 | 1.645162605 | 9.579430841 | 8.590771778 | 8.175061096 | 9.376537567 | 9.40011621 | 9.862742364 | 10.22886814 |
| ENSG00000090989 | EXOC1 | 1.401182696 | 0.001537778 | 0.242985399 | 0.642930217 | 1.644168095 | 8.138771685 | 7.572420273 | 7.285536013 | 9.00785664 | 9.17201613 | 8.069864557 | 7.81253023 |
| ENSG00000174996 | KLC2 | 0.765435081 | 0.292274211 | 0.876682379 | 0.211608759 | 1.64211746 | 9.33149386 | 8.433041974 | 7.646393253 | 9.391694063 | 9.39049411 | 9.539738196 | 9.46262491 |
| ENSG00000188313 | PLSCR1 | 3.552397159 | 0.002181398 | -1.913169018 | 0.120834165 | 1.639228141 | 9.435834703 | 5.946638857 | 6.958202216 | 11.7198617 | 11.7308771 | 6.433059984 | 6.222348126 |
| ENSG00000165175 | MID1IP1 | 0.072099296 | 0.935263493 | 1.566971447 | 0.02674158 | 1.639070742 | 6.96351253 | 7.874700259 | 8.721235202 | 8.076429088 | 8.11025689 | 9.811219795 | 9.291382799 |
| ENSG00000135945 | REV1 | 0.74852836 | 0.632487313 | 0.890002794 | 0.557451959 | 1.638531155 | 4.00722531 | 8.376385542 | 6.073378139 | 7.772969103 | 7.93600799 | 7.934143724 | 8.055192982 |
| ENSG00000126970 | ZC4H2 | 0.531303922 | 0.714622915 | 1.107225595 | 0.410036586 | 1.638529518 | 5.330594134 | 7.572420273 | 4.058998049 | 7.030886362 | 6.74548212 | 7.302260672 | 7.593325136 |
| ENSG00000138080 | EMILIN1 | 1.307326783 | 0.021422337 | 0.329362623 | 0.61033641 | 1.636689406 | 11.21173326 | 10.20420974 | 10.0085654 | 11.86363439 | 11.8983971 | 10.96297653 | 10.84383043 |
| ENSG00000116062 | MSH6 | 0.090956424 | 0.921327192 | 1.544274286 | 0.035456159 | 1.63523071 | 7.096806373 | 8.876041558 | 8.922968322 | 8.540455898 | 8.65720817 | 10.10054783 | 9.984841938 |
| ENSG00000163428 | LRRC58 | 0.600075182 | 0.628451704 | 1.034973928 | 0.372265366 | 1.63504911 | 6.138360429 | 9.540933215 | 8.486635645 | 9.236404433 | 9.18523054 | 9.834322366 | 9.423285228 |
| ENSG00000083520 | DIS3 | 0.921912354 | 0.36126721 | 0.712269046 | 0.483045217 | 1.6341814 | 6.366709671 | 9.055168103 | 8.608698767 | 9.366644887 | 9.24080977 | 9.174915459 | 9.014359877 |
| ENSG00000125356 | NDUFA1 | 0.959406731 | 0.196836305 | 0.673670303 | 0.372076473 | 1.633077034 | 9.097018025 | 7.572420273 | 7.734844356 | 9.204447245 | 9.30367823 | 8.862639958 | 9.07227736 |
| ENSG00000153140 | CETN3 | 1.270834451 | 0.354199679 | 0.362180615 | 0.808803514 | 1.633015066 | 6.009025997 | 5.72254323 | 2 | 6.628980152 | 6.43421531 | 5.948096914 | 5.349527629 |
| ENSG00000113407 | TARS | 1.224644322 | 0.00192427 | 0.408114017 | 0.353971254 | 1.632758339 | 9.966689162 | 9.558016677 | 9.165043284 | 10.82354685 | 10.8180948 | 10.10214984 | 9.904473108 |
| ENSG00000119979 | FAM45A | 2.078249598 | 0.230676825 | -0.445502583 | 0.819020478 | 1.632747015 | 2 | 7.232742069 | 8.296287027 | 9.394352287 | 9.27639101 | 6.814608286 | 6.89352185 |
| ENSG00000159905 | ZNF221 | 0.897070195 | 0.34242241 | 0.73424395 | 0.435748117 | 1.631314146 | 4.00722531 | 5.72254323 | 4.874587708 | 5.920568914 | 5.85535975 | 6.004008792 | 5.408276315 |
| ENSG00000127527 | EPS15L1 | 0.713174283 | 0.549476534 | 0.917114158 | 0.424715914 | 1.630288441 | 8.468948322 | 7.393647515 | 5.156626401 | 8.276040626 | 8.20512425 | 8.360229299 | 8.520227711 |
| ENSG00000118007 | STAG1 | 1.368359545 | 0.425434611 | 0.259833587 | 0.891849925 | 1.628193132 | 3.327711234 | 8.717821467 | 6.073378139 | 8.726347845 | 8.73530306 | 7.49992804 | 7.763868281 |
| ENSG00000122707 | RECK | 0.798318628 | 0.162633024 | 0.82970276 | 0.137078393 | 1.628021388 | 8.777722034 | 9.225224669 | 9.553445883 | 9.989990706 | 10.0367996 | 9.514829317 | 10.43169893 |
| ENSG00000077458 | FAM76B | 1.271183012 | 0.291215116 | 0.356267888 | 0.787590496 | 1.6274509 | 3.327711234 | 6.535001748 | 5.772716771 | 6.851641821 | 7.02417148 | 6.417131325 | 5.601925227 |
| ENSG00000184863 | RBM33 | 0.804947935 | 0.293290194 | 0.822348746 | 0.271724882 | 1.627296681 | 9.423197387 | 7.761270542 | 8.175061096 | 9.443949663 | 9.41572435 | 9.443604635 | 9.450795495 |
| ENSG00000059758 | CDK17 | 1.863858309 | 0.287529501 | -0.237163828 | 0.906201712 | 1.626694481 | 3.327711234 | 7.669933291 | 9.757248847 | 10.32001706 | 10.3626704 | 8.436147258 | 8.047821014 |
| ENSG00000151623 | NR3C2 | 2.237766043 | 0.21912951 | -0.611497525 | 0.762269886 | 1.626268518 | 2 | 4.93638933 | 2 | 5.731694 | 5.49945122 | 3.429858444 | 3.287118176 |
| ENSG00000141968 | VAV1 | -0.727833888 | 0.863750456 | 2.3533402 | NA | 1.625506312 | 2 | 2 | 2 | 2.141737246 | 2.2680111 | 3.648325314 | 2.551896972 |
| ENSG00000145002 | FAM86B2 | 3.643594916 | 0.392859546 | -2.018322498 | 0.658188823 | 1.625272418 | 2 | 2 | 2 | 4.070463721 | 4.05806467 | 2 | 2.172608095 |
| ENSG00000103248 | MTHFSD | 0.679015328 | 0.7690856 | 0.945471571 | 0.673457912 | 1.624486899 | 3.327711234 | 7.60566235 | 2 | 6.722207358 | 6.84106088 | 7.134272047 | 6.940057611 |
| ENSG00000120256 | LRP11 | 2.290879515 | 5.30E-06 | -0.666830253 | 0.243487701 | 1.624049262 | 6.89194353 | 6.228547981 | 7.342949392 | 9.119171929 | 9.12531607 | 6.387074688 | 6.059610627 |
| ENSG00000103769 | RAB11A | 0.813024953 | 0.171886783 | 0.809866696 | 0.16559668 | 1.622891649 | 10.13887448 | 9.608085847 | 8.805342542 | 10.45261097 | 10.4010763 | 10.34909821 | 10.49507022 |
| ENSG00000155506 | LARP1 | 0.584608584 | 0.167456107 | 1.037951569 | 0.00811335 | 1.622560153 | 10.39976609 | 11.19358635 | 10.93284392 | 11.42564506 | 11.5004268 | 11.8938265 | 11.93845516 |
| ENSG00000182117 | NOP10 | 0.460003357 | 0.705592343 | 1.162494387 | 0.291093097 | 1.622497744 | 10.36714848 | 7.790474954 | 8.007629717 | 9.690252889 | 9.67835667 | 10.25510417 | 10.50250454 |
| ENSG00000024048 | UBR2 | 1.11325866 | 0.211811753 | 0.508393903 | 0.590415354 | 1.621652562 | 6.89194353 | 9.125613817 | 7.896971941 | 9.309580714 | 9.41658653 | 8.890924823 | 8.624257514 |
| ENSG00000184368 | MAP7D2 | 0.312651914 | 0.912931516 | 1.307560401 | 0.605429695 | 1.620212314 | 2 | 3.076665209 | 2 | 2.695456494 | 2.68924543 | 2.885263063 | 3.383972333 |
| ENSG00000177885 | GRB2 | 0.483849626 | 0.223040801 | 1.135222266 | 0.001455406 | 1.619071892 | 9.509418609 | 10.18234215 | 10.04361983 | 10.40726916 | 10.440607 | 11.00668325 | 11.13810331 |
| ENSG00000066230 | SLC9A3 | 0.65994633 | 0.638989421 | 0.958903786 | 0.472427727 | 1.618850116 | 5.330594134 | 4.11352228 | 2 | 4.783303513 | 5.0259382 | 5.233158834 | 5.108633494 |
| ENSG00000168591 | TMUB2 | 0.635565825 | 0.685365166 | 0.982613336 | 0.507844439 | 1.618179162 | 9.371511179 | 7.979858146 | 4.523675391 | 8.828161587 | 9.00091592 | 9.341208071 | 9.177355133 |
| ENSG00000127022 | CANX | 1.19703719 | 0.091109722 | 0.419435358 | 0.586489016 | 1.616472548 | 11.08508709 | 12.53629536 | 12.65268348 | 13.43492421 | 13.4459705 | 12.35521156 | 12.91691141 |
| ENSG00000166130 | IKBIP | 1.104195523 | 0.000183037 | 0.512156221 | 0.10784488 | 1.616351744 | 9.247966931 | 9.114109157 | 8.763901704 | 10.19294689 | 10.1196108 | 9.581426916 | 9.552946519 |
| ENSG00000149131 | SERPING1 | 0.79024395 | 0.58832515 | 0.825797951 | 0.56390268 | 1.616041901 | 10.08373159 | 10.12043894 | 5.772716771 | 10.28987826 | 10.3897054 | 10.52368776 | 10.211653 |
| ENSG00000187944 | C2orf66 | 4.484493801 | 0.16845319 | -2.869577966 | 0.429144636 | 1.614915836 | 2 | 2 | 2 | 4.846787657 | 4.63360314 | 2.094598369 | 2 |
| ENSG00000129521 | EGLN3 | -2.272307319 | 0.358886773 | 3.887075281 | 0.044173979 | 1.614767962 | 2 | 3.076665209 | 2 | 2.141737246 | 2.14022021 | 5.180212365 | 4.927687581 |
| ENSG00000128294 | TPST2 | 0.405448059 | 0.825869978 | 1.208008421 | 0.465968326 | 1.61345648 | 3.327711234 | 7.954278496 | 9.180874355 | 8.551612617 | 8.50013797 | 9.124657997 | 9.49551136 |
| ENSG00000181045 | SLC26A11 | 1.154385238 | 0.311127936 | 0.458184811 | 0.706713381 | 1.612570049 | 4.815609471 | 7.790474954 | 8.042706615 | 8.550024073 | 8.56222967 | 7.957931324 | 7.775072826 |
| ENSG00000078687 | TNRC6C | 0.334909021 | 0.876735407 | 1.277429889 | 0.501916413 | 1.61233891 | 5.095959458 | 7.874700259 | 2 | 6.815074525 | 6.86652414 | 7.88535852 | 7.620910859 |
| ENSG00000173726 | TOMM20 | 0.546546066 | 0.571724149 | 1.065753203 | 0.233224134 | 1.612299269 | 10.43166252 | 9.814097497 | 8.007629717 | 10.28606625 | 10.2535095 | 10.77166745 | 10.80349856 |
| ENSG00000135245 | HILPDA | 0.605664469 | 0.633945061 | 1.00573159 | 0.400315488 | 1.611396059 | 4.467311984 | 7.979858146 | 7.451345288 | 7.849869274 | 7.80253898 | 8.071319244 | 8.355338318 |
| ENSG00000183808 | RBM12B | 0.637575315 | 0.727420094 | 0.973184423 | 0.576686766 | 1.610759737 | 7.485143553 | 7.60566235 | 2 | 7.659688863 | 7.54308677 | 8.234726135 | 7.548321824 |
| ENSG00000160613 | PCSK7 | 0.509564468 | 0.647667448 | 1.099565519 | 0.282627158 | 1.609129987 | 6.96351253 | 7.572420273 | 9.478569125 | 8.890998469 | 8.95298909 | 9.544459308 | 9.471261544 |
| ENSG00000151892 | GFRA1 | -0.909223561 | 0.639879212 | 2.517321104 | 0.146527599 | 1.608097543 | 4.00722531 | 3.686303414 | 8.35328246 | 5.987823101 | 6.04571775 | 8.737205015 | 9.794249791 |
| ENSG00000143382 | ADAMTSL4 | -0.588907582 | 0.482526112 | 2.195689955 | 0.002195337 | 1.606782373 | 6.89194353 | 8.414402626 | 8.408111533 | 7.450827758 | 7.51437845 | 9.664732333 | 10.64977092 |

| | | | | | | | | | | | | | |
|---|---|---|---|---|---|---|---|---|---|---|---|---|---|
| ENSG00000168538 | TRAPPC11 | 0.97372773 | 0.549783955 | 0.632783393 | 0.704091061 | 1.606511123 | 3.327711234 | 8.077869079 | 8.742726175 | 8.875861021 | 8.81462916 | 8.598808264 | 8.410985412 |
| ENSG00000204104 | TRAF3IP1 | 1.283137273 | 0.090973094 | 0.321597152 | 0.705349408 | 1.604734425 | 7.790810999 | 6.464272742 | 6.073378139 | 8.201097719 | 8.24644267 | 7.283556699 | 7.277588383 |
| ENSG00000110048 | OSBP | 0.538649209 | 0.510049398 | 1.065961154 | 0.154093392 | 1.604610363 | 8.716893051 | 8.522788974 | 10.14394518 | 9.832293064 | 9.87684923 | 10.4654111 | 10.28950788 |
| ENSG00000081665 | ZNF506 | 0.980130288 | 0.083005001 | 0.623601875 | 0.282259948 | 1.603732163 | 5.866944937 | 5.946638857 | 6.629488387 | 7.165617905 | 7.06433609 | 6.932953913 | 6.587749895 |
| ENSG00000168813 | ZNF507 | 0.445453963 | 0.825681195 | 1.157777407 | 0.528988964 | 1.60323137 | 8.468948322 | 7.274681896 | 2 | 7.932663244 | 7.77322015 | 8.692059118 | 8.409548182 |
| ENSG00000063015 | SEZ6 | 0.318951311 | 0.925027909 | 1.282946641 | 0.676479319 | 1.601897952 | 2 | 3.076665209 | 2 | 3.09461492 | 2.14022021 | 2.885263063 | 3.36036153 |
| ENSG00000176055 | MBLAC2 | 0.46497279 | 0.767183028 | 1.133719986 | 0.430383523 | 1.598692775 | 2 | 5.303523747 | 6.203069917 | 5.697663076 | 5.71663161 | 6.615624322 | 5.984375752 |
| ENSG00000147145 | LPAR4 | 0.051036888 | 0.985675142 | 1.545605011 | 0.538360258 | 1.5966419 | 2 | 3.076665209 | 2 | 2.695456494 | 2.49394783 | 2.962551659 | 3.560117069 |
| ENSG00000163257 | DCAF16 | 0.5975618 | 0.270310495 | 0.996454348 | 0.048281896 | 1.594016147 | 7.53297911 | 7.819099893 | 8.511888463 | 8.657201865 | 8.54024645 | 9.157900392 | 8.81165749 |
| ENSG00000159069 | FBXW5 | 0.359202117 | 0.265297898 | 1.232024834 | 9.85E-06 | 1.591226952 | 9.90440318 | 9.479495904 | 9.577565424 | 9.966365455 | 10.0727955 | 10.86939366 | 10.91202892 |
| ENSG00000175197 | DDIT3 | 2.025519724 | 0.03133681 | -0.434883932 | 0.687792014 | 1.590635792 | 7.27616306 | 7.503546079 | 8.721235202 | 10.01035013 | 9.96317668 | 8.224302521 | 6.251520369 |
| ENSG00000132388 | UBE2G1 | 0.655842194 | 0.480535674 | 0.934026411 | 0.288509468 | 1.589868605 | 8.716893051 | 9.203675681 | 6.882775709 | 9.253098082 | 9.16587611 | 9.475267633 | 9.49788026 |
| ENSG00000174326 | SLC16A11 | -1.303393354 | 0.803778942 | 2.891486596 | NA | 1.588093242 | 2 | 2 | 2 | 2.270786764 | 2 | 3.663836244 | 3.383972333 |
| ENSG00000138081 | FBXO11 | 1.60413458 | 0.023977132 | -0.01696545 | 0.984890159 | 1.58716913 | 8.340423424 | 8.395519306 | 6.629488387 | 9.591782342 | 9.56705484 | 8.029274808 | 7.916365698 |
| ENSG00000011451 | WIZ | 0.21696598 | 0.876259116 | 1.370023574 | 0.242288367 | 1.586989553 | 10.09173937 | 8.671479029 | 6.534109959 | 9.226493643 | 9.30554121 | 10.35074632 | 10.47545674 |
| ENSG00000005436 | GCFC2 | 0.32914893 | 0.590895562 | 1.255710828 | 0.016427091 | 1.584859758 | 5.532353208 | 6.389896633 | 6.073378139 | 6.429169081 | 6.31350454 | 7.204892513 | 7.324100729 |
| ENSG00000136816 | TOR1B | 0.303325723 | 0.743829336 | 1.27893809 | 0.11156057 | 1.582263813 | 9.646202137 | 8.029695761 | 7.858115942 | 9.060130342 | 9.04395769 | 10.08496939 | 9.956825691 |
| ENSG00000168496 | FEN1 | 0.157505675 | 0.879872883 | 1.422331745 | 0.091732358 | 1.579837421 | 4.815609471 | 6.78762695 | 6.629488387 | 6.422239259 | 6.51372389 | 7.265878131 | 8.025476363 |
| ENSG00000224383 | PRR29 | 0.045282321 | 0.988285735 | 1.533819392 | 0.562631262 | 1.579101714 | 2 | 3.076665209 | 2 | 2.270786764 | 2.86123014 | 3.256555576 | 3.311948089 |
| ENSG00000102753 | KPNA3 | 1.031479043 | 0.396224049 | 0.547518559 | 0.666185912 | 1.578997602 | 8.631580145 | 9.34816733 | 5.772716771 | 9.58946437 | 9.50841342 | 9.168813127 | 8.960904642 |
| ENSG00000139626 | ITGB7 | 2.395407291 | 0.520119153 | -0.817698537 | NA | 1.577708754 | 2 | 2 | 2 | 3.709785322 | 2.49394783 | 2.267001569 | 2.117335275 |
| ENSG00000171163 | ZNF692 | 0.078437621 | 0.94475044 | 1.498258008 | 0.107289325 | 1.576695628 | 6.563806279 | 6.311476754 | 4.058998049 | 6.155987605 | 5.99120031 | 7.700314972 | 7.124815549 |
| ENSG00000177663 | IL17RA | 1.12509812 | 0.005668424 | 0.451099935 | 0.308943128 | 1.576198055 | 8.107517749 | 8.101365653 | 8.784770914 | 9.458417697 | 9.49623046 | 8.885560124 | 8.726491728 |
| ENSG00000072401 | UBE2D1 | 2.041656551 | 0.092065517 | -0.466008713 | 0.732916452 | 1.575647838 | 3.327711234 | 7.189546432 | 6.718950348 | 8.521127045 | 8.36552655 | 6.134985075 | 5.88031069 |
| ENSG00000204963 | PCDHA7 | 1.031511683 | 0.686593442 | 0.5428848 | 0.836490254 | 1.574396483 | 2 | 3.686303414 | 2 | 3.840289928 | 2.86123014 | 2.858551861 | 3.383972333 |
| | C19ORF68 | 6.763667823 | 0.007296304 | -5.191140549 | 0.097775866 | 1.572527274 | 2 | 2 | 2 | 6.86703417 | 6.81514011 | 2 | 2 |
| ENSG00000135164 | DMTF1 | 1.290761951 | 0.229716518 | 0.278269063 | 0.81770874 | 1.569031013 | 8.586954094 | 9.018611864 | 5.930865648 | 9.584041242 | 9.64048078 | 8.648381562 | 8.565456872 |
| ENSG00000129038 | LOXL1 | -1.042748507 | 0.1406021 | 2.61159446 | 3.49E-05 | 1.568845953 | 10.1311248 | 8.834582582 | 9.703463497 | 8.668958796 | 8.55598266 | 11.62494827 | 12.69103003 |
| ENSG00000148840 | PPRC1 | 0.375594481 | 0.796912783 | 1.192751377 | 0.35900228 | 1.568345858 | 4.815609471 | 8.623598428 | 8.825624954 | 8.559529208 | 8.56534297 | 9.422673149 | 9.321499526 |
| ENSG00000187624 | C17orf97 | 1.223030104 | 0.362279345 | 0.345126821 | 0.81390919 | 1.568156925 | 6.009025997 | 5.303523747 | 2 | 6.172600389 | 6.42738748 | 5.357600875 | 5.606950214 |
| ENSG00000087152 | ATXN7L3 | 0.801371311 | 0.115157031 | 0.766626984 | 0.126902787 | 1.567998295 | 9.371511179 | 9.664384941 | 8.631919287 | 10.07473944 | 10.0971999 | 10.18343032 | 9.906002844 |
| ENSG00000143119 | CD53 | -3.75307254 | 0.427813453 | 5.320513139 | 0.191567177 | 1.567440599 | 2 | 2 | 2 | 2 | 2 | 6.075339068 | 4.456980017 |
| ENSG00000144366 | GULP1 | 1.020822872 | 0.558047538 | 0.546556523 | 0.762852238 | 1.567379395 | 2 | 7.538394143 | 7.398165132 | 7.982979299 | 7.81305436 | 7.396955563 | 7.473262559 |
| ENSG00000100426 | ZBED4 | 0.309073886 | 0.882971353 | 1.256306518 | 0.49597846 | 1.565380404 | 2 | 7.847167909 | 8.561105505 | 7.93509946 | 8.01555453 | 9.010625386 | 8.806206367 |
| ENSG00000241553 | ARPC4 | 0.964088954 | 0.066974888 | 0.598861386 | 0.271400583 | 1.56295034 | 8.836089686 | 9.479495904 | 9.565556058 | 10.25617662 | 10.3209133 | 9.471140938 | 10.26984055 |
| ENSG00000041802 | LSG1 | 1.095908181 | 0.379885407 | 0.466881519 | 0.723926237 | 1.562789699 | 4.815609471 | 8.191702463 | 8.460932916 | 8.861844103 | 8.9229643 | 8.162177243 | 8.37093032 |
| ENSG00000135387 | CAPRIN1 | 0.981449287 | 0.036333097 | 0.580933636 | 0.235600774 | 1.562382923 | 10.21416435 | 11.16599533 | 10.50627919 | 11.68934031 | 11.6446463 | 11.30065805 | 11.23297706 |
| ENSG00000176953 | NFATC2IP | 0.91172266 | 0.073765025 | 0.650196725 | 0.210659124 | 1.561919385 | 8.042902253 | 7.790474954 | 8.805342542 | 9.140440229 | 9.20831741 | 8.90322929 | 8.927151259 |
| ZNF664-FAM101A | | 6.75243517 | 0.007655523 | -5.191140397 | 0.098726147 | 1.561294773 | 2 | 2 | 2 | 6.772115916 | 6.88657603 | 2 | 2 |
| ENSG00000111450 | STX2 | 0.128420195 | 0.83752462 | 1.429972387 | 0.003580998 | 1.558392581 | 8.009475818 | 8.451443573 | 9.099922433 | 8.723531969 | 8.70598173 | 9.874038788 | 10.13088292 |
| ENSG00000107929 | LARP4B | 0.505761843 | 0.348682275 | 1.052587342 | 0.032404634 | 1.558349185 | 8.777722034 | 9.608085847 | 8.65477198 | 9.530270581 | 9.64342987 | 10.12018263 | 10.14455455 |
| ENSG00000213625 | LEPROT | 0.541514584 | 0.309909983 | 1.01485488 | 0.038648151 | 1.556396464 | 9.949165863 | 9.479495904 | 10.49998282 | 10.57051624 | 10.576422 | 10.87669185 | 11.19708549 |
| ENSG00000106299 | WASL | 0.738975847 | 0.127670293 | 0.817110483 | 0.083336867 | 1.55608633 | 8.716893051 | 8.862353853 | 8.143093325 | 9.40405742 | 9.28113268 | 9.673379144 | 9.116112551 |
| ENSG00000167378 | IRGQ | 1.101504635 | 0.018749668 | 0.453086382 | 0.372076473 | 1.554591017 | 9.602033055 | 8.590771778 | 9.116480286 | 10.22863675 | 10.2769623 | 9.58550484 | 9.628710015 |
| ENSG00000104894 | CD37 | -3.753078007 | 0.418387377 | 5.305433271 | 0.182610051 | 1.552355264 | 2 | 2 | 2 | 2 | 2 | 5.906324469 | 4.836507146 |
| ENSG00000156983 | BRPF1 | 0.562716433 | 0.71541993 | 0.987426887 | 0.495779443 | 1.55014332 | 7.867441304 | 7.819099893 | 3.369317972 | 7.810605345 | 7.87959733 | 8.287622092 | 8.239587639 |
| ENSG00000203778 | FAM229B | 0.222195793 | 0.841373243 | 1.325329055 | 0.153932236 | 1.547524848 | 7.159102577 | 5.595927442 | 4.874587708 | 6.470063213 | 6.33555364 | 7.452712791 | 7.488317792 |
| ENSG00000185963 | BICD2 | -0.029761675 | 0.974752037 | 1.577000746 | 0.035746075 | 1.547239071 | 7.82963488 | 8.655694736 | 9.778213818 | 8.970341944 | 8.91444416 | 10.54098392 | 10.54206659 |
| ENSG00000067992 | PDK3 | -0.925825098 | 0.133293863 | 2.47273291 | 4.64E-06 | 1.546907812 | 6.96351253 | 6.78762695 | 6.322058544 | 5.808110275 | 5.87556634 | 8.651302479 | 9.515022972 |
| ENSG00000125304 | TM9SF2 | 1.328431105 | 0.003598276 | 0.217617475 | 0.688299482 | 1.546048581 | 10.49341471 | 10.11467741 | 9.516493218 | 11.40228438 | 11.4404381 | 10.15592861 | 10.45487637 |
| ENSG00000107959 | PITRM1 | 1.155638531 | 0.2265035 | 0.389246848 | 0.708157776 | 1.544885378 | 7.485143553 | 8.297207175 | 9.868959282 | 10.00340207 | 10.0798882 | 9.172206498 | 9.379543083 |
| ENSG00000135900 | MRPL44 | 0.980969658 | 0.126065169 | 0.563066174 | 0.397324398 | 1.544035832 | 8.285626337 | 7.467835385 | 6.80318718 | 8.557949361 | 8.66303256 | 8.219062336 | 8.177610383 |
| ENSG00000179918 | SEPHS2 | 0.785183514 | 0.359261467 | 0.757725853 | 0.368129487 | 1.542909367 | 9.435834703 | 8.077869079 | 7.502634626 | 9.302050417 | 9.40098777 | 9.185700723 | 9.452192236 |
| ENSG00000105472 | CLEC11A | 1.083810115 | 0.336744609 | 0.458185513 | 0.702091517 | 1.541995627 | 10.21416435 | 7.431218269 | 8.110401118 | 10.15373922 | 10.1822948 | 9.571181392 | 9.518693865 |
| ENSG00000145817 | YIPF5 | 1.527811672 | 7.90E-06 | 0.013063092 | 0.976832019 | 1.540874764 | 9.567995545 | 10.14890644 | 10.21486071 | 11.55640578 | 11.5034733 | 9.997579827 | 10.04022229 |
| ENSG00000139613 | SMARCC2 | 1.214634889 | 0.000674918 | 0.325667974 | 0.42448075 | 1.540302864 | 9.262228249 | 8.623598428 | 8.805342542 | 10.091763 | 10.1598144 | 9.166092673 | 9.313452824 |
| ENSG00000152465 | NMT2 | 0.402296678 | 0.624897619 | 1.13765861 | 0.120205411 | 1.539955288 | 6.009025997 | 7.572420273 | 7.858115942 | 7.797276258 | 7.67575955 | 8.401292507 | 8.520893436 |
| ENSG00000273686 | | 3.02803197 | 0.000112893 | -1.490559479 | 0.076831332 | 1.537472492 | 14.00909496 | 12.13658528 | 12.19766108 | 16.12211306 | 16.0595297 | 11.70438214 | 11.43146841 |
| ENSG00000169994 | MYO7B | -2.306377453 | 0.589234072 | 3.843806445 | 0.2779606 | 1.537428992 | 2 | 2 | 2 | 2.141737246 | 2 | 4.406883583 | 4.002845082 |
| ENSG00000170837 | GPR27 | -2.306294028 | 0.594639663 | 3.843667276 | 0.286305677 | 1.537373247 | 2 | 2 | 2 | 2.141737246 | 2 | 3.709392871 | 4.594883674 |
| ENSG00000166794 | PPIB | 2.624040511 | 2.15E-08 | -1.086913849 | 0.032897253 | 1.537126662 | 11.01523161 | 9.975384048 | 10.30429804 | 13.09939874 | 13.1366789 | 9.22511553 | 9.582902386 |
| ENSG00000160951 | PTGER1 | 0.67480488 | 0.785196727 | 0.861961693 | 0.719120052 | 1.536766573 | 2 | 6.140560021 | 2 | 5.461211713 | 5.19627118 | 5.673303969 | 5.313094696 |
| ENSG00000166887 | VPS39 | 0.779667697 | 0.582172352 | 0.756969232 | 0.587973908 | 1.536636929 | 6.009025997 | 9.943237334 | 10.28245311 | 10.31300956 | 10.3971503 | 10.31864645 | 10.34732599 |
| ENSG00000116017 | ARID3A | 0.938628264 | 0.034657972 | 0.597281455 | 0.19277085 | 1.535909719 | 6.96351253 | 6.899080544 | 7.451345288 | 7.959237999 | 8.10598728 | 7.842553218 | 7.539150216 |
| ENSG00000083750 | RRAGB | 0.545912091 | 0.749224767 | 0.989565837 | 0.537860646 | 1.535477929 | 2 | 7.051611315 | 6.629488387 | 6.820355514 | 6.83074839 | 7.20223931 | 7.307219812 |
| ENSG00000259956 | RBM15B | 0.357682257 | 0.723107864 | 1.176838784 | 0.187681315 | 1.534521041 | 9.521325812 | 9.910357957 | 7.502634626 | 9.603316663 | 9.69266358 | 10.36252773 | 10.56015326 |
| ENSG00000254087 | LYN | 0.669938087 | 0.663958104 | 0.864290813 | 0.560286335 | 1.5342289 | 6.257048221 | 7.819099893 | 3.369317972 | 7.363085426 | 7.36873214 | 7.272216802 | 7.792821043 |
| ENSG00000198768 | APCDD1L | -3.753070016 | 0.432702068 | 5.284632105 | 0.200448801 | 1.531562089 | 2 | 2 | 2 | 2 | 2 | 6.057834398 | 4.376876162 |
| ENSG00000060718 | COL11A1 | 0.062045834 | 0.917858995 | 1.467812658 | 0.001387224 | 1.529858492 | 8.493334308 | 9.067150531 | 8.721235202 | 8.959612762 | 8.72141482 | 10.56053732 | 9.836041609 |
| ENSG00000118689 | FOXO3 | 0.982886009 | 0.028297569 | 0.546902502 | 0.246468033 | 1.529788512 | 8.39321512 | 9.328390744 | 9.06622548 | 9.941135381 | 9.98252 | 9.535001584 | 9.521025034 |
| ENSG00000164081 | TEX264 | 0.81612179 | 0.560680027 | 0.712256235 | 0.61097572 | 1.528378025 | 4.467311984 | 8.732943944 | 7.896971941 | 8.600009476 | 8.69180693 | 8.504505158 | 8.582138547 |
| ENSG00000048740 | CELF2 | -1.675075969 | 0.033290369 | 3.200383366 | 8.95E-06 | 1.525307397 | 8.009475818 | 6.535001748 | 6.43197674 | 5.662809983 | 5.51241389 | 9.915528427 | 10.67451395 |
| ENSG00000159335 | PTMS | -0.108964465 | 0.858427987 | 1.633216959 | 0.000478561 | 1.524252493 | 10.34051281 | 9.599860728 | 10.47451889 | 10.06030508 | 10.0901921 | 11.50728966 | 12.06640492 |
| ENSG00000115145 | STAM2 | 0.368938665 | 0.745829169 | 1.155285084 | 0.256540578 | 1.524223749 | 9.710019321 | 7.002537973 | 8.143093325 | 9.083378928 | 9.04395769 | 9.849026448 | 9.842180628 |
| ENSG00000054118 | THRAP3 | 0.580642444 | 0.42279064 | 0.943353434 | 0.162472292 | 1.523995877 | 9.730679447 | 9.386925327 | 10.68342405 | 10.60955021 | 10.6290713 | 10.52089571 | 11.32970929 |
| ENSG00000113621 | TXNDC15 | 0.964678134 | 0.085477895 | 0.557316731 | 0.339494409 | 1.521994865 | 9.957954117 | 9.125613817 | 8.763901704 | 10.30172628 | 10.3608797 | 9.891848754 | 9.958792353 |
| ENSG00000129474 | AJUBA | -0.935412269 | 0.220826448 | 2.457132537 | 0.000280937 | 1.521720268 | 7.624139768 | 9.079034257 | 7.552162999 | 7.29643414 | 7.42122542 | 10.81887028 | 10.61511909 |
| ENSG00000123297 | TSFM | 0.669041231 | 0.748404234 | 0.852589884 | 0.675028301 | 1.521631115 | 3.327711234 | 8.101365653 | 3.369317972 | 7.222570549 | 7.33264456 | 7.433611594 | 7.481493974 |
| ENSG00000163728 | TTC14 | 1.424467638 | 0.323982764 | 0.097074046 | 0.953777463 | 1.521541683 | 4.00722531 | 7.60566235 | 8.742726175 | 9.144656522 | 9.13791797 | 8.05229293 | 7.565201648 |
| ENSG00000107862 | GBF1 | 0.504258577 | 0.565891391 | 1.015438168 | 0.209646083 | 1.519696746 | 9.966689162 | 10.66082477 | 8.561105505 | 10.45261097 | 10.4856715 | 10.87565151 | 11.07581394 |
| ENSG00000168827 | GFM1 | 0.841601312 | 0.527890193 | 0.675526665 | 0.611774286 | 1.517127977 | 8.444143041 | 8.004992147 | 4.523675391 | 8.532433657 | 8.55127993 | 8.281350892 | 8.468021799 |
| ENSG00000071189 | SNX13 | 1.079685743 | 0.20811252 | 0.437307577 | 0.634083172 | 1.516993321 | 6.563806279 | 7.819099893 | 8.784770914 | 9.076774616 | 9.0282492 | 8.445151776 | 8.387086051 |
| ENSG00000101844 | ATG4A | 0.861666882 | 0.106568751 | 0.654927626 | 0.222928155 | 1.516594508 | 6.468621203 | 6.311476754 | 6.073378139 | 7.148924055 | 7.0991228 | 7.134272047 | 6.676467423 |
| ENSG00000116198 | CEP104 | 0.727198684 | 0.306713831 | 0.789290831 | 0.25426536 | 1.516489515 | 8.737455672 | 7.232742069 | 7.896971941 | 8.813671111 | 8.78422202 | 9.072586388 | 8.611812925 |
| ENSG00000165915 | SLC39A13 | 0.460323398 | 0.628146304 | 1.05502017 | 0.222891601 | 1.515343568 | 10.6695979 | 9.540933215 | 8.408111533 | 10.23260335 | 10.3245927 | 10.86845263 | 10.87644816 |
| ENSG00000066933 | MYO9A | 0.050370042 | 0.941473373 | 1.463826739 | 0.006147699 | 1.514196781 | 8.653385977 | 7.431218269 | 7.896971941 | 8.109160159 | 8.14188248 | 9.505174684 | 9.546749232 |
| ENSG00000127220 | ABHD8 | 0.207251615 | 0.906926658 | 1.306877806 | 0.388725469 | 1.514129421 | 8.418903792 | 6.78762695 | 3.369317972 | 7.497728447 | 7.4450836 | 8.634179825 | 8.468712049 |
| ENSG00000114948 | ADAM23 | -0.04237584 | 0.981811904 | 1.554516217 | 0.308354984 | 1.512140377 | 3.327711234 | 7.099070191 | 4.058998049 | 5.839660368 | 5.67111583 | 7.094763493 | 7.446882182 |
| ENSG00000131408 | NR1H2 | 0.714729851 | 0.312262093 | 0.791290565 | 0.250438486 | 1.506020416 | 9.460782017 | 8.124485672 | 9.703463497 | 9.886796832 | 10.0096294 | 10.04351195 | 10.00781983 |
| ENSG00000100739 | BDKRB1 | 6.957728462 | 5.66E-09 | -5.452162361 | 0.000459097 | 1.5055661 | 2 | 3.076665209 | 4.874587708 | 10.20914183 | 10.2074225 | 2.139668783 | 2 |
| ENSG00000137673 | MMP7 | 0.380498449 | 0.932860886 | 1.124670754 | 0.794654458 | 1.505169203 | 2 | 2 | 4.058998049 | 3.349643765 | 3.08597292 | 4.45225155 | 2 |
| ENSG00000178338 | ZNF354B | 0.71697161 | 0.660186751 | 0.785602946 | 0.622327372 | 1.502574556 | 4.467311984 | 3.076665209 | 6.80318718 | 6.18902405 | 6.23747437 | 6.656902456 | 5.76365862 |
| ENSG00000129245 | FXR2 | 0.692227303 | 0.726949143 | 0.809704735 | 0.677249145 | 1.501932037 | 10.28571918 | 8.004992147 | 3.369317972 | 9.620825434 | 9.7138614 | 9.73525344 | 9.833096297 |
| ENSG00000226173 | | 9.031355207 | 7.09E-25 | -7.531023029 | 2.36E-05 | 1.500332178 | 4.00722531 | 4.93638933 | 2 | 12.78877383 | 12.8690299 | 2.048074405 | 2 |
| ENSG00000188566 | NDOR1 | 1.252038139 | 0.036509535 | 0.247697023 | 0.720064769 | 1.499735162 | 6.009025997 | 7.232742069 | 6.322058544 | 7.836899669 | 7.88708154 | 6.85908071 | 6.905807071 |
| ENSG00000150990 | DHX37 | 0.796898186 | 0.486515971 | 0.702551526 | 0.536069347 | 1.499449713 | 8.16936288 | 6.728495673 | 5.156626401 | 7.877998516 | 7.99953197 | 7.782541424 | 7.909264332 |
| ENSG00000076351 | SLC46A1 | 0.414187895 | 0.704833394 | 1.085019226 | 0.269817196 | 1.499207121 | 4.815609471 | 7.051611315 | 5.59507254 | 6.592405944 | 6.49425375 | 7.463831776 | 6.856025745 |
| ENSG00000151702 | FLI1 | 0.980274673 | 0.185790369 | 0.518417262 | 0.502693319 | 1.498691935 | 6.73718411 | 8.297207175 | 7.163363831 | 8.503178519 | 8.56534297 | 7.844256003 | 8.280260977 |
| ENSG00000175467 | SART1 | 0.786270634 | 0.514396922 | 0.711786367 | 0.549060202 | 1.498057001 | 5.866944937 | 7.638155693 | 9.116480286 | 8.817637515 | 8.89354047 | 8.641543003 | 8.910481888 |
| ENSG00000174206 | C12orf66 | 0.769230005 | 0.765179552 | 0.728610609 | 0.776060045 | 1.497840614 | 2 | 2 | 6.203069917 | 5.304221918 | 5.53799497 | 5.65789504 | 5.0581604 |
| ENSG00000161642 | ZNF385A | 0.606651925 | 0.506977228 | 0.890561685 | 0.300692047 | 1.49721361 | 5.70932728 | 7.669933291 | 7.896971941 | 7.885575888 | 8.03365075 | 8.204552906 | 8.277112026 |
| ENSG00000069956 | MAPK6 | 0.886924672 | 0.447339181 | 0.609444354 | 0.60616609 | 1.496869026 | 6.89194353 | 10.17128279 | 9.565556058 | 10.32281055 | 10.2496441 | 10.12894058 | 9.881848605 |
| ENSG00000131435 | PDLIM4 | 1.009381612 | 0.37278455 | 0.486476779 | 0.683005276 | 1.495858391 | 6.816637977 | 8.557180776 | 9.926432745 | 9.886796832 | 9.97434539 | 9.173900195 | 9.614433792 |
| ENSG00000106771 | TMEM245 | 0.066977524 | 0.954792088 | 1.42703289 | 0.149298501 | 1.494010414 | 8.418903792 | 9.277729677 | 11.03218278 | 10.11236021 | 10.0027561 | 11.51592898 | 11.30721324 |
| ENSG00000236236 | | 6.684376293 | 0.010013464 | -5.191139549 | 0.104453095 | 1.493236744 | 2 | 2 | 2 | 6.872128674 | 6.64900057 | 2 | 2 |
| ENSG00000125826 | RBCK1 | 1.19135046 | 0.299070637 | 0.301613139 | 0.811551833 | 1.492963599 | 10.84091191 | 9.067150531 | 7.734844356 | 10.91339224 | 10.9608743 | 9.847329288 | 10.22784916 |
| ENSG00000163040 | CCDC74A | 1.36350568 | 0.322678131 | 0.128655245 | 0.935446753 | 1.492160925 | 6.138360429 | 5.457120953 | 2 | 6.592405944 | 6.57060569 | 5.680947153 | 5.115701921 |

| | | | | | | | | | | | | | |
|---|---|---|---|---|---|---|---|---|---|---|---|---|---|
| ENSG00000186908 | ZDHHC17 | 0.626810959 | 0.701399082 | 0.864952357 | 0.580654321 | 1.491763316 | 4.00722531 | 7.35507212 | 9.031722608 | 8.535647908 | 8.44034872 | 8.846588786 | 8.591041727 |
| ENSG00000162599 | NFIA | -1.304498935 | 0.421581455 | 2.795585088 | 0.056539999 | 1.491086152 | 2 | 6.602424674 | 7.225742779 | 5.22679461 | 5.13053105 | 8.677327975 | 9.460197359 |
| ENSG00000164144 | ARFIP1 | 0.812716076 | 0.023841784 | 0.674852841 | 0.062801111 | 1.487568917 | 8.075571721 | 7.60566235 | 7.818184403 | 8.595396228 | 8.68323472 | 8.412813334 | 8.589138522 |
| ENSG00000197472 | ZNF695 | 2.595814869 | 0.43323806 | -1.108884633 | NA | 1.486930235 | 2 | 2 | 2 | 3.289987342 | 3.39680241 | 2.094598369 | 2.225841198 |
| ENSG00000091317 | CMTM6 | 1.005606206 | 0.346801992 | 0.480108719 | 0.670289164 | 1.485714926 | 6.89194353 | 9.806969599 | 9.491321578 | 10.17399201 | 10.1807731 | 9.696187102 | 9.61068621 |
| ENSG00000234636 | MED14OS | -0.344263529 | 0.88827382 | 1.82541239 | 0.372590042 | 1.481148861 | 2 | 3.686303414 | 2 | 2.784612485 | 2.68924543 | 3.968654369 | 4.033026654 |
| ENSG00000127241 | MASP1 | 0.195588784 | 0.945987763 | 1.285484574 | 0.62811405 | 1.481073358 | 2 | 7.669933291 | 2 | 6.321555392 | 6.35006802 | 7.110413993 | 7.619668339 |
| ENSG00000145476 | CYP4V2 | 2.19020448 | 0.001757739 | -0.710133785 | 0.364514502 | 1.480070694 | 8.138771685 | 6.602424674 | 6.629488387 | 9.446513412 | 9.49459826 | 6.710164669 | 6.528072775 |
| ENSG00000166181 | API5 | 0.877757332 | 0.681482862 | 0.597668722 | 0.783429907 | 1.475426053 | 2 | 8.820493869 | 9.746650996 | 9.618553696 | 9.67619834 | 9.396799738 | 9.340856676 |
| ENSG00000120685 | PROSER1 | 0.805055032 | 0.667251957 | 0.669610608 | 0.72102077 | 1.474665639 | 8.19931886 | 7.60566235 | 2 | 8.188900477 | 8.11238697 | 8.046387837 | 7.988411589 |
| ENSG00000057757 | PITHD1 | 0.369681713 | 0.829611423 | 1.104469726 | 0.473241998 | 1.47415144 | 8.564112515 | 7.538394143 | 3.369317972 | 7.966401501 | 7.92393105 | 8.64301114 | 8.697929554 |
| ENSG00000144228 | SPOPL | 0.796430225 | 0.667396303 | 0.676273006 | 0.715242599 | 1.472703231 | 8.075571721 | 7.232742069 | 2 | 7.994705029 | 7.84926609 | 7.966556098 | 7.62463201 |
| ENSG00000088325 | TPX2 | -0.691421764 | 0.69860994 | 2.163872253 | 0.170653805 | 1.472450489 | 6.257048221 | 8.777382925 | 3.369317972 | 6.856790872 | 6.71776877 | 9.639702688 | 9.556197588 |
| ENSG00000065978 | YBX1 | 0.338371471 | 0.574012887 | 1.128820778 | 0.03233724 | 1.467192249 | 11.55650736 | 10.73780697 | 10.47451889 | 11.35143572 | 11.3225709 | 11.95568405 | 12.27937268 |
| ENSG00000089041 | P2RX7 | -3.753058783 | 0.451325678 | 5.21975834 | 0.22882416 | 1.466699556 | 2 | 2 | 2 | 2 | 2 | 6.109723842 | 3.8919413 |
| ENSG00000120802 | TMPO | 0.213656314 | 0.789422857 | 1.252627734 | 0.06088768 | 1.466284048 | 6.816637977 | 7.467835385 | 8.175061096 | 7.917958914 | 7.66709116 | 9.157215598 | 8.388545823 |
| ENSG00000174799 | CEP135 | 1.356151837 | 0.02780363 | 0.10995017 | 0.885109991 | 1.466102007 | 5.095959458 | 5.72254323 | 5.930865648 | 7.106324235 | 6.68379438 | 5.783905948 | 5.670730563 |
| | C10ORF10 | 6.655558276 | 0.010887704 | -5.191139318 | 0.106137285 | 1.464418958 | 2 | 2 | 2 | 6.804454198 | 6.66650236 | 2 | 2 |
| ENSG00000179412 | HNRNPCL4 | 6.655363271 | 0.010826309 | -5.191139354 | 0.105933593 | 1.464223917 | 2 | 2 | 2 | 6.772115916 | 6.70088158 | 2 | 2 |
| ENSG00000277058 | HNRNPCL3 | 6.655363271 | 0.010826309 | -5.191139354 | 0.105933593 | 1.464223917 | 2 | 2 | 2 | 6.772115916 | 6.70088158 | 2 | 2 |
| ENSG00000167986 | DDB1 | 0.239612301 | 0.722229354 | 1.224195326 | 0.032155546 | 1.463807627 | 10.85509492 | 9.923599762 | 11.25043551 | 11.00087894 | 11.0255982 | 11.85871828 | 12.12225468 |
| ENSG00000256222 | MTRNR2L3 | 5.591831631 | 4.78E-09 | -4.128620665 | 0.000386199 | 1.463210966 | 3.327711234 | 4.11352228 | 3.369317972 | 8.812346549 | 8.72280968 | 2.267001569 | 2.059860178 |
| ENSG00000137073 | UBAP2 | 0.416869038 | 0.771061294 | 1.045910893 | 0.426580825 | 1.462779931 | 7.096806373 | 8.968383114 | 5.156626401 | 8.145379268 | 8.30721295 | 8.736746727 | 8.956482366 |
| ENSG00000161920 | MED11 | 0.171049298 | 0.91899196 | 1.291651781 | 0.372076473 | 1.462701079 | 6.009025997 | 6.535001748 | 2 | 5.94010643 | 5.90535527 | 6.700796638 | 7.232780814 |
| ENSG00000102189 | EEA1 | 0.404893583 | 0.22951741 | 1.05704694 | 0.000380145 | 1.461940523 | 7.904282243 | 7.979858146 | 8.143093325 | 8.420396336 | 8.39906692 | 9.131995505 | 8.975062684 |
| ENSG00000100461 | RBM23 | 0.336502472 | 0.769854407 | 1.124791274 | 0.271460899 | 1.461293746 | 7.031698319 | 9.479495904 | 7.552162999 | 8.697938662 | 8.83153649 | 9.349025111 | 9.724584499 |
| ENSG00000132326 | PER2 | 0.371507834 | 0.743918027 | 1.089700854 | 0.283648588 | 1.461208688 | 5.532353208 | 3.686303414 | 6.203069917 | 5.742861563 | 5.80356929 | 6.33715871 | 6.554323192 |
| ENSG00000157796 | WDR19 | 0.91895177 | 0.54826185 | 0.54209662 | 0.730737838 | 1.46104839 | 3.327711234 | 7.572420273 | 8.175061096 | 8.211183901 | 8.29225867 | 7.972017867 | 7.783974227 |
| ENSG00000241973 | PI4KA | 0.914313599 | 0.363354318 | 0.545084021 | 0.599502335 | 1.45939762 | 6.96351253 | 9.488434476 | 9.25752426 | 9.833599446 | 9.8316678 | 9.116226322 | 9.746095808 |
| ENSG00000104299 | INTS9 | 0.727600993 | 0.483194833 | 0.730902673 | 0.471266152 | 1.458503667 | 5.866944937 | 6.666836837 | 4.058998049 | 6.622948477 | 6.57060569 | 6.437578914 | 6.746446439 |
| ENSG00000181830 | SLC35C1 | 0.365252948 | 0.632952844 | 1.092214246 | 0.107174752 | 1.457467194 | 7.435667355 | 8.487557258 | 9.132850258 | 8.847689662 | 8.89230135 | 9.549947816 | 9.633334927 |
| ENSG00000138769 | CDKL2 | -0.272308266 | 0.92422517 | 1.72943474 | 0.477216955 | 1.457126474 | 2 | 3.076665209 | 2 | 2.498691189 | 2.49394783 | 3.632645806 | 3.128433649 |
| ENSG00000214029 | ZNF891 | -0.113213865 | 0.96332191 | 1.568644749 | 0.475128388 | 1.455430883 | 2 | 5.72254323 | 2 | 4.268917409 | 4.31639476 | 6.167993056 | 5.177806074 |
| ENSG00000139194 | RBP5 | -2.312508989 | 0.59184685 | 3.766473241 | 0.294086803 | 1.453964252 | 2 | 2 | 2 | 2 | 2.14022021 | 4.04171018 | 4.26688425 |
| ENSG00000154265 | ABCA5 | 1.275954619 | 0.261985421 | 0.177233642 | 0.892291608 | 1.453188261 | 3.327711234 | 6.389896633 | 5.39244166 | 6.755670824 | 6.70653264 | 5.489669427 | 5.871985934 |
| ENSG00000174989 | FBXW8 | 0.336225709 | 0.803737808 | 1.116765599 | 0.35248148 | 1.452991308 | 8.16936288 | 8.433041974 | 4.874587708 | 8.219202381 | 8.02009991 | 8.932738969 | 8.856124078 |
| ENSG00000164932 | CTHRC1 | 2.817461463 | 0.183470484 | -1.366018491 | 0.540695624 | 1.451442972 | 10.01801709 | 6.844429755 | 2 | 11.36888555 | 11.41801 | 7.754744914 | 6.454714037 |
| ENSG00000163191 | S100A11 | 0.976670055 | 0.053673848 | 0.474464559 | 0.376596657 | 1.451134614 | 11.76115866 | 10.88044608 | 11.2690689 | 12.30537237 | 12.3415581 | 11.46717251 | 12.10631899 |
| ENSG00000013561 | RNF14 | 0.721743853 | 0.289703301 | 0.728135391 | 0.274614244 | 1.449879244 | 8.696033109 | 7.35507212 | 8.805342542 | 9.160358775 | 9.08900105 | 9.166773267 | 9.095255687 |
| ENSG00000077097 | TOP2B | 0.726971824 | 0.156616494 | 0.720822273 | 0.152087349 | 1.447794097 | 8.653385977 | 9.616264339 | 8.941672936 | 9.847245515 | 9.8667903 | 9.696894073 | 9.990140122 |
| ENSG00000270011 | ZNF559-ZNF | 2.704834238 | 0.205847326 | -1.257083543 | 0.588778065 | 1.447750695 | 2 | 3.076665209 | 2 | 4.070463721 | 4.0930011 | 2.267001569 | 2.277179783 |
| ENSG00000009307 | CSDE1 | 0.595187581 | 0.252535231 | 0.851944878 | 0.083938238 | 1.447132458 | 11.27639669 | 12.26437678 | 11.50743625 | 12.34816175 | 12.3371251 | 12.62183347 | 12.57626311 |
| ENSG00000131375 | CAPN7 | 0.664464164 | 0.410513137 | 0.780941405 | 0.315074518 | 1.445405569 | 7.159102577 | 8.862353853 | 8.434763955 | 8.986874783 | 8.9648262 | 9.426088142 | 8.654907296 |
| ENSG00000129682 | FGF13 | 3.442674196 | 0.051696866 | -1.997418794 | 0.289718205 | 1.445255402 | 2 | 4.442710129 | 2 | 6.205262847 | 6.25300433 | 2.717032197 | 2.225841198 |
| ENSG00000197746 | PSAP | 0.699686461 | 0.312262093 | 0.745101598 | 0.270301865 | 1.444788059 | 13.57805376 | 12.02889368 | 12.68078978 | 13.59239489 | 13.6113618 | 13.66815092 | 13.62615866 |
| ENSG00000138600 | SPPL2A | 1.090226872 | 0.014945429 | 0.352482535 | 0.478845029 | 1.442709407 | 8.716893051 | 8.376385542 | 9.272377397 | 9.922839878 | 9.90476967 | 9.066399259 | 9.285518343 |
| ENSG00000106609 | TMEM248 | 0.570622247 | 0.038260026 | 0.871744277 | 0.000791829 | 1.442366524 | 10.19198706 | 9.975384048 | 10.08627303 | 10.65902291 | 10.6501178 | 10.8087678 | 11.08788243 |
| ENSG00000104368 | PLAT | 0.933909 | 0.375381504 | 0.507954471 | 0.642633186 | 1.441863471 | 9.913467515 | 9.386925327 | 7.029880454 | 10.12041043 | 10.1401137 | 9.643130384 | 9.76644922 |
| ENSG00000180198 | RCC1 | 0.173498941 | 0.776910404 | 1.268196754 | 0.011308372 | 1.441695696 | 8.042902253 | 7.467835385 | 7.646393253 | 7.915493556 | 7.89452713 | 9.319337188 | 8.553793978 |
| ENSG00000174325 | DIRC1 | 2.777382728 | 0.415953852 | -1.335735674 | NA | 1.441647055 | 2 | 2 | 2 | 3.515101962 | 3.39680241 | 2.048074405 | 2.225841198 |
| ENSG00000159674 | SPON2 | 4.432245561 | 7.02E-14 | -2.992057847 | 8.05E-07 | 1.440187714 | 9.579430841 | 8.077869079 | 8.608698767 | 13.28530217 | 13.333318 | 6.189586148 | 5.73181021 |
| ENSG00000143933 | CALM2 | 0.079029916 | 0.925549897 | 1.361058578 | 0.044255311 | 1.440088494 | 12.06557695 | 10.71514053 | 10.6093157 | 11.38412223 | 11.3595809 | 12.63532708 | 12.66976376 |
| ENSG00000151689 | INPP1 | 0.585704816 | 0.753072123 | 0.853645868 | 0.633525288 | 1.439350684 | 2 | 7.669933291 | 7.691296561 | 7.638916932 | 7.72670378 | 7.934942996 | 7.958255135 |
| ENSG00000106263 | EIF3B | 0.659177633 | 0.335863227 | 0.779400216 | 0.239064344 | 1.438577849 | 9.27634997 | 10.75272247 | 9.935792933 | 10.72738111 | 10.8187472 | 10.70950083 | 11.05711825 |
| ENSG00000264230 | ANXA8L1 | 0.728621256 | 0.79405451 | 0.708340691 | 0.794934423 | 1.436961948 | 2 | 2 | 3.369317972 | 2.947924294 | 2.77779907 | 3.10571689 | 2.551896972 |
| ENSG00000139908 | TSSK4 | -2.312515245 | 0.59365851 | 3.748797708 | 0.299320086 | 1.436282463 | 2 | 2 | 2 | 2 | 2.14022021 | 4.311629396 | 3.956354764 |
| ENSG00000091128 | LAMB4 | 2.389898298 | 0.472233499 | -0.954775133 | NA | 1.435123165 | 2 | 2 | 2 | 3.227757282 | 3.2184422 | 2.183373728 | 2.172608095 |
| ENSG00000107890 | ANKRD26 | 1.192835034 | 0.617138584 | 0.239975319 | 0.926858085 | 1.432810353 | 3.327711234 | 8.255926468 | 2 | 7.905589775 | 7.90685167 | 7.229857522 | 6.666875073 |
| ENSG00000113712 | CSNK1A1 | 0.827855572 | 0.026772451 | 0.603795783 | 0.114660271 | 1.431651355 | 10.31337612 | 9.664384941 | 10.26027233 | 10.91647777 | 10.9436312 | 10.74502644 | 10.66701521 |
| ENSG00000166263 | STXBP4 | 0.869195315 | 0.687246529 | 0.561050262 | 0.798439725 | 1.430245577 | 3.327711234 | 7.099070191 | 2 | 6.529309835 | 6.45450701 | 6.162543904 | 6.235386446 |
| ENSG00000064687 | ABCA7 | 0.461555301 | 0.793086452 | 0.967668431 | 0.551426737 | 1.429223732 | 2 | 6.389896633 | 6.958202216 | 6.483440992 | 6.67805337 | 7.414254101 | 6.622902297 |
| ENSG00000055211 | GINM1 | 0.873207979 | 0.192577616 | 0.54849531 | 0.426645605 | 1.421703288 | 9.219014269 | 7.954278496 | 9.413059136 | 9.851765671 | 9.8547535 | 9.685540753 | 9.355859082 |
| ENSG00000048140 | TSPAN17 | 0.333952478 | 0.691746353 | 1.085725779 | 0.143504721 | 1.419678257 | 7.82963488 | 8.255926468 | 9.528916236 | 8.996237745 | 9.11260313 | 9.695243935 | 9.903197088 |
| ENSG00000154640 | BTG3 | 0.861246365 | 0.052183296 | 0.55799092 | 0.222263411 | 1.419237286 | 8.855032352 | 8.522788974 | 8.076950877 | 9.397888985 | 9.35135081 | 9.256785236 | 8.863491214 |
| ENSG00000151611 | MMAA | 1.082634973 | 0.364486576 | 0.334003092 | 0.796772666 | 1.416638065 | 6.366709671 | 6.666836837 | 3.369317972 | 7.080147179 | 7.05992818 | 6.609629839 | 6.033722345 |
| ENSG00000146197 | SCUBE3 | -1.401374117 | 0.003403655 | 2.816668851 | 3.17E-10 | 1.415294733 | 10.98545464 | 10.73405387 | 9.907528187 | 9.176921588 | 9.25437635 | 13.42909294 | 13.42090046 |
| ENSG00000104343 | UBE2W | 0.812306953 | 0.152981087 | 0.60172012 | 0.294411384 | 1.414027073 | 7.159102577 | 8.191702463 | 7.818184403 | 8.645348336 | 8.53866332 | 8.628756176 | 8.090601559 |
| ENSG00000235718 | MFRP | 6.604703856 | 0.013055734 | -5.1911387 | 0.110027396 | 1.413565156 | 2 | 2 | 2 | 6.592405944 | 6.77805036 | 2 | 2 |
| ENSG00000035928 | RFC1 | 0.422075165 | 0.469673243 | 0.989968242 | 0.058838964 | 1.412043407 | 8.228665467 | 8.929529568 | 9.426401271 | 9.336553206 | 9.38345569 | 9.934236219 | 9.916159841 |
| ENSG00000157985 | AGAP1 | -0.354943038 | 0.765528102 | 1.766455666 | 0.081972081 | 1.411512628 | 9.081016759 | 6.78762695 | 6.718950348 | 7.659688863 | 7.59580267 | 9.624919616 | 9.822065034 |
| ENSG00000007516 | BAIAP3 | 0.960589052 | 0.653881546 | 0.447916702 | 0.840565911 | 1.408505754 | 2 | 4.11352228 | 2 | 3.79808662 | 3.74316441 | 3.616794016 | 3.209956562 |
| ENSG00000068400 | GRIPAP1 | 1.278317088 | 0.426620169 | 0.129649256 | 0.942926896 | 1.407966344 | 3.327711234 | 7.901717006 | 8.784770914 | 9.120242823 | 9.09762836 | 7.785203203 | 8.139104389 |
| ENSG00000197021 | CXorf40B | 0.961075455 | 0.421979907 | 0.44687078 | 0.721847383 | 1.407946235 | 7.867441304 | 6.602424674 | 4.523675391 | 7.751013478 | 7.88209639 | 7.199581219 | 7.421437279 |
| ENSG00000184588 | PDE4B | 3.658029975 | 0.027591082 | -2.251839286 | 0.192951753 | 1.406190688 | 8.009475818 | 8.540087354 | 2 | 11.33538698 | 11.3820459 | 5.916072173 | 5.122735884 |
| ENSG00000048028 | USP28 | 1.374769857 | 0.361961292 | 0.031180275 | 0.985627735 | 1.405950132 | 4.815609471 | 9.055168103 | 6.203069917 | 9.069031178 | 9.10727282 | 7.817634286 | 7.701812481 |
| ENSG00000166839 | ANKDD1A | 0.325350326 | 0.763884658 | 1.079851944 | 0.259132418 | 1.405202269 | 5.866944937 | 3.686303414 | 5.39244166 | 5.501219927 | 5.48637102 | 6.325014994 | 6.052261256 |
| ENSG00000102934 | PLLP | -2.677433158 | 0.311541535 | 4.081321192 | 0.072160727 | 1.403888034 | 2 | 2 | 4.058998049 | 2.3892346 | 2 | 6.444330878 | 5.777095613 |
| ENSG00000075213 | SEMA3A | 4.434025939 | 0.002427568 | -3.031905249 | 0.046484399 | 1.40212069 | 7.82963488 | 6.899080544 | 2 | 11.25064042 | 11.2638957 | 3.878466556 | 4.388595494 |
| ENSG00000100997 | ABHD12 | 0.974046616 | 0.023504727 | 0.426059232 | 0.358251684 | 1.400105848 | 9.689059027 | 9.102512015 | 8.845626171 | 10.24099634 | 10.2094139 | 9.746906545 | 9.608808761 |
| ENSG00000175602 | CCDC85B | 0.43807655 | 0.290292991 | 0.961423588 | 0.010103248 | 1.399500137 | 8.468948322 | 8.395519306 | 9.031722608 | 9.052296807 | 9.12531607 | 9.588555735 | 9.629944775 |
| ENSG00000132470 | ITGB4 | -0.24792847 | 0.882573218 | 1.646719013 | 0.23817122 | 1.398790543 | 2 | 4.710564154 | 5.39244166 | 4.804776789 | 3.7864959 | 5.72966459 | 6.329589667 |
| ENSG00000176076 | KCNE5 | 1.818453729 | 0.39609411 | -0.419947643 | 0.859331521 | 1.398506086 | 2 | 3.686303414 | 2 | 4.237680128 | 3.69849104 | 2.775316226 | 2.632958241 |
| ENSG00000166888 | STAT6 | 1.251382143 | 0.15370945 | 0.146509939 | 0.886854075 | 1.397892082 | 10.58135316 | 10.57531312 | 8.408111533 | 11.38946169 | 11.400249 | 10.19365153 | 10.3849277 |
| ENSG00000176720 | BOK | 0.118171837 | 0.907693271 | 1.278553681 | 0.122162715 | 1.396725518 | 9.820154628 | 8.88960062 | 7.646393253 | 9.146760056 | 9.16690126 | 10.08514941 | 10.50738357 |
| ENSG00000139324 | TMTC3 | 0.838286751 | 0.099683072 | 0.556713676 | 0.285220143 | 1.395000426 | 9.031919741 | 8.101365653 | 8.742726175 | 9.566077486 | 9.43372295 | 9.424097045 | 8.98571134 |
| ENSG00000095319 | NUP188 | 0.717488889 | 0.625823305 | 0.677333671 | 0.642345888 | 1.394822561 | 5.330594134 | 9.74864196 | 8.325066185 | 9.418058443 | 9.34954609 | 9.05871952 | 9.582583566 |
| ENSG00000025772 | TOMM34 | 0.656889175 | 0.43568023 | 0.737673219 | 0.364791599 | 1.394562393 | 9.128497779 | 8.540087354 | 7.098165388 | 9.064587625 | 9.19128891 | 9.089555166 | 9.320352741 |
| ENSG00000006747 | SCIN | -0.349116 | 0.895265695 | 1.742330461 | 0.444850169 | 1.393214461 | 2 | 2 | 4.523675391 | 2.947924294 | 3.28037825 | 5.158477451 | 3.640730416 |
| ENSG00000112578 | BYSL | 0.35607883 | 0.818460344 | 1.035923597 | 0.457596529 | 1.392002427 | 7.750913381 | 6.666836837 | 3.369317972 | 7.035434774 | 7.19067478 | 7.747483829 | 7.81035357 |
| ENSG00000236287 | ZBED5 | 0.567393814 | 0.613157373 | 0.824596383 | 0.439516998 | 1.391990197 | 7.53297911 | 8.414402626 | 5.59507254 | 8.235106869 | 8.05595642 | 8.700544163 | 8.025476363 |
| ENSG00000137871 | ZNF280D | 1.251533933 | 0.457673604 | 0.137462375 | 0.942145445 | 1.388996307 | 7.485143553 | 7.189546432 | 2 | 8.058664058 | 7.94320592 | 6.959905775 | 6.866539459 |
| ENSG00000143811 | PYCR2 | 0.483678808 | 0.738128035 | 0.905115067 | 0.503902094 | 1.388793875 | 8.609439662 | 8.469613413 | 4.523675391 | 8.43079185 | 8.52753255 | 8.902003555 | 8.892589222 |
| ENSG00000111832 | RWDD1 | 0.491798737 | 0.606852863 | 0.895662475 | 0.311515104 | 1.387461212 | 5.70932728 | 7.979858146 | 7.734844356 | 7.900612272 | 7.93841129 | 8.327733606 | 8.307525005 |
| ENSG00000163798 | SLC4A1AP | 0.839573721 | 0.39058013 | 0.546619362 | 0.583098537 | 1.386193083 | 5.70932728 | 8.101365653 | 8.110401118 | 8.509731121 | 8.45385092 | 8.164905085 | 8.220043091 |
| ENSG00000162377 | COA7 | 0.147282294 | 0.905710932 | 1.23790996 | 0.231007098 | 1.385192254 | 5.532353208 | 5.595927442 | 7.777116083 | 6.83085985 | 6.84618961 | 7.948463298 | 7.854331622 |
| ENSG00000155846 | PPARGC1B | -0.907058336 | 0.649972951 | 2.291975461 | 0.193576 | 1.384917125 | 2 | 4.710564154 | 2 | 2.947924294 | 3.15372745 | 5.502666283 | 5.385063124 |
| ENSG00000159348 | CYB5R1 | 1.018129065 | 0.563160425 | 0.364768853 | 0.846218002 | 1.382897918 | 9.602033055 | 8.376385542 | 3.369317972 | 9.566077486 | 9.5427783 | 8.804787505 | 8.99962918 |
| ENSG00000089505 | CMTM1 | 0.459261904 | 0.851584865 | 0.923120377 | 0.687191321 | 1.382382282 | 2 | 5.72254323 | 2 | 4.846787657 | 4.68029874 | 5.371837532 | 4.935697922 |
| ENSG00000088930 | XRN2 | 1.38655917 | 0.00265282 | -0.004594978 | 0.993931149 | 1.381964192 | 8.468948322 | 9.34816733 | 8.536706852 | 10.21867227 | 10.2394476 | 8.779646757 | 8.908956896 |
| ENSG00000116990 | MYCL | 1.068673975 | 0.63692076 | 0.313032287 | 0.896941641 | 1.381706261 | 4.00722531 | 2 | 2 | 3.663521627 | 3.50452879 | 3.12825509 | 3.128433649 |
| ENSG00000138138 | ATAD1 | 0.777713839 | 0.131645633 | 0.603921854 | 0.246132276 | 1.381635693 | 8.87372952 | 9.308339287 | 8.296287027 | 9.645581273 | 9.67979376 | 9.661597094 | 9.294890064 |
| ENSG00000023839 | ABCC2 | 3.708523217 | 0.018902587 | -2.327696826 | 0.168373716 | 1.380826391 | 2 | 2 | 4.874587708 | 6.710878132 | 6.78340763 | 2.492221014 | 2.466008509 |
| ENSG00000170322 | NFRKB | 0.580937352 | 0.757563401 | 0.799136928 | 0.661209052 | 1.38007428 | 7.384433967 | 8.004992147 | 2 | 7.694616532 | 7.76240929 | 7.778094182 | 8.091498142 |
| ENSG00000119661 | DNAL1 | 0.741131022 | 0.254759301 | 0.638513901 | 0.323686171 | 1.379644923 | 6.73718411 | 6.140560021 | 7.398165132 | 7.64488235 | 7.45182838 | 7.247980242 | 7.629578661 |
| ENSG00000180979 | LRRC57 | 0.384070624 | 0.468786454 | 0.993677881 | 0.034406558 | 1.377748505 | 6.96351253 | 7.35507212 | 7.029880454 | 7.626911583 | 7.38291816 | 8.337438499 | 7.836259222 |
| ENSG00000163320 | CGGBP1 | 0.701783468 | 0.340527707 | 0.67505467 | 0.352263729 | 1.376838138 | 8.075571721 | 9.680074956 | 9.601288346 | 10.04910261 | 9.9102895 | 10.02786376 | 9.877696666 |
| ENSG00000184270 | HIST2H2AB | 5.156998032 | 0.047659578 | -3.780590748 | 0.238854949 | 1.376407284 | 2 | 2 | 2 | 5.334065616 | 5.33407135 | 2.048074405 | 2 |

| | | | | | | | | | | | | | |
|---|---|---|---|---|---|---|---|---|---|---|---|---|---|
| ENSG00000138646 | HERC5 | 4.993410556 | 0.006399324 | -3.617901426 | 0.056674464 | 1.37550913 | 7.384433967 | 3.076665209 | 2 | 10.79901337 | 10.7874282 | 3.296906311 | 2.851942824 |
| ENSG00000088367 | EPB41L1 | 0.218500927 | 0.781202672 | 1.156329651 | 0.082638 | 1.374830577 | 9.081016759 | 7.60566235 | 7.934808822 | 8.543652346 | 8.58541725 | 9.426940628 | 9.560413054 |
| ENSG00000179021 | C3orf38 | 1.194016373 | 0.001824265 | 0.179909193 | 0.694591795 | 1.373925566 | 7.485143553 | 8.053983481 | 7.97167868 | 9.129845271 | 8.95180002 | 8.158075786 | 7.912312056 |
| ENSG00000130528 | HRC | -2.306155035 | 0.603854595 | 3.679349392 | 0.324412375 | 1.373194356 | 2 | 2 | 2 | 2.141737246 | 2 | 4.434275199 | 3.640730416 |
| ENSG00000147905 | ZCCHC7 | 0.839636411 | 0.550029831 | 0.532686065 | 0.712026005 | 1.372322477 | 7.33131396 | 8.297207175 | 4.058998049 | 8.221200057 | 8.15847021 | 8.069136662 | 7.680519528 |
| ENSG00000017260 | ATP2C1 | 1.403532336 | 0.003430894 | -0.031767183 | 0.959231573 | 1.371765153 | 8.981092835 | 9.574900212 | 10.08627303 | 10.99096084 | 11.0379776 | 9.558011928 | 9.609747791 |
| ENSG00000198860 | TSEN15 | 0.017155117 | 0.974752037 | 1.354100349 | 0.00095192 | 1.371255467 | 7.096806373 | 6.535001748 | 6.80318718 | 6.907295206 | 6.75642009 | 8.189896068 | 8.087908463 |
| ENSG00000158710 | TAGLN2 | 0.047091251 | 0.941979106 | 1.32017456 | 0.009454427 | 1.367265811 | 11.92025849 | 11.21259083 | 11.95457585 | 11.73269717 | 11.8262615 | 12.5783741 | 13.4085426 |
| ENSG00000007237 | GAS7 | 0.49187094 | 0.715717569 | 0.874968264 | 0.489336405 | 1.366839204 | 4.00722531 | 7.431218269 | 6.073378139 | 6.882264028 | 6.92586261 | 7.586913431 | 6.879055607 |
| ENSG00000198000 | NOL8 | 1.005251372 | 0.391746117 | 0.361145665 | 0.774275034 | 1.366397037 | 5.095959458 | 8.433041974 | 7.398165132 | 8.548433777 | 8.47387021 | 7.845956781 | 7.906210156 |
| ENSG00000198932 | GPRASP1 | 0.423226861 | 0.820270883 | 0.942728832 | 0.582327511 | 1.365955693 | 7.667647223 | 6.899080544 | 2 | 7.206526483 | 7.14977896 | 7.6993745 | 7.672153316 |
| ENSG00000107099 | DOCK8 | -1.685494736 | 0.547559684 | 3.051139857 | 0.200422954 | 1.36564512 | 2 | 3.076665209 | 2 | 2.270786764 | 2.14022021 | 4.927401733 | 3.383972333 |
| ENSG00000136738 | STAM | 0.8045575 | 0.39960716 | 0.559438241 | 0.563920281 | 1.363995741 | 7.031698319 | 7.761270542 | 9.227351203 | 9.130908275 | 9.05949698 | 8.563064723 | 9.093466633 |
| ENSG00000114062 | UBE3A | 0.440406244 | 0.270653422 | 0.923065399 | 0.01114802 | 1.363471643 | 9.159305284 | 9.514922632 | 8.960138144 | 9.712435568 | 9.63233954 | 10.30804289 | 9.980492541 |
| ENSG00000133246 | PRAM1 | -1.317630003 | 0.811052763 | 2.679408937 | NA | 1.361778934 | 2 | 2 | 2 | 2 | 2.2680111 | 3.753554863 | 2.918126211 |
| ENSG00000140577 | CRTC3 | -0.066473927 | 0.972436025 | 1.428044416 | 0.3943541 | 1.36157049 | 3.327711234 | 8.376385542 | 9.516493218 | 8.447952961 | 8.38855922 | 10.01865665 | 9.775403823 |
| ENSG00000119487 | MAPKAP1 | 0.538496117 | 0.266338693 | 0.822240598 | 0.071322601 | 1.360736714 | 8.631580145 | 9.28800521 | 9.528916236 | 9.682298904 | 9.78101395 | 9.875912956 | 10.1419604 |
| ENSG00000001460 | STPG1 | -0.019137042 | 0.990169856 | 1.379863925 | 0.290178986 | 1.360726883 | 3.327711234 | 5.838937808 | 7.285536013 | 6.069927018 | 6.30607931 | 7.5848779 | 7.497817236 |
| ENSG00000170581 | STAT2 | 1.910923717 | 0.011653721 | -0.550643497 | 0.518870344 | 1.36028022 | 9.064836028 | 10.57111193 | 11.0793996 | 12.37330564 | 12.3590431 | 9.645573761 | 10.1326262 |
| ENSG00000078967 | UBE2D4 | -0.046581936 | 0.982078574 | 1.403401411 | 0.429681573 | 1.356819475 | 3.327711234 | 8.234834549 | 4.523675391 | 6.687949235 | 6.84106088 | 8.00436125 | 8.34108458 |
| ENSG00000213030 | CGB8 | 4.227093562 | 0.222198443 | -2.873147706 | 0.450687102 | 1.353945856 | 2 | 2 | 2 | 4.299492648 | 4.72553021 | 2.094598369 | 2 |
| ENSG00000165861 | ZFYVE1 | 0.647958669 | 0.406678307 | 0.705658678 | 0.352303034 | 1.353617347 | 6.96351253 | 8.717821467 | 8.408111533 | 8.843805136 | 8.85843657 | 9.089196199 | 8.701457496 |
| ENSG00000197818 | SLC9A8 | 1.289202533 | 0.479814084 | 0.06325251 | 0.974738711 | 1.352455043 | 2 | 7.315436877 | 8.175061096 | 8.43768081 | 8.56378716 | 6.981729486 | 7.55613706 |
| ENSG00000119912 | IDE | 1.079395607 | 0.321826745 | 0.271408808 | 0.821189708 | 1.350804415 | 8.696033109 | 8.639735838 | 5.772716771 | 9.29542901 | 9.20133 | 8.331380596 | 8.556393893 |
| ENSG00000184574 | LPAR5 | -3.753073107 | 0.429843212 | 5.103853444 | 0.215047296 | 1.350780337 | 2 | 2 | 2 | 2 | 2 | 5.152992171 | 5.408276315 |
| ENSG00000186532 | SMYD4 | 0.554583697 | 0.467615355 | 0.796010886 | 0.268035588 | 1.350594583 | 6.73718411 | 6.78762695 | 5.59507254 | 7.097651217 | 6.92586261 | 7.583859056 | 6.811011855 |
| ENSG00000150667 | FSIP1 | 2.330116428 | 0.19061882 | -0.982692834 | 0.609511386 | 1.347423595 | 2 | 4.11352228 | 2 | 5.022023783 | 4.70309173 | 2.746468522 | 2.671842496 |
| ENSG00000100650 | SRSF5 | 1.070491019 | 0.066558504 | 0.276300998 | 0.674724562 | 1.346792017 | 10.12333326 | 9.664384941 | 9.31604076 | 10.79800992 | 10.8145014 | 9.415246149 | 10.43664657 |
| ENSG00000136003 | ISCU | 0.705866774 | 0.108937139 | 0.639944088 | 0.142663282 | 1.345810862 | 8.444143041 | 9.079034257 | 8.805342542 | 9.495204222 | 9.51406378 | 9.66376837 | 9.172702781 |
| ENSG00000119541 | VPS4B | 0.829602835 | 0.194826666 | 0.514919321 | 0.434435201 | 1.344522156 | 8.19931886 | 9.13702746 | 7.777116083 | 9.354865403 | 9.28018559 | 9.056150495 | 8.953033422 |
| ENSG00000149260 | CAPN5 | 0.976454458 | 0.144865788 | 0.367810783 | 0.610962735 | 1.344265241 | 8.777722034 | 8.191702463 | 7.163363831 | 9.106258804 | 9.20332986 | 8.568225482 | 8.536120927 |
| ENSG00000171408 | PDE7B | 1.749236283 | 0.18647032 | -0.405396365 | 0.785325587 | 1.343839918 | 3.327711234 | 6.602424674 | 7.342949392 | 8.199071993 | 8.185025 | 6.518722958 | 5.519049808 |
| ENSG00000102096 | PIM2 | 0.689218553 | 0.50100135 | 0.651567895 | 0.518473647 | 1.340786448 | 6.009025997 | 7.051611315 | 4.523675391 | 6.946463121 | 6.81514011 | 6.892381078 | 6.797888492 |
| ENSG00000103168 | TAF1C | 0.58195562 | 0.758330669 | 0.758707361 | 0.680152864 | 1.340662981 | 2 | 7.979858146 | 7.451345288 | 7.762033056 | 7.71833702 | 7.862014947 | 7.964141273 |
| ENSG00000186416 | NKRF | 0.515584257 | 0.717276749 | 0.823598215 | 0.541656025 | 1.339182472 | 3.327711234 | 7.232742069 | 7.098165388 | 7.230526165 | 7.03319408 | 7.399274054 | 7.470508284 |
| ENSG00000132950 | ZMYM5 | 1.047587954 | 0.338720894 | 0.291281792 | 0.807912915 | 1.338869747 | 4.815609471 | 7.393647515 | 5.772716771 | 7.437142666 | 7.40393901 | 6.952030889 | 6.362645675 |
| ENSG00000263091 | | 6.529182539 | 0.016750741 | -5.191137785 | 0.116242229 | 1.338044753 | 2 | 2 | 2 | 6.716553866 | 6.507263 | 2 | 2 |
| ENSG00000128563 | PRKRIP1 | 0.708902101 | 0.432702068 | 0.629050326 | 0.481058497 | 1.337952427 | 6.257048221 | 7.051611315 | 8.35328246 | 8.192977693 | 8.15020018 | 8.035250217 | 8.148612304 |
| ENSG00000139182 | CLSTN3 | 0.807572614 | 0.311127936 | 0.529393998 | 0.516505072 | 1.336966612 | 8.16936288 | 7.099070191 | 9.014156722 | 9.060130342 | 9.11366683 | 8.789402187 | 8.832722224 |
| ENSG00000138698 | RAP1GDS1 | 0.373397181 | 0.374982962 | 0.961892657 | 0.010215226 | 1.335289838 | 8.696033109 | 8.717821467 | 8.536706852 | 8.989221226 | 9.05839257 | 9.860850997 | 9.303427654 |
| ENSG00000130711 | PRDM12 | -0.727834839 | NA | 2.062642664 | NA | 1.334807825 | 2 | 2 | 2 | 2.141737246 | 2.2680111 | 3.316666208 | 2.70970617 |
| ENSG00000121406 | ZNF549 | 0.201396428 | 0.892276535 | 1.132121948 | 0.375035076 | 1.333518376 | 4.467311984 | 7.232742069 | 4.874587708 | 6.29903474 | 6.27599024 | 7.5848779 | 6.622902297 |
| ENSG00000174718 | KIAA1551 | 1.59802648 | 0.216138361 | -0.265113279 | 0.856832448 | 1.3329132 | 4.467311984 | 8.317412869 | 8.175061096 | 9.299216392 | 9.29432703 | 7.81936672 | 6.973513842 |
| ENSG00000100439 | ABHD4 | -0.302406377 | 0.645053038 | 1.634518405 | 0.002808849 | 1.332112028 | 9.423197387 | 8.717821467 | 9.897982195 | 9.091045942 | 9.14937381 | 11.31477425 | 10.72478611 |
| ENSG00000198792 | TMEM184B | 0.204291317 | 0.755058981 | 1.127428756 | 0.041403012 | 1.331720073 | 11.24442723 | 10.20962523 | 10.26027233 | 10.83337521 | 10.8819159 | 11.56276827 | 11.96792586 |
| ENSG00000196141 | SPATS2L | 1.003066049 | 0.009469798 | 0.328010493 | 0.445736555 | 1.331076542 | 10.41898845 | 10.66082477 | 9.91701143 | 11.34549558 | 11.3895304 | 10.59558475 | 10.78602048 |
| ENSG00000131849 | ZNF132 | 0.730629218 | 0.734036455 | 0.599560837 | 0.781762153 | 1.330190055 | 2 | 3.686303414 | 7.225742779 | 6.449761039 | 6.49425375 | 6.589466407 | 6.055940621 |
| ENSG00000197024 | ZNF398 | 0.311820074 | 0.863136627 | 1.017904937 | 0.52813899 | 1.329725011 | 6.653098118 | 8.57407404 | 3.369317972 | 7.662632018 | 7.67863747 | 8.448514034 | 8.277899908 |
| ENSG00000214338 | SOGA3 | 4.704047971 | 7.47E-06 | -3.374993898 | 0.003522037 | 1.329054073 | 3.327711234 | 5.131603625 | 2 | 8.446246007 | 8.49850989 | 2.34604626 | 2.466008509 |
| ENSG00000197385 | ZNF860 | 1.123722657 | 0.624036401 | 0.204073846 | 0.934813535 | 1.327796503 | 2 | 4.11352228 | 2 | 4.205751501 | 3.45167086 | 2.911488689 | 3.560117069 |
| ENSG00000123496 | IL13RA2 | 6.518686237 | 0.017140458 | -5.191137724 | 0.11669998 | 1.327548512 | 2 | 2 | 2 | 6.535745081 | 6.67228943 | 2 | 2 |
| ENSG00000010017 | RANBP9 | 0.66743219 | 0.689816154 | 0.658822683 | 0.689674098 | 1.326254873 | 9.48530526 | 9.018611864 | 4.058998049 | 9.352133347 | 9.38433737 | 9.332135462 | 9.386865301 |
| ENSG00000102984 | ZNF821 | 1.276527354 | 0.394463454 | 0.047459591 | 0.97742548 | 1.323986465 | 4.00722531 | 5.72254323 | 2 | 5.764940652 | 5.81407719 | 4.742549839 | 4.644600858 |
| ENSG00000134186 | PRPF38B | 0.517293757 | 0.573772614 | 0.806683812 | 0.353054225 | 1.323977569 | 8.444143041 | 8.702538795 | 6.629488387 | 8.705093576 | 8.66738547 | 9.242652114 | 8.641501544 |
| ENSG00000112739 | PRPF4B | 1.027931166 | 0.443895169 | 0.295110562 | 0.839017584 | 1.323041728 | 5.532353208 | 9.461450882 | 9.413059136 | 9.925905286 | 9.91944253 | 9.133737044 | 9.251427401 |
| ENSG00000198028 | ZNF560 | 2.64071853 | 0.619792462 | -1.317696067 | NA | 1.323022463 | 2 | 2 | 2 | 3.023133209 | 3.65239003 | 2.267001569 | 2 |
| ENSG00000049759 | NEDD4L | 0.897021324 | 0.585414444 | 0.424957284 | 0.804379534 | 1.321978608 | 3.327711234 | 8.147241016 | 5.930865648 | 7.759286043 | 7.78127547 | 7.153627862 | 7.448282715 |
| ENSG00000097096 | SYDE2 | -0.625910628 | 0.561312002 | 1.94297005 | 0.037743848 | 1.317059421 | 3.327711234 | 5.457120953 | 5.59507254 | 4.387512679 | 4.72553021 | 7.198250335 | 6.567271433 |
| ENSG00000128989 | ARPP19 | 0.165278394 | 0.693142866 | 1.150624343 | 0.000727777 | 1.315902737 | 10.32700827 | 9.770792087 | 9.897982195 | 10.23309841 | 10.1280575 | 11.28232575 | 11.03596654 |
| ENSG00000107864 | CPEB3 | 1.193248963 | 0.616474704 | 0.122408887 | 0.963128357 | 1.31565785 | 2 | 6.228547981 | 2 | 5.997179931 | 5.78232107 | 4.971696071 | 4.878668439 |
| ENSG00000050730 | TNIP3 | 5.278658201 | 3.43E-08 | -3.963986266 | 0.000493944 | 1.314671935 | 4.467311984 | 2 | 4.058998049 | 8.562683721 | 8.50824913 | 2.183373728 | 2.225841198 |
| ENSG00000175931 | UBE2O | 0.767217738 | 0.251320848 | 0.547414142 | 0.420661534 | 1.314631879 | 8.998234871 | 7.928237109 | 7.600047325 | 9.066811112 | 9.05839257 | 8.879345195 | 8.808934503 |
| ENSG00000025293 | PHF20 | -0.338398805 | 0.546723851 | 1.65214476 | 0.000435758 | 1.313745954 | 9.384607364 | 8.414402626 | 8.699419242 | 8.551612617 | 8.55441692 | 10.69494761 | 10.34826401 |
| ENSG00000196502 | SULT1A1 | 0.173054131 | 0.91483416 | 1.138493556 | 0.417064406 | 1.311547686 | 2 | 5.303523747 | 5.59507254 | 5.003571641 | 5.18011485 | 5.381251181 | 6.394961217 |
| ENSG00000171634 | BPTF | 0.549226425 | 0.249172977 | 0.760810055 | 0.09315428 | 1.31003648 | 8.981092835 | 9.792607274 | 9.798878489 | 10.09557938 | 10.1442836 | 10.43310438 | 10.22110577 |
| ENSG00000150961 | SEC24D | 0.684985163 | 0.568972239 | 0.620813126 | 0.604252415 | 1.305798289 | 8.107517749 | 11.47791526 | 10.46161637 | 11.22249509 | 11.2720171 | 11.24050906 | 11.12393198 |
| ENSG00000197785 | ATAD3A | 0.875196511 | 0.035423229 | 0.430458309 | 0.329501828 | 1.30565482 | 7.867441304 | 7.979858146 | 7.285536013 | 8.625876185 | 8.6021877 | 8.05817395 | 8.282618188 |
| ENSG00000188629 | ZNF177 | 3.116275225 | 0.307253487 | -1.81071869 | 0.582427016 | 1.305556535 | 2 | 2 | 2 | 3.754611435 | 3.60476711 | 2.139668783 | 2.059860178 |
| ENSG00000136628 | EPRS | 0.310691232 | 0.848581742 | 0.992799923 | 0.495779443 | 1.303491155 | 6.009025997 | 10.54992047 | 10.67785662 | 10.36226743 | 10.3769158 | 11.13151297 | 10.96374572 |
| ENSG00000266967 | AARSD1 | 2.288202724 | 0.0437622 | -0.985071943 | 0.424233954 | 1.303130781 | 4.815609471 | 4.710564154 | 2 | 6.358325776 | 6.34282908 | 3.851628042 | 3.261853426 |
| ENSG00000157350 | ST3GAL2 | 0.59452867 | 0.561228414 | 0.707605596 | 0.476093865 | 1.302134266 | 7.27616306 | 9.04308532 | 9.91701143 | 9.697445926 | 9.68838639 | 9.750316091 | 9.858600332 |
| ENSG00000177125 | ZBTB34 | 0.625263369 | 0.495809834 | 0.676569455 | 0.448338304 | 1.301832825 | 7.82963488 | 5.838937808 | 7.734844356 | 7.915493556 | 8.03365075 | 8.261735106 | 7.744618542 |
| ENSG00000134884 | ARGLU1 | 0.426615264 | 0.262230813 | 0.874912052 | 0.011674378 | 1.301527317 | 8.609439662 | 9.13702746 | 8.677268317 | 9.215512556 | 9.29994502 | 9.76229903 | 9.643457842 |
| ENSG00000104081 | BMF | 1.206977699 | 0.066640853 | 0.094242073 | 0.906201712 | 1.301219772 | 7.709880939 | 7.099070191 | 6.322058544 | 8.300810597 | 8.37087451 | 7.525629979 | 6.89352185 |
| ENSG00000138942 | RNF185 | 0.753552529 | 0.362156704 | 0.547631922 | 0.513597151 | 1.301184451 | 7.27616306 | 9.256956408 | 8.922968322 | 9.431062134 | 9.48147379 | 9.139295883 | 9.35623215 |
| ENSG00000140941 | MAP1LC3B | 0.674155513 | 0.014015823 | 0.626349588 | 0.022878027 | 1.300505101 | 10.88304938 | 10.54564457 | 10.93284392 | 11.44991178 | 11.4855078 | 11.43709348 | 11.40286138 |
| ENSG00000037757 | MRI1 | 0.98618286 | 0.026148248 | 0.313945787 | 0.523269622 | 1.300128647 | 6.96351253 | 6.602424674 | 7.225742779 | 7.900612272 | 7.89946959 | 7.366471135 | 7.105417192 |
| ENSG00000130717 | UCK1 | 0.410851794 | 0.832844524 | 0.887329259 | 0.621823406 | 1.298181053 | 5.330594134 | 9.00621767 | 4.058998049 | 7.985332082 | 7.9809999 | 8.586144131 | 8.30675314 |
| ENSG00000118495 | PLAGL1 | -0.256252962 | 0.610659792 | 1.553473465 | 0.000161735 | 1.297220504 | 8.586954094 | 9.396454291 | 9.031722608 | 8.814994458 | 8.77079897 | 10.76043889 | 10.40454019 |
| ENSG00000188895 | MSL1 | 0.163067369 | 0.832844524 | 1.133431982 | 0.075265997 | 1.296499352 | 8.418903792 | 9.770792087 | 9.849281147 | 9.596407142 | 9.68338024 | 10.73840307 | 10.46321306 |
| ENSG00000158315 | RHBDL2 | 1.567223455 | 0.079559698 | -0.270954938 | 0.793999085 | 1.296268518 | 4.467311984 | 4.11352228 | 5.772716771 | 6.172600389 | 6.53928196 | 4.667592615 | 4.729989799 |
| ENSG00000132970 | WASF3 | 0.748993669 | 0.631428733 | 0.543059568 | 0.730619275 | 1.292053237 | 3.327711234 | 8.124485672 | 7.451345288 | 8.03841338 | 7.98565528 | 7.561260509 | 8.021718375 |
| ENSG00000139433 | GLTP | 0.094509813 | 0.92127192 | 1.194993879 | 0.127181705 | 1.289503692 | 9.473095744 | 9.225224669 | 7.502634626 | 9.025111392 | 9.07923325 | 9.956251889 | 10.3156405 |
| ENSG00000137558 | PI15 | 3.187522367 | 0.520929186 | -1.898269048 | 0.714892595 | 1.289253319 | 2 | 2 | 2 | 4.358765249 | 2.59489879 | 2.183373728 | 2 |
| ENSG00000188191 | PRKAR1B | -0.623322183 | 0.738744016 | 1.912289698 | 0.252976607 | 1.288967515 | 7.82963488 | 7.954278496 | 2 | 6.682159616 | 6.75642009 | 9.203388342 | 9.215277965 |
| ENSG00000064932 | SBNO2 | 4.87138545 | 2.78E-06 | -3.582558676 | 0.001115074 | 1.288826774 | 6.257048221 | 5.303523747 | 2 | 10.05078854 | 10.0529704 | 2.803598391 | 2.817676617 |
| ENSG00000160094 | ZNF362 | -0.115443843 | 0.853173474 | 1.403617473 | 0.003902794 | 1.28817363 | 7.218819781 | 7.819099893 | 8.143093325 | 7.677258395 | 7.64666005 | 9.39534863 | 8.884335219 |
| ENSG00000135365 | PHF21A | 0.706991528 | 0.444373895 | 0.578851208 | 0.529459204 | 1.285842737 | 7.218819781 | 8.955547846 | 9.528916236 | 9.506715015 | 9.59550527 | 9.537898028 | 9.301104226 |
| ENSG00000108349 | CASC3 | 0.063208646 | 0.887804971 | 1.220828874 | 0.000290492 | 1.28403752 | 9.081016759 | 9.632483534 | 9.658943166 | 9.496029474 | 9.59321927 | 10.75343304 | 10.64198913 |
| ENSG00000153786 | ZDHHC7 | 0.201159824 | 0.677597403 | 1.082744095 | 0.00728342 | 1.283903919 | 9.048471759 | 9.763446433 | 9.196533591 | 9.539906319 | 9.60989979 | 10.32155264 | 10.57433282 |
| ENSG00000114416 | FXR1 | 0.616203858 | 0.437089015 | 0.666183631 | 0.385866049 | 1.282387489 | 9.358295023 | 10.26789142 | 8.460932916 | 10.15269291 | 10.1721198 | 10.34594659 | 10.06485442 |
| ENSG00000181220 | ZNF746 | 0.608965037 | 0.628451704 | 0.673350024 | 0.583522797 | 1.282315061 | 8.107517749 | 8.255926468 | 4.874587708 | 8.274117511 | 8.27334517 | 8.459665302 | 8.203551241 |
| ENSG00000153956 | CACNA2D1 | 1.315333476 | 0.09651388 | -0.036280733 | 0.969875015 | 1.279052743 | 6.653098118 | 8.414402626 | 8.536706852 | 9.404936476 | 9.36927498 | 7.765568476 | 8.285755157 |
| ENSG00000023228 | NDUFS1 | 0.333020735 | 0.80400762 | 0.945803147 | 0.434676981 | 1.278823882 | 10.43166252 | 8.317412869 | 7.225742779 | 9.573133444 | 9.62340548 | 10.14267068 | 10.27181939 |
| ENSG00000213689 | TREX1 | 7.431041385 | 1.07E-06 | -6.152947209 | 0.007313014 | 1.278094176 | 3.327711234 | 2 | 2 | 8.347282003 | 8.51952909 | 2 | 2 |
| ENSG00000107362 | ABHD17B | 0.888628652 | 0.472233499 | 0.386905027 | 0.765722308 | 1.275533678 | 7.579279349 | 6.899080544 | 4.058998049 | 7.626911583 | 7.64666005 | 7.309674888 | 6.963754056 |
| ENSG00000180104 | EXOC3 | 0.464319523 | 0.632226513 | 0.8108536 | 0.370531531 | 1.275173123 | 6.653098118 | 8.234834549 | 9.031722608 | 8.736160353 | 8.71722214 | 8.99461439 | 9.142307541 |
| ENSG00000114739 | ACVR2B | 0.587447774 | 0.835281211 | 0.687415517 | 0.80197737 | 1.274863291 | 2 | 7.431218269 | 2 | 6.476767609 | 6.48112613 | 6.797131715 | 6.311234697 |
| ENSG00000185482 | STAC3 | 0.270502426 | 0.910353949 | 1.003483677 | 0.640556462 | 1.273986103 | 2 | 4.442710129 | 2 | 3.566290607 | 3.55551834 | 4.434275199 | 3.620995423 |
| ENSG00000137563 | GGH | 0.289102994 | 0.897634669 | 0.982428951 | 0.630693854 | 1.271531944 | 8.998234871 | 6.728495673 | 2 | 7.823812411 | 8.11663771 | 8.518969664 | 8.792487762 |
| ENSG00000213024 | NUP62 | 0.585000677 | 0.041502924 | 0.686417874 | 0.013709533 | 1.271418551 | 8.716893051 | 8.806266215 | 8.960138144 | 9.381905119 | 9.43882443 | 9.425519537 | 9.592751029 |
| ENSG00000244693 | CTAGE8 | 6.461139415 | 0.020346777 | -5.191137047 | 0.12122066 | 1.270002368 | 2 | 2 | 2 | 6.598566398 | 6.50077304 | 2 | 2 |
| ENSG00000140326 | CDAN1 | 0.85835598 | 0.579168793 | 0.41117404 | 0.79903783 | 1.26953002 | 4.467311984 | 7.790474954 | 4.523675391 | 7.315259362 | 7.32531744 | 7.131485609 | 6.590289384 |
| ENSG00000214212 | C19orf38 | 0.161627712 | 0.954276444 | 1.106667846 | 0.669090392 | 1.268295557 | 2 | 3.686303414 | 2 | 3.09461492 | 2.77779907 | 3.956110469 | 2.746601469 |
| ENSG00000253797 | UTP14C | 2.009682769 | 0.028016568 | -0.741648019 | 0.459674606 | 1.268034751 | 6.96351253 | 8.522788974 | 6.073378139 | 9.539105795 | 9.54672161 | 6.892381078 | 6.775747403 |
| ENSG00000166923 | GREM1 | 9.892013099 | 9.06E-05 | -8.626227425 | 0.002061106 | 1.265785674 | 2 | 2 | 7.646393253 | 15.91267495 | 15.9260404 | 2 | 2.117335275 |
| ENSG00000187257 | RSBN1L | 0.499191645 | 0.355599712 | 0.766280887 | 0.129182035 | 1.265472532 | 6.96351253 | 6.602424674 | 7.098165388 | 7.315259362 | 7.43490694 | 7.877896341 | 7.348307474 |

| | | | | | | | | | | | | |
|---|---|---|---|---|---|---|---|---|---|---|---|---|
| ENSG00000106608 | URGCP | 0.122473556 | 0.755348516 | 1.142525267 | 0.000248821 | 1.264998823 | 8.468948322 | 8.929529568 | 8.608698767 | 8.74312865 | 8.87484109 | 9.776414628 | 9.870140686 |
| ENSG00000132256 | TRIM5 | 0.714338675 | 0.056070605 | 0.55038726 | 0.145899169 | 1.264725935 | 9.590776211 | 9.608085847 | 10.16000817 | 10.52717074 | 10.5141456 | 10.26324148 | 10.44543965 |
| ENSG00000162976 | PQLC3 | 0.12828678 | 0.936431891 | 1.135486397 | 0.41910268 | 1.263773177 | 3.327711234 | 7.761270542 | 7.163363831 | 7.026323566 | 7.12888795 | 8.203889907 | 7.909264332 |
| | C8ORF4 | 6.454709124 | 0.020858537 | -5.191136926 | 0.122048565 | 1.263572198 | 2 | 2 | 2 | 6.622948477 | 6.46120797 | 2 | 2 |
| ENSG00000188811 | NHLRC3 | 0.887478925 | 0.141958529 | 0.375875142 | 0.559807045 | 1.263354067 | 6.468621203 | 6.728495673 | 7.163363831 | 7.64488235 | 7.70428369 | 7.512833218 | 6.732719555 |
| ENSG00000172037 | LAMB2 | 0.12525297 | 0.872778119 | 1.137554932 | 0.073580128 | 1.262807902 | 11.45462681 | 12.18137127 | 12.98461505 | 12.43032753 | 12.4993288 | 13.44378564 | 13.5095186 |
| ENSG00000118777 | ABCG2 | 0.994336921 | 0.679435427 | 0.267847808 | 0.916676409 | 1.262184729 | 2 | 2 | 4.058998049 | 3.566290607 | 3.55551834 | 3.374372158 | 2.817676617 |
| ENSG00000165046 | LETM2 | 0.950913582 | 0.540002704 | 0.309563122 | 0.852566533 | 1.260476704 | 2 | 6.04685525 | 4.523675391 | 5.829220145 | 5.87556634 | 5.269098128 | 5.263040529 |
| ENSG00000248871 | TNFSF12-TN | 6.45062145 | 0.020985554 | -5.191136917 | 0.12214799 | 1.259484533 | 2 | 2 | 2 | 6.490083648 | 6.5890786 | 2 | 2 |
| ENSG00000185900 | POMK | 0.315173632 | 0.896905654 | 0.942240171 | 0.673457912 | 1.257413803 | 7.218819781 | 2 | 3.369317972 | 6.024891728 | 6.08095082 | 6.508020621 | 6.777976866 |
| ENSG00000120709 | FAM53C | 0.438291171 | 0.444381183 | 0.818634548 | 0.119894702 | 1.256925718 | 8.39321512 | 9.479495904 | 8.742726175 | 9.347568393 | 9.42432312 | 9.811002128 | 9.717051741 |
| ENSG00000186230 | ZNF749 | 0.815707374 | 0.672951696 | 0.438549691 | 0.825605649 | 1.254257065 | 4.00722531 | 2 | 6.322058544 | 5.764940652 | 5.78232107 | 5.744329515 | 5.020995508 |
| ENSG00000177156 | TALDO1 | 1.428315628 | 0.084560698 | -0.17414904 | 0.857985777 | 1.254166588 | 10.87842776 | 9.506147172 | 8.884815469 | 11.4071185 | 11.4507864 | 9.577081455 | 10.04670882 |
| ENSG00000117724 | CENPF | 0.902162979 | 0.479099956 | 0.351611266 | 0.794825276 | 1.253774244 | 5.70932728 | 6.389896633 | 3.369317972 | 6.761173376 | 6.1322378 | 5.376552034 | 6.374480702 |
| ENSG00000120688 | WBP4 | 0.749823083 | 0.182445823 | 0.503108989 | 0.379197781 | 1.252932072 | 6.468621203 | 7.60566235 | 7.163363831 | 7.922877029 | 7.87709394 | 7.552954291 | 7.755972999 |
| ENSG00000142733 | MAP3K6 | -0.22635506 | 0.780859156 | 1.478136525 | 0.028009055 | 1.251781464 | 6.96351253 | 6.228547981 | 7.818184403 | 6.739036112 | 7.06873058 | 8.647894167 | 8.517561736 |
| ENSG00000112992 | NNT | 0.287057887 | 0.473839036 | 0.964128369 | 0.005328635 | 1.251186256 | 8.418903792 | 8.955547846 | 8.904018013 | 9.063474594 | 9.07049489 | 9.627394005 | 9.844575759 |
| ENSG00000178950 | GAK | 0.907126694 | 0.405114539 | 0.344048874 | 0.767139907 | 1.251175569 | 7.667647223 | 8.414402626 | 10.31867969 | 10.07473944 | 10.2153718 | 9.581682124 | 9.590533012 |
| ENSG00000059769 | DNAJC25 | 0.833172492 | 0.266620128 | 0.417428792 | 0.595040124 | 1.250601284 | 6.89194353 | 5.457120953 | 6.958202216 | 7.426792977 | 7.29937545 | 7.141214687 | 6.753261212 |
| ENSG00000074800 | ENO1 | 0.381845511 | 0.62849957 | 0.86860756 | 0.227933253 | 1.250453071 | 13.50647618 | 12.46336633 | 12.12587352 | 13.1870944 | 13.2239193 | 13.18080919 | 14.06901306 |
| ENSG00000129484 | PARP2 | 0.72513945 | 0.706120595 | 0.525041335 | 0.787590496 | 1.250180785 | 6.009025997 | 7.928237109 | 2 | 7.391747748 | 7.43490694 | 7.382965824 | 7.032593922 |
| ENSG00000214087 | ARL16 | 0.071420768 | 0.981947437 | 1.178401278 | 0.677268519 | 1.249822045 | 2 | 4.442710129 | 2 | 4.034562184 | 2.2680111 | 4.652122235 | 3.496556178 |
| ENSG00000104331 | IMPAD1 | 0.52689232 | 0.558026599 | 0.722033677 | 0.401569414 | 1.248925997 | 9.771132748 | 11.22333918 | 9.301632639 | 10.88783887 | 10.8525899 | 10.86928913 | 11.23743934 |
| ENSG00000107021 | TBC1D13 | 0.454709684 | 0.596370107 | 0.79310466 | 0.321893604 | 1.247814344 | 6.89194353 | 8.414402626 | 8.904018013 | 8.700804887 | 8.77752611 | 9.05908615 | 9.091227191 |
| ENSG00000119599 | DCAF4 | 0.306014985 | 0.682009332 | 0.936932636 | 0.153226253 | 1.242947621 | 5.866944937 | 6.78762695 | 5.59507254 | 6.567497652 | 6.42738748 | 6.950450741 | 7.24894374 |
| ENSG00000136636 | KCTD3 | 0.565623692 | 0.479092457 | 0.676703693 | 0.377779769 | 1.242327385 | 6.89194353 | 8.255926468 | 8.721235202 | 8.729158235 | 8.66883353 | 8.632702669 | 8.966780003 |
| ENSG00000135148 | TRAFD1 | 0.604752538 | 0.625382427 | 0.637568344 | 0.599804676 | 1.242320882 | 6.366709671 | 9.792607274 | 9.149036566 | 9.565291356 | 9.64195608 | 9.506788283 | 9.756025697 |
| ENSG00000171160 | MORN4 | 0.294653693 | 0.820270883 | 0.947535653 | 0.411135188 | 1.242189346 | 5.532353208 | 7.538394143 | 4.874587708 | 6.766655021 | 6.73446058 | 7.607112326 | 7.119550903 |
| ENSG00000147144 | CCDC120 | 0.045756452 | 0.972563056 | 1.193750536 | 0.292297008 | 1.239506988 | 4.467311984 | 6.78762695 | 4.523675391 | 5.910700032 | 5.62411716 | 6.879153024 | 6.847559257 |
| ENSG00000126264 | HCST | 1.583146848 | 0.705820174 | -0.344491346 | 0.938709627 | 1.238655501 | 2 | 3.076665209 | 2 | 3.023133209 | 3.55551834 | 2.831336763 | 2 |
| ENSG00000128585 | MKLN1 | 1.593595479 | 1.48E-05 | -0.355172744 | 0.405823254 | 1.238422735 | 9.097018025 | 9.13702746 | 9.703463497 | 10.9050282 | 10.9439303 | 9.1257085 | 8.827350262 |
| ENSG00000187808 | SOWAHD | -3.753052045 | 0.464100906 | 4.990667867 | 0.267778932 | 1.237615822 | 2 | 2 | 2 | 2 | 2 | 3.943456545 | 5.842465082 |
| ENSG00000261852 | | 6.428088805 | 0.022632059 | -5.191136546 | 0.124325213 | 1.236952259 | 2 | 2 | 2 | 6.622948477 | 6.40670796 | 2 | 2 |
| ENSG00000135094 | SDS | -3.753041652 | 0.477619809 | 4.989068695 | 0.285220143 | 1.236027043 | 2 | 2 | 2 | 2 | 2 | 5.948096914 | 3.45257121 |
| ENSG00000198890 | PRMT6 | 0.491246344 | 0.580694192 | 0.744570475 | 0.373980691 | 1.235816819 | 5.095959458 | 6.899080544 | 6.80318718 | 6.946463121 | 6.94988431 | 6.845542321 | 7.474637727 |
| ENSG00000172534 | HCFC1 | 0.382702769 | 0.497766732 | 0.851706864 | 0.095670604 | 1.234409633 | 10.09173937 | 9.102512015 | 9.287079172 | 9.898079272 | 9.97960581 | 10.29362669 | 10.51174389 |
| ENSG00000197162 | ZNF785 | 0.534270216 | 0.7112679 | 0.699645659 | 0.614845785 | 1.233915875 | 7.27616306 | 3.686303414 | 6.073378139 | 6.809774134 | 6.77267312 | 7.349785676 | 6.397863414 |
| ENSG00000108671 | PSMD11 | 1.070589285 | 0.410778027 | 0.162315296 | 0.911555747 | 1.23290458 | 8.963744667 | 10.13758676 | 6.203069917 | 10.17141133 | 10.2549564 | 9.132692373 | 9.467467793 |
| ENSG00000134897 | BIVM | 1.404561463 | 0.001667272 | -0.172041986 | 0.751485356 | 1.232519476 | 6.653098118 | 7.467835385 | 7.029880454 | 8.509731121 | 8.45552985 | 7.046766788 | 6.81969491 |
| ENSG00000135931 | ARMC9 | 0.774999784 | 0.468786454 | 0.457500279 | 0.677816167 | 1.232500063 | 7.159102577 | 8.147241016 | 9.788583153 | 9.504256136 | 9.57170671 | 9.32818621 | 9.109046142 |
| ENSG00000196793 | ZNF239 | 0.864176831 | 0.732095163 | 0.367321646 | 0.889682343 | 1.231498477 | 2 | 6.389896633 | 2 | 5.720439317 | 5.76075522 | 5.650128416 | 4.80186789 |
| ENSG00000156973 | PDE6D | 0.277087405 | 0.905443737 | 0.952157694 | 0.652626737 | 1.229245099 | 2 | 5.838937808 | 8.805342542 | 7.659688863 | 7.68723697 | 8.222339682 | 8.44505627 |
| ENSG00000197928 | ZNF677 | -0.998317719 | 0.34076194 | 2.227299702 | 0.017280596 | 1.228981984 | 4.467311984 | 6.464272742 | 4.523675391 | 4.670864177 | 4.60967607 | 7.780764172 | 7.491038307 |
| ENSG00000177239 | MAN1B1 | 0.538537864 | 0.398531703 | 0.689184858 | 0.259166961 | 1.227722723 | 10.36053552 | 9.616264339 | 9.031722608 | 10.25422683 | 10.3604317 | 10.57522267 | 10.33147536 |
| ENSG00000108639 | SYNGR2 | -0.921760439 | 0.099215121 | 2.148909569 | 1.87E-05 | 1.22714913 | 8.564112515 | 9.814097497 | 9.344432377 | 8.434240442 | 8.40255255 | 11.50163612 | 11.44626306 |
| ENSG00000071655 | MBD3 | 0.412606487 | 0.544427162 | 0.813229475 | 0.194894738 | 1.225835962 | 9.143983763 | 8.929529568 | 10.22252928 | 9.934453939 | 9.97960581 | 10.40871423 | 10.30214935 |
| ENSG00000062282 | DGAT2 | 2.226664638 | 0.328037547 | -1.001644031 | 0.679226319 | 1.225020607 | 2 | 5.457120953 | 2 | 6.006476466 | 6.19790195 | 3.981090143 | 2.592996859 |
| ENSG00000110711 | AIP | 0.343487372 | 0.557439662 | 0.879481937 | 0.094200705 | 1.222969309 | 8.285626337 | 8.395519306 | 7.285536013 | 8.444537031 | 8.38327653 | 8.985384832 | 8.903351429 |
| ENSG00000156162 | DPY19L4 | 0.932702663 | 0.355475457 | 0.289899631 | 0.792195215 | 1.222602294 | 5.70932728 | 8.356994601 | 7.502634626 | 8.532433657 | 8.41468635 | 8.026277797 | 7.628343587 |
| ENSG00000112655 | PTK7 | 0.379374176 | 0.717288969 | 0.84276378 | 0.381294572 | 1.222137956 | 8.586954094 | 9.181799937 | 10.80076732 | 10.19141925 | 10.2409086 | 10.98311289 | 10.29165501 |
| ENSG00000131724 | IL13RA1 | 0.190993908 | 0.879872883 | 1.030687472 | 0.340611737 | 1.221681381 | 7.867441304 | 9.114109157 | 10.78012988 | 9.9332358 | 9.90292502 | 10.85056146 | 10.65311485 |
| ENSG00000181704 | YIPF6 | 0.548785042 | 0.653114396 | 0.671885752 | 0.570388322 | 1.220670794 | 9.867565369 | 8.762721501 | 6.534109959 | 9.504256136 | 9.44136842 | 9.676966742 | 9.509331338 |
| ENSG00000197119 | SLC25A29 | 0.011884056 | 0.996696604 | 1.208047455 | 0.555606024 | 1.219931511 | 3.327711234 | 8.981105197 | 4.058998049 | 7.514115926 | 7.46188679 | 8.628756176 | 8.702631562 |
| ENSG00000053747 | LAMA3 | 2.092431071 | 0.357081274 | -0.8736922 | 0.718279305 | 1.218738871 | 2 | 2 | 6.073378139 | 6.401247806 | 6.64900057 | 3.782262616 | 4.077147165 |
| ENSG00000125630 | POLR1B | 0.483880942 | 0.811736938 | 0.734529546 | 0.704080508 | 1.218410488 | 6.563806279 | 8.317412869 | 2 | 7.471115032 | 7.71833702 | 7.875400346 | 7.815789075 |
| ENSG00000174652 | ZNF266 | 0.226337215 | 0.894519627 | 0.990930822 | 0.509248144 | 1.217268037 | 3.327711234 | 8.124485672 | 7.777116083 | 7.714607209 | 7.53992502 | 8.419110791 | 8.344849238 |
| ENSG00000205659 | LIN52 | 1.057953528 | 0.484134885 | 0.158201208 | 0.925792761 | 1.216154736 | 3.327711234 | 7.145017454 | 8.143093325 | 8.22518713 | 8.20112674 | 7.373563392 | 7.285445324 |
| ENSG00000205060 | SLC35B4 | 0.42468515 | 0.64079571 | 0.791317758 | 0.350276926 | 1.216002908 | 8.285626337 | 8.557180776 | 6.534109959 | 8.461536372 | 8.43865202 | 8.52641307 | 9.052154518 |
| ENSG00000187026 | KRTAP21-2 | 6.406676273 | 0.023816364 | -5.191136358 | 0.125696822 | 1.215539915 | 2 | 2 | 2 | 6.436065776 | 6.55815769 | 2 | 2 |
| ENSG00000118965 | WDR35 | 0.513544122 | 0.706615573 | 0.701785344 | 0.591567059 | 1.215329466 | 3.327711234 | 6.728495673 | 6.80318718 | 6.799114572 | 6.68379438 | 6.552446484 | 7.219718817 |
| ENSG00000185215 | TNFAIP2 | 1.607488718 | 0.001272822 | -0.393426114 | 0.492233435 | 1.214062604 | 9.204317094 | 9.949724236 | 9.624627483 | 11.20241092 | 11.2557283 | 8.747705648 | 9.599699825 |
| ENSG00000117262 | GPR89A | 2.125124191 | 0.01309428 | -0.911151284 | 0.327334994 | 1.213972907 | 7.096806373 | 5.303523747 | 5.156626401 | 8.157948442 | 8.22494733 | 5.217477917 | 5.361470313 |
| ENSG00000063761 | ADCK1 | 0.132299465 | 0.94581868 | 1.081572881 | 0.535270007 | 1.213872346 | 4.00722531 | 6.464272742 | 2 | 5.289065324 | 5.36298494 | 6.075339068 | 6.323497254 |
| ENSG00000124596 | OARD1 | 0.842331764 | 0.561188023 | 0.370004329 | 0.808290096 | 1.212336093 | 4.467311984 | 7.232742069 | 4.058998049 | 6.766655021 | 6.84618961 | 6.442083733 | 6.261114822 |
| ENSG00000140939 | NOL3 | 0.510329087 | 0.625823305 | 0.699772728 | 0.484010617 | 1.210101816 | 8.009475818 | 5.457120953 | 7.552162999 | 7.823812411 | 7.85436579 | 8.084345843 | 7.964141273 |
| ENSG00000185973 | TMLHE | 0.510790208 | 0.785196727 | 0.698778403 | 0.698000726 | 1.20956861 | 3.327711234 | 7.60566235 | 4.058998049 | 6.744602355 | 6.67805337 | 6.850634117 | 6.934072182 |
| ENSG00000144746 | ARL6IP5 | 0.148820374 | 0.699491428 | 1.060461035 | 0.000697154 | 1.20928141 | 10.36053552 | 9.962611189 | 10.06936278 | 10.31581664 | 10.2588077 | 11.01180608 | 11.36127648 |
| ENSG00000144635 | DYNC1LI1 | 0.453441296 | 0.352000159 | 0.755263567 | 0.095600702 | 1.208704864 | 8.564112515 | 8.029695761 | 7.691296561 | 8.616799622 | 8.5575471 | 8.782757919 | 8.983298084 |
| ENSG00000166860 | ZBTB39 | 0.743440654 | 0.547789118 | 0.464155309 | 0.715036782 | 1.207595963 | 4.00722531 | 6.602424674 | 7.451345288 | 7.242377933 | 7.36158648 | 7.157741966 | 6.89352185 |
| ENSG00000063245 | EPN1 | 0.206286405 | 0.678236128 | 1.000474967 | 0.017257913 | 1.206761372 | 10.16946352 | 10.02537381 | 9.358421373 | 10.06698504 | 10.1301615 | 10.77032462 | 11.00215671 |
| ENSG00000214518 | KRTAP2-2 | 6.39723104 | 0.024565732 | -5.191136201 | 0.126534939 | 1.206094839 | 2 | 2 | 2 | 6.573765157 | 6.39974839 | 2 | 2 |
| ENSG00000145868 | FBXO38 | 0.616711396 | 0.735527035 | 0.588203517 | 0.744872588 | 1.204914913 | 3.327711234 | 7.393647515 | 9.25752426 | 8.542055007 | 8.74768919 | 8.72431761 | 8.508191835 |
| ENSG00000013573 | DDX11 | 0.395491876 | 0.817474674 | 0.809328711 | 0.610398832 | 1.204820587 | 6.89194353 | 5.946638857 | 2 | 6.306580752 | 6.32824132 | 6.757921389 | 6.666875073 |
| ENSG00000164754 | RAD21 | 0.906442989 | 0.421979907 | 0.297367984 | 0.806378761 | 1.203810974 | 7.096806373 | 10.019219 | 9.981704053 | 10.46530827 | 10.3577408 | 10.04073005 | 9.524680706 |
| ENSG00000135185 | TMEM243 | 0.935304702 | 0.108937139 | 0.267982346 | 0.678808079 | 1.203287047 | 6.563806279 | 7.002537973 | 5.930865648 | 7.52061915 | 7.46188679 | 7.018390248 | 6.6426114 |
| ENSG00000159261 | CLDN14 | 3.013777299 | 0.340147427 | -1.811414342 | 0.593306846 | 1.202362957 | 2 | 2 | 2 | 3.881293667 | 3.28037825 | 2.139668783 | 2.059860178 |
| ENSG00000136002 | ARHGEF4 | 0.319437888 | 0.809107998 | 0.881907257 | 0.460053783 | 1.201345145 | 3.327711234 | 6.535001748 | 6.534109959 | 6.260701031 | 6.40670796 | 7.007793418 | 6.728114749 |
| ENSG00000137309 | HMGA1 | 1.457476422 | 0.02162451 | -0.257309585 | 0.728468477 | 1.200166836 | 12.23783517 | 10.89059638 | 10.9968459 | 12.95434551 | 12.9845144 | 11.20292911 | 11.30788843 |
| ENSG00000177614 | PGBD5 | 0.058134433 | 0.979836893 | 1.14146624 | 0.568016472 | 1.199600673 | 2 | 3.076665209 | 4.874587708 | 3.566290607 | 4.02226117 | 5.31402805 | 3.430065404 |
| ENSG00000125534 | PPDPF | 0.317595519 | 0.528648153 | 0.881134787 | 0.048142698 | 1.198730306 | 9.848787621 | 9.687856435 | 10.53735535 | 10.39449089 | 10.3808971 | 11.00144677 | 10.89633773 |
| ENSG00000185527 | PDE6G | -0.689500438 | 0.806161404 | 1.887553383 | 0.436601068 | 1.198052945 | 2 | 2 | 3.369317972 | 2.270786764 | 2.49394783 | 3.709392871 | 3.261853426 |
| ENSG00000166987 | MBD6 | 0.39437071 | 0.310604101 | 0.802948057 | 0.023498087 | 1.197318767 | 7.709880939 | 8.234834549 | 8.206335842 | 8.47833774 | 8.44880239 | 8.917453109 | 8.817088095 |
| ENSG00000110324 | IL10RA | -3.753061303 | 0.45109867 | 4.950042938 | 0.25649001 | 1.196981634 | 2 | 2 | 2 | 2 | 2 | 5.532544554 | 4.624919101 |
| ENSG00000175893 | ZDHHC21 | 0.802867199 | 0.673891173 | 0.393379168 | 0.842165885 | 1.196246367 | 2 | 6.951736442 | 8.206335842 | 7.990026167 | 7.85436579 | 7.401588825 | 7.633277551 |
| ENSG00000204071 | TCEAL6 | 6.386627013 | 0.025128901 | -5.191136117 | 0.127181705 | 1.195490896 | 2 | 2 | 2 | 6.529309835 | 6.42738748 | 2 | 2 |
| ENSG00000143753 | DEGS1 | 0.487663813 | 0.296880612 | 0.707280041 | 0.108707597 | 1.194943853 | 11.24801458 | 10.51971956 | 10.48092713 | 11.26741277 | 11.2940008 | 11.46066561 | 11.53840508 |
| ENSG00000091009 | RBM27 | 1.532065852 | 0.322947052 | -0.337211671 | 0.8452659 | 1.194854181 | 3.327711234 | 8.487557258 | 6.882775709 | 8.894758143 | 8.82115545 | 7.127295835 | 6.907844475 |
| ENSG00000100889 | PCK2 | -0.909509019 | 0.622871051 | 2.10366349 | 0.21055748 | 1.194154471 | 7.096806373 | 8.147241016 | 2 | 6.252910529 | 6.29861567 | 9.49599671 | 8.884335219 |
| ENSG00000107938 | EDRF1 | 1.071471277 | 0.124202911 | 0.122568935 | 0.882372947 | 1.194040212 | 5.532353208 | 6.951736442 | 6.80318718 | 7.574744049 | 7.6318872 | 6.924930115 | 6.389139248 |
| ENSG00000108588 | CCDC47 | 1.046741446 | 0.118180147 | 0.146944645 | 0.850843239 | 1.193686092 | 8.855032352 | 8.834582582 | 10.14394518 | 10.44792714 | 10.4653998 | 9.445288547 | 9.667106106 |
| ENSG00000168229 | PTGDR | 3.498201202 | 0.410778027 | -2.305529472 | 0.608101226 | 1.19267173 | 2 | 2 | 2 | 4.139692846 | 3.74316441 | 2.139668783 | 2 |
| ENSG00000122068 | FYTTD1 | 0.643295307 | 0.433957193 | 0.549214404 | 0.500662365 | 1.192509711 | 8.19931886 | 9.828248506 | 10.11951029 | 10.28176567 | 10.1828016 | 10.13590886 | 10.14282564 |
| ENSG00000109111 | SUPT6H | 0.994403086 | 0.200056839 | 0.197560017 | 0.821567233 | 1.191963103 | 9.304185368 | 9.988044813 | 11.14543241 | 11.32775878 | 11.3528401 | 10.50900237 | 10.58041299 |
| ENSG00000131043 | AAR2 | 0.207705645 | 0.925454021 | 0.983888918 | 0.625198138 | 1.191594563 | 9.317904228 | 8.732943944 | 2 | 8.598473365 | 8.75996988 | 9.352020423 | 9.545113933 |
| ENSG00000103249 | CLCN7 | 1.852015065 | 0.044720075 | -0.660486039 | 0.513707623 | 1.191529025 | 9.720386367 | 9.114109157 | 7.029880454 | 10.81464371 | 10.8602152 | 8.353668905 | 8.325926932 |
| ENSG00000178502 | KLHL11 | 1.107943816 | 0.365386306 | 0.083289212 | 0.952652139 | 1.191233028 | 5.330594134 | 7.431218269 | 4.523675391 | 7.370304568 | 7.4074129 | 6.575182432 | 6.199243628 |
| ENSG00000160233 | LRRC3 | -1.262213643 | 0.343731455 | 2.450841418 | 0.042225848 | 1.188627775 | 3.327711234 | 6.311476754 | 4.058998049 | 4.070463721 | 4.1930041 | 7.710619897 | 7.264927592 |
| ENSG00000236761 | CTAGE9 | 6.379500933 | 0.025566866 | -5.191136046 | 0.127639854 | 1.188364887 | 2 | 2 | 2 | 6.496695859 | 6.44777477 | 2 | 2 |
| ENSG00000140450 | ARRDC4 | 0.676292883 | 0.392311627 | 0.508704795 | 0.523029563 | 1.184997678 | 8.39321512 | 8.862353853 | 9.809100877 | 9.874791085 | 9.7546632 | 10.07738813 | 9.037341395 |
| ENSG00000176396 | EID2 | 0.145612922 | 0.942460221 | 1.038942724 | 0.568673191 | 1.184555647 | 8.138771685 | 6.728495673 | 2 | 7.2696592 | 7.06433609 | 7.813294077 | 8.245238611 |
| ENSG00000128915 | ICE2 | 0.442601512 | 0.72366688 | 0.741005176 | 0.531002733 | 1.183606687 | 4.815609471 | 7.761270542 | 8.110401118 | 7.93509946 | 7.81567127 | 8.269993575 | 8.063441666 |
| ENSG00000128510 | CPA4 | -0.198874274 | 0.925672996 | 1.381966079 | 0.458652814 | 1.183091805 | 5.70932728 | 2 | 6.073378139 | 5.110902178 | 5.22805098 | 4.841348851 | 7.417152486 |
| ENSG00000168517 | HEXIM2 | 1.608810586 | 0.465557972 | -0.42577379 | 0.858491391 | 1.183036797 | 2 | 5.72254323 | 2 | 5.742861563 | 5.84514933 | 4.271694079 | 3.789585079 |
| ENSG00000171045 | TSNARE1 | 0.563391967 | 0.726445618 | 0.619359922 | 0.694591795 | 1.182751889 | 3.327711234 | 3.686303414 | 6.534109959 | 5.731694 | 5.78232107 | 5.818966034 | 5.799216178 |
| ENSG00000136950 | ARPC5L | 0.686902497 | 0.164314918 | 0.495501931 | 0.321838434 | 1.182404428 | 7.790810999 | 8.469613413 | 7.600047325 | 8.622857004 | 8.76268478 | 8.437839892 | 8.569323645 |
| ENSG00000077721 | UBE2A | 0.665370224 | 0.130840883 | 0.51702798 | 0.244791857 | 1.182398204 | 9.544849385 | 8.717821467 | 9.31604076 | 9.932626344 | 9.84581957 | 9.692647041 | 9.790113704 |
| ENSG00000133116 | KL | -1.3104834 | 0.740521619 | 2.49193891 | NA | 1.18145551 | 2 | 2 | 2 | 2.141737246 | 2.14022021 | 3.632645806 | 2.817676617 |

| | | | | | | | | | | | | | |
|---|---|---|---|---|---|---|---|---|---|---|---|---|---|
| ENSG00000106460 | TMEM106B | 0.33118287 | 0.471798515 | 0.849959936 | 0.03766925 | 1.181142807 | 8.674867122 | 8.317412869 | 8.110401118 | 8.795015659 | 8.62772819 | 9.423527655 | 9.002492046 |
| ENSG00000151789 | ZNF385D | 1.104154447 | 0.628529867 | 0.07684531 | 0.974738711 | 1.180999758 | 4.00722531 | 2 | 2 | 3.709785322 | 3.50452879 | 2.937246078 | 3.071407821 |
| ENSG00000198964 | SGMS1 | 0.354144603 | 0.639949127 | 0.826787776 | 0.229804148 | 1.180932379 | 8.042902253 | 6.899080544 | 8.511888463 | 8.312101014 | 8.29037844 | 8.685424964 | 8.847658168 |
| ENSG00000277796 | | 6.37079232 | 0.028382077 | -5.191135276 | 0.132037535 | 1.179657044 | 2 | 2 | 2 | 6.172600389 | 6.70653264 | 2 | 2 |
| ENSG00000133226 | SRRM1 | 0.650664961 | 0.398806121 | 0.528569389 | 0.492624208 | 1.179234349 | 9.751047882 | 8.451443573 | 8.175061096 | 9.643348214 | 9.57865651 | 9.376644377 | 9.594965643 |
| ENSG00000095970 | TREM2 | -3.753036251 | 0.484606942 | 4.932028529 | 0.300671384 | 1.178992278 | 2 | 2 | 2 | 2 | 2 | 5.912830251 | 3.311948089 |
| ENSG00000213918 | DNASE1 | -0.317453789 | 0.770049657 | 1.495745194 | 0.106971668 | 1.178291406 | 5.095959458 | 6.728495673 | 5.930865648 | 5.720439317 | 5.85535975 | 8.066950774 | 6.649933445 |
| ENSG00000100934 | SEC23A | 0.400990749 | 0.693814531 | 0.776537474 | 0.412307433 | 1.177528223 | 8.653385977 | 11.13502303 | 10.62096771 | 10.8418393 | 10.8803543 | 11.2572217 | 11.21457718 |
| ENSG00000106993 | CDC37L1 | 0.388413332 | 0.851528683 | 0.787357733 | 0.684823328 | 1.175771065 | 2 | 6.602424674 | 8.296287027 | 7.517371202 | 7.46188679 | 8.123435792 | 7.589522245 |
| ENSG00000129353 | SLC44A2 | 0.381112134 | 0.48510951 | 0.794557118 | 0.110465713 | 1.175669252 | 11.09707093 | 10.30378101 | 10.21486071 | 10.9418361 | 11.0082583 | 11.20209938 | 11.55296168 |
| ENSG00000140750 | ARHGAP17 | 0.223195816 | 0.52733186 | 0.952321044 | 0.001347185 | 1.17551686 | 9.015175615 | 9.090820894 | 9.439621147 | 9.385472425 | 9.43967292 | 10.06739966 | 10.20648952 |
| ENSG00000175582 | RAB6A | 0.648597485 | 0.237056501 | 0.526640176 | 0.339165925 | 1.175237661 | 9.740899611 | 9.883504052 | 10.76969939 | 10.87774275 | 10.8262286 | 10.76561488 | 10.69456799 |
| ENSG00000162735 | PEX19 | 0.58017549 | 0.406294093 | 0.594422415 | 0.382921621 | 1.174597905 | 8.418903792 | 9.433952963 | 9.972637969 | 9.987058621 | 9.98368402 | 9.865889142 | 10.1219154 |
| ENSG00000090104 | RGS1 | -2.312741879 | 0.674378575 | 3.487252386 | 0.473060066 | 1.174510507 | 2 | 2 | 2 | 2 | 2.14022021 | 4.727866721 | 2.117335275 |
| ENSG00000157259 | GATAD1 | 0.228276904 | 0.712244219 | 0.946091631 | 0.07587755 | 1.174368534 | 8.39321512 | 8.981105197 | 8.325066185 | 8.83470036 | 8.82375772 | 9.894727355 | 9.074092868 |
| ENSG00000115687 | PASK | 0.936025938 | 0.598708746 | 0.237651304 | 0.901746027 | 1.173677242 | 3.327711234 | 5.595927442 | 2 | 5.178261742 | 5.28958453 | 4.933813368 | 4.241259475 |
| ENSG00000100744 | GSKIP | 0.846414387 | 0.686262329 | 0.326729498 | 0.883002588 | 1.173143885 | 2 | 6.389896633 | 8.784770914 | 8.332573384 | 8.248381 | 7.942116583 | 7.595854839 |
| ENSG00000007168 | PAFAH1B1 | 0.295041452 | 0.820993582 | 0.877746485 | 0.453149799 | 1.172787937 | 7.867441304 | 10.51535307 | 11.2690689 | 10.78523872 | 10.6795076 | 11.16263905 | 11.4526447 |
| ENSG00000221869 | CEBPD | 0.366906266 | 0.530881055 | 0.804922155 | 0.128777677 | 1.171828422 | 8.107517749 | 9.018611864 | 8.408111533 | 8.887228971 | 8.97656697 | 9.621696553 | 9.058587737 |
| ENSG00000127616 | SMARCA4 | 0.410396067 | 0.52243871 | 0.75762372 | 0.199517168 | 1.168019786 | 9.174465797 | 10.3778065 | 9.503962296 | 10.17296029 | 10.2079206 | 10.37096889 | 10.68539259 |
| ENSG00000156030 | ELMSAN1 | 0.254378258 | 0.67218509 | 0.913302252 | 0.080754483 | 1.16768051 | 8.313285031 | 7.819099893 | 8.825624954 | 8.593855195 | 8.64842725 | 9.023081974 | 9.489401982 |
| ENSG00000144485 | HES6 | -0.358795464 | 0.820253704 | 1.525429812 | 0.260131361 | 1.166634348 | 4.467311984 | 5.72254323 | 2 | 4.470469939 | 4.45712836 | 6.353989995 | 5.896817464 |
| ENSG00000040275 | SPDL1 | 0.91932035 | 0.476505377 | 0.245471105 | 0.860801801 | 1.164791455 | 8.586954094 | 8.376385542 | 4.874587708 | 8.84120963 | 8.89725143 | 8.242494717 | 8.162331091 |
| ENSG00000186132 | C2orf76 | 0.392198987 | 0.841622412 | 0.77191035 | 0.674363565 | 1.164109337 | 5.532353208 | 2 | 3.369317972 | 4.739373666 | 4.50979031 | 5.002522427 | 4.903386193 |
| ENSG00000173039 | RELA | 0.366588791 | 0.528469036 | 0.79751928 | 0.12999296 | 1.164108071 | 9.384607364 | 10.07368926 | 10.48730704 | 10.43937195 | 10.3914606 | 10.65972326 | 11.01000224 |
| ENSG00000108001 | EBF3 | -0.681559707 | 0.809107998 | 1.843010066 | 0.459209794 | 1.161450359 | 2 | 3.686303414 | 2 | 2.868578001 | 2.2680111 | 4.443291373 | 3.311948089 |
| ENSG00000147649 | MTDH | 0.491231989 | 0.207531949 | 0.670161882 | 0.070946784 | 1.161393871 | 10.05943807 | 10.40166101 | 10.75391088 | 10.92814295 | 10.9161551 | 11.07299152 | 11.12797955 |
| ENSG00000173409 | ARV1 | 0.539420588 | 0.78336925 | 0.621013239 | 0.744740559 | 1.160433827 | 2 | 6.602424674 | 8.076950877 | 7.523859802 | 7.42122564 | 7.340163764 | 7.738907626 |
| ENSG00000102910 | LONP2 | 0.088296307 | 0.931617375 | 1.071865002 | 0.215498263 | 1.160161309 | 8.716893051 | 8.540087354 | 10.45512158 | 9.601017164 | 9.60234163 | 10.62621766 | 10.53549289 |
| ENSG00000185803 | SLC52A2 | 0.609850721 | 0.451559843 | 0.548922807 | 0.492084442 | 1.158773528 | 9.384607364 | 9.506147172 | 7.691296561 | 9.653739712 | 9.68624302 | 9.646549955 | 9.571380235 |
| ENSG00000102890 | ELMO3 | -0.100042052 | 0.961244994 | 1.258662839 | 0.488001567 | 1.158620787 | 4.00722531 | 3.076665209 | 2 | 2.868578001 | 3.45167086 | 4.434275199 | 3.660199087 |
| ENSG00000100522 | GNPNAT1 | 1.07264829 | 0.275627669 | 0.085785265 | 0.940230037 | 1.158433555 | 6.73718411 | 7.638155693 | 9.196533591 | 9.32080297 | 9.22907886 | 8.237320313 | 8.356084628 |
| ENSG00000105649 | RAB3A | 1.011556307 | 0.535687826 | 0.146444276 | 0.935496579 | 1.158000582 | 7.031698319 | 5.72254323 | 2 | 6.941624891 | 6.9210099 | 6.109723842 | 6.099376274 |
| ENSG00000004399 | PLXND1 | -0.843742838 | 0.137345288 | 2.001327315 | 8.35E-05 | 1.157584477 | 9.966689162 | 9.256956408 | 8.784770914 | 8.499891024 | 8.65866647 | 11.5508907 | 11.26163861 |
| ENSG00000151876 | FBXO4 | 0.645412024 | 0.734810023 | 0.511751494 | 0.789905257 | 1.157163518 | 3.327711234 | 5.946638857 | 2 | 5.093559143 | 5.41912973 | 5.348031117 | 4.911532223 |
| ENSG00000012048 | BRCA1 | 1.170332113 | 0.235251572 | -0.013544381 | 0.990611315 | 1.156787732 | 3.327711234 | 5.595927442 | 4.874587708 | 6.061031724 | 5.87556634 | 4.854926465 | 4.943664032 |
| ENSG00000151502 | VPS26B | -0.352645484 | 0.174638808 | 1.509076506 | 7.66E-12 | 1.156431022 | 9.128497779 | 9.170736405 | 9.083172337 | 8.820275739 | 8.73806474 | 10.62163296 | 10.64214212 |
| ENSG00000136451 | VEZF1 | 0.645775755 | 0.725609862 | 0.5091375 | 0.784027326 | 1.154913255 | 3.327711234 | 9.566483142 | 9.099922433 | 9.479434065 | 9.35854723 | 9.395058233 | 9.164631393 |
| | CCL3L3 | 6.345960029 | 0.030507988 | -5.191134925 | 0.134274864 | 1.154825104 | 2 | 2 | 2 | 6.139181293 | 6.68951263 | 2 | 2 |
| ENSG00000052841 | TTC17 | 0.589023168 | 0.704833394 | 0.565402969 | 0.71335944 | 1.154426137 | 5.532353208 | 10.26269067 | 8.865353883 | 9.785808044 | 9.75124983 | 9.970549666 | 9.477790979 |
| ENSG00000136436 | CALCOCO2 | 0.679720163 | 0.329996554 | 0.474513853 | 0.503157092 | 1.154234016 | 10.19198706 | 9.828248506 | 8.608698767 | 10.37802153 | 10.3478309 | 10.24273081 | 10.06895712 |
| ENSG00000118292 | C1orf54 | 0.801092904 | 0.536848085 | 0.352581418 | 0.796677889 | 1.153674322 | 7.27616306 | 5.303523747 | 4.058998049 | 6.892328675 | 6.90143415 | 6.401024833 | 6.528072775 |
| ENSG00000089682 | RBM41 | 1.159527669 | 0.156709059 | -0.006060647 | 0.995062288 | 1.153467022 | 7.031698319 | 7.847167909 | 5.930865648 | 8.338106704 | 8.23671175 | 7.427945125 | 6.793487375 |
| ENSG00000080546 | SESN1 | 0.144001584 | 0.943253539 | 1.009117024 | 0.581437534 | 1.153118608 | 2 | 7.761270542 | 6.958202216 | 7.080147179 | 6.89649843 | 8.359038717 | 7.006211772 |
| ENSG00000214530 | STARD10 | 0.030227345 | 0.982666154 | 1.122817542 | 0.340932973 | 1.153044887 | 4.815609471 | 8.191702463 | 8.007629717 | 7.52061915 | 7.71833702 | 8.879345195 | 8.494699808 |
| ENSG00000160318 | CLDND2 | 0.728617391 | 0.790784372 | 0.42284701 | 0.879570438 | 1.151464401 | 2 | 3.076665209 | 2 | 2.947924294 | 2.77779907 | 2.68698273 | 2.782576693 |
| ENSG00000128699 | ORMDL1 | 0.662936493 | 0.551800049 | 0.488234036 | 0.666867266 | 1.151170529 | 7.975256505 | 9.523645035 | 6.718950348 | 9.211498619 | 9.13163078 | 9.123606729 | 8.861915706 |
| ENSG00000164168 | TMEM184C | 0.528923682 | 0.446900339 | 0.621561566 | 0.353466558 | 1.150075248 | 7.435667355 | 8.297207175 | 8.978369997 | 8.874592369 | 8.90710092 | 9.002642099 | 8.961885532 |
| ENSG00000124479 | NDP | 3.047145665 | 0.523661392 | -1.897326629 | 0.702841297 | 1.149819036 | 2 | 2 | 2 | 3.663521627 | 3.60476711 | 2.183373728 | 2 |
| ENSG00000181004 | BBS12 | 0.720736108 | 0.670452104 | 0.428957693 | 0.804379534 | 1.149693801 | 2 | 5.595927442 | 6.882775709 | 6.573765157 | 6.42738748 | 6.407949645 | 6.014942101 |
| ENSG00000121653 | MAPK8IP1 | 0.490442097 | 0.699694748 | 0.659100455 | 0.588785095 | 1.149542552 | 3.327711234 | 6.389896633 | 6.43197674 | 6.415275989 | 6.34282908 | 6.237020416 | 6.791281771 |
| ENSG00000130649 | CYP2E1 | 0.44687083 | 0.77994267 | 0.701307938 | 0.64228189 | 1.148178767 | 2 | 4.11352228 | 4.058998049 | 4.17310023 | 3.82856379 | 4.45225155 | 3.908316285 |
| ENSG00000177045 | SIX5 | 0.702989977 | 0.73397139 | 0.445167031 | 0.833553126 | 1.148157008 | 5.866944937 | 8.395519306 | 2 | 7.691737974 | 7.80517502 | 7.628021136 | 7.352801367 |
| ENSG00000168297 | PXK | 0.443718202 | 0.796967396 | 0.704354423 | 0.668486483 | 1.148072625 | 7.384433967 | 8.451443573 | 3.369317972 | 7.922877029 | 7.8745862 | 8.589193677 | 7.532563185 |
| ENSG00000129292 | PHF20L1 | 0.898744777 | 0.306048059 | 0.249111812 | 0.795874263 | 1.147856589 | 8.16936288 | 7.73146271 | 9.69246155 | 9.711725296 | 9.64416621 | 9.123606729 | 8.935663927 |
| ENSG00000138686 | BBS7 | 0.982018923 | 0.531143362 | 0.164758624 | 0.925295809 | 1.146777547 | 3.327711234 | 6.899080544 | 8.266922072 | 8.136938566 | 8.15847021 | 7.557113378 | 7.09829807 |
| ENSG00000068079 | IFI35 | 2.555363014 | 0.238266302 | -1.409111794 | 0.532346835 | 1.14625122 | 10.12333326 | 6.464272742 | 2 | 11.18792842 | 11.2037318 | 7.05561307 | 7.451079708 |
| ENSG00000162368 | CMPK1 | 0.429866735 | 0.623107483 | 0.716331193 | 0.382493233 | 1.146197928 | 8.816894989 | 9.159587375 | 10.57976717 | 10.23853288 | 10.0760734 | 10.61877565 | 10.24769118 |
| ENSG00000196911 | KPNA5 | 0.44367403 | 0.811172496 | 0.701474344 | 0.690278517 | 1.145148375 | 2 | 6.78762695 | 4.874587708 | 6.069927018 | 5.96314905 | 6.45105139 | 6.041166521 |
| ENSG00000009780 | FAM76A | 0.659856564 | 0.706615573 | 0.485200421 | 0.784357456 | 1.145056985 | 2 | 6.602424674 | 7.342949392 | 7.093295078 | 7.09050444 | 6.857395343 | 6.985139097 |
| ENSG00000172817 | CYP7B1 | 0.212015851 | 0.927391636 | 0.932843185 | 0.658380896 | 1.144859036 | 2 | 4.11352228 | 2 | 3.227757282 | 3.33976443 | 3.694366604 | 3.807156736 |
| ENSG00000116035 | VAX2 | 0.560041001 | 0.807722186 | 0.584741022 | 0.795268255 | 1.144782022 | 2 | 4.93638933 | 2 | 4.329433326 | 3.98554648 | 3.904814898 | 4.4231929 |
| ENSG00000022840 | RNF10 | 0.416942322 | 0.478107313 | 0.726754355 | 0.181447155 | 1.143696677 | 9.761125267 | 10.51535307 | 9.589475645 | 10.4457931 | 10.4188272 | 10.42288506 | 11.00215671 |
| ENSG00000197362 | ZNF786 | 0.398418583 | 0.760990111 | 0.74505019 | 0.543096343 | 1.143468773 | 4.815609471 | 6.311476754 | 3.369317972 | 5.627094009 | 5.77157844 | 6.140539117 | 5.896817464 |
| ENSG00000243244 | STON1 | 0.384134807 | 0.847860259 | 0.757507382 | 0.686224112 | 1.141642189 | 2 | 8.124485672 | 7.163363831 | 7.555872146 | 7.50468064 | 8.11499695 | 7.63942137 |
| ENSG00000198001 | IRAK4 | 0.711978449 | 0.654474154 | 0.429571327 | 0.793476131 | 1.141549776 | 4.467311984 | 5.457120953 | 8.408111533 | 7.799951945 | 7.73502229 | 7.717139459 | 7.22299542 |
| ENSG00000100441 | KHNYN | 0.504755617 | 0.671131573 | 0.636303123 | 0.578462503 | 1.14105874 | 9.27634997 | 9.599860728 | 6.534109959 | 9.381905119 | 9.53167935 | 9.77507622 | 9.376603759 |
| ENSG00000130347 | RTN4IP1 | 0.166216354 | 0.91483416 | 0.974699122 | 0.473386442 | 1.140915475 | 3.327711234 | 5.72254323 | 3.369317972 | 4.623314311 | 4.97065356 | 5.4091291 | 5.631815919 |
| ENSG00000176125 | UFSP1 | -0.075266601 | 0.973704667 | 1.216068003 | 0.549060202 | 1.140801402 | 2 | 4.710564154 | 2 | 3.663521627 | 3.39680241 | 4.620674722 | 4.365060851 |
| ENSG00000104852 | SNRNP70 | 0.12276665 | 0.903601303 | 1.016934303 | 0.229671798 | 1.139700954 | 10.43166252 | 8.234834549 | 9.91701143 | 9.971714969 | 9.83811764 | 10.6925865 | 10.89595288 |
| ENSG00000256223 | ZNF10 | 0.765890814 | 0.767238548 | 0.373765404 | 0.889574473 | 1.139656218 | 2 | 6.666836837 | 2 | 5.94010643 | 5.88556448 | 5.758846871 | 5.306932076 |
| ENSG00000133561 | GIMAP6 | -0.733152585 | NA | 1.872209133 | NA | 1.139056548 | 2 | 2 | 2 | 2 | 2.38539787 | 3.336159116 | 2.421067459 |
| ENSG00000260230 | FRRS1L | -2.801422478 | 0.166629297 | 3.940160475 | 0.03232554 | 1.138737997 | 2 | 2 | 5.59507254 | 2.695456494 | 2.49394783 | 8.139473554 | 7.373589743 |
| ENSG00000168505 | GBX2 | -3.753052554 | 0.464636768 | 4.891395323 | 0.278925497 | 1.138342769 | 2 | 2 | 2 | 2 | 2 | 5.650128416 | 4.188602707 |
| ENSG00000169306 | IL1RAPL1 | 3.443389262 | 0.422792655 | -2.305804935 | 0.610398832 | 1.137584327 | 2 | 2 | 2 | 3.79808662 | 4.02226117 | 2.139668783 | 2 |
| ENSG00000087053 | MTMR2 | 0.18195346 | 0.856036852 | 0.95560691 | 0.265190311 | 1.13756037 | 7.624139768 | 9.814097497 | 9.478569125 | 9.432787137 | 9.40794139 | 10.31542752 | 10.05385647 |
| ENSG00000183814 | LIN9 | 1.028460086 | 0.669545698 | 0.108968658 | 0.967122742 | 1.137428744 | 2 | 5.838937808 | 2 | 5.433909581 | 5.30456641 | 4.984105765 | 3.987514168 |
| ENSG00000172840 | PDP2 | 0.652630915 | 0.441837321 | 0.484326975 | 0.569975156 | 1.136957889 | 8.39321512 | 6.602424674 | 7.029880454 | 8.211183901 | 8.16669309 | 8.017249226 | 8.029224588 |
| ENSG00000163959 | SLC51A | -3.753058285 | 0.456587162 | 4.889008004 | 0.268957569 | 1.135949719 | 2 | 2 | 2 | 2 | 2 | 4.596630906 | 5.464725981 |
| ENSG00000137876 | RSL24D1 | 0.665345072 | 0.484223993 | 0.469559815 | 0.626190802 | 1.134904887 | 9.159305284 | 8.297207175 | 6.882775709 | 9.053418491 | 9.03948698 | 9.14414236 | 8.485861839 |
| ENSG00000144711 | IQSEC1 | -0.524042989 | 0.620673635 | 1.658540477 | 0.075265997 | 1.134497488 | 9.761125267 | 8.929529568 | 7.029880454 | 8.422134132 | 8.44542686 | 10.64526705 | 10.56015326 |
| ENSG00000223609 | HBD | -3.753046562 | 0.472493119 | 4.885780587 | 0.289935311 | 1.132734025 | 2 | 2 | 2 | 2 | 2 | 3.865109708 | 5.745545708 |
| ENSG00000182993 | C12orf60 | 0.649701868 | 0.698007365 | 0.482729901 | 0.773825851 | 1.132431769 | 3.327711234 | 4.11352228 | 2 | 3.997744386 | 3.7864959 | 3.930690651 | 3.600986722 |
| ENSG00000117151 | CTBS | 2.007010237 | 0.000140956 | -0.874802202 | 0.127639854 | 1.132208035 | 7.667647223 | 6.728495673 | 7.97167868 | 9.56056556 | 9.46656462 | 6.781219718 | 6.574984961 |
| ENSG00000122729 | ACO1 | -0.364205092 | 0.506977228 | 1.495666997 | 0.001282361 | 1.130861904 | 9.384607364 | 8.981105197 | 8.699419242 | 8.643859774 | 8.73115061 | 10.12806717 | 10.85575293 |
| ENSG00000168273 | SMIM4 | 1.223491477 | 0.293606106 | -0.09417951 | 0.944101725 | 1.129311966 | 6.257048221 | 4.11352228 | 4.058998049 | 6.29144905 | 6.33555364 | 4.868377487 | 5.300743019 |
| ENSG00000177084 | POLE | 0.201821637 | 0.85332146 | 0.927330666 | 0.325557697 | 1.129152302 | 5.095959458 | 7.572420273 | 6.718950348 | 7.003290579 | 6.97819215 | 7.799317009 | 7.589522245 |
| ENSG00000129083 | COPB1 | 0.99160389 | 0.026999437 | 0.136787676 | 0.797729416 | 1.128391566 | 10.91047246 | 10.31889309 | 10.01740919 | 11.45776963 | 11.4495233 | 10.70165563 | 10.49167828 |
| ENSG00000186187 | ZNRF1 | -0.136334662 | 0.85249814 | 1.262536412 | 0.032526136 | 1.12620175 | 8.313285031 | 8.004992147 | 6.882775709 | 7.706073672 | 7.7377845 | 9.041753004 | 9.163352825 |
| ENSG00000187866 | FAM122A | 0.421154344 | 0.832844524 | 0.701539516 | 0.710275609 | 1.12269386 | 2 | 7.503546079 | 8.206335842 | 7.783822874 | 7.68723697 | 8.061837451 | 7.96021985 |
| ENSG00000109436 | TBC1D9 | 1.593312096 | 0.007712736 | -0.471224391 | 0.483213396 | 1.122087705 | 8.586954094 | 7.503546079 | 7.734844356 | 9.587916984 | 9.60006644 | 7.880388025 | 7.128314671 |
| ENSG00000108821 | COL1A1 | -0.05553266 | 0.878292377 | 1.177415735 | 1.22E-05 | 1.121883075 | 16.50604222 | 16.62872997 | 16.91596224 | 16.60604717 | 16.6700869 | 17.77225622 | 17.96405002 |
| ENSG00000113732 | ATP6V0E1 | 1.007120342 | 0.010191064 | 0.114701849 | 0.809379135 | 1.121822191 | 10.25752168 | 10.24169813 | 10.87563042 | 11.50833956 | 11.4778868 | 10.4659642 | 10.72723989 |
| ENSG00000170088 | TMEM192 | 0.2460735 | 0.687336686 | 0.875200914 | 0.099247094 | 1.121274414 | 6.653098118 | 7.60566235 | 6.882775709 | 7.437142666 | 7.28811304 | 7.86621124 | 8.087908463 |
| ENSG00000155827 | RNF20 | 0.72042312 | 0.021458981 | 0.400271475 | 0.22346137 | 1.120694595 | 8.313285031 | 8.004992147 | 8.076950877 | 8.786946023 | 8.90218458 | 8.520567915 | 8.53677936 |
| ENSG00000257218 | GATC | 0.773533602 | 0.227425425 | 0.346420599 | 0.610398832 | 1.119954201 | 7.867441304 | 8.234834549 | 6.80318718 | 8.538855014 | 8.50986596 | 8.201899078 | 7.996092644 |
| ENSG00000006715 | VPS41 | 0.469161983 | 0.726949143 | 0.648859687 | 0.616400319 | 1.118021671 | 5.70932728 | 8.623598428 | 9.465702942 | 9.005540335 | 9.09655277 | 9.035431928 | 9.401760658 |
| ENSG00000125875 | TBC1D20 | 0.219383541 | 0.733353042 | 0.898453203 | 0.107495597 | 1.117836744 | 10.08373159 | 9.632483534 | 8.805342542 | 9.820482079 | 9.81541592 | 10.53229574 | 10.45592109 |
| ENSG00000160785 | SLC25A44 | 0.552797788 | 0.807307128 | 0.564434798 | 0.799962733 | 1.117232587 | 9.771132748 | 7.701025996 | 2 | 9.05677832 | 9.03724641 | 8.846588786 | 9.243382473 |
| ENSG00000122490 | PQLC1 | 0.341916036 | 0.588043549 | 0.774042542 | 0.176369212 | 1.115958578 | 7.579279349 | 8.234834549 | 8.865353883 | 8.679168098 | 8.63218877 | 9.128855427 | 9.038271684 |
| ENSG00000188368 | PRR19 | 0.765281136 | 0.727489842 | 0.347897664 | 0.879564373 | 1.1131788 | 2 | 4.11352228 | 2 | 3.709785322 | 3.55551834 | 3.483289517 | 3.236138336 |
| ENSG00000169239 | CA5B | -0.393660247 | 0.739140058 | 1.50648611 | 0.143980561 | 1.112825863 | 5.095959458 | 6.228547981 | 7.777116083 | 6.358325776 | 6.42052719 | 8.546948034 | 7.845855157 |
| ENSG00000205268 | PDE7A | 0.701456821 | 0.463053095 | 0.411257407 | 0.675028301 | 1.112714227 | 4.00722531 | 6.140560021 | 6.073378139 | 6.29144905 | 6.41363412 | 6.247352872 | 5.888587685 |
| ENSG00000011376 | LARS2 | 0.695737617 | 0.730413605 | 0.416976322 | 0.840253981 | 1.112713939 | 8.737455672 | 7.145017454 | 2 | 8.145379268 | 8.36552655 | 7.860333006 | 8.099542409 |
| ENSG00000168461 | RAB31 | 0.687629817 | 0.052939325 | 0.424785026 | 0.249077972 | 1.112414843 | 10.27168932 | 10.57950211 | 9.972637969 | 10.99271605 | 10.9764528 | 10.80702258 | 10.63231802 |
| ENSG00000184160 | ADRA2C | -1.232188041 | 0.604386987 | 2.343096719 | 0.261724935 | 1.110908678 | 2 | 3.686303414 | 2 | 2.270786764 | 2.59489879 | 4.530483275 | 4.254128754 |
| ENSG00000109736 | MFSD10 | 0.565104143 | 0.560004695 | 0.543823767 | 0.570388322 | 1.10892791 | 9.886101849 | 8.777382925 | 7.552162999 | 9.567648462 | 9.62265847 | 9.330617798 | 9.782638733 |

| | | | | | | | | | | | | | |
|---|---|---|---|---|---|---|---|---|---|---|---|---|---|
| ENSG00000152380 | FAM151B | 0.706214314 | 0.758269479 | 0.399613243 | 0.864992366 | 1.105827557 | 2 | 5.303523747 | 2 | 4.415698455 | 4.76938658 | 4.487545924 | 4.201948172 |
| ENSG00000109572 | CLCN3 | 0.527413376 | 0.704195879 | 0.574941694 | 0.673640399 | 1.10235507 | 5.70932728 | 8.732943944 | 9.714382178 | 9.286870873 | 9.3166689 | 9.394767778 | 9.301879118 |
| ENSG00000130813 | C19orf66 | 1.941426824 | 0.247157846 | -0.839151685 | 0.638604205 | 1.102275139 | 10.37373127 | 5.303523747 | 6.629488387 | 10.85444304 | 10.8725209 | 8.094396944 | 8.107542072 |
| ENSG00000187231 | SESTD1 | 0.422866345 | 0.84453415 | 0.678015251 | 0.740101564 | 1.100881596 | 8.313285031 | 9.181799937 | 2 | 8.660150093 | 8.64695856 | 9.113404777 | 8.664579282 |
| ENSG00000000419 | DPM1 | 1.475511389 | 0.163819362 | -0.375704265 | 0.749808479 | 1.099807124 | 8.367060731 | 6.04685525 | 6.073378139 | 8.744518279 | 8.69038176 | 7.006273211 | 6.78907279 |
| ENSG00000170961 | HAS2 | 3.818563938 | 0.005459329 | -2.718923171 | 0.055260188 | 1.099640767 | 6.563806279 | 8.834582582 | 9.670202709 | 12.64894221 | 12.6252174 | 7.046766788 | 3.67940853 |
| ENSG00000138764 | CCNG2 | 1.539674367 | 0.191786205 | -0.443022055 | 0.732581748 | 1.096652312 | 5.330594134 | 8.004992147 | 8.960138144 | 9.587916984 | 9.56082893 | 7.690882553 | 7.532563185 |
| ENSG00000106261 | ZKSCAN1 | 0.213179912 | 0.73312176 | 0.883200121 | 0.102696941 | 1.096380033 | 9.317904228 | 9.806969599 | 10.40208303 | 10.16675444 | 10.0755276 | 11.01398252 | 10.52458037 |
| ENSG00000109046 | WSB1 | 0.800023724 | 0.120735975 | 0.296161657 | 0.595909021 | 1.096185381 | 9.646202137 | 9.975384048 | 10.67785662 | 10.95239719 | 10.972354 | 10.56209021 | 10.34957621 |
| ENSG00000133433 | GSTT2B | 6.285527061 | 0.032987346 | -5.191134849 | 0.135323833 | 1.094392212 | 2 | 2 | 2 | 6.379946408 | 6.3857277 | 2 | 2 |
| ENSG00000176834 | VSIG10 | 1.154333199 | 0.488856774 | -0.064278552 | 0.972248849 | 1.090054647 | 2 | 6.535001748 | 7.285536013 | 7.43025113 | 7.58969985 | 6.475428841 | 6.179140936 |
| ENSG00000169598 | DFFB | 0.559215493 | 0.508392192 | 0.530616028 | 0.521153279 | 1.08983152 | 4.467311984 | 5.72254323 | 4.874587708 | 5.565529336 | 5.76075522 | 5.696113106 | 5.581648407 |
| ENSG00000014138 | POLA2 | 1.612032121 | 0.170202551 | -0.52241666 | 0.684865709 | 1.089615461 | 3.327711234 | 6.602424674 | 5.39244166 | 7.303993732 | 7.08183429 | 4.996409624 | 5.361470313 |
| ENSG00000069020 | MAST4 | 1.183144606 | 0.550957372 | -0.094351207 | 0.966440292 | 1.088793398 | 2 | 7.051611315 | 8.825624954 | 8.76109026 | 8.80938679 | 7.808940772 | 7.174740504 |
| ENSG00000111911 | HINT3 | 0.378883407 | 0.33193825 | 0.709185207 | 0.048045799 | 1.088068614 | 8.228665467 | 8.414402626 | 8.076950877 | 8.643859774 | 8.60974666 | 9.121501891 | 8.764652886 |
| ENSG00000100883 | SRP54 | 1.058306802 | 0.001669747 | 0.028985164 | 0.94688402 | 1.087291966 | 8.517314938 | 8.732943944 | 9.116480286 | 9.853698554 | 9.86489641 | 8.794695657 | 8.877074023 |
| ENSG00000129625 | REEP5 | 1.097955562 | 0.002123735 | -0.01101708 | 0.981056901 | 1.086938482 | 10.46286899 | 9.792607274 | 9.999667074 | 11.21699976 | 11.1947131 | 10.06976682 | 10.12913753 |
| ENSG00000136770 | DNAJC1 | 1.545376082 | 0.179452048 | -0.460795479 | 0.716145684 | 1.084580603 | 9.410448395 | 8.276714465 | 6.073378139 | 10.06420548 | 9.92611804 | 8.364981826 | 7.531242162 |
| ENSG00000155096 | AZIN1 | 0.868543828 | 0.049761083 | 0.216007871 | 0.66704421 | 1.084551659 | 9.790941832 | 9.726146444 | 10.43545992 | 10.89848915 | 10.871578 | 10.02617706 | 10.41621675 |
| ENSG00000115827 | DCAF17 | 0.415496067 | 0.535201232 | 0.668042874 | 0.288186427 | 1.083538941 | 7.750913381 | 6.535001748 | 7.098165388 | 7.662632018 | 7.54938955 | 8.059640467 | 7.615934347 |
| ENSG00000128815 | WDFY4 | -3.753037904 | 0.483854018 | 4.836149506 | 0.31009769 | 1.083111602 | 2 | 2 | 2 | 2 | 2 | 5.769639817 | 3.539239711 |
| ENSG00000177302 | TOP3A | 0.489107526 | 0.71007843 | 0.593939031 | 0.641832906 | 1.083046557 | 5.095959458 | 7.503546079 | 8.677268317 | 8.178656739 | 8.185025 | 8.443467705 | 8.107542072 |
| ENSG00000174796 | THAP6 | 0.751217359 | 0.39960716 | 0.331225818 | 0.724423786 | 1.082443177 | 5.330594134 | 6.311476754 | 7.163363831 | 7.157295126 | 7.18663697 | 7.125896536 | 6.283256815 |
| ENSG00000153113 | CAST | 0.803328538 | 0.007872825 | 0.278649428 | 0.404998648 | 1.081977966 | 10.52923584 | 10.12617756 | 10.29705308 | 11.10597219 | 11.1469365 | 10.45945184 | 10.73371512 |
| ENSG00000148400 | NOTCH1 | -0.526973024 | 0.706963346 | 1.608762051 | 0.195995925 | 1.081789027 | 10.00110988 | 7.189546432 | 6.882775709 | 8.244958867 | 8.20312688 | 10.56040784 | 10.0915399 |
| ENSG00000197894 | ADH5 | 0.575321858 | 0.148344449 | 0.503610428 | 0.203432268 | 1.078932286 | 10.11549941 | 9.558016677 | 9.452720984 | 10.32559865 | 10.2962973 | 10.23236508 | 10.24668541 |
| ENSG00000167716 | WDR81 | 2.949186931 | 0.096199 | -1.870553838 | 0.305382004 | 1.078633093 | 4.815609471 | 7.790474954 | 2 | 9.253098082 | 9.35225233 | 4.914492341 | 4.479073019 |
| ENSG00000197283 | SYNGAP1 | 3.826808083 | 9.25E-05 | -2.748504671 | 0.007503255 | 1.078303412 | 5.095959458 | 6.464272742 | 3.369317972 | 9.211498619 | 9.21229508 | 3.534812207 | 3.128433649 |
| ENSG00000163516 | ANKZF1 | 0.416287819 | 0.811431778 | 0.661789553 | 0.689452881 | 1.078077372 | 3.327711234 | 8.337339476 | 8.608698767 | 8.374462892 | 8.27524772 | 8.624303429 | 8.5102048 |
| ENSG00000139842 | CUL4A | 0.555931936 | 0.494662338 | 0.521035324 | 0.516558603 | 1.07696726 | 10.01801709 | 9.377333006 | 8.110401118 | 9.883647099 | 9.95310979 | 9.787077426 | 9.974915827 |
| ENSG00000143418 | CERS2 | 1.186806401 | 1.75E-05 | -0.110086832 | 0.74998849 | 1.076719568 | 10.13887448 | 10.3138733 | 10.07784268 | 11.34755451 | 11.384692 | 9.888345571 | 10.23598093 |
| ENSG00000183864 | TOB2 | 0.356721654 | 0.611774065 | 0.717345209 | 0.267661148 | 1.074066863 | 9.33149386 | 8.777382925 | 7.818184403 | 9.115954467 | 9.13163078 | 9.491928968 | 9.472294474 |
| ENSG00000131236 | CAP1 | 0.929941166 | 0.283881375 | 0.142956637 | 0.885544215 | 1.072897803 | 11.70484245 | 11.00090285 | 9.478569125 | 11.89635181 | 11.9405179 | 11.05867027 | 11.20311249 |
| ENSG00000145592 | RPL37 | 0.428416973 | 0.715571124 | 0.644315383 | 0.565593677 | 1.072732356 | 11.31110672 | 9.114109157 | 8.845626171 | 10.61183616 | 10.6621638 | 11.06379481 | 10.60510859 |
| ENSG00000138376 | BARD1 | 0.026003698 | 0.987001945 | 1.044337043 | 0.453525552 | 1.070340741 | 5.70932728 | 6.140560021 | 2 | 5.514313187 | 5.37722731 | 6.529346487 | 6.251520369 |
| ENSG00000110917 | MLEC | -0.012444626 | 0.98350331 | 1.082670239 | 0.024735408 | 1.070225613 | 10.19198706 | 10.83911531 | 11.16147894 | 10.74553862 | 10.7983847 | 11.63457429 | 12.06463678 |
| ENSG00000155906 | RMND1 | -0.079283928 | 0.947432314 | 1.14894686 | 0.268972233 | 1.069662933 | 8.075571721 | 6.951736442 | 5.156626401 | 7.157295126 | 7.00595521 | 8.4151065 | 8.146888246 |
| ENSG00000105784 | RUNDC3B | -2.305941104 | 0.617291379 | 3.371333613 | 0.392847555 | 1.065392509 | 2 | 2 | 2 | 2.141737246 | 2 | 3.930690651 | 3.789585079 |
| ENSG00000102144 | PGK1 | 0.412534376 | 0.716492539 | 0.65255626 | 0.544753443 | 1.065090635 | 12.91048752 | 11.64711661 | 10.23774519 | 12.3799161 | 12.4031015 | 12.09610914 | 13.02096146 |
| ENSG00000148843 | PDCD11 | 0.463596082 | 0.693529621 | 0.601348771 | 0.596070489 | 1.064944853 | 6.009025997 | 8.169643011 | 9.099922433 | 8.687861792 | 8.68752719 | 8.896269648 | 8.74825912 |
| ENSG00000136463 | TACO1 | 0.279991015 | 0.848343006 | 0.784530242 | 0.554021903 | 1.064251258 | 7.096806373 | 8.004992147 | 4.058998049 | 7.348537858 | 7.4074129 | 7.755649985 | 7.980689421 |
| ENSG00000138696 | BMPR1B | 3.743685578 | 0.008473525 | -2.680087471 | 0.079084492 | 1.063598108 | 2 | 4.710564154 | 2 | 6.841288255 | 6.83591385 | 2.384001718 | 2.421067459 |
| ENSG00000169019 | COMMD8 | 0.71883224 | 0.575783958 | 0.344344166 | 0.797495402 | 1.063176406 | 6.89194353 | 6.728495673 | 3.369317972 | 7.088925745 | 6.88657603 | 6.685680098 | 6.572418365 |
| ENSG00000166974 | MAPRE2 | -0.043076625 | 0.961267251 | 1.106247473 | 0.136956979 | 1.063170848 | 9.87686338 | 8.414402626 | 8.325066185 | 8.971529165 | 9.05949698 | 10.00711202 | 10.29710605 |
| ENSG00000080822 | CLDND1 | 0.791530259 | 0.115289102 | 0.271623456 | 0.620529852 | 1.063153715 | 9.219014269 | 8.981105197 | 8.206335842 | 9.706743577 | 9.59398167 | 9.246842239 | 9.012940881 |
| ENSG00000104825 | NFKBIB | 0.841733084 | 0.501949395 | 0.219443847 | 0.872155485 | 1.061176932 | 8.75772933 | 6.140560021 | 6.073378139 | 8.447952961 | 8.34212018 | 7.674704537 | 7.883613091 |
| ENSG00000181634 | TNFSF15 | 2.274944218 | 0.006236051 | -1.21466539 | 0.172169048 | 1.060278828 | 5.330594134 | 4.93638933 | 5.39244166 | 7.384635376 | 7.35080119 | 4.644324387 | 3.539239711 |
| ENSG00000072201 | LNX1 | -0.686353273 | 0.774715444 | 1.746433287 | 0.423520139 | 1.060080014 | 2 | 5.946638857 | 2 | 4.205751501 | 3.82856379 | 6.054896192 | 6.222348126 |
| ENSG00000126351 | THRA | -0.175274449 | 0.855650806 | 1.235243857 | 0.122176759 | 1.059969408 | 7.27616306 | 8.124485672 | 9.301632639 | 8.325162455 | 8.2657098 | 9.759594615 | 9.626855891 |
| ENSG00000065833 | ME1 | 0.898469613 | 0.031533344 | 0.160345423 | 0.740394508 | 1.058815037 | 7.975256505 | 8.687092496 | 8.325066185 | 9.26185831 | 9.25051318 | 8.35963413 | 8.672390305 |
| ENSG00000132549 | VPS13B | 0.218176471 | 0.841552857 | 0.838384585 | 0.381507263 | 1.056561057 | 7.031698319 | 9.470501606 | 9.31604076 | 9.135152476 | 9.18927228 | 9.619709545 | 9.922723748 |
| ENSG00000203879 | GDI1 | 0.025887369 | 0.94878661 | 1.029112279 | 0.001098499 | 1.054999648 | 10.98115017 | 10.60026656 | 10.56174313 | 10.7438026 | 10.760856 | 11.58483423 | 11.90525006 |
| ENSG00000143632 | ACTA1 | 2.668055973 | 0.128993961 | -1.613897053 | 0.376931525 | 1.05415892 | 2 | 5.303523747 | 4.058998049 | 6.622948477 | 6.90143415 | 3.782262616 | 2.326754077 |
| ENSG00000203668 | CHML | -0.798688733 | 0.46309526 | 1.851376473 | 0.056913062 | 1.05268774 | 7.579279349 | 5.595927442 | 5.39244166 | 5.987823101 | 5.56313034 | 8.552166664 | 8.116378869 |
| ENSG00000131188 | PRR7 | -0.184287874 | 0.864318577 | 1.235244229 | 0.166131291 | 1.050956355 | 4.467311984 | 5.303523747 | 6.534109959 | 5.304221918 | 5.67111583 | 6.704551153 | 6.957866334 |
| ENSG00000102870 | ZNF629 | 0.077305014 | 0.881572243 | 0.972960291 | 0.016733148 | 1.050265305 | 8.981092835 | 8.234834549 | 8.585098385 | 8.658676732 | 8.74631819 | 9.470589819 | 9.704799739 |
| ENSG00000243444 | PALM2 | 2.958923197 | 0.000162583 | -1.909910494 | 0.022627193 | 1.049012702 | 4.815609471 | 5.595927442 | 4.523675391 | 7.855024645 | 7.92150349 | 3.904814898 | 3.15612297 |
| ENSG00000122482 | ZNF644 | 0.736589359 | 0.130096658 | 0.311249551 | 0.548805363 | 1.047838911 | 8.042902253 | 8.356994601 | 8.486635645 | 9.035366048 | 9.03836713 | 8.93593651 | 8.20189164 |
| ENSG00000111445 | RFC5 | -0.050085257 | 0.982507576 | 1.096457513 | 0.587973908 | 1.046372255 | 3.327711234 | 6.844429755 | 2 | 5.258265913 | 5.53799497 | 6.475428841 | 6.474094059 |
| ENSG00000196357 | ZNF565 | 0.187934195 | 0.880488774 | 0.856953985 | 0.427508497 | 1.044888181 | 4.00722531 | 5.946638857 | 4.523675391 | 5.178261742 | 5.37722731 | 6.210860813 | 5.486701579 |
| ENSG00000198355 | PIM3 | 0.648521463 | 0.30880261 | 0.395842158 | 0.544751469 | 1.044363621 | 8.540903468 | 8.004992147 | 9.272377397 | 9.286870873 | 9.38697919 | 8.843186716 | 9.295279234 |
| ENSG00000113569 | NUP155 | 0.286692032 | 0.88899497 | 0.757349417 | 0.688288292 | 1.044041449 | 8.340423424 | 7.315436877 | 2 | 7.635924947 | 7.61395775 | 8.171702207 | 7.998006535 |
| ENSG00000018625 | ATP1A2 | -1.499673253 | 0.356981191 | 2.541713422 | 0.085269823 | 1.042040169 | 4.00722531 | 3.686303414 | 4.874587708 | 3.79808662 | 2.2680111 | 5.991769846 | 6.91798856 |
| ENSG00000132964 | CDK8 | 0.730076069 | 0.096173427 | 0.311295071 | 0.505195354 | 1.04137114 | 7.435667355 | 7.189546432 | 7.451345288 | 8.15586117 | 7.98565528 | 7.858649102 | 7.432801564 |
| ENSG00000140859 | KIFC3 | 0.466035741 | 0.717903042 | 0.575134688 | 0.646513979 | 1.041170428 | 10.29268328 | 7.051611315 | 8.434763955 | 9.629876784 | 9.64121862 | 9.524419769 | 9.934763826 |
| ENSG00000198668 | CALM1 | 0.394443059 | 0.507120384 | 0.646417246 | 0.244987518 | 1.040860304 | 11.62983336 | 11.30149645 | 10.61515347 | 11.64928415 | 11.6210902 | 11.53917738 | 12.16696017 |
| ENSG00000176472 | ZNF575 | -0.082248565 | 0.95389048 | 1.122473112 | 0.355912095 | 1.040224547 | 4.467311984 | 4.442710129 | 2 | 3.997744386 | 3.94787299 | 4.977914261 | 4.927687581 |
| ENSG00000100060 | MFNG | -2.312593073 | 0.61703084 | 3.352633357 | 0.397192742 | 1.040040284 | 2 | 2 | 2 | 2 | 2.14022021 | 3.838019203 | 3.858622973 |
| ENSG00000179152 | TCAIM | -0.039175195 | 0.957413691 | 1.078840541 | 0.070856581 | 1.039665346 | 6.89194353 | 6.666836837 | 7.896971941 | 7.222570549 | 7.18663697 | 8.268726114 | 8.333525663 |
| ENSG00000086712 | TXLNG | 0.717213474 | 0.347836352 | 0.322251568 | 0.68935002 | 1.039465042 | 6.96351253 | 6.78762695 | 8.236946975 | 8.237082658 | 8.11663771 | 7.992893101 | 7.552234734 |
| ENSG00000123159 | GIPC1 | -0.199251919 | 0.80934297 | 1.238003 | 0.072624316 | 1.038751081 | 8.444143041 | 9.763446433 | 10.17589429 | 9.414570905 | 9.44559847 | 10.776023 | 10.94340508 |
| ENSG00000125629 | INSIG2 | 0.676805393 | 0.262504755 | 0.361636402 | 0.56576762 | 1.038441794 | 9.048471759 | 8.147241016 | 7.818184403 | 9.128781483 | 9.07923325 | 8.786305336 | 8.796891925 |
| ENSG00000198947 | DMD | -1.690087007 | 0.215891254 | 2.728010258 | 0.032479705 | 1.037923251 | 4.467311984 | 8.639735838 | 6.629488387 | 6.006476466 | 5.62411716 | 10.13920704 | 10.15230914 |
| ENSG00000162385 | MAGOH | 1.532787592 | 0.254759301 | -0.495026377 | 0.735074986 | 1.037761215 | 2 | 5.72254323 | 6.073378139 | 6.687949235 | 6.93069905 | 4.952879058 | 4.919632516 |
| ENSG00000016402 | IL20RA | -2.305744058 | 0.629279802 | 3.343053519 | 0.419331922 | 1.03730946 | 2 | 2 | 2 | 2.141737246 | 2 | 3.12825509 | 4.316807515 |
| ENSG00000064651 | SLC12A2 | -0.286900474 | 0.608680785 | 1.324134251 | 0.00487588 | 1.037233776 | 7.750913381 | 7.954278496 | 8.236946975 | 7.739909082 | 7.68150967 | 8.877267588 | 9.632103066 |
| ENSG00000105538 | RASIP1 | -2.141005764 | 0.314789729 | 3.177135768 | 0.077959309 | 1.036130003 | 2 | 3.076665209 | 3.369317972 | 2.270786764 | 2.2680111 | 5.747972572 | 4.30448772 |
| ENSG00000139641 | ESYT1 | 0.166129026 | 0.802782927 | 0.869843956 | 0.12255258 | 1.035972983 | 9.829762239 | 10.54135596 | 9.465702942 | 10.15216947 | 10.2168574 | 10.57483812 | 11.14297651 |
| ENSG00000140564 | FURIN | 0.288205309 | 0.700066046 | 0.74686252 | 0.268928351 | 1.035067829 | 10.43166252 | 10.69982928 | 9.149036566 | 10.47329248 | 10.5706238 | 10.94899883 | 11.01106879 |
| ENSG00000143157 | POGK | -0.267435407 | 0.73397139 | 1.302277072 | 0.051791194 | 1.034841665 | 7.579279349 | 9.018611864 | 9.149036566 | 8.468280417 | 8.47387021 | 10.19981619 | 9.837378403 |
| ENSG00000170049 | KCNAB3 | -0.998593852 | 0.650789466 | 2.033042986 | 0.312242938 | 1.034449134 | 2 | 5.457120953 | 2 | 3.615725078 | 3.28037825 | 6.146071859 | 5.685057129 |
| ENSG00000205476 | CCDC85C | 0.062450639 | 0.97026695 | 0.971383889 | 0.518789265 | 1.033834528 | 8.257427002 | 7.315436877 | 3.369317972 | 7.261917051 | 7.47187556 | 8.221684808 | 8.303661547 |
| ENSG00000145241 | CENPC | 0.737335375 | 0.665819919 | 0.295352372 | 0.868784979 | 1.032687747 | 2 | 6.951736442 | 7.029880454 | 7.062428156 | 7.21862736 | 6.723178551 | 6.707209278 |
| ENSG00000143028 | SYPL2 | -0.725021169 | 0.73917724 | 1.756600624 | 0.370804133 | 1.031579455 | 2 | 4.93638933 | 2 | 3.462030052 | 3.08597292 | 5.30910438 | 5.028505473 |
| ENSG00000139636 | LMBR1L | 1.105290613 | 0.227729767 | -0.07456722 | 0.944363944 | 1.030723393 | 8.285626337 | 6.228547981 | 6.718950348 | 8.379838154 | 8.4936155 | 7.461614824 | 7.069464998 |
| ENSG00000132600 | PRMT7 | 0.51675037 | 0.730963227 | 0.511140876 | 0.730619275 | 1.027891246 | 7.031698319 | 7.73146271 | 3.369317972 | 7.381065997 | 7.40393901 | 7.182183216 | 7.566491953 |
| ENSG00000215421 | ZNF407 | 0.5686966 | 0.625452298 | 0.458940139 | 0.694164121 | 1.027636738 | 8.285626337 | 6.04685525 | 5.930865648 | 7.68596357 | 7.79989812 | 7.474865722 | 7.78175402 |
| ENSG00000101343 | CRNKL1 | 0.813860906 | 0.484988029 | 0.213246242 | 0.866182611 | 1.027107149 | 5.095959458 | 7.73146271 | 8.325066185 | 8.408172799 | 8.3183278 | 7.833151805 | 7.708840864 |
| ENSG00000198356 | ASNA1 | 0.386143043 | 0.815821608 | 0.640815325 | 0.683411628 | 1.026958368 | 4.467311984 | 9.13702746 | 9.372276026 | 9.034230235 | 9.13163078 | 9.281189863 | 9.390148213 |
| ENSG00000167552 | TUBA1A | 0.090832195 | 0.912408896 | 0.935642388 | 0.16984745 | 1.026474583 | 12.00538348 | 11.37081044 | 11.11279376 | 11.66716327 | 11.6066186 | 11.6643988 | 12.99956744 |
| ENSG00000028839 | TBPL1 | 0.23083103 | 0.803843283 | 0.79361221 | 0.333586444 | 1.02444324 | 5.866944937 | 7.572420273 | 7.896971941 | 7.629222299 | 7.50792049 | 8.024025946 | 8.215937769 |
| ENSG00000158292 | GPR153 | 0.172262748 | 0.93751235 | 0.851972567 | 0.674981326 | 1.024235315 | 4.00722531 | 7.189546432 | 2 | 5.94010643 | 6.00043082 | 6.913621294 | 6.238627697 |
| ENSG00000101193 | GID8 | 0.212815897 | 0.854355329 | 0.811271147 | 0.428528777 | 1.024087045 | 9.048471759 | 9.214490408 | 6.534109959 | 8.864402784 | 8.88981993 | 9.379582949 | 9.557820379 |
| ENSG00000154359 | LONRF1 | -0.214675993 | 0.756822448 | 1.237237166 | 0.032014466 | 1.022561174 | 6.009025997 | 6.311476754 | 6.958202216 | 6.29144905 | 6.24526024 | 7.914016653 | 7.380942161 |
| ENSG00000145220 | LYAR | 0.777686828 | 0.583250623 | 0.243726873 | 0.872485883 | 1.021413701 | 7.435667355 | 6.04685525 | 3.369317972 | 7.106324235 | 7.14562489 | 6.554528258 | 6.664466987 |
| ENSG00000181007 | ZFP82 | 0.326092604 | 0.86184497 | 0.694844839 | 0.689153776 | 1.020937443 | 7.435667355 | 6.728495673 | 2 | 6.856790872 | 6.89649843 | 7.464938976 | 6.959831579 |
| ENSG00000184939 | ZFP90 | -0.368934056 | 0.812768397 | 1.386547575 | 0.309101085 | 1.017613519 | 4.467311984 | 8.762721501 | 8.699419242 | 7.748245381 | 7.89204953 | 9.775522494 | 9.304974529 |
| ENSG00000196664 | TLR7 | -1.048995563 | 0.625524857 | 2.064716866 | 0.274873919 | 1.015721302 | 3.327711234 | 2 | 3.369317972 | 2.498691189 | 2.59489879 | 4.73522696 | 3.735548302 |
| ENSG00000170144 | HNRNPA3 | 1.073278195 | 0.009101188 | -0.057819775 | 0.911555747 | 1.015458421 | 8.963744667 | 9.599860728 | 9.788583153 | 10.56737883 | 10.5550472 | 9.357098224 | 9.508324596 |
| ENSG00000144230 | GPR17 | -3.753054793 | 0.463231524 | 4.766481437 | 0.290456069 | 1.013426644 | 2 | 2 | 2 | 2 | 2 | 4.888321757 | 5.080008298 |
| ENSG00000160049 | DFFA | -0.511979825 | 0.466937891 | 1.525252154 | 0.013281894 | 1.013272329 | 8.517314938 | 8.747909548 | 7.285536013 | 7.893113669 | 7.72670378 | 9.984514132 | 9.656845513 |
| ENSG00000079841 | RIMS1 | -0.47657268 | 0.849039274 | 1.489662828 | 0.50032005 | 1.013090148 | 2 | 2 | 4.523675391 | 2.695456494 | 3.33976443 | 4.416072028 | 4.341135913 |
| ENSG00000151612 | ZNF827 | 0.521782981 | 0.730447087 | 0.491289602 | 0.743325344 | 1.013072582 | 7.82963488 | 5.131603625 | 9.439621147 | 8.814994458 | 8.84440862 | 8.457441933 | 9.075906094 |
| ENSG00000138767 | CNOT6L | 0.865882135 | 0.649796974 | 0.141301366 | 0.944972617 | 1.007183501 | 8.19931886 | 8.234834549 | 2 | 8.545247918 | 8.44711562 | 7.953205078 | 7.589522245 |
| ENSG00000132004 | FBXW9 | 0.470400488 | 0.864688463 | 0.534936422 | 0.841812696 | 1.00533691 | 2 | 6.844429755 | 2 | 5.900763176 | 5.70538658 | 5.515547099 | 6.148451724 |
| ENSG00000154144 | TBRG1 | 1.072404723 | 0.009121679 | -0.068444653 | 0.894874854 | 1.00396007 | 7.904282243 | 8.191702463 | 8.721235202 | 9.382797773 | 9.36034073 | 8.266822833 | 8.215115301 |
| ENSG00000088833 | NSFL1C | 0.391879512 | 0.638674782 | 0.611911371 | 0.438092003 | 1.003790883 | 7.384433967 | 9.181799937 | 9.049077186 | 9.123450746 | 9.12742405 | 9.240067514 | 9.440980262 |

| | | | | | | | | | | | | | |
|---|---|---|---|---|---|---|---|---|---|---|---|---|---|
| ENSG00000100867 | DHRS2 | 2.39111736 | 0.48535932 | -1.391306034 | NA | 0.999811326 | 2 | 2 | 2 | 3.349643765 | 3.08597292 | 2.094598369 | 2.172608095 |
| ENSG00000132912 | DCTN4 | 0.758615305 | 0.499380069 | 0.240884264 | 0.841697129 | 0.999499569 | 9.064836028 | 10.019219 | 6.958202216 | 9.927741409 | 9.87559569 | 9.481025247 | 9.285910049 |
| ENSG00000115446 | UNC50 | 0.787275219 | 0.498492641 | 0.212068028 | 0.866223157 | 0.999343246 | 8.586954094 | 7.979858146 | 5.39244166 | 8.537252353 | 8.66157866 | 8.193240267 | 7.85221717 |
| ENSG00000147044 | CASK | 0.739810008 | 0.490247308 | 0.25931473 | 0.82055968 | 0.999124738 | 7.435667355 | 8.540087354 | 5.772716771 | 8.435961651 | 8.32569041 | 8.070592084 | 7.725108479 |
| ENSG00000186862 | PDZD7 | -1.455601688 | 0.465603893 | 2.454510686 | 0.172295853 | 0.998908998 | 2 | 4.93638933 | 2 | 2.784612485 | 3.01487888 | 5.630526941 | 5.984375752 |
| ENSG00000144935 | TRPC1 | -0.225534374 | 0.87294376 | 1.224308878 | 0.308986373 | 0.998774504 | 5.532353208 | 6.311476754 | 8.585098385 | 7.210554257 | 7.15392111 | 8.802599622 | 8.380498741 |
| ENSG00000197620 | CXorf40A | 1.057486238 | 0.214681289 | -0.063347435 | 0.949476741 | 0.994138803 | 7.750913381 | 6.389896633 | 5.772716771 | 7.855024645 | 7.96222667 | 6.791847192 | 6.841176624 |
| ENSG00000112210 | RAB23 | 0.155173961 | 0.898417957 | 0.83714063 | 0.429336127 | 0.992314591 | 7.096806373 | 9.835272298 | 8.677268317 | 9.095408867 | 9.07268445 | 10.10392776 | 9.313068526 |
| ENSG00000158417 | EIF5B | 0.775800356 | 0.001397551 | 0.215239376 | 0.43458157 | 0.991039732 | 9.98400217 | 9.71856931 | 9.888372618 | 10.58687666 | 10.6877448 | 10.10570349 | 10.05247582 |
| ENSG00000115816 | CEBPZ | 0.913897797 | 0.524696221 | 0.076785035 | 0.962239953 | 0.990682833 | 9.304185368 | 8.702538795 | 4.874587708 | 9.380118153 | 9.41313469 | 8.741779902 | 8.366492631 |
| ENSG00000110077 | MS4A6A | -3.753048705 | 0.471586224 | 4.743349317 | 0.30371251 | 0.990300612 | 2 | 2 | 2 | 2 | 2 | 5.390603804 | 4.365060851 |
| ENSG00000102054 | RBBP7 | 0.505658307 | 0.282799793 | 0.484102298 | 0.295471026 | 0.989760606 | 9.081016759 | 9.396454291 | 8.511888463 | 9.569217728 | 9.52928986 | 9.448650487 | 9.603161713 |
| ENSG00000147853 | AK3 | 0.196565549 | 0.821777402 | 0.793094323 | 0.295788824 | 0.989659873 | 9.771132748 | 8.255926468 | 8.296287027 | 9.136211578 | 9.16894939 | 9.662803762 | 9.824224109 |
| ENSG00000148180 | GSN | -0.341803699 | 0.222084596 | 1.329588194 | 3.17E-08 | 0.987784495 | 11.50945836 | 11.42768197 | 11.4724895 | 11.0877485 | 11.1685936 | 12.61732815 | 12.95921509 |
| ENSG00000116212 | LRRC42 | 0.206151623 | 0.89994774 | 0.781614332 | 0.596648352 | 0.987765955 | 4.467311984 | 9.067150531 | 7.777116083 | 8.23312837 | 8.21902896 | 8.599816608 | 8.967268538 |
| ENSG00000014641 | MDH1 | 0.454799739 | 0.252546786 | 0.532893651 | 0.164653895 | 0.98769339 | 9.128497779 | 9.608085847 | 9.767769415 | 9.978816937 | 9.97960581 | 9.937430448 | 10.16748181 |
| ENSG00000006125 | AP2B1 | -0.059340782 | 0.906842758 | 1.046305156 | 0.007498635 | 0.986964374 | 10.70134604 | 10.51971956 | 11.19694409 | 10.75866452 | 10.7900919 | 11.69591971 | 12.04029115 |
| ENSG00000141219 | C17orf80 | 0.342885744 | 0.812947514 | 0.643422318 | 0.635522997 | 0.986308062 | 7.579279349 | 7.901717006 | 4.058998049 | 7.574744049 | 7.5462416 | 8.113585664 | 7.540464021 |
| ENSG00000128191 | DGCR8 | 0.408678617 | 0.774885822 | 0.576114862 | 0.676798727 | 0.984793479 | 8.928409675 | 8.356994601 | 4.874587708 | 8.543652346 | 8.54024645 | 8.660513226 | 8.753366804 |
| ENSG00000156858 | PRR14 | 0.358733407 | 0.35676743 | 0.624980194 | 0.082216561 | 0.9837136 | 7.709880939 | 8.169643011 | 7.858115942 | 8.217201935 | 8.35656898 | 8.588177878 | 8.514222325 |
| ENSG00000126777 | KTN1 | 1.232746047 | 0.301721022 | -0.249059101 | 0.852211698 | 0.983686946 | 7.940205782 | 9.583268459 | 11.32712828 | 11.44115415 | 11.4604341 | 10.03738467 | 9.9049832 |
| ENSG00000174428 | GTF2IRD2B | 2.16333317 | 0.004958412 | -1.1859433 | 0.148125821 | 0.97738987 | 6.138360429 | 6.140560021 | 4.058998049 | 7.842101511 | 7.7866208 | 4.555645384 | 4.80186789 |
| ENSG00000138190 | EXOC6 | 0.211431143 | 0.92098972 | 0.765865694 | 0.69597295 | 0.977296837 | 7.750913381 | 5.131603625 | 2 | 6.470063213 | 6.72891807 | 7.142599214 | 7.138761401 |
| ENSG00000165102 | HGSNAT | 0.205670872 | 0.571578307 | 0.770397839 | 0.014292331 | 0.976068711 | 10.1770105 | 10.05576005 | 9.714382178 | 10.19904132 | 10.2039308 | 10.63902691 | 10.88021462 |
| ENSG00000186314 | PRELID2 | -0.145847117 | 0.861743464 | 1.120327607 | 0.097911265 | 0.97448049 | 5.532353208 | 5.838937808 | 4.874587708 | 5.128039203 | 5.56313034 | 6.597565628 | 6.487779256 |
| ENSG00000197965 | MPZL1 | 0.562287718 | 0.213853811 | 0.412004017 | 0.36789886 | 0.974291735 | 10.70134604 | 11.30149645 | 11.57188978 | 11.79739298 | 11.796598 | 11.61238761 | 11.68058894 |
| ENSG00000167525 | PROCA1 | -0.437619502 | 0.801824404 | 1.409791463 | 0.350123893 | 0.972171962 | 3.327711234 | 4.11352228 | 2 | 3.023133209 | 3.33976443 | 4.487545924 | 4.489993931 |
| ENSG00000139990 | DCAF5 | 0.293798487 | 0.740549771 | 0.678148369 | 0.406804715 | 0.971946856 | 8.418903792 | 9.203675681 | 10.34701993 | 9.757180673 | 9.90230962 | 10.18712501 | 10.24124211 |
| ENSG00000141985 | SH3GL1 | 0.251616342 | 0.685162585 | 0.719270908 | 0.192814682 | 0.97088725 | 9.497412313 | 10.63690565 | 9.888372618 | 10.31488156 | 10.3657986 | 10.692232 | 10.91329715 |
| ENSG00000006740 | ARHGAP44 | -2.000977692 | 0.160968748 | 2.971857812 | 0.023498087 | 0.970880121 | 5.866944937 | 4.710564154 | 2 | 3.754611435 | 2.86123014 | 7.652523812 | 7.697107799 |
| ENSG00000184056 | VPS33B | 1.409591725 | 0.38839201 | -0.4405703 | 0.803056929 | 0.969021425 | 5.532353208 | 7.002537973 | 2 | 7.292639438 | 7.23044321 | 5.180212365 | 5.772630506 |
| ENSG00000175746 | C15orf54 | 3.275372785 | 0.463403238 | -2.306869944 | 0.621054729 | 0.968502842 | 2 | 2 | 2 | 3.921164147 | 3.65239003 | 2.139668783 | 2 |
| ENSG00000153107 | ANAPC1 | 1.46230928 | 0.438477481 | -0.494509678 | 0.806845941 | 0.967799602 | 7.384433967 | 8.717821467 | 2 | 9.097585392 | 9.04730166 | 7.067324363 | 7.221358049 |
| ENSG00000103199 | ZNF500 | 0.368055054 | 0.841373243 | 0.59875438 | 0.730981666 | 0.966809434 | 2 | 7.051611315 | 7.225742779 | 6.89733479 | 6.96881778 | 7.267148095 | 7.038185015 |
| ENSG00000096746 | HNRNPH3 | 0.640198224 | 0.283681909 | 0.326526354 | 0.600974746 | 0.966724578 | 9.848787621 | 8.94259736 | 8.742726175 | 9.936887136 | 9.85793083 | 9.753944082 | 9.395240163 |
| ENSG00000184465 | WDR27 | -0.232320908 | 0.866469882 | 1.198849994 | 0.309292345 | 0.966529086 | 3.327711234 | 6.535001748 | 6.534109959 | 5.829220145 | 5.80356929 | 7.447120996 | 6.856025745 |
| ENSG00000179172 | HNRNPCL1 | 5.693619691 | 0.000723945 | -4.72751968 | 0.050406299 | 0.966100011 | 2 | 3.076665209 | 2 | 6.772115916 | 6.70088158 | 2.048074405 | 2 |
| ENSG00000065154 | OAT | 0.287525884 | 0.576906213 | 0.676414691 | 0.14670132 | 0.963940574 | 10.08373159 | 10.21502048 | 9.385998894 | 10.2136641 | 10.2399348 | 10.40756301 | 10.79568025 |
| ENSG00000151746 | BICD1 | -0.318323161 | 0.841373243 | 1.281139929 | 0.357318652 | 0.962816768 | 4.815609471 | 6.535001748 | 8.865353883 | 7.230526165 | 7.36158648 | 8.973764228 | 8.762965592 |
| ENSG00000151687 | ANKAR | 0.826485518 | 0.566726644 | 0.13592821 | 0.932067558 | 0.962413728 | 4.815609471 | 4.11352228 | 2 | 4.783303513 | 4.65713986 | 4.45225155 | 3.789585079 |
| ENSG00000157916 | RER1 | 0.785540321 | 0.052158739 | 0.176086749 | 0.701998223 | 0.96162707 | 10.03472846 | 9.975384048 | 9.399592459 | 10.63487332 | 10.594821 | 9.816217077 | 10.17638265 |
| ENSG00000112246 | SIM1 | 6.152526882 | 0.045644094 | -5.191133054 | 0.147030332 | 0.961393828 | 2 | 2 | 2 | 6.197166296 | 6.31350454 | 2 | 2 |
| ENSG00000130985 | UBA1 | 0.440666976 | 0.136814806 | 0.517710141 | 0.070779599 | 0.958377117 | 11.26227543 | 11.41367288 | 11.47570169 | 11.78038858 | 11.8717671 | 11.71605378 | 12.06987785 |
| ENSG00000239732 | TLR9 | 2.215435294 | NA | -1.257443073 | NA | 0.957992221 | 2 | 2 | 2 | 3.349643765 | 2.86123014 | 2.225793514 | 2.059860178 |
| ENSG00000164031 | DNAJB14 | 0.823251859 | 0.342311654 | 0.134148093 | 0.89140859 | 0.957399952 | 7.867441304 | 7.503546079 | 9.31604076 | 9.363935069 | 9.15970984 | 8.828207602 | 8.284971554 |
| ENSG00000127080 | IPPK | 1.797416607 | 0.340701465 | -0.840604118 | 0.673761372 | 0.956812489 | 4.467311984 | 7.503546079 | 2 | 7.831679003 | 7.89946959 | 5.085488617 | 5.576534358 |
| ENSG00000132382 | MYBBP1A | 0.630073531 | 0.498588661 | 0.326632109 | 0.734711071 | 0.956705639 | 7.031698319 | 8.395519306 | 9.31604076 | 9.100844036 | 9.19330273 | 8.702422971 | 8.977974666 |
| ENSG00000163788 | SNRK | 0.676046577 | 0.738698334 | 0.280070079 | 0.895191045 | 0.956116656 | 8.855032352 | 7.538394143 | 2 | 8.413424144 | 8.4504872 | 8.195243072 | 7.869046644 |
| ENSG00000111077 | TNS2 | -0.007060232 | 0.988487935 | 0.9625769 | 0.010218512 | 0.955516669 | 7.485143553 | 7.819099893 | 7.858115942 | 7.608714138 | 7.83385794 | 8.544855285 | 8.807298241 |
| ENSG00000168795 | ZBTB5 | -0.302071613 | 0.845142817 | 1.257001835 | 0.352660759 | 0.954930223 | 9.015175615 | 5.946638857 | 5.772716771 | 7.423326516 | 7.42465798 | 8.985770579 | 8.93516457 |
| ENSG00000181610 | MRPS23 | 0.662266074 | 0.563764353 | 0.286075545 | 0.812905528 | 0.948341619 | 5.330594134 | 7.538394143 | 8.434763955 | 8.184811706 | 8.26379462 | 7.908330388 | 7.79832288 |
| ENSG00000113163 | COL4A3BP | 0.78690186 | 0.315977132 | 0.161120301 | 0.854072961 | 0.948022162 | 6.816637977 | 8.255926468 | 8.677268317 | 8.859280876 | 8.91322288 | 8.425380879 | 8.089704419 |
| ENSG00000108474 | PIGL | -0.328546146 | 0.908151255 | 1.276095261 | 0.617024792 | 0.947549115 | 2 | 7.099070191 | 2 | 5.461211713 | 5.14724905 | 7.094763493 | 6.471341372 |
| ENSG00000113391 | FAM172A | -0.632176171 | 0.77110538 | 1.577981272 | 0.428900076 | 0.945805101 | 2 | 4.93638933 | 8.206335842 | 6.069927018 | 6.26836889 | 8.138086022 | 8.479026367 |
| ENSG00000270276 | HIST2H4B | 6.136220417 | 0.047200445 | -5.19113285 | 0.148400754 | 0.945087567 | 2 | 2 | 2 | 6.205262847 | 6.27599024 | 2 | 2 |
| ENSG00000270882 | HIST2H4A | 6.136220417 | 0.047200445 | -5.19113285 | 0.148400754 | 0.945087567 | 2 | 2 | 2 | 6.205262847 | 6.27599024 | 2 | 2 |
| ENSG00000197054 | ZNF763 | 1.93776038 | 0.124543846 | -0.993223815 | 0.459077723 | 0.944536565 | 4.467311984 | 3.076665209 | 3.369317972 | 5.48800675 | 5.11361705 | 3.316666208 | 2.885414003 |
| ENSG00000204604 | ZNF468 | 0.39481804 | 0.641177228 | 0.54794382 | 0.498062551 | 0.942761861 | 7.975256505 | 7.35507212 | 6.203069917 | 7.683067677 | 7.78928606 | 8.173735115 | 7.529919928 |
| ENSG00000144504 | ANKMY1 | 0.006628273 | 0.997720641 | 0.934353219 | 0.715181868 | 0.940981492 | 2 | 6.728495673 | 2 | 5.37770462 | 5.16377554 | 6.444330878 | 5.727202496 |
| ENSG00000106526 | ACTR3C | 4.722420473 | 0.104495453 | -3.783183945 | 0.271243655 | 0.939236529 | 2 | 2 | 2 | 4.887609717 | 5.00774437 | 2.048074405 | 2 |
| ENSG00000182704 | TSKU | 1.333894466 | 0.101156806 | -0.395120135 | 0.66187124 | 0.93877433 | 10.4253394 | 9.267380431 | 8.35328246 | 10.88185282 | 10.9629415 | 9.168473351 | 9.232450166 |
| ENSG00000182022 | CHST15 | -0.357797314 | 0.896917828 | 1.295610925 | 0.603099089 | 0.937813611 | 2 | 3.076665209 | 8.65477198 | 6.722207358 | 6.80463998 | 7.96967964 | 8.700282474 |
| ENSG00000137124 | ALDH1B1 | -1.736201203 | 0.15408844 | 2.67070397 | 0.019811774 | 0.934502768 | 9.556468883 | 9.338312924 | 5.772716771 | 7.182120794 | 7.24604864 | 11.48917421 | 11.67267253 |
| ENSG00000176058 | TPRN | -0.198993166 | 0.897826717 | 1.133271685 | 0.396880252 | 0.934278519 | 7.53297911 | 7.099070191 | 3.369317972 | 6.5612028 | 6.64900057 | 8.001311979 | 7.795024296 |
| ENSG00000137040 | RANBP6 | 0.447299193 | 0.832297095 | 0.48576192 | 0.813013269 | 0.933061113 | 2 | 8.623598428 | 8.631919287 | 8.469961514 | 8.51470562 | 8.663409709 | 8.38416207 |
| ENSG00000167658 | EEF2 | -0.458414385 | 0.092448935 | 1.389973675 | 7.63E-09 | 0.93155929 | 13.98403793 | 13.59338974 | 13.78504471 | 13.32578388 | 13.349718 | 15.15818234 | 15.21287184 |
| ENSG00000136709 | WDR33 | 0.504663202 | 0.348531182 | 0.426873457 | 0.426694614 | 0.931536659 | 7.667647223 | 8.671479029 | 8.631919287 | 8.896009193 | 8.89477851 | 8.800408416 | 8.835935827 |
| ENSG00000214491 | SEC14L6 | -3.753044207 | 0.478107313 | 4.683926509 | 0.319695652 | 0.930882302 | 2 | 2 | 2 | 2 | 2 | 4.240999983 | 5.373314945 |
| ENSG00000054611 | TBC1D22A | 0.092945177 | 0.95203552 | 0.837260472 | 0.549261882 | 0.930205649 | 5.330594134 | 7.467835385 | 9.344432377 | 8.197043419 | 8.32935768 | 8.582578161 | 9.325696621 |
| ENSG00000160209 | PDXK | 0.391451624 | 0.586677368 | 0.538272345 | 0.43412023 | 0.929723969 | 10.60951384 | 9.903691196 | 9.049077186 | 10.36723728 | 10.3923374 | 10.50200781 | 10.55007877 |
| ENSG00000166579 | NDEL1 | 0.169276525 | 0.58791184 | 0.760162197 | 0.003916559 | 0.929438721 | 9.790941832 | 9.624396729 | 9.452720984 | 9.755803193 | 9.83940415 | 10.43832796 | 10.33356098 |
| ENSG00000021574 | SPAST | 0.403711937 | 0.553397759 | 0.525586259 | 0.421989779 | 0.929298196 | 6.009025997 | 6.899080544 | 7.398165132 | 7.292639438 | 7.23826702 | 7.52030827 | 7.234405279 |
| ENSG00000174697 | LEP | -3.273501426 | 0.257133882 | 4.202783823 | 0.04664104 | 0.929282398 | 2 | 2 | 3.369317972 | 2 | 2.14022021 | 4.952879058 | 5.646532192 |
| ENSG00000171467 | ZNF318 | 0.160372977 | 0.807381293 | 0.767802743 | 0.172094144 | 0.928175721 | 8.257427002 | 8.317412869 | 9.301632639 | 8.868232317 | 8.8597051 | 9.574262696 | 9.351374718 |
| ENSG00000135069 | PSAT1 | -0.843829148 | 0.330982566 | 1.770155812 | 0.0252943 | 0.926326664 | 8.107517749 | 9.514922632 | 7.691296561 | 7.880528731 | 7.7866208 | 10.72217346 | 10.05546557 |
| ENSG00000196323 | ZBTB44 | 0.564131808 | 0.139917893 | 0.361776331 | 0.35529554 | 0.925908139 | 7.940205782 | 7.847167909 | 8.076950877 | 8.595396228 | 8.41985547 | 8.465761957 | 8.135631371 |
| ENSG00000134900 | TPP2 | 1.279534093 | 0.019532885 | -0.354108996 | 0.566151826 | 0.925425096 | 7.867441304 | 8.834582582 | 7.777116083 | 9.516508835 | 9.52849248 | 7.775419242 | 8.024537783 |
| ENSG00000162994 | CLHC1 | 0.428936699 | 0.869954492 | 0.495908778 | 0.845529489 | 0.924845477 | 2 | 6.228547981 | 2 | 5.076005082 | 5.30456641 | 5.357600875 | 5.14363432 |
| ENSG00000148606 | POLR3A | 0.923141841 | 0.546911592 | 0.001096196 | 0.999379962 | 0.924238036 | 4.467311984 | 8.147241016 | 9.413059136 | 9.263797825 | 9.28868707 | 8.419110791 | 8.303661547 |
| ENSG00000141873 | SLC39A3 | 0.261054917 | 0.804796579 | 0.661390601 | 0.491172843 | 0.922445518 | 7.904282243 | 8.213429689 | 5.772716771 | 7.810605345 | 7.95749494 | 8.21774931 | 8.340330468 |
| ENSG00000081059 | TCF7 | 1.735139078 | 0.174638808 | -0.812848508 | 0.547277519 | 0.922290569 | 4.00722531 | 7.002537973 | 4.523675391 | 7.366699513 | 7.61094772 | 4.933813368 | 5.23734678 |
| ENSG00000174013 | FBXO45 | 0.190028402 | 0.845522872 | 0.732212602 | 0.392103714 | 0.922241004 | 6.138360429 | 6.311476754 | 7.691296561 | 7.161462517 | 6.96410766 | 7.936540212 | 7.145684098 |
| ENSG00000118257 | NRP2 | 1.286851975 | 0.18345602 | -0.365231606 | 0.731701381 | 0.921620369 | 7.904282243 | 9.943237334 | 10.57976717 | 11.10272512 | 11.1482347 | 9.622689032 | 9.319587711 |
| ENSG00000104142 | VPS18 | 0.336098373 | 0.673891173 | 0.585371264 | 0.434090596 | 0.921469637 | 7.579279349 | 8.88960062 | 9.330306409 | 9.089953145 | 9.11579188 | 9.258063226 | 9.437458576 |
| ENSG00000136108 | CKAP2 | 0.322096409 | 0.886772235 | 0.598836776 | 0.779280283 | 0.920933185 | 2 | 8.671479029 | 6.073378139 | 7.64488235 | 7.63485389 | 7.801071566 | 8.014172904 |
| ENSG00000176393 | RNPEP | 0.298304148 | 0.516317205 | 0.622613483 | 0.13571439 | 0.920917631 | 9.689059027 | 9.00621767 | 9.031722608 | 9.538304826 | 9.60612566 | 9.848602345 | 9.940746393 |
| ENSG00000182718 | ANXA2 | 0.572992579 | 0.339969687 | 0.347057397 | 0.574245941 | 0.920049975 | 12.53220074 | 12.90092286 | 11.97287769 | 13.08541585 | 13.0958833 | 12.36465981 | 13.23554421 |
| ENSG00000108551 | RASD1 | 2.279770452 | 0.154170396 | -1.361551386 | 0.411359472 | 0.918219065 | 6.563806279 | 7.315436877 | 2 | 8.655725489 | 8.66593596 | 5.646229369 | 4.468068808 |
| ENSG00000077549 | CAPZB | 0.124576649 | 0.78138675 | 0.793394116 | 0.033633667 | 0.917970765 | 10.85038273 | 10.60848937 | 11.07092887 | 10.93576629 | 11.0202196 | 11.36604996 | 11.88101829 |
| ENSG00000071564 | TCF3 | 0.049191117 | 0.922182516 | 0.868381052 | 0.02966485 | 0.917572169 | 9.371511179 | 9.591588447 | 8.845626171 | 9.331013912 | 9.37992356 | 10.27544091 | 10.05844914 |
| ENSG00000177943 | MAMDC4 | -0.271815082 | 0.82182886 | 1.187726095 | 0.251562038 | 0.915911014 | 4.815609471 | 4.11352228 | 6.322058544 | 4.846787657 | 5.33407135 | 6.437578914 | 6.482320745 |
| ENSG00000121764 | HCRTR1 | -2.312619827 | 0.625382427 | 3.227469533 | 0.431879804 | 0.914849705 | 2 | 2 | 2 | 2 | 2.14022021 | 3.5515855 | 3.940519005 |
| ENSG00000168702 | LRP1B | -1.221676017 | 0.650879996 | 2.136415202 | 0.390307972 | 0.914739185 | 2 | 2 | 4.874587708 | 3.162721474 | 2.59489879 | 3.483289517 | 6.099376274 |
| ENSG00000011426 | ANLN | -0.852311962 | 0.39960716 | 1.766858142 | 0.054043826 | 0.91454618 | 4.815609471 | 6.899080544 | 7.342949392 | 5.968925393 | 5.7929843 | 8.065491673 | 8.702044649 |
| ENSG00000151366 | NDUFC2 | 1.998035816 | 1.22E-07 | -1.084724888 | 0.006842904 | 0.913310928 | 7.940205782 | 8.124485672 | 7.342949392 | 9.825086737 | 9.83360578 | 6.750676314 | 6.834765628 |
| ENSG00000166510 | CCDC68 | 2.579521469 | 0.148604754 | -1.666536728 | 0.36789886 | 0.912984741 | 2 | 4.93638933 | 2 | 6.006476466 | 5.86549842 | 2.937246078 | 2.671842496 |
| ENSG00000198791 | CNOT7 | 0.563324495 | 0.382177977 | 0.347609504 | 0.599062079 | 0.910933999 | 9.128497779 | 9.277729677 | 7.934808822 | 9.481935579 | 9.42860336 | 9.284642781 | 9.19623278 |
| ENSG00000177000 | MTHFR | 0.217303306 | 0.858812103 | 0.692522158 | 0.527241133 | 0.909825464 | 5.866944937 | 8.806266215 | 8.35328246 | 8.339946438 | 8.33118782 | 8.948260474 | 8.647610514 |
| ENSG00000115421 | PAPOLG | 0.740952464 | 0.47470898 | 0.168631681 | 0.882958039 | 0.909584146 | 7.750913381 | 6.666836837 | 5.39244166 | 7.68016596 | 7.56502709 | 7.450478673 | 6.538630327 |
| ENSG00000148384 | INPP5E | 0.018377555 | 0.993323827 | 0.890652875 | 0.62915305 | 0.909030429 | 8.910412251 | 3.686303414 | 5.39244166 | 7.464384256 | 7.52720828 | 8.359038717 | 8.348604099 |
| ENSG00000073803 | MAP3K13 | 0.28406475 | 0.789639588 | 0.623017578 | 0.525386697 | 0.907082329 | 5.532353208 | 7.60566235 | 7.97167868 | 7.632926745 | 7.64076905 | 7.715279732 | 8.186029473 |
| ENSG00000121933 | TMIGD3 | -1.303156549 | NA | 2.209632429 | NA | 0.906475879 | 2 | 2 | 2 | 2.270786764 | 2 | 3.374372158 | 2.817676617 |
| ENSG00000180785 | OR51E1 | -3.753019292 | 0.508933699 | 4.658420497 | 0.362166499 | 0.905401205 | 2 | 2 | 2 | 2 | 2 | 5.673303969 | 3.071407821 |
| ENSG00000153885 | KCTD15 | -0.762435854 | 0.705011053 | 1.667707654 | 0.366065794 | 0.905271799 | 6.366709671 | 8.433041974 | 2 | 6.496695859 | 6.3857277 | 9.095643988 | 8.472158355 |
| ENSG00000101417 | PXMP4 | -0.490532569 | 0.699868245 | 1.393516287 | 0.219792859 | 0.902983718 | 6.138360429 | 4.710564154 | 7.777116083 | 6.078767801 | 6.37866593 | 8.100821584 | 8.025476363 |
| ENSG00000173145 | NOC3L | 0.778773877 | 0.561910399 | 0.123856322 | 0.933434132 | 0.902630199 | 4.467311984 | 7.393647515 | 8.408111533 | 8.27796118 | 8.17690623 | 7.6993745 | 7.459438299 |
| ENSG00000181315 | ZNF322 | 1.923333267 | 0.401419262 | -1.020850463 | 0.669668955 | 0.902482805 | 3.327711234 | 8.169643011 | 2 | 8.556367783 | 8.52912793 | 6.167993056 | 4.998228167 |

| | | | | | | | | | | | | | |
|---|---|---|---|---|---|---|---|---|---|---|---|---|---|
| ENSG00000145331 | TRMT10A | 0.292390824 | 0.848343006 | 0.609645708 | 0.668751914 | 0.902036532 | 5.866944937 | 5.131603625 | 2 | 5.22679461 | 5.34860058 | 5.4091291 | 5.727202496 |
| ENSG00000197702 | PARVA | -0.273296636 | 0.617138584 | 1.175091892 | 0.011742973 | 0.901795256 | 11.53317465 | 11.00402286 | 10.53735535 | 10.78388778 | 10.8343167 | 12.12445714 | 12.37594932 |
| ENSG00000169299 | PGM2 | 0.560930906 | 0.679562529 | 0.339875403 | 0.806377308 | 0.900806309 | 5.095959458 | 8.929529568 | 8.35328246 | 8.70652031 | 8.69749358 | 8.360824221 | 8.596104698 |
| ENSG00000233276 | GPX1 | 0.638104076 | 0.096396156 | 0.262432478 | 0.523669368 | 0.900536554 | 10.33377634 | 9.741182361 | 10.38167985 | 10.8110007 | 10.8154823 | 10.41287971 | 10.4637325 |
| ENSG00000108381 | ASPA | -2.312620553 | 0.625452298 | 3.21211454 | 0.434435201 | 0.899493987 | 2 | 2 | 2 | 2 | 2.14022021 | 3.865109708 | 3.620995423 |
| ENSG00000275774 | HNRNPCL2 | 6.090511484 | 0.052262733 | -5.191132222 | 0.152607456 | 0.899379261 | 2 | 2 | 2 | 6.24507773 | 6.14893615 | 2 | 2 |
| ENSG00000180730 | SHISA2 | 0.12113734 | 0.895301283 | 0.777710723 | 0.314028074 | 0.898848063 | 5.095959458 | 4.93638933 | 4.874587708 | 4.946755474 | 5.19627118 | 5.463319095 | 5.85096143 |
| ENSG00000164604 | GPR85 | 2.158465299 | 0.258233759 | -1.259768128 | 0.525988168 | 0.898697171 | 2 | 5.131603625 | 2 | 5.808110275 | 5.65950892 | 3.193831101 | 3.012034821 |
| ENSG00000168939 | SPRY3 | 6.089355567 | 0.053105198 | -5.191132041 | 0.153638105 | 0.898223526 | 2 | 2 | 2 | 6.043074888 | 6.33555364 | 2 | 2 |
| ENSG00000085978 | ATG16L1 | 0.167973519 | 0.925390417 | 0.72743341 | 0.657735862 | 0.895406929 | 9.048471759 | 6.844429755 | 4.058998049 | 7.93753157 | 7.95749494 | 8.639092784 | 8.344849238 |
| ENSG00000277445 | | 6.086068487 | 0.052698164 | -5.191132177 | 0.152915343 | 0.89493631 | 2 | 2 | 2 | 6.213314212 | 6.17362678 | 2 | 2 |
| ENSG00000112234 | FBXL4 | 0.022881158 | 0.988321837 | 0.870956549 | 0.523326453 | 0.893837707 | 7.709880939 | 7.874700259 | 4.058998049 | 7.315259362 | 7.26532099 | 8.545902039 | 7.519298205 |
| ENSG00000176105 | YES1 | 1.020375881 | 0.45109867 | -0.129095132 | 0.932122597 | 0.891280749 | 6.138360429 | 8.702538795 | 10.21486071 | 10.17656808 | 10.1014956 | 9.035431928 | 8.955990164 |
| ENSG00000253729 | PRKDC | 0.130856207 | 0.881476754 | 0.759696252 | 0.306456109 | 0.890552459 | 10.25752168 | 10.76380876 | 8.960138144 | 10.33577583 | 10.2722069 | 10.85827134 | 11.0013221 |
| ENSG00000170540 | ARL6IP1 | 0.921247843 | 0.013687742 | -0.030974428 | 0.947678709 | 0.890273414 | 9.867565369 | 9.828248506 | 9.25752426 | 10.6209439 | 10.5725591 | 9.772842777 | 9.511677672 |
| ENSG00000116785 | CFHR3 | 6.081321087 | 0.053285833 | -5.191132085 | 0.153445957 | 0.890189002 | 2 | 2 | 2 | 6.130704144 | 6.24526024 | 2 | 2 |
| ENSG00000105323 | HNRNPUL1 | 0.258289549 | 0.614578038 | 0.630428437 | 0.173219998 | 0.888717986 | 10.09970294 | 10.96932597 | 10.33998689 | 10.78253558 | 10.7730255 | 11.05931184 | 11.23378941 |
| ENSG00000134283 | PPHLN1 | 0.568997041 | 0.256686981 | 0.319564774 | 0.538569152 | 0.888561816 | 8.042902253 | 8.777382925 | 7.934808822 | 8.890998469 | 8.86097251 | 8.536980328 | 8.71548362 |
| ENSG00000041357 | PSMA4 | 1.089045304 | 0.051734578 | -0.203152552 | 0.753039803 | 0.885892752 | 9.895281535 | 10.00059544 | 8.784770914 | 10.75659993 | 10.7278927 | 9.439949402 | 9.469883152 |
| ENSG00000142751 | GPN2 | -0.060073814 | 0.903601303 | 0.945689283 | 0.014292331 | 0.885615469 | 8.16936288 | 7.572420273 | 7.600047325 | 7.748245381 | 7.7377845 | 8.751793877 | 8.718388644 |
| ENSG00000107882 | SUFU | 0.306686545 | 0.624119761 | 0.578772212 | 0.317915266 | 0.885458757 | 8.075571721 | 8.337339476 | 7.098165388 | 8.244958867 | 8.23280093 | 8.496406213 | 8.51889534 |
| ENSG00000100299 | ARSA | 0.760705594 | 0.462743191 | 0.123745651 | 0.914220926 | 0.884451246 | 9.810482605 | 9.540933215 | 7.163363831 | 9.896204982 | 10.0513059 | 9.280561171 | 9.401037612 |
| ENSG00000144028 | SNRNP200 | 0.267618875 | 0.627929301 | 0.615638492 | 0.220561335 | 0.883257367 | 9.800745303 | 10.69982928 | 9.878698603 | 10.47538626 | 10.437637 | 10.73931845 | 10.86548261 |
| ENSG00000055483 | USP36 | 0.456804579 | 0.837724165 | 0.425238463 | 0.846443289 | 0.882043042 | 9.646202137 | 8.522788974 | 2 | 9.044420506 | 9.08358266 | 9.081998027 | 8.98184819 |
| ENSG00000153147 | SMARCA5 | 1.461034258 | 0.230018039 | -0.582050769 | 0.656825095 | 0.87898349 | 6.73718411 | 10.42046463 | 10.03493575 | 11.18281219 | 11.1680817 | 9.368088514 | 8.869253401 |
| ENSG00000100209 | HSCB | 1.285765749 | 0.122091309 | -0.407201284 | 0.657203666 | 0.878564465 | 6.468621203 | 6.666836837 | 4.523675391 | 7.423326516 | 7.39346687 | 5.83618166 | 5.740981724 |
| ENSG00000089472 | HEPH | 0.897327111 | 0.344828673 | -0.019938651 | 0.985591183 | 0.87738846 | 7.159102577 | 8.955547846 | 9.647595056 | 9.782429661 | 9.81672282 | 8.752700801 | 9.011994108 |
| ENSG00000138834 | MAPK8IP3 | 0.149487957 | 0.941569674 | 0.727848987 | 0.700873168 | 0.877336943 | 3.327711234 | 9.357954881 | 10.11951029 | 9.318005579 | 9.40098777 | 10.23155211 | 9.562677828 |
| ENSG00000177685 | CRACR2B | -1.277880903 | 0.525895193 | 2.155135907 | 0.229495532 | 0.877255004 | 4.00722531 | 3.076665209 | 2 | 2.947924294 | 2.2680111 | 4.530483275 | 5.080008298 |
| ENSG00000164885 | CDK5 | 1.173767623 | 0.357945396 | -0.298390699 | 0.832869996 | 0.875376925 | 5.866944937 | 7.051611315 | 3.369317972 | 7.144720234 | 7.28433928 | 5.853194273 | 5.768151538 |
| ENSG00000166912 | MTMR10 | 2.061851837 | 0.196711636 | -1.186910784 | 0.474542652 | 0.874941053 | 5.095959458 | 7.002537973 | 2 | 7.72026832 | 7.86955761 | 4.513461373 | 4.959465495 |
| ENSG00000103494 | RPGRIP1L | 0.776031139 | 0.449317736 | 0.098056787 | 0.932122597 | 0.874087926 | 5.095959458 | 7.431218269 | 5.930865648 | 7.234487582 | 7.26914476 | 6.637393426 | 6.554323192 |
| ENSG00000185728 | YTHDF3 | 0.565985618 | 0.490472675 | 0.308067801 | 0.717164009 | 0.874053418 | 8.493334308 | 10.34373336 | 10.16797309 | 10.47287335 | 10.3997688 | 10.20612015 | 10.15273873 |
| ENSG00000100036 | SLC35E4 | 0.050659701 | 0.975506192 | 0.823303735 | 0.577037638 | 0.873963436 | 3.327711234 | 7.145017454 | 7.934808822 | 7.057964187 | 7.12888795 | 7.786974998 | 7.908246991 |
| ENSG00000138279 | ANXA7 | 1.163196814 | 0.001734804 | -0.290430333 | 0.494470997 | 0.872766481 | 9.509418609 | 9.203675681 | 8.784770914 | 10.33393077 | 10.3751428 | 8.818278819 | 8.993886349 |
| ENSG00000167904 | TMEM68 | 0.273580879 | 0.726499281 | 0.597706096 | 0.406622399 | 0.871286976 | 7.485143553 | 6.602424674 | 6.534109959 | 7.222570549 | 7.18258783 | 7.904255004 | 6.990916783 |
| ENSG00000101266 | CSNK2A1 | 0.017037327 | 0.982192754 | 0.853062979 | 0.169861871 | 0.870100306 | 9.87686338 | 8.487557258 | 9.681375057 | 9.464333207 | 9.4921465 | 10.31204759 | 10.30948877 |
| ENSG00000278608 | | -1.051907038 | 0.54826185 | 1.921945112 | 0.23044932 | 0.870038074 | 5.532353208 | 3.076665209 | 2 | 3.406930997 | 3.60476711 | 5.846413283 | 6.055940621 |
| ENSG00000184988 | TMEM106A | 1.122529167 | 0.215763131 | -0.252667569 | 0.803497966 | 0.869861598 | 6.009025997 | 7.928237109 | 6.203069917 | 8.136938566 | 8.07792246 | 6.732402829 | 6.797888492 |
| ENSG00000108556 | CHRNE | 1.164843359 | 0.641266479 | -0.295981584 | 0.912467872 | 0.868861775 | 2 | 2 | 4.058998049 | 3.754611435 | 3.60476711 | 3.1504466 | 2.466008509 |
| ENSG00000103061 | SLC7A6OS | 0.581916242 | 0.75774249 | 0.286431994 | 0.884256906 | 0.868348236 | 2 | 5.457120953 | 7.398165132 | 6.682159616 | 6.76185814 | 6.533574091 | 6.338680316 |
| ENSG00000140612 | SEC11A | 0.694762107 | 0.002963985 | 0.173078253 | 0.517968728 | 0.86784036 | 9.820154628 | 9.792607274 | 9.589475645 | 10.43593562 | 10.4274075 | 9.90822107 | 9.915400542 |
| ENSG00000015475 | BID | 0.692624333 | 0.315574141 | 0.174894954 | 0.818501743 | 0.867519287 | 8.340423424 | 8.655694736 | 7.098165388 | 8.808365552 | 8.91566441 | 8.390261842 | 8.309838123 |
| ENSG00000169413 | RNASE6 | -3.753043198 | 0.479762551 | 4.619939118 | 0.329937069 | 0.86689592 | 2 | 2 | 2 | 2 | 2 | 5.201624686 | 4.411752372 |
| ENSG00000130270 | ATP8B3 | 0.303113923 | 0.879863666 | 0.563319946 | 0.763737563 | 0.866433869 | 5.330594134 | 2 | 3.369317972 | 4.139692846 | 4.60967607 | 4.73522696 | 4.468068808 |
| ENSG00000065911 | MTHFD2 | 0.170554272 | 0.870694345 | 0.695207239 | 0.450551283 | 0.865761511 | 9.740899611 | 9.277729677 | 7.646393253 | 9.319871109 | 9.27353851 | 10.22666466 | 9.247812734 |
| ENSG00000196616 | ADH1B | -3.753029746 | 0.496719668 | 4.618022397 | 0.352068654 | 0.864992652 | 2 | 2 | 2 | 2 | 2 | 3.584558147 | 5.524371359 |
| ENSG00000186567 | CEACAM19 | 0.14689704 | 0.888172933 | 0.716469029 | 0.429217146 | 0.863366069 | 7.27616306 | 5.303523747 | 7.398165132 | 7.123515401 | 6.96410766 | 7.68709225 | 7.509939612 |
| ENSG00000196187 | TMEM63A | 0.902111331 | 0.263458099 | -0.040304867 | 0.966513683 | 0.861806463 | 8.963744667 | 7.145017454 | 8.922968322 | 9.384581425 | 9.49623046 | 8.63565547 | 8.365010363 |
| ENSG00000163251 | FZD5 | 0.696367696 | 0.777603672 | 0.165303749 | 0.948615522 | 0.861671446 | 2 | 2 | 5.39244166 | 4.76150579 | 4.37434574 | 4.301748724 | 3.987514168 |
| ENSG00000164951 | PDP1 | 1.137783553 | 0.309598975 | -0.276432394 | 0.823246972 | 0.861351159 | 6.73718411 | 7.315436877 | 9.426401271 | 9.483600848 | 9.38873772 | 7.851893762 | 8.200230127 |
| ENSG00000067955 | CBFB | 0.750150736 | 0.586657904 | 0.105633508 | 0.944101725 | 0.855784244 | 4.815609471 | 8.469613413 | 8.865353883 | 8.868232317 | 8.90587341 | 8.293242958 | 8.204380326 |
| ENSG00000185869 | ZNF829 | 0.289262986 | 0.741924158 | 0.566170355 | 0.487033897 | 0.855433341 | 6.009025997 | 5.131603625 | 5.156626401 | 5.797438367 | 5.69405321 | 6.315225883 | 5.606950214 |
| ENSG00000184162 | NR2C2AP | 0.296627234 | 0.851528683 | 0.558524888 | 0.706382819 | 0.855152122 | 2 | 5.838937808 | 5.59507254 | 5.406080789 | 5.53799497 | 5.957567336 | 5.419744333 |
| ENSG00000110435 | PDHX | 0.810141002 | 0.530053668 | 0.044886282 | 0.974562729 | 0.855027284 | 5.095959458 | 8.124485672 | 8.825624954 | 8.790986483 | 8.80544247 | 8.146391268 | 7.929461924 |
| ENSG00000196739 | COL27A1 | 0.414369583 | 0.832844524 | 0.44059289 | 0.818406267 | 0.854962473 | 4.00722531 | 6.602424674 | 9.69246155 | 8.612239836 | 8.78288532 | 8.620830655 | 8.824117457 |
| ENSG00000254004 | ZNF260 | 0.428945275 | 0.703668984 | 0.425073265 | 0.701904883 | 0.85401854 | 5.866944937 | 8.487557258 | 8.236946975 | 8.386974157 | 8.27334517 | 8.671105351 | 7.875307424 |
| ENSG00000106479 | ZNF862 | 0.484074915 | 0.711397752 | 0.365834806 | 0.781762153 | 0.849909721 | 5.095959458 | 6.666836837 | 8.266922072 | 7.723090567 | 7.62891439 | 7.935741825 | 7.052989523 |
| ENSG00000198876 | DCAF12 | 0.045116198 | 0.984043262 | 0.804645547 | 0.702841297 | 0.849761745 | 2 | 9.225224669 | 9.601288346 | 8.866956935 | 8.91200057 | 9.601198809 | 9.685927623 |
| ENSG00000164111 | ANXA5 | 1.327146299 | 0.007634516 | -0.477581154 | 0.383233513 | 0.849565144 | 12.53220074 | 11.53079123 | 11.86432331 | 13.36165586 | 13.3648174 | 11.31669201 | 11.76793522 |
| ENSG00000124216 | SNAI1 | 1.457601135 | 0.092161915 | -0.60805415 | 0.513107667 | 0.849546984 | 6.89194353 | 4.93638933 | 7.098165388 | 7.994705029 | 7.95512324 | 5.909581027 | 6.048572483 |
| ENSG00000116906 | GNPAT | -0.054544392 | 0.913792484 | 0.903966368 | 0.022591207 | 0.849421975 | 8.340423424 | 8.655694736 | 7.896971941 | 8.289431124 | 8.28283292 | 9.200732369 | 9.26976224 |
| ENSG00000166035 | LIPC | -3.753045051 | 0.478107313 | 4.60227063 | 0.329251531 | 0.849225578 | 2 | 2 | 2 | 2 | 2 | 4.894909044 | 4.793076545 |
| ENSG00000028528 | SNX1 | 0.543793405 | 0.389803748 | 0.302797257 | 0.643384314 | 0.846590662 | 10.1770105 | 9.599860728 | 8.825624954 | 10.16675444 | 10.1893748 | 9.830461743 | 10.03813112 |
| ENSG00000136938 | ANP32B | 0.176496191 | 0.85461878 | 0.66970055 | 0.431645387 | 0.846196741 | 9.820154628 | 9.624396729 | 7.777116083 | 9.53348965 | 9.45234084 | 9.921992929 | 10.04416401 |
| ENSG00000198589 | LRBA | 0.220023917 | 0.903601303 | 0.624642391 | 0.709548861 | 0.844666308 | 4.467311984 | 7.761270542 | 9.746650996 | 8.713632895 | 8.75042728 | 8.942707845 | 9.301879118 |
| ENSG00000139974 | SLC38A6 | 1.775907192 | 0.170726471 | -0.931480008 | 0.492815255 | 0.844427184 | 6.138360429 | 5.457120953 | 2 | 7.035434774 | 6.9210099 | 4.530483275 | 4.479073019 |
| ENSG00000111412 | C12orf49 | 0.058810721 | 0.962673456 | 0.785016795 | 0.483687127 | 0.843827515 | 9.567995545 | 6.464272742 | 8.460932916 | 8.692189072 | 8.7045705 | 9.439386236 | 9.398504094 |
| ENSG00000145979 | TBC1D7 | 0.95345227 | 0.483591352 | -0.109864133 | 0.942375924 | 0.843588137 | 4.00722531 | 6.728495673 | 8.007629717 | 7.826439377 | 7.99029569 | 6.816344356 | 6.928061817 |
| ENSG00000137942 | FNBP1L | 0.243594988 | 0.86389717 | 0.598431476 | 0.646493605 | 0.842026464 | 4.815609471 | 8.376385542 | 6.882775709 | 7.549526251 | 7.58049694 | 8.007404089 | 7.81253023 |
| ENSG00000115602 | IL1RL1 | 1.050376516 | 0.706793237 | -0.208700024 | 0.944101725 | 0.841676493 | 3.327711234 | 2 | 2 | 3.09461492 | 2.94009902 | 2.717032197 | 2.277179783 |
| ENSG00000123353 | ORMDL2 | 0.69694514 | 0.461193236 | 0.143006378 | 0.89090828 | 0.839951518 | 5.70932728 | 7.847167909 | 7.552162999 | 7.997038782 | 7.96694292 | 7.039353222 | 7.751441934 |
| ENSG00000123505 | AMD1 | 0.653863444 | 0.190053842 | 0.183977645 | 0.737229272 | 0.837841089 | 8.517314938 | 9.28800521 | 8.960138144 | 9.634381252 | 9.58403885 | 8.825624065 | 9.402122045 |
| ENSG00000106070 | GRB10 | 0.139451949 | 0.877879848 | 0.698114301 | 0.369714606 | 0.83756625 | 7.33131396 | 8.834582582 | 9.099922433 | 8.697938662 | 8.78422202 | 9.528926528 | 9.018137056 |
| ENSG00000163703 | CRELD1 | 0.523771572 | 0.540424469 | 0.312389379 | 0.723575557 | 0.836160951 | 9.189468652 | 9.415325457 | 7.451345288 | 9.388142127 | 9.47486651 | 9.126758239 | 9.311530309 |
| ENSG00000125810 | CD93 | -3.240232983 | 0.241642985 | 4.074703051 | 0.080865234 | 0.834470068 | 4.00722531 | 2 | 2 | 2 | 2.2680111 | 6.750676314 | 4.935697922 |
| ENSG00000197927 | C2orf27A | 6.024854329 | 0.059854337 | -5.191131311 | 0.158650282 | 0.833723018 | 2 | 2 | 2 | 6.155987605 | 6.11534392 | 2 | 2 |
| ENSG00000108055 | SMC3 | 0.613629925 | 0.528710992 | 0.219942769 | 0.832583267 | 0.833572694 | 6.653098118 | 8.487557258 | 9.083172337 | 8.983347946 | 8.99861459 | 8.727088905 | 8.462487876 |
| ENSG00000157837 | SPPL3 | 0.60113162 | 0.055580779 | 0.230012424 | 0.498291593 | 0.831144043 | 8.674867122 | 8.732943944 | 8.631919287 | 9.155143657 | 9.39224836 | 8.807408592 | 9.006300384 |
| ENSG00000204991 | SPIRE2 | 0.768574185 | 0.518554054 | 0.062538131 | 0.962746693 | 0.831112316 | 3.327711234 | 4.11352228 | 5.59507254 | 5.258265913 | 5.33407135 | 4.813804156 | 4.532869193 |
| ENSG00000124067 | SLC12A4 | -0.308948409 | 0.64165426 | 1.139991917 | 0.048141929 | 0.831043507 | 10.64259676 | 9.277729677 | 10.06936278 | 9.780398828 | 9.80096123 | 11.24663807 | 11.22431746 |
| ENSG00000004455 | AK2 | 0.33594499 | 0.588546732 | 0.494004961 | 0.401725514 | 0.829949951 | 9.509418609 | 9.067150531 | 10.22252928 | 10.0010786 | 10.0204455 | 9.97191327 | 10.34111971 |
| ENSG00000143390 | RFX5 | 0.601621043 | 0.632030586 | 0.226184039 | 0.864675088 | 0.827805082 | 9.233563228 | 9.308339287 | 6.073378139 | 9.384581425 | 9.33957954 | 8.590716036 | 9.301491724 |
| ENSG00000151729 | SLC25A4 | -1.148721657 | 0.021949338 | 1.976256754 | 2.29E-05 | 0.827535097 | 8.19931886 | 8.276714465 | 9.116480286 | 7.501020868 | 7.41433569 | 10.39163988 | 10.69986899 |
| ENSG00000164902 | PHAX | 0.16016597 | 0.891428469 | 0.666155953 | 0.521375485 | 0.826321923 | 7.031698319 | 8.607278467 | 9.757248847 | 8.99156386 | 9.02712059 | 9.390403917 | 9.624380013 |
| ENSG00000127311 | HELB | -3.75303753 | 0.487478481 | 4.578340914 | 0.345156474 | 0.825303385 | 2 | 2 | 2 | 2 | 2 | 5.289239922 | 4.133950982 |
| ENSG00000176261 | ZBTB8OS | 0.66195109 | 0.290102247 | 0.163284069 | 0.81284055 | 0.825235159 | 6.96351253 | 6.666836837 | 6.073378139 | 7.277360022 | 7.23826702 | 6.430795201 | 7.052989523 |
| ENSG00000162885 | B3GALNT2 | 1.274167524 | 0.03553166 | -0.449649254 | 0.498871333 | 0.82451827 | 6.366709671 | 7.761270542 | 7.098165388 | 8.399377854 | 8.50176361 | 6.833591108 | 6.693102083 |
| ENSG00000103489 | XYLT1 | 1.241140879 | 0.33935585 | -0.419215645 | 0.765333198 | 0.821925234 | 5.532353208 | 8.968383114 | 8.784770914 | 9.562930393 | 9.62936766 | 8.512558892 | 7.023227011 |
| ENSG00000144893 | MED12L | 1.491947248 | 0.466289371 | -0.67080781 | 0.755643195 | 0.821139438 | 2 | 4.11352228 | 2 | 4.299492648 | 4.02226117 | 2.911488689 | 2.746601469 |
| ENSG00000090581 | GNPTG | 1.456902272 | 0.000157844 | -0.63649638 | 0.128956336 | 0.820405892 | 8.696033109 | 7.928237109 | 8.434763955 | 9.777686595 | 9.86994125 | 7.709686122 | 7.792821043 |
| ENSG00000059145 | UNKL | 0.497601167 | 0.819371474 | 0.322797511 | 0.883858724 | 0.820398677 | 2 | 6.464272742 | 8.784770914 | 7.954442493 | 7.96458672 | 7.764669607 | 7.81035357 |
| ENSG00000127324 | TSPAN8 | -3.753041074 | 0.483277973 | 4.573335546 | 0.340038826 | 0.820294472 | 2 | 2 | 2 | 2 | 2 | 4.425202323 | 5.12973572 |
| ENSG00000170876 | TMEM43 | 0.53320276 | 0.264473942 | 0.287043709 | 0.564270352 | 0.820246469 | 9.839306291 | 9.479495904 | 10.38851302 | 10.48414712 | 10.4753644 | 10.18963871 | 10.27773968 |
| ENSG00000157500 | APPL1 | -0.140283637 | 0.89458451 | 0.957485929 | 0.28290231 | 0.817202292 | 6.366709671 | 8.053983481 | 8.721235202 | 7.982979299 | 7.7377845 | 9.023833473 | 8.872386739 |
| ENSG00000126243 | LRFN3 | -0.184121648 | 0.930075716 | 1.000651291 | 0.595347869 | 0.816529643 | 2 | 7.73146271 | 8.511888463 | 7.366699513 | 7.46855365 | 8.522696167 | 8.643336952 |
| ENSG00000134970 | TMED7 | 1.174301886 | 0.01542766 | -0.35892372 | 0.507672408 | 0.815378166 | 9.081016759 | 10.13189344 | 9.636156974 | 10.90440671 | 10.7993766 | 9.226421872 | 9.418291476 |
| ENSG00000159713 | TPPP3 | -0.633996037 | 0.776246706 | 1.446420481 | 0.466865178 | 0.812424444 | 3.327711234 | 2 | 3.369317972 | 2.784612485 | 2.59489879 | 4.188345803 | 3.407202941 |
| ENSG00000113645 | WWC1 | -0.578770667 | 0.745616581 | 1.390428178 | 0.393412229 | 0.811657511 | 3.327711234 | 4.442710129 | 7.451345288 | 5.540148523 | 5.5755357 | 6.737909218 | 7.900082338 |
| ENSG00000078403 | MLLT10 | 0.563691884 | 0.727489842 | 0.246867335 | 0.884612514 | 0.810559219 | 3.327711234 | 7.874700259 | 8.175061096 | 8.060896687 | 7.99722844 | 7.914016653 | 7.491038307 |
| ENSG00000179943 | FIZ1 | -0.216174217 | 0.910642404 | 1.025997529 | 0.544751469 | 0.809823312 | 7.33131396 | 7.467835385 | 2 | 6.640968371 | 6.61935067 | 7.900986387 | 7.772838868 |
| ENSG00000158717 | RNF166 | 0.266590644 | 0.621265967 | 0.542873113 | 0.271232177 | 0.809463757 | 7.159102577 | 6.535001748 | 6.958202216 | 7.178012736 | 7.1246732 | 7.577730839 | 7.244113847 |
| ENSG00000162688 | AGL | 0.188936235 | 0.67705568 | 0.619438698 | 0.117356112 | 0.808374933 | 7.709880939 | 7.315436877 | 7.163363831 | 7.54315232 | 7.64959655 | 8.034504643 | 8.005636809 |
| ENSG00000103194 | USP10 | 0.670683225 | 0.286739789 | 0.136285877 | 0.846777728 | 0.806969101 | 9.358295023 | 9.514922632 | 8.206335842 | 9.772927884 | 9.8316678 | 9.139642601 | 9.39160489 |
| ENSG00000168785 | TSPAN5 | 0.102453402 | 0.868745346 | 0.70167226 | 0.173666446 | 0.804125662 | 9.87686338 | 8.929529568 | 9.272377397 | 9.50916971 | 9.51647861 | 9.861061271 | 10.32119294 |
| ENSG00000135469 | COQ10A | 0.554811035 | 0.812947514 | 0.249196039 | 0.91758425 | 0.804007074 | 2 | 7.431218269 | 3.369317972 | 6.503277904 | 6.50077304 | 6.159811591 | 6.261114822 |
| ENSG00000160695 | VPS11 | 0.717707119 | 0.098800583 | 0.086214535 | 0.865952727 | 0.803921654 | 9.015175615 | 8.540087354 | 8.175061096 | 9.32080297 | 9.33867006 | 8.734453103 | 8.671191358 |

| | | | | | | | | | | | | | |
|---|---|---|---|---|---|---|---|---|---|---|---|---|---|
| ENSG00000124571 | XPO5 | 0.537074534 | 0.807640508 | 0.266513584 | 0.906201712 | 0.803588118 | 2 | 9.608085847 | 8.784770914 | 9.232448283 | 9.18320541 | 9.117986989 | 8.735697007 |
| ENSG00000168566 | SNRNP48 | 0.027250279 | 0.98895591 | 0.775861933 | 0.665712192 | 0.803112211 | 2 | 6.602424674 | 7.691296561 | 6.658765962 | 6.75096147 | 7.484724631 | 7.376535208 |
| ENSG00000113161 | HMGCR | 0.657821859 | 0.642184909 | 0.145106523 | 0.92531104 | 0.802928382 | 6.257048221 | 10.30883598 | 9.849281147 | 10.19396441 | 10.2413953 | 9.19106327 | 10.08660193 |
| ENSG00000170153 | RNF150 | -0.515288328 | 0.710155984 | 1.317215291 | 0.291682761 | 0.801926963 | 7.159102577 | 6.666836837 | 3.369317972 | 5.900763176 | 5.9536761 | 7.944499874 | 7.380942161 |
| ENSG00000139973 | SYT16 | -5.525988803 | 0.016319666 | 6.327562674 | 2.26E-05 | 0.801573871 | 2 | 3.686303414 | 2 | 2 | 2 | 8.037484631 | 8.249261511 |
| ENSG00000086102 | NFX1 | 0.268601402 | 0.596790354 | 0.532570646 | 0.255070887 | 0.801172048 | 8.737455672 | 9.00621767 | 8.110401118 | 8.935480545 | 8.9398551 | 9.113757772 | 9.280809554 |
| ENSG00000118508 | RAB32 | -0.372588738 | 0.718561383 | 1.173755194 | 0.201042704 | 0.801166455 | 9.689059027 | 8.487557258 | 7.398165132 | 8.48998335 | 8.40429221 | 9.531835181 | 10.32653371 |
| ENSG00000166167 | BTRC | 0.286855192 | 0.487512694 | 0.512933018 | 0.178705011 | 0.79978821 | 8.39321512 | 8.702538795 | 9.014156722 | 8.995070693 | 9.02146427 | 9.336073933 | 9.121389737 |
| ENSG00000008869 | HEATR5B | 0.805403442 | 0.075112275 | -0.005946302 | 0.991443049 | 0.799457139 | 7.867441304 | 8.433041974 | 8.076950877 | 8.95000826 | 8.94703391 | 8.387344963 | 7.855387687 |
| ENSG00000102743 | SLC25A15 | 0.789582192 | 0.175685887 | 0.00939077 | 0.988988478 | 0.798972961 | 5.70932728 | 6.535001748 | 6.80318718 | 7.140504127 | 7.22257674 | 6.310306306 | 6.541257688 |
| ENSG00000133112 | TPT1 | 0.54815232 | 0.396129098 | 0.250393898 | 0.713019003 | 0.798546218 | 13.2480265 | 12.32176563 | 11.95227171 | 13.16475722 | 13.159391 | 12.98073122 | 12.73798674 |
| ENSG00000170852 | KBTBD2 | 0.67606666 | 0.24500653 | 0.122340718 | 0.852603315 | 0.798407378 | 9.556468883 | 8.57407404 | 8.486635645 | 9.665524776 | 9.59016561 | 9.145869306 | 9.00677572 |
| ENSG00000106404 | CLDN15 | 0.336920065 | 0.800325517 | 0.459912606 | 0.718917407 | 0.796832671 | 6.468621203 | 5.457120953 | 4.523675391 | 5.930370745 | 6.08095082 | 6.765130262 | 4.935697922 |
| ENSG00000108094 | CUL2 | 0.951809761 | 0.031800415 | -0.155778524 | 0.764528151 | 0.796031238 | 8.340423424 | 8.124485672 | 8.960138144 | 9.508351943 | 9.40359931 | 8.413960373 | 8.301338496 |
| ENSG00000143641 | GALNT2 | 0.330995207 | 0.599159126 | 0.462746739 | 0.441227897 | 0.793741946 | 10.70657014 | 9.876711741 | 10.48730704 | 10.73509104 | 10.7174353 | 10.32552007 | 11.24571875 |
| ENSG00000114853 | ZBTB47 | -0.107273677 | 0.95872661 | 0.899071624 | 0.638141963 | 0.791797947 | 5.095959458 | 7.538394143 | 11.2124308 | 9.587916984 | 9.70541961 | 10.62175706 | 10.67884542 |
| ENSG00000197647 | ZNF433 | 0.048819949 | 0.968309509 | 0.739149791 | 0.495902901 | 0.78796974 | 4.00722531 | 6.140560021 | 4.523675391 | 5.144975053 | 5.40529704 | 5.889930493 | 5.937276556 |
| ENSG00000266338 | NBPF15 | 2.826620958 | 1.89E-12 | -2.042125544 | 6.31E-07 | 0.784495415 | 9.509418609 | 8.820493869 | 8.65477198 | 11.81237837 | 11.9057847 | 7.000176314 | 7.067643653 |
| ENSG00000132773 | TOE1 | 0.544250285 | 0.475730142 | 0.239700084 | 0.763860964 | 0.783950369 | 5.095959458 | 5.946638857 | 6.322058544 | 6.336376195 | 6.43421531 | 5.787450547 | 6.353705253 |
| ENSG00000151923 | TIAL1 | 0.480944899 | 0.45218119 | 0.302842651 | 0.64238228 | 0.78378755 | 8.981092835 | 9.497318007 | 8.143093325 | 9.405814997 | 9.51083771 | 9.277413594 | 9.28708453 |
| ENSG00000144224 | UBXN4 | 0.576083489 | 0.247133389 | 0.205879857 | 0.70178553 | 0.781963347 | 9.913467515 | 10.29870826 | 10.89014791 | 11.01130667 | 10.9843247 | 10.67666663 | 10.57865558 |
| ENSG00000181513 | ACBD4 | -0.007765027 | 0.9947462 | 0.789450554 | 0.388123757 | 0.781685527 | 5.70932728 | 4.710564154 | 6.80318718 | 5.930370745 | 5.9725602 | 6.728720192 | 6.688368862 |
| ENSG00000161980 | POLR3K | 0.635042208 | 0.583878202 | 0.144760974 | 0.908703812 | 0.779803182 | 4.467311984 | 6.228547981 | 7.451345288 | 7.044488801 | 7.17037177 | 6.645228724 | 6.62538071 |
| ENSG00000006007 | GDE1 | 0.482887973 | 0.424587993 | 0.296662533 | 0.632283385 | 0.779550506 | 9.344956677 | 9.415325457 | 10.42220168 | 10.32187999 | 10.2683912 | 10.13868678 | 10.0802927 |
| ENSG00000056972 | TRAF3IP2 | 0.762673903 | 0.494744255 | 0.016384721 | 0.989360801 | 0.779058624 | 7.159102577 | 6.951736442 | 9.344432377 | 9.006698952 | 9.00436103 | 8.102245405 | 8.414572237 |
| ENSG00000068697 | LAPTM4A | 0.838295897 | 0.25593159 | -0.059916177 | 0.944157185 | 0.778379719 | 11.82259453 | 10.18234215 | 10.66665689 | 11.88757367 | 11.9071656 | 10.94354834 | 11.05528124 |
| ENSG00000165113 | GKAP1 | 0.623902657 | 0.707242233 | 0.153894679 | 0.930784885 | 0.777797336 | 3.327711234 | 4.442710129 | 2 | 4.237680128 | 3.94787299 | 3.891700877 | 3.600986722 |
| ENSG00000164284 | GRPEL2 | 0.413878932 | 0.807722186 | 0.363722849 | 0.830158879 | 0.77760178 | 3.327711234 | 8.169643011 | 6.629488387 | 7.419851705 | 7.49817903 | 7.709686122 | 7.032593922 |
| ENSG00000085840 | ORC1 | 0.917154 | 0.579813548 | -0.14080448 | 0.938141666 | 0.77634952 | 2 | 4.11352228 | 3.369317972 | 4.205751501 | 3.98554648 | 3.172301926 | 3.539239711 |
| ENSG00000089327 | FXYD5 | 1.232159408 | 0.064313177 | -0.457350024 | 0.527954208 | 0.774809384 | 9.189468652 | 10.00683004 | 10.60345421 | 11.29470341 | 11.2513856 | 9.143105199 | 9.929007159 |
| ENSG00000136603 | SKIL | 0.849416769 | 0.459780125 | -0.074910149 | 0.953777463 | 0.77450662 | 6.96351253 | 9.896993485 | 9.165043284 | 9.960996031 | 9.94357048 | 9.471967222 | 8.399447278 |
| ENSG00000188868 | ZNF563 | 0.187425615 | 0.83752462 | 0.58662152 | 0.466293248 | 0.774047135 | 4.467311984 | 5.131603625 | 4.874587708 | 4.927308402 | 5.13053105 | 5.4091291 | 5.367404784 |
| ENSG00000147475 | ERLIN2 | 0.044203307 | 0.954430302 | 0.728217684 | 0.27098548 | 0.772420992 | 7.867441304 | 8.687092496 | 9.330306409 | 8.855427477 | 8.72141482 | 9.239097094 | 9.667406786 |
| ENSG00000160818 | GPATCH4 | 0.397662508 | 0.788243206 | 0.374427002 | 0.798199695 | 0.77208951 | 8.19931886 | 7.761270542 | 4.058998049 | 7.818544084 | 7.88209639 | 7.664138901 | 7.974870548 |
| ENSG00000211456 | SACM1L | 0.849545211 | 0.330348729 | -0.077669145 | 0.938363512 | 0.771876067 | 6.816637977 | 8.732943944 | 8.960138144 | 9.294480609 | 9.25630406 | 8.401292507 | 8.310608339 |
| ENSG00000132031 | MATN3 | -0.285803555 | 0.865218379 | 1.05687192 | 0.475546641 | 0.771068364 | 2 | 5.72254323 | 4.523675391 | 4.739373666 | 4.2865223 | 5.47655442 | 5.917188835 |
| ENSG00000196313 | POM121 | 1.571856908 | 1.15E-05 | -0.801139597 | 0.036686534 | 0.770717311 | 9.509418609 | 8.88960062 | 8.922968322 | 10.68711744 | 10.7104214 | 8.263645107 | 8.413138577 |
| ENSG00000244462 | RBM12 | 0.072046434 | 0.915618176 | 0.697658473 | 0.220786883 | 0.769704907 | 9.015175615 | 9.357954881 | 9.798878489 | 9.482768454 | 9.5116449 | 9.593626268 | 10.50503024 |
| ENSG00000163214 | DHX57 | 0.506592458 | 0.765893437 | 0.262037193 | 0.882673743 | 0.768629651 | 6.009025997 | 8.981105197 | 4.523675391 | 8.115618198 | 8.14812525 | 7.86872317 | 7.913326535 |
| ENSG00000104805 | NUCB1 | 0.105258976 | 0.865039257 | 0.662599467 | 0.202582154 | 0.767858443 | 11.87225278 | 10.80015962 | 11.38294124 | 11.50362834 | 11.5383508 | 11.93825096 | 12.20505008 |
| ENSG00000187553 | CYP26C1 | -1.317823125 | NA | 2.084595296 | NA | 0.76677217 | 2 | 2 | 2 | 2 | 2.2680111 | 2.42098416 | 3.496556178 |
| ENSG00000006756 | ARSD | 0.344409115 | 0.832844524 | 0.421956775 | 0.789486933 | 0.76636589 | 3.327711234 | 7.928237109 | 7.225742779 | 7.454228844 | 7.36873214 | 7.260787065 | 7.68409025 |
| ENSG00000123213 | NLN | -0.286283385 | 0.705469615 | 1.052568411 | 0.109470992 | 0.766285025 | 7.159102577 | 8.234834549 | 6.882775709 | 7.348537858 | 7.21862736 | 8.437839892 | 8.74825912 |
| ENSG00000213281 | NRAS | 0.165587793 | 0.926059063 | 0.600027601 | 0.71788604 | 0.765615395 | 4.467311984 | 9.506147172 | 9.788583153 | 9.260887574 | 9.24663965 | 9.455629465 | 9.88417883 |
| ENSG00000185049 | NELFA | -0.152023692 | 0.926636636 | 0.916796689 | 0.531002733 | 0.764772997 | 7.218819781 | 8.004992147 | 3.369317972 | 6.917187326 | 7.03319408 | 8.042684843 | 7.998962529 |
| ENSG00000010810 | FYN | 0.615393721 | 0.016642205 | 0.148936832 | 0.611272575 | 0.764330553 | 9.98400217 | 9.969011754 | 10.2528023 | 10.64829839 | 10.7226735 | 10.18913632 | 10.25190776 |
| ENSG00000122406 | RPL5 | -0.014555117 | 0.989000921 | 0.777628505 | 0.391227832 | 0.763073388 | 10.96815916 | 10.01303782 | 12.24925065 | 11.31401914 | 11.3818252 | 11.96059845 | 12.2987658 |
| ENSG00000063978 | RNF4 | 0.267067696 | 0.814508396 | 0.493563436 | 0.643682096 | 0.760631132 | 10.46903009 | 9.318399851 | 7.734844356 | 9.787831283 | 9.86805151 | 9.931235194 | 10.16726922 |
| ENSG00000114126 | TFDP2 | 0.282717026 | 0.525364769 | 0.477172796 | 0.248228307 | 0.759889822 | 8.564112515 | 8.147241016 | 8.865353883 | 8.845101141 | 8.80807321 | 9.079470158 | 8.9569744 |
| | PGBD3 | 5.950976265 | 0.06938652 | -5.191130213 | 0.165757831 | 0.759846051 | 2 | 2 | 2 | 5.987823101 | 6.14061114 | 2 | 2 |
| ENSG00000137185 | ZSCAN9 | 0.217735002 | 0.838100556 | 0.542095224 | 0.576450932 | 0.759830227 | 6.89194353 | 6.464272742 | 4.523675391 | 6.573765157 | 6.37156942 | 6.956760977 | 6.595354992 |
| ENSG00000067334 | DNTTIP2 | 0.663881989 | 0.065602331 | 0.092408771 | 0.827527224 | 0.756290761 | 9.646202137 | 9.090820894 | 9.132850258 | 9.981765804 | 9.95963168 | 9.383101353 | 9.420790513 |
| ENSG00000204950 | LRRC10B | 0.729575348 | 0.803737808 | 0.026450843 | NA | 0.756026191 | 2 | 3.076665209 | 2 | 3.023133209 | 2.68924543 | 2.775316226 | 2.374681325 |
| ENSG00000172336 | POP7 | -0.108509041 | 0.931219161 | 0.864421771 | 0.431786601 | 0.75591273 | 5.330594134 | 7.099070191 | 8.325066185 | 7.292639438 | 7.23826702 | 8.083625221 | 8.344849238 |
| ENSG00000131351 | HAUS8 | 0.659419991 | 0.714367242 | 0.096052417 | 0.960499071 | 0.755472408 | 2 | 5.595927442 | 3.369317972 | 4.965943888 | 5.0259382 | 4.478802948 | 4.553838275 |
| ENSG00000277276 | | 5.946400417 | 0.069869263 | -5.19113017 | 0.165966488 | 0.755270247 | 2 | 2 | 2 | 6.006476466 | 6.11534392 | 2 | 2 |
| ENSG00000092847 | AGO1 | -0.139571875 | 0.897105982 | 0.89358936 | 0.332631208 | 0.754017485 | 7.096806373 | 8.834582582 | 9.478569125 | 8.624367384 | 8.63218877 | 9.354412203 | 9.901919938 |
| ENSG00000012983 | MAP4K5 | 0.353933401 | 0.573001609 | 0.399742189 | 0.513742625 | 0.75367559 | 8.653385977 | 9.876711741 | 9.099922433 | 9.720931668 | 9.59016561 | 9.685065753 | 9.719953612 |
| ENSG00000113194 | FAF2 | 0.333332237 | 0.682283472 | 0.420100791 | 0.592465838 | 0.753433028 | 8.892187472 | 10.32888058 | 8.904018013 | 9.814540128 | 9.94297217 | 9.826590762 | 10.09422624 |
| ENSG00000155366 | RHOC | -0.057131599 | 0.914640194 | 0.808680632 | 0.058348141 | 0.751549033 | 11.68645261 | 10.8668003 | 11.23155829 | 11.20997529 | 11.2746349 | 11.9737357 | 12.2297432 |
| ENSG00000164683 | HEY1 | -0.629543348 | 0.74150751 | 1.377393305 | 0.429740549 | 0.747849957 | 2 | 4.442710129 | 4.523675391 | 3.709785322 | 3.45167086 | 5.880003936 | 3.753786374 |
| ENSG00000138107 | ACTR1A | 0.025256241 | 0.957413691 | 0.722520133 | 0.060082802 | 0.747776374 | 10.94624409 | 10.3138733 | 10.37481416 | 10.5472158 | 10.6475497 | 11.22220981 | 11.3625769 |
| ENSG00000197279 | ZNF165 | 0.434421618 | 0.816209881 | 0.313267318 | 0.865495099 | 0.747688936 | 2 | 3.686303414 | 3.369317972 | 3.462030052 | 3.50452879 | 3.336159116 | 3.474731315 |
| ENSG00000129535 | NRL | 0.833898886 | 0.704833394 | -0.087372175 | 0.97020102 | 0.74652671 | 2 | 4.11352228 | 2 | 3.615725078 | 3.74316441 | 3.12825509 | 3.100202476 |
| ENSG00000185009 | AP3M1 | 0.030722918 | 0.917958722 | 0.715207469 | 0.00165041 | 0.745930388 | 9.435834703 | 9.298208073 | 9.330306409 | 9.398771806 | 9.37105521 | 10.06867477 | 10.0637127 |
| ENSG00000154310 | TNIK | 1.827411195 | 0.154010606 | -1.083654903 | 0.415216121 | 0.743756292 | 5.532353208 | 7.274681896 | 4.058998049 | 8.001694993 | 7.94320592 | 5.707383854 | 4.4231929 |
| ENSG00000185238 | PRMT3 | 0.119594784 | 0.940829302 | 0.62262279 | 0.675301063 | 0.742217574 | 5.330594134 | 3.686303414 | 7.398165132 | 6.155987605 | 6.44101097 | 6.748859346 | 6.824016918 |
| ENSG00000136986 | DERL1 | 0.453663783 | 0.252999676 | 0.28779831 | 0.479315748 | 0.741462093 | 10.20680973 | 9.616264339 | 9.624627483 | 10.32001706 | 10.2669578 | 10.10747704 | 10.14908313 |
| ENSG00000183378 | OVCH2 | -2.312654496 | 0.63692076 | 3.053538922 | 0.477568485 | 0.740884426 | 2 | 2 | 2 | 2 | 2.14022021 | 3.904814898 | 3.311948089 |
| ENSG00000083937 | CHMP2B | 1.080304031 | 0.610039424 | -0.340077117 | 0.881655142 | 0.740226915 | 9.533135543 | 7.393647515 | 2 | 9.358500114 | 9.27923788 | 7.910770103 | 7.917377332 |
| ENSG00000137210 | TMEM14B | 0.040664041 | 0.970129688 | 0.699451551 | 0.467490985 | 0.740115592 | 8.468948322 | 8.147241016 | 6.073378139 | 7.992367495 | 7.8441483 | 8.298842011 | 8.798540026 |
| ENSG00000062725 | APPBP2 | 0.471913563 | 0.370967351 | 0.266728981 | 0.62541695 | 0.738642544 | 8.340423424 | 9.267380431 | 9.049077186 | 9.414570905 | 9.40359931 | 9.390112523 | 8.992926984 |
| ENSG00000148925 | BTBD10 | -0.048702723 | 0.950599592 | 0.786600747 | 0.243287647 | 0.737898023 | 8.257427002 | 7.232742069 | 8.763901704 | 8.139053376 | 8.17690623 | 9.12745764 | 8.825734765 |
| ENSG00000148803 | FUOM | -0.068902512 | 0.973095237 | 0.806287644 | 0.668910133 | 0.737385132 | 2 | 7.503546079 | 5.39244166 | 6.034011955 | 6.33555364 | 6.812870125 | 7.221358049 |
| ENSG00000166435 | XRRA1 | 1.264325494 | 0.012324317 | -0.527008228 | 0.337799565 | 0.737317266 | 7.096806373 | 7.051611315 | 6.073378139 | 8.033874364 | 8.08227582 | 6.277913523 | 6.359671687 |
| ENSG00000187837 | HIST1H1C | 1.252085452 | 0.361789525 | -0.515315037 | 0.72430187 | 0.736770414 | 7.159102577 | 4.93638933 | 8.825624954 | 8.874592369 | 8.99861459 | 6.601598251 | 7.619668339 |
| ENSG00000134874 | DZIP1 | 0.693639681 | 0.040664998 | 0.042969742 | 0.918154036 | 0.736609423 | 8.609439662 | 9.055168103 | 8.825624954 | 9.56056556 | 9.51003006 | 9.007589341 | 8.760149043 |
| ENSG00000176723 | ZNF843 | -1.451194262 | 0.548895656 | 2.187793737 | 0.329251531 | 0.736599475 | 2 | 4.710564154 | 2 | 3.162721474 | 2.2680111 | 5.680947153 | 4.836507146 |
| ENSG00000269743 | SLC25A53 | -0.623718491 | 0.70144113 | 1.360015682 | 0.350712355 | 0.736297191 | 2 | 3.686303414 | 4.523675391 | 3.566290607 | 3.01487888 | 4.742549839 | 4.827924991 |
| ENSG00000121579 | NAA50 | 0.710934866 | 0.241676601 | 0.024755818 | 0.972083517 | 0.735690684 | 9.128497779 | 9.532315021 | 10.39531398 | 10.56580756 | 10.4140861 | 9.813611959 | 9.802760311 |
| ENSG00000167380 | ZNF226 | 0.705278586 | 0.717276749 | 0.029693219 | 0.988584182 | 0.734971805 | 6.816637977 | 8.297207175 | 2 | 7.860161659 | 7.86703672 | 7.382965824 | 6.996671423 |
| ENSG00000108950 | FAM20A | -1.296538697 | 0.468067566 | 2.03148493 | 0.212636396 | 0.734946233 | 2 | 4.93638933 | 3.369317972 | 2.947924294 | 3.28037825 | 4.749835737 | 6.245088435 |
| ENSG00000079337 | RAPGEF3 | -0.675175632 | 0.706047591 | 1.409615799 | 0.38737291 | 0.734440168 | 2 | 5.595927442 | 3.369317972 | 3.79808662 | 4.02226117 | 5.650128416 | 5.694529741 |
| ENSG00000136758 | YME1L1 | 0.766372421 | 0.011999587 | -0.03256478 | 0.933173996 | 0.733807641 | 10.9638027 | 10.54135596 | 10.50627919 | 11.44607351 | 11.4539395 | 10.63927213 | 10.66581178 |
| ENSG00000186310 | NAP1L3 | -0.677658251 | 0.673764708 | 1.409891167 | 0.339618327 | 0.732232916 | 6.73718411 | 6.228547981 | 2 | 5.406080789 | 5.2591458 | 7.099048677 | 7.491038307 |
| ENSG00000130479 | MAP1S | 0.094107393 | 0.962673456 | 0.63755838 | 0.737112257 | 0.731665773 | 10.22148166 | 9.267380431 | 3.369317972 | 9.32173423 | 9.35585278 | 9.843290512 | 9.91489412 |
| ENSG00000178568 | ERBB4 | 1.897171808 | 0.438149869 | -1.166234078 | 0.647618864 | 0.730937729 | 2 | 3.076665209 | 2 | 3.462030052 | 3.55551834 | 2.34604626 | 2.225841198 |
| ENSG00000120696 | KBTBD7 | 0.499927771 | 0.827964997 | 0.228281734 | 0.922471927 | 0.728209505 | 2 | 4.710564154 | 2 | 3.95996233 | 3.90918929 | 3.865109708 | 3.600986722 |
| ENSG00000141570 | CBX8 | 0.209034854 | 0.913321769 | 0.518237765 | 0.774562861 | 0.72727262 | 2 | 5.457120953 | 7.098165388 | 6.139181293 | 6.1322378 | 6.639356243 | 6.182510897 |
| ENSG00000100395 | L3MBTL2 | 0.391586259 | 0.751627932 | 0.334853349 | 0.787590496 | 0.726439608 | 7.867441304 | 8.607278467 | 5.39244166 | 8.102673083 | 8.26379462 | 7.904255004 | 8.324402372 |
| ENSG00000153443 | UBALD1 | 0.139357912 | 0.784445644 | 0.587065145 | 0.181113049 | 0.726423058 | 7.790810999 | 8.255926468 | 8.631919287 | 8.354580458 | 8.44880239 | 8.79381476 | 8.895672339 |
| ENSG00000136279 | DBNL | 0.052238189 | 0.945987763 | 0.670688704 | 0.311659052 | 0.722926893 | 8.87372952 | 10.00059544 | 10.31867969 | 9.894954101 | 9.91151326 | 10.29113189 | 10.71681868 |
| ENSG00000170873 | MTSS1 | -0.486431102 | 0.821446892 | 1.20828405 | 0.537860646 | 0.721852948 | 7.867441304 | 4.93638933 | 2 | 6.139181293 | 5.88556448 | 7.929338774 | 7.302581437 |
| ENSG00000156876 | SASS6 | 0.69682043 | 0.535083376 | 0.024989665 | 0.984153231 | 0.721810095 | 4.467311984 | 5.457120953 | 4.058998049 | 5.565529336 | 5.30456641 | 5.243518884 | 4.292061816 |
| ENSG00000110060 | PUS3 | 0.144755353 | 0.925454021 | 0.576270283 | 0.685289111 | 0.721025636 | 4.467311984 | 7.503546079 | 4.874587708 | 6.394182228 | 6.48112613 | 7.0614806 | 6.607941737 |
| ENSG00000126246 | IGFLR1 | 1.877711305 | 0.412886226 | -1.156777166 | 0.627060885 | 0.720934139 | 2 | 3.686303414 | 2 | 3.79808662 | 4.22485516 | 2.267001569 | 2.632958241 |
| ENSG00000241878 | PISD | 0.530568303 | 0.682219847 | 0.190109916 | 0.889682343 | 0.720678219 | 5.330594134 | 8.522788974 | 8.845626171 | 8.703665429 | 8.70032852 | 8.421393978 | 8.30675314 |
| ENSG00000089639 | GMIP | 0.763540462 | 0.521751449 | -0.043423958 | 0.97326848 | 0.720116503 | 5.866944937 | 5.72254323 | 8.206335842 | 7.762033056 | 7.8720741 | 6.932953913 | 7.12130792 |
| ENSG00000265491 | RNF115 | -0.052792507 | 0.961244994 | 0.772648205 | 0.422682091 | 0.719855698 | 9.031919741 | 7.099070191 | 9.61300511 | 8.817637515 | 8.88981993 | 9.584231726 | 9.7588503 |
| ENSG00000183856 | IQGAP3 | -0.005465152 | 0.99798804 | 0.722433311 | 0.778850367 | 0.716968159 | 2 | 5.946638857 | 2 | 4.237680128 | 4.83276841 | 4.301748724 | 5.740981724 |
| ENSG00000156650 | KAT6B | 0.372152156 | 0.704032536 | 0.344744086 | 0.722618004 | 0.716896242 | 6.73718411 | 8.671479029 | 9.049077186 | 8.795015659 | 8.82894823 | 8.782313877 | 8.787516955 |
| ENSG00000101181 | MTG2 | 0.081296255 | 0.951240273 | 0.635405798 | 0.603099089 | 0.716702053 | 8.653385977 | 6.78762695 | 5.59507254 | 7.433701013 | 7.79460185 | 8.072046038 | 8.262059922 |
| ENSG00000110713 | NUP98 | 0.252390497 | 0.675311646 | 0.463754551 | 0.408562547 | 0.716145047 | 9.371511179 | 9.726146444 | 10.48730704 | 10.20056091 | 10.1772163 | 10.41631794 | 10.38273198 |
| ENSG00000037637 | FBXO42 | -0.081679049 | 0.966682851 | 0.796978917 | 0.660328198 | 0.715299868 | 4.467311984 | 6.602424674 | 9.888372618 | 8.441112993 | 8.34030387 | 9.172206498 | 9.34725177 |
| ENSG00000109689 | STIM2 | 1.221595148 | 0.064886519 | -0.509159716 | 0.475128388 | 0.712435431 | 6.366709671 | 7.503546079 | 7.285536013 | 8.270263574 | 8.39732094 | 7.000176314 | 6.172377306 |
| ENSG00000087338 | GMCL1 | 0.234219105 | 0.878292377 | 0.476682733 | 0.7369322 | 0.710901837 | 4.00722531 | 7.002537973 | 8.076950877 | 7.402350992 | 7.26532099 | 7.626042826 | 7.51663051 |
| ENSG00000129450 | SIGLEC9 | -2.3053361 | 0.655168404 | 3.015558758 | 0.511065006 | 0.710222658 | 2 | 2 | 2 | 2.141737246 | 2 | 4.177580368 | 2.671842496 |

| | | | | | | | | | | | | | |
|---|---|---|---|---|---|---|---|---|---|---|---|---|---|
| ENSG00000159140 | SON | 0.209791795 | 0.632755739 | 0.498061208 | 0.211429442 | 0.707853002 | 12.19201182 | 11.60869109 | 11.55072361 | 12.02642465 | 12.0201065 | 12.24198271 | 12.3774204 |
| ENSG00000142669 | SH3BGRL3 | 0.213325757 | 0.767351465 | 0.494310673 | 0.457548655 | 0.70763643 | 12.15618594 | 11.54371287 | 11.28013451 | 11.89463174 | 11.9469477 | 11.52908949 | 12.65872837 |
| ENSG00000176340 | COX8A | 0.065830708 | 0.959731105 | 0.640122079 | 0.591055458 | 0.705952787 | 10.71696191 | 8.505280662 | 7.896971941 | 9.596407142 | 9.67763759 | 10.04998226 | 10.35313191 |
| ENSG00000163482 | STK36 | 0.384555724 | 0.806161404 | 0.32109905 | 0.836918007 | 0.705654774 | 4.00722531 | 8.451443573 | 7.285536013 | 7.883054517 | 7.76782485 | 7.551912644 | 7.949380652 |
| ENSG00000113211 | PCDHB6 | 0.765917475 | 0.729728652 | -0.060708141 | 0.979401422 | 0.705209334 | 2 | 4.11352228 | 2 | 3.566290607 | 3.69849104 | 3.12825509 | 3.128433649 |
| ENSG00000126562 | WNK4 | -0.046117571 | 0.98350331 | 0.749927954 | 0.71854685 | 0.703810383 | 2 | 5.946638857 | 8.536706852 | 7.057964187 | 7.21466714 | 8.028526139 | 7.796124663 |
| ENSG00000126524 | SBDS | 0.237945637 | 0.759076344 | 0.465716875 | 0.52135424 | 0.703662511 | 10.27168932 | 8.702538795 | 10.08627303 | 10.05135008 | 10.0869461 | 10.28267996 | 10.30968141 |
| ENSG00000084234 | APLP2 | 0.138867788 | 0.888548519 | 0.564754808 | 0.519747981 | 0.703622596 | 12.76283692 | 12.46223249 | 10.67226762 | 12.35897559 | 12.3412176 | 12.61180942 | 12.92290535 |
| ENSG00000083642 | PDS5B | 0.961684702 | 0.116820694 | -0.258548017 | 0.704939784 | 0.703136686 | 8.16936288 | 8.747909548 | 7.342949392 | 9.162439556 | 9.15351711 | 8.016494289 | 7.882577494 |
| ENSG00000186448 | ZNF197 | 0.762802852 | 0.45354552 | -0.060158447 | 0.958081539 | 0.702644405 | 5.866944937 | 8.004992147 | 8.460932916 | 8.52598359 | 8.57464274 | 7.918064594 | 7.535201607 |
| ENSG00000089351 | GRAMD1A | 0.065293134 | 0.957587914 | 0.635887649 | 0.567222525 | 0.701180783 | 10.87842776 | 8.522788974 | 8.65477198 | 9.793884031 | 9.90906469 | 10.66105227 | 10.13327938 |
| ENSG00000139154 | AEBP2 | 0.19165712 | 0.898282189 | 0.508418955 | 0.714850336 | 0.700076076 | 5.330594134 | 6.728495673 | 8.884815469 | 7.880528731 | 7.86451142 | 8.331380596 | 8.020777347 |
| ENSG00000105053 | VRK3 | 0.84807364 | 0.085388044 | -0.148377441 | 0.794186359 | 0.699696199 | 7.53297911 | 6.666836837 | 7.451345288 | 8.124184191 | 8.0358969 | 7.14536429 | 7.063994051 |
| ENSG00000196136 | SERPINA3 | 2.900846406 | 0.361569449 | -2.201624459 | NA | 0.699221947 | 2 | 2 | 2 | 3.754611435 | 3.28037825 | 2.094598369 | 2.059860178 |
| ENSG00000165410 | CFL2 | -0.526830228 | 0.237527811 | 1.225690773 | 0.002219876 | 0.698860545 | 10.00110988 | 9.806969599 | 9.196533591 | 9.269600773 | 9.09439916 | 10.8677203 | 10.99006746 |
| ENSG00000137507 | LRRC32 | -1.801641984 | 0.000721583 | 2.496108099 | 1.02E-06 | 0.694466116 | 7.867441304 | 8.903033433 | 9.149036566 | 6.922108067 | 7.0150921 | 11.25322188 | 11.19155596 |
| ENSG00000119953 | SMNDC1 | 0.407490713 | 0.500733183 | 0.28692416 | 0.639193832 | 0.694414873 | 8.285626337 | 7.467835385 | 7.163363831 | 8.184811706 | 8.04707549 | 8.045647998 | 7.950369405 |
| ENSG00000138095 | LRPPRC | 0.689403621 | 0.373352069 | 0.004536763 | 0.996064263 | 0.693940384 | 8.075571721 | 9.903691196 | 9.132850258 | 9.903687576 | 9.91517834 | 9.208355151 | 9.247812734 |
| ENSG00000170365 | SMAD1 | 1.163739471 | 0.44522898 | -0.471075957 | 0.770200762 | 0.692663514 | 3.327711234 | 7.761270542 | 5.772716771 | 7.731524332 | 7.66128317 | 6.146071859 | 6.092227206 |
| ENSG00000198858 | R3HDM4 | -0.092017401 | 0.908769676 | 0.782991588 | 0.241623158 | 0.690974187 | 9.371511179 | 9.13702746 | 7.896971941 | 8.830780654 | 8.85207713 | 9.490026756 | 9.902175458 |
| ENSG00000153130 | SCOC | 0.431171138 | 0.414030475 | 0.259400849 | 0.632223509 | 0.690571987 | 9.112843761 | 8.777382925 | 8.266922072 | 9.269600773 | 9.10299435 | 9.244909847 | 8.748827534 |
| ENSG00000105723 | GSK3A | 0.346300297 | 0.753326748 | 0.342935954 | 0.753191906 | 0.689236251 | 7.384433967 | 8.981105197 | 10.10298738 | 9.519758723 | 9.57865651 | 9.589063591 | 9.501932203 |
| ENSG00000185630 | PBX1 | 0.291979726 | 0.76909522 | 0.396506292 | 0.681431908 | 0.688486017 | 8.716893051 | 8.655694736 | 6.534109959 | 8.575233246 | 8.51952909 | 8.477329966 | 8.806206367 |
| ENSG00000196581 | AJAP1 | -2.312721121 | 0.663582024 | 3.000648273 | 0.525943482 | 0.687927152 | 2 | 2 | 2 | 2 | 2.14022021 | 4.230621763 | 2.466008509 |
| ENSG00000168763 | CNNM3 | 0.191214924 | 0.798284713 | 0.495398142 | 0.464069117 | 0.686613065 | 6.468621203 | 7.35507212 | 6.629488387 | 7.066878356 | 7.0774796 | 7.697491716 | 6.940057611 |
| ENSG00000175727 | MLXIP | -0.106435288 | 0.932860886 | 0.792855718 | 0.478845029 | 0.68642043 | 7.384433967 | 10.35355093 | 10.26027233 | 9.707456306 | 9.71175558 | 10.80232183 | 10.37796311 |
| ENSG00000105088 | OLFM2 | -1.421783508 | 0.032828609 | 2.106590658 | 0.000537851 | 0.68480715 | 5.095959458 | 6.04685525 | 6.073378139 | 4.623314311 | 4.4836996 | 8.086505553 | 7.54177663 |
| ENSG00000164347 | GFM2 | 0.822242668 | 0.114309143 | -0.138557445 | 0.817289897 | 0.683685223 | 7.33131396 | 7.790474954 | 8.408111533 | 8.674801527 | 8.76132797 | 7.832294085 | 7.706501872 |
| ENSG00000158545 | ZC3H18 | 0.185246201 | 0.86184497 | 0.495939515 | 0.609008368 | 0.681185716 | 9.497412313 | 7.790474954 | 7.451345288 | 8.731963162 | 8.71722214 | 9.172206498 | 8.878113577 |
| ENSG00000164830 | OXR1 | -0.058745524 | 0.919646442 | 0.739480662 | 0.120443179 | 0.680735138 | 8.138771685 | 7.638155693 | 8.296287027 | 8.022463952 | 7.94559728 | 9.019318595 | 8.475596448 |
| ENSG00000157869 | RAB28 | 0.331510088 | 0.710348924 | 0.348229093 | 0.690278517 | 0.679739181 | 7.384433967 | 6.602424674 | 5.39244166 | 6.931899482 | 7.05550676 | 7.107581047 | 6.907844475 |
| ENSG00000090621 | PABPC4 | -0.134142475 | 0.863478741 | 0.813553039 | 0.214752155 | 0.679410564 | 9.097018025 | 10.20420974 | 8.845626171 | 9.381011913 | 9.3808074 | 10.21453767 | 10.42655664 |
| ENSG00000169105 | CHST14 | -0.213981495 | 0.857034032 | 0.89204791 | 0.389087222 | 0.678066415 | 9.219014269 | 6.535001748 | 7.734844356 | 7.966401501 | 8.06478301 | 9.089196199 | 9.131450704 |
| ENSG00000023318 | ERP44 | 1.082973146 | 0.140291258 | -0.406234004 | 0.609008368 | 0.676739142 | 10.27168932 | 9.377333006 | 8.486635645 | 10.64606954 | 10.6257199 | 9.041010784 | 9.257431793 |
| ENSG00000140688 | C16orf58 | 0.864123103 | 0.030796809 | -0.187500877 | 0.683460163 | 0.676622227 | 8.075571721 | 8.717821467 | 8.175061096 | 9.210493388 | 9.22416269 | 8.256629373 | 8.090601559 |
| ENSG00000196074 | SYCP2 | 0.925280289 | 0.53071634 | -0.248706415 | 0.876525531 | 0.676573875 | 4.00722531 | 3.076665209 | 4.058998049 | 4.299492648 | 4.58534546 | 3.943456545 | 3.071407821 |
| ENSG00000169814 | BTD | -0.089574366 | 0.968773308 | 0.765027889 | 0.721493405 | 0.675453522 | 9.358295023 | 7.051611315 | 2 | 7.997038782 | 7.91663606 | 8.59173005 | 8.984746522 |
| ENSG00000135250 | SRPK2 | 0.236762685 | 0.745616581 | 0.437842352 | 0.523326453 | 0.674605037 | 8.042902253 | 8.607278467 | 9.503962296 | 9.089953145 | 9.06500642 | 9.256145816 | 9.298000491 |
| ENSG00000080007 | DDX43 | -3.753029217 | 0.499533315 | 4.427034123 | 0.376931525 | 0.674004906 | 2 | 2 | 2 | 2 | 2 | 4.099885429 | 5.115701921 |
| ENSG00000128000 | ZNF780B | 1.466298839 | 0.185150547 | -0.79240235 | 0.492708893 | 0.673896488 | 4.00722531 | 7.145017454 | 6.882775709 | 7.987681034 | 7.92393105 | 5.951260634 | 5.59182244 |
| ENSG00000091651 | ORC6 | 0.722352048 | 0.358652871 | -0.048806546 | 0.956860836 | 0.673545502 | 4.815609471 | 5.131603625 | 4.874587708 | 5.639097847 | 5.5755357 | 5.068107405 | 4.74829783 |
| ENSG00000077713 | SLC25A43 | 0.30946912 | 0.834827076 | 0.36331945 | 0.800774229 | 0.67278857 | 9.344956677 | 6.78762695 | 5.930865648 | 8.401141138 | 8.40081078 | 8.39955642 | 8.506848296 |
| ENSG00000100372 | SLC25A17 | 0.390579384 | 0.690529475 | 0.279042295 | 0.778789835 | 0.669621679 | 6.563806279 | 7.503546079 | 8.763901704 | 8.270263574 | 8.26762243 | 7.98365249 | 8.31522102 |
| ENSG00000133619 | KRBA1 | -0.139560346 | 0.929222345 | 0.808649548 | 0.56354884 | 0.669089202 | 6.96351253 | 7.60566235 | 3.369317972 | 6.598566398 | 6.71216164 | 7.544599974 | 7.608437243 |
| ENSG00000185163 | DDX51 | -0.222412329 | 0.744473447 | 0.891354026 | 0.129961155 | 0.668941697 | 6.257048221 | 7.274681896 | 6.43197674 | 6.664650034 | 6.39275508 | 7.52137298 | 7.702986259 |
| ENSG00000130176 | CNN1 | -2.587342088 | 1.24E-07 | 3.255981994 | 6.95E-12 | 0.668639906 | 9.33149386 | 8.169643011 | 9.116480286 | 6.401247806 | 6.43421531 | 12.18290317 | 12.20354771 |
| ENSG00000108946 | PRKAR1A | 0.473396567 | 0.025286919 | 0.195233305 | 0.393866241 | 0.668629872 | 11.25159303 | 11.26034006 | 11.31641863 | 11.76026481 | 11.7374476 | 11.37858912 | 11.55807875 |
| ENSG00000111907 | TPD52L1 | -2.80785372 | 0.056626744 | 3.476135259 | 0.011014824 | 0.668281539 | 2 | 6.140560021 | 4.058998049 | 2.947924294 | 3.01487888 | 8.504505158 | 7.980689421 |
| ENSG00000171928 | TVP23B | 1.26126292 | 0.008469405 | -0.595567931 | 0.247751753 | 0.665694989 | 8.586954094 | 7.874700259 | 8.784770914 | 9.749588203 | 9.6675326 | 8.062569037 | 7.647978997 |
| ENSG00000172340 | SUCLG2 | 0.621465982 | 0.098154133 | 0.043870756 | 0.923063983 | 0.665336738 | 7.82963488 | 8.053983481 | 7.934808822 | 8.600009476 | 8.51792306 | 7.804574291 | 8.153772149 |
| ENSG00000196182 | STK40 | -0.108538451 | 0.955340357 | 0.773123121 | 0.668910133 | 0.66458467 | 5.330594134 | 9.943237334 | 5.772716771 | 8.36726458 | 8.40776525 | 9.321478134 | 9.198731239 |
| ENSG00000071205 | ARHGAP10 | -0.378429518 | 0.74176673 | 1.040811095 | 0.315894652 | 0.662381577 | 9.174465797 | 6.844429755 | 7.029880454 | 7.739909082 | 7.70992146 | 9.136171672 | 9.117873759 |
| ENSG00000184897 | H1FX | 0.061003135 | 0.925672996 | 0.600957565 | 0.277409224 | 0.6619607 | 9.048471759 | 10.14890644 | 10.01740919 | 9.841412998 | 9.91151326 | 10.28769449 | 10.53186455 |
| ENSG00000204138 | PHACTR4 | 0.439862739 | 0.346404872 | 0.221978032 | 0.649746012 | 0.66184077 | 8.674867122 | 9.067150531 | 9.491321578 | 9.517321994 | 9.58557298 | 9.189724501 | 9.467467793 |
| ENSG00000173272 | MZT2A | 0.635991565 | 0.239055913 | 0.023355279 | 0.97050839 | 0.659346843 | 8.674867122 | 8.077869079 | 7.646393253 | 8.865680425 | 8.7802082 | 8.376795097 | 8.038552764 |
| ENSG00000114812 | VIPR1 | -3.753032715 | 0.495884604 | 4.411319177 | 0.373787431 | 0.658286463 | 2 | 2 | 2 | 2 | 2 | 4.572179576 | 4.784231301 |
| ENSG00000126003 | PLAGL2 | 0.173951396 | 0.939485429 | 0.484127414 | 0.827299212 | 0.65807881 | 4.467311984 | 9.298208073 | 3.369317972 | 7.920420067 | 8.00183183 | 8.290747499 | 8.243626305 |
| ENSG00000135407 | AVIL | 1.147763814 | 0.149004691 | -0.490124537 | 0.564099325 | 0.657639277 | 5.330594134 | 5.72254323 | 4.058998049 | 6.221320894 | 6.35727082 | 4.799832052 | 4.80186789 |
| ENSG00000110057 | UNC93B1 | 1.216890579 | 0.03681088 | -0.559451855 | 0.369090445 | 0.657438723 | 8.493334308 | 7.538394143 | 8.865353883 | 9.615519136 | 9.58019637 | 7.716209895 | 7.953331607 |
| ENSG00000117477 | CCDC181 | -0.687342928 | 0.818202168 | 1.3443249 | 0.615729155 | 0.656981971 | 2 | 3.076665209 | 2 | 2.3892346 | 2.38539787 | 3.483289517 | 2.782576693 |
| ENSG00000147654 | EBAG9 | 0.671124345 | 0.29397165 | -0.01471447 | 0.984191003 | 0.656409875 | 5.532353208 | 6.602424674 | 6.629488387 | 6.907295206 | 7.06873058 | 6.466611899 | 6.182510897 |
| ENSG00000084070 | SMAP2 | 0.574032486 | 0.058669633 | 0.080469439 | 0.820851865 | 0.654501925 | 8.609439662 | 8.623598428 | 8.561105505 | 9.193296408 | 9.14313645 | 8.549559709 | 8.797990868 |
| ENSG00000078699 | CBFA2T2 | 0.717445664 | 0.509306243 | -0.063398539 | 0.958174026 | 0.654047125 | 9.262228249 | 8.791896851 | 6.43197674 | 9.234427714 | 9.33867006 | 8.708512253 | 8.287321088 |
| ENSG00000137845 | ADAM10 | 0.380986816 | 0.580787928 | 0.271718197 | 0.696976163 | 0.652705013 | 9.371511179 | 10.53705456 | 10.77492406 | 10.69863405 | 10.7502937 | 10.47451025 | 10.7443005 |
| ENSG00000169490 | TM2D2 | 0.459646722 | 0.729210347 | 0.192338625 | 0.890313426 | 0.651985347 | 10.07567912 | 8.297207175 | 6.629488387 | 9.39612172 | 9.43287094 | 8.904453984 | 9.358096047 |
| ENSG00000274286 | ADRA2B | -1.303057274 | NA | 1.955008429 | NA | 0.651951155 | 2 | 2 | 2 | 2.270786764 | 2 | 3.374372158 | 2.466008509 |
| ENSG00000166557 | TMED3 | 0.098887683 | 0.886208951 | 0.551950234 | 0.351453916 | 0.650837918 | 11.32815351 | 10.56689847 | 10.13584611 | 10.84605279 | 10.8765996 | 11.47344391 | 11.13386651 |
| ENSG00000198816 | ZNF358 | -0.167727553 | 0.912094181 | 0.8181049 | 0.542439157 | 0.650377348 | 5.532353208 | 9.648522413 | 8.845626171 | 8.568972129 | 8.64106878 | 9.63011094 | 9.536251143 |
| ENSG00000142959 | BEST4 | 0.368676804 | 0.881476754 | 0.281032589 | 0.90832745 | 0.649709394 | 2 | 4.710564154 | 2 | 3.709785322 | 3.94787299 | 3.336159116 | 4.105828626 |
| ENSG00000100225 | FBXO7 | 0.192228706 | 0.491293064 | 0.457157417 | 0.068317498 | 0.649386123 | 9.98400217 | 9.936721131 | 9.735974718 | 10.06086293 | 10.1036386 | 10.29767155 | 10.39367729 |
| ENSG00000006638 | TBXA2R | -0.672448741 | 0.612543393 | 1.321420778 | 0.278961643 | 0.648972037 | 7.485143553 | 4.11352228 | 5.930865648 | 5.686138681 | 5.86549842 | 7.765568476 | 7.613439639 |
| ENSG00000140025 | EFCAB11 | 0.720938107 | 0.328225936 | -0.072180271 | 0.932122597 | 0.648757837 | 5.532353208 | 5.595927442 | 4.523675391 | 5.949776857 | 6.00960265 | 5.169385838 | 5.331426166 |
| ENSG00000111711 | GOLT1B | 0.734388726 | 0.357059066 | -0.086199066 | 0.92531104 | 0.64818966 | 7.867441304 | 9.672251278 | 8.960138144 | 9.793884031 | 9.69337522 | 9.18804929 | 8.613062226 |
| ENSG00000100354 | TNRC6B | 0.468445227 | 0.751472272 | 0.179582185 | 0.908291211 | 0.648027413 | 5.532353208 | 9.159587375 | 9.897982195 | 9.411074916 | 9.58173459 | 9.159610958 | 9.262217379 |
| ENSG00000240682 | ISY1 | 0.831790376 | 0.046962517 | -0.18386986 | 0.699954446 | 0.647920516 | 7.53297911 | 7.60566235 | 7.285536013 | 8.184811706 | 8.41468635 | 7.113241388 | 7.478755384 |
| ENSG00000105011 | ASF1B | 0.201402563 | 0.941409766 | 0.445454879 | 0.867353663 | 0.646857442 | 2 | 5.303523747 | 2 | 4.299492648 | 4.05806467 | 3.215043717 | 5.013446246 |
| ENSG00000180616 | SSTR2 | 0.439292997 | 0.883067069 | 0.205720839 | 0.944462782 | 0.645013836 | 2 | 3.686303414 | 4.523675391 | 4.268917409 | 3.7864959 | 4.652122235 | 2.059860178 |
| ENSG00000145332 | KLHL8 | 0.466896889 | 0.739729032 | 0.177570652 | 0.904611962 | 0.644467541 | 4.00722531 | 7.73146271 | 7.451345288 | 7.517371202 | 7.5462416 | 7.57363085 | 6.830475749 |
| ENSG00000115109 | EPB41L5 | 0.228223992 | 0.895400127 | 0.416046231 | 0.800593168 | 0.644270223 | 7.904282243 | 8.376385542 | 3.369317972 | 7.786523606 | 7.86955761 | 7.895248429 | 8.122532579 |
| ENSG00000149218 | ENDOD1 | -0.590608988 | 0.698378828 | 1.234270431 | 0.37875047 | 0.643661443 | 6.009025997 | 9.443177293 | 10.67785662 | 9.108419034 | 9.00091592 | 11.02039807 | 10.7082226 |
| ENSG00000105953 | OGDH | -0.333603187 | 0.757468139 | 0.976930842 | 0.314242081 | 0.643327655 | 9.015175615 | 9.470501606 | 11.23913884 | 9.870342608 | 9.95429779 | 10.85172587 | 11.51294866 |
| ENSG00000128311 | TST | -0.077009948 | 0.970008162 | 0.71953003 | 0.706265396 | 0.642520083 | 8.367060731 | 7.189546432 | 2 | 7.169761359 | 7.32163988 | 8.042684843 | 8.013226939 |
| ENSG00000078061 | ARAF | 0.185215922 | 0.878396152 | 0.457245194 | 0.681460704 | 0.642461117 | 10.1311248 | 8.433041974 | 7.502634626 | 9.288777073 | 9.28113268 | 9.351421858 | 9.734089014 |
| ENSG00000047249 | ATP6V1H | 0.731288694 | 0.410778027 | -0.089555611 | 0.92948232 | 0.641733082 | 9.590776211 | 8.234834549 | 7.552162999 | 9.451627281 | 9.42603674 | 8.52322774 | 8.718968947 |
| ENSG00000143751 | SDE2 | 0.385301785 | 0.770265142 | 0.256361497 | 0.848992364 | 0.641663282 | 6.138360429 | 7.638155693 | 9.465702942 | 8.722121967 | 8.72837565 | 8.676372393 | 8.514222325 |
| ENSG00000118855 | MFSD1 | 1.338847245 | 0.090297135 | -0.698589273 | 0.399585878 | 0.640257972 | 9.473095744 | 9.497318007 | 7.600047325 | 10.35546265 | 10.4786707 | 8.247650629 | 8.526871175 |
| ENSG00000167766 | ZNF83 | 0.250508946 | 0.822912318 | 0.388757271 | 0.715258735 | 0.639266217 | 7.384433967 | 8.053983481 | 9.714382178 | 8.948803189 | 8.99976571 | 9.366608223 | 8.80183059 |
| ENSG00000137497 | NUMA1 | 0.764556484 | 0.074018972 | -0.126678652 | 0.798498793 | 0.637877832 | 9.957954117 | 10.73780697 | 10.13584611 | 11.05899696 | 11.1061 | 10.23447665 | 10.15187942 |
| ENSG00000144118 | RALB | 0.65815223 | 0.419141353 | -0.022869885 | 0.980129329 | 0.635282345 | 7.790810999 | 9.405920729 | 9.658943166 | 9.756492097 | 9.85856545 | 8.987312532 | 9.262217379 |
| ENSG00000120370 | GORAB | 0.737608778 | 0.618090751 | -0.103277143 | 0.948615522 | 0.634331635 | 3.327711234 | 7.701025996 | 7.163363831 | 7.739909082 | 7.52720828 | 6.898949899 | 6.735016458 |
| ENSG00000177479 | ARIH2 | 0.42940172 | 0.643876123 | 0.204891525 | 0.833193204 | 0.634293245 | 7.904282243 | 10.00683004 | 8.763901704 | 9.545497589 | 9.61366407 | 9.366904403 | 9.346500876 |
| ENSG00000171067 | C11orf24 | -0.465032786 | 0.412187433 | 1.098216319 | 0.031149835 | 0.633183533 | 9.751047882 | 9.308339287 | 10.38167985 | 9.397888985 | 9.43372295 | 10.80297867 | 11.12612127 |
| ENSG00000110108 | TMEM109 | -0.369019317 | 0.456750419 | 1.001937238 | 0.022450111 | 0.63291792 | 10.21416435 | 9.656475478 | 9.385998894 | 9.422406043 | 9.43116542 | 10.61939729 | 10.9472486 |
| ENSG00000166716 | ZNF592 | -0.159860838 | 0.814508396 | 0.792373589 | 0.172021653 | 0.632512752 | 9.90440318 | 8.747909548 | 8.941672936 | 9.04103174 | 9.21427981 | 10.07358251 | 10.08142138 |
| ENSG00000101346 | POFUT1 | -0.093734502 | 0.934842791 | 0.725920886 | 0.472907496 | 0.632186384 | 7.667647223 | 10.34373336 | 9.725218844 | 9.484432763 | 9.55770587 | 10.15524287 | 10.49980559 |
| ENSG00000173281 | PPP1R3B | -0.274112122 | 0.832844524 | 0.906257737 | 0.432001677 | 0.632145615 | 5.532353208 | 7.002537973 | 8.608698767 | 7.292639438 | 7.27676203 | 8.620333861 | 8.243626305 |
| ENSG00000139946 | PELI2 | 0.408936114 | 0.820262885 | 0.223102144 | 0.902593423 | 0.632038259 | 4.00722531 | 7.669933291 | 4.058998049 | 6.825617242 | 6.58294721 | 6.583362013 | 6.476841503 |
| ENSG00000067221 | STOML1 | -0.333755093 | 0.864010109 | 0.965409337 | 0.583360046 | 0.631654245 | 7.096806373 | 7.847167909 | 2 | 6.509830055 | 6.75096147 | 7.923712697 | 7.875307424 |
| ENSG00000182700 | IGIP | -0.207628746 | 0.935007649 | 0.838608868 | 0.72430187 | 0.630980123 | 2 | 4.442710129 | 8.904018013 | 7.29643414 | 7.07311172 | 8.367352231 | 8.044120856 |
| ENSG00000215883 | CYB5RL | -0.586558378 | 0.829400292 | 1.217266624 | 0.62887288 | 0.630708246 | 2 | 2 | 6.43197674 | 4.299492648 | 4.60967607 | 6.231826305 | 5.88031069 |
| ENSG00000110719 | TCIRG1 | 0.850235308 | 0.47919324 | -0.219575757 | 0.866382985 | 0.630659551 | 8.468948322 | 8.213429689 | 10.93284392 | 10.55277887 | 10.6618002 | 9.549165021 | 9.535921843 |
| ENSG00000159873 | CCDC117 | 0.30079432 | 0.834110208 | 0.327368933 | 0.815505549 | 0.628163253 | 5.532353208 | 9.125613817 | 9.287079172 | 8.946390023 | 9.00894176 | 9.197405512 | 8.781418172 |
| ENSG00000145990 | GFOD1 | -0.137542462 | 0.946595613 | 0.765410802 | 0.690305992 | 0.62786834 | 2 | 6.140560021 | 3.369317972 | 4.574143677 | 4.89348248 | 5.614651724 | 5.464725981 |
| ENSG00000163393 | SLC22A15 | 1.716002374 | 0.261861084 | -1.088309073 | 0.488633039 | 0.627693301 | 7.218819781 | 4.93638933 | 4.058998049 | 7.617841552 | 7.7377845 | 5.73334482 | 3.620995423 |
| ENSG00000093217 | XYLB | 0.326800842 | 0.898630945 | 0.30068208 | 0.906004994 | 0.627482922 | 2 | 5.72254323 | 2 | 4.523237872 | 4.76938658 | 4.340871764 | 4.870334174 |
| ENSG00000185088 | RPS27L | 0.142703709 | 0.866763421 | 0.484684871 | 0.523602241 | 0.62738858 | 10.56418906 | 9.055168103 | 9.272377397 | 9.899327447 | 9.97024066 | 10.38434358 | 10.15873961 |
| ENSG00000159322 | ADPGK | 1.101298396 | 0.021693999 | -0.474522217 | 0.360306411 | 0.626776179 | 8.493334308 | 9.532315021 | 8.99637432 | 10.17605323 | 10.1644413 | 8.616851507 | 8.591675571 |

| | | | | | | | | | | | | | |
|---|---|---|---|---|---|---|---|---|---|---|---|---|---|
| ENSG00000072756 | TRNT1 | 0.554107726 | 0.638803946 | 0.072107368 | 0.955191984 | 0.626215094 | 4.467311984 | 7.431218269 | 7.098165388 | 7.514115926 | 7.12888795 | 7.045287119 | 6.662054875 |
| ENSG00000147416 | ATP6V1B2 | 0.342690976 | 0.620548015 | 0.283011522 | 0.683637326 | 0.625702498 | 10.86913986 | 9.514922632 | 10.01740919 | 10.54562241 | 10.621987 | 10.34293862 | 10.68628308 |
| ENSG00000102158 | MAGT1 | 0.405366028 | 0.794639251 | 0.220282841 | 0.890244223 | 0.625648869 | 6.009025997 | 9.903691196 | 10.60345421 | 10.19498122 | 10.1052438 | 9.745312661 | 10.15809785 |
| ENSG00000089177 | KIF16B | 0.121472673 | 0.946330866 | 0.504131598 | 0.770708651 | 0.62560427 | 3.327711234 | 6.464272742 | 8.486635645 | 7.355829978 | 7.37584259 | 7.629009275 | 7.843728236 |
| ENSG00000168724 | DNAJC21 | 0.054615641 | 0.965988399 | 0.570827326 | 0.625700973 | 0.625442967 | 6.257048221 | 9.415325457 | 8.904018013 | 8.828161587 | 8.65866647 | 9.38982107 | 9.113025255 |
| ENSG00000184058 | TBX1 | 0.426355313 | 0.904107254 | 0.198309074 | NA | 0.624664387 | 2 | 3.076665209 | 2 | 2.600425781 | 2.86123014 | 3.035909414 | 2.117335275 |
| ENSG00000141644 | MBD1 | 0.17860786 | 0.725609862 | 0.445869422 | 0.335637906 | 0.624477282 | 9.667789716 | 8.791896851 | 9.372276026 | 9.476928206 | 9.5116449 | 9.793481551 | 9.726317257 |
| ENSG00000119707 | RBM25 | 0.758418743 | 0.082371227 | -0.134187154 | 0.790197999 | 0.624231589 | 9.533135543 | 9.00621767 | 9.849281147 | 10.25178588 | 10.2564019 | 9.285583053 | 9.441331959 |
| ENSG00000124574 | ABCC10 | 0.538200022 | 0.794520103 | 0.084581124 | 0.968806022 | 0.622781146 | 8.855032352 | 8.169643011 | 2 | 8.483340231 | 8.53072154 | 7.959503304 | 8.152913455 |
| ENSG00000162441 | LZIC | 0.383633885 | 0.540633385 | 0.238930513 | 0.710603737 | 0.622564398 | 8.075571721 | 6.899080544 | 7.502634626 | 7.968781453 | 7.90439517 | 7.935741825 | 7.638194697 |
| ENSG00000116337 | AMPD2 | 0.287346392 | 0.783350857 | 0.334864048 | 0.741073451 | 0.62221044 | 10.27872131 | 9.497318007 | 7.777116083 | 9.829676744 | 9.75806852 | 9.783975576 | 9.897056431 |
| | MB21D1 | 5.812770451 | 0.089797897 | -5.191128106 | 0.178878036 | 0.621642346 | 2 | 2 | 2 | 5.818703821 | 6.04571775 | 2 | 2 |
| ENSG00000143110 | C1orf162 | -0.991610824 | 0.747288938 | 1.612814046 | 0.570849073 | 0.621203222 | 2 | 3.686303414 | 2 | 2.141737246 | 2.77779907 | 4.369533167 | 2.851942824 |
| ENSG00000060138 | YBX3 | -0.175481614 | 0.556828814 | 0.795601632 | 0.00161096 | 0.620120019 | 10.76282024 | 10.47993704 | 10.75391088 | 10.51622947 | 10.4720505 | 11.37888285 | 11.54332823 |
| ENSG00000101199 | ARFGAP1 | 0.080262096 | 0.930786574 | 0.539802381 | 0.509619901 | 0.620064478 | 9.174465797 | 9.170736405 | 10.74328839 | 9.901820563 | 10.0524158 | 10.50025386 | 10.37096473 |
| ENSG00000137714 | FDX1 | 0.566631682 | 0.606376439 | 0.053311991 | 0.965290088 | 0.619943672 | 5.095959458 | 7.701025996 | 7.818184403 | 7.836899669 | 7.84926609 | 7.328045423 | 7.352801367 |
| ENSG00000036054 | TBC1D23 | 0.946620385 | 0.517222971 | -0.32700503 | 0.833980846 | 0.619615355 | 5.330594134 | 9.876711741 | 8.922968322 | 9.921611889 | 9.82778401 | 8.888863837 | 8.265241863 |
| ENSG00000100991 | TRPC4AP | 0.477177862 | 0.131912603 | 0.142027702 | 0.684760578 | 0.619205565 | 10.25752168 | 9.785372135 | 10.10298738 | 10.52838133 | 10.5400919 | 10.16293862 | 10.23638632 |
| ENSG00000111727 | HCFC2 | 0.051103744 | 0.943613626 | 0.567683783 | 0.365809256 | 0.618787527 | 7.975256505 | 7.35507212 | 7.342949392 | 7.762033056 | 7.49817903 | 8.522696167 | 7.628343587 |
| ENSG00000266028 | SRGAP2 | 0.813129394 | 0.185725588 | -0.194938161 | 0.776974137 | 0.618191233 | 8.257427002 | 9.540933215 | 9.452720984 | 9.965769838 | 10.0345549 | 9.097429939 | 8.89156005 |
| ENSG00000197933 | ZNF823 | 0.448309168 | 0.680957539 | 0.168672028 | 0.883235347 | 0.616981195 | 4.815609471 | 5.457120953 | 3.369317972 | 5.110902178 | 5.3193943 | 5.073924441 | 4.861951484 |
| ENSG00000142002 | DPP9 | 0.324779296 | 0.712692238 | 0.291650235 | 0.738292369 | 0.616429531 | 9.913467515 | 7.979858146 | 9.647595056 | 9.731481912 | 9.69692814 | 9.611983935 | 9.747518555 |
| ENSG00000127804 | METTL16 | -0.375412501 | 0.845522872 | 0.990627978 | 0.572896533 | 0.615215477 | 3.327711234 | 7.35507212 | 9.344432377 | 7.691737974 | 7.76782485 | 9.067857443 | 9.091227191 |
| ENSG00000111206 | FOXM1 | -0.035906969 | 0.982223298 | 0.650248729 | 0.653855654 | 0.61434176 | 4.00722531 | 7.002537973 | 5.59507254 | 5.94010643 | 6.03677338 | 5.399896186 | 7.297928101 |
| ENSG00000116273 | PHF13 | -0.190946053 | 0.876278112 | 0.805142932 | 0.452613409 | 0.614196879 | 8.009475818 | 7.189546432 | 9.819251342 | 8.609191952 | 8.53866332 | 9.456464678 | 9.661984931 |
| ENSG00000177200 | CHD9 | 0.493037439 | 0.531143362 | 0.119799193 | 0.889682343 | 0.612836632 | 9.730679447 | 10.16572116 | 8.434763955 | 10.08464853 | 10.1222557 | 9.942606032 | 9.484973508 |
| ENSG00000106948 | AKNA | 0.41894905 | 0.812190473 | 0.191667889 | 0.915522323 | 0.61061694 | 4.815609471 | 4.442710129 | 8.236946975 | 7.214570819 | 7.32531744 | 7.398115274 | 6.590289384 |
| ENSG00000138193 | PLCE1 | -2.822940798 | 0.076976946 | 3.432529709 | 0.024210868 | 0.609588911 | 2 | 6.04685525 | 7.163363831 | 4.205751501 | 3.08597292 | 9.229031012 | 9.739818999 |
| ENSG00000165527 | ARF6 | -0.303321537 | 0.56715642 | 0.911886078 | 0.051766586 | 0.608564541 | 10.02639697 | 10.55418374 | 10.89974571 | 10.22266632 | 10.2457682 | 11.18498388 | 11.66824733 |
| ENSG00000005302 | MSL3 | 0.817945855 | 0.166721726 | -0.209403614 | 0.749808479 | 0.608542241 | 7.53297911 | 7.503546079 | 8.561105505 | 8.699372486 | 8.80807321 | 7.626042826 | 7.847978946 |
| ENSG00000158195 | WASF2 | -0.411521236 | 0.701800878 | 1.019934752 | 0.294703553 | 0.608413517 | 8.631580145 | 11.13786633 | 11.1494608 | 10.25958244 | 10.2802818 | 11.67532131 | 11.72127105 |
| ENSG00000067248 | DHX29 | 0.807474287 | 0.586183061 | -0.199260153 | 0.90169025 | 0.608214134 | 9.521325812 | 8.806266215 | 4.874587708 | 9.483600848 | 9.43627594 | 8.389678938 | 8.53018147 |
| ENSG00000143862 | ARL8A | 0.366700699 | 0.606992001 | 0.2413599 | 0.738195992 | 0.608060599 | 9.966689162 | 9.443177293 | 8.434763955 | 9.81982307 | 9.73267673 | 9.618217491 | 9.685333873 |
| ENSG00000103449 | SALL1 | -0.865482026 | 0.601928679 | 1.473122051 | 0.340932973 | 0.607640026 | 2 | 6.140560021 | 6.80318718 | 5.501219927 | 4.72553021 | 7.525623979 | 7.191563076 |
| ENSG00000129422 | MTUS1 | -1.747217422 | 0.300402677 | 2.353336895 | 0.133579211 | 0.606119473 | 2 | 5.838937808 | 3.369317972 | 3.406930997 | 3.28037825 | 7.02591204 | 6.476841503 |
| ENSG00000164251 | F2RL1 | 1.173668482 | 0.298019815 | -0.568401734 | 0.631319446 | 0.605266748 | 4.815609471 | 6.464272742 | 7.858115942 | 8.027038952 | 7.97164382 | 6.109723842 | 6.485052582 |
| ENSG00000131591 | C1orf159 | 0.21922025 | 0.910380971 | 0.384813594 | 0.836009948 | 0.604033844 | 7.579279349 | 6.389896633 | 2 | 6.788375664 | 6.71216164 | 6.978631953 | 6.839042789 |
| ENSG00000120805 | ARL1 | 0.189446954 | 0.474064658 | 0.41428372 | 0.083172408 | 0.603730674 | 9.87686338 | 9.969011754 | 10.05225195 | 10.15582956 | 10.1567216 | 10.44437493 | 10.31333667 |
| ENSG00000168621 | GDNF | 1.837270373 | 0.361789525 | -1.234699506 | 0.547274868 | 0.602570867 | 8.107517749 | 3.076665209 | 5.156626401 | 8.554784469 | 8.50501001 | 6.412547789 | 3.287118176 |
| ENSG00000152443 | ZNF776 | 0.530347231 | 0.751472272 | 0.071069699 | 0.968477418 | 0.60141693 | 3.327711234 | 8.077869079 | 8.486635645 | 8.22518713 | 8.28849576 | 7.731011524 | 7.875307424 |
| ENSG00000126653 | NSRP1 | 0.515759029 | 0.190607653 | 0.083378073 | 0.853024401 | 0.599137102 | 9.031919741 | 8.376385542 | 8.631919287 | 9.231457548 | 9.19128891 | 8.774744193 | 8.790280628 |
| ENSG00000197816 | CCDC180 | 0.856490318 | 0.713785889 | -0.257825018 | 0.917242457 | 0.5986653 | 4.815609471 | 2 | 2 | 4.17310023 | 4.1930041 | 3.584558147 | 3.183290849 |
| ENSG00000184867 | ARMCX2 | 0.090214716 | 0.968342042 | 0.507342978 | 0.817314365 | 0.597557695 | 9.304185368 | 9.74864196 | 2 | 9.104095335 | 8.99976571 | 9.577081455 | 9.34837738 |
| ENSG00000138964 | PARVG | -2.305380043 | 0.65272748 | 2.901291844 | NA | 0.595911802 | 2 | 2 | 2 | 2.141737246 | 2 | 3.968654369 | 2.918126211 |
| ENSG00000116213 | WRAP73 | 0.035702978 | 0.98413673 | 0.559810573 | 0.744740559 | 0.595513551 | 6.89194353 | 7.145017454 | 2 | 6.604700657 | 6.39275508 | 7.137053113 | 6.864442837 |
| ENSG00000166689 | PLEKHA7 | 1.134436764 | 0.64718241 | -0.539503272 | 0.835180221 | 0.594933492 | 2 | 4.710564154 | 2 | 4.804776789 | 3.86943969 | 2.592885947 | 3.67940853 |
| ENSG00000108924 | HLF | -3.753025965 | 0.504957686 | 4.347911412 | 0.392715269 | 0.594885447 | 2 | 2 | 2 | 2 | 2 | 4.199031499 | 4.959465495 |
| ENSG00000177842 | ZNF620 | -0.77334746 | 0.51181261 | 1.367432796 | 0.199468909 | 0.594085335 | 3.327711234 | 4.710564154 | 5.156626401 | 3.754611435 | 4.16043394 | 6.146071859 | 5.331426166 |
| ENSG00000170323 | FABP4 | -3.274019969 | 0.353103921 | 3.864473174 | 0.176769831 | 0.590453205 | 3.327711234 | 2 | 2 | 2 | 2.14022021 | 5.86330637 | 2.918126211 |
| ENSG00000106245 | BUD31 | 1.279279043 | 0.055604743 | -0.688930562 | 0.327012123 | 0.590348481 | 8.75772933 | 7.790474954 | 7.163363831 | 9.324524407 | 9.31482024 | 7.347386207 | 7.407104868 |
| ENSG00000120948 | TARDBP | 0.737973686 | 0.453160638 | -0.149007239 | 0.891004263 | 0.588966447 | 9.544849385 | 9.792607274 | 7.342949392 | 9.991161874 | 9.93276281 | 8.972597003 | 9.180307928 |
| ENSG00000144579 | CTDSP1 | 0.054622793 | 0.920437233 | 0.531945668 | 0.247703343 | 0.586568462 | 9.262228249 | 10.02537381 | 10.05225195 | 9.878593178 | 9.87935305 | 10.22388773 | 10.47442608 |
| ENSG00000164609 | SLU7 | 0.967330404 | 0.47627622 | -0.381197367 | 0.791764439 | 0.586133037 | 9.613202746 | 9.45234302 | 5.59507254 | 9.959201773 | 9.95963168 | 8.541185654 | 8.694392964 |
| ENSG00000152082 | MZT2B | 0.389739051 | 0.344601719 | 0.196109417 | 0.64966917 | 0.585848467 | 8.855032352 | 8.955547846 | 9.503962296 | 9.535096495 | 9.50030288 | 9.323005438 | 9.327600363 |
| ENSG00000107679 | PLEKHA1 | -0.88111134 | 0.066039055 | 1.466108281 | 0.000983739 | 0.58499694 | 8.928409675 | 7.954278496 | 8.585098385 | 7.717440541 | 7.61696151 | 10.02598953 | 9.957809357 |
| ENSG00000130303 | BST2 | 1.87506185 | 0.031250396 | -1.290836001 | 0.149212315 | 0.584225849 | 7.904282243 | 8.903033433 | 10.17589429 | 11.14332431 | 11.165776 | 8.306273771 | 7.633277551 |
| ENSG00000198039 | ZNF273 | 0.511518232 | 0.667273997 | 0.071813148 | 0.955158571 | 0.58333138 | 4.00722531 | 4.710564154 | 3.369317972 | 4.76150579 | 4.40247105 | 4.281782022 | 4.188602707 |
| ENSG00000123739 | PLA2G12A | -0.881958413 | 0.405292907 | 1.464475475 | 0.136351095 | 0.582517062 | 5.330594134 | 8.077869079 | 7.934808822 | 6.658765962 | 6.70088158 | 9.03282106 | 8.926146458 |
| ENSG00000126247 | CAPNS1 | 0.232691374 | 0.76542366 | 0.349232324 | 0.640632886 | 0.581923698 | 10.06758145 | 11.35379164 | 11.53848775 | 11.31939084 | 11.3798372 | 11.143491 | 11.72993391 |
| ENSG00000133243 | BTBD2 | -0.12748489 | 0.941619909 | 0.708836412 | 0.659176557 | 0.581351522 | 5.095959458 | 10.36815316 | 9.809100877 | 9.418929011 | 9.43457445 | 10.34052773 | 10.1734218 |
| ENSG00000099992 | TBC1D10A | 0.254166249 | 0.891376437 | 0.327035955 | 0.855655091 | 0.581202204 | 9.081016759 | 6.464272742 | 4.058998049 | 7.855024645 | 8.13141746 | 7.821097076 | 8.284971554 |
| ENSG00000103064 | SLC7A6 | 0.333915797 | 0.573636123 | 0.247257103 | 0.681162733 | 0.5811729 | 8.963744667 | 8.671479029 | 7.934808822 | 8.899755849 | 8.9350493 | 8.558404203 | 9.061336073 |
| ENSG00000204335 | SP5 | -1.310474982 | NA | 1.890539153 | NA | 0.58006417 | 2 | 2 | 2 | 2.141737246 | 2.14022021 | 2.3070652 | 3.407202941 |
| ENSG00000090924 | PLEKHG2 | -0.787861386 | 0.330789334 | 1.366938184 | 0.069363787 | 0.579076798 | 8.928409675 | 8.004992147 | 9.926432745 | 8.420396336 | 8.33118782 | 10.43593029 | 10.5951862 |
| LCMT1 | | 0.473654576 | 0.806161404 | 0.105131983 | 0.958081539 | 0.578786559 | 7.624139768 | 7.901717006 | 2 | 7.577865504 | 7.74603955 | 7.108998216 | 7.476011586 |
| ENSG00000158321 | AUTS2 | 0.740931805 | 0.476344102 | -0.162682632 | 0.886854075 | 0.578249173 | 4.467311984 | 7.002537973 | 6.80318718 | 7.265793319 | 7.07311172 | 6.471027105 | 6.113568999 |
| ENSG00000141665 | FBXO15 | 0.189126244 | 0.950359078 | 0.388294319 | NA | 0.577420563 | 2 | 3.076665209 | 2 | 2.784612485 | 2.49394783 | 2.937246078 | 2.466008509 |
| ENSG00000175356 | SCUBE2 | 1.134152537 | 0.292494247 | -0.556991643 | 0.622741559 | 0.577160893 | 3.327711234 | 5.131603625 | 4.058998049 | 5.48800675 | 5.30456641 | 4.04171018 | 3.924507493 |
| ENSG00000159917 | ZNF235 | 0.877181161 | 0.661258435 | -0.300595956 | 0.887741643 | 0.576585205 | 6.138360429 | 3.076665209 | 2 | 5.686138681 | 5.40529704 | 4.563936166 | 4.479073019 |
| ENSG00000164307 | ERAP1 | 0.842849817 | 0.323812042 | -0.266336103 | 0.774665597 | 0.576513714 | 8.257427002 | 9.506147172 | 10.40208303 | 10.45897359 | 10.4897738 | 9.359183899 | 9.380643788 |
| ENSG00000170222 | ADPRM | 0.504669156 | 0.737405845 | 0.071284027 | 0.965258777 | 0.575953183 | 3.327711234 | 4.93638933 | 6.958202216 | 6.139181293 | 6.33555364 | 5.960710379 | 5.694529741 |
| ENSG00000126062 | TMEM115 | 0.024336174 | 0.973778114 | 0.551335334 | 0.392649764 | 0.575671507 | 9.848787621 | 10.08551947 | 8.699419242 | 9.611716938 | 9.75534491 | 10.1231663 | 10.29224003 |
| ENSG00000000457 | SCYL3 | 0.693695718 | 0.709351398 | -0.11844719 | 0.952589029 | 0.575248529 | 5.70932728 | 7.503546079 | 2 | 6.89733479 | 7.07311172 | 6.302895343 | 6.110033886 |
| ENSG00000032219 | ARID4A | 0.866051575 | 0.221769007 | -0.295466468 | 0.701056475 | 0.570585107 | 7.33131396 | 6.951736442 | 8.408111533 | 8.567402595 | 8.5227358 | 7.577730839 | 7.203223239 |
| ENSG00000215440 | NPEPL1 | 1.04014288 | 0.024327615 | -0.470247838 | 0.344285984 | 0.569895042 | 7.940205782 | 7.145017454 | 7.398165132 | 8.425603456 | 8.66157866 | 6.970858896 | 7.156006254 |
| ENSG00000066557 | LRRC40 | 0.688715983 | 0.226161058 | -0.119765167 | 0.852959423 | 0.568950817 | 6.257048221 | 6.602424674 | 7.225742779 | 7.381065997 | 7.44169935 | 6.782996409 | 6.454714037 |
| ENSG00000141736 | ERBB2 | 0.003098851 | 0.997041173 | 0.565529491 | 0.400315488 | 0.568628341 | 9.204317094 | 10.14890644 | 8.721235202 | 9.471064221 | 9.51003006 | 9.95290183 | 10.1419604 |
| ENSG00000197265 | GTF2E2 | 0.089094941 | 0.962823674 | 0.47952356 | 0.795268255 | 0.568618502 | 3.327711234 | 8.029695761 | 9.439621147 | 8.439397922 | 8.39557285 | 8.934738262 | 8.660355764 |
| ENSG00000105948 | TTC26 | 0.463794743 | 0.709997456 | 0.104107428 | 0.937449278 | 0.567902171 | 4.467311984 | 6.844429755 | 4.874587708 | 6.252910529 | 6.26836889 | 5.614651724 | 6.175763085 |
| ENSG00000100599 | RIN3 | -0.091464764 | 0.916568051 | 0.65852947 | 0.380792862 | 0.567064706 | 8.16936288 | 6.464272742 | 7.646393253 | 7.494428494 | 7.47849652 | 8.050818921 | 8.377561365 |
| ENSG00000139517 | LNX2 | -0.35865738 | 0.62100324 | 0.925075168 | 0.150484813 | 0.566417788 | 5.70932728 | 6.311476754 | 6.322058544 | 5.839660368 | 5.78232107 | 7.354572674 | 6.617932662 |
| ENSG00000198912 | C1orf174 | -0.335229607 | 0.714367242 | 0.901503383 | 0.273716572 | 0.566273777 | 7.750913381 | 7.73146271 | 5.772716771 | 7.021746293 | 7.00136498 | 8.271259924 | 8.17168765 |
| ENSG00000124588 | NQO2 | -0.043265075 | 0.970415686 | 0.608621201 | 0.565677502 | 0.565356125 | 7.940205782 | 8.297207175 | 5.59507254 | 7.45762193 | 7.78127547 | 8.086505553 | 8.432373803 |
| ENSG00000228716 | DHFR | 1.574295665 | 0.262309738 | -1.01262014 | 0.482463036 | 0.561675525 | 3.327711234 | 7.638155693 | 6.80318718 | 8.254744042 | 8.32017198 | 5.238348159 | 6.189227296 |
| ENSG00000142867 | BCL10 | 0.688522688 | 0.465992837 | -0.128078692 | 0.902389555 | 0.560443996 | 8.998234871 | 6.728495673 | 7.858115942 | 8.84120963 | 8.80016655 | 7.966556098 | 8.059781405 |
| ENSG00000183775 | KCTD16 | -1.32373524 | 0.651415542 | 1.884035983 | 0.500719988 | 0.560300743 | 2 | 2 | 8.608698767 | 5.880681751 | 5.69405321 | 9.08127623 | 8.697340724 |
| ENSG00000140993 | TIGD7 | 1.383448926 | 0.114419311 | -0.823289016 | 0.366065794 | 0.56015991 | 4.815609471 | 5.131603625 | 5.156626401 | 6.351046291 | 6.28357155 | 4.742549839 | 3.807156736 |
| ENSG00000107341 | UBE2R2 | 0.099049751 | 0.833540602 | 0.460653339 | 0.256328984 | 0.55970309 | 10.29268328 | 10.09139839 | 9.589475645 | 10.09992864 | 10.1401137 | 10.58136152 | 10.37225646 |
| ENSG00000213901 | SLC23A3 | 1.041495701 | 0.653324928 | -0.482809942 | 0.842082048 | 0.558685759 | 2 | 2 | 4.874587708 | 4.358765249 | 4.37434574 | 3.500668998 | 3.042026731 |
| ENSG00000107262 | BAG1 | 1.269388211 | 0.193841835 | -0.710886391 | 0.48356437 | 0.55850182 | 6.653098118 | 9.170736405 | 7.734844356 | 9.437090644 | 9.51728265 | 7.35696023 | 7.66855289 |
| ENSG00000018869 | ZNF582 | 0.40975959 | 0.804532535 | 0.148707015 | 0.930981294 | 0.558466606 | 2 | 5.457120953 | 5.59507254 | 5.406080789 | 5.34860058 | 5.606648123 | 4.456980017 |
| ENSG00000113360 | DROSHA | 0.190244139 | 0.895265695 | 0.367452602 | 0.788187265 | 0.557696741 | 8.737455672 | 9.148351514 | 5.39244166 | 8.583021623 | 8.66303256 | 8.729394257 | 8.865589219 |
| ENSG00000148943 | LIN7C | 0.611774396 | 0.697744171 | -0.056832455 | 0.97269175 | 0.554941941 | 4.00722531 | 8.414402626 | 8.805342542 | 8.729158235 | 8.61275919 | 8.062569037 | 7.959237827 |
| ENSG00000181016 | LSMEM1 | 1.319856956 | 0.552095038 | -0.764960798 | 0.738292369 | 0.554896158 | 2 | 2 | 4.874587708 | 4.739373666 | 4.43005853 | 3.235948948 | 3.012034821 |
| ENSG00000169359 | SLC33A1 | 0.577512865 | 0.436535627 | -0.02271399 | 0.977998169 | 0.554798875 | 7.975256505 | 7.002537973 | 8.65477198 | 8.551612617 | 8.62474679 | 8.021770574 | 7.970003494 |
| ENSG00000140092 | FBLN5 | -0.028871809 | 0.96053419 | 0.583595048 | 0.245225364 | 0.554723238 | 10.56993315 | 9.549500232 | 10.24529339 | 10.19090967 | 10.1079152 | 10.71590919 | 10.80418236 |
| ENSG00000114857 | NKTR | 0.551877239 | 0.549066587 | 0.001733458 | 0.998969048 | 0.553610697 | 8.16936288 | 10.23640192 | 8.825624954 | 9.908654498 | 9.88247671 | 9.413813486 | 9.279629951 |
| ENSG00000158856 | DMTN | -0.182885058 | 0.911458513 | 0.736069965 | 0.613476865 | 0.553184907 | 2 | 4.11352228 | 4.874587708 | 3.840289928 | 4.02226117 | 4.933813368 | 4.329022995 |
| ENSG00000165704 | HPRT1 | 0.530899193 | 0.331331826 | 0.022080833 | 0.971808785 | 0.552980026 | 7.33131396 | 6.602424674 | 6.718950348 | 7.523859802 | 7.32531744 | 6.763331418 | 7.085754446 |
| ENSG00000085415 | SEH1L | 0.156729678 | 0.915012803 | 0.396143222 | 0.775777561 | 0.552872901 | 4.815609471 | 8.639735838 | 7.858115942 | 7.959237999 | 7.91663606 | 7.853585565 | 8.435907908 |
| ENSG00000162302 | RPS6KA4 | -0.19404378 | 0.81054847 | 0.746547465 | 0.290479733 | 0.552503685 | 7.218819781 | 8.234834549 | 8.825624954 | 8.015574133 | 8.06917614 | 8.721540982 | 9.186615069 |
| ENSG00000123600 | METTL8 | 0.344325292 | 0.768318166 | 0.207860119 | 0.861870179 | 0.552185411 | 5.330594134 | 8.213429689 | 7.163363831 | 7.778406195 | 7.55253065 | 7.494516625 | 7.576773052 |
| ENSG00000077152 | UBE2T | 0.464710236 | 0.83197366 | 0.086301493 | 0.969279885 | 0.551011729 | 3.327711234 | 6.602424674 | 2 | 5.890757403 | 5.39133044 | 5.445480974 | 5.177806074 |
| ENSG00000099949 | LZTR1 | 0.578189271 | 0.714821112 | -0.027280425 | 0.987221496 | 0.550908846 | 9.204317094 | 9.28800521 | 4.523675391 | 9.215512556 | 9.31111579 | 8.638111529 | 8.688479312 |
| ENSG00000135838 | NPL | -1.39040001 | 0.570724655 | 1.93920775 | 0.403634871 | 0.54880774 | 2 | 5.838937808 | 2 | 3.349643765 | 3.55551834 | 6.898949899 | 4.853519801 |
| ENSG00000112964 | GHR | -1.571678786 | 0.303447732 | 2.120125507 | 0.137639917 | 0.548446721 | 2 | 5.131603625 | 5.930865648 | 3.79808662 | 3.7864959 | 6.092633892 | 7.603417442 |
| ENSG00000161904 | LEMD2 | -0.029171313 | 0.974752037 | 0.577385874 | 0.476093865 | 0.548214561 | 10.03472846 | 8.234834549 | 9.165043284 | 9.298270478 | 9.28868707 | 10.17076319 | 9.558468984 |

| | | | | | | | | | | | | | |
|---|---|---|---|---|---|---|---|---|---|---|---|---|---|
| ENSG00000171357 | LURAP1 | 0.480006047 | 0.641227131 | 0.064939631 | 0.953719218 | 0.544945678 | 4.815609471 | 4.93638933 | 4.058998049 | 5.37770462 | 4.72553021 | 4.572179576 | 4.870334174 |
| ENSG00000072274 | TFRC | 0.373316877 | 0.801833271 | 0.171429488 | 0.91083145 | 0.544746365 | 6.89194353 | 11.05905903 | 10.62675862 | 10.70863622 | 10.6759114 | 10.207113 | 10.72810493 |
| ENSG00000140367 | UBE2Q2 | 0.528758618 | 0.371475952 | 0.015911812 | 0.981190943 | 0.54467043 | 8.107517749 | 8.376385542 | 9.242516611 | 9.215512556 | 9.14521857 | 8.733534631 | 8.60805852 |
| ENSG00000090612 | ZNF268 | 0.273809934 | 0.899665024 | 0.270207096 | 0.900092939 | 0.54401703 | 4.467311984 | 9.203675681 | 4.058998049 | 7.975897842 | 7.99492122 | 7.938932725 | 8.023598593 |
| ENSG00000008735 | MAPK8IP2 | 1.762634467 | 0.435930907 | -1.22196406 | 0.594394464 | 0.540670407 | 2 | 6.04685525 | 2 | 6.372775472 | 6.12381559 | 3.904814898 | 3.496556178 |
| ENSG00000114405 | C3orf14 | 1.588040315 | 0.212730396 | -1.047578356 | 0.424853745 | 0.540461959 | 5.866944937 | 3.686303414 | 3.369317972 | 6.252910529 | 6.1068222 | 4.122514202 | 3.660199087 |
| ENSG00000137509 | PRCP | 0.444470304 | 0.528239125 | 0.095580686 | 0.901587015 | 0.54005099 | 10.78274301 | 9.549500232 | 9.624627483 | 10.51093171 | 10.5791198 | 9.972107965 | 10.39258649 |
| ENSG00000132958 | TPTE2 | 3.417477183 | 0.412304476 | -2.881694072 | 0.511130806 | 0.53578311 | 2 | 2 | 2 | 3.921164147 | 3.86943969 | 2.094598369 | 2 |
| ENSG00000134684 | YARS | -0.150292897 | 0.773531565 | 0.686064919 | 0.1272864 | 0.535772022 | 10.46903009 | 10.61666558 | 9.746650996 | 10.1342586 | 10.217847 | 11.06635027 | 10.95305641 |
| ENSG00000124209 | RAB22A | 0.21428859 | 0.512728968 | 0.321004834 | 0.29554872 | 0.535293424 | 9.097018025 | 9.308339287 | 9.478569125 | 9.55265467 | 9.47734778 | 9.564482569 | 9.676696925 |
| ENSG00000180773 | SLC36A4 | -0.050617735 | 0.977133743 | 0.585837449 | 0.721968109 | 0.535219714 | 3.327711234 | 7.431218269 | 8.486635645 | 7.491120977 | 7.39696604 | 7.862855183 | 8.250064747 |
| ENSG00000114933 | INO80D | 0.588163966 | 0.08123451 | -0.053524017 | 0.894789442 | 0.534639949 | 8.340423424 | 8.337339476 | 8.042706615 | 8.869506572 | 8.79487126 | 8.296978073 | 8.091498142 |
| ENSG00000004961 | HCCS | 0.047907451 | 0.981829576 | 0.484544516 | 0.80343258 | 0.532451967 | 3.327711234 | 6.899080544 | 9.491321578 | 8.197043419 | 8.18299958 | 8.518969664 | 8.717227336 |
| ENSG00000162873 | KLHDC8A | 3.425341355 | 0.548693531 | -2.893582639 | 0.624498271 | 0.531758716 | 2 | 2 | 2 | 4.694064037 | 2 | 2.094598369 | 2 |
| ENSG00000165792 | METTL17 | 0.674370657 | 0.747288938 | -0.143426893 | 0.948326217 | 0.530943764 | 5.095959458 | 8.004992147 | 2 | 7.348537858 | 7.21466714 | 6.435321218 | 6.549111181 |
| ENSG00000101945 | SUV39H1 | 0.060868933 | 0.966162087 | 0.469384177 | 0.725041248 | 0.530253109 | 3.327711234 | 5.838937808 | 4.523675391 | 4.907595604 | 5.09650239 | 4.841348851 | 5.75463059 |
| ENSG00000109606 | DHX15 | 0.224522547 | 0.625309666 | 0.305222215 | 0.487297152 | 0.529744761 | 9.460782017 | 10.10308486 | 9.478569125 | 9.961593621 | 9.92005068 | 9.927424911 | 10.10957632 |
| ENSG00000093144 | ECHDC1 | 0.566803269 | 0.255069337 | -0.037263695 | 0.948615522 | 0.529539575 | 8.777722034 | 8.469613413 | 7.934808822 | 9.066811112 | 8.92538942 | 8.175765162 | 8.593575433 |
| ENSG00000071246 | VASH1 | -0.090076681 | 0.965522332 | 0.619212773 | 0.752411321 | 0.529136091 | 7.031698319 | 8.101365653 | 2 | 6.912249745 | 7.10341274 | 8.229523742 | 6.864442837 |
| ENSG00000085788 | DDHD2 | -0.263089682 | 0.666522361 | 0.792115073 | 0.142327697 | 0.529025391 | 9.048471759 | 8.077869079 | 9.180874355 | 8.540455898 | 8.61576544 | 9.703944794 | 9.543476778 |
| ENSG00000002919 | SNX11 | 0.145580757 | 0.950922102 | 0.383146536 | 0.870919566 | 0.528727293 | 9.48530526 | 6.389896633 | 2 | 8.139053376 | 8.27714776 | 8.304419417 | 8.571895748 |
| ENSG00000158828 | PINK1 | 0.823447028 | 0.368407723 | -0.295601233 | 0.763865232 | 0.527845795 | 7.579279349 | 9.896993485 | 9.491321578 | 10.06531795 | 10.1359316 | 9.018187664 | 8.957466266 |
| ENSG00000156050 | FAM161B | 0.035179293 | 0.981829576 | 0.492030307 | 0.724837675 | 0.5272096 | 7.096806373 | 6.951736442 | 3.369317972 | 6.64692533 | 6.42052719 | 7.246693284 | 6.64749689 |
| ENSG00000171813 | PWWP2B | -0.200322101 | 0.905447484 | 0.726596858 | 0.632965021 | 0.526274757 | 3.327711234 | 7.274681896 | 7.896971941 | 6.841288255 | 6.91125523 | 8.098683215 | 7.370638252 |
| ENSG00000256806 | C17orf100 | 0.315682678 | 0.853531206 | 0.209968464 | 0.902389591 | 0.525651142 | 2 | 6.140560021 | 4.874587708 | 5.363304456 | 5.48637102 | 5.196301326 | 5.453611367 |
| ENSG00000109971 | HSPA8 | -0.02427492 | 0.945043724 | 0.549373331 | 0.054901185 | 0.525098411 | 13.57305719 | 13.45717518 | 13.13440312 | 13.36470886 | 13.3867068 | 13.98526566 | 13.91224726 |
| ENSG00000184277 | TM2D3 | -0.156150533 | 0.885189733 | 0.681139646 | 0.471649635 | 0.524989113 | 6.138360429 | 8.356994601 | 8.460932916 | 7.829061569 | 7.80253898 | 8.634179825 | 8.649438172 |
| ENSG00000168438 | CDC40 | 1.042076293 | 0.305827583 | -0.519108036 | 0.626596436 | 0.522968257 | 8.517314938 | 7.431218269 | 5.772716771 | 8.627383409 | 8.68323472 | 7.192914486 | 7.04745562 |
| ENSG00000187513 | GJA4 | -3.75301868 | 0.514704475 | 4.275376839 | 0.414211825 | 0.522358159 | 2 | 2 | 2 | 2 | 2 | 4.996409624 | 3.956354764 |
| ENSG00000140968 | IRF8 | -3.753012457 | 0.521753772 | 4.274127284 | 0.423211461 | 0.521114827 | 2 | 2 | 2 | 2 | 2 | 5.147485955 | 3.580696614 |
| ENSG00000100722 | ZC3H14 | 0.715911054 | 0.519228435 | -0.195016238 | 0.870966304 | 0.520894817 | 8.75772933 | 9.763446433 | 6.80318718 | 9.615519136 | 9.57479964 | 8.568225482 | 8.804020138 |
| ENSG00000080815 | PSEN1 | 0.446225931 | 0.313361774 | 0.074576157 | 0.882758816 | 0.520802088 | 10.34051281 | 9.632483534 | 9.69246155 | 10.38025813 | 10.3568427 | 10.05256229 | 9.941741079 |
| ENSG00000156232 | WHAMM | 0.191056695 | 0.88968542 | 0.329369043 | 0.800584348 | 0.520425738 | 5.866944937 | 6.844429755 | 8.904018013 | 7.917958914 | 7.97632944 | 8.249579355 | 7.895982618 |
| ENSG00000197329 | PELI1 | 0.613614272 | 0.510049398 | -0.096171248 | 0.926209474 | 0.517443024 | 6.73718411 | 7.901717006 | 5.772716771 | 7.659688863 | 7.69294162 | 7.086154752 | 6.874895611 |
| ENSG00000083844 | ZNF264 | 0.186591168 | 0.880334493 | 0.330656153 | 0.777382971 | 0.517247321 | 6.563806279 | 7.761270542 | 9.330306409 | 8.559529208 | 8.43013848 | 8.82044307 | 8.430957733 |
| ENSG00000105792 | CFAP69 | 1.882285193 | 0.28497373 | -1.365756055 | 0.445853185 | 0.516529138 | 2 | 5.457120953 | 3.369317972 | 5.968925393 | 6.00043082 | 3.782262616 | 2.746601469 |
| ENSG00000138346 | DNA2 | 0.302394651 | 0.911302045 | 0.210220908 | 0.937572507 | 0.512615559 | 2 | 5.72254323 | 2 | 4.470469939 | 4.76938658 | 5.068107405 | 3.735548302 |
| ENSG00000198483 | ANKRD35 | 0.100337558 | 0.957413691 | 0.411815082 | 0.820281623 | 0.51215264 | 5.866944937 | 2 | 5.156626401 | 5.128039203 | 5.06165168 | 4.675266036 | 5.842465082 |
| ENSG00000127423 | AUNIP | -1.13735904 | 0.478467424 | 1.649047666 | 0.258489631 | 0.511688626 | 2 | 4.442710129 | 3.369317972 | 3.023133209 | 2.94009902 | 4.690491514 | 5.171036168 |
| ENSG00000125895 | TMEM74B | 0.055935097 | 0.97695465 | 0.455345742 | 0.802702997 | 0.511280839 | 2 | 4.710564154 | 3.369317972 | 3.709785322 | 3.98554648 | 4.530483275 | 3.640730416 |
| ENSG00000188997 | KCTD21 | 0.037948177 | 0.983064817 | 0.472154082 | 0.778107221 | 0.51010226 | 8.540903468 | 7.232742069 | 3.369317972 | 7.464384256 | 7.55566492 | 8.008923105 | 7.860656447 |
| ENSG00000274320 | | 4.557005953 | 1.43E-09 | -4.049428319 | 5.31E-06 | 0.507577634 | 4.467311984 | 3.686303414 | 4.523675391 | 8.514626115 | 8.40429221 | 2.34604626 | 2.225841198 |
| ENSG00000135414 | GDF11 | -0.045325292 | 0.98350331 | 0.552011935 | 0.79391096 | 0.506686643 | 7.867441304 | 9.246456519 | 2 | 8.109160159 | 8.08227582 | 8.744517886 | 8.624876935 |
| ENSG00000135953 | MFSD9 | 0.18708487 | 0.917958722 | 0.319080795 | 0.85594901 | 0.506165665 | 6.816637977 | 7.002537973 | 2 | 6.580005553 | 6.49425375 | 6.732402829 | 6.592824411 |
| ENSG00000213672 | NCKIPSD | -0.22307134 | 0.92127192 | 0.727033664 | 0.728633609 | 0.503962324 | 9.448362284 | 7.954278496 | 2 | 8.013270196 | 8.15847021 | 8.919069795 | 9.128396091 |
| ENSG00000086758 | HUWE1 | 0.006466878 | 0.979997472 | 0.495012427 | 0.012586703 | 0.501479304 | 11.55070946 | 11.55651981 | 11.51057165 | 11.49601699 | 11.5944788 | 11.96871957 | 12.09663891 |
| ENSG00000135723 | FHOD1 | -0.269855881 | 0.719407817 | 0.771109912 | 0.252830324 | 0.501254031 | 7.709880939 | 8.848535039 | 8.536706852 | 8.076429088 | 8.27144011 | 8.659546439 | 9.604418524 |
| ENSG00000064703 | DDX20 | 0.250560262 | 0.903601303 | 0.250565645 | 0.90231922 | 0.501125906 | 2 | 8.169643011 | 7.97167868 | 7.756533788 | 7.7377845 | 7.852739912 | 7.633277551 |
| ENSG00000137955 | RABGGTB | 0.267875162 | 0.90624054 | 0.229811202 | 0.918425911 | 0.497686363 | 9.371511179 | 7.701025996 | 2 | 8.423869837 | 8.47883193 | 8.447394152 | 8.379764957 |
| ENSG00000167895 | TMC8 | -2.305372515 | 0.653114396 | 2.801478826 | NA | 0.496106312 | 2 | 2 | 2 | 2.141737246 | 2 | 3.767980144 | 3.100202476 |
| ENSG00000047315 | POLR2B | 0.49489604 | 0.506870806 | 0.000940844 | 0.999263508 | 0.495836884 | 8.075571721 | 9.616264339 | 9.577565424 | 9.715981692 | 9.75874862 | 9.306733157 | 9.183254692 |
| ENSG00000072609 | CHFR | -0.013742937 | 0.990573498 | 0.509208692 | 0.625351493 | 0.495465755 | 9.2903348 | 8.191702463 | 6.629488387 | 8.402902269 | 8.38151133 | 8.880175401 | 8.939154596 |
| ENSG00000138182 | KIF20B | 0.215590503 | 0.937471333 | 0.279467763 | 0.919037965 | 0.495058266 | 2 | 6.666836837 | 2 | 5.474671438 | 5.33407135 | 5.650128416 | 5.250250853 |
| ENSG00000183397 | C19orf71 | -3.2611602 | 0.222140997 | 3.756160483 | 0.095497186 | 0.495000284 | 2 | 2 | 4.058998049 | 2.270786764 | 2 | 6.192262702 | 5.379200993 |
| ENSG00000142065 | ZFP14 | 0.160028995 | 0.876399317 | 0.333876736 | 0.725625206 | 0.493905732 | 5.532353208 | 6.04685525 | 6.073378139 | 6.113598934 | 6.00043082 | 6.7543034 | 5.367404784 |
| ENSG00000072694 | FCGR2B | -3.753019347 | 0.514396922 | 4.246589535 | 0.417384315 | 0.493570189 | 2 | 2 | 2 | 2 | 2 | 4.854926465 | 4.147809227 |
| ENSG00000122884 | P4HA1 | 0.708014246 | 0.209654079 | -0.215031078 | 0.727736031 | 0.492983169 | 12.17515252 | 11.22065959 | 11.34128524 | 12.38036311 | 12.3222493 | 11.13046669 | 11.67715885 |
| ENSG00000138674 | SEC31A | 0.071369211 | 0.931247849 | 0.419753429 | 0.573211571 | 0.49112264 | 11.55650736 | 12.49586584 | 10.86096535 | 11.8525616 | 11.878197 | 12.26512815 | 12.15992538 |
| ENSG00000134851 | TMEM165 | 1.061274942 | 0.089945888 | -0.572890886 | 0.380616152 | 0.488384057 | 10.91047246 | 9.763446433 | 9.658943166 | 11.2913796 | 11.2838779 | 9.56938092 | 9.742104636 |
| ENSG00000143575 | HAX1 | -0.053316308 | 0.966718151 | 0.541674471 | 0.64649554 | 0.488358163 | 9.751047882 | 9.246456519 | 6.534109959 | 8.941551548 | 9.0020652 | 9.573749602 | 9.552621009 |
| ENSG00000170234 | PWWP2A | 0.203743946 | 0.883143303 | 0.282244147 | 0.832998352 | 0.485988093 | 5.330594134 | 7.503546079 | 8.763901704 | 7.971157485 | 7.97164382 | 8.292619498 | 7.754841567 |
| ENSG00000163738 | MTHFD2L | 1.26578566 | 0.190366567 | -0.784705442 | 0.432381295 | 0.481080218 | 5.70932728 | 5.457120953 | 3.369317972 | 6.268449691 | 6.40670796 | 4.675266036 | 4.292061816 |
| ENSG00000188986 | NELFB | 0.047152593 | 0.956792211 | 0.43391207 | 0.581663843 | 0.481064662 | 8.138771685 | 8.169643011 | 9.61300511 | 8.875861021 | 8.84312657 | 9.201728932 | 9.283950453 |
| ENSG00000178904 | DPY19L3 | -0.154814657 | 0.926810059 | 0.634270891 | 0.683005276 | 0.479456235 | 4.00722531 | 8.433041974 | 8.742726175 | 7.971157485 | 7.80253898 | 8.437839892 | 8.866637078 |
| ENSG00000131171 | SH3BGRL | 0.363167157 | 0.681613489 | 0.116176724 | 0.901737951 | 0.479343881 | 9.635285994 | 10.05576005 | 8.042706615 | 9.869705991 | 9.79168681 | 9.645817872 | 9.522355441 |
| ENSG00000166164 | BRD7 | 0.658072595 | 0.54335456 | -0.179318203 | 0.879564373 | 0.478754392 | 8.797441463 | 8.169643011 | 5.930865648 | 8.648320862 | 8.74905888 | 7.782541424 | 7.958255135 |
| ENSG00000143502 | SUSD4 | 2.146708035 | NA | -1.67157831 | NA | 0.475129725 | 2 | 2 | 2 | 2.868578001 | 3.28037825 | 2.048074405 | 2.172608095 |
| ENSG00000157600 | TMEM164 | 0.357308632 | 0.863599094 | 0.116466122 | 0.955852858 | 0.473774754 | 8.009475818 | 8.522788974 | 2 | 8.065351606 | 8.06917614 | 7.694662924 | 7.954317658 |
| ENSG00000135870 | RC3H1 | 0.668102033 | 0.533299715 | -0.194518901 | 0.866671393 | 0.473583132 | 9.435834703 | 8.029695761 | 6.718950348 | 9.163478821 | 9.08575246 | 8.213803048 | 8.325926932 |
| ENSG00000183648 | NDUFB1 | 0.630682073 | 0.522626682 | -0.158023864 | 0.883361661 | 0.472658208 | 8.39321512 | 6.602424674 | 6.43197674 | 8.058664058 | 8.04261444 | 7.093332266 | 7.441266415 |
| ENSG00000162337 | LRP5 | -0.992399322 | 0.479583688 | 1.464617861 | 0.268612714 | 0.472218538 | 10.31337612 | 6.464272742 | 7.97167868 | 8.109160159 | 8.06258142 | 10.27622952 | 10.738159 |
| ENSG00000050393 | MCUR1 | 0.025756811 | 0.970415686 | 0.446241172 | 0.471041116 | 0.471997983 | 7.33131396 | 7.819099893 | 8.585098385 | 8.130575473 | 7.91419618 | 8.427654172 | 8.454843905 |
| ENSG00000123219 | CENPK | 0.52621539 | 0.832844524 | -0.054631528 | 0.983038213 | 0.471583862 | 5.095959458 | 2 | 2 | 4.237680128 | 4.12711147 | 3.904814898 | 3.580696614 |
| ENSG00000159164 | SV2A | -0.198618799 | 0.859388318 | 0.66990954 | 0.502317168 | 0.471290741 | 7.031698319 | 9.235879652 | 9.601288346 | 8.754208387 | 8.8041253 | 9.475816969 | 9.79094187 |
| ENSG00000284770 | TBCE | -0.206672183 | 0.891546154 | 0.677390394 | 0.621584245 | 0.470718211 | 4.00722531 | 7.979858146 | 7.646393253 | 7.035434774 | 7.14562489 | 8.000548654 | 7.910280955 |
| ENSG00000139722 | VPS37B | 0.276163281 | 0.832844524 | 0.193652363 | 0.883160796 | 0.469815644 | 6.138360429 | 8.147241016 | 9.439621147 | 8.75144642 | 8.68895519 | 8.555288804 | 8.717808107 |
| ENSG00000110076 | NRXN2 | -2.494587274 | 0.24868502 | 2.963059295 | 0.142448826 | 0.468472022 | 2 | 5.457120953 | 2 | 2.947924294 | 2.38539787 | 6.106889539 | 7.287011591 |
| ENSG00000105556 | MIER2 | 0.494932959 | 0.538867025 | -0.027148661 | 0.97520757 | 0.467784298 | 9.410448395 | 7.761270542 | 8.143093325 | 9.128781483 | 9.08466797 | 8.676372393 | 8.496731673 |
| ENSG00000134371 | CDC73 | 0.515606155 | 0.618837592 | -0.048949378 | 0.966283503 | 0.466656778 | 9.886101849 | 8.029695761 | 7.691296561 | 9.407570435 | 9.37460909 | 8.933938877 | 8.719549018 |
| ENSG00000103197 | TSC2 | -0.004368852 | 0.997041173 | 0.470884716 | 0.587320792 | 0.466515864 | 7.82963488 | 9.916994052 | 9.272377397 | 9.219515356 | 9.25534053 | 9.6668989 | 9.752061939 |
| ENSG00000117616 | RSRP1 | 0.620521451 | 0.463760408 | -0.154387581 | 0.866824598 | 0.46613387 | 9.579430841 | 8.255926468 | 7.97167868 | 9.381011913 | 9.41399842 | 8.958123037 | 8.201061123 |
| ENSG00000129351 | ILF3 | -0.026050793 | 0.970543625 | 0.491765087 | 0.432815399 | 0.465714294 | 9.761125267 | 10.92721483 | 11.01019887 | 10.61487847 | 10.6676062 | 11.02725374 | 11.27827087 |
| ENSG00000101265 | RASSF2 | -1.099073218 | 0.313653871 | 1.563904084 | 0.120494972 | 0.464830866 | 4.467311984 | 4.442710129 | 5.39244166 | 4.329433326 | 3.39680241 | 6.537789343 | 5.867805474 |
| ENSG00000162695 | SLC30A7 | 0.957095799 | 0.033936427 | -0.492281157 | 0.302337918 | 0.464814642 | 8.696033109 | 9.235879652 | 8.325066185 | 9.773608662 | 9.74577157 | 8.287622092 | 8.355338318 |
| ENSG00000196866 | HIST1H2AD | 5.655350111 | 0.114206862 | -5.191125726 | 0.193537693 | 0.464224385 | 2 | 2 | 2 | 5.829220145 | 5.74985018 | 2 | 2 |
| ENSG00000197043 | ANXA6 | 0.595260866 | 0.562304924 | -0.131396613 | 0.90737308 | 0.463864253 | 12.07772327 | 9.506147172 | 10.89495479 | 11.74177109 | 11.8025467 | 10.84589444 | 11.22329526 |
| ENSG00000164366 | CCDC127 | -0.267947256 | 0.663217573 | 0.728343034 | 0.183892387 | 0.460395778 | 8.138771685 | 7.979858146 | 7.029880454 | 7.546342806 | 7.5240115 | 8.651788724 | 8.365751687 |
| ENSG00000177853 | ZNF518A | 0.548347441 | 0.779920665 | -0.08830035 | 0.966407561 | 0.460047091 | 7.159102577 | 7.954278496 | 2 | 7.536750104 | 7.62295033 | 7.433611594 | 6.251520369 |
| ENSG00000167670 | CHAF1A | 0.323744655 | 0.709501472 | 0.134427913 | 0.883002588 | 0.458172568 | 5.330594134 | 6.04685525 | 7.029880454 | 6.449761039 | 6.73998187 | 6.277913523 | 6.55171954 |
| ENSG00000157578 | LCA5L | 0.456777446 | 0.846875777 | -0.001124696 | 0.99945274 | 0.45565275 | 2 | 4.11352228 | 2 | 3.406930997 | 3.45167086 | 3.215043717 | 3.100202476 |
| ENSG00000090006 | LTBP4 | -0.961029126 | 0.16672399 | 1.414780339 | 0.031022179 | 0.453751213 | 10.1076228 | 8.702538795 | 9.577565424 | 8.573670512 | 8.65720817 | 10.51529534 | 11.33084866 |
| ENSG00000141027 | NCOR1 | 0.224337312 | 0.67689906 | 0.229122074 | 0.666121968 | 0.453459386 | 10.19941743 | 10.36815316 | 11.08782089 | 10.81365108 | 10.8423598 | 10.91840948 | 10.74187559 |
| ENSG00000122778 | KIAA1549 | 1.208045071 | 0.492183068 | -0.755393369 | 0.675750281 | 0.452651702 | 3.327711234 | 8.124485672 | 4.874587708 | 7.860161659 | 7.97632944 | 6.184218099 | 5.842465082 |
| ENSG00000122958 | VPS26A | 0.714145897 | 0.138731984 | -0.261790378 | 0.616400319 | 0.452355519 | 8.493334308 | 9.235879652 | 9.426401271 | 9.814540128 | 9.81606952 | 8.728933481 | 8.953526633 |
| ENSG00000133028 | SCO1 | 0.035348309 | 0.946040014 | 0.415821794 | 0.360288866 | 0.451170102 | 7.750913381 | 8.395519306 | 8.585098385 | 8.330724217 | 8.31278108 | 8.707577112 | 8.688479312 |
| ENSG00000174130 | TLR6 | 0.232326039 | 0.82001794 | 0.218165907 | 0.826511578 | 0.450491946 | 4.467311984 | 5.131603625 | 4.058998049 | 4.867343068 | 4.85329164 | 4.799832052 | 4.895193906 |
| ENSG00000161996 | WDR90 | 0.371630407 | 0.802275213 | 0.077282386 | 0.960122226 | 0.448912794 | 4.00722531 | 6.228547981 | 7.858115942 | 7.021746293 | 7.18258783 | 6.67615119 | 6.949978473 |
| ENSG00000197857 | ZNF44 | 0.396165672 | 0.531143362 | 0.052628058 | 0.939798462 | 0.44879373 | 6.96351253 | 6.535001748 | 5.772716771 | 6.841288255 | 6.93551934 | 6.613628927 | 6.501335855 |
| ENSG00000137601 | NEK1 | 0.446082949 | 0.821801475 | 0.001545863 | 0.999378208 | 0.447628812 | 2 | 7.701025996 | 8.042706615 | 7.857595438 | 7.62593544 | 7.399274054 | 7.218077721 |
| ENSG00000168872 | DDX19A | 0.924886113 | 0.114918885 | -0.477598324 | 0.438543047 | 0.447287789 | 8.836089686 | 8.169643011 | 7.552162999 | 9.263797825 | 9.12531607 | 7.688988646 | 7.921416787 |
| ENSG00000159958 | TNFRSF13C | -2.305383504 | 0.652705192 | 2.752312442 | NA | 0.446928938 | 2 | 2 | 2 | 2.141737246 | 2 | 3.429858444 | 3.45257121 |
| ENSG00000166135 | HIF1AN | -0.303943487 | 0.720184992 | 0.750860814 | 0.331246196 | 0.446917327 | 8.797441463 | 9.225224669 | 10.51254821 | 9.389919188 | 9.41227044 | 10.38463615 | 10.51542313 |
| ENSG00000122678 | POLM | -0.469530625 | 0.711000931 | 0.914884548 | 0.435748117 | 0.445353923 | 4.467311984 | 6.464272742 | 7.734844356 | 6.18902405 | 6.39974839 | 7.672789259 | 7.599641096 |
| ENSG00000058262 | SEC61A1 | 0.104146997 | 0.758250661 | 0.339996795 | 0.259132418 | 0.444143792 | 12.36881891 | 12.36244576 | 12.5724427 | 12.5306927 | 12.5526578 | 12.57786247 | 12.95272298 |
| ENSG00000008130 | NADK | 0.308607842 | 0.715896274 | 0.135087304 | 0.879564373 | 0.443695146 | 10.16187685 | 8.505280662 | 8.805342542 | 9.607141034 | 9.69905569 | 9.435720286 | 9.525344379 |
| ENSG00000145833 | DDX46 | 0.384107681 | 0.658522427 | 0.059490682 | 0.949302416 | 0.443598364 | 8.910412251 | 9.34816733 | 7.398165132 | 9.161399541 | 9.13163078 | 8.637620652 | 8.990045056 |
| ENSG00000113810 | SMC4 | 1.027342969 | 0.531143362 | -0.58387539 | 0.728705652 | 0.443467578 | 3.327711234 | 8.639735838 | 8.143093325 | 8.888486565 | 8.84312657 | 7.571576479 | 6.913939481 |

| | | | | | | | | | | | | | |
|---|---|---|---|---|---|---|---|---|---|---|---|---|---|
| ENSG00000108107 | RPL28 | 0.256014654 | 0.722734802 | 0.186905236 | 0.79812555 | 0.44291989 | 11.99474663 | 11.41601719 | 10.48730704 | 11.65889637 | 11.7025256 | 11.59074223 | 11.63265648 |
| ENSG00000167601 | AXL | -0.17954499 | 0.64928304 | 0.620514132 | 0.072784469 | 0.440969141 | 11.59081378 | 12.01658788 | 12.05247881 | 11.70101821 | 11.7431293 | 12.33714807 | 12.68506543 |
| ENSG00000035499 | DEPDC1B | -0.815681997 | 0.717844348 | 1.255985836 | 0.543323761 | 0.44030384 | 2 | 3.076665209 | 3.369317972 | 2.784612485 | 2.38539787 | 3.891700877 | 3.383972333 |
| ENSG00000185480 | PARPBP | 0.621515017 | 0.731956618 | -0.181546781 | 0.925393616 | 0.439968237 | 2 | 5.457120953 | 3.369317972 | 5.058234798 | 4.60967607 | 4.281782022 | 4.09155917 |
| ENSG00000119616 | FCF1 | 0.15164245 | 0.915100867 | 0.288319839 | 0.833878439 | 0.439962288 | 6.138360429 | 7.145017454 | 9.372276026 | 8.343618883 | 8.32385328 | 8.483352582 | 8.454146986 |
| ENSG00000171502 | COL24A1 | 4.098040732 | 0.015484621 | -3.658411256 | 0.050033331 | 0.439629476 | 2 | 4.11352228 | 2 | 6.490083648 | 6.48770488 | 2.048074405 | 2.225841198 |
| ENSG00000124214 | STAU1 | 0.21797436 | 0.66172885 | 0.221460203 | 0.651770088 | 0.439434563 | 10.87842776 | 10.019219 | 10.59756881 | 10.73648843 | 10.776052 | 10.67331779 | 10.8415673 |
| ENSG00000113282 | CLINT1 | 0.155826179 | 0.833643937 | 0.28232663 | 0.686013943 | 0.438152809 | 8.836089686 | 9.664384941 | 10.30429804 | 9.852410253 | 9.89922864 | 10.06575858 | 9.935263322 |
| ENSG00000135404 | CD63 | 0.851768641 | 0.126838044 | -0.414006577 | 0.482146792 | 0.437762064 | 13.5666075 | 12.46789278 | 12.83381799 | 13.87862076 | 13.8845443 | 12.38044183 | 12.81904935 |
| ENSG00000189376 | C8orf76 | 1.596256949 | 0.077375879 | -1.158858494 | 0.208960532 | 0.437398454 | 7.33131396 | 5.595927442 | 5.39244166 | 7.829061569 | 8.02236724 | 5.48093935 | 5.101530266 |
| ENSG00000099940 | SNAP29 | -0.071602366 | 0.897105982 | 0.505247226 | 0.279927595 | 0.43364486 | 8.797441463 | 9.125613817 | 8.266922072 | 8.719297823 | 8.69607402 | 9.142413344 | 9.405730946 |
| ENSG00000144895 | EIF2A | 0.417555453 | 0.463231524 | 0.015729636 | 0.980204682 | 0.433285089 | 9.410448395 | 8.356994601 | 9.31604076 | 9.491070865 | 9.5268964 | 9.096358634 | 9.123144518 |
| ENSG00000176986 | SEC24C | 0.564020013 | 0.108473389 | -0.131324007 | 0.738341738 | 0.432696006 | 10.00110988 | 9.549500232 | 10.02619908 | 10.39714397 | 10.468729 | 9.620951746 | 9.851208114 |
| ENSG00000147443 | DOK2 | -1.310474018 | NA | 1.74257277 | NA | 0.432098752 | 2 | 2 | 2 | 2.141737246 | 2.14022021 | 3.316666208 | 2.277179783 |
| ENSG00000167550 | RHEBL1 | 2.025768347 | 0.170091125 | -1.594720972 | 0.290456069 | 0.431047375 | 2 | 4.442710129 | 3.369317972 | 5.37770462 | 5.18011485 | 2.962551659 | 2.551896972 |
| ENSG00000136783 | NIPSNAP3A | 0.650220131 | 0.568853498 | -0.219976 | 0.856534247 | 0.430244132 | 6.009025997 | 7.315436877 | 8.865353883 | 8.503178519 | 8.46222608 | 7.435871964 | 7.79392309 |
| ENSG00000172831 | CES2 | 0.548635003 | 0.196836305 | -0.1197069 | 0.801853641 | 0.428928104 | 9.143983763 | 8.687092496 | 8.742726175 | 9.314267267 | 9.50922197 | 8.504505158 | 8.962865756 |
| ENSG00000125257 | ABCC4 | 1.278618505 | 0.065445182 | -0.849779024 | 0.23478853 | 0.428839481 | 7.940205782 | 8.590771778 | 6.882775709 | 9.192278409 | 9.28397021 | 7.263334844 | 7.010010317 |
| ENSG00000155636 | RBM45 | 0.536246302 | 0.798939121 | -0.108021726 | 0.961024044 | 0.428224576 | 2 | 7.431218269 | 4.523675391 | 6.490083648 | 6.66650236 | 5.963846589 | 5.976631893 |
| ENSG00000158716 | DUSP23 | -0.214008818 | 0.912931516 | 0.641126259 | 0.723919381 | 0.427117441 | 2 | 7.002537973 | 7.646393253 | 6.456560224 | 6.70653264 | 7.244115921 | 7.567781104 |
| ENSG00000144736 | SHQ1 | 0.055629245 | 0.918975348 | 0.371427376 | 0.433240072 | 0.427056621 | 7.031698319 | 7.315436877 | 7.098165388 | 7.230526165 | 7.19871665 | 7.296052918 | 7.719319651 |
| ENSG00000005812 | FBXL3 | 0.114730942 | 0.934796779 | 0.3121848 | 0.816103107 | 0.426915741 | 6.257048221 | 8.101365653 | 9.746650996 | 8.851563757 | 8.67892944 | 9.022330083 | 8.902329908 |
| ENSG00000068323 | TFE3 | -0.045164025 | 0.943520433 | 0.47177711 | 0.397331772 | 0.426613085 | 10.04301211 | 9.71095217 | 10.61515347 | 10.06476182 | 10.1833083 | 10.2814236 | 10.92805327 |
| ENSG00000204084 | INPP5B | 0.282747339 | 0.805210364 | 0.143413153 | 0.902554091 | 0.426160492 | 5.095959458 | 7.538394143 | 7.818184403 | 7.527093191 | 7.45854177 | 7.4096614 | 7.302581437 |
| ENSG00000104129 | DNAJC17 | 0.611591022 | 0.707094181 | -0.186533355 | 0.914029999 | 0.425057667 | 2 | 5.303523747 | 6.43197674 | 6.01571348 | 5.9725602 | 5.119635168 | 5.390901532 |
| ENSG00000067560 | RHOA | -0.531493973 | 0.413879953 | 0.955686714 | 0.111723144 | 0.424192741 | 13.24354095 | 12.11221489 | 12.12791662 | 12.06350133 | 12.0702787 | 13.36457616 | 13.72000673 |
| ENSG00000060982 | BCAT1 | 0.484623668 | 0.410126067 | -0.060656194 | 0.927796772 | 0.423967475 | 9.579430841 | 10.76748534 | 10.09465441 | 10.75004251 | 10.6795076 | 10.21634679 | 10.12520267 |
| ENSG00000082212 | ME2 | -0.005589821 | 0.997041173 | 0.429390169 | 0.710106162 | 0.423800348 | 6.257048221 | 7.538394143 | 9.083172337 | 8.093978021 | 8.02236724 | 8.641543003 | 8.315988368 |
| ENSG00000080298 | RFX3 | 0.430323887 | 0.85603014 | -0.006743294 | 0.998049427 | 0.423580593 | 2 | 7.315436877 | 3.369317972 | 6.283823264 | 6.26836889 | 5.979426183 | 5.740981724 |
| ENSG00000117682 | DHDDS | -0.169865697 | 0.82182886 | 0.592892691 | 0.372337042 | 0.423026994 | 7.940205782 | 9.256956408 | 9.330306409 | 8.762462704 | 8.83669913 | 9.427224679 | 9.676995613 |
| ENSG00000136478 | TEX2 | -0.796491874 | 0.258819968 | 1.218655036 | 0.066009623 | 0.422163162 | 8.19931886 | 9.192779271 | 9.859153765 | 8.411675819 | 8.47717992 | 10.48669015 | 10.40706314 |
| ENSG00000131653 | TRAF7 | 0.05815062 | 0.891376437 | 0.36319483 | 0.314431097 | 0.42134545 | 9.931427102 | 9.34816733 | 9.636156974 | 9.723752619 | 9.70118012 | 9.98393497 | 10.04786406 |
| ENSG00000089820 | ARHGAP4 | -0.105338751 | 0.959724553 | 0.526570637 | 0.793118059 | 0.421231886 | 3.327711234 | 5.72254323 | 2 | 4.387512679 | 4.4836996 | 5.463319095 | 4.215171317 |
| ENSG00000164442 | CITED2 | -0.740184813 | 0.145213802 | 1.157965778 | 0.015354606 | 0.417780966 | 12.16759578 | 11.29386853 | 11.51370025 | 10.97299388 | 10.9635316 | 12.59272163 | 13.09279326 |
| ENSG00000055070 | SZRD1 | 0.110964005 | 0.87974571 | 0.306461429 | 0.645978013 | 0.417425434 | 10.87379128 | 9.910357957 | 11.1930463 | 10.86406385 | 10.8617987 | 10.79364543 | 11.2814173 |
| ENSG00000100227 | POLDIP3 | -0.193262414 | 0.588054914 | 0.60926883 | 0.051318907 | 0.416006416 | 9.317904228 | 9.799806309 | 9.714382178 | 9.44052423 | 9.43116542 | 10.23609884 | 10.23354621 |
| ENSG00000151304 | SRFBP1 | 0.672006078 | 0.500544612 | -0.257276447 | 0.807781951 | 0.414729631 | 6.96351253 | 4.93638933 | 7.285536013 | 7.322721205 | 7.33993465 | 6.520853931 | 6.341697848 |
| ENSG00000158985 | CDC42SE2 | 0.16112983 | 0.890185466 | 0.253462113 | 0.819999326 | 0.414591942 | 6.257048221 | 8.955547846 | 7.934808822 | 8.291333947 | 8.21902896 | 8.382079673 | 8.310608339 |
| ENSG00000132664 | POLR3F | 0.226318052 | 0.908151255 | 0.18763624 | 0.923213606 | 0.413954293 | 2 | 7.35507212 | 7.502634626 | 7.144720234 | 7.01963893 | 6.956760977 | 7.130061054 |
| ENSG00000130939 | UBE4B | -0.446981021 | 0.665847367 | 0.860753237 | 0.36864007 | 0.413772217 | 8.340423424 | 9.672251278 | 7.225742779 | 8.235106869 | 8.38327653 | 9.600191431 | 9.615680827 |
| ENSG00000169188 | APEX2 | -0.417885488 | 0.778992908 | 0.826262323 | 0.551011557 | 0.408376835 | 8.257427002 | 6.951736442 | 4.058998049 | 6.809774134 | 6.81514011 | 8.001311979 | 8.051511707 |
| ENSG00000005075 | POLR2J | 0.831128975 | 0.162897673 | -0.423333156 | 0.497326437 | 0.407795819 | 8.540903468 | 8.101365653 | 7.225742779 | 8.811020771 | 8.9398551 | 7.535143176 | 7.736616914 |
| ENSG00000108433 | GOSR2 | 0.213126221 | 0.858165041 | 0.194536156 | 0.8682894 | 0.407662377 | 9.397585735 | 10.19331737 | 7.285536013 | 9.59332559 | 9.59930725 | 9.557233497 | 9.59812349 |
| ENSG00000172382 | PRSS27 | -0.114106222 | 0.964071163 | 0.520997366 | 0.832967479 | 0.406891144 | 2 | 5.131603625 | 2 | 4.034562184 | 3.55551834 | 4.698044417 | 3.735548302 |
| ENSG00000164244 | PRRC1 | -0.016544213 | 0.984277478 | 0.423070971 | 0.586489016 | 0.406526758 | 8.517314938 | 10.26269067 | 9.945092785 | 9.740563462 | 9.72712735 | 10.10286127 | 10.23739929 |
| ENSG00000285077 | ARHGAP11B | 5.596258368 | 0.124401402 | -5.191124779 | 0.199257862 | 0.40513359 | 2 | 2 | 2 | 5.742861563 | 5.72778967 | 2 | 2 |
| ENSG00000240849 | TMEM189 | 0.080489471 | 0.891774766 | 0.324585425 | 0.536695429 | 0.405074896 | 9.867565369 | 9.148351514 | 10.13584611 | 9.823772619 | 9.87684923 | 10.00976979 | 10.17299833 |
| ENSG00000180190 | TDRP | 0.17629205 | 0.950011373 | 0.228437564 | 0.936599654 | 0.404729614 | 2 | 7.189546432 | 2 | 6.069927018 | 5.6121243 | 6.060766632 | 5.736403255 |
| ENSG00000119041 | GTF3C3 | 0.214441688 | 0.711059078 | 0.189559746 | 0.740394508 | 0.404001435 | 8.257427002 | 8.590771778 | 7.646393253 | 8.321442665 | 8.54182783 | 8.57541967 | 8.225770986 |
| ENSG00000128654 | MTX2 | 0.497158934 | 0.765943216 | -0.093678129 | 0.957844617 | 0.403480805 | 7.82963488 | 8.505280662 | 3.369317972 | 8.054188409 | 8.22100445 | 7.726402307 | 7.370638252 |
| ENSG00000115459 | ELMOD3 | 0.491898806 | 0.818202168 | -0.088960767 | 0.968233365 | 0.402938038 | 2 | 6.535001748 | 8.561105505 | 7.783822874 | 7.78395061 | 7.527744791 | 6.81752904 |
| ENSG00000115042 | FAHD2A | 0.104866349 | 0.957853023 | 0.298003595 | 0.880432293 | 0.402869943 | 2 | 7.189546432 | 7.97167868 | 7.178012736 | 7.14977896 | 7.27600668 | 7.424286753 |
| ENSG00000141258 | SGSM2 | -0.231899655 | 0.784771092 | 0.634643574 | 0.408028698 | 0.402743919 | 9.602033055 | 7.847167909 | 8.65477198 | 8.697938662 | 8.58694992 | 9.389237988 | 9.609434849 |
| ENSG00000095261 | PSMD5 | 0.052562094 | 0.969963817 | 0.350106186 | 0.792798504 | 0.40266828 | 6.138360429 | 9.308339287 | 9.735974718 | 9.019382707 | 9.12320501 | 9.412953205 | 9.321117366 |
| ENSG00000159055 | MIS18A | 0.281576951 | 0.829527562 | 0.120894106 | 0.927960061 | 0.402471057 | 3.327711234 | 6.228547981 | 5.156626401 | 5.662809983 | 5.5755357 | 5.524070859 | 5.419744333 |
| ENSG00000070501 | POLB | 0.131048924 | 0.945043724 | 0.271410444 | 0.886197127 | 0.402459369 | 2 | 7.145017454 | 5.59507254 | 6.104969671 | 6.19790195 | 6.412547789 | 6.145001162 |
| ENSG00000155016 | CYP2U1 | -1.948421925 | 0.21164903 | 2.350350764 | 0.117096798 | 0.401928839 | 3.327711234 | 6.844429755 | 8.699419242 | 5.578054239 | 5.6826301 | 9.896164504 | 9.729200569 |
| ENSG00000134001 | EIF2S1 | 0.036417628 | 0.97026695 | 0.364506672 | 0.689674098 | 0.4009243 | 10.41898845 | 10.00059544 | 8.266922072 | 9.854342273 | 9.84901661 | 10.1988183 | 10.15788386 |
| ENSG00000170949 | ZNF160 | 0.046840041 | 0.980534836 | 0.354070062 | 0.84516091 | 0.400910103 | 4.00722531 | 8.57407404 | 9.839340503 | 8.7747563 | 8.86350401 | 9.343316829 | 8.866637078 |
| ENSG00000090686 | USP48 | 0.139316038 | 0.785196727 | 0.260691772 | 0.585262994 | 0.40000781 | 9.27634997 | 8.903033433 | 8.408111533 | 9.014783325 | 9.07377798 | 9.207032339 | 9.121828633 |
| ENSG00000140105 | WARS | 1.735181766 | 6.81E-20 | -1.335924481 | 2.85E-12 | 0.399257285 | 11.98187803 | 11.89916567 | 12.03298648 | 13.68671956 | 13.7250429 | 10.72066788 | 10.55040485 |
| ENSG00000180667 | YOD1 | 0.752673724 | 0.528794428 | -0.35453126 | 0.776319002 | 0.398142464 | 5.095959458 | 7.099070191 | 8.325066185 | 8.098332103 | 8.08661608 | 7.033394818 | 6.985139097 |
| ENSG00000105856 | HBP1 | 0.893583194 | 0.095791495 | -0.496250216 | 0.374026188 | 0.397332978 | 10.3868074 | 9.246456519 | 9.703463497 | 10.73613921 | 10.7523441 | 9.380463355 | 9.338970358 |
| ENSG00000173334 | TRIB1 | 1.233596475 | 0.437642291 | -0.837219839 | 0.603961617 | 0.396376636 | 3.327711234 | 7.669933291 | 8.511888463 | 8.811020771 | 8.8041253 | 7.106162485 | 6.350712758 |
| ENSG00000122741 | DCAF10 | -0.372568876 | 0.499134136 | 0.767791807 | 0.12649658 | 0.395222932 | 9.544849385 | 9.876711741 | 9.25752426 | 9.246247604 | 9.18117744 | 10.69848203 | 9.887021781 |
| ENSG00000144566 | RAB5A | 0.314636547 | 0.761583294 | 0.079166474 | 0.942375924 | 0.393803021 | 10.25038534 | 10.19877383 | 7.896971941 | 10.1262855 | 10.0557402 | 9.870700911 | 9.842979447 |
| ENSG00000109929 | SC5D | 0.507420964 | 0.630303294 | -0.115193585 | 0.919715042 | 0.392227378 | 7.27616306 | 9.506147172 | 9.926432745 | 9.83881318 | 9.7131598 | 9.103841116 | 9.210746501 |
| ENSG00000168228 | ZCCHC4 | 0.168438463 | 0.909988721 | 0.221517673 | 0.878629811 | 0.389956136 | 3.327711234 | 6.535001748 | 6.958202216 | 6.456560224 | 6.35006802 | 6.712030998 | 6.141542327 |
| ENSG00000129636 | ITFG1 | 1.08655283 | 0.000528881 | -0.696690155 | 0.033633667 | 0.389862675 | 9.87686338 | 9.506147172 | 9.330306409 | 10.6527458 | 10.689886 | 8.918261679 | 8.874992666 |
| ENSG00000103043 | VAC14 | 0.870900054 | 0.190847995 | -0.481365673 | 0.486954891 | 0.389534381 | 9.751047882 | 8.395519306 | 8.608698767 | 9.870978946 | 9.95192082 | 8.513629334 | 8.622397655 |
| ENSG00000146243 | IRAK1BP1 | -0.460686808 | 0.598708746 | 0.846798209 | 0.285556212 | 0.386111401 | 4.815609471 | 5.303523747 | 4.523675391 | 4.497095146 | 4.60967607 | 5.54934408 | 5.85096143 |
| ENSG00000108669 | CYTH1 | 1.157143121 | 0.577257973 | -0.774706838 | 0.714892595 | 0.382436284 | 9.344956677 | 7.274681896 | 2 | 9.170732795 | 9.26781654 | 7.506395059 | 7.094725293 |
| ENSG00000147162 | OGT | 0.250308661 | 0.842102843 | 0.131638517 | 0.91798278 | 0.381947177 | 8.009475818 | 11.09461697 | 10.87075861 | 10.7217479 | 10.7601769 | 10.70903366 | 10.53087342 |
| ENSG00000165568 | AKR1E2 | -3.753012982 | 0.522811731 | 4.134909296 | 0.440692188 | 0.381896314 | 2 | 2 | 2 | 2 | 2 | 4.230621763 | 4.644600858 |
| ENSG00000108312 | UBTF | -0.019561082 | 0.958728989 | 0.401453196 | 0.209143226 | 0.381892114 | 8.910412251 | 9.338312924 | 9.212024683 | 9.09649754 | 9.19932737 | 9.531835181 | 9.601903806 |
| ENSG00000169884 | WNT10B | 0.323788626 | 0.912780559 | 0.057816098 | 0.984296564 | 0.381604724 | 2 | 3.686303414 | 2 | 3.406930997 | 2.49394783 | 3.082820997 | 2.671842496 |
| ENSG00000090674 | MCOLN1 | 0.869931401 | 0.462190956 | -0.488709593 | 0.688288292 | 0.381221808 | 5.866944937 | 7.431218269 | 9.014156722 | 8.811020771 | 8.8250571 | 7.426809156 | 7.529919928 |
| ENSG00000173269 | MMRN2 | -0.795478245 | 0.543644578 | 1.176369084 | 0.329610478 | 0.38089084 | 3.327711234 | 4.442710129 | 4.058998049 | 3.406930997 | 3.55551834 | 5.381251181 | 4.500832793 |
| ENSG00000113583 | C5orf15 | 0.760159909 | 0.044804086 | -0.379432904 | 0.345156474 | 0.380727005 | 10.32020829 | 9.821190352 | 10.28245311 | 10.97151121 | 10.8541818 | 9.563449215 | 9.964920949 |
| ENSG00000170500 | LONRF2 | -0.550063596 | 0.847129001 | 0.928944661 | 0.729041892 | 0.378881066 | 2 | 2 | 5.39244166 | 4.17310023 | 2.77779907 | 4.133696865 | 5.230851202 |
| ENSG00000166192 | SENP8 | -0.319481101 | 0.875544825 | 0.698032268 | 0.710952496 | 0.378551167 | 3.327711234 | 5.946638857 | 2 | 4.387512679 | 4.4836996 | 5.29920634 | 5.32534152 |
| ENSG00000221887 | HMSD | -2.305309923 | 0.657619829 | 2.682918479 | NA | 0.377608557 | 2 | 2 | 2 | 2.141737246 | 2 | 3.215043717 | 3.560117069 |
| ENSG00000106052 | TAX1BP1 | 0.542143813 | 0.242184305 | -0.167632904 | 0.739634542 | 0.374510909 | 9.931427102 | 10.82153847 | 10.55568472 | 11.0564817 | 10.9924432 | 10.30294393 | 10.33052636 |
| ENSG00000166986 | MARS | -0.176045435 | 0.830362923 | 0.550451877 | 0.453090834 | 0.374406442 | 10.74769552 | 10.44362967 | 9.099922433 | 10.06309216 | 10.0934307 | 10.93288529 | 10.65962843 |
| ENSG00000177191 | B3GNT8 | 0.748484633 | 0.679837753 | -0.374229653 | 0.841697129 | 0.374254981 | 3.327711234 | 5.595927442 | 2 | 5.040242897 | 5.09650239 | 4.281782022 | 3.972018587 |
| ENSG00000175215 | CTDSP2 | 0.157315925 | 0.714313079 | 0.216483673 | 0.600319397 | 0.373799599 | 10.6855593 | 11.35134391 | 11.14139273 | 11.20443196 | 11.2817502 | 11.23321719 | 11.36896909 |
| ENSG00000178381 | ZFAND2A | 1.057111836 | 0.405184227 | -0.683463692 | 0.59841801 | 0.373648145 | 8.228665467 | 6.951736442 | 4.523675391 | 8.190940526 | 8.31648126 | 6.556607032 | 6.559516442 |
| ENSG00000002549 | LAP3 | 1.68013421 | 0.089428115 | -1.307618354 | 0.189945688 | 0.372515856 | 10.57565445 | 9.318399851 | 7.818184403 | 11.30956651 | 11.3163477 | 8.038972319 | 8.590407604 |
| ENSG00000158050 | DUSP2 | 0.672976564 | 0.692285797 | -0.302025746 | 0.864452193 | 0.370950818 | 3.327711234 | 5.131603625 | 2 | 4.71689672 | 4.56059747 | 3.838019203 | 3.908316285 |
| ENSG00000104859 | CLASRP | 0.111502162 | 0.945987763 | 0.258247961 | 0.874266884 | 0.369750123 | 7.867441304 | 7.874700259 | 3.369317972 | 7.370304568 | 7.48837134 | 7.514972902 | 7.630812678 |
| ENSG00000103942 | HOMER2 | -1.135905223 | 0.554188019 | 1.505294743 | 0.402484163 | 0.36938952 | 4.467311984 | 3.686303414 | 2 | 3.406930997 | 2.49394783 | 5.385935071 | 4.133950982 |
| ENSG00000115652 | UXS1 | 0.776340888 | 0.300991733 | -0.410023098 | 0.59982426 | 0.36631779 | 7.940205782 | 7.954278496 | 9.301632639 | 9.347568393 | 9.28302499 | 7.854430723 | 8.379764957 |
| ENSG00000089876 | DHX32 | 0.643468041 | 0.262230813 | -0.27731805 | 0.650030799 | 0.366149991 | 9.143983763 | 8.820493869 | 7.934808822 | 9.392580682 | 9.32220073 | 8.351275892 | 8.532824248 |
| ENSG00000183696 | UPP1 | 1.266044689 | 0.379217697 | -0.902218522 | 0.536069347 | 0.363826167 | 8.653385977 | 8.356994601 | 4.058998049 | 9.12771691 | 9.29713876 | 7.081831041 | 7.078537388 |
| ENSG00000171855 | IFNB1 | 5.553656509 | 0.131910476 | -5.191124081 | 0.203845577 | 0.362532428 | 2 | 2 | 2 | 5.720439317 | 5.67111583 | 2 | 2 |
| ENSG00000146592 | CREB5 | -0.692286799 | 0.706018071 | 1.05452867 | 0.544666992 | 0.362241871 | 2 | 7.431218269 | 7.502634626 | 6.351046291 | 6.11534392 | 7.634923938 | 8.175075046 |
| ENSG00000149761 | NUDT22 | 0.02636921 | 0.969963817 | 0.33551982 | 0.594804601 | 0.361889029 | 7.790810999 | 8.255926468 | 7.098165388 | 7.748245381 | 7.90193448 | 7.898530056 | 8.332009115 |
| ENSG00000092330 | TINF2 | -0.083206249 | 0.919060361 | 0.443207042 | 0.54430284 | 0.360000793 | 9.886101849 | 9.030900487 | 8.325066185 | 9.099758639 | 9.17507637 | 9.452841924 | 9.843245622 |
| ENSG00000168395 | ING5 | -0.114428817 | 0.89152261 | 0.473846762 | 0.522984627 | 0.359417945 | 8.009475818 | 7.928237109 | 6.534109959 | 7.440576128 | 7.60187978 | 8.327124877 | 7.83304634 |
| ENSG00000186020 | ZNF529 | -0.20706616 | 0.7690856 | 0.566385978 | 0.372265366 | 0.359319819 | 6.257048221 | 7.051611315 | 7.285536013 | 6.887305128 | 6.55189318 | 7.724554489 | 7.184857568 |
| ENSG00000018236 | CNTN1 | -2.305113648 | 0.671272366 | 2.664419918 | NA | 0.35930627 | 2 | 2 | 2 | 2.141737246 | 2 | 3.878466556 | 2.632958241 |
| ENSG00000168096 | ANKS3 | -0.223217395 | 0.89133285 | 0.581224057 | 0.698780079 | 0.358006662 | 7.159102577 | 7.874700259 | 3.369317972 | 6.79375511 | 6.82036165 | 7.652523812 | 7.523290521 |
| ENSG00000116016 | EPAS1 | -0.150314401 | 0.816128663 | 0.508155904 | 0.374026188 | 0.357841503 | 11.30078111 | 11.24195824 | 12.26789926 | 11.51181191 | 11.5558697 | 12.09526031 | 12.28222225 |
| ENSG00000198142 | SOWAHC | -0.032330226 | 0.988321837 | 0.389880862 | 0.853341364 | 0.357550636 | 3.327711234 | 5.457120953 | 9.301632639 | 7.867833045 | 7.72948195 | 8.307508685 | 8.118139752 |
| ENSG00000128606 | LRRC17 | 0.144974069 | 0.954132822 | 0.21106439 | 0.933991771 | 0.356038459 | 2 | 4.93638933 | 8.763901704 | 7.474468681 | 7.37938471 | 6.231826305 | 8.154630332 |
| ENSG00000125124 | BBS2 | 0.577133677 | 0.442359404 | -0.221605734 | 0.78167297 | 0.355527944 | 8.444143041 | 7.847167909 | 9.439621147 | 9.302050417 | 9.29056951 | 8.645454719 | 8.347853908 |
| ENSG00000182107 | TMEM30B | -4.714951099 | 0.096084829 | 5.069828543 | 0.017532061 | 0.354877444 | 2 | 2 | 3.369317972 | 2 | 2 | 6.315225883 | 5.945233978 |

| | | | | | | | | | | | | | |
|---|---|---|---|---|---|---|---|---|---|---|---|---|---|
| ENSG00000171574 | ZNF584 | -0.368117364 | 0.829692566 | 0.721958874 | 0.650570904 | 0.35384151 | 4.467311984 | 5.131603625 | 8.296287027 | 6.496695859 | 6.67805337 | 7.522436904 | 7.763868281 |
| ENSG00000214063 | TSPAN4 | 0.665183565 | 0.367097793 | -0.313957455 | 0.686181755 | 0.35122611 | 10.27168932 | 8.655694736 | 9.990713521 | 10.4041951 | 10.4959054 | 9.275521747 | 9.651383746 |
| ENSG00000253305 | PCDHGB6 | 2.673746455 | 2.54E-11 | -2.322627031 | 1.33E-08 | 0.351119424 | 7.975256505 | 7.189546432 | 7.342949392 | 10.1568736 | 10.2084185 | 5.4091291 | 5.250250853 |
| ENSG00000102302 | FGD1 | -0.270392058 | 0.845986004 | 0.621020103 | 0.629551284 | 0.350628046 | 8.609439662 | 8.702538795 | 5.156626401 | 7.867833045 | 7.8745862 | 8.802599622 | 8.697929554 |
| ENSG00000113140 | SPARC | -0.065144413 | 0.945987763 | 0.414536766 | 0.640863647 | 0.349392353 | 17.20663094 | 15.68411684 | 15.37962849 | 16.25522284 | 16.2674118 | 16.88548476 | 16.58038783 |
| ENSG00000105289 | TJP3 | -2.312716832 | 0.660533364 | 2.660540267 | NA | 0.347823435 | 2 | 2 | 2 | 2 | 2.14022021 | 3.679182183 | 3.012034821 |
| ENSG00000158882 | TOMM40L | 0.589226498 | 0.391687888 | -0.242092853 | 0.738292369 | 0.347133645 | 7.485143553 | 7.669933291 | 6.203069917 | 7.862723323 | 7.81305436 | 7.078941352 | 6.97156716 |
| ENSG00000105085 | MED26 | 0.112380464 | 0.956619854 | 0.234703581 | 0.909725832 | 0.347084045 | 2 | 5.72254323 | 7.818184403 | 6.693715713 | 6.6253296 | 6.797131715 | 6.757786576 |
| ENSG00000173473 | SMARCC1 | 0.123426302 | 0.937032541 | 0.22223712 | 0.886082448 | 0.345663421 | 6.009025997 | 10.19877383 | 10.19940021 | 9.746817367 | 9.80557618 | 9.744173094 | 9.99470022 |
| ENSG00000198074 | AKR1B10 | 2.407764739 | 0.033569383 | -2.062283764 | 0.080940119 | 0.345480975 | 3.327711234 | 3.076665209 | 4.523675391 | 5.639097847 | 5.84514933 | 2.68698273 | 2.592996859 |
| ENSG00000044090 | CUL7 | -0.40056868 | 0.387701261 | 0.745963881 | 0.079886981 | 0.345395201 | 8.816894989 | 9.277729677 | 9.330306409 | 8.734762645 | 8.790887 | 10.15695659 | 9.592751029 |
| ENSG00000139890 | REM2 | 0.35164789 | 0.892338755 | -0.00634432 | 0.99833637 | 0.34530357 | 2 | 4.710564154 | 2 | 4.17310023 | 3.33976443 | 3.904814898 | 3.128433649 |
| ENSG00000109066 | TMEM104 | -0.470113089 | 0.841054398 | 0.814877255 | 0.712935639 | 0.344764166 | 2 | 8.376385542 | 10.15199903 | 8.483340231 | 8.46556258 | 9.678637907 | 9.817737166 |
| ENSG00000064933 | PMS1 | -0.331265394 | 0.554864335 | 0.675575647 | 0.184184495 | 0.344310254 | 6.96351253 | 6.389896633 | 6.43197674 | 6.351046291 | 6.23747437 | 7.405054033 | 7.108963622 |
| ENSG00000136238 | RAC1 | 0.052881929 | 0.908151255 | 0.291207553 | 0.463927375 | 0.344089482 | 11.6325805 | 11.0102427 | 11.1169142 | 11.32636749 | 11.3369882 | 11.55492576 | 11.58427143 |
| ENSG00000220201 | ZGLP1 | 1.112468172 | 0.610039424 | -0.768776294 | 0.728258838 | 0.343691879 | 2 | 4.710564154 | 2 | 4.358765249 | 4.45712836 | 3.316666208 | 2.885414003 |
| ENSG00000078246 | TULP3 | -0.127027878 | 0.906452235 | 0.469862921 | 0.629414332 | 0.342835043 | 7.031698319 | 8.732943944 | 9.465702942 | 8.551612617 | 8.63070345 | 9.157900392 | 9.208682031 |
| ENSG00000008282 | SYPL1 | 0.541407413 | 0.47627622 | -0.199393519 | 0.804937917 | 0.342013894 | 8.444143041 | 9.733683991 | 10.16000817 | 10.15948039 | 10.1380242 | 9.493286157 | 9.324553169 |
| ENSG00000086570 | FAT2 | -0.873554179 | 0.719674266 | 1.214937869 | 0.5943601 | 0.34138369 | 2 | 4.11352228 | 2 | 2.868578001 | 2.59489879 | 3.411599105 | 4.329022995 |
| ENSG00000108395 | TRIM37 | -0.567127415 | 0.61090653 | 0.908329073 | 0.385615654 | 0.341201658 | 7.33131396 | 8.806266215 | 10.16797309 | 8.616799622 | 8.64695856 | 10.00197246 | 10.18816565 |
| ENSG00000164104 | HMGB2 | -0.031902352 | 0.961244994 | 0.372654252 | 0.527923969 | 0.3407519 | 7.096806373 | 6.666836837 | 6.203069917 | 6.739036112 | 6.60126376 | 7.244115921 | 6.853913776 |
| ENSG00000184381 | PLA2G6 | -0.358225706 | 0.738128035 | 0.698703989 | 0.480246049 | 0.340478282 | 6.138360429 | 4.11352228 | 5.156626401 | 5.076005082 | 4.95174433 | 5.812021848 | 6.124122661 |
| ENSG00000134871 | COL4A2 | -0.950016152 | 0.009719754 | 1.288787746 | 0.000263742 | 0.338771594 | 14.59904125 | 14.8709666 | 14.86603163 | 13.80219439 | 13.8658061 | 16.3592535 | 15.7151211 |
| ENSG00000212916 | MAP10 | 0.496611675 | 0.844549702 | -0.161624375 | 0.950047826 | 0.3349873 | 2 | 5.838937808 | 2 | 4.804776789 | 4.98931816 | 4.271694079 | 4.4231929 |
| ENSG00000175697 | GPR156 | -0.599159954 | 0.807354269 | 0.934047778 | 0.686224112 | 0.334887824 | 2 | 2 | 4.874587708 | 3.09461492 | 3.33976443 | 4.378961816 | 4.175132639 |
| ENSG00000198818 | SFT2D1 | 1.190548665 | 0.277947155 | -0.856570415 | 0.441358531 | 0.33397825 | 8.285626337 | 8.376385542 | 5.39244166 | 9.027396512 | 9.01008467 | 7.138441638 | 6.860240429 |
| ENSG00000244274 | DBNDD2 | 0.537577342 | 0.78255046 | -0.20454159 | 0.919589888 | 0.333035751 | 7.096806373 | 4.710564154 | 2 | 6.415275989 | 6.17362678 | 5.747972572 | 5.448021784 |
| ENSG00000134202 | GSTM3 | 0.73469941 | 0.237601833 | -0.401963468 | 0.535268829 | 0.332735942 | 9.590776211 | 8.297207175 | 8.941672936 | 9.75855684 | 9.77093593 | 8.419110791 | 8.824656761 |
| ENSG00000101161 | PRPF6 | 0.030666537 | 0.978813376 | 0.298991975 | 0.78199021 | 0.329658513 | 8.75772933 | 10.32888058 | 7.896971941 | 9.354865403 | 9.41227044 | 9.531835181 | 9.760542411 |
| ENSG00000101544 | ADNP2 | 0.093728418 | 0.957587914 | 0.235562774 | 0.893536663 | 0.329291192 | 4.00722531 | 8.848535039 | 9.301632639 | 8.693628619 | 8.54656162 | 8.540135471 | 8.955005256 |
| ENSG00000127334 | DYRK2 | -0.651156 | 0.47726999 | 0.979451989 | 0.255860639 | 0.328295989 | 9.689059027 | 7.60566235 | 8.325066185 | 8.153770873 | 8.16053033 | 9.838172685 | 9.706848979 |
| ENSG00000162517 | PEF1 | -0.376728753 | 0.761228536 | 0.70446934 | 0.544666992 | 0.327740588 | 10.19198706 | 9.656475478 | 6.958202216 | 9.055659246 | 9.10085035 | 9.936433011 | 10.34300322 |
| ENSG00000197763 | TXNRD3 | -0.455317158 | 0.876694325 | 0.781094657 | 0.778128936 | 0.325777499 | 2 | 7.538394143 | 2 | 5.753943344 | 5.43283104 | 6.649130473 | 6.879055607 |
| ENSG00000183826 | BTBD9 | -0.264336653 | 0.79719998 | 0.587326095 | 0.534923556 | 0.322989442 | 8.777722034 | 6.464272742 | 8.266922072 | 7.839502934 | 7.86955761 | 8.512558892 | 8.857178832 |
| ENSG00000100422 | CERK | -0.980822534 | 0.159032929 | 1.302578883 | 0.049414607 | 0.321756349 | 8.107517749 | 9.386925327 | 9.735974718 | 8.192977693 | 8.31648126 | 10.64282315 | 10.39531195 |
| ENSG00000185761 | ADAMTSL5 | -1.163383471 | 0.36509383 | 1.485029319 | 0.226994232 | 0.321645848 | 4.815609471 | 7.002537973 | 8.536706852 | 6.283823264 | 6.37866593 | 8.707577112 | 9.103279088 |
| ENSG00000147400 | CETN2 | 0.191259023 | 0.681917972 | 0.129658052 | 0.783874152 | 0.320917075 | 8.39321512 | 8.004992147 | 7.896971941 | 8.327018759 | 8.27524772 | 8.037484631 | 8.418864688 |
| ENSG00000164077 | MON1A | -0.391601305 | 0.798773623 | 0.710683882 | 0.621538139 | 0.319082577 | 3.327711234 | 6.78762695 | 7.600047325 | 6.436065776 | 6.22964624 | 7.473766117 | 7.322574236 |
| ENSG00000125046 | SSUH2 | -2.26658333 | 0.496719668 | 2.583924385 | 0.378260987 | 0.317341055 | 2 | 3.076665209 | 2 | 2.270786764 | 2 | 3.724264246 | 4.215171317 |
| ENSG00000048342 | CC2D2A | -0.526428713 | 0.434235922 | 0.84254902 | 0.17668291 | 0.316120307 | 6.257048221 | 7.35507212 | 7.451345288 | 6.61689148 | 6.60126376 | 7.669911566 | 8.169990965 |
| ENSG00000124164 | VAPB | -0.450168724 | 0.386935633 | 0.765766047 | 0.112309718 | 0.315597323 | 9.913467515 | 8.94259736 | 9.31604076 | 8.956018563 | 9.03612482 | 10.13382191 | 10.2749799 |
| ENSG00000148339 | SLC25A25 | 0.105362775 | 0.960095276 | 0.210213871 | 0.922180661 | 0.315576646 | 3.327711234 | 6.389896633 | 9.670202709 | 8.31397419 | 8.37442884 | 8.487716929 | 8.407389648 |
| ENSG00000173890 | GPR160 | -0.949099384 | 0.739704558 | 1.264018438 | 0.631802652 | 0.314919054 | 3.327711234 | 2 | 2 | 2.3892346 | 2.2680111 | 3.316666208 | 2.918126211 |
| ENSG00000127463 | EMC1 | 0.135579942 | 0.748334322 | 0.17742303 | 0.666121968 | 0.313002972 | 10.48735708 | 10.09725346 | 9.954333071 | 10.30502636 | 10.3590869 | 10.20578905 | 10.52474633 |
| ENSG00000115840 | SLC25A12 | 0.58092257 | 0.704833394 | -0.268690395 | 0.865952727 | 0.312232175 | 4.467311984 | 6.602424674 | 8.677268317 | 8.04293816 | 8.01099479 | 7.027411703 | 7.34530375 |
| ENSG00000224420 | ADM5 | 0.442104509 | 0.870799344 | -0.130902488 | 0.962322454 | 0.311202021 | 2 | 3.686303414 | 2 | 3.162721474 | 3.01487888 | 3.035909414 | 2.551896972 |
| ENSG00000101811 | CSTF2 | 0.444066547 | 0.841155419 | -0.133016263 | 0.953164556 | 0.311050284 | 5.095959458 | 8.124485672 | 2 | 7.178012736 | 7.14977896 | 6.448814696 | 6.748721609 |
| ENSG00000165264 | NDUFB6 | 0.20267941 | 0.663582024 | 0.108311867 | 0.82058396 | 0.310991278 | 7.790810999 | 7.431218269 | 7.451345288 | 7.751013478 | 7.77322015 | 7.485815918 | 7.83304634 |
| ENSG00000169740 | ZNF32 | 0.167444393 | 0.925360182 | 0.143165409 | 0.935870665 | 0.310609802 | 8.653385977 | 7.847167909 | 3.369317972 | 7.890605449 | 7.92877397 | 7.752026287 | 8.008487789 |
| ENSG00000143786 | CNIH3 | -3.753002793 | 0.531143362 | 4.063478256 | 0.463438411 | 0.310475463 | 2 | 2 | 2 | 2 | 2 | 4.894909044 | 3.620995423 |
| ENSG00000166394 | CYB5R2 | -0.788680511 | 0.679907921 | 1.098833864 | 0.546009747 | 0.310153353 | 4.00722531 | 7.928237109 | 3.369317972 | 5.753943344 | 5.77157844 | 7.54878318 | 7.592058619 |
| ENSG00000143850 | PLEKHA6 | -2.472504455 | 0.206338371 | 2.779670271 | 0.12649658 | 0.307165816 | 2 | 5.303523747 | 2 | 2.695456494 | 2.59489879 | 6.623578417 | 6.350712758 |
| ENSG00000150768 | DLAT | 0.342651512 | 0.696339473 | -0.035963831 | 0.969519452 | 0.306687681 | 6.96351253 | 8.557180776 | 8.941672936 | 8.696503411 | 8.72420319 | 8.275052313 | 8.394370181 |
| ENSG00000136699 | SMPD4 | 0.57626911 | 0.474064658 | -0.27033677 | 0.747340082 | 0.30593234 | 8.19931886 | 8.671479029 | 9.907528187 | 9.660380652 | 9.70753469 | 8.738579005 | 8.940648015 |
| ENSG00000125447 | GGA3 | -0.049967095 | 0.978749512 | 0.355575727 | 0.841817241 | 0.305608632 | 9.473095744 | 7.431218269 | 4.058998049 | 8.172475401 | 8.185025 | 8.730775702 | 8.410266976 |
| ENSG00000149532 | CPSF7 | -0.253588145 | 0.79810609 | 0.559056811 | 0.54005296 | 0.305468666 | 7.904282243 | 10.17682307 | 9.589475645 | 9.256998085 | 9.23691006 | 9.978712022 | 10.12848246 |
| ENSG00000204970 | PCDHA1 | 1.31715843 | 0.63692076 | -1.0137313 | 0.721493405 | 0.30342713 | 2 | 2 | 3.369317972 | 3.566290607 | 2.59489879 | 2.3070652 | 2.326754077 |
| ENSG00000137575 | SDCBP | 0.747351224 | 0.062366663 | -0.444074916 | 0.286844962 | 0.303276308 | 10.23600589 | 10.99464252 | 10.56174313 | 11.41302923 | 11.3444829 | 10.19181371 | 10.18921304 |
| ENSG00000163867 | ZMYM6 | 0.929586919 | 0.083215069 | -0.62762161 | 0.254347507 | 0.301965309 | 6.73718411 | 6.666836837 | 7.552162999 | 7.942383523 | 7.93600799 | 6.554528258 | 6.286392397 |
| ENSG00000120686 | UFM1 | 0.697907139 | 0.008153101 | -0.39596759 | 0.153721088 | 0.301939548 | 9.678463567 | 9.616264339 | 9.439621147 | 10.30502636 | 10.2501278 | 9.282760396 | 9.088984267 |
| ENSG00000001036 | FUCA2 | 0.978976395 | 0.318498522 | -0.677681152 | 0.497375263 | 0.301295243 | 11.12465208 | 9.849218119 | 8.585098385 | 11.17819202 | 11.1831065 | 9.328794492 | 9.705385533 |
| ENSG00000109519 | GRPEL1 | 0.402574787 | 0.322710742 | -0.101501492 | 0.8207453 | 0.301073295 | 8.777722034 | 8.955547846 | 8.763901704 | 9.220514323 | 9.25341153 | 8.445151776 | 8.979912728 |
| ENSG00000115364 | MRPL19 | 0.065861987 | 0.930416944 | 0.234392513 | 0.738292369 | 0.3002545 | 8.609439662 | 7.954278496 | 9.372276026 | 8.838609445 | 8.80148734 | 9.030579407 | 8.942636839 |
| ENSG00000129071 | MBD4 | 0.371026554 | 0.396920108 | -0.073248655 | 0.881261401 | 0.297777899 | 7.667647223 | 8.213429689 | 8.206335842 | 8.372666678 | 8.46722793 | 8.11499695 | 7.836259222 |
| ENSG00000154978 | VOPP1 | 0.532657661 | 0.442135948 | -0.235524605 | 0.745524236 | 0.297133056 | 9.533135543 | 8.732943944 | 10.23774519 | 10.14060551 | 10.1695649 | 9.313509983 | 9.46366404 |
| ENSG00000149480 | MTA2 | 0.158961698 | 0.938309757 | 0.138005201 | 0.946447056 | 0.296966899 | 4.00722531 | 8.317412869 | 10.68897008 | 9.475255213 | 9.57865651 | 9.404033465 | 9.603161713 |
| ENSG00000110328 | GALNT18 | -1.279411486 | 0.644633417 | 1.576326301 | 0.556063931 | 0.296914815 | 7.709880939 | 2 | 2 | 5.022023783 | 4.98931816 | 7.940525532 | 7.405663765 |
| ENSG00000188735 | TMEM120B | -0.212749574 | 0.747695986 | 0.509480363 | 0.398667475 | 0.29673079 | 8.39321512 | 7.393647515 | 7.552162999 | 7.617841552 | 7.65252709 | 8.57182706 | 8.077990544 |
| ENSG00000056097 | ZFR | 0.218235773 | 0.831805706 | 0.078155638 | 0.940405661 | 0.29639141 | 8.586954094 | 10.76012279 | 10.72180585 | 10.54003162 | 10.5157558 | 10.5244845 | 10.23780428 |
| ENSG00000164306 | PRIMPOL | 0.191009417 | 0.934841508 | 0.101577588 | 0.966283503 | 0.292587005 | 2 | 7.051611315 | 3.369317972 | 5.85002558 | 5.77157844 | 5.853194273 | 5.59182244 |
| ENSG00000134153 | EMC7 | 0.809839434 | 0.361201385 | -0.518364081 | 0.568052013 | 0.291475353 | 11.14403498 | 9.532315021 | 9.212024683 | 11.05732061 | 11.0114006 | 9.635775323 | 9.781528029 |
| ENSG00000134086 | VHL | 0.716140297 | 0.361166693 | -0.426450434 | 0.597996964 | 0.289689863 | 9.730679447 | 7.954278496 | 9.528916236 | 9.95800436 | 9.97493083 | 8.929133243 | 8.725336927 |
| ENSG00000117748 | RPA2 | 0.195379655 | 0.892020352 | 0.094149931 | 0.947678709 | 0.289529586 | 4.815609471 | 8.540087354 | 7.502634626 | 7.834291698 | 7.75969386 | 7.700314972 | 7.695929227 |
| ENSG00000162437 | RAVER2 | -2.523347567 | 0.115204374 | 2.812791749 | 0.069547827 | 0.289444182 | 7.031698319 | 7.701025996 | 2 | 4.329433326 | 4.72553021 | 9.907610449 | 9.241364212 |
| ENSG00000102401 | ARMCX3 | -0.222727419 | 0.589615937 | 0.511671467 | 0.171340702 | 0.288944047 | 9.751047882 | 10.01303782 | 10.37481416 | 9.918537335 | 9.77026155 | 10.65645596 | 10.49540898 |
| ENSG00000082516 | GEMIN5 | -0.025150469 | 0.988931662 | 0.311747426 | 0.857059231 | 0.286596957 | 3.327711234 | 7.35507212 | 8.585098385 | 7.494428494 | 7.53039799 | 7.86621124 | 7.819040576 |
| ENSG00000147394 | ZNF185 | -1.44889044 | 0.361913214 | 1.734753922 | 0.256088094 | 0.285863483 | 7.218819781 | 6.602424674 | 2 | 5.040242897 | 5.06165168 | 8.087943566 | 8.053353518 |
| ENSG00000164252 | AGGF1 | 0.417207795 | 0.386972978 | -0.133190519 | 0.797784693 | 0.284017276 | 7.940205782 | 8.029695761 | 7.225742779 | 8.18685754 | 8.20112674 | 7.586913431 | 7.715835172 |
| ENSG00000221983 | UBA52 | 0.28194718 | 0.466152501 | 0.001097915 | 0.998296152 | 0.283045096 | 10.85979178 | 10.28338212 | 10.75391088 | 10.91431859 | 10.950196 | 10.57970149 | 10.72117 |
| ENSG00000138443 | ABI2 | -0.114947602 | 0.914640194 | 0.397429098 | 0.686301227 | 0.282481496 | 10.57565445 | 8.955547846 | 8.434763955 | 9.523810814 | 9.49541459 | 10.08622907 | 9.949921184 |
| ENSG00000219438 | FAM19A5 | -5.493632287 | 0.008781104 | 5.775082705 | 0.000612377 | 0.281450418 | 2 | 5.838937808 | 2 | 2.141737246 | 2.14022021 | 9.791276442 | 10.16514154 |
| ENSG00000014824 | SLC30A9 | 0.26606654 | 0.628420441 | 0.015114582 | 0.979817504 | 0.281181122 | 8.910412251 | 9.640525262 | 8.699419242 | 9.439666599 | 9.38521851 | 9.130600759 | 9.195398998 |
| ENSG00000101412 | E2F1 | 0.262198119 | 0.895675696 | 0.01857733 | 0.992804891 | 0.280775449 | 4.00722531 | 6.311476754 | 2 | 5.304221918 | 5.33407135 | 4.713032626 | 5.408276315 |
| ENSG00000173175 | ADCY5 | -2.573709635 | 0.040691338 | 2.853263347 | 0.012415381 | 0.279553712 | 4.467311984 | 5.131603625 | 2 | 2.947924294 | 2.59489879 | 6.737909218 | 7.218077721 |
| ENSG00000090932 | DLL3 | -0.211479066 | 0.937710599 | 0.489182567 | 0.850325768 | 0.277703501 | 2 | 3.686303414 | 2 | 2.947924294 | 2.59489879 | 3.215043717 | 3.012034821 |
| ENSG00000082213 | C5orf22 | 0.262785018 | 0.52833407 | 0.01412643 | 0.974934399 | 0.276911448 | 8.138771685 | 8.376385542 | 8.677268317 | 8.615281294 | 8.72698617 | 8.544855285 | 8.296681143 |
| ENSG00000117625 | RCOR3 | 0.497717397 | 0.489413619 | -0.221749375 | 0.769071028 | 0.275968023 | 8.228665467 | 8.590771778 | 7.029880454 | 8.579911316 | 8.59458893 | 7.979784698 | 7.767238791 |
| ENSG00000198089 | SFI1 | 0.45442313 | 0.8426245 | -0.179186615 | 0.938715125 | 0.275236515 | 8.367060731 | 2 | 5.59507254 | 7.487805859 | 7.37584259 | 6.911998488 | 6.716538017 |
| ENSG00000111875 | ASF1A | 0.816556944 | 0.606992001 | -0.541506321 | 0.736592879 | 0.275050623 | 8.39321512 | 7.431218269 | 3.369317972 | 8.262524677 | 8.2130862 | 6.978631953 | 6.836905794 |
| ENSG00000102172 | SMS | 0.885170732 | 0.042971153 | -0.610740832 | 0.173383246 | 0.2744299 | 9.304185368 | 8.540087354 | 8.742726175 | 9.780398828 | 9.77026155 | 8.135306947 | 8.432373803 |
| ENSG00000100234 | TIMP3 | -0.70024055 | 0.172180866 | 0.973040539 | 0.045739026 | 0.272799989 | 11.34834729 | 11.47116739 | 10.57976717 | 10.46446525 | 10.5064723 | 11.88836944 | 12.38191627 |
| ENSG00000189007 | ADAT2 | 0.407146053 | 0.767234656 | -0.135750206 | 0.925711712 | 0.271395847 | 3.327711234 | 6.389896633 | 4.874587708 | 5.686138681 | 5.84514933 | 5.413723492 | 5.115701921 |
| ENSG00000164466 | SFXN1 | -0.192671668 | 0.84124654 | 0.463650998 | 0.599737667 | 0.27097933 | 8.075571721 | 9.792607274 | 10.18377222 | 9.406692983 | 9.40011621 | 10.05293048 | 10.06028212 |
| ENSG00000116205 | TCEANC2 | -0.050928545 | 0.975272867 | 0.320654386 | 0.839520202 | 0.269725841 | 8.285626337 | 5.303523747 | 4.874587708 | 7.035434774 | 6.82556437 | 7.517109417 | 7.032593922 |
| ENSG00000066322 | ELOVL1 | 0.169251729 | 0.837999 | 0.099893994 | 0.90483434 | 0.269145723 | 10.04301211 | 10.35843478 | 8.721235202 | 9.997003493 | 10.0656679 | 9.762974342 | 10.13806041 |
| ENSG00000184203 | PPP1R2 | 0.066073126 | 0.93880631 | 0.202839418 | 0.805557931 | 0.268912544 | 8.797441463 | 7.099070191 | 8.460932916 | 8.36726458 | 8.31463236 | 8.589193677 | 8.353844539 |
| ENSG00000125449 | ARMC7 | -0.452982853 | 0.806791315 | 0.721305296 | 0.68221623 | 0.268322443 | 2 | 7.315436877 | 6.958202216 | 6.237202172 | 6.0722229 | 7.544599974 | 6.969617847 |
| ENSG00000136928 | GABBR2 | 3.152133138 | 0.484134885 | -2.884875253 | 0.537048771 | 0.267257884 | 2 | 2 | 2 | 4.070463721 | 3.2184422 | 2.094598369 | 2 |
| ENSG00000124356 | STAMBP | 0.016671501 | 0.97361691 | 0.249722254 | 0.582427016 | 0.266393755 | 9.371511179 | 8.639735838 | 9.014156722 | 9.059013867 | 9.04284131 | 9.154473165 | 9.399952364 |
| ENSG00000118181 | RPS25 | -0.065555729 | 0.96006048 | 0.331674606 | 0.793834795 | 0.266118877 | 11.41687936 | 9.067150531 | 8.631919287 | 10.1868266 | 10.1923984 | 10.58976003 | 10.58153023 |
| ENSG00000013583 | HEBP1 | -0.299050976 | 0.636158103 | 0.564339135 | 0.336343976 | 0.265288159 | 9.667789716 | 8.487557258 | 8.960138144 | 8.825537758 | 8.81593678 | 9.513492253 | 9.826379958 |
| ENSG00000161542 | PRPSAP1 | -0.617934684 | 0.248274186 | 0.883129936 | 0.080480446 | 0.265195253 | 8.910412251 | 7.928237109 | 8.65477198 | 7.908072101 | 7.96694292 | 9.594132343 | 9.230009473 |
| ENSG00000120253 | NUP43 | -0.226177785 | 0.761391811 | 0.490744732 | 0.475879402 | 0.264566947 | 7.33131396 | 7.60566235 | 8.561105505 | 7.617841552 | 7.78928606 | 8.670625574 | 8.096865967 |
| ENSG00000166848 | TERF2IP | -0.04866475 | 0.968106553 | 0.313151181 | 0.787590496 | 0.264486431 | 11.12854954 | 9.799806309 | 8.236946975 | 10.1096668 | 10.1127112 | 10.58772851 | 10.34582388 |
| ENSG00000127328 | RAB3IP | -1.091698095 | 0.181178512 | 1.355327483 | 0.079746219 | 0.263629388 | 6.653098118 | 4.93638933 | 6.718950348 | 5.273747807 | 5.2591458 | 7.593002849 | 7.578053054 |
| ENSG00000061794 | MRPS35 | 0.402569088 | 0.453915688 | -0.139826853 | 0.80714548 | 0.262742235 | 7.867441304 | 8.655694736 | 7.818184403 | 8.667494409 | 8.47883193 | 8.060373167 | 8.020777347 |
| ENSG00000135045 | C9orf40 | -0.513069073 | 0.77088407 | 0.77264167 | 0.647304758 | 0.259572597 | 6.816637977 | 6.140560021 | 2 | 5.501219927 | 5.48637102 | 7.168655895 | 6.007361012 |

| | | | | | | | | | | | | | |
|---|---|---|---|---|---|---|---|---|---|---|---|---|---|
| ENSG00000120341 | SEC16B | 2.531520275 | 0.160878275 | -2.272819584 | 0.220299195 | 0.258700691 | 2 | 4.11352228 | 2 | 5.210797711 | 4.85329164 | 2.384001718 | 2.277179783 |
| ENSG00000101557 | USP14 | 0.19342017 | 0.749436229 | 0.064542482 | 0.919580542 | 0.257962652 | 10.29268328 | 9.170736405 | 9.647595056 | 9.966365455 | 9.9684779 | 9.799418565 | 9.877696666 |
| ENSG00000107819 | SFXN3 | -0.137527033 | 0.89442148 | 0.394948587 | 0.677551443 | 0.257421554 | 8.609439662 | 9.225224669 | 10.63827116 | 9.60637697 | 9.61666844 | 9.84030732 | 10.39094874 |
| ENSG00000132842 | AP3B1 | 0.459559235 | 0.346404872 | -0.202249687 | 0.694809381 | 0.257309547 | 9.262228249 | 9.648522413 | 8.825624954 | 9.730780958 | 9.760787 | 8.878514511 | 9.270949935 |
| ENSG00000026559 | KCNG1 | 0.006543717 | 0.997264364 | 0.250432588 | 0.90231922 | 0.256976305 | 7.53297911 | 7.431218269 | 2 | 6.892328675 | 6.95464102 | 7.754744914 | 6.131115787 |
| ENSG00000103496 | STX4 | 0.621890845 | 0.6972049 | -0.365387503 | 0.82326635 | 0.256503342 | 9.699577239 | 6.728495673 | 5.59507254 | 8.943972814 | 9.00436103 | 7.842553218 | 8.139104389 |
| ENSG00000169764 | UGP2 | -0.040407387 | 0.969622978 | 0.295513021 | 0.768207195 | 0.255105634 | 8.87372952 | 9.386925327 | 11.01019887 | 10.04403298 | 9.9912274 | 10.33841489 | 10.36671232 |
| ENSG00000137522 | RNF121 | -0.076870303 | 0.945987763 | 0.331845763 | 0.758720739 | 0.25497546 | 9.829762239 | 7.954278496 | 7.552162999 | 8.705093576 | 8.75724985 | 9.203056613 | 9.067272902 |
| ENSG00000124104 | SNX21 | 0.199529729 | 0.908645221 | 0.054798054 | 0.974300795 | 0.254327784 | 9.344956677 | 7.467835385 | 4.523675391 | 8.385193462 | 8.29975519 | 8.00283742 | 8.373881215 |
| ENSG00000197576 | HOXA4 | 5.444097041 | 0.152989085 | -5.19112223 | 0.214752155 | 0.252974811 | 2 | 2 | 2 | 5.578054239 | 5.6121243 | 2 | 2 |
| ENSG00000130749 | ZC3H4 | 0.104946368 | 0.932700573 | 0.148008473 | 0.90481909 | 0.252954841 | 6.257048221 | 8.213429689 | 9.272377397 | 8.468280417 | 8.47387021 | 8.598808264 | 8.423144406 |
| ENSG00000131584 | ACAP3 | -0.49295913 | 0.704833394 | 0.744710054 | 0.546275271 | 0.251750925 | 9.521325812 | 8.487557258 | 5.930865648 | 8.054188409 | 8.14812525 | 9.274574892 | 9.378441538 |
| ENSG00000158092 | NCK1 | 0.643947459 | 0.09566079 | -0.392486426 | 0.324893899 | 0.251461032 | 8.107517749 | 8.433041974 | 8.076950877 | 8.869506572 | 8.84824797 | 8.006643981 | 7.640647 |
| ENSG00000144840 | RABL3 | 0.346167148 | 0.815788738 | -0.095228085 | 0.950246079 | 0.250939062 | 7.218819781 | 4.710564154 | 8.65477198 | 7.920420067 | 7.92635454 | 7.560224843 | 7.417152486 |
| ENSG00000144115 | THNSL2 | 0.19459412 | 0.853531206 | 0.055877649 | 0.958419113 | 0.250471768 | 6.009025997 | 7.503546079 | 7.896971941 | 7.471115032 | 7.57741621 | 7.856119555 | 6.695462882 |
| ENSG00000126107 | HECTD3 | -0.12059712 | 0.946341641 | 0.370318808 | 0.83342673 | 0.249721687 | 4.00722531 | 9.055168103 | 9.577565424 | 8.634896 | 8.67172529 | 9.117986989 | 9.162073123 |
| ENSG00000280618 | | 0.295294585 | 0.789557312 | -0.047576572 | 0.967612474 | 0.247718013 | 7.709880939 | 9.916994052 | 10.36791565 | 9.986471488 | 10.0044775 | 9.504367208 | 9.789837542 |
| ENSG00000086065 | CHMP5 | 1.274788156 | 0.059166153 | -1.027419376 | 0.131477233 | 0.24736878 | 10.06758145 | 8.395519306 | 9.465702942 | 10.75935206 | 10.6994819 | 8.431623835 | 8.449958388 |
| ENSG00000119686 | FLVCR2 | -0.705082321 | 0.770289193 | 0.951386928 | 0.682959333 | 0.246304607 | 2 | 5.595927442 | 2 | 3.515101962 | 3.94787299 | 5.020706951 | 5.12973572 |
| ENSG00000117643 | MAN1C1 | -3.073944738 | 0.000984598 | 3.315699931 | 5.62E-05 | 0.241755192 | 5.095959458 | 4.710564154 | 5.156626401 | 3.227757282 | 2.38539787 | 7.990588491 | 8.256474583 |
| ENSG00000132313 | MRPL35 | -0.048694147 | 0.975624098 | 0.290292955 | 0.850843239 | 0.241598808 | 8.963744667 | 8.395519306 | 4.523675391 | 8.098332103 | 8.13141746 | 8.440093652 | 8.459018364 |
| ENSG00000057252 | SOAT1 | 1.217252998 | 0.120812437 | -0.977811135 | 0.215617166 | 0.239441864 | 9.678463567 | 7.928237109 | 9.809100877 | 10.55277887 | 10.568298 | 8.513629334 | 8.230662576 |
| ENSG00000100078 | PLA2G3 | -3.753003693 | 0.531143362 | 3.991383105 | 0.472178974 | 0.238379412 | 2 | 2 | 2 | 2 | 2 | 3.663836244 | 4.793076545 |
| ENSG00000117360 | PRPF3 | 0.395945225 | 0.446038292 | -0.157611675 | 0.774520328 | 0.23833355 | 8.631580145 | 7.701025996 | 8.076950877 | 8.516254096 | 8.62474679 | 8.040458475 | 8.009436866 |
| ENSG00000204174 | NPY4R | -2.312727365 | 0.665128529 | 2.549864217 | NA | 0.237136852 | 2 | 2 | 2 | 2 | 2.14022021 | 3.082820997 | 3.518055793 |
| ENSG00000001497 | LAS1L | -0.139063308 | 0.865194998 | 0.375201627 | 0.614795534 | 0.236138319 | 7.031698319 | 8.451443573 | 7.502634626 | 7.739909082 | 7.57741621 | 7.919680595 | 8.377561365 |
| ENSG00000197959 | DNM3 | 0.88044281 | 0.690529475 | -0.644857767 | 0.773343718 | 0.235585043 | 2 | 4.442710129 | 2 | 3.997744386 | 3.98554648 | 3.1504466 | 2.885414003 |
| ENSG00000165195 | PIGA | 0.223188316 | 0.885649929 | 0.011876821 | 0.993931149 | 0.235065137 | 4.00722531 | 7.954278496 | 6.43197674 | 7.048994596 | 7.14145882 | 6.768721236 | 7.004308743 |
| ENSG00000174109 | C16orf91 | -0.154432088 | 0.895265695 | 0.386779044 | 0.721563265 | 0.232346957 | 6.009025997 | 4.93638933 | 7.342949392 | 6.252910529 | 6.26070706 | 6.697032326 | 6.847559257 |
| ENSG00000160991 | ORAI2 | -0.365112075 | 0.646065476 | 0.595823027 | 0.42719519 | 0.230710953 | 8.257427002 | 9.514922632 | 9.990713521 | 9.085573664 | 9.0282492 | 9.842864719 | 10.1649286 |
| ENSG00000185825 | BCAP31 | 0.73125249 | 0.08771873 | -0.501155301 | 0.25275751 | 0.230097189 | 10.28571918 | 10.36815316 | 10.96062172 | 11.28423127 | 11.314036 | 9.882764149 | 10.23557543 |
| ENSG00000167695 | FAM57A | -0.187076095 | 0.851584865 | 0.416758582 | 0.653326175 | 0.229682487 | 7.096806373 | 8.505280662 | 9.330306409 | 8.392303091 | 8.38503958 | 8.901185819 | 9.067272902 |
| ENSG00000122550 | KLHL7 | 0.243521005 | 0.634071993 | -0.014690366 | 0.978701383 | 0.22883064 | 8.340423424 | 7.979858146 | 8.721235202 | 8.553199415 | 8.66883353 | 8.541185654 | 8.145162126 |
| ENSG00000213865 | C8orf44 | 0.231672835 | 0.895723155 | -0.005176996 | 0.997969492 | 0.22649584 | 2 | 4.11352228 | 5.772716771 | 4.887609717 | 4.79082474 | 4.727866721 | 4.543391831 |
| ENSG00000123989 | CHPF | -0.383315135 | 0.579274794 | 0.608588335 | 0.350450714 | 0.2252732 | 12.67495574 | 11.56710608 | 11.59570544 | 11.64594186 | 11.6755879 | 12.93232161 | 12.30336808 |
| ENSG00000125454 | SLC25A19 | -0.226539289 | 0.895062834 | 0.447909933 | 0.781250208 | 0.221370644 | 4.815609471 | 6.04685525 | 2 | 4.927308402 | 4.811949 | 5.610655473 | 5.313094696 |
| ENSG00000110321 | EIF4G2 | 0.093160006 | 0.923175097 | 0.128043827 | 0.893295691 | 0.221203832 | 11.54488817 | 13.1166165 | 13.63150382 | 13.10089132 | 13.0834501 | 13.092274 | 13.16103071 |
| ENSG00000105401 | CDC37 | 0.159106845 | 0.660816284 | 0.06073039 | 0.873992896 | 0.219837236 | 10.82178169 | 10.58784379 | 10.9788475 | 10.9293654 | 10.9941769 | 10.69565518 | 11.01473643 |
| ENSG00000281300 | | 0.272858106 | 0.74150751 | -0.053355396 | 0.951751695 | 0.219502709 | 8.855032352 | 10.1878402 | 8.742726175 | 9.677214349 | 9.71736428 | 9.08740002 | 9.611311483 |
| ENSG00000189144 | ZNF573 | -0.256129893 | 0.903607792 | 0.475063712 | 0.812849666 | 0.218933819 | 2 | 3.686303414 | 6.203069917 | 4.984880435 | 4.2865223 | 5.665620077 | 4.853519801 |
| ENSG00000088205 | DDX18 | 0.255611721 | 0.620673635 | -0.039254371 | 0.944101725 | 0.216357349 | 8.855032352 | 9.470501606 | 8.585098385 | 9.288777073 | 9.27068036 | 8.985384832 | 8.98860193 |
| ENSG00000196504 | PRPF40A | 0.600319555 | 0.328511143 | -0.384220452 | 0.541584801 | 0.216099103 | 9.189468652 | 9.540933215 | 10.43545992 | 10.42427719 | 10.4106283 | 9.472517815 | 9.400675953 |
| ENSG00000153187 | HNRNPU | 0.372226925 | 0.423319756 | -0.156146007 | 0.74998849 | 0.216080918 | 11.99688029 | 11.23931304 | 11.37961538 | 11.95574745 | 11.9417163 | 11.31077711 | 11.52345097 |
| ENSG00000084073 | ZMPSTE24 | 0.480550405 | 0.40310156 | -0.264608533 | 0.657720059 | 0.215941872 | 8.674867122 | 9.055168103 | 9.757248847 | 9.730780958 | 9.68767229 | 8.797335126 | 9.122267395 |
| ENSG00000179364 | PACS2 | -0.496102049 | 0.644633417 | 0.710089012 | 0.488684721 | 0.213986963 | 10.18451821 | 8.433041974 | 7.734844356 | 8.610716699 | 8.73115061 | 9.873413524 | 9.86726424 |
| ENSG00000119699 | TGFB3 | -1.349845927 | 0.395836393 | 1.56355649 | 0.307397922 | 0.213710562 | 4.467311984 | 7.189546432 | 9.465702942 | 6.846474326 | 6.88657603 | 10.05477007 | 9.327600363 |
| ENSG00000106723 | SPIN1 | 0.008439274 | 0.993765388 | 0.203721466 | 0.833507886 | 0.21216074 | 8.107517749 | 10.27823701 | 9.798878489 | 9.704603278 | 9.61215953 | 9.994516174 | 9.697162691 |
| ENSG00000187522 | HSPA14 | 0.276097347 | 0.899484135 | -0.064831352 | 0.975799274 | 0.211265996 | 9.048471759 | 8.557180776 | 2 | 8.509731121 | 8.52753255 | 8.319187939 | 8.030160124 |
| ENSG00000157445 | CACNA2D3 | 1.001545229 | 0.680089245 | -0.791204884 | 0.744740559 | 0.210340345 | 2 | 2 | 4.874587708 | 4.329433326 | 4.3456612 | 3.411599105 | 2.746601469 |
| ENSG00000119402 | FBXW2 | -0.390109076 | 0.612785008 | 0.600269925 | 0.410154943 | 0.210160849 | 10.29961392 | 8.655694736 | 10.01740919 | 9.37922384 | 9.4657318 | 10.49389479 | 10.31947205 |
| ENSG00000242247 | ARFGAP3 | 0.47797379 | 0.307672642 | -0.268193506 | 0.580066685 | 0.209780283 | 9.922475255 | 10.07961649 | 9.227351203 | 10.28320062 | 10.2506114 | 9.587285313 | 9.457765716 |
| ENSG00000091483 | FH | -0.649045267 | 0.536848085 | 0.857196104 | 0.394793944 | 0.208150837 | 9.657036304 | 7.467835385 | 7.646393253 | 8.015574133 | 7.94798467 | 9.291523922 | 9.635180748 |
| ENSG00000111860 | CEP85L | 0.391709223 | 0.813408453 | -0.183771725 | 0.914029999 | 0.207937498 | 8.075571721 | 6.844429755 | 3.369317972 | 7.437142666 | 7.4074129 | 7.17948788 | 6.457498602 |
| ENSG00000163539 | CLASP2 | -0.133490794 | 0.905447484 | 0.339418097 | 0.744205289 | 0.205927303 | 6.89194353 | 8.762721501 | 9.439621147 | 8.596935617 | 8.53707846 | 9.200400028 | 8.850309106 |
| ENSG00000171314 | PGAM1 | 1.273560737 | 0.003434646 | -1.067809076 | 0.015942135 | 0.205751661 | 11.8768947 | 11.30403016 | 11.2764554 | 12.76859676 | 12.8021511 | 10.09805229 | 10.72853726 |
| ENSG00000182568 | SATB1 | 0.920447446 | 0.392191303 | -0.715587924 | 0.507672408 | 0.204859521 | 6.653098118 | 7.60566235 | 4.874587708 | 7.691737974 | 7.64666005 | 6.370627179 | 5.745545708 |
| ENSG00000276380 | UBE2NL | -3.753004132 | 0.531143362 | 3.956244847 | 0.476306746 | 0.203240715 | 2 | 2 | 2 | 2 | 2 | 4.155805465 | 4.445805334 |
| ENSG00000102362 | SYTL4 | 0.760695124 | 0.525220439 | -0.558049914 | 0.643328101 | 0.20264521 | 3.327711234 | 5.838937808 | 6.322058544 | 6.12217689 | 6.51372389 | 5.294231737 | 4.911532223 |
| ENSG00000051620 | HEBP2 | -0.037507482 | 0.954566832 | 0.239962107 | 0.69597295 | 0.202454624 | 8.418903792 | 8.395519306 | 9.372276026 | 8.740345368 | 8.78555748 | 9.177281635 | 8.884335219 |
| ENSG00000196678 | ERI2 | 0.143281824 | 0.941473373 | 0.058767787 | 0.975626005 | 0.202049611 | 5.095959458 | 7.051611315 | 2 | 6.034011955 | 5.93454154 | 6.043083161 | 5.75463059 |
| ENSG00000153037 | SRP19 | 0.732947727 | 0.051734578 | -0.532399491 | 0.16559668 | 0.200548236 | 7.750913381 | 8.276714465 | 7.858115942 | 8.756965077 | 8.67172529 | 7.511762185 | 7.424286753 |
| ENSG00000133597 | ADCK2 | -1.357163776 | 0.474361527 | 1.557209677 | 0.393866241 | 0.200045901 | 2 | 6.228547981 | 8.236946975 | 5.797438367 | 5.58783529 | 8.596789458 | 8.419578856 |
| ENSG00000211584 | SLC48A1 | -1.063214715 | 0.164591992 | 1.262818251 | 0.085639381 | 0.199603537 | 6.73718411 | 8.376385542 | 7.225742779 | 6.61689148 | 6.57678965 | 9.08127623 | 8.632905293 |
| ENSG00000197563 | PIGN | -0.120212127 | 0.802441015 | 0.31880469 | 0.460415275 | 0.198592563 | 7.667647223 | 7.819099893 | 7.858115942 | 7.737119576 | 7.59275449 | 7.86872317 | 8.296681143 |
| ENSG00000116171 | SCP2 | -0.084683343 | 0.893473749 | 0.281711832 | 0.622327372 | 0.197028489 | 9.635285994 | 9.04308532 | 10.13584611 | 9.59332559 | 9.57557184 | 9.835820934 | 10.0543164 |
| ENSG00000149269 | PAK1 | -0.127222321 | 0.943520433 | 0.319518619 | 0.855882497 | 0.192296298 | 8.313285031 | 8.791896851 | 3.369317972 | 7.815902688 | 7.95036812 | 8.438967212 | 8.201061123 |
| ENSG00000172733 | PURG | -3.003890167 | 0.115773136 | 3.193670324 | 0.05188662 | 0.189780157 | 2 | 4.442710129 | 2 | 2.270786764 | 2.2680111 | 6.18690462 | 5.799216178 |
| ENSG00000169733 | RFNG | -0.367144045 | 0.551369059 | 0.555981394 | 0.340755867 | 0.188837349 | 9.521325812 | 8.317412869 | 9.099922433 | 8.679168098 | 8.70598173 | 9.745540467 | 9.458460889 |
| ENSG00000198933 | TBKBP1 | -0.216249597 | 0.892276535 | 0.404160465 | 0.789905257 | 0.187910869 | 4.815609471 | 7.315436877 | 9.049077186 | 7.620871236 | 7.74878077 | 8.464101787 | 8.107542072 |
| ENSG00000183691 | NOG | 0.738148119 | 0.771089389 | -0.550454572 | 0.83120048 | 0.187693547 | 2 | 2 | 5.156626401 | 4.443344104 | 4.3456612 | 2.962551659 | 3.824516949 |
| ENSG00000131669 | NINJ1 | 1.31380199 | 0.191634596 | -1.126147825 | 0.262168393 | 0.187654165 | 11.11289609 | 8.806266215 | 9.014156722 | 11.32961175 | 11.3882124 | 9.01441148 | 8.838074258 |
| ENSG00000129472 | RAB2B | 0.253788403 | 0.680957539 | -0.066311077 | 0.919852079 | 0.187477326 | 8.998234871 | 7.954278496 | 8.677268317 | 8.851563757 | 8.85080188 | 8.752700801 | 8.276323713 |
| ENSG00000184182 | UBE2F | 0.395241817 | 0.634203222 | -0.208270778 | 0.808778437 | 0.186971039 | 8.716893051 | 7.979858146 | 6.882775709 | 8.399377854 | 8.46722793 | 7.636890116 | 8.016062975 |
| ENSG00000140459 | CYP11A1 | 1.053106589 | 0.435021415 | -0.867161651 | 0.519967781 | 0.185944938 | 5.70932728 | 5.131603625 | 2 | 5.860316854 | 5.92487824 | 4.350488971 | 4.077147165 |
| ENSG00000145536 | ADAMTS16 | 1.504784136 | NA | -1.318858121 | NA | 0.185926015 | 2 | 2 | 2 | 3.289987342 | 2 | 2.267001569 | 2 |
| ENSG00000168802 | CHTF8 | -0.113819961 | 0.913307146 | 0.298836189 | 0.760366765 | 0.185016228 | 10.69610297 | 8.540087354 | 9.272377397 | 9.635130631 | 9.70753469 | 10.06100716 | 10.10403354 |
| ENSG00000169446 | MMGT1 | -0.22720727 | 0.4795951 | 0.412001116 | 0.161146297 | 0.184793846 | 8.609439662 | 8.395519306 | 8.561105505 | 8.327018759 | 8.26379462 | 8.902820827 | 8.953526633 |
| ENSG00000175470 | PPP2R2D | -0.109462885 | 0.918662967 | 0.294045875 | 0.771060512 | 0.18458299 | 9.304185368 | 8.147241016 | 6.958202216 | 8.276040626 | 8.37974396 | 8.835930569 | 8.611187869 |
| ENSG00000198453 | ZNF568 | -0.226029368 | 0.855033687 | 0.410526425 | 0.723926237 | 0.184497057 | 6.96351253 | 6.728495673 | 4.058998049 | 6.237202172 | 6.08095082 | 6.848938847 | 6.683620062 |
| ENSG00000156515 | HK1 | -0.271967927 | 0.648950726 | 0.456326486 | 0.417520898 | 0.184358559 | 10.86913986 | 10.66871049 | 11.59570544 | 10.77948848 | 10.8775392 | 11.24446364 | 11.8127295 |
| ENSG00000104884 | ERCC2 | -0.986691777 | 0.224578964 | 1.170487408 | 0.134337451 | 0.183795631 | 8.910412251 | 6.951736442 | 8.608698767 | 7.464384256 | 7.32898565 | 9.455350953 | 9.598438895 |
| ENSG00000167625 | ZNF526 | -0.118078139 | 0.942994458 | 0.300557579 | 0.852305768 | 0.18247944 | 4.00722531 | 8.147241016 | 8.721235202 | 7.739909082 | 7.84670947 | 8.350677018 | 8.045972122 |
| ENSG00000080839 | RBL1 | 0.234662717 | 0.810928754 | -0.053752286 | 0.957763507 | 0.180910431 | 4.467311984 | 5.457120953 | 6.322058544 | 5.870535235 | 5.74985018 | 5.703636713 | 5.367404784 |
| ENSG00000167987 | VPS37C | -0.498392697 | 0.735274416 | 0.679163131 | 0.633185132 | 0.180770434 | 5.532353208 | 8.702538795 | 9.868959282 | 8.401141138 | 8.33848526 | 9.382222556 | 9.678488127 |
| ENSG00000131732 | ZCCHC9 | 0.307819285 | 0.880694572 | -0.127610649 | 0.950246079 | 0.180208636 | 8.19931886 | 7.761270542 | 2 | 7.72026832 | 7.73225478 | 7.485815918 | 7.087553083 |
| ENSG00000179010 | MRFAP1 | -0.179507098 | 0.750738031 | 0.358723106 | 0.495887113 | 0.179216007 | 12.20129367 | 11.34889202 | 11.55983287 | 11.56959791 | 11.5718246 | 11.84150468 | 12.33347507 |
| ENSG00000116750 | UCHL5 | 0.077581319 | 0.943613626 | 0.101066463 | 0.927960061 | 0.178647782 | 8.39321512 | 6.844429755 | 6.203069917 | 7.423326516 | 7.61696151 | 7.754744914 | 7.302581437 |
| ENSG00000174500 | GCSAM | -3.753004463 | 0.531143362 | 3.929715656 | 0.479315748 | 0.176711193 | 2 | 2 | 2 | 2 | 2 | 4.065262523 | 4.479073019 |
| ENSG00000174807 | CD248 | 0.040614105 | 0.95872661 | 0.136045666 | 0.860323788 | 0.176659771 | 11.90670479 | 11.14918383 | 11.90777898 | 11.73793909 | 11.7341661 | 11.03547504 | 12.34136406 |
| ENSG00000139826 | ABHD13 | 0.314585124 | 0.882991782 | -0.138449871 | 0.948273373 | 0.176135253 | 8.716893051 | 7.538394143 | 2 | 8.004017475 | 7.95274764 | 7.69371876 | 7.354296226 |
| ENSG00000038358 | EDC4 | 0.121530913 | 0.760985399 | 0.054255525 | 0.895713612 | 0.175786438 | 8.816894989 | 8.671479029 | 8.296287027 | 8.71221318 | 8.75588792 | 8.715971642 | 8.617426278 |
| ENSG00000125912 | NCLN | 0.318492111 | 0.765322807 | -0.142750166 | 0.897608659 | 0.175741945 | 10.9638027 | 9.523645035 | 8.511888463 | 10.26201025 | 10.3936516 | 9.752131227 | 9.978555258 |
| ENSG00000181638 | ZFP41 | 0.652521426 | 0.628255701 | -0.477551362 | 0.725841069 | 0.174970065 | 5.095959458 | 5.595927442 | 8.076950877 | 7.341208691 | 7.64666005 | 6.503717356 | 6.308152703 |
| ENSG00000175110 | MRPS22 | 0.465645869 | 0.671105744 | -0.291276648 | 0.794825276 | 0.174369222 | 8.75772933 | 7.467835385 | 6.073378139 | 8.219202381 | 8.33118782 | 7.54878318 | 7.515294811 |
| ENSG00000143457 | GOLPH3L | 0.089045969 | 0.945455549 | 0.084909936 | 0.948481065 | 0.173955905 | 9.081016759 | 7.638155693 | 5.930865648 | 8.205140657 | 8.09094333 | 8.292619498 | 7.980689421 |
| ENSG00000114626 | ABTB1 | -0.566277698 | 0.442942747 | 0.740038173 | 0.291119632 | 0.173760475 | 8.228665467 | 7.538394143 | 9.116480286 | 7.862723323 | 7.88459112 | 9.284329221 | 9.035479014 |
| ENSG00000132676 | DAP3 | 0.696580944 | 0.071684427 | -0.526069333 | 0.179352679 | 0.170511611 | 8.564112515 | 9.148351514 | 8.960138144 | 9.626112287 | 9.58787113 | 8.2308261 | 8.541379994 |
| ENSG00000126067 | PSMB2 | 0.284453727 | 0.590007772 | -0.114039899 | 0.837844551 | 0.170413828 | 10.61508061 | 9.763446433 | 10.40882047 | 10.57520961 | 10.5990046 | 9.95546434 | 10.39149486 |
| ENSG00000143183 | TMCO1 | 0.198478225 | 0.580892944 | -0.028086121 | 0.943162724 | 0.170392105 | 9.848787621 | 9.591588447 | 10.03493575 | 10.10264028 | 9.95489142 | 9.713525389 | 9.891147007 |
| ENSG00000197614 | MFAP5 | -6.293764667 | 0.009458344 | 6.464054075 | 0.000144114 | 0.170289408 | 2 | 2 | 4.523675391 | 2 | 2 | 8.858854473 | 9.204544205 |
| ENSG00000185800 | DMWD | -0.232771492 | 0.495143159 | 0.402562575 | 0.20216279 | 0.169791083 | 9.509418609 | 9.328390744 | 9.681375057 | 9.255049401 | 9.30181283 | 10.02879997 | 9.781805785 |
| ENSG00000198919 | DZIP3 | -0.66365869 | 0.596379651 | 0.833434142 | 0.487153954 | 0.169775452 | 4.00722531 | 7.051611315 | 5.59507254 | 5.527288686 | 5.36298494 | 6.987904673 | 6.711881188 |
| ENSG00000119414 | PPP6C | -0.132259033 | 0.839051597 | 0.301094124 | 0.616273347 | 0.168835091 | 8.609439662 | 9.396454291 | 9.767769415 | 9.134092596 | 9.26781654 | 9.471967222 | 9.778469175 |
| ENSG00000166913 | YWHAB | 0.029253429 | 0.932390851 | 0.137995077 | 0.664743846 | 0.167248506 | 11.65707219 | 11.51118914 | 11.56587392 | 11.60502731 | 11.6113331 | 11.51539471 | 11.89353851 |
| ENSG00000244734 | HBB | -3.753004582 | 0.531143362 | 3.920236544 | 0.480467544 | 0.167231962 | 2 | 2 | 2 | 2 | 2 | 3.517841605 | 4.77533149 |
| ENSG00000166477 | LEO1 | 0.627154582 | 0.768745599 | -0.461226456 | 0.831750977 | 0.165928126 | 9.204317094 | 7.315436877 | 2 | 8.610716699 | 8.57000035 | 7.543552275 | 7.493753702 |
| ENSG00000174574 | AKIRIN1 | 0.248849046 | 0.660813516 | -0.085059258 | 0.887886595 | 0.163789788 | 10.51143645 | 9.479495904 | 9.888372618 | 10.30502636 | 10.2316302 | 10.00844152 | 9.859916395 |

| | | | | | | | | | | | | | |
|---|---|---|---|---|---|---|---|---|---|---|---|---|---|
| ENSG00000035664 | DAPK2 | 0.146921773 | 0.933881218 | 0.016239749 | 0.992804891 | 0.163161522 | 2 | 3.686303414 | 4.058998049 | 3.663521627 | 3.50452879 | 3.393105702 | 3.600986722 |
| ENSG00000164219 | PGGT1B | -0.38930468 | 0.770303179 | 0.552167114 | 0.668751914 | 0.162862434 | 4.467311984 | 7.638155693 | 8.007629717 | 6.960881212 | 6.9210099 | 7.906701615 | 7.811442311 |
| ENSG00000160256 | FAM207A | -0.25706012 | 0.869968338 | 0.418807778 | 0.779317373 | 0.161747657 | 7.096806373 | 7.572420273 | 3.369317972 | 6.610808945 | 6.53928196 | 7.030406362 | 7.402777233 |
| ENSG00000144061 | NPHP1 | -0.301402955 | 0.855036819 | 0.462054644 | 0.767357706 | 0.160651688 | 2 | 4.442710129 | 5.59507254 | 4.329433326 | 4.40247105 | 4.749835737 | 5.230851202 |
| ENSG00000189266 | PNRC2 | 1.061078172 | 2.12E-05 | -0.901125083 | 0.000363392 | 0.15995309 | 9.344956677 | 9.079034257 | 9.212024683 | 10.28128703 | 10.2559202 | 8.268091966 | 8.370191653 |
| ENSG00000167685 | ZNF444 | -0.415774581 | 0.445961619 | 0.575678697 | 0.263479222 | 0.159904116 | 8.75772933 | 7.73146271 | 8.266922072 | 7.857595438 | 7.93118934 | 8.917048655 | 8.830038743 |
| ENSG00000122481 | RWDD3 | 1.279523504 | 0.09807061 | -1.122201444 | 0.147783571 | 0.157322061 | 5.095959458 | 5.457120953 | 4.523675391 | 6.276156955 | 6.28357155 | 4.369533167 | 3.924507493 |
| ENSG00000171475 | WIPF2 | -0.185978238 | 0.756699054 | 0.340746183 | 0.54493451 | 0.154767944 | 9.895281535 | 8.955547846 | 9.990713521 | 9.474417988 | 9.51406378 | 9.924409668 | 10.10714012 |
| ENSG00000122359 | ANXA11 | -0.845036612 | 0.107655331 | 0.999517029 | 0.048821317 | 0.154480417 | 10.71696191 | 9.672251278 | 10.23774519 | 9.425008328 | 9.43031191 | 11.05011945 | 11.45403965 |
| ENSG00000168936 | TMEM129 | -0.497495127 | 0.418653107 | 0.651530748 | 0.266310577 | 0.154035621 | 10.05943807 | 9.102512015 | 8.960138144 | 8.976268302 | 8.95536429 | 10.2523815 | 9.948684752 |
| ENSG00000164338 | UTP15 | 0.323112187 | 0.8692818 | -0.16947804 | 0.932012683 | 0.153634146 | 2 | 7.099070191 | 7.777116083 | 7.136275664 | 7.30311013 | 6.768721236 | 6.721179863 |
| ENSG00000196267 | ZNF836 | 0.501921846 | 0.584830919 | -0.350950647 | 0.705751145 | 0.150971199 | 4.467311984 | 5.72254323 | 6.203069917 | 6.024891728 | 6.16544343 | 5.565950232 | 4.998228167 |
| ENSG00000100285 | NEFH | 0.954626963 | 0.638803946 | -0.80423798 | 0.692520331 | 0.150388983 | 3.327711234 | 2 | 5.59507254 | 5.178261742 | 5.13053105 | 4.177580368 | 3.100202476 |
| ENSG00000180329 | CCDC43 | 0.173580732 | 0.890295464 | -0.023258728 | 0.9854015 | 0.150322004 | 8.87372952 | 7.189546432 | 6.073378139 | 8.015574133 | 7.9809999 | 7.763770178 | 7.844792088 |
| ENSG00000138744 | NAAA | -0.093691901 | 0.895265695 | 0.243558866 | 0.709981742 | 0.149866965 | 6.89194353 | 7.572420273 | 6.629488387 | 6.956091174 | 7.06873058 | 7.57363085 | 7.063994051 |
| ENSG00000134453 | RBM17 | 0.263449376 | 0.761694883 | -0.113682288 | 0.900290106 | 0.149767088 | 9.262228249 | 8.414402626 | 7.342949392 | 8.796356221 | 8.80807321 | 8.407640344 | 8.449958388 |
| ENSG00000142544 | CTU1 | -0.429305922 | 0.671131573 | 0.578706802 | 0.546392953 | 0.14940088 | 4.467311984 | 5.946638857 | 6.534109959 | 5.527288686 | 5.43283104 | 6.723178551 | 6.041166521 |
| ENSG00000119421 | NDUFA8 | -0.367486831 | 0.52974812 | 0.516355792 | 0.350276926 | 0.148868961 | 8.674867122 | 8.395519306 | 7.552162999 | 7.885575888 | 7.95749494 | 8.687797808 | 8.894645365 |
| ENSG00000171421 | MRPL36 | -0.046960713 | 0.943613626 | 0.195658162 | 0.756508981 | 0.148697449 | 8.816894989 | 7.790474954 | 7.934808822 | 8.151677543 | 8.2561084 | 8.296978073 | 8.578306014 |
| ENSG00000160360 | GPSM1 | -0.309862264 | 0.687785088 | 0.458019465 | 0.531674005 | 0.148157201 | 10.22876205 | 9.018611864 | 8.784770914 | 9.242318391 | 9.12003262 | 9.849450427 | 10.03696805 |
| ENSG00000148019 | CEP78 | -1.324193242 | 0.432204028 | 1.471684378 | 0.365364036 | 0.147491136 | 7.384433967 | 7.315436877 | 2 | 5.662809983 | 5.40529704 | 8.175088797 | 8.273956186 |
| ENSG00000162600 | OMA1 | -0.203073784 | 0.909988721 | 0.350531029 | 0.837543189 | 0.147457245 | 2 | 6.311476754 | 6.534109959 | 5.818703821 | 5.55061739 | 6.453284622 | 5.913137508 |
| ENSG00000165983 | PTER | -0.243882774 | 0.897905614 | 0.391057722 | 0.830862053 | 0.147174948 | 6.816637977 | 5.457120953 | 2 | 5.242616075 | 5.74985018 | 6.495072127 | 5.576534358 |
| ENSG00000145907 | G3BP1 | -0.100208559 | 0.903052048 | 0.247307787 | 0.746005167 | 0.147099228 | 9.509418609 | 10.25223258 | 11.0793996 | 10.3559173 | 10.2835937 | 10.4654111 | 10.84343131 |
| ENSG00000112167 | SAYSD1 | -0.04034197 | 0.966682851 | 0.187381742 | 0.841697129 | 0.147039773 | 6.257048221 | 6.04685525 | 7.734844356 | 6.841288255 | 6.83074839 | 7.125896536 | 6.983208048 |
| ENSG00000116729 | WLS | 0.684697395 | 0.309576645 | -0.539594781 | 0.42570287 | 0.145102614 | 10.78274301 | 9.328390744 | 9.888372618 | 10.82025579 | 10.7937466 | 9.4949131 | 9.673706637 |
| ENSG00000100591 | AHSA1 | 0.218997933 | 0.711659309 | -0.073956398 | 0.906192676 | 0.145041536 | 9.858207046 | 8.806266215 | 9.725218844 | 9.76541805 | 9.72782219 | 9.390112523 | 9.517693638 |
| ENSG00000167657 | DAPK3 | -0.5182187 | 0.702916163 | 0.662815882 | 0.613365058 | 0.144597182 | 9.678463567 | 6.666836837 | 10.50627919 | 9.129845271 | 9.09439916 | 10.1379928 | 10.42282144 |
| ENSG00000165092 | ALDH1A1 | -2.312736954 | 0.66900125 | 2.456900924 | NA | 0.14416397 | 2 | 2 | 2 | 2 | 2.14022021 | 3.336159116 | 3.183290849 |
| ENSG00000164197 | RNF180 | -1.638219227 | 0.477431438 | 1.781086424 | 0.420543923 | 0.142867197 | 2 | 5.595927442 | 2 | 3.462030052 | 2.77779907 | 6.101204172 | 5.492143571 |
| ENSG00000148335 | NTMT1 | 0.097148178 | 0.945043724 | 0.045199248 | 0.974393699 | 0.142347426 | 8.836089686 | 7.638155693 | 5.156626401 | 7.920420067 | 7.96929529 | 7.780764172 | 7.998962529 |
| ENSG00000181924 | COA4 | -0.112712638 | 0.871671212 | 0.254522634 | 0.693056959 | 0.141809996 | 8.777722034 | 7.819099893 | 8.206335842 | 8.166267464 | 8.24450173 | 8.166267073 | 8.88330014 |
| ENSG00000138029 | HADHB | 0.842504196 | 0.326142247 | -0.700704906 | 0.415037148 | 0.14179929 | 10.79750658 | 9.821190352 | 8.65477198 | 10.81728741 | 10.8765996 | 9.287148813 | 9.332538367 |
| ENSG00000179151 | EDC3 | -0.214651712 | 0.887194753 | 0.356447452 | 0.804092949 | 0.14179574 | 8.39321512 | 9.664384941 | 5.59507254 | 8.429064458 | 8.43184521 | 8.978035992 | 9.013887033 |
| ENSG00000141522 | ARHGDIA | -0.048854471 | 0.919059559 | 0.189998088 | 0.668446136 | 0.141143616 | 11.63805915 | 11.38763081 | 11.44653121 | 11.40843409 | 11.4820112 | 11.30553333 | 11.98394814 |
| ENSG00000104964 | AES | 0.606176984 | 0.113076042 | -0.465694334 | 0.228436335 | 0.14048265 | 11.20436621 | 10.98835491 | 11.65641302 | 11.86666856 | 11.9609044 | 10.83416038 | 10.85443307 |
| ENSG00000134291 | TMEM106C | -1.051733303 | 0.606090823 | 1.191434973 | 0.546604763 | 0.13970167 | 9.317904228 | 8.101365653 | 2 | 7.144720234 | 7.29937545 | 9.472517815 | 9.399590433 |
| ENSG00000174446 | SNAPC5 | -0.259148328 | 0.751436954 | 0.398086351 | 0.609008368 | 0.138938023 | 7.53297911 | 6.311476754 | 6.073378139 | 6.483440992 | 6.58294721 | 7.20223931 | 7.133547492 |
| ENSG00000165259 | HDX | 0.992676878 | 0.554602856 | -0.853784441 | 0.611576383 | 0.138892437 | 7.159102577 | 5.838937808 | 2 | 6.998639523 | 7.0774796 | 5.233158834 | 5.367404784 |
| ENSG00000104325 | DECR1 | 1.50997339 | 0.008952796 | -1.371345329 | 0.018674882 | 0.13862806 | 7.709880939 | 8.590771778 | 7.342949392 | 9.511620236 | 9.46072474 | 6.484192226 | 6.808832901 |
| ENSG00000156831 | NSMCE2 | -0.014728869 | 0.988861116 | 0.152158107 | 0.882006656 | 0.137429238 | 7.27616306 | 7.638155693 | 5.39244166 | 6.975156635 | 7.10768997 | 7.183528998 | 7.226264598 |
| ENSG00000196268 | ZNF493 | 0.135484548 | 0.912378494 | 0.001261307 | 0.999263508 | 0.136745855 | 6.366709671 | 5.131603625 | 4.874587708 | 5.870535235 | 5.5755357 | 6.112552588 | 4.819291477 |
| ENSG00000169609 | C15orf40 | -0.480584539 | 0.483180449 | 0.615096175 | 0.345033356 | 0.134511636 | 7.384433967 | 6.389896633 | 7.029880454 | 6.693715713 | 6.31350454 | 7.849352333 | 7.242500282 |
| ENSG00000183908 | LRRC55 | -3.753005002 | 0.531143362 | 3.886550564 | 0.484010617 | 0.133545562 | 2 | 2 | 2 | 2 | 2 | 4.301748724 | 4.201948172 |
| ENSG00000123595 | RAB9A | 0.58969212 | 0.665129218 | -0.456633599 | 0.737989924 | 0.133058521 | 9.410448395 | 7.274681896 | 5.930865648 | 8.758341449 | 8.84952549 | 7.672789259 | 7.864857644 |
| ENSG00000182150 | ERCC6L2 | 0.270108633 | 0.78979414 | -0.139155702 | 0.893479206 | 0.130952931 | 6.563806279 | 7.503546079 | 8.763901704 | 8.190940526 | 8.10812366 | 7.580798201 | 7.897008641 |
| ENSG00000138031 | ADCY3 | -0.38226598 | 0.832844524 | 0.511157275 | 0.768563703 | 0.128891295 | 9.957954117 | 7.232742069 | 4.874587708 | 8.227176542 | 8.240612 | 9.206039434 | 9.025192723 |
| ENSG00000186283 | TOR3A | -0.363589973 | 0.654620374 | 0.4922311 | 0.527634025 | 0.128641127 | 8.16936288 | 7.819099893 | 6.43197674 | 7.366699513 | 7.21466714 | 8.300083299 | 7.941446131 |
| ENSG00000155755 | TMEM237 | -0.089267388 | 0.898841046 | 0.215086404 | 0.741726039 | 0.125819016 | 8.777722034 | 8.655694736 | 8.042706615 | 8.447952961 | 8.43184521 | 8.200570331 | 9.133628617 |
| ENSG00000054654 | SYNE2 | -0.661766456 | 0.431378597 | 0.786675412 | 0.326487945 | 0.124908956 | 6.96351253 | 5.838937808 | 6.322058544 | 6.229283386 | 5.22805098 | 7.380620955 | 6.983208048 |
| ENSG00000159267 | HLCS | -0.69383182 | 0.690326721 | 0.818105889 | 0.62915305 | 0.124274069 | 4.00722531 | 7.572420273 | 9.553445883 | 7.700356475 | 7.56502709 | 9.155502188 | 9.098381203 |
| ENSG00000105221 | AKT2 | -0.213532524 | 0.686696702 | 0.337675408 | 0.498871333 | 0.124142884 | 10.56418906 | 9.785372135 | 10.63252637 | 10.13213675 | 10.1868502 | 10.57445347 | 10.83274721 |
| ENSG00000182983 | ZNF662 | 0.067634408 | 0.966177728 | 0.05519012 | 0.972248923 | 0.122824527 | 4.00722531 | 3.686303414 | 6.534109959 | 5.258265913 | 5.48637102 | 5.536762811 | 5.171036168 |
| ENSG00000134917 | ADAMTS8 | -2.312742802 | 0.671489931 | 2.434718438 | NA | 0.121975636 | 2 | 2 | 2 | 2 | 2.14022021 | 3.011868911 | 3.45257121 |
| ENSG00000184209 | SNRNP35 | -0.309438561 | 0.885805368 | 0.430852016 | 0.835405544 | 0.121413456 | 8.444143041 | 6.899080544 | 2 | 7.044488801 | 6.94511186 | 7.093332266 | 8.15205425 |
| ENSG00000140632 | GLYR1 | 0.172051055 | 0.87178855 | -0.051638902 | 0.962322454 | 0.120412153 | 8.444143041 | 9.170736405 | 10.67226762 | 9.900574544 | 9.90906469 | 9.601450544 | 9.758568089 |
| ENSG00000133110 | POSTN | 1.124764394 | 0.189674145 | -1.006611135 | 0.237882611 | 0.118153258 | 12.82540395 | 12.83762542 | 12.94228516 | 13.99473163 | 13.9927475 | 12.58710778 | 10.34375594 |
| ENSG00000139800 | ZIC5 | -5.650660466 | 0.021326273 | 5.768595101 | 0.000779484 | 0.117934635 | 4.00722531 | 2 | 2 | 2 | 2 | 7.54878318 | 7.873223515 |
| ENSG00000063241 | ISOC2 | 0.062234787 | 0.962823674 | 0.054655989 | 0.968477418 | 0.116890776 | 9.771132748 | 6.728495673 | 7.777116083 | 8.700804887 | 8.70315789 | 8.499651249 | 8.866113243 |
| ENSG00000086232 | EIF2AK1 | -0.139632416 | 0.857804601 | 0.256205743 | 0.726349422 | 0.116573327 | 11.15554094 | 10.35843478 | 9.624627483 | 10.35500785 | 10.3918991 | 10.69730485 | 10.8360945 |
| ENSG00000130766 | SESN2 | -0.026807827 | 0.985675142 | 0.142593536 | 0.925786782 | 0.11578571 | 5.866944937 | 8.623598428 | 9.61300511 | 8.618316353 | 8.69891174 | 9.365719319 | 7.957271773 |
| ENSG00000164061 | BSN | -0.588057193 | 0.749240001 | 0.703259459 | 0.690278517 | 0.115202266 | 2 | 3.076665209 | 4.874587708 | 3.515101962 | 3.2184422 | 4.388329244 | 4.188602707 |
| ENSG00000164332 | UBLCP1 | 0.364238938 | 0.548198478 | -0.251122869 | 0.68474076 | 0.113116069 | 8.075571721 | 7.572420273 | 8.742726175 | 8.616799622 | 8.50338773 | 7.927733561 | 7.979721237 |
| ENSG00000101247 | NDUFAF5 | 0.104170899 | 0.960095276 | 0.00888354 | 0.996970186 | 0.113054439 | 2 | 5.131603625 | 7.600047325 | 6.29903474 | 6.44101097 | 6.394066622 | 6.158754168 |
| ENSG00000221926 | TRIM16 | 0.695012933 | 0.155535752 | -0.582855663 | 0.233677793 | 0.11215727 | 8.075571721 | 7.538394143 | 7.934808822 | 8.469961514 | 8.61726622 | 6.970858896 | 7.55093161 |
| ENSG00000160124 | CCDC58 | 0.925705657 | 0.212930755 | -0.813581975 | 0.272130443 | 0.112123683 | 6.257048221 | 4.93638933 | 5.59507254 | 6.586219071 | 6.49425375 | 4.785723309 | 5.080008298 |
| ENSG00000105229 | PIAS4 | 0.001619009 | 0.997903682 | 0.110454042 | 0.879418834 | 0.11207305 | 7.27616306 | 7.503546079 | 8.434763955 | 7.815902688 | 7.83385794 | 8.135306947 | 7.694749691 |
| ENSG00000198680 | TUSC1 | -0.523961365 | 0.623796324 | 0.634836873 | 0.538227094 | 0.110875507 | 5.70932728 | 7.189546432 | 8.35328246 | 6.965655398 | 6.91125523 | 8.054501123 | 8.087908463 |
| ENSG00000153814 | JAZF1 | -0.51078519 | 0.716843352 | 0.621204661 | 0.650182334 | 0.110419471 | 4.815609471 | 8.356994601 | 8.805342542 | 7.605658789 | 7.52080763 | 8.622816121 | 8.733401188 |
| ENSG00000087263 | OGFOD1 | 0.243155379 | 0.647252339 | -0.133170437 | 0.808262771 | 0.109984942 | 9.751047882 | 8.916342327 | 9.735974718 | 9.739866907 | 9.76891184 | 9.240714099 | 9.507317152 |
| ENSG00000156502 | SUPV3L1 | 0.403968942 | 0.383159098 | -0.297355898 | 0.523553954 | 0.106613044 | 7.667647223 | 7.232742069 | 7.934808822 | 8.100504225 | 7.94320592 | 7.384136831 | 7.287011591 |
| ENSG00000130669 | PAK4 | -0.069464078 | 0.935745276 | 0.174666789 | 0.834220385 | 0.105202712 | 8.228665467 | 8.540087354 | 9.798878489 | 8.936696791 | 8.96953398 | 9.110223929 | 9.276085345 |
| ENSG00000101367 | MAPRE1 | -0.241070389 | 0.803176349 | 0.34573858 | 0.708647078 | 0.104668191 | 9.031919741 | 10.47993704 | 11.20470823 | 10.20863848 | 10.2873694 | 10.66238005 | 10.98693891 |
| ENSG00000155189 | AGPAT5 | -0.227270954 | 0.897554944 | 0.33192247 | 0.845529489 | 0.104651517 | 3.327711234 | 7.145017454 | 8.408111533 | 7.084543139 | 7.17852729 | 7.635907362 | 7.719319651 |
| ENSG00000158773 | USF1 | 0.735214758 | 0.253424754 | -0.631272195 | 0.325173109 | 0.103942563 | 8.418903792 | 8.623598428 | 7.225742779 | 8.901002576 | 8.98007065 | 7.526684775 | 7.656486163 |
| ENSG00000154945 | ANKRD40 | -0.110773417 | 0.879824258 | 0.21457261 | 0.75370958 | 0.103799193 | 8.87372952 | 9.405920729 | 10.21486071 | 9.539906319 | 9.44475345 | 9.737087614 | 9.892433725 |
| ENSG00000213886 | UBD | -0.230175068 | 0.490900748 | 0.332986909 | 0.291605818 | 0.102811841 | 12.10369464 | 12.08742575 | 12.30989775 | 11.97318846 | 11.9067054 | 12.28739969 | 12.69058631 |
| ENSG00000139998 | RAB15 | -0.905376566 | 0.497088189 | 1.007976617 | 0.435748117 | 0.102600051 | 8.257427002 | 5.838937808 | 9.69246155 | 7.799951945 | 7.66998641 | 9.795023124 | 9.443089155 |
| ENSG00000054267 | ARID4B | 0.663278599 | 0.106608823 | -0.561032867 | 0.172094144 | 0.102245732 | 8.963744667 | 9.415325457 | 9.478569125 | 9.990576409 | 9.93757622 | 8.965573751 | 8.490627472 |
| ENSG00000143420 | ENSA | -0.311081418 | 0.638989421 | 0.411916465 | 0.518445971 | 0.100835047 | 10.12333326 | 8.903033433 | 9.990713521 | 9.449924671 | 9.45653888 | 9.963712247 | 10.3557463 |
| ENSG00000114378 | HYAL1 | -2.576747169 | 0.023043738 | 2.677500958 | 0.005400389 | 0.100753789 | 4.00722531 | 3.686303414 | 4.058998049 | 2.498691189 | 2.59489879 | 6.148830293 | 6.286392397 |
| ENSG00000118407 | FILIP1 | -0.134945661 | 0.95389048 | 0.235444579 | 0.918692085 | 0.100498918 | 3.327711234 | 2 | 3.369317972 | 2.947924294 | 2.86123014 | 3.316666208 | 2.817676617 |
| ENSG00000107518 | ATRNL1 | -2.619049532 | 0.06531578 | 2.719353566 | 0.036821565 | 0.100304034 | 3.327711234 | 5.131603625 | 3.369317972 | 3.023133209 | 2.2680111 | 6.723178551 | 6.676467423 |
| ENSG00000165169 | DYNLT3 | -0.047117529 | 0.926779701 | 0.147358141 | 0.758720739 | 0.100240612 | 8.228665467 | 8.639735838 | 8.408111533 | 8.427334995 | 8.35836495 | 8.824761857 | 8.298235265 |
| ENSG00000120318 | ARAP3 | 0.428592663 | 0.807381293 | -0.328708492 | 0.85201813 | 0.09988417 | 3.327711234 | 8.053983481 | 5.772716771 | 7.165617905 | 7.2537885 | 6.645228724 | 6.29264319 |
| ENSG00000114744 | COMMD2 | -0.31713075 | 0.501509841 | 0.416627805 | 0.353451393 | 0.099497056 | 8.797441463 | 8.101365653 | 8.825624954 | 8.264463296 | 8.31463236 | 9.076575731 | 8.953033422 |
| ENSG00000106105 | GARS | -0.135522495 | 0.770162624 | 0.232052925 | 0.597996964 | 0.09653043 | 10.80726569 | 11.31160468 | 11.51057165 | 11.09457533 | 11.1130312 | 11.59940158 | 11.3296143 |
| ENSG00000169379 | ARL13B | 0.184058571 | 0.91483416 | -0.088085159 | 0.959372651 | 0.095973411 | 7.82963488 | 5.838937808 | 3.369317972 | 6.872128674 | 6.71216164 | 6.697032326 | 6.347714043 |
| ENSG00000196466 | ZNF799 | 0.32154846 | 0.878583032 | -0.226725516 | 0.913644152 | 0.094822944 | 6.366709671 | 3.686303414 | 2 | 5.319220934 | 5.33407135 | 5.174809258 | 4.411752372 |
| ENSG00000197321 | SVIL | -0.408748845 | 0.435404672 | 0.502584803 | 0.316308322 | 0.093835958 | 9.567995545 | 9.687856435 | 10.46808205 | 9.604082348 | 9.50841342 | 10.47890139 | 10.45051519 |
| ENSG00000129187 | DCTD | 0.141752681 | 0.790629813 | -0.049052226 | 0.929877038 | 0.092700455 | 10.41260941 | 9.583268459 | 10.12770132 | 10.22615207 | 10.2128923 | 9.831105898 | 10.20379719 |
| ENSG00000171403 | KRT9 | -3.753005516 | 0.531143362 | 3.845397614 | 0.489327719 | 0.092392098 | 2 | 2 | 2 | 2 | 2 | 3.355392153 | 4.757365452 |
| ENSG00000139133 | ALG10 | 0.730302692 | 0.610603213 | -0.638154406 | 0.655811705 | 0.092148286 | 5.095959458 | 4.93638933 | 2 | 5.304221918 | 5.0259382 | 4.321442857 | 3.67940853 |
| ENSG00000160058 | BSDC1 | 0.239466113 | 0.755240998 | -0.147510376 | 0.851219083 | 0.091955737 | 11.20066853 | 9.608085847 | 10.36791565 | 10.7284349 | 10.8180948 | 10.38082821 | 10.39513041 |
| ENSG00000042493 | CAPG | 0.138321124 | 0.922673812 | -0.046934828 | 0.97326848 | 0.091386296 | 10.05943807 | 8.522788974 | 6.43197674 | 9.061245954 | 9.18219179 | 8.904861985 | 8.972631533 |
| ENSG00000088320 | REM1 | -3.753005537 | 0.531143362 | 3.843727009 | 0.48942971 | 0.090721472 | 2 | 2 | 2 | 2 | 2 | 3.5515855 | 4.673629022 |
| ENSG00000135108 | FBXO21 | -0.500315198 | 0.606861222 | 0.590937067 | 0.529397088 | 0.090621869 | 7.159102577 | 9.214490408 | 9.478569125 | 8.408172799 | 8.43695333 | 9.532891421 | 9.476418815 |
| ENSG00000159166 | LAD1 | -2.272292985 | 0.446344169 | 2.361568347 | 0.360634167 | 0.089275362 | 2 | 2 | 3.369317972 | 2.141737246 | 2.14022021 | 2.962551659 | 4.365060851 |
| ENSG00000167977 | KCTD5 | 0.202930992 | 0.843549372 | -0.116704837 | 0.910950051 | 0.086226155 | 9.567995545 | 7.35507212 | 9.426401271 | 9.221512599 | 9.31018819 | 8.971428833 | 8.926648946 |
| ENSG00000088451 | TGDS | 0.585439624 | 0.441837321 | -0.499295236 | 0.508321704 | 0.086144388 | 5.866944937 | 6.389896633 | 6.43197674 | 6.815074525 | 6.81514011 | 6.192262702 | 5.217771644 |
| ENSG00000177426 | TGIF1 | 0.715227142 | 0.649501644 | -0.629684135 | 0.688459235 | 0.085543007 | 11.03613077 | 7.538394143 | 7.285536013 | 10.36768823 | 10.3932137 | 9.095286533 | 8.987157359 |
| ENSG00000137055 | PLAA | 0.313375341 | 0.594131176 | -0.228329459 | 0.700693286 | 0.085045882 | 9.397585735 | 8.469613413 | 9.528916236 | 9.489414201 | 9.52849248 | 8.852100097 | 9.080880746 |
| ENSG00000113205 | PCDHB3 | 0.252871434 | 0.901083982 | -0.168269121 | 0.93349971 | 0.084602314 | 2 | 4.11352228 | 3.369317972 | 3.921164147 | 3.2184422 | 3.059555873 | 3.580696614 |
| ENSG00000116809 | ZBTB17 | 0.224745907 | 0.780077895 | -0.140358189 | 0.863604833 | 0.084387717 | 8.009475818 | 6.899080544 | 8.561105505 | 8.14116509 | 8.23084153 | 7.793159202 | 7.860656447 |
| ENSG00000198538 | ZNF28 | -0.051772415 | 0.959782419 | 0.136008115 | 0.895080873 | 0.0842357 | 6.563806279 | 7.572420273 | 8.699419242 | 7.810605345 | 7.81567127 | 8.259184498 | 7.678134128 |
| ENSG00000084093 | REST | 0.043534826 | 0.981829576 | 0.040471047 | 0.982913396 | 0.084005872 | 10.37373127 | 9.13702746 | 4.523675391 | 9.436230969 | 9.27923788 | 9.357396362 | 9.35623215 |
| ENSG00000106462 | EZH2 | 1.754839289 | 0.168980059 | -1.67232205 | 0.184671758 | 0.08251724 | 5.095959458 | 7.669933291 | 4.523675391 | 8.195011988 | 8.14604733 | 4.996409624 | 4.853519801 |

| | | | | | | | | | | | | | |
|---|---|---|---|---|---|---|---|---|---|---|---|---|---|
| ENSG00000169220 | RGS14 | -0.192835627 | 0.796574107 | 0.273048131 | 0.699954446 | 0.080212504 | 6.257048221 | 5.838937808 | 6.073378139 | 5.651002631 | 6.0722229 | 6.560755617 | 6.018717758 |
| ENSG00000181026 | AEN | -0.379036406 | 0.751922716 | 0.458997684 | 0.694619921 | 0.079961278 | 10.46903009 | 9.159587375 | 7.451345288 | 9.103012382 | 9.13268055 | 10.16889595 | 9.695099637 |
| ENSG00000013503 | POLR3B | 0.015723098 | 0.990899695 | 0.063929416 | 0.963128357 | 0.079652514 | 5.095959458 | 8.004992147 | 6.073378139 | 6.941624891 | 6.9210099 | 6.898949899 | 7.052989523 |
| ENSG00000146386 | ABRACL | 1.343802342 | 0.104122789 | -1.265648571 | 0.122892685 | 0.078153771 | 7.579279349 | 6.899080544 | 5.39244166 | 8.124184191 | 8.25995663 | 5.574182174 | 5.781546943 |
| ENSG00000109654 | TRIM2 | 1.111586373 | 0.067047287 | -1.033894582 | 0.087579018 | 0.077691791 | 7.384433967 | 7.572420273 | 6.203069917 | 8.321442665 | 8.21705076 | 6.151583464 | 6.222348126 |
| ENSG00000101003 | GINS1 | -0.119721355 | 0.956634366 | 0.195676188 | 0.930031781 | 0.075954834 | 3.327711234 | 6.535001748 | 2 | 4.946755474 | 5.19627118 | 5.294231737 | 5.419744333 |
| ENSG00000196214 | ZNF766 | -0.835896695 | 0.538896186 | 0.911815618 | 0.491920975 | 0.075918923 | 4.00722531 | 7.467835385 | 7.858115942 | 6.415275989 | 6.26836889 | 8.32468739 | 7.68646583 |
| ENSG00000188690 | UROS | -0.012446354 | 0.991325463 | 0.085741807 | 0.939676905 | 0.073295453 | 8.777722034 | 8.94259736 | 6.43197674 | 8.312101014 | 8.47717992 | 8.411091064 | 8.572538058 |
| ENSG00000198586 | TLK1 | 0.587415981 | 0.090299556 | -0.516934659 | 0.133943241 | 0.070481322 | 9.473095744 | 9.785372135 | 9.287079172 | 10.13796435 | 10.1004228 | 8.902412249 | 9.130578617 |
| ENSG00000179218 | CALR | 0.47090534 | 0.488092913 | -0.401092241 | 0.552503225 | 0.069813099 | 14.00486055 | 13.04571907 | 13.08372738 | 13.88981009 | 13.9512905 | 12.51624569 | 13.43750115 |
| ENSG00000076242 | MLH1 | 0.164459775 | 0.928883922 | -0.097245773 | 0.958016332 | 0.067214001 | 7.485143553 | 8.834582582 | 3.369317972 | 7.895617537 | 7.93841129 | 7.474865722 | 7.824443511 |
| ENSG00000169813 | HNRNPF | -0.316547493 | 0.511669208 | 0.382947352 | 0.408802956 | 0.066399859 | 10.6695979 | 10.51097332 | 9.819251342 | 10.08080308 | 10.0457438 | 10.73943283 | 10.7818607 |
| ENSG00000257923 | CUX1 | -0.120181579 | 0.80412149 | 0.185576553 | 0.687726485 | 0.065394974 | 9.992581382 | 10.68045884 | 10.09465441 | 10.14850007 | 10.1954158 | 10.38302632 | 10.56516425 |
| ENSG00000175387 | SMAD2 | -0.008842393 | 0.993765388 | 0.074070093 | 0.943575121 | 0.065227699 | 10.3933013 | 9.523645035 | 8.042706615 | 9.587916984 | 9.61516703 | 9.673618595 | 9.694509651 |
| ENSG00000158604 | TMED4 | -0.069277019 | 0.889181201 | 0.131845981 | 0.778721241 | 0.062568961 | 10.72212986 | 10.22575081 | 9.963514551 | 10.24492915 | 10.2934839 | 10.47230966 | 10.46753608 |
| ENSG00000198934 | MAGEE1 | -0.470179312 | 0.821622214 | 0.530848573 | 0.79391096 | 0.060669262 | 7.53297911 | 2 | 5.59507254 | 5.829220145 | 5.88556448 | 6.710164669 | 6.899677537 |
| ENSG00000116863 | ADPRHL2 | -0.269339926 | 0.709351398 | 0.329566777 | 0.63788893 | 0.060226851 | 9.371511179 | 8.101365653 | 8.486635645 | 8.474993083 | 8.4936155 | 8.776082909 | 9.327219816 |
| ENSG00000198156 | NPIPB6 | 3.412484602 | 0.002364079 | -3.352561433 | 0.0045879 | 0.059923168 | 4.00722531 | 3.076665209 | 5.156626401 | 7.412876874 | 7.37229174 | 2.656294034 | 2.225841198 |
| ENSG00000189057 | FAM111B | 0.061777076 | 0.982251788 | -0.004232548 | 0.998980237 | 0.057544528 | 2 | 2 | 4.058998049 | 3.227757282 | 2.77779907 | 2.831336763 | 3.128433649 |
| ENSG00000135535 | CD164 | 0.801733694 | 0.187195342 | -0.744753344 | 0.21592192 | 0.05698035 | 10.57565445 | 11.8974889 | 11.39316998 | 12.2479868 | 12.1255035 | 10.79309453 | 10.47751587 |
| ENSG00000153823 | PID1 | 2.844507483 | 1.38E-11 | -2.788501435 | 4.74E-11 | 0.056006048 | 8.444143041 | 8.451443573 | 8.865353883 | 11.43428179 | 11.4250965 | 6.210860813 | 5.486701579 |
| ENSG00000152683 | SLC30A6 | 0.133804407 | 0.798798132 | -0.077891002 | 0.884519325 | 0.055913405 | 9.344956677 | 8.505280662 | 8.784770914 | 9.060130342 | 9.03948698 | 8.792933324 | 8.885369555 |
| ENSG00000125818 | PSMF1 | -0.301844421 | 0.550280394 | 0.356520575 | 0.467088383 | 0.054676154 | 10.16946352 | 9.930175364 | 9.301632639 | 9.551067273 | 9.53961588 | 10.03328532 | 10.35107441 |
| ENSG00000008083 | JARID2 | 0.678435393 | 0.246800072 | -0.62386462 | 0.282300357 | 0.054570773 | 8.418903792 | 8.234834549 | 9.344432377 | 9.39612172 | 9.44390794 | 8.038228667 | 8.216759768 |
| ENSG00000104671 | DCTN6 | 0.572252845 | 0.585414444 | -0.517816635 | 0.619379532 | 0.054436209 | 8.998234871 | 7.315436877 | 6.629488387 | 8.593855195 | 8.53707846 | 7.38764416 | 7.59079099 |
| ENSG00000104660 | LEPROTL1 | -0.294151256 | 0.653912395 | 0.348545652 | 0.58421129 | 0.054394396 | 9.48530526 | 8.169643011 | 8.99637432 | 8.707945635 | 8.6601233 | 9.373994535 | 9.267780572 |
| ENSG00000205358 | MT1H | 5.244689346 | 0.254666268 | -5.191106557 | 0.288611054 | 0.053582789 | 2 | 2 | 2 | 3.349643765 | 6.22964624 | 2 | 2 |
| ENSG00000109270 | LAMTOR3 | 0.226051414 | 0.788297603 | -0.173338053 | 0.837335975 | 0.052713361 | 7.031698319 | 8.732943944 | 8.561105505 | 8.461536372 | 8.55598286 | 8.237320313 | 7.981656955 |
| ENSG00000180340 | FZD2 | -0.941041324 | 0.39960716 | 0.993614281 | 0.361984045 | 0.052572957 | 9.966689162 | 8.717821467 | 6.882775709 | 8.036145657 | 8.10812366 | 9.985285987 | 9.997333704 |
| ENSG00000256771 | ZNF253 | 0.60617024 | 0.655168404 | -0.553716899 | 0.682815119 | 0.052453342 | 4.00722531 | 7.274681896 | 5.59507254 | 6.841288255 | 6.73998187 | 5.976323695 | 5.343519008 |
| ENSG00000089775 | ZBTB25 | -0.224623135 | 0.916883733 | 0.276393186 | 0.896486957 | 0.051770051 | 2 | 5.946638857 | 8.143093325 | 6.682159616 | 6.59518404 | 7.324389982 | 6.889403418 |
| ENSG00000167981 | ZNF597 | 0.235758526 | 0.914584038 | -0.184395296 | 0.933063871 | 0.05136323 | 4.815609471 | 7.669933291 | 2 | 6.610808945 | 6.45450701 | 6.249924465 | 6.011156536 |
| ENSG00000143612 | C1orf43 | -0.15964596 | 0.897978564 | 0.210815293 | 0.861704178 | 0.051169393 | 12.07166289 | 10.17682307 | 9.344432377 | 10.8123265 | 10.8532269 | 11.15913757 | 11.24491309 |
| ENSG00000164182 | NDUFAF2 | 0.228877906 | 0.841054398 | -0.17892196 | 0.874995212 | 0.049955946 | 4.815609471 | 7.051611315 | 7.451345288 | 6.989292186 | 7.06873058 | 6.725028131 | 6.541257688 |
| ENSG00000173327 | MAP3K11 | -0.278785305 | 0.778392611 | 0.327819439 | 0.733748666 | 0.049034134 | 8.16936288 | 9.018611864 | 10.31150679 | 9.171766106 | 9.14209426 | 9.531042494 | 9.957809357 |
| ENSG00000196923 | PDLIM7 | -0.488793225 | 0.373373057 | 0.537153918 | 0.314029219 | 0.048360693 | 11.89987989 | 10.8599287 | 11.47570169 | 10.96228447 | 11.0056822 | 11.80075995 | 12.19032173 |
| ENSG00000100258 | LMF2 | 0.247878395 | 0.574576953 | -0.199926028 | 0.652472336 | 0.047952367 | 11.01943573 | 10.43902632 | 10.35401885 | 10.86374419 | 10.9005244 | 10.36223065 | 10.50536667 |
| ENSG00000108518 | PFN1 | -0.296046398 | 0.735002613 | 0.343934277 | 0.687843562 | 0.047887879 | 13.52950438 | 11.87039211 | 12.15221073 | 12.40121794 | 12.4293961 | 12.75624689 | 13.30233737 |
| ENSG00000105246 | EBI3 | -3.236747458 | 0.116637236 | 3.283447776 | 0.088390629 | 0.046700318 | 2 | 5.595927442 | 2 | 2.270786764 | 2.68924543 | 7.184873526 | 7.351304958 |
| ENSG00000196867 | ZFP28 | -0.681603878 | 0.417942235 | 0.727482504 | 0.3708938 | 0.045878627 | 7.790810999 | 5.946638857 | 7.225742779 | 6.634986713 | 6.35727082 | 8.04860508 | 7.669754031 |
| ENSG00000138375 | SMARCAL1 | 0.096570903 | 0.93008201 | -0.051458881 | 0.963377583 | 0.045112022 | 9.143983763 | 7.503546079 | 6.958202216 | 8.260583448 | 8.29037844 | 8.006643981 | 8.242819476 |
| ENSG00000186326 | RGS9BP | -1.269142941 | 0.673171288 | 1.314192505 | 0.638181881 | 0.045049564 | 2 | 3.076665209 | 2 | 2.3892346 | 2.14022021 | 3.316666208 | 2.981406154 |
| ENSG00000100105 | PATZ1 | -0.192719933 | 0.887105191 | 0.23763546 | 0.85594901 | 0.044915527 | 4.815609471 | 8.101365653 | 7.858115942 | 7.273514749 | 7.31425651 | 7.840848421 | 7.576773052 |
| ENSG00000138271 | GPR87 | 5.234806949 | 0.196836305 | -5.191118168 | 0.237913947 | 0.043688782 | 2 | 2 | 2 | 5.161714397 | 5.6121243 | 2 | 2 |
| ENSG00000079739 | PGM1 | -0.142188312 | 0.819577124 | 0.183949838 | 0.759271438 | 0.041761526 | 10.45668146 | 9.45234302 | 10.40208303 | 9.948991943 | 10.1009593 | 10.20380084 | 10.48640496 |
| ENSG00000100612 | DHRS7 | 1.36032436 | 0.021528722 | -1.319098561 | 0.025554781 | 0.041225799 | 8.836089686 | 9.055168103 | 10.08627303 | 10.80601808 | 10.7686428 | 8.264281212 | 7.971952287 |
| ENSG00000156531 | PHF6 | 0.02668943 | 0.983064817 | 0.014312686 | 0.99072569 | 0.041002116 | 6.009025997 | 8.747909548 | 8.742726175 | 8.343618883 | 8.240612 | 8.539610093 | 7.965119966 |
| ENSG00000122390 | NAA60 | -0.245891688 | 0.748322371 | 0.286443035 | 0.702091517 | 0.040551346 | 9.886101849 | 9.506147172 | 8.825066185 | 9.155143657 | 9.11153864 | 9.465620252 | 9.836041609 |
| ENSG00000163399 | ATP1A1 | 0.494224993 | 0.256360353 | -0.453699805 | 0.292481588 | 0.040525187 | 11.18203661 | 11.53295289 | 11.95457585 | 12.06794823 | 12.1009069 | 11.25282129 | 11.01284462 |
| ENSG00000213965 | NUDT19 | -0.271711096 | 0.770883416 | 0.309651774 | 0.73428898 | 0.037940678 | 7.53297911 | 8.57407404 | 6.534109959 | 7.484483106 | 7.56191311 | 8.047866378 | 8.139971339 |
| ENSG00000197150 | ABCB8 | 0.055154152 | 0.98047943 | -0.017271214 | 0.993931149 | 0.037882938 | 2 | 7.761270542 | 8.960138144 | 7.971157485 | 7.94559728 | 7.529862489 | 8.173382342 |
| ENSG00000116791 | CRYZ | -0.350658531 | 0.58220518 | 0.386734344 | 0.531194792 | 0.036075813 | 7.667647223 | 6.602424674 | 7.691296561 | 6.979883891 | 7.10768997 | 7.684242973 | 7.841598174 |
| ENSG00000138735 | PDE5A | -2.077168152 | 0.072123063 | 2.112488153 | 0.062801111 | 0.03532 | 7.33131396 | 10.24697497 | 7.691296561 | 7.030886362 | 6.97819215 | 11.36070623 | 10.90630802 |
| ENSG00000136826 | KLF4 | -0.899844595 | 0.309576645 | 0.934911781 | 0.278767895 | 0.035067186 | 7.940205782 | 7.761270542 | 9.624627483 | 7.778406195 | 7.84670947 | 9.674815254 | 9.581945714 |
| ENSG00000186575 | NF2 | -0.921481395 | 0.053285833 | 0.955799625 | 0.041273153 | 0.03431823 | 9.771132748 | 9.540933215 | 10.32581712 | 8.930605282 | 9.06500642 | 10.58238213 | 11.10707312 |
| ENSG00000163898 | LIPH | -2.968225821 | 0.197278637 | 3.001843828 | 0.167372108 | 0.033618007 | 2 | 2 | 5.59507254 | 2.270786764 | 2.77779907 | 6.104049656 | 7.411419562 |
| ENSG00000165533 | TTC8 | 0.920318672 | 0.629335817 | -0.886811352 | 0.638784658 | 0.03350732 | 8.16936288 | 8.337339476 | 2 | 8.607665593 | 8.57154947 | 6.812870125 | 6.826173076 |
| ENSG00000173264 | GPR137 | -0.607812403 | 0.072172046 | 0.639840174 | 0.051748984 | 0.032027771 | 8.797441463 | 9.055168103 | 8.742726175 | 8.178656739 | 8.36552655 | 9.385442197 | 9.62840116 |
| ENSG00000114904 | NEK4 | 0.398671978 | 0.739013584 | -0.367078515 | 0.758322907 | 0.031593463 | 5.70932728 | 6.228547981 | 8.236946975 | 7.626911583 | 7.44845993 | 6.940933331 | 6.635252003 |
| ENSG00000134759 | ELP2 | 0.001695854 | 0.997481474 | 0.027815915 | 0.963863059 | 0.029511769 | 9.233563228 | 9.821190352 | 8.865353883 | 9.342988948 | 9.39224836 | 9.536845448 | 9.234886737 |
| ENSG00000188725 | SMIM15 | 0.266076458 | 0.534490579 | -0.236746999 | 0.57774123 | 0.029329459 | 8.39321512 | 8.747909548 | 8.110401118 | 8.780186674 | 8.64254348 | 8.257907501 | 8.172535245 |
| ENSG00000244687 | UBE2V1 | 2.853630014 | 7.05E-08 | -2.824695714 | 1.20E-07 | 0.028934299 | 7.904282243 | 7.002537973 | 8.35328246 | 10.63786749 | 10.7125292 | 5.073924441 | 5.250250853 |
| ENSG00000196950 | SLC39A10 | -1.387512202 | 0.01604719 | 1.415518104 | 0.012677767 | 0.028005902 | 8.609439662 | 8.820493869 | 7.691296561 | 7.182120794 | 7.00136498 | 9.515096581 | 10.13979504 |
| ENSG00000038210 | PI4K2B | 1.006379072 | 0.030828564 | -0.97962898 | 0.035010376 | 0.026750091 | 6.816637977 | 7.274681896 | 7.734844356 | 8.310225403 | 8.30535212 | 6.302895343 | 6.443521721 |
| ENSG00000113300 | CNOT6 | 0.219908419 | 0.63056237 | -0.193411407 | 0.672104764 | 0.026497012 | 8.609439662 | 8.469613413 | 9.165043284 | 9.022822647 | 8.96128525 | 8.596789458 | 8.568679902 |
| ENSG00000134533 | RERG | -3.224023371 | 0.182596921 | 3.249411075 | 0.154661864 | 0.025387704 | 2 | 5.595927442 | 2 | 2.141737246 | 2.77779907 | 5.790986459 | 7.927454842 |
| ENSG00000092201 | SUPT16H | -0.035021336 | 0.972180135 | 0.060210061 | 0.952633805 | 0.025188725 | 10.29961392 | 10.28338212 | 8.236946975 | 9.864602909 | 9.80886355 | 9.868192425 | 9.992781936 |
| ENSG00000070718 | AP3M2 | 0.187464546 | 0.676328124 | -0.162463892 | 0.716074072 | 0.025000654 | 8.468948322 | 8.717821467 | 8.845626171 | 8.88597028 | 8.85716692 | 8.749978316 | 8.258072602 |
| ENSG00000125651 | GTF2F1 | 0.008618337 | 0.994208627 | 0.016231548 | 0.988291799 | 0.024849884 | 10.51739401 | 9.497318007 | 8.110401118 | 9.650037049 | 9.73059821 | 9.428927807 | 9.925491821 |
| ENSG00000134986 | NREP | -0.920685924 | 0.233103112 | 0.944815089 | 0.211911316 | 0.024129165 | 9.317904228 | 10.09725346 | 10.84117713 | 9.300161685 | 9.29807478 | 11.5927062 | 10.53268998 |
| ENSG00000185008 | ROBO2 | -1.915387485 | 0.346794529 | 1.939502735 | 0.321893604 | 0.02411525 | 3.327711234 | 5.457120953 | 3.369317972 | 3.709785322 | 2.14022021 | 4.914492341 | 6.847559257 |
| ENSG00000105419 | MEIS3 | 0.03782926 | 0.968309509 | -0.013764803 | 0.988584182 | 0.024064457 | 9.423197387 | 9.090820894 | 7.398165132 | 8.939126212 | 8.87232943 | 8.773404234 | 8.932164792 |
| ENSG00000171159 | C9orf16 | -0.411472194 | 0.616466414 | 0.435524494 | 0.587446222 | 0.0240523 | 9.128497779 | 7.572420273 | 9.287079172 | 8.392303091 | 8.46222608 | 9.428644092 | 9.085388321 |
| ENSG00000107223 | EDF1 | 0.247038642 | 0.603540731 | -0.223519937 | 0.635737899 | 0.023518705 | 10.83615308 | 10.07368926 | 10.5916593 | 10.7730345 | 10.7844255 | 10.14560825 | 10.45661716 |
| ENSG00000205250 | E2F4 | -0.09503982 | 0.941409766 | 0.113864925 | 0.930293595 | 0.018825105 | 8.928409675 | 9.656475478 | 6.43197674 | 8.658676732 | 8.84568954 | 8.927527801 | 8.996281974 |
| ENSG00000161533 | ACOX1 | -0.085763735 | 0.890295464 | 0.10450308 | 0.862199682 | 0.018739345 | 9.384607364 | 8.337339476 | 8.884815469 | 8.929383887 | 8.74768919 | 9.088477995 | 8.969221026 |
| ENSG00000175854 | SWI5 | -0.173691914 | 0.91115308 | 0.190284861 | 0.900816595 | 0.016592947 | 8.228665467 | 7.538394143 | 4.058998049 | 7.25413313 | 7.18663697 | 7.566427714 | 7.585709303 |
| ENSG00000073578 | SDHA | 0.128196666 | 0.871671212 | -0.113589152 | 0.885335383 | 0.014607514 | 10.3868074 | 9.102512015 | 9.116480286 | 9.773608662 | 9.81802855 | 9.649231086 | 9.454981667 |
| ENSG00000176024 | ZNF613 | -0.152765746 | 0.95872661 | 0.16674461 | 0.955794916 | 0.013978864 | 2 | 7.761270542 | 2 | 6.164317909 | 6.01871653 | 6.569017168 | 6.199243628 |
| ENSG00000164002 | EXO5 | 0.476387863 | 0.654150801 | -0.462585389 | 0.661465236 | 0.013802473 | 5.70932728 | 6.535001748 | 4.058998049 | 6.268449691 | 6.18985553 | 5.381251181 | 5.337485258 |
| ENSG00000140995 | DEF8 | -0.271183895 | 0.691499043 | 0.280227773 | 0.677268519 | 0.009043879 | 8.313285031 | 9.13702746 | 9.69246155 | 8.902248225 | 8.86476809 | 9.53894984 | 9.314605104 |
| ENSG00000164096 | C4orf3 | 0.247453559 | 0.800804384 | -0.238800439 | 0.804586396 | 0.008653119 | 11.02362764 | 9.055168103 | 10.75919294 | 10.77541564 | 10.7167355 | 9.763874267 | 10.63077694 |
| ENSG00000239789 | MRPS17 | 0.156115041 | 0.861781723 | -0.148134601 | 0.865271077 | 0.00798044 | 5.70932728 | 4.93638933 | 6.43197674 | 6.01571348 | 5.84514933 | 5.610655473 | 5.689801209 |
| ENSG00000261652 | C15orf65 | 0.059447946 | 0.982666154 | -0.051798017 | 0.9847789 | 0.007649928 | 2 | 2 | 5.772716771 | 4.358765249 | 4.45712836 | 4.369533167 | 4.26688425 |
| ENSG00000180694 | TMEM64 | -0.4232975 | 0.800895544 | 0.430174925 | 0.794249133 | 0.006877425 | 3.327711234 | 8.234834549 | 7.818184403 | 7.136275664 | 7.01053089 | 8.072046038 | 7.725108479 |
| ENSG00000166619 | BLCAP | -0.458332466 | 0.633070574 | 0.46472442 | 0.623383229 | 0.006391955 | 10.36053552 | 8.147241016 | 9.778213818 | 9.219515356 | 9.24275568 | 9.799418565 | 10.43134488 |
| ENSG00000156110 | ADK | -0.392679346 | 0.554575573 | 0.398472379 | 0.541021035 | 0.005793033 | 8.609439662 | 7.572420273 | 7.646393253 | 7.739909082 | 7.52080763 | 8.145010376 | 8.639663797 |
| ENSG00000183617 | MRPL54 | 0.139716208 | 0.829668812 | -0.133926852 | 0.833626336 | 0.005789356 | 7.709880939 | 8.234834549 | 7.225742779 | 7.949630994 | 7.91175216 | 7.422256313 | 7.868000534 |
| ENSG00000162928 | PEX13 | -0.110719781 | 0.92127192 | 0.115621079 | 0.917565744 | 0.004901298 | 7.867441304 | 7.002537973 | 9.358421373 | 8.350935846 | 8.23084153 | 8.595273498 | 8.431665941 |
| ENSG00000159692 | CTBP1 | 0.227414581 | 0.785042442 | -0.223888236 | 0.785920862 | 0.003526345 | 10.58702944 | 9.148351514 | 9.132850258 | 9.99525349 | 10.046858 | 9.585759327 | 9.558144718 |
| ENSG00000134697 | GNL2 | 0.107636163 | 0.915320238 | -0.104464283 | 0.917689829 | 0.003171881 | 10.34721797 | 8.903033433 | 8.325066185 | 9.571568433 | 9.54593381 | 9.230659332 | 9.456374363 |
| ENSG00000129219 | PLD2 | -0.409619152 | 0.686593442 | 0.412054733 | 0.680326628 | 0.002435581 | 8.075571721 | 8.029695761 | 5.772716771 | 7.25803034 | 7.17037177 | 7.922907177 | 8.110198799 |
| ENSG00000206418 | RAB12 | -0.12038533 | 0.807798742 | 0.122667827 | 0.800369309 | 0.002282497 | 9.410448395 | 8.876041558 | 8.699419242 | 8.924487942 | 8.88981993 | 9.263482089 | 9.023314583 |
| ENSG00000177150 | FAM210A | 0.09577006 | 0.853531206 | -0.094616071 | 0.85221514 | 0.001153988 | 7.485143553 | 7.954278496 | 7.285536013 | 7.703217906 | 7.71833702 | 7.595026956 | 7.449681889 |
| ENSG00000187741 | FANCA | 0.48459663 | 0.739704558 | -0.483929471 | 0.736888343 | 0.00066716 | 5.095959458 | 5.595927442 | 2 | 5.194621444 | 5.41912973 | 4.369533167 | 4.574506939 |
| ENSG00000090520 | DNAJB11 | 0.824184447 | 0.129332125 | -0.827672174 | 0.121602573 | -0.003487727 | 9.931427102 | 9.981728319 | 9.031722608 | 10.5182619 | 10.5464078 | 8.594767824 | 9.133193298 |
| ENSG00000151806 | GUF1 | -0.021509767 | 0.98350331 | 0.017711322 | 0.986497228 | -0.003798445 | 6.563806279 | 8.469613413 | 6.958202216 | 7.53353831 | 7.60490877 | 7.815031728 | 7.366199662 |
| ENSG00000077147 | TM9SF3 | 0.248699125 | 0.824376979 | -0.25564107 | 0.815972748 | -0.006941945 | 9.533135543 | 11.14353618 | 12.21698408 | 11.61835275 | 11.5523437 | 11.12496121 | 11.03806084 |
| ENSG00000104415 | WISP1 | 0.306557854 | 0.879440766 | -0.313755922 | 0.873635383 | -0.007198068 | 7.82963488 | 6.78762695 | 2 | 7.136275664 | 7.1246732 | 6.019162896 | 6.909879006 |
| ENSG00000164751 | PEX2 | -0.365502806 | 0.428026621 | 0.357342936 | 0.427508497 | -0.00815987 | 8.564112515 | 8.487557258 | 7.818184403 | 7.956842238 | 7.98565528 | 8.615356492 | 8.751666251 |
| ENSG00000159592 | GPBP1L1 | 0.174020261 | 0.852628537 | -0.18249688 | 0.842165885 | -0.008476619 | 8.586954094 | 9.672251278 | 10.60345421 | 10.06865021 | 9.96140527 | 9.71002767 | 9.610373471 |
| ENSG00000115677 | HDLBP | -0.400608993 | 0.490606244 | 0.391938251 | 0.492233435 | -0.008670742 | 12.46905053 | 13.35763551 | 13.55612433 | 12.77838142 | 12.8186338 | 13.41539496 | 13.74726846 |
| ENSG00000083535 | PIBF1 | 0.203374661 | 0.831652978 | -0.213591031 | 0.818512537 | -0.01021637 | 7.27616306 | 7.232742069 | 5.59507254 | 7.114945424 | 7.05992818 | 7.015370516 | 6.254725615 |
| ENSG00000164163 | ABCE1 | 0.087257416 | 0.93880631 | -0.097800389 | 0.932122597 | -0.010542973 | 7.27616306 | 9.549500232 | 7.734844356 | 8.639384849 | 8.62772819 | 8.441219213 | 8.459712934 |
| ENSG00000081154 | PCNP | -0.114738178 | 0.887076711 | 0.103904364 | 0.895904824 | -0.010833813 | 10.29961392 | 10.019219 | 8.742726175 | 9.751662842 | 9.67691814 | 10.03403152 | 9.823144976 |
| ENSG00000095627 | TDRD1 | -3.753006835 | 0.531143362 | 3.739779159 | 0.502368069 | -0.013227677 | 2 | 2 | 2 | 2 | 2 | 4.73522696 | 3.100202476 |
| ENSG00000134909 | ARHGAP32 | 0.25841915 | 0.772399916 | -0.272656027 | 0.757050516 | -0.014236878 | 6.563806279 | 8.124485672 | 6.629488387 | 7.590284224 | 7.5462416 | 7.083273719 | 7.017577533 |

| | | | | | | | | | | | | | |
|---|---|---|---|---|---|---|---|---|---|---|---|---|---|
| ENSG00000187240 | DYNC2H1 | 0.747023097 | 0.622552387 | -0.763066631 | 0.609510346 | -0.016043533 | 5.532353208 | 8.234834549 | 4.523675391 | 7.778406195 | 7.59580267 | 6.034159474 | 6.403650353 |
| ENSG00000247596 | TWF2 | 0.229905916 | 0.858507416 | -0.246811198 | 0.844695037 | -0.016905281 | 10.02639697 | 7.790474954 | 7.342949392 | 9.115954467 | 9.12426093 | 8.679237241 | 8.614934151 |
| ENSG00000159788 | RGS12 | 0.025756057 | 0.978514936 | -0.043143322 | 0.964338803 | -0.017387265 | 7.031698319 | 6.728495673 | 8.408111533 | 7.584088227 | 7.61995902 | 7.669911566 | 7.385335693 |
| ENSG00000108829 | LRRC59 | -0.18812856 | 0.744523835 | 0.168125379 | 0.770703322 | -0.020003181 | 11.58798587 | 10.55418374 | 11.22775303 | 10.98773739 | 11.0025273 | 11.20375836 | 11.48422157 |
| ENSG00000111247 | RAD51AP1 | 0.129754643 | 0.95872661 | -0.151641092 | 0.952529433 | -0.021886449 | 2 | 5.457120953 | 2 | 4.329433326 | 4.1930041 | 3.981090143 | 4.105828626 |
| ENSG00000157388 | CACNA1D | -2.312760619 | NA | 2.290327653 | NA | -0.022432966 | 2 | 2 | 2 | 2 | 2.14022021 | 3.355392153 | 2.950113106 |
| ENSG00000185418 | TARSL2 | -0.706561556 | 0.05996039 | 0.683960643 | 0.06220751 | -0.022600912 | 7.750913381 | 8.029695761 | 7.734844356 | 7.110641269 | 7.21069602 | 8.389678938 | 8.652479133 |
| ENSG00000281763 | | 0.968011833 | 0.184484665 | -0.991503157 | 0.165663949 | -0.023491325 | 7.384433967 | 7.954278496 | 9.049077186 | 9.294480609 | 9.21229508 | 7.346184975 | 7.296373648 |
| ENSG00000080854 | IGSF9B | -2.632918738 | 0.274396109 | 2.608273723 | 0.252282956 | -0.024645015 | 2 | 2 | 5.156626401 | 2.784612485 | 2.14022021 | 5.944926241 | 6.151894053 |
| ENSG00000183762 | KREMEN1 | -1.813818676 | 0.151077965 | 1.788647565 | 0.146121333 | -0.02517111 | 4.815609471 | 4.93638933 | 7.691296561 | 4.71689672 | 4.87352701 | 8.261097876 | 8.126912071 |
| ENSG00000143093 | STRIP1 | 0.125951833 | 0.832844524 | -0.151303335 | 0.79391096 | -0.025351503 | 8.540903468 | 8.791896851 | 9.503962296 | 9.088859521 | 9.16485023 | 8.734912119 | 8.960904642 |
| ENSG00000278312 | | 5.165535877 | 0.209831388 | -5.191117205 | 0.243536295 | -0.025581327 | 2 | 2 | 2 | 5.420062284 | 5.2591458 | 2 | 2 |
| ENSG00000111707 | SUDS3 | -0.167811788 | 0.831652978 | 0.141553566 | 0.855655091 | -0.026258222 | 8.16936288 | 8.806266215 | 9.714382178 | 8.830780654 | 8.90464485 | 9.201064633 | 9.148782641 |
| ENSG00000197006 | METTL9 | 0.393840225 | 0.316169441 | -0.420217078 | 0.273363547 | -0.026376853 | 10.54098126 | 10.47544827 | 9.907528187 | 10.71680073 | 10.7410309 | 9.900262774 | 9.934514013 |
| ENSG00000178229 | ZNF543 | 0.140462711 | 0.906603349 | -0.167352161 | 0.886082448 | -0.02688945 | 4.815609471 | 4.93638933 | 6.80318718 | 5.949776857 | 5.92487824 | 5.699879814 | 5.601925227 |
| ENSG00000213463 | SYNJ2BP | -0.473686445 | 0.595799399 | 0.445396054 | 0.614013611 | -0.028290391 | 7.435667355 | 9.079034257 | 9.528916236 | 8.545247918 | 8.34936267 | 9.365719319 | 9.358096047 |
| ENSG00000115289 | PCGF1 | -0.11974355 | 0.903607792 | 0.090966367 | 0.926534356 | -0.028777182 | 6.009025997 | 6.844429755 | 5.156626401 | 6.139181293 | 6.00960265 | 6.550361701 | 5.933281326 |
| ENSG00000186715 | MST1L | 2.625926345 | 0.180152411 | -2.656164129 | 0.189944554 | -0.030237784 | 3.327711234 | 3.076665209 | 2 | 4.647285135 | 4.76938658 | 2.3070652 | 2.059860178 |
| ENSG00000114850 | SSR3 | -0.135197821 | 0.767206549 | 0.104060828 | 0.820771665 | -0.031136993 | 11.98618033 | 11.72292201 | 12.41570295 | 11.95199324 | 11.9166432 | 12.0510638 | 12.28629031 |
| ENSG00000126934 | MAP2K2 | -0.259423584 | 0.539543786 | 0.227592017 | 0.588973243 | -0.031831568 | 10.5468183 | 9.962611189 | 10.11127249 | 9.922839878 | 10.0130538 | 10.28941421 | 10.6018087 |
| ENSG00000112214 | FHL5 | -2.31280349 | NA | 2.280676918 | NA | -0.032126572 | 2 | 2 | 2 | 2 | 2.14022021 | 3.724264246 | 2.225841198 |
| ENSG00000215784 | FAM72D | 5.157088758 | 0.211381873 | -5.191117084 | 0.244323998 | -0.034028325 | 2 | 2 | 2 | 5.363304456 | 5.30456641 | 2 | 2 |
| ENSG00000101400 | SNTA1 | -0.929869578 | 0.679377579 | 0.895198777 | 0.686960124 | -0.034670802 | 2 | 7.35507212 | 9.888372618 | 7.54315232 | 7.6957855 | 9.441918755 | 9.408611581 |
| ENSG00000117691 | NENF | 0.042761786 | 0.930551849 | -0.077769508 | 0.871169659 | -0.035007722 | 9.304185368 | 8.590771778 | 9.149036566 | 9.04103174 | 9.12214832 | 8.969089651 | 8.955990164 |
| ENSG00000197037 | ZSCAN25 | -0.255123004 | 0.870868026 | 0.219695388 | 0.887741643 | -0.035427616 | 4.00722531 | 7.051611315 | 8.266922072 | 7.035434774 | 6.96410766 | 7.401588825 | 7.520630205 |
| ENSG00000108651 | UTP6 | 0.153781289 | 0.850026272 | -0.190546327 | 0.808588735 | -0.036765038 | 9.031919741 | 8.101365653 | 7.451345288 | 8.481674661 | 8.50338773 | 8.043426202 | 8.25407423 |
| ENSG00000100554 | ATP6V1D | 0.240566546 | 0.875544825 | -0.278305839 | 0.851903823 | -0.037739293 | 10.60951384 | 7.954278496 | 7.098165388 | 9.60255057 | 9.55848727 | 8.924311548 | 9.193312432 |
| ENSG00000197451 | HNRNPAB | -0.038120033 | 0.970008162 | -0.00130432 | 0.99924195 | -0.039424353 | 10.78274301 | 9.225224669 | 8.805342542 | 9.847892119 | 9.81476202 | 9.723505443 | 9.999484803 |
| ENSG00000040608 | RTN4R | -2.113096077 | 0.213019329 | 2.073560684 | 0.192445639 | -0.039535393 | 3.327711234 | 4.93638933 | 2 | 2.784612485 | 2.68924543 | 6.247352872 | 4.870334174 |
| ENSG00000167136 | ENDOG | 0.129521566 | 0.946720822 | -0.171400072 | 0.931410087 | -0.041878505 | 2 | 6.464272742 | 7.691296561 | 6.70517998 | 6.81514011 | 6.525106458 | 6.418016952 |
| ENSG00000129680 | MAP7D3 | -0.559199719 | 0.362105114 | 0.516598619 | 0.391760019 | -0.0426011 | 8.257427002 | 7.099070191 | 8.110401118 | 7.288834728 | 7.40393901 | 8.271892681 | 8.526208204 |
| ENSG00000100749 | VRK1 | 0.339892743 | 0.764551483 | -0.383369704 | 0.729041892 | -0.043476961 | 5.866944937 | 5.838937808 | 3.369317972 | 5.742861563 | 5.70538658 | 5.008609438 | 5.12973572 |
| ENSG00000099337 | KCNK6 | -1.644887264 | 0.000817299 | 1.601178555 | 0.001008927 | -0.043708709 | 9.410448395 | 8.687092496 | 8.721235202 | 7.381065997 | 7.33264456 | 10.18174778 | 10.87774805 |
| ENSG00000079931 | MOXD1 | 1.347517964 | 0.150716856 | -1.393053101 | 0.129597634 | -0.045535138 | 7.218819781 | 5.838937808 | 7.934808822 | 8.590768181 | 8.49688027 | 6.329884754 | 5.250250853 |
| ENSG00000106049 | HIBADH | -0.161514578 | 0.867570088 | 0.115975185 | 0.904611962 | -0.045539394 | 9.204317094 | 9.267380431 | 7.398165132 | 8.615281294 | 8.74768919 | 8.630730772 | 9.224707098 |
| ENSG00000113638 | TTC33 | 0.204126551 | 0.890526182 | -0.251110802 | 0.861440521 | -0.046984251 | 4.00722531 | 7.232742069 | 7.691296561 | 7.161462517 | 7.15392111 | 7.031901364 | 6.314310122 |
| ENSG00000100938 | GMPR2 | -0.452442272 | 0.837958929 | 0.405388112 | 0.852566533 | -0.04705416 | 9.448362284 | 8.993716073 | 2 | 8.207157886 | 8.21705076 | 8.908121842 | 9.199563099 |
| ENSG00000007923 | DNAJC11 | 0.590114964 | 0.304803313 | -0.637567816 | 0.255124481 | -0.047452852 | 8.777722034 | 8.077869079 | 7.734844356 | 8.772033429 | 8.91688363 | 7.422256313 | 7.821204179 |
| ENSG00000128626 | MRPS12 | -0.282336713 | 0.568853498 | 0.234569271 | 0.635038067 | -0.047767441 | 7.435667355 | 8.124485672 | 8.206335842 | 7.723090567 | 7.64959655 | 8.094396944 | 8.289666801 |
| ENSG00000175224 | ATG13 | -0.016703666 | 0.977090174 | -0.031829675 | 0.956622971 | -0.048533341 | 10.72727936 | 10.23640192 | 9.735974718 | 10.22266632 | 10.3213737 | 10.28691212 | 10.22805302 |
| ENSG00000131459 | GFPT2 | 2.066295679 | 1.75E-08 | -2.114839439 | 7.94E-09 | -0.048543761 | 8.981092835 | 9.540933215 | 9.465702942 | 11.38256115 | 11.4395901 | 7.475964489 | 7.019463152 |
| ENSG00000170604 | IRF2BP1 | -0.587530217 | 0.256335494 | 0.537509834 | 0.292216612 | -0.050020383 | 8.257427002 | 8.337339476 | 9.099922433 | 8.015574133 | 8.04930085 | 9.301474629 | 8.970683659 |
| ENSG00000039068 | CDH1 | -2.266386616 | 0.514865066 | 2.215885979 | 0.479004141 | -0.050500637 | 2 | 2 | 3.369317972 | 2.270786764 | 2 | 3.256555576 | 4.077147165 |
| ENSG00000143839 | REN | 0.532894795 | 0.86129847 | -0.583481377 | NA | -0.050586582 | 2 | 3.076665209 | 2 | 2.600425781 | 2.94009902 | 2.492221014 | 2.326754077 |
| ENSG00000083544 | TDRD3 | -0.215708993 | 0.838446668 | 0.165115452 | 0.875759484 | -0.050593541 | 6.009025997 | 7.819099893 | 6.073378139 | 6.598566398 | 6.80989959 | 7.203566521 | 6.936070085 |
| ENSG00000139726 | DENR | 0.048379803 | 0.953297746 | -0.099039498 | 0.904744852 | -0.050659695 | 7.940205782 | 9.125613817 | 9.61300511 | 9.138327453 | 9.05063789 | 8.844037986 | 9.046617405 |
| ENSG00000197429 | IPP | -0.620632979 | 0.786839864 | 0.569907744 | 0.80197737 | -0.050725235 | 2 | 4.93638933 | 8.511888463 | 6.542151749 | 6.36443784 | 7.655436363 | 7.561323797 |
| ZASP | | 5.139299599 | 0.216431863 | -5.19111649 | 0.247207591 | -0.051816891 | 2 | 2 | 2 | 5.110902178 | 5.49945122 | 2 | 2 |
| ENSG00000131508 | UBE2D2 | -0.001591732 | 0.99707802 | -0.050689183 | 0.900190897 | -0.052280915 | 10.19941743 | 9.849218119 | 9.670202709 | 9.95740528 | 9.88434767 | 9.918562269 | 9.824763374 |
| ENSG00000104863 | LIN7B | -0.604037908 | 0.823403844 | 0.551374774 | 0.836490254 | -0.052663133 | 2 | 6.666836837 | 2 | 4.623314311 | 4.74762503 | 5.812021848 | 5.606950214 |
| ENSG00000132837 | DMGDH | -3.753007336 | 0.531143362 | 3.699693042 | 0.507814827 | -0.053314294 | 2 | 2 | 2 | 2 | 2 | 4.177580368 | 4.033026654 |
| ENSG00000103490 | PYCARD | 1.128472467 | 0.042046869 | -1.181974984 | 0.031574569 | -0.053502517 | 6.009025997 | 6.311476754 | 6.80318718 | 7.484483106 | 7.50792049 | 5.50697268 | 5.072762365 |
| ENSG00000140259 | MFAP1 | 0.099816723 | 0.812768397 | -0.153390516 | 0.701056475 | -0.053573793 | 8.816894989 | 8.607278467 | 9.149036566 | 8.95000826 | 8.98589128 | 8.640073371 | 8.790280628 |
| ENSG00000133315 | MACROD1 | -1.333178413 | 0.378770423 | 1.27920325 | 0.385805831 | -0.053975163 | 5.532353208 | 6.464272742 | 2 | 4.299492648 | 4.4836996 | 6.950450741 | 6.533361209 |
| ENSG00000122643 | NT5C3A | 2.346825624 | 1.72E-05 | -2.401265782 | 1.16E-05 | -0.054440159 | 7.790810999 | 8.004992147 | 6.629488387 | 9.976453497 | 9.86489641 | 5.385935071 | 5.250250853 |
| ENSG00000109618 | SEPSECS | 0.455770577 | 0.825685372 | -0.510241723 | 0.800237157 | -0.054471145 | 7.867441304 | 5.72254323 | 2 | 7.017154451 | 7.05107174 | 6.473229652 | 5.621921006 |
| ENSG00000180304 | OAZ2 | -0.846487328 | 0.295027628 | 0.789785345 | 0.321533624 | -0.056701983 | 10.73241055 | 10.04368224 | 8.784770914 | 9.240349764 | 9.19128891 | 10.62881333 | 11.03421896 |
| ENSG00000205133 | TRIQK | -0.606508367 | 0.75782421 | 0.548925663 | 0.779935476 | -0.057582704 | 8.418903792 | 7.790474954 | 2 | 6.993973425 | 6.95464102 | 8.195910056 | 8.006587762 |
| ENSG00000125037 | EMC3 | 0.653337446 | 0.357081274 | -0.711368549 | 0.302366413 | -0.058031103 | 10.05943807 | 8.590771778 | 9.06622548 | 10.02414665 | 10.0164701 | 8.458554046 | 8.8444706 |
| ENSG00000181896 | ZNF101 | 0.072479198 | 0.936390504 | -0.131175177 | 0.883777495 | -0.058695978 | 7.27616306 | 5.72254323 | 6.073378139 | 6.529309835 | 6.61935067 | 6.412547789 | 6.347114043 |
| ENSG00000156011 | PSD3 | -0.608683441 | 0.770602177 | 0.549577902 | 0.792987538 | -0.059105539 | 9.112843761 | 8.639735838 | 2 | 7.783822874 | 7.64959655 | 9.001879477 | 8.705562554 |
| ENSG00000183454 | GRIN2A | 3.499811463 | 0.072123063 | -3.561167325 | 0.069213816 | -0.061355862 | 2 | 5.72254323 | 2 | 7.584088227 | 7.6378145 | 2.225793514 | 2.632958241 |
| ENSG00000103018 | CYB5B | 0.116503589 | 0.873933481 | -0.178752917 | 0.798439725 | -0.062249328 | 9.646202137 | 10.13189344 | 8.904018013 | 9.793884011 | 9.73406074 | 9.116578627 | 9.754611319 |
| ENSG00000136878 | USP20 | -0.513326422 | 0.845989871 | 0.450274894 | 0.863507758 | -0.063051528 | 2 | 5.303523747 | 10.03493575 | 8.013270196 | 7.98797735 | 9.104551714 | 8.784193542 |
| ENSG00000197756 | RPL37A | -0.305413301 | 0.729728652 | 0.241634951 | 0.786775559 | -0.06377835 | 12.02014453 | 10.25223258 | 10.59756881 | 10.81431291 | 10.9173739 | 11.34169508 | 11.4806329 |
| ENSG00000169857 | AVEN | -0.543240672 | 0.728541941 | 0.47892818 | 0.759498144 | -0.064312492 | 5.095959458 | 8.053983481 | 9.767769415 | 8.040677544 | 8.10170499 | 9.075126337 | 9.09033044 |
| ENSG00000119147 | C2orf40 | -3.753007415 | 0.531143362 | 3.688559532 | 0.509267049 | -0.064447943 | 2 | 2 | 2 | 2 | 2 | 3.679182183 | 4.4231929 |
| ENSG00000137764 | MAP2K5 | -0.458516318 | 0.820634582 | 0.393897343 | 0.843924153 | -0.064618975 | 6.257048221 | 8.029695761 | 2 | 6.470063213 | 6.32824132 | 7.32316944 | 7.114266974 |
| ENSG00000108852 | MPP2 | -0.704735147 | 0.581619703 | 0.638581505 | 0.614736828 | -0.066153641 | 8.737455672 | 6.728495673 | 5.59507254 | 6.941624891 | 6.87658492 | 7.947671484 | 8.454843905 |
| ENSG00000254535 | PABPC4L | -0.764698938 | 0.749679501 | 0.697931827 | 0.767692243 | -0.066767111 | 2 | 4.93638933 | 2 | 3.349643765 | 3.15372745 | 4.628601098 | 3.789585079 |
| ENSG00000164941 | INTS8 | 0.13935577 | 0.91483416 | -0.207388302 | 0.870629355 | -0.068032532 | 7.384433967 | 6.311476754 | 9.149036566 | 8.215198712 | 8.21705076 | 8.145700987 | 7.540464021 |
| ENSG00000124535 | WRNIP1 | -0.404514607 | 0.466549268 | 0.334573672 | 0.544273144 | -0.069940935 | 9.646202137 | 8.671479029 | 9.399592459 | 8.902248225 | 8.87609528 | 9.808387568 | 9.411127426 |
| ENSG00000146278 | PNRC1 | 0.150142013 | 0.903601303 | -0.220966631 | 0.853313931 | -0.070824618 | 10.91499272 | 10.02537381 | 7.896971941 | 10.22017133 | 10.2074225 | 9.80073459 | 9.886246986 |
| ENSG00000169570 | DTWD2 | -0.572296878 | 0.479814084 | 0.500615318 | 0.527954208 | -0.07168156 | 5.095959458 | 5.595927442 | 6.629488387 | 5.348759107 | 5.40529704 | 6.358763086 | 6.392053169 |
| ENSG00000134028 | ADAMDEC1 | -3.753007591 | 0.531143362 | 3.679246468 | 0.51059361 | -0.073761123 | 2 | 2 | 2 | 2 | 2 | 4.792794928 | 2.671842496 |
| ENSG00000240583 | AQP1 | -2.57392445 | 0.015963161 | 2.498910616 | 0.009732169 | -0.075013834 | 3.327711234 | 4.442710129 | 5.156626401 | 2.868578001 | 2.77779907 | 7.091899617 | 6.301968891 |
| AP5S1 | AP5S1 | -0.591292888 | 0.621621222 | 0.512578007 | 0.668751914 | -0.078714881 | 8.257427002 | 7.099070191 | 5.156626401 | 6.761173376 | 6.72891807 | 7.557113378 | 8.042267212 |
| ENSG00000125846 | ZNF133 | 0.076002756 | 0.965988399 | -0.155705956 | 0.931082222 | -0.0797032 | 3.327711234 | 8.505280662 | 8.175061096 | 7.880528731 | 7.84670947 | 7.812424466 | 7.435628703 |
| ENSG00000162946 | DISC1 | -0.308759446 | 0.803672304 | 0.228982632 | 0.853313931 | -0.079776814 | 4.467311984 | 7.189546432 | 5.930865648 | 5.900763176 | 6.06344184 | 6.173421703 | 6.755525668 |
| ENSG00000139746 | RBM26 | 0.210812497 | 0.785196727 | -0.290969555 | 0.695197299 | -0.080157058 | 7.624139768 | 8.414402626 | 9.180874355 | 8.727753724 | 8.77079897 | 8.3832514 | 8.110198799 |
| ENSG00000134901 | KDELC1 | -0.200735699 | 0.88827382 | 0.119665379 | 0.933175306 | -0.08107032 | 9.233563228 | 7.467835385 | 5.772716771 | 7.90310317 | 7.93600799 | 8.407640344 | 8.039482273 |
| ENSG00000162736 | NCSTN | 0.303259433 | 0.760220023 | -0.384710651 | 0.690174836 | -0.081451218 | 11.77862605 | 10.30378101 | 9.725218844 | 11.1637228 | 11.1856376 | 10.15661401 | 10.75777696 |
| ENSG00000164088 | PPM1M | -1.037700595 | 0.602587452 | 0.956219249 | 0.628501113 | -0.081481346 | 8.998234871 | 7.819099893 | 2 | 6.851641821 | 7.0150921 | 9.012898235 | 8.764090674 |
| ENSG00000148719 | DNAJB12 | 0.286929743 | 0.658476741 | -0.369110368 | 0.554801672 | -0.082180625 | 8.468948322 | 9.7032946 | 9.491321578 | 9.596407142 | 9.60461324 | 8.797335126 | 9.085838304 |
| ENSG00000198055 | GRK6 | -0.438010624 | 0.813586965 | 0.355269764 | 0.847870601 | -0.08274086 | 7.624139768 | 9.235879652 | 3.369317972 | 7.602596956 | 7.6986238 | 8.458554046 | 8.405948829 |
| ENSG00000175505 | CLCF1 | 0.689730479 | 0.702820767 | -0.775036065 | 0.661877164 | -0.085305586 | 10.19198706 | 6.464272742 | 5.39244166 | 9.418058443 | 9.46739696 | 7.794921259 | 8.166591596 |
| ENSG00000196417 | ZNF765 | 0.221614833 | 0.898841046 | -0.308023976 | 0.855423548 | -0.086409143 | 4.00722531 | 5.838937808 | 8.35328246 | 7.315259362 | 7.22257674 | 6.880813179 | 6.620419619 |
| ENSG00000133131 | MORC4 | -1.099311311 | 0.581310291 | 1.010371243 | 0.609366282 | -0.088940068 | 2 | 8.607278467 | 8.978369997 | 7.101994243 | 7.19470132 | 9.277728661 | 9.174819346 |
| ENSG00000143971 | ETAA1 | -0.25784823 | 0.722734802 | 0.16888198 | 0.818792684 | -0.08896625 | 6.89194353 | 6.844429755 | 7.97167868 | 7.080147179 | 7.06873058 | 7.682340325 | 7.268103222 |
| ENSG00000069329 | VPS35 | 0.12725414 | 0.723898621 | -0.216267428 | 0.524504404 | -0.089013289 | 10.1076228 | 10.5284131 | 10.47451889 | 10.55079452 | 10.4683133 | 10.25622378 | 10.07327514 |
| ENSG00000014216 | CAPN1 | 0.058958566 | 0.931034311 | -0.149087597 | 0.81957996 | -0.090129031 | 10.50545418 | 9.969011754 | 11.10036136 | 10.65089438 | 10.6603451 | 10.16770645 | 10.68286652 |
| ENSG00000263513 | FAM72C | 5.09949797 | 0.224142044 | -5.19111587 | 0.250703287 | -0.0916179 | 2 | 2 | 2 | 5.406080789 | 5.14724905 | 2 | 2 |
| ENSG00000129696 | TTI2 | -0.098629479 | 0.932710628 | 0.006621213 | 0.995596508 | -0.092008266 | 4.815609471 | 7.431218269 | 7.225742779 | 6.733448311 | 6.82556437 | 6.857395343 | 6.905807071 |
| ENSG00000164237 | CMBL | -0.777166665 | 0.148518474 | 0.684913234 | 0.196709806 | -0.092253432 | 8.285626337 | 7.315436877 | 7.858115942 | 7.066878356 | 7.14145882 | 8.725241966 | 8.326688609 |
| ENSG00000185340 | GAS2L1 | -0.738784008 | 0.511176757 | 0.645891117 | 0.562211 | -0.092892891 | 9.761125267 | 8.414402626 | 6.80318718 | 8.029321023 | 8.07355593 | 9.468935198 | 9.375131848 |
| ENSG00000164172 | MOCS2 | -0.91551534 | 0.02176637 | 0.820962917 | 0.038452755 | -0.094552423 | 8.009475818 | 8.276714465 | 8.742726175 | 7.474468681 | 7.46855365 | 9.148628127 | 9.220615045 |
| ENSG00000176978 | DPP7 | 0.248337997 | 0.55246072 | -0.344003024 | 0.388543168 | -0.095665027 | 9.657036304 | 9.090820894 | 9.658943165 | 9.74820345 | 9.72155655 | 9.242974864 | 9.041986855 |
| ENSG00000163293 | NIPAL1 | -0.552528815 | 0.838100556 | 0.455035109 | 0.863507947 | -0.097493706 | 2 | 2 | 4.523675391 | 3.162721474 | 2.86123014 | 3.059556873 | 4.018014792 |
| ENSG00000089012 | SIRPG | 3.688652407 | 0.324700963 | -3.788106481 | 0.356167588 | -0.099454074 | 2 | 2 | 2 | 3.997744386 | 4.1930041 | 2.048074405 | 2 |
| LOC391322 | | 5.090365617 | 0.225518037 | -5.191115802 | 0.251137486 | -0.100750185 | 2 | 2 | 2 | 5.304221918 | 5.24368216 | 2 | 2 |
| ENSG00000079332 | SAR1A | 0.037020759 | 0.946595613 | -0.138302831 | 0.796658198 | -0.101282072 | 11.52728203 | 10.72273565 | 10.86096535 | 11.10164114 | 11.1336271 | 10.76122774 | 11.1039664 |
| ENSG00000102893 | PHKB | -0.337609616 | 0.734347184 | 0.235865202 | 0.815278413 | -0.101744414 | 7.435667355 | 8.848535039 | 9.647595056 | 8.540455898 | 8.59458893 | 8.710380719 | 9.464702422 |
| ENSG00000108848 | LUC7L3 | 0.280348145 | 0.688878059 | -0.38289239 | 0.568416252 | -0.102544245 | 9.097018025 | 10.47544827 | 10.24529339 | 10.31347779 | 10.3523439 | 9.798979623 | 9.533614641 |
| ENSG00000140650 | PMM2 | 1.229317439 | 0.457169403 | -1.332963721 | 0.406143499 | -0.103646281 | 4.815609471 | 9.159587375 | 5.772716771 | 8.914645924 | 9.06280517 | 6.057834398 | 6.811011855 |
| ENSG00000173786 | CNP | 0.536250022 | 0.166335577 | -0.639987179 | 0.087003979 | -0.103737156 | 11.44840376 | 10.81088865 | 10.92346452 | 11.61683606 | 11.629295 | 10.43987727 | 10.45835585 |
| ENSG00000172936 | MYD88 | 0.667055738 | 0.524245263 | -0.771631147 | 0.445282505 | -0.104575409 | 10.87842776 | 8.317412869 | 9.385998894 | 10.58648927 | 10.544437 | 9.070041959 | 9.194147421 |
| ENSG00000144214 | LYG1 | 1.128863324 | 0.636158103 | -1.236214608 | 0.599062079 | -0.107351284 | 2 | 4.11352228 | 2 | 3.95996233 | 3.7864959 | 2.831336763 | 2.326754077 |

| | | | | | | | | | | | | | |
|---|---|---|---|---|---|---|---|---|---|---|---|---|---|
| ENSG00000172262 | ZNF131 | -0.167723303 | 0.88199664 | 0.060084184 | 0.957826958 | -0.107639119 | 6.468621203 | 7.979858146 | 8.99637432 | 7.999368766 | 7.96458672 | 8.402448739 | 7.979721237 |
| ENSG00000102858 | MGRN1 | -0.164980448 | 0.837724165 | 0.057127718 | 0.944157185 | -0.10785273 | 10.58135316 | 8.981105197 | 9.859153765 | 9.739866907 | 9.825189 | 9.850086161 | 10.14369036 |
| ENSG00000147202 | DIAPH2 | -0.009902157 | 0.98668938 | -0.09815555 | 0.86473808 | -0.108057707 | 7.667647223 | 7.638155693 | 8.408111533 | 7.947219213 | 7.91907183 | 7.992125306 | 7.68409025 |
| ENSG00000100106 | TRIOBP | -0.393773771 | 0.282435386 | 0.285201224 | 0.441391676 | -0.108572547 | 10.84091191 | 11.0620559 | 10.48730704 | 10.4186301 | 10.4312521 | 11.00896231 | 11.19009178 |
| ENSG00000135506 | OS9 | 0.196765202 | 0.803469715 | -0.305510217 | 0.684964093 | -0.108745015 | 12.12726019 | 11.32164258 | 10.5187901 | 11.6625764 | 11.6670996 | 11.04522364 | 11.2724516 |
| ENSG00000136100 | VPS36 | 0.032194515 | 0.984117158 | -0.141023285 | 0.932122597 | -0.108828771 | 4.467311984 | 9.079034257 | 8.845626171 | 8.509731121 | 8.38151133 | 8.185201128 | 8.361298015 |
| ENSG00000142655 | PEX14 | 0.048766082 | 0.912239853 | -0.158729418 | 0.694164121 | -0.109963336 | 8.340423424 | 8.147241016 | 8.236946975 | 8.31397419 | 8.26379462 | 7.8927823 | 8.25407423 |
| ENSG00000164434 | FABP7 | 5.081056982 | 0.357361029 | -5.191085614 | 0.364514502 | -0.110028632 | 2 | 2 | 2 | 6.18902405 | 2 | 2 | 2 |
| ENSG00000104853 | CLPTM1 | 0.136944888 | 0.88199664 | -0.247599815 | 0.776294535 | -0.110654927 | 11.65707219 | 9.71856931 | 11.00131064 | 11.12343458 | 11.1375544 | 10.6268361 | 10.85746696 |
| ENSG00000161021 | MAML1 | -0.142232871 | 0.798768209 | 0.031456365 | 0.956825769 | -0.110776506 | 9.397585735 | 10.27307349 | 9.681375057 | 9.668456061 | 9.71736428 | 10.01733654 | 9.69686815 |
| ENSG00000100023 | PPIL2 | -0.146465024 | 0.84752019 | 0.034517058 | 0.965258777 | -0.111947967 | 9.2903348 | 8.029695761 | 8.175061096 | 8.486665614 | 8.44711562 | 8.489350161 | 8.787516955 |
| ENSG00000128383 | APOBEC3A | 3.546549242 | 0.37096152 | -3.659109335 | 0.391759399 | -0.112560093 | 2 | 2 | 2 | 3.881293667 | 4.0930011 | 2 | 2.059860178 |
| ENSG00000162408 | NOL9 | -0.198992558 | 0.739382058 | 0.082234266 | 0.89539314 | -0.116758292 | 7.667647223 | 7.669933291 | 8.561105505 | 7.849869274 | 7.80517502 | 8.199240359 | 8.002780183 |
| ENSG00000160959 | LRRC14 | -0.421822364 | 0.780343245 | 0.304887908 | 0.841557303 | -0.116934456 | 9.33149386 | 7.503546079 | 5.156626401 | 7.72026832 | 7.77859536 | 8.52322774 | 8.405948829 |
| ENSG00000104723 | TUSC3 | 0.743984443 | 0.336346335 | -0.861453029 | 0.250810834 | -0.117468586 | 11.06895221 | 9.583268459 | 9.589475645 | 10.99738621 | 11.0105443 | 9.271730593 | 9.525676102 |
| ENSG00000138756 | BMP2K | 0.823005348 | 0.691466494 | -0.94085223 | 0.641992641 | -0.117846882 | 8.313285031 | 5.595927442 | 2 | 7.772969103 | 7.72392024 | 6.057834398 | 6.037449234 |
| ENSG00000138750 | NUP54 | 0.395870355 | 0.727489842 | -0.51437404 | 0.640632886 | -0.118503685 | 5.866944937 | 7.73146271 | 8.677268317 | 8.178656739 | 8.24644267 | 7.539353852 | 7.060335194 |
| ENSG00000188295 | ZNF669 | -0.056530125 | 0.93987322 | -0.062357442 | 0.933434132 | -0.118887567 | 5.866944937 | 5.595927442 | 5.772716771 | 5.527288686 | 5.83486613 | 5.822425633 | 5.529673353 |
| ENSG00000105518 | TMEM205 | 0.27273544 | 0.762344209 | -0.392492194 | 0.651136675 | -0.119756754 | 9.112843761 | 8.671479029 | 7.163363831 | 8.693628619 | 8.89230135 | 7.903438544 | 8.34108458 |
| ENSG00000138326 | RPS24 | -0.056224848 | 0.960095276 | -0.063679825 | 0.955859616 | -0.119904673 | 12.1882823 | 10.04368224 | 10.11951029 | 11.08473443 | 11.1095698 | 11.09800658 | 11.08146599 |
| ENSG00000120162 | MOB3B | 0.294559198 | 0.886587309 | -0.414509753 | 0.834220385 | -0.119950555 | 2 | 5.131603625 | 3.369317972 | 4.443344104 | 4.1930041 | 4.209638631 | 3.183290849 |
| ENSG00000107745 | MICU1 | 0.16171994 | 0.838100556 | -0.281821586 | 0.705599521 | -0.120101646 | 8.468948322 | 8.903033433 | 9.907528187 | 9.346653666 | 9.41658653 | 8.722467118 | 9.13188655 |
| ENSG00000122435 | TRMT13 | 0.226016381 | 0.869626608 | -0.347389185 | 0.792195215 | -0.121372804 | 3.327711234 | 6.602424674 | 6.203069917 | 6.139181293 | 6.14061114 | 5.72966459 | 5.470251342 |
| ENSG00000103671 | TRIP4 | 0.250201204 | 0.827911815 | -0.372890939 | 0.73312167 | -0.122689735 | 8.517314938 | 7.60566235 | 5.772716771 | 7.877998516 | 7.97632944 | 7.200910877 | 7.425709383 |
| ENSG00000183963 | SMTN | 0.139738834 | 0.865062033 | -0.263940627 | 0.731726336 | -0.124201794 | 10.81695923 | 9.235879652 | 9.809100877 | 10.16416077 | 10.3172245 | 9.839240397 | 9.841914256 |
| ENSG00000198300 | PEG3 | -3.104077876 | 0.151263538 | 2.979204955 | 0.14497505 | -0.124872922 | 5.70932728 | 2 | 2 | 2.3892346 | 2.68924543 | 7.281044413 | 6.681239787 |
| ENSG00000189046 | ALKBH2 | -0.661534528 | 0.362647422 | 0.536247735 | 0.454637787 | -0.125286793 | 5.095959458 | 6.140560021 | 6.322058544 | 5.461211713 | 5.21224859 | 6.599583349 | 6.298867018 |
| ENSG00000206535 | LNP1 | -0.692243041 | 0.764891131 | 0.566756099 | 0.804586396 | -0.125486943 | 2 | 4.710564154 | 2 | 3.162721474 | 3.15372745 | 3.943456545 | 4.018014792 |
| ENSG00000164749 | HNF4G | 5.06526085 | 0.231765386 | -5.19111517 | 0.254503115 | -0.12585432 | 2 | 2 | 2 | 5.391962469 | 5.09650239 | 2 | 2 |
| ENSG00000128534 | LSM8 | 0.250241119 | 0.613329998 | -0.376290351 | 0.422595617 | -0.126049232 | 8.19931886 | 7.431218269 | 7.691296561 | 8.093978021 | 7.99953197 | 7.44824109 | 7.417152486 |
| ENSG00000166197 | NOLC1 | -0.351775852 | 0.747677329 | 0.225612128 | 0.839662804 | -0.126163724 | 9.781071289 | 9.74864196 | 7.225742779 | 8.946390023 | 8.95655043 | 9.527337514 | 9.522687851 |
| ENSG00000110436 | SLC1A2 | 1.22415269 | 0.262567938 | -1.35061094 | 0.206789707 | -0.12645825 | 4.00722531 | 4.710564154 | 5.772716771 | 6.24507773 | 5.9725602 | 4.281782022 | 3.383972333 |
| ENSG00000204065 | TCEAL5 | 5.064183952 | 0.231292431 | -5.191115294 | 0.253975361 | -0.126931342 | 2 | 2 | 2 | 5.22679461 | 5.27444544 | 2 | 2 |
| ENSG00000119401 | TRIM32 | -0.617886692 | 0.387577122 | 0.490577215 | 0.49246368 | -0.127309477 | 9.358295023 | 8.356994601 | 9.788583153 | 8.687861792 | 8.64401667 | 10.06283648 | 9.394150543 |
| ENSG00000102119 | EMD | -0.660502642 | 0.156708581 | 0.532685955 | 0.253721513 | -0.127816687 | 9.384607364 | 8.834582582 | 8.536706852 | 8.300810597 | 8.31463236 | 9.362752343 | 9.608182402 |
| ENSG00000254402 | LRRC24 | 5.063203033 | 0.232506713 | -5.191115056 | 0.255070887 | -0.127912023 | 2 | 2 | 2 | 5.058234798 | 5.41912973 | 2 | 2 |
| ENSG00000139631 | CSAD | -0.19896804 | 0.822767511 | 0.069869286 | 0.93862683 | -0.129098754 | 5.70932728 | 7.503546079 | 7.098165388 | 6.664650034 | 6.86652414 | 7.130090369 | 6.920008846 |
| ENSG00000186104 | CYP2R1 | -0.868671148 | 0.609720423 | 0.739294327 | 0.662784961 | -0.129376822 | 6.563806279 | 5.131603625 | 2 | 4.887609717 | 4.50979031 | 6.499401217 | 5.722580018 |
| ENSG00000109103 | UNC119 | -0.560922838 | 0.132777783 | 0.431278991 | 0.249078729 | -0.129643848 | 8.564112515 | 8.687092496 | 8.143093325 | 7.852449262 | 8.01555453 | 8.921491433 | 8.912005269 |
| ENSG00000154781 | CCDC174 | -0.028383074 | 0.981947437 | -0.102869885 | 0.933434132 | -0.131252958 | 6.96351253 | 8.469613413 | 5.772716771 | 7.395290829 | 7.52080763 | 7.551912644 | 7.199901369 |
| ENSG00000106868 | SUSD1 | 0.078161509 | 0.972415697 | -0.212139513 | 0.926209474 | -0.133978004 | 9.48530855 | 8.004992147 | 2 | 8.276040626 | 8.5606705 | 7.853585565 | 8.376826085 |
| ENSG00000168116 | KIAA1586 | -0.111413518 | 0.865906878 | -0.028200703 | 0.966627129 | -0.139614221 | 7.435667355 | 6.78762695 | 6.43197674 | 6.836083475 | 6.83591385 | 7.034886728 | 6.786860422 |
| ENSG00000148834 | GSTO1 | 1.180001383 | 0.145540049 | -1.319626219 | 0.094172652 | -0.139624835 | 10.87379128 | 9.090820894 | 9.372276026 | 11.17226686 | 11.1906866 | 8.512023373 | 8.854540499 |
| ENSG00000068028 | RASSF1 | -0.462387907 | 0.814176064 | 0.322546449 | 0.870457209 | -0.139841457 | 9.771132748 | 7.73146271 | 3.369317972 | 8.006336225 | 8.10598728 | 8.815677424 | 8.849779308 |
| ENSG00000105127 | AKAP8 | 0.056492744 | 0.976731083 | -0.197035528 | 0.918257682 | -0.140542784 | 9.358295023 | 7.901717006 | 3.369317972 | 8.312101014 | 8.27524772 | 7.832294085 | 8.228218854 |
| ENSG00000228300 | C19orf24 | 0.103553021 | 0.911517345 | -0.244497292 | 0.779935476 | -0.140944272 | 8.910412251 | 7.274681896 | 7.502634626 | 8.100504225 | 8.26953254 | 7.802823992 | 7.885682058 |
| ENSG00000188428 | BLOC1S5 | -0.775764479 | 0.588826811 | 0.634144483 | 0.660050259 | -0.141619996 | 8.540903468 | 6.78762695 | 4.523675391 | 6.634986713 | 6.65485813 | 8.266822833 | 7.718159092 |
| ENSG00000077514 | POLD3 | 0.244068877 | 0.888744026 | -0.385763569 | 0.81669225 | -0.141694692 | 3.327711234 | 7.051611315 | 8.175061096 | 7.444001439 | 7.36516374 | 7.100474247 | 6.403650353 |
| ENSG00000196693 | ZNF33B | 0.732167398 | 0.582172352 | -0.875271313 | 0.496317147 | -0.143103915 | 4.467311984 | 7.790474954 | 6.073378139 | 7.391747748 | 7.44845993 | 6.17883 | 5.502966356 |
| ENSG00000152518 | ZFP36L2 | -0.054832111 | 0.943520433 | -0.088403082 | 0.909584743 | -0.143235193 | 9.667789716 | 10.28850895 | 11.15748401 | 10.42297598 | 10.4683133 | 10.32536768 | 10.49439247 |
| ENSG00000168806 | LCMT2 | -0.347810519 | 0.871671212 | 0.204451173 | 0.92531104 | -0.143359346 | 5.866944937 | 8.191702463 | 2 | 6.542151749 | 6.57678965 | 6.973973146 | 7.20156326 |
| ENSG00000137815 | RTF1 | -0.032809666 | 0.957978247 | -0.110780833 | 0.855882497 | -0.143590498 | 9.33149386 | 10.35843478 | 9.798878489 | 9.858840283 | 9.86300004 | 9.579383624 | 9.961982472 |
| ENSG00000160961 | ZNF333 | -0.83138998 | 0.556481113 | 0.686038205 | 0.628179702 | -0.145351776 | 3.327711234 | 7.35507212 | 7.029880454 | 5.959382894 | 5.80356929 | 7.49992804 | 7.150854407 |
| ENSG00000163930 | BAP1 | -0.402677717 | 0.821801475 | 0.256859692 | 0.886860792 | -0.145818025 | 10.62615013 | 9.814097497 | 4.874587708 | 9.279220784 | 9.33684939 | 9.99413276 | 9.934264157 |
| ENSG00000079112 | CDH17 | 1.191350439 | 0.691746353 | -1.339554992 | 0.65246542 | -0.148204553 | 2 | 2 | 3.369317972 | 3.462030052 | 2.59489879 | 2.384001718 | 2.117335275 |
| ENSG00000133872 | SARAF | 0.190770447 | 0.453919948 | -0.339878612 | 0.14757638 | -0.149108165 | 11.40085503 | 11.35867467 | 11.337759 | 11.58267051 | 11.5298098 | 10.90501116 | 11.13897086 |
| ENSG00000130175 | PRKCSH | -0.015770433 | 0.982399308 | -0.133680416 | 0.84468287 | -0.149450849 | 11.91575472 | 10.67655334 | 11.07517045 | 11.25795672 | 11.3433499 | 11.07135679 | 11.28768964 |
| ENSG00000189298 | ZKSCAN3 | -0.051563396 | 0.954030819 | -0.098965724 | 0.911774385 | -0.15052912 | 5.532353208 | 6.951736442 | 6.534109959 | 6.442929658 | 6.39275508 | 6.68757835 | 5.968846243 |
| ENSG00000007080 | CCDC124 | -0.704147357 | 0.218319116 | 0.551781169 | 0.33664227 | -0.152366188 | 8.444143041 | 9.298208073 | 8.236946975 | 8.022463952 | 8.07355593 | 9.154129994 | 9.41686151 |
| ENSG00000089195 | TRMT6 | -0.054259863 | 0.953443586 | -0.098450355 | 0.915858934 | -0.152710219 | 7.579279349 | 5.946638857 | 7.600047325 | 6.998639523 | 7.29187696 | 6.895669227 | 7.296373648 |
| ENSG00000099377 | HSD3B7 | 0.118799896 | 0.946375129 | -0.271594625 | 0.878225373 | -0.152794729 | 3.327711234 | 7.145017454 | 8.536706852 | 7.419851705 | 7.68723697 | 7.14536429 | 7.211494604 |
| ENSG00000114209 | PDCD10 | 0.724515808 | 0.09063243 | -0.877630512 | 0.033859643 | -0.153114704 | 8.586954094 | 7.874700259 | 8.486635645 | 9.085573664 | 9.0282492 | 7.565395753 | 7.382408159 |
| ENSG00000113318 | MSH3 | -0.226755965 | 0.866211938 | 0.073611841 | 0.956916625 | -0.153144124 | 8.540903468 | 7.572420273 | 5.156626401 | 7.467753569 | 7.36873214 | 7.538302335 | 7.869046644 |
| ENSG00000101126 | ADNP | -0.388176227 | 0.262259514 | 0.234373629 | 0.510878389 | -0.153802599 | 10.16187685 | 10.17682307 | 9.963514551 | 9.718812329 | 9.71806384 | 10.54229576 | 10.10114286 |
| ENSG00000141646 | SMAD4 | -0.16174422 | 0.798465488 | 0.007556273 | 0.99072569 | -0.154187947 | 8.418903792 | 9.497318007 | 9.358421373 | 9.067921572 | 8.94225201 | 9.314738721 | 9.019079808 |
| ENSG00000136522 | MRPL47 | 0.170765628 | 0.857034032 | -0.325413179 | 0.714828755 | -0.154647551 | 8.564112515 | 6.602424674 | 8.110401118 | 8.168339746 | 8.09094333 | 7.486906381 | 7.782864551 |
| ENSG00000124116 | WFDC3 | 2.298319033 | 0.309909983 | -2.453493777 | 0.278679003 | -0.155174743 | 2 | 4.11352228 | 2 | 4.497095146 | 5.06165168 | 2.094598369 | 2.466008509 |
| ENSG00000171847 | FAM90A1 | -3.753008627 | 0.531143362 | 3.596315706 | 0.521375485 | -0.15669292 | 2 | 2 | 2 | 2 | 2 | 3.648325314 | 4.329022995 |
| ENSG00000159023 | EPB41 | -0.411845891 | 0.775566336 | 0.254314532 | 0.862994354 | -0.157531358 | 5.532353208 | 7.538394143 | 9.358421373 | 7.786523606 | 7.81828345 | 8.66581901 | 8.2175813 |
| ENSG00000090861 | AARS | -0.549024616 | 0.434146426 | 0.391243777 | 0.580587983 | -0.157780839 | 11.04027452 | 10.47993704 | 9.69246155 | 9.930185944 | 9.98658999 | 11.22245533 | 10.47373855 |
| ENSG00000004487 | KDM1A | -0.247815143 | 0.555167478 | 0.086833785 | 0.846443289 | -0.160981358 | 9.219014269 | 9.785372135 | 9.859153765 | 9.376537567 | 9.42860336 | 9.67529364 | 9.793698997 |
| ENSG00000062194 | GPBP1 | 0.365033286 | 0.302694048 | -0.526044964 | 0.115721683 | -0.161011678 | 9.922475255 | 9.923599762 | 9.809100877 | 10.26878646 | 10.2316302 | 9.581682124 | 9.105499915 |
| ENSG00000100504 | PYGL | -0.346029015 | 0.887450188 | 0.184317181 | 0.939787567 | -0.161711833 | 10.07567912 | 7.431218269 | 2 | 8.404661252 | 8.32752521 | 8.850830124 | 8.931163478 |
| ENSG00000110042 | DTX4 | 0.012807243 | 0.996622494 | -0.17466616 | 0.942414968 | -0.161858917 | 3.327711234 | 7.232742069 | 2 | 5.808110275 | 5.80356929 | 5.206928476 | 5.96101835 |
| ENSG00000176890 | TYMS | 0.413780593 | 0.785196727 | -0.576516307 | 0.692374562 | -0.162735714 | 4.00722531 | 6.78762695 | 4.058998049 | 6.024891728 | 5.96314905 | 5.136410075 | 5.04340913 |
| ENSG00000152455 | SUV39H2 | -0.115440766 | 0.912408896 | -0.048301005 | 0.963748226 | -0.163741771 | 4.467311984 | 6.464272742 | 5.59507254 | 5.651002631 | 5.64780787 | 5.737015686 | 5.685057129 |
| ENSG00000261456 | TUBB8 | 2.904451352 | 0.182906893 | -3.068766981 | 0.176029831 | -0.164315629 | 2 | 3.686303414 | 2 | 5.076005082 | 4.60967607 | 2.048074405 | 2.225841198 |
| ENSG00000137824 | RMDN3 | -0.285236357 | 0.859119352 | 0.11949187 | 0.941500505 | -0.165744487 | 9.423197387 | 9.055168103 | 4.874587708 | 8.409925372 | 8.42842973 | 8.724779862 | 8.908956896 |
| ENSG00000135372 | NAT10 | -0.015488382 | 0.983101649 | -0.150577523 | 0.830158879 | -0.166065905 | 7.624139768 | 8.147241016 | 8.941672936 | 8.341783829 | 8.30162324 | 8.091532339 | 8.279474384 |
| ENSG00000102393 | GLA | 0.915968831 | 0.261009892 | -1.082848018 | 0.169103768 | -0.166879186 | 9.233563228 | 8.916342327 | 7.225742779 | 9.586367937 | 9.60688128 | 7.624061799 | 7.615934347 |
| ENSG00000001631 | KRIT1 | 1.112915557 | 0.009363387 | -1.281493089 | 0.002330612 | -0.168577532 | 7.031698319 | 7.847167909 | 7.646393253 | 8.624367384 | 8.68323472 | 6.339575235 | 6.298867018 |
| ENSG00000124251 | TP53TG5 | -3.753008776 | 0.531143362 | 3.584371155 | 0.52282666 | -0.168637621 | 2 | 2 | 2 | 2 | 2 | 4.065262523 | 3.972018587 |
| ENSG00000134882 | UBAC2 | -0.639763313 | 0.293257156 | 0.470414923 | 0.443446023 | -0.16934839 | 9.87686338 | 8.702538795 | 9.859153765 | 8.957217624 | 8.91077722 | 10.12998797 | 9.939751019 |
| ENSG00000166024 | R3HCC1L | 0.716930081 | 0.731288333 | -0.887331173 | 0.663750141 | -0.170401093 | 7.159102577 | 8.993716073 | 2 | 8.399377854 | 8.56534297 | 7.08038692 | 6.723495198 |
| ENSG00000010322 | NISCH | -0.410757721 | 0.630961153 | 0.239934587 | 0.785559516 | -0.170823134 | 11.29385609 | 9.77810053 | 9.647595056 | 10.00165982 | 10.0733424 | 10.61790491 | 10.75225974 |
| ENSG00000164344 | KLKB1 | 5.019901363 | 0.241359928 | -5.191114409 | 0.258315695 | -0.171213046 | 2 | 2 | 2 | 5.161714397 | 5.2591458 | 2 | 2 |
| ENSG00000140157 | NIPA2 | 0.368795086 | 0.607262407 | -0.540103428 | 0.427903866 | -0.171308342 | 9.771132748 | 9.656475478 | 8.325066185 | 9.763363111 | 9.73820484 | 8.766685705 | 8.924637944 |
| ENSG00000188785 | ZNF548 | -0.487132927 | 0.539576503 | 0.315570148 | 0.696257748 | -0.171562779 | 7.750913381 | 6.899080544 | 6.203069917 | 6.529309835 | 6.70088158 | 7.655436363 | 7.076727469 |
| ENSG00000112685 | EXOC2 | 0.070767565 | 0.925390417 | -0.24296372 | 0.728500949 | -0.172196155 | 8.075571721 | 8.395519306 | 9.399592459 | 8.814994458 | 8.79619691 | 8.548515606 | 8.439433377 |
| ENSG00000005020 | SKAP2 | 1.322376003 | 0.146554142 | -1.496790599 | 0.091551895 | -0.174414596 | 5.095959458 | 6.951736442 | 7.502634626 | 8.096156705 | 8.12931531 | 5.60262961 | 5.197927245 |
| ENSG00000198732 | SMOC1 | -0.897966126 | 0.642184909 | 0.722898539 | 0.708681786 | -0.175067588 | 2 | 6.951736442 | 4.874587708 | 5.22679461 | 4.53541753 | 6.877490957 | 5.708623051 |
| ENSG00000119655 | NPC2 | 0.469571185 | 0.653653157 | -0.645079769 | 0.521361073 | -0.175508584 | 10.73752355 | 9.170736405 | 8.296287027 | 10.21566945 | 10.2350555 | 9.082719462 | 9.149213283 |
| ENSG00000164597 | COG5 | -0.407417581 | 0.282232621 | 0.22971684 | 0.555749022 | -0.177700741 | 8.777722034 | 9.00621767 | 8.805342542 | 8.540455898 | 8.38855922 | 9.286209561 | 8.880709189 |
| ENSG00000030110 | BAK1 | 0.311070522 | 0.694579859 | -0.489644477 | 0.5142789 | -0.178573955 | 8.16936288 | 8.590771778 | 6.882775709 | 8.361842177 | 8.35477078 | 7.57465694 | 7.566491953 |
| ENSG00000117676 | RPS6KA1 | -0.172718641 | 0.92551187 | -0.006050293 | 0.997764007 | -0.178768934 | 3.327711234 | 7.503546079 | 4.874587708 | 5.880681751 | 6.21386141 | 6.587434478 | 5.708623051 |
| ENSG00000178685 | PARP10 | -6.445436812 | 0.00487994 | 6.266656504 | 4.62E-05 | -0.178780308 | 2 | 4.442710129 | 2 | 2 | 2 | 9.024959988 | 8.967756908 |
| ENSG00000130338 | TULP4 | -0.104915675 | 0.807865742 | -0.074460146 | 0.863495873 | -0.179375821 | 8.910412251 | 9.488434476 | 9.099922433 | 9.126651551 | 9.04841459 | 9.051735782 | 9.18199253 |
| ENSG00000160113 | NR2F6 | -0.588957269 | 0.708284536 | 0.408052604 | 0.798191812 | -0.180904665 | 9.556468883 | 10.02537381 | 5.156626401 | 8.709369553 | 8.63070345 | 9.650204809 | 9.670710139 |
| ENSG00000121236 | TRIM6 | -0.697825074 | 0.682343107 | 0.514098307 | 0.764313677 | -0.183726767 | 3.327711234 | 4.710564154 | 7.502634626 | 5.651002631 | 5.37722731 | 6.678062013 | 6.640162436 |
| ENSG00000159885 | ZNF222 | 0.382712088 | 0.826909189 | -0.56657688 | 0.733788685 | -0.183864792 | 2 | 6.535001748 | 6.534109959 | 6.401247806 | 6.29111322 | 5.680947153 | 5.20457242 |
| ENSG00000168818 | STX18 | -0.138402314 | 0.828815955 | -0.045520675 | 0.944101725 | -0.183922989 | 7.579279349 | 8.356994601 | 7.502634626 | 7.734324667 | 7.7515168 | 7.600074836 | 8.035760639 |
| ENSG00000187824 | TMEM220 | -0.012661715 | 0.993765388 | -0.171668597 | 0.906716859 | -0.184330313 | 4.467311984 | 6.666836837 | 4.058998049 | 5.674521485 | 5.45984938 | 5.096960836 | 5.694529741 |
| ENSG00000124541 | RRP36 | -0.059332324 | 0.960501625 | -0.127534968 | 0.916470603 | -0.186867292 | 9.699577239 | 8.88960062 | 6.80318718 | 8.814994458 | 8.83669913 | 8.501810558 | 8.976033998 |
| ENSG00000100532 | CGRRF1 | -0.213919403 | 0.911376403 | 0.026289302 | 0.988820823 | -0.187630101 | 2 | 7.189546432 | 7.098165388 | 6.372775472 | 6.3857277 | 6.752490997 | 6.449128733 |
| ENSG00000172465 | TCEAL1 | -0.533120785 | 0.377368801 | 0.343869441 | 0.574881251 | -0.189251344 | 7.27616306 | 7.002537973 | 8.076950877 | 7.101994243 | 6.88158913 | 7.77631144 | 7.924438973 |
| ENSG00000181929 | PRKAG1 | -0.409735831 | 0.273150492 | 0.220311444 | 0.569975156 | -0.189424388 | 9.064836028 | 9.616264339 | 9.272377397 | 8.923261356 | 8.94464494 | 9.643863831 | 9.470572512 |
| ENSG00000143486 | EIF2D | -0.420515895 | 0.436019907 | 0.230238162 | 0.679605718 | -0.190277733 | 8.367060731 | 7.761270542 | 8.763901704 | 7.978262192 | 7.88209639 | 8.498029643 | 8.644559262 |

| | | | | | | | | | | | | | |
|---|---|---|---|---|---|---|---|---|---|---|---|---|---|
| ENSG00000130726 | TRIM28 | -0.318138124 | 0.721422916 | 0.12564137 | 0.893463941 | -0.192496754 | 10.38028415 | 11.54799449 | 9.61300511 | 10.41208663 | 10.4248388 | 10.96542343 | 10.74871193 |
| ENSG00000115685 | PPP1R7 | 0.245583982 | 0.713873535 | -0.438890974 | 0.484010617 | -0.193306992 | 9.262228249 | 8.433041974 | 8.007629717 | 8.887228971 | 8.92538942 | 8.110758945 | 8.33882106 |
| ENSG00000181035 | SLC25A42 | -0.909432098 | 0.282224272 | 0.715033388 | 0.397171615 | -0.19439871 | 8.285626337 | 6.899080544 | 6.43197674 | 6.5612028 | 6.53293476 | 8.187215117 | 8.068914703 |
| ENSG00000142046 | TMEM91 | 0.359423836 | 0.803441373 | -0.55426411 | 0.686301227 | -0.194840274 | 6.366709671 | 4.11352228 | 4.058998049 | 5.709096143 | 5.45984938 | 5.119635168 | 4.329022995 |
| ENSG00000160679 | CHTOP | -0.237742771 | 0.818399248 | 0.042260431 | 0.968580178 | -0.195482341 | 7.579279349 | 9.916994052 | 9.031722608 | 8.892252782 | 8.90341524 | 9.366311982 | 8.955990164 |
| ENSG00000170075 | GPR37L1 | -0.934829896 | 0.666836354 | 0.736927311 | 0.728110981 | -0.197902584 | 3.327711234 | 2 | 4.058998049 | 2.695456494 | 2.94009902 | 3.035909414 | 4.279527956 |
| ENSG00000205456 | TP53TG3D | 2.689899487 | 0.584143941 | -2.887860292 | NA | -0.197960805 | 2 | 2 | 2 | 3.462030052 | 3.33976443 | 2.094598369 | 2 |
| ENSG00000172508 | CARNS1 | -1.877744509 | 0.239916541 | 1.679545043 | 0.27708601 | -0.198199466 | 5.70932728 | 3.686303414 | 2 | 3.289987342 | 3.08597292 | 6.249924465 | 5.722580018 |
| ENSG00000180884 | ZNF792 | 0.844886219 | 0.634552412 | -1.046156816 | 0.543191241 | -0.201270597 | 3.327711234 | 4.442710129 | 7.734844356 | 7.182120794 | 7.1246732 | 5.381251181 | 5.355511328 |
| ENSG00000106804 | C5 | 0.05731432 | 0.98350331 | -0.260451927 | 0.926209474 | -0.203137607 | 2 | 2 | 6.203069917 | 4.867343068 | 4.72553021 | 4.5389195 | 4.522269241 |
| ENSG00000109339 | MAPK10 | -1.79975479 | 0.19577655 | 1.596489171 | 0.239770681 | -0.203265618 | 6.009025997 | 5.303523747 | 2 | 3.663521627 | 3.7864959 | 6.247352872 | 6.983208048 |
| ENSG00000175106 | TVP23C | 2.293573275 | 0.0383061 | -2.496989556 | 0.022406102 | -0.203416281 | 6.366709671 | 4.93638933 | 8.143093325 | 9.305820479 | 9.2754408 | 4.742549839 | 4.711446447 |
| ENSG00000161714 | PLCD3 | -1.192254961 | 0.362599152 | 0.988693842 | 0.450551283 | -0.203561119 | 10.38028415 | 9.470501606 | 6.322058544 | 8.239055745 | 8.32935768 | 10.226175 | 10.64611413 |
| ENSG00000278129 | ZNF8 | -0.717489598 | 0.772450402 | 0.511136166 | 0.839017584 | -0.206353432 | 8.540903468 | 3.686303414 | 2 | 6.29144905 | 6.36443784 | 7.646681011 | 7.370638252 |
| ENSG00000197019 | SERTAD1 | 0.417988553 | 0.645473237 | -0.625087887 | 0.469777313 | -0.207099334 | 7.709880939 | 7.274681896 | 9.149036566 | 8.699372486 | 8.673169 | 7.710619897 | 7.603417442 |
| ENSG00000111339 | ART4 | -3.753009283 | 0.531143362 | 3.543772668 | 0.52769064 | -0.209236615 | 2 | 2 | 2 | 2 | 2 | 3.584558147 | 4.30448772 |
| ENSG00000108039 | XPNPEP1 | -0.178410127 | 0.725609862 | -0.030901903 | 0.954650147 | -0.20931203 | 9.384607364 | 10.03150247 | 9.413059136 | 9.393466757 | 9.53961588 | 9.406340618 | 9.797550144 |
| ENSG00000101132 | PFDN4 | 0.359400783 | 0.806299763 | -0.570489539 | 0.682815119 | -0.211088756 | 7.904282243 | 6.04685525 | 4.523675391 | 7.214570819 | 7.01963893 | 6.635427934 | 5.6562603 |
| ENSG00000111291 | GPRC5D | -2.312787877 | NA | 2.101534343 | NA | -0.211253534 | 2 | 2 | 2 | 2 | 2.14022021 | 2.656294034 | 3.383972333 |
| ENSG00000087269 | NOP14 | -0.017273546 | 0.983773789 | -0.195530034 | 0.811305502 | -0.21280358 | 8.19931886 | 8.053983481 | 9.528916236 | 8.7747563 | 8.69607402 | 8.531177917 | 8.587232804 |
| ENSG00000069399 | BCL3 | -0.217029673 | 0.829171998 | 0.003519938 | 0.99765513 | -0.213509735 | 7.33131396 | 7.669933291 | 9.180874355 | 7.975897842 | 8.17894021 | 7.968899389 | 8.566746948 |
| ENSG00000163444 | TMEM183A | 0.14796585 | 0.770265142 | -0.361744671 | 0.43609424 | -0.213778821 | 8.946185348 | 8.213429689 | 8.978369997 | 8.898508045 | 8.88608977 | 8.406488267 | 8.368713182 |
| ENSG00000174903 | RAB1B | -0.050492189 | 0.911631829 | -0.163441364 | 0.694750517 | -0.213933553 | 10.16946352 | 10.54564457 | 10.73794767 | 10.43679547 | 10.468729 | 10.16430247 | 10.49710157 |
| ENSG00000178449 | COX14 | -0.373011824 | 0.364106676 | 0.158041123 | 0.715368383 | -0.214970701 | 8.009475818 | 8.337339476 | 7.818184403 | 7.778406195 | 7.64666005 | 8.14846013 | 8.319819 |
| ENSG00000169508 | GPR183 | 0.451986539 | 0.772510709 | -0.667748816 | 0.657862568 | -0.215762277 | 2 | 5.303523747 | 4.874587708 | 5.022023783 | 5.00774437 | 4.397636241 | 3.698365557 |
| ENSG00000244754 | N4BP2L2 | -0.284859376 | 0.561250332 | 0.067541064 | 0.899650959 | -0.217318312 | 10.03472846 | 9.308339287 | 9.725218844 | 9.466018907 | 9.40011621 | 9.995091105 | 9.536580367 |
| ENSG00000112651 | MRPL2 | -0.194087928 | 0.695551109 | -0.023571064 | 0.965258777 | -0.217658992 | 7.667647223 | 7.701025996 | 7.163363831 | 7.178012736 | 7.5014335 | 7.491260006 | 7.539150216 |
| ENSG00000196517 | SLC6A9 | 0.15427454 | 0.922635322 | -0.373485386 | 0.804586396 | -0.219210847 | 9.886101849 | 6.78762695 | 6.629488387 | 8.636393836 | 8.83798691 | 8.464655389 | 7.922424886 |
| ENSG00000121931 | LRIF1 | 1.010778063 | 0.21521423 | -1.230076257 | 0.117872694 | -0.219298193 | 8.418903792 | 7.274681896 | 7.225742779 | 8.756965077 | 8.73530306 | 7.037865927 | 5.821002687 |
| ENSG00000103356 | EARS2 | -0.715844376 | 0.651333637 | 0.495853842 | 0.758228816 | -0.219990534 | 9.064836028 | 8.451443573 | 4.058998049 | 7.454228844 | 7.60187978 | 8.750886382 | 8.696162343 |
| ENSG00000164056 | SPRY1 | -0.672693996 | 0.490423307 | 0.450711093 | 0.645796163 | -0.221982903 | 4.00722531 | 6.140560021 | 5.930865648 | 5.076005082 | 4.98931816 | 6.148830293 | 5.96101835 |
| ENSG00000170631 | ZNF16 | -0.686798113 | 0.424587993 | 0.463814822 | 0.592366323 | -0.222983291 | 5.095959458 | 7.051611315 | 6.718950348 | 5.920568914 | 5.81407719 | 6.913621294 | 7.011905845 |
| ENSG00000130312 | MRPL34 | -0.538931517 | 0.233859725 | 0.315316476 | 0.498844659 | -0.223615041 | 9.081016759 | 8.433041974 | 8.325066185 | 8.134820652 | 8.10170499 | 8.972986183 | 8.953526633 |
| ENSG00000169241 | SLC50A1 | -0.136979759 | 0.918959987 | -0.086712743 | 0.948481065 | -0.223692502 | 6.366709671 | 9.695596167 | 8.35328246 | 8.47833774 | 8.63367258 | 8.55268749 | 8.660959882 |
| ENSG00000165487 | MICU2 | 0.565512876 | 0.113360191 | -0.790357671 | 0.020227058 | -0.224844795 | 7.750913381 | 7.638155693 | 7.691296561 | 8.190940526 | 8.30348889 | 6.828438696 | 7.010010317 |
| ENSG00000112599 | GUCA1B | 0.543759983 | 0.786662631 | -0.769331037 | 0.689804433 | -0.225571054 | 3.327711234 | 2 | 4.523675391 | 4.17310023 | 3.69849104 | 3.374372158 | 2.782576693 |
| ENSG00000173898 | SPTBN2 | 0.053083641 | 0.969053019 | -0.279742031 | 0.83342673 | -0.226658391 | 6.73718411 | 8.57407404 | 10.15199903 | 9.174861605 | 9.09008229 | 8.779201757 | 8.826273465 |
| ENSG00000111229 | ARPC3 | 0.243193657 | 0.766736242 | -0.469982289 | 0.540654844 | -0.226788633 | 10.83137851 | 9.34816733 | 9.465702942 | 10.24689154 | 10.334663 | 9.493286157 | 9.664095852 |
| ENSG00000129009 | ISLR | -1.873353696 | 0.108536572 | 1.64613969 | 0.156194904 | -0.227214005 | 7.27616306 | 9.386925327 | 10.58572549 | 7.68596357 | 7.84158259 | 10.46000723 | 11.77297383 |
| ENSG00000112576 | CCND3 | 0.209151165 | 0.739140058 | -0.437017365 | 0.454500807 | -0.2278662 | 9.940323746 | 10.19331737 | 8.99637432 | 9.975862031 | 10.0345549 | 9.349025111 | 9.375867991 |
| ENSG00000113648 | H2AFY | -0.323209233 | 0.507503623 | 0.093821546 | 0.857692232 | -0.229387687 | 10.00958825 | 9.930175364 | 10.70548138 | 9.95800436 | 9.91151326 | 10.2789076 | 10.41871938 |
| ENSG00000164086 | DUSP7 | -1.235104683 | 0.011460822 | 1.005296744 | 0.041611142 | -0.229807939 | 9.304185368 | 8.276714465 | 8.805342542 | 7.605658789 | 7.65837038 | 9.972302634 | 9.715307813 |
| ENSG00000119965 | C10orf88 | -0.251712227 | 0.908179413 | 0.021177901 | 0.992406225 | -0.230534327 | 2 | 7.099070191 | 8.585098385 | 7.206526483 | 7.20272083 | 7.775419242 | 7.083953563 |
| ENSG00000178734 | LMO7DN | -2.304491744 | NA | 2.073147501 | NA | -0.231344243 | 2 | 2 | 2 | 2.141737246 | 2 | 2.094598369 | 3.620995423 |
| ENSG00000154240 | CEP112 | 0.255500264 | 0.898841046 | -0.487606773 | 0.798598135 | -0.232106509 | 2 | 6.228547981 | 7.691296561 | 6.912249745 | 6.71216164 | 5.893224226 | 6.29264319 |
| ENSG00000164037 | SLC9B1 | -0.065088061 | 0.975808161 | -0.168287498 | 0.936648304 | -0.233375559 | 4.00722531 | 2 | 3.369317972 | 3.162721474 | 3.33976443 | 3.44788957 | 2.885414003 |
| ENSG00000138668 | HNRNPD | 0.064090442 | 0.930344224 | -0.297657877 | 0.658096973 | -0.233567435 | 10.26462289 | 9.415325457 | 9.788583153 | 9.933844998 | 9.92065856 | 8.953791814 | 9.99637663 |
| ENSG00000096060 | FKBP5 | -0.207604329 | 0.720126758 | -0.026482445 | 0.966296927 | -0.234086773 | 6.653098118 | 7.232742069 | 6.534109959 | 6.535745081 | 6.77267312 | 6.795372357 | 6.870723585 |
| ENSG00000169375 | SIN3A | -0.083808217 | 0.936411459 | -0.150403035 | 0.885109991 | -0.234211252 | 7.750913381 | 9.763446433 | 10.03493575 | 9.351221511 | 9.43031191 | 9.182002376 | 9.455329967 |
| ENSG00000165006 | UBAP1 | 0.003456296 | 0.997041173 | -0.238403072 | 0.765637652 | -0.234946776 | 10.75275473 | 9.055168103 | 9.878698603 | 10.02643329 | 10.0928914 | 9.852414792 | 9.785134696 |
| ENSG00000261992 | | 2.570211224 | 0.020861927 | -2.805997252 | 0.011198174 | -0.235786027 | 4.00722531 | 7.099070191 | 5.156626401 | 8.454760653 | 8.53866332 | 3.5515855 | 3.753786374 |
| ENSG00000143869 | GDF7 | -3.75300962 | 0.531143362 | 3.516785402 | 0.530948199 | -0.236224218 | 2 | 2 | 2 | 2 | 2 | 4.099885429 | 3.824516949 |
| ENSG00000223547 | ZNF844 | 0.1068001 | 0.967125696 | -0.344782889 | 0.893909641 | -0.237982788 | 9.112843761 | 4.442710129 | 2 | 7.688853661 | 7.70428369 | 7.593002849 | 6.813187523 |
| ENSG00000163162 | RNF149 | 1.004383823 | 0.150830829 | -1.242581671 | 0.064830738 | -0.238197848 | 9.031919741 | 8.702538795 | 10.21486071 | 10.47329248 | 10.4624805 | 8.284489899 | 8.186029473 |
| ENSG00000160208 | RRP1B | -0.608056259 | 0.465935157 | 0.369851433 | 0.665677302 | -0.238204826 | 7.33131396 | 8.433041974 | 9.287079172 | 7.952038749 | 7.96222667 | 8.97687222 | 8.868207441 |
| ENSG00000128708 | HAT1 | -0.124151001 | 0.765831172 | -0.114623698 | 0.78199021 | -0.238774699 | 8.493334308 | 8.557180776 | 8.922968322 | 8.508095759 | 8.57927024 | 8.680667534 | 8.413855585 |
| ENSG00000266964 | FXYD1 | -0.682576334 | 0.844391096 | 0.443615594 | NA | -0.238960741 | 2 | 3.076665209 | 2 | 2.600425781 | 2.14022021 | 3.035909414 | 2.374681325 |
| ENSG00000077232 | DNAJC10 | 0.048098393 | 0.945987763 | -0.2888739 | 0.661912042 | -0.240775507 | 9.820154628 | 10.70750522 | 11.16943577 | 10.71219168 | 10.7184845 | 10.2683239 | 10.48230911 |
| ENSG00000154719 | MRPL39 | 0.00963404 | 0.991315828 | -0.251874214 | 0.757701514 | -0.242240174 | 7.159102577 | 7.145017454 | 5.59507254 | 6.687949235 | 6.93069905 | 6.668482487 | 6.449128733 |
| ENSG00000144199 | FAHD2B | 0.28701399 | 0.843262224 | -0.530019742 | 0.697571096 | -0.243005751 | 7.218819781 | 5.131603625 | 4.058998049 | 6.29144905 | 6.36443784 | 5.553513572 | 5.576534358 |
| ENSG00000196072 | BLOC1S2 | 0.402554446 | 0.643136765 | -0.646919484 | 0.431126874 | -0.244365038 | 9.448362284 | 7.761270542 | 7.97167868 | 8.964391141 | 9.03275482 | 7.893604812 | 8.026414333 |
| ENSG00000163114 | PDHA2 | 2.49907863 | 0.47708865 | -2.743628696 | NA | -0.244550067 | 2 | 2 | 2 | 3.515101962 | 3.01487888 | 2.048074405 | 2.059860178 |
| ENSG00000170275 | CRTAP | -0.436085469 | 0.329070084 | 0.191107576 | 0.686807452 | -0.244977894 | 12.05742222 | 11.2524908 | 11.87895445 | 11.31705777 | 11.3474248 | 11.97640709 | 11.94082017 |
| ENSG00000120458 | MSANTD2 | 0.088026536 | 0.977429905 | -0.333130707 | 0.914220926 | -0.245104171 | 2 | 8.255926468 | 2 | 6.772115916 | 6.82556437 | 6.809387508 | 5.812327524 |
| ENSG00000115112 | TFCP2L1 | -2.272298728 | 0.421397646 | 2.02689433 | 0.408802956 | -0.245404399 | 3.327711234 | 2 | 2 | 2.141737246 | 2.14022021 | 3.724264246 | 3.45257121 |
| ENSG00000165699 | TSC1 | 0.103088394 | 0.941409766 | -0.349713884 | 0.794078028 | -0.24662549 | 6.468621203 | 10.13758676 | 9.491321578 | 9.430198858 | 9.44390794 | 9.052472506 | 8.91909319 |
| ENSG00000119650 | IFT43 | -0.16868943 | 0.825992773 | -0.077948508 | 0.920635926 | -0.246637938 | 6.89194353 | 8.169643011 | 7.098165388 | 7.43025113 | 7.26914476 | 7.402744819 | 7.477384137 |
| ENSG00000142731 | PLK4 | 0.060189533 | 0.981829576 | -0.308364472 | 0.903920298 | -0.248174939 | 2 | 5.595927442 | 2 | 4.329433326 | 4.31639476 | 4.111244182 | 3.956354764 |
| ENSG00000103995 | CEP152 | 0.032510219 | 0.987070736 | -0.281992007 | 0.886770342 | -0.249481788 | 7.159102577 | 5.303523747 | 2 | 6.104969671 | 5.85535975 | 6.075339068 | 5.171036168 |
| ENSG00000006704 | GTF2IRD1 | 0.214170463 | 0.908850677 | -0.464133024 | 0.792727651 | -0.249962561 | 8.910412251 | 8.147241016 | 3.369317972 | 8.229163214 | 8.22100445 | 7.887011553 | 7.131805326 |
| ENSG00000185024 | BRF1 | -0.270858864 | 0.593368805 | 0.019857579 | 0.971342629 | -0.251001284 | 7.485143553 | 8.297207175 | 8.206335842 | 7.748245381 | 7.80780625 | 8.135306947 | 7.989373961 |
| ENSG00000106344 | RBM28 | -0.548783563 | 0.287337376 | 0.29731032 | 0.576448639 | -0.251473242 | 8.285626337 | 7.393647515 | 8.206335842 | 7.477814553 | 7.45182838 | 8.223648538 | 8.365751687 |
| ENSG00000157349 | DDX19B | 0.607799866 | 0.356507018 | -0.859663141 | 0.168742422 | -0.251863275 | 8.39321512 | 7.761270542 | 7.029880454 | 8.578353644 | 8.26953254 | 6.860764111 | 7.116030435 |
| ENSG00000131778 | CHD1L | 0.209072136 | 0.765491992 | -0.461261219 | 0.476851144 | -0.252189083 | 8.042902253 | 9.00621767 | 9.212024683 | 8.946390023 | 9.13582529 | 8.033758684 | 8.659751394 |
| ENSG00000130714 | POMT1 | -0.200339662 | 0.895675696 | -0.053302122 | 0.972083517 | -0.253641784 | 6.468621203 | 9.821190352 | 7.098165388 | 8.29892024 | 8.42671895 | 8.586652837 | 8.428121416 |
| ENSG00000164024 | METAP1 | -0.002955704 | 0.997204246 | -0.254087282 | 0.748418682 | -0.257042986 | 9.410448395 | 8.687092496 | 7.691296561 | 8.696503411 | 8.81332035 | 8.551124447 | 8.462487876 |
| ENSG00000131174 | COX7B | -0.148225362 | 0.877236903 | -0.109803382 | 0.908314446 | -0.258028745 | 10.00110988 | 8.029695761 | 8.941672936 | 9.077877436 | 9.04395769 | 8.986927198 | 9.20330054 |
| ENSG00000196505 | GDAP2 | 0.060586962 | 0.929816715 | -0.319688245 | 0.605467095 | -0.259101283 | 7.159102577 | 7.701025996 | 8.296287027 | 7.770242853 | 7.92393105 | 7.644728141 | 7.285445324 |
| ENSG00000109390 | NDUFC1 | 0.593314518 | 0.27254373 | -0.852761818 | 0.095706512 | -0.2594473 | 7.867441304 | 7.099070191 | 6.958202216 | 7.913023977 | 7.97164382 | 6.442083733 | 6.62538071 |
| ENSG00000043462 | LCP2 | -3.75300991 | 0.531143362 | 3.49355199 | 0.534071787 | -0.25945792 | 2 | 2 | 2 | 2 | 2 | 4.416072028 | 3.261853426 |
| ENSG00000077684 | JADE1 | -1.447264575 | 0.000515432 | 1.185162936 | 0.004574546 | -0.262101638 | 8.517314938 | 8.101365653 | 8.110401118 | 6.907295206 | 6.76727577 | 9.678399287 | 9.125772694 |
| ENSG00000103187 | COTL1 | -0.031438266 | 0.962823674 | -0.231211623 | 0.716517864 | -0.262649889 | 10.30651143 | 9.608085847 | 10.81097638 | 10.26394956 | 10.3149142 | 9.889995187 | 10.26666875 |
| ENSG00000171302 | CANT1 | -0.116940358 | 0.911517345 | -0.1472908 | 0.886197127 | -0.264231159 | 9.730679447 | 8.791896851 | 11.0057616 | 9.910512694 | 10.096124 | 9.891025241 | 10.05638425 |
| ENSG00000139921 | TMX1 | 0.636548941 | 0.357278052 | -0.900960221 | 0.169365312 | -0.26441128 | 9.233563228 | 9.7032946 | 8.143093325 | 9.842062221 | 9.7546632 | 8.242494717 | 8.304435067 |
| ENSG00000132005 | RFX1 | -0.215216665 | 0.885711022 | -0.049397118 | 0.97326848 | -0.264613783 | 8.313285031 | 7.232742069 | 4.523675391 | 7.114945424 | 7.17852729 | 7.648631242 | 6.862343163 |
| ENSG00000171103 | TRMT61B | -0.254571096 | 0.82798291 | -0.01036649 | 0.993322572 | -0.264937586 | 5.095959458 | 6.535001748 | 7.777116083 | 6.598566398 | 6.60126376 | 6.845542321 | 6.824016918 |
| ENSG00000104738 | MCM4 | 0.50220602 | 0.76909522 | -0.767152561 | 0.640432051 | -0.264946541 | 7.975256505 | 8.747909548 | 3.369317972 | 8.336264621 | 8.36015869 | 7.021403672 | 7.18149309 |
| ENSG00000132432 | SEC61G | 0.840272721 | 0.482249983 | -1.106697088 | 0.33238218 | -0.266424367 | 9.624286623 | 7.145017454 | 7.029880454 | 9.296376788 | 9.2990102 | 7.366471135 | 7.386797235 |
| ENSG00000103591 | AAGAB | -0.099901681 | 0.957170231 | -0.16841465 | 0.929020974 | -0.26831633 | 9.975371636 | 8.929529568 | 4.058998049 | 8.908460392 | 8.84568954 | 8.648868792 | 8.954019676 |
| ENSG00000213762 | ZNF134 | -0.433201056 | 0.598164094 | 0.164876005 | 0.849628112 | -0.268325051 | 8.138771685 | 7.503546079 | 9.242516611 | 7.987681034 | 8.09740995 | 8.691586261 | 8.574463273 |
| ENSG00000186716 | BCR | -0.422862257 | 0.58752019 | 0.152757637 | 0.853263424 | -0.270104621 | 9.90440318 | 9.298208073 | 8.206335842 | 8.799033614 | 8.94703391 | 9.258063226 | 9.61318568 |
| ENSG00000164062 | APEH | -0.486505286 | 0.342227985 | 0.213843628 | 0.692464563 | -0.272661657 | 9.317904228 | 9.664384941 | 8.721235202 | 8.813671111 | 8.80016655 | 9.338794282 | 9.649254128 |
| ENSG00000204366 | ZBTB12 | 4.918259755 | 0.264205073 | -5.19111239 | 0.268308947 | -0.272852634 | 2 | 2 | 2 | 5.144975053 | 5.09650239 | 2 | 2 |
| ENSG00000162341 | TPCN2 | -0.149570362 | 0.92127192 | -0.12342748 | 0.935063964 | -0.272997842 | 4.467311984 | 7.145017454 | 8.380957451 | 7.114945424 | 7.32531744 | 7.024410815 | 7.444077031 |
| ENSG00000143303 | RRNAD1 | -0.389838099 | 0.850552412 | 0.116069491 | 0.956087085 | -0.273768608 | 8.444143041 | 7.145017454 | 2 | 7.03996889 | 6.92586261 | 7.57363085 | 7.372114752 |
| ENSG00000103653 | CSK | -0.249590421 | 0.818844433 | -0.024695377 | 0.982605796 | -0.274285798 | 7.709880939 | 8.376385542 | 10.01740919 | 8.707945635 | 8.86603107 | 9.033940582 | 8.990045056 |
| ENSG00000122122 | SASH3 | -3.753010097 | 0.531143362 | 3.478557533 | 0.53589394 | -0.274452564 | 2 | 2 | 2 | 2 | 2 | 4.388329244 | 3.287118176 |
| ENSG00000167535 | CACNB3 | -0.535222119 | 0.462870065 | 0.259076474 | 0.731726336 | -0.276145645 | 8.540903468 | 7.901717006 | 9.426401271 | 8.266399315 | 8.18097132 | 9.145869306 | 8.86086441 |
| ENSG00000085998 | POMGNT1 | -0.14809326 | 0.929222345 | -0.1288043 | 0.938226757 | -0.27689756 | 6.138360429 | 9.814097497 | 10.9743126 | 9.788505066 | 9.83037437 | 9.63945754 | 9.996137262 |
| ENSG00000131748 | STARD3 | 0.077710168 | 0.918662967 | -0.355023574 | 0.605467095 | -0.277313406 | 9.344956677 | 8.820493869 | 7.896971941 | 8.817637515 | 8.94344897 | 8.576956626 | 8.318287967 |
| ENSG00000050426 | LETMD1 | 0.028909185 | 0.981338611 | -0.306490148 | 0.78794815 | -0.277580962 | 6.563806279 | 8.191702463 | 9.330306409 | 8.361842177 | 8.53549186 | 8.311822576 | 7.8990585 |
| ENSG00000198833 | UBE2J1 | -0.631823285 | 0.018342641 | 0.353588945 | 0.208031037 | -0.27823434 | 10.03472846 | 10.40166101 | 10.23774519 | 9.575477782 | 9.63530531 | 10.58174434 | 10.5931302 |
| ENSG00000160062 | ZBTB8A | 0.06366546 | 0.921327192 | -0.342856776 | 0.548805363 | -0.279191316 | 6.563806279 | 7.099070191 | 7.600047325 | 7.206526483 | 7.20671394 | 6.916861442 | 6.700172923 |
| ENSG00000127337 | YEATS4 | -0.294045454 | 0.778918686 | 0.014299537 | 0.989360801 | -0.279745917 | 6.366709671 | 6.04685525 | 4.058998049 | 5.461211713 | 5.58783529 | 5.776790516 | 5.842465082 |
| ENSG00000178913 | TAF7 | 0.050843435 | 0.94878661 | -0.331012808 | 0.656735751 | -0.280169373 | 11.32815351 | 10.46189757 | 9.757248847 | 10.71183653 | 10.7051385 | 10.45207267 | 10.19151462 |
| ENSG00000165650 | PDZD8 | -0.742302471 | 0.025103126 | 0.461880925 | 0.177835207 | -0.280421546 | 9.423197387 | 9.079034257 | 9.553445883 | 8.695066731 | 8.54970888 | 9.898624863 | 9.736956851 |
| ENSG00000197417 | SHPK | 0.531164676 | 0.711436164 | -0.813510386 | 0.550336725 | -0.28234571 | 7.218819781 | 6.04685525 | 3.369317972 | 6.699459234 | 6.77805036 | 5.217477917 | 5.694529741 |

| | | | | | | | | | | | | | |
|---|---|---|---|---|---|---|---|---|---|---|---|---|---|
| ENSG00000075975 | MKRN2 | -0.506629535 | 0.182413091 | 0.22241316 | 0.579833528 | -0.284216375 | 9.128497779 | 8.671479029 | 9.099922433 | 8.394075037 | 8.54970888 | 9.32239471 | 9.058587737 |
| ENSG00000136527 | TRA2B | 0.195146288 | 0.899681269 | -0.479737451 | 0.73847616 | -0.284591162 | 10.31337612 | 9.214490408 | 6.073378139 | 9.579376566 | 9.47155148 | 8.567710236 | 9.096596023 |
| ENSG00000124486 | USP9X | -0.274949579 | 0.732066351 | -0.010297947 | 0.990275412 | -0.285247525 | 10.37373127 | 12.01813185 | 11.91252756 | 11.34022038 | 11.3165787 | 11.58011589 | 11.6052181 |
| ENSG00000117614 | SYF2 | 0.826514757 | 0.35107637 | -1.112132153 | 0.188112133 | -0.285617395 | 9.174465797 | 7.393647515 | 7.451345288 | 9.095408867 | 9.05839257 | 7.241533944 | 7.091143645 |
| ENSG00000170345 | FOS | 0.801000683 | 0.531497817 | -1.087323117 | 0.374771224 | -0.286322434 | 7.709880939 | 6.311476754 | 5.59507254 | 7.632926745 | 7.54308677 | 6.473229652 | 4.388595494 |
| ENSG00000104870 | FCGRT | -0.708525534 | 0.01449096 | 0.421708717 | 0.162812823 | -0.286816817 | 10.45668146 | 10.01303782 | 10.28245311 | 9.535899247 | 9.57170671 | 10.65512272 | 10.70441818 |
| ENSG00000197380 | DACT3 | -1.069680083 | 0.461552688 | 0.782834549 | 0.591158809 | -0.286845535 | 5.532353208 | 8.505280662 | 5.156626401 | 6.155987605 | 6.23747437 | 8.123435792 | 7.846917442 |
| ENSG00000138640 | FAM13A | 0.366463059 | 0.651205609 | -0.653475363 | 0.388123757 | -0.287012304 | 6.96351253 | 8.505280662 | 7.777116083 | 8.323303759 | 8.17486938 | 6.854018691 | 7.561323797 |
| ENSG00000135373 | EHF | 4.344324678 | 0.119484573 | -4.631915205 | 0.096576281 | -0.287590527 | 2 | 2 | 5.930865648 | 8.512996295 | 8.66303256 | 2 | 2.466008509 |
| ENSG00000180929 | GPR62 | 0.312648954 | 0.920079197 | -0.600961736 | NA | -0.288312782 | 2 | 3.076665209 | 2 | 2.695456494 | 2.68924543 | 2.526562374 | 2.277179783 |
| ENSG00000197405 | C5AR1 | -2.127526704 | 0.415078879 | 1.838185114 | 0.475101652 | -0.28934159 | 2 | 2 | 6.629488387 | 3.289987342 | 3.65239003 | 7.552954291 | 5.475755622 |
| ENSG00000099995 | SF3A1 | -0.473480464 | 0.332121646 | 0.180953862 | 0.728484312 | -0.292526602 | 10.03472846 | 9.532315021 | 10.46808205 | 9.58946437 | 9.58480612 | 10.19698705 | 10.27951104 |
| ENSG00000105993 | DNAJB6 | -0.080799685 | 0.91483416 | -0.214222638 | 0.762852238 | -0.295022323 | 9.761125267 | 9.328390744 | 10.66665689 | 9.972901061 | 9.92065856 | 9.970159828 | 9.639171944 |
| ENSG00000144063 | MALL | 2.329003389 | 0.210602238 | -2.624472601 | 0.169780322 | -0.295469212 | 3.327711234 | 3.076665209 | 2 | 4.598938466 | 4.31639476 | 2.267001569 | 2.117335275 |
| ENSG00000099875 | MKNK2 | -0.434975865 | 0.506870806 | 0.139355994 | 0.842279911 | -0.295619871 | 9.128497779 | 9.956182102 | 10.27509734 | 9.363030664 | 9.49296422 | 10.30943039 | 9.608182402 |
| ENSG00000095066 | HOOK2 | -0.740354847 | 0.485315834 | 0.441378876 | 0.684334335 | -0.298975971 | 5.095959458 | 7.274681896 | 5.59507254 | 5.731694 | 5.53799497 | 6.633459761 | 6.864442837 |
| ENSG00000105755 | ETHE1 | 1.014608558 | 0.588479045 | -1.313687866 | 0.466340006 | -0.299079309 | 9.931427102 | 7.819099893 | 3.369317972 | 9.685196338 | 9.64637295 | 7.160478198 | 7.545707309 |
| ENSG00000140943 | MBTPS1 | -0.236671143 | 0.758253496 | -0.063604667 | 0.937496618 | -0.300275809 | 9.473095744 | 10.72651826 | 11.04521411 | 10.32930776 | 10.3121369 | 10.35149484 | 10.62242656 |
| ENSG00000101439 | CST3 | -0.317544985 | 0.382104545 | 0.014747639 | 0.971501984 | -0.302797346 | 12.06760845 | 11.65309132 | 11.74160414 | 11.47105152 | 11.5564566 | 11.65521291 | 12.01499609 |
| ENSG00000184517 | ZFP1 | -0.579695524 | 0.280922232 | 0.276018159 | 0.621070373 | -0.303677365 | 6.73718411 | 6.951736442 | 6.80318718 | 6.237202172 | 6.32824132 | 7.350983915 | 6.808832901 |
| ENSG00000081148 | IMPG2 | 0.592108463 | 0.82454488 | -0.896362502 | 0.724892691 | -0.304254039 | 2 | 2 | 4.058998049 | 2.947924294 | 3.60476711 | 2.592885947 | 2.632958241 |
| ENSG00000148468 | FAM171A1 | -0.958628982 | 0.391821726 | 0.654318564 | 0.564387793 | -0.304310418 | 6.366709671 | 9.470501606 | 8.631919287 | 7.794595599 | 7.57432889 | 9.179643937 | 9.378808813 |
| ENSG00000196363 | WDR5 | -0.519330302 | 0.300898696 | 0.211592223 | 0.690410177 | -0.307738079 | 8.946185348 | 8.101365653 | 8.206335842 | 7.985332082 | 7.91907183 | 8.694421086 | 8.653693725 |
| ENSG00000188677 | PARVB | -0.706777786 | 0.33594872 | 0.397773984 | 0.600319397 | -0.309003802 | 8.816894989 | 7.928237109 | 7.163363831 | 7.507583254 | 7.35080119 | 8.538032809 | 8.497408326 |
| | C16ORF52 | 6.662861068 | 3.49E-06 | -6.971965631 | 0.00142934 | -0.309104563 | 2 | 3.686303414 | 2 | 8.521127045 | 8.4504872 | 2 | 2 |
| ENSG00000122176 | FMOD | -2.626720108 | 0.09807061 | 2.317276872 | 0.141678745 | -0.309443236 | 4.00722531 | 8.124485672 | 9.954333071 | 6.18083559 | 6.18985553 | 10.74946305 | 11.29947428 |
| ENSG00000163866 | SMIM12 | -0.579138185 | 0.611410824 | 0.269322701 | 0.82055968 | -0.309815484 | 8.928409675 | 8.916342327 | 6.203069917 | 7.836899669 | 7.91175216 | 8.355461064 | 8.998195614 |
| ENSG00000131620 | ANO1 | -5.694929374 | 0.031180503 | 5.384931636 | 0.006312157 | -0.309992338 | 2 | 2 | 4.058998049 | 2 | 2 | 7.761068522 | 6.883203642 |
| ENSG00000113758 | DBN1 | -0.768604753 | 0.176804511 | 0.458586172 | 0.435298592 | -0.310018581 | 12.14083108 | 11.13786633 | 11.33422412 | 10.83272206 | 10.8436425 | 11.75696777 | 12.31632069 |
| ENSG00000143341 | HMCN1 | -1.155513076 | 0.359275511 | 0.845295301 | 0.506163186 | -0.310217775 | 5.095959458 | 7.847167909 | 8.742726175 | 6.682159616 | 6.76727577 | 8.263008721 | 9.010572782 |
| ENSG00000116128 | BCL9 | -1.108634351 | 0.437795312 | 0.794932144 | 0.58065443 | -0.313702207 | 4.467311984 | 7.928237109 | 8.941672936 | 6.846474326 | 6.94511186 | 8.652274805 | 8.868207441 |
| ENSG00000151655 | ITIH2 | -3.753010588 | 0.531143362 | 3.439230986 | 0.540908493 | -0.313779602 | 2 | 2 | 2 | 2 | 2 | 3.648325314 | 4.133950982 |
| ENSG00000164889 | SLC4A2 | 0.292226397 | 0.632716815 | -0.606782476 | 0.280587322 | -0.31455608 | 9.957954117 | 10.24169813 | 10.91875185 | 10.67405041 | 10.7672916 | 9.474992887 | 10.10714012 |
| ENSG00000103811 | CTSH | -2.245863179 | 0.340614989 | 1.930768819 | 0.406108709 | -0.315094361 | 2 | 6.535001748 | 2 | 3.09461492 | 3.60476711 | 7.078941352 | 6.681239787 |
| ENSG00000115053 | NCL | -0.2026247 | 0.804551315 | -0.113806381 | 0.891733131 | -0.316431081 | 12.60813735 | 11.81817171 | 10.9743126 | 11.7358446 | 11.7585122 | 11.56380211 | 12.06480799 |
| ENSG00000066027 | PPP2R5A | -0.546959254 | 0.337437807 | 0.230348902 | 0.702388158 | -0.316610352 | 7.867441304 | 8.395519306 | 8.434763955 | 7.751013478 | 7.68437616 | 8.840629888 | 8.008487789 |
| ENSG00000164253 | WDR41 | -0.912043258 | 0.015510476 | 0.594863974 | 0.125418384 | -0.317179284 | 9.358295023 | 8.687092496 | 8.960138144 | 8.128448191 | 8.11451391 | 9.675771867 | 9.551318233 |
| ENSG00000163563 | MNDA | -3.753010643 | 0.531143362 | 3.434835446 | 0.541339971 | -0.318175197 | 2 | 2 | 2 | 2 | 2 | 4.291799915 | 3.383972333 |
| ENSG00000140718 | FTO | -0.592341579 | 0.473769274 | 0.273867478 | 0.750952725 | -0.3184741 | 8.39321512 | 9.962611189 | 10.26027233 | 9.160358775 | 9.13687201 | 9.809259615 | 10.18669803 |
| ENSG00000188352 | FOCAD | -0.663288356 | 0.478725082 | 0.344685142 | 0.72255273 | -0.318603214 | 9.27634997 | 8.702538795 | 6.958202216 | 7.971157485 | 7.91419618 | 8.816978708 | 9.051693903 |
| ENSG00000109881 | CCDC34 | -0.026566918 | 0.984043262 | -0.292568167 | 0.82055968 | -0.319135085 | 4.467311984 | 6.389896633 | 4.058998049 | 5.406080789 | 5.33407135 | 5.22272385 | 5.035976547 |
| ENSG00000154096 | THY1 | 1.429765059 | 0.329061986 | -1.75058436 | 0.214752155 | -0.320819301 | 11.83218596 | 10.09725346 | 6.958202216 | 12.06377966 | 12.1177047 | 8.817412208 | 9.018137056 |
| ENSG00000166387 | PPFIBP2 | -3.151962709 | 9.48E-10 | 2.831000483 | 7.94E-09 | -0.320962226 | 6.96351253 | 7.979858146 | 7.163363831 | 4.387512679 | 4.68029874 | 10.22568517 | 10.28813986 |
| ENSG00000130706 | ADRM1 | 0.234740894 | 0.854286984 | -0.557743108 | 0.635985655 | -0.323002214 | 10.60951384 | 9.936721131 | 7.398165132 | 10.01956245 | 10.0874876 | 9.068221759 | 9.438868282 |
| ENSG00000181038 | METTL23 | -0.198545844 | 0.927391636 | -0.126193953 | 0.954172164 | -0.324739797 | 8.75772933 | 7.002537973 | 2 | 7.454228844 | 7.26532099 | 7.402744819 | 7.463599504 |
| ENSG00000178719 | GRINA | 0.193606804 | 0.713585581 | -0.518453854 | 0.275620191 | -0.32484705 | 11.57090097 | 12.00728915 | 12.48428878 | 12.24651637 | 12.2775093 | 11.42398765 | 11.66734563 |
| ENSG00000154370 | TRIM11 | -0.481847093 | 0.366470827 | 0.155438733 | 0.787507734 | -0.326408359 | 8.444143041 | 7.503546079 | 7.858115942 | 7.514115926 | 7.5014335 | 8.243140215 | 8.015118249 |
| ENSG00000100528 | CNIH1 | 0.011953501 | 0.981947437 | -0.338522703 | 0.456684488 | -0.326569202 | 9.689059027 | 10.40638494 | 9.897982195 | 10.02471865 | 10.0651181 | 9.849450427 | 9.524348754 |
| ENSG00000137770 | CTDSPL2 | -0.180827354 | 0.917096582 | -0.145945653 | 0.933175306 | -0.326773007 | 4.815609471 | 7.232742069 | 9.516493218 | 8.164192202 | 7.95986274 | 8.171702207 | 8.024537783 |
| ENSG00000068615 | REEP1 | -2.266295273 | 0.523661392 | 1.939415207 | 0.549471093 | -0.326880066 | 2 | 3.076665209 | 2 | 2.270786764 | 2 | 3.616794016 | 3.45257121 |
| ENSG00000175348 | TMEM9B | 0.211873006 | 0.707193196 | -0.539138228 | 0.290417779 | -0.327265222 | 8.517314938 | 8.590771778 | 9.372276026 | 9.110576034 | 9.06390621 | 8.191903518 | 8.481080411 |
| ENSG00000232838 | PET117 | 1.204012107 | 0.213514288 | -1.532923502 | 0.101181698 | -0.328911395 | 5.532353208 | 3.686303414 | 5.772716771 | 6.372775472 | 6.26070706 | 3.878466556 | 4.002845082 |
| ENSG00000157514 | TSC22D3 | -0.801445298 | 0.519400194 | 0.472389047 | 0.711063784 | -0.329056251 | 9.800745303 | 9.367676479 | 6.534109959 | 8.227176542 | 8.38151133 | 9.998344725 | 8.956482366 |
| ENSG00000075618 | FSCN1 | -1.219378269 | 0.042276369 | 0.889851411 | 0.145971903 | -0.329526858 | 12.26818393 | 11.34889202 | 11.21627664 | 10.46993609 | 10.4778448 | 12.1473965 | 12.9123837 |
| ENSG00000175806 | MSRA | 0.20145032 | 0.869968338 | -0.531157651 | 0.637573045 | -0.329707332 | 6.138360429 | 5.946638857 | 8.007629717 | 7.106324235 | 7.29937545 | 5.886629224 | 6.928061817 |
| ENSG00000145293 | ENOPH1 | -0.56886352 | 0.658741986 | 0.237697997 | 0.8601928 | -0.331165522 | 8.816894989 | 8.732943944 | 5.39244166 | 7.703217906 | 7.6986238 | 8.446833885 | 8.542691771 |
| ENSG00000112799 | LY86 | -3.75301081 | 0.531143362 | 3.421462616 | 0.543166952 | -0.331548194 | 2 | 2 | 2 | 2 | 2 | 4.350488971 | 3.236138336 |
| ENSG00000065621 | GSTO2 | 1.027162119 | 0.583738181 | -1.360193934 | 0.450687102 | -0.333031815 | 4.467311984 | 5.131603625 | 2 | 5.22679461 | 5.28958453 | 3.943456545 | 2.509591799 |
| ENSG00000087087 | SRRT | -0.214681381 | 0.666056796 | -0.118733646 | 0.816271471 | -0.333415027 | 9.689059027 | 9.148351514 | 9.049077186 | 9.103012382 | 9.11579188 | 8.960480054 | 9.413638893 |
| ENSG00000132763 | MMACHC | -0.724796769 | 0.694034631 | 0.391321272 | 0.836584165 | -0.333475497 | 2 | 5.946638857 | 7.451345288 | 5.639097847 | 5.64780787 | 6.601598251 | 6.784644656 |
| ENSG00000152454 | ZNF256 | -0.482964696 | 0.789639588 | 0.149212982 | 0.936599654 | -0.333751714 | 2 | 5.946638857 | 7.098165388 | 5.614989456 | 5.63601114 | 6.244776687 | 6.185873005 |
| ENSG00000049769 | PPP1R3F | -0.791654401 | 0.722734802 | 0.456518602 | 0.84189993 | -0.3351358 | 8.631580145 | 5.303523747 | 2 | 6.422239259 | 6.44101097 | 7.976682981 | 7.20156326 |
| ENSG00000026652 | AGPAT4 | -0.016225951 | 0.994366162 | -0.319234495 | 0.878328838 | -0.335460446 | 2 | 8.540087354 | 7.896971941 | 7.703217906 | 7.62593544 | 7.329261848 | 7.407104868 |
| ENSG00000170037 | CNTROB | -0.690427135 | 0.712692238 | 0.352016513 | 0.855655091 | -0.338410622 | 3.327711234 | 7.393647515 | 9.478569125 | 7.52061915 | 7.54938955 | 8.554768917 | 8.568035872 |
| ENSG00000183255 | PTTG1IP | -0.561400578 | 0.26397465 | 0.222048046 | 0.68121881 | -0.339352532 | 12.60953239 | 11.91084881 | 12.89867565 | 11.96531395 | 11.9706237 | 12.66522651 | 12.83161361 |
| ENSG00000168653 | NDUFS5 | -0.374840834 | 0.429456377 | 0.0350944 | 0.947397878 | -0.339746434 | 10.43166252 | 9.616264339 | 10.2528023 | 9.74820345 | 9.78034427 | 10.06484606 | 10.2730054 |
| ENSG00000183570 | PCBP3 | -2.53387251 | 0.159913355 | 2.1936141 | 0.212256603 | -0.34025841 | 3.327711234 | 6.464272742 | 2 | 3.227757282 | 3.2184422 | 7.263334844 | 7.100081147 |
| ENSG00000138613 | APH1B | -0.097602976 | 0.855515488 | -0.243235275 | 0.619827156 | -0.340838251 | 7.709880939 | 7.503546079 | 8.236946975 | 7.826439377 | 7.65837038 | 7.545646914 | 7.656486163 |
| ENSG00000282019 | | 0.314889248 | 0.567016969 | -0.655966684 | 0.193744404 | -0.341077436 | 11.27287932 | 10.67263725 | 10.9968459 | 11.30745256 | 11.3225709 | 9.86021999 | 10.70939118 |
| ENSG00000156931 | VPS8 | 0.439489774 | 0.686884062 | -0.780852722 | 0.443851376 | -0.341362949 | 8.042902253 | 7.189546432 | 9.681375057 | 9.084476714 | 9.13896318 | 7.890311949 | 7.920407983 |
| ENSG00000204314 | PRRT1 | 4.848574462 | 0.281547017 | -5.191110886 | 0.275561942 | -0.342536424 | 2 | 2 | 2 | 4.965943888 | 5.14724905 | 2 | 2 |
| ENSG00000143196 | DPT | 1.066229779 | 0.665819919 | -1.409802585 | 0.555970778 | -0.343572806 | 2 | 3.686303414 | 2 | 3.663521627 | 3.2184422 | 2.457042231 | 2.326754077 |
| ENSG00000023330 | ALAS1 | 0.03383807 | 0.96176107 | -0.378811465 | 0.554788482 | -0.344973395 | 10.25038534 | 8.916342327 | 9.926432745 | 9.791869268 | 9.8686817 | 9.39882884 | 9.441331959 |
| ENSG00000184900 | SUMO3 | -0.269288051 | 0.641252598 | -0.07577366 | 0.902642273 | -0.345061712 | 10.67493801 | 9.680074956 | 10.63827116 | 10.1305433 | 10.1248958 | 10.21552475 | 10.41854076 |
| ENSG00000213339 | QTRT1 | -0.177225064 | 0.83752462 | -0.167949222 | 0.842413632 | -0.345174286 | 5.866944937 | 7.35507212 | 7.451345288 | 6.820355514 | 6.94032357 | 6.672321934 | 7.08034504 |
| ENSG00000152315 | KCNK13 | -2.312813426 | NA | 1.96738736 | NA | -0.345426066 | 2 | 2 | 2 | 2 | 2.14022021 | 3.429858444 | 2.374681325 |
| ENSG00000137500 | CCDC90B | 0.137049903 | 0.824880011 | -0.482771905 | 0.376596657 | -0.345722003 | 7.624139768 | 8.356994601 | 8.721235202 | 8.474993083 | 8.39906692 | 7.925322387 | 7.720479276 |
| ENSG00000115318 | LOXL3 | -0.845395728 | 0.128214828 | 0.497340444 | 0.386914635 | -0.348055284 | 11.13243651 | 10.16572116 | 10.23015729 | 9.713855064 | 9.760787 | 10.83544511 | 11.28239915 |
| ENSG00000090061 | CCNK | 1.63949277 | 0.005747675 | -1.989313961 | 0.000598834 | -0.349821191 | 8.340423424 | 7.638155693 | 9.083172337 | 10.07418693 | 10.1132431 | 6.459963649 | 6.585205928 |
| ENSG00000104450 | SPAG1 | -0.332171113 | 0.719366965 | -0.019045175 | 0.984485378 | -0.351216288 | 5.866944937 | 6.389896633 | 4.523675391 | 5.501219927 | 5.51241389 | 5.699879814 | 5.884455123 |
| ENSG00000119559 | C19orf25 | -0.180412213 | 0.7329405 | -0.171833291 | 0.741322616 | -0.352245504 | 8.16936288 | 8.487557258 | 7.600047325 | 7.910550164 | 8.01327646 | 7.934143724 | 8.007538088 |
| ENSG00000162687 | KCNT2 | -2.292481394 | 0.020855853 | 1.940176582 | 0.042725112 | -0.352304812 | 5.70932728 | 4.93638933 | 5.156626401 | 3.79808662 | 3.08597292 | 6.287958221 | 7.647978997 |
| ENSG00000153914 | SREK1 | 0.082394618 | 0.922706766 | -0.437203124 | 0.564260714 | -0.354808505 | 9.2903348 | 9.949724236 | 8.236946975 | 9.435370781 | 9.37460909 | 8.975319061 | 8.797990868 |
| ENSG00000142864 | SERBP1 | -0.347526853 | 0.713335141 | -0.007736128 | 0.993931149 | -0.355262981 | 11.61879218 | 11.04700893 | 9.399592459 | 10.6042022 | 10.6035548 | 10.85003188 | 11.03013303 |
| ENSG00000123485 | HJURP | 0.134682881 | 0.930008622 | -0.492002843 | 0.728500949 | -0.357319962 | 4.00722531 | 6.311476754 | 4.058998049 | 5.304221918 | 5.45984938 | 4.301748724 | 5.217771644 |
| ENSG00000136982 | DSCC1 | -0.38728097 | 0.824427896 | 0.028613038 | 0.987221496 | -0.358667932 | 4.00722531 | 5.131603625 | 2 | 3.997744386 | 3.90918929 | 4.604690117 | 3.841670744 |
| ENSG00000129559 | NEDD8 | -0.12730055 | 0.841925584 | -0.231538879 | 0.699954446 | -0.358839429 | 10.59831544 | 9.549500232 | 9.798878489 | 9.940529254 | 9.91151326 | 9.674576002 | 9.956333606 |
| ENSG00000133142 | TCEAL4 | -0.875346126 | 0.174064658 | 0.514688406 | 0.439219521 | -0.36065772 | 10.54098126 | 9.102512015 | 9.878698603 | 9.076774616 | 9.09116272 | 10.31389217 | 10.60620688 |
| ENSG00000174808 | BTC | 4.829973619 | 0.286761062 | -5.191110333 | 0.278346547 | -0.361136714 | 2 | 2 | 2 | 5.210797711 | 4.85329164 | 2 | 2 |
| ENSG00000138433 | CIR1 | 0.291220428 | 0.662232293 | -0.65292133 | 0.283396195 | -0.361700902 | 9.567995545 | 8.623598428 | 8.35328246 | 9.274418797 | 9.19330273 | 8.39955642 | 8.191057445 |
| ENSG00000132612 | VPS4A | -0.652060826 | 0.133069745 | 0.286115089 | 0.53558846 | -0.365945736 | 10.01801709 | 9.566483142 | 10.37481416 | 9.306761457 | 9.43372295 | 10.37686315 | 10.2307005 |
| ENSG00000131437 | KIF3A | -0.431843689 | 0.679228019 | 0.064281625 | 0.954416752 | -0.367562064 | 8.009475818 | 6.140560021 | 8.460932916 | 7.484483106 | 7.29937545 | 8.195910056 | 7.477384137 |
| ENSG00000161036 | LRWD1 | 0.006316062 | 0.995663772 | -0.374833258 | 0.687802707 | -0.368517197 | 8.075571721 | 7.051611315 | 5.930865648 | 7.230526165 | 7.32163988 | 6.887434761 | 6.926052784 |
| ENSG00000107554 | DNMBP | 0.359179529 | 0.680321745 | -0.72974284 | 0.364350502 | -0.370563311 | 9.097018025 | 8.993716073 | 10.63827116 | 10.12148041 | 10.1644413 | 9.042865619 | 9.076811853 |
| ENSG00000198171 | DDRGK1 | 0.4683342435 | 0.318114195 | -0.839688279 | 0.05360207 | -0.371345844 | 9.720386367 | 8.862353853 | 9.242516611 | 9.763363111 | 9.79633147 | 8.544855285 | 8.415288533 |
| ENSG00000188130 | MAPK12 | -0.276521267 | 0.824387358 | -0.095317955 | 0.940405661 | -0.371839222 | 8.313285031 | 8.297207175 | 5.39244166 | 7.514115926 | 7.57741621 | 7.583859056 | 7.851158781 |
| ENSG00000187109 | NAP1L1 | 0.046752507 | 0.954376589 | -0.419373972 | 0.570849073 | -0.372621464 | 11.37484003 | 11.00713613 | 9.725218844 | 10.93180718 | 10.8737771 | 10.45025691 | 10.42584592 |
| ENSG00000151498 | ACAD8 | -0.012714951 | 0.989340687 | -0.36149967 | 0.676479319 | -0.374214621 | 7.031698319 | 7.232742069 | 8.65477198 | 7.805288479 | 7.82349365 | 7.262061517 | 7.656486163 |
| ENSG00000130349 | C6orf203 | -0.281215463 | 0.832852241 | -0.093454231 | 0.944699256 | -0.374669694 | 7.709880939 | 5.72254323 | 4.874587708 | 6.34372989 | 6.31350454 | 6.589466407 | 6.420873184 |
| ENSG00000037042 | TUBG2 | -0.024159918 | 0.9732242 | -0.351339828 | 0.588182865 | -0.375499746 | 8.928409675 | 7.669933291 | 8.486635645 | 8.458152491 | 8.38151133 | 7.890311949 | 8.278687361 |
| ENSG00000119535 | CSF3R | -3.753011378 | 0.531143362 | 3.375955232 | 0.548441916 | -0.377056147 | 2 | 2 | 2 | 2 | 2 | 4.350488971 | 3.128433649 |
| ENSG00000181544 | FANCB | -1.085443007 | 0.646818108 | 0.707299006 | 0.767139907 | -0.378144001 | 2 | 5.131603625 | 2 | 3.406930997 | 2.94009902 | 4.675266036 | 4.175132639 |
| ENSG00000166246 | C16orf71 | -1.465964689 | 0.514678379 | 1.087083002 | 0.62219346 | -0.378881688 | 2 | 4.710564154 | 2 | 2.947924294 | 2.59489879 | 4.369533167 | 4.400220394 |
| ENSG00000165526 | RPUSD4 | -0.28898036 | 0.598988441 | -0.089920304 | 0.878637249 | -0.378900664 | 7.218819781 | 7.701025996 | 8.206335842 | 7.471115032 | 7.48508724 | 7.730090858 | 7.613439639 |
| ENSG00000168288 | MMADHC | 0.341018218 | 0.739039557 | -0.723078583 | 0.446930583 | -0.382060365 | 11.03197509 | 9.256956408 | 8.904018013 | 10.42037002 | 10.3613276 | 9.343015767 | 9.320352741 |
| ENSG00000107829 | FBXW4 | -0.489265943 | 0.733300911 | 0.105977197 | 0.944157185 | -0.383288746 | 5.095959458 | 8.762721501 | 9.180874355 | 7.915493556 | 8.01782901 | 8.637620652 | 8.467331218 |
| ENSG00000162512 | SDC3 | -0.820574106 | 0.328511143 | 0.436623076 | 0.618028749 | -0.383951029 | 10.27872131 | 10.96932597 | 8.904018013 | 9.44822005 | 9.47817393 | 10.54203348 | 10.87071469 |

| | | | | | | | | | | | | | |
|---|---|---|---|---|---|---|---|---|---|---|---|---|---|
| ENSG00000101363 | MANBAL | -0.630226369 | 0.280938145 | 0.246234635 | 0.692848923 | -0.383991734 | 9.761125267 | 9.203675681 | 8.536706852 | 8.660150093 | 8.59306436 | 9.386026816 | 9.599699825 |
| ENSG00000168454 | TXNDC2 | -0.467978772 | 0.884927669 | 0.083148957 | NA | -0.384829815 | 2 | 3.076665209 | 2 | 2.270786764 | 2.59489879 | 2.62493832 | 2.592996859 |
| ENSG00000177674 | AGTRAP | 0.471036861 | 0.461552688 | -0.855961766 | 0.146023342 | -0.384924905 | 9.635285994 | 8.337339476 | 9.242516611 | 9.524619869 | 9.72990471 | 8.397238383 | 8.22740336 |
| ENSG00000114784 | EIF1B | 0.161973187 | 0.842086376 | -0.547382956 | 0.450077395 | -0.38540977 | 9.410448395 | 7.979858146 | 8.585098385 | 8.976268302 | 8.89230135 | 8.52588267 | 7.864857644 |
| ENSG00000167680 | SEMA6B | -1.434806159 | 0.531386353 | 1.049038569 | 0.64532651 | -0.38576759 | 2 | 4.93638933 | 2 | 2.784612485 | 3.01487888 | 4.920961476 | 4.062589736 |
| ENSG00000168993 | CPLX1 | -0.951757503 | 0.77464352 | 0.565839125 | NA | -0.385918378 | 2 | 3.076665209 | 2 | 2.270786764 | 2.38539787 | 3.172301926 | 2.277179783 |
| ENSG00000221883 | ARIH2OS | 0.016977224 | 0.993765388 | -0.404928644 | 0.832938355 | -0.38795142 | 3.327711234 | 2 | 4.874587708 | 3.709785322 | 3.90918929 | 3.465698119 | 3.580696614 |
| | LOC400927-( | 7.696521343 | 1.01E-12 | -8.08496528 | 2.68E-05 | -0.388443937 | 4.467311984 | 2 | 3.369317972 | 10.62924246 | 10.6114832 | 2 | 2 |
| ENSG00000269190 | FBXO17 | -0.59840767 | 0.274205421 | 0.209305397 | 0.722120418 | -0.389102273 | 8.468948322 | 8.213429689 | 7.502634626 | 7.474468681 | 7.58663873 | 8.525352075 | 8.094184551 |
| ENSG00000130300 | PLVAP | 0.312648823 | 0.918611539 | -0.702564565 | NA | -0.389915742 | 2 | 3.076665209 | 2 | 2.695456494 | 2.68924543 | 2.34604626 | 2.421067459 |
| ENSG00000163909 | HEYL | -3.817131392 | 0.089224711 | 3.426737723 | 0.062305464 | -0.390393669 | 4.467311984 | 2 | 2 | 2.141737246 | 2.14022021 | 6.430795201 | 5.529673353 |
| ENSG00000115325 | DOK1 | -0.953509061 | 0.478310707 | 0.563101614 | 0.683905619 | -0.390407447 | 10.09173937 | 6.228547981 | 8.408111533 | 7.978262192 | 8.07355593 | 9.35888613 | 9.680872942 |
| ENSG00000165832 | TRUB1 | -0.239131436 | 0.910161622 | -0.151618213 | 0.942375924 | -0.390749649 | 2 | 7.979858146 | 8.380957451 | 7.437142666 | 7.33264456 | 7.485815918 | 7.456657479 |
| ENSG00000107731 | UNC5B | -0.166573799 | 0.932700573 | -0.224271105 | 0.909359225 | -0.390844903 | 5.095959458 | 8.717821467 | 10.20715117 | 8.846395982 | 9.0020652 | 9.693119552 | 6.746446439 |
| ENSG00000213512 | GBP7 | 4.799379942 | 0.295753807 | -5.191109406 | 0.2826457 | -0.391729465 | 2 | 2 | 2 | 5.242616075 | 4.74762503 | 2 | 2 |
| ENSG00000126216 | TUBGCP3 | -0.29237324 | 0.88199664 | -0.101714477 | 0.958899411 | -0.394087718 | 6.138360429 | 8.820493869 | 3.369317972 | 7.226553841 | 7.16216989 | 7.417689057 | 7.342293759 |
| ENSG00000174276 | ZNHIT2 | -0.420402802 | 0.793600622 | 0.024263172 | 0.988451404 | -0.39613963 | 7.667647223 | 5.946638857 | 3.369317972 | 6.147608921 | 6.08962627 | 6.625560108 | 6.451924088 |
| ENSG00000074410 | CA12 | -7.177270833 | 0.004413569 | 6.780332 | 0.00027703 | -0.396938833 | 2 | 5.131603625 | 2 | 2 | 2 | 10.2041324 | 10.26647028 |
| ENSG00000069431 | ABCC9 | 2.651333088 | 0.000213646 | -3.050071296 | 1.90E-05 | -0.398738208 | 7.218819781 | 6.389896633 | 7.342949392 | 9.610955295 | 9.67979376 | 4.742549839 | 3.560117069 |
| ENSG00000104976 | SNAPC2 | -0.411184583 | 0.58474982 | 0.01215568 | 0.988303878 | -0.399028903 | 9.247966931 | 8.213429689 | 7.734844356 | 8.117764469 | 8.15020018 | 8.411091064 | 8.680755147 |
| ENSG00000102738 | MRPS31 | 0.04882451 | 0.972268825 | -0.447956678 | 0.732916452 | -0.399132167 | 8.138771685 | 5.946638857 | 5.156626401 | 6.941624891 | 7.10341274 | 6.649130473 | 6.435070131 |
| ENSG00000217128 | FNIP1 | 0.649110465 | 0.367412771 | -1.049329733 | 0.119894702 | -0.400219268 | 8.009475818 | 9.640525262 | 9.180874355 | 9.769519169 | 9.70964668 | 8.155334989 | 7.950369405 |
| ENSG00000136868 | SLC31A1 | -0.403211133 | 0.126527284 | -0.000501601 | 0.998980237 | -0.403712735 | 10.81212061 | 10.63690565 | 10.49998282 | 10.20813496 | 10.2972338 | 10.68714125 | 10.62196123 |
| ENSG00000126814 | TRMT5 | 0.113467258 | 0.891428469 | -0.51741566 | 0.477751115 | -0.403948402 | 8.493334308 | 6.899080544 | 8.007629717 | 8.03159949 | 8.05595642 | 7.536196998 | 7.30412922 |
| ENSG00000163660 | CCNL1 | 0.509099059 | 0.507120384 | -0.914021907 | 0.197757479 | -0.404922848 | 10.16946352 | 8.469613413 | 9.330306409 | 10.03327157 | 9.94357048 | 8.513094212 | 8.635366595 |
| ENSG00000129480 | DTD2 | -0.853764353 | 0.092249336 | 0.447679973 | 0.392537339 | -0.406084381 | 7.667647223 | 6.78762695 | 7.098165388 | 6.436065776 | 6.35727082 | 7.599066672 | 7.713507501 |
| ENSG00000149548 | CCDC15 | -0.359428662 | 0.890529219 | -0.046850069 | 0.985627735 | -0.406278731 | 2 | 5.303523747 | 2 | 3.754611435 | 3.7864959 | 4.321442857 | 3.580696614 |
| ENSG00000185252 | ZNF74 | -0.760161024 | 0.670378341 | 0.353783967 | 0.848992364 | -0.406377057 | 4.467311984 | 8.124485672 | 4.058998049 | 6.006476466 | 6.02777321 | 7.389977651 | 6.690737414 |
| ENSG00000119509 | INVS | 0.05438548 | 0.951349347 | -0.461378071 | 0.570028066 | -0.40699259 | 6.563806279 | 7.538394143 | 8.408111533 | 7.723090567 | 7.75969386 | 7.139828829 | 7.32867054 |
| ENSG00000135541 | AHI1 | 0.113969824 | 0.8692818 | -0.522714425 | 0.383915641 | -0.408744601 | 8.892187472 | 8.820493869 | 7.818184403 | 8.684969711 | 8.72001862 | 8.307508685 | 7.79392309 |
| ENSG00000134324 | LPIN1 | -0.029593605 | 0.981338611 | -0.380126411 | 0.738292369 | -0.409720016 | 7.33131396 | 9.949724236 | 10.0085654 | 9.438808458 | 9.51325794 | 8.712712904 | 9.450446099 |
| ENSG00000165023 | DIRAS2 | -3.753011788 | 0.531143362 | 3.343185637 | 0.552967424 | -0.40982615 | 2 | 2 | 2 | 2 | 2 | 3.891700877 | 3.789585079 |
| ENSG00000119638 | NEK9 | -0.559096197 | 0.211593503 | 0.149047176 | 0.763378865 | -0.410049021 | 10.75275473 | 9.969011754 | 10.16000817 | 9.704603278 | 9.84261543 | 10.52222593 | 10.43823328 |
| ENSG00000187172 | BAGE2 | 4.776907368 | 0.298608151 | -5.191109433 | 0.282784279 | -0.414202065 | 2 | 2 | 2 | 5.022023783 | 4.97065356 | 2 | 2 |
| | BAGE3 | 4.776907368 | 0.298608151 | -5.191109433 | 0.282784279 | -0.414202065 | 2 | 2 | 2 | 5.022023783 | 4.97065356 | 2 | 2 |
| ENSG00000163466 | ARPC2 | -0.016043333 | 0.979923499 | -0.400831512 | 0.464862114 | -0.416874845 | 12.54683767 | 11.64711661 | 11.60453593 | 12.00455363 | 11.9665083 | 11.53983427 | 11.66026272 |
| ENSG00000263002 | ZNF234 | 0.130780385 | 0.906858652 | -0.547873817 | 0.581597072 | -0.417093432 | 5.70932728 | 7.979858146 | 7.163363831 | 7.391747748 | 7.32898565 | 7.027411703 | 6.283256815 |
| ENSG00000104973 | MED25 | 0.056064753 | 0.94475044 | -0.47328207 | 0.513742625 | -0.417217318 | 8.075571721 | 9.566483142 | 8.561105505 | 8.94033939 | 8.92660044 | 8.252146986 | 8.55118937 |
| ENSG00000123130 | ACOT9 | -0.3305405 | 0.719060419 | -0.08688559 | 0.929627216 | -0.417426091 | 10.19198706 | 8.124485672 | 9.330306409 | 9.088859521 | 9.14105132 | 9.177956973 | 9.516692718 |
| ENSG00000144802 | NFKBIZ | 1.575569171 | 0.000562684 | -1.993060148 | 7.93E-06 | -0.417490977 | 8.674867122 | 9.235879652 | 9.725218844 | 10.83010652 | 10.8554541 | 7.269684675 | 7.34530375 |
| ENSG00000272602 | ZNF595 | -0.088069461 | 0.918975348 | -0.329808738 | 0.678175142 | -0.417878199 | 5.532353208 | 6.535001748 | 6.322058544 | 6.096288482 | 6.14061114 | 6.226613426 | 5.453611367 |
| ENSG00000130703 | OSBPL2 | -0.418607639 | 0.379355575 | 0.000545772 | 0.999263508 | -0.418061868 | 9.820154628 | 9.479495904 | 8.960138144 | 8.960808841 | 9.12847688 | 9.539475459 | 9.383208837 |
| ENSG00000154217 | PITPNC1 | 1.219595666 | 0.513480912 | -1.638544022 | 0.356668405 | -0.418948356 | 8.075571721 | 7.572420273 | 2 | 8.422134132 | 8.5227358 | 6.165271053 | 5.050803619 |
| ENSG00000143333 | RGS16 | -2.95109697 | 0.112520621 | 2.530101149 | 0.158865702 | -0.420995821 | 4.00722531 | 5.946638857 | 2 | 3.09461492 | 2.49394783 | 7.903438544 | 5.470251342 |
| ENSG00000113368 | LMNB1 | -0.336411325 | 0.902572075 | -0.084804685 | 0.974738711 | -0.421216009 | 2 | 6.311476754 | 2 | 4.739373666 | 4.45712836 | 4.799832052 | 4.836507146 |
| ENSG00000175115 | PACS1 | -0.46145765 | 0.38975761 | 0.038960289 | 0.948687403 | -0.422497361 | 10.21416435 | 10.48441188 | 11.2124308 | 10.2335933 | 10.2457682 | 10.67463433 | 10.80007469 |
| ENSG00000168246 | UBTD2 | -0.081745029 | 0.841645125 | -0.341503902 | 0.336902641 | -0.423248931 | 8.493334308 | 8.077869079 | 8.460932916 | 8.264463296 | 8.26953254 | 7.979009894 | 8.042267212 |
| ENSG00000123562 | MORF4L2 | -0.581917382 | 0.372864851 | 0.157670578 | 0.825446679 | -0.424246804 | 12.78399552 | 11.83228301 | 11.44325334 | 11.55026089 | 11.5548912 | 12.2230071 | 12.35668755 |
| ENSG00000087365 | SF3B2 | -0.410632528 | 0.228220774 | -0.013883577 | 0.972127213 | -0.424516105 | 11.21540273 | 11.22868348 | 10.75391088 | 10.6575484 | 10.6895293 | 10.99886917 | 11.13690957 |
| ENSG00000127191 | TRAF2 | -0.188672857 | 0.91485625 | -0.236242463 | 0.892291608 | -0.42491532 | 9.33149386 | 7.002537973 | 4.523675391 | 7.910550164 | 7.81567127 | 7.894426854 | 7.735470193 |
| ENSG00000130816 | DNMT1 | 0.037177735 | 0.942177035 | -0.462618249 | 0.290479733 | -0.425440515 | 9.189468652 | 8.747909548 | 9.553445883 | 9.196346103 | 9.26686067 | 8.704299335 | 8.767460665 |
| ENSG00000067842 | ATP2B3 | -0.949020201 | 0.764636496 | 0.523208041 | NA | -0.42581216 | 2 | 3.076665209 | 2 | 2.3892346 | 2.2680111 | 3.059555873 | 2.421067459 |
| ENSG00000282608 | ADORA3 | -3.753011995 | 0.531143362 | 3.326572216 | 0.555044393 | -0.426439779 | 2 | 2 | 2 | 2 | 2 | 4.311629396 | 3.100202476 |
| ENSG00000154059 | IMPACT | -0.428872594 | 0.544237044 | 0.001789586 | 0.998392474 | -0.427083009 | 8.418903792 | 9.018611864 | 7.502634626 | 8.045195239 | 8.00412803 | 8.510415622 | 8.386355612 |
| ENSG00000223496 | EXOSC6 | -0.614809495 | 0.350255547 | 0.186066726 | 0.794799105 | -0.428742769 | 8.609439662 | 7.315436877 | 8.175061096 | 7.530319349 | 7.5014335 | 8.522164399 | 8.045046786 |
| ENSG00000117984 | CTSD | 0.231031255 | 0.739013584 | -0.660426985 | 0.290456069 | -0.429395729 | 13.1138925 | 12.07561123 | 12.00879923 | 12.71608724 | 12.7327132 | 12.08025667 | 11.53610185 |
| ENSG00000184207 | PGP | -0.698538837 | 0.485453797 | 0.268935019 | 0.799962733 | -0.429603818 | 9.262228249 | 7.051611315 | 7.600047325 | 7.54315232 | 7.65837038 | 8.39955642 | 8.694982998 |
| ENSG00000164300 | SERINC5 | -1.225885445 | 0.000133455 | 0.794739261 | 0.016927551 | -0.431146185 | 9.556468883 | 9.203675681 | 9.452720984 | 8.209172298 | 8.17894021 | 10.37745126 | 9.997094495 |
| ENSG00000088035 | ALG6 | 0.336731388 | 0.802652992 | -0.768002364 | 0.53460268 | -0.431270976 | 5.866944937 | 6.602424674 | 3.369317972 | 6.104969671 | 6.14893615 | 5.328699035 | 4.861951484 |
| ENSG00000124767 | GLO1 | -0.311041572 | 0.723978556 | -0.121499654 | 0.895505527 | -0.432541226 | 11.18951828 | 10.13758676 | 9.242516611 | 10.1096668 | 10.0798882 | 10.32445299 | 10.24184794 |
| ENSG00000188452 | CERKL | -3.753012085 | 0.531143362 | 3.319361679 | 0.556010306 | -0.433650406 | 2 | 2 | 2 | 2 | 2 | 4.133696865 | 3.430065404 |
| ENSG00000108424 | KPNB1 | -0.365434296 | 0.448739712 | -0.068951794 | 0.897115004 | -0.43438609 | 11.24801458 | 12.01039539 | 12.06319515 | 11.43600295 | 11.4718853 | 11.68200198 | 11.81524183 |
| ENSG00000237440 | ZNF737 | 0.482665512 | 0.538070994 | -0.917548007 | 0.206491773 | -0.434882495 | 6.468621203 | 5.595927442 | 6.958202216 | 6.907295206 | 6.86652414 | 5.825876956 | 5.243813244 |
| ENSG00000129562 | DAD1 | -0.781454762 | 0.204362279 | 0.346305835 | 0.596208763 | -0.435148928 | 11.0444064 | 9.785372135 | 10.11951029 | 9.617795654 | 9.66027117 | 10.87544335 | 10.64107086 |
| | BAGE4 | 4.755655615 | 0.303181423 | -5.191108963 | 0.284871836 | -0.435453347 | 2 | 2 | 2 | 5.022023783 | 4.93258395 | 2 | 2 |
| | BAGE5 | 4.755655615 | 0.303181423 | -5.191108963 | 0.284871836 | -0.435453347 | 2 | 2 | 2 | 5.022023783 | 4.93258395 | 2 | 2 |
| ENSG00000168152 | THAP9 | -0.399282679 | 0.732697705 | -0.036597239 | 0.975799274 | -0.435879919 | 4.00722531 | 5.72254323 | 5.59507254 | 4.846787657 | 5.04390545 | 5.707383854 | 4.634793542 |
| ENSG00000160791 | CCR5 | -3.753012115 | 0.531143362 | 3.316981831 | 0.556282123 | -0.436030283 | 2 | 2 | 2 | 2 | 2 | 4.397636241 | 2.851942824 |
| ENSG00000156469 | MTERF3 | -0.088321698 | 0.970209755 | -0.348475655 | 0.883002588 | -0.436797353 | 8.493334308 | 4.93638933 | 2 | 6.936770382 | 6.96881778 | 6.940933331 | 6.415155055 |
| ENSG00000150540 | HNMT | -0.008814267 | 0.997041173 | -0.428312676 | 0.814923166 | -0.437126943 | 2 | 6.04685525 | 4.523675391 | 5.058234798 | 4.89348248 | 5.026717879 | 4.047883918 |
| ENSG00000163629 | PTPN13 | -0.782776319 | 0.323240837 | 0.345199017 | 0.681431908 | -0.437577301 | 8.737455672 | 9.488434476 | 7.600047325 | 8.063125866 | 8.00412803 | 9.030953258 | 9.25982657 |
| ENSG00000147457 | CHMP7 | -0.471442728 | 0.566726644 | 0.033270424 | 0.970602478 | -0.438172304 | 10.21416435 | 8.356994601 | 9.272377397 | 8.983347946 | 9.02372946 | 9.494370989 | 9.515691102 |
| ENSG00000131931 | THAP1 | 0.106782665 | 0.931219161 | -0.545002917 | 0.62909049 | -0.438220252 | 5.532353208 | 6.228547981 | 7.818184403 | 7.021746293 | 6.8715633 | 6.743394671 | 5.727202496 |
| ENSG00000166821 | PEX11A | -0.700804628 | 0.763884658 | 0.262423149 | 0.914220926 | -0.43838148 | 2 | 8.101365653 | 4.058998049 | 5.910700032 | 6.01871653 | 6.944112782 | 6.81969491 |
| ENSG00000148248 | SURF4 | -0.456817869 | 0.254887129 | 0.017476406 | 0.97020102 | -0.439341463 | 12.4934348 | 12.26697646 | 12.92005579 | 12.12173968 | 12.1362731 | 12.4719126 | 12.72309441 |
| ENSG00000130119 | GNL3L | 0.249124511 | 0.616043469 | -0.688558682 | 0.119746169 | -0.439434171 | 7.485143553 | 8.004992147 | 8.007629717 | 8.126317766 | 8.07574085 | 6.953609309 | 7.380942161 |
| ENSG00000137710 | RDX | -0.3761076 | 0.470515237 | -0.06420754 | 0.911555747 | -0.44031514 | 9.800745303 | 10.7784592 | 10.45512158 | 10.06253517 | 9.98949012 | 10.4114447 | 10.25910765 |
| ENSG00000150893 | FREM2 | 3.189270163 | 0.579885898 | -3.630743412 | 0.538029429 | -0.441473249 | 2 | 2 | 2 | 4.497095146 | 2 | 2 | 2.059860178 |
| ENSG00000118705 | RPN2 | 0.492700388 | 0.491521702 | -0.936852603 | 0.15499647 | -0.444152216 | 12.44893028 | 12.38655348 | 11.11279376 | 12.58338056 | 12.5997269 | 10.81040203 | 11.44713892 |
| ENSG00000174343 | CHRNA9 | 2.085402755 | 0.549630367 | -2.530047934 | 0.467835004 | -0.444645179 | 2 | 3.076665209 | 2 | 3.663521627 | 3.60476711 | 2.225793514 | 2 |
| ENSG00000272047 | GTF2H5 | -0.88040055 | 0.135121389 | 0.435588432 | 0.480467544 | -0.444812118 | 8.737455672 | 7.538394143 | 7.97167868 | 7.25413313 | 7.34356593 | 8.479522918 | 8.699106494 |
| ENSG00000204370 | SDHD | 0.118472909 | 0.911517345 | -0.563774258 | 0.54895459 | -0.445301349 | 9.624286623 | 8.414402626 | 7.342949392 | 8.931825643 | 8.80148734 | 8.255989884 | 8.127786376 |
| ENSG00000198937 | CCDC167 | -1.032466979 | 0.269034612 | 0.58641793 | 0.541021035 | -0.446049049 | 5.532353208 | 7.274681896 | 5.772716771 | 5.273747807 | 5.64780787 | 6.702675117 | 7.24894374 |
| ENSG00000163071 | SPATA18 | -1.810630226 | 0.36589956 | 1.362507773 | 0.498878282 | -0.448122453 | 8.75772933 | 6.04685525 | 2 | 5.674521485 | 5.64780787 | 9.069314155 | 8.28340307 |
| ENSG00000130560 | UBAC1 | -0.521811316 | 0.754977416 | 0.073193703 | 0.967559124 | -0.448617613 | 5.330594134 | 6.728495673 | 9.714382178 | 7.797276258 | 7.88956768 | 8.529591383 | 8.324402372 |
| ENSG00000130304 | SLC27A1 | -1.968299204 | 0.090738947 | 1.518811452 | 0.194468033 | -0.449487752 | 9.579430841 | 8.004992147 | 6.073378139 | 6.476767609 | 6.67805337 | 9.809259615 | 10.18984111 |
| ENSG00000146828 | SLC12A9 | -0.489036958 | 0.690575637 | 0.039124964 | 0.975799274 | -0.449911994 | 7.159102577 | 6.78762695 | 9.426401271 | 7.767511442 | 7.84670947 | 8.214461509 | 8.433081317 |
| ENSG00000181817 | LSM10 | -0.699910125 | 0.395581532 | 0.249140629 | 0.77791101 | -0.450769496 | 8.855032352 | 8.777382925 | 7.029880454 | 7.641902724 | 7.83385794 | 8.535927079 | 8.80566012 |
| ENSG00000184432 | COPB2 | -0.068203533 | 0.8177838 | -0.38304489 | 0.123771525 | -0.451248423 | 11.84171405 | 11.77133279 | 11.53541251 | 11.630805 | 11.6775673 | 11.31807122 | 11.36155524 |
| ENSG00000197497 | ZNF665 | -0.879137244 | 0.548487869 | 0.426486725 | 0.780515702 | -0.452650519 | 5.866944937 | 3.076665209 | 6.958202216 | 5.210797711 | 5.13053105 | 6.508020621 | 6.235386446 |
| ENSG00000179094 | PER1 | -1.019970907 | 0.283678293 | 0.567049685 | 0.564587637 | -0.452921222 | 6.468621203 | 8.433041974 | 8.922968322 | 7.165617905 | 7.34718809 | 8.626284124 | 8.992447063 |
| ENSG00000106524 | ANKMY2 | -0.232083296 | 0.859987098 | -0.222028629 | 0.863495873 | -0.454111924 | 5.70932728 | 8.124485672 | 9.031722608 | 7.975897842 | 7.8720741 | 7.922101208 | 7.947401109 |
| ENSG00000124126 | PREX1 | 0.30164994 | 0.851414911 | -0.755802734 | 0.608695439 | -0.454152794 | 4.00722531 | 7.274681896 | 8.511888463 | 7.805288479 | 7.74329312 | 6.775876478 | 6.714211481 |
| ENSG00000104915 | STX10 | -0.449984679 | 0.651645795 | -0.00536009 | 0.996190831 | -0.455344769 | 9.317904228 | 7.572420273 | 7.285536013 | 7.920420067 | 7.90439517 | 8.268091966 | 8.428831018 |
| ENSG00000132604 | TERF2 | -0.382436626 | 0.448617153 | -0.074427984 | 0.893566028 | -0.456864609 | 8.285626337 | 8.337339476 | 9.06622548 | 8.244958867 | 8.20711886 | 8.622816121 | 8.431665941 |
| ENSG00000278588 | HIST1H2BI | 3.332290267 | 0.416320096 | -3.789453408 | 0.385135236 | -0.457163141 | 2 | 2 | 2 | 3.709785322 | 3.94787299 | 2.048074405 | 2 |
| ENSG00000183186 | C2CD4C | -2.417001423 | 0.380422171 | 1.958170196 | 0.438543047 | -0.458831227 | 2 | 3.686303414 | 2 | 2.270786764 | 2.14022021 | 4.644324387 | 3.100202476 |
| ENSG00000130283 | GDF1 | 4.731989459 | 0.310556285 | -5.191107995 | 0.28942322 | -0.459118536 | 2 | 2 | 2 | 4.670864177 | 5.19627118 | 2 | 2 |
| ENSG00000221955 | SLC12A8 | 0.31515015 | 0.743030025 | -0.775047219 | 0.376725096 | -0.459897069 | 5.532353208 | 6.951736442 | 5.772716771 | 6.658765962 | 6.43421531 | 5.873348082 | 5.072762365 |
| ENSG00000147419 | CCDC25 | -0.360204808 | 0.307768638 | -0.100127129 | 0.794947828 | -0.460331937 | 8.674867122 | 8.522788974 | 8.941672936 | 8.291333947 | 8.42842973 | 8.563064723 | 8.672390305 |
| ENSG00000182533 | CAV3 | -3.753012419 | 0.531143362 | 3.292621821 | 0.560073901 | -0.460390598 | 2 | 2 | 2 | 2 | 2 | 3.534812207 | 4.033026654 |
| ENSG00000105793 | GTPBP10 | 0.4004161 | 0.613073282 | -0.861440078 | 0.234784916 | -0.461023978 | 7.975256505 | 6.899080544 | 6.718950348 | 7.700356475 | 7.6957855 | 6.793610851 | 6.06327132 |
| ENSG00000088854 | C20orf194 | -0.507358909 | 0.419141353 | 0.045878387 | 0.94830231 | -0.461480522 | 9.602033055 | 9.102512015 | 8.266922072 | 8.568972129 | 8.60672783 | 9.172206498 | 9.09971864 |
| ENSG00000174804 | FZD4 | -0.832506003 | 0.252767379 | 0.369133192 | 0.632354165 | -0.46337281 | 8.16936288 | 9.00621767 | 9.868959282 | 8.401141138 | 8.29975519 | 9.571952335 | 9.513351291 |
| ENSG00000142230 | SAE1 | -0.502129464 | 0.293107873 | 0.037464402 | 0.945470349 | -0.464665062 | 9.33149386 | 9.910357957 | 10.14394518 | 9.279220784 | 9.38785872 | 9.676010922 | 10.0434692 |
| ENSG00000179593 | ALOX15B | -2.304510325 | NA | 1.839071568 | NA | -0.465438757 | 2 | 2 | 2 | 2.141737246 | 2 | 3.336159116 | 2.374681325 |
| ENSG00000070087 | PFN2 | -0.22401762 | 0.725681209 | -0.241832042 | 0.699954446 | -0.465849663 | 11.93813426 | 10.83211025 | 10.9833682 | 11.14542979 | 11.0834766 | 11.04836626 | 11.14405721 |

| | | | | | | | | | | | | | |
|---|---|---|---|---|---|---|---|---|---|---|---|---|---|
| ENSG00000137106 | GRHPR | -0.677157603 | 0.351836711 | 0.209944735 | 0.790974913 | -0.467212868 | 10.31337612 | 8.929529568 | 9.031722608 | 8.879660299 | 8.91566441 | 9.610733985 | 9.930260564 |
| ENSG00000196646 | ZNF136 | -0.37988894 | 0.838336958 | -0.087559394 | 0.963967825 | -0.467448334 | 3.327711234 | 6.464272742 | 8.677268317 | 7.030886362 | 7.02868983 | 7.398115274 | 7.224630935 |
| ENSG00000117593 | DARS2 | -0.489935222 | 0.762843325 | 0.022263962 | 0.989426938 | -0.46767126 | 8.228665467 | 7.467835385 | 3.369317972 | 6.936770382 | 6.79936111 | 7.142599214 | 7.561323797 |
| ENSG00000185650 | ZFP36L1 | -0.185027 | 0.747940998 | -0.282660775 | 0.608206076 | -0.467687775 | 12.58421296 | 11.76398989 | 11.67888398 | 11.86539179 | 11.9037876 | 11.91539999 | 11.64644938 |
| ENSG00000164654 | MIOS | -0.692195506 | 0.667332365 | 0.222633871 | 0.896455571 | -0.469561635 | 3.327711234 | 8.101365653 | 7.777116083 | 6.503277904 | 6.91614081 | 7.846806418 | 7.334741191 |
| ENSG00000106789 | CORO2A | -0.929541459 | 0.472536264 | 0.459826948 | 0.731494326 | -0.469714511 | 5.532353208 | 5.303523747 | 8.076950877 | 5.900763176 | 6.08095082 | 7.446000032 | 7.194904178 |
| ENSG00000155659 | VSIG4 | -3.753012542 | 0.531143362 | 3.28277859 | 0.5613132 | -0.470233952 | 2 | 2 | 2 | 2 | 2 | 3.336159116 | 4.147809227 |
| ENSG00000122507 | BBS9 | -0.421924028 | 0.51002601 | -0.050958512 | 0.942419363 | -0.47288254 | 7.096806373 | 7.790474954 | 6.80318718 | 6.907295206 | 6.88657603 | 6.989444349 | 7.482861322 |
| ENSG00000103024 | NME3 | -0.748282811 | 0.517016221 | 0.275130362 | 0.821586671 | -0.473152449 | 5.70932728 | 8.124485672 | 8.784770914 | 7.226553841 | 7.32163988 | 8.480618146 | 8.045046786 |
| ENSG00000132376 | INPP5K | -0.499913102 | 0.472081543 | 0.024911465 | 0.973993243 | -0.475001637 | 9.460782017 | 8.213429689 | 8.380957451 | 8.239055745 | 8.35836495 | 8.607356811 | 9.003921351 |
| ENSG00000130830 | MPP1 | 0.147431068 | 0.92573854 | -0.622734117 | 0.668910133 | -0.475303048 | 6.73718411 | 7.431218269 | 3.369317972 | 6.861921612 | 6.61935067 | 6.307840213 | 5.641643428 |
| ENSG00000157379 | DHRS1 | -0.530381467 | 0.560175881 | 0.053914202 | 0.957558622 | -0.476467264 | 8.609439662 | 6.728495673 | 7.342949392 | 7.333842102 | 7.17037177 | 7.539353852 | 8.065268319 |
| ENSG00000151893 | CACUL1 | -0.100145265 | 0.910744811 | -0.376728453 | 0.639943914 | -0.476873718 | 10.6316532 | 9.656475478 | 8.845626171 | 9.833599446 | 9.75261615 | 9.567062722 | 9.468158308 |
| ENSG00000143493 | INTS7 | -0.431947436 | 0.745616581 | -0.046376079 | 0.973327949 | -0.478323515 | 8.75772933 | 7.901717006 | 5.156626401 | 7.416368504 | 7.5014335 | 8.047866378 | 7.59079099 |
| ENSG00000140153 | WDR20 | -0.494254005 | 0.784685584 | 0.01549131 | 0.993715232 | -0.478762694 | 3.327711234 | 8.639735838 | 9.083172337 | 7.807949361 | 7.83643742 | 8.304419417 | 8.344097092 |
| ENSG00000142453 | CARM1 | -0.219233946 | 0.628892599 | -0.259631295 | 0.554505321 | -0.478865241 | 9.097018025 | 9.672251278 | 9.829330889 | 9.36754703 | 9.32862783 | 9.237802185 | 9.375131848 |
| ENSG00000196511 | TPK1 | -0.481094844 | 0.827605625 | 0.001487897 | 0.999379962 | -0.479606947 | 6.96351253 | 3.686303414 | 2 | 5.178261742 | 5.06165168 | 5.740677235 | 5.343519008 |
| ENSG00000011638 | TMEM159 | 0.02503224 | 0.981947437 | -0.50720192 | 0.605467095 | -0.48216968 | 9.460782017 | 7.572420273 | 7.552162999 | 8.468280417 | 8.56534297 | 8.147081218 | 7.819040576 |
| ENSG00000196557 | CACNA1H | -4.446332599 | 0.001101634 | 3.96396397 | 0.002319776 | -0.482368629 | 4.815609471 | 5.457120953 | 7.163363831 | 3.289987342 | 2.14022021 | 10.56415812 | 9.224298414 |
| ENSG00000185013 | NT5C1B | -3.753012696 | 0.531143362 | 3.270429411 | 0.562988004 | -0.482583285 | 2 | 2 | 2 | 2 | 2 | 3.824280769 | 3.753786374 |
| ENSG00000197748 | CFAP43 | -0.270544793 | 0.930223809 | -0.212138476 | NA | -0.482683269 | 2 | 2 | 3.369317972 | 2.600425781 | 2.38539787 | 2.560105256 | 2.466008509 |
| ENSG00000103005 | USB1 | -0.622056241 | 0.313384316 | 0.138069322 | 0.840370813 | -0.483986919 | 10.14658276 | 9.018611864 | 9.439621147 | 8.99156386 | 8.99053093 | 9.415532511 | 10.01563471 |
| ENSG00000141086 | CTRL | 3.303288798 | 0.423427099 | -3.789547534 | 0.387476301 | -0.486258735 | 2 | 2 | 2 | 3.709785322 | 3.90918929 | 2.048074405 | 2 |
| ENSG00000160345 | C9orf116 | -0.373825054 | 0.767978648 | -0.112498978 | 0.932122597 | -0.486324032 | 5.095959458 | 3.076665209 | 5.156626401 | 4.415698455 | 4.2865223 | 4.628601098 | 4.500832793 |
| ENSG00000197467 | COL13A1 | -0.191519993 | 0.882732465 | -0.295501866 | 0.811305502 | -0.487021859 | 10.01801709 | 7.431218269 | 7.818184403 | 8.673343061 | 8.74631819 | 8.3234671 | 8.8444706 |
| ENSG00000156928 | MALSU1 | -0.522516611 | 0.531143362 | 0.031597085 | 0.972248923 | -0.490919527 | 7.096806373 | 7.002537973 | 8.561105505 | 7.182120794 | 7.26532099 | 7.799317009 | 7.73202455 |
| ENSG00000010278 | CD9 | -0.305378411 | 0.62163201 | -0.186655633 | 0.768414094 | -0.492034044 | 10.39976609 | 9.540933215 | 9.778213818 | 9.6388717 | 9.65662671 | 9.320255125 | 10.1060314 |
| ENSG00000247746 | USP51 | 0.538522384 | 0.835029603 | -1.032345622 | 0.671068215 | -0.493823238 | 2 | 5.946638857 | 2 | 5.022023783 | 5.04390545 | 3.663836244 | 3.875378318 |
| ENSG00000143793 | C1orf35 | -0.425337119 | 0.605487532 | -0.06944062 | 0.938021298 | -0.494777739 | 5.866944937 | 7.538394143 | 6.629488387 | 6.394182228 | 6.50077304 | 6.857395343 | 6.718860807 |
| ENSG00000249459 | ZNF286B | 1.703577931 | 0.009229661 | -2.19850669 | 0.000640345 | -0.494928759 | 5.866944937 | 5.303523747 | 6.322058544 | 7.454228844 | 7.54308677 | 4.220168345 | 3.698365557 |
| ENSG00000126698 | DNAJC8 | -0.512052302 | 0.085098867 | 0.016137809 | 0.965171887 | -0.495914494 | 9.922475255 | 9.988044813 | 10.14394518 | 9.4935523 | 9.52769466 | 9.870283133 | 10.18564881 |
| ENSG00000094963 | FMO2 | -3.753012883 | 0.531143362 | 3.255441713 | 0.564749006 | -0.497571171 | 2 | 2 | 2 | 2 | 2 | 3.968654369 | 3.560117069 |
| ENSG00000165304 | MELK | -0.42804397 | 0.72338932 | -0.070368526 | 0.956587684 | -0.498412496 | 5.095959458 | 7.315436877 | 5.156626401 | 5.860316854 | 5.89549381 | 6.205571486 | 6.222348126 |
| ENSG00000175906 | ARL4D | -1.308546862 | 0.001161121 | 0.810087604 | 0.051735128 | -0.498459258 | 8.313285031 | 8.487557258 | 7.734844356 | 6.89733479 | 6.98285659 | 9.094571356 | 8.947101695 |
| ENSG00000128590 | DNAJB9 | 0.529489348 | 0.202610543 | -1.028694667 | 0.007054497 | -0.499205319 | 9.602033055 | 9.386925327 | 9.399592459 | 10.07086745 | 9.90967722 | 8.720150661 | 8.104880443 |
| ENSG00000184445 | KNTC1 | -0.039938889 | 0.98345284 | -0.459357011 | 0.800241379 | -0.4992959 | 6.563806279 | 7.393647515 | 2 | 6.503277904 | 6.3857277 | 6.165271053 | 5.921228818 |
| ENSG00000101901 | ALG13 | -0.087670616 | 0.95389048 | -0.412575306 | 0.775706431 | -0.500245922 | 4.467311984 | 8.505280662 | 8.110401118 | 7.759286043 | 7.64371756 | 7.210184325 | 7.540464021 |
| ENSG00000126091 | ST3GAL3 | 1.094643044 | 0.409632583 | -1.595111975 | 0.203815171 | -0.500468932 | 3.327711234 | 7.189546432 | 6.629488387 | 7.444001439 | 7.51115308 | 5.056402493 | 4.845038551 |
| ENSG00000198034 | RPS4X | -1.008285021 | 0.09741056 | 0.50708177 | 0.428827907 | -0.501203251 | 12.96817362 | 12.07857394 | 11.63358652 | 11.31822478 | 11.3401725 | 12.76832102 | 12.91428517 |
| ENSG00000139644 | TMBIM6 | -0.183409291 | 0.704833394 | -0.318053632 | 0.48356437 | -0.501462924 | 13.55579352 | 13.03584485 | 12.75140232 | 12.94621622 | 12.9937154 | 12.77798604 | 12.89097863 |
| ENSG00000120913 | PDLIM2 | -1.167899459 | 0.329735621 | 0.666307116 | 0.591168456 | -0.501592344 | 10.81212061 | 9.181799937 | 7.398165132 | 8.537252353 | 8.6021877 | 10.2787502 | 10.49946786 |
| ENSG00000104327 | CALB1 | 1.604315142 | NA | -2.106159237 | NA | -0.501844094 | 2 | 2 | 2 | 3.349643765 | 2 | 2.048074405 | 2.117335275 |
| ENSG00000136696 | IL36B | 2.3867222 | 0.653114396 | -2.890379571 | NA | -0.50365737 | 2 | 2 | 2 | 2.868578001 | 3.50452879 | 2.094598369 | 2 |
| ENSG00000268738 | HSFX2 | 3.148095335 | 0.471298431 | -3.65198425 | 0.426694614 | -0.503888915 | 2 | 2 | 2 | 3.615725078 | 3.7864959 | 2 | 2.059860178 |
| ENSG00000136859 | ANGPTL2 | 0.958441931 | 0.359313288 | -1.463660174 | 0.136560497 | -0.505218242 | 7.750913381 | 10.02537381 | 10.57976717 | 10.77881047 | 10.8513151 | 8.585126182 | 8.204380326 |
| ENSG00000117877 | CD3EAP | -0.420975567 | 0.696640403 | -0.084354597 | 0.941762599 | -0.505330164 | 4.467311984 | 6.602424674 | 7.029880454 | 5.94010643 | 6.02777321 | 6.329884754 | 6.27380897 |
| ENSG00000143919 | CAMKMT | -0.74495469 | 0.654365619 | 0.239599592 | 0.891849925 | -0.505355097 | 4.00722531 | 5.595927442 | 2 | 3.881293667 | 4.05806467 | 4.713032626 | 4.810605986 |
| ENSG00000117410 | ATP6V0B | 0.523872413 | 0.446344169 | -1.02938217 | 0.101902753 | -0.505509757 | 10.96815916 | 10.01303782 | 9.541233192 | 10.78692561 | 10.8560898 | 9.325140951 | 9.222662522 |
| ENSG00000156675 | RAB11FIP1 | -1.46971541 | 0.001242812 | 0.964048559 | 0.039698009 | -0.505666851 | 8.564112515 | 8.053983481 | 8.110401118 | 7.066878356 | 6.52655952 | 9.393605372 | 9.0072509 |
| ENSG00000111725 | PRKAB1 | -0.540637368 | 0.503519889 | 0.034765456 | 0.969279885 | -0.505871912 | 8.855032352 | 7.35507212 | 7.502634626 | 7.523859802 | 7.55253065 | 8.286996197 | 7.892900165 |
| ENSG00000189171 | S100A13 | 0.372984484 | 0.628218242 | -0.879686532 | 0.208642569 | -0.506702048 | 10.41898845 | 9.00621767 | 9.132850258 | 10.02070987 | 10.0596091 | 8.644477784 | 8.931664222 |
| ENSG00000147127 | RAB41 | -3.753013009 | 0.531143362 | 3.245340444 | 0.565942887 | -0.507672566 | 2 | 2 | 2 | 2 | 2 | 3.824280769 | 3.717076715 |
| ENSG00000136937 | NCBP1 | -0.162628686 | 0.625452298 | -0.345394911 | 0.25649001 | -0.508023597 | 8.87372952 | 8.820493869 | 8.631919287 | 8.600009476 | 8.63959257 | 8.360229299 | 8.514222325 |
| ENSG00000108469 | RECQL5 | -0.27876941 | 0.883143303 | -0.229519543 | 0.902568424 | -0.508288954 | 8.836089686 | 6.78762695 | 3.369317972 | 7.218576229 | 7.4074129 | 7.534088584 | 7.171352322 |
| ENSG00000107789 | MINPP1 | 0.397894023 | 0.460218392 | -0.906631667 | 0.061688239 | -0.508737644 | 8.107517749 | 9.13702746 | 8.742726175 | 9.140440229 | 9.10513517 | 7.799317009 | 7.866953664 |
| ENSG00000135341 | MAP3K7 | -0.346529224 | 0.253811546 | -0.162273743 | 0.61128327 | -0.508802966 | 9.710019321 | 9.632483534 | 9.788583153 | 9.410199593 | 9.31943747 | 9.686964814 | 9.395603187 |
| ENSG00000198369 | SPRED2 | -0.043555355 | 0.97688117 | -0.465886487 | 0.738195992 | -0.509441842 | 10.27872131 | 9.256956408 | 6.203069917 | 9.236404433 | 9.33137356 | 8.901594745 | 8.828426256 |
| ENSG00000196275 | GTF2IRD2 | 2.239858855 | 0.000123128 | -2.750189176 | 3.60E-06 | -0.51033032 | 6.009025997 | 4.93638933 | 5.930865648 | 7.842101511 | 7.7866208 | 3.411599105 | 3.496556178 |
| ENSG00000086061 | DNAJA1 | 0.317525555 | 0.427142195 | -0.834362317 | 0.019134157 | -0.516836762 | 11.75612888 | 11.1062775 | 11.60453593 | 11.82786535 | 11.8332199 | 10.71265031 | 10.64779131 |
| ENSG00000140350 | ANP32A | -0.470714904 | 0.080529035 | -0.046638469 | 0.88481517 | -0.517353373 | 9.544849385 | 9.45234302 | 9.399592459 | 9.004380786 | 8.99400091 | 9.319337188 | 9.515022972 |
| ENSG00000168803 | ADAL | -0.202746277 | 0.849223293 | -0.314714979 | 0.756352835 | -0.517461255 | 5.095959458 | 6.602424674 | 7.451345288 | 6.548530093 | 6.39974839 | 6.361143723 | 6.37742435 |
| ENSG00000145604 | SKP2 | -0.843026096 | 0.230169878 | 0.324642737 | 0.667089979 | -0.51838336 | 6.816637977 | 8.276714465 | 7.451345288 | 6.64692533 | 6.99214046 | 8.091532339 | 7.835189056 |
| ENSG00000008988 | RPS20 | 0.063171163 | 0.947611632 | -0.581620626 | 0.504030629 | -0.518449463 | 11.80565454 | 9.726146444 | 10.64969256 | 11.01936533 | 11.0524721 | 10.41459985 | 10.37059545 |
| ENSG00000158793 | NIT1 | -0.415679516 | 0.56929975 | -0.103074638 | 0.896486957 | -0.518754153 | 8.609439662 | 7.315436877 | 7.451345288 | 7.366699513 | 7.62891439 | 7.723629692 | 7.891871215 |
| ENSG00000186642 | PDE2A | 2.154636385 | 0.521613346 | -2.673457098 | 0.416362316 | -0.518820713 | 4.00722531 | 2 | 2 | 4.598938466 | 4.22485516 | 2.384001718 | 2 |
| ENSG00000101868 | POLA1 | -0.719855632 | 0.699169627 | 0.201024206 | 0.91912988 | -0.518831426 | 2 | 5.595927442 | 7.342949392 | 5.48800675 | 5.48637102 | 6.295446113 | 6.394961217 |
| ENSG00000105865 | DUS4L | -0.353117369 | 0.842102843 | -0.165840021 | 0.927015754 | -0.51895739 | 2 | 6.228547981 | 4.874587708 | 4.965943888 | 4.811949 | 5.136410075 | 4.967301796 |
| ENSG00000161249 | DMKN | 1.411713556 | 0.336817477 | -1.931655142 | 0.177744904 | -0.519941586 | 4.00722531 | 3.686303414 | 2 | 4.523237872 | 4.53541753 | 2.560105256 | 2.551896972 |
| ENSG00000135502 | SLC26A10 | -0.021894558 | 0.994204236 | -0.498211204 | 0.851505238 | -0.520105762 | 2 | 6.228547981 | 2 | 4.927308402 | 4.65713986 | 4.065262523 | 4.673629022 |
| ENSG00000087274 | ADD1 | -0.661402979 | 0.144346027 | 0.140422638 | 0.784356511 | -0.520980341 | 11.44840376 | 11.39952642 | 12.14415806 | 11.02366403 | 11.0657237 | 11.71465816 | 11.96548264 |
| ENSG00000198700 | IPO9 | -0.466302111 | 0.44286374 | -0.056268332 | 0.934216456 | -0.522570443 | 10.64259676 | 10.21502048 | 9.426401271 | 9.685919789 | 9.74164918 | 9.951520129 | 10.27517721 |
| ENSG00000100239 | PPP6R2 | -0.339710906 | 0.622552387 | -0.184626096 | 0.795874263 | -0.524337002 | 10.21416435 | 8.862353853 | 10.09465441 | 9.470224559 | 9.52449896 | 9.594132343 | 9.707433942 |
| ENSG00000076685 | NT5C2 | 0.619875524 | 0.414405236 | -1.144497961 | 0.102021981 | -0.524622436 | 7.82963488 | 9.566483142 | 8.922968322 | 9.585592789 | 9.52928986 | 7.711553069 | 7.905190659 |
| ENSG00000177889 | UBE2N | -0.205279718 | 0.564402109 | -0.319371875 | 0.342242445 | -0.524651593 | 9.710019321 | 9.74864196 | 9.399592459 | 9.425874714 | 9.42603674 | 9.162002347 | 9.449047665 |
| ENSG00000059573 | ALDH18A1 | -0.760203741 | 0.420363638 | 0.235122235 | 0.817323895 | -0.525081507 | 8.75772933 | 10.14890644 | 11.1494608 | 9.570001722 | 9.55848727 | 10.51809824 | 10.592497 |
| ENSG00000143379 | SETDB1 | -0.356940962 | 0.642211794 | -0.168947463 | 0.83342673 | -0.525888425 | 7.975256505 | 7.393647515 | 8.941672936 | 7.770242853 | 7.99722844 | 8.104378504 | 8.041339497 |
| ENSG00000106605 | BLVRA | -0.4705541 | 0.688047343 | -0.055775819 | 0.965290088 | -0.526329919 | 6.89194353 | 7.638155693 | 9.553445883 | 8.045195239 | 7.97398853 | 8.313667445 | 8.517561736 |
| ENSG00000143393 | PI4KB | -0.271247402 | 0.657619829 | -0.255724868 | 0.674724562 | -0.52697227 | 10.11549941 | 9.994333773 | 11.04521411 | 10.19090967 | 10.1964202 | 10.14836757 | 10.26746235 |
| ENSG00000168397 | ATG4B | -0.444372317 | 0.679837753 | -0.083905149 | 0.942222441 | -0.528277466 | 10.37373127 | 8.791896851 | 7.896971941 | 8.94033939 | 8.94941893 | 9.3448212 | 9.259427717 |
| ENSG00000183258 | DDX41 | -0.215415878 | 0.719874208 | -0.313318145 | 0.58526661 | -0.528734023 | 10.08373159 | 9.055168103 | 9.344432377 | 9.296376788 | 9.39224836 | 9.10206309 | 9.38027698 |
| ENSG00000167920 | TMEM99 | -0.559619717 | 0.790662721 | 0.030546701 | 0.988721265 | -0.529073016 | 7.579279349 | 4.710564154 | 2 | 5.720439317 | 5.63601114 | 6.189586148 | 6.270645889 |
| ENSG00000053372 | MRTO4 | -0.264381851 | 0.844391096 | -0.266448056 | 0.840299173 | -0.530829906 | 9.247966931 | 8.395519306 | 5.772716771 | 8.017874397 | 8.21506984 | 7.915637193 | 8.295125345 |
| ENSG00000041988 | THAP3 | -0.314487505 | 0.884847189 | -0.216732526 | 0.920142601 | -0.531220032 | 8.797441463 | 7.538394143 | 2 | 7.504305792 | 7.32531744 | 7.755649985 | 7.221358049 |
| ENSG00000104131 | EIF3J | -0.422834632 | 0.341899875 | -0.108802272 | 0.823280183 | -0.531636904 | 9.509418609 | 8.747909548 | 9.272377397 | 8.838609445 | 8.73115061 | 9.159610958 | 9.032214052 |
| ENSG00000111863 | ADTRP | 1.162316178 | 0.63692076 | -1.693955742 | 0.472178974 | -0.531639564 | 5.70932728 | 2 | 2 | 5.627094009 | 4.89348248 | 3.011868911 | 3.287118176 |
| ENSG00000124160 | NCOA5 | -0.187429141 | 0.721693747 | -0.344220826 | 0.483542133 | -0.531649967 | 8.009475818 | 8.077869079 | 8.784770914 | 8.091796043 | 8.19510963 | 8.047866378 | 7.929461924 |
| ENSG00000172586 | CHCHD1 | -0.396916874 | 0.665128529 | -0.135108393 | 0.889682343 | -0.532025267 | 8.696033109 | 7.467835385 | 6.718950348 | 7.402350992 | 7.53358066 | 7.639834368 | 7.80708241 |
| ENSG00000154473 | BUB3 | 0.073428397 | 0.930787681 | -0.605660809 | 0.410371427 | -0.532232412 | 10.38028415 | 9.849218119 | 8.677268317 | 9.904309377 | 9.8316678 | 9.04656815 | 9.326077571 |
| ENSG00000099849 | RASSF7 | 0.311097401 | 0.851362698 | -0.843711819 | 0.575697563 | -0.532614419 | 5.70932728 | 6.140560021 | 2 | 5.590471342 | 5.81407719 | 4.875056259 | 4.456980017 |
| ENSG00000072954 | TMEM38A | -2.560121127 | 0.021321155 | 2.026401413 | 0.059608695 | -0.533719714 | 5.70932728 | 3.076665209 | 5.59507254 | 3.162721474 | 3.28037825 | 7.263334844 | 6.791281771 |
| ENSG00000177556 | ATOX1 | 0.239745556 | 0.654620374 | -0.774646829 | 0.10141086 | -0.534901273 | 8.540903468 | 8.356994601 | 9.212024683 | 8.964391141 | 9.00321357 | 7.850199974 | 8.101323949 |
| ENSG00000018189 | RUFY3 | 0.062613515 | 0.956151149 | -0.598454477 | 0.56087963 | -0.535840962 | 6.563806279 | 9.192779271 | 8.845626171 | 8.663092307 | 8.60521604 | 8.021018 | 7.941446131 |
| ENSG00000182670 | TTC3 | -1.082736415 | 0.002249586 | 0.546269475 | 0.148400754 | -0.536466941 | 11.17451594 | 11.86009815 | 11.54461865 | 10.52353286 | 10.4222654 | 12.09105348 | 12.10897797 |
| ENSG00000125459 | MSTO1 | 0.766343328 | 0.385890403 | -1.302905354 | 0.113061522 | -0.536562026 | 7.709880939 | 6.728495673 | 5.772716771 | 7.753776273 | 7.62891439 | 5.371837532 | 5.984375752 |
| ENSG00000069248 | NUP133 | -0.323248605 | 0.68974372 | -0.213389073 | 0.795568436 | -0.536637678 | 9.384607364 | 8.777382925 | 7.600047325 | 8.468280417 | 8.41468635 | 8.52588267 | 8.574463273 |
| ENSG00000147533 | GOLGA7 | 0.085117712 | 0.91483416 | -0.622487469 | 0.362015414 | -0.537369757 | 10.44422623 | 9.13702746 | 9.212024683 | 9.786482773 | 9.84133177 | 9.185364902 | 9.031747025 |
| ENSG00000176927 | EFCAB5 | 0.625196449 | 0.705896131 | -1.16377545 | 0.45324474 | -0.538579001 | 4.00722531 | 4.442710129 | 2 | 4.387512679 | 4.2560182 | 3.374372158 | 2.782576693 |
| ENSG00000103091 | WDR59 | -0.179933155 | 0.933928913 | -0.360426775 | 0.864992366 | -0.54035993 | 9.015175615 | 8.376385542 | 2 | 7.985332082 | 7.96694292 | 7.737439787 | 7.8574975 |
| ENSG00000107521 | HPS1 | -0.274982011 | 0.7400867 | -0.266443695 | 0.745570382 | -0.541425706 | 9.678463567 | 8.337339476 | 10.0085654 | 9.164517339 | 9.28113268 | 8.928330746 | 9.483608161 |
| ENSG00000119801 | YPEL5 | -0.203512359 | 0.678957281 | -0.33814267 | 0.464069117 | -0.541655028 | 11.26227543 | 10.49775373 | 10.99236731 | 10.7658674 | 10.7271979 | 10.41960523 | 10.78241603 |
| ENSG00000185808 | PIGP | 0.547465598 | 0.60735938 | -1.089923456 | 0.266749214 | -0.542457857 | 7.82963488 | 6.389896633 | 5.39244166 | 7.477814553 | 7.33629421 | 5.711121288 | 5.96101835 |
| ENSG00000232859 | LYRM9 | -0.359549707 | 0.823564926 | -0.18335188 | 0.910950051 | -0.542901587 | 2 | 5.457120953 | 5.930865648 | 4.739373666 | 4.98931816 | 5.002522427 | 5.04340913 |
| ENSG00000220205 | VAMP2 | -0.747394758 | 0.29492176 | 0.203976061 | 0.794797351 | -0.543418697 | 10.83137851 | 9.806969599 | 9.330306409 | 9.352133347 | 9.41830936 | 10.11137118 | 10.52474633 |
| ENSG00000167562 | ZNF701 | 0.288281909 | 0.894855735 | -0.831847131 | 0.677551443 | -0.543565222 | 2 | 5.946638857 | 8.206335842 | 7.242377933 | 7.12888795 | 6.200262696 | 6.018717758 |
| ENSG00000166444 | ST5 | -0.05940787 | 0.973280525 | -0.484269041 | 0.773865459 | -0.54367691 | 4.467311984 | 8.671479029 | 9.829330889 | 8.719297823 | 8.76674758 | 8.115702075 | 8.504157459 |
| ENSG00000205838 | TTC23L | -0.470220855 | 0.898417957 | -0.073536776 | NA | -0.543757631 | 2 | 2 | 3.369317972 | 2.141737246 | 2.68924543 | 2.858551861 | 2.172608095 |
| ENSG00000126947 | ARMCX1 | 0.312282624 | 0.88983543 | -0.856851867 | 0.679613476 | -0.544569244 | 2 | 9.018611864 | 9.287079172 | 8.837307593 | 8.93865515 | 7.851047117 | 7.608437243 |

| | | | | | | | | | | | | | |
|---|---|---|---|---|---|---|---|---|---|---|---|---|---|
| ENSG00000183598 | HIST2H3D | 2.22040346 | 0.366576641 | -2.765139141 | 0.274695513 | -0.544735681 | 3.327711234 | 2 | 2 | 3.840289928 | 3.60476711 | 2.139668783 | 2.059860178 |
| ENSG00000119574 | ZBTB45 | -0.835758564 | 0.40909689 | 0.289863466 | 0.789642976 | -0.545895098 | 6.009025997 | 8.053983481 | 8.65477198 | 7.101994243 | 7.12888795 | 8.218405973 | 8.218402364 |
| ENSG00000033050 | ABCF2 | -0.526179896 | 0.320074535 | -0.021044968 | 0.972248923 | -0.547224863 | 9.031919741 | 9.591588447 | 10.07784268 | 9.04780133 | 9.16073939 | 9.513759765 | 9.69539454 |
| ENSG00000173705 | SUSD5 | -2.988949723 | 0.167659551 | 2.439457996 | 0.259652636 | -0.549491727 | 4.00722531 | 8.747909548 | 2 | 4.387512679 | 4.58534546 | 9.346023568 | 9.888570123 |
| ENSG00000090013 | BLVRB | -0.726777038 | 0.34076194 | 0.177212571 | 0.833507886 | -0.549564467 | 8.910412251 | 7.669933291 | 7.342949392 | 7.384635376 | 7.46522407 | 8.411091064 | 8.209344857 |
| ENSG00000140307 | GTF2A2 | -0.127542247 | 0.836817752 | -0.422375949 | 0.442025836 | -0.549918196 | 8.737455672 | 7.790474954 | 8.143093325 | 8.151677543 | 8.13979554 | 7.648631242 | 8.036691948 |
| ENSG00000272886 | DCP1A | 0.176031795 | 0.664619951 | -0.726465044 | 0.037142991 | -0.550433249 | 9.423197387 | 9.969011754 | 9.491321578 | 9.792541168 | 9.86426457 | 8.943502388 | 8.920102913 |
| ENSG00000118420 | UBE3D | -0.838008382 | 0.748522947 | 0.28585686 | 0.917287256 | -0.552151523 | 2 | 6.464272742 | 2 | 4.358765249 | 4.2865223 | 5.108342547 | 5.436777225 |
| ENSG00000169217 | CD2BP2 | -0.458906254 | 0.110464035 | -0.093606868 | 0.774164954 | -0.552513122 | 9.740899611 | 9.741182361 | 10.05225195 | 9.419799055 | 9.36660051 | 9.803144189 | 9.708895312 |
| ENSG00000196199 | MPHOSPH8 | -0.040488267 | 0.958996611 | -0.514265926 | 0.463376829 | -0.554754193 | 10.13887448 | 8.762721501 | 8.960138144 | 9.333786218 | 9.42689279 | 8.890924823 | 8.930161469 |
| ENSG00000115486 | GGCX | -0.752394173 | 0.255434807 | 0.196724388 | 0.788486582 | -0.555669784 | 9.33149386 | 9.71095217 | 10.67785662 | 9.290680758 | 9.25437635 | 9.998918132 | 10.40688307 |
| ENSG00000133103 | COG6 | -0.577561003 | 0.545365418 | 0.02124989 | 0.984153231 | -0.556311113 | 7.485143553 | 9.338312924 | 9.647595056 | 8.570539957 | 8.45217004 | 9.432893972 | 8.682541321 |
| ENSG00000176946 | THAP4 | -0.796610869 | 0.134337633 | 0.238329579 | 0.684431323 | -0.55828129 | 9.839306291 | 9.090820894 | 8.763901704 | 8.511364632 | 8.51470562 | 9.51135037 | 9.567841145 |
| ENSG00000074842 | MYDGF | 0.459085104 | 0.546357577 | -1.017728065 | 0.142297927 | -0.558642961 | 12.02642475 | 10.69982928 | 10.57378414 | 11.67082229 | 11.7644504 | 10.1958205 | 10.29185004 |
| ENSG00000131165 | CHMP1A | -0.540643499 | 0.113304945 | -0.018040217 | 0.965819335 | -0.558683716 | 9.931427102 | 10.27823701 | 10.40882047 | 9.594867189 | 9.76350037 | 10.09537369 | 10.30156833 |
| ENSG00000168286 | THAP11 | -0.664409581 | 0.747940998 | 0.105637722 | 0.962309739 | -0.558771859 | 9.048471759 | 8.433041974 | 2 | 7.474468681 | 7.60793141 | 8.424811996 | 8.1631842 |
| ENSG00000164294 | GPX8 | -0.411764502 | 0.466152501 | -0.147843567 | 0.805630147 | -0.559608068 | 10.64259676 | 11.26034006 | 11.65074017 | 10.82486117 | 10.8372174 | 11.31967864 | 10.82737534 |
| ENSG00000164985 | PSIP1 | -0.982323736 | 0.053105198 | 0.422389996 | 0.436296276 | -0.55993374 | 8.107517749 | 9.067150531 | 8.536706852 | 7.723090567 | 7.59884443 | 9.2270746 | 8.846065265 |
| ENSG00000148153 | INIP | -0.495582726 | 0.540424469 | -0.064728386 | 0.942414968 | -0.560311112 | 8.674867122 | 6.899080544 | 7.858115942 | 7.555872146 | 7.42465798 | 8.014983227 | 7.805990373 |
| ENSG00000135040 | NAA35 | 0.482691334 | 0.465166199 | -1.043440822 | 0.081548025 | -0.560749488 | 7.218819781 | 8.487557258 | 7.646393253 | 8.260583448 | 8.47221251 | 6.831875681 | 6.920008846 |
| ENSG00000119812 | FAM98A | -0.309421518 | 0.844353006 | -0.252607028 | 0.872155485 | -0.562028546 | 11.58798587 | 8.687092496 | 7.858115942 | 9.979997207 | 9.95667084 | 9.922798958 | 10.12037877 |
| ENSG00000134255 | CEPT1 | -0.238369693 | 0.866444985 | -0.324153776 | 0.811836432 | -0.562523469 | 9.015175615 | 8.101365653 | 5.39244166 | 7.88809286 | 7.88209639 | 8.093681326 | 7.432801564 |
| ENSG00000069345 | DNAJA2 | -0.110424631 | 0.880685896 | -0.452231511 | 0.483213396 | -0.562656143 | 9.460782017 | 9.298208073 | 10.5187901 | 9.766786386 | 9.74233706 | 9.473068198 | 9.353618642 |
| ENSG00000108883 | EFTUD2 | -0.09124473 | 0.931237974 | -0.47196316 | 0.624555723 | -0.56320789 | 8.19931886 | 10.5327403 | 10.19940021 | 9.814540128 | 9.88932507 | 9.355308099 | 9.582902386 |
| ENSG00000076344 | RGS11 | -1.12951116 | 0.499226274 | 0.564918274 | 0.742760586 | -0.564592886 | 6.468621203 | 4.710564154 | 2 | 4.268917409 | 4.40247105 | 5.769639817 | 5.867805474 |
| ENSG00000188917 | TRMT2B | -0.185200018 | 0.866763421 | -0.379463232 | 0.712935639 | -0.564663249 | 8.493334308 | 7.099070191 | 6.322058544 | 7.491120977 | 7.31425651 | 6.831875681 | 7.519298205 |
| ENSG00000121440 | PDZRN3 | -1.027568175 | 0.010924161 | 0.461775773 | 0.285670744 | -0.565792401 | 8.696033109 | 9.125613817 | 9.503962296 | 8.096156705 | 8.15847021 | 9.574006172 | 9.632719128 |
| ENSG00000103066 | PLA2G15 | -0.557409656 | 0.667251957 | -0.008450378 | 0.995181121 | -0.565860034 | 9.397585725 | 6.899080544 | 10.46808205 | 8.967964567 | 8.97070853 | 9.711427775 | 9.286693143 |
| ENSG00000011523 | CEP68 | -1.456681457 | 0.128993961 | 0.890648401 | 0.366387369 | -0.566033056 | 8.609439662 | 6.04685525 | 8.042706615 | 6.529309835 | 6.44101097 | 8.971428833 | 8.560285 |
| ENSG00000122873 | CISD1 | 0.0310295 | 0.984020078 | -0.598546204 | 0.67741085 | -0.567516704 | 9.344956677 | 5.595927442 | 7.029880454 | 8.134820652 | 8.14188248 | 7.497765909 | 7.54177663 |
| ENSG00000158435 | CNOT11 | -0.620183109 | 0.031572147 | 0.052347231 | 0.882077604 | -0.567835878 | 8.836089686 | 8.717821467 | 8.960138144 | 8.188900477 | 8.2580338 | 8.82044307 | 8.955005256 |
| ENSG00000148429 | USP6NL | -0.317917381 | 0.7192138 | -0.25019443 | 0.778789835 | -0.568111811 | 8.87372952 | 7.002537973 | 7.777116083 | 7.742693204 | 7.79725241 | 7.842553218 | 7.830900438 |
| ENSG00000071626 | DAZAP1 | -0.45589922 | 0.421603569 | -0.112334874 | 0.85594901 | -0.568234094 | 10.23600589 | 9.181799937 | 10.16797309 | 9.496029474 | 9.46072474 | 9.801172998 | 9.839247834 |
| ENSG00000178988 | MRFAP1L1 | -0.477158994 | 0.285057543 | -0.091376166 | 0.854928523 | -0.56853516 | 9.27634997 | 9.443177293 | 8.677268317 | 8.717883675 | 8.68036596 | 8.97997353 | 9.176932811 |
| ENSG00000189091 | SF3B3 | -0.463282559 | 0.306087183 | -0.106286769 | 0.833139147 | -0.569569328 | 10.29961392 | 11.0102427 | 10.3329194 | 10.10372352 | 10.1499977 | 10.40871423 | 10.55349898 |
| ENSG00000161970 | RPL26 | -0.261757328 | 0.723145763 | -0.308586551 | 0.669799484 | -0.570343879 | 11.87225278 | 10.76012279 | 10.47451889 | 10.90160668 | 10.9124925 | 10.89908444 | 10.82049707 |
| ENSG00000148848 | ADAM12 | -0.527621407 | 0.552447214 | -0.044945214 | 0.964355081 | -0.572566621 | 9.90440318 | 11.73423878 | 10.19160738 | 10.30737895 | 10.341489 | 10.65609247 | 10.94166594 |
| ENSG00000134152 | KATNBL1 | -0.431824705 | 0.688569122 | -0.141406967 | 0.901623508 | -0.573231672 | 8.797441463 | 7.051611315 | 6.629488387 | 7.481152682 | 7.29563107 | 7.950836136 | 7.334741191 |
| ENSG00000090971 | NAT14 | -0.596194885 | 0.338188336 | 0.022244885 | 0.974689151 | -0.57395 | 7.904282243 | 7.60566235 | 8.805342542 | 7.593372273 | 7.61696151 | 8.149149092 | 8.274745794 |
| ENSG00000174599 | TRAM1L1 | 0.20669036 | 0.938887874 | -0.781544753 | 0.759271438 | -0.574854393 | 2 | 6.228547981 | 2 | 5.093559143 | 4.89348248 | 4.230621763 | 4.119958326 |
| ENSG00000104957 | CCDC130 | -0.129824559 | 0.939161162 | -0.446127603 | 0.783596985 | -0.575952163 | 9.317904228 | 7.538394143 | 4.523675391 | 7.956842238 | 8.06698125 | 7.852739912 | 7.533882999 |
| ENSG00000108774 | RAB5C | 0.419406787 | 0.609297559 | -0.996402484 | 0.179070824 | -0.576995697 | 11.13631304 | 9.277729677 | 10.44859742 | 10.89160678 | 10.8918703 | 9.406340618 | 9.551969768 |
| ENSG00000169372 | CRADD | -0.473285668 | 0.660274906 | -0.104213486 | 0.928829011 | -0.577499154 | 8.075571721 | 6.389896633 | 5.772716771 | 6.670510205 | 6.58294721 | 6.920094328 | 7.043754524 |
| ENSG00000079313 | REXO1 | -0.249060334 | 0.881733734 | -0.328745469 | 0.838156295 | -0.577805803 | 5.532353208 | 8.356994601 | 10.19160738 | 8.729158235 | 8.7882247 | 8.637620652 | 8.72128783 |
| ENSG00000154736 | ADAMTS5 | -1.68048304 | 0.019792117 | 1.102482692 | 0.138594698 | -0.578000348 | 9.509418609 | 8.806266215 | 9.116480286 | 7.577865504 | 7.4450836 | 9.210337096 | 10.87267185 |
| ENSG00000196792 | STRN3 | -0.166749335 | 0.795687476 | -0.411447473 | 0.48317362 | -0.578196808 | 8.631580145 | 9.424669273 | 8.677268317 | 8.813671111 | 8.78422202 | 8.860117405 | 8.17168765 |
| ENSG00000156697 | UTP14A | 0.057620144 | 0.931706466 | -0.637483115 | 0.263817749 | -0.579862972 | 7.867441304 | 7.538394143 | 7.818184403 | 7.77569021 | 7.82609171 | 6.593521701 | 7.5085977 |
| ENSG00000183011 | NAA38 | -0.573129708 | 0.322499342 | -0.006834356 | 0.991913393 | -0.579964064 | 7.624139768 | 8.077869079 | 8.784770914 | 7.626911583 | 7.72113133 | 8.254069715 | 8.209344857 |
| ENSG00000198920 | KIAA0753 | -0.207053354 | 0.911517345 | -0.373879291 | 0.834048189 | -0.580932644 | 4.00722531 | 6.899080544 | 9.212024683 | 7.694616532 | 7.74054144 | 7.626042826 | 7.478755384 |
| ENSG00000165609 | NUDT5 | -0.087885325 | 0.875335246 | -0.49317686 | 0.298915693 | -0.581062186 | 9.344956677 | 8.876041558 | 8.460932916 | 8.780186674 | 8.92053511 | 8.547470747 | 8.347103327 |
| ENSG00000272968 | RBAK-RBAKI | 4.607708252 | 0.33707801 | -5.191105638 | 0.300167308 | -0.583397386 | 2 | 2 | 2 | 4.783303513 | 4.91316568 | 2 | 2 |
| ENSG00000115257 | PCSK4 | -0.826258821 | 0.575921504 | 0.2421454 | 0.878301109 | -0.584113421 | 2 | 5.303523747 | 4.874587708 | 3.754611435 | 4.16043394 | 4.742549839 | 4.919632516 |
| ENSG00000144848 | ATG3 | 0.214629708 | 0.8177838 | -0.798844229 | 0.328235173 | -0.58421452 | 9.48530526 | 8.590771778 | 7.451345288 | 8.956018563 | 8.9374542 | 7.926930284 | 7.963161916 |
| ENSG00000226479 | TMEM185B | -0.26784716 | 0.833109956 | -0.31773546 | 0.796772666 | -0.58558262 | 6.96351253 | 10.02537381 | 8.511888463 | 8.726347845 | 8.74219737 | 8.511487655 | 8.839676004 |
| ENSG00000160294 | MCM3AP | -0.430802875 | 0.694540471 | -0.156820535 | 0.892291608 | -0.587623409 | 8.517314938 | 10.00059544 | 11.24667977 | 9.925905286 | 9.84261543 | 10.16089043 | 10.15467031 |
| ENSG00000258289 | CHURC1 | -0.156335845 | 0.867371688 | -0.43163824 | 0.612133959 | -0.587974085 | 7.940205782 | 9.125613817 | 9.935792933 | 9.093229054 | 9.02486072 | 8.859275573 | 8.709655965 |
| ENSG00000172725 | CORO1B | -0.274850826 | 0.860190935 | -0.313898764 | 0.836381584 | -0.58874959 | 10.96815916 | 9.497318007 | 6.629488387 | 9.597945453 | 9.612912 | 9.364533261 | 9.74381649 |
| ENSG00000170954 | ZNF415 | -0.700866648 | 0.299070637 | 0.111111273 | 0.884519325 | -0.589755375 | 7.435667355 | 6.228547981 | 7.451345288 | 6.429169081 | 6.46787796 | 7.380620955 | 7.065820006 |
| ENSG00000064652 | SNX24 | -0.346768386 | 0.715717569 | -0.243482787 | 0.800241379 | -0.590251173 | 7.82963488 | 6.666836837 | 5.772716771 | 6.604700657 | 6.71776877 | 6.763331418 | 6.760043946 |
| ENSG00000046647 | GEMIN8 | -0.329660131 | 0.839097157 | -0.261833768 | 0.871406162 | -0.591493899 | 4.467311784 | 6.311476754 | 8.536706852 | 6.699459234 | 7.20671394 | 6.86244555 | 7.199901369 |
| ENSG00000163655 | GMPS | -0.855966987 | 0.476162581 | 0.264437503 | 0.837824475 | -0.591529484 | 6.73718411 | 9.479495904 | 10.13584611 | 8.562683721 | 8.41813449 | 9.629617332 | 9.577792788 |
| ENSG00000100714 | MTHFD1 | -0.243431884 | 0.659761977 | -0.352723843 | 0.501916413 | -0.596155726 | 8.493334308 | 8.053983481 | 8.941672936 | 8.339946438 | 8.24644267 | 7.985968197 | 8.362041249 |
| ENSG00000196586 | MYO6 | -0.94671452 | 0.25067706 | 0.350377906 | 0.692464563 | -0.596336613 | 10.46903009 | 9.318399851 | 8.486635445 | 8.74312865 | 8.68036596 | 9.928428593 | 10.0700947 |
| ENSG00000171016 | PYGO1 | -0.623689633 | 0.285611782 | 0.027066836 | 0.968381186 | -0.596622796 | 6.563806279 | 7.503546079 | 7.163363831 | 6.490083648 | 6.58294721 | 7.364099281 | 6.934072182 |
| ENSG00000151135 | TMEM263 | -0.346761724 | 0.549118904 | -0.250189977 | 0.671298904 | -0.596951701 | 9.820154628 | 10.93377402 | 10.50627919 | 10.18324441 | 10.1079152 | 10.36845651 | 10.10403354 |
| ENSG00000099194 | SCD | -1.294475896 | 0.116761103 | 0.696769724 | 0.420713272 | -0.597706172 | 11.43901855 | 12.88662532 | 13.02902175 | 11.33354142 | 11.2928274 | 12.49302884 | 13.81844695 |
| ENSG00000186777 | ZNF732 | -0.140686901 | 0.956087463 | -0.460748492 | 0.853015332 | -0.601435393 | 2 | 4.710564154 | 2 | 3.162721474 | 3.74316441 | 3.44788957 | 3.100202476 |
| ENSG00000172409 | CLP1 | -0.380791427 | 0.792847991 | -0.222153235 | 0.881261401 | -0.602944661 | 5.532353208 | 6.140560021 | 8.721235202 | 7.190301983 | 7.02868983 | 7.267148095 | 7.264927592 |
| ENSG00000112200 | ZNF451 | -0.091354897 | 0.947190051 | -0.511854195 | 0.69384525 | -0.603209091 | 6.653098118 | 9.903691196 | 10.22252928 | 9.502614552 | 9.42088976 | 9.260296988 | 8.790832728 |
| ENSG00000169131 | ZNF354A | -0.641524676 | 0.471535675 | 0.038077764 | 0.969515502 | -0.603446912 | 8.340423424 | 6.844429755 | 7.225742779 | 6.979883891 | 6.99214046 | 8.097255872 | 6.992837548 |
| ENSG00000198218 | QRICH1 | -0.38646303 | 0.507060161 | -0.217189213 | 0.717711114 | -0.603652243 | 10.78274301 | 10.04368224 | 9.681375057 | 9.794554993 | 9.92005068 | 9.96488668 | 10.08705154 |
| ENSG00000100664 | EIF5 | 0.238934899 | 0.528830242 | -0.843680275 | 0.010052906 | -0.604745376 | 11.20066853 | 11.41367288 | 11.77848812 | 11.71721007 | 11.7275807 | 10.58937934 | 10.69220572 |
| ENSG00000174442 | ZWILCH | -0.302881186 | 0.877766587 | -0.301874933 | 0.875414673 | -0.604756119 | 7.709880939 | 7.099070191 | 2 | 6.682159616 | 6.46787796 | 6.60762613 | 6.55171954 |
| ENSG00000117632 | STMN1 | -1.053974223 | 0.144179754 | 0.446131448 | 0.562234526 | -0.607842775 | 10.06758145 | 9.256956408 | 8.561105505 | 8.447952961 | 8.30907138 | 9.421818136 | 10.2100027 |
| ENSG00000155959 | VBP1 | -0.330010766 | 0.350112527 | -0.2804866 | 0.423741616 | -0.610497366 | 8.367060731 | 8.505280662 | 8.805342542 | 8.260583448 | 8.22297724 | 8.250863742 | 8.3297313 |
| ENSG00000067365 | METTL22 | -0.844323078 | 0.596631897 | 0.232884774 | 0.892291608 | -0.611438304 | 3.327711234 | 7.928237109 | 8.236946975 | 6.548530093 | 6.86146731 | 7.786974998 | 7.735470193 |
| ENSG00000171681 | ATF7IP | -0.13875923 | 0.931483663 | -0.4732533 | 0.755970837 | -0.61201253 | 6.138360429 | 10.59613751 | 10.46808205 | 9.873521491 | 9.81541592 | 9.737316722 | 9.244592076 |
| ENSG00000110934 | BIN2 | -3.753014319 | 0.531143362 | 3.140454489 | 0.579831678 | -0.61255983 | 2 | 2 | 2 | 2 | 2 | 4.340871764 | 2.509591799 |
| ENSG00000138078 | PREPL | -1.044563781 | 0.115443406 | 0.431895709 | 0.543191241 | -0.612668072 | 9.097018025 | 9.148351514 | 10.38167985 | 8.679168098 | 8.58694992 | 9.938028579 | 10.24728895 |
| ENSG00000079134 | THOC1 | 0.045561935 | 0.937917387 | -0.658375081 | 0.175083229 | -0.612813146 | 6.653098118 | 6.602424674 | 6.958202216 | 6.716553866 | 6.84618961 | 6.262714117 | 5.941260754 |
| ENSG00000171094 | ALK | -0.300867655 | 0.918975348 | -0.313933711 | 0.914220926 | -0.614801366 | 2 | 3.686303414 | 2 | 2.868578001 | 2.59489879 | 2.987421008 | 2.421067459 |
| ENSG00000215186 | GOLGA6B | 2.275697383 | NA | -2.890808174 | NA | -0.61511079 | 2 | 2 | 2 | 3.289987342 | 3.01487888 | 2.094598369 | 2 |
| ENSG00000163749 | CCDC158 | -3.753014354 | 0.531143362 | 3.137666994 | 0.580041481 | -0.61534736 | 2 | 2 | 2 | 2 | 2 | 3.904814898 | 3.45257121 |
| ENSG00000099866 | MADCAM1 | -3.753014354 | 0.531143362 | 3.137634335 | 0.580041481 | -0.615380019 | 2 | 2 | 2 | 2 | 2 | 3.616794016 | 3.771796761 |
| ENSG00000116704 | SLC35D1 | 0.36739613 | 0.736046082 | -0.983389439 | 0.31933214 | -0.615993309 | 8.042902253 | 7.669933291 | 9.935792933 | 9.29163166 | 9.2582292 | 8.069136662 | 7.795024296 |
| ENSG00000250571 | GLI4 | -0.540873287 | 0.695743402 | -0.0758065 | 0.959231573 | -0.616679787 | 4.00722531 | 6.728495673 | 7.777116083 | 6.306580752 | 6.31350454 | 6.833591108 | 6.671679221 |
| ENSG00000235750 | KIAA0040 | -4.596837097 | 0.067812259 | 3.979743478 | 0.026072796 | -0.617093619 | 2 | 4.11352228 | 2 | 2.141737246 | 2 | 6.162543904 | 6.543880272 |
| ENSG00000113312 | TTC1 | -0.402800155 | 0.498492641 | -0.214504071 | 0.726968373 | -0.617304226 | 9.761125267 | 8.820493869 | 8.784770914 | 8.773395507 | 8.81724322 | 8.838922815 | 9.112583673 |
| ENSG00000147592 | LACTB2 | 0.298046019 | 0.885491359 | -0.91703026 | 0.624817493 | -0.618984241 | 2 | 5.303523747 | 7.451345288 | 6.586219071 | 6.32824132 | 5.238348159 | 5.419744333 |
| ENSG00000066654 | THUMPD1 | -0.15355873 | 0.652929267 | -0.465701477 | 0.123069403 | -0.619260208 | 9.128497779 | 9.277729677 | 9.049077186 | 9.055659246 | 8.95417718 | 8.809589199 | 8.573180082 |
| ENSG00000119718 | EIF2B2 | -0.483427235 | 0.780664977 | -0.136127533 | 0.940230037 | -0.619554768 | 9.602033055 | 8.702538795 | 4.058998049 | 8.219202381 | 8.13770558 | 8.603340274 | 8.435907908 |
| ENSG00000146576 | C7orf26 | -0.82866958 | 0.713873535 | 0.207217801 | 0.931938712 | -0.621451778 | 9.646202137 | 6.951736442 | 2 | 7.461007055 | 7.45854177 | 8.816111316 | 8.037622656 |
| ENSG00000244509 | APOBEC3C | -1.652682766 | 0.000161722 | 1.030492447 | 0.024397067 | -0.622190319 | 10.25752168 | 9.308339287 | 9.69246155 | 8.184811706 | 8.14396641 | 10.76235392 | 10.89582457 |
| ENSG00000141098 | GFOD2 | -0.462355817 | 0.571137034 | -0.160963313 | 0.853015332 | -0.62331913 | 7.667647223 | 7.051611315 | 8.763901704 | 7.527093191 | 7.5587924 | 7.919680595 | 7.752576035 |
| ENSG00000079335 | CDC14A | 1.151836057 | 0.643838836 | -1.775317014 | 0.453489046 | -0.623480957 | 2 | 6.464272742 | 2 | 6.164317909 | 5.99120031 | 3.616794016 | 3.640730416 |
| ENSG00000198925 | ATG9A | -0.235863056 | 0.924580052 | -0.3888638 | 0.872875203 | -0.624726857 | 11.07704221 | 10.3729879 | 2 | 9.91915277 | 9.97902226 | 9.760045703 | 9.833096297 |
| ENSG00000135930 | EIF4E2 | -0.616459212 | 0.224421978 | -0.009576999 | 0.987221496 | -0.626036211 | 10.51143645 | 9.540933215 | 10.21486071 | 9.49932577 | 9.55457603 | 9.979293284 | 10.26904825 |
| ENSG00000108106 | UBE2S | -0.068856502 | 0.946330866 | -0.558281911 | 0.546275271 | -0.627138413 | 7.867441304 | 9.890264534 | 8.380957451 | 8.902248225 | 8.93264039 | 8.242494717 | 8.601149963 |
| ENSG00000143155 | TIPRL | -0.079630242 | 0.892338755 | -0.547834415 | 0.270301865 | -0.627464657 | 9.358295023 | 9.298208073 | 8.536706852 | 9.119171929 | 8.94225201 | 8.749978316 | 8.362784101 |
| ENSG00000102900 | NUP93 | -0.231503024 | 0.687937839 | -0.396995356 | 0.46213389 | -0.62849838 | 9.064836028 | 8.337339476 | 8.143093325 | 8.356399318 | 8.32569041 | 8.015738956 | 8.323639487 |
| ENSG00000205765 | C5orf51 | -0.229423351 | 0.872289176 | -0.399519184 | 0.767357706 | -0.628942534 | 6.653098118 | 9.298208073 | 10.42884603 | 9.283050899 | 9.17201613 | 9.13790818 | 8.977004658 |
| ENSG00000134222 | PSRC1 | -0.65543864 | 0.579599436 | 0.02550263 | 0.984296564 | -0.629936011 | 4.00722531 | 5.303523747 | 6.718950348 | 5.093559143 | 5.18011485 | 5.769639817 | 5.740981724 |
| ENSG00000020256 | ZFP64 | -0.316366794 | 0.470185896 | -0.313588187 | 0.462949352 | -0.629954981 | 8.586954094 | 8.124485672 | 8.380957451 | 8.172475401 | 7.93600799 | 8.218405973 | 7.885682058 |
| ENSG00000167842 | MIS12 | -0.245522546 | 0.798798132 | -0.384761482 | 0.674981326 | -0.630284028 | 7.709880939 | 8.639735838 | 6.534109959 | 7.626911583 | 7.64666005 | 7.576706933 | 7.420010428 |
| ENSG00000105821 | DNAJC2 | 0.623718823 | 0.424331051 | -1.255204448 | 0.078779916 | -0.631485624 | 8.493334308 | 7.099070191 | 7.098165388 | 8.325162455 | 8.34212018 | 6.66270409 | 6.320441373 |
| ENSG00000100865 | CINP | -0.794845561 | 0.188055699 | 0.162058285 | 0.811488554 | -0.632787276 | 8.87372952 | 7.979858146 | 7.691296561 | 7.536750104 | 7.44169935 | 8.30132352 | 8.550537483 |
| ENSG00000152439 | ZNF773 | -0.597365944 | 0.657839596 | -0.03632003 | 0.980129329 | -0.633685974 | 4.815609471 | 6.78762695 | 8.296287027 | 6.592405944 | 6.71776877 | 7.514972902 | 6.793487375 |

| | | | | | | | | | | | | | |
|---|---|---|---|---|---|---|---|---|---|---|---|---|---|
| ENSG00000077463 | SIRT6 | -0.176298047 | 0.9141407 | -0.457637205 | 0.765367121 | -0.633935251 | 8.468948322 | 8.356994601 | 4.058998049 | 7.641902724 | 7.74329312 | 7.382965824 | 7.452476174 |
| ENSG00000139168 | ZCRB1 | -0.447824897 | 0.547023674 | -0.187298609 | 0.811488554 | -0.635123506 | 9.204317094 | 7.901717006 | 7.896971941 | 7.939959586 | 8.11451391 | 8.206540078 | 8.362784101 |
| ENSG00000063322 | MED29 | -0.702671303 | 0.042012415 | 0.067318098 | 0.87167541 | -0.635353205 | 9.533135543 | 9.842262059 | 9.344432377 | 8.912174893 | 8.87609528 | 9.526011998 | 9.781250219 |
| ENSG00000196704 | AMZ2 | -0.563813816 | 0.52931101 | -0.071750342 | 0.942331353 | -0.635564159 | 9.886101849 | 9.04308532 | 7.818184403 | 8.561107327 | 8.61275919 | 9.313202635 | 8.789175794 |
| ENSG00000120675 | DNAJC15 | -0.013034994 | 0.988750252 | -0.622853573 | 0.43187425 | -0.635888567 | 9.646202137 | 7.847167909 | 8.805342542 | 8.979812467 | 8.87609528 | 8.093681326 | 8.524881346 |
| ENSG00000153132 | CLGN | 3.896739606 | 0.059474197 | -4.532649953 | 0.04575727 | -0.635910348 | 2 | 4.11352228 | 2 | 6.29144905 | 6.23747437 | 2.139668783 | 2 |
| ENSG00000102763 | VWA8 | -0.079360543 | 0.912239853 | -0.560797814 | 0.362015414 | -0.640158357 | 7.579279349 | 8.077869079 | 8.865353883 | 8.134820652 | 8.24450173 | 7.721778317 | 7.712342256 |
| ENSG00000189045 | ANKDD1B | 0.79519526 | 0.764636496 | -1.435993395 | 0.569190163 | -0.640798135 | 2 | 3.686303414 | 2 | 3.349643765 | 3.2184422 | 2.526562374 | 2.225841198 |
| ENSG00000173545 | ZNF622 | 0.163787964 | 0.870709566 | -0.805306437 | 0.354345163 | -0.641518473 | 10.29961392 | 8.981105197 | 8.236946975 | 9.571568433 | 9.60989979 | 8.46742022 | 8.774177123 |
| ENSG00000005243 | COPZ2 | -0.397874657 | 0.644424353 | -0.24441318 | 0.782285213 | -0.642287837 | 10.00958825 | 8.702538795 | 8.266922072 | 8.773395507 | 8.81985255 | 8.741779902 | 9.130142376 |
| ENSG00000183137 | CEP57L1 | -0.200779158 | 0.84860975 | -0.441595986 | 0.650306201 | -0.642375143 | 6.468621203 | 5.595927442 | 7.451345288 | 6.529309835 | 6.45450701 | 6.635427934 | 5.759151667 |
| ENSG00000133706 | LARS | -0.716962803 | 0.112779646 | 0.073946123 | 0.890497536 | -0.643016681 | 9.567995545 | 10.3778065 | 10.34701993 | 9.4022977 | 9.45821469 | 10.15507139 | 10.27813351 |
| ENSG00000149091 | DGKZ | -0.59966218 | 0.323956816 | -0.043488757 | 0.950328841 | -0.643150937 | 9.048471759 | 8.317412869 | 7.818184403 | 7.783822874 | 7.99261031 | 8.371491092 | 8.506176057 |
| ENSG00000111783 | RFX4 | -0.264565931 | 0.954430302 | -0.379713394 | NA | -0.644279325 | 2 | 3.076665209 | 2 | 2.868578001 | 2 | 2.42098416 | 2.509591799 |
| ENSG00000162623 | TYW3 | -0.200143982 | 0.901917502 | -0.445664958 | 0.770017372 | -0.64580894 | 5.095959458 | 8.101365653 | 9.577565424 | 8.283707551 | 8.28283292 | 8.128335751 | 7.94839122 |
| ENSG00000085644 | ZNF213 | -0.736683411 | 0.720923534 | 0.089920732 | 0.967776844 | -0.646762679 | 2 | 7.761270542 | 8.978369997 | 7.084543139 | 7.29937545 | 8.002074901 | 8.008487789 |
| ENSG00000158796 | DEDD | -0.275835888 | 0.751627932 | -0.372347239 | 0.659557507 | -0.648183127 | 8.19931886 | 9.591588447 | 7.934808822 | 8.459845423 | 8.54656162 | 8.331987532 | 8.481080411 |
| ENSG00000128789 | PSMG2 | 0.079183001 | 0.906842758 | -0.728389395 | 0.191278435 | -0.649206394 | 9.048471759 | 8.004992147 | 8.110401118 | 8.565831351 | 8.51952909 | 7.754744914 | 7.738907626 |
| ENSG00000197562 | RAB40C | -0.86797583 | 0.314840892 | 0.217937121 | 0.818792684 | -0.650038709 | 6.563806279 | 8.169643011 | 8.721235202 | 7.19843704 | 7.23436042 | 8.329558253 | 8.232289427 |
| ENSG00000033800 | PIAS1 | -0.204423255 | 0.725318811 | -0.446377864 | 0.405169521 | -0.650801119 | 9.720386367 | 9.256956408 | 8.631919287 | 9.078979414 | 9.0561812 | 8.880175401 | 8.773059882 |
| ENSG00000155508 | CNOT8 | -0.098443205 | 0.749513987 | -0.55332789 | 0.032897253 | -0.651771095 | 9.848787621 | 9.856140801 | 9.577565424 | 9.677941812 | 9.66390646 | 9.248772045 | 9.187873195 |
| ENSG00000116679 | IVNS1ABP | -0.719299668 | 0.004354518 | 0.066866122 | 0.83065631 | -0.652433546 | 10.79260216 | 10.92392403 | 10.9788475 | 10.14007766 | 10.2252472 | 11.09344985 | 10.82912339 |
| ENSG00000157036 | EXOG | -0.433054609 | 0.8339564 | -0.220803647 | 0.916320081 | -0.653858256 | 8.517314938 | 8.147241016 | 2 | 7.265793319 | 7.42122542 | 7.601082296 | 7.503217537 |
| ENSG00000042753 | AP2S1 | -0.228640559 | 0.716867438 | -0.425333629 | 0.470865845 | -0.653974188 | 11.11682541 | 10.24169813 | 10.16797309 | 10.33023354 | 10.3657986 | 9.86021999 | 10.39476727 |
| ENSG00000165633 | VSTM4 | 0.096833233 | 0.935203319 | -0.751832131 | 0.47448492 | -0.654998899 | 9.810482605 | 8.124485672 | 7.225742779 | 8.88471049 | 8.88234995 | 8.151214006 | 7.929461924 |
| ENSG00000187189 | TSPYL4 | -0.679907357 | 0.506977228 | 0.024308167 | 0.983104465 | -0.65559919 | 10.19198706 | 7.819099893 | 8.536706852 | 8.537252353 | 8.51470562 | 9.183011957 | 9.264206697 |
| ENSG00000120725 | SIL1 | 0.110702037 | 0.88608019 | -0.767838136 | 0.236344937 | -0.657136099 | 10.52923584 | 9.308339287 | 9.344432377 | 9.924679904 | 9.98542831 | 8.927527801 | 9.221024773 |
| ENSG00000213551 | DNAJC9 | 0.072534587 | 0.96300973 | -0.733644664 | 0.610519493 | -0.661110077 | 4.467311984 | 8.451453573 | 6.629488387 | 7.419851705 | 7.32898565 | 6.85908071 | 6.270645889 |
| ENSG00000163644 | PPM1K | 0.219184769 | 0.676973067 | -0.882163295 | 0.051399479 | -0.662978526 | 9.790941832 | 8.94259736 | 9.636156974 | 9.717397705 | 9.71245786 | 8.759935742 | 8.468021799 |
| ENSG00000108823 | SGCA | -6.969609488 | 0.004926903 | 6.305707469 | 0.000505952 | -0.663902019 | 2 | 4.93638933 | 2 | 2 | 2 | 9.503020408 | 9.607555771 |
| ENSG00000156170 | NDUFAF6 | -0.420383412 | 0.883547191 | -0.243761687 | 0.932122597 | -0.664145099 | 2 | 6.78762695 | 2 | 4.965943888 | 4.93258395 | 5.463319095 | 4.624919101 |
| ENSG00000160570 | DEDD2 | 0.138738393 | 0.870775814 | -0.804406717 | 0.267163049 | -0.665668324 | 8.468948322 | 8.522788974 | 9.849281147 | 9.303936679 | 9.15867957 | 8.502349881 | 8.061612696 |
| ENSG00000100814 | CCNB1IP1 | -1.076106611 | 0.375913231 | 0.410403987 | 0.751265003 | -0.665702623 | 9.097018025 | 8.623598428 | 5.59507254 | 7.402350992 | 7.21466714 | 8.922700729 | 8.609311074 |
| ENSG00000010803 | SCMH1 | -0.277061528 | 0.879733274 | -0.390452834 | 0.82326635 | -0.667514361 | 4.00722531 | 8.147241016 | 9.491321578 | 8.111316053 | 8.15640714 | 7.770949957 | 8.236348546 |
| ENSG00000132024 | CC2D1A | -0.757746568 | 0.395924587 | 0.08933408 | 0.930031445 | -0.668412488 | 8.367060731 | 7.638155693 | 9.670202709 | 8.113468729 | 7.99261031 | 9.038038079 | 8.72879856 |
| ENSG00000243612 | | 4.522200632 | 0.357169987 | -5.191103674 | 0.309101085 | -0.668903042 | 2 | 2 | 2 | 4.783303513 | 4.76938658 | 2 | 2 |
| ENSG00000161999 | JMJD8 | 0.55359923 | 0.591037548 | -1.222796329 | 0.19277085 | -0.669197099 | 8.228665467 | 9.125613817 | 6.534109959 | 8.860563059 | 8.86476809 | 7.043805932 | 7.184857568 |
| ENSG00000125814 | NAPB | 0.12000917 | 0.946420592 | -0.790736576 | 0.632809239 | -0.670727406 | 3.327711234 | 8.623598428 | 7.225742779 | 7.731524332 | 7.56813436 | 6.797131715 | 6.735016458 |
| ENSG00000134575 | ACP2 | -0.03312885 | 0.965522332 | -0.643597838 | 0.329793525 | -0.676726688 | 10.38028415 | 9.159587375 | 9.809100877 | 9.826399658 | 9.83811764 | 8.760386723 | 9.57619232 |
| ENSG00000145882 | PCYOX1L | -0.925209778 | 0.633232538 | 0.248125467 | 0.905614752 | -0.677084311 | 8.517314938 | 7.467835385 | 2 | 6.604700657 | 6.6253296 | 7.92612656 | 7.561323797 |
| ENSG00000197050 | ZNF420 | -0.637484772 | 0.736035171 | -0.040096268 | 0.984191003 | -0.67758104 | 2 | 7.315436877 | 6.882775709 | 6.024891728 | 5.87556634 | 7.021403672 | 5.736403255 |
| ENSG00000204673 | AKT1S1 | -0.75201632 | 0.194034006 | 0.073894597 | 0.913418819 | -0.678121723 | 10.63713535 | 9.540933215 | 9.935792933 | 9.318938646 | 9.40098777 | 9.908424552 | 10.41299269 |
| ENSG00000081479 | LRP2 | 0.903630838 | 0.780723424 | -1.582436706 | NA | -0.678805868 | 3.327711234 | 2 | 2 | 3.406930997 | 2.2680111 | 2.267001569 | 2.172608095 |
| ENSG00000102241 | HTATSF1 | -0.533255052 | 0.615433095 | -0.146526195 | 0.897819577 | -0.679781247 | 9.159305284 | 10.22575081 | 7.552162999 | 8.80304041 | 8.84312657 | 9.126058498 | 9.283558215 |
| ENSG00000204954 | C12orf73 | -0.939665952 | 0.603040858 | 0.25804034 | 0.894354862 | -0.681625612 | 7.485143553 | 5.946638857 | 2 | 5.474671438 | 5.47317115 | 6.666558925 | 6.522764885 |
| ENSG00000174720 | LARP7 | 0.571068433 | 0.595406493 | -1.252768244 | 0.201673674 | -0.681699811 | 9.646202137 | 8.57407404 | 6.882775709 | 9.323594948 | 9.32954365 | 7.496683627 | 7.553536683 |
| ENSG00000137574 | TGS1 | 0.199041584 | 0.644240767 | -0.880848614 | 0.016663213 | -0.68180703 | 8.444143041 | 7.979858146 | 7.896971941 | 8.372666678 | 8.26762243 | 7.287316943 | 7.234405279 |
| ENSG00000136156 | ITM2B | -0.063490797 | 0.930223809 | -0.620145547 | 0.314835008 | -0.683636345 | 13.36497316 | 12.09476099 | 12.43892907 | 12.67459172 | 12.6714587 | 12.21628363 | 12.00990713 |
| ENSG00000139725 | RHOF | 1.298616881 | 0.319708718 | -1.982595458 | 0.111042614 | -0.683978577 | 2 | 4.710564154 | 5.39244166 | 5.786686927 | 5.6826301 | 3.256555576 | 3.100202476 |
| ENSG00000117713 | ARID1A | -0.747495615 | 0.104588075 | 0.063481634 | 0.908444479 | -0.684013981 | 10.51143645 | 10.88044608 | 9.999667074 | 9.740563462 | 9.78969165 | 10.42715191 | 10.70632165 |
| ENSG00000106853 | PTGR1 | 0.198460197 | 0.922914669 | -0.884632563 | 0.635038067 | -0.686172366 | 10.05124847 | 8.276714465 | 3.369317972 | 9.054539304 | 9.03275482 | 8.00436125 | 7.940451241 |
| ENSG00000010539 | ZNF200 | -0.423210266 | 0.832296582 | -0.263529404 | 0.895505527 | -0.68673967 | 8.042902253 | 6.844429755 | 2 | 6.548530093 | 6.6253296 | 6.793610851 | 6.685996416 |
| ENSG00000136897 | MRPL50 | -0.181051149 | 0.918272566 | -0.505789887 | 0.760322555 | -0.686841035 | 9.204317094 | 7.701025996 | 4.058998049 | 7.980622673 | 7.8286851 | 7.53725005 | 7.636966981 |
| ENSG00000185627 | PSMD13 | 0.097628454 | 0.942177035 | -0.784822686 | 0.51368513 | -0.687194231 | 10.45046729 | 10.42046463 | 7.163363831 | 9.987645515 | 10.0546329 | 9.096001355 | 9.189967637 |
| ENSG00000003056 | M6PR | 0.506130278 | 0.527988504 | -1.195215789 | 0.09771017 | -0.689085511 | 10.87842776 | 9.648522413 | 9.196530712 | 10.59035845 | 10.5993844 | 8.841482668 | 8.959923085 |
| ENSG00000160803 | UBQLN4 | -0.541871481 | 0.729601901 | -0.147853209 | 0.92969272 | -0.68972469 | 9.567995545 | 9.386925327 | 4.874587708 | 8.356399318 | 8.4215744 | 8.85040655 | 8.704390872 |
| ENSG00000109472 | CPE | -1.552499928 | 0.010454649 | 0.861916476 | 0.174415528 | -0.690583452 | 7.435667355 | 8.101365653 | 8.721235202 | 6.716553866 | 6.60731796 | 9.306424362 | 8.688479312 |
| ENSG00000198590 | C3orf35 | 0.049413734 | 0.984043262 | -0.740485644 | 0.752160737 | -0.69107191 | 2 | 4.710564154 | 2 | 3.462030052 | 3.74316441 | 3.235948948 | 3.012034821 |
| ENSG00000185885 | IFITM1 | 2.734619264 | 1.48E-05 | -3.42800425 | 2.95E-08 | -0.693384986 | 11.06489013 | 11.97107825 | 12.69464022 | 14.73480042 | 14.8444446 | 8.878929913 | 8.359066013 |
| ENSG00000163510 | CWC22 | 0.108078671 | 0.914220207 | -0.801476359 | 0.350702316 | -0.693397688 | 6.468621203 | 8.414402626 | 8.408111533 | 8.160032699 | 8.06258142 | 7.418832227 | 6.990916783 |
| ENSG00000143252 | SDHC | -0.513034651 | 0.664949829 | -0.181221083 | 0.885544215 | -0.694255734 | 10.97683282 | 8.191702463 | 9.014156722 | 9.351221511 | 9.38345569 | 9.670982446 | 9.72342817 |
| ENSG00000143748 | NVL | 0.49240785 | 0.768037744 | -1.186696552 | 0.439027681 | -0.694288701 | 8.468948322 | 7.503546079 | 3.369317972 | 7.942383523 | 8.04037874 | 6.194934299 | 6.512090157 |
| ENSG00000169951 | ZNF764 | -0.513841021 | 0.570100628 | -0.180942344 | 0.85102559 | -0.694783365 | 6.366709671 | 7.954278496 | 6.322058544 | 6.580005552 | 6.65485813 | 6.840432491 | 7.030725399 |
| ENSG00000258429 | PDF | 0.922452662 | 0.277913797 | -1.617327019 | 0.041273153 | -0.694874356 | 5.095959458 | 6.78762695 | 5.59507254 | 6.861921612 | 6.97351258 | 4.705557986 | 4.468068808 |
| ENSG00000179262 | RAD23A | -0.38832239 | 0.516281765 | -0.306556222 | 0.608622814 | -0.694878612 | 10.65345793 | 9.514922632 | 9.868959282 | 9.682298904 | 9.72712735 | 9.770158073 | 9.802212758 |
| ENSG00000171202 | TMEM126A | 0.18854378 | 0.812790339 | -0.88366746 | 0.194894738 | -0.695123679 | 7.750913381 | 6.951736442 | 6.322058544 | 7.250225364 | 7.36516374 | 6.040114727 | 6.479583725 |
| ENSG00000186625 | KATNA1 | -0.575170916 | 0.762607571 | -0.119969386 | 0.952575081 | -0.695140301 | 7.790810999 | 7.761270542 | 2 | 6.70517998 | 6.59518404 | 7.146744843 | 7.032593922 |
| ENSG00000023041 | ZDHHC6 | 0.014674799 | 0.989593043 | -0.709856172 | 0.465968326 | -0.695181373 | 9.33149386 | 8.356994601 | 6.80318718 | 8.551612617 | 8.4705529 | 7.81936672 | 7.775072826 |
| ENSG00000100523 | DDHD1 | -0.86623261 | 0.442343803 | 0.167694651 | 0.893305945 | -0.698537959 | 8.928409675 | 6.951736442 | 9.963514551 | 8.166267464 | 8.24644267 | 9.462574845 | 8.95451255 |
| ENSG00000109220 | CHIC2 | 0.272855103 | 0.838658392 | -0.971937213 | 0.413080568 | -0.699082109 | 6.468621203 | 7.701025996 | 9.624627483 | 8.765203682 | 8.76944977 | 7.441507444 | 7.633277551 |
| ENSG00000160131 | VMA21 | -0.384977691 | 0.699347285 | -0.314640192 | 0.752090127 | -0.699617883 | 7.096806373 | 8.968383114 | 9.465702942 | 8.474993083 | 8.37797443 | 8.390261842 | 8.596104698 |
| ENSG00000039650 | PNKP | 0.112777733 | 0.945043724 | -0.81309559 | 0.584265208 | -0.700317857 | 9.064836028 | 7.503546079 | 4.523675391 | 8.006336225 | 8.10598728 | 6.882471426 | 7.376535208 |
| ENSG00000131187 | F12 | -1.573884004 | 0.322289819 | 0.873426203 | 0.581597072 | -0.700457801 | 4.00722531 | 4.442710129 | 2 | 2.784612485 | 3.01487888 | 4.881704255 | 4.047883918 |
| ENSG00000038274 | MAT2B | 0.209731134 | 0.531497817 | -0.910221558 | 0.001387469 | -0.700490424 | 9.33149386 | 9.74864196 | 9.491321578 | 9.75786892 | 9.73613428 | 8.679237241 | 8.589138522 |
| ENSG00000068796 | KIF2A | -0.096176004 | 0.91483416 | -0.605494076 | 0.439526418 | -0.70167008 | 9.613202746 | 8.234834549 | 8.076950877 | 8.75144642 | 8.69038176 | 8.387928811 | 8.024537783 |
| ENSG00000189180 | ZNF33A | -0.007768121 | 0.990196005 | -0.694142437 | 0.183892387 | -0.701910558 | 8.340423424 | 8.876041558 | 8.784770914 | 8.686416476 | 8.673169 | 8.356654601 | 7.529919928 |
| ENSG00000104714 | ERICH1 | -0.355390101 | 0.716902068 | -0.347574936 | 0.719914201 | -0.702965037 | 8.138771685 | 7.051611315 | 5.930865648 | 6.917187326 | 7.00595521 | 6.894026089 | 7.041900409 |
| ENSG00000178498 | DTX3 | -0.566066236 | 0.560330239 | -0.139268549 | 0.89539314 | -0.705334785 | 9.262228249 | 6.951736442 | 8.884815469 | 7.971157485 | 8.20512425 | 8.361418899 | 8.652479133 |
| ENSG00000148814 | LRRC27 | -0.373731148 | 0.667273997 | -0.331702542 | 0.699954446 | -0.705433691 | 5.866944937 | 6.228547981 | 7.451345288 | 6.456560224 | 6.1572134 | 6.475428841 | 6.222348126 |
| ENSG00000183628 | DGCR6 | 3.483451168 | 0.001734804 | -4.189830944 | 0.0005163 | -0.706379776 | 4.467311984 | 2 | 4.874587708 | 7.214570819 | 7.36516374 | 2.267001569 | 2.225841198 |
| ENSG00000177981 | ASB8 | -0.904564328 | 0.342300881 | 0.197703669 | 0.851990553 | -0.70686066 | 9.112843761 | 8.903033433 | 6.718950348 | 7.691737974 | 7.66419009 | 8.673501845 | 8.851368118 |
| ENSG00000143742 | SRP9 | 0.311437891 | 0.339794204 | -1.018435596 | 0.000322111 | -0.706997705 | 9.602033055 | 9.532315021 | 9.735974718 | 9.971714969 | 9.89552277 | 8.768928693 | 8.438024223 |
| ENSG00000117153 | KLHL12 | -0.761696312 | 0.208501379 | 0.05435047 | 0.939029498 | -0.707345842 | 9.317904228 | 8.777382925 | 8.007629717 | 8.045195239 | 8.04484669 | 8.97493051 | 8.716646332 |
| ENSG00000166598 | HSP90B1 | 0.184182085 | 0.768318166 | -0.893262949 | 0.095461226 | -0.709080864 | 13.77366519 | 13.31367773 | 13.11599373 | 13.60545465 | 13.6187793 | 11.9668157 | 12.94188924 |
| ENSG00000112062 | MAPK14 | -0.459810925 | 0.5253082 | -0.250831467 | 0.735959689 | -0.710642392 | 8.928409675 | 9.856140801 | 10.50627919 | 9.428470756 | 9.45737703 | 9.733646623 | 9.563001078 |
| ENSG00000067064 | IDI1 | -0.249191734 | 0.609720423 | -0.462582903 | 0.305744318 | -0.711774637 | 10.31337612 | 10.3778065 | 10.37481416 | 10.14218788 | 10.0717013 | 9.429778618 | 10.24608161 |
| ENSG00000230989 | HSBP1 | 0.05615337 | 0.916366826 | -0.768137428 | 0.076093247 | -0.711984057 | 10.86447339 | 10.13758676 | 10.16797309 | 10.45981982 | 10.5113233 | 9.558011928 | 9.763920691 |
| ENSG00000258947 | TUBB3 | 2.731258988 | 0.103095955 | -3.443750539 | 0.033898917 | -0.712491551 | 4.815609471 | 7.393647515 | 2 | 8.737556707 | 8.73392024 | 3.235948948 | 3.36036153 |
| ENSG00000134330 | IAH1 | -0.855988712 | 0.354286428 | 0.143106249 | 0.89090828 | -0.712882463 | 7.031698319 | 8.820493869 | 9.399592459 | 7.836899669 | 7.89700047 | 8.855481238 | 8.852426354 |
| ENSG00000086189 | DIMT1 | -0.463414708 | 0.501025025 | -0.25058415 | 0.72430187 | -0.713998858 | 8.564112515 | 7.847167909 | 7.163363831 | 7.641902724 | 7.37584259 | 7.807195765 | 7.634508409 |
| ENSG00000148672 | GLUD1 | -0.554142247 | 0.172055101 | -0.159942323 | 0.72045523 | -0.71408457 | 10.34051281 | 9.770792087 | 10.40882047 | 9.641857585 | 9.64931004 | 9.909238196 | 10.15638509 |
| ENSG00000101290 | CDS2 | -1.053419015 | 0.005630034 | 0.338205045 | 0.425433438 | -0.71521397 | 9.810482605 | 10.32888058 | 10.55568472 | 9.205456697 | 9.22613117 | 10.54687782 | 10.65402549 |
| ENSG00000145247 | OCIAD2 | -4.072265994 | 0.010739295 | 3.356105552 | 0.029365887 | -0.716160442 | 6.73718411 | 4.710564154 | 2 | 2.784612485 | 2.59489879 | 7.804574291 | 9.308063313 |
| ENSG00000133055 | MYBPH | 0.493741669 | 0.908850677 | -1.210616419 | 0.763520469 | -0.71687475 | 2 | 4.442710129 | 2 | 3.709785322 | 3.60476711 | 3.215043717 | 2 |
| ENSG00000167637 | ZNF283 | -0.55273791 | 0.663582024 | -0.1696176 | 0.900290106 | -0.72235551 | 5.330594134 | 5.72254323 | 7.934808822 | 6.24507773 | 6.30607931 | 6.90385696 | 6.320441373 |
| ENSG00000254656 | RTL1 | -3.753015698 | 0.531143362 | 3.030052067 | 0.594469008 | -0.722963631 | 2 | 2 | 2 | 2 | 2 | 3.215043717 | 3.940519005 |
| ENSG00000183655 | KLHL25 | -1.033074505 | 0.509300208 | 0.30855418 | 0.853872436 | -0.724520325 | 3.327711234 | 6.04685525 | 7.818184403 | 5.742861563 | 5.60003091 | 7.227249878 | 6.592824411 |
| ENSG00000066405 | CLDN18 | -3.753015734 | 0.531143362 | 3.027155226 | 0.594837647 | -0.725860508 | 2 | 2 | 2 | 2 | 2 | 3.600766115 | 3.640730416 |
| ENSG00000142173 | COL6A2 | -0.23343907 | 0.561212599 | -0.492559317 | 0.18084322 | -0.725998387 | 14.83498075 | 15.27905033 | 15.4147785 | 14.94766741 | 14.9789182 | 14.83726528 | 14.55780273 |
| ENSG00000129451 | KLK10 | 1.048071971 | 0.713118332 | -1.774722936 | NA | -0.726650966 | 3.327711234 | 2 | 2 | 2.868578001 | 3.15372745 | 2.267001569 | 2.117335275 |
| ENSG00000120533 | ENY2 | 0.389468241 | 0.410440873 | -1.118428901 | 0.00730204 | -0.72896066 | 8.946185348 | 8.317412869 | 8.236946975 | 8.941551548 | 8.89725143 | 7.31213785 | 7.545707309 |
| ENSG00000115484 | CCT4 | -0.516912688 | 0.509536538 | -0.214574049 | 0.795143796 | -0.731486738 | 11.46391134 | 10.57950211 | 9.703463497 | 10.23507695 | 10.2462532 | 10.62559896 | 10.45348223 |
| ENSG00000131941 | RHPN2 | -1.395983159 | 0.409632583 | 0.663470545 | 0.707028482 | -0.732512615 | 6.138360429 | 7.315436877 | 2 | 5.058234798 | 4.97065356 | 6.775876478 | 7.073100803 |
| ENSG00000013441 | CLK1 | 0.227551906 | 0.759044086 | -0.960181772 | 0.135257068 | -0.732629866 | 8.586954094 | 8.981105197 | 9.91701143 | 9.537503411 | 9.45653888 | 8.557885437 | 8.034828728 |
| ENSG00000114054 | PCCB | -1.065878069 | 0.533389071 | 0.332987587 | 0.856270394 | -0.732890482 | 9.27634997 | 8.623598428 | 3.369317972 | 7.363085426 | 7.37584259 | 8.617349501 | 8.865065003 |
| ENSG00000162804 | SNED1 | -0.56809851 | 0.852516729 | -0.165843844 | 0.957730283 | -0.733942354 | 2 | 2 | 7.818184403 | 5.614989456 | 5.86549842 | 6.223999906 | 6.011156536 |
| ENSG00000198754 | OXCT2 | 1.403792126 | 0.527945306 | -2.139465314 | 0.314064542 | -0.735673188 | 2 | 2 | 4.523675391 | 4.237680128 | 4.37434574 | 2.42098416 | 2.421067459 |

| | | | | | | | | | | | | | |
|---|---|---|---|---|---|---|---|---|---|---|---|---|---|
| ENSG00000007866 | TEAD3 | -1.182569814 | 5.52E-06 | 0.4462193 | 0.115808265 | -0.736350515 | 9.829762239 | 9.608085847 | 9.478569125 | 8.425603456 | 8.51792306 | 10.06045791 | 10.11949997 |
| ENSG00000171311 | EXOSC1 | -0.25209171 | 0.701991975 | -0.485566511 | 0.42719519 | -0.737658221 | 7.53297911 | 8.451443573 | 7.398165132 | 7.536750104 | 7.73502229 | 7.507470083 | 7.307219812 |
| ENSG00000166938 | DIS3L | -0.478369298 | 0.644658664 | -0.260510374 | 0.807781951 | -0.738879672 | 6.257048221 | 8.806266215 | 8.175061096 | 7.659688863 | 7.57432889 | 7.722704302 | 7.932467322 |
| ENSG00000175591 | P2RY2 | -0.549092175 | 0.770226986 | -0.191406919 | 0.921796599 | -0.740499094 | 4.00722531 | 2 | 4.058998049 | 2.947924294 | 3.45167086 | 3.465698119 | 3.407202941 |
| ENSG00000176715 | ACSF3 | -0.741623488 | 0.654365619 | -0.000964374 | 0.999439688 | -0.742587862 | 4.00722531 | 7.467835385 | 9.180874355 | 7.093295078 | 7.44845993 | 8.066950774 | 7.945418846 |
| ENSG00000196227 | FAM217B | -0.20961905 | 0.892020352 | -0.53360601 | 0.708830978 | -0.74322506 | 4.467311984 | 7.145017454 | 8.486635645 | 7.322721205 | 7.14562489 | 7.113241388 | 6.70252219 |
| ENSG00000176909 | MAMSTR | -1.042388583 | 0.45032434 | 0.298624224 | 0.838351035 | -0.743764358 | 5.095959458 | 4.710564154 | 2 | 3.615725078 | 3.69849104 | 4.612704558 | 4.766376438 |
| ENSG00000164542 | KIAA0895 | -2.051046402 | 0.076794225 | 1.304574033 | 0.2660773 | -0.746472369 | 3.327711234 | 6.602424674 | 5.930865648 | 4.205751501 | 3.90918929 | 7.238947338 | 6.879055607 |
| ENSG00000113248 | PCDHB15 | -0.482761088 | 0.848659281 | -0.263829319 | 0.91758425 | -0.746590406 | 2 | 5.131603625 | 2 | 3.566290607 | 3.55551834 | 3.500668998 | 3.8919413 |
| ENSG00000105849 | TWISTNB | -0.472807519 | 0.735527035 | -0.275049971 | 0.848412544 | -0.747857489 | 4.815609471 | 8.469613413 | 8.608698767 | 7.61176303 | 7.47849652 | 7.709686122 | 7.770601445 |
| ENSG00000164938 | TP53INP1 | -0.721682378 | 0.226126239 | -0.027389064 | 0.968966553 | -0.749071441 | 8.340423424 | 9.583268459 | 9.344432377 | 8.480007165 | 8.45720682 | 9.270464658 | 9.033147652 |
| ENSG00000132603 | NIP7 | -0.240170675 | 0.492183068 | -0.508943782 | 0.10794541 | -0.749114458 | 8.517314938 | 8.57407404 | 8.408111533 | 8.256693138 | 8.27714776 | 7.876232824 | 8.118139752 |
| ENSG00000172247 | C1QTNF4 | -0.83569486 | 0.671489931 | 0.08650516 | 0.967162683 | -0.749189701 | 4.00722531 | 3.076665209 | 2 | 2.695456494 | 2.94009902 | 3.355392153 | 3.236138336 |
| ENSG00000131233 | GJA9 | 1.895849372 | 0.478107313 | -2.645114166 | 0.323165276 | -0.749264794 | 2 | 2 | 3.369317972 | 3.289987342 | 3.69849104 | 2.048074405 | 2.172608095 |
| ENSG00000167543 | TP53I13 | -0.447114158 | 0.216533153 | -0.304285684 | 0.408760989 | -0.751399842 | 9.533135543 | 9.148351514 | 9.703463497 | 9.052296807 | 9.00894176 | 9.22478876 | 9.118313725 |
| ENSG00000067182 | TNFRSF1A | -0.443807411 | 0.357081274 | -0.308681762 | 0.527954208 | -0.752489173 | 11.27990552 | 10.45735223 | 10.9651998 | 10.49161431 | 10.499165 | 10.42217269 | 10.81195485 |
| ENSG00000164405 | UQCRQ | -0.690048273 | 0.331987296 | -0.062563889 | 0.938941066 | -0.752612162 | 10.43795805 | 9.00621767 | 9.31604076 | 9.025111392 | 9.04952667 | 9.472793033 | 9.827456676 |
| ENSG00000113734 | BNIP1 | -0.413104475 | 0.841034276 | -0.339603665 | 0.867480863 | -0.75270814 | 8.138771685 | 6.389896633 | 2 | 6.509830055 | 6.58294721 | 6.631488899 | 6.602920209 |
| ENSG00000163374 | YY1AP1 | 0.007381116 | 0.994991731 | -0.760288974 | 0.408562547 | -0.752907859 | 7.867441304 | 9.00621767 | 10.18377222 | 9.316137634 | 9.32587686 | 8.502889002 | 8.614310446 |
| ENSG00000168906 | MAT2A | 0.297915708 | 0.392191303 | -1.050952283 | 0.000451663 | -0.753036575 | 10.71177538 | 10.79296235 | 11.17734896 | 11.19073463 | 11.2191332 | 9.94340063 | 9.771213272 |
| ENSG00000139405 | RITA1 | -0.897382372 | 0.254887129 | 0.144282138 | 0.872268569 | -0.753100233 | 8.674867122 | 7.051611315 | 8.65477198 | 7.395290829 | 7.42465798 | 8.108635266 | 8.699694604 |
| ENSG00000143553 | SNAPIN | -0.700203091 | 0.315751098 | -0.054334941 | 0.945727701 | -0.754538031 | 9.204317094 | 8.433041974 | 7.646393253 | 7.893113669 | 7.85436579 | 8.405911884 | 8.608684933 |
| ENSG00000100568 | VTI1B | -1.015884291 | 0.129065162 | 0.260397382 | 0.72665212 | -0.755486909 | 10.34721797 | 9.030900487 | 9.099922433 | 8.592312513 | 8.64401667 | 9.984900111 | 9.779860367 |
| ENSG00000134061 | CD180 | -3.753016105 | 0.531143362 | 2.997444774 | 0.599074031 | -0.755571331 | 2 | 2 | 2 | 2 | 2 | 3.824280769 | 3.33635788 |
| ENSG00000130803 | ZNF317 | -0.34059655 | 0.768745599 | -0.415147762 | 0.713898124 | -0.755744312 | 7.384433967 | 8.451443573 | 10.09465441 | 8.727753724 | 8.7297638 | 8.747705648 | 8.555744353 |
| ENSG00000170017 | ALCAM | -0.818554266 | 0.380566315 | 0.061354536 | 0.954328634 | -0.757199729 | 10.19941743 | 12.56192683 | 11.91252756 | 11.03846779 | 11.0284211 | 11.67585929 | 12.11505304 |
| ENSG00000020129 | NCDN | -0.709516206 | 0.247942337 | -0.049204948 | 0.9448613 | -0.758721154 | 9.624286623 | 9.34816733 | 10.5187901 | 9.19431369 | 9.23104065 | 9.641172698 | 10.06667928 |
| ENSG00000186184 | POLR1D | -0.697000015 | 0.141109656 | -0.062201313 | 0.911626073 | -0.759201328 | 9.867565369 | 9.018611864 | 9.165043284 | 8.745906571 | 8.66448499 | 9.360076838 | 9.310760585 |
| ENSG00000265681 | RPL17 | 2.243812984 | 0.001788016 | -3.003222343 | 1.68E-05 | -0.759409359 | 9.371511179 | 7.819099893 | 7.896971941 | 10.84767009 | 10.7121781 | 5.367107572 | 5.888587685 |
| ENSG00000102030 | NAA10 | -0.238628141 | 0.483277973 | -0.521244258 | 0.090068933 | -0.759872399 | 8.696033109 | 8.671479029 | 8.805342542 | 8.43079185 | 8.54182783 | 8.085066106 | 8.322112506 |
| ENSG00000180658 | OR2A4 | 4.430848314 | 0.379335003 | -5.191101477 | 0.319243495 | -0.760253163 | 2 | 2 | 2 | 4.71689672 | 4.68029874 | 2 | 2 |
| ENSG00000186141 | POLR3C | 0.188434243 | 0.878396152 | -0.948967804 | 0.369406917 | -0.760533561 | 9.27634997 | 8.702538795 | 6.322058544 | 8.703665429 | 8.75179438 | 7.517109417 | 7.691205288 |
| | LOC1002870 | 4.42963167 | 0.380422171 | -5.191101266 | 0.320161353 | -0.761469596 | 2 | 2 | 2 | 4.523237872 | 4.85329164 | 2 | 2 |
| ENSG00000143228 | NUF2 | -0.305415355 | 0.863478741 | -0.457459847 | 0.784860167 | -0.762875202 | 3.327711234 | 3.686303414 | 4.058998049 | 3.79808662 | 3.15372745 | 2.911488689 | 3.789585079 |
| ENSG00000132305 | IMMT | -0.624315133 | 0.119466705 | -0.138685612 | 0.759164528 | -0.763000745 | 10.51143645 | 9.930175364 | 9.888372618 | 9.514066606 | 9.5188894 | 9.904553464 | 10.08929747 |
| ENSG00000100211 | CBY1 | -0.844188375 | 0.618625373 | 0.080715812 | 0.965981107 | -0.763472563 | 3.327711234 | 7.60566235 | 8.845626171 | 6.946463121 | 6.98750599 | 7.863694929 | 7.876348251 |
| ENSG00000148090 | AUH | 0.086325479 | 0.910324107 | -0.850604442 | 0.174826446 | -0.764278963 | 6.653098118 | 6.728495673 | 5.59507254 | 6.401247806 | 6.60731796 | 5.511266259 | 5.736403255 |
| ENSG00000163811 | WDR43 | -0.205956893 | 0.546145518 | -0.558474945 | 0.065974684 | -0.764431839 | 8.963744667 | 9.079034257 | 8.677268317 | 8.720710586 | 8.71442024 | 8.363795161 | 8.373881215 |
| ENSG00000172732 | MUS81 | -0.870252996 | 0.236297945 | 0.10543871 | 0.901615613 | -0.764814286 | 8.928409675 | 8.077869079 | 7.225742779 | 7.359462263 | 7.41087845 | 8.449073649 | 8.230662576 |
| ENSG00000110075 | PPP6R3 | -0.255308503 | 0.743393183 | -0.51062498 | 0.482463036 | -0.765933482 | 9.219014269 | 9.842262059 | 10.79563557 | 9.828366803 | 9.85983387 | 9.439667847 | 9.726317257 |
| ENSG00000100297 | MCM5 | -0.633613974 | 0.744136523 | -0.132717644 | 0.948456219 | -0.766331618 | 6.257048221 | 7.901717006 | 2 | 6.172600389 | 6.1068222 | 6.479817188 | 6.741885308 |
| ENSG00000275111 | ZNF2 | -0.751537883 | 0.318559047 | -0.015535465 | 0.985627735 | -0.767073348 | 5.532353208 | 7.145017454 | 6.80318718 | 5.978405189 | 5.90535527 | 6.739740021 | 6.533361209 |
| ENSG00000146457 | WTAP | 1.104051464 | 0.056526916 | -1.871312113 | 0.00053196 | -0.767260649 | 11.09308737 | 10.07961649 | 11.35530464 | 12.0672543 | 12.010716 | 8.917857451 | 9.213631806 |
| ENSG00000178386 | ZNF223 | 1.03095917 | 0.689206562 | -1.79855974 | 0.457954601 | -0.76760057 | 2 | 6.666836837 | 2 | 6.276156955 | 6.03677338 | 3.878466556 | 3.620995423 |
| ENSG00000040487 | PQLC2 | 0.048525777 | 0.97419422 | -0.816136983 | 0.543323761 | -0.767611206 | 8.928409675 | 8.077869079 | 4.874587708 | 8.040677544 | 8.12510181 | 7.318276922 | 7.149133029 |
| ENSG00000169599 | NFU1 | 0.098102766 | 0.948315617 | -0.866485877 | 0.52982455 | -0.768383111 | 8.653385977 | 7.274681896 | 4.523675391 | 7.64488235 | 7.7377845 | 6.85908071 | 6.640162436 |
| ENSG00000146701 | MDH2 | -0.401839214 | 0.649861077 | -0.366925034 | 0.677816167 | -0.768764247 | 11.26227543 | 10.73029099 | 9.242516611 | 10.20511011 | 10.2578459 | 10.05697448 | 10.44946654 |
| ENSG00000189184 | PCDH18 | -0.971954753 | 0.273930661 | 0.20287422 | 0.838475663 | -0.769080533 | 8.910412251 | 11.05605593 | 10.91875185 | 9.585592789 | 9.60989979 | 10.95797194 | 10.55024182 |
| ENSG00000143437 | ARNT | -0.514955601 | 0.473719789 | -0.254417888 | 0.732916452 | -0.769373488 | 8.963744667 | 9.549500232 | 10.48730704 | 9.238378447 | 9.34321171 | 9.488938651 | 9.610060665 |
| ENSG00000163714 | U2SURP | 0.082294193 | 0.788320904 | -0.853303927 | 0.00040416 | -0.771009734 | 9.895281535 | 9.672251278 | 9.658943166 | 9.873521491 | 9.77900397 | 8.910968188 | 8.88330014 |
| ENSG00000148737 | TCF7L2 | 0.111250568 | 0.912378494 | -0.884793352 | 0.300068725 | -0.773542784 | 7.624139768 | 9.030900487 | 7.163363831 | 8.262524677 | 8.31278108 | 7.052670333 | 7.528596482 |
| ENSG00000257137 | C12orf80 | -3.753016332 | 0.531143362 | 2.979279574 | 0.601502996 | -0.773736758 | 2 | 2 | 2 | 2 | 2 | 3.694366604 | 3.474731315 |
| ENSG00000103160 | HSDL1 | -0.460190638 | 0.368923829 | -0.314881125 | 0.54430284 | -0.775071763 | 8.39321512 | 8.297207175 | 9.083172337 | 8.119907552 | 8.22887947 | 8.475683062 | 8.142569067 |
| ENSG00000141562 | NARF | -0.803470841 | 0.528830242 | 0.028345377 | 0.984153231 | -0.775125464 | 9.730679447 | 9.170736405 | 6.073378139 | 8.149581172 | 8.18097132 | 9.031700669 | 8.943630224 |
| ENSG00000174891 | RSRC1 | -0.09600776 | 0.912378494 | -0.679615697 | 0.362166499 | -0.775623457 | 6.366709671 | 7.145017454 | 8.042706615 | 7.246306984 | 7.24604864 | 6.775876478 | 6.580104496 |
| ENSG00000161203 | AP2M1 | -0.711537532 | 0.194855335 | -0.064464673 | 0.920627804 | -0.776002205 | 12.92624669 | 11.92408613 | 12.80346116 | 11.90507685 | 11.9019417 | 12.28442381 | 12.77427944 |
| ENSG00000187742 | SECISBP2 | 0.271740744 | 0.682283472 | -1.049402849 | 0.067723467 | -0.777662105 | 8.696033109 | 7.60566235 | 7.646393253 | 8.310225403 | 8.36909406 | 7.046766788 | 7.043754524 |
| ENSG00000083123 | BCKDHB | -0.578093281 | 0.372026795 | -0.202920521 | 0.769672064 | -0.781013802 | 7.096806373 | 7.979858146 | 7.225742779 | 6.917187326 | 6.95938211 | 6.915242278 | 7.608437243 |
| ENSG00000227500 | SCAMP4 | -0.299617992 | 0.612075895 | -0.483490541 | 0.383684848 | -0.783108533 | 10.3868074 | 9.71095217 | 10.76969939 | 10.00861625 | 10.0939697 | 9.63945754 | 10.06667928 |
| ENSG00000172594 | SMPDL3A | 0.817711623 | 0.639687692 | -1.600829744 | 0.322890646 | -0.78311812 | 6.468621203 | 7.35507212 | 2 | 7.311513918 | 7.11195456 | 5.102662916 | 4.793076545 |
| ENSG00000132199 | ENOSF1 | 0.93093729 | 0.301027058 | -1.714233072 | 0.039100734 | -0.783295782 | 6.73718411 | 7.393647515 | 8.825624954 | 8.76109026 | 8.91200057 | 6.305369896 | 6.209190898 |
| ENSG00000186001 | LRCH3 | -1.213156515 | 0.018412016 | 0.429814514 | 0.446243463 | -0.783342001 | 9.423197387 | 8.297207175 | 8.865353883 | 7.717440541 | 7.74603955 | 9.285896341 | 9.423285228 |
| ENSG00000174165 | ZDHHC24 | -0.887370916 | 0.326333176 | 0.102972339 | 0.920980141 | -0.784398577 | 6.816637977 | 8.297207175 | 9.083172337 | 7.426792977 | 7.49817903 | 8.244430344 | 8.606804877 |
| ENSG00000104133 | SPG11 | -0.182243705 | 0.751812007 | -0.602994111 | 0.239452646 | -0.785237816 | 10.21416435 | 9.170736405 | 9.935792933 | 9.672111811 | 9.63233954 | 9.254546025 | 9.213219973 |
| ENSG00000157429 | ZNF19 | -0.706972789 | 0.715896274 | -0.078355275 | 0.969731551 | -0.785328065 | 2 | 5.72254323 | 3.369317972 | 3.840289928 | 4.0930011 | 4.588526421 | 4.329022995 |
| ENSG00000100412 | ACO2 | -0.070239013 | 0.930680662 | -0.716289889 | 0.296655486 | -0.786528902 | 10.26462289 | 9.079034257 | 8.884815469 | 9.429335066 | 9.52049436 | 8.702892291 | 8.95451255 |
| ENSG00000135046 | ANXA1 | 0.148279409 | 0.838100556 | -0.935432899 | 0.122999873 | -0.787153489 | 12.2287862 | 11.12072186 | 11.18521898 | 11.7618096 | 11.7467334 | 10.37347689 | 10.92000025 |
| ENSG00000170298 | LGALS9B | 4.403162088 | 0.386463861 | -5.191100782 | 0.322255484 | -0.787938695 | 2 | 2 | 2 | 4.623314311 | 4.72553021 | 2 | 2 |
| ENSG00000087303 | NID2 | -1.784652458 | 0.04311869 | 0.996697504 | 0.281004483 | -0.787954954 | 11.06489013 | 12.86533873 | 12.38869272 | 10.48912954 | 10.5165603 | 12.24081159 | 13.87892942 |
| ENSG00000121578 | B4GALT4 | -0.628803586 | 0.323982764 | -0.160058835 | 0.819657623 | -0.788862421 | 9.720386367 | 8.671479029 | 8.561105505 | 8.564258394 | 8.34755546 | 8.803037464 | 9.035944835 |
| ENSG00000120093 | HOXB3 | 3.440116023 | 2.27E-06 | -4.229033288 | 5.01E-08 | -0.788917265 | 6.257048221 | 4.93638933 | 4.523675391 | 8.773395507 | 8.7017439 | 2.68698273 | 2.509591799 |
| ENSG00000170545 | SMAGP | -0.377645643 | 0.720482866 | -0.411818067 | 0.690405965 | -0.78946371 | 8.16936288 | 5.946638857 | 8.325066185 | 7.416368504 | 7.43490694 | 7.259511486 | 7.513957873 |
| ENSG00000136940 | PDCL | -0.498403694 | 0.315955574 | -0.291951572 | 0.568416252 | -0.790355266 | 8.493334308 | 8.337339476 | 9.180874355 | 8.27796118 | 8.16053033 | 8.360824221 | 8.485179747 |
| ENSG00000180479 | ZNF571 | -1.18262253 | 0.51002601 | 0.390717577 | 0.837824475 | -0.791904953 | 2 | 6.535001748 | 4.874587708 | 4.358765249 | 4.43005853 | 6.239610476 | 5.080008298 |
| ENSG00000129515 | SNX6 | 0.012819438 | 0.982918686 | -0.8071713 | 0.093525732 | -0.794351862 | 10.47516499 | 9.756063186 | 10.75919294 | 10.42297598 | 10.3742555 | 9.549686932 | 9.613809871 |
| ENSG00000070756 | PABPC1 | -1.41941659 | 0.011400228 | 0.624991304 | 0.301662931 | -0.794425285 | 12.09176592 | 13.33984242 | 12.62546556 | 11.38211482 | 11.3356213 | 13.17422418 | 13.59831928 |
| ENSG00000173809 | TDRD12 | -3.753016599 | 0.531143362 | 2.95789763 | NA | -0.795118969 | 2 | 2 | 2 | 2 | 2 | 4.111244182 | 2.70970617 |
| ENSG00000151694 | ADAM17 | 0.604207722 | 0.599217549 | -1.400269442 | 0.17932719 | -0.796061721 | 7.579279349 | 9.608085847 | 10.65536953 | 10.39138945 | 10.3116735 | 8.308125745 | 8.411703491 |
| ENSG00000101040 | ZMYND8 | 0.621322073 | 0.613329998 | -1.417530458 | 0.205343649 | -0.796208385 | 6.653098118 | 8.57407404 | 9.972637969 | 9.524619869 | 9.61140668 | 7.52137298 | 7.58443608 |
| ENSG00000101928 | MOSPD1 | 0.636176159 | 0.510282114 | -1.432618851 | 0.101255863 | -0.796442692 | 9.371511179 | 7.002537973 | 8.585098385 | 9.26185831 | 9.21725179 | 7.341370018 | 7.045606259 |
| ENSG00000125779 | PANK2 | 0.204708302 | 0.91483416 | -1.001686296 | 0.558314161 | -0.796977994 | 4.815609471 | 7.874700259 | 10.44859742 | 9.293531584 | 9.33502642 | 8.102956788 | 8.13650041 |
| ENSG00000126214 | KLC1 | -0.078543467 | 0.903601303 | -0.719650802 | 0.17883612 | -0.798194269 | 10.81212061 | 9.672251278 | 10.19160738 | 10.19447291 | 10.2457682 | 9.614730018 | 9.54838268 |
| ENSG00000186767 | SPIN4 | -0.507340314 | 0.825685372 | -0.291486472 | 0.900758644 | -0.798826787 | 3.327711234 | 6.899080544 | 2 | 5.040242897 | 5.04390545 | 5.50697268 | 4.895193906 |
| ENSG00000003989 | SLC7A2 | -2.822325533 | 0.000804913 | 2.022984424 | 0.018492461 | -0.799341108 | 7.53297911 | 6.602424674 | 8.325066185 | 4.946755474 | 5.0259382 | 10.20860099 | 8.706733286 |
| ENSG00000170430 | MGMT | -0.911892944 | 0.636476225 | 0.11214637 | 0.957713961 | -0.799746574 | 8.138771685 | 8.297207175 | 2 | 6.580005553 | 6.9210099 | 7.564363052 | 7.927454842 |
| ENSG00000107669 | ATE1 | -0.713252281 | 0.467358725 | -0.087686419 | 0.936069485 | -0.8009387 | 7.159102577 | 9.532315021 | 8.266922072 | 8.017874397 | 7.84670947 | 8.541185654 | 8.562873254 |
| ENSG00000140854 | KATNB1 | -0.401060072 | 0.816824789 | -0.404474633 | 0.811558361 | -0.805534705 | 8.998234871 | 7.145017454 | 4.058998049 | 7.437142666 | 7.37938471 | 6.995586672 | 7.72395257 |
| ENSG00000011105 | TSPAN9 | -1.258169275 | 0.059358025 | 0.451020717 | 0.535037046 | -0.807148558 | 10.1076228 | 8.540087354 | 9.272377397 | 8.119907552 | 8.27144011 | 9.76634617 | 10.00971824 |
| ENSG00000075239 | ACAT1 | -0.452618404 | 0.839051597 | -0.355207014 | 0.872932235 | -0.807825418 | 9.48530526 | 8.717821467 | 2 | 8.087422161 | 8.16258751 | 7.813294077 | 8.53941009 |
| ENSG00000008300 | CELSR3 | -0.362106571 | 0.872289176 | -0.44614836 | 0.837960391 | -0.808254931 | 2 | 5.595927442 | 8.296287027 | 6.456560224 | 6.6952083 | 6.747040087 | 6.202567011 |
| ENSG00000147130 | ZMYM3 | -0.86215367 | 0.045918436 | 0.053690538 | 0.919423567 | -0.808463132 | 9.509418609 | 9.726146444 | 8.922968322 | 8.491639362 | 8.64989445 | 9.378702006 | 9.577792788 |
| ENSG00000103852 | TTC23 | -0.390353436 | 0.618255555 | -0.418111507 | 0.584589142 | -0.808464943 | 7.790810999 | 8.029695761 | 9.287079172 | 8.124184191 | 8.15227214 | 8.17644121 | 8.042267212 |
| ENSG00000125285 | SOX21 | 1.504773636 | NA | -2.31353874 | NA | -0.808765105 | 2 | 2 | 2 | 3.289987342 | 2 | 2.139668783 | 2 |
| ENSG00000142686 | C1orf216 | -0.424151114 | 0.856794892 | -0.387222396 | 0.866925741 | -0.81137351 | 10.14658276 | 8.469613413 | 2 | 8.50645854 | 8.56844956 | 8.446833885 | 8.691439168 |
| ENSG00000164815 | ORC5 | 0.196939654 | 0.903601303 | -1.00927417 | 0.478299157 | -0.812334516 | 7.031698319 | 7.701025996 | 3.369317972 | 6.931899482 | 7.18258783 | 6.075339068 | 5.727202496 |
| ENSG00000174876 | AMY1B | 4.378401314 | 0.396344882 | -5.191099045 | 0.329252781 | -0.812697731 | 2 | 2 | 2 | 5.003571641 | 4.1930041 | 2 | 2 |
| ENSG00000187733 | AMY1C | 4.378401314 | 0.396344882 | -5.191099045 | 0.329252781 | -0.812697731 | 2 | 2 | 2 | 5.003571641 | 4.1930041 | 2 | 2 |
| ENSG00000237763 | AMY1A | 4.378401314 | 0.396344882 | -5.191099045 | 0.329252781 | -0.812697731 | 2 | 2 | 2 | 5.003571641 | 4.1930041 | 2 | 2 |
| ENSG00000174871 | CNIH2 | -1.317562515 | 0.484134885 | 0.504847856 | 0.798479518 | -0.812714659 | 3.327711234 | 5.946638857 | 2 | 3.615725078 | 3.74316441 | 5.152992171 | 5.122735884 |
| ENSG00000113013 | HSPA9 | 0.035813284 | 0.922635322 | -0.849958115 | 0.002681332 | -0.814144831 | 11.61323974 | 11.14918383 | 11.31999735 | 11.41040523 | 11.4063384 | 10.52620926 | 10.52258738 |
| ENSG00000165502 | RPL36AL | -0.756669681 | 0.175327851 | -0.058776214 | 0.929596218 | -0.815445895 | 10.93293345 | 9.890264534 | 10.43545992 | 9.70102904 | 9.75056618 | 10.11225475 | 10.67571014 |
| ENSG00000141385 | AFG3L2 | -0.394929577 | 0.781110464 | -0.421431842 | 0.762852238 | -0.816361419 | 6.563806279 | 8.623598428 | 10.37481416 | 8.821593044 | 8.87609528 | 8.717365994 | 8.921111931 |
| ENSG00000145740 | SLC30A5 | -0.188498162 | 0.747712189 | -0.628940987 | 0.226495503 | -0.817439149 | 9.533135543 | 9.994333773 | 8.978369997 | 9.389919188 | 9.36392108 | 9.106681409 | 8.750531435 |
| ENSG00000112679 | DUSP22 | -1.517154802 | 0.399795702 | 0.698289329 | 0.712878225 | -0.818865473 | 3.327711234 | 7.431218269 | 9.681375057 | 6.877205252 | 6.9210099 | 9.204383073 | 8.935663927 |
| ENSG00000068912 | ERLEC1 | 0.131133123 | 0.784300093 | -0.950665033 | 0.01565797 | -0.81953191 | 10.48735708 | 11.22601381 | 10.7904855 | 11.02423622 | 10.9741121 | 10.02786376 | 9.804948449 |
| ENSG00000101856 | PGRMC1 | 0.063372714 | 0.910745594 | -0.883821062 | 0.048909865 | -0.820448348 | 11.23361136 | 10.33384851 | 10.62096771 | 10.85508643 | 10.8278498 | 9.914515727 | 9.879774129 |
| ENSG00000139832 | RAB20 | -0.69209745 | 0.687848199 | -0.130509183 | 0.943446144 | -0.822606633 | 2 | 6.389896633 | 6.882775709 | 5.461211713 | 5.45984938 | 6.132200015 | 5.807970301 |

| | | | | | | | | | | | | | |
|---|---|---|---|---|---|---|---|---|---|---|---|---|---|
| ENSG00000133895 | MEN1 | -0.4141856 | 0.258819968 | -0.410037778 | 0.253726618 | -0.824223378 | 8.910412251 | 8.732943944 | 8.408111533 | 8.283707551 | 8.29788471 | 8.382079673 | 8.20189164 |
| ENSG00000088986 | DYNLL1 | -0.747653177 | 0.168573196 | -0.078885901 | 0.901746027 | -0.826539078 | 11.59645304 | 10.47993704 | 11.20470823 | 10.41208663 | 10.4222654 | 10.92667044 | 11.22656376 |
| ENSG00000171843 | MLLT3 | -1.00193187 | 0.020898129 | 0.174727391 | 0.728258838 | -0.827204479 | 7.904282243 | 8.469613413 | 8.677268317 | 7.437142666 | 7.36516374 | 8.69913345 | 8.403784899 |
| ENSG00000111665 | CDCA3 | -1.099058047 | 0.461217823 | 0.271075497 | 0.865035836 | -0.82798255 | 3.327711234 | 5.595927442 | 3.369317972 | 3.615725078 | 3.82856379 | 4.443291373 | 5.08721802 |
| ENSG00000123836 | PFKFB2 | -1.124398403 | 0.417004646 | 0.295614931 | 0.844695037 | -0.828783472 | 5.532353208 | 5.595927442 | 8.561105505 | 6.29903474 | 6.11534392 | 7.800194554 | 7.321046127 |
| ENSG00000144021 | CIAO1 | -0.556662669 | 0.535148173 | -0.272381801 | 0.772041492 | -0.829044471 | 8.468948322 | 9.890264534 | 10.65536953 | 9.360314042 | 9.3808074 | 9.661597094 | 9.64498544 |
| ENSG00000123352 | SPATS2 | 0.08295674 | 0.888897361 | -0.912174398 | 0.055208449 | -0.829217658 | 9.460782017 | 9.424669273 | 8.631919287 | 9.239364442 | 9.36749255 | 8.503427922 | 8.106655408 |
| ENSG00000206073 | SERPINB4 | 4.361726697 | 0.395913674 | -5.19109977 | 0.326816217 | -0.829373073 | 2 | 2 | 2 | 4.647285135 | 4.63360314 | 2 | 2 |
| ENSG00000163106 | HPGDS | -3.753017028 | 0.531143362 | 2.923522658 | NA | -0.829494371 | 2 | 2 | 2 | 2 | 2 | 4.122514202 | 2.592996859 |
| ENSG00000111775 | COX6A1 | -0.153334045 | 0.888333431 | -0.67660597 | 0.480204433 | -0.829940015 | 10.95065381 | 8.806266215 | 9.212024683 | 9.808573603 | 9.8342512 | 9.283701895 | 9.318056434 |
| ENSG00000156256 | USP16 | -0.121597233 | 0.86184497 | -0.70848483 | 0.230307325 | -0.830082063 | 9.848787621 | 8.806266215 | 8.742726175 | 9.115954467 | 9.09332116 | 8.469076579 | 8.575104441 |
| ENSG00000051009 | FAM160A2 | -0.803891693 | 0.711497595 | -0.026200495 | 0.990893061 | -0.830092188 | 2 | 7.35507212 | 9.426401271 | 7.377487766 | 7.35440525 | 8.085066106 | 8.17168765 |
| ENSG00000174226 | SNX31 | -0.689658684 | 0.830335275 | -0.140441925 | NA | -0.830100609 | 2 | 2 | 3.369317972 | 2.270786764 | 2.49394783 | 2.717032197 | 2.326754077 |
| ENSG00000107164 | FUBP3 | -0.211167459 | 0.573001609 | -0.619987338 | 0.061569555 | -0.831154798 | 9.358295023 | 9.424669273 | 9.735974718 | 9.347568393 | 9.2591908 | 8.737663157 | 9.042914155 |
| ENSG00000008056 | SYN1 | -0.366213846 | 0.86184497 | -0.465427973 | 0.818406267 | -0.83164182 | 4.00722531 | 6.666836837 | 2 | 5.076005082 | 5.00774437 | 5.136410075 | 4.74829783 |
| ENSG00000123416 | TUBA1B | 0.555260257 | 0.446354696 | -1.387157924 | 0.033296326 | -0.831897667 | 11.87457535 | 11.31160468 | 10.81097638 | 11.97200261 | 11.9319504 | 9.288712876 | 10.49422299 |
| ENSG00000161277 | THAP8 | -0.783428623 | 0.79343476 | -0.049023761 | 0.987416408 | -0.832452384 | 2 | 8.255926468 | 2 | 5.829220145 | 6.09824986 | 6.653021698 | 6.681239787 |
| ENSG00000187609 | EXD3 | -0.907111839 | 0.628146304 | 0.074333621 | 0.97068214 | -0.832778218 | 7.975256505 | 6.666836837 | 2 | 5.997179931 | 6.06344184 | 7.11465301 | 6.804465095 |
| ENSG00000096093 | EFHC1 | -0.862562624 | 0.657619829 | 0.02643922 | 0.989740833 | -0.836123405 | 2 | 8.004992147 | 8.380957451 | 6.856790872 | 6.72335419 | 7.631969638 | 7.680519528 |
| ENSG00000105810 | CDK6 | -0.613054718 | 0.222830111 | -0.223549996 | 0.681431908 | -0.836604713 | 9.895281535 | 10.4709455 | 10.63252637 | 9.799243015 | 9.71034999 | 9.788404752 | 10.42815449 |
| ENSG00000112249 | ASCC3 | -0.666142661 | 0.413198702 | -0.171414284 | 0.847380153 | -0.837556945 | 9.143983763 | 10.48887288 | 11.04953186 | 9.778365132 | 9.74233706 | 10.42644164 | 10.05638425 |
| ENSG00000003509 | NDUFAF7 | -0.500746205 | 0.685162585 | -0.337368504 | 0.78794815 | -0.838114709 | 6.009025997 | 6.728495673 | 8.784770914 | 7.25803034 | 7.09050444 | 7.341370018 | 7.330190599 |
| ENSG00000174606 | ANGEL2 | -0.053196802 | 0.948155695 | -0.786841734 | 0.266613509 | -0.840038536 | 7.485143553 | 7.538394143 | 8.825624954 | 8.013270196 | 8.05374131 | 7.462723726 | 7.149133029 |
| ENSG00000092841 | MYL6 | -0.924475435 | 0.082516252 | 0.083995036 | 0.89539314 | -0.840480399 | 13.99047696 | 12.86619626 | 13.43923225 | 12.56285085 | 12.5963066 | 13.42632465 | 13.73277744 |
| ENSG00000108788 | MLX | -0.366266811 | 0.846600015 | -0.474284399 | 0.795688254 | -0.84055121 | 10.73241055 | 7.60566235 | 5.39244166 | 8.973900679 | 8.96835847 | 8.999589189 | 8.714319971 |
| ENSG00000149554 | CHEK1 | 0.074734476 | 0.95872661 | -0.915626936 | 0.476899959 | -0.84089246 | 8.257427002 | 7.002537973 | 4.523675391 | 7.423326516 | 7.21862736 | 6.442083733 | 6.29264319 |
| ENSG00000124155 | PIGT | -0.722037561 | 0.21973507 | -0.119196381 | 0.858897472 | -0.841233943 | 11.69698977 | 10.5284131 | 10.81605395 | 10.38070504 | 10.3835453 | 10.85732303 | 11.09951659 |
| ENSG00000112320 | SOBP | -2.600639488 | 0.12809176 | 1.758051981 | 0.301787266 | -0.842587507 | 2 | 6.228547981 | 4.058998049 | 2.947924294 | 3.28037825 | 7.123093858 | 5.957088417 |
| ENSG00000182208 | MOB2 | -0.370245189 | 0.835111495 | -0.473034216 | 0.783596985 | -0.843279404 | 4.00722531 | 8.053983481 | 9.385998894 | 7.847284664 | 8.0314011 | 8.115702075 | 7.501869356 |
| ENSG00000100592 | DAAM1 | -0.132274982 | 0.91483416 | -0.711760248 | 0.51563255 | -0.844035231 | 6.563806279 | 9.34816733 | 9.358421373 | 8.750063451 | 8.72837565 | 8.203226602 | 8.128660152 |
| ENSG00000148572 | NRBF2 | -0.336507198 | 0.539576503 | -0.508095075 | 0.327879239 | -0.844602272 | 9.262228249 | 8.255926468 | 9.049077186 | 8.540455898 | 8.61275919 | 8.431623835 | 8.382697855 |
| ENSG00000149243 | KLHL35 | -1.37566174 | 0.396224049 | 0.530893528 | 0.757048961 | -0.844768212 | 2 | 5.595927442 | 6.534109959 | 4.54891529 | 4.22485516 | 6.430795201 | 5.631815919 |
| ENSG00000167397 | VKORC1 | -0.257793347 | 0.634741192 | -0.587732727 | 0.23570663 | -0.845526074 | 10.62615013 | 10.09139839 | 11.0961933 | 10.37846912 | 10.4269797 | 10.01071781 | 10.13480234 |
| ENSG00000188706 | ZDHHC9 | -0.281238257 | 0.718010015 | -0.565042876 | 0.435161964 | -0.846281133 | 9.699577239 | 9.842262059 | 11.12920538 | 10.09721189 | 10.0934307 | 9.765223103 | 9.858600332 |
| ENSG00000099308 | MAST3 | -0.731270766 | 0.587266932 | -0.115025835 | 0.937781729 | -0.846296601 | 7.384433967 | 8.993716073 | 5.39244166 | 7.093295078 | 7.29187696 | 8.108635266 | 7.402777233 |
| ENSG00000078487 | ZCWPW1 | -0.305189258 | 0.912501485 | -0.541184491 | 0.839285386 | -0.846373749 | 2 | 6.389896633 | 2 | 4.739373666 | 4.65713986 | 4.555645384 | 4.445805334 |
| ENSG00000087206 | UIMC1 | -0.461885165 | 0.57369092 | -0.385043515 | 0.639193832 | -0.846928681 | 8.946185348 | 7.315436877 | 7.646393253 | 7.694616532 | 7.69009211 | 7.759264603 | 7.776188509 |
| ENSG00000127955 | GNAI1 | 0.90491794 | 0.445130278 | -1.751923622 | 0.10692029 | -0.847005682 | 9.590776211 | 7.954278496 | 6.43197674 | 9.452477833 | 9.3913715 | 6.932953913 | 6.674075308 |
| ENSG00000144034 | TPRKB | 0.020338142 | 0.981338611 | -0.867481653 | 0.218585328 | -0.847143511 | 6.468621203 | 5.303523747 | 5.772716771 | 5.987823101 | 5.86549842 | 4.959178705 | 5.263040529 |
| ENSG00000102974 | CTCF | -0.589671174 | 0.696183014 | -0.258014716 | 0.869968927 | -0.84768589 | 5.330594134 | 9.225224669 | 9.868959282 | 8.461536372 | 8.44034872 | 8.966355803 | 8.564165641 |
| ENSG00000241945 | PWP2 | 2.380060642 | 0.015303223 | -3.228098505 | 0.000786588 | -0.848037863 | 6.366709671 | 4.11352228 | 4.523675391 | 7.587189552 | 7.59884443 | 2.911488689 | 3.042026731 |
| ENSG00000124357 | NAGK | 0.028049655 | 0.969963817 | -0.876392798 | 0.155720174 | -0.848343143 | 10.05124847 | 9.863030423 | 8.699419242 | 9.604082348 | 9.7485133 | 8.63860224 | 8.908448207 |
| ENSG00000135775 | COG2 | -0.810240914 | 0.394385857 | -0.039564334 | 0.970602478 | -0.849805248 | 6.257048221 | 8.590771778 | 8.325066185 | 7.119236776 | 7.34718809 | 7.993660487 | 7.991296779 |
| ENSG00000147439 | BIN3 | 0.512694328 | 0.380422171 | -1.362681562 | 0.0088054 | -0.849987234 | 6.89194353 | 7.901717006 | 7.97167868 | 8.117764469 | 8.22691474 | 6.257611854 | 6.437892833 |
| ENSG00000111554 | MDM1 | -0.253497527 | 0.895265695 | -0.59688528 | 0.737997178 | -0.850382807 | 7.27616306 | 6.535001748 | 2 | 6.252910529 | 6.04571775 | 6.126613714 | 5.459179378 |
| ENSG00000087258 | GNAO1 | -1.646812564 | 0.438555858 | 0.795900003 | 0.711952414 | -0.850912561 | 2 | 4.442710129 | 3.369317972 | 3.09461492 | 2.2680111 | 4.652122235 | 3.580696614 |
| ENSG00000215218 | UBE2QL1 | -0.634066049 | 0.805573299 | -0.217756881 | 0.934077269 | -0.85122293 | 2 | 4.442710129 | 2 | 3.289987342 | 2.68924543 | 3.355392153 | 3.128433649 |
| ENSG00000175104 | TRAF6 | -0.214395175 | 0.913299181 | -0.639036511 | 0.726284264 | -0.853431687 | 9.48530526 | 7.928237109 | 3.369317972 | 8.203120604 | 8.04484669 | 7.954782214 | 7.411419562 |
| ENSG00000132846 | ZBED3 | -3.430389364 | 0.00368158 | 2.576928351 | 0.019796165 | -0.853461013 | 3.327711234 | 5.946638857 | 4.058998049 | 2.695456494 | 2.68924543 | 7.310906894 | 7.376535208 |
| ENSG00000165511 | C10orf25 | -4.71493145 | 0.159913355 | 3.86095831 | 0.168877692 | -0.85397314 | 2 | 3.076665209 | 2 | 2 | 2 | 5.238348159 | 4.80186789 |
| ENSG00000157191 | NECAP2 | -0.608128088 | 0.349396138 | -0.246852991 | 0.721502431 | -0.854981078 | 10.98974631 | 9.664384941 | 10.7325871 | 9.878593178 | 10.032869 | 10.11190139 | 10.49794712 |
| ENSG00000159882 | ZNF230 | -0.283145771 | 0.890436443 | -0.572832591 | 0.766433306 | -0.855978362 | 2 | 5.838937808 | 7.646393253 | 6.113598934 | 6.22964624 | 5.991769846 | 5.803599879 |
| ENSG00000122218 | COPA | -0.280481638 | 0.822518774 | -0.576360313 | 0.620145964 | -0.856841951 | 9.613202746 | 12.42197616 | 12.73541118 | 11.78647256 | 11.835317 | 11.44398809 | 11.58379362 |
| ENSG00000172748 | ZNF596 | 0.032164838 | 0.99113768 | -0.889151603 | 0.736787432 | -0.856986765 | 2 | 6.728495673 | 2 | 5.22679461 | 5.36298494 | 4.757085024 | 4.188602707 |
| ENSG00000221821 | C6orf226 | -0.299607266 | 0.848265166 | -0.557922518 | 0.70356954 | -0.857529784 | 6.816637977 | 4.11352228 | 4.058998049 | 5.348759107 | 5.30456641 | 5.147485955 | 5.04340913 |
| ENSG00000164221 | CCDC112 | -0.411690879 | 0.793896601 | -0.446247973 | 0.772041492 | -0.857938851 | 3.327711234 | 5.72254323 | 7.285536013 | 5.949776857 | 5.62411716 | 6.019162896 | 5.453611367 |
| ENSG00000184937 | WT1 | 4.333155878 | 0.402624908 | -5.191099051 | 0.329804519 | -0.857943173 | 2 | 2 | 2 | 4.623314311 | 4.60967607 | 2 | 2 |
| ENSG00000176371 | ZSCAN2 | 0.148764131 | 0.926779701 | -1.007828347 | 0.478389629 | -0.859064216 | 3.327711234 | 7.431218269 | 7.502634626 | 7.044488801 | 7.09482005 | 5.825876956 | 6.092227206 |
| ENSG00000117174 | ZNHIT6 | -0.628005157 | 0.185239825 | -0.231210621 | 0.6505088 | -0.859215778 | 7.867441304 | 8.623598428 | 8.631919287 | 7.860161659 | 7.7377845 | 8.064031095 | 8.307525005 |
| ENSG00000170456 | DENND5B | -0.549889132 | 0.599957698 | -0.309392507 | 0.774853287 | -0.859281638 | 6.816637977 | 9.090820894 | 9.372276026 | 8.272191829 | 8.20512425 | 8.654217494 | 8.273956186 |
| ENSG00000102471 | NDFIP2 | -0.717865055 | 0.412901143 | -0.142063649 | 0.884519325 | -0.859928703 | 7.624139768 | 9.415325457 | 9.670202709 | 8.576794289 | 8.26953254 | 8.868927112 | 9.123144518 |
| ENSG00000203995 | ZYG11A | -1.231491347 | 0.362148669 | 0.369762653 | 0.797269964 | -0.861728694 | 5.532353208 | 3.076665209 | 4.058998049 | 3.515101962 | 3.69849104 | 4.68289886 | 5.005857272 |
| ENSG00000153721 | CNKSR3 | -1.45282579 | 0.222084596 | 0.589632032 | 0.642308336 | -0.863193758 | 5.330594134 | 6.228547981 | 8.325066185 | 5.949776857 | 5.6121243 | 7.703132714 | 7.799420735 |
| ENSG00000123395 | ATG101 | -0.111301012 | 0.960285623 | -0.751979112 | 0.722957397 | -0.863280124 | 9.602033055 | 9.170736405 | 2 | 8.731963162 | 8.69180693 | 8.061837451 | 8.095972716 |
| ENSG00000135211 | TMEM60 | 0.08684711 | 0.863599094 | -0.950444913 | 0.017855055 | -0.863597803 | 7.667647223 | 7.954278496 | 8.380957451 | 8.109160159 | 8.11663771 | 7.165935143 | 7.025105265 |
| ENSG00000149483 | TMEM138 | -0.533020446 | 0.431896348 | -0.330604232 | 0.633167305 | -0.863624678 | 9.015175615 | 7.669933291 | 8.922968322 | 8.18685754 | 8.04707549 | 8.294489071 | 8.341838298 |
| ENSG00000182318 | ZSCAN22 | -0.71392501 | 0.227604787 | -0.150860318 | 0.818089356 | -0.864785327 | 6.257048221 | 6.602424674 | 6.322058544 | 5.578054239 | 5.88556448 | 6.471027105 | 6.029985803 |
| | SQRDL | 7.931448953 | 2.55E-05 | -8.797640877 | 0.000397994 | -0.866191925 | 2 | 5.303523747 | 2 | 11.5430919 | 11.5891353 | 2 | 2 |
| ENSG00000030582 | GRN | 0.084565189 | 0.89152261 | -0.952636948 | 0.055557322 | -0.868071759 | 13.24264217 | 12.43127714 | 12.26418878 | 12.78278409 | 12.810703 | 11.66933017 | 11.84618268 |
| ENSG00000135316 | SYNCRIP | -0.024408915 | 0.957978247 | -0.844214721 | 0.023138941 | -0.868623636 | 10.91499272 | 10.67655334 | 10.29705308 | 10.65570314 | 10.5990046 | 9.63331528 | 9.968585649 |
| ENSG00000240654 | C1QTNF9 | 2.775429691 | 0.558941971 | -3.645545009 | NA | -0.870115319 | 2 | 2 | 2 | 3.289987342 | 3.60476711 | 2 | 2.059860178 |
| ENSG00000118096 | IFT46 | -0.353280875 | 0.841155419 | -0.517607621 | 0.758720739 | -0.870888496 | 3.327711234 | 7.467835385 | 8.561105505 | 7.210554257 | 7.21069602 | 6.977080688 | 7.12130792 |
| ENSG00000198799 | LRIG2 | -0.625671101 | 0.778610361 | -0.246398313 | 0.915259822 | -0.872069413 | 2 | 6.311476754 | 8.978369997 | 6.960881212 | 7.04216061 | 7.538302335 | 7.18149309 |
| ENSG00000117707 | PROX1 | 1.729358872 | 0.39960716 | -2.6023719 | 0.193600095 | -0.873013028 | 2 | 4.11352228 | 2 | 4.358765249 | 4.3456612 | 2.3070652 | 2.225841198 |
| ENSG00000001167 | NFYA | 0.196518321 | 0.922962824 | -1.069884866 | 0.554691707 | -0.873366545 | 3.327711234 | 8.395519306 | 9.849281147 | 8.922033725 | 8.91566441 | 7.988280194 | 7.258555286 |
| ENSG00000089163 | SIRT4 | -0.770774745 | 0.762599354 | -0.103624975 | 0.969279885 | -0.874399721 | 2 | 5.72254323 | 2 | 3.709785322 | 3.86943969 | 4.580376151 | 3.940519005 |
| ENSG00000242802 | AP5Z1 | 0.151358295 | 0.897065049 | -1.025859389 | 0.301496071 | -0.874501094 | 10.14658276 | 7.538394143 | 8.742726175 | 9.260887574 | 9.40446877 | 8.246363378 | 8.086110269 |
| ENSG00000168259 | DNAJC7 | 0.431926879 | 0.59184685 | -1.307191403 | 0.066431499 | -0.875264524 | 9.081016759 | 8.732943944 | 10.34701993 | 9.956206375 | 10.0238444 | 8.147081218 | 8.373144057 |
| ENSG00000173889 | PHC3 | -0.523221719 | 0.605635278 | -0.352201142 | 0.731545698 | -0.875422861 | 9.031919741 | 9.405920729 | 11.08361639 | 9.672841851 | 9.54908243 | 10.12491852 | 9.332538367 |
| ENSG00000138069 | RAB1A | -0.362273235 | 0.611600602 | -0.515207353 | 0.448615444 | -0.877480588 | 11.99046984 | 10.71514053 | 10.76445573 | 10.93819728 | 10.9088206 | 10.74627917 | 10.79526758 |
| ENSG00000174429 | ABRA | -3.753017631 | 0.531143362 | 2.875241323 | NA | -0.877776308 | 2 | 2 | 2 | 2 | 2 | 3.943456545 | 2.918126211 |
| ENSG00000018699 | TTC27 | -0.424746247 | 0.694658127 | -0.454282726 | 0.669330321 | -0.879028973 | 6.73718411 | 6.602424674 | 8.699419242 | 7.303993732 | 7.23044321 | 7.118879607 | 7.349806994 |
| ENSG00000116161 | CACYBP | -0.513470559 | 0.199526015 | -0.366654998 | 0.364637341 | -0.880125557 | 8.653385977 | 9.256956408 | 8.825624954 | 8.381625466 | 8.48213028 | 8.492068113 | 8.659146771 |
| ENSG00000175463 | TBC1D10C | -4.714929056 | 0.166109341 | 3.834444079 | 0.180853 | -0.880484976 | 2 | 3.076665209 | 2 | 2 | 2 | 5.343222328 | 4.584731282 |
| ENSG00000112972 | HMGCS1 | -0.548976929 | 0.520760111 | -0.332844435 | 0.702841297 | -0.881821364 | 10.4253394 | 10.9629268 | 11.04521411 | 10.31066614 | 10.2664796 | 9.284329221 | 11.15567751 |
| ENSG00000156976 | EIF4A2 | -0.898575935 | 0.021970728 | 0.016739917 | 0.972606493 | -0.881836018 | 12.07772327 | 11.34889202 | 11.87652621 | 10.90191807 | 10.9002162 | 11.92100809 | 11.70048584 |
| ENSG00000141198 | TOM1L1 | 0.043629322 | 0.98350331 | -0.925990102 | 0.632223509 | -0.882360779 | 2 | 6.04685525 | 8.042706615 | 6.872128674 | 6.79936111 | 5.747972572 | 6.06327132 |
| ENSG00000115468 | EFHD1 | -5.20880374 | 0.001784151 | 4.325195715 | 0.007589468 | -0.883608025 | 2 | 5.838937808 | 7.818184403 | 2.947924294 | 2.38539787 | 10.96796381 | 10.70295227 |
| ENSG00000052850 | ALX4 | -4.644542973 | 0.032131199 | 3.760189132 | 0.052358157 | -0.884353841 | 2 | 5.595927442 | 2 | 2.270786764 | 2.14022021 | 6.855708005 | 8.277112026 |
| ENSG00000137473 | TTC29 | -3.753017715 | 0.531143362 | 2.868543381 | NA | -0.884474334 | 2 | 2 | 2 | 2 | 2 | 3.1504466 | 3.807156736 |
| ENSG00000152760 | TCTEX1D1 | -2.16178391 | 0.34076194 | 1.276455785 | 0.56576762 | -0.885328125 | 2 | 4.442710129 | 2 | 2.3892346 | 2.49394783 | 3.838019203 | 4.634793542 |
| ENSG00000146085 | MUT | -0.80078433 | 0.002905845 | -0.087459749 | 0.788768394 | -0.888244079 | 8.609439662 | 8.777382925 | 8.721235202 | 7.947219213 | 7.88459112 | 8.594261972 | 8.645780537 |
| ENSG00000152527 | PLEKHH2 | -1.702469625 | 0.193080814 | 0.813509091 | 0.553184301 | -0.888960533 | 4.00722531 | 7.099070191 | 8.110401118 | 5.590471342 | 5.49945122 | 7.891135869 | 8.017950573 |
| ENSG00000148459 | PDSS1 | -0.348869546 | 0.869037712 | -0.541603384 | 0.78794815 | -0.89047293 | 3.327711234 | 5.946638857 | 2 | 4.497095146 | 4.31639476 | 4.065262523 | 4.4231929 |
| ENSG00000159784 | FAM131B | -2.353108564 | 0.171142026 | 1.462005178 | 0.404583432 | -0.891103386 | 4.00722531 | 3.076665209 | 7.502634626 | 4.205751501 | 3.86943969 | 7.616110375 | 7.363232998 |
| ENSG00000160200 | CBS | 2.06449838 | 0.095138308 | -2.956868203 | 0.01263727 | -0.892369824 | 7.031698319 | 5.303523747 | 3.369317972 | 7.885575888 | 7.909304 | 3.500668998 | 3.474731315 |
| ENSG00000136492 | BRIP1 | 0.050874581 | 0.96670681 | -0.943921186 | 0.372909708 | -0.893046605 | 6.009025997 | 5.131603625 | 7.600047325 | 6.710878132 | 6.57678965 | 5.843010798 | 5.555894927 |
| ENSG00000119446 | RBM18 | -0.350174233 | 0.471757988 | -0.545589037 | 0.230518351 | -0.89576327 | 9.371511179 | 8.57407404 | 8.805342542 | 8.524366558 | 8.68323472 | 8.474034275 | 8.348604099 |
| ENSG00000168010 | ATG16L2 | -0.80248032 | 0.688658118 | -0.094194107 | 0.965819335 | -0.896674427 | 2 | 7.274681896 | 8.561105505 | 6.658765962 | 6.74548212 | 7.583859056 | 7.164551992 |
| ENSG00000111087 | GLI1 | -1.463685389 | 0.38727917 | 0.567007642 | 0.749352532 | -0.896777747 | 2 | 5.838937808 | 4.058998049 | 3.840289928 | 3.33976443 | 5.253805068 | 5.211187127 |
| ENSG00000172115 | CYCS | -0.465498577 | 0.304264796 | -0.431974514 | 0.333367096 | -0.897473092 | 9.128497779 | 9.821190352 | 9.714382178 | 9.175891964 | 9.07159009 | 8.96830908 | 9.325696621 |
| ENSG00000204965 | PCDHA5 | 1.375664444 | 0.734590946 | -2.273389914 | NA | -0.89772547 | 2 | 3.076665209 | 2 | 3.566290607 | 2.68924543 | 2.267001569 | 2 |
| ENSG00000107937 | GTPBP4 | -0.269704631 | 0.617709569 | -0.628716929 | 0.199257862 | -0.898421561 | 10.21416435 | 9.267380431 | 9.746650996 | 9.51325161 | 9.53247497 | 9.010625386 | 9.306519748 |
| ENSG00000015479 | MATR3 | -0.598687344 | 0.315977132 | -0.300077566 | 0.632223509 | -0.898764911 | 10.59268348 | 11.60869109 | 11.79401137 | 10.84151467 | 10.8026781 | 11.25410279 | 10.97265421 |
| ENSG00000170260 | ZNF212 | -0.317940103 | 0.846875777 | -0.58159067 | 0.707642617 | -0.899530773 | 4.00722531 | 7.145017454 | 8.608698767 | 7.178012736 | 7.22257674 | 6.779440836 | 7.09293558 |
| ENSG00000169692 | AGPAT2 | -0.476854282 | 0.303980957 | -0.422825724 | 0.357318652 | -0.899680006 | 10.07567912 | 9.28800521 | 9.439621147 | 9.155143657 | 9.17813013 | 9.269197611 | 9.169734363 |
| ENSG00000163931 | TKT | -0.262807629 | 0.764891131 | -0.637000343 | 0.42991362 | -0.899807973 | 12.54246216 | 11.12645935 | 10.87075861 | 11.44585997 | 11.4554086 | 10.87814707 | 11.25184716 |

| | | | | | | | | | | | | | |
|---|---|---|---|---|---|---|---|---|---|---|---|---|---|
| ENSG00000189306 | RRP7A | -0.521224982 | 0.600951592 | -0.379137277 | 0.706382819 | -0.900362259 | 7.218819781 | 9.433952963 | 9.577565424 | 8.559529208 | 8.53549186 | 8.713178889 | 8.662770717 |
| ENSG00000072736 | NFATC3 | -0.335498078 | 0.699202151 | -0.565903784 | 0.487891953 | -0.901401862 | 8.564112515 | 8.147241016 | 9.888372618 | 8.676258519 | 8.78422202 | 8.571313099 | 8.431665941 |
| ENSG00000263012 | | 4.289461303 | 0.413324098 | -5.191097924 | 0.334615401 | -0.901636622 | 2 | 2 | 2 | 4.623314311 | 4.53541753 | 2 | 2 |
| ENSG00000126903 | SLC10A3 | -0.595105891 | 0.406922957 | -0.306929649 | 0.681460704 | -0.902035539 | 8.228665467 | 9.256956408 | 9.849281147 | 8.62888906 | 8.70032852 | 8.973764228 | 8.927151259 |
| ENSG00000176435 | CLEC14A | 0.411845342 | 0.872658363 | -1.313930786 | 0.57016511 | -0.902085444 | 2 | 4.93638933 | 2 | 3.79808662 | 4.2560182 | 3.172301926 | 2.70970617 |
| ENSG00000188243 | COMMD6 | -0.609410265 | 0.543796126 | -0.292932725 | 0.781011291 | -0.902342989 | 9.344956677 | 6.951736442 | 8.076950877 | 7.917958914 | 7.75424764 | 8.260460366 | 8.030160124 |
| ENSG00000122566 | HNRNPA2B1 | 0.129803125 | 0.859640066 | -1.033409471 | 0.086189352 | -0.903606346 | 12.53953777 | 11.56288089 | 11.58978828 | 12.12053612 | 12.0843183 | 10.48450807 | 11.28720059 |
| ENSG00000170348 | TMED10 | -0.43056494 | 0.405040536 | -0.473581285 | 0.34601665 | -0.904146225 | 11.6462383 | 12.26046845 | 12.60507648 | 11.78883165 | 11.7967636 | 11.58578877 | 11.89713248 |
| ENSG00000185000 | DGAT1 | -0.497127047 | 0.173242934 | -0.409888911 | 0.258351845 | -0.907015959 | 8.19931886 | 8.317412869 | 7.896971941 | 7.731524332 | 7.59884443 | 7.793159202 | 7.710008939 |
| ENSG00000196352 | CD55 | 0.14904633 | 0.843023983 | -1.056777029 | 0.090455282 | -0.907730699 | 9.460782017 | 9.04308532 | 10.41552659 | 9.893075744 | 9.91456814 | 8.827346937 | 8.573180082 |
| ENSG00000111726 | CMAS | 0.076956106 | 0.932714208 | -0.986120532 | 0.19694971 | -0.909164426 | 9.810482605 | 8.848535039 | 7.934808822 | 9.073461088 | 9.20032903 | 8.18049086 | 7.985520618 |
| ENSG00000105270 | CLIP3 | -1.243946507 | 0.000578198 | 0.334741296 | 0.415479043 | -0.909205211 | 10.70657014 | 10.86336859 | 10.31150679 | 9.420668575 | 9.39487575 | 10.75037144 | 11.17958994 |
| ENSG00000117523 | PRRC2C | -0.368748207 | 0.599701246 | -0.540509066 | 0.418666323 | -0.909257273 | 12.99617415 | 11.77499027 | 11.7548855 | 11.91590947 | 11.9470968 | 11.75034236 | 11.76961671 |
| ENSG00000143633 | C1orf131 | -0.42514176 | 0.657619829 | -0.484306584 | 0.60340423 | -0.909448344 | 6.816637977 | 6.602424674 | 4.874587708 | 5.764940652 | 6.0722229 | 5.606648123 | 6.095806168 |
| ENSG00000248587 | GDNF-AS1 | -3.753018043 | 0.531143362 | 2.842223862 | NA | -0.910794181 | 2 | 2 | 2 | 2 | 2 | 4.177580368 | 2.172608095 |
| ENSG00000101444 | AHCY | -1.491125563 | 0.100898224 | 0.579714891 | 0.553494044 | -0.911410673 | 8.19931886 | 10.71514053 | 9.972637969 | 8.488325436 | 8.46556258 | 10.45039667 | 10.61574244 |
| ENSG00000006625 | GGCT | -0.436415669 | 0.75303653 | -0.475268398 | 0.726905737 | -0.911684066 | 4.467311984 | 7.503546079 | 8.266922072 | 7.062428156 | 6.91125523 | 6.942523933 | 6.959831579 |
| ENSG00000131067 | GGT7 | -0.991296372 | 0.32983595 | 0.078176498 | 0.946354454 | -0.913119874 | 9.128497779 | 6.78762695 | 9.358421373 | 7.734324667 | 7.8745862 | 8.60534994 | 9.079977539 |
| ENSG00000132485 | ZRANB2 | 0.267577005 | 0.793086452 | -1.181043444 | 0.17880563 | -0.913466439 | 8.468948322 | 9.45234302 | 10.7325871 | 10.15112202 | 10.0656679 | 8.792492404 | 8.534803159 |
| ENSG00000123342 | MMP19 | -0.343216791 | 0.599957698 | -0.57266796 | 0.350332806 | -0.91588475 | 8.540903468 | 9.00621767 | 8.076950877 | 8.18685754 | 8.32385328 | 7.567458938 | 8.381232151 |
| ENSG00000141965 | FEM1A | 0.958753507 | 0.440574326 | -1.874788236 | 0.099916399 | -0.916034729 | 4.815609471 | 7.60566235 | 8.380957471 | 8.447952961 | 8.50663048 | 5.870008604 | 5.596882677 |
| ENSG00000215712 | TMEM242 | -0.255475968 | 0.822912318 | -0.660815641 | 0.524163889 | -0.916291609 | 9.064836028 | 7.790474954 | 6.43197674 | 7.913023977 | 7.85181819 | 7.594015257 | 7.36768071 |
| ENSG00000108773 | KAT2A | -0.562527749 | 0.611381475 | -0.353843886 | 0.754015748 | -0.916371634 | 6.96351253 | 8.607278467 | 9.788583153 | 8.244958867 | 8.36731141 | 8.560477404 | 8.465257491 |
| ENSG00000135249 | RINT1 | -0.210061135 | 0.672900335 | -0.707298361 | 0.103932144 | -0.917359496 | 7.750913381 | 8.317412869 | 8.408111533 | 7.952038749 | 8.01099479 | 7.656405908 | 7.310303798 |
| ENSG00000101017 | CD40 | -0.841773245 | 0.545297456 | -0.075731858 | 0.961240785 | -0.917505103 | 7.579279349 | 4.11352228 | 5.772716771 | 5.552894743 | 5.74985018 | 6.137764768 | 6.574984961 |
| ENSG00000245680 | ZNF585B | -0.171704304 | 0.931077597 | -0.746305578 | 0.684505063 | -0.918009882 | 4.00722531 | 6.899080544 | 9.746650996 | 8.244958867 | 8.15640714 | 7.727325329 | 7.539150216 |
| ENSG00000123349 | PFDN5 | -0.739749089 | 0.203773754 | -0.178534888 | 0.783874152 | -0.918283977 | 11.31794963 | 10.05576005 | 10.94216274 | 10.13160579 | 10.1148377 | 10.71171785 | 10.65326667 |
| ENSG00000107625 | DDX50 | -0.707737923 | 0.115125804 | -0.212073884 | 0.668986346 | -0.919811807 | 8.564112515 | 9.308339287 | 9.25752426 | 8.369067528 | 8.39557285 | 9.012519675 | 8.718388644 |
| ENSG00000136319 | TTC5 | -0.939069025 | 0.623053677 | 0.019237992 | 0.992804891 | -0.919831033 | 2 | 8.147241016 | 8.007629717 | 6.496695859 | 6.68951263 | 7.507470083 | 7.543088047 |
| ENSG00000137364 | TPMT | -0.084788095 | 0.923175097 | -0.835473645 | 0.259652636 | -0.92026174 | 8.493334308 | 6.78762695 | 7.934808822 | 7.781117077 | 7.83127384 | 6.902223127 | 7.226264598 |
| ENSG00000163155 | LYSMD1 | -0.899142457 | 0.264205073 | -0.026008944 | 0.977454517 | -0.925151401 | 8.39321512 | 6.666836837 | 7.163363831 | 6.744602355 | 6.68951263 | 7.674704537 | 7.446882182 |
| ENSG00000095794 | CREM | 0.078155987 | 0.965486841 | -1.004719295 | 0.5332435 | -0.926563307 | 7.624139768 | 8.747909548 | 3.369317972 | 7.786523606 | 7.8390123 | 6.840432491 | 6.671679221 |
| ENSG00000127586 | CHTF18 | -1.245844645 | 0.422347566 | 0.31901841 | 0.850046117 | -0.926826235 | 6.366709671 | 6.228547981 | 2 | 4.867343068 | 4.40247105 | 6.106889539 | 6.007361012 |
| ENSG00000211460 | TSN | -0.607665902 | 0.272669235 | -0.319435694 | 0.579833528 | -0.927101596 | 9.858207046 | 8.848535039 | 9.859153765 | 9.027396512 | 8.94344897 | 9.183011957 | 9.35623215 |
| ENSG00000243137 | PSG4 | -0.689618725 | 0.809318998 | -0.238668087 | 0.935195648 | -0.928286812 | 2 | 2 | 5.156626401 | 3.462030052 | 3.28037825 | 2.775316226 | 4.215171317 |
| ENSG00000100330 | MTMR3 | 0.89786758 | 0.258164018 | -1.826764319 | 0.012003587 | -0.928896739 | 8.653385977 | 9.102512015 | 7.163363831 | 9.340234287 | 9.47403846 | 6.670403488 | 6.78907279 |
| ENSG00000197170 | PSMD12 | -0.053115656 | 0.96053419 | -0.876505282 | 0.352564766 | -0.929620937 | 10.66423795 | 9.246456519 | 8.296287027 | 9.735680502 | 9.61140668 | 8.884319278 | 8.828963952 |
| ENSG00000182916 | TCEAL7 | -0.533750205 | 0.821911287 | -0.39608692 | 0.866925741 | -0.929837125 | 8.836089686 | 5.303523747 | 2 | 6.956091174 | 6.76185814 | 6.619606851 | 7.293259708 |
| ENSG00000113649 | TCERG1 | -0.560341252 | 0.338203518 | -0.36957139 | 0.53558846 | -0.929912641 | 8.540903468 | 9.74864196 | 9.452720984 | 8.795015659 | 8.76268478 | 8.962049262 | 8.974090715 |
| ENSG00000166171 | DPCD | -0.747925753 | 0.367406838 | -0.182842706 | 0.840762669 | -0.930768458 | 7.82963488 | 7.189546432 | 6.073378139 | 6.542151749 | 6.41363412 | 6.643273885 | 7.324100729 |
| ENSG00000100593 | ISM2 | 0.035142853 | 0.983064817 | -0.965922648 | 0.503743146 | -0.930779795 | 4.815609471 | 4.11352228 | 2 | 3.997744386 | 4.16043394 | 3.500668998 | 3.261853426 |
| | FLJ46635 | 6.687811334 | 3.17E-09 | -7.618732524 | 0.000102845 | -0.93092119 | 4.00722531 | 3.076665209 | 2 | 9.191259691 | 9.11153864 | 2 | 2 |
| ENSG00000140497 | SCAMP2 | -0.608625479 | 0.175108303 | -0.322887944 | 0.490052989 | -0.931513424 | 10.65885801 | 9.849218119 | 10.43545992 | 9.731481912 | 9.7573881 | 9.899648775 | 10.1477907 |
| ENSG00000140479 | PCSK6 | -3.255253103 | 0.197230313 | 2.323465978 | 0.289491881 | -0.931787124 | 2 | 2 | 4.058998049 | 2.141737246 | 2.14022021 | 4.875056259 | 4.161535617 |
| ENSG00000167434 | CA4 | 4.259065389 | 0.420476172 | -5.191097152 | 0.337980836 | -0.932031763 | 2 | 2 | 2 | 4.54891529 | 4.56059747 | 2 | 2 |
| ENSG00000131238 | PPT1 | -0.901741279 | 0.370378063 | -0.030767515 | 0.978413041 | -0.932508794 | 10.65345793 | 10.65289571 | 8.143093325 | 9.310519243 | 9.2754408 | 10.11877641 | 10.19985319 |
| ENSG00000089063 | TMEM230 | -0.56149375 | 0.036303182 | -0.37228396 | 0.176769831 | -0.93377771 | 10.16946352 | 10.40166101 | 10.49365885 | 9.798574229 | 9.80557618 | 9.916742729 | 10.06074001 |
| ENSG00000111642 | CHD4 | 0.938064047 | 0.156773734 | -1.872429896 | 0.001983235 | -0.934365849 | 9.635285994 | 11.03485733 | 9.981704053 | 11.23934953 | 11.3276219 | 8.350077895 | 8.613062226 |
| ENSG00000243955 | GSTA1 | -3.753018341 | 0.531143362 | 2.81840284 | NA | -0.934615501 | 2 | 2 | 2 | 2 | 2 | 3.256555576 | 3.67940853 |
| ENSG00000145375 | SPATA5 | -0.755197651 | 0.301889468 | -0.17955702 | 0.82326635 | -0.934754671 | 6.009025997 | 7.051611315 | 7.646393253 | 6.387081877 | 6.25300433 | 6.892381078 | 6.853913776 |
| ENSG00000148450 | MSRB2 | 0.023547138 | 0.961717317 | -0.959821522 | 0.014490584 | -0.936274383 | 8.228665467 | 8.337339476 | 7.734844356 | 8.260583448 | 8.03813956 | 7.11465301 | 7.261744957 |
| ENSG00000166451 | CENPN | 0.871378771 | 0.654593217 | -1.807853543 | 0.314782677 | -0.936474771 | 3.327711234 | 8.124485672 | 4.058998049 | 7.577865504 | 7.47849652 | 4.959178705 | 5.0581604 |
| ENSG00000196262 | PPIA | 0.006893371 | 0.995663772 | -0.943532934 | 0.317937692 | -0.936639563 | 13.42879239 | 11.43000365 | 11.4724895 | 12.47653022 | 12.4088539 | 11.23248597 | 11.71466685 |
| ENSG00000147804 | SLC39A4 | -1.079156221 | 0.554764081 | 0.140968958 | 0.944101725 | -0.938187263 | 7.435667355 | 5.595927442 | 2 | 5.003571641 | 5.43283104 | 6.307840213 | 6.418016952 |
| ENSG00000115806 | GORASP2 | -0.912975714 | 0.070246253 | -0.0260772 | 0.96694554 | -0.939052914 | 11.38790617 | 10.55418374 | 10.42220168 | 9.95860319 | 9.92914219 | 10.84536314 | 10.8107304 |
| ENSG00000112304 | ACOT13 | -0.712216627 | 0.242017731 | -0.230413521 | 0.726496359 | -0.942630147 | 7.485143553 | 8.276714465 | 7.163363831 | 6.941624891 | 7.12888795 | 7.644728141 | 7.355789538 |
| ENSG00000006194 | ZNF263 | -0.797105313 | 0.267233798 | -0.146127865 | 0.856534247 | -0.943233179 | 9.613202746 | 7.979858146 | 9.242516611 | 8.323303759 | 8.28094035 | 9.041010784 | 8.845533906 |
| ENSG00000176101 | SSNA1 | -0.819374387 | 0.068942241 | -0.125584727 | 0.810523076 | -0.944959114 | 9.384607364 | 8.590771778 | 8.699419242 | 8.122047456 | 8.12299044 | 8.785862385 | 8.830575839 |
| ENSG00000197885 | NKIRAS1 | -0.45643516 | 0.356981191 | -0.488536759 | 0.30872573 | -0.944971919 | 8.16936288 | 7.467835385 | 8.236946975 | 7.568480804 | 7.50792049 | 7.396955563 | 7.609689469 |
| ENSG00000181827 | RFX7 | -0.597535707 | 0.43635695 | -0.347679369 | 0.660564083 | -0.945215076 | 8.928409675 | 8.317412869 | 9.935792933 | 8.657201865 | 8.58388295 | 9.053576888 | 8.656119846 |
| ENSG00000120910 | PPP3CC | 0.162394592 | 0.720965573 | -1.108131193 | 0.002976633 | -0.945736601 | 9.128497779 | 9.640525262 | 9.839340503 | 9.718812329 | 9.73682479 | 8.376795097 | 8.551840963 |
| ENSG00000135346 | CGA | 4.245265296 | 0.424867908 | -5.191096489 | 0.340621206 | -0.945831193 | 2 | 2 | 2 | 4.739373666 | 4.31639476 | 2 | 2 |
| ENSG00000113761 | ZNF346 | -0.479184796 | 0.704032536 | -0.466673393 | 0.707642617 | -0.945858189 | 6.563806279 | 9.04308532 | 6.534109959 | 7.311513918 | 7.54938955 | 7.55607473 | 7.330190599 |
| ENSG00000172954 | LCLAT1 | -1.231221128 | 0.000206873 | 0.284296702 | 0.452298819 | -0.946924426 | 8.138771685 | 8.414402626 | 8.608698767 | 7.288834728 | 7.08617588 | 8.657126631 | 8.703804674 |
| ENSG00000076382 | SPAG5 | -0.27188904 | 0.873236533 | -0.675112666 | 0.667404344 | -0.947001706 | 4.467311984 | 6.899080544 | 3.369317972 | 5.540148523 | 5.33407125 | 4.927401733 | 5.217771644 |
| ENSG00000109919 | MTCH2 | -0.373815969 | 0.531143362 | -0.573223648 | 0.306515269 | -0.947039616 | 9.317904228 | 10.17682307 | 9.25752426 | 9.287824288 | 9.27353851 | 8.824761857 | 9.301491724 |
| ENSG00000137338 | PGBD1 | -0.596942659 | 0.755058981 | -0.352401867 | 0.856791105 | -0.949344526 | 7.975256505 | 7.099070191 | 2 | 6.503277904 | 6.41363412 | 6.843841054 | 6.525421271 |
| ENSG00000154957 | ZNF18 | -1.301488248 | 0.327520219 | 0.351220096 | 0.808868142 | -0.950268152 | 3.327711234 | 6.951736442 | 7.029880454 | 5.058234798 | 5.43283104 | 6.91847879 | 6.678855578 |
| ENSG00000079215 | SLC1A3 | 0.251770411 | 0.875544825 | -1.203224152 | 0.385615654 | -0.951453741 | 4.00722531 | 7.232742069 | 7.342949392 | 7.311513918 | 6.66069199 | 6.239610476 | 4.604965121 |
| ENSG00000182923 | CEP63 | -0.699336718 | 0.542375976 | -0.254048322 | 0.835954015 | -0.95338504 | 7.790810999 | 6.311476754 | 9.242516611 | 7.593372273 | 7.49817903 | 8.055236437 | 7.908246991 |
| ENSG00000139567 | ACVRL1 | -0.603130636 | 0.709997456 | -0.350465232 | 0.833427256 | -0.953595869 | 8.228665467 | 7.73146271 | 3.369317972 | 6.788375664 | 6.93069905 | 6.890734189 | 7.293259708 |
| ENSG00000154889 | MPPE1 | 0.504189461 | 0.761525791 | -1.458355537 | 0.330230765 | -0.954166076 | 4.467311984 | 5.838937808 | 8.677268317 | 7.767511442 | 7.90193448 | 6.037140174 | 5.846719511 |
| ENSG00000096872 | IFT74 | -0.070208941 | 0.906452235 | -0.884714224 | 0.063233483 | -0.954923165 | 7.750913381 | 7.145017454 | 7.552162999 | 7.546342806 | 7.29563107 | 6.414841377 | 6.830475749 |
| ENSG00000125821 | DTD1 | -0.333105627 | 0.687168755 | -0.621840679 | 0.419204582 | -0.954946306 | 7.33131396 | 7.503546079 | 8.65477198 | 7.584088227 | 7.66419009 | 6.892381078 | 7.685278529 |
| ENSG00000087245 | MMP2 | 2.283428878 | 2.70E-09 | -3.238428522 | 7.49E-18 | -0.954999643 | 13.28952258 | 13.04041058 | 12.62979741 | 15.27638247 | 15.31259 | 10.02448838 | 9.483266623 |
| ENSG00000112759 | SLC29A1 | -1.966626334 | 7.67E-05 | 1.011534546 | 0.054367856 | -0.955091788 | 9.064836028 | 7.928237109 | 8.763901704 | 6.716553866 | 6.73998187 | 9.582192405 | 9.730352282 |
| ENSG00000133997 | MED6 | -0.20250219 | 0.897514654 | -0.754787767 | 0.593195109 | -0.957289957 | 9.435834703 | 7.145017454 | 5.59507254 | 7.994705029 | 8.00412803 | 7.538302335 | 7.372114752 |
| ENSG00000124228 | DDX27 | -0.205559941 | 0.868087762 | -0.75192653 | 0.492905469 | -0.957486471 | 6.366709671 | 9.308339287 | 8.978369997 | 8.456457569 | 8.47883193 | 8.008163797 | 7.844792088 |
| ENSG00000169519 | METTL15 | -0.580748814 | 0.218893333 | -0.377609872 | 0.431879804 | -0.958358686 | 7.750913381 | 7.315436877 | 7.342949392 | 7.012547947 | 6.80989959 | 7.284811203 | 6.913939481 |
| ENSG00000105808 | RASA4 | 2.02716002 | 0.116189409 | -2.985539079 | 0.015451866 | -0.958379059 | 7.27616306 | 5.838937808 | 3.369317972 | 8.058664058 | 8.29037844 | 3.374372158 | 3.940519005 |
| ENSG00000162971 | TYW5 | -0.193148398 | 0.885491359 | -0.76531685 | 0.518870344 | -0.958465248 | 5.095959458 | 7.189546432 | 8.296287027 | 7.173892946 | 7.1744553 | 6.848938847 | 6.350712758 |
| ENSG00000072042 | RDH11 | -0.421287761 | 0.383589636 | -0.537467239 | 0.245497099 | -0.958755 | 11.27990552 | 10.62073637 | 10.99236731 | 10.57129953 | 10.5644133 | 10.11543113 | 10.72478611 |
| ENSG00000237649 | KIFC1 | 4.229413014 | 0.428189845 | -5.191096228 | 0.341748605 | -0.961683214 | 2 | 2 | 2 | 4.670864177 | 4.37434574 | 2 | 2 |
| ENSG00000178234 | GALNT11 | -1.273314428 | 0.008246269 | 0.311412266 | 0.569500316 | -0.961902162 | 9.219014269 | 8.191702463 | 8.825624954 | 7.497728447 | 7.58969985 | 9.030953258 | 9.18199253 |
| ENSG00000006432 | MAP3K9 | -2.006749345 | 0.136243503 | 1.042989566 | 0.451766731 | -0.963759778 | 3.327711234 | 4.93638933 | 6.43197674 | 3.840289928 | 3.65239003 | 5.532544554 | 6.885273196 |
| ENSG00000177105 | RHOG | -1.035739137 | 0.225157865 | 0.071270443 | 0.94333501 | -0.964468694 | 10.48735708 | 8.607278467 | 8.99637432 | 8.527598812 | 8.62175922 | 9.438259243 | 9.877176832 |
| ENSG00000109062 | SLC9A3R1 | -2.516670841 | 2.20E-06 | 1.552125614 | 0.004741368 | -0.964545227 | 9.064836028 | 8.451443573 | 9.132850258 | 6.542151749 | 6.35006802 | 9.873621976 | 10.86770856 |
| ENSG00000173598 | NUDT4 | 1.04488855 | 0.408736505 | -2.011825795 | 0.083380799 | -0.966937245 | 5.866944937 | 9.433952963 | 9.399592459 | 9.96278806 | 9.90046182 | 7.052670333 | 6.769038269 |
| ENSG00000138741 | TRPC3 | 0.052615046 | 0.988129802 | -1.02040173 | NA | -0.967786683 | 2 | 2 | 3.369317972 | 2.784612485 | 2.38539787 | 2.094598369 | 2.509591799 |
| ENSG00000081913 | PHLPP1 | -0.779068799 | 0.119536182 | -0.189136776 | 0.733835313 | -0.968205575 | 8.418903792 | 7.572420273 | 7.691296561 | 7.261917051 | 7.08617588 | 7.726402307 | 7.779530392 |
| ENSG00000102098 | SCML2 | -0.680484809 | 0.79686468 | -0.28805225 | 0.915553996 | -0.968537058 | 2 | 6.140560021 | 2 | 4.299492648 | 4.05806467 | 4.580376151 | 4.411752372 |
| ENSG00000118579 | MED28 | -0.783544033 | 0.410160907 | -0.185189464 | 0.858897472 | -0.968733496 | 9.839306291 | 8.317412869 | 7.646393253 | 8.033874364 | 8.2130862 | 8.747250685 | 8.687293668 |
| ENSG00000060762 | MPC1 | -0.551078529 | 0.673480955 | -0.417786053 | 0.74998849 | -0.968864581 | 8.285626337 | 6.899080544 | 4.874587708 | 6.733448311 | 6.72335419 | 6.656902456 | 7.032593922 |
| ENSG00000050628 | PTGER3 | -3.753018769 | 0.531143362 | 2.784113098 | NA | -0.968905671 | 2 | 2 | 2 | 2 | 2 | 3.904814898 | 2.817676617 |
| ENSG00000146247 | PHIP | -0.155588464 | 0.82307937 | -0.814032451 | 0.167996939 | -0.969620914 | 9.635285994 | 9.181799937 | 8.536706852 | 8.993902695 | 9.06939887 | 8.68637457 | 7.99033569 |
| ENSG00000104691 | UBXN8 | 0.202230594 | 0.910912588 | -1.172812323 | 0.454394312 | -0.970581729 | 6.366709671 | 6.728495673 | 2 | 6.306580752 | 6.09824986 | 4.868377487 | 5.028505473 |
| ENSG00000084774 | CAD | -1.171714305 | 0.468098579 | 0.200641015 | 0.910973565 | -0.97107329 | 4.467311984 | 9.192779271 | 9.981704053 | 7.847284664 | 7.98332947 | 9.375172844 | 9.166334389 |
| ENSG00000108344 | PSMD3 | -0.386729933 | 0.415575369 | -0.584645633 | 0.190011047 | -0.971375566 | 11.12854954 | 10.27307349 | 10.7904855 | 10.39404824 | 10.3760296 | 10.08784705 | 10.28128023 |
| ENSG00000124380 | SNRNP27 | 0.088089724 | 0.876599527 | -1.060003043 | 0.017748375 | -0.971913318 | 8.609439662 | 7.790474954 | 8.511888463 | 8.469961514 | 8.37974396 | 7.38764416 | 7.20156326 |
| ENSG00000146263 | MMS22L | -0.863653788 | 0.548900135 | -0.110075421 | 0.944363944 | -0.973729209 | 5.330594134 | 7.35507212 | 4.058998049 | 5.391962469 | 5.44640346 | 6.556607032 | 5.481238982 |
| ENSG00000181467 | RAP2B | -0.520543922 | 0.177849888 | -0.453607577 | 0.236728689 | -0.9741515 | 9.128497779 | 8.671479029 | 9.049077186 | 8.469961514 | 8.41468635 | 8.670145638 | 8.326688609 |
| ENSG00000113522 | RAD50 | 1.313332074 | 6.08E-05 | -2.287581648 | 6.20E-13 | -0.974249574 | 7.940205782 | 8.169643011 | 8.296287027 | 9.439666599 | 9.44813055 | 6.063692919 | 5.785984581 |
| ENSG00000173200 | PARP15 | -0.951404396 | 0.729390378 | -0.023090082 | 0.993837021 | -0.974494478 | 2 | 2 | 4.058998049 | 2.3892346 | 2.77779907 | 3.12825509 | 2.817676617 |
| ENSG00000143256 | PFDN2 | -0.124203324 | 0.878583032 | -0.850698724 | 0.209877866 | -0.974902048 | 9.410448395 | 7.847167909 | 8.742726175 | 8.646835365 | 8.7017439 | 7.824551575 | 8.078894996 |
| ENSG00000111011 | RSRC2 | -0.152667876 | 0.73835556 | -0.822752251 | 0.032989268 | -0.975420127 | 10.03472846 | 9.367676479 | 9.541233192 | 9.483600848 | 9.55926825 | 8.944693381 | 8.762402722 |
| ENSG00000095321 | CRAT | -0.992565275 | 0.015303223 | 0.016177903 | 0.974668275 | -0.976387372 | 10.25038534 | 9.506147172 | 10.17589429 | 9.012478123 | 9.03500236 | 9.937829229 | 10.11090342 |
| ENSG00000276219 | | 4.213560972 | 0.431878529 | -5.191095877 | 0.343240742 | -0.977534905 | 2 | 2 | 2 | 4.623314311 | 4.40247105 | 2 | 2 |

| | | | | | | | | | | | | | |
|---|---|---|---|---|---|---|---|---|---|---|---|---|---|
| ENSG00000138381 | ASNSD1 | -0.531450811 | 0.46101922 | -0.448468429 | 0.530188487 | -0.979919241 | 9.992581382 | 8.469613413 | 9.116480286 | 8.837307593 | 8.75724985 | 8.975319061 | 8.778080665 |
| ENSG00000138867 | GUCD1 | -0.670311904 | 0.344773612 | -0.309765391 | 0.679613476 | -0.980077295 | 9.435834703 | 10.76012279 | 11.01019887 | 9.872886274 | 9.88621621 | 10.15112163 | 10.32138402 |
| ENSG00000163017 | ACTG2 | -4.788328933 | 4.43E-06 | 3.808237203 | 0.000275121 | -0.98009173 | 9.031919741 | 9.608085847 | 7.934808822 | 5.22679461 | 2.94009902 | 12.73669842 | 12.88226211 |
| ENSG00000147677 | EIF3H | -0.462542674 | 0.509773777 | -0.518460546 | 0.445705675 | -0.981003219 | 11.22635546 | 10.1878402 | 9.829330889 | 10.15373922 | 10.0027561 | 9.982775948 | 10.06508265 |
| ENSG00000148399 | DPH7 | -0.501680551 | 0.628146304 | -0.480194257 | 0.639193832 | -0.981874808 | 8.228665467 | 8.376385542 | 5.930865648 | 7.450827758 | 7.27676203 | 7.460505069 | 7.310303798 |
| ENSG00000065135 | GNAI3 | -0.33786649 | 0.665819919 | -0.64579722 | 0.372471975 | -0.98366371 | 9.975371636 | 10.00683004 | 11.37951548 | 10.32141448 | 10.2277056 | 9.968404253 | 9.968585649 |
| ENSG00000141738 | GRB7 | -1.910181048 | 0.484988029 | 0.926275767 | 0.732364476 | -0.983905282 | 4.00722531 | 2 | 2 | 2.498691189 | 2.14022021 | 3.355392153 | 3.640730416 |
| ENSG00000147118 | ZNF182 | -0.349476832 | 0.863750456 | -0.636204604 | 0.738292369 | -0.985681436 | 2 | 6.728495673 | 7.858115942 | 6.580055553 | 6.40670796 | 6.539892359 | 5.816671626 |
| ENSG00000139372 | TDG | -0.405174861 | 0.42301813 | -0.580744317 | 0.223260545 | -0.985919178 | 8.418903792 | 9.328390744 | 8.884815469 | 8.483340231 | 8.57309694 | 8.440093652 | 8.265241863 |
| ENSG00000124786 | SLC35B3 | -0.450749114 | 0.468547999 | -0.536315886 | 0.368689059 | -0.987065 | 8.674867122 | 8.806266215 | 9.735974718 | 8.797695539 | 8.60370266 | 8.777865933 | 8.440841157 |
| ENSG00000106080 | FKBP14 | -0.982017415 | 0.000876231 | -0.005425955 | 0.988654005 | -0.987443371 | 9.667789716 | 10.02537381 | 10.02619908 | 9.003220305 | 8.87609528 | 9.806423533 | 10.01019245 |
| ENSG00000136045 | PWP1 | -0.3446276 | 0.802652992 | -0.643404871 | 0.618163882 | -0.988032471 | 10.56418906 | 8.505280662 | 7.225742779 | 9.094319372 | 9.01919552 | 8.506119506 | 8.977489744 |
| ENSG00000092140 | G2E3 | -0.499155588 | 0.70056887 | -0.489111855 | 0.702388158 | -0.988267443 | 8.777722034 | 6.844429755 | 5.772716771 | 7.226553841 | 7.1204461 | 7.502086936 | 6.775747403 |
| ENSG00000189157 | FAM47E | -0.692198405 | NA | -0.297892091 | NA | -0.990090496 | 2 | 3.076665209 | 2 | 2.141737246 | 2.59489879 | 2.746468522 | 2.172608095 |
| ENSG00000060656 | PTPRU | -1.398671931 | 0.438437558 | 0.40685429 | 0.83510727 | -0.991817642 | 7.485143553 | 8.053983481 | 2 | 5.818703821 | 5.96314905 | 7.496683627 | 7.746896594 |
| ENSG00000115944 | COX7A2L | -0.727388631 | 0.288657299 | -0.266017491 | 0.718208015 | -0.993406122 | 10.93738399 | 9.470501606 | 9.954333071 | 9.488585154 | 9.56627807 | 10.06265365 | 9.90753096 |
| ENSG00000189164 | ZNF527 | -0.700395514 | 0.298583304 | -0.293225455 | 0.679226319 | -0.993620969 | 5.70932728 | 6.535001748 | 6.80318718 | 5.578054239 | 5.92487824 | 6.287958221 | 5.980509018 |
| ENSG00000106305 | AIMP2 | -0.592516099 | 0.121962743 | -0.401257829 | 0.300265097 | -0.993773928 | 7.790810999 | 7.638155693 | 8.110401118 | 7.29643414 | 7.24604864 | 7.407359556 | 7.507254539 |
| ENSG00000181090 | EHMT1 | -0.681066656 | 0.047519069 | -0.313772281 | 0.389671888 | -0.994838937 | 9.624286623 | 9.114109157 | 9.301632639 | 8.624367384 | 8.7394436 | 9.123256136 | 8.966780003 |
| ENSG00000117592 | PRDX6 | -0.239475942 | 0.798171474 | -0.75592572 | 0.364514502 | -0.995401662 | 12.19014827 | 11.0102427 | 10.31150679 | 11.13619556 | 11.1492725 | 10.26514949 | 10.91671583 |
| ENSG00000134461 | ANKRD16 | -0.761361661 | 0.497204407 | -0.234572277 | 0.845108741 | -0.995933938 | 4.467311984 | 5.72254323 | 7.098165388 | 5.44762523 | 5.40529704 | 5.906324469 | 5.917188835 |
| ENSG00000104946 | TBC1D17 | -0.651769769 | 0.11481363 | -0.344612634 | 0.425827846 | -0.996382403 | 9.204317094 | 9.277729677 | 9.788583153 | 8.663092307 | 8.92175022 | 8.967528086 | 9.22674878 |
| ENSG00000169228 | RAB24 | 0.505905412 | 0.639874207 | -1.503429378 | 0.117817538 | -0.997518566 | 7.27616306 | 9.256956408 | 7.029880454 | 8.71221318 | 8.75316018 | 6.850634117 | 6.671679221 |
| ENSG00000117143 | UAP1 | -0.347035768 | 0.715844099 | -0.652017568 | 0.464454793 | -0.999053336 | 11.50945836 | 9.506147172 | 10.06936278 | 10.24689154 | 10.2962973 | 9.757337057 | 10.15166451 |
| ENSG00000143889 | HNRNPLL | -0.454001731 | 0.471675519 | -0.545794425 | 0.366665941 | -0.999796155 | 9.27634997 | 9.405920729 | 8.175061096 | 8.655725489 | 8.54498542 | 8.584107514 | 8.433081317 |
| ENSG00000205795 | CYS1 | -1.146892964 | 0.031446236 | 0.146651379 | 0.812849666 | -1.000241584 | 6.73718411 | 6.389896633 | 5.930865648 | 5.334065616 | 5.3193943 | 6.593521701 | 6.468583424 |
| ENSG00000049192 | ADAMTS6 | -6.966291077 | 0.005644199 | 5.964844852 | 0.001443968 | -1.001446225 | 2 | 4.93638933 | 2 | 2 | 2 | 9.311664913 | 9.109488808 |
| ENSG00000124140 | SLC12A5 | -1.694515975 | 0.696746436 | 0.692075095 | NA | -1.00237688 | 2 | 3.076665209 | 2 | 2 | 2.38539787 | 3.256555576 | 2.277179783 |
| ENSG00000133026 | MYH10 | -1.0751562 | 0.017562521 | 0.071417679 | 0.899395052 | -1.003738521 | 9.689059027 | 10.36330215 | 10.22252928 | 9.052296807 | 9.05063789 | 9.846904685 | 10.47046835 |
| ENSG00000077063 | CTTNBP2 | 0.671947796 | 0.722734802 | -1.676061712 | 0.332277495 | -1.004113916 | 2 | 6.228547981 | 4.058998049 | 5.614989456 | 5.64780787 | 3.981090143 | 3.33635788 |
| ENSG00000178252 | WDR6 | -0.851252735 | 0.101553845 | -0.153453706 | 0.79696914 | -1.004706441 | 10.60951384 | 11.07398149 | 9.981704053 | 9.729378027 | 9.82128771 | 10.36712466 | 10.56854893 |
| ENSG00000173581 | CCDC106 | -1.173624922 | 0.085637804 | 0.168611979 | 0.832583267 | -1.005012943 | 8.042902253 | 6.951736442 | 8.486635645 | 6.777556219 | 6.83074839 | 7.91644678 | 8.289666801 |
| ENSG00000158161 | EYA3 | -0.430936556 | 0.641177228 | -0.575457645 | 0.517359347 | -1.006394201 | 9.533135543 | 7.431218269 | 9.031722608 | 8.542055007 | 8.40081078 | 8.517903178 | 8.114615835 |
| ENSG00000095209 | TMEM38B | 0.61478547 | 0.617393398 | -1.623248815 | 0.141216873 | -1.008463345 | 5.70932728 | 7.467835385 | 8.721235202 | 8.339946438 | 8.39031583 | 6.60762613 | 5.601925227 |
| ENSG00000167900 | TK1 | -1.703695037 | 0.416993309 | 0.694324972 | 0.753823132 | -1.009370065 | 4.467311984 | 8.004992147 | 2 | 5.161714397 | 4.811949 | 6.826717129 | 7.562617572 |
| ENSG00000102882 | MAPK3 | -1.270453101 | 0.018021461 | 0.260401752 | 0.675928124 | -1.010051349 | 10.40620204 | 9.624396729 | 10.77492406 | 9.061245954 | 9.09332116 | 10.36963935 | 10.80076011 |
| ENSG00000115263 | GCG | 4.180601725 | 0.439229935 | -5.191095048 | 0.346533496 | -1.010493323 | 2 | 2 | 2 | 4.415698455 | 4.56059747 | 2 | 2 |
| ENSG00000130177 | CDC16 | -0.359285126 | 0.386972978 | -0.652036102 | 0.088880848 | -1.011321228 | 10.16946352 | 9.994333773 | 9.491321578 | 9.468543769 | 9.63900396 | 9.240714099 | 9.289430628 |
| ENSG00000066827 | ZFAT | -0.917367075 | 0.600101312 | -0.094170699 | 0.961240785 | -1.011537774 | 8.87372952 | 5.72254323 | 4.058998049 | 6.548530093 | 6.64311914 | 7.564363052 | 7.199901369 |
| ENSG00000150316 | CWC15 | -0.383108875 | 0.688362385 | -0.628825453 | 0.483865109 | -1.011934329 | 9.358295023 | 8.702538795 | 7.163363831 | 8.339946438 | 8.22887947 | 7.807195765 | 8.246044089 |
| ENSG00000134077 | THUMPD3 | -0.69790279 | 0.493931013 | -0.316128509 | 0.767357706 | -1.014031299 | 9.886101849 | 7.761270542 | 7.934808822 | 8.207157886 | 8.15640714 | 8.512558892 | 8.604294318 |
| ENSG00000197406 | DIO3 | -4.714921645 | 0.185413931 | 3.700594983 | 0.227412477 | -1.014326662 | 2 | 2 | 3.369317972 | 2 | 2 | 5.427419529 | 4.047883918 |
| ENSG00000186073 | C15orf41 | -0.771909234 | 0.703942894 | -0.243109286 | 0.909865545 | -1.015018521 | 2 | 7.189546432 | 4.523675391 | 5.289065324 | 5.0259382 | 5.755231204 | 5.519049808 |
| ENSG00000112996 | MRPS30 | -0.772328214 | 0.454478461 | -0.242935465 | 0.826586773 | -1.015263679 | 9.048471759 | 7.979858146 | 6.534109959 | 7.416368504 | 7.44169935 | 8.25278818 | 7.561323797 |
| ENSG00000134440 | NARS | -0.312058155 | 0.456600383 | -0.703601755 | 0.06253927 | -1.015659911 | 11.18951828 | 10.69597594 | 10.56174313 | 10.5227232 | 10.53732 | 10.24788588 | 10.02270259 |
| ENSG00000135617 | PRADC1 | -1.489308489 | 0.42315744 | 0.47306558 | 0.812849666 | -1.016242908 | 2 | 7.954278496 | 6.322058544 | 5.514313187 | 5.27444544 | 6.877490957 | 7.55613706 |
| ENSG00000196405 | EVL | -0.955602265 | 0.099823785 | -0.061350795 | 0.930293595 | -1.01695306 | 8.653385977 | 8.213429689 | 7.451345288 | 7.222570549 | 7.28055563 | 7.917255914 | 8.312916523 |
| ENSG00000076321 | KLHL20 | -0.653300232 | 0.535619507 | -0.364521826 | 0.73782085 | -1.018122059 | 7.096806373 | 8.590771778 | 9.778213818 | 8.211183901 | 8.22691474 | 8.360229299 | 8.635366595 |
| ENSG00000206047 | YBX2 | -3.753019391 | 0.531143362 | 2.734339999 | NA | -1.018679391 | 2 | 2 | 2 | 2 | 2 | 3.235948948 | 3.600986722 |
| ENSG00000119772 | DNMT3A | -1.285555143 | 0.000994715 | 0.266627837 | 0.553713666 | -1.018927307 | 8.75772933 | 8.848535039 | 8.143093325 | 7.285019958 | 7.42465798 | 8.933539019 | 8.835400723 |
| ENSG00000105677 | TMEM147 | -0.218083986 | 0.586581206 | -0.801759187 | 0.020991008 | -1.019843173 | 9.567995545 | 9.608085847 | 10.09465441 | 9.56056556 | 9.55535913 | 8.888863837 | 9.062251023 |
| ENSG00000106415 | GLCCI1 | -0.145434447 | 0.935309765 | -0.874493388 | 0.589352188 | -1.019927835 | 3.327711234 | 8.297207175 | 6.718950348 | 6.975156635 | 7.08183429 | 6.609629839 | 5.980509018 |
| ENSG00000111666 | CHPT1 | -0.335904197 | 0.725609862 | -0.686877691 | 0.439455057 | -1.022781887 | 7.218819781 | 9.067150531 | 9.399592459 | 8.451360823 | 8.5339035 | 8.25278818 | 8.032029379 |
| ENSG00000114902 | SPCS1 | -0.155654521 | 0.689206562 | -0.867327956 | 0.007526564 | -1.022982477 | 10.3933013 | 9.863030423 | 9.972637969 | 9.902443169 | 9.97258765 | 9.24458753 | 9.215277965 |
| ENSG00000137727 | ARHGAP20 | -1.379135718 | 0.496719668 | 0.355077963 | 0.872155485 | -1.024057755 | 2 | 6.311476754 | 8.561105505 | 5.753943344 | 6.1068222 | 8.012713664 | 7.043754524 |
| ENSG00000152620 | NADK2 | -0.33567604 | 0.596166767 | -0.688667967 | 0.236074775 | -1.024344007 | 7.53297911 | 8.639735838 | 8.434763955 | 8.004017475 | 7.89452713 | 7.80632247 | 7.36768071 |
| ENSG00000151233 | GXYLT1 | 0.04000535 | 0.984309875 | -1.065317209 | 0.571694567 | -1.02531186 | 8.444143041 | 7.928237109 | 2 | 7.813256448 | 7.52080763 | 6.788313392 | 6.38621943 |
| ENSG00000138385 | SSB | -0.298071409 | 0.716902068 | -0.728020609 | 0.33116471 | -1.026092018 | 10.5468183 | 9.298208073 | 8.978369997 | 9.479434065 | 9.47734778 | 8.848286818 | 9.22919499 |
| ENSG00000265354 | TIMM23 | -0.363394063 | 0.451037592 | -0.662721366 | 0.13467376 | -1.026115428 | 9.358295023 | 10.06773758 | 10.0085654 | 9.446513412 | 9.52289845 | 8.97493051 | 9.373289845 |
| ENSG00000066185 | ZMYND12 | -1.643982387 | 0.530264818 | 0.617123023 | 0.816030298 | -1.026859363 | 4.00722531 | 2 | 2 | 2.498691189 | 2.2680111 | 3.355392153 | 3.261853426 |
| | C9ORF172 | 4.164200733 | 0.443453067 | -5.191094556 | 0.348435042 | -1.026893823 | 2 | 2 | 2 | 4.358765249 | 4.58534546 | 2 | 2 |
| ENSG00000117395 | EBNA1BP2 | -0.489329829 | 0.483350437 | -0.537659561 | 0.42790044 | -1.02698939 | 9.410448395 | 8.376385542 | 8.296287027 | 8.297027403 | 8.30907138 | 7.88204676 | 8.551840963 |
| ENSG00000164109 | MAD2L1 | -0.484360616 | 0.869037712 | -0.543011972 | 0.849422471 | -1.027372588 | 2 | 7.35507212 | 2 | 5.461211713 | 5.34860058 | 5.4091291 | 5.294527297 |
| ENSG00000013523 | ANGEL1 | -1.42340446 | 0.027864804 | 0.396018716 | 0.58348958 | -1.027385745 | 7.031698319 | 8.317412869 | 8.585098385 | 6.70517998 | 6.75642009 | 8.547470747 | 8.473534575 |
| ENSG00000197982 | C1orf122 | -0.741883924 | 0.003365499 | -0.286509314 | 0.298467142 | -1.028393238 | 9.720386367 | 9.549500232 | 9.788583153 | 8.875861021 | 9.01919552 | 9.409219381 | 9.393787153 |
| ENSG00000131711 | MAP1B | -1.300081414 | 1.21E-06 | 0.271456028 | 0.38415025 | -1.028625386 | 14.04768507 | 14.1025305 | 13.70724472 | 12.6623295 | 12.6644892 | 14.13023131 | 14.33147456 |
| ENSG00000253719 | ATXN7L3B | -0.545349227 | 0.601702841 | -0.483428727 | 0.641992641 | -1.028777954 | 8.609439662 | 10.39692156 | 11.28380426 | 9.929575199 | 9.90722553 | 9.990100744 | 9.970048927 |
| ENSG00000169139 | UBE2V2 | -0.641315379 | 0.122576344 | -0.38799175 | 0.36482448 | -1.029307129 | 10.04301211 | 9.770792087 | 9.31604076 | 9.212503151 | 8.98589128 | 9.422103197 | 9.28512653 |
| ENSG00000166411 | IDH3A | 0.318612567 | 0.698007365 | -1.347964944 | 0.056148162 | -1.029352377 | 9.435834703 | 8.191702463 | 7.818184403 | 8.984524517 | 8.95655043 | 7.354572674 | 7.302581437 |
| ENSG00000215114 | UBXN2B | -1.2637439 | 0.274972023 | 0.23425562 | 0.856791105 | -1.02948828 | 8.716893051 | 7.979858146 | 5.39244166 | 6.652857793 | 6.68379438 | 8.327733606 | 7.8990585 |
| ENSG00000006744 | ELAC2 | -0.583073264 | 0.128141904 | -0.446478064 | 0.24798411 | -1.029551327 | 9.2903348 | 9.308339287 | 9.859153765 | 8.953617446 | 8.90341524 | 9.012898235 | 9.113908014 |
| ENSG00000078902 | TOLLIP | -0.78474762 | 0.246264529 | -0.245429321 | 0.738292369 | -1.030176941 | 10.64259676 | 9.170736405 | 10.50627919 | 9.476091952 | 9.43797543 | 9.993365625 | 9.994460574 |
| ENSG00000145217 | SLC26A1 | -2.29935171 | 0.321145767 | 1.26867813 | 0.582452573 | -1.03067358 | 2 | 2 | 4.874587708 | 2.600425781 | 2.38539787 | 4.636484163 | 4.468068808 |
| ENSG00000137947 | GTF2B | -0.045147061 | 0.942460221 | -0.985546951 | 0.051573713 | -1.030694011 | 9.27634997 | 8.395519306 | 8.585098385 | 8.7747563 | 8.73668456 | 8.045647998 | 7.570355956 |
| ENSG00000160691 | SHC1 | -0.2916018 | 0.677691017 | -0.740252767 | 0.242732554 | -1.031854567 | 13.23633495 | 12.21407135 | 12.01984373 | 12.28032204 | 12.3252367 | 11.59118594 | 12.07747769 |
| ENSG00000084090 | STARD7 | -0.562764547 | 0.080556335 | -0.469765291 | 0.144905041 | -1.032529838 | 11.0567315 | 10.82507101 | 10.66102425 | 10.26346497 | 10.3268876 | 10.22290636 | 10.53664546 |
| ENSG00000197530 | MIB2 | -0.786602906 | 0.395924587 | -0.246410108 | 0.805557931 | -1.033013014 | 9.204317094 | 8.317412869 | 10.48730704 | 8.776115811 | 8.87484109 | 9.582192405 | 9.103279088 |
| ENSG00000092010 | PSME1 | 0.395657733 | 0.372520317 | -1.428897717 | 0.000181365 | -1.033239984 | 10.58135316 | 9.863030423 | 10.36791565 | 10.68422387 | 10.702313 | 8.712712904 | 9.025661876 |
| ENSG00000182872 | RBM10 | -0.512027161 | 0.129332125 | -0.522102101 | 0.113430713 | -1.034129263 | 9.384607364 | 8.929529568 | 9.031722608 | 8.610716699 | 8.62474679 | 8.550081477 | 8.663373824 |
| ENSG00000184634 | MED12 | -0.583817841 | 0.793896601 | -0.450504846 | 0.840192213 | -1.034322687 | 9.048471759 | 9.896993485 | 2 | 8.394075037 | 8.35836495 | 8.479522918 | 8.536120927 |
| ENSG00000124217 | MOCS3 | -0.717670031 | 0.145361929 | -0.318336489 | 0.538306766 | -1.03600652 | 6.73718411 | 7.051611315 | 7.502634626 | 6.436065776 | 6.43421531 | 6.713894915 | 6.915965441 |
| ENSG00000153391 | INO80C | -1.187439341 | 0.254030721 | 0.149543086 | 0.900996869 | -1.037896255 | 8.16936288 | 5.946638857 | 8.585098385 | 6.772115916 | 6.76727577 | 7.65057884 | 8.401617719 |
| ENSG00000138231 | DBR1 | -0.432770584 | 0.664549192 | -0.605328164 | 0.52471944 | -1.038098748 | 7.940205782 | 5.838937808 | 7.934808822 | 6.922108067 | 7.22257674 | 7.013858277 | 6.804465095 |
| | MFSD7 | 5.981813646 | 1.49E-05 | -7.019994838 | 0.001016559 | -1.038181192 | 3.327711234 | 3.076665209 | 2 | 7.807949361 | 7.91419618 | 2 | 2 |
| ENSG00000148296 | SURF6 | -0.815192985 | 0.463646059 | -0.223295935 | 0.85266974 | -1.03848892 | 9.886101849 | 9.235879652 | 6.882775709 | 8.343618883 | 8.2790453 | 8.756774919 | 9.025661876 |
| ENSG00000105438 | KDELR1 | -0.815192346 | 0.161771312 | -0.223711006 | 0.728068232 | -1.038903353 | 12.7234361 | 11.54799449 | 11.82708281 | 11.3298432 | 11.2885951 | 11.79620367 | 11.99698602 |
| ENSG00000130313 | PGLS | -0.549436069 | 0.429519861 | -0.489800131 | 0.476243419 | -1.0392362 | 10.16946352 | 9.318399851 | 8.677268317 | 8.998569022 | 8.94464494 | 9.077661821 | 8.982814948 |
| ENSG00000134627 | PIWIL4 | 0.013562299 | 0.994867193 | -1.052862191 | 0.533469206 | -1.039299892 | 6.89194353 | 5.303523747 | 2 | 5.742861563 | 5.7929843 | 4.940196634 | 4.673629022 |
| STRN4 | | -0.799781446 | 0.240478094 | -0.242290328 | 0.744693168 | -1.042071773 | 10.04301211 | 9.936721131 | 11.27276688 | 9.725161029 | 9.78502553 | 10.23236508 | 10.38511052 |
| ENSG00000122966 | CIT | 0.198267028 | 0.917583206 | -1.240349219 | 0.460189911 | -1.042082192 | 2 | 6.728495673 | 7.163363831 | 6.628980152 | 6.55189318 | 4.881704255 | 5.545463418 |
| | FAM26F | 4.14761108 | 0.447517543 | -5.191094031 | 0.350450714 | -1.043482951 | 2 | 2 | 2 | 4.299492648 | 4.60967607 | 2 | 2 |
| ENSG00000215305 | VPS16 | -0.362172227 | 0.504589952 | -0.6820602 | 0.170920391 | -1.044232426 | 8.653385977 | 8.451443573 | 7.646393253 | 7.944803394 | 7.97164382 | 7.632955077 | 7.65527392 |
| ENSG00000125691 | RPL23 | -1.232762946 | 0.076121912 | 0.188510057 | 0.815972748 | -1.044252889 | 12.39494173 | 10.85302421 | 11.16147894 | 10.43162866 | 10.3715902 | 11.78472189 | 11.85544917 |
| ENSG00000077044 | DGKD | -1.561494534 | 0.006399185 | 0.516659845 | 0.4152616 | -1.044834689 | 7.790810999 | 8.903033433 | 9.132850258 | 7.119236776 | 7.24216308 | 9.199402547 | 9.259028753 |
| ENSG00000268916 | CSAG3 | 2.596819353 | 0.600260828 | -3.642853624 | NA | -1.046034272 | 2 | 2 | 2 | 3.406930997 | 3.28037825 | 2 | 2.059860178 |
| ENSG00000075413 | MARK3 | -0.598099596 | 0.428026621 | -0.448678689 | 0.553628122 | -1.046778286 | 9.397585735 | 9.923599762 | 11.01903267 | 9.692414568 | 9.6668081 | 9.886487495 | 9.767852046 |
| ENSG00000148229 | POLE3 | -0.613991752 | 0.141601992 | -0.433642759 | 0.305685651 | -1.047563451 | 9.771132748 | 9.102512015 | 9.180874355 | 8.81895723 | 8.72280968 | 8.892982869 | 9.005824891 |
| ENSG00000162068 | NTN3 | -3.753019755 | 0.531143362 | 2.705187516 | NA | -1.047832239 | 2 | 2 | 2 | 2 | 2 | 3.193831101 | 3.600986722 |
| ENSG00000091073 | DTX2 | 0.285678018 | 0.895265695 | -1.33476464 | 0.48296181 | -1.049086622 | 9.384607364 | 5.72254323 | 3.369317972 | 8.139053376 | 8.27714776 | 6.595545082 | 6.662054875 |
| ENSG00000115750 | TAF1B | -0.165805705 | 0.931725033 | -0.883506509 | 0.616621341 | -1.049312214 | 7.485143553 | 7.35507212 | 2 | 6.722207358 | 6.67228943 | 5.957567336 | 6.070564958 |
| ENSG00000196296 | ATP2A1 | -0.847983713 | 0.528630833 | -0.202008767 | 0.888119558 | -1.049992479 | 4.00722531 | 4.710564154 | 3.369317972 | 3.289987342 | 3.7864959 | 4.251304078 | 3.735548302 |
| ENSG00000144649 | FAM198A | -3.753019798 | 0.531143362 | 2.70169638 | NA | -1.05132516 | 2 | 2 | 2 | 2 | 2 | 3.943456545 | 2.551896972 |
| ENSG00000196123 | KIAA0895L | -0.604667641 | 0.758269479 | -0.447136853 | 0.821473491 | -1.051804494 | 8.228665467 | 7.638155693 | 2 | 6.804454198 | 6.80463998 | 6.647180917 | 7.211494604 |
| ENSG00000050130 | JKAMP | -0.252360647 | 0.413845402 | -0.803564711 | 0.002720068 | -1.055925358 | 10.19198706 | 9.930175364 | 9.849281147 | 9.7700774 | 9.71456265 | 9.261571873 | 9.131450704 |
| ENSG00000130254 | SAFB2 | 0.014483507 | 0.988129802 | -1.070905702 | 0.17572907 | -1.056422195 | 9.667789716 | 7.669933291 | 9.149036566 | 9.026254405 | 9.08466797 | 8.089380147 | 7.866953664 |
| ENSG00000134690 | CDCA8 | -0.998788202 | 0.701852817 | -0.061954108 | 0.982373817 | -1.06074231 | 2 | 5.946638857 | 2 | 4.17310023 | 3.28037825 | 4.331190017 | 4.683176719 |
| ENSG00000113552 | GNPDA1 | -0.352412221 | 0.291168491 | -0.71048918 | 0.019469114 | -1.062901401 | 9.015175615 | 9.079034257 | 9.165043284 | 8.727753724 | 8.74768919 | 8.539084523 | 8.208518621 |

| | | | | | | | | | | | | | |
|---|---|---|---|---|---|---|---|---|---|---|---|---|---|
| ENSG00000076604 | TRAF4 | -0.810166681 | 0.339253229 | -0.255295834 | 0.781935426 | -1.065462514 | 10.66423795 | 8.993716073 | 8.978369997 | 8.939126212 | 9.00665321 | 9.526277199 | 9.522687851 |
| ENSG00000183807 | FAM162B | -3.753019983 | 0.531143362 | 2.686904797 | NA | -1.066115186 | 2 | 2 | 2 | 2 | 2 | 4.053534413 | 2.172608095 |
| ENSG00000105290 | APLP1 | -1.692367215 | 0.000150601 | 0.625353647 | 0.198968723 | -1.067013568 | 8.696033109 | 7.928237109 | 8.585098385 | 6.79375511 | 6.76185814 | 8.816978708 | 9.255833064 |
| ENSG00000138796 | HADH | -1.954646762 | 0.0205779 | 0.887014627 | 0.324714138 | -1.067632135 | 8.540903468 | 7.954278496 | 6.203069917 | 6.096288482 | 5.83486613 | 8.667743558 | 8.793039018 |
| ENSG00000143631 | FLG | -5.678913119 | 0.051473882 | 4.609797636 | 0.048277535 | -1.069115484 | 4.00722531 | 2 | 2 | 2 | 2 | 5.843010798 | 7.124815549 |
| ENSG00000100344 | PNPLA3 | 0.450219626 | 0.748958558 | -1.520636012 | 0.223084039 | -1.070416386 | 4.00722531 | 7.761270542 | 7.342949392 | 7.497728447 | 7.47518984 | 5.343222328 | 5.829625997 |
| ENSG00000025796 | SEC63 | -0.215568052 | 0.736896378 | -0.856732574 | 0.125144676 | -1.072300626 | 9.613202746 | 10.83561703 | 10.48092713 | 10.23111715 | 10.130687 | 9.652393285 | 9.425420127 |
| ENSG00000114650 | SCAP | -0.723306477 | 0.472007774 | -0.349084178 | 0.738195992 | -1.072390656 | 11.07300288 | 9.916994052 | 8.511888463 | 9.417187348 | 9.50598504 | 9.855795196 | 9.813124536 |
| ENSG00000254636 | ARMS2 | -3.753020081 | 0.531143362 | 2.679094865 | NA | -1.073925216 | 2 | 2 | 2 | 2 | 2 | 2.592885947 | 3.908316285 |
| ENSG00000139173 | TMEM117 | -1.0307138 | 0.249172977 | -0.043443591 | 0.967312407 | -1.074157392 | 7.031698319 | 6.228547981 | 8.296287027 | 6.328984825 | 6.52655952 | 7.462723726 | 7.302581437 |
| C8ORF46 | C8ORF46 | 4.115551581 | 0.454568815 | -5.191093324 | 0.353041048 | -1.075541743 | 2 | 2 | 2 | 4.443344104 | 4.43005853 | 2 | 2 |
| ENSG00000080802 | CNOT4 | -0.291036276 | 0.561250332 | -0.784555497 | 0.079746219 | -1.075591772 | 8.444143041 | 9.203675681 | 9.014156722 | 8.616799622 | 8.65720817 | 8.332594213 | 7.939455664 |
| ENSG00000058600 | POLR3E | -0.704409354 | 0.691987391 | -0.371700361 | 0.839520202 | -1.076109715 | 3.327711234 | 8.687092496 | 9.180874355 | 7.745471964 | 7.63485389 | 8.082904238 | 7.950369405 |
| ENSG00000130935 | NOL11 | -0.101461579 | 0.92127192 | -0.975199093 | 0.260131361 | -1.076660672 | 8.777722034 | 8.077869079 | 6.534109959 | 8.022463952 | 7.90193448 | 7.192914486 | 7.019463152 |
| ENSG00000197903 | HIST1H2BK | 1.985308756 | 0.000911216 | -3.061996054 | 1.08E-07 | -1.076687298 | 8.418903792 | 8.234834549 | 9.503962296 | 10.78827371 | 10.8233057 | 5.610655473 | 6.070564958 |
| ENSG00000123737 | EXOSC9 | 0.135971339 | 0.823963146 | -1.214230678 | 0.013699636 | -1.078259339 | 7.904282243 | 7.538394143 | 8.561105505 | 8.190940526 | 8.18704757 | 6.875826972 | 6.862343163 |
| ENSG00000150459 | SAP18 | -0.379302959 | 0.572669004 | -0.699836155 | 0.262859223 | -1.079139114 | 11.10500521 | 9.71856931 | 10.37481416 | 10.1390214 | 10.1206693 | 9.729967164 | 9.887021781 |
| ENSG00000124191 | TOX2 | 3.135425563 | 0.030198621 | -4.214677163 | 0.004583447 | -1.0792516 | 2 | 4.442710129 | 4.874587708 | 7.057964187 | 7.00136498 | 2.42098416 | 2.059860178 |
| ENSG00000173418 | NAA20 | -0.71582275 | 0.077472236 | -0.364804997 | 0.391016933 | -1.080627747 | 9.87686338 | 9.277729677 | 9.399592459 | 8.756965077 | 8.89725143 | 9.00682933 | 9.329501596 |
| ENSG00000101474 | APMAP | -0.837960835 | 0.003812454 | -0.24436069 | 0.453575633 | -1.082321526 | 10.58135316 | 10.32888058 | 10.79563557 | 9.715981692 | 9.77093593 | 10.35791611 | 10.31179874 |
| ENSG00000164663 | USP49 | -0.366608227 | 0.395924587 | -0.716294607 | 0.068345365 | -1.082902834 | 7.218819781 | 7.35507212 | 7.646393253 | 7.106324235 | 7.00595521 | 6.784770914 | 6.649933445 |
| ENSG00000172878 | METAP1D | -1.333634932 | 0.122761253 | 0.25034146 | 0.796537901 | -1.083293472 | 6.009025997 | 4.710564154 | 6.43197674 | 4.825935136 | 4.4836996 | 5.893224226 | 6.261114822 |
| ENSG00000116918 | TSNAX | -0.23161751 | 0.59184685 | -0.853405132 | 0.022521058 | -1.085022642 | 8.946185348 | 8.791896851 | 8.845626171 | 8.630393142 | 8.63367258 | 7.718068425 | 8.266830206 |
| ENSG00000100228 | RAB36 | -0.067777108 | 0.954792088 | -1.017444033 | 0.326936342 | -1.085221141 | 7.940205782 | 5.838937808 | 6.322058544 | 6.693715713 | 7.1204461 | 5.47655442 | 6.409414172 |
| ENSG00000138398 | PPIG | -0.835046209 | 0.299078862 | -0.250252344 | 0.775592511 | -1.085298553 | 10.61508061 | 8.981105197 | 9.149036566 | 8.948803189 | 8.95298909 | 9.680545441 | 9.365898224 |
| ENSG00000144524 | COPS7B | -0.899273878 | 0.197832048 | -0.188602195 | 0.810298197 | -1.087876073 | 8.75772933 | 8.557180776 | 9.981704053 | 8.302698479 | 8.39732094 | 9.021201511 | 9.087187414 |
| ENSG00000177854 | TMEM187 | -1.511734095 | 0.372864851 | 0.42356458 | 0.818512537 | -1.088169514 | 2 | 6.535001748 | 7.285536013 | 5.058234798 | 4.95174433 | 7.10332117 | 6.420873184 |
| ENSG00000168434 | COG7 | -0.541646215 | 0.807156873 | -0.546691561 | 0.801957964 | -1.088337775 | 9.544849385 | 7.901717006 | 2 | 7.799951945 | 7.86451142 | 7.754744914 | 7.897008641 |
| ENSG00000113384 | GOLPH3 | -0.829911318 | 0.016111406 | -0.258601544 | 0.500300419 | -1.088512862 | 10.86447339 | 10.48887288 | 11.10451741 | 10.04628835 | 9.976101 | 10.62868983 | 10.53186455 |
| ENSG00000235711 | ANKRD34C | -3.753020266 | 0.531143362 | 2.664244902 | NA | -1.088775364 | 2 | 2 | 2 | 2 | 2 | 3.663836244 | 3.042026731 |
| ENSG00000189042 | ZNF567 | -0.999306923 | 0.164064972 | -0.090437257 | 0.914220926 | -1.08974418 | 7.33131396 | 6.04685525 | 6.958202216 | 5.85002558 | 5.96314905 | 7.062943761 | 6.403650353 |
| ENSG00000156735 | BAG4 | -0.755886478 | 0.572651771 | -0.335536356 | 0.811551833 | -1.091422834 | 5.532353208 | 9.055168103 | 9.31604076 | 7.880528731 | 7.95274764 | 8.404758425 | 8.25407423 |
| ENSG00000136866 | ZFP37 | -0.192600727 | 0.918975348 | -0.899117159 | 0.59841801 | -1.091717887 | 2 | 6.666836837 | 6.203069917 | 5.949776857 | 5.47317115 | 5.47655442 | 4.56420962 |
| ENSG00000186825 | C2orf27B | 4.09781558 | 0.459397746 | -5.191092741 | 0.35529554 | -1.093277161 | 2 | 2 | 2 | 4.299492648 | 4.53541753 | 2 | 2 |
| ENSG00000090889 | KIF4A | -0.92370137 | 0.493535102 | -0.170060956 | 0.908291211 | -1.093762326 | 4.00722531 | 5.457120953 | 3.369317972 | 4.105493497 | 3.60476711 | 4.199031499 | 4.683176719 |
| ENSG00000119523 | ALG2 | -0.429174892 | 0.463676051 | -0.664721244 | 0.225285675 | -1.093896136 | 10.19941743 | 9.090820894 | 9.426401271 | 9.198375657 | 9.24178305 | 8.958909138 | 9.013414035 |
| ENSG00000254685 | FPGT | -0.032255367 | 0.986960349 | -1.06169026 | 0.544751469 | -1.093945627 | 2 | 7.35507212 | 7.225742779 | 6.727838783 | 6.66650236 | 6.022174708 | 5.349527629 |
| ENSG00000126010 | GRPR | -2.510834149 | 0.304711188 | 1.416197542 | 0.5723508 | -1.094636607 | 2 | 6.844429755 | 2 | 3.615725078 | 3.15372745 | 6.959905775 | 6.38621943 |
| ENSG00000175166 | PSMD2 | -0.122160074 | 0.815788738 | -0.975768286 | 0.022364812 | -1.097928359 | 12.29444214 | 11.72102722 | 11.60159843 | 11.76318136 | 11.8025467 | 10.68536117 | 11.14037952 |
| ENSG00000135052 | GOLM1 | -0.191672842 | 0.776256344 | -0.907158598 | 0.117096798 | -1.09883144 | 10.19198706 | 11.24195824 | 11.28380426 | 10.7993477 | 10.7910896 | 9.771724757 | 10.33696731 |
| ENSG00000123454 | DBH | -3.75302041 | 0.531143362 | 2.652696705 | NA | -1.100323705 | 2 | 2 | 2 | 2 | 2 | 3.059555873 | 3.640730416 |
| ENSG00000163754 | GYG1 | -0.371503323 | 0.566726644 | -0.729814426 | 0.223025633 | -1.101317749 | 10.3868074 | 9.055168103 | 9.981704053 | 9.486925632 | 9.58250309 | 9.170850107 | 9.187873195 |
| ENSG00000031698 | SARS | -0.841891493 | 0.020700446 | -0.260311713 | 0.521153279 | -1.102203206 | 10.75779627 | 11.273328 | 11.42675132 | 10.35364261 | 10.3264289 | 10.89918683 | 10.94091995 |
| ENSG00000064012 | CASP8 | 0.361723117 | 0.782156761 | -1.464349169 | 0.1981584 | -1.102626051 | 5.70932728 | 6.535001748 | 8.631919287 | 7.751013478 | 7.93360068 | 6.049001768 | 6.113568999 |
| ENSG00000109107 | ALDOC | -1.631666292 | 0.014506629 | 0.529005985 | 0.474487232 | -1.102660307 | 9.247966931 | 8.004992147 | 9.330306409 | 7.318995108 | 7.40393901 | 9.026085624 | 9.848823977 |
| ENSG00000065717 | TLE2 | -2.202525568 | 0.104055955 | 1.098151797 | 0.435639036 | -1.104373771 | 5.330594134 | 7.099070191 | 3.369317972 | 4.17310023 | 3.98554648 | 7.380620955 | 6.590289384 |
| ENSG00000116747 | TROVE2 | -0.810171124 | 0.324359661 | -0.294591559 | 0.737997178 | -1.104762683 | 8.107517749 | 9.28800521 | 9.945092785 | 8.534041678 | 8.4504872 | 9.332742081 | 8.576385923 |
| ENSG00000101189 | MRGBP | -0.526649021 | 0.073702888 | -0.578131923 | 0.042592362 | -1.104780944 | 8.540903468 | 8.590771778 | 8.742726175 | 8.080836374 | 8.12931531 | 8.007404089 | 8.100433454 |
| ENSG00000157110 | RBPMS | -1.236102294 | 0.023538321 | 0.13095559 | 0.842587044 | -1.105146704 | 11.14017917 | 9.981728319 | 10.26027233 | 9.244284335 | 9.38609912 | 10.70435357 | 10.65205172 |
| ENSG00000276340 | | 3.48264496 | 0.06001882 | -4.58784196 | 0.012795456 | -1.105197 | 2 | 6.04685525 | 3.369317972 | 8.049698831 | 8.06258142 | 2.526562374 | 2.059860178 |
| ENSG00000152944 | MED21 | -0.671823575 | 0.386556156 | -0.433764128 | 0.582952722 | -1.105587703 | 9.27634997 | 7.503546079 | 8.65477198 | 8.036145657 | 7.92393105 | 8.290747499 | 8.13650041 |
| ENSG00000092096 | SLC22A17 | -1.388560145 | 0.008760885 | 0.282655365 | 0.645132043 | -1.105904779 | 8.910412251 | 9.34816733 | 9.963514551 | 8.109160159 | 8.08227582 | 10.00483003 | 9.446246726 |
| ENSG00000068489 | PRR11 | -0.99505996 | 0.148587465 | -0.111784064 | 0.889135814 | -1.106844025 | 6.257048221 | 6.311476754 | 6.958202216 | 5.775854763 | 5.40529704 | 6.007052374 | 6.757786576 |
| ENSG00000137825 | ITPKA | 1.211040879 | 0.619674661 | -2.320849903 | 0.311632672 | -1.109809025 | 2 | 4.442710129 | 2 | 4.034562184 | 4.40247105 | 2.592885947 | 2.172608095 |
| ENSG00000154134 | ROBO3 | -1.112312767 | 0.594639301 | 0.002295035 | 0.999263508 | -1.110017732 | 8.998234871 | 6.666836837 | 2 | 6.640968371 | 6.56439512 | 7.388811377 | 7.931466218 |
| ENSG00000090054 | SPTLC1 | -0.271876088 | 0.64079571 | -0.838929137 | 0.103603737 | -1.110805225 | 10.47516499 | 9.849218119 | 9.426401271 | 9.720225568 | 9.70188757 | 8.979198827 | 9.298000491 |
| ENSG00000164850 | GPER1 | -1.515881534 | 0.472437308 | 0.403493614 | 0.859659709 | -1.11238792 | 2 | 6.140560021 | 8.904018013 | 6.01571348 | 6.11534392 | 7.34858644 | 8.3297313 |
| ENSG00000126106 | TMEM53 | -0.881957126 | 0.633758183 | -0.230431598 | 0.908314446 | -1.112388724 | 2 | 7.572420273 | 6.203069917 | 5.742861563 | 5.5755357 | 6.104049656 | 6.420873184 |
| ENSG00000186300 | ZNF555 | -0.987253966 | 0.409036401 | -0.127031685 | 0.92548706 | -1.114285651 | 8.313285031 | 5.946638857 | 5.930865648 | 6.18083559 | 6.30607931 | 7.385306888 | 6.666875073 |
| ENSG00000172809 | RPL38 | 0.147168197 | 0.911517345 | -1.262702636 | 0.255721676 | -1.115534439 | 11.08508709 | 8.732943944 | 8.408111533 | 10.00861625 | 10.150516 | 8.73123589 | 8.62735196 |
| ENSG00000171723 | GPHN | -0.718806105 | 0.76909522 | -0.396997115 | 0.875239371 | -1.11580322 | 3.327711234 | 7.928237109 | 2 | 5.870535235 | 5.63601114 | 5.985611216 | 6.127623461 |
| ENSG00000144659 | SLC25A38 | -0.261422 | 0.691119414 | -0.857927577 | 0.137924768 | -1.119349577 | 8.418903792 | 8.255926468 | 9.385998894 | 8.43079185 | 8.59306436 | 7.877064822 | 7.972925696 |
| ENSG00000174173 | TRMT10C | -0.274382254 | 0.489001987 | -0.845369057 | 0.014719532 | -1.11975131 | 8.009475818 | 8.213429689 | 8.536706852 | 8.04293816 | 7.94798467 | 7.438128798 | 7.434215826 |
| ENSG00000181915 | ADO | -0.664592586 | 0.613081998 | -0.456440194 | 0.731701381 | -1.121032779 | 7.975256505 | 7.145017454 | 10.28245311 | 8.474993083 | 8.38855922 | 8.909749013 | 8.303661547 |
| ENSG00000070413 | DGCR2 | -1.024565366 | 0.326833114 | -0.096642018 | 0.935870665 | -1.121207384 | 8.107517749 | 10.09139839 | 11.0057616 | 9.111653326 | 9.16073939 | 10.06594101 | 10.05293618 |
| ENSG00000090487 | SPG21 | -0.379978563 | 0.790662721 | -0.742082637 | 0.57806806 | -1.1220612 | 11.36826231 | 8.623598428 | 8.143093325 | 9.702459798 | 9.76350037 | 9.381049994 | 9.365156975 |
| ENSG00000164897 | TMUB1 | -1.011003586 | 0.013877267 | -0.111369997 | 0.822143119 | -1.122373583 | 9.448362284 | 9.04308532 | 8.631919287 | 8.109160159 | 8.04930085 | 8.930736901 | 9.00677572 |
| ENSG00000101337 | TM9SF4 | -0.51424953 | 0.174135326 | -0.608248361 | 0.095497186 | -1.122497891 | 11.0567315 | 10.61258327 | 11.23155829 | 10.49945465 | 10.4499017 | 10.28346463 | 10.47270666 |
| ENSG00000147434 | CHRNA6 | -3.753020699 | 0.531143362 | 2.629570255 | NA | -1.123450444 | 2 | 2 | 2 | 2 | 2 | 2.831336763 | 3.753786374 |
| ENSG00000112335 | SNX3 | -0.453307125 | 0.221900832 | -0.672384849 | 0.054916044 | -1.125691974 | 11.40085503 | 10.81088865 | 10.94216274 | 10.62924246 | 10.612236 | 10.43381138 | 10.37004136 |
| ENSG00000066926 | FECH | -0.628884337 | 0.15622962 | -0.496992381 | 0.263184603 | -1.125876718 | 8.855032352 | 8.820493869 | 8.325066185 | 8.03841338 | 8.09740995 | 7.934143724 | 8.421719243 |
| ENSG00000198625 | MDM4 | -0.135381601 | 0.891546154 | -0.990914893 | 0.234649837 | -1.126296495 | 7.031698319 | 8.639735838 | 9.180874355 | 8.397612413 | 8.39732094 | 7.559188434 | 7.552234734 |
| ENSG00000180053 | NKX2-6 | 4.064225256 | 0.466248007 | -5.191091905 | 0.358489009 | -1.126866649 | 2 | 2 | 2 | 4.387512679 | 4.40247105 | 2 | 2 |
| ENSG00000158863 | FAM160B2 | -1.037349073 | 0.115878835 | -0.08951956 | 0.909689184 | -1.126868633 | 8.946185348 | 9.532315021 | 10.45512158 | 8.752828065 | 8.74357228 | 9.722117015 | 9.65623967 |
| ENSG00000132938 | MTUS2 | 0.048126111 | NA | -1.17731756 | NA | -1.129191449 | 2 | 3.076665209 | 2 | 2.498691189 | 2.68924543 | 2.139668783 | 2.421067459 |
| ENSG00000065328 | MCM10 | 0.596241688 | 0.712287044 | -1.72789612 | 0.23639043 | -1.131654432 | 4.467311984 | 3.076665209 | 4.058998049 | 4.268917409 | 4.53541753 | 2.457042231 | 3.236138336 |
| ENSG00000079974 | RABL2B | -0.290694919 | 0.816824789 | -0.841246714 | 0.456128443 | -1.131941633 | 8.586954094 | 5.595927442 | 7.342949392 | 7.25803034 | 7.42122542 | 6.86244555 | 6.750993196 |
| ENSG00000099256 | PRTFDC1 | 0.014689275 | 0.995306348 | -1.146930482 | 0.562234526 | -1.132241207 | 8.653385977 | 6.535001748 | 2 | 7.363085426 | 7.42122542 | 6.060766632 | 6.457498602 |
| ENSG00000221914 | PPP2R2A | -0.466955826 | 0.419187861 | -0.665678892 | 0.223069519 | -1.132634718 | 10.44422623 | 9.7032946 | 9.399592459 | 9.44052423 | 9.46489849 | 9.431761894 | 9.053995509 |
| ENSG00000250317 | SMIM20 | -0.861004115 | 0.162308072 | -0.271714808 | 0.685920699 | -1.132718924 | 8.586954094 | 7.467835385 | 7.600047325 | 7.226553841 | 7.02417148 | 7.5848779 | 7.814703611 |
| ENSG00000177830 | CHID1 | -0.278496529 | 0.653630592 | -0.855157639 | 0.120415896 | -1.133654168 | 9.521325812 | 9.672251278 | 10.6093157 | 9.717397705 | 9.76214432 | 9.156187796 | 9.176932811 |
| ENSG00000179348 | GATA2 | 0.576961711 | 0.73580717 | -1.71095258 | 0.265569714 | -1.133990868 | 5.70932728 | 6.728495673 | 2 | 6.394182228 | 6.23747437 | 4.065262523 | 4.479073019 |
| ENSG00000068438 | FTSJ1 | -0.497724693 | 0.550603127 | -0.636382395 | 0.428827907 | -1.134107088 | 10.16187685 | 8.968383114 | 8.380957451 | 8.830780654 | 8.91077722 | 8.561512889 | 8.887952157 |
| ENSG00000121022 | COPS5 | -0.30013257 | 0.652929267 | -0.834525836 | 0.162167432 | -1.134658406 | 9.98400217 | 8.94259736 | 8.805342542 | 9.010169232 | 9.08140959 | 8.469076579 | 8.562873254 |
| ENSG00000163806 | SPDYA | -0.308445516 | 0.912408896 | -0.826984552 | 0.74998849 | -1.135430067 | 2 | 4.11352228 | 2 | 3.289987342 | 2.59489879 | 2.858551861 | 2.632958241 |
| ENSG00000178802 | MPI | -1.324798667 | 0.296385835 | 0.189180733 | 0.89578964 | -1.135617935 | 9.90440318 | 9.159587375 | 6.073378139 | 7.781117077 | 7.71553728 | 8.897090174 | 9.525344379 |
| ENSG00000101310 | SEC23B | -0.44579734 | 0.570041492 | -0.690645682 | 0.352432367 | -1.136443022 | 10.5351205 | 9.170736405 | 9.099922433 | 9.325453269 | 9.31389502 | 8.886799903 | 9.244188987 |
| ENSG00000172354 | GNB2 | -0.803057664 | 0.001780256 | -0.33350613 | 0.233678267 | -1.136563793 | 11.47313651 | 11.66496702 | 11.85201631 | 10.85926146 | 10.8800418 | 11.28460067 | 11.39025627 |
| ENSG00000172215 | CXCR6 | -3.753020892 | 0.531143362 | 2.61409108 | NA | -1.138929812 | 2 | 2 | 2 | 2 | 2 | 3.374372158 | 3.33635788 |
| ENSG00000183018 | SPNS2 | -0.785316419 | 0.760985399 | -0.354349952 | 0.893566028 | -1.139666371 | 2 | 2 | 4.523675391 | 3.023133209 | 2.77779907 | 3.059555873 | 3.183290849 |
| ENSG00000137103 | TMEM8B | -2.187153245 | 0.006538839 | 1.046967506 | 0.222377009 | -1.140185738 | 7.159102577 | 8.337339476 | 9.272377397 | 6.365568715 | 6.36443784 | 9.828312481 | 9.166759824 |
| ENSG00000089248 | ERP29 | -0.726412325 | 0.184252877 | -0.4152589 | 0.464454793 | -1.141671225 | 10.22148166 | 10.04368224 | 11.08782089 | 9.829021922 | 9.77295717 | 10.06265365 | 10.15809785 |
| ENSG00000187735 | TCEA1 | -0.325764207 | 0.609032994 | -0.816283644 | 0.15331931 | -1.142047851 | 9.810482605 | 9.04308532 | 9.301632639 | 9.209487455 | 8.97188212 | 8.98885284 | 8.092394167 |
| ENSG00000172081 | MOB3A | -1.055308018 | 0.025780112 | -0.086853857 | 0.88223205 | -1.142161875 | 9.895281535 | 9.591588447 | 10.54349105 | 9.009013399 | 9.01919552 | 9.872787989 | 10.07440932 |
| ENSG00000072571 | HMMR | 0.418119138 | 0.753072123 | -1.560339307 | 0.184116925 | -1.142220169 | 3.327711234 | 4.11352228 | 4.523675391 | 4.523237872 | 4.2560182 | 2.858551861 | 3.236138336 |
| ENSG00000162929 | KIAA1841 | -0.113369153 | 0.930344224 | -1.030176514 | 0.354864291 | -1.143545666 | 8.009475818 | 6.140560021 | 5.39244166 | 6.861921612 | 6.80463998 | 6.17883 | 5.703940561 |
| ENSG00000111605 | CPSF6 | -0.710023136 | 0.036085138 | -0.433878178 | 0.216308317 | -1.143901314 | 9.174465797 | 9.71095217 | 9.553445883 | 8.799033614 | 8.790887 | 8.99920712 | 9.135368584 |
| ENSG00000154721 | JAM2 | -2.779269962 | 0.000125058 | 1.63518117 | 0.030981543 | -1.144088793 | 7.435667355 | 8.395519306 | 9.399592459 | 5.987823101 | 5.86549842 | 10.40597858 | 10.0730482 |
| ENSG00000213462 | ERV3-1 | -0.615699963 | 0.758753417 | -0.528469189 | 0.792987538 | -1.144169153 | 2 | 6.899080544 | 8.266922072 | 6.483440992 | 6.65485813 | 6.384736493 | 6.883203642 |
| ENSG00000253831 | ETV3L | 4.046834707 | 0.470055465 | -5.191091418 | 0.360419662 | -1.144256711 | 2 | 2 | 2 | 4.387512679 | 4.37434574 | 2 | 2 |
| ENSG00000166471 | TMEM41B | -0.107699193 | 0.749240001 | -1.036899714 | 0.00010592 | -1.144598907 | 9.112843761 | 9.267380431 | 9.439621147 | 9.204447245 | 9.13791797 | 8.316736994 | 8.182667731 |
| ENSG00000236438 | FAM157A | 4.046438119 | 0.470185896 | -5.191091389 | 0.360479076 | -1.14465327 | 2 | 2 | 2 | 4.329433326 | 4.43005853 | 2 | 2 |
| ENSG00000198283 | OR5B21 | 4.046239732 | 0.470185896 | -5.191091358 | 0.360546357 | -1.144851626 | 2 | 2 | 2 | 4.299492648 | 4.45712836 | 2 | 2 |
| ENSG00000187475 | HIST1H1T | 2.496167913 | 0.622552387 | -3.641286977 | NA | -1.145119063 | 2 | 2 | 2 | 3.227757282 | 3.33976443 | 2 | 2.059860178 |
| ENSG00000100632 | ERH | -0.155987479 | 0.815259329 | -0.989849544 | 0.074729474 | -1.145837023 | 9.657036304 | 8.607278467 | 9.788583153 | 9.224503288 | 9.33228764 | 8.413960373 | 8.490627472 |
| ENSG00000277157 | HIST1H4D | 2.495013423 | 0.623809291 | -3.641029376 | NA | -1.146015953 | 2 | 2 | 2 | 3.09461492 | 3.45167086 | 2 | 2.059860178 |
| ENSG00000196236 | XPNPEP3 | -0.654425037 | 0.27254373 | -0.492208669 | 0.41217935 | -1.146633705 | 7.867441304 | 7.572420273 | 8.608698767 | 7.426792977 | 7.44169935 | 7.807195765 | 7.340786405 |

| | | | | | | | | | | | | | |
|---|---|---|---|---|---|---|---|---|---|---|---|---|---|
| ENSG00000159792 | PSKH1 | -1.024090349 | 0.088708728 | -0.123403334 | 0.861696561 | -1.147493683 | 10.60392551 | 9.225224669 | 10.06083273 | 9.012478123 | 9.08358266 | 9.942208568 | 9.945960865 |
| ENSG00000158987 | RAPGEF6 | 0.824149344 | 0.225654811 | -1.972827325 | 0.001286666 | -1.148677981 | 7.579279349 | 8.053983481 | 6.629488387 | 8.471640654 | 8.23084153 | 5.853194273 | 5.436777225 |
| ENSG00000027847 | B4GALT7 | -0.809951852 | 0.35543975 | -0.339107353 | 0.715680486 | -1.149059205 | 9.371511179 | 9.549500232 | 7.451345288 | 8.302698479 | 8.19108423 | 8.701014095 | 8.725914443 |
| ENSG00000111602 | TIMELESS | -0.507191113 | 0.734475554 | -0.642076828 | 0.659411517 | -1.149267941 | 4.00722531 | 7.954278496 | 8.206335842 | 7.053486362 | 7.04662306 | 6.7543034 | 7.067643653 |
| ENSG00000197448 | GSTK1 | -0.321106308 | 0.714968044 | -0.828641851 | 0.297804511 | -1.149748159 | 10.80726569 | 8.993716073 | 9.516493218 | 9.645581273 | 9.67619834 | 9.046937881 | 9.258230494 |
| ENSG00000205744 | DENND1C | -3.753021038 | 0.531143362 | 2.602399887 | NA | -1.150621152 | 2 | 2 | 2 | 2 | 2 | 3.753554863 | 2.782576693 |
| ENSG00000187699 | C2orf88 | 0.992527916 | 0.688047343 | -2.14343518 | 0.347127935 | -1.150907263 | 2 | 6.04685525 | 2 | 5.651002631 | 5.40529704 | 3.256555576 | 3.012034821 |
| ENSG00000100150 | DEPDC5 | -0.367554978 | 0.759077898 | -0.783438795 | 0.480467544 | -1.150993774 | 6.563806279 | 7.73146271 | 4.874587708 | 6.470063213 | 6.47451726 | 5.941748584 | 6.202567011 |
| ENSG00000171865 | RNASEH1 | -0.826204556 | 0.610039424 | -0.325089976 | 0.849222745 | -1.151294533 | 9.710019321 | 7.393647515 | 4.874587708 | 7.68016596 | 7.54938955 | 8.041943102 | 8.172535245 |
| ENSG00000109917 | ZPR1 | 0.001821929 | 0.999189606 | -1.15327064 | 0.597939463 | -1.151448711 | 10.45668146 | 8.717821467 | 2 | 9.248208205 | 9.2611121 | 7.964208996 | 8.244432684 |
| ENSG00000169249 | ZRSR2 | -0.412763623 | 0.589874122 | -0.739491901 | 0.297771937 | -1.152255525 | 6.816637977 | 7.002537973 | 8.175061096 | 7.035434774 | 7.07311172 | 6.895669227 | 6.562106074 |
| ENSG00000156603 | MED19 | -0.652145459 | 0.671489931 | -0.501849913 | 0.744573138 | -1.153995372 | 8.564112515 | 8.433041974 | 4.058998049 | 7.388195945 | 7.23044321 | 7.430214384 | 7.485592137 |
| ENSG00000115514 | TXNDC9 | 0.076419501 | 0.966538801 | -1.230486828 | 0.443133621 | -1.154067328 | 9.556468883 | 7.874700259 | 4.058998049 | 8.535647908 | 8.38503958 | 7.168655895 | 7.189889619 |
| ENSG00000171148 | TADA3 | -0.747342758 | 0.043381631 | -0.406810223 | 0.293835258 | -1.154152982 | 10.51739401 | 10.04368224 | 9.926432745 | 9.469384409 | 9.41227044 | 9.671462104 | 9.88081174 |
| ENSG00000167749 | KLK4 | -3.753021086 | 0.531143362 | 2.598550145 | NA | -1.154470941 | 2 | 2 | 2 | 2 | 2 | 4.029788235 | 2 |
| ENSG00000047597 | XK | -3.753021089 | 0.531143362 | 2.598372343 | NA | -1.154648746 | 2 | 2 | 2 | 2 | 2 | 3.534812207 | 3.128433649 |
| ENSG00000142168 | SOD1 | -0.16787414 | 0.842541719 | -0.988260321 | 0.165667977 | -1.156134462 | 11.62432333 | 10.27307349 | 10.17589429 | 10.67405041 | 10.6970002 | 9.672420943 | 10.03999009 |
| ENSG00000180992 | MRPL14 | -0.714221574 | 0.298900033 | -0.442153267 | 0.529629876 | -1.156374841 | 9.613202746 | 9.367676479 | 8.143093325 | 8.458152491 | 8.46722793 | 8.537506665 | 8.903351429 |
| ENSG00000170464 | DNAJC18 | -1.025453499 | 0.19054829 | -0.131282963 | 0.885564891 | -1.156736462 | 9.015175615 | 7.60566235 | 9.491321578 | 7.842101571 | 7.91907183 | 8.60585192 | 8.906411656 |
| ENSG00000153531 | ADPRHL1 | -2.238699462 | 0.019769712 | 1.079610291 | 0.276208632 | -1.159089171 | 5.532353208 | 4.442710129 | 6.43197674 | 3.754611435 | 3.82856379 | 6.176128386 | 7.04745562 |
| ENSG00000082068 | WDR70 | -0.192375771 | 0.895675696 | -0.967872766 | 0.450687102 | -1.160248538 | 4.815609471 | 7.819099893 | 8.631919287 | 7.608714138 | 7.53358066 | 6.674237832 | 6.947999751 |
| ENSG00000120693 | SMAD9 | -0.680608217 | 0.720923534 | -0.479783777 | 0.80363944 | -1.160391994 | 2 | 7.35507212 | 7.777116083 | 6.456560224 | 6.25300433 | 6.840432491 | 6.185873005 |
| ENSG00000182473 | EXOC7 | 0.299855149 | 0.804920943 | -1.460851325 | 0.158887679 | -1.160996175 | 8.928409675 | 10.00059544 | 11.760164 | 10.96972999 | 11.0210702 | 9.114110681 | 9.358840932 |
| ENSG00000106006 | HOXA6 | 4.029633139 | 0.473893575 | -5.191090891 | 0.362261771 | -1.161457752 | 2 | 2 | 2 | 4.443344104 | 4.2865223 | 2 | 2 |
| ENSG00000161677 | JOSD2 | -0.492820998 | 0.463053095 | -0.668822922 | 0.292072257 | -1.161643919 | 8.138771685 | 7.315436877 | 8.65477198 | 7.688853661 | 7.59580267 | 7.309674888 | 7.615934347 |
| ENSG00000130764 | LRRC47 | -1.09320368 | 0.019638737 | -0.069009013 | 0.906104302 | -1.162212693 | 9.886101849 | 8.993716073 | 9.859153765 | 8.522747711 | 8.56844956 | 9.640192857 | 9.479504351 |
| ENSG00000154767 | XPC | -0.068601836 | 0.928929217 | -1.095030209 | 0.077869736 | -1.163632045 | 8.737455672 | 8.862353853 | 9.972637969 | 9.209487455 | 9.25726695 | 8.377971121 | 8.035760639 |
| | DOPEY1 | 5.787338782 | 0.000322116 | -6.951186767 | 0.002485079 | -1.163847985 | 2 | 3.686303414 | 2 | 7.59952861 | 7.60793141 | 2 | 2 |
| ENSG00000196642 | RABL6 | -1.067807383 | 0.129332125 | -0.096933607 | 0.908291211 | -1.164740989 | 9.533135543 | 9.936721131 | 11.10036136 | 9.272493518 | 9.29994502 | 10.30912217 | 10.19381254 |
| ENSG00000149743 | TRPT1 | -0.681622314 | 0.396175926 | -0.483223535 | 0.55075414 | -1.164845849 | 8.696033109 | 7.189546432 | 7.342949392 | 7.017154451 | 7.43490694 | 7.381793866 | 7.485592137 |
| ENSG00000134248 | LAMTOR5 | -0.439608041 | 0.367412771 | -0.725821236 | 0.110555942 | -1.165429278 | 9.678463567 | 8.903033433 | 8.978369997 | 8.756965077 | 8.82635531 | 8.403604044 | 8.606804877 |
| ENSG00000140682 | TGFB1I1 | -1.721461203 | 0.000159191 | 0.55499921 | 0.275628117 | -1.166461993 | 10.77281601 | 9.785372135 | 10.64969256 | 8.723531969 | 8.77618319 | 10.99934685 | 11.03223581 |
| ENSG00000111653 | ING4 | -0.269533606 | 0.810373984 | -0.89821696 | 0.366670893 | -1.167750566 | 9.48530526 | 8.968383114 | 6.80318718 | 8.517880242 | 8.5339035 | 8.018003769 | 7.783974227 |
| ENSG00000237155 | | 8.868271774 | 3.94E-07 | -10.03617909 | 2.17E-05 | -1.167907314 | 3.327711234 | 6.389896633 | 2 | 13.72712598 | 13.7531302 | 2 | 2 |
| ENSG00000156042 | CFAP70 | 0.705110877 | 0.648413066 | -1.873537101 | 0.184133611 | -1.168426224 | 4.00722531 | 4.11352228 | 2 | 4.268917409 | 4.12711147 | 2.68698273 | 2.671842496 |
| ENSG00000168301 | KCTD6 | -1.131126886 | 0.162604083 | -0.037654054 | 0.968812919 | -1.16878094 | 7.709880939 | 6.389896633 | 6.203069917 | 5.808110275 | 5.89549381 | 7.156371901 | 6.567271433 |
| ENSG00000130921 | C12orf65 | -1.642240652 | 0.172958175 | 0.473136163 | 0.721484546 | -1.169104489 | 6.009025997 | 7.051611315 | 9.227351203 | 6.408278948 | 6.49425375 | 8.407640344 | 8.616180753 |
| ENSG00000265808 | SEC22B | 0.098604492 | 0.887804971 | -1.269110638 | 0.021623829 | -1.170056146 | 10.905938 | 10.62073637 | 11.80936937 | 11.31682426 | 11.2878885 | 10.0293614 | 9.841114847 |
| ENSG00000112796 | ENPP5 | -3.753021288 | 0.531143362 | 2.582403787 | NA | -1.170617501 | 2 | 2 | 2 | 2 | 2 | 3.429858444 | 3.236138336 |
| ENSG00000172113 | NME6 | -0.771283877 | 0.042332258 | -0.401246888 | 0.312303334 | -1.172530766 | 8.444143041 | 8.337339476 | 8.042706615 | 7.494428494 | 7.56191311 | 8.055971376 | 7.705330953 |
| ENSG00000161558 | TMEM143 | -1.038307391 | 0.600231188 | -0.137510473 | 0.948992109 | -1.175817864 | 2 | 7.189546432 | 4.523675391 | 4.825935136 | 5.0259382 | 5.73334482 | 5.745545708 |
| ENSG00000120727 | PAIP2 | 0.162892612 | 0.818265844 | -1.338872718 | 0.020011794 | -1.175980105 | 10.40620204 | 9.308339287 | 9.272377397 | 9.921611889 | 9.92914219 | 8.310591351 | 8.548580051 |
| ENSG00000139323 | POC1B | -0.601141023 | 0.52931101 | -0.57627557 | 0.538360258 | -1.177416593 | 8.138771685 | 6.04685525 | 8.007629717 | 7.062428156 | 7.05992818 | 7.279786628 | 6.860240429 |
| ENSG00000130724 | CHMP2A | -0.892985259 | 0.003805818 | -0.284446477 | 0.407829503 | -1.177431736 | 9.646202137 | 9.835272298 | 9.344432377 | 8.74173768 | 8.73806474 | 9.286209561 | 9.399228411 |
| ENSG00000157502 | MUM1L1 | 2.874762525 | 0.340693523 | -4.052563301 | 0.164968351 | -1.177800775 | 2 | 2 | 5.39244166 | 6.586219071 | 6.56439512 | 2.457042231 | 2 |
| ENSG00000180902 | D2HGDH | -0.842005764 | 0.718118984 | -0.338593179 | 0.890313426 | -1.180598943 | 8.998234871 | 5.131603625 | 2 | 6.772115916 | 6.6253296 | 7.071691703 | 7.294817518 |
| ENSG00000170448 | NFXL1 | -0.469905016 | 0.825160194 | -0.710841106 | 0.725592339 | -1.180746122 | 2 | 7.954278496 | 5.930865648 | 6.29144905 | 6.23747437 | 6.471027105 | 5.414021719 |
| ENSG00000143436 | MRPL9 | -0.24747599 | 0.525575922 | -0.933784237 | 0.00498916 | -1.181260227 | 8.586954094 | 8.101365653 | 8.266922072 | 8.078634414 | 8.08661608 | 7.402744819 | 7.415721389 |
| ENSG00000166501 | PRKCB | -1.54516659 | 0.585723622 | 0.362551568 | 0.90481909 | -1.182615021 | 2 | 4.710564154 | 2 | 2.600425781 | 2.86123014 | 4.478802948 | 2.509591799 |
| ENSG00000182901 | RGS7 | -4.045406193 | 0.078250731 | 2.860868663 | 0.170814432 | -1.18453753 | 2 | 2 | 5.156626401 | 2.270786764 | 2.14022021 | 5.582367412 | 6.755525668 |
| ENSG00000149100 | EIF3M | -0.150033946 | 0.878396152 | -1.035201712 | 0.205129503 | -1.185235658 | 11.02362764 | 9.443177293 | 9.227351203 | 10.02357442 | 9.94834802 | 9.046937881 | 9.162926384 |
| ENSG00000143761 | ARF1 | -0.787357312 | 0.219396158 | -0.398410952 | 0.553274344 | -1.185768264 | 13.70485113 | 12.52221914 | 12.496991 | 12.21628798 | 12.2605626 | 12.55201409 | 12.69877307 |
| ENSG00000114391 | RPL24 | -1.028670926 | 0.274935376 | -0.157167075 | 0.884519325 | -1.185838001 | 11.93591186 | 9.7032946 | 10.27509734 | 9.893702135 | 9.97960581 | 10.78801637 | 10.82400773 |
| ENSG00000150787 | PTS | -0.223914523 | 0.713961855 | -0.962593332 | 0.066744424 | -1.186507855 | 7.384433967 | 8.053983481 | 8.536706852 | 7.839502934 | 7.84926609 | 7.149501993 | 7.094725293 |
| ENSG00000151665 | PIGF | 0.098292346 | 0.946981783 | -1.285788624 | 0.318708894 | -1.187496279 | 8.16936288 | 5.838937808 | 4.874587708 | 7.123515401 | 6.98285659 | 5.843010798 | 5.626876946 |
| ENSG00000158156 | XKR8 | -0.196746492 | 0.911302045 | -0.991066311 | 0.52301169 | -1.187812803 | 8.19931886 | 9.192779271 | 4.058998049 | 8.060896687 | 8.00183183 | 7.393470826 | 7.09829807 |
| ENSG00000119878 | CRIPT | -0.153951358 | 0.865062033 | -1.033998291 | 0.171003885 | -1.18794965 | 8.493334308 | 7.315436877 | 6.80318718 | 7.61176303 | 7.51437845 | 6.840432491 | 6.564691065 |
| ENSG00000106799 | TGFBR1 | -1.012134609 | 0.236430115 | -0.176837368 | 0.855445922 | -1.188971977 | 8.75772933 | 10.59199661 | 10.89974571 | 9.40053583 | 9.26015177 | 10.10818585 | 10.21659261 |
| ENSG00000168685 | IL7R | -4.714927092 | 0.174626306 | 3.525365365 | 0.236344937 | -1.189561727 | 2 | 2 | 3.369317972 | 2 | 2 | 4.771475226 | 4.720747915 |
| ENSG00000178445 | GLDC | -1.244783879 | 0.571216016 | 0.054998358 | 0.981401003 | -1.18978552 | 4.00722531 | 2 | 2 | 2.868578001 | 2.49394783 | 2.775316226 | 3.735548302 |
| ENSG00000152147 | GEMIN6 | -1.264558124 | 0.433185867 | 0.074449815 | 0.967718357 | -1.190108309 | 8.228665467 | 4.710564154 | 4.523675391 | 5.731694 | 5.58783529 | 7.060015953 | 6.786860422 |
| ENSG00000149084 | HSD17B12 | -0.060175527 | 0.908645221 | -1.130435371 | 0.004943283 | -1.190610898 | 10.04301211 | 9.433952963 | 10.13584611 | 9.85562885 | 9.82323967 | 8.603340274 | 8.93066256 |
| ENSG00000197943 | PLCG2 | -1.007344125 | 0.725609862 | -0.183544572 | 0.951929397 | -1.190888697 | 2 | 5.131603625 | 2 | 3.615725078 | 2.68924543 | 4.350488971 | 2.70970617 |
| ENSG00000198478 | SH3BGRL2 | -2.543519618 | 0.12806343 | 1.351654715 | 0.43784645 | -1.191864904 | 7.904282243 | 7.73146271 | 2 | 4.71689672 | 5.0259382 | 8.621327277 | 8.536120927 |
| ENSG00000100413 | POLR3H | -0.717518303 | 0.293044241 | -0.476806375 | 0.492177662 | -1.194324678 | 9.435834703 | 8.522788974 | 9.798878489 | 8.637890117 | 8.62175922 | 8.429356794 | 9.204544205 |
| ENSG00000185127 | C6orf120 | -0.161189784 | 0.7690856 | -1.033164346 | 0.024076465 | -1.194354131 | 9.097018025 | 8.607278467 | 9.491321578 | 8.986874783 | 8.90095288 | 8.237320313 | 7.908246991 |
| ENSG00000099804 | CDC34 | -1.098214769 | 0.003251348 | -0.096852425 | 0.834220385 | -1.195067193 | 9.613202746 | 9.799806309 | 10.07784268 | 8.667494409 | 8.83024294 | 9.968209057 | 9.482583303 |
| ENSG00000158234 | FAIM | -1.188685845 | 0.111035703 | -0.00691277 | 0.993931149 | -1.195598615 | 5.532353208 | 5.946638857 | 6.882775709 | 5.22679461 | 5.0259382 | 5.982522014 | 6.418016952 |
| ENSG00000124406 | ATP8A1 | -0.326281744 | 0.908151255 | -0.870210468 | 0.737997178 | -1.196492213 | 2 | 5.838937808 | 2 | 3.881293667 | 4.4836996 | 4.199031499 | 3.261853426 |
| ENSG00000164684 | ZNF704 | -4.681164486 | 0.120133693 | 3.48449402 | 0.244635807 | -1.196670467 | 2 | 7.189546432 | 2 | 2.947924294 | 2 | 8.864319213 | 9.249822 |
| ENSG00000155329 | ZCCHC10 | -1.125852032 | 0.392041264 | -0.071672076 | 0.962347915 | -1.197524108 | 8.631580145 | 7.232742069 | 4.874587708 | 6.34372989 | 6.63128385 | 7.754744914 | 7.227896413 |
| ENSG00000124279 | FASTKD3 | -0.412984984 | 0.736587839 | -0.785155802 | 0.493181118 | -1.198140786 | 7.435667355 | 5.595927442 | 4.874587708 | 5.94010643 | 6.03677338 | 5.798032389 | 5.470251342 |
| ENSG00000261986 | | 3.992752324 | 0.480588858 | -5.191090144 | 0.364514502 | -1.19833782 | 2 | 2 | 2 | 4.299492648 | 4.37434574 | 2 | 2 |
| ENSG00000123131 | PRDX4 | -0.167805768 | 0.812887832 | -1.03078147 | 0.081818603 | -1.198587237 | 11.98187803 | 11.00713613 | 10.69999858 | 11.16683559 | 11.1737025 | 10.29019523 | 10.3282462 |
| ENSG00000258890 | CEP95 | -0.230072844 | 0.72758145 | -0.970619971 | 0.089151012 | -1.200692815 | 8.009475818 | 7.145017454 | 7.896971941 | 7.617841552 | 7.36516374 | 7.05561307 | 6.432241896 |
| ENSG00000081307 | UBA5 | -0.075360792 | 0.948180321 | -1.125461762 | 0.26264969 | -1.200822554 | 9.423197387 | 8.903033433 | 6.629488387 | 8.71221318 | 8.57618689 | 7.698433415 | 7.517964974 |
| ENSG00000131467 | PSME3 | -0.644123202 | 0.216076248 | -0.557707131 | 0.282374528 | -1.201830332 | 10.59831544 | 10.70750522 | 11.46604358 | 10.35864221 | 10.3102824 | 10.18813102 | 10.62118535 |
| ENSG00000196683 | TOMM7 | -0.458227466 | 0.624036401 | -0.744900827 | 0.397192742 | -1.203128294 | 10.85509492 | 8.806266215 | 9.426401271 | 9.486925632 | 9.51808625 | 9.259659123 | 9.174396282 |
| ENSG00000110344 | UBE4A | 0.003342091 | 0.997041173 | -1.206904309 | 0.069457906 | -1.203562217 | 8.716893051 | 10.3138733 | 10.0085654 | 9.856914282 | 9.80557618 | 8.508269163 | 8.741420621 |
| ENSG00000184436 | THAP7 | -1.127729573 | 0.177620155 | -0.080646188 | 0.935063964 | -1.208375761 | 6.653098118 | 7.790474954 | 8.721235202 | 6.820355514 | 6.88158913 | 8.009682014 | 7.720479276 |
| ENSG00000145725 | PPIP5K2 | -0.435021466 | 0.724360006 | -0.774457851 | 0.503743146 | -1.209479317 | 6.468621203 | 9.71856931 | 9.099922433 | 8.606137616 | 8.42329129 | 8.263008721 | 8.096865967 |
| ENSG00000108840 | HDAC5 | -1.516003589 | 0.001826988 | 0.305239027 | 0.590415354 | -1.210764562 | 10.73752355 | 9.632483534 | 10.21486071 | 8.736160353 | 8.77349359 | 10.49619812 | 10.63370358 |
| ENSG00000128928 | IVD | -0.91586399 | 0.357081274 | -0.295565704 | 0.784238084 | -1.211429695 | 8.418903792 | 9.405920729 | 10.88048583 | 9.055659246 | 8.95891978 | 9.434872973 | 9.792045353 |
| ENSG00000227471 | AKR1B15 | 0.931753887 | 0.646472565 | -2.143240536 | 0.258155626 | -1.211486649 | 2 | 3.686303414 | 3.369317972 | 3.79808662 | 3.82856379 | 2.183373728 | 2.551896972 |
| ENSG00000085760 | MTIF2 | -0.340799619 | 0.778610361 | -0.872892019 | 0.429681573 | -1.213691638 | 6.563806279 | 7.638155693 | 9.372276026 | 8.029321023 | 7.92393105 | 7.49992804 | 7.409982764 |
| ENSG00000105697 | HAMP | -3.753021845 | 0.531143362 | 2.537792347 | NA | -1.215229498 | 2 | 2 | 2 | 2 | 2 | 3.838019203 | 2.466008509 |
| ENSG00000163746 | PLSCR2 | 3.522153188 | 0.132111686 | -4.737651392 | 0.093843117 | -1.215498205 | 2 | 2 | 3.369317972 | 4.76150579 | 4.72553021 | 2.048074405 | 2 |
| ENSG00000167642 | SPINT2 | -2.692463962 | 0.033287513 | 1.476863431 | 0.264448359 | -1.215600532 | 8.87372952 | 6.951736442 | 4.874587708 | 5.319220934 | 4.95174433 | 9.024959988 | 9.272136653 |
| ENSG00000198910 | L1CAM | -2.303493203 | 0.355101172 | 1.087122409 | 0.676785453 | -1.216370793 | 2 | 6.728495673 | 2 | 3.754611435 | 3.08597292 | 5.916072173 | 6.559516442 |
| ENSG00000180822 | PSMG4 | -0.849991972 | 0.469998704 | -0.366789801 | 0.766202864 | -1.216781773 | 6.89194353 | 4.93638933 | 7.97167868 | 6.229283386 | 6.23747437 | 6.66270409 | 6.725806822 |
| ENSG00000185947 | ZNF267 | 0.228934351 | 0.873367801 | -1.446044454 | 0.232591517 | -1.217110103 | 6.468621203 | 5.946638857 | 8.941672936 | 8.076429088 | 7.83385794 | 6.322573937 | 6.347714043 |
| ENSG00000138032 | PPM1B | -0.213538106 | 0.805756178 | -1.004572637 | 0.176769831 | -1.218110743 | 7.218819781 | 7.73146271 | 8.904018013 | 8.010962573 | 7.81305436 | 7.120285726 | 7.162846889 |
| ENSG00000120896 | SORBS3 | -1.304273883 | 0.000812533 | 0.085866456 | 0.862065438 | -1.218407427 | 11.01943573 | 10.38739569 | 11.0408834 | 9.486095154 | 9.60082524 | 10.7588599 | 11.0829319 |
| ENSG00000179912 | R3HDM2 | -0.287119244 | 0.313894643 | -0.931749324 | 0.000162448 | -1.218868568 | 9.262228249 | 9.318399851 | 9.516493218 | 9.05677832 | 9.10834046 | 8.41853943 | 8.469401969 |
| ENSG00000151332 | MBIP | -0.275493029 | 0.761078417 | -0.943405153 | 0.240015951 | -1.218898182 | 8.418903792 | 7.847167909 | 6.43197674 | 7.517371202 | 7.50468064 | 7.0047514 | 6.70252219 |
| ENSG00000150907 | FOXO1 | -1.11039219 | 0.496139185 | -0.110031993 | 0.951839566 | -1.220424184 | 8.075571721 | 8.732943944 | 3.369317972 | 6.761173376 | 6.83591385 | 7.942116583 | 7.581886257 |
| ENSG00000115207 | GTF3C2 | -0.747664724 | 0.584419874 | -0.473468838 | 0.733879184 | -1.221133562 | 6.009025997 | 9.599860728 | 9.990713521 | 8.418656444 | 8.63959257 | 8.75451294 | 8.853483814 |
| ENSG00000070081 | NUCB2 | -0.083879657 | 0.895420772 | -1.137713491 | 0.024456784 | -1.221593147 | 11.15935596 | 10.06176125 | 10.7904855 | 10.67441498 | 10.632786 | 9.73685847 | 9.457070207 |
| ENSG00000172301 | COPRS | -0.984533662 | 0.10165634 | -0.237692245 | 0.724423786 | -1.222225907 | 8.716893051 | 9.415325457 | 9.69246155 | 8.31397419 | 8.39382264 | 8.590716036 | 9.466776947 |
| ENSG00000221838 | AP4M1 | -0.238673657 | 0.907686866 | -0.984903803 | 0.591055458 | -1.22357746 | 8.075571721 | 7.60566235 | 2 | 7.088925745 | 7.02417148 | 6.394066622 | 6.283256815 |
| ENSG00000124006 | OBSL1 | -1.744648452 | 1.68E-06 | 0.520617436 | 0.202653423 | -1.224031016 | 10.65345793 | 10.00683004 | 10.41552659 | 8.655725489 | 8.63663561 | 10.70294658 | 11.07343347 |
| ENSG00000025423 | HSD17B6 | -2.271957134 | 0.312908582 | 1.047460096 | 0.64966917 | -1.224497038 | 2 | 5.131603625 | 2 | 2.695456494 | 2.59489879 | 5.04460184 | 4.329022995 |
| ENSG00000144724 | PTPRG | -0.733891142 | 0.356968286 | -0.491677397 | 0.54430284 | -1.225568539 | 8.16936288 | 9.869887299 | 9.714382178 | 8.720710586 | 8.67749148 | 8.604847786 | 9.210746501 |
| ENSG00000105701 | FKBP8 | -0.880137726 | 0.125555633 | -0.345709765 | 0.575949683 | -1.22584749 | 11.75612888 | 11.45985033 | 12.62546556 | 11.1619039 | 11.1500503 | 11.55082553 | 11.81625909 |
| ENSG00000071553 | ATP6AP1 | -0.466164649 | 0.279628699 | -0.759984737 | 0.058901421 | -1.226149386 | 12.09176592 | 11.53295289 | 11.46280979 | 11.27800726 | 11.238766 | 10.7784742 | 11.13059894 |
| ENSG00000173535 | TNFRSF10C | -2.635772979 | 0.150647216 | 1.409381717 | 0.456223157 | -1.226391261 | 6.96351253 | 4.442710129 | 2 | 3.754611435 | 3.2184422 | 7.528804029 | 6.081436739 |
| ENSG00000175264 | CHST1 | -1.465913847 | 0.150126699 | 0.234915292 | 0.835954015 | -1.230998555 | 4.815609471 | 4.710564154 | 4.874587708 | 3.515101962 | 3.7864959 | 5.404520029 | 4.434543419 |

| | | | | | | | | | | | | | |
|---|---|---|---|---|---|---|---|---|---|---|---|---|---|
| ENSG00000167112 | TRUB2 | -0.926787122 | 0.124764984 | -0.304438165 | 0.64383682 | -1.231225288 | 8.777722034 | 8.848535039 | 7.600047325 | 7.668500375 | 7.53358066 | 8.138779954 | 8.286538335 |
| ENSG00000169062 | UPF3A | -0.358895314 | 0.278511335 | -0.875134355 | 0.002981664 | -1.234029669 | 8.716893051 | 8.791896851 | 8.434763955 | 8.345451606 | 8.2657098 | 7.790512074 | 7.804897508 |
| ENSG00000105325 | FZR1 | -0.759358114 | 0.289112399 | -0.47489108 | 0.517968728 | -1.234249195 | 9.233563228 | 8.834582582 | 10.33998689 | 8.814994458 | 8.90218458 | 9.029083034 | 9.245800665 |
| ENSG00000189052 | CGB5 | 3.955962672 | 0.484983469 | -5.191090298 | 0.364514502 | -1.235127626 | 2 | 2 | 2 | 4.299492648 | 4.31639476 | 2 | 2 |
| ENSG00000129103 | SUMF2 | -0.268284522 | 0.430332959 | -0.967577995 | 0.000907642 | -1.235862517 | 11.41687936 | 10.95004268 | 11.18128933 | 10.87615882 | 10.9746976 | 10.30356295 | 10.1499441 |
| ENSG00000096092 | TMEM14A | -0.596384742 | 0.30261211 | -0.640430354 | 0.252589462 | -1.236815097 | 7.435667355 | 7.393647515 | 6.718950348 | 6.670510205 | 6.6253296 | 6.297933466 | 6.860240429 |
| ENSG00000100567 | PSMA3 | 1.073845132 | 0.008831138 | -2.312505628 | 1.51E-09 | -1.238660496 | 8.367060731 | 9.114109157 | 8.608698767 | 9.777686595 | 9.8316678 | 6.375345632 | 6.574984961 |
| ENSG00000065548 | ZC3H15 | -0.251925311 | 0.619188542 | -0.987075196 | 0.024026933 | -1.239000506 | 9.975371636 | 9.090820894 | 9.714382178 | 9.442237963 | 9.32495871 | 8.681620276 | 8.628587883 |
| ENSG00000143569 | UBAP2L | -0.610243518 | 0.054358412 | -0.630170278 | 0.043410967 | -1.240413796 | 11.17074085 | 11.26554928 | 11.65357938 | 10.73124123 | 10.805643 | 10.72402432 | 10.77364464 |
| ENSG00000062598 | ELMO2 | -0.117790785 | 0.902927943 | -1.122848425 | 0.156984236 | -1.24063921 | 9.27634997 | 9.318399851 | 10.95142175 | 9.954406146 | 9.96022311 | 8.996529805 | 8.918588063 |
| ENSG00000161057 | PSMC2 | -0.163562982 | 0.876384005 | -1.07813224 | 0.220472648 | -1.241695222 | 11.09707093 | 9.988044813 | 8.784770914 | 10.09339983 | 10.0755276 | 9.10597186 | 9.240151898 |
| ENSG00000183513 | COA5 | -0.556485408 | 0.476505377 | -0.685699244 | 0.359691834 | -1.242184653 | 8.367060731 | 7.819099893 | 6.629488387 | 7.218576229 | 7.23826702 | 7.231159579 | 6.96179415 |
| ENSG00000170584 | NUDCD2 | -0.861353401 | 0.247792329 | -0.381528184 | 0.628768646 | -1.242881585 | 8.340423424 | 6.666836837 | 7.97167868 | 7.057964187 | 6.8715633 | 7.446000032 | 7.422862719 |
| ENSG00000171530 | TBCA | 0.714065427 | 0.119484573 | -1.957436161 | 1.67E-06 | -1.243370735 | 9.128497779 | 8.276714465 | 8.922968322 | 9.557406408 | 9.48558803 | 6.814608286 | 6.965711302 |
| ENSG00000198258 | UBL5 | -0.511104089 | 0.50195138 | -0.732565587 | 0.308715893 | -1.243669676 | 10.1076228 | 8.505280662 | 9.06622548 | 8.883449599 | 8.86223882 | 8.563064723 | 8.738561653 |
| ENSG00000170619 | COMMD5 | -0.501499429 | 0.517483563 | -0.742427492 | 0.308005991 | -1.243926922 | 8.87372952 | 7.315436877 | 8.865353883 | 8.056427969 | 7.96222667 | 7.742013931 | 7.803803816 |
| ENSG00000122705 | CLTA | -0.411708981 | 0.534043967 | -0.834256808 | 0.170526163 | -1.245965789 | 11.43901855 | 10.17128279 | 10.48092713 | 10.37578145 | 10.4062943 | 9.8349648 | 10.09355512 |
| ENSG00000142534 | RPS11 | -1.050142282 | 0.001272822 | -0.196791982 | 0.608318434 | -1.246934263 | 12.5084683 | 12.0319538 | 12.60800691 | 11.32032301 | 11.3871132 | 12.17607424 | 12.23624376 |
| ENSG00000182871 | COL18A1 | -2.163111779 | 4.34E-11 | 0.914191499 | 0.009411948 | -1.24892028 | 11.57946871 | 11.28620006 | 11.86677216 | 9.431062134 | 9.44728702 | 12.33642988 | 12.66340894 |
| ENSG00000139410 | SDSL | -0.417631209 | 0.694698997 | -0.8326945 | 0.396311091 | -1.250325709 | 4.467311984 | 6.535001748 | 6.882775709 | 5.920568914 | 5.85535975 | 5.390603804 | 5.611957759 |
| ENSG00000184840 | TMED9 | -0.812155317 | 0.45032434 | -0.438352168 | 0.693056959 | -1.250507485 | 12.79016004 | 9.883504052 | 12.06958695 | 11.14910699 | 11.2355938 | 11.55082553 | 11.58148201 |
| ENSG00000149136 | SSRP1 | -0.758585444 | 0.227473008 | -0.492313634 | 0.444980407 | -1.250899078 | 10.59268348 | 10.52407288 | 9.301632639 | 9.520570052 | 9.46656462 | 9.634053733 | 9.873532744 |
| ENSG00000196562 | SULF2 | -2.107320846 | 8.35E-06 | 0.855476969 | 0.094172652 | -1.251843877 | 8.564112515 | 8.777382925 | 8.904018013 | 6.652857793 | 6.72891807 | 9.067857443 | 9.995179393 |
| ENSG00000012660 | ELOVL5 | -0.818189104 | 0.00026044 | -0.43432858 | 0.06810855 | -1.252517684 | 10.787681 | 10.79656548 | 10.85604368 | 10.07142123 | 9.92005068 | 10.28628591 | 10.46857168 |
| ENSG00000236443 | | 6.92426161 | 1.47E-05 | -8.177102486 | 0.000313985 | -1.252840876 | 2 | 4.710564154 | 2 | 9.914221922 | 9.97024066 | 2 | 2 |
| ENSG00000176293 | ZNF135 | -2.177791623 | 0.234266721 | 0.922612291 | 0.63300769 | -1.255179332 | 2 | 6.728495673 | 4.058998049 | 3.406930997 | 3.86943969 | 6.428526856 | 6.120613345 |
| ENSG00000136840 | ST6GALNAC4 | -1.072634979 | 0.007453569 | -0.183628146 | 0.695554779 | -1.256263125 | 9.189468652 | 8.732943944 | 8.721235202 | 7.893113669 | 7.77859536 | 8.463547972 | 8.92815536 |
| ENSG00000198301 | SDAD1 | 0.04320208 | 0.964905581 | -1.300226088 | 0.108113626 | -1.257024008 | 9.435834703 | 9.079034257 | 7.342949392 | 8.887228971 | 8.92538942 | 7.726402307 | 7.420010428 |
| ENSG00000103152 | MPG | -0.134148305 | 0.861594641 | -1.123753562 | 0.072934705 | -1.257901867 | 10.41260941 | 9.203675681 | 9.165043284 | 9.511620236 | 9.65150894 | 8.597799214 | 8.604294318 |
| ENSG00000177570 | SAMD12 | -6.314953619 | 0.018254181 | 5.055363165 | 0.014763919 | -1.259590454 | 2 | 2 | 4.523675391 | 2 | 2 | 7.550870243 | 7.783974227 |
| ENSG00000115350 | POLE4 | -0.631675074 | 0.66635587 | -0.628356846 | 0.664743846 | -1.26003192 | 8.19931886 | 7.874700259 | 4.058998049 | 6.89733479 | 6.89154576 | 6.605619634 | 7.140495192 |
| ENSG00000125970 | RALY | -1.140700254 | 0.001373482 | -0.121985554 | 0.781250208 | -1.262685808 | 11.0444064 | 10.59199661 | 10.88048583 | 9.707456306 | 9.71806384 | 10.45472238 | 10.95773505 |
| ENSG00000132635 | PCED1A | -0.995816471 | 0.645580573 | -0.267691322 | 0.908686871 | -1.263507793 | 9.957954117 | 8.376385542 | 2 | 7.753776273 | 7.86451142 | 8.554248841 | 8.501461594 |
| ENSG00000052802 | MSMO1 | -0.589940449 | 0.40395265 | -0.674786311 | 0.323536602 | -1.26472676 | 10.59268348 | 11.09168706 | 11.95227171 | 10.73718662 | 10.7317082 | 10.25846039 | 10.95736623 |
| ENSG00000125944 | HNRNPR | -0.679071953 | 0.426810018 | -0.586089117 | 0.489050844 | -1.265161069 | 11.18578229 | 10.72273565 | 9.301632639 | 9.922226014 | 9.92126619 | 9.564740792 | 10.35667886 |
| ENSG00000125703 | ATG4C | -0.808719727 | 0.531143362 | -0.456761313 | 0.730288533 | -1.26548104 | 7.096806373 | 4.11352228 | 7.225742779 | 5.94010643 | 5.8245091 | 6.514451546 | 5.8339184 |
| ENSG00000134265 | NAPG | -0.902204743 | 0.2695484 | -0.363557123 | 0.677816167 | -1.265761867 | 9.829762239 | 7.847167909 | 8.941672936 | 8.23312837 | 8.14812525 | 8.784976076 | 8.659751394 |
| ENSG00000008853 | RHOBTB2 | -1.643593783 | 0.000799417 | 0.377695109 | 0.501916413 | -1.265898674 | 8.631580145 | 8.862353853 | 9.658943166 | 7.504305792 | 7.48837134 | 9.581682124 | 9.399228411 |
| ENSG00000172775 | FAM192A | -0.52491216 | 0.490581765 | -0.741048207 | 0.303875747 | -1.265960367 | 8.87372952 | 9.377333006 | 10.44859742 | 9.213506984 | 9.18117744 | 8.79381476 | 9.150074181 |
| ENSG00000169976 | SF3B5 | -1.249628785 | 0.107655331 | -0.016603094 | 0.985856566 | -1.266231878 | 10.71177538 | 9.102512015 | 9.196533591 | 8.550024073 | 8.70739158 | 9.676249936 | 10.01492601 |
| ENSG00000168476 | REEP4 | -0.981920557 | 0.642855534 | -0.28447624 | 0.900496697 | -1.266396797 | 9.448362284 | 7.572420273 | 2 | 7.265793319 | 7.24216308 | 7.988280194 | 7.883613091 |
| ENSG00000253304 | TMEM200B | -0.649873396 | 0.398651947 | -0.617174701 | 0.415812209 | -1.267048096 | 9.143983763 | 7.790474954 | 7.777116083 | 7.723090567 | 7.77052502 | 7.991357103 | 7.529919928 |
| ENSG00000180957 | PITPNB | -0.800552055 | 0.018276173 | -0.467707399 | 0.186976308 | -1.268259455 | 10.33377634 | 10.019219 | 9.767769415 | 9.260887574 | 9.26398924 | 9.499242667 | 9.681766233 |
| ENSG00000087157 | PGS1 | -0.871035955 | 0.11367876 | -0.39735864 | 0.495466824 | -1.268394595 | 8.042902253 | 8.777382925 | 9.242516611 | 7.905589775 | 7.909304 | 8.375618113 | 8.373144057 |
| ENSG00000146918 | NCAPG2 | -1.317319372 | 0.310604101 | 0.048669962 | 0.973559952 | -1.26864941 | 4.00722531 | 7.051611315 | 7.858115942 | 5.818703821 | 5.64780787 | 6.948868859 | 7.114266974 |
| ENSG00000094804 | CDC6 | -0.893111324 | 0.528693205 | -0.377976936 | 0.798498793 | -1.27108826 | 5.532353208 | 6.666836837 | 3.369317972 | 5.076005082 | 4.79082474 | 4.771475226 | 5.8339184 |
| ENSG00000153902 | LGI4 | -2.435450369 | 0.179526399 | 1.164155868 | 0.52507775 | -1.271294501 | 4.00722531 | 4.93638933 | 2 | 3.023133209 | 2.2680111 | 5.489669427 | 4.522269241 |
| ENSG00000182552 | RWDD4 | -0.417995771 | 0.751812007 | -0.85606099 | 0.486233434 | -1.274056761 | 9.262228249 | 6.78762695 | 6.43197674 | 7.691737974 | 7.6378145 | 7.31213785 | 7.157719456 |
| ENSG00000198951 | NAGA | -1.444442656 | 0.015168155 | 0.170092649 | 0.812274339 | -1.274350007 | 9.097018025 | 10.06773758 | 8.763901704 | 8.001694993 | 7.98797735 | 9.483758913 | 9.696573548 |
| ENSG00000132434 | LANCL2 | -0.612725678 | 0.702917775 | -0.661993304 | 0.674981326 | -1.274718982 | 4.00722531 | 8.687092496 | 8.677268317 | 7.370304568 | 7.66419009 | 7.25183424 | 7.670954172 |
| ENSG00000187531 | SIRT7 | -0.801713481 | 0.374121952 | -0.47327325 | 0.610378131 | -1.274986731 | 8.540903468 | 6.389896633 | 8.076950877 | 7.080147179 | 7.1744553 | 7.308441828 | 7.575491915 |
| ENSG00000077380 | DYNC1I2 | -0.078164343 | 0.943520433 | -1.197581085 | 0.197561623 | -1.275745428 | 10.26462289 | 10.54992047 | 8.042706615 | 9.911131561 | 9.86046766 | 8.707577112 | 8.837005439 |
| ENSG00000177879 | AP3S1 | -0.595367349 | 0.350878999 | -0.6811832 | 0.270175158 | -1.276550548 | 10.46286899 | 9.470501606 | 9.212024683 | 9.290680758 | 9.16279627 | 9.173900195 | 9.111258115 |
| ENSG00000145348 | TBCK | -0.621014685 | 0.426810018 | -0.656106944 | 0.387074442 | -1.277121629 | 7.709880939 | 9.018611864 | 8.960138144 | 7.966401501 | 8.15434112 | 7.461614824 | 8.435201779 |
| ENSG00000197912 | SPG7 | -0.893655615 | 0.105206276 | -0.383484141 | 0.515470896 | -1.277139756 | 10.84091191 | 9.648522413 | 10.26770388 | 9.357592295 | 9.52049436 | 9.930233463 | 9.967609306 |
| ENSG00000243709 | LEFTY1 | 0.637088348 | 0.832852241 | -1.916287705 | NA | -1.279199356 | 3.327711234 | 2 | 2 | 3.023133209 | 2.59489879 | 2.183373728 | 2.172608095 |
| ENSG00000144231 | POLR2D | -0.970429022 | 0.002036692 | -0.311235857 | 0.371207868 | -1.281664878 | 9.174465797 | 8.88960062 | 8.677268317 | 7.990026167 | 7.94798467 | 8.619836896 | 8.619914108 |
| ENSG00000135070 | ISCA1 | -0.682192379 | 0.284613367 | -0.600352267 | 0.34281457 | -1.282544646 | 9.710019321 | 8.414402626 | 8.784770914 | 8.43768081 | 8.35656898 | 8.471282103 | 8.485861839 |
| ENSG00000138587 | MNS1 | -2.27056554 | 0.313384316 | 0.987578811 | 0.670910636 | -1.28298673 | 2 | 5.131603625 | 2 | 2.784612485 | 2.49394783 | 4.588526421 | 4.766376438 |
| ENSG00000167004 | PDIA3 | 0.237438585 | 0.699202151 | -1.520696938 | 0.00262317 | -1.283258353 | 13.1430848 | 12.32301536 | 12.37843207 | 12.88916971 | 12.9165825 | 10.74479855 | 11.46056057 |
| ENSG00000182220 | ATP6AP2 | -0.212260129 | 0.652346084 | -1.072729571 | 0.006872527 | -1.2849897 | 11.9089726 | 11.13786633 | 11.34128524 | 11.30297958 | 11.2729696 | 10.44029952 | 10.41854076 |
| ENSG00000102178 | UBL4A | -0.84219375 | 0.153980097 | -0.442886933 | 0.473146336 | -1.285080683 | 8.586954094 | 9.799806309 | 9.61300511 | 8.551612617 | 8.62623826 | 9.135476483 | 8.817088095 |
| ENSG00000089220 | PEBP1 | -1.074419213 | 0.012676892 | -0.21108279 | 0.674981326 | -1.285502002 | 12.37539409 | 11.95020188 | 11.6648807 | 10.9673516 | 10.9400378 | 11.51999629 | 12.0611512 |
| ENSG00000132879 | FBXO44 | -0.938157072 | 0.586967447 | -0.350715008 | 0.84812366 | -1.288872079 | 9.657036304 | 7.538394143 | 4.058998049 | 7.536750104 | 7.40393901 | 8.126237806 | 7.966097995 |
| ENSG00000141456 | PELP1 | -1.39756533 | 0.004858833 | 0.107548874 | 0.861820805 | -1.290016456 | 8.87372952 | 9.28800521 | 9.963514551 | 7.994705029 | 8.12299044 | 9.584486439 | 9.517360075 |
| ENSG00000095485 | CWF19L1 | -0.47484668 | 0.565600882 | -0.815636972 | 0.290853649 | -1.290483652 | 8.998234871 | 7.60566235 | 7.398165132 | 7.700356475 | 7.73502229 | 7.358152527 | 7.409982764 |
| ENSG00000198131 | ZNF544 | -0.639659198 | 0.48167816 | -0.652572391 | 0.462949352 | -1.292231589 | 8.946185348 | 7.189546432 | 9.196533591 | 8.004017475 | 8.0713677 | 7.680435163 | 8.303661547 |
| ENSG00000099960 | SLC7A4 | -4.714923371 | 0.18460402 | 3.422223338 | 0.26685414 | -1.292700033 | 2 | 3.076665209 | 2 | 2 | 2 | 4.644324387 | 4.673629022 |
| ENSG00000184924 | PTRHD1 | -0.874313282 | 0.358348477 | -0.419297372 | 0.675137901 | -1.293610653 | 8.340423424 | 7.274681896 | 6.073378139 | 6.70517998 | 6.61334686 | 6.807643042 | 7.342293759 |
| ENSG00000082515 | MRPL22 | -0.257195395 | 0.734460813 | -1.037921361 | 0.114849347 | -1.295116756 | 7.384433967 | 8.687092496 | 7.502634626 | 7.751013478 | 7.7377845 | 7.016881172 | 6.947999751 |
| ENSG00000111696 | NT5DC3 | -1.851588158 | 0.056230678 | 0.556211282 | 0.604593035 | -1.295376876 | 6.366709671 | 8.433041974 | 9.180874355 | 6.658765962 | 6.507263 | 8.920684672 | 8.965313405 |
| ENSG00000164265 | SCGB3A2 | -4.229205342 | 0.146700672 | 2.931093823 | 0.205667088 | -1.298111519 | 4.00722531 | 2 | 2 | 2.141737246 | 2 | 4.920961476 | 5.191251321 |
| ENSG00000151445 | VIPAS39 | -0.497351743 | 0.821777402 | -0.800862822 | 0.701757981 | -1.298214564 | 9.497412313 | 8.414402626 | 2 | 7.93509946 | 8.03365075 | 7.675661224 | 7.698285408 |
| ENSG00000171490 | RSL1D1 | -0.592088588 | 0.07325688 | -0.706578269 | 0.027268332 | -1.298666856 | 11.04852648 | 10.50217373 | 10.81097638 | 10.22664935 | 10.1974239 | 10.12211394 | 10.07350205 |
| ENSG00000005022 | SLC25A5 | -1.11220466 | 0.000728334 | -0.18849253 | 0.629414332 | -1.300697189 | 10.65885801 | 10.24169813 | 10.22252928 | 9.298270478 | 9.26303083 | 10.02917428 | 10.35294498 |
| ENSG00000113594 | LIFR | -1.194965702 | 0.01989306 | -0.109672298 | 0.859331521 | -1.304638 | 6.89194353 | 7.901717006 | 7.342949392 | 6.328984825 | 6.26836889 | 7.35218116 | 7.337766965 |
| ENSG00000136875 | PRPF4 | -0.603775814 | 0.163887287 | -0.701113654 | 0.093533047 | -1.304889468 | 8.367060731 | 8.505280662 | 9.083172337 | 8.040677544 | 8.12931531 | 8.030023089 | 7.94839122 |
| ENSG00000173153 | ESRRA | -0.610315052 | 0.491805453 | -0.696387182 | 0.418338064 | -1.306702235 | 9.509418609 | 8.356994601 | 7.552162999 | 8.096156705 | 8.08878133 | 8.247007147 | 7.720479276 |
| ENSG00000277462 | ZNF670 | -0.243709916 | 0.857421507 | -1.065214007 | 0.3650389 | -1.308923923 | 4.00722531 | 4.710564154 | 6.534109959 | 5.258265913 | 5.24368216 | 4.620674722 | 4.456980017 |
| ENSG00000163812 | ZDHHC3 | -0.830420795 | 0.190607653 | -0.479726951 | 0.465741681 | -1.310147746 | 10.9638027 | 9.664384941 | 9.981704053 | 9.466861018 | 9.50517467 | 9.700423741 | 9.958300939 |
| ENSG00000170633 | RNF34 | -0.573360651 | 0.751812007 | -0.737374828 | 0.676516223 | -1.310735478 | 4.00722531 | 8.124485672 | 9.926432745 | 8.111316053 | 8.19912382 | 8.032265607 | 7.955303035 |
| ENSG00000072135 | PTPN18 | -1.154923281 | 0.349969585 | -0.158500863 | 0.910302233 | -1.313424144 | 5.70932728 | 8.414402626 | 9.344432377 | 7.238438152 | 7.37229174 | 8.090815298 | 8.456236734 |
| ENSG00000092203 | TOX4 | -0.977112121 | 0.06635276 | -0.338413585 | 0.561013956 | -1.315525706 | 9.699577239 | 10.36815316 | 10.88048583 | 9.41544358 | 9.42689279 | 10.06648818 | 10.04717102 |
| ENSG00000101916 | TLR8 | -3.753023129 | 0.531143362 | 2.434989022 | NA | -1.318034107 | 2 | 2 | 2 | 2 | 2 | 3.679182183 | 2.632958241 |
| ENSG00000164114 | MAP9 | -0.392110266 | 0.568384743 | -0.925943256 | 0.136351095 | -1.318053522 | 8.285626337 | 7.503546079 | 8.884815469 | 7.971157485 | 7.90193448 | 7.540404604 | 7.274433587 |
| ENSG00000147669 | POLR2K | -0.751971497 | 0.315751098 | -0.569520156 | 0.451396209 | -1.321491653 | 10.11549941 | 8.981105197 | 8.486635645 | 8.595396228 | 8.63367258 | 8.889688585 | 8.694392964 |
| ENSG00000132670 | PTPRA | -0.220863738 | 0.794969986 | -1.10072301 | 0.127918786 | -1.321586748 | 9.646202137 | 10.69982928 | 11.46604358 | 10.54043168 | 10.5879489 | 9.613232802 | 9.758568089 |
| ENSG00000164331 | ANKRA2 | -0.626260046 | 0.335559831 | -0.695385451 | 0.269861386 | -1.321645497 | 8.39321512 | 8.655694736 | 7.502634626 | 7.691737974 | 7.61094772 | 7.874567387 | 7.219718817 |
| ENSG00000204237 | OXLD1 | -0.792549038 | 0.235847119 | -0.529336823 | 0.437014882 | -1.321885861 | 8.19931886 | 6.899080544 | 8.076950877 | 6.970413838 | 7.1204461 | 7.47706242 | 7.105417192 |
| ENSG00000183765 | CHEK2 | 0.761990294 | 0.668927862 | -2.085063831 | 0.191114239 | -1.323073537 | 8.816894989 | 6.464272742 | 3.369317972 | 8.213192703 | 8.31463236 | 5.545162502 | 5.513708555 |
| ENSG00000186806 | VSIG10L | -0.027965848 | 0.989663573 | -1.297571982 | 0.491100919 | -1.32553783 | 3.327711234 | 8.169643011 | 4.058998049 | 6.658765962 | 6.73446058 | 5.557671049 | 5.448021784 |
| ENSG00000139697 | SBNO1 | -0.566935356 | 0.205078218 | -0.759074013 | 0.07517382 | -1.326009369 | 10.00958825 | 10.019219 | 10.62096771 | 9.652999939 | 9.70612498 | 9.68720202 | 9.258230494 |
| ENSG00000152700 | SAR1B | -0.756204641 | 0.154010606 | -0.570300222 | 0.286633227 | -1.326504863 | 10.58135316 | 9.591588447 | 10.60345421 | 9.598713994 | 9.54829592 | 9.721422299 | 9.795075587 |
| ENSG00000101452 | DHX35 | -0.89625539 | 0.565047493 | -0.431302369 | 0.792001716 | -1.327557758 | 3.327711234 | 7.467835385 | 8.110401118 | 6.283823264 | 6.52015598 | 6.944112782 | 6.755525668 |
| ENSG00000172819 | RARG | -0.529522998 | 0.428683492 | -0.798079675 | 0.204147015 | -1.327602672 | 9.219014269 | 9.930175364 | 10.65536953 | 9.477763976 | 9.56860713 | 9.220861732 | 9.292162928 |
| ENSG00000105186 | ANKRD27 | -0.628122517 | 0.212797514 | -0.70082079 | 0.15123158 | -1.328943307 | 9.579430841 | 8.687092496 | 9.049077186 | 8.537252353 | 8.51470562 | 8.627767864 | 8.256474583 |
| ENSG00000127580 | WDR24 | -0.724908677 | 0.728237469 | -0.60446095 | 0.773321468 | -1.329369626 | 2 | 7.431218269 | 8.941672936 | 7.165617905 | 7.01053089 | 7.403899888 | 6.979338179 |
| ENSG00000204514 | ZNF814 | -0.93519648 | 0.200334149 | -0.394779781 | 0.610752069 | -1.329976261 | 6.563806279 | 7.954278496 | 7.398165132 | 6.470063213 | 6.56439512 | 7.360534171 | 6.622902297 |
| ENSG00000168528 | SERINC2 | -0.157578465 | 0.88438939 | -1.172409374 | 0.19277085 | -1.329987839 | 10.56993315 | 8.876041558 | 8.511888463 | 9.404936476 | 9.51083771 | 8.195243072 | 8.670591511 |
| ENSG00000185946 | RNPC3 | 0.239382337 | 0.881720015 | -1.569991233 | 0.245468671 | -1.330608897 | 7.384433967 | 7.503546079 | 3.369317972 | 7.093295078 | 7.18663697 | 5.22272385 | 5.611957759 |
| ENSG00000054179 | ENTPD2 | -1.269030326 | 0.690257999 | -0.062199576 | NA | -1.331229902 | 2 | 2 | 3.369317972 | 2.3892346 | 2.14022021 | 2.68698273 | 2.421067459 |
| ENSG00000143851 | PTPN7 | -3.753023295 | 0.531143362 | 2.421672991 | NA | -1.331350304 | 2 | 2 | 2 | 2 | 2 | 3.796405078 | 2.326754077 |
| ENSG00000104731 | KLHDC4 | -1.295929675 | 0.534439893 | -0.036624464 | 0.987416408 | -1.332554139 | 9.015175615 | 6.389896633 | 2 | 6.408278948 | 6.39275508 | 7.618102344 | 7.620910859 |
| ENSG00000132823 | OSER1 | -0.397262233 | 0.790951603 | -0.936139736 | 0.495682443 | -1.333401969 | 10.59268348 | 8.820493869 | 6.43197674 | 9.067921572 | 9.02033034 | 8.466314923 | 8.553143267 |
| ENSG00000235708 | | 3.857403838 | 0.496953626 | -5.191090708 | 0.364514502 | -1.333686869 | 2 | 2 | 2 | 3.95996233 | 4.45712836 | 2 | 2 |
| | C15ORF52 | 6.757112528 | 2.69E-09 | -8.090970261 | 3.49E-05 | -1.333857732 | 4.467311984 | 2 | 3.369317972 | 9.677214349 | 9.70330142 | 2 | 2 |
| ENSG00000010244 | ZNF207 | -0.29019246 | 0.619015165 | -1.043678015 | 0.039749923 | -1.333870475 | 11.55939759 | 10.54992047 | 10.77492406 | 10.69648168 | 10.7797423 | 9.842864719 | 10.11906036 |

| | | | | | | | | | | | | | |
|---|---|---|---|---|---|---|---|---|---|---|---|---|---|
| ENSG00000172292 | CERS6 | -1.129786443 | 0.580580155 | -0.204186762 | 0.927897448 | -1.333973204 | 8.313285031 | 9.338312924 | 2 | 7.366699513 | 7.0774796 | 8.263008721 | 7.998006535 |
| ENSG00000149257 | SERPINH1 | -1.197349674 | 0.048926942 | -0.137559493 | 0.850018602 | -1.334909167 | 14.21556964 | 13.04192929 | 13.01031565 | 12.31942734 | 12.3646385 | 13.33642019 | 13.46361928 |
| ENSG00000168995 | SIGLEC7 | -3.753023353 | 0.531143362 | 2.417054371 | NA | -1.335968981 | 2 | 2 | 2 | 2 | 2 | 3.500668998 | 2.918126211 |
| ENSG00000106511 | MEOX2 | -3.753023356 | 0.531143362 | 2.416770992 | NA | -1.336252364 | 2 | 2 | 2 | 2 | 2 | 2.560105256 | 3.698365557 |
| ENSG00000136213 | CHST12 | -0.96881867 | 0.686456446 | -0.367501968 | 0.884851957 | -1.336320637 | 10.59268348 | 7.35507212 | 2 | 8.166267464 | 8.22691474 | 8.798213877 | 8.784747976 |
| ENSG00000168427 | KLHL30 | -6.103015833 | 0.006359008 | 4.765727346 | 0.001345688 | -1.337288487 | 3.327711234 | 2 | 5.39244166 | 2 | 2.14022021 | 8.737205015 | 8.44505627 |
| ENSG00000143552 | NUP210L | 0.311380123 | 0.927479541 | -1.650689564 | NA | -1.339309442 | 2 | 2 | 3.369317972 | 2.600425781 | 2.77779907 | 2.34604626 | 2.059860178 |
| ENSG00000168374 | ARF4 | -0.711093699 | 0.138848227 | -0.63010723 | 0.187257885 | -1.341200929 | 13.00468149 | 12.17723115 | 12.27897381 | 11.84965902 | 11.8008967 | 11.74630693 | 12.05047073 |
| ENSG00000189369 | GSPT2 | -0.584219674 | 0.313991908 | -0.757985921 | 0.170908786 | -1.342205595 | 8.444143041 | 7.73146271 | 8.904018013 | 7.829061569 | 7.87709394 | 7.623070264 | 7.740051619 |
| ENSG00000118816 | CCNI | -1.489329495 | 6.29E-05 | 0.146842973 | 0.750755237 | -1.342486521 | 11.91800836 | 11.58596781 | 12.33487604 | 10.46657187 | 10.5157558 | 12.21344546 | 12.03038993 |
| ENSG00000173674 | EIF1AX | 0.201137383 | 0.592806105 | -1.544819259 | 3.41E-07 | -1.343681876 | 9.189468652 | 9.608085847 | 9.301632639 | 9.650778342 | 9.50922197 | 7.95714469 | 7.741194706 |
| ENSG00000071282 | LMCD1 | -1.910770364 | 3.75E-06 | 0.56549001 | 0.22056224 | -1.345280354 | 8.75772933 | 9.114109157 | 9.491321578 | 7.341208691 | 7.19067478 | 9.471691847 | 9.919950354 |
| ENSG00000159761 | C16orf86 | -1.155511044 | 0.525813654 | -0.190008756 | 0.925215975 | -1.3455198 | 6.653098118 | 4.442710129 | 2 | 4.415698455 | 4.37434574 | 5.196301326 | 5.243813244 |
| ENSG00000169045 | HNRNPH1 | -0.210304565 | 0.695936302 | -1.135357644 | 0.01186525 | -1.345662209 | 11.51244431 | 10.69982928 | 10.71094342 | 10.86022322 | 10.7716784 | 9.816867623 | 9.96809756 |
| ENSG00000180879 | SSR4 | 0.060099163 | 0.930223809 | -1.405854382 | 0.008676361 | -1.34575522 | 11.48534568 | 10.60438382 | 10.31867969 | 10.95779865 | 10.9454246 | 9.404033465 | 9.572665028 |
| ENSG00000151575 | TEX9 | -0.032780982 | 0.990154914 | -1.313383998 | 0.576534502 | -1.34616498 | 2 | 5.457120953 | 2 | 4.070463721 | 4.1930041 | 3.355392153 | 3.15612297 |
| ENSG00000101407 | TTI1 | -0.790665793 | 0.629335817 | -0.556755248 | 0.736888343 | -1.347421041 | 4.00722531 | 9.00621767 | 9.180874355 | 7.683067677 | 7.8208909 | 8.04860508 | 7.915353355 |
| ENSG00000163159 | VPS72 | -1.345433715 | 0.031713694 | -0.002754463 | 0.997599173 | -1.348188178 | 9.761125267 | 8.255926468 | 9.132850258 | 7.831679003 | 7.85181819 | 9.154473165 | 9.183675168 |
| ENSG00000125170 | DOK4 | -0.84824569 | 0.02494348 | -0.499999872 | 0.205129503 | -1.348245562 | 9.097018025 | 9.687856435 | 9.735974718 | 8.682071821 | 8.70598173 | 8.967137431 | 9.108160402 |
| ENSG00000132017 | DCAF15 | -0.545980619 | 0.427719039 | -0.803827796 | 0.214346841 | -1.349808414 | 7.975256505 | 7.638155693 | 8.978369997 | 7.734324667 | 7.80517502 | 7.446000032 | 7.585709303 |
| ENSG00000164638 | SLC29A4 | -2.19838324 | 0.154820857 | 0.848179658 | 0.608615553 | -1.350203582 | 5.70932728 | 8.101365653 | 3.369317972 | 4.946755474 | 4.63360314 | 7.777203086 | 7.511280276 |
| ENSG00000147121 | KRBOX4 | -0.255405013 | 0.711497595 | -1.096616753 | 0.064293553 | -1.352021765 | 7.031698319 | 7.002537973 | 8.042706615 | 7.114945424 | 7.2537885 | 6.216130818 | 6.520103598 |
| ENSG00000100483 | VCPKMT | -0.782080185 | 0.567927475 | -0.570127742 | 0.681431908 | -1.352207927 | 4.00722531 | 6.535001748 | 7.646393253 | 5.870535235 | 6.00960265 | 6.475428841 | 5.703940561 |
| ENSG00000145022 | TCTA | -0.681381135 | 0.463231524 | -0.670834195 | 0.460415275 | -1.35221533 | 9.957954117 | 9.267380431 | 8.236946975 | 8.579911316 | 8.69891174 | 7.795801481 | 9.184515751 |
| ENSG00000167995 | BEST1 | -5.447844384 | 0.060484839 | 4.095447447 | 0.078793483 | -1.352396937 | 2 | 3.686303414 | 2 | 2 | 2 | 6.134985075 | 5.6562603 |
| ENSG00000213413 | PVRIG | 1.199646954 | 0.594033397 | -2.553241732 | 0.220592048 | -1.353594778 | 2 | 5.303523747 | 2 | 5.110902178 | 4.93258395 | 2.592885947 | 2.632958241 |
| ENSG00000070961 | ATP2B1 | 0.849295143 | 0.258493876 | -2.20534949 | 0.00097064 | -1.356054347 | 9.820154628 | 11.631062 | 11.16147894 | 11.90445538 | 11.8959261 | 8.858011903 | 8.855068552 |
| ENSG00000113971 | NPHP3 | 0.901872832 | 0.366585534 | -2.258151107 | 0.011804798 | -1.356278275 | 6.816637977 | 9.00621767 | 7.163363831 | 8.915879855 | 8.89354047 | 5.935372196 | 5.73181021 |
| ENSG00000141380 | SS18 | -0.351722409 | 0.588327109 | -1.004943761 | 0.081826473 | -1.35666617 | 10.41898845 | 10.92721483 | 9.589475645 | 10.06420548 | 10.0623662 | 9.357098224 | 9.467122411 |
| ENSG00000198093 | ZNF649 | -0.720133918 | 0.59691102 | -0.636600467 | 0.638247848 | -1.356734385 | 8.16936288 | 6.04685525 | 4.874587708 | 6.358325776 | 6.24526024 | 6.57313027 | 6.162172032 |
| ENSG00000155256 | ZFYVE27 | -0.259602376 | 0.795427588 | -1.097196177 | 0.205667088 | -1.356798553 | 10.30651143 | 8.395519306 | 8.677268317 | 9.135152476 | 9.12531607 | 8.515767839 | 8.047821014 |
| ENSG00000178307 | TMEM11 | -1.094985281 | 3.05E-05 | -0.262572416 | 0.37875047 | -1.357557697 | 9.247966931 | 9.067150531 | 9.049077186 | 8.115618198 | 7.95986274 | 8.899958346 | 8.824117457 |
| ENSG00000144713 | RPL32 | -1.157783461 | 0.168223251 | -0.202174821 | 0.833553126 | -1.359958282 | 12.97143233 | 11.07398149 | 11.39656351 | 10.92967084 | 10.8937291 | 11.79433308 | 11.93421333 |
| ENSG00000136819 | C9orf78 | -0.500868214 | 0.362508916 | -0.859655308 | 0.09315428 | -1.360523522 | 10.22148166 | 9.367676479 | 9.242516611 | 9.147810675 | 9.21229508 | 8.79205135 | 8.856124078 |
| ENSG00000144182 | LIPT1 | -0.698661558 | 0.676600545 | -0.663404235 | 0.687792014 | -1.362065793 | 3.327711234 | 6.78762695 | 4.058998049 | 5.040242897 | 4.79082474 | 5.108342547 | 4.77533149 |
| ENSG00000150753 | CCT5 | -0.876278972 | 0.000559668 | -0.488974869 | 0.068678166 | -1.365253841 | 11.13243651 | 11.19085087 | 11.2124308 | 10.33669748 | 10.2722069 | 10.51007546 | 10.85192199 |
| | APITD1-COR | 4.786738953 | 0.038248139 | -6.152940704 | 0.029610048 | -1.366201751 | 3.327711234 | 2 | 2 | 5.590471342 | 6.11534392 | 2 | 2 |
| ENSG00000148730 | EIF4EBP2 | -1.005701782 | 0.113324776 | -0.361215687 | 0.600600207 | -1.366917469 | 10.19941743 | 10.95971654 | 11.51370025 | 9.98823217 | 9.98077219 | 10.25141933 | 10.92466139 |
| ENSG00000173588 | CEP83 | -0.718490093 | 0.43340631 | -0.648721108 | 0.47051548 | -1.3672112 | 6.257048221 | 6.228547981 | 4.874587708 | 5.590471342 | 4.85329164 | 5.528313928 | 5.101530266 |
| ENSG00000198825 | INPP5F | -1.100725665 | 0.201926325 | -0.267641644 | 0.781305953 | -1.368367309 | 8.16936288 | 10.28338212 | 9.196533591 | 8.374462892 | 8.37974396 | 9.422673149 | 8.942636839 |
| ENSG00000163904 | SENP2 | -0.786173651 | 0.231765386 | -0.582662911 | 0.379991231 | -1.368836562 | 8.609439662 | 9.308339287 | 10.08627303 | 8.684969711 | 8.67028013 | 8.851676897 | 8.906411656 |
| ENSG00000181222 | POLR2A | -0.721324321 | 0.290454869 | -0.647694924 | 0.338044189 | -1.369019245 | 11.15554094 | 12.70874416 | 12.17214792 | 11.42152442 | 11.4291609 | 11.40918076 | 11.58331566 |
| ENSG00000117450 | PRDX1 | 0.221158612 | 0.693063949 | -1.590613978 | 0.000467379 | -1.369455366 | 12.03890377 | 11.07694755 | 11.57488832 | 11.83750166 | 11.8354782 | 9.820764745 | 10.21041545 |
| ENSG00000198298 | ZNF485 | -1.22731531 | 0.624769744 | -0.144134019 | 0.957898398 | -1.371449329 | 2 | 5.838937808 | 2 | 3.663521627 | 3.45167086 | 4.596630906 | 4.077147165 |
| ENSG00000148337 | CIZ1 | -0.908724213 | 0.126427717 | -0.465406477 | 0.456352327 | -1.37413069 | 11.25516263 | 10.20962523 | 10.09465441 | 9.734282329 | 9.69337522 | 10.1959872 | 10.11377463 |
| ENSG00000183943 | PRKX | -1.317039212 | 0.385680172 | -0.058183581 | 0.97269175 | -1.375222793 | 4.00722531 | 7.393647515 | 8.536706852 | 6.18083559 | 6.33555364 | 7.992125306 | 6.659638722 |
| ENSG00000127554 | GFER | -0.436483398 | 0.609544685 | -0.938771847 | 0.228844962 | -1.375255245 | 8.586954094 | 6.78762695 | 8.460932916 | 7.700356475 | 7.70992146 | 7.279786628 | 7.145684098 |
| ENSG00000102978 | POLR2C | -0.886016599 | 0.00307497 | -0.489529851 | 0.12122066 | -1.37554645 | 10.04301211 | 10.00683004 | 10.44204362 | 9.253098082 | 9.33320115 | 9.679353528 | 9.696573548 |
| ENSG00000108590 | MED31 | 1.10063578 | 0.076688241 | -2.476579785 | 1.87E-05 | -1.375944005 | 5.095959458 | 6.04685525 | 5.930865648 | 6.782976084 | 6.83074839 | 3.694366604 | 3.717076715 |
| ENSG00000170312 | CDK1 | -0.770369935 | 0.652705192 | -0.606385745 | 0.723776062 | -1.37675568 | 3.327711234 | 6.311476754 | 3.369317972 | 4.670864177 | 4.16043394 | 4.350488971 | 4.766376438 |
| ENSG00000172922 | RNASEH2C | -0.505808342 | 0.680089245 | -0.872869338 | 0.447209151 | -1.37867768 | 9.544849385 | 8.607278467 | 6.322058544 | 8.207157886 | 8.12087597 | 7.54878318 | 8.019835704 |
| ENSG00000102595 | UGGT2 | -0.538138459 | 0.471279327 | -0.842516768 | 0.228217127 | -1.380655227 | 8.653385977 | 9.883504052 | 8.434763955 | 8.587674547 | 8.63663561 | 8.291371768 | 8.331250243 |
| ENSG00000134308 | YWHAQ | -0.331323461 | 0.581201394 | -1.050065408 | 0.046784923 | -1.381388869 | 12.90368053 | 11.83753936 | 12.01322718 | 12.00411861 | 11.9950723 | 11.18792045 | 11.36989314 |
| ENSG00000177951 | BET1L | -0.97979521 | 0.07251956 | -0.401610275 | 0.493153856 | -1.381405485 | 9.710019321 | 10.86336859 | 10.28245311 | 9.333786218 | 9.43967292 | 9.799418565 | 10.11001882 |
| ENSG00000165389 | SPTSSA | -1.119292087 | 0.20765518 | -0.262341327 | 0.792037041 | -1.381633415 | 9.015175615 | 8.557180776 | 6.718950348 | 7.288834728 | 7.27676203 | 8.061837451 | 8.18182607 |
| ENSG00000177732 | SOX12 | -1.169108991 | 0.304573038 | -0.212911558 | 0.867658115 | -1.382020549 | 7.435667355 | 9.903691196 | 10.74860941 | 8.71221318 | 8.74905888 | 9.689335129 | 9.671309937 |
| ENSG00000150455 | TIRAP | -0.34830253 | 0.881863329 | -1.035345443 | 0.625695912 | -1.383647973 | 8.367060731 | 5.131603625 | 2 | 6.516352584 | 6.68951263 | 5.982522014 | 5.925257518 |
| ENSG00000186687 | LYRM7 | -1.366298137 | 0.004556921 | -0.017419314 | 0.976832019 | -1.383717451 | 8.19931886 | 7.503546079 | 7.398165132 | 6.415275989 | 6.42052719 | 7.884531292 | 7.545707309 |
| ENSG00000136732 | GYPC | 0.032080908 | 0.981681548 | -1.416062718 | 0.210179769 | -1.38398181 | 10.03472846 | 9.574900212 | 6.629488387 | 9.339314897 | 9.35585278 | 8.015738956 | 7.805990373 |
| ENSG00000250021 | C15orf38-AP | 6.661749443 | 1.04E-09 | -8.046248376 | 3.01E-05 | -1.384498933 | 4.467311984 | 3.076665209 | 2 | 9.565291356 | 9.5364465 | 2 | 2 |
| ENSG00000143653 | SCCPDH | -0.739918221 | 0.213670455 | -0.646803359 | 0.274221768 | -1.386721579 | 9.771132748 | 8.702538795 | 9.819251342 | 8.736160353 | 8.81462916 | 8.658095041 | 9.051693903 |
| ENSG00000065427 | KARS | -0.567409088 | 0.518602353 | -0.819986079 | 0.323125242 | -1.387395168 | 11.48534568 | 10.28850895 | 9.491321578 | 10.10480594 | 10.0733424 | 9.714922106 | 9.950662535 |
| ENSG00000100109 | TFIP11 | -0.222675115 | 0.825480361 | -1.16498424 | 0.174300946 | -1.387659355 | 9.810482605 | 8.639735838 | 7.600047325 | 8.736160353 | 8.73530306 | 7.88535852 | 7.72163797 |
| ENSG00000145919 | BOD1 | -0.412434843 | 0.359833218 | -0.977563354 | 0.016000913 | -1.389998197 | 9.384607364 | 9.949724236 | 10.16797309 | 9.466861018 | 9.45318143 | 8.997295259 | 8.793039018 |
| ENSG00000167377 | ZNF23 | 1.152120315 | 0.506977228 | -2.542587672 | 0.107289325 | -1.390467357 | 7.384433967 | 5.838937808 | 2 | 7.398825229 | 7.31795292 | 4.017766949 | 4.047883918 |
| ENSG00000116661 | FBXO2 | -1.097844092 | 0.585772669 | -0.293384722 | 0.892702877 | -1.391228814 | 7.667647223 | 4.93638933 | 2 | 5.258265913 | 5.3193943 | 6.275391383 | 5.73181021 |
| ENSG00000121864 | ZNF639 | -0.774832779 | 0.535148173 | -0.618292881 | 0.623218139 | -1.39312566 | 7.33131396 | 7.431218269 | 10.04361983 | 8.134820652 | 8.0358969 | 8.360824221 | 8.109313767 |
| ENSG00000115541 | HSPE1 | 0.204671961 | 0.826701316 | -1.598596648 | 0.036351322 | -1.393924687 | 8.998234871 | 7.393647515 | 7.552162999 | 8.370868225 | 8.37620272 | 6.855708005 | 6.320441373 |
| ENSG00000116985 | BMP8B | -1.849749257 | 0.485018665 | 0.455290059 | 0.872155485 | -1.394459198 | 2 | 2 | 4.874587708 | 2.868578001 | 2.38539787 | 4.388329244 | 3.236138336 |
| ENSG00000277586 | NEFL | 0.882350118 | 0.469356107 | -2.27697561 | 0.037729049 | -1.394625492 | 4.815609471 | 6.228547981 | 4.523675391 | 6.5612028 | 5.83486613 | 3.981090143 | 3.042026731 |
| ENSG00000172663 | TMEM134 | -1.233917966 | 0.041949021 | -0.162672449 | 0.820142868 | -1.396590415 | 8.963744667 | 8.276714465 | 7.552162999 | 7.202487431 | 7.12888795 | 8.209845962 | 8.222500688 |
| ENSG00000137168 | PPIL1 | -0.948615344 | 0.065184591 | -0.44863372 | 0.410449693 | -1.397249065 | 9.064836028 | 8.557180776 | 8.175061096 | 7.791909949 | 7.62295033 | 7.924517766 | 8.434495305 |
| ENSG00000163882 | POLR2H | -0.617911612 | 0.425471375 | -0.780737085 | 0.291836129 | -1.398648697 | 9.771132748 | 8.557180776 | 8.206335842 | 8.266399315 | 8.51470562 | 8.102245405 | 8.356830552 |
| ENSG00000130244 | FAM98C | -1.222438416 | 0.536668932 | -0.17783759 | 0.935430183 | -1.400276006 | 8.138771685 | 5.72254323 | 2 | 5.639097847 | 5.6826301 | 6.842137779 | 6.418016952 |
| ENSG00000244005 | NFS1 | -1.084533436 | 0.154269477 | -0.315831952 | 0.705751145 | -1.400365387 | 7.940205782 | 7.145017454 | 8.884815469 | 6.993973425 | 7.18663697 | 7.992893101 | 7.679327321 |
| ENSG00000188529 | SRSF10 | -0.541982045 | 0.087151155 | -0.859008322 | 0.00372404 | -1.400990367 | 9.931427102 | 10.19331737 | 10.40208303 | 9.719519122 | 9.57325401 | 9.319337188 | 9.346500876 |
| ENSG00000149089 | APIP | -0.187606438 | 0.917511462 | -1.213563819 | 0.443133621 | -1.401170257 | 8.564112515 | 7.002537973 | 3.369317972 | 7.178012736 | 7.31055061 | 6.170709933 | 6.341697848 |
| ENSG00000171824 | EXOSC10 | -0.189222728 | 0.874589067 | -1.213366406 | 0.228416028 | -1.402589134 | 10.58702944 | 9.102512015 | 7.858115942 | 9.382797773 | 9.42945789 | 8.342266573 | 8.438024223 |
| ENSG00000116815 | CD58 | 0.406033062 | 0.730567296 | -1.809396083 | 0.074893479 | -1.403363021 | 8.16936288 | 5.595927442 | 8.175061096 | 8.024753265 | 8.16258751 | 6.297933466 | 5.502966356 |
| ENSG00000151148 | UBE3B | -0.946776905 | 0.408383735 | -0.457013817 | 0.702456266 | -1.403790722 | 8.285626337 | 8.806266215 | 10.9968459 | 8.920805049 | 8.93022744 | 9.375172844 | 9.447647875 |
| ENSG00000138780 | GSTCD | -1.027438739 | 0.085059099 | -0.376369956 | 0.555749022 | -1.403808695 | 7.33131396 | 7.051611315 | 6.43197674 | 5.997179931 | 6.01871653 | 6.894026089 | 6.29575846 |
| ENSG00000011304 | PTBP1 | -0.734396537 | 0.021914674 | -0.672570304 | 0.036138686 | -1.40696684 | 11.99046984 | 11.55651981 | 12.04601058 | 11.14385097 | 11.1479752 | 11.11829161 | 11.29179109 |
| ENSG00000118707 | TGIF2 | -1.332572964 | 0.380936196 | -0.074711323 | 0.966296927 | -1.407284287 | 9.112843761 | 8.451443573 | 4.058998049 | 6.965655398 | 6.94988431 | 8.17644121 | 8.200230127 |
| ENSG00000125520 | SLC2A4RG | -1.56405937 | 1.61E-10 | 0.15548627 | 0.604131365 | -1.4085731 | 9.624286623 | 9.785372135 | 9.589475645 | 8.098332103 | 8.14396641 | 9.917551697 | 9.728912497 |
| ENSG00000110768 | GTF2H1 | -0.346117262 | 0.549552855 | -1.062491844 | 0.03743516 | -1.408609106 | 9.081016759 | 9.246456519 | 10.10298738 | 9.209487455 | 9.19732195 | 8.511487655 | 8.474909484 |
| ENSG00000198898 | CAPZA2 | -0.388805723 | 0.565888439 | -1.020156226 | 0.093459108 | -1.408961949 | 9.839306291 | 10.08551947 | 11.10866152 | 10.09230881 | 10.0396007 | 9.561639062 | 9.302266408 |
| ENSG00000031823 | RANBP3 | -0.686172248 | 0.360070875 | -0.723068617 | 0.323118682 | -1.409240865 | 10.62062599 | 9.203675681 | 9.25752426 | 9.15618819 | 9.18421833 | 8.96830908 | 9.281988193 |
| ENSG00000132394 | EEFSEC | -1.382070308 | 0.475887749 | -0.02749862 | 0.989709187 | -1.409568927 | 2 | 8.469613413 | 8.65477198 | 6.716553866 | 6.56439512 | 7.886185273 | 8.032963099 |
| ENSG00000137834 | SMAD6 | -1.63141754 | 0.239845221 | 0.221532425 | 0.890149172 | -1.409885115 | 6.73718411 | 4.93638933 | 8.941672936 | 6.172600389 | 6.08962627 | 7.768261726 | 8.064355282 |
| ENSG00000172179 | PRL | 3.778423042 | 0.507060161 | -5.191091037 | 0.364514502 | -1.412667994 | 2 | 2 | 2 | 4.387512679 | 3.90918929 | 2 | 2 |
| ENSG00000213928 | IRF9 | 2.337281161 | 0.138263648 | -3.75071641 | 0.01144673 | -1.41343525 | 9.081016759 | 5.595927442 | 4.523675391 | 9.909893561 | 10.0596091 | 3.930690651 | 4.489993931 |
| ENSG00000160917 | CPSF4 | -0.879546085 | 0.490472675 | -0.534525116 | 0.683123451 | -1.414071201 | 5.330594134 | 7.954278496 | 8.941672936 | 7.234487582 | 7.08183429 | 7.688988646 | 7.274433587 |
| ENSG00000243927 | MRPS6 | -0.772125796 | 0.20465032 | -0.6430478 | 0.290375858 | -1.415173595 | 9.371511179 | 9.079034257 | 10.22252928 | 8.870779703 | 8.87484109 | 9.228705128 | 8.729950594 |
| ENSG00000161298 | ZNF382 | -2.012945014 | 0.285993957 | 0.597456042 | 0.770772698 | -1.415488972 | 2 | 5.131603625 | 7.552162999 | 4.358765249 | 4.50979031 | 7.256956941 | 6.141542327 |
| ENSG00000186272 | ZNF17 | -0.636105664 | 0.750725045 | -0.780816649 | 0.688756148 | -1.416922313 | 2 | 6.228547981 | 8.007629717 | 6.237202172 | 6.1572134 | 6.282944614 | 5.799216178 |
| ENSG00000100196 | KDELR3 | -1.396370439 | 2.29E-06 | -0.020553286 | 0.958967393 | -1.416923724 | 11.44215374 | 11.07100932 | 11.5927499 | 9.976453497 | 10.0044775 | 11.41320151 | 11.31068228 |
| ENSG00000105281 | SLC1A5 | -0.92707703 | 0.009556581 | -0.491065545 | 0.194623596 | -1.418142575 | 11.13243651 | 11.07990752 | 10.9743126 | 10.07142123 | 10.2039308 | 10.85013781 | 10.23171747 |
| ENSG00000008405 | CRY1 | -1.433307673 | 3.66E-07 | 0.014893458 | 0.968553627 | -1.418414215 | 8.444143041 | 8.234834549 | 8.296287027 | 6.936770382 | 6.90635305 | 8.351874517 | 8.326688609 |
| ENSG00000261371 | PECAM1 | -1.067670142 | 0.558688888 | -0.351251475 | 0.856791105 | -1.418921617 | 7.579279349 | 3.076665209 | 5.39244166 | 5.304221918 | 5.36298494 | 6.660772804 | 4.711446447 |
| ENSG00000102780 | DGKH | -0.620610512 | 0.318938708 | -0.798759634 | 0.180023668 | -1.419370145 | 8.631580145 | 9.479495904 | 9.819251342 | 8.778830996 | 8.77079897 | 8.859275573 | 8.279474384 |
| ENSG00000129204 | USP6 | 2.038085675 | 0.103636128 | -3.457762684 | 0.003716817 | -1.419677009 | 3.327711234 | 4.442710129 | 6.534109959 | 7.2696592 | 7.29187696 | 2.937246078 | 2.817676617 |
| ENSG00000144029 | MRPS5 | -0.926735356 | 0.271482843 | -0.493608252 | 0.573477412 | -1.420343609 | 9.521325812 | 9.214490408 | 7.451345288 | 8.029321023 | 8.08010078 | 8.478974993 | 8.485861839 |
| ENSG00000173950 | XXYLT1 | -1.318582962 | 0.113829415 | -0.102655292 | 0.918231554 | -1.421238254 | 10.5468183 | 8.395519306 | 9.647595056 | 8.471640654 | 8.46389529 | 9.505174684 | 9.825841298 |
| ENSG00000187726 | DNAJB13 | -2.278728721 | 0.566415954 | 0.857377365 | NA | -1.421351356 | 2 | 3.076665209 | 2 | 2 | 2.2680111 | 3.193831101 | 2.592996859 |
| ENSG00000243943 | ZNF512 | -0.721375787 | 0.739034243 | -0.70164277 | 0.743381156 | -1.423018557 | 9.751047882 | 8.762721501 | 2 | 7.975897842 | 8.11451391 | 8.005883472 | 8.124285971 |
| ENSG00000180318 | ALX1 | -3.753024452 | NA | 2.32901662 | NA | -1.424007832 | 2 | 2 | 2 | 2 | 2 | 3.393105702 | 2.950113106 |
| ENSG00000007520 | TSR3 | -0.719501244 | 0.48167816 | -0.706964579 | 0.480981934 | -1.426465823 | 7.667647223 | 8.607278467 | 10.12770132 | 8.468280417 | 8.41468635 | 8.388512422 | 8.516894472 |

| | | | | | | | | | | | | | |
|---|---|---|---|---|---|---|---|---|---|---|---|---|---|
| ENSG00000153250 | RBMS1 | -0.324072554 | 0.626363416 | -1.102754054 | 0.058325459 | -1.426826607 | 10.19941743 | 9.235879652 | 10.58572549 | 9.845303961 | 9.72155655 | 9.050629989 | 8.969708735 |
| ENSG00000239672 | NME1 | -0.40652937 | 0.548500399 | -1.020583053 | 0.094571914 | -1.427112423 | 9.304185368 | 8.993716073 | 10.09465441 | 9.078979414 | 9.1882629 | 8.099396357 | 8.855068552 |
| ENSG00000122557 | HERPUD2 | -1.177713482 | 0.019477382 | -0.249556202 | 0.66811141 | -1.427269684 | 9.646202137 | 8.903033433 | 8.699419242 | 8.054188409 | 7.89452713 | 9.058352796 | 8.707903068 |
| ENSG00000067191 | CACNB1 | -1.300005714 | 0.184030119 | -0.127446876 | 0.911555747 | -1.42745259 | 8.737455672 | 6.228547981 | 7.600047325 | 6.604700567 | 6.57060569 | 7.494516625 | 7.926450253 |
| ENSG00000114354 | TFG | -0.678566375 | 0.013719923 | -0.749424836 | 0.005580838 | -1.427991211 | 11.59363617 | 11.56499503 | 11.26536143 | 10.82288924 | 10.7877614 | 10.62175706 | 10.83956746 |
| ENSG00000055950 | MRPL43 | -1.186486767 | 0.005498029 | -0.241706463 | 0.626190802 | -1.42819323 | 9.848787621 | 9.367676479 | 8.99637432 | 8.23312837 | 8.30721295 | 9.093139938 | 9.311530309 |
| ENSG00000072422 | RHOBTB1 | -1.947510101 | 7.69E-06 | 0.519113486 | 0.288589938 | -1.428396615 | 8.998234871 | 8.356994601 | 8.175061096 | 6.79375511 | 6.48770488 | 9.068221759 | 9.066817089 |
| ENSG00000103495 | MAZ | 0.308644541 | 0.754977416 | -1.737572729 | 0.03707375 | -1.428928188 | 10.1076228 | 9.514922632 | 7.777116083 | 9.727271068 | 9.72712735 | 7.778984728 | 7.620910859 |
| ENSG00000187151 | ANGPTL5 | -0.461622215 | NA | -0.968100316 | NA | -1.429722531 | 2 | 3.076665209 | 2 | 2.600425781 | 2.2680111 | 2 | 2.592996859 |
| ENSG00000135378 | PRRG4 | -0.197249917 | 0.860437532 | -1.232726573 | 0.188401866 | -1.42997649 | 4.815609471 | 4.11352228 | 5.39244166 | 4.647285135 | 4.68029874 | 4.029788235 | 3.640730416 |
| ENSG00000169189 | NSMCE1 | -1.285333831 | 0.164037813 | -0.144794174 | 0.893463707 | -1.430128005 | 8.816894989 | 6.535001748 | 7.646393253 | 6.548530093 | 6.83074839 | 7.618102344 | 7.972925696 |
| ENSG00000204969 | PCDHA2 | 0.921247835 | 0.755643383 | -2.351531034 | 0.392854305 | -1.4302832 | 2 | 3.686303414 | 2 | 3.79808662 | 2.68924543 | 2.094598369 | 2.326754077 |
| ENSG00000130635 | COL5A1 | -1.482334768 | 7.19E-08 | 0.051939177 | 0.888119558 | -1.43039559 | 14.59270693 | 14.92196164 | 14.44962988 | 13.14048624 | 13.231656 | 14.71263075 | 14.72864471 |
| ENSG00000130309 | COLGALT1 | -1.00600095 | 0.073563931 | -0.424715337 | 0.481274312 | -1.430716287 | 11.55939759 | 11.29386853 | 12.38698767 | 10.79734056 | 10.8423598 | 11.61288715 | 11.15073956 |
| ENSG00000133193 | FAM104A | -0.102568997 | 0.885376787 | -1.328475655 | 0.017716114 | -1.431044652 | 7.975256505 | 8.862353853 | 9.301632639 | 8.693628619 | 8.72559535 | 7.53303322 | 7.470508284 |
| ENSG00000176597 | B3GNT5 | 0.242569915 | 0.89066231 | -1.675116994 | 0.260870186 | -1.432547079 | 2 | 6.311476754 | 5.59507254 | 5.686138681 | 5.70538658 | 4.220168345 | 3.875378318 |
| | LOC441155 | 6.629128683 | 2.58E-09 | -8.062274789 | 3.27E-05 | -1.433146106 | 2 | 3.076665209 | 4.523675391 | 9.50916971 | 9.55926825 | 2 | 2 |
| ENSG00000182400 | TRAPPC6B | 0.532177133 | 0.39788284 | -1.966817605 | 0.000275322 | -1.434640472 | 8.963744667 | 8.234834549 | 7.691296561 | 8.902248225 | 8.93264039 | 6.595545082 | 6.338680316 |
| C2ORF82 | C2ORF82 | 3.754287162 | 0.510049398 | -5.191091137 | 0.364514502 | -1.436803975 | 2 | 2 | 2 | 3.997744386 | 4.2865223 | 2 | 2 |
| ENSG00000181481 | RNF135 | -1.060973779 | 0.159914848 | -0.3770057 | 0.644088306 | -1.437979479 | 9.435834703 | 8.169643011 | 7.858115942 | 7.638916932 | 7.5835711 | 8.478426859 | 8.059781405 |
| ENSG00000125247 | TMTC4 | -0.829745383 | 0.737269626 | -0.608647798 | 0.807781951 | -1.438393181 | 3.327711234 | 8.053983481 | 2 | 6.006476466 | 5.48637102 | 6.197600957 | 5.713290392 |
| ENSG00000183155 | RABIF | -0.774001053 | 0.727489842 | -0.664679952 | 0.765135376 | -1.438681005 | 9.761125267 | 7.60566235 | 2 | 7.708923796 | 7.7127321 | 7.692773977 | 7.933467731 |
| ENSG00000153574 | RPIA | -0.508310456 | 0.83752462 | -0.931078713 | 0.688544443 | -1.439389169 | 2 | 4.442710129 | 8.608698767 | 6.634986713 | 6.60731796 | 6.361143723 | 6.059610627 |
| ENSG00000146733 | PSPH | -1.106136133 | 0.034068642 | -0.333539415 | 0.562413083 | -1.439675548 | 8.009475818 | 7.638155693 | 7.163363831 | 6.634986713 | 6.507263 | 7.55607473 | 7.036323724 |
| ENSG00000076248 | UNG | -0.711430401 | 0.458940186 | -0.729159452 | 0.435504299 | -1.440589853 | 7.218819781 | 7.315436877 | 9.116480286 | 7.474468681 | 7.45518898 | 7.307207714 | 7.575491915 |
| ENSG00000198551 | ZNF627 | -0.736992151 | 0.259012717 | -0.703752297 | 0.272931073 | -1.440744448 | 7.33131396 | 8.213429689 | 8.585098385 | 7.527093191 | 7.28055563 | 7.688988646 | 7.142226902 |
| ENSG00000185730 | ZNF696 | -1.206417215 | 0.150100354 | -0.236414309 | 0.801869817 | -1.442831524 | 7.384433967 | 6.389896633 | 5.59507254 | 5.304221918 | 5.6826301 | 6.617616961 | 6.172377306 |
| ENSG00000167100 | SAMD14 | -1.703804007 | 0.164180702 | 0.25929675 | 0.853657176 | -1.444507257 | 9.174465797 | 5.595927442 | 9.180874355 | 6.998639523 | 6.95464102 | 8.88183438 | 8.928657149 |
| ENSG00000102103 | PQBP1 | -0.581297924 | 0.242935896 | -0.863633598 | 0.06554197 | -1.444931523 | 8.696033109 | 8.101365653 | 8.922968322 | 7.956842238 | 8.10384772 | 7.561260509 | 7.925444964 |
| ENSG00000111652 | COPS7A | -1.207752034 | 0.012083417 | -0.237246178 | 0.672805615 | -1.444998213 | 10.52923584 | 9.849218119 | 10.70548138 | 9.161399541 | 9.24080977 | 9.862322274 | 10.41907654 |
| ENSG00000160972 | PPP1R16A | -0.431265645 | 0.298315448 | -1.013771771 | 0.006260318 | -1.445037416 | 8.540903468 | 8.234834549 | 8.742726175 | 8.008651253 | 8.16464176 | 7.358152527 | 7.66011679 |
| ENSG00000135678 | CPM | -2.119571761 | 0.141954442 | 0.673008447 | 0.669265476 | -1.446563314 | 6.366709671 | 4.93638933 | 7.225742779 | 4.783303513 | 4.22485516 | 7.944499874 | 4.729989799 |
| ENSG00000099246 | RAB18 | -0.545024483 | 0.401480288 | -0.90154789 | 0.13626007 | -1.446572373 | 11.31110672 | 10.43440824 | 9.981704053 | 10.19294689 | 10.0858626 | 9.868610809 | 9.69686815 |
| ENSG00000138468 | SENP7 | -0.892532448 | 0.14099447 | -0.554276852 | 0.370202745 | -1.446809299 | 7.031698319 | 7.669933291 | 6.718950348 | 6.442929658 | 6.25300433 | 6.90385696 | 6.400759785 |
| ENSG00000132122 | SPATA6 | -0.678603675 | 0.735978187 | -0.769194256 | 0.695554779 | -1.447797931 | 2 | 7.051611315 | 4.523675391 | 5.058234798 | 5.21224859 | 5.085488617 | 5.028505473 |
| ENSG00000164924 | YWHAZ | -0.580887586 | 0.032660886 | -0.867471222 | 0.000762931 | -1.448358808 | 13.26936508 | 12.87899828 | 13.07845487 | 12.54471931 | 12.4609778 | 12.16436478 | 12.26796737 |
| ENSG00000164117 | FBXO8 | -0.57968563 | 0.154817843 | -0.868708389 | 0.023427954 | -1.448394019 | 8.716893051 | 8.732943944 | 8.408111533 | 8.03841338 | 8.07574085 | 7.992893101 | 7.512619696 |
| ENSG00000121075 | TBX4 | -0.422645388 | 0.878583032 | -1.027103859 | 0.684198796 | -1.449749246 | 3.327711234 | 3.076665209 | 2 | 2.600425781 | 2.86123014 | 2.803598391 | 2.172608095 |
| ENSG00000180834 | MAP6D1 | -0.031101789 | 0.989593043 | -1.419041335 | 0.494816556 | -1.450143124 | 2 | 3.076665209 | 6.534109959 | 5.128039203 | 5.04390545 | 4.065262523 | 3.824516949 |
| ENSG00000167720 | SRR | -0.290034751 | 0.611381475 | -1.161143055 | 0.017728145 | -1.451177805 | 7.031698319 | 7.701025996 | 6.958202216 | 7.044888801 | 6.96410766 | 6.034159474 | 6.29575846 |
| ENSG00000126787 | DLGAP5 | -0.769622348 | 0.659371222 | -0.682174821 | 0.692464563 | -1.451797168 | 3.327711234 | 5.595927442 | 3.369317972 | 4.070463721 | 3.82856379 | 3.193831101 | 4.543391831 |
| ENSG00000101246 | ARFRP1 | -0.989362484 | 0.140578614 | -0.46414343 | 0.513597151 | -1.453505915 | 9.771132748 | 8.234834549 | 9.426401271 | 8.279879181 | 8.30907138 | 8.72431761 | 8.898748881 |
| ENSG00000140386 | SCAPER | -1.333650035 | 0.392352229 | -0.119913384 | 0.945540243 | -1.453563419 | 8.444143041 | 7.274681896 | 3.369317972 | 6.237202172 | 6.02777321 | 7.224637511 | 7.376535208 |
| ENSG00000256683 | ZNF350 | -0.05048753 | 0.962823674 | -1.404177641 | 0.120415896 | -1.45466517 | 6.366709671 | 7.051611315 | 8.585098385 | 7.552702688 | 7.61995902 | 6.239610476 | 6.317379004 |
| ENSG00000106399 | RPA3 | -0.339597249 | 0.845967999 | -1.116215091 | 0.474892185 | -1.45581234 | 6.563806279 | 5.946638857 | 2 | 5.474671438 | 5.39133044 | 4.888321757 | 4.594883674 |
| ENSG00000103876 | FAH | 0.211613337 | 0.882991782 | -1.667700467 | 0.16021677 | -1.45608713 | 8.737455672 | 7.393647515 | 5.156626401 | 7.893113669 | 7.95274764 | 5.829320042 | 6.338680316 |
| ENSG00000093134 | VNN3 | 3.7349147 | 0.512728968 | -5.191091218 | 0.364514502 | -1.456176518 | 2 | 2 | 2 | 4.299492648 | 3.94787299 | 2 | 2 |
| ENSG00000243480 | AMY2A | 3.7349147 | 0.512728968 | -5.191091218 | 0.364514502 | -1.456176518 | 2 | 2 | 2 | 4.299492648 | 3.94787299 | 2 | 2 |
| ENSG00000148671 | ADIRF | 5.484290174 | 0.001294782 | -6.94073151 | 0.0032699 | -1.456441336 | 2 | 3.686303414 | 2 | 7.206526483 | 7.38291816 | 2 | 2 |
| ENSG00000114923 | SLC4A3 | -1.953576601 | 0.064794697 | 0.496994794 | 0.675190643 | -1.456581806 | 8.19931886 | 6.951736442 | 5.156626401 | 5.334065616 | 5.44640346 | 7.79139499 | 7.65527392 |
| ENSG00000196290 | NIF3L1 | -0.981880931 | 0.634672806 | -0.475018853 | 0.825446679 | -1.456899784 | 9.189468652 | 7.538394143 | 2 | 7.101994243 | 6.99676009 | 7.598057803 | 7.488317792 |
| ENSG00000165416 | SUGT1 | -0.503973745 | 0.555006373 | -0.95304876 | 0.228768931 | -1.457022504 | 9.635285994 | 8.147241016 | 7.97167868 | 8.23312837 | 8.34755546 | 7.948463298 | 7.742336888 |
| ENSG00000197008 | ZNF138 | -0.202166378 | 0.912234522 | -1.255968314 | 0.428993571 | -1.458134692 | 6.73718411 | 6.228547981 | 2 | 5.839660368 | 5.69405321 | 5.04460184 | 4.56420962 |
| ENSG00000100028 | SNRPD3 | -0.517787483 | 0.303181423 | -0.941101226 | 0.042409585 | -1.458888708 | 10.29268328 | 9.488434476 | 10.11127249 | 9.453327884 | 9.51647861 | 8.799092094 | 9.289039876 |
| ENSG00000145685 | LHFPL2 | -0.217390226 | 0.738608861 | -1.242864596 | 0.023412451 | -1.460254822 | 10.82178169 | 11.2913169 | 11.88621715 | 11.20190521 | 11.1611534 | 9.780867042 | 10.45800828 |
| ENSG00000105699 | LSR | -0.720705481 | 0.610159623 | -0.739801544 | 0.591655943 | -1.460507025 | 4.815609471 | 6.464272742 | 3.369317972 | 4.907595604 | 4.68029874 | 4.5389195 | 4.990558504 |
| | DOPEY1 | 5.654690303 | 0.001390334 | -7.116254018 | 0.003086571 | -1.461563716 | 2 | 2 | 4.058998049 | 7.656739692 | 7.61395775 | 2 | 2 |
| ENSG00000116030 | SUMO1 | -0.400867737 | 0.516230128 | -1.061288523 | 0.053149142 | -1.46215626 | 10.9638027 | 9.806969599 | 10.10298738 | 10.01956245 | 9.93396767 | 9.241683432 | 9.394877047 |
| ENSG00000115524 | SF3B1 | -0.476136794 | 0.512666787 | -0.986575183 | 0.136166444 | -1.462711977 | 11.25159303 | 12.86276311 | 12.26789926 | 11.78444742 | 11.8053473 | 11.31623198 | 11.25345031 |
| ENSG00000149923 | PPP4C | -0.739347384 | 0.556147109 | -0.723812487 | 0.559807045 | -1.463159871 | 11.65437133 | 9.648522413 | 8.380957451 | 9.783105971 | 9.75261615 | 9.646061941 | 9.908803154 |
| ENSG00000198929 | NOS1AP | -2.326272487 | 0.374953074 | 0.862118573 | 0.751485356 | -1.464153914 | 2 | 4.93638933 | 2 | 2.868578001 | 2.14022021 | 4.5389195 | 4.215171317 |
| ENSG00000105968 | H2AFV | -1.49748547 | 2.86E-09 | 0.032261687 | 0.92532897 | -1.465223783 | 10.01801709 | 9.71095217 | 9.963514551 | 8.388752656 | 8.43865202 | 9.864840978 | 9.997333704 |
| ENSG00000109184 | DCUN1D4 | -0.290397939 | 0.711436164 | -1.17512651 | 0.083368021 | -1.465524449 | 8.228665467 | 9.672251278 | 9.839340503 | 9.143603603 | 9.087919 | 8.25342909 | 8.226587404 |
| ENSG00000109814 | UGDH | -0.949179364 | 0.182358952 | -0.517074894 | 0.485029668 | -1.466254258 | 11.74347718 | 11.29641566 | 10.29705308 | 10.29273067 | 10.2722069 | 10.21173728 | 11.08495917 |
| ENSG00000164610 | RP9 | -0.255713334 | 0.813586965 | -1.212553702 | 0.191291972 | -1.468267036 | 8.042902253 | 6.140560021 | 6.203069917 | 6.79375511 | 6.8715633 | 6.05195199 | 5.781546943 |
| ENSG00000181458 | TMEM45A | -0.581751928 | 0.458819694 | -0.886880763 | 0.228318728 | -1.468632691 | 8.87372952 | 9.890264534 | 10.67226762 | 9.425008328 | 9.39487575 | 9.023457772 | 9.185775709 |
| ENSG00000134253 | TRIM45 | -1.477220988 | 0.133069745 | 0.007867964 | 0.994638415 | -1.469353024 | 7.031698319 | 5.303523747 | 5.39244166 | 4.623314311 | 4.95174433 | 6.468821191 | 5.717942682 |
| ENSG00000234465 | PINLYP | 0.353675172 | 0.898417957 | -1.823203018 | 0.452416189 | -1.469527846 | 8.631580145 | 3.076665209 | 2 | 7.281195074 | 7.57123495 | 5.274160001 | 5.459179378 |
| ENSG00000243678 | NME2 | 2.996174115 | 2.62E-08 | -4.46594169 | 2.08E-16 | -1.469767575 | 8.257427002 | 6.899080544 | 7.646393253 | 10.6638047 | 10.6679683 | 3.632645806 | 3.807156736 |
| ENSG00000198015 | MRPL42 | -0.813537967 | 0.211593503 | -0.658880454 | 0.312110723 | -1.472418421 | 8.418903792 | 9.298208073 | 9.897982195 | 8.538855014 | 8.49850989 | 8.740408958 | 8.5998903 |
| ENSG00000186074 | CD300LF | -3.753025058 | NA | 2.280545605 | NA | -1.472479453 | 2 | 2 | 2 | 2 | 2 | 3.56816602 | 2.592996859 |
| ENSG00000150337 | FCGR1A | -3.753025059 | NA | 2.280455726 | NA | -1.472569333 | 2 | 2 | 2 | 2 | 2 | 3.393105702 | 2.885414003 |
| ENSG00000100379 | KCTD17 | -0.901036234 | 0.184561268 | -0.573055838 | 0.410673825 | -1.474092072 | 8.674867122 | 7.638155693 | 9.165043284 | 7.656739692 | 7.79460185 | 7.935741825 | 8.155488004 |
| ENSG00000164520 | RAET1E | 0.744135918 | 0.704833394 | -2.218980077 | 0.210711023 | -1.474844159 | 3.327711234 | 3.076665209 | 4.523675391 | 4.358765249 | 4.2865223 | 2.139668783 | 2.950113106 |
| ENSG00000136098 | NEK3 | -1.132339128 | 0.514610732 | -0.342709505 | 0.853576247 | -1.475048633 | 7.33131396 | 6.844429755 | 2 | 5.433909581 | 5.53799497 | 6.398709156 | 6.018717758 |
| ENSG00000150756 | FAM173B | -1.212035397 | 0.33586874 | -0.26369942 | 0.850222943 | -1.475734816 | 7.579279349 | 5.595927442 | 4.523675391 | 5.334065616 | 5.30456641 | 6.000958775 | 6.368575325 |
| ENSG00000141428 | C18orf21 | -0.501521848 | 0.579491849 | -0.974293965 | 0.243352449 | -1.475815813 | 8.19931886 | 6.535001748 | 6.718950348 | 6.86703417 | 6.86146731 | 6.609629839 | 6.179140936 |
| ENSG00000118898 | PPL | -3.359310002 | 0.069933875 | 1.882931046 | 0.267661148 | -1.476378956 | 3.327711234 | 4.442710129 | 2 | 2.270786764 | 2.2680111 | 4.291799915 | 5.661099865 |
| ENSG00000163902 | RPN1 | -0.336938743 | 0.464545264 | -1.140550693 | 0.004152549 | -1.477489436 | 13.07503738 | 12.29271883 | 12.51583706 | 12.31382177 | 12.3448457 | 11.46661987 | 11.58427143 |
| ENSG00000007038 | PRSS21 | -3.75302512 | NA | 2.275517957 | NA | -1.477507164 | 2 | 2 | 2 | 2 | 2 | 3.411599105 | 2.851942824 |
| ENSG00000136014 | USP44 | -3.524175504 | 0.099060337 | 2.044396167 | 0.2908151 | -1.479779337 | 4.00722531 | 3.686303414 | 2 | 2.3892346 | 2 | 5.338397457 | 4.711446447 |
| ENSG00000166226 | CCT2 | -0.495386419 | 0.434185114 | -0.985971139 | 0.088712251 | -1.481357558 | 11.19324461 | 9.869887299 | 10.68897008 | 10.15895941 | 10.2133886 | 9.598679043 | 9.790665867 |
| ENSG00000197062 | ZSCAN26 | -0.939520837 | 0.643088643 | -0.541975076 | 0.795268255 | -1.481495912 | 8.963744667 | 8.077869079 | 2 | 7.053486362 | 7.12888795 | 7.775419242 | 7.105417192 |
| ENSG00000136695 | IL36RN | 3.709533236 | 0.51607452 | -5.191091323 | 0.364514502 | -1.481558088 | 2 | 2 | 2 | 3.79808662 | 4.37434574 | 2 | 2 |
| ENSG00000198719 | DLL1 | 4.671220287 | 0.046826482 | -6.1529404 | 0.031054949 | -1.481720113 | 3.327711234 | 2 | 2 | 5.94010643 | 5.5755357 | 2 | 2 |
| ENSG00000115128 | SF3B6 | -1.022274183 | 0.074775117 | -0.462745207 | 0.4484575 | -1.48501939 | 10.22876205 | 9.079034257 | 9.287079172 | 8.567402595 | 8.64401667 | 9.236830239 | 9.080880746 |
| ENSG00000100142 | POLR2F | -0.487974879 | 0.487504654 | -0.997945978 | 0.11985978 | -1.485920857 | 8.998234871 | 7.979858146 | 7.691296561 | 7.849869274 | 7.85944752 | 7.155000534 | 7.531242162 |
| ENSG00000116396 | KCNC4 | -0.983417547 | 0.657356622 | -0.504081051 | 0.826586773 | -1.487498598 | 2 | 6.535001748 | 9.227351203 | 6.917187326 | 6.87658492 | 7.789628618 | 6.753261212 |
| ENSG00000100711 | ZFYVE21 | -0.990488931 | 0.002467749 | -0.499913486 | 0.148716503 | -1.490402417 | 8.631580145 | 9.030900487 | 8.677268317 | 7.834291698 | 7.80517502 | 8.199905499 | 8.399447278 |
| ENSG00000121940 | CLCC1 | -0.512170113 | 0.725856392 | -0.978992381 | 0.473680232 | -1.491162494 | 6.563806279 | 8.862353853 | 10.68342405 | 8.948803189 | 9.07159009 | 8.53487306 | 8.562873254 |
| ENSG00000171566 | PLRG1 | -0.281395807 | 0.85537103 | -1.211740181 | 0.367297237 | -1.493135989 | 5.70932728 | 9.806969599 | 9.868959282 | 8.999733249 | 9.03050377 | 8.162859687 | 8.023598593 |
| ENSG00000100365 | NCF4 | -1.689351372 | 0.585157799 | 0.196190376 | NA | -1.493160996 | 2 | 2 | 3.369317972 | 2.141737246 | 2.2680111 | 2.775316226 | 2.509591799 |
| ENSG00000181690 | PLAG1 | -0.94318561 | 0.632299403 | -0.550935149 | 0.786091427 | -1.494120759 | 2 | 8.077869079 | 7.285536013 | 6.336376195 | 6.18176397 | 7.157741966 | 5.807970301 |
| ENSG00000161544 | CYGB | 1.790440909 | 0.121386739 | -3.285133478 | 0.002227172 | -1.494692569 | 5.330594134 | 7.901717006 | 7.163363831 | 8.873322599 | 8.9398551 | 4.742549839 | 3.236138336 |
| ENSG00000180376 | CCDC66 | -0.214800383 | 0.832111917 | -1.280732508 | 0.130070296 | -1.495532891 | 6.009025997 | 7.669933291 | 7.398165132 | 6.989292186 | 6.97819215 | 6.405645065 | 5.331426166 |
| ENSG00000141378 | PTRH2 | -0.827347491 | 0.475887749 | -0.668735638 | 0.56354884 | -1.49608313 | 9.810482605 | 7.35507212 | 7.451345288 | 7.870381136 | 7.86703672 | 7.852739912 | 8.178454506 |
| ENSG00000173992 | CCS | -1.06872093 | 0.080136816 | -0.427404128 | 0.513742625 | -1.496125058 | 7.485143553 | 8.317412869 | 7.098165388 | 6.610808945 | 6.78340763 | 7.421115854 | 7.213143202 |
| ENSG00000148985 | PGAP2 | -1.08788745 | 0.41446225 | -0.408473555 | 0.773867441 | -1.496361005 | 4.467311984 | 8.147241016 | 8.266922072 | 6.70517998 | 6.53293476 | 7.393470826 | 7.154291015 |
| ENSG00000204519 | ZNF551 | -1.143274248 | 0.484587476 | -0.354255952 | 0.840156112 | -1.4975302 | 3.327711234 | 5.946638857 | 8.110401118 | 5.85002558 | 5.6826301 | 6.562825444 | 6.460277802 |
| ENSG00000160446 | ZDHHC12 | -0.752761006 | 0.391821726 | -0.746948385 | 0.384717028 | -1.499709391 | 9.678463567 | 8.557180776 | 7.600047325 | 8.004017475 | 8.21109983 | 8.046387837 | 8.184349581 |
| ENSG00000158966 | CACHD1 | 0.614358526 | 0.762344209 | -2.114546499 | 0.240936153 | -1.500187973 | 2 | 8.004992147 | 8.699419242 | 8.404661252 | 8.44034872 | 5.815498119 | 5.75463059 |
| ENSG00000088682 | COQ9 | -0.844264512 | 0.710858197 | -0.655956933 | 0.774684899 | -1.500221445 | 10.00110988 | 7.503546079 | 2 | 7.895617537 | 7.74603955 | 8.085066106 | 7.926450253 |
| ENSG00000106009 | BRAT1 | -0.882232392 | 0.172591839 | -0.618282231 | 0.34684373 | -1.500514623 | 7.975256505 | 8.88960062 | 9.413059136 | 7.956842238 | 8.04484669 | 8.408791501 | 8.097758666 |
| ENSG00000100290 | BIK | -0.46395321 | 0.884340692 | -1.036757009 | NA | -1.500710219 | 2 | 2 | 3.369317972 | 2.498691189 | 2.38539787 | 2.34604626 | 2.277179783 |
| ENSG00000167487 | KLHL26 | -0.683026678 | 0.749679501 | -0.818284099 | 0.695197299 | -1.501310777 | 2 | 7.467835385 | 9.180874355 | 7.398825229 | 7.23044321 | 7.009312024 | 7.342293759 |
| ENSG00000110717 | NDUFS8 | -0.886082181 | 0.192239251 | -0.615312951 | 0.37239741 | -1.501395132 | 10.20680973 | 9.792607274 | 8.784770914 | 8.770670063 | 8.88484424 | 8.742693141 | 9.382110089 |
| ENSG00000127125 | PPCS | -1.050015005 | 0.13450058 | -0.451884141 | 0.545019112 | -1.501899146 | 9.174465797 | 8.124485672 | 9.798878489 | 8.143273718 | 8.14188248 | 8.787633372 | 8.677773268 |

| | | | | | | | | | | | | | |
|---|---|---|---|---|---|---|---|---|---|---|---|---|---|
| ENSG00000084754 | HADHA | -0.496525205 | 0.192239251 | -1.005578357 | 0.003786604 | -1.502103562 | 11.29385609 | 11.15761406 | 11.73357624 | 10.91339224 | 10.9261795 | 10.27622952 | 10.53615161 |
| ENSG00000140839 | CLEC18B | 1.705091093 | 0.505493848 | -3.207345203 | 0.193871288 | -1.502254109 | 2 | 2 | 4.058998049 | 3.663521627 | 4.40247105 | 2.225793514 | 2.059860178 |
| ENSG00000275499 | | 3.688784418 | 0.518691015 | -5.19109141 | 0.364514502 | -1.502306992 | 2 | 2 | 2 | 4.034562184 | 4.16043394 | 2 | 2 |
| ENSG00000141759 | TXNL4A | -1.271934671 | 0.002781499 | -0.230696277 | 0.644509054 | -1.502630949 | 9.895281535 | 9.695596167 | 10.44204362 | 8.846395982 | 8.71020716 | 9.606476037 | 9.996137262 |
| ENSG00000131013 | PPIL4 | -0.453831438 | 0.713282111 | -1.050125704 | 0.352922604 | -1.503957142 | 9.544849385 | 9.102512015 | 6.322058544 | 8.409925372 | 8.37797443 | 7.968899389 | 7.623392692 |
| ENSG00000125166 | GOT2 | -0.906739182 | 0.29492176 | -0.597219871 | 0.498114559 | -1.503959053 | 11.13631304 | 10.3729879 | 8.941672936 | 9.486925632 | 9.51808625 | 9.735024004 | 9.882366758 |
| ENSG00000153561 | RMND5A | -0.716052345 | 0.274396109 | -0.788300626 | 0.215146032 | -1.504352971 | 8.836089686 | 9.741182361 | 8.536706852 | 8.369067528 | 8.48542112 | 8.611362196 | 8.047821014 |
| ENSG00000125962 | ARMCX5 | -0.659193896 | 0.681796824 | -0.845337078 | 0.584335308 | -1.504530975 | 4.00722531 | 6.78762695 | 8.65477198 | 6.777556219 | 6.86146731 | 6.697032326 | 6.582657467 |
| ENSG00000141447 | OSBPL1A | -1.238172128 | 0.000876231 | -0.269517126 | 0.529270693 | -1.507689254 | 8.816894989 | 8.540087354 | 9.165043284 | 7.497728447 | 7.76240929 | 8.5805365 | 8.600520269 |
| ENSG00000103522 | IL21R | -0.27649774 | 0.881480919 | -1.23149971 | 0.444246609 | -1.50799745 | 4.467311984 | 2 | 4.874587708 | 3.79808662 | 4.0930011 | 3.12825509 | 3.474731315 |
| ENSG00000196177 | ACADSB | -1.029774972 | 0.311246056 | -0.478242461 | 0.656134931 | -1.508017434 | 8.75772933 | 8.004992147 | 6.322058544 | 7.003290579 | 6.99214046 | 6.939340974 | 7.955303035 |
| ENSG00000125734 | GPR108 | -0.22548368 | 0.658522427 | -1.2831875 | 0.002463508 | -1.50867118 | 10.20680973 | 9.338312924 | 9.788583153 | 9.544700162 | 9.64121862 | 8.435582605 | 8.645170029 |
| ENSG00000177042 | TMEM80 | -1.858223319 | 0.063207012 | 0.349021993 | 0.761747773 | -1.509201326 | 6.96351253 | 8.540087354 | 9.767769415 | 7.003290579 | 7.01053089 | 9.444727462 | 8.851897333 |
| ENSG00000105193 | RPS16 | -1.281089022 | 0.01776689 | -0.229430473 | 0.717035549 | -1.510519496 | 11.98187803 | 11.0620559 | 10.85110517 | 10.04346859 | 10.1695649 | 11.11952254 | 11.19322749 |
| ENSG00000113369 | ARRDC3 | 0.835550045 | 0.367367297 | -2.346301935 | 0.004207821 | -1.51075189 | 8.16936288 | 10.2574711 | 10.40208303 | 10.75866452 | 10.7065491 | 7.354572674 | 7.77172059 |
| ENSG00000079482 | OPHN1 | -1.342147925 | 0.041012176 | -0.169669794 | 0.827324651 | -1.511817718 | 8.75772933 | 7.232742069 | 7.97167868 | 6.861921612 | 6.73998187 | 7.938932725 | 7.952344882 |
| ENSG00000106603 | COA1 | -1.098712843 | 0.172176659 | -0.413178828 | 0.633693854 | -1.511891671 | 9.761125267 | 8.469613413 | 7.934808822 | 7.875463856 | 7.81828345 | 8.350077895 | 8.677773268 |
| ENSG00000142039 | CCDC97 | -1.037416179 | 0.268404698 | -0.474955777 | 0.631857848 | -1.512371957 | 6.96351253 | 8.213429689 | 9.344432377 | 7.461007055 | 7.45518898 | 7.858649102 | 8.146888246 |
| ENSG00000167748 | KLK1 | -3.753025559 | NA | 2.240376398 | NA | -1.512649161 | 2 | 2 | 2 | 2 | 2 | 3.355392153 | 2.885414003 |
| ENSG00000121390 | PSPC1 | -0.268697088 | 0.561781402 | -1.244586815 | 0.001446259 | -1.513283903 | 9.204317094 | 8.522788974 | 8.585098385 | 8.496596021 | 8.57309694 | 7.569519177 | 7.57933192 |
| ENSG00000052126 | PLEKHA5 | -0.378322239 | 0.784875307 | -1.135866233 | 0.360441295 | -1.514188472 | 6.009025997 | 9.225224669 | 9.798878489 | 8.616799622 | 8.66593596 | 7.912394292 | 7.875307424 |
| ENSG00000060339 | CCAR1 | -0.070807413 | 0.932860886 | -1.444113969 | 0.03239564 | -1.514921383 | 8.564112515 | 10.07368926 | 10.16000817 | 9.656695015 | 9.72225407 | 8.433321781 | 8.213468956 |
| ENSG00000115884 | SDC1 | -1.108062972 | 0.125657681 | -0.406904112 | 0.603099089 | -1.514967085 | 9.317904228 | 8.169643011 | 7.734844356 | 7.501020868 | 7.45182838 | 8.301943231 | 8.014172904 |
| ENSG00000109787 | KLF3 | -0.802655777 | 0.113360191 | -0.712517264 | 0.156916815 | -1.515173041 | 8.87372952 | 9.591588447 | 9.926432745 | 8.76109026 | 8.69891174 | 8.877267588 | 8.759585072 |
| ENSG00000101871 | MID1 | -0.466563266 | 0.535988719 | -1.050102332 | 0.125781284 | -1.516665597 | 9.159305284 | 8.191702463 | 9.227351203 | 8.501535708 | 8.42671895 | 7.223329552 | 8.343344553 |
| ENSG00000154277 | UCHL1 | 1.197562714 | 0.11571852 | -2.714450969 | 7.98E-05 | -1.516888254 | 12.2287862 | 10.52407288 | 10.98787477 | 12.60218578 | 12.6580329 | 8.450751194 | 8.965802436 |
| ENSG00000068308 | OTUD5 | -1.206015961 | 0.145426901 | -0.31108441 | 0.735231663 | -1.517100371 | 10.46903009 | 9.246456519 | 11.38635888 | 9.36574218 | 9.45570025 | 10.30927629 | 10.29555071 |
| ENSG00000057608 | GDI2 | -0.810529571 | 0.033905673 | -0.707352429 | 0.065229023 | -1.517882001 | 12.08576437 | 11.86868155 | 11.36226358 | 10.97949959 | 11.0082583 | 11.06132633 | 11.13168895 |
| ENSG00000196724 | ZNF418 | -0.925398621 | 0.531161556 | -0.593149232 | 0.694164121 | -1.518547853 | 3.327711234 | 7.145017454 | 5.59507254 | 5.003571641 | 5.41912973 | 5.762453498 | 5.243813244 |
| ENSG00000083290 | ULK2 | -1.438061174 | 0.014508855 | -0.081431294 | 0.911833873 | -1.519492468 | 8.107517749 | 8.747909548 | 9.491321578 | 7.412876874 | 7.53039799 | 8.789844052 | 8.828963952 |
| ENSG00000144535 | DIS3L2 | -0.677730176 | 0.466156888 | -0.84237625 | 0.34391267 | -1.520106426 | 8.696033109 | 7.701025996 | 6.534109959 | 7.218576229 | 7.24992376 | 6.824993504 | 7.287011591 |
| ENSG00000131747 | TOP2A | -0.870834295 | 0.691466494 | -0.649960397 | 0.768665969 | -1.520794692 | 5.532353208 | 8.469613413 | 2 | 6.542151749 | 5.87556634 | 5.963846589 | 6.824016918 |
| ENSG00000138160 | KIF11 | -0.826752313 | 0.629335817 | -0.694261571 | 0.685433873 | -1.521013883 | 4.815609471 | 7.60566235 | 3.369317972 | 5.709096143 | 5.3193943 | 5.646229369 | 5.651404445 |
| ENSG00000117461 | PIK3R3 | 0.438651746 | 0.616855364 | -1.959697256 | 0.008669082 | -1.52104551 | 6.257048221 | 7.847167909 | 7.896971941 | 7.956842238 | 7.92150349 | 5.22272385 | 5.96101835 |
| ENSG00000089234 | BRAP | -0.313432194 | 0.807472212 | -1.208002059 | 0.283292274 | -1.521434253 | 6.816637977 | 8.903033433 | 10.11951029 | 8.85414072 | 8.82115545 | 8.050818921 | 7.850099615 |
| ENSG00000184678 | HIST2H2BE | 1.251191287 | 0.322921865 | -2.773344459 | 0.016029651 | -1.522153172 | 9.829762239 | 8.469613413 | 6.629488387 | 10.05695346 | 10.0864045 | 5.085488617 | 6.725806822 |
| ENSG00000129691 | ASH2L | -0.809502457 | 0.725609862 | -0.712720593 | 0.757049561 | -1.522223051 | 10.5468183 | 8.540087354 | 2 | 8.496596021 | 8.46889137 | 8.512023373 | 8.642113606 |
| ENSG00000113558 | SKP1 | -0.203023344 | 0.591297391 | -1.31971002 | 1.88E-05 | -1.522733364 | 11.25872341 | 11.65110249 | 11.78885549 | 11.38768406 | 11.3734127 | 10.15695659 | 10.36819284 |
| ENSG00000213639 | PPP1CB | -0.406891588 | 0.40149941 | -1.11598792 | 0.009399663 | -1.522879508 | 10.71696191 | 11.32663546 | 11.5927499 | 10.91524435 | 10.7827547 | 10.24289218 | 10.03650256 |
| ENSG00000136930 | PSMB7 | -0.213376017 | 0.665819919 | -1.310253853 | 0.001254829 | -1.523629869 | 11.22271378 | 10.60026656 | 10.48092713 | 10.54402725 | 10.6398176 | 9.361563842 | 9.626546639 |
| ENSG00000108963 | DPH1 | 0.068801649 | 0.909699221 | -1.59333673 | 0.000463383 | -1.524535081 | 9.015175615 | 8.297207175 | 9.301632639 | 8.998569022 | 8.98472903 | 7.377096489 | 7.330190599 |
| ENSG00000100083 | GGA1 | -0.783253417 | 0.282216889 | -0.741732146 | 0.300571978 | -1.524985563 | 9.48530526 | 8.57407404 | 10.26027233 | 8.782894215 | 8.84952549 | 8.837641184 | 8.877593894 |
| ENSG00000034713 | GABARAPL2 | -0.715646633 | 0.515922264 | -0.810088591 | 0.447054694 | -1.525735224 | 11.51244431 | 8.929529568 | 9.601288346 | 9.750280081 | 9.71456265 | 9.693591907 | 9.58130758 |
| ENSG00000160087 | UBE2J2 | -1.090020813 | 0.219962273 | -0.43575758 | 0.647499233 | -1.525778393 | 10.72727936 | 9.067150531 | 8.805342542 | 8.70652031 | 8.73115061 | 9.100282869 | 9.594016939 |
| ENSG00000160888 | IER2 | -0.061436621 | 0.939941719 | -1.466000871 | 0.024374582 | -1.527437492 | 9.2903348 | 9.461450882 | 10.53119344 | 9.822457304 | 9.79566886 | 8.751793877 | 7.974870548 |
| ENSG00000143774 | GUK1 | -0.763172071 | 0.08383018 | -0.765000872 | 0.078125458 | -1.528172943 | 11.38138789 | 10.58784379 | 11.34831197 | 10.36045596 | 10.4119259 | 10.34308917 | 10.42495703 |
| ENSG00000160447 | PKN3 | -1.882949173 | 0.229349511 | 0.354401046 | 0.840156112 | -1.528548127 | 2 | 5.838937808 | 6.718950348 | 3.997744386 | 4.31639476 | 6.146071859 | 6.102937566 |
| ENSG00000145439 | CBR4 | -0.737664447 | 0.096461716 | -0.79118843 | 0.065559433 | -1.528852877 | 7.579279349 | 7.572420273 | 7.97167868 | 6.984595707 | 7.00595521 | 7.137053113 | 6.718860807 |
| ENSG00000029639 | TFB1M | -0.4082241 | 0.818265925 | -1.120703993 | 0.484174515 | -1.528928094 | 8.716893051 | 4.11352228 | 6.073378139 | 6.979883891 | 7.01053089 | 6.423979439 | 6.189227296 |
| ENSG00000130520 | LSM4 | -1.210506601 | 0.043308828 | -0.31888359 | 0.635737899 | -1.529390191 | 10.3933013 | 9.00621767 | 9.735974718 | 8.627383409 | 8.60370266 | 9.410081891 | 9.585450415 |
| ENSG00000125319 | C17orf53 | -0.458279944 | 0.796259474 | -1.07111521 | 0.509775204 | -1.529395154 | 6.563806279 | 5.946638857 | 2 | 5.194621444 | 5.44640346 | 4.350488971 | 5.122735884 |
| ENSG00000127184 | COX7C | -1.095890149 | 0.172471782 | -0.434308518 | 0.614417433 | -1.530198667 | 10.94624409 | 9.298208073 | 9.330306409 | 9.009013399 | 8.96835847 | 9.679115027 | 9.612561218 |
| ENSG00000120798 | NR2C1 | -0.40937927 | 0.550957372 | -1.120862181 | 0.067400343 | -1.530241451 | 7.435667355 | 8.451443573 | 8.784770914 | 8.008651253 | 7.83643742 | 7.4096614 | 7.021346309 |
| ENSG00000133398 | MED10 | -0.540072041 | 0.670923419 | -0.990279311 | 0.402364322 | -1.530351352 | 9.781071289 | 8.639735838 | 6.322058544 | 8.268332738 | 8.30907138 | 7.896069536 | 7.792821043 |
| ENSG00000164548 | TRA2A | -0.289058478 | 0.785196727 | -1.241436909 | 0.175908868 | -1.530495388 | 10.30651143 | 8.213429689 | 8.511888463 | 9.084476714 | 8.99284518 | 7.8927823 | 8.276323713 |
| ENSG00000011465 | DCN | -1.300564051 | 0.000196745 | -0.230972622 | 0.57665812 | -1.531536673 | 15.07579791 | 14.52772935 | 14.69777459 | 13.48165944 | 13.4889246 | 14.33331165 | 14.74646283 |
| ENSG00000143499 | SMYD2 | -1.20383463 | 0.000526024 | -0.329051721 | 0.400109235 | -1.532886352 | 9.097018025 | 8.820493869 | 8.784770914 | 7.77569021 | 7.65837038 | 8.360824221 | 8.771382398 |
| ENSG00000115073 | ACTR1B | -1.724873193 | 0.006218119 | 0.191814705 | 0.802148981 | -1.533058487 | 9.820154628 | 8.57407404 | 10.01740919 | 7.862723323 | 7.90685167 | 9.524685263 | 10.00306286 |
| ENSG00000101361 | NOP56 | -0.458244792 | 0.498614729 | -1.07491066 | 0.078357385 | -1.533155452 | 10.49341471 | 9.125613817 | 9.577565424 | 9.401417034 | 9.37815424 | 8.741323065 | 8.813288819 |
| ENSG00000105726 | ATP13A1 | -0.241074372 | 0.712294566 | -1.293141582 | 0.018856875 | -1.534215953 | 10.70657014 | 10.11467741 | 9.426401271 | 9.889939704 | 9.97843848 | 8.793374109 | 8.978944023 |
| ENSG00000151287 | TEX30 | -0.685556728 | 0.822912318 | -0.850474766 | 0.774493502 | -1.536031493 | 7.82963488 | 2 | 2 | 5.775854763 | 5.51241389 | 5.870008604 | 4.998228167 |
| ENSG00000125977 | EIF2S2 | -0.664128936 | 0.182039968 | -0.872007296 | 0.066689274 | -1.536136232 | 11.6462383 | 10.74528399 | 10.87563042 | 10.49120048 | 10.4749506 | 10.33765955 | 10.2112406 |
| ENSG00000085733 | CTTN | -0.575420157 | 0.225627059 | -0.961289951 | 0.030397832 | -1.536710109 | 12.31510994 | 12.10496808 | 12.71516972 | 11.83244654 | 11.8189434 | 11.05646846 | 11.74325939 |
| ENSG00000008382 | MPND | -1.099671906 | 0.528056559 | -0.437169822 | 0.811488554 | -1.536841728 | 7.159102577 | 6.140560021 | 2 | 4.927308402 | 5.37722731 | 5.528313928 | 5.980509018 |
| ENSG00000105185 | PDCD5 | -0.147994077 | 0.762607571 | -1.389288388 | 0.000378379 | -1.537282465 | 8.910412251 | 8.234834549 | 8.941672936 | 8.538855014 | 8.61275919 | 7.346184975 | 7.364717093 |
| ENSG00000165280 | VCP | -0.78814434 | 0.069078369 | -0.751508703 | 0.080413293 | -1.539653043 | 13.09907181 | 12.65418243 | 12.49382591 | 11.97939826 | 11.9900129 | 11.70033342 | 12.28374298 |
| ENSG00000172007 | RAB33B | -1.406044698 | 0.04814826 | -0.136530118 | 0.872932235 | -1.542574816 | 6.816637977 | 8.234834549 | 8.076950877 | 6.496695859 | 6.44777477 | 8.042684843 | 7.258555286 |
| ENSG00000105700 | KXD1 | -0.61331871 | 0.240254037 | -0.929314931 | 0.058802978 | -1.542633641 | 10.15425007 | 9.842262059 | 9.132850258 | 9.137269904 | 9.18421833 | 8.701014095 | 8.980396837 |
| ENSG00000166428 | PLD4 | -3.753025943 | NA | 2.209644739 | NA | -1.543381204 | 2 | 2 | 2 | 2 | 2 | 3.465698119 | 2.671842496 |
| ENSG00000120837 | NFYB | -1.305834681 | 0.335280148 | -0.240727325 | 0.874489793 | -1.546562006 | 8.963744667 | 8.053983481 | 4.874587708 | 6.820355514 | 6.72891807 | 8.171023934 | 7.334741191 |
| ENSG00000126267 | COX6B1 | -0.771415386 | 0.189075569 | -0.775325282 | 0.178213485 | -1.546740669 | 10.79750658 | 9.558016677 | 10.11127249 | 9.435370781 | 9.51325794 | 9.35261874 | 9.580349849 |
| ENSG00000076043 | REXO2 | -0.663142779 | 0.417114533 | -0.88385748 | 0.25649001 | -1.547000258 | 11.05263482 | 9.695596167 | 9.31604076 | 9.549478128 | 9.57865651 | 9.399697583 | 9.287475811 |
| ENSG00000176731 | C8orf59 | -0.289203395 | 0.654150801 | -1.257925748 | 0.023011611 | -1.547129143 | 7.940205782 | 6.78762695 | 7.646393253 | 7.2696592 | 7.21069602 | 6.205571486 | 6.403650353 |
| C4ORF22 | C4ORF22 | 3.64349023 | 0.524450905 | -5.191091598 | 0.364514502 | -1.547601368 | 2 | 2 | 2 | 4.070463721 | 4.05806467 | 2 | 2 |
| ENSG00000163013 | FBXO41 | -0.664050218 | 0.733353042 | -0.883593821 | 0.639142933 | -1.547644039 | 2 | 6.78762695 | 7.97167868 | 6.164317909 | 6.40670796 | 6.320128742 | 5.799216178 |
| ENSG00000167595 | PROSER3 | -1.504591278 | 0.284673982 | -0.04387781 | 0.978224387 | -1.548469089 | 5.330594134 | 5.457120953 | 8.561105505 | 5.968925393 | 5.65950892 | 7.20886319 | 7.224630935 |
| ENSG00000175084 | DES | -4.343113373 | 0.000228835 | 2.793790816 | 0.017043752 | -1.549322557 | 4.00722531 | 6.389896633 | 7.342949392 | 2.868578001 | 3.15372745 | 8.285116882 | 9.732652955 |
| ENSG00000125485 | DDX31 | -0.611843019 | 0.766447439 | -0.939274038 | 0.633808927 | -1.551117057 | 2 | 7.051611315 | 8.561105505 | 6.887305128 | 6.76185814 | 6.708295923 | 6.283256815 |
| ENSG00000136273 | HUS1 | -0.479326724 | 0.721229196 | -1.073205031 | 0.384149032 | -1.552531755 | 9.556468883 | 6.899080544 | 6.718950348 | 7.865280446 | 7.88209639 | 7.138441638 | 7.431385914 |
| ENSG00000139910 | NOVA1 | -2.320477225 | 0.005058045 | 0.766312867 | 0.379558392 | -1.554164358 | 6.138360429 | 4.93638933 | 5.156626401 | 3.840289928 | 3.33976443 | 6.028179536 | 6.365613546 |
| ENSG00000141741 | MIEN1 | -1.034735058 | 0.198785891 | -0.519481132 | 0.538306766 | -1.55421619 | 10.07567912 | 8.213429689 | 8.960138144 | 8.283707551 | 8.23475766 | 8.530649266 | 8.973604485 |
| ENSG00000118690 | ARMC2 | -0.916760786 | 0.649972951 | -0.638191759 | 0.754844937 | -1.554952545 | 6.563806279 | 3.686303414 | 2 | 4.387512679 | 4.45712836 | 4.727866721 | 4.574506939 |
| ENSG00000132321 | IQCA1 | -2.976211788 | 0.367097793 | 1.419422184 | 0.655105561 | -1.556789604 | 2 | 3.686303414 | 2 | 2 | 2.2680111 | 3.709392871 | 3.600986722 |
| ENSG00000005156 | LIG3 | -1.17583695 | 0.002482135 | -0.380992218 | 0.37369315 | -1.556829168 | 8.444143041 | 7.790474954 | 8.236946975 | 6.946463121 | 7.09482005 | 7.797560316 | 7.799420735 |
| ENSG00000112308 | C6orf62 | -0.262200101 | 0.537440375 | -1.294785324 | 0.000280988 | -1.556985426 | 11.34834729 | 11.72481431 | 12.03516535 | 11.50956605 | 11.424668 | 10.32567245 | 10.54091836 |
| ENSG00000161929 | SCIMP | -1.476497549 | 0.433973734 | -0.081541656 | 0.968570087 | -1.558039206 | 3.327711234 | 4.11352228 | 2 | 2.695456494 | 2.68924543 | 3.648325314 | 3.071407821 |
| ENSG00000186684 | CYP27C1 | -3.538553439 | 0.002397074 | 1.980122694 | 0.05519677 | -1.558430745 | 3.327711234 | 4.11352228 | 5.156626401 | 2.3892346 | 2.49394783 | 5.985611216 | 6.254725615 |
| ENSG00000141499 | WRAP53 | -0.773911016 | 0.691746353 | -0.786555731 | 0.683005276 | -1.560466748 | 2 | 6.78762695 | 8.110401118 | 6.449761039 | 6.08962627 | 6.192262702 | 6.341697848 |
| ENSG00000103021 | CCDC113 | -1.676310401 | 0.065038902 | 0.115720952 | 0.914753421 | -1.560589449 | 4.815609471 | 6.228547981 | 5.59507254 | 4.237680128 | 4.22485516 | 6.192262702 | 5.217771644 |
| ENSG00000175203 | DCTN2 | -0.944028635 | 0.000672752 | -0.618412546 | 0.032293519 | -1.562441181 | 10.7426185 | 10.77115258 | 10.41552659 | 9.678668908 | 9.74577157 | 9.934835676 | 10.13219057 |
| ENSG00000151882 | CCL28 | -1.893647614 | 0.121412754 | 0.330317647 | 0.807192459 | -1.563329967 | 5.095959458 | 3.076665209 | 5.156626401 | 3.462030052 | 3.08597292 | 4.705557986 | 5.1574003 |
| ENSG00000135457 | TFCP2 | -0.757422699 | 0.724832375 | -0.80631297 | 0.702841297 | -1.56373567 | 2 | 8.806266215 | 9.589475645 | 7.959237999 | 7.88459112 | 7.815899769 | 7.930464419 |
| ENSG00000111669 | TPI1 | -1.127406666 | 0.002169292 | -0.437410043 | 0.278589089 | -1.56481671 | 13.67917225 | 13.35152472 | 13.71677814 | 12.44482528 | 12.4839871 | 12.81250915 | 13.43018847 |
| ENSG00000145191 | EIF2B5 | -0.804637358 | 0.201926325 | -0.760673233 | 0.221569388 | -1.565310591 | 8.367060731 | 9.079034257 | 9.757248847 | 8.442826028 | 8.31092742 | 8.336833852 | 8.502136032 |
| ENSG00000070423 | RNF126 | -1.041649651 | 0.219156141 | -0.524957525 | 0.554505321 | -1.566607176 | 10.58135316 | 9.04308532 | 8.825624954 | 8.616799622 | 8.73115061 | 8.860117405 | 9.454284815 |
| ENSG00000117009 | KMO | -1.689389614 | 0.594639663 | 0.119757502 | NA | -1.569632112 | 2 | 2 | 3.369317972 | 2.141737246 | 2.2680111 | 2.831336763 | 2.374681325 |
| ENSG00000109113 | RAB34 | -0.989161904 | 0.172780449 | -0.580837262 | 0.439051343 | -1.569999166 | 11.32476023 | 11.13502303 | 9.681375057 | 9.905552176 | 9.87371334 | 10.08262707 | 10.48282173 |
| ENSG00000132170 | PPARG | -3.627929422 | 0.014454932 | 2.057192496 | 0.133507552 | -1.570736925 | 2 | 5.131603625 | 4.058998049 | 2.270786764 | 2.49394783 | 6.019162896 | 6.124122661 |
| ENSG00000137171 | KLC4 | -1.079102505 | 0.605635767 | -0.493149075 | 0.8207453 | -1.572251581 | 9.460782017 | 7.954278496 | 2 | 7.242377933 | 7.27295842 | 7.943705881 | 7.708840864 |
| ENSG00000182393 | IFNL1 | 3.618567051 | 0.52739221 | -5.191091702 | 0.364514502 | -1.572524651 | 2 | 2 | 2 | 3.840289928 | 4.22485516 | 2 | 2 |
| ENSG00000106346 | USP42 | -0.646391046 | 0.459554196 | -0.928964617 | 0.259642722 | -1.575355663 | 7.384433967 | 8.862353853 | 9.439621147 | 8.078634414 | 8.21705076 | 8.035995406 | 7.685278529 |
| ENSG00000144744 | UBA3 | -0.46264031 | 0.066039024 | -1.11279563 | 8.97E-07 | -1.57543594 | 9.740899611 | 9.532315021 | 9.636156974 | 9.203437087 | 9.14937381 | 8.548515606 | 8.515559017 |
| ENSG00000162241 | SLC25A45 | -0.501689755 | 0.784300093 | -1.074075288 | 0.525858337 | -1.575765043 | 3.327711234 | 7.73146271 | 4.874587708 | 6.096288482 | 5.74985018 | 5.125248504 | 5.641643428 |
| ENSG00000177432 | NAP1L5 | -1.099016961 | 0.2546055 | -0.477191524 | 0.640791054 | -1.576208485 | 7.82963488 | 5.946638857 | 8.35328246 | 6.670510205 | 6.53928196 | 6.975527754 | 7.407104868 |
| ENSG00000170445 | HARS | -0.340930011 | 0.509715065 | -1.235474309 | 0.005486623 | -1.57640432 | 10.34721797 | 9.71095217 | 9.926432745 | 9.689531608 | 9.6675326 | 8.42082352 | 9.084037526 |
| ENSG00000197905 | TEAD4 | -3.045366003 | 0.085712737 | 1.46837253 | 0.431244286 | -1.576993474 | 8.313285031 | 6.666836837 | 2 | 4.205751501 | 4.50979031 | 7.907516231 | 9.041986855 |
| ENSG00000006453 | BAIAP2L1 | -2.240292079 | 0.000642562 | 0.662773446 | 0.364514502 | -1.577518634 | 8.540903468 | 7.761270542 | 6.882775709 | 5.720439317 | 5.73886209 | 8.681620276 | 8.372406522 |

| | | | | | | | | | | | | | |
|---|---|---|---|---|---|---|---|---|---|---|---|---|---|
| ENSG00000132300 | PTCD3 | -0.435290779 | 0.17053846 | -1.142801875 | 4.85E-05 | -1.578092653 | 9.689059027 | 9.591588447 | 9.272377397 | 9.109497937 | 9.08575246 | 8.335018389 | 8.456236734 |
| ENSG00000123268 | ATF1 | -0.432699073 | 0.70815223 | -1.146098997 | 0.271141542 | -1.57879807 | 6.816637977 | 7.189546432 | 9.227351203 | 7.614805492 | 7.83643742 | 7.107581047 | 6.955898408 |
| ENSG00000123545 | NDUFAF4 | -0.622473606 | 0.459212556 | -0.958363308 | 0.222598244 | -1.580836914 | 7.904282243 | 6.228547981 | 7.896971941 | 6.912249745 | 6.90635305 | 6.763331418 | 6.38329369 |
| ENSG00000172239 | PAIP1 | 0.208741726 | 0.688378388 | -1.790387884 | 1.65E-05 | -1.581646158 | 9.730679447 | 9.599860728 | 8.941672936 | 9.654479106 | 9.69052657 | 7.874567387 | 7.488317792 |
| ENSG00000183597 | TANGO2 | -0.028212384 | 0.989759352 | -1.554498541 | 0.412109699 | -1.582710925 | 10.16187685 | 3.686303414 | 6.718950348 | 8.622857004 | 8.75316018 | 7.24282551 | 7.133547492 |
| ENSG00000160075 | SSU72 | -0.96190806 | 0.128533462 | -0.622094426 | 0.338976671 | -1.584002485 | 11.31794963 | 10.27823701 | 9.981704053 | 9.657432896 | 9.71526357 | 10.05624005 | 9.993021861 |
| ENSG00000136240 | KDELR2 | -0.990308687 | 0.021365645 | -0.593811641 | 0.186486326 | -1.584120328 | 13.33577055 | 12.72774099 | 12.5282657 | 11.92359756 | 11.9094643 | 12.18307138 | 12.43161868 |
| ENSG00000165506 | DNAAF2 | -0.995616896 | 0.371190997 | -0.588779591 | 0.607699942 | -1.584396488 | 8.418903792 | 6.899080544 | 5.59507254 | 6.422239259 | 6.46787796 | 6.961475607 | 6.697819825 |
| ENSG00000134824 | FADS2 | -1.864043699 | 0.038357022 | 0.278389614 | 0.79369186 | -1.585654084 | 13.04132235 | 10.81444733 | 12.64414369 | 10.5575302 | 10.5925339 | 12.01712251 | 13.1828323 |
| ENSG00000103502 | CDIPT | -0.887887772 | 0.232682693 | -0.698227105 | 0.351780632 | -1.586114878 | 11.59645304 | 10.03760521 | 10.27509734 | 9.840113674 | 10.0039039 | 10.05164138 | 10.17236288 |
| ENSG00000153291 | SLC25A27 | -0.758270085 | 0.416637582 | -0.82840854 | 0.353010925 | -1.586678625 | 4.467311984 | 4.710564154 | 5.59507254 | 4.387512679 | 4.31639476 | 4.406883583 | 4.175132639 |
| ENSG00000186638 | KIF24 | -1.095436761 | 0.617500397 | -0.49370262 | 0.829885014 | -1.589139381 | 3.327711234 | 6.78762695 | 2 | 4.470469939 | 4.43005853 | 4.434275199 | 5.343519008 |
| ENSG00000101546 | RBFA | -1.08429857 | 0.218893333 | -0.507955978 | 0.583582174 | -1.592254548 | 6.366709671 | 8.169643011 | 6.629488387 | 6.268449691 | 6.25300433 | 6.645228724 | 6.96179415 |
| ENSG00000145741 | BTF3 | -0.968552646 | 0.205806676 | -0.62502701 | 0.426712094 | -1.593579655 | 11.98832668 | 10.13189344 | 11.05383672 | 10.27841188 | 10.2892535 | 10.56905757 | 10.6815274 |
| ENSG00000178537 | SLC25A20 | -0.848717697 | 0.647476405 | -0.74512894 | 0.687731208 | -1.593846637 | 3.327711234 | 7.761270542 | 9.61300511 | 7.571615825 | 7.5462416 | 7.505319234 | 7.800517754 |
| ENSG00000130005 | GAMT | -1.371363454 | 0.17057763 | -0.22318916 | 0.845529489 | -1.594552615 | 8.981092835 | 8.862353853 | 6.322058544 | 7.157295126 | 7.05992818 | 8.275052313 | 8.192729549 |
| ENSG00000085872 | CHERP | -0.821368824 | 0.142652965 | -0.773903291 | 0.161318642 | -1.595272114 | 9.081016759 | 8.791896851 | 9.868959282 | 8.409925372 | 8.59153817 | 8.680667534 | 8.407389648 |
| ENSG00000160051 | IQCC | -1.85580017 | 0.023144808 | 0.260385785 | 0.778780813 | -1.595414385 | 5.095959458 | 5.595927442 | 4.523675391 | 3.515101962 | 3.82856379 | 5.449961224 | 5.349527629 |
| ENSG00000185875 | THNSL1 | -0.857589202 | 0.254248233 | -0.738251224 | 0.318663122 | -1.595840426 | 5.532353208 | 6.535001748 | 5.930865648 | 5.258265913 | 5.3193943 | 5.669467138 | 5.0581604 |
| ENSG00000114779 | ABHD14B | -0.863713487 | 0.259714186 | -0.733137779 | 0.337851136 | -1.596851266 | 9.460782017 | 8.234834549 | 9.963514551 | 8.446246007 | 8.59763324 | 8.357847151 | 8.89823658 |
| ENSG00000168556 | ING2 | -1.084290375 | 0.098414809 | -0.513204273 | 0.457693718 | -1.597494648 | 8.285626337 | 7.538394143 | 6.958202216 | 6.693715713 | 6.5890786 | 6.915242278 | 7.428550441 |
| ENSG00000139291 | TMEM19 | -0.790567012 | 0.443668589 | -0.80840849 | 0.422686222 | -1.598975502 | 9.544849385 | 6.899080544 | 8.677268317 | 7.990026167 | 7.90439517 | 7.79139499 | 8.057030103 |
| ENSG00000132963 | POMP | 0.408542176 | 0.501949395 | -2.008784389 | 7.65E-05 | -1.600242213 | 10.16946352 | 9.267380431 | 10.49365885 | 10.46951599 | 10.4658164 | 8.04860508 | 8.087009646 |
| ENSG00000221978 | CCNL2 | -0.991542303 | 0.004574811 | -0.608812607 | 0.096019822 | -1.60035491 | 10.80239439 | 11.14636276 | 11.40332671 | 10.1289481 | 10.1685416 | 10.66298318 | 10.38437908 |
| ENSG00000159346 | ADIPOR1 | -0.974731373 | 0.009276725 | -0.626763721 | 0.107612604 | -1.601495094 | 11.55650736 | 11.20447669 | 11.91489601 | 10.55555238 | 10.6694157 | 10.89014846 | 11.02814425 |
| ENSG00000011260 | UTP18 | -0.35636865 | 0.596166767 | -1.24636268 | 0.033143403 | -1.60273133 | 8.946185348 | 8.147241016 | 7.646393253 | 8.036145657 | 7.95274764 | 7.236356087 | 7.0024032 |
| ENSG00000169251 | NMD3 | -0.682558965 | 0.35660897 | -0.921119735 | 0.190829323 | -1.6036787 | 10.22148166 | 9.785372135 | 8.585098385 | 9.031955922 | 8.97188212 | 8.977260248 | 8.517561736 |
| ENSG00000176087 | SLC35A4 | -1.19061362 | 0.531143362 | -0.4159707 | 0.837702779 | -1.60658432 | 4.00722531 | 10.93049815 | 11.18913795 | 9.248208205 | 9.35135081 | 9.73685847 | 10.34300322 |
| ENSG00000213699 | SLC35F6 | -1.072914023 | 0.095993057 | -0.536000703 | 0.42991362 | -1.608914726 | 10.1076228 | 9.308339287 | 10.81605395 | 9.12451847 | 9.14521857 | 9.620703391 | 9.715016953 |
| ENSG00000161547 | SRSF2 | -0.510673857 | 0.396866142 | -1.099227674 | 0.044653042 | -1.609901531 | 10.84565509 | 10.61258327 | 10.09465441 | 10.05303338 | 10.0306182 | 8.890924823 | 9.861231258 |
| ENSG00000163634 | THOC7 | -0.63158217 | 0.601636131 | -0.979785688 | 0.39010814 | -1.611367858 | 9.858207046 | 8.590771778 | 6.629488387 | 8.223194971 | 8.284723 | 7.727325329 | 8.074367056 |
| ENSG00000151500 | THYN1 | -0.499212016 | 0.561438629 | -1.112709714 | 0.155175621 | -1.61192173 | 7.940205782 | 8.671479029 | 6.718950348 | 7.497728447 | 7.50468064 | 7.031901364 | 6.764548117 |
| ENSG00000152457 | DCLRE1C | -0.319364436 | 0.530705838 | -1.293177938 | 0.00277408 | -1.612542375 | 7.867441304 | 7.393647515 | 7.896971941 | 7.450827758 | 7.37938471 | 6.679970309 | 6.232137896 |
| ENSG00000071894 | CPSF1 | -1.014369994 | 0.127598657 | -0.598733174 | 0.38415025 | -1.613103168 | 10.68025843 | 10.39692156 | 9.132850258 | 9.196346103 | 9.20432875 | 9.588047701 | 9.638558632 |
| ENSG00000188878 | FBF1 | -0.403293552 | 0.834689821 | -1.210594242 | 0.484697884 | -1.613887794 | 7.384433967 | 5.946638857 | 2 | 5.860316854 | 5.93454154 | 5.289239922 | 5.020995508 |
| ENSG00000141627 | DYM | -1.369262251 | 0.00015596 | -0.245000521 | 0.565942887 | -1.614262772 | 9.751047882 | 9.591588447 | 10.16797309 | 8.444537031 | 8.54498542 | 9.772619242 | 9.429325923 |
| ENSG00000011132 | APBA3 | -1.018981023 | 0.321495834 | -0.595565021 | 0.5745905 | -1.614546044 | 7.096806373 | 7.761270542 | 9.541233192 | 7.464384256 | 7.55566492 | 7.975129619 | 7.875307424 |
| ENSG00000148308 | GTF3C5 | -0.906934479 | 0.007452299 | -0.708208404 | 0.039648512 | -1.615142884 | 9.189468652 | 9.030900487 | 9.589475645 | 8.379838154 | 8.3920703 | 8.566163393 | 8.598629536 |
| ENSG00000173933 | RBM4 | 0.476986874 | 0.671489931 | -2.093092726 | 0.029674646 | -1.616105851 | 9.613202746 | 8.747909548 | 6.718950348 | 9.288777073 | 9.21725179 | 6.683779344 | 6.77351449 |
| ENSG00000116489 | CAPZA1 | -0.53108163 | 0.216092164 | -1.0873797 | 0.00553577 | -1.618461329 | 11.35168561 | 11.51338035 | 12.09904656 | 11.17175047 | 11.1497911 | 10.63300595 | 10.57785604 |
| ENSG00000166902 | MRPL16 | -1.140996869 | 0.146175165 | -0.477928167 | 0.568416252 | -1.618926835 | 9.810482605 | 8.623598428 | 8.007629717 | 7.745471964 | 8.01099479 | 8.385592001 | 8.676578788 |
| ENSG00000072958 | AP1M1 | -1.179511609 | 0.016308157 | -0.44127264 | 0.412109699 | -1.62078425 | 10.51739401 | 9.828248506 | 10.87075861 | 9.331013912 | 9.24954577 | 9.880069077 | 10.1574558 |
| ENSG00000172009 | THOP1 | -1.049925723 | 0.646239362 | -0.572169259 | 0.808778437 | -1.622094982 | 10.35389212 | 7.874700259 | 2 | 7.927778434 | 8.01555453 | 8.066950774 | 8.741991736 |
| ENSG00000187601 | MAGEH1 | -1.475539763 | 0.000257387 | -0.146569589 | 0.768207195 | -1.622109352 | 8.87372952 | 9.00621767 | 8.236946975 | 7.352188525 | 7.23826702 | 8.678282924 | 8.523553267 |
| ENSG00000113593 | PPWD1 | -0.452728914 | 0.736046082 | -1.170912284 | 0.337870848 | -1.623641198 | 9.613202746 | 8.029695761 | 6.073378139 | 8.139053376 | 8.0314011 | 7.55607473 | 7.18317631 |
| ENSG00000136908 | DPM2 | -1.143887217 | 0.000621727 | -0.479820398 | 0.184008652 | -1.623707615 | 8.998234871 | 8.806266215 | 9.287079172 | 7.900612272 | 7.91663606 | 8.427654172 | 8.688479312 |
| ENSG00000197980 | LEKR1 | -0.045903241 | 0.986149479 | -1.580136215 | 0.503376289 | -1.626039456 | 2 | 4.11352228 | 2 | 2.947924294 | 3.28037825 | 2.560105256 | 2.421067459 |
| ENSG00000198026 | ZNF335 | -0.699916836 | 0.404128024 | -0.927196939 | 0.246468033 | -1.627113775 | 9.497412313 | 8.655694736 | 7.502634626 | 8.078634414 | 8.07355593 | 7.752026287 | 7.946410318 |
| ENSG00000115266 | APC2 | -1.383856991 | 0.019668123 | -0.243780579 | 0.719764201 | -1.627637571 | 6.257048221 | 6.140560021 | 6.80318718 | 5.319220934 | 4.93258395 | 6.353989995 | 5.995914157 |
| ENSG00000173614 | NMNAT1 | -0.949663814 | 0.251382472 | -0.678120911 | 0.419378285 | -1.627784725 | 8.963744667 | 7.274681896 | 7.451345288 | 7.165617905 | 7.18663697 | 7.511762185 | 7.366199662 |
| ENSG00000149547 | EI24 | -1.172465159 | 0.032927043 | -0.456108007 | 0.445567406 | -1.628573165 | 11.60207034 | 10.93704246 | 10.37481416 | 9.906793906 | 9.87119971 | 10.47807907 | 10.71812545 |
| ENSG00000108798 | ABI3 | -2.999620341 | 0.29492176 | 1.370981987 | 0.608991096 | -1.628638354 | 2 | 3.686303414 | 2 | 2.141737246 | 2.14022021 | 4.122514202 | 2.918126211 |
| ENSG00000156482 | RPL30 | -0.528807095 | 0.554920781 | -1.09989818 | 0.180853 | -1.628705275 | 12.12920672 | 10.26789142 | 10.54960077 | 10.6534857 | 10.7506356 | 10.11437312 | 10.15230914 |
| ENSG00000182199 | SHMT2 | -1.435263653 | 1.76E-05 | -0.193900156 | 0.632771785 | -1.629163809 | 11.20066853 | 10.65686569 | 11.25791777 | 9.61323902 | 9.64710779 | 10.9086776 | 10.82495146 |
| ENSG00000185361 | TNFAIP8L1 | -2.079156997 | 0.080087979 | 0.449921476 | 0.736888343 | -1.629235521 | 5.532353208 | 6.666836837 | 8.784770914 | 5.514313187 | 5.72778967 | 7.978234673 | 8.126037235 |
| ENSG00000185842 | DNAH14 | -0.561511866 | 0.536732782 | -1.068651791 | 0.200135445 | -1.630163657 | 4.815609471 | 5.595927442 | 5.59507254 | 4.804776789 | 4.97065356 | 4.764298067 | 4.077147165 |
| ENSG00000213626 | LBH | -2.985600854 | 1.31E-32 | 1.354928113 | 1.32E-07 | -1.63067274 | 12.05127578 | 11.68259927 | 11.80936937 | 8.887228971 | 8.87358581 | 13.30687429 | 13.10455459 |
| ENSG00000116898 | MRPS15 | -0.857248483 | 0.226279403 | -0.774491692 | 0.270404914 | -1.631740174 | 9.657036304 | 9.114109157 | 8.042706615 | 8.178656739 | 8.28661061 | 8.11499695 | 8.491306993 |
| ENSG00000168256 | NKIRAS2 | -1.206616729 | 0.059908089 | -0.425231973 | 0.543799327 | -1.631848702 | 11.50046327 | 10.14890644 | 10.34701993 | 9.569217728 | 9.61666844 | 10.29003906 | 10.44841713 |
| ENSG00000115368 | WDR75 | -0.359640461 | 0.497802375 | -1.274060212 | 0.005383097 | -1.633700673 | 9.233563228 | 8.590771778 | 9.180874355 | 8.670421699 | 8.66738547 | 8.044167181 | 7.422862719 |
| ENSG00000150054 | MPP7 | -4.742671606 | 4.12E-06 | 3.108839248 | 4.41E-05 | -1.633832358 | 4.00722531 | 4.93638933 | 5.59507254 | 2.3892346 | 2.2680111 | 7.743839536 | 8.06618078 |
| ENSG00000184698 | OR51M1 | -3.753027084 | NA | 2.118273668 | NA | -1.634753416 | 2 | 2 | 2 | 2 | 2 | 2.717032197 | 3.36036153 |
| ENSG00000273780 | | 4.40134359 | 0.045693057 | -6.036288545 | 0.005795713 | -1.634944954 | 2 | 7.002537973 | 2 | 9.715273165 | 9.87810168 | 2.094598369 | 2.326754077 |
| ENSG00000168040 | FADD | 0.230984751 | 0.80683022 | -1.866914004 | 0.015246955 | -1.635929252 | 9.33149386 | 7.315436877 | 8.408111533 | 8.805705438 | 8.790887 | 6.571075185 | 6.895576665 |
| ENSG00000105355 | PLIN3 | -0.931847766 | 0.022017331 | -0.70420287 | 0.090276485 | -1.636050636 | 11.40085503 | 11.30655942 | 11.91962127 | 10.63224832 | 10.642768 | 10.62126059 | 11.07286611 |
| ENSG00000177025 | C19orf18 | -0.23625302 | 0.91483416 | -1.400102605 | 0.468311342 | -1.636355625 | 4.00722531 | 4.11352228 | 2 | 3.462030052 | 3.55551834 | 3.256555576 | 2.326754077 |
| ENSG00000172269 | DPAGT1 | -1.095534861 | 0.042310832 | -0.540845681 | 0.344232516 | -1.636380542 | 9.848787621 | 9.549500232 | 8.699419242 | 8.388752656 | 8.32569041 | 8.738579005 | 9.057211603 |
| ENSG00000091536 | MYO15A | -1.248981788 | 0.635835342 | -0.388235429 | 0.890313426 | -1.637217217 | 6.138360429 | 2 | 2 | 3.709785322 | 3.74316441 | 4.521997428 | 4.215171317 |
| ENSG00000028116 | VRK2 | 0.486062392 | 0.417669822 | -2.123516636 | 2.79E-05 | -1.637454244 | 8.777722034 | 8.356994601 | 7.552162999 | 8.807036108 | 8.78555748 | 6.332313483 | 6.165581817 |
| ENSG00000133983 | COX16 | 0.908258014 | 0.077275225 | -2.545911432 | 4.05E-08 | -1.637653418 | 7.384433967 | 6.899080544 | 7.691296561 | 8.235106869 | 8.24450173 | 4.73522696 | 5.164234344 |
| ENSG00000168924 | LETM1 | -1.26761458 | 0.000702835 | -0.370414385 | 0.377920795 | -1.638028965 | 9.790941832 | 9.599860728 | 10.17589429 | 8.643859774 | 8.58388295 | 9.284329221 | 9.695689382 |
| ENSG00000136044 | APPL2 | -1.57493892 | 0.025870272 | -0.06350371 | 0.94272587 | -1.63844263 | 10.30651143 | 8.717821467 | 9.116480286 | 7.982979299 | 7.98565528 | 9.22478876 | 9.698340256 |
| ENSG00000104763 | ASAH1 | -0.183684071 | 0.826100734 | -1.454933106 | 0.032980835 | -1.638617177 | 11.04027452 | 9.672251278 | 11.36918911 | 10.70364381 | 10.642031 | 9.52813224 | 9.274507165 |
| ENSG00000104897 | SF3A2 | -1.163909042 | 5.66E-05 | -0.47541993 | 0.12973302 | -1.639328973 | 9.992581382 | 9.574900212 | 9.990713521 | 8.699372486 | 8.71161289 | 9.379289361 | 9.3981418 |
| ENSG00000076003 | MCM6 | -1.745902078 | 0.002944485 | 0.106152008 | 0.886197127 | -1.63975007 | 7.485143553 | 8.671479029 | 7.734844356 | 6.456560224 | 6.28357155 | 8.341061068 | 7.98359008 |
| ENSG00000100767 | PAPLN | -3.019452803 | 7.07E-05 | 1.379151089 | 0.069529485 | -1.640301713 | 5.532353208 | 5.72254323 | 6.203069917 | 3.615725078 | 3.15372745 | 6.651077397 | 7.531242162 |
| ENSG00000128683 | GAD1 | 2.152475678 | NA | -3.793052289 | NA | -1.640576611 | 2 | 2 | 2 | 3.406930997 | 2.68924543 | 2.048074405 | 2 |
| ENSG00000013375 | PGM3 | -0.467614483 | 0.337673274 | -1.1739301 | 0.007115471 | -1.641544584 | 10.84091191 | 11.37322539 | 11.78108696 | 10.91431859 | 10.9155453 | 10.26324148 | 10.15574229 |
| ENSG00000168036 | CTNNB1 | -0.573688045 | 0.076570922 | -1.068714112 | 0.000343749 | -1.642402157 | 12.47671498 | 12.06519355 | 12.56341295 | 11.7822503 | 11.8378939 | 11.25618281 | 11.37339918 |
| ENSG00000144550 | CPNE9 | -0.272311451 | 0.93880631 | -1.37034212 | NA | -1.642653571 | 2 | 2 | 3.369317972 | 2.498691189 | 2.49394783 | 2.42098416 | 2.059860178 |
| ENSG00000075218 | GTSE1 | -1.214188438 | 0.566306066 | -0.42863272 | 0.849170035 | -1.642821158 | 3.327711234 | 6.666836837 | 2 | 4.17310023 | 4.3456612 | 4.644324387 | 5.14363432 |
| C1QTNF9B-AS1 | C1QTNF9B-A | 3.546158349 | 0.533465331 | -5.191092003 | 0.364514502 | -1.644933655 | 2 | 2 | 2 | 3.840289928 | 4.12711147 | 2 | 2 |
| ENSG00000196235 | SUPT5H | -0.549656807 | 0.262050542 | -1.097783477 | 0.014619621 | -1.647440284 | 11.42640922 | 10.62884366 | 11.54155646 | 10.69755827 | 10.7065491 | 10.11666451 | 10.19339501 |
| ENSG00000187051 | RPS19BP1 | -0.817011244 | 0.386453724 | -0.83084243 | 0.366751756 | -1.647853673 | 10.499447 | 8.955547846 | 8.325066185 | 8.752828065 | 8.74357228 | 8.836358413 | 8.624876935 |
| ENSG00000138303 | ASCC1 | -0.950919044 | 0.001032412 | -0.697042141 | 0.018248047 | -1.647961185 | 8.540903468 | 8.639735838 | 8.408111533 | 7.549526251 | 7.64959655 | 7.802823992 | 7.894955865 |
| ENSG00000079246 | XRCC5 | -0.71954502 | 0.020700446 | -0.928790617 | 0.001956856 | -1.648335637 | 11.52136524 | 11.94371713 | 11.760164 | 11.06651655 | 11.0013784 | 10.66815974 | 10.9674134 |
| ENSG00000154258 | ABCA9 | 1.136848049 | 0.409917847 | -2.789465146 | 0.029770746 | -1.652617097 | 4.00722531 | 3.076665209 | 4.523675391 | 4.825935136 | 4.89348248 | 2.3070652 | 2.671842496 |
| ENSG00000132341 | RAN | -0.88339077 | 0.110591109 | -0.769276209 | 0.163161971 | -1.652666979 | 12.24323744 | 11.26814685 | 11.34128524 | 10.8245327 | 10.7897592 | 10.67988853 | 11.12776105 |
| ENSG00000175826 | CTDNEP1 | -0.819918827 | 0.121301732 | -0.833882783 | 0.108401159 | -1.653801609 | 10.51143645 | 10.32389548 | 9.452720984 | 9.298270478 | 9.39837151 | 9.194070966 | 9.464010251 |
| ENSG00000167257 | RNF214 | -0.758673162 | 0.72338932 | -0.898179109 | 0.668849935 | -1.656852271 | 9.397585735 | 7.669933291 | 2 | 7.409376773 | 7.49817903 | 7.371203179 | 7.263337153 |
| ENSG00000088179 | PTPN4 | -0.67020026 | 0.107207965 | -0.986972721 | 0.011684435 | -1.65717298 | 7.940205782 | 8.004992147 | 8.561105505 | 7.549526251 | 7.51437845 | 7.178138321 | 7.266516281 |
| ENSG00000213190 | MLLT11 | 0.362709659 | 0.26497168 | -2.020085307 | 1.53E-13 | -1.657375648 | 10.31337612 | 10.3387994 | 10.32581712 | 10.70292919 | 10.6726672 | 8.085786009 | 8.521558854 |
| ENSG00000157800 | SLC37A3 | -1.165995899 | 0.002229097 | -0.491522291 | 0.235579015 | -1.65751819 | 9.602033055 | 10.32888058 | 10.10298738 | 8.852852814 | 8.91688363 | 9.458689553 | 9.645596027 |
| ENSG00000078668 | VDAC3 | -1.13255362 | 0.002449695 | -0.525754295 | 0.1906313 | -1.658307916 | 10.1076228 | 10.10308486 | 10.68897008 | 9.164517339 | 9.23104065 | 9.671222295 | 9.921211649 |
| ENSG00000185624 | P4HB | -0.255405536 | 0.684986885 | -1.40319177 | 0.007503255 | -1.658597306 | 15.03873922 | 13.88920025 | 14.24271598 | 14.1891151 | 14.2452844 | 12.93237165 | 13.19573601 |
| ENSG00000169964 | TMEM42 | -1.620193879 | 0.057338722 | -0.03919371 | 0.970049089 | -1.659387589 | 6.366709671 | 7.467835385 | 8.536706852 | 6.18902405 | 6.1068222 | 7.852739912 | 7.470508284 |
| ENSG00000162073 | PAQR4 | -1.400612513 | 0.633071268 | -0.259187572 | 0.935195648 | -1.659800085 | 2 | 2 | 8.110401118 | 5.273747807 | 5.24368216 | 6.68757835 | 5.812327524 |
| ENSG00000198908 | BHLHB9 | -0.953071876 | 0.644603834 | -0.707044466 | 0.7339259 | -1.660116342 | 2 | 5.946638857 | 8.325066185 | 6.221320894 | 5.9536761 | 6.577231679 | 6.02248356 |
| ENSG00000177613 | CSTF2T | -0.940116861 | 0.524284348 | -0.720048381 | 0.62682033 | -1.660165242 | 5.095959458 | 9.159587375 | 9.681375057 | 7.898117066 | 8.02914793 | 8.278834759 | 8.077990544 |
| ENSG00000282684 | | -0.56064183 | 0.704833394 | -1.100631355 | 0.423483563 | -1.661273185 | 4.815609471 | 8.655694736 | 9.116480286 | 7.818544084 | 7.79989812 | 7.249266052 | 7.311843322 |
| ENSG00000214300 | SPDYE3 | -1.779759788 | 0.200017793 | 0.11793418 | 0.942222441 | -1.661825608 | 6.563806279 | 3.076665209 | 5.39244166 | 4.17310023 | 3.94787299 | 5.31402805 | 5.980509018 |
| ENSG00000070814 | TCOF1 | -1.20654221 | 5.45E-05 | -0.45531789 | 0.163428103 | -1.6618601 | 9.262228249 | 9.506147172 | 9.553445883 | 8.16211395 | 8.33484115 | 9.126408411 | 8.849249315 |
| ENSG00000129467 | ADCY4 | -3.923771053 | 0.064138813 | 2.261508784 | 0.2596534 | -1.662262269 | 2 | 5.303523747 | 2 | 2.270786764 | 2.2680111 | 6.031172603 | 5.957088417 |
| ENSG00000105383 | CD33 | -3.753027428 | NA | 2.090678645 | NA | -1.662348783 | 2 | 2 | 2 | 2 | 2 | 3.316666208 | 2.746601469 |
| ENSG00000140280 | LYSMD2 | 0.364222112 | 0.867562397 | -2.026640189 | 0.276521038 | -1.662418077 | 8.564112515 | 6.389896633 | 2 | 7.590284224 | 7.67863747 | 5.30910438 | 5.414021719 |
| ENSG00000107175 | CREB3 | -0.59862801 | 0.509478857 | -1.064724746 | 0.204926957 | -1.663352756 | 11.00254523 | 10.08551947 | 8.825624954 | 9.665524776 | 9.59245646 | 9.010625386 | 9.306519748 |
| ENSG00000120156 | TEK | -0.73527338 | 0.588183879 | -0.928368419 | 0.477216955 | -1.6636418 | 4.467311984 | 8.077869079 | 7.818184403 | 6.64692533 | 6.79406287 | 6.109723842 | 6.866539459 |

| | | | | | | | | | | | | | |
|---|---|---|---|---|---|---|---|---|---|---|---|---|---|
| ENSG00000164896 | FASTK | -1.579612167 | 0.001157054 | -0.085267373 | 0.89140859 | -1.66487954 | 9.112843761 | 9.733683991 | 10.23774519 | 8.18685754 | 8.21109983 | 9.706989438 | 9.653813751 |
| ENSG00000186918 | ZNF395 | -0.661608553 | 0.390208914 | -1.003416452 | 0.165231608 | -1.665025006 | 8.910412251 | 9.648522413 | 10.49998282 | 9.076774616 | 9.2611121 | 8.444590638 | 9.138408485 |
| ENSG00000183401 | CCDC159 | -0.643879931 | 0.734506972 | -1.021307844 | 0.571333187 | -1.665187775 | 7.384433967 | 5.72254323 | 2 | 5.602782483 | 5.63601114 | 4.977914261 | 5.524371359 |
| ENSG00000177542 | SLC25A22 | -1.183600466 | 0.16737365 | -0.482143441 | 0.599074031 | -1.665743907 | 10.29961392 | 9.102512015 | 8.434763955 | 8.323303759 | 8.30721295 | 8.465761957 | 9.403927624 |
| ENSG00000116157 | GPX7 | -1.419725076 | 5.57E-07 | -0.246570324 | 0.457548655 | -1.6662954 | 8.737455672 | 8.955547846 | 9.149036566 | 7.565338955 | 7.5462416 | 8.688745854 | 8.734549554 |
| ENSG00000111667 | USP5 | -0.906122188 | 0.071713825 | -0.76141596 | 0.130742576 | -1.667538149 | 11.00254523 | 10.55418374 | 10.03493575 | 9.634381252 | 9.72643218 | 9.583976969 | 10.03183938 |
| ENSG00000160117 | ANKLE1 | -0.577098276 | 0.672951696 | -1.091582506 | 0.38391229 | -1.668680782 | 3.327711234 | 4.442710129 | 4.523675391 | 3.840289928 | 3.69849104 | 3.709392871 | 3.071407821 |
| ENSG00000086589 | RBM22 | -0.496515901 | 0.163759565 | -1.172718904 | 0.00023687 | -1.669234804 | 9.097018025 | 9.386925327 | 9.439621147 | 8.861844103 | 8.78422202 | 8.322245777 | 7.965119966 |
| ENSG00000204923 | FBXO48 | -0.750522552 | 0.612898467 | -0.918923507 | 0.519595531 | -1.669446059 | 3.327711234 | 4.442710129 | 6.322058544 | 4.358765249 | 4.72553021 | 4.698044417 | 4.062589736 |
| ENSG00000166261 | ZNF202 | -0.63444433 | 0.744523835 | -1.035313942 | 0.575633447 | -1.669758271 | 3.327711234 | 5.946638857 | 8.99637432 | 7.057964187 | 6.89649843 | 6.800644007 | 6.350712758 |
| ENSG00000134955 | SLC37A2 | -1.167834343 | 0.524450905 | -0.503083062 | 0.79391096 | -1.670917405 | 7.82963488 | 3.686303414 | 4.523675391 | 5.194621444 | 5.51241389 | 6.569017168 | 4.943664032 |
| ENSG00000147383 | NSDHL | -0.798609859 | 0.276678479 | -0.872861557 | 0.220822202 | -1.671471416 | 8.631580145 | 8.451443573 | 9.577565424 | 8.180711314 | 8.185025 | 7.491260006 | 8.541379994 |
| ENSG00000121067 | SPOP | -0.402862358 | 0.361569449 | -1.26921843 | 0.000964865 | -1.672080788 | 9.992581382 | 9.256956408 | 9.528916236 | 9.221512599 | 9.22219152 | 8.455215133 | 8.265241863 |
| ENSG00000165555 | NOXRED1 | -0.554290629 | 0.837802275 | -1.121803927 | 0.655320771 | -1.676094556 | 2 | 4.710564154 | 2 | 3.566290607 | 2.77779907 | 3.059555873 | 2.782576693 |
| ENSG00000136718 | IMP4 | -0.8271799 | 0.243498812 | -0.851034041 | 0.220441583 | -1.678213941 | 9.751047882 | 9.318399851 | 8.110401118 | 8.408172799 | 8.37442884 | 8.47458408 | 8.252471774 |
| ENSG00000204070 | SYS1 | -0.82929883 | 0.272167028 | -0.849117501 | 0.250993106 | -1.678416331 | 9.790941832 | 8.687092496 | 8.110401118 | 8.178656739 | 8.248381 | 8.109343506 | 8.274745794 |
| ENSG00000160783 | PMF1 | -0.973629399 | 0.058967988 | -0.706376544 | 0.176639946 | -1.680005943 | 8.910412251 | 8.004992147 | 8.236946975 | 7.467753569 | 7.48179565 | 7.523500045 | 7.922424886 |
| ENSG00000179163 | FUCA1 | -0.91088221 | 0.059073176 | -0.769819761 | 0.110655463 | -1.680701971 | 9.27634997 | 8.297207175 | 8.922968322 | 8.008651253 | 7.95274764 | 8.098683215 | 8.140837768 |
| ENSG00000140830 | TXNL4B | -0.061577044 | 0.96857017 | -1.619344579 | 0.220279485 | -1.680921622 | 7.709880939 | 8.213429689 | 4.058998049 | 7.388195945 | 7.39346687 | 6.15433139 | 5.586744392 |
| ENSG00000138785 | INTS12 | -0.859407716 | 0.707528996 | -0.822117432 | 0.717366226 | -1.681525148 | 9.790941832 | 6.899080544 | 2 | 7.552702688 | 7.54308677 | 7.70407074 | 7.453871289 |
| ENSG00000198839 | ZNF277 | -0.456424362 | 0.582172352 | -1.227926763 | 0.097270762 | -1.684351125 | 7.667647223 | 6.535001748 | 8.296287027 | 7.144720234 | 7.27676203 | 6.621594001 | 6.29575846 |
| ENSG00000092208 | GEMIN2 | -1.163895752 | 0.296880612 | -0.521028965 | 0.655988002 | -1.684924717 | 4.00722531 | 5.595927442 | 6.629488387 | 4.927308402 | 4.4836996 | 5.371837532 | 5.191251321 |
| ENSG00000162231 | NXF1 | -0.250476754 | 0.70357284 | -1.434540897 | 0.008847264 | -1.685017651 | 10.29268328 | 8.993716073 | 9.746650996 | 9.534293296 | 9.50598504 | 8.315509958 | 8.376090431 |
| ENSG00000136754 | ABI1 | -0.302496756 | 0.713785889 | -1.383100844 | 0.049810851 | -1.685597599 | 10.87379128 | 9.067150531 | 10.36098398 | 9.99525349 | 9.95667084 | 8.871434321 | 8.931664222 |
| ENSG00000142186 | SCYL1 | -0.875594879 | 0.099672705 | -0.812896239 | 0.122835348 | -1.688491117 | 11.88613247 | 10.78935021 | 11.15347798 | 10.44280012 | 10.5080911 | 10.47574661 | 10.59834356 |
| ENSG00000085274 | MYNN | -0.620357133 | 0.200056839 | -1.071745961 | 0.01704757 | -1.692013095 | 7.904282243 | 8.124485672 | 8.35328246 | 7.484483106 | 7.57123495 | 7.364099281 | 6.746446439 |
| ENSG00000155363 | MOV10 | 0.088830676 | 0.823806404 | -1.781015875 | 2.32E-09 | -1.692185199 | 10.46903009 | 10.61258327 | 10.68342405 | 10.6531158 | 10.705844 | 8.559959383 | 9.039201374 |
| ENSG00000089006 | SNX5 | -1.149570905 | 0.105639361 | -0.542732043 | 0.470337112 | -1.692302948 | 8.444143041 | 9.523645035 | 8.110401118 | 7.717440541 | 7.67575955 | 8.582068017 | 7.933467731 |
| ENSG00000064666 | CNN2 | -1.92583233 | 1.07E-06 | 0.231867908 | 0.632223509 | -1.693964421 | 10.9550501 | 11.23666298 | 11.68722117 | 9.368448609 | 9.44052092 | 11.32281177 | 11.75674553 |
| ENSG00000261509 | TP53TG3B | 3.49665459 | 0.539543786 | -5.19109221 | 0.364514502 | -1.694437619 | 2 | 2 | 2 | 3.95996233 | 3.94787299 | 2 | 2 |
| ENSG00000104969 | SGTA | -1.006142379 | 0.122841764 | -0.688622659 | 0.300383908 | -1.694765038 | 10.92846914 | 9.799806309 | 9.577565424 | 9.180006057 | 9.2754408 | 9.47746372 | 9.60786912 |
| ENSG00000160410 | SHKBP1 | -0.805313112 | 0.087707776 | -0.88963815 | 0.052243716 | -1.694951262 | 10.02639697 | 9.424669273 | 9.503962296 | 8.83470036 | 8.91566441 | 8.447394152 | 9.070459565 |
| ENSG00000100304 | TTLL12 | -1.72350201 | 0.003221255 | 0.028321413 | 0.970049089 | -1.695180597 | 9.2903348 | 8.834582582 | 10.19160738 | 7.815902688 | 7.86955761 | 9.664732333 | 9.483608161 |
| ENSG00000070610 | GBA2 | -1.098720387 | 0.061878981 | -0.597117846 | 0.332654942 | -1.695838233 | 10.15425007 | 8.876041558 | 9.31604076 | 8.503178519 | 8.40776525 | 9.012898235 | 8.890530144 |
| ENSG00000023697 | DERA | -0.822996441 | 0.052698164 | -0.877385031 | 0.03408921 | -1.700381471 | 7.750913381 | 8.053983481 | 8.325066185 | 7.330144653 | 7.1744553 | 7.019897747 | 7.363232998 |
| ENSG00000119048 | UBE2B | -0.22311307 | 0.768318166 | -1.478495115 | 0.018944745 | -1.701608185 | 10.03472846 | 8.487557258 | 9.553445883 | 9.29542901 | 9.23691006 | 8.102956788 | 7.940451241 |
| ENSG00000128694 | OSGEPL1 | -1.261765594 | 0.333096085 | -0.441502877 | 0.751814452 | -1.70326847 | 4.00722531 | 6.04685525 | 7.398165132 | 5.273747807 | 5.14724905 | 6.320128742 | 5.486701579 |
| ENSG00000160199 | PKNOX1 | -0.583609933 | 0.444231654 | -1.119795332 | 0.109134859 | -1.703405264 | 7.904282243 | 8.94259736 | 9.636156974 | 8.352759303 | 8.4705529 | 7.86621124 | 7.902127837 |
| ENSG00000115239 | ASB3 | 1.604573695 | 0.105639361 | -3.309429154 | 0.000302537 | -1.704855459 | 5.866944937 | 5.457120953 | 7.451345288 | 7.944803394 | 8.19912382 | 4.017766949 | 3.236138336 |
| ENSG00000100219 | XBP1 | -0.927139927 | 0.023535526 | -0.779682169 | 0.059210657 | -1.706822096 | 12.34389158 | 11.70001862 | 11.60746745 | 10.99183871 | 11.0010911 | 11.18195716 | 11.10429958 |
| ENSG00000115282 | TTC31 | -1.085566035 | 0.003498455 | -0.621711953 | 0.110225496 | -1.707277988 | 8.910412251 | 8.337339476 | 8.922968322 | 7.605658789 | 7.73225478 | 8.127636775 | 8.125161869 |
| ENSG00000105662 | CRTC1 | -0.635790466 | 0.302788797 | -1.071513371 | 0.061577604 | -1.707303836 | 9.740899611 | 8.522788974 | 9.031722608 | 8.501535708 | 8.60067114 | 8.292619498 | 7.927454842 |
| ENSG00000172375 | C2CD2L | -1.005923373 | 0.650131252 | -0.702852776 | 0.755000321 | -1.708776149 | 9.781071289 | 2 | 8.65477198 | 7.728718552 | 7.77322015 | 8.28637003 | 7.764992659 |
| ENSG00000189114 | BLOC1S3 | -0.785214634 | 0.507120384 | -0.923594453 | 0.41925397 | -1.708809087 | 6.009025997 | 8.469613413 | 9.25752426 | 7.555872146 | 7.74054144 | 7.489084836 | 7.54177663 |
| ENSG00000055147 | FAM114A2 | -0.94425923 | 0.330789334 | -0.766679321 | 0.431024961 | -1.710938551 | 7.624139768 | 7.232742069 | 9.385998894 | 7.440576128 | 7.5014335 | 7.644728141 | 7.645539136 |
| ENSG00000101460 | MAP1LC3A | -1.33262446 | 0.214603975 | -0.3794546 | 0.747588951 | -1.712079059 | 10.72727936 | 8.777382925 | 8.007629717 | 8.336264621 | 8.2960118 | 9.135476483 | 9.375867991 |
| ENSG00000205309 | NT5M | -2.253563752 | 0.25952816 | 0.540829163 | 0.8046279 | -1.712734589 | 2 | 6.228547981 | 3.369317972 | 3.162721474 | 3.33976443 | 5.951260634 | 4.522269241 |
| ENSG00000132286 | TIMM10B | -0.80307707 | 0.244724681 | -0.909869845 | 0.173666446 | -1.712946915 | 8.468948322 | 9.632483534 | 10.06083273 | 8.734762645 | 8.72837565 | 8.571313099 | 8.678370137 |
| ENSG00000128944 | KNSTRN | -1.374195032 | 0.02660565 | -0.339538951 | 0.627355322 | -1.713733983 | 8.138771685 | 7.761270542 | 6.718950348 | 6.213314212 | 6.43421531 | 7.25183424 | 7.395535532 |
| ENSG00000167723 | TRPV3 | 0.80838034 | 0.786688692 | -2.523361146 | 0.345358207 | -1.714980806 | 2 | 2 | 7.896719741 | 7.144720234 | 7.1204461 | 4.443291373 | 3.735548302 |
| ENSG00000143882 | ATP6V1C2 | -1.562550723 | 0.533990735 | -0.152570063 | 0.955158571 | -1.715120786 | 2 | 4.11352228 | 2 | 2.600425781 | 2.38539787 | 2.937246078 | 3.183290849 |
| ENSG00000113387 | SUB1 | -0.603790046 | 0.34984013 | -1.112500619 | 0.062080537 | -1.716290665 | 10.65885801 | 9.34816733 | 9.714382178 | 9.451627281 | 9.37638276 | 8.911780399 | 8.906921063 |
| ENSG00000275221 | HIST1H2AK | 3.47228305 | 0.542787593 | -5.191092311 | 0.364514502 | -1.718809261 | 2 | 2 | 2 | 4.139692846 | 3.69849104 | 2 | 2 |
| ENSG00000007392 | LUC7L | -0.384197583 | 0.728252779 | -1.334843017 | 0.170852952 | -1.7190406 | 9.858207046 | 8.433041974 | 7.342949392 | 8.496596021 | 8.54656162 | 7.876232824 | 7.221358049 |
| ENSG00000168765 | GSTM4 | -1.430444599 | 0.006866454 | -0.29018578 | 0.633808927 | -1.720630379 | 7.218819781 | 7.60566235 | 8.266922072 | 6.436065776 | 6.30607931 | 7.303499023 | 7.627107456 |
| ENSG00000162664 | ZNF326 | -0.238381594 | 0.68556684 | -1.482536287 | 0.002183465 | -1.720917882 | 8.816894989 | 8.451443573 | 7.734844356 | 8.190940526 | 8.14396641 | 6.864125031 | 7.032593922 |
| ENSG00000090432 | MUL1 | -0.834000644 | 0.024480315 | -0.887427738 | 0.01510731 | -1.721428382 | 10.06758145 | 9.433952963 | 9.829330889 | 8.924487942 | 9.01008467 | 8.808281231 | 9.014359877 |
| ENSG00000136891 | TEX10 | -0.608772049 | 0.180706341 | -1.113790138 | 0.007855038 | -1.722562187 | 9.219014269 | 8.590771778 | 9.413059136 | 8.485003881 | 8.52593541 | 7.950836136 | 8.061612696 |
| ENSG00000179335 | CLK3 | -0.841309968 | 0.047154938 | -0.88633467 | 0.03310095 | -1.727644637 | 9.624286623 | 9.114109157 | 9.839340503 | 8.707945635 | 8.72698617 | 8.832932104 | 8.492665077 |
| ENSG00000170271 | FAXDC2 | -1.273023646 | 0.289625434 | -0.455235384 | 0.72430187 | -1.72825903 | 5.330594134 | 7.790474954 | 8.825624954 | 6.687949235 | 6.6253296 | 7.512833218 | 7.383872668 |
| ENSG00000120992 | LYPLA1 | -0.64617318 | 0.140124762 | -1.082377142 | 0.007862246 | -1.728550322 | 9.262228249 | 9.00621767 | 8.511888463 | 8.401141138 | 8.23671175 | 8.007404089 | 7.764992659 |
| ENSG00000155066 | PROM2 | -0.701839933 | 0.69608789 | -1.026980046 | 0.544850149 | -1.72881998 | 5.095959458 | 3.076665209 | 3.369317972 | 3.921164147 | 3.2184422 | 3.796405078 | 2.851942824 |
| ENSG00000068001 | HYAL2 | -0.81823896 | 0.135737919 | -0.911462058 | 0.087275793 | -1.729701018 | 8.777722034 | 9.308339287 | 9.935792933 | 8.592312513 | 8.61726622 | 8.581557692 | 8.440137439 |
| ENSG00000135392 | DNAJC14 | -0.73349074 | 0.182358952 | -0.996864893 | 0.056539999 | -1.730355633 | 9.751047882 | 8.791896851 | 8.865353883 | 8.50645854 | 8.44542686 | 8.071319244 | 8.347853908 |
| ENSG00000125149 | C16orf70 | -0.708356003 | 0.283441752 | -1.022345889 | 0.100649724 | -1.730701892 | 8.816894989 | 7.847167909 | 7.502634626 | 7.447418636 | 7.49164798 | 7.258234779 | 7.062165783 |
| ENSG00000189129 | PLAC9 | -3.360469219 | 6.63E-07 | 1.629205074 | 0.017958187 | -1.731264145 | 6.563806279 | 7.189546432 | 7.029880454 | 3.921164147 | 4.02226117 | 7.778984728 | 9.054915125 |
| ENSG00000135829 | DHX9 | -0.603325679 | 0.433957193 | -1.129454314 | 0.110981627 | -1.732779992 | 9.657036304 | 10.94030351 | 11.36918911 | 10.225157 | 10.2084185 | 9.457299408 | 9.895260472 |
| ENSG00000127540 | UQCR11 | -0.571339478 | 0.181227887 | -1.16154079 | 0.002838737 | -1.732880269 | 10.27872131 | 9.842262059 | 9.491321578 | 9.32080297 | 9.35585278 | 8.713644723 | 8.790832728 |
| ENSG00000162604 | TM2D1 | -0.524645985 | 0.627093532 | -1.212612674 | 0.217050348 | -1.737258659 | 8.418903792 | 6.389896633 | 9.06622548 | 7.857595438 | 7.74603955 | 7.253116621 | 6.996671423 |
| ENSG00000131061 | ZNF341 | -1.187749495 | 0.585838677 | -0.549831424 | 0.808868217 | -1.737580919 | 8.009475818 | 4.442710129 | 2 | 5.461211713 | 5.43283104 | 6.285453595 | 5.736403255 |
| ENSG00000120053 | GOT1 | -1.27971996 | 8.61E-05 | -0.457955749 | 0.199124017 | -1.737675708 | 8.564112515 | 9.018611864 | 8.699419242 | 7.423326516 | 7.60793141 | 8.288873067 | 8.36352657 |
| ENSG00000167186 | COQ7 | -1.498338725 | 0.039474306 | -0.240213869 | 0.777011185 | -1.738552594 | 7.031698319 | 7.189546432 | 8.408111533 | 6.043074888 | 6.39275508 | 7.123093858 | 7.701812481 |
| ENSG00000244045 | TMEM199 | 0.199713957 | 0.738698334 | -1.938695275 | 4.72E-05 | -1.738981317 | 8.653385977 | 8.029695761 | 7.646393253 | 8.415170354 | 8.31648126 | 6.132200015 | 6.426568739 |
| ENSG00000144668 | ITGA9 | -3.268949188 | 0.370967351 | 1.529795759 | 0.642128837 | -1.739153429 | 2 | 3.076665209 | 2 | 2.141737246 | 2 | 3.374372158 | 3.183290849 |
| ENSG00000106144 | CASP2 | -1.080647744 | 0.0034093 | -0.658897142 | 0.086198005 | -1.739544886 | 8.836089686 | 8.234834549 | 8.742726175 | 7.565338955 | 7.5462416 | 7.909957323 | 8.027351693 |
| ENSG00000054219 | LY75 | -2.433843174 | 0.123996487 | 0.693116674 | 0.667996442 | -1.7407265 | 2 | 4.11352228 | 4.058998049 | 2.498691189 | 2.49394783 | 4.053534413 | 4.329022995 |
| ENSG00000146731 | CCT6A | -0.805963363 | 0.044014587 | -0.934907752 | 0.016521712 | -1.740871114 | 10.29961392 | 11.00402286 | 10.96062172 | 9.955006472 | 10.0164701 | 9.900058137 | 9.813939602 |
| ENSG00000155876 | RRAGA | -1.030804312 | 0.189102784 | -0.711938895 | 0.372465764 | -1.742743207 | 11.86992646 | 10.1878402 | 11.87895445 | 10.43765481 | 10.4983508 | 10.43140618 | 11.07104905 |
| ENSG00000063046 | EIF4B | -1.181224725 | 0.012942716 | -0.561561847 | 0.270007586 | -1.742786573 | 11.23722567 | 12.27732859 | 11.73357624 | 10.66012781 | 10.6058245 | 11.22089964 | 11.28171193 |
| ENSG00000072682 | P4HA2 | -0.462151904 | 0.568285409 | -1.281306023 | 0.076411855 | -1.743457927 | 12.76534233 | 11.35134391 | 11.19694409 | 11.46811054 | 11.5206165 | 10.54583177 | 10.79719235 |
| ENSG00000139880 | CDH24 | -1.050174708 | 0.599957698 | -0.693477864 | 0.732916452 | -1.743652573 | 8.313285031 | 6.140560021 | 2 | 6.155987605 | 5.88556448 | 6.446574529 | 6.276965131 |
| ENSG00000203710 | CR1 | -0.678242903 | 0.774549338 | -1.065777093 | 0.631689024 | -1.744019997 | 2 | 3.076665209 | 3.369317972 | 2.784612485 | 2.49394783 | 2.526562374 | 2.509591799 |
| ENSG00000157216 | SSBP3 | -0.878512015 | 0.010660073 | -0.866073188 | 0.011165872 | -1.744585203 | 9.567995545 | 9.055168103 | 9.491321578 | 8.579911316 | 8.44204342 | 8.444590638 | 8.601149963 |
| ENSG00000135473 | PAN2 | -1.0495667 | 0.615039451 | -0.69627695 | 0.742149828 | -1.745843651 | 9.460782017 | 8.053983481 | 2 | 7.288834728 | 7.33629421 | 7.751118939 | 7.558732759 |
| ENSG00000106330 | MOSPD3 | -1.457889951 | 0.146764584 | -0.288171706 | 0.799963632 | -1.746061656 | 8.910412251 | 6.311476754 | 7.818184403 | 6.610808945 | 6.60731796 | 7.362911889 | 8.047821014 |
| ENSG00000167468 | GPX4 | -0.776421644 | 0.280259419 | -0.969858885 | 0.15923574 | -1.746280529 | 12.02224098 | 10.41578665 | 11.01019887 | 10.49657106 | 10.5609081 | 10.28189487 | 10.38857981 |
| ENSG00000213029 | SPHAR | 3.444489832 | 0.546436072 | -5.191092427 | 0.364514502 | -1.746602595 | 2 | 2 | 2 | 3.95996233 | 3.86943969 | 2 | 2 |
| ENSG00000059378 | PARP12 | -7.139314663 | 0.000113553 | 5.392448157 | 4.05E-09 | -1.746866506 | 3.327711234 | 4.442710129 | 3.369317972 | 2 | 2 | 8.878929913 | 8.75393321 |
| ENSG00000115661 | STK16 | -1.175643234 | 0.392717095 | -0.5714471 | 0.6908855 | -1.747090334 | 8.107517749 | 4.93638933 | 9.083172337 | 6.820355514 | 7.14145882 | 7.450478673 | 7.695929227 |
| ENSG00000147601 | TERF1 | -0.66729278 | 0.115572829 | -1.081859076 | 0.006258366 | -1.749151856 | 8.39321512 | 8.732943944 | 9.049077186 | 8.074220386 | 8.10384772 | 7.496683627 | 7.845855157 |
| ENSG00000176714 | CCDC121 | -1.68853666 | 0.309909983 | -0.061884741 | 0.973199511 | -1.750421402 | 4.00722531 | 4.710564154 | 2 | 2.947924294 | 2.94009902 | 4.311629396 | 3.45257121 |
| ENSG00000133316 | WDR74 | -0.123108401 | 0.780669035 | -1.631865051 | 1.78E-06 | -1.754973452 | 7.667647223 | 7.847167909 | 8.110401118 | 7.656739892 | 7.86198169 | 6.315225883 | 6.289521179 |
| ENSG00000070018 | LRP6 | -1.567855721 | 2.60E-06 | -0.187360505 | 0.646525284 | -1.755216226 | 10.90138923 | 10.30378101 | 10.5250051 | 9.077877436 | 8.99284518 | 10.29953459 | 10.51207876 |
| ENSG00000276043 | UHRF1 | -0.816974489 | 0.493227914 | -0.939594658 | 0.413811187 | -1.756569347 | 4.467311984 | 6.899080544 | 5.930865648 | 5.242616075 | 5.43283104 | 4.596630906 | 5.670730563 |
| ENSG00000082153 | BZW1 | 0.881558018 | 5.06E-05 | -2.638888745 | 8.51E-38 | -1.757330727 | 11.07704221 | 11.07990752 | 11.18913795 | 12.03638639 | 11.9548335 | 8.385007206 | 8.589138522 |
| ENSG00000169230 | PRELID1 | -0.51133154 | 0.210210865 | -1.246118336 | 0.000633362 | -1.757449877 | 10.29268328 | 9.656475478 | 10.24529339 | 9.584817224 | 9.57557184 | 8.744061916 | 8.949576212 |
| ENSG00000185298 | CCDC137 | -1.244707294 | 0.234625812 | -0.513865031 | 0.646098069 | -1.758572325 | 9.579430841 | 6.666836837 | 8.742726175 | 7.536750104 | 7.51115308 | 8.189226297 | 8.293567867 |
| ENSG00000058729 | RIOK2 | -0.542847744 | 0.253506089 | -1.215767922 | 0.004575334 | -1.758615666 | 8.540903468 | 8.732943944 | 9.301632639 | 8.334420183 | 8.37620272 | 7.851893762 | 7.513957873 |
| ENSG00000155368 | DBI | -0.809067216 | 0.137065313 | -0.950084653 | 0.070975318 | -1.759151869 | 10.1770105 | 9.591588447 | 10.44859729 | 9.324524407 | 9.29056951 | 8.749524069 | 9.491441317 |
| ENSG00000138336 | TET1 | -2.820769075 | 0.107655331 | 1.060360439 | 0.572841244 | -1.760408636 | 4.00722531 | 8.053983481 | 3.369317972 | 4.034562184 | 4.1930041 | 7.86872317 | 7.386797235 |
| ENSG00000243335 | KCTD7 | -1.456930929 | 0.432787266 | -0.30496971 | 0.882758816 | -1.76190064 | 3.327711234 | 7.874700259 | 9.954333071 | 7.206526483 | 7.29937545 | 8.693476763 | 7.991296779 |
| ENSG00000179044 | EXOC3L1 | -2.265410344 | 0.340614989 | 0.50332903 | 0.842603619 | -1.762081313 | 2 | 4.710564154 | 2 | 2.3892346 | 2.59489879 | 3.517841605 | 4.241259475 |
| ENSG00000100983 | GSS | -0.727673359 | 0.573772614 | -1.03774661 | 0.399735181 | -1.765419969 | 10.39976609 | 10.17128279 | 6.80318718 | 9.073461088 | 9.0169232 | 8.500191379 | 8.94213989 |
| ENSG00000173275 | ZNF449 | -1.261453385 | 0.451753365 | -0.505040778 | 0.775706431 | -1.766494163 | 3.327711234 | 7.874700259 | 8.35328246 | 6.496695859 | 6.19790195 | 7.669911566 | 6.074197996 |
| ENSG00000115204 | MPV17 | -1.076987109 | 0.000427769 | -0.689672307 | 0.030115812 | -1.766659417 | 9.781071289 | 9.540933215 | 9.344432377 | 8.456457569 | 8.53866332 | 8.767583318 | 8.987639044 |
| ENSG00000102225 | CDK16 | -1.168614058 | 0.013435581 | -0.59862171 | 0.233839673 | -1.767235768 | 11.43272766 | 10.55843445 | 11.2764554 | 9.953204745 | 9.98600927 | 10.27764795 | 10.75820049 |
| ENSG00000128524 | ATP6V1F | -0.834221369 | 0.00864231 | -0.933661072 | 0.002696789 | -1.767882441 | 10.83137851 | 10.63288029 | 10.37481416 | 9.777686595 | 9.81017641 | 9.832607809 | 9.542821395 |

| | | | | | | | | | | | | | |
|---|---|---|---|---|---|---|---|---|---|---|---|---|---|
| ENSG00000089289 | IGBP1 | -0.883057606 | 0.282118771 | -0.886134988 | 0.270404914 | -1.769192594 | 9.886101849 | 9.148351514 | 7.858115942 | 8.27796118 | 8.34212018 | 8.264917037 | 8.348604099 |
| ENSG00000101464 | PIGU | -0.788337454 | 0.653305306 | -0.981822215 | 0.56354884 | -1.770159669 | 9.143983763 | 7.701025996 | 3.369317972 | 7.210554257 | 7.30311013 | 7.006273211 | 7.130061054 |
| ENSG00000147684 | NDUFB9 | -1.085676338 | 0.128503333 | -0.684578009 | 0.352068654 | -1.770254347 | 10.39976609 | 9.672251278 | 8.677268317 | 8.66162195 | 8.67461127 | 8.929935295 | 9.189967637 |
| ENSG00000125772 | GPCPD1 | -0.006350983 | 0.997041173 | -1.764073338 | 0.146403239 | -1.770424321 | 8.042902253 | 8.806266215 | 5.156626401 | 8.045195239 | 7.86703672 | 6.653021698 | 5.703940561 |
| ENSG00000155893 | PXYLP1 | -2.627399147 | 0.11276309 | 0.856078881 | 0.635038067 | -1.771320266 | 7.750913381 | 6.602424674 | 2 | 4.523237872 | 4.16043394 | 7.651551654 | 7.448282715 |
| ENSG00000090372 | STRN4 | 0.503234931 | 0.722734802 | -2.276067687 | 0.061919051 | -1.772832755 | 7.667647223 | 4.11352228 | 7.285536013 | 7.370304568 | 7.53039799 | 5.185595313 | 4.4231929 |
| ENSG00000155463 | OXA1L | -0.854357095 | 0.166802524 | -0.918773302 | 0.12650708 | -1.773130398 | 10.76282024 | 9.770792087 | 9.491321578 | 9.269600773 | 9.26207178 | 9.210337096 | 9.193312432 |
| APOBEC3A_B | APOBEC3A_E | 3.417070077 | 0.549630367 | -5.191092541 | 0.364514502 | -1.774022464 | 2 | 2 | 2 | 3.881293667 | 3.90918929 | 2 | 2 |
| ENSG00000115241 | PPM1G | -1.177677465 | 0.068908006 | -0.596451758 | 0.381507263 | -1.774129223 | 9.720386367 | 9.856140801 | 11.04521411 | 9.085573664 | 9.23983583 | 9.656518038 | 9.823954402 |
| ENSG00000107874 | CUEDC2 | -1.066450857 | 0.020143473 | -0.7077506 | 0.134444561 | -1.774201457 | 9.839306291 | 9.13702746 | 9.014156722 | 8.272191829 | 8.36552655 | 8.541185654 | 8.796342139 |
| ENSG00000130159 | ECSIT | -1.539720937 | 0.453495317 | -0.234597927 | 0.918342152 | -1.774318864 | 9.497412313 | 7.73146271 | 2 | 6.670510205 | 6.89154576 | 8.00283742 | 8.112850643 |
| ENSG00000153896 | ZNF599 | -1.444145213 | 0.391746117 | -0.332390111 | 0.856791105 | -1.776535324 | 2 | 6.728495673 | 6.322058544 | 4.387512679 | 4.95174433 | 6.013120342 | 5.256659864 |
| ENSG00000124383 | MPHOSPH10 | -0.462618842 | 0.444641873 | -1.31489722 | 0.013709627 | -1.777516062 | 9.064836028 | 8.317412869 | 9.528916236 | 8.610716699 | 8.5668971 | 7.824551575 | 7.66855289 |
| ENSG00000136159 | NUDT15 | -1.11502814 | 0.114654417 | -0.663096892 | 0.363476242 | -1.778125032 | 7.867441304 | 6.951736442 | 8.585098385 | 6.917187326 | 6.78340763 | 7.249266052 | 7.330190599 |
| ENSG00000188315 | C3orf62 | -1.221862388 | 0.34076194 | -0.556557952 | 0.681335361 | -1.77842034 | 5.330594134 | 6.311476754 | 8.536706852 | 6.197166296 | 6.14893615 | 6.984820384 | 6.607941737 |
| ENSG00000136631 | VPS45 | -0.799752177 | 0.021412208 | -0.978927477 | 0.003559969 | -1.778679654 | 8.855032352 | 9.415325457 | 9.049077186 | 8.31397419 | 8.36373948 | 8.162177243 | 8.162331091 |
| ENSG00000214866 | DCDC2C | 2.01372804 | NA | -3.793256136 | NA | -1.779528096 | 2 | 2 | 2 | 3.227757282 | 2.77779907 | 2.048074405 | 2 |
| ENSG00000054938 | CHRDL2 | -0.474665523 | 0.863478741 | -1.305465833 | 0.59982426 | -1.780131356 | 2 | 4.11352228 | 2 | 2.868578001 | 2.94009902 | 2.384001718 | 2.746601469 |
| ENSG00000254986 | DPP3 | -0.442120541 | 0.69646163 | -1.341220527 | 0.182472647 | -1.783341068 | 10.37373127 | 7.60566235 | 9.165043284 | 8.986874783 | 9.02712059 | 7.815899769 | 8.367233194 |
| ENSG00000182628 | SKA2 | -1.405351301 | 0.006781203 | -0.381722421 | 0.514884694 | -1.787073722 | 8.444143041 | 8.876041558 | 9.478569125 | 7.66556918 | 7.5462416 | 8.357250999 | 8.834865421 |
| ENSG00000124098 | FAM210B | -1.170804543 | 0.157179445 | -0.620297409 | 0.473386442 | -1.791101952 | 9.957954117 | 9.308339287 | 11.28013451 | 9.284006841 | 9.22514727 | 9.674097378 | 9.920202701 |
| ENSG00000132792 | CTNNBL1 | 0.054045573 | 0.970008162 | -1.84536911 | 0.1204601 | -1.791323537 | 10.19941743 | 8.540087354 | 6.534109959 | 9.110576034 | 9.18523054 | 7.111828383 | 7.424286753 |
| ENSG00000054983 | GALC | -0.813797303 | 0.618562037 | -0.97813634 | 0.536324873 | -1.791933643 | 4.00722531 | 9.235879652 | 8.825624954 | 7.647855834 | 7.71553728 | 7.482539577 | 7.558732759 |
| ENSG00000230124 | ACBD6 | -1.245699672 | 0.134484147 | -0.547787683 | 0.536069347 | -1.793487355 | 8.564112515 | 8.124485672 | 10.02619908 | 7.978262192 | 7.84158259 | 8.505581591 | 8.68907177 |
| ENSG00000083720 | OXCT1 | -1.654833767 | 0.192239251 | -0.1389763 | 0.926455594 | -1.793810068 | 6.138360429 | 7.538394143 | 9.61300511 | 6.727838783 | 6.89649843 | 7.81936672 | 8.654907296 |
| ENSG00000151657 | KIN | -0.313383138 | 0.501522547 | -1.480710014 | 0.000144114 | -1.794093153 | 7.384433967 | 7.60566235 | 7.691296561 | 7.437142666 | 7.05992818 | 6.208218574 | 6.066922748 |
| ENSG00000105672 | ETV2 | -2.128048467 | 0.18838102 | 0.332250817 | 0.852228272 | -1.795797651 | 4.00722531 | 4.93638933 | 2 | 2.868578001 | 2.77779907 | 3.878466556 | 4.766376438 |
| ENSG00000166347 | CYB5A | -1.443442522 | 0.011601873 | -0.352763341 | 0.586489016 | -1.796205864 | 8.946185348 | 7.790474954 | 8.042706615 | 6.941624891 | 6.9210099 | 7.827137031 | 8.148612304 |
| ENSG00000144285 | SCN1A | 3.394729588 | 0.552383336 | -5.191092634 | 0.364514502 | -1.796363046 | 2 | 2 | 2 | 4.415698455 | 3.01487888 | 2 | 2 |
| ENSG00000167740 | CYB5D2 | -0.862273979 | 0.361913214 | -0.934191509 | 0.309378866 | -1.796465488 | 8.16936288 | 7.790474954 | 9.849281147 | 8.006336225 | 8.07792246 | 7.795801481 | 8.128660152 |
| ENSG00000083807 | SLC27A5 | -2.820952644 | 0.094280197 | 1.023609331 | 0.565006761 | -1.797343314 | 6.366709671 | 4.11352228 | 2 | 2.868578001 | 3.2184422 | 5.85994354 | 6.202567011 |
| ENSG00000162415 | ZSWIM5 | -1.468499382 | 0.491724432 | -0.329676106 | 0.888119558 | -1.798175489 | 2 | 4.710564154 | 8.143093325 | 5.128039203 | 5.49945122 | 6.499401217 | 6.257923755 |
| ENSG00000170502 | NUDT9 | -0.690830579 | 0.061299815 | -1.109018462 | 0.001305492 | -1.799849041 | 9.031919741 | 8.903033433 | 9.149036566 | 8.390528966 | 8.29788471 | 7.679481638 | 8.148612304 |
| ENSG00000165275 | TRMT10B | -0.832699964 | 0.105432147 | -0.967245941 | 0.050027637 | -1.799945906 | 6.96351253 | 7.538394143 | 6.718950348 | 6.351046291 | 6.31350454 | 6.239610476 | 6.168983563 |
| ENSG00000169087 | HSPBAP1 | -0.837565425 | 0.40748464 | -0.962602854 | 0.322244282 | -1.800168279 | 7.27616306 | 4.93638933 | 6.629488387 | 5.686138681 | 5.83486613 | 5.755231204 | 5.529673353 |
| ENSG00000143498 | TAF1A | 0.0412019 | 0.98108975 | -1.841725037 | 0.189314051 | -1.800523137 | 3.327711234 | 7.051611315 | 5.59507254 | 6.087554739 | 6.00960265 | 4.82764224 | 3.735548302 |
| ENSG00000100324 | TAB1 | -1.611230071 | 0.047807202 | -0.189870024 | 0.844740034 | -1.801100095 | 8.946185348 | 7.73146271 | 9.878698603 | 7.423326516 | 7.58663873 | 8.907307568 | 8.909973736 |
| ENSG00000119778 | ATAD2B | -0.942424713 | 0.592997115 | -0.859171093 | 0.62358047 | -1.801595807 | 5.330594134 | 5.595927442 | 9.589475645 | 7.261917051 | 7.20671394 | 7.64179387 | 6.89352185 |
| ENSG00000235173 | HGH1 | -0.684259048 | 0.120319176 | -1.117699968 | 0.00651667 | -1.801959016 | 8.340423424 | 9.125613817 | 9.014156722 | 8.16211395 | 8.22297724 | 7.797560316 | 7.734322559 |
| ENSG00000187713 | TMEM203 | -1.214164078 | 0.005219399 | -0.593637129 | 0.200464134 | -1.807801207 | 9.771132748 | 9.599860728 | 8.884815469 | 8.260583448 | 8.26953254 | 8.81307133 | 8.94213989 |
| ENSG00000180611 | MB21D2 | -1.002166037 | 0.209002923 | -0.806671869 | 0.311846327 | -1.808837906 | 7.218819781 | 7.315436877 | 8.805342542 | 7.021746293 | 6.95938211 | 7.108998216 | 7.24894374 |
| ENSG00000150433 | TMEM218 | -1.147782609 | 0.22267971 | -0.661120463 | 0.496156922 | -1.808903073 | 8.19931886 | 7.467835385 | 5.772716771 | 6.237202172 | 6.44777477 | 6.679970309 | 6.934072182 |
| ENSG00000115286 | NDUFS7 | -0.565256653 | 0.328225936 | -1.244312109 | 0.017698699 | -1.809568763 | 8.816894989 | 8.791896851 | 7.734844356 | 7.90310317 | 8.04261444 | 7.309674888 | 7.308762629 |
| ENSG00000186298 | PPP1CC | -0.994512735 | 0.048767966 | -0.815141233 | 0.108876124 | -1.809653968 | 10.77778805 | 11.1798568 | 10.08627303 | 9.787831283 | 9.73059821 | 9.881520895 | 9.99254197 |
| ENSG00000213066 | FGFR1OP | -0.202638848 | 0.917642114 | -1.607149943 | 0.338976671 | -1.809788791 | 3.327711234 | 7.099070191 | 9.132850258 | 7.745471964 | 7.61696151 | 6.522981762 | 6.092227206 |
| ENSG00000171853 | TRAPPC12 | -0.837503368 | 0.243555525 | -0.972978461 | 0.160951705 | -1.810481828 | 9.219014269 | 8.414402626 | 10.06083273 | 8.485003881 | 8.61275919 | 8.367944224 | 8.463180778 |
| ENSG00000230778 | ANKRD63 | -3.753029279 | NA | 1.942520064 | NA | -1.810509214 | 2 | 2 | 2 | 2 | 2 | 3.316666208 | 2.551896972 |
| ENSG00000065183 | WDR3 | -0.480735168 | 0.749500183 | -1.331216074 | 0.328380953 | -1.811951243 | 4.815609471 | 8.834582582 | 9.149036566 | 7.994705029 | 7.96694292 | 7.20886319 | 7.085754446 |
| ENSG00000137285 | TUBB2B | 0.626752522 | 0.478306674 | -2.439446584 | 0.00124221 | -1.812694061 | 8.797441463 | 7.669933291 | 6.958202216 | 8.744518279 | 8.49850989 | 5.304163849 | 5.953147751 |
| ENSG00000042088 | TDP1 | -1.054720866 | 0.619674661 | -0.758931177 | 0.72430187 | -1.813652043 | 9.423197387 | 7.393647515 | 2 | 6.960881212 | 7.27676203 | 7.380620955 | 7.449681889 |
| ENSG00000205710 | C17orf107 | -0.387943423 | 0.833035958 | -1.426404567 | 0.376931525 | -1.814347989 | 3.327711234 | 5.595927442 | 3.369317972 | 4.268917409 | 4.22485516 | 3.956110469 | 2.817676617 |
| ENSG00000167850 | CD300C | -3.753029357 | NA | 1.936281607 | NA | -1.816747749 | 2 | 2 | 2 | 2 | 2 | 3.336159116 | 2.509591799 |
| ENSG00000135297 | MTO1 | -1.378484603 | 0.309156734 | -0.438427681 | 0.765406629 | -1.816912284 | 9.304185368 | 7.761270542 | 5.156626401 | 6.809774134 | 6.90143415 | 7.875400346 | 7.661324972 |
| ENSG00000168612 | ZSWIM1 | -1.365921434 | 0.421293524 | -0.454337938 | 0.802964883 | -1.820259372 | 5.095959458 | 5.303523747 | 9.287079172 | 6.664650034 | 6.3857277 | 7.427945125 | 7.399884913 |
| ENSG00000016864 | GLT8D1 | -1.245262453 | 1.94E-08 | -0.575269375 | 0.014854455 | -1.820531827 | 10.22148166 | 10.03760521 | 10.06083273 | 8.88597028 | 8.85335127 | 9.592106979 | 9.475732243 |
| ENSG00000134779 | TPGS2 | -1.338374893 | 0.007266635 | -0.483255265 | 0.377517471 | -1.821630158 | 10.61508061 | 9.994333773 | 10.95602907 | 9.260887574 | 9.21922971 | 9.779310259 | 10.34732599 |
| ENSG00000130643 | CALY | -2.165107267 | 0.363586907 | 0.343271464 | 0.896486957 | -1.821835803 | 2 | 5.457120953 | 2 | 3.023133209 | 2.59489879 | 4.281782022 | 4.56420962 |
| ENSG00000079459 | FDFT1 | -1.18789947 | 0.018486008 | -0.635458807 | 0.233690382 | -1.823358276 | 12.15999925 | 11.49353862 | 11.33422412 | 10.5227232 | 10.5229798 | 10.69695151 | 11.37275397 |
| ENSG00000260001 | TGFBR3L | -0.514941899 | 0.796317099 | -1.309072932 | 0.470609658 | -1.824014831 | 7.031698319 | 4.93638933 | 2 | 5.194621444 | 5.41912973 | 4.927401733 | 4.228274364 |
| ENSG00000146842 | TMEM209 | -0.411593042 | 0.573609291 | -1.412455249 | 0.027000298 | -1.82404829 | 7.159102577 | 8.687092496 | 8.042706615 | 7.781117077 | 7.60490877 | 6.748859346 | 6.695462882 |
| ENSG00000106803 | SEC61B | -0.72445178 | 0.220626804 | -1.099943045 | 0.048384016 | -1.824394826 | 10.75779627 | 9.656475478 | 9.757248847 | 9.36574218 | 9.48312089 | 8.981908469 | 9.120072249 |
| ENSG00000169403 | PTAFR | -2.151126908 | 0.22333671 | 0.324202717 | 0.87135418 | -1.826924191 | 2 | 4.93638933 | 6.203069917 | 3.566290607 | 3.39680241 | 6.098353065 | 4.279527956 |
| ENSG00000171169 | NAIF1 | -1.049166523 | 0.292000423 | -0.778471903 | 0.436962292 | -1.827638426 | 8.586954094 | 7.099070191 | 6.203069917 | 6.604700657 | 6.61935067 | 6.732402829 | 7.000495137 |
| ENSG00000167969 | ECI1 | -1.429353167 | 0.113104573 | -0.399015997 | 0.689674098 | -1.828369164 | 9.2903348 | 8.88960062 | 6.882775709 | 7.326437704 | 7.19470132 | 8.350677018 | 8.191057445 |
| ENSG00000092098 | RNF31 | 0.007444297 | 0.997041173 | -1.835826586 | 0.188561332 | -1.828382289 | 4.815609471 | 9.298208073 | 9.565556058 | 8.809693772 | 8.96600458 | 6.997118176 | 7.154291015 |
| ENSG00000152219 | ARL14EP | -1.221563461 | 0.117596105 | -0.606885809 | 0.460026214 | -1.828449269 | 9.317904228 | 7.503546079 | 8.408111533 | 7.539954763 | 7.21862736 | 8.191903518 | 7.757103546 |
| ENSG00000160161 | CILP2 | -1.837861712 | 0.339484846 | 0.008464701 | 0.997302572 | -1.829397011 | 2 | 4.93638933 | 7.600047325 | 4.329433326 | 4.811949 | 6.295446113 | 6.209190898 |
| ENSG00000101940 | WDR13 | -1.342543079 | 0.004333387 | -0.487120177 | 0.347421792 | -1.829663256 | 9.048471759 | 10.02537381 | 9.589475645 | 8.31771326 | 8.24255817 | 8.960480054 | 9.277661801 |
| ENSG00000179933 | C14orf119 | -0.807449987 | 0.17161955 | -1.023492492 | 0.070716745 | -1.830942479 | 10.787681 | 9.488434476 | 10.33998689 | 9.484432763 | 9.50355257 | 9.313509983 | 9.244188987 |
| ENSG00000133627 | ACTR3B | -1.178210712 | 0.468786454 | -0.65333065 | 0.695048367 | -1.831541363 | 6.366709671 | 5.838937808 | 2 | 4.574143677 | 4.53541753 | 5.338397457 | 4.56420962 |
| ENSG00000082269 | FAM135A | -0.87090656 | 0.346696909 | -0.964609556 | 0.281743965 | -1.835516116 | 7.435667355 | 7.60566235 | 5.59507254 | 6.442929658 | 6.11534392 | 6.468821191 | 5.867805474 |
| ENSG00000117133 | RPF1 | -0.424422243 | 0.463796219 | -1.411674819 | 0.004833789 | -1.836097062 | 8.981092835 | 8.029695761 | 8.110401118 | 7.994705029 | 8.04261444 | 7.142599214 | 6.957866334 |
| ENSG00000128581 | IFT22 | -0.9229347 | 0.098318255 | -0.914520143 | 0.094766332 | -1.837454843 | 7.667647223 | 8.762721501 | 8.380957451 | 7.398825229 | 7.47518984 | 7.240241221 | 7.625870264 |
| ENSG00000162642 | C1orf52 | -0.576915297 | 0.323982764 | -1.261414655 | 0.017661584 | -1.838329952 | 9.48530526 | 8.317412869 | 9.06622548 | 8.50645854 | 8.40602977 | 7.903438544 | 7.650414739 |
| ENSG00000125676 | THOC2 | -0.5500445 | 0.275627669 | -1.288292303 | 0.004493423 | -1.838336803 | 9.613202746 | 10.40166101 | 10.47451889 | 9.670650622 | 9.65954302 | 9.101707221 | 8.738561653 |
| ENSG00000136715 | SAP130 | -0.912909408 | 0.232251697 | -0.925610486 | 0.217050348 | -1.838519894 | 9.423197387 | 8.169643011 | 9.999667074 | 8.413424144 | 8.52593541 | 8.356654601 | 8.553143267 |
| ENSG00000136888 | ATP6V1G1 | -0.462434135 | 0.148344449 | -1.376465231 | 9.97E-07 | -1.838899366 | 11.05263482 | 10.91068521 | 11.39316998 | 10.6795094 | 10.6610729 | 9.751223945 | 9.76757159 |
| ENSG00000156261 | CCT8 | -1.03424512 | 0.024637077 | -0.805336308 | 0.085735367 | -1.839581428 | 11.55070946 | 10.96932597 | 11.91252756 | 10.49945465 | 10.4901834 | 10.85352355 | 10.57977418 |
| ENSG00000256053 | APOPT1 | -0.911483305 | 0.221454877 | -0.9285398 | 0.203560175 | -1.840023105 | 9.384607364 | 8.124485672 | 7.818184403 | 7.694616532 | 7.72670378 | 7.764669607 | 7.619668339 |
| ENSG00000172590 | MRPL52 | -0.702207078 | 0.328545024 | -1.138804281 | 0.090404285 | -1.841011359 | 9.448362284 | 8.297207175 | 8.007629717 | 8.102673083 | 7.95512324 | 7.493431902 | 7.702986259 |
| ENSG00000117859 | OSBPL9 | -0.711873687 | 0.160563675 | -1.131309045 | 0.017373749 | -1.843182732 | 10.87379128 | 10.40638494 | 9.999667074 | 9.728676049 | 9.79367921 | 9.588047701 | 9.050772231 |
| ENSG00000118242 | MREG | -0.357053667 | 0.844353006 | -1.486456233 | 0.350123893 | -1.8435099 | 5.866944937 | 3.686303414 | 7.97167868 | 6.268449691 | 6.51372389 | 5.899789217 | 4.434543419 |
| ENSG00000242372 | EIF6 | -0.621665761 | 0.392541278 | -1.222385348 | 0.066638754 | -1.84405111 | 9.460782017 | 10.49775373 | 11.07092887 | 9.80724435 | 9.92248069 | 9.067128536 | 9.44519497 |
| ENSG00000224051 | CPTP | -1.539097829 | 9.08E-09 | -0.306105539 | 0.320201587 | -1.845203368 | 9.174465797 | 9.246456519 | 9.385998894 | 7.626911583 | 7.86198169 | 8.918665794 | 9.013414035 |
| ENSG00000130772 | MED18 | -0.706958615 | 0.582810301 | -1.141095955 | 0.346063415 | -1.84805457 | 9.567995545 | 7.145017454 | 6.629488387 | 7.674344958 | 7.69009211 | 7.389977651 | 7.112501355 |
| ENSG00000138085 | ATRAID | -0.336040531 | 0.466780097 | -1.512253766 | 9.42E-05 | -1.848294297 | 10.78274301 | 10.03760521 | 10.26027233 | 10.02529042 | 10.0907323 | 9.003785277 | 8.764090674 |
| ENSG00000103150 | MLYCD | -1.037113817 | 0.620052795 | -0.814047595 | 0.698497879 | -1.851161412 | 9.189468652 | 7.189546432 | 2 | 6.882264028 | 6.95938211 | 7.118879607 | 7.156006254 |
| ENSG00000175283 | DOLK | -1.123616904 | 0.310065927 | -0.728583383 | 0.519747981 | -1.852200287 | 7.27616306 | 8.777382925 | 10.23774519 | 8.124184191 | 8.10812366 | 8.234076861 | 8.735123394 |
| ENSG00000221909 | FAM200A | -0.841016854 | 0.653037922 | -1.012805232 | 0.576450932 | -1.853822086 | 7.750913381 | 6.535001748 | 2 | 6.01571348 | 5.78232107 | 5.954417431 | 5.497565112 |
| ENSG00000168067 | MAP4K2 | -1.000983521 | 0.239153967 | -0.852857501 | 0.314114017 | -1.853841022 | 8.075571721 | 7.002537973 | 9.031722608 | 7.388195945 | 7.14977896 | 7.264607048 | 7.557435494 |
| ENSG00000141012 | GALNS | -0.494248942 | 0.468067566 | -1.359892289 | 0.024203174 | -1.854141231 | 10.22148166 | 9.433952963 | 8.845626171 | 9.065699797 | 9.17099461 | 8.064761569 | 8.435907908 |
| ENSG00000074582 | BCS1L | -1.69898779 | 0.282799717 | -0.155843521 | 0.932122597 | -1.854831312 | 8.797441463 | 7.60566235 | 3.369317972 | 6.221320894 | 6.00960265 | 7.583859056 | 7.629578661 |
| ENSG00000083812 | ZNF324 | -1.243644273 | 0.252882834 | -0.61123652 | 0.591752395 | -1.854880793 | 7.27616306 | 6.228547981 | 8.978369997 | 6.755670824 | 6.67805337 | 7.402744819 | 7.253757517 |
| ENSG00000115233 | PSMD14 | 0.031967913 | 0.959724553 | -1.8871424 | 8.98E-05 | -1.855174487 | 9.397585735 | 10.49332014 | 10.0085654 | 10.03951164 | 10.0971999 | 8.00283742 | 8.309838123 |
| ENSG00000133313 | CNDP2 | -0.590511677 | 0.262050542 | -1.264931992 | 0.008241509 | -1.855443669 | 10.1770105 | 9.806969599 | 10.7325871 | 9.703174645 | 9.69550802 | 8.770720575 | 9.248214811 |
| ENSG00000134910 | STT3A | -0.751468199 | 0.004499435 | -1.104750285 | 1.27E-05 | -1.856218484 | 11.85353634 | 12.2341388 | 12.05891816 | 11.3321557 | 11.2822233 | 10.8829182 | 11.02321901 |
| ENSG00000138162 | TACC2 | -1.598173952 | 0.05437861 | -0.258072438 | 0.790020846 | -1.85624639 | 8.228665467 | 7.315436877 | 9.413059136 | 7.101994243 | 6.88657603 | 8.242494717 | 8.376090431 |
| ENSG00000104435 | STMN2 | 3.334135715 | 0.560525946 | -5.191092886 | 0.364514502 | -1.856957171 | 2 | 2 | 2 | 3.95996233 | 3.69849104 | 2 | 2 |
| ENSG00000267631 | CGB1 | 3.333486812 | 0.560547313 | -5.191092889 | 0.364514502 | -1.857606077 | 2 | 2 | 2 | 3.881293667 | 3.7864959 | 2 | 2 |
| ENSG00000196531 | NACA | -0.733498575 | 0.01701725 | -1.124448786 | 0.000113299 | -1.857947361 | 12.08776766 | 11.6550774 | 11.65357938 | 11.03478106 | 11.1254767 | 10.65645596 | 10.72420814 |
| ENSG00000126254 | RBM42 | -1.152010977 | 0.007139137 | -0.706835124 | 0.113430713 | -1.858846101 | 10.15425007 | 10.10308486 | 9.399592459 | 8.842507967 | 8.71722214 | 9.052840727 | 9.375131848 |
| ENSG00000011243 | AKAP8L | -0.827286911 | 0.365077256 | -1.031794375 | 0.240278931 | -1.859081286 | 10.6316532 | 8.317412869 | 10.02619908 | 9.067921572 | 9.16279627 | 8.967137431 | 8.857178832 |
| ENSG00000160471 | COX6B2 | -2.71967371 | 0.053462108 | 0.859391647 | 0.53844608 | -1.860282063 | 3.327711234 | 4.11352228 | 4.058998049 | 2.270786764 | 2.68924543 | 4.749835737 | 4.341135913 |
| ENSG00000130702 | LAMA5 | -3.853562229 | 3.52E-10 | 1.992053188 | 0.001927913 | -1.861509041 | 10.65885801 | 9.267380431 | 10.45512158 | 6.799114572 | 5.99120031 | 12.28712585 | 12.170995 |
| ENSG00000149357 | LAMTOR1 | -0.76984554 | 0.077518916 | -1.09352983 | 0.008257259 | -1.863375371 | 10.11549941 | 10.31889309 | 9.788583153 | 9.277301907 | 9.37372144 | 8.661962194 | 9.281988193 |
| ENSG00000176428 | VPS37D | -1.854760975 | 0.163796432 | -0.008709311 | 0.995654398 | -1.863470286 | 4.00722531 | 5.595927442 | 7.552162999 | 4.887609717 | 4.4836996 | 6.566956211 | 6.155328188 |
| ENSG00000172244 | C5orf34 | -0.082691058 | 0.959439303 | -1.785125131 | 0.194134168 | -1.867816189 | 5.330594134 | 5.72254323 | 2 | 4.984880435 | 4.95174433 | 3.753554863 | 3.496556178 |

| | | | | | | | | | | | | | |
|---|---|---|---|---|---|---|---|---|---|---|---|---|---|
| ENSG00000183549 | ACSM5 | -1.066539995 | 0.683688476 | -0.803253127 | 0.758322907 | -1.869793122 | 5.095959458 | 2 | 2 | 2.868578001 | 3.33976443 | 3.1504466 | 3.383972333 |
| ENSG00000213380 | COG8 | -0.882028892 | 0.216791462 | -0.988367085 | 0.153226253 | -1.870395977 | 9.720386367 | 8.213429689 | 8.721235202 | 8.15586117 | 8.14188248 | 8.248293824 | 7.805990373 |
| ENSG00000081721 | DUSP12 | -0.323234139 | 0.539294232 | -1.547424126 | 0.00046995 | -1.870658265 | 8.075571721 | 7.467835385 | 8.35328246 | 7.714607209 | 7.64959655 | 6.417131325 | 6.569847194 |
| ENSG00000169371 | SNUPN | -0.437578912 | 0.7863872 | -1.433249597 | 0.317845862 | -1.870828509 | 7.709880939 | 8.806266215 | 4.058998049 | 7.447418636 | 7.32531744 | 6.372988335 | 6.476841503 |
| ENSG00000100029 | PES1 | -0.856364746 | 0.130846679 | -1.016250272 | 0.063605295 | -1.872615018 | 9.556468883 | 9.566483142 | 10.49365885 | 9.085573664 | 9.09332116 | 8.656642182 | 9.161219358 |
| ENSG00000111640 | GAPDH | -0.392590719 | 0.588718348 | -1.480409062 | 0.01853175 | -1.87299978 | 15.9392534 | 14.99021973 | 14.54216152 | 14.85807235 | 14.9133335 | 13.50068265 | 14.04500215 |
| ENSG00000157212 | PAXIP1 | -0.767301793 | 0.633945061 | -1.106271385 | 0.471669087 | -1.873573178 | 4.00722531 | 6.389896633 | 8.561105505 | 6.476767609 | 6.65485813 | 6.375345632 | 6.102937566 |
| ENSG00000103343 | ZNF174 | -1.375675597 | 0.02259857 | -0.498241116 | 0.448152433 | -1.873916713 | 6.89194353 | 7.538394143 | 8.206335842 | 6.29903474 | 6.32089174 | 7.310906894 | 6.977399345 |
| ENSG00000108733 | PEX12 | -0.778350223 | 0.68311016 | -1.096247004 | 0.546129722 | -1.874597227 | 7.667647223 | 7.847167909 | 2 | 6.387081877 | 6.49425375 | 6.401024833 | 5.821002687 |
| ENSG00000066056 | TIE1 | -2.119341245 | 0.430098143 | 0.244499856 | 0.932012683 | -1.874841389 | 3.327711234 | 3.076665209 | 2 | 2.270786764 | 2.2680111 | 3.534812207 | 2.277179783 |
| ENSG00000160908 | ZNF394 | -0.79673613 | 0.468098579 | -1.084585415 | 0.297503104 | -1.881321545 | 8.009475818 | 6.844429755 | 9.565556058 | 7.77569021 | 7.75969386 | 7.430214384 | 7.539150216 |
| ENSG00000188186 | LAMTOR4 | -1.081721227 | 0.109717472 | -0.800664525 | 0.242561306 | -1.882385752 | 8.696033109 | 8.124485672 | 9.636156974 | 7.826439377 | 7.93600799 | 8.061837451 | 8.250064747 |
| ENSG00000167208 | SNX20 | -3.274145399 | 0.384580714 | 1.391574176 | 0.686186 | -1.882571223 | 3.327711234 | 2 | 2 | 2 | 2.14022021 | 3.56816602 | 2.70970617 |
| ENSG00000070540 | WIPI1 | -0.15575854 | 0.878154523 | -1.727995187 | 0.032363747 | -1.883753727 | 11.08909278 | 10.23108619 | 8.904018013 | 10.17037776 | 10.1777249 | 8.380906995 | 8.810568912 |
| ENSG00000239264 | TXNDC5 | 1.05421316 | 0.236638612 | -2.938135257 | 0.000190879 | -1.883922097 | 9.922475255 | 11.29641566 | 9.099922433 | 11.39964074 | 11.4928844 | 7.522436904 | 7.446882182 |
| ENSG00000169193 | CCDC126 | -1.045011227 | 0.050710673 | -0.838962517 | 0.114203801 | -1.883973744 | 6.257048221 | 6.728495673 | 6.958202216 | 5.742861563 | 5.63601114 | 6.031172603 | 5.713290392 |
| ENSG00000004478 | FKBP4 | -1.43220214 | 7.83E-05 | -0.452418607 | 0.264775609 | -1.884620746 | 10.62062599 | 10.019219 | 10.31150679 | 8.807036108 | 9.00894176 | 10.04184345 | 9.709771424 |
| ENSG00000149636 | DSN1 | -0.871960099 | 0.680832734 | -1.012728243 | 0.623122945 | -1.884688342 | 2 | 6.728495673 | 8.865353883 | 6.739036112 | 6.73446058 | 6.830158212 | 6.329589667 |
| ENSG00000109794 | FAM149A | -1.215980497 | 0.444480686 | -0.669010756 | 0.681460704 | -1.884991253 | 3.327711234 | 3.686303414 | 6.203069917 | 3.921164147 | 4.05806467 | 4.749835737 | 3.987514168 |
| ENSG00000124688 | MAD2L1BP | -0.719692421 | 0.594639663 | -1.165901893 | 0.359547024 | -1.885594315 | 10.25752168 | 7.099070191 | 7.600047325 | 8.27796118 | 8.32935768 | 7.964991788 | 7.755972999 |
| ENSG00000120314 | WDR55 | -0.969961853 | 0.119633373 | -0.918549985 | 0.135132661 | -1.888511838 | 9.667789716 | 8.732943944 | 8.35328246 | 8.069792811 | 8.06478301 | 8.015738956 | 8.213468956 |
| ENSG00000138175 | ARL3 | -1.239095719 | 0.000196923 | -0.650625843 | 0.064617229 | -1.889721562 | 8.928409675 | 8.862353853 | 9.116480286 | 7.739909082 | 7.75424764 | 8.10153367 | 8.521558854 |
| ENSG00000004534 | RBM6 | -0.806477441 | 0.209654079 | -1.083807814 | 0.077031166 | -1.890285255 | 9.751047882 | 9.549500232 | 10.79563557 | 9.374743937 | 9.29807478 | 8.870181261 | 9.230416542 |
| ENSG00000105771 | SMG9 | -1.395311639 | 0.018810941 | -0.496851551 | 0.446195114 | -1.89216319 | 8.946185348 | 8.848535039 | 10.0085654 | 7.944803394 | 8.02463101 | 9.054312673 | 8.665783734 |
| ENSG00000119977 | TCTN3 | -0.913533668 | 0.037504648 | -0.978636098 | 0.023427954 | -1.892169765 | 10.5468183 | 9.77810053 | 10.11951029 | 9.254074072 | 9.28868707 | 8.987312532 | 9.397416937 |
| ENSG00000068745 | IP6K2 | -0.647920548 | 0.322658513 | -1.244833036 | 0.038886962 | -1.892753584 | 11.00254523 | 9.956182102 | 9.69246155 | 9.650037049 | 9.72504082 | 9.135128763 | 9.053075307 |
| ENSG00000173465 | SSSCA1 | -0.955502843 | 0.134496548 | -0.9398925 | 0.132964024 | -1.895395343 | 8.468948322 | 8.029695761 | 7.029880454 | 6.989292186 | 7.06433609 | 6.986363353 | 7.096512788 |
| ENSG00000139269 | INHBE | -1.116606922 | 0.340882919 | -0.779590624 | 0.501137387 | -1.896197546 | 4.00722531 | 4.710564154 | 4.058998049 | 3.406930997 | 3.60476711 | 3.956110469 | 3.474731315 |
| ENSG00000094841 | UPRT | -0.745168249 | 0.725625854 | -1.151607611 | 0.569190163 | -1.89677586 | 9.143983763 | 7.35507212 | 2 | 7.084543139 | 7.31425651 | 6.603610344 | 6.990916783 |
| ENSG00000132016 | C19orf57 | -2.94504289 | 0.251382472 | 1.047610924 | 0.686181755 | -1.897431967 | 2 | 4.442710129 | 2 | 2.3892346 | 2.14022021 | 4.580376151 | 3.36036153 |
| ENSG00000104388 | RAB2A | -0.886813196 | 0.040278645 | -1.010802116 | 0.016731085 | -1.897615312 | 10.40620204 | 10.92062569 | 11.28380426 | 10.05191139 | 10.0067695 | 9.945980056 | 9.864906512 |
| ENSG00000172766 | NAA16 | -0.526577523 | 0.743333936 | -1.371565114 | 0.348530091 | -1.898142637 | 4.815609471 | 5.595927442 | 8.511888463 | 6.682159616 | 6.68951263 | 6.162543904 | 5.540219242 |
| ENSG00000075188 | NUP37 | -0.777194817 | 0.06885972 | -1.121106679 | 0.005233988 | -1.898301496 | 7.485143553 | 7.274681896 | 7.451345288 | 6.772115916 | 6.51372389 | 6.442083733 | 6.185873005 |
| ENSG00000177946 | CENPBD1 | -1.084430391 | 0.200330183 | -0.814021348 | 0.340884972 | -1.89845174 | 8.16936288 | 6.535001748 | 8.585098385 | 6.936770382 | 6.91614081 | 7.309674888 | 7.056667034 |
| ENSG00000065665 | SEC61A2 | -0.747878712 | 0.182596921 | -1.15063918 | 0.028720091 | -1.898517892 | 7.975256505 | 7.73146271 | 6.882775709 | 6.975156635 | 6.77267312 | 6.525106458 | 6.454714037 |
| ENSG00000141696 | P3H4 | -1.669941213 | 0.000673737 | -0.228679604 | 0.699954446 | -1.898620817 | 11.43901855 | 11.08286144 | 10.32581712 | 9.362125691 | 9.35044873 | 10.93158439 | 10.63739192 |
| ENSG00000104047 | DTWD1 | -0.00138869 | 0.998610961 | -1.897269266 | 0.021362205 | -1.898657956 | 8.981092835 | 7.051611315 | 8.65477198 | 8.504819462 | 8.35656898 | 6.977080688 | 6.033722345 |
| ENSG00000188613 | NANOS1 | -2.831437171 | 0.110015664 | 0.932321797 | 0.616643045 | -1.899115374 | 3.327711234 | 5.72254323 | 2 | 2.868578001 | 2.59489879 | 5.404520029 | 5.256659864 |
| ENSG00000100387 | RBX1 | -0.493920466 | 0.421603569 | -1.405330405 | 0.009454429 | -1.899250871 | 9.820154628 | 8.687092496 | 9.031722608 | 8.747293529 | 8.78689171 | 7.669911566 | 8.041339497 |
| ENSG00000166704 | ZNF606 | -0.985956364 | 0.524696221 | -0.913941865 | 0.548117081 | -1.899898228 | 3.327711234 | 7.701025996 | 7.600047325 | 5.968925393 | 6.32824132 | 6.57313027 | 5.772630506 |
| ENSG00000213638 | ADAT3 | -0.58257636 | 0.735527035 | -1.319123538 | 0.405619867 | -1.901699898 | 6.653098118 | 5.72254323 | 2 | 5.22679461 | 5.14724905 | 4.727866721 | 4.353147976 |
| ENSG00000139620 | KANSL2 | -0.795248293 | 0.244358561 | -1.108355667 | 0.087576727 | -1.90360396 | 9.460782017 | 8.213429689 | 8.206335842 | 7.952038749 | 7.98332947 | 7.758361797 | 7.554837457 |
| ENSG00000119777 | TMEM214 | -1.401362596 | 0.00362723 | -0.502360353 | 0.346019765 | -1.903722949 | 12.14852894 | 11.26554928 | 11.21627664 | 10.1842688 | 10.2389602 | 11.03863895 | 11.17536771 |
| ENSG00000166341 | DCHS1 | -0.943679579 | 0.498655693 | -0.960605511 | 0.481615811 | -1.90428509 | 8.716893051 | 10.34373336 | 6.203069917 | 8.287525788 | 8.2960118 | 8.102956788 | 8.428831018 |
| ENSG00000166402 | TUB | -2.144466826 | 3.26E-09 | 0.239949868 | 0.584589142 | -1.904516958 | 8.892187472 | 9.357954881 | 8.904018013 | 6.975156635 | 6.95464102 | 9.501402587 | 9.092571273 |
| ENSG00000136271 | DDX56 | -0.736825346 | 0.017838528 | -1.170161114 | 6.48E-05 | -1.906986459 | 9.435834703 | 9.806969599 | 9.452720984 | 8.861844103 | 8.83412012 | 8.322856568 | 8.509534124 |
| ENSG00000114956 | DGUOK | -1.070024587 | 0.174638808 | -0.837455896 | 0.29039103 | -1.907480482 | 10.56993315 | 9.04308532 | 8.941672936 | 8.655725489 | 8.66738547 | 8.806098644 | 8.973118091 |
| ENSG00000175198 | PCCA | -0.779731748 | 0.680833278 | -1.127848164 | 0.531540246 | -1.907579912 | 7.27616306 | 7.928237109 | 2 | 6.147608921 | 6.48112613 | 6.084012396 | 5.900914815 |
| ENSG00000053900 | ANAPC4 | -0.492183521 | 0.415575369 | -1.415558465 | 0.007718915 | -1.907741986 | 7.485143553 | 7.274681896 | 8.325066185 | 7.190301983 | 7.36158648 | 6.363520437 | 6.412287469 |
| ENSG00000163520 | FBLN2 | -3.418532982 | 1.39E-11 | 1.510735424 | 0.004914775 | -1.907797558 | 10.29268328 | 10.50217373 | 11.45632039 | 7.450827758 | 7.45854177 | 12.37599757 | 12.32716121 |
| ENSG00000108219 | TSPAN14 | -1.773409223 | 0.006990516 | -0.135497237 | 0.868228993 | -1.908906459 | 8.716893051 | 9.488434476 | 10.36791565 | 7.917958914 | 7.92635454 | 9.450887435 | 9.631178479 |
| ENSG00000138777 | PPA2 | -0.555544611 | 0.597540846 | -1.355171217 | 0.153080856 | -1.910715828 | 9.497412313 | 7.572420273 | 7.342949392 | 7.939959586 | 7.93118934 | 7.283556699 | 7.010010317 |
| ENSG00000166224 | SGPL1 | -1.794137607 | 2.30E-07 | -0.11821586 | 0.790777785 | -1.912353467 | 10.65885801 | 10.17128279 | 10.02619908 | 8.463225341 | 8.59001038 | 10.26657884 | 10.11575905 |
| ENSG00000221878 | PSG7 | 0.603075583 | 0.838765858 | -2.516796132 | 0.336805988 | -1.913720548 | 2 | 2 | 4.523675391 | 3.566290607 | 3.86943969 | 2.094598369 | 2.551896972 |
| ENSG00000161243 | FBXO27 | -0.589699445 | 0.417837871 | -1.325270232 | 0.042999625 | -1.914969677 | 5.70932728 | 6.78762695 | 7.098165388 | 5.997179931 | 6.16544343 | 5.253805068 | 5.540219242 |
| ENSG00000146414 | SHPRH | -0.880006645 | 0.166330801 | -1.035084258 | 0.091257767 | -1.915090904 | 9.048471759 | 7.790474954 | 8.143093325 | 7.656739692 | 7.45182838 | 7.536196998 | 7.263337153 |
| ENSG00000169750 | RAC3 | -2.500657914 | 0.009223953 | 0.585138567 | 0.59025161 | -1.915519347 | 7.940205782 | 6.464272742 | 5.772716771 | 4.17310023 | 5.09650239 | 7.779874724 | 7.357281305 |
| ENSG00000148300 | REXO4 | -1.09268405 | 0.265307052 | -0.823981991 | 0.405601197 | -1.916666042 | 8.493334308 | 7.790474954 | 10.11127249 | 8.083034979 | 8.02009991 | 8.547470747 | 8.042267212 |
| ENSG00000143401 | ANP32E | -0.797447389 | 0.152575674 | -1.119685736 | 0.034280104 | -1.917133125 | 8.946185348 | 10.13758676 | 9.809100877 | 8.925713487 | 8.91931898 | 8.610361892 | 8.596104698 |
| ENSG00000168214 | RBPJ | -0.590515183 | 0.181720064 | -1.328608244 | 0.000859069 | -1.919123427 | 10.40620204 | 11.03790484 | 10.42220168 | 10.04177408 | 10.0912724 | 9.508668537 | 9.131014727 |
| ENSG00000073605 | GSDMB | -1.830044684 | 0.396964364 | -0.090831287 | 0.9703912 | -1.920875971 | 2 | 6.04685525 | 9.031722608 | 5.959382894 | 5.77157844 | 8.11499695 | 6.556922154 |
| ENSG00000116586 | LAMTOR2 | -0.618696954 | 0.474824893 | -1.303975027 | 0.097545528 | -1.922671981 | 7.904282243 | 8.029695761 | 9.577565424 | 7.985332082 | 8.21506984 | 7.31213785 | 7.543088047 |
| ENSG00000124570 | SERPINB6 | -1.074575172 | 0.033835146 | -0.848266152 | 0.098450928 | -1.922841324 | 10.55263183 | 10.50217373 | 11.4530647 | 9.796566011 | 9.86931161 | 10.00616163 | 10.10957632 |
| ENSG00000142235 | LMTK3 | -2.121366872 | 0.407875143 | 0.197112577 | 0.944880505 | -1.924254295 | 2 | 2 | 6.322058544 | 3.349643765 | 3.2184422 | 4.875056259 | 5.171036168 |
| ENSG00000135093 | USP30 | -1.130211699 | 0.001506743 | -0.794161396 | 0.029318923 | -1.924373095 | 8.468948322 | 8.053983481 | 8.110401118 | 7.202487431 | 7.0150921 | 7.52030827 | 7.352801367 |
| ENSG00000130844 | ZNF331 | -1.416840574 | 0.203718914 | -0.507976455 | 0.674475323 | -1.924817029 | 5.70932728 | 8.762721501 | 8.486635645 | 6.658765962 | 6.85130017 | 7.89195932 | 7.339277475 |
| ENSG00000099139 | PCSK5 | -1.593256233 | 0.373200768 | -0.331608266 | 0.866860327 | -1.924864498 | 2 | 7.431218269 | 5.772716771 | 4.825935136 | 4.83276841 | 6.118193494 | 5.799216178 |
| ENSG00000175354 | PTPN2 | -0.947531645 | 0.046063656 | -0.979592082 | 0.035498785 | -1.927123727 | 8.075571721 | 8.147241016 | 8.742726175 | 7.344877929 | 7.48837134 | 7.138441638 | 7.599641096 |
| ENSG00000155530 | LRGUK | 0.636138544 | 0.887753545 | -2.563665455 | NA | -1.927526911 | 2 | 3.076665209 | 2 | 2.947924294 | 2.68924543 | 2 | 2.225841198 |
| ENSG00000125835 | SNRPB | -0.813527426 | 0.322678131 | -1.114675047 | 0.154104664 | -1.928202473 | 11.56228205 | 10.00683004 | 9.868959282 | 9.843359791 | 9.93095363 | 9.558011928 | 9.617549359 |
| ENSG00000102057 | KCND1 | -2.657562356 | 0.038689773 | 0.728421195 | 0.608329799 | -1.92914116 | 3.327711234 | 6.140560021 | 7.225742779 | 3.997744386 | 3.90918929 | 6.566956211 | 7.263337153 |
| ENSG00000068903 | SIRT2 | -1.313469899 | 0.046693782 | -0.617051441 | 0.37875047 | -1.93052134 | 10.20680973 | 8.820493869 | 8.99637432 | 8.246921218 | 8.10598728 | 8.809589199 | 8.922120243 |
| ENSG00000100104 | SRRD | -1.203941151 | 0.005734682 | -0.727502136 | 0.107979874 | -1.931443287 | 7.975256505 | 7.874700259 | 8.536706852 | 7.053486362 | 6.89154576 | 7.27600668 | 7.58443608 |
| ENSG00000178184 | PARD6G | -2.664378177 | 0.000246448 | 0.731394398 | 0.370750556 | -1.932983778 | 8.517314938 | 6.899080544 | 8.845626171 | 5.720439317 | 5.72778967 | 9.047307518 | 8.987157359 |
| ENSG00000077585 | GPR137B | -0.800891219 | 0.39960716 | -1.133445654 | 0.207913863 | -1.934336873 | 9.410448395 | 7.274681896 | 7.858115942 | 7.731524332 | 7.6318872 | 7.36172352 | 7.352801367 |
| ENSG00000173456 | RNF26 | -1.264579801 | 0.0026249 | -0.670356175 | 0.132316489 | -1.934935976 | 10.02639697 | 10.27307349 | 9.426401271 | 8.686416476 | 8.70739158 | 9.170171434 | 9.394150543 |
| ENSG00000106400 | ZNHIT1 | -1.20987264 | 8.45E-05 | -0.725385471 | 0.024667973 | -1.935258112 | 10.24321353 | 9.962611189 | 10.40208303 | 8.902248225 | 9.10620439 | 9.378702006 | 9.591800868 |
| ENSG00000187954 | CYHR1 | -1.508137045 | 0.075046884 | -0.427647392 | 0.650098837 | -1.935784437 | 10.75779627 | 8.916342327 | 8.978369997 | 8.325162455 | 8.32752521 | 9.457577544 | 9.334433107 |
| ENSG00000165695 | AK8 | -2.37126688 | 0.525394621 | 0.435132908 | 0.911626073 | -1.936133972 | 2 | 3.686303414 | 2 | 2 | 2.38539787 | 3.27687201 | 2.782576693 |
| ENSG00000253250 | C8orf88 | -0.3318592 | 0.731338839 | -1.604558181 | 0.051506371 | -1.936417381 | 6.73718411 | 5.131603625 | 5.39244166 | 5.590471342 | 5.63601114 | 4.5389195 | 4.456980017 |
| ENSG00000111581 | NUP107 | -0.554285875 | 0.461962192 | -1.384164641 | 0.039291523 | -1.938450516 | 8.586954094 | 7.051611315 | 8.486635645 | 7.608714138 | 7.66419009 | 6.78654324 | 6.870723585 |
| ENSG00000204175 | GPRIN2 | -5.272940451 | 0.045443019 | 3.330987016 | 0.100713788 | -1.941953435 | 2 | 4.710564154 | 2 | 2 | 2.14022021 | 6.295446113 | 6.476841503 |
| ENSG00000018280 | SLC11A1 | -4.24482746 | 0.149345693 | 2.299669348 | 0.404282769 | -1.945158112 | 2 | 4.710564154 | 2 | 2 | 2.2680111 | 6.013120342 | 4.30448772 |
| ENSG00000089335 | ZNF302 | -0.434034974 | 0.537614614 | -1.513098076 | 0.013738586 | -1.947133051 | 9.448362284 | 8.356994601 | 8.486635645 | 8.486665614 | 8.34755546 | 7.679481638 | 6.944034147 |
| ENSG00000103275 | UBE2I | -0.902037226 | 0.183896625 | -1.047486668 | 0.110655039 | -1.949523894 | 11.0110152 | 9.470501606 | 10.76969939 | 9.623093601 | 9.68910013 | 9.52123003 | 9.502269352 |
| ENSG00000174791 | RIN1 | -0.533806087 | 0.820701528 | -1.418456857 | 0.506072054 | -1.952262944 | 10.05943807 | 7.847167909 | 2 | 8.174538791 | 8.28849576 | 7.107581047 | 7.581886257 |
| ENSG00000127837 | AAMP | -0.82104526 | 0.145540049 | -1.132305779 | 0.03494019 | -1.953351039 | 10.98115017 | 9.923599762 | 10.05225195 | 9.524619869 | 9.6360458 | 9.157900392 | 9.377706707 |
| ENSG00000176410 | DNAJC30 | -0.88937043 | 0.601416718 | -1.064226363 | 0.518736842 | -1.953596793 | 9.667789716 | 8.234834549 | 4.058998049 | 7.697489359 | 7.66709116 | 7.172727426 | 7.78508305 |
| ENSG00000149541 | B3GAT3 | -0.791011956 | 0.335922911 | -1.164846019 | 0.132957972 | -1.955857975 | 9.98400217 | 8.487557258 | 10.45512158 | 9.038768132 | 9.09008229 | 8.821740064 | 8.553793978 |
| ENSG00000148288 | GBGT1 | -1.303343492 | 0.399649306 | -0.653388538 | 0.685289111 | -1.95673203 | 5.095959458 | 7.393647515 | 3.369317972 | 4.927308402 | 5.0259382 | 5.638399518 | 5.475755622 |
| ENSG00000174744 | BRMS1 | -0.675775574 | 0.530106276 | -1.281779516 | 0.19506376 | -1.957555089 | 7.218819781 | 8.169643011 | 9.767769415 | 8.040677544 | 8.15434112 | 7.315824427 | 7.669754031 |
| ENSG00000196670 | ZFP62 | -1.226416917 | 0.230742948 | -0.732799523 | 0.487153954 | -1.95921644 | 7.867441304 | 6.389896633 | 9.014156722 | 7.026323566 | 6.79406287 | 7.648631242 | 7.085754446 |
| ENSG00000104613 | INTS10 | -0.76503763 | 0.185014751 | -1.195009111 | 0.027565187 | -1.960046741 | 9.579430841 | 9.71095217 | 8.561105505 | 8.633396609 | 8.58848096 | 8.321634728 | 8.036691948 |
| ENSG00000198873 | GRK5 | -1.871723471 | 0.022553657 | -0.088691394 | 0.93161544 | -1.960414865 | 8.39321512 | 7.393647515 | 9.553445883 | 6.856790872 | 6.88158913 | 8.568740544 | 8.663373824 |
| ENSG00000122417 | ODF2L | -0.596781482 | 0.416081529 | -1.365221429 | 0.038915612 | -1.96200291 | 7.867441304 | 7.467835385 | 8.941672936 | 7.650823203 | 7.62593544 | 6.812870125 | 6.963754056 |
| ENSG00000179476 | C14orf28 | -1.253880973 | 0.34242241 | -0.708223712 | 0.604288014 | -1.962104685 | 4.00722531 | 7.431218269 | 7.818184403 | 6.061031724 | 5.74985018 | 6.548273902 | 6.298867018 |
| ENSG00000214113 | LYRM4 | -1.365768996 | 7.59E-05 | -0.596789914 | 0.107088128 | -1.96255891 | 8.367060731 | 8.671479029 | 8.865353883 | 7.292639438 | 7.31795292 | 7.912394292 | 8.192729549 |
| ENSG00000100968 | NFATC4 | -1.005263412 | 0.003443909 | -0.957327373 | 0.005335787 | -1.962590785 | 11.30078111 | 11.03180337 | 10.65536953 | 9.968745466 | 10.0651181 | 10.08064212 | 10.05017178 |
| ENSG00000173436 | MINOS1 | -0.212613779 | 0.845989871 | -1.750320965 | 0.050410355 | -1.962934744 | 9.27634997 | 6.899080544 | 8.143093325 | 8.244958867 | 8.15020018 | 6.426254939 | 6.926052784 |
| ENSG00000086598 | TMED2 | -1.012217447 | 2.18E-05 | -0.950805551 | 6.72E-05 | -1.963022999 | 12.57852557 | 12.38176406 | 12.74476087 | 11.56900979 | 11.5589968 | 11.61644135 | 11.63419556 |
| ENSG00000123992 | DNPEP | -0.480229876 | 0.398651947 | -1.483936858 | 0.002818955 | -1.964166734 | 9.521325812 | 9.835272298 | 10.57378414 | 9.573915313 | 9.5600488 | 8.411091064 | 8.714901913 |
| ENSG00000185261 | KIAA0825 | 0.05272122 | 0.976237425 | -2.017397166 | 0.167574724 | -1.964675947 | 6.009025997 | 4.710564154 | 2 | 5.040242897 | 5.00774437 | 3.500668998 | 3.36036153 |
| ENSG00000168077 | SCARA3 | -3.153472278 | 0.027752249 | 1.188099738 | 0.447054694 | -1.965372541 | 4.467311984 | 9.424669273 | 9.746650996 | 6.024891728 | 5.89549381 | 9.843290512 | 10.50603928 |
| ENSG00000114757 | PEX5L | 0.610599278 | 0.812967163 | -2.576535868 | 0.255841947 | -1.96593659 | 2 | 2 | 5.156626401 | 4.205751501 | 4.37434574 | 2.560105256 | 2.466008509 |
| ENSG00000171204 | TMEM126B | -0.586816629 | 0.582509943 | -1.379614378 | 0.153073981 | -1.966431007 | 9.384607364 | 8.004992147 | 6.80318718 | 7.947219213 | 7.75424764 | 7.186216802 | 6.963754056 |
| ENSG00000115649 | CNPPD1 | -1.415707947 | 0.175994512 | -0.551669443 | 0.623961728 | -1.96737739 | 10.71177538 | 8.077869079 | 8.784770914 | 8.207157886 | 8.2561084 | 9.130600759 | 9.042450579 |
| ENSG00000100603 | SNW1 | -0.718507627 | 0.161771312 | -1.249073198 | 0.008693809 | -1.967580825 | 10.58135316 | 10.03150247 | 9.565556058 | 9.353955293 | 9.45234084 | 8.998824951 | 8.744273936 |

| | | | | | | | | | | | | | |
|---|---|---|---|---|---|---|---|---|---|---|---|---|---|
| ENSG00000151131 | C12orf45 | -1.695968875 | 0.150440253 | -0.272034196 | 0.839792588 | -1.968003071 | 7.27616306 | 4.11352228 | 6.80318718 | 5.003571641 | 4.97065356 | 6.462183138 | 6.099376274 |
| ENSG00000127452 | FBXL12 | -1.1462805 | 0.000234475 | -0.821864184 | 0.00991729 | -1.968144683 | 8.696033109 | 8.981105197 | 8.585098385 | 7.656739692 | 7.62295033 | 8.035995406 | 7.876348251 |
| ENSG00000215009 | ACSM4 | -3.753031254 | NA | 1.784370277 | NA | -1.968660976 | 2 | 2 | 2 | 2 | 2 | 3.296906311 | 2.374681325 |
| ENSG00000239382 | ALKBH6 | 0.412414357 | 0.777645763 | -2.381867968 | 0.052961956 | -1.969453611 | 7.975256505 | 6.78762695 | 4.058998049 | 7.311513918 | 7.44845993 | 4.521997428 | 4.998228167 |
| ENSG00000168646 | AXIN2 | -2.414081402 | 0.132833397 | 0.44444572 | 0.80714548 | -1.969635682 | 7.27616306 | 6.311476754 | 2 | 4.358765249 | 3.98554648 | 6.823267818 | 6.649933445 |
| ENSG00000051128 | HOMER3 | -0.369714932 | 0.736493595 | -1.600684686 | 0.091962344 | -1.970399618 | 10.36053552 | 7.638155693 | 9.69246155 | 9.231457548 | 9.24566964 | 7.919680595 | 8.114615835 |
| ENSG00000167130 | DOLPP1 | -1.236898213 | 0.002958911 | -0.735048593 | 0.09054815 | -1.971946806 | 7.904282243 | 8.655694736 | 8.561105505 | 7.194375246 | 7.20671394 | 7.604100461 | 7.772838868 |
| ENSG00000137513 | NARS2 | -1.341770162 | 0.218538145 | -0.630804851 | 0.582427016 | -1.972575013 | 8.39321512 | 7.189546432 | 5.39244166 | 6.18083559 | 6.1322378 | 6.802396951 | 6.868633039 |
| ENSG00000100353 | EIF3D | -1.061939404 | 0.009987576 | -0.91115098 | 0.028762582 | -1.973090384 | 11.12854954 | 10.58784379 | 11.39994907 | 10.00745917 | 10.0249756 | 10.05219399 | 10.27241252 |
| ENSG00000135736 | CCDC102A | -2.186562086 | 9.68E-11 | 0.211994081 | 0.60541273 | -1.974568005 | 8.418903792 | 8.451443573 | 8.865353883 | 6.314087501 | 6.58294721 | 8.776974697 | 8.820877391 |
| ENSG00000139292 | LGR5 | -8.636823166 | 0.006111607 | 6.661981911 | 0.013890006 | -1.974841255 | 6.563806279 | 2 | 2 | 2 | 2 | 10.36904805 | 12.22132013 |
| ENSG00000178567 | EPM2AIP1 | -0.398020988 | 0.58791184 | -1.5775621 | 0.012430339 | -1.975583088 | 8.19931886 | 9.443177293 | 9.725218844 | 8.842507967 | 8.88359764 | 7.551912644 | 7.83304634 |
| ENSG00000197747 | S100A10 | -1.159039485 | 0.074921525 | -0.81711527 | 0.218712264 | -1.976154755 | 12.07772327 | 11.08580931 | 11.37608156 | 10.43033406 | 10.4058602 | 10.08262707 | 11.2179681 |
| ENSG00000057149 | SERPINB3 | 3.212872721 | 0.576613399 | -5.191093391 | 0.364514502 | -1.97822067 | 2 | 2 | 2 | 3.754611435 | 3.74316441 | 2 | 2 |
| ENSG00000142676 | RPL11 | -0.809746002 | 0.196949817 | -1.168610324 | 0.049227778 | -1.978356326 | 12.74263541 | 11.65309132 | 11.52304554 | 11.25454706 | 11.2935316 | 10.80407274 | 11.01980841 |
| ENSG00000109016 | DHRS7B | -1.202074434 | 0.00733148 | -0.77661153 | 0.093798515 | -1.978685964 | 8.042902253 | 8.414402626 | 8.941672936 | 7.29643414 | 7.36158648 | 7.808940772 | 7.679327321 |
| ENSG00000124657 | OR2B6 | 3.212167157 | 0.576690781 | -5.191093394 | 0.364514502 | -1.978926237 | 2 | 2 | 2 | 3.663521627 | 3.82856379 | 2 | 2 |
| ENSG00000181789 | COPG1 | -0.780587023 | 0.038727587 | -1.200326262 | 0.000760041 | -1.980913285 | 12.45670208 | 11.94046379 | 12.39039575 | 11.47251976 | 11.5264191 | 10.84185164 | 11.2855366 |
| ENSG00000108961 | RANGRF | 0.480266899 | 0.465481438 | -2.462844644 | 8.00E-06 | -1.982577745 | 6.257048221 | 6.951736442 | 7.163363831 | 7.300218887 | 7.31425651 | 4.841348851 | 4.279527956 |
| ENSG00000134640 | MTNR1B | -3.75303144 | NA | 1.769476796 | NA | -1.983554643 | 2 | 2 | 2 | 2 | 2 | 2 | 3.45257121 |
| ENSG00000087077 | TRIP6 | -1.476012375 | 0.016330246 | -0.508032625 | 0.454394312 | -1.984045001 | 11.30767305 | 9.863030423 | 10.48730704 | 9.162439556 | 9.24275568 | 10.0143146 | 10.30389101 |
| ENSG00000121769 | FABP3 | -3.298042147 | 0.070855582 | 1.313496339 | 0.501916413 | -1.984545808 | 8.228665467 | 6.899080544 | 2 | 3.840289928 | 4.43005853 | 7.128693779 | 9.102389799 |
| ENSG00000141959 | PFKL | -1.267501046 | 0.0402643 | -0.717352625 | 0.267777028 | -1.984853671 | 11.90215845 | 11.0102427 | 12.3188683 | 10.55990001 | 10.5883315 | 10.74069043 | 11.42517001 |
| ENSG00000122483 | CCDC18 | 0.158810538 | 0.884722082 | -2.144114391 | 0.013198781 | -1.985303853 | 4.467311984 | 4.93638933 | 6.203069917 | 5.602782483 | 5.43283104 | 3.663836244 | 3.620995423 |
| ENSG00000129460 | NGDN | -0.906034206 | 0.542476242 | -1.081746589 | 0.453341798 | -1.987780795 | 8.493334308 | 8.414402626 | 4.058998049 | 7.035434774 | 7.00595521 | 6.895669227 | 6.804465095 |
| ENSG00000099341 | PSMD8 | -0.82996723 | 0.022676194 | -1.157943885 | 0.000884741 | -1.987911115 | 11.68380625 | 11.08286144 | 11.6648807 | 10.63112186 | 10.7139326 | 10.28393523 | 10.40562199 |
| ENSG00000178814 | OPLAH | -3.009742941 | 0.04127093 | 1.021064353 | 0.524825533 | -1.988678589 | 3.327711234 | 8.124485672 | 6.073378139 | 4.070463721 | 4.31639476 | 7.608114885 | 8.141703677 |
| ENSG00000213563 | C8orf82 | -1.290134976 | 0.066092631 | -0.699127347 | 0.342120369 | -1.989262323 | 8.609439662 | 8.356994601 | 9.868959282 | 7.842101511 | 7.81305436 | 8.330165956 | 8.485861839 |
| ENSG00000178531 | CTXN1 | -2.198409884 | 0.015090069 | 0.208358998 | 0.850369177 | -1.990050886 | 7.53297911 | 7.901717006 | 9.703463497 | 6.463327516 | 6.64900057 | 8.64301114 | 9.144469138 |
| ENSG00000197128 | ZNF772 | -1.092459106 | 0.182231831 | -0.897596521 | 0.271454037 | -1.990055627 | 8.042902253 | 7.35507212 | 6.073378139 | 6.351046291 | 6.27599024 | 6.649130473 | 6.332626253 |
| ENSG00000072818 | ACAP1 | -2.736342537 | 0.025398486 | 0.745383558 | 0.575961295 | -1.990958978 | 6.563806279 | 4.93638933 | 3.369317972 | 3.227757282 | 3.45167086 | 6.262714117 | 6.102937566 |
| ENSG00000109670 | FBXW7 | -1.00719304 | 0.163278075 | -0.984068378 | 0.165345082 | -1.991261418 | 7.975256505 | 8.557180776 | 9.553445883 | 7.834291698 | 7.86198169 | 8.075674526 | 7.632045641 |
| ENSG00000125450 | NUP85 | -1.000862645 | 0.030910658 | -0.991082597 | 0.030536043 | -1.991945242 | 8.87372952 | 8.147241016 | 8.076950877 | 7.474468681 | 7.37584259 | 7.373563392 | 7.495109486 |
| ENSG00000168758 | SEMA4C | -0.883692972 | 0.47314822 | -1.108498065 | 0.347715703 | -1.992191037 | 9.204317094 | 6.311476754 | 9.636156974 | 8.063125866 | 8.04707549 | 8.124837479 | 7.469129172 |
| ENSG00000173611 | SCAI | -1.444114499 | 0.11039739 | -0.548114876 | 0.572896533 | -1.992229375 | 6.653098118 | 7.002537973 | 8.585098385 | 6.276156955 | 6.26836889 | 7.422256313 | 6.771278116 |
| ENSG00000171960 | PPIH | -0.38878686 | 0.790951603 | -1.604295043 | 0.207863251 | -1.993081903 | 7.159102577 | 6.140560021 | 3.369317972 | 5.829220145 | 5.88556448 | 4.813804156 | 4.720747915 |
| ENSG00000100147 | CCDC134 | -0.892296184 | 0.639595086 | -1.101677267 | 0.548817763 | -1.993973451 | 7.53297911 | 5.595927442 | 2 | 5.527288686 | 5.41912973 | 5.163941955 | 5.396716407 |
| ENSG00000175087 | PDIK1L | -1.148355845 | 0.203854133 | -0.845996768 | 0.352922604 | -1.994352612 | 5.330594134 | 7.002537973 | 7.552162999 | 5.808110275 | 5.7929843 | 6.231826305 | 5.917188835 |
| ENSG00000167306 | MYO5B | -1.479266756 | 0.429843212 | -0.517023685 | 0.793470518 | -1.996290441 | 3.327711234 | 3.686303414 | 4.874587708 | 3.663521627 | 2.2680111 | 4.017766949 | 3.36036153 |
| ENSG00000010270 | STARD3NL | -0.509100469 | 0.545365418 | -1.487630084 | 0.046421006 | -1.996730552 | 10.55842202 | 9.030900487 | 8.960138144 | 9.244284335 | 9.17914662 | 8.100821584 | 8.372406522 |
| ENSG00000030066 | NUP160 | -0.882853817 | 0.175376463 | -1.114449466 | 0.074468007 | -1.997303283 | 9.751047882 | 8.687092496 | 10.16797309 | 8.792330793 | 8.76404032 | 8.576444489 | 8.520893436 |
| ENSG00000109133 | TMEM33 | -0.765805955 | 0.159553366 | -1.23214525 | 0.015610198 | -1.997951205 | 9.556468883 | 9.835272298 | 10.63252637 | 9.364838907 | 9.27258642 | 8.899548956 | 8.812201472 |
| ENSG00000181191 | PJA1 | -1.358119156 | 0.024706507 | -0.639844499 | 0.321669264 | -1.997963655 | 9.940323746 | 9.090820894 | 8.536706852 | 7.927778434 | 7.99492122 | 8.677805528 | 8.663976679 |
| ENSG00000159720 | ATP6V0D1 | -1.006531095 | 0.106307854 | -0.991447946 | 0.106512505 | -1.997979041 | 11.4883819 | 10.81799725 | 10.17589429 | 9.867156706 | 9.97493083 | 9.632330087 | 10.1885847 |
| ENSG00000167645 | YIF1B | -1.607826157 | 0.000819269 | -0.390250763 | 0.479041989 | -1.99807692 | 10.99402524 | 10.04973379 | 10.30429804 | 8.94033939 | 8.86981338 | 9.877784692 | 10.3208107 |
| ENSG00000068354 | TBC1D25 | -1.473829905 | 0.080169719 | -0.524547096 | 0.566749731 | -1.998377002 | 7.904282243 | 8.655694736 | 9.972637969 | 7.574744049 | 7.70992146 | 8.397238383 | 8.740849281 |
| ENSG00000142945 | KIF2C | -1.200981895 | 0.579599436 | -0.799393626 | 0.716536102 | -2.000375521 | 3.327711234 | 6.389896633 | 2 | 4.139692846 | 3.98554648 | 3.632645806 | 4.870334174 |
| ENSG00000197872 | FAM49A | -1.509076804 | 0.269781523 | -0.493173211 | 0.736281057 | -2.002250015 | 5.866944937 | 5.72254323 | 3.369317972 | 4.237680128 | 3.90918929 | 5.485310992 | 3.956354764 |
| ENSG00000105983 | LMBR1 | -1.261044467 | 0.012401427 | -0.741569248 | 0.159452507 | -2.002613715 | 9.751047882 | 9.624396729 | 8.721235202 | 8.235106869 | 8.13351654 | 8.546425131 | 8.836470732 |
| ENSG00000157315 | TMED6 | -1.727636454 | 0.551856542 | -0.276348822 | 0.930293595 | -2.003985276 | 2 | 2 | 4.523675391 | 2.141737246 | 2.86123014 | 3.193831101 | 3.15612297 |
| ENSG00000102805 | CLN5 | 0.250672925 | 0.81413118 | -2.255507857 | 0.008495313 | -2.004834932 | 7.485143553 | 8.522788974 | 9.859153765 | 9.246247604 | 9.12952895 | 6.774090992 | 6.681239787 |
| ENSG00000168439 | STIP1 | -1.100515019 | 0.064248308 | -0.905100299 | 0.130055652 | -2.005615318 | 11.81051487 | 10.63690565 | 10.72720655 | 10.04910261 | 10.0804323 | 10.15764151 | 10.35481313 |
| ENSG00000181061 | HIGD1A | -0.379279531 | 0.138339617 | -1.626920767 | 1.55E-13 | -2.006200298 | 10.27872131 | 10.27823701 | 10.28245311 | 9.979407193 | 9.81998493 | 8.748615145 | 8.57061027 |
| ENSG00000130305 | NSUN5 | -0.078779429 | 0.927016036 | -1.928094709 | 0.003735066 | -2.006874138 | 8.928409675 | 7.701025996 | 7.451345288 | 8.098332103 | 8.10170499 | 6.442083733 | 6.151894053 |
| ENSG00000138755 | CXCL9 | 3.182066078 | 0.580892944 | -5.191093519 | 0.364514502 | -2.009027441 | 2 | 2 | 2 | 3.840289928 | 3.60476711 | 2 | 2 |
| ENSG00000149489 | ROM1 | -1.111319894 | 0.543001285 | -0.897762871 | 0.624398897 | -2.009082765 | 2 | 5.303523747 | 7.098165388 | 4.887609717 | 4.91316568 | 5.294231737 | 4.845038551 |
| ENSG00000198721 | ECI2 | -0.627232681 | 0.237182775 | -1.383775969 | 0.003956363 | -2.01100865 | 9.790941832 | 8.848535039 | 9.344432377 | 8.716468139 | 8.7882247 | 7.754744914 | 8.21922296 |
| ENSG00000183763 | TRAIP | -1.472449467 | 0.34408635 | -0.539348743 | 0.738292369 | -2.01179821 | 4.00722531 | 4.710564154 | 2 | 3.09461492 | 3.01487888 | 3.584558147 | 3.640730416 |
| ENSG00000125089 | SH3TC1 | -0.698766396 | 0.666953311 | -1.31327117 | 0.382062502 | -2.012037566 | 3.327711234 | 4.442710129 | 6.80318718 | 4.825935136 | 5.04390545 | 4.698044417 | 4.077147165 |
| ENSG00000175220 | ARHGAP1 | -1.531481834 | 1.13E-07 | -0.483019712 | 0.132806015 | -2.014501546 | 11.63532243 | 11.11207261 | 11.34831197 | 9.851120801 | 9.85411719 | 10.9107105 | 10.88487658 |
| ENSG00000131828 | PDHA1 | -1.467941458 | 0.149386626 | -0.549512242 | 0.617024792 | -2.0174537 | 7.624139768 | 9.34816733 | 10.49365885 | 8.087422161 | 8.15434112 | 8.908528807 | 9.141874832 |
| ENSG00000137692 | DCUN1D5 | -0.471660928 | 0.556356511 | -1.546181213 | 0.028158563 | -2.017842142 | 8.892187472 | 8.540087354 | 10.16797309 | 8.919575326 | 8.8960155 | 7.86621124 | 7.826599033 |
| ENSG00000178974 | FBXO34 | -1.241719352 | 0.112781159 | -0.779776816 | 0.333347859 | -2.021496168 | 8.816894989 | 8.433041974 | 10.19940021 | 8.157948442 | 8.09740995 | 8.627767864 | 8.53941009 |
| ENSG00000104472 | CHRAC1 | -1.282479556 | 0.208331946 | -0.740011308 | 0.483045217 | -2.022490864 | 8.19931886 | 7.315436877 | 9.839340503 | 7.467753569 | 7.65837038 | 7.871230734 | 8.295903454 |
| ENSG00000072210 | ALDH3A2 | -1.548096601 | 6.14E-06 | -0.475277218 | 0.213131367 | -2.023373819 | 9.262228249 | 9.181799937 | 9.735974718 | 7.949630994 | 7.80517502 | 8.861379233 | 9.012940881 |
| ENSG00000126581 | BECN1 | -0.363209869 | 0.535148173 | -1.661106577 | 0.000789378 | -2.024316446 | 10.46286899 | 9.624396729 | 9.601288346 | 9.608667949 | 9.57557184 | 8.017249226 | 8.545311752 |
| ENSG00000183431 | SF3A3 | -0.820497454 | 0.112331583 | -1.204477417 | 0.013717681 | -2.024974871 | 10.15425007 | 9.318399851 | 9.242516611 | 8.744518279 | 8.88733423 | 8.325906648 | 8.541379994 |
| ENSG00000106077 | ABHD11 | -1.31921711 | 0.299548911 | -0.705984112 | 0.592201451 | -2.025201222 | 5.532353208 | 8.317412869 | 5.930865648 | 5.870535235 | 5.93454154 | 5.988693817 | 6.849680541 |
| ENSG00000128609 | NDUFA5 | -1.111050976 | 0.030289936 | -0.914488795 | 0.076831332 | -2.025539772 | 9.27634997 | 9.102512015 | 8.206335842 | 7.920420067 | 7.74878077 | 8.071319244 | 7.988411589 |
| ENSG00000160993 | ALKBH4 | -1.361948653 | 0.169717282 | -0.663618232 | 0.523865784 | -2.025566885 | 5.866944937 | 7.701025996 | 8.561105505 | 6.34372989 | 6.50077304 | 7.086154752 | 7.107191496 |
| ENSG00000170909 | OSCAR | 0.990917799 | 0.661870947 | -3.016653633 | 0.142957636 | -2.025735834 | 2 | 4.710564154 | 2 | 4.205751501 | 4.40247105 | 2.3070652 | 2.326754077 |
| ENSG00000136942 | RPL35 | -0.661416994 | 0.198770347 | -1.367240027 | 0.003541393 | -2.028657021 | 11.83218596 | 10.79296235 | 11.5927499 | 10.81860745 | 10.7983847 | 10.12000692 | 10.08929747 |
| ENSG00000112812 | PRSS16 | -1.268935106 | NA | -0.760475137 | NA | -2.029410243 | 2 | 3.076665209 | 2 | 2.3892346 | 2.14022021 | 2.457042231 | 2.277179783 |
| ENSG00000141376 | BCAS3 | -1.237405045 | 0.018249043 | -0.793043121 | 0.142869248 | -2.030448166 | 8.009475818 | 8.297207175 | 8.978369997 | 7.03996889 | 7.46522407 | 7.594015257 | 7.803803816 |
| ENSG00000142528 | ZNF473 | -1.294700994 | 0.347668061 | -0.736398972 | 0.605488001 | -2.031099966 | 4.467311984 | 7.467835385 | 8.631919287 | 6.351046291 | 6.40670796 | 7.164572841 | 6.612945848 |
| ENSG00000075624 | ACTB | -2.561100716 | 1.41E-11 | 0.529340817 | 0.227054772 | -2.031759899 | 16.68863805 | 16.31538107 | 16.69915842 | 13.99575309 | 14.0391125 | 16.69559387 | 17.42791496 |
| ENSG00000003756 | RBM5 | -0.641277527 | 0.370429303 | -1.390936088 | 0.032616532 | -2.032213616 | 9.410448395 | 10.42046463 | 10.93751085 | 9.781753035 | 9.70964668 | 9.249735981 | 8.700282474 |
| ENSG00000170522 | ELOVL6 | -1.117788119 | 0.010240552 | -0.915758966 | 0.037527109 | -2.033547085 | 9.473095744 | 9.04308532 | 8.677268317 | 7.949630994 | 8.04037874 | 8.009682014 | 8.359066013 |
| ENSG00000102100 | SLC35A2 | -1.07130572 | 0.024397609 | -0.962953911 | 0.043346526 | -2.034259631 | 10.05943807 | 9.055168103 | 9.735974718 | 8.559529208 | 8.65428713 | 8.708044758 | 8.721866969 |
| ENSG00000120820 | GLT8D2 | -0.315322219 | 0.756395306 | -1.719243 | 0.045247931 | -2.034565219 | 8.340423424 | 6.899080544 | 9.149036566 | 8.020170999 | 8.13770558 | 6.387074688 | 6.975457902 |
| ENSG00000183734 | ASCL2 | 0.428702331 | 0.895301283 | -2.464206963 | NA | -2.035504632 | 2 | 2 | 3.369317972 | 2.784612485 | 2.68924543 | 2.183373728 | 2.059860178 |
| ENSG00000128917 | DLL4 | -0.319243902 | 0.899437889 | -1.717342599 | 0.435253955 | -2.036586501 | 3.327711234 | 3.076665209 | 2 | 2.695456494 | 2.86123014 | 2.457042231 | 2.225841198 |
| ENSG00000148143 | ZNF462 | -0.600667567 | 0.274064931 | -1.437648543 | 0.003474236 | -2.03831611 | 8.586954094 | 9.34816733 | 8.486635645 | 8.164192202 | 8.37442884 | 7.238947338 | 7.635738219 |
| ENSG00000203705 | TATDN3 | -1.083216496 | 0.007764718 | -0.957891001 | 0.017643691 | -2.041107497 | 7.435667355 | 7.002537973 | 7.285536013 | 6.147608921 | 6.25300433 | 6.346800618 | 6.29264319 |
| ENSG00000182816 | KRTAP13-2 | 3.149850445 | 0.585258297 | -5.191093653 | 0.364514502 | -2.041243208 | 2 | 2 | 2 | 3.840289928 | 3.55551834 | 2 | 2 |
| ENSG00000221826 | PSG3 | 3.148744888 | 0.585407002 | -5.191093658 | 0.364514502 | -2.042348769 | 2 | 2 | 2 | 3.709785322 | 3.69849104 | 2 | 2 |
| ENSG00000167565 | SERTAD3 | -1.42809821 | 0.019551277 | -0.616855707 | 0.349687984 | -2.044953918 | 9.544849385 | 8.169643011 | 9.439621147 | 7.748245381 | 7.76240929 | 8.508806077 | 8.5998903 |
| ENSG00000147140 | NONO | -1.291049643 | 4.44E-07 | -0.756305654 | 0.004768505 | -2.047355297 | 12.32703082 | 12.0991443 | 12.40057196 | 10.97654609 | 11.0056822 | 11.37866256 | 11.65799488 |
| ENSG00000188107 | EYS | -0.664692424 | 0.700085754 | -1.382727378 | 0.380887317 | -2.047419802 | 2 | 4.710564154 | 4.058998049 | 3.406930997 | 3.65239003 | 3.296906311 | 2.885414003 |
| ENSG00000198939 | ZFP2 | -2.167560754 | 0.377617138 | 0.118927886 | 0.966296927 | -2.048632868 | 2 | 5.457120953 | 2 | 2.600425781 | 3.01487888 | 4.604690117 | 3.771796761 |
| ENSG00000174611 | KY | -1.727915059 | 0.516230128 | -0.32113025 | 0.912100129 | -2.049045309 | 2 | 2 | 6.534109959 | 3.754611435 | 3.60476711 | 4.778616857 | 4.77533149 |
| ENSG00000156970 | BUB1B | -1.071492119 | 0.552001887 | -0.977751614 | 0.582327511 | -2.049243734 | 4.467311984 | 5.72254323 | 2 | 4.237680128 | 3.45167086 | 3.483289517 | 4.329022995 |
| ENSG00000114098 | ARMC8 | -0.774435765 | 0.117793766 | -1.275050434 | 0.005763848 | -2.049486199 | 9.358295023 | 9.181799937 | 8.380957451 | 8.260583448 | 8.27524772 | 7.829717863 | 7.718159092 |
| ENSG00000169635 | HIC2 | -1.176507782 | 0.479746549 | -0.874612071 | 0.601817251 | -2.051119853 | 3.327711234 | 8.717821467 | 7.225742779 | 6.503277904 | 6.42052719 | 6.872493235 | 6.620419619 |
| ENSG00000092199 | HNRNPC | -0.558270541 | 0.266356869 | -1.493041497 | 0.000802417 | -2.051312038 | 12.70528324 | 11.70386119 | 12.28996399 | 11.72268474 | 11.7403773 | 10.85595215 | 10.73830212 |
| ENSG00000103365 | GGA2 | -0.90240295 | 0.059528826 | -1.1499592 | 0.012605537 | -2.05236215 | 10.65885801 | 9.856140801 | 9.907528187 | 9.265734736 | 9.31482024 | 8.857590434 | 9.210333843 |
| ENSG00000111596 | CNOT2 | -0.634966291 | 0.347067822 | -1.41750044 | 0.020707999 | -2.05246673 | 10.41898845 | 10.51535307 | 9.099922433 | 9.504256136 | 9.5116449 | 8.765787532 | 8.695572791 |
| ENSG00000108654 | DDX5 | -0.655566717 | 0.054833942 | -1.398865024 | 7.33E-06 | -2.054431741 | 13.38629018 | 13.41853031 | 13.71813493 | 12.86525712 | 12.8550256 | 11.86229406 | 12.33409073 |
| ENSG00000183607 | GKN2 | -4.714890868 | 0.267434798 | 2.66031661 | 0.501315475 | -2.054574258 | 2 | 2 | 3.369317972 | 2 | 2 | 4.68289886 | 2.885414003 |
| ENSG00000168495 | POLR3D | -1.717609067 | 0.018003832 | -0.338580569 | 0.687802707 | -2.056189636 | 10.96815916 | 9.267380431 | 9.767769415 | 8.499891024 | 8.45385092 | 10.07285649 | 9.57683272 |
| ENSG00000161179 | YDJC | -1.041348544 | 0.105982801 | -1.015195056 | 0.108572393 | -2.0565436 | 7.750913381 | 8.451443573 | 7.029880454 | 6.86703417 | 6.84106088 | 6.934553331 | 6.821857532 |
| ENSG00000109265 | KIAA1211 | -2.116334345 | 0.364034842 | 0.059636281 | 0.982250031 | -2.056698064 | 4.467311984 | 6.140560021 | 2 | 4.070463721 | 2.14022021 | 5.454427604 | 4.574506939 |
| ENSG00000106682 | EIF4H | -1.015186601 | 0.000770868 | -1.041931615 | 0.000500148 | -2.057118215 | 12.15618594 | 12.47802568 | 12.42404098 | 11.34526663 | 11.3460677 | 11.09943318 | 11.50951469 |
| ENSG00000104356 | POP1 | -0.992270248 | 0.452776776 | -1.065792066 | 0.405644179 | -2.058062315 | 4.467311984 | 7.928237109 | 8.042706615 | 6.463327516 | 6.54560135 | 6.123812431 | 6.690737414 |
| ENSG00000166260 | COX11 | -1.127625251 | 0.163112784 | -0.93309944 | 0.248020572 | -2.060724691 | 9.064836028 | 7.145017454 | 8.076950877 | 7.322721205 | 7.04662306 | 7.415399995 | 7.34530375 |
| ENSG00000177058 | SLC38A9 | -0.349687697 | 0.623572187 | -1.711284141 | 0.004335271 | -2.060971838 | 7.750913381 | 7.099070191 | 8.143093325 | 7.409376773 | 7.33993465 | 6.405645065 | 5.621921006 |
| ENSG00000113068 | PFDN1 | -0.971378036 | 0.195462641 | -1.089744722 | 0.133624198 | -2.061122758 | 10.27872131 | 9.975384048 | 8.561105505 | 8.7747563 | 8.84440862 | 8.435582605 | 8.91098986 |

| | | | | | | | | | | | | | |
|---|---|---|---|---|---|---|---|---|---|---|---|---|---|
| ENSG00000113575 | PPP2CA | -0.665488824 | 0.254264315 | -1.395727582 | 0.008663163 | -2.061216406 | 11.63532243 | 11.19631665 | 10.37481416 | 10.52191308 | 10.4662328 | 9.770381989 | 9.763076863 |
| ENSG00000160352 | ZNF714 | -0.975262235 | 0.226076026 | -1.088082269 | 0.161988102 | -2.063344504 | 5.532353208 | 6.899080544 | 7.342949392 | 5.85002558 | 5.84514933 | 5.899789217 | 5.566251551 |
| ENSG00000143590 | EFNA3 | -1.587282335 | 0.454602329 | -0.47716361 | 0.834398737 | -2.064445945 | 7.27616306 | 3.686303414 | 2 | 4.497095146 | 4.37434574 | 5.294231737 | 5.492143571 |
| ENSG00000116752 | BCAS2 | -0.586929969 | 0.212730396 | -1.478319932 | 0.000422063 | -2.065249901 | 8.998234871 | 8.317412869 | 8.266922072 | 8.006336225 | 7.96458672 | 7.21150425 | 7.008112295 |
| ENSG00000149557 | FEZ1 | -2.034280737 | 9.55E-08 | -0.031277067 | 0.951751695 | -2.065557804 | 9.317904228 | 9.214490408 | 9.878698603 | 7.447418636 | 7.52720828 | 9.312280199 | 9.605988 |
| ENSG00000198755 | RPL10A | -1.661080764 | 0.031352923 | -0.405469398 | 0.643682096 | -2.066550162 | 12.38357115 | 11.11207261 | 10.57976717 | 9.893075744 | 9.92187357 | 10.79562693 | 11.4501129 |
| ENSG00000112592 | TBP | -0.79419611 | 0.705469615 | -1.272418549 | 0.522285759 | -2.06661466 | 9.262228249 | 7.979858146 | 2 | 7.363085426 | 7.43830714 | 6.973973146 | 6.897628558 |
| ENSG00000121152 | NCAPH | -0.893962105 | 0.479054624 | -1.173404366 | 0.324488998 | -2.06736647 | 5.095959458 | 4.442710129 | 4.523675391 | 4.299492648 | 3.60476711 | 3.215043717 | 4.201948172 |
| ENSG00000087237 | CETP | -1.990290946 | 0.491841862 | -0.077762142 | 0.979817504 | -2.068053088 | 2 | 3.686303414 | 2 | 2.3892346 | 2.14022021 | 2.656294034 | 2.981406154 |
| ENSG00000173156 | RHOD | -1.890666282 | 0.000155088 | -0.179074568 | 0.772803831 | -2.06974085 | 8.564112515 | 8.255926468 | 9.272377397 | 6.877205252 | 6.92586261 | 8.758581951 | 8.374617996 |
| ENSG00000100124 | ANKRD54 | -1.418971662 | 9.11E-07 | -0.650911841 | 0.033763115 | -2.069883503 | 8.674867122 | 8.487557258 | 8.845626171 | 7.311513918 | 7.23826702 | 8.028526139 | 8.027351693 |
| ENSG00000049167 | ERCC8 | -0.762166898 | 0.354286428 | -1.309121026 | 0.086247207 | -2.071287924 | 5.70932728 | 7.145017454 | 7.552162999 | 6.229283386 | 6.29111322 | 5.880003936 | 5.601925227 |
| ENSG00000116221 | MRPL37 | -1.325856072 | 7.64E-06 | -0.747485644 | 0.016746626 | -2.073341716 | 10.06758145 | 10.5327403 | 10.46808205 | 9.021676912 | 9.08249654 | 9.549165021 | 9.701279971 |
| ENSG00000136404 | TM6SF1 | -2.961693707 | 0.292274211 | 0.887956545 | 0.753691694 | -2.073737162 | 2 | 4.11352228 | 2 | 2.141737246 | 2.2680111 | 4.261535101 | 2.70970617 |
| ENSG00000116133 | DHCR24 | -2.070992053 | 0.008138998 | -0.003053784 | 0.997910957 | -2.074045837 | 10.85509492 | 11.67869957 | 12.47629272 | 9.772927884 | 9.72990471 | 10.95954387 | 12.34756929 |
| ENSG00000140264 | SERF2 | -1.216026267 | 0.024616474 | -0.860464805 | 0.121912821 | -2.076491072 | 13.48990769 | 12.30162123 | 12.79448541 | 11.70155491 | 11.7573216 | 11.92764893 | 12.22668595 |
| C1ORF204 | C1ORF204 | 4.076422354 | 0.127676681 | -6.152937149 | 0.045223643 | -2.076514795 | 2 | 2 | 3.369317972 | 5.242616075 | 5.21224859 | 2 | 2 |
| ENSG00000127084 | FGD3 | -3.753032607 | NA | 1.675958486 | NA | -2.077074121 | 2 | 2 | 2 | 2 | 2 | 3.296906311 | 2.225841198 |
| ENSG00000148835 | TAF5 | -0.912185141 | 0.627312688 | -1.165594607 | 0.519747981 | -2.077779747 | 7.485143553 | 6.464272742 | 2 | 5.686138681 | 5.60003091 | 5.873348082 | 4.729989799 |
| ENSG00000074071 | MRPS34 | -1.516297959 | 0.011057942 | -0.561526914 | 0.39030005 | -2.077824873 | 10.52923584 | 9.114109157 | 9.746650996 | 8.369067528 | 8.44034872 | 9.278673449 | 9.421147165 |
| ENSG00000180964 | TCEAL8 | -1.541239587 | 0.068056382 | -0.536616216 | 0.561193746 | -2.077855804 | 10.91499272 | 9.090820894 | 9.132850258 | 8.545247918 | 8.35297033 | 9.352020423 | 9.53921146 |
| ENSG00000164054 | SHISA5 | -0.959436447 | 0.11734453 | -1.118874571 | 0.0593663 | -2.078311018 | 13.0669361 | 11.78952816 | 12.18004629 | 11.45755782 | 11.5200149 | 11.12136697 | 11.51211186 |
| ENSG00000177683 | THAP5 | -0.927878178 | 0.062310616 | -1.151133898 | 0.01654823 | -2.079012076 | 8.540903468 | 8.94259736 | 9.25752426 | 8.08961076 | 7.95749494 | 8.096541671 | 7.439859051 |
| ENSG00000250486 | FAM218A | -1.607501491 | 0.502887909 | -0.473536614 | 0.85221514 | -2.081038105 | 2 | 4.93638933 | 2 | 2.868578001 | 2.77779907 | 3.44788957 | 3.407202941 |
| ENSG00000255423 | EBLN2 | -0.942267394 | 0.544448 | -1.13905094 | 0.447209151 | -2.081318334 | 6.009025997 | 5.72254323 | 2 | 4.694064037 | 4.3456612 | 4.301748724 | 4.334543419 |
| ENSG00000205269 | TMEM170B | -2.576016677 | 0.18345602 | 0.493845173 | 0.82055968 | -2.082171504 | 5.095959458 | 2 | 7.502634626 | 4.329433326 | 3.45167086 | 6.503717356 | 6.791281771 |
| ENSG00000174915 | PTDSS2 | 0.36738411 | 0.750452376 | -2.449982092 | 0.01054493 | -2.082597982 | 6.468621203 | 9.181799937 | 7.646393253 | 8.565831351 | 8.5323134 | 5.747972572 | 5.892708443 |
| ENSG00000162869 | PPP1R21 | -0.782631205 | 0.343443781 | -1.300247477 | 0.091607485 | -2.082878682 | 7.904282243 | 7.35507212 | 9.132850258 | 7.641902724 | 7.45854177 | 6.900587441 | 7.18317631 |
| ENSG00000183283 | DAZAP2 | -0.989226305 | 0.008368835 | -1.094293246 | 0.002981509 | -2.083519551 | 12.13696669 | 11.5394185 | 12.0674595 | 10.93880439 | 10.9602832 | 10.65463759 | 11.0127263 |
| ENSG00000087111 | PIGS | -1.589815922 | 6.22E-05 | -0.494181501 | 0.267250193 | -2.083997423 | 11.03613077 | 10.63690565 | 11.47570169 | 9.546294576 | 9.46072474 | 10.53756755 | 10.65129185 |
| ENSG00000134717 | BTF3L4 | -1.173861854 | 0.036724367 | -0.910173715 | 0.109945258 | -2.084035569 | 9.533135543 | 8.655694736 | 9.897982195 | 8.31397419 | 8.25031674 | 8.658579002 | 8.417435289 |
| ENSG00000150967 | ABCB9 | -1.502092645 | 0.262309738 | -0.582234456 | 0.685116055 | -2.084327101 | 4.00722531 | 6.899080544 | 8.007629717 | 5.501219927 | 5.70538658 | 6.656902456 | 6.254725615 |
| ENSG00000109065 | NAT9 | -1.377833883 | 5.90E-06 | -0.709982111 | 0.027377975 | -2.087815993 | 9.219014269 | 9.214490408 | 9.287079172 | 7.831679003 | 7.92150349 | 8.722467118 | 8.321348409 |
| ENSG00000204920 | ZNF155 | -0.400395083 | 0.843607025 | -1.69070491 | 0.340409749 | -2.091099993 | 8.19931886 | 7.315436877 | 2 | 6.892328675 | 6.83591385 | 5.758846871 | 5.519049808 |
| ENSG00000066117 | SMARCD1 | -1.109896217 | 0.014298064 | -0.982413282 | 0.031125527 | -2.092309499 | 10.09970294 | 9.632483534 | 10.5187901 | 8.979812467 | 9.06280517 | 8.969089651 | 9.308063313 |
| ENSG00000091164 | TXNL1 | -0.298097527 | 0.345376097 | -1.794255988 | 7.57E-12 | -2.092353516 | 10.08373159 | 10.07961649 | 9.778213818 | 9.739170016 | 9.64416621 | 8.305655918 | 8.104880443 |
| ENSG00000121766 | ZCCHC17 | -0.762343363 | 0.524515475 | -1.330022907 | 0.230984425 | -2.092366271 | 7.624139768 | 9.159587375 | 10.78531698 | 9.003220305 | 8.9229643 | 8.308125745 | 8.489267468 |
| ENSG00000143801 | PSEN2 | -0.687317921 | 0.731439684 | -1.405395592 | 0.450551283 | -2.092713513 | 3.327711234 | 7.393647515 | 9.981704053 | 7.93509946 | 7.96694292 | 7.045287119 | 7.425709383 |
| ENSG00000262473 | | -0.933462667 | 0.052647422 | -1.162633663 | 0.012352962 | -2.09609633 | 10.32020829 | 9.640525262 | 9.358421373 | 8.926937992 | 8.87860041 | 8.659546439 | 8.692030411 |
| ENSG00000109534 | GAR1 | -0.600033088 | 0.484940021 | -1.496291752 | 0.051515981 | -2.09632484 | 7.53297911 | 6.04685525 | 7.934808822 | 6.693715713 | 6.85130017 | 5.9289675 | 5.917188835 |
| ENSG00000066044 | ELAVL1 | -1.369092423 | 0.137045092 | -0.728332207 | 0.449734118 | -2.09742463 | 11.56516075 | 10.78209868 | 9.06622548 | 9.372948073 | 9.48558803 | 9.859588707 | 10.2503029 |
| ENSG00000089053 | ANAPC5 | -1.207990897 | 0.000112284 | -0.88961613 | 0.005607507 | -2.097607027 | 10.83615308 | 10.69597594 | 11.24291424 | 9.734282329 | 9.74233706 | 10.0712216 | 10.03929326 |
| ENSG00000130958 | SLC35D2 | -0.172280399 | 0.924945325 | -1.927062991 | 0.205394441 | -2.09934339 | 3.327711234 | 7.099070191 | 8.561105505 | 7.226553841 | 7.33264456 | 5.707383854 | 5.508347454 |
| ENSG00000235961 | PNMA6A | -1.639652215 | 0.377308259 | -0.460154278 | 0.818792684 | -2.099806493 | 4.00722531 | 6.389896633 | 2 | 3.997744386 | 3.60476711 | 4.920961476 | 4.543391831 |
| ENSG00000112855 | HARS2 | -0.548037367 | 0.759077898 | -1.552375832 | 0.336472054 | -2.100413199 | 10.22148166 | 8.213429689 | 4.523675391 | 8.469961514 | 8.40081078 | 7.551912644 | 7.336254871 |
| ENSG00000102760 | RGCC | -1.434307064 | 0.092448935 | -0.667325699 | 0.457819373 | -2.101632764 | 5.70932728 | 7.701025996 | 7.552162999 | 5.978405189 | 5.72778967 | 6.768721236 | 6.371531035 |
| STUB1 | | -0.76082784 | 0.292677052 | -1.341444289 | 0.044959455 | -2.102272129 | 9.344956677 | 9.267380431 | 10.66665689 | 9.222510184 | 9.08249654 | 8.582068017 | 8.575104441 |
| ENSG00000181264 | TMEM136 | -1.056033762 | 0.23823472 | -1.04675281 | 0.234600074 | -2.102786571 | 9.112843761 | 8.029695761 | 7.029880454 | 7.19843704 | 7.32531744 | 7.580798201 | 6.879055607 |
| ENSG00000117054 | ACADM | -0.646657581 | 0.085171677 | -1.456389229 | 1.70E-05 | -2.103046809 | 8.797441463 | 8.916342327 | 8.380957451 | 8.047448792 | 8.11663771 | 7.182183216 | 7.388257299 |
| ENSG00000110171 | TRIM3 | -1.82921755 | 5.58E-06 | -0.274608862 | 0.566108671 | -2.103826412 | 8.892187472 | 8.213429689 | 8.922968322 | 6.989292186 | 6.82556437 | 8.509342792 | 8.350852333 |
| ENSG00000242173 | ARHGDIG | -3.981249662 | 0.222165985 | 1.876685604 | 0.508679168 | -2.104564058 | 2 | 3.686303414 | 2 | 2.141737246 | 2 | 4.350488971 | 3.518055793 |
| ENSG00000242689 | CNTF | 0.083289726 | 0.976290713 | -2.188022253 | 0.364514502 | -2.104732527 | 5.70932728 | 2 | 2 | 4.623314311 | 4.0930011 | 2.987421008 | 2.817676617 |
| ENSG00000205084 | TMEM231 | -0.809742842 | 0.353886558 | -1.295976176 | 0.112305841 | -2.105719018 | 8.540903468 | 6.899080544 | 6.882775709 | 6.83085985 | 6.91614081 | 6.334738131 | 6.476841503 |
| ENSG00000182931 | WFDC10B | 0.975198196 | 0.745371929 | -3.081618455 | NA | -2.10642026 | 2 | 3.076665209 | 2 | 2.947924294 | 3.01487888 | 2.048074405 | 2.117335275 |
| ENSG00000215021 | PHB2 | -1.287840505 | 0.00289076 | -0.818717585 | 0.069605713 | -2.10655809 | 11.21173326 | 10.70367235 | 10.30429804 | 9.504256136 | 9.50517467 | 9.879031168 | 10.05867839 |
| ENSG00000123810 | B9D2 | -0.253584399 | 0.833633118 | -1.856740859 | 0.061919051 | -2.110325258 | 4.467311984 | 5.838937808 | 6.882775709 | 5.870535235 | 5.71663161 | 4.053534413 | 4.664017718 |
| ENSG00000127314 | RAP1B | 0.466421923 | 0.210210321 | -2.578299024 | 2.69E-16 | -2.111877102 | 10.76282024 | 10.32389548 | 10.26027233 | 10.95839757 | 10.9039101 | 8.040458475 | 7.761616892 |
| ENSG00000104880 | ARHGEF18 | 0.463870764 | 0.790720929 | -2.576151125 | 0.080743277 | -2.112280361 | 9.521325812 | 8.053983481 | 4.058998049 | 8.873322599 | 8.85589616 | 6.004008792 | 5.812327524 |
| ENSG00000138801 | PAPSS1 | -1.387634426 | 0.028439901 | -0.72504218 | 0.278961643 | -2.112676607 | 9.448362284 | 10.66477302 | 10.89014791 | 9.044420506 | 9.10940731 | 9.504636417 | 9.935263322 |
| ENSG00000247315 | ZCCHC3 | -1.376204999 | 0.05194085 | -0.737066541 | 0.321260238 | -2.11327154 | 10.1076228 | 8.702538795 | 8.677268317 | 7.930222906 | 7.99953197 | 8.715506558 | 8.465949064 |
| ENSG00000101134 | DOK5 | -1.754704169 | 0.370378063 | -0.359515339 | 0.868498866 | -2.114219509 | 2 | 8.004992147 | 5.772716771 | 5.003571641 | 5.18011485 | 5.692336538 | 6.845434848 |
| ENSG00000205220 | PSMB10 | 1.054921317 | 0.208618256 | -3.171075496 | 1.82E-05 | -2.116154179 | 8.981092835 | 8.077869079 | 6.958202216 | 9.20242622 | 9.34139676 | 4.927401733 | 5.436777225 |
| ENSG00000149600 | COMMD7 | -1.718294834 | 4.31E-06 | -0.397969727 | 0.35517853 | -2.116264561 | 10.12333429 | 9.835272298 | 10.55568472 | 8.461536372 | 8.5227358 | 9.673139654 | 9.922471842 |
| ENSG00000135698 | MPHOSPH6 | -0.18216611 | 0.834344332 | -1.934319378 | 0.005059275 | -2.116485488 | 8.313285031 | 8.169643011 | 6.629488387 | 7.745471964 | 7.65837038 | 5.846413283 | 6.165581817 |
| ENSG00000162576 | MXRA8 | -1.176189101 | 0.063034875 | -0.941350049 | 0.140177518 | -2.11753915 | 12.38520102 | 11.02260251 | 11.83960343 | 10.64271979 | 10.7121781 | 10.55326835 | 11.19937456 |
| ENSG00000158062 | UBXN11 | -1.349043937 | 0.351836711 | -0.769002611 | 0.607673137 | -2.118046548 | 3.327711234 | 7.35507212 | 7.691296561 | 5.786686927 | 5.62411716 | 6.129409568 | 6.362645675 |
| ENSG00000142661 | MYOM3 | 0.177060991 | 0.925027909 | -2.295675093 | 0.140636706 | -2.118614102 | 3.327711234 | 3.076665209 | 5.156626401 | 4.237680128 | 4.31639476 | 3.059555873 | 2.374681325 |
| ENSG00000110011 | DNAJC4 | -1.322214058 | 0.029718519 | -0.796436742 | 0.207938624 | -2.118650799 | 9.410448395 | 8.834582582 | 7.97167868 | 7.539954763 | 7.56191311 | 8.156706039 | 7.971952287 |
| ENSG00000154832 | CXXC1 | -0.913939358 | 0.508212783 | -1.205838075 | 0.360288866 | -2.119777433 | 10.09970294 | 7.761270542 | 6.43197674 | 7.90310317 | 8.02009991 | 7.710619897 | 7.640647 |
| ENSG00000040633 | PHF23 | -1.10030544 | 0.082476467 | -1.022478313 | 0.103484196 | -2.122783753 | 10.9550501 | 9.558016677 | 10.04361983 | 9.170732795 | 9.24469897 | 9.153099991 | 9.406811858 |
| ENSG00000254858 | MPV17L2 | -1.525100242 | 0.496398079 | -0.599510505 | 0.800446891 | -2.124610747 | 9.895281535 | 6.666836837 | 2 | 6.861921612 | 7.06433609 | 7.840848421 | 7.8990585 |
| ENSG00000168884 | TNIP2 | -1.21430188 | 0.210376805 | -0.912053614 | 0.352068654 | -2.126355493 | 8.19931886 | 9.090820894 | 10.64399316 | 8.454760653 | 8.46722793 | 8.810024926 | 8.708487604 |
| ENSG00000034677 | RNF19A | -0.967481232 | 0.592445571 | -1.158976143 | 0.507424606 | -2.126457374 | 5.095959458 | 8.916342327 | 11.27276688 | 9.005540335 | 8.99284518 | 8.977648172 | 8.618670729 |
| ENSG00000197157 | SND1 | -1.233975571 | 0.003720538 | -0.892787473 | 0.041810498 | -2.126763044 | 11.81293891 | 12.0227539 | 12.60360903 | 10.88941003 | 11.01539 | 11.11468069 | 11.45412679 |
| ENSG00000148297 | MED22 | -1.249475396 | 0.167187365 | -0.877479567 | 0.340137275 | -2.126954964 | 10.49341471 | 8.607278467 | 8.608698767 | 8.260583448 | 8.32385328 | 8.500191379 | 8.804567006 |
| ENSG00000164484 | TMEM200A | -0.241635679 | 0.841373243 | -1.885785413 | 0.056612408 | -2.127421092 | 8.042902253 | 10.73780697 | 10.48092713 | 9.948991943 | 9.86300004 | 7.734688324 | 8.667588528 |
| ENSG00000161526 | SAP30BP | -0.677381007 | 0.194794952 | -1.452407411 | 0.002155456 | -2.129788418 | 10.36053552 | 9.599860728 | 9.465702942 | 9.129845271 | 9.24857772 | 8.234076861 | 8.589138522 |
| ENSG00000128408 | RIBC2 | -0.95464234 | NA | -1.175904597 | NA | -2.130546937 | 2 | 3.076665209 | 2 | 2.141737246 | 2.49394783 | 2.139668783 | 2.421067459 |
| ENSG00000171443 | ZNF524 | -1.467514393 | 0.494956066 | -0.66327148 | 0.768414094 | -2.130785873 | 9.233563228 | 6.311476754 | 2 | 6.456560224 | 6.36443784 | 7.399274054 | 6.936070085 |
| ENSG00000099821 | POLRMT | -1.749952438 | 0.000101449 | -0.381087311 | 0.46434877 | -2.13103975 | 9.751047882 | 8.834582582 | 9.358421373 | 7.632926745 | 7.62593544 | 9.154129994 | 8.782528961 |
| ENSG00000096384 | HSP90AB1 | -1.222249147 | 0.015366572 | -0.909472281 | 0.078322326 | -2.131721428 | 14.27259948 | 13.49148113 | 13.23355721 | 12.49274548 | 12.5359925 | 12.69454834 | 12.94852295 |
| ENSG00000162244 | RPL29 | -1.350051289 | 0.003642609 | -0.783522785 | 0.109386471 | -2.133574074 | 13.06388636 | 12.15494759 | 12.24549183 | 11.14279746 | 11.2554874 | 11.78466647 | 11.74703401 |
| ENSG00000062485 | CS | -1.569986492 | 1.29E-06 | -0.563878443 | 0.114873438 | -2.133864935 | 11.41687936 | 10.91068521 | 11.12511995 | 9.613999459 | 9.58557298 | 10.41187535 | 10.77042905 |
| ENSG00000103507 | BCKDK | -1.484445057 | 0.016810729 | -0.650323288 | 0.331375959 | -2.134768345 | 10.92846914 | 10.09725346 | 9.541233192 | 8.777474042 | 8.87358581 | 9.375172844 | 9.889085869 |
| ENSG00000213366 | GSTM2 | -0.111145106 | 0.907490514 | -2.023794363 | 0.005993894 | -2.134939468 | 6.73718411 | 8.213429689 | 8.677268317 | 7.964017617 | 7.9809999 | 5.944926241 | 6.280114403 |
| ENSG00000049089 | COL9A2 | -1.897603495 | 0.314701066 | -0.237350424 | 0.911000084 | -2.134953919 | 2 | 5.457120953 | 3.369317972 | 2.868578001 | 3.2184422 | 4.45225155 | 3.735548302 |
| ENSG00000119231 | SENP5 | -1.098278025 | 0.099672705 | -1.037220261 | 0.115817438 | -2.135498287 | 9.992581382 | 10.19331737 | 11.37608156 | 9.587142668 | 9.53803206 | 9.743945072 | 9.492120456 |
| ENSG00000112306 | RPS12 | -0.604811495 | 0.391805366 | -1.532989412 | 0.015161888 | -2.137800906 | 11.88152073 | 10.60848937 | 10.55568472 | 10.54322901 | 10.558176 | 9.584995729 | 9.665902758 |
| ENSG00000196150 | ZNF250 | -1.519515677 | 0.065184591 | -0.621080581 | 0.482146792 | -2.140596258 | 8.468948322 | 7.538394143 | 6.43197674 | 6.260701031 | 6.21386141 | 7.314596615 | 6.847559257 |
| ENSG00000164414 | SLC35A1 | -0.840710877 | 0.689206562 | -1.300547699 | 0.514552506 | -2.141258576 | 9.081016759 | 8.077869079 | 2 | 7.222570549 | 7.30311013 | 7.269684675 | 6.155328188 |
| ENSG00000065029 | ZNF76 | -1.300075503 | 0.479762551 | -0.841719437 | 0.654382498 | -2.14179494 | 9.839306291 | 7.73146271 | 3.369317972 | 7.226553841 | 7.35080119 | 7.597048228 | 7.866953664 |
| ENSG00000105254 | TBCB | -1.371426893 | 0.001908478 | -0.770442772 | 0.098635116 | -2.141869665 | 10.47516499 | 9.916994052 | 10.48092713 | 8.878394985 | 9.01464729 | 9.169152824 | 9.843245622 |
| ENSG00000171307 | ZDHHC16 | -1.241774201 | 0.025049412 | -0.900945411 | 0.112157708 | -2.142719611 | 9.031919741 | 9.770792087 | 10.30429804 | 8.596935617 | 8.52113334 | 8.808281231 | 8.980880783 |
| ENSG00000166313 | APBB1 | -1.353908123 | 0.119684351 | -0.790396134 | 0.380413475 | -2.144304256 | 9.602033055 | 9.192779271 | 11.18521898 | 8.866956935 | 8.9648262 | 9.215609012 | 9.697162691 |
| ENSG00000151640 | DPYSL4 | -1.24937098 | 0.3854131 | -0.89508524 | 0.538134884 | -2.14445622 | 9.740899611 | 8.590771778 | 5.156626401 | 7.494428494 | 7.51115308 | 7.585896024 | 8.07345976 |
| ENSG00000035141 | FAM136A | -1.302371541 | 0.318470503 | -0.842336971 | 0.528809949 | -2.144708512 | 10.52332707 | 9.424669273 | 6.43197674 | 8.323303759 | 8.18704757 | 8.685424964 | 8.737416477 |
| ENSG00000117543 | DPH5 | -1.556251877 | 0.003880482 | -0.588969591 | 0.313919617 | -2.145221468 | 7.867441304 | 7.701025996 | 6.958202216 | 5.949776857 | 6.17362678 | 7.228554289 | 6.704867638 |
| ENSG00000007376 | RPUSD1 | -1.274345095 | 0.103167922 | -0.870968929 | 0.274579713 | -2.145314024 | 9.48530526 | 8.806266215 | 7.502634626 | 7.474468681 | 7.62891439 | 8.008923105 | 7.887748062 |
| ENSG00000166343 | MSS51 | -1.183391095 | 0.215393228 | -0.962760677 | 0.305318653 | -2.146151773 | 4.467311984 | 5.838937808 | 4.874587708 | 4.268917409 | 4.16043394 | 4.588526421 | 4.161535617 |
| ENSG00000149150 | SLC43A1 | -0.858469304 | 0.64998964 | -1.289995795 | 0.473584699 | -2.148465099 | 6.468621203 | 7.790474954 | 2 | 5.949776857 | 5.85535975 | 5.744329515 | 5.217771644 |
| ENSG00000115758 | ODC1 | -0.1931383 | 0.754387465 | -1.955509775 | 6.32E-05 | -2.148648075 | 11.13631304 | 10.45735223 | 9.981704053 | 10.40463465 | 10.4171049 | 8.641053292 | 8.675981178 |
| ENSG00000115866 | DARS | -0.575199767 | 0.146453455 | -1.573767851 | 7.33E-06 | -2.148967618 | 10.29268328 | 10.15453315 | 10.79563557 | 9.896204982 | 9.83554116 | 8.958516141 | 8.783638895 |
| ENSG00000099250 | NRP1 | 0.958570297 | 0.326219261 | -3.107835333 | 0.000254856 | -2.149265036 | 9.949165863 | 11.13502303 | 12.32065573 | 12.39824131 | 12.4025576 | 8.72385521 | 7.845855157 |
| ENSG00000147536 | GINS4 | -0.626037056 | 0.667279239 | -1.524142648 | 0.247374227 | -2.150179704 | 4.467311984 | 6.04685525 | 3.369317972 | 4.299492648 | 4.74762503 | 3.500668998 | 4.119958326 |
| ENSG00000167461 | RAB8A | 0.088773946 | 0.88983543 | -2.240286826 | 2.25E-06 | -2.151512881 | 9.189468652 | 9.981728319 | 10.18377222 | 9.928352931 | 9.93817677 | 7.371203179 | 7.845855157 |
| ENSG00000165392 | WRN | -0.774610683 | 0.262822363 | -1.378502184 | 0.031613028 | -2.153112867 | 8.855032352 | 7.35507212 | 8.206335842 | 7.471115032 | 7.52080763 | 7.037865927 | 6.76679494 |
| ENSG00000159111 | MRPL10 | -1.186228999 | 0.069674535 | -0.966967751 | 0.140757366 | -2.153196751 | 9.751047882 | 8.862353853 | 8.236946975 | 7.954442493 | 7.86703672 | 8.131128273 | 8.123409541 |

| | | | | | | | | | | | | | |
|---|---|---|---|---|---|---|---|---|---|---|---|---|---|
| ENSG00000134480 | CCNH | -0.718925031 | 0.235186197 | -1.435721066 | 0.009577248 | -2.154646097 | 7.579279349 | 8.557180776 | 8.865353883 | 7.66556918 | 7.77591026 | 7.0047514 | 7.040043908 |
| ENSG00000142657 | PGD | -0.571217333 | 0.282839894 | -1.583470171 | 0.000737309 | -2.154687504 | 11.94035323 | 11.46212086 | 11.83710798 | 11.15721618 | 11.2218486 | 9.63035768 | 10.57737611 |
| ENSG00000282291 | | -1.044444823 | 0.227473008 | -1.110945777 | 0.187955191 | -2.1553906 | 8.564112515 | 6.78762695 | 8.904018013 | 7.377487766 | 7.22257674 | 7.240241221 | 7.234405279 |
| ENSG00000113119 | TMCO6 | -1.573620583 | 0.469993356 | -0.583918864 | 0.799962733 | -2.157539447 | 3.327711234 | 7.099070191 | 2 | 4.237680128 | 4.37434574 | 5.258920789 | 5.020995508 |
| ENSG00000170801 | HTRA3 | -1.243059234 | 0.534490579 | -0.914742312 | 0.65200599 | -2.157801546 | 2 | 8.540087354 | 6.629488387 | 6.078767801 | 6.1572134 | 6.710164669 | 6.081436739 |
| ENSG00000105426 | PTPRS | -1.232478259 | 0.021674606 | -0.926066976 | 0.091407 | -2.158545235 | 11.23361136 | 10.05576005 | 11.06240811 | 9.621581886 | 9.65662671 | 9.993749244 | 9.892948091 |
| ENSG00000163521 | GLB1L | -1.654111368 | 0.11657228 | -0.50465664 | 0.664628785 | -2.158768009 | 6.257048221 | 7.701025996 | 8.742726175 | 6.351046291 | 6.22177541 | 6.544089217 | 7.928458732 |
| ENSG00000168209 | DDIT4 | -0.079418835 | 0.946595613 | -2.079863715 | 0.030994241 | -2.159282551 | 9.975371636 | 10.18234215 | 8.677268317 | 9.662587523 | 9.67547818 | 8.413960373 | 6.165581817 |
| ENSG00000116251 | RPL22 | -0.566414944 | 0.43599311 | -1.593151683 | 0.013079078 | -2.159566628 | 11.37155492 | 10.42046463 | 9.798878489 | 10.1342586 | 10.0891109 | 9.029831415 | 9.148782641 |
| ENSG00000176473 | WDR25 | -1.056627003 | 0.237601833 | -1.104091482 | 0.206132822 | -2.160718485 | 8.517314938 | 7.393647515 | 6.322058544 | 6.682159616 | 6.61935067 | 6.645228724 | 6.564691065 |
| ENSG00000121064 | SCPEP1 | -0.088973988 | 0.866613986 | -2.072256591 | 1.22E-07 | -2.161230579 | 10.66423795 | 10.08551947 | 10.55568472 | 10.35409783 | 10.3773587 | 8.072772466 | 8.659146771 |
| ENSG00000175764 | TTLL11 | -0.907893093 | 0.495809834 | -1.254964516 | 0.320805825 | -2.162857609 | 4.00722531 | 7.35507212 | 7.646393253 | 6.130704144 | 6.1068222 | 5.610655473 | 5.957088417 |
| ENSG00000063176 | SPHK2 | -1.447991278 | 0.000620294 | -0.714922697 | 0.112037873 | -2.162913975 | 8.653385977 | 8.356994601 | 8.941672936 | 7.127781374 | 7.34718809 | 7.689935911 | 8.186868687 |
| ENSG00000147862 | NFIB | -4.361180572 | 0.003309136 | 2.19708251 | 0.15962782 | -2.164098062 | 6.138360429 | 7.099070191 | 8.511888463 | 4.443344104 | 2.14022021 | 9.475816969 | 9.976372702 |
| ENSG00000163512 | AZI2 | 0.054463863 | 0.954731956 | -2.218968904 | 0.003108279 | -2.164505042 | 10.83137851 | 9.04308532 | 9.301632639 | 10.02414665 | 9.9946957 | 7.633939843 | 7.871136591 |
| ENSG00000101751 | POLI | -1.218655126 | 0.171886783 | -0.946001939 | 0.291093097 | -2.164657065 | 7.709880939 | 7.928237109 | 9.589475645 | 7.477814553 | 7.45854177 | 7.989819469 | 7.425709383 |
| ENSG00000215271 | HOMEZ | -1.327430678 | 0.13908327 | -0.837653451 | 0.363335037 | -2.165084129 | 6.468621203 | 8.191702463 | 8.825624954 | 6.841288255 | 6.80463998 | 7.413107295 | 7.173047408 |
| ENSG00000172425 | TTC36 | -1.412364034 | 0.617411375 | -0.752734138 | NA | -2.165098172 | 2 | 3.686303414 | 2 | 2.498691189 | 2.2680111 | 2.592885947 | 2.551896972 |
| ENSG00000185513 | L3MBTL1 | -1.738756165 | 0.178468688 | -0.427415494 | 0.765367121 | -2.166171659 | 6.257048221 | 4.11352228 | 7.502634626 | 4.443344104 | 5.2591458 | 6.265258498 | 5.913137508 |
| ENSG00000166575 | TMEM135 | -0.532039691 | 0.283678293 | -1.635024255 | 0.000157738 | -2.167063946 | 8.257427002 | 8.929529568 | 8.266922072 | 8.051945366 | 7.95274764 | 6.730562686 | 7.09829807 |
| ENSG00000064309 | CDON | -1.555266634 | 0.261480902 | -0.613220349 | 0.677268519 | -2.168486983 | 3.327711234 | 5.457120953 | 6.322058544 | 4.647285135 | 3.39680241 | 5.026717879 | 4.827924991 |
| ENSG00000122882 | ECD | -1.018024708 | 0.148427006 | -1.15249115 | 0.091654931 | -2.170515859 | 9.247966931 | 8.255926468 | 9.849281147 | 8.16211395 | 8.32385328 | 7.877064822 | 8.31522102 |
| ENSG00000142694 | EVA1B | -0.559275813 | 0.387605511 | -1.611769558 | 0.004649784 | -2.171045371 | 10.45668146 | 9.192779271 | 10.41552659 | 9.556615538 | 9.58019637 | 8.417967842 | 8.623017875 |
| ENSG00000141933 | TPGS1 | -1.054944753 | 0.309968682 | -1.116114634 | 0.269817196 | -2.171059387 | 6.138360429 | 7.503546079 | 8.805342542 | 6.766655021 | 6.89649843 | 6.875826972 | 6.664466987 |
| ENSG00000162191 | UBXN1 | -1.057926716 | 0.118824374 | -1.113533801 | 0.093533047 | -2.171460516 | 10.49341471 | 9.405920729 | 9.014156722 | 8.738951712 | 8.71722214 | 8.568225482 | 8.770822803 |
| ENSG00000145916 | RMND5B | -2.11045593 | 0.212730396 | -0.061225755 | 0.974934399 | -2.171681685 | 9.790941832 | 8.57407404 | 3.369317972 | 6.554880361 | 6.77267312 | 8.771615683 | 8.568035872 |
| ENSG00000140030 | GPR65 | -1.293666759 | 0.379713551 | -0.878455955 | 0.547274868 | -2.172122714 | 4.00722531 | 3.686303414 | 4.523675391 | 3.162721474 | 3.28037825 | 3.904814898 | 2.851942824 |
| ENSG00000181991 | MRPS11 | -0.680881651 | 0.487705929 | -1.49432213 | 0.093525732 | -2.175203781 | 9.497412313 | 8.777382925 | 7.098165388 | 8.033874364 | 8.13561258 | 7.001702955 | 7.52727182 |
| ENSG00000164118 | CEP44 | -1.345880238 | 0.200330183 | -0.829846219 | 0.442768139 | -2.175726457 | 7.27616306 | 8.806266215 | 6.322058544 | 6.554880361 | 6.507263 | 7.406207255 | 6.514766253 |
| ENSG00000204524 | ZNF805 | -0.664555185 | 0.319838395 | -1.513398195 | 0.012183874 | -2.17795338 | 8.340423424 | 7.874700259 | 7.029880454 | 7.173892946 | 7.21862736 | 6.609629839 | 6.102937566 |
| ENSG00000196411 | EPHB4 | -1.5359722 | 0.007257564 | -0.642205806 | 0.299826316 | -2.178178006 | 9.761125267 | 8.981105197 | 10.34701993 | 8.300810597 | 8.24644267 | 9.224134997 | 9.090778885 |
| ENSG00000124467 | PSG8 | 3.010841338 | 0.603581513 | -5.191094232 | 0.364514502 | -2.180252894 | 2 | 2 | 2 | 3.566290607 | 3.65239003 | 2 | 2 |
| ENSG00000103051 | COG4 | -1.312708597 | 0.133069745 | -0.868260095 | 0.332318898 | -2.180968692 | 10.82658808 | 9.599860728 | 8.585098385 | 8.649804831 | 8.64989445 | 8.967137431 | 9.204129769 |
| ENSG00000103145 | HCFC1R1 | -1.994206651 | 8.69E-06 | -0.188420544 | 0.733788685 | -2.182627196 | 9.304185368 | 8.356994601 | 8.65477198 | 6.841288255 | 6.89154576 | 8.645942939 | 8.62735196 |
| ENSG00000178057 | NDUFAF3 | -1.131240099 | 0.005333784 | -1.052346434 | 0.009656574 | -2.183586533 | 10.38028415 | 9.842262059 | 9.809100877 | 8.890998469 | 8.92781046 | 8.73399394 | 9.203715214 |
| ENSG00000166562 | SEC11C | 0.378295764 | 0.642216602 | -2.564401426 | 0.000116198 | -2.186105662 | 9.143983763 | 7.819099893 | 8.206335842 | 8.888486565 | 8.85589616 | 5.467744375 | 6.403650353 |
| ENSG00000206562 | METTL6 | -0.976484232 | 0.310604101 | -1.210496329 | 0.189811454 | -2.186980561 | 7.096806373 | 7.928237109 | 9.372276026 | 7.501020868 | 7.45518898 | 7.452712791 | 7.013798886 |
| ENSG00000137720 | C11orf1 | -0.740990951 | 0.467848857 | -1.447401486 | 0.120494972 | -2.188392437 | 6.009025997 | 5.303523747 | 7.398165132 | 5.818703821 | 5.76075522 | 4.946551782 | 5.32534152 |
| ENSG00000122965 | RBM19 | -0.570828713 | 0.733731931 | -1.618292414 | 0.283648588 | -2.189121127 | 10.29268328 | 9.308339287 | 4.874587708 | 8.677714042 | 8.82635531 | 7.634923938 | 7.80161394 |
| ENSG00000274229 | | 0.214527813 | 0.881720015 | -2.403755015 | 0.036373066 | -2.189227202 | 5.532353208 | 5.838937808 | 8.35328246 | 7.377487766 | 7.35440525 | 5.008609438 | 4.819291477 |
| ENSG00000160194 | NDUFV3 | -1.999244895 | 0.002312454 | -0.190030931 | 0.814643023 | -2.189275827 | 10.13887448 | 8.451443573 | 9.330306409 | 7.487805859 | 7.48837134 | 9.258701797 | 9.289430628 |
| ENSG00000155275 | TRMT44 | -1.943957136 | 0.248222521 | -0.246613369 | 0.899287217 | -2.190570505 | 4.00722531 | 6.140560021 | 9.227351203 | 5.764940652 | 6.1322378 | 7.66606569 | 7.513957873 |
| ENSG00000141179 | PCTP | -0.939198182 | 0.259291944 | -1.251524989 | 0.113966986 | -2.190723172 | 6.366709671 | 6.951736442 | 8.143093325 | 6.429169081 | 6.44101097 | 6.327451929 | 5.921228818 |
| ENSG00000189056 | RELN | 1.263108008 | 0.42279064 | -3.454829089 | 0.01363715 | -2.191721081 | 9.143983763 | 5.303523747 | 5.930865648 | 8.984524517 | 9.09332116 | 5.014670875 | 3.940519005 |
| ENSG00000129055 | ANAPC13 | -1.203518796 | 0.036049949 | -0.988459454 | 0.088195247 | -2.191978251 | 11.18578229 | 10.01303782 | 10.19160738 | 9.404936476 | 9.3166689 | 9.480751596 | 9.664095852 |
| ENSG00000100429 | HDAC10 | -0.161779528 | 0.88968542 | -2.031065003 | 0.028205751 | -2.19284453 | 7.867441304 | 5.946638857 | 8.511888463 | 7.494428494 | 7.72670378 | 5.634468599 | 5.98823215 |
| ENSG00000006282 | SPATA20 | -1.329637576 | 0.053270494 | -0.864457185 | 0.222891601 | -2.194094761 | 9.790941832 | 9.828248506 | 11.15347798 | 9.069031178 | 9.09547637 | 9.304879394 | 9.750075969 |
| ENSG00000182103 | FAM181B | -3.756860774 | 0.095646206 | 1.559671051 | 0.467249874 | -2.197189723 | 2 | 4.442710129 | 3.369317972 | 2.3892346 | 2 | 4.996409624 | 4.664017718 |
| ENSG00000124659 | TBCC | -0.903545994 | 0.343732241 | -1.29566471 | 0.151114596 | -2.199210703 | 8.228665467 | 8.297207175 | 10.18377222 | 8.270263574 | 8.35477078 | 7.733770002 | 8.096865967 |
| ENSG00000122386 | ZNF205 | -1.579350541 | 0.470709201 | -0.620189124 | 0.78961933 | -2.199539665 | 9.304185368 | 6.04685525 | 2 | 6.172600389 | 6.48770488 | 7.446000032 | 7.052989523 |
| ENSG00000110799 | VWF | -1.337647337 | 0.496719668 | -0.864052557 | 0.663337914 | -2.201699894 | 4.815609471 | 2 | 4.058998049 | 2.600425781 | 3.50452879 | 3.215043717 | 3.560117069 |
| ENSG00000232258 | TMEM114 | -3.753034173 | NA | 1.550617918 | NA | -2.202416255 | 2 | 2 | 2 | 2 | 2 | 3.316666208 | 2 |
| ENSG00000164615 | CAMLG | -1.394804046 | 0.000720759 | -0.808152988 | 0.062311706 | -2.202957033 | 9.839306291 | 9.214490408 | 9.385998894 | 8.143273718 | 8.09740995 | 8.930336153 | 8.425990514 |
| ENSG00000163961 | RNF168 | -1.413858365 | 0.252244014 | -0.789123127 | 0.538029429 | -2.202981491 | 7.82963488 | 7.928237109 | 10.71094342 | 7.992367495 | 8.16464176 | 8.636147016 | 8.757891838 |
| ENSG00000128254 | C22orf24 | -2.272289522 | NA | 0.068762586 | NA | -2.203526936 | 2 | 3.076665209 | 2 | 2.141737246 | 2.14022021 | 2.3070652 | 2.851942824 |
| C4ORF32 | C4ORF32 | 3.946973061 | 0.156616494 | -6.152935894 | 0.050696253 | -2.205962832 | 2 | 3.076665209 | 2 | 5.144975053 | 5.07918227 | 2 | 2 |
| ENSG00000112237 | CCNC | -0.986314438 | 0.013722176 | -1.220837991 | 0.001608419 | -2.20715243 | 9.613202746 | 9.114109157 | 8.884815469 | 8.22518713 | 8.29225867 | 8.102245405 | 7.949380652 |
| ENSG00000145494 | NDUFS6 | -1.03437552 | 0.115157031 | -1.172807772 | 0.065682577 | -2.207183292 | 9.820154628 | 8.671479029 | 8.608698767 | 8.136938566 | 8.10812366 | 7.728247761 | 8.205208936 |
| ENSG00000125648 | SLC25A23 | -2.797117806 | 0.000187015 | 0.589446427 | 0.498291593 | -2.207671379 | 9.159305284 | 9.488434476 | 10.94679967 | 7.307758725 | 7.33264456 | 10.32871658 | 10.95206949 |
| ENSG00000185085 | INTS5 | -1.568545663 | 0.465126545 | -0.639656873 | 0.778107221 | -2.208202537 | 2 | 8.820493869 | 10.26027233 | 7.647855834 | 7.51437845 | 8.483352582 | 8.510875165 |
| ENSG00000188807 | TMEM201 | -1.547629512 | 0.34368375 | -0.660791586 | 0.702769774 | -2.208421098 | 4.00722531 | 7.761270542 | 9.577565424 | 6.658765962 | 7.03768431 | 7.759264603 | 7.688837505 |
| ENSG00000172270 | BSG | -1.379684693 | 0.000410517 | -0.829604782 | 0.043393159 | -2.209289475 | 14.26110617 | 13.45717518 | 13.73632778 | 12.47506605 | 12.4814142 | 13.03518302 | 13.02078497 |
| ENSG00000129255 | MPDU1 | -0.948520699 | 0.446716354 | -1.2626001 | 0.286667458 | -2.211120799 | 10.905938 | 8.848535039 | 7.777116083 | 8.778830996 | 8.85207713 | 8.068408401 | 8.839676004 |
| ENSG00000205464 | ATP6AP1L | -1.38509727 | 0.281735266 | -0.826566376 | 0.527954208 | -2.211663647 | 4.00722531 | 6.04685525 | 5.156626401 | 4.034562184 | 4.2560182 | 5.141958644 | 3.717076715 |
| ENSG00000167515 | TRAPPC2L | -1.027432373 | 0.001958103 | -1.184721503 | 0.000243834 | -2.212153877 | 8.981092835 | 8.655694736 | 8.631919287 | 7.703217906 | 7.79460185 | 7.722704302 | 7.455265057 |
| ENSG00000168175 | MAPK1IP1L | -1.420197724 | 0.002089486 | -0.792517791 | 0.104511913 | -2.212715515 | 11.4974524 | 10.76380876 | 10.68342405 | 9.618553696 | 9.60989979 | 9.997388539 | 10.44649123 |
| ENSG00000198270 | TMEM116 | -0.819256458 | 0.551171197 | -1.393506958 | 0.278176873 | -2.212763416 | 6.009025997 | 3.686303414 | 7.029880454 | 5.319220934 | 5.36298494 | 5.119635168 | 4.479073019 |
| ENSG00000024526 | DEPDC1 | -0.917973963 | 0.646818108 | -1.296329576 | 0.49588682 | -2.214303538 | 3.327711234 | 5.595927442 | 2 | 3.840289928 | 3.60476711 | 3.12825509 | 3.753786374 |
| ENSG00000127838 | PNKD | -1.362401838 | 0.000182564 | -0.852269228 | 0.024727678 | -2.214671066 | 9.384607364 | 9.318399851 | 8.941672936 | 7.807949361 | 7.95274764 | 8.151901655 | 8.585960925 |
| ENSG00000177917 | ARL6IP6 | -1.023225645 | 0.162619057 | -1.191610779 | 0.092106832 | -2.214836424 | 8.928409675 | 7.431218269 | 8.721235202 | 7.555872146 | 7.40393901 | 7.606109069 | 6.955898408 |
| ENSG00000126088 | UROD | -1.017517835 | 0.311127936 | -1.19774905 | 0.217196921 | -2.215266885 | 11.25159303 | 9.386925327 | 8.922968322 | 9.206465443 | 9.21922971 | 8.881419814 | 9.171855284 |
| ENSG00000275824 | | 2.975761116 | 0.60852038 | -5.191094378 | 0.364514502 | -2.215333262 | 2 | 2 | 2 | 3.709785322 | 3.45167086 | 2 | 2 |
| ENSG00000175029 | CTBP2 | -1.309103948 | 0.053925622 | -0.9063733 | 0.192038633 | -2.215477247 | 10.81695923 | 9.936721131 | 9.180874355 | 8.767939462 | 8.88857762 | 9.315659588 | 9.137106455 |
| ENSG00000007541 | PIGQ | -1.362080496 | 0.250618358 | -0.854056733 | 0.48296181 | -2.216137229 | 7.790810999 | 8.487557258 | 10.80588088 | 8.310225403 | 8.23671175 | 8.794695657 | 8.757891838 |
| ENSG00000065882 | TBC1D1 | -2.290429903 | 1.87E-09 | 0.073217966 | 0.886475445 | -2.217211937 | 10.43795805 | 11.04094592 | 11.18913795 | 8.645348336 | 8.64548836 | 10.83619401 | 11.14243586 |
| ENSG00000276780 | | 2.973682718 | 0.608680785 | -5.191094386 | 0.364514502 | -2.217411669 | 2 | 2 | 2 | 3.462030052 | 3.69849104 | 2 | 2 |
| ENSG00000120742 | SERP1 | -1.302941071 | 0.00562177 | -0.9151278 | 0.0593663 | -2.218068871 | 10.80239439 | 11.1990418 | 11.79914889 | 10.01841412 | 10.032869 | 10.53493405 | 10.27793661 |
| ENSG00000188385 | JAKMIP3 | -1.680836372 | 0.503822785 | -0.538918214 | 0.839017584 | -2.219754586 | 5.095959458 | 2 | 2 | 2.784612485 | 2.86123014 | 3.316666208 | 3.539239711 |
| ENSG00000105479 | CCDC114 | -1.971754019 | 0.445453858 | -0.250433898 | 0.928829011 | -2.222187917 | 2 | 4.11352228 | 2 | 2.498691189 | 2.2680111 | 3.12825509 | 2.885414003 |
| ENSG00000105568 | PPP2R1A | -1.51544718 | 0.013543185 | -0.708219232 | 0.282259948 | -2.223666412 | 13.00891643 | 11.68842912 | 11.95227171 | 10.78388778 | 10.8589471 | 11.40838965 | 11.81815607 |
| ENSG00000161513 | FDXR | -1.780141894 | 0.001111835 | -0.445244223 | 0.470583967 | -2.225386117 | 8.107517749 | 7.002537973 | 7.600047325 | 6.069927018 | 5.74985018 | 7.296052918 | 7.085754446 |
| ENSG00000167118 | URM1 | -1.176792429 | 0.003796745 | -1.049811712 | 0.010307848 | -2.22660414 | 9.667789716 | 9.799806309 | 10.28977157 | 8.716468139 | 8.82894823 | 8.65615757 | 9.108160402 |
| ENSG00000165802 | NSMF | -2.005869944 | 3.74E-05 | -0.221445723 | 0.712407637 | -2.227315667 | 10.60392551 | 9.849218119 | 10.9743126 | 8.550024073 | 8.55284928 | 10.44941809 | 10.1885847 |
| ENSG00000143036 | SLC44A3 | -2.100027672 | 0.198785891 | -0.128750671 | 0.945755737 | -2.228778343 | 5.330594134 | 6.535001748 | 2 | 4.070463721 | 3.45167086 | 5.650128416 | 5.136701757 |
| ENSG00000104824 | HNRNPL | -0.997737671 | 0.07806983 | -1.231300657 | 0.024328003 | -2.229038327 | 10.60951384 | 11.01334259 | 11.57188978 | 10.13319806 | 10.1143063 | 9.439667847 | 10.23456118 |
| ENSG00000166526 | ZNF3 | -1.076189529 | 0.065058738 | -1.153990213 | 0.043110722 | -2.230179741 | 9.730679447 | 8.916342327 | 8.460932916 | 8.024753265 | 8.11025689 | 8.020265032 | 7.962181894 |
| ENSG00000010219 | DYRK4 | -1.16411001 | 0.024656108 | -1.068064396 | 0.03854522 | -2.232174406 | 8.87322763 | 8.077869079 | 7.858115942 | 7.165617905 | 7.22257674 | 7.223329552 | 7.349806994 |
| ENSG00000106278 | PTPRZ1 | 0.919798563 | 0.795095011 | -3.152347578 | 0.316133759 | -2.232549015 | 4.00722531 | 5.72254323 | 2 | 6.351046291 | 2 | 2.987421008 | 2.225841198 |
| ENSG00000005206 | SPPL2B | -1.239553907 | 0.000599658 | -0.993243478 | 0.006770897 | -2.232797385 | 9.128497779 | 9.497318007 | 9.829330889 | 8.315844936 | 8.25031674 | 8.538032809 | 8.515559017 |
| ENSG00000131788 | PIAS3 | -1.479479638 | 4.06E-06 | -0.753659176 | 0.02758368 | -2.233138813 | 10.67493801 | 10.2574711 | 10.67226762 | 9.09649754 | 9.04952667 | 9.605472338 | 9.962227574 |
| ENSG00000179029 | TMEM107 | -1.283971544 | 0.467608963 | -0.951543788 | 0.591655943 | -2.235515332 | 7.579279349 | 6.899080544 | 2 | 5.602782483 | 5.40529704 | 5.853194273 | 5.772630506 |
| ENSG00000146066 | HIGD2A | -0.877542401 | 0.305220054 | -1.358628375 | 0.090878546 | -2.236170776 | 10.60951384 | 9.018611864 | 8.904018013 | 8.880924505 | 8.84696933 | 8.108635266 | 8.621777169 |
| ENSG00000178952 | TUFM | -1.445346686 | 0.001031241 | -0.791781021 | 0.09006775 | -2.237126707 | 11.34164739 | 11.01952247 | 10.39531398 | 9.514881142 | 9.54514558 | 10.06192211 | 10.29067944 |
| ENSG00000090263 | MRPS33 | -1.116087612 | 0.13727197 | -1.12118299 | 0.127460764 | -2.237270601 | 8.716893051 | 7.051611315 | 8.561105505 | 7.238438152 | 7.11195456 | 6.987904673 | 7.334741191 |
| ENSG00000009844 | VTA1 | -0.702279529 | 0.210105576 | -1.535491697 | 0.002467188 | -2.237771226 | 9.358295023 | 10.12043894 | 10.55568472 | 9.401417034 | 9.38433737 | 8.538032809 | 8.594208165 |
| ENSG00000086544 | ITPKC | -1.359983844 | 0.224142044 | -0.878592479 | 0.443890552 | -2.238576323 | 10.32020829 | 8.191702463 | 7.552162999 | 7.839502934 | 7.85944752 | 8.486081846 | 8.140837768 |
| ENSG00000143603 | KCNN3 | 0.409005538 | 0.825160194 | -2.647728319 | 0.088351175 | -2.238722781 | 5.70932728 | 2 | 5.59507254 | 5.662809983 | 5.24368216 | 2.858551861 | 3.383972333 |
| ENSG00000180185 | FAHD1 | -0.980068427 | 0.018147366 | -1.258906888 | 0.001610393 | -2.238975315 | 9.384607364 | 8.671479029 | 9.083172337 | 8.134820652 | 8.06698125 | 7.661243884 | 7.975841991 |
| ENSG00000145354 | CISD2 | -0.49047917 | 0.444744248 | -1.748753509 | 0.001504602 | -2.239232678 | 8.16936288 | 9.338312924 | 8.434763955 | 8.235106869 | 8.284723 | 6.879153024 | 7.162846889 |
| ENSG00000182154 | MRPL41 | -1.588057746 | 6.91E-05 | -0.651709368 | 0.132806015 | -2.239767115 | 9.761125267 | 9.549500232 | 10.2528023 | 8.285617933 | 8.32752521 | 9.028334265 | 9.414713898 |
| ENSG00000116754 | SRSF11 | -0.817823715 | 0.10052636 | -1.422645765 | 0.002070266 | -2.24046948 | 11.69173081 | 11.04700893 | 10.72720655 | 10.44622016 | 10.3446634 | 9.924007159 | 9.650471437 |
| ENSG00000184205 | TSPYL2 | -0.211559498 | 0.709501472 | -2.030127664 | 6.23E-06 | -2.241687162 | 10.03472846 | 9.74864196 | 10.69449487 | 10.00861625 | 9.99584995 | 8.031518488 | 8.347103327 |
| ENSG00000155660 | PDIA4 | -0.444636972 | 0.449163778 | -1.797227554 | 0.000305645 | -2.241864527 | 12.21051618 | 12.12373461 | 12.3988809 | 11.79320259 | 11.8160057 | 9.765897048 | 10.91886419 |
| ENSG00000253958 | CLDN23 | 0.449065549 | 0.838658392 | -2.691832659 | 0.148381845 | -2.242767109 | 2 | 5.595927442 | 5.39244166 | 5.258265913 | 5.49945122 | 3.56816602 | 2.225841198 |
| ENSG00000137166 | FOXP4 | -1.360343882 | 0.091805172 | -0.883772828 | 0.285220143 | -2.24411671 | 8.674867122 | 10.46189757 | 8.978369997 | 8.287525788 | 8.21506984 | 8.888863837 | 8.535462193 |
| ENSG00000091844 | RGS17 | 0.913290076 | 0.756463845 | -3.158151206 | 0.226910457 | -2.24486113 | 2 | 2 | 7.225742779 | 6.548530093 | 6.5890786 | 3.694366604 | 2.551896972 |
| ENSG00000113048 | MRPS27 | -1.901084501 | 4.21E-05 | -0.343923282 | 0.529270693 | -2.245007782 | 9.790941832 | 10.15453315 | 9.116480286 | 7.920420067 | 7.81567127 | 9.2896505 | 9.52069224 |

| | | | | | | | | | | | | | |
|---|---|---|---|---|---|---|---|---|---|---|---|---|---|
| ENSG00000177595 | PIDD1 | -2.148390332 | 0.000432249 | -0.096843529 | 0.901842972 | -2.24523386 | 8.313285031 | 7.431218269 | 7.818184403 | 5.900763176 | 5.74985018 | 8.219062336 | 7.20488131 |
| ENSG00000164675 | IQUB | -0.172435443 | 0.948671972 | -2.072847484 | 0.382824967 | -2.245282926 | 2 | 4.710564154 | 2 | 3.289987342 | 3.60476711 | 2.746468522 | 2.326754077 |
| ENSG00000147509 | RGS20 | -0.724283143 | 0.774707448 | -1.521369687 | 0.518400462 | -2.24565283 | 7.435667355 | 3.076665209 | 2 | 5.242616075 | 5.3193943 | 5.136410075 | 3.698365557 |
| ENSG00000167196 | FBXO22 | -1.348318061 | 0.03918001 | -0.898215753 | 0.181113049 | -2.246533814 | 9.720386367 | 8.356994601 | 9.465702942 | 8.03841338 | 7.86198169 | 8.689219644 | 8.029224588 |
| ENSG00000124733 | MEA1 | -1.069733664 | 0.000423512 | -1.177265764 | 8.02E-05 | -2.246999428 | 10.60951384 | 10.26269067 | 10.62675862 | 9.420668575 | 9.46239569 | 9.171189325 | 9.481899659 |
| ENSG00000111785 | RIC8B | -1.150532879 | 0.256719105 | -1.096538557 | 0.272887212 | -2.247071436 | 9.423197387 | 9.125613817 | 6.718950348 | 7.706073672 | 7.65545169 | 7.836577593 | 7.623392692 |
| ENSG00000146670 | CDCA5 | -1.410718714 | 0.209225902 | -0.836757696 | 0.46727486 | -2.24747641 | 4.00722531 | 5.946638857 | 6.80318718 | 4.783303513 | 4.63360314 | 4.933813368 | 5.44241046 |
| ENSG00000279566 | | 0.111479647 | 0.935007649 | -2.35988275 | 0.031285282 | -2.248403103 | 4.00722531 | 7.099070191 | 7.098165388 | 6.628980152 | 6.78340763 | 4.604690117 | 4.316807515 |
| ENSG00000104979 | C19orf53 | -1.237939981 | 0.003686449 | -1.013896994 | 0.019496236 | -2.251836975 | 10.70134604 | 9.916994052 | 10.05225195 | 9.014783325 | 9.05063789 | 9.122905457 | 9.376603759 |
| ENSG00000006652 | IFRD1 | -0.392406963 | 0.659880399 | -1.860096626 | 0.014001426 | -2.25250359 | 10.43166252 | 9.433952963 | 10.41552659 | 9.788505066 | 9.75261615 | 8.917048655 | 7.260151003 |
| ENSG00000162521 | RBBP4 | -1.146900147 | 1.72E-05 | -1.106953526 | 3.11E-05 | -2.253853673 | 11.15935596 | 10.72651826 | 10.9788475 | 9.834904647 | 9.80623425 | 9.863792053 | 9.857019471 |
| ENSG00000169955 | ZNF747 | -0.794819935 | 0.41162884 | -1.459179416 | 0.101692496 | -2.25399935 | 9.081016759 | 6.844429755 | 7.691296561 | 7.409376773 | 7.35800033 | 6.615624322 | 6.853913776 |
| ENSG00000173875 | ZNF791 | -1.611222687 | 1.37E-05 | -0.643153255 | 0.109831184 | -2.254375942 | 9.602033055 | 9.04308532 | 8.922968322 | 7.605658789 | 7.64959655 | 8.59173005 | 8.571253152 |
| ENSG00000186193 | SAPCD2 | -1.762136861 | 0.256571989 | -0.494518798 | 0.76594204 | -2.256655659 | 5.330594134 | 3.076665209 | 3.369317972 | 3.227757282 | 2.94009902 | 4.166733998 | 3.560117069 |
| ENSG00000124299 | PEPD | -1.041253114 | 0.046045679 | -1.215856711 | 0.016706175 | -2.257109825 | 10.1076228 | 9.125613817 | 10.06083273 | 8.755587391 | 8.82765235 | 8.426517973 | 8.789175794 |
| ENSG00000157778 | PSMG3 | -1.661810848 | 0.000636258 | -0.596359836 | 0.265494962 | -2.258206683 | 9.159305284 | 9.192779271 | 8.175061096 | 7.250225364 | 7.31055061 | 8.350677018 | 8.302113262 |
| ENSG00000100296 | THOC5 | -0.99690658 | 0.408476816 | -1.261410345 | 0.27417388 | -2.258316925 | 10.41898845 | 8.469613413 | 7.285536013 | 8.404661252 | 8.19309833 | 7.919680595 | 8.154630332 |
| ENSG00000131143 | COX4I1 | -1.35501981 | 0.037220596 | -0.905044725 | 0.176711799 | -2.260064535 | 12.56276862 | 11.17709514 | 11.34831197 | 10.46277773 | 10.5044462 | 10.82664326 | 11.03141843 |
| ENSG00000214253 | FIS1 | -1.330651364 | 0.047054927 | -0.930690492 | 0.174415528 | -2.261341856 | 10.46903009 | 8.88960062 | 9.963514551 | 8.600009476 | 8.57618689 | 8.749978316 | 9.18619545 |
| ENSG00000196419 | XRCC6 | -1.204224519 | 0.003463152 | -1.05718026 | 0.011096104 | -2.261404779 | 12.2324126 | 11.69616579 | 11.57188978 | 10.64941153 | 10.6715842 | 10.54700852 | 11.0276759 |
| ENSG00000114646 | CSPG5 | -2.76125645 | 0.240049702 | 0.499502519 | 0.847549471 | -2.26175393 | 2 | 5.595927442 | 2 | 2.695456494 | 2.59489879 | 5.002522427 | 4.26688425 |
| ENSG00000006025 | OSBPL7 | -1.510331624 | 0.362508675 | -0.751588907 | 0.665769105 | -2.261920531 | 7.031698319 | 6.844429755 | 2 | 5.058234798 | 4.93258395 | 5.528313928 | 5.816671626 |
| ENSG00000136521 | NDUFB5 | -0.819325113 | 0.296385835 | -1.44326904 | 0.046743116 | -2.262594153 | 10.24321353 | 9.00621767 | 8.511888463 | 8.637890117 | 8.62474679 | 8.017249226 | 8.013226939 |
| ENSG00000198492 | YTHDF2 | -1.067228125 | 0.02109997 | -1.197228368 | 0.008293918 | -2.264456494 | 10.83615308 | 11.05905903 | 10.10298738 | 9.660380652 | 9.65662671 | 9.438259243 | 9.614745652 |
| ENSG00000104064 | GABPB1 | -0.848373848 | 0.159914848 | -1.417666854 | 0.011593579 | -2.266040702 | 9.317904228 | 8.169643011 | 8.325066185 | 7.831679003 | 7.88459112 | 7.355766946 | 7.244113847 |
| ENSG00000134809 | TIMM10 | -1.266065996 | 0.253395005 | -1.000227401 | 0.367386473 | -2.266293398 | 8.928409675 | 6.666836837 | 6.534109959 | 6.542151749 | 6.64311914 | 6.649130473 | 7.025105265 |
| ENSG00000170855 | TRIAP1 | -1.315411071 | 0.001935204 | -0.951260472 | 0.029190238 | -2.266671543 | 9.579430841 | 8.762721501 | 9.196533591 | 7.942383523 | 7.88209639 | 8.40533527 | 8.123409541 |
| ENSG00000203943 | SAMD13 | -0.954618247 | NA | -1.312738577 | NA | -2.267356824 | 2 | 3.076665209 | 2 | 2.141737246 | 2.49394783 | 2.34604626 | 2.172608095 |
| ENSG00000099385 | BCL7C | -1.095522159 | 0.055040826 | -1.172004681 | 0.036112934 | -2.26752684 | 9.473095744 | 8.317412869 | 9.083172337 | 7.905589775 | 7.98565528 | 7.612118169 | 8.090601559 |
| ENSG00000107020 | PLGRKT | -0.617146737 | 0.712294566 | -1.650467387 | 0.273834457 | -2.267614124 | 8.928409675 | 5.72254323 | 4.874587708 | 6.902323594 | 7.02417148 | 5.85994354 | 6.088639344 |
| ENSG00000119760 | SUPT7L | -1.071609232 | 2.41E-05 | -1.196421287 | 1.59E-06 | -2.26803052 | 10.28571918 | 10.08551947 | 10.26770388 | 9.16866395 | 9.12637045 | 8.893394126 | 9.143604888 |
| ENSG00000091140 | DLD | -1.10968401 | 0.041566879 | -1.158951777 | 0.03070475 | -2.268635787 | 9.521325812 | 10.12043894 | 10.67226762 | 9.081180848 | 9.06939887 | 8.806972075 | 9.216922247 |
| ENSG00000100842 | EFS | -2.024661756 | 0.016912578 | -0.246026092 | 0.808868217 | -2.270687848 | 7.624139768 | 9.405920729 | 10.01740919 | 7.348537858 | 7.28811304 | 9.151037776 | 8.987157359 |
| ENSG00000198342 | ZNF442 | -0.12462535 | 0.966716761 | -2.147352734 | 0.414376291 | -2.271978083 | 6.96351253 | 2 | 2 | 5.514313187 | 5.16377554 | 4.029788235 | 3.287118176 |
| ENSG00000005483 | KMT2E | -1.147279858 | 2.18E-05 | -1.125339535 | 2.86E-05 | -2.272619393 | 11.17451594 | 11.03180337 | 11.46604358 | 10.11987514 | 10.059057 | 10.10907137 | 10.11421585 |
| ENSG00000140365 | COMMD4 | -0.709513067 | 0.370174297 | -1.563926135 | 0.029540521 | -2.273439202 | 9.710019321 | 8.147241016 | 8.434763955 | 8.252792309 | 8.20512425 | 7.124495877 | 7.614687532 |
| ENSG00000214046 | SMIM7 | -0.932868345 | 9.38E-05 | -1.34220685 | 5.38E-09 | -2.275075195 | 9.810482605 | 9.806969599 | 9.788583153 | 8.845101141 | 8.90464485 | 8.362013331 | 8.57061027 |
| ENSG00000004142 | POLDIP2 | -1.639080064 | 0.015702631 | -0.636468649 | 0.389304796 | -2.275548712 | 11.67050115 | 10.12617756 | 10.40208303 | 9.214510118 | 9.31574487 | 10.23382726 | 10.29340937 |
| ENSG00000189067 | LITAF | -2.341892435 | 0.001497712 | 0.066313198 | 0.944363944 | -2.275579237 | 11.32135895 | 9.357954881 | 10.16797309 | 8.227176542 | 8.13141746 | 10.49619812 | 10.64412949 |
| ENSG00000124422 | USP22 | -1.525446723 | 7.50E-09 | -0.753247842 | 0.007273542 | -2.278694565 | 12.20498975 | 12.43011779 | 12.5724427 | 10.82584613 | 10.9454246 | 11.53219589 | 11.77339293 |
| ENSG00000140391 | TSPAN3 | -1.354506888 | 0.000454418 | -0.924781457 | 0.021329954 | -2.279288344 | 12.45359838 | 12.03348143 | 12.54671079 | 11.02423622 | 10.9915755 | 11.10893748 | 11.70415409 |
| ENSG00000242715 | CCDC169 | -2.262767666 | 0.229708761 | -0.017991203 | 0.993459837 | -2.280758869 | 4.467311984 | 2 | 4.058998049 | 2.784612485 | 2.38539787 | 4.133696865 | 3.311948089 |
| ENSG00000132661 | NXT1 | -1.689938669 | 0.000161722 | -0.591232458 | 0.228666677 | -2.281171127 | 8.540903468 | 7.847167909 | 7.818184403 | 6.29903474 | 6.60731796 | 7.479255779 | 7.569069104 |
| ENSG00000176102 | CSTF3 | -0.24509708 | 0.913189302 | -2.03661714 | 0.283648588 | -2.281714219 | 2 | 8.834582582 | 9.196533591 | 8.128448191 | 8.2790453 | 6.514451546 | 6.406535141 |
| ENSG00000185359 | HGS | -0.43566131 | 0.558317273 | -1.84624715 | 0.003646438 | -2.281908459 | 11.67583792 | 10.47544827 | 10.23774519 | 10.49161431 | 10.5225794 | 8.967528086 | 9.22715677 |
| ENSG00000169714 | CNBP | -1.179373957 | 0.007634516 | -1.104069052 | 0.012705607 | -2.283443009 | 12.74263541 | 11.89244687 | 12.08228623 | 11.09020988 | 11.1267944 | 11.30784912 | 11.0480252 |
| ENSG00000088766 | CRLS1 | -1.220196657 | 0.003161248 | -1.063632302 | 0.010607918 | -2.283828959 | 8.716893051 | 9.308339287 | 9.385998894 | 7.966401501 | 7.95512324 | 8.300083299 | 7.902127837 |
| ENSG00000086200 | IPO11 | -1.005402617 | 0.561371291 | -1.280360574 | 0.443230611 | -2.285763191 | 4.815609471 | 7.099070191 | 10.05225195 | 7.791909949 | 7.56502709 | 7.388811377 | 7.438450313 |
| ENSG00000176623 | RMDN1 | -1.379470876 | 0.07497303 | -0.906482068 | 0.255070887 | -2.285952944 | 10.07567912 | 8.101365653 | 9.242516611 | 7.990026167 | 7.97632944 | 8.409941741 | 8.488586985 |
| ENSG00000170166 | HOXD4 | 4.622411758 | 0.024200025 | -6.908755836 | 0.007633479 | -2.286344079 | 2 | 3.686303414 | 2 | 6.358325776 | 6.507263 | 2 | 2 |
| ENSG00000005486 | RHBDD2 | -1.142763801 | 0.005164205 | -1.144675825 | 0.004822148 | -2.287439627 | 11.10895605 | 10.50217373 | 10.31867969 | 9.520570052 | 9.56938265 | 9.548642921 | 9.537896513 |
| ENSG00000169282 | KCNAB1 | -2.866026939 | 0.057963449 | 0.578470371 | 0.732581748 | -2.287556569 | 2 | 5.72254323 | 6.629488387 | 3.227757282 | 3.55551834 | 6.290458506 | 6.182510897 |
| ENSG00000198856 | OSTC | -1.207263016 | 0.122079915 | -1.08172518 | 0.163562907 | -2.288988195 | 12.16380252 | 10.89059638 | 10.30429804 | 10.14902484 | 10.1121791 | 10.29128794 | 10.21946626 |
| ENSG00000134597 | RBMX2 | -1.335131993 | 0.537027555 | -0.954261094 | 0.665712192 | -2.289393087 | 9.800745303 | 7.35507212 | 2 | 7.136275664 | 7.17037177 | 7.610117916 | 7.434215826 |
| ENSG00000282604 | | 2.901620803 | 0.617500397 | -5.191094686 | NA | -2.289473883 | 2 | 2 | 2 | 3.840289928 | 3.15372745 | 2 | 2 |
| ENSG00000160796 | NBEAL2 | -2.320581108 | 0.119776467 | 0.028915421 | 0.987221496 | -2.291665687 | 3.327711234 | 8.433041974 | 8.175061096 | 5.501219927 | 5.56313034 | 7.809812485 | 7.745758018 |
| ENSG00000169255 | B3GALNT1 | -0.53422772 | 0.341329168 | -1.757762809 | 0.000306618 | -2.291990529 | 7.667647223 | 8.717821467 | 8.325066185 | 7.821180652 | 7.7377845 | 6.448814696 | 6.721179863 |
| ENSG00000183395 | PMCH | 2.898995389 | 0.617831455 | -5.191094697 | NA | -2.292099308 | 2 | 2 | 2 | 3.566290607 | 3.50452879 | 2 | 2 |
| ENSG00000104818 | CGB2 | 2.898557361 | 0.617843763 | -5.191094699 | NA | -2.292537338 | 2 | 2 | 2 | 3.515101962 | 3.55551834 | 2 | 2 |
| ENSG00000111786 | SRSF9 | -0.72130009 | 0.191457657 | -1.571477772 | 0.00166238 | -2.292777862 | 11.04027452 | 10.12617756 | 10.55568472 | 9.971121557 | 9.82972721 | 8.676372393 | 9.35772346 |
| ENSG00000161016 | RPL8 | -1.746583118 | 0.008869853 | -0.547352454 | 0.463132585 | -2.293935572 | 14.02224794 | 12.74089149 | 12.49857094 | 11.4920928 | 11.5216186 | 12.62892023 | 12.77734718 |
| ENSG00000157335 | CLEC18C | 2.895487483 | 0.618153694 | -5.191094712 | NA | -2.295607229 | 2 | 2 | 2 | 3.09461492 | 3.86943969 | 2 | 2 |
| ENSG00000106615 | RHEB | -0.387692371 | 0.563160425 | -1.909258143 | 0.000698186 | -2.296950514 | 10.5468183 | 9.203675681 | 9.681375057 | 9.568433308 | 9.49541459 | 8.056705941 | 7.993217039 |
| ENSG00000137996 | RTCA | -0.436392366 | 0.36394741 | -1.862264393 | 4.91E-06 | -2.298656759 | 9.624286623 | 9.246456519 | 8.721235202 | 8.804373539 | 8.81854848 | 7.425672291 | 7.388257299 |
| ENSG00000242259 | C22orf39 | -2.029238061 | 0.20665794 | -0.270183223 | 0.885032716 | -2.299421284 | 9.174465797 | 8.505280662 | 3.369317972 | 6.283823264 | 6.3857277 | 8.152588976 | 7.925444964 |
| ENSG00000165669 | FAM204A | -1.102087491 | 0.02474593 | -1.197803088 | 0.012880352 | -2.299890579 | 9.233563228 | 8.213429689 | 8.904018013 | 7.731524332 | 7.76782485 | 7.590975897 | 7.718159092 |
| ENSG00000178921 | PFAS | -1.882318281 | 0.089428115 | -0.418773429 | 0.736888343 | -2.301091709 | 5.70932728 | 8.94259736 | 8.434763955 | 6.221320894 | 6.53293476 | 7.580798201 | 7.998006535 |
| ENSG00000164039 | BDH2 | -0.200662073 | 0.72600899 | -2.102873539 | 3.32E-06 | -2.303535612 | 9.081016759 | 8.557180776 | 8.042706615 | 8.456457569 | 8.39031583 | 6.558682816 | 6.574984961 |
| ENSG00000180035 | ZNF48 | -2.117473647 | 0.364034842 | -0.186474716 | 0.944101725 | -2.303948363 | 2 | 5.457120953 | 9.926432745 | 6.449761039 | 6.23747437 | 8.437839892 | 7.975841991 |
| ENSG00000172167 | MTBP | -0.655140304 | 0.713187979 | -1.649914328 | 0.306062327 | -2.305054632 | 2 | 4.93638933 | 6.322058544 | 4.907595604 | 4.40247105 | 4.029788235 | 3.735548302 |
| ENSG00000181523 | SGSH | -0.876697944 | 0.290539585 | -1.428754501 | 0.06486798 | -2.305452444 | 11.41049097 | 9.506147172 | 10.17589429 | 9.682298904 | 9.73682479 | 8.99192853 | 9.312299623 |
| ENSG00000092068 | SLC7A8 | -1.50688755 | 0.352079152 | -0.800853777 | 0.634425125 | -2.307741327 | 3.327711234 | 6.602424674 | 8.007629717 | 5.527288686 | 5.47317115 | 6.736076089 | 5.171036168 |
| ENSG00000276045 | ORAI1 | -1.324521159 | 4.93E-06 | -0.984825126 | 0.000794685 | -2.309346285 | 8.777722034 | 8.88960062 | 9.116480286 | 7.574744049 | 7.67575955 | 8.034504643 | 7.880504067 |
| ENSG00000114125 | RNF7 | -0.99127646 | 0.000114327 | -1.318449761 | 1.21E-07 | -2.309726221 | 10.59268348 | 10.23640192 | 10.37481416 | 9.408447353 | 9.43116542 | 9.043977376 | 9.144037078 |
| ENSG00000081760 | AACS | -1.37912481 | 0.176302078 | -0.930629026 | 0.370342728 | -2.309753836 | 8.285626337 | 8.623598428 | 10.62675862 | 8.16211395 | 8.248381 | 8.340457938 | 8.903351429 |
| ENSG00000076555 | ACACB | -0.941847677 | 0.644285833 | -1.370048322 | 0.480467544 | -2.311895999 | 2 | 5.131603625 | 7.734844356 | 5.514313187 | 5.49945122 | 5.141958644 | 5.101530266 |
| ENSG00000144867 | SRPRB | -1.061129909 | 9.86E-08 | -1.25137751 | 1.53E-10 | -2.31250742 | 10.56418096 | 10.59613751 | 10.45512158 | 9.453327884 | 9.51406378 | 9.313817266 | 9.275690962 |
| ENSG00000109445 | ZNF330 | -1.091404952 | 0.626071633 | -1.222427621 | 0.575697563 | -2.313832573 | 10.1076228 | 7.761270542 | 2 | 7.626911583 | 7.7866208 | 7.816767288 | 7.297928101 |
| ENSG00000262327 | | 0.495253758 | 0.754341134 | -2.80927219 | 0.035591756 | -2.314018432 | 3.327711234 | 7.901717006 | 7.285536013 | 7.697489359 | 7.42122542 | 4.496236235 | 4.522269241 |
| ENSG00000072778 | ACADVL | -0.765741747 | 0.220214001 | -1.548822257 | 0.006729172 | -2.314564004 | 12.74390633 | 11.45757623 | 12.68078978 | 11.61151504 | 11.6612917 | 10.75286657 | 10.95009364 |
| ENSG00000131966 | ACTR10 | -0.933681522 | 0.167317726 | -1.381272582 | 0.030563837 | -2.314954104 | 9.097018025 | 10.22575081 | 10.63827116 | 9.26766905 | 9.11047337 | 8.885973502 | 8.593575433 |
| ENSG00000046651 | OFD1 | -1.104535205 | 0.210990211 | -1.210980993 | 0.157715974 | -2.315516198 | 7.159102577 | 7.928237109 | 9.272377397 | 7.288834728 | 7.31055061 | 7.228554289 | 7.162846889 |
| ENSG00000204152 | TIMM23B | 0.738379625 | 0.184268122 | -3.054916354 | 2.33E-10 | -2.31653673 | 7.82963488 | 8.414402626 | 8.677268317 | 9.061245954 | 9.09870316 | 5.056402493 | 5.708623051 |
| ENSG00000139354 | GAS2L3 | -1.313177251 | 0.203115525 | -1.003511173 | 0.332515383 | -2.316688424 | 6.138360429 | 5.303523747 | 7.691296561 | 5.433909581 | 5.51241389 | 5.711121288 | 5.803599879 |
| ENSG00000180592 | SKIDA1 | -2.122610663 | 0.150705666 | -0.196133708 | 0.908941926 | -2.318744371 | 3.327711234 | 6.04685525 | 4.058998049 | 3.566290607 | 3.15372745 | 4.188345803 | 5.256659864 |
| ENSG00000159648 | TEPP | -4.714896419 | 0.259738629 | 2.395429686 | 0.541103884 | -2.319466734 | 2 | 3.076665209 | 2 | 2 | 2 | 3.500668998 | 4.133950982 |
| ENSG00000044459 | CNTLN | -0.501100764 | 0.749056274 | -1.820480322 | 0.186484076 | -2.321581086 | 8.653385977 | 7.572420273 | 4.058998049 | 7.140504127 | 7.21466714 | 6.05195199 | 5.768151538 |
| ENSG00000073464 | CLCN4 | -2.546933067 | 0.304297653 | 0.225115946 | 0.935446753 | -2.32181712 | 2 | 5.131603625 | 2 | 2.784612485 | 2.2680111 | 4.053534413 | 4.062589736 |
| ENSG00000183049 | CAMK1D | -3.456064464 | 0.010024408 | 1.13402239 | 0.445282505 | -2.322042074 | 6.009025997 | 6.78762695 | 9.849281147 | 5.552894743 | 4.74762503 | 9.636512519 | 9.638558632 |
| ENSG00000110104 | CCDC86 | -1.2902047 | 0.040174325 | -1.032673905 | 0.103916882 | -2.322878605 | 9.922475255 | 8.487557258 | 9.541233192 | 8.164192202 | 8.13979554 | 8.211825862 | 8.577666268 |
| ENSG00000205592 | MUC19 | -0.69218051 | NA | -1.630843029 | NA | -2.323023539 | 2 | 2 | 3.369317972 | 2.141737246 | 2.59489879 | 2.094598369 | 2.326754077 |
| ENSG00000106638 | TBL2 | -1.345717549 | 0.00058344 | -0.977856437 | 0.015303411 | -2.323573987 | 10.71696191 | 9.936721131 | 10.56174313 | 9.088859521 | 9.11260313 | 9.492471997 | 9.439572619 |
| ENSG00000089847 | ANKRD24 | -1.702281039 | 0.515393703 | -0.621498504 | 0.821109617 | -2.323779543 | 2 | 2 | 5.930865648 | 3.162721474 | 3.39680241 | 4.166733998 | 3.841670744 |
| ENSG00000084453 | SLCO1A2 | -4.13918349 | 0.143847237 | 1.815238694 | 0.433240072 | -2.323944795 | 2 | 3.076665209 | 3.369317972 | 2 | 2.14022021 | 3.411599105 | 4.511590831 |
| ENSG00000100601 | ALKBH1 | -0.98210296 | 0.169054392 | -1.342381546 | 0.04761818 | -2.324484506 | 7.709880939 | 7.051611315 | 8.536706852 | 6.89733479 | 6.95464102 | 6.800644007 | 6.317379004 |
| ENSG00000170745 | KCNS3 | -1.549251646 | 0.346696909 | -0.77832167 | 0.651103372 | -2.327573316 | 3.327711234 | 7.051611315 | 8.076950877 | 5.753943344 | 5.49945122 | 6.981729486 | 5.217771644 |
| ENSG00000157734 | SNX22 | -0.043751437 | 0.96929411 | -2.28445244 | 0.011166291 | -2.328203877 | 4.815609471 | 5.131603625 | 6.718950348 | 5.753943344 | 5.74985018 | 3.851628042 | 3.875378318 |
| ENSG00000204852 | TCTN1 | -0.806251977 | 0.09337333 | -1.522199296 | 0.000549691 | -2.328451272 | 9.317904228 | 8.623598428 | 9.196533591 | 8.268332738 | 8.2790453 | 7.267148095 | 7.820122783 |
| ENSG00000112984 | KIF20A | -1.87813222 | 0.346404872 | -0.4505043 | 0.836821477 | -2.328636521 | 4.467311984 | 6.389896633 | 2 | 3.881293667 | 3.50452879 | 3.500668998 | 5.502966356 |
| ENSG00000132463 | GRSF1 | -1.049428755 | 0.040810687 | -1.279562264 | 0.010048431 | -2.328991019 | 11.92474824 | 10.79656548 | 11.35181254 | 10.38872576 | 10.3760296 | 10.16225622 | 10.14369036 |
| ENSG00000112081 | SRSF3 | -0.970041883 | 0.034923735 | -1.360447729 | 0.001927913 | -2.330489613 | 11.1478805 | 10.68045884 | 11.35181254 | 10.1698607 | 10.064018 | 9.363643077 | 10.02058586 |
| ENSG00000186907 | RTN4RL2 | -1.334223468 | 0.576805834 | -0.996591445 | 0.670031027 | -2.330814913 | 2 | 3.076665209 | 3.369317972 | 2.270786764 | 2.59489879 | 2.526562374 | 2.551896972 |
| ENSG00000134014 | ELP3 | -0.953354461 | 0.226126239 | -1.377977717 | 0.064751221 | -2.331332178 | 10.02639697 | 8.147241016 | 9.031722608 | 8.327018759 | 8.31463236 | 7.910770103 | 7.893928381 |
| ENSG00000105996 | HOXA2 | 2.859699628 | 0.622818782 | -5.191094861 | NA | -2.331395233 | 2 | 2 | 2 | 3.566290607 | 3.45167086 | 2 | 2 |
| ENSG00000269556 | TMEM185A | -0.287996468 | 0.61090653 | -2.043692893 | 8.40E-06 | -2.331689361 | 8.716893051 | 8.469613413 | 9.452720984 | 8.621345041 | 8.68180105 | 6.921708059 | 6.938065226 |
| ENSG00000059377 | TBXAS1 | -1.810122381 | 0.383862151 | -0.521865964 | 0.814923166 | -2.331988344 | 2 | 3.076665209 | 5.772716771 | 3.289987342 | 3.08597292 | 4.478802948 | 3.430065404 |
| ENSG00000167554 | ZNF610 | -0.680012292 | 0.72338932 | -1.652451174 | 0.346462637 | -2.332463466 | 7.82963488 | 6.728495673 | 2 | 6.213314212 | 6.11534392 | 5.553513572 | 4.911532223 |

| | | | | | | | | | | | | | |
|---|---|---|---|---|---|---|---|---|---|---|---|---|---|
| ENSG00000189186 | DCAF8L2 | 2.858349037 | 0.622871051 | -5.191094867 | NA | -2.332745829 | 2 | 2 | 2 | 3.406930997 | 3.60476711 | 2 | 2 |
| ENSG00000205457 | TP53TG3C | 2.858349037 | 0.622871051 | -5.191094867 | NA | -2.332745829 | 2 | 2 | 2 | 3.406930997 | 3.60476711 | 2 | 2 |
| ENSG00000204843 | DCTN1 | -1.526318982 | 7.80E-05 | -0.807587227 | 0.049586809 | -2.333906209 | 12.38845524 | 11.752905 | 11.92668027 | 10.48788555 | 10.5593475 | 10.98908978 | 11.45586848 |
| ENSG00000174177 | CTU2 | -1.671168656 | 0.000201031 | -0.665481145 | 0.169565436 | -2.336649802 | 7.790810999 | 7.35507212 | 8.143093325 | 6.069927018 | 6.26836889 | 7.254397864 | 7.010010317 |
| ENSG00000197568 | HHLA3 | -2.29886036 | 0.091749244 | -0.03900373 | 0.981056901 | -2.33786409 | 4.815609471 | 5.838937808 | 8.511888463 | 5.110902178 | 5.0259382 | 7.106162485 | 7.261744957 |
| ENSG00000179222 | MAGED1 | -1.891845568 | 1.02E-07 | -0.446306585 | 0.274614244 | -2.338152154 | 13.66850366 | 13.11877686 | 13.67337272 | 11.58558106 | 11.6507048 | 12.84207702 | 13.25435596 |
| ENSG00000154822 | PLCL2 | -1.402842217 | 0.336743265 | -0.937496201 | 0.527954208 | -2.340338417 | 3.327711234 | 7.002537973 | 7.818184403 | 5.578054239 | 5.6121243 | 6.14330814 | 5.900914815 |
| ENSG00000166166 | TRMT61A | -1.325764294 | 0.541643419 | -1.014846099 | 0.643682096 | -2.340610393 | 9.858207046 | 7.431218269 | 2 | 7.062428156 | 7.36516374 | 7.577730839 | 7.470508284 |
| ENSG00000080603 | SRCAP | 0.732592001 | 0.076242146 | -3.074283342 | 2.18E-17 | -2.341691342 | 9.839306291 | 9.267380431 | 9.990713521 | 10.40683041 | 10.5056622 | 6.603610344 | 6.795689611 |
| ENSG00000162999 | DUSP19 | -2.017941933 | 0.064138247 | -0.324818287 | 0.794186359 | -2.34276022 | 4.00722531 | 5.595927442 | 6.322058544 | 3.709785322 | 4.05806467 | 5.669467138 | 4.77533149 |
| ENSG00000135845 | PIGC | -1.309772255 | 0.051823689 | -1.033061409 | 0.127753231 | -2.342833664 | 7.624139768 | 8.968383114 | 7.858115942 | 7.003290579 | 7.00136498 | 7.043805932 | 7.467748741 |
| ENSG00000204815 | TTC25 | -4.714896876 | 0.259172295 | 2.371842652 | 0.544666992 | -2.343054224 | 2 | 3.076665209 | 2 | 2 | 2 | 3.584558147 | 4.047883918 |
| ENSG00000086730 | LAT2 | -1.096798019 | 0.495803512 | -1.246960318 | 0.421398024 | -2.343758338 | 2 | 5.131603625 | 5.772716771 | 4.17310023 | 3.94787299 | 4.220168345 | 3.620995423 |
| ENSG00000125871 | MGME1 | -1.01947662 | 0.590423475 | -1.324883836 | 0.466737112 | -2.344360456 | 9.839306291 | 7.638155693 | 3.369317972 | 7.433701013 | 7.65252709 | 7.528804029 | 6.90172361 |
| ENSG00000171208 | NETO2 | 0.070757617 | 0.957153132 | -2.415742104 | 0.021928563 | -2.344984486 | 5.095959458 | 7.819099893 | 7.896971941 | 7.454228844 | 7.44169935 | 5.498346994 | 4.56420962 |
| ENSG00000204131 | NHSL2 | -5.041827371 | 2.82E-11 | 2.695341458 | 4.13E-05 | -2.346485913 | 6.653098118 | 6.602424674 | 6.629488387 | 3.162721474 | 2.2680111 | 9.371930177 | 9.166759824 |
| ENSG00000143198 | MGST3 | -0.875260002 | 0.03287065 | -1.471529449 | 0.000125997 | -2.346789451 | 10.6695979 | 9.916994052 | 10.09465441 | 9.377433547 | 9.40446877 | 8.780980934 | 8.817088095 |
| ENSG00000167699 | GLOD4 | -1.30548518 | 0.003515112 | -1.041406181 | 0.022449102 | -2.346891362 | 9.895281535 | 8.94259736 | 9.503962296 | 8.237082658 | 8.16464176 | 8.44290591 | 8.481080411 |
| ENSG00000103253 | HAGHL | -1.913144243 | 0.253947704 | -0.434737665 | 0.815972748 | -2.347881908 | 3.327711234 | 6.140560021 | 8.561105505 | 5.242616075 | 5.6121243 | 7.170014349 | 6.38329369 |
| ENSG00000171735 | CAMTA1 | -1.108577622 | 0.000797451 | -1.239908348 | 0.000123316 | -2.34848597 | 8.737455672 | 9.102512015 | 8.784770914 | 7.839502934 | 7.74878077 | 7.523500045 | 7.796124663 |
| ENSG00000149564 | ESAM | -3.553350736 | 0.191751955 | 1.204423273 | 0.682126239 | -2.348927463 | 7.53297911 | 2 | 2 | 3.162721474 | 3.2184422 | 7.954782214 | 5.230851202 |
| ENSG00000138382 | METTL5 | -1.195804692 | 0.057338722 | -1.155569744 | 0.062814356 | -2.351374436 | 8.892187472 | 8.124485672 | 7.451345288 | 7.088925745 | 7.11620658 | 7.127295835 | 7.156006254 |
| ENSG00000261236 | BOP1 | -1.765817308 | 0.315574141 | -0.586349597 | 0.758401903 | -2.352166905 | 10.05943807 | 8.968383114 | 3.369317972 | 7.273514749 | 7.32531744 | 8.432756021 | 8.483814596 |
| ENSG00000094880 | CDC23 | -1.544722217 | 0.002664855 | -0.808647298 | 0.138024336 | -2.353369515 | 9.112843761 | 8.791896851 | 9.849281147 | 7.697489359 | 7.87709394 | 8.667743558 | 8.345600992 |
| ENSG00000131196 | NFATC1 | -0.537446258 | 0.603040858 | -1.817241866 | 0.044024163 | -2.354688124 | 6.366709671 | 7.847167909 | 8.884815469 | 7.481152682 | 7.5240115 | 6.341987719 | 6.189227296 |
| ENSG00000188312 | CENPP | -2.391385183 | 0.105206276 | 0.035740098 | 0.983983084 | -2.355645086 | 5.866944937 | 5.303523747 | 2 | 3.289987342 | 3.28037825 | 4.68289886 | 5.464725981 |
| ENSG00000122033 | MTIF3 | -1.093634091 | 0.277913797 | -1.263630306 | 0.19380488 | -2.357264397 | 9.957954117 | 8.191702463 | 7.552162999 | 7.821180652 | 7.89700047 | 7.653495316 | 7.730874172 |
| ENSG00000179604 | CDC42EP4 | -1.662896303 | 0.013325886 | -0.694414606 | 0.34077058 | -2.357310909 | 10.84565509 | 9.159587375 | 9.945092785 | 8.512996295 | 8.4705529 | 9.43006211 | 9.474701772 |
| ENSG00000165516 | KLHDC2 | -1.577325208 | 2.50E-05 | -0.780342398 | 0.050340543 | -2.357667606 | 9.966689162 | 9.405920729 | 9.25752426 | 8.022463952 | 8.00412803 | 8.809589199 | 8.791936295 |
| ENSG00000137207 | YIPF3 | -1.002572688 | 0.100199697 | -1.355829046 | 0.020018787 | -2.358401735 | 12.24682774 | 10.87022386 | 11.73893313 | 10.68241243 | 10.761874 | 10.21716837 | 10.50889441 |
| ENSG00000143891 | GALM | -0.365255376 | 0.678162682 | -1.993707682 | 0.006618212 | -2.358963058 | 8.928409675 | 8.929529568 | 7.163363831 | 8.136938566 | 8.23084153 | 6.791847192 | 6.371531035 |
| ENSG00000086827 | ZW10 | -0.708521021 | 0.643838836 | -1.650567202 | 0.23711641 | -2.359088223 | 9.602033055 | 8.395519306 | 4.874587708 | 7.893113669 | 7.85690889 | 6.931352719 | 6.979338179 |
| ENSG00000089091 | DZANK1 | -0.487562922 | 0.596116556 | -1.872776187 | 0.018343763 | -2.360339109 | 4.815609471 | 5.457120953 | 5.59507254 | 4.783303513 | 5.00774437 | 4.04171018 | 3.496556178 |
| ENSG00000138442 | WDR12 | -1.309718157 | 0.00730559 | -1.050653936 | 0.03362727 | -2.360372094 | 7.975256505 | 8.981105197 | 8.608698767 | 7.178012736 | 7.4074129 | 7.617106703 | 7.480125329 |
| ENSG00000102901 | CENPT | -1.586388482 | 0.015795913 | -0.774368531 | 0.268917609 | -2.360757014 | 8.631580145 | 9.067150531 | 10.11951029 | 7.826439377 | 7.85690889 | 8.437839892 | 8.822498333 |
| ENSG00000165525 | NEMF | -0.581379031 | 0.292697717 | -1.78044759 | 0.000219832 | -2.361826621 | 10.31337612 | 9.664384941 | 9.212024683 | 9.252121432 | 9.19330273 | 7.962642137 | 8.106655408 |
| ENSG00000189241 | TSPYL1 | -1.242898683 | 0.026251815 | -1.119507998 | 0.045942033 | -2.362406682 | 11.03197509 | 11.06803102 | 12.09069073 | 10.26540233 | 10.2237702 | 10.18023182 | 10.53367986 |
| ENSG00000196937 | FAM3C | -0.523693312 | 0.41446225 | -1.83878744 | 0.000872643 | -2.362480752 | 9.358295023 | 9.192779271 | 9.963514551 | 9.097585392 | 8.94225201 | 7.148124077 | 8.133891722 |
| ENSG00000115317 | HTRA2 | -0.768904034 | 0.420476172 | -1.593941173 | 0.066662083 | -2.362845206 | 8.586954094 | 6.899080544 | 9.212024683 | 7.748245381 | 7.75424764 | 7.153627862 | 6.704867638 |
| ENSG00000138101 | DTNB | -1.927012004 | 0.236297945 | -0.436282743 | 0.808868142 | -2.363294747 | 3.327711234 | 7.051611315 | 4.058998049 | 4.329433326 | 3.82856379 | 5.233158834 | 5.470251342 |
| ENSG00000154914 | USP43 | -3.260477536 | 0.180554595 | 0.896722638 | 0.719815802 | -2.363754897 | 2 | 4.710564154 | 2 | 2.270786764 | 2.2680111 | 3.517841605 | 4.692661645 |
| ENSG00000160271 | RALGDS | 0.378887015 | 0.822240381 | -2.743139454 | 0.04807007 | -2.364252439 | 4.467311984 | 8.169643011 | 9.503962296 | 8.772033429 | 8.83927354 | 5.873348082 | 5.680297397 |
| ENSG00000196655 | TRAPPC4 | -1.211055401 | 0.239632794 | -1.154136623 | 0.257017948 | -2.365192024 | 10.76782679 | 9.405920729 | 7.97167868 | 8.58612524 | 8.60672783 | 8.596789458 | 8.706733286 |
| ENSG00000128973 | CLN6 | -0.798504988 | 0.65512513 | -1.567601677 | 0.345156474 | -2.366106665 | 3.327711234 | 7.002537973 | 8.904018013 | 6.965655398 | 6.84106088 | 5.773219597 | 6.479583725 |
| ENSG00000173812 | EIF1 | -1.464644325 | 0.001638082 | -0.904660897 | 0.063520955 | -2.369305223 | 13.61880139 | 12.90092286 | 12.62979741 | 11.59003251 | 11.7051743 | 12.24770346 | 12.16749172 |
| ENSG00000140990 | NDUFB10 | -1.249015401 | 0.024854983 | -1.120870943 | 0.044632457 | -2.369886344 | 10.72727936 | 9.656475478 | 9.714382178 | 8.894758143 | 8.86223882 | 8.86850882 | 9.131014727 |
| ENSG00000158815 | FGF17 | -1.996104488 | 0.484805583 | -0.373850616 | NA | -2.369955103 | 2 | 3.686303414 | 2 | 2.270786764 | 2.2680111 | 2.911488689 | 2.466008509 |
| ENSG00000092978 | GPATCH2 | -0.866295714 | 0.314797955 | -1.505481679 | 0.059927165 | -2.371777393 | 7.867441304 | 7.901717006 | 9.541233192 | 7.789219291 | 7.82349365 | 7.367655602 | 6.967665897 |
| ENSG00000135917 | SLC19A3 | -0.277912652 | 0.929574989 | -2.093957699 | 0.446373473 | -2.371870351 | 2 | 2 | 4.058998049 | 3.162721474 | 2.49394783 | 2.42098416 | 2.172608095 |
| ENSG00000114395 | CYB561D2 | -0.781741296 | 0.335570391 | -1.590245828 | 0.032165651 | -2.371987124 | 7.218819781 | 8.029695761 | 9.014156722 | 7.416368504 | 7.58049694 | 6.473229652 | 6.92404095 |
| ENSG00000198457 | | 2.816988068 | 0.628255701 | -5.191095039 | NA | -2.374106971 | 2 | 2 | 2 | 3.289987342 | 3.65239003 | 2 | 2 |
| ENSG00000230048 | | 2.816988068 | 0.628255701 | -5.191095039 | NA | -2.374106971 | 2 | 2 | 2 | 3.289987342 | 3.65239003 | 2 | 2 |
| ENSG00000135437 | RDH5 | -1.963667222 | 0.273150492 | -0.411518528 | 0.836906381 | -2.37518575 | 7.485143553 | 5.838937808 | 2 | 4.574143677 | 4.53541753 | 5.338397457 | 6.356691554 |
| ENSG00000111142 | METAP2 | -1.107829734 | 0.375381504 | -1.267459319 | 0.294668948 | -2.375289052 | 11.61879218 | 10.28850895 | 7.896971941 | 9.481935579 | 9.49704586 | 9.394477264 | 9.264206697 |
| ENSG00000103528 | SYT17 | -2.006951012 | 0.464078443 | -0.368596857 | 0.9017845 | -2.37554787 | 2 | 5.131603625 | 2 | 3.09461492 | 2.2680111 | 3.993419637 | 3.128433649 |
| ENSG00000129347 | KRI1 | -0.892533283 | 0.357725763 | -1.483067682 | 0.101181698 | -2.375600965 | 9.247966931 | 6.78762695 | 8.296287027 | 7.565338955 | 7.5240115 | 6.91847879 | 7.019463152 |
| ENSG00000162236 | STX5 | -1.258619398 | 3.89E-05 | -1.118255254 | 0.000269416 | -2.376874652 | 10.54098126 | 10.20962523 | 10.06083273 | 8.995070693 | 9.06830201 | 9.270148001 | 9.065448786 |
| ENSG00000160310 | PRMT2 | -1.542002651 | 0.028640021 | -0.835007287 | 0.261300134 | -2.377009938 | 11.86526254 | 10.06176125 | 11.03653965 | 9.656695015 | 9.60158363 | 10.27085842 | 10.39367729 |
| ENSG00000064225 | ST3GAL6 | -0.776786731 | 0.542758789 | -1.600543405 | 0.171406992 | -2.377330137 | 5.532353208 | 5.595927442 | 7.858115942 | 6.069927018 | 5.96314905 | 5.699879814 | 4.624919101 |
| ENSG00000115694 | STK25 | -1.856753364 | 2.35E-05 | -0.521281042 | 0.294091971 | -2.378034407 | 11.75612888 | 10.87363932 | 11.01462253 | 9.349396108 | 9.48312089 | 10.65609247 | 10.83475651 |
| ENSG00000129910 | CDH15 | -2.839931294 | 0.278879149 | 0.461880553 | 0.8682894 | -2.378050741 | 4.467311984 | 2 | 2 | 2.141737246 | 2.38539787 | 3.393105702 | 3.771796761 |
| ENSG00000115268 | RPS15 | -1.518581679 | 0.045000662 | -0.859539528 | 0.278339715 | -2.378121207 | 12.8217977 | 11.42768197 | 11.04521411 | 10.42210785 | 10.5020111 | 11.06251538 | 11.17557911 |
| ENSG00000089280 | FUS | -1.138176776 | 0.115157031 | -1.242427765 | 0.078025727 | -2.380604541 | 11.75864596 | 10.28338212 | 10.67226762 | 9.919767943 | 9.90107802 | 9.383101353 | 10.13371468 |
| ENSG00000124532 | MRS2 | -1.098632886 | 0.044976341 | -1.282667678 | 0.016076822 | -2.381300563 | 9.533135543 | 8.414402626 | 8.742726175 | 7.944803394 | 7.82609171 | 7.621085148 | 7.786191022 |
| ENSG00000129518 | EAPP | -1.125236709 | 0.23034597 | -1.256491817 | 0.167672785 | -2.381728526 | 7.667647223 | 8.968383114 | 10.11951029 | 8.14116509 | 8.11238697 | 7.968118715 | 8.026414333 |
| ENSG00000092853 | CLSPN | -0.298340196 | 0.795427588 | -2.083461977 | 0.028515268 | -2.381802172 | 4.00722531 | 6.04685525 | 6.073378139 | 5.363304456 | 5.39133044 | 3.616794016 | 4.133950982 |
| ENSG00000163950 | SLBP | -1.051182924 | 0.123980108 | -1.334033596 | 0.042011267 | -2.38521652 | 10.62062599 | 9.225224669 | 9.452720984 | 8.85414072 | 8.86097251 | 8.75224741 | 8.37829627 |
| ENSG00000091622 | PITPNM3 | -2.76033306 | 0.217903135 | 0.373749869 | 0.885335383 | -2.38658319 | 8.418903792 | 3.686303414 | 2 | 4.470469939 | 4.31639476 | 6.710164669 | 7.662532142 |
| ENSG00000109332 | UBE2D3 | -0.396707135 | 0.530264818 | -1.990650103 | 0.000153586 | -2.387357238 | 12.78523054 | 11.5838842 | 11.87409387 | 11.82063682 | 11.7367574 | 10.23350246 | 10.14239308 |
| ENSG00000171612 | SLC25A33 | -1.418797749 | 0.11191982 | -0.969552463 | 0.284322026 | -2.388350213 | 7.82963488 | 7.051611315 | 5.59507254 | 5.786686927 | 5.69405321 | 5.951260634 | 6.344709081 |
| ENSG00000165775 | FUNDC2 | -0.939485128 | 0.050150442 | -1.44927295 | 0.001318425 | -2.388758079 | 9.858207046 | 9.181799937 | 8.978369997 | 8.461536372 | 8.45385092 | 7.801948045 | 8.093289637 |
| ENSG00000115761 | NOL10 | -0.801389379 | 0.032486453 | -1.587656244 | 4.20E-06 | -2.389045622 | 9.064836028 | 8.505280662 | 8.561105505 | 7.975897842 | 7.89946959 | 7.191577434 | 7.145684098 |
| ENSG00000100897 | DCAF11 | -1.654492672 | 0.075396159 | -0.735273417 | 0.459209794 | -2.389766089 | 8.928409675 | 9.055168103 | 10.9833682 | 8.349110081 | 8.33666436 | 8.91583461 | 9.52733357 |
| ENSG00000130811 | EIF3G | -0.945135751 | 0.055248319 | -1.445048181 | 0.001888215 | -2.390183932 | 11.20805444 | 10.23640192 | 10.49998282 | 9.759931695 | 9.77093593 | 9.16404896 | 9.365527647 |
| ENSG00000112339 | HBS1L | -1.489994623 | 0.020769965 | -0.900969844 | 0.179539773 | -2.390964467 | 10.58702944 | 9.04308532 | 10.15199903 | 8.595396228 | 8.5606705 | 9.333651532 | 8.967268538 |
| ENSG00000100284 | TOM1 | -0.886990838 | 0.234289292 | -1.504192442 | 0.030748092 | -2.391183279 | 11.39439513 | 9.608085847 | 10.95602907 | 9.898079272 | 9.98833077 | 9.335468714 | 9.324171817 |
| ENSG00000221923 | ZNF880 | -2.00199728 | 1.69E-06 | -0.389639827 | 0.414555096 | -2.391637107 | 7.867441304 | 7.393647515 | 7.451345288 | 5.808110275 | 5.51241389 | 7.320725255 | 7.071284046 |
| ENSG00000125743 | SNRPD2 | -1.698215173 | 0.012684784 | -0.695023133 | 0.348375224 | -2.393238306 | 11.59363617 | 10.11467741 | 10.31867969 | 9.121312924 | 9.16279627 | 9.895138115 | 10.34976357 |
| ENSG00000090534 | THPO | -1.514149752 | 0.557788547 | -0.879231858 | 0.735231663 | -2.39338161 | 4.467311984 | 2 | 2 | 2.498691189 | 2.68924543 | 2.803598391 | 2.885414003 |
| ENSG00000136518 | ACTL6A | -0.760550136 | 0.577531912 | -1.633217922 | 0.19059348 | -2.393768059 | 9.624286623 | 8.747909548 | 5.59507254 | 7.905589775 | 8.0358969 | 7.14536429 | 7.091143645 |
| ENSG00000163933 | RFT1 | -1.071573789 | 0.149609361 | -1.322315687 | 0.064234196 | -2.393889476 | 7.485143553 | 8.820493869 | 9.242516611 | 7.737119576 | 7.51759662 | 7.247980242 | 7.512619696 |
| ENSG00000143799 | PARP1 | -2.104812025 | 0.016039408 | -0.289180365 | 0.78196159 | -2.39399239 | 8.075571721 | 9.7032946 | 10.5916593 | 7.728718552 | 7.67287587 | 9.404322061 | 9.583539815 |
| ENSG00000186792 | HYAL3 | -1.145310131 | 0.410023637 | -1.248914557 | 0.354839138 | -2.394224688 | 9.384607364 | 5.838937808 | 7.163363831 | 6.979883891 | 7.1246732 | 6.320128742 | 7.392628644 |
| ENSG00000124702 | KLHDC3 | -1.529057663 | 0.008765708 | -0.865309946 | 0.158640059 | -2.394367609 | 11.34500123 | 9.943237334 | 10.69449487 | 9.189220094 | 9.30274583 | 9.827882244 | 9.982668878 |
| ENSG00000088876 | ZNF343 | -1.168952059 | 0.015631681 | -1.226111886 | 0.009845724 | -2.395063945 | 7.709880939 | 8.004992147 | 8.65477198 | 7.066878356 | 6.99214046 | 7.036377096 | 6.909879006 |
| ENSG00000120333 | MRPS14 | -1.650476833 | 0.00097459 | -0.745411943 | 0.165549719 | -2.395888775 | 8.444143041 | 8.395519306 | 9.272377397 | 7.062428156 | 7.20671394 | 7.801948045 | 8.213468956 |
| ENSG00000168216 | LMBRD1 | -0.996266532 | 0.093060766 | -1.401943262 | 0.012771461 | -2.398209794 | 9.509418609 | 9.949724236 | 8.631919287 | 8.572106084 | 8.37442884 | 8.130430649 | 8.019835704 |
| ENSG00000149499 | EML3 | -1.262960871 | 0.127215703 | -1.135338823 | 0.167152492 | -2.398299695 | 10.59831544 | 8.834582582 | 8.941672936 | 8.408172799 | 8.48542112 | 8.643989069 | 8.499436383 |
| ENSG00000135912 | TTLL4 | -1.135490385 | 0.115204374 | -1.264480641 | 0.071132677 | -2.399971026 | 7.53297911 | 8.762721501 | 8.904018013 | 7.337530098 | 7.46188679 | 7.637872201 | 6.78907279 |
| ENSG00000135336 | ORC3 | -0.818107797 | 0.114510897 | -1.581917597 | 0.000867841 | -2.400025394 | 8.39321512 | 9.424669273 | 9.287079172 | 8.266399315 | 8.3220138 | 7.526684775 | 7.56002886 |
| ENSG00000133884 | DPF2 | -1.333195971 | 2.62E-06 | -1.068290536 | 0.000198102 | -2.401486507 | 10.43166252 | 10.08551947 | 10.12770132 | 8.94033939 | 8.85080188 | 9.276783254 | 9.030812517 |
| ENSG00000185122 | HSF1 | -1.316295693 | 0.084119732 | -1.085853344 | 0.154872063 | -2.402149037 | 10.06758145 | 10.03150247 | 11.62203638 | 9.454177434 | 9.4789996 | 9.748725972 | 9.641010317 |
| ENSG00000148120 | C9orf3 | -2.207939752 | 0.002472293 | -0.194521891 | 0.83065631 | -2.402461643 | 10.30651143 | 8.590771778 | 8.845626171 | 7.307758725 | 7.24992376 | 9.253264914 | 9.275296471 |
| ENSG00000129194 | SOX15 | -1.692273371 | 0.513268686 | -0.710400391 | 0.792987538 | -2.402673763 | 5.70932728 | 2 | 2 | 3.09461492 | 3.2184422 | 3.968654369 | 3.560117069 |
| ENSG00000170266 | GLB1 | -0.774713475 | 0.312204876 | -1.62798819 | 0.020041931 | -2.402701665 | 12.04303957 | 10.54992047 | 10.56174313 | 10.44237204 | 10.480321 | 9.599435435 | 9.623450463 |
| ENSG00000156500 | FAM122C | -0.921006404 | 0.185725588 | -1.482598634 | 0.022298533 | -2.403605039 | 5.532353208 | 6.78762695 | 6.718950348 | 5.639097847 | 5.55061739 | 5.201624686 | 4.967301796 |
| ENSG00000128602 | SMO | -2.447779376 | 0.125723688 | 0.042814771 | 0.982373817 | -2.404964606 | 8.586954094 | 9.338312924 | 3.369317972 | 6.130704144 | 6.00043082 | 8.600320517 | 8.359810397 |
| ENSG00000171953 | ATPAF2 | -1.3360803 | 0.362647422 | -1.069861039 | 0.467461223 | -2.405941338 | 8.696033109 | 8.376385542 | 4.058998049 | 6.733448311 | 6.64900057 | 6.770513376 | 7.107191496 |
| ENSG00000166819 | PLIN1 | -2.917714303 | 0.414321642 | 0.5103855 | 0.893463941 | -2.407328802 | 2 | 4.11352228 | 2 | 2.3892346 | 2 | 3.851628042 | 2.782576693 |
| ENSG00000179115 | FARSA | -1.131057795 | 0.079842495 | -1.276697175 | 0.042070567 | -2.40775497 | 9.761125267 | 9.443177293 | 10.82615578 | 8.992733751 | 9.0282492 | 8.758581951 | 8.966291302 |
| ENSG00000137713 | PPP2R1B | -0.617945508 | 0.555396013 | -1.791555442 | 0.054407162 | -2.40950095 | 8.075571721 | 9.849218119 | 10.79563557 | 9.343906001 | 9.32771142 | 8.151214006 | 8.195234077 |
| ENSG00000100325 | ASCC2 | -1.310325259 | 0.141361699 | -1.100083477 | 0.217009028 | -2.410408736 | 10.36714848 | 10.40166101 | 8.175061096 | 8.567402595 | 8.72280968 | 8.788960186 | 8.91909319 |
| ENSG00000002330 | BAD | -1.316928886 | 0.010257088 | -1.093999654 | 0.035245998 | -2.41092854 | 10.19941743 | 9.680074956 | 9.06622548 | 8.378048624 | 8.4504872 | 8.498570381 | 8.760149043 |
| ENSG00000228474 | OST4 | -0.950055588 | 0.255415577 | -1.461186493 | 0.063181453 | -2.411242081 | 11.93368604 | 10.24697497 | 10.28245311 | 10.14060551 | 10.0678647 | 9.421247846 | 9.751494798 |
| TXNRD3NB | | 2.779646541 | 0.633071268 | -5.191095194 | NA | -2.411448653 | 2 | 2 | 2 | 3.754611435 | 3.08597292 | 2 | 2 |
| ENSG00000125991 | ERGIC3 | -0.622827783 | 0.227473008 | -1.788743136 | 8.35E-05 | -2.411570919 | 12.34389158 | 11.5089946 | 11.58086679 | 11.22647859 | 11.2550054 | 9.838386291 | 10.28324346 |
| CCL4L1 | CCL4L1 | 2.77917106 | 0.633108144 | -5.191095196 | NA | -2.411924136 | 2 | 2 | 2 | 3.709785322 | 3.15372745 | 2 | 2 |

| | | | | | | | | | | | | | |
|---|---|---|---|---|---|---|---|---|---|---|---|---|---|
| ENSG00000187147 | RNF220 | -1.614846515 | 7.06E-11 | -0.797787146 | 0.00214576 | -2.412633661 | 10.15425007 | 10.34373336 | 10.40882047 | 8.654247601 | 8.74631819 | 9.400276454 | 9.615992417 |
| ENSG00000183527 | PSMG1 | -1.483496717 | 0.065152211 | -0.929834309 | 0.262970634 | -2.413331026 | 9.613202746 | 7.60566235 | 8.585098385 | 7.348537858 | 7.37584259 | 7.726402307 | 8.063441666 |
| ENSG00000100216 | TOMM22 | -1.21282245 | 0.16375923 | -1.201348877 | 0.159642087 | -2.414171327 | 10.26462289 | 9.962611189 | 8.143093325 | 8.425603456 | 8.58234702 | 8.16558624 | 8.800186246 |
| ENSG00000161204 | ABCF3 | -0.697375118 | 0.144179754 | -1.717773543 | 5.72E-05 | -2.415148662 | 10.35389212 | 9.828248506 | 9.503962296 | 9.16555511 | 9.31943747 | 8.054501123 | 8.393643421 |
| ENSG00000233013 | FAM157B | 2.775837411 | 0.63346707 | -5.19109521 | NA | -2.415257799 | 2 | 2 | 2 | 3.349643765 | 3.55551834 | 2 | 2 |
| ENSG00000142156 | COL6A1 | -1.080159768 | 0.001097079 | -1.335120741 | 3.45E-05 | -2.415280509 | 15.47506374 | 15.88236009 | 16.10171637 | 14.74860836 | 14.7761579 | 14.53459064 | 14.47994585 |
| ENSG00000142408 | CACNG8 | 1.031173708 | 0.164591992 | -3.447823315 | 1.75E-07 | -2.416649607 | 7.031698319 | 6.04685525 | 7.098165388 | 7.731524332 | 7.8441483 | 4.099885429 | 3.36036153 |
| ENSG00000103510 | KAT8 | -1.405839974 | 0.202605116 | -1.011063154 | 0.365582948 | -2.416903127 | 7.579279349 | 6.899080544 | 9.624627483 | 7.053486362 | 7.21069602 | 7.562295432 | 7.474637727 |
| ARL16 | ARL16 | -1.689851255 | 0.059194088 | -0.727098403 | 0.44897681 | -2.416949658 | 10.3933013 | 8.820493869 | 8.143093325 | 7.767511442 | 7.76240929 | 8.585635246 | 8.833258322 |
| ENSG00000109084 | TMEM97 | -3.994036968 | 0.000161536 | 1.576876245 | 0.17143346 | -2.417160723 | 6.563806279 | 9.890264534 | 9.287079172 | 5.258265913 | 5.28958453 | 10.00444935 | 11.15428371 |
| ENSG00000176142 | TMEM39A | -1.061123227 | 0.025464282 | -1.356942512 | 0.003062764 | -2.418065739 | 10.06758145 | 10.62479571 | 9.681375057 | 9.088859521 | 9.15867957 | 8.975707507 | 8.669391068 |
| ENSG00000168303 | MPLKIP | -1.446987185 | 0.003251505 | -0.974791901 | 0.05485101 | -2.421779085 | 8.631580145 | 8.356994601 | 9.372276026 | 7.373900638 | 7.46855365 | 7.857806412 | 7.911296863 |
| ENSG00000196981 | WDR5B | -1.227634633 | 0.305395568 | -1.194966814 | 0.308265999 | -2.422601447 | 8.313285031 | 6.464272742 | 5.39244166 | 6.006476466 | 6.08095082 | 6.363520437 | 5.713290392 |
| ENSG00000100926 | TM9SF1 | 0.744337927 | 0.120540527 | -3.168215297 | 2.92E-14 | -2.423877371 | 9.730679447 | 9.680074956 | 8.845626171 | 10.17296029 | 10.2564019 | 6.285453595 | 6.449128733 |
| ENSG00000154928 | EPHB1 | 0.156378889 | 0.958503495 | -2.580359002 | 0.320401275 | -2.423980113 | 2 | 2 | 7.342949392 | 5.949776857 | 5.98191036 | 3.709392871 | 3.640730416 |
| ENSG00000165338 | HECTD2 | -1.130992805 | 0.497739811 | -1.293126407 | 0.423874761 | -2.424119212 | 3.327711234 | 8.395519306 | 8.805342642 | 6.965655398 | 6.9210099 | 6.920094328 | 6.640162436 |
| ENSG00000100380 | ST13 | -1.073394837 | 0.000288085 | -1.35292882 | 2.77E-06 | -2.426323656 | 11.71783599 | 11.21797502 | 11.41005835 | 10.41732378 | 10.3649055 | 10.05770853 | 10.16578017 |
| ENSG00000160014 | CALM3 | -1.507094011 | 8.30E-07 | -0.91924387 | 0.003997578 | -2.426337881 | 12.27171276 | 11.92243809 | 12.25299971 | 10.58958544 | 10.7125292 | 11.02190347 | 11.42774592 |
| ENSG00000244038 | DDOST | -0.53102967 | 0.196009997 | -1.895794324 | 9.77E-08 | -2.426823993 | 12.49946702 | 11.86696895 | 12.46016642 | 11.73985636 | 11.8032062 | 10.25766199 | 10.54730407 |
| ENSG00000198417 | MT1F | -0.457299918 | 0.864318577 | -1.972431597 | 0.397984664 | -2.429731514 | 5.866944937 | 3.076665209 | 2 | 4.139692846 | 4.22485516 | 3.724264246 | 2.225841198 |
| ENSG00000066427 | ATXN3 | -0.84829045 | 0.241359928 | -1.582866209 | 0.017961959 | -2.431156659 | 9.174465797 | 7.954278496 | 9.553445883 | 8.219202381 | 8.16874151 | 7.670871435 | 7.242500282 |
| ENSG00000120158 | RCL1 | -1.160191438 | 0.086279573 | -1.271025631 | 0.052104808 | -2.431217069 | 6.653098118 | 6.602424674 | 7.777116083 | 5.920568914 | 6.0722229 | 5.737015686 | 6.037449234 |
| ENSG00000037241 | RPL26L1 | -1.112851175 | 0.217853755 | -1.318925339 | 0.129597634 | -2.431776514 | 9.128497779 | 7.701025996 | 7.029880454 | 7.017154451 | 7.24216308 | 6.800644007 | 7.056667034 |
| ENSG00000164008 | C1orf50 | -0.972882603 | 0.362218166 | -1.459371044 | 0.146023342 | -2.432253647 | 4.815609471 | 7.393647515 | 7.398165132 | 6.024891728 | 5.98191036 | 5.630526941 | 5.470251342 |
| ENSG00000089199 | CHGB | 0.72863364 | 0.815259329 | -3.161053607 | NA | -2.432419967 | 2 | 3.076665209 | 2 | 2.947924294 | 2.77779907 | 2.094598369 | 2.059860178 |
| ENSG00000173137 | ADCK5 | -0.894661572 | 0.679561713 | -1.538575323 | 0.447991703 | -2.433236895 | 4.815609471 | 7.979858146 | 2 | 5.602782483 | 5.85535975 | 5.163941955 | 5.136701757 |
| ENSG00000071994 | PDCD2 | -1.345766343 | 0.085059099 | -1.087555679 | 0.165533397 | -2.433322022 | 10.46286899 | 9.488434476 | 8.486635645 | 8.328872678 | 8.3920703 | 8.694893015 | 8.532824248 |
| ENSG00000169288 | MRPL1 | -0.4523484 | 0.819093952 | -1.981316235 | 0.246641741 | -2.433664635 | 7.904282243 | 7.145017454 | 2 | 6.687949235 | 6.44101097 | 4.984105765 | 5.300743019 |
| ENSG00000126261 | UBA2 | -1.065176416 | 0.003556198 | -1.368805479 | 0.000108944 | -2.433981895 | 10.64803757 | 10.31889309 | 10.06936278 | 9.364838907 | 9.24178305 | 9.171528462 | 8.812201472 |
| ENSG00000166441 | RPL27A | -1.077582217 | 0.216892178 | -1.357541586 | 0.105389356 | -2.435123803 | 13.20898726 | 11.54156729 | 11.25418149 | 11.19022482 | 11.2114531 | 10.88757038 | 10.95453551 |
| ENSG00000188060 | RAB42 | -2.535197459 | 0.009053015 | 0.098780736 | 0.935481323 | -2.436416724 | 5.70932728 | 6.464272742 | 7.934808822 | 4.76150579 | 4.56059747 | 6.479817188 | 7.523290521 |
| ENSG00000092108 | SCFD1 | -0.311607502 | 0.200056839 | -2.125006438 | 3.30E-25 | -2.43661394 | 9.678463567 | 9.71095217 | 9.69246155 | 9.398771806 | 9.37016537 | 7.628021136 | 7.55613706 |
| ENSG00000092445 | TYRO3 | -1.4334481 | 0.466616843 | -1.00397755 | 0.614013611 | -2.437425651 | 8.696033109 | 8.762721501 | 2 | 6.820355514 | 6.68379438 | 7.450478673 | 6.81969491 |
| ENSG00000183066 | WBP2NL | -3.274201019 | 0.39960716 | 0.835741962 | NA | -2.438459057 | 2 | 3.076665209 | 2 | 2 | 2.14022021 | 2.775316226 | 3.042026731 |
| ENSG00000170791 | CHCHD7 | -0.927506885 | 0.219104153 | -1.511349067 | 0.032176413 | -2.438855952 | 8.963744667 | 7.638155693 | 7.451345288 | 7.238438152 | 7.30683517 | 6.621594001 | 6.78907279 |
| ENSG00000100154 | TTC28 | -1.434657469 | 0.044804086 | -1.005168975 | 0.169441906 | -2.439826444 | 9.128497779 | 10.24697497 | 10.92816184 | 8.924487942 | 8.77214691 | 9.133040681 | 9.407171982 |
| ENSG00000147912 | FBXO10 | -0.32217918 | 0.707966552 | -2.118278493 | 0.002715313 | -2.440457673 | 6.89194353 | 7.638155693 | 8.65477198 | 7.638916932 | 7.53675633 | 6.013120342 | 5.699242824 |
| ENSG00000189043 | NDUFA4 | -1.408724358 | 0.036457816 | -1.032224967 | 0.13261638 | -2.440949324 | 11.25516263 | 9.910357957 | 9.868959282 | 9.12771691 | 9.06390621 | 9.544459308 | 9.39196883 |
| ENSG00000011009 | LYPLA2 | -1.167205954 | 0.007244134 | -1.275329222 | 0.002785432 | -2.442535176 | 9.940323746 | 9.386925327 | 9.099922433 | 8.332573384 | 8.38855922 | 8.129732687 | 8.368713182 |
| ENSG00000107833 | NPM3 | -1.401625857 | 0.500480698 | -1.046764268 | 0.617051806 | -2.448390125 | 3.327711234 | 3.686303414 | 8.536706852 | 5.775854763 | 5.62411716 | 6.132200015 | 5.925257518 |
| ENSG00000125967 | NECAB3 | -0.968328445 | 0.249938781 | -1.483047119 | 0.06093067 | -2.451375565 | 9.317904228 | 7.431218269 | 9.344432377 | 7.915493556 | 8.01555453 | 7.576706933 | 7.336254871 |
| ENSG00000112149 | CD83 | -1.17879237 | 0.522626682 | -1.273926492 | 0.476851144 | -2.452718862 | 9.410448395 | 6.899080544 | 3.369317972 | 6.951285179 | 6.89649843 | 6.706424753 | 6.947999751 |
| ENSG00000165644 | COMTD1 | -1.441929243 | 0.538564148 | -1.010838243 | 0.672486215 | -2.452767486 | 2 | 3.076665209 | 7.502636426 | 4.670864177 | 4.74762503 | 4.965450964 | 5.191251321 |
| ENSG00000071539 | TRIP13 | -1.57508851 | 0.033658406 | -0.878145479 | 0.250261337 | -2.453233989 | 5.532353208 | 6.311476754 | 6.322058544 | 4.739373666 | 4.63360314 | 4.834511823 | 5.646532192 |
| ENSG00000206530 | CFAP44 | 0.064394262 | 0.976724726 | -2.517690141 | 0.165966488 | -2.453295879 | 2 | 5.303523747 | 7.777116083 | 6.573765157 | 6.44101097 | 4.45225155 | 3.924507493 |
| ENSG00000102710 | SUPT20H | -0.700168705 | 0.104334281 | -1.753769447 | 5.48E-06 | -2.453938151 | 10.04301211 | 10.00683004 | 9.399592459 | 9.240349764 | 9.05507424 | 8.27442094 | 7.919398473 |
| ENSG00000083457 | ITGAE | -1.732153583 | 0.204801787 | -0.721804029 | 0.62045962 | -2.453957612 | 8.816894989 | 7.393647515 | 4.523675391 | 5.987823101 | 6.14893615 | 6.855708005 | 7.198237561 |
| ENSG00000165572 | KBTBD6 | -1.715287526 | 0.016480221 | -0.739159184 | 0.337319235 | -2.45444671 | 7.33131396 | 8.57407404 | 9.031722608 | 6.744602355 | 6.84106088 | 8.015738956 | 7.408544534 |
| ENSG00000115539 | PDCL3 | -0.747051005 | 0.417929197 | -1.707457056 | 0.04003872 | -2.454508061 | 8.963744667 | 7.431218269 | 7.225742779 | 7.250225364 | 7.46522407 | 6.028179536 | 6.750993196 |
| ENSG00000164972 | C9orf24 | 0.180799142 | NA | -2.635600556 | NA | -2.454801415 | 2 | 3.076665209 | 2 | 2.141737246 | 3.01487888 | 2.048074405 | 2.172608095 |
| ENSG00000255071 | SAA2-SAA4 | 2.735440379 | 0.638989421 | -5.191095378 | NA | -2.455654999 | 2 | 2 | 2 | 3.615725078 | 3.2184422 | 2 | 2 |
| ENSG00000141448 | GATA6 | -1.774247105 | 0.217082182 | -0.681643909 | 0.660486629 | -2.455891014 | 5.095959458 | 9.461450882 | 9.963514551 | 7.481152682 | 7.37229174 | 8.644477784 | 8.344849238 |
| ENSG00000100351 | GRAP2 | 2.734950075 | 0.638989421 | -5.19109538 | NA | -2.456145305 | 2 | 2 | 2 | 3.566290607 | 3.28037825 | 2 | 2 |
| ENSG00000143452 | HORMAD1 | 2.734950075 | 0.638989421 | -5.19109538 | NA | -2.456145305 | 2 | 2 | 2 | 3.566290607 | 3.28037825 | 2 | 2 |
| ENSG00000168028 | RPSA | -0.660448224 | 0.125657681 | -1.795976725 | 2.78E-06 | -2.456424949 | 12.12920672 | 11.81817171 | 11.29839064 | 11.09892761 | 11.1580635 | 9.950137104 | 10.04115072 |
| ENSG00000145248 | SLC10A4 | -0.654581947 | 0.834689821 | -1.802813409 | 0.524456316 | -2.457395356 | 4.00722531 | 2 | 2 | 2.947924294 | 2.38539787 | 2.183373728 | 2.509591799 |
| ENSG00000128266 | GNAZ | -3.495302408 | 0.078054688 | 1.037158048 | 0.631197419 | -2.458144361 | 7.218819781 | 3.686303414 | 2 | 3.406930997 | 2.68924543 | 6.639356243 | 6.881131115 |
| ENSG00000115657 | ABCB6 | 0.23366888 | 0.897633629 | -2.694035618 | 0.066291657 | -2.460366738 | 3.327711234 | 7.538394143 | 8.65477198 | 7.810605345 | 7.92635454 | 5.147485955 | 5.04340913 |
| ENSG00000171291 | ZNF439 | -1.476717063 | 0.097724298 | -0.985806346 | 0.278635374 | -2.462523409 | 8.777722034 | 6.464272742 | 8.266922072 | 6.761173376 | 6.58294721 | 7.190239142 | 7.107191496 |
| ENSG00000157911 | PEX10 | -1.438990687 | 0.010939322 | -1.024994481 | 0.077118013 | -2.463985168 | 9.174465797 | 7.928237109 | 8.511888463 | 7.136275664 | 7.27676203 | 7.489084836 | 7.726263462 |
| ENSG00000145388 | METTL14 | -1.005462991 | 0.326315684 | -1.459239239 | 0.131842802 | -2.46470223 | 9.613202746 | 7.315436877 | 9.981704053 | 8.395844809 | 8.29788471 | 8.092965353 | 7.678134128 |
| ENSG00000120833 | SOCS2 | -1.703404432 | 0.007905366 | -0.764833759 | 0.26685414 | -2.468238191 | 9.048471759 | 8.590771778 | 7.451345288 | 6.902323594 | 6.77267312 | 7.744751472 | 7.761616892 |
| ENSG00000078177 | N4BP2 | -0.742726027 | 0.674378575 | -1.72719082 | 0.282968343 | -2.469916847 | 4.815609471 | 9.386925327 | 5.930865648 | 7.373900638 | 7.13728069 | 6.484192226 | 6.117095472 |
| ENSG00000108010 | GLRX3 | -0.927438342 | 0.186621771 | -1.543306874 | 0.01873645 | -2.470745216 | 10.47516499 | 8.820493869 | 9.788583153 | 8.951212326 | 8.89106117 | 8.281350892 | 8.341838298 |
| ENSG00000176973 | FAM89B | -0.650222403 | 0.135425564 | -1.820824546 | 2.32E-06 | -2.471046949 | 9.579430841 | 8.955547846 | 8.978369997 | 8.508095759 | 8.59915299 | 7.546693093 | 7.242500282 |
| ENSG00000141577 | CEP131 | -1.503146073 | 0.349375154 | -0.968407835 | 0.554691707 | -2.471553908 | 4.00722531 | 6.602424674 | 8.978369997 | 6.29903474 | 6.1572134 | 6.768721236 | 6.711881188 |
| ENSG00000156171 | DRAM2 | -1.378331148 | 0.05355269 | -1.097398895 | 0.127604991 | -2.475730043 | 9.810482605 | 8.505280662 | 9.963514551 | 8.205140657 | 8.17078703 | 8.828637742 | 7.978752403 |
| ENSG00000113240 | CLK4 | -1.282797327 | 0.459733797 | -1.193154164 | 0.483761199 | -2.475951491 | 3.327711234 | 8.255926468 | 9.272377397 | 7.080147179 | 6.96881778 | 7.402744819 | 6.748721609 |
| ENSG00000163694 | RBM47 | 0.221322555 | 0.912449258 | -2.698238287 | 0.099707517 | -2.476915732 | 3.327711234 | 8.124485672 | 5.930865648 | 7.148924055 | 7.02868983 | 5.04460184 | 3.311948089 |
| ENSG00000163479 | SSR2 | -1.240945569 | 7.78E-06 | -1.236669843 | 7.69E-06 | -2.477615413 | 12.31339889 | 11.87380718 | 11.94533713 | 10.80101819 | 10.8343167 | 10.78391875 | 10.85917895 |
| ENSG00000090273 | NUDC | -1.318668884 | 2.14E-06 | -1.158976386 | 3.35E-05 | -2.47764527 | 10.81695923 | 10.57950211 | 10.78531698 | 9.449072612 | 9.38257346 | 9.382222556 | 9.745241486 |
| ENSG00000111885 | MAN1A1 | -0.972724244 | 0.410778027 | -1.505252507 | 0.174488324 | -2.477976751 | 8.138771685 | 11.40661699 | 10.94679967 | 9.718812329 | 9.73198422 | 8.817412208 | 9.49551136 |
| ENSG00000092931 | MFSD11 | -0.497911016 | 0.506773631 | -1.980318906 | 0.001927913 | -2.478229923 | 9.590776211 | 8.053983481 | 8.585098385 | 8.385193462 | 8.39732094 | 7.042323222 | 6.828326017 |
| ENSG00000064490 | RFXANK | -0.550785144 | 0.41863397 | -1.92864787 | 0.001004097 | -2.479433014 | 8.716893051 | 8.732943944 | 7.398165132 | 7.942383523 | 7.78127547 | 6.535683256 | 6.514766253 |
| ENSG00000276429 | | -0.289992535 | 0.926059063 | -2.190312314 | NA | -2.480304848 | 2 | 3.686303414 | 2 | 2.868578001 | 2.59489879 | 2.34604626 | 2.117335275 |
| ENSG00000274675 | | 5.533347298 | 3.06E-07 | -8.014388639 | 2.83E-05 | -2.481041342 | 4.00722531 | 3.686303414 | 2 | 8.408172799 | 8.3920703 | 2 | 2 |
| ENSG00000047579 | DTNBP1 | -1.541893308 | 0.103759715 | -0.940463524 | 0.334815084 | -2.482356831 | 5.532353208 | 7.979858146 | 7.646393253 | 5.720439317 | 6.0722229 | 6.482006365 | 6.468583424 |
| ENSG00000240230 | COX19 | -1.191135624 | 0.481054906 | -1.292312467 | 0.432545818 | -2.483448091 | 4.467311984 | 8.433041974 | 10.11127249 | 7.734324667 | 7.7919464 | 8.060373167 | 7.116030435 |
| ENSG00000205138 | SDHAF1 | -1.912580883 | 1.14E-06 | -0.572103521 | 0.187327641 | -2.484684404 | 8.517314938 | 7.819099893 | 8.266922072 | 6.306580752 | 6.41363412 | 7.740186014 | 7.571641661 |
| ENSG00000132589 | FLOT2 | -1.791795935 | 0.001903145 | -0.693013459 | 0.273571221 | -2.484809394 | 11.59363617 | 10.19877383 | 10.76969399 | 9.134092596 | 9.23006009 | 10.27180772 | 10.28088727 |
| ENSG00000114503 | NCBP2 | -1.259509244 | 0.038369791 | -1.225589541 | 0.042265942 | -2.485098785 | 11.4295719 | 10.019219 | 10.59756881 | 9.56371781 | 9.52209753 | 9.542100683 | 9.61068621 |
| ENSG00000136270 | TBRG4 | -1.692754155 | 0.016213051 | -0.794770971 | 0.291201113 | -2.487525126 | 10.41260941 | 8.590771778 | 9.809100877 | 8.033874364 | 8.17078703 | 8.980360725 | 9.003921351 |
| ENSG00000204967 | PCDHA4 | -0.797552948 | 0.773531565 | -1.691651574 | 0.510353967 | -2.489204521 | 2 | 4.93638933 | 2 | 3.663521627 | 2.59489879 | 2.885263063 | 2.671842496 |
| ENSG00000172803 | SNX32 | -3.590444403 | 0.117068598 | 1.09847501 | 0.64228189 | -2.491969392 | 2 | 5.303523747 | 2 | 2.3892346 | 2.2680111 | 5.169385838 | 4.644600858 |
| ENSG00000101391 | CDK5RAP1 | -1.086453648 | 0.551434318 | -1.405731389 | 0.424102084 | -2.492185037 | 3.327711234 | 8.234834549 | 9.778213818 | 7.477814553 | 7.6378145 | 7.190239142 | 7.30412922 |
| ENSG00000104408 | EIF3E | -1.162283456 | 0.018126811 | -1.330553564 | 0.005669552 | -2.49283702 | 11.44528213 | 10.65289571 | 10.47451889 | 9.767470067 | 9.75874862 | 9.700188698 | 9.482925003 |
| ENSG00000197223 | C1D | -0.743765488 | 0.261139503 | -1.749680458 | 0.003253036 | -2.493445945 | 8.39321512 | 7.002537973 | 8.042706615 | 7.119236776 | 7.24992376 | 6.213498222 | 6.22561877 |
| ENSG00000138617 | PARP16 | -1.684196757 | 0.307444336 | -0.809832031 | 0.64013824 | -2.494028788 | 3.327711234 | 7.60566235 | 8.978369997 | 6.205262847 | 6.29861567 | 7.354572674 | 6.771278116 |
| ENSG00000164904 | ALDH7A1 | -1.379845873 | 0.169072016 | -1.115312513 | 0.267163049 | -2.495158386 | 6.366709671 | 9.125613817 | 8.460932916 | 7.057964187 | 6.98750599 | 7.17543541 | 7.378005688 |
| ENSG00000173486 | FKBP2 | -0.74082676 | 0.089224711 | -1.755905645 | 7.55E-06 | -2.496732405 | 10.09970294 | 9.532315021 | 9.61300511 | 9.033093526 | 9.03275482 | 7.749302529 | 8.26365177 |
| ENSG00000072195 | SPEG | -2.369885796 | 1.98E-27 | -0.126913143 | 0.64228189 | -2.496798939 | 10.34051281 | 10.20420974 | 10.16000817 | 7.930222906 | 7.84158259 | 10.16430247 | 10.05385647 |
| ENSG00000175376 | EIF1AD | -1.373307353 | 0.084538754 | -1.126137762 | 0.157821042 | -2.499445115 | 8.963744667 | 8.356994601 | 10.26027233 | 8.040677544 | 8.0713677 | 8.269359984 | 8.3297313 |
| ENSG00000136305 | CIDEB | 2.69135397 | 0.644717004 | -5.191095562 | NA | -2.499741591 | 2 | 2 | 2 | 3.615725078 | 3.15372745 | 2 | 2 |
| ENSG00000084072 | PPIE | -1.183533564 | 0.000170819 | -1.319550952 | 1.92E-05 | -2.503084516 | 9.031919741 | 9.506147172 | 9.385998894 | 8.083034979 | 8.21902896 | 8.080016699 | 7.954317658 |
| ENSG00000084623 | EIF3I | -1.258944205 | 0.016890353 | -1.244518511 | 0.017714167 | -2.503462716 | 12.02014453 | 11.31412071 | 10.89495479 | 10.21466712 | 10.244312 | 10.06210503 | 10.40544175 |
| ENSG00000166448 | TMEM130 | -3.306543218 | 0.015891709 | 0.80250793 | 0.600600207 | -2.504035288 | 5.095959458 | 4.11352228 | 7.600047325 | 3.406930997 | 3.69849104 | 6.484192226 | 7.55093161 |
| ENSG00000170638 | TRABD | -1.214990905 | 0.023974609 | -1.291229392 | 0.014940374 | -2.506220297 | 9.204317094 | 8.848535039 | 9.999667074 | 8.209172298 | 8.24255817 | 8.137391755 | 8.164036804 |
| ENSG00000026297 | RNASET2 | -0.604166158 | 0.39960716 | -1.90405453 | 0.002334223 | -2.508220688 | 9.344956677 | 7.901717006 | 9.344432377 | 8.409925372 | 8.39031583 | 7.156371901 | 7.09293558 |
| ENSG00000114316 | USP4 | -1.358396914 | 0.210105576 | -1.149886507 | 0.287769738 | -2.508283421 | 8.418903792 | 10.01303782 | 11.4724895 | 9.091045942 | 9.11791381 | 9.270781246 | 9.351374718 |
| ENSG00000132254 | ARFIP2 | -1.688715308 | 4.33E-05 | -0.820000426 | 0.064008656 | -2.508715734 | 10.76782679 | 9.981728319 | 10.20715117 | 8.637890117 | 8.71582187 | 9.363049315 | 9.697162691 |
| ENSG00000120438 | TCP1 | -1.343045428 | 0.014834268 | -1.167558676 | 0.035608878 | -2.510604104 | 11.69961208 | 10.56267266 | 10.70548138 | 9.730079663 | 9.75670736 | 9.880276569 | 9.955102659 |
| ENSG00000056998 | GYG2 | -1.275670648 | 0.542790331 | -1.235176457 | 0.549961603 | -2.510847105 | 5.095959458 | 8.101365653 | 2 | 5.627094009 | 5.39133044 | 5.185595313 | 5.842465082 |
| ENSG00000149929 | HIRIP3 | -1.920024056 | 0.049155716 | -0.591005282 | 0.58225381 | -2.511029338 | 7.975256505 | 7.669933291 | 5.39244166 | 5.363304456 | 5.71663161 | 7.027411703 | 6.538630327 |
| ENSG00000087884 | AAMDC | -0.508865694 | 0.361961292 | -2.002402428 | 2.64E-05 | -2.511268122 | 7.790810799 | 7.145017454 | 7.163363831 | 6.89733479 | 6.90143415 | 5.243518884 | 5.703940561 |
| ENSG00000206452 | | 7.351477013 | 1.74E-28 | -9.863379109 | 4.03E-19 | -2.511902097 | 8.39321512 | 6.899080544 | 7.451345288 | 15.03289325 | 15.033612 | 2.139668783 | 2 |
| ENSG00000189077 | TMEM120A | -1.013818542 | 0.218893333 | -1.500562585 | 0.053864969 | -2.514381127 | 8.107517749 | 8.234834549 | 9.788583153 | 7.88809286 | 7.95749494 | 7.277267764 | 7.597118029 |
| ENSG00000132383 | RPA1 | -1.800693141 | 1.50E-10 | -0.713940931 | 0.018611632 | -2.514634072 | 10.41260941 | 10.06176125 | 10.53119344 | 8.517880242 | 8.59306436 | 9.626899466 | 9.642234599 |

| | | | | | | | | | | | | | |
|---|---|---|---|---|---|---|---|---|---|---|---|---|---|
| ENSG00000162755 | KLHDC9 | -4.714897443 | 0.260096091 | 2.194821195 | 0.579335184 | -2.520076248 | 3.327711234 | 2 | 2 | 2 | 2 | 3.663836244 | 3.753786374 |
| ENSG00000023734 | STRAP | -1.585814145 | 4.55E-19 | -0.934286384 | 2.32E-07 | -2.52010053 | 11.53611196 | 11.59220064 | 11.46604358 | 9.97822644 | 9.92369416 | 10.57355555 | 10.6272262 |
| ENSG00000255302 | EID1 | -1.345382648 | 0.010846156 | -1.175038423 | 0.027547688 | -2.520421071 | 12.5114563 | 11.40897279 | 11.9030147 | 10.73544052 | 10.5971045 | 10.57624762 | 11.05929667 |
| ENSG00000025770 | NCAPH2 | -0.965401479 | 0.522626682 | -1.556370322 | 0.269833543 | -2.521771802 | 9.262228249 | 8.213429689 | 4.523675391 | 7.261917051 | 7.40393901 | 6.7543034 | 6.769038269 |
| ENSG00000155957 | TMBIM4 | 0.482971231 | 0.616184132 | -3.004908379 | 0.000143403 | -2.521937148 | 10.65885801 | 9.214490408 | 8.561105505 | 10.24001146 | 10.2267227 | 6.541992314 | 7.008112295 |
| ENSG00000186395 | KRT10 | -1.874933779 | 0.000458914 | -0.647948757 | 0.271141542 | -2.522882535 | 8.517314938 | 7.669933291 | 8.486635645 | 6.379946408 | 6.50077304 | 7.249266052 | 7.931466218 |
| ENSG00000179409 | GEMIN4 | -1.948956231 | 0.036163998 | -0.576433476 | 0.577440741 | -2.525389706 | 10.4253394 | 8.655694736 | 8.076950877 | 7.437142666 | 7.53992502 | 8.758581951 | 8.921111931 |
| ENSG00000133250 | ZNF414 | -1.439957176 | 0.158024792 | -1.090019896 | 0.288238766 | -2.529977072 | 6.138360429 | 7.572420273 | 8.865353883 | 6.554880361 | 6.48112613 | 6.759726988 | 6.944034147 |
| ENSG00000095370 | SH2D3C | -4.136307932 | 0.13805413 | 1.605422752 | 0.482371468 | -2.53088518 | 2 | 3.076665209 | 3.369317972 | 2.141737246 | 2 | 4.199031499 | 3.539239711 |
| ENSG00000064961 | HMG20B | -1.917185419 | 0.011775752 | -0.614019779 | 0.468899775 | -2.531205197 | 11.45151864 | 9.443177293 | 10.40208303 | 8.660150093 | 8.8354102 | 10.14940096 | 9.933764315 |
| ENSG00000187800 | PEAR1 | -4.029083255 | 0.000130593 | 1.497766824 | 0.191871601 | -2.53131643 | 9.48530526 | 7.928237109 | 6.073378139 | 4.54891529 | 4.70309173 | 9.944988515 | 9.866740633 |
| ENSG00000140382 | HMG20A | -1.101509743 | 0.259792504 | -1.430309444 | 0.125991152 | -2.531819187 | 8.107517749 | 9.856140801 | 10.75919294 | 8.80037045 | 8.87734839 | 8.501271034 | 8.526208204 |
| ENSG00000131475 | VPS25 | -1.716982965 | 3.83E-11 | -0.816160463 | 0.002803772 | -2.533143428 | 10.29961392 | 10.11467741 | 9.972637969 | 8.501535708 | 8.35477078 | 9.239097094 | 9.402483342 |
| ENSG00000249915 | PDCD6 | -1.418723484 | 0.007048618 | -1.114702138 | 0.037743848 | -2.533425621 | 10.98115017 | 9.994333773 | 9.954333071 | 8.957217624 | 9.00321357 | 9.201396821 | 9.358840932 |
| ENSG00000140740 | UQCRC2 | -1.164271916 | 0.084747221 | -1.369347333 | 0.036253153 | -2.533619249 | 11.78851328 | 10.75272247 | 10.23774519 | 9.914221922 | 9.91273599 | 9.670982446 | 9.746665075 |
| ENSG00000105875 | WDR91 | -1.142941207 | 0.348943914 | -1.391322395 | 0.236277861 | -2.534263602 | 6.816637977 | 6.602424674 | 9.287079172 | 6.941624891 | 7.05107174 | 7.039353222 | 6.406535141 |
| ENSG00000138439 | FAM117B | -1.603435067 | 0.095126919 | -0.931159035 | 0.348561445 | -2.534594102 | 6.009025997 | 8.376385542 | 7.163363831 | 6.12217689 | 5.78232107 | 6.835304498 | 6.317379004 |
| ENSG00000162676 | GFI1 | -1.149162646 | 0.64718241 | -1.386186799 | 0.564260714 | -2.535349444 | 2 | 4.442710129 | 2 | 2.784612485 | 2.77779907 | 2.62493832 | 2.746601469 |
| ENSG00000104823 | ECH1 | -1.064238361 | 0.065953631 | -1.471430642 | 0.007688776 | -2.535669004 | 10.66423795 | 9.558016677 | 9.735974718 | 8.986874783 | 9.03724641 | 8.381493453 | 8.805113667 |
| ENSG00000163319 | MRPS18C | 0.736875933 | 0.336111036 | -3.275694411 | 6.73E-07 | -2.538818478 | 7.384433967 | 6.844429755 | 5.772716771 | 7.433701013 | 7.61995902 | 3.724264246 | 4.105828626 |
| ENSG00000115163 | CENPA | -1.034967767 | 0.594008398 | -1.505836243 | 0.408557169 | -2.54080401 | 3.327711234 | 4.442710129 | 2 | 2.868578001 | 3.15372745 | 2.592885947 | 2.981406154 |
| ENSG00000099817 | POLR2E | -1.462215913 | 6.63E-08 | -1.078956419 | 9.13E-05 | -2.541172332 | 11.28689767 | 10.93049815 | 11.19694409 | 9.691694368 | 9.68194672 | 9.929030467 | 10.19568993 |
| ENSG00000057935 | MTA3 | -1.638743131 | 0.045222505 | -0.903196302 | 0.291783001 | -2.541939433 | 8.340423424 | 9.055168103 | 10.39531398 | 7.910550164 | 7.88209639 | 8.557366484 | 8.684325286 |
| ENSG00000166845 | C18orf54 | -2.010427205 | 0.249656226 | -0.533196495 | 0.78196159 | -2.543623701 | 6.73718411 | 7.572420273 | 2 | 4.984880435 | 4.60967607 | 6.546183077 | 5.596882677 |
| ENSG00000152217 | SETBP1 | -2.307417779 | 0.399970556 | -0.236594169 | 0.939029732 | -2.544011949 | 2 | 7.467835385 | 2 | 4.497095146 | 3.15372745 | 5.790986459 | 5.680297397 |
| ENSG00000152556 | PFKM | -1.161165901 | 0.041111897 | -1.385061597 | 0.012102831 | -2.546227498 | 10.905938 | 10.36815316 | 9.757248847 | 9.30110636 | 9.22317745 | 8.710380719 | 9.308834477 |
| LOC643802 | | 2.644834365 | 0.650789466 | -5.191095755 | NA | -2.54626139 | 2 | 2 | 2 | 3.515101962 | 3.2184422 | 2 | 2 |
| ENSG00000173421 | CCDC36 | -4.302539298 | 0.080939257 | 1.754977963 | 0.440478859 | -2.547561335 | 2 | 2 | 4.874587708 | 2.141737246 | 2.14022021 | 5.113999907 | 4.810605986 |
| ENSG00000173068 | BNC2 | -1.906033142 | 0.192994348 | -0.641585281 | 0.686960124 | -2.547618423 | 4.467311984 | 9.461450882 | 8.585098385 | 6.64692533 | 6.6952083 | 7.625052652 | 8.134761809 |
| ENSG00000079785 | DDX1 | -0.730365011 | 0.152640264 | -1.817621152 | 6.70E-05 | -2.547986163 | 11.50346787 | 10.60848937 | 10.64399316 | 10.27505029 | 10.2286877 | 9.099213681 | 9.237724211 |
| ENSG00000267680 | ZNF224 | 0.115567252 | 0.851541104 | -2.663737628 | 6.59E-09 | -2.548170376 | 8.564112515 | 7.928237109 | 8.884815469 | 8.607665593 | 8.62921659 | 6.101204172 | 5.717942682 |
| ENSG00000025156 | HSF2 | -1.404530659 | 0.00103094 | -1.145778391 | 0.00795665 | -2.55030905 | 7.975256505 | 8.124485672 | 8.608698767 | 6.907295206 | 6.86146731 | 7.328045423 | 6.911910672 |
| ENSG00000176531 | PHLDB3 | -1.342017454 | 0.047187606 | -1.208568546 | 0.071752409 | -2.550586 | 7.53297911 | 6.844429755 | 7.691296561 | 6.12217689 | 6.08095082 | 6.647180917 | 5.631815919 |
| ENSG00000162065 | TBC1D24 | -1.021005853 | 0.165981382 | -1.529770818 | 0.027684956 | -2.550776671 | 9.015175615 | 7.954278496 | 9.624627483 | 7.925329813 | 8.07792246 | 7.729169604 | 7.237648733 |
| ENSG00000087086 | FTL | -0.595571128 | 0.474577536 | -1.955843277 | 0.006881281 | -2.551414404 | 16.60962533 | 15.07629484 | 15.31092757 | 15.14447115 | 15.3260317 | 14.24616937 | 13.38261223 |
| ENSG00000187994 | RINL | -2.043352044 | 0.077472236 | -0.50816533 | 0.691742513 | -2.551517375 | 4.00722531 | 5.72254323 | 6.958202216 | 4.139692846 | 4.2865223 | 5.805044075 | 5.191251321 |
| ENSG00000142937 | RPS8 | -1.806521396 | 0.005229301 | -0.746404159 | 0.289477544 | -2.552925555 | 13.4577386 | 12.25916332 | 11.94765236 | 10.89848915 | 10.9127981 | 11.96837804 | 11.96003159 |
| ENSG00000165895 | ARHGAP42 | -1.860656202 | 0.02322092 | -0.693177086 | 0.437976776 | -2.553833287 | 8.674867122 | 8.414402626 | 6.534109959 | 6.422239259 | 6.22964624 | 7.54878318 | 7.357281305 |
| ENSG00000137700 | SLC37A4 | -2.068533792 | 0.133069745 | -0.486992724 | 0.752096732 | -2.555526516 | 9.174465797 | 8.276714465 | 4.523675391 | 6.321555392 | 6.14893615 | 7.747483829 | 7.787298144 |
| ENSG00000196470 | SIAH1 | -1.035211346 | 0.61318036 | -1.520450822 | 0.434760049 | -2.555662169 | 2 | 7.761270542 | 9.06622548 | 6.927012081 | 7.00595521 | 6.564892306 | 6.432241896 |
| ENSG00000116001 | TIA1 | -0.734038039 | 0.241411697 | -1.821743751 | 0.001138858 | -2.555781791 | 10.31337612 | 10.28338212 | 9.083172337 | 9.32080297 | 9.20931286 | 8.344674564 | 8.014172904 |
| ENSG00000086848 | ALG9 | -0.650027665 | 0.386972978 | -1.906734159 | 0.003824718 | -2.556761824 | 7.790810999 | 9.415325457 | 8.486635645 | 8.020170999 | 8.13979554 | 6.953609309 | 6.746446439 |
| ENSG00000163636 | PSMD6 | -0.910745534 | 0.248988278 | -1.646048421 | 0.024540431 | -2.556793955 | 11.25516263 | 9.785372135 | 9.624627483 | 9.567648462 | 9.45905186 | 8.669665542 | 8.890530144 |
| ENSG00000108592 | FTSJ3 | -0.960510941 | 0.099701083 | -1.596632906 | 0.003306461 | -2.557143846 | 10.54098126 | 9.338312924 | 9.859153765 | 9.000896538 | 9.07814384 | 8.187215117 | 8.603037398 |
| ENSG00000170779 | CDCA4 | -1.506535558 | 0.446883723 | -1.050964245 | 0.599804676 | -2.557499803 | 9.143983763 | 8.276714465 | 2 | 6.777556219 | 6.67228943 | 7.010829034 | 7.307219812 |
| | GRAMD3 | 5.84523908 | 0.002088916 | -8.403369954 | 0.000643868 | -2.558130875 | 2 | 4.93638933 | 2 | 9.115954467 | 9.07268445 | 2 | 2 |
| ENSG00000138073 | PREB | -1.843309437 | 3.28E-13 | -0.717064659 | 0.008040575 | -2.560374097 | 10.43795805 | 10.1432577 | 10.19160738 | 8.36726458 | 8.49198035 | 9.625662378 | 9.46608577 |
| ENSG00000055044 | NOP58 | -0.871078877 | 0.391678983 | -1.689929696 | 0.070032223 | -2.561008573 | 10.48127392 | 8.955547846 | 8.007629717 | 8.601543953 | 8.68180165 | 7.667989909 | 7.98359008 |
| ENSG00000006283 | CACNA1G | -0.195789567 | 0.946040014 | -2.365925276 | 0.350450714 | -2.561714843 | 5.095959458 | 2 | 2 | 3.921164147 | 3.33976443 | 2.803598391 | 2.277179783 |
| ENSG00000260027 | HOXB7 | 2.225840194 | 0.17177866 | -4.788035147 | 0.00161096 | -2.562194953 | 6.257048221 | 6.140560021 | 2 | 7.778406195 | 7.81305436 | 2.775316226 | 2.172608095 |
| ENSG00000175426 | PCSK1 | -0.042376226 | 0.985464453 | -2.520318135 | 0.209322464 | -2.562694362 | 4.00722531 | 3.076665209 | 2 | 3.162721474 | 3.28037825 | 2.139668783 | 2.466008509 |
| ENSG00000100478 | AP4S1 | -0.948620416 | 0.1187722 | -1.614094853 | 0.00411419 | -2.562715269 | 7.435667355 | 6.602424674 | 6.718950348 | 6.221320894 | 5.86549842 | 5.243518884 | 5.611957759 |
| ENSG00000137154 | RPS6 | -2.117565557 | 0.000754913 | -0.446278858 | 0.542075443 | -2.563844414 | 13.91124438 | 12.65219511 | 12.48588269 | 11.06512699 | 11.0362957 | 12.64097106 | 12.79530838 |
| ENSG00000164919 | COX6C | -1.323853676 | 0.181146687 | -1.241042578 | 0.204538137 | -2.564896254 | 11.19324461 | 8.623598428 | 9.61300511 | 8.873322599 | 8.88359764 | 8.996146925 | 8.924134755 |
| ENSG00000128283 | CDC42EP1 | -2.462144322 | 1.03E-05 | -0.104904689 | 0.886770342 | -2.56704901 | 11.65166541 | 10.42977533 | 11.78627063 | 8.941551548 | 8.95891978 | 11.24494713 | 11.34933218 |
| ENSG00000088832 | FKBP1A | -0.911099104 | 0.046672139 | -1.659362251 | 9.35E-05 | -2.570461355 | 12.78892924 | 12.15073083 | 12.82880267 | 11.71011514 | 11.7106329 | 10.63583525 | 11.23002844 |
| ENSG00000173692 | PSMD1 | -0.977842178 | 0.00912284 | -1.593006423 | 7.08E-06 | -2.570848601 | 11.83695787 | 11.2524908 | 11.63070764 | 10.57442844 | 10.6570657 | 9.778197243 | 10.19777306 |
| ENSG00000174684 | B4GAT1 | -1.932399052 | 0.028063103 | -0.639924708 | 0.510549382 | -2.57232376 | 8.564112515 | 9.756063186 | 11.00131064 | 8.170409056 | 8.19309833 | 9.404322061 | 9.518360533 |
| ENSG00000110442 | COMMD9 | -1.260664045 | 0.01578088 | -1.311695765 | 0.010977411 | -2.572359809 | 9.931427102 | 9.030900487 | 9.180874355 | 8.246921218 | 8.12299044 | 7.833151805 | 8.386355612 |
| ENSG00000197858 | GPAA1 | -1.523865517 | 0.023379435 | -1.048609463 | 0.129694026 | -2.57247498 | 11.90443341 | 11.00713613 | 10.2528023 | 9.659644277 | 9.72295127 | 10.01544857 | 10.30040559 |
| ENSG00000137288 | UQCC2 | -1.251444048 | 5.20E-05 | -1.323561371 | 1.28E-05 | -2.575005419 | 8.609439662 | 8.607278467 | 8.699419242 | 7.409376773 | 7.40393901 | 7.16729616 | 7.488317792 |
| ENSG00000117385 | P3H1 | -1.259894582 | 0.000302512 | -1.31686521 | 0.000139067 | -2.576759792 | 12.79261849 | 12.0991443 | 12.47147373 | 11.24500602 | 11.2004816 | 11.17706738 | 11.15492717 |
| ENSG00000197915 | HRNR | -1.713156076 | 0.536668932 | -0.864937987 | 0.76003079 | -2.578094063 | 2 | 2 | 4.523675391 | 2.784612485 | 2.2680111 | 2.68698273 | 3.042026731 |
| ENSG00000197557 | TTC30A | -0.525682284 | 0.51591099 | -2.054473486 | 0.00290832 | -2.58015577 | 7.159102577 | 6.899080544 | 8.325066185 | 7.044488801 | 7.11620658 | 5.328699035 | 5.88031069 |
| ENSG00000166900 | STX3 | -0.403990471 | 0.693142866 | -2.176662563 | 0.011271266 | -2.580653034 | 7.218819781 | 8.834582582 | 9.465702942 | 8.293234263 | 8.44880239 | 7.012344451 | 6.141542327 |
| ENSG00000123136 | DDX39A | -0.747397278 | 0.461018278 | -1.835200875 | 0.04279221 | -2.582598153 | 10.30651143 | 8.124485672 | 8.511888463 | 8.573670512 | 8.57000035 | 7.264607048 | 7.704159082 |
| ENSG00000155868 | MED7 | -1.354877831 | 0.187957925 | -1.228695089 | 0.22807891 | -2.58357292 | 9.048471759 | 6.464272742 | 7.451345288 | 6.777556219 | 6.66069199 | 6.923319986 | 6.755525668 |
| ENSG00000013275 | PSMC4 | -0.86350316 | 0.009557939 | -1.720831861 | 2.87E-08 | -2.584335021 | 10.96815916 | 10.67655334 | 11.02781271 | 9.991161874 | 10.0804323 | 8.950238388 | 9.384672535 |
| ENSG00000185774 | KCNIP4 | -3.084074005 | 0.145965965 | 0.499547159 | 0.814954199 | -2.584526847 | 3.327711234 | 3.686303414 | 2 | 2.270786764 | 2.14022021 | 3.738983888 | 3.236138336 |
| ENSG00000163517 | HDAC11 | -1.74837232 | 0.207275017 | -0.838609577 | 0.565593677 | -2.586981897 | 9.423197387 | 4.93638933 | 8.460932916 | 6.799114572 | 6.72891807 | 7.846806418 | 7.420010428 |
| ENSG00000118620 | ZNF430 | -0.98702689 | 0.329061986 | -1.600737849 | 0.090736393 | -2.587764739 | 8.716893051 | 7.189546432 | 6.322058544 | 6.887305128 | 6.68379438 | 6.329884754 | 6.06327132 |
| ENSG00000104549 | SQLE | -1.383773033 | 3.96E-05 | -1.204073959 | 0.000386271 | -2.587846992 | 11.38790617 | 11.31412071 | 11.25043551 | 9.943557345 | 9.93336537 | 9.772172069 | 10.39603787 |
| ENSG00000150995 | ITPR1 | -1.620199993 | 0.003298939 | -0.969746244 | 0.093130425 | -2.589946237 | 8.564112515 | 9.090820894 | 9.399592459 | 7.494428494 | 7.42465798 | 8.483898848 | 7.570355956 |
| ENSG00000221829 | FANCG | -1.545302846 | 0.018712352 | -1.045057988 | 0.120202987 | -2.590360834 | 8.009475818 | 7.051611315 | 6.629488387 | 5.870535235 | 5.86549842 | 6.490730008 | 6.168983563 |
| ENSG00000168904 | LRRC28 | -1.769551214 | 0.313384316 | -0.821419288 | 0.658193363 | -2.590970502 | 3.327711234 | 7.189546432 | 9.426401271 | 6.237202172 | 6.57060569 | 7.27600668 | 7.379474672 |
| ENSG00000149313 | AASDHPPT | -1.0191591 | 0.000526024 | -1.572086721 | 2.19E-08 | -2.591245821 | 9.579430841 | 9.443177293 | 9.61300511 | 8.527598812 | 8.54024645 | 8.154648976 | 7.799420735 |
| ENSG00000197798 | FAM118B | -1.309153718 | 0.028899691 | -1.282157518 | 0.030994241 | -2.591311236 | 9.397585735 | 8.191702463 | 8.325066185 | 7.426792977 | 7.47849652 | 7.449360315 | 7.5085977 |
| ENSG00000168306 | ACOX2 | -1.320073357 | 0.514104281 | -1.271863345 | 0.521952482 | -2.591936702 | 8.631580145 | 6.535001748 | 2 | 5.94010643 | 6.22964624 | 5.849807762 | 6.37742435 |
| ENSG00000064995 | TAF11 | -1.485313498 | 0.008096943 | -1.109078045 | 0.053144626 | -2.594391543 | 9.740899611 | 8.451443573 | 9.344432377 | 7.872924734 | 7.72392024 | 8.249579355 | 8.087009646 |
| ENSG00000117595 | IRF6 | -1.559325673 | 0.088235741 | -1.036215921 | 0.257062588 | -2.595541594 | 4.467311984 | 5.131603625 | 5.930865648 | 3.95996233 | 4.02226117 | 4.612704558 | 4.147809227 |
| ENSG00000153982 | GDPD1 | -1.173957089 | 0.329606836 | -1.422559614 | 0.218679367 | -2.596516702 | 6.653098118 | 4.442710129 | 6.958202216 | 5.348759107 | 5.16377554 | 5.511266259 | 4.329022995 |
| ENSG00000164053 | ATRIP | -1.74446742 | 0.35107637 | -0.85312825 | 0.664743846 | -2.597595671 | 8.444143041 | 7.002537973 | 2 | 5.674521485 | 5.64780787 | 6.633459761 | 6.350712758 |
| ENSG00000147852 | VLDLR | -2.14589275 | 0.121803593 | -0.452699259 | 0.773865459 | -2.598592009 | 5.866944937 | 10.03150247 | 10.71094342 | 7.706073672 | 7.75697332 | 9.499512834 | 9.306133598 |
| ENSG00000167315 | ACAA2 | -0.895151351 | 0.366470827 | -1.70450735 | 0.061919368 | -2.5996587 | 10.67493801 | 10.45279253 | 8.143093325 | 9.20040236 | 9.24275568 | 8.385007206 | 8.45344973 |
| ENSG00000198612 | COPS8 | -1.208971145 | 0.022546075 | -1.392596627 | 0.007169426 | -2.601567771 | 10.89224845 | 9.71095217 | 10.2528023 | 9.140440229 | 9.18117744 | 9.039153561 | 8.915553582 |
| ENSG00000132507 | EIF5A | -1.170452283 | 0.095852222 | -1.432166163 | 0.03456438 | -2.602618446 | 13.1801726 | 11.60869109 | 12.27160023 | 11.30085596 | 11.3548657 | 10.73519466 | 11.33634299 |
| ENSG00000110801 | PSMD9 | -0.426797024 | 0.27007879 | -2.176125681 | 2.65E-11 | -2.602922705 | 9.497412313 | 9.181799937 | 9.06622548 | 8.76657222 | 8.90218458 | 6.931352719 | 7.280736296 |
| ENSG00000099800 | TIMM13 | -1.639907209 | 4.27E-05 | -0.964477411 | 0.02217593 | -2.60438462 | 10.27168932 | 9.856140801 | 9.735974718 | 8.241026137 | 8.44204342 | 8.708044758 | 9.266193276 |
| ENSG00000184117 | NIPSNAP1 | -1.59298922 | 0.098030868 | -1.01203173 | 0.305936987 | -2.60502095 | 11.17074085 | 9.756063186 | 8.585098385 | 8.630393142 | 8.62325378 | 9.01024623 | 9.372921162 |
| ENSG00000128463 | EMC4 | -1.172058568 | 0.000671618 | -1.436448016 | 1.90E-05 | -2.608506584 | 10.27168932 | 10.65289571 | 10.90452077 | 9.460955886 | 9.46906021 | 9.100282869 | 9.297612054 |
| ENSG00000140319 | SRP14 | -1.111684397 | 0.026613498 | -1.497065242 | 0.001875691 | -2.60874964 | 12.39655881 | 11.30403016 | 11.77848812 | 10.77677453 | 10.7940784 | 10.34218565 | 10.45766062 |
| ENSG00000122692 | SMU1 | -1.139544928 | 0.035422866 | -1.469888805 | 0.004813198 | -2.609433733 | 10.64803757 | 10.17682307 | 9.426401271 | 8.998569022 | 9.06939887 | 8.627767864 | 8.781973674 |
| ENSG00000179292 | TMEM151A | -3.642713847 | 0.022267907 | 1.032728626 | 0.538703409 | -2.60998522 | 3.327711234 | 6.04685525 | 3.369317972 | 2.498691189 | 2.68924543 | 6.384736493 | 4.870334174 |
| ENSG00000166199 | ALKBH3 | -1.44687535 | 0.392541278 | -1.16352346 | 0.491522021 | -2.61039881 | 4.467311984 | 6.899080544 | 9.735974718 | 6.993973425 | 6.91125523 | 7.010829034 | 7.417152486 |
| ENSG00000130511 | SSBP4 | -1.451039229 | 5.97E-06 | -1.159742067 | 0.000327357 | -2.610781296 | 8.892187472 | 9.298208073 | 8.904018013 | 7.641902724 | 7.59275449 | 7.794921259 | 8.004685229 |
| ENSG00000186222 | BLOC1S4 | -1.364973807 | 0.10052636 | -1.246545702 | 0.129711398 | -2.611519509 | 9.435834703 | 8.540087354 | 7.285536013 | 7.359462263 | 7.29187696 | 7.381793866 | 7.499169207 |
| ENSG00000124802 | EEF1E1 | -1.758670335 | 0.171238625 | -0.853210539 | 0.527954208 | -2.611880874 | 9.613202746 | 7.467835385 | 5.930865648 | 6.604700657 | 6.77267312 | 7.486906381 | 7.649197382 |
| ENSG00000271601 | LIX1L | -1.682859333 | 0.000127603 | -0.933408299 | 0.044614383 | -2.616267632 | 10.00958825 | 9.994333773 | 10.80076732 | 8.612239836 | 8.67749148 | 9.271730593 | 9.497203828 |
| ENSG00000239306 | RBM14 | -1.73607159 | 0.00477741 | -0.881478543 | 0.177968622 | -2.617550133 | 8.696033109 | 8.916342327 | 10.03493575 | 7.734324667 | 7.50468064 | 8.501271034 | 8.428831018 |
| ENSG00000138036 | DYNC2LI1 | -1.06835703 | 0.183576111 | -1.549586184 | 0.041032339 | -2.617943213 | 7.031698319 | 6.666836837 | 8.35328246 | 6.476767609 | 6.52015598 | 5.99483933 | 6.088639344 |
| ENSG00000172171 | TEFM | -0.865238974 | 0.496953626 | -1.753483537 | 0.13163848 | -2.618722511 | 8.340423424 | 5.946638857 | 5.59507254 | 6.379946408 | 6.29861567 | 5.665620077 | 5.337485258 |
| ENSG00000204764 | RANBP17 | -0.079418282 | 0.968054519 | -2.539757575 | 0.120755432 | -2.619175857 | 5.095959458 | 6.951736442 | 2 | 5.590471342 | 5.80356929 | 3.600766115 | 3.735548302 |
| ENSG00000262814 | MRPL12 | -0.986216212 | 0.010979795 | -1.633428064 | 7.73E-06 | -2.619644275 | 9.015175615 | 9.558016677 | 9.399592459 | 8.458152491 | 8.26953254 | 7.523500045 | 7.910280955 |
| ENSG00000157326 | DHRS4 | -0.548370075 | 0.552095038 | -2.071467958 | 0.009438355 | -2.619838033 | 7.435667355 | 6.844429755 | 8.784770914 | 7.359462263 | 7.40045674 | 5.744329515 | 6.077821908 |
| ENSG00000136261 | BZW2 | -1.700552031 | 0.002045073 | -0.920038741 | 0.114871963 | -2.620590772 | 8.609439662 | 9.13702746 | 9.897982195 | 7.577865504 | 7.68150967 | 8.375029261 | 8.422432 |
| ENSG00000111325 | OGFOD2 | -0.098276887 | 0.939485429 | -2.523066777 | 0.014049355 | -2.621343664 | 6.89194353 | 7.790474954 | 4.874587708 | 6.892328675 | 6.83074839 | 4.291799915 | 4.919632516 |
| ENSG00000139163 | ETNK1 | -0.681545763 | 0.433017846 | -1.940020778 | 0.011321064 | -2.621566541 | 8.444143041 | 9.969011754 | 10.46808205 | 9.253098082 | 9.08466797 | 8.153275969 | 7.657697387 |

| | | | | | | | | | | | | | |
|---|---|---|---|---|---|---|---|---|---|---|---|---|---|
| ENSG00000166188 | ZNF319 | -1.813418683 | 0.095496255 | -0.809202493 | 0.483542133 | -2.622621176 | 8.517314938 | 6.535001748 | 9.725218844 | 6.998639523 | 6.96410766 | 7.937338157 | 7.987448576 |
| ENSG00000139684 | ESD | -1.15286446 | 0.001617597 | -1.470493779 | 3.33E-05 | -2.623358239 | 10.69084077 | 10.20962523 | 10.07784268 | 9.195330254 | 9.21030762 | 8.745429394 | 9.017193688 |
| ENSG00000172000 | ZNF556 | -1.689559495 | NA | -0.934455208 | NA | -2.624014703 | 2 | 2 | 3.369317972 | 2.141737246 | 2.2680111 | 2.560105256 | 2.059860178 |
| ENSG00000139537 | CCDC65 | -4.55116024 | 0.129548839 | 1.926937082 | 0.452105431 | -2.624223159 | 2 | 4.11352228 | 2 | 2 | 2.14022021 | 4.487545924 | 4.468068808 |
| ENSG00000135929 | CYP27A1 | -1.016436552 | 0.428291678 | -1.609317978 | 0.179430199 | -2.625754529 | 10.32700827 | 7.35507212 | 8.236946975 | 8.080836374 | 8.27334517 | 8.122032742 | 6.764548117 |
| ENSG00000198246 | SLC29A3 | -1.701685904 | 0.426363818 | -0.924296957 | 0.677074052 | -2.625982862 | 8.797441463 | 5.595927442 | 2 | 5.709096143 | 5.78232107 | 6.830158212 | 6.011156536 |
| ENSG00000163832 | ELP6 | -1.391521201 | 0.000834812 | -1.238942864 | 0.002969599 | -2.630464064 | 8.468948322 | 8.607278467 | 9.132850258 | 7.330144653 | 7.45182838 | 7.344982741 | 7.714671806 |
| ENSG00000172243 | CLEC7A | -2.910230673 | 0.281095622 | 0.279452138 | 0.926160637 | -2.630778536 | 2 | 3.076665209 | 4.523675391 | 2 | 2.59489879 | 4.076896059 | 3.287118176 |
| ENSG00000083635 | NUFIP1 | -1.310248101 | 0.476911496 | -1.322119671 | 0.462769314 | -2.632367772 | 8.009475818 | 7.232742069 | 2 | 6.034011955 | 5.64780787 | 5.825876956 | 5.859408034 |
| ENSG00000130429 | ARPC1B | 1.170717704 | 0.030750279 | -3.803621366 | 6.75E-15 | -2.632903662 | 9.751047882 | 9.055168103 | 10.20715117 | 10.86086404 | 10.956731 | 5.812021848 | 6.209190898 |
| ENSG00000105048 | TNNT1 | -2.791980982 | 0.167654826 | 0.1586039 | 0.944157185 | -2.633377082 | 2 | 4.710564154 | 3.369317972 | 2.270786764 | 2.59489879 | 4.397636241 | 3.183290849 |
| ENSG00000104872 | PIH1D1 | -1.464657731 | 0.073359589 | -1.169373468 | 0.155406001 | -2.634031198 | 10.54098126 | 9.125613817 | 8.631919287 | 8.14748175 | 8.284723 | 8.295734109 | 8.695572791 |
| ENSG00000168778 | TCTN2 | -1.321132538 | 0.544796958 | -1.315088558 | 0.540695624 | -2.636221096 | 2 | 6.04685525 | 9.132850258 | 6.372775472 | 6.49425375 | 6.320128742 | 6.55171954 |
| ENSG00000103549 | RNF40 | -0.68179176 | 0.290429859 | -1.954624818 | 0.000553999 | -2.636416579 | 10.36714848 | 10.73405387 | 11.7010103 | 10.33623673 | 10.3980237 | 9.077661821 | 9.12358288 |
| ENSG00000166025 | AMOTL1 | -1.826509718 | 0.00023389 | -0.811316219 | 0.131351033 | -2.637825937 | 11.29385609 | 12.35391197 | 11.48210469 | 9.911750163 | 10.0187431 | 10.90816893 | 11.04559838 |
| ENSG00000107902 | LHPP | -1.974592171 | 0.011022574 | -0.66563123 | 0.437688449 | -2.640223401 | 7.53297911 | 7.874700259 | 9.344432377 | 6.456560224 | 6.63128385 | 7.873733947 | 7.760489878 |
| ENSG00000176896 | TCEANC | -1.204541873 | 0.142431446 | -1.436028821 | 0.066886602 | -2.640570694 | 5.095959458 | 6.311476754 | 5.930865648 | 4.804776789 | 4.79082474 | 4.888321757 | 4.241259475 |
| ENSG00000188582 | PAQR9 | -0.979601134 | 0.729390378 | -1.661422481 | 0.531906196 | -2.641023615 | 2 | 2 | 4.523675391 | 2.784612485 | 2.86123014 | 2.803598391 | 2.277179783 |
| ENSG00000167775 | CD320 | -1.728474749 | 5.26E-05 | -0.913289515 | 0.044041567 | -2.641764265 | 9.839306291 | 10.13758676 | 9.31604076 | 8.080836374 | 8.10384772 | 8.670625574 | 9.095255687 |
| ENSG00000167792 | NDUFV1 | -1.883932015 | 0.106436871 | -0.758473301 | 0.544775711 | -2.642405317 | 11.49141174 | 9.624396729 | 8.076950877 | 8.481674661 | 8.49034333 | 9.418393009 | 9.766168491 |
| ENSG00000165028 | NIPSNAP3B | -0.040677459 | 0.984108348 | -2.601824053 | 0.129124437 | -2.642501511 | 6.89194353 | 5.303523747 | 2 | 5.662809983 | 5.77157844 | 4.065262523 | 2.950113106 |
| ENSG00000162694 | EXTL2 | -1.120129499 | 0.063823695 | -1.522784402 | 0.008354549 | -2.642913901 | 8.737455672 | 9.532315021 | 10.14394518 | 8.491639362 | 8.44711562 | 8.091532339 | 8.052432906 |
| ENSG00000160323 | ADAMTS13 | -1.620127722 | 0.409685056 | -1.022997444 | 0.610287918 | -2.643125166 | 8.19931886 | 5.838937808 | 2 | 5.406080789 | 5.3193943 | 6.14330814 | 5.636738041 |
| ENSG00000100442 | FKBP3 | -0.365381286 | 0.708408469 | -2.278207921 | 0.00476195 | -2.643589206 | 10.31337612 | 8.169643011 | 9.287079172 | 9.230466133 | 9.05063789 | 6.911998488 | 7.537835213 |
| ENSG00000143947 | RPS27A | -1.591494026 | 0.022379195 | -1.052614599 | 0.144035339 | -2.644108625 | 12.91500773 | 11.22868348 | 11.74958762 | 10.55198546 | 10.5550472 | 11.06935627 | 11.11271466 |
| ENSG00000119705 | SLIRP | -0.520564664 | 0.417361165 | -2.123563626 | 9.65E-05 | -2.64412829 | 8.75772933 | 7.467835385 | 8.143093325 | 7.641902724 | 7.7515168 | 6.121005698 | 6.179140936 |
| ENSG00000144031 | ANKRD53 | -1.384688194 | 0.606861222 | -1.259577141 | 0.635737899 | -2.644265335 | 2 | 6.535001748 | 2 | 4.17310023 | 3.69849104 | 3.663836244 | 4.353147976 |
| ENSG00000135916 | ITM2C | -1.833493593 | 0.079559698 | -0.813208684 | 0.4666278 | -2.646702278 | 13.49066495 | 11.08580931 | 11.03218278 | 10.52353286 | 10.5361304 | 11.51552829 | 11.58004535 |
| ENSG00000131370 | SH3BP5 | -0.35044294 | 0.673891173 | -2.296590822 | 0.000751772 | -2.647033762 | 8.981092835 | 8.029695761 | 7.225742779 | 7.947219213 | 7.86955761 | 5.912830251 | 6.13807518 |
| ENSG00000123933 | MXD4 | -2.105516965 | 2.39E-07 | -0.542337742 | 0.243516721 | -2.647854707 | 11.99474663 | 11.1687783 | 11.37608156 | 9.457570641 | 9.45737703 | 10.87804317 | 11.14048782 |
| ENSG00000169021 | UQCRFS1 | -1.353037861 | 1.46E-05 | -1.29507286 | 3.12E-05 | -2.648110721 | 10.55263183 | 10.63288029 | 10.23015729 | 9.100844036 | 9.17201613 | 9.005308106 | 9.361816632 |
| ENSG00000088812 | ATRN | -1.751682897 | 7.29E-06 | -0.897632548 | 0.031375886 | -2.649315446 | 12.07772327 | 11.23134822 | 11.67049834 | 9.920382853 | 9.98368402 | 10.80974856 | 10.79787914 |
| ENSG00000136143 | SUCLA2 | -1.435463591 | 3.53E-05 | -1.213918803 | 0.000505533 | -2.649382394 | 9.579430841 | 9.806969599 | 9.165043284 | 8.174538791 | 8.06917614 | 8.288873067 | 8.392188801 |
| ENSG00000164418 | GRIK2 | 1.094154401 | 0.686593442 | -3.745897463 | 0.117040598 | -2.651743062 | 2 | 7.145017454 | 2 | 6.622948477 | 6.74548212 | 2.62493832 | 3.128433649 |
| ENSG00000102580 | DNAJC3 | -1.164267877 | 0.048513855 | -1.487678899 | 0.008884136 | -2.651946776 | 11.13631304 | 10.87704672 | 12.02204252 | 10.2793709 | 10.2602493 | 9.628876603 | 10.20855713 |
| ENSG00000101997 | CCDC22 | -1.146499301 | 0.573001609 | -1.507415987 | 0.440486964 | -2.653915288 | 9.27634997 | 8.522788974 | 2 | 7.226553841 | 7.26148706 | 6.798888929 | 6.983208048 |
| ENSG00000122034 | GTF3A | -1.401091958 | 0.007345799 | -1.253461593 | 0.017178874 | -2.654553552 | 10.5468183 | 9.566483142 | 9.528916236 | 8.573670512 | 8.56534297 | 8.69064008 | 8.740277714 |
| ENSG00000072858 | SIDT1 | -2.555456211 | 0.290311321 | -0.099517732 | 0.968964721 | -2.654973944 | 2 | 3.076665209 | 3.369317972 | 2.141737246 | 2.2680111 | 3.035909414 | 2.70970617 |
| ENSG00000185101 | ANO9 | -3.296402287 | 0.34076194 | 0.641386778 | 0.857444981 | -2.65501551 | 2 | 2 | 4.058998049 | 2.270786764 | 2 | 3.27687201 | 3.45257121 |
| ENSG00000100908 | EMC9 | -1.815677544 | 0.140439423 | -0.840523447 | 0.518736842 | -2.656200991 | 8.586954094 | 7.393647515 | 4.874587708 | 5.797438367 | 5.91514978 | 6.591495479 | 6.951954485 |
| ENSG00000128050 | PAICS | -2.018958335 | 5.10E-06 | -0.637409796 | 0.197561623 | -2.656368131 | 11.52136524 | 10.74528399 | 10.67226762 | 9.04780133 | 8.99630959 | 10.23447665 | 10.54305007 |
| ENSG00000171863 | RPS7 | -1.127041599 | 0.141109656 | -1.531983301 | 0.035956933 | -2.6590249 | 12.09972935 | 10.50217373 | 10.5916593 | 10.16260233 | 10.1174913 | 9.788404752 | 9.683551156 |
| ENSG00000131779 | PEX11B | -1.736000205 | 1.09E-05 | -0.923501813 | 0.026953886 | -2.659502018 | 9.810482605 | 9.170736405 | 9.06622548 | 7.662632018 | 7.67863747 | 8.496947559 | 8.444354606 |
| ENSG00000162188 | GNG3 | -1.180843895 | 0.691648862 | -1.478851842 | NA | -2.659695737 | 2 | 3.686303414 | 2 | 2.600425781 | 2.2680111 | 2.457042231 | 2.277179783 |
| ENSG00000254999 | BRK1 | -1.246820537 | 0.00326405 | -1.413981193 | 0.000672535 | -2.66080173 | 11.88382844 | 11.29895829 | 11.95227171 | 10.46993609 | 10.5189709 | 10.02242176 | 10.58009362 |
| ENSG00000171791 | BCL2 | -3.151414997 | 0.211336072 | 0.489153135 | 0.863199064 | -2.662261862 | 2 | 6.311476754 | 2 | 3.09461492 | 2.38539787 | 5.598599872 | 4.975095763 |
| ENSG00000162086 | ZNF75A | -0.443137041 | 0.452334443 | -2.219924126 | 6.05E-06 | -2.663061168 | 7.904282243 | 8.004992147 | 8.677268317 | 7.839502934 | 7.75424764 | 6.353989995 | 5.745545708 |
| ENSG00000178028 | DMAP1 | -1.360529697 | 0.089666875 | -1.302702256 | 0.099409613 | -2.663231953 | 9.761125267 | 7.928237109 | 8.486635645 | 7.626911583 | 7.55566492 | 7.4096614 | 7.853274783 |
| ENSG00000109771 | LRP2BP | -2.665273284 | 0.028200875 | 0.001134234 | 0.999378208 | -2.664139049 | 5.095959458 | 3.076665209 | 5.39244166 | 3.09461492 | 2.77779907 | 4.785723309 | 4.77533149 |
| ENSG00000169885 | CALML6 | -1.825849609 | 0.486921279 | -0.838741498 | 0.757654618 | -2.664591106 | 2 | 2 | 5.39244166 | 2.868578001 | 2.94009902 | 3.782262616 | 3.012034821 |
| ENSG00000166546 | BEAN1 | -2.495415615 | 0.206479362 | -0.169455254 | 0.940405661 | -2.664870869 | 5.70932728 | 3.076665209 | 2 | 2.947924294 | 2.68924543 | 4.628601098 | 3.824516949 |
| ENSG00000091640 | SPAG7 | -1.368037655 | 0.041735857 | -1.300409776 | 0.051271517 | -2.668447431 | 10.80726569 | 9.799806309 | 9.242516611 | 8.703665429 | 8.77214691 | 8.66870487 | 8.930161469 |
| ENSG00000177646 | ACAD9 | -1.023946107 | 0.002766788 | -1.645473103 | 4.02E-07 | -2.669419211 | 9.497412313 | 9.34816733 | 9.227351203 | 8.287525788 | 8.40429221 | 7.492346363 | 7.943433856 |
| ENSG00000121104 | FAM117A | -1.499318005 | 0.001565551 | -1.170555059 | 0.014545829 | -2.669873064 | 7.624139768 | 7.232742069 | 7.858115942 | 6.069927018 | 6.20590375 | 6.189586148 | 6.674075308 |
| ENSG00000152464 | RPP38 | -1.45418264 | 0.016061522 | -1.215698565 | 0.04533349 | -2.669881206 | 7.667647223 | 7.35507212 | 8.608698767 | 6.604700657 | 6.507263 | 6.879153024 | 6.685996416 |
| ENSG00000100583 | SAMD15 | -0.151461032 | 0.957050083 | -2.519037623 | 0.296825224 | -2.670498655 | 2 | 5.838937808 | 2 | 4.299492648 | 4.40247105 | 2.492221014 | 3.128433649 |
| ENSG00000225697 | SLC26A6 | -1.123590379 | 0.02438057 | -1.546987375 | 0.001186636 | -2.670577755 | 10.04301211 | 9.159587375 | 9.116480286 | 8.402902269 | 8.37797443 | 7.952415862 | 7.993217039 |
| ENSG00000118402 | ELOVL4 | -0.607791358 | 0.70562308 | -2.062866833 | 0.144465782 | -2.670658191 | 9.2903348 | 7.051611315 | 4.874587708 | 7.501020868 | 7.37584259 | 6.353989995 | 5.641643428 |
| ENSG00000187514 | PTMA | -1.117788414 | 0.000160157 | -1.553492864 | 6.15E-08 | -2.671281278 | 12.84803656 | 12.60879193 | 12.81745404 | 11.66367858 | 11.6257578 | 10.96258464 | 11.42080713 |
| ENSG00000157322 | CLEC18A | 1.60707571 | 0.620052795 | -4.278823407 | 0.16716103 | -2.671747697 | 2 | 2 | 4.058998049 | 3.709785322 | 4.2865223 | 2.139668783 | 2 |
| ENSG00000105404 | RABAC1 | -0.916965935 | 6.72E-05 | -1.755355504 | 1.12E-15 | -2.672321439 | 11.18203661 | 10.94030351 | 11.08361639 | 10.10372352 | 10.2069242 | 9.250699274 | 9.389783813 |
| ENSG00000113851 | CRBN | -1.101261666 | 0.16737365 | -1.57191523 | 0.037364726 | -2.673176895 | 9.949165863 | 9.328390744 | 8.007629717 | 8.203120604 | 8.2130862 | 7.958717528 | 7.493753702 |
| ENSG00000171766 | GATM | -2.133495613 | 0.271773169 | -0.539741233 | 0.796690808 | -2.673236845 | 3.327711234 | 5.303523747 | 2 | 3.09461492 | 2.59489879 | 4.122514202 | 3.383972333 |
| ENSG00000163348 | PYGO2 | -2.057662074 | 0.030750279 | -0.617013918 | 0.560994904 | -2.674675992 | 8.777722034 | 8.655694736 | 10.89495479 | 7.842101511 | 7.75424764 | 9.304879394 | 9.136237781 |
| ENSG00000160633 | SAFB | -0.823829629 | 0.419016636 | -1.851373685 | 0.04446759 | -2.675203314 | 11.35168561 | 9.148351514 | 9.359998894 | 9.460955886 | 9.54672161 | 8.619836896 | 8.336553984 |
| ENSG00000152763 | WDR78 | -1.104732095 | 0.621262845 | -1.572104225 | 0.460375683 | -2.676836321 | 2 | 3.686303414 | 7.163363831 | 4.846787657 | 4.63360314 | 4.771475226 | 3.771796761 |
| ENSG00000173818 | ENDOV | -1.887932946 | 0.035043531 | -0.789542026 | 0.412508059 | -2.677474971 | 7.384433967 | 6.728495673 | 8.845626171 | 5.987823101 | 6.20590375 | 7.417689057 | 6.830475749 |
| ENSG00000157557 | ETS2 | -0.778494503 | 0.034731167 | -1.900266644 | 1.39E-08 | -2.678761147 | 10.58135316 | 10.19877383 | 10.61515347 | 9.630628506 | 9.76553205 | 8.344674564 | 8.793039018 |
| ENSG00000276600 | RAB7B | -6.383706317 | 0.02747405 | 3.700763167 | 0.125991152 | -2.68294315 | 2 | 4.442710129 | 2 | 2 | 2 | 6.535683256 | 6.308152703 |
| ENSG00000197147 | LRRC8B | -2.976643232 | 0.067576467 | 0.292435438 | 0.87857982 | -2.684207794 | 2 | 4.93638933 | 6.322058544 | 3.289987342 | 2.77779907 | 5.822425633 | 5.115701921 |
| ENSG00000214367 | HAUS3 | -1.186130103 | 0.539576503 | -1.499491715 | 0.422327856 | -2.685621818 | 9.371511179 | 6.04685525 | 3.369317972 | 6.841288255 | 6.72891807 | 6.774090992 | 6.124122661 |
| ENSG00000059691 | GATB | -1.208054723 | 0.450332207 | -1.478455479 | 0.335253459 | -2.686510202 | 9.544849385 | 6.228547981 | 5.39244166 | 6.975156635 | 6.99676009 | 6.613628927 | 6.828326017 |
| ENSG00000163719 | MTMR14 | -1.143716011 | 0.061530373 | -1.543248341 | 0.008306252 | -2.686964353 | 9.657036304 | 8.276714465 | 9.272377397 | 8.069792811 | 8.01099479 | 7.538302335 | 7.750306941 |
| ENSG00000109956 | B3GAT1 | -0.679672253 | NA | -2.007887435 | NA | -2.687559687 | 2 | 2 | 3.369317972 | 2.695456494 | 2 | 2.267001569 | 2.059860178 |
| ENSG00000104953 | TLE6 | -1.015917121 | 0.506649307 | -1.672557516 | 0.235344728 | -2.688474637 | 3.327711234 | 4.11352228 | 4.058998049 | 3.162721474 | 3.28037825 | 3.1504466 | 2.551896972 |
| ENSG00000112305 | SMAP1 | -0.405347018 | 0.531143362 | -2.283931628 | 1.91E-05 | -2.689278646 | 7.667647223 | 8.916342327 | 8.175061096 | 7.966401501 | 7.9408106 | 6.081127077 | 6.189227296 |
| ENSG00000165209 | STRBP | -2.513067601 | 0.113748903 | -0.179643173 | 0.92531104 | -2.692710774 | 7.031698319 | 7.051611315 | 2 | 4.205751501 | 4.2865223 | 6.327451929 | 6.298867018 |
| ENSG00000136243 | NUPL2 | -0.644162096 | 0.466937891 | -2.049959787 | 0.007873748 | -2.694121883 | 8.716893051 | 7.35507212 | 6.882775709 | 7.25803034 | 7.21862736 | 6.040114727 | 5.73181021 |
| ENSG00000255221 | CARD17 | 2.49669361 | 0.671489931 | -5.191096372 | NA | -2.694402762 | 2 | 2 | 2 | 3.289987342 | 3.28037825 | 2 | 2 |
| ENSG00000260458 | KCNJ18 | 2.494380499 | 0.67181002 | -5.191096382 | NA | -2.696715883 | 2 | 2 | 2 | 3.023133209 | 3.50452879 | 2 | 2 |
| ENSG00000113946 | CLDN16 | -0.330168266 | 0.8694764 | -2.367401932 | 0.16091081 | -2.697570199 | 2 | 3.686303414 | 4.058998049 | 3.227757282 | 3.33976443 | 2.3070652 | 2.551896972 |
| ENSG00000246705 | H2AFJ | -2.270478075 | 0.001819985 | -0.427980748 | 0.61684613 | -2.698458823 | 11.33491623 | 9.726146444 | 9.849281147 | 8.227176542 | 8.2657098 | 9.773736569 | 10.32462856 |
| ENSG00000205629 | LCMT1 | -1.270145134 | 6.34E-05 | -1.429840715 | 4.83E-06 | -2.699985849 | 11.57090097 | 11.72481431 | 12.10735427 | 10.60534987 | 10.4938644 | 10.48300598 | 10.29360417 |
| ENSG00000137074 | APTX | -1.316596536 | 0.001277928 | -1.38396252 | 0.000591536 | -2.700559056 | 9.473095744 | 8.702538795 | 9.399592459 | 7.915493556 | 7.93118934 | 7.86621124 | 7.847978946 |
| ENSG00000075142 | SRI | -1.061104853 | 0.118143225 | -1.640685473 | 0.010224534 | -2.701790326 | 10.36714848 | 9.763446433 | 8.742726175 | 8.676258519 | 8.75452469 | 8.194575779 | 8.089704419 |
| ENSG00000197124 | ZNF682 | -1.138388056 | 0.587855308 | -1.564424519 | 0.435048513 | -2.702812575 | 2 | 4.442710129 | 7.502634626 | 5.144975053 | 4.97065356 | 4.894909044 | 4.456980017 |
| ENSG00000115946 | PNO1 | -1.00033789 | 0.144467139 | -1.703465995 | 0.007523544 | -2.703803885 | 9.613202746 | 8.276714465 | 9.809100877 | 8.29513208 | 8.45720682 | 7.633939843 | 7.736616914 |
| ENSG00000160207 | HSF2BP | -1.464562995 | 0.580207026 | -1.240486161 | 0.637526628 | -2.705049156 | 2 | 2 | 5.930865648 | 3.406930997 | 3.45167086 | 3.483289517 | 3.660199087 |
| ENSG00000266472 | MRPS21 | -1.65245985 | 0.000151748 | -1.05408672 | 0.020815791 | -2.70654657 | 10.80239439 | 9.876711741 | 10.42220168 | 8.713632895 | 8.8250571 | 9.229031012 | 9.488040822 |
| ENSG00000184281 | TSSC4 | -1.877719365 | 0.000108969 | -0.829011922 | 0.113843563 | -2.706731287 | 10.27168932 | 9.298208073 | 9.385998894 | 7.920420067 | 7.79989812 | 8.844037986 | 8.948586914 |
| ENSG00000103707 | MTFMT | -0.913380507 | 0.089385355 | -1.793702475 | 0.000260301 | -2.707082982 | 8.39321512 | 7.669933291 | 8.763901704 | 7.370304568 | 7.5014335 | 6.569017168 | 6.600402875 |
| ENSG00000128059 | PPAT | -1.344522704 | 0.210236498 | -1.36320072 | 0.194132356 | -2.707723424 | 5.866944937 | 8.820493869 | 8.175061096 | 6.799114572 | 6.70088158 | 6.928144991 | 6.506723027 |
| ENSG00000129244 | ATP1B2 | -1.917192296 | 0.201282385 | -0.790878288 | 0.615603284 | -2.708070584 | 3.327711234 | 5.595927442 | 4.874587708 | 3.95996233 | 2.59489879 | 4.281782022 | 4.188602707 |
| ENSG00000197114 | ZGPAT | -0.167499847 | 0.938052303 | -2.540675104 | 0.155898571 | -2.708174951 | 2 | 7.874700259 | 8.65477198 | 7.653784481 | 7.4949172 | 4.996409624 | 5.59182244 |
| ENSG00000140006 | WDR89 | -1.468634891 | 0.462772837 | -1.239945807 | 0.531674005 | -2.708580698 | 2 | 7.145017454 | 8.825624954 | 6.321555392 | 6.11534392 | 6.645228724 | 6.199243628 |
| ENSG00000130733 | YIPF2 | -1.364333532 | 0.000422235 | -1.347033319 | 0.000461516 | -2.711366851 | 10.18451821 | 9.821190352 | 10.55568472 | 8.85414072 | 8.86350401 | 9.009866973 | 8.728222198 |
| ENSG00000171885 | AQP4 | -1.9755469 | 0.503635732 | -0.736238562 | 0.811558361 | -2.711785461 | 2 | 4.710564154 | 3.369317972 | 3.162721474 | 2 | 3.517841605 | 3.042026731 |
| ENSG00000082397 | EPB41L3 | -1.188354604 | 0.185185634 | -1.523546298 | 0.076272108 | -2.711900902 | 8.540903468 | 10.26789142 | 8.865353883 | 8.195011988 | 8.31648126 | 8.396077967 | 7.226264598 |
| ENSG00000143222 | UFC1 | -1.15750876 | 0.00359668 | -1.554421305 | 4.97E-05 | -2.711930065 | 10.43795805 | 10.10889278 | 10.12770132 | 9.070139391 | 9.09116272 | 8.313052751 | 8.984263871 |
| ENSG00000099864 | PALM | -6.191029247 | 2.32E-11 | 3.478268537 | 7.32E-05 | -2.712760709 | 8.444143041 | 8.255926468 | 6.958202216 | 3.349643765 | 2.2680111 | 11.50493671 | 11.45325516 |
| ENSG00000101353 | MROH8 | -1.928752576 | 0.253424754 | -0.7842146 | 0.660736901 | -2.712967176 | 4.467311984 | 6.140560021 | 2 | 3.663521627 | 3.39680241 | 4.166733998 | 4.56420962 |
| ENSG00000008513 | ST3GAL1 | -0.463592125 | 0.585792769 | -2.249594884 | 0.001633879 | -2.713187008 | 9.699577239 | 10.64091981 | 11.63645966 | 10.37891658 | 10.4333835 | 8.69913345 | 8.55898913 |
| ENSG00000113504 | SLC12A7 | -0.648809293 | 0.417929197 | -2.065040465 | 0.003051234 | -2.713849759 | 5.330594134 | 6.951736442 | 6.718950348 | 5.85002558 | 5.92487824 | 4.636484163 | 4.614976609 |
| ENSG00000105696 | TMEM59L | -0.992596598 | 0.280090671 | -1.721736235 | 0.043186443 | -2.714332834 | 7.435667355 | 6.535001748 | 8.460932916 | 6.622948477 | 6.79936111 | 6.394066622 | 5.513708555 |
| ENSG00000077312 | SNRPA | -1.510383335 | 0.17350281 | -1.203969365 | 0.2795154 | -2.7143527 | 10.00110988 | 9.04308532 | 6.958202216 | 7.608714138 | 7.66128317 | 7.347386207 | 8.353097069 |
| ENSG00000076053 | RBM7 | -1.028807136 | 0.216615116 | -1.685952615 | 0.030507451 | -2.714759751 | 9.966689162 | 9.067150531 | 7.896971941 | 8.246921218 | 8.13141746 | 7.660277586 | 7.418582165 |
| ENSG00000178665 | ZNF713 | -1.310774944 | 0.229253543 | -1.404459345 | 0.180731158 | -2.715234289 | 5.532353208 | 4.11352228 | 5.772716771 | 4.358765249 | 3.94787299 | 4.416072028 | 3.67940853 |

| | | | | | | | | | | | | | |
|---|---|---|---|---|---|---|---|---|---|---|---|---|---|
| ENSG00000127952 | STYXL1 | -1.276951996 | 0.000482403 | -1.441764877 | 5.35E-05 | -2.718716873 | 8.418903792 | 8.395519306 | 8.865353883 | 7.277360022 | 7.35440525 | 7.291067412 | 7.006211772 |
| ENSG00000183780 | SLC35F3 | 3.655394797 | 0.061530373 | -6.375815045 | 0.009454427 | -2.720420248 | 2 | 4.442710129 | 2 | 6.456560224 | 6.37156942 | 2 | 2.059860178 |
| ENSG00000160948 | VPS28 | -1.532671859 | 2.60E-08 | -1.187853066 | 1.93E-05 | -2.720524924 | 9.886101849 | 9.863030423 | 10.19160738 | 8.420396336 | 8.50663048 | 8.697250351 | 8.906411656 |
| ENSG00000075856 | SART3 | -1.099774519 | 0.07662408 | -1.621826317 | 0.00575638 | -2.721600835 | 10.19941743 | 9.246456519 | 10.69449487 | 9.060130342 | 9.07377798 | 8.596789458 | 8.500786839 |
| ENSG00000182791 | CCDC87 | -0.845544851 | 0.593517792 | -1.880467942 | 0.19011257 | -2.726012793 | 2 | 4.710564154 | 4.523675391 | 3.663521627 | 3.45167086 | 2.885263063 | 3.042026731 |
| ENSG00000274750 | HIST1H3E | -1.235446277 | 0.478344278 | -1.490698374 | 0.3708938 | -2.726144651 | 6.138360429 | 2 | 5.930865648 | 4.443344104 | 4.37434574 | 4.04171018 | 4.353147976 |
| ENSG00000181666 | HKR1 | -1.626451258 | 0.159712355 | -1.101278066 | 0.350712355 | -2.727729323 | 6.89194353 | 9.13702746 | 10.37481416 | 7.756533788 | 7.79989812 | 8.344072943 | 8.245238611 |
| ENSG00000100347 | SAMM50 | -1.264982956 | 0.048902337 | -1.464195165 | 0.019274967 | -2.729178121 | 9.940323746 | 9.192779271 | 8.434763955 | 8.029321023 | 8.09955907 | 7.737439787 | 7.989373961 |
| ENSG00000162631 | NTNG1 | 2.014492482 | 0.519228435 | -4.744437094 | 0.151238336 | -2.729944612 | 2 | 3.076665209 | 2 | 3.754611435 | 3.39680241 | 2.048074405 | 2 |
| ENSG00000105202 | FBL | -1.477345348 | 7.66E-05 | -1.257094294 | 0.000823875 | -2.734439641 | 9.730679447 | 9.170736405 | 9.06622548 | 7.872924734 | 7.90685167 | 8.042684843 | 8.168292282 |
| ENSG00000010379 | SLC6A13 | -1.402290725 | 0.649972951 | -1.33398239 | NA | -2.736273115 | 2 | 3.686303414 | 2 | 2.141737246 | 2.59489879 | 2.384001718 | 2.421067459 |
| ENSG00000182749 | PAQR7 | -1.514977072 | 0.140239643 | -1.223385197 | 0.235763044 | -2.738362269 | 9.886101849 | 7.572420273 | 10.51254821 | 8.217201935 | 8.28849576 | 8.521632434 | 8.560932499 |
| ENSG00000116478 | HDAC1 | -1.037816453 | 0.075639549 | -1.700839492 | 0.001834875 | -2.738655944 | 10.56418906 | 9.377333006 | 10.30429804 | 9.103012382 | 9.1566168 | 8.199240359 | 8.703218237 |
| ENSG00000122970 | IFT81 | -0.709468198 | 0.648413066 | -2.029614387 | 0.143651762 | -2.739082585 | 4.467311984 | 7.701025996 | 9.165043284 | 7.292639438 | 7.44169935 | 6.013120342 | 6.182510897 |
| ENSG00000149932 | TMEM219 | -1.259822914 | 0.007788736 | -1.48259412 | 0.001323898 | -2.742417034 | 11.09707093 | 10.16572116 | 11.10866152 | 9.570001722 | 9.62041512 | 9.388071115 | 9.359585433 |
| ENSG00000178053 | MLF1 | -1.619840107 | 0.164250542 | -1.124035266 | 0.341075041 | -2.743875373 | 4.467311984 | 7.73146271 | 6.80318718 | 5.334065616 | 5.34860058 | 5.876679847 | 5.708623051 |
| ENSG00000161381 | PLXDC1 | -3.326036282 | 0.003441205 | 0.581518973 | 0.633084435 | -2.744517309 | 4.00722531 | 5.303523747 | 4.874587708 | 2.695456494 | 2.68924543 | 5.843010798 | 4.614976609 |
| ENSG00000168958 | MFF | -1.393334224 | 0.04626209 | -1.351597989 | 0.051017141 | -2.744932213 | 10.905938 | 9.148351514 | 10.31150679 | 8.909699622 | 8.8960155 | 8.99844268 | 8.887952157 |
| ENSG00000172421 | EFCAB3 | 2.445126554 | 0.678957281 | -5.191096587 | NA | -2.745970033 | 2 | 2 | 2 | 3.349643765 | 3.15372745 | 2 | 2 |
| ENSG00000114383 | TUSC2 | -1.525617039 | 0.000408074 | -1.220504723 | 0.00539006 | -2.746121761 | 9.371511179 | 9.102512015 | 9.981704053 | 7.954442493 | 8.08010078 | 8.331380596 | 8.307525005 |
| ENSG00000130414 | NDUFA10 | -1.596793132 | 0.001128625 | -1.149438745 | 0.023237554 | -2.746231876 | 11.11289609 | 9.988044813 | 10.55568472 | 9.013631184 | 9.05285777 | 9.421533019 | 9.531964377 |
| ENSG00000140478 | GOLGA6D | 2.444527733 | 0.679016801 | -5.191096589 | NA | -2.746568856 | 2 | 2 | 2 | 3.289987342 | 3.2184422 | 2 | 2 |
| ENSG00000100027 | YPEL1 | -1.544692282 | 0.078250731 | -1.201969897 | 0.165909543 | -2.74666218 | 5.70932728 | 4.710564154 | 6.073378139 | 4.299492648 | 4.1930041 | 4.785723309 | 4.201948172 |
| ENSG00000143376 | SNX27 | -1.705306929 | 7.72E-05 | -1.04150704 | 0.021475934 | -2.746813969 | 10.41898845 | 10.03760521 | 10.95142175 | 8.80304041 | 8.83669913 | 9.593120016 | 9.355112655 |
| ENSG00000244607 | CCDC13 | 2.44332935 | 0.67918823 | -5.191096594 | NA | -2.747767244 | 2 | 2 | 2 | 3.162721474 | 3.33976443 | 2 | 2 |
| ENSG00000147123 | NDUFB11 | -1.31226377 | 0.004606222 | -1.436315885 | 0.001618861 | -2.748579655 | 10.72727936 | 9.856140801 | 10.6093157 | 9.140440229 | 9.1347778 | 8.790727376 | 9.208268782 |
| ENSG00000141349 | G6PC3 | -1.281104061 | 0.000257308 | -1.468030187 | 2.01E-05 | -2.749134248 | 10.95943304 | 10.94355721 | 10.85110517 | 9.617795654 | 9.66608324 | 9.095643988 | 9.744671657 |
| ENSG00000277789 | | 2.44153019 | 0.679377579 | -5.191096602 | NA | -2.749566411 | 2 | 2 | 2 | 2.947924294 | 3.50452879 | 2 | 2 |
| ENSG00000198746 | GPATCH3 | -0.980049618 | 0.029202849 | -1.770795086 | 2.20E-05 | -2.750844703 | 8.418903792 | 7.928237109 | 8.763901704 | 7.527093191 | 7.34356593 | 6.69514648 | 6.64749689 |
| ENSG00000110721 | CHKA | -0.682091869 | 0.055943651 | -2.072538672 | 7.22E-11 | -2.754630541 | 7.940205782 | 8.077869079 | 8.143093325 | 7.398825229 | 7.37584259 | 6.173421703 | 5.917188835 |
| ENSG00000150510 | FAM124A | -2.589072218 | 0.023121834 | -0.16562593 | 0.904524117 | -2.754698148 | 4.815609471 | 6.602424674 | 4.058998049 | 3.515101962 | 3.50452879 | 5.4588802 | 5.436777225 |
| ENSG00000156873 | PHKG2 | -1.36942671 | 0.33304377 | -1.386463661 | 0.317269735 | -2.755890371 | 5.532353208 | 8.395519306 | 9.981704053 | 7.471115032 | 7.5462416 | 7.268416942 | 7.68646583 |
| ENSG00000169223 | LMAN2 | -1.072799535 | 0.002515173 | -1.683256345 | 6.68E-07 | -2.75605588 | 11.99260982 | 11.50459547 | 12.10320639 | 10.79667089 | 10.8375393 | 10.05880891 | 10.34413215 |
| ENSG00000158850 | B4GALT3 | -1.170095797 | 0.00375557 | -1.586909881 | 4.46E-05 | -2.757005678 | 10.27872131 | 10.00683004 | 9.491321578 | 8.748679155 | 8.84824797 | 8.492068113 | 8.273166147 |
| ENSG00000109083 | IFT20 | -1.267985809 | 0.039760769 | -1.491716215 | 0.012813892 | -2.759702024 | 9.304185368 | 8.414402626 | 9.859153765 | 7.97352961 | 8.12087597 | 7.792277365 | 7.864857644 |
| ENSG00000134049 | IER3IP1 | -0.301466294 | 0.686253129 | -2.461428383 | 3.77E-05 | -2.762894677 | 9.940323746 | 10.06773758 | 8.65477198 | 9.411949708 | 9.34954609 | 7.458282995 | 7.008112295 |
| ENSG00000178896 | EXOSC4 | -1.351040043 | 0.028339952 | -1.412956388 | 0.019994373 | -2.763996432 | 8.963744667 | 8.433041974 | 7.502634626 | 7.148924055 | 7.03768431 | 7.037865927 | 7.030725399 |
| ENSG00000165819 | METTL3 | -1.630582495 | 0.003661907 | -1.134993189 | 0.049999298 | -2.765575685 | 9.931427102 | 8.623598428 | 9.301632639 | 7.797276258 | 7.73502229 | 8.363201462 | 8.13650041 |
| ENSG00000120509 | PDZD11 | -1.622688606 | 0.061601819 | -1.143049473 | 0.196946345 | -2.76573808 | 10.3868074 | 8.903033433 | 8.206335842 | 7.883054517 | 7.82349365 | 8.299462788 | 8.352349212 |
| ENSG00000137054 | POLR1E | -2.15193064 | 6.41E-05 | -0.61494156 | 0.305318653 | -2.7668722 | 8.493334308 | 7.874700259 | 7.285536013 | 5.920568914 | 5.86549842 | 7.219398545 | 7.497817236 |
| ENSG00000087301 | TXNDC16 | -2.485832536 | 0.226635372 | -0.281790676 | 0.906201712 | -2.767623212 | 2 | 8.57407404 | 5.156626401 | 4.907595604 | 4.74762503 | 6.870823472 | 6.860240429 |
| ENSG00000103067 | ESRP2 | -2.178340454 | 0.18239431 | -0.589632131 | 0.725309613 | -2.767972585 | 4.00722531 | 3.076665209 | 3.369317972 | 2.600425781 | 2.38539787 | 3.082820997 | 3.236138336 |
| ENSG00000185909 | KLHDC8B | -2.386750533 | 2.68E-05 | -0.38244151 | 0.571407617 | -2.769192043 | 9.922475255 | 9.203675681 | 8.511888463 | 6.946463121 | 7.00595521 | 9.04915428 | 8.834865421 |
| ENSG00000144455 | SUMF1 | -2.064266434 | 4.41E-07 | -0.705835076 | 0.117409111 | -2.77010151 | 9.922475255 | 9.461450882 | 9.878698603 | 7.656739692 | 7.7866208 | 8.687797808 | 9.363673333 |
| ENSG00000173914 | RBM4B | -0.955215117 | 0.609360413 | -1.815702999 | 0.293236581 | -2.770918116 | 10.04301211 | 8.762721501 | 3.369317972 | 7.994705029 | 8.04707549 | 7.039353222 | 7.305675343 |
| ENSG00000205544 | TMEM256 | -1.496500894 | 0.114732583 | -1.274568724 | 0.177686131 | -2.771069617 | 8.963744667 | 6.899080544 | 7.098165388 | 6.379946408 | 6.64311914 | 6.635427934 | 6.81969491 |
| ENSG00000213015 | ZNF580 | -1.922105601 | 4.02E-08 | -0.850018367 | 0.023768024 | -2.772123968 | 10.29268328 | 9.962611189 | 9.735974718 | 8.113468729 | 8.10384772 | 8.955762164 | 9.35623215 |
| ENSG00000134375 | TIMM17A | -1.252383117 | 0.019565207 | -1.524953306 | 0.003407034 | -2.777336423 | 10.86447339 | 9.7032946 | 10.73794767 | 9.312394472 | 9.22907886 | 8.99461439 | 9.00677572 |
| ENSG00000204977 | TRIM13 | -1.195801459 | 0.129734214 | -1.583231116 | 0.03600172 | -2.779032575 | 10.787681 | 9.090820894 | 9.372276026 | 8.745906571 | 8.78288532 | 8.58766971 | 8.139971339 |
| ENSG00000185515 | BRCC3 | -1.146149607 | 0.028178148 | -1.63393744 | 0.001006735 | -2.780087047 | 8.228665467 | 9.159587375 | 9.242516611 | 7.895617537 | 7.72670378 | 7.178138321 | 7.478755384 |
| ENSG00000049768 | FOXP3 | -1.303391381 | 0.647700657 | -1.479173632 | 0.590766167 | -2.782565013 | 2 | 2 | 4.523675391 | 2.695456494 | 2.68924543 | 2.267001569 | 2.918126211 |
| ENSG00000166483 | WEE1 | -2.005488868 | 1.07E-11 | -0.778345209 | 0.01361034 | -2.783834077 | 9.448362284 | 9.506147172 | 9.385998894 | 7.440576128 | 7.4949172 | 8.864319213 | 8.459712934 |
| ENSG00000125656 | CLPP | -1.681753265 | 0.000354648 | -1.10316997 | 0.024438946 | -2.784923235 | 10.30651143 | 9.357954881 | 10.13584611 | 8.392303091 | 8.23671175 | 8.692059118 | 9.062708281 |
| ENSG00000165124 | SVEP1 | -1.626227311 | 0.006210955 | -1.159050227 | 0.057959502 | -2.785277538 | 9.081016759 | 9.648522413 | 10.49998282 | 8.254744042 | 8.240612 | 8.708979596 | 8.710239791 |
| ENSG00000149806 | FAU | -1.648659952 | 0.009564066 | -1.138680571 | 0.083207596 | -2.787340524 | 11.98832668 | 10.81444733 | 10.54960077 | 9.585592789 | 9.64784224 | 10.11349084 | 10.13610645 |
| ENSG00000139579 | NABP2 | -2.068636208 | 0.011100643 | -0.720515516 | 0.423520139 | -2.789151723 | 8.981092835 | 8.101365653 | 10.2528023 | 7.307758725 | 7.36516374 | 8.587161363 | 8.735123394 |
| ENSG00000182149 | IST1 | -1.325006323 | 0.000658244 | -1.465679894 | 0.000129752 | -2.790686217 | 11.59081378 | 11.08286144 | 11.89103474 | 10.2136641 | 10.2573647 | 10.12614381 | 10.06416949 |
| ENSG00000163795 | ZNF513 | -1.386915597 | 0.528732117 | -1.406266698 | 0.513903548 | -2.793182296 | 10.24321353 | 8.101365653 | 2 | 7.552702688 | 7.62295033 | 7.501007892 | 7.634508409 |
| ENSG00000106089 | STX1A | -1.47088652 | 0.423427099 | -1.322374525 | 0.467016155 | -2.793261045 | 9.371511179 | 6.535001748 | 3.369317972 | 6.535745081 | 6.5890786 | 6.800644007 | 6.602920209 |
| ENSG00000048544 | MRPS10 | -1.493452259 | 0.013191227 | -1.302169748 | 0.032017989 | -2.795622007 | 9.435834703 | 9.159587375 | 10.46161637 | 8.429064458 | 8.19309833 | 8.444590638 | 8.562873254 |
| ENSG00000255408 | PCDHA3 | -0.578115245 | 0.807827682 | -2.217525796 | 0.288238766 | -2.795641041 | 3.327711234 | 4.442710129 | 2 | 3.709785322 | 2.59489879 | 2.746468522 | 2.277179783 |
| ENSG00000109180 | OCIAD1 | -0.67248825 | 0.157050937 | -2.123699882 | 3.31E-07 | -2.796188132 | 11.59081378 | 10.75272247 | 10.85110517 | 10.47496775 | 10.4123582 | 9.045828404 | 8.952540041 |
| ENSG00000142621 | FHAD1 | 1.545639723 | 0.441837321 | -4.342347618 | 0.017198932 | -2.796707895 | 3.327711234 | 2 | 6.322058544 | 6.379946408 | 6.35006802 | 2.492221014 | 2.326754077 |
| ENSG00000173660 | UQCRH | -1.173193291 | 0.016661881 | -1.62382569 | 0.000529876 | -2.797018982 | 10.40620204 | 9.405920729 | 9.658943166 | 8.722121967 | 8.72001862 | 8.254710055 | 8.295903454 |
| ENSG00000111971 | | 2.393509416 | NA | -5.191096802 | NA | -2.797587386 | 2 | 2 | 2 | 3.566290607 | 2.77779907 | 2 | 2 |
| ENSG00000128039 | SRD5A3 | -0.37854259 | 0.529979238 | -2.41928596 | 8.50E-07 | -2.797828551 | 8.892187472 | 8.834582582 | 9.778213818 | 8.85414072 | 8.86097251 | 7.045287119 | 6.635252003 |
| ENSG00000124102 | PI3 | 2.391647173 | NA | -5.191096809 | NA | -2.799449637 | 2 | 2 | 2 | 3.406930997 | 3.01487888 | 2 | 2 |
| ENSG00000173113 | TRMT112 | -1.11611536 | 0.197898228 | -1.68351778 | 0.039331005 | -2.799633139 | 10.79260216 | 8.806266215 | 9.287079172 | 8.772033429 | 8.78689171 | 8.066221408 | 8.356084628 |
| ENSG00000186081 | KRT5 | 2.391025901 | NA | -5.191096812 | NA | -2.800070911 | 2 | 2 | 2 | 3.349643765 | 3.08597292 | 2 | 2 |
| ENSG00000167195 | GOLGA6C | 2.390404361 | NA | -5.191096814 | NA | -2.800692453 | 2 | 2 | 2 | 3.289987342 | 3.15372745 | 2 | 2 |
| ENSG00000097046 | CDC7 | -1.287739307 | 0.395924587 | -1.51398348 | 0.298583938 | -2.801722786 | 4.00722531 | 7.232742069 | 4.523675391 | 4.907595604 | 4.76938658 | 4.940196634 | 4.279527956 |
| ENSG00000135637 | CCDC142 | -1.258851613 | 0.030150603 | -1.54328255 | 0.005900513 | -2.802134162 | 8.418903792 | 8.297207175 | 7.225742779 | 6.782976084 | 6.90635305 | 6.713894915 | 6.415155055 |
| ENSG00000175756 | AURKAIP1 | -1.647923258 | 0.011477015 | -1.154481014 | 0.085841182 | -2.802404272 | 11.27639669 | 10.03760521 | 9.829330889 | 8.873322599 | 8.90955283 | 9.331225056 | 9.430034933 |
| ENSG00000008226 | DLEC1 | -2.682186671 | 0.470185896 | -0.124467237 | 0.974678148 | -2.806653909 | 4.00722531 | 2 | 2 | 2 | 2.38539787 | 2.937246078 | 2.918126211 |
| ENSG00000053371 | AKR7A2 | -1.915836314 | 0.009469798 | -0.890877431 | 0.261047618 | -2.806713746 | 10.51739401 | 8.777382925 | 9.132850258 | 7.717440541 | 7.84926609 | 8.705236603 | 8.882264318 |
| ENSG00000132517 | SLC52A1 | -4.295368089 | 0.204890893 | 1.487155305 | 0.630693854 | -2.808212784 | 2 | 2 | 4.058998049 | 2 | 2.14022021 | 4.378961816 | 3.287118176 |
| ENSG00000128604 | IRF5 | -2.70196544 | 0.307069196 | -0.10628783 | 0.9716719 | -2.80825327 | 2 | 2 | 5.930865648 | 2.868578001 | 2.68924543 | 4.984105765 | 3.474731315 |
| ENSG00000185236 | RAB11B | -1.788554232 | 0.000255505 | -1.020626731 | 0.048365141 | -2.809180963 | 11.79588449 | 10.68823825 | 11.38976845 | 9.5407064 | 9.61140668 | 10.17195017 | 10.49286637 |
| ENSG00000172671 | ZFAND4 | -1.739592812 | 0.000482793 | -1.06971743 | 0.036844987 | -2.809310243 | 7.096806373 | 7.002537973 | 7.600047325 | 5.709096143 | 5.47317115 | 6.428526856 | 5.976631893 |
| ENSG00000140691 | ARMC5 | -1.74852013 | 0.015778095 | -1.061783584 | 0.158766578 | -2.810303714 | 8.564112515 | 7.538394143 | 9.372276026 | 6.922108067 | 6.98750599 | 7.517109417 | 7.72395257 |
| ENSG00000111850 | SMIM8 | -1.73633672 | 0.042566554 | -1.0743204 | 0.221353293 | -2.81065712 | 5.532353208 | 7.467835385 | 6.322058544 | 4.927308402 | 5.19627118 | 5.725974948 | 5.601925227 |
| ENSG00000076554 | TPD52 | -5.696119424 | 0.097440073 | 2.884803238 | 0.348299029 | -2.811316186 | 4.00722531 | 2 | 2 | 2 | 2 | 5.125248504 | 4.951586397 |
| ENSG00000126215 | XRCC3 | -2.387137916 | 0.023103998 | -0.424858383 | 0.726496359 | -2.811996298 | 5.095959458 | 8.053983481 | 6.80318718 | 5.003571641 | 4.76938658 | 6.577231679 | 6.815359915 |
| ENSG00000132781 | MUTYH | -1.392388392 | 0.486454906 | -1.42093167 | 0.467550744 | -2.813320062 | 8.042902253 | 5.457120953 | 2 | 5.433909581 | 5.34860058 | 5.399896186 | 5.331426166 |
| | ATP5L2 | 4.1986317 | 0.01486761 | -7.012000278 | 0.002624999 | -2.813368578 | 3.327711234 | 3.076665209 | 2 | 5.978405189 | 6.26836889 | 2 | 2 |
| ENSG00000188493 | C19orf54 | -1.872842394 | 0.004587427 | -0.941691474 | 0.179051068 | -2.814533868 | 8.367060731 | 8.806266215 | 7.163363831 | 6.456560224 | 6.43421531 | 7.447120996 | 7.234405279 |
| ENSG00000130713 | EXOSC2 | -1.545021592 | 0.001154449 | -1.269739242 | 0.008252225 | -2.814760834 | 8.586954094 | 7.928237109 | 8.941672936 | 7.044488801 | 6.99676009 | 7.232460462 | 7.342293759 |
| ENSG00000116649 | SRM | -1.700989737 | 4.24E-05 | -1.115484574 | 0.009830805 | -2.81647431 | 11.09707093 | 10.92392403 | 10.32581712 | 9.061245954 | 9.18624204 | 9.476915012 | 9.905748 |
| ENSG00000091947 | TMEM101 | -1.683820391 | 0.001858408 | -1.133031403 | 0.042948413 | -2.816851795 | 9.48530526 | 8.234834549 | 9.180874355 | 7.398825229 | 7.38644297 | 7.908330388 | 7.95628774 |
| ENSG00000099899 | TRMT2A | -1.615725765 | 0.238681925 | -1.202504385 | 0.385194227 | -2.818230151 | 10.72212986 | 8.57407404 | 6.534109959 | 7.939959586 | 7.84670947 | 8.394916616 | 8.201061123 |
| ENSG00000172361 | CFAP53 | -1.266585231 | 0.479204668 | -1.55370456 | 0.361514946 | -2.82028979 | 2 | 3.686303414 | 5.39244166 | 3.515101962 | 3.2184422 | 3.27687201 | 3.128433649 |
| ENSG00000119969 | HELLS | -1.368220969 | 0.315594945 | -1.454353246 | 0.271500407 | -2.822574215 | 3.327711234 | 5.457120953 | 6.80318718 | 4.670864177 | 4.45712836 | 4.713032626 | 4.241259475 |
| ENSG00000084652 | TXLNA | -1.708759619 | 0.00731379 | -1.114026819 | 0.092695101 | -2.822786438 | 11.22998797 | 11.46891107 | 12.66680507 | 10.19396441 | 10.2616895 | 10.86970719 | 10.77126859 |
| ENSG00000169727 | GPS1 | -1.53398694 | 1.93E-07 | -1.288878354 | 1.42E-05 | -2.822865294 | 11.04852648 | 10.58367898 | 10.94216274 | 9.303936679 | 9.37815424 | 9.47746372 | 9.685630779 |
| ENSG00000144642 | RBMS3 | -1.301597805 | 8.19E-06 | -1.522154119 | 1.02E-07 | -2.823751924 | 10.11549941 | 9.695596167 | 9.935792933 | 8.693628619 | 8.56378716 | 8.499110916 | 8.319053686 |
| ENSG00000036672 | USP2 | -3.586497444 | 0.131232486 | 0.760600863 | 0.764012718 | -2.825896581 | 2 | 5.303523747 | 2 | 2.3892346 | 2.2680111 | 5.026717879 | 4.105828626 |
| ENSG00000049449 | RCN1 | -0.569253257 | 0.037200815 | -2.258619376 | 1.15E-20 | -2.827872634 | 12.81818241 | 12.56720988 | 12.41904395 | 12.06975087 | 12.0139954 | 10.32993244 | 10.38126632 |
| ENSG00000197299 | BLM | -1.678591066 | 0.502067192 | -1.15163983 | 0.646513979 | -2.830230896 | 2 | 5.457120953 | 2 | 3.162721474 | 2.94009902 | 3.193831101 | 3.496556178 |
| ENSG00000213741 | RPS29 | -0.939736459 | 0.367712724 | -1.890544924 | 0.047947959 | -2.830281383 | 11.47006801 | 9.256956408 | 9.287079172 | 9.408447353 | 9.5300868 | 8.631223999 | 8.415288533 |
| ENSG00000112312 | GMNN | -1.230192995 | 0.246051772 | -1.600121881 | 0.113988623 | -2.830314877 | 8.257427002 | 7.35507212 | 5.39244166 | 6.306580752 | 6.14893615 | 5.944926241 | 5.816671626 |
| ENSG00000182899 | RPL35A | -1.368613317 | 0.054635828 | -1.463156328 | 0.036332662 | -2.831769646 | 12.38030589 | 10.60438382 | 11.4530647 | 10.2522744 | 10.3291788 | 10.17482878 | 10.21885096 |
| ENSG00000196408 | NOXO1 | 0.427506011 | 0.918975348 | -3.261183263 | NA | -2.833677252 | 3.327711234 | 2 | 2 | 2.695456494 | 2.77779907 | 2.139668783 | 2 |
| ENSG00000183161 | FANCF | -1.567136812 | 0.054791031 | -1.266645596 | 0.122000983 | -2.833782408 | 8.19931886 | 6.311476754 | 7.552162999 | 6.034011955 | 6.02777321 | 6.595545082 | 5.964937606 |
| ENSG00000179256 | SMCO3 | -4.714884954 | 0.291724472 | 1.876909989 | 0.666037299 | -2.837974965 | 2 | 3.076665209 | 2 | 2 | 2 | 3.429858444 | 3.560117069 |
| ENSG00000140463 | BBS4 | -1.496333213 | 0.001579876 | -1.342422527 | 0.004656627 | -2.83875574 | 9.143983763 | 8.820493869 | 8.110401118 | 7.238438152 | 7.32531744 | 7.487996019 | 7.375063227 |
| ENSG00000086666 | ZFAND6 | -0.772040178 | 0.42301813 | -2.066836511 | 0.01584134 | -2.83887669 | 11.26935334 | 8.903033433 | 10.02619908 | 9.689531608 | 9.5196921 | 8.404758425 | 8.236348546 |
| ENSG00000269343 | ZNF587B | -1.453356309 | 0.361244176 | -1.385990349 | 0.376070469 | -2.839346657 | 4.467311984 | 6.899080544 | 9.301632639 | 6.628980152 | 6.55189318 | 6.272864826 | 6.957866334 |

| | | | | | | | | | | | | | |
|---|---|---|---|---|---|---|---|---|---|---|---|---|---|
| ENSG00000178149 | DALRD3 | -1.755367373 | 0.254759301 | -1.085673002 | 0.492815255 | -2.841040375 | 4.00722531 | 7.538394143 | 9.180874355 | 6.34372989 | 6.29861567 | 7.019897747 | 6.905807071 |
| ENSG00000198440 | ZNF583 | -1.428157938 | 0.466937891 | -1.413738675 | 0.461896088 | -2.841896613 | 2 | 6.666836837 | 8.434763955 | 5.709096143 | 6.00960265 | 6.049001768 | 5.689801209 |
| ENSG00000167103 | PIP5KL1 | -2.625472152 | 0.164314918 | -0.217613221 | 0.922180661 | -2.843085373 | 7.384433967 | 2 | 5.156626401 | 3.709785322 | 3.98554648 | 5.696113106 | 6.070564958 |
| ENSG00000087502 | ERGIC2 | -1.373127721 | 0.023590666 | -1.469970646 | 0.013945355 | -2.843098367 | 10.16187685 | 8.862353853 | 10.12770132 | 8.488325436 | 8.43865202 | 8.508806077 | 8.211820731 |
| ENSG00000135077 | HAVCR2 | -3.283783577 | 0.0122849 | 0.437812053 | 0.768665969 | -2.845971524 | 5.095959458 | 4.710564154 | 5.59507254 | 2.784612485 | 2.94009902 | 6.320128742 | 3.875378318 |
| ENSG00000198185 | ZNF334 | -1.287598273 | 0.481054906 | -1.559342235 | 0.373139559 | -2.846940508 | 5.330594134 | 7.145017454 | 2 | 4.783303513 | 4.83276841 | 4.727866721 | 4.411752372 |
| ENSG00000144591 | GMPPA | -1.322859122 | 6.53E-06 | -1.524522455 | 1.19E-07 | -2.847381577 | 10.05943807 | 9.680074956 | 10.04361983 | 8.624367384 | 8.61726622 | 8.303181855 | 8.530842618 |
| ENSG00000131269 | ABCB7 | -1.495433426 | 0.000142273 | -1.353171442 | 0.000546278 | -2.848604869 | 8.493334308 | 8.029695761 | 8.763901704 | 6.936770382 | 7.03319408 | 7.124495877 | 7.123062801 |
| ENSG00000100726 | TELO2 | -1.537748665 | 0.075134033 | -1.310959432 | 0.128768928 | -2.848708096 | 9.344956677 | 7.145017454 | 9.165043284 | 7.288834728 | 7.33629421 | 7.578754018 | 7.488317792 |
| ENSG00000164051 | CCDC51 | -2.121352662 | 0.061450217 | -0.727491856 | 0.556696605 | -2.848844518 | 9.2903348 | 6.464272742 | 6.958202216 | 5.949776857 | 6.18176397 | 7.262061517 | 7.544398272 |
| ENSG00000147041 | SYTL5 | -3.322470231 | 0.318527649 | 0.472854102 | 0.895713612 | -2.849616129 | 2 | 4.442710129 | 2 | 2.3892346 | 2 | 3.865109708 | 3.407202941 |
| ENSG00000091137 | SLC26A4 | -1.378080213 | 0.329342401 | -1.471898841 | 0.280146209 | -2.849979054 | 3.327711234 | 4.93638933 | 5.59507254 | 3.754611435 | 3.82856379 | 4.199031499 | 3.012034821 |
| ENSG00000132002 | DNAJB1 | -1.961071125 | 1.94E-05 | -0.889587149 | 0.072934705 | -2.850658274 | 11.9425688 | 10.95327452 | 11.82959564 | 9.641857585 | 9.71876306 | 10.85384055 | 10.63508781 |
| ENSG00000120697 | ALG5 | -0.543338838 | 0.594639663 | -2.30757328 | 0.008111825 | -2.850882118 | 10.33377634 | 7.874700259 | 9.049077186 | 8.868232317 | 8.88857762 | 7.107581047 | 7.178120748 |
| ENSG00000174695 | TMEM167A | -1.458430501 | 0.002590338 | -1.393133705 | 0.003999086 | -2.851564207 | 10.56418906 | 11.54371287 | 11.55983287 | 9.856914282 | 9.81737584 | 9.975607996 | 9.82503293 |
| ENSG00000072506 | HSD17B10 | -1.642857519 | 0.008043844 | -1.211128127 | 0.056459449 | -2.853985646 | 9.98400217 | 8.791896851 | 9.767769415 | 7.922877029 | 8.01555453 | 7.912394292 | 8.757326984 |
| ENSG00000143368 | SF3B4 | -1.590690912 | 0.005010769 | -1.263385199 | 0.029032528 | -2.854076111 | 11.30078111 | 10.67263725 | 10.16000817 | 9.206465443 | 9.19832501 | 9.122554694 | 9.843777824 |
| ENSG00000169919 | GUSB | -2.026631842 | 4.84E-14 | -0.829392316 | 0.003408467 | -2.856024158 | 9.720386367 | 9.497318007 | 9.746650996 | 7.596453725 | 7.70428369 | 8.946676188 | 8.707903068 |
| ENSG00000169925 | BRD3 | -1.559884908 | 3.61E-06 | -1.296705458 | 0.000135243 | -2.856590366 | 10.31337612 | 10.42977533 | 9.888372618 | 8.622857004 | 8.73530306 | 9.082719462 | 8.783638895 |
| ENSG00000274671 | | -1.8727386 | 0.390227632 | -0.98436935 | 0.649493169 | -2.85710795 | 2 | 3.686303414 | 3.369317972 | 2.270786764 | 2.59489879 | 2.526562374 | 2.918126211 |
| ENSG00000141449 | GREB1L | 1.996700168 | 0.195586526 | -4.854346519 | 0.000723382 | -2.857646351 | 2 | 6.535001748 | 4.874587708 | 7.25803034 | 7.36158648 | 2.457042231 | 2.374681325 |
| ENSG00000130950 | NUTM2F | 2.330936339 | NA | -5.191097062 | NA | -2.860160723 | 2 | 2 | 2 | 2.947924294 | 3.39680241 | 2 | 2 |
| ENSG00000184792 | OSBP2 | -3.658561394 | 8.02E-08 | 0.798298348 | 0.292247026 | -2.860263046 | 8.107517749 | 7.002537973 | 7.342949392 | 4.237680128 | 4.1930041 | 7.667028121 | 8.797990868 |
| ENSG00000111361 | EIF2B1 | -1.609522312 | 0.003680018 | -1.25129718 | 0.027315593 | -2.860819492 | 10.23600589 | 9.159587375 | 10.38167985 | 8.319579168 | 8.50986596 | 8.640073371 | 8.89361766 |
| ENSG00000168569 | TMEM223 | -1.790798158 | 0.270288869 | -1.074110395 | 0.521409207 | -2.864908553 | 9.112843761 | 8.337339476 | 3.369317972 | 6.470063213 | 6.46120797 | 7.315824427 | 6.977399345 |
| ENSG00000127989 | MTERF1 | -1.158270159 | 0.237918279 | -1.707244474 | 0.065523007 | -2.865514633 | 8.696033109 | 6.389896633 | 7.225742779 | 6.658765962 | 6.5890786 | 6.287958221 | 5.888587685 |
| ENSG00000107949 | BCCIP | -1.073710119 | 0.000513088 | -1.791945454 | 1.17E-09 | -2.865655573 | 10.02639697 | 9.741182361 | 10.17589429 | 9.010169232 | 8.82765235 | 8.110051399 | 8.305980863 |
| ENSG00000172728 | FUT10 | -1.636585398 | 0.000131823 | -1.229108657 | 0.004594153 | -2.865694054 | 7.624139768 | 8.395519306 | 7.934808822 | 6.516352584 | 6.35727082 | 6.884127769 | 6.77351449 |
| ENSG00000159596 | TMEM69 | -1.428699774 | 0.112341523 | -1.446821659 | 0.101201855 | -2.875521433 | 10.04301211 | 8.955547846 | 7.691296561 | 7.791909949 | 7.77322015 | 7.541454591 | 7.958255135 |
| ENSG00000054690 | PLEKHH1 | -2.493628445 | 0.00938604 | -0.382286219 | 0.723692046 | -2.875914664 | 4.00722531 | 5.457120953 | 5.59507254 | 3.023133209 | 3.50452879 | 4.868377487 | 4.827924991 |
| ENSG00000095303 | PTGS1 | -0.782840344 | 0.46185912 | -2.093187639 | 0.026578616 | -2.876027983 | 8.493334308 | 6.535001748 | 8.884815469 | 7.559034656 | 7.4280824 | 6.672321934 | 5.59182244 |
| ENSG00000118162 | KPTN | -1.869266624 | 0.001675231 | -1.006952166 | 0.104207046 | -2.876218791 | 7.384433967 | 7.099070191 | 6.322058544 | 5.194621444 | 5.30456641 | 5.751606453 | 6.286392397 |
| ENSG00000182810 | DDX28 | -1.597583075 | 0.000547787 | -1.278718494 | 0.006075791 | -2.876301569 | 8.39321512 | 7.503546079 | 7.896971941 | 6.387081877 | 6.44777477 | 6.62753908 | 6.813187523 |
| ENSG00000104998 | IL27RA | -2.382374605 | 0.207400524 | -0.494350545 | 0.815972748 | -2.87672515 | 8.737455672 | 6.951736442 | 2 | 5.144975053 | 5.39133044 | 7.332904981 | 6.693102083 |
| ENSG00000185303 | SFTPA2 | -3.274362978 | 0.456607119 | 0.397169276 | NA | -2.877193702 | 2 | 2 | 3.369317972 | 2 | 2.14022021 | 3.1504466 | 2.117335275 |
| ENSG00000184154 | LRTOMT | -1.941909325 | 0.007283609 | -0.93674479 | 0.215049853 | -2.878654115 | 5.532353208 | 6.389896633 | 6.629488387 | 4.237680128 | 4.74762503 | 5.102662916 | 5.626876946 |
| ENSG00000106268 | NUDT1 | -2.238690145 | 0.164314918 | -0.640278309 | 0.717839521 | -2.878968455 | 8.367060731 | 5.946638857 | 3.369317972 | 5.003571641 | 4.95174433 | 6.363520437 | 6.528072775 |
| ENSG00000101004 | NINL | -0.522048186 | 0.546426658 | -2.357065361 | 0.001300235 | -2.879113547 | 7.709880939 | 7.538394143 | 9.180874355 | 7.700356475 | 7.9408106 | 6.189586148 | 5.905000563 |
| ENSG00000146109 | ABT1 | -1.642894336 | 4.76E-05 | -1.237528247 | 0.002625096 | -2.880422584 | 8.609439662 | 9.214490408 | 9.439621147 | 7.481152682 | 7.53358066 | 7.850199974 | 7.958255135 |
| ENSG00000182307 | C8orf33 | -2.060294166 | 1.17E-06 | -0.821146659 | 0.075265997 | -2.881440825 | 10.15425007 | 9.415325457 | 10.29705308 | 7.944803394 | 7.97164382 | 9.286835797 | 9.0731854 |
| ENSG00000196465 | MYL6B | -1.000167292 | 0.112165694 | -1.881954767 | 0.001141619 | -2.882122059 | 10.25038534 | 9.04308532 | 9.624627483 | 8.724940594 | 8.7297638 | 7.451596164 | 8.173382342 |
| ENSG00000140374 | ETFA | -0.90830538 | 0.448379821 | -1.974905765 | 0.068863117 | -2.883211145 | 10.98974631 | 9.377333006 | 7.734844356 | 9.011324139 | 9.02712059 | 7.882875413 | 8.042267212 |
| ENSG00000182518 | FAM104B | -1.958390638 | 0.008742661 | -0.926577288 | 0.238534046 | -2.884967927 | 6.563806279 | 7.051611315 | 5.39244166 | 4.647285135 | 4.79082474 | 5.4588802 | 5.794819115 |
| ACRC | ACRC | 3.266878519 | 0.310604101 | -6.152931129 | 0.075017773 | -2.886052609 | 3.327711234 | 2 | 2 | 4.54891529 | 4.50979031 | 2 | 2 |
| ENSG00000196407 | THEM5 | -4.71488067 | 0.300402677 | 1.825880697 | 0.681431908 | -2.888999974 | 2 | 3.076665209 | 2 | 2 | 2 | 3.753554863 | 3.100202476 |
| ENSG00000067533 | RRP15 | -1.519848591 | 0.134992306 | -1.36934732 | 0.174527088 | -2.889195911 | 8.285626337 | 7.538394143 | 10.10298738 | 7.53353831 | 7.57432889 | 7.68709225 | 7.715835172 |
| ENSG00000172346 | CSDC2 | -3.313112567 | 0.098913801 | 0.423502963 | 0.856270394 | -2.889609604 | 8.816894989 | 6.389896633 | 2 | 4.443344104 | 4.43005853 | 6.380048704 | 8.62735196 |
| ENSG00000133328 | HRASLS2 | 0.55292621 | 0.832844524 | -3.442660783 | 0.126117296 | -2.889734573 | 2 | 2 | 4.874587708 | 3.95996233 | 4.02226117 | 2.267001569 | 2.225841198 |
| ENSG00000117155 | SSX2IP | -1.381858822 | 0.262822363 | -1.508804617 | 0.208845264 | -2.890663439 | 7.384433967 | 8.655694736 | 5.156626401 | 6.422239259 | 6.22964624 | 6.439833081 | 5.933281326 |
| ENSG00000105443 | CYTH2 | -0.887013828 | 0.103494474 | -2.004361861 | 4.61E-05 | -2.891375688 | 10.92846914 | 9.821190352 | 10.81605395 | 9.730780958 | 9.69763767 | 8.673022865 | 8.533484187 |
| DUS2 | DUS2 | -1.703530555 | 0.041956419 | -1.187910495 | 0.163887111 | -2.891441051 | 6.138360429 | 8.124485672 | 7.777116083 | 5.753943344 | 6.09824986 | 6.109723842 | 6.681239787 |
| ENSG00000184708 | EIF4ENIF1 | -1.552376744 | 0.527194576 | -1.340369743 | 0.581686453 | -2.892746486 | 2 | 6.899080544 | 10.94679967 | 7.870381136 | 7.95274764 | 8.224302521 | 8.009436866 |
| ENSG00000069509 | FUNDC1 | -1.340440469 | 0.02968828 | -1.55354509 | 0.009522887 | -2.893985559 | 8.674867122 | 7.35507212 | 7.818184403 | 6.634986713 | 6.84106088 | 6.546183077 | 6.528072775 |
| ENSG00000143156 | NME7 | -0.635275875 | 0.236508204 | -2.258822944 | 1.17E-06 | -2.894098818 | 8.285626337 | 7.701025996 | 7.285536013 | 7.157295126 | 7.23044321 | 5.590506472 | 5.708623051 |
| ENSG00000178573 | MAF | -4.055114762 | 5.66E-10 | 1.158654855 | 0.095534894 | -2.896459907 | 7.435667355 | 7.874700259 | 7.552162999 | 4.443344104 | 3.28037825 | 8.601831186 | 8.937160962 |
| ENSG00000163006 | CCDC138 | -1.038489915 | 0.530264818 | -1.857977125 | 0.223744982 | -2.89646704 | 2 | 5.946638857 | 4.874587708 | 4.17310023 | 4.12711147 | 3.600766115 | 3.518055793 |
| ENSG00000071859 | FAM50A | -1.267594533 | 0.058083878 | -1.629664137 | 0.011313199 | -2.89725867 | 11.16695586 | 9.648522413 | 10.11127249 | 9.135152476 | 9.24469897 | 8.916644086 | 8.741420621 |
| ENSG00000186871 | ERCC6L | -1.828039726 | 0.479583688 | -1.069803142 | 0.684964093 | -2.897842868 | 2 | 5.72254323 | 2 | 2.947924294 | 3.28037825 | 3.10571689 | 3.940519005 |
| ENSG00000110881 | ASIC1 | -2.822067849 | 0.154010606 | -0.077982186 | 0.973251353 | -2.900050035 | 4.00722531 | 9.055168103 | 3.369317972 | 4.804776789 | 4.98931816 | 7.619097297 | 7.302581437 |
| ENSG00000178343 | SHISA3 | -3.917262418 | 0.211000866 | 1.01686808 | 0.747201125 | -2.900394338 | 2 | 4.442710129 | 2 | 2.270786764 | 2 | 3.838019203 | 4.241259475 |
| ENSG00000049323 | LTBP1 | -2.232952584 | 0.019076624 | -0.668287599 | 0.530366429 | -2.901240184 | 10.95943304 | 13.78286584 | 12.98235842 | 10.74032429 | 10.7523441 | 12.56045519 | 12.00372897 |
| ENSG00000159593 | NAE1 | -0.284064734 | 0.801824404 | -2.617375397 | 0.003770117 | -2.901440131 | 10.14658276 | 8.077869079 | 8.042706615 | 8.83470036 | 8.8354102 | 6.739740021 | 6.33565646 |
| ENSG00000182508 | LHFPL1 | 0.991084307 | 0.713785889 | -3.893577725 | 0.126945765 | -2.902493418 | 2 | 3.686303414 | 2 | 3.09461492 | 3.65239003 | 2.094598369 | 2.059860178 |
| ENSG00000022976 | ZNF839 | -1.191892872 | 0.324504083 | -1.713020334 | 0.134216355 | -2.904913206 | 8.816894989 | 7.538394143 | 5.39244166 | 6.604700657 | 6.71776877 | 6.22922222 | 6.099376274 |
| ENSG00000157020 | SEC13 | -1.700063959 | 0.002924743 | -1.205998832 | 0.041029321 | -2.906062791 | 12.55410083 | 11.50459547 | 11.32712828 | 10.16571753 | 10.245283 | 10.56905757 | 10.81765528 |
| ENSG00000169905 | TOR1AIP2 | -1.568062107 | 0.001457494 | -1.33900646 | 0.007299395 | -2.907068567 | 11.06081658 | 11.57970793 | 12.18594181 | 10.09829921 | 10.135408 | 10.25574405 | 10.42957331 |
| ENSG00000186787 | SPIN2B | -0.891140938 | 0.305697747 | -2.016692222 | 0.010048431 | -2.90783316 | 6.653098118 | 4.93638933 | 6.43197674 | 5.304221918 | 5.36298494 | 4.311629396 | 4.4231929 |
| ENSG00000188342 | GTF2F2 | -1.255585176 | 0.230926542 | -1.65398868 | 0.098757188 | -2.909573856 | 10.56993315 | 8.916342327 | 7.818184403 | 8.285617933 | 8.29788471 | 7.87872738 | 7.919398473 |
| ENSG00000140332 | TLE3 | -0.957558593 | 0.091400188 | -1.952837437 | 0.000158283 | -2.910396029 | 9.521325812 | 9.102512015 | 9.385998894 | 8.341783829 | 8.44880239 | 7.85696323 | 6.782425482 |
| ENSG00000177600 | RPLP2 | -1.233378285 | 0.05970955 | -1.677421517 | 0.007405859 | -2.910799802 | 12.53660743 | 11.2524908 | 11.23155829 | 10.55871559 | 10.6016606 | 10.02279773 | 10.24406712 |
| ENSG00000162384 | C1orf123 | -1.680155279 | 3.10E-11 | -1.231056894 | 1.11E-06 | -2.911212173 | 9.097018025 | 9.067150531 | 8.922968322 | 7.341208691 | 7.41087845 | 7.767364534 | 7.863808491 |
| ENSG00000117862 | TXNDC12 | 0.126603713 | 0.850047138 | -3.038859502 | 8.79E-10 | -2.91225579 | 9.448362284 | 9.267380431 | 10.28977157 | 9.817184017 | 9.90967722 | 6.956760977 | 6.501335855 |
| ENSG00000125901 | MRPS26 | -1.84124714 | 0.053531997 | -1.071311069 | 0.280385375 | -2.912558209 | 9.931427102 | 8.191702463 | 7.502634626 | 7.157295126 | 7.05107174 | 7.87872738 | 7.836259222 |
| ENSG00000157343 | ARMC12 | -2.209184749 | 0.361916636 | -0.70364882 | 0.784923737 | -2.912833568 | 2 | 5.303523747 | 2 | 2.695456494 | 2.77779907 | 3.648325314 | 3.407202941 |
| ENSG00000128645 | HOXD1 | 2.276328435 | NA | -5.191097289 | NA | -2.914768854 | 2 | 2 | 2 | 3.349643765 | 2.94009902 | 2 | 2 |
| ENSG00000159377 | PSMB4 | -1.27920546 | 0.028200875 | -1.636480699 | 0.003631339 | -2.915686159 | 12.32363489 | 11.03180337 | 11.38976845 | 10.38694723 | 10.4355118 | 10.02129326 | 10.08862406 |
| C10ORF35 | C10ORF35 | 2.274983007 | NA | -5.191097295 | NA | -2.916114288 | 2 | 2 | 2 | 3.227757282 | 3.08597292 | 2 | 2 |
| ENSG00000146267 | FAXC | -0.590465287 | 0.764636496 | -2.327173673 | 0.178036985 | -2.917638959 | 3.327711234 | 5.131603625 | 2 | 3.881293667 | 3.39680241 | 2.656294034 | 2.782576693 |
| ENSG00000241394 | | 3.234980731 | 0.324107683 | -6.15293023 | 0.079108348 | -2.917949498 | 2 | 3.076665209 | 2 | 4.443344104 | 4.56059747 | 2 | 2 |
| ENSG00000177301 | KCNA2 | -3.267594726 | 0.479080051 | 0.349492645 | NA | -2.918102081 | 3.327711234 | 2 | 2 | 2.141737246 | 2 | 3.1504466 | 2.059860178 |
| ENSG00000188321 | ZNF559 | -1.433094729 | 0.232537767 | -1.485994487 | 0.205709574 | -2.919089216 | 5.532353208 | 7.60566235 | 8.99637432 | 6.61689148 | 6.52655952 | 6.444330878 | 6.595354992 |
| ENSG00000145908 | ZNF300 | -1.031027365 | 0.528056559 | -1.891275903 | 0.209481138 | -2.922303268 | 3.327711234 | 6.951736442 | 8.380957451 | 6.336376195 | 6.23747437 | 5.390603804 | 5.576534358 |
| ENSG00000159147 | DONSON | -1.633613125 | 0.324734879 | -1.291013947 | 0.438543047 | -2.924627072 | 4.00722531 | 7.431218269 | 9.553445883 | 6.710878132 | 6.67805337 | 7.305972543 | 6.676467423 |
| ENSG00000197111 | PCBP2 | -1.140136896 | 1.43E-09 | -1.786060833 | 2.69E-22 | -2.926197728 | 12.13890018 | 12.13942551 | 12.14011481 | 11.00812841 | 10.9935992 | 10.25558411 | 10.45103923 |
| ENSG00000125775 | SDCBP2 | -0.716892038 | 0.683648319 | -2.211161536 | 0.154998812 | -2.928053574 | 2 | 6.228547981 | 6.43197674 | 4.984880435 | 5.27444544 | 4.209638631 | 3.539239711 |
| ENSG00000101638 | ST8SIA5 | 0.047842333 | 0.98350331 | -2.977151451 | 0.121757443 | -2.929309118 | 3.327711234 | 4.442710129 | 2 | 3.663521627 | 3.65239003 | 2.526562374 | 2.117335275 |
| ENSG00000173715 | C11orf80 | -1.32983744 | 0.35107637 | -1.600036379 | 0.243971969 | -2.929873819 | 6.816637977 | 7.60566235 | 3.369317972 | 5.461211713 | 5.51241389 | 5.371837532 | 5.101530266 |
| ENSG00000141510 | TP53 | -2.052360401 | 4.07E-05 | -0.881162947 | 0.104308168 | -2.933523348 | 9.689059027 | 9.981728319 | 10.70548138 | 8.10700104 | 8.19711812 | 9.03878183 | 9.542821395 |
| ENSG00000154743 | TSEN2 | -2.000499156 | 0.160777328 | -0.936142833 | 0.531906196 | -2.936641989 | 3.327711234 | 6.311476754 | 7.691296561 | 4.71689672 | 4.85329164 | 5.747972572 | 5.727202496 |
| ENSG00000136717 | BIN1 | -1.541385316 | 0.114575748 | -1.39724306 | 0.149336017 | -2.938628376 | 11.40728614 | 9.726146444 | 8.865353883 | 8.896009193 | 8.81854848 | 9.017055847 | 8.984263871 |
| ENSG00000162961 | DPY30 | -1.035164775 | 0.196944427 | -1.903752319 | 0.010141651 | -2.938917093 | 9.590776211 | 8.356994601 | 7.777116083 | 7.786523606 | 7.72670378 | 7.04972158 | 6.755525668 |
| ENSG00000166979 | EVA1C | -5.394579411 | 0.144043985 | 2.453381047 | 0.471404763 | -2.941198364 | 2 | 3.686303414 | 2 | 2 | 2 | 4.742549839 | 4.002845082 |
| ENSG00000139531 | SUOX | -1.803511166 | 1.68E-06 | -1.14178967 | 0.003249295 | -2.945300835 | 8.716893051 | 9.125613817 | 8.434763955 | 7.003290579 | 7.04216061 | 7.578754018 | 7.752576035 |
| ENSG00000068784 | SRBD1 | -1.852376092 | 0.000396333 | -1.093829276 | 0.044809508 | -2.946205368 | 8.586954094 | 7.572420273 | 8.042706615 | 6.29903474 | 6.34282908 | 6.795372357 | 7.266516281 |
| ENSG00000172172 | MRPL13 | -1.058127543 | 0.001405808 | -1.888266535 | 1.63E-09 | -2.946394078 | 9.435834703 | 8.929529568 | 9.014156722 | 8.132699624 | 8.05374131 | 7.282301103 | 7.277588383 |
| ENSG00000075826 | SEC31B | -1.244659541 | 0.251199372 | -1.702351617 | 0.098657948 | -2.947011158 | 8.797441463 | 5.838937808 | 8.076950877 | 6.733448311 | 6.84106088 | 6.275391383 | 6.420873184 |
| ENSG00000145386 | CCNA2 | -0.72920507 | 0.613825095 | -2.218995266 | 0.082655654 | -2.948200336 | 3.327711234 | 7.232742069 | 6.629488387 | 5.797438367 | 5.71663161 | 4.311629396 | 4.574506939 |
| ENSG00000069188 | SDK2 | -4.335476493 | 0.008885084 | 1.387217689 | 0.446296881 | -2.948258805 | 2 | 8.191702463 | 7.451345288 | 3.709785322 | 3.33976443 | 8.243140215 | 8.989083133 |
| ENSG00000105851 | PIK3CG | -4.714876406 | 0.309004906 | 1.76567576 | 0.696976163 | -2.949200647 | 2 | 2 | 3.369317972 | 2 | 2 | 3.917810786 | 2.671842496 |
| ENSG00000128578 | STRIP2 | 0.405287939 | 0.717477947 | -3.355798427 | 0.000222971 | -2.950510487 | 4.815609471 | 7.002537973 | 7.502634626 | 7.097651217 | 7.31055061 | 3.943456545 | 3.789585079 |
| ENSG00000188976 | NOC2L | -1.29646654 | 0.020861927 | -1.654986545 | 0.002249766 | -2.951453085 | 10.25752168 | 9.470501606 | 10.76969939 | 8.894758143 | 9.03836713 | 8.566163393 | 8.658541894 |
| ENSG00000108578 | BLMH | -2.158021755 | 1.58E-08 | -0.796506842 | 0.055158384 | -2.954519597 | 9.646202137 | 9.994333773 | 9.227351203 | 7.43025113 | 7.61696151 | 8.792933324 | 8.941145478 |
| ENSG00000161395 | PGAP3 | -2.027646649 | 0.052880994 | -0.926955633 | 0.406108709 | -2.954602282 | 9.204317094 | 8.505280662 | 6.073378139 | 6.394182228 | 6.47451726 | 7.568489426 | 7.427130611 |
| ENSG00000275763 | C18orf65 | -1.340359273 | 0.60992929 | -1.616798567 | 0.521209652 | -2.95715784 | 5.095959458 | 2 | 2 | 3.023133209 | 2.94009902 | 3.011868911 | 2.671842496 |
| ENSG00000138131 | LOXL4 | -3.398085076 | 0.044129227 | 0.437475729 | 0.827299212 | -2.960609347 | 11.22998797 | 9.680074956 | 4.058998049 | 6.722207358 | 6.73446058 | 10.59545838 | 10.42424552 |
| ENSG00000126749 | EMG1 | -1.411132853 | 0.126534396 | -1.550709512 | 0.084711655 | -2.961842365 | 10.06758145 | 8.255926468 | 7.97167868 | 7.614805492 | 7.76240929 | 7.473766117 | 7.629578661 |
| ENSG00000160145 | KALRN | -2.068350804 | 0.055017038 | -0.895801648 | 0.432665781 | -2.964152452 | 4.467311984 | 5.946638857 | 7.029880454 | 4.358765249 | 4.2865223 | 5.684753614 | 4.845038551 |

| | | | | | | | | | | | | | |
|---|---|---|---|---|---|---|---|---|---|---|---|---|---|
| ENSG00000272674 | PCDHB16 | -0.332466583 | 0.854355329 | -2.632021705 | 0.077889486 | -2.964488287 | 2 | 5.838937808 | 6.629488387 | 5.406080789 | 5.45984938 | 3.600766115 | 3.560117069 |
| ENSG00000086300 | SNX10 | -3.328513255 | 0.235745334 | 0.363994978 | 0.908329356 | -2.964518277 | 2 | 4.442710129 | 4.523675391 | 2 | 2.68924543 | 5.079718116 | 2.551896972 |
| ENSG00000277823 | | 0.86753033 | 0.547023674 | -3.832853984 | 0.002956694 | -2.965323654 | 3.327711234 | 5.303523747 | 3.369317972 | 4.984880435 | 5.22805098 | 2.457042231 | 2.277179783 |
| ENSG00000145982 | FARS2 | -1.567643794 | 0.001787884 | -1.398134492 | 0.005270287 | -2.965778287 | 8.16936288 | 7.393647515 | 8.076950877 | 6.221320894 | 6.53928196 | 6.322573937 | 6.748721609 |
| ENSG00000064545 | TMEM161A | -1.737631161 | 4.68E-10 | -1.229004447 | 1.21E-05 | -2.966635607 | 9.27634997 | 9.125613817 | 8.884815469 | 7.348537858 | 7.43490694 | 7.884531292 | 7.897008641 |
| ENSG00000134815 | DHX34 | -1.142415944 | 0.004955448 | -1.82446991 | 2.11E-06 | -2.966885855 | 8.631580145 | 8.607278467 | 8.110401118 | 7.366699513 | 7.33264456 | 6.879153024 | 6.468583424 |
| ENSG00000099783 | HNRNPM | -1.049065596 | 0.034081755 | -1.920477646 | 2.97E-05 | -2.969543242 | 11.55650736 | 10.64091981 | 10.92346452 | 10.04177408 | 10.0490837 | 8.919069795 | 9.398504094 |
| ENSG00000180287 | PLD5 | 2.220966612 | NA | -5.19109752 | NA | -2.970130908 | 2 | 2 | 2 | 3.754611435 | 2 | 2 | 2 |
| ENSG00000141026 | MED9 | -1.817004559 | 0.017562521 | -1.153465527 | 0.145326684 | -2.970470086 | 7.27616306 | 8.469613413 | 9.272377397 | 6.622948477 | 6.91125523 | 7.405054033 | 7.431385914 |
| ENSG00000125445 | MRPS7 | -1.677018674 | 3.25E-06 | -1.294360838 | 0.000405269 | -2.971379511 | 10.33377634 | 9.936721131 | 9.819251342 | 8.374462892 | 8.38855922 | 8.517903178 | 8.967756908 |
| ENSG00000167881 | SRP68 | -1.72116603 | 6.13E-07 | -1.251265239 | 0.000394133 | -2.972431269 | 11.44840376 | 10.76748534 | 11.24291424 | 9.449924671 | 9.47652116 | 9.863162331 | 9.995658407 |
| ENSG00000213402 | PTPRCAP | 2.215367921 | NA | -5.191097543 | NA | -2.975729622 | 2 | 2 | 2 | 3.349643765 | 2.86123014 | 2 | 2 |
| ENSG00000197261 | C6orf141 | 2.214666508 | NA | -5.191097546 | NA | -2.976431038 | 2 | 2 | 2 | 3.289987342 | 2.94009902 | 2 | 2 |
| ENSG00000131746 | TNS4 | -1.14431883 | 0.671525544 | -1.832828218 | 0.46940984 | -2.977147047 | 2 | 2 | 4.874587708 | 2.695456494 | 3.15372745 | 2.746468522 | 2.551896972 |
| ENSG00000087299 | L2HGDH | -1.504187788 | 0.024718288 | -1.473096048 | 0.026400043 | -2.977283836 | 8.313285031 | 6.844429755 | 8.007629717 | 6.260701031 | 6.48770488 | 6.575182432 | 6.219070051 |
| ENSG00000213186 | TRIM59 | -1.931935821 | 0.029396517 | -1.046826589 | 0.261292425 | -2.97876241 | 7.218819781 | 7.232742069 | 9.116480286 | 6.372775472 | 6.16544343 | 7.262061517 | 6.969617847 |
| ENSG00000137709 | POU2F3 | 0.281501418 | 0.900708025 | -3.261256887 | 0.087927265 | -2.979755469 | 2 | 3.076665209 | 4.058998049 | 3.566290607 | 3.28037825 | 2.267001569 | 2.117335275 |
| ENSG00000104341 | LAPTM4B | -1.995784548 | 1.17E-07 | -0.98408491 | 0.014322982 | -2.979869458 | 11.52432667 | 11.07100932 | 10.72720655 | 9.166592134 | 9.1462585 | 10.09769543 | 10.22560487 |
| ENSG00000173085 | COQ2 | -1.336501493 | 0.067838142 | -1.644320155 | 0.019496236 | -2.980821648 | 7.384433967 | 7.145017454 | 6.203069917 | 5.627094009 | 5.8245091 | 4.959178705 | 5.812327524 |
| ENSG00000179583 | CIITA | -3.146555903 | 0.010592892 | 0.162685833 | 0.91151969 | -2.98387007 | 4.467311984 | 3.686303414 | 5.930865648 | 3.023133209 | 2.59489879 | 5.29920634 | 4.895193906 |
| ENSG00000109911 | ELP4 | -1.215831692 | 0.002366651 | -1.768477708 | 3.24E-06 | -2.9843094 | 7.159102577 | 7.638155693 | 7.646393253 | 6.29903474 | 6.35727082 | 5.856572853 | 5.759151667 |
| ENSG00000198105 | ZNF248 | -1.565474758 | 0.025250552 | -1.41921743 | 0.042401898 | -2.984692188 | 9.015175615 | 8.687092496 | 7.552162999 | 7.062428156 | 6.94988431 | 7.504242605 | 6.676467423 |
| ENSG00000213465 | ARL2 | -1.690329851 | 1.46E-05 | -1.296727026 | 0.001089398 | -2.987056876 | 10.27872131 | 9.869887299 | 10.65536953 | 8.584574266 | 8.65574839 | 8.888863837 | 9.123144518 |
| ENSG00000101843 | PSMD10 | -1.617939451 | 2.37E-07 | -1.369262656 | 1.34E-05 | -2.987202106 | 10.29268328 | 9.821190352 | 9.788583153 | 8.395844809 | 8.36195019 | 8.653732067 | 8.595472797 |
| ENSG00000105819 | PMPCB | -1.052010169 | 0.647922415 | -1.93524808 | 0.36717983 | -2.987258249 | 10.95943304 | 9.256956408 | 2 | 8.723531969 | 8.71301725 | 7.668951058 | 8.004685229 |
| ENSG00000011295 | TTC19 | -1.917075085 | 0.000527243 | -1.07133348 | 0.066638754 | -2.988408565 | 9.143983763 | 9.357954881 | 10.34701993 | 7.72026832 | 7.90685167 | 8.673501845 | 8.627970054 |
| ENSG00000185787 | MORF4L1 | -1.201776164 | 0.004971461 | -1.786707463 | 1.25E-05 | -2.988483628 | 12.38357115 | 11.7323588 | 11.6249326 | 10.77982736 | 10.7278927 | 9.988176769 | 10.33299247 |
| ENSG00000105676 | ARMC6 | -1.693392669 | 0.415754832 | -1.297003875 | 0.534817139 | -2.990396544 | 10.00958825 | 8.955547846 | 2 | 7.265793319 | 7.38644297 | 7.52030827 | 7.88671543 |
| ENSG00000198522 | GPN1 | -1.70032859 | 0.017807933 | -1.290971179 | 0.07779425 | -2.991299769 | 10.24321353 | 8.791896851 | 8.699419242 | 7.66556918 | 7.82609171 | 8.158075786 | 8.143433938 |
| ENSG00000065559 | MAP2K4 | -1.46252136 | 9.99E-09 | -1.528963963 | 1.40E-09 | -2.991485323 | 10.35389212 | 10.39692156 | 10.07784268 | 8.821593044 | 8.83927354 | 8.677327975 | 8.847127396 |
| ENSG00000101220 | C20orf27 | -2.351245525 | 8.15E-08 | -0.640652616 | 0.191295729 | -2.991898141 | 8.540903468 | 9.090820894 | 9.301632639 | 6.750147204 | 6.66069199 | 8.084345843 | 8.621777169 |
| ENSG00000141505 | ASGR1 | -0.768297517 | 0.770303179 | -2.224033478 | 0.350313231 | -2.992330996 | 2 | 5.72254323 | 2 | 3.709785322 | 3.86943969 | 2.831336763 | 3.012034821 |
| ENSG00000186889 | TMEM17 | -1.16375963 | 0.223356242 | -1.828593342 | 0.040825791 | -2.992352972 | 5.330594134 | 7.232742069 | 5.772716771 | 5.319220934 | 5.27444544 | 4.563936166 | 4.853519801 |
| ENSG00000170035 | UBE2E3 | -1.320305758 | 0.018442639 | -1.675947628 | 0.001960035 | -2.996253386 | 9.886101849 | 9.680074956 | 10.85604368 | 8.894758143 | 8.94822691 | 8.581047186 | 8.557692095 |
| ENSG00000134490 | TMEM241 | -2.052992793 | 0.000110521 | -0.943470288 | 0.095306826 | -2.996463081 | 7.159102577 | 8.213429689 | 8.143093325 | 5.829220145 | 6.03677338 | 6.835304498 | 7.138761401 |
| AIM1 | AIM1 | 4.694414354 | 0.001071039 | -7.69206992 | 0.000303606 | -2.997655566 | 3.327711234 | 2 | 4.058998049 | 7.277360022 | 7.24216308 | 2 | 2 |
| ENSG00000132128 | LRRC41 | -1.703656495 | 1.82E-09 | -1.294097729 | 6.85E-06 | -2.997754224 | 12.13890018 | 11.63710343 | 11.99095026 | 10.18886958 | 10.2817021 | 10.65548645 | 10.63324187 |
| ENSG00000198624 | CCDC69 | -2.227708961 | 0.220329561 | -0.771266985 | 0.69448539 | -2.998975946 | 7.867441304 | 6.602424674 | 2 | 4.497095146 | 4.97065356 | 6.546183077 | 5.343519008 |
| ENSG00000130165 | ELOF1 | -1.350172552 | 0.002910526 | -1.649905434 | 0.000184124 | -3.000077986 | 10.3933013 | 9.599860728 | 10.09465441 | 8.648320862 | 8.790887 | 8.1339154 | 8.666987181 |
| ENSG00000025039 | RRAGD | -2.093950469 | 0.326199902 | -0.906757807 | 0.687882518 | -3.000708276 | 7.940205782 | 4.710564154 | 2 | 4.846787657 | 4.3456612 | 6.277913523 | 4.584731282 |
| ENSG00000115234 | SNX17 | -1.725074421 | 3.94E-06 | -1.278790639 | 0.000828683 | -3.00386506 | 11.55361133 | 10.87363932 | 11.37951548 | 9.52946469 | 9.62041512 | 9.830891212 | 10.1871175 |
| ENSG00000113460 | BRIX1 | -0.909553973 | 0.392859546 | -2.095417921 | 0.029300298 | -3.004971895 | 9.975371636 | 7.928237109 | 7.646393253 | 8.033874364 | 7.99953197 | 7.000176314 | 6.70252219 |
| ENSG00000101294 | HM13 | -0.707625289 | 0.328352926 | -2.299306145 | 0.000259672 | -3.006931434 | 13.29992879 | 12.01813185 | 11.84209456 | 11.81386838 | 11.8575514 | 10.21552475 | 10.27852722 |
| ENSG00000094631 | HDAC6 | -1.895789628 | 0.013182156 | -1.11217494 | 0.165176225 | -3.007964568 | 9.2903348 | 9.479495904 | 11.02781271 | 8.266399315 | 8.28283292 | 8.891336668 | 9.193729987 |
| ENSG00000160214 | RRP1 | -1.575924681 | 3.00E-05 | -1.433199538 | 0.000139649 | -3.009124219 | 8.910412251 | 8.623598428 | 9.014156722 | 7.25413313 | 7.35080119 | 7.168655895 | 7.673351463 |
| ENSG00000185246 | PRPF39 | -1.347598948 | 0.093898169 | -1.662724662 | 0.031770277 | -3.01032361 | 7.867441304 | 9.433952963 | 9.829330889 | 7.971157485 | 7.8745862 | 7.862014947 | 7.314917451 |
| ENSG00000188938 | FAM120AOS | -1.636032674 | 0.013857003 | -1.374766813 | 0.040881571 | -3.010799488 | 10.72727936 | 9.102512015 | 10.5187901 | 8.687861792 | 8.60067114 | 8.870181261 | 8.936662123 |
| ENSG00000125503 | PPP1R12C | -0.364745252 | 0.823079233 | -2.649364472 | 0.048314828 | -3.014109724 | 4.467311984 | 8.834582582 | 9.132850258 | 8.047448792 | 8.10812366 | 5.758846871 | 5.980509018 |
| ENSG00000128185 | DGCR6L | -1.200828209 | 0.018747898 | -1.814125993 | 0.00017855 | -3.014954203 | 9.579430841 | 8.57407404 | 9.553445883 | 8.024753265 | 8.19309833 | 7.49884738 | 7.519298205 |
| ENSG00000134905 | CARS2 | -1.324335305 | 0.041201278 | -1.690756831 | 0.006799071 | -3.015092136 | 9.112843761 | 9.616264339 | 10.44204362 | 8.464912335 | 8.5606705 | 7.752026287 | 8.464565585 |
| ENSG00000121481 | RNF2 | -1.081161613 | 0.047693588 | -1.934479411 | 0.000132575 | -3.015641024 | 8.075571721 | 9.055168103 | 8.236946975 | 7.514115926 | 7.41778467 | 6.800644007 | 6.457498602 |
| ENSG00000134748 | PRPF38A | -1.744481102 | 0.00491763 | -1.271806202 | 0.045894998 | -3.016287304 | 8.564112515 | 9.424669273 | 10.07784268 | 7.683067677 | 7.82609171 | 8.366167515 | 8.060697341 |
| ENSG00000109586 | GALNT7 | -1.201541154 | 0.312073196 | -1.815891835 | 0.104923324 | -3.01743299 | 6.89194353 | 7.669933291 | 9.859153765 | 7.571615825 | 7.46188679 | 6.959905775 | 6.879055607 |
| ENSG00000174371 | EXO1 | -1.582113614 | 0.445687918 | -1.438287276 | 0.478933744 | -3.020400889 | 3.327711234 | 6.04685525 | 2 | 3.566290607 | 3.55551834 | 3.1504466 | 4.033026654 |
| ENSG00000138074 | SLC5A6 | -1.475567486 | 0.004333061 | -1.545630712 | 0.002429239 | -3.021198198 | 8.716893051 | 9.030900487 | 7.896971941 | 7.25803034 | 7.09482005 | 7.172727426 | 7.045606259 |
| ENSG00000151247 | EIF4E | -0.397285183 | 0.127670293 | -2.624192266 | 6.26E-32 | -3.021477449 | 9.48530526 | 9.45234302 | 9.399592459 | 9.023967474 | 9.07923325 | 6.923319986 | 6.802276223 |
| ENSG00000248496 | | 0.465467901 | 0.882596592 | -3.487248838 | 0.204625268 | -3.021780938 | 2 | 3.686303414 | 2 | 2.600425781 | 3.45167086 | 2.139668783 | 2.059860178 |
| ENSG00000102898 | NUTF2 | -1.90002317 | 0.000227045 | -1.124964274 | 0.038160787 | -3.024987444 | 11.63532243 | 10.56689847 | 10.83118029 | 9.160358775 | 9.22317745 | 9.710961224 | 10.17764973 |
| ENSG00000172500 | FIBP | -1.369063414 | 0.124157787 | -1.65626892 | 0.053857397 | -3.025332334 | 10.73241055 | 9.828248506 | 8.408111533 | 8.575233246 | 8.58848096 | 8.001311979 | 8.544002356 |
| ENSG00000155621 | C9orf85 | -1.886320148 | 0.102696843 | -1.139576989 | 0.339445471 | -3.025897138 | 8.87372952 | 6.535001748 | 6.203069917 | 6.104969671 | 5.6826301 | 6.864125031 | 6.317379004 |
| ENSG00000243414 | TICAM2 | 2.938094859 | 0.042732218 | -5.965951271 | 4.21E-05 | -3.027856411 | 6.009025997 | 2 | 5.772716771 | 8.197043419 | 8.09094333 | 2.267001569 | 2.117335275 |
| ENSG00000182093 | WRB | -2.171185878 | 0.179023799 | -0.856725455 | 0.621252089 | -3.027911334 | 8.836089686 | 9.203675681 | 3.369317972 | 6.476767609 | 6.20590375 | 7.570548195 | 7.666147605 |
| ENSG00000108298 | RPL19 | -1.714035409 | 0.007676211 | -1.317618211 | 0.045052759 | -3.031653619 | 14.0671076 | 12.60879193 | 12.86966683 | 11.60598322 | 11.628365 | 11.91230876 | 12.1074829 |
| ENSG00000113845 | TIMMDC1 | -1.708321996 | 6.38E-05 | -1.324068613 | 0.002374237 | -3.03239061 | 10.85509492 | 9.910357957 | 10.40208303 | 8.750063451 | 8.72837565 | 9.133737044 | 9.106830771 |
| ENSG00000198556 | ZNF789 | -2.049804939 | 0.052744177 | -0.98303222 | 0.376725096 | -3.032837159 | 7.53297911 | 4.710564154 | 6.80318718 | 4.694064037 | 4.98931816 | 5.866661379 | 5.745545708 |
| C8ORF44-SGK3 | C8ORF44-SG | 4.084720255 | 0.094840682 | -7.118346281 | 0.012419506 | -3.033626025 | 4.00722531 | 2 | 2 | 6.221320894 | 6.01871653 | 2 | 2 |
| ENSG00000168071 | CCDC88B | -1.207917327 | 0.496209542 | -1.828206954 | 0.272924984 | -3.036124281 | 2 | 5.72254323 | 7.029880454 | 4.783303513 | 4.95174433 | 4.530483275 | 4.147809227 |
| ENSG00000146038 | DCDC2 | -5.988849441 | 0.004694342 | 2.949836413 | 0.032616532 | -3.039013027 | 4.467311984 | 3.686303414 | 4.058998049 | 2.141737246 | 2 | 5.264018433 | 7.420010428 |
| ENSG00000173638 | SLC19A1 | -3.088606608 | 0.101839732 | 0.049378201 | 0.982938888 | -3.039228407 | 9.473095744 | 8.029695761 | 2 | 5.37770462 | 5.39133044 | 8.480070636 | 8.303661547 |
| ENSG00000163938 | GNL3 | -1.409152506 | 0.029278378 | -1.630409413 | 0.009785674 | -3.039561919 | 11.51542408 | 10.03150247 | 10.45512158 | 9.352133347 | 9.45234084 | 9.221844492 | 9.143604888 |
| ENSG00000158423 | RIBC1 | -2.341431519 | 0.335233605 | -0.700496564 | 0.785640197 | -3.041928082 | 2 | 4.93638933 | 2 | 2.498691189 | 2.59489879 | 3.12825509 | 3.430065404 |
| ENSG00000105058 | FAM32A | -1.439768434 | 0.000211037 | -1.602982406 | 2.82E-05 | -3.04275084 | 11.0567315 | 11.35134391 | 11.85201631 | 9.987058621 | 10.0546329 | 9.810348932 | 9.905493112 |
| ENSG00000165140 | FBP1 | -3.110850661 | 0.143276504 | 0.066987241 | 0.976621338 | -3.04386342 | 4.00722531 | 3.686303414 | 2 | 2.3892346 | 2.14022021 | 3.968654369 | 2.851942824 |
| ENSG00000081051 | AFP | 2.146580184 | NA | -5.191097829 | NA | -3.044517646 | 2 | 2 | 2 | 2.868578001 | 3.28037825 | 2 | 2 |
| ENSG00000276416 | | 2.146580184 | NA | -5.191097829 | NA | -3.044517646 | 2 | 2 | 2 | 2.868578001 | 3.28037825 | 2 | 2 |
| ENSG00000164169 | PRMT9 | -1.531876955 | 0.022263838 | -1.513729795 | 0.022702853 | -3.04560675 | 8.716893051 | 7.232742069 | 7.896971941 | 6.503277904 | 6.64900057 | 6.78654324 | 6.374480702 |
| ENSG00000123472 | ATPAF1 | -2.037699642 | 1.12E-05 | -1.009792196 | 0.041403012 | -3.047491837 | 9.317904228 | 10.24169813 | 10.3329194 | 8.040677544 | 7.9809999 | 9.027210383 | 9.028005357 |
| ENSG00000144792 | ZNF660 | -3.277040795 | 0.138590087 | 0.22787428 | 0.931173501 | -3.049166515 | 7.750913381 | 3.076665209 | 2 | 3.709785322 | 3.28037825 | 6.361143723 | 6.55171954 |
| ENSG00000011485 | PPP5C | -1.218742544 | 0.274233692 | -1.83316658 | 0.080940119 | -3.051909123 | 11.09308737 | 9.396454291 | 8.076950877 | 8.875861021 | 8.75042728 | 8.181838221 | 8.23147623 |
| ENSG00000100263 | RHBDD3 | -0.946572914 | 0.02294794 | -2.105599802 | 3.51E-08 | -3.052172716 | 9.048471759 | 8.848535039 | 8.380957451 | 7.770242853 | 7.93118934 | 6.875826972 | 6.554323192 |
| ENSG00000101558 | VAPA | -1.424759949 | 0.004733672 | -1.62809317 | 0.000988461 | -3.052853119 | 12.01804503 | 10.86336859 | 11.62203638 | 10.22764339 | 10.076619 | 9.985093062 | 9.917171619 |
| ENSG00000108523 | RNF167 | -1.530576527 | 0.083356641 | -1.523426899 | 0.080069004 | -3.054003426 | 8.75772933 | 10.01303782 | 11.13734171 | 8.786946023 | 8.71862106 | 8.811331315 | 8.707318295 |
| ENSG00000068120 | COASY | -1.998771622 | 0.003237238 | -1.055358205 | 0.143232507 | -3.054129827 | 11.3042312 | 9.969011754 | 9.746650996 | 8.511364632 | 8.54340749 | 9.380756704 | 9.541509736 |
| ENSG00000183751 | TBL3 | -1.68640805 | 0.00564355 | -1.367743222 | 0.027293207 | -3.054151272 | 10.27872131 | 8.848535039 | 9.372276026 | 7.862723323 | 8.03365075 | 8.238615655 | 8.290447859 |
| ENSG00000144381 | HSPD1 | -1.528959734 | 0.002781572 | -1.525231758 | 0.002713948 | -3.054191492 | 12.05127578 | 11.51338035 | 10.85110517 | 10.05583453 | 9.99930717 | 10.07793097 | 9.983635087 |
| ENSG00000074319 | TSG101 | -1.314516695 | 0.046353446 | -1.740049976 | 0.005978172 | -3.054566672 | 11.33828573 | 9.756063186 | 10.47451889 | 9.364838907 | 9.3477391 | 8.864319213 | 9.000106719 |
| ENSG00000196172 | ZNF681 | -0.602852828 | 0.534490579 | -2.452311566 | 0.003295058 | -3.055164394 | 5.330594134 | 5.303523747 | 6.322058544 | 5.420062284 | 4.87352701 | 3.968654369 | 3.33635788 |
| ENSG00000136504 | KAT7 | -1.698164695 | 0.016178162 | -1.357928828 | 0.058095053 | -3.056093523 | 10.6695979 | 9.616264339 | 8.922968322 | 8.270263574 | 8.19108423 | 8.439530542 | 8.684919451 |
| ENSG00000137571 | SLCO5A1 | -0.285692137 | 0.923175097 | -2.770547903 | 0.26979692 | -3.05624004 | 2 | 2 | 6.534109959 | 4.647285135 | 4.95174433 | 3.172301926 | 2.918126211 |
| ENSG00000015520 | NPC1L1 | -0.361943862 | 0.902303961 | -2.695735001 | 0.28326297 | -3.057678862 | 2 | 2 | 4.523675391 | 2.947924294 | 3.28037825 | 2.42098416 | 2.172608095 |
| ENSG00000153015 | CWC27 | -1.356298752 | 0.537670546 | -1.703163671 | 0.423520139 | -3.059462424 | 10.38028415 | 8.337339476 | 2 | 7.805288479 | 7.7377845 | 7.496683627 | 7.364717093 |
| ENSG00000157823 | AP3S2 | -7.829077115 | 2.22E-05 | 4.769480262 | 4.48E-07 | -3.059596854 | 5.095959458 | 3.076665209 | 4.523675391 | 2 | 2 | 8.949447548 | 8.815461059 |
| ENSG00000080189 | SLC35C2 | -1.618928447 | 0.001327229 | -1.441265184 | 0.004549414 | -3.060193631 | 11.00254523 | 9.890264534 | 10.89014791 | 9.045548328 | 9.07377798 | 9.228053138 | 9.244188987 |
| ENSG00000141504 | SAT2 | -1.539639543 | 0.009141757 | -1.521372547 | 0.009785845 | -3.06101209 | 9.064836028 | 9.256956408 | 10.29705308 | 8.072008297 | 8.16464176 | 7.978234673 | 8.280260977 |
| ENSG00000183978 | COA3 | -1.771829753 | 4.37E-14 | -1.290994807 | 3.66E-08 | -3.06282456 | 9.646202137 | 9.591588447 | 9.624627483 | 7.852449262 | 7.87959733 | 8.239909835 | 8.433788484 |
| ENSG00000163684 | RPP14 | -1.116571654 | 0.022400263 | -1.947309363 | 2.05E-05 | -3.063881017 | 9.590776211 | 8.687092496 | 9.541233192 | 8.128448191 | 8.30162324 | 7.263334844 | 7.528596482 |
| ENSG00000105989 | WNT2 | -0.469788082 | 0.869037712 | -2.59442452 | 0.290390881 | -3.064212602 | 2 | 4.11352228 | 2 | 2.784612485 | 3.01487888 | 2.34604626 | 2.172608095 |
| ENSG00000129235 | TXNDC17 | -0.713868446 | 0.400402339 | -2.352775361 | 0.001407955 | -3.066643807 | 9.730679447 | 8.191702463 | 8.076950877 | 8.172475401 | 8.16874151 | 6.39173974 | 6.735016458 |
| ENSG00000171649 | ZIK1 | -1.960514747 | 0.231268956 | -1.106775706 | 0.515169146 | -3.067290454 | 7.904282243 | 3.076665209 | 8.631919287 | 6.024891728 | 5.67111583 | 6.651077397 | 6.683620062 |
| ENSG00000054356 | PTPRN | 1.690624157 | 0.293493132 | -4.758295453 | 0.000933854 | -3.067671297 | 7.218819781 | 7.099070191 | 2 | 8.197043419 | 8.29975519 | 2.803598391 | 2.981406154 |
| ENSG00000103254 | FAM173A | -2.048969371 | 0.012059009 | -1.018824376 | 0.240215699 | -3.067793747 | 9.031919741 | 6.951736442 | 7.858115942 | 6.283823264 | 6.11534392 | 7.307207714 | 7.065820006 |
| ENSG00000224501 | | 5.880166147 | 1.31E-06 | -8.948903545 | 4.41E-06 | -3.068737398 | 5.532353208 | 6.140560021 | 2 | 11.06373609 | 11.1154227 | 2.048074405 | 2 |
| ENSG00000044574 | HSPA5 | -1.35386169 | 0.004629196 | -1.714988044 | 0.000217575 | -3.068849734 | 14.35016946 | 14.38095284 | 14.45655588 | 13.03264551 | 13.053554 | 12.0433481 | 13.12347144 |
| ENSG00000171135 | JAGN1 | -1.499148661 | 0.05137125 | -1.570046472 | 0.037822688 | -3.069195133 | 10.81695923 | 8.94259736 | 9.714382178 | 8.474993083 | 8.60067114 | 8.247007147 | 8.661563746 |
| ENSG00000144741 | SLC25A26 | -1.15363733 | 0.113003183 | -1.915789108 | 0.004724491 | -3.069426438 | 8.009475818 | 7.189546432 | 8.865353883 | 6.984595707 | 7.0991228 | 6.21875862 | 6.397863414 |
| ENSG00000091129 | NRCAM | -0.96034012 | 0.636480383 | -2.110669155 | 0.256957702 | -3.071009275 | 7.485143553 | 5.457120953 | 2 | 5.614989456 | 5.0259382 | 4.841348851 | 3.640730416 |
| ENSG00000156966 | B3GNT7 | -2.323965597 | 0.103167922 | -0.747926458 | 0.626190802 | -3.071892055 | 6.563806279 | 3.686303414 | 4.058998049 | 3.289987342 | 3.69849104 | 4.727866721 | 4.673629022 |

| | | | | | | | | | | | | | |
|---|---|---|---|---|---|---|---|---|---|---|---|---|---|
| ENSG00000162980 | ARL5A | -1.309748859 | 9.35E-07 | -1.762697496 | 1.12E-11 | -3.072446356 | 9.678463567 | 9.461450882 | 9.31604076 | 8.192977693 | 8.19711812 | 7.794040499 | 7.702986259 |
| ENSG00000125352 | RNF113A | -1.609162483 | 0.205222597 | -1.465545733 | 0.244978858 | -3.074708217 | 9.90440318 | 6.311476754 | 7.818184403 | 7.140504127 | 7.12888795 | 7.184873526 | 7.358771532 |
| ENSG00000185420 | SMYD3 | -1.759250575 | 0.087690677 | -1.316832353 | 0.205699167 | -3.076082928 | 9.219014269 | 6.311476754 | 8.460932916 | 6.567497652 | 6.81514011 | 6.987904673 | 7.24894374 |
| ENSG00000065057 | NTHL1 | -1.977519851 | 0.045765337 | -1.100794008 | 0.28541815 | -3.078313858 | 5.532353208 | 7.145017454 | 8.206335842 | 5.210797711 | 5.65950892 | 6.339575235 | 6.192573806 |
| ENSG00000042429 | MED17 | -0.00959098 | 0.98413673 | -3.069187804 | 1.33E-17 | -3.078778783 | 8.468948322 | 8.557180776 | 9.083172337 | 8.677714042 | 8.75316018 | 5.856572853 | 5.641643428 |
| ENSG00000105447 | GRWD1 | -1.886902234 | 3.57E-05 | -1.192481162 | 0.012195095 | -3.079383396 | 10.32020829 | 9.318399851 | 9.981704053 | 7.999368766 | 8.11238697 | 8.624303429 | 8.852955181 |
| ENSG00000268350 | FAM156A | 2.4913919 | 0.00247249 | -5.571144053 | 3.39E-10 | -3.079752153 | 5.095959458 | 5.303523747 | 6.203069917 | 7.932663244 | 8.05595642 | 2.457042231 | 2.117335275 |
| ENSG00000136811 | ODF2 | -1.750700067 | 0.001137082 | -1.329818597 | 0.015985371 | -3.080518664 | 9.975371636 | 9.018611864 | 10.26027233 | 8.10700104 | 8.09740995 | 8.394916616 | 8.631056556 |
| ENSG00000160688 | FLAD1 | -1.62245189 | 0.031612654 | -1.458666202 | 0.053383916 | -3.081118093 | 8.19931886 | 9.616264339 | 10.18377222 | 7.875463856 | 7.99953197 | 8.090815298 | 8.109313767 |
| ENSG00000102312 | PORCN | -1.025004207 | 0.246800072 | -2.056530739 | 0.011165872 | -3.081534946 | 9.174465797 | 7.431218269 | 9.658943166 | 7.959237999 | 8.04707549 | 7.074595936 | 6.911910672 |
| ENSG00000151117 | TMEM86A | -1.929448734 | 0.338160854 | -1.158714592 | 0.575727591 | -3.088163326 | 8.696033109 | 6.228547981 | 2 | 5.697663076 | 5.33407135 | 6.595545082 | 5.781546943 |
| ENSG00000106348 | IMPDH1 | -1.817148954 | 0.001196996 | -1.272281492 | 0.028326696 | -3.089430445 | 10.6855593 | 9.792607274 | 9.703463497 | 8.372666678 | 8.28094035 | 9.188719608 | 8.451355939 |
| ENSG00000175309 | PHYKPL | -2.023682526 | 0.082586793 | -1.066124659 | 0.3818783 | -3.089807184 | 10.21416435 | 8.57407404 | 6.718950348 | 6.960881212 | 7.28433928 | 8.031518488 | 8.105768199 |
| ENSG00000106537 | TSPAN13 | -1.396448384 | 0.000123302 | -1.693739403 | 1.70E-06 | -3.090187788 | 8.855032352 | 8.557180776 | 9.06622548 | 7.471115032 | 7.44845993 | 7.337748225 | 6.979338179 |
| ENSG00000136197 | C7orf25 | 1.241453084 | 0.013063165 | -4.332176982 | 1.30E-18 | -3.090723898 | 6.563806279 | 6.844429755 | 6.073738139 | 7.767511442 | 7.73502229 | 3.082820997 | 3.071407821 |
| ENSG00000174444 | RPL4 | -2.024695643 | 2.44E-07 | -1.067197314 | 0.01038035 | -3.091892957 | 14.03891066 | 13.65124938 | 13.1675554 | 11.62194852 | 11.6561905 | 12.54875431 | 12.64134588 |
| ENSG00000146070 | PLA2G7 | -5.757650451 | 0.143394379 | 2.665343097 | 0.470583967 | -3.092307354 | 2 | 2 | 4.058998049 | 2 | 2 | 5.665620077 | 3.183290849 |
| ENSG00000074621 | SLC24A1 | -1.683266519 | 0.142466415 | -1.409808209 | 0.218735712 | -3.093074728 | 8.16936288 | 5.457120953 | 8.763901704 | 6.314087501 | 6.39974839 | 6.770513376 | 6.449128733 |
| ENSG00000156958 | GALK2 | -1.286934305 | 0.029405943 | -1.808631341 | 0.001305534 | -3.095565646 | 7.709880939 | 8.414402626 | 9.06622548 | 7.246306984 | 7.22257674 | 6.775876478 | 6.681239787 |
| ENSG00000115216 | NRBP1 | -1.070846204 | 0.000303054 | -2.025137944 | 6.50E-13 | -3.095984148 | 12.59129083 | 12.130888 | 12.496991 | 11.35234742 | 11.346294 | 10.28346463 | 10.50250454 |
| ENSG00000169302 | STK32A | -2.931582097 | 0.063038247 | -0.165438755 | 0.925215975 | -3.097020852 | 4.467311984 | 3.076665209 | 3.369317972 | 2.270786764 | 2.49394783 | 3.616794016 | 3.640730416 |
| ENSG00000146281 | PM20D2 | -2.925863372 | 0.006769346 | -0.171226502 | 0.899648072 | -3.097089874 | 5.095959458 | 8.255926468 | 7.97167868 | 4.927308402 | 4.83276841 | 7.019897747 | 7.79392309 |
| ENSG00000278828 | HIST1H3H | -1.827585588 | 0.465624329 | -1.270861932 | 0.611925414 | -3.09844752 | 2 | 5.595927442 | 2 | 2.947924294 | 3.15372745 | 3.355392153 | 3.383972333 |
| ENSG00000161653 | NAGS | -1.215152529 | 0.549630367 | -1.885657988 | 0.325814666 | -3.100810516 | 2 | 5.303523747 | 7.896971941 | 5.406080789 | 5.41912973 | 5.04460184 | 4.553838275 |
| ENSG00000116014 | KISS1R | -2.139640728 | 0.413888829 | -0.96170965 | 0.718413093 | -3.101350378 | 2 | 4.442710129 | 2 | 2.3892346 | 2.49394783 | 2.526562374 | 3.128433649 |
| ENSG00000106633 | GCK | -6.511738061 | 0.004400767 | 3.4098477 | 0.037799318 | -3.101890361 | 2 | 4.11352228 | 3.369317972 | 2 | 2 | 6.539892359 | 5.941260754 |
| ENSG00000165637 | VDAC2 | -0.809731587 | 0.010720102 | -2.294168443 | 2.36E-15 | -3.103900029 | 10.76282024 | 10.74155032 | 10.44204362 | 9.880490474 | 9.81868097 | 8.174412115 | 8.554444396 |
| ENSG00000065618 | COL17A1 | -0.792792949 | 0.777561305 | -2.313494587 | 0.360490023 | -3.106287536 | 2 | 2 | 4.523675391 | 3.162721474 | 2.59489879 | 2.34604626 | 2.421067459 |
| ENSG00000096996 | IL12RB1 | -1.755034746 | 0.112122648 | -1.351872643 | 0.211980478 | -3.106907389 | 4.815609471 | 4.442710129 | 5.156626401 | 3.09461492 | 3.7864959 | 4.029788235 | 3.407202941 |
| ENSG00000262347 | | 2.083592507 | NA | -5.191098092 | NA | -3.107505584 | 2 | 2 | 2 | 3.227757282 | 2.86123014 | 2 | 2 |
| ENSG00000262635 | | 2.083592507 | NA | -5.191098092 | NA | -3.107505584 | 2 | 2 | 2 | 3.227757282 | 2.86123014 | 2 | 2 |
| ENSG00000005194 | CIAPIN1 | -1.825269606 | 7.55E-05 | -1.282776324 | 0.006953081 | -3.10804593 | 10.41898845 | 9.461450882 | 9.809100877 | 8.113468729 | 8.16464176 | 8.497488703 | 8.833794221 |
| ENSG00000143344 | RGL1 | -1.160359622 | 0.02195944 | -1.947705552 | 4.18E-05 | -3.108065174 | 9.48530526 | 10.3387994 | 10.08627303 | 8.85414072 | 8.86729294 | 7.707816755 | 8.379764957 |
| ENSG00000158552 | ZFAND2B | -1.843885282 | 0.00308587 | -1.264503022 | 0.048849278 | -3.108388304 | 7.384433967 | 8.590771778 | 8.865353883 | 6.463327516 | 6.73998187 | 7.191577434 | 7.143956535 |
| ENSG00000102043 | MTMR8 | -3.973125072 | 0.250010976 | 0.864339097 | 0.79275994 | -3.108785975 | 2 | 3.686303414 | 2 | 2 | 2.14022021 | 3.316666208 | 3.287118176 |
| ENSG00000140395 | WDR61 | -1.507089239 | 0.481509008 | -1.602452967 | 0.442728081 | -3.109542205 | 10.29961392 | 9.214490408 | 2 | 7.742693204 | 7.82609171 | 7.522436904 | 7.842663598 |
| ENSG00000224902 | GAGE12H | 2.078979719 | NA | -5.191098111 | NA | -3.112118392 | 2 | 2 | 2 | 2.784612485 | 3.28037825 | 2 | 2 |
| ENSG00000227488 | GAGE12D | 2.078979719 | NA | -5.191098111 | NA | -3.112118392 | 2 | 2 | 2 | 2.784612485 | 3.28037825 | 2 | 2 |
| ENSG00000237671 | GAGE12C | 2.078979719 | NA | -5.191098111 | NA | -3.112118392 | 2 | 2 | 2 | 2.784612485 | 3.28037825 | 2 | 2 |
| ENSG00000157895 | C12orf43 | -1.698405274 | 0.002362021 | -1.415722985 | 0.012368406 | -3.114128259 | 9.657036304 | 8.356994601 | 9.099922433 | 7.497728447 | 7.4074129 | 7.613117257 | 7.838397174 |
| ENSG00000167985 | SDHAF2 | -0.704891918 | 0.45794139 | -2.4121831 | 0.00322719 | -3.117075019 | 7.709880939 | 8.414402626 | 9.849281147 | 8.211183901 | 8.26762243 | 6.401024833 | 6.730418989 |
| ENSG00000138430 | OLA1 | -1.498887563 | 0.000504609 | -1.618342512 | 0.000138943 | -3.117230075 | 10.73241055 | 9.821190352 | 10.23015729 | 8.850273548 | 8.78288532 | 8.615855003 | 8.776966445 |
| ENSG00000163879 | DNALI1 | -1.79398759 | 0.001441426 | -1.324907294 | 0.021329954 | -3.118894884 | 8.564112515 | 7.315436877 | 7.896971941 | 6.205262847 | 6.32824132 | 6.779440836 | 6.649933445 |
| ENSG00000136371 | MTHFS | 0.753496723 | 0.328133599 | -3.872875558 | 7.48E-09 | -3.119378836 | 5.532353208 | 6.728495673 | 6.534109959 | 7.062428156 | 7.11195456 | 3.44788957 | 2.950113106 |
| ENSG00000135597 | REPS1 | -1.403156936 | 0.006685849 | -1.716738756 | 0.000620507 | -3.119895692 | 9.848787621 | 8.94259736 | 8.825624954 | 7.890605449 | 7.89452713 | 7.52030827 | 7.646759583 |
| ENSG00000156709 | AIFM1 | -1.449152112 | 0.094412709 | -1.670754756 | 0.046421006 | -3.119906867 | 10.1311248 | 7.847167909 | 9.287079172 | 7.942383523 | 7.91175216 | 7.570548195 | 7.836259222 |
| ENSG00000083814 | ZNF671 | -1.851825747 | 0.137320752 | -1.26979095 | 0.316737831 | -3.121616696 | 4.815609471 | 7.954278496 | 8.677268317 | 5.968925393 | 6.12381559 | 6.569017168 | 6.632790505 |
| ENSG00000100220 | RTCB | -1.651679818 | 8.64E-12 | -1.470220619 | 1.18E-09 | -3.121900437 | 10.89224845 | 11.19631665 | 11.10036136 | 9.381905119 | 9.46406471 | 9.51135037 | 9.691260406 |
| ENSG00000105364 | MRPL4 | -1.35782012 | 0.025804104 | -1.766170701 | 0.002612948 | -3.123990821 | 10.19198706 | 8.762721501 | 9.439621147 | 8.205140657 | 8.25995663 | 7.782541424 | 7.877388328 |
| ENSG00000013306 | SLC25A39 | -2.207377073 | 1.45E-08 | -0.916984594 | 0.029610048 | -3.124361667 | 11.17074085 | 10.3387994 | 10.92346452 | 8.664561167 | 8.64254348 | 9.853049221 | 10.01350757 |
| ENSG00000151552 | QDPR | -1.628010996 | 0.00011228 | -1.496640055 | 0.000373663 | -3.124651051 | 9.460782017 | 8.876041558 | 8.763901704 | 7.504305792 | 7.41433569 | 7.380620955 | 7.770601445 |
| ENSG00000005448 | WDR54 | -2.169291662 | 0.012923624 | -0.956445294 | 0.307150063 | -3.125736956 | 7.667647223 | 7.274681896 | 9.372276026 | 6.24507773 | 6.34282908 | 7.371203179 | 7.553536683 |
| ENSG00000161847 | RAVER1 | -0.934008443 | 0.194014185 | -2.192569818 | 0.000682045 | -3.126578261 | 10.73752355 | 9.159587375 | 9.636156974 | 9.087765067 | 9.05396643 | 7.706881163 | 7.938459399 |
| ENSG00000160953 | MUM1 | -1.854738532 | 0.029129406 | -1.27223163 | 0.145413199 | -3.126970162 | 8.674867122 | 9.192779271 | 10.78531698 | 7.959237999 | 8.04930085 | 8.578491945 | 8.581500498 |
| ENSG00000068137 | PLEKHH3 | -7.777679205 | 0.000179038 | 4.650527711 | 0.000501743 | -3.127151494 | 4.00722531 | 5.303523747 | 2 | 2 | 2 | 8.601831186 | 8.820336672 |
| ENSG00000130734 | ATG4D | -1.700156812 | 0.018749668 | -1.428257696 | 0.050135255 | -3.128414508 | 8.16936288 | 8.376385542 | 9.767769415 | 7.226553841 | 7.33264456 | 7.599066672 | 7.491038307 |
| ENSG00000101052 | IFT52 | -1.432294335 | 0.023304992 | -1.696142413 | 0.00573183 | -3.128436748 | 7.82963488 | 8.929529568 | 9.242516611 | 7.250225364 | 7.48508724 | 6.898949899 | 7.30412922 |
| ENSG00000196387 | ZNF140 | -1.426029986 | 0.21980537 | -1.702581488 | 0.128777677 | -3.128611474 | 9.90440318 | 7.979858146 | 6.80318718 | 7.467753569 | 7.28433928 | 7.245405178 | 6.959831579 |
| ENSG00000156384 | SFR1 | -1.390682524 | 0.212937379 | -1.738716993 | 0.103755371 | -3.129399517 | 4.815609471 | 6.228547981 | 7.552162999 | 5.040242897 | 5.51241389 | 4.785723309 | 5.164234344 |
| ENSG00000149262 | INTS4 | -1.076726332 | 0.009386453 | -2.053368782 | 1.07E-07 | -3.130095114 | 9.031919741 | 8.702538795 | 8.266922072 | 7.700356475 | 7.57741621 | 6.782996409 | 6.592824411 |
| ENSG00000174917 | C19orf70 | -1.953625519 | 0.001999158 | -1.178164612 | 0.075316635 | -3.131790131 | 10.29268328 | 9.225224669 | 8.784770914 | 7.638916932 | 7.65252709 | 8.391426944 | 8.426701165 |
| ENSG00000140876 | NUDT7 | -0.644663743 | 0.53650408 | -2.487330361 | 0.005903997 | -3.131994104 | 6.009025997 | 4.442710129 | 5.156626401 | 4.927308402 | 4.56059747 | 3.5515855 | 3.183290849 |
| ENSG00000143545 | RAB13 | -0.4034801 | 0.603581513 | -2.728861368 | 1.55E-05 | -3.132341468 | 11.72814722 | 10.68823825 | 10.23015729 | 10.59344634 | 10.646815 | 7.977459036 | 8.579584657 |
| ENSG00000138772 | ANXA3 | -1.289688989 | 0.377120449 | -1.845199683 | 0.180357273 | -3.134888672 | 3.327711234 | 7.099070191 | 6.203069917 | 4.887609717 | 5.16377554 | 3.968654369 | 4.975095763 |
| ENSG00000213398 | LCAT | -2.322764489 | 0.180251842 | -0.813066887 | 0.666185912 | -3.135831376 | 4.00722531 | 6.04685525 | 9.491321578 | 5.697663076 | 5.93454154 | 7.20886319 | 7.314917451 |
| ENSG00000111671 | SPSB2 | -1.69106728 | 0.361961292 | -1.445202012 | 0.434158696 | -3.136269293 | 8.009475818 | 6.464272742 | 2 | 5.144975053 | 5.40529704 | 5.773219597 | 5.171036168 |
| ENSG00000171033 | PKIA | -0.349575223 | 0.810928754 | -2.788967961 | 0.019419395 | -3.138543184 | 7.435667355 | 7.901717006 | 4.058998049 | 6.846474326 | 6.80989959 | 4.5389195 | 4.654341953 |
| ENSG00000158526 | TSR2 | -1.58673438 | 0.10165634 | -1.553735039 | 0.103797879 | -3.140469419 | 8.16936288 | 9.246456519 | 10.7325871 | 8.199071993 | 8.16669309 | 8.018003769 | 8.389275155 |
| ENSG00000111203 | ITFG2 | -1.438225729 | 0.079815728 | -1.704231608 | 0.031808421 | -3.142457338 | 9.015175615 | 7.638155693 | 7.163363831 | 6.902323594 | 6.59518404 | 6.398709156 | 6.597881141 |
| ENSG00000086475 | SEPHS1 | -1.815115834 | 0.004366202 | -1.327605728 | 0.042293147 | -3.142721562 | 9.957954117 | 9.181799937 | 8.380957451 | 7.565338955 | 7.47518984 | 7.871230734 | 8.117259579 |
| ENSG00000014919 | COX15 | -2.080549762 | 9.22E-14 | -1.06287066 | 0.000205276 | -3.143420421 | 9.384607364 | 9.04308532 | 9.358421373 | 7.285019958 | 7.14145882 | 8.203226602 | 8.222500688 |
| ENSG00000197769 | MAP1LC3C | -3.667236413 | 0.106547308 | 0.523724053 | 0.8125854 | -3.14351236 | 3.327711234 | 3.686303414 | 2 | 2.141737246 | 2.14022021 | 3.059555873 | 3.875378318 |
| ENSG00000131051 | RBM39 | -1.346221476 | 0.000486999 | -1.79851751 | 1.53E-06 | -3.144738986 | 11.94698976 | 11.46438781 | 12.00213172 | 10.49037246 | 10.4683133 | 10.27528313 | 9.730640067 |
| ENSG00000167191 | GPRC5B | -2.221440644 | 0.152949994 | -0.924596489 | 0.576450932 | -3.146037133 | 8.367060731 | 8.053983481 | 3.369317972 | 5.742861563 | 5.30456641 | 6.013120342 | 7.252154709 |
| ENSG00000092094 | OSGEP | -1.981780603 | 0.078887275 | -1.165062339 | 0.318479973 | -3.146842942 | 9.189468652 | 7.73146271 | 6.073738139 | 6.213314212 | 6.27599024 | 6.516588833 | 7.405663765 |
| ENSG00000066136 | NFYC | -1.592715612 | 0.053699169 | -1.554239616 | 0.056973044 | -3.146955228 | 10.41260941 | 8.337339476 | 9.242516611 | 7.956842238 | 8.03813956 | 7.805448645 | 8.234726269 |
| ENSG00000006695 | COX10 | -1.3254945 | 0.002318605 | -1.822562878 | 1.32E-05 | -3.148057378 | 8.998234871 | 8.433041974 | 9.31604076 | 7.706073672 | 7.5835711 | 7.187558829 | 7.133547492 |
| ENSG00000198918 | RPL39 | 0.31552691 | 0.794914423 | -3.464563105 | 0.000271376 | -3.149036194 | 11.36165447 | 8.671479029 | 9.439621147 | 10.5720824 | 10.6178696 | 7.05561307 | 6.637709309 |
| ENSG00000188580 | NKAIN2 | -4.714867831 | 0.326061292 | 1.563730225 | 0.743102056 | -3.151137606 | 3.327711234 | 2 | 2 | 2 | 2 | 3.810410249 | 2.509591799 |
| ENSG00000172426 | RSPH9 | -2.437210686 | 0.239611233 | -0.715755875 | 0.74998849 | -3.15296656 | 2 | 6.728495673 | 3.369317972 | 3.709785322 | 3.08597292 | 4.675266036 | 4.766376438 |
| ENSG00000183741 | CBX6 | -2.837658697 | 4.09E-10 | -0.317659578 | 0.568601621 | -3.155318275 | 12.00326238 | 11.83753936 | 12.8250297 | 9.446513412 | 9.47072153 | 11.90217794 | 12.03837446 |
| ENSG00000121900 | TMEM54 | -1.745989375 | 3.30E-07 | -1.409647446 | 3.87E-05 | -3.155636821 | 9.081016759 | 8.732943944 | 8.511888463 | 7.017154451 | 7.13728069 | 7.474865722 | 7.333225922 |
| ENSG00000169071 | ROR2 | -2.404322397 | 2.22E-07 | -0.751911471 | 0.141701619 | -3.156233868 | 8.981092835 | 8.702538795 | 8.296287027 | 6.147608921 | 6.51372389 | 7.615113359 | 8.214292363 |
| ENSG00000137547 | MRPL15 | -1.392095001 | 0.070384687 | -1.767787106 | 0.017112266 | -3.159882106 | 10.62062599 | 9.13702746 | 8.941672936 | 8.378048624 | 8.40776525 | 7.926930284 | 8.111967236 |
| ENSG00000135519 | KCNH3 | -2.866879578 | 0.085388044 | -0.293203334 | 0.876355454 | -3.160082912 | 4.00722531 | 5.131603625 | 2 | 2.498691189 | 2.68924543 | 3.663836244 | 4.30448772 |
| ENSG00000134215 | VAV3 | -3.26480331 | 0.214781105 | 0.104693083 | 0.9716719 | -3.160110227 | 5.095959458 | 2 | 2 | 2.498691189 | 2.14022021 | 3.709392871 | 4.018014792 |
| ENSG00000125995 | ROMO1 | -1.758207273 | 0.011008907 | -1.404003345 | 0.045772843 | -3.162210618 | 10.27872131 | 8.522788974 | 9.849281147 | 7.895617537 | 8.05374131 | 8.230175068 | 8.416004473 |
| ENSG00000111639 | MRPL51 | -1.957583815 | 3.74E-05 | -1.205467005 | 0.015413637 | -3.16305082 | 10.76782679 | 10.00059544 | 11.00131064 | 8.624367384 | 8.77214691 | 9.230984775 | 9.634258132 |
| ENSG00000176809 | LRRC37A3 | -0.43855229 | 0.738258264 | -2.724767398 | 0.012793396 | -3.163319688 | 7.485143553 | 5.838937808 | 9.049077186 | 7.536750104 | 7.5835711 | 5.665620077 | 5.04340913 |
| ENSG00000141543 | EIF4A3 | -1.153710363 | 0.020935936 | -2.009690129 | 1.73E-05 | -3.163400492 | 10.85509492 | 9.77810053 | 10.31150679 | 9.189220954 | 9.27163371 | 8.287622092 | 8.472158355 |
| ENSG00000089022 | MAPKAPK5 | -1.544805806 | 0.032240446 | -1.619202788 | 0.022948755 | -3.164008594 | 10.15425007 | 8.337339476 | 9.413059136 | 7.872924734 | 8.02236724 | 7.98365249 | 7.760489878 |
| ENSG00000241343 | RPL36A | 0.355396136 | 0.798171474 | -3.520346329 | 0.001414241 | -3.164950193 | 9.435834703 | 7.189546432 | 6.322058544 | 8.578353644 | 8.63663561 | 5.180212365 | 4.604965121 |
| ENSG00000185033 | SEMA4B | -0.708437271 | 0.739704558 | -2.459500234 | 0.19190395 | -3.167937505 | 3.327711234 | 7.002537973 | 10.36098398 | 8.223194971 | 8.2130862 | 6.802396951 | 6.148451724 |
| ENSG00000176209 | SMIM19 | -1.728844326 | 0.048492007 | -1.440034936 | 0.101374362 | -3.168879262 | 8.340423424 | 6.602424674 | 8.904018013 | 6.652857793 | 6.41363412 | 6.942523933 | 6.676467423 |
| ENSG00000234545 | FAM133B | 0.00462502 | 0.993750338 | -3.177468018 | 5.76E-16 | -3.172842997 | 7.159102577 | 7.638155693 | 7.398165132 | 7.562190249 | 7.27295842 | 4.513461373 | 4.468068808 |
| ENSG00000138663 | COPS4 | -1.837488763 | 0.226126239 | -1.335920794 | 0.384717028 | -3.173409557 | 11.04852648 | 9.842262059 | 5.59507254 | 8.15586117 | 8.20911072 | 8.596284315 | 8.756196613 |
| ENSG00000215915 | ATAD3C | 0.863287917 | 0.423402777 | -4.038169391 | 1.75E-05 | -3.174818474 | 5.70932728 | 4.710564154 | 6.203069917 | 6.205262847 | 6.66650236 | 3.035909414 | 2.421067459 |
| ENSG00000111271 | ACAD10 | -2.320346671 | 1.82E-07 | -0.856166051 | 0.075069882 | -3.176512722 | 8.631580145 | 7.790474954 | 8.631919287 | 6.164317909 | 6.11534392 | 7.596037946 | 7.50456446 |
| ENSG00000165185 | KIAA1958 | -2.883892779 | 0.000634997 | -0.294516668 | 0.774877964 | -3.178409448 | 8.39321512 | 6.666836837 | 6.629488387 | 4.907595604 | 4.65713986 | 7.454943454 | 6.868633039 |
| ENSG00000111678 | C12orf57 | -1.232119611 | 0.002939392 | -1.947354638 | 7.85E-07 | -3.179474249 | 10.43166252 | 9.77810053 | 9.809100877 | 8.724940594 | 8.8960155 | 8.266187847 | 7.925444964 |
| ENSG00000003096 | KLHL13 | -1.436834464 | 0.485444732 | -1.743785096 | 0.377312046 | -3.18061956 | 7.53297911 | 4.442710129 | 2 | 4.739373666 | 4.91316568 | 4.68289886 | 4.445805334 |
| ENSG00000075945 | KIFAP3 | -0.877263934 | 0.204923899 | -2.306017727 | 0.000176526 | -3.183281662 | 9.159305284 | 9.308339287 | 10.54349105 | 8.957217624 | 8.92053511 | 7.463831776 | 7.593325136 |
| ENSG00000135521 | LTV1 | -1.136777379 | 0.105267278 | -2.048136311 | 0.001583285 | -3.18491369 | 9.957954117 | 8.623598428 | 8.65477198 | 8.03159949 | 8.15640714 | 6.981729486 | 7.39408282 |
| ENSG00000075415 | SLC25A3 | -1.967623414 | 2.45E-06 | -1.218694756 | 0.005232053 | -3.18631817 | 13.48155154 | 12.85154862 | 12.65834871 | 11.08473443 | 11.0668225 | 11.63795282 | 11.9881668 |
| ENSG00000159445 | THEM4 | -1.176196487 | 0.195586526 | -2.016097737 | 0.017084929 | -3.192294224 | 8.075571721 | 7.979858146 | 5.930865648 | 6.436065776 | 6.50077304 | 5.463319095 | 5.867805474 |
| ENSG00000110921 | MVK | -2.478584938 | 1.51E-06 | -0.714421304 | 0.213651438 | -3.193006241 | 8.609439662 | 8.777382925 | 8.007629717 | 6.096288482 | 6.09824986 | 7.3389565 | 8.150334303 |
| ENSG00000213023 | SYT3 | -2.596859319 | 0.310866 | -0.596519797 | 0.83342673 | -3.193379116 | 2 | 6.78762695 | 2 | 3.09461492 | 3.50452879 | 4.90146639 | 4.683176719 |

| | | | | | | | | | | | | | |
|---|---|---|---|---|---|---|---|---|---|---|---|---|---|
| ENSG00000174939 | ASPHD1 | -0.016198647 | 0.994366162 | -3.179527011 | 0.060577873 | -3.195725659 | 3.327711234 | 7.274681896 | 9.735974718 | 8.356399318 | 8.41985547 | 5.699879814 | 4.903386193 |
| ENSG00000171017 | LRRC8E | 0.809193459 | 0.647852448 | -4.009332601 | 0.008063128 | -3.200139142 | 4.00722531 | 3.686303414 | 7.451345288 | 6.851641821 | 6.84106088 | 3.193831101 | 2.746601469 |
| ENSG00000196345 | ZKSCAN7 | -1.230370475 | 0.497739811 | -1.969885509 | 0.246828935 | -3.200255984 | 3.327711234 | 5.457120953 | 8.460932916 | 5.808110275 | 5.99120031 | 5.467744375 | 4.935697922 |
| ENSG00000173065 | FAM222B | -1.806429878 | 0.052665973 | -1.394522655 | 0.139393652 | -3.200952533 | 10.85509492 | 9.433952963 | 8.325066185 | 8.083034979 | 8.13141746 | 8.617847323 | 8.402340474 |
| ENSG00000184451 | CCR10 | -0.786298074 | 0.568853498 | -2.415219258 | 0.047712153 | -3.201517333 | 6.816637977 | 4.11352228 | 7.818184403 | 6.213314212 | 6.03677338 | 4.659878161 | 4.664017718 |
| ENSG00000180228 | PRKRA | -1.410014827 | 0.020506295 | -1.794635629 | 0.002248156 | -3.204650456 | 10.09173937 | 8.732943944 | 9.491321578 | 8.237082658 | 8.04037874 | 7.966556098 | 7.52727182 |
| ENSG00000075234 | TTC38 | -1.521916634 | 0.000559325 | -1.684161406 | 9.95E-05 | -3.20607804 | 8.910412251 | 9.624396729 | 8.845626171 | 7.546342806 | 7.7919464 | 7.473766117 | 7.55613706 |
| ENSG00000105427 | CNFN | -2.22207455 | 0.433973734 | -0.984512121 | 0.730619275 | -3.206586671 | 2 | 4.11352228 | 2 | 2.270786764 | 2.38539787 | 2.3070652 | 2.981406154 |
| ENSG00000124615 | MOCS1 | -2.156727084 | 0.029463335 | -1.051284839 | 0.318173125 | -3.208011923 | 10.21416435 | 7.431218269 | 8.825624954 | 7.214570819 | 7.00595521 | 8.422534217 | 7.929461924 |
| ENSG00000108405 | P2RX1 | -2.317421058 | 0.393143611 | -0.890887438 | 0.751167749 | -3.208308496 | 2 | 2 | 4.874587708 | 2.3892346 | 2.59489879 | 3.429858444 | 2.592996859 |
| ENSG00000139719 | VPS33A | -1.437397486 | 0.000590286 | -1.771753812 | 1.32E-05 | -3.209151299 | 8.716893051 | 9.125613817 | 9.565556058 | 7.797276258 | 7.7127321 | 7.511762185 | 7.339277475 |
| ENSG00000127948 | POR | -1.175092022 | 0.213853811 | -2.037069712 | 0.020727627 | -3.212161733 | 11.71265261 | 10.22039562 | 9.31604076 | 9.594867189 | 9.57402703 | 8.705236603 | 8.751666251 |
| ENSG00000063177 | RPL18 | -1.528241052 | 0.037063786 | -1.684153415 | 0.019287415 | -3.212394467 | 13.33745529 | 12.04564457 | 11.58682058 | 10.98215259 | 11.0102587 | 10.86216341 | 10.81887387 |
| ENSG00000135451 | TROAP | -1.351648363 | 0.40455867 | -1.860943279 | 0.222909496 | -3.212591642 | 4.467311984 | 5.303523747 | 3.369317972 | 3.663521627 | 3.50452879 | 2.492221014 | 3.771796761 |
| ENSG00000103351 | CLUAP1 | -1.415619842 | 0.001569431 | -1.798028146 | 3.33E-05 | -3.213647989 | 9.189468652 | 8.414402626 | 9.301632639 | 7.571615825 | 7.65837038 | 7.214140484 | 7.269688419 |
| ENSG00000133401 | PDZD2 | -2.726302003 | 0.015969093 | -0.487456875 | 0.708194244 | -3.213758877 | 7.435667355 | 4.710564154 | 6.073378139 | 4.034562184 | 4.0930011 | 5.338397457 | 6.454714037 |
| ENSG00000178031 | ADAMTSL1 | -1.448227096 | 0.050246433 | -1.765915442 | 0.013607938 | -3.214142538 | 9.2903348 | 7.60566235 | 9.25752426 | 7.405868159 | 7.52720828 | 7.292315404 | 7.010010317 |
| ENSG00000108641 | B9D1 | -1.69507273 | 0.313384316 | -1.520417151 | 0.362166499 | -3.215489881 | 3.327711234 | 9.00621767 | 7.225742779 | 6.172600389 | 6.18176397 | 6.054896192 | 6.582657467 |
| ENSG00000131148 | EMC8 | -1.876555051 | 0.022721789 | -1.339187896 | 0.113061522 | -3.215742946 | 10.71696191 | 8.469613413 | 10.26770388 | 8.241026137 | 8.21109983 | 8.644966335 | 8.861915706 |
| ENSG00000185339 | TCN2 | -0.967461608 | 0.66635587 | -2.248793758 | 0.27100504 | -3.216255366 | 10.22148166 | 8.356994601 | 2 | 7.997038782 | 8.06917614 | 6.514451546 | 7.010010317 |
| ENSG00000058799 | YIPF1 | -1.26138459 | 0.19884911 | -1.956547474 | 0.034086028 | -3.217932063 | 8.493334308 | 8.395519306 | 10.54349105 | 8.205140657 | 8.31092742 | 7.535143176 | 7.614687532 |
| ENSG00000140455 | USP3 | -1.356328573 | 0.057486219 | -1.861671367 | 0.006279821 | -3.21799994 | 10.01801709 | 8.433041974 | 8.763901704 | 7.885575888 | 7.91175216 | 7.414254101 | 7.392628644 |
| ENSG00000198346 | ZNF813 | -1.355565707 | 0.552447214 | -1.862964168 | 0.393071188 | -3.218529875 | 2 | 5.946638857 | 9.478569125 | 6.727838783 | 6.66650236 | 6.529346487 | 5.807970301 |
| ENSG00000132768 | DPH2 | -1.769524639 | 1.38E-05 | -1.45050121 | 0.000405269 | -3.220025849 | 9.473095744 | 8.732943944 | 9.465702942 | 7.507583254 | 7.51437845 | 7.914827151 | 7.726263462 |
| ENSG00000205155 | PSENEN | -1.354352576 | 0.001192861 | -1.865856029 | 3.61E-06 | -3.220208605 | 10.03472846 | 9.225224669 | 9.61300511 | 8.258639605 | 8.37087451 | 7.673747216 | 7.938459399 |
| ENSG00000280759 | | 0.276321907 | 0.80351594 | -3.499018556 | 4.78E-05 | -3.222696649 | 7.218819781 | 9.599860728 | 9.788583153 | 9.445659335 | 9.57479964 | 5.832754931 | 5.8339184 |
| ENSG00000111845 | PAK1IP1 | -1.346332111 | 0.25067706 | -1.878808464 | 0.091455421 | -3.225140575 | 9.533135543 | 7.232742069 | 6.718950348 | 7.017154451 | 7.09050444 | 6.426254939 | 6.64749689 |
| ENSG00000120498 | TEX11 | -0.689897333 | NA | -2.536410019 | NA | -3.226307352 | 3.327711234 | 2 | 2 | 2.270786764 | 2.49394783 | 2.225793514 | 2 |
| ENSG00000172977 | KAT5 | -0.456014166 | 0.494444165 | -2.770814605 | 4.29E-07 | -3.226828771 | 9.031919741 | 7.761270542 | 8.904018013 | 8.211183901 | 8.20911072 | 5.843010798 | 6.088639344 |
| ENSG00000144712 | CAND2 | -4.31403024 | 0.001659143 | 1.086478296 | 0.481274312 | -3.227551944 | 8.468948322 | 7.538394143 | 3.369317972 | 3.663521627 | 3.74316441 | 8.653246476 | 8.524881346 |
| ENSG00000175536 | LIPT2 | -1.429348958 | 0.588043549 | -1.798366985 | 0.477852963 | -3.227715943 | 2 | 6.389896633 | 2 | 3.921164147 | 3.69849104 | 3.517841605 | 3.600986722 |
| ENSG00000184831 | APOO | -0.866874323 | 0.26760945 | -2.361070556 | 0.000646284 | -3.227944878 | 7.384433967 | 7.954278496 | 6.203069917 | 6.652857793 | 6.37866593 | 5.096960836 | 5.177806074 |
| ENSG00000109475 | RPL34 | -1.453430509 | 0.0347897 | -1.774572927 | 0.007862246 | -3.228003436 | 12.02642475 | 10.49332014 | 10.73794767 | 9.782429661 | 9.82323967 | 9.514027227 | 9.452192236 |
| ENSG00000159915 | ZNF233 | -3.070505084 | 0.200961249 | -0.157541105 | 0.954416752 | -3.228046189 | 2 | 5.457120953 | 2 | 2.600425781 | 2.38539787 | 4.005644654 | 4.062589736 |
| ENSG00000143314 | MRPL24 | -1.946766933 | 0.047029973 | -1.281641272 | 0.20389337 | -3.228408205 | 9.87686338 | 8.848535039 | 7.029880454 | 7.153115662 | 7.00595521 | 7.662209535 | 7.796124663 |
| ENSG00000182287 | AP1S2 | -1.818970173 | 5.14E-05 | -1.409864214 | 0.002070948 | -3.228834387 | 9.521325812 | 10.07368926 | 10.45512158 | 8.291333947 | 8.22887947 | 8.455772155 | 8.847658168 |
| ENSG00000116288 | PARK7 | -1.813736751 | 0.00200525 | -1.415175479 | 0.018674367 | -3.22891223 | 12.20314289 | 11.00090285 | 11.16546284 | 9.764048416 | 9.73820484 | 9.853683372 | 10.39258649 |
| ENSG00000104412 | EMC2 | -1.143510177 | 0.020547487 | -2.086449586 | 5.75E-06 | -3.229959763 | 9.219014269 | 8.376385542 | 9.212024683 | 7.885575888 | 7.81567127 | 7.118879607 | 6.714211481 |
| ENSG00000052344 | PRSS8 | -1.272306145 | NA | -1.957980749 | NA | -3.230286895 | 2 | 2 | 3.369317972 | 2.270786764 | 2.2680111 | 2.225793514 | 2.117335275 |
| ENSG00000205978 | NYNRIN | -3.190992818 | 0.000382069 | -0.041259188 | 0.971943653 | -3.232252006 | 7.579279349 | 9.246456519 | 10.3329194 | 6.283823264 | 6.35006802 | 9.418964428 | 9.383574901 |
| ENSG00000075131 | TIPIN | -1.590095484 | 0.195468305 | -1.645722453 | 0.169009791 | -3.235817936 | 6.89194353 | 4.442710129 | 7.858115942 | 5.348759107 | 5.53799497 | 5.485310992 | 5.300743019 |
| ENSG00000165119 | HNRNPK | -1.269156098 | 0.007300149 | -1.967329379 | 1.22E-05 | -3.236485477 | 13.19511653 | 12.21272354 | 13.06678708 | 11.65132285 | 11.5824281 | 10.81822064 | 11.01556332 |
| ENSG00000160949 | TONSL | -2.35116929 | 0.003304802 | -0.888820301 | 0.306338229 | -3.239989591 | 8.285626337 | 6.228547981 | 7.285536013 | 5.258265913 | 5.27444544 | 6.589466407 | 6.669279147 |
| ENSG00000076356 | PLXNA2 | -0.941906806 | 0.141052921 | -2.299141956 | 6.03E-05 | -3.241048762 | 9.521325812 | 8.433041974 | 8.699419242 | 8.04293816 | 8.01327646 | 7.030406362 | 6.283256815 |
| ENSG00000163002 | NUP35 | -1.142491144 | 0.561228414 | -2.099137768 | 0.251602132 | -3.241628912 | 2 | 6.951736442 | 8.434763955 | 6.372775472 | 6.00960265 | 5.174809258 | 5.448021784 |
| ENSG00000160298 | C21orf58 | -2.116359363 | 0.207952625 | -1.126628622 | 0.519967781 | -3.242987985 | 3.327711234 | 5.595927442 | 8.408111533 | 5.144975053 | 4.98931816 | 5.843010798 | 6.088639344 |
| ENSG00000101680 | LAMA1 | -0.59559462 | 0.725223616 | -2.648600573 | 0.069125054 | -3.244195194 | 4.467311984 | 9.799806309 | 9.516493218 | 8.459845423 | 8.55910965 | 6.706424753 | 6.27380897 |
| ENSG00000114767 | RRP9 | -1.17200548 | 0.560973962 | -2.073811899 | 0.270301865 | -3.245817379 | 2 | 7.954278496 | 9.031722608 | 6.820355514 | 6.91614081 | 5.893224226 | 6.117095472 |
| ENSG00000227993 | | 1.944484639 | NA | -5.191098671 | NA | -3.246614031 | 2 | 2 | 2 | 3.515101962 | 2.14022021 | 2 | 2 |
| LOC388436 | | 1.942794459 | NA | -5.191098678 | NA | -3.248304219 | 2 | 2 | 2 | 3.406930997 | 2.38539787 | 2 | 2 |
| LOC79999 | | 1.942794459 | NA | -5.191098678 | NA | -3.248304219 | 2 | 2 | 2 | 3.406930997 | 2.38539787 | 2 | 2 |
| ENSG00000040341 | STAU2 | -1.945790322 | 0.00914796 | -1.302947795 | 0.091865326 | -3.248738118 | 10.29268328 | 8.297207175 | 9.725218844 | 7.767511442 | 7.68150967 | 8.283862644 | 8.428831018 |
| ENSG00000145781 | COMMD10 | -1.35439217 | 0.355101172 | -1.898309764 | 0.171406992 | -3.252701934 | 9.304185368 | 6.844429755 | 5.156626401 | 6.652857793 | 6.75642009 | 6.287958221 | 6.077821908 |
| ENSG00000165678 | GHITM | -1.256683603 | 0.001101852 | -1.997404454 | 5.61E-08 | -3.254088057 | 12.42852545 | 11.64511952 | 12.12382753 | 10.85540801 | 10.8352843 | 10.17584339 | 10.03417286 |
| ENSG00000132423 | COQ3 | -1.82036736 | 0.003124535 | -1.436891553 | 0.020274693 | -3.257258913 | 6.73718411 | 6.464272742 | 7.163363831 | 5.022023783 | 5.19627118 | 5.04460184 | 5.777095613 |
| ENSG00000166927 | MS4A7 | -5.813753075 | 0.008410532 | 2.553012312 | 0.094049143 | -3.260740763 | 3.327711234 | 3.076665209 | 4.874587708 | 2.141737246 | 2 | 6.736076089 | 5.211187127 |
| ENSG00000144048 | DUSP11 | -1.517327996 | 0.128187989 | -1.743539808 | 0.071471553 | -3.260867803 | 9.957954117 | 8.639735838 | 7.225742779 | 7.571615825 | 7.4450836 | 7.292315404 | 7.285445324 |
| ENSG00000135124 | P2RX4 | -1.53880268 | 0.047549457 | -1.722651715 | 0.023355089 | -3.261454395 | 9.384607364 | 8.053983481 | 10.06083273 | 7.772969103 | 7.93600799 | 7.768261726 | 7.578053054 |
| ENSG00000163467 | TSACC | -2.954451323 | 0.433930421 | -0.307969437 | NA | -3.262420759 | 2 | 3.686303414 | 2 | 2 | 2.2680111 | 2.858551861 | 2.551896972 |
| ENSG00000151092 | NGLY1 | -1.696170591 | 0.028034076 | -1.567024633 | 0.042177264 | -3.263195224 | 10.52923584 | 9.28800521 | 8.65477198 | 8.051945366 | 7.99722844 | 8.350077895 | 7.922424886 |
| ENSG00000134146 | DPH6 | -1.506328392 | 0.011645431 | -1.75785231 | 0.002051968 | -3.264180701 | 6.257048221 | 6.04685525 | 5.772716771 | 4.804776789 | 4.56059747 | 4.388329244 | 4.56420962 |
| ENSG00000130255 | RPL36 | -1.542744708 | 0.025691449 | -1.724252857 | 0.0110979 | -3.266997565 | 12.84685418 | 11.30908426 | 11.50429402 | 10.49037246 | 10.5479824 | 10.38712053 | 10.28813986 |
| ENSG00000181218 | HIST3H2A | -1.233958818 | 0.162328517 | -2.033097102 | 0.013453741 | -3.267055921 | 7.904282243 | 5.838937808 | 6.882775709 | 5.930370745 | 5.91514978 | 5.152992171 | 5.230851202 |
| ENSG00000147099 | HDAC8 | -1.337292166 | 0.003152689 | -1.932920181 | 8.03E-06 | -3.270212347 | 8.981092835 | 8.487557258 | 8.076950877 | 7.300218887 | 7.19470132 | 6.571075185 | 6.76679494 |
| ENSG00000172339 | ALG14 | -2.29010037 | 0.001529373 | -0.980683561 | 0.203900623 | -3.270783931 | 6.816637977 | 6.389896633 | 7.777116083 | 4.965943888 | 4.97065356 | 5.740677235 | 6.490500786 |
| ENSG00000117906 | RCN2 | -1.771190389 | 4.12E-07 | -1.502090286 | 2.02E-05 | -3.273280674 | 10.83137851 | 10.38739569 | 11.03218278 | 9.104095335 | 8.9083274 | 9.174577117 | 9.372183513 |
| ENSG00000135447 | PPP1R1A | -3.274332991 | NA | 0.000667656 | NA | -3.273665335 | 2 | 3.076665209 | 2 | 2 | 2.14022021 | 2.831336763 | 2.277179783 |
| ENSG00000249158 | PCDHA11 | -0.115394523 | 0.971678075 | -3.159633292 | 0.260567982 | -3.275027815 | 2 | 4.442710129 | 2 | 3.754611435 | 2.59489879 | 2.384001718 | 2.059860178 |
| ENSG00000111196 | MAGOHB | -1.186691458 | 0.149439706 | -2.090713858 | 0.006027795 | -3.277405316 | 6.009025997 | 7.790474954 | 7.896971941 | 6.422239259 | 6.18176397 | 5.454427604 | 5.475755622 |
| ENSG00000182944 | EWSR1 | -1.609530875 | 0.001194252 | -1.66855669 | 0.000700711 | -3.278087565 | 12.44424702 | 11.84799496 | 11.70922096 | 10.40155496 | 10.4582997 | 9.89698509 | 10.72781664 |
| ENSG00000146006 | LRRTM2 | -1.509514994 | 0.619188542 | -1.769458033 | 0.544666992 | -3.278973027 | 2 | 2 | 4.874587708 | 3.162721474 | 2.2680111 | 2.962551659 | 2.326754077 |
| ENSG00000144730 | IL17RD | -2.572184652 | 0.034056966 | -0.706945026 | 0.600178142 | -3.279129678 | 6.653098118 | 4.442710129 | 7.646393253 | 4.739373666 | 3.98554648 | 6.428526856 | 5.576534358 |
| ENSG00000102387 | TAF7L | -0.167871197 | 0.949370102 | -3.111850687 | 0.17053521 | -3.279721885 | 2 | 5.131603625 | 2 | 3.997744386 | 3.50452879 | 2.457042231 | 2.326754077 |
| ENSG00000101158 | NELFCD | -1.839211277 | 0.001901239 | -1.440975095 | 0.017396937 | -3.280186372 | 10.86447339 | 9.785372135 | 9.601288346 | 8.302698479 | 8.43525263 | 8.58766971 | 8.922120243 |
| ENSG00000121058 | COIL | -1.881312812 | 8.27E-06 | -1.398990737 | 0.001105945 | -3.280303549 | 9.317904228 | 9.055168103 | 8.408111533 | 7.140504127 | 7.11195456 | 7.599066672 | 7.594590542 |
| ENSG00000115998 | C2orf42 | -1.204042864 | 0.371555413 | -2.077542733 | 0.097255856 | -3.281585597 | 5.70932728 | 7.790474954 | 9.601288346 | 7.202487431 | 7.32898565 | 6.601598251 | 6.222348126 |
| ENSG00000117640 | MTFR1L | -2.042220691 | 4.16E-07 | -1.240531521 | 0.003164856 | -3.282752212 | 10.92399098 | 10.17128279 | 10.90928008 | 8.570539957 | 8.77214691 | 9.356203439 | 9.576512555 |
| ENSG00000100462 | PRMT5 | -1.089872826 | 0.103636128 | -2.196808914 | 0.000327125 | -3.28668174 | 10.76782679 | 10.39216648 | 9.196533591 | 9.154098367 | 9.19430858 | 7.960288653 | 8.18854565 |
| ENSG00000075336 | TIMM21 | -2.210798707 | 0.00980268 | -1.076794988 | 0.237465262 | -3.287593695 | 8.910412251 | 8.077869079 | 6.534109959 | 5.968925393 | 6.02777321 | 6.939340974 | 7.213143202 |
| ENSG00000162643 | WDR63 | -2.371507072 | 0.340882919 | -0.917168223 | 0.725298877 | -3.288675295 | 2 | 5.457120953 | 2 | 2.600425781 | 2.86123014 | 3.738983888 | 3.209956562 |
| ENSG00000134830 | C5AR2 | -4.888871402 | 0.119616016 | 1.595282587 | 0.570917019 | -3.293588815 | 2 | 2 | 4.523675391 | 2.141737246 | 2 | 4.504874511 | 4.468068808 |
| ENSG00000144451 | SPAG16 | -1.561398972 | 0.092788449 | -1.736664611 | 0.053870623 | -3.298063583 | 8.609439662 | 6.899080544 | 6.534109959 | 6.221320894 | 6.05460702 | 5.870008604 | 6.074197996 |
| ENSG00000160179 | ABCG1 | -3.86670838 | 0.031195752 | 0.568434488 | 0.760016149 | -3.298273893 | 2 | 4.710564154 | 4.058998049 | 2.270786764 | 2.2680111 | 4.907994066 | 3.753786374 |
| ENSG00000149485 | FADS1 | -1.789411877 | 0.001617597 | -1.510907928 | 0.008692652 | -3.300319806 | 11.9710658 | 12.57771829 | 13.02134859 | 10.77779286 | 10.8190733 | 10.54072141 | 11.4663611 |
| ENSG00000108561 | C1QBP | -1.625068872 | 1.26E-05 | -1.675589445 | 5.59E-06 | -3.300658317 | 10.80239439 | 10.24697497 | 10.70548138 | 8.925713487 | 9.04284131 | 8.684474732 | 9.149213283 |
| ENSG00000255346 | NOX5 | -1.685204822 | NA | -1.616714281 | NA | -3.301919103 | 2 | 2 | 3.369317972 | 2.270786764 | 2.14022021 | 2.3070652 | 2.117335275 |
| ENSG00000123415 | SMUG1 | -1.661081805 | 0.298816194 | -1.643258582 | 0.293628603 | -3.304340387 | 10.06758145 | 8.356994601 | 4.523675391 | 7.182120794 | 7.32531744 | 7.178138321 | 7.361747375 |
| ENSG00000170315 | UBB | -0.215382625 | 0.589880261 | -3.08949696 | 4.27E-23 | -3.304879585 | 13.49896863 | 12.99094813 | 13.12010506 | 13.01072495 | 12.997898 | 9.946970917 | 10.30079327 |
| ENSG00000168894 | RNF181 | -1.378317163 | 0.015510476 | -1.92835502 | 0.000397713 | -3.306672183 | 10.64259676 | 9.608085847 | 9.503962296 | 8.705093576 | 8.58234702 | 7.988280194 | 8.206864727 |
| ENSG00000111319 | SCNN1A | -2.243686076 | 0.284911179 | -1.063435121 | 0.609433523 | -3.307121197 | 4.00722531 | 2 | 3.369317972 | 2.3892346 | 2.38539787 | 2.592885947 | 2.918126211 |
| ENSG00000105204 | DYRK1B | -1.408068361 | 0.061452906 | -1.900656496 | 0.008312099 | -3.308724856 | 10.27168932 | 8.607278467 | 9.491321578 | 8.188900477 | 8.23671175 | 7.238947338 | 8.093289637 |
| C9ORF47 | C9ORF47 | 3.55872953 | 0.181478081 | -6.868095241 | 0.022211412 | -3.309365711 | 2 | 3.686303414 | 2 | 5.304221918 | 5.49945122 | 2 | 2 |
| ENSG00000166851 | PLK1 | -1.860537634 | 0.233747682 | -1.453217456 | 0.354062737 | -3.31375509 | 5.70932728 | 7.538394143 | 3.369317972 | 4.443344104 | 4.91316568 | 4.311629396 | 5.534955934 |
| ENSG00000125266 | EFNB2 | -2.504929968 | 0.000463831 | -0.81200393 | 0.306556213 | -3.316933898 | 8.367060731 | 9.540933215 | 8.007629717 | 6.442929658 | 6.26070706 | 7.538302335 | 8.346352355 |
| ENSG00000266173 | STRADA | -0.585227325 | 0.607759609 | -2.73212405 | 0.004514133 | -3.317351375 | 7.031698319 | 7.928237109 | 9.703463497 | 8.08961076 | 8.05595642 | 5.960710379 | 6.033722345 |
| ENSG00000188153 | COL4A5 | -4.128755702 | 5.15E-14 | 0.811034428 | 0.195723468 | -3.317721274 | 10.62062599 | 11.09754095 | 10.20715117 | 6.856790872 | 6.32824132 | 10.94325044 | 11.89815768 |
| ENSG00000205643 | CDPF1 | -2.413974421 | 0.038304889 | -0.905419131 | 0.473584699 | -3.319393552 | 7.790810999 | 8.029695761 | 4.523675391 | 5.194621444 | 5.06165168 | 6.623578417 | 6.423723772 |
| ENSG00000125351 | UPF3B | -1.388872626 | 0.054916895 | -1.931289596 | 0.005050688 | -3.320162222 | 9.233563228 | 7.847167909 | 7.858115942 | 7.140504127 | 7.06873058 | 6.784770914 | 6.344709081 |
| ENSG00000064199 | SPA17 | -0.662119866 | 0.739570871 | -2.659264029 | 0.12597772 | -3.321383896 | 6.366709671 | 2 | 7.502634626 | 5.797438367 | 5.89549381 | 4.111244182 | 4.133950982 |
| ENSG00000179988 | PSTK | -1.936403817 | 0.232124999 | -1.386296887 | 0.397152832 | -3.322700704 | 6.73718411 | 6.899080544 | 2 | 4.54891529 | 4.50979031 | 5.050514232 | 4.943664032 |
| ENSG00000170291 | ELP5 | -1.898609238 | 0.001127253 | -1.42481004 | 0.017257913 | -3.323419279 | 10.19198706 | 8.834582582 | 9.25752426 | 7.641902724 | 7.68437616 | 8.112879504 | 8.145162126 |
| ENSG00000006634 | DBF4 | -0.97996363 | 0.483277973 | -2.345011727 | 0.062128736 | -3.324975357 | 4.00722531 | 6.844429755 | 7.97167868 | 6.087554739 | 6.01871653 | 4.959178705 | 4.664017718 |
| ENSG00000163710 | PCOLCE2 | -2.327202144 | 0.013562863 | -0.998330999 | 0.327109241 | -3.325533144 | 10.39976609 | 9.181799937 | 7.691296561 | 7.148924055 | 7.21466714 | 8.214461509 | 8.716065093 |
| ENSG00000004779 | NDUFAB1 | -1.312994211 | 0.277913797 | -2.013300759 | 0.076665265 | -3.32629497 | 10.95065381 | 8.053983481 | 8.380957451 | 8.427334995 | 8.45888185 | 7.717139459 | 7.788404416 |
| ENSG00000099326 | MZF1 | -1.779172387 | 0.046591905 | -1.548344576 | 0.083814249 | -3.327516963 | 10.00958825 | 7.538394143 | 9.196533591 | 7.433701013 | 7.51115308 | 7.807195765 | 7.581886257 |
| ENSG00000119203 | CPSF3 | -1.352066671 | 0.000484199 | -1.975622689 | 1.12E-07 | -3.32768936 | 9.992581382 | 9.308339287 | 9.839340503 | 8.293234263 | 8.4936155 | 7.662209535 | 7.893928381 |
| ENSG00000092470 | WDR76 | -2.489599971 | 0.160982131 | -0.838932864 | 0.664628785 | -3.328532835 | 4.467311984 | 3.686303414 | 8.460932916 | 4.783303513 | 4.63360314 | 6.320128742 | 6.074197996 |

| | | | | | | | | | | | | | |
|---|---|---|---|---|---|---|---|---|---|---|---|---|---|
| ENSG00000120942 | UBIAD1 | -2.483932806 | 2.42E-05 | -0.844719255 | 0.193664118 | -3.328652061 | 9.174465797 | 8.317412869 | 9.778213818 | 6.83085985 | 6.6952083 | 8.226915494 | 8.495377414 |
| ENSG00000196787 | HIST1H2AG | -1.8110768 | 0.303863659 | -1.518313141 | 0.38169184 | -3.329389941 | 6.009025997 | 2 | 5.156626401 | 3.462030052 | 3.82856379 | 3.5515855 | 4.119958326 |
| ENSG00000119333 | WDR34 | -2.720110838 | 6.47E-10 | -0.609408649 | 0.221852238 | -3.329519488 | 8.586954094 | 9.34816733 | 9.149036566 | 6.470063213 | 6.33555364 | 8.167627778 | 8.706148039 |
| ENSG00000105829 | BET1 | -0.965490684 | 0.199905975 | -2.366255847 | 0.000442304 | -3.331746531 | 10.31337612 | 8.94259736 | 9.25752426 | 8.663092307 | 8.67605209 | 7.685193358 | 6.748721609 |
| ENSG00000183386 | FHL3 | -1.98969218 | 0.002660765 | -1.342123932 | 0.050500882 | -3.331816112 | 10.40620204 | 8.687092496 | 9.935792933 | 7.849869274 | 7.88209639 | 8.504505158 | 8.504830639 |
| ENSG00000135111 | TBX3 | 0.741908334 | 0.215420072 | -4.079739943 | 8.90E-16 | -3.337831609 | 9.848787621 | 10.85302421 | 10.62675862 | 11.25235087 | 11.2358381 | 6.818078339 | 6.052261256 |
| ENSG00000150779 | TIMM8B | -1.878362543 | 0.000246664 | -1.460639075 | 0.005222606 | -3.339001618 | 9.031919741 | 9.74864196 | 9.972637969 | 7.700356475 | 7.85181819 | 7.808068532 | 8.490627472 |
| ENSG00000112667 | DNPH1 | -1.855826818 | 0.003101681 | -1.483672163 | 0.020210784 | -3.339498981 | 8.564112515 | 8.029695761 | 9.465702942 | 6.851641821 | 7.10341274 | 7.195584877 | 7.480125329 |
| ENSG00000114805 | PLCH1 | 0.94653885 | 0.791030588 | -4.287170024 | 0.187701731 | -3.340631174 | 2 | 2 | 4.058998049 | 3.754611435 | 3.33976443 | 2.139668783 | 2 |
| ENSG00000196152 | ZNF79 | -2.034200824 | 0.035999029 | -1.306828658 | 0.190459822 | -3.341029482 | 8.75772933 | 6.899080544 | 6.629488387 | 5.614989456 | 5.9725602 | 6.784770914 | 6.120613345 |
| ENSG00000180817 | PPA1 | -1.267028349 | 0.000270729 | -2.075065917 | 4.67E-10 | -3.342094267 | 9.966689162 | 9.583268459 | 10.17589429 | 8.716468139 | 8.61576544 | 7.743839536 | 7.985520618 |
| ENSG00000166997 | CNPY4 | -1.621830367 | 0.175251415 | -1.720375596 | 0.14015005 | -3.342205962 | 10.05124847 | 6.464272742 | 8.865353883 | 7.405868159 | 7.53358066 | 7.172727426 | 7.552234734 |
| ENSG00000176387 | HSD11B2 | -2.279075096 | NA | -1.065945952 | NA | -3.345021049 | 2 | 3.076665209 | 2 | 2 | 2.2680111 | 2.183373728 | 2.421067459 |
| ENSG00000196998 | WDR45 | -1.16091898 | 0.002141811 | -2.187365409 | 8.46E-10 | -3.348284389 | 10.48735708 | 9.916994052 | 10.08627303 | 8.954818504 | 9.09655277 | 7.800194554 | 8.197734265 |
| ENSG00000159479 | MED8 | -1.61785831 | 0.001260177 | -1.731718434 | 0.00046483 | -3.349576744 | 9.98400217 | 9.664384941 | 10.74328839 | 8.540455898 | 8.64548836 | 8.371491092 | 8.583413798 |
| ENSG00000118246 | FASTKD2 | -1.416906597 | 0.00174339 | -1.933215755 | 9.32E-06 | -3.350122352 | 9.800745303 | 9.079034257 | 9.981704053 | 8.237082658 | 8.284723 | 7.887011553 | 7.604674031 |
| ENSG00000135940 | COX5B | -1.736609351 | 1.46E-05 | -1.615316898 | 5.52E-05 | -3.351926248 | 11.0110152 | 10.16572116 | 10.80588088 | 8.996237745 | 8.94941893 | 9.05798598 | 9.12795919 |
| ENSG00000135976 | ANKRD36 | 0.588414133 | 0.447517543 | -3.941525298 | 8.89E-10 | -3.353111166 | 7.435667355 | 8.029695761 | 6.43197674 | 8.004017475 | 8.05374131 | 3.709392871 | 4.105828626 |
| ENSG00000100031 | GGT1 | -3.998070797 | 0.000344331 | 0.644699248 | 0.624398897 | -3.353371548 | 5.095959458 | 7.761270542 | 8.845626171 | 4.139692846 | 4.2865223 | 8.524290298 | 8.522888769 |
| ENSG00000141295 | SCRN2 | -1.55589932 | 0.18345602 | -1.801370682 | 0.111449285 | -3.357270002 | 10.37373127 | 8.820493869 | 6.958202216 | 7.807949361 | 7.73225478 | 7.418832227 | 7.633277551 |
| ENSG00000111846 | GCNT2 | -3.847749128 | 0.277191183 | 0.489848761 | 0.897052185 | -3.357900367 | 4.467311984 | 2 | 2 | 2.270786764 | 2 | 4.166733998 | 2.746601469 |
| ENSG00000168002 | POLR2G | -1.357680491 | 3.07E-05 | -2.002302217 | 1.97E-10 | -3.359982708 | 10.02639697 | 9.842262059 | 9.452720984 | 8.453061739 | 8.44034872 | 7.731011524 | 7.887748062 |
| ENSG00000166352 | C11orf74 | -2.099013205 | 0.053811959 | -1.263527514 | 0.262299105 | -3.362540719 | 8.797441463 | 6.140560021 | 6.718950348 | 5.697663076 | 5.65950892 | 6.679970309 | 6.209190898 |
| ENSG00000156508 | EEF1A1 | -0.978216924 | 0.193269017 | -2.384937671 | 0.000390307 | -3.363154595 | 17.08591914 | 15.97550326 | 15.3216779 | 15.32423008 | 15.3484 | 13.88227622 | 13.9759469 |
| ENSG00000165898 | ISCA2 | -1.762167361 | 0.284397666 | -1.601633275 | 0.325226668 | -3.363800636 | 9.810482605 | 7.099070191 | 4.523675391 | 6.902323594 | 6.55189318 | 6.952030889 | 6.830475749 |
| ENSG00000117569 | PTBP2 | -1.027728329 | 0.067297628 | -2.336303923 | 4.48E-06 | -3.364032252 | 8.009475818 | 7.431218269 | 7.029880454 | 6.509830055 | 6.5890786 | 5.519815274 | 5.108633494 |
| ENSG00000141994 | DUS3L | -1.970023159 | 0.012565918 | -1.394425228 | 0.083889202 | -3.364448387 | 7.624139768 | 6.311476754 | 8.076950877 | 5.651002631 | 5.58783529 | 5.725974948 | 6.487779256 |
| ENSG00000276618 | | 0.528139611 | 0.536685094 | -3.893143644 | 3.28E-08 | -3.365004032 | 6.366709671 | 7.099070191 | 8.143093325 | 8.010962573 | 7.77859536 | 4.065262523 | 3.717076715 |
| ENSG00000176108 | CHMP6 | -1.403727195 | 0.026683521 | -1.962897564 | 0.001164068 | -3.36662476 | 8.138771685 | 8.876041558 | 9.565556058 | 7.593372273 | 7.5835711 | 6.78654324 | 7.261744957 |
| ENSG00000132323 | ILKAP | -1.292117676 | 0.058026394 | -2.075725193 | 0.001141619 | -3.367842868 | 9.159305284 | 7.572420273 | 8.408111533 | 7.238438152 | 7.2537885 | 6.372988335 | 6.597881141 |
| ENSG00000131069 | ACSS2 | -2.205732901 | 0.002297097 | -1.162942825 | 0.130070296 | -3.368675727 | 9.27634997 | 9.949724236 | 10.91875185 | 7.97352961 | 8.06037647 | 8.502889002 | 9.444142447 |
| ENSG00000124194 | GDAP1L1 | -3.268197946 | NA | -0.101351706 | NA | -3.369549652 | 3.327711234 | 2 | 2 | 2.141737246 | 2 | 2.775316226 | 2.277179783 |
| ENSG00000130638 | ATXN10 | -1.116534054 | 0.025368878 | -2.254640707 | 1.04E-06 | -3.371174761 | 12.0533275 | 11.15761406 | 11.67329897 | 10.57910912 | 10.5377163 | 9.073312544 | 9.707726334 |
| ENSG00000138161 | CUZD1 | -4.714874885 | 0.316503916 | 1.343344311 | 0.777296986 | -3.371530574 | 2 | 2 | 3.369317972 | 2 | 2 | 3.483289517 | 2.782576693 |
| ENSG00000170175 | CHRNB1 | -2.347219242 | 0.011059186 | -1.025394169 | 0.301108761 | -3.372613411 | 8.855032352 | 7.847167909 | 6.203069917 | 5.686138681 | 5.7929843 | 6.987904673 | 7.004308743 |
| ENSG00000165688 | PMPCA | -1.860888377 | 0.00022855 | -1.512424816 | 0.003213124 | -3.373313193 | 10.905938 | 10.20962523 | 9.757248847 | 8.550024073 | 8.48870446 | 8.738579005 | 8.980880783 |
| ENSG00000008018 | PSMB1 | -1.386223632 | 0.008722986 | -1.988515687 | 8.31E-05 | -3.374739319 | 12.12140479 | 11.38044608 | 10.92816184 | 10.17862562 | 10.1787417 | 9.514829317 | 9.640397786 |
| ENSG00000187187 | ZNF546 | -1.26332498 | 0.145734001 | -2.111429528 | 0.008953574 | -3.374754508 | 7.790810999 | 5.838937808 | 7.342949392 | 5.870535235 | 6.08095082 | 5.427419529 | 4.935697922 |
| ENSG00000114770 | ABCC5 | -1.703988231 | 0.046022764 | -1.671367178 | 0.048008107 | -3.375355409 | 10.56993315 | 9.067150531 | 8.486635645 | 7.968781453 | 7.95986274 | 8.179816707 | 7.786191022 |
| ENSG00000102309 | PIN4 | -1.509577716 | 0.074080876 | -1.867600188 | 0.022061204 | -3.377177904 | 9.344956677 | 7.274681896 | 8.110401118 | 7.093295078 | 6.9210099 | 6.546183077 | 6.77351449 |
| ENSG00000186952 | TMEM232 | -1.136270537 | 0.672530091 | -2.240949443 | 0.364984948 | -3.37721998 | 2 | 4.442710129 | 2 | 3.023133209 | 2.49394783 | 2.457042231 | 2.374681325 |
| ENSG00000092820 | EZR | -1.763154728 | 7.61E-10 | -1.614209238 | 1.78E-08 | -3.377363966 | 11.59363617 | 11.29641566 | 11.80426818 | 9.849184459 | 9.78969165 | 9.904961437 | 10.02856616 |
| ENSG00000106976 | DNM1 | -1.119002369 | 0.059087638 | -2.260478879 | 3.17E-05 | -3.379481248 | 9.048471759 | 8.848535039 | 9.926432745 | 8.192977693 | 8.29037844 | 7.381793866 | 6.808832901 |
| ENSG00000100319 | ZMAT5 | -0.862376441 | 0.206736001 | -2.517625799 | 2.84E-05 | -3.38000224 | 7.579279349 | 6.535001748 | 7.934808822 | 6.70517998 | 6.51372389 | 4.996409624 | 5.1574003 |
| ENSG00000184752 | NDUFA12 | -1.7497014 | 2.54E-06 | -1.631001354 | 1.13E-05 | -3.380702754 | 10.22876205 | 9.523645035 | 10.11951029 | 8.207157886 | 8.29037844 | 8.287622092 | 8.441544531 |
| ENSG00000174007 | CEP19 | -1.987976371 | 0.121367017 | -1.394544244 | 0.284420805 | -3.382520615 | 8.586954094 | 5.72254323 | 5.772716771 | 5.334065616 | 5.58783529 | 6.292954465 | 5.670730563 |
| ENSG00000156374 | PCGF6 | -1.223537341 | 0.299078862 | -2.160129433 | 0.04761818 | -3.383666774 | 7.867441304 | 5.595927442 | 8.884815469 | 6.856790872 | 6.68951263 | 5.812021848 | 5.957088417 |
| ENSG00000103035 | PSMD7 | -0.30868507 | 0.593059711 | -3.075145486 | 1.98E-11 | -3.383830556 | 10.56993315 | 9.975384048 | 9.528916236 | 9.759931695 | 9.80096123 | 6.872493235 | 7.211494604 |
| ENSG00000197275 | RAD54B | 0.165785468 | 0.8692818 | -3.551278188 | 3.27E-06 | -3.385492719 | 7.384433967 | 5.72254323 | 5.930865648 | 6.676346669 | 6.71776877 | 3.465698119 | 3.600986722 |
| ENSG00000116670 | MAD2L2 | -1.987838337 | 0.000301908 | -1.398950146 | 0.01372949 | -3.386788483 | 8.981092835 | 8.702538795 | 9.907528187 | 7.210554257 | 7.43830714 | 7.900168075 | 7.911296863 |
| ENSG00000125931 | CITED1 | 0.189234629 | NA | -3.576521674 | NA | -3.387287045 | 2 | 3.076665209 | 2 | 2.784612485 | 2.49394783 | 2 | 2.117335275 |
| ENSG00000254726 | MEX3A | -1.359962849 | 0.017662579 | -2.028932363 | 0.000185282 | -3.388895212 | 7.53297911 | 7.901717006 | 8.380957451 | 6.628980152 | 6.67805337 | 6.351597515 | 5.581648407 |
| ENSG00000133119 | RFC3 | -1.824072844 | 0.26060449 | -1.565543326 | 0.33100194 | -3.38961617 | 6.816637977 | 6.535001748 | 2 | 4.329433326 | 4.65713986 | 4.698044417 | 4.739172856 |
| ENSG00000171320 | ESCO2 | -1.415944649 | 0.632952844 | -1.974019541 | 0.483986068 | -3.389964191 | 2 | 5.457120953 | 2 | 3.162721474 | 3.2184422 | 2.183373728 | 3.383972333 |
| ENSG00000089225 | TBX5 | 1.971842902 | 0.454426917 | -5.364103587 | 0.024326118 | -3.392260686 | 7.096806373 | 2 | 2 | 7.433701013 | 7.5014335 | 2.492221014 | 2.172608095 |
| ENSG00000130544 | ZNF557 | -1.286843367 | 0.034522208 | -2.10683189 | 0.000220962 | -3.393675257 | 8.138771685 | 7.847167909 | 9.116480286 | 7.246306984 | 7.16216989 | 6.464399218 | 6.362645675 |
| ENSG00000143476 | DTL | -1.053401382 | 0.425471375 | -2.340552915 | 0.050033331 | -3.393954297 | 5.866944937 | 6.311476754 | 3.369317972 | 4.783303513 | 4.63360314 | 3.296906311 | 4.033026654 |
| ENSG00000239697 | TNFSF12 | -7.588163977 | 0.012519549 | 4.193389218 | 0.112167652 | -3.394774758 | 2 | 2 | 5.59507254 | 2 | 2 | 8.200570331 | 7.940451241 |
| ENSG00000101138 | CSTF1 | -1.877395837 | 0.097789727 | -1.51782923 | 0.181655674 | -3.395225067 | 10.85038273 | 8.655694736 | 7.818184403 | 7.831679003 | 7.82609171 | 8.190565529 | 8.175075046 |
| ENSG00000260238 | PMF1-BGLAF | 4.302478968 | 0.00486902 | -7.699596909 | 0.000411973 | -3.397117941 | 3.327711234 | 2 | 4.058998049 | 6.882264028 | 6.89154576 | 2 | 2 |
| ENSG00000239497 | | 3.46729034 | 0.201926325 | -6.865008823 | 0.02391519 | -3.397718483 | 2 | 3.686303414 | 2 | 5.406080789 | 5.22805098 | 2 | 2 |
| ENSG00000232119 | MCTS1 | -1.103235195 | 5.29E-06 | -2.296845116 | 3.47E-23 | -3.400080311 | 9.112843761 | 9.030900487 | 9.099922433 | 7.992367495 | 7.98565528 | 6.798888929 | 6.849680541 |
| ENSG00000184986 | TMEM121 | -2.417528864 | 0.254666268 | -0.985719619 | 0.664743846 | -3.403248483 | 2 | 5.457120953 | 8.978369997 | 5.058234798 | 5.39133044 | 6.728720192 | 6.38329369 |
| ENSG00000167131 | CCDC103 | 3.459773362 | 0.204544536 | -6.864493493 | 0.024210868 | -3.404720131 | 2 | 3.686303414 | 2 | 5.433909581 | 5.18011485 | 2 | 2 |
| ENSG00000213694 | S1PR3 | -1.341429455 | 0.035353509 | -2.063645981 | 0.000605536 | -3.405075436 | 10.34721797 | 11.70001862 | 10.55568472 | 9.673571522 | 9.65077635 | 9.070405724 | 8.806752407 |
| ENSG00000079150 | FKBP7 | -1.576649607 | 4.43E-06 | -1.828928819 | 5.90E-08 | -3.405578425 | 10.3868074 | 9.726146444 | 10.08627303 | 8.48998335 | 8.55441692 | 8.226915494 | 8.318287967 |
| ENSG00000103148 | NPRL3 | -2.012679758 | 0.016182913 | -1.393911544 | 0.105991636 | -3.406591302 | 10.18451821 | 7.979858146 | 10.08627303 | 7.653784481 | 7.76511961 | 8.27442094 | 8.36352657 |
| ENSG00000140043 | PTGR2 | -0.508132946 | 0.632193989 | -2.898503948 | 0.001039403 | -3.406636893 | 5.095959458 | 6.228547981 | 7.451345288 | 6.061031724 | 6.08962627 | 4.029788235 | 3.987514168 |
| ENSG00000197479 | PCDHB11 | -1.720783467 | 0.323876103 | -1.690868758 | 0.321893604 | -3.411652225 | 3.327711234 | 4.11352228 | 6.958202216 | 4.470469939 | 3.7864959 | 3.465698119 | 4.673629022 |
| ENSG00000130299 | GTPBP3 | -2.57362459 | 0.011022574 | -0.838443753 | 0.455138882 | -3.412068343 | 6.89194353 | 7.73146271 | 9.61300511 | 6.069927018 | 6.00043082 | 7.903438544 | 7.481493974 |
| ENSG00000171700 | RGS19 | -1.86459538 | 0.393969236 | -1.550181566 | 0.477543645 | -3.414776946 | 10.27168932 | 7.669933291 | 2 | 7.161462517 | 6.98750599 | 7.566427714 | 7.173047408 |
| ENSG00000189050 | RNFT1 | -0.708793704 | 0.161952785 | -2.706210751 | 7.95E-10 | -3.415004455 | 7.485143553 | 7.274681896 | 8.076950877 | 6.872128674 | 7.02868983 | 5.158477451 | 5.013446246 |
| ENSG00000140987 | ZSCAN32 | 0.355659839 | 0.857804601 | -3.771434895 | 0.017740157 | -3.415775056 | 2 | 7.73146271 | 7.451345288 | 7.461007055 | 7.28811304 | 3.709392871 | 3.753786374 |
| ENSG00000083817 | ZNF416 | -2.122778502 | 0.342855793 | -1.293374475 | 0.573911706 | -3.416152977 | 9.473095744 | 5.457120953 | 2 | 6.034011955 | 5.8245091 | 6.726875343 | 6.714211481 |
| ENSG00000111231 | GPN3 | -1.5584113 | 0.033644986 | -1.858367505 | 0.008995102 | -3.416778805 | 8.493334308 | 7.145017454 | 8.960138144 | 6.872128674 | 6.82556437 | 6.482006365 | 6.635252003 |
| ENSG00000214114 | MYCBP | -1.718682334 | 0.027069469 | -1.698854443 | 0.027684956 | -3.417536777 | 9.460782017 | 7.467835385 | 8.805342542 | 7.044888801 | 7.14977896 | 7.286064617 | 6.926052784 |
| ENSG00000153283 | CD96 | -0.512076852 | 0.684436474 | -2.905864967 | 0.006426945 | -3.417941819 | 3.327711234 | 4.93638933 | 4.874587708 | 4.139692846 | 4.16043394 | 2.592885947 | 2.851942824 |
| ENSG00000164758 | MED30 | -1.35382089 | 0.316685768 | -2.064226672 | 0.104988652 | -3.418047562 | 8.340423424 | 5.457120953 | 5.39244166 | 5.808110275 | 5.7929843 | 5.238348159 | 5.065479856 |
| ENSG00000119865 | CNRIP1 | -1.213219989 | 0.020787524 | -2.206039432 | 6.54E-06 | -3.419259421 | 9.567995545 | 8.607278467 | 9.516493218 | 8.145379268 | 8.02689124 | 7.319501608 | 6.876977108 |
| ENSG00000196209 | SIRPB2 | -3.2549399 | 0.223929761 | -0.164944228 | 0.955098488 | -3.419884127 | 4.467311984 | 2 | 2 | 2.141737246 | 2.2680111 | 3.355392153 | 3.042026731 |
| ENSG00000116353 | MECR | -2.260600449 | 2.59E-08 | -1.163360375 | 0.006050817 | -3.423960824 | 8.928409675 | 8.762721501 | 8.143093325 | 6.321555392 | 6.55815769 | 7.483632518 | 7.528596482 |
| ENSG00000112640 | PPP2R5D | -1.30993595 | 0.002199247 | -2.114237164 | 2.07E-07 | -3.424173113 | 10.47516499 | 10.81444733 | 9.897982195 | 9.123450746 | 9.16073939 | 8.350677018 | 8.340330468 |
| ENSG00000166037 | CEP57 | -1.700066037 | 0.000102258 | -1.725732502 | 6.95E-05 | -3.425798539 | 9.992581382 | 9.988044813 | 9.61300511 | 8.160032699 | 8.21902896 | 8.512558892 | 7.704159082 |
| ENSG00000241685 | ARPC1A | -1.137514228 | 0.071563439 | -2.288614783 | 7.56E-05 | -3.426129011 | 11.08508709 | 9.672251278 | 10.59756881 | 9.405814997 | 9.4506582 | 8.041943102 | 8.497408326 |
| ENSG00000198736 | MSRB1 | -2.697469927 | 6.62E-06 | -0.730038392 | 0.28128563 | -3.42750832 | 10.22876205 | 8.88960062 | 9.465702942 | 7.044888801 | 6.89649843 | 8.52588267 | 9.20537272 |
| ENSG00000026103 | FAS | -7.50185366 | 0.013123487 | 4.074321781 | 0.12071938 | -3.427531878 | 2 | 5.457120953 | 2 | 2 | 2 | 8.431623835 | 6.953927794 |
| ENSG00000141971 | MVB12A | -1.916201373 | 2.91E-09 | -1.513042822 | 3.20E-06 | -3.429244195 | 9.667789716 | 9.566483142 | 10.10298738 | 7.807949361 | 7.9809999 | 8.282607314 | 8.305980863 |
| ENSG00000068078 | FGFR3 | -4.118918135 | 0.002790444 | 0.688108591 | 0.669750001 | -3.430809543 | 4.00722531 | 8.395519306 | 6.882775709 | 4.034562184 | 3.2184422 | 7.343779505 | 8.413138577 |
| ENSG00000108423 | TUBD1 | -1.424955426 | 0.377098982 | -2.00662184 | 0.189170169 | -3.431577266 | 4.00722531 | 7.099070191 | 8.922968322 | 6.503277904 | 6.18176397 | 6.265258498 | 5.122735884 |
| ENSG00000135972 | MRPS9 | -1.680578336 | 3.47E-05 | -1.753245255 | 1.12E-05 | -3.433823591 | 8.609439662 | 8.702538795 | 7.934808822 | 6.825617242 | 6.80463998 | 6.741568503 | 6.748721609 |
| ENSG00000125864 | BFSP1 | -2.782011048 | 0.114937899 | -0.651984954 | 0.738292369 | -3.433996002 | 7.159102577 | 4.93638933 | 2 | 3.462030052 | 3.69849104 | 5.404520029 | 5.1574003 |
| ENSG00000114735 | HEMK1 | -2.05696178 | 0.011629766 | -1.377653262 | 0.101809263 | -3.434615042 | 8.075571721 | 8.414402626 | 9.935792933 | 6.931899482 | 7.11195456 | 7.959503304 | 7.351304958 |
| ENSG00000146802 | TMEM168 | -2.724427729 | 0.038918148 | -0.710952009 | 0.632473342 | -3.435379738 | 5.866944937 | 8.671479029 | 10.38167985 | 6.567497652 | 6.54560135 | 8.524821284 | 8.521558854 |
| ENSG00000104848 | KCNA7 | -3.303461926 | 0.138710156 | -0.132117258 | 0.958081539 | -3.435579184 | 3.327711234 | 2 | 4.874587708 | 2.141737246 | 2.49394783 | 4.199031499 | 3.012034821 |
| ENSG00000077454 | LRCH4 | -1.002720144 | 0.151162081 | -2.43305922 | 0.000100317 | -3.435779364 | 9.646202137 | 8.317412869 | 9.798878489 | 8.361842177 | 8.41985547 | 6.732402829 | 7.208191747 |
| ENSG00000160973 | FOXH1 | -1.065674404 | 0.609297559 | -2.374996797 | 0.210029949 | -3.440671201 | 3.327711234 | 4.442710129 | 3.369317972 | 3.09461492 | 3.2184422 | 2.094598369 | 2.950113106 |
| ENSG00000178741 | COX5A | -1.425748602 | 0.012388648 | -2.015558622 | 0.000217912 | -3.441307224 | 10.85038273 | 9.506147172 | 10.31150679 | 8.915879855 | 8.89106117 | 8.228220207 | 8.405948829 |
| ENSG00000136111 | TBC1D4 | -1.243695041 | 0.53875552 | -2.200340519 | 0.244585846 | -3.44403556 | 2 | 8.732943944 | 8.904018013 | 7.003290579 | 7.04662306 | 6.462183138 | 5.641643428 |
| ENSG00000184990 | SIVA1 | -1.854457368 | 0.000192559 | -1.589986836 | 0.001505204 | -3.444444203 | 9.790941832 | 8.88960062 | 8.805342542 | 7.45762193 | 7.34718809 | 7.588946094 | 7.73202455 |
| ENSG00000121073 | SLC35B1 | -1.130091452 | 0.352172376 | -2.315406009 | 0.037018249 | -3.445497461 | 11.10500521 | 9.648522413 | 7.646393253 | 8.919575326 | 8.95536429 | 7.470462265 | 8.013226939 |
| ENSG00000177971 | IMP3 | -1.238621773 | 0.104398641 | -2.210240979 | 0.001732243 | -3.448862751 | 10.36053552 | 9.318399851 | 8.742726175 | 8.379838154 | 8.42671895 | 6.990982383 | 7.795024296 |
| ENSG00000148950 | IMMP1L | -2.929895877 | 0.025122781 | -0.519643155 | 0.727097723 | -3.449539032 | 6.96351253 | 3.686303414 | 5.156626401 | 3.566290607 | 3.33976443 | 5.582367412 | 5.101530266 |
| ENSG00000105523 | FAM83E | -0.682514016 | NA | -2.77322601 | NA | -3.455740026 | 3.327711234 | 2 | 2 | 2.600425781 | 2.14022021 | 2.139668783 | 2.059860178 |
| ENSG00000112936 | C7 | -4.468944188 | 0.058148984 | 1.012406089 | 0.688459235 | -3.456538099 | 2 | 6.140560021 | 2 | 2.270786764 | 2.38539787 | 5.744329515 | 5.524371359 |
| ENSG00000117266 | CDK18 | -3.940775204 | 9.00E-05 | 0.484186722 | 0.674724562 | -3.456588482 | 4.00722531 | 6.602424674 | 6.43197674 | 2.947924294 | 3.08597292 | 6.631488899 | 6.412287469 |
| ENSG00000168701 | TMEM208 | -1.271461383 | 0.023102077 | -2.186548244 | 3.02E-05 | -3.458009627 | 10.01801709 | 8.848535039 | 9.132850258 | 8.078634414 | 8.23866319 | 7.31213785 | 7.213143202 |
| ENSG00000197183 | NOL4L | -1.708083455 | 0.003882515 | -1.751315223 | 0.002836757 | -3.459398678 | 9.790941832 | 10.48887288 | 11.24667977 | 8.923261356 | 8.9350493 | 8.911780399 | 8.860338475 |

| | | | | | | | | | | | | | |
|---|---|---|---|---|---|---|---|---|---|---|---|---|---|
| ENSG00000149823 | VPS51 | -2.503100841 | 1.27E-08 | -0.956954704 | 0.045706083 | -3.460055545 | 10.86913986 | 10.26269067 | 9.868959282 | 7.860161659 | 7.95749494 | 9.402878503 | 9.47607557 |
| ENSG00000091732 | ZC3HC1 | -1.692458134 | 0.015080556 | -1.767779418 | 0.010097543 | -3.460237552 | 8.468948322 | 8.356994601 | 9.788583153 | 7.381065997 | 7.33264456 | 7.037865927 | 7.493753702 |
| ENSG00000174963 | ZIC4 | -5.848028296 | 0.02131845 | 2.387399116 | 0.371277929 | -3.46062918 | 2 | 2 | 8.175061096 | 2.600425781 | 2.38539787 | 9.167113444 | 8.729374692 |
| ENSG00000198393 | ZNF26 | -1.832099588 | 3.11E-09 | -1.630578709 | 8.72E-08 | -3.462678298 | 8.493334308 | 8.469613413 | 8.380957451 | 6.5612028 | 6.75096147 | 6.959905775 | 6.73730971 |
| ENSG00000130348 | QRSL1 | -1.476009883 | 0.09269647 | -1.987266103 | 0.017733439 | -3.463275985 | 8.340423424 | 7.638155693 | 9.778213818 | 7.322721205 | 7.49164798 | 6.905488945 | 6.920008846 |
| ENSG00000114686 | MRPL3 | -2.008763579 | 0.000154681 | -1.4566835 | 0.00769046 | -3.46544708 | 10.90138923 | 9.74864196 | 9.926432745 | 8.325162455 | 8.25418043 | 8.858433249 | 8.813832186 |
| ENSG00000111224 | PARP11 | -1.631307484 | 0.245194354 | -1.835175436 | 0.177686131 | -3.46648292 | 5.330594134 | 7.928237109 | 9.746650996 | 6.984595707 | 6.95938211 | 6.670403488 | 6.872811106 |
| ENSG00000167173 | C15orf39 | -2.403937999 | 0.000497524 | -1.065341432 | 0.153080856 | -3.469279431 | 8.609439662 | 10.46189757 | 10.01740919 | 7.461007055 | 7.55566492 | 8.921088109 | 8.729374692 |
| ENSG00000126432 | PRDX5 | -2.104367014 | 1.08E-10 | -1.367922295 | 4.38E-05 | -3.472289308 | 12.27171276 | 11.67674577 | 11.97968146 | 9.866518681 | 9.92490661 | 10.48396204 | 10.76271041 |
| ENSG00000171806 | METTL18 | -1.558340727 | 0.283520814 | -1.914806233 | 0.168954524 | -3.47314696 | 3.327711234 | 6.844429755 | 7.777116083 | 5.391962469 | 5.36298494 | 5.038665118 | 5.080008298 |
| ENSG00000213619 | NDUFS3 | -1.803785018 | 0.003882515 | -1.669976944 | 0.007710626 | -3.473761962 | 10.63713535 | 9.34816733 | 9.358421373 | 8.174538791 | 8.07792246 | 8.193908178 | 8.321348409 |
| ENSG00000119328 | FAM206A | -1.469701851 | 0.011808534 | -2.005465282 | 0.000337252 | -3.475167133 | 8.586954094 | 8.762721501 | 9.439621147 | 7.471115032 | 7.58049694 | 7.388811377 | 6.479583725 |
| ENSG00000114107 | CEP70 | -2.036398766 | 0.113145409 | -1.439053219 | 0.267993165 | -3.475451985 | 7.435667355 | 5.72254323 | 4.058998049 | 4.647285135 | 4.3456612 | 5.248671144 | 4.729989799 |
| ENSG00000178821 | TMEM52 | -3.235623428 | 0.213844269 | -0.241178707 | 0.934216456 | -3.476802135 | 3.327711234 | 4.710564154 | 2 | 2.600425781 | 2 | 3.824280769 | 3.36036153 |
| ENSG00000168542 | COL3A1 | -1.839042531 | 0.000470613 | -1.63922295 | 0.00198533 | -3.478265481 | 16.75364453 | 17.94737345 | 17.6457676 | 15.67013347 | 15.7143961 | 15.65953104 | 16.09257205 |
| ENSG00000090097 | PCBP4 | -1.793889858 | 2.65E-05 | -1.685585717 | 7.79E-05 | -3.479475576 | 10.12333326 | 9.869887299 | 10.67226762 | 8.534041678 | 8.41813449 | 8.366759995 | 8.773618611 |
| ENSG00000115274 | INO80B | 0.630479521 | 0.765831172 | -4.112528841 | 0.020434196 | -3.48204932 | 8.042902253 | 5.303523747 | 2 | 7.182120794 | 7.38291816 | 3.429858444 | 3.100202476 |
| ENSG00000185813 | PCYT2 | -2.252014583 | 0.00492344 | -1.23314764 | 0.14497505 | -3.485162223 | 9.2903348 | 8.590771778 | 10.48092713 | 7.315259362 | 7.55566492 | 7.918064594 | 8.826273465 |
| ENSG00000135083 | CCNJL | -2.57372519 | 0.065557037 | -0.911958345 | 0.547700965 | -3.485683535 | 8.609439662 | 6.844429755 | 4.058998049 | 5.076005082 | 4.97065356 | 6.62753908 | 6.487779256 |
| ENSG00000113231 | PDE8B | -4.417789674 | 0.059087638 | 0.930571828 | 0.718590706 | -3.487217846 | 2 | 4.442710129 | 6.534109959 | 2.868578001 | 2 | 6.167993056 | 6.141542327 |
| ENSG00000177839 | PCDHB9 | -0.622813787 | 0.560547313 | -2.867135503 | 0.001736775 | -3.48994929 | 4.467311984 | 5.131603625 | 5.930865648 | 4.946755474 | 4.50979031 | 2.885263063 | 3.36036153 |
| ENSG00000069812 | HES2 | -5.003615516 | 1.29E-07 | 1.513504272 | 0.049489333 | -3.490111244 | 5.70932728 | 4.710564154 | 5.772716771 | 2.498691189 | 2.2680111 | 6.916861442 | 6.81969491 |
| ENSG00000131116 | ZNF428 | -2.000503166 | 0.014874866 | -1.490182016 | 0.075998868 | -3.490685182 | 10.29268328 | 8.053983481 | 9.503962296 | 7.602596956 | 7.53358066 | 7.872065621 | 8.243626305 |
| ENSG00000185736 | ADARB2 | -3.099915293 | 0.235251572 | -0.391499871 | 0.89090828 | -3.491415164 | 4.00722531 | 3.686303414 | 2 | 2.498691189 | 2 | 3.5515855 | 2.817676617 |
| ENSG00000167193 | CRK | 0.213058791 | 0.869626608 | -3.705207236 | 0.000170129 | -3.492148445 | 7.53297911 | 10.40638494 | 10.73794767 | 10.26878646 | 10.3186089 | 6.368262153 | 6.506723027 |
| ENSG00000073050 | XRCC1 | -0.862364579 | 0.531143362 | -2.630032084 | 0.030786735 | -3.492396662 | 6.563806279 | 9.770792087 | 10.75919294 | 8.996237745 | 8.91688363 | 7.117472117 | 7.310303798 |
| ENSG00000100360 | IFT27 | -1.993732473 | 0.001266304 | -1.498939392 | 0.018037816 | -3.492671866 | 8.540903468 | 8.317412869 | 9.658943166 | 6.927012081 | 7.07311172 | 7.607112326 | 7.348307474 |
| ENSG00000175193 | PARL | -1.022735918 | 0.222583351 | -2.470271493 | 0.001016114 | -3.493007411 | 9.33149386 | 9.328390744 | 7.451345288 | 7.910550164 | 7.93118934 | 6.262714117 | 6.728114749 |
| ENSG00000115875 | SRSF7 | -1.59404917 | 1.29E-07 | -1.900831537 | 1.51E-10 | -3.494880707 | 10.80239439 | 10.34373336 | 10.63252637 | 9.071247833 | 8.960103 | 8.57541967 | 8.838074258 |
| ENSG00000158352 | SHROOM4 | -1.816222161 | 0.065340768 | -1.680168269 | 0.086198005 | -3.49639043 | 8.418903792 | 6.602424674 | 9.372276026 | 6.722207358 | 6.75642009 | 6.99251878 | 6.73730971 |
| ENSG00000119514 | GALNT12 | 0.155928047 | 0.959439303 | -3.653666704 | 0.156449615 | -3.497738657 | 2 | 6.951736442 | 2 | 5.651002631 | 5.5755357 | 3.256555576 | 2.277179783 |
| ENSG00000183770 | FOXL2 | -2.102840854 | 0.318751901 | -1.396904785 | 0.506531437 | -3.49974564 | 2 | 4.93638933 | 3.369317972 | 2.270786764 | 3.08597292 | 3.10571689 | 3.012034821 |
| ENSG00000173163 | COMMD1 | -2.016185542 | 0.000261061 | -1.486662178 | 0.008669082 | -3.50284772 | 8.653385977 | 8.862353853 | 9.681375057 | 7.165617905 | 7.13309042 | 7.296052918 | 7.961201206 |
| ENSG00000135119 | RNFT2 | -3.863695921 | 0.120642255 | 0.359767725 | 0.899395052 | -3.503928196 | 5.866944937 | 2 | 2 | 2.3892346 | 2.38539787 | 5.119635168 | 4.26688425 |
| ENSG00000170836 | PPM1D | -2.060402138 | 0.010529822 | -1.444456981 | 0.081300083 | -3.50485912 | 8.892187472 | 7.638155693 | 9.565556058 | 6.993973425 | 6.73998187 | 7.855275386 | 6.946018311 |
| ENSG00000104147 | OIP5 | -2.47867519 | 0.219945377 | -1.029188944 | 0.611176359 | -3.507864134 | 2 | 4.11352228 | 3.369317972 | 2.270786764 | 2.49394783 | 3.059555873 | 2.671842496 |
| ENSG00000188051 | TMEM221 | -4.236390062 | 0.181775686 | 0.723535319 | 0.827324651 | -3.512854743 | 2 | 4.710564154 | 2 | 2 | 2.2680111 | 4.378961816 | 3.660199087 |
| ENSG00000151465 | CDC123 | -1.350463592 | 0.013140909 | -2.163343968 | 2.62E-05 | -3.51380756 | 10.24321353 | 9.591588447 | 10.8211137 | 9.027396512 | 8.88608977 | 8.031518488 | 8.267623722 |
| ENSG00000196510 | ANAPC7 | -1.943938208 | 0.002029323 | -1.576112334 | 0.014055845 | -3.520050543 | 9.509418609 | 9.566483142 | 10.85604368 | 8.188900477 | 8.19108423 | 8.501271034 | 8.603665995 |
| ENSG00000166321 | NUDT13 | -5.561781257 | 0.053222658 | 2.041649252 | 0.409820322 | -3.520132005 | 3.327711234 | 3.076665209 | 2 | 2 | 2 | 3.993419637 | 4.4231929 |
| ENSG00000181873 | IBA57 | -2.329720303 | 0.001871365 | -1.191009952 | 0.133346754 | -3.520730255 | 7.82963488 | 7.315436877 | 9.099922433 | 6.024891728 | 6.01871653 | 6.907119087 | 7.290139032 |
| ENSG00000148444 | COMMD3 | -1.258682611 | 0.041439227 | -2.266565931 | 7.62E-05 | -3.525248541 | 9.219014269 | 8.004992147 | 9.330306409 | 7.641902724 | 7.78395061 | 6.631488899 | 6.830475749 |
| ENSG00000171703 | TCEA2 | -2.489921442 | 7.80E-14 | -1.035827321 | 0.003055989 | -3.525748763 | 9.015175615 | 9.523645035 | 9.516493218 | 7.030886362 | 6.80463998 | 8.396658292 | 8.292788497 |
| ENSG00000164330 | EBF1 | -3.552344476 | 0.035568009 | 0.026404939 | 0.989740833 | -3.525939537 | 4.00722531 | 5.946638857 | 9.636156974 | 4.867343068 | 4.79082474 | 8.335623797 | 8.070734444 |
| ENSG00000163093 | BBS5 | -1.904669714 | 0.022559831 | -1.624845548 | 0.052961956 | -3.529515262 | 8.075571721 | 6.951736442 | 9.132850258 | 6.542151749 | 6.35727082 | 6.788313392 | 6.649933445 |
| ENSG00000166484 | MAPK7 | -1.985393183 | 0.014133546 | -1.54596813 | 0.060804392 | -3.531361313 | 10.06758145 | 9.415325457 | 11.46927014 | 8.606137616 | 8.6021877 | 8.856747125 | 9.202055802 |
| ENSG00000012061 | ERCC1 | -1.718976823 | 0.000636626 | -1.819734831 | 0.000254518 | -3.538711654 | 10.59268348 | 10.45279253 | 11.40332671 | 9.129845271 | 9.20332986 | 8.789844052 | 9.299553193 |
| ENSG00000205352 | PRR13 | -1.120577092 | 0.004077242 | -2.419188168 | 3.03E-11 | -3.53976526 | 11.1478805 | 10.53705456 | 10.81605395 | 9.726568065 | 9.74782835 | 8.194575779 | 8.664579282 |
| ENSG00000172530 | BANP | -1.423067436 | 0.065440948 | -2.119550532 | 0.00367982 | -3.542617969 | 8.340423424 | 7.638155693 | 9.083172337 | 7.030886362 | 7.11195456 | 6.850634117 | 5.745545708 |
| ENSG00000071073 | MGAT4A | -3.936130652 | 0.025901628 | 0.391523554 | 0.839662804 | -3.544607098 | 4.00722531 | 3.076665209 | 5.156626401 | 2.141737246 | 2.49394783 | 5.253805068 | 3.45257121 |
| ENSG00000168421 | RHOH | -0.689891946 | NA | -2.85482501 | NA | -3.544716956 | 2 | 2 | 3.369317972 | 2.270786764 | 2.49394783 | 2.183373728 | 2 |
| ENSG00000079999 | KEAP1 | -1.735876694 | 0.000654354 | -1.808845576 | 0.000335973 | -3.544722269 | 11.78604782 | 10.63690565 | 11.06240811 | 9.522191343 | 9.49786081 | 9.55463569 | 9.309990451 |
| ENSG00000184349 | EFNA5 | -1.827711024 | 0.017388264 | -1.717578649 | 0.024438946 | -3.545289673 | 8.009475818 | 7.002537973 | 6.629488387 | 5.501219927 | 5.6826301 | 5.163941955 | 6.085042537 |
| ENSG00000184508 | HDDC3 | -2.031994048 | 0.300898696 | -1.517319492 | 0.442768139 | -3.54931354 | 8.493334308 | 6.04685525 | 2 | 5.210797711 | 5.28958453 | 5.783905948 | 5.651404445 |
| ENSG00000088356 | PDRG1 | -1.618651544 | 0.444705393 | -1.93200564 | 0.343240742 | -3.550657184 | 2 | 9.214490408 | 10.11951029 | 7.549526251 | 7.56502709 | 7.396955563 | 7.089349481 |
| ENSG00000009830 | POMT2 | -1.576620121 | 0.079474352 | -1.976283048 | 0.022630598 | -3.552903168 | 10.71696191 | 8.297207175 | 9.714382178 | 8.399377854 | 8.23475766 | 8.084345843 | 7.746896594 |
| ENSG00000010626 | LRRC23 | -2.406293704 | 0.213291302 | -1.14802207 | 0.572525943 | -3.554315775 | 8.75772933 | 6.464272742 | 2 | 5.258265913 | 5.09650239 | 6.410250549 | 6.270645889 |
| ENSG00000117697 | NSL1 | -1.623972346 | 0.004014422 | -1.931005171 | 0.000443279 | -3.554977517 | 9.567995545 | 8.255926468 | 9.083172337 | 7.381065997 | 7.53358066 | 7.081831041 | 7.234405279 |
| ENSG00000164022 | AIMP1 | -2.204028677 | 1.78E-07 | -1.353027116 | 0.002032168 | -3.557055793 | 10.86913986 | 10.35355093 | 10.11127249 | 8.358215887 | 8.21902896 | 8.859696551 | 9.36293094 |
| ENSG00000130332 | LSM7 | -2.026120884 | 1.06E-08 | -1.531479688 | 1.77E-05 | -3.557600573 | 9.544849385 | 9.00621767 | 8.941672936 | 7.214570819 | 7.17037177 | 7.731931603 | 7.617180085 |
| ENSG00000179965 | ZNF771 | -2.166040682 | 0.272317502 | -1.395287289 | 0.489011537 | -3.561327971 | 9.174465797 | 7.393647515 | 2 | 5.808110275 | 5.94414054 | 6.184218099 | 6.934072182 |
| ENSG00000225921 | NOL7 | -1.32099438 | 0.188330316 | -2.242890692 | 0.016663213 | -3.563885072 | 11.14017917 | 8.687092496 | 9.31604076 | 8.856713087 | 8.72420319 | 7.720851738 | 8.026414333 |
| ENSG00000172893 | DHCR7 | -2.269616933 | 0.000618475 | -1.296104 | 0.064397771 | -3.565720933 | 12.43483465 | 11.08875118 | 11.45632039 | 9.506715015 | 9.5188894 | 9.895754036 | 10.89838855 |
| ENSG00000066583 | ISOC1 | -1.165115742 | 0.007365759 | -2.401628783 | 2.89E-09 | -3.566744525 | 9.174465797 | 8.395519306 | 8.805342542 | 7.668500375 | 7.67575955 | 6.623578417 | 6.30506411 |
| ENSG00000120068 | HOXB8 | 0.198584326 | 0.959782419 | -3.766014049 | 0.278767895 | -3.567429723 | 2 | 4.11352228 | 2 | 3.566290607 | 2.77779907 | 2.225793514 | 2 |
| ENSG00000099957 | P2RX6 | -1.205102171 | 0.478725082 | -2.363274878 | 0.128944742 | -3.568377049 | 2 | 5.946638857 | 6.534109959 | 4.867343068 | 4.40247105 | 3.517841605 | 3.956354764 |
| ENSG00000179846 | NKPD1 | -3.288868317 | 0.144551494 | -0.279795812 | 0.909865545 | -3.568664129 | 3.327711234 | 2 | 4.523675391 | 2.141737246 | 2.38539787 | 3.838019203 | 2.782576693 |
| ENSG00000154814 | OXNAD1 | -1.96648695 | 0.003507653 | -1.602285032 | 0.018502076 | -3.568771981 | 7.904282243 | 6.389896633 | 7.734844356 | 5.674521485 | 5.51241389 | 5.880003936 | 5.984375752 |
| ENSG00000163607 | GTPBP8 | -1.944303288 | 0.011645431 | -1.626133316 | 0.036276935 | -3.570436604 | 7.709880939 | 6.78762695 | 8.65477198 | 6.18083559 | 5.85535975 | 6.151583464 | 6.485052582 |
| ENSG00000099840 | IZUMO4 | -2.663207901 | 0.306087183 | -0.907459002 | 0.742986506 | -3.570666903 | 6.366709671 | 2 | 2 | 2.947924294 | 3.08597292 | 3.917810786 | 4.365060851 |
| ENSG00000186364 | NUDT17 | -2.245903227 | 0.204923899 | -1.325733839 | 0.465968326 | -3.571637067 | 2 | 4.93638933 | 6.958202216 | 3.754611435 | 3.90918929 | 4.470006665 | 4.624919101 |
| ENSG00000169607 | CKAP2L | -1.166087187 | 0.440854209 | -2.409638558 | 0.080217537 | -3.575725746 | 3.327711234 | 6.844429755 | 7.502634626 | 5.48800675 | 5.65950892 | 3.956110469 | 4.878668439 |
| ENSG00000184908 | CLCNKB | -2.622284719 | 0.332592522 | -0.954247419 | 0.730981666 | -3.576532138 | 2 | 4.442710129 | 2 | 2.141737246 | 2.49394783 | 2.831336763 | 2.885414003 |
| ENSG00000148225 | WDR31 | -1.373364343 | 0.499737617 | -2.205670585 | 0.24759592 | -3.579034928 | 8.009475818 | 5.303523747 | 2 | 5.242616075 | 5.47317115 | 4.785723309 | 4.468068808 |
| ENSG00000212123 | PRR22 | -0.857323989 | 0.606318567 | -2.721738442 | 0.063742291 | -3.57906243 | 5.330594134 | 4.93638933 | 2 | 3.840289928 | 4.0930011 | 2.592885947 | 3.071407821 |
| ENSG00000168393 | DTYMK | -2.41094093 | 7.18E-05 | -1.169223096 | 0.07118462 | -3.580164026 | 9.371511179 | 7.847167909 | 8.865353883 | 6.379946408 | 6.53928196 | 7.643750713 | 7.68646583 |
| ENSG00000274233 | | 5.932951618 | 1.39E-11 | -9.514223057 | 4.86E-08 | -3.581271438 | 5.532353208 | 4.442710129 | 6.80318718 | 11.68446157 | 11.7403773 | 2.048074405 | 2 |
| ENSG00000101911 | PRPS2 | -1.843158571 | 0.00066186 | -1.73979524 | 0.001265798 | -3.58295381 | 8.228665467 | 8.955547846 | 9.503962296 | 7.182120794 | 7.16216989 | 7.374742053 | 7.162846889 |
| ENSG00000099795 | NDUFB7 | -1.864024363 | 2.38E-05 | -1.719818498 | 9.78E-05 | -3.583842861 | 10.40620204 | 9.799806309 | 10.74860941 | 8.512996295 | 8.51631523 | 8.505043475 | 8.795241939 |
| ENSG00000170515 | PA2G4 | -1.984151225 | 6.57E-12 | -1.601455801 | 3.42E-08 | -3.585607026 | 10.59831544 | 10.88383745 | 10.94679967 | 8.875861021 | 8.80938679 | 9.036921734 | 9.387595483 |
| ENSG00000131650 | KREMEN2 | -4.135256377 | 0.152528849 | 0.549381073 | 0.835379021 | -3.585875304 | 2 | 3.076665209 | 3.369317972 | 2.141737246 | 2 | 3.296906311 | 3.128433649 |
| ENSG00000139352 | ASCL1 | 1.604282604 | NA | -5.191100087 | NA | -3.586817483 | 2 | 2 | 2 | 3.349643765 | 2 | 2 | 2 |
| ENSG00000164651 | SP8 | 1.604282604 | NA | -5.191100087 | NA | -3.586817483 | 2 | 2 | 2 | 3.349643765 | 2 | 2 | 2 |
| ENSG00000205085 | FAM71F2 | 0.244943963 | 0.733731931 | -3.831861407 | 3.45E-11 | -3.586917444 | 6.009025997 | 5.946638857 | 6.882775709 | 6.634986713 | 6.48770488 | 3.193831101 | 3.261853426 |
| ENSG00000164867 | NOS3 | -1.775766957 | 0.514867285 | -1.812264705 | 0.487850169 | -3.588031662 | 2 | 4.442710129 | 2 | 2.270786764 | 2.77779907 | 2.560105256 | 2.509591799 |
| ENSG00000182195 | LDOC1 | -2.204884541 | 0.008885084 | -1.384408492 | 0.114946876 | -3.589293033 | 11.22998797 | 9.114109157 | 10.23015729 | 8.207157886 | 8.28283292 | 8.464101787 | 9.47710506 |
| ENSG00000163528 | CHCHD4 | -1.790898925 | 1.25E-07 | -1.798784779 | 6.41E-08 | -3.589683754 | 8.716893051 | 8.234834549 | 8.35328246 | 6.722207358 | 6.66650236 | 6.615624322 | 6.755525668 |
| ENSG00000122986 | HVCN1 | -1.535146258 | 0.464599186 | -2.055849578 | 0.301670224 | -3.590995835 | 8.631580145 | 5.72254323 | 2 | 5.775854763 | 5.77157844 | 5.431956133 | 5.150533729 |
| ENSG00000172273 | HINFP | -1.726454085 | 0.032879246 | -1.866353287 | 0.018961223 | -3.592807372 | 9.689059027 | 7.638155693 | 9.227351203 | 7.409376773 | 7.33993465 | 7.485815918 | 6.940057611 |
| ENSG00000151164 | RAD9B | -2.096425631 | 0.080529035 | -1.497352903 | 0.195903487 | -3.593778534 | 4.815609471 | 3.686303414 | 4.058998049 | 2.868578001 | 2.94009902 | 3.193831101 | 3.236138336 |
| ENSG00000100982 | PCIF1 | -2.426363993 | 0.000144714 | -1.174741061 | 0.085873925 | -3.601105055 | 8.540903468 | 8.981105197 | 10.09465441 | 6.931899482 | 7.00595521 | 8.181164698 | 8.206864727 |
| ENSG00000159063 | ALG8 | -1.198407589 | 0.013588226 | -2.402727677 | 9.90E-08 | -3.601135266 | 10.16946352 | 9.148351514 | 9.819251342 | 8.589222193 | 8.56844956 | 7.268416942 | 7.509939612 |
| ENSG00000112290 | WASF1 | -2.07867103 | 0.005775587 | -1.524367933 | 0.048366399 | -3.603038963 | 10.03472846 | 8.255926468 | 8.608698767 | 7.173892946 | 7.09482005 | 7.57363085 | 7.77172059 |
| ENSG00000205758 | CRYZL1 | -2.08027555 | 0.00311339 | -1.523380335 | 0.034894638 | -3.603655886 | 8.107517749 | 8.004992147 | 9.528916236 | 6.739036112 | 6.63128385 | 7.304736313 | 7.140495192 |
| ENSG00000100445 | SDR39U1 | -0.544588524 | 0.4795951 | -3.059082512 | 1.70E-06 | -3.603671035 | 8.075571721 | 6.535001748 | 7.285536013 | 6.887305128 | 6.90143415 | 4.45225155 | 4.720747915 |
| ENSG00000168779 | SHOX2 | 2.466187679 | 0.136952876 | -6.072150278 | 7.55E-05 | -3.605962599 | 7.53297911 | 4.93638933 | 6.203069917 | 8.996237745 | 9.01122668 | 2.746468522 | 2 |
| ENSG00000155926 | SLA | -3.482756978 | 0.180987334 | -0.124709963 | 0.967122742 | -3.607466941 | 2 | 5.457120953 | 2 | 2.270786764 | 2.49394783 | 4.612704558 | 3.128433649 |
| ENSG00000103550 | KNOP1 | -2.213052118 | 0.000484789 | -1.394472418 | 0.034772335 | -3.607524536 | 9.247966931 | 7.701025996 | 9.014156722 | 6.610808945 | 6.63721363 | 7.325609491 | 7.503217537 |
| ENSG00000163900 | TMEM41A | -2.073855138 | 0.015143652 | -1.534582337 | 0.079105356 | -3.608437475 | 10.5468183 | 8.147241016 | 10.05225195 | 7.753776273 | 7.90439571 | 8.359038717 | 8.365751687 |
| ENSG00000116455 | WDR77 | -1.100581639 | 0.352465423 | -2.510340481 | 0.019191632 | -3.61092212 | 9.678463567 | 9.785372135 | 6.534109959 | 8.172475401 | 8.09955907 | 6.57313027 | 6.926052784 |
| ENSG00000095637 | SORBS1 | -4.525387627 | 2.17E-05 | 0.914023915 | 0.450687102 | -3.611363711 | 6.89194353 | 7.638155693 | 8.042706615 | 4.205751501 | 2.59489879 | 9.0649396 | 7.553536683 |
| ENSG00000105379 | ETFB | -0.909211624 | 0.138339617 | -2.703734565 | 5.85E-07 | -3.612946189 | 9.521325812 | 9.680074956 | 8.434763955 | 8.385193462 | 8.43354993 | 6.442083733 | 6.843307307 |
| ENSG00000166133 | RPUSD2 | -2.100088569 | 0.055248319 | -1.513652937 | 0.174415528 | -3.613741506 | 9.473095744 | 6.666836837 | 7.342949392 | 6.172600389 | 6.41363412 | 6.915242278 | 6.802276223 |
| ENSG00000205089 | CCNI2 | -2.988493921 | 0.246112995 | -0.625650407 | 0.820069606 | -3.614144328 | 2 | 2 | 4.874587708 | 2.3892346 | 2.2680111 | 3.082820997 | 3.33635788 |
| ENSG00000112294 | ALDH5A1 | -3.539901984 | 0.014471998 | -0.074971641 | 0.967718357 | -3.614873625 | 9.204317094 | 5.131603625 | 5.772716771 | 4.574143677 | 4.4836996 | 7.49884738 | 7.959237827 |
| ENSG00000087266 | SH3BP2 | -2.723402345 | 0.000192819 | -0.894924827 | 0.271739141 | -3.618327172 | 10.18451821 | 8.791896851 | 10.85110517 | 7.501020868 | 7.44169935 | 9.331225056 | 9.219795239 |
| ENSG00000100320 | RBFOX2 | 0.296182991 | 0.646648662 | -3.916679819 | 8.18E-15 | -3.620496828 | 10.36053552 | 11.22601381 | 9.999667074 | 10.90999052 | 10.932824 | 6.741568503 | 6.777976866 |

| | | | | | | | | | | | | | |
|---|---|---|---|---|---|---|---|---|---|---|---|---|---|
| ENSG00000150672 | DLG2 | -2.432653974 | 0.17115254 | -1.188122844 | 0.519747981 | -3.620776817 | 3.327711234 | 6.535001748 | 3.369317972 | 3.754611435 | 2.86123014 | 4.628601098 | 3.753786374 |
| ENSG00000187607 | ZNF286A | -0.877660646 | 0.118145526 | -2.745032915 | 2.95E-08 | -3.622693561 | 8.16936288 | 8.916342327 | 9.330306409 | 8.040677544 | 7.98565528 | 6.346800618 | 6.044874254 |
| ENSG00000142530 | FAM71E1 | -2.524122909 | 0.130894481 | -1.099762113 | 0.525717368 | -3.623885021 | 5.866944937 | 4.11352228 | 2 | 3.023133209 | 2.94009902 | 3.917810786 | 3.789585079 |
| ENSG00000168061 | SAC3D1 | -2.713385481 | 2.34E-06 | -0.913523648 | 0.14682989 | -3.62690913 | 9.015175615 | 7.73146271 | 8.143093325 | 5.651002631 | 5.88556448 | 7.613117257 | 7.36768071 |
| ENSG00000176222 | ZNF404 | -2.203347059 | 0.047516317 | -1.424089185 | 0.212221448 | -3.627436244 | 8.107517749 | 5.131603625 | 7.451345288 | 5.474671438 | 4.98931816 | 6.307840213 | 5.519049808 |
| ENSG00000165283 | STOML2 | -1.755447 | 5.57E-11 | -1.872453879 | 1.66E-12 | -3.627900879 | 10.82658808 | 10.64091981 | 11.04953136 | 9.038768132 | 9.1566168 | 8.908528807 | 9.053535482 |
| ENSG00000181449 | SOX2 | -0.329057479 | 0.929222345 | -3.298958478 | 0.294154458 | -3.628015957 | 6.009025997 | 3.076665209 | 2 | 5.258265913 | 2 | 2.775316226 | 2.466008509 |
| ENSG00000184428 | TOP1MT | -1.65209828 | 0.40383052 | -1.977299366 | 0.298583938 | -3.629397646 | 9.143983763 | 7.819099893 | 2 | 6.328984825 | 6.55189318 | 6.192262702 | 6.077821908 |
| ENSG00000105393 | BABAM1 | -1.38007444 | 0.001374548 | -2.249683942 | 4.31E-08 | -3.629758382 | 10.90138923 | 10.08551947 | 10.30429804 | 9.070139931 | 9.12531607 | 8.066221408 | 8.390004118 |
| ENSG00000167526 | RPL13 | -2.129557167 | 2.45E-05 | -1.502061633 | 0.003920794 | -3.6316188 | 14.1971304 | 13.09483299 | 13.25240846 | 11.44714072 | 11.4992064 | 12.10283739 | 12.09753175 |
| ENSG00000125870 | SNRPB2 | -1.247286924 | 0.003814226 | -2.384530105 | 3.95E-09 | -3.631817029 | 10.46903009 | 9.785372135 | 9.703463497 | 8.842507967 | 8.73115061 | 7.538302335 | 7.783974227 |
| ENSG00000115947 | ORC4 | -1.200777561 | 0.080140412 | -2.431837291 | 0.000105608 | -3.632614852 | 9.966689162 | 8.522788974 | 8.784770914 | 8.164192202 | 7.91663606 | 6.833591108 | 6.853913776 |
| ENSG00000174886 | NDUFA11 | -1.146968652 | 0.018093579 | -2.489666033 | 2.61E-08 | -3.636634685 | 10.81212061 | 9.835272298 | 10.39531398 | 9.254074072 | 9.26207178 | 7.722704302 | 8.111083289 |
| ENSG00000120963 | ZNF706 | -1.726215063 | 0.000129797 | -1.912726101 | 1.61E-05 | -3.638941164 | 9.423197387 | 8.834582582 | 9.746650996 | 7.593372273 | 7.74603955 | 7.64179387 | 7.317985043 |
| ENSG00000143977 | SNRPG | -0.840799687 | 0.355105968 | -2.798354263 | 0.000388373 | -3.63915395 | 9.646202137 | 7.954278496 | 7.777116083 | 7.802622679 | 7.97632944 | 5.916072173 | 6.085042537 |
| ENSG00000173540 | GMPPB | -2.110869745 | 0.001518244 | -1.530489917 | 0.025520755 | -3.641359661 | 10.18451821 | 8.903033433 | 8.784770914 | 7.326437704 | 7.38291816 | 7.648631242 | 8.154630332 |
| ENSG00000088970 | KIZ | -2.151385061 | 0.258818497 | -1.491238907 | 0.441681135 | -3.642623968 | 8.998234871 | 8.395519306 | 2 | 6.221320894 | 5.89549381 | 6.934553331 | 6.409414172 |
| ENSG00000124302 | CHST8 | -1.685210108 | NA | -1.958538237 | NA | -3.643748345 | 3.327711234 | 2 | 2 | 2.270786764 | 2.14022021 | 2.225793514 | 2.117335275 |
| ENSG00000198189 | HSD17B11 | -0.869358171 | 0.259379662 | -2.775096155 | 3.64E-05 | -3.644454326 | 9.602033055 | 8.124485672 | 8.699419242 | 8.060896687 | 8.09094333 | 6.595545082 | 5.73181021 |
| ENSG00000175575 | PAAF1 | -2.030884785 | 0.006372593 | -1.615524157 | 0.032727299 | -3.646408942 | 7.27616306 | 9.235879652 | 8.784770914 | 6.586219071 | 6.72891807 | 7.02890981 | 7.091143645 |
| ENSG00000124784 | RIOK1 | -2.005750877 | 0.013980351 | -1.641433741 | 0.046918064 | -3.647184618 | 8.042902253 | 8.053983481 | 9.819251342 | 6.975156635 | 6.87658492 | 7.214140484 | 7.343799539 |
| ENSG00000166295 | ANAPC16 | -2.195258404 | 8.86E-11 | -1.451939527 | 2.70E-05 | -3.64719793 | 10.94624409 | 10.30378101 | 10.46161637 | 8.369067528 | 8.45720682 | 9.166773267 | 9.133628617 |
| ENSG00000146205 | ANO7 | -2.05665234 | 0.211593503 | -1.590628748 | 0.336327263 | -3.647281088 | 4.00722531 | 4.710564154 | 8.110401118 | 4.846787657 | 4.811949 | 4.813804156 | 5.566251551 |
| ENSG00000168268 | NT5DC2 | -2.740937459 | 7.81E-05 | -0.908465151 | 0.24090402 | -3.64940261 | 9.544849385 | 11.35867467 | 11.23535355 | 8.258639605 | 8.11663771 | 9.886487495 | 10.11928018 |
| ENSG00000198826 | ARHGAP11A | 1.363812056 | 0.096765227 | -5.013355815 | 2.79E-11 | -3.649543759 | 7.384433967 | 5.457120953 | 7.098165388 | 8.237082658 | 8.11025689 | 2.962551659 | 2.817676617 |
| ENSG00000167394 | ZNF668 | -1.984227211 | 0.059087638 | -1.667507514 | 0.113881928 | -3.651734725 | 9.533135543 | 6.844429755 | 9.888372618 | 7.230526165 | 7.31425651 | 7.35696023 | 7.77730333 |
| ENSG00000100376 | FAM118A | -2.656045639 | 3.73E-08 | -0.995711638 | 0.057483341 | -3.651757277 | 8.737455672 | 9.148351514 | 9.849281147 | 6.693715713 | 6.72335419 | 8.380906995 | 8.278687361 |
| ENSG00000186446 | ZNF501 | -1.975499903 | 0.182550229 | -1.676448045 | 0.256069609 | -3.651947948 | 7.940205782 | 5.595927442 | 4.058998049 | 4.846787657 | 4.89348248 | 5.553513572 | 4.532869193 |
| ENSG00000115919 | KYNU | 2.195338677 | 0.183470484 | -5.851181933 | 8.00E-05 | -3.655843256 | 8.946185348 | 6.899080544 | 5.39244166 | 9.894954101 | 9.98949012 | 3.465698119 | 2.117335275 |
| ENSG00000134072 | CAMK1 | -0.877574953 | 0.328352926 | -2.779129522 | 0.00039065 | -3.656704476 | 8.517314938 | 8.124485672 | 6.322058544 | 6.998639523 | 7.1246732 | 5.29920634 | 5.250250853 |
| ENSG00000197409 | HIST1H3D | 4.174455099 | 0.0614041 | -7.832784015 | 0.0032699 | -3.658328916 | 2 | 4.442710129 | 2 | 6.892328675 | 6.89154576 | 2 | 2 |
| ENSG00000100577 | GSTZ1 | -2.024617176 | 0.010351963 | -1.637205551 | 0.040450017 | -3.661822727 | 6.73718411 | 8.053983481 | 8.845626171 | 6.18083559 | 6.12381559 | 6.581321461 | 6.457498602 |
| ENSG00000164181 | ELOVL7 | 0.427419808 | 0.801141925 | -4.093743296 | 0.003861307 | -3.666323487 | 3.327711234 | 3.076665209 | 5.59507254 | 4.804776789 | 4.79082474 | 2.384001718 | 2.277179783 |
| ENSG00000137269 | LRRC1 | -3.132524052 | 1.05E-08 | -0.534226561 | 0.391911003 | -3.666750613 | 7.031698319 | 8.169643011 | 7.600047325 | 4.867343068 | 4.63360314 | 7.301021256 | 7.010010317 |
| ENSG00000051108 | HERPUD1 | -2.042365688 | 1.10E-13 | -1.625683424 | 4.19E-09 | -3.668049112 | 11.68115503 | 11.69809349 | 11.80171081 | 9.688809967 | 9.69266358 | 10.31849333 | 9.855700762 |
| ENSG00000143621 | ILF2 | -1.63125322 | 1.06E-07 | -2.036817822 | 1.42E-11 | -3.668071042 | 11.90670479 | 11.49797153 | 11.85201631 | 10.103182 | 10.1654674 | 9.532099314 | 9.905493112 |
| ENSG00000182544 | MFSD5 | -1.796561723 | 0.006464219 | -1.874362668 | 0.004058514 | -3.670924391 | 9.720386367 | 9.821190352 | 11.12920538 | 8.538855014 | 8.63811484 | 8.357250999 | 8.652479133 |
| ENSG00000155115 | GTF3C6 | -1.551766794 | 0.007945031 | -2.119538983 | 0.000159314 | -3.671305778 | 10.16187685 | 9.125613817 | 9.132850258 | 7.980622673 | 8.06258142 | 7.141214687 | 7.729722875 |
| ENSG00000196872 | KIAA1211L | -2.26484723 | NA | -1.408755192 | NA | -3.673602423 | 3.327711234 | 2 | 2 | 2.270786764 | 2 | 2.457042231 | 2 |
| ENSG00000115129 | TP53I3 | -0.770867164 | 0.082840433 | -2.903618983 | 9.23E-14 | -3.674486147 | 9.810482605 | 8.981105197 | 9.516493218 | 8.612239836 | 8.79487126 | 6.601598251 | 6.637709309 |
| ENSG00000084463 | WBP11 | -1.961738666 | 5.50E-07 | -1.713267422 | 1.38E-05 | -3.675006088 | 10.85979178 | 10.74900803 | 11.46280979 | 9.12771691 | 9.08032183 | 9.179643937 | 9.504290592 |
| ENSG00000162076 | FLYWCH2 | -2.742507628 | 5.93E-05 | -0.933162719 | 0.218284684 | -3.675670347 | 10.67493801 | 8.94259736 | 9.491321578 | 7.161462517 | 7.19871665 | 8.842760893 | 9.070004758 |
| ENSG00000102390 | PBDC1 | -1.741456241 | 0.003736544 | -1.934682727 | 0.001035679 | -3.676138968 | 9.839306291 | 8.540087354 | 9.61300511 | 7.62389457 | 7.78395061 | 7.220710071 | 7.762743025 |
| ENSG00000125652 | ALKBH7 | -1.948869505 | 0.001101338 | -1.728560455 | 0.004000831 | -3.677429961 | 8.540903468 | 9.77810053 | 9.945092785 | 7.587189552 | 7.64371756 | 7.680435163 | 7.96902811 |
| ENSG00000129214 | SHBG | -1.920262068 | 0.227286772 | -1.758381905 | 0.259652636 | -3.678643974 | 2 | 5.946638857 | 5.156626401 | 3.349643765 | 3.7864959 | 3.709392871 | 3.67940853 |
| ENSG00000249115 | HAUS5 | -2.670192805 | 0.005703498 | -1.010247387 | 0.336805988 | -3.680440192 | 8.418903792 | 7.503546079 | 5.59507254 | 4.965943888 | 5.14724905 | 6.670403488 | 6.517437394 |
| ENSG00000242616 | GNG10 | 1.574616214 | 0.107655331 | -5.255339221 | 4.08E-09 | -3.680723007 | 7.82963488 | 5.595927442 | 6.43197674 | 8.519504557 | 8.37442884 | 2.592885947 | 3.012034821 |
| ENSG00000105705 | SUGP1 | -1.643490761 | 0.125913128 | -2.039594976 | 0.048332079 | -3.683085736 | 10.73752355 | 8.147241016 | 8.486635645 | 7.944803394 | 8.02236724 | 7.535143176 | 7.652846376 |
| ENSG00000182362 | YBEY | -1.86129031 | 0.055248429 | -1.822084201 | 0.057302601 | -3.683374511 | 9.219014269 | 6.951736442 | 7.342949392 | 6.314087501 | 6.45450701 | 6.599583349 | 6.222348126 |
| ENSG00000145996 | CDKAL1 | -1.349326115 | 0.000260005 | -2.334720798 | 3.27E-11 | -3.684046912 | 8.228665467 | 8.414402626 | 8.721235202 | 7.075737784 | 7.20671394 | 6.037140174 | 6.341697848 |
| ENSG00000274636 | | -1.093179795 | 0.20769014 | -2.591232987 | 0.000878719 | -3.684412782 | 8.19931886 | 8.029695761 | 9.839340503 | 7.842101511 | 7.86703672 | 6.210860813 | 6.569847194 |
| ENSG00000179091 | CYC1 | -2.451965793 | 1.00E-09 | -1.232550347 | 0.003618457 | -3.68451614 | 11.55650736 | 10.89732378 | 10.71638486 | 8.654247601 | 8.67172529 | 9.894932749 | 9.853324044 |
| ENSG00000148704 | VAX1 | 1.504743261 | NA | -5.191100501 | NA | -3.686357241 | 2 | 2 | 2 | 3.289987342 | 2 | 2 | 2 |
| ENSG00000137642 | SORL1 | -2.814773015 | 0.099672705 | -0.871650503 | 0.63881429 | -3.686423518 | 6.73718411 | 5.838937808 | 2 | 3.709785322 | 3.28037825 | 5.54934408 | 4.175132639 |
| ENSG00000112699 | GMDS | -3.651057349 | 0.00122169 | -0.035418248 | 0.98058233 | -3.686475497 | 9.143983763 | 6.844429755 | 6.073378139 | 4.497095146 | 4.60967607 | 7.513903457 | 8.244432684 |
| ENSG00000106554 | CHCHD3 | -1.668324508 | 0.047516317 | -2.018951945 | 0.013432092 | -3.687276453 | 11.25159303 | 9.680074956 | 9.287079172 | 8.651287275 | 8.70598173 | 8.064761569 | 8.557692095 |
| ENSG00000183114 | FAM43B | -4.135560015 | 0.094524523 | 0.446816255 | 0.866824598 | -3.68874376 | 2 | 5.131603625 | 2 | 2.270786764 | 2.14022021 | 4.369533167 | 4.077147165 |
| ENSG00000152433 | ZNF547 | -1.208308489 | 0.517222971 | -2.480988959 | 0.144498426 | -3.689297447 | 2 | 6.78762695 | 7.777116083 | 5.614989456 | 5.69405321 | 4.771475226 | 4.254128754 |
| ENSG00000165060 | FXN | -1.760695092 | 0.000118649 | -1.929712313 | 1.52E-05 | -3.690407405 | 7.218819781 | 8.053983481 | 7.777116083 | 5.920568914 | 6.14061114 | 5.83960027 | 5.913137508 |
| ENSG00000118894 | EEF2KMT | -1.347257272 | 0.000891154 | -2.343722092 | 9.99E-10 | -3.690979364 | 7.940205782 | 7.503546079 | 7.734844356 | 6.422239259 | 6.42738748 | 5.318934974 | 5.651404445 |
| ENSG00000171219 | CDC42BPG | -2.44717364 | 0.317860928 | -1.24414551 | 0.620242104 | -3.69131915 | 2 | 5.303523747 | 2 | 2.695456494 | 2.59489879 | 3.193831101 | 3.209956562 |
| ENSG00000165091 | TMC1 | -2.934152049 | 0.094962877 | -0.760569431 | 0.673665014 | -3.69472148 | 4.467311984 | 3.076665209 | 3.369317972 | 2.498691189 | 2.2680111 | 2.775316226 | 3.620995423 |
| ENSG00000204856 | FAM216A | -2.360376501 | 2.96E-06 | -1.334668957 | 0.010747713 | -3.695045458 | 8.228665467 | 7.189546432 | 8.007629717 | 5.627094009 | 5.5755357 | 6.660772804 | 6.463051659 |
| ENSG00000069696 | DRD4 | 2.456607006 | 0.531143362 | -6.152922849 | 0.118418414 | -3.696315843 | 2 | 3.076665209 | 2 | 4.034562184 | 3.74316441 | 2 | 2 |
| ENSG00000167965 | MLST8 | -1.900589226 | 1.51E-13 | -1.796825922 | 1.67E-12 | -3.697415147 | 9.966689162 | 9.632483534 | 9.897982195 | 8.008651253 | 7.89452573 | 8.054501123 | 8.055192982 |
| ENSG00000136146 | MED4 | -1.307354572 | 0.000365107 | -2.39089293 | 6.83E-12 | -3.698247502 | 9.384607364 | 9.532315021 | 8.904018013 | 8.047448792 | 7.96694292 | 7.060015953 | 6.828326017 |
| ENSG00000155438 | NIFK | -1.44242687 | 0.392191303 | -2.259039051 | 0.152446886 | -3.701465921 | 10.78274301 | 8.234834549 | 5.156626401 | 8.060896687 | 7.9809999 | 7.127295835 | 7.310303798 |
| ENSG00000128218 | VPREB3 | -2.196040437 | 0.192819624 | -1.506706287 | 0.357318652 | -3.702746724 | 4.467311984 | 3.076665209 | 3.369317972 | 2.695456494 | 2.49394783 | 3.12825509 | 2.551896972 |
| ENSG00000108666 | C17orf75 | -1.564510963 | 0.010010073 | -2.139677821 | 0.000234844 | -3.704188783 | 8.946185348 | 7.572420273 | 8.486635645 | 6.956091174 | 6.84618961 | 6.495072127 | 6.189227296 |
| ENSG00000269335 | IKBKG | -0.09599818 | 0.931706466 | -3.610649544 | 1.84E-05 | -3.706647723 | 9.509418609 | 7.431218269 | 7.646393253 | 8.473317843 | 8.37087451 | 4.914492341 | 5.217771644 |
| ENSG00000176974 | SHMT1 | -2.516160992 | 0.052939325 | -1.19389386 | 0.38415025 | -3.710054852 | 4.00722531 | 7.60566235 | 7.934808822 | 4.907595604 | 4.89348248 | 5.614651724 | 6.465820193 |
| ENSG00000070785 | EIF2B3 | -1.834498545 | 0.108894003 | -1.877093828 | 0.094510534 | -3.711592372 | 10.60951384 | 7.538394143 | 8.460932916 | 7.620871236 | 7.65252709 | 7.50961773 | 7.675744778 |
| ENSG00000277114 | | 0.183107515 | 0.88199664 | -3.895019836 | 2.43E-05 | -3.711912321 | 9.112843761 | 8.451443573 | 6.203069917 | 8.543652346 | 8.51631523 | 4.496236235 | 4.845038551 |
| ENSG00000138413 | IDH1 | -1.273351737 | 8.49E-07 | -2.440293228 | 1.28E-22 | -3.713644966 | 11.18203661 | 11.02567599 | 11.1494608 | 9.883016326 | 9.81672282 | 8.504505158 | 8.856651551 |
| ENSG00000106927 | AMBP | -1.689547294 | NA | -2.024447659 | NA | -3.713994953 | 3.327711234 | 2 | 2 | 2.141737246 | 2.2680111 | 2.048074405 | 2.277179783 |
| ENSG00000000460 | C1orf112 | -1.557433747 | 0.42536274 | -2.156634796 | 0.244329036 | -3.714068543 | 2 | 7.538394143 | 5.156626401 | 4.623314311 | 5.00774437 | 4.281782022 | 4.376876162 |
| KBTBD7 | | -2.383550139 | 0.227664887 | -1.331741485 | 0.51714034 | -3.715291624 | 2 | 8.124485672 | 9.528916236 | 6.147608921 | 6.04571775 | 7.370021622 | 6.780202889 |
| ENSG00000182224 | CYB5D1 | 1.080766552 | 0.614749417 | -4.799212743 | 0.009476259 | -3.718446191 | 4.00722531 | 7.638155693 | 2 | 7.281195074 | 7.20671394 | 2.656294034 | 2.746601469 |
| ENSG00000160606 | TLCD1 | -1.667180138 | 0.045709955 | -2.052820317 | 0.010696797 | -3.720000455 | 8.367060731 | 6.602424674 | 7.029880454 | 5.880681751 | 5.98191036 | 5.83960027 | 5.250250853 |
| ENSG00000118137 | APOA1 | -2.279096386 | NA | -1.443110015 | NA | -3.722206401 | 2 | 3.076665209 | 2 | 2 | 2.2680111 | 2.3070652 | 2.172608095 |
| ENSG00000170369 | CST2 | -1.798987805 | 0.506870806 | -1.925158545 | 0.463894914 | -3.72414635 | 2 | 2 | 6.43197674 | 3.615725078 | 3.50452879 | 3.483289517 | 3.474731315 |
| ENSG00000178585 | CTNNBIP1 | -2.269518934 | 2.03E-08 | -1.455009623 | 0.00046003 | -3.724528557 | 9.87686338 | 9.267380431 | 9.647595056 | 7.29643414 | 7.44845993 | 7.86872317 | 8.425990514 |
| ENSG00000115211 | EIF2B4 | -1.979559514 | 7.83E-05 | -1.747106666 | 0.000520893 | -3.72666618 | 9.064836028 | 10.15453315 | 10.09465441 | 7.90310317 | 7.87709394 | 8.209185391 | 8.02265879 |
| ENSG00000092051 | JPH4 | -2.34391398 | 0.571578307 | -1.382752923 | NA | -3.726666903 | 2 | 3.686303414 | 2 | 2.3892346 | 2 | 2.560105256 | 2.172608095 |
| ENSG00000110696 | C11orf58 | -1.372936263 | 2.65E-05 | -2.35600139 | 6.03E-14 | -3.728937654 | 11.75108151 | 11.35623522 | 11.98194227 | 10.37308875 | 10.321834 | 9.334257513 | 9.402844548 |
| ENSG00000141506 | PIK3R5 | -6.294901401 | 0.069759872 | 2.565570935 | 0.424499453 | -3.729330466 | 4.467311984 | 2 | 2 | 2 | 2 | 5.638399518 | 4.827924991 |
| ENSG00000164167 | LSM6 | -1.698939168 | 0.051267618 | -2.031814542 | 0.016000913 | -3.73075371 | 8.444143041 | 6.602424674 | 6.882775709 | 5.651002631 | 6.14061114 | 5.532544554 | 5.680297397 |
| ENSG00000074201 | CLNS1A | -1.836815808 | 0.000228835 | -1.894546371 | 0.000126518 | -3.73136218 | 10.4253394 | 9.74864196 | 10.92346452 | 8.551612617 | 8.67605209 | 8.550603056 | 8.565456872 |
| ENSG00000128652 | HOXD3 | 3.695879336 | 0.133069745 | -7.428917731 | 0.00861456 | -3.733038395 | 2 | 4.11352228 | 2 | 6.221320894 | 5.85535975 | 2 | 2 |
| ENSG00000121851 | POLR3GL | -2.125469189 | 0.249844477 | -1.609191597 | 0.387074442 | -3.734660786 | 10.56993315 | 8.469613413 | 3.369317972 | 7.123515401 | 7.27295842 | 7.557113378 | 7.838397174 |
| ENSG00000155229 | MMS19 | -2.068986494 | 7.69E-05 | -1.667034708 | 0.001725967 | -3.736021202 | 10.27872131 | 10.29870826 | 11.35181254 | 8.689305661 | 8.66303256 | 9.068585983 | 9.080880746 |
| ENSG00000088727 | KIF9 | -1.870325148 | 0.094962877 | -1.866364722 | 0.087552565 | -3.73668987 | 7.218819781 | 4.710564154 | 5.59507254 | 4.670864177 | 4.37434574 | 4.331190017 | 4.711446447 |
| ENSG00000019485 | PRDM11 | -3.209696514 | 1.69E-08 | -0.529931981 | 0.419909413 | -3.739628496 | 8.418903792 | 7.145017454 | 7.896971941 | 4.694064037 | 5.04390545 | 7.392307374 | 7.375063227 |
| ENSG00000143320 | CRABP2 | -2.708688939 | 0.059985568 | -1.032380413 | 0.508404219 | -3.741069352 | 10.62062599 | 8.169643011 | 6.073378139 | 6.727838783 | 6.60731796 | 8.790285782 | 7.570355956 |
| ENSG00000119630 | PGF | -2.201802798 | 0.051696866 | -1.546702877 | 0.180993471 | -3.748505675 | 11.36165447 | 9.181799937 | 10.57976717 | 8.423869837 | 8.4504872 | 9.870283133 | 7.239267729 |
| ENSG00000175792 | RUVBL1 | -2.068213016 | 0.028160569 | -1.680753662 | 0.078046587 | -3.748966678 | 10.72212986 | 8.671479029 | 8.561105505 | 7.580980221 | 7.6986238 | 7.754744914 | 8.246849118 |
| ENSG00000172006 | ZNF554 | -2.530654306 | 0.03867891 | -1.218803815 | 0.347291869 | -3.749458121 | 5.532353208 | 6.228547981 | 8.699419242 | 4.946755474 | 5.22805098 | 6.405645065 | 6.189227296 |
| ENSG00000163389 | POGLUT1 | -1.605053931 | 0.053222658 | -2.145677328 | 0.006975264 | -3.750731259 | 8.653385977 | 9.71856931 | 10.8211137 | 8.372666678 | 8.4215744 | 7.697491716 | 8.014172904 |
| ENSG00000149179 | C11orf49 | -2.129848935 | 0.000301351 | -1.621374021 | 0.007196293 | -3.751222956 | 9.2903348 | 9.181799937 | 10.30429804 | 7.559034656 | 7.59580267 | 7.698433415 | 8.376826085 |
| ENSG00000196263 | ZNF471 | -1.692384577 | 0.138710156 | -2.059115306 | 0.061458022 | -3.751499882 | 6.89194353 | 7.431218269 | 9.703463497 | 6.882264028 | 6.89649843 | 6.726875343 | 6.317379004 |
| ENSG00000092531 | SNAP23 | -0.975103395 | 0.356756632 | -2.778314828 | 0.002890079 | -3.753418223 | 10.87842776 | 9.030900487 | 8.35328246 | 8.977450658 | 8.73530306 | 6.940933331 | 7.22299542 |
| ENSG00000108559 | NUP88 | -2.178480526 | 0.002960272 | -1.576631777 | 0.036695546 | -3.755112303 | 10.499447 | 8.522788974 | 10.01740919 | 7.756533788 | 7.70992146 | 8.234076861 | 8.412421212 |
| ENSG00000090776 | EFNB1 | -3.216916072 | 6.36E-16 | -0.540155787 | 0.237178826 | -3.757071859 | 10.1311248 | 9.396454291 | 9.426401271 | 6.554880361 | 6.507263 | 9.080554073 | 9.224707098 |
| ENSG00000205639 | MFSD2B | -2.882831177 | 0.274413253 | -0.87434742 | 0.746319043 | -3.757178597 | 2 | 2 | 4.523675391 | 2.270786764 | 2.2680111 | 2.885263063 | 2.851942824 |
| ENSG00000143870 | PDIA6 | -0.827110247 | 0.110962 | -2.930398561 | 1.05E-10 | -3.757508808 | 13.4024762 | 12.68949384 | 13.26731409 | 12.31674406 | 12.3321074 | 9.760045703 | 10.57593535 |
| ENSG00000109576 | AADAT | -0.965864931 | 0.606417049 | -2.794841251 | 0.093132318 | -3.760706183 | 2 | 6.535001748 | 7.691296561 | 5.662809983 | 5.8245091 | 4.188345803 | 4.147809227 |
| ENSG00000143294 | PRCC | -2.14398563 | 0.00010839 | -1.617292711 | 0.004386566 | -3.761278342 | 11.55361133 | 10.2574711 | 11.39656351 | 9.022822647 | 9.04952667 | 9.407492813 | 9.697162691 |

| | | | | | | | | | | | | | |
|---|---|---|---|---|---|---|---|---|---|---|---|---|---|
| ENSG00000111906 | HDDC2 | -2.140180457 | 0.077791464 | -1.621108577 | 0.186670425 | -3.761289034 | 10.78274301 | 8.376385542 | 7.451345288 | 7.416368504 | 7.47849652 | 7.703132714 | 8.172535245 |
| ENSG00000130413 | STK33 | -2.748468738 | 0.156896095 | -1.013527895 | 0.621070373 | -3.761996633 | 2 | 6.04685525 | 3.369317972 | 2.947924294 | 2.86123014 | 3.738983888 | 4.161535617 |
| ENSG00000015133 | CCDC88C | -4.61149939 | 0.000157859 | 0.848445785 | 0.51600333 | -3.763053604 | 3.327711234 | 6.04685525 | 6.43197674 | 2.784612485 | 2.38539787 | 6.643273885 | 6.476841503 |
| ENSG00000168887 | C2orf68 | -2.474760944 | 0.000105278 | -1.288706324 | 0.057525246 | -3.763467268 | 10.57565445 | 9.203675681 | 9.212024683 | 7.377487766 | 7.36516374 | 8.448514034 | 8.621777169 |
| ENSG00000171497 | PPID | -1.54393187 | 0.000579367 | -2.221469966 | 2.62E-07 | -3.765401836 | 9.304185368 | 9.114109157 | 9.999667074 | 7.990026167 | 7.99492122 | 7.41654498 | 7.234405279 |
| ENSG00000267060 | PTGES3L | -2.4230083 | 0.082092336 | -1.343816018 | 0.34518468 | -3.766824317 | 4.815609471 | 3.076665209 | 6.322058544 | 3.462030052 | 3.28037825 | 4.144793515 | 4.105828626 |
| ENSG00000074219 | TEAD2 | -1.905161836 | 2.94E-11 | -1.862645313 | 5.55E-11 | -3.767807149 | 10.22876205 | 9.849218119 | 9.990713521 | 8.14748175 | 8.13141746 | 8.038972319 | 8.311378144 |
| ENSG00000171161 | ZNF672 | -2.257698958 | 0.019726538 | -1.51134006 | 0.130419453 | -3.769039018 | 10.09173937 | 7.819099893 | 10.71638486 | 7.751013478 | 7.70428369 | 8.558922783 | 8.360554398 |
| ENSG00000115526 | CHST10 | -2.150768359 | 0.264068306 | -1.619360501 | 0.405240029 | -3.77012886 | 9.233563228 | 8.639735838 | 2 | 6.408278948 | 6.17362678 | 6.723178551 | 6.881131115 |
| ENSG00000130768 | SMPDL3B | -2.280655953 | 0.439157025 | -1.490207222 | 0.60383961 | -3.770863176 | 2 | 2 | 4.058998049 | 2.3892346 | 2.14022021 | 2.457042231 | 2.421067459 |
| ENSG00000113966 | ARL6 | -6.676995518 | 0.034423594 | 2.905179202 | 0.302589526 | -3.771816316 | 2 | 4.710564154 | 2 | 2 | 2 | 5.938563912 | 5.964937606 |
| ENSG00000105072 | C19orf44 | -2.201841002 | 0.06478162 | -1.571274513 | 0.19477976 | -3.773115515 | 5.532353208 | 7.145017454 | 8.941672936 | 5.709096143 | 5.70538658 | 6.518722958 | 6.037449234 |
| ENSG00000159352 | PSMD4 | -1.554560929 | 0.014617921 | -2.219036896 | 0.000262877 | -3.773597825 | 11.73838527 | 10.75272247 | 10.23015729 | 9.496029474 | 9.49704586 | 8.740866084 | 8.926648946 |
| ENSG00000196584 | XRCC2 | -2.512037998 | 0.00175424 | -1.262393448 | 0.1204601 | -3.774431446 | 5.095959458 | 5.457120953 | 5.930865648 | 3.663521627 | 3.33976443 | 4.667592615 | 4.147809227 |
| ENSG00000103740 | ACSBG1 | -2.265402063 | NA | -1.509106822 | NA | -3.774508885 | 3.327711234 | 2 | 2 | 2.270786764 | 2 | 2.384001718 | 2.059860178 |
| ENSG00000171105 | INSR | -2.700704443 | 2.76E-08 | -1.073920519 | 0.04093731 | -3.774624962 | 8.87372952 | 9.377333006 | 9.972637969 | 6.676346669 | 6.94988431 | 8.510951738 | 8.304435067 |
| ENSG00000167600 | CYP2S1 | 0.353678511 | 0.805509199 | -4.128604467 | 0.000243363 | -3.774925956 | 9.031919741 | 8.029695761 | 5.930865648 | 8.52598359 | 8.4504872 | 4.881704255 | 3.36036153 |
| ENSG00000126953 | TIMM8A | -2.042459879 | 0.318672895 | -1.734587782 | 0.395470363 | -3.777047661 | 8.631580145 | 5.595927442 | 2 | 5.319220934 | 5.27444544 | 5.606648123 | 5.545463418 |
| ENSG00000204682 | CASC10 | -3.818023117 | 0.018991992 | 0.040669336 | 0.984153231 | -3.777353782 | 2 | 6.389896633 | 7.600047325 | 3.462030052 | 3.28037825 | 6.653021698 | 6.525421271 |
| ENSG00000100348 | TXN2 | -2.289469922 | 3.47E-11 | -1.488550109 | 2.51E-05 | -3.778020031 | 10.85979178 | 10.61258327 | 10.16797309 | 8.279879181 | 8.32017198 | 9.004546892 | 9.176932811 |
| ENSG00000168275 | COA6 | -0.681563848 | 0.599977946 | -3.097949075 | 0.004921372 | -3.779512923 | 9.533135543 | 6.951736442 | 6.718950348 | 7.756533788 | 7.56502709 | 5.279204176 | 5.448021784 |
| ENSG00000137821 | LRRC49 | -1.377398925 | 0.360439432 | -2.406835401 | 0.085406775 | -3.784234326 | 5.095959458 | 6.78762695 | 9.358421373 | 6.750147204 | 6.67805337 | 5.498346994 | 5.949196291 |
| ENSG00000178761 | FAM219B | -1.594091223 | 0.006709768 | -2.194994331 | 0.000101151 | -3.789085554 | 9.613202746 | 10.45735223 | 10.95602907 | 8.839910123 | 8.87358581 | 8.490981547 | 7.99033569 |
| ENSG00000180043 | FAM71E2 | 0.009748828 | 0.997264364 | -3.801467427 | NA | -3.791718599 | 2 | 3.686303414 | 2 | 3.023133209 | 2.68924543 | 2.048074405 | 2.117335275 |
| ENSG00000182180 | MRPS16 | -2.0761821 | 0.001705352 | -1.720429479 | 0.010504502 | -3.796611579 | 11.33491623 | 9.640525262 | 10.89974571 | 8.738951712 | 8.69607402 | 8.876019586 | 9.24176809 |
| ENSG00000141552 | ANAPC11 | -2.293521235 | 0.00203162 | -1.504057384 | 0.051696684 | -3.797578619 | 11.0567315 | 9.396454291 | 9.439621147 | 7.898117066 | 7.92635454 | 8.806535425 | 8.566746948 |
| ENSG00000140848 | CPNE2 | -2.358579249 | 0.00115848 | -1.440440541 | 0.057729712 | -3.799019791 | 10.04301211 | 8.451443573 | 8.511888463 | 6.825617242 | 6.94032357 | 7.91644678 | 7.627107456 |
| ENSG00000185090 | MANEAL | -1.580474379 | 0.148482497 | -2.21921459 | 0.031377287 | -3.799688969 | 4.815609471 | 6.464272742 | 4.523675391 | 4.237680128 | 4.1930041 | 3.5515855 | 3.908316285 |
| ENSG00000133111 | RFXAP | -2.161269038 | 0.120812437 | -1.639075269 | 0.242857126 | -3.800344307 | 3.327711234 | 6.728495673 | 7.600047325 | 4.907595604 | 4.4836996 | 5.238348159 | 5.08721802 |
| ENSG00000278718 | | 5.184904515 | 5.55E-07 | -8.985401462 | 1.52E-06 | -3.800496947 | 2 | 4.710564154 | 4.523675391 | 8.965583267 | 9.06610578 | 2 | 2 |
| ENSG00000163435 | ELF3 | -0.891381543 | 0.756699054 | -2.910152584 | 0.258406162 | -3.801534127 | 2 | 2 | 5.156626401 | 3.289987342 | 3.2184422 | 2.183373728 | 2.632958241 |
| ENSG00000120832 | MTERF2 | -2.313825135 | 3.57E-08 | -1.488565641 | 0.000365718 | -3.802390775 | 7.435667355 | 7.051611315 | 7.502634626 | 5.22679461 | 5.07918227 | 6.004008792 | 5.803599879 |
| ENSG00000174021 | GNG5 | -0.340513131 | 0.73397139 | -3.463378622 | 1.24E-05 | -3.803891753 | 11.18951828 | 9.214490408 | 9.385998894 | 9.903687576 | 9.87245708 | 6.790081374 | 6.832622283 |
| ENSG00000100151 | PICK1 | -2.160547903 | 0.001152009 | -1.644218291 | 0.015445588 | -3.804766194 | 8.946185348 | 7.315436877 | 8.536706852 | 6.252910529 | 6.35727082 | 6.643273885 | 6.942047249 |
| ENSG00000167384 | ZNF180 | -1.802952037 | 0.045495366 | -2.001879898 | 0.023062272 | -3.804831934 | 8.16936288 | 6.389896633 | 8.742726175 | 6.276156955 | 6.32824132 | 6.327451929 | 5.863612864 |
| ENSG00000072849 | DERL2 | -1.688716022 | 0.003767538 | -2.118724889 | 0.000181601 | -3.807440911 | 11.02362764 | 9.664384941 | 10.19160738 | 8.748679155 | 8.69607402 | 8.158760173 | 8.423856459 |
| ENSG00000136997 | MYC | -2.783743917 | 1.78E-07 | -1.025106821 | 0.079657735 | -3.808850738 | 10.33377634 | 10.30883598 | 10.85110517 | 7.64488235 | 7.86451142 | 9.959004933 | 8.818171767 |
| ENSG00000282813 | | -0.823281207 | 0.008466761 | -2.989698467 | 1.25E-25 | -3.812979673 | 9.015175615 | 8.777382925 | 9.212024683 | 8.209172298 | 8.17486938 | 6.118193494 | 6.066922748 |
| ENSG00000169605 | GKN1 | -1.657973751 | 0.598164094 | -2.159638551 | NA | -3.817612302 | 2 | 3.686303414 | 2 | 2.270786764 | 2.38539787 | 2.094598369 | 2.374681325 |
| ENSG00000198157 | HMGN5 | -2.089526425 | 0.005993307 | -1.728932469 | 0.022948166 | -3.818458894 | 7.33131396 | 5.946638857 | 5.930865648 | 4.623314311 | 4.68029874 | 5.008609438 | 4.911532223 |
| ENSG00000132530 | XAF1 | -6.80400418 | 0.003278093 | 2.985144362 | 0.082953796 | -3.818859818 | 2 | 4.442710129 | 3.369317972 | 2 | 2 | 6.098353065 | 6.192573806 |
| ENSG00000148690 | FRA10AC1 | -1.743454482 | 0.021100389 | -2.075994104 | 0.004667079 | -3.819448586 | 8.75772933 | 7.051611315 | 8.408111533 | 6.687949235 | 6.35727082 | 6.495072127 | 5.867805474 |
| ENSG00000159388 | BTG2 | -3.709146373 | 6.38E-08 | -0.110342388 | 0.903960775 | -3.81948876 | 9.800745303 | 7.979858146 | 8.941672936 | 5.697663076 | 5.27444544 | 8.986541761 | 8.970196279 |
| ENSG00000067829 | IDH3G | -1.679391555 | 0.000132612 | -2.140906339 | 5.70E-07 | -3.820297894 | 10.65345793 | 9.828248506 | 10.01740919 | 8.511364632 | 8.57000035 | 7.88204676 | 8.26365177 |
| ENSG00000227268 | KLLN | -2.042693191 | 0.040034859 | -1.778918593 | 0.064804976 | -3.821611784 | 4.815609471 | 5.131603625 | 4.874587708 | 3.162721474 | 3.60476711 | 3.838019203 | 3.236138336 |
| ENSG00000169902 | TPST1 | -0.985218914 | 0.051337047 | -2.836408395 | 3.75E-10 | -3.821627309 | 11.57661845 | 11.38044608 | 10.51254821 | 10.20914183 | 10.2755373 | 8.372671446 | 8.434495305 |
| ENSG00000071889 | FAM3A | -2.597372674 | 8.38E-05 | -1.22427697 | 0.085129079 | -3.821649644 | 10.72727936 | 9.443177293 | 11.23535355 | 8.027038952 | 8.09955907 | 9.442199872 | 9.404649223 |
| ENSG00000180998 | GPR137C | -1.822721076 | 0.051085048 | -1.999888047 | 0.025986563 | -3.822609124 | 5.330594134 | 4.93638933 | 6.073378139 | 4.268917409 | 3.60476711 | 4.065262523 | 3.580696614 |
| ENSG00000070182 | SPTB | -4.696921001 | 0.003484322 | 0.871082879 | 0.610378131 | -3.825838123 | 2 | 4.93638933 | 6.203069917 | 2.270786764 | 2.49394783 | 6.194934299 | 5.685057129 |
| ENSG00000174255 | ZNF80 | -0.407315917 | 0.898417957 | -3.421105488 | NA | -3.828421406 | 2 | 3.686303414 | 2 | 2.3892346 | 2.94009902 | 2.094598369 | 2.117335275 |
| ENSG00000178695 | KCTD12 | -2.856359724 | 0.000330537 | -0.972758352 | 0.27169027 | -3.829118076 | 10.09970294 | 12.1280309 | 11.12102291 | 8.521127045 | 8.48377664 | 10.79903314 | 9.770373699 |
| ENSG00000095059 | DHPS | 0.346167383 | 0.803450392 | -4.175769161 | 0.000117891 | -3.829601778 | 5.70932728 | 8.004992147 | 9.358421373 | 8.655725489 | 8.68609779 | 4.271694079 | 4.574506939 |
| ENSG00000129932 | DOHH | -2.054542028 | 0.025764885 | -1.775917993 | 0.055140779 | -3.83046002 | 9.820154628 | 8.029695761 | 7.600047325 | 6.83085985 | 6.78340763 | 6.826717129 | 7.290139032 |
| ENSG00000133640 | LRRIQ1 | -0.606697389 | 0.89458451 | -3.224061659 | NA | -3.830759048 | 2 | 3.686303414 | 2 | 2.868578001 | 2.2680111 | 2 | 2.225841198 |
| ENSG00000107951 | MTPAP | -1.78081899 | 0.002316839 | -2.051553129 | 0.00033142 | -3.832372119 | 8.257427002 | 9.338312924 | 9.624627483 | 7.433701013 | 7.41778467 | 7.308441828 | 6.998584547 |
| ENSG00000242485 | MRPL20 | -1.745749803 | 0.053757037 | -2.086687652 | 0.017626246 | -3.832437455 | 11.08909278 | 8.747909548 | 9.681375057 | 8.453061739 | 8.39382264 | 7.813294077 | 8.317521841 |
| ENSG00000099901 | RANBP1 | -1.764400934 | 0.000559325 | -2.069334105 | 3.52E-05 | -3.833735039 | 10.85038273 | 9.695596167 | 10.13584611 | 8.493293474 | 8.60823803 | 8.160127972 | 8.335797499 |
| ENSG00000198203 | SULT1C2 | -2.971855773 | 0.156934326 | -0.86237888 | 0.699250106 | -3.834234653 | 4.815609471 | 2 | 4.874587708 | 2.868578001 | 2.2680111 | 4.251304078 | 2.746601469 |
| ENSG00000087842 | PIR | -0.639701428 | 0.770739289 | -3.194763885 | 0.08973642 | -3.834465313 | 8.892187472 | 6.389896633 | 2 | 6.825617242 | 7.02417148 | 4.406883583 | 4.74829783 |
| ENSG00000111816 | FRK | -0.859330552 | 0.753351211 | -2.978867578 | 0.220679476 | -3.83819813 | 2 | 5.595927442 | 2 | 3.462030052 | 3.7864959 | 2.775316226 | 2.326754077 |
| ENSG00000275035 | | -0.310285079 | 0.591636403 | -3.529626893 | 1.21E-14 | -3.839911972 | 9.448362284 | 9.558016677 | 8.631919287 | 8.959612762 | 8.96600458 | 5.594558848 | 6.044874254 |
| ENSG00000105197 | TIMM50 | -2.029766069 | 0.060971292 | -1.810753622 | 0.094172652 | -3.840519691 | 11.72814722 | 8.747909548 | 9.69246155 | 8.615281294 | 8.54182783 | 8.798213877 | 8.793590064 |
| ENSG00000185721 | DRG1 | -1.93347528 | 3.56E-05 | -1.907166471 | 4.14E-05 | -3.840641751 | 10.87842776 | 10.47993704 | 9.809100877 | 8.532433657 | 8.5323134 | 8.438967212 | 8.668790472 |
| ENSG00000148290 | SURF1 | -2.072046376 | 0.013140909 | -1.771313451 | 0.035664619 | -3.843359827 | 10.00110988 | 9.583268459 | 7.691296561 | 7.218576229 | 7.4450836 | 7.580798201 | 7.678134128 |
| ENSG00000186106 | ANKRD46 | -2.231422692 | 0.001370226 | -1.61369549 | 0.024210868 | -3.845118182 | 9.204317094 | 7.790474954 | 8.143093325 | 6.476767609 | 6.17362678 | 7.260787065 | 6.487779256 |
| ENSG00000006757 | PNPLA4 | -1.62020862 | 0.037309591 | -2.225324907 | 0.002657411 | -3.845533526 | 7.53297911 | 6.04685525 | 7.777116083 | 5.839660368 | 5.62411716 | 5.357600875 | 4.990558504 |
| ENSG00000241852 | C8orf58 | -2.690439847 | 3.01E-09 | -1.155395502 | 0.016699027 | -3.845835349 | 8.19931886 | 9.067150531 | 9.149036566 | 6.283823264 | 6.19790195 | 7.799317009 | 7.647978997 |
| ENSG00000277868 | | 0.771339671 | 0.474821199 | -4.617859537 | 1.52E-06 | -3.846519866 | 5.70932728 | 4.11352228 | 5.39244166 | 5.987823101 | 5.7929843 | 2.267001569 | 2.551896972 |
| ENSG00000169564 | PCBP1 | -1.680726547 | 0.000480633 | -2.166161773 | 3.72E-06 | -3.84688832 | 11.96017193 | 12.60776686 | 13.00587874 | 10.89692786 | 10.9185917 | 10.19915101 | 10.61776649 |
| ENSG00000112333 | NR2E1 | -2.265463264 | NA | -1.584277594 | NA | -3.849740857 | 2 | 3.076665209 | 2 | 2.270786764 | 2 | 2.267001569 | 2.172608095 |
| ENSG00000161180 | CCDC116 | -1.707015647 | 0.503519889 | -2.142973562 | 0.377510751 | -3.849989209 | 2 | 5.72254323 | 2 | 3.227757282 | 3.15372745 | 2.911488689 | 3.012034821 |
| ENSG00000141480 | ARRB2 | -3.452887662 | 9.74E-08 | -0.397214844 | 0.605488001 | -3.850102506 | 8.285626337 | 7.099070191 | 6.958202216 | 4.387512679 | 4.40247105 | 7.349785676 | 7.0024032 |
| ENSG00000129437 | KLK14 | -0.804999641 | 0.851584865 | -3.046248923 | 0.42731388 | -3.851248564 | 4.00722531 | 2 | 2 | 2.695456494 | 2.59489879 | 2.3070652 | 2 |
| ENSG00000168078 | PBK | -2.221140542 | 0.404611241 | -1.631007353 | 0.541689634 | -3.852147895 | 2 | 6.728495673 | 2 | 3.566290607 | 3.45167086 | 3.393105702 | 4.30448772 |
| ENSG00000131876 | SNRPA1 | -1.816873126 | 4.44E-10 | -2.035949497 | 9.47E-13 | -3.852822623 | 8.75772933 | 9.00621767 | 9.196533591 | 7.210554257 | 7.20671394 | 6.997118176 | 6.994755759 |
| ENSG00000004975 | DVL2 | -2.276711233 | 1.87E-10 | -1.577401833 | 1.45E-05 | -3.854113066 | 10.19198706 | 10.3138733 | 10.77492406 | 8.076429088 | 8.29037844 | 9.03282106 | 8.706733286 |
| ENSG00000173141 | MRPL57 | -2.277933507 | 6.66E-06 | -1.578224869 | 0.002429355 | -3.856158375 | 10.48127392 | 9.256956408 | 10.05225195 | 7.772969103 | 7.7377845 | 8.435582605 | 8.454146986 |
| ENSG00000136856 | SLC2A8 | -2.68938828 | 2.13E-05 | -1.166934836 | 0.086492666 | -3.856323116 | 7.790810999 | 8.317412869 | 9.344432377 | 6.01571348 | 6.01871653 | 7.501007892 | 7.459438299 |
| ENSG00000214309 | MBLAC1 | -2.313935402 | 0.330996743 | -1.54514167 | 0.523941689 | -3.859077071 | 2 | 3.076665209 | 8.110401118 | 4.598938466 | 4.37434574 | 5.056402493 | 5.23734678 |
| ENSG00000258405 | ZNF578 | -0.123055046 | 0.882608621 | -3.73691165 | 3.08E-09 | -3.859966696 | 6.009025997 | 5.457120953 | 6.534109959 | 6.052081244 | 5.7929843 | 3.1504466 | 3.071407821 |
| ENSG00000182185 | RAD51B | -1.262039147 | 0.097603025 | -2.598036428 | 0.000178603 | -3.860075574 | 6.73718411 | 6.140560021 | 7.600047325 | 5.501219927 | 5.94414054 | 4.720468739 | 4.376876162 |
| ENSG00000127993 | RBM48 | -1.536561637 | 0.019027101 | -2.324107618 | 0.000182351 | -3.860669255 | 9.667789716 | 8.101365653 | 8.99637432 | 7.467753569 | 7.60490877 | 6.902223127 | 6.630324799 |
| ENSG00000276016 | | 1.801725424 | 0.183376109 | -5.662747663 | 2.35E-06 | -3.861022239 | 4.467311984 | 8.903033433 | 8.561105505 | 9.973493741 | 9.99411823 | 3.296906311 | 3.261853426 |
| ENSG00000184076 | UQCR10 | -2.212507752 | 2.03E-07 | -1.64981629 | 0.000142527 | -3.862324042 | 11.22998797 | 10.34373336 | 10.5677762 | 8.529212228 | 8.59611189 | 9.011762258 | 9.222253259 |
| ENSG00000197498 | RPF2 | -1.974792092 | 0.044706135 | -1.88767679 | 0.052955478 | -3.862468882 | 9.931427102 | 8.433041974 | 7.225742779 | 6.86703417 | 7.11620658 | 7.142599214 | 7.017577533 |
| ENSG00000179271 | GADD45GIP1 | -2.446650585 | 7.04E-09 | -1.41598486 | 0.001295182 | -3.862635445 | 11.28689767 | 10.43902632 | 11.25043551 | 8.564258394 | 8.64548836 | 9.489754807 | 9.755177237 |
| ENSG00000146830 | GIGYF1 | -1.718931088 | 0.000942464 | -2.144185925 | 2.25E-05 | -3.863117013 | 11.08508709 | 10.68045884 | 11.86432331 | 9.591010098 | 9.57093244 | 9.235857639 | 9.076359045 |
| ENSG00000114279 | FGF12 | -1.079259586 | 0.685162585 | -2.786061371 | 0.249055682 | -3.865320957 | 3.327711234 | 2 | 3.369317972 | 2.498691189 | 2.59489879 | 2.3070652 | 2.059860178 |
| ENSG00000129749 | CHRNA10 | -1.453422557 | 0.420476172 | -2.417494272 | 0.146465505 | -3.870916828 | 2 | 3.686303414 | 4.874587708 | 2.784612485 | 3.2184422 | 2.62493832 | 2.592996859 |
| ENSG00000198931 | APRT | -2.264612082 | 1.58E-09 | -1.607893212 | 2.58E-05 | -3.872505294 | 11.05263482 | 10.41578665 | 10.81605395 | 8.481674661 | 8.57927024 | 8.931137537 | 9.396328961 |
| ENSG00000172053 | QARS | -1.887774779 | 9.17E-10 | -1.98958197 | 7.62E-11 | -3.877356749 | 10.75779627 | 11.14636276 | 11.2690689 | 9.199389365 | 9.18725282 | 8.954974347 | 9.217743686 |
| ENSG00000166337 | TAF10 | -0.975071819 | 0.059920834 | -2.903984911 | 3.52E-10 | -3.87905673 | 9.730679447 | 8.806266215 | 9.624627483 | 8.48998335 | 8.45217004 | 6.344396176 | 6.795689611 |
| ENSG00000132010 | ZNF20 | 1.006776474 | 0.64718241 | -4.886194747 | 0.009972043 | -3.879418273 | 2 | 7.145017454 | 3.369317972 | 6.64692533 | 6.68951263 | 2.526562374 | 2.466008509 |
| ENSG00000110900 | TSPAN11 | -2.85905115 | 0.336346335 | -1.022008984 | 0.746521334 | -3.881060135 | 2 | 6.535001748 | 2 | 3.566290607 | 2.14022021 | 4.563936166 | 3.789585079 |
| ENSG00000181163 | NPM1 | -1.253882606 | 0.006082261 | -2.629183428 | 6.73E-10 | -3.883066034 | 13.18766392 | 12.40319236 | 12.30268094 | 11.4306175 | 11.4400142 | 10.00254443 | 10.12169598 |
| ENSG00000267041 | ZNF850 | -2.455002732 | 7.55E-05 | -1.428279617 | 0.024512417 | -3.883282349 | 6.138360429 | 6.78762695 | 7.285536013 | 4.598938466 | 4.53541753 | 5.688550058 | 5.197927245 |
| ENSG00000187049 | TMEM216 | -1.965100049 | 0.010246566 | -1.919038988 | 0.011668792 | -3.884139037 | 7.435667355 | 8.004992147 | 9.272377397 | 6.496695859 | 6.55815769 | 6.726875343 | 6.397863414 |
| ENSG00000196993 | NPIPB9 | 4.310627574 | 0.078977124 | -8.196111779 | 0.003680239 | -3.885484205 | 2 | 2 | 4.874587708 | 7.464384256 | 7.28433928 | 2 | 2 |
| ENSG00000167508 | MVD | -2.630594423 | 4.37E-05 | -1.256031582 | 0.069461561 | -3.886626005 | 11.29385609 | 10.18234215 | 10.01740919 | 8.017874397 | 7.99029569 | 8.849559031 | 9.74381649 |
| ENSG00000103175 | WFDC1 | -6.347586576 | 0.088075356 | 2.458955323 | 0.48356437 | -3.888631253 | 2 | 4.442710129 | 2 | 2 | 2 | 6.004008792 | 3.518055793 |
| ENSG00000149925 | ALDOA | 0.754807922 | 0.138897807 | -4.644005666 | 1.17E-26 | -3.889197745 | 13.2480265 | 12.70011329 | 12.47147373 | 13.57404708 | 13.6230809 | 7.838287442 | 8.520227711 |
| ENSG00000167107 | ACSF2 | -2.006363815 | 0.083215069 | -1.884840404 | 0.099932882 | -3.891204219 | 8.138771685 | 6.728495673 | 10.02619908 | 6.970413838 | 6.87658492 | 7.123093858 | 6.955898408 |
| ENSG00000113328 | CCNG1 | -2.29788585 | 1.79E-08 | -1.594349121 | 0.000138385 | -3.892234971 | 12.13503061 | 12.10787117 | 11.42009729 | 9.671381402 | 9.59092963 | 10.55053302 | 10.07531602 |
| ENSG00000105372 | RPS19 | -2.202437969 | 0.002511597 | -1.690129418 | 0.023933333 | -3.892567387 | 14.21465325 | 12.58815072 | 12.63123848 | 11.06929166 | 11.2419313 | 11.8028959 | 11.52303567 |
| ENSG00000079102 | RUNX1T1 | -2.867529076 | 0.000512189 | -1.025139082 | 0.258080827 | -3.892668159 | 8.228665467 | 8.876041558 | 6.534109959 | 5.501219927 | 5.34860058 | 7.222020406 | 7.114266974 |
| ENSG00000153774 | CFDP1 | -2.039603955 | 1.47E-06 | -1.853309167 | 1.22E-05 | -3.892913122 | 9.966689162 | 9.148351514 | 9.963514551 | 7.778406195 | 7.65545169 | 7.832294085 | 7.967075361 |
| ENSG00000132475 | H3F3B | -1.214826516 | 8.03E-05 | -2.67958432 | 4.79E-20 | -3.894410836 | 13.8494174 | 13.36796481 | 13.44661752 | 12.31872784 | 12.3926233 | 10.7763575 | 11.00156061 |

| | | | | | | | | | | | | | |
|---|---|---|---|---|---|---|---|---|---|---|---|---|---|
| ENSG00000130741 | EIF2S3 | -2.110548708 | 1.84E-05 | -1.787692098 | 0.000329966 | -3.898240806 | 12.62202739 | 11.54799449 | 11.79143574 | 9.989990706 | 9.92369416 | 10.14543562 | 10.40110911 |
| ENSG00000229833 | PET100 | -2.11327185 | 0.009616999 | -1.787693837 | 0.029998862 | -3.900965688 | 9.233563228 | 7.901717006 | 7.225742779 | 6.268449691 | 6.35006802 | 6.387074688 | 6.821857532 |
| ENSG00000162878 | PKDCC | -3.694071816 | 2.82E-11 | -0.207944937 | 0.762652241 | -3.902016753 | 7.435667355 | 8.414402626 | 7.600047325 | 4.387512679 | 4.53541753 | 7.927733561 | 7.402777233 |
| ENSG00000004059 | ARF5 | -1.693558566 | 0.007453224 | -2.20955286 | 0.000297907 | -3.903111426 | 11.25872341 | 10.39216648 | 9.703463497 | 8.887228971 | 8.92417737 | 8.298842011 | 8.485179747 |
| ENSG00000104679 | R3HCC1 | -2.389451038 | 7.79E-08 | -1.513970806 | 0.000947533 | -3.903421844 | 9.031919741 | 9.461450882 | 9.935792933 | 7.148924055 | 7.18258783 | 7.794921259 | 8.220043091 |
| ENSG00000114446 | IFT57 | -2.035599235 | 0.000166301 | -1.867999732 | 0.000564093 | -3.903598967 | 10.65345793 | 9.763446433 | 9.528916236 | 8.096156705 | 7.99722844 | 7.966556098 | 8.423144406 |
| ENSG00000136247 | ZDHHC4 | -1.642937886 | 1.89E-05 | -2.262337109 | 1.30E-09 | -3.905274995 | 9.48530526 | 10.24697497 | 9.809100877 | 8.244958867 | 8.26379462 | 7.726402307 | 7.558732759 |
| ENSG00000078549 | ADCYAP1R1 | -2.857267657 | 0.30511035 | -1.049781032 | 0.714833968 | -3.907048689 | 4.467311984 | 3.076665209 | 2 | 2.600425781 | 2 | 3.059555873 | 2.782576693 |
| ENSG00000112118 | MCM3 | -2.022542096 | 1.27E-14 | -1.884657509 | 3.15E-13 | -3.907199605 | 9.699577239 | 9.532315021 | 9.426401271 | 7.650823203 | 7.45854177 | 7.645704907 | 7.738907626 |
| ENSG00000181409 | AATK | -2.992329908 | 0.04707563 | -0.915282495 | 0.574881251 | -3.907612403 | 2 | 6.04685525 | 6.203069917 | 3.227757282 | 3.28037825 | 4.927401733 | 4.604965121 |
| ENSG00000126226 | PCID2 | -2.390825551 | 0.000325357 | -1.517985768 | 0.029053832 | -3.908811318 | 10.43166252 | 8.993716073 | 10.7325871 | 7.829061569 | 7.87709394 | 8.787190829 | 8.638437331 |
| ENSG00000174851 | YIF1A | -2.328154763 | 0.004901099 | -1.581150502 | 0.065337615 | -3.909305265 | 12.43010534 | 10.54992047 | 10.54349105 | 9.159317259 | 9.14000763 | 9.770158073 | 10.0054433 |
| ENSG00000100823 | APEX1 | -2.088626512 | 0.000646898 | -1.820919803 | 0.003232449 | -3.909546315 | 12.32703082 | 11.1798568 | 11.03218278 | 9.555824233 | 9.55143939 | 9.532363398 | 10.0598241 |
| ENSG00000071243 | ING3 | -1.515418789 | 0.340882919 | -2.395273179 | 0.108394135 | -3.910691967 | 9.174465797 | 7.232742069 | 4.058998049 | 6.490083648 | 6.47451726 | 5.883320383 | 5.385063124 |
| ENSG00000176912 | TYMSOS | -2.032935026 | 0.457936387 | -1.881014387 | 0.474487232 | -3.913949413 | 4.467311984 | 2 | 2 | 2.3892346 | 2.49394783 | 2.62493832 | 2.326754077 |
| ENSG00000064201 | TSPAN32 | -5.378269045 | 0.193918683 | 1.462625117 | 0.721847383 | -3.915643928 | 2 | 3.686303414 | 2 | 2 | 2 | 3.810410249 | 3.496556178 |
| ENSG00000163611 | SPICE1 | -0.183942839 | 0.930416944 | -3.731927815 | 0.026364101 | -3.915870654 | 2 | 6.951736442 | 8.206335842 | 6.975156635 | 6.94511186 | 3.981090143 | 3.67940853 |
| ENSG00000213218 | CSH2 | -2.02005436 | 1.72E-08 | -1.898575248 | 1.17E-07 | -3.918629607 | 12.61092608 | 12.35391197 | 11.94069548 | 10.39581804 | 10.2213052 | 10.58097861 | 10.26647028 |
| ENSG00000102897 | LYRM1 | -2.356941602 | 0.001097653 | -1.563509017 | 0.03672502 | -3.92045062 | 9.624286623 | 7.73146271 | 9.132850258 | 6.872128674 | 6.52655952 | 7.523500045 | 7.434215826 |
| ENSG00000166788 | SAAL1 | -1.951194139 | 2.04E-05 | -1.971598675 | 1.09E-05 | -3.922792814 | 7.624139768 | 7.467835385 | 7.646393253 | 5.37770462 | 5.98191036 | 5.578280598 | 5.799216178 |
| ENSG00000170860 | LSM3 | -2.327033938 | 0.000247921 | -1.600595461 | 0.014861287 | -3.927629399 | 9.90440318 | 8.487557258 | 8.805342542 | 6.989292186 | 6.82556437 | 7.385306888 | 7.816873723 |
| ENSG00000154734 | ADAMTS1 | -2.988339337 | 1.88E-09 | -0.941731744 | 0.089211519 | -3.930071081 | 13.36414695 | 14.33244562 | 14.1682348 | 11.05732061 | 10.9956201 | 12.65313458 | 13.39512515 |
| ENSG00000162572 | SCNN1D | -6.674484607 | 0.038139545 | 2.743766215 | 0.346792247 | -3.930718392 | 2 | 4.710564154 | 2 | 2 | 2 | 5.960710379 | 5.616947984 |
| ENSG00000154917 | RAB6B | -4.378820634 | 3.60E-05 | 0.447536513 | 0.727840402 | -3.931284121 | 8.998234871 | 6.140560021 | 6.80318718 | 4.070463721 | 3.69849104 | 8.362607519 | 8.213468956 |
| ENSG00000070190 | DAPP1 | -2.69666285 | 0.461270807 | -1.235246424 | 0.735593063 | -3.931909274 | 3.327711234 | 2 | 3.369317972 | 2.270786764 | 2.14022021 | 2.885263063 | 2 |
| ENSG00000169891 | REPS2 | -3.909409499 | 0.035924136 | -0.022701537 | 0.991906099 | -3.932111036 | 4.00722531 | 2 | 6.073378139 | 2.3892346 | 2.59489879 | 5.068107405 | 4.522269241 |
| ENSG00000113643 | RARS | -1.603996342 | 1.72E-05 | -2.328429289 | 1.24E-10 | -3.932425631 | 11.29732276 | 11.1062775 | 11.83710798 | 9.796566011 | 9.89552277 | 9.167793558 | 9.084037526 |
| ENSG00000153303 | FRMD1 | -6.349006909 | 0.069096826 | 2.414250931 | 0.458681676 | -3.934755977 | 2 | 2 | 4.523675391 | 2 | 2 | 5.233158834 | 5.171036168 |
| ENSG00000269113 | TRABD2B | -4.656984455 | 0.085124351 | 0.720073454 | 0.762230217 | -3.936911002 | 3.327711234 | 3.686303414 | 2 | 2.141737246 | 2 | 3.968654369 | 3.236138336 |
| ENSG00000148303 | RPL7A | -2.357877165 | 4.83E-05 | -1.579481534 | 0.008812882 | -3.937358699 | 13.72622802 | 12.49142644 | 12.49857094 | 10.63074617 | 10.7181349 | 11.36441926 | 11.53461925 |
| ENSG00000164587 | RPS14 | -2.1375451 | 0.000306209 | -1.802800252 | 0.002681406 | -3.940345353 | 13.61046475 | 12.28504423 | 12.43563386 | 10.75418751 | 10.7887607 | 11.04735028 | 11.16166619 |
| ENSG00000168672 | FAM84B | -3.418437295 | 0.013205398 | -0.52504514 | 0.744221494 | -3.943482436 | 6.563806279 | 7.638155693 | 6.073378139 | 4.598938466 | 2.49394783 | 7.033394818 | 5.300743019 |
| ENSG00000203724 | C1orf53 | -1.678364247 | 0.452192995 | -2.268098977 | 0.28290231 | -3.946463224 | 6.73718411 | 3.076665209 | 2 | 3.95996233 | 3.86943969 | 3.235948948 | 3.735548302 |
| ENSG00000171720 | HDAC3 | -2.052798017 | 0.000167361 | -1.89434553 | 0.000529876 | -3.947143547 | 10.99829153 | 9.890264534 | 11.01903267 | 8.692189072 | 8.66593596 | 8.558922783 | 9.068639477 |
| ENSG00000196353 | CPNE4 | -0.682261839 | NA | -3.26627238 | NA | -3.94853422 | 2 | 2 | 3.369317972 | 2.600425781 | 2.14022021 | 2.139668783 | 2 |
| ENSG00000273889 | | 3.326218139 | 0.023775024 | -7.277302715 | 0.000509009 | -3.951084576 | 3.327711234 | 4.93638933 | 2 | 6.951285179 | 6.86652414 | 2.048074405 | 2 |
| ENSG00000282712 | | -2.644574848 | 3.77E-06 | -1.308509555 | 0.032293519 | -3.953084403 | 10.62062599 | 9.298208073 | 10.5677762 | 7.62389457 | 7.68437616 | 9.129204663 | 8.797990868 |
| ENSG00000166377 | ATP9B | -2.048193411 | 0.219718753 | -1.906749228 | 0.248813053 | -3.954942639 | 4.00722531 | 7.819099893 | 9.926432745 | 6.573765157 | 6.73446058 | 6.544089217 | 7.004308743 |
| ENSG00000125434 | SLC25A35 | -2.530254191 | 0.080615128 | -1.426739883 | 0.342972522 | -3.956994075 | 8.540903468 | 4.710564154 | 5.59507254 | 4.598938466 | 5.07918227 | 5.783905948 | 5.917188835 |
| ENSG00000116668 | SWT1 | -0.836177203 | 0.34242241 | -3.121738262 | 3.85E-05 | -3.957915465 | 8.367060731 | 7.189546432 | 6.43197674 | 6.727838783 | 6.74548212 | 4.813804156 | 4.456980017 |
| ENSG00000136425 | CIB2 | -2.910146109 | 0.000120788 | -1.049430483 | 0.202663689 | -3.959576593 | 6.138360429 | 8.029695761 | 8.007629717 | 4.907595604 | 4.87352701 | 6.625560108 | 6.580104496 |
| ENSG00000162722 | TRIM58 | -2.282268355 | 0.000942464 | -1.679119516 | 0.016000913 | -3.961387871 | 6.366709671 | 6.228547981 | 7.502634626 | 4.804776789 | 4.60967607 | 5.376552034 | 5.080008298 |
| ENSG00000150867 | PIP4K2A | -2.097563948 | 0.000133101 | -1.863896758 | 0.000738183 | -3.961460706 | 9.829762239 | 9.916994052 | 10.98787477 | 8.323303759 | 8.19912382 | 8.560995239 | 8.422432 |
| ENSG00000100038 | TOP3B | 1.178871387 | 0.520431515 | -5.140376719 | 0.001103414 | -3.961505332 | 7.579279349 | 5.595927442 | 2 | 7.517371202 | 7.44169935 | 2.62493832 | 2.632958241 |
| ENSG00000178187 | ZNF454 | -1.684439742 | 0.465624329 | -2.277636103 | 0.297700044 | -3.962075844 | 7.750913381 | 3.686303414 | 2 | 4.670864177 | 4.811949 | 4.713032626 | 3.580696614 |
| ENSG00000188612 | SUMO2 | -0.677781373 | 0.112733934 | -3.284831852 | 5.07E-19 | -3.962613225 | 11.35501623 | 10.53705456 | 10.94216274 | 10.29320552 | 10.3181476 | 7.726402307 | 7.7169976 |
| ENSG00000144554 | FANCD2 | -1.662381983 | 0.01538745 | -2.303103448 | 0.00039122 | -3.965485431 | 6.563806279 | 6.602424674 | 5.39244166 | 4.71689672 | 4.85329164 | 4.144793515 | 4.365060851 |
| ENSG00000152520 | PAN3 | -1.416425746 | 1.46E-05 | -2.550269735 | 3.81E-16 | -3.966695481 | 10.26462289 | 9.726146444 | 10.18377222 | 8.726347845 | 8.60370266 | 7.47706242 | 7.618424749 |
| ENSG00000171303 | KCNK3 | -3.585325644 | 0.170442388 | -0.38265554 | 0.897819577 | -3.967981184 | 2 | 5.946638857 | 2 | 2.270786764 | 2.68924543 | 4.757085024 | 3.474731315 |
| ENSG00000204922 | UQCC3 | -2.058449394 | 0.002456458 | -1.90968772 | 0.004838638 | -3.968137114 | 7.624139768 | 7.315436877 | 8.805342542 | 6.18083559 | 5.94414054 | 6.121005698 | 6.289521179 |
| ENSG00000146966 | DENND2A | -4.626790015 | 0.007864549 | 0.657661307 | 0.746440634 | -3.969128708 | 4.467311984 | 8.356994601 | 3.369317972 | 3.406930997 | 2.77779907 | 6.947285242 | 7.985520618 |
| ENSG00000004776 | HSPB6 | -2.93652211 | 7.80E-14 | -1.03544146 | 0.015286773 | -3.971963571 | 11.98618033 | 11.60869109 | 11.23913884 | 8.71221318 | 8.72559535 | 10.35239254 | 10.82845131 |
| ENSG00000165349 | SLC7A3 | -6.111975319 | 0.022595921 | 2.136030136 | 0.343763347 | -3.975945184 | 2 | 3.686303414 | 3.369317972 | 2 | 2 | 5.163941955 | 4.175132639 |
| ENSG00000278615 | C11orf98 | 2.32441749 | 0.017885637 | -6.302472513 | 3.91E-09 | -3.978055024 | 4.00722531 | 6.389896633 | 5.156626401 | 7.823812411 | 7.67863747 | 2.139668783 | 2.225841198 |
| ENSG00000188511 | C22orf34 | -4.714861447 | 0.349875999 | 0.736465838 | NA | -3.978395609 | 3.327711234 | 2 | 2 | 2 | 2 | 3.12825509 | 2.551896972 |
| ENSG00000136444 | RSAD1 | -2.35799345 | 0.021848633 | -1.623926875 | 0.125080013 | -3.981920326 | 10.16187685 | 7.467835385 | 10.56174313 | 7.477814553 | 7.61696151 | 8.431623835 | 8.089704419 |
| ENSG00000117481 | NSUN4 | -2.072487319 | 0.001588398 | -1.910888006 | 0.00367301 | -3.983375325 | 9.624286623 | 9.00621767 | 10.62675862 | 7.865280446 | 7.84158259 | 8.133219123 | 7.880504067 |
| ENSG00000197077 | KIAA1671 | -4.494077293 | 0.000258322 | 0.509837767 | 0.732916452 | -3.984239526 | 5.095959458 | 9.125613817 | 9.014156722 | 4.739373666 | 3.7864959 | 8.862639958 | 9.193729987 |
| ENSG00000168282 | MGAT2 | 0.017811008 | 0.97365672 | -4.005556737 | 1.06E-24 | -3.987745729 | 9.646202137 | 9.461450882 | 10.02619908 | 9.726568065 | 9.76553205 | 5.502666283 | 6.085042537 |
| ENSG00000189403 | HMGB1 | -1.013176637 | 0.000102258 | -2.975369944 | 6.22E-34 | -3.988546582 | 11.73583256 | 11.56921405 | 11.32712828 | 10.56659341 | 10.5185694 | 8.626778874 | 8.555744353 |
| ENSG00000149182 | ARFGAP2 | -1.96124918 | 1.15E-06 | -2.028499237 | 4.02E-07 | -3.989748418 | 10.81212061 | 10.60438382 | 11.3657305 | 8.99156386 | 9.0282492 | 8.745429394 | 9.117433659 |
| ENSG00000105409 | ATP1A3 | -0.814787487 | 0.799141442 | -3.175181128 | NA | -3.989968614 | 2 | 3.686303414 | 2 | 2.3892346 | 2.68924543 | 2.183373728 | 2.059860178 |
| ENSG00000203780 | FANK1 | -1.8533706 | 0.52209421 | -2.137700197 | 0.443907894 | -3.991070797 | 7.975256505 | 2 | 2 | 4.867343068 | 4.60967607 | 4.434275199 | 4.574506939 |
| ENSG00000156463 | SH3RF2 | -3.828005727 | 1.68E-05 | -0.164834954 | 0.886529537 | -3.992840682 | 8.16936288 | 6.602424674 | 9.031722608 | 4.825935136 | 4.43005853 | 8.424242888 | 7.610940609 |
| ENSG00000101049 | SGK2 | -1.96427339 | 0.371250007 | -2.029262534 | 0.340738496 | -3.993535924 | 4.00722531 | 2 | 5.59507254 | 3.162721474 | 3.08597292 | 3.584558147 | 2.326754077 |
| ENSG00000145779 | TNFAIP8 | -1.76671239 | 0.000311823 | -2.228631861 | 3.04E-06 | -3.995344251 | 10.60392551 | 9.648522413 | 10.10298738 | 8.50645854 | 8.31648126 | 7.661243884 | 8.206037069 |
| ENSG00000232112 | TMA7 | -0.436472147 | 0.53856935 | -3.559289503 | 6.02E-10 | -3.99576165 | 9.751047882 | 8.505280662 | 8.561105505 | 8.565831351 | 8.68466696 | 5.498346994 | 5.717942682 |
| ENSG00000140678 | ITGAX | -5.313768404 | 0.000214851 | 1.315936527 | 0.365712823 | -3.997831877 | 5.532353208 | 4.442710129 | 6.534109959 | 2.498691189 | 2.2680111 | 7.779874724 | 5.020995508 |
| ENSG00000170085 | SIMC1 | -2.08239117 | 0.063652249 | -1.916490618 | 0.086034904 | -3.998881788 | 6.653098118 | 7.819099893 | 9.767769415 | 6.598566398 | 6.60731796 | 6.854018691 | 6.664466987 |
| ENSG00000134317 | GRHL1 | -3.141792377 | 0.231366574 | -0.857519077 | 0.760773185 | -3.999311455 | 2 | 2 | 5.59507254 | 2.498691189 | 2.49394783 | 3.993419637 | 3.012034821 |
| ENSG00000173599 | PC | -3.452268164 | 7.91E-09 | -0.549258392 | 0.437595013 | -4.001526556 | 10.03472846 | 8.791896851 | 10.11951029 | 6.283823264 | 6.45450701 | 8.858433249 | 9.497542084 |
| ENSG00000113272 | THG1L | -1.542808826 | 0.098785844 | -2.458966743 | 0.004927313 | -4.001775569 | 9.448362284 | 8.607278467 | 6.882775709 | 7.123515401 | 7.14977896 | 6.22922222 | 6.283256815 |
| ENSG00000095002 | MSH2 | -1.940580305 | 0.166634602 | -2.063738248 | 0.130907295 | -4.004318554 | 5.095959458 | 9.067150531 | 9.819251342 | 7.165617905 | 6.88657603 | 7.054142452 | 6.757786576 |
| ENSG00000111859 | NEDD9 | -4.664254769 | 3.58E-09 | 0.659504155 | 0.483694039 | -4.004750614 | 10.84565509 | 9.648522413 | 8.65477198 | 5.720439317 | 5.13053105 | 10.42644164 | 10.83448877 |
| ENSG00000157404 | KIT | -6.585958835 | 2.35E-06 | 2.580644073 | 0.038115632 | -4.005314762 | 3.327711234 | 6.899080544 | 6.882775709 | 2.141737246 | 2.38539787 | 8.774297678 | 9.001538388 |
| ENSG00000154358 | OBSCN | -2.864076839 | 0.002641029 | -1.141405662 | 0.271435731 | -4.005482501 | 7.384433967 | 8.029695761 | 9.897982195 | 6.12217689 | 5.99120031 | 7.901804236 | 7.509939612 |
| ENSG00000140983 | RHOT2 | -2.247528706 | 0.001122662 | -1.758374401 | 0.012725186 | -4.005903107 | 10.96815916 | 9.687856435 | 11.54767436 | 8.649804831 | 8.71862106 | 9.068221759 | 9.264604232 |
| ENSG00000214941 | ZSWIM7 | -1.657717807 | 0.029528379 | -2.349512031 | 0.001185546 | -4.007229837 | 7.031698319 | 7.701025996 | 8.845626171 | 6.321555392 | 6.55189318 | 5.798032389 | 5.781546943 |
| ENSG00000118640 | VAMP8 | -5.20613658 | 4.93E-11 | 1.197480927 | 0.112264981 | -4.008655653 | 5.70932728 | 7.145017454 | 6.80318718 | 2.947924294 | 2.49394783 | 7.871230734 | 7.822284765 |
| ENSG00000095380 | NANS | -2.302644131 | 5.49E-06 | -1.70672696 | 0.000987139 | -4.009371091 | 11.21906289 | 10.25223258 | 10.35401885 | 8.347282003 | 8.42671895 | 8.68304821 | 9.222253259 |
| ENSG00000132780 | NASP | -1.562459675 | 4.23E-05 | -2.447728359 | 2.84E-11 | -4.010188033 | 10.19198706 | 9.988044813 | 10.67226762 | 8.833394974 | 8.67749148 | 7.735606062 | 8.017950573 |
| ENSG00000178038 | ALS2CL | -3.23045781 | 9.46E-06 | -0.782590397 | 0.346522734 | -4.013048208 | 8.313285031 | 7.761270542 | 9.553445883 | 5.363304456 | 5.83486613 | 7.860333006 | 8.06618078 |
| ENSG00000118514 | ALDH8A1 | -0.995986601 | 0.647922415 | -3.017406365 | 0.121270625 | -4.013392966 | 3.327711234 | 2 | 5.59507254 | 3.462030052 | 3.74316441 | 2.775316226 | 2.374681325 |
| ENSG00000107872 | FBXL15 | -2.749073429 | 0.001201769 | -1.264573534 | 0.165497653 | -4.013646964 | 9.781071289 | 7.538394143 | 8.35328246 | 6.043074888 | 6.29861567 | 7.765568476 | 7.432801564 |
| ENSG00000196812 | ZSCAN16 | -2.09293164 | 0.128681564 | -1.921146846 | 0.156413245 | -4.014078486 | 7.27616306 | 5.838937808 | 3.369317972 | 4.54891529 | 4.12711147 | 4.45225155 | 4.532869193 |
| ENSG00000258366 | RTEL1 | 0.201185467 | 0.929816715 | -4.217600115 | 0.01949823 | -4.016414648 | 8.836089686 | 6.389896633 | 2 | 7.674344958 | 7.72670378 | 3.782262616 | 3.735548302 |
| ENSG00000163624 | CDS1 | -4.544913561 | 0.001261372 | 0.527691741 | 0.678808079 | -4.01722182 | 4.00722531 | 4.442710129 | 4.058998049 | 2.141737246 | 2.2680111 | 4.813804156 | 4.411752372 |
| ENSG00000107672 | NSMCE4A | -1.776068587 | 0.000414663 | -2.243182897 | 4.38E-06 | -4.019251484 | 9.410448395 | 8.376385542 | 8.784770914 | 7.035434774 | 7.29187696 | 6.848938847 | 6.572418365 |
| ENSG00000276537 | | 2.127586277 | 0.675664186 | -6.15290235 | 0.207247817 | -4.025316073 | 3.327711234 | 2 | 2 | 4.387512679 | 2.14022021 | 2 | 2 |
| ENSG00000111012 | CYP27B1 | -1.535223147 | 0.586969261 | -2.491456204 | 0.348235366 | -4.02667935 | 2 | 2 | 6.958202216 | 3.997744386 | 4.2865223 | 3.709392871 | 3.183290849 |
| ENSG00000084092 | NOA1 | -2.290291142 | 0.042563867 | -1.738638103 | 0.129312271 | -4.028929244 | 10.88304938 | 7.274681896 | 10.06083273 | 7.734324667 | 7.76240929 | 8.368535975 | 8.210170621 |
| ENSG00000134460 | IL2RA | -4.714863752 | 0.346842553 | 0.683679441 | NA | -4.03118431 | 2 | 2 | 3.369317972 | 2 | 2 | 3.059555873 | 2.592996859 |
| ENSG00000106538 | RARRES2 | -2.408903693 | 0.00100424 | -1.622696675 | 0.030216428 | -4.031600367 | 6.563806279 | 7.315436877 | 7.225742779 | 4.907595604 | 4.76938658 | 4.792794928 | 6.029985803 |
| ENSG00000004777 | ARHGAP33 | -2.623391007 | 0.097604871 | -1.416296178 | 0.388467305 | -4.039687185 | 3.327711234 | 5.838937808 | 8.206335842 | 4.623314311 | 4.40247105 | 5.318934974 | 5.79040861 |
| ENSG00000071082 | RPL31 | -2.104525544 | 0.008305601 | -1.940907626 | 0.015419052 | -4.04543317 | 13.4089001 | 11.54799449 | 11.75752717 | 10.16260233 | 10.594821 | 10.57342723 | 10.54239449 |
| ENSG00000105514 | RAB3D | -1.975549148 | 3.61E-05 | -2.072133493 | 1.12E-05 | -4.047682641 | 8.313285031 | 9.067150531 | 9.06622548 | 6.86703417 | 6.96881778 | 7.087593115 | 6.504031956 |
| ENSG00000265241 | RBM8A | -1.710194665 | 0.001772867 | -2.338617076 | 9.46E-06 | -4.048811741 | 11.63805915 | 10.43902632 | 10.69999858 | 9.331013912 | 9.30367823 | 8.590716036 | 8.790280628 |
| ENSG00000116991 | SIPA1L2 | -2.23045264 | 2.44E-09 | -1.818502483 | 1.15E-06 | -4.048955123 | 8.716893051 | 9.298208073 | 9.385998894 | 7.080147179 | 6.85130017 | 7.318276922 | 7.421437279 |
| ENSG00000163947 | ARHGEF3 | -2.277131909 | 1.80E-08 | -1.773408352 | 9.79E-06 | -4.050540261 | 7.867441304 | 8.124485672 | 7.734844356 | 5.85002558 | 5.6121243 | 6.384736493 | 6.003555476 |
| ENSG00000188001 | TPRG1 | -2.319502576 | 0.394151052 | -1.734428368 | 0.517691224 | -4.053930944 | 2 | 2 | 4.874587708 | 2.270786764 | 2.68924543 | 2.746468522 | 2.632958241 |
| ENSG00000125735 | TNFSF14 | -2.008647228 | 0.439449514 | -2.047251027 | 0.416586577 | -4.055898255 | 4.00722531 | 3.686303414 | 5.156626401 | 3.406930997 | 2.59489879 | 3.632645806 | 2 |
| ENSG00000060688 | SNRNP40 | -1.835455387 | 0.006474078 | -2.220840585 | 0.000702882 | -4.056295972 | 10.3933013 | 8.848535039 | 9.149036566 | 7.831679003 | 7.7866208 | 7.439255893 | 7.424286753 |
| ENSG00000134825 | TMEM258 | -1.579212179 | 0.005110816 | -2.478577493 | 3.83E-06 | -4.057789672 | 10.68025843 | 9.367676479 | 9.888372618 | 8.483340231 | 8.53707846 | 7.630983525 | 7.618424749 |
| ENSG00000124608 | AARS2 | -2.454335495 | 1.09E-05 | -1.604343016 | 0.005416742 | -4.058678511 | 9.159305284 | 8.557180776 | 9.868959282 | 6.841288255 | 6.91125523 | 7.86872317 | 7.519298205 |
| ENSG00000108591 | DRG2 | -1.62289858 | 0.00099561 | -2.43628185 | 2.47E-07 | -4.05918043 | 9.761125267 | 8.717821467 | 9.658943166 | 7.860161659 | 7.81828345 | 7.093332266 | 6.994755759 |
| ENSG00000169688 | MT1B | 3.102662925 | 0.349375154 | -7.162250279 | 0.038778125 | -4.059587353 | 3.327711234 | 2 | 3.369317972 | 2.270786764 | 6.16544343 | 2 | 2 |
| ENSG00000143258 | USP21 | -2.261615406 | 0.000351568 | -1.798032375 | 0.005237084 | -4.059647781 | 9.710019321 | 8.297207175 | 9.798878489 | 7.148924055 | 7.20671394 | 7.760166845 | 7.488317792 |

| | | | | | | | | | | | | | |
|---|---|---|---|---|---|---|---|---|---|---|---|---|---|
| ENSG00000206052 | DOK6 | 1.405407403 | 0.644450822 | -5.469115644 | 0.05396136 | -4.063708241 | 2 | 4.93638933 | 2 | 4.76150579 | 4.89348248 | 2.139668783 | 2 |
| ENSG00000158864 | NDUFS2 | -1.606048985 | 0.034923735 | -2.458644412 | 0.000632311 | -4.064693397 | 12.12140479 | 10.59613751 | 10.46808205 | 9.677941812 | 9.66318013 | 8.6083592 | 9.011046713 |
| ENSG00000046604 | DSG2 | -1.603137284 | 0.21912951 | -2.465422082 | 0.044648908 | -4.068559365 | 6.96351253 | 11.07694755 | 10.36098398 | 8.595396228 | 8.67605209 | 8.105088837 | 7.375063227 |
| ENSG00000112299 | VNN1 | 0.447035016 | 0.908850677 | -4.515773881 | 0.166491674 | -4.068738865 | 2 | 2 | 5.39244166 | 4.329433326 | 4.45712836 | 2.34604626 | 2 |
| ENSG00000174547 | MRPL11 | -2.559584611 | 4.10E-08 | -1.511395897 | 0.00174195 | -4.070980508 | 8.836089686 | 9.04308532 | 9.624627483 | 6.456560224 | 6.89649843 | 7.473766117 | 7.917377332 |
| ENSG00000119711 | ALDH6A1 | -2.361555662 | 9.30E-07 | -1.71107761 | 0.000496194 | -4.072633272 | 10.22148166 | 9.246456519 | 9.935792933 | 7.565338955 | 7.46855365 | 7.86621124 | 8.398723071 |
| ENSG00000143149 | ALDH9A1 | -1.876795081 | 0.000107715 | -2.1962839 | 3.90E-06 | -4.073078981 | 11.52432667 | 11.2017618 | 10.48730704 | 9.227487791 | 9.29807478 | 8.696779192 | 9.158655025 |
| ENSG00000161981 | SNRNP25 | -2.106658025 | 2.43E-05 | -1.971925492 | 7.31E-05 | -4.078583517 | 9.579430841 | 8.94259736 | 8.585098385 | 6.989292186 | 7.05550676 | 6.884127769 | 7.380942161 |
| ENSG00000152582 | SPEF2 | -1.021502616 | 0.358192878 | -3.057134067 | 0.001780787 | -4.078636683 | 7.579279349 | 6.311476754 | 5.156626401 | 5.808110275 | 5.60003091 | 4.301748724 | 3.560117069 |
| ENSG00000183117 | CSMD1 | -3.274287034 | NA | -0.805382712 | NA | -4.079669746 | 2 | 2 | 3.369317972 | 2 | 2.14022021 | 2.34604626 | 2.374681325 |
| ENSG00000188010 | MORN2 | -2.013108095 | 0.059053709 | -2.06872979 | 0.048121028 | -4.081837885 | 8.963744667 | 6.535001748 | 6.718950348 | 5.900763176 | 5.93454154 | 5.680947153 | 6.029985803 |
| ENSG00000123444 | KBTBD4 | -1.491339268 | 0.164155144 | -2.591119054 | 0.009186901 | -4.082458322 | 9.829762239 | 7.701025996 | 7.285536013 | 7.277360022 | 7.24604864 | 6.121005698 | 6.283256815 |
| ENSG00000171316 | CHD7 | -3.267178547 | 0.072641384 | -0.816510901 | 0.688732389 | -4.083689448 | 2 | 8.623598428 | 7.552162999 | 4.647285135 | 4.50979031 | 7.249266052 | 6.202567011 |
| ENSG00000100804 | PSMB5 | -2.287111303 | 2.00E-11 | -1.798618707 | 1.80E-07 | -4.08573001 | 12.75277152 | 12.12373461 | 12.47147373 | 10.18529247 | 10.1913912 | 10.49633349 | 10.83475651 |
| ENSG00000273899 | NOL12 | -1.454610926 | 0.024278747 | -2.631214326 | 1.24E-05 | -4.085825253 | 8.285626337 | 7.051611315 | 8.408111533 | 6.436065776 | 6.75642009 | 5.578280598 | 5.419744333 |
| ENSG00000125877 | ITPA | -1.745642414 | 0.001757739 | -2.340218575 | 1.40E-05 | -4.085860988 | 10.26462289 | 8.955547846 | 9.788583153 | 8.040677544 | 8.02463101 | 7.297296608 | 7.586981404 |
| ENSG00000165512 | ZNF22 | -1.872028989 | 0.000217137 | -2.213857035 | 7.73E-06 | -4.085886024 | 9.397585735 | 8.623598428 | 9.212024683 | 7.303993732 | 7.22651534 | 7.267148095 | 6.498634707 |
| ENSG00000101445 | PPP1R16B | -4.714862249 | 0.350187134 | 0.628868624 | NA | -4.085993625 | 3.327711234 | 2 | 2 | 2 | 2 | 2.987421008 | 2.632958241 |
| ENSG00000206561 | COLQ | -1.092926828 | 0.52243871 | -2.993115709 | 0.048466418 | -4.086042537 | 2 | 6.535001748 | 6.322058544 | 5.003571641 | 4.79082474 | 3.616794016 | 3.261853426 |
| ENSG00000101082 | SLA2 | -2.272291722 | NA | -1.817334692 | NA | -4.089626414 | 2 | 2 | 3.369317972 | 2.141737246 | 2.14022021 | 2.3070652 | 2.059860178 |
| ENSG00000167272 | POP5 | -2.432244185 | 0.002164759 | -1.6595707 | 0.042927735 | -4.091814885 | 9.533135543 | 7.467835385 | 9.165043284 | 6.61689148 | 6.52655952 | 6.995586672 | 7.58443608 |
| ENSG00000157456 | CCNB2 | -3.117936322 | 0.008633495 | -0.974865279 | 0.456415864 | -4.092801601 | 5.532353208 | 8.147241016 | 5.39244166 | 4.105493497 | 4.22485516 | 6.237020416 | 5.803599879 |
| ENSG00000213337 | ANKRD39 | -1.882584646 | 0.237679353 | -2.21124893 | 0.150762161 | -4.093833576 | 3.327711234 | 7.819099893 | 8.845626171 | 6.034011955 | 6.04571775 | 5.650128416 | 5.812327524 |
| ENSG00000108528 | SLC25A11 | -2.521856677 | 0.000352701 | -1.57818069 | 0.032735676 | -4.100037367 | 11.61045549 | 10.00059544 | 10.07784268 | 8.270263574 | 8.25224987 | 9.112345273 | 9.274901872 |
| ENSG00000148688 | RPP30 | -1.432975224 | 4.29E-05 | -2.668084107 | 1.46E-15 | -4.101059331 | 9.081016759 | 8.505280662 | 8.884815469 | 7.402350992 | 7.44845993 | 6.310306306 | 6.155328188 |
| ENSG00000142082 | SIRT3 | -2.461833108 | 0.005262242 | -1.640730414 | 0.072675764 | -4.102563522 | 10.01801709 | 8.557180776 | 7.600047325 | 6.64692533 | 6.65485813 | 7.431347676 | 7.462213768 |
| ENSG00000204310 | AGPAT1 | 0.410878677 | 0.638989421 | -4.514703695 | 7.08E-11 | -4.103825018 | 9.886101849 | 9.962611189 | 8.585098385 | 10.00630116 | 10.0147629 | 4.547306681 | 5.708623051 |
| ENSG00000104960 | PTOV1 | -1.841406574 | 0.09807061 | -2.264710257 | 0.034772335 | -4.106116831 | 8.39321512 | 11.41132475 | 11.74958762 | 9.26766905 | 9.22711441 | 8.721540982 | 8.926146458 |
| ENSG00000228439 | TSTD3 | -1.799717749 | 0.195586526 | -2.307060351 | 0.08238425 | -4.1067781 | 5.866944937 | 4.11352228 | 7.734844356 | 4.825935136 | 5.00774437 | 4.122514202 | 4.784231301 |
| ENSG00000143033 | MTF2 | -1.220617891 | 0.077120195 | -2.887568314 | 3.65E-06 | -4.108186205 | 9.624286623 | 9.277729677 | 8.042706615 | 7.883054517 | 7.95274764 | 6.501560901 | 6.070564958 |
| ENSG00000146143 | PRIM2 | -1.393397885 | 0.144179754 | -2.716230668 | 0.001888215 | -4.109628553 | 8.16936288 | 6.951736442 | 9.426401271 | 7.140504127 | 7.14977896 | 5.9289675 | 5.8339184 |
| ENSG00000158792 | SPATA2L | -2.870413146 | 3.34E-06 | -1.241553185 | 0.06185232 | -4.111966331 | 9.781071289 | 8.433041974 | 8.511888463 | 6.18083559 | 6.30607931 | 7.811554331 | 7.830900438 |
| ENSG00000150782 | IL18 | -4.138416799 | 0.160411864 | 0.025762837 | 0.992845825 | -4.112653962 | 2 | 3.076665209 | 3.369317972 | 2 | 2.14022021 | 2.962551659 | 2.918126211 |
| ENSG00000182512 | GLRX5 | -2.687893699 | 6.96E-11 | -1.428580325 | 0.00085782 | -4.116474024 | 9.371511179 | 9.74864196 | 10.26027233 | 7.226553841 | 7.14145882 | 8.322245777 | 8.512884393 |
| ENSG00000197958 | RPL12 | -1.622928267 | 0.041502924 | -2.496824919 | 0.000889083 | -4.119753186 | 12.96272604 | 11.1062775 | 11.41341243 | 10.4770591 | 10.4252672 | 9.479382558 | 9.675800489 |
| ENSG00000135439 | AGAP2 | -3.24997781 | 0.000184497 | -0.872061786 | 0.331999289 | -4.122039596 | 6.009025997 | 4.93638933 | 4.874587708 | 2.947924294 | 3.01487888 | 4.713032626 | 4.479073019 |
| ENSG00000114738 | MAPKAPK3 | -2.032169671 | 0.000520688 | -2.092126696 | 0.00030778 | -4.124296366 | 8.855032352 | 10.18234215 | 10.04361983 | 7.867833045 | 7.7127321 | 7.52030827 | 7.919398473 |
| ENSG00000213420 | GPC2 | -2.510815813 | 0.368000274 | -1.615743826 | 0.570028066 | -4.126559639 | 2 | 2 | 7.552162999 | 4.205751501 | 3.45167086 | 4.612704558 | 4.543391831 |
| ENSG00000139637 | C12orf10 | -2.29291377 | 0.021786137 | -1.834413619 | 0.070636438 | -4.127327388 | 7.940205782 | 9.479495904 | 10.88048583 | 7.617841552 | 7.61995902 | 7.904255004 | 8.2175813 |
| ENSG00000137265 | IRF4 | -2.738963038 | 0.494607899 | -1.390369092 | 0.728964147 | -4.12933213 | 2 | 2 | 4.058998049 | 2 | 2.38539787 | 2.68698273 | 2.225841198 |
| ENSG00000197635 | DPP4 | 1.418582456 | 0.22891277 | -5.549139117 | 4.72E-08 | -4.130556661 | 9.966689162 | 11.08875118 | 10.5187901 | 12.01019697 | 12.0165567 | 6.004008792 | 3.287118176 |
| ENSG00000124641 | MED20 | -2.152940447 | 0.021869406 | -1.979443369 | 0.035200747 | -4.132383816 | 10.32700827 | 8.848535039 | 7.734844356 | 7.352188525 | 7.09050444 | 7.395794919 | 7.396986783 |
| ENSG00000173258 | ZNF483 | -3.286393055 | 0.005315748 | -0.846721205 | 0.516558603 | -4.13311426 | 5.70932178 | 4.11352228 | 7.163363831 | 3.515101962 | 3.33976443 | 5.565950232 | 5.094391891 |
| ENSG00000182481 | KPNA2 | -2.17564536 | 1.97E-07 | -1.958855476 | 3.04E-06 | -4.134500836 | 11.32135895 | 11.15761406 | 11.06667478 | 9.03081742 | 9.00665321 | 8.721540982 | 9.611624018 |
| ENSG00000254415 | SIGLEC14 | -2.272269617 | NA | -1.864114193 | NA | -4.13638381 | 3.327711234 | 2 | 2 | 2.141737246 | 2.14022021 | 2.34604626 | 2 |
| ENSG00000088538 | DOCK3 | -2.907730496 | 0.004463515 | -1.233812293 | 0.247215086 | -4.141542789 | 4.00722531 | 6.04685525 | 6.073378139 | 3.09461492 | 3.55551834 | 4.73522696 | 4.365060851 |
| ENSG00000112039 | FANCE | -3.202899781 | 0.05874363 | -0.940038522 | 0.616621341 | -4.142938303 | 3.327711234 | 6.464272742 | 9.272377397 | 4.946755474 | 4.811949 | 6.85908071 | 7.091143645 |
| ENSG00000134262 | AP4B1 | -1.829161762 | 0.012981199 | -2.316394632 | 0.001112159 | -4.145556394 | 7.904282243 | 7.60566235 | 9.25752426 | 6.586219071 | 6.71776877 | 6.231826305 | 6.145001162 |
| ENSG00000136450 | SRSF1 | -0.281801498 | 0.625823305 | -3.866489444 | 1.06E-17 | -4.148290942 | 11.0278074 | 10.55843445 | 10.47451889 | 10.43636561 | 10.4171049 | 6.382394503 | 7.261744957 |
| ENSG00000138363 | ATIC | -2.130942498 | 0.007823935 | -2.019524015 | 0.011776447 | -4.150466513 | 11.79097454 | 9.806969599 | 10.2528023 | 8.696503411 | 8.82115545 | 8.738121153 | 8.991966981 |
| ENSG00000090447 | TFAP4 | -1.50117788 | 0.156963258 | -2.649612687 | 0.0069133 | -4.150790567 | 5.095959458 | 6.464272742 | 7.225742779 | 5.003571641 | 5.21224859 | 4.580376151 | 3.539239711 |
| ENSG00000136161 | RCBTB2 | -1.464145166 | 0.004451468 | -2.686754113 | 2.84E-08 | -4.150899279 | 8.16936288 | 8.191702463 | 9.099922433 | 7.110641269 | 7.11620658 | 6.05195199 | 5.829625997 |
| ENSG00000249853 | HS3ST5 | -4.714850357 | NA | 0.563186642 | NA | -4.151663714 | 2 | 3.076665209 | 2 | 2 | 2 | 2.656294034 | 2.918126211 |
| ENSG00000173653 | RCE1 | -2.107610669 | 2.97E-11 | -2.045098163 | 5.42E-11 | -4.152708832 | 8.998234871 | 8.522788974 | 8.904018013 | 6.739036112 | 6.76185814 | 6.840432491 | 6.780202889 |
| ENSG00000206432 | TMEM200C | -4.104486027 | 0.208966396 | -0.054440508 | 0.988291799 | -4.158926535 | 2 | 5.131603625 | 2 | 2.3892346 | 2 | 3.824280769 | 3.841670744 |
| ENSG00000100526 | CDKN3 | -3.394864229 | 0.020261568 | -0.764979749 | 0.638543357 | -4.159843978 | 4.467311984 | 6.899080544 | 3.369317972 | 3.162721474 | 3.01487888 | 4.675266036 | 5.263040529 |
| ENSG00000140807 | NKD1 | -2.996124858 | 0.007885856 | -1.163746285 | 0.324775694 | -4.159871142 | 5.532353208 | 5.131603625 | 5.772716771 | 3.615725078 | 2.59489879 | 4.920961476 | 3.841670744 |
| ENSG00000167088 | SNRPD1 | -1.824356305 | 0.002127716 | -2.337195356 | 4.87E-05 | -4.161551662 | 9.613202746 | 9.055168103 | 10.42884603 | 8.024753265 | 7.97398853 | 7.294808156 | 7.673351463 |
| ENSG00000143815 | LBR | -1.641265328 | 0.117116078 | -2.520486414 | 0.010618344 | -4.161751743 | 6.73718411 | 9.770792087 | 8.922968322 | 7.366699513 | 7.26914476 | 6.516588833 | 6.420873184 |
| SGK223 | | 3.789280321 | 0.010861394 | -7.958771341 | 0.00018896 | -4.16949102 | 2 | 4.11352228 | 3.369317972 | 6.676346669 | 6.60731796 | 2 | 2 |
| ENSG00000237724 | | 6.032035861 | 3.82E-06 | -10.20292155 | 4.30E-07 | -4.170885689 | 2 | 5.838937808 | 5.59507254 | 11.05284078 | 11.0926765 | 2 | 2 |
| ENSG00000155970 | MICU3 | -1.483454347 | 0.010160299 | -2.688001103 | 6.59E-07 | -4.171455449 | 8.39321512 | 7.790474954 | 8.325066185 | 6.936770382 | 6.51372389 | 5.963846589 | 5.122735884 |
| ENSG00000196976 | LAGE3 | -2.560581659 | 1.10E-07 | -1.613563454 | 0.001058529 | -4.174145113 | 8.946185348 | 8.101365653 | 8.042706615 | 5.949776857 | 5.93454154 | 6.953609309 | 6.723495198 |
| ENSG00000145216 | FIP1L1 | -1.343521195 | 0.001400964 | -2.832846833 | 7.33E-13 | -4.176368028 | 10.43166252 | 9.664384941 | 10.26027233 | 8.709369553 | 8.91566441 | 7.49884738 | 7.18317631 |
| ENSG00000176563 | CNTD1 | -3.907643128 | 0.247434707 | -0.274604462 | 0.940403768 | -4.18224759 | 2 | 4.442710129 | 2 | 2.270786764 | 2 | 3.082820997 | 3.287118176 |
| ENSG00000149273 | RPS3 | -2.703850462 | 0.00015819 | -1.480131722 | 0.051545976 | -4.183982184 | 14.5778178 | 13.33179442 | 12.66820966 | 11.02794996 | 11.064624 | 12.23258705 | 12.30365825 |
| ENSG00000088305 | DNMT3B | -2.785777254 | 0.03769062 | -1.400246955 | 0.321669264 | -4.18602421 | 7.435667355 | 8.451443573 | 4.058998049 | 4.574143677 | 5.14724905 | 6.265258498 | 6.02248356 |
| ENSG00000125122 | LRRC29 | -2.105138126 | 0.290270981 | -2.082801127 | 0.284366681 | -4.187939254 | 2 | 5.303523747 | 7.934808822 | 4.299492648 | 4.95174433 | 4.406883583 | 4.911532223 |
| ENSG00000221845 | C7orf65 | -1.654777715 | 0.611381475 | -2.535197629 | NA | -4.189975344 | 2 | 3.686303414 | 2 | 2.141737246 | 2.49394783 | 2.094598369 | 2.277179783 |
| ENSG00000082014 | SMARCD3 | -2.863010627 | 4.09E-15 | -1.333828897 | 0.000426182 | -4.196839525 | 10.24321353 | 9.672251278 | 9.999667074 | 7.288834728 | 7.01963893 | 8.458554046 | 8.842341636 |
| ENSG00000107562 | CXCL12 | -2.167232486 | 0.140578614 | -2.030401863 | 0.162319088 | -4.197634348 | 5.095959458 | 9.77810053 | 9.888372618 | 7.226553841 | 7.05107174 | 7.753839274 | 6.55171954 |
| ENSG00000114942 | EEF1B2 | -2.101515578 | 1.14E-06 | -2.097563343 | 1.08E-06 | -4.199078921 | 12.0285121 | 11.20989118 | 11.18913795 | 9.397005623 | 9.47403846 | 9.401144325 | 9.477790979 |
| ENSG00000101098 | RIMS4 | -2.925573511 | 0.525963919 | -1.273544341 | NA | -4.199117853 | 2 | 3.686303414 | 2 | 2.141737246 | 2.14022021 | 2.717032197 | 2 |
| ENSG00000141576 | RNF157 | -1.004421908 | 0.226266075 | -3.195654948 | 1.32E-05 | -4.200076856 | 6.257048221 | 5.131603625 | 6.534109959 | 5.058234798 | 5.22805098 | 3.600766115 | 3.261853426 |
| ENSG00000146540 | C7orf50 | -2.243535041 | 2.89E-05 | -1.957199505 | 0.000292529 | -4.200734546 | 11.24801458 | 10.03150247 | 11.22011226 | 8.639384849 | 8.75588792 | 8.946279845 | 9.018137056 |
| ENSG00000165480 | SKA3 | -1.663892767 | 0.424434971 | -2.541593581 | 0.192038633 | -4.205486348 | 3.327711234 | 6.389896633 | 2 | 3.840289928 | 3.60476711 | 3.082820997 | 3.261853426 |
| ENSG00000154309 | DISP1 | -2.025321698 | 0.169999755 | -2.181003845 | 0.128777677 | -4.206325543 | 8.981092835 | 9.532315021 | 4.523675391 | 6.739036112 | 6.74548212 | 6.913621294 | 6.179140936 |
| ENSG00000075651 | PLD1 | -0.527281898 | 0.466093816 | -3.682987251 | 4.40E-10 | -4.210269149 | 8.609439662 | 9.338312924 | 8.631919287 | 8.302698479 | 8.45888185 | 5.805044075 | 4.702084619 |
| ENSG00000171425 | ZNF581 | -2.662847745 | 2.29E-05 | -1.54782397 | 0.019121696 | -4.210671715 | 10.05124847 | 8.671479029 | 8.784770914 | 6.652857793 | 6.73998187 | 7.896069536 | 7.657697387 |
| ENSG00000002822 | MAD1L1 | -1.913063352 | 0.000238058 | -2.29844177 | 6.24E-06 | -4.211505122 | 9.233563228 | 8.590771778 | 9.809100877 | 7.355829978 | 7.44845993 | 6.998648055 | 7.056667034 |
| ENSG00000141030 | COPS3 | -2.050817656 | 4.13E-13 | -2.162543657 | 1.01E-14 | -4.213361313 | 10.6316532 | 10.16572116 | 10.49365885 | 8.466597359 | 8.33666436 | 8.311207095 | 8.273956186 |
| ENSG00000277149 | TYW1B | 0.605383334 | 0.738608861 | -4.820048804 | 0.001245628 | -4.21466547 | 7.709880939 | 4.93638933 | 3.369317972 | 6.861921612 | 7.05107174 | 3.011868911 | 2.466008509 |
| ENSG00000094916 | CBX5 | -3.034570118 | 1.60E-07 | -1.180556748 | 0.062327547 | -4.215126866 | 11.78110422 | 12.03957582 | 13.12215634 | 9.425008328 | 9.39575048 | 11.04531616 | 11.44389555 |
| ENSG00000021488 | SLC7A9 | -0.951896666 | NA | -3.266656142 | NA | -4.218552808 | 2 | 3.076665209 | 2 | 2.270786764 | 2.38539787 | 2.139668783 | 2 |
| ENSG00000187045 | TMPRSS6 | -4.714849728 | NA | 0.495045834 | NA | -4.219803894 | 2 | 3.076665209 | 2 | 2 | 2 | 2.746468522 | 2.782576693 |
| ENSG00000141002 | TCF25 | -2.080722321 | 5.11E-05 | -2.13995874 | 2.71E-05 | -4.220681061 | 12.36386778 | 11.14636276 | 12.05462845 | 9.819163759 | 9.90722553 | 9.678399287 | 9.921211649 |
| ENSG00000104907 | TRMT1 | -2.079582955 | 0.045753442 | -2.141467495 | 0.036785154 | -4.22105045 | 10.7426185 | 9.159587375 | 7.734844356 | 7.590284224 | 7.6957855 | 7.517109417 | 7.646759583 |
| ENSG00000002016 | RAD52 | -2.200725991 | 8.36E-05 | -2.022707819 | 0.000278998 | -4.22343381 | 8.586954094 | 7.761270542 | 8.408111533 | 6.130704144 | 6.17362678 | 6.685680098 | 5.829625997 |
| ENSG00000184903 | IMMP2L | -2.609422897 | 0.131120567 | -1.614882952 | 0.363318728 | -4.224305849 | 2 | 7.099070191 | 7.858115942 | 4.694064037 | 4.43005853 | 5.371837532 | 5.497565112 |
| ENSG00000123179 | EBPL | -2.537193051 | 3.63E-10 | -1.689200497 | 3.32E-05 | -4.226393548 | 8.816894989 | 8.433041974 | 8.076950877 | 6.069927018 | 5.9536761 | 6.947285242 | 6.681239787 |
| ENSG00000004864 | SLC25A13 | -1.448274795 | 0.018609883 | -2.778295586 | 1.19E-06 | -4.226570381 | 8.418903792 | 8.255926468 | 9.478569125 | 7.510853288 | 7.26914476 | 5.966975997 | 6.251520369 |
| ENSG00000153395 | LPCAT1 | 0.624144657 | 0.534490579 | -4.85178306 | 4.21E-09 | -4.227638402 | 8.285626337 | 8.414402626 | 6.073378139 | 8.47833774 | 8.57772939 | 3.767980144 | 3.430065404 |
| ENSG00000185475 | TMEM179B | -0.902091175 | 0.007572761 | -3.326251409 | 3.23E-27 | -4.228342584 | 10.56993315 | 10.29361762 | 9.990713521 | 9.363935069 | 9.44728702 | 6.940933331 | 7.09293558 |
| ENSG00000155897 | ADCY8 | -0.406115909 | 0.921327192 | -3.822229845 | 0.276521038 | -4.228344404 | 2 | 2 | 5.39244166 | 3.754611435 | 3.74316441 | 2.526562374 | 2 |
| ENSG00000128951 | DUT | -1.808386179 | 0.000125098 | -2.420206445 | 1.22E-07 | -4.228592623 | 10.32020829 | 9.806969599 | 9.242516611 | 8.128448191 | 7.99492122 | 7.47156439 | 7.453871289 |
| ENSG00000121897 | LIAS | -2.03075785 | 0.182549331 | -2.203312785 | 0.136300946 | -4.234070635 | 3.327711234 | 6.844429755 | 8.266922072 | 5.22679461 | 5.28958453 | 5.217477917 | 4.982847849 |
| ENSG00000100564 | PIGH | -2.240029562 | 1.57E-08 | -1.99477978 | 3.58E-07 | -4.234809341 | 8.737455672 | 8.029695761 | 8.511888463 | 6.328984825 | 6.20590375 | 6.591495479 | 6.406535141 |
| ENSG00000117448 | AKR1A1 | -2.51463696 | 3.01E-09 | -1.720565617 | 7.33E-05 | -4.235202577 | 10.72212986 | 10.67655334 | 11.4530647 | 8.464912335 | 8.5227358 | 9.08235879 | 9.456374363 |
| ENSG00000123999 | INHA | -2.760519025 | 0.117331848 | -1.475249467 | 0.420838949 | -4.235768492 | 7.435667355 | 5.457120953 | 2 | 3.881293667 | 3.7864959 | 4.705557986 | 4.998228167 |
| ENSG00000099889 | ARVCF | -3.484293365 | 0.059528826 | -0.757307043 | 0.719120052 | -4.241600409 | 3.327711234 | 6.228547981 | 9.990713521 | 5.210797711 | 5.16377554 | 7.838287442 | 7.705330953 |
| ENSG00000147789 | ZNF7 | -2.514604218 | 0.126534396 | -1.727603416 | 0.302902594 | -4.242207634 | 10.48735708 | 9.256956408 | 4.058998049 | 6.979883891 | 6.9210099 | 7.76915836 | 7.664943458 |
| ENSG00000115041 | KCNIP3 | -3.263888985 | 0.195586526 | -0.979170449 | 0.719746226 | -4.243059434 | 2 | 6.602424674 | 2 | 2.784612485 | 2.94009902 | 4.443291373 | 4.188602707 |
| ENSG00000138821 | SLC39A8 | 0.187305977 | 0.896917828 | -4.430753744 | 4.25E-05 | -4.243447767 | 10.46903009 | 9.664384941 | 6.629488387 | 9.818504147 | 9.75398117 | 5.304163849 | 5.32534152 |
| ENSG00000075886 | TUBA3D | -4.714857915 | 0.359799649 | 0.46899834 | NA | -4.245859575 | 3.327711234 | 2 | 2 | 2 | 2 | 2.962551659 | 2.509591799 |
| ENSG00000182511 | FES | -1.986383755 | 0.007076225 | -2.260698028 | 0.001669835 | -4.247081783 | 8.716893051 | 7.669933291 | 7.029880454 | 6.213314212 | 5.87556634 | 6.000958775 | 5.561082532 |
| ENSG00000179314 | WSCD1 | -2.820838506 | 0.193784218 | -1.431608823 | 0.523327331 | -4.252447329 | 3.327711234 | 3.076665209 | 6.43197674 | 3.566290607 | 2.2680111 | 4.572179576 | 2.782576693 |
| ENSG00000282228 | | 3.869341033 | 0.00912284 | -8.124752487 | 0.000134379 | -4.255411454 | 4.00722531 | 2 | 4.058998049 | 6.841288255 | 6.91614081 | 2 | 2 |

| | | | | | | | | | | | | | |
|---|---|---|---|---|---|---|---|---|---|---|---|---|---|
| ENSG00000164032 | H2AFZ | -2.002628403 | 0.002807298 | -2.254037866 | 0.000612879 | -4.256666269 | 11.75864596 | 10.46189757 | 10.19940021 | 9.004380786 | 8.96246653 | 8.77875662 | 8.688479312 |
| ENSG00000164483 | SAMD3 | 0.079213662 | 0.978795893 | -4.338678352 | 0.072675764 | -4.25946469 | 2 | 6.04685525 | 2 | 4.574143677 | 4.85329164 | 2.139668783 | 2.509591799 |
| ENSG00000174748 | RPL15 | -2.536511692 | 1.81E-06 | -1.724169186 | 0.001703637 | -4.260680878 | 15.07731105 | 13.94139794 | 13.96781386 | 11.88662999 | 11.907319 | 12.69917208 | 12.7181685 |
| ENSG00000116285 | ERRFI1 | -3.445193644 | 1.21E-10 | -0.818722001 | 0.178997619 | -4.263915645 | 12.22696957 | 11.76214832 | 10.94216274 | 8.315844936 | 8.29975519 | 11.16604696 | 10.62056434 |
| ENSG00000101146 | RAE1 | -1.945865425 | 0.007764718 | -2.321646823 | 0.001101245 | -4.267512248 | 10.33377634 | 9.566483142 | 8.380957451 | 7.714607209 | 7.70145652 | 7.272216802 | 7.405663765 |
| ENSG00000162148 | PPP1R32 | -2.12461538 | 0.455923164 | -2.144788336 | 0.439942697 | -4.269403716 | 7.485143553 | 2 | 2 | 4.237680128 | 3.98554648 | 4.504874511 | 3.539239711 |
| ENSG00000106443 | PHF14 | -1.748733837 | 0.012863597 | -2.520859829 | 0.000171403 | -4.269593666 | 8.737455672 | 9.386925327 | 10.43545992 | 7.913023977 | 8.00183183 | 7.380620955 | 6.998584547 |
| ENSG00000180011 | ZADH2 | -2.425169201 | 0.000561169 | -1.845898062 | 0.010373989 | -4.271067263 | 9.112843761 | 10.80732116 | 9.565556058 | 7.659688863 | 7.57123495 | 8.391426944 | 7.945418846 |
| ENSG00000183196 | CHST6 | -3.476319743 | 0.000392437 | -0.795446046 | 0.459125626 | -4.271765788 | 5.70932728 | 5.131603625 | 6.718950348 | 3.515101962 | 2.86123014 | 5.614651724 | 4.784231301 |
| ENSG00000186868 | MAPT | -0.841642191 | 0.785196727 | -3.430392708 | 0.211625477 | -4.272034899 | 2 | 4.11352228 | 2 | 2.270786764 | 3.08597292 | 2.183373728 | 2.117335275 |
| ENSG00000119335 | SET | -2.037288311 | 6.98E-17 | -2.235151912 | 2.36E-20 | -4.272440222 | 13.10501837 | 12.73245137 | 12.99808116 | 10.95209651 | 10.883164 | 10.65245252 | 10.78574353 |
| ENSG00000175581 | MRPL48 | -2.733656528 | 0.000192513 | -1.538810554 | 0.045885615 | -4.272467082 | 9.33149386 | 7.572420273 | 8.236946975 | 5.651002631 | 6.1322378 | 6.833591108 | 7.240884911 |
| ENSG00000196378 | ZNF34 | -2.463933152 | 0.029667098 | -1.809079256 | 0.116235371 | -4.273012408 | 8.540903468 | 6.04685525 | 5.930865648 | 5.144975053 | 4.97065356 | 5.650128416 | 5.651404445 |
| ENSG00000180767 | CHST13 | -4.71485825 | 0.359699571 | 0.441164482 | NA | -4.273693768 | 2 | 2 | 3.369317972 | 2 | 2 | 2.457042231 | 2.981406154 |
| ENSG00000198838 | RYR3 | -5.597645813 | 0.020793632 | 1.323282552 | 0.49597846 | -4.27436326 | 2 | 4.710564154 | 3.369317972 | 2.141737246 | 2 | 4.927401733 | 4.77533149 |
| ENSG00000173083 | HPSE | -2.918327577 | 0.004432859 | -1.357134314 | 0.196524797 | -4.275461891 | 5.70932728 | 4.442710129 | 5.930865648 | 3.09461492 | 3.33976443 | 3.753554863 | 4.702084619 |
| ENSG00000072133 | RPS6KA6 | 0.806248133 | 0.245194354 | -5.082589645 | 5.16E-14 | -4.276341511 | 5.330594134 | 5.838937808 | 5.930865648 | 6.586219071 | 6.37866593 | 2.384001718 | 2.509591799 |
| ENSG00000258818 | RNASE4 | 0.969838731 | 0.479607778 | -5.246697738 | 6.73E-06 | -4.276859007 | 4.467311984 | 5.838937808 | 7.97167868 | 7.662632018 | 7.77322015 | 2.775316226 | 2.746601469 |
| SGK494 | | -1.994962249 | 0.160878275 | -2.282714022 | 0.09629221 | -4.27767627 | 4.00722531 | 5.838937808 | 7.934808822 | 4.965943888 | 4.79082474 | 4.854926465 | 4.376876162 |
| ENSG00000165916 | PSMC3 | -1.974811426 | 3.49E-06 | -2.305593611 | 3.76E-08 | -4.280405037 | 12.2521965 | 11.65904139 | 11.62203638 | 9.906173175 | 9.90169395 | 9.22478876 | 9.85622839 |
| ENSG00000106852 | LHX6 | -4.714844908 | NA | 0.432161051 | NA | -4.282683858 | 2 | 3.076665209 | 2 | 2 | 2 | 2.492221014 | 2.950113106 |
| ENSG00000241058 | NSUN6 | -1.651485406 | 0.28126265 | -2.631642351 | 0.066527636 | -4.283127757 | 3.327711234 | 8.213429689 | 7.858115942 | 5.808110275 | 6.00043082 | 5.158477451 | 4.861951484 |
| ENSG00000151353 | TMEM18 | -2.46283301 | 1.98E-05 | -1.822344153 | 0.001998187 | -4.285177163 | 9.317904228 | 8.88960062 | 10.2528023 | 7.173892946 | 7.16627666 | 7.679481638 | 7.90417044 |
| ENSG00000178035 | IMPDH2 | -2.853552045 | 8.15E-15 | -1.432135897 | 0.000173991 | -4.285687942 | 11.22271378 | 10.50658023 | 11.0793996 | 8.047448792 | 8.20911072 | 9.646061941 | 9.423285228 |
| ENSG00000111644 | ACRBP | -4.523349016 | 0.191972185 | 0.237613751 | 0.945777268 | -4.285735265 | 2 | 4.11352228 | 2 | 2 | 2.14022021 | 3.5515855 | 2.885414003 |
| ENSG00000213782 | DDX47 | 0.825515056 | 0.058148984 | -5.114871015 | 1.24E-36 | -4.289355959 | 7.904282243 | 7.847167909 | 8.460932916 | 8.904736303 | 8.91077722 | 3.44788957 | 3.660199087 |
| ENSG00000149922 | TBX6 | -2.461324868 | 0.351613795 | -1.828385169 | 0.489050844 | -4.289710038 | 6.366709671 | 2 | 2 | 3.09461492 | 3.15372745 | 3.648325314 | 3.33635788 |
| ENSG00000133678 | TMEM254 | -2.426997614 | 0.000470613 | -1.863250393 | 0.008592718 | -4.290248008 | 9.87686338 | 8.053983481 | 9.399592459 | 6.960881212 | 6.84106088 | 7.577730839 | 7.310303798 |
| ENSG00000082146 | STRADB | -1.45499528 | 0.093501828 | -2.835328769 | 0.000357492 | -4.290324049 | 10.51739401 | 8.848535039 | 8.631919287 | 8.132699624 | 8.17486938 | 7.043805932 | 6.520103598 |
| ENSG00000146072 | TNFRSF21 | -2.104851197 | 1.26E-20 | -2.185732952 | 1.52E-22 | -4.29058415 | 11.39115429 | 11.49575678 | 11.71466889 | 9.419799055 | 9.46239569 | 9.339699927 | 9.381743653 |
| ENSG00000130684 | ZNF337 | -0.730734877 | 0.726676555 | -3.560128799 | 0.046040945 | -4.290863676 | 8.963744667 | 8.540087354 | 2 | 7.593372273 | 7.33993465 | 5.130840083 | 4.445805334 |
| ENSG00000135587 | SMPD2 | -1.727000154 | 0.389329355 | -2.564534088 | 0.174243611 | -4.291534242 | 2 | 7.467835385 | 9.132850258 | 6.24507773 | 6.30607931 | 5.598599872 | 5.385063124 |
| ENSG00000160752 | FDPS | -1.839542505 | 0.001710311 | -2.453048621 | 1.52E-05 | -4.292591126 | 11.9425688 | 11.21259083 | 10.9743126 | 9.611716938 | 9.59474367 | 8.401292507 | 9.412563085 |
| ENSG00000137409 | MTCH1 | -2.362103678 | 2.44E-28 | -1.933194682 | 1.51E-19 | -4.29529836 | 13.38791698 | 13.15009613 | 13.13338652 | 10.84346133 | 10.8937291 | 11.2301273 | 11.36081175 |
| ENSG00000142546 | NOSIP | -1.698915675 | 0.010564109 | -2.597004276 | 3.88E-05 | -4.295919951 | 10.33377634 | 9.318399851 | 11.01903267 | 8.686416476 | 8.69607402 | 7.792277365 | 7.817957556 |
| ENSG00000171659 | GPR34 | -5.943512128 | 0.124766724 | 1.646750108 | 0.666185912 | -4.29676202 | 2 | 4.11352228 | 2 | 2 | 2 | 4.652122235 | 3.580696614 |
| ENSG00000244187 | TMEM141 | -2.386398369 | 0.000315137 | -1.915459779 | 0.00450069 | -4.301858147 | 10.72212986 | 9.7032946 | 9.014156722 | 7.638916932 | 7.60490877 | 8.033012338 | 8.134761809 |
| ENSG00000159685 | CHCHD6 | -2.568306076 | 0.154641078 | -1.737033835 | 0.345156474 | -4.305339911 | 8.540903468 | 7.73146271 | 2 | 5.242616075 | 5.13053105 | 5.985611216 | 5.909074772 |
| ENSG00000132026 | RTBDN | -3.267921895 | NA | -1.037693814 | NA | -4.305615709 | 2 | 3.076665209 | 2 | 2.141737246 | 2 | 2.34604626 | 2.277179783 |
| ENSG00000099953 | MMP11 | 2.697238703 | 0.225340974 | -7.003620934 | 0.00729012 | -4.306382231 | 3.327711234 | 3.076665209 | 2 | 4.783303513 | 4.74762503 | 2 | 2 |
| ENSG00000102977 | ACD | -2.149438427 | 0.061614336 | -2.157117255 | 0.056913062 | -4.306555682 | 6.138360429 | 8.337339476 | 9.647595056 | 6.470063213 | 6.60126376 | 6.681876084 | 6.359671687 |
| ENSG00000141127 | PRPSAP2 | -2.278195301 | 0.02238434 | -2.030130933 | 0.04257856 | -4.308326234 | 10.73241055 | 8.777382925 | 8.110401118 | 7.52061915 | 7.27295842 | 7.504242605 | 7.772838868 |
| ENSG00000160883 | HK3 | -4.891135293 | 0.019121386 | 0.582088643 | 0.776890378 | -4.30904625 | 4.467311984 | 4.442710129 | 2 | 2.141737246 | 2.14022021 | 5.102662916 | 3.33635788 |
| ENSG00000189227 | C15orf61 | -2.3055924 | 0.049214138 | -2.008241882 | 0.085942697 | -4.313834283 | 8.493334308 | 5.595927442 | 6.203069917 | 5.040242897 | 5.2591458 | 5.357600875 | 5.481238982 |
| ENSG00000134602 | STK26 | -0.766769206 | 0.472853254 | -3.548345304 | 7.94E-05 | -4.31511451 | 7.384433967 | 8.053983481 | 6.882775709 | 6.882264028 | 6.67228943 | 4.875056259 | 3.236138336 |
| ENSG00000166002 | SMCO4 | -2.916793333 | 1.88E-08 | -1.40350695 | 0.009028137 | -4.320300283 | 8.340423424 | 7.503546079 | 7.398165132 | 5.076005082 | 5.0259382 | 6.595545082 | 6.280114403 |
| ENSG00000105854 | PON2 | -1.307627094 | 0.006236051 | -3.013930513 | 1.02E-11 | -4.321557608 | 12.73754052 | 11.86525432 | 11.84954227 | 10.89755258 | 10.9176784 | 9.099926561 | 9.308834477 |
| ENSG00000137270 | GCM1 | -1.685160218 | NA | -2.637185881 | NA | -4.322346099 | 2 | 2 | 3.369317972 | 2.270786764 | 2.14022021 | 2.048074405 | 2.172608095 |
| ENSG00000110200 | ANAPC15 | -0.570346951 | 0.39460075 | -3.754088505 | 2.49E-11 | -4.324435456 | 7.579279349 | 7.002537973 | 6.322058544 | 6.509830055 | 6.507263 | 3.663836244 | 3.858622973 |
| ENSG00000178982 | EIF3K | -1.646317268 | 0.006115397 | -2.682792963 | 2.43E-06 | -4.329110232 | 11.40407417 | 10.3778065 | 10.26770388 | 9.137269904 | 9.14209426 | 7.742013931 | 8.409548182 |
| ENSG00000140398 | NEIL1 | -2.551524043 | 0.043880019 | -1.78039183 | 0.167056356 | -4.331915873 | 5.330594134 | 5.72254323 | 8.408111533 | 4.647285135 | 4.93258395 | 5.598599872 | 5.349527629 |
| ENSG00000183742 | MACC1 | -1.454222696 | 0.396614281 | -2.877976511 | 0.067194295 | -4.332199208 | 4.00722531 | 4.442710129 | 4.523675391 | 3.227757282 | 3.39680241 | 3.011868911 | 2.117335275 |
| ENSG00000157184 | CPT2 | -2.106928155 | 0.042833181 | -2.227990579 | 0.029472075 | -4.334918734 | 10.52332707 | 7.467835385 | 9.503962296 | 7.416368504 | 7.66128317 | 7.116063252 | 7.680519528 |
| ENSG00000105173 | CCNE1 | -2.436148741 | 0.017542283 | -1.899789931 | 0.067889203 | -4.335938672 | 9.097018025 | 6.666836837 | 6.958202216 | 5.590471342 | 5.73886209 | 6.368262153 | 5.933281326 |
| ENSG00000110074 | FOXRED1 | -2.479105376 | 2.36E-05 | -1.858223982 | 0.001847366 | -4.337329358 | 9.344956677 | 7.954278496 | 9.049077186 | 6.442929658 | 6.48770488 | 6.892381078 | 7.216434756 |
| ENSG00000140519 | RHCG | 0.426466622 | 0.883547191 | -4.765419947 | 0.046152919 | -4.338953325 | 2 | 5.946638857 | 2 | 4.804776789 | 5.04390545 | 2.094598369 | 2.374681325 |
| ENSG00000117305 | HMGCL | -2.332109314 | 8.13E-05 | -2.007174944 | 0.000776127 | -4.339284258 | 11.00254523 | 9.479495904 | 10.46161637 | 8.130575473 | 8.12087597 | 8.382079673 | 8.508191835 |
| ENSG00000267796 | LIN37 | -1.203561142 | 0.02230134 | -3.136026983 | 8.28E-11 | -4.339588125 | 7.750913381 | 6.951736442 | 7.163363831 | 6.043074888 | 6.27599024 | 4.311629396 | 4.56420962 |
| ENSG00000243279 | PRAF2 | -1.963555623 | 5.91E-06 | -2.376243765 | 2.33E-08 | -4.339799388 | 11.60207034 | 10.74900803 | 11.13734171 | 9.280179267 | 9.21427981 | 8.586144131 | 9.052614986 |
| ENSG00000224186 | C5orf66 | -3.184715625 | 0.207275017 | -1.162347279 | 0.654429096 | -4.347062903 | 2 | 4.93638933 | 2 | 2.270786764 | 2.38539787 | 2.858551861 | 3.183290849 |
| ENSG00000183569 | SERHL2 | -2.498143801 | 0.007772622 | -1.850439926 | 0.049827446 | -4.348583727 | 6.96351253 | 6.140560021 | 5.156626401 | 3.921164147 | 4.22485516 | 4.996409624 | 4.047883918 |
| ENSG00000128656 | CHN1 | -0.127433417 | 0.768745599 | -4.221767959 | 4.76E-38 | -4.349201375 | 9.678463567 | 9.497318007 | 9.735974718 | 9.481935579 | 9.54198834 | 5.773219597 | 5.256659864 |
| ENSG00000167513 | CDT1 | -2.647622022 | 0.047576493 | -1.702714908 | 0.210457673 | -4.35033693 | 4.815609471 | 7.145017454 | 4.058998049 | 3.709785322 | 3.7864959 | 4.209638631 | 4.692661645 |
| ENSG00000116885 | OSCP1 | -1.68733769 | 0.104928256 | -2.663011099 | 0.006338722 | -4.350348789 | 8.816894989 | 6.464272742 | 7.029880454 | 5.920568914 | 6.37866593 | 4.907994066 | 5.555894927 |
| ENSG00000084110 | HAL | -1.239517263 | 0.651205609 | -3.110867832 | 0.213537395 | -4.350385095 | 2 | 4.442710129 | 2 | 2.947924294 | 2.49394783 | 2.3070652 | 2.172608095 |
| ENSG00000107614 | TRDMT1 | -2.297502033 | 0.040383064 | -2.052952093 | 0.065986003 | -4.350454126 | 8.586954094 | 7.002537973 | 5.59507254 | 5.565529336 | 5.13053105 | 5.185595313 | 5.900914815 |
| ENSG00000184500 | PROS1 | -2.013366448 | 0.021931339 | -2.339087599 | 0.006324904 | -4.352454048 | 7.159102577 | 8.955547846 | 9.491321578 | 6.727838783 | 6.95938211 | 6.184218099 | 6.81752904 |
| ENSG00000130287 | NCAN | -0.691402429 | 0.877879848 | -3.662884712 | 0.344285984 | -4.354287141 | 5.330594134 | 2 | 2 | 4.237680128 | 2 | 2.183373728 | 2.421067459 |
| ENSG00000144908 | ALDH1L1 | -5.370789922 | 0.223978244 | 1.016112502 | 0.82326635 | -4.35467742 | 2 | 3.686303414 | 2 | 2 | 2 | 3.709392871 | 2.918126211 |
| ENSG00000093009 | CDC45 | -1.599969163 | 0.384530388 | -2.755635887 | 0.105831447 | -4.35560505 | 4.815609471 | 5.457120953 | 2 | 3.566290607 | 3.39680241 | 2.384001718 | 3.209956562 |
| ENSG00000105171 | POP4 | -2.134359894 | 0.035018296 | -2.224672916 | 0.025955343 | -4.35903281 | 10.1770105 | 7.701025996 | 10.71638486 | 7.77569021 | 7.96222667 | 7.637872201 | 7.915353355 |
| ENSG00000154328 | NEIL2 | -3.028369118 | 9.83E-08 | -1.3308437 | 0.028788809 | -4.359212818 | 8.418903792 | 9.13702746 | 9.849281147 | 6.283823264 | 6.29111322 | 7.828858098 | 8.030160124 |
| ENSG00000016391 | CHDH | -2.78577691 | 0.200963739 | -1.57474248 | 0.483865109 | -4.36051939 | 2 | 3.686303414 | 7.896971941 | 3.997744386 | 3.94787299 | 5.113999907 | 4.784231301 |
| ENSG00000054148 | PHPT1 | -2.796281836 | 1.26E-12 | -1.567438839 | 0.0001198 | -4.363720674 | 11.62156041 | 10.73780697 | 11.18521898 | 8.408172799 | 8.47717992 | 9.676010922 | 9.648645089 |
| ENSG00000198498 | TMA16 | -1.942734809 | 0.000281341 | -2.421108857 | 3.29E-06 | -4.363843666 | 9.317904228 | 8.191702463 | 8.408111533 | 6.856790872 | 6.77805036 | 6.380048704 | 6.338680316 |
| ENSG00000100321 | SYNGR1 | -3.325444423 | 0.00013616 | -1.0390358 | 0.287184913 | -4.364480223 | 8.418903792 | 10.160138 | 11.05812878 | 6.965655398 | 6.94032357 | 8.947864566 | 9.417934118 |
| FAM196A | | 3.047043755 | 0.282799717 | -7.41184454 | 0.016000913 | -4.364800785 | 2 | 4.11352228 | 2 | 5.578054239 | 5.27444544 | 2 | 2 |
| ENSG00000166153 | DEPDC4 | -4.714846108 | NA | 0.348050893 | NA | -4.366795215 | 2 | 3.076665209 | 2 | 2 | 2 | 2.775316226 | 2.632958241 |
| ENSG00000121716 | PILRB | -1.40749318 | 0.000820237 | -2.959316772 | 8.37E-14 | -4.366809952 | 7.667647223 | 8.101365653 | 8.380957451 | 6.687949235 | 6.72335419 | 5.125248504 | 5.379200993 |
| ENSG00000171222 | SCAND1 | -2.270396896 | 0.00464393 | -2.105492452 | 0.00889203 | -4.375889347 | 11.57661845 | 9.470501606 | 10.23774519 | 8.43079185 | 8.44204342 | 8.405911884 | 8.77026299 |
| ENSG00000168918 | INPP5D | -5.710311396 | 0.063965155 | 1.33187271 | 0.641832906 | -4.378438686 | 2 | 5.131603625 | 2 | 2 | 2.14022021 | 5.318934974 | 4.468068808 |
| ENSG00000166165 | CKB | -2.553831851 | 0.000979681 | -1.825040907 | 0.022751919 | -4.378872758 | 11.89987989 | 9.849218119 | 10.94216274 | 8.654247601 | 8.5130942 | 8.904861985 | 9.623140479 |
| ENSG00000119522 | DENND1A | -2.29484459 | 7.63E-10 | -2.085758425 | 2.19E-08 | -4.380603015 | 11.06489013 | 10.42512749 | 10.46808205 | 8.310225403 | 8.48706372 | 8.47458408 | 8.730526266 |
| ENSG00000281775 | | -2.321325986 | 0.140439423 | -2.061992265 | 0.175908868 | -4.383318251 | 2 | 4.442710129 | 5.39244166 | 2.947924294 | 2.94009902 | 3.10571689 | 3.042026731 |
| ENSG00000128271 | ADORA2A | -0.210848303 | 0.93643841 | -4.176360917 | 0.050866589 | -4.387209221 | 8.468948322 | 3.686303414 | 2 | 6.772115916 | 6.71776877 | 3.679182183 | 3.071407821 |
| ENSG00000150593 | PDCD4 | -2.078674416 | 2.09E-09 | -2.3146147 | 1.49E-11 | -4.393289115 | 10.98974631 | 10.93377402 | 10.36098398 | 8.705093576 | 8.73668456 | 8.373850835 | 8.592942424 |
| ENSG00000127884 | ECHS1 | -2.117503775 | 0.000600374 | -2.277492832 | 0.000185631 | -4.394996607 | 11.9688936 | 10.47544827 | 11.40332671 | 9.288777073 | 9.30087923 | 8.823467578 | 9.392696434 |
| ENSG00000188542 | DUSP28 | -2.58938675 | 0.206839295 | -1.808417952 | 0.383915641 | -4.397804702 | 2 | 4.93638933 | 8.266922072 | 4.237680128 | 4.68029874 | 5.185595313 | 5.101530266 |
| ENSG00000100911 | PSME2 | -1.125518655 | 0.001649914 | -3.272665835 | 6.16E-23 | -4.39818449 | 11.06081658 | 10.76012279 | 11.05383672 | 9.813216364 | 9.86994125 | 7.401588825 | 7.974870548 |
| ENSG00000145075 | CCDC39 | -4.138691062 | 0.08619446 | -0.260855499 | 0.917615645 | -4.399546561 | 2 | 3.686303414 | 4.058998049 | 2 | 2.2680111 | 3.429858444 | 3.209956562 |
| ENSG00000257591 | ZNF625 | -0.494132803 | 0.69646163 | -3.907194904 | 0.000163346 | -4.401327706 | 4.00722531 | 6.666836837 | 6.43197674 | 5.639097847 | 5.63601114 | 3.27687201 | 2.782576693 |
| ENSG00000169972 | PUSL1 | -2.487064709 | 0.003076004 | -1.915493088 | 0.025713804 | -4.402557797 | 9.839306291 | 8.433041974 | 7.600047325 | 6.394182228 | 6.57678965 | 7.046766788 | 7.030725399 |
| ENSG00000102317 | RBM3 | -2.053825982 | 8.26E-05 | -2.352706225 | 4.59E-06 | -4.406532207 | 12.2287862 | 11.5089946 | 12.58588223 | 10.07584382 | 10.1695649 | 9.450048986 | 10.12366954 |
| ENSG00000120149 | MSX2 | -4.01613522 | 0.028092908 | -0.390908642 | 0.856832448 | -4.407043862 | 2 | 7.538394143 | 4.874587708 | 3.023133209 | 3.08597292 | 6.084012396 | 5.540219242 |
| ENSG00000171872 | KLF17 | -0.368871079 | 0.903690956 | -4.039825047 | 0.110909474 | -4.408696126 | 5.330594134 | 2 | 2 | 3.462030052 | 3.94787299 | 2.094598369 | 2.374681325 |
| ENSG00000184232 | OAF | -0.703391006 | 0.013540843 | -3.708025421 | 3.86E-48 | -4.411416427 | 12.50547411 | 12.28376116 | 12.32244095 | 11.64798529 | 11.6934836 | 8.476781207 | 8.856124078 |
| ENSG00000048540 | LMO3 | -2.910507749 | 0.443895169 | -1.502667696 | 0.691938174 | -4.413175445 | 2 | 4.11352228 | 2 | 2.3892346 | 2 | 2.34604626 | 2.632958241 |
| ENSG00000138629 | UBL7 | -2.079491442 | 4.06E-07 | -2.339675536 | 7.27E-09 | -4.419166978 | 9.790941832 | 9.591588447 | 10.41552659 | 7.818544084 | 8.00183183 | 7.646681011 | 7.66855289 |
| ENSG00000089009 | RPL6 | -1.897938492 | 0.000107449 | -2.52368001 | 1.19E-07 | -4.421618502 | 13.60557937 | 12.48474162 | 12.84130843 | 11.15408254 | 11.1637233 | 10.52182699 | 10.54714069 |
| ENSG00000168734 | PKIG | -2.603770309 | 1.51E-06 | -1.818697293 | 0.0010959 | -4.422467602 | 12.15236252 | 10.82507101 | 11.29839064 | 8.929383887 | 8.94583992 | 9.76049665 | 9.673706637 |
| ENSG00000079616 | KIF22 | -2.266195314 | 0.054418754 | -2.158459119 | 0.064499754 | -4.424654433 | 5.330594134 | 8.623598428 | 8.978369997 | 6.12217689 | 6.06344184 | 6.159811591 | 6.228882015 |
| ENSG00000148187 | MRRF | -2.091001587 | 0.006764239 | -2.334725638 | 0.002043428 | -4.425727225 | 9.143983763 | 8.124485672 | 10.16000817 | 7.222570549 | 7.38644297 | 6.940933331 | 7.188214218 |
| ENSG00000106355 | LSM5 | -1.940396692 | 0.184268122 | -2.48891469 | 0.075415372 | -4.429311382 | 9.790941832 | 6.389896633 | 6.073378139 | 6.604700657 | 6.46787796 | 6.115375798 | 5.913137508 |
| ENSG00000165912 | PACSIN3 | -4.016791514 | 8.05E-20 | -0.412853954 | 0.432665696 | -4.429645468 | 10.09970294 | 9.640525262 | 10.44204362 | 6.104969671 | 6.20590375 | 9.35351575 | 9.952884305 |
| ENSG00000114993 | RTKN | -2.375172972 | 1.68E-07 | -2.055476077 | 5.85E-06 | -4.430649049 | 8.39321512 | 9.367676479 | 9.165043284 | 6.727838783 | 6.68379438 | 6.911998488 | 7.110733573 |
| ENSG00000066651 | TRMT11 | -1.790438497 | 0.00837844 | -2.642272647 | 4.40E-05 | -4.432711145 | 9.473095744 | 8.077869079 | 8.266922072 | 7.057964187 | 6.91125523 | 6.39173974 | 5.909074772 |
| ENSG00000198901 | PRC1 | -0.380881241 | 0.882991782 | -4.058008017 | 0.05160469 | -4.438889258 | 3.327711234 | 7.638155693 | 2 | 5.978405189 | 5.6121243 | 2.746468522 | 3.236138336 |

| | | | | | | | | | | | | | |
|---|---|---|---|---|---|---|---|---|---|---|---|---|---|
| ENSG00000142748 | FCN3 | -3.267967567 | NA | -1.173994584 | NA | -4.441962152 | 3.327711234 | 2 | 2 | 2.141737246 | 2 | 2.139668783 | 2.421067459 |
| ENSG00000174628 | IQCK | -1.051467089 | 0.267665409 | -3.390614321 | 4.50E-05 | -4.44208141 | 5.532353208 | 6.899080544 | 7.600047325 | 5.829220145 | 5.9725602 | 3.56816602 | 4.188602707 |
| ENSG00000095539 | SEMA4G | -4.327543174 | 1.82E-10 | -0.116703132 | 0.895080873 | -4.444246306 | 7.940205782 | 7.002537973 | 8.721235202 | 4.17310023 | 3.98554648 | 7.988280194 | 7.865906035 |
| ENSG00000153551 | CMTM7 | -1.93894812 | 0.224460923 | -2.50676097 | 0.100917556 | -4.445709091 | 8.797441463 | 8.777382925 | 3.369317972 | 6.321555392 | 6.35006802 | 5.714849065 | 5.884455123 |
| ENSG00000152463 | OLAH | -2.48161252 | 0.171886783 | -1.964716804 | 0.262927366 | -4.446329324 | 4.467311984 | 3.076665209 | 4.058998049 | 2.784612485 | 2.38539787 | 3.172301926 | 2.277179783 |
| ENSG00000178175 | ZNF366 | -2.833911415 | 0.210179323 | -1.613634509 | 0.458154419 | -4.447545923 | 4.00722531 | 2 | 3.369317972 | 2.3892346 | 2.14022021 | 2.656294034 | 2.466008509 |
| ENSG00000124256 | ZBP1 | 0.889944644 | 0.794914423 | -5.339607229 | 0.108157469 | -4.449662585 | 2 | 3.686303414 | 2 | 3.227757282 | 3.39680241 | 2 | 2.059860178 |
| ENSG00000125675 | GRIA3 | -0.766696868 | 0.101694894 | -3.687056554 | 3.03E-19 | -4.453753422 | 7.790810999 | 8.213429689 | 8.110401118 | 7.19843704 | 7.39696604 | 4.820739789 | 4.341135913 |
| ENSG00000198960 | ARMCX6 | -0.963538377 | 0.632341559 | -3.491125796 | 0.046421006 | -4.454664173 | 11.07300288 | 9.624396729 | 3.369317972 | 9.03081742 | 8.93865515 | 6.653021698 | 6.353705253 |
| ENSG00000112208 | BAG2 | -2.748083716 | 4.18E-10 | -1.708623079 | 0.000160068 | -4.456706795 | 10.70134604 | 10.00683004 | 10.66665689 | 7.711768301 | 7.81567127 | 9.079108672 | 8.428831018 |
| ENSG00000262795 | | 1.185567138 | 0.106775299 | -5.642589474 | 1.30E-18 | -4.457022336 | 10.75779627 | 9.549500232 | 9.06622548 | 11.14332431 | 11.1678257 | 4.5389195 | 4.634793542 |
| ENSG00000124257 | NEURL2 | -0.564756938 | 0.547445831 | -3.893061904 | 1.07E-06 | -4.457818842 | 4.467311984 | 5.946638857 | 5.59507254 | 4.927308402 | 5.0259382 | 2.68698273 | 2.817676617 |
| ENSG00000185019 | UBOX5 | -2.534605911 | 0.154872672 | -1.924989195 | 0.284006025 | -4.459595106 | 4.00722531 | 7.572420273 | 10.53119344 | 6.535745081 | 6.75096147 | 7.318276922 | 7.150854407 |
| ENSG00000117519 | CNN3 | -3.028740839 | 9.53E-18 | -1.433299637 | 9.81E-05 | -4.462040476 | 13.44216416 | 13.08602666 | 13.81388507 | 10.47580465 | 10.4286902 | 11.92065507 | 12.15971167 |
| ENSG00000107537 | PHYH | -1.601330271 | 0.006908109 | -2.862902456 | 2.76E-07 | -4.464232727 | 9.247966931 | 8.671479029 | 10.03493575 | 7.781117077 | 7.89452713 | 6.728720192 | 6.485052582 |
| ENSG00000282092 | | -2.052632319 | 0.027195324 | -2.414057901 | 0.007716323 | -4.46669022 | 11.61879218 | 9.549500232 | 9.426401271 | 8.532433657 | 8.53866332 | 8.157391075 | 8.199398653 |
| ENSG00000019549 | SNAI2 | -1.414370394 | 0.000431872 | -3.059285977 | 6.84E-16 | -4.473656371 | 10.46286899 | 10.72651826 | 11.28013451 | 9.490242771 | 9.41744821 | 7.849352333 | 7.803803816 |
| ENSG00000162227 | TAF6L | -1.952789477 | 0.003175086 | -2.522113292 | 8.10E-05 | -4.474902769 | 9.174465797 | 7.701025996 | 9.227351203 | 6.902323594 | 6.95464102 | 6.252491482 | 6.501335855 |
| ENSG00000186854 | TRABD2A | -0.395997655 | 0.900708025 | -4.080171036 | 0.117000847 | -4.476168691 | 2 | 2 | 7.934808822 | 6.024891728 | 6.00960265 | 3.1504466 | 3.100202476 |
| ENSG00000163918 | RFC4 | -2.724782038 | 0.000177796 | -1.752277051 | 0.019191632 | -4.477059088 | 8.138771685 | 6.535001748 | 7.225742779 | 5.093559143 | 4.65713986 | 5.963846589 | 5.534955934 |
| ENSG00000162426 | SLC45A1 | -3.748899505 | 4.57E-09 | -0.728342206 | 0.297854277 | -4.477241711 | 7.031698319 | 6.389896633 | 7.691296561 | 3.615725078 | 3.98554648 | 6.394066622 | 6.443521721 |
| ENSG00000186185 | KIF18B | -1.97731369 | 0.081326682 | -2.501128109 | 0.020270597 | -4.478441799 | 5.095959458 | 4.710564154 | 5.772716771 | 3.566290607 | 3.74316441 | 2.746468522 | 3.735548302 |
| ENSG00000151413 | NUBPL | -2.598241543 | 0.059358025 | -1.880472475 | 0.179070824 | -4.478714018 | 4.467311984 | 7.572420273 | 9.083172337 | 5.391962469 | 5.5755357 | 6.054896192 | 6.245088435 |
| ENSG00000262771 | | -0.397414504 | 0.713282111 | -4.082060635 | 1.77E-06 | -4.479475139 | 10.07567912 | 8.623598428 | 7.646393253 | 8.812346549 | 8.64106878 | 4.927401733 | 5.425444337 |
| ENSG00000163913 | IFT122 | -2.521055087 | 5.60E-06 | -1.959369974 | 0.000476276 | -4.480425061 | 8.19931886 | 8.276714465 | 9.287079172 | 6.34372989 | 6.08095082 | 6.591495479 | 6.90172361 |
| ENSG00000135446 | CDK4 | -1.530959234 | 0.001413507 | -2.950779787 | 7.14E-11 | -4.481739022 | 11.43587654 | 10.95971654 | 10.3329194 | 9.449072612 | 9.45653888 | 8.019511671 | 8.07345976 |
| ENSG00000105732 | ZNF574 | -2.619166143 | 1.36E-05 | -1.865690947 | 0.002532488 | -4.48485709 | 10.55263183 | 9.235879652 | 9.358421373 | 7.202487431 | 7.31055061 | 8.124136806 | 7.854331622 |
| ENSG00000211445 | GPX3 | -3.541138623 | 0.099087773 | -0.943820715 | 0.692261286 | -4.484959338 | 9.435834703 | 5.303523747 | 2 | 4.415698455 | 4.811949 | 7.039353222 | 6.990916783 |
| ENSG00000116874 | WARS2 | -2.669164388 | 0.025103126 | -1.821076511 | 0.136586769 | -4.4902409 | 9.204317094 | 8.004992147 | 5.39244166 | 5.44762523 | 5.81407719 | 6.315225883 | 6.55171954 |
| ENSG00000125458 | NT5C | -1.846700504 | 0.012348836 | -2.643915503 | 0.000175425 | -4.490616006 | 9.90440318 | 8.414402626 | 8.35328246 | 7.238438152 | 7.28433928 | 6.292954465 | 6.666875073 |
| ENSG00000137818 | RPLP1 | -2.550695091 | 0.000263662 | -1.942278445 | 0.006817972 | -4.492973536 | 14.19387916 | 12.50910266 | 12.73541118 | 10.78321184 | 10.8262286 | 11.377928 | 11.44564963 |
| ENSG00000175390 | EIF3F | -2.878719451 | 8.10E-05 | -1.614696736 | 0.036755663 | -4.493416186 | 12.3737531 | 11.29895829 | 10.33998689 | 8.727753724 | 8.67892944 | 9.855584152 | 10.05638425 |
| ENSG00000206237 | | -7.140990513 | 0.030869987 | 2.647339749 | 0.387344103 | -4.493650763 | 2 | 5.131603625 | 2 | 2 | 2 | 6.643273885 | 5.379200993 |
| ENSG00000170264 | FAM161A | -1.901868268 | 0.125866269 | -2.592122795 | 0.028494013 | -4.493991063 | 8.257427002 | 5.838937808 | 5.59507254 | 5.391962469 | 5.21224859 | 5.085488617 | 4.175132639 |
| ENSG00000184347 | SLIT3 | -2.891323511 | 0.000877941 | -1.605140827 | 0.080940119 | -4.496464339 | 8.340423424 | 10.90735644 | 10.54960077 | 7.454228844 | 7.39346687 | 8.583088126 | 8.790280628 |
| ENSG00000149016 | TUT1 | -2.390036566 | 0.123445724 | -2.106998351 | 0.172021653 | -4.497034917 | 4.00722531 | 8.639735838 | 9.670202709 | 6.394182228 | 6.29861567 | 6.797131715 | 6.412287469 |
| ENSG00000102003 | SYP | -2.266043021 | 0.405931256 | -2.232655683 | 0.399776674 | -4.498698704 | 2 | 2 | 5.930865648 | 2.695456494 | 3.2184422 | 3.235948948 | 2.70970617 |
| ENSG00000101457 | DNTTIP1 | -1.458540022 | 3.28E-06 | -3.040551381 | 5.84E-24 | -4.499091404 | 9.730679447 | 9.214490408 | 9.601288346 | 8.049698831 | 8.11451391 | 6.514451546 | 6.569847194 |
| ENSG00000154710 | RABGEF1 | -0.062118211 | 0.923323112 | -4.438404128 | 1.42E-21 | -4.500522339 | 9.839306291 | 8.791896851 | 9.746650996 | 9.470224559 | 9.45905186 | 5.31402805 | 5.164234344 |
| ENSG00000114854 | TNNC1 | -4.938308435 | 0.055017038 | 0.435118632 | 0.884384341 | -4.503189804 | 2 | 6.602424674 | 2 | 2.3892346 | 2.2680111 | 4.301748724 | 6.199243628 |
| ENSG00000242265 | PEG10 | -2.921424252 | 0.000136045 | -1.582276947 | 0.051489483 | -4.503701199 | 10.86447339 | 10.79656548 | 9.301632639 | 7.802622679 | 7.32163988 | 9.39882884 | 8.140837768 |
| ENSG00000256269 | HMBS | -2.274377351 | 2.48E-08 | -2.231882164 | 3.19E-08 | -4.506259514 | 9.262228249 | 9.148351514 | 8.805342542 | 6.907295206 | 6.79406287 | 6.546183077 | 7.173047408 |
| ENSG00000127561 | SYNGR3 | -4.308922776 | 0.012336386 | -0.19827647 | 0.927674983 | -4.507199247 | 9.699577239 | 5.457120953 | 4.058998049 | 4.299492648 | 4.16043394 | 8.138086022 | 7.889811111 |
| ENSG00000147155 | EBP | -2.521266248 | 1.90E-07 | -1.986311599 | 5.05E-05 | -4.507577847 | 10.499447 | 10.42046463 | 10.34701993 | 7.925329813 | 7.91907183 | 7.785203203 | 8.903351429 |
| ENSG00000187764 | SEMA4D | -4.003595085 | 0.038376006 | -0.505833584 | 0.817094045 | -4.509428669 | 3.327711234 | 6.228547981 | 2 | 2.498691189 | 2.49394783 | 4.478802948 | 4.511590831 |
| ENSG00000175093 | SPSB4 | -4.914508033 | 0.153618856 | 0.404333163 | 0.90832745 | -4.51017487 | 2 | 4.442710129 | 2 | 2.141737246 | 2 | 2.885263063 | 4.09155917 |
| ENSG00000205403 | CFI | 1.640931884 | 0.739034243 | -6.152908519 | 0.188634646 | -4.511976635 | 3.327711234 | 2 | 2 | 3.921164147 | 2.38539787 | 2 | 2 |
| ENSG00000108384 | RAD51C | -3.23751219 | 4.78E-08 | -1.276012855 | 0.044054943 | -4.513525045 | 8.313285031 | 7.35507212 | 8.805342542 | 5.210797711 | 5.14724905 | 6.929749747 | 7.100081147 |
| ENSG00000196418 | ZNF124 | -1.064139952 | 0.265307052 | -3.449667956 | 3.87E-05 | -4.513807909 | 5.095959458 | 7.315436877 | 6.629488387 | 5.697663076 | 5.53799497 | 3.753554863 | 3.560117069 |
| ENSG00000083845 | RPS5 | -2.605916067 | 7.43E-06 | -1.91167943 | 0.001355764 | -4.517595497 | 13.44607356 | 12.4880879 | 11.99095026 | 10.16208248 | 10.1772163 | 10.79661666 | 10.92453561 |
| ENSG00000088448 | ANKRD10 | -1.981012621 | 0.000345401 | -2.537199661 | 2.52E-06 | -4.518212282 | 10.9638027 | 11.11784453 | 12.07171128 | 9.539105795 | 9.45318143 | 9.196072618 | 8.639050694 |
| ENSG00000106571 | GLI3 | -1.729083462 | 0.329061986 | -2.792303668 | 0.092655285 | -4.52138713 | 3.327711234 | 8.806266215 | 9.972637969 | 7.292639438 | 7.15805139 | 6.157074092 | 6.251520369 |
| ENSG00000184730 | APOBR | -4.852502712 | 0.163759565 | 0.328916851 | 0.927015754 | -4.523585862 | 4.467311984 | 2 | 2 | 2 | 2.14022021 | 3.904814898 | 2.981406154 |
| ENSG00000128833 | MYO5C | -1.568108721 | 0.408062848 | -2.955699573 | 0.090151257 | -4.523808294 | 4.467311984 | 2 | 5.39244166 | 3.023133209 | 3.60476711 | 2.457042231 | 2.851942824 |
| ENSG00000116544 | DLGAP3 | -3.17403894 | 0.090685651 | -1.350306946 | 0.47372156 | -4.524345886 | 3.327711234 | 5.131603625 | 2 | 2.3892346 | 2.49394783 | 3.235948948 | 3.128433649 |
| ENSG00000133265 | HSPBP1 | -2.642216434 | 0.000117983 | -1.886088433 | 0.007647248 | -4.528304867 | 10.85038273 | 9.114109157 | 9.681375057 | 7.419851705 | 7.48508724 | 8.05302937 | 8.325164853 |
| ENSG00000205213 | LGR4 | -2.193175539 | 0.011385558 | -2.335426669 | 0.006294563 | -4.528602208 | 8.009475818 | 10.17682307 | 10.20715117 | 7.68596357 | 7.48508724 | 7.749302529 | 7.071284046 |
| ENSG00000148057 | IDNK | -2.097834973 | 0.000427594 | -2.433421112 | 2.08E-05 | -4.531256085 | 6.138360429 | 6.228547981 | 6.958202216 | 4.523237872 | 4.63360314 | 4.166733998 | 4.434543419 |
| ENSG00000081803 | CADPS2 | -1.936834138 | 0.100236935 | -2.59519686 | 0.021232762 | -4.532030997 | 5.70932728 | 9.018611864 | 8.042706615 | 6.213314212 | 6.27599024 | 6.049001768 | 5.035976547 |
| ENSG00000183010 | PYCR1 | -3.323062067 | 1.38E-15 | -1.211535802 | 0.006709168 | -4.534597869 | 10.85509492 | 11.45529853 | 11.79914889 | 8.151677543 | 8.08010078 | 10.32063553 | 10.09400257 |
| ENSG00000179776 | CDH5 | -3.705167669 | 0.223171036 | -0.834169893 | 0.794825276 | -4.539337562 | 4.467311984 | 2 | 3.369317972 | 2 | 2.38539787 | 2.225793514 | 3.620995423 |
| ENSG00000134057 | CCNB1 | -2.08730944 | 0.217193372 | -2.456657974 | 0.132407854 | -4.543967414 | 10.33377634 | 8.522788974 | 4.058998049 | 7.186217188 | 6.94511186 | 6.45105139 | 6.936070085 |
| ENSG00000275835 | TUBGCP5 | 0.19331591 | 0.795707131 | -4.737769604 | 1.79E-16 | -4.544453695 | 8.138771685 | 6.78762695 | 7.451345288 | 7.728718552 | 7.75697332 | 3.393105702 | 3.496556178 |
| ENSG00000185250 | PPIL6 | -2.351325797 | 0.087667241 | -2.193347549 | 0.102264609 | -4.544673346 | 5.095959458 | 3.686303414 | 6.43197674 | 3.566290607 | 3.55551834 | 4.099885429 | 3.042026731 |
| ENSG00000219626 | FAM228B | -2.957940194 | 0.005993307 | -1.58923962 | 0.15962782 | -4.547179815 | 6.96351253 | 5.595927442 | 8.631919287 | 4.694064037 | 4.89348248 | 5.982522014 | 6.081436739 |
| ENSG00000173762 | CD7 | -1.243602627 | 0.543429761 | -3.304483212 | 0.073251024 | -4.548085839 | 4.00722531 | 5.838937808 | 2 | 3.566290607 | 3.90918929 | 2.937246078 | 2.277179783 |
| ENSG00000100075 | SLC25A1 | -2.926677736 | 3.04E-08 | -1.621437631 | 0.003509963 | -4.548115367 | 12.37211025 | 11.7361163 | 12.79319859 | 9.410199593 | 9.47652116 | 10.26642009 | 11.10229933 |
| ENSG00000180875 | GREM2 | 1.846634285 | 0.204890893 | -6.400736252 | 1.54E-06 | -4.554101967 | 7.667647223 | 6.311476754 | 3.369317972 | 8.429064458 | 8.39732094 | 2.183373728 | 2.509591799 |
| ENSG00000163344 | PMVK | -2.79980626 | 1.97E-11 | -1.754692418 | 3.48E-05 | -4.554498679 | 9.48530526 | 8.791896851 | 9.399592459 | 6.463327516 | 6.55189318 | 7.269684675 | 7.734322559 |
| ENSG00000107371 | EXOSC3 | -1.957311072 | 0.010509078 | -2.600123291 | 0.000408599 | -4.557434363 | 9.544849385 | 7.60566235 | 9.212024683 | 7.062428156 | 7.0774796 | 6.643273885 | 6.228882015 |
| ENSG00000146233 | CYP39A1 | -2.525297988 | 0.378746936 | -2.033022451 | 0.464069117 | -4.558320439 | 4.467311984 | 2 | 2 | 2.141737246 | 2.49394783 | 2.42098416 | 2.466008509 |
| ENSG00000130826 | DKC1 | -2.412927436 | 1.60E-05 | -2.148828569 | 0.000128848 | -4.561756005 | 10.81695923 | 9.506147172 | 9.91701143 | 7.711768301 | 7.8720741 | 7.947671484 | 8.152913455 |
| ENSG00000114026 | OGG1 | -2.776802482 | 0.000233444 | -1.785046595 | 0.023283239 | -4.561849077 | 9.975371636 | 8.916342327 | 7.934808822 | 6.456560224 | 6.45450701 | 7.231159579 | 7.576773052 |
| ENSG00000205832 | C16orf96 | -1.655751365 | 0.603581513 | -2.918808134 | NA | -4.574559499 | 2 | 3.686303414 | 2 | 2.270786764 | 2.38539787 | 2.225793514 | 2.059860178 |
| ENSG00000171858 | RPS21 | 0.042190627 | 0.975050331 | -4.616901626 | 4.29E-06 | -4.574710999 | 10.66423795 | 7.669933291 | 8.561105505 | 9.590237441 | 9.53247497 | 4.984105765 | 5.164234344 |
| ENSG00000126249 | PDCD2L | -1.899913539 | 0.24142045 | -2.676926263 | 0.08152866 | -4.576839802 | 6.653098118 | 6.666836837 | 2 | 4.387512679 | 4.45712836 | 3.738983888 | 3.908316285 |
| ENSG00000139546 | TARBP2 | -2.316788808 | 0.006786037 | -2.260066244 | 0.008020131 | -4.576855051 | 10.4253394 | 8.469613413 | 8.511888463 | 7.182120794 | 7.14145882 | 7.225944286 | 7.208191747 |
| ENSG00000127995 | CASD1 | -2.035455771 | 0.000900255 | -2.541739229 | 1.93E-05 | -4.577195001 | 7.790810999 | 8.317412869 | 9.212024683 | 6.70517998 | 6.42052719 | 6.106889539 | 6.070564958 |
| ENSG00000132196 | HSD17B7 | -2.008452393 | 0.000987743 | -2.570076508 | 1.32E-05 | -4.578528901 | 8.892187472 | 8.101365653 | 7.600047325 | 6.197166296 | 6.47451726 | 5.561816579 | 6.029985803 |
| ENSG00000242110 | AMACR | -0.752507816 | 0.467358725 | -3.826557069 | 1.00E-05 | -4.579064885 | 6.89194353 | 6.140560021 | 8.434763955 | 6.788375664 | 6.68951263 | 4.005644654 | 4.033026654 |
| ENSG00000073792 | IGF2BP2 | -2.285877773 | 0.000232057 | -2.305139112 | 0.000190063 | -4.591016885 | 10.76782679 | 11.7859074 | 12.23415624 | 9.449072612 | 9.42432312 | 9.726278296 | 9.024253959 |
| ENSG00000203667 | COX20 | 0.20935167 | 0.691159366 | -4.801777492 | 7.99E-31 | -4.592425822 | 8.19931886 | 8.029695761 | 8.825624954 | 8.624367384 | 8.56222967 | 3.956110469 | 4.018014792 |
| ENSG00000116771 | AGMAT | -1.528571505 | 0.18711518 | -3.065600115 | 0.003824718 | -4.59417162 | 7.485143553 | 4.710564154 | 5.930865648 | 5.022023783 | 5.07918227 | 3.709392871 | 3.908316285 |
| ENSG00000167536 | DHRS13 | -1.593935651 | 0.403432532 | -3.000991273 | 0.087671418 | -4.594926924 | 8.540903468 | 8.101365653 | 2 | 6.252910529 | 6.18985553 | 4.667592615 | 5.164234344 |
| ENSG00000128394 | APOBEC3F | -1.257146228 | 0.103636128 | -3.338166463 | 1.28E-06 | -4.595312691 | 8.313285031 | 8.414402626 | 9.91701143 | 7.844695415 | 7.8286851 | 5.818966034 | 5.85096143 |
| ENSG00000112706 | IMPG1 | -4.714853507 | NA | 0.118009987 | NA | -4.59684352 | 2 | 2 | 3.369317972 | 2 | 2 | 2.831336763 | 2.374681325 |
| ENSG00000104886 | PLEKHJ1 | -2.24715343 | 0.008799219 | -2.350296206 | 0.005580838 | -4.597449636 | 11.10895605 | 9.814097497 | 8.763901704 | 7.890605449 | 8.05816814 | 8.029274808 | 7.702986259 |
| ENSG00000101019 | UQCC1 | -2.218331982 | 7.03E-05 | -2.379217993 | 1.60E-05 | -4.597549974 | 10.05124847 | 10.31889309 | 11.32712828 | 8.451360823 | 8.48706372 | 8.259184498 | 8.359810397 |
| ENSG00000038382 | TRIO | -10.17036391 | 1.66E-10 | 5.572729617 | 4.43E-40 | -4.597634297 | 6.468621203 | 6.602424674 | 6.43197674 | 2 | 2 | 11.91858564 | 12.11041621 |
| ENSG00000103227 | LMF1 | -2.802250581 | 2.65E-07 | -1.800394129 | 0.001377103 | -4.60264471 | 10.93293345 | 9.632483534 | 10.0085654 | 7.507583254 | 7.53675633 | 8.513094212 | 8.503483965 |
| ENSG00000197345 | MRPL21 | -2.11786704 | 0.000242984 | -2.489908409 | 1.06E-05 | -4.607775449 | 10.00958825 | 8.747909548 | 9.014156722 | 7.330144653 | 7.21466714 | 6.761530329 | 7.049302614 |
| ENSG00000042317 | SPATA7 | -1.837055166 | 0.349953578 | -2.771112346 | 0.133937533 | -4.608167512 | 9.064836028 | 7.503546079 | 2 | 6.252910529 | 6.00043082 | 5.328699035 | 5.211187127 |
| ENSG00000148334 | PTGES2 | -2.809290381 | 6.57E-07 | -1.798909455 | 0.002132231 | -4.608199837 | 10.5351205 | 9.943237334 | 11.37263945 | 7.93753157 | 7.95512324 | 8.932738969 | 8.9569744 |
| ENSG00000160999 | SH2B2 | -2.467693687 | 0.156306514 | -2.141008569 | 0.214813421 | -4.608702256 | 2 | 6.666836837 | 7.691296561 | 4.598938466 | 4.31639476 | 4.875056259 | 4.584731282 |
| ENSG00000162910 | MRPL55 | -2.744818422 | 0.001690934 | -1.864646484 | 0.039416005 | -4.609464906 | 10.69084077 | 9.267380431 | 8.236946975 | 6.887305128 | 7.17037177 | 7.930942203 | 7.859604233 |
| ENSG00000048162 | NOP16 | -2.630688127 | 2.60E-05 | -1.980606238 | 0.001917231 | -4.611294365 | 9.975371636 | 8.414402626 | 9.809100877 | 6.998639523 | 6.90635305 | 7.602089053 | 7.570355956 |
| ENSG00000101182 | PSMA7 | -1.947630913 | 2.54E-09 | -2.664506542 | 9.01E-17 | -4.612137455 | 12.50097114 | 11.98535617 | 12.49540932 | 10.35909586 | 10.4427247 | 9.529720379 | 9.829607703 |
| ENSG00000129084 | PSMA1 | -0.777816678 | 0.231292431 | -3.837025637 | 3.47E-12 | -4.614842315 | 11.72042071 | 10.5284131 | 10.51254821 | 10.29936444 | 10.2247551 | 7.328045423 | 7.140495192 |
| ENSG00000143942 | CHAC2 | -2.122832336 | 0.281057743 | -2.492205525 | 0.18752152 | -4.615037861 | 7.031698319 | 4.11352228 | 2 | 3.95996233 | 3.7864959 | 3.5515855 | 3.67940853 |
| ENSG00000139343 | SNRPF | -1.822118949 | 0.061746837 | -2.796656721 | 0.002350467 | -4.61877567 | 9.800745303 | 7.901717006 | 7.646393253 | 7.062428156 | 6.94032357 | 5.618636935 | 6.415155055 |
| ENSG00000215788 | TNFRSF25 | -3.74087585 | 5.06E-05 | -0.87857808 | 0.399807732 | -4.61945393 | 6.366709671 | 6.899080544 | 8.721235202 | 4.299492648 | 4.22485516 | 6.879153024 | 6.797888492 |
| ENSG00000250120 | PCDHA10 | -1.924367209 | 0.309909983 | -2.703232216 | 0.129124437 | -4.627599425 | 2 | 5.595927442 | 4.058998049 | 3.566290607 | 2.68924543 | 2.831336763 | 2.782576693 |
| ENSG00000092036 | HAUS4 | -1.956120163 | 0.013801401 | -2.671822571 | 0.000442844 | -4.627942734 | 8.716893051 | 6.844429755 | 8.143093325 | 6.18902405 | 6.18985553 | 5.83618166 | 5.122735884 |
| C11ORF63 | C11ORF63 | 3.425839387 | 0.007764718 | -8.062198824 | 5.46E-05 | -4.636359437 | 3.327711234 | 3.076665209 | 4.058998049 | 6.415275989 | 6.37156942 | 2 | 2 |
| ENSG00000166669 | ATF7IP2 | -2.142888627 | 0.255328097 | -2.495491578 | 0.16984745 | -4.638380205 | 7.904282243 | 6.140560021 | 2 | 4.739373666 | 4.74762503 | 4.875056259 | 3.841670744 |
| ENSG00000143434 | SEMA6C | -3.213550721 | 0.009447578 | -1.425483299 | 0.284063725 | -4.63903402 | 8.257427002 | 5.457120953 | 9.636156974 | 5.44762523 | 5.51241389 | 7.214140484 | 7.131805326 |
| ENSG00000249715 | FER1L5 | -3.69764103 | 0.296352897 | -0.943637677 | 0.799962733 | -4.641278707 | 4.815609471 | 2 | 2 | 2 | 2.38539787 | 3.1504466 | 2.885414003 |

| | | | | | | | | | | | | | |
|---|---|---|---|---|---|---|---|---|---|---|---|---|---|
| ENSG00000185608 | MRPL40 | -2.386890139 | 6.78E-15 | -2.25801557 | 9.13E-14 | -4.64490571 | 9.90440318 | 9.821190352 | 10.19160738 | 7.700356475 | 7.52720828 | 7.575682301 | 7.892900165 |
| ENSG00000037897 | METTL1 | -2.170052174 | 2.70E-06 | -2.476143993 | 4.64E-08 | -4.646196168 | 9.128497779 | 8.623598428 | 8.143093325 | 6.535745081 | 6.59518404 | 6.146071859 | 6.392053169 |
| ENSG00000033011 | ALG1 | -1.370514765 | 0.050222651 | -3.276002079 | 2.43E-07 | -4.646516844 | 9.848787621 | 8.451443573 | 10.06936278 | 8.237082658 | 8.2618769 | 6.18690462 | 6.577546999 |
| ENSG00000176386 | CDC26 | 0.026528123 | 0.98350331 | -4.673803218 | 1.36E-06 | -4.647275095 | 8.313285031 | 5.595927442 | 6.718950348 | 7.210554257 | 7.41433569 | 3.172301926 | 3.45257121 |
| ENSG00000168101 | NUDT16L1 | -2.912962579 | 6.16E-12 | -1.735096289 | 6.34E-05 | -4.648058868 | 9.689059027 | 9.470501606 | 10.32581712 | 6.922108067 | 7.07311172 | 8.024776954 | 8.269209446 |
| ENSG00000147586 | MRPS28 | -1.184708729 | 0.259704244 | -3.465200367 | 0.000181365 | -4.649909097 | 8.228665467 | 6.04685525 | 8.721235202 | 6.846474326 | 6.90143415 | 4.461156421 | 5.028505473 |
| ENSG00000154263 | ABCA10 | -0.010440377 | 0.997204246 | -4.640254224 | 0.065538486 | -4.650694601 | 5.095959458 | 2 | 3.369317972 | 3.95996233 | 4.02226117 | 2.3070652 | 2 |
| ENSG00000166582 | CENPV | -2.539074624 | 9.53E-09 | -2.112156974 | 1.41E-06 | -4.651231599 | 7.624139768 | 8.337339476 | 8.35328246 | 5.686138681 | 5.71663161 | 6.257611854 | 5.921228818 |
| ENSG00000100092 | SH3BP1 | -1.459548697 | 0.038232977 | -3.193249005 | 7.36E-07 | -4.652797701 | 7.940205782 | 8.777382925 | 9.503962296 | 7.395290829 | 7.47849652 | 6.069527745 | 5.414021719 |
| ENSG00000157150 | TIMP4 | -5.710104213 | 0.175377978 | 1.055328099 | 0.80714548 | -4.654776114 | 4.00722531 | 2 | 2 | 2 | 2 | 3.517841605 | 3.698365557 |
| ENSG00000145088 | EAF2 | -3.642744331 | 0.067543282 | -1.012293539 | 0.626957756 | -4.655037869 | 5.532353208 | 3.076665209 | 2 | 2.3892346 | 2.38539787 | 3.753554863 | 3.287118176 |
| ENSG00000102359 | SRPX2 | -0.674175041 | 0.366343821 | -3.983257264 | 1.32E-10 | -4.657432305 | 10.56993315 | 9.506147172 | 11.21627664 | 9.925905286 | 9.9084519 | 6.52722803 | 6.782425482 |
| ENSG00000179455 | MKRN3 | -2.295974599 | 0.467226103 | -2.361537829 | NA | -4.657512428 | 2 | 2 | 4.058998049 | 2.141737246 | 2.38539787 | 2.384001718 | 2.117335275 |
| ENSG00000204859 | ZBTB48 | -2.818448397 | 0.029091374 | -1.840110264 | 0.167292121 | -4.65855866 | 7.218819781 | 6.389896633 | 9.999667074 | 5.890757403 | 6.04571775 | 7.096193303 | 6.678855578 |
| ENSG00000179528 | LBX2 | -2.081937252 | 0.046816693 | -2.576871601 | 0.010481522 | -4.658808854 | 7.435667355 | 6.228547981 | 5.59507254 | 4.443344104 | 4.95174433 | 4.792794928 | 3.580696614 |
| ENSG00000114988 | LMAN2L | -2.360586636 | 2.84E-14 | -2.300966777 | 7.29E-14 | -4.661553414 | 10.70657014 | 10.18234215 | 10.40208303 | 8.104838684 | 8.09740995 | 8.034504643 | 8.275534969 |
| ENSG00000186280 | KDM4D | -2.381522771 | 0.108329575 | -2.280721305 | 0.117290762 | -4.662244076 | 7.750913381 | 4.442710129 | 4.523675391 | 4.17310023 | 4.43005853 | 4.240999983 | 4.522269241 |
| ENSG00000168491 | CCDC110 | -5.228467136 | 0.114708096 | 0.566028953 | 0.864047058 | -4.662438183 | 2 | 4.710564154 | 2 | 2.141737246 | 2 | 4.397636241 | 3.287118176 |
| ENSG00000243646 | IL10RB | 0.882504731 | 0.035933276 | -5.545334816 | 4.84E-45 | -4.662830085 | 8.998234871 | 8.469613413 | 8.266922072 | 9.445659335 | 9.52609769 | 3.616794016 | 3.620995423 |
| ENSG00000186166 | CCDC84 | -7.16191671 | 0.001267977 | 2.494755485 | 0.126314768 | -4.667161226 | 2 | 4.11352228 | 4.523675391 | 2 | 2 | 5.919306827 | 6.117095472 |
| ENSG00000213221 | DNLZ | -0.643122044 | 0.738608861 | -4.028629473 | 0.012596707 | -4.671571517 | 2 | 6.602424674 | 7.398165132 | 5.85002558 | 5.91514978 | 2.858551861 | 3.496556178 |
| ENSG00000085741 | WNT11 | -1.865907254 | 0.451437436 | -2.808233216 | 0.223758017 | -4.674140471 | 3.327711234 | 4.442710129 | 2 | 2.784612485 | 2.49394783 | 2.62493832 | 2.059860178 |
| ENSG00000033122 | LRRC7 | -5.368611553 | 0.233789503 | 0.6895371 | 0.885698578 | -4.679074452 | 2 | 3.686303414 | 2 | 2 | 2 | 3.393105702 | 2.918126211 |
| ENSG00000167178 | ISLR2 | -4.737640468 | 0.063719227 | 0.056502107 | 0.984481872 | -4.681138361 | 3.327711234 | 2 | 5.59507254 | 2.3892346 | 2 | 3.981090143 | 4.729989799 |
| ENSG00000185664 | PMEL | -2.445670188 | 0.159914848 | -2.237854931 | 0.190022549 | -4.683525119 | 4.00722531 | 3.076665209 | 7.163363831 | 3.566290607 | 3.90918929 | 4.04171018 | 3.735548302 |
| ENSG00000197360 | ZNF98 | 2.502419524 | 0.463096875 | -7.186499402 | 0.0384565 | -4.684079878 | 2 | 2 | 4.058998049 | 4.907595604 | 4.58534546 | 2 | 2 |
| ENSG00000163596 | ICA1L | -1.7015804 | 0.064757647 | -2.984220001 | 0.000486561 | -4.685800401 | 7.709880939 | 6.602424674 | 8.978369997 | 6.394182228 | 6.44101097 | 5.233158834 | 5.224326245 |
| ENSG00000216937 | CCDC7 | -5.368128939 | 0.236297945 | 0.68031239 | 0.887741643 | -4.687816549 | 2 | 3.686303414 | 2 | 2 | 2 | 3.429858444 | 2.851942824 |
| ENSG00000136104 | RNASEH2B | -1.982636363 | 0.041735857 | -2.70560579 | 0.003589152 | -4.688242154 | 8.836089686 | 7.638155693 | 6.203069917 | 5.870535235 | 6.14061114 | 5.50697268 | 5.164234344 |
| ENSG00000126583 | PRKCG | -1.165973858 | 0.525963919 | -3.522638245 | 0.033795077 | -4.688612103 | 4.00722531 | 4.11352228 | 2 | 2.784612485 | 3.15372745 | 2.225793514 | 2.277179783 |
| ENSG00000100246 | DNAL4 | -2.619723939 | 0.133013379 | -2.069751855 | 0.238769141 | -4.689475794 | 10.19941743 | 8.395519306 | 3.369317972 | 6.463327516 | 6.3857277 | 7.001702955 | 6.903766785 |
| ENSG00000147996 | CBWD5 | 0.088618796 | 0.936463188 | -4.779206221 | 8.18E-09 | -4.690587424 | 6.468621203 | 7.189546432 | 8.721235202 | 7.815902688 | 7.89452713 | 3.600766115 | 3.518055793 |
| ENSG00000126602 | TRAP1 | -2.499874855 | 3.35E-18 | -2.192842944 | 1.32E-14 | -4.6927178 | 10.09173937 | 10.45735223 | 10.45512158 | 7.872924734 | 7.85944752 | 8.298220966 | 8.025476363 |
| ENSG00000146409 | SLC18B1 | -0.792451644 | 0.38329635 | -3.901297752 | 3.57E-07 | -4.693749395 | 7.940205782 | 8.169643011 | 6.073378139 | 6.820355514 | 6.94032357 | 4.199031499 | 4.002845082 |
| ENSG00000137261 | KIAA0319 | -2.841916096 | 0.335597482 | -1.856828377 | 0.527954208 | -4.698744473 | 2 | 2 | 5.156626401 | 2.600425781 | 2.2680111 | 3.172301926 | 2.225841198 |
| ENSG00000105486 | LIG1 | -2.289156198 | 0.015510476 | -2.41006366 | 0.009656574 | -4.699219858 | 9.064836028 | 7.274681896 | 6.718950348 | 5.764940652 | 5.91514978 | 5.790986459 | 5.66592325 |
| ENSG00000102409 | BEX4 | -3.007776614 | 2.48E-05 | -1.694539017 | 0.024328003 | -4.702315631 | 10.39976609 | 9.415325457 | 8.460932916 | 6.777556219 | 6.57060569 | 8.065491673 | 7.839464964 |
| ENSG00000139131 | YARS2 | -2.470013347 | 2.21E-05 | -2.232359425 | 0.000128473 | -4.702372771 | 9.895281535 | 8.747909548 | 10.17589429 | 7.261917051 | 7.30311013 | 7.389977651 | 7.629578661 |
| ENSG00000136936 | XPA | -2.504884803 | 1.83E-08 | -2.198284261 | 6.32E-07 | -4.703169064 | 8.87372952 | 8.124485672 | 8.380957451 | 6.043074888 | 6.0722229 | 6.104049656 | 6.556922154 |
| ENSG00000162300 | ZFPL1 | -1.155398194 | 0.047405404 | -3.555933641 | 1.01E-11 | -4.711331835 | 8.836089686 | 8.276714465 | 9.478569125 | 7.742693204 | 7.85690889 | 5.29920634 | 5.689801209 |
| ENSG00000205581 | HMGN1 | -2.166960092 | 1.87E-06 | -2.545842288 | 1.29E-08 | -4.71280238 | 12.58563132 | 11.631062 | 11.83210411 | 9.952002342 | 9.8762226 | 9.39882884 | 9.664095852 |
| ENSG00000196917 | HCAR1 | -3.210806623 | 0.017635531 | -1.503699076 | 0.262013998 | -4.714505699 | 4.815609471 | 3.686303414 | 4.874587708 | 2.868578001 | 2.2680111 | 3.193831101 | 3.620995423 |
| ENSG00000118004 | COLEC11 | -3.118107787 | 0.060051979 | -1.599393784 | 0.339979696 | -4.717501571 | 5.866944937 | 3.076665209 | 3.369317972 | 2.695456494 | 2.68924543 | 3.082820997 | 3.771796761 |
| ENSG00000099622 | CIRBP | -2.522868492 | 9.07E-05 | -2.195792329 | 0.000737133 | -4.718660821 | 12.79629833 | 11.26554928 | 11.74693137 | 9.548682898 | 9.58250309 | 9.561639062 | 10.15938108 |
| ENSG00000086504 | MRPL28 | -2.582010875 | 0.000229156 | -2.137594344 | 0.002665419 | -4.719605219 | 11.89302256 | 10.28850895 | 10.41552659 | 8.476666381 | 8.51148098 | 8.745884933 | 9.101054838 |
| ENSG00000196597 | ZNF782 | -2.301156941 | 0.210296787 | -2.424349569 | 0.175083229 | -4.72550651 | 7.82963488 | 5.838937808 | 2 | 4.647285135 | 4.3456612 | 4.496236235 | 4.30448772 |
| ENSG00000137274 | BPHL | -2.082339909 | 0.000152306 | -2.643564675 | 7.01E-07 | -4.725904584 | 8.444143041 | 8.029695761 | 7.225742779 | 5.920568914 | 6.02777321 | 5.30910438 | 5.59182244 |
| C1ORF101 | C1ORF101 | 2.097297326 | 0.569007348 | -6.82617952 | 0.063908241 | -4.728822194 | 2 | 3.686303414 | 2 | 3.921164147 | 4.31639476 | 2 | 2 |
| ENSG00000102125 | TAZ | -3.057516353 | 0.01540171 | -1.67201104 | 0.208915771 | -4.729527393 | 10.80239439 | 7.847167909 | 7.285536013 | 6.516352584 | 6.48112613 | 7.947671484 | 7.733174012 |
| ENSG00000179133 | C10orf67 | -1.867638499 | 0.412971533 | -2.868200202 | 0.174243611 | -4.735838701 | 2 | 3.686303414 | 3.369317972 | 2.3892346 | 2.49394783 | 2.34604626 | 2.117335275 |
| ENSG00000186603 | HPDL | -3.688973308 | 0.138339617 | -1.047325201 | 0.694164121 | -4.73629851 | 2 | 6.04685525 | 2 | 2.498691189 | 2.49394783 | 3.600766115 | 4.033026654 |
| ENSG00000128045 | RASL11B | -1.685120655 | 0.465481438 | -3.053275297 | 0.150568629 | -4.738395952 | 4.815609471 | 2 | 3.369317972 | 2.868578001 | 2.77779907 | 2.592885947 | 2.117335275 |
| ENSG00000156345 | CDK20 | -2.137216206 | 1.92E-05 | -2.601458078 | 9.03E-08 | -4.738674284 | 8.418903792 | 7.638155693 | 8.721235202 | 6.314087501 | 6.16544343 | 5.80853718 | 5.803599879 |
| ENSG00000183091 | NEB | -1.575604775 | 0.464545264 | -3.165236125 | 0.107674894 | -4.7408409 | 2 | 3.686303414 | 6.80318718 | 3.709785322 | 4.37434574 | 3.27687201 | 2.782576693 |
| ENSG00000142632 | ARHGEF19 | -1.951778298 | 0.00464393 | -2.789113156 | 2.29E-05 | -4.740891454 | 7.904282243 | 8.147241016 | 6.534109959 | 5.686138681 | 5.93454154 | 5.147485955 | 4.959465495 |
| ENSG00000116667 | C1orf21 | -0.610190794 | 0.626320647 | -4.132478441 | 5.41E-05 | -4.742669235 | 9.579430841 | 8.169643011 | 7.398165132 | 8.080836374 | 8.05152279 | 5.390603804 | 3.580696614 |
| ENSG00000105708 | ZNF14 | -2.67941796 | 0.065321338 | -2.065120807 | 0.15956158 | -4.744538766 | 9.064836028 | 4.11352228 | 8.380957451 | 5.742861563 | 5.49945122 | 6.512311087 | 5.799216178 |
| ENSG00000125611 | CHCHD5 | -2.267518028 | 1.16E-06 | -2.477118383 | 5.74E-08 | -4.744636411 | 8.009475818 | 7.761270542 | 8.608698767 | 6.087554739 | 5.84514933 | 5.650128416 | 5.892708443 |
| ENSG00000175489 | LRRC25 | -4.289541304 | 0.053258105 | -0.456165881 | 0.852228272 | -4.745707185 | 5.330594134 | 2 | 4.058998049 | 2.141737246 | 2.38539787 | 4.612704558 | 2.782576693 |
| ENSG00000000003 | TSPAN6 | -1.838782479 | 0.06234238 | -2.909639032 | 0.001700357 | -4.748421512 | 11.14017917 | 10.28850895 | 8.236946975 | 8.463225341 | 8.50338773 | 7.371203179 | 7.486955608 |
| ENSG00000127419 | TMEM175 | -2.791061697 | 0.00273558 | -1.962836675 | 0.041307496 | -4.753898373 | 10.5351205 | 7.701025996 | 9.972637969 | 7.066878356 | 7.04662306 | 7.850199974 | 7.881541153 |
| ENSG00000203727 | SAMD5 | -7.108078761 | 6.88E-06 | 2.354019353 | 0.11939466 | -4.754059408 | 5.330594134 | 8.101365653 | 5.156626401 | 2.498691189 | 2 | 8.803475173 | 9.499569953 |
| ENSG00000172531 | PPP1CA | -2.396594435 | 2.89E-06 | -2.363423642 | 3.76E-06 | -4.760018077 | 12.44893208 | 11.35379164 | 12.18004629 | 9.651519255 | 9.6940865 | 9.370453824 | 9.978070531 |
| ENSG00000167994 | RAB3IL1 | -2.736607533 | 0.002135362 | -2.029726251 | 0.026370078 | -4.766333784 | 9.751047882 | 7.819099893 | 7.691296561 | 6.043074888 | 6.1322378 | 6.986363353 | 6.495928492 |
| ENSG00000105223 | PLD3 | -2.585429367 | 1.30E-09 | -2.184286322 | 3.67E-07 | -4.769715689 | 14.53844446 | 13.54172576 | 14.13052015 | 11.49622323 | 11.5874136 | 11.99889247 | 11.88528995 |
| ENSG00000155542 | SETD9 | -2.242750623 | 0.003398446 | -2.527241574 | 0.000696708 | -4.769992197 | 6.653098118 | 6.464272742 | 8.042706615 | 5.161714397 | 5.07918227 | 5.008609438 | 4.720747915 |
| ENSG00000122188 | LAX1 | -0.817456483 | 0.750090129 | -3.953094314 | NA | -4.770550797 | 2 | 3.076665209 | 3.369317972 | 2.498691189 | 2.68924543 | 2.048074405 | 2.117335275 |
| ENSG00000167703 | SLC43A2 | 1.038180231 | 0.338195259 | -5.810959195 | 1.98E-09 | -4.772778964 | 4.467311984 | 6.464272742 | 7.285536013 | 7.366699513 | 7.57432889 | 2.457042231 | 2.466008509 |
| ENSG00000198691 | ABCA4 | -5.796113525 | 0.004986977 | 1.022643632 | 0.633776471 | -4.773469893 | 2 | 3.686303414 | 6.629488387 | 2.270786764 | 2.14022021 | 5.348031117 | 6.735016458 |
| ENSG00000165985 | C1QL3 | -5.230329694 | 0.110001901 | 0.450647488 | 0.89140859 | -4.779682205 | 2 | 4.710564154 | 2 | 2.141737246 | 2 | 4.155805465 | 3.496556178 |
| ENSG00000119688 | ABCD4 | -1.965648098 | 9.30E-08 | -2.815897603 | 4.04E-15 | -4.781545701 | 10.02639697 | 9.377333006 | 10.06083273 | 7.908072101 | 7.89700047 | 7.093332266 | 7.049302614 |
| ENSG00000108465 | CDK5RAP3 | -1.482476571 | 0.005652183 | -3.302411436 | 3.40E-11 | -4.784888007 | 10.63713535 | 9.514922632 | 9.714382178 | 8.52598359 | 8.60974666 | 6.805896464 | 6.76679494 |
| ENSG00000187778 | MCRS1 | -2.397691655 | 0.001665412 | -2.388161872 | 0.001640149 | -4.785853526 | 11.39439513 | 9.267380431 | 10.72720655 | 8.272191829 | 8.36552655 | 8.181164698 | 8.463180778 |
| ENSG00000242028 | HYPK | -0.739806885 | 0.374603554 | -4.047064153 | 6.49E-09 | -4.786871038 | 9.112843761 | 7.274681896 | 8.922968322 | 7.900612272 | 7.90193448 | 4.628601098 | 4.943664032 |
| ENSG00000176422 | SPRYD4 | -2.643677087 | 0.044211426 | -2.145041935 | 0.105430104 | -4.788719022 | 5.330594134 | 8.777382925 | 9.788583153 | 6.252910529 | 6.24526024 | 6.382394503 | 7.0024032 |
| ENSG00000162769 | FLVCR1 | -2.151280351 | 0.002764986 | -2.641178057 | 0.000153674 | -4.792458408 | 7.218819781 | 8.395519306 | 9.083172337 | 6.379946408 | 6.26836889 | 5.73334482 | 5.984375752 |
| ENSG00000128228 | SDF2L1 | -2.082877571 | 0.004233667 | -2.710093588 | 0.000117999 | -4.792971159 | 8.493334308 | 8.981105197 | 10.06936278 | 7.311513918 | 7.24992376 | 6.176128386 | 7.043754524 |
| ENSG00000148053 | NTRK2 | -6.883116315 | 3.31E-17 | 2.089972671 | 0.00812552 | -4.793143644 | 9.460782017 | 7.979858146 | 8.206335842 | 3.227757282 | 2.49394783 | 10.76381664 | 10.80130822 |
| ENSG00000167588 | GPD1 | -3.644448457 | 0.163221965 | -1.150443461 | 0.670105815 | -4.794891918 | 2 | 2 | 5.156626401 | 2.270786764 | 2.2680111 | 3.172301926 | 3.042026731 |
| ENSG00000113555 | PCDH12 | -3.400722338 | 0.055734443 | -1.394530153 | 0.454313899 | -4.79525249 | 3.327711234 | 3.076665209 | 6.629488387 | 2.784612485 | 2.94009902 | 4.496236235 | 3.560117069 |
| ENSG00000171848 | RRM2 | -1.877847766 | 0.197515156 | -2.919869946 | 0.03297173 | -4.797717712 | 6.009025997 | 8.607278467 | 5.59507254 | 5.697663076 | 5.51241389 | 3.534812207 | 5.319231103 |
| ENSG00000105552 | BCAT2 | -4.481655373 | 1.64E-19 | -0.316261374 | 0.603832507 | -4.797916747 | 9.922475255 | 9.235879652 | 10.20715117 | 5.501219927 | 5.47317115 | 9.156873079 | 9.819361638 |
| ENSG00000152422 | XRCC4 | -1.938188199 | 0.323812042 | -2.860795288 | 0.123001355 | -4.798983487 | 8.998234871 | 7.189546432 | 2 | 5.949776857 | 5.88556448 | 5.174809258 | 4.975095763 |
| ENSG00000112319 | EYA4 | -1.911169269 | 0.56213732 | -2.894023354 | 0.352263729 | -4.805192623 | 2 | 4.442710129 | 2 | 2.784612485 | 2.14022021 | 2.457042231 | 2.059860178 |
| ENSG00000161265 | U2AF1L4 | -2.370767589 | 0.018178424 | -2.435574735 | 0.013782273 | -4.806342324 | 7.096806373 | 8.169643011 | 5.39244166 | 4.927308402 | 5.19627118 | 5.096960836 | 4.919632516 |
| ENSG00000105655 | ISYNA1 | -4.29448041 | 2.38E-10 | -0.517854588 | 0.529356411 | -4.812334998 | 11.82499837 | 10.29870826 | 10.53119344 | 6.634986713 | 6.95938211 | 10.1044607 | 10.86508943 |
| ENSG00000087191 | PSMC5 | -1.883214555 | 6.93E-08 | -2.930609003 | 6.43E-18 | -4.813823558 | 11.69698977 | 11.21797502 | 11.50114494 | 9.553447714 | 9.6580856 | 8.288873067 | 8.800734569 |
| ENSG00000284308 | C2orf81 | -0.307347548 | 0.919403818 | -4.507017051 | 0.067088938 | -4.814364599 | 2 | 3.076665209 | 9.972637969 | 8.072008297 | 8.12931531 | 3.968654369 | 4.445805334 |
| ENSG00000094914 | AAAS | -2.703633752 | 4.39E-09 | -2.110732377 | 5.75E-06 | -4.814366129 | 10.58135316 | 9.514922632 | 10.28977157 | 7.450827758 | 7.57741621 | 8.047866378 | 8.146025444 |
| ENSG00000168876 | ANKRD49 | -2.412092037 | 0.000284738 | -2.405463285 | 0.000265718 | -4.817555322 | 9.544849385 | 7.979858146 | 9.212024683 | 6.598566398 | 6.76185814 | 6.958334233 | 6.356691554 |
| ENSG00000104427 | ZC2HC1A | -1.922268931 | 0.000361141 | -2.895721584 | 2.14E-08 | -4.817990514 | 8.564112515 | 8.376385542 | 8.076950877 | 6.628980152 | 6.31350454 | 5.944926241 | 5.050803619 |
| ENSG00000143512 | HHIPL2 | -0.461738822 | 0.877879848 | -4.356975009 | 0.093525732 | -4.818713831 | 2 | 4.11352228 | 2 | 2.784612485 | 3.01487888 | 2.048074405 | 2.117335275 |
| ENSG00000082175 | PGR | -7.568426039 | 5.42E-05 | 2.744655933 | 0.008689971 | -4.823770106 | 4.467311984 | 4.442710129 | 3.369317972 | 2 | 2 | 6.564892306 | 6.721179863 |
| ENSG00000196912 | ANKRD36B | 0.087548172 | 0.937842924 | -4.911357586 | 4.72E-09 | -4.823809414 | 8.493334308 | 7.431218269 | 6.073378139 | 7.68016596 | 7.77859536 | 3.235948948 | 3.539239711 |
| ENSG00000184164 | CRELD2 | -1.851429071 | 0.000497308 | -2.973893501 | 5.08E-09 | -4.825322571 | 11.09308737 | 10.04973379 | 10.31867969 | 8.658676732 | 8.76944977 | 7.283556699 | 7.874265846 |
| ENSG00000173320 | STOX2 | -8.592463118 | 0.000879422 | 3.762512432 | 0.078680066 | -4.829950685 | 3.327711234 | 6.389896633 | 2 | 2 | 2 | 7.88783736 | 9.135803248 |
| ENSG00000170190 | SLC16A5 | -1.148947855 | 0.516317205 | -3.681776913 | 0.017536365 | -4.830724769 | 3.327711234 | 6.228547981 | 8.631919287 | 6.229283386 | 6.20590375 | 4.209638631 | 3.789585079 |
| ENSG00000137872 | SEMA6D | -3.044131671 | 3.01E-06 | -1.788563609 | 0.008697305 | -4.83269528 | 9.751047882 | 9.903691196 | 8.408111533 | 6.682159616 | 6.30607931 | 7.372383768 | 8.007538088 |
| ENSG00000180739 | S1PR5 | -3.588976041 | 0.012263291 | -1.24516644 | 0.422686222 | -4.834142481 | 3.327711234 | 5.72254323 | 7.552162999 | 3.566290607 | 3.2184422 | 5.524070859 | 4.845038551 |
| ENSG00000178217 | SH2D4B | -2.279673765 | NA | -2.556080522 | NA | -4.835754287 | 2 | 3.076665209 | 2 | 2 | 2.2680111 | 2 | 2.225841198 |
| ENSG00000158516 | CPA2 | -0.211660454 | 0.957159975 | -4.624675754 | 0.176199638 | -4.836336209 | 2 | 2 | 4.523675391 | 3.09461492 | 3.33976443 | 2 | 2.172608095 |
| ENSG00000152348 | ATG10 | -2.863694405 | 0.013710187 | -1.972689632 | 0.098632015 | -4.836384037 | 6.009025997 | 7.35507212 | 9.358421373 | 5.697663076 | 5.16377554 | 6.221381643 | 6.344709081 |
| ENSG00000169418 | NPR1 | -3.13976362 | 0.362647422 | -1.704408617 | NA | -4.844172237 | 2 | 3.076665209 | 3.369317972 | 2 | 2.2680111 | 2.592885947 | 2.059860178 |
| ENSG00000274137 | | 0.631595084 | 0.851491754 | -5.476662478 | 0.070384971 | -4.845067394 | 2 | 4.442710129 | 2 | 3.663521627 | 3.86943969 | 2.094598369 | 2 |
| ENSG00000173928 | SWSAP1 | -2.604010526 | 0.003687423 | -2.249697 | 0.011459401 | -4.853707526 | 5.095959458 | 7.099070191 | 6.203069917 | 4.205751501 | 3.94787299 | 4.111244182 | 4.574506939 |
| ENSG00000103274 | NUBP1 | -1.447555273 | 0.409695646 | -3.407027323 | 0.031257426 | -4.854582596 | 3.327711234 | 8.213429689 | 9.528916236 | 7.017154451 | 7.03319408 | 4.984105765 | 5.367404784 |
| ENSG00000132205 | EMILIN2 | -5.534893574 | 3.52E-37 | 0.679199574 | 0.122835348 | -4.855694001 | 8.963744667 | 8.903033433 | 8.941672936 | 4.034562184 | 3.65239003 | 9.291211854 | 9.87222905 |

| | | | | | | | | | | | | |
|---|---|---|---|---|---|---|---|---|---|---|---|---|
| ENSG00000064547 | LPAR2 | -2.319969023 | 0.196401085 | -2.539184703 | 0.145238766 | -4.859153726 | 2 | 7.979858146 | 7.646393253 | 4.907595604 | 5.22805098 | 5.147485955 | 4.56420962 |
| ENSG00000143543 | JTB | -1.696342669 | 6.34E-06 | -3.172310127 | 1.50E-18 | -4.868652796 | 11.39115429 | 10.60848937 | 11.06667478 | 9.306761457 | 9.42174886 | 7.918872821 | 7.890841531 |
| ENSG00000159210 | SNF8 | -2.276531891 | 1.33E-08 | -2.594708053 | 5.39E-11 | -4.871239944 | 10.95065381 | 10.20420974 | 10.80076732 | 8.315844936 | 8.51952909 | 7.914016653 | 8.277899908 |
| ENSG00000143061 | IGSF3 | -5.079826241 | 0.000608515 | 0.208038412 | 0.912467872 | -4.87178783 | 8.836089686 | 5.946638857 | 7.163363831 | 4.034562184 | 2.14022021 | 7.900168075 | 8.070734444 |
| ENSG00000136541 | ERMN | -1.781096991 | 0.519400194 | -3.091102541 | 0.227974555 | -4.872199532 | 6.257048221 | 2 | 2 | 3.462030052 | 3.45167086 | 2.987421008 | 2.509591799 |
| ENSG00000121413 | ZSCAN18 | -3.927726104 | 3.67E-10 | -0.945249452 | 0.178782858 | -4.872975556 | 9.790941832 | 9.125613817 | 8.143093325 | 5.433909581 | 5.3193943 | 8.382079673 | 8.068003971 |
| ENSG00000132330 | SCLY | -1.623165215 | 0.170583392 | -3.252494044 | 0.002658477 | -4.875659259 | 4.815609471 | 8.004992147 | 8.110401118 | 6.01571348 | 5.96314905 | 4.470006665 | 4.604965121 |
| ENSG00000183831 | ANKRD45 | -1.033641353 | 0.671332479 | -3.850136164 | 0.071834001 | -4.883777517 | 5.095959458 | 3.076665209 | 5.156626401 | 3.921164147 | 3.82856379 | 2 | 2.817676617 |
| ENSG00000179057 | IGSF22 | -4.327890363 | 0.104928256 | -0.562376437 | 0.845791525 | -4.890266799 | 2 | 2 | 5.39244166 | 2.270786764 | 2.14022021 | 3.824280769 | 3.407202941 |
| ENSG00000051180 | RAD51 | -1.194502498 | 0.544293669 | -3.69835901 | 0.034894396 | -4.892861507 | 4.00722531 | 6.535001748 | 2 | 4.299492648 | 4.1930041 | 2.717032197 | 2.746601469 |
| ENSG00000076770 | MBNL3 | -3.59743331 | 0.20811252 | -1.295698263 | 0.671246939 | -4.893131573 | 2 | 6.535001748 | 2 | 3.09461492 | 2.14022021 | 4.388329244 | 3.518055793 |
| ENSG00000025434 | NR1H3 | -3.860107629 | 5.50E-05 | -1.03639301 | 0.331498803 | -4.896500639 | 8.564112515 | 7.051611315 | 6.203069917 | 3.95996233 | 4.22485516 | 7.006273211 | 6.029985803 |
| ENSG00000163072 | NOSTRIN | -0.27232781 | NA | -4.624687993 | NA | -4.897015802 | 2 | 3.076665209 | 2 | 2.498691189 | 2.49394783 | 2 | 2.059860178 |
| ENSG00000204779 | FOXD4L5 | -0.27232781 | NA | -4.624687993 | NA | -4.897015802 | 2 | 3.076665209 | 2 | 2.498691189 | 2.49394783 | 2 | 2.059860178 |
| ENSG00000134138 | MEIS2 | 0.151546153 | 0.946324665 | -5.049667028 | 0.004461713 | -4.898120876 | 9.304185368 | 7.790474954 | 2 | 8.291333947 | 8.32569041 | 3.930690651 | 3.287118176 |
| ENSG00000184838 | PRR16 | -2.776953546 | 7.75E-07 | -2.121523889 | 0.000211346 | -4.898477435 | 12.17703553 | 10.78209868 | 11.27276688 | 8.820275739 | 8.70315789 | 9.372225266 | 9.453587626 |
| ENSG00000183828 | NUDT14 | -2.493900685 | 1.08E-05 | -2.406450547 | 1.70E-05 | -4.900351232 | 8.257427002 | 8.255926468 | 7.451345288 | 5.590471342 | 5.69405321 | 5.269098128 | 6.070564958 |
| ENSG00000231179 | | 4.64729485 | 4.53E-08 | -9.547908867 | 5.74E-08 | -4.900614016 | 4.467311984 | 3.686303414 | 5.39244166 | 9.048926513 | 9.03387903 | 2 | 2 |
| ENSG00000102226 | USP11 | -2.33509028 | 3.20E-07 | -2.5700453 | 1.30E-08 | -4.90513558 | 12.31339889 | 11.77681554 | 11.53848775 | 9.582488026 | 9.58633944 | 8.950238388 | 9.664095852 |
| ENSG00000055163 | CYFIP2 | -5.051257881 | 2.46E-11 | 0.145927746 | 0.883149421 | -4.905330134 | 9.247966931 | 8.337339476 | 7.691296561 | 4.358765249 | 3.33976443 | 8.787633372 | 8.631056556 |
| ENSG00000120656 | TAF12 | -2.120680056 | 0.01532262 | -2.784881166 | 0.000942469 | -4.905561222 | 10.50545418 | 8.029695761 | 9.647595056 | 7.61176303 | 7.61395775 | 7.101898411 | 6.815359915 |
| ENSG00000186281 | GPAT2 | -0.03847003 | 0.971851048 | -4.867139066 | 7.50E-09 | -4.905609095 | 5.095959458 | 5.946638857 | 7.029880454 | 6.113598934 | 6.26836889 | 2.803598391 | 2.551896972 |
| ENSG00000163041 | H3F3A | -0.099378217 | 0.918552138 | -4.80771073 | 9.68E-12 | -4.907088946 | 10.70134604 | 8.94259736 | 9.149036566 | 9.771565364 | 9.68051177 | 5.113999907 | 5.243813244 |
| ENSG00000149926 | FAM57B | -4.314913587 | 0.098465408 | -0.593591055 | 0.83342673 | -4.908504642 | 2 | 5.303523747 | 2 | 2.270786764 | 2.14022021 | 3.891700877 | 3.236138336 |
| ENSG00000142507 | PSMB6 | -2.42672637 | 4.65E-08 | -2.482719083 | 1.88E-08 | -4.909445453 | 12.04303957 | 11.22333918 | 11.37608156 | 9.115954467 | 9.23006009 | 8.83721372 | 9.353992291 |
| ENSG00000129007 | CALML4 | -0.244148483 | 0.911302045 | -4.666230137 | 0.008139262 | -4.91037862 | 3.327711234 | 5.72254323 | 2 | 4.237680128 | 4.40247105 | 2.267001569 | 2.225841198 |
| ENSG00000182986 | ZNF320 | -0.492395181 | 0.767793449 | -4.422283937 | 0.000984041 | -4.914679117 | 4.467311984 | 8.077869079 | 9.541233192 | 8.051945366 | 7.83385794 | 4.388329244 | 4.241259475 |
| ENSG00000099338 | CATSPERG | -3.469601242 | 0.185378637 | -1.44511084 | 0.59846514 | -4.914712082 | 2 | 2 | 6.322058544 | 2.784612485 | 2.49394783 | 3.851628042 | 3.580696614 |
| ENSG00000122971 | ACADS | -2.349711083 | 0.043170473 | -2.570977484 | 0.023974857 | -4.920688567 | 10.1076228 | 6.899080544 | 7.858115942 | 6.586219071 | 6.64900057 | 6.448814696 | 6.362645675 |
| ENSG00000186409 | CCDC30 | -3.594627814 | 0.009670067 | -1.329764501 | 0.356359551 | -4.924392316 | 3.327711234 | 4.710564154 | 6.203069917 | 2.947924294 | 2.49394783 | 4.360042493 | 3.717076715 |
| ENSG00000103154 | NECAB2 | -1.262196899 | 0.63899539 | -3.662522468 | 0.12986224 | -4.924719367 | 2 | 5.457120953 | 2 | 3.023133209 | 3.50452879 | 2.457042231 | 2.225841198 |
| ENSG00000139117 | CPNE8 | -1.536160579 | 1.61E-06 | -3.391810096 | 3.85E-28 | -4.927970675 | 9.233563228 | 8.955547846 | 9.452720984 | 7.739909082 | 7.66998641 | 6.000958775 | 5.8339184 |
| ENSG00000143842 | SOX13 | -3.048887989 | 0.025673766 | -1.883698464 | 0.184734803 | -4.932586452 | 7.159102577 | 6.844429755 | 10.54960077 | 6.104969671 | 6.31350454 | 7.526684775 | 7.112501355 |
| ENSG00000177181 | RIMKLA | -3.038277886 | 0.039262977 | -1.898934576 | 0.177577912 | -4.937212462 | 4.00722531 | 3.076665209 | 4.874587708 | 2.3892346 | 2.59489879 | 3.059555873 | 2.782576693 |
| ENSG00000132329 | RAMP1 | -4.379358957 | 8.43E-12 | -0.557913638 | 0.469381607 | -4.937272595 | 11.53317465 | 9.828248506 | 10.56174313 | 6.387081877 | 6.5890786 | 10.07829275 | 10.4074232 |
| ENSG00000243449 | C4orf48 | -1.524021493 | 0.006372593 | -3.413716111 | 5.01E-11 | -4.937737604 | 9.509418609 | 8.234834549 | 8.922968322 | 7.546342806 | 7.39696604 | 5.677130623 | 5.661099865 |
| ENSG00000102796 | DHRS12 | -2.46731552 | 0.005841121 | -2.475122787 | 0.005037402 | -4.942438308 | 8.19931886 | 6.140560021 | 6.629488387 | 4.804776789 | 5.09650239 | 4.82764224 | 5.065479856 |
| ENSG00000188338 | SLC38A3 | -4.217645728 | 0.211077851 | -0.72479927 | 0.838764132 | -4.942444997 | 2 | 4.710564154 | 2 | 2.270786764 | 2 | 3.1504466 | 3.071407821 |
| ENSG00000112742 | TTK | -1.821698296 | 0.18207078 | -3.121269706 | 0.013968105 | -4.942968003 | 3.327711234 | 5.303523747 | 6.80318718 | 4.299492648 | 4.02226117 | 3.235948948 | 3.311948089 |
| ENSG00000184949 | FAM227A | -1.813640266 | 0.04670103 | -3.132531481 | 0.000226183 | -4.946171747 | 8.107517749 | 5.946638857 | 7.342949392 | 5.614989456 | 5.67111583 | 4.778616857 | 4.119958326 |
| ENSG00000172031 | EPHX4 | -1.52434525 | 0.625452298 | -3.423937053 | 0.234784916 | -4.948282303 | 2 | 4.11352228 | 2 | 2.784612485 | 2.14022021 | 2.183373728 | 2.117335275 |
| ENSG00000138621 | PPCDC | -2.676228311 | 0.000137549 | -2.274086759 | 0.001181762 | -4.950315071 | 7.867441304 | 6.464272742 | 8.175061096 | 5.128039203 | 5.13053105 | 5.399896186 | 5.576534358 |
| ENSG00000139767 | SRRM4 | -1.684927862 | NA | -3.266544435 | NA | -4.951472297 | 3.327711234 | 2 | 2 | 2.270786764 | 2.14022021 | 2.139668783 | 2 |
| ENSG00000115392 | FANCL | -1.50538167 | 0.369747208 | -3.448592596 | 0.023364532 | -4.953974267 | 6.96351253 | 8.791896851 | 3.369317972 | 6.336376195 | 5.92487824 | 4.188345803 | 4.614976609 |
| ENSG00000162639 | HENMT1 | -3.156811168 | 0.0823951 | -1.799592305 | 0.339683272 | -4.956403473 | 7.940205782 | 2 | 7.777116083 | 4.268917409 | 4.50979031 | 5.413723492 | 5.717942682 |
| ENSG00000075914 | EXOSC7 | -2.527333704 | 1.65E-05 | -2.429432185 | 3.20E-05 | -4.956765888 | 10.25752168 | 8.806266215 | 9.757248847 | 7.190301983 | 7.26148706 | 7.137053113 | 7.485592137 |
| ENSG00000184220 | CMSS1 | -1.838012178 | 0.033603888 | -3.119808442 | 0.000115924 | -4.95782062 | 9.931427102 | 7.847167909 | 8.380957451 | 7.136275664 | 7.24604864 | 5.783905948 | 6.127623461 |
| ENSG00000168237 | GLYCTK | -2.252040527 | 0.081994807 | -2.706384443 | 0.030137313 | -4.95842497 | 7.867441304 | 6.899080544 | 4.058998049 | 4.804776789 | 4.91316568 | 4.088436536 | 4.784231301 |
| ENSG00000102996 | MMP15 | -2.161754699 | 0.145889895 | -2.797876162 | 0.049160384 | -4.959630861 | 9.128497779 | 8.747909548 | 4.058998049 | 6.351046291 | 6.22177541 | 5.912830251 | 5.431121909 |
| ENSG00000263465 | SRSF8 | -3.035408903 | 1.58E-08 | -1.924827851 | 0.000509252 | -4.960236753 | 10.09970294 | 10.26789142 | 11.23913884 | 7.641902724 | 7.59275449 | 8.456328963 | 8.929660203 |
| ENSG00000117308 | GALE | -2.583324944 | 0.000237364 | -2.377639692 | 0.000732419 | -4.960964636 | 9.895281535 | 8.191702463 | 9.839340503 | 6.917187326 | 6.97351258 | 6.884127769 | 7.366199662 |
| ENSG00000281289 | | -0.466588576 | 0.501076699 | -4.497286059 | 4.97E-16 | -4.963874635 | 11.75108151 | 11.12072186 | 10.81097638 | 10.80468647 | 10.8275257 | 6.197600957 | 7.274433587 |
| ENSG00000163554 | SPTA1 | 1.18850625 | 0.815821608 | -6.15290972 | NA | -4.96440347 | 2 | 3.076665209 | 2 | 3.227757282 | 2.94009902 | 2 | 2 |
| ENSG00000167733 | HSD11B1L | -3.258044978 | 0.014463959 | -1.707431402 | 0.225712688 | -4.96547638 | 9.90440318 | 5.303523747 | 8.206335842 | 5.742861563 | 5.44640346 | 7.034886728 | 7.108963622 |
| ENSG00000080986 | NDC80 | -2.29615253 | 0.25952816 | -2.676152219 | 0.172121765 | -4.972704749 | 4.00722531 | 7.232742069 | 2 | 4.070463721 | 3.69849104 | 3.534812207 | 3.717076715 |
| ENSG00000185985 | SLITRK2 | -3.946936777 | 0.304282027 | -1.025576405 | NA | -4.972513182 | 2 | 3.686303414 | 2 | 2.141737246 | 2 | 2.526562374 | 2.421067459 |
| ENSG00000173801 | JUP | -4.046316203 | 4.83E-14 | -0.928317511 | 0.124173275 | -4.974633714 | 10.79260216 | 9.792607274 | 10.63252637 | 6.401247806 | 6.55815769 | 9.046937881 | 9.909311718 |
| ENSG00000160404 | TOR2A | -2.546747528 | 9.16E-10 | -2.429635244 | 1.94E-09 | -4.976382772 | 7.709880939 | 7.928237109 | 7.646393253 | 5.076005082 | 5.5755357 | 5.4588802 | 5.448021784 |
| ENSG00000103202 | NME4 | -3.090903811 | 6.87E-21 | -1.88586447 | 1.79E-08 | -4.976768281 | 12.70789053 | 12.13942551 | 12.78286249 | 9.472742079 | 9.49948932 | 10.70235993 | 10.67076945 |
| ENSG00000049249 | TNFRSF9 | -1.980787242 | 0.283681909 | -2.996196676 | 0.084214225 | -4.976983918 | 5.70932728 | 2 | 6.203069917 | 3.566290607 | 3.98554648 | 3.483289517 | 2.70970617 |
| ENSG00000115271 | GCA | -1.019730776 | 0.61367293 | -3.960954879 | 0.023453844 | -4.980685654 | 9.699577239 | 6.228547981 | 3.369317972 | 7.25803034 | 7.24992376 | 4.778616857 | 4.26688425 |
| ENSG00000187446 | CHP1 | -3.075621196 | 1.79E-10 | -1.906476596 | 0.000128861 | -4.982097791 | 12.2450337 | 12.130888 | 13.15658907 | 9.518946937 | 9.5188894 | 10.52859399 | 10.82373799 |
| ENSG00000185338 | SOCS1 | -1.735097523 | 0.214129738 | -3.24711639 | 0.011787753 | -4.982213913 | 3.327711234 | 7.051611315 | 7.342949392 | 5.273747807 | 4.811949 | 3.917810786 | 3.753786374 |
| ENSG00000165923 | AGBL2 | -0.924566665 | 0.683136708 | -4.059394064 | 0.037830739 | -4.983960729 | 2 | 3.686303414 | 6.629488387 | 4.415698455 | 4.50979031 | 2.858551861 | 2.277179783 |
| ENSG00000149292 | TTC12 | -2.459036696 | 0.010207854 | -2.525778024 | 0.00747349 | -4.984814719 | 8.075571721 | 8.234834549 | 5.59507254 | 5.420062284 | 5.28958453 | 5.253805068 | 5.337485258 |
| ENSG00000120327 | PCDHB14 | -2.706685541 | 0.065184591 | -2.285147173 | 0.119716833 | -4.991832714 | 8.946185348 | 4.93638933 | 5.930865648 | 5.273747807 | 4.79082474 | 5.125248504 | 5.680297397 |
| ENSG00000126768 | TIMM17B | -2.801422942 | 7.06E-07 | -2.190577088 | 0.000131177 | -4.99200003 | 9.521325812 | 9.785372135 | 10.83618737 | 7.352188525 | 7.4314987 | 8.006643981 | 7.975841991 |
| ENSG00000101624 | CEP76 | -7.356310008 | 0.000213254 | 2.364097994 | 0.061577604 | -4.992212014 | 4.00722531 | 4.442710129 | 3.369317972 | 2 | 2 | 6.403336799 | 5.66592325 |
| ENSG00000128886 | ELL3 | -1.934155753 | 0.329010237 | -3.062946824 | 0.098395659 | -4.997102576 | 3.327711234 | 5.838937808 | 2 | 3.289987342 | 3.15372745 | 2.775316226 | 2.592996859 |
| ENSG00000084207 | GSTP1 | -2.434793316 | 2.51E-13 | -2.565520677 | 8.21E-15 | -5.000313993 | 13.99742029 | 13.37521182 | 13.63868632 | 11.27488516 | 11.2450896 | 10.97342027 | 11.27047363 |
| ENSG00000248712 | CCDC153 | -2.978832861 | 0.097774135 | -2.023677177 | 0.245643493 | -5.002510038 | 5.095959458 | 3.686303414 | 2 | 2.600425781 | 2.38539787 | 2.937246078 | 2.746601469 |
| ENSG00000176887 | SOX11 | -2.91512532 | 0.165445345 | -2.087830201 | 0.325446617 | -5.002955521 | 5.330594134 | 6.899080544 | 7.285536013 | 4.946755474 | 2 | 5.294231737 | 4.033026654 |
| ENSG00000144362 | PHOSPHO2 | -2.826173316 | 0.087079822 | -2.18134046 | 0.179432455 | -5.007513776 | 5.532353208 | 5.457120953 | 2 | 3.162721474 | 2.77779907 | 3.679182183 | 2.885414003 |
| ENSG00000251247 | ZNF345 | -2.441085205 | 0.000709823 | -2.568125024 | 0.000273989 | -5.009210229 | 7.096806373 | 6.389896633 | 8.076950877 | 4.965943888 | 5.13053105 | 4.927401733 | 4.951586397 |
| ENSG00000171453 | POLR1C | -2.478408851 | 0.000577828 | -2.531226706 | 0.000383385 | -5.009635557 | 9.931427102 | 9.159587375 | 7.97167868 | 6.727838783 | 6.8563927 | 6.797131715 | 6.685996416 |
| ENSG00000118156 | ZNF541 | -3.942941507 | 0.314561941 | -1.068275731 | NA | -5.011217238 | 2 | 3.686303414 | 2 | 2.141737246 | 2 | 2.62493832 | 2.277179783 |
| ENSG00000151062 | CACNA2D4 | -1.772056201 | 0.349594603 | -3.240774395 | 0.064469289 | -5.012830596 | 2 | 5.72254323 | 4.058998049 | 3.406930997 | 3.28037825 | 2.858551861 | 2.374681325 |
| ENSG00000118939 | UCHL3 | -1.07159716 | 0.067726512 | -3.943142416 | 3.45E-14 | -5.014739576 | 9.247966931 | 8.590771778 | 8.175061096 | 7.734324667 | 7.62891439 | 4.675266036 | 5.230851202 |
| ENSG00000198673 | FAM19A2 | -2.973704527 | 0.359143464 | -2.042479493 | NA | -5.01618402 | 2 | 3.686303414 | 2 | 2.141737246 | 2.14022021 | 2.384001718 | 2.117335275 |
| ENSG00000166664 | CHRFAM7A | 0.297969523 | 0.938141609 | -5.31764086 | 0.135257068 | -5.019671338 | 2 | 3.686303414 | 2 | 3.289987342 | 2.59489879 | 2 | 2.059860178 |
| ENSG00000165495 | PKNOX2 | -5.810936506 | 6.78E-05 | 0.789932568 | 0.606490645 | -5.021003938 | 6.468621203 | 6.311476754 | 4.058998049 | 2.600425781 | 2 | 6.934553331 | 6.420873184 |
| ENSG00000137135 | ARHGEF39 | -3.972937789 | 2.04E-10 | -1.048901537 | 0.082588601 | -5.021839326 | 6.468621203 | 5.946638857 | 6.43197674 | 3.09461492 | 3.15372745 | 5.440986767 | 5.177806074 |
| ENSG00000123612 | ACVR1C | -1.160338721 | 0.605487532 | -3.862183134 | 0.05062544 | -5.022521856 | 4.00722531 | 2 | 6.80318718 | 4.623314311 | 4.22485516 | 3.011868911 | 2.421067459 |
| ENSG00000196689 | TRPV1 | -1.202187086 | 0.130759852 | -3.826654609 | 5.48E-08 | -5.028841695 | 7.82963488 | 6.78762695 | 7.97167868 | 6.422239259 | 6.46787796 | 4.530483275 | 3.560117069 |
| ENSG00000183484 | GPR132 | -3.420455736 | 0.191214633 | -1.610136931 | 0.536100907 | -5.030592667 | 2 | 4.11352228 | 4.058998049 | 2 | 2.49394783 | 3.035909414 | 2.466008509 |
| ENSG00000110063 | DCPS | -2.526364674 | 0.004029857 | -2.506681562 | 0.004060884 | -5.033046237 | 7.904282243 | 7.538394143 | 9.681375057 | 6.213314212 | 6.25300433 | 6.132200015 | 6.362645675 |
| ENSG00000167113 | COQ4 | -2.270793906 | 0.000548094 | -2.762742767 | 1.64E-05 | -5.033536673 | 10.06758145 | 9.090820894 | 10.74860941 | 7.791909949 | 7.93841129 | 7.138441638 | 7.595854839 |
| ENSG00000010256 | UQCRC1 | -2.648252308 | 7.96E-08 | -2.38878366 | 1.40E-06 | -5.037035968 | 12.87959993 | 11.65706076 | 12.32778341 | 9.741955564 | 9.71666438 | 9.930233463 | 10.0430058 |
| ENSG00000112218 | GPR63 | -3.942178547 | 0.046457783 | -1.095669087 | 0.607590138 | -5.037847634 | 2 | 4.11352228 | 6.43197674 | 2.695456494 | 2.49394783 | 4.813804156 | 3.236138336 |
| ENSG00000197181 | PIWIL2 | -2.373744051 | 0.197017964 | -2.668821168 | 0.131223185 | -5.042565219 | 2 | 4.93638933 | 6.203069917 | 3.462030052 | 3.2184422 | 3.584558147 | 2.592996859 |
| ENSG00000112983 | BRD8 | -2.194553477 | 2.32E-09 | -2.848928872 | 2.64E-15 | -5.043482348 | 10.30651143 | 10.40166101 | 10.9743126 | 8.394075037 | 8.42500614 | 7.877064822 | 7.643095142 |
| ENSG00000177990 | DPY19L2 | -1.333273022 | 0.490247308 | -3.712520981 | 0.031552246 | -5.045794004 | 2 | 4.442710129 | 6.534109959 | 4.070463721 | 4.2560182 | 2.937246078 | 2.509591799 |
| ENSG00000250506 | CDK3 | -0.713893604 | 0.627296544 | -4.33402832 | 0.000436182 | -5.047921924 | 4.00722531 | 5.131603625 | 7.163363831 | 5.433909581 | 5.2591458 | 2.987421008 | 2.592996859 |
| ENSG00000162746 | FCRLB | -2.262911454 | 0.42327046 | -2.785121485 | 0.302902594 | -5.048032939 | 2 | 2 | 7.600047325 | 4.139692846 | 4.05806467 | 3.891700877 | 3.518055793 |
| ENSG00000007968 | E2F2 | -2.541016252 | 0.168998459 | -2.507853962 | 0.143612995 | -5.048870213 | 3.327711234 | 3.686303414 | 3.369317972 | 2.498691189 | 2.2680111 | 2.225793514 | 2.551896972 |
| ENSG00000196459 | TRAPPC2 | -2.579135975 | 0.02856454 | -2.471100475 | 0.035163709 | -5.05023645 | 6.257048221 | 9.566483142 | 10.11127249 | 6.846474326 | 6.75642009 | 6.770513376 | 7.030725399 |
| ENSG00000105737 | GRIK5 | -3.865905063 | 0.070362082 | -1.184604528 | 0.610420234 | -5.050509591 | 7.384433967 | 4.442710129 | 2 | 3.515101962 | 2.2680111 | 5.158477451 | 4.654341953 |
| ENSG00000205277 | MUC12 | -0.64418138 | 0.824467159 | -4.412483878 | 0.080287228 | -5.056665259 | 4.00722531 | 3.686303414 | 2 | 3.289987342 | 2.86123014 | 2 | 2.225841198 |
| ENSG00000123329 | ARHGAP9 | -2.692630895 | 0.30607807 | -2.369375652 | 0.351780632 | -5.062006547 | 2 | 4.93638933 | 2 | 2.498691189 | 2.38539787 | 2.717032197 | 2.326754077 |
| ENSG00000205981 | DNAJC19 | -2.201621971 | 0.004323654 | -2.86384485 | 0.00012132 | -5.065466821 | 9.602033055 | 7.790474954 | 8.143093325 | 6.640968371 | 6.52015598 | 6.043083161 | 5.859408034 |
| ENSG00000162814 | SPATA17 | -1.086829034 | 0.700085754 | -3.981006543 | 0.120755432 | -5.067835577 | 2 | 4.11352228 | 2 | 2.784612485 | 2.49394783 | 2.094598369 | 2.117335275 |
| ENSG00000100129 | EIF3L | -2.547723959 | 5.90E-12 | -2.523438871 | 7.67E-12 | -5.071162831 | 12.67495574 | 12.31800992 | 11.88379871 | 9.790524529 | 9.78101395 | 9.705350813 | 9.90753096 |
| ENSG00000100417 | PMM1 | -2.617658014 | 0.004211074 | -2.459127358 | 0.007309398 | -5.076785372 | 11.22998797 | 8.639735838 | 11.22011226 | 8.134820652 | 8.17282964 | 8.330773404 | 8.289666801 |
| ENSG00000122952 | ZWINT | -1.621264387 | 0.279328335 | -3.460930205 | 0.011144177 | -5.082194592 | 4.467311984 | 8.607278467 | 6.629488387 | 5.709096143 | 6.00960265 | 3.648325314 | 4.692661645 |
| ENSG00000117480 | FAAH | -3.059847662 | 0.251780872 | -2.022827376 | 0.456415864 | -5.082675038 | 2 | 7.467835385 | 2 | 3.162721474 | 3.69849104 | 4.144793515 | 4.241259475 |
| ENSG00000187068 | C3orf70 | -4.935165763 | 0.00012717 | -0.148058114 | 0.927617649 | -5.083223878 | 5.330594134 | 5.457120953 | 7.734844356 | 3.162721474 | 2.38539787 | 6.96617488 | 5.740981724 |
| ENSG00000278677 | HIST1H2AM | 1.734386574 | 0.661931364 | -6.820010762 | 0.076887988 | -5.085624188 | 2 | 3.686303414 | 2 | 4.034562184 | 3.65239003 | 2 | 2 |

| | | | | | | | | | | | | | |
|---|---|---|---|---|---|---|---|---|---|---|---|---|---|
| ENSG00000171551 | ECEL1 | -2.260954841 | 0.473662292 | -2.824844769 | 0.343361584 | -5.08579961 | 4.467311984 | 2 | 2 | 2.498691189 | 2.2680111 | 2.457042231 | 2.059860178 |
| ENSG00000070808 | CAMK2A | -2.686641803 | 0.309385265 | -2.404205556 | 0.353451393 | -5.090847359 | 2 | 6.04685525 | 2 | 2.868578001 | 2.86123014 | 2.560105256 | 3.33635788 |
| ENSG00000196388 | INCA1 | -4.094173561 | 1.39E-09 | -1.00085468 | 0.150262362 | -5.095028241 | 7.27616306 | 6.04685525 | 6.629488387 | 3.462030052 | 3.15372745 | 5.773219597 | 5.781546943 |
| ENSG00000070669 | ASNS | -1.197890674 | 0.260096091 | -3.903232262 | 2.61E-05 | -5.101122936 | 8.042902253 | 8.487557258 | 9.099922433 | 7.474468681 | 7.38291816 | 5.598599872 | 3.474731315 |
| ENSG00000179168 | GGN | -3.359632588 | 0.230602145 | -1.742075653 | 0.550704482 | -5.101708241 | 2 | 2 | 8.434763955 | 3.840289928 | 3.98554648 | 5.333556395 | 5.1574003 |
| ENSG00000272803 | | 1.050345858 | 0.841054398 | -6.152908521 | NA | -5.102562663 | 2 | 3.076665209 | 2 | 3.09461492 | 2.94009902 | 2 | 2 |
| ENSG00000203734 | ECT2L | -2.992645648 | 0.299078862 | -2.113796894 | 0.453489046 | -5.106442542 | 3.327711234 | 2 | 4.523675391 | 2 | 2.59489879 | 2.384001718 | 2.70970617 |
| ENSG00000166136 | NDUFB8 | -2.24562039 | 2.29E-06 | -2.861025607 | 7.99E-10 | -5.106645997 | 10.80726569 | 10.40638494 | 11.43337479 | 8.70652031 | 8.71161289 | 7.88204676 | 8.291228494 |
| ENSG00000113141 | IK | -2.116105516 | 7.08E-06 | -2.994103991 | 6.55E-11 | -5.110209507 | 11.96017193 | 11.04700893 | 11.07517045 | 9.333786218 | 9.29994502 | 8.290747499 | 8.587232804 |
| ENSG00000126457 | PRMT1 | -2.38357581 | 2.76E-13 | -2.730361015 | 2.54E-17 | -5.113936825 | 12.1807942 | 11.80749691 | 11.98870353 | 9.518946937 | 9.71876306 | 9.005308106 | 9.506645132 |
| ENSG00000108479 | GALK1 | -2.929900287 | 1.38E-07 | -2.185214482 | 0.000112171 | -5.115114769 | 10.57565445 | 9.28800521 | 10.58572549 | 7.373900638 | 7.35080119 | 8.1339154 | 8.050589918 |
| ENSG00000077522 | ACTN2 | -0.609883562 | 0.786065276 | -4.505958315 | 0.015805954 | -5.115841877 | 2 | 7.051611315 | 4.523675391 | 5.194621444 | 5.19627118 | 2.911488689 | 2.277179783 |
| ENSG00000232196 | MTRNR2L4 | -2.895933461 | 0.31597318 | -2.222115566 | 0.426131057 | -5.118049027 | 2 | 2 | 4.523675391 | 2.141737246 | 2.38539787 | 2.526562374 | 2.277179783 |
| ENSG00000139174 | PRICKLE1 | -3.427465026 | 7.08E-15 | -1.692541117 | 0.000184508 | -5.120006142 | 9.358295023 | 10.00683004 | 9.149036566 | 6.328984825 | 6.06344184 | 7.751118939 | 8.000872619 |
| ENSG00000100802 | C14orf93 | -3.107121795 | 0.028072086 | -2.013438278 | 0.167056356 | -5.120560073 | 8.892187472 | 7.145017454 | 4.058998049 | 4.846787657 | 4.76938658 | 5.755231204 | 5.821002687 |
| ENSG00000184857 | TMEM186 | -2.870692168 | 0.009037926 | -2.251233438 | 0.043960412 | -5.121925606 | 7.384433967 | 6.666836837 | 9.636156974 | 5.686138681 | 5.70538658 | 6.252491482 | 6.298867018 |
| ENSG00000122779 | TRIM24 | -2.836267939 | 6.87E-17 | -2.288108095 | 1.09E-11 | -5.124376034 | 9.384607364 | 9.726146444 | 9.972637969 | 6.946463121 | 6.89154576 | 7.5848779 | 7.305675343 |
| ENSG00000198885 | ITPRIPL1 | -6.279917631 | 0.011943408 | 1.149097583 | 0.590927058 | -5.130820048 | 2 | 5.457120953 | 3.369317972 | 2 | 2.14022021 | 5.561816579 | 4.975095763 |
| ENSG00000196284 | SUPT3H | -2.363348556 | 0.002165672 | -2.76779845 | 0.000221057 | -5.131147006 | 8.468948322 | 6.728495673 | 7.098165388 | 5.304221918 | 5.45984938 | 4.965450964 | 5.080008298 |
| ENSG00000158106 | RHPN1 | -4.45974661 | 0.001582119 | -0.671918282 | 0.680497021 | -5.131664892 | 3.327711234 | 5.72254323 | 7.600047325 | 2.784612485 | 3.08597292 | 5.866661379 | 5.689801209 |
| ENSG00000139187 | KLRG1 | -3.369245073 | 0.103759715 | -1.765127294 | 0.395075292 | -5.134372367 | 3.327711234 | 4.11352228 | 5.156626401 | 2 | 2.86123014 | 3.465698119 | 2.782576693 |
| ENSG00000177483 | RBM44 | -2.478776718 | 0.355466174 | -2.656363428 | 0.306441895 | -5.135140145 | 6.653098118 | 2 | 2 | 3.289987342 | 3.28037825 | 3.12825509 | 3.236138336 |
| ENSG00000152382 | TADA1 | -3.048057322 | 2.50E-18 | -2.087816244 | 9.26E-10 | -5.135873566 | 8.586954094 | 8.487557258 | 8.721235202 | 5.651002631 | 5.65950892 | 6.750676314 | 6.33565646 |
| ENSG00000147642 | SYBU | -2.447423646 | 0.300270694 | -2.689387289 | 0.236344937 | -5.136810934 | 2 | 5.131603625 | 4.874587708 | 2.141737246 | 3.39680241 | 3.172301926 | 2.277179783 |
| ENSG00000131044 | TTLL9 | -3.289599331 | 0.291724472 | -1.848588755 | NA | -5.138188086 | 2 | 2 | 4.058998049 | 2.141737246 | 2.14022021 | 2.183373728 | 2.509591799 |
| ENSG00000164934 | DCAF13 | -2.176343636 | 8.35E-06 | -2.96230526 | 4.82E-10 | -5.138648897 | 10.87379128 | 9.71856931 | 10.50627919 | 8.250837933 | 8.30535212 | 7.562295432 | 7.446882182 |
| ENSG00000091879 | ANGPT2 | -3.493527871 | 0.033910747 | -1.648604047 | 0.31486887 | -5.142131918 | 2 | 4.442710129 | 5.39244166 | 2.3892346 | 2.59489879 | 3.172301926 | 3.430065404 |
| ENSG00000034533 | ASTE1 | -2.74063204 | 0.089597823 | -2.402252871 | 0.134152882 | -5.142884912 | 3.327711234 | 8.004992147 | 9.272377397 | 5.602782483 | 5.53799497 | 6.101204172 | 5.626876946 |
| ENSG00000171695 | LKAAEAR1 | -4.71484831 | NA | -0.428764264 | NA | -5.143612575 | 3.327711234 | 2 | 2 | 2 | 2 | 2.34604626 | 2.551896972 |
| ENSG00000183850 | ZNF730 | -1.529785684 | 0.40900069 | -3.61439882 | 0.031808421 | -5.144184503 | 2 | 5.595927442 | 4.058998049 | 3.515101962 | 3.33976443 | 2.62493832 | 2.326754077 |
| ENSG00000008323 | PLEKHG6 | -4.953190056 | 0.00193985 | -0.192624187 | 0.893463707 | -5.145814243 | 4.00722531 | 4.11352228 | 4.058998049 | 2.141737246 | 2.14022021 | 3.865109708 | 3.972018587 |
| ENSG00000114388 | NPRL2 | -3.04462513 | 1.06E-09 | -2.102320052 | 3.26E-05 | -5.146945182 | 9.219014269 | 8.791896851 | 9.945092785 | 6.429169081 | 6.39275508 | 7.389977651 | 7.247335571 |
| ENSG00000188846 | RPL14 | -2.804206368 | 0.000258981 | -2.349592439 | 0.00257688 | -5.153798807 | 14.1140585 | 12.18687301 | 12.48428878 | 10.36543204 | 10.4218361 | 10.81442522 | 10.87956593 |
| ENSG00000181826 | RELL1 | -1.673472769 | 0.017236601 | -3.480381397 | 8.73E-08 | -5.153854167 | 9.189468652 | 8.522788974 | 10.14394518 | 7.77569021 | 7.7866208 | 6.290458506 | 5.73181021 |
| ENSG00000198400 | NTRK1 | -2.340407617 | 0.587097377 | -2.814009894 | 0.495558428 | -5.154417511 | 2 | 2 | 4.523675391 | 2.141737246 | 2.59489879 | 2.526562374 | 2 |
| ENSG00000152782 | PANK1 | -2.59362437 | 2.08E-09 | -2.56163968 | 1.62E-09 | -5.15526405 | 8.493334308 | 8.213429689 | 8.175061096 | 5.890757403 | 5.69405321 | 6.101204172 | 5.481238982 |
| ENSG00000112394 | SLC16A10 | -5.105098758 | 0.072402975 | -0.050572864 | 0.985591183 | -5.155671622 | 4.00722531 | 3.686303414 | 2 | 2.141737246 | 2 | 3.981090143 | 2.551896972 |
| ENSG00000172458 | IL17D | -3.170783934 | 5.25E-08 | -1.985722877 | 0.000708612 | -5.156506811 | 7.750913381 | 6.899080544 | 8.076950877 | 4.846787657 | 4.50979031 | 5.574182174 | 5.892708443 |
| ENSG00000164011 | ZNF691 | -2.544404862 | 0.193269017 | -2.614825389 | 0.17143346 | -5.159230251 | 9.497412313 | 8.053983481 | 2 | 5.818703821 | 5.99120031 | 6.025180246 | 5.631815919 |
| ENSG00000178201 | VN1R1 | -2.260479029 | 0.022144103 | -2.902359696 | 0.00197542 | -5.162838725 | 5.095959458 | 5.72254323 | 5.39244166 | 3.349643765 | 3.82856379 | 3.534812207 | 2.782576693 |
| ENSG00000113838 | TBCCD1 | -2.968950515 | 1.11E-06 | -2.193956319 | 0.000399783 | -5.162906835 | 9.533135543 | 8.255926468 | 9.725218844 | 6.401247806 | 6.37866593 | 7.314596615 | 6.932071508 |
| ENSG00000110536 | PTPMT1 | -1.636914751 | 0.000911216 | -3.526775072 | 3.10E-14 | -5.163689823 | 10.44422623 | 9.488434476 | 9.516493218 | 8.262524677 | 8.26379462 | 6.405645065 | 6.440710022 |
| ENSG00000138798 | EGF | -1.094951156 | 0.614705994 | -4.069851924 | 0.031811212 | -5.16480308 | 7.667647223 | 2 | 4.874587708 | 5.37770462 | 5.18011485 | 3.193831101 | 2.950113106 |
| ENSG00000159640 | ACE | -3.159779869 | 0.070689945 | -2.005683369 | 0.265563294 | -5.165463239 | 4.00722531 | 6.228547981 | 9.735974718 | 5.319220934 | 5.22805098 | 5.673303969 | 6.800084026 |
| ENSG00000137822 | TUBGCP4 | -3.173533864 | 7.42E-07 | -1.997614731 | 0.002624999 | -5.171148596 | 10.24321353 | 8.557180776 | 9.849281147 | 6.542151749 | 6.63128385 | 7.861174222 | 7.58443608 |
| ENSG00000128309 | MPST | -3.604988768 | 3.69E-29 | -1.568191441 | 2.02E-06 | -5.173180209 | 11.68645261 | 11.14918383 | 11.14543241 | 7.794595599 | 7.74329312 | 9.849026448 | 9.719663687 |
| ENSG00000182551 | ADI1 | -1.086974998 | 0.08673281 | -4.087695236 | 1.86E-13 | -5.174670234 | 10.92399098 | 9.770792087 | 10.94216274 | 9.526236618 | 9.58250309 | 6.226613426 | 6.907844475 |
| ENSG00000084710 | EFR3B | -2.171832571 | 0.000246655 | -3.003380532 | 1.06E-07 | -5.175213103 | 6.366709671 | 6.464272742 | 6.073378139 | 4.17310023 | 4.56059747 | 3.865109708 | 3.620995423 |
| ENSG00000127252 | HRASLS | -4.714827672 | NA | -0.461211974 | NA | -5.176039646 | 2 | 3.076665209 | 2 | 2 | 2 | 2.592885947 | 2.277179783 |
| ENSG00000174938 | SEZ6L2 | -2.439477113 | 0.003647879 | -2.742995672 | 0.00087746 | -5.182472785 | 9.521325812 | 10.95649913 | 11.58682058 | 8.451360823 | 8.52753255 | 8.666781604 | 7.473262559 |
| ENSG00000178562 | CD28 | -5.364087556 | 0.256719105 | 0.179652726 | NA | -5.18443483 | 2 | 3.686303414 | 2 | 2 | 2 | 2.656294034 | 3.128433649 |
| ENSG00000179981 | TSHZ1 | -2.746361989 | 4.24E-07 | -2.43840777 | 7.48E-06 | -5.184769759 | 9.624286623 | 10.56689847 | 9.503962296 | 7.348537858 | 7.18663697 | 7.824551575 | 7.261744957 |
| ENSG00000105669 | COPE | -2.118843219 | 0.000132612 | -3.066563094 | 1.06E-08 | -5.185406313 | 12.65887593 | 11.39001779 | 11.67049834 | 9.865880373 | 9.93276281 | 8.878929913 | 9.032214052 |
| ENSG00000186468 | RPS23 | -2.236162052 | 0.003447259 | -2.949803335 | 6.64E-05 | -5.185965387 | 13.58728744 | 11.87721417 | 11.91489601 | 10.45431041 | 10.4749506 | 9.533683094 | 9.944472938 |
| ENSG00000130307 | USHBP1 | -4.714849208 | NA | -0.478836164 | NA | -5.193685372 | 3.327711234 | 2 | 2 | 2 | 2 | 2.457042231 | 2.421067459 |
| ENSG00000204291 | COL15A1 | -1.820620713 | 0.390209318 | -3.373264286 | 0.084214225 | -5.193884999 | 9.112843761 | 10.07368926 | 2 | 7.19843704 | 7.3899592 | 6.272864826 | 5.136701757 |
| ENSG00000147403 | RPL10 | -2.676284864 | 3.42E-09 | -2.522244157 | 2.61E-08 | -5.198529021 | 14.59235421 | 13.63725175 | 13.71949044 | 11.36663123 | 11.3868933 | 11.52875862 | 11.53247504 |
| ENSG00000151090 | THRB | -8.016323189 | 0.012863597 | 2.814737749 | 0.349246545 | -5.20158544 | 2 | 5.946638857 | 2 | 2 | 2 | 7.231159579 | 7.05482945 |
| ENSG00000087589 | CASS4 | -3.515538608 | 0.097462907 | -1.688046001 | 0.436077057 | -5.203584609 | 6.138360429 | 3.076665209 | 2 | 2.3892346 | 2.77779907 | 3.767980144 | 3.15612297 |
| ENSG00000139428 | MMAB | -3.388666524 | 0.004883137 | -1.816971306 | 0.153651094 | -5.20563783 | 10.60392551 | 8.903033433 | 6.629488387 | 6.087554739 | 6.22964624 | 7.288568183 | 7.98359008 |
| ENSG00000108826 | MRPL27 | -2.425275503 | 1.61E-05 | -2.782103156 | 4.91E-07 | -5.207378659 | 10.77778805 | 9.71856931 | 11.0961933 | 8.192977693 | 8.26379462 | 7.726402307 | 8.014172904 |
| ENSG00000133835 | HSD17B4 | -2.298584966 | 1.16E-07 | -2.910274492 | 8.30E-12 | -5.208859459 | 11.78604782 | 11.15761406 | 11.98419955 | 9.432787137 | 9.3477391 | 8.495864663 | 9.023784348 |
| ENSG00000081248 | CACNA1S | -4.901640115 | 0.168638386 | -0.309021422 | 0.933175306 | -5.210661537 | 2 | 2 | 4.523675391 | 2.141737246 | 2 | 3.465698119 | 2.782576693 |
| ENSG00000078795 | PKD2L2 | -4.714830514 | NA | -0.495948699 | NA | -5.210779213 | 2 | 3.076665209 | 2 | 2 | 2 | 2.492221014 | 2.374681325 |
| ENSG00000187550 | SBK2 | -4.714828882 | NA | -0.496724947 | NA | -5.211553829 | 2 | 3.076665209 | 2 | 2 | 2 | 2.3070652 | 2.551896972 |
| ENSG00000235692 | | 3.876577836 | 0.144656154 | -9.088813613 | 0.001863963 | -5.212235778 | 2 | 5.595927442 | 2 | 7.593372273 | 8.02689124 | 2 | 2 |
| ENSG00000171049 | FPR2 | 0.095658118 | 0.979884745 | -5.316861428 | 0.095340194 | -5.22120331 | 2 | 2 | 4.874587708 | 3.709785322 | 3.65239003 | 2.139668783 | 2 |
| ENSG00000110700 | RPS13 | -2.269050489 | 0.000449511 | -2.952338869 | 2.64E-06 | -5.221389359 | 12.86682403 | 11.39001779 | 11.54767436 | 9.869069092 | 9.79633147 | 9.22217193 | 9.081783389 |
| ENSG00000119004 | CYP20A1 | -1.860206358 | 1.11E-06 | -3.361865339 | 7.99E-20 | -5.222071697 | 9.761125267 | 9.308339287 | 10.09465441 | 7.947219213 | 7.8720741 | 6.448814696 | 6.454714037 |
| ENSG00000188227 | ZNF793 | -2.469841783 | 0.00490488 | -2.754892907 | 0.001287325 | -5.22473469 | 6.816637977 | 6.04685525 | 8.076950877 | 5.040242897 | 4.76938658 | 4.907994066 | 4.376876162 |
| ENSG00000090382 | LYZ | -4.834325205 | 0.000258152 | -0.390907799 | 0.799372311 | -5.225233004 | 6.009025997 | 4.710564154 | 6.629488387 | 2.784612485 | 2.38539787 | 6.332313483 | 4.077147165 |
| ENSG00000175634 | RPS6KB2 | -2.16212771 | 6.61E-05 | -3.063280767 | 5.35E-09 | -5.225408477 | 10.93293345 | 9.687856435 | 10.10298738 | 8.117764469 | 8.2561084 | 7.174082053 | 7.424286753 |
| ENSG00000132141 | CCT6B | -1.934775756 | 0.289625434 | -3.303844079 | 0.051418119 | -5.238619836 | 7.031698319 | 4.93638933 | 2 | 4.139692846 | 4.0930011 | 3.12825509 | 3.261853426 |
| ENSG00000197165 | SULT1A2 | -0.401881372 | 0.865062033 | -4.839313456 | 0.012183874 | -5.241194828 | 2 | 3.076665209 | 6.073378139 | 4.17310023 | 4.56059747 | 2.183373728 | 2.326754077 |
| ENSG00000186469 | GNG2 | 0.793673617 | 0.719144616 | -6.036972871 | 0.000856346 | -5.243299253 | 2 | 6.666836837 | 8.380951741 | 8.352759303 | 8.3220138 | 2.384001718 | 3.071407821 |
| ENSG00000117650 | NEK2 | -2.654295033 | 0.0437622 | -2.58990333 | 0.042567233 | -5.244198363 | 5.095959458 | 4.710564154 | 5.930865648 | 2.868578001 | 3.60476711 | 2.592885947 | 3.807156736 |
| ENSG00000177383 | MAGEF1 | -3.00055542 | 6.64E-07 | -2.245195527 | 0.000267013 | -5.245750947 | 11.61323974 | 10.06773758 | 11.38294124 | 8.115618198 | 8.23866319 | 8.803037464 | 9.038736604 |
| ENSG00000239857 | GET4 | -0.924331123 | 0.014691493 | -4.32229185 | 5.02E-36 | -5.246622974 | 9.358295023 | 8.876041558 | 9.344432377 | 8.401141138 | 8.16874151 | 5.174809258 | 4.927687581 |
| ENSG00000137474 | MYO7A | -2.804837511 | 0.317447824 | -2.444102606 | 0.377779769 | -5.248940118 | 2 | 6.728495673 | 2 | 3.09461492 | 3.2184422 | 3.917810786 | 2.466008509 |
| ENSG00000072041 | SLC6A15 | 0.623019225 | 0.613825095 | -5.872427975 | 3.28E-09 | -5.24940875 | 6.96351253 | 9.377333006 | 9.132850258 | 9.389919188 | 9.4789996 | 4.029788235 | 2.782576693 |
| ENSG00000170537 | TMC7 | -2.867085001 | 1.19E-05 | -2.383490046 | 0.000194399 | -5.250575047 | 7.031698319 | 5.838937808 | 6.958202216 | 4.205751501 | 4.05806467 | 4.620674722 | 4.411752372 |
| ENSG00000205593 | DENND6B | -2.752343336 | 0.073577427 | -2.500603183 | 0.100794035 | -5.252946519 | 6.138360429 | 3.686303414 | 8.380957451 | 4.647285135 | 4.50979031 | 4.478802948 | 5.050803619 |
| ENSG00000196549 | MME | 1.42053898 | 0.310604101 | -6.675785682 | 2.01E-08 | -5.255246703 | 9.521325812 | 10.66871049 | 9.714382178 | 11.44991178 | 11.5052982 | 4.580376151 | 2.277179783 |
| ENSG00000258724 | AC105001.2 | -2.894412166 | 0.042121045 | -2.361940131 | 0.099493727 | -5.256352298 | 9.800745303 | 5.72254323 | 6.629488387 | 5.639097847 | 5.65950892 | 6.14330814 | 6.148451724 |
| ENSG00000129226 | CD68 | 4.33784884 | 9.67E-12 | -9.599103572 | 8.57E-26 | -5.261254733 | 7.867441304 | 7.467835385 | 8.941672936 | 12.53882471 | 12.5522656 | 2.094598369 | 2.172608095 |
| ENSG00000100519 | PSMC6 | -2.133221765 | 2.99E-06 | -3.129799853 | 1.76E-12 | -5.263021619 | 10.88765625 | 9.890264534 | 10.57378414 | 8.390528966 | 8.38151133 | 7.561260509 | 7.232780814 |
| ENSG00000104980 | TIMM44 | -2.838813397 | 4.29E-05 | -2.427207651 | 0.000524836 | -5.266021048 | 11.1517158 | 9.235879652 | 10.31150679 | 7.61176303 | 7.62593544 | 8.087943566 | 7.955303035 |
| ENSG00000112029 | FBXO5 | -2.464408836 | 0.012174173 | -2.806608673 | 0.003453677 | -5.271017508 | 7.579279349 | 6.728495673 | 9.242516611 | 6.01571348 | 5.65950892 | 5.773219597 | 5.243813244 |
| ENSG00000139329 | LUM | -2.824466728 | 1.41E-11 | -2.449557666 | 5.55E-09 | -5.274024394 | 16.17280313 | 15.36952225 | 15.37683927 | 12.86908439 | 12.8662715 | 13.36227546 | 13.1116706 |
| ENSG00000142227 | EMP3 | -2.069818727 | 0.00012366 | -3.204755769 | 6.95E-10 | -5.274574496 | 13.01629778 | 12.74835258 | 13.77334518 | 11.15408254 | 11.2034821 | 9.611983935 | 10.38089967 |
| ENSG00000132561 | MATN2 | -4.756069867 | 2.32E-11 | -0.518588354 | 0.547274868 | -5.274658222 | 10.5351205 | 10.01303782 | 9.116480286 | 5.258265913 | 5.49945122 | 8.624798858 | 10.01917298 |
| ENSG00000153898 | MCOLN2 | -3.267747582 | NA | -2.007121799 | NA | -5.274869381 | 2 | 2 | 3.369317972 | 2.141737246 | 2 | 2.267001569 | 2.059860178 |
| ENSG00000125246 | CLYBL | -6.864212226 | 0.004631619 | 1.588686189 | 0.428703065 | -5.275526036 | 2 | 4.11352228 | 4.058998049 | 2 | 2 | 4.946551782 | 4.911532223 |
| ENSG00000178199 | ZC3H12D | -2.464325013 | 0.136166712 | -2.811970161 | 0.074118334 | -5.276295174 | 4.00722531 | 3.686303414 | 5.39244166 | 3.227757282 | 2.49394783 | 2.3070652 | 3.100202476 |
| ENSG00000137968 | SLC44A5 | -1.216513667 | 0.704230942 | -4.061054085 | 0.160285094 | -5.277567752 | 2 | 4.442710129 | 2 | 3.162721474 | 2.14022021 | 2.139668783 | 2.117335275 |
| ENSG00000197168 | NEK5 | -3.479276254 | 0.022625554 | -1.798941204 | 0.244978858 | -5.278217458 | 3.327711234 | 3.686303414 | 6.322058544 | 2.784612485 | 2.68924543 | 3.679182183 | 3.620995423 |
| ENSG00000174586 | ZNF497 | -4.169334373 | 0.017841169 | -1.110335883 | 0.567222525 | -5.279670256 | 7.485143553 | 5.303523747 | 2 | 2.784612485 | 3.15372745 | 5.212218839 | 5.164234344 |
| ENSG00000135314 | KHDC1 | -2.307355796 | 0.048902337 | -2.973446363 | 0.008292896 | -5.280802159 | 7.975256505 | 5.595927442 | 9.180874355 | 5.880681751 | 6.03677338 | 5.253805068 | 5.436777225 |
| ENSG00000180481 | GLIPR1L2 | -3.197439316 | 0.009787285 | -2.085084525 | 0.091103426 | -5.282523841 | 5.095959458 | 3.686303414 | 6.322058544 | 2.784612485 | 3.2184422 | 3.782262616 | 3.580696614 |
| ENSG00000178026 | LRRC75B | -3.367271956 | 0.002459978 | -1.919341381 | 0.100435749 | -5.286613337 | 9.790941832 | 7.669933291 | 6.534109959 | 5.334065616 | 5.43283104 | 6.664632795 | 6.815359915 |
| ENSG00000164404 | GDF9 | -1.557206713 | 0.341930387 | -3.730634867 | 0.012400861 | -5.28784158 | 5.70932728 | 4.710564154 | 2 | 3.566290607 | 3.60476711 | 2.42098416 | 2.632958241 |
| ENSG00000142252 | GEMIN7 | -2.244382866 | 0.003103036 | -3.045636933 | 3.12E-05 | -5.290019799 | 10.32020829 | 8.376385542 | 9.196533591 | 7.234487582 | 7.36158648 | 6.307840213 | 6.714211481 |
| ENSG00000007264 | MATK | -4.895470583 | 0.221769007 | -0.399847035 | 0.92531104 | -5.295317618 | 2 | 4.442710129 | 2 | 2.141737246 | 2 | 3.694366604 | 2.117335275 |
| ENSG00000130294 | KIF1A | -3.086666879 | 0.093055209 | -2.209060781 | 0.23233796 | -5.29572766 | 5.70932728 | 7.099070191 | 3.369317972 | 4.17310023 | 2.2680111 | 4.563936166 | 3.539239711 |
| ENSG00000182858 | ALG12 | -2.429881004 | 7.59E-05 | -2.866707318 | 1.90E-06 | -5.296588322 | 9.423197387 | 9.396454291 | 10.71638486 | 7.596453725 | 7.56191311 | 7.1328795 | 7.173047408 |
| ENSG00000038295 | TLL1 | -6.778467247 | 0.070466756 | 1.475628484 | 0.694164121 | -5.302838763 | 2 | 2 | 4.874587708 | 2 | 2 | 4.984105765 | 4.489993931 |
| ENSG00000214078 | CPNE1 | -2.436759782 | 6.67E-12 | -2.878902 | 2.08E-16 | -5.315661782 | 12.37703322 | 11.69036718 | 11.92902561 | 9.566077486 | 9.62564419 | 9.284956273 | 9.015777478 |

| | | | | | | | | | | | | | |
|---|---|---|---|---|---|---|---|---|---|---|---|---|---|
| ENSG00000079257 | LXN | -1.6952289 | 0.421185937 | -3.624861961 | 0.058124528 | -5.320090861 | 2 | 9.558016677 | 10.08627303 | 7.602596956 | 7.57741621 | 5.72966459 | 5.75463059 |
| ENSG00000182247 | UBE2E2 | -3.766639373 | 6.77E-08 | -1.556196738 | 0.039587574 | -5.32283611 | 10.905938 | 9.608085847 | 11.62203638 | 7.202487431 | 7.19470132 | 9.318112363 | 9.43534143 |
| ENSG00000055955 | ITIH4 | -0.415138926 | 0.812035173 | -4.912195342 | 0.00050089 | -5.327334268 | 3.327711234 | 4.710564154 | 6.534109959 | 4.984880435 | 5.09650239 | 2.592885947 | 2.172608095 |
| ENSG00000163481 | RNF25 | -2.140639062 | 0.001573471 | -3.186962192 | 9.15E-07 | -5.327601254 | 10.66423795 | 9.599860728 | 11.39656351 | 8.551612617 | 8.64548836 | 7.489084836 | 7.644317657 |
| ENSG00000106588 | PSMA2 | -0.338558935 | 0.684355643 | -4.989928093 | 8.86E-15 | -5.328487027 | 10.91047246 | 10.17682307 | 9.06622548 | 9.856271709 | 9.94057646 | 5.352823931 | 5.419744333 |
| ENSG00000147408 | CSGALNACT1 | 0.596561273 | 0.609297559 | -5.925193933 | 1.76E-10 | -5.32863266 | 7.624139768 | 10.06176125 | 10.68897008 | 10.508071 | 10.5313622 | 4.521997428 | 4.077147165 |
| ENSG00000137100 | DCTN3 | -2.533085322 | 0.000136488 | -2.795985443 | 1.95E-05 | -5.329070765 | 11.21906289 | 9.514922632 | 10.60345421 | 8.058664058 | 8.11663771 | 7.534088584 | 8.074367056 |
| ENSG00000171174 | RBKS | -2.098681182 | 0.276988468 | -3.23527053 | 0.074313018 | -5.333951712 | 7.82963488 | 5.303523747 | 2 | 4.574143677 | 4.60967607 | 3.753554863 | 3.67940853 |
| ENSG00000100316 | RPL3 | -3.447394931 | 1.00E-09 | -1.886969902 | 0.00142758 | -5.334364833 | 15.10304219 | 14.24449601 | 13.70997499 | 11.02051291 | 11.0264456 | 12.33499242 | 12.79296778 |
| ENSG00000122194 | PLG | 0.815550983 | 0.88199664 | -6.152906239 | NA | -5.337355257 | 2 | 3.076665209 | 2 | 2.947924294 | 2.86123014 | 2 | 2 |
| ENSG00000067836 | ROGDI | -2.851375896 | 0.05996039 | -2.49201118 | 0.100049958 | -5.343387076 | 10.26462289 | 9.055168103 | 4.523675391 | 6.387081877 | 6.46120797 | 6.615624322 | 6.907844475 |
| ENSG00000274853 | | 0.278272838 | 0.941333105 | -5.622324578 | 0.072262076 | -5.34405174 | 2 | 6.389896633 | 2 | 5.144975053 | 5.24368216 | 2.34604626 | 2 |
| ENSG00000029993 | HMGB3 | -2.248983658 | 0.001093029 | -3.099134895 | 3.10E-06 | -5.348118554 | 9.740899611 | 8.124485672 | 9.798878489 | 7.226553841 | 7.12888795 | 6.252491482 | 6.460277802 |
| ENSG00000182389 | CACNB4 | -7.099535728 | 4.34E-06 | 1.745786128 | 0.216304915 | -5.3537496 | 3.327711234 | 6.228547981 | 7.502634626 | 2.270786764 | 2.14022021 | 8.374440168 | 7.926450253 |
| ENSG00000110723 | EXPH5 | -4.714241971 | 3.68E-12 | -0.646097219 | 0.370178383 | -5.360339191 | 6.89194353 | 6.728495673 | 7.600047325 | 3.289987342 | 3.08597292 | 6.835304498 | 6.044874254 |
| ENSG00000105255 | FSD1 | -3.628417327 | 0.001262337 | -1.736281814 | 0.147650336 | -5.364699141 | 8.39321512 | 5.595927442 | 9.242516611 | 5.040242897 | 4.79082474 | 6.448814696 | 6.856025745 |
| ENSG00000196503 | ARL9 | -6.302008823 | 0.116790251 | 0.936073716 | 0.822442961 | -5.365935108 | 4.467311984 | 2 | 2 | 2 | 2 | 3.917810786 | 3.972018587 |
| ENSG00000188732 | FAM221A | -3.64173839 | 0.024113792 | -1.727049321 | 0.297867673 | -5.368787711 | 2 | 5.595927442 | 6.073378139 | 3.023133209 | 2.49394783 | 3.810410249 | 3.972018587 |
| ENSG00000121417 | ZNF211 | -2.521089314 | 2.03E-10 | -2.849840721 | 2.94E-13 | -5.370930035 | 9.781071289 | 9.433952963 | 10.08627303 | 7.352188525 | 7.24216308 | 7.204892513 | 6.709547124 |
| ENSG00000139180 | NDUFA9 | -1.959092392 | 3.91E-09 | -3.412358903 | 6.12E-26 | -5.371451295 | 10.73241055 | 10.18234215 | 10.43545992 | 8.496596021 | 8.54024645 | 6.921708059 | 7.245725608 |
| ENSG00000077942 | FBLN1 | -4.258088042 | 7.83E-33 | -1.114098082 | 0.003857671 | -5.372186123 | 12.60534322 | 12.66703369 | 13.29303402 | 8.631895657 | 8.65866647 | 11.71663489 | 11.83397961 |
| ENSG00000146005 | PSD2 | -4.882661078 | 0.007379211 | -0.495359572 | 0.807725848 | -5.37802065 | 5.866944937 | 2 | 5.39244166 | 2.498691189 | 2.14022021 | 4.749835737 | 4.594883674 |
| ENSG00000006042 | TMEM98 | -2.250767307 | 8.02E-12 | -3.130721761 | 3.65E-22 | -5.381489068 | 11.99260982 | 11.41132475 | 11.64504491 | 9.390806898 | 9.52209753 | 8.437275901 | 8.718968947 |
| ENSG00000120699 | EXOSC8 | -2.243692539 | 0.000292848 | -3.140319568 | 1.54E-07 | -5.384012107 | 9.646202137 | 8.337339476 | 8.408111533 | 6.766655021 | 6.68951263 | 5.896510455 | 5.859408034 |
| ENSG00000161618 | ALDH16A1 | -1.977700577 | 0.005775587 | -3.408165493 | 4.70E-07 | -5.38586607 | 10.19198706 | 8.522788974 | 10.09465441 | 7.77569021 | 7.85436579 | 6.106889539 | 6.690737414 |
| ENSG00000110871 | COQ5 | -2.88607071 | 1.96E-08 | -2.500785343 | 1.21E-06 | -5.386856053 | 10.7426185 | 9.664384941 | 9.725218844 | 7.341208691 | 7.21466714 | 7.708751741 | 7.600900977 |
| ENSG00000185670 | ZBTB3 | -2.94853045 | 0.033002523 | -2.444866944 | 0.079075233 | -5.393397394 | 9.521325812 | 6.311476754 | 5.772716771 | 5.348759107 | 5.36298494 | 5.922534244 | 5.703940561 |
| ENSG00000176236 | C10orf111 | -2.691103879 | 0.215094303 | -2.702319198 | 0.186685975 | -5.393423078 | 4.00722531 | 2 | 4.058998049 | 2.141737246 | 2.59489879 | 2.34604626 | 2.421067459 |
| ENSG00000182272 | B4GALNT4 | -4.28047 | 1.94E-11 | -1.115048908 | 0.093533047 | -5.395518907 | 7.384433967 | 7.232742069 | 7.451345288 | 3.997744386 | 3.08597292 | 6.490730008 | 6.037449234 |
| ENSG00000104221 | BRF2 | -2.010010171 | 0.128591494 | -3.388528225 | 0.005996167 | -5.398538397 | 10.28571918 | 8.276714465 | 6.203069917 | 7.030886362 | 7.18258783 | 5.938563912 | 5.636738041 |
| ENSG00000111752 | PHC1 | -1.704861308 | 0.022800644 | -3.695767723 | 8.62E-08 | -5.400629031 | 7.709880939 | 9.45234302 | 9.553445883 | 7.467753569 | 7.39696604 | 5.54096877 | 5.534955934 |
| ENSG00000135486 | HNRNPA1 | -1.533780238 | 0.036573046 | -3.867184397 | 6.12E-09 | -5.400964634 | 13.34081888 | 12.14791279 | 11.66769225 | 11.04862828 | 11.0151054 | 8.379146188 | 8.978459426 |
| ENSG00000163682 | RPL9 | -2.063789586 | 0.000258152 | -3.338067229 | 7.40E-10 | -5.401856814 | 13.22727653 | 12.10351434 | 12.00213172 | 10.488715 | 10.5056622 | 9.285896341 | 9.16888512 |
| ENSG00000177710 | SLC35G5 | -3.310912207 | 0.25952816 | -2.096891849 | 0.459893021 | -5.407804056 | 2 | 2 | 4.523675391 | 2.141737246 | 2.2680111 | 2.62493832 | 2.225841198 |
| ENSG00000105519 | CAPS | -3.073598565 | 0.000252864 | -2.334865857 | 0.006381105 | -5.408464422 | 8.696033109 | 6.78762695 | 9.083172337 | 5.319220934 | 5.6826301 | 6.353989995 | 6.026239558 |
| ENSG00000065268 | WDR18 | -2.756752835 | 3.26E-29 | -2.653322968 | 5.48E-28 | -5.410075803 | 10.62615013 | 10.52407288 | 10.5916593 | 7.789219291 | 7.89946959 | 7.789628618 | 8.089704419 |
| ENSG00000167815 | PRDX2 | -2.380441212 | 1.24E-05 | -3.029880795 | 1.28E-08 | -5.410322007 | 12.40300905 | 11.52428675 | 12.82880267 | 9.939922873 | 10.0021819 | 9.083440537 | 9.532294581 |
| ENSG00000145491 | ROPN1L | -1.400327048 | 0.64165426 | -4.013904333 | 0.14113949 | -5.414231381 | 2 | 4.93638933 | 2 | 3.162721474 | 2.59489879 | 2.048074405 | 2.326754077 |
| ENSG00000150048 | CLEC1A | -4.714845832 | NA | -0.70039391 | NA | -5.415239742 | 2 | 2 | 3.369317972 | 2 | 2 | 2.526562374 | 2.225841198 |
| ENSG00000143443 | C1orf56 | -2.939989567 | 0.008914562 | -2.476389297 | 0.028992424 | -5.416378863 | 6.96351253 | 6.140560021 | 9.180874355 | 5.210797711 | 5.21224859 | 5.646229369 | 5.616947984 |
| ENSG00000261594 | TPBGL | -3.98888454 | 0.04048095 | -1.427744792 | 0.492084442 | -5.416629332 | 2 | 7.189546432 | 4.058998049 | 2.947924294 | 2.77779907 | 4.340871764 | 4.729989799 |
| ENSG00000167220 | HDHD2 | -0.444075163 | 0.604439857 | -4.973517294 | 3.77E-13 | -5.417592457 | 8.816894989 | 8.671479029 | 7.029880454 | 7.975897842 | 7.88459112 | 3.694366604 | 3.987514168 |
| ENSG00000126522 | ASL | -2.052524417 | 7.15E-05 | -3.365163885 | 1.24E-11 | -5.417688302 | 10.00110988 | 9.656475478 | 10.08627303 | 7.852449262 | 7.92877397 | 5.982522014 | 7.052989523 |
| ENSG00000136634 | IL10 | -2.299951907 | 0.135958069 | -3.119619501 | 0.03159523 | -5.419571407 | 4.00722531 | 3.686303414 | 5.156626401 | 3.023133209 | 2.77779907 | 2.858551861 | 2.225841198 |
| ENSG00000181392 | SYNE4 | -2.732794853 | 0.310115955 | -2.688666072 | 0.299826316 | -5.421460926 | 2 | 5.303523747 | 2 | 2.270786764 | 2.77779907 | 2.68698273 | 2.421067459 |
| ENSG00000136842 | TMOD1 | -2.169481631 | 0.01952524 | -3.254418203 | 0.000222254 | -5.423899834 | 7.384433967 | 7.002537973 | 9.212024683 | 6.197166296 | 6.00043082 | 5.079718116 | 5.136701757 |
| ENSG00000282017 | | 4.452552565 | 1.97E-07 | -9.879054415 | 1.89E-08 | -5.426501851 | 5.095959458 | 5.457120953 | 3.369317972 | 9.144656522 | 9.20831741 | 2 | 2 |
| ENSG00000175866 | BAIAP2 | -2.594709551 | 0.001044702 | -2.83182366 | 0.000279638 | -5.426533211 | 10.73752355 | 8.540087354 | 10.13840817 | 7.536750104 | 7.4450836 | 7.0047514 | 7.477384137 |
| ENSG00000236609 | ZNF853 | -5.186081431 | 0.004381278 | -0.241601976 | 0.915858934 | -5.427683406 | 2 | 6.389896633 | 8.631919287 | 3.289987342 | 2.77779907 | 7.220710071 | 6.963754056 |
| ENSG00000141642 | ELAC1 | -2.4638844 | 2.93E-11 | -2.969968439 | 9.95E-17 | -5.433852838 | 7.709880939 | 7.928237109 | 7.552162999 | 5.406080789 | 5.39133044 | 5.014670875 | 4.878668439 |
| ENSG00000185272 | RBM11 | -3.928668535 | 0.367319832 | -1.506125853 | NA | -5.434794388 | 2 | 3.686303414 | 2 | 2.141737246 | 2 | 2.592885947 | 2.059860178 |
| ENSG00000183783 | KCTD8 | -2.272296594 | NA | -3.163834183 | NA | -5.436130777 | 2 | 3.076665209 | 2 | 2.141737246 | 2.14022021 | 2.094598369 | 2.059860178 |
| ENSG00000157152 | SYN2 | -4.714827224 | NA | -0.722507286 | NA | -5.43733451 | 2 | 3.076665209 | 2 | 2 | 2 | 2.183373728 | 2.551896972 |
| ENSG00000139679 | LPAR6 | -2.553110641 | 6.75E-05 | -2.884887074 | 4.37E-06 | -5.437997715 | 7.709880939 | 9.214490408 | 8.763901704 | 6.328984825 | 6.0722229 | 6.126613714 | 5.616947984 |
| ENSG00000168158 | OR2C1 | -3.274419209 | NA | -2.16845216 | NA | -5.442871369 | 2 | 3.076665209 | 2 | 2 | 2.14022021 | 2.183373728 | 2.117335275 |
| ENSG00000134419 | RPS15A | -1.308702615 | 0.104928256 | -4.134382029 | 6.17E-09 | -5.443084645 | 12.36881891 | 10.50217373 | 10.89974571 | 10.18324441 | 10.1913912 | 7.457170674 | 7.322574236 |
| ENSG00000196632 | WNK3 | -4.19005784 | 0.000268707 | -1.254198291 | 0.303073664 | -5.444256131 | 7.096806373 | 4.710564154 | 5.39244166 | 3.162721474 | 2.59489879 | 5.085488617 | 4.784231301 |
| ENSG00000183160 | TMEM119 | 0.177052961 | 0.935262104 | -5.621263283 | 0.000779055 | -5.444610323 | 3.327711234 | 7.503546079 | 10.12770182 | 8.860563059 | 9.02486072 | 4.04171018 | 3.183290849 |
| ENSG00000101365 | IDH3B | -2.747553956 | 2.33E-13 | -2.698083585 | 4.47E-13 | -5.44563754 | 11.17828117 | 10.68435379 | 11.4724895 | 8.332573384 | 8.48870446 | 8.380320298 | 8.538752857 |
| ENSG00000183208 | GDPGP1 | -3.249548955 | 0.067089529 | -2.20273728 | 0.219193182 | -5.452286235 | 2 | 4.93638933 | 7.029880454 | 3.289987342 | 3.15372745 | 4.065262523 | 3.735548302 |
| ENSG00000137491 | SLCO2B1 | -5.510419188 | 4.62E-13 | 0.057372666 | 0.948539975 | -5.453046522 | 6.816637977 | 7.274681896 | 7.098165388 | 3.023133209 | 2.49394783 | 7.535143176 | 6.577546999 |
| ENSG00000090266 | NDUFB2 | -1.947482514 | 6.54E-05 | -3.508728283 | 5.48E-14 | -5.456210797 | 10.55263183 | 10.32389548 | 11.2690689 | 8.792330793 | 8.87484109 | 7.016881172 | 7.532563185 |
| ENSG00000160050 | CCDC28B | -3.372157953 | 1.10E-06 | -2.084140776 | 0.003524935 | -5.456298728 | 7.384433967 | 9.055168103 | 9.120850258 | 5.44762523 | 5.48637102 | 6.537789343 | 6.802276223 |
| ENSG00000183048 | SLC25A10 | -2.457916224 | 0.111165454 | -3.001427158 | 0.043204146 | -5.459343382 | 3.327711234 | 7.232742069 | 8.608698767 | 5.242616075 | 5.14724905 | 4.778616857 | 4.664017718 |
| ENSG00000170542 | SERPINB9 | -2.288658808 | 4.28E-07 | -3.177396975 | 5.80E-13 | -5.466055784 | 8.87372952 | 9.338312924 | 9.809100877 | 6.975156635 | 7.27676203 | 6.151583464 | 6.397863414 |
| ENSG00000011332 | DPF1 | -3.535453246 | 0.055846839 | -1.939049735 | 0.314029219 | -5.474502981 | 8.540903468 | 6.535001748 | 2 | 4.034562184 | 4.16043394 | 5.798032389 | 4.990558504 |
| ENSG00000077935 | SMC1B | -4.714845442 | NA | -0.762122318 | NA | -5.476967761 | 3.327711234 | 2 | 2 | 2 | 2 | 2.457042231 | 2.277179783 |
| ENSG00000124713 | GNMT | -5.361287587 | 0.272396482 | -0.116288376 | NA | -5.477575964 | 2 | 3.686303414 | 2 | 2 | 2 | 3.035909414 | 2.466008509 |
| ENSG00000106565 | TMEM176B | -7.559657967 | 0.004968903 | 2.08049792 | 0.37875047 | -5.479160047 | 3.327711234 | 5.303523747 | 2 | 2 | 2 | 6.633459761 | 4.870334174 |
| ENSG00000198546 | ZNF511 | -2.584153856 | 4.79E-08 | -2.898037807 | 5.29E-10 | -5.482191663 | 10.08373159 | 9.470501606 | 9.06622548 | 6.917187326 | 7.17852729 | 6.609629839 | 6.879055607 |
| ENSG00000125744 | RTN2 | -4.144309417 | 9.03E-05 | -1.338824382 | 0.255323155 | -5.483133799 | 6.563806279 | 8.557180776 | 9.999667074 | 5.076005082 | 4.91316568 | 7.542503814 | 7.728570659 |
| ENSG00000111728 | ST8SIA1 | -5.038737353 | 0.002086628 | -0.446042366 | 0.78845613 | -5.484779719 | 4.467311984 | 4.442710129 | 4.874587708 | 2.3892346 | 2 | 4.727866721 | 3.496556178 |
| ENSG00000162711 | NLRP3 | -1.182299042 | 0.503132794 | -4.317336004 | 0.00511008 | -5.499635046 | 3.327711234 | 4.11352228 | 6.958202216 | 4.329433326 | 4.83276841 | 2.858551861 | 2.421067459 |
| ENSG00000154102 | C16orf74 | -1.080885267 | 0.207420969 | -4.420442568 | 6.64E-09 | -5.501327836 | 5.095959458 | 6.389896633 | 6.882775709 | 5.161714397 | 5.41912973 | 2.858551861 | 2.950113106 |
| ENSG00000153404 | PLEKHG4B | -0.680571742 | 0.47367927 | -4.824630167 | 1.93E-09 | -5.505201908 | 5.330594134 | 6.666836837 | 7.029880454 | 5.987823101 | 5.72778967 | 3.011868911 | 2.592996859 |
| ENSG00000089169 | RPH3A | -3.274632245 | NA | -2.232807753 | NA | -5.507439999 | 2 | 2 | 3.369317972 | 2 | 2.14022021 | 2 | 2.277179783 |
| ENSG00000180628 | PCGF5 | -9.862163256 | 3.35E-08 | 4.352433529 | 1.39E-06 | -5.509729727 | 7.159102577 | 5.303523747 | 5.39244166 | 2 | 2 | 10.55391886 | 10.42691187 |
| ENSG00000049541 | RFC2 | -2.481982974 | 4.43E-09 | -3.028193633 | 2.80E-13 | -5.510176607 | 9.048471759 | 8.903033433 | 9.670202709 | 6.772115916 | 6.83591385 | 6.104049656 | 6.440710022 |
| ENSG00000188322 | SBK1 | -2.486941789 | 0.123764005 | -3.024770589 | 0.047712153 | -5.511712378 | 4.815609471 | 4.93638933 | 2 | 2.947924294 | 2.68924543 | 2.746468522 | 2.466008509 |
| ENSG00000007341 | ST7L | -1.89430328 | 0.011777196 | -3.62015317 | 2.52E-07 | -5.51445645 | 9.344956677 | 8.191702463 | 10.09465441 | 7.571615825 | 7.49164798 | 5.598599872 | 6.110033886 |
| ENSG00000156194 | PPEF2 | -2.285847504 | 0.307789068 | -3.230039261 | 0.123018585 | -5.515886764 | 3.327711234 | 2 | 4.523675391 | 2.498691189 | 2.49394783 | 2.42098416 | 2.117335275 |
| ENSG00000251287 | ALG1L2 | -0.968874539 | 0.617906824 | -4.547523412 | 0.00836966 | -5.516397952 | 2 | 4.11352228 | 5.156626401 | 3.615725078 | 3.45167086 | 2.094598369 | 2.326754077 |
| ENSG00000145757 | SPATA9 | -4.714828243 | NA | -0.804067876 | NA | -5.518896119 | 2 | 3.076665209 | 2 | 2 | 2 | 2.526562374 | 2.172608095 |
| ENSG00000232357 | | 0.632905779 | NA | -6.152904184 | NA | -5.519998405 | 2 | 3.076665209 | 2 | 2.695456494 | 2.94009902 | 2 | 2 |
| ENSG00000168658 | VWA3B | -4.714849816 | NA | -0.80525164 | NA | -5.520101456 | 2 | 2 | 3.369317972 | 2 | 2 | 2.34604626 | 2.374681325 |
| ENSG00000167085 | PHB | -2.103109834 | 2.20E-07 | -3.418832795 | 4.03E-18 | -5.521942629 | 11.73071355 | 10.90735644 | 11.27276688 | 9.233438338 | 9.2591908 | 7.759264603 | 8.109313767 |
| ENSG00000111684 | LPCAT3 | -2.253867145 | 4.33E-05 | -3.275776191 | 8.30E-10 | -5.529643336 | 9.98400217 | 9.814097497 | 8.805342542 | 7.461007055 | 7.32531744 | 6.121005698 | 6.64749689 |
| ENSG00000204025 | TRPC5OS | -1.689725496 | NA | -3.84188224 | NA | -5.531607736 | 2 | 2 | 3.369317972 | 2.141737246 | 2.2680111 | 2.094598369 | 2 |
| ENSG00000124614 | RPS10 | 0.701967592 | 0.208187941 | -6.239551535 | 6.69E-38 | -5.537583943 | 9.839306291 | 8.88960062 | 8.99637432 | 9.991747101 | 10.0159013 | 3.753554863 | 3.474731315 |
| ENSG00000148291 | SURF2 | -2.902998451 | 8.05E-06 | -2.635249522 | 4.68E-05 | -5.538247974 | 7.384433967 | 8.607278467 | 9.083172337 | 5.731694 | 5.69405321 | 5.919306827 | 6.003555476 |
| ENSG00000169744 | LDB2 | -2.660481288 | 0.002374404 | -2.878828424 | 0.000829997 | -5.539309713 | 7.159102577 | 8.505280662 | 9.541233192 | 6.078767801 | 6.14893615 | 5.536762811 | 6.205882756 |
| ENSG00000141401 | IMPA2 | -4.500645908 | 2.16E-23 | -1.042359386 | 0.016160066 | -5.543005294 | 7.904282243 | 7.467835385 | 7.818184403 | 3.566290607 | 3.86943969 | 6.797131715 | 6.640162436 |
| ENSG00000160712 | IL6R | -3.783080999 | 1.02E-06 | -1.760361385 | 0.032913606 | -5.543442385 | 8.228665467 | 8.191702463 | 9.954333071 | 5.474671438 | 5.30456641 | 7.107581047 | 7.482861322 |
| ENSG00000222036 | POTEM | 2.197137771 | 0.359275511 | -7.740757829 | 0.001039403 | -5.543620058 | 2 | 3.076665209 | 6.718950348 | 7.261917051 | 7.45182838 | 2 | 2.117335275 |
| ENSG00000142149 | HUNK | -5.223451517 | 0.000177532 | -0.321425915 | 0.848992364 | -5.544877432 | 5.532353208 | 7.761270542 | 4.523675391 | 3.023133209 | 2.2680111 | 6.106889539 | 6.440710022 |
| ENSG00000173531 | MST1 | -2.56486526 | 0.009601079 | -2.98002712 | 0.002011167 | -5.544892379 | 7.218819781 | 7.002537973 | 9.014156722 | 5.514313187 | 5.64780787 | 5.692336538 | 4.468068808 |
| ENSG00000131697 | NPHP4 | -8.218910365 | 0.011791352 | 2.665519571 | 0.386563583 | -5.553390794 | 2 | 6.140560021 | 2 | 2 | 2 | 7.107581047 | 7.282307681 |
| ENSG00000032742 | IFT88 | -1.603579855 | 0.004849167 | -3.9555963 | 1.00E-13 | -5.559176156 | 8.138771685 | 7.189546432 | 8.35328246 | 6.516352584 | 6.29111322 | 4.388329244 | 4.228274364 |
| ENSG00000148219 | ASTN2 | -3.67143832 | 3.43E-09 | -1.888921999 | 0.002753906 | -5.560360319 | 7.159102577 | 8.169643011 | 7.502634626 | 4.139692846 | 4.45712836 | 6.189586148 | 5.448021784 |
| ENSG00000111348 | ARHGDIB | -5.455175358 | 9.58E-09 | -0.110211668 | 0.928701424 | -5.565387026 | 6.89194353 | 6.666836837 | 8.699419242 | 3.09461492 | 3.15372745 | 7.000176314 | 8.043194332 |
| ENSG00000142185 | TRPM2 | -5.052655269 | 0.066277575 | -0.521310982 | 0.867018813 | -5.573966251 | 2 | 2 | 6.43197674 | 2.141737246 | 2.38539787 | 5.06226682 | 3.640730416 |
| ENSG00000243811 | APOBEC3D | -0.763600369 | 0.519891195 | -4.811206526 | 8.56E-07 | -5.574806895 | 8.836089686 | 6.140560021 | 7.225742779 | 7.127781374 | 7.00595521 | 3.838019203 | 3.236138336 |
| ENSG00000243364 | EFNA4 | -2.734643466 | 0.002564821 | -2.840751602 | 0.001501013 | -5.575395068 | 6.366709671 | 8.623598428 | 8.960138144 | 5.742861563 | 5.67111583 | 5.519815274 | 5.694529741 |
| ENSG00000204822 | MRPL53 | -1.499673725 | 0.000513088 | -4.078517812 | 1.96E-23 | -5.578191537 | 9.015175615 | 8.451443573 | 8.206335842 | 7.206526483 | 7.03768431 | 4.727866721 | 4.74829783 |
| ENSG00000175691 | ZNF77 | -3.862738353 | 4.62E-07 | -1.727247177 | 0.030676858 | -5.58998553 | 7.940205782 | 6.140560021 | 7.777116083 | 4.034562184 | 3.94787299 | 5.570072074 | 6.026239558 |
| ENSG00000050438 | SLC4A8 | -2.729947229 | 0.000381151 | -2.860384315 | 0.000144965 | -5.590331544 | 7.384433967 | 6.311476754 | 8.110401118 | 5.058234798 | 4.68029874 | 4.971696071 | 4.532869193 |
| ENSG00000267534 | S1PR2 | -2.811936332 | 1.54E-07 | -2.779924211 | 1.89E-07 | -5.591860543 | 11.38138789 | 10.07961649 | 11.21627664 | 8.209172298 | 8.19108423 | 8.286996197 | 8.174228942 |

| | | | | | | | | | | | | | |
|---|---|---|---|---|---|---|---|---|---|---|---|---|---|
| ENSG00000162105 | SHANK2 | -5.356243215 | 0.3034352 | -0.235705581 | NA | -5.591948797 | 2 | 3.686303414 | 2 | 2 | 2 | 3.12825509 | 2.172608095 |
| ENSG00000159217 | IGF2BP1 | -3.000094819 | 5.97E-08 | -2.593141327 | 3.01E-06 | -5.593236146 | 10.27168932 | 10.17682307 | 8.960138144 | 6.970413838 | 6.94032357 | 7.227249878 | 7.464983909 |
| ENSG00000144161 | ZC3H8 | -2.765446723 | 0.132233055 | -2.830764878 | 0.115974173 | -5.596211602 | 9.567995545 | 4.710564154 | 4.523675391 | 5.304221918 | 5.53799497 | 5.610655473 | 5.072762365 |
| ENSG00000130204 | TOMM40 | -3.140777063 | 8.01E-07 | -2.459363326 | 0.000143403 | -5.600140389 | 11.90670479 | 10.21502048 | 11.35530464 | 8.172475401 | 8.21109983 | 8.631223999 | 9.067728571 |
| ENSG00000151322 | NPAS3 | -2.641850386 | 0.498888183 | -2.959466395 | 0.433240072 | -5.601316781 | 2 | 2 | 5.772716771 | 3.227757282 | 2 | 2.831336763 | 2.374681325 |
| ENSG00000257727 | CNPY2 | -0.623078445 | 0.437949596 | -4.978724279 | 1.98E-14 | -5.601802724 | 11.88843282 | 10.19331737 | 10.68342405 | 10.47329248 | 10.4914114 | 5.997902297 | 6.38329369 |
| ENSG00000123154 | WDR83 | -2.776310286 | 0.001441038 | -2.827382446 | 0.001008927 | -5.603692732 | 8.910412251 | 6.535001748 | 7.734844356 | 5.348759107 | 5.40529704 | 5.294231737 | 5.367404784 |
| ENSG00000166455 | C16orf46 | 0.993798497 | 0.620673635 | -6.597513509 | 0.004363792 | -5.603715012 | 2 | 3.686303414 | 4.058998049 | 4.237680128 | 4.16043394 | 2.048074405 | 2 |
| ENSG00000100916 | BRMS1L | -2.981783755 | 1.25E-09 | -2.628759094 | 6.91E-08 | -5.610542849 | 8.910412251 | 8.337339476 | 9.465702942 | 6.130704144 | 6.00043082 | 6.433059984 | 6.368575325 |
| ENSG00000125384 | PTGER2 | -2.088313092 | 0.043683252 | -3.523522495 | 0.000269768 | -5.611835587 | 7.485143553 | 5.838937808 | 5.156626401 | 4.443344104 | 4.74762503 | 3.632645806 | 3.407202941 |
| BAGE | BAGE | 0.536121113 | 0.921779028 | -6.152907829 | NA | -5.616786716 | 3.327711234 | 2 | 2 | 2.868578001 | 2.68924543 | 2 | 2 |
| ENSG00000276559 | | 0.533830534 | NA | -6.152903416 | NA | -5.619072882 | 2 | 3.076665209 | 2 | 2.695456494 | 2.86123014 | 2 | 2 |
| C1ORF228 | C1ORF228 | 0.532684864 | NA | -6.15290287 | NA | -5.620218006 | 2 | 3.076665209 | 2 | 2.600425781 | 2.94009902 | 2 | 2 |
| ENSG00000197238 | HIST1H4J | 1.121302967 | 0.716222677 | -6.741748624 | 0.028630588 | -5.620445657 | 4.815609471 | 2 | 2 | 4.034562184 | 4.70309173 | 2.048074405 | 2 |
| ENSG00000172602 | RND1 | -1.810557502 | 0.525987563 | -3.81002244 | 0.143109331 | -5.620579942 | 2 | 2 | 7.285356013 | 4.17310023 | 4.22485516 | 3.082820997 | 2.746601469 |
| ENSG00000156968 | MPV17L | -2.445403252 | 0.068833104 | -3.175379454 | 0.011825656 | -5.620782706 | 3.327711234 | 4.11352228 | 5.156626401 | 2.784612485 | 2.86123014 | 2.42098416 | 2.671842496 |
| ENSG00000164742 | ADCY1 | -3.171556926 | 0.000605979 | -2.450544804 | 0.007241092 | -5.62210173 | 5.330594134 | 6.602424674 | 6.073378139 | 3.754611435 | 3.15372745 | 4.350488971 | 3.496556178 |
| ENSG00000105650 | PDE4C | -0.409337839 | 0.744383084 | -5.213053439 | 2.82E-07 | -5.622391278 | 6.138360429 | 5.131603625 | 7.285356013 | 6.069927018 | 6.01871653 | 2.885263063 | 2.326754077 |
| ENSG00000127564 | PKMYT1 | -3.30417846 | 0.097197518 | -2.320903004 | 0.245873893 | -5.625081464 | 4.00722531 | 6.951736442 | 2 | 3.023133209 | 3.15372745 | 3.193831101 | 4.062589736 |
| ENSG00000073536 | NLE1 | -3.423939609 | 0.013427184 | -2.204805387 | 0.124686589 | -5.628744996 | 5.532353208 | 8.414402626 | 10.45512158 | 5.860316854 | 5.91514978 | 7.170014349 | 6.920008846 |
| ENSG00000073598 | FNDC8 | -4.71482967 | NA | -0.918350298 | NA | -5.633179968 | 2 | 3.076665209 | 2 | 2 | 2 | 2.139668783 | 2.509591799 |
| ENSG00000145198 | VWA5B2 | -5.770533712 | 0.210545477 | 0.128695385 | 0.979494354 | -5.641838327 | 2 | 2 | 4.058998049 | 2 | 2 | 3.411599105 | 2.671842496 |
| ENSG00000182173 | TSEN54 | -3.513727833 | 5.73E-07 | -2.12875232 | 0.003405686 | -5.642480152 | 8.716893051 | 7.928237109 | 9.670202709 | 5.501219927 | 5.58783529 | 7.138441638 | 6.493217192 |
| ENSG00000136895 | GARNL3 | -4.435800805 | 0.004574811 | -1.209030371 | 0.480515064 | -5.644831176 | 7.096806373 | 6.602424674 | 2 | 2.947924294 | 2.86123014 | 5.113999907 | 5.275717819 |
| ENSG00000089157 | RPLP0 | -2.983586948 | 3.25E-07 | -2.663640485 | 5.80E-06 | -5.647227432 | 15.2059963 | 13.96680524 | 13.87841176 | 11.50874851 | 11.5006301 | 11.85381349 | 11.7940561 |
| ENSG00000007402 | CACNA2D2 | -4.220297015 | 0.044533055 | -1.437575908 | 0.519532223 | -5.657872923 | 2 | 6.78762695 | 3.369317972 | 2.3892346 | 2.77779907 | 4.461156421 | 3.824516949 |
| ENSG00000196684 | HSH2D | 2.978536753 | 0.010013464 | -8.636686184 | 5.78E-06 | -5.658149431 | 3.327711234 | 3.686303414 | 4.523675391 | 6.503277904 | 6.52655952 | 2 | 2 |
| ENSG00000119888 | EPCAM | -1.988740547 | 0.099708601 | -3.669409432 | 0.001060079 | -5.658149979 | 5.532353208 | 4.442710129 | 4.523675391 | 3.615725078 | 3.15372745 | 2.34604626 | 2.817676617 |
| ENSG00000101213 | PTK6 | -1.774560473 | 0.226187947 | -3.885318875 | 0.003695718 | -5.659879348 | 5.866944937 | 3.076665209 | 6.718950348 | 4.299492648 | 4.2560182 | 3.12825509 | 2.671842496 |
| ENSG00000137142 | IGFBPL1 | -4.007562267 | 0.044313047 | -1.654678115 | 0.435161964 | -5.662240382 | 3.327711234 | 4.710564154 | 7.818184403 | 3.79808662 | 2.14022021 | 5.19095825 | 4.594883674 |
| ENSG00000131470 | PSMC3IP | -6.915579233 | 0.060051979 | 1.252248085 | 0.736382011 | -5.663331148 | 2 | 4.93638933 | 2 | 2 | 2 | 4.848153631 | 4.500832793 |
| ENSG00000171914 | TLN2 | -2.656584905 | 5.47E-07 | -3.012056512 | 8.45E-09 | -5.668641417 | 9.473095744 | 10.7784592 | 10.49998282 | 7.734324667 | 7.70145652 | 7.317051196 | 7.421437279 |
| ENSG00000185133 | INPP5J | -4.312730425 | 0.025597813 | -1.356678302 | 0.523941689 | -5.669408727 | 8.797441463 | 5.72254323 | 2 | 3.462030052 | 3.74316441 | 5.919306827 | 6.22561877 |
| ENSG00000110002 | VWA5A | -2.959473434 | 0.111422752 | -2.712008516 | 0.139882696 | -5.67148195 | 2 | 7.099070191 | 8.434763955 | 4.783303513 | 4.40247105 | 5.217477917 | 4.254128754 |
| ENSG00000166508 | MCM7 | -2.381611287 | 0.00285755 | -3.291856891 | 1.87E-05 | -5.673468178 | 11.46082313 | 9.540933215 | 9.788583153 | 8.054188409 | 8.27334517 | 7.263334844 | 7.287011591 |
| ENSG00000161692 | DBF4B | -7.186616285 | 0.003507653 | 1.511986395 | 0.469777313 | -5.67462989 | 4.815609471 | 3.686303414 | 2 | 2 | 2 | 5.233158834 | 5.050803619 |
| ENSG00000172554 | SNTG2 | -3.267987542 | NA | -2.410565385 | NA | -5.678552927 | 2 | 2 | 3.369317972 | 2.141737246 | 2 | 2.139668783 | 2.117335275 |
| FAM109B | | 3.542226018 | 0.202033543 | -9.221259714 | 0.002040977 | -5.679033696 | 2 | 5.72254323 | 2 | 7.688853661 | 7.56502709 | 2 | 2 |
| ENSG00000250067 | YJEFN3 | -2.867718649 | 0.060644377 | -2.812220476 | 0.061661317 | -5.679939125 | 6.138360429 | 4.11352228 | 8.266922072 | 4.268917409 | 4.56059747 | 5.026717879 | 3.539239711 |
| ENSG00000027644 | INSRR | -4.714846806 | NA | -0.965171305 | NA | -5.68001811 | 2 | 2 | 3.369317972 | 2 | 2 | 2.225793514 | 2.421067459 |
| ENSG00000179889 | PDXDC1 | 0.285280584 | 0.658379805 | -5.967628985 | 1.10E-32 | -5.6823484 | 9.556468883 | 10.50217373 | 10.82615578 | 10.6575484 | 10.6866731 | 4.604690117 | 4.720747915 |
| ENSG00000277594 | | -0.215819269 | 0.521983587 | -5.471254644 | 6.06E-88 | -5.687073913 | 9.790941832 | 9.470501606 | 9.809100877 | 9.460955886 | 9.49867529 | 4.45225155 | 4.543391831 |
| ENSG00000168938 | PPIC | -1.980615237 | 6.40E-06 | -3.707325905 | 1.46E-18 | -5.687941143 | 12.87844315 | 12.11942549 | 11.95917313 | 10.38917005 | 10.4101955 | 8.646430994 | 8.718388644 |
| ENSG00000160602 | NEK8 | -2.631537481 | 0.184561268 | -3.057253159 | 0.110806839 | -5.68879064 | 9.081016759 | 6.844429755 | 2 | 5.022023783 | 5.49945122 | 4.547306681 | 5.191251321 |
| ENSG00000127774 | EMC6 | -1.184143175 | 0.236859596 | -4.507265645 | 1.91E-07 | -5.69140882 | 9.940323746 | 8.255926468 | 7.502634626 | 7.68596357 | 7.84158259 | 4.604690117 | 4.720747915 |
| ENSG00000221944 | TIGD1 | -2.348640043 | 0.09997999 | -3.345154001 | 0.013555764 | -5.693794044 | 8.946185348 | 5.946638857 | 5.156626401 | 5.501219927 | 5.2591458 | 4.496236235 | 4.543391831 |
| ENSG00000226887 | ERVMER34-1 | -6.087705029 | 0.03936298 | 0.392695896 | 0.894957398 | -5.695009133 | 3.327711234 | 3.686303414 | 2 | 2 | 2 | 3.465698119 | 3.383972333 |
| ENSG00000179841 | AKAP5 | -5.138841843 | 2.69E-09 | -0.559698596 | 0.570917019 | -5.69854044 | 8.107517749 | 7.60566235 | 5.930865648 | 3.09461492 | 3.2184422 | 7.222020406 | 6.556922154 |
| ENSG00000223355 | | 1.112841442 | 0.803400294 | -6.811852417 | 0.098632015 | -5.699010975 | 2 | 3.686303414 | 2 | 3.406930997 | 3.45167086 | 2 | 2 |
| ENSG00000010319 | SEMA3G | -4.714843179 | NA | -0.98739121 | NA | -5.702234389 | 2 | 2 | 3.369317972 | 2 | 2 | 2.457042231 | 2.172608095 |
| ENSG00000088881 | EBF4 | -3.923443132 | 0.03077866 | -1.781414597 | 0.357240472 | -5.704857729 | 8.609439662 | 7.393647515 | 2 | 3.79808662 | 4.1930041 | 6.339575235 | 5.080008298 |
| ENSG00000092621 | PHGDH | -4.146330811 | 5.71E-50 | -1.559550837 | 3.68E-08 | -5.705881647 | 10.95943304 | 11.22333918 | 11.33068056 | 7.026323566 | 7.1204461 | 9.765672435 | 9.468503442 |
| ENSG00000177721 | ANXA2R | -2.358113072 | 0.390208914 | -3.348594186 | 0.196027747 | -5.706707258 | 7.031698319 | 2 | 2 | 3.663521627 | 3.55551834 | 3.035909414 | 3.012034821 |
| ENSG00000152785 | BMP3 | -0.770910103 | 0.685014501 | -4.93596478 | 0.003079009 | -5.706874884 | 4.467311984 | 2 | 5.156626401 | 3.462030052 | 3.98554648 | 2.183373728 | 2.172608095 |
| ENSG00000183248 | PRR36 | -4.645835368 | 0.022086075 | -1.062080228 | 0.635771871 | -5.707915595 | 7.031698319 | 4.710564154 | 3.369317972 | 3.023133209 | 2 | 5.108342547 | 4.522269241 |
| ENSG00000066294 | CD84 | -7.290646322 | 1.86E-05 | 1.581931318 | 0.277833834 | -5.708715004 | 5.532353208 | 5.131603625 | 6.958202216 | 2 | 2.2680111 | 8.392009143 | 5.807970301 |
| ENSG00000092758 | COL9A3 | -2.387101439 | 0.196009997 | -3.326789827 | 0.056372767 | -5.713891266 | 4.00722531 | 6.140560021 | 2 | 3.349643765 | 3.01487888 | 2.803598391 | 2.671842496 |
| ENSG00000120457 | KCNJ5 | -6.422109074 | 1.33E-06 | 0.699428731 | 0.552462172 | -5.722680343 | 5.866944937 | 5.303523747 | 6.203069917 | 2.141737246 | 2.2680111 | 7.13566325 | 5.275717819 |
| ENSG00000164530 | PI16 | -4.552787018 | 0.009264454 | -1.169924018 | 0.536839311 | -5.722711036 | 6.96351253 | 4.442710129 | 3.369317972 | 2.3892346 | 2.77779907 | 5.362362054 | 3.261853426 |
| ENSG00000100122 | CRYBB1 | -4.714830856 | NA | -1.013335316 | NA | -5.728166172 | 2 | 3.076665209 | 2 | 2 | 2 | 2.3070652 | 2.326754077 |
| ENSG00000205302 | SNX2 | -2.396139511 | 1.40E-12 | -3.339601419 | 9.69E-24 | -5.73574093 | 11.5419687 | 11.02874293 | 10.90452077 | 8.811020771 | 8.7882247 | 7.920487917 | 7.81361733 |
| ENSG00000174327 | SLC16A13 | -3.024040813 | 0.002356861 | -2.721696654 | 0.006299526 | -5.745737467 | 7.750913381 | 6.389896633 | 9.212024683 | 5.406080789 | 5.22805098 | 5.385935071 | 5.777095613 |
| ENSG00000184113 | CLDN5 | -3.676275169 | 0.107251275 | -2.076621361 | 0.332960127 | -5.75289653 | 4.467311984 | 2 | 3.369317972 | 2.270786764 | 2.14022021 | 2.746468522 | 2.326754077 |
| ENSG00000197191 | CYSRT1 | -2.710608345 | 0.246577339 | -3.04372318 | 0.17862881 | -5.754331524 | 7.709880939 | 3.076665209 | 2 | 3.406930997 | 4.22485516 | 3.838019203 | 3.407202941 |
| ENSG00000175322 | ZNF519 | -7.073928332 | 0.007237601 | 1.318824782 | 0.575697563 | -5.75510355 | 4.00722531 | 4.442710129 | 2 | 2 | 2 | 5.445480974 | 3.924507493 |
| ENSG00000104067 | TJP1 | -0.313870404 | 0.609297559 | -5.448495208 | 4.91E-30 | -5.762365613 | 9.957954117 | 11.07694755 | 10.89974571 | 10.40770778 | 10.411061 | 5.248671144 | 5.561082532 |
| ENSG00000172828 | CES3 | -2.52177591 | 0.070700526 | -3.247478581 | 0.014653769 | -5.769254491 | 5.70932728 | 3.076665209 | 6.203069917 | 3.663521627 | 3.2184422 | 3.256555576 | 2.782576693 |
| ENSG00000037965 | HOXC8 | 2.78602211 | 0.161394558 | -8.564311002 | 3.44E-05 | -5.778288892 | 2 | 7.274681896 | 5.156626401 | 8.726347845 | 8.75179438 | 2 | 2.117335275 |
| ENSG00000117399 | CDC20 | -3.019043807 | 0.06933392 | -2.76032911 | 0.093560552 | -5.779372917 | 5.532353208 | 8.191702463 | 3.369317972 | 4.139692846 | 4.1930041 | 3.663836244 | 4.845038551 |
| ENSG00000011143 | MKS1 | -3.181567559 | 0.03191533 | -2.599036831 | 0.082396059 | -5.780604389 | 5.532353208 | 6.228547981 | 9.746650996 | 5.242616075 | 5.36298494 | 5.997902297 | 5.6562603 |
| ENSG00000198286 | CARD11 | -2.009925884 | 0.179023799 | -3.770860079 | 0.006264056 | -5.780785962 | 4.00722531 | 7.874700259 | 6.43197674 | 4.846787657 | 5.06165168 | 4.065262523 | 2.851942824 |
| ENSG00000053501 | USE1 | -3.12698751 | 9.46E-09 | -2.654195332 | 1.02E-06 | -5.781182842 | 8.716893051 | 8.356994601 | 9.565556058 | 5.797438367 | 6.04571775 | 6.25505394 | 6.482320745 |
| ENSG00000166126 | AMN | -3.313644142 | 0.05663139 | -2.470706278 | 0.147985348 | -5.78435042 | 5.70932728 | 5.131603625 | 2 | 3.023133209 | 2.38539787 | 3.374372158 | 2.851942824 |
| KIAA1024 | | 3.075736712 | 0.097826657 | -8.86289843 | 0.000128163 | -5.787161719 | 2 | 5.131603625 | 3.369317972 | 6.841288255 | 6.80989959 | 2 | 2 |
| ENSG00000161958 | FGF11 | -7.506857536 | 0.001043297 | 1.715172966 | 0.346462637 | -5.79168457 | 4.467311984 | 4.710564154 | 2 | 2 | 2 | 5.714849065 | 5.513708555 |
| ENSG00000165443 | PHYHIPL | -1.161820149 | 0.85127879 | -4.629885346 | 0.404031117 | -5.791705495 | 2 | 2 | 4.058998049 | 2.947924294 | 2 | 2 | 2.117335275 |
| ENSG00000269526 | ERVV-1 | -2.679910807 | 0.080019873 | -3.112889967 | 0.031407323 | -5.792800774 | 3.327711234 | 5.131603625 | 4.058998049 | 2.3892346 | 3.01487888 | 2.717032197 | 2.421067459 |
| ENSG00000217555 | CKLF | 1.915844149 | 0.024710822 | -7.710405559 | 6.16E-09 | -5.79456141 | 4.467311984 | 5.457120953 | 5.39244166 | 6.936770382 | 7.0150921 | 2 | 2.117335275 |
| ENSG00000100902 | PSMA6 | -0.707202112 | 0.177877384 | -5.090712338 | 1.82E-30 | -5.79791445 | 10.79750658 | 10.62479571 | 11.50429402 | 10.3445078 | 10.2977019 | 5.766051132 | 6.245088435 |
| ENSG00000132182 | NUP210 | -5.072029644 | 0.069723472 | -0.726312722 | 0.82055968 | -5.798342366 | 2 | 3.076665209 | 7.552162999 | 2.947924294 | 2 | 5.4183033 | 5.349527629 |
| ENSG00000102316 | MAGED2 | -3.371957411 | 2.15E-18 | -2.428852309 | 4.42E-10 | -5.80080972 | 14.55193054 | 13.77512507 | 14.47913858 | 10.92262909 | 10.9555451 | 11.73344351 | 12.01464163 |
| ENSG00000274897 | PANO1 | -2.707054243 | 0.000257308 | -3.094294375 | 1.67E-05 | -5.801348618 | 6.89194353 | 6.140560021 | 7.646393253 | 4.647285135 | 4.40247105 | 3.917810786 | 4.468068808 |
| SDCCAG3 | | 3.760323096 | 0.178862879 | -9.562122974 | 0.001453943 | -5.801799878 | 2 | 6.04685525 | 2 | 8.188900477 | 8.16669309 | 2 | 2 |
| ENSG00000176894 | PXMP2 | -3.113939749 | 0.019148526 | -2.693978359 | 0.04323921 | -5.807918109 | 5.330594134 | 8.929529568 | 6.629488387 | 4.54891529 | 5.00774437 | 4.443291373 | 5.641643428 |
| ENSG00000172992 | DCAKD | -3.629918095 | 3.68E-15 | -2.178006002 | 3.78E-06 | -5.807924097 | 11.32476023 | 10.2574711 | 11.0408834 | 7.326437704 | 7.35800033 | 8.908528807 | 8.619914108 |
| ENSG00000155974 | GRIP1 | -4.756701131 | 3.43E-07 | -1.052603666 | 0.222734185 | -5.809304797 | 5.70932728 | 5.946638857 | 4.874587708 | 2.3892346 | 2.59489879 | 4.496236235 | 4.819291477 |
| ENSG00000186510 | CLCNKA | -3.120185185 | 0.123586937 | -2.690859731 | 0.177601555 | -5.811044916 | 6.89194353 | 3.686303414 | 2 | 2.947924294 | 3.28037825 | 3.12825509 | 3.580696614 |
| ENSG00000141956 | PRDM15 | -8.396704274 | 0.010714621 | 2.583251146 | 0.410543204 | -5.813453127 | 2 | 6.311476754 | 2 | 2 | 2 | 7.393470826 | 7.179807904 |
| ENSG00000005882 | PDK2 | -2.483494776 | 4.82E-11 | -3.330788151 | 2.60E-19 | -5.814282927 | 10.71177538 | 10.019219 | 10.19160738 | 7.893113669 | 7.85436579 | 6.880813179 | 7.194904178 |
| ENSG00000163808 | KIF15 | -2.864249596 | 0.053270494 | -2.951393618 | 0.040825791 | -5.815643213 | 3.327711234 | 4.93638933 | 7.163363831 | 3.754611435 | 3.33976443 | 3.44788957 | 3.560117069 |
| ENSG00000152580 | IGSF10 | -1.323460311 | 0.376950306 | -4.499959406 | 0.000566042 | -5.823419718 | 3.327711234 | 7.189546432 | 7.858115942 | 5.697663076 | 5.81407719 | 3.193831101 | 3.311948089 |
| LACE1 | | 2.408813911 | 0.218303953 | -8.233753636 | 0.000491458 | -5.824939725 | 2 | 4.442710129 | 3.369317972 | 5.674521485 | 5.52526112 | 2 | 2 |
| ENSG00000119943 | PYROXD2 | -3.227629349 | 0.000979119 | -2.599635942 | 0.00888012 | -5.827265292 | 9.771132748 | 7.60566235 | 7.398165132 | 5.686138681 | 5.44640346 | 5.873348082 | 6.392053169 |
| ENSG00000144645 | OSBPL10 | -3.905305431 | 7.66E-05 | -1.922421354 | 0.06883389 | -5.827726785 | 7.750913381 | 9.890264534 | 10.9833682 | 6.328984825 | 6.1068222 | 8.115702075 | 8.175920653 |
| ENSG00000198130 | HIBCH | -2.972399659 | 4.27E-05 | -2.858956838 | 7.66E-05 | -5.831356497 | 8.816894989 | 8.337339476 | 10.06083273 | 6.276156955 | 6.42052719 | 6.112552588 | 6.73730971 |
| ENSG00000128655 | PDE11A | -0.902708467 | 0.623515815 | -4.928843276 | 0.002241819 | -5.831551743 | 3.327711234 | 3.686303414 | 5.772716771 | 3.881293667 | 4.02226117 | 2.384001718 | 2.059860178 |
| ENSG00000174928 | C3orf33 | -2.511600261 | 0.01328895 | -3.321702374 | 0.000647301 | -5.833302634 | 7.975256505 | 5.303523747 | 7.97167868 | 5.22679461 | 5.00774437 | 4.461156421 | 4.400220394 |
| ENSG00000120437 | ACAT2 | -3.685225265 | 2.73E-07 | -2.150815102 | 0.004116939 | -5.836040367 | 10.64259676 | 10.45735223 | 11.35878831 | 7.206526483 | 7.24216308 | 7.589961352 | 9.363302184 |
| ENSG00000124334 | IL9R | 0.315244232 | NA | -6.152900572 | NA | -5.837656339 | 2 | 3.076665209 | 2 | 2.868578001 | 2.49394783 | 2 | 2 |
| ENSG00000149050 | ZNF214 | -2.006714445 | 0.298900033 | -3.831290932 | 0.031405303 | -5.838005377 | 6.816637977 | 4.442710129 | 2 | 3.462030052 | 4.16043394 | 2.911488689 | 2.671842496 |
| ENSG00000159167 | STC1 | -0.667936062 | 0.251211523 | -5.170477927 | 1.12E-26 | -5.838413988 | 14.17657127 | 14.52311524 | 14.25678827 | 13.66436424 | 13.6532952 | 8.449073649 | 9.64284635 |
| ENSG00000138722 | MMRN1 | 0.311262088 | 0.954792088 | -6.152906275 | NA | -5.841644187 | 3.327711234 | 2 | 2 | 2.600425781 | 2.77779907 | 2 | 2 |
| ENSG00000151773 | CCDC122 | -2.631347155 | 0.03724796 | -3.214736424 | 0.008475154 | -5.84608358 | 8.367060731 | 4.442710129 | 7.646393253 | 5.040242897 | 5.04390545 | 4.68289886 | 4.388595494 |
| ENSG00000214706 | IFRD2 | -2.876584283 | 0.000236891 | -2.972734099 | 0.000128272 | -5.849318382 | 11.54488817 | 9.523645035 | 10.0085654 | 7.723090567 | 7.82609171 | 7.580798201 | 7.775072826 |
| ENSG00000198879 | SFMBT2 | -3.305167628 | 0.003841732 | -2.544234824 | 0.024727678 | -5.849402452 | 4.467311984 | 5.457120953 | 6.718950348 | 3.227757282 | 3.2184422 | 4.099885429 | 3.15612297 |
| ENSG00000198108 | CHSY3 | -3.077877634 | 0.015709434 | -2.774409449 | 0.030010873 | -5.852287083 | 6.563806279 | 10.21502048 | 7.734844356 | 6.01571348 | 5.92487824 | 6.672321934 | 5.670730563 |
| ENSG00000183977 | PP2D1 | -5.929102931 | 0.17848626 | 0.069123494 | 0.988547953 | -5.859979436 | 2 | 4.11352228 | 2 | 2 | 2 | 2.885263063 | 3.36036153 |

| | | | | | | | | | | | | | |
|---|---|---|---|---|---|---|---|---|---|---|---|---|---|
| ENSG00000105135 | ILVBL | -2.897677338 | 0.00019891 | -2.962783188 | 0.000128383 | -5.860460526 | 12.35224864 | 10.3729879 | 10.7325871 | 8.494945692 | 8.58694992 | 8.519502611 | 8.433788484 |
| ENSG00000105258 | POLR2I | -1.914721447 | 6.07E-05 | -3.947419325 | 5.73E-18 | -5.862140772 | 8.075571721 | 8.639735838 | 8.904018013 | 6.727838783 | 6.68379438 | 4.563936166 | 5.065479856 |
| ENSG00000186153 | WWOX | -3.380763447 | 5.17E-10 | -2.485862198 | 3.51E-06 | -5.866625645 | 7.975256505 | 7.099070191 | 8.206335842 | 4.670864177 | 4.65713986 | 5.29920634 | 5.586744392 |
| ENSG00000012171 | SEMA3B | -4.323920682 | 0.000207528 | -1.544743911 | 0.230479854 | -5.868664593 | 7.159102577 | 9.367676479 | 11.14139273 | 5.94010643 | 5.58783529 | 8.271892681 | 8.627970054 |
| ENSG00000277848 | | -3.342118036 | 0.377695639 | -2.530003629 | 0.501628498 | -5.872121665 | 2 | 2 | 5.772716771 | 2.868578001 | 2 | 3.10571689 | 2.374681325 |
| ENSG00000213977 | TAX1BP3 | -1.183021211 | 0.128900041 | -4.691199573 | 4.05E-12 | -5.874220784 | 12.18266987 | 10.86336859 | 10.46161637 | 10.15895941 | 10.2128923 | 6.464399218 | 6.951954485 |
| ENSG00000136933 | RABEPK | -3.308791196 | 7.66E-09 | -2.569561703 | 9.03E-06 | -5.878352899 | 10.73241055 | 9.338312924 | 9.735974718 | 6.799114572 | 6.79406287 | 7.490172831 | 7.539150216 |
| ENSG00000175894 | TSPEAR | -4.907623604 | 0.169783339 | -0.973104849 | 0.781385428 | -5.880728453 | 2 | 2 | 4.523675391 | 2 | 2.14022021 | 2.885263063 | 2.782576693 |
| ENSG00000159588 | CCDC17 | -2.560252802 | 0.183917549 | -3.320505948 | 0.070571408 | -5.88075875 | 2 | 4.11352228 | 6.80318718 | 3.462030052 | 3.39680241 | 3.10571689 | 2.885414003 |
| ENSG00000078295 | ADCY2 | -3.005952788 | 0.049756502 | -2.87671747 | 0.050493994 | -5.882670258 | 3.327711234 | 5.72254323 | 4.058998049 | 3.09461492 | 2.38539787 | 2.858551861 | 2.817676617 |
| ENSG00000139574 | NPFF | 1.497320901 | 0.697720524 | -7.382468173 | 0.046085237 | -5.885147272 | 2 | 4.11352228 | 2 | 3.997744386 | 4.1930041 | 2 | 2 |
| ENSG00000113249 | HAVCR1 | -2.523435341 | 0.452856856 | -3.366042298 | NA | -5.889477639 | 3.327711234 | 3.076665209 | 2 | 2.270786764 | 2.14022021 | 2.225793514 | 2 |
| ENSG00000221953 | C1orf229 | -2.96669094 | 0.001787884 | -2.923521076 | 0.001111552 | -5.890212016 | 5.532353208 | 4.442710129 | 5.39244166 | 2.868578001 | 3.15372745 | 3.1504466 | 2.918126211 |
| ENSG00000188739 | RBM34 | 0.035973197 | 0.946943756 | -5.929690793 | 8.03E-46 | -5.893717596 | 9.128497779 | 8.639735838 | 8.380957451 | 8.850273548 | 8.71722214 | 3.355392153 | 3.560117069 |
| KIAA0754 | | 3.759412481 | 5.01E-06 | -9.656991958 | 2.84E-08 | -5.897579477 | 4.467311984 | 4.11352228 | 5.39244166 | 8.36365191 | 8.17087703 | 2 | 2 |
| ENSG00000083838 | ZNF446 | -2.773027074 | 0.005296826 | -3.128008354 | 0.001317742 | -5.901035428 | 9.858207046 | 7.60566235 | 7.451345288 | 6.113598934 | 6.00043082 | 5.794513725 | 5.6562603 |
| ENSG00000129946 | SHC2 | -5.299612986 | 9.97E-07 | -0.615007634 | 0.631219007 | -5.91462062 | 7.750913381 | 8.671479029 | 5.39244166 | 3.515101962 | 2.94009902 | 7.229857522 | 7.164551992 |
| ENSG00000166507 | NDST2 | 0.065957187 | 0.965522332 | -5.98426236 | 1.79E-07 | -5.918305173 | 5.095959458 | 8.356994601 | 8.978369997 | 8.279879181 | 8.19711812 | 3.336159116 | 2.782576693 |
| ENSG00000133134 | BEX2 | -2.985852403 | 0.262549653 | -2.933962993 | 0.258145418 | -5.919815396 | 2 | 6.04685525 | 2 | 2.695456494 | 2.77779907 | 3.059555873 | 2.374681325 |
| ENSG00000083828 | ZNF586 | -2.928077257 | 1.02E-05 | -2.991831371 | 4.97E-06 | -5.919908628 | 8.540903468 | 7.638155693 | 9.344432377 | 5.900763176 | 5.74985018 | 5.829320042 | 5.703940561 |
| ENSG00000184047 | DIABLO | -1.290437361 | 0.026279618 | -4.631518648 | 9.20E-19 | -5.92195601 | 8.836089686 | 10.13189344 | 9.859153765 | 8.435961651 | 8.4215744 | 5.352823931 | 5.101530266 |
| KIAA1524 | | 2.301178906 | 0.246621847 | -8.223203132 | 0.000537577 | -5.922024226 | 3.327711234 | 4.442710129 | 2 | 5.48800675 | 5.49945122 | 2 | 2 |
| ENSG00000187840 | EIF4EBP1 | -3.570848582 | 5.84E-30 | -2.352735747 | 7.47E-14 | -5.923584329 | 11.4295719 | 11.70577865 | 11.97287769 | 8.10700104 | 8.22494733 | 9.512153949 | 9.220615045 |
| ENSG00000204323 | SMIM5 | -3.554071922 | 0.068922496 | -2.372714846 | 0.201906734 | -5.926786768 | 4.00722531 | 2 | 4.874587708 | 2.270786764 | 2.38539787 | 2.831336763 | 2.466008509 |
| ENSG00000079385 | CEACAM1 | -0.501327831 | 0.711000931 | -5.429034288 | 2.17E-06 | -5.930362119 | 6.366709671 | 5.131603625 | 4.058998049 | 4.783303513 | 5.24368216 | 2.42098416 | 2.172608095 |
| ENSG00000099284 | H2AFY2 | -2.778943325 | 2.58E-05 | -3.152442269 | 1.22E-06 | -5.931385594 | 10.77778805 | 9.04308532 | 10.0085654 | 7.370304568 | 7.34718809 | 6.793610851 | 7.173047408 |
| ENSG00000170477 | KRT4 | -3.307512029 | 0.244213659 | -2.631699019 | 0.328032674 | -5.939211048 | 2 | 2 | 4.523675391 | 2.141737246 | 2.2680111 | 2.3070652 | 2.326754077 |
| ENSG00000141404 | GNAL | -2.547272335 | 0.097789727 | -3.392118984 | 0.020661164 | -5.939391319 | 4.00722531 | 7.232742069 | 4.523675391 | 3.95996233 | 3.69849104 | 3.648325314 | 2.782576693 |
| ENSG00000175183 | CSRP2 | -3.688230353 | 2.28E-17 | -2.252176169 | 3.65E-07 | -5.940406522 | 11.74347718 | 11.51775281 | 11.93370489 | 8.091796043 | 8.05374131 | 9.888964398 | 8.952046492 |
| ENSG00000196368 | NUDT11 | -3.093195233 | 2.77E-14 | -2.861498629 | 7.19E-13 | -5.954693862 | 9.317904228 | 8.702538795 | 8.561105505 | 5.839660368 | 5.94414054 | 6.063692919 | 6.155328188 |
| ENSG00000154783 | FGD5 | -5.642938303 | 0.023928454 | -0.314833554 | 0.912648061 | -5.957771857 | 2 | 6.464272742 | 3.369317972 | 2 | 2.38539787 | 4.727866721 | 4.943664032 |
| ENSG00000129354 | AP1M2 | -2.5393452 | 0.565149117 | -3.420879855 | NA | -5.960225055 | 2 | 3.076665209 | 3.369317972 | 2.3892346 | 2 | 2 | 2.225841198 |
| ENSG00000198242 | RPL23A | -1.301075153 | 0.238103591 | -4.663514451 | 1.01E-06 | -5.964589604 | 13.27201035 | 10.95971654 | 10.66102425 | 10.86725658 | 10.8053139 | 7.343779505 | 7.646759583 |
| ENSG00000138615 | CILP | -3.542395856 | 0.116580901 | -2.423633665 | 0.270671739 | -5.966029521 | 3.327711234 | 2 | 5.39244166 | 2.270786764 | 2.49394783 | 2.3070652 | 3.071407821 |
| ENSG00000089685 | BIRC5 | -3.267426643 | 0.09831254 | -2.701304581 | 0.171340702 | -5.968731223 | 8.777722034 | 8.276714465 | 2 | 4.927308402 | 4.85329164 | 3.917810786 | 6.106490089 |
| ENSG00000137841 | PLCB2 | -5.819285924 | 0.003349323 | -0.151771079 | 0.945901868 | -5.971057003 | 4.467311984 | 6.464272742 | 2 | 2.141737246 | 2.2680111 | 5.684753614 | 4.254128754 |
| ENSG00000183145 | RIPPLY3 | -4.129595859 | 0.056157333 | -1.843894367 | 0.336472054 | -5.973490226 | 3.327711234 | 3.686303414 | 3.369317972 | 2 | 2.2680111 | 2.492221014 | 2.671842496 |
| ENSG00000206557 | TRIM71 | -2.272300684 | NA | -3.706147444 | NA | -5.978448128 | 2 | 2 | 3.369317972 | 2.141737246 | 2.14022021 | 2.048074405 | 2.059860178 |
| ENSG00000156299 | TIAM1 | -4.598898952 | 0.001047492 | -1.379959447 | 0.377007831 | -5.978858399 | 4.815609471 | 9.687856435 | 7.552162999 | 4.443344104 | 3.86943969 | 7.421115854 | 6.654794245 |
| ENSG00000229253 | | 2.809731662 | 0.125649219 | -8.791761509 | 0.000102415 | -5.982029847 | 2 | 6.464272742 | 4.058998049 | 7.862723323 | 7.89204953 | 2.048074405 | 2 |
| ENSG00000164045 | CDC25A | -2.918103606 | 0.132130317 | -3.066932137 | 0.105736164 | -5.985035743 | 2 | 7.60566235 | 9.358421373 | 5.348759107 | 5.37722731 | 5.174809258 | 5.282014932 |
| ENSG00000213930 | GALT | -2.805115736 | 2.09E-08 | -3.180194488 | 1.15E-10 | -5.985310224 | 9.810482605 | 9.148351514 | 10.28977157 | 7.106324235 | 6.99676009 | 6.492902701 | 6.864442837 |
| ENSG00000132692 | BCAN | -2.930383489 | 0.329877736 | -3.058722372 | 0.29402979 | -5.989105861 | 6.138360429 | 2 | 2 | 3.227757282 | 2.14022021 | 2.560105256 | 2.885414003 |
| ENSG00000183979 | NPB | 1.205208052 | 0.779312814 | -7.2009967 | 0.072675764 | -5.995788648 | 2 | 2 | 4.058998049 | 3.997744386 | 3.45167086 | 2 | 2 |
| ENSG00000091106 | NLRC4 | -6.052267331 | 0.061260107 | 0.054409274 | 0.987221496 | -5.997858057 | 2 | 5.457120953 | 2 | 2.141737246 | 2 | 4.321442857 | 4.033026654 |
| ENSG00000107736 | CDH23 | -5.068745434 | 0.019562832 | -0.929551962 | 0.704904787 | -5.998297396 | 7.33131396 | 2 | 5.772716771 | 2.695456494 | 2.49394783 | 6.18690462 | 2.918126211 |
| ENSG00000188152 | NUTM2G | -2.589737387 | 0.078050197 | -3.411508216 | 0.014587726 | -6.001245602 | 6.366709671 | 4.11352228 | 4.058998049 | 2.868578001 | 3.60476711 | 3.10571689 | 2.551896972 |
| ENSG00000155465 | SLC7A7 | -7.48915279 | 0.036927252 | 1.487607952 | 0.681460704 | -6.001544837 | 2 | 5.457120953 | 2 | 2 | 2 | 5.85994354 | 4.702084619 |
| ENSG00000275003 | | 3.668199943 | 7.10E-05 | -9.670202421 | 5.99E-08 | -6.002002478 | 4.815609471 | 5.303523747 | 3.369317972 | 8.087422161 | 8.29225867 | 2 | 2 |
| ENSG00000101447 | FAM83D | -3.858745153 | 0.040290943 | -2.147899311 | 0.272931073 | -6.006644465 | 8.716893051 | 6.311476754 | 2 | 4.070463721 | 3.7864959 | 5.594558848 | 5.072762365 |
| ENSG00000224320 | | 6.352557529 | 4.47E-16 | -12.35985957 | 1.17E-13 | -6.007302044 | 9.435834703 | 8.487557258 | 7.342949392 | 14.97000283 | 15.0270464 | 2 | 2.059860178 |
| ENSG00000221946 | FXYD7 | -3.541739941 | 0.175994512 | -2.466424933 | 0.318708894 | -6.008164874 | 2 | 3.686303414 | 4.058998049 | 2.3892346 | 2 | 2.34604626 | 2.466008509 |
| FAM212B | | 1.77877644 | 0.616682216 | -7.788822028 | 0.025732112 | -6.010045588 | 2 | 4.442710129 | 2 | 4.739373666 | 4.56059747 | 2 | 2 |
| ENSG00000163827 | LRRC2 | -3.955281323 | 1.06E-23 | -2.057402515 | 1.31E-07 | -6.012683838 | 9.397585735 | 9.583268459 | 8.805342542 | 5.514313187 | 5.43283104 | 7.229857522 | 7.316452062 |
| ENSG00000129295 | LRRC6 | -2.515558894 | 0.099584363 | -3.500911581 | 0.015939794 | -6.016470475 | 7.667647223 | 3.076665209 | 6.80318718 | 4.358765249 | 4.56059747 | 3.981090143 | 3.383972333 |
| ENSG00000111305 | GSG1 | -5.305962037 | 0.000519886 | -0.712295942 | 0.662699449 | -6.018257979 | 5.330594134 | 5.303523747 | 5.772716771 | 2.498691189 | 2.14022021 | 3.082820997 | 5.621921006 |
| ENSG00000185100 | ADSSL1 | -3.888036874 | 0.031289994 | -2.132566732 | 0.257569188 | -6.020603606 | 8.19931886 | 6.602424674 | 2 | 3.663521627 | 3.65239003 | 5.532544554 | 4.316807515 |
| ENSG00000100162 | CENPM | -3.607001569 | 0.055550539 | -2.415477461 | 0.19591446 | -6.02247903 | 2 | 6.04685525 | 4.523675391 | 2.695456494 | 2.59489879 | 2.592885947 | 3.620995423 |
| ENSG00000152092 | ASTN1 | -3.257830586 | 0.415495349 | -2.772249815 | NA | -6.030080401 | 4.00722531 | 2 | 2 | 2.270786764 | 2 | 2.267001569 | 2.117335275 |
| ENSG00000124920 | MYRF | -5.006120668 | 0.001020099 | -1.03109984 | 0.557541861 | -6.037220509 | 3.327711234 | 9.488434476 | 8.536706852 | 4.034562184 | 3.82856379 | 7.530920174 | 7.478755384 |
| ENSG00000167157 | PRRX2 | -1.545585785 | 0.008829144 | -4.494458167 | 9.48E-17 | -6.040043953 | 9.922475255 | 10.60848937 | 10.51254821 | 8.81895723 | 8.86350401 | 6.430795201 | 5.300743019 |
| ENSG00000124787 | RPP40 | -2.958596051 | 0.000111259 | -3.087876448 | 3.73E-05 | -6.046472499 | 8.228665467 | 6.666836837 | 6.718950348 | 4.574143677 | 4.72553021 | 4.425202323 | 4.654341953 |
| ENSG00000169594 | BNC1 | -0.502532108 | 0.826909189 | -5.544105587 | 0.00236338 | -6.046637696 | 8.75772933 | 5.838937808 | 2 | 6.841288255 | 6.90635305 | 2.937246078 | 2.851942824 |
| ENSG00000179361 | ARID3B | -2.567134284 | 4.35E-06 | -3.4836795 | 1.48E-10 | -6.050813784 | 7.904282243 | 9.246456519 | 8.763901704 | 6.268449691 | 6.21386141 | 5.48093935 | 5.294527297 |
| ENSG00000185522 | LMNTD2 | 1.736570041 | 0.627146058 | -7.788319215 | 0.026370027 | -6.051749174 | 2 | 4.442710129 | 2 | 4.694064037 | 4.53541753 | 2 | 2 |
| ENSG00000105366 | SIGLEC8 | -7.292000576 | 0.009532048 | 1.225977991 | 0.642633186 | -6.066022584 | 2 | 3.076665209 | 5.156626401 | 2 | 2 | 5.4183033 | 4.365060851 |
| ENSG00000170293 | CMTM8 | -4.702817756 | 0.008961131 | -1.365143398 | 0.471189091 | -6.067961154 | 6.563806279 | 4.710564154 | 2 | 2.600425781 | 2.2680111 | 4.443291373 | 3.940519005 |
| ENSG00000239779 | WBP1 | -2.29091872 | 2.50E-07 | -3.780245939 | 1.81E-18 | -6.071164659 | 10.41260941 | 10.09725346 | 9.528916236 | 7.844695415 | 7.72948195 | 6.554528258 | 6.099376274 |
| ENSG00000143845 | ETNK2 | -5.090913068 | 1.88E-08 | -0.982155136 | 0.345256613 | -6.073068204 | 9.886101849 | 9.159587375 | 7.029880454 | 4.358765249 | 4.2865223 | 8.10153367 | 8.159768736 |
| ENSG00000132639 | SNAP25 | -2.332867806 | 0.374118489 | -3.740650567 | 0.127716243 | -6.073518374 | 2 | 5.946638857 | 2 | 3.162721474 | 2.77779907 | 2.34604626 | 2.551896972 |
| ENSG00000175567 | UCP2 | -6.607932831 | 1.01E-12 | 0.534129128 | 0.621093779 | -6.073803703 | 9.2903348 | 8.820493869 | 7.098165388 | 3.349643765 | 2.59489879 | 8.843612414 | 9.474014382 |
| ENSG00000165025 | SYK | -6.194459543 | 0.00015131 | 0.116304882 | 0.94131204 | -6.078154661 | 4.467311984 | 5.457120953 | 5.156626401 | 2.270786764 | 2 | 5.769639817 | 4.228274364 |
| ENSG00000166473 | PKD1L2 | -3.340116522 | 0.303359251 | -2.741232677 | 0.388123757 | -6.081349199 | 2 | 2 | 5.59507254 | 2.600425781 | 2.2680111 | 2.048074405 | 3.042026731 |
| ENSG00000179750 | APOBEC3B | -1.441623918 | 0.442550676 | -4.644766334 | 0.004851999 | -6.086390252 | 7.096806373 | 6.228547981 | 2 | 4.927308402 | 4.79082474 | 2.384001718 | 3.042026731 |
| ENSG00000004848 | ARX | 0.057512093 | NA | -6.152891209 | NA | -6.095379116 | 2 | 3.076665209 | 2 | 3.023133209 | 2 | 2 | 2 |
| ENSG00000156150 | ALX3 | -5.208991304 | 0.015381668 | -0.88962232 | 0.718049597 | -6.098613624 | 2 | 3.686303414 | 8.076950877 | 2.868578001 | 2.49394783 | 5.227950776 | 6.110033886 |
| ENSG00000135525 | MAP7 | -2.174014894 | 0.325432015 | -3.926481863 | 0.055557322 | -6.100496757 | 2 | 4.710564154 | 6.958202216 | 3.921164147 | 3.7864959 | 3.27687201 | 2.172608095 |
| ENSG00000255690 | TRIL | -1.206558954 | 0.744917483 | -4.895462432 | 0.14640672 | -6.102021386 | 2 | 4.442710129 | 2 | 2.784612485 | 2.68924543 | 2.139668783 | 2 |
| ENSG00000164967 | RPP25L | -3.145081331 | 1.74E-07 | -2.958225501 | 7.63E-07 | -6.103306832 | 9.895281535 | 8.557180776 | 8.742726175 | 5.987823101 | 6.24526024 | 6.329884754 | 6.267475858 |
| ENSG00000167578 | RAB4B | -0.357529845 | 0.651333637 | -5.752443527 | 1.13E-19 | -6.109973372 | 8.609439662 | 7.35507212 | 8.763901704 | 7.895617537 | 8.10812366 | 3.10571689 | 3.518055793 |
| ENSG00000255823 | MTRNR2L8 | 2.137933062 | 0.000124982 | -8.255775274 | 7.50E-54 | -6.117842212 | 12.04716355 | 11.40425734 | 11.07517045 | 13.69231094 | 13.7137319 | 3.393105702 | 4.119958326 |
| ENSG00000213937 | CLDN9 | -4.714835794 | NA | -1.408760762 | NA | -6.123596556 | 3.327711234 | 2 | 2 | 2 | 2 | 2.457042231 | 2 |
| ENSG00000099282 | TSPAN15 | -4.500605437 | 9.17E-11 | -1.624493525 | 0.012879978 | -6.125098962 | 6.96351253 | 6.311476754 | 6.203069917 | 3.09461492 | 2.86123014 | 5.243518884 | 4.77533149 |
| ENSG00000162174 | ASRGL1 | -4.203388606 | 0.001834437 | -1.926657598 | 0.162098142 | -6.130046204 | 3.327711234 | 5.457120953 | 6.882775709 | 2.947924294 | 2.59489879 | 4.261535101 | 4.062589736 |
| ENSG00000106004 | HOXA5 | 3.152518844 | 0.109012781 | -9.282679047 | 9.67E-05 | -6.130160203 | 3.327711234 | 5.595927442 | 2 | 7.318995108 | 7.29563107 | 2 | 2 |
| ENSG00000230510 | PPP5D1 | -2.859398144 | 0.296800324 | -3.271303146 | 0.21491217 | -6.13070129 | 7.159102577 | 2 | 2 | 3.227757282 | 3.50452879 | 3.296906311 | 2.950113106 |
| ENSG00000141750 | STAC2 | -4.234243163 | 0.000402124 | -1.896874215 | 0.118732434 | -6.131117378 | 4.467311984 | 5.131603625 | 7.029880454 | 3.023133209 | 2.59489879 | 4.240999983 | 4.341135913 |
| ENSG00000275713 | HIST1H2BH | -2.35973833 | 0.102509199 | -3.771424259 | 0.005334947 | -6.13116259 | 5.095959458 | 3.686303414 | 5.156626401 | 2.947924294 | 3.2184422 | 2.183373728 | 2.782576693 |
| ENSG00000036530 | CYP46A1 | -5.702854155 | 0.085411097 | -0.429617921 | 0.899648072 | -6.132472075 | 2 | 5.131603625 | 2 | 2 | 2.14022021 | 3.5515855 | 3.600986722 |
| ENSG00000277273 | | 1.447353505 | 0.38484122 | -7.589402908 | 3.59E-07 | -6.142049402 | 8.042902253 | 6.728495673 | 3.369317972 | 8.392303091 | 8.41468635 | 2.34604626 | 2.059860178 |
| ENSG00000101057 | MYBL2 | -3.441584444 | 0.08215819 | -2.703068684 | 0.17344883 | -6.144653128 | 7.53297911 | 8.077869079 | 2 | 4.17310023 | 4.12711147 | 3.12825509 | 5.497565112 |
| ENSG00000275758 | | 3.054860948 | 0.065314973 | -9.202745486 | 2.44E-05 | -6.147884538 | 4.00722531 | 5.303523747 | 2 | 7.106324235 | 7.16216989 | 2 | 2 |
| ENSG00000007944 | MYLIP | -4.549262982 | 3.12E-21 | -1.598794858 | 0.000483841 | -6.148057841 | 7.53297911 | 8.255926468 | 7.896971941 | 3.881293667 | 3.7864959 | 6.21875862 | 6.528072775 |
| ENSG00000143224 | PPOX | -3.298352811 | 1.41E-05 | -2.855760288 | 0.000180075 | -6.154113099 | 10.32020829 | 8.639735838 | 8.631919287 | 6.197166296 | 6.21386141 | 6.370627179 | 6.845434848 |
| ENSG00000274538 | | 4.487177927 | 0.022529066 | -10.64180081 | 1.04E-05 | -6.15462288 | 6.96351253 | 4.11352228 | 2 | 9.960398193 | 9.97960581 | 2 | 2 |
| ENSG00000127533 | F2RL3 | -1.548941724 | 0.421603569 | -4.610711786 | 0.008733743 | -6.159653511 | 4.00722531 | 2 | 5.39244166 | 3.227757282 | 3.28037825 | 2.267001569 | 2.172608095 |
| ENSG00000237651 | C2orf74 | -1.106922622 | 0.293404788 | -5.053462323 | 3.54E-08 | -6.160384945 | 7.096806373 | 4.93638933 | 7.398165132 | 5.674521485 | 5.81407719 | 2.885263063 | 2.817676617 |
| ENSG00000204947 | ZNF425 | -3.340504203 | 0.041942911 | -2.822116768 | 0.087160749 | -6.162620971 | 10.16187685 | 7.274681896 | 4.523675391 | 5.662809983 | 5.44640346 | 6.522981762 | 5.306932076 |
| ENSG00000162493 | PDPN | -3.039980349 | 8.67E-05 | -3.12858548 | 4.66E-05 | -6.168565829 | 11.82739822 | 10.26789142 | 9.935792933 | 7.789219291 | 8.02009991 | 7.529862489 | 8.065268319 |
| ENSG00000168090 | COPS6 | -3.111546986 | 2.50E-21 | -3.059026213 | 6.21E-21 | -6.170573199 | 12.55554908 | 12.10787117 | 12.69739443 | 9.385472425 | 9.35495351 | 9.260615815 | 9.568163239 |
| ENSG00000172349 | IL16 | -4.008885746 | 0.062028799 | -2.162400456 | 0.328297809 | -6.171286202 | 3.327711234 | 7.274681896 | 2 | 2.868578001 | 2.86123014 | 4.369533167 | 3.45257121 |
| ENSG00000214960 | ISPD | -3.321706049 | 0.108852636 | -2.850155126 | 0.16687418 | -6.171861175 | 2 | 5.72254323 | 8.99637432 | 4.598938466 | 4.37434574 | 4.360042493 | 5.275717819 |
| ENSG00000103512 | NOMO1 | 5.214018352 | 2.24E-09 | -11.38734235 | 3.58E-11 | -6.173324 | 6.138360429 | 8.540087354 | 7.398165132 | 12.84248342 | 12.8881928 | 2.048074405 | 2 |
| ENSG00000236802 | | 2.58790661 | 0.39960716 | -8.768137527 | 0.005507299 | -6.180230917 | 2 | 5.303523747 | 2 | 6.237202172 | 6.29861567 | 2 | 2 |
| ENSG00000140488 | CELF6 | 1.605126409 | 0.665554125 | -7.785613641 | 0.030137313 | -6.180487232 | 2 | 4.442710129 | 2 | 4.825935136 | 4.0930011 | 2 | 2 |
| ENSG00000122644 | ARL4A | -3.094286203 | 0.022602116 | -3.094014217 | 0.021646679 | -6.18830042 | 9.800745303 | 7.232742069 | 5.59507254 | 5.406080789 | 5.63601114 | 5.673303969 | 5.361470313 |
| ENSG00000115255 | REEP6 | -4.264089253 | 1.37E-17 | -1.925906465 | 7.02E-05 | -6.189995718 | 8.418903792 | 7.669933291 | 7.818184403 | 3.997744386 | 4.1930041 | 5.925754458 | 6.323497254 |

| | | | | | | | | | | | | | |
|---|---|---|---|---|---|---|---|---|---|---|---|---|---|
| RPL5 | | -3.426502808 | 3.05E-10 | -2.767241072 | 4.92E-07 | -6.193743881 | 14.65348042 | 13.77101014 | 13.26731409 | 10.58338644 | 10.5971045 | 11.25778079 | 11.23845157 |
| ENSG00000276880 | | 3.722894776 | 0.197881096 | -9.918217824 | 0.001201322 | -6.195323048 | 2 | 6.389896633 | 2 | 8.480007165 | 8.50501001 | 2 | 2 |
| ENSG00000154330 | PGM5 | -6.306243667 | 8.41E-22 | 0.108657833 | 0.886350236 | -6.197585834 | 8.285626337 | 7.431218269 | 8.677268317 | 2.947924294 | 2.94009902 | 8.11429148 | 8.498084662 |
| ENSG00000131652 | THOC6 | -2.932491451 | 0.002493632 | -3.267232643 | 0.00059676 | -6.199724093 | 10.08373159 | 8.590771778 | 7.285536013 | 6.415275989 | 5.99120031 | 6.013120342 | 5.785984581 |
| ENSG00000117419 | ERI3 | -3.462016095 | 5.94E-16 | -2.73894972 | 1.81E-10 | -6.200965815 | 11.84881904 | 11.10917796 | 12.08228623 | 8.30646685 | 8.27334517 | 8.898729828 | 9.105499915 |
| ENSG00000107719 | PALD1 | -7.213411651 | 3.71E-07 | 1.011747829 | 0.331819555 | -6.201663822 | 4.467311984 | 6.602424674 | 6.203069917 | 2.141737246 | 2.14022021 | 6.672321934 | 7.245725608 |
| ENSG00000170899 | GSTA4 | -3.905737588 | 2.36E-06 | -2.299161018 | 0.007985652 | -6.204898606 | 11.1517158 | 9.45234302 | 8.99637432 | 6.483440992 | 6.18985553 | 7.707816755 | 8.07345976 |
| ENSG00000128849 | CGNL1 | -4.788827804 | 0.001963782 | -1.420598506 | 0.412081701 | -6.209426309 | 3.327711234 | 9.125613817 | 9.385989894 | 4.268917409 | 4.1930041 | 6.777659758 | 7.669754031 |
| ENSG00000113212 | PCDHB7 | -3.567519177 | 0.11963969 | -2.642761968 | 0.25426536 | -6.210281145 | 2 | 3.686303414 | 8.608698767 | 3.881293667 | 3.94787299 | 4.053534413 | 5.065479856 |
| ENSG00000142675 | CNKSR1 | -2.628268342 | 0.167222535 | -3.58421651 | 0.046360366 | -6.212484852 | 4.00722531 | 6.04685525 | 2 | 2.784612485 | 3.2184422 | 2.42098416 | 2.782576693 |
| ENSG00000250298 | GIMD1 | -5.356545357 | 0.301633714 | -0.870082842 | NA | -6.226628199 | 2 | 3.686303414 | 2 | 2 | 2 | 2.225793514 | 2.746601469 |
| ENSG00000156990 | RPUSD3 | -3.77738056 | 9.44E-12 | -2.452256692 | 1.38E-05 | -6.229637251 | 9.48530526 | 9.770792087 | 10.79563557 | 6.394182228 | 6.44101097 | 7.718068425 | 7.685278529 |
| ENSG00000189375 | TBC1D28 | -3.266347321 | NA | -2.973323727 | NA | -6.239671048 | 2 | 2 | 3.369317972 | 2.141737246 | 2 | 2 | 2.172608095 |
| ENSG00000116299 | KIAA1324 | -2.761101002 | 0.000140922 | -3.481058872 | 4.83E-07 | -6.242159875 | 6.468621203 | 5.457120953 | 5.930865648 | 3.615725078 | 3.74316441 | 3.082820997 | 3.36036153 |
| ENSG00000160325 | CACFD1 | -4.000452667 | 2.75E-22 | -2.242980036 | 5.74E-08 | -6.243432703 | 10.52332707 | 9.640525262 | 10.12770132 | 6.221320894 | 6.20590375 | 8.027027634 | 7.79392309 |
| ENSG00000058673 | ZC3H11A | 1.632961161 | 0.161589056 | -7.878798843 | 1.70E-14 | -6.245837682 | 7.096806373 | 10.3729879 | 9.06622548 | 11.01361374 | 11.0053956 | 2.526562374 | 2.981406154 |
| ENSG00000124587 | PEX6 | -9.582991334 | 7.09E-05 | 3.335438201 | 0.089784827 | -6.247553133 | 5.095959458 | 7.189546432 | 2 | 2 | 2 | 9.264754164 | 9.135368584 |
| ENSG00000038427 | VCAN | -2.276547037 | 0.005458837 | -3.980417822 | 2.59E-07 | -6.256964859 | 14.09725211 | 15.20082333 | 14.71112305 | 12.46182124 | 12.4639002 | 11.44980174 | 9.396691711 |
| ENSG00000109099 | PMP22 | -4.158591292 | 8.52E-08 | -2.101219001 | 0.010976293 | -6.259810293 | 13.04545123 | 11.83928723 | 10.75919294 | 8.067573917 | 8.00183183 | 9.933236571 | 10.20524753 |
| ENSG00000121410 | A1BG | -1.981672481 | 0.141050239 | -4.279421188 | 0.000488691 | -6.261093669 | 7.82963488 | 6.311476754 | 4.058998049 | 4.887609717 | 4.95174433 | 3.429858444 | 2.981406154 |
| ENSG00000206075 | SERPINB5 | -0.10866337 | NA | -6.1528919 | NA | -6.26155527 | 2 | 3.076665209 | 2 | 2.141737246 | 2.86123014 | 2 | 2 |
| C10ORF54 | C10ORF54 | 3.271702513 | 0.061207735 | -9.54965 | 2.00E-05 | -6.277947487 | 4.00722531 | 5.72254323 | 2 | 7.641902724 | 7.72670378 | 2 | 2 |
| RSG1 | | 1.736097025 | 0.409961505 | -8.015221648 | 0.000920864 | -6.279124623 | 2 | 3.686303414 | 4.058998049 | 4.867343068 | 4.74762503 | 2 | 2 |
| ENSG00000189366 | ALG1L | -0.054961239 | 0.983101649 | -6.224415426 | 0.006603439 | -6.279376666 | 4.00722531 | 2 | 4.874587708 | 3.881293667 | 4.05806467 | 2 | 2.117335275 |
| ENSG00000116761 | CTH | -2.614442803 | 0.105550682 | -3.667273491 | 0.017079544 | -6.281716294 | 3.327711234 | 7.315436877 | 8.99637432 | 5.194621444 | 5.45984938 | 4.620674722 | 4.201948172 |
| ENSG00000140368 | PSTPIP1 | -2.916410313 | 0.113077307 | -3.369053257 | 0.054407162 | -6.28546357 | 5.095959458 | 2 | 5.156626401 | 2.947924294 | 2.49394783 | 2.746468522 | 2.374681325 |
| ENSG00000147687 | TATDN1 | -2.36559657 | 0.016877493 | -3.920997154 | 2.62E-05 | -6.286593724 | 9.159305284 | 6.535001748 | 7.600047325 | 5.829220145 | 5.91514978 | 4.675266036 | 4.292061816 |
| WASH1 | | 3.778839716 | 4.28E-07 | -10.06616877 | 3.39E-09 | -6.287329051 | 5.70932728 | 4.442710129 | 4.874587708 | 8.709369553 | 8.67892944 | 2 | 2 |
| ENSG00000105639 | JAK3 | -5.185796118 | 2.42E-29 | -1.105656333 | 0.016663848 | -6.29145245 | 8.631580145 | 8.433041974 | 9.165043284 | 3.921164147 | 4.05806467 | 7.845956781 | 7.496463996 |
| ENSG00000143153 | ATP1B1 | -4.173132512 | 2.33E-16 | -2.123586716 | 4.86E-05 | -6.296719229 | 9.751047882 | 10.5327403 | 11.00131064 | 6.548530093 | 6.25300433 | 8.436147258 | 8.373881215 |
| ENSG00000064042 | LIMCH1 | -3.821456848 | 3.80E-06 | -2.475559969 | 0.003767414 | -6.297016817 | 8.87372952 | 9.159587375 | 10.22252928 | 5.997179931 | 5.6121243 | 5.957567336 | 7.729722875 |
| ENSG00000158169 | FANCC | -3.584807492 | 6.85E-08 | -2.717058924 | 5.01E-05 | -6.301866415 | 8.855032352 | 8.169643011 | 9.868959282 | 5.578054239 | 5.72778967 | 6.587434478 | 6.341697848 |
| ENSG00000095739 | BAMBI | -3.113241934 | 7.09E-25 | -3.194772901 | 1.36E-26 | -6.308014835 | 11.97540042 | 11.9979301 | 12.22845487 | 8.939126212 | 8.99746255 | 8.612361808 | 9.119193255 |
| ENSG00000166317 | SYNPO2L | -2.812696988 | 0.006359606 | -3.497211817 | 0.000344212 | -6.309908806 | 5.532353208 | 4.11352228 | 5.772716771 | 3.349643765 | 2.94009902 | 2.858551861 | 2.782576693 |
| ENSG00000073584 | SMARCE1 | -1.326736072 | 0.02238434 | -4.984090895 | 1.65E-21 | -6.310826967 | 11.28689767 | 10.06773758 | 10.34701993 | 9.356683903 | 9.3304589 | 5.622611168 | 5.933281326 |
| ENSG00000116691 | MIIP | -3.490545862 | 2.64E-05 | -2.820605041 | 0.000805329 | -6.311150903 | 10.76282024 | 9.74864196 | 8.35328246 | 6.358325776 | 6.6253296 | 7.367655602 | 6.879055607 |
| ENSG00000175334 | BANF1 | -3.175588299 | 3.35E-11 | -3.136048036 | 5.11E-11 | -6.311636334 | 11.86992646 | 11.67869957 | 12.70835896 | 9.020530266 | 8.96364685 | 8.841056341 | 9.199978848 |
| ENSG00000085563 | ABCB1 | -3.989131511 | 0.137045092 | -2.325489115 | 0.391460007 | -6.314620626 | 2 | 2 | 6.534109959 | 2.3892346 | 2.68924543 | 3.011868911 | 3.539239711 |
| ENSG00000131203 | IDO1 | 4.335844248 | 5.41E-12 | -10.65097162 | 1.14E-10 | -6.315127375 | 5.330594134 | 5.131603625 | 6.203069917 | 9.828366803 | 9.82843204 | 2 | 2 |
| ENSG00000104899 | AMH | -2.993312612 | 0.162903923 | -3.327957938 | 0.1084754 | -6.32127055 | 2 | 3.686303414 | 7.552162999 | 3.754611435 | 3.39680241 | 3.517841605 | 3.209956562 |
| ENSG00000196839 | ADA | -4.0295066 | 3.38E-08 | -2.300507259 | 0.002441344 | -6.330013859 | 10.56418906 | 9.308339287 | 8.742726175 | 5.674521485 | 5.94414054 | 7.754744914 | 7.117791742 |
| ENSG00000229676 | ZNF492 | -0.43242731 | 0.753072123 | -5.899819064 | 1.29E-06 | -6.332246374 | 3.327711234 | 5.303523747 | 5.772716771 | 4.887609717 | 4.58534546 | 2.225793514 | 2.117335275 |
| ENSG00000133027 | PEMT | -3.052477033 | 5.28E-06 | -3.281999785 | 7.28E-07 | -6.334476819 | 10.65345793 | 8.88960062 | 9.945092785 | 6.922108067 | 7.04662306 | 6.516588833 | 6.975457902 |
| ENSG00000159189 | C1QC | -8.819166189 | 0.009987576 | 2.480872075 | 0.457914331 | -6.338294114 | 6.73718411 | 2 | 2 | 2 | 2 | 7.961073574 | 7.126566171 |
| ENSG00000139835 | GRTP1 | -4.238769752 | 0.000109757 | -2.101920099 | 0.065664664 | -6.34068985 | 5.866944937 | 6.04685525 | 8.325066185 | 3.79808662 | 3.2184422 | 4.990270811 | 5.431121909 |
| ENSG00000157613 | CREB3L1 | -2.636167296 | 3.61E-30 | -3.711752071 | 3.25E-60 | -6.347919367 | 13.53171353 | 13.28510734 | 13.37100627 | 10.75728846 | 10.7747077 | 9.824651369 | 9.550340379 |
| ENSG00000197013 | ZNF429 | -3.165809985 | 3.68E-07 | -3.184693586 | 2.21E-07 | -6.350503571 | 9.2903348 | 8.053983481 | 8.561105505 | 5.651002631 | 5.67111583 | 5.997902297 | 5.171036168 |
| ENSG00000241644 | INMT | -6.562504686 | 4.16E-12 | 0.212000844 | 0.806099073 | -6.350503843 | 6.468621203 | 5.946638857 | 7.285536013 | 2.270786764 | 2.38539787 | 6.880813179 | 6.843307307 |
| ENSG00000171130 | ATP6V0E2 | -9.019076261 | 7.51E-05 | 2.666571585 | 0.135257068 | -6.352504676 | 5.330594134 | 6.389896633 | 2 | 2 | 2 | 7.934942996 | 8.027351693 |
| ENSG00000170915 | PAQR8 | -4.465783421 | 2.21E-10 | -1.886952923 | 0.011313199 | -6.352736344 | 8.631580145 | 8.414402626 | 10.05225195 | 5.076005082 | 4.811949 | 7.527744791 | 7.179807904 |
| ENSG00000123405 | NFE2 | -3.149433291 | 0.292538621 | -3.211457726 | 0.265028723 | -6.360891017 | 2 | 2 | 5.59507254 | 2.3892346 | 2.59489879 | 2.094598369 | 2.782576693 |
| ENSG00000162738 | VANGL2 | -4.246640559 | 0.001661187 | -2.118709255 | 0.133193465 | -6.365349814 | 4.467311984 | 6.844429755 | 7.97167868 | 3.921164147 | 2.59489879 | 4.854926465 | 5.197927245 |
| ENSG00000120324 | PCDHB10 | -2.812253919 | 0.004232874 | -3.55507942 | 0.000177803 | -6.367333339 | 7.750913381 | 5.131603625 | 7.225742779 | 4.623314311 | 4.31639476 | 3.824280769 | 3.972018587 |
| ENSG00000269858 | EGLN2 | -1.529765116 | 0.038412766 | -4.837610565 | 2.84E-13 | -6.367375681 | 9.678463567 | 10.73029099 | 11.2764554 | 9.170732795 | 9.18725282 | 5.404520029 | 6.356691554 |
| ENSG00000183072 | NKX2-5 | -2.928039123 | NA | -3.43957706 | NA | -6.367616183 | 2 | 3.686303414 | 2 | 2.270786764 | 2 | 2.139668783 | 2.059860178 |
| ENSG00000134817 | APLNR | -5.861641128 | 0.009292487 | -0.512247179 | 0.783438027 | -6.373888307 | 3.327711234 | 4.710564154 | 3.369317972 | 2 | 2.14022021 | 3.256555576 | 3.924507493 |
| ENSG00000165828 | PRAP1 | -3.903256076 | 0.236880364 | -2.47418632 | 0.432546542 | -6.377442396 | 2 | 2 | 4.523675391 | 2.141737246 | 2.14022021 | 2.592885947 | 2.059860178 |
| ENSG00000128536 | CDHR3 | -4.533747038 | 0.023289156 | -1.848209808 | 0.357295649 | -6.381956845 | 3.327711234 | 3.686303414 | 5.930865648 | 2 | 2.59489879 | 3.465698119 | 3.33635788 |
| ENSG00000121775 | TMEM39B | -3.10718328 | 3.85E-07 | -3.276451299 | 6.38E-08 | -6.383634579 | 10.77281601 | 9.192779271 | 10.04361983 | 6.979883891 | 7.15805139 | 6.748859346 | 7.051147246 |
| ENSG00000136490 | LIMD2 | -2.999263815 | 6.45E-06 | -3.384930428 | 2.38E-07 | -6.384194243 | 10.41898845 | 9.203675681 | 10.98787477 | 7.307758725 | 7.49817903 | 7.184873526 | 6.858134626 |
| ENSG00000180539 | C9orf139 | -4.723388046 | 0.019086098 | -1.662938928 | 0.39030005 | -6.386326974 | 4.00722531 | 2 | 5.39244166 | 2.141737246 | 2.2680111 | 3.059555873 | 3.311948089 |
| ENSG00000005102 | MEOX1 | -1.956653747 | 0.494684323 | -4.430990863 | 0.100649724 | -6.38764461 | 2 | 4.11352228 | 2 | 2.3892346 | 2.38539787 | 2.094598369 | 2.059860178 |
| ENSG00000112293 | GPLD1 | -4.925186787 | 1.90E-07 | -1.462586071 | 0.137179363 | -6.387772858 | 6.366709671 | 5.946638857 | 7.934808822 | 2.947924294 | 3.08597292 | 5.212212839 | 5.941260754 |
| ENSG00000143315 | PIGM | -3.449328599 | 3.03E-08 | -2.940188759 | 2.58E-06 | -6.389517358 | 10.91047246 | 9.71095217 | 9.439621147 | 6.699459234 | 6.83591385 | 7.107581047 | 7.402777233 |
| ENSG00000124243 | BCAS4 | -3.414952104 | 0.012746065 | -2.977651473 | 0.03118692 | -6.392603577 | 10.58702944 | 9.04308532 | 5.772716771 | 6.069927018 | 6.18176397 | 6.99251878 | 5.884455123 |
| ENSG00000168743 | NPNT | -5.13042098 | 0.00593236 | -1.270790473 | 0.503651726 | -6.401211453 | 4.467311984 | 6.140560021 | 2 | 2.3892346 | 2.14022021 | 3.956110469 | 4.033026654 |
| ENSG00000104903 | LYL1 | -2.91622395 | 0.310429853 | -3.49225438 | 0.206767732 | -6.40847873 | 8.609439662 | 2 | 2 | 4.497095146 | 4.2865223 | 3.956110469 | 3.940519005 |
| ENSG00000131142 | CCL25 | -3.90849343 | NA | -2.500887928 | NA | -6.409381358 | 2 | 3.686303414 | 2 | 2 | 2.14022021 | 2.34604626 | 2 |
| ENSG00000109705 | NKX3-2 | -3.166723115 | 0.293193512 | -3.246396184 | 0.258826994 | -6.413119299 | 2 | 4.93638933 | 2 | 2.270786764 | 2.38539787 | 2.526562374 | 2.059860178 |
| ENSG00000100116 | GCAT | -4.421529978 | 6.04E-07 | -1.991650102 | 0.035807788 | -6.413180081 | 10.46286899 | 7.819099893 | 9.180874355 | 5.110902178 | 5.39133044 | 7.360534171 | 7.734322259 |
| ENSG00000152078 | TMEM56 | -4.468742746 | 0.003502679 | -1.947417712 | 0.225514494 | -6.416160458 | 7.750913381 | 4.710564154 | 4.058998049 | 2.868578001 | 3.01487888 | 5.158477451 | 3.875378318 |
| ENSG00000156466 | GDF6 | -4.470043918 | 0.008892695 | -1.953734636 | 0.280587322 | -6.423778554 | 2 | 6.78762695 | 7.818184403 | 3.289987342 | 3.01487888 | 5.367107572 | 4.532869193 |
| ENSG00000135423 | GLS2 | -3.219145402 | 0.110411514 | -3.214532542 | 0.090248812 | -6.433677944 | 2 | 4.93638933 | 3.369317972 | 2.270786764 | 2.49394783 | 2.3070652 | 2.466008509 |
| ENSG00000105549 | THEG | -1.689719683 | NA | -4.750701575 | NA | -6.440421258 | 3.327711234 | 2 | 2 | 2.141737246 | 2.2680111 | 2.048074405 | 2 |
| ENSG00000187325 | TAF9B | -4.295020596 | 7.15E-10 | -2.147935182 | 0.003244189 | -6.442955778 | 8.586954094 | 9.785372135 | 10.40882047 | 5.797438367 | 5.36298494 | 7.335328635 | 7.905190659 |
| ENSG00000144401 | METTL21A | -4.012655142 | 2.10E-06 | -2.431354917 | 0.005702324 | -6.444010059 | 10.09970294 | 7.60566235 | 9.61300511 | 5.742861563 | 5.30456641 | 7.042323222 | 7.030725399 |
| AAED1 | AAED1 | 2.9970704 | 0.316606609 | -9.454960587 | 0.002374237 | -6.457890186 | 2 | 5.946638857 | 2 | 7.288834728 | 7.35800033 | 2 | 2 |
| ENSG00000188958 | UTS2B | -3.412134 | 0.050115737 | -3.045900128 | 0.076041644 | -6.458034128 | 6.816637977 | 2 | 6.203069917 | 3.289987342 | 3.2184422 | 3.193831101 | 3.717076715 |
| ENSG00000079462 | PAFAH1B3 | -2.785292415 | 9.03E-05 | -3.673638726 | 1.06E-07 | -6.458931141 | 9.699577239 | 8.053983481 | 8.460932916 | 6.172600389 | 6.22177541 | 5.09123613 | 5.611957759 |
| ENSG00000221886 | ZBED8 | -3.105307493 | 0.003088289 | -3.358663345 | 0.001128494 | -6.463970838 | 9.410448395 | 7.669933291 | 7.398165132 | 5.662809983 | 5.24368216 | 5.853194273 | 4.147809227 |
| ENSG00000101441 | CST4 | -2.015987669 | 0.463053095 | -4.448751409 | 0.076272108 | -6.464739078 | 6.009025997 | 2 | 2 | 3.406930997 | 2.86123014 | 2.267001569 | 2.326754077 |
| ENSG00000133056 | PIK3C2B | -3.976611236 | 0.00014002 | -2.48819672 | 0.020094762 | -6.464807956 | 5.330594134 | 6.844429755 | 8.296287027 | 3.921164147 | 3.60476711 | 5.026717879 | 4.886954834 |
| ENSG00000239900 | ADSL | -1.952488125 | 0.089992166 | -4.512511118 | 1.43E-05 | -6.464999243 | 11.22998797 | 9.856140801 | 7.777116083 | 8.323303759 | 8.2130862 | 5.445480974 | 6.081436739 |
| ENSG00000067082 | KLF6 | -10.85985302 | 3.03E-07 | 4.394545817 | 0.003667911 | -6.465307207 | 3.327711234 | 8.053983481 | 7.285536013 | 2 | 2 | 11.74305903 | 11.27895974 |
| ENSG00000221818 | EBF2 | -2.883592822 | 0.259714186 | -3.582009403 | 0.1415706 | -6.465602225 | 2 | 5.946638857 | 2 | 2.695456494 | 2.77779907 | 2.526562374 | 2.466008509 |
| ENSG00000250510 | GPR162 | -4.942838189 | 0.003933311 | -1.527145835 | 0.422022879 | -6.469984024 | 2 | 8.255926468 | 8.585098385 | 3.515101962 | 3.50452879 | 6.295446113 | 6.446327951 |
| ENSG00000178662 | CSRNP3 | -2.447602326 | 0.204890893 | -4.035612278 | 0.025703955 | -6.483214605 | 5.532353208 | 6.666836837 | 2 | 3.881293667 | 3.39680241 | 3.1504466 | 2.277179783 |
| ENSG00000132854 | KANK4 | -3.418114719 | 0.0513415 | -3.065889717 | 0.067996482 | -6.484004436 | 4.467311984 | 4.710564154 | 4.523675391 | 2.868578001 | 2.14022021 | 3.035909414 | 2.172608095 |
| ENSG00000166450 | PRTG | -4.097347604 | 0.016979993 | -2.38901161 | 0.179503712 | -6.486359214 | 2 | 7.467835385 | 8.206335842 | 3.881293667 | 3.50452879 | 4.933813368 | 5.191251321 |
| ENSG00000152779 | SLC16A12 | -8.515481866 | 1.96E-05 | 2.0291173 | 0.130742576 | -6.486364566 | 5.532353208 | 3.076665209 | 5.39244166 | 2 | 2 | 6.956760977 | 6.77351449 |
| ENSG00000197558 | SSPO | -4.382770748 | 4.84E-06 | -2.106257067 | 0.025223108 | -6.489027815 | 4.815609471 | 6.389896633 | 6.718950348 | 2.947924294 | 2.77779907 | 4.612704558 | 3.940519005 |
| ENSG00000127903 | ZNF835 | -4.953460517 | 0.000578232 | -1.536197752 | 0.248020572 | -6.489658269 | 4.815609471 | 4.93638933 | 4.874587708 | 2.141737246 | 2.38539787 | 4.177580368 | 2.885414003 |
| ENSG00000163406 | SLC15A2 | -3.073477486 | 0.097604871 | -3.418153178 | 0.05364723 | -6.491630664 | 2 | 4.710564154 | 5.772716771 | 2.695456494 | 2.86123014 | 2.937246078 | 2.277179783 |
| ENSG00000239388 | ASB14 | -6.334381965 | 0.137320752 | -0.159959538 | 0.972248923 | -6.494341503 | 2 | 4.442710129 | 2 | 2 | 2 | 3.082820997 | 3.383972333 |
| ENSG00000169738 | DCXR | -3.758129096 | 1.69E-08 | -2.73658218 | 5.40E-05 | -6.494711276 | 11.31453223 | 10.17682307 | 9.565556058 | 6.782976084 | 6.8715633 | 7.673747216 | 7.959237827 |
| ENSG00000189051 | RNF222 | -3.379654187 | 0.014912024 | -3.115294339 | 0.014975157 | -6.494948526 | 4.815609471 | 3.686303414 | 4.058998049 | 2.600425781 | 2.2680111 | 2.526562374 | 2.509591799 |
| ENSG00000157423 | HYDIN | 1.720136134 | 0.63772496 | -8.215131356 | 0.020693942 | -6.494995222 | 2 | 2 | 4.874587708 | 4.887609717 | 5.04390545 | 2 | 2 |
| ENSG00000214357 | NEURL1B | -5.089017723 | 0.007426265 | -1.421661869 | 0.499776567 | -6.510679592 | 8.609439662 | 6.311476754 | 2 | 3.227757282 | 2.94009902 | 6.546183077 | 4.878668439 |
| ENSG00000106628 | POLD2 | -3.843833185 | 7.52E-17 | -2.669949508 | 9.75E-09 | -6.513782693 | 12.10171335 | 11.10337119 | 11.95227171 | 7.932663244 | 7.98332947 | 8.890512862 | 9.316523529 |
| ENSG00000175063 | UBE2C | -2.960011455 | 0.136473583 | -3.554960259 | 0.063416684 | -6.514971714 | 8.16936288 | 6.04685525 | 2 | 4.574143677 | 3.7864959 | 3.035909414 | 4.279527956 |
| ENSG00000179403 | VWA1 | -4.807593177 | 3.89E-11 | -1.708331177 | 0.030035111 | -6.515924354 | 8.855032352 | 9.45234302 | 10.80076732 | 5.334065616 | 5.22805098 | 8.14846013 | 8.333525663 |
| ENSG00000161654 | LSM12 | -1.117665073 | 0.001509359 | -5.403025079 | 1.96E-60 | -6.520690152 | 9.810482605 | 10.3778065 | 10.11951029 | 8.993902695 | 9.02937692 | 4.778616857 | 5.050803619 |
| ENSG00000143375 | CGN | -5.295554524 | 0.000476048 | -1.230088134 | 0.402910626 | -6.525642658 | 3.327711234 | 4.710564154 | 6.203069917 | 2.3892346 | 2.14022021 | 3.981090143 | 4.30448772 |
| ENSG00000221866 | PLXNA4 | -2.14060325 | 0.043735108 | -4.385446608 | 7.65E-06 | -6.526049858 | 5.70932728 | 8.505280662 | 7.029880454 | 5.578054239 | 5.37722731 | 3.810410249 | 3.474731315 |
| ENSG00000203666 | EFCAB2 | -2.180393508 | 0.039380876 | -4.348576621 | 8.63E-06 | -6.528970129 | 9.867565369 | 7.315436877 | 7.600047325 | 6.727838783 | 6.48112613 | 4.563936166 | 4.702084619 |
| ENSG00000096080 | MRPS18A | -2.174962168 | 1.11E-21 | -4.354308223 | 3.29E-84 | -6.529270391 | 10.08373159 | 10.18234215 | 10.19940021 | 7.925329813 | 8.06917614 | 5.83618166 | 5.957088417 |
| ENSG00000148483 | TMEM236 | -2.043029682 | 0.04359401 | -4.489856859 | 1.95E-06 | -6.532886541 | 5.866944937 | 5.457120953 | 6.629488387 | 4.070463721 | 4.43005853 | 3.059555873 | 2.374681325 |

| | | | | | | | | | | | | | |
|---|---|---|---|---|---|---|---|---|---|---|---|---|---|
| ENSG00000276836 | | 3.302544663 | 0.267308677 | -9.836235702 | 0.001533521 | -6.533691039 | 2 | 6.311476754 | 2 | 7.952038749 | 8.04037874 | 2 | 2 |
| ENSG00000155269 | GPR78 | -7.15566821 | 0.000888659 | 0.619763878 | 0.721968109 | -6.535904332 | 3.327711234 | 4.11352228 | 4.058998049 | 2 | 2 | 3.993419637 | 4.654341953 |
| ENSG00000197580 | BCO2 | -2.157754861 | 0.049611259 | -4.378591365 | 2.08E-05 | -6.536346227 | 5.095959458 | 6.389896633 | 4.523675391 | 3.709785322 | 3.86943969 | 2.775316226 | 2.421067459 |
| ENSG00000145681 | HAPLN1 | -2.902655537 | 2.84E-09 | -3.63950248 | 3.75E-14 | -6.542158017 | 12.33210977 | 12.88493391 | 12.79705561 | 9.787157185 | 9.80096123 | 8.438403662 | 9.49551136 |
| ENSG00000148600 | CDHR1 | -4.864277365 | 0.147262659 | -1.679838981 | 0.62915305 | -6.544116346 | 2 | 5.838937808 | 2 | 2.3892346 | 2 | 3.483289517 | 3.012034821 |
| ENSG00000221916 | C19orf73 | -4.220546286 | 0.028174937 | -2.325332533 | 0.219716772 | -6.545878819 | 6.009025997 | 3.686303414 | 2 | 2.270786764 | 2.49394783 | 3.12825509 | 3.071407821 |
| ENSG00000184434 | LRRC19 | -5.331954657 | 0.129527338 | -1.214832331 | 0.722001338 | -6.546786989 | 2 | 2 | 4.874587708 | 2 | 2.14022021 | 2.885263063 | 2.950113106 |
| ENSG00000181444 | ZNF467 | -4.623637704 | 0.005365654 | -1.923521997 | 0.275529427 | -6.547159701 | 6.816637977 | 7.73146271 | 2 | 3.023133209 | 3.08597292 | 5.279204176 | 4.654341953 |
| ENSG00000133477 | FAM83F | -3.274641452 | NA | -3.275338026 | NA | -6.549979478 | 3.327711234 | 2 | 2 | 2 | 2.14022021 | 2.139668783 | 2 |
| ENSG00000186665 | C17orf58 | -3.552255389 | 1.60E-11 | -2.997859944 | 1.06E-08 | -6.550115333 | 9.895281535 | 9.00621767 | 8.805342542 | 5.870535235 | 5.84514933 | 6.148830293 | 6.580104496 |
| ENSG00000177888 | ZBTB41 | -9.132791786 | 0.000878889 | 2.582621841 | 0.294969622 | -6.550169945 | 3.327711234 | 6.951736442 | 2 | 2 | 2 | 8.454657895 | 7.36768071 |
| ENSG00000172345 | STARD5 | -2.63738959 | 0.089597823 | -3.914282488 | 0.00758297 | -6.551672078 | 3.327711234 | 7.874700259 | 5.59507254 | 4.299492648 | 4.22485516 | 3.429858444 | 3.287118176 |
| ENSG00000135821 | GLUL | -3.380924697 | 5.17E-27 | -3.174973938 | 3.22E-24 | -6.555898635 | 13.54926601 | 13.27868079 | 13.63150382 | 10.14639907 | 10.08857 | 10.56105514 | 10.03766601 |
| ENSG00000111664 | GNB3 | -3.81100689 | 4.13E-08 | -2.746330616 | 7.98E-05 | -6.557337506 | 7.031698319 | 8.376385542 | 8.805342542 | 4.54891529 | 4.76938658 | 5.762453498 | 5.425444337 |
| ENSG00000103742 | IGDCC4 | -4.849807418 | 1.09E-21 | -1.708836738 | 0.001300235 | -6.558644156 | 9.761125267 | 10.89732378 | 10.89495479 | 5.775854763 | 5.93454154 | 8.84531395 | 8.966780003 |
| ENSG00000196476 | C20orf96 | -3.160081867 | 0.000123283 | -3.400183862 | 2.69E-05 | -6.560265729 | 8.855032352 | 6.951736442 | 9.272377397 | 5.614989456 | 5.5755357 | 5.4091291 | 5.343519008 |
| ENSG00000198870 | STKLD1 | -6.567421389 | 0.006598327 | 0.00521396 | 0.998392474 | -6.562207429 | 3.327711234 | 3.686303414 | 3.369317972 | 2 | 2 | 3.767980144 | 3.128433649 |
| ENSG00000164175 | SLC45A2 | -1.141466304 | 0.767351465 | -5.423888063 | NA | -6.565354367 | 2 | 3.686303414 | 2 | 2.3892346 | 2.49394783 | 2.048074405 | 2 |
| ENSG00000249437 | NAIP | 0.777128134 | 0.688362385 | -7.345186234 | 0.00088495 | -6.5680581 | 4.815609471 | 4.11352228 | 2 | 4.694064037 | 4.65713986 | 2 | 2.059860178 |
| ENSG00000181029 | TRAPPC5 | -1.317505784 | 0.020153341 | -5.251573957 | 2.37E-24 | -6.56907974 | 10.00958825 | 8.88960062 | 9.31604076 | 8.087422161 | 8.25224987 | 4.705557986 | 4.254128754 |
| ENSG00000143507 | DUSP10 | -8.476851957 | 0.012899357 | 1.907127198 | 0.572810622 | -6.56972476 | 2 | 6.389896633 | 2 | 2 | 2 | 6.700796638 | 6.725806822 |
| ENSG00000189334 | S100A14 | -2.702368988 | 0.340614989 | -3.86791188 | 0.148181182 | -6.570280867 | 2 | 2 | 5.930865648 | 3.023133209 | 2.49394783 | 2.183373728 | 2.592996859 |
| ENSG00000138814 | PPP3CA | -3.036802417 | 5.56E-16 | -3.542851721 | 1.23E-21 | -6.579654138 | 12.6398868 | 11.82171249 | 12.18790165 | 9.271529914 | 9.18016239 | 8.769824912 | 8.677773268 |
| ENSG00000104369 | JPH1 | -2.346464006 | NA | -4.236840948 | NA | -6.583304954 | 2 | 3.686303414 | 2 | 2.270786764 | 2.14022021 | 2 | 2.117335275 |
| ENSG00000067177 | PHKA1 | -3.843156158 | 0.001102522 | -2.74035435 | 0.023941231 | -6.583510507 | 6.009025997 | 9.079034257 | 10.05225195 | 5.614989456 | 5.16377554 | 6.216130818 | 6.62538071 |
| ENSG00000204334 | ERICH2 | -5.357235097 | 0.297702978 | -1.227700922 | NA | -6.584936019 | 2 | 3.686303414 | 2 | 2 | 2 | 2.492221014 | 2.326754077 |
| ENSG00000278570 | NR2E3 | -4.081491178 | 0.138334654 | -2.504566288 | 0.353971254 | -6.586057467 | 5.532353208 | 2 | 2 | 2.3892346 | 2.14022021 | 2.592885947 | 2.782576693 |
| ENSG00000141542 | RAB40B | -4.689197675 | 2.29E-13 | -1.897949097 | 0.004530905 | -6.587146772 | 9.397585735 | 8.147241016 | 9.735974718 | 4.54891529 | 4.95174433 | 7.245405178 | 7.469129172 |
| ENSG00000137486 | ARRB1 | -4.747506592 | 3.76E-64 | -1.844559914 | 5.15E-12 | -6.592066507 | 9.781071289 | 9.975384048 | 10.15199903 | 5.391962469 | 5.34860058 | 8.193908178 | 8.098650814 |
| ENSG00000243130 | PSG11 | -1.961750566 | 0.612906092 | -4.630858748 | 0.193576 | -6.592609314 | 2 | 2 | 5.156626401 | 2.868578001 | 2.59489879 | 2 | 2.277179783 |
| ENSG00000151023 | ENKUR | -6.722679782 | 0.103499918 | 0.129425954 | 0.976784704 | -6.593253829 | 4.815609471 | 2 | 2 | 2 | 2 | 3.824280769 | 3.496556178 |
| ENSG00000180316 | PNPLA1 | -2.315078565 | 0.277730712 | -4.279888926 | 0.032357779 | -6.594967491 | 2 | 3.686303414 | 4.874587708 | 2.947924294 | 2.2680111 | 2.267001569 | 2.117335275 |
| ENSG00000274349 | ZNF658 | -7.226503039 | 0.008722986 | 0.625708348 | 0.816103107 | -6.600794691 | 2 | 4.93638933 | 3.369317972 | 2 | 2 | 4.604690117 | 4.215171317 |
| ENSG00000067445 | TRO | -4.118843864 | 7.56E-18 | -2.485194238 | 3.15E-07 | -6.604038102 | 10.70657014 | 11.02260251 | 11.79401137 | 7.178012736 | 7.15805139 | 8.615855003 | 8.917071621 |
| ENSG00000134339 | SAA2 | -0.095954528 | 0.95806358 | -6.511955713 | 0.00029754 | -6.607910241 | 3.327711234 | 3.686303414 | 4.874587708 | 4.237680128 | 3.7864959 | 2.094598369 | 2 |
| ENSG00000197471 | SPN | -4.572547566 | 0.000585277 | -2.037754811 | 0.13837407 | -6.610302377 | 6.009025997 | 5.595927442 | 7.225742779 | 2.947924294 | 2.86123014 | 5.352823931 | 2.918126211 |
| ENSG00000136883 | KIF12 | -6.327055199 | 0.009371257 | -0.2863352 | 0.899397925 | -6.613390399 | 2 | 5.303523747 | 4.058998049 | 2 | 2.14022021 | 4.053534413 | 4.201948172 |
| ENSG00000166257 | SCN3B | -2.479588393 | 0.248222521 | -4.135649491 | 0.039885498 | -6.615237884 | 2 | 4.93638933 | 5.156626401 | 3.162721474 | 2.59489879 | 2.592885947 | 2.059860178 |
| ENSG00000276231 | PIK3R6 | -5.92259887 | 0.210036317 | -0.694632556 | 0.891629211 | -6.617231426 | 2 | 4.11352228 | 2 | 2 | 2 | 3.082820997 | 2.374681325 |
| ENSG00000179083 | FAM133A | -4.461464499 | 0.4401696 | -2.157208443 | NA | -6.618672942 | 2 | 4.11352228 | 2 | 2.141737246 | 2 | 2.592885947 | 2 |
| ENSG00000175130 | MARCKSL1 | -3.038407205 | 1.66E-06 | -3.581482148 | 9.80E-09 | -6.619889353 | 12.84803656 | 11.60046943 | 11.39994907 | 9.170732795 | 8.96600458 | 8.750432421 | 8.278687361 |
| ENSG00000110777 | POU2AF1 | -3.216485679 | 0.180445785 | -3.403949267 | 0.128471526 | -6.620434946 | 2 | 3.076665209 | 4.523675391 | 2.3892346 | 2.14022021 | 2.094598369 | 2.374681325 |
| ENSG00000106258 | CYP3A5 | -6.333519186 | 0.141463415 | -0.290047912 | 0.950961155 | -6.623567098 | 2 | 4.442710129 | 2 | 2 | 2 | 3.355392153 | 2.950113106 |
| ENSG00000136274 | NACAD | -4.893455574 | 5.05E-52 | -1.731384416 | 6.74E-08 | -6.62483999 | 10.79260216 | 10.47993704 | 10.27509734 | 5.674521485 | 5.81407719 | 8.683523874 | 8.925140957 |
| ENSG00000259494 | MRPL46 | -2.36496909 | 0.000693009 | -4.260677995 | 1.37E-10 | -6.625647085 | 9.913467515 | 8.124485672 | 9.180874355 | 6.861921612 | 6.97351258 | 5.318934974 | 4.951586397 |
| ENSG00000113108 | APBB3 | -3.186491972 | 4.75E-07 | -3.441968785 | 3.68E-08 | -6.628460758 | 9.174465797 | 9.415325457 | 10.67226762 | 6.825617242 | 6.71776877 | 6.550361701 | 6.504031956 |
| ENSG00000170425 | ADORA2B | -4.351601387 | 1.72E-07 | -2.277218075 | 0.009126321 | -6.628819462 | 10.1770105 | 7.790474954 | 8.884815469 | 5.22679461 | 4.93258395 | 7.045287119 | 7.006211772 |
| ENSG00000214140 | PRCD | -2.345048585 | 0.288045778 | -4.285541223 | 0.036191293 | -6.630589808 | 2 | 3.686303414 | 6.958202216 | 3.515101962 | 3.74316441 | 2.911488689 | 2.277179783 |
| ENSG00000153976 | HS3ST3A1 | 1.662782493 | 0.067806127 | -8.294650994 | 1.05E-19 | -6.6318685 | 8.777722034 | 7.393647515 | 7.734844356 | 9.729378027 | 9.75056618 | 2.457042231 | 2.059860178 |
| ENSG00000118520 | ARG1 | -5.356083266 | 0.304345203 | -1.284695939 | NA | -6.640779206 | 2 | 3.686303414 | 2 | 2 | 2 | 2.225793514 | 2.551896972 |
| ENSG00000106992 | AK1 | -2.727037821 | 6.53E-18 | -3.914018734 | 1.74E-36 | -6.641056555 | 11.40728614 | 10.92392403 | 11.31283101 | 8.494945692 | 8.5339035 | 7.510690356 | 7.162846889 |
| ENSG00000167772 | ANGPTL4 | -9.207749523 | 0.00619284 | 2.564334236 | 0.434435201 | -6.643415287 | 2 | 7.099070191 | 2 | 2 | 2 | 8.295111724 | 7.796124663 |
| ENSG00000163590 | PPM1L | -3.040874197 | 0.000224661 | -3.603165473 | 7.62E-06 | -6.64403967 | 9.219014269 | 7.467835385 | 7.600047325 | 5.44762523 | 5.37722731 | 5.233158834 | 4.500832793 |
| ENSG00000111181 | SLC6A12 | -4.621291534 | 0.099215121 | -2.024506914 | 0.467805568 | -6.645798447 | 2 | 5.595927442 | 2 | 2.270786764 | 2.14022021 | 3.336159116 | 2.421067459 |
| ENSG00000162490 | DRAXIN | -4.767006397 | 0.091611673 | -1.881579943 | 0.522888638 | -6.64858634 | 2 | 2 | 6.718950348 | 2.3892346 | 2.38539787 | 4.321442857 | 2.632958241 |
| ENSG00000272398 | CD24 | -1.986477718 | 0.258310503 | -4.665096546 | 0.003362603 | -6.651574264 | 7.579279349 | 6.464272742 | 4.874587708 | 5.709096143 | 2.49394783 | 3.12825509 | 2.817676617 |
| ENSG00000154920 | EME1 | -3.720648415 | 0.172696309 | -2.931252995 | 0.282001843 | -6.65190141 | 7.750913381 | 2 | 2 | 3.09461492 | 3.33976443 | 3.838019203 | 3.600986722 |
| ENSG00000233927 | RPS28 | -2.152344809 | 0.007826211 | -4.500652286 | 2.32E-09 | -6.652997094 | 12.23602991 | 10.30378101 | 10.57378414 | 9.162439556 | 9.16587611 | 6.689474108 | 7.006211772 |
| ENSG00000213203 | GIMAP1 | -4.652947049 | 0.125363988 | -2.001644348 | 0.497432421 | -6.654591397 | 2 | 2 | 5.156626401 | 2.141737246 | 2.14022021 | 3.082820997 | 2.225841198 |
| ENSG00000108018 | SORCS1 | -1.678203375 | 0.463760408 | -4.984926131 | 0.020501539 | -6.663129507 | 2 | 3.686303414 | 4.523675391 | 3.162721474 | 2.2680111 | 2.139668783 | 2.059860178 |
| ENSG00000049283 | EPN3 | -5.329909916 | 0.047154938 | -1.338105606 | 0.562965652 | -6.668015521 | 3.327711234 | 4.442710129 | 2 | 2.141737246 | 2 | 2.937246078 | 2.782576693 |
| ENSG00000180425 | C11orf71 | -2.673517027 | 0.004642582 | -3.998858779 | 9.20E-06 | -6.672375806 | 5.095959458 | 6.844429755 | 7.502634626 | 4.299492648 | 4.43005853 | 3.584558147 | 3.209956562 |
| ENSG00000160588 | MPZL3 | -2.491739414 | 0.031747165 | -4.183536376 | 0.000122298 | -6.675275781 | 6.009025997 | 6.311476754 | 8.763901704 | 5.022023783 | 5.40529704 | 3.738983888 | 3.940519005 |
| ENSG00000138606 | SHF | -3.490543888 | 0.000842579 | -3.185233067 | 0.002274706 | -6.675776955 | 9.219014269 | 6.666836837 | 7.502634626 | 4.804776789 | 4.97065356 | 5.646229369 | 4.411752372 |
| ENSG00000081692 | JMJD4 | -3.103009048 | 4.95E-09 | -3.573323601 | 8.34E-12 | -6.676332649 | 10.03472846 | 9.298208073 | 10.59756881 | 6.861921612 | 7.13728069 | 6.501560901 | 6.602920209 |
| ENSG00000137804 | NUSAP1 | -2.186868798 | 0.312962462 | -4.491847258 | 0.023862255 | -6.678716056 | 5.095959458 | 8.791896851 | 2 | 5.348759107 | 5.18011485 | 3.56816602 | 3.33635788 |
| ENSG00000120280 | CXorf21 | -6.079476156 | 0.061292986 | -0.605175485 | 0.854029073 | -6.684651641 | 3.327711234 | 3.686303414 | 2 | 2 | 2 | 3.172301926 | 2.509591799 |
| ENSG00000104490 | NCALD | -3.129249586 | 0.029030546 | -3.557325377 | 0.01090815 | -6.686574963 | 3.327711234 | 7.790474954 | 8.110401118 | 4.670864177 | 4.3456612 | 4.301748724 | 4.033026654 |
| ENSG00000005073 | HOXA11 | 0.401930175 | 0.910682148 | -7.088987141 | 0.018823615 | -6.687056966 | 2 | 6.140560021 | 2 | 4.867343068 | 5.2591458 | 2 | 2.117335275 |
| ENSG00000147955 | SIGMAR1 | -3.910680253 | 4.62E-15 | -2.777520299 | 3.89E-08 | -6.688200551 | 12.23783517 | 13.06750563 | 13.48935805 | 9.157231967 | 9.07923325 | 10.37421371 | 10.10580955 |
| ENSG00000198336 | MYL4 | -5.074940461 | 0.004232874 | -1.616193909 | 0.331819555 | -6.69113437 | 5.095959458 | 5.131603625 | 2 | 2.141737246 | 2.2680111 | 3.616794016 | 3.209956562 |
| ENSG00000275723 | | 0.555788798 | 0.74129049 | -7.247413427 | 2.49E-06 | -6.691624629 | 4.467311984 | 6.78762695 | 3.369317972 | 6.104969671 | 6.1322378 | 2.139668783 | 2.059860178 |
| ENSG00000140988 | RPS2 | -3.514315121 | 1.46E-23 | -3.179254443 | 1.09E-19 | -6.693569564 | 15.52119227 | 14.86497311 | 14.88548414 | 11.61170542 | 11.6111448 | 11.99501747 | 11.89546496 |
| ENSG00000175538 | KCNE3 | -6.336073589 | 1.71E-05 | -0.360398763 | 0.768057473 | -6.696472352 | 5.095959458 | 5.72254323 | 4.523675391 | 2.270786764 | 2 | 5.096960836 | 4.673629022 |
| ENSG00000171970 | ZNF57 | -3.499984826 | 0.076937101 | -3.19778614 | 0.103567515 | -6.697770966 | 8.138771685 | 2 | 9.541233192 | 5.161714397 | 5.00774437 | 5.152992171 | 5.534955934 |
| ENSG00000162494 | LRRC38 | -6.722832954 | 0.105206276 | 0.021597822 | 0.996386969 | -6.701235133 | 4.815609471 | 2 | 2 | 2 | 2 | 3.465698119 | 3.717076715 |
| ENSG00000213123 | TCTEX1D2 | -2.317190026 | 0.00351465 | -4.387592864 | 5.47E-09 | -6.70478289 | 8.285626337 | 6.389896633 | 7.342949392 | 5.334065616 | 5.33407135 | 3.824280769 | 3.474731315 |
| ENSG00000175311 | ANKS4B | -2.288519456 | 0.303863659 | -4.421217157 | 0.035456159 | -6.709736613 | 3.327711234 | 2 | 4.523675391 | 2.270786764 | 2.68924543 | 2.139668783 | 2.117335275 |
| ENSG00000248383 | PCDHAC1 | -1.559085901 | 0.597182199 | -5.151333222 | 0.053429704 | -6.710419123 | 2 | 5.595927442 | 2 | 3.615725078 | 2.59489879 | 2.225793514 | 2.059860178 |
| ENSG00000113209 | PCDHB5 | -3.865067198 | 0.009778948 | -2.846371561 | 0.059149315 | -6.711438759 | 4.467311984 | 4.442710129 | 8.007629717 | 3.515101962 | 3.28037825 | 4.230621763 | 3.987514168 |
| ENSG00000174844 | DNAH12 | -1.992070871 | 0.332298792 | -4.72375139 | 0.013961124 | -6.715822261 | 5.330594134 | 2 | 3.369317972 | 2.947924294 | 2.86123014 | 2.183373728 | 2.172608095 |
| ENSG00000164695 | CHMP4C | -4.812011232 | 0.019668123 | -1.904081558 | 0.298022192 | -6.71609279 | 4.00722531 | 3.076665209 | 4.523675391 | 2.270786764 | 2 | 3.082820997 | 2.509591799 |
| ENSG00000276130 | | 0.134273143 | 0.941709668 | -6.856800162 | 1.13E-06 | -6.722527019 | 3.327711234 | 7.189546432 | 8.511888463 | 7.574744049 | 7.56502709 | 2.560105256 | 2.326754077 |
| ENSG00000204128 | C2orf72 | -4.147324099 | 0.029155797 | -2.580907751 | 0.188229922 | -6.72823185 | 2 | 6.535001748 | 8.784770914 | 4.139692846 | 3.33976443 | 5.264018433 | 4.810605986 |
| ENSG00000166152 | C16orf78 | -4.714843385 | NA | -2.016634707 | NA | -6.731478091 | 2 | 2 | 3.369317972 | 2 | 2 | 2.048074405 | 2.277179783 |
| ENSG00000196338 | NLGN3 | -3.378105244 | 0.019026761 | -3.359701518 | 0.018122416 | -6.737806762 | 4.00722531 | 6.535001748 | 8.007629717 | 4.470469939 | 3.08597292 | 4.111244182 | 3.771796761 |
| ENSG00000158769 | F11R | -5.363410182 | 2.71E-05 | -1.376745334 | 0.331999289 | -6.740155516 | 5.330594134 | 6.78762695 | 9.099922433 | 3.515101962 | 2.94009902 | 6.804147769 | 6.175763085 |
| ENSG00000259224 | SLC35G6 | -4.909479384 | 0.178354575 | -1.833983107 | 0.591752395 | -6.743462492 | 2 | 2 | 4.523675391 | 2 | 2.14022021 | 2.526562374 | 2.509591799 |
| ENSG00000163463 | KRTCAP2 | -1.017729437 | 0.022336639 | -5.729301564 | 3.95E-46 | -6.747031001 | 10.60951384 | 9.883504052 | 10.29705308 | 9.18103275 | 9.37016537 | 4.547306681 | 4.990558504 |
| ENSG00000172938 | MRGPRD | -5.35066353 | 0.341002462 | -1.396803887 | NA | -6.747467417 | 2 | 3.686303414 | 2 | 2 | 2 | 2.62493832 | 2.059860178 |
| ENSG00000151025 | GPR158 | -4.164332465 | 0.014678051 | -2.585527336 | 0.127281504 | -6.749859801 | 2 | 6.228547981 | 5.39244166 | 2.3892346 | 2.77779907 | 3.584558147 | 3.042026731 |
| ENSG00000106333 | PCOLCE | -2.194943715 | 0.006082629 | -4.555405434 | 1.03E-09 | -6.750349149 | 13.90444097 | 12.06817771 | 12.10735427 | 10.77405547 | 10.777395 | 8.408791501 | 8.448559482 |
| ENSG00000164659 | KIAA1324L | -3.034931185 | 0.000894709 | -3.715869858 | 2.93E-05 | -6.750801044 | 7.27616306 | 8.147241016 | 9.778213818 | 5.829220145 | 5.83486613 | 5.136410075 | 5.282014932 |
| ENSG00000013293 | SLC7A14 | -5.301545438 | 3.17E-06 | -1.450227645 | 0.148588423 | -6.751773083 | 4.815609471 | 5.946638857 | 5.39244166 | 2.270786764 | 2.38539787 | 4.580376151 | 3.771796761 |
| ENSG00000270647 | TAF15 | 4.01823386 | 7.78E-05 | -10.78197588 | 3.36E-09 | -6.763742023 | 6.73718411 | 4.93638933 | 4.523675391 | 9.58171079 | 9.70118012 | 2 | 2 |
| ENSG00000222046 | DCDC2B | -5.70095682 | 0.092053435 | -1.064541299 | 0.746296139 | -6.765498119 | 2 | 5.131603625 | 2 | 2 | 2.14022021 | 3.10571689 | 3.261853426 |
| ENSG00000172322 | CLEC12A | -4.714843915 | NA | -2.059063613 | NA | -6.773907528 | 2 | 3.076665209 | 2 | 2 | 2 | 2.3070652 | 2 |
| ENSG00000136861 | CDK5RAP2 | -10.008403 | 6.36E-07 | 3.233647726 | 0.017427038 | -6.774755274 | 3.327711234 | 6.78762695 | 6.958202216 | 2 | 2 | 9.374289203 | 9.658056436 |
| ENSG00000163737 | PF4 | 0.430442642 | 0.929073514 | -7.205877804 | 0.091283544 | -6.775435162 | 2 | 2 | 4.058998049 | 3.349643765 | 3.15372745 | 2 | 2 |
| ENSG00000102174 | PHEX | -2.045665446 | 0.470185896 | -4.729864647 | 0.066331445 | -6.775530093 | 2 | 2 | 6.629488387 | 3.754611435 | 3.28037825 | 2.225793514 | 2.509591799 |
| ENSG00000274332 | | 2.972797449 | 0.005904618 | -9.749750899 | 1.33E-07 | -6.776953451 | 3.327711234 | 4.710564154 | 5.59507254 | 7.52061915 | 7.65252709 | 2 | 2 |
| ENSG00000123901 | GPR83 | 0.373106406 | 0.912408896 | -7.151034639 | 0.022791391 | -6.777928233 | 3.327711234 | 2 | 3.369317972 | 3.162721474 | 3.2184422 | 2 | 2 |
| ENSG00000162373 | BEND5 | -4.550327648 | 0.18345602 | -2.230395892 | 0.52422773 | -6.78072354 | 2 | 6.228547981 | 2 | 2 | 2.59489879 | 2.987421008 | 3.36036153 |
| ENSG00000167528 | ZNF641 | -3.696739741 | 3.53E-13 | -3.088325889 | 1.26E-09 | -6.78506563 | 9.895281535 | 10.57111193 | 11.13327928 | 7.101994243 | 6.81514011 | 7.64179387 | 7.469129172 |
| ENSG00000280109 | PLAC4 | -6.98570882 | 0.00256088 | 0.197761667 | 0.926910758 | -6.787947153 | 3.327711234 | 3.076665209 | 4.523675391 | 2 | 2 | 3.851628042 | 3.956354764 |
| ENSG00000168026 | TTC21A | -2.841564868 | 0.010605213 | -3.951326211 | 0.000206862 | -6.792891079 | 8.540903468 | 5.303523747 | 8.408111533 | 5.144975053 | 5.36298494 | 4.416072028 | 4.201948172 |
| ENSG00000282076 | | -5.918632626 | 0.231845171 | -0.87445001 | 0.868498866 | -6.793082636 | 2 | 4.11352228 | 2 | 2 | 2 | 3.082820997 | 2.172608095 |

| | | | | | | | | | | | | | |
|---|---|---|---|---|---|---|---|---|---|---|---|---|---|
| ENSG00000162931 | TRIM17 | -5.887024764 | 0.008715217 | -0.910980693 | 0.686224112 | -6.798005457 | 4.467311984 | 5.838937808 | 2 | 2 | 2.2680111 | 4.291799915 | 3.824516949 |
| ENSG00000171763 | SPATA5L1 | -8.350058768 | 0.002045695 | 1.550249701 | 0.535543384 | -6.799809067 | 3.327711234 | 6.140560021 | 2 | 2 | 2 | 6.459963649 | 6.007361012 |
| ENSG00000128918 | ALDH1A2 | -8.763441633 | 0.010230123 | 1.954961215 | 0.565006761 | -6.808480418 | 2 | 6.666836837 | 2 | 2 | 2 | 6.870823472 | 7.184857568 |
| ENSG00000157103 | SLC6A1 | -6.154617278 | 0.073455594 | -0.654145229 | 0.864027588 | -6.808762508 | 2 | 2 | 6.534109959 | 2.270786764 | 2 | 5.206928476 | 3.071407821 |
| ENSG00000196781 | TLE1 | -3.501483099 | 6.54E-10 | -3.311047677 | 4.33E-09 | -6.812530775 | 9.189468652 | 10.28338212 | 10.38851302 | 6.604700657 | 6.59518404 | 7.086154752 | 6.397863414 |
| ENSG00000172476 | RAB40A | -4.548920596 | 0.023677801 | -2.264288582 | 0.268354747 | -6.813209178 | 2 | 3.686303414 | 6.80318718 | 2.600425781 | 2.38539787 | 3.374372158 | 3.771796761 |
| ENSG00000178732 | GP5 | -1.59423393 | 0.680095757 | -5.219127408 | 0.143374446 | -6.813361338 | 2 | 2 | 4.523675391 | 2.3892346 | 2.77779907 | 2 | 2.117335275 |
| ENSG00000183032 | SLC25A21 | -2.720602046 | 0.007588734 | -4.09281006 | 2.41E-05 | -6.813412107 | 4.00722531 | 5.946638857 | 5.772716771 | 3.227757282 | 3.45167086 | 2.746468522 | 2.592996859 |
| ENSG00000152705 | CATSPER3 | -0.014430392 | 0.997481474 | -6.801334936 | NA | -6.815765328 | 2 | 3.686303414 | 2 | 2.784612485 | 2.86123014 | 2 | 2 |
| ENSG00000151150 | ANK3 | -6.605399503 | 0.000274049 | -0.214587226 | 0.922485577 | -6.819986729 | 6.563806279 | 7.73146271 | 2 | 2.498691189 | 2.14022021 | 6.265258498 | 6.697819825 |
| ENSG00000008311 | AASS | -3.157717325 | 2.30E-07 | -3.668858462 | 1.05E-09 | -6.826575787 | 10.40620204 | 9.367676479 | 10.61515347 | 7.273514749 | 6.90635305 | 6.129409568 | 6.967665897 |
| ENSG00000100359 | SGSM3 | -1.110924551 | 0.425388379 | -5.71728328 | 8.73E-07 | -6.828207832 | 10.80239439 | 9.687856435 | 6.43197674 | 8.652768197 | 8.76268478 | 4.443291373 | 4.341135913 |
| ENSG00000169851 | PCDH7 | -0.18975763 | 0.891428469 | -6.638706158 | 9.95E-11 | -6.828463788 | 7.096806373 | 9.640525262 | 8.325066185 | 8.474993083 | 8.57309694 | 3.429858444 | 2.466008509 |
| ENSG00000255561 | FDXACB1 | -2.621183218 | 0.000209397 | -4.209545196 | 6.55E-10 | -6.830728414 | 6.009025997 | 7.189546432 | 7.398165132 | 4.670864177 | 4.4836996 | 3.600766115 | 3.183290849 |
| ENSG00000169515 | CCDC8 | -6.348721498 | 8.90E-57 | -0.482273415 | 0.217451672 | -6.830994913 | 9.579430841 | 9.45234302 | 9.358421373 | 3.921164147 | 3.33976443 | 8.949843022 | 9.023784348 |
| ENSG00000053524 | MCF2L2 | -4.898926261 | 2.95E-05 | -1.932687402 | 0.079412418 | -6.831613663 | 4.815609471 | 5.838937808 | 5.772716771 | 2.498691189 | 2.38539787 | 4.416072028 | 3.183290849 |
| ZCCHC16 | | -0.682150113 | NA | -6.152897122 | NA | -6.835047235 | 2 | 2 | 3.369317972 | 2.600425781 | 2.14022021 | 2 | 2 |
| ENSG00000167889 | MGAT5B | -1.57445286 | 0.574255931 | -5.264750371 | 0.040597375 | -6.83920323 | 2 | 5.303523747 | 2 | 3.162721474 | 2.86123014 | 2.048074405 | 2.172608095 |
| ENSG00000011600 | TYROBP | -8.543007558 | 0.001045597 | 1.696453442 | 0.469777313 | -6.846554116 | 2 | 5.457120953 | 5.59507254 | 2 | 2 | 7.370021622 | 4.654341953 |
| ENSG00000162989 | KCNJ3 | -0.695296316 | NA | -6.152894342 | NA | -6.848190658 | 3.327711234 | 2 | 2 | 2 | 2.68924543 | 2 | 2 |
| ENSG00000132554 | RGS22 | -4.706542691 | 0.173138308 | -2.148470731 | 0.538164917 | -6.855013422 | 5.70932728 | 2 | 2 | 2 | 2.38539787 | 3.059555873 | 2.782576693 |
| ENSG00000117148 | ACTL8 | -2.264883678 | NA | -4.592592627 | NA | -6.857476305 | 3.327711234 | 2 | 2 | 2.270786764 | 2 | 2 | 2.059860178 |
| ENSG00000104883 | PEX11G | -3.896840758 | 0.028846345 | -2.964103099 | 0.098174415 | -6.860943857 | 2 | 5.72254323 | 7.734844356 | 3.349643765 | 3.2184422 | 3.709392871 | 4.077147165 |
| ENSG00000188316 | ENO4 | -3.02360277 | 0.244954821 | -3.840065885 | 0.120439864 | -6.863668655 | 2 | 5.72254323 | 2 | 2.695456494 | 2.49394783 | 2.267001569 | 2.466008509 |
| ENSG00000174483 | BBS1 | -1.245597302 | 0.004306044 | -5.623913513 | 2.21E-41 | -6.869510815 | 8.39321512 | 9.079034257 | 8.511888463 | 7.555872146 | 7.38644297 | 3.534812207 | 3.717076715 |
| ENSG00000091972 | CD200 | -2.860506944 | 0.123445724 | -4.012332126 | 0.023237554 | -6.87283907 | 2 | 7.35507212 | 8.143093325 | 4.574143677 | 4.60967607 | 4.144793515 | 3.071407821 |
| ENSG00000164929 | BAALC | -1.575386997 | 0.458262387 | -5.298471226 | 0.004094667 | -6.873858223 | 8.009475818 | 8.101365653 | 2 | 6.034011955 | 5.90535527 | 3.56816602 | 2.326754077 |
| ENSG00000132570 | PCBD2 | -2.870715279 | 0.001070151 | -4.009950858 | 2.12E-06 | -6.880666138 | 9.533135543 | 7.431218269 | 7.734844356 | 5.709096143 | 5.83486613 | 4.907994066 | 4.584731282 |
| ENSG00000171056 | SOX7 | -3.821924634 | 0.042638051 | -3.059800957 | 0.091166431 | -6.88172559 | 4.467311984 | 2 | 5.59507254 | 2.600425781 | 2.2680111 | 2.775316226 | 2.592996859 |
| ENSG00000047648 | ARHGAP6 | -8.699215263 | 0.000183235 | 1.816459899 | 0.345156474 | -6.882755364 | 2 | 5.131603625 | 6.073378139 | 2 | 2 | 6.854018691 | 6.826173076 |
| ENSG00000170390 | DCLK2 | -9.039883501 | 0.000133459 | 2.156113255 | 0.272249094 | -6.883770245 | 2 | 5.131603625 | 6.534109959 | 2 | 2 | 7.620091565 | 7.372114752 |
| ENSG00000002746 | HECW1 | -5.51247322 | 0.110888355 | -1.372732343 | 0.702980293 | -6.885205563 | 2 | 2 | 5.930865648 | 2.270786764 | 2 | 3.968654369 | 2.782576693 |
| ENSG00000102935 | ZNF423 | -6.341994898 | 6.31E-07 | -0.546304068 | 0.721493405 | -6.888298966 | 7.82963488 | 8.337339476 | 8.296287027 | 3.462030052 | 2 | 7.235058713 | 7.94839122 |
| ENSG00000229117 | RPL41 | -0.261886328 | 0.778918686 | -6.629029513 | 2.22E-21 | -6.89091584 | 13.57662794 | 11.93393502 | 11.8245656 | 12.38638425 | 12.4592006 | 6.167993056 | 6.110033886 |
| ENSG00000280789 | PAGR1 | -2.528066735 | 0.000510004 | -4.364485862 | 3.30E-10 | -6.892552597 | 8.586954094 | 8.747909548 | 10.23774519 | 6.836083475 | 6.97351258 | 5.31402805 | 5.050803619 |
| ENSG00000268182 | SMIM17 | -2.954997614 | 0.042927754 | -3.945297768 | 0.004306061 | -6.900295382 | 4.00722531 | 3.686303414 | 5.930865648 | 2.868578001 | 2.86123014 | 2.68698273 | 2.277179783 |
| ENSG00000160219 | GAB3 | -3.074736351 | 0.297702978 | -3.8265522 | 0.17543064 | -6.901288551 | 2 | 2 | 8.960138144 | 4.71689672 | 4.37434574 | 4.350488971 | 3.430065404 |
| ENSG00000085511 | MAP3K4 | -9.193262886 | 8.41E-08 | 2.290101112 | 0.003182712 | -6.903161773 | 5.095959458 | 5.457120953 | 6.073378139 | 2 | 2 | 8.055236437 | 7.446882182 |
| ENSG00000186517 | ARHGAP30 | -5.656725498 | 0.003872181 | -1.256663467 | 0.553184301 | -6.913388966 | 6.009025997 | 2 | 7.029880454 | 2.141737246 | 2.59489879 | 5.390603804 | 4.161535617 |
| ENSG00000234370 | | 2.53827299 | 0.42279064 | -9.453460001 | 0.003089496 | -6.91518701 | 2 | 5.946638857 | 2 | 6.89733479 | 6.8563927 | 2 | 2 |
| ENSG00000171596 | NMUR1 | -5.096485169 | 0.000446568 | -1.822064017 | 0.13626007 | -6.918549187 | 4.815609471 | 3.686303414 | 5.156626401 | 2.141737246 | 2.2680111 | 3.500668998 | 3.100202476 |
| ENSG00000263324 | | 1.84449953 | 0.58949285 | -8.763419456 | 0.008958685 | -6.918919926 | 2 | 5.303523747 | 2 | 5.48800675 | 5.64780787 | 2 | 2 |
| ENSG00000117983 | MUC5B | -6.365294729 | 0.15396863 | -0.556177131 | 0.908964084 | -6.92147186 | 2 | 2 | 4.523675391 | 2 | 2 | 2.746468522 | 3.287118176 |
| ENSG00000136011 | STAB2 | -5.77466907 | 0.240025405 | -1.148955839 | 0.823280183 | -6.923624909 | 2 | 2 | 4.058998049 | 2 | 2 | 2.746468522 | 2.326754077 |
| ENSG00000165685 | TMEM52B | -4.806729887 | 1.68E-06 | -2.118733091 | 0.011322831 | -6.925462978 | 5.532353208 | 5.303523747 | 4.874587708 | 2.3892346 | 2.38539787 | 3.44788957 | 3.698365557 |
| ENSG00000167747 | C19orf48 | -3.35843027 | 2.10E-13 | -3.569330345 | 2.97E-15 | -6.927760616 | 11.11682541 | 10.22575081 | 10.18377222 | 7.234487582 | 7.26914476 | 7.084714955 | 7.008112295 |
| FAM63A | | 2.405452298 | 0.453804781 | -9.339539276 | 0.003688051 | -6.934086977 | 2 | 5.838937808 | 2 | 6.693715713 | 6.58294721 | 2 | 2 |
| KIRREL | | 5.158291542 | 0.007775879 | -12.09743171 | 4.23E-07 | -6.93914017 | 8.138771685 | 6.535001748 | 2 | 12.08984274 | 12.09486 | 2 | 2 |
| ENSG00000182108 | DEXI | -1.686075257 | 0.032137123 | -5.253424417 | 1.42E-13 | -6.939499674 | 9.015175615 | 8.968383114 | 10.66102425 | 8.098332103 | 8.11451391 | 4.813804156 | 4.683176719 |
| ENSG00000112245 | PTP4A1 | 1.578118641 | 9.13E-06 | -8.526874136 | 3.10E-94 | -6.948755494 | 10.00110988 | 10.17682307 | 9.636156974 | 11.59292826 | 11.4681476 | 2.831336763 | 2.632958241 |
| ENSG00000118596 | SLC16A7 | -9.47604362 | 8.87E-09 | 2.526053271 | 2.14E-05 | -6.949990349 | 6.138360429 | 5.457120953 | 5.930865648 | 2 | 2 | 8.283862644 | 8.303661547 |
| ENSG00000243789 | JMJD7 | -1.875520376 | 0.087812072 | -5.078823983 | 4.39E-07 | -6.954344359 | 7.485143553 | 5.457120953 | 5.772716771 | 4.647285135 | 4.95174433 | 2.937246078 | 2.466008509 |
| ENSG00000111716 | LDHB | -1.920607782 | 0.000159152 | -5.03562715 | 8.57E-26 | -6.956234932 | 13.27377117 | 12.34163262 | 12.23415624 | 10.79834448 | 10.7574572 | 7.52030827 | 7.838397174 |
| ENSG00000198795 | ZNF521 | -2.650621421 | 1.63E-05 | -4.30934473 | 3.85E-13 | -6.959966151 | 9.189468652 | 9.994333773 | 8.845626171 | 6.710878132 | 6.92586261 | 5.630526941 | 4.784231301 |
| ENSG00000174842 | GLMN | -2.62118417 | 2.05E-07 | -4.346827941 | 1.02E-18 | -6.968012111 | 8.444143041 | 8.004992147 | 7.818184403 | 5.552894743 | 5.63601114 | 4.397636241 | 3.771796761 |
| ENSG00000196381 | ZNF781 | -2.633529135 | 0.231366574 | -4.337045341 | 0.035498586 | -6.970574476 | 2 | 3.686303414 | 7.691296561 | 4.17310023 | 3.65239003 | 2.746468522 | 3.042026731 |
| ENSG00000188033 | ZNF490 | -2.123872662 | 0.010511796 | -4.852995767 | 3.16E-10 | -6.976868429 | 8.836089686 | 7.315436877 | 9.491321578 | 6.567497652 | 6.84618961 | 4.45225155 | 4.047883918 |
| ENSG00000151617 | EDNRA | -10.10846269 | 2.18E-05 | 3.125221379 | 0.108834364 | -6.983241314 | 2 | 7.467835385 | 6.322058544 | 2 | 2 | 9.660147759 | 9.353244897 |
| ENSG00000110169 | HPX | -3.234946827 | 0.104928256 | -3.757255872 | 0.050341437 | -6.992202699 | 2 | 4.442710129 | 7.225742779 | 3.349643765 | 3.2184422 | 3.316666208 | 2.592996859 |
| ENSG00000144559 | TAMM41 | -2.647638335 | 0.001742638 | -4.348741952 | 6.87E-08 | -6.996380287 | 8.737455672 | 6.535001748 | 8.585098385 | 5.742861563 | 5.6121243 | 4.321442857 | 4.09155917 |
| ENSG00000248487 | ABHD14A | -2.921066707 | 5.43E-11 | -4.080501577 | 1.01E-20 | -7.001568284 | 10.16946352 | 9.461450882 | 10.09465441 | 6.975156635 | 7.13309042 | 5.614651724 | 6.222348126 |
| ENSG00000141485 | SLC13A5 | -5.353356344 | 0.321299399 | -1.650007447 | NA | -7.003363792 | 2 | 3.686303414 | 2 | 2 | 2 | 2.492221014 | 2.117335275 |
| ENSG00000125510 | OPRL1 | -3.10342041 | 0.111760424 | -3.900196091 | 0.037278996 | -7.003616502 | 2 | 6.389896633 | 8.608698767 | 4.268917409 | 4.63360314 | 4.209638631 | 3.36036153 |
| ENSG00000187950 | OVCH1 | -4.714846799 | NA | -2.290034007 | NA | -7.004880806 | 3.327711234 | 2 | 2 | 2 | 2 | 2.048074405 | 2.225841198 |
| ENSG00000212993 | POU5F1B | -4.714846799 | NA | -2.290034007 | NA | -7.004880806 | 2 | 2 | 3.369317972 | 2 | 2 | 2.048074405 | 2.225841198 |
| ENSG00000233670 | PIRT | -4.714846799 | NA | -2.290034007 | NA | -7.004880806 | 2 | 2 | 3.369317972 | 2 | 2 | 2.048074405 | 2.225841198 |
| ENSG00000182054 | IDH2 | -5.479192863 | 3.25E-12 | -1.525935501 | 0.081867095 | -7.005128364 | 12.96599706 | 11.11496145 | 10.94216274 | 6.610808945 | 6.52655952 | 10.39976816 | 10.52657062 |
| FAM35A | | 3.311894226 | 0.104743804 | -10.32039177 | 1.93E-05 | -7.008497545 | 4.00722531 | 6.602424674 | 2 | 8.521127045 | 8.44542686 | 2 | 2 |
| ENSG00000173198 | CYSLTR1 | -5.356213821 | 0.303739764 | -1.654712232 | NA | -7.010926053 | 2 | 3.686303414 | 2 | 2 | 2 | 2.3070652 | 2.326754077 |
| ENSG00000186376 | ZNF75D | -8.826762869 | 0.010358674 | 1.813054923 | 0.600860223 | -7.013707947 | 2 | 6.728495673 | 2 | 2 | 2 | 7.215456797 | 6.649933445 |
| ENSG00000230341 | | 4.652755098 | 0.000455587 | -11.66680928 | 4.87E-09 | -7.014054182 | 3.327711234 | 6.951736442 | 7.225742779 | 11.12849931 | 11.1856376 | 2 | 2 |
| ENSG00000282725 | | 2.490622278 | 0.257707314 | -9.505995517 | 0.00014105 | -7.015373239 | 3.327711234 | 5.838937808 | 2 | 6.836083475 | 6.92586261 | 2 | 2 |
| ENSG00000273814 | | -2.54495056 | 0.007426265 | -4.479319824 | 6.57E-07 | -7.024270384 | 7.218819781 | 7.274681896 | 6.203069917 | 4.694064037 | 4.58534546 | 2.592885947 | 3.698365557 |
| ENSG00000167769 | ACER1 | -3.258919674 | 0.42533738 | -3.770356692 | NA | -7.029276366 | 4.00722531 | 2 | 2 | 2.270786764 | 2 | 2.139668783 | 2.059860178 |
| ENSG00000158402 | CDC25C | -4.057995483 | 0.017176325 | -2.973161929 | 0.073368879 | -7.031157411 | 5.866944937 | 4.11352228 | 4.874587708 | 2.498691189 | 2.59489879 | 2.139668783 | 3.518055793 |
| ENSG00000162063 | CCNF | -3.744447104 | 0.000715365 | -3.287378557 | 0.0031038 | -7.031825661 | 9.966689162 | 6.899080544 | 7.691296561 | 5.258265913 | 5.13053105 | 5.880003936 | 5.275717819 |
| ENSG00000108786 | HSD17B1 | -4.975355424 | 1.32E-07 | -2.059772089 | 0.036346775 | -7.035127513 | 8.586954094 | 7.928237109 | 5.930865648 | 3.406930997 | 3.55551834 | 6.157074092 | 5.492143571 |
| ENSG00000101180 | HRH3 | -2.279698943 | NA | -4.755634545 | NA | -7.035333488 | 3.327711234 | 2 | 2 | 2 | 2.2680111 | 2.048074405 | 2 |
| ENSG00000164035 | EMCN | -6.220805942 | 0.048800801 | -0.822700439 | 0.793235017 | -7.043506382 | 2 | 3.076665209 | 4.058998049 | 2 | 2 | 2.803598391 | 2.885414003 |
| ENSG00000118418 | HMGN3 | -3.101489362 | 1.51E-11 | -3.943026506 | 2.60E-18 | -7.044515869 | 10.39976609 | 9.479495904 | 10.46161637 | 7.071314871 | 7.14977896 | 6.200262696 | 6.397863414 |
| ENSG00000150627 | WDR17 | -3.438114238 | 0.061530373 | -3.610434087 | 0.034202855 | -7.048548325 | 4.00722531 | 3.686303414 | 4.058998049 | 2.270786764 | 2.38539787 | 2.492221014 | 2.059860178 |
| ENSG00000112182 | BACH2 | -3.75424811 | 0.001787884 | -3.306294099 | 0.00487318 | -7.06054221 | 5.095959458 | 7.099070191 | 4.874587708 | 3.462030052 | 2.68924543 | 3.411599105 | 3.36036153 |
| ENSG00000113262 | GRM6 | -4.346470638 | 0.002731351 | -2.723306076 | 0.04476747 | -7.069776714 | 4.00722531 | 5.131603625 | 5.156626401 | 2.3892346 | 2.38539787 | 3.374372158 | 2.374681325 |
| ENSG00000128908 | INO80 | -9.781593735 | 9.76E-07 | 2.709802537 | 0.045524045 | -7.071791198 | 3.327711234 | 6.666836837 | 6.629488387 | 2 | 2 | 8.56358163 | 8.964334843 |
| ENSG00000144648 | ACKR2 | -5.412912389 | 0.001841909 | -1.661048135 | 0.250598944 | -7.073960524 | 4.815609471 | 3.076665209 | 4.874587708 | 2.141737246 | 2.14022021 | 3.500668998 | 2.950113106 |
| ENSG00000155761 | SPAG17 | -3.939960696 | NA | -3.137124751 | NA | -7.077085448 | 2 | 3.686303414 | 2 | 2 | 2.14022021 | 2.183373728 | 2.059860178 |
| ENSG00000070748 | CHAT | -7.374391673 | 0.063677965 | 0.297288468 | 0.945727701 | -7.077103205 | 2 | 2 | 5.39244166 | 2 | 2 | 3.694366604 | 4.692661645 |
| ENSG00000131771 | PPP1R1B | -8.381054853 | 0.0240574 | 1.303754532 | 0.736281057 | -7.077300321 | 2 | 2 | 6.322058544 | 2 | 2 | 4.764298067 | 6.716538017 |
| ENSG00000151093 | OXSM | -8.626633086 | 0.013163141 | 1.544505659 | 0.664743846 | -7.082127427 | 2 | 6.535001748 | 2 | 2 | 2 | 6.723178551 | 6.257923755 |
| ENSG00000166800 | LDHAL6A | -5.717190871 | 0.245408313 | -1.366196483 | NA | -7.083387354 | 4.00722531 | 2 | 2 | 2 | 2 | 2.384001718 | 2.551896972 |
| ENSG00000242866 | STRC | -2.203021367 | 0.268445403 | -4.881805824 | 0.009775788 | -7.084827191 | 3.327711234 | 4.442710129 | 4.058998049 | 2.868578001 | 2.59489879 | 2.267001569 | 2 |
| ENSG00000150625 | GPM6A | -1.723001656 | 0.673891173 | -5.371237693 | 0.143595385 | -7.094239349 | 2 | 3.686303414 | 5.772716771 | 3.997744386 | 2 | 2.3070652 | 2 |
| ENSG00000277322 | GOLGA6L6 | -0.948849888 | NA | -6.152895398 | NA | -7.101745286 | 2 | 3.076665209 | 2 | 2.3892346 | 2.2680111 | 2 | 2 |
| ENSG00000141527 | CARD14 | -4.139825693 | 0.014871185 | -2.964134098 | 0.075781139 | -7.103959792 | 5.866944937 | 5.838937808 | 2 | 2.498691189 | 2.68924543 | 3.465698119 | 2.632958241 |
| ENSG00000262164 | | -0.951975516 | NA | -6.152900175 | NA | -7.104875692 | 3.327711234 | 2 | 2 | 2.270786764 | 2.38539787 | 2 | 2 |
| ENSG00000149090 | PAMR1 | -3.403897696 | 1.40E-12 | -3.702244944 | 7.52E-15 | -7.106142641 | 12.7464448 | 12.011946 | 11.58978828 | 8.74173768 | 8.86476809 | 8.545902039 | 8.471469752 |
| ENSG00000103196 | CRISPLD2 | -4.594103397 | 4.70E-18 | -2.513817703 | 3.04E-06 | -7.1079211 | 9.730679447 | 10.70750522 | 10.5677762 | 5.818703821 | 5.9725602 | 8.255350111 | 7.432801564 |
| ENSG00000147689 | FAM83A | -0.955123359 | NA | -6.152895424 | NA | -7.108018783 | 2 | 3.076665209 | 2 | 2.141737246 | 2.49394783 | 2 | 2 |
| ENSG00000281618 | | -0.394987181 | 0.791895481 | -6.722049756 | 2.19E-08 | -7.117036938 | 8.16936288 | 5.838937808 | 5.156626401 | 6.658765962 | 6.54560135 | 2.492221014 | 2.225841198 |
| ENSG00000203880 | PCMTD2 | -2.366075884 | 0.09480535 | -4.751033479 | 0.000240677 | -7.117109363 | 6.009025997 | 10.29361762 | 11.06667478 | 7.88809286 | 7.76511961 | 5.677130623 | 5.402507939 |
| ENSG00000187266 | EPOR | -3.768379058 | 3.05E-08 | -3.351974948 | 7.97E-07 | -7.120354006 | 10.19198706 | 9.566483142 | 8.511888463 | 5.764940652 | 6.02777321 | 6.617616961 | 5.876154317 |
| KIAA1468 | | 3.400312508 | 0.141109656 | -10.52116369 | 5.18E-05 | -7.120851181 | 3.327711234 | 6.899080544 | 2 | 8.769305409 | 8.77079897 | 2 | 2 |
| ENSG00000115155 | OTOF | -2.901569052 | 0.118359009 | -4.219995301 | 0.01651049 | -7.121564353 | 4.467311984 | 2 | 5.930865648 | 2.947924294 | 2.77779907 | 2.3070652 | 2.509591799 |
| ENSG00000267467 | APOC4 | -3.266366214 | NA | -3.85538923 | NA | -7.121755443 | 2 | 2 | 3.369317972 | 2.141737246 | 2 | 2.094598369 | 2 |
| ENSG00000076944 | STXBP2 | -8.940725521 | 1.83E-05 | 1.817349165 | 0.237913947 | -7.123376357 | 4.00722531 | 6.464272742 | 4.058998049 | 2 | 2 | 6.60762613 | 7.427130611 |
| ENSG00000140263 | SORD | -2.751401586 | 0.000444894 | -4.373655282 | 5.78E-09 | -7.125056868 | 11.30767305 | 9.090820894 | 10.55568472 | 7.810605345 | 7.88209639 | 6.060766632 | 6.463051659 |
| ENSG00000117407 | ARTN | -3.291869925 | 0.228339937 | -3.833746182 | 0.145444579 | -7.125616107 | 7.33131396 | 2 | 2 | 2.947924294 | 3.45167086 | 2.987421008 | 2.885414003 |

| | | | | | | | | | | | | | |
|---|---|---|---|---|---|---|---|---|---|---|---|---|---|
| ENSG00000175267 | VWA3A | -3.735287495 | 0.205850144 | -3.391593819 | 0.223815362 | -7.126881315 | 2 | 2 | 4.874587708 | 2.270786764 | 2.14022021 | 2.384001718 | 2.117335275 |
| ENSG00000257949 | TEN1 | 0.285901302 | 0.815735054 | -7.422753287 | 3.61E-13 | -7.136851985 | 5.532353208 | 7.099070191 | 7.896971941 | 7.409376773 | 7.41433569 | 2.384001718 | 2.117335275 |
| ENSG00000001630 | CYP51A1 | -1.512557732 | 8.90E-05 | -5.626604663 | 1.27E-54 | -7.139162395 | 12.03475608 | 12.29526798 | 12.5064448 | 10.81166375 | 10.7499517 | 6.363520437 | 7.006211772 |
| ENSG00000263247 | | 1.138688392 | 0.611390057 | -8.279959 | 0.000717338 | -7.141270608 | 4.00722531 | 4.11352228 | 2 | 4.358765249 | 4.68029874 | 2 | 2 |
| ENSG00000204052 | LRRC73 | -3.476430874 | 0.156635009 | -3.673518055 | 0.124283687 | -7.149948929 | 2 | 3.076665209 | 8.884815469 | 3.79808662 | 4.4836996 | 4.209638631 | 3.824516949 |
| ENSG00000275331 | | -2.361071219 | 0.292493649 | -4.803703344 | 0.020998348 | -7.164774563 | 2 | 4.11352228 | 6.073378139 | 3.289987342 | 3.01487888 | 2.048074405 | 2.509591799 |
| ENSG00000146674 | IGFBP3 | -3.68734947 | 1.66E-10 | -3.479648448 | 1.71E-09 | -7.166997918 | 18.91474819 | 17.97982537 | 17.67771703 | 14.61266148 | 14.5955599 | 15.15895754 | 14.35371509 |
| ENSG00000101670 | LIPG | -3.371234418 | 0.022265894 | -3.802369345 | 0.008354549 | -7.173603764 | 4.815609471 | 8.848535039 | 8.042706615 | 5.501219927 | 3.39680241 | 3.930690651 | 4.819291477 |
| ENSG00000163060 | TEKT4 | -0.788086037 | 0.835945164 | -6.385655728 | 0.048778416 | -7.173741765 | 2 | 2 | 6.073378139 | 4.205751501 | 3.7864959 | 2 | 2.172608095 |
| ENSG00000185883 | ATP6V0C | 0.919084527 | 0.312710697 | -8.093079573 | 6.01E-26 | -7.173995046 | 11.29385609 | 10.43440824 | 9.227351203 | 11.44093988 | 11.4908391 | 2.911488689 | 3.518055793 |
| ENSG00000125966 | MMP24 | -3.748860756 | 0.001718833 | -3.427985143 | 0.003544773 | -7.176845899 | 4.00722531 | 6.464272742 | 7.502634626 | 3.349643765 | 3.50452879 | 3.767980144 | 3.496556178 |
| ENSG00000144354 | CDCA7 | -4.208562527 | 0.0051495 | -2.978850774 | 0.043361049 | -7.187413301 | 5.532353208 | 6.311476754 | 3.369317972 | 2.947924294 | 2.2680111 | 3.5515855 | 2.782576693 |
| ENSG00000132386 | SERPINF1 | -2.285983518 | 0.00129592 | -4.904121042 | 2.05E-13 | -7.19010456 | 12.20867639 | 11.49353862 | 10.62096771 | 9.316137634 | 9.28868707 | 7.15911073 | 6.117095472 |
| ENSG00000113073 | SLC4A9 | -4.714849116 | NA | -2.4755493 | NA | -7.190398416 | 3.327711234 | 2 | 2 | 2 | 2 | 2.183373728 | 2.059860178 |
| ENSG00000143867 | OSR1 | -4.466187301 | 1.63E-29 | -2.724380871 | 5.97E-12 | -7.190568172 | 11.46082313 | 11.30403016 | 10.6093157 | 6.693715713 | 6.81514011 | 8.4151065 | 8.502810156 |
| ENSG00000152467 | ZSCAN1 | -5.339116445 | 0.049220147 | -1.852179755 | 0.4196504 | -7.1912962 | 2 | 4.442710129 | 3.369317972 | 2.141737246 | 2 | 2.717032197 | 2.592996859 |
| ENSG00000102001 | CACNA1F | -3.844975143 | 0.201485907 | -3.350093036 | 0.228416028 | -7.195068179 | 4.467311984 | 2 | 2 | 2.141737246 | 2.14022021 | 2.267001569 | 2.117335275 |
| ENSG00000116833 | NR5A2 | 0.427144668 | 0.889250843 | -7.624265618 | 0.008231431 | -7.19712095 | 4.00722531 | 3.076665209 | 2 | 3.615725078 | 3.39680241 | 2 | 2 |
| ENSG00000185379 | RAD51D | -8.994560501 | 0.000545577 | 1.795291149 | 0.444326413 | -7.199269352 | 4.00722531 | 6.728495673 | 2 | 2 | 2 | 6.91847879 | 7.271271876 |
| ENSG00000138185 | ENTPD1 | -7.14516145 | 3.43E-12 | -0.055052448 | 0.963369049 | -7.200213897 | 6.468621203 | 7.051611315 | 8.65477198 | 2.3892346 | 2.49394783 | 7.713417604 | 7.566491953 |
| MICALCL | | 1.279490899 | 0.739013584 | -8.482857592 | 0.019310208 | -7.203366692 | 5.095959458 | 2 | 2 | 4.694064037 | 4.93258395 | 2 | 2 |
| ENSG00000255974 | CYP2A6 | -5.921966286 | 0.213235791 | -1.283293243 | 0.795527614 | -7.205259529 | 2 | 4.11352228 | 2 | 2 | 2 | 2.717032197 | 2.374681325 |
| ENSG00000131831 | RAI2 | -3.584719693 | 0.169023265 | -3.622880325 | 0.151262675 | -7.207600017 | 2 | 6.728495673 | 2 | 2.695456494 | 2.86123014 | 2.885263063 | 2.632958241 |
| ENSG00000235169 | SMIM1 | -3.493803484 | 0.011522367 | -3.716215776 | 0.005905716 | -7.21001926 | 6.138360429 | 7.638155693 | 3.369317972 | 3.566290607 | 3.60476711 | 3.429858444 | 3.45257121 |
| ENSG00000163541 | SUCLG1 | -3.61603909 | 1.62E-24 | -3.599096157 | 1.02E-24 | -7.215135247 | 12.2979074 | 11.55226345 | 12.0110149 | 8.31771326 | 8.44880239 | 8.350677018 | 8.450657333 |
| ENSG00000111641 | NOP2 | -1.060293117 | 0.148044571 | -6.162148017 | 5.22E-22 | -7.222441134 | 8.653385977 | 8.607278467 | 9.999667074 | 8.192977693 | 8.185025 | 3.767980144 | 3.474731315 |
| ENSG00000169918 | OTUD7A | -1.57398932 | 0.665819919 | -5.649699528 | 0.08996128 | -7.223688849 | 2 | 2 | 5.156626401 | 2.868578001 | 2.94009902 | 2.139668783 | 2 |
| ENSG00000168334 | XIRP1 | -6.351386854 | 0.00128458 | -0.8756172 | 0.706090029 | -7.227004054 | 6.468621203 | 2 | 9.287079172 | 2.784612485 | 2.77779907 | 6.599583349 | 7.382408159 |
| ENSG00000108839 | ALOX12 | -7.812000502 | 0.003103036 | 0.584387153 | 0.820330666 | -7.227613348 | 5.532353208 | 3.686303414 | 2 | 2 | 2 | 4.977914261 | 4.77533149 |
| ENSG00000278016 | | 1.477988164 | 0.273697678 | -8.70994383 | 5.47E-06 | -7.231955667 | 4.00722531 | 6.140560021 | 4.058998049 | 6.529309835 | 6.49425375 | 2.048074405 | 2 |
| ENSG00000236251 | | 2.219081045 | 0.499226274 | -9.452439843 | 0.003709986 | -7.233358798 | 2 | 5.946638857 | 2 | 6.548530093 | 6.5890786 | 2 | 2 |
| ENSG00000196247 | ZNF107 | -8.06118447 | 0.003497481 | 0.827591063 | 0.759279214 | -7.233593408 | 3.327711234 | 5.838937808 | 2 | 2 | 2 | 5.489669427 | 5.115701921 |
| ENSG00000048545 | GUCA1A | -4.497800463 | 0.106714989 | -2.740822152 | 0.290479733 | -7.238622615 | 2 | 4.93638933 | 2 | 2.141737246 | 2.14022021 | 2.34604626 | 2.509591799 |
| ENSG00000173171 | MTX1 | -2.329923526 | 8.13E-11 | -4.910879187 | 2.32E-44 | -7.240802713 | 10.30651143 | 9.763446433 | 9.647595056 | 7.671425626 | 7.5835711 | 5.180212365 | 5.191251321 |
| ENSG00000114115 | RBP1 | -5.426888414 | 2.00E-06 | -1.814534877 | 0.132538191 | -7.241423291 | 7.579279349 | 8.147241016 | 6.073378139 | 3.515101962 | 2.2680111 | 5.912830251 | 5.601925227 |
| ENSG00000130383 | FUT5 | -2.180897957 | 0.560280703 | -5.068594121 | NA | -7.249492078 | 2 | 4.11352228 | 2 | 2.3892346 | 2.2680111 | 2.094598369 | 2 |
| ENSG00000165061 | ZMAT4 | -0.009001438 | 0.997041173 | -7.245577349 | 0.0017986 | -7.254578788 | 4.00722531 | 4.710564154 | 2 | 4.105493497 | 3.82856379 | 2 | 2.059860178 |
| ENSG00000275591 | XKR5 | -4.099733326 | 0.046591905 | -3.157391475 | 0.101262724 | -7.257124801 | 4.467311984 | 2 | 4.874587708 | 2.3892346 | 2.14022021 | 2.656294034 | 2.277179783 |
| ENSG00000163281 | GNPDA2 | -9.12979646 | 8.28E-05 | 1.868548558 | 0.33137707 | -7.261247902 | 2 | 6.311476754 | 5.772716771 | 2 | 2 | 7.370021622 | 7.244113847 |
| ENSG00000213983 | AP1G2 | -3.424076578 | 2.48E-08 | -3.840861813 | 2.33E-10 | -7.26493839 | 9.858207046 | 8.903033433 | 10.54349105 | 6.535745081 | 6.55815769 | 6.173421703 | 6.127623461 |
| ENSG00000183230 | CTNNA3 | -4.313181095 | 0.268695832 | -2.955567277 | NA | -7.268748372 | 2 | 2 | 4.058998049 | 2 | 2.14022021 | 2.183373728 | 2.172608095 |
| ENSG00000157554 | ERG | -7.415118608 | 0.000447336 | 0.145421179 | 0.955098488 | -7.269697429 | 7.904282243 | 7.572420273 | 2 | 2.498691189 | 2 | 7.138441638 | 7.480125329 |
| ENSG00000176920 | FUT2 | -4.076037969 | 0.003539212 | -3.195766352 | 0.023388267 | -7.271804321 | 5.330594134 | 5.131603625 | 8.380957451 | 3.566290607 | 3.55551834 | 4.667592615 | 3.496556178 |
| ENSG00000123191 | ATP7B | -4.771899015 | 0.000111745 | -2.504504303 | 0.055960012 | -7.276403318 | 6.138360429 | 10.65289571 | 10.26027233 | 5.461211713 | 5.11361705 | 7.636890116 | 7.221358049 |
| ENSG00000130208 | APOC1 | -7.199897653 | 2.59E-05 | -0.08049147 | 0.969279885 | -7.280389123 | 8.564112515 | 5.838937808 | 4.874587708 | 2.270786764 | 2.38539787 | 8.069864557 | 4.614976609 |
| ENSG00000044524 | EPHA3 | -4.528441877 | 0.021694583 | -2.753822464 | 0.167853283 | -7.282264341 | 6.96351253 | 5.595927442 | 3.369317972 | 3.162721474 | 2 | 3.679182183 | 3.600986722 |
| ENSG00000077943 | ITGA8 | -9.740964361 | 8.18E-12 | 2.456847263 | 0.014767163 | -7.284117098 | 6.73718411 | 9.308339287 | 8.325066185 | 2 | 2.2680111 | 10.91760099 | 10.91075957 |
| ENSG00000154548 | SRSF12 | -5.616779291 | 1.34E-07 | -1.671065452 | 0.091257767 | -7.287844743 | 4.815609471 | 6.389896633 | 6.80318718 | 2.600425781 | 2.2680111 | 4.572179576 | 4.836507146 |
| ENSG00000198535 | C2CD4A | -0.927164516 | 0.593368805 | -6.360693908 | 0.000181365 | -7.287858424 | 4.815609471 | 3.076665209 | 4.874587708 | 3.754611435 | 3.69849104 | 2.139668783 | 2 |
| ENSG00000112130 | RNF8 | -9.309382629 | 1.85E-06 | 2.021162054 | 0.119389862 | -7.288220575 | 6.009025997 | 6.311476754 | 3.369317972 | 2 | 2 | 7.608114885 | 7.657697387 |
| ENSG00000187955 | COL14A1 | -6.398570552 | 2.51E-09 | -0.899267399 | 0.469467139 | -7.297837951 | 7.975256505 | 9.876711741 | 8.110401118 | 3.709785322 | 2.68924543 | 6.984820384 | 8.654300638 |
| ENSG00000172296 | SPTLC3 | -5.334385335 | 7.32E-05 | -1.964118453 | 0.173373971 | -7.298503788 | 5.532353208 | 9.114109157 | 5.772716771 | 3.289987342 | 3.15372745 | 5.899789217 | 5.876154317 |
| ENSG00000154274 | C4orf19 | -3.780242667 | 0.033496988 | -3.518278354 | 0.035333386 | -7.298521022 | 2 | 4.11352228 | 5.39244166 | 2.3892346 | 2.38539787 | 2.526562374 | 2.374681325 |
| ENSG00000074964 | ARHGEF10L | -10.02313825 | 7.67E-05 | 2.721253357 | 0.215756786 | -7.301884893 | 5.095959458 | 7.701025996 | 2 | 2 | 2 | 8.898320089 | 9.148351871 |
| ENSG00000178718 | RPP25 | -3.47470936 | 1.67E-06 | -3.829713848 | 9.17E-08 | -7.304423208 | 11.59363617 | 9.799806309 | 10.17589429 | 7.416368504 | 7.1744553 | 7.16729616 | 6.707209278 |
| ENSG00000196433 | ASMT | 1.102825182 | 0.530264818 | -8.40915886 | 8.76E-05 | -7.306333679 | 4.00722531 | 3.076665209 | 4.058998049 | 4.887609717 | 4.2560182 | 2 | 2 |
| ENSG00000112499 | SLC22A2 | -2.941116692 | NA | -4.366530099 | NA | -7.30764679 | 2 | 3.686303414 | 2 | 2 | 2.2680111 | 2.048074405 | 2.059860178 |
| ENSG00000146904 | EPHA1 | -6.366579175 | 0.003151939 | -0.949929746 | 0.56354884 | -7.316508921 | 5.330594134 | 3.686303414 | 3.369317972 | 2.141737246 | 2 | 3.709392871 | 3.640730416 |
| ENSG00000017483 | SLC38A5 | -1.938841357 | 0.232267301 | -5.379119682 | 0.000181791 | -7.317961039 | 5.532353208 | 10.33384851 | 11.48529557 | 8.378048624 | 8.65866647 | 5.622611168 | 4.692661645 |
| ENSG00000198553 | KCNRG | -1.116800039 | 0.445130278 | -6.202649388 | 8.07E-07 | -7.319449427 | 6.816637977 | 3.686303414 | 7.691296561 | 5.775854763 | 5.67111583 | 2.42098416 | 2.466008509 |
| ENSG00000116774 | OLFML3 | -2.026624482 | 0.000112999 | -5.298063024 | 1.42E-26 | -7.324687505 | 11.92025849 | 11.13502303 | 11.3657305 | 9.518946937 | 9.46323044 | 5.751606453 | 6.678855578 |
| ENSG00000155719 | OTOA | -4.60799897 | 0.198940934 | -2.723055667 | 0.434305685 | -7.331054637 | 5.095959458 | 2 | 2 | 2 | 2.2680111 | 2.62493832 | 2.277179783 |
| ENSG00000002933 | TMEM176A | -7.893702893 | 0.043881335 | 0.559083642 | 0.89666844 | -7.334619251 | 2 | 5.838937808 | 2 | 2 | 2 | 5.65789504 | 3.383972333 |
| ENSG00000007255 | TRAPPC6A | -5.028145953 | 8.88E-11 | -2.308998493 | 0.003531764 | -7.337144446 | 8.586954094 | 7.002537973 | 8.865353883 | 3.462030052 | 4.05806467 | 6.221381643 | 5.976631893 |
| ENSG00000131969 | ABHD12B | -7.016177485 | 0.014880133 | -0.330164151 | 0.911348375 | -7.346341636 | 2 | 4.710564154 | 3.369317972 | 2 | 2 | 3.534812207 | 3.580696614 |
| ENSG00000090659 | CD209 | -5.822339823 | 0.011601873 | -1.524888586 | 0.391227832 | -7.347228409 | 3.327711234 | 3.076665209 | 4.874587708 | 2 | 2.14022021 | 3.059555873 | 2.950113106 |
| ENSG00000171119 | NRTN | -4.762463435 | 0.076570922 | -2.585778126 | 0.32269585 | -7.348241562 | 2 | 5.72254323 | 2 | 2.141737246 | 2.2680111 | 2.911488689 | 2.592996859 |
| ENSG00000161040 | FBXL13 | -3.042775762 | 0.00311339 | -4.310516251 | 1.21E-05 | -7.353292013 | 4.00722531 | 6.311476754 | 6.073378139 | 3.406930997 | 3.28037825 | 2.717032197 | 2.70970617 |
| ENSG00000125337 | KIF25 | -3.277360534 | 0.307672642 | -4.07701041 | NA | -7.354370944 | 3.327711234 | 2 | 3.369317972 | 2 | 2.2680111 | 2.048074405 | 2.117335275 |
| ENSG00000181856 | SLC2A4 | -5.422764569 | 0.000189892 | -1.931970692 | 0.190983933 | -7.354735261 | 4.467311984 | 4.442710129 | 7.502634626 | 2.498691189 | 2.49394783 | 4.133696865 | 4.77533149 |
| ENSG00000143318 | CASQ1 | -7.155722656 | 0.031793498 | -0.200078167 | 0.955794916 | -7.355800823 | 2 | 2 | 6.534109959 | 2.141737246 | 2 | 5.279204176 | 4.341135913 |
| ENSG00000064300 | NGFR | -3.145990044 | 0.30948285 | -4.212838646 | 0.152092183 | -7.35882869 | 8.16936288 | 2 | 2 | 3.881293667 | 3.86943969 | 2.183373728 | 3.771796761 |
| ENSG00000153093 | ACOXL | -5.717458901 | 0.248966881 | -1.644857412 | NA | -7.362316314 | 4.00722531 | 2 | 2 | 2 | 2 | 2.42098416 | 2.374681325 |
| ENSG00000145912 | NHP2 | -3.293430786 | 1.02E-26 | -4.070683353 | 2.57E-41 | -7.364114139 | 11.24801458 | 10.85648059 | 11.23155829 | 7.865280446 | 7.83643742 | 6.889085418 | 7.269688419 |
| ENSG00000188833 | ENTPD8 | -3.934371945 | NA | -3.442966189 | NA | -7.377338134 | 2 | 3.686303414 | 2 | 2.141737246 | 2 | 2.139668783 | 2.059860178 |
| | TMEM2 | 4.464384879 | 0.016810729 | -11.85656107 | 3.38E-07 | -7.392176194 | 2 | 7.638155693 | 6.882775709 | 11.15094209 | 11.1665449 | 2 | 2 |
| ENSG00000148357 | HMCN2 | -4.980558743 | 0.023814605 | -2.415407066 | 0.252589462 | -7.395965809 | 5.866944937 | 3.076665209 | 2 | 2.141737246 | 2.2680111 | 2.746468522 | 3.100202476 |
| ENSG00000027869 | SH2D2A | -4.610475974 | 0.011054842 | -2.789299276 | 0.130582534 | -7.399775249 | 7.384433967 | 2 | 6.073378139 | 2.600425781 | 3.01487888 | 4.122514202 | 3.620995423 |
| ENSG00000183873 | SCN5A | -8.731545354 | 5.66E-17 | 1.327858406 | 0.034579718 | -7.403686948 | 8.285626337 | 7.232742069 | 8.408111533 | 2.270786764 | 2.14022021 | 9.348125304 | 9.39051252 |
| ENSG00000083782 | EPYC | -4.714851945 | NA | -2.702126509 | NA | -7.416978454 | 2 | 3.076665209 | 2 | 2 | 2 | 2.094598369 | 2.117335275 |
| ENSG00000163075 | CFAP221 | -4.714851945 | NA | -2.702126509 | NA | -7.416978454 | 2 | 3.076665209 | 2 | 2 | 2 | 2.094598369 | 2.117335275 |
| ENSG00000187416 | LHFPL3 | -1.264590424 | NA | -6.152896712 | NA | -7.417487136 | 3.327711234 | 2 | 2 | 2.498691189 | 2 | 2 | 2 |
| ENSG00000076826 | CAMSAP3 | -4.110428619 | 0.215385243 | -3.307109735 | NA | -7.417538354 | 3.327711234 | 3.076665209 | 2 | 2.141737246 | 2 | 2.183373728 | 2.059860178 |
| ENSG00000206450 | | 6.82787963 | 3.49E-09 | -14.25098675 | 5.44E-14 | -7.423107115 | 10.28571918 | 7.35507212 | 7.858115942 | 15.91381407 | 15.914652 | 2 | 2 |
| ENSG00000169989 | TIGD4 | -3.370163206 | 0.150595623 | -4.056211772 | 0.069988141 | -7.426374978 | 6.257048221 | 3.076665209 | 2 | 2.695456494 | 2.68924543 | 2.746468522 | 2.117335275 |
| ENSG00000122824 | NUDT10 | -2.463081335 | 0.006270092 | -4.970281116 | 8.37E-09 | -7.433362451 | 5.532353208 | 6.04685525 | 7.285536013 | 4.237680128 | 4.31639476 | 2.911488689 | 2.551896972 |
| ENSG00000100288 | CHKB | -1.251876575 | 0.286739789 | -6.18222984 | 6.12E-10 | -7.434106415 | 10.47516499 | 8.147241016 | 7.646393253 | 8.115618198 | 8.02689124 | 3.709392871 | 3.620995423 |
| ENSG00000162592 | CCDC27 | -5.354606606 | NA | -2.083569118 | NA | -7.438175723 | 2 | 3.686303414 | 2 | 2 | 2 | 2.3070652 | 2.172608095 |
| ENSG00000177669 | MBOAT4 | -5.354606606 | NA | -2.083569118 | NA | -7.438175723 | 2 | 3.686303414 | 2 | 2 | 2 | 2.3070652 | 2.172608095 |
| ENSG00000275066 | SYNRG | 2.096089733 | 4.03E-08 | -9.535470391 | 3.75E-24 | -7.439380658 | 7.384433967 | 7.538394143 | 7.342949392 | 9.481102223 | 9.51728265 | 2.048074405 | 2.117335275 |
| ENSG00000077348 | EXOSC5 | -3.774743395 | 2.05E-27 | -3.667449417 | 1.36E-27 | -7.442192812 | 9.27634997 | 9.125613817 | 9.647595056 | 5.651002631 | 5.73886209 | 5.916072173 | 5.661099865 |
| ENSG00000134548 | SPX | -4.214016713 | 0.044421992 | -3.233175701 | 0.100117636 | -7.447192414 | 4.467311984 | 4.93638933 | 2 | 2.141737246 | 2.38539787 | 2.746468522 | 2.172608095 |
| ENSG00000169291 | SHE | -4.277328697 | 0.03787204 | -3.170841277 | 0.091135376 | -7.448169974 | 4.815609471 | 2 | 4.058998049 | 2.141737246 | 2.2680111 | 2.492221014 | 2.326754077 |
| ENSG00000259803 | SLC22A31 | -3.266144789 | 0.236109579 | -4.182088972 | 0.112822788 | -7.448233761 | 2 | 2 | 7.646393253 | 3.462030052 | 3.39680241 | 2.831336763 | 3.012034821 |
| ENSG00000242114 | MTFP1 | -2.299003718 | 0.037305226 | -5.149943633 | 4.03E-07 | -7.448947352 | 5.866944937 | 7.847167909 | 9.099922433 | 5.85002558 | 5.9536761 | 3.393105702 | 3.698365557 |
| ENSG00000114631 | PODXL2 | -4.195090941 | 0.005822636 | -3.25538067 | 0.035456159 | -7.450471612 | 9.771132748 | 6.899080544 | 4.523675391 | 4.329433326 | 4.60967607 | 5.449961224 | 5.101530266 |
| ENSG00000162650 | ATXN7L2 | -3.685579341 | 0.092779568 | -3.775324588 | 0.07900979 | -7.460903929 | 10.19198706 | 6.228547981 | 2 | 5.161714397 | 5.18001485 | 5.385935071 | 4.720747915 |
| ENSG00000112493 | | 1.019555333 | 0.178423108 | -8.484508813 | 4.04E-38 | -7.464953481 | 9.90440318 | 10.71514053 | 11.698263 | 11.94339693 | 12.0045649 | 3.193831101 | 3.311948089 |
| FLJ45513 | | -2.157159825 | 0.186455215 | -5.308015409 | 0.000470077 | -7.465175234 | 4.00722531 | 3.686303414 | 6.43197674 | 3.754611435 | 3.28037825 | 2.42098416 | 2.117335275 |
| ENSG00000171004 | HS6ST2 | -4.407404017 | 0.012745818 | -3.068524083 | 0.066070408 | -7.4759281 | 4.815609471 | 5.457120953 | 2 | 2.3892346 | 2.2680111 | 2.937246078 | 2.466008509 |
| ENSG00000175938 | ORAI3 | -2.893394265 | 8.76E-05 | -4.587678293 | 1.07E-10 | -7.481072559 | 9.966689162 | 10.84955954 | 11.95917313 | 8.256693138 | 8.29225867 | 6.823267818 | 6.38621943 |
| ENSG00000188687 | SLC4A5 | -2.94702531 | 0.004653794 | -4.54774615 | 4.83E-06 | -7.49477146 | 5.330594134 | 8.004992147 | 7.225742779 | 4.647285135 | 4.37434574 | 3.648325314 | 2.981406154 |
| ENSG00000130035 | GALNT8 | -3.496890138 | 0.200007734 | -4.001562792 | 0.119659197 | -7.49845293 | 2 | 4.93638933 | 2 | 2.270786764 | 2.2680111 | 2.267001569 | 2.117335275 |
| ENSG00000053770 | AP5M1 | -10.05192841 | 1.07E-06 | 2.549097282 | 0.085204406 | -7.502831124 | 6.257048221 | 7.35507212 | 3.369317972 | 2 | 2 | 8.971818328 | 8.795792144 |
| ENSG00000244694 | PTCHD4 | -3.620098403 | 0.007552518 | -3.889756118 | 0.003544156 | -7.509854521 | 10.16946352 | 8.747909548 | 6.203069917 | 5.565529336 | 5.63601114 | 6.05195199 | 3.956354764 |
| ENSG00000159496 | RGL4 | -7.210263709 | 0.014740108 | -0.308147262 | 0.920627804 | -7.518410971 | 3.327711234 | 4.93638933 | 2 | 2 | 2 | 4.099885429 | 3.15612297 |
| ENSG00000099954 | CECR2 | -4.469420821 | 0.01227951 | -3.054120226 | 0.089598569 | -7.523541047 | 3.327711234 | 4.442710129 | 7.646393253 | 3.162721474 | 2.49394783 | 4.155805465 | 2.950113106 |

| | | | | | | | | | | | | | |
|---|---|---|---|---|---|---|---|---|---|---|---|---|---|
| ENSG00000147180 | ZNF711 | -3.601214357 | 0.021326273 | -3.926202568 | 0.01054493 | -7.527416925 | 9.304185368 | 7.928237109 | 3.369317972 | 4.694064037 | 4.91316568 | 4.713032626 | 4.329022995 |
| ENSG00000119946 | CNNM1 | -1.988513466 | 0.338460636 | -5.540646221 | 0.003325449 | -7.529159687 | 2 | 6.666836837 | 4.874587708 | 3.406930997 | 4.1930041 | 2.094598369 | 2.466008509 |
| ENSG00000261984 | | 0.235311236 | 0.957988784 | -7.77474648 | 0.051766586 | -7.539435244 | 2 | 4.442710129 | 2 | 3.515101962 | 3.45167086 | 2 | 2 |
| ENSG00000277641 | | 0.8765805 | 0.510230014 | -8.417217731 | 7.17E-06 | -7.540637231 | 3.327711234 | 4.93638933 | 5.59507254 | 5.627094009 | 5.71663161 | 2.048074405 | 2 |
| ENSG00000119725 | ZNF410 | -0.323873996 | 0.567927475 | -7.218000242 | 6.72E-52 | -7.541874238 | 9.886101849 | 9.079034257 | 9.385998894 | 9.147810675 | 9.18219179 | 3.316666208 | 2.918126211 |
| ENSG00000276234 | TADA2A | -0.006494088 | 0.997264364 | -7.535964045 | 1.13E-05 | -7.542458133 | 3.327711234 | 5.131603625 | 6.958202216 | 5.860316854 | 5.74985018 | 2.183373728 | 2 |
| ENSG00000166323 | C11orf65 | -5.461781934 | 0.035685956 | -2.082922548 | 0.327076201 | -7.544704481 | 4.467311984 | 3.686303414 | 2 | 2 | 2.14022021 | 2.560105256 | 2.671842496 |
| ENSG00000119640 | ACYP1 | -8.464097617 | 0.000432249 | 0.917829853 | 0.672805615 | -7.546267764 | 5.095959458 | 5.72254323 | 2 | 2 | 2 | 6.069527745 | 5.379200993 |
| ENSG00000014257 | ACPP | -1.429974544 | 0.607921054 | -6.124774846 | 0.013724207 | -7.55474939 | 2 | 6.389896633 | 2 | 3.515101962 | 4.05806467 | 2.094598369 | 2.172608095 |
| ENSG00000149256 | TENM4 | -2.327187128 | 0.021090306 | -5.228116143 | 1.97E-08 | -7.555303271 | 6.89194353 | 9.74864196 | 9.624627483 | 6.809774134 | 7.0150921 | 4.099885429 | 4.468068808 |
| ENSG00000233904 | | 4.552894217 | 1.58E-27 | -12.10997428 | 7.79E-37 | -7.557080066 | 9.590776211 | 9.828248506 | 10.36791565 | 14.52150018 | 14.5038897 | 2.094598369 | 2.059860178 |
| | FAM21A | 4.308104878 | 0.001718115 | -11.86966308 | 3.68E-09 | -7.561558198 | 4.00722531 | 7.901717006 | 6.073378139 | 10.96824395 | 11.0618712 | 2 | 2 |
| ENSG00000163803 | PLB1 | -3.055748027 | 0.092426088 | -4.508534932 | 0.008640454 | -7.564282958 | 4.815609471 | 6.728495673 | 2 | 3.289987342 | 3.08597292 | 2.384001718 | 2.70970617 |
| ENSG00000186654 | PRR5 | -5.178183329 | 1.44E-16 | -2.387719552 | 0.000156432 | -7.565902882 | 8.257427002 | 9.338312924 | 9.636156974 | 4.034562184 | 4.56059747 | 6.593521701 | 7.052989523 |
| ENSG00000102967 | DHODH | -4.677106342 | 5.06E-07 | -2.899655912 | 0.002409889 | -7.576762254 | 8.340423424 | 7.145017454 | 9.819251342 | 4.205751501 | 4.63360314 | 5.849807762 | 6.155328188 |
| ENSG00000130595 | TNNT3 | -4.714853953 | NA | -2.862908728 | NA | -7.577762681 | 2 | 2 | 3.369317972 | 2 | 2 | 2.183373728 | 2 |
| ENSG00000134571 | MYBPC3 | -4.714853953 | NA | -2.862908728 | NA | -7.577762681 | 2 | 2 | 3.369317972 | 2 | 2 | 2.183373728 | 2 |
| ENSG00000182118 | FAM89A | -4.414205491 | 3.93E-24 | -3.164941884 | 1.53E-13 | -7.579147375 | 10.40620204 | 9.470501606 | 10.05225195 | 5.602782483 | 5.8245091 | 6.814608286 | 6.985139097 |
| ENSG00000166426 | CRABP1 | -2.09703255 | 0.59764617 | -5.48294404 | 0.131551868 | -7.57997659 | 5.330594134 | 3.686303414 | 2 | 3.462030052 | 2 | 2 | 2.225841198 |
| ENSG00000160190 | SLC37A1 | -4.608508766 | 5.23E-11 | -2.989985959 | 2.82E-05 | -7.598494725 | 9.48530526 | 9.067150531 | 10.82615578 | 5.709096143 | 5.28958453 | 7.0614806 | 7.028854452 |
| ENSG00000164879 | CA3 | -3.258811503 | 0.279279457 | -4.346135018 | 0.124359227 | -7.60494652 | 4.467311984 | 2 | 2 | 2.270786764 | 2.14022021 | 2.139668783 | 2.059860178 |
| ENSG00000178301 | AQP11 | -3.836393823 | 0.094108586 | -3.770951581 | 0.093533047 | -7.607345403 | 8.674867122 | 3.686303414 | 2 | 3.515101962 | 3.98554648 | 4.133696865 | 3.407202941 |
| ENSG00000104213 | PDGFRL | -4.454901444 | 0.002654125 | -3.158708133 | 0.036579278 | -7.613609577 | 8.609439662 | 7.232742069 | 3.369317972 | 3.227757282 | 3.90918929 | 4.620674722 | 4.56420962 |
| ENSG00000182985 | CADM1 | -2.572201978 | 0.000362367 | -5.049887956 | 2.39E-13 | -7.622089935 | 8.816894989 | 8.57407404 | 8.585098385 | 6.580005553 | 5.56313034 | 4.45225155 | 3.36036153 |
| ENSG00000213213 | CCDC183 | -3.935253116 | 0.002161564 | -3.692392833 | 0.003339591 | -7.62764595 | 7.667647223 | 4.710564154 | 5.59507254 | 3.615725078 | 2.86123014 | 3.648325314 | 3.183290849 |
| ENSG00000184786 | TCTE3 | 0.462511581 | 0.915012803 | -8.091247274 | 0.035305431 | -7.628735693 | 2 | 4.710564154 | 2 | 3.95996233 | 3.74316441 | 2 | 2 |
| ENSG00000187122 | SLIT1 | -4.488759494 | 0.111077329 | -3.149151524 | 0.255596226 | -7.637911018 | 2 | 2 | 6.203069917 | 2.141737246 | 2.49394783 | 2.492221014 | 2.918126211 |
| ENSG00000101842 | VSIG1 | -3.041902923 | 0.018452657 | -4.597833015 | 0.000187375 | -7.639735938 | 4.00722531 | 4.710564154 | 6.073378139 | 3.227757282 | 2.68924543 | 2.267001569 | 2.551896972 |
| ENSG00000124374 | PAIP2B | -5.887672326 | 0.001065014 | -1.768704212 | 0.346462637 | -7.656376539 | 2 | 5.72254323 | 7.451345288 | 2.498691189 | 2.2680111 | 4.82764224 | 4.489993931 |
| ENSG00000140832 | MARVELD3 | -2.942951558 | 0.047394637 | -4.71647036 | 0.000808652 | -7.659421918 | 5.095959458 | 3.686303414 | 5.930865648 | 2.784612485 | 3.2184422 | 2.592885947 | 2.117335275 |
| SPTBN2 | | -5.05438082 | 5.25E-10 | -2.611195864 | 0.001592231 | -7.665576684 | 7.384433967 | 9.235879652 | 8.143093325 | 3.997744386 | 3.69849104 | 6.339575235 | 5.355511328 |
| ENSG00000178425 | NT5DC1 | -3.258837153 | 0.000175511 | -4.40692796 | 1.74E-07 | -7.665765114 | 11.49443523 | 10.05576005 | 9.014156722 | 7.461007055 | 7.14977896 | 6.356378514 | 6.044874254 |
| ENSG00000121905 | HPCA | -6.330136512 | 0.159956387 | -1.336745393 | 0.774562861 | -7.666881905 | 2 | 4.442710129 | 2 | 2 | 2 | 2.68698273 | 2.671842496 |
| ENSG00000104814 | MAP4K1 | -3.079721128 | 0.135425564 | -4.587247123 | 0.019298939 | -7.666968251 | 4.00722531 | 4.11352228 | 2 | 2.3892346 | 2.2680111 | 2.183373728 | 2.059860178 |
| ENSG00000103313 | MEFV | -2.804933335 | 0.293493132 | -4.863747603 | 0.054080042 | -7.668680937 | 4.467311984 | 2 | 4.058998049 | 2.498691189 | 2.38539787 | 2.225793514 | 2 |
| LOC101927322 | | -0.316051399 | 0.947611632 | -7.364747287 | 0.090083987 | -7.680798687 | 2 | 4.11352228 | 2 | 2.784612485 | 3.08597292 | 2 | 2 |
| ENSG00000076650 | GPATCH1 | -8.727619887 | 0.001787802 | 1.043819807 | 0.702427566 | -7.68380008 | 3.327711234 | 6.535001748 | 2 | 2 | 2 | 6.134985075 | 6.124122661 |
| ENSG00000137504 | CREBZF | -10.4230331 | 2.14E-10 | 2.73573816 | 5.18E-06 | -7.687294943 | 6.257048221 | 7.232742069 | 6.534109959 | 2 | 2 | 9.58830174 | 9.275296471 |
| ENSG00000162779 | AXDND1 | -2.467268746 | 0.518587294 | -5.221725268 | 0.139816214 | -7.688994014 | 2 | 2 | 5.39244166 | 2.868578001 | 2.38539787 | 2 | 2.225841198 |
| ENSG00000215845 | TSTD1 | -6.31141716 | 0.260096091 | -1.37951911 | 0.816073612 | -7.69093627 | 4.467311984 | 2 | 2 | 2 | 2 | 3.10571689 | 2 |
| ENSG00000183914 | DNAH2 | -4.32004127 | 0.057973134 | -3.3790224 | 0.117356112 | -7.69906367 | 3.327711234 | 2 | 5.59507254 | 2.3892346 | 2.14022021 | 2.656294034 | 2.277179783 |
| ENSG00000188766 | SPRED3 | -9.705983476 | 8.07E-05 | 2.003805329 | 0.351170929 | -7.702178147 | 7.031698319 | 2 | 6.073378139 | 2 | 2 | 7.999020789 | 8.012280354 |
| ENSG00000174992 | ZG16 | -3.341702017 | 0.390304325 | -4.366498708 | 0.237112054 | -7.708200725 | 2 | 2 | 5.156626401 | 2.3892346 | 2.2680111 | 2 | 2.326754077 |
| ENSG00000177599 | ZNF491 | -4.321437394 | 0.002076371 | -3.391161278 | 0.017563259 | -7.712598672 | 6.73718411 | 5.457120953 | 9.809100877 | 4.237680128 | 4.56059747 | 5.279204176 | 5.122735884 |
| ENSG00000152213 | ARL11 | -5.282571443 | 0.005767867 | -2.431271419 | 0.172175696 | -7.713842862 | 4.467311984 | 3.076665209 | 5.772716771 | 2.270786764 | 2.14022021 | 3.56816602 | 2.326754077 |
| ENSG00000174951 | FUT1 | -6.085683707 | 0.000171376 | -1.628846418 | 0.305874629 | -7.714530125 | 4.00722531 | 4.442710129 | 7.163363831 | 2.3892346 | 2.14022021 | 4.785723309 | 4.047883918 |
| ENSG00000135951 | TSGA10 | -5.173927284 | 0.000269848 | -2.551172099 | 0.064276738 | -7.725099383 | 4.815609471 | 4.11352228 | 6.958202216 | 2.600425781 | 2.2680111 | 3.917810786 | 3.407202941 |
| ENSG00000117281 | CD160 | -6.513226326 | 0.056070605 | -1.212032821 | 0.719815802 | -7.725259147 | 2 | 2 | 5.930865648 | 2.141737246 | 2 | 3.767980144 | 3.407202941 |
| ENSG00000142765 | SYTL1 | -4.974597533 | 0.012493831 | -2.76043213 | 0.156315531 | -7.735029663 | 4.815609471 | 6.228547981 | 2 | 2.141737246 | 2.49394783 | 3.517841605 | 2.592996859 |
| ENSG00000064270 | ATP2C2 | -6.64699832 | 0.1248043 | -1.096217164 | 0.809379135 | -7.743215484 | 2 | 4.710564154 | 2 | 2 | 2 | 2.775316226 | 3.042026731 |
| GSG2 | | 0.457419818 | 0.872186145 | -8.203524697 | 0.002583929 | -7.746104878 | 3.327711234 | 4.442710129 | 2 | 3.79808662 | 4.05806467 | 2 | 2 |
| ENSG00000135643 | KCNMB4 | -4.063898862 | 6.73E-06 | -3.69059255 | 2.96E-05 | -7.754491412 | 6.257048221 | 7.002537973 | 8.143093325 | 3.79808662 | 3.69849104 | 4.425202323 | 3.45257121 |
| ENSG00000053702 | NRIP2 | -7.388744978 | 0.000873217 | -0.372661787 | 0.847683815 | -7.761406764 | 4.467311984 | 3.076665209 | 6.43197674 | 2.141737246 | 2 | 5.19095825 | 4.624919101 |
| ENSG00000100207 | TCF20 | 3.191959513 | 0.067543282 | -10.96090608 | 7.15E-07 | -7.768946563 | 6.563806279 | 6.311476754 | 2 | 8.945181925 | 9.05174826 | 2 | 2 |
| ENSG00000105610 | KLF1 | -4.714856513 | NA | -3.067962815 | NA | -7.782819328 | 3.327711234 | 2 | 2 | 2 | 2 | 2.048074405 | 2.117335275 |
| ENSG00000184313 | MROH7 | -4.714856513 | NA | -3.067962815 | NA | -7.782819328 | 2 | 2 | 3.369317972 | 2 | 2 | 2.048074405 | 2.117335275 |
| ENSG00000187905 | LRRC74B | -2.322118113 | 0.384425629 | -5.461106365 | 0.035786107 | -7.783224478 | 2 | 3.686303414 | 4.058998049 | 2.600425781 | 2.2680111 | 2 | 2.117335275 |
| ENSG00000145687 | SSBP2 | -4.093157877 | 7.28E-11 | -3.703338296 | 3.67E-09 | -7.796496173 | 11.55650736 | 9.930175364 | 10.42884603 | 6.777556219 | 6.74548212 | 7.143982415 | 7.133547492 |
| ENSG00000139116 | KIF21A | -3.560088831 | 0.00022 | -4.241726641 | 6.92E-06 | -7.801815473 | 10.26462289 | 9.616264339 | 7.342949392 | 6.172600389 | 5.86549842 | 5.696113106 | 5.020995508 |
| ENSG00000160097 | FNDC5 | -5.545922568 | 0.001837562 | -2.25956146 | 0.232527319 | -7.805484028 | 8.517314938 | 6.666836837 | 2 | 2.695456494 | 3.01487888 | 5.274160001 | 5.0581604 |
| ENSG00000205730 | ITPRIPL2 | -10.84048959 | 2.77E-11 | 3.030981561 | 8.41E-08 | -7.809508031 | 7.579279349 | 7.099070191 | 6.629488387 | 2 | 2 | 10.36875231 | 9.891404443 |
| ENSG00000169783 | LINGO1 | -4.766183544 | 0.004706762 | -3.0480724 | 0.059798852 | -7.814255944 | 5.095959458 | 5.595927442 | 4.874587708 | 2.270786764 | 2.49394783 | 3.5515855 | 2.117335275 |
| ENSG00000102683 | SGCG | -7.634646801 | 0.048826572 | -0.181852135 | 0.96694554 | -7.816498937 | 2 | 5.595927442 | 2 | 2 | 2 | 4.122514202 | 4.09155917 |
| | FAM21C | 4.144919415 | 0.004043945 | -11.96204842 | 5.20E-09 | -7.817129001 | 3.327711234 | 7.669933291 | 7.029880454 | 10.89380019 | 10.9947543 | 2 | 2 |
| ENSG00000112041 | TULP1 | -6.460314077 | 0.036314274 | -1.359626892 | 0.643682096 | -7.819940969 | 3.327711234 | 4.11352228 | 2 | 2 | 2 | 2.656294034 | 2.782576693 |
| ENSG00000101230 | ISM1 | -4.546711419 | 0.121082311 | -3.276598737 | 0.256356989 | -7.823310156 | 2 | 6.228547981 | 2 | 2.3892346 | 2.2680111 | 3.10571689 | 2.117335275 |
| ENSG00000168243 | GNG4 | -2.668237937 | 0.000392466 | -5.160114391 | 2.87E-12 | -7.828352328 | 6.468621203 | 5.72254323 | 7.285536013 | 4.268917409 | 4.1930041 | 2.746468522 | 2.70970617 |
| ENSG00000086696 | HSD17B2 | -1.67973259 | NA | -6.15289844 | NA | -7.83263103 | 3.327711234 | 2 | 2 | 2.3892346 | 2 | 2 | 2 |
| ENSG00000177511 | ST8SIA3 | -1.67973259 | NA | -6.15289844 | NA | -7.83263103 | 2 | 3.076665209 | 2 | 2.3892346 | 2 | 2 | 2 |
| ENSG00000179674 | ARL14 | -1.67973259 | NA | -6.15289844 | NA | -7.83263103 | 2 | 2 | 3.369317972 | 2.3892346 | 2 | 2 | 2 |
| ENSG00000163794 | UCN | -4.163513252 | 4.79E-05 | -3.669333235 | 0.000246207 | -7.832846487 | 7.27616306 | 5.457120953 | 7.398165132 | 3.406930997 | 3.39680241 | 4.188345803 | 3.042026731 |
| FAM159B | | -1.684809032 | NA | -6.152898461 | NA | -7.837707494 | 3.327711234 | 2 | 2 | 2.270786764 | 2.14022021 | 2 | 2 |
| ENSG00000126016 | AMOT | -3.706884221 | 1.26E-06 | -4.131098309 | 4.06E-08 | -7.837982531 | 7.750913381 | 9.030900487 | 9.888372618 | 5.590471342 | 5.49945122 | 5.352823931 | 4.943664032 |
| ENSG00000118849 | RARRES1 | -3.09077255 | 0.007968904 | -4.749255884 | 1.89E-05 | -7.840028434 | 8.19931886 | 5.131603625 | 7.398165132 | 4.825935136 | 4.12711147 | 3.632645806 | 2.918126211 |
| ENSG00000273045 | C2orf15 | -5.645298628 | 0.024143857 | -2.19620941 | 0.272132584 | -7.841508039 | 3.327711234 | 4.442710129 | 3.369317972 | 2.141737246 | 2 | 2.962551659 | 2.225841198 |
| FAM69A | | 3.058138885 | 0.021240574 | -10.90091899 | 2.89E-08 | -7.842780108 | 4.00722531 | 6.951736442 | 4.874587708 | 8.866956935 | 8.74494589 | 2 | 2 |
| ENSG00000101342 | TLDC2 | -1.689899584 | NA | -6.152899015 | NA | -7.842798598 | 2 | 2 | 3.369317972 | 2.141737246 | 2.2680111 | 2 | 2 |
| ENSG00000148513 | ANKRD30A | -1.689899584 | NA | -6.152899015 | NA | -7.842798598 | 2 | 2 | 3.369317972 | 2.141737246 | 2.2680111 | 2 | 2 |
| ENSG00000185155 | MIXL1 | -3.777051334 | 0.088711396 | -4.071345042 | 0.050890944 | -7.848396376 | 4.815609471 | 2 | 4.523675391 | 2.3892346 | 2.2680111 | 2.457042231 | 2.059860178 |
| ENSG00000073111 | MCM2 | -8.973330894 | 5.99E-05 | 1.116216388 | 0.55075414 | -7.857114506 | 3.327711234 | 6.602424674 | 4.058998049 | 2 | 2 | 6.421700344 | 6.440710022 |
| ENSG00000149970 | CNKSR2 | -0.084977929 | 0.985057747 | -7.772879773 | 0.056973044 | -7.857857701 | 2 | 4.442710129 | 2 | 3.289987342 | 3.28037825 | 2 | 2 |
| ENSG00000197444 | OGDHL | -6.864517935 | 2.90E-07 | -0.996339845 | 0.258826994 | -7.860857781 | 5.095959458 | 5.595927442 | 6.203069917 | 2.141737246 | 2.14022021 | 4.959178705 | 4.634793542 |
| ENSG00000273936 | | 2.766180259 | 0.412532734 | -10.6292347 | 0.001426759 | -7.86305444 | 2 | 2 | 7.098165388 | 8.29513208 | 8.20312688 | 2 | 2 |
| ENSG00000165181 | C9orf84 | -5.920850288 | 0.219396158 | -1.947551005 | NA | -7.868401293 | 2 | 4.11352228 | 2 | 2 | 2 | 2.42098416 | 2.326754077 |
| ENSG00000182636 | NDN | -2.654672902 | 8.34E-07 | -5.215169693 | 1.24E-23 | -7.869842596 | 11.09707093 | 11.72670414 | 11.9774171 | 8.930605282 | 9.06720431 | 6.902223127 | 5.921228818 |
| ENSG00000011021 | CLCN6 | -9.916922455 | 2.65E-05 | 2.040016697 | 0.306126091 | -7.876905758 | 2 | 6.844429755 | 6.80318718 | 2 | 2 | 8.095112207 | 8.388545823 |
| ENSG00000152256 | PDK1 | -10.33366013 | 2.75E-08 | 2.454367982 | 0.028914977 | -7.879292152 | 4.815609471 | 6.666836837 | 7.451345288 | 2 | 2 | 8.619836896 | 9.415788104 |
| ENSG00000141524 | TMC6 | -7.96295697 | 3.34E-05 | 0.083521766 | 0.968477418 | -7.879435204 | 7.435667355 | 7.145017454 | 2 | 2.141737246 | 2.14022021 | 7.06440544 | 6.512090157 |
| ENSG00000244752 | CRYBB2 | 0.334995649 | 0.857831154 | -8.215329601 | 7.91E-05 | -7.880333951 | 5.866944937 | 3.686303414 | 3.369317972 | 4.965943888 | 5.11361705 | 2 | 2.059860178 |
| ENSG00000118363 | SPCS2 | -1.979504154 | 3.28E-06 | -5.902301931 | 1.30E-47 | -7.881806085 | 11.40085503 | 10.76380876 | 11.55376643 | 9.340234287 | 9.26686067 | 5.264018433 | 5.713290392 |
| ENSG00000214946 | TBC1D26 | -4.319667212 | 0.137326507 | -3.563979717 | 0.182639149 | -7.883646928 | 2 | 2 | 4.874587708 | 2.141737246 | 2.14022021 | 2.183373728 | 2.277179783 |
| ENSG00000163584 | RPL22L1 | -9.584813518 | 5.13E-08 | 1.697345864 | 0.066316548 | -7.887467654 | 5.866944937 | 6.389896633 | 5.39244166 | 2 | 2 | 6.910373854 | 8.044120856 |
| ENSG00000167332 | OR51E2 | -5.548334952 | 0.002908324 | -2.345206938 | 0.136426732 | -7.893541891 | 4.815609471 | 3.686303414 | 4.874587708 | 2 | 2.2680111 | 3.296906311 | 2.509591799 |
| ENSG00000155011 | DKK2 | -9.220171088 | 1.15E-16 | 1.318894326 | 0.26264969 | -7.901276761 | 10.05124847 | 10.43902632 | 10.26770388 | 2.498691189 | 2.68924543 | 9.035059236 | 12.44811907 |
| ENSG00000128268 | MGAT3 | -7.987489068 | 1.04E-18 | 0.084417422 | 0.93891146 | -7.903071646 | 8.367060731 | 9.125613817 | 9.972637969 | 3.023133209 | 2.2680111 | 9.018941717 | 9.676696925 |
| ENSG00000130948 | HSD17B3 | -1.109788004 | 0.842086376 | -6.794331781 | NA | -7.904119785 | 2 | 3.686303414 | 2 | 2.270786764 | 2.59489879 | 2 | 2 |
| ENSG00000172765 | TMCC1 | -10.289763 | 5.66E-08 | 2.38251872 | 0.046132155 | -7.907244284 | 6.468621203 | 7.503546079 | 4.523675391 | 2 | 2 | 9.067128536 | 8.838074258 |
| ENSG00000120075 | HOXB5 | 1.645717256 | 0.639879212 | -9.556172559 | 0.004627208 | -7.910455303 | 2 | 6.04685525 | 2 | 6.034011955 | 6.20590375 | 2 | 2 |
| ENSG00000278644 | | 3.355728593 | 0.030567914 | -11.27408828 | 7.49E-08 | -7.918359687 | 4.467311984 | 4.11352228 | 7.502634626 | 9.441381351 | 9.50436385 | 2 | 2 |
| ENSG00000151725 | CENPU | -8.431826025 | 0.002799889 | 0.505271417 | 0.861696561 | -7.926554608 | 3.327711234 | 6.228547981 | 2 | 2 | 2 | 5.48093935 | 5.224326245 |
| ENSG00000072864 | NDE1 | 1.748893796 | 0.019764706 | -9.67824636 | 4.03E-09 | -7.929352564 | 6.563806279 | 6.04685525 | 5.156626401 | 7.72026832 | 7.72392024 | 2.048074405 | 2 |
| ENSG00000263028 | | -0.576718982 | 0.908905358 | -7.363631838 | 0.093855546 | -7.94035082 | 2 | 4.11352228 | 2 | 2.784612485 | 2.86123014 | 2 | 2 |
| ENSG00000105926 | MPP6 | -3.316647476 | 2.59E-18 | -4.632790306 | 2.14E-35 | -7.949437782 | 10.31337612 | 9.532315021 | 9.963514551 | 6.628980152 | 6.77267312 | 5.463319095 | 5.470251342 |
| ENSG00000145861 | C1QTNF2 | -3.462139231 | 0.085759206 | -4.490621973 | 0.020067196 | -7.952761204 | 2 | 5.595927442 | 8.35328246 | 3.881293667 | 3.94787299 | 2.717032197 | 3.620995423 |
| CXORF57 | CXORF57 | 0.389956872 | 0.927635884 | -8.348843051 | 0.027892955 | -7.958886179 | 2 | 4.93638933 | 2 | 4.034562184 | 3.94787299 | 2 | 2 |
| ENSG00000175643 | RMI2 | -4.016140565 | 1.10E-05 | -3.948932939 | 1.17E-05 | -7.965073504 | 8.963744667 | 6.311476754 | 8.380957451 | 4.523237872 | 4.45712836 | 4.360042493 | 4.711446447 |
| ENSG00000160408 | ST6GALNAC6 | -1.037050583 | 0.05128447 | -6.932428654 | 5.85E-47 | -7.969479237 | 9.410448395 | 9.806969599 | 10.50627919 | 8.889743063 | 9.00436103 | 3.584558147 | 3.640730416 |

| | | | | | | | | | | | | | |
|---|---|---|---|---|---|---|---|---|---|---|---|---|---|
| ENSG00000281216 | | 0.41515615 | 0.858540161 | -8.388242103 | 6.86E-05 | -7.973085953 | 5.095959458 | 6.844429755 | 2 | 6.01571348 | 6.1322378 | 2.094598369 | 2 |
| ENSG00000091428 | RAPGEF4 | -3.873370245 | 0.111902276 | -4.104928123 | 0.084137979 | -7.978298369 | 2 | 3.076665209 | 8.904018013 | 3.95996233 | 3.82856379 | 4.053534413 | 3.311948089 |
| ENSG00000163793 | DNAJC5G | -4.714859103 | NA | -3.275341825 | NA | -7.990200928 | 2 | 2 | 3.369317972 | 2 | 2 | 2.139668783 | 2 |
| ENSG00000114541 | FRMD4B | -5.208588659 | 0.000989884 | -2.7844008 | 0.081866442 | -7.992989459 | 4.00722531 | 7.790474954 | 4.523675391 | 2.868578001 | 2.38539787 | 3.930690651 | 4.09155917 |
| ENSG00000189221 | MAOA | -5.168266186 | 2.82E-07 | -2.826675469 | 0.006640877 | -7.994941655 | 9.710019321 | 6.535001748 | 8.434763955 | 3.881293667 | 4.05806467 | 6.025180246 | 5.941260754 |
| ENSG00000126733 | DACH2 | -5.351054262 | NA | -2.646154545 | NA | -7.997208807 | 2 | 3.686303414 | 2 | 2 | 2 | 2.267001569 | 2.059860178 |
| ENSG00000181585 | TMIE | -4.787825534 | 7.29E-05 | -3.218019891 | 0.005716042 | -8.005845425 | 4.467311984 | 6.140560021 | 6.958202216 | 2.947924294 | 2.38539787 | 3.796405078 | 3.128433649 |
| | PTRF | 4.944056095 | 0.000481753 | -12.95200717 | 2.15E-10 | -8.007951071 | 8.493334308 | 8.213429689 | 4.058998049 | 12.72227161 | 12.7420294 | 2 | 2 |
| ENSG00000150773 | PIH1D2 | -3.591269091 | 0.081497157 | -4.418196197 | 0.025954595 | -8.009465288 | 8.042902253 | 4.710564154 | 2 | 3.023133209 | 3.94787299 | 2.911488689 | 3.209956562 |
| ENSG00000273092 | | 0.445113535 | 0.875544825 | -8.45773136 | 0.001770226 | -8.012617825 | 2 | 4.710564154 | 3.369317972 | 4.105493497 | 4.12711147 | 2 | 2 |
| ENSG00000167011 | NAT16 | -4.275099753 | 0.230926542 | -3.737763775 | 0.276037949 | -8.012863528 | 5.330594134 | 2 | 2 | 2.3892346 | 2 | 2.3070652 | 2.277179783 |
| ENSG00000122026 | RPL21 | -1.446465991 | 0.012491705 | -6.56714669 | 4.13E-36 | -8.013612681 | 12.3116858 | 11.07990752 | 11.28013451 | 10.20863848 | 10.2316302 | 5.180212365 | 5.32534152 |
| ENSG00000105640 | RPL18A | -3.020363654 | 2.65E-13 | -4.999920094 | 3.70E-35 | -8.020283747 | 14.41861704 | 13.51299452 | 13.73900361 | 10.88311511 | 10.9667728 | 8.97881132 | 8.930161469 |
| ENSG00000206457 | | -2.458159791 | 0.003324438 | -5.562509536 | 2.20E-12 | -8.020669327 | 9.081016759 | 6.844429755 | 8.175061096 | 5.818703821 | 6.00960265 | 3.483289517 | 3.311948089 |
| ENSG00000165182 | CXorf58 | -3.266386349 | NA | -4.755635337 | NA | -8.022021685 | 2 | 3.076665209 | 2 | 2.141737246 | 2 | 2.048074405 | 2 |
| ENSG00000140015 | KCNH5 | -4.50160502 | 0.263585024 | -3.522093654 | NA | -8.023698674 | 2 | 4.11352228 | 2 | 2 | 2.14022021 | 2.048074405 | 2.225841198 |
| ENSG00000273562 | | -0.718326707 | 0.045753442 | -7.309677838 | 1.07E-101 | -8.028004546 | 9.992581382 | 10.42512749 | 10.38167985 | 9.538304826 | 9.59092963 | 3.694366604 | 3.407202941 |
| ENSG00000135622 | SEMA4F | -10.0415489 | 3.52E-09 | 2.01032082 | 0.009503324 | -8.031228085 | 5.532353208 | 6.228547981 | 7.029880454 | 2 | 2 | 8.250863742 | 8.429540272 |
| ENSG00000136457 | CHAD | -6.905808951 | 0.103461762 | -1.131294053 | 0.79925824 | -8.037103004 | 2 | 4.93638933 | 2 | 2 | 2 | 3.1504466 | 2.885414003 |
| ENSG00000176244 | ACBD7 | -3.487789473 | 0.017030773 | -4.555998402 | 0.001137782 | -8.043787876 | 4.467311984 | 4.442710129 | 7.225742779 | 3.406930997 | 3.01487888 | 2.384001718 | 2.981406154 |
| ENSG00000225351 | | 0.520975498 | 0.902061374 | -8.567700748 | 0.021646679 | -8.04672525 | 2 | 5.131603625 | 2 | 4.070463721 | 4.43005853 | 2 | 2 |
| ENSG00000017427 | IGF1 | -2.299783352 | 0.063606895 | -5.747529047 | 2.08E-06 | -8.047312399 | 3.327711234 | 5.457120953 | 6.073378139 | 3.615725078 | 3.39680241 | 2.225793514 | 2.225841198 |
| ENSG00000178997 | EXD1 | -1.723016183 | 0.331987296 | -6.325135913 | 0.000900485 | -8.048152096 | 4.00722531 | 3.076665209 | 4.523675391 | 3.023133209 | 2.77779907 | 2.094598369 | 2 |
| ENSG00000147485 | PXDNL | -2.577708703 | 0.360667996 | -5.474779316 | 0.038705514 | -8.052488019 | 5.70932728 | 2 | 2 | 2.868578001 | 2.59489879 | 2.183373728 | 2.059860178 |
| ENSG00000258315 | C17orf49 | -1.035685505 | 0.138710156 | -7.033797317 | 8.01E-29 | -8.069482821 | 9.448362284 | 8.916342327 | 7.858115942 | 7.852449262 | 7.86451142 | 2.858551861 | 2.981406154 |
| ENSG00000151715 | TMEM45B | -6.634665615 | 0.007712736 | -1.43484687 | 0.499362489 | -8.069512485 | 4.00722531 | 3.076665209 | 3.369317972 | 2 | 2 | 2.62493832 | 2.885414003 |
| ENSG00000217236 | SP9 | -1.869239158 | 0.736046082 | -6.201410801 | 0.232022674 | -8.070649959 | 2 | 4.442710129 | 2 | 2.868578001 | 2 | 2 | 2.059860178 |
| ENSG00000204316 | MRPL38 | -2.168937464 | 0.090597396 | -5.905314633 | 2.49E-07 | -8.074252097 | 6.96351253 | 9.532315021 | 11.13734171 | 7.857595438 | 7.88459112 | 4.443291373 | 4.376876162 |
| ENSG00000198547 | C20orf203 | -4.354314252 | 0.135212277 | -3.721407008 | 0.190983933 | -8.07572126 | 2 | 2 | 6.534109959 | 2.270786764 | 2.59489879 | 2.987421008 | 2.172608095 |
| ENSG00000145335 | SNCA | -3.503130056 | 0.011808471 | -4.575243457 | 0.000616002 | -8.078373513 | 7.435667355 | 5.946638857 | 5.156626401 | 3.289987342 | 3.74316441 | 3.393105702 | 2.225841198 |
| ENSG00000052723 | SIKE1 | -10.25432616 | 3.38E-07 | 2.168972494 | 0.126596199 | -8.085353669 | 6.138360429 | 7.638155693 | 4.058998049 | 2 | 2 | 8.856325286 | 8.547926983 |
| ENSG00000102383 | ZDHHC15 | -6.629529493 | 7.68E-05 | -1.456487376 | 0.303132265 | -8.086016869 | 6.257048221 | 5.131603625 | 4.523675391 | 2.270786764 | 2 | 3.483289517 | 4.729989799 |
| ENSG00000251664 | PCDHA12 | -1.748071823 | 0.375025567 | -6.33831043 | 0.001016114 | -8.086382253 | 2 | 4.710564154 | 5.156626401 | 3.663521627 | 2.59489879 | 2.094598369 | 2.059860178 |
| ENSG00000028203 | VEZT | -10.6396868 | 1.40E-07 | 2.542088366 | 0.074814288 | -8.097598431 | 4.815609471 | 8.169643011 | 5.772716771 | 2 | 2 | 9.364533261 | 9.55554796 |
| ENSG00000147697 | GSDMC | -4.713003623 | 0.012324317 | -3.393966221 | 0.02899481 | -8.106969844 | 3.327711234 | 4.11352228 | 4.058998049 | 2.141737246 | 2.14022021 | 2.183373728 | 2.466008509 |
| ENSG00000204130 | RUFY2 | -9.784738061 | 5.33E-07 | 1.676781731 | 0.209143226 | -8.10795633 | 4.00722531 | 7.051611315 | 5.930865648 | 2 | 2 | 7.821097076 | 7.699462058 |
| ENSG00000130812 | ANGPTL6 | -3.896567784 | 0.041733892 | -4.21165994 | 0.02393307 | -8.108227724 | 2 | 5.303523747 | 7.818184403 | 2.868578001 | 3.60476711 | 3.193831101 | 3.012034821 |
| ENSG00000156521 | TYSND1 | -9.634272822 | 0.000253258 | 1.517285455 | 0.537280787 | -8.116987366 | 2 | 7.35507212 | 4.523675391 | 2 | 2 | 7.62703232 | 7.266516281 |
| ENSG00000078596 | ITM2A | -9.12072594 | 0.000230097 | 0.990904139 | 0.667030736 | -8.129821801 | 5.866944937 | 6.228547981 | 2 | 2 | 2 | 5.449961224 | 7.038185015 |
| ENSG00000203730 | TEDDM1 | -4.310708038 | 0.287734175 | -3.819179155 | NA | -8.129887193 | 2 | 2 | 4.058998049 | 2.141737246 | 2 | 2.139668783 | 2.059860178 |
| ENSG00000175879 | HOXD8 | 0.163765202 | 0.954704452 | -8.296826198 | 0.000379643 | -8.133060996 | 2 | 4.710564154 | 8.266922072 | 6.941624891 | 7.00136498 | 2.225793514 | 2 |
| ENSG00000134940 | ACRV1 | -4.316940784 | 0.286739789 | -3.819117152 | NA | -8.136057936 | 2 | 2 | 4.058998049 | 2 | 2.14022021 | 2.139668783 | 2.059860178 |
| ENSG00000171777 | RASGRP4 | -5.921468764 | 0.000261061 | -2.215733224 | 0.079481787 | -8.137201988 | 4.00722531 | 4.11352228 | 5.772716771 | 2.141737246 | 2.14022021 | 3.316666208 | 3.128433649 |
| ENSG00000144868 | TMEM108 | -5.921610411 | 0.00263431 | -2.233406937 | 0.271739141 | -8.155017349 | 2 | 5.303523747 | 7.600047325 | 2.600425781 | 2.14022021 | 4.388329244 | 4.241259475 |
| ENSG00000185669 | SNAI3 | -8.108471055 | 0.002287795 | -0.049258881 | 0.985627735 | -8.157729936 | 2 | 5.72254323 | 4.058998049 | 2 | 2 | 4.572179576 | 4.614976609 |
| ENSG00000149212 | SESN3 | -6.043683458 | 0.000183037 | -2.127725201 | 0.234784916 | -8.171408659 | 9.720386367 | 11.83928723 | 11.25791777 | 6.205262847 | 2.14022021 | 9.205045846 | 8.898748881 |
| ENSG00000176732 | PFN4 | -4.184584311 | 0.045774818 | -3.987666235 | 0.051151516 | -8.172250546 | 4.00722531 | 2 | 7.225742779 | 2.600425781 | 2.94009902 | 3.035909414 | 2.671842496 |
| ENSG00000164744 | SUN3 | -4.055992744 | 0.0002601 | -4.122741157 | 6.67E-05 | -8.178733901 | 4.815609471 | 4.93638933 | 6.073378139 | 2.868578001 | 2.38539787 | 2.492221014 | 2.746601469 |
| ENSG00000183785 | TUBA8 | -1.414553994 | 0.626912572 | -6.768220511 | 0.008084243 | -8.182774506 | 7.218819781 | 2 | 2 | 4.470469939 | 4.45712836 | 2.183373728 | 2.117335275 |
| ENSG00000112280 | COL9A1 | -3.908994771 | 0.318446459 | -4.275385014 | 0.248264907 | -8.184379785 | 2 | 2 | 4.523675391 | 2.270786764 | 2 | 2.048074405 | 2.172608095 |
| ENSG00000125731 | SH2D3A | -6.684344949 | 0.04311869 | -1.503649251 | 0.668127909 | -8.1879942 | 2 | 2 | 7.029880454 | 2 | 2.2680111 | 4.563936166 | 3.807156736 |
| ENSG00000130748 | TMEM160 | -4.451027118 | 3.55E-22 | -3.741185745 | 1.54E-16 | -8.192212863 | 10.59268348 | 9.883504052 | 10.83618737 | 6.229283386 | 6.00043082 | 6.575182432 | 6.988993457 |
| ENSG00000105707 | HPN | -4.497524415 | 0.256095992 | -3.700534463 | NA | -8.198058878 | 2 | 4.11352228 | 2 | 2.141737246 | 2 | 2.183373728 | 2.059860178 |
| ENSG00000177455 | CD19 | -4.921981918 | 0.169028813 | -3.277524651 | 0.340755867 | -8.199506569 | 2 | 2 | 5.39244166 | 2.270786764 | 2 | 2.560105256 | 2.225841198 |
| ENSG00000149043 | SYT8 | -4.463407562 | 0.426810018 | -3.738925722 | NA | -8.202333284 | 2 | 4.11352228 | 2 | 2.141737246 | 2 | 2.225793514 | 2 |
| ENSG00000120915 | EPHX2 | -3.920430492 | 0.071694155 | -4.288045428 | 0.041704347 | -8.20847592 | 2 | 7.274681896 | 3.369317972 | 3.162721474 | 2.59489879 | 2.746468522 | 2.746601469 |
| ENSG00000174521 | TTC9B | -7.599762989 | 0.062938344 | -0.616736153 | 0.89090828 | -8.216499142 | 2 | 2 | 5.59507254 | 2 | 2 | 3.256555576 | 4.133950982 |
| ENSG00000104835 | SARS2 | -2.577917676 | 0.002299809 | -5.639041225 | 1.67E-12 | -8.216958902 | 8.631580145 | 7.60566235 | 9.829330889 | 6.394182228 | 6.48770488 | 3.56816602 | 4.002845082 |
| ENSG00000162981 | FAM84A | -3.816477598 | 0.026840833 | -4.401833525 | 0.008734147 | -8.218311123 | 8.777722034 | 3.076665209 | 5.930865648 | 4.205751501 | 3.69849104 | 3.411599105 | 3.698365557 |
| ENSG00000172578 | KLHL6 | -8.880091066 | 3.61E-05 | 0.654724851 | 0.717384422 | -8.225366215 | 5.330594134 | 4.93638933 | 5.59507254 | 2 | 2 | 6.730562686 | 3.789585079 |
| ENSG00000156239 | N6AMT1 | -9.304525542 | 4.54E-06 | 1.078830949 | 0.480374581 | -8.225694593 | 3.327711234 | 5.72254323 | 6.534109959 | 2 | 2 | 6.752490997 | 6.671679221 |
| ENSG00000154001 | PPP2R5E | -10.6193326 | 5.61E-07 | 2.390194103 | 0.137100827 | -8.229138502 | 6.816637977 | 7.928237109 | 3.369317972 | 2 | 2 | 9.372225266 | 9.207028324 |
| ENSG00000254505 | CHMP4A | -1.485925425 | 0.031197086 | -6.747618894 | 3.83E-26 | -8.233544318 | 9.371511179 | 8.234834549 | 9.277397397 | 7.568480804 | 7.57741621 | 3.429858444 | 2.782576693 |
| ENSG00000143494 | VASH2 | -3.387758571 | 0.001996857 | -4.850219398 | 4.30E-06 | -8.237977968 | 4.815609471 | 5.595927442 | 7.029880454 | 3.663521627 | 2.94009902 | 2.68698273 | 2.592996859 |
| ENSG00000117122 | MFAP2 | -2.216041035 | 1.54E-07 | -6.022403265 | 4.66E-50 | -8.2384443 | 12.94404887 | 12.01658788 | 12.7906215 | 10.39758568 | 10.4494805 | 6.631488899 | 6.709547124 |
| ENSG00000070019 | GUCY2C | -7.315427985 | 0.072562696 | -0.92498604 | 0.832164708 | -8.240414025 | 2 | 5.303523747 | 2 | 2 | 2 | 3.256555576 | 3.474731315 |
| ENSG00000173114 | LRRN3 | 0.232349922 | 0.871079042 | -8.480854046 | 8.69E-16 | -8.248504124 | 8.493334308 | 11.66890396 | 9.196533591 | 10.71361142 | 10.6567008 | 2.987421008 | 2.981406154 |
| ENSG00000075340 | ADD2 | -4.686251096 | 0.044143813 | -3.565292167 | 0.12312733 | -8.251543262 | 5.70932728 | 6.951736442 | 3.369317972 | 3.09461492 | 2 | 3.738983888 | 2.172608095 |
| ENSG00000102931 | ARL2BP | -1.683562084 | 1.31E-05 | -6.568866985 | 3.06E-70 | -8.252429068 | 10.64259676 | 11.34889202 | 11.14139273 | 9.425874714 | 9.36749255 | 4.563936166 | 4.895193906 |
| ENSG00000153707 | PTPRD | -3.985384569 | 2.00E-07 | -4.271238749 | 1.34E-08 | -8.256623318 | 8.418903792 | 8.540087354 | 8.460932916 | 5.022023783 | 4.31639476 | 5.032703865 | 3.539239711 |
| ENSG00000174527 | MYO1H | -6.147200853 | 0.078819008 | -2.112993703 | 0.521878465 | -8.260194556 | 2 | 2 | 5.59507254 | 2.141737246 | 2 | 3.035909414 | 2.70970617 |
| ENSG00000265107 | GJA5 | -6.200667092 | 0.076916982 | -2.061980352 | 0.534101559 | -8.262647444 | 2 | 5.595927442 | 2 | 2 | 2.14022021 | 3.27687201 | 2.466008509 |
| ENSG00000169682 | SPNS1 | -0.936467442 | 0.260860892 | -7.326832695 | 3.01E-25 | -8.263300138 | 11.1478805 | 10.24697497 | 9.099922433 | 9.411074916 | 9.50760441 | 3.5515855 | 3.698365557 |
| ENSG00000057468 | MSH4 | -4.701379156 | 0.12992063 | -3.565502008 | 0.219849101 | -8.266881164 | 2 | 5.131603625 | 2 | 2.141737246 | 2.14022021 | 2.492221014 | 2.059860178 |
| ENSG00000174672 | BRSK2 | -4.269003447 | 4.07E-06 | -4.002006617 | 1.00E-05 | -8.271010064 | 7.750913381 | 6.140560021 | 7.97167868 | 3.615725078 | 3.7864959 | 4.340871764 | 3.236138336 |
| ENSG00000176020 | AMIGO3 | -0.511628837 | 0.738820346 | -7.764333107 | 1.64E-07 | -8.275961944 | 4.00722531 | 6.844429755 | 4.874587708 | 5.363304456 | 5.22805098 | 2.048074405 | 2.117335275 |
| ENSG00000274588 | DGKK | -4.714862689 | NA | -3.562536068 | NA | -8.277398758 | 2 | 2 | 3.369317972 | 2 | 2 | 2 | 2.117335275 |
| ENSG00000150687 | PRSS23 | -12.52688681 | 4.17E-09 | 4.239490501 | 0.006951324 | -8.287396309 | 4.815609471 | 8.337339476 | 9.963514551 | 2 | 2 | 13.13637395 | 12.9374082 |
| ENSG00000281991 | TMEM265 | 1.357342268 | 0.614181381 | -9.646566628 | 0.000418144 | -8.289224361 | 2 | 3.076665209 | 6.073378139 | 5.808110275 | 6.05460702 | 2 | 2 |
| ENSG00000137880 | GCHFR | -3.001264997 | 0.169644746 | -5.28933233 | 0.009371668 | -8.290597326 | 8.138771685 | 4.11352228 | 2 | 3.709785322 | 4.2560182 | 2.858551861 | 2.509591799 |
| ENSG00000144847 | IGSF11 | -4.294481686 | 0.308981021 | -4.00289349 | NA | -8.297375176 | 3.327711234 | 2 | 3.369317972 | 2 | 2.14022021 | 2 | 2.172608095 |
| ENSG00000166342 | NETO1 | -1.611209185 | 0.267451577 | -6.692112733 | 4.58E-05 | -8.303321918 | 4.00722531 | 4.442710129 | 4.523675391 | 3.289987342 | 3.15372745 | 2 | 2.117335275 |
| ENSG00000156398 | SFXN2 | -4.932658877 | 0.000145787 | -3.377560998 | 0.010590595 | -8.310219875 | 8.285626337 | 8.487557258 | 4.058998049 | 3.515101962 | 3.50452879 | 4.461156421 | 4.878668439 |
| ENSG00000196218 | RYR1 | -6.147374274 | 0.001876889 | -2.184792088 | 0.270921627 | -8.332166362 | 5.330594134 | 2 | 7.163363831 | 2.141737246 | 2.38539787 | 4.144793515 | 3.987514168 |
| ENSG00000167371 | PRRT2 | -5.725564472 | 4.87E-08 | -2.607300211 | 0.017146254 | -8.332864683 | 9.751047882 | 7.099070191 | 7.285536013 | 3.566290607 | 3.39680241 | 6.167993056 | 5.929275 |
| ENSG00000102755 | FLT1 | -2.842433666 | 0.002647799 | -5.492280748 | 8.75E-10 | -8.334714414 | 6.96351253 | 9.461450882 | 9.180874355 | 6.052081244 | 6.16544343 | 4.177580368 | 3.407202941 |
| ENSG00000169684 | CHRNA5 | -2.152418644 | 0.17464992 | -6.183284048 | 7.48E-05 | -8.335702692 | 4.00722531 | 5.595927442 | 6.534109959 | 3.349643765 | 3.2184422 | 2.225793514 | 2 |
| ENSG00000137860 | SLC28A2 | -4.535087516 | 0.000118113 | -3.805951425 | 0.000130252 | -8.341038941 | 4.815609471 | 4.710564154 | 4.874587708 | 2.3892346 | 2.2680111 | 2.592885947 | 2.421067459 |
| ENSG00000121297 | TSHZ3 | -11.19889992 | 8.95E-06 | 2.855829252 | 0.195006512 | -8.343070671 | 2 | 8.607278467 | 7.225742779 | 2 | 2 | 9.930433865 | 10.64657174 |
| C18ORF8 | C18ORF8 | 2.575875378 | 0.201291754 | -10.91944443 | 3.48E-06 | -8.343569049 | 5.330594134 | 7.002537973 | 2 | 8.192977693 | 8.48870446 | 2 | 2 |
| ENSG00000158528 | PPP1R9A | -6.909648872 | 1.04E-09 | -1.435742136 | 0.160592365 | -8.345391008 | 7.667647223 | 6.464272742 | 6.534109959 | 2.600425781 | 2 | 5.703636713 | 5.566251551 |
| ENSG00000258839 | MC1R | -3.926095421 | 7.06E-09 | -4.42829409 | 3.41E-11 | -8.354389511 | 9.015175615 | 8.876041558 | 10.34701993 | 5.651002631 | 5.84514933 | 5.557671049 | 4.967301796 |
| MURC | | 0.613013414 | 0.708408469 | -8.96820504 | 9.87E-06 | -8.355191626 | 3.327711234 | 4.710564154 | 4.058998049 | 4.694064037 | 4.63360314 | 2 | 2 |
| ENSG00000101695 | RNF125 | -4.883842015 | 3.65E-05 | -3.471846155 | 0.002150339 | -8.35568817 | 5.866944937 | 4.93638933 | 7.163363831 | 2.695456494 | 2.68924543 | 3.810410249 | 2.817676617 |
| ENSG00000134539 | KLRD1 | -5.211543383 | 1.42E-06 | -3.151391331 | 0.001491164 | -8.362934714 | 5.532353208 | 6.04685525 | 6.718950348 | 2.3892346 | 2.68924543 | 3.981090143 | 2.918126211 |
| ENSG00000150722 | PPP1R1C | -3.868344267 | 0.179078755 | -4.497393972 | 0.102444617 | -8.365738239 | 6.257048221 | 2 | 2 | 2.600425781 | 2.38539787 | 2.094598369 | 2.551896972 |
| ENSG00000121989 | ACVR2A | -3.7571674 | 2.37E-07 | -4.609495127 | 1.09E-10 | -8.366662527 | 8.674867122 | 10.43440824 | 10.49365885 | 6.442929658 | 6.32089174 | 5.873348082 | 5.224326245 |
| ENSG00000134042 | MRO | -5.077531456 | 0.002498829 | -3.30055484 | 0.024375228 | -8.378086296 | 4.00722531 | 4.11352228 | 5.39244166 | 2.3892346 | 2 | 2.592885947 | 2.632958241 |
| ENSG00000276053 | | -2.94131502 | 0.096461716 | -5.443491129 | 0.001584572 | -8.384806149 | 3.327711234 | 3.076665209 | 5.39244166 | 2.498691189 | 2.68924543 | 2.183373728 | 2.059860178 |
| ENSG00000175768 | TOMM5 | -1.430502724 | 0.009834683 | -6.954310385 | 3.52E-41 | -8.384813109 | 10.1770105 | 9.067150531 | 10.19940021 | 8.48998335 | 8.46722793 | 3.465698119 | 3.620995423 |
| ENSG00000111275 | ALDH2 | -3.79531481 | 5.31E-14 | -4.590057941 | 3.19E-20 | -8.385372752 | 12.34221434 | 11.71342308 | 12.67800366 | 8.425603456 | 8.6021877 | 7.270951294 | 8.083408769 |
| ENSG00000013725 | CD6 | -5.090677785 | 0.084498748 | -3.298873329 | 0.256352368 | -8.389551114 | 6.468621203 | 2 | 2 | 2.3892346 | 2.14022021 | 3.172301926 | 2.225841198 |
| ENSG00000144481 | TRPM8 | -3.308311549 | 0.142581803 | -5.084609126 | 0.018557201 | -8.392920675 | 3.327711234 | 2 | 5.156626401 | 2.498691189 | 2.2680111 | 2.183373728 | 2.059860178 |
| ENSG00000019991 | HGF | -2.610368696 | 4.44E-05 | -5.786433723 | 2.04E-21 | -8.39680242 | 12.95725781 | 12.29271883 | 11.22775303 | 9.730780958 | 9.70541961 | 6.505870593 | 6.681239787 |
| ENSG00000262860 | | -0.566630431 | 0.17595559 | -7.830526086 | 5.23E-87 | -8.397156517 | 9.98400217 | 10.09139839 | 9.778213818 | 9.44822005 | 9.33867006 | 3.256555576 | 2.817676617 |
| ENSG00000187554 | TLR5 | -4.974476058 | 0.01813334 | -3.43040959 | 0.106524826 | -8.404885648 | 8.009475818 | 4.442710129 | 2 | 2.695456494 | 2.86123014 | 4.144793515 | 2.817676617 |
| ENSG00000103534 | TMC5 | -6.855645088 | 0.018661592 | -1.551401759 | 0.566749731 | -8.407046847 | 2 | 4.11352228 | 4.058998049 | 2 | 2 | 2.987421008 | 2.592996859 |

| | | | | | | | | | | | | | |
|---|---|---|---|---|---|---|---|---|---|---|---|---|---|
| ENSG00000110887 | DAO | -4.714864442 | NA | -3.702891357 | NA | -8.417755799 | 2 | 2 | 3.369317972 | 2 | 2 | 2.048074405 | 2.059860178 |
| ENSG00000163630 | SYNPR | -2.264901538 | NA | -6.152900876 | NA | -8.417802414 | 3.327711234 | 2 | 2 | 2.270786764 | 2 | 2 | 2 |
| ENSG00000182162 | P2RY8 | -2.264901538 | NA | -6.152900876 | NA | -8.417802414 | 2 | 2 | 3.369317972 | 2.270786764 | 2 | 2 | 2 |
| ENSG00000213088 | ACKR1 | -2.264901538 | NA | -6.152900876 | NA | -8.417802414 | 2 | 3.076665209 | 2 | 2.270786764 | 2 | 2 | 2 |
| ENSG00000081181 | ARG2 | -4.978877912 | 4.09E-07 | -3.439577014 | 0.000543061 | -8.418454926 | 9.913467515 | 7.701025996 | 7.342949392 | 4.268917409 | 4.05806467 | 5.431956133 | 5.524371359 |
| ENSG00000143570 | SLC39A1 | -1.015058581 | 0.128900041 | -7.404132485 | 2.78E-37 | -8.419191066 | 11.40407417 | 10.50217373 | 10.06083273 | 9.763363111 | 9.74714308 | 3.56816602 | 4.062589736 |
| ENSG00000185436 | IFNLR1 | -4.907122666 | 0.003483277 | -3.51321328 | 0.038609931 | -8.420335946 | 8.19931886 | 3.076665209 | 6.882775709 | 3.289987342 | 2.86123014 | 4.596630906 | 2.918126211 |
| ENSG00000187838 | PLSCR3 | -0.819140721 | 0.153204144 | -7.601578383 | 1.04E-48 | -8.420719103 | 9.87686338 | 9.770792087 | 9.465702942 | 8.852852814 | 8.94822691 | 2.68698273 | 3.36036153 |
| ENSG00000164818 | DNAAF5 | -10.39606276 | 1.06E-07 | 1.973105738 | 0.13968042 | -8.422957017 | 7.384433967 | 7.051611315 | 4.058998049 | 2 | 2 | 8.508269163 | 8.791384617 |
| ENSG00000112195 | TREML2 | -2.272292639 | NA | -6.152900907 | NA | -8.425193546 | 3.327711234 | 2 | 2 | 2.141737246 | 2.14022021 | 2 | 2 |
| ENSG00000127954 | STEAP4 | -2.272292639 | NA | -6.152900907 | NA | -8.425193546 | 3.327711234 | 2 | 2 | 2.141737246 | 2.14022021 | 2 | 2 |
| ENSG00000154162 | CDH12 | -2.272292639 | NA | -6.152900907 | NA | -8.425193546 | 2 | 3.076665209 | 2 | 2.141737246 | 2.14022021 | 2 | 2 |
| ENSG00000171903 | CYP4F11 | -2.272292639 | NA | -6.152900907 | NA | -8.425193546 | 2 | 2 | 3.369317972 | 2.141737246 | 2.14022021 | 2 | 2 |
| ENSG00000187682 | ERAS | -2.272292639 | NA | -6.152900907 | NA | -8.425193546 | 2 | 3.076665209 | 2 | 2.141737246 | 2.14022021 | 2 | 2 |
| ENSG00000188760 | TMEM198 | -3.994860196 | 0.045036382 | -4.437476016 | 0.023011611 | -8.432336213 | 3.327711234 | 6.140560021 | 10.79563557 | 5.258265913 | 5.55061739 | 5.050514232 | 4.982847849 |
| ENSG00000232125 | DYTN | -2.279714938 | NA | -6.152900938 | NA | -8.432615876 | 2 | 2 | 3.369317972 | 2 | 2.2680111 | 2 | 2 |
| ENSG00000164591 | MYOZ3 | -3.711368126 | 0.007972283 | -4.734956983 | 0.000431486 | -8.446325109 | 3.327711234 | 5.946638857 | 7.451345288 | 3.227757282 | 3.39680241 | 2.656294034 | 2.918126211 |
| ENSG00000178460 | MCMDC2 | -5.231075808 | 0.003364094 | -3.215814514 | 0.059657664 | -8.446890321 | 6.138360429 | 2 | 5.772716771 | 2.3892346 | 2.2680111 | 3.12825509 | 2.918126211 |
| ENSG00000177954 | RPS27 | -1.857961243 | 0.009302257 | -6.589312106 | 4.00E-24 | -8.447273348 | 13.26583049 | 11.77133279 | 11.88621475 | 10.6389887 | 10.608468 | 6.267798399 | 5.626876946 |
| ENSG00000100336 | APOL4 | -2.241829077 | 0.181893587 | -6.207118945 | 9.70E-05 | -8.448948023 | 2 | 6.04685525 | 5.772716771 | 3.615725078 | 3.55551834 | 2.225793514 | 2.117335275 |
| ENSG00000181778 | TMEM252 | -6.904904597 | 0.108874129 | -1.545885393 | 0.726496359 | -8.450789991 | 2 | 4.93638933 | 2 | 2 | 2 | 2.656294034 | 2.981406154 |
| ENSG00000205560 | CPT1B | -1.040155465 | 0.374504554 | -7.41399312 | 1.68E-13 | -8.454148586 | 9.128497779 | 8.337339476 | 7.098165388 | 7.426792977 | 7.34718809 | 2.885263063 | 2.172608095 |
| ENSG00000154639 | CXADR | -5.268802638 | 8.96E-07 | -3.192510234 | 0.002994187 | -8.461312872 | 7.904282243 | 6.951736442 | 8.076950877 | 3.79808662 | 2.2680111 | 4.799832052 | 4.664017718 |
| ENSG00000110660 | SLC35F2 | -4.49227673 | 2.53E-22 | -3.975508115 | 2.70E-18 | -8.467784845 | 10.67493801 | 10.3486505 | 10.89014791 | 6.096288482 | 6.36443784 | 7.100474247 | 6.228882015 |
| ENSG00000147604 | RPL7 | -2.520368497 | 2.37E-05 | -5.948317011 | 1.06E-25 | -8.468685508 | 13.83696579 | 12.50910266 | 12.67101473 | 10.59383186 | 10.6405557 | 7.02591204 | 7.399884913 |
| ENSG00000139193 | CD27 | -5.800223441 | 0.012783133 | -2.669379989 | 0.124418769 | -8.46960343 | 4.815609471 | 3.076665209 | 3.369317972 | 2 | 2.14022021 | 2.560105256 | 2.509591799 |
| ENSG00000213973 | ZNF99 | -3.103299821 | 0.028626428 | -5.366845874 | 0.000114785 | -8.470145695 | 4.467311984 | 3.686303414 | 5.39244166 | 2.947924294 | 2.38539787 | 2.225793514 | 2.117335275 |
| ENSG00000166482 | MFAP4 | -5.19366202 | 2.22E-43 | -3.297175868 | 3.20E-18 | -8.490837888 | 13.98833381 | 13.22554292 | 13.33431742 | 8.29513208 | 8.46055493 | 10.28283693 | 10.24487326 |
| ENSG00000169495 | HTRA4 | -5.589013049 | 0.05900662 | -2.905619054 | 0.305537213 | -8.494632103 | 5.095959458 | 3.076665209 | 4.874587708 | 2.270786764 | 2 | 3.215043717 | 2 |
| ENSG00000130203 | APOE | -7.661671916 | 9.77E-19 | -0.839493052 | 0.404588211 | -8.501164967 | 11.55361133 | 10.54992047 | 9.541233192 | 3.663521627 | 3.65239003 | 10.52700461 | 8.918588063 |
| C21ORF33 | C21ORF33 | 2.764328772 | 1.75E-05 | -11.26783267 | 7.28E-12 | -8.503503899 | 5.532353208 | 6.666836837 | 6.073378139 | 8.866956935 | 8.89230135 | 2 | 2 |
| ENSG00000154655 | L3MBTL4 | -7.123833766 | 0.094901214 | -1.391077815 | 0.752794932 | -8.514911582 | 2 | 5.131603625 | 2 | 2 | 2 | 3.296906311 | 2.632958241 |
| ENSG00000138670 | RASGEF1B | -6.714249292 | 0.001224561 | -1.801392401 | 0.420881674 | -8.515641693 | 2 | 5.303523747 | 8.486635645 | 2.600425781 | 2.14022021 | 5.515547099 | 5.197927245 |
| ENSG00000120658 | ENOX1 | -3.646558176 | 0.008394964 | -4.87327213 | 0.000249325 | -8.519830306 | 8.340423424 | 4.11352228 | 7.225742779 | 3.663521627 | 4.37434574 | 3.517841605 | 2.885414003 |
| ENSG00000262599 | | 2.300716665 | 0.21839016 | -10.82527777 | 1.46E-06 | -8.524561103 | 6.138360429 | 6.464272742 | 2 | 7.966401501 | 8.00183183 | 2 | 2 |
| ENSG00000165156 | ZHX1 | -11.1628576 | 5.93E-09 | 2.637275244 | 0.034537867 | -8.525582353 | 4.815609471 | 8.077869079 | 7.858115942 | 2 | 2 | 10.1584972 | 9.994220888 |
| ENSG00000121749 | TBC1D15 | -10.20629097 | 3.17E-09 | 1.679187787 | 0.04896152 | -8.527103185 | 5.70932728 | 7.232742069 | 6.203069917 | 2 | 2 | 8.317962987 | 8.024537783 |
| ENSG00000147082 | CCNB3 | -4.181746413 | 0.117474983 | -4.349476455 | 0.091962344 | -8.531222868 | 2 | 2 | 6.718950348 | 2.498691189 | 2.59489879 | 2.526562374 | 2.466008509 |
| ENSG00000145431 | PDGFC | -10.16274107 | 2.81E-10 | 1.631054027 | 0.001817437 | -8.531687047 | 6.468621203 | 6.666836837 | 6.322058544 | 2 | 2 | 8.025527571 | 8.148612304 |
| ENSG00000253313 | C1orf210 | -3.542120474 | 0.117068598 | -4.99666063 | 0.020430732 | -8.538781103 | 5.095959458 | 2 | 3.369317972 | 2.141737246 | 2.49394783 | 2.139668783 | 2.117335275 |
| ENSG00000136160 | EDNRB | -2.142512811 | 0.135211488 | -6.396994331 | 7.48E-07 | -8.539507142 | 5.330594134 | 7.393647515 | 8.608698767 | 6.078767801 | 4.85329164 | 2.987421008 | 2.225841198 |
| PAPD7 | | 2.69890454 | 0.243995258 | -11.24052974 | 1.07E-05 | -8.541625201 | 2 | 7.538394143 | 4.523675391 | 8.71221318 | 8.8597051 | 2 | 2 |
| ENSG00000165816 | VWA2 | -4.034281966 | 0.075317795 | -4.512544053 | 0.031407323 | -8.546826019 | 3.327711234 | 2 | 4.874587708 | 2.270786764 | 2.14022021 | 2.183373728 | 2.117335275 |
| ENSG00000176533 | GNG7 | -7.083173098 | 0.000597845 | -1.472374717 | 0.52560346 | -8.555547815 | 9.473095744 | 4.442710129 | 5.156626401 | 2.947924294 | 2 | 6.329884754 | 6.769038269 |
| ENSG00000198830 | HMGN2 | -2.491651151 | 3.13E-08 | -6.07371094 | 1.53E-44 | -8.565362092 | 13.12856249 | 12.23944331 | 12.26604521 | 10.11558569 | 10.1233123 | 6.475428841 | 6.70252219 |
| ENSG00000164764 | SBSPON | -9.831310988 | 1.11E-07 | 1.265358485 | 0.28061232 | -8.565952503 | 7.579279349 | 8.101365653 | 6.718950348 | 2 | 2.14022021 | 7.204892513 | 9.568485262 |
| FAM132A | | -0.021682735 | 0.997023429 | -8.547044429 | 0.029864907 | -8.568727164 | 2 | 2 | 5.156626401 | 3.95996233 | 3.69849104 | 2 | 2 |
| ENSG00000182534 | MXRA7 | -12.35399936 | 7.96E-11 | 3.785104003 | 0.001419617 | -8.568895359 | 8.696033109 | 9.479495904 | 6.073378139 | 2 | 2 | 12.20004322 | 12.59921797 |
| ENSG00000273111 | LYPD4 | -4.714866347 | NA | -3.855404194 | NA | -8.570270541 | 2 | 2 | 3.369317972 | 2 | 2 | 2.094598369 | 2 |
| ENSG00000126217 | MCF2L | -4.857180072 | 0.006460277 | -3.72545735 | 0.02899481 | -8.582637422 | 3.327711234 | 5.595927442 | 5.930865648 | 2.695456494 | 2 | 2.746468522 | 2.746601469 |
| ENSG00000249709 | ZNF564 | -1.656407914 | 0.126664884 | -6.937301915 | 1.35E-12 | -8.593709829 | 7.667647223 | 6.389896633 | 9.049077186 | 6.483440992 | 6.46120797 | 2.803598391 | 2.421067459 |
| ENSG00000183798 | EMILIN3 | -7.921652248 | 0.000223811 | -0.675473578 | 0.76623592 | -8.597125826 | 3.327711234 | 3.686303414 | 8.175061096 | 2.141737246 | 2.14022021 | 6.68757835 | 4.886954834 |
| ENSG00000102962 | CCL22 | -6.746059063 | 0.004659743 | -1.862006037 | 0.331905931 | -8.608065099 | 4.00722531 | 3.076665209 | 5.772716771 | 2 | 2.14022021 | 3.796405078 | 2.632958241 |
| ENSG00000263956 | NBPF11 | -8.588621875 | 3.56E-06 | -0.019871992 | 0.988303878 | -8.608493867 | 4.467311984 | 5.131603625 | 5.39244166 | 2 | 2 | 5.333556395 | 4.644600858 |
| ENSG00000203872 | C6orf163 | -4.642164344 | 6.61E-06 | -3.966733813 | 1.26E-05 | -8.608898157 | 5.330594134 | 5.131603625 | 5.930865648 | 2.600425781 | 2.38539787 | 2.911488689 | 2.509591799 |
| ENSG00000171476 | HOPX | -8.150395409 | 7.74E-10 | -0.474627519 | 0.765722308 | -8.625022928 | 8.910412251 | 9.018611864 | 9.589475645 | 3.023133209 | 2 | 9.475267633 | 7.124815549 |
| ENSG00000035115 | SH3YL1 | -9.973094041 | 1.95E-07 | 1.341404725 | 0.301388454 | -8.631689316 | 6.73718411 | 6.844429755 | 4.058998049 | 2 | 2 | 7.711553069 | 7.51663051 |
| ENSG00000129173 | E2F8 | -4.463457689 | 0.426591614 | -4.171107697 | NA | -8.634565387 | 2 | 4.11352228 | 2 | 2.141737246 | 2 | 2 | 2.172608095 |
| ENSG00000185920 | PTCH1 | -3.441483434 | 0.000299697 | -5.200367555 | 1.44E-08 | -8.641850989 | 7.940205782 | 8.968383114 | 10.5916593 | 6.276156955 | 6.1322378 | 4.914492341 | 4.254128754 |
| DFNA5 | DFNA5 | 2.947447778 | 0.00310241 | -11.58996188 | 9.47E-11 | -8.642514104 | 7.33131396 | 6.311476754 | 4.874587708 | 9.375641031 | 9.38697919 | 2 | 2 |
| ENSG00000188603 | CLN3 | -1.447655537 | 0.151335244 | -7.194948335 | 3.35E-16 | -8.642603872 | 8.816894989 | 7.954278496 | 10.49998282 | 8.027038952 | 8.06698125 | 3.215043717 | 3.071407821 |
| ENSG00000132746 | ALDH3B2 | -0.43456371 | 0.925925983 | -8.222534378 | 0.04466989 | -8.657098087 | 2 | 2 | 4.874587708 | 3.406930997 | 3.28037825 | 2 | 2 |
| ENSG00000186377 | CYP4X1 | -7.122038384 | 0.106975585 | -1.536315939 | 0.736592879 | -8.658354323 | 2 | 5.131603625 | 2 | 2 | 2 | 3.355392153 | 2.326754077 |
| ENSG00000175189 | INHBC | -4.698902984 | 0.178468905 | -3.96947095 | 0.227933253 | -8.668373933 | 2 | 5.131603625 | 2 | 2 | 2.2680111 | 2.225793514 | 2.225841198 |
| ENSG00000185442 | FAM174B | -6.11173065 | 0.000539749 | -2.56057086 | 0.167093515 | -8.67230151 | 8.609439662 | 7.002537973 | 2 | 2.600425781 | 2.77779907 | 5.032703865 | 5.04340913 |
| ENSG00000082293 | COL19A1 | -9.351515316 | 2.71E-05 | 0.675566399 | 0.729761219 | -8.675948917 | 6.89194353 | 3.686303414 | 4.874587708 | 2 | 2 | 6.998648055 | 5.23734678 |
| ENSG00000186472 | PCLO | -3.699056986 | 0.000154408 | -4.998623253 | 1.24E-07 | -8.697680238 | 5.330594134 | 5.595927442 | 7.225742779 | 3.462030052 | 3.08597292 | 2.656294034 | 2.671842496 |
| ENSG00000183206 | POTEC | -5.722021535 | 0.315977132 | -2.976059234 | NA | -8.698080769 | 4.00722531 | 2 | 2 | 2 | 2 | 2 | 2.326754077 |
| ENSG00000082805 | ERC1 | -11.79844799 | 5.30E-12 | 3.093965264 | 0.000109562 | -8.70448273 | 8.910412251 | 7.467835385 | 7.342949392 | 2 | 2 | 10.96307448 | 11.34764456 |
| ENSG00000132821 | VSTM2L | -4.836547722 | 1.03E-08 | -3.868123206 | 4.34E-06 | -8.704670928 | 8.107517749 | 8.077869079 | 9.963514551 | 4.205751501 | 4.65713986 | 5.524070859 | 4.990558504 |
| ENSG00000169435 | RASSF6 | -1.912217925 | NA | -6.792803666 | NA | -8.705021591 | 2 | 3.686303414 | 2 | 2.270786764 | 2.2680111 | 2 | 2 |
| ENSG00000175040 | CHST2 | -3.622808264 | 3.21E-24 | -5.083572829 | 1.15E-47 | -8.706381093 | 12.01383685 | 11.86868155 | 11.27276688 | 8.128448191 | 8.16874151 | 6.752490997 | 6.693102083 |
| ENSG00000005513 | SOX8 | -5.456174556 | 0.002847427 | -3.263745065 | 0.079040984 | -8.719919621 | 2 | 6.666836837 | 7.858115942 | 2.947924294 | 2.38539787 | 4.331190017 | 3.383972333 |
| ENSG00000122696 | SLC25A51 | -9.930833826 | 1.35E-09 | 1.210898415 | 0.053870623 | -8.71993541 | 6.563806279 | 6.04685525 | 6.203069917 | 2 | 2 | 7.576706933 | 7.307219812 |
| ENSG00000160963 | COL26A1 | -6.903232752 | 0.006218119 | -1.819726119 | 0.515470896 | -8.722958872 | 2 | 5.838937808 | 9.272377397 | 2.947924294 | 2 | 6.387074688 | 5.651404445 |
| ENSG00000155657 | TTN | -7.298688991 | 8.09E-20 | -1.429552842 | 0.031032608 | -8.728241833 | 7.485143553 | 7.701025996 | 8.65477198 | 2.498691189 | 2.49394783 | 6.838725184 | 6.418016952 |
| ENSG00000161647 | MPP3 | -9.235880668 | 0.00017318 | 0.502616922 | 0.833626336 | -8.733263746 | 6.653098118 | 5.457120953 | 2 | 2 | 2 | 6.043083161 | 6.151894053 |
| ENSG00000196730 | DAPK1 | -7.932083006 | 1.64E-17 | -0.805470164 | 0.423841772 | -8.73755317 | 8.257427002 | 9.514922632 | 9.868959282 | 3.09461492 | 2.2680111 | 8.102245405 | 8.910481888 |
| ENSG00000197757 | HOXC6 | 2.781743257 | 0.000109273 | -11.52015758 | 5.31E-12 | -8.738414322 | 5.532353208 | 7.002537973 | 6.322058544 | 9.108419034 | 9.18523054 | 2 | 2 |
| ENSG00000162461 | SLC25A34 | -3.486057466 | 0.002696999 | -5.258017198 | 2.43E-06 | -8.744074664 | 7.750913381 | 5.131603625 | 8.805342542 | 4.574143677 | 4.63360314 | 3.411599105 | 3.209956562 |
| ENSG00000105141 | CASP14 | -1.767982629 | 0.757468139 | -6.978261313 | NA | -8.746243943 | 3.327711234 | 3.076665209 | 2 | 2.600425781 | 2 | 2 | 2 |
| ENSG00000275176 | | -1.427090698 | 0.036933389 | -7.323155226 | 1.80E-31 | -8.750245923 | 10.25038534 | 9.799806309 | 8.677268317 | 8.356399318 | 8.23866319 | 3.035909414 | 3.36036153 |
| ENSG00000282013 | | -2.011336843 | 0.09200362 | -6.743923212 | 1.55E-08 | -8.755260055 | 5.330594134 | 6.844429755 | 4.523675391 | 4.470469939 | 3.7864959 | 2.183373728 | 2.172608095 |
| ENSG00000196167 | COLCA1 | -5.224251378 | 0.069118897 | -3.532172815 | 0.139445727 | -8.756424192 | 2 | 3.076665209 | 4.523675391 | 2 | 2.14022021 | 2.267001569 | 2.172608095 |
| ENSG00000131400 | NAPSA | -4.855420362 | 0.08813762 | -3.903612913 | 0.165909543 | -8.759033275 | 2 | 2 | 7.646393253 | 2.3892346 | 2.86123014 | 3.483289517 | 2.466008509 |
| ENSG00000168309 | FAM107A | -3.611816348 | 0.333096085 | -5.148755537 | 0.145619319 | -8.760571885 | 2 | 2 | 6.882775709 | 3.349643765 | 2 | 2.3070652 | 2.374681325 |
| ENSG00000274287 | | 2.509532951 | 0.191385897 | -11.27269051 | 7.81E-07 | -8.763157561 | 6.563806279 | 6.899080544 | 2 | 8.643859774 | 8.6202631 | 2 | 2 |
| ENSG00000184408 | KCND2 | -1.622495207 | 0.475123305 | -7.15374341 | 0.000355261 | -8.776238616 | 3.327711234 | 7.73146271 | 4.523675391 | 4.927308402 | 4.83276841 | 2.34604626 | 2 |
| ENSG00000146700 | SSC4D | -5.061922099 | 0.003918069 | -3.715892052 | 0.03672502 | -8.777814151 | 2 | 7.315436877 | 8.585098385 | 3.227757282 | 3.2184422 | 3.878466556 | 4.353147976 |
| ENSG00000124743 | KLHL31 | -4.25542698 | 0.121505956 | -4.527758492 | 0.084356716 | -8.783185471 | 2 | 5.946638857 | 2 | 2.3892346 | 2.2680111 | 2.42098416 | 2.117335275 |
| ENSG00000049247 | UTS2 | -5.722022661 | 0.315977132 | -3.066254937 | NA | -8.788277598 | 4.00722531 | 2 | 2 | 2 | 2 | 2.3070652 | 2 |
| ENSG00000262102 | | 1.838633944 | 0.040274925 | -10.62863604 | 7.19E-20 | -8.790002097 | 7.579279349 | 8.213429689 | 9.330306409 | 10.40770778 | 10.3617753 | 2 | 2.172608095 |
| ENSG00000133067 | LGR6 | -7.048940524 | 0.112812992 | -1.746561326 | 0.701839324 | -8.79550185 | 5.095959458 | 2 | 2 | 2 | 2 | 2.384001718 | 3.128433649 |
| ENSG00000056487 | PHF21B | -2.951105923 | 0.4329 | -5.844412848 | 0.104679282 | -8.79551877 | 4.815609471 | 2 | 2 | 2.498691189 | 2.14022021 | 2.094598369 | 2 |
| ENSG00000198780 | FAM169A | -8.880227701 | 0.000417585 | 0.083657378 | 0.973405153 | -8.796570323 | 2 | 5.131603625 | 6.322058544 | 2 | 2 | 5.130840083 | 5.596882677 |
| DFNB59 | DFNB59 | 0.173907023 | 0.967784468 | -8.973094126 | 0.018179008 | -8.799187103 | 5.532353208 | 2 | 2 | 4.268917409 | 4.3456612 | 2 | 2 |
| LOC100505549 | | -0.375850398 | 0.866106266 | -8.425968206 | 0.000263926 | -8.801818604 | 3.327711234 | 3.076665209 | 4.523675391 | 3.462030052 | 3.55551834 | 2 | 2 |
| ENSG00000178127 | NDUFV2 | -1.012998663 | 0.018486008 | -7.793799238 | 1.71E-80 | -8.806797901 | 10.82658808 | 10.30883598 | 10.19160738 | 9.431924894 | 9.48805095 | 3.534812207 | 3.183290849 |
| ENSG00000185332 | TMEM105 | -4.605225129 | 0.037892098 | -4.211631868 | 0.030705557 | -8.816856997 | 4.00722531 | 2 | 4.523675391 | 2.141737246 | 2.14022021 | 2.139668783 | 2.225841198 |
| ENSG00000110925 | CSRNP2 | -11.15200943 | 3.83E-11 | 2.333273607 | 0.001960387 | -8.818735828 | 6.468621203 | 7.35507212 | 8.110401118 | 2 | 2 | 9.862952363 | 9.660173108 |
| ZCCHC6 | | 3.048727327 | 0.136445865 | -11.87199321 | 6.12E-07 | -8.823265879 | 2 | 7.790474954 | 6.629488387 | 9.747510575 | 9.77766243 | 2 | 2 |
| ENSG00000239389 | PCDHA13 | -1.807756167 | 0.606861222 | -7.019857217 | 0.032017989 | -8.827613384 | 2 | 5.838937808 | 2 | 3.615725078 | 2.59489879 | 2.094598369 | 2 |
| ENSG00000126231 | PROZ | -6.327750635 | 0.173924989 | -2.507194626 | 0.586489016 | -8.834945261 | 2 | 4.442710129 | 2 | 2 | 2 | 2.3070652 | 2.374681325 |
| ENSG00000149328 | GLB1L2 | -8.212637917 | 0.032740713 | -0.62254555 | 0.883777495 | -8.835183467 | 2 | 6.140560021 | 2 | 2 | 2 | 4.029788235 | 4.376876162 |
| ENSG00000135436 | FAM186B | -7.296346575 | 0.016923525 | -1.539847577 | 0.598500734 | -8.836194152 | 2 | 3.076665209 | 5.156626401 | 2 | 2 | 2.987421008 | 3.042026731 |
| ENSG00000185432 | METTL7A | -4.571068776 | 4.89E-12 | -4.270936279 | 7.74E-11 | -8.842005055 | 9.602033055 | 9.386925327 | 10.9833682 | 5.720439317 | 5.70538658 | 5.941748584 | 6.041166521 |
| ENSG00000137133 | HINT2 | -1.471265027 | 0.015072011 | -7.372557783 | 4.04E-32 | -8.843822811 | 8.138771685 | 7.393647515 | 8.631919287 | 6.722207358 | 6.67228943 | 2.457042231 | 2.551896972 |

| | | | | | | | | | | | | | |
|---|---|---|---|---|---|---|---|---|---|---|---|---|---|
| FAM46C | | 1.323203032 | 0.522107672 | -10.17235291 | 1.06E-05 | -8.849149876 | 2 | 5.946638857 | 5.39244166 | 6.449761039 | 6.35006802 | 2 | 2 |
| ENSG00000100784 | RPS6KA5 | -3.651067716 | 5.78E-09 | -5.200817337 | 2.84E-17 | -8.851885053 | 7.435667355 | 8.687092496 | 7.552162999 | 4.694064037 | 4.50979031 | 3.429858444 | 3.474731315 |
| ENSG00000166148 | AVPR1A | -3.848235967 | 0.063373552 | -5.004206108 | 0.011122193 | -8.852442075 | 4.815609471 | 3.076665209 | 4.874587708 | 2.270786764 | 2.49394783 | 2.34604626 | 2 |
| ENSG00000065675 | PRKCQ | -4.894035037 | 0.092604558 | -3.96315447 | 0.139450354 | -8.857189508 | 2 | 5.303523747 | 2 | 2.141737246 | 2.14022021 | 2.384001718 | 2.117335275 |
| ENSG00000229685 | | -0.669681593 | 0.827069926 | -8.196348046 | 0.004848864 | -8.86602964 | 3.327711234 | 4.442710129 | 2 | 3.406930997 | 2.94009902 | 2 | 2 |
| ENSG00000205790 | DPP9-AS1 | -5.15581557 | 0.061765197 | -3.71312009 | 0.159274764 | -8.86893566 | 6.138360429 | 2 | 2 | 2.270786764 | 2.14022021 | 2.457042231 | 2.551896972 |
| ENSG00000111057 | KRT18 | -7.205494005 | 5.15E-45 | -1.666620892 | 0.002334223 | -8.872114898 | 13.06795125 | 12.58294393 | 12.96874405 | 5.775854763 | 5.80356929 | 10.48723515 | 11.70766686 |
| ENSG00000070831 | CDC42 | -0.254236562 | 0.647611714 | -8.631855866 | 1.97E-83 | -8.886092428 | 11.84408627 | 10.88722087 | 11.66769225 | 11.27271974 | 11.2619781 | 3.584558147 | 3.430065404 |
| ENSG00000275199 | | 2.615342728 | 1.08E-13 | -11.50336603 | 2.88E-14 | -8.888023304 | 7.667647223 | 7.874700259 | 7.858115942 | 10.42037002 | 10.3769158 | 2 | 2.059860178 |
| ENSG00000107295 | SH3GL2 | -4.903350562 | 0.190366567 | -3.987312017 | 0.23639043 | -8.890662579 | 2 | 4.442710129 | 2 | 2.141737246 | 2 | 2.094598369 | 2.172608095 |
| ENSG00000106336 | FBXO24 | -4.790967994 | 4.51E-07 | -4.103746911 | 3.07E-06 | -8.894714905 | 7.218819781 | 5.595927442 | 5.772716771 | 2.600425781 | 2.94009902 | 3.059555873 | 3.15612297 |
| ENSG00000115423 | DNAH6 | -4.909396728 | 0.189533864 | -3.987386504 | 0.236114884 | -8.896783232 | 2 | 4.442710129 | 2 | 2 | 2.14022021 | 2.094598369 | 2.172608095 |
| ENSG00000005844 | ITGAL | -6.787403344 | 0.000653446 | -2.11833373 | 0.241720341 | -8.905737073 | 5.095959458 | 4.442710129 | 6.534109959 | 2 | 2.2680111 | 4.530483275 | 2.509591799 |
| ENSG00000170381 | SEMA3E | -2.220503513 | 0.381987044 | -6.691343857 | 0.014208037 | -8.911847371 | 2 | 3.076665209 | 4.523675391 | 2.600425781 | 2.38539787 | 2.048074405 | 2 |
| ENSG00000253293 | HOXA10 | -0.232066183 | 0.77088407 | -8.679860892 | 5.79E-28 | -8.911927075 | 7.33131396 | 8.414402626 | 8.076950877 | 7.66556918 | 7.89452713 | 2.094598369 | 2.326754077 |
| ENSG00000260456 | C16orf95 | -3.854147484 | 6.98E-05 | -5.06806292 | 7.33E-08 | -8.922210404 | 7.82963488 | 5.946638857 | 8.805342542 | 4.299492648 | 4.43005853 | 3.465698119 | 3.496556178 |
| ENSG00000197728 | RPS26 | -1.862533858 | 0.010140946 | -7.059909052 | 9.83E-27 | -8.922442911 | 12.76659341 | 11.14918383 | 11.2616444 | 10.02586197 | 10.1121791 | 5.096960836 | 5.005857272 |
| ENSG00000120885 | CLU | -12.09598686 | 8.61E-12 | 3.172273717 | 0.000751755 | -8.923713145 | 7.096806373 | 8.317412869 | 9.099922433 | 2 | 2 | 10.80516598 | 12.02939606 |
| ENSG00000118997 | DNAH7 | -3.306188189 | 0.071924384 | -5.62482031 | 0.001128494 | -8.931008499 | 7.940205782 | 3.686303414 | 4.058998049 | 3.881293667 | 3.45167086 | 2.225793514 | 2.782576693 |
| ENSG00000181350 | LRRC75A | -6.247230689 | 1.06E-24 | -2.685065441 | 5.27E-07 | -8.93229613 | 8.540903468 | 8.317412869 | 8.460932916 | 3.406930997 | 2.68924543 | 6.104049656 | 5.524371359 |
| ENSG00000183682 | BMP8A | -2.897290428 | 0.047389207 | -6.036255466 | 2.84E-05 | -8.933545894 | 4.467311984 | 5.457120953 | 5.39244166 | 3.162721474 | 2.94009902 | 2.3070652 | 2 |
| ENSG00000183476 | SH2D7 | -7.312914106 | 0.087631985 | -1.623463913 | 0.713040563 | -8.936378019 | 5.330594134 | 2 | 2 | 2 | 2 | 3.235948948 | 2.671842496 |
| ENSG00000176971 | FIBIN | -2.819308032 | 0.019422499 | -6.120299217 | 4.32E-08 | -8.939607248 | 8.16936288 | 7.874700259 | 10.65536953 | 6.664650034 | 6.73998187 | 4.301748724 | 3.100202476 |
| ENSG00000163666 | HESX1 | -4.739421164 | 0.005361039 | -4.203594486 | 0.006521275 | -8.94301565 | 4.00722531 | 3.076665209 | 5.772716771 | 2.141737246 | 2.38539787 | 2.42098416 | 2.326754077 |
| ENSG00000008516 | MMP25 | -5.588913485 | 1.69E-06 | -3.354392925 | 0.002719265 | -8.94330641 | 7.750913381 | 4.710564154 | 6.629488387 | 2.784612485 | 2.49394783 | 3.851628042 | 3.8919413 |
| ENSG00000114638 | UPK1B | -3.623415465 | 0.008955678 | -5.320746297 | 7.54E-05 | -8.944161762 | 4.815609471 | 3.686303414 | 5.59507254 | 2.498691189 | 2.68924543 | 2.094598369 | 2.326754077 |
| ENSG00000243509 | TNFRSF6B | 2.11717415 | 0.35702693 | -11.06242038 | 1.01E-05 | -8.94524623 | 7.33131396 | 4.710564154 | 2 | 8.013270196 | 8.06037647 | 2 | 2 |
| ENSG00000198739 | LRRTM3 | -5.348717897 | NA | -3.607198229 | NA | -8.955916126 | 2 | 3.686303414 | 2 | 2 | 2 | 2 | 2.172608095 |
| ENSG00000073350 | LLGL2 | -6.393050103 | 1.15E-05 | -2.563584407 | 0.088390629 | -8.956634511 | 8.816894989 | 6.844429755 | 4.058998049 | 2.784612485 | 2.49394783 | 5.494014735 | 4.766376438 |
| ENSG00000108309 | RUNDC3A | -5.576260859 | 0.101847838 | -3.385964266 | 0.301898338 | -8.962225124 | 2 | 5.946638857 | 2 | 2 | 2.2680111 | 2.746468522 | 2.326754077 |
| ENSG00000114491 | UMPS | -10.35150231 | 8.73E-07 | 1.388521435 | 0.403003915 | -8.962980876 | 4.815609471 | 7.954278496 | 4.874587708 | 2 | 2 | 8.065491673 | 8.004685229 |
| ENSG00000078114 | NEBL | -4.800292958 | 0.000717672 | -4.168090963 | 0.002733705 | -8.968383921 | 5.095959458 | 7.232742069 | 7.098165388 | 3.462030052 | 2.14022021 | 2.885263063 | 3.600986722 |
| ENSG00000147676 | MAL2 | -2.819832398 | 0.236944877 | -6.168775167 | 0.010855766 | -8.988607564 | 4.00722531 | 4.442710129 | 2 | 2.600425781 | 2.2680111 | 2.094598369 | 2 |
| ENSG00000110025 | SNX15 | -2.489448759 | 0.000175762 | -6.502247605 | 2.71E-24 | -8.991696364 | 9.031919741 | 8.820493869 | 10.26770388 | 7.026323566 | 7.10768997 | 3.753554863 | 3.407202941 |
| ENSG00000154025 | SLC5A10 | -4.027467709 | 0.076121912 | -4.967523868 | 0.022614967 | -8.994991577 | 6.89194353 | 2 | 3.369317972 | 2.600425781 | 2.77779907 | 2.183373728 | 2.592996859 |
| ENSG00000159409 | CELF3 | -5.287422401 | 0.316965083 | -3.709799834 | 0.464069117 | -8.997222236 | 4.815609471 | 2 | 2 | 2 | 2.14022021 | 2.384001718 | 2 |
| ENSG00000132522 | GPS2 | -2.495879482 | 0.000130459 | -6.51422681 | 1.35E-25 | -9.010106292 | 9.820154628 | 9.7032946 | 11.16546284 | 7.847284664 | 7.9809999 | 4.188345803 | 4.254128754 |
| ENSG00000161328 | LRRC56 | 0.415154999 | 0.80351594 | -9.430936084 | 2.69E-06 | -9.015781085 | 4.00722531 | 3.686303414 | 5.39244166 | 4.846787657 | 4.93258395 | 2 | 2 |
| FAM46B | | -1.275027707 | 0.803231804 | -7.745841124 | 0.092628209 | -9.020868831 | 4.467311984 | 2 | 2 | 3.023133209 | 2.2680111 | 2 | 2 |
| ENSG00000112530 | PACRG | -4.920635672 | 0.101317769 | -4.100315804 | 0.140205611 | -9.020951476 | 2 | 2 | 5.39244166 | 2.141737246 | 2.14022021 | 2.094598369 | 2.374681325 |
| ENSG00000186714 | CCDC73 | -3.534066641 | 0.110888355 | -5.491448592 | 0.011099437 | -9.025515233 | 4.467311984 | 3.076665209 | 3.369317972 | 2.141737246 | 2.38539787 | 2.139668783 | 2 |
| ENSG00000121316 | PLBD1 | -6.249731075 | 9.63E-07 | -2.780238627 | 0.036333082 | -9.029969702 | 9.521325812 | 7.790474954 | 5.39244166 | 3.289987342 | 2.77779907 | 6.013120342 | 5.28828468 |
| ENSG00000113532 | ST8SIA4 | -4.40086726 | 2.85E-05 | -4.631446485 | 4.23E-06 | -9.032313745 | 6.009025997 | 5.131603625 | 7.451345288 | 2.784612485 | 3.2184422 | 2.987421008 | 2.817676617 |
| ENSG00000131409 | LRRC4B | -6.150814966 | 3.19E-08 | -2.886955554 | 0.008426519 | -9.037770519 | 8.257427002 | 7.002537973 | 7.225742779 | 3.162721474 | 2.14022021 | 5.279204176 | 4.365060851 |
| ENSG00000146250 | PRSS35 | -5.732426358 | 1.31E-12 | -3.307462401 | 3.12E-05 | -9.039888758 | 9.358295023 | 8.540087354 | 7.342949392 | 3.566290607 | 3.45167086 | 4.996409624 | 5.794819115 |
| ENSG00000103485 | QPRT | -5.411317114 | 1.59E-15 | -3.631332846 | 1.30E-07 | -9.04264996 | 12.68953976 | 10.73405387 | 12.16619564 | 6.580005553 | 6.83591385 | 8.353071024 | 8.548580051 |
| KIAA1683 | | -0.819455867 | 0.862549976 | -8.22321805 | 0.048204701 | -9.042673917 | 2 | 2 | 4.874587708 | 3.09461492 | 3.15372745 | 2 | 2 |
| ENSG00000130377 | ACSBG2 | -6.030379454 | 0.032965099 | -3.014697064 | 0.207305039 | -9.045076518 | 5.330594134 | 3.076665209 | 2 | 2.141737246 | 2 | 2.656294034 | 2.326754077 |
| ENSG00000116151 | MORN1 | -8.938552657 | 0.001473428 | -0.106899518 | 0.971501984 | -9.045452175 | 6.73718411 | 3.686303414 | 2 | 2 | 2 | 5.385935071 | 5.12973572 |
| ENSG00000076864 | RAP1GAP | -3.936320765 | 0.033732905 | -5.109282795 | 0.004137709 | -9.04560356 | 8.340423424 | 6.140560021 | 2 | 3.515101962 | 3.74316441 | 3.12825509 | 2.746601469 |
| ENSG00000203952 | CCDC160 | -5.351245399 | NA | -3.704699042 | NA | -9.055944441 | 2 | 3.686303414 | 2 | 2 | 2 | 2.048074405 | 2.117335275 |
| ENSG00000185818 | NAT8L | -5.557439942 | 0.060113247 | -3.502538417 | 0.246468033 | -9.05997836 | 3.327711234 | 2 | 8.561105505 | 3.227757282 | 2 | 4.209638631 | 3.560117069 |
| ENSG00000111729 | CLEC4A | -3.093615941 | 0.246202736 | -5.969359255 | 0.017180311 | -9.062975196 | 2 | 6.728495673 | 4.058998049 | 3.227757282 | 2.94009902 | 2.384001718 | 2 |
| ENSG00000188729 | OSTN | -7.626437245 | 0.00603454 | -1.437757604 | 0.577689617 | -9.06419485 | 2 | 4.11352228 | 5.156626401 | 2 | 2 | 3.517841605 | 2.918126211 |
| ENSG00000239887 | C1orf226 | -6.228130235 | 1.21E-13 | -2.838663224 | 0.00060039 | -9.066793459 | 7.384433967 | 8.540087354 | 9.399592459 | 3.406930997 | 3.01487888 | 6.247352872 | 5.459179378 |
| ENSG00000184697 | CLDN6 | -5.490354241 | 7.26E-05 | -3.577119961 | 0.00396904 | -9.067474202 | 6.653098118 | 4.442710129 | 4.874587708 | 2.3892346 | 2.2680111 | 2.62493832 | 3.261853426 |
| ENSG00000182580 | EPHB3 | -3.33661751 | 3.01E-05 | -5.73347956 | 1.27E-13 | -9.07009707 | 9.410448395 | 8.169643011 | 9.413059136 | 5.709096143 | 5.99120031 | 4.311629396 | 3.100202476 |
| SSFA2 | | 3.420202164 | 0.110962 | -12.50890703 | 3.39E-07 | -9.088704865 | 2 | 7.099070191 | 8.486635645 | 10.78928396 | 10.7458412 | 2 | 2 |
| ENSG00000185313 | SCN10A | -6.787714655 | 0.141312369 | -2.3014087 | 0.620241915 | -9.089123354 | 2 | 2 | 4.874587708 | 2 | 2 | 2.225793514 | 2.746601469 |
| ENSG00000188818 | ZDHHC11 | -3.910597957 | 7.50E-09 | -5.179499429 | 4.87E-15 | -9.090097386 | 8.107517749 | 7.189546432 | 7.734844356 | 4.139692846 | 4.16043394 | 3.600766115 | 2.851942824 |
| ENSG00000164616 | FBXL21 | -2.313590056 | NA | -6.786762896 | NA | -9.100352952 | 2 | 3.686303414 | 2 | 2.3892346 | 2 | 2 | 2 |
| ENSG00000073067 | CYP2W1 | -6.64826247 | 0.0100797 | -2.460348368 | 0.257500244 | -9.108610838 | 3.327711234 | 3.076665209 | 4.058998049 | 2 | 2 | 2.560105256 | 2.277179783 |
| ENSG00000153157 | SYCP2L | -5.344862832 | 0.053302303 | -3.774755322 | 0.139882696 | -9.119618154 | 2 | 5.72254323 | 2 | 2.141737246 | 2.14022021 | 2.34604626 | 2.421067459 |
| ENSG00000275004 | ZNF280B | 0.867006497 | 0.82454488 | -9.990466838 | 0.0043482 | -9.123460341 | 2 | 6.464272742 | 2 | 5.753943344 | 5.84514933 | 2 | 2 |
| ENSG00000272573 | MUSTN1 | -3.025209809 | 0.209836547 | -6.101335096 | 0.006951324 | -9.126544905 | 2 | 3.076665209 | 7.029880454 | 3.023133209 | 3.39680241 | 2.34604626 | 2.059860178 |
| ENSG00000002587 | HS3ST1 | -7.479676011 | 0.088683678 | -1.652494525 | 0.716686548 | -9.132170537 | 2 | 5.457120953 | 2 | 2 | 2 | 3.5515855 | 2.277179783 |
| ENSG00000170439 | METTL7B | -7.744282548 | 0.022810377 | -1.390272372 | 0.685308246 | -9.13455492 | 2 | 2 | 7.098165388 | 2.141737246 | 2 | 3.767980144 | 4.810605986 |
| ENSG00000135406 | PRPH | -2.884993976 | 0.39960716 | -6.256309328 | 0.058578003 | -9.141303304 | 2 | 5.131603625 | 2 | 2.498691189 | 2.38539787 | 2.094598369 | 2 |
| ENSG00000085276 | MECOM | 1.096223441 | 0.255069337 | -10.24003077 | 3.86E-09 | -9.143807332 | 5.330594134 | 5.595927442 | 4.523675391 | 6.306580752 | 6.18985553 | 2 | 2 |
| ENSG00000144120 | TMEM177 | -5.373406159 | 0.008781104 | -3.775402278 | 0.073502138 | -9.148808437 | 11.09308737 | 8.337339476 | 2 | 4.358765249 | 4.79082474 | 6.121005698 | 5.909074772 |
| ENSG00000270885 | RASL10B | -6.641779007 | 4.57E-35 | -2.513294687 | 1.93E-08 | -9.155073694 | 8.928409675 | 8.732943944 | 8.825624954 | 3.162721474 | 3.01487888 | 6.693258166 | 5.964937606 |
| ENSG00000133488 | SEC14L4 | -4.558139254 | 0.000995812 | -4.623065075 | 0.000264451 | -9.181204329 | 4.467311984 | 4.11352228 | 5.930865648 | 2.270786764 | 2.49394783 | 2.225793514 | 2.509591799 |
| ENSG00000134569 | LRP4 | -5.551129981 | 1.12E-23 | -3.632478162 | 4.19E-11 | -9.183608144 | 10.39976609 | 11.60252924 | 10.54960077 | 5.720439317 | 5.3193943 | 7.38764416 | 7.327148877 |
| ENSG00000118298 | CA14 | -5.914670816 | 0.001428136 | -3.270750099 | 0.080069004 | -9.185420915 | 7.435667355 | 7.051611315 | 2 | 2.784612485 | 2.14022021 | 3.917810786 | 3.753786374 |
| ENSG00000113430 | IRX4 | -1.110796052 | 0.642697566 | -8.074886486 | 0.001390996 | -9.185682538 | 4.00722531 | 3.076665209 | 3.369317972 | 2.498691189 | 3.2184422 | 2 | 2 |
| ENSG00000174721 | FGFBP3 | -4.079700049 | 4.53E-07 | -5.120148678 | 7.87E-11 | -9.199848727 | 7.975256505 | 7.572420273 | 6.322058544 | 4.070463721 | 3.50452879 | 2.885263063 | 3.383972333 |
| ENSG00000149809 | TM7SF2 | -5.225748195 | 2.09E-10 | -3.985138795 | 1.55E-06 | -9.21088699 | 11.7536074 | 9.664384941 | 9.868959282 | 5.639097847 | 5.65950892 | 6.593521701 | 7.019463152 |
| CXORF23 | CXORF23 | 1.289983191 | 0.317766838 | -10.50325948 | 2.54E-08 | -9.213276289 | 4.00722531 | 6.311476754 | 5.156626401 | 6.64692533 | 6.72335419 | 2 | 2 |
| ENSG00000055732 | MCOLN3 | -5.965033708 | 0.012863941 | -3.249557376 | 0.170653805 | -9.214591085 | 3.327711234 | 7.232742069 | 2 | 2.270786764 | 2.2680111 | 3.782262616 | 2.326754077 |
| ENSG00000165588 | OTX2 | -1.873823373 | 0.75247423 | -7.347203331 | NA | -9.221026705 | 2 | 4.11352228 | 2 | 2.695456494 | 2 | 2 | 2 |
| ENSG00000101162 | TUBB1 | -6.623571243 | 0.017686038 | -2.598380446 | 0.280327402 | -9.22195169 | 2 | 3.076665209 | 5.930865648 | 2 | 2.14022021 | 2.962551659 | 2.782576693 |
| C15ORF57 | C15ORF57 | 1.957197659 | 0.477372356 | -11.1792536 | 6.19E-05 | -9.222055944 | 2 | 7.572420273 | 2 | 7.949630994 | 8.03813956 | 2 | 2 |
| C11ORF70 | C11ORF70 | -0.067243687 | 0.978926263 | -9.15795506 | 0.000205725 | -9.225198747 | 2 | 4.710564154 | 4.874587708 | 4.358765249 | 4.16043394 | 2 | 2 |
| ENSG00000186010 | NDUFA13 | -1.754794694 | 0.000341826 | -7.472595551 | 6.42E-58 | -9.227390246 | 11.57090097 | 10.82859492 | 10.9788875 | 9.375641031 | 9.4506582 | 4.360042493 | 3.717076715 |
| ENSG00000179950 | PUF60 | -1.832139293 | 0.001073603 | -7.39702634 | 3.47E-45 | -9.229165633 | 11.87225278 | 10.60438382 | 11.34128524 | 9.535096495 | 9.53565307 | 4.487545924 | 4.062589736 |
| ENSG00000037280 | FLT4 | -3.819333165 | 0.024113792 | -5.4127327 | 0.000879192 | -9.232065864 | 2 | 5.595927442 | 5.772716771 | 2.3892346 | 2.86123014 | 2.267001569 | 2.225841198 |
| ENSG00000179008 | C14orf39 | -4.472698561 | 0.417669822 | -4.762285414 | NA | -9.234983975 | 2 | 4.11352228 | 2 | 2.14022021 | 2 | 2 | 2.117335275 |
| ENSG00000165995 | CACNB2 | -4.438402305 | 1.70E-09 | -4.79814636 | 1.38E-11 | -9.236548665 | 7.159102577 | 6.728495673 | 8.296287027 | 3.754611435 | 3.50452879 | 3.483289517 | 3.311948089 |
| ENSG00000100099 | HPS4 | -10.57878833 | 5.28E-10 | 1.339830436 | 0.106515827 | -9.23895789 | 5.866944937 | 6.899080544 | 7.502634626 | 2 | 2 | 8.257268579 | 8.164036804 |
| ENSG00000087085 | ACHE | -5.915666262 | 0.006363593 | -3.323911874 | 0.132570043 | -9.239578136 | 8.367060731 | 3.686303414 | 2 | 2.498691189 | 2.59489879 | 3.993419637 | 3.841670744 |
| ENSG00000166333 | ILK | -1.787852532 | 7.19E-05 | -7.453077686 | 9.30E-69 | -9.240930219 | 12.63852082 | 11.71912994 | 12.19376522 | 10.42514402 | 10.4678974 | 4.757085024 | 5.164234344 |
| ENSG00000152954 | NRSN1 | -4.869901954 | 0.112165694 | -4.371267003 | 0.095595607 | -9.241168957 | 3.327711234 | 2 | 4.058998049 | 2.141737246 | 2 | 2.139668783 | 2.059860178 |
| ENSG00000164007 | CLDN19 | -3.278233873 | 0.100124896 | -5.971927857 | 0.002694218 | -9.25016173 | 5.095959458 | 2 | 4.058998049 | 2.498691189 | 2.38539787 | 2.094598369 | 2.059860178 |
| ENSG00000156273 | BACH1 | -12.18596686 | 5.48E-14 | 2.935344994 | 2.42E-07 | -9.250621871 | 9.048471759 | 8.077869079 | 8.076950877 | 2 | 2 | 11.35100741 | 11.44090872 |
| ENSG00000138449 | SLC40A1 | -4.630644566 | 1.45E-06 | -4.623012999 | 1.32E-06 | -9.253657565 | 8.855032352 | 10.61666558 | 11.5927499 | 6.078767801 | 6.28357155 | 6.717615541 | 5.355511328 |
| ENSG00000159625 | DRC7 | -5.348721669 | NA | -3.909212506 | NA | -9.257934175 | 2 | 3.686303414 | 2 | 2 | 2 | 2.139668783 | 2 |
| ENSG00000103316 | CRYM | -1.622278315 | 0.416358129 | -7.637386406 | 4.01E-05 | -9.259664721 | 3.327711234 | 7.35507212 | 3.369317972 | 4.299492648 | 4.70309173 | 2.139668783 | 2.059860178 |
| ENSG00000257108 | NHLRC4 | -3.633539461 | 0.218319116 | -5.626321793 | 0.042927735 | -9.259861254 | 9.497412313 | 2 | 2 | 4.497095146 | 4.58534546 | 3.411599105 | 2.817676617 |
| ENSG00000276776 | | -2.310147786 | 0.491805453 | -6.954903563 | 0.038868875 | -9.26505135 | 2 | 4.93638933 | 2 | 2.498691189 | 2.59489879 | 2.048074405 | 2 |
| ENSG00000100979 | PLTP | -4.953066125 | 1.90E-33 | -4.320555636 | 3.62E-26 | -9.273621761 | 12.70658747 | 12.08595421 | 12.07171128 | 7.447418636 | 7.35080119 | 7.692773977 | 8.287321088 |
| ENSG00000120907 | ADRA1A | -3.156612311 | 0.260350896 | -6.12596684 | 0.035505946 | -9.282579151 | 4.00722531 | 3.076665209 | 2 | 2.270786764 | 2.14022021 | 2 | 2.059860178 |
| ENSG00000136010 | ALDH1L2 | -12.02795897 | 1.92E-08 | 2.737644662 | 0.096828736 | -9.290314309 | 4.00722531 | 9.055168103 | 8.677268317 | 2 | 2 | 11.14002933 | 10.9359368 |
| ENSG00000163141 | BNIPL | -4.371243619 | 0.000335735 | -4.930409759 | 2.62E-05 | -9.301653378 | 6.73718411 | 4.93638933 | 6.882775709 | 2.498691189 | 3.33976443 | 3.011868911 | 2.374681325 |
| ENSG00000122592 | HOXA7 | -0.753268983 | 0.712692238 | -8.549181414 | 1.70E-06 | -9.302450397 | 2 | 6.535001748 | 8.007629717 | 6.155987605 | 6.16544343 | 2.094598369 | 2.117335275 |
| ENSG00000185015 | CA13 | -3.49491669 | 0.000216249 | -5.807792767 | 2.55E-10 | -9.302709457 | 6.96351253 | 6.728495673 | 8.699419242 | 4.523237872 | 4.4836996 | 3.215043717 | 2.632958241 |
| ENSG00000007372 | PAX6 | -3.494807205 | 0.343130213 | -5.810803188 | 0.095461226 | -9.305610392 | 2 | 6.04685525 | 2 | 2.947924294 | 2 | 2.139668783 | 2.117335275 |
| ENSG00000012504 | NR1H4 | -4.714875553 | NA | -4.59262614 | NA | -9.307501693 | 2 | 3.076665209 | 2 | 2 | 2 | 2 | 2.059860178 |

| | | | | | | | | | | | | | |
|---|---|---|---|---|---|---|---|---|---|---|---|---|---|
| ENSG00000110148 | CCKBR | -4.714875553 | NA | -4.59262614 | NA | -9.307501693 | 2 | 2 | 3.369317972 | 2 | 2 | 2 | 2.059860178 |
| ENSG00000143536 | CRNN | -4.714875553 | NA | -4.59262614 | NA | -9.307501693 | 2 | 2 | 3.369317972 | 2 | 2 | 2 | 2.059860178 |
| ENSG00000158764 | ITLN2 | -4.714875553 | NA | -4.59262614 | NA | -9.307501693 | 3.327711234 | 2 | 2 | 2 | 2 | 2 | 2.059860178 |
| ENSG00000164485 | IL22RA2 | -4.714875553 | NA | -4.59262614 | NA | -9.307501693 | 3.327711234 | 2 | 2 | 2 | 2 | 2 | 2.059860178 |
| DIEXF | DIEXF | 2.618381995 | 0.118775449 | -11.93512057 | 2.19E-08 | -9.316738578 | 4.00722531 | 8.124485672 | 5.39244166 | 9.413697701 | 9.3808074 | 2 | 2 |
| ENSG00000242852 | ZNF709 | -2.873862561 | 0.000409777 | -6.445494044 | 9.57E-16 | -9.319356606 | 7.790810999 | 7.274681896 | 8.076950877 | 5.22679461 | 4.811949 | 2.987421008 | 2.277179783 |
| ENSG00000160111 | CPAMD8 | -6.309756955 | 1.99E-07 | -3.011519001 | 0.012430339 | -9.321275956 | 8.777722034 | 6.728495673 | 5.772716771 | 2.784612485 | 2.59489879 | 4.166733998 | 5.269393099 |
| ENSG00000125386 | FAM193A | -10.87123541 | 1.99E-05 | 1.54592997 | 0.514884694 | -9.325305445 | 2 | 8.376385542 | 6.629488387 | 2 | 2 | 8.834217923 | 8.561579708 |
| ENSG00000281935 | | -3.72128549 | 0.310601177 | -5.605302867 | 0.110476291 | -9.326588358 | 3.327711234 | 4.442710129 | 2 | 2 | 2.38539787 | 2 | 2.117335275 |
| ENSG00000115942 | ORC2 | -10.19018438 | 4.80E-08 | 0.859331952 | 0.49395309 | -9.330852423 | 6.257048221 | 7.393647515 | 4.874587708 | 2 | 2 | 7.468255486 | 7.239267729 |
| C5ORF42 | C5ORF42 | 2.365488182 | 0.012923624 | -11.70105769 | 2.78E-11 | -9.335569506 | 5.095959458 | 7.274681896 | 6.629488387 | 8.888486565 | 8.93865515 | 2 | 2 |
| ENSG00000162739 | SLAMF6 | -6.756425581 | 0.041283325 | -2.588067254 | 0.400363179 | -9.344492835 | 3.327711234 | 2 | 4.523675391 | 2 | 2 | 2.592885947 | 2.225841198 |
| ENSG00000111837 | MAK | -6.394656053 | 0.002696708 | -2.952516259 | 0.123771208 | -9.347172312 | 4.00722531 | 3.076665209 | 6.534109959 | 2.270786764 | 2 | 3.355392153 | 2.746601469 |
| ENSG00000165501 | LRR1 | -9.501166508 | 1.09E-06 | 0.146007581 | 0.927015754 | -9.355158927 | 4.00722531 | 6.311476754 | 6.322058544 | 2 | 2 | 6.358763086 | 5.555894927 |
| ENSG00000172139 | SLC9C1 | -6.076796326 | 0.07115111 | -3.278432416 | 0.285782968 | -9.355228742 | 3.327711234 | 3.686303414 | 2 | 2 | 2 | 2.183373728 | 2.172608095 |
| ENSG00000168878 | SFTPB | -4.552996939 | 0.0022619 | -4.80666582 | 0.000575272 | -9.359662759 | 6.138360429 | 3.076665209 | 5.156626401 | 2.270786764 | 2.59489879 | 2.384001718 | 2.374681325 |
| ENSG00000178177 | LCORL | -9.669388924 | 9.78E-05 | 0.308402375 | 0.902248935 | -9.360986549 | 6.816637977 | 6.311476754 | 2 | 2 | 2 | 6.759726988 | 5.699242824 |
| ENSG00000173894 | CBX2 | -5.650263698 | 0.000181905 | -3.716396113 | 0.01661716 | -9.366659811 | 4.467311984 | 7.145017454 | 9.819251342 | 3.754611435 | 3.08597292 | 4.977914261 | 4.903386193 |
| ENSG00000225968 | ELFN1 | -0.245586573 | 0.916562039 | -9.128284118 | 1.01E-06 | -9.373870691 | 9.048471759 | 6.728495673 | 2 | 7.477814553 | 7.51115308 | 2.139668783 | 2.117335275 |
| ENSG00000169122 | FAM110B | -4.529856819 | 1.20E-32 | -4.851467383 | 1.65E-38 | -9.381324202 | 11.23722567 | 10.73780697 | 10.95142175 | 6.394182228 | 6.63721363 | 6.471027105 | 5.900914815 |
| LOC100289561 | | 1.14969477 | 0.693364598 | -10.53450164 | 0.000205959 | -9.384806869 | 2 | 3.076665209 | 6.958202216 | 6.503277904 | 6.65485813 | 2 | 2 |
| ENSG00000167851 | CD300A | -8.981285315 | 0.024264145 | -0.409803841 | 0.928509131 | -9.391089156 | 6.89194353 | 2 | 2 | 2 | 2 | 5.790986459 | 3.33635788 |
| ENSG00000085999 | RAD54L | -3.837015852 | 0.008868951 | -5.556783634 | 7.71E-05 | -9.393799485 | 8.631580145 | 6.04685525 | 4.058998049 | 3.95996233 | 3.82856379 | 2.885263063 | 2.851942824 |
| ENSG00000172197 | MBOAT1 | -5.089870948 | 5.85E-08 | -4.304512596 | 3.26E-06 | -9.394383544 | 9.371511179 | 7.35507212 | 7.600047325 | 3.709785322 | 3.86943969 | 4.820739789 | 3.771796761 |
| ENSG00000185352 | HS6ST3 | -1.461938528 | 0.66293501 | -7.934470917 | 0.015041283 | -9.396409446 | 2 | 5.838937808 | 2 | 3.023133209 | 3.69849104 | 2.048074405 | 2 |
| ENSG00000117154 | IGSF21 | -8.342939498 | 0.000720316 | -1.065699173 | 0.633257223 | -9.40863867 | 3.327711234 | 5.303523747 | 5.156626401 | 2 | 2 | 4.675266036 | 2.551896972 |
| ENSG00000197594 | ENPP1 | -3.613263977 | 9.94E-10 | -5.797674171 | 1.38E-23 | -9.410938148 | 10.59831544 | 10.69597594 | 11.85448613 | 7.587189552 | 7.58049694 | 5.747972572 | 5.20457242 |
| ENSG00000241935 | HOGA1 | -5.977961501 | 0.000176081 | -3.434575397 | 0.00390558 | -9.412536898 | 4.467311984 | 4.710564154 | 5.39244166 | 2.270786764 | 2 | 2.526562374 | 2.817676617 |
| ENSG00000101203 | COL20A1 | -3.266415942 | NA | -6.152905045 | NA | -9.419320987 | 2 | 2 | 3.369317972 | 2.141737246 | 2 | 2 | 2 |
| ENSG00000105251 | SHD | -3.266415942 | NA | -6.152905045 | NA | -9.419320987 | 2 | 3.076665209 | 2 | 2.141737246 | 2 | 2 | 2 |
| ENSG00000105507 | CABP5 | -3.266415942 | NA | -6.152905045 | NA | -9.419320987 | 2 | 2 | 3.369317972 | 2.141737246 | 2 | 2 | 2 |
| ENSG00000111218 | PRMT8 | -3.266415942 | NA | -6.152905045 | NA | -9.419320987 | 3.327711234 | 2 | 2 | 2.141737246 | 2 | 2 | 2 |
| ENSG00000144119 | C1QL2 | -3.266415942 | NA | -6.152905045 | NA | -9.419320987 | 3.327711234 | 2 | 2 | 2.141737246 | 2 | 2 | 2 |
| ENSG00000154227 | CERS3 | -3.266415942 | NA | -6.152905045 | NA | -9.419320987 | 3.327711234 | 2 | 2 | 2.141737246 | 2 | 2 | 2 |
| ENSG00000176979 | TRIM60 | -3.266415942 | NA | -6.152905045 | NA | -9.419320987 | 2 | 2 | 3.369317972 | 2.141737246 | 2 | 2 | 2 |
| ENSG00000177238 | TRIM72 | -3.266415942 | NA | -6.152905045 | NA | -9.419320987 | 2 | 2 | 3.369317972 | 2.141737246 | 2 | 2 | 2 |
| TCEB3B | | -3.266415942 | NA | -6.152905045 | NA | -9.419320987 | 2 | 2 | 3.369317972 | 2.141737246 | 2 | 2 | 2 |
| ENSG00000166105 | GLB1L3 | -5.961174586 | 0.047911602 | -3.4626934 | 0.18059707 | -9.423867987 | 3.327711234 | 4.11352228 | 4.523675391 | 2 | 2.14022021 | 2.656294034 | 2 |
| ENSG00000134709 | HOOK1 | -3.274667334 | NA | -6.152905079 | NA | -9.427572413 | 3.327711234 | 2 | 2 | 2 | 2.14022021 | 2 | 2 |
| ENSG00000188467 | SLC24A5 | -3.274667334 | NA | -6.152905079 | NA | -9.427572413 | 3.327711234 | 2 | 2 | 2 | 2.14022021 | 2 | 2 |
| ENSG00000215910 | C1orf167 | -3.274667334 | NA | -6.152905079 | NA | -9.427572413 | 2 | 2 | 3.369317972 | 2 | 2.14022021 | 2 | 2 |
| ENSG00000224982 | TMEM233 | -3.274667334 | NA | -6.152905079 | NA | -9.427572413 | 3.327711234 | 2 | 2 | 2 | 2.14022021 | 2 | 2 |
| ENSG00000034053 | APBA2 | -2.298892941 | 0.120909818 | -7.1411304 | 6.45E-08 | -9.44002334 | 5.330594134 | 8.451443573 | 10.40208303 | 6.877205252 | 6.96410766 | 3.10571689 | 2.918126211 |
| ENSG00000221947 | XKR9 | -4.633164255 | 0.000141512 | -4.811316835 | 1.89E-05 | -9.44448109 | 6.138360429 | 4.11352228 | 5.59507254 | 2.498691189 | 2.49394783 | 2.592885947 | 2.277179783 |
| ENSG00000186523 | FAM86B1 | -8.195174119 | 0.002889631 | -1.250864602 | 0.632223509 | -9.446038721 | 4.00722531 | 5.838937808 | 2 | 2 | 2 | 3.694366604 | 3.771796761 |
| ENSG00000082556 | OPRK1 | -2.282004473 | 0.539424899 | -7.166215364 | NA | -9.448219837 | 3.327711234 | 2 | 3.369317972 | 2.3892346 | 2.14022021 | 2 | 2 |
| ENSG00000155792 | DEPTOR | -5.635020626 | 1.74E-33 | -3.817558779 | 9.25E-20 | -9.452579405 | 9.097018025 | 8.414402626 | 9.06622548 | 3.709785322 | 3.7864959 | 5.352823931 | 5.072762365 |
| ENSG00000167799 | NUDT8 | -6.078207805 | 1.67E-09 | -3.378882334 | 0.000676782 | -9.45709014 | 9.015175615 | 6.535001748 | 8.110401118 | 2.695456494 | 3.33976443 | 4.521997428 | 5.355511328 |
| ENSG00000100433 | KCNK10 | -4.871880039 | 0.214919497 | -4.588365892 | 0.197851694 | -9.460245932 | 4.00722531 | 2 | 3.369317972 | 2.141737246 | 2 | 2 | 2.172608095 |
| ENSG00000256043 | CTSO | 2.518209193 | 0.22821255 | -11.98509122 | 4.79E-07 | -9.466882026 | 7.82963488 | 6.899080544 | 2 | 9.314267267 | 9.37992356 | 2 | 2 |
| ENSG00000118231 | CRYGD | -4.714877588 | NA | -4.755636499 | NA | -9.470514087 | 3.327711234 | 2 | 2 | 2 | 2 | 2.048074405 | 2 |
| ENSG00000120057 | SFRP5 | -4.714877588 | NA | -4.755636499 | NA | -9.470514087 | 2 | 3.076665209 | 2 | 2 | 2 | 2.048074405 | 2 |
| ENSG00000120659 | TNFSF11 | -4.714877588 | NA | -4.755636499 | NA | -9.470514087 | 3.327711234 | 2 | 2 | 2 | 2 | 2.048074405 | 2 |
| ENSG00000167656 | LY6D | -4.714877588 | NA | -4.755636499 | NA | -9.470514087 | 3.327711234 | 2 | 2 | 2 | 2 | 2.048074405 | 2 |
| ENSG00000135374 | ELF5 | -4.714877588 | NA | -4.755636499 | NA | -9.470514087 | 2 | 2 | 3.369317972 | 2 | 2 | 2.048074405 | 2 |
| ENSG00000166828 | SCNN1G | -4.714877588 | NA | -4.755636499 | NA | -9.470514087 | 2 | 2 | 3.369317972 | 2 | 2 | 2.048074405 | 2 |
| ENSG00000138829 | FBN2 | -1.841293539 | 0.187271236 | -7.637621359 | 2.73E-10 | -9.478914898 | 9.886101849 | 11.13217411 | 10.22252928 | 8.717883675 | 8.64695856 | 4.177580368 | 2.172608095 |
| ENSG00000171798 | KNDC1 | -3.393955771 | 0.058360787 | -6.086479205 | 0.00036219 | -9.480434976 | 4.00722531 | 5.72254323 | 6.80318718 | 3.09461492 | 3.33976443 | 2.492221014 | 2 |
| DYX1C1 | DYX1C1 | -0.313157584 | 0.92048073 | -9.170133465 | 0.001470898 | -9.483291049 | 2 | 3.076665209 | 5.59507254 | 3.95996233 | 4.1930041 | 2 | 2 |
| ENSG00000134532 | SOX5 | -2.61075546 | 0.02564158 | -6.875338351 | 3.02E-10 | -9.486093811 | 9.460782017 | 7.232742069 | 6.629488387 | 5.900763176 | 5.67111583 | 2.962551659 | 2.466008509 |
| ENSG00000140470 | ADAMTS17 | -5.480176461 | 0.010326783 | -4.0266686 | 0.041653228 | -9.506845061 | 2 | 6.04685525 | 4.523675391 | 2.141737246 | 2.2680111 | 2.775316226 | 2.172608095 |
| ENSG00000128011 | LRFN1 | -7.146894066 | 3.42E-05 | -2.374468542 | 0.180259128 | -9.521362609 | 8.797441463 | 6.311476754 | 3.369317972 | 2.141737246 | 2.59489879 | 5.385935071 | 5.0581604 |
| ENSG00000262993 | | -0.48425623 | 0.9141407 | -9.037376823 | 0.020804985 | -9.521633053 | 2 | 2 | 5.59507254 | 3.881293667 | 3.82856379 | 2 | 2 |
| ENSG00000243989 | ACY1 | -2.763972119 | 3.85E-06 | -6.759634347 | 4.25E-30 | -9.523606466 | 8.836089686 | 9.726146444 | 8.99637432 | 6.429169081 | 6.63128385 | 2.885263063 | 3.560117069 |
| ENSG00000169116 | PARM1 | -5.823709821 | 5.46E-07 | -3.70131653 | 0.001429245 | -9.525026351 | 7.159102577 | 8.169643011 | 8.585098385 | 3.227757282 | 3.01487888 | 5.376552034 | 2.918126211 |
| ENSG00000153086 | ACMSD | -6.308503199 | 0.198785891 | -3.219698056 | 0.502996981 | -9.528201255 | 4.467311984 | 2 | 2 | 2 | 2 | 2.094598369 | 2.326754077 |
| ENSG00000157851 | DPYSL5 | -2.090876387 | 0.429299106 | -7.440091821 | 0.003569866 | -9.530968209 | 4.00722531 | 3.076665209 | 6.073378139 | 4.034562184 | 2 | 2.048074405 | 2.059860178 |
| ENSG00000182771 | GRID1 | -4.389913195 | 0.108587351 | -5.143489803 | 0.050004237 | -9.533402998 | 2 | 6.899080544 | 2 | 2.784612485 | 2.2680111 | 2.225793514 | 2.466008509 |
| ENSG00000108576 | SLC6A4 | -4.673346195 | 0.00707651 | -4.86148271 | 0.00262317 | -9.534828905 | 4.00722531 | 3.076665209 | 6.073378139 | 2.141737246 | 2.49394783 | 2.3070652 | 2.277179783 |
| ENSG00000171724 | VAT1L | -0.273508912 | 0.898841046 | -9.261922775 | 3.73E-07 | -9.535431687 | 2 | 7.431218269 | 7.934808822 | 6.841288255 | 6.89649843 | 2.094598369 | 2.059860178 |
| ENSG00000120519 | SLC10A7 | -9.227681347 | 0.001746789 | -0.312552328 | 0.922262647 | -9.540233675 | 3.327711234 | 7.051611315 | 2 | 2 | 2 | 5.557671049 | 5.080008298 |
| ENSG00000172717 | FAM71D | -5.777463822 | 0.263300675 | -3.764916273 | NA | -9.542380095 | 2 | 2 | 4.058998049 | 2 | 2 | 2.094598369 | 2.117335275 |
| ENSG00000171811 | CFAP46 | -4.853611866 | 0.000158354 | -4.693609436 | 9.42E-05 | -9.547221301 | 4.815609471 | 4.442710129 | 6.958202216 | 2.498691189 | 2.59489879 | 2.526562374 | 2.671842496 |
| ENSG00000086205 | FOLH1 | -2.53786421 | 0.647487207 | -7.010980844 | NA | -9.548845054 | 2 | 3.076665209 | 3.369317972 | 2.3892346 | 2 | 2 | 2 |
| ENSG00000135426 | TESPA1 | -2.331937113 | 0.693189026 | -7.220255073 | NA | -9.552192186 | 2 | 2 | 4.058998049 | 2.498691189 | 2 | 2 | 2 |
| ENSG00000074603 | DPP8 | -11.58238691 | 1.03E-06 | 2.022651988 | 0.327334994 | -9.559734917 | 3.327711234 | 6.311476754 | 9.272377397 | 2 | 2 | 9.734335478 | 10.02035048 |
| ENSG00000133466 | C1QTNF6 | -4.679833671 | 2.18E-15 | -4.887376452 | 6.17E-17 | -9.567210124 | 11.18951828 | 11.54156729 | 12.48906523 | 7.214570819 | 7.19871665 | 7.340163764 | 6.567271433 |
| ENSG00000120949 | TNFRSF8 | -0.684143448 | 0.798171474 | -8.885886846 | 0.000746214 | -9.570030293 | 4.00722531 | 4.93638933 | 2 | 3.515101962 | 3.69849104 | 2 | 2 |
| ENSG00000133661 | SFTPD | -6.770737513 | 0.005229614 | -2.801594952 | 0.142275223 | -9.572332465 | 3.327711234 | 5.838937808 | 3.369317972 | 2 | 2.14022021 | 2.42098416 | 3.183290849 |
| ENSG00000151065 | DCP1B | -10.24604317 | 1.76E-08 | 0.671007165 | 0.568416252 | -9.575036004 | 5.70932728 | 7.467835385 | 5.772716771 | 2 | 2 | 7.410810947 | 7.021346309 |
| ENSG00000120215 | MLANA | -7.957891014 | 9.47E-05 | -1.619075469 | 0.26362544 | -9.576966483 | 4.00722531 | 4.11352228 | 5.156626401 | 2 | 2 | 3.215043717 | 3.45257121 |
| ENSG00000174469 | CNTNAP2 | -0.340280901 | 0.861961559 | -9.245574325 | 1.55E-05 | -9.585855226 | 4.00722531 | 5.131603625 | 3.369317972 | 4.415698455 | 3.74316441 | 2 | 2 |
| ENSG00000278693 | | -1.504648649 | 0.744265156 | -8.083819497 | 0.057485677 | -9.588468146 | 2 | 4.710564154 | 2 | 2.600425781 | 2.86123014 | 2 | 2 |
| ENSG00000077616 | NAALAD2 | -6.290110121 | 1.81E-07 | -3.303450617 | 0.001415339 | -9.593560738 | 6.366709671 | 6.666836837 | 6.073378139 | 2 | 2.59489879 | 3.917810786 | 3.183290849 |
| ENSG00000183876 | ARSI | -5.167447291 | 1.57E-07 | -4.430999993 | 7.73E-06 | -9.598447285 | 9.699577239 | 9.7032946 | 12.18594181 | 6.024891728 | 5.90535527 | 6.257611854 | 6.983208048 |
| ENSG00000092421 | SEMA6A | -3.69221611 | 0.000874942 | -5.911542084 | 3.32E-08 | -9.603758194 | 6.468621203 | 8.955547846 | 6.882775709 | 4.946755474 | 3.69849104 | 2.962551659 | 2.981406154 |
| KIAA1644 | | 0.458675229 | 0.838765858 | -10.06489549 | 1.41E-05 | -9.606220256 | 3.327711234 | 6.311476754 | 3.369317972 | 5.348759107 | 5.62411716 | 2 | 2 |
| ENSG00000111732 | AICDA | -5.301181447 | 0.029843903 | -4.307425203 | 0.049504709 | -9.608606651 | 5.095959458 | 2 | 4.523675391 | 2 | 2.2680111 | 2.3070652 | 2.225841198 |
| ENSG00000167244 | IGF2 | -7.106405473 | 2.72E-26 | -2.50878481 | 0.000269482 | -9.615190283 | 11.43272766 | 10.35843478 | 11.90539881 | 4.647285135 | 4.40247105 | 8.295111724 | 9.273717427 |
| ENSG00000116141 | MARK1 | -6.570694032 | 0.002109106 | -3.049560204 | 0.17183963 | -9.620254236 | 8.946185348 | 6.602424674 | 3.369317972 | 3.023133209 | 2 | 5.445480974 | 3.560117069 |
| ENSG00000185739 | SRL | -4.901492851 | 0.043683252 | -4.725376344 | 0.036686779 | -9.626869195 | 4.815609471 | 2 | 5.156626401 | 2 | 2.38539787 | 2.183373728 | 2.277179783 |
| ENSG00000087076 | HSD17B14 | -5.165150556 | 0.000229321 | -4.463813486 | 0.001491443 | -9.628964042 | 10.07567912 | 7.979858146 | 4.874587708 | 4.105493497 | 3.98554648 | 4.778616857 | 4.411752372 |
| ENSG00000174607 | UGT8 | -4.285884686 | 0.00723449 | -5.34432116 | 0.000429856 | -9.630205846 | 4.467311984 | 3.076665209 | 6.073378139 | 2.600425781 | 2.2680111 | 2.183373728 | 2.277179783 |
| ENSG00000197734 | C14orf178 | -1.976390974 | 0.550280394 | -7.661164955 | 0.019537987 | -9.637555929 | 2 | 3.076665209 | 4.058998049 | 2.270786764 | 2.59489879 | 2 | 2 |
| ENSG00000205126 | ACCSL | -5.659310487 | 0.146668283 | -3.98341597 | 0.263275845 | -9.642726457 | 2 | 2 | 5.156626401 | 2.141737246 | 2 | 2.048074405 | 2.374681325 |
| ENSG00000105997 | HOXA3 | -1.159561272 | 0.632487313 | -8.492420972 | 8.35E-05 | -9.651982244 | 2 | 6.228547981 | 7.646393253 | 5.514313187 | 5.39133044 | 2 | 2.172608095 |
| ENSG00000164611 | PTTG1 | -5.000651862 | 4.37E-08 | -4.655083815 | 3.23E-07 | -9.655735677 | 11.60207034 | 10.17682307 | 8.784770914 | 5.818703821 | 5.62411716 | 5.790986459 | 6.264298846 |
| ENSG00000136286 | MYO1G | -6.154365316 | 0.068005802 | -3.502697552 | 0.285172239 | -9.657062868 | 2 | 2 | 6.534109959 | 2.270786764 | 2 | 2.656294034 | 2.782576693 |
| ENSG00000274749 | KRTAP7-1 | -1.437172553 | 0.756466007 | -8.22409231 | 0.053208271 | -9.661264862 | 2 | 2 | 4.874587708 | 2.784612485 | 2.86123014 | 2 | 2 |
| ENSG00000188038 | NRN1L | -1.602219675 | 0.601592548 | -8.067794794 | 0.008182544 | -9.670014469 | 4.467311984 | 3.076665209 | 2 | 2.695456494 | 2.68924543 | 2 | 2 |
| ENSG00000050555 | LAMC3 | -9.484313036 | 1.29E-06 | -0.185824603 | 0.907728859 | -9.670137638 | 6.009025997 | 8.317412869 | 6.073378139 | 2.141737246 | 2 | 7.64179387 | 6.066922748 |
| ENSG00000146090 | RASGEF1C | -6.056270527 | 0.239187141 | -3.620866478 | NA | -9.677136735 | 3.327711234 | 3.686303414 | 2 | 2 | 2 | 2.267001569 | 2 |
| KIAA0907 | | 2.155858833 | 0.046288831 | -11.83324546 | 5.92E-11 | -9.677386622 | 6.816637977 | 7.431218269 | 4.874587708 | 8.838609445 | 8.8354102 | 2 | 2 |
| KIAA0430 | | 2.71020169 | 0.146925888 | -12.38847703 | 3.38E-08 | -9.678275338 | 3.327711234 | 8.57407404 | 6.073378139 | 9.920382853 | 9.95903999 | 2 | 2 |
| ENSG00000243725 | TTC4 | -1.857927988 | 0.087151155 | -7.824078055 | 4.49E-16 | -9.682006043 | 11.16695586 | 9.079034257 | 8.486635645 | 8.188900477 | 8.23866319 | 3.316666208 | 2.885414003 |
| ENSG00000070031 | SCT | -2.8987599 | NA | -6.786766493 | NA | -9.685526394 | 2 | 3.686303414 | 2 | 2.270786764 | 2 | 2 | 2 |
| ENSG00000176769 | TCERG1L | -5.868593177 | 0.112267143 | -3.817621297 | 0.255124481 | -9.686214474 | 5.330594134 | 2 | 2 | 2 | 2.14022021 | 2.42098416 | 2.117335275 |

| | | | | | | | | | | | | | |
|---|---|---|---|---|---|---|---|---|---|---|---|---|---|
| ENSG00000204572 | KRTAP5-10 | -5.351139648 | NA | -4.339290733 | NA | -9.690430382 | 2 | 3.686303414 | 2 | 2 | 2 | 2.048074405 | 2.059860178 |
| FAM188B | | 0.190163045 | 0.940171576 | -9.881256754 | 6.53E-05 | -9.691093709 | 2 | 4.710564154 | 5.930865648 | 5.242616075 | 4.91316568 | 2 | 2 |
| C14ORF37 | C14ORF37 | 0.687530997 | 0.864318577 | -10.38308252 | 0.003179488 | -9.69555152 | 2 | 6.844429755 | 2 | 6.052081244 | 5.94414054 | 2 | 2 |
| METTL13 | | 1.721117104 | 0.415430055 | -11.41995173 | 1.16E-06 | -9.698834623 | 3.327711234 | 7.73146271 | 4.058998049 | 7.930222906 | 8.06478301 | 2 | 2 |
| ENSG00000139053 | PDE6H | -2.909528447 | NA | -6.790147504 | NA | -9.699675951 | 2 | 3.686303414 | 2 | 2.141737246 | 2.14022021 | 2 | 2 |
| ENSG00000278466 | | -1.619499588 | 0.725192089 | -8.083708447 | 0.057916885 | -9.703208034 | 2 | 4.710564154 | 2 | 2.600425781 | 2.77779907 | 2 | 2 |
| LINC01272 | | -0.667248722 | 0.882991782 | -9.037663623 | 0.022211493 | -9.704912344 | 2 | 2 | 5.59507254 | 3.462030052 | 3.94787299 | 2 | 2 |
| ENSG00000241863 | | 1.010020098 | 0.343732241 | -10.716063 | 1.47E-09 | -9.706042906 | 4.815609471 | 5.303523747 | 6.43197674 | 6.622948477 | 6.61935067 | 2 | 2 |
| ENSG00000253710 | ALG11 | -10.25887356 | 2.35E-08 | 0.532510279 | 0.670289164 | -9.726363286 | 7.218819781 | 6.78762695 | 4.874587708 | 2 | 2 | 7.296052918 | 6.889403418 |
| ENSG00000077327 | SPAG6 | -5.231284555 | 0.170583392 | -4.496896198 | 0.190985267 | -9.728180753 | 2 | 3.076665209 | 4.523675391 | 2.141737246 | 2 | 2.225793514 | 2 |
| ENSG00000161911 | TREML1 | -6.374193244 | 0.254759301 | -3.356945819 | 0.544850149 | -9.731139063 | 2 | 2 | 4.523675391 | 2 | 2 | 2.384001718 | 2 |
| C14ORF79 | C14ORF79 | 0.625514659 | 0.800325517 | -10.35798053 | 2.76E-05 | -9.732465876 | 4.815609471 | 6.464272742 | 2 | 5.860316854 | 5.9725602 | 2 | 2 |
| ENSG00000100281 | HMGXB4 | -11.17041226 | 2.38E-10 | 1.435472407 | 0.146347806 | -9.734939854 | 5.866944937 | 7.847167909 | 7.934808822 | 2 | 2 | 8.954974347 | 8.824656761 |
| ENSG00000157570 | TSPAN18 | -8.367380174 | 7.23E-06 | -1.367718486 | 0.515962661 | -9.73509866 | 2 | 9.04308532 | 9.819251342 | 2.3892346 | 2.49394783 | 7.669911566 | 7.425709383 |
| ENSG00000187024 | PTRH1 | -2.260094196 | 6.91E-05 | -7.477363189 | 4.99E-35 | -9.737457385 | 8.910412251 | 7.73146271 | 8.408111533 | 6.130704144 | 6.31350454 | 2.492221014 | 2.632958241 |
| ENSG00000177791 | MYOZ1 | -7.774987661 | 0.002472458 | -1.964362713 | 0.377325083 | -9.739350374 | 6.816637977 | 4.93638933 | 2 | 2.141737246 | 2 | 4.470006665 | 3.15612297 |
| ENSG00000167984 | NLRC3 | -9.032350952 | 1.51E-05 | -0.712800784 | 0.674004992 | -9.745151736 | 4.467311984 | 4.11352228 | 6.534109959 | 2 | 2 | 4.933813368 | 4.683176719 |
| ENSG00000129465 | RIPK3 | -7.573099357 | 0.006745422 | -2.181097942 | 0.384504677 | -9.754197299 | 2 | 4.442710129 | 4.874587708 | 2 | 2 | 3.082820997 | 2.551896972 |
| ENSG00000162510 | MATN1 | -5.614675221 | 0.038677883 | -4.139573421 | 0.117012619 | -9.754248642 | 2 | 6.951736442 | 2 | 2.3892346 | 2.14022021 | 2.746468522 | 2.551896972 |
| ENSG00000205323 | SARNP | -1.519439032 | 0.091109722 | -8.24281081 | 1.38E-23 | -9.762249842 | 10.29961392 | 8.191702463 | 8.884815469 | 7.870381136 | 7.91663606 | 2.746468522 | 2.509591799 |
| ENSG00000183098 | GPC6 | -4.165323448 | 9.44E-10 | -5.60667 | 4.95E-17 | -9.771993448 | 11.72300081 | 11.39001779 | 11.46280979 | 7.409376773 | 7.3899592 | 4.861667652 | 6.649933445 |
| ENSG00000196208 | GREB1 | -3.365859624 | 0.010093341 | -6.407622793 | 2.68E-07 | -9.773482416 | 5.330594134 | 7.002537973 | 8.721235202 | 4.299492648 | 4.68029874 | 2.937246078 | 2.277179783 |
| ENSG00000046653 | GPM6B | -5.209935754 | 0.000797434 | -4.565503077 | 0.003524935 | -9.775438831 | 10.51143645 | 10.19331737 | 10.03493575 | 6.12217689 | 2.14022021 | 6.060766632 | 5.475755622 |
| ENSG00000149609 | C20orf144 | -7.781583353 | 0.000139777 | -2.00000822 | 0.160186683 | -9.781591573 | 4.815609471 | 4.11352228 | 4.058998049 | 2 | 2 | 3.12825509 | 2.918126211 |
| ENSG00000037474 | NSUN2 | -11.25314853 | 1.26E-12 | 1.468568827 | 0.00139739 | -9.7845797 | 7.485143553 | 7.538394143 | 7.646393253 | 2 | 2 | 8.784089222 | 9.198731239 |
| ENSG00000137078 | SIT1 | -4.503632515 | 0.122517925 | -5.281950837 | 0.055158384 | -9.785583352 | 5.532353208 | 2 | 2 | 2.141737246 | 2.2680111 | 2.183373728 | 2.059860178 |
| DOPEY2 | DOPEY2 | 0.803976364 | 0.470185896 | -10.60017072 | 2.60E-09 | -9.796194361 | 4.815609471 | 5.131603625 | 6.322058544 | 6.314087501 | 6.31350454 | 2 | 2 |
| ENSG00000174123 | TLR10 | -4.322785776 | 0.029045946 | -5.475724756 | 0.003778129 | -9.798510531 | 4.00722531 | 4.442710129 | 5.59507254 | 2.141737246 | 2.59489879 | 2.34604626 | 2 |
| ENSG00000215277 | RNF212B | -6.223749529 | 0.071713825 | -3.578886724 | 0.258978492 | -9.802636253 | 2 | 3.076665209 | 4.058998049 | 2 | 2 | 2.094598369 | 2.225841198 |
| ENSG00000143416 | SELENBP1 | -5.658491069 | 5.98E-38 | -4.144467625 | 1.88E-22 | -9.802958693 | 10.37373127 | 10.56689847 | 10.48092713 | 5.003571641 | 5.00774437 | 5.883320383 | 6.777976866 |
| ENSG00000196132 | MYT1 | -2.458887501 | 0.676103313 | -7.34720617 | NA | -9.806093671 | 2 | 4.11352228 | 2 | 2.498691189 | 2 | 2 | 2 |
| ENSG00000148200 | NR6A1 | -5.491447637 | 3.92E-14 | -4.325676052 | 4.18E-10 | -9.817123688 | 8.696033109 | 8.702538795 | 7.451345288 | 3.289987342 | 3.69849104 | 3.904814898 | 4.711446447 |
| ENSG00000257704 | INAFM1 | -10.6503249 | 6.41E-11 | 0.831907094 | 0.198260297 | -9.818417806 | 6.468621203 | 7.274681896 | 7.029880454 | 2 | 2 | 7.936540212 | 7.608437243 |
| ENSG00000078399 | HOXA9 | 0.866762487 | 0.692130582 | -10.68863965 | 3.73E-06 | -9.821877162 | 8.075571721 | 6.728495673 | 2 | 7.849869274 | 7.81043269 | 2.048074405 | 2 |
| ENSG00000278458 | | 0.864173997 | 0.357366207 | -10.68961091 | 4.60E-16 | -9.825436908 | 8.631580145 | 8.077869079 | 6.534109959 | 8.828161587 | 8.85080188 | 2.048074405 | 2.059860178 |
| ENSG00000067606 | PRKCZ | -5.811239629 | 9.01E-16 | -4.016966336 | 3.37E-09 | -9.828205966 | 8.928409675 | 7.467835385 | 8.460932916 | 2.947924294 | 3.60476711 | 4.547306681 | 4.692661645 |
| ENSG00000140022 | STON2 | -3.668013738 | 0.028758071 | -6.161769618 | 0.000171403 | -9.829783356 | 6.366709671 | 4.11352228 | 4.523675391 | 3.227757282 | 2.14022021 | 2.225793514 | 2.117335275 |
| ENSG00000103241 | FOXF1 | 0.627910111 | 0.429614324 | -10.46037607 | 1.73E-43 | -9.832465955 | 9.448362284 | 10.3729879 | 10.9743126 | 10.99534488 | 11.0444066 | 2.42098416 | 2.172608095 |
| ENSG00000166348 | USP54 | -10.57228383 | 4.92E-09 | 0.736721984 | 0.521821801 | -9.835561849 | 5.330594134 | 7.572420273 | 6.958202216 | 2 | 2 | 7.374742053 | 7.815789075 |
| ENSG00000139874 | SSTR1 | -1.720495514 | 0.100358479 | -8.121028913 | 2.11E-18 | -9.841524427 | 9.159305284 | 9.479495904 | 10.98787477 | 8.425603456 | 8.37620272 | 3.393105702 | 2.421067459 |
| ENSG00000275851 | | 1.390386206 | 0.00491763 | -11.23263323 | 2.27E-22 | -9.842247027 | 8.493334308 | 8.053983481 | 8.904018013 | 9.856271709 | 9.94177481 | 2.048074405 | 2.059860178 |
| ENSG00000124491 | F13A1 | -7.937952649 | 0.023977132 | -1.908350303 | 0.584074945 | -9.846302953 | 2 | 2 | 7.285536013 | 2.141737246 | 2 | 2.911488689 | 4.77533149 |
| ENSG00000166106 | ADAMTS15 | -7.764846826 | 1.33E-22 | -2.091370697 | 0.008689336 | -9.856217524 | 10.84565509 | 9.181799937 | 9.658943166 | 3.162721474 | 3.15372745 | 8.440656542 | 7.352801367 |
| ENSG00000110693 | SOX6 | -5.130332885 | 0.000206644 | -4.727415802 | 0.000527491 | -9.857748687 | 4.467311984 | 8.88960062 | 7.342949392 | 3.754611435 | 2.77779907 | 3.56816602 | 3.640730416 |
| ENSG00000163138 | PACRGL | -9.438025288 | 0.00015151 | -0.42501945 | 0.863495873 | -9.863044739 | 2 | 6.602424674 | 6.073378139 | 2 | 2 | 5.574182174 | 5.263040529 |
| ENSG00000273494 | | 1.477871687 | 0.199198205 | -11.34247695 | 4.70E-10 | -9.864605265 | 7.218819781 | 5.303523747 | 5.39244166 | 7.717440541 | 7.64959655 | 2 | 2 |
| ENSG00000164398 | ACSL6 | -5.573414439 | 0.065318905 | -4.300367991 | 0.094621254 | -9.87378243 | 2 | 3.076665209 | 4.874587708 | 2 | 2.14022021 | 2.267001569 | 2.059860178 |
| ENSG00000164303 | ENPP6 | -6.936203223 | 0.019358115 | -2.9426348 | 0.265970284 | -9.878838022 | 2 | 3.686303414 | 4.523675391 | 2 | 2 | 2.384001718 | 2.374681325 |
| ENSG00000249242 | TMEM150C | -7.390115514 | 0.000146126 | -2.489272896 | 0.188560075 | -9.87938841 | 7.53297911 | 7.051611315 | 2 | 2.270786764 | 2.14022021 | 5.038665118 | 3.580696614 |
| ENSG00000134873 | CLDN10 | -5.719642876 | 0.279078313 | -4.172783063 | NA | -9.892425939 | 4.00722531 | 2 | 2 | 2 | 2 | 2.094598369 | 2.059860178 |
| ENSG00000180806 | HOXC9 | 0.647196807 | 0.87294376 | -10.54547715 | 0.002764101 | -9.898280346 | 2 | 7.002537973 | 2 | 5.997179931 | 6.22177541 | 2 | 2 |
| ENSG00000159495 | TGM7 | -7.376832792 | 0.088220418 | -2.522513956 | 0.557806634 | -9.899346748 | 2 | 2 | 5.39244166 | 2 | 2 | 2.457042231 | 2.782576693 |
| ENSG00000123453 | SARDH | -6.926164686 | 0.000531042 | -2.975417907 | 0.159315752 | -9.901582593 | 9.740899611 | 6.951736442 | 2 | 2.3892346 | 3.01487888 | 5.113999907 | 5.799216178 |
| ENSG00000275544 | | 0.320344354 | 0.816209881 | -10.22447944 | 3.48E-08 | -9.90413509 | 4.00722531 | 5.72254323 | 5.39244166 | 5.319220934 | 5.6826301 | 2 | 2 |
| FASTKD1 | | -9.916450527 | 1.53E-07 | 0.010827395 | 0.994111049 | -9.905623132 | 4.467311984 | 6.844429755 | 6.534109959 | 2 | 2 | 6.514451546 | 5.992078266 |
| ENSG00000164509 | IL31RA | -4.195212263 | 0.05396828 | -5.718234092 | 0.006194322 | -9.913446354 | 4.00722531 | 3.076665209 | 6.073378139 | 2.3892346 | 2.49394783 | 2 | 2.326754077 |
| ENSG00000117501 | MROH9 | -2.404784033 | 0.074138279 | -7.50953086 | 5.26E-08 | -9.914314893 | 5.70932728 | 4.11352228 | 7.163363831 | 3.921164147 | 4.16043394 | 2.183373728 | 2.059860178 |
| ENSG00000096063 | SRPK1 | -11.86597221 | 1.08E-07 | 1.949722911 | 0.293464301 | -9.916249295 | 3.327711234 | 8.57407404 | 8.865353883 | 2 | 2 | 9.942009796 | 10.23192078 |
| ENSG00000101104 | PABPC1L | -9.545242738 | 0.009875401 | -0.380860639 | 0.928476068 | -9.926103377 | 2 | 7.431218269 | 2 | 2 | 2 | 5.553513572 | 5.576534358 |
| FAM105A | | 0.230333488 | 0.931034311 | -10.15849807 | 6.58E-05 | -9.928164583 | 2 | 6.311476754 | 4.523675391 | 5.44762523 | 5.28958453 | 2 | 2 |
| ENSG00000131153 | GINS2 | -4.784546605 | 0.000809213 | -5.148481356 | 0.000241045 | -9.93302796 | 4.467311984 | 7.73146271 | 9.577565424 | 4.205751501 | 3.74316441 | 3.393105702 | 4.002845082 |
| ENSG00000160472 | TMEM190 | -4.249515682 | 0.020143473 | -5.685169427 | 0.001220103 | -9.934685109 | 2 | 5.303523747 | 6.43197674 | 2.868578001 | 2.2680111 | 2.225793514 | 2.277179783 |
| LHFP | | 2.351478829 | 0.270743954 | -12.30670196 | 2.71E-07 | -9.955223134 | 7.82963488 | 7.669933291 | 2 | 9.500971099 | 9.50192864 | 2 | 2 |
| ENSG00000129673 | AANAT | -4.565203177 | 0.001530486 | -5.39050916 | 8.91E-05 | -9.955712337 | 6.89194353 | 4.11352228 | 4.523675391 | 2.695456494 | 2.49394783 | 2.492221014 | 2.225841198 |
| ENSG00000266524 | GDF10 | -7.631186755 | 0.070544451 | -2.327471391 | 0.582819507 | -9.958658146 | 2 | 5.595927442 | 2 | 2 | 2 | 3.10571689 | 2.421067459 |
| ENSG00000177689 | MAGEB10 | -5.115961052 | 0.090138531 | -4.847316292 | 0.061222989 | -9.963277344 | 3.327711234 | 3.686303414 | 3.369317972 | 2.141737246 | 2 | 2 | 2.172608095 |
| ENSG00000149599 | DUSP15 | -7.304799952 | 6.68E-08 | -2.659121213 | 0.001534373 | -9.963921164 | 6.366709671 | 6.228547981 | 5.59507254 | 2 | 2.2680111 | 4.076896059 | 3.539239711 |
| ENSG00000107954 | NEURL1 | -5.242875089 | 6.94E-06 | -4.72866621 | 2.20E-05 | -9.971541299 | 6.468621203 | 4.93638933 | 7.934808822 | 2.600425781 | 3.01487888 | 3.1504466 | 2.981406154 |
| ENSG00000175711 | B3GNTL1 | -3.238793576 | 1.85E-08 | -6.734145207 | 7.00E-26 | -9.972938782 | 7.031698319 | 7.431218269 | 7.029880454 | 4.070463721 | 4.40247105 | 2.526562374 | 2.277179783 |
| C10ORF126 | C10ORF126 | -2.762355664 | 0.637132746 | -7.220257012 | NA | -9.982612676 | 2 | 2 | 4.058998049 | 2 | 2.38539787 | 2 | 2 |
| ENSG00000137747 | TMPRSS13 | -5.497590506 | 0.131120567 | -4.495270677 | 0.167372108 | -9.992861183 | 2 | 3.686303414 | 4.523675391 | 2 | 2.14022021 | 2.267001569 | 2 |
| ENSG00000114982 | KANSL3 | -11.59646661 | 5.60E-06 | 1.602754801 | 0.502872336 | -9.993711812 | 2 | 8.916342327 | 7.818184403 | 2 | 2 | 9.572466068 | 9.382476431 |
| ENSG00000225526 | MKRN2OS | -6.386696582 | 0.002486363 | -3.60847078 | 0.011199128 | -9.995167362 | 4.00722531 | 3.686303414 | 5.156626401 | 2.141737246 | 2 | 2.492221014 | 2.374681325 |
| ENSG00000162032 | SPSB3 | -1.774829325 | 0.004401152 | -8.225041028 | 1.11E-38 | -9.999870353 | 9.867565369 | 8.955547846 | 8.742726175 | 7.447418636 | 7.59275449 | 2.746468522 | 2.421067459 |
| ENSG00000172782 | FADS6 | -4.338830398 | 0.430594842 | -5.661712092 | NA | -10.00054249 | 2 | 2 | 4.058998049 | 2 | 2.14022021 | 2 | 2.059860178 |
| ENSG00000204120 | GIGYF2 | -11.7635322 | 3.04E-08 | 1.754036813 | 0.298525733 | -10.00949538 | 8.963744667 | 8.147241016 | 4.058998049 | 2 | 2 | 9.764548842 | 9.829339 |
| ENSG00000137806 | NDUFAF1 | -10.57929367 | 5.15E-05 | 0.566146416 | 0.832608099 | -10.01314725 | 3.327711234 | 8.395519306 | 3.369317972 | 2 | 2 | 7.377096489 | 7.523290521 |
| ENSG00000134765 | DSC1 | -4.025123429 | 0.116793916 | -5.992076208 | 0.016541828 | -10.01719964 | 3.327711234 | 3.686303414 | 4.874587708 | 2 | 2.49394783 | 2.139668783 | 2 |
| ENSG00000179921 | GPBAR1 | -5.0926111 | 0.006994946 | -4.927784355 | 0.00641103 | -10.02039545 | 2 | 6.951736442 | 4.874587708 | 2.270786764 | 2.59489879 | 2.42098416 | 2.551896972 |
| ENSG00000197859 | ADAMTSL2 | -7.151771138 | 0.000312209 | -2.869834418 | 0.021969175 | -10.02160556 | 4.467311984 | 5.72254323 | 4.523675391 | 2 | 2.14022021 | 3.235948948 | 2.70970617 |
| ENSG00000138622 | HCN4 | -4.86948051 | 0.358760146 | -5.159771245 | 0.300546566 | -10.02925176 | 4.467311984 | 2 | 2 | 2 | 2.14022021 | 2 | 2.117335275 |
| ENSG00000147481 | SNTG1 | -2.422318011 | 0.408537877 | -7.607671445 | 0.011593579 | -10.02998946 | 5.532353208 | 3.076665209 | 2 | 3.227757282 | 2.14022021 | 2 | 2.059860178 |
| ENSG00000173376 | NDNF | -3.292459913 | 0.096340918 | -6.742230536 | 0.000204916 | -10.03469045 | 9.810482605 | 8.414402626 | 2 | 5.540148523 | 5.49945122 | 3.215043717 | 2.671842496 |
| ENSG00000145287 | PLAC8 | -4.665353723 | 0.355780597 | -5.375532347 | 0.265033519 | -10.04088607 | 2 | 2 | 5.156626401 | 2.270786764 | 2 | 2 | 2.172608095 |
| ENSG00000160838 | LRRC71 | -7.120925614 | 0.115550873 | -2.921490749 | 0.513903548 | -10.04241636 | 2 | 5.131603625 | 2 | 2 | 2 | 2.139668783 | 2.671842496 |
| ENSG00000122787 | AKR1D1 | -5.85125065 | 0.044023974 | -4.193069055 | 0.085631007 | -10.04431971 | 2 | 3.076665209 | 5.156626401 | 2 | 2.14022021 | 2.3070652 | 2.117335275 |
| ENSG00000077279 | DCX | -0.866394391 | 0.867562397 | -9.181236409 | 0.041403012 | -10.0476308 | 2 | 3.076665209 | 5.59507254 | 4.443344104 | 2 | 2 | 2 |
| ENSG00000214435 | AS3MT | -1.871216429 | 0.230405007 | -8.181422145 | 3.44E-08 | -10.05263857 | 4.00722531 | 6.389896633 | 8.175061096 | 5.022023783 | 5.45984938 | 2.225793514 | 2.059860178 |
| ENSG00000167914 | GSDMA | -5.500376863 | 5.50E-08 | -4.555965921 | 9.56E-09 | -10.05634278 | 5.70932728 | 5.303523747 | 5.930865648 | 2.3892346 | 2.2680111 | 2.526562374 | 2.632958241 |
| ENSG00000188050 | RNF133 | -2.302199829 | 0.646239362 | -7.762536398 | 0.098562133 | -10.06473623 | 2 | 4.442710129 | 2 | 2.141737246 | 2.59489879 | 2 | 2 |
| ENSG00000129993 | CBFA2T3 | -7.31463298 | 0.12645146 | -2.755315109 | 0.56576762 | -10.06994809 | 5.330594134 | 2 | 2 | 2 | 2 | 2.885263063 | 2.059860178 |
| ENSG00000107831 | FGF8 | -5.130252105 | 0.075259489 | -4.942420879 | 0.04383944 | -10.07267298 | 2 | 3.686303414 | 4.058998049 | 2.141737246 | 2 | 2.048074405 | 2.117335275 |
| ENSG00000113597 | TRAPPC13 | -10.64442529 | 1.79E-10 | 0.569981245 | 0.477216955 | -10.07444405 | 7.27616306 | 7.232742069 | 6.073378139 | 2 | 2 | 7.507470083 | 7.531242162 |
| ENSG00000169347 | GP2 | -3.10536534 | 0.569289533 | -6.978678352 | NA | -10.08404369 | 3.327711234 | 3.076665209 | 2 | 2 | 2.2680111 | 2 | 2 |
| ENSG00000157399 | ARSE | -6.314006236 | 3.18E-06 | -3.77267318 | 0.00483723 | -10.08667942 | 5.532353208 | 5.838937808 | 8.960138144 | 2.600425781 | 2.77779907 | 3.943456545 | 4.411752372 |
| C11ORF84 | C11ORF84 | 1.687867633 | 0.574946111 | -11.78481732 | 6.09E-05 | -10.09694968 | 2 | 3.076665209 | 8.206335842 | 8.281794636 | 8.36909406 | 2 | 2 |
| ENSG00000133863 | TEX15 | -5.976355109 | 0.01740322 | -4.12065394 | 0.072993837 | -10.09700905 | 2 | 3.686303414 | 6.203069917 | 2.270786764 | 2 | 2.492221014 | 2.421067459 |
| ENSG00000100479 | POLE2 | -5.090979276 | 0.000152287 | -5.007783194 | 0.000138854 | -10.09876247 | 5.095959458 | 5.838937808 | 8.805342542 | 3.227757282 | 3.15372745 | 3.215043717 | 3.261853426 |
| FAM175A | | 0.933414558 | 0.685014501 | -11.03370827 | 4.30E-06 | -10.10029371 | 5.866944937 | 6.951736442 | 2 | 6.693715713 | 6.99676009 | 2 | 2 |
| ENSG00000134757 | DSG3 | -3.464711159 | 0.024857174 | -6.646224924 | 2.14E-05 | -10.11093608 | 5.095959458 | 3.076665209 | 6.322058544 | 3.023133209 | 2.68924543 | 2.183373728 | 2.059860178 |
| ENSG00000163492 | CCDC141 | -6.148302798 | 0.088683678 | -3.967054424 | 0.223166355 | -10.11535722 | 2 | 2 | 5.59507254 | 2.141737246 | 2 | 2.42098416 | 2.172608095 |
| ENSG00000166532 | RIMKLB | -11.12736895 | 3.54E-11 | 1.007157273 | 0.209686284 | -10.12021168 | 6.468621203 | 7.979858146 | 7.451345288 | 2 | 2 | 8.390844511 | 8.457628219 |
| ENSG00000100416 | TRMU | -10.86540298 | 4.06E-07 | 0.735053436 | 0.690305992 | -10.13034954 | 7.750913381 | 7.701025996 | 3.369317972 | 2 | 2 | 7.815899769 | 7.975841991 |
| ENSG00000183784 | C9orf66 | -2.223661317 | 0.483832498 | -7.908453632 | 0.012742463 | -10.13211495 | 2 | 4.11352228 | 3.369317972 | 2.270786764 | 2.59489879 | 2 | 2 |
| ENSG00000100764 | PSMC1 | -1.155571272 | 0.030953326 | -8.987064166 | 7.28E-71 | -10.14263544 | 11.673172 | 10.63690565 | 10.92346452 | 9.985884116 | 10.0004577 | 3.193831101 | 2.950113106 |
| ENSG00000182809 | CRIP2 | -11.71385361 | 1.69E-09 | 1.571040194 | 0.254071608 | -10.14281342 | 8.981092835 | 5.457120953 | 7.818184403 | 2 | 2 | 9.481572395 | 9.645290766 |
| ENSG00000116138 | DNAJC16 | -11.01813008 | 1.11E-11 | 0.872909047 | 0.16091081 | -10.14522104 | 7.096806373 | 7.35507212 | 7.502634626 | 2 | 2 | 8.488805956 | 7.79722419 |

| | | | | | | | | | | | | | |
|---|---|---|---|---|---|---|---|---|---|---|---|---|---|
| ENSG00000171864 | PRND | -2.133685994 | 0.469910803 | -8.012454714 | 0.007417387 | -10.14614071 | 2 | 3.686303414 | 4.058998049 | 2.3892346 | 2.59489879 | 2 | 2 |
| ENSG00000072832 | CRMP1 | -7.581284676 | 7.82E-14 | -2.565593357 | 0.016796686 | -10.14687803 | 10.68025843 | 10.17682307 | 10.53119344 | 4.17310023 | 2.2680111 | 7.821961477 | 8.033896215 |
| ENSG00000259511 | UBE2Q2L | -1.717639196 | 0.348810364 | -8.43060126 | 0.000122942 | -10.14824046 | 3.327711234 | 3.686303414 | 6.073378139 | 3.79808662 | 3.28037825 | 2.048074405 | 2 |
| ENSG00000188886 | ASTL | -4.330366975 | 0.426589417 | -5.818463937 | NA | -10.14883091 | 2 | 2 | 4.058998049 | 2.141737246 | 2 | 2.048074405 | 2 |
| ENSG00000160055 | TMEM234 | -10.04860468 | 9.06E-05 | -0.105918353 | 0.968806022 | -10.15452304 | 2 | 7.538394143 | 5.930865648 | 2 | 2 | 6.159811591 | 6.412287469 |
| ENSG00000171241 | SHCBP1 | -9.18142449 | 4.50E-05 | -0.9850953 | 0.610378131 | -10.16651979 | 4.815609471 | 6.602424674 | 4.058998049 | 2 | 2 | 3.465698119 | 5.367404784 |
| ENSG00000167393 | PPP2R3B | 0.472389098 | 0.843094589 | -10.6422298 | 8.41E-06 | -10.1698407 | 5.70932728 | 6.464272742 | 2 | 6.113598934 | 5.96314905 | 2 | 2 |
| ENSG00000100335 | MIEF1 | -11.91069952 | 9.14E-13 | 1.734347106 | 0.018944745 | -10.17635241 | 8.367060731 | 8.671479029 | 7.163363831 | 2 | 2 | 9.816867623 | 10.02411203 |
| ENSG00000117400 | MPL | -5.591133322 | 0.000344238 | -4.592356244 | 0.00050057 | -10.18348957 | 3.327711234 | 4.93638933 | 5.930865648 | 2.270786764 | 2.14022021 | 2.3070652 | 2.466008509 |
| ENSG00000204661 | C5orf60 | -6.316738481 | 0.259792504 | -3.88257138 | 0.480467544 | -10.19930986 | 2 | 4.442710129 | 2 | 2 | 2 | 2.267001569 | 2 |
| ENSG00000007314 | SCN4A | -6.518892884 | 0.052743577 | -3.681947206 | 0.26164227 | -10.20084009 | 2 | 2 | 6.882775709 | 2.270786764 | 2 | 2.62493832 | 2.950113106 |
| ENSG00000278259 | MYO19 | 1.339806523 | 0.020322886 | -11.54177324 | 6.54E-13 | -10.20196671 | 6.138360429 | 6.602424674 | 6.534109959 | 7.668500375 | 7.81567127 | 2 | 2 |
| ENSG00000274437 | | -1.876980624 | 0.676338863 | -8.342515455 | 0.046438008 | -10.21949608 | 2 | 4.93638933 | 2 | 2.695456494 | 2.68924543 | 2 | 2 |
| ENSG00000158805 | ZNF276 | -10.35975872 | 4.18E-08 | 0.138143076 | 0.926209474 | -10.22161564 | 5.095959458 | 7.638155693 | 6.203069917 | 2 | 2 | 6.828438696 | 6.81752904 |
| ENSG00000197565 | COL4A6 | -7.432898579 | 7.60E-20 | -2.789664753 | 0.00089073 | -10.22256333 | 10.93738399 | 10.27823701 | 11.60159843 | 4.497095146 | 3.28037825 | 7.511762185 | 8.756761909 |
| FAM175B | | 1.612099843 | 0.271602878 | -11.83607762 | 1.98E-09 | -10.22397777 | 4.467311984 | 7.819099893 | 6.073378139 | 8.241026137 | 8.36015869 | 2 | 2 |
| ENSG00000112852 | PCDHB2 | -3.494243948 | 4.00E-07 | -6.732391012 | 9.18E-22 | -10.22663496 | 8.285626337 | 7.232742069 | 8.460932916 | 5.110902178 | 4.37434574 | 2.831336763 | 2.551896972 |
| ENSG00000204961 | PCDHA9 | -2.466974621 | 0.072787538 | -7.769108173 | 1.97E-06 | -10.23608279 | 5.095959458 | 4.11352228 | 5.930865648 | 3.754611435 | 2.68924543 | 2.048074405 | 2.059860178 |
| ENSG00000110492 | MDK | -3.503319687 | 6.86E-17 | -6.733303407 | 6.68E-61 | -10.23662309 | 15.12369229 | 14.28706125 | 14.46263956 | 11.1619039 | 11.178285 | 8.153962636 | 7.737762724 |
| ENSG00000189152 | GRAPL | -2.887118327 | 0.574423555 | -7.355188093 | NA | -10.24230642 | 2 | 4.11352228 | 2 | 2.270786764 | 2.14022021 | 2 | 2 |
| ENSG00000110675 | ELMOD1 | -2.708942111 | 0.055113052 | -7.544524075 | 1.03E-06 | -10.25346619 | 3.327711234 | 5.131603625 | 6.534109959 | 3.462030052 | 3.28037825 | 2.094598369 | 2.059860178 |
| ENSG00000161960 | EIF4A1 | -1.576981298 | 0.00333081 | -8.680529921 | 3.06E-70 | -10.25751122 | 14.3380183 | 13.0897025 | 13.68660132 | 12.19827324 | 12.2368395 | 5.147485955 | 5.385063124 |
| ENSG00000187091 | PLCD1 | -11.10713881 | 2.88E-10 | 0.84476993 | 0.415980405 | -10.26236888 | 8.19931886 | 6.140560021 | 7.225742779 | 2 | 2 | 8.173057797 | 8.312147538 |
| ENSG00000152766 | ANKRD22 | -3.898358967 | 0.280551441 | -6.36855056 | 0.075446691 | -10.26690953 | 3.327711234 | 2 | 4.058998049 | 2 | 2.2680111 | 2.048074405 | 2 |
| ENSG00000273645 | | -0.632000939 | 0.821711576 | -9.644177121 | 0.000333873 | -10.27617806 | 2 | 5.838937808 | 4.058998049 | 4.497095146 | 3.82856379 | 2 | 2 |
| ENSG00000114737 | CISH | -6.175383589 | 8.54E-15 | -4.10291182 | 2.50E-08 | -10.27829541 | 8.228665467 | 7.73146271 | 8.608698767 | 2.695456494 | 3.28037825 | 4.854926465 | 3.753786374 |
| ENSG00000188124 | OR2AG2 | -5.34207996 | 0.297547855 | -4.936335774 | 0.303359822 | -10.27841573 | 2 | 2 | 4.874587708 | 2.141737246 | 2 | 2.183373728 | 2 |
| ENSG00000140538 | NTRK3 | -6.415887319 | 2.30E-14 | -3.864391854 | 3.32E-06 | -10.28027917 | 8.107517749 | 9.733683991 | 10.16797309 | 4.034562184 | 3.2184422 | 5.47655442 | 6.070564958 |
| ENSG00000198515 | CNGA1 | -5.549252842 | 0.026538199 | -4.733297017 | 0.017063108 | -10.28254986 | 3.327711234 | 3.076665209 | 4.523675391 | 2 | 2.14022021 | 2.183373728 | 2.059860178 |
| ENSG00000204653 | ASPDH | -7.986976854 | 0.018770404 | -2.302142949 | 0.483351919 | -10.2891198 | 7.33131396 | 2 | 2 | 2.141737246 | 2 | 4.133696865 | 3.539239711 |
| ENSG00000148824 | MTG1 | -3.780933361 | 5.14E-10 | -6.512839766 | 1.14E-27 | -10.29377313 | 11.59081378 | 10.11467741 | 10.53735535 | 7.157295126 | 7.12888795 | 4.425202323 | 4.80186789 |
| ENSG00000283632 | EXOC3L2 | -6.214447173 | 8.23E-05 | -4.081406202 | 0.000455111 | -10.29585338 | 4.815609471 | 4.442710129 | 5.772716771 | 2 | 2.2680111 | 2.560105256 | 2.509591799 |
| ENSG00000070047 | PHRF1 | 2.113697121 | 0.016075341 | -12.41507756 | 4.57E-13 | -10.30138044 | 5.866944937 | 7.572420273 | 7.777116083 | 9.423273993 | 9.32035914 | 2 | 2 |
| ENSG00000149294 | NCAM1 | -3.761369168 | 1.95E-26 | -6.541255686 | 1.72E-73 | -10.30262485 | 9.590776211 | 9.849218119 | 9.91701143 | 6.229283386 | 5.98191036 | 3.930690651 | 3.539239711 |
| ENSG00000104059 | FAM189A1 | -4.955370987 | 0.072436276 | -5.352941038 | 0.041455988 | -10.30831202 | 2 | 6.311476754 | 2 | 2.3892346 | 2.14022021 | 2.139668783 | 2.277179783 |
| ENSG00000170777 | TPD52L3 | -1.75939606 | 0.699135667 | -8.549505251 | 0.042633274 | -10.30890131 | 2 | 2 | 5.156626401 | 3.023133209 | 2.59489879 | 2 | 2 |
| ENSG00000168772 | CXXC4 | -2.555548494 | 0.608677328 | -7.762696922 | 0.097737025 | -10.31824542 | 2 | 4.442710129 | 2 | 2.498691189 | 2.14022021 | 2 | 2 |
| ENSG00000133169 | BEX1 | -3.664095194 | 0.000450648 | -6.654759639 | 7.40E-11 | -10.31885483 | 8.517314938 | 7.002537973 | 7.398165132 | 4.965943888 | 3.45167086 | 2.858551861 | 2.326754077 |
| ENSG00000165449 | SLC16A9 | -4.245789734 | 0.021230506 | -6.073534284 | 0.000610079 | -10.31932402 | 3.327711234 | 7.503546079 | 4.874587708 | 3.162721474 | 2.68924543 | 2.560105256 | 2.059860178 |
| ENSG00000243566 | UPK3B | -0.076996901 | 0.946943756 | -10.24253115 | 5.61E-09 | -10.31952805 | 5.330594134 | 4.710564154 | 5.59507254 | 5.093559143 | 5.24368216 | 2 | 2 |
| ENSG00000161896 | IP6K3 | -5.214833571 | 0.161801288 | -5.111754636 | 0.127753231 | -10.32658821 | 2 | 4.710564154 | 2 | 2.141737246 | 2 | 2.094598369 | 2.059860178 |
| ENSG00000108830 | RND2 | -6.782419063 | 9.78E-10 | -3.545650612 | 0.001925481 | -10.32806968 | 8.042902253 | 8.687092496 | 11.2239377 | 3.881293667 | 3.55551834 | 6.770513376 | 6.192573806 |
| ENSG00000115604 | IL18R1 | -4.892537721 | 0.002904929 | -5.437773105 | 0.000537219 | -10.33031083 | 6.138360429 | 3.076665209 | 6.629488387 | 2.3892346 | 2.68924543 | 2.62493832 | 2.117335275 |
| ENSG00000156427 | FGF18 | -2.323571535 | 0.095672957 | -8.009567465 | 1.04E-06 | -10.333139 | 3.327711234 | 5.303523747 | 6.322058544 | 3.349643765 | 3.74316441 | 2.048074405 | 2.059860178 |
| ENSG00000160224 | AIRE | -6.667745254 | 0.060038494 | -3.66909289 | 0.257630995 | -10.33683814 | 2 | 2 | 6.073378139 | 2.141737246 | 2 | 2.68698273 | 2.277179783 |
| ENSG00000119703 | ZC2HC1C | -8.152558374 | 0.000302608 | -2.188967794 | 0.216878928 | -10.34152617 | 3.327711234 | 5.595927442 | 4.058998049 | 2 | 2 | 3.27687201 | 2.950113106 |
| ENSG00000165566 | AMER2 | -1.939665717 | 0.536258443 | -8.409089082 | 0.00728342 | -10.3487548 | 4.815609471 | 3.076665209 | 2 | 2.947924294 | 2.38539787 | 2 | 2 |
| MRE11A | | 1.277049117 | 0.455679163 | -11.62825593 | 2.28E-08 | -10.35120682 | 3.327711234 | 7.431218269 | 6.534109959 | 7.697489359 | 7.83385794 | 2 | 2 |
| ENSG00000132801 | ZSWIM3 | -9.97172802 | 0.000140391 | -0.380279518 | 0.888924545 | -10.35200754 | 2 | 5.595927442 | 7.552162999 | 2 | 2 | 6.208218574 | 5.661099865 |
| C20ORF24 | C20ORF24 | 2.596321998 | 1.06E-07 | -12.96308008 | 2.57E-16 | -10.36675808 | 7.384433967 | 7.73146271 | 8.236946975 | 10.35954937 | 10.4372122 | 2 | 2 |
| ENSG00000240563 | L1TD1 | -4.976668327 | 1.62E-06 | -5.395666678 | 1.70E-08 | -10.372335 | 6.257048221 | 4.710564154 | 6.43197674 | 2.498691189 | 2.59489879 | 2.34604626 | 2.509591799 |
| ENSG00000151812 | SLC35F4 | -5.909156026 | 0.299070637 | -4.469653862 | NA | -10.37880989 | 2 | 4.11352228 | 2 | 2 | 2 | 2.139668783 | 2 |
| ENSG00000104808 | DHDH | -4.636138097 | 0.004337502 | -5.745634189 | 0.000248367 | -10.38177229 | 6.366709671 | 3.076665209 | 7.098165388 | 2.784612485 | 2.77779907 | 2.656294034 | 2.117335275 |
| ENSG00000156413 | FUT6 | -4.42090499 | 0.033708167 | -5.972479219 | 0.002979942 | -10.39338421 | 5.330594134 | 3.076665209 | 4.058998049 | 2.141737246 | 2.38539787 | 2.183373728 | 2 |
| ENSG00000182463 | TSHZ2 | 0.135752291 | 0.866211938 | -10.53479726 | 1.86E-28 | -10.39904497 | 9.219014269 | 8.124485672 | 8.805342542 | 8.882187604 | 8.94703391 | 2.183373728 | 2 |
| ENSG00000187867 | PALM3 | -7.006105345 | 0.003634803 | -3.393880019 | 0.151324285 | -10.39998536 | 2 | 3.076665209 | 7.896971941 | 2.270786764 | 2.14022021 | 3.891700877 | 3.042026731 |
| ENSG00000125430 | HS3ST3B1 | -0.692138198 | 0.404366557 | -9.709717307 | 9.73E-42 | -10.40185551 | 10.3933013 | 10.93377402 | 11.13734171 | 10.16571753 | 10.1623867 | 2.937246078 | 2.225841198 |
| ENSG00000179958 | DCTPP1 | -10.91505623 | 1.24E-05 | 0.508156642 | 0.839388001 | -10.40689958 | 8.042902253 | 7.503546079 | 2 | 2 | 2 | 7.698433415 | 7.749171054 |
| ENSG00000122121 | XPNPEP2 | -6.324145281 | 0.197561323 | -4.083758392 | 0.388256033 | -10.40790367 | 2 | 4.442710129 | 2 | 2 | 2 | 2.183373728 | 2.059860178 |
| KIAA1107 | | -0.175914858 | 0.943520433 | -10.23614636 | 2.71E-05 | -10.41206122 | 2 | 6.04685525 | 5.39244166 | 5.040242897 | 5.11361705 | 2 | 2 |
| ENSG00000165702 | GFI1B | -5.782208905 | 0.310604101 | -4.629901785 | NA | -10.41211069 | 2 | 2 | 4.058998049 | 2 | 2 | 2 | 2.117335275 |
| ENSG00000173947 | PIFO | -7.308042304 | 0.000329894 | -3.105516049 | 0.113001279 | -10.41355835 | 8.009475818 | 5.457120953 | 2 | 2.270786764 | 2.14022021 | 4.188345803 | 3.640730416 |
| ENSG00000206487 | | 0.36933921 | 0.903336546 | -10.78329731 | 9.67E-05 | -10.4139581 | 7.159102577 | 3.686303414 | 2 | 6.18083559 | 5.96314905 | 2 | 2 |
| ENSG00000273633 | | 0.623191972 | 0.535148173 | -11.04121752 | 1.65E-10 | -10.41802555 | 5.532353208 | 6.602424674 | 5.39244166 | 6.415275989 | 6.6952083 | 2 | 2 |
| ENSG00000179178 | TMEM125 | -7.122792137 | 0.101694894 | -3.302267961 | 0.435161964 | -10.4250601 | 2 | 5.131603625 | 2 | 2 | 2 | 2.3070652 | 2.374681325 |
| ENSG00000099797 | TECR | -3.972898074 | 1.08E-14 | -6.452238029 | 1.60E-37 | -10.4251361 | 12.40622339 | 12.09036432 | 13.15258056 | 8.666028533 | 8.65574839 | 5.870008604 | 6.543880272 |
| ENSG00000187546 | AGMO | -6.045235962 | 0.008560993 | -4.380409777 | 0.009711649 | -10.42564574 | 4.467311984 | 3.076665209 | 4.523675391 | 2 | 2.14022021 | 2.3070652 | 2.117335275 |
| ENSG00000152270 | PDE3B | -7.52462084 | 0.003384073 | -2.905915258 | 0.173291381 | -10.4305361 | 2 | 6.04685525 | 5.772716771 | 2 | 2.14022021 | 3.648325314 | 2.466008509 |
| ENSG00000157927 | RADIL | -6.304711784 | 0.000654271 | -4.129020388 | 0.029853681 | -10.43373217 | 9.771132748 | 8.234834549 | 2 | 2.947924294 | 3.33976443 | 4.946551782 | 4.434543419 |
| ENSG00000143858 | SYT2 | -6.462986461 | 1.50E-07 | -3.973184496 | 6.73E-06 | -10.43617096 | 6.563806279 | 5.303523747 | 5.39244166 | 2.141737246 | 2.2680111 | 2.911488689 | 2.885414003 |
| ENSG00000170509 | HSD17B13 | -3.600633972 | 0.058720914 | -6.838053044 | 0.000264771 | -10.43868702 | 6.563806279 | 3.076665209 | 4.874587708 | 2.784612485 | 2.94009902 | 2.225793514 | 2 |
| ENSG00000188056 | TREML4 | -4.583584441 | 0.350743351 | -5.864202859 | 0.211555168 | -10.4477873 | 2 | 2 | 5.59507254 | 2 | 2.38539787 | 2 | 2.172608095 |
| ENSG00000167992 | VWCE | -5.15114554 | 1.39E-19 | -5.316757111 | 1.60E-22 | -10.46790265 | 8.674867122 | 8.487557258 | 9.014156722 | 4.205751501 | 3.74316441 | 4.188345803 | 3.45257121 |
| ENSG00000137817 | PARP6 | -11.0111514 | 2.02E-05 | 0.540602513 | 0.837702779 | -10.47054889 | 6.73718411 | 8.522788974 | 2 | 2 | 2 | 7.979009894 | 7.708840864 |
| ENSG00000276725 | | 3.334281347 | 0.007899101 | -13.80601618 | 4.57E-13 | -10.47173484 | 6.366709671 | 8.004992147 | 9.767769415 | 11.9782175 | 11.9757514 | 2 | 2 |
| ENSG00000160336 | ZNF761 | -10.80890899 | 3.58E-08 | 0.335295972 | 0.83322656 | -10.47361302 | 4.815609471 | 8.077869079 | 6.882775709 | 2 | 2 | 7.672789259 | 7.188214218 |
| ENSG00000189283 | FHIT | -6.411620904 | 0.000831714 | -4.068860445 | 0.028762582 | -10.48048135 | 7.709880939 | 5.946638857 | 2 | 2.498691189 | 2.14022021 | 3.374372158 | 3.012034821 |
| ENSG00000273608 | | -2.883971028 | 0.184286434 | -7.602854752 | 0.002034082 | -10.48682578 | 4.467311984 | 4.93638933 | 2 | 2.784612485 | 2.38539787 | 2 | 2.059860178 |
| ENSG00000158164 | TMSB15A | -4.748581971 | 0.002024004 | -5.740473276 | 0.000112767 | -10.48905525 | 7.867441304 | 4.442710129 | 4.523675391 | 3.023133209 | 2.68924543 | 2.3070652 | 2.671842496 |
| ENSG00000183837 | PNMA3 | -7.577960677 | 9.33E-05 | -2.924274164 | 0.087889252 | -10.50223484 | 4.467311984 | 7.73146271 | 3.369317972 | 2.141737246 | 2.14022021 | 3.824280769 | 3.858622973 |
| ENSG00000151779 | NBAS | -11.98454683 | 1.20E-08 | 1.472737666 | 0.381333928 | -10.51180916 | 8.107517749 | 9.298208073 | 4.523675391 | 2 | 2 | 9.845205026 | 9.620658212 |
| ENSG00000155886 | SLC24A2 | -7.376877829 | 0.088683678 | -3.140808401 | 0.458690094 | -10.51768623 | 2 | 2 | 5.39244166 | 2 | 2 | 2.457042231 | 2.421067459 |
| ENSG00000105679 | GAPDHS | -7.387306758 | 0.000897252 | -3.135157803 | 0.058838964 | -10.52246456 | 4.467311984 | 4.11352228 | 3.369317972 | 2 | 2 | 2.42098416 | 2.466008509 |
| ENSG00000114455 | HHLA2 | -4.274928745 | 0.116793916 | -6.259681219 | 0.018312083 | -10.53460996 | 3.327711234 | 3.076665209 | 5.39244166 | 2.498691189 | 2 | 2.139668783 | 2 |
| ENSG00000136297 | MMD2 | -4.925449014 | 0.174608987 | -5.631099488 | 0.099807208 | -10.5565485 | 3.327711234 | 2 | 5.156626401 | 2 | 2.2680111 | 2 | 2.172608095 |
| ENSG00000162896 | PIGR | -5.375702787 | 0.003354987 | -5.182876358 | 0.001604999 | -10.55857915 | 4.00722531 | 3.686303414 | 5.930865648 | 2.270786764 | 2.14022021 | 2.384001718 | 2.059860178 |
| GLTSCR1L | | 1.107596301 | 0.24800863 | -11.67515994 | 1.30E-11 | -10.56756364 | 6.563806279 | 7.189546432 | 5.39244166 | 7.61176303 | 7.68150967 | 2 | 2 |
| ENSG00000162551 | ALPL | -9.794040328 | 1.18E-11 | -0.776454747 | 0.583004866 | -10.57049507 | 10.24321353 | 6.464272742 | 8.460932916 | 2.141737246 | 2.2680111 | 8.307508685 | 8.362041249 |
| ENSG00000278268 | | 1.353750717 | 0.446344169 | -11.92431011 | 1.83E-08 | -10.5705594 | 6.563806279 | 7.847167909 | 3.369317972 | 8.104838684 | 8.16258751 | 2 | 2 |
| ENSG00000166960 | CCDC178 | -5.348738396 | NA | -5.226496393 | NA | -10.57523479 | 2 | 3.686303414 | 2 | 2 | 2 | 2 | 2.059860178 |
| ENSG00000070729 | CNGB1 | -5.367408608 | 0.033441444 | -5.209246101 | 0.031846812 | -10.57665471 | 6.89194353 | 4.11352228 | 2 | 2.600425781 | 2 | 2.3070652 | 2.421067459 |
| C17ORF96 | C17ORF96 | 0.639767216 | 0.746656687 | -11.2235719 | 2.74E-07 | -10.58380469 | 7.384433967 | 5.131603625 | 3.369317972 | 6.716553866 | 6.78874509 | 2 | 2 |
| ENSG00000165548 | TMEM63C | -6.661343021 | 5.93E-05 | -3.922664511 | 0.006869845 | -10.58400753 | 4.00722531 | 6.844429755 | 6.073378139 | 2.3892346 | 2 | 3.27687201 | 2.70970617 |
| ENSG00000036565 | SLC18A1 | -5.722047765 | 0.315977132 | -4.86259819 | NA | -10.58464595 | 4.00722531 | 2 | 2 | 2 | 2 | 2.094598369 | 2 |
| ENSG00000185899 | TAS2R60 | -5.722047765 | 0.315977132 | -4.86259819 | NA | -10.58464595 | 4.00722531 | 2 | 2 | 2 | 2 | 2.094598369 | 2 |
| ENSG00000115970 | THADA | -11.42710371 | 3.55E-12 | 0.8373566 | 0.236344937 | -10.58974711 | 7.031698319 | 8.234834549 | 7.646393253 | 2 | 2 | 8.484444907 | 8.618670729 |
| ENSG00000142512 | SIGLEC10 | -6.237910941 | 6.31E-07 | -4.352306371 | 2.03E-05 | -10.59021731 | 6.73718411 | 4.710564154 | 6.073378139 | 2.3892346 | 2.14022021 | 2.911488689 | 2.70970617 |
| ENSG00000104290 | FZD3 | -4.891462303 | 8.97E-06 | -5.723230863 | 1.18E-07 | -10.61469317 | 8.39321512 | 9.079034257 | 10.12770132 | 5.461211713 | 3.08597292 | 4.311629396 | 3.717076715 |
| ENSG00000166111 | SVOP | -5.693854256 | 0.004148594 | -4.925266221 | 0.010530438 | -10.61912048 | 5.095959458 | 5.303523747 | 7.342949392 | 2.498691189 | 2.38539787 | 3.1504466 | 2 |
| ENSG00000152056 | AP1S3 | -10.15173285 | 6.50E-06 | -0.475984082 | 0.818792684 | -10.62771693 | 4.815609471 | 4.11352228 | 7.818184403 | 2 | 2 | 5.60262961 | 6.374480702 |
| ENSG00000136870 | ZNF189 | -11.02127394 | 2.34E-08 | 0.393112666 | 0.805557931 | -10.62816127 | 8.493334308 | 5.457120953 | 6.322058544 | 2 | 2 | 7.731011524 | 7.699462058 |
| ENSG00000158865 | SLC5A11 | -4.913887185 | 0.052698164 | -5.718060012 | 0.015355655 | -10.6319472 | 2 | 4.442710129 | 4.523675391 | 2 | 2.2680111 | 2.048074405 | 2.117335275 |
| KIAA1033 | | 2.485909118 | 0.121382642 | -13.1181187 | 2.55E-10 | -10.63220959 | 8.696033109 | 8.337339476 | 4.058998049 | 10.45048384 | 10.4363623 | 2 | 2 |
| ENSG00000075790 | BCAP29 | -11.57003314 | 2.55E-12 | 0.913422699 | 0.21425456 | -10.65661044 | 6.96351253 | 8.029695761 | 8.296287027 | 2 | 2 | 8.784089222 | 8.755631095 |
| ENSG00000187714 | SLC18A3 | -11.64662828 | 3.59E-07 | 0.979926503 | 0.636488065 | -10.66670178 | 3.327711234 | 7.638155693 | 9.049077186 | 2 | 2 | 8.327733606 | 9.32645842 |
| ENSG00000140285 | FGF7 | 1.211330454 | 0.026902025 | -11.88065779 | 3.77E-53 | -10.66932733 | 11.25159303 | 10.4709455 | 10.26770388 | 11.94702268 | 11.9280254 | 2.183373728 | 2.117335275 |
| ENSG00000227046 | | -0.137945454 | 0.887194753 | -10.53853819 | 7.39E-31 | -10.67648365 | 8.716893051 | 9.148351514 | 10.12770132 | 9.284006841 | 9.35044873 | 2.048074405 | 2.277179783 |

| | | | | | | | | | | | | | |
|---|---|---|---|---|---|---|---|---|---|---|---|---|---|
| ENSG00000108506 | INTS2 | -10.5333326 | 7.46E-09 | -0.1474957 | 0.911348375 | -10.6808283 | 5.70932728 | 7.73146271 | 6.322058544 | 2 | 2 | 6.915242278 | 6.482320745 |
| ORAOV1 | | 0.904997511 | 0.584585811 | -11.58869739 | 1.11E-08 | -10.68369988 | 4.00722531 | 7.572420273 | 5.930865648 | 7.409376773 | 7.31795292 | 2 | 2 |
| ENSG00000151834 | GABRA2 | -3.900269968 | NA | -6.786772651 | NA | -10.68704262 | 2 | 3.686303414 | 2 | 2.141737246 | 2 | 2 | 2 |
| LOC101928841 | | -1.404710281 | 0.472233499 | -9.283404836 | 3.64E-05 | -10.68811512 | 3.327711234 | 3.686303414 | 5.39244166 | 3.162721474 | 3.60476711 | 2 | 2 |
| ENSG00000171804 | WDR87 | -4.761724232 | 0.190366567 | -5.93050971 | 0.087552565 | -10.69223394 | 2 | 2 | 5.772716771 | 2.270786764 | 2.14022021 | 2.183373728 | 2 |
| ENSG00000151849 | CENPJ | -10.67400755 | 2.03E-05 | -0.018571533 | 0.994326215 | -10.69257909 | 7.82963488 | 7.232742069 | 2 | 2 | 2 | 7.096193303 | 6.843307307 |
| ENSG00000009724 | MASP2 | -8.42033613 | 2.25E-05 | -2.277203894 | 0.084945224 | -10.69754002 | 4.00722531 | 6.140560021 | 7.098165388 | 2.141737246 | 2 | 4.271694079 | 4.147809227 |
| ENSG00000231115 | | 1.434197884 | 0.353802368 | -12.14499354 | 1.32E-09 | -10.71079565 | 7.940205782 | 7.002537973 | 4.058998049 | 8.418656444 | 8.44373613 | 2 | 2 |
| ENSG00000178150 | ZNF114 | -3.65550041 | 3.67E-05 | -7.055629045 | 7.96E-12 | -10.71112946 | 5.70932728 | 5.595927442 | 6.073378139 | 3.023133209 | 3.01487888 | 2.048074405 | 2.225841198 |
| ENSG00000103310 | ZP2 | -3.83322114 | 0.117474983 | -6.884495864 | 0.009186901 | -10.717717 | 2 | 4.11352228 | 4.058998049 | 2.141737246 | 2.2680111 | 2.048074405 | 2 |
| ENSG00000167711 | SERPINF2 | -1.96683547 | 0.652705192 | -8.751637799 | 0.032897253 | -10.71847327 | 2 | 5.303523747 | 2 | 2.600425781 | 3.01487888 | 2 | 2 |
| ENSG00000278220 | | -0.563704798 | 0.589153227 | -10.15577884 | 5.50E-20 | -10.71948364 | 9.397585735 | 7.503546079 | 7.600047325 | 7.93509946 | 7.84926609 | 2.094598369 | 2.117335275 |
| ENSG00000249693 | THEGL | -6.261510552 | 0.009864327 | -4.463785803 | 0.042370211 | -10.72529636 | 6.563806279 | 3.076665209 | 2 | 2.141737246 | 2.14022021 | 2.62493832 | 2.225841198 |
| ENSG00000227345 | PARG | -11.54707696 | 2.37E-07 | 0.817143763 | 0.68474076 | -10.7299332 | 3.327711234 | 7.901717006 | 8.784770914 | 2 | 2 | 8.772510235 | 8.520227711 |
| ENSG00000106113 | CRHR2 | -6.787711414 | 0.141109656 | -3.945633085 | 0.374026188 | -10.7333445 | 2 | 2 | 4.874587708 | 2 | 2 | 2.139668783 | 2.225841198 |
| ENSG00000100033 | PRODH | -2.923944469 | 0.611410824 | -7.812268902 | 0.146527599 | -10.73621337 | 2 | 2 | 4.523675391 | 2.498691189 | 2 | 2 | 2 |
| ENSG00000161649 | CD300LG | -6.214815037 | 0.005938317 | -4.522604353 | 0.005913437 | -10.73741939 | 3.327711234 | 3.686303414 | 5.156626401 | 2.141737246 | 2 | 2.267001569 | 2.172608095 |
| ENSG00000213920 | MDP1 | -0.86155592 | 0.421069804 | -9.884272757 | 1.22E-08 | -10.74582868 | 5.70932728 | 5.457120953 | 7.029880454 | 5.552894743 | 5.28958453 | 2.048074405 | 2 |
| SLC35E2 | | 1.870313829 | 0.079842495 | -12.62060784 | 1.59E-12 | -10.75029402 | 6.96351253 | 6.140560021 | 8.434763955 | 9.297323944 | 9.3719445 | 2 | 2 |
| ENSG00000102468 | HTR2A | -4.013841674 | 0.016302514 | -6.73677473 | 2.73E-05 | -10.7506164 | 7.624139768 | 7.393647515 | 2 | 3.515101962 | 3.50452879 | 2.384001718 | 2.326754077 |
| ENSG00000154118 | JPH3 | -7.312907584 | 0.089666875 | -3.446106902 | 0.409348476 | -10.75901449 | 2 | 5.303523747 | 2 | 2 | 2 | 2.42098416 | 2.277179783 |
| ENSG00000145425 | RPS3A | -3.01940258 | 3.49E-06 | -7.743628534 | 1.17E-35 | -10.76303111 | 14.59129552 | 13.01511834 | 13.26731409 | 10.81133226 | 10.7571169 | 6.234425698 | 6.041166521 |
| ENSG00000109758 | HGFAC | -5.224695425 | 0.076258148 | -5.538727963 | 0.047007566 | -10.76342339 | 2 | 2 | 6.203069917 | 2.141737246 | 2.2680111 | 2.267001569 | 2.059860178 |
| ENSG00000164933 | SLC25A32 | -1.011859062 | 0.101317769 | -9.758843052 | 4.68E-47 | -10.77070211 | 8.87372952 | 9.583268459 | 10.12770132 | 8.697938662 | 8.51952909 | 2.3070652 | 2.277179783 |
| ENSG00000136485 | DCAF7 | -12.77309566 | 6.45E-16 | 2.001898355 | 1.17E-05 | -10.77119731 | 8.797441463 | 9.45234302 | 8.805342542 | 2 | 2 | 11.07117504 | 11.03036682 |
| ENSG00000170412 | GPRC5C | -8.841100487 | 8.71E-10 | -1.962081699 | 0.207075162 | -10.80318219 | 10.95065381 | 9.599860728 | 6.203069917 | 2.270786764 | 2.86123014 | 6.934553331 | 8.522223965 |
| ENSG00000004846 | ABCB5 | -3.636086403 | 0.055580779 | -7.167115288 | 0.0004491 | -10.80320169 | 3.327711234 | 4.442710129 | 5.39244166 | 2.784612485 | 2.14022021 | 2.094598369 | 2 |
| ENSG00000136099 | PCDH8 | -3.459198015 | 0.549630367 | -7.347211024 | NA | -10.80640904 | 2 | 4.11352228 | 2 | 2.270786764 | 2 | 2 | 2 |
| ENSG00000240021 | TEX35 | -6.224365591 | 0.078050197 | -4.584238494 | 0.158852787 | -10.80860408 | 2 | 3.076665209 | 4.058998049 | 2 | 2 | 2.048074405 | 2.117335275 |
| ENSG00000166831 | RBPMS2 | -6.579278571 | 3.89E-27 | -4.232504956 | 1.03E-12 | -10.81178353 | 10.72212986 | 9.664384941 | 11.15748401 | 4.237680128 | 4.4836996 | 6.668482487 | 6.235386446 |
| ENSG00000166159 | LRTM2 | -4.010229511 | 0.267665409 | -6.803922986 | 0.05016305 | -10.8141525 | 2 | 2 | 6.203069917 | 2.141737246 | 2.68924543 | 2.139668783 | 2 |
| ENSG00000173559 | NABP1 | -11.4858725 | 2.78E-12 | 0.667140311 | 0.358514023 | -10.81873219 | 8.16936288 | 7.051611315 | 7.934808822 | 2 | 2 | 8.620333861 | 8.240396277 |
| ENSG00000174227 | PIGG | -11.79669995 | 1.61E-11 | 0.973864058 | 0.33413809 | -10.8228359 | 8.340423424 | 8.639735838 | 6.534109959 | 2 | 2 | 8.968699418 | 9.135803248 |
| ENSG00000065491 | TBC1D22B | -11.60503449 | 5.89E-07 | 0.765113596 | 0.724892691 | -10.8399209 | 3.327711234 | 7.002537973 | 9.180874355 | 2 | 2 | 8.839349773 | 8.449958388 |
| ENSG00000275397 | | -1.851766344 | 0.241676601 | -8.990533159 | 2.36E-07 | -10.8422995 | 3.327711234 | 5.72254323 | 7.502634626 | 4.647285135 | 4.68029874 | 2.048074405 | 2.059860178 |
| ENSG00000135472 | FAIM2 | -6.026586202 | 7.61E-20 | -4.81985232 | 4.06E-15 | -10.84643852 | 9.015175615 | 7.761270542 | 9.149036566 | 3.515101962 | 3.28037825 | 4.122514202 | 4.388595494 |
| ENSG00000236353 | | 1.685755569 | 0.465280682 | -12.53473824 | 3.04E-07 | -10.84898267 | 2 | 7.790474954 | 8.143093325 | 9.088859521 | 9.04395769 | 2 | 2 |
| TMEM110 | | 1.572753159 | 0.490581765 | -12.4238707 | 3.09E-07 | -10.85111755 | 7.940205782 | 7.790474954 | 2 | 8.801706047 | 8.88608977 | 2 | 2 |
| ENSG00000141431 | ASXL3 | -6.148180098 | 0.000186547 | -4.703411087 | 0.000900485 | -10.85159119 | 4.815609471 | 6.389896633 | 4.874587708 | 2.141737246 | 2.2680111 | 2.803598391 | 2.117335275 |
| ENSG00000149646 | CNBD2 | -4.355569236 | 0.057769744 | -6.496632179 | 0.003752234 | -10.85220142 | 4.467311984 | 4.11352228 | 5.59507254 | 2 | 2.68924543 | 2.183373728 | 2 |
| ENSG00000182489 | XKRX | -3.588488467 | 0.038100108 | -7.271258537 | 3.24E-05 | -10.859742 | 2 | 5.457120953 | 6.718950348 | 3.023133209 | 2.94009902 | 2.094598369 | 2.117335275 |
| ENSG00000196131 | VN1R2 | -3.198602686 | 0.344365344 | -7.661936541 | 0.022824771 | -10.86053923 | 2 | 3.076665209 | 4.058998049 | 2.141737246 | 2.2680111 | 2 | 2 |
| ENSG00000140795 | MYLK3 | -5.312639057 | 0.000106612 | -5.548052449 | 2.49E-05 | -10.86069151 | 6.009025997 | 4.11352228 | 7.97167868 | 2.868578001 | 2.59489879 | 2.62493832 | 2.671842496 |
| C14ORF183 | C14ORF183 | -4.714895037 | NA | -6.152911075 | NA | -10.86780611 | 3.327711234 | 2 | 2 | 2 | 2 | 2 | 2 |
| C16ORF93 | C16ORF93 | -4.714895037 | NA | -6.152911075 | NA | -10.86780611 | 2 | 3.076665209 | 2 | 2 | 2 | 2 | 2 |
| CCRN4L | CCRN4L | -4.714895037 | NA | -6.152911075 | NA | -10.86780611 | 3.327711234 | 2 | 2 | 2 | 2 | 2 | 2 |
| ENSG00000006071 | ABCC8 | -4.714895037 | NA | -6.152911075 | NA | -10.86780611 | 2 | 2 | 3.369317972 | 2 | 2 | 2 | 2 |
| ENSG00000007908 | SELE | -4.714895037 | NA | -6.152911075 | NA | -10.86780611 | 2 | 2 | 3.369317972 | 2 | 2 | 2 | 2 |
| ENSG00000069206 | ADAM7 | -4.714895037 | NA | -6.152911075 | NA | -10.86780611 | 2 | 2 | 3.369317972 | 2 | 2 | 2 | 2 |
| ENSG00000083307 | GRHL2 | -4.714895037 | NA | -6.152911075 | NA | -10.86780611 | 2 | 3.076665209 | 2 | 2 | 2 | 2 | 2 |
| ENSG00000101276 | SLC52A3 | -4.714895037 | NA | -6.152911075 | NA | -10.86780611 | 3.327711234 | 2 | 2 | 2 | 2 | 2 | 2 |
| ENSG00000103089 | FA2H | -4.714895037 | NA | -6.152911075 | NA | -10.86780611 | 3.327711234 | 2 | 2 | 2 | 2 | 2 | 2 |
| ENSG00000106069 | CHN2 | -4.714895037 | NA | -6.152911075 | NA | -10.86780611 | 3.327711234 | 2 | 2 | 2 | 2 | 2 | 2 |
| ENSG00000107014 | RLN2 | -4.714895037 | NA | -6.152911075 | NA | -10.86780611 | 3.327711234 | 2 | 2 | 2 | 2 | 2 | 2 |
| ENSG00000111432 | FZD10 | -4.714895037 | NA | -6.152911075 | NA | -10.86780611 | 3.327711234 | 2 | 2 | 2 | 2 | 2 | 2 |
| ENSG00000124429 | POF1B | -4.714895037 | NA | -6.152911075 | NA | -10.86780611 | 2 | 2 | 3.369317972 | 2 | 2 | 2 | 2 |
| ENSG00000124564 | SLC17A3 | -4.714895037 | NA | -6.152911075 | NA | -10.86780611 | 2 | 2 | 3.369317972 | 2 | 2 | 2 | 2 |
| ENSG00000125571 | IL37 | -4.714895037 | NA | -6.152911075 | NA | -10.86780611 | 2 | 3.076665209 | 2 | 2 | 2 | 2 | 2 |
| ENSG00000127152 | BCL11B | -4.714895037 | NA | -6.152911075 | NA | -10.86780611 | 2 | 3.076665209 | 2 | 2 | 2 | 2 | 2 |
| ENSG00000128276 | RFPL3 | -4.714895037 | NA | -6.152911075 | NA | -10.86780611 | 2 | 3.076665209 | 2 | 2 | 2 | 2 | 2 |
| ENSG00000131864 | USP29 | -4.714895037 | NA | -6.152911075 | NA | -10.86780611 | 2 | 2 | 3.369317972 | 2 | 2 | 2 | 2 |
| ENSG00000145384 | FABP2 | -4.714895037 | NA | -6.152911075 | NA | -10.86780611 | 3.327711234 | 2 | 2 | 2 | 2 | 2 | 2 |
| ENSG00000146285 | SCML4 | -4.714895037 | NA | -6.152911075 | NA | -10.86780611 | 2 | 3.076665209 | 2 | 2 | 2 | 2 | 2 |
| ENSG00000148386 | LCN9 | -4.714895037 | NA | -6.152911075 | NA | -10.86780611 | 2 | 3.076665209 | 2 | 2 | 2 | 2 | 2 |
| ENSG00000149742 | SLC22A9 | -4.714895037 | NA | -6.152911075 | NA | -10.86780611 | 2 | 3.076665209 | 2 | 2 | 2 | 2 | 2 |
| ENSG00000153779 | TGIF2LX | -4.714895037 | NA | -6.152911075 | NA | -10.86780611 | 2 | 3.076665209 | 2 | 2 | 2 | 2 | 2 |
| ENSG00000158816 | VWA5B1 | -4.714895037 | NA | -6.152911075 | NA | -10.86780611 | 2 | 3.076665209 | 2 | 2 | 2 | 2 | 2 |
| ENSG00000162670 | BRINP3 | -4.714895037 | NA | -6.152911075 | NA | -10.86780611 | 2 | 3.076665209 | 2 | 2 | 2 | 2 | 2 |
| ENSG00000162951 | LRRTM1 | -4.714895037 | NA | -6.152911075 | NA | -10.86780611 | 2 | 3.076665209 | 2 | 2 | 2 | 2 | 2 |
| ENSG00000163440 | PDCL2 | -4.714895037 | NA | -6.152911075 | NA | -10.86780611 | 2 | 2 | 3.369317972 | 2 | 2 | 2 | 2 |
| ENSG00000165192 | ASB11 | -4.714895037 | NA | -6.152911075 | NA | -10.86780611 | 2 | 3.076665209 | 2 | 2 | 2 | 2 | 2 |
| ENSG00000165643 | SOHLH1 | -4.714895037 | NA | -6.152911075 | NA | -10.86780611 | 2 | 3.076665209 | 2 | 2 | 2 | 2 | 2 |
| ENSG00000171772 | SYCE1 | -4.714895037 | NA | -6.152911075 | NA | -10.86780611 | 3.327711234 | 2 | 2 | 2 | 2 | 2 | 2 |
| ENSG00000171954 | CYP4F22 | -4.714895037 | NA | -6.152911075 | NA | -10.86780611 | 2 | 2 | 3.369317972 | 2 | 2 | 2 | 2 |
| ENSG00000173612 | GPRC6A | -4.714895037 | NA | -6.152911075 | NA | -10.86780611 | 2 | 2 | 3.369317972 | 2 | 2 | 2 | 2 |
| ENSG00000173673 | HES3 | -4.714895037 | NA | -6.152911075 | NA | -10.86780611 | 3.327711234 | 2 | 2 | 2 | 2 | 2 | 2 |
| ENSG00000173769 | TOPAZ1 | -4.714895037 | NA | -6.152911075 | NA | -10.86780611 | 2 | 3.076665209 | 2 | 2 | 2 | 2 | 2 |
| ENSG00000174667 | OR7D4 | -4.714895037 | NA | -6.152911075 | NA | -10.86780611 | 3.327711234 | 2 | 2 | 2 | 2 | 2 | 2 |
| ENSG00000175766 | EIF4E1B | -4.714895037 | NA | -6.152911075 | NA | -10.86780611 | 3.327711234 | 2 | 2 | 2 | 2 | 2 | 2 |
| ENSG00000176742 | OR51V1 | -4.714895037 | NA | -6.152911075 | NA | -10.86780611 | 3.327711234 | 2 | 2 | 2 | 2 | 2 | 2 |
| ENSG00000177910 | SPATA31C2 | -4.714895037 | NA | -6.152911075 | NA | -10.86780611 | 2 | 2 | 3.369317972 | 2 | 2 | 2 | 2 |
| ENSG00000178243 | C9orf62 | -4.714895037 | NA | -6.152911075 | NA | -10.86780611 | 2 | 2 | 3.369317972 | 2 | 2 | 2 | 2 |
| ENSG00000178522 | AMBN | -4.714895037 | NA | -6.152911075 | NA | -10.86780611 | 3.327711234 | 2 | 2 | 2 | 2 | 2 | 2 |
| ENSG00000183035 | CYLC1 | -4.714895037 | NA | -6.152911075 | NA | -10.86780611 | 2 | 2 | 3.369317972 | 2 | 2 | 2 | 2 |
| ENSG00000183695 | MRGPRX2 | -4.714895037 | NA | -6.152911075 | NA | -10.86780611 | 3.327711234 | 2 | 2 | 2 | 2 | 2 | 2 |
| ENSG00000187833 | C2orf78 | -4.714895037 | NA | -6.152911075 | NA | -10.86780611 | 3.327711234 | 2 | 2 | 2 | 2 | 2 | 2 |
| ENSG00000197322 | C17orf102 | -4.714895037 | NA | -6.152911075 | NA | -10.86780611 | 2 | 3.076665209 | 2 | 2 | 2 | 2 | 2 |
| ENSG00000198178 | CLEC4C | -4.714895037 | NA | -6.152911075 | NA | -10.86780611 | 2 | 2 | 3.369317972 | 2 | 2 | 2 | 2 |
| ENSG00000203857 | HSD3B1 | -4.714895037 | NA | -6.152911075 | NA | -10.86780611 | 2 | 2 | 3.369317972 | 2 | 2 | 2 | 2 |
| ENSG00000204450 | TRIM64 | -4.714895037 | NA | -6.152911075 | NA | -10.86780611 | 3.327711234 | 2 | 2 | 2 | 2 | 2 | 2 |
| ENSG00000214891 | TRIM64C | -4.714895037 | NA | -6.152911075 | NA | -10.86780611 | 3.327711234 | 2 | 2 | 2 | 2 | 2 | 2 |
| ENSG00000225550 | | -4.714895037 | NA | -6.152911075 | NA | -10.86780611 | 2 | 2 | 3.369317972 | 2 | 2 | 2 | 2 |
| ENSG00000242289 | | -4.714895037 | NA | -6.152911075 | NA | -10.86780611 | 3.327711234 | 2 | 2 | 2 | 2 | 2 | 2 |
| ENSG00000261972 | | -4.714895037 | NA | -6.152911075 | NA | -10.86780611 | 2 | 2 | 3.369317972 | 2 | 2 | 2 | 2 |
| ENSG00000274571 | | -4.714895037 | NA | -6.152911075 | NA | -10.86780611 | 2 | 3.076665209 | 2 | 2 | 2 | 2 | 2 |
| ENSG00000275164 | | -4.714895037 | NA | -6.152911075 | NA | -10.86780611 | 2 | 2 | 3.369317972 | 2 | 2 | 2 | 2 |
| ENSG00000277132 | | -4.714895037 | NA | -6.152911075 | NA | -10.86780611 | 2 | 2 | 3.369317972 | 2 | 2 | 2 | 2 |
| ENSG00000278369 | | -4.714895037 | NA | -6.152911075 | NA | -10.86780611 | 2 | 2 | 3.369317972 | 2 | 2 | 2 | 2 |
| ENSG00000278767 | | -4.714895037 | NA | -6.152911075 | NA | -10.86780611 | 3.327711234 | 2 | 2 | 2 | 2 | 2 | 2 |
| ENSG00000280585 | | -4.714895037 | NA | -6.152911075 | NA | -10.86780611 | 2 | 3.076665209 | 2 | 2 | 2 | 2 | 2 |
| GPR110 | | -4.714895037 | NA | -6.152911075 | NA | -10.86780611 | 2 | 3.076665209 | 2 | 2 | 2 | 2 | 2 |
| GPR123 | | -4.714895037 | NA | -6.152911075 | NA | -10.86780611 | 3.327711234 | 2 | 2 | 2 | 2 | 2 | 2 |
| GPR64 | | -4.714895037 | NA | -6.152911075 | NA | -10.86780611 | 3.327711234 | 2 | 2 | 2 | 2 | 2 | 2 |
| IGJ | | -4.714895037 | NA | -6.152911075 | NA | -10.86780611 | 3.327711234 | 2 | 2 | 2 | 2 | 2 | 2 |
| KIAA0226L | | -4.714895037 | NA | -6.152911075 | NA | -10.86780611 | 2 | 3.076665209 | 2 | 2 | 2 | 2 | 2 |
| LOC100144595 | | -4.714895037 | NA | -6.152911075 | NA | -10.86780611 | 2 | 3.076665209 | 2 | 2 | 2 | 2 | 2 |
| MTL5 | | -4.714895037 | NA | -6.152911075 | NA | -10.86780611 | 3.327711234 | 2 | 2 | 2 | 2 | 2 | 2 |

| | | | | | | | | | | | | | |
|---|---|---|---|---|---|---|---|---|---|---|---|---|---|
| OR8G2 | | -4.714895037 | NA | -6.152911075 | NA | -10.86780611 | 3.327711234 | 2 | 2 | 2 | 2 | 2 | 2 |
| PPAPDC1A | | -4.714895037 | NA | -6.152911075 | NA | -10.86780611 | 2 | 3.076665209 | 2 | 2 | 2 | 2 | 2 |
| PTPLAD2 | | -4.714895037 | NA | -6.152911075 | NA | -10.86780611 | 2 | 3.076665209 | 2 | 2 | 2 | 2 | 2 |
| TBC1D29 | | -4.714895037 | NA | -6.152911075 | NA | -10.86780611 | 2 | 2 | 3.369317972 | 2 | 2 | 2 | 2 |
| ENSG00000148948 | LRRC4C | -5.235279469 | 0.052566108 | -5.632609557 | 0.032176413 | -10.86788903 | 2 | 8.356994601 | 2 | 2.868578001 | 2.68924543 | 2.457042231 | 2.782576693 |
| ENSG00000188064 | WNT7B | 0.093289743 | 0.942460221 | -10.96145999 | 1.15E-09 | -10.86817025 | 5.330594134 | 5.131603625 | 6.718950348 | 5.85002558 | 6.1068222 | 2 | 2 |
| ENSG00000278224 | PRICKLE4 | -1.998134819 | 0.203739617 | -8.8800782 | 1.55E-09 | -10.87821302 | 7.096806373 | 4.442710129 | 9.180874355 | 5.709096143 | 6.26070706 | 2.225793514 | 2.117335275 |
| ENSG00000088386 | SLC15A1 | -5.913859286 | 0.009557939 | -4.966615816 | 0.014415881 | -10.8804751 | 4.00722531 | 2 | 6.073378139 | 2.141737246 | 2.14022021 | 2.139668783 | 2.374681325 |
| ENSG00000134755 | DSC2 | -7.589917958 | 8.27E-07 | -3.306643811 | 0.025387429 | -10.89656177 | 5.330594134 | 8.655694736 | 7.342949392 | 2.600425781 | 2 | 4.933813368 | 4.133950982 |
| ENSG00000206486 | | 1.55874316 | 0.375485391 | -12.46711933 | 4.09E-09 | -10.90837617 | 7.904282243 | 7.874700259 | 3.369317972 | 8.877128559 | 8.86981338 | 2 | 2 |
| ENSG00000274619 | | -3.332946884 | 0.103095955 | -7.576489386 | 0.000402763 | -10.90943627 | 6.366709671 | 3.686303414 | 2 | 2.784612485 | 2.77779907 | 2.048074405 | 2.059860178 |
| ENSG00000085721 | RRN3 | 2.04455034 | 0.215393228 | -12.96288571 | 4.79E-10 | -10.91833537 | 4.00722531 | 7.954278496 | 8.699419242 | 9.825086737 | 9.87245708 | 2 | 2 |
| ENSG00000145020 | AMT | -4.115265624 | 5.47E-15 | -6.803378979 | 2.48E-36 | -10.9186446 | 8.418903792 | 8.356994601 | 9.212024683 | 4.574143677 | 5.00774437 | 3.035909414 | 2.851942824 |
| ENSG00000274205 | | -2.168774972 | 0.618130061 | -8.751619065 | 0.032955861 | -10.92039404 | 2 | 5.303523747 | 2 | 2.868578001 | 2.59489879 | 2 | 2 |
| ENSG00000177380 | PPFIA3 | -10.42715365 | 5.02E-05 | -0.493375238 | 0.850432248 | -10.92052888 | 2 | 6.464272742 | 7.858115942 | 2 | 2 | 6.471027105 | 6.066922748 |
| ENSG00000206384 | COL6A6 | -7.112101964 | 0.110527921 | -3.808595292 | 0.374771224 | -10.92069726 | 2 | 2 | 5.156626401 | 2 | 2 | 2.267001569 | 2.225841198 |
| CRIPAK | CRIPAK | 0.282907584 | 0.891679753 | -11.20486194 | 2.47E-07 | -10.92195436 | 4.00722531 | 7.393647515 | 4.523675391 | 6.372775472 | 6.41363412 | 2 | 2 |
| ENSG00000131477 | RAMP2 | -5.785507797 | 4.21E-13 | -5.138277419 | 1.76E-12 | -10.92378522 | 8.19931886 | 7.002537973 | 7.398165132 | 2.695456494 | 3.08597292 | 3.500668998 | 2.918126211 |
| ENSG00000166452 | AKIP1 | -11.22121889 | 1.36E-12 | 0.293466232 | 0.579833528 | -10.92775266 | 7.667647223 | 7.35507212 | 7.552162999 | 2 | 2 | 7.908330388 | 7.712342256 |
| ENSG00000127377 | CRYGN | -6.792926016 | 0.219632896 | -4.137171692 | 0.446373473 | -10.93009771 | 2 | 2 | 4.874587708 | 2 | 2 | 2.3070652 | 2 |
| ENSG00000064787 | BCAS1 | -4.811022476 | 0.003211642 | -6.12144584 | 0.000101067 | -10.93246832 | 5.330594134 | 4.710564154 | 6.073378139 | 2.3892346 | 2.49394783 | 2 | 2.374681325 |
| C14ORF80 | C14ORF80 | -0.985161742 | 0.691415318 | -9.952522714 | 8.72E-05 | -10.93768446 | 4.815609471 | 5.946638857 | 2 | 4.070463721 | 4.2560182 | 2 | 2 |
| ENSG00000140905 | GCSH | -3.340065298 | 1.25E-06 | -7.598430924 | 2.58E-27 | -10.93849622 | 9.761125267 | 8.762721501 | 8.076950877 | 5.839660368 | 5.73886209 | 2.62493832 | 2.851942824 |
| ENSG00000145428 | RNF175 | -6.89373913 | 0.211937735 | -4.045947259 | 0.456684488 | -10.93968639 | 2 | 4.93638933 | 2 | 2 | 2 | 2.34604626 | 2 |
| ENSG00000092871 | RFFL | -10.75752715 | 7.26E-07 | -0.183787871 | 0.929020974 | -10.94131502 | 7.218819781 | 7.928237109 | 3.369317972 | 2 | 2 | 7.054142452 | 6.718860807 |
| ENSG00000108064 | TFAM | -10.73799738 | 1.95E-05 | -0.204555746 | 0.937781729 | -10.94255313 | 7.27616306 | 7.901717006 | 2 | 2 | 2 | 6.926538448 | 6.784644656 |
| ENSG00000184984 | CHRM5 | -6.342649843 | 0.058965253 | -4.603252611 | 0.159680036 | -10.94590245 | 7.27616306 | 2 | 2 | 2.3892346 | 2 | 2.526562374 | 2.671842496 |
| ENSG00000106635 | BCL7B | -11.98197576 | 9.99E-09 | 1.035936717 | 0.546449437 | -10.94603904 | 4.815609471 | 7.819099893 | 9.399592459 | 2 | 2 | 9.208685664 | 9.386500071 |
| ENSG00000100490 | CDKL1 | -6.110087161 | 1.19E-34 | -4.839609684 | 2.70E-24 | -10.94969685 | 10.6695979 | 9.566483142 | 10.3329194 | 4.497095146 | 4.37434574 | 5.436478516 | 5.641643428 |
| ENSG00000225766 | DHRS4L1 | -6.726323115 | 0.161320596 | -4.234444293 | 0.360286316 | -10.96076741 | 4.815609471 | 2 | 2 | 2 | 2 | 2.225793514 | 2.059860178 |
| ENSG00000101197 | BIRC7 | -6.902545864 | 0.124626428 | -4.060540354 | 0.347421792 | -10.96308622 | 2 | 4.93638933 | 2 | 2 | 2 | 2.139668783 | 2.225841198 |
| ENSG00000155714 | PDZD9 | -5.540811277 | NA | -5.424506616 | NA | -10.96531789 | 3.327711234 | 3.076665209 | 2 | 2 | 2 | 2 | 2.059860178 |
| C6ORF25 | C6ORF25 | -2.24713036 | 0.484134885 | -8.719140106 | 0.006385211 | -10.96627047 | 2 | 4.710564154 | 4.058998049 | 3.09461492 | 2.14022021 | 2 | 2 |
| ENSG00000103723 | AP3B2 | -5.928696813 | 0.003412541 | -5.041333296 | 0.009098098 | -10.97003011 | 2 | 4.442710129 | 7.451345288 | 2.3892346 | 2.2680111 | 2.68698273 | 2.421067459 |
| ENSG00000090020 | SLC9A1 | -12.37663178 | 2.27E-06 | 1.4039576 | 0.580976488 | -10.97267418 | 9.730679447 | 8.505280662 | 2 | 2 | 2 | 10.00483003 | 10.11178748 |
| ENSG00000214021 | TTLL3 | -11.18209087 | 1.56E-11 | 0.209039688 | 0.805203765 | -10.97305119 | 7.867441304 | 6.666836837 | 7.691296561 | 2 | 2 | 7.794040499 | 7.583161732 |
| ENSG00000074706 | IPCEF1 | -5.436732231 | 0.001665744 | -5.537811219 | 0.000600322 | -10.97454345 | 4.00722531 | 3.686303414 | 6.882775709 | 2.3892346 | 2.2680111 | 2.139668783 | 2.466008509 |
| ENSG00000173930 | SLCO4C1 | -5.341844051 | 0.292553183 | -5.640950222 | 0.235899221 | -10.98279427 | 2 | 2 | 4.874587708 | 2.141737246 | 2 | 2 | 2.117335275 |
| ENSG00000197457 | STMN3 | -10.83812774 | 4.44E-08 | -0.147747222 | 0.930565059 | -10.98587496 | 4.815609471 | 6.78762695 | 8.175061096 | 2 | 2 | 7.164572841 | 6.834765628 |
| ENSG00000123388 | HOXC11 | -1.966603157 | 0.446354696 | -9.01964157 | 0.000772226 | -10.98624473 | 3.327711234 | 6.899080544 | 2 | 3.881293667 | 3.82856379 | 2 | 2.059860178 |
| ENSG00000262655 | SPON1 | -6.454288152 | 4.77E-24 | -4.536199583 | 4.99E-14 | -10.99048774 | 9.031919741 | 10.18234215 | 9.330306409 | 3.615725078 | 3.74316441 | 4.757085024 | 5.576534358 |
| ENSG00000189401 | OTUD6A | -2.473213181 | 0.416432783 | -8.522914684 | 0.005690978 | -10.99612787 | 3.327711234 | 2 | 4.874587708 | 2.498691189 | 2.59489879 | 2 | 2 |
| ENSG00000139190 | VAMP1 | -10.92567912 | 3.98E-05 | -0.070967109 | 0.980129329 | -10.99664622 | 8.564112515 | 6.140560021 | 2 | 2 | 2 | 6.977080688 | 7.337766965 |
| ENSG00000163472 | TMEM79 | -10.35301359 | 1.51E-06 | -0.653624853 | 0.725795325 | -11.00663845 | 4.815609471 | 4.93638933 | 7.97167868 | 2 | 2 | 6.329884754 | 5.73181021 |
| ENSG00000143537 | ADAM15 | -12.23583408 | 1.15E-11 | 1.227804143 | 0.271312302 | -11.00802994 | 9.521325812 | 7.790474954 | 7.225742779 | 2 | 2 | 9.709327107 | 9.77707664 |
| ENSG00000104892 | KLC3 | -6.824269236 | 3.85E-05 | -4.190487225 | 0.005913437 | -11.01475646 | 5.532353208 | 5.303523747 | 7.691296561 | 2.270786764 | 2.2680111 | 3.753554863 | 2.225841198 |
| PAPD4 | | 1.715743281 | 0.178633142 | -12.74051721 | 1.07E-11 | -11.02477393 | 5.330594134 | 7.538394143 | 8.486635645 | 9.32823627 | 9.27258642 | 2 | 2 |
| ENSG00000204271 | SPIN3 | -10.65100279 | 2.79E-08 | -0.374414553 | 0.799779812 | -11.02541734 | 7.579279349 | 7.274681896 | 4.874587708 | 2 | 2 | 6.978631953 | 6.113568999 |
| ENSG00000100197 | CYP2D6 | -4.169268349 | 0.024179818 | -6.857519279 | 0.000132405 | -11.02678763 | 2 | 5.946638857 | 7.398165132 | 3.162721474 | 2.86123014 | 2.094598369 | 2.326754077 |
| ENSG00000169860 | P2RY1 | -5.354374524 | 0.000357896 | -5.67301759 | 0.000101069 | -11.02739211 | 8.564112515 | 6.389896633 | 4.874587708 | 3.09461492 | 2.86123014 | 3.256555576 | 2.225841198 |
| ENSG00000214456 | PLIN5 | -3.281624021 | 0.566095204 | -7.754807607 | 0.150262362 | -11.03643163 | 2 | 4.442710129 | 2 | 2.3892346 | 2 | 2 | 2 |
| ENSG00000151650 | VENTX | -6.655997315 | 0.001848041 | -4.381272043 | 0.002806858 | -11.03726936 | 4.815609471 | 4.11352228 | 4.874587708 | 2.141737246 | 2 | 2.492221014 | 2.117335275 |
| ENSG00000142856 | ITGB3BP | -9.848917295 | 5.86E-06 | -1.190506176 | 0.506141411 | -11.03942347 | 7.218819781 | 5.946638857 | 3.369317972 | 2 | 2 | 5.362362054 | 4.80186789 |
| ENSG00000185507 | IRF7 | 2.286326652 | 0.364372706 | -13.32952463 | 3.88E-07 | -11.04319798 | 2 | 7.099070191 | 9.516493218 | 10.45939676 | 10.4511645 | 2 | 2 |
| ENSG00000183971 | NPW | -6.771120983 | 1.09E-23 | -4.279092802 | 9.86E-15 | -11.05021379 | 8.39321512 | 8.101365653 | 8.434763955 | 3.09461492 | 2.38539787 | 4.111244182 | 4.532869193 |
| ENSG00000123572 | NRK | -3.295087203 | 0.501376348 | -7.763149484 | 0.095340194 | -11.05823669 | 2 | 4.442710129 | 2 | 2.270786764 | 2.14022021 | 2 | 2 |
| ENSG00000134962 | KLB | -6.546162767 | 2.97E-06 | -4.519800024 | 8.28E-07 | -11.06596279 | 5.866944937 | 5.303523747 | 4.874587708 | 2.141737246 | 2.14022021 | 2.457042231 | 2.551896972 |
| ENSG00000141946 | ZIM3 | -4.093048618 | 0.447737947 | -6.979174365 | NA | -11.07222298 | 3.327711234 | 3.076665209 | 2 | 2.141737246 | 2 | 2 | 2 |
| ZZZ3 | | 1.72023297 | 0.462658193 | -12.79272949 | 2.16E-07 | -11.07249652 | 2 | 8.255926468 | 8.206335842 | 9.341153092 | 9.37283325 | 2 | 2 |
| ENSG00000184716 | SERINC4 | -2.512882636 | 0.570041492 | -8.560768256 | 0.042020739 | -11.07365089 | 2 | 5.131603625 | 2 | 2.3892346 | 2.68924543 | 2 | 2 |
| ENSG00000042980 | ADAM28 | -5.117629559 | 0.001609982 | -5.957004487 | 0.000170129 | -11.07463405 | 3.327711234 | 9.102512015 | 9.385998894 | 3.663521627 | 4.22485516 | 3.824280769 | 2.782576693 |
| ENSG00000188761 | BCL2L15 | -3.575929364 | 0.000884837 | -7.499833173 | 7.94E-09 | -11.07576254 | 5.095959458 | 4.93638933 | 6.203069917 | 3.09461492 | 2.68924543 | 2.048074405 | 2.117335275 |
| ENSG00000100206 | DMC1 | -5.093182261 | 2.90E-05 | -5.990510056 | 1.85E-07 | -11.08369232 | 6.138360429 | 4.11352228 | 6.203069917 | 2.498691189 | 2.38539787 | 2.183373728 | 2.326754077 |
| ENSG00000156475 | PPP2R2B | -6.037970237 | 0.000628711 | -5.046777834 | 0.001833742 | -11.08474807 | 6.468621203 | 6.228547981 | 3.369317972 | 2.498691189 | 2 | 2.34604626 | 2.632958241 |
| ENSG00000221874 | ZNF816-ZNF | -0.553597633 | 0.615612823 | -10.53129492 | 2.22E-09 | -11.08489255 | 6.009025997 | 4.93638933 | 5.39244166 | 4.846787657 | 5.14724905 | 2 | 2 |
| ENSG00000149792 | MRPL49 | -12.20477645 | 1.07E-13 | 1.105015236 | 0.114203801 | -11.09976122 | 8.696033109 | 8.876041558 | 7.600047325 | 2 | 2 | 9.677205599 | 9.498218357 |
| ENSG00000009694 | TENM1 | -4.114751231 | 0.094457417 | -6.98511165 | 0.005140501 | -11.09986288 | 2 | 5.457120953 | 4.058998049 | 2.498691189 | 2.14022021 | 2.094598369 | 2 |
| ENSG00000275482 | | -8.605878214 | 0.032120021 | -2.500363523 | 0.536372468 | -11.10624174 | 2 | 2 | 6.534109959 | 2 | 2 | 2.962551659 | 3.407202941 |
| ENSG00000149418 | ST14 | -7.858164537 | 0.008902973 | -3.252010353 | 0.302590335 | -11.11017489 | 9.48530526 | 2 | 2 | 2.141737246 | 2.49394783 | 5.622611168 | 3.100202476 |
| ENSG00000129596 | CDO1 | -5.227528608 | 2.42E-13 | -5.886102358 | 1.62E-17 | -11.11363097 | 7.867441304 | 8.004992147 | 9.014156722 | 3.881293667 | 3.45167086 | 3.534812207 | 2.918126211 |
| ENSG00000175606 | TMEM70 | -11.079488 | 9.40E-10 | -0.037885657 | 0.976631077 | -11.11737366 | 8.042902253 | 7.467835385 | 5.930865648 | 2 | 2 | 7.759264603 | 6.777976866 |
| ENSG00000186998 | EMID1 | -9.29456353 | 1.57E-06 | -1.824524713 | 0.155516119 | -11.11908824 | 4.467311984 | 6.464272742 | 5.39244166 | 2 | 2 | 4.340871764 | 3.858622973 |
| ENSG00000127249 | ATP13A4 | -4.863570684 | 0.003100255 | -6.26157906 | 8.27E-05 | -11.12514974 | 4.467311984 | 3.686303414 | 6.43197674 | 2.600425781 | 2.14022021 | 2.094598369 | 2.225841198 |
| ENSG00000126259 | KIRREL2 | -7.377446451 | 0.095756389 | -3.750711391 | 0.380084058 | -11.12815784 | 2 | 2 | 5.39244166 | 2 | 2 | 2.42098416 | 2.172608095 |
| ENSG00000187736 | NHEJ1 | -2.533923061 | 0.243498812 | -8.599806627 | 8.74E-06 | -11.13372969 | 10.1076228 | 7.002537973 | 2 | 6.139181293 | 6.29111322 | 2.34604626 | 2.326754077 |
| PDDC1 | | 1.329037417 | 0.146925888 | -12.46282641 | 3.08E-13 | -11.13378899 | 6.468621203 | 6.844429755 | 8.143093325 | 8.622857004 | 8.6601233 | 2 | 2 |
| ENSG00000168824 | NSG1 | -5.817086771 | 0.002194986 | -5.317226639 | 0.004511358 | -11.13431341 | 8.855032352 | 6.04685525 | 2 | 2.784612485 | 2.86123014 | 2.962551659 | 3.15612297 |
| LDOC1L | | 1.802432738 | 0.262282921 | -12.9402382 | 2.62E-10 | -11.13780546 | 4.467311984 | 7.503546079 | 8.865353883 | 9.562930393 | 9.6076365 | 2 | 2 |
| NOTCH2NL | | 2.090751501 | 1.65E-06 | -13.23367905 | 2.75E-17 | -11.14292755 | 8.107517749 | 7.73146271 | 8.380957451 | 10.14113316 | 10.1878605 | 2 | 2 |
| ENSG00000100271 | TTLL1 | -10.47728339 | 7.69E-09 | -0.671010393 | 0.567464659 | -11.14829378 | 7.667647223 | 6.389896633 | 5.59507254 | 2 | 2 | 6.202919533 | 6.120613345 |
| ENSG00000182712 | CMC4 | 0.142958714 | 0.932714208 | -11.31423997 | 1.03E-08 | -11.17128125 | 4.00722531 | 6.04685525 | 7.163363831 | 6.221320894 | 6.49425375 | 2 | 2 |
| ENSG00000005801 | ZNF195 | -10.79490579 | 2.62E-10 | -0.3783951 | 0.692464563 | -11.17330089 | 7.579279349 | 7.274681896 | 6.073378139 | 2 | 2 | 6.910373854 | 6.556922154 |
| ENSG00000205307 | SAP25 | -2.103462308 | 0.620548015 | -9.070085737 | 0.023691722 | -11.17354805 | 2 | 5.595927442 | 2 | 2.784612485 | 3.01487888 | 2 | 2 |
| FAM65C | | 0.932914255 | 0.699694748 | -12.10821409 | 8.32E-07 | -11.17529984 | 2 | 7.051611315 | 7.934808822 | 7.805288479 | 7.99029569 | 2 | 2 |
| ENSG00000133574 | GIMAP4 | -8.639596161 | 0.033496988 | -2.538246193 | 0.53558846 | -11.17784235 | 6.563806279 | 2 | 2 | 2 | 2 | 3.584558147 | 2.671842496 |
| ENSG00000188404 | SELL | -6.36955191 | 0.194034006 | -4.822383156 | 0.305382004 | -11.19193507 | 2 | 2 | 4.523675391 | 2 | 2 | 2.094598369 | 2.059860178 |
| ENSG00000136280 | CCM2 | -12.0228659 | 3.23E-06 | 0.825348472 | 0.74998849 | -11.19751743 | 9.262228249 | 8.395519306 | 2 | 2 | 2 | 9.059819132 | 9.200394478 |
| ENSG00000173406 | DAB1 | -5.608686014 | 0.029904856 | -5.594748368 | 0.021753975 | -11.20343438 | 2 | 3.686303414 | 6.43197674 | 2.3892346 | 2 | 2.139668783 | 2.277179783 |
| ENSG00000262446 | | 2.786059108 | 0.057449191 | -13.99400566 | 2.70E-12 | -11.20794655 | 6.257048221 | 7.701025996 | 10.09465441 | 11.56232762 | 11.6699949 | 2 | 2 |
| ENSG00000155749 | ALS2CR12 | -7.480660753 | 0.080529035 | -3.728880907 | 0.366065794 | -11.20954166 | 2 | 5.457120953 | 2 | 2 | 2 | 2.42098416 | 2.225841198 |
| ENSG00000180098 | TRNAU1AP | -10.92360151 | 1.44E-05 | -0.296321874 | 0.909854675 | -11.21992338 | 7.904282243 | 7.701025996 | 2 | 2 | 2 | 6.811129868 | 7.073100803 |
| ENSG00000187486 | KCNJ11 | -5.863425867 | 0.001566427 | -5.359886174 | 0.000548532 | -11.22331204 | 4.467311984 | 3.076665209 | 5.772716771 | 2.141737246 | 2.14022021 | 2.267001569 | 2.117335275 |
| ENSG00000136206 | SPDYE1 | -4.351104907 | 1.77E-06 | -6.878282717 | 3.02E-14 | -11.22938762 | 7.53297911 | 6.78762695 | 9.165043284 | 4.105493497 | 4.22485516 | 2.656294034 | 2.70970617 |
| ENSG00000135097 | MSI1 | -4.654383493 | 0.00837844 | -6.580063627 | 0.00011376 | -11.23444712 | 3.327711234 | 6.228547981 | 8.236946975 | 3.709785322 | 2.2680111 | 2.34604626 | 2.466008509 |
| ENSG00000175161 | CADM2 | -5.940672672 | 0.024598953 | -5.296986907 | 0.034422353 | -11.23765958 | 2 | 6.78762695 | 3.369317972 | 2.3892346 | 2 | 2.34604626 | 2.277179783 |
| ENSG00000173714 | WFIKKN2 | -6.514511259 | 0.067812259 | -4.728779335 | 0.139393652 | -11.24329059 | 2 | 2 | 5.930865648 | 2.141737246 | 2 | 2.139668783 | 2.326754077 |
| ENSG00000262664 | OVCA2 | 1.261478724 | 0.077972459 | -12.50861499 | 2.24E-14 | -11.24713627 | 7.624139768 | 6.535001748 | 7.734844356 | 8.621345041 | 8.61876544 | 2 | 2 |
| ENSG00000140986 | RPL3L | -7.767575773 | 0.061554206 | -3.480473755 | 0.387383324 | -11.24804953 | 2 | 5.72254323 | 2 | 2 | 2 | 2.526562374 | 2.374681325 |
| ENSG00000123119 | NECAB1 | -8.543920042 | 3.20E-09 | -2.70819072 | 0.007742505 | -11.25211076 | 5.70932728 | 7.73146271 | 7.691296561 | 2.270786764 | 2 | 5.014670875 | 4.500832793 |
| ENSG00000090661 | CERS4 | -9.07733102 | 7.47E-16 | -2.177463576 | 0.036658542 | -11.2547946 | 8.653385977 | 8.732943944 | 10.21486071 | 2.695456494 | 2 | 7.35696023 | 7.12130792 |
| ENSG00000175544 | CABP4 | -4.470351502 | 0.000803779 | -6.799951317 | 3.88E-07 | -11.27030282 | 5.70932728 | 4.442710129 | 7.029880454 | 2.600425781 | 2.94009902 | 2.3070652 | 2.059860178 |
| ENSG00000235568 | NFAM1 | -8.453291088 | 0.001037349 | -2.819880321 | 0.268955903 | -11.27317141 | 8.631580145 | 2 | 5.39244166 | 2 | 2.2680111 | 5.201624686 | 3.560117069 |
| ENSG00000146223 | RPL7L1 | -12.74650484 | 4.37E-15 | 1.463685777 | 0.02080403 | -11.28281907 | 9.567995545 | 8.916342327 | 8.35328246 | 2 | 2 | 10.44086233 | 10.53252493 |
| ENSG00000101115 | SALL4 | -5.483366009 | 0.000214633 | -5.804171324 | 6.33E-05 | -11.28753733 | 8.998234871 | 8.169643011 | 3.369317972 | 3.09461492 | 3.50452879 | 3.10571689 | 3.15612297 |
| ENSG00000180815 | MAP3K15 | -5.604732726 | 0.002905853 | -5.689621497 | 0.001144452 | -11.29435422 | 6.468621203 | 5.303523747 | 2 | 2.3892346 | 2.14022021 | 2.183373728 | 2.326754077 |
| ENSG00000278311 | GGNBP2 | -0.043529066 | 0.946720822 | -11.26402862 | 1.45E-37 | -11.30755769 | 10.02639697 | 9.328390744 | 9.116480286 | 9.453327884 | 9.54672161 | 2.094598369 | 2.117335275 |
| ENSG00000161551 | ZNF577 | -10.42537512 | 9.35E-08 | -0.885450561 | 0.538569152 | -11.31082568 | 7.031698319 | 7.393647515 | 4.523675391 | 2 | 2 | 6.210860813 | 5.534955934 |

| | | | | | | | | | | | | | |
|---|---|---|---|---|---|---|---|---|---|---|---|---|---|
| ENSG00000130054 | FAM155B | -5.17807393 | 0.041322629 | -6.137040139 | 0.010904086 | -11.31511407 | 5.330594134 | 2 | 5.156626401 | 2.3892346 | 2 | 2.048074405 | 2.172608095 |
| ENSG00000166535 | A2ML1 | -5.718323644 | 0.033296552 | -5.604410975 | 0.013216683 | -11.32273462 | 2 | 4.11352228 | 4.523675391 | 2 | 2.14022021 | 2.094598369 | 2.059860178 |
| ENSG00000270231 | NBPF8 | -10.28913583 | 6.46E-09 | -1.040161614 | 0.309292345 | -11.32929745 | 5.330594134 | 6.951736442 | 7.029880454 | 2 | 2 | 5.751606453 | 5.524371359 |
| ENSG00000138759 | FRAS1 | -5.35030153 | 0.000342454 | -5.983026886 | 3.91E-05 | -11.33332842 | 4.467311984 | 6.04685525 | 8.978369997 | 3.406930997 | 2.77779907 | 2.885263063 | 2.746601469 |
| ENSG00000139915 | MDGA2 | -9.812290361 | 0.00012717 | -1.524768009 | 0.518789265 | -11.33705837 | 2 | 6.728495673 | 6.718950348 | 2 | 2 | 4.261535101 | 5.171036168 |
| ENSG00000147573 | TRIM55 | -5.666131956 | 0.248162973 | -5.671296701 | 0.217009028 | -11.33742866 | 2 | 2 | 5.156626401 | 2.141737246 | 2 | 2.139668783 | 2 |
| ENSG00000101144 | BMP7 | -4.553851332 | 0.000609233 | -6.783658355 | 1.99E-07 | -11.33750969 | 6.257048221 | 7.901717006 | 6.43197674 | 3.663521627 | 2.68924543 | 2.62493832 | 2.059860178 |
| ENSG00000059915 | PSD | -9.823442231 | 9.55E-05 | -1.517182165 | 0.508771779 | -11.3406244 | 6.73718411 | 6.728495673 | 2 | 2 | 2 | 5.014670875 | 4.553838275 |
| ENSG00000274746 | | 0.08068852 | 0.915816833 | -11.42531807 | 1.51E-12 | -11.34462955 | 6.257048221 | 6.311476754 | 6.43197674 | 6.358325776 | 6.46120797 | 2 | 2 |
| ENSG00000179915 | NRXN1 | -6.742771104 | 0.003824257 | -4.607439907 | 0.03720716 | -11.35021101 | 2 | 7.393647515 | 5.156626401 | 2 | 2.38539787 | 2.775316226 | 2.70970617 |
| ENSG00000132874 | SLC14A2 | -4.753223019 | 0.001232604 | -6.602537614 | 5.95E-06 | -11.35576063 | 3.327711234 | 4.93638933 | 6.718950348 | 2.498691189 | 2.49394783 | 2.139668783 | 2.172608095 |
| ENSG00000151239 | TWF1 | -12.02871329 | 1.07E-14 | 0.669833705 | 0.072935017 | -11.35887959 | 8.367060731 | 8.297207175 | 8.296287027 | 2 | 2 | 8.990775914 | 8.975548422 |
| ENSG00000151136 | BTBD11 | -4.448857177 | 5.91E-07 | -6.916005359 | 4.18E-15 | -11.36486254 | 7.975256505 | 7.538394143 | 9.69246155 | 4.329433326 | 4.70309173 | 3.059555873 | 2.70970617 |
| ENSG00000105357 | MYH14 | -6.875270634 | 0.017873155 | -4.490486649 | 0.068307911 | -11.36575728 | 6.138360429 | 2 | 3.369317972 | 2 | 2.14022021 | 2.492221014 | 2.172608095 |
| GLTSCR1 | | 0.619961047 | 0.549337318 | -11.98647271 | 4.43E-12 | -11.36651166 | 6.257048221 | 6.140560021 | 7.691296561 | 7.409376773 | 7.53675633 | 2 | 2 |
| ENSG00000102078 | SLC25A14 | -10.63734492 | 2.43E-06 | -0.733306816 | 0.719764201 | -11.37065174 | 8.16936288 | 6.228547981 | 3.369317972 | 2 | 2 | 6.134985075 | 6.365613546 |
| ENSG00000141338 | ABCA8 | -5.933387283 | 1.18E-13 | -5.441707717 | 2.20E-12 | -11.375095 | 7.667647223 | 9.148351514 | 9.868959282 | 3.663521627 | 3.7864959 | 3.865109708 | 4.279527956 |
| ENSG00000149798 | CDC42EP2 | -0.576516072 | 0.47052531 | -10.80576308 | 8.17E-32 | -11.38227915 | 9.800745303 | 9.687856435 | 8.266922072 | 8.825537758 | 8.83024294 | 2.139668783 | 2.117335275 |
| ENSG00000160191 | PDE9A | -7.771446495 | 1.07E-14 | -3.611989388 | 2.44E-06 | -11.38343588 | 7.940205782 | 7.145017454 | 8.236946975 | 2.600425781 | 2 | 4.521997428 | 4.445805334 |
| ENSG00000188517 | COL25A1 | -7.311788406 | 0.098313191 | -4.073182681 | 0.338044189 | -11.38497109 | 2 | 5.303523747 | 2 | 2 | 2 | 2.34604626 | 2.117335275 |
| FAM58A | | 0.463312622 | 0.832568498 | -11.85395809 | 1.38E-07 | -11.39064546 | 3.327711234 | 8.029695761 | 5.59507254 | 7.194375246 | 7.19067478 | 2 | 2 |
| ENSG00000076662 | ICAM3 | -2.196742263 | 0.071313222 | -9.200291526 | 3.96E-13 | -11.39703379 | 5.330594134 | 7.538394143 | 8.805342542 | 5.651002631 | 5.74985018 | 2.139668783 | 2.117335275 |
| ENSG00000185917 | SETD4 | -10.50780014 | 2.40E-07 | -0.896311266 | 0.575748624 | -11.4041114 | 4.00722531 | 7.503546079 | 7.163363831 | 2 | 2 | 6.031172603 | 5.925257518 |
| ENSG00000146555 | SDK1 | -6.70105056 | 1.44E-22 | -4.705243813 | 6.37E-14 | -11.40629437 | 8.777722034 | 9.256956408 | 9.301632639 | 3.663521627 | 2.59489879 | 4.563936166 | 4.757365452 |
| ENSG00000064419 | TNPO3 | -12.19101027 | 1.86E-07 | 0.783328922 | 0.723171334 | -11.40768135 | 3.327711234 | 7.874700259 | 9.69246155 | 2 | 2 | 9.357396362 | 9.149213283 |
| ENSG00000165959 | CLMN | -8.004046152 | 2.09E-17 | -3.422128161 | 5.33E-05 | -11.42617431 | 8.998234871 | 9.71856931 | 7.896971941 | 2.784612485 | 2.38539787 | 6.17883 | 5.08721802 |
| ENSG00000233838 | DPH3P1 | -2.272785914 | 0.371549126 | -9.156282842 | 0.000688863 | -11.42906876 | 2 | 4.710564154 | 4.874587708 | 3.09461492 | 2.59489879 | 2 | 2 |
| PAPD5 | | 0.81902333 | 0.552458672 | -12.25029884 | 8.17E-11 | -11.43127551 | 5.532353208 | 6.311476754 | 8.206335842 | 7.956842238 | 7.89946959 | 2 | 2 |
| ENSG00000183067 | IGSF5 | -4.273585166 | 0.468786454 | -7.160095846 | NA | -11.43368101 | 4.00722531 | 2 | 2 | 2.141737246 | 2 | 2 | 2 |
| ENSG00000057294 | PKP2 | -9.500787323 | 1.68E-40 | -1.940136956 | 0.001145858 | -11.44092428 | 9.761125267 | 11.06803102 | 10.89495479 | 2.695456494 | 2.59489879 | 8.662927365 | 8.831649431 |
| ENSG00000103546 | SLC6A2 | -5.782221289 | 0.310604101 | -5.659984698 | NA | -11.44220599 | 2 | 2 | 4.058998049 | 2 | 2 | 2 | 2.059860178 |
| ENSG00000151962 | RBM46 | -5.782221289 | 0.310604101 | -5.659984698 | NA | -11.44220599 | 2 | 2 | 4.058998049 | 2 | 2 | 2 | 2.059860178 |
| ENSG00000187260 | WDR86 | -5.060410446 | 6.20E-06 | -6.382223529 | 5.35E-09 | -11.44263397 | 5.095959458 | 8.053983481 | 7.342949392 | 3.162721474 | 3.01487888 | 2.717032197 | 2.326754077 |
| ENSG00000169242 | EFNA1 | -7.771296355 | 4.06E-26 | -3.678922528 | 5.66E-19 | -11.45021888 | 8.042902253 | 8.297207175 | 7.896971941 | 2.498691189 | 2.2680111 | 4.596630906 | 4.692661645 |
| ENSG00000198276 | UCKL1 | -11.41892874 | 3.58E-11 | -0.032199547 | 0.976216195 | -11.45112829 | 7.940205782 | 6.464272742 | 8.236946975 | 2 | 2 | 7.55607473 | 7.809264007 |
| ENSG00000134333 | LDHA | -15.49828348 | 2.83E-20 | 4.04501214 | 6.15E-08 | -11.45327134 | 12.14468514 | 11.87039211 | 11.12102291 | 2 | 2 | 14.89971348 | 16.37279217 |
| ENSG00000164346 | NSA2 | -11.85877711 | 9.97E-11 | 0.40245299 | 0.757887688 | -11.45632412 | 8.468948322 | 8.747909548 | 6.073378139 | 2 | 2 | 8.46188525 | 8.632905293 |
| ENSG00000134489 | HRH4 | -5.444563695 | 0.032971338 | -6.015127923 | 0.011004133 | -11.45969162 | 4.00722531 | 3.076665209 | 5.39244166 | 2.270786764 | 2 | 2.183373728 | 2 |
| ENSG00000142606 | MMEL1 | -3.795265784 | 0.438840927 | -7.668728463 | 0.099180241 | -11.46399425 | 2 | 3.076665209 | 4.058998049 | 2 | 2.2680111 | 2 | 2 |
| ENSG00000222011 | FAM185A | -9.722379955 | 1.19E-05 | -1.747001284 | 0.331745601 | -11.46938124 | 4.00722531 | 4.710564154 | 7.342949392 | 2 | 2 | 4.271694079 | 4.739172856 |
| ENSG00000170092 | SPDYE5 | -5.112677079 | 3.83E-12 | -6.364127774 | 1.46E-18 | -11.47680485 | 8.75772933 | 8.029695761 | 9.703463497 | 4.105493497 | 4.31639476 | 3.600766115 | 3.012034821 |
| ENSG00000196376 | SLC35F1 | -4.486605005 | 0.002131393 | -6.992973131 | 7.61E-07 | -11.47957814 | 8.716893051 | 7.002537973 | 7.734844356 | 4.647285135 | 2.38539787 | 2.885263063 | 2.172608095 |
| ENSG00000179603 | GRM8 | -6.316754489 | 0.259792504 | -5.164454097 | 0.343240742 | -11.48120859 | 2 | 4.442710129 | 2 | 2 | 2 | 2 | 2.117335275 |
| ENSG00000197977 | ELOVL2 | -6.46885598 | 7.55E-32 | -5.012423032 | 7.98E-24 | -11.48127901 | 9.189468652 | 8.993716073 | 9.703463497 | 3.79808662 | 3.08597292 | 4.644324387 | 4.489993931 |
| ENSG00000186628 | FSD2 | -5.457089203 | 0.014432388 | -6.028480779 | 0.004360709 | -11.48556998 | 4.467311984 | 4.93638933 | 6.073378139 | 2 | 2.49394783 | 2.34604626 | 2 |
| ENSG00000261210 | CLEC19A | -4.2997745 | 0.413845402 | -7.185847532 | NA | -11.48562203 | 3.327711234 | 2 | 3.369317972 | 2.141737246 | 2 | 2 | 2 |
| ENSG00000197245 | FAM110D | -5.452161757 | 0.259887271 | -6.035288495 | 0.184040613 | -11.48745025 | 2 | 4.93638933 | 2 | 2.141737246 | 2 | 2.094598369 | 2 |
| ENSG00000188710 | QRFP | -8.256524309 | 0.044161456 | -3.23326795 | 0.423067088 | -11.48979226 | 2 | 2 | 6.203069917 | 2 | 2 | 2.592885947 | 2.782576693 |
| ENSG00000275947 | | 0.003331231 | 0.998610961 | -11.49595175 | 2.67E-06 | -11.49262052 | 7.33131396 | 6.464272742 | 2 | 6.463327516 | 6.34282908 | 2 | 2 |
| ENSG00000109805 | NCAPG | -9.656309494 | 0.000123773 | -1.838191629 | 0.414043461 | -11.49450112 | 6.468621203 | 6.666836837 | 2 | 2 | 2 | 4.425202323 | 4.365060851 |
| ENSG00000142549 | IGLON5 | -6.142972663 | 0.008570711 | -5.352510618 | 0.016521712 | -11.49548328 | 2 | 3.076665209 | 7.451345288 | 2.3892346 | 2.14022021 | 2.457042231 | 2.421067459 |
| ENSG00000140557 | ST8SIA2 | -8.67163755 | 2.01E-09 | -2.827434814 | 0.034961639 | -11.49907236 | 9.384607364 | 6.140560021 | 8.143093325 | 2.498691189 | 2 | 5.485310992 | 5.855190914 |
| ENSG00000122085 | MTERF4 | -11.58578581 | 2.00E-07 | 0.07717287 | 0.971645188 | -11.50861294 | 3.327711234 | 8.433041974 | 8.460932916 | 2 | 2 | 7.953993861 | 7.965119966 |
| ENSG00000133256 | PDE6B | -9.198676589 | 5.04E-06 | -2.316255556 | 0.09744279 | -11.51493215 | 4.00722531 | 7.35507212 | 7.600047325 | 2.141737246 | 2 | 4.713032626 | 4.919632516 |
| ENSG00000135702 | CHST5 | -4.51175308 | 0.034034402 | -7.005963514 | 0.000867089 | -11.51771659 | 4.00722531 | 2 | 6.718950348 | 2.3892346 | 2.59489879 | 2.139668783 | 2.059860178 |
| ENSG00000115966 | ATF2 | -11.86021747 | 1.39E-07 | 0.338376042 | 0.876848851 | -11.52184143 | 8.998234871 | 8.376385542 | 3.369317972 | 2 | 2 | 8.677327975 | 8.270001654 |
| ENSG00000163009 | C2orf48 | -6.20536066 | 0.073013981 | -5.321767667 | 0.100973768 | -11.52712833 | 3.327711234 | 3.076665209 | 3.369317972 | 2 | 2 | 2.094598369 | 2 |
| ENSG00000146374 | RSPO3 | 0.052626333 | 0.935309765 | -11.58402482 | 4.27E-70 | -11.53139848 | 11.16316091 | 11.56288089 | 10.5916593 | 11.2472134 | 11.1780308 | 2.3070652 | 2.172608095 |
| ENSG00000164756 | SLC30A8 | -6.120968334 | 0.079559698 | -5.420132068 | 0.080856643 | -11.5411004 | 4.467311984 | 2 | 4.874587708 | 2.141737246 | 2 | 2 | 2.225841198 |
| ENSG00000054116 | TRAPPC3 | -12.47447469 | 1.71E-13 | 0.931273086 | 0.283758637 | -11.54320161 | 9.233563228 | 7.538394143 | 9.014156722 | 2 | 2 | 9.589063591 | 9.776797972 |
| ENSG00000203963 | C1orf141 | -4.342016007 | 0.461270807 | -7.20026365 | NA | -11.56227966 | 2 | 2 | 4.058998049 | 2 | 2.14022021 | 2 | 2 |
| ENSG00000167797 | CDK2AP2 | -12.09496163 | 1.69E-07 | 0.525115653 | 0.81343336 | -11.56984597 | 9.497412313 | 8.077869079 | 3.369317972 | 2 | 2 | 8.48662708 | 9.229602289 |
| CCDC94 | CCDC94 | 0.408555274 | 0.687246529 | -11.98362655 | 2.42E-12 | -11.57507128 | 6.009025997 | 7.572420273 | 6.534109959 | 7.238438152 | 7.29187696 | 2 | 2 |
| FAM65B | | -1.085365989 | 0.699672674 | -10.49021982 | 0.00014179 | -11.57558581 | 2 | 6.78762695 | 4.058998049 | 4.623314311 | 4.40247105 | 2 | 2 |
| ENSG00000066382 | MPPED2 | -4.01554557 | 0.087551534 | -7.564140281 | 0.001771437 | -11.57968585 | 2 | 5.303523747 | 5.59507254 | 2.784612485 | 2.14022021 | 2.094598369 | 2 |
| ENSG00000198354 | DCAF12L2 | -4.555422923 | 0.171807065 | -7.024773705 | 0.036177376 | -11.58019663 | 3.327711234 | 4.710564154 | 2 | 2 | 2.2680111 | 2.048074405 | 2 |
| ENSG00000100427 | MLC1 | -5.054391892 | 0.06096044 | -6.531306957 | 0.011880654 | -11.58569885 | 8.797441463 | 5.838937808 | 2 | 3.79808662 | 2 | 2.592885947 | 2.466008509 |
| ENSG00000234539 | | 0.980459012 | 0.34242241 | -12.57150599 | 1.75E-13 | -11.59104698 | 8.540903468 | 7.431218269 | 9.714382178 | 9.794554993 | 9.84837777 | 2.048074405 | 2 |
| ENSG00000250423 | KIAA1210 | -4.525369495 | 0.054013417 | -7.072794876 | 0.003131786 | -11.59816437 | 3.327711234 | 4.11352228 | 5.39244166 | 2.498691189 | 2 | 2.094598369 | 2 |
| ENSG00000144057 | ST6GAL2 | -8.428903445 | 2.74E-05 | -3.173722932 | 0.017988009 | -11.60262638 | 4.00722531 | 4.710564154 | 5.59507254 | 2 | 2 | 2.885263063 | 2.671842496 |
| ENSG00000118971 | CCND2 | -7.170469912 | 1.35E-17 | -4.43400073 | 1.70E-07 | -11.60447064 | 11.27287932 | 12.52221914 | 12.23794467 | 5.753943344 | 3.90918929 | 8.07712337 | 7.173047408 |
| ENSG00000177272 | KCNA3 | -5.782223324 | 0.310604101 | -5.822995563 | NA | -11.60521889 | 2 | 2 | 4.058998049 | 2 | 2 | 2.048074405 | 2 |
| ENSG00000181433 | SAGE1 | -5.782223324 | 0.310604101 | -5.822995563 | NA | -11.60521889 | 2 | 2 | 4.058998049 | 2 | 2 | 2.048074405 | 2 |
| ENSG00000221855 | TAS2R41 | -5.782223324 | 0.310604101 | -5.822995563 | NA | -11.60521889 | 2 | 2 | 4.058998049 | 2 | 2 | 2.048074405 | 2 |
| ENSG00000283597 | FAM169B | -5.782223324 | 0.310604101 | -5.822995563 | NA | -11.60521889 | 2 | 2 | 4.058998049 | 2 | 2 | 2.048074405 | 2 |
| ENSG00000164816 | DEFA5 | -6.308772637 | 0.203115525 | -5.297225996 | 0.267757907 | -11.60599863 | 4.467311984 | 2 | 2 | 2 | 2 | 2.048074405 | 2.059860178 |
| ENSG00000150977 | RILPL2 | -11.37511313 | 2.22E-05 | -0.232117745 | 0.936069485 | -11.60723087 | 9.015175615 | 6.535001748 | 2 | 2 | 2 | 7.414254101 | 7.484227376 |
| ENSG00000183214 | | 1.701104417 | 0.229879807 | -13.31969473 | 6.54E-12 | -11.61859031 | 9.233563228 | 7.761270542 | 5.59507254 | 9.871615002 | 9.85284374 | 2 | 2 |
| ENSG00000161405 | IKZF3 | -4.049556505 | 2.70E-05 | -7.573352037 | 7.04E-13 | -11.62290854 | 5.866944937 | 5.946638857 | 7.646393253 | 3.462030052 | 3.2184422 | 2.139668783 | 2.225841198 |
| C9ORF69 | C9ORF69 | 1.102549783 | 0.651333637 | -12.72628115 | 3.01E-07 | -11.62373137 | 8.517314938 | 7.701025996 | 2 | 8.668958796 | 8.68752719 | 2 | 2 |
| ENSG00000166292 | TMEM100 | -6.800269969 | 0.002935656 | -4.823763123 | 0.018611532 | -11.62403309 | 2 | 4.442710129 | 6.958202216 | 2.141737246 | 2.14022021 | 2.746468522 | 2.172608095 |
| ENSG00000174307 | PHLDA3 | -13.15099308 | 8.42E-15 | 1.525807187 | 0.068954301 | -11.62518589 | 9.233563228 | 8.395519306 | 10.15199903 | 2 | 2 | 11.14219385 | 10.73500669 |
| ENSG00000111817 | DSE | -12.78042044 | 6.69E-14 | 1.147655919 | 0.196144269 | -11.63276452 | 7.709880939 | 9.540933215 | 9.344432377 | 2 | 2 | 10.25270208 | 10.15809785 |
| ENSG00000057663 | ATG5 | -11.80084196 | 3.29E-11 | 0.167720517 | 0.892291608 | -11.63312144 | 7.218819781 | 7.232742069 | 9.031722608 | 2 | 2 | 8.446833885 | 8.046896864 |
| ENSG00000131495 | NDUFA2 | -11.63950783 | 1.01E-12 | 0.002460274 | 0.997910957 | -11.63704755 | 8.444143041 | 7.35507212 | 7.818184403 | 2 | 2 | 7.863694929 | 8.010385318 |
| ENSG00000123364 | HOXC13 | -6.324312712 | 0.196499175 | -5.312812131 | 0.2601381 | -11.63712484 | 2 | 4.442710129 | 2 | 2 | 2 | 2.048074405 | 2.059860178 |
| ENSG00000198874 | TYW1 | -11.03805944 | 7.94E-12 | -0.60798078 | 0.322726336 | -11.64604022 | 7.33131396 | 6.951736442 | 7.691296561 | 2 | 2 | 6.741568503 | 6.77351449 |
| ENSG00000114520 | SNX4 | -11.61088299 | 8.12E-10 | -0.035966521 | 0.981056901 | -11.64684951 | 5.532353208 | 8.487557258 | 8.296287027 | 2 | 2 | 7.796681167 | 7.945418846 |
| ENSG00000158270 | COLEC12 | -4.07531155 | 1.52E-26 | -7.572224813 | 2.21E-88 | -11.64753636 | 12.45359838 | 11.96628737 | 12.11972681 | 8.115618198 | 8.16053033 | 5.113999907 | 4.489993931 |
| FAM64A | | -1.934293694 | 0.478094553 | -9.724957265 | 0.000490945 | -11.65925096 | 4.00722531 | 5.946638857 | 2 | 3.663521627 | 2.94009902 | 2 | 2 |
| ENSG00000117971 | CHRNB4 | -7.377636148 | 0.09809518 | -4.283479977 | 0.316388957 | -11.66111612 | 2 | 2 | 5.39244166 | 2 | 2 | 2.3070652 | 2.117335275 |
| ENSG00000119673 | ACOT2 | -11.50897888 | 3.36E-09 | -0.174678538 | 0.914220926 | -11.68365742 | 5.866944937 | 6.951736442 | 8.941672936 | 2 | 2 | 7.514972902 | 7.749171054 |
| ENSG00000187010 | RHD | -6.360899818 | 1.17E-05 | -5.323353569 | 6.69E-05 | -11.68425339 | 5.70932728 | 4.710564154 | 7.896971941 | 2.141737246 | 2.59489879 | 2.62493832 | 2.782576693 |
| ENSG00000163312 | HELQ | -10.89444951 | 5.69E-06 | -0.795745677 | 0.726349422 | -11.69019518 | 3.327711234 | 5.457120953 | 8.608698767 | 2 | 2 | 6.464399218 | 6.415155055 |
| ENSG00000116793 | PHTF1 | -11.53533369 | 2.77E-11 | -0.158909943 | 0.886082448 | -11.69424363 | 7.485143553 | 8.607278467 | 6.80318718 | 2 | 2 | 7.581819208 | 7.768360547 |
| ENSG00000128573 | FOXP2 | -10.24819411 | 1.08E-06 | -1.450926419 | 0.377312046 | -11.69912053 | 4.815609471 | 7.790474954 | 5.156626401 | 2 | 2 | 5.422868615 | 5.013446246 |
| ENSG00000185561 | TLCD2 | -1.669140352 | 0.088873 | -10.03719629 | 2.93E-20 | -11.70633664 | 9.112843761 | 6.899080544 | 8.960138144 | 7.030886362 | 6.90143415 | 2.183373728 | 2.059860178 |
| ENSG00000179826 | MRGPRX3 | -2.328133005 | 0.395924587 | -9.378670545 | 0.000955525 | -11.70680355 | 4.00722531 | 2 | 5.59507254 | 2.947924294 | 2.94009902 | 2 | 2 |
| ENSG00000138400 | MDH1B | -8.302056391 | 0.00072502 | -3.415389947 | 0.088031079 | -11.71744634 | 3.327711234 | 5.946638857 | 3.369317972 | 2 | 2 | 2.526562374 | 2.746601469 |
| ENSG00000204842 | ATXN2 | -12.44782983 | 1.65E-15 | 0.719980652 | 0.085133627 | -11.72784918 | 8.981092835 | 8.607278467 | 8.585098385 | 2 | 2 | 9.507057041 | 9.389054739 |
| ENSG00000258873 | DUXA | -3.13323377 | 0.253527674 | -8.596440643 | 0.002795201 | -11.72967441 | 2 | 4.11352228 | 4.523675391 | 2.270786764 | 2.49394783 | 2 | 2 |
| ENSG00000213246 | SUPT4H1 | -0.258818229 | 0.705820174 | -11.47531105 | 1.41E-30 | -11.73412928 | 9.317904228 | 8.862353853 | 9.788583153 | 9.130908275 | 9.09332116 | 2 | 2.172608095 |
| ENSG00000156574 | NODAL | -7.731852348 | 0.013721829 | -4.006964958 | 0.155852109 | -11.73881731 | 5.532353208 | 2 | 3.369317972 | 2 | 2 | 2.267001569 | 2.374681325 |
| ENSG00000205784 | ARRDC5 | -7.377218045 | 0.09269647 | -4.364318018 | 0.297469212 | -11.74153606 | 2 | 2 | 5.39244166 | 2 | 2 | 2.183373728 | 2.225841198 |
| ENSG00000198488 | B3GNT6 | -4.639784195 | 0.152526608 | -7.108696838 | 0.030044768 | -11.74848103 | 4.00722531 | 2 | 4.523675391 | 2 | 2.2680111 | 2.048074405 | 2 |
| ENSG00000188800 | TMCO2 | -3.939068832 | 0.490247308 | -7.812273827 | 0.146527599 | -11.75134266 | 2 | 2 | 4.523675391 | 2 | 2.2680111 | 2 | 2 |
| ENSG00000215182 | MUC5AC | -6.311472343 | 0.260096091 | -5.452019587 | 0.316598154 | -11.76349193 | 4.467311984 | 2 | 2 | 2 | 2 | 2.094598369 | 2 |

| | | | | | | | | | | | | | |
|---|---|---|---|---|---|---|---|---|---|---|---|---|---|
| ENSG00000180901 | KCTD2 | -12.35318797 | 1.64E-13 | 0.583530244 | 0.493973015 | -11.76965773 | 8.653385977 | 7.638155693 | 9.227351203 | 2 | 2 | 9.330314074 | 9.09971864 |
| C1ORF186 | C1ORF186 | -2.861428734 | 0.303181423 | -8.90948017 | 0.002100994 | -11.7709089 | 2 | 3.686303414 | 5.156626401 | 2.3892346 | 2.68924543 | 2 | 2 |
| ENSG00000187672 | ERC2 | -3.325711167 | 0.248162973 | -8.452557288 | 0.001960035 | -11.77826846 | 7.975256505 | 2 | 2 | 3.754611435 | 3.45167086 | 2.048074405 | 2.117335275 |
| ENSG00000055118 | KCNH2 | -6.623621744 | 0.00113016 | -5.155938531 | 0.01045232 | -11.77956028 | 8.855032352 | 6.464272742 | 3.369317972 | 2.947924294 | 2 | 3.374372158 | 2.981406154 |
| ENSG00000176049 | JAKMIP2 | -5.316216112 | 0.000101449 | -6.464337693 | 1.04E-06 | -11.7805538 | 4.815609471 | 6.535001748 | 7.342949392 | 3.023133209 | 2.14022021 | 2.42098416 | 2.225841198 |
| ENSG00000008128 | CDK11A | -9.810091451 | 1.80E-05 | -1.970506485 | 0.296532366 | -11.78059794 | 5.70932728 | 3.076665209 | 7.285536013 | 2 | 2 | 4.555645384 | 4.254128754 |
| ENSG00000279247 | | 0.582874456 | 0.536668932 | -12.3730144 | 2.78E-13 | -11.79013994 | 7.435667355 | 7.73146271 | 6.073378139 | 7.786523606 | 7.84670947 | 2 | 2 |
| ENSG00000082258 | CCNT2 | -12.08444572 | 1.18E-09 | 0.285544726 | 0.864992366 | -11.79890099 | 5.330594134 | 9.090820894 | 8.677268317 | 2 | 2 | 8.915429701 | 8.343344553 |
| ENSG00000085365 | SCAMP1 | -12.21476515 | 1.27E-12 | 0.407843852 | 0.688358595 | -11.8069213 | 7.33131396 | 8.337339476 | 9.242516611 | 2 | 2 | 9.054312673 | 8.744843922 |
| ENSG00000147869 | CER1 | -4.460698837 | 0.446344169 | -7.347214041 | NA | -11.80791288 | 2 | 4.11352228 | 2 | 2.141737246 | 2 | 2 | 2 |
| ENSG00000125482 | TTF1 | -11.51465582 | 4.31E-11 | -0.300244921 | 0.785640197 | -11.81490074 | 7.031698319 | 7.099070191 | 8.677268317 | 2 | 2 | 7.570548195 | 7.466366987 |
| ENSG00000086159 | AQP6 | -6.103938211 | 0.003797562 | -5.714546501 | 0.002015098 | -11.81848471 | 4.00722531 | 3.076665209 | 6.203069917 | 2.270786764 | 2 | 2.139668783 | 2.225841198 |
| ENSG00000148832 | PAOX | -10.73491832 | 4.08E-05 | -1.085510539 | 0.67150607 | -11.82042886 | 8.257427002 | 6.464272742 | 2 | 2 | 2 | 6.123812431 | 5.896817464 |
| ENSG00000057704 | TMCC3 | -7.210969533 | 4.61E-05 | -4.614307755 | 0.008435628 | -11.82527729 | 3.327711234 | 8.876041558 | 8.042706615 | 2.784612485 | 2.14022021 | 4.360042493 | 2.885414003 |
| ENSG00000196428 | TSC22D2 | -12.2094344 | 1.73E-11 | 0.383709879 | 0.762484783 | -11.82572452 | 6.563806279 | 8.806266215 | 9.083172337 | 2 | 2 | 9.120096958 | 8.588503563 |
| ENSG00000274803 | | 1.549641848 | 0.001926435 | -13.37758773 | 2.18E-17 | -11.82794588 | 8.493334308 | 7.73146271 | 8.380957451 | 9.744735743 | 9.79301538 | 2 | 2 |
| C4ORF26 | C4ORF26 | -2.473695048 | 0.082516252 | -9.356020495 | 2.26E-06 | -11.82971554 | 4.467311984 | 4.442710129 | 4.523675391 | 3.023133209 | 2.68924543 | 2 | 2 |
| ENSG00000137411 | VARS2 | 0.086766998 | 0.97361691 | -11.9199787 | 1.73E-06 | -11.8332117 | 2 | 7.819099893 | 6.718950348 | 6.804454198 | 6.97351258 | 2 | 2 |
| KIAA1429 | | 1.423251986 | 0.561312002 | -13.25720292 | 1.37E-07 | -11.83395093 | 2 | 8.590771778 | 8.784770914 | 9.498502402 | 9.54829592 | 2 | 2 |
| AHSA2 | AHSA2 | 1.169167687 | 0.468786454 | -13.01076319 | 9.71E-11 | -11.8415955 | 8.609439662 | 8.169643011 | 4.523675391 | 9.055659246 | 8.99630959 | 2 | 2 |
| ENSG00000204315 | FKBPL | -0.320113913 | 0.901083982 | -11.52237777 | 2.79E-06 | -11.84249169 | 7.33131396 | 6.535001748 | 2 | 6.087554739 | 6.1572134 | 2 | 2 |
| ENSG00000198216 | CACNA1E | -6.581324925 | 0.015327439 | -5.264524835 | 0.018578065 | -11.84584976 | 2 | 3.686303414 | 5.772716771 | 2 | 2.14022021 | 2.225793514 | 2.117335275 |
| ENSG00000144619 | CNTN4 | -7.288076049 | 0.003506502 | -4.558299952 | 0.020804985 | -11.846376 | 6.009025997 | 5.303523747 | 2 | 2 | 2.14022021 | 2.592885947 | 2.225841198 |
| ENSG00000167705 | RILP | -0.320900533 | 0.810928754 | -11.53234378 | 2.79E-10 | -11.85324431 | 4.815609471 | 7.099070191 | 6.534109959 | 6.12217689 | 6.14061114 | 2 | 2 |
| ENSG00000079277 | MKNK1 | -11.61818679 | 3.30E-09 | -0.235813485 | 0.886197127 | -11.85400028 | 8.737455672 | 8.029695761 | 5.156626401 | 2 | 2 | 7.593002849 | 7.769481431 |
| ENSG00000182612 | TSPAN10 | -9.760773894 | 0.000196845 | -2.09375021 | 0.377546443 | -11.8545241 | 6.816637977 | 2 | 6.534109959 | 2 | 2 | 4.230621763 | 4.316807515 |
| ENSG00000274352 | | 0.54407073 | 0.721937953 | -12.40094308 | 1.33E-10 | -11.85687235 | 5.095959458 | 6.78762695 | 8.325066185 | 7.753776273 | 7.85690889 | 2 | 2 |
| ENSG00000146955 | RAB19 | -5.909172668 | 0.299070637 | -5.949946493 | NA | -11.85911916 | 2 | 4.11352228 | 2 | 2 | 2 | 2.048074405 | 2 |
| ENSG00000135720 | DYNC1LI2 | -13.35792816 | 2.14E-13 | 1.492916392 | 0.193664118 | -11.86501177 | 7.53297911 | 9.77810053 | 10.37481416 | 2 | 2 | 11.08807051 | 11.16422519 |
| ENSG00000102230 | PCYT1B | -6.734923166 | 0.001035933 | -5.141820047 | 0.011593579 | -11.87674321 | 4.00722531 | 6.311476754 | 8.99637432 | 2.947924294 | 2 | 2.560105256 | 3.735548302 |
| ENSG00000166206 | GABRB3 | -6.837100448 | 2.67E-06 | -5.04358904 | 0.000388995 | -11.88068949 | 5.095959458 | 8.487557258 | 8.865353883 | 3.023133209 | 2.2680111 | 4.188345803 | 2.782576693 |
| ENSG00000064692 | SNCAIP | -8.006285685 | 0.0515939 | -3.87654536 | 0.32803778 | -11.88283104 | 2 | 5.946638857 | 2 | 2 | 2 | 2.492221014 | 2.326754077 |
| ENSG00000163357 | DCST1 | -7.048924995 | 0.0437622 | -4.842740951 | 0.122820268 | -11.89166595 | 2 | 2 | 6.43197674 | 2.141737246 | 2 | 2.384001718 | 2.225841198 |
| ENSG00000105675 | ATP4A | -7.795680548 | 0.078688092 | -4.096652357 | 0.339717672 | -11.8923329 | 2 | 2 | 5.772716771 | 2 | 2 | 2.094598369 | 2.509591799 |
| ENSG00000140505 | CYP1A2 | -4.759514134 | 0.002299086 | -7.134187263 | 2.29E-05 | -11.8937014 | 4.815609471 | 4.442710129 | 4.874587708 | 2.3892346 | 2.14022021 | 2 | 2.117335275 |
| ENSG00000188039 | NWD1 | -5.698437886 | 0.000620065 | -6.19559333 | 5.79E-05 | -11.89403122 | 5.095959458 | 3.076665209 | 6.629488387 | 2.270786764 | 2.2680111 | 2.267001569 | 2.117335275 |
| ENSG00000274113 | | -3.346245925 | 0.459630918 | -8.550310407 | 0.048142698 | -11.89655633 | 2 | 2 | 5.156626401 | 2.3892346 | 2.2680111 | 2 | 2 |
| ENSG00000196440 | ARMCX4 | -11.93437 | 1.77E-09 | 0.036926389 | 0.983041858 | -11.89744361 | 5.330594134 | 8.276714465 | 9.099922433 | 2 | 2 | 7.997491306 | 8.487906181 |
| ENSG00000141540 | TTYH2 | -6.924481069 | 3.88E-17 | -4.985870582 | 1.20E-10 | -11.91035165 | 9.635285994 | 8.433041974 | 9.385998894 | 3.162721474 | 3.15372745 | 5.06226682 | 3.620995423 |
| ENSG00000088827 | SIGLEC1 | -8.691716762 | 0.008946207 | -3.220806611 | 0.344285984 | -11.91252337 | 8.998234871 | 2 | 2 | 2 | 2.2680111 | 4.017766949 | 4.810605986 |
| ASUN | ASUN | 0.481836265 | 0.63848867 | -12.39464432 | 5.13E-13 | -11.91280806 | 7.709880939 | 7.572420273 | 5.930865648 | 7.762033056 | 7.71553728 | 2 | 2 |
| CECR5 | CECR5 | 0.366054764 | 0.856283733 | -12.2857366 | 9.75E-09 | -11.91968184 | 3.327711234 | 7.928237109 | 7.451345288 | 7.474468681 | 7.56191311 | 2 | 2 |
| ENSG00000170743 | SYT9 | -4.979317108 | 0.0367628 | -6.970414547 | 0.002588125 | -11.94973166 | 5.532353208 | 2 | 6.882775709 | 2.498691189 | 2.49394783 | 2.267001569 | 2 |
| C20ORF196 | C20ORF196 | -1.392742392 | 0.479076764 | -10.55998943 | 2.54E-06 | -11.95273182 | 6.73718411 | 4.442710129 | 3.369317972 | 4.497095146 | 4.12711147 | 2 | 2 |
| ENSG00000183605 | SFXN4 | -10.72430324 | 3.32E-07 | -1.230430845 | 0.465241848 | -11.95473409 | 4.815609471 | 8.276714465 | 5.772716771 | 2 | 2 | 5.948096914 | 5.785984581 |
| ENSG00000213397 | HAUS7 | -10.98864145 | 8.48E-12 | -0.970840759 | 0.087948816 | -11.95948221 | 7.218819781 | 7.431218269 | 7.225742779 | 2 | 2 | 6.566956211 | 6.124122661 |
| ENSG00000129595 | EPB41L4A | -6.811214176 | 0.00048963 | -5.14870361 | 0.005004527 | -11.95991779 | 2 | 6.602424674 | 7.552162999 | 2.270786764 | 2.2680111 | 3.059555873 | 2.277179783 |
| ENSG00000114745 | GORASP1 | -12.7729576 | 1.53E-14 | 0.804637332 | 0.309292345 | -11.96832027 | 8.981092835 | 8.169643011 | 9.658943166 | 2 | 2 | 10.01280126 | 9.682361453 |
| ENSG00000188996 | HUS1B | -6.167729364 | 0.034288024 | -5.804404078 | 0.038665093 | -11.97213344 | 2 | 2 | 7.502634626 | 2.141737246 | 2.38539787 | 2.526562374 | 2.117335275 |
| ENSG00000198734 | F5 | -5.627186626 | 0.073376458 | -6.350758056 | 0.032176413 | -11.97794468 | 2 | 5.303523747 | 4.874587708 | 2.270786764 | 2 | 2 | 2.172608095 |
| ENSG00000214717 | ZBED1 | 0.356187885 | 0.409421012 | -12.36780755 | 2.36E-16 | -12.01161967 | 8.517314938 | 8.57407404 | 8.865353883 | 9.003220305 | 9.01919552 | 2 | 2.059860178 |
| ENSG00000198650 | TAT | -5.673167515 | 0.003210494 | -6.343828219 | 0.000386994 | -12.01699573 | 2 | 5.131603625 | 6.073378139 | 2.141737246 | 2.2680111 | 2.094598369 | 2.172608095 |
| ENSG00000163743 | RCHY1 | -11.05331973 | 3.80E-12 | -0.968840643 | 0.054375467 | -12.02216037 | 7.27616306 | 7.35507212 | 7.451345288 | 2 | 2 | 6.537789343 | 6.30506411 |
| ENSG00000152936 | LMNTD1 | -7.907439722 | 0.000297995 | -4.129035841 | 0.009975921 | -12.03647556 | 5.095959458 | 4.442710129 | 3.369317972 | 2 | 2 | 2.384001718 | 2.277179783 |
| ENSG00000180818 | HOXC10 | -1.768889645 | 0.007425994 | -10.27258538 | 3.85E-22 | -12.04147503 | 8.009475818 | 8.607278467 | 7.502634626 | 6.449761039 | 6.34282908 | 2.139668783 | 2 |
| ENSG00000178538 | CA8 | -5.278263139 | 0.28002852 | -6.765143029 | 0.14640672 | -12.04340617 | 4.815609471 | 2 | 2 | 2.141737246 | 2 | 2.048074405 | 2 |
| ENSG00000112232 | KHDRBS2 | -6.08167052 | 0.206479362 | -5.971598528 | NA | -12.05326905 | 2 | 3.686303414 | 3.369317972 | 2 | 2 | 2 | 2.059860178 |
| ENSG00000172689 | MS4A10 | -2.505016233 | 0.204752691 | -9.558068051 | 3.68E-05 | -12.06308428 | 5.70932728 | 3.686303414 | 3.369317972 | 3.162721474 | 2.68924543 | 2 | 2 |
| ENSG00000227314 | | -1.157799709 | 0.632341559 | -10.90855769 | 1.31E-05 | -12.0663574 | 2 | 5.838937808 | 6.80318718 | 4.783303513 | 4.83276841 | 2 | 2 |
| KIAA1161 | | -0.848878913 | 0.39383571 | -11.22197659 | 8.65E-11 | -12.0708555 | 6.653098118 | 5.303523747 | 6.203069917 | 5.461211713 | 5.24368216 | 2 | 2 |
| ENSG00000170310 | STX8 | -11.30542105 | 1.03E-05 | -0.770458239 | 0.762852238 | -12.07587929 | 2 | 7.954278496 | 8.380957451 | 2 | 2 | 6.890734189 | 6.826173076 |
| ENSG00000277111 | | 1.280704712 | 0.531291719 | -13.37175277 | 3.22E-09 | -12.09104806 | 4.00722531 | 6.899080544 | 9.577565424 | 9.425874714 | 9.56238793 | 2 | 2 |
| ENSG00000168487 | BMP1 | -13.84759372 | 3.87E-18 | 1.756474322 | 0.000911929 | -12.0911194 | 9.940323746 | 10.59199661 | 9.681375057 | 2 | 2 | 11.75249754 | 11.99495099 |
| ENSG00000173848 | NET1 | -10.83603939 | 2.56E-05 | -1.257243643 | 0.609260112 | -12.09328303 | 8.19931886 | 7.002537973 | 2 | 2 | 2 | 5.899789217 | 5.995914157 |
| ENSG00000125841 | NRSN2 | -13.07703593 | 2.09E-16 | 0.977255282 | 0.070847274 | -12.09978064 | 9.790941832 | 9.181799937 | 8.978369997 | 2 | 2 | 10.19181371 | 10.45974529 |
| ENSG00000145147 | SLIT2 | -4.30094119 | 1.22E-77 | -7.799068754 | 5.15E-236 | -12.10000994 | 12.36552004 | 12.44396886 | 12.56943906 | 8.221200057 | 8.13770558 | 4.742549839 | 4.998228167 |
| ENSG00000007174 | DNAH9 | -7.377636719 | 0.09807061 | -4.729825173 | 0.266306871 | -12.10746189 | 2 | 2 | 5.39244166 | 2 | 2 | 2.094598369 | 2.225841198 |
| ENSG00000156509 | FBXO43 | -5.757281393 | 0.094654603 | -6.351317244 | 0.054350719 | -12.10859864 | 7.096806373 | 2 | 2 | 2 | 2.49394783 | 2.139668783 | 2.225841198 |
| ENSG00000174669 | SLC29A2 | -11.04671382 | 1.93E-09 | -1.063496548 | 0.374026188 | -12.11021037 | 9.800745303 | 7.145017454 | 8.143093325 | 2 | 2.14022021 | 7.400431904 | 7.991296779 |
| ENSG00000205857 | NANOGNB | -3.824774706 | 0.098666313 | -8.288504572 | 0.000931021 | -12.11327928 | 3.327711234 | 4.11352228 | 3.369317972 | 2.141737246 | 2.2680111 | 2 | 2 |
| ENSG00000185565 | LSAMP | -4.24241185 | 1.07E-17 | -7.871442763 | 2.53E-50 | -12.11385461 | 9.848787621 | 9.318399851 | 8.978369997 | 5.433909581 | 5.21224859 | 2.62493832 | 2.950113106 |
| TMEM57 | | 1.194007149 | 0.147447249 | -13.31226247 | 1.43E-15 | -12.11825532 | 8.716893051 | 7.099070191 | 8.266922072 | 9.425874714 | 9.27068036 | 2 | 2 |
| ENSG00000172795 | DCP2 | -11.98760188 | 3.92E-13 | -0.131780348 | 0.880025861 | -12.11938223 | 7.82963488 | 8.297207175 | 8.608698767 | 2 | 2 | 8.567194806 | 7.558732759 |
| C17ORF104 | C17ORF104 | -5.348757964 | NA | -6.786781917 | NA | -12.13553988 | 2 | 3.686303414 | 2 | 2 | 2 | 2 | 2 |
| EMR2 | EMR2 | -5.348757964 | NA | -6.786781917 | NA | -12.13553988 | 2 | 3.686303414 | 2 | 2 | 2 | 2 | 2 |
| ENSG00000125533 | BHLHE23 | -5.348757964 | NA | -6.786781917 | NA | -12.13553988 | 2 | 3.686303414 | 2 | 2 | 2 | 2 | 2 |
| ENSG00000125831 | CST11 | -5.348757964 | NA | -6.786781917 | NA | -12.13553988 | 2 | 3.686303414 | 2 | 2 | 2 | 2 | 2 |
| ENSG00000129152 | MYOD1 | -5.348757964 | NA | -6.786781917 | NA | -12.13553988 | 2 | 3.686303414 | 2 | 2 | 2 | 2 | 2 |
| ENSG00000138136 | LBX1 | -5.348757964 | NA | -6.786781917 | NA | -12.13553988 | 2 | 3.686303414 | 2 | 2 | 2 | 2 | 2 |
| ENSG00000160182 | TFF1 | -5.348757964 | NA | -6.786781917 | NA | -12.13553988 | 2 | 3.686303414 | 2 | 2 | 2 | 2 | 2 |
| ENSG00000166884 | OR4D6 | -5.348757964 | NA | -6.786781917 | NA | -12.13553988 | 2 | 3.686303414 | 2 | 2 | 2 | 2 | 2 |
| ENSG00000177627 | C12orf54 | -5.348757964 | NA | -6.786781917 | NA | -12.13553988 | 2 | 3.686303414 | 2 | 2 | 2 | 2 | 2 |
| ENSG00000184895 | SRY | -5.348757964 | NA | -6.786781917 | NA | -12.13553988 | 2 | 3.686303414 | 2 | 2 | 2 | 2 | 2 |
| ENSG00000187021 | PNLIPRP1 | -5.348757964 | NA | -6.786781917 | NA | -12.13553988 | 2 | 3.686303414 | 2 | 2 | 2 | 2 | 2 |
| ENSG00000203972 | GLYATL3 | -5.348757964 | NA | -6.786781917 | NA | -12.13553988 | 2 | 3.686303414 | 2 | 2 | 2 | 2 | 2 |
| LOC158434 | | -5.348757964 | NA | -6.786781917 | NA | -12.13553988 | 2 | 3.686303414 | 2 | 2 | 2 | 2 | 2 |
| LOC554223 | | -5.348757964 | NA | -6.786781917 | NA | -12.13553988 | 2 | 3.686303414 | 2 | 2 | 2 | 2 | 2 |
| ENSG00000153208 | MERTK | -8.330002951 | 4.25E-13 | -3.810448963 | 1.96E-05 | -12.14045191 | 7.218819781 | 7.979858146 | 8.65477198 | 2.141737246 | 2.38539787 | 5.026717879 | 3.698365557 |
| ENSG00000088992 | TESC | -6.645796621 | 0.008820478 | -5.495306889 | 0.006698657 | -12.14110351 | 2 | 4.442710129 | 5.59507254 | 2.141737246 | 2 | 2.139668783 | 2.172608095 |
| ENSG00000156218 | ADAMTSL3 | -9.551059078 | 1.19E-05 | -2.591763349 | 0.119536754 | -12.14282243 | 7.624139768 | 8.053983481 | 3.369317972 | 2.141737246 | 2 | 4.240999983 | 5.331426166 |
| ENSG00000121351 | IAPP | -4.810459005 | 0.01037751 | -7.337001848 | 0.000224682 | -12.14746085 | 3.327711234 | 4.11352228 | 5.772716771 | 2.3892346 | 2.14022021 | 2.094598369 | 2 |
| ENSG00000131368 | MRPS25 | -11.91988663 | 1.03E-10 | -0.240130228 | 0.861691876 | -12.16001686 | 6.138360429 | 8.356994601 | 8.941672936 | 2 | 2 | 7.989819469 | 7.962181894 |
| ENSG00000198169 | ZNF251 | 0.37991282 | 0.731653953 | -12.55306612 | 5.23E-13 | -12.17315331 | 7.904282243 | 7.73146271 | 5.930865648 | 7.764774849 | 7.82349365 | 2 | 2 |
| ENSG00000196166 | C8orf86 | -6.688981317 | 0.011629766 | -5.489659405 | 0.011176913 | -12.17864072 | 2 | 4.11352228 | 5.772716771 | 2 | 2.14022021 | 2.094598369 | 2.225841198 |
| ENSG00000101892 | ATP1B4 | -6.012530641 | 0.002612888 | -6.167822536 | 0.000780217 | -12.18035318 | 4.467311984 | 4.11352228 | 6.534109959 | 2.141737246 | 2.2680111 | 2.34604626 | 2 |
| ENSG00000177108 | ZDHHC22 | -5.439089556 | 0.235789306 | -6.741445825 | 0.125265786 | -12.18053538 | 2 | 5.838937808 | 2 | 2.270786764 | 2 | 2 | 2.117335275 |
| ENSG00000275145 | | 0.674640756 | 0.534490579 | -12.85781365 | 1.85E-13 | -12.18317289 | 7.82963488 | 6.228547981 | 8.380957451 | 8.372666678 | 8.39732094 | 2 | 2 |
| ENSG00000164687 | FABP5 | -3.618495289 | 2.77E-08 | -8.568745409 | 5.04E-31 | -12.1872407 | 7.940205782 | 8.487557258 | 9.165043284 | 5.22679461 | 5.07918227 | 2.457042231 | 2.172608095 |
| ENSG00000262139 | | -1.811725244 | 0.363985262 | -10.38284826 | 5.34E-08 | -12.1945735 | 4.00722531 | 6.602424674 | 9.541233192 | 6.34372989 | 6.44777477 | 2.139668783 | 2 |
| ENSG00000198690 | FAN1 | -0.256025795 | 0.751438346 | -11.94621334 | 7.63E-14 | -12.20223914 | 8.609439662 | 7.35507212 | 8.460932916 | 8.008651253 | 7.95274764 | 2.048074405 | 2 |
| ENSG00000172209 | GPR22 | -6.082025372 | 0.200172026 | -6.120648555 | NA | -12.20267393 | 2 | 3.686303414 | 3.369317972 | 2 | 2 | 2.048074405 | 2 |
| ENSG00000273611 | ZNHIT3 | 0.401786876 | 0.632693884 | -12.63806389 | 1.81E-14 | -12.23627701 | 7.096806373 | 7.002537973 | 8.143093325 | 7.789219291 | 8.00183183 | 2 | 2 |
| ENSG00000198911 | SREBF2 | -14.02624043 | 1.57E-16 | 1.774688958 | 0.036922742 | -12.25155147 | 8.998234871 | 10.79296235 | 10.55568472 | 2 | 2 | 11.78809862 | 12.31502585 |
| ENSG00000146350 | TBC1D32 | -10.49706159 | 7.45E-10 | -1.75785863 | 0.030972925 | -12.25492022 | 6.138360429 | 6.899080544 | 7.225742779 | 2 | 2 | 5.463319095 | 4.827924991 |
| ENSG00000235925 | | -4.184022211 | 0.461270807 | -8.072043517 | 0.131367278 | -12.25606573 | 2 | 4.710564154 | 2 | 2.270786764 | 2 | 2 | 2 |
| ENSG00000040731 | CDH10 | -7.013778021 | 0.013496022 | -5.247450512 | 0.0277865 | -12.26122853 | 4.00722531 | 6.140560021 | 2 | 2.141737246 | 2 | 2.384001718 | 2.059860178 |
| ENSG00000188037 | CLCN1 | -7.313594147 | 0.10027796 | -4.949137279 | 0.243333363 | -12.26273143 | 5.330594134 | 2 | 2 | 2 | 2 | 2.094598369 | 2.172608095 |

| | | | | | | | | | | | | | |
|---|---|---|---|---|---|---|---|---|---|---|---|---|---|
| ENSG00000137221 | TJAP1 | -12.11503824 | 4.24E-11 | -0.150285432 | 0.914220926 | -12.26532368 | 6.653098118 | 8.029695761 | 9.358421373 | 2 | 2 | 8.2308261 | 8.282618188 |
| ENSG00000166278 | C2 | -2.264092894 | 0.118462931 | -10.00245017 | 5.21E-07 | -12.26654307 | 5.866944937 | 4.11352228 | 4.523675391 | 3.289987342 | 3.33976443 | 2 | 2 |
| ENSG00000275272 | | 1.215906566 | 0.479762551 | -13.48264086 | 6.86E-11 | -12.26673429 | 4.467311984 | 8.981105197 | 8.784770914 | 9.50916971 | 9.57325401 | 2 | 2 |
| LOC101929372 | | -2.847117428 | 0.067812259 | -9.428713219 | 4.82E-06 | -12.27583065 | 4.467311984 | 3.686303414 | 5.156626401 | 2.784612485 | 2.68924543 | 2 | 2 |
| ENSG00000136731 | UGGT1 | -13.26115653 | 4.03E-13 | 0.980725115 | 0.420726478 | -12.28043141 | 7.709880939 | 9.256956408 | 10.46808205 | 2 | 2 | 10.17922029 | 10.79389117 |
| ENSG00000196372 | ASB13 | -11.80891404 | 9.60E-09 | -0.482544917 | 0.784973872 | -12.29145895 | 5.330594134 | 7.274681896 | 9.301632639 | 2 | 2 | 7.824551575 | 7.402777233 |
| ENSG00000101850 | GPR143 | -6.409514792 | 0.012298147 | -5.882397703 | 0.011978839 | -12.2919125 | 6.653098118 | 3.686303414 | 2 | 2.270786764 | 2 | 2.225793514 | 2.172608095 |
| ENSG00000168356 | SCN11A | -4.995017113 | 3.91E-07 | -7.30394485 | 3.61E-12 | -12.29896196 | 5.70932728 | 6.464272742 | 6.43197674 | 2.270786764 | 2.94009902 | 2.139668783 | 2.172608095 |
| ENSG00000243232 | PCDHAC2 | -4.919142453 | 0.00744278 | -7.382113747 | 3.27E-05 | -12.3012562 | 2 | 7.572420273 | 8.631919287 | 3.663521627 | 3.01487888 | 2.492221014 | 2.225841198 |
| ENSG00000135549 | PKIB | -7.074772514 | 3.85E-11 | -5.23854643 | 1.80E-07 | -12.31331894 | 9.064836028 | 7.701025996 | 8.460932916 | 3.09461492 | 3.2680111 | 4.261535101 | 2.981406154 |
| ENSG00000104237 | RP1 | -6.162182594 | 0.191888665 | -6.158915791 | 0.161791931 | -12.32109839 | 2 | 2 | 5.59507254 | 2 | 2.14022021 | 2.139668783 | 2 |
| ENSG00000006712 | PAF1 | -12.41922437 | 2.49E-15 | 0.09114618 | 0.864675088 | -12.32807819 | 8.631580145 | 8.876041558 | 8.585098385 | 2 | 2 | 8.593755943 | 8.974576781 |
| ENSG00000189325 | C6orf222 | -6.741933956 | 0.054872489 | -5.591441052 | 0.07449588 | -12.33337501 | 6.138360429 | 2 | 2 | 2.141737246 | 2 | 2.139668783 | 2.172608095 |
| ENSG00000099910 | KLHL22 | -12.30375428 | 1.10E-07 | -0.032973816 | 0.988654005 | -12.33672809 | 9.678463567 | 8.356994601 | 3.369317972 | 2 | 2 | 8.547470747 | 8.571253152 |
| ENSG00000230678 | | 2.218160985 | 0.005379744 | -14.55708655 | 3.52E-18 | -12.33892556 | 9.839306291 | 9.7032946 | 8.110401118 | 11.58732457 | 11.6368986 | 2 | 2 |
| ENSG00000198892 | SHISA4 | -12.32519976 | 1.02E-13 | -0.018554049 | 0.984013064 | -12.34375381 | 8.564112515 | 7.819099893 | 9.165043284 | 2 | 2 | 8.355461064 | 8.800186246 |
| ENSG00000165300 | SLITRK5 | -3.45488058 | 0.081376146 | -8.894318415 | 2.61E-05 | -12.34919899 | 6.816637977 | 6.602424674 | 2 | 3.227757282 | 3.45167086 | 2.094598369 | 2 |
| KIAA0368 | | 1.659883296 | 0.045222786 | -14.01058717 | 6.61E-17 | -12.35070387 | 8.586954094 | 9.591588447 | 7.858115942 | 10.53762892 | 10.4811454 | 2 | 2 |
| GATSL2 | | 0.256129708 | 0.85890231 | -12.60724115 | 9.68E-12 | -12.35111144 | 5.330594134 | 8.124485672 | 7.734844356 | 7.691737974 | 7.75969386 | 2 | 2 |
| ENSG00000277222 | | 0.649253099 | 0.638882507 | -13.00187982 | 3.34E-12 | -12.35262672 | 7.904282243 | 8.671479029 | 5.59507254 | 8.485003881 | 8.51952909 | 2 | 2 |
| ENSG00000166676 | TVP23A | -8.666674186 | 1.14E-05 | -3.686436205 | 0.003736867 | -12.35311039 | 4.00722531 | 5.303523747 | 5.59507254 | 2 | 2 | 2.560105256 | 2.782576693 |
| ENSG00000125863 | MKKS | -11.59708441 | 2.03E-13 | -0.759915527 | 0.109215771 | -12.35699993 | 7.867441304 | 7.73146271 | 8.076950877 | 2 | 2 | 7.253116621 | 7.041900409 |
| ENSG00000278191 | | -0.319686122 | 0.609297559 | -12.03768856 | 2.74E-42 | -12.35737468 | 9.657036304 | 10.42512749 | 10.67785662 | 9.941741253 | 10.0485276 | 2.094598369 | 2.117335275 |
| ENSG00000135968 | GCC2 | -12.29040957 | 1.02E-14 | -0.073386875 | 0.90832745 | -12.36379644 | 8.340423424 | 8.623598428 | 8.742726175 | 2 | 2 | 8.766685705 | 8.187707412 |
| C14ORF169 | C14ORF169 | -0.155423582 | 0.951713702 | -12.21256669 | 7.89E-07 | -12.36799028 | 7.82963488 | 7.467835385 | 2 | 6.79375511 | 7.07311172 | 2 | 2 |
| ENSG00000148204 | CRB2 | -5.661369677 | 2.68E-05 | -6.708658738 | 3.27E-07 | -12.37002841 | 5.866944937 | 8.356994601 | 7.97167868 | 3.566290607 | 2.14022021 | 2.492221014 | 2.671842496 |
| UPK3BL | | 0.76253004 | 0.52243871 | -13.13491161 | 2.10E-13 | -12.37238157 | 8.042902253 | 6.228547981 | 8.742726175 | 8.756965077 | 8.73530306 | 2 | 2 |
| ENSG00000268104 | SLC6A14 | -3.954065089 | 0.077815532 | -8.421495402 | 0.000590127 | -12.37556049 | 4.00722531 | 3.076665209 | 4.058998049 | 2.270786764 | 2.14022021 | 2 | 2 |
| FAM179B | | 0.508899109 | 0.669953753 | -12.88542673 | 4.12E-13 | -12.37652762 | 6.138360429 | 7.669933291 | 8.536706852 | 8.252792309 | 8.24450173 | 2 | 2 |
| ENSG00000165821 | SALL2 | -12.20845016 | 2.81E-08 | -0.173872419 | 0.935195648 | -12.38232257 | 4.00722531 | 8.522788974 | 9.452720984 | 2 | 2 | 8.359038717 | 8.292008706 |
| ENSG00000100575 | TIMM9 | -11.42002461 | 5.47E-11 | -0.971051924 | 0.322038974 | -12.39107654 | 6.96351253 | 7.051611315 | 8.561105505 | 2 | 2 | 6.793610851 | 6.757786576 |
| ENSG00000180953 | ST20 | -10.07904516 | 3.76E-08 | -2.32019752 | 0.030674362 | -12.39924268 | 6.89194353 | 5.131603625 | 6.718950348 | 2 | 2 | 4.604690117 | 4.033026654 |
| ENSG00000158022 | TRIM63 | -6.6339874 | 0.232251697 | -5.774538775 | 0.284291641 | -12.40852618 | 2 | 4.710564154 | 2 | 2 | 2 | 2.094598369 | 2 |
| ENSG00000167355 | OR51B5 | -6.6339874 | 0.232251697 | -5.774538775 | 0.284291641 | -12.40852618 | 2 | 4.710564154 | 2 | 2 | 2 | 2.094598369 | 2 |
| ENSG00000183166 | CALN1 | -4.336732455 | 0.006051819 | -8.076182533 | 7.28E-05 | -12.41291499 | 4.467311984 | 4.11352228 | 5.156626401 | 2.498691189 | 2.14022021 | 2 | 2.059860178 |
| LOC100129940 | | -3.179564059 | 0.071563439 | -9.233816296 | 2.39E-05 | -12.41338036 | 4.00722531 | 4.11352228 | 4.874587708 | 2.784612485 | 2.2680111 | 2 | 2 |
| ENSG00000242950 | ERVW-1 | -8.976030159 | 0.000142843 | -3.44155507 | 0.067503656 | -12.41758523 | 5.866944937 | 4.710564154 | 5.39244166 | 2 | 2 | 3.411599105 | 2.117335275 |
| ENSG00000179909 | ZNF154 | -11.62394365 | 1.16E-07 | -0.795310609 | 0.682815119 | -12.41925426 | 4.467311984 | 6.899080544 | 9.196533591 | 2 | 2 | 7.413107295 | 6.811011855 |
| ENSG00000274336 | | -1.592850842 | 0.060764772 | -10.82774001 | 1.22E-29 | -12.42059085 | 8.257427002 | 9.763446433 | 10.36098398 | 8.078634414 | 8.15640714 | 2.139668783 | 2.172608095 |
| ENSG00000106236 | NPTX2 | -4.752847069 | 0.008060333 | -7.669087203 | 1.11E-05 | -12.42193427 | 8.418903792 | 7.503546079 | 2 | 3.709785322 | 2.94009902 | 2.225793514 | 2.326754077 |
| ENSG00000071909 | MYO3B | -6.256016886 | 0.000309039 | -6.17206197 | 0.000109005 | -12.42807886 | 4.815609471 | 4.93638933 | 7.163363831 | 2.498691189 | 2 | 2.139668783 | 2.421067459 |
| ENSG00000100191 | SLC5A4 | -3.614339817 | 0.412400906 | -8.815340497 | 0.038171018 | -12.42968031 | 2 | 2 | 5.39244166 | 2.270786764 | 2.38539787 | 2 | 2 |
| ENSG00000143520 | FLG2 | -6.341839545 | 0.028112563 | -6.100208285 | 0.02348089 | -12.44204783 | 3.327711234 | 2 | 6.629488387 | 2.270786764 | 2 | 2.048074405 | 2.277179783 |
| ENSG00000105991 | HOXA1 | -3.127467243 | 0.283185204 | -9.315728061 | 0.000895747 | -12.4431953 | 2 | 2 | 8.206335842 | 3.921164147 | 3.90918929 | 2.048074405 | 2.059860178 |
| ENSG00000101443 | WFDC2 | -4.278873395 | 0.449348839 | -8.166895787 | 0.126314768 | -12.44576918 | 4.815609471 | 2 | 2 | 2.270786764 | 2 | 2 | 2 |
| ENSG00000092020 | PPP2R3C | -11.63257136 | 1.98E-07 | -0.820045937 | 0.684386831 | -12.4526173 | 8.367060731 | 8.607278467 | 3.369317972 | 2 | 2 | 7.229857522 | 7.017577533 |
| ENSG00000105879 | CBLL1 | -12.21098105 | 1.84E-13 | -0.248627088 | 0.773784823 | -12.45960814 | 9.064836028 | 8.451443573 | 7.646393253 | 2 | 2 | 8.387928811 | 8.107542072 |
| ENSG00000276458 | | 1.961193339 | 0.00103094 | -14.42447698 | 2.49E-19 | -12.46328364 | 9.761125267 | 8.57407404 | 9.242516611 | 11.22796957 | 11.2186389 | 2 | 2 |
| ENSG00000138794 | CASP6 | -10.92136727 | 9.47E-08 | -1.550150621 | 0.318191025 | -12.47151789 | 4.467311984 | 7.002537973 | 8.236946975 | 2 | 2 | 5.873348082 | 5.626876946 |
| ENSG00000104889 | RNASEH2A | -3.249360569 | 0.001989926 | -9.226462722 | 5.88E-17 | -12.47582329 | 9.317904228 | 7.874700259 | 8.110401118 | 5.44762523 | 5.45984938 | 2.384001718 | 2 |
| ENSG00000100294 | MCAT | -11.6482657 | 1.45E-11 | -0.832668297 | 0.38169184 | -12.480934 | 8.716893051 | 7.60566235 | 6.958202216 | 2 | 2 | 7.046766788 | 7.20984412 |
| ENSG00000178999 | AURKB | -9.562667003 | 0.000666479 | -2.921969106 | 0.252840018 | -12.48463611 | 7.218819781 | 4.93638933 | 2 | 2 | 2 | 2.885263063 | 3.972018587 |
| ENSG00000170893 | TRH | -4.669555896 | 0.150100354 | -7.81792621 | 0.014967451 | -12.48748211 | 2 | 6.602424674 | 2 | 2.3892346 | 2.38539787 | 2.094598369 | 2 |
| ENSG00000277292 | | 1.371981594 | 0.38950809 | -13.86451467 | 4.85E-12 | -12.49253308 | 8.609439662 | 9.687856435 | 5.39244166 | 10.03611131 | 10.116961 | 2 | 2 |
| ENSG00000188501 | LCTL | -8.120379301 | 0.000660114 | -4.375139629 | 0.020100516 | -12.49551893 | 5.70932728 | 3.686303414 | 3.369317972 | 2 | 2 | 2.225793514 | 2.421067459 |
| GUCY1B3 | | -2.152768715 | 0.444097907 | -10.35569575 | 0.000288228 | -12.50846447 | 2 | 3.686303414 | 6.718950348 | 3.663521627 | 3.55551834 | 2 | 2 |
| ENSG00000172673 | THEMIS | -5.539445712 | NA | -6.977472046 | NA | -12.51691776 | 3.327711234 | 3.076665209 | 2 | 2 | 2 | 2 | 2 |
| ENSG00000173610 | UGT2A1 | -5.539445712 | NA | -6.977472046 | NA | -12.51691776 | 3.327711234 | 3.076665209 | 2 | 2 | 2 | 2 | 2 |
| GYLTL1B | | -5.539445712 | NA | -6.977472046 | NA | -12.51691776 | 3.327711234 | 3.076665209 | 2 | 2 | 2 | 2 | 2 |
| ENSG00000081870 | HSPB11 | -11.03172181 | 2.80E-10 | -1.491084716 | 0.115989891 | -12.52280653 | 8.138771685 | 6.464272742 | 6.882775709 | 2 | 2 | 5.846413283 | 5.976631893 |
| IKBKAP | | 0.520361125 | 0.860774154 | -13.06568038 | 1.30E-06 | -12.54531926 | 6.257048221 | 9.338312924 | 2 | 8.422134132 | 8.45385092 | 2 | 2 |
| ENSG00000001626 | CFTR | -6.327834002 | 0.180258969 | -6.218732251 | 0.177528888 | -12.54656625 | 4.00722531 | 2 | 3.369317972 | 2 | 2 | 2 | 2.059860178 |
| ENSG00000274334 | | 1.821147738 | 2.36E-05 | -14.37319675 | 2.97E-20 | -12.55204901 | 9.521325812 | 8.834582582 | 9.227351203 | 11.01246067 | 11.0519173 | 2 | 2 |
| ENSG00000124333 | VAMP7 | 0.797738087 | 0.711497595 | -13.36705364 | 4.06E-09 | -12.56931556 | 9.384607364 | 7.761270542 | 3.369317972 | 9.013631184 | 9.00894176 | 2 | 2 |
| ENSG00000176390 | CRLF3 | -11.79658333 | 2.54E-10 | -0.774038384 | 0.552503225 | -12.57062171 | 6.563806279 | 7.35507212 | 9.132850258 | 2 | 2 | 7.543552275 | 7.083953563 |
| ENSG00000113396 | SLC27A6 | -4.342983683 | 0.441471424 | -8.231006809 | 0.122968358 | -12.57399049 | 2 | 2 | 4.874587708 | 2.270786764 | 2 | 2 | 2 |
| ENSG00000197061 | HIST1H4C | -5.09783059 | 4.94E-11 | -7.481114574 | 8.58E-21 | -12.57894516 | 7.485143553 | 8.623598428 | 7.600047325 | 3.406930997 | 3.60476711 | 2.225793514 | 2.632958241 |
| ENSG00000147246 | HTR2C | -5.572711136 | 0.331951645 | -7.010737885 | NA | -12.58344902 | 2 | 3.076665209 | 3.369317972 | 2 | 2 | 2 | 2 |
| ENSG00000215183 | MSMP | -5.572711136 | 0.331951645 | -7.010737885 | NA | -12.58344902 | 2 | 3.076665209 | 3.369317972 | 2 | 2 | 2 | 2 |
| ENSG00000215612 | HMX1 | -5.572711136 | 0.331951645 | -7.010737885 | NA | -12.58344902 | 2 | 3.076665209 | 3.369317972 | 2 | 2 | 2 | 2 |
| ENSG00000274148 | | -5.572711136 | 0.331951645 | -7.010737885 | NA | -12.58344902 | 2 | 3.076665209 | 3.369317972 | 2 | 2 | 2 | 2 |
| ENSG00000124469 | CEACAM8 | -4.060706074 | 0.073252828 | -8.524440808 | 0.000564093 | -12.58514688 | 3.327711234 | 4.442710129 | 3.369317972 | 2.141737246 | 2.2680111 | 2 | 2 |
| ENSG00000261272 | MUC22 | -4.357797095 | 0.439805367 | -8.231006883 | 0.122968358 | -12.58880398 | 2 | 2 | 4.874587708 | 2 | 2.2680111 | 2 | 2 |
| ENSG00000162627 | SNX7 | -12.23530431 | 3.92E-15 | -0.355662032 | 0.385661678 | -12.59096634 | 8.444143041 | 8.469613413 | 8.65477198 | 2 | 2 | 8.175088797 | 8.170839557 |
| ENSG00000170382 | LRRN2 | -8.814088647 | 9.47E-07 | -3.777054503 | 0.024032158 | -12.59114315 | 7.33131396 | 9.367676479 | 4.523675391 | 2.3892346 | 2 | 4.209638631 | 4.911532223 |
| ENSG00000274679 | | -0.438209956 | 0.8692818 | -12.15418301 | 1.32E-06 | -12.59239297 | 8.075571721 | 6.899080544 | 2 | 6.503277904 | 6.71216164 | 2 | 2 |
| ENSG00000010295 | IFFO1 | -12.57019922 | 1.12E-12 | -0.025345352 | 0.983983084 | -12.59554457 | 9.710019321 | 7.467835385 | 8.561105505 | 2 | 2 | 8.687797808 | 8.960904642 |
| ENSG00000224994 | | 0.097669107 | 0.970415686 | -12.69857939 | 3.87E-07 | -12.60091029 | 8.418903792 | 7.790474954 | 2 | 7.632926745 | 7.68723697 | 2 | 2 |
| ENSG00000156853 | ZNF689 | -11.86761683 | 1.10E-09 | -0.738942385 | 0.619706292 | -12.60655921 | 9.064836028 | 8.101365653 | 5.59507254 | 2 | 2 | 7.631969638 | 7.208191747 |
| ENSG00000274726 | | 0.513373421 | 0.848265166 | -13.12113115 | 2.15E-07 | -12.60775772 | 2 | 8.255926468 | 8.805342542 | 8.474993083 | 8.49688027 | 2 | 2 |
| ENSG00000067646 | ZFY | -1.411414029 | 0.548376437 | -11.19904027 | 6.38E-06 | -12.61045429 | 2 | 6.535001748 | 6.80318718 | 4.867343068 | 4.811949 | 2 | 2 |
| ENSG00000175482 | POLD4 | -11.56930387 | 4.16E-12 | -1.041334753 | 0.17880563 | -12.61063862 | 7.096806373 | 7.73146271 | 8.460932916 | 2 | 2 | 6.737909218 | 6.957866334 |
| ENSG00000112562 | SMOC2 | -12.14982802 | 4.80E-13 | -0.461975105 | 0.605291131 | -12.61180313 | 9.646202137 | 10.5284131 | 9.049077186 | 2.141737246 | 2 | 8.922700729 | 9.777912323 |
| ENSG00000278165 | | -1.126322424 | 0.671489931 | -11.49022653 | 1.33E-05 | -12.61654896 | 7.709880939 | 5.303523747 | 2 | 5.273747807 | 5.41912973 | 2 | 2 |
| WBSCR27 | | -1.803144906 | 0.255045672 | -10.813955 | 1.29E-07 | -12.61709991 | 5.70932728 | 3.686303414 | 6.629488387 | 4.329433326 | 4.05806467 | 2 | 2 |
| ENSG00000113263 | ITK | -4.871273372 | 0.39788284 | -7.749525689 | 0.15053059 | -12.62079906 | 4.467311984 | 2 | 2 | 2 | 2.14022021 | 2 | 2 |
| ENSG00000126453 | BCL2L12 | -11.30045417 | 3.13E-11 | -1.321620218 | 0.117817538 | -12.62207439 | 8.138771685 | 7.701025996 | 6.534109959 | 2 | 2 | 6.384736493 | 6.264298846 |
| ENSG00000103184 | SEC14L5 | -6.619953715 | 0.004736683 | -6.00328339 | 0.007273889 | -12.62323711 | 2 | 3.686303414 | 7.896971941 | 2.270786764 | 2.2680111 | 2.62493832 | 2.117335275 |
| ENSG00000143125 | PROK1 | -6.893773833 | 0.211937735 | -5.741480647 | 0.28326297 | -12.63525448 | 2 | 4.93638933 | 2 | 2 | 2 | 2 | 2.117335275 |
| ENSG00000161270 | NPHS1 | -5.276243131 | 0.000778431 | -7.361277387 | 4.93E-06 | -12.63752052 | 4.00722531 | 6.04685525 | 5.59507254 | 2.3892346 | 2.2680111 | 2 | 2.172608095 |
| ENSG00000162819 | BROX | -12.37233166 | 3.40E-14 | -0.267832444 | 0.72536009 | -12.6401641 | 8.517314938 | 9.192779271 | 8.007629717 | 2 | 2 | 8.441219213 | 8.347103327 |
| RGAG4 | | -0.855146204 | 0.741924158 | -11.79670064 | 4.41E-06 | -12.65184684 | 7.904282243 | 6.04685525 | 2 | 5.930370745 | 5.8245091 | 2 | 2 |
| ENSG00000021461 | CYP3A43 | -4.886905564 | 0.3060178 | -7.765291355 | 0.08248437 | -12.65219692 | 4.00722531 | 2 | 3.369317972 | 2 | 2.14022021 | 2 | 2 |
| ENSG00000121005 | CRISPLD1 | -5.253162432 | 1.53E-08 | -7.399935151 | 2.49E-15 | -12.65309758 | 7.33131396 | 8.540087354 | 7.691296561 | 3.79808662 | 2.77779907 | 2.62493832 | 2.225841198 |
| ENSG00000273993 | | 1.768344456 | 0.039535444 | -14.42460227 | 1.33E-17 | -12.65625782 | 9.751047882 | 9.549500232 | 7.896971941 | 11.04157998 | 11.0202196 | 2 | 2 |
| MKL1 | | 0.638085541 | 0.312941507 | -13.2954831 | 8.05E-17 | -12.65739756 | 7.975256505 | 8.607278467 | 7.691296561 | 8.734762645 | 8.82765235 | 2 | 2 |
| ENSG00000088298 | EDEM2 | -11.97601654 | 8.83E-11 | -0.68575406 | 0.592260592 | -12.6617706 | 8.468948322 | 6.228547981 | 8.960138144 | 2 | 2 | 7.375919751 | 7.782864551 |
| ENSG00000148680 | HTR7 | -3.74896538 | 1.29E-15 | -8.913719586 | 1.39E-67 | -12.66268497 | 10.29268328 | 10.11467741 | 10.6093157 | 6.788375664 | 6.51372389 | 2.962551659 | 2.466008509 |
| ENSG00000168314 | MOBP | -4.819003224 | 0.000517884 | -7.847488171 | 1.21E-06 | -12.66649139 | 5.70932728 | 4.11352228 | 5.59507254 | 2.600425781 | 2.14022021 | 2.048074405 | 2.059860178 |
| ENSG00000117834 | SLC5A9 | -7.378166581 | 0.105084134 | -5.292584825 | 0.222721235 | -12.67075141 | 2 | 2 | 5.39244166 | 2 | 2 | 2.048074405 | 2.172608095 |
| ENSG00000091704 | CPA1 | -6.316771709 | 0.259792504 | -6.357550623 | 0.247715412 | -12.67432233 | 2 | 4.442710129 | 2 | 2 | 2 | 2.048074405 | 2 |
| ENSG00000078900 | TP73 | -8.38562036 | 0.118145526 | -4.292443785 | 0.421230824 | -12.67806415 | 2 | 2 | 6.322058544 | 2 | 2 | 2.717032197 | 2 |
| ENSG00000278787 | | 0.203389326 | 0.900945558 | -12.88259594 | 2.39E-11 | -12.67920661 | 8.609439662 | 7.761270542 | 5.156626401 | 7.877998516 | 8.0087095 | 2 | 2 |
| ENSG00000113456 | RAD1 | -12.55161576 | 7.23E-11 | -0.130094179 | 0.935525414 | -12.68170994 | 10.00958825 | 7.638155693 | 7.163363831 | 2 | 2 | 8.956155911 | 8.409548182 |
| ENSG00000105750 | ZNF85 | -1.101529146 | 0.649152876 | -11.58253749 | 3.24E-06 | -12.68406664 | 7.384433967 | 6.602424674 | 2 | 5.540148523 | 5.36298494 | 2 | 2 |
| ENSG00000169035 | KLK7 | -4.89956118 | 0.304282027 | -7.785332211 | 0.081408101 | -12.68489339 | 3.327711234 | 2 | 4.058998049 | 2.141737246 | 2 | 2 | 2 |
| ENSG00000158901 | WFDC8 | -4.906920562 | 0.3034352 | -7.785307147 | 0.081225915 | -12.69222771 | 3.327711234 | 2 | 4.058998049 | 2 | 2.14022021 | 2 | 2 |

| | | | | | | | | | | | | | |
|---|---|---|---|---|---|---|---|---|---|---|---|---|---|
| ENSG00000136267 | DGKB | -7.071610953 | 2.05E-05 | -5.621341267 | 1.33E-05 | -12.69295222 | 5.70932728 | 4.442710129 | 6.718950348 | 2.141737246 | 2.14022021 | 2.139668783 | 2.551896972 |
| ENSG00000125998 | FAM83C | -7.263208274 | 0.042408891 | -5.438757272 | 0.091397761 | -12.70196555 | 2 | 2 | 6.629488387 | 2 | 2.14022021 | 2.094598369 | 2.374681325 |
| ENSG00000251692 | PTX4 | -6.984341156 | 0.054419619 | -5.722322335 | 0.079932646 | -12.70666349 | 6.366709671 | 2 | 2 | 2.141737246 | 2 | 2.267001569 | 2.059860178 |
| ENSG00000168280 | KIF5C | -8.796695681 | 0.000119329 | -3.915658607 | 0.02587428 | -12.71235429 | 6.257048221 | 4.710564154 | 3.369317972 | 2 | 2 | 2.858551861 | 2.374681325 |
| ENSG00000164953 | TMEM67 | -10.74109327 | 1.20E-06 | -1.974810314 | 0.271491721 | -12.71590359 | 3.327711234 | 7.051611315 | 8.007629717 | 2 | 2 | 5.212212839 | 5.197927245 |
| ENSG00000078725 | BRINP1 | -8.307126965 | 1.54E-21 | -4.411862449 | 2.10E-12 | -12.71898941 | 7.940205782 | 8.297207175 | 9.287079172 | 2.3892346 | 2.38539787 | 4.416072028 | 4.543391831 |
| ENSG00000103429 | BFAR | 1.018108613 | 0.129065162 | -13.75081432 | 1.54E-17 | -12.73270571 | 8.540903468 | 9.102512015 | 7.896971941 | 9.607904693 | 9.61516703 | 2 | 2 |
| ENSG00000152240 | HAUS1 | -11.06536989 | 4.41E-11 | -1.668398597 | 0.028762582 | -12.73376848 | 7.384433967 | 7.874700259 | 6.534109959 | 2 | 2 | 5.846413283 | 5.708623051 |
| ENSG00000164362 | TERT | -4.925748858 | 0.391949897 | -7.812274021 | 0.146527599 | -12.73802288 | 2 | 2 | 4.523675391 | 2.141737246 | 2 | 2 | 2 |
| ENSG00000178690 | DYNAP | -4.925748858 | 0.391949897 | -7.812274021 | 0.146527599 | -12.73802288 | 2 | 2 | 4.523675391 | 2.141737246 | 2 | 2 | 2 |
| LOC100505478 | | -4.925748858 | 0.391949897 | -7.812274021 | 0.146527599 | -12.73802288 | 2 | 2 | 4.523675391 | 2.141737246 | 2 | 2 | 2 |
| LOC101927844 | | -4.925748858 | 0.391949897 | -7.812274021 | 0.146527599 | -12.73802288 | 2 | 2 | 4.523675391 | 2.141737246 | 2 | 2 | 2 |
| ENSG00000131242 | RAB11FIP4 | -7.729505549 | 2.12E-07 | -5.008899594 | 0.000399783 | -12.73840514 | 9.2903348 | 8.147241016 | 4.058998049 | 2.3892346 | 2.49394783 | 3.44788957 | 4.002845082 |
| ENSG00000109182 | CWH43 | -4.934021191 | 0.391478048 | -7.812274062 | 0.146527599 | -12.74629525 | 2 | 2 | 4.523675391 | 2 | 2.14022021 | 2 | 2 |
| ENSG00000179873 | NLRP11 | -4.934021191 | 0.391478048 | -7.812274062 | 0.146527599 | -12.74629525 | 2 | 2 | 4.523675391 | 2 | 2.14022021 | 2 | 2 |
| LOC401052 | | -2.255686448 | 0.096199279 | -10.49442464 | 6.32E-08 | -12.75011109 | 4.815609471 | 6.311476754 | 4.523675391 | 3.709785322 | 3.55551834 | 2 | 2 |
| ENSG00000179299 | NSUN7 | -9.438832349 | 0.000473619 | -3.318704751 | 0.161118254 | -12.7575371 | 2 | 5.131603625 | 7.029880454 | 2 | 2 | 3.336159116 | 3.071407821 |
| ENSG00000158691 | ZSCAN12 | -11.4926839 | 3.32E-10 | -1.265781202 | 0.275530102 | -12.7584651 | 8.444143041 | 5.946638857 | 8.007629717 | 2 | 2 | 6.653021698 | 6.465820193 |
| ENSG00000145029 | NICN1 | -0.232343267 | 0.890583775 | -12.53307148 | 1.50E-10 | -12.76541475 | 4.815609471 | 7.315436877 | 8.325066185 | 7.210554257 | 7.14145882 | 2 | 2 |
| ENSG00000118094 | TREH | -7.35669725 | 0.038214495 | -5.425947201 | 0.089458136 | -12.78264445 | 2 | 2 | 6.718950348 | 2 | 2.14022021 | 2.384001718 | 2.117335275 |
| FDX1L | | -0.086779889 | 0.964831294 | -12.69828054 | 1.64E-09 | -12.78506043 | 4.00722531 | 7.393647515 | 8.585098385 | 7.412876874 | 7.54308677 | 2 | 2 |
| ENSG00000170128 | GPR25 | -6.374226846 | 0.254759301 | -6.415006478 | 0.242863138 | -12.78923332 | 2 | 2 | 4.523675391 | 2 | 2 | 2.048074405 | 2 |
| ENSG00000167281 | RBFOX3 | -7.688375697 | 0.000100149 | -5.1046219 | 3.46E-05 | -12.7929976 | 6.257048221 | 4.442710129 | 5.39244166 | 2 | 2.14022021 | 2.526562374 | 2.225841198 |
| ENSG00000057593 | F7 | -7.853237434 | 0.017646781 | -4.945726844 | 0.122999873 | -12.79896428 | 8.16936288 | 2 | 2 | 2 | 2.2680111 | 2.858551861 | 2.782576693 |
| ENSG00000277781 | | -1.64984114 | 0.59764617 | -11.15952631 | 0.000224386 | -12.80936745 | 7.579279349 | 3.076665209 | 2 | 4.574143677 | 4.63360314 | 2 | 2 |
| ENSG00000141551 | CSNK1D | -13.86570067 | 3.85E-17 | 1.046242571 | 0.156611002 | -12.8194581 | 10.83137851 | 9.616264339 | 9.624627483 | 2 | 2 | 11.12950693 | 11.24350211 |
| ENSG00000236250 | | -0.004419361 | 0.997264364 | -12.83033452 | 2.25E-12 | -12.83475389 | 8.716893051 | 6.389896633 | 6.882775709 | 7.671425626 | 7.70710533 | 2 | 2 |
| C9ORF142 | C9ORF142 | -0.980392997 | 0.435112068 | -11.85540488 | 1.09E-10 | -12.83579787 | 7.709880939 | 5.72254323 | 5.930865648 | 5.829220145 | 5.80356929 | 2 | 2 |
| ENSG00000122012 | SV2C | -6.993216357 | 0.002648354 | -5.843072697 | 0.0009597 | -12.83628905 | 4.00722531 | 3.076665209 | 6.073378139 | 2.141737246 | 2 | 2.139668783 | 2.172608095 |
| ENSG00000157045 | NTAN1 | 0.490899228 | 0.328495839 | -13.33051845 | 1.44E-17 | -12.83961922 | 8.009475818 | 8.469613413 | 8.007629717 | 8.631895657 | 8.70880006 | 2 | 2 |
| ENSG00000095596 | CYP26A1 | -6.936177672 | 0.137415848 | -5.904203418 | 0.175522875 | -12.84038109 | 2 | 2 | 6.322058544 | 2.141737246 | 2 | 2 | 2.277179783 |
| KIAA0922 | | -0.82949807 | 0.806463377 | -12.01372263 | 8.79E-05 | -12.8432207 | 2 | 3.076665209 | 8.434763955 | 6.096288482 | 6.11534392 | 2 | 2 |
| ENSG00000168955 | TM4SF20 | -4.898782875 | 0.038349318 | -7.954746123 | 0.002144187 | -12.853529 | 4.467311984 | 2 | 5.39244166 | 2.270786764 | 2.14022021 | 2.048074405 | 2 |
| ENSG00000042781 | USH2A | -4.853111882 | 0.160424416 | -8.009923621 | 0.01873645 | -12.8630355 | 2 | 2 | 6.80318718 | 2.600425781 | 2.14022021 | 2.094598369 | 2 |
| ENSG00000147465 | STAR | -6.260696622 | 6.45E-06 | -6.605962556 | 6.38E-07 | -12.86665918 | 4.815609471 | 7.503546079 | 8.110401118 | 2.947924294 | 2.14022021 | 2.384001718 | 2.592996859 |
| ENSG00000142794 | NBPF3 | -12.12338782 | 3.69E-09 | -0.744905701 | 0.664543864 | -12.86829352 | 5.095959458 | 8.234834549 | 9.426401271 | 2 | 2 | 7.564363052 | 7.787298144 |
| ENSG00000188000 | OR7D2 | -5.297874984 | 0.00490603 | -7.573754977 | 8.11E-05 | -12.87162996 | 4.467311984 | 3.686303414 | 6.629488387 | 2.141737246 | 2.49394783 | 2.139668783 | 2 |
| CXORF30 | CXORF30 | -5.722076455 | 0.315977132 | -7.16010507 | NA | -12.88218153 | 4.00722531 | 2 | 2 | 2 | 2 | 2 | 2 |
| ENSG00000112494 | UNC93A | -5.722076455 | 0.315977132 | -7.16010507 | NA | -12.88218153 | 4.00722531 | 2 | 2 | 2 | 2 | 2 | 2 |
| ENSG00000122859 | NEUROG3 | -5.722076455 | 0.315977132 | -7.16010507 | NA | -12.88218153 | 4.00722531 | 2 | 2 | 2 | 2 | 2 | 2 |
| ENSG00000147378 | FATE1 | -5.722076455 | 0.315977132 | -7.16010507 | NA | -12.88218153 | 4.00722531 | 2 | 2 | 2 | 2 | 2 | 2 |
| ENSG00000147596 | PRDM14 | -5.722076455 | 0.315977132 | -7.16010507 | NA | -12.88218153 | 4.00722531 | 2 | 2 | 2 | 2 | 2 | 2 |
| ENSG00000186766 | FOXI2 | -5.722076455 | 0.315977132 | -7.16010507 | NA | -12.88218153 | 4.00722531 | 2 | 2 | 2 | 2 | 2 | 2 |
| ENSG00000187140 | FOXD3 | -5.722076455 | 0.315977132 | -7.16010507 | NA | -12.88218153 | 4.00722531 | 2 | 2 | 2 | 2 | 2 | 2 |
| ENSG00000188984 | AADACL3 | -5.722076455 | 0.315977132 | -7.16010507 | NA | -12.88218153 | 4.00722531 | 2 | 2 | 2 | 2 | 2 | 2 |
| ENSG00000221864 | KRTAP12-2 | -5.722076455 | 0.315977132 | -7.16010507 | NA | -12.88218153 | 4.00722531 | 2 | 2 | 2 | 2 | 2 | 2 |
| ENSG00000236311 | TLX1NB | -5.722076455 | 0.315977132 | -7.16010507 | NA | -12.88218153 | 4.00722531 | 2 | 2 | 2 | 2 | 2 | 2 |
| ENSG00000262851 | | -5.722076455 | 0.315977132 | -7.16010507 | NA | -12.88218153 | 4.00722531 | 2 | 2 | 2 | 2 | 2 | 2 |
| ENSG00000274247 | | -5.722076455 | 0.315977132 | -7.16010507 | NA | -12.88218153 | 4.00722531 | 2 | 2 | 2 | 2 | 2 | 2 |
| PVRL4 | | -5.722076455 | 0.315977132 | -7.16010507 | NA | -12.88218153 | 4.00722531 | 2 | 2 | 2 | 2 | 2 | 2 |
| ENSG00000115091 | ACTR3 | -13.96049305 | 1.98E-18 | 1.069713911 | 0.05471533 | -12.89077914 | 9.740899611 | 10.12043894 | 10.69449487 | 2 | 2 | 11.21433092 | 11.39162185 |
| ENSG00000135919 | SERPINE2 | -13.56397427 | 4.16E-17 | 0.672532644 | 0.301003469 | -12.89144162 | 10.1770105 | 10.00683004 | 9.149036566 | 2 | 2 | 10.70716344 | 10.29009378 |
| ENSG00000120539 | MASTL | -11.45088282 | 2.14E-10 | -1.440673571 | 0.185080874 | -12.89155639 | 7.435667355 | 8.607278467 | 6.322058544 | 2 | 2 | 6.398709156 | 6.311234697 |
| ENSG00000185674 | LYG2 | -4.7126954 | 0.071179771 | -8.18123764 | 0.003695138 | -12.89393304 | 6.009025997 | 3.076665209 | 2 | 2.141737246 | 2.38539787 | 2.048074405 | 2 |
| ENSG00000173349 | SFT2D3 | -1.213253737 | 0.16475399 | -11.68638544 | 5.10E-12 | -12.89963918 | 5.70932728 | 7.002537973 | 6.718950348 | 5.461211713 | 5.43283104 | 2 | 2 |
| KIAA0196 | | 0.914338792 | 0.049220147 | -13.81829273 | 8.16E-19 | -12.90395394 | 8.910412251 | 8.297207175 | 8.742726175 | 9.558986849 | 9.59169325 | 2 | 2 |
| ENSG00000158486 | DNAH3 | -8.488497044 | 0.006549526 | -4.415598232 | 0.116476087 | -12.90409528 | 3.327711234 | 2 | 6.322058544 | 2 | 2 | 2.42098416 | 2.374681325 |
| ENSG00000166925 | TSC22D4 | -13.0996434 | 2.98E-16 | 0.189920614 | 0.779791018 | -12.90972278 | 9.678463567 | 8.762721501 | 9.553445883 | 2 | 2 | 9.42466621 | 9.702747585 |
| PNMAL1 | | -0.445604814 | 0.829870645 | -12.4710654 | 1.04E-08 | -12.91667022 | 3.327711234 | 7.503546079 | 8.206335842 | 6.951285179 | 6.86652414 | 2 | 2 |
| ENSG00000164406 | LEAP2 | -8.68298256 | 0.010953704 | -4.241079016 | 0.179051068 | -12.92406158 | 2 | 2 | 8.007629717 | 2 | 2.14022021 | 3.059555873 | 3.100202476 |
| ENSG00000095464 | PDE6C | -8.11853734 | 0.051793985 | -4.814988926 | 0.225799741 | -12.93352627 | 2 | 2 | 6.073378139 | 2 | 2 | 2.267001569 | 2.225841198 |
| ENSG00000113492 | AGXT2 | -5.752387615 | 0.313419222 | -7.190416608 | NA | -12.94280422 | 3.327711234 | 2 | 3.369317972 | 2 | 2 | 2 | 2 |
| ENSG00000276111 | | 0.357623164 | 0.765831172 | -13.31564053 | 4.78E-14 | -12.95801737 | 9.128497779 | 7.232742069 | 7.285536013 | 8.50645854 | 8.54182783 | 2 | 2 |
| ENSG00000180251 | SLC9A4 | -5.151078209 | 0.004731125 | -7.821644382 | 5.40E-05 | -12.97272259 | 5.095959458 | 3.076665209 | 6.322058544 | 2.270786764 | 2.38539787 | 2 | 2.117335275 |
| ENSG00000140694 | PARN | 0.255817641 | 0.8694764 | -13.22894554 | 2.94E-12 | -12.9731279 | 5.70932728 | 7.979858146 | 8.99637432 | 8.274117511 | 8.39906692 | 2 | 2 |
| ENSG00000276761 | | 0.100855 | 0.948149753 | -13.08718321 | 6.55E-12 | -12.98632821 | 5.330594134 | 8.487557258 | 8.408111533 | 8.091796043 | 7.99722844 | 2 | 2 |
| ENSG00000106809 | OGN | -6.031065761 | 8.21E-05 | -6.956506522 | 1.37E-06 | -12.98757228 | 6.96351253 | 4.710564154 | 4.523675391 | 2.270786764 | 2.2680111 | 2.225793514 | 2.059860178 |
| ENSG00000158813 | EDA | -7.896228145 | 0.000424952 | -5.093311737 | 0.021646679 | -12.98953988 | 9.48530526 | 4.710564154 | 2 | 2.3892346 | 2.2680111 | 3.851628042 | 2.981406154 |
| ENSG00000182601 | HS3ST4 | -7.697051212 | 0.02779988 | -5.294788322 | 0.093601225 | -12.99183953 | 4.00722531 | 6.899080544 | 2 | 2.141737246 | 2 | 2.62493832 | 2 |
| LOC403312 | | -3.896927747 | 0.16105271 | -9.103711155 | 0.001804299 | -13.0006389 | 2 | 4.93638933 | 4.523675391 | 2.498691189 | 2.14022021 | 2 | 2 |
| CCDC64 | CCDC64 | -5.782241252 | 0.310604101 | -7.220270619 | NA | -13.00251187 | 2 | 2 | 4.058998049 | 2 | 2 | 2 | 2 |
| ENSG00000062096 | ARSF | -5.782241252 | 0.310604101 | -7.220270619 | NA | -13.00251187 | 2 | 2 | 4.058998049 | 2 | 2 | 2 | 2 |
| ENSG00000092054 | MYH7 | -5.782241252 | 0.310604101 | -7.220270619 | NA | -13.00251187 | 2 | 2 | 4.058998049 | 2 | 2 | 2 | 2 |
| ENSG00000095981 | KCNK16 | -5.782241252 | 0.310604101 | -7.220270619 | NA | -13.00251187 | 2 | 2 | 4.058998049 | 2 | 2 | 2 | 2 |
| ENSG00000102010 | BMX | -5.782241252 | 0.310604101 | -7.220270619 | NA | -13.00251187 | 2 | 2 | 4.058998049 | 2 | 2 | 2 | 2 |
| ENSG00000124237 | C20orf85 | -5.782241252 | 0.310604101 | -7.220270619 | NA | -13.00251187 | 2 | 2 | 4.058998049 | 2 | 2 | 2 | 2 |
| ENSG00000142025 | DMRTC2 | -5.782241252 | 0.310604101 | -7.220270619 | NA | -13.00251187 | 2 | 2 | 4.058998049 | 2 | 2 | 2 | 2 |
| ENSG00000149305 | HTR3B | -5.782241252 | 0.310604101 | -7.220270619 | NA | -13.00251187 | 2 | 2 | 4.058998049 | 2 | 2 | 2 | 2 |
| ENSG00000164729 | SLC35G3 | -5.782241252 | 0.310604101 | -7.220270619 | NA | -13.00251187 | 2 | 2 | 4.058998049 | 2 | 2 | 2 | 2 |
| ENSG00000166948 | TGM6 | -5.782241252 | 0.310604101 | -7.220270619 | NA | -13.00251187 | 2 | 2 | 4.058998049 | 2 | 2 | 2 | 2 |
| ENSG00000176571 | CNBD1 | -5.782241252 | 0.310604101 | -7.220270619 | NA | -13.00251187 | 2 | 2 | 4.058998049 | 2 | 2 | 2 | 2 |
| ENSG00000186007 | LEMD1 | -5.782241252 | 0.310604101 | -7.220270619 | NA | -13.00251187 | 2 | 2 | 4.058998049 | 2 | 2 | 2 | 2 |
| ENSG00000186115 | CYP4F2 | -5.782241252 | 0.310604101 | -7.220270619 | NA | -13.00251187 | 2 | 2 | 4.058998049 | 2 | 2 | 2 | 2 |
| ENSG00000188389 | PDCD1 | -5.782241252 | 0.310604101 | -7.220270619 | NA | -13.00251187 | 2 | 2 | 4.058998049 | 2 | 2 | 2 | 2 |
| ENSG00000197364 | S100A7L2 | -5.782241252 | 0.310604101 | -7.220270619 | NA | -13.00251187 | 2 | 2 | 4.058998049 | 2 | 2 | 2 | 2 |
| ENSG00000243073 | PRAMEF4 | -5.782241252 | 0.310604101 | -7.220270619 | NA | -13.00251187 | 2 | 2 | 4.058998049 | 2 | 2 | 2 | 2 |
| ENSG00000275290 | | -5.782241252 | 0.310604101 | -7.220270619 | NA | -13.00251187 | 2 | 2 | 4.058998049 | 2 | 2 | 2 | 2 |
| ENSG00000167654 | ATCAY | -4.452907503 | 0.000108779 | -8.550016843 | 6.97E-10 | -13.00292435 | 5.70932728 | 5.595927442 | 7.285536013 | 3.09461492 | 2.77779907 | 2.139668783 | 2 |
| DCDC5 | DCDC5 | -3.773338772 | 0.395836393 | -9.233723885 | 0.031648006 | -13.00706266 | 2 | 2 | 5.772716771 | 2.141737246 | 2.59489879 | 2 | 2 |
| SHFM1 | | 0.35673098 | 0.663997039 | -13.37571625 | 3.03E-16 | -13.01898527 | 8.87372952 | 7.701025996 | 7.818184403 | 8.6030768 | 8.56222967 | 2 | 2 |
| ENSG00000134762 | DSC3 | -8.373471517 | 1.35E-22 | -4.648868319 | 8.89E-12 | -13.02233984 | 8.107517749 | 9.71095217 | 9.049077186 | 2.600425781 | 2.38539787 | 4.45225155 | 4.886954834 |
| ENSG00000136877 | FPGS | -12.79956306 | 3.61E-13 | -0.224375912 | 0.849874923 | -13.02393897 | 10.00110988 | 8.487557258 | 7.934808822 | 2 | 2 | 8.745429394 | 8.967756908 |
| ENSG00000171298 | GAA | -13.33152331 | 5.46E-17 | 0.300230051 | 0.622741559 | -13.03129326 | 9.810482605 | 9.13702746 | 9.798878489 | 2 | 2 | 10.13451789 | 9.646206355 |
| ENSG00000179059 | ZFP42 | -3.282875429 | 0.019241842 | -9.748471479 | 8.05E-07 | -13.03134691 | 4.00722531 | 4.710564154 | 5.39244166 | 2.695456494 | 2.68924543 | 2 | 2 |
| ENSG00000104413 | ESRP1 | -3.913696628 | 0.095454435 | -9.120252558 | 0.000408188 | -13.03394919 | 5.330594134 | 3.076665209 | 3.369317972 | 2.498691189 | 2.14022021 | 2 | 2 |
| ENSG00000095574 | IKZF5 | -11.91929528 | 5.82E-10 | -1.116480837 | 0.421230824 | -13.03577611 | 9.2903348 | 7.572420273 | 6.203069917 | 2 | 2 | 7.310906894 | 6.895576665 |
| ENSG00000181626 | ANKRD62 | -3.579583143 | 0.048902337 | -9.461874563 | 2.12E-05 | -13.04145771 | 5.095959458 | 3.076665209 | 4.874587708 | 2.600425781 | 2.38539787 | 2 | 2 |
| ENSG00000141560 | FN3KRP | -12.35966775 | 8.26E-15 | -0.687893824 | 0.210610076 | -13.04756158 | 8.981092835 | 8.276714465 | 8.608698767 | 2 | 2 | 7.792277365 | 8.125161869 |
| ENSG00000105641 | SLC5A5 | -4.454697808 | 0.001222537 | -8.59390351 | 2.23E-07 | -13.04860132 | 5.70932728 | 3.686303414 | 6.958202216 | 2.695456494 | 2.77779907 | 2.048074405 | 2.059860178 |
| YAE1D1 | | 0.066412391 | 0.952060394 | -13.12149659 | 2.03E-14 | -13.05508419 | 8.39321512 | 6.602424674 | 8.35328246 | 8.067573917 | 8.02236724 | 2 | 2 |
| ENSG00000105619 | TFPT | -0.198527604 | 0.927391636 | -12.85907318 | 5.89E-09 | -13.05760078 | 8.696033109 | 7.701025996 | 3.369317972 | 7.574744049 | 7.47849652 | 2 | 2 |
| ENSG00000182973 | CNOT10 | -11.9416969 | 4.07E-08 | -1.117160033 | 0.547700965 | -13.05885694 | 4.00722531 | 8.376385542 | 9.116480286 | 2 | 2 | 7.200910877 | 7.074915275 |
| ENSG00000277936 | | -1.219553478 | 0.036653216 | -11.8412522 | 5.76E-24 | -13.06080568 | 9.624286623 | 8.806266215 | 8.845626171 | 8.022463952 | 7.8441483 | 2 | 2.117335275 |
| USMG5 | | 0.592707849 | 0.558267032 | -13.65915509 | 1.42E-15 | -13.06644724 | 9.358295023 | 7.60566235 | 7.934808822 | 9.107339324 | 9.087919 | 2 | 2 |
| ENSG00000241258 | CRCP | -12.75265474 | 1.12E-13 | -0.319059538 | 0.757887688 | -13.07171428 | 9.858207046 | 8.053983481 | 8.585098385 | 2 | 2 | 8.693476763 | 8.747121619 |
| ENSG00000115474 | KCNJ13 | -7.890320323 | 0.069608997 | -5.183913337 | 0.210403187 | -13.07423366 | 2 | 5.838937808 | 2 | 2 | 2 | 2.267001569 | 2.059860178 |
| ENSG00000151208 | DLG5 | 0.946619861 | 0.544427162 | -14.02422289 | 9.81E-13 | -13.07760303 | 8.892187472 | 9.770792087 | 5.772716771 | 9.785133 | 9.83940415 | 2 | 2 |

| | | | | | | | | | | | | | |
|---|---|---|---|---|---|---|---|---|---|---|---|---|---|
| METTL21B | | -0.164614766 | 0.867562397 | -12.92011309 | 1.11E-14 | -13.08472786 | 7.031698319 | 7.431218269 | 8.486635645 | 7.608714138 | 7.6318872 | 2 | 2 |
| ENSG00000170296 | GABARAP | -1.611393801 | 0.001986931 | -11.47611189 | 3.43E-89 | -13.08750569 | 12.52334676 | 11.631062 | 12.02204252 | 10.48331504 | 10.5088999 | 2.3070652 | 2.632958241 |
| FAM63B | | 0.104299992 | 0.886731176 | -13.19308775 | 1.42E-16 | -13.08878775 | 8.517314938 | 7.638155693 | 7.858115942 | 8.153770873 | 8.15227214 | 2 | 2 |
| TCEB3 | | 1.207145953 | 0.029205956 | -14.29825298 | 1.74E-19 | -13.09110702 | 9.344956677 | 9.405920729 | 8.511888463 | 10.34908243 | 10.3410349 | 2 | 2 |
| ENSG00000179520 | SLC17A8 | -4.606992163 | 0.317035976 | -8.487631592 | 0.055828012 | -13.09462376 | 5.095959458 | 2 | 2 | 2.141737246 | 2.14022021 | 2 | 2 |
| ENSG00000178171 | AMER3 | -4.619434376 | 0.132570607 | -8.500075522 | 0.005867842 | -13.1195099 | 4.467311984 | 2 | 4.058998049 | 2.141737246 | 2.14022021 | 2 | 2 |
| ENSG00000145526 | CDH18 | -2.538642158 | 0.180807042 | -10.5887568 | 2.93E-06 | -13.12739896 | 3.327711234 | 6.728495673 | 4.523675391 | 3.462030052 | 3.55551834 | 2 | 2 |
| WHSC1 | | 0.410874856 | 0.851584865 | -13.53932924 | 1.31E-09 | -13.12845438 | 4.00722531 | 7.572420273 | 9.647595056 | 8.836004565 | 8.75996988 | 2 | 2 |
| ENSG00000123178 | SPRYD7 | -11.90346413 | 2.62E-11 | -1.235825273 | 0.249275363 | -13.1392894 | 9.048471759 | 7.901717006 | 6.80318718 | 2 | 2 | 6.793610851 | 7.157719456 |
| ENSG00000112110 | MRPL18 | -12.86028827 | 2.44E-12 | -0.28044676 | 0.836549691 | -13.14073503 | 10.20680973 | 8.213429689 | 7.777116083 | 2 | 2 | 8.791168836 | 8.936163111 |
| ENSG00000089818 | NECAP1 | -12.64187708 | 2.82E-11 | -0.500405728 | 0.728705652 | -13.14228281 | 6.468621203 | 8.929529568 | 9.788583153 | 2 | 2 | 8.493696429 | 8.366492631 |
| ENSG00000132975 | GPR12 | -4.62776989 | 0.26661026 | -8.515304106 | 0.035498586 | -13.143074 | 2 | 4.442710129 | 4.058998049 | 2.270786764 | 2 | 2 | 2 |
| LOC730183 | | -3.23303716 | 0.237182775 | -9.919903902 | 0.000573235 | -13.15294106 | 4.00722531 | 2 | 6.203069917 | 2.695456494 | 2.86123014 | 2 | 2 |
| ENSG00000146938 | NLGN4X | -5.743979669 | 0.000186547 | -7.415074417 | 8.19E-07 | -13.15905409 | 7.33131396 | 8.451443573 | 8.631919287 | 3.921164147 | 2 | 2.656294034 | 2.374681325 |
| ENSG00000198176 | TFDP1 | -13.36288166 | 2.38E-16 | 0.203186476 | 0.795874263 | -13.15969519 | 9.410448395 | 9.04308532 | 10.22252928 | 2 | 2 | 9.682212468 | 9.992301965 |
| ENSG00000181396 | OGFOD3 | -12.73140247 | 6.60E-13 | -0.429351748 | 0.711063784 | -13.16075422 | 9.567995545 | 7.315436877 | 9.31604076 | 2 | 2 | 8.389095798 | 8.766899546 |
| ENSG00000114030 | KPNA1 | -13.04867229 | 2.65E-13 | -0.118548274 | 0.927047173 | -13.16722057 | 9.646202137 | 9.896993485 | 7.451345288 | 2 | 2 | 9.316273172 | 9.101944949 |
| ENSG00000166265 | CYYR1 | -8.254333141 | 5.24E-14 | -4.917134783 | 8.31E-07 | -13.17146792 | 9.064836028 | 7.669933291 | 9.926432745 | 2.868578001 | 2.14022021 | 4.757085024 | 4.201948172 |
| ENSG00000215301 | DDX3X | -14.60166651 | 7.61E-19 | 1.427485843 | 0.048671297 | -13.17418067 | 9.87686338 | 11.37322539 | 11.00131064 | 2 | 2 | 12.25835913 | 12.34728781 |
| ENSG00000177875 | CCDC184 | -5.977339088 | 2.17E-19 | -7.206894801 | 2.45E-28 | -13.18423389 | 8.910412251 | 9.338312924 | 10.49998282 | 4.205751501 | 4.05806467 | 3.316666208 | 3.261853426 |
| HDGFRP3 | | 0.415373786 | 0.54644592 | -13.60506532 | 2.07E-17 | -13.18969153 | 8.910412251 | 7.847167909 | 8.434763955 | 8.920805049 | 8.81332035 | 2 | 2 |
| ENSG00000173826 | KCNH6 | -7.311189766 | 0.00159118 | -5.879111206 | 0.000746591 | -13.19030097 | 4.815609471 | 5.303523747 | 5.39244166 | 2 | 2.14022021 | 2.34604626 | 2 |
| ENSG00000172932 | ANKRD13D | -12.63145204 | 1.25E-10 | -0.562257512 | 0.720810194 | -13.19370955 | 9.886101849 | 6.140560021 | 8.763901704 | 2 | 2 | 8.503966641 | 8.200230127 |
| ENSG00000147206 | NXF3 | -7.377794219 | 0.099954534 | -5.830216784 | 0.1713815 | -13.208011 | 2 | 2 | 5.39244166 | 2 | 2 | 2.094598369 | 2.059860178 |
| ENSG00000189409 | MMP23B | -5.655699077 | 1.54E-11 | -7.556847067 | 4.86E-19 | -13.21254614 | 8.797441463 | 7.35507212 | 7.225742779 | 3.09461492 | 3.2184422 | 2.526562374 | 2.277179783 |
| ENSG00000224859 | | -1.116934851 | 0.307515243 | -12.11087282 | 8.47E-12 | -13.22780767 | 7.790810999 | 6.728495673 | 5.772716771 | 5.860316854 | 5.99120031 | 2 | 2 |
| ENSG00000183347 | GBP6 | -3.134380253 | 0.162341892 | -10.10111709 | 5.93E-05 | -13.23549735 | 3.327711234 | 3.076665209 | 6.43197674 | 2.784612485 | 3.01487888 | 2 | 2 |
| ENSG00000164258 | NDUFS4 | -12.57619298 | 1.59E-12 | -0.66051748 | 0.560120837 | -13.23671046 | 9.771132748 | 8.414402626 | 7.502634626 | 2 | 2 | 8.328950294 | 8.077085525 |
| ENSG00000206240 | | -4.678493711 | 0.301217705 | -8.559133998 | 0.049513463 | -13.23762771 | 2 | 5.131603625 | 2 | 2.141737246 | 2.14022021 | 2 | 2 |
| ENSG00000092200 | RPGRIP1 | -7.871301313 | 0.022319086 | -5.367362785 | 0.083573371 | -13.2386641 | 7.218819781 | 2 | 2 | 2.141737246 | 2 | 2.267001569 | 2.466008509 |
| ENSG00000162669 | HFM1 | -5.0895306 | 0.132434625 | -8.149185267 | 0.018662618 | -13.23871587 | 2 | 2 | 6.073378139 | 2.141737246 | 2.2680111 | 2.048074405 | 2 |
| ENSG00000084693 | AGBL5 | -12.84303213 | 2.06E-15 | -0.412541922 | 0.550336725 | -13.25557405 | 8.517314938 | 9.624396729 | 9.014156722 | 2 | 2 | 8.659062802 | 8.773618611 |
| KIAA1462 | | 1.670783585 | 0.008361244 | -14.92697467 | 1.89E-20 | -13.25619108 | 10.28571918 | 9.733683991 | 9.014156722 | 11.44243908 | 11.4276648 | 2 | 2 |
| CCDC64B | CCDC64B | -5.90919196 | 0.299070637 | -7.347222912 | NA | -13.25641487 | 2 | 4.11352228 | 2 | 2 | 2 | 2 | 2 |
| ENSG00000095777 | MYO3A | -5.90919196 | 0.299070637 | -7.347222912 | NA | -13.25641487 | 2 | 4.11352228 | 2 | 2 | 2 | 2 | 2 |
| ENSG00000187135 | VSTM2B | -5.90919196 | 0.299070637 | -7.347222912 | NA | -13.25641487 | 2 | 4.11352228 | 2 | 2 | 2 | 2 | 2 |
| ENSG00000198502 | HLA-DRB5 | -5.90919196 | 0.299070637 | -7.347222912 | NA | -13.25641487 | 2 | 4.11352228 | 2 | 2 | 2 | 2 | 2 |
| ENSG00000198889 | DCAF12L1 | -5.90919196 | 0.299070637 | -7.347222912 | NA | -13.25641487 | 2 | 4.11352228 | 2 | 2 | 2 | 2 | 2 |
| ENSG00000261701 | HPR | -5.90919196 | 0.299070637 | -7.347222912 | NA | -13.25641487 | 2 | 4.11352228 | 2 | 2 | 2 | 2 | 2 |
| FAM132B | | -5.90919196 | 0.299070637 | -7.347222912 | NA | -13.25641487 | 2 | 4.11352228 | 2 | 2 | 2 | 2 | 2 |
| MGC57346-CRHR1 | | -5.90919196 | 0.299070637 | -7.347222912 | NA | -13.25641487 | 2 | 4.11352228 | 2 | 2 | 2 | 2 | 2 |
| RLTPR | | -5.90919196 | 0.299070637 | -7.347222912 | NA | -13.25641487 | 2 | 4.11352228 | 2 | 2 | 2 | 2 | 2 |
| ENSG00000105671 | DDX49 | -12.36623839 | 2.66E-06 | -0.903753128 | 0.733879184 | -13.26999152 | 9.613202746 | 8.717821467 | 2 | 2 | 2 | 7.718996793 | 7.804897508 |
| ENSG00000143374 | TARS2 | -12.34114446 | 1.63E-12 | -0.934580394 | 0.356359551 | -13.27572485 | 7.485143553 | 8.213429689 | 9.478569125 | 2 | 2 | 7.821097076 | 7.58443608 |
| ENSG00000108947 | EFNB3 | -7.526421059 | 4.32E-60 | -5.751249783 | 2.00E-44 | -13.27767084 | 10.87379128 | 10.42046463 | 10.88048583 | 4.034562184 | 3.33976443 | 5.227950776 | 5.080008298 |
| ENSG00000197408 | CYP2B6 | -5.19778721 | 0.105639361 | -8.081641931 | 0.007517278 | -13.27942914 | 4.00722531 | 3.076665209 | 3.369317972 | 2.141737246 | 2 | 2 | 2 |
| ENSG00000173212 | MAB21L3 | -4.201127538 | 3.85E-05 | -9.079281669 | 2.86E-10 | -13.28040921 | 5.866944937 | 5.946638857 | 7.029880454 | 2.695456494 | 3.33976443 | 2.094598369 | 2 |
| ENSG00000089558 | KCNH4 | -8.606301984 | 0.038063431 | -4.694710271 | 0.238242877 | -13.30101226 | 2 | 6.534109959 | 2 | 2 | 2 | 2.526562374 | 2.172608095 |
| ENSG00000171564 | FGB | -5.211748248 | 0.104948235 | -8.09560952 | 0.007458119 | -13.30735777 | 3.327711234 | 3.076665209 | 4.058998049 | 2.141737246 | 2 | 2 | 2 |
| ENSG00000234012 | | -0.087025891 | 0.953443586 | -13.22132257 | 1.49E-12 | -13.30834846 | 8.855032352 | 8.191702463 | 5.772716771 | 7.971157485 | 8.01327646 | 2 | 2 |
| DGCR14 | DGCR14 | -0.185604848 | 0.948155695 | -13.12340523 | 7.80E-07 | -13.30901008 | 2 | 6.951736442 | 9.301632639 | 7.671425626 | 7.91663606 | 2 | 2 |
| ENSG00000172404 | DNAJB7 | -7.217886898 | 0.016301152 | -6.0961952 | 0.026809473 | -13.3140821 | 3.327711234 | 3.686303414 | 4.523675391 | 2 | 2 | 2 | 2.117335275 |
| ENSG00000205086 | C2orf91 | -7.049850941 | 1.37E-05 | -6.274028666 | 7.98E-07 | -13.32387961 | 4.815609471 | 4.93638933 | 6.882775709 | 2.141737246 | 2.14022021 | 2.139668783 | 2.326754077 |
| ENSG00000177551 | NHLH2 | -7.815564298 | 0.006677491 | -5.510274084 | 0.028934847 | -13.32583838 | 2 | 4.11352228 | 5.39244166 | 2 | 2 | 2.139668783 | 2.117335275 |
| ENSG00000107485 | GATA3 | -2.172403245 | 6.75E-06 | -11.17879028 | 4.36E-30 | -13.35119353 | 8.892187472 | 8.717821467 | 9.385998894 | 6.998639523 | 6.77267312 | 2.139668783 | 2 |
| ENSG00000262826 | | -0.345465442 | 0.802167143 | -13.00684046 | 4.09E-13 | -13.3523059 | 7.031698319 | 9.328390744 | 10.14394518 | 8.83470036 | 9.08900105 | 2 | 2.059860178 |
| ENSG00000204576 | PRR3 | -0.959633238 | 0.438991892 | -12.40569542 | 1.05E-11 | -13.36532866 | 8.107517749 | 7.145017454 | 5.59507254 | 6.252910529 | 6.44777477 | 2 | 2 |
| ENSG00000237515 | SHISA9 | -5.361255648 | 1.54E-08 | -8.010819823 | 2.34E-12 | -13.37207547 | 5.866944937 | 6.228547981 | 5.772716771 | 2.3892346 | 2.49394783 | 2.048074405 | 2.117335275 |
| ENSG00000133665 | DYDC2 | -7.117149644 | 0.195586526 | -6.257707052 | 0.241041529 | -13.3748567 | 2 | 2 | 5.156626401 | 2 | 2 | 2.094598369 | 2 |
| ENSG00000145700 | ANKRD31 | -6.33890392 | 0.027618711 | -7.037515692 | 0.011808867 | -13.37641961 | 2 | 2 | 8.460932916 | 2.141737246 | 2.68924543 | 2.34604626 | 2.225841198 |
| ENSG00000163875 | MEAF6 | -12.68723021 | 8.83E-15 | -0.694371216 | 0.333586444 | -13.38160142 | 9.219014269 | 9.28800521 | 8.143093325 | 2 | 2 | 8.43671169 | 8.114615835 |
| ENSG00000106025 | TSPAN12 | -6.339285328 | 2.52E-21 | -7.043211472 | 5.34E-27 | -13.3824968 | 10.51143645 | 10.08551947 | 10.64969256 | 4.387512679 | 4.40247105 | 4.340871764 | 3.100202476 |
| ENSG00000132581 | SDF2 | -13.05714281 | 2.70E-14 | -0.327688651 | 0.74998849 | -13.38483146 | 10.18451821 | 8.655694736 | 8.561105505 | 2 | 2 | 8.898729828 | 9.120511546 |
| ENSG00000232569 | | 0.081057017 | 0.94947311 | -13.47923331 | 4.12E-14 | -13.39817629 | 8.517314938 | 6.535001748 | 8.99637432 | 8.29513208 | 8.52113334 | 2 | 2 |
| ENSG00000161664 | ASB16 | -8.371875973 | 1.07E-11 | -5.03478882 | 4.50E-16 | -13.40666479 | 6.653098118 | 7.393647515 | 7.225742779 | 2.141737246 | 2.14022021 | 3.1504466 | 2.885414003 |
| TMEM55A | | -0.197454611 | 0.942460221 | -13.21116938 | 2.27E-07 | -13.40862399 | 9.015175615 | 8.147241016 | 2 | 7.883054517 | 7.86451142 | 2 | 2 |
| ENSG00000185651 | UBE2L3 | -13.60797543 | 4.25E-16 | 0.192094079 | 0.838840227 | -13.41588135 | 10.32700827 | 10.13758676 | 8.742726175 | 2 | 2 | 9.91066097 | 10.22907185 |
| WBSCR17 | | -2.65998083 | 0.635961632 | -10.75799121 | 0.036435838 | -13.41797204 | 7.218819781 | 2 | 2 | 4.268917409 | 2 | 2 | 2 |
| ENSG00000273274 | ZBTB8B | -8.444355487 | 0.00194296 | -4.986197436 | 0.031633742 | -13.43055292 | 2 | 4.710564154 | 5.930865648 | 2 | 2 | 2.267001569 | 2.277179783 |
| PARK2 | | -3.013797638 | 0.001914878 | -10.41715801 | 3.25E-09 | -13.43095564 | 5.330594134 | 5.457120953 | 5.39244166 | 3.023133209 | 3.2184422 | 2 | 2 |
| ENSG00000129158 | SERGEF | -11.76330276 | 7.42E-06 | -1.673202893 | 0.502368069 | -13.43650565 | 9.097018025 | 7.954278496 | 2 | 2 | 2 | 6.15433139 | 6.664466987 |
| ENSG00000160753 | RUSC1 | -13.22329334 | 2.35E-15 | -0.21432229 | 0.815853642 | -13.43761563 | 9.800745303 | 8.395519306 | 9.897982195 | 2 | 2 | 9.373699808 | 9.204129769 |
| ENSG00000256453 | DND1 | -2.678863452 | 0.275620582 | -10.76656258 | 4.54E-05 | -13.44542604 | 6.89194353 | 5.131603625 | 2 | 3.227757282 | 3.7864959 | 2 | 2 |
| ENSG00000111237 | VPS29 | -13.15756618 | 5.95E-17 | -0.294410882 | 0.571387495 | -13.45197706 | 9.635285994 | 9.04308532 | 9.577565424 | 2 | 2 | 9.039153561 | 9.246605829 |
| ENSG00000185043 | CIB1 | -12.8352089 | 1.36E-16 | -0.619119918 | 0.099357865 | -13.45432881 | 9.204317094 | 9.067150531 | 9.083172337 | 2 | 2 | 8.383836906 | 8.617426278 |
| ENSG00000076258 | FMO4 | -7.213680525 | 5.19E-18 | -6.243966311 | 3.06E-15 | -13.45764684 | 10.58702944 | 8.655694736 | 9.301632639 | 3.162721474 | 3.39680241 | 3.517841605 | 4.254128754 |
| CCDC53 | CCDC53 | -0.18209407 | 0.726199435 | -13.27636273 | 1.54E-17 | -13.4584568 | 7.940205782 | 8.169643011 | 8.266922072 | 7.834291698 | 8.06258142 | 2 | 2 |
| ENSG00000077800 | FKBP6 | -5.29251548 | 0.242094498 | -8.178131881 | 0.05484371 | -13.47064736 | 4.467311984 | 2 | 3.369317972 | 2.141737246 | 2 | 2 | 2 |
| ENSG00000133606 | MKRN1 | -13.66668064 | 1.01E-16 | 0.195163271 | 0.818699747 | -13.47151737 | 9.740899611 | 9.170736405 | 10.57976717 | 2 | 2 | 10.15198119 | 10.12826404 |
| ENSG00000137776 | SLTM | -13.17535572 | 1.08E-14 | -0.299169983 | 0.76623592 | -13.47452571 | 8.674867122 | 8.834582582 | 10.28245311 | 2 | 2 | 9.123606729 | 9.194564734 |
| ENSG00000132622 | HSPA12B | -9.586962431 | 0.000361141 | -3.897195551 | 0.094043218 | -13.48415798 | 7.159102577 | 5.303523747 | 2 | 2 | 2 | 3.12825509 | 2.817676617 |
| C21ORF2 | C21ORF2 | -1.244363451 | 0.610039424 | -12.24822983 | 9.55E-07 | -13.49259329 | 7.624139768 | 7.761270542 | 2 | 5.797438367 | 6.06344184 | 2 | 2 |
| ENSG00000132424 | PNISR | -13.62782197 | 2.21E-17 | 0.127188939 | 0.859499684 | -13.50063304 | 10.24321353 | 10.06773758 | 9.212024683 | 2 | 2 | 10.1332997 | 9.926748282 |
| ENSG00000233348 | | -0.026271299 | 0.979836893 | -13.47875154 | 7.40E-26 | -13.50502284 | 9.448362284 | 10.77481051 | 11.41675873 | 10.6993508 | 10.7574572 | 2.048074405 | 2.059860178 |
| ENSG00000178971 | CTC1 | -12.60110027 | 1.12E-11 | -0.90555008 | 0.478299157 | -13.50665035 | 8.444143041 | 7.051611315 | 9.878698603 | 2 | 2 | 8.135306947 | 7.831973788 |
| METTL10 | | -0.670044914 | 0.511816142 | -12.84182284 | 8.12E-14 | -13.51186775 | 6.468621203 | 7.761270542 | 8.325066185 | 6.841288255 | 7.23436042 | 2 | 2 |
| ENSG00000204469 | PRRC2A | 1.47440345 | 0.22593371 | -14.99175078 | 2.07E-16 | -13.51734733 | 10.29268328 | 10.34373336 | 7.502634626 | 11.2911419 | 11.3161167 | 2 | 2 |
| ENSG00000140090 | SLC24A4 | -6.770615967 | 9.87E-06 | -6.752907173 | 1.04E-06 | -13.52352314 | 6.563806279 | 3.686303414 | 7.451345288 | 2.141737246 | 2.38539787 | 2.267001569 | 2.277179783 |
| ENSG00000028310 | BRD9 | -12.73727835 | 6.54E-12 | -0.789303848 | 0.541300866 | -13.5265822 | 7.159102577 | 8.57407404 | 10.01740919 | 2 | 2 | 8.102245405 | 8.366492631 |
| ENSG00000171657 | GPR82 | -5.367180514 | 1.95E-06 | -8.167944363 | 8.64E-11 | -13.53512488 | 6.468621203 | 4.93638933 | 7.225742779 | 2.695456494 | 2.49394783 | 2.139668783 | 2.059860178 |
| PRKCDBP | | 0.625518912 | 0.357361029 | -14.16216082 | 1.09E-18 | -13.53664191 | 9.031919741 | 8.317412869 | 9.465702942 | 9.600249849 | 9.65954302 | 2 | 2 |
| ENSG00000105398 | SULT2A1 | -6.053707398 | 0.284753947 | -7.491740154 | NA | -13.54544755 | 3.327711234 | 3.686303414 | 2 | 2 | 2 | 2 | 2 |
| ENSG00000185291 | IL3RA | -6.053707398 | 0.284753947 | -7.491740154 | NA | -13.54544755 | 3.327711234 | 3.686303414 | 2 | 2 | 2 | 2 | 2 |
| ENSG00000188931 | CFAP126 | -6.053707398 | 0.284753947 | -7.491740154 | NA | -13.54544755 | 3.327711234 | 3.686303414 | 2 | 2 | 2 | 2 | 2 |
| FAM134A | | 0.751093792 | 0.050385816 | -14.29987726 | 1.69E-20 | -13.54878347 | 8.910412251 | 9.267380431 | 9.227351203 | 9.86268459 | 9.92126619 | 2 | 2 |
| ENSG00000203618 | GP1BB | -3.605323692 | 0.140291258 | -9.947425181 | 0.000198824 | -13.55274887 | 6.009025997 | 4.710564154 | 2 | 2.784612485 | 2.49394783 | 2 | 2 |
| MESDC1 | | -0.195814437 | 0.80683022 | -13.35923513 | 1.68E-16 | -13.55504957 | 8.107517749 | 7.572420273 | 8.742726175 | 7.975897842 | 8.06478301 | 2 | 2 |
| ENSG00000277361 | | -5.767422297 | 0.012348082 | -7.789661105 | 0.000788654 | -13.5570834 | 3.327711234 | 6.602424674 | 2 | 2.270786764 | 2.14022021 | 2.048074405 | 2.059860178 |
| ENSG00000228605 | | -0.403432102 | 0.479099956 | -13.15747584 | 6.53E-17 | -13.56090794 | 8.257427002 | 8.101365653 | 7.600047325 | 7.620871236 | 7.61696151 | 2 | 2 |
| ENSG00000205212 | CCDC144NL | -7.714817204 | 0.052218839 | -5.855722478 | 0.116476087 | -13.57053968 | 2 | 4.710564154 | 4.874587708 | 2 | 2 | 2.183373728 | 2 |
| ENSG00000039537 | C6 | -7.795128177 | 0.069954594 | -5.782872663 | 0.155516119 | -13.57800084 | 2 | 2 | 5.772716771 | 2 | 2 | 2.094598369 | 2.117335275 |
| ENSG00000166573 | GALR1 | -5.352754032 | 0.345459763 | -8.231010601 | 0.122968358 | -13.58376463 | 2 | 2 | 4.874587708 | 2 | 2.14022021 | 2 | 2 |
| ENSG00000197079 | KRT35 | -4.352464363 | 0.306644136 | -9.233082652 | 0.026495931 | -13.58554702 | 2 | 2 | 5.772716771 | 2.270786764 | 2.2680111 | 2 | 2 |
| ENSG00000198797 | BRINP2 | -4.352464363 | 0.306644136 | -9.233082652 | 0.026495931 | -13.58554702 | 2 | 2 | 5.772716771 | 2.270786764 | 2.2680111 | 2 | 2 |
| ENSG00000145782 | ATG12 | -13.15173497 | 6.96E-16 | -0.440586495 | 0.547884639 | -13.59232146 | 9.233563228 | 8.820493869 | 9.999667074 | 2 | 2 | 9.128156703 | 8.849779308 |
| ENSG00000135253 | KCP | -7.526784484 | 1.02E-11 | -6.070520547 | 4.10E-09 | -13.59730503 | 8.075571721 | 7.431218269 | 9.868959282 | 2.695456494 | 2.68924543 | 3.851628042 | 2.817676617 |
| ENSG00000170289 | CNGB3 | -6.792958407 | 0.219632896 | -6.833743268 | 0.207921012 | -13.62670167 | 2 | 2 | 4.874587708 | 2 | 2 | 2.048074405 | 2 |

| | | | | | | | | | | | | | |
|---|---|---|---|---|---|---|---|---|---|---|---|---|---|
| ZUFSP | | -0.680129093 | 0.819250325 | -12.94904445 | 2.35E-06 | -13.62917354 | 9.247966931 | 5.946638857 | 2 | 7.210554257 | 7.0774796 | 2 | 2 |
| ZAK | | 0.306681779 | 0.476871248 | -13.93942295 | 1.68E-19 | -13.63274117 | 8.797441463 | 8.929529568 | 8.608698767 | 9.142549916 | 9.03948698 | 2 | 2 |
| ENSG00000267795 | SMIM22 | -5.88365399 | 0.18239431 | -7.756818372 | 0.068882273 | -13.64047236 | 6.816637977 | 2 | 2 | 2 | 2.38539787 | 2 | 2.117335275 |
| C1ORF27 | C1ORF27 | 0.28962736 | 0.622871051 | -13.93042441 | 8.43E-19 | -13.64079705 | 9.189468652 | 8.557180776 | 8.486635645 | 9.09649754 | 9.03500236 | 2 | 2 |
| C22ORF29 | C22ORF29 | -0.676253082 | 0.67805098 | -12.97061253 | 6.04E-11 | -13.64686561 | 4.815609471 | 8.147241016 | 8.536706852 | 7.19843704 | 7.14145882 | 2 | 2 |
| ENSG00000099617 | EFNA2 | -7.139318474 | 3.27E-05 | -6.507953873 | 1.74E-05 | -13.64727235 | 4.467311984 | 6.140560021 | 7.502634626 | 2.3892346 | 2 | 2.139668783 | 2.466008509 |
| ENSG00000186188 | FFAR4 | -8.576603099 | 0.003221255 | -5.082835142 | 0.045885615 | -13.65943824 | 5.095959458 | 2 | 5.930865648 | 2 | 2 | 2.42098416 | 2.117335275 |
| ENSG00000170989 | S1PR1 | -4.815036411 | 1.35E-22 | -8.846644732 | 3.33E-69 | -13.66168114 | 11.79588449 | 10.95649913 | 11.52614724 | 6.652857793 | 6.75642009 | 3.600766115 | 3.012034821 |
| C15ORF59 | C15ORF59 | -2.416554496 | 0.000657371 | -11.25138614 | 1.01E-11 | -13.66794064 | 6.138360429 | 6.04685525 | 6.322058544 | 4.070463721 | 4.05806467 | 2 | 2 |
| C6ORF1 | C6ORF1 | -0.206284109 | 0.924421511 | -13.46278677 | 9.23E-10 | -13.66907087 | 9.497412313 | 7.761270542 | 4.058998049 | 8.083034979 | 8.14188248 | 2 | 2 |
| ENSG00000185324 | CDK10 | -13.15852617 | 5.37E-14 | -0.515032055 | 0.635985655 | -13.67355822 | 10.32020829 | 8.213429689 | 8.99637432 | 2 | 2 | 8.909342392 | 8.947596938 |
| ZCCHC11 | | -0.027731978 | 0.988931662 | -13.64703244 | 7.15E-11 | -13.67476442 | 4.467311984 | 8.968383114 | 9.132850258 | 8.43768081 | 8.50338773 | 2 | 2 |
| ENSG00000185231 | MC2R | -4.105025039 | 0.113850795 | -9.570617064 | 0.000597178 | -13.6756421 | 2 | 4.11352228 | 5.772716771 | 2.3892346 | 2.38539787 | 2 | 2 |
| ENSG00000008838 | MED24 | -13.21901299 | 1.60E-16 | -0.46341329 | 0.462478098 | -13.68242629 | 9.966689162 | 9.256956408 | 9.132850258 | 2 | 2 | 8.834217923 | 9.219795239 |
| ENSG00000184302 | SIX6 | -5.401822757 | 0.079965857 | -8.280733647 | 0.004388858 | -13.6825564 | 3.327711234 | 4.11352228 | 3.369317972 | 2 | 2.14022021 | 2 | 2 |
| ENSG00000142494 | SLC47A1 | -9.741728272 | 1.39E-19 | -3.941742978 | 7.96E-10 | -13.68347125 | 9.112843761 | 8.376385542 | 9.478569125 | 2.3892346 | 2 | 5.494014735 | 4.982847849 |
| ENSG00000274768 | | -0.735204362 | 0.507489288 | -12.95154607 | 1.61E-13 | -13.68675043 | 7.940205782 | 6.311476754 | 8.460932916 | 7.157295126 | 7.02868983 | 2 | 2 |
| ENSG00000134398 | ERN2 | -7.88169826 | 0.003560394 | -5.815713758 | 0.00911236 | -13.69741202 | 2 | 4.11352228 | 7.098165388 | 2.141737246 | 2 | 2.183373728 | 2.374681325 |
| ENSG00000142920 | AZIN2 | -11.44686907 | 5.76E-11 | -2.253045802 | 0.014208037 | -13.69991487 | 8.042902253 | 6.464272742 | 8.206335842 | 2 | 2 | 5.829320042 | 5.306932076 |
| C17ORF62 | C17ORF62 | 0.367019755 | 0.677535675 | -14.06991694 | 1.63E-17 | -13.70289718 | 9.410448395 | 9.067150531 | 7.858115942 | 9.255049401 | 9.3074018 | 2 | 2 |
| ENSG00000166897 | ELFN2 | -8.507976653 | 0.002889025 | -5.203716627 | 0.035760415 | -13.71169328 | 2 | 4.11352228 | 6.203069917 | 2 | 2 | 2.267001569 | 2.225841198 |
| ENSG00000276879 | | -0.210278977 | 0.720334587 | -13.50623021 | 8.34E-18 | -13.71650919 | 8.653385977 | 7.979858146 | 8.380957451 | 8.083034979 | 8.21705076 | 2 | 2 |
| ENSG00000236759 | | 0.08354393 | 0.970415686 | -13.80462737 | 1.12E-09 | -13.72108344 | 9.031919741 | 9.396454291 | 3.369317972 | 8.720710586 | 8.75316018 | 2 | 2 |
| ENSG00000185829 | ARL17A | -0.133839918 | 0.936681124 | -13.60578406 | 3.22E-12 | -13.73962398 | 5.532353208 | 8.607278467 | 9.272377397 | 8.458152491 | 8.17894021 | 2 | 2 |
| ENSG00000261893 | | -0.678638453 | 0.52775508 | -13.06457567 | 5.60E-14 | -13.74321412 | 6.468621203 | 8.522788974 | 8.076950877 | 7.178012736 | 7.33629421 | 2 | 2 |
| ENSG00000164107 | HAND2 | -3.486107573 | 1.15E-17 | -10.25967088 | 5.90E-56 | -13.74577845 | 9.602033055 | 9.799806309 | 9.132850258 | 6.147608921 | 6.11534392 | 2.183373728 | 2.225841198 |
| ENSG00000253457 | SMIM18 | -6.52614098 | 0.066974888 | -7.23731629 | 0.032955861 | -13.76345727 | 2 | 2 | 6.882775709 | 2.141737246 | 2.14022021 | 2 | 2.172608095 |
| ENSG00000131471 | AOC3 | -10.70248582 | 4.66E-05 | -3.063981332 | 0.188337529 | -13.76646716 | 3.327711234 | 4.442710129 | 8.486635645 | 2 | 2 | 4.875056259 | 3.128433649 |
| ENSG00000163702 | IL17RC | -12.75921795 | 2.00E-16 | -1.009208698 | 0.004268256 | -13.76842665 | 9.015175615 | 8.929529568 | 9.180874355 | 2 | 2 | 8.008923105 | 8.0807022 |
| ENSG00000175287 | PHYHD1 | -5.801580116 | 1.57E-19 | -7.966901167 | 2.58E-35 | -13.76848128 | 11.00678643 | 9.726146444 | 9.553445883 | 4.739373666 | 4.63360314 | 3.215043717 | 3.071407821 |
| ENSG00000168883 | USP39 | -13.2934196 | 1.37E-17 | -0.477008462 | 0.257956922 | -13.77042806 | 9.771132748 | 9.308339287 | 9.61300511 | 2 | 2 | 9.048046506 | 9.150504437 |
| ENSG00000144045 | DQX1 | -8.579162176 | 0.013324274 | -5.197452812 | 0.100093385 | -13.77661499 | 7.904282243 | 2 | 2 | 2 | 2.14022021 | 2.858551861 | 2.326754077 |
| FAM109A | | -0.451586965 | 0.830558266 | -13.33791714 | 1.14E-09 | -13.7895041 | 8.836089686 | 3.686303414 | 8.677268317 | 7.77569021 | 7.72113133 | 2 | 2 |
| MINA | | -0.755327424 | 0.144739115 | -13.03495125 | 1.07E-16 | -13.79027868 | 7.904282243 | 7.60566235 | 8.143093325 | 7.206526483 | 7.10341274 | 2 | 2 |
| HDGFRP2 | | 0.186631764 | 0.877879848 | -13.98008848 | 1.06E-15 | -13.79345671 | 9.751047882 | 8.147241016 | 7.777116083 | 9.042162213 | 8.98356585 | 2 | 2 |
| ENSG00000277101 | | -2.055750797 | 0.3418564 | -11.74414397 | 9.52E-07 | -13.79989477 | 3.327711234 | 4.710564154 | 8.042706615 | 4.825935136 | 4.68029874 | 2 | 2 |
| ENSG00000106631 | MYL7 | -8.50956081 | 0.006112518 | -5.300232775 | 0.055557322 | -13.80979359 | 2 | 4.11352228 | 6.203069917 | 2 | 2 | 2.384001718 | 2.059860178 |
| ENSG00000099715 | PCDH11Y | -3.340271952 | 0.001558837 | -10.46984689 | 5.98E-09 | -13.81011884 | 5.70932728 | 5.595927442 | 4.874587708 | 3.09461492 | 2.86123014 | 2 | 2 |
| ENSG00000171956 | FOXB1 | -6.187415475 | 0.271909254 | -7.625449901 | 0.158721188 | -13.81286538 | 4.00722531 | 3.076665209 | 2 | 2 | 2 | 2 | 2 |
| ENSG00000229859 | PGA3 | -6.187415475 | 0.271909254 | -7.625449901 | 0.158721188 | -13.81286538 | 4.00722531 | 3.076665209 | 2 | 2 | 2 | 2 | 2 |
| DSCR3 | DSCR3 | 0.406119041 | 0.641103815 | -14.22110952 | 7.15E-18 | -13.81499048 | 8.367060731 | 8.655694736 | 9.778213818 | 9.494378497 | 9.44644299 | 2 | 2 |
| ENSG00000157765 | SLC34A2 | -7.156853465 | 0.044631453 | -6.664841554 | 0.037819082 | -13.82169502 | 2 | 2 | 6.534109959 | 2.141737246 | 2 | 2.139668783 | 2.059860178 |
| ENSG00000124207 | CSE1L | -13.47961673 | 1.01E-17 | -0.356600542 | 0.487618515 | -13.83621727 | 9.966689162 | 9.896993485 | 9.330306409 | 2 | 2 | 9.490570501 | 9.313452824 |
| ENSG00000145703 | IQGAP2 | -8.28118198 | 2.66E-14 | -5.559895014 | 4.22E-08 | -13.84107699 | 8.138771685 | 10.51535307 | 9.149036566 | 2.947924294 | 2.38539787 | 4.82764224 | 3.600986722 |
| ENSG00000130182 | ZSCAN10 | -8.257009427 | 0.007711404 | -5.590550153 | 0.043076236 | -13.84755958 | 5.866944937 | 3.686303414 | 3.369317972 | 2 | 2 | 2.3070652 | 2 |
| ENSG00000124171 | PARD6B | -7.579293108 | 1.54E-35 | -6.27728074 | 2.19E-31 | -13.85657385 | 9.247966931 | 10.42512749 | 9.735974718 | 3.289987342 | 3.01487888 | 3.917810786 | 4.105828626 |
| ENSG00000278243 | | -2.4971412 | 1.55E-05 | -11.36077855 | 7.04E-40 | -13.85791975 | 10.45046729 | 9.181799937 | 9.963514551 | 7.467753569 | 7.49817903 | 2.139668783 | 2.117335275 |
| ENSG00000277808 | | -0.894630207 | 0.453256583 | -12.96772705 | 5.28E-13 | -13.86235726 | 8.797441463 | 6.666836837 | 7.098165388 | 6.993973425 | 6.91614081 | 2 | 2 |
| ENSG00000225950 | NTF4 | -8.354143543 | 1.54E-07 | -5.511148677 | 4.66E-05 | -13.86529222 | 8.892187472 | 4.93638933 | 6.80318718 | 2.141737246 | 2.2680111 | 2.717032197 | 3.36036153 |
| ENSG00000069702 | TGFBR3 | -7.669132351 | 1.05E-49 | -6.204385754 | 1.02E-33 | -13.87351811 | 12.90140438 | 13.71594413 | 13.89650122 | 5.930370745 | 6.04571775 | 6.908747388 | 7.758233206 |
| ENSG00000009765 | IYD | -5.738930757 | 0.012035229 | -8.136794873 | 0.000893267 | -13.87572563 | 4.467311984 | 3.076665209 | 5.59507254 | 2.270786764 | 2 | 2 | 2.059860178 |
| ENSG00000224399 | | 1.198598149 | 0.064643779 | -15.09365288 | 5.76E-21 | -13.89505473 | 9.344956677 | 9.7032946 | 10.50627919 | 11.06373609 | 11.193456 | 2 | 2 |
| ENSG00000126856 | PRDM7 | -7.009217066 | 0.000505431 | -6.887504838 | 4.20E-06 | -13.8967219 | 4.467311984 | 4.442710129 | 5.59507254 | 2 | 2.14022021 | 2.094598369 | 2.059860178 |
| ENSG00000198626 | RYR2 | -6.232557318 | 0.267451577 | -7.670592308 | 0.155664455 | -13.90314963 | 2 | 3.076665209 | 4.058998049 | 2 | 2 | 2 | 2 |
| ENSG00000178722 | C5orf64 | -4.513156642 | 0.145868949 | -9.390247654 | 0.003187497 | -13.9034043 | 3.327711234 | 2 | 5.772716771 | 2.141737246 | 2.38539787 | 2 | 2 |
| ENSG00000170965 | PLAC1 | -7.152339744 | 0.000737017 | -6.757816737 | 0.000295983 | -13.91015648 | 2 | 7.189546432 | 5.156626401 | 2.141737246 | 2.14022021 | 2.139668783 | 2.225841198 |
| ENSG00000188738 | FSIP2 | -6.66840998 | 0.048767966 | -7.254927537 | 0.024153443 | -13.92333752 | 3.327711234 | 2 | 6.958202216 | 2.270786764 | 2 | 2.183373728 | 2 |
| ENSG00000262911 | | -2.902286552 | 0.46731457 | -11.03034587 | 0.003707085 | -13.93263242 | 7.485143553 | 2 | 2 | 3.515101962 | 3.60476711 | 2 | 2 |
| ENSG00000146839 | ZAN | -8.118681973 | 0.053640085 | -5.814677517 | 0.143469849 | -13.93335949 | 2 | 2 | 6.073378139 | 2 | 2 | 2.139668783 | 2.117335275 |
| ENSG00000134352 | IL6ST | -14.45905833 | 1.03E-17 | 0.525578583 | 0.569078707 | -13.93347975 | 9.829762239 | 11.51993408 | 10.30429804 | 2 | 2 | 11.30104558 | 11.21817336 |
| ENSG00000109072 | VTN | -7.325258166 | 6.91E-46 | -6.614351664 | 4.48E-50 | -13.93960983 | 9.957954117 | 9.883504052 | 9.31604076 | 3.09461492 | 3.33976443 | 3.483289517 | 3.841670744 |
| ENSG00000142684 | ZNF593 | -1.051825233 | 0.400917152 | -12.89800631 | 1.94E-12 | -13.94983154 | 8.737455672 | 7.232742069 | 6.203069917 | 6.70517998 | 6.76727577 | 2 | 2 |
| TLDC1 | | 0.268551794 | 0.717276749 | -14.22165853 | 9.12E-19 | -13.95310674 | 9.590776211 | 8.451443573 | 8.941276636 | 9.248208205 | 9.41572435 | 2 | 2 |
| ENSG00000074276 | CDHR2 | -9.257380034 | 0.000203506 | -4.699283745 | 0.019733629 | -13.95666378 | 7.53297911 | 2 | 7.646393253 | 2 | 2.14022021 | 3.336159116 | 2.918126211 |
| FAM65A | | 0.649355228 | 0.375015921 | -14.60720526 | 1.80E-19 | -13.95785003 | 9.624286623 | 8.590771778 | 9.859153765 | 10.07142123 | 10.1222557 | 2 | 2 |
| ENSG00000167037 | SGSM1 | -6.857044899 | 0.000357712 | -7.11254174 | 8.00E-05 | -13.96958664 | 5.095959458 | 4.710564154 | 7.896971941 | 2.270786764 | 2.2680111 | 2.42098416 | 2 |
| ENSG00000187533 | PRR27 | -4.753989863 | 0.125204611 | -9.217338062 | 0.0036002 | -13.97132793 | 3.327711234 | 2 | 5.59507254 | 2.141737246 | 2.2680111 | 2 | 2 |
| ENSG00000164076 | CAMKV | -5.798736043 | 0.033319681 | -8.175347995 | 0.003846635 | -13.97408404 | 3.327711234 | 2 | 6.073378139 | 2.141737246 | 2.14022021 | 2 | 2.059860178 |
| ENSG00000106484 | MEST | -8.372547702 | 1.25E-68 | -5.609638722 | 4.67E-32 | -13.98218642 | 14.10617399 | 14.13775586 | 14.74413016 | 6.130704144 | 6.01871653 | 8.209845962 | 9.163352825 |
| ENSG00000168066 | SF1 | -14.23770904 | 3.99E-20 | 0.24790574 | 0.55564101 | -13.9898033 | 10.64259676 | 10.40638494 | 10.48730704 | 2 | 2 | 10.56183151 | 10.93805674 |
| ENSG00000105371 | ICAM4 | -1.808584946 | 0.325172387 | -12.18882026 | 2.28E-08 | -13.99740521 | 8.042902253 | 7.002537973 | 3.369317972 | 5.334065616 | 5.39133044 | 2 | 2 |
| ENSG00000119185 | ITGB1BP1 | -13.4066322 | 6.44E-16 | -0.59921296 | 0.46274242 | -14.00584516 | 9.143983763 | 9.13702746 | 10.39531398 | 2 | 2 | 9.018187664 | 9.160792285 |
| ENSG00000138115 | CYP2C8 | -4.566504753 | 0.278762742 | -9.443357308 | 0.022493273 | -14.00986206 | 2 | 5.946638857 | 2 | 2.141737246 | 2.38539787 | 2 | 2 |
| ENSG00000169231 | THBS3 | -12.52439286 | 1.16E-13 | -1.487778717 | 0.068525886 | -14.01217157 | 9.567995545 | 8.169643011 | 8.236946975 | 2 | 2 | 7.303499023 | 7.386797235 |
| ENSG00000134207 | SYT6 | -8.780631461 | 0.002190909 | -5.234758886 | 0.033320023 | -14.01539035 | 7.790810999 | 5.838937808 | 2 | 2 | 2.14022021 | 3.10571689 | 2.059860178 |
| ENSG00000168546 | GFRA2 | -8.635947176 | 0.002952151 | -5.380343607 | 0.031427993 | -14.01629078 | 7.940205782 | 3.076665209 | 2 | 2.141737246 | 2 | 2.560105256 | 2.592996859 |
| ENSG00000081985 | IL12RB2 | -8.591904186 | 1.75E-05 | -5.426242068 | 5.40E-05 | -14.01814625 | 4.00722531 | 5.131603625 | 5.59507254 | 2 | 2 | 2.225793514 | 2.225841198 |
| ENSG00000070761 | CFAP20 | -12.73816904 | 1.09E-15 | -1.28020112 | 0.012470755 | -14.01837016 | 9.33149386 | 9.018611864 | 8.631919287 | 2 | 2 | 7.550870243 | 7.939455664 |
| ENSG00000162344 | FGF19 | -7.061235183 | 0.103636128 | -6.957483202 | 0.10104293 | -14.01871839 | 4.815609471 | 2 | 3.369317972 | 2 | 2 | 2 | 2.059860178 |
| ENSG00000234906 | APOC2 | -7.732461207 | 0.154760847 | -6.292981812 | 0.230555284 | -14.02544302 | 5.70932728 | 2 | 2 | 2 | 2 | 2.139668783 | 2 |
| ENSG00000152904 | GGPS1 | -12.60083552 | 1.41E-15 | -1.424849411 | 0.001960035 | -14.02568493 | 8.981092835 | 8.505280662 | 9.116480286 | 2 | 2 | 7.569519177 | 7.388257299 |
| ENSG00000106078 | COBL | -6.875053582 | 0.002972167 | -7.151272395 | 0.001015228 | -14.02632598 | 7.435667355 | 5.72254323 | 2 | 2.3892346 | 2 | 2.225793514 | 2.117335275 |
| ENSG00000248098 | BCKDHA | -3.256806157 | 1.52E-21 | -10.76982474 | 1.30E-97 | -14.0266309 | 10.76282024 | 10.62073637 | 11.10036136 | 7.527093191 | 7.69009211 | 2.384001718 | 2.277179783 |
| ENSG00000137948 | BRDT | -5.07440467 | 0.088829476 | -8.961032753 | 0.002934332 | -14.03543742 | 3.327711234 | 3.076665209 | 5.156626401 | 2.270786764 | 2 | 2 | 2 |
| ENSG00000153406 | NMRAL1 | -0.739014975 | 0.78336925 | -13.30942351 | 2.38E-07 | -14.04843848 | 9.174465797 | 8.124485672 | 2 | 7.419851705 | 7.45854177 | 2 | 2 |
| ENSG00000181092 | ADIPOQ | -4.424162868 | 0.030240619 | -9.627995708 | 4.88E-05 | -14.05215858 | 4.00722531 | 3.076665209 | 5.772716771 | 2.3892346 | 2.2680111 | 2 | 2 |
| ENSG00000152430 | BOLL | -7.533907016 | 0.02131845 | -6.524496432 | 0.032648562 | -14.05840345 | 2 | 3.076665209 | 5.39244166 | 2 | 2 | 2.048074405 | 2.059860178 |
| ENSG00000106331 | PAX4 | -6.311501034 | 0.260096091 | -7.749537009 | 0.15053059 | -14.06103804 | 4.467311984 | 2 | 2 | 2 | 2 | 2 | 2 |
| ENSG00000150275 | PCDH15 | -6.311501034 | 0.260096091 | -7.749537009 | 0.15053059 | -14.06103804 | 4.467311984 | 2 | 2 | 2 | 2 | 2 | 2 |
| ENSG00000173578 | XCR1 | -6.311501034 | 0.260096091 | -7.749537009 | 0.15053059 | -14.06103804 | 4.467311984 | 2 | 2 | 2 | 2 | 2 | 2 |
| ENSG00000262068 | | -6.311501034 | 0.260096091 | -7.749537009 | 0.15053059 | -14.06103804 | 4.467311984 | 2 | 2 | 2 | 2 | 2 | 2 |
| ENSG00000135100 | HNF1A | -6.295107047 | 0.060284857 | -7.768395628 | 0.016033391 | -14.06350268 | 2 | 3.076665209 | 5.59507254 | 2 | 2.14022021 | 2.048074405 | 2 |
| ENSG00000133454 | MYO18B | -6.296088423 | 0.027564901 | -7.770488809 | 0.005130285 | -14.06657723 | 2 | 2 | 7.225742779 | 2.141737246 | 2.2680111 | 2.094598369 | 2.059860178 |
| ENSG00000112511 | PHF1 | 0.405436725 | 0.784005445 | -14.47485979 | 1.17E-14 | -14.06942306 | 10.47516499 | 8.101365653 | 7.777116083 | 9.737775223 | 9.70683001 | 2 | 2 |
| BAI3 | BAI3 | -6.316786837 | 0.259792504 | -7.754822878 | 0.150262362 | -14.07160971 | 2 | 4.442710129 | 2 | 2 | 2 | 2 | 2 |
| ENSG00000170178 | HOXD12 | -6.316786837 | 0.259792504 | -7.754822878 | 0.150262362 | -14.07160971 | 2 | 4.442710129 | 2 | 2 | 2 | 2 | 2 |
| ENSG00000171557 | FGG | -6.316786837 | 0.259792504 | -7.754822878 | 0.150262362 | -14.07160971 | 2 | 4.442710129 | 2 | 2 | 2 | 2 | 2 |
| ENSG00000185038 | MROH2A | -9.609100087 | 0.016756486 | -4.462630594 | 0.253035563 | -14.07173068 | 2 | 2 | 7.502634626 | 2 | 2 | 2.3070652 | 3.071407821 |
| ENSG00000158941 | CCAR2 | -13.43243078 | 3.92E-13 | -0.64361168 | 0.626190802 | -14.07604246 | 10.81212061 | 8.687092496 | 8.266922072 | 2 | 2 | 9.096715824 | 9.048465472 |
| ENSG00000139219 | COL2A1 | -6.100985492 | 0.069715937 | -7.978140109 | 0.014446357 | -14.0791256 | 2 | 8.004992147 | 2 | 2.3892346 | 2.38539787 | 2 | 2.225841198 |
| FAM60A | | -0.551873472 | 0.437489824 | -13.52910957 | 4.75E-17 | -14.08098304 | 7.624139768 | 8.639735838 | 8.65477198 | 7.826439377 | 7.84926609 | 2 | 2 |
| ENSG00000132591 | ERAL1 | -13.13949605 | 1.38E-10 | -0.943410876 | 0.573181246 | -14.08290693 | 5.70932728 | 9.936721131 | 9.999667074 | 2 | 2 | 8.260460366 | 8.680159264 |
| ENSG00000109205 | ODAM | -6.332665187 | 0.258590042 | -7.770701427 | 0.149193138 | -14.10336661 | 4.00722531 | 2 | 3.369317972 | 2 | 2 | 2 | 2 |
| ENSG00000185873 | TMPRSS11B | -6.332665187 | 0.258590042 | -7.770701427 | 0.149193138 | -14.10336661 | 4.00722531 | 2 | 3.369317972 | 2 | 2 | 2 | 2 |
| ENSG00000269404 | SPIB | -4.320593116 | 0.012187376 | -9.784150252 | 5.93E-06 | -14.10474337 | 3.327711234 | 5.303523747 | 5.156626401 | 2.270786764 | 2.49394783 | 2 | 2 |
| HN1L | | -0.012484548 | 0.989907129 | -14.09403604 | 2.22E-17 | -14.10652059 | 7.790810999 | 9.277729677 | 9.301632639 | 8.941551548 | 8.91566441 | 2 | 2 |
| ENSG00000124772 | CPNE5 | -7.423746325 | 0.002373085 | -6.68507495 | 0.004716908 | -14.10882128 | 2 | 3.076665209 | 9.049077186 | 2.3892346 | 2.2680111 | 2.717032197 | 2.277179783 |

| | | | | | | | | | | | | | |
|---|---|---|---|---|---|---|---|---|---|---|---|---|---|
| ENSG00000164796 | CSMD3 | -7.117158698 | 0.195586526 | -6.994938778 | 0.193494022 | -14.11209748 | 2 | 2 | 5.156626401 | 2 | 2 | 2 | 2.059860178 |
| ENSG00000143740 | SNAP47 | -13.15471154 | 1.54E-15 | -0.973651914 | 0.191295729 | -14.12836345 | 10.08373159 | 9.159587375 | 8.721235202 | 2 | 2 | 8.485536406 | 8.455540488 |
| ENSG00000179284 | DAND5 | -7.335211515 | 4.32E-05 | -6.795975234 | 5.38E-06 | -14.13118675 | 5.095959458 | 4.442710129 | 7.285536013 | 2 | 2.2680111 | 2.225793514 | 2.172608095 |
| ENSG00000184571 | PIWIL3 | -5.631910168 | 0.180807042 | -8.51039148 | 0.032038249 | -14.14230165 | 4.467311984 | 2 | 4.058998049 | 2 | 2.14022021 | 2 | 2 |
| ENSG00000237693 | IRGM | -5.136413417 | 0.195269534 | -9.010219612 | 0.020431994 | -14.14663303 | 4.467311984 | 2 | 4.874587708 | 2 | 2.2680111 | 2 | 2 |
| ENSG00000129221 | AIPL1 | -3.973038858 | 0.009111513 | -10.17691385 | 7.16E-07 | -14.1499527 | 4.00722531 | 4.710564154 | 6.073378139 | 2.695456494 | 2.49394783 | 2 | 2 |
| ENSG00000172461 | FUT9 | -5.132190322 | 0.203546074 | -9.01969419 | 0.022575955 | -14.15188451 | 4.00722531 | 2 | 5.156626401 | 2.270786764 | 2 | 2 | 2 |
| ENSG00000186335 | SLC36A2 | -4.965132895 | 0.000153054 | -9.191413818 | 1.11E-06 | -14.15654671 | 5.330594134 | 4.442710129 | 6.43197674 | 2.3892346 | 2.49394783 | 2 | 2.059860178 |
| ENSG00000174231 | PRPF8 | 0.903723139 | 0.128591494 | -15.06668007 | 2.17E-21 | -14.16295693 | 9.556468883 | 10.41578665 | 9.565556058 | 10.78692561 | 10.8304401 | 2 | 2 |
| ENSG00000072182 | ASIC4 | -8.821317541 | 0.000718231 | -5.351264156 | 0.012057282 | -14.1725817 | 7.975256505 | 5.131603625 | 2 | 2 | 2.14022021 | 2.775316226 | 2.509591799 |
| ENSG00000204574 | ABCF1 | 0.62818542 | 0.45794139 | -14.80598736 | 2.83E-19 | -14.17780194 | 9.940323746 | 10.03760521 | 8.536706852 | 10.2683035 | 10.2798081 | 2 | 2 |
| ENSG00000122694 | GLIPR2 | -13.02488182 | 1.16E-13 | -1.158805807 | 0.256746052 | -14.18368763 | 10.22876205 | 8.255926468 | 8.631919287 | 2 | 2 | 8.156020677 | 8.1631842 |
| ENSG00000163817 | SLC6A20 | -6.359240203 | 0.00830029 | -7.825703885 | 0.00072327 | -14.18494409 | 3.327711234 | 4.11352228 | 5.156626401 | 2.141737246 | 2 | 2.048074405 | 2 |
| BAI1 | BAI1 | -6.374248881 | 0.254759301 | -7.81228564 | 0.146527599 | -14.18653452 | 2 | 2 | 4.523675391 | 2 | 2 | 2 | 2 |
| ENSG00000090402 | SI | -6.374248881 | 0.254759301 | -7.81228564 | 0.146527599 | -14.18653452 | 2 | 2 | 4.523675391 | 2 | 2 | 2 | 2 |
| ENSG00000173976 | RAX2 | -6.374248881 | 0.254759301 | -7.81228564 | 0.146527599 | -14.18653452 | 2 | 2 | 4.523675391 | 2 | 2 | 2 | 2 |
| ENSG00000177535 | OR2B11 | -6.374248881 | 0.254759301 | -7.81228564 | 0.146527599 | -14.18653452 | 2 | 2 | 4.523675391 | 2 | 2 | 2 | 2 |
| ENSG00000196169 | KIF19 | -6.374248881 | 0.254759301 | -7.81228564 | 0.146527599 | -14.18653452 | 2 | 2 | 4.523675391 | 2 | 2 | 2 | 2 |
| ENSG00000197849 | OR8G1 | -6.374248881 | 0.254759301 | -7.81228564 | 0.146527599 | -14.18653452 | 2 | 2 | 4.523675391 | 2 | 2 | 2 | 2 |
| ENSG00000204278 | TMEM235 | -6.374248881 | 0.254759301 | -7.81228564 | 0.146527599 | -14.18653452 | 2 | 2 | 4.523675391 | 2 | 2 | 2 | 2 |
| KIAA1024L | | -6.374248881 | 0.254759301 | -7.81228564 | 0.146527599 | -14.18653452 | 2 | 2 | 4.523675391 | 2 | 2 | 2 | 2 |
| ENSG00000146215 | CRIP3 | -7.963749969 | 0.041788425 | -6.230397844 | 0.089897497 | -14.19414781 | 2 | 5.131603625 | 4.874587708 | 2 | 2 | 2 | 2.172608095 |
| ENSG00000103460 | TOX3 | -5.153483515 | 0.004642354 | -9.056703084 | 6.26E-06 | -14.2101866 | 4.467311984 | 7.35507212 | 6.073378139 | 3.09461492 | 2 | 2.048074405 | 2.059860178 |
| MKL2 | | -0.729808455 | 0.786127952 | -13.48817505 | 1.59E-07 | -14.2179835 | 2 | 9.04308532 | 8.784770914 | 7.587189552 | 7.65837038 | 2 | 2 |
| ENSG00000169084 | DHRSX | 0.023989968 | 0.97625994 | -14.24874417 | 9.15E-19 | -14.22475421 | 9.678463567 | 8.732943944 | 8.631919287 | 9.088859521 | 9.14729768 | 2 | 2 |
| ENSG00000119121 | TRPM6 | -6.872347801 | 1.57E-05 | -7.353033792 | 6.74E-08 | -14.22538159 | 4.815609471 | 4.93638933 | 6.629488387 | 2.141737246 | 2.14022021 | 2.139668783 | 2.059860178 |
| C19ORF43 | C19ORF43 | 0.31116038 | 0.620673635 | -14.54257451 | 3.69E-20 | -14.23141413 | 8.87372952 | 9.328390744 | 9.809100877 | 9.662587523 | 9.7285167 | 2 | 2 |
| ENSG00000274348 | | -5.676957393 | 0.312833791 | -8.555216825 | 0.106808076 | -14.23217422 | 2 | 2 | 5.156626401 | 2 | 2.14022021 | 2 | 2 |
| ENSG00000273761 | | 1.110839328 | 0.389718246 | -15.34707919 | 6.74E-17 | -14.23623986 | 11.04027452 | 10.17128279 | 7.896971941 | 11.28351448 | 11.307311 | 2 | 2 |
| ENSG00000228896 | | -0.222099596 | 0.757468139 | -14.01845808 | 1.94E-18 | -14.24055768 | 8.631580145 | 9.377333006 | 8.380957451 | 8.627383409 | 8.66448499 | 2 | 2 |
| MGEA5 | | 0.897085534 | 0.227537839 | -15.14055467 | 1.36E-20 | -14.24346914 | 9.015175615 | 10.22039562 | 10.36791565 | 10.90937116 | 10.841718 | 2 | 2 |
| ENSG00000138795 | LEF1 | -12.22464565 | 6.96E-13 | -2.02647135 | 0.014389895 | -14.251117 | 9.064836028 | 7.538394143 | 8.561105505 | 2 | 2 | 6.86244555 | 6.110033886 |
| ENSG00000175416 | CLTB | -14.2012623 | 1.06E-15 | -0.052809054 | 0.967312407 | -14.25407135 | 10.82658808 | 8.747909548 | 10.99236731 | 2 | 2 | 10.0143146 | 10.74572502 |
| ENSG00000148377 | IDI2 | -5.187613553 | 0.230018039 | -9.068258036 | 0.031238942 | -14.25587159 | 2 | 5.595927442 | 2 | 2.141737246 | 2.14022021 | 2 | 2 |
| ENSG00000169894 | MUC3A | -8.200514991 | 0.007915488 | -6.05667295 | 0.028736671 | -14.25718794 | 4.00722531 | 3.076665209 | 5.772716771 | 2 | 2 | 2 | 2.225841198 |
| ENSG00000144460 | NYAP2 | -5.607446158 | 0.001761963 | -8.652867946 | 5.24E-05 | -14.2603141 | 5.330594134 | 3.076665209 | 5.772716771 | 2.141737246 | 2.2680111 | 2.048074405 | 2 |
| C10ORF76 | C10ORF76 | -0.322824657 | 0.646065476 | -13.93903468 | 3.17E-18 | -14.26185934 | 9.189468652 | 8.124485672 | 8.865353883 | 8.480007165 | 8.45552985 | 2 | 2 |
| ENSG00000276023 | DUSP14 | -0.123610629 | 0.916429292 | -14.14462222 | 2.89E-16 | -14.26823285 | 9.895281535 | 8.255926468 | 8.076950877 | 8.797695539 | 8.93625225 | 2 | 2 |
| ENSG00000276262 | | 0.229416734 | 0.556481113 | -14.5047901 | 3.18E-21 | -14.27537337 | 9.233563228 | 9.479495904 | 9.31604076 | 9.562930393 | 9.59092963 | 2 | 2 |
| ENSG00000121211 | MND1 | -4.697602833 | 0.052698164 | -9.584753606 | 0.000289395 | -14.28235644 | 4.815609471 | 3.686303414 | 5.156626401 | 2.498691189 | 2 | 2 | 2 |
| LOC100506127 | | -2.298672601 | 0.026904691 | -11.98847475 | 1.90E-11 | -14.28714735 | 7.031698319 | 5.595927442 | 7.451345288 | 5.022023783 | 4.43005853 | 2 | 2 |
| ENSG00000004939 | SLC4A1 | -4.704554992 | 0.005114776 | -9.587761432 | 4.75E-06 | -14.29231642 | 4.00722531 | 4.93638933 | 4.874587708 | 2.3892346 | 2.14022021 | 2 | 2 |
| XRCC6BP1 | | -6.42963078 | 0.250146839 | -7.86766823 | 0.143441027 | -14.29729901 | 3.327711234 | 4.11352228 | 2 | 2 | 2 | 2 | 2 |
| ENSG00000174775 | HRAS | -0.665222545 | 0.762165237 | -13.63297395 | 2.24E-09 | -14.2981965 | 9.497412313 | 8.414402626 | 3.369317972 | 7.783822874 | 7.8720741 | 2 | 2 |
| ENSG00000083099 | LYRM2 | -13.1824408 | 1.37E-16 | -1.12160777 | 0.040313458 | -14.30404857 | 9.473095744 | 9.842262059 | 8.922968322 | 2 | 2 | 8.317962987 | 8.384893621 |
| ATP5H | ATP5H | 0.134093222 | 0.900045869 | -14.44356965 | 1.23E-17 | -14.30947643 | 10.1076228 | 8.717821467 | 8.460932916 | 9.442237963 | 9.40011621 | 2 | 2 |
| ENSG00000112031 | MTRF1L | -11.76326854 | 6.24E-06 | -2.55384678 | 0.280522158 | -14.31711532 | 8.855032352 | 8.376385542 | 2 | 2 | 2 | 5.4183033 | 5.772630506 |
| ENSG00000129197 | RPAIN | -12.7561083 | 5.75E-15 | -1.573851337 | 0.016521712 | -14.32995997 | 9.624286623 | 8.557180776 | 8.699419242 | 2 | 2 | 7.340163764 | 7.622152309 |
| ENSG00000225452 | | -0.577759306 | 0.518953771 | -13.7617705 | 1.52E-16 | -14.3395298 | 9.064836028 | 7.503546079 | 8.845626171 | 7.978262192 | 8.10170499 | 2 | 2 |
| ENSG00000165496 | RPL10L | -4.941079555 | 0.246947394 | -9.404217843 | 0.024667973 | -14.3452974 | 2 | 2 | 5.930865648 | 2.141737246 | 2.2680111 | 2 | 2 |
| ENSG00000168913 | ENHO | -5.06033404 | 2.33E-05 | -9.29082057 | 3.03E-07 | -14.35115461 | 5.095959458 | 6.389896633 | 5.156626401 | 2.3892346 | 2.49394783 | 2 | 2.059860178 |
| FAM127C | | -0.418679626 | 0.74561264 | -13.93501074 | 1.11E-14 | -14.35369037 | 9.820154628 | 7.73146271 | 7.691296561 | 8.376256872 | 8.36195019 | 2 | 2 |
| ATP5C1 | ATP5C1 | 0.234071156 | 0.616184132 | -14.59952796 | 3.56E-21 | -14.3654568 | 9.128497779 | 9.574900212 | 9.577565424 | 9.659644277 | 9.6919516 | 2 | 2 |
| ENSG00000255633 | MTRNR2L9 | 1.260937129 | 0.050999384 | -15.63460297 | 2.10E-22 | -14.37366584 | 11.05263482 | 10.23108619 | 9.878698603 | 11.71437628 | 11.7503284 | 2 | 2 |
| C19ORF60 | C19ORF60 | -0.697619923 | 0.52966791 | -13.68580611 | 5.14E-15 | -14.38342603 | 8.946185348 | 6.899080544 | 8.978369997 | 7.915493556 | 7.77859536 | 2 | 2 |
| ENSG00000146147 | MLIP | -8.010943983 | 0.005097116 | -6.374823614 | 0.0121807 | -14.3857676 | 2 | 4.93638933 | 5.156626401 | 2 | 2 | 2.048074405 | 2.117335275 |
| ENSG00000095906 | NUBP2 | -13.37098513 | 2.87E-16 | -1.02330037 | 0.141867727 | -14.3942855 | 10.15425007 | 8.88960062 | 9.647595056 | 2 | 2 | 8.803037464 | 8.445757593 |
| ENSG00000007062 | PROM1 | -4.820861753 | 0.024772728 | -9.574125249 | 7.64E-06 | -14.394987 | 8.009475818 | 4.11352228 | 8.65477198 | 4.299492648 | 2 | 2.139668783 | 2.059860178 |
| ENSG00000101489 | CELF4 | -9.098000185 | 0.003622044 | -5.299899812 | 0.058994081 | -14.3979 | 2 | 3.076665209 | 6.958202216 | 2 | 2 | 2.34604626 | 2.326754077 |
| ENSG00000069011 | PITX1 | -9.530622271 | 1.53E-49 | -4.875014526 | 6.54E-52 | -14.4056368 | 10.05943807 | 9.969011754 | 10.14394518 | 2.141737246 | 2.68924543 | 5.284230775 | 5.373314945 |
| ENSG00000175728 | C11orf44 | -6.540806075 | 0.005639608 | -7.878213232 | 0.000556851 | -14.41901931 | 4.00722531 | 3.686303414 | 6.629488387 | 2.270786764 | 2 | 2 | 2.117335275 |
| ENSG00000111344 | RASAL1 | -8.541398506 | 0.110944274 | -5.885666017 | 0.257880462 | -14.42706452 | 6.468621203 | 2 | 2 | 2 | 2 | 2.3070652 | 2 |
| C14ORF159 | C14ORF159 | -0.637267069 | 0.412535896 | -13.79621605 | 2.90E-17 | -14.43348312 | 9.27634997 | 8.317412869 | 8.042706615 | 8.001694993 | 8.0314011 | 2 | 2 |
| ENSG00000120055 | C10orf95 | -7.63341989 | 0.000128216 | -6.806241439 | 7.29E-07 | -14.43966133 | 4.815609471 | 6.228547981 | 4.874587708 | 2 | 2.14022021 | 2.183373728 | 2.059860178 |
| ENSG00000130540 | SULT4A1 | -7.918588658 | 9.89E-05 | -6.528364208 | 0.000440485 | -14.44695287 | 2 | 6.389896633 | 8.536706852 | 2.141737246 | 2.2680111 | 2.592885947 | 2.374681325 |
| ENSG00000139624 | CERS5 | -13.87944916 | 7.09E-19 | -0.572959182 | 0.211244657 | -14.45240834 | 10.45046729 | 9.863030423 | 10.10298738 | 2 | 2 | 9.48539663 | 9.682361453 |
| ENSG00000172638 | EFEMP2 | -15.19076238 | 3.64E-22 | 0.733895414 | 0.133545445 | -14.45686696 | 11.05263482 | 11.45757623 | 11.79401137 | 2 | 2 | 12.08332398 | 12.30626723 |
| ENSG00000142615 | CELA2A | -6.941485794 | 0.110411514 | -7.521344054 | 0.064830738 | -14.46282985 | 2 | 6.311476754 | 2 | 2.141737246 | 2 | 2.094598369 | 2 |
| FAM46A | | 1.077822517 | 0.099949746 | -15.54368658 | 3.52E-22 | -14.46586406 | 10.97683282 | 9.994333773 | 9.926432745 | 11.47607929 | 11.4412857 | 2 | 2 |
| ENSG00000149534 | MS4A2 | -5.293072206 | 0.093600232 | -9.173716631 | 0.003771401 | -14.46678884 | 2 | 3.076665209 | 5.59507254 | 2.141737246 | 2.14022021 | 2 | 2 |
| ENSG00000132915 | PDE6A | -6.229649612 | 7.67E-08 | -8.251493144 | 1.49E-11 | -14.48114276 | 5.70932728 | 5.838937808 | 7.502634626 | 2.270786764 | 2.49394783 | 2.094598369 | 2.117335275 |
| ENSG00000065485 | PDIA5 | -13.20482224 | 2.48E-14 | -1.276820651 | 0.18084322 | -14.48164289 | 10.26462289 | 8.395519306 | 9.196533591 | 2 | 2 | 7.800194554 | 8.545966004 |
| ENSG00000106066 | CPVL | -9.458097166 | 2.26E-28 | -5.027260586 | 9.12E-15 | -14.48535775 | 10.20680973 | 10.30378101 | 9.212024683 | 2.270786764 | 2.59489879 | 5.498346994 | 4.624919101 |
| ENSG00000169567 | HINT1 | -14.15117912 | 6.77E-18 | -0.341170456 | 0.674002095 | -14.49234958 | 11.08107026 | 9.756063186 | 10.11127249 | 2 | 2 | 10.15489989 | 10.01893736 |
| C10ORF11 | C10ORF11 | -5.803758376 | 0.176505928 | -8.689208628 | 0.032138031 | -14.492967 | 5.095959458 | 3.076665209 | 2 | 2.141737246 | 2 | 2 | 2 |
| ENSG00000109625 | CPZ | -7.974450263 | 2.09E-48 | -6.51859453 | 1.64E-38 | -14.49304479 | 11.50046327 | 10.85302421 | 10.78531698 | 3.349643765 | 3.90918929 | 4.470006665 | 5.050803619 |
| ENSG00000112559 | MDFI | -9.57413003 | 1.09E-17 | -4.921879054 | 6.18E-08 | -14.49600908 | 8.816849489 | 8.317412869 | 10.63827116 | 2.3892346 | 2.2680111 | 5.056402493 | 4.720747915 |
| ENSG00000185823 | NPAP1 | -4.01805327 | 0.009929837 | -10.4825017 | 4.78E-07 | -14.50055497 | 5.330594134 | 3.686303414 | 6.322058544 | 2.600425781 | 2.77779907 | 2 | 2 |
| ENSG00000169436 | COL22A1 | -10.46128628 | 0.000161309 | -4.042909604 | 0.099862782 | -14.50419589 | 2 | 7.189546432 | 7.502634626 | 2 | 2 | 3.943456545 | 2.466008509 |
| ENSG00000010361 | FUZ | -11.79714539 | 3.49E-08 | -2.709716671 | 0.099493727 | -14.50686206 | 9.384607364 | 6.844429755 | 5.156626401 | 2 | 2 | 5.367107572 | 5.611957759 |
| ENSG00000123728 | RAP2C | -13.15654538 | 8.50E-16 | -1.351437201 | 0.045513428 | -14.50798258 | 9.992581382 | 9.308339287 | 8.742726175 | 2 | 2 | 8.238615655 | 7.945418846 |
| ENSG00000167874 | TMEM88 | -6.823950656 | 5.91E-20 | -7.696517034 | 2.91E-26 | -14.52046769 | 10.48127392 | 9.030900487 | 9.478569125 | 3.566290607 | 3.55551834 | 3.336159116 | 2.671842496 |
| ENSG00000135778 | NTPCR | -13.10609433 | 2.48E-16 | -1.42092595 | 0.000770859 | -14.52702028 | 9.810482605 | 8.862353853 | 9.344432377 | 2 | 2 | 7.938932725 | 8.02265879 |
| ENSG00000107105 | ELAVL2 | -6.483105542 | 0.003077009 | -8.045766637 | 0.00017766 | -14.52887218 | 7.031698319 | 6.602424674 | 2 | 2.270786764 | 2.2680111 | 2.183373728 | 2 |
| ENSG00000160801 | PTH1R | -10.64483158 | 5.54E-22 | -3.895065173 | 2.97E-06 | -14.53989675 | 11.19324461 | 10.16572116 | 8.865353883 | 2.270786764 | 2.2680111 | 6.446574529 | 6.607941737 |
| FAM134B | | -3.938877033 | 0.329259892 | -10.63135102 | 0.007111387 | -14.57022806 | 7.096806373 | 2 | 2 | 3.09461492 | 2.38539787 | 2 | 2 |
| ENSG00000146477 | SLC22A3 | -6.788645509 | 0.000650953 | -7.782599483 | 6.24E-05 | -14.57124499 | 9.699577239 | 7.638155693 | 2 | 2.947924294 | 2.68924543 | 2.717032197 | 2.172608095 |
| FBXO18 | | 0.068479378 | 0.966538801 | -14.64133722 | 2.44E-14 | -14.57285784 | 10.12333326 | 9.876711741 | 6.534109959 | 9.5407064 | 9.56472327 | 2 | 2 |
| ENSG00000105388 | CEACAM5 | -4.056571036 | 0.000112827 | -10.52310257 | 3.86E-09 | -14.57967361 | 5.532353208 | 5.131603625 | 5.772716771 | 2.784612485 | 2.59489879 | 2 | 2 |
| ENSG00000188869 | TMC3 | -8.156454012 | 0.033350515 | -6.423648687 | 0.073787407 | -14.5801027 | 2 | 4.93638933 | 5.39244166 | 2 | 2 | 2 | 2.172608095 |
| PPAPDC3 | | -6.571764211 | 0.237552448 | -8.009803437 | 0.134877558 | -14.58156765 | 2 | 3.686303414 | 4.058998049 | 2 | 2 | 2 | 2 |
| ENSG00000112695 | COX7A2 | -13.94715525 | 1.67E-18 | -0.656257344 | 0.245121437 | -14.60341259 | 10.66423795 | 9.77810053 | 10.09465441 | 2 | 2 | 9.615976512 | 9.526007748 |
| ENSG00000019186 | CYP24A1 | -5.864440266 | 0.296385835 | -8.742701355 | 0.09817373 | -14.60714162 | 2 | 5.303523747 | 2 | 2 | 2.14022021 | 2 | 2 |
| ENSG00000204655 | MOG | -5.070852807 | 0.002572364 | -9.537908922 | 3.48E-06 | -14.60876173 | 4.815609471 | 4.11352228 | 4.874587708 | 2.270786764 | 2.14022021 | 2 | 2 |
| ENSG00000236826 | | -1.367998144 | 0.003213414 | -13.24280334 | 2.71E-17 | -14.61080148 | 8.228665467 | 8.234834549 | 7.777116083 | 6.739036112 | 6.78874509 | 2 | 2 |
| ENSG00000153310 | FAM49B | -12.7603331 | 3.18E-15 | -1.857089521 | 0.001845751 | -14.61742262 | 9.317904228 | 8.317412869 | 9.301632639 | 2 | 2 | 7.229857522 | 7.20156326 |
| ENSG00000104381 | GDAP1 | -10.78915608 | 1.89E-06 | -3.832578405 | 0.030281135 | -14.62173448 | 3.327711234 | 7.467835385 | 7.818184403 | 2 | 2 | 3.1504466 | 4.161535617 |
| ENSG00000168291 | PDHB | -14.11610866 | 7.15E-20 | -0.508877974 | 0.174047291 | -14.62498663 | 10.38028415 | 10.28850895 | 10.50627919 | 2 | 2 | 9.718407995 | 10.03720074 |
| ENSG00000119866 | BCL11A | -6.798860851 | 1.02E-13 | -7.839891856 | 2.62E-18 | -14.63875271 | 9.159305284 | 9.443177293 | 10.9968459 | 4.105493497 | 3.39680241 | 3.483289517 | 2.671842496 |
| ENSG00000034239 | EFCAB1 | -7.530200908 | 0.088259606 | -7.114480642 | 0.082535125 | -14.64468155 | 2 | 2 | 6.882775709 | 2 | 2.14022021 | 2.183373728 | 2 |
| ENSG00000187173 | LCE2A | -5.088927893 | 0.228197547 | -9.556970578 | 0.021560447 | -14.64589847 | 2 | 2 | 6.073378139 | 2.270786764 | 2.14022021 | 2 | 2 |
| ENSG00000142405 | NLRP12 | -6.568462167 | 0.000391481 | -8.081158265 | 1.00E-05 | -14.64962043 | 5.095959458 | 4.442710129 | 6.203069917 | 2 | 2.2680111 | 2.094598369 | 2 |
| ENSG00000276583 | | -1.764321195 | 0.37459299 | -12.89210488 | 1.11E-08 | -14.65642607 | 9.174465797 | 5.131603625 | 5.156626401 | 5.930370745 | 6.16544343 | 2 | 2 |
| ENSG00000188817 | SNTN | -5.885624626 | 0.150440253 | -8.770864238 | 0.023476231 | -14.65648886 | 4.815609471 | 2 | 4.058998049 | 2.141737246 | 2 | 2 | 2 |
| ENSG00000064601 | CTSA | -14.14126232 | 4.52E-19 | -0.517346091 | 0.357318652 | -14.65860841 | 10.36053552 | 9.956182102 | 10.81605395 | 2 | 2 | 9.808169474 | 9.992301965 |
| ENSG00000254585 | MAGEL2 | -6.954278537 | 0.000178451 | -7.716531771 | 1.94E-05 | -14.67081031 | 9.015175615 | 8.376385542 | 2 | 2.868578001 | 2.38539787 | 2.267001569 | 2.551896972 |
| ENSG00000104231 | ZFAND1 | -12.82274637 | 5.44E-15 | -1.852267733 | 0.005439997 | -14.6750141 | 9.304185368 | 8.255926468 | 9.491321578 | 2 | 2 | 7.393470826 | 7.159430626 |

| | | | | | | | | | | | | | |
|---|---|---|---|---|---|---|---|---|---|---|---|---|---|
| TCEB1 | | -0.329902758 | 0.728428075 | -14.3626911 | 8.41E-18 | -14.69259386 | 9.957954117 | 8.356994601 | 8.825624954 | 8.866956935 | 8.89354047 | 2 | 2 |
| ENSG00000171462 | DLK2 | -8.226103791 | 9.52E-14 | -6.476392256 | 4.63E-11 | -14.70249605 | 9.771132748 | 6.78762695 | 8.978369997 | 2.3892346 | 2.59489879 | 3.10571689 | 3.383972333 |
| C6ORF7 | C6ORF7 | -6.634016091 | 0.232251697 | -8.072056094 | 0.131367278 | -14.70607218 | 2 | 4.710564154 | 2 | 2 | 2 | 2 | 2 |
| ENSG00000147081 | AKAP4 | -6.634016091 | 0.232251697 | -8.072056094 | 0.131367278 | -14.70607218 | 2 | 4.710564154 | 2 | 2 | 2 | 2 | 2 |
| ENSG00000234224 | TMEM229A | -6.634016091 | 0.232251697 | -8.072056094 | 0.131367278 | -14.70607218 | 2 | 4.710564154 | 2 | 2 | 2 | 2 | 2 |
| ENSG00000166091 | CMTM5 | -8.903303657 | 0.029202849 | -5.808833203 | 0.132363963 | -14.71213686 | 6.816637977 | 2 | 2 | 2 | 2 | 2.3070652 | 2.117335275 |
| ENSG00000170043 | TRAPPC1 | -13.89984743 | 3.70E-19 | -0.814544884 | 0.037552988 | -14.71439232 | 10.12333326 | 10.13189344 | 10.27509734 | 2 | 2 | 9.136171672 | 9.566552047 |
| PROSC | | -0.804143832 | 0.387134584 | -13.91440642 | 1.65E-16 | -14.71855025 | 9.497412313 | 8.522788974 | 7.734844356 | 7.908072101 | 8.02689124 | 2 | 2 |
| ENSG00000106436 | MYL10 | -6.649135118 | 0.104612178 | -8.087137268 | 0.040810128 | -14.73627239 | 4.00722531 | 3.076665209 | 3.369317972 | 2 | 2 | 2 | 2 |
| ENSG00000177138 | FAM9B | -6.649135118 | 0.104612178 | -8.087137268 | 0.040810128 | -14.73627239 | 4.00722531 | 3.076665209 | 3.369317972 | 2 | 2 | 2 | 2 |
| ENSG00000165194 | PCDH19 | -6.405811888 | 3.84E-09 | -8.336691452 | 1.01E-14 | -14.74250334 | 7.435667355 | 10.12617756 | 10.59756881 | 4.415698455 | 3.15372745 | 2.831336763 | 2.746601469 |
| ENSG00000169696 | ASPSCR1 | -12.6882173 | 5.24E-16 | -2.06364047 | 1.44E-07 | -14.75185777 | 9.128497779 | 9.067150531 | 8.677268317 | 2 | 2 | 6.892381078 | 6.996671423 |
| FAM188A | | -0.502377633 | 0.633147923 | -14.25351273 | 1.00E-16 | -14.75589036 | 8.609439662 | 8.053983481 | 9.954333071 | 8.634896 | 8.5668971 | 2 | 2 |
| HEXDC | | -1.319385132 | 0.29269712 | -13.44565284 | 2.81E-13 | -14.76503797 | 7.159102577 | 7.145017454 | 9.372276026 | 7.030886362 | 6.98285659 | 2 | 2 |
| ENSG00000182111 | ZNF716 | -4.214659108 | 0.008633495 | -10.55160311 | 5.84E-07 | -14.76626222 | 5.70932728 | 3.686303414 | 6.203069917 | 2.3892346 | 2.86123014 | 2 | 2 |
| ENSG00000130640 | TUBGCP2 | -13.98388753 | 1.64E-16 | -0.782485775 | 0.389710269 | -14.76637331 | 11.01943573 | 9.969011754 | 9.242516611 | 2 | 2 | 9.300234537 | 9.64468005 |
| ENSG00000116120 | FARSB | -13.07157525 | 8.53E-16 | -1.697115563 | 0.00668594 | -14.76869081 | 9.761125267 | 9.470501606 | 8.585098385 | 2 | 2 | 7.593002849 | 7.754841567 |
| ENSG00000171792 | RHNO1 | -13.23516909 | 6.39E-16 | -1.534986162 | 0.023485564 | -14.77015225 | 10.08373159 | 8.916342327 | 9.301632639 | 2 | 2 | 7.734688324 | 8.2175813 |
| ENSG00000197976 | AKAP17A | -0.334493911 | 0.608546899 | -14.44325679 | 9.21E-20 | -14.7777507 | 8.737455672 | 9.235879652 | 9.725218844 | 8.948803189 | 8.96246653 | 2 | 2 |
| ENSG00000168118 | RAB4A | -13.41316603 | 7.19E-16 | -1.368629529 | 0.072863536 | -14.78179555 | 9.397585735 | 8.903033433 | 10.38167985 | 2 | 2 | 8.344674564 | 8.326688609 |
| ENSG00000215252 | GOLGA8B | -12.35454731 | 9.91E-12 | -2.431538537 | 0.023328609 | -14.78608585 | 7.031698319 | 8.522788974 | 9.452720984 | 2 | 2 | 6.613628927 | 5.825320785 |
| ENSG00000110786 | PTPN5 | -8.435085551 | 0.00980946 | -6.354874854 | 0.032404634 | -14.78996041 | 6.257048221 | 2 | 3.369317972 | 2 | 2 | 2.048074405 | 2.172608095 |
| ENSG00000183077 | AFMID | -13.22570334 | 2.53E-17 | -1.569613398 | 7.04E-05 | -14.79531674 | 9.710019321 | 9.192779271 | 9.577565424 | 2 | 2 | 7.941321277 | 7.964141273 |
| ENSG00000120500 | ARR3 | -10.05036302 | 0.010013464 | -4.746903055 | 0.207737426 | -14.79726608 | 2 | 2 | 7.934808822 | 2 | 2 | 2.987421008 | 2.592996859 |
| ENSG00000121774 | KHDRBS1 | -14.15596877 | 1.46E-18 | -0.641715032 | 0.323609074 | -14.7976838 | 10.29268328 | 10.82153847 | 10.07784268 | 2 | 2 | 9.337585872 | 10.14022837 |
| ENSG00000106290 | TAF6 | -13.42732077 | 7.36E-09 | -1.370481282 | 0.515079411 | -14.79780205 | 10.83137851 | 9.386925327 | 4.058998049 | 2 | 2 | 8.334412727 | 8.361298015 |
| RFWD2 | | -0.67174415 | 0.202157273 | -14.13160836 | 2.07E-19 | -14.80335251 | 8.892187472 | 9.328390744 | 8.608698767 | 8.30646685 | 8.32017198 | 2 | 2 |
| ENSG00000149575 | SCN2B | -6.686111317 | 0.228106188 | -8.124151971 | 0.128611101 | -14.81026329 | 2 | 3.076665209 | 4.523675391 | 2 | 2 | 2 | 2 |
| ENSG00000164068 | RNF123 | -13.31657807 | 2.35E-16 | -1.500732546 | 0.017993014 | -14.81731061 | 9.949165863 | 8.848535039 | 9.778213818 | 2 | 2 | 7.925322387 | 8.273956186 |
| ENSG00000106477 | CEP41 | -12.54716139 | 6.18E-13 | -2.273830802 | 0.013724207 | -14.82099219 | 8.444143041 | 7.701025996 | 9.670202709 | 2 | 2 | 6.466611899 | 6.735016458 |
| ENSG00000131097 | HIGD1B | -8.844192859 | 0.000361969 | -6.006402815 | 0.002878524 | -14.85059567 | 4.00722531 | 3.076665209 | 6.534109959 | 2 | 2 | 2.139668783 | 2.225841198 |
| ENSG00000196305 | IARS | -14.79715466 | 2.16E-17 | -0.054606411 | 0.964424173 | -14.85176107 | 11.0444064 | 11.81994319 | 9.516493218 | 2 | 2 | 11.10946858 | 10.92151362 |
| ENSG00000132153 | DHX30 | -14.21754694 | 4.45E-19 | -0.635823066 | 0.277366793 | -14.85337 | 10.60392551 | 9.890264534 | 10.83618737 | 2 | 2 | 9.789951757 | 9.929007159 |
| ENSG00000205937 | RNPS1 | -14.18909178 | 8.38E-18 | -0.670528225 | 0.394793944 | -14.85962001 | 10.86447339 | 10.72273565 | 9.426401271 | 2 | 2 | 9.774629808 | 9.82152476 |
| ENSG00000206488 | | -1.384774084 | 0.112473946 | -13.4773873 | 1.40E-15 | -14.86216138 | 9.015175615 | 8.191702463 | 7.285536013 | 6.998639523 | 6.94988431 | 2 | 2 |
| ENSG00000165646 | SLC18A2 | -10.15970545 | 9.87E-07 | -4.702979697 | 0.001061118 | -14.86268514 | 5.532353208 | 4.710564154 | 7.646393253 | 2 | 2 | 2.717032197 | 3.012034821 |
| ENSG00000113494 | PRLR | -4.974023368 | 1.57E-05 | -9.894679504 | 7.28E-17 | -14.86870287 | 9.189468652 | 10.44821838 | 8.677268317 | 4.825935136 | 4.91316568 | 2.492221014 | 2 |
| C19ORF35 | C19ORF35 | -5.494384846 | 0.050150442 | -9.375028706 | 0.001103783 | -14.86941355 | 5.532353208 | 4.11352228 | 2 | 2.141737246 | 2.14022021 | 2 | 2 |
| ENSG00000050767 | COL23A1 | -10.04471992 | 1.72E-05 | -4.830673043 | 0.008020131 | -14.87539296 | 3.327711234 | 7.145017454 | 6.629488387 | 2 | 2 | 3.12825509 | 2.277179783 |
| ENSG00000175344 | CHRNA7 | -4.267047451 | 0.00687121 | -10.61145584 | 4.11E-07 | -14.87850329 | 4.467311984 | 5.131603625 | 6.43197674 | 2.947924294 | 2.2680111 | 2 | 2 |
| ENSG00000111704 | NANOG | -4.709498309 | 0.017236601 | -10.17069549 | 1.43E-05 | -14.8801938 | 5.532353208 | 3.076665209 | 5.772716771 | 2.141737246 | 2.59489879 | 2 | 2 |
| TMEM55B | | -0.619344319 | 0.312833791 | -14.26461476 | 2.54E-19 | -14.88395908 | 9.128497779 | 9.497318007 | 8.536706852 | 8.463225341 | 8.52912793 | 2 | 2 |
| ENSG00000206260 | PRR23A | -6.726646522 | 0.22487948 | -8.164687682 | 0.1264478 | -14.8913342 | 3.327711234 | 4.442710129 | 2 | 2 | 2 | 2 | 2 |
| ENSG00000116574 | RHOU | -11.86960172 | 4.34E-07 | -3.023121758 | 0.122820268 | -14.89272348 | 3.327711234 | 7.503546079 | 9.385998894 | 2 | 2 | 5.206928476 | 5.343519008 |
| ENSG00000125815 | CST8 | -6.728866918 | 0.224699399 | -8.166908105 | 0.126314768 | -14.89577502 | 4.815609471 | 2 | 2 | 2 | 2 | 2 | 2 |
| ENSG00000166006 | KCNC2 | -6.728866918 | 0.224699399 | -8.166908105 | 0.126314768 | -14.89577502 | 4.815609471 | 2 | 2 | 2 | 2 | 2 | 2 |
| ENSG00000173302 | GPR148 | -6.728866918 | 0.224699399 | -8.166908105 | 0.126314768 | -14.89577502 | 4.815609471 | 2 | 2 | 2 | 2 | 2 | 2 |
| ENSG00000187537 | POTEG | -6.728866918 | 0.224699399 | -8.166908105 | 0.126314768 | -14.89577502 | 4.815609471 | 2 | 2 | 2 | 2 | 2 | 2 |
| ENSG00000250799 | PRODH2 | -6.728866918 | 0.224699399 | -8.166908105 | 0.126314768 | -14.89577502 | 4.815609471 | 2 | 2 | 2 | 2 | 2 | 2 |
| GPR126 | | -6.728866918 | 0.224699399 | -8.166908105 | 0.126314768 | -14.89577502 | 4.815609471 | 2 | 2 | 2 | 2 | 2 | 2 |
| ENSG00000157017 | GHRL | -9.458575507 | 5.43E-05 | -5.439157772 | 0.002753691 | -14.89773328 | 7.031698319 | 4.93638933 | 3.369317972 | 2 | 2 | 2.526562374 | 2.225841198 |
| ENSG00000163870 | TPRA1 | -13.73012926 | 3.50E-17 | -1.189440521 | 0.076250044 | -14.91956978 | 10.60392551 | 9.405920729 | 9.746650996 | 2 | 2 | 8.904045868 | 8.744843922 |
| ENSG00000147257 | GPC3 | -8.568828859 | 4.36E-07 | -6.353539263 | 1.66E-05 | -14.92236812 | 4.815609471 | 9.267380431 | 6.073378139 | 2.270786764 | 2.14022021 | 2.775316226 | 2.782576693 |
| ENSG00000164920 | OSR2 | -9.019658285 | 9.03E-14 | -5.90511781 | 2.35E-07 | -14.9247761 | 11.39115429 | 8.522788974 | 8.865353883 | 2.270786764 | 2.94009902 | 4.675266036 | 4.4231929 |
| ENSG00000106302 | HYAL4 | -6.745125132 | 0.223475046 | -8.183166522 | 0.12555257 | -14.92829165 | 4.467311984 | 2 | 3.369317972 | 2 | 2 | 2 | 2 |
| ENSG00000160799 | CCDC12 | -12.68067124 | 1.59E-14 | -2.248848554 | 0.000969832 | -14.9295198 | 9.521325812 | 8.234834549 | 8.865353883 | 2 | 2 | 6.529346487 | 6.957866334 |
| ENSG00000137868 | STRA6 | -9.263080828 | 1.17E-127 | -5.669527633 | 8.93E-82 | -14.93260846 | 12.22332945 | 12.08889578 | 11.86921686 | 3.515101962 | 3.39680241 | 6.305369896 | 6.607941737 |
| SKIV2L2 | | -0.492965462 | 0.590107068 | -14.45670328 | 4.45E-18 | -14.94966875 | 8.367060731 | 9.04308532 | 10.01740919 | 8.754208387 | 8.86729294 | 2 | 2 |
| ENSG00000213593 | TMX2 | -13.77837956 | 2.52E-17 | -1.17245226 | 0.079075233 | -14.95083182 | 10.60392551 | 9.386925327 | 9.926432745 | 2 | 2 | 8.733534631 | 9.033614226 |
| ENSG00000185518 | SV2B | -7.670631414 | 0.000153646 | -7.281248782 | 3.91E-05 | -14.9518802 | 4.467311984 | 4.11352228 | 7.818184403 | 2.270786764 | 2 | 2.139668783 | 2.225841198 |
| ENSG00000180919 | OR56B4 | -7.539301946 | 0.167050868 | -7.417087298 | 0.164554173 | -14.95638924 | 5.532353208 | 2 | 2 | 2 | 2 | 2 | 2.059860178 |
| ENSG00000186487 | MYT1L | -7.539301946 | 0.167050868 | -7.417087298 | 0.164554173 | -14.95638924 | 5.532353208 | 2 | 2 | 2 | 2 | 2 | 2.059860178 |
| ENSG00000160932 | LY6E | 1.793329995 | 0.004405565 | -16.75486538 | 1.72E-25 | -14.96153539 | 12.20867639 | 11.14070404 | 11.1494608 | 13.35183121 | 13.4154034 | 2 | 2 |
| ENSG00000198740 | ZNF652 | -13.67492962 | 6.43E-17 | -1.287404341 | 0.06185232 | -14.96233396 | 9.358295023 | 9.616264339 | 10.58572549 | 2 | 2 | 8.615855003 | 8.731676922 |
| ENSG00000276726 | | -0.658723751 | 0.284645724 | -14.30880379 | 2.10E-19 | -14.96752754 | 8.928409675 | 8.747909548 | 9.636156974 | 8.439397922 | 8.5606705 | 2 | 2 |
| ENSG00000139344 | AMDHD1 | -7.468678914 | 4.67E-07 | -7.501704868 | 1.99E-08 | -14.97038378 | 8.340423424 | 6.666836837 | 4.523675391 | 2.270786764 | 2.2680111 | 2.34604626 | 2.172608095 |
| ENSG00000223680 | | -0.37160466 | 0.376209416 | -14.60099574 | 2.64E-21 | -14.9726004 | 9.2903348 | 9.34816733 | 9.670202709 | 9.088859521 | 9.06170328 | 2 | 2 |
| ENSG00000162613 | FUBP1 | -14.01498891 | 7.93E-20 | -0.958247214 | 0.001183427 | -14.97323612 | 10.23600589 | 10.3486505 | 10.28977157 | 2 | 2 | 9.260934571 | 9.412921777 |
| ENSG00000136960 | ENPP2 | -14.07643467 | 4.80E-16 | -0.904918884 | 0.366230344 | -14.98135355 | 10.69610297 | 10.81799725 | 8.845626171 | 2 | 2 | 9.115521453 | 9.726317257 |
| ENSG00000112514 | CUTA | 0.08436588 | 0.917096582 | -15.06924913 | 8.36E-21 | -14.98488325 | 10.51143645 | 9.405920729 | 9.553445883 | 9.96278806 | 10.0255408 | 2 | 2 |
| ENSG00000160188 | RSPH1 | -9.578082527 | 0.002195057 | -5.412479764 | 0.054430004 | -14.99056229 | 7.435667355 | 3.076665209 | 2 | 2 | 2 | 2.42098416 | 2.421067459 |
| ENSG00000275095 | | -6.777177849 | 0.220845539 | -8.21521964 | 0.123777974 | -14.99239749 | 3.327711234 | 2 | 4.523675391 | 2 | 2 | 2 | 2 |
| GPR111 | | -6.777177849 | 0.220845539 | -8.21521964 | 0.123777974 | -14.99239749 | 3.327711234 | 2 | 4.523675391 | 2 | 2 | 2 | 2 |
| ENSG00000170653 | ATF7 | -13.73562594 | 5.07E-16 | -1.260681139 | 0.142209608 | -14.99630708 | 10.62062599 | 10.08551947 | 8.784770914 | 2 | 2 | 8.838495731 | 8.680159264 |
| ENSG00000227315 | | -0.070225361 | 0.951527415 | -14.92659877 | 5.17E-18 | -14.99682413 | 10.65345793 | 9.267380431 | 8.608698767 | 9.690252889 | 9.70683001 | 2 | 2 |
| FAM127B | | -0.786482898 | 0.693382761 | -14.21600609 | 6.19E-11 | -15.00248898 | 10.09970294 | 8.916342327 | 4.523675391 | 8.310225403 | 8.2561084 | 2 | 2 |
| ENSG00000180871 | CXCR2 | -6.767945083 | 0.023174508 | -8.243652478 | 0.004151263 | -15.01159756 | 5.330594134 | 2 | 5.156626401 | 2.141737246 | 2 | 2.048074405 | 2 |
| KIAA1456 | | -3.389014803 | 0.00362723 | -11.62811412 | 4.56E-10 | -15.01712892 | 7.218819781 | 4.93638933 | 6.629488387 | 3.754611435 | 3.50452879 | 2 | 2 |
| ENSG00000142224 | IL19 | -4.279424535 | 0.089858968 | -10.74374022 | 9.33E-05 | -15.02316475 | 5.095959458 | 2 | 6.882775709 | 2.600425781 | 2.77779907 | 2 | 2 |
| ENSG00000171595 | DNAI2 | -6.792977138 | 0.219632896 | -8.231019126 | 0.122968358 | -15.02399626 | 2 | 2 | 4.874587708 | 2 | 2 | 2 | 2 |
| ENSG00000174957 | OR5J2 | -6.792977138 | 0.219632896 | -8.231019126 | 0.122968358 | -15.02399626 | 2 | 2 | 4.874587708 | 2 | 2 | 2 | 2 |
| ENSG00000185247 | MAGEA11 | -6.792977138 | 0.219632896 | -8.231019126 | 0.122968358 | -15.02399626 | 2 | 2 | 4.874587708 | 2 | 2 | 2 | 2 |
| ENSG00000203985 | LDLRAD1 | -6.792977138 | 0.219632896 | -8.231019126 | 0.122968358 | -15.02399626 | 2 | 2 | 4.874587708 | 2 | 2 | 2 | 2 |
| ENSG00000204979 | MS4A13 | -6.792977138 | 0.219632896 | -8.231019126 | 0.122968358 | -15.02399626 | 2 | 2 | 4.874587708 | 2 | 2 | 2 | 2 |
| ENSG00000214842 | RAD51AP2 | -6.792977138 | 0.219632896 | -8.231019126 | 0.122968358 | -15.02399626 | 2 | 2 | 4.874587708 | 2 | 2 | 2 | 2 |
| ENSG00000224586 | GPX5 | -6.792977138 | 0.219632896 | -8.231019126 | 0.122968358 | -15.02399626 | 2 | 2 | 4.874587708 | 2 | 2 | 2 | 2 |
| KIAA1407 | | -6.792977138 | 0.219632896 | -8.231019126 | 0.122968358 | -15.02399626 | 2 | 2 | 4.874587708 | 2 | 2 | 2 | 2 |
| LOC400736 | | -6.792977138 | 0.219632896 | -8.231019126 | 0.122968358 | -15.02399626 | 2 | 2 | 4.874587708 | 2 | 2 | 2 | 2 |
| TSSC1 | | -1.130376201 | 0.455839475 | -13.8936314 | 1.04E-12 | -15.0240076 | 9.839306291 | 6.228547981 | 8.206335842 | 7.635924947 | 7.61995902 | 2 | 2 |
| ENSG00000119782 | FKBP1B | -11.97726192 | 4.11E-13 | -3.055129513 | 3.53E-06 | -15.03239143 | 8.75772933 | 7.503546079 | 8.296287027 | 2 | 2 | 5.449961224 | 5.230851202 |
| ENSG00000168003 | SLC3A2 | -14.38795367 | 4.02E-18 | -0.651225656 | 0.425726443 | -15.03917932 | 11.27287932 | 10.28338212 | 10.16797309 | 2 | 2 | 10.39962341 | 9.49042201 |
| ENSG00000185414 | MRPL30 | -13.32041696 | 1.34E-15 | -1.721708713 | 0.023047207 | -15.04212567 | 9.751047882 | 8.540087354 | 10.10298738 | 2 | 2 | 7.905886539 | 7.88671543 |
| LOC100130705 | | -2.821619435 | 0.161350886 | -12.24130383 | 1.59E-07 | -15.06292327 | 4.00722531 | 4.93638933 | 8.536706852 | 4.523237872 | 4.53541753 | 2 | 2 |
| ENSG00000226103 | | -1.093798183 | 0.21164903 | -13.97168477 | 8.63E-17 | -15.06548295 | 9.189468652 | 7.60566235 | 9.180874355 | 7.762033056 | 7.71833702 | 2 | 2 |
| ENSG00000165730 | STOX1 | -7.442097548 | 0.000249977 | -7.624778317 | 8.46E-05 | -15.06687587 | 2 | 7.761270542 | 8.35328246 | 2.3892346 | 2.2680111 | 2.492221014 | 2.059860178 |
| ENSG00000259431 | THTPA | -12.31089089 | 2.15E-14 | -2.757091209 | 4.58E-07 | -15.0679821 | 8.737455672 | 8.029695761 | 8.904018013 | 2 | 2 | 5.773219597 | 6.06327132 |
| ENSG00000183207 | RUVBL2 | -13.63357715 | 5.76E-17 | -1.435023108 | 0.029186087 | -15.06860026 | 9.667789716 | 10.47544827 | 9.344432377 | 2 | 2 | 8.269359984 | 8.677773268 |
| ENSG00000231479 | | -6.109289301 | 0.022800644 | -8.988407294 | 0.00043275 | -15.0976966 | 4.467311984 | 4.442710129 | 3.369317972 | 2 | 2.14022021 | 2 | 2 |
| ENSG00000146707 | POMZP3 | -1.217813529 | 0.004679711 | -13.88020313 | 4.51E-19 | -15.09801666 | 8.946185348 | 8.469613413 | 8.742726175 | 7.605658789 | 7.44845993 | 2 | 2 |
| ENSG00000276935 | | 0.211062032 | 0.672102316 | -15.31040795 | 4.84E-23 | -15.09934592 | 10.25752168 | 9.785372135 | 10.35401885 | 10.31394586 | 10.4067283 | 2 | 2 |
| ENSG00000132716 | DCAF8 | -13.92335577 | 6.89E-19 | -1.182365804 | 0.007963312 | -15.10572157 | 10.52923584 | 9.856140801 | 10.14394518 | 2 | 2 | 9.004166135 | 9.046155018 |
| ENSG00000177082 | WDR73 | -11.78635855 | 3.65E-12 | -3.326584015 | 1.33E-05 | -15.11294257 | 7.096806373 | 8.124485672 | 8.631919287 | 2 | 2 | 4.720468739 | 5.122735884 |
| ENSG00000135441 | BLOC1S1 | -12.90212148 | 2.39E-15 | -2.219834748 | 0.000347201 | -15.12195623 | 9.771132748 | 8.717821467 | 8.825624954 | 2 | 2 | 6.929749747 | 7.069464998 |
| ENSG00000256713 | PGA5 | -8.494447398 | 0.020941472 | -6.633428607 | 0.052849619 | -15.127876 | 2 | 5.131603625 | 5.772716771 | 2 | 2 | 2.183373728 | 2 |
| ENSG00000197587 | DMBX1 | -7.559141922 | 0.00244 | -7.58307887 | 0.002021405 | -15.14222079 | 4.00722531 | 4.442710129 | 4.058998049 | 2 | 2 | 2.048074405 | 2 |
| ENSG00000183560 | IZUMO1R | -8.501434221 | 0.112649338 | -6.649536383 | 0.198160911 | -15.1509706 | 2 | 2 | 6.43197674 | 2 | 2 | 2.183373728 | 2 |
| ENSG00000163638 | ADAMTS9 | -6.446454156 | 6.47E-21 | -8.71783705 | 2.37E-37 | -15.16429121 | 10.4253394 | 12.28376116 | 12.15221073 | 5.501219927 | 5.52526112 | 3.767980144 | 3.539239711 |

| | | | | | | | | | | | | | |
|---|---|---|---|---|---|---|---|---|---|---|---|---|---|
| ENSG00000112486 | CCR6 | -5.3535989 | 0.017029927 | -9.813362902 | 7.80E-05 | -15.1669618 | 4.467311984 | 4.11352228 | 5.59507254 | 2 | 2.38539787 | 2 | 2 |
| ENSG00000107821 | KAZALD1 | -12.13963394 | 5.37E-11 | -3.03291559 | 0.006692734 | -15.17254953 | 9.015175615 | 6.602424674 | 8.721235202 | 2 | 2 | 5.922534244 | 4.935697922 |
| FYB | | -2.964008061 | 0.01313323 | -12.22015017 | 6.86E-11 | -15.18415823 | 7.709880939 | 5.131603625 | 7.398165132 | 4.470469939 | 4.31639476 | 2 | 2 |
| ENSG00000166946 | CCNDBP1 | -13.73523604 | 1.17E-16 | -1.45722189 | 0.050939912 | -15.19245793 | 10.62615013 | 9.090820894 | 9.935792933 | 2 | 2 | 8.557885437 | 8.575104441 |
| ENSG00000062822 | POLD1 | -12.74634957 | 5.77E-16 | -2.452134595 | 9.90E-09 | -15.19848416 | 8.946185348 | 8.806266215 | 9.301632639 | 2 | 2 | 6.775876478 | 6.460277802 |
| ENSG00000105607 | GCDH | -12.93997429 | 3.67E-13 | -2.263840759 | 0.025515228 | -15.20381505 | 9.657036304 | 7.538394143 | 9.647595056 | 2 | 2 | 6.599583349 | 7.305675343 |
| ENSG00000234058 | | -1.226728429 | 0.058415094 | -13.97911518 | 5.01E-19 | -15.20584361 | 10.79260216 | 10.19331737 | 9.541233192 | 9.000896538 | 9.08575246 | 2 | 2.059860178 |
| CXORF36 | CXORF36 | -5.166788697 | 0.00280218 | -10.04472805 | 2.54E-06 | -15.21151674 | 5.330594134 | 3.686303414 | 5.59507254 | 2.141737246 | 2.38539787 | 2 | 2 |
| C16ORF59 | C16ORF59 | -3.67031966 | 0.133069745 | -11.55192205 | 1.54E-05 | -15.22224171 | 7.750913381 | 5.457120953 | 2 | 3.515101962 | 3.28037825 | 2 | 2 |
| BZRAP1 | BZRAP1 | -6.89380618 | 0.211937735 | -8.331849428 | 0.117817538 | -15.22565561 | 2 | 4.93638933 | 2 | 2 | 2 | 2 | 2 |
| ENSG00000107807 | TLX1 | -6.89380618 | 0.211937735 | -8.331849428 | 0.117817538 | -15.22565561 | 2 | 4.93638933 | 2 | 2 | 2 | 2 | 2 |
| ENSG00000215644 | GCGR | -6.89380618 | 0.211937735 | -8.331849428 | 0.117817538 | -15.22565561 | 2 | 4.93638933 | 2 | 2 | 2 | 2 | 2 |
| ENSG00000164512 | ANKRD55 | -9.192421077 | 0.002250083 | -6.038444766 | 0.023409169 | -15.23086584 | 6.563806279 | 3.076665209 | 5.39244166 | 2 | 2 | 2 | 2.421067459 |
| ENSG00000112038 | OPRM1 | -4.889419966 | 0.001551816 | -10.35113795 | 4.22E-07 | -15.24055792 | 5.866944937 | 4.442710129 | 5.39244166 | 2.141737246 | 2.59489879 | 2 | 2 |
| ENSG00000277161 | PIGW | -2.014849662 | 0.021504527 | -13.24445448 | 8.04E-15 | -15.25930414 | 7.667647223 | 7.232742069 | 8.884815469 | 6.197166296 | 6.09824986 | 2 | 2 |
| ENSG00000262938 | | -3.584520074 | 0.021123194 | -11.67731288 | 2.14E-08 | -15.26183295 | 6.366709671 | 4.11352228 | 7.552162999 | 3.840289928 | 3.15372745 | 2 | 2 |
| ENSG00000162594 | IL23R | -6.912501382 | 0.065184591 | -8.350498504 | 0.021735943 | -15.26299989 | 4.00722531 | 3.686303414 | 3.369317972 | 2 | 2 | 2 | 2 |
| TMEM133 | | -2.662084253 | 0.254759301 | -12.60780517 | 6.68E-07 | -15.26988943 | 8.042902253 | 8.053983481 | 2 | 4.825935136 | 5.11361705 | 2 | 2 |
| ENSG00000106018 | VIPR2 | -7.779674245 | 8.42E-05 | -7.500122301 | 1.35E-05 | -15.27979655 | 4.467311984 | 4.11352228 | 7.934808822 | 2.141737246 | 2.14022021 | 2.267001569 | 2.059860178 |
| ENSG00000109775 | UFSP2 | -13.08518192 | 5.93E-16 | -2.19675268 | 0.000174596 | -15.2819346 | 9.48530526 | 8.655694736 | 9.757248847 | 2 | 2 | 7.15911073 | 7.242500282 |
| ENSG00000124092 | CTCFL | -6.925668757 | 0.064637739 | -8.363665992 | 0.021520027 | -15.28933475 | 3.327711234 | 3.686303414 | 4.058998049 | 2 | 2 | 2 | 2 |
| ENSG00000188089 | PLA2G4E | -6.925668757 | 0.064637739 | -8.363665992 | 0.021520027 | -15.28933475 | 3.327711234 | 3.686303414 | 4.058998049 | 2 | 2 | 2 | 2 |
| NAT6 | | -2.220188079 | 0.398597718 | -13.07227825 | 1.21E-06 | -15.29246633 | 2 | 6.78762695 | 9.272377397 | 5.797438367 | 5.7929843 | 2 | 2 |
| ENSG00000130023 | ERMARD | -1.872917371 | 0.021456873 | -13.43314638 | 1.20E-15 | -15.30606375 | 9.015175615 | 7.901717006 | 7.502634626 | 6.490083648 | 6.43421531 | 2 | 2 |
| ENSG00000262666 | | -0.366907173 | 0.45354552 | -14.94026092 | 6.31E-22 | -15.30716809 | 9.397585735 | 9.962611189 | 9.91701143 | 9.443949663 | 9.38961618 | 2 | 2 |
| SPG20 | | -0.005682111 | 0.991458614 | -15.31574177 | 4.13E-23 | -15.32142388 | 10.36053552 | 10.24697497 | 9.798878489 | 10.16260233 | 10.1380242 | 2 | 2 |
| ENSG00000163032 | VSNL1 | -6.539056825 | 7.49E-08 | -8.788117477 | 1.16E-12 | -15.3271743 | 5.330594134 | 9.192779271 | 8.486635645 | 2.947924294 | 2.86123014 | 2.3070652 | 2.172608095 |
| ENSG00000132429 | POPDC3 | -10.86832945 | 1.34E-07 | -4.459484791 | 0.001777936 | -15.32781424 | 7.33131396 | 5.457120953 | 7.858115942 | 2 | 2 | 3.917810786 | 2.509591799 |
| C16ORF62 | C16ORF62 | -0.464588229 | 0.325432015 | -14.86325685 | 9.86E-22 | -15.32784507 | 9.90440318 | 9.814097497 | 9.330306409 | 9.229474035 | 9.25726695 | 2 | 2 |
| ENSG00000214128 | TMEM213 | -4.930732845 | 0.002086464 | -10.39831696 | 6.19E-07 | -15.3290498 | 5.70932728 | 3.686303414 | 5.930865648 | 2.498691189 | 2.2680111 | 2 | 2 |
| ENSG00000003147 | ICA1 | -9.188238992 | 8.14E-07 | -6.142037264 | 8.12E-09 | -15.33027626 | 6.73718411 | 7.467835385 | 6.43197674 | 2.141737246 | 2 | 2.775316226 | 2.172608095 |
| WBSCR22 | | -0.741501843 | 0.050767575 | -14.58986963 | 2.54E-21 | -15.33137147 | 9.317904228 | 9.328390744 | 9.636156974 | 8.667494409 | 8.72698617 | 2 | 2 |
| ENSG00000131126 | TEX101 | -6.912408562 | 4.90E-05 | -8.420364274 | 6.04E-07 | -15.33277284 | 5.866944937 | 4.11352228 | 6.43197674 | 2.141737246 | 2.14022021 | 2.094598369 | 2 |
| ENSG00000134183 | GNAT2 | -5.066557127 | 0.066916943 | -10.2705426 | 0.000470077 | -15.33709973 | 2 | 3.686303414 | 6.629488387 | 2.3892346 | 2.2680111 | 2 | 2 |
| ENSG00000228736 | | -1.883056232 | 0.00091871 | -13.45940308 | 5.25E-18 | -15.34245931 | 9.097018025 | 9.77810053 | 10.16797309 | 7.759286043 | 7.99261031 | 2 | 2.059860178 |
| ENSG00000171345 | KRT19 | -9.162663296 | 9.42E-58 | -6.181341334 | 1.24E-28 | -15.34400463 | 13.81717923 | 12.97195359 | 13.23923849 | 4.523237872 | 4.4836996 | 6.535683256 | 7.7169976 |
| ENSG00000140474 | ULK3 | -13.20756868 | 8.10E-14 | -2.139245613 | 0.031375886 | -15.34681429 | 10.44422623 | 8.469613413 | 8.677268317 | 2 | 2 | 7.441507444 | 7.308762629 |
| ENSG00000104112 | SCG3 | -8.418985146 | 0.000222305 | -6.930586026 | 0.000233074 | -15.34957117 | 5.095959458 | 4.710564154 | 4.874587708 | 2 | 2 | 2.139668783 | 2 |
| ENSG00000066735 | KIF26A | -9.826830654 | 3.24E-17 | -5.523091891 | 1.42E-12 | -15.34992254 | 9.317904228 | 8.124485672 | 9.601288346 | 2.270786764 | 2.14022021 | 4.416072028 | 3.45257121 |
| ENSG00000132677 | RHBG | -4.830358739 | 0.001254131 | -10.5209238 | 2.08E-07 | -15.35128254 | 6.138360429 | 4.11352228 | 5.59507254 | 2.600425781 | 2.2680111 | 2 | 2 |
| ENSG00000096088 | PGC | -8.499123044 | 0.043548782 | -6.854680034 | 0.085882266 | -15.35380308 | 2 | 2 | 6.43197674 | 2 | 2 | 2.048074405 | 2.117335275 |
| ENSG00000236279 | CLEC2L | -6.442563403 | 0.055836832 | -8.919681762 | 0.009166093 | -15.36224517 | 2 | 2 | 6.80318718 | 2.141737246 | 2.14022021 | 2.048074405 | 2 |
| ENSG00000249860 | MTRNR2L5 | -5.462490201 | 0.145397751 | -9.920895953 | 0.008020131 | -15.38338615 | 4.467311984 | 2 | 6.073378139 | 2 | 2.38539787 | 2 | 2 |
| ENSG00000110958 | PTGES3 | -14.51915393 | 1.21E-20 | -0.866564387 | 0.040753925 | -15.38571831 | 11.09308737 | 10.68823825 | 10.54960077 | 2 | 2 | 10.02373723 | 9.833364301 |
| ENSG00000235744 | | -6.252432749 | 0.022930009 | -9.135703281 | 0.000485822 | -15.38813603 | 4.815609471 | 3.076665209 | 4.523675391 | 2.141737246 | 2 | 2 | 2 |
| WHSC1L1 | | -0.35432536 | 0.645266607 | -15.04490678 | 1.35E-20 | -15.39923214 | 9.771132748 | 9.159587375 | 10.44859742 | 9.503435578 | 9.5631668 | 2 | 2 |
| ENSG00000179526 | SHARPIN | -13.90787624 | 1.94E-16 | -1.494133143 | 0.075316635 | -15.40200938 | 9.509418609 | 9.514922632 | 10.99236731 | 2 | 2 | 8.715506558 | 8.686107048 |
| ENSG00000148408 | CACNA1B | -7.674766543 | 0.000411445 | -7.729518925 | 0.000176184 | -15.40428547 | 8.87372952 | 5.457120953 | 2 | 2.3892346 | 2.14022021 | 2.139668783 | 2.374681325 |
| ENSG00000112584 | FAM120B | -0.539235531 | 0.478107313 | -14.87283797 | 4.59E-20 | -15.4120735 | 10.19941743 | 9.77810053 | 8.865353883 | 9.134092596 | 9.22219152 | 2 | 2 |
| ENSG00000132793 | LPIN3 | -13.25025861 | 8.61E-15 | -2.177751608 | 0.009956274 | -15.42801022 | 9.097018025 | 8.590771778 | 10.3329194 | 2 | 2 | 7.264607048 | 7.488317792 |
| ENSG00000121207 | LRAT | -7.448122727 | 0.002296166 | -7.982813349 | 0.000175842 | -15.43093608 | 6.257048221 | 3.076665209 | 5.156626401 | 2.141737246 | 2 | 2.094598369 | 2 |
| ENSG00000116147 | TNR | -8.801683798 | 0.099210255 | -6.629479945 | 0.19771526 | -15.43116374 | 2 | 2 | 6.718950348 | 2 | 2 | 2.225793514 | 2 |
| ENSG00000165186 | PTCHD1 | -8.332942952 | 0.005854733 | -7.112067552 | 0.006569922 | -15.4450105 | 6.653098118 | 6.728495673 | 2 | 2.141737246 | 2 | 2.3070652 | 2 |
| ENSG00000127445 | PIN1 | -14.40589026 | 2.15E-19 | -1.049874247 | 0.072769585 | -15.45576451 | 10.24321353 | 10.44362967 | 11.18521898 | 2 | 2 | 9.509205303 | 9.753478818 |
| ENSG00000070985 | TRPM5 | -8.971940699 | 0.092116635 | -6.489959127 | 0.207305039 | -15.46189983 | 2 | 2 | 6.882775709 | 2 | 2 | 2 | 2.277179783 |
| ENSG00000233701 | PRR23C | -7.012551534 | 0.072225893 | -8.450552993 | 0.025405862 | -15.46310453 | 3.327711234 | 3.076665209 | 4.523675391 | 2 | 2 | 2 | 2 |
| ENSG00000278501 | | -1.165614616 | 0.133025275 | -14.29769285 | 3.75E-18 | -15.46330746 | 9.556468883 | 9.377333006 | 8.110401118 | 7.992367495 | 7.98797735 | 2 | 2 |
| ENSG00000116560 | SFPQ | -14.89649027 | 2.08E-20 | -0.568705322 | 0.390730653 | -15.4651956 | 11.21906289 | 10.50658023 | 11.5927499 | 2 | 2 | 10.38419728 | 10.79595529 |
| ENSG00000060971 | ACAA1 | -13.27852987 | 2.04E-14 | -2.187496504 | 0.017043752 | -15.46602638 | 10.3933013 | 8.356994601 | 9.212024683 | 2 | 2 | 7.328045423 | 7.466366987 |
| ENSG00000186501 | TMEM222 | -13.8302112 | 4.22E-17 | -1.641114526 | 0.019407296 | -15.47132573 | 10.70134604 | 9.34816733 | 9.963514551 | 2 | 2 | 8.27378929 | 8.657936763 |
| ENSG00000261308 | FIGNL2 | -5.413771978 | 2.29E-07 | -10.06137279 | 1.11E-15 | -15.47514477 | 7.904282243 | 6.899080544 | 9.165043284 | 3.406930997 | 3.55551834 | 2.183373728 | 2 |
| ENSG00000089123 | TASP1 | -12.17459535 | 8.19E-11 | -3.304480918 | 0.004191574 | -15.47907627 | 9.544849385 | 7.051611315 | 7.502634626 | 2 | 2 | 5.130840083 | 5.448021784 |
| ENSG00000186103 | ARGFX | -4.896554062 | 0.000372472 | -10.58535628 | 5.55E-08 | -15.48191034 | 5.330594134 | 4.442710129 | 6.322058544 | 2.498691189 | 2.38539787 | 2 | 2 |
| ENSG00000111481 | COPZ1 | -14.86133795 | 8.95E-19 | -0.623884695 | 0.483654059 | -15.48522265 | 11.92474824 | 10.44821838 | 10.5250051 | 2 | 2 | 10.3948386 | 10.62521539 |
| ENSG00000174944 | P2RY14 | -6.304305732 | 0.110411514 | -9.182835103 | 0.014179007 | -15.48714084 | 5.330594134 | 2 | 4.058998049 | 2 | 2.14022021 | 2 | 2 |
| ENSG00000213614 | HEXA | -14.05819155 | 5.66E-18 | -1.432596893 | 0.029339151 | -15.49078845 | 9.533135543 | 10.40166101 | 10.80076732 | 2 | 2 | 9.027210383 | 8.784193542 |
| ENSG00000089250 | NOS1 | -7.039193397 | 0.000599658 | -8.457358393 | 1.71E-05 | -15.49655179 | 4.467311984 | 5.131603625 | 5.156626401 | 2.141737246 | 2 | 2 | 2.059860178 |
| ENSG00000256045 | MTRNR2L10 | -1.1593458 | 0.015370181 | -14.34910788 | 4.61E-20 | -15.50845368 | 9.48530526 | 9.225224669 | 8.784770914 | 7.992367495 | 8.09955907 | 2 | 2 |
| ENSG00000213585 | VDAC1 | -15.17577068 | 1.02E-21 | -0.334233443 | 0.578130043 | -15.51000412 | 11.69698977 | 10.91068521 | 11.6249326 | 2 | 2 | 10.90276596 | 11.30412264 |
| ENSG00000101076 | HNF4A | -8.192913294 | 0.12809176 | -7.333484136 | 0.159405443 | -15.52639743 | 6.138360429 | 2 | 2 | 2 | 2 | 2.094598369 | 2 |
| ENSG00000175324 | LSM1 | -13.08563544 | 7.17E-14 | -2.450177896 | 0.008494375 | -15.53581334 | 10.27872131 | 8.505280662 | 8.561105505 | 2 | 2 | 6.845542321 | 7.05850228 |
| ENSG00000016490 | CLCA1 | -7.052226778 | 0.200056839 | -8.490272004 | 0.109819312 | -15.54249878 | 5.095959458 | 2 | 2 | 2 | 2 | 2 | 2 |
| ENSG00000111700 | SLCO1B3 | -7.052226778 | 0.200056839 | -8.490272004 | 0.109819312 | -15.54249878 | 5.095959458 | 2 | 2 | 2 | 2 | 2 | 2 |
| ENSG00000135903 | PAX3 | -7.052226778 | 0.200056839 | -8.490272004 | 0.109819312 | -15.54249878 | 5.095959458 | 2 | 2 | 2 | 2 | 2 | 2 |
| ENSG00000169344 | UMOD | -7.052226778 | 0.200056839 | -8.490272004 | 0.109819312 | -15.54249878 | 5.095959458 | 2 | 2 | 2 | 2 | 2 | 2 |
| ENSG00000176204 | LRRTM4 | -7.052226778 | 0.200056839 | -8.490272004 | 0.109819312 | -15.54249878 | 5.095959458 | 2 | 2 | 2 | 2 | 2 | 2 |
| GPR116 | | -7.052226778 | 0.200056839 | -8.490272004 | 0.109819312 | -15.54249878 | 5.095959458 | 2 | 2 | 2 | 2 | 2 | 2 |
| ENSG00000277439 | | -5.53570051 | 0.010136565 | -10.00699945 | 3.45E-05 | -15.54269996 | 4.467311984 | 4.442710129 | 5.772716771 | 2.3892346 | 2 | 2 | 2 |
| ENSG00000105392 | CRX | -4.748011745 | 0.000224082 | -10.79658532 | 1.56E-08 | -15.54459706 | 5.095959458 | 5.131603625 | 6.534109959 | 2.3892346 | 2.68924543 | 2 | 2 |
| ENSG00000168522 | FNTA | -14.01407159 | 3.59E-16 | -1.53209832 | 0.093567925 | -15.54616991 | 11.10500521 | 9.828248506 | 9.344432377 | 2 | 2 | 9.071132978 | 8.384893621 |
| ENSG00000164100 | NDST3 | -4.833562552 | 6.51E-05 | -10.71419516 | 7.79E-09 | -15.54775771 | 6.366709671 | 5.131603625 | 5.156626401 | 2.498691189 | 2.49394783 | 2 | 2 |
| ENSG00000213523 | SRA1 | -14.32276614 | 2.87E-19 | -1.236780175 | 0.02758368 | -15.55954631 | 11.03197509 | 10.04973379 | 10.55568472 | 2 | 2 | 9.257104839 | 9.470917069 |
| ENSG00000170906 | NDUFA3 | -1.395467487 | 0.099672705 | -14.17223311 | 2.69E-17 | -15.5677006 | 9.720386367 | 8.029695761 | 8.825624954 | 7.614805492 | 7.66709116 | 2 | 2 |
| ENSG00000140009 | ESR2 | -7.073513292 | 0.198716426 | -8.511558784 | 0.108834364 | -15.58507208 | 2 | 4.442710129 | 4.058998049 | 2 | 2 | 2 | 2 |
| ENSG00000163283 | ALPP | -7.073513292 | 0.198716426 | -8.511558784 | 0.108834364 | -15.58507208 | 2 | 4.442710129 | 4.058998049 | 2 | 2 | 2 | 2 |
| ENSG00000278624 | | -4.769906078 | 0.230213505 | -10.81606893 | 0.006042694 | -15.585975 | 7.27616306 | 2 | 2 | 2.270786764 | 2.77779907 | 2 | 2 |
| ENSG00000101448 | EPPIN | -6.350235027 | 0.251780872 | -9.236790792 | 0.078206869 | -15.58702582 | 2 | 2 | 5.772716771 | 2.141737246 | 2 | 2 | 2 |
| C11ORF57 | C11ORF57 | -0.978915344 | 0.031601758 | -14.61937116 | 5.85E-21 | -15.5982865 | 9.423197387 | 9.13702746 | 9.767769415 | 8.488325436 | 8.49524881 | 2 | 2 |
| ENSG00000204590 | GNL1 | -0.510261238 | 0.52563759 | -15.08820336 | 2.33E-20 | -15.5984646 | 10.45668146 | 9.018611864 | 9.972637969 | 9.414570905 | 9.42860336 | 2 | 2 |
| ENSG00000230514 | | -3.109853895 | 0.264849717 | -12.49832319 | 1.41E-05 | -15.60817709 | 2 | 4.710564154 | 8.865353883 | 4.523237872 | 4.4836996 | 2 | 2 |
| ENSG00000187715 | KBTBD12 | -7.051324359 | 5.20E-05 | -8.558765592 | 7.17E-07 | -15.61008995 | 4.815609471 | 5.595927442 | 6.629488387 | 2 | 2.2680111 | 2.094598369 | 2 |
| ENSG00000061455 | PRDM6 | -8.105223695 | 3.04E-16 | -7.505796291 | 5.96E-18 | -15.61101999 | 9.521325812 | 8.876041558 | 7.342949392 | 2.3892346 | 2.59489879 | 2.858551861 | 2.509591799 |
| ENSG00000262634 | | -3.766816078 | 7.03E-05 | -11.85594244 | 1.52E-11 | -15.62275852 | 5.70932728 | 6.666836837 | 7.398165132 | 3.406930997 | 3.65239003 | 2 | 2 |
| TMEM5 | | -1.982739612 | 0.007860301 | -13.64404128 | 1.64E-16 | -15.6267809 | 8.816894989 | 7.503546079 | 8.825624954 | 6.442929658 | 6.66650236 | 2 | 2 |
| ENSG00000214050 | FBXO16 | -10.77380502 | 3.09E-05 | -4.853576213 | 0.026057893 | -15.62738124 | 2 | 7.431218269 | 7.858115942 | 2 | 2 | 3.011868911 | 3.183290849 |
| ENSG00000104901 | DKKL1 | -10.10801817 | 8.75E-07 | -5.537657909 | 5.74E-05 | -15.64567607 | 8.674867122 | 4.442710129 | 8.042706615 | 2.141737246 | 2 | 3.27687201 | 3.012034821 |
| ENSG00000182352 | C17orf77 | -7.10439383 | 0.19645832 | -8.542439707 | 0.107319759 | -15.64683354 | 3.327711234 | 2 | 4.874587708 | 2 | 2 | 2 | 2 |
| ENSG00000186867 | QRFPR | -7.10439383 | 0.19645832 | -8.542439707 | 0.107319759 | -15.64683354 | 3.327711234 | 2 | 4.874587708 | 2 | 2 | 2 | 2 |
| ENSG00000090316 | MAEA | -14.38497159 | 1.21E-17 | -1.26219615 | 0.129694026 | -15.64716774 | 11.10895605 | 9.424669273 | 10.94216274 | 2 | 2 | 9.299303769 | 9.502606422 |
| ENSG00000143578 | CREB3L4 | -11.84905781 | 1.37E-10 | -3.798963071 | 0.000427063 | -15.64802089 | 8.009475818 | 6.464272742 | 8.978369997 | 2 | 2 | 4.888321757 | 4.201948172 |
| ATP5J | ATP5J | -0.948312734 | 0.133144408 | -14.70922721 | 3.38E-20 | -15.65753994 | 10.09173937 | 9.246456519 | 9.116480286 | 8.607665593 | 8.6142631 | 2 | 2 |
| ENSG00000080644 | CHRNA3 | -5.597564427 | 0.165082329 | -10.06072829 | 0.011703886 | -15.65829271 | 2 | 6.535001748 | 2 | 2.141737246 | 2.2680111 | 2 | 2 |
| ENSG00000176406 | RIMS2 | -5.77108761 | 9.28E-14 | -9.887696775 | 1.49E-15 | -15.65878438 | 6.73718411 | 7.274681896 | 7.552162999 | 2.695456494 | 2.77779907 | 2 | 2.117335275 |
| ENSG00000183463 | URAD | -7.113826926 | 0.19577655 | -8.55187292 | 0.10692029 | -15.66569985 | 2 | 5.131603625 | 2 | 2 | 2 | 2 | 2 |
| ENSG00000280165 | PCDH20 | -7.113826926 | 0.19577655 | -8.55187292 | 0.10692029 | -15.66569985 | 2 | 5.131603625 | 2 | 2 | 2 | 2 | 2 |
| KIAA1045 | | -7.113826926 | 0.19577655 | -8.55187292 | 0.10692029 | -15.66569985 | 2 | 5.131603625 | 2 | 2 | 2 | 2 | 2 |
| ATP5E | ATP5E | -0.190455801 | 0.8426245 | -15.4787687 | 9.76E-21 | -15.6692245 | 10.86447339 | 9.192779271 | 10.43545992 | 10.11773201 | 10.1395916 | 2 | 2 |

| | | | | | | | | | | | | | |
|---|---|---|---|---|---|---|---|---|---|---|---|---|---|
| ENSG00000157703 | SVOPL | -7.117178334 | 0.195586526 | -8.555224371 | 0.106808076 | -15.67240271 | 2 | 2 | 5.156626401 | 2 | 2 | 2 | 2 |
| ENSG00000163530 | DPPA2 | -7.117178334 | 0.195586526 | -8.555224371 | 0.106808076 | -15.67240271 | 2 | 2 | 5.156626401 | 2 | 2 | 2 | 2 |
| ENSG00000172680 | MOS | -7.117178334 | 0.195586526 | -8.555224371 | 0.106808076 | -15.67240271 | 2 | 2 | 5.156626401 | 2 | 2 | 2 | 2 |
| ENSG00000178279 | TNP2 | -7.117178334 | 0.195586526 | -8.555224371 | 0.106808076 | -15.67240271 | 2 | 2 | 5.156626401 | 2 | 2 | 2 | 2 |
| ENSG00000184945 | AQP12A | -7.117178334 | 0.195586526 | -8.555224371 | 0.106808076 | -15.67240271 | 2 | 2 | 5.156626401 | 2 | 2 | 2 | 2 |
| ENSG00000226182 | | -7.117178334 | 0.195586526 | -8.555224371 | 0.106808076 | -15.67240271 | 2 | 2 | 5.156626401 | 2 | 2 | 2 | 2 |
| ENSG00000274131 | | -7.117178334 | 0.195586526 | -8.555224371 | 0.106808076 | -15.67240271 | 2 | 2 | 5.156626401 | 2 | 2 | 2 | 2 |
| LOC93432 | | -7.117178334 | 0.195586526 | -8.555224371 | 0.106808076 | -15.67240271 | 2 | 2 | 5.156626401 | 2 | 2 | 2 | 2 |
| PAK7 | | -7.117178334 | 0.195586526 | -8.555224371 | 0.106808076 | -15.67240271 | 2 | 2 | 5.156626401 | 2 | 2 | 2 | 2 |
| ENSG00000135409 | AMHR2 | -7.118982796 | 0.195586526 | -8.557028856 | 0.106808076 | -15.67601165 | 4.467311984 | 4.11352228 | 2 | 2 | 2 | 2 | 2 |
| ENSG00000187871 | GFRAL | -7.118982796 | 0.195586526 | -8.557028856 | 0.106808076 | -15.67601165 | 4.467311984 | 4.11352228 | 2 | 2 | 2 | 2 | 2 |
| KIAA0101 | | -3.590580565 | 0.0269659 | -12.08703751 | 1.30E-08 | -15.67761808 | 8.257427002 | 5.131603625 | 5.156626401 | 3.95996233 | 3.65239003 | 2 | 2 |
| ENSG00000205923 | CEMP1 | -4.814037773 | 0.101317769 | -10.86373652 | 0.00042153 | -15.67777429 | 2 | 3.076665209 | 7.285536013 | 2.498691189 | 2.59489879 | 2 | 2 |
| ENSG00000234495 | | -0.772670762 | 0.407875143 | -14.91721321 | 8.10E-19 | -15.68988397 | 10.35389212 | 9.863030423 | 8.486635645 | 9.010169232 | 8.97188212 | 2 | 2 |
| ENSG00000152253 | SPC25 | -5.229708605 | 1.25E-07 | -10.46583678 | 1.54E-09 | -15.69554539 | 5.866944937 | 6.602424674 | 7.502634626 | 2.947924294 | 2.59489879 | 2 | 2.059860178 |
| PNMAL2 | | -4.913682134 | 0.051116463 | -10.79246418 | 9.20E-05 | -15.70614631 | 2 | 4.710564154 | 7.029880454 | 2.3892346 | 2.59489879 | 2 | 2 |
| ENSG00000278605 | | -1.008427944 | 0.058610905 | -14.69818553 | 9.76E-21 | -15.70661348 | 9.48530526 | 9.923599762 | 9.083172337 | 8.576794289 | 8.50338773 | 2 | 2 |
| ENSG00000198570 | RD3 | -7.134863608 | 0.045685903 | -8.57285883 | 0.01343349 | -15.70772244 | 4.00722531 | 4.11352228 | 3.369317972 | 2 | 2 | 2 | 2 |
| ENSG00000196990 | FAM163B | -9.319182413 | 0.02522648 | -6.393070585 | 0.105745608 | -15.712253 | 7.218819781 | 2 | 2 | 2 | 2 | 2.048074405 | 2.326754077 |
| ENSG00000175164 | ABO | -9.618288215 | 0.013898856 | -6.100210188 | 0.098152435 | -15.7184984 | 2 | 7.503546079 | 2 | 2 | 2 | 2.384001718 | 2.172608095 |
| ENSG00000149972 | CNTN5 | -6.377230614 | 0.007453224 | -9.343005079 | 0.000345154 | -15.72023569 | 2 | 7.099070191 | 4.523675391 | 2.141737246 | 2.2680111 | 2 | 2.059860178 |
| ENSG00000100314 | CABP7 | -9.013755098 | 0.00103094 | -6.70836408 | 0.004532191 | -15.72211918 | 6.563806279 | 4.93638933 | 2 | 2 | 2 | 2.139668783 | 2.117335275 |
| C7ORF73 | C7ORF73 | -1.474003947 | 0.016470777 | -14.25123002 | 6.20E-19 | -15.72523397 | 9.635285994 | 8.820493869 | 8.631919287 | 7.694616532 | 7.58663873 | 2 | 2 |
| ENSG00000254440 | PBOV1 | -7.146204142 | 0.04538649 | -8.584199478 | 0.013325953 | -15.73040362 | 3.327711234 | 4.11352228 | 4.058998049 | 2 | 2 | 2 | 2 |
| ENSG00000090863 | GLG1 | -15.66086299 | 1.96E-18 | -0.074831209 | 0.955098488 | -15.73569419 | 9.940323746 | 12.16194825 | 12.59477286 | 2 | 2 | 11.83531067 | 11.88587147 |
| ENSG00000062038 | CDH3 | -9.285646132 | 5.92E-17 | -6.457447048 | 5.93E-20 | -15.74309318 | 8.963744667 | 7.761270542 | 8.825624954 | 2.3892346 | 2 | 3.172301926 | 2.950113106 |
| ENSG00000197753 | LHFPL5 | -6.4366656 | 0.017181187 | -9.315746314 | 0.000301187 | -15.75241191 | 4.815609471 | 3.076665209 | 4.874587708 | 2 | 2.14022021 | 2 | 2 |
| ENSG00000143158 | MPC2 | -14.10392285 | 3.77E-17 | -1.652946779 | 0.036294563 | -15.75686962 | 10.96815916 | 9.298208073 | 10.42220168 | 2 | 2 | 8.568225482 | 8.889499502 |
| ENSG00000166869 | CHP2 | -6.443879943 | 0.018338429 | -9.322937604 | 0.000346886 | -15.76681755 | 4.467311984 | 3.076665209 | 5.156626401 | 2 | 2.14022021 | 2 | 2 |
| RTFDC1 | | -0.550458491 | 0.459812839 | -15.22821567 | 4.35E-21 | -15.77867416 | 9.810482605 | 9.443177293 | 10.67226762 | 9.506715015 | 9.5348592 | 2 | 2 |
| ENSG00000206403 | | -0.802709566 | 0.499134136 | -14.98336311 | 3.81E-17 | -15.78607267 | 10.75275473 | 9.45234302 | 8.206335842 | 8.978632046 | 9.07377798 | 2 | 2 |
| ENSG00000165379 | LRFN5 | -6.625870382 | 3.47E-24 | -9.167241308 | 6.52E-43 | -15.79311169 | 9.867565369 | 10.32888058 | 11.27276688 | 4.139692846 | 4.45712836 | 2.911488689 | 2.551896972 |
| ENSG00000183172 | SMDT1 | -2.064171506 | 0.031754273 | -13.72989142 | 2.77E-15 | -15.79406292 | 9.460782017 | 8.004992147 | 7.552162999 | 6.64692533 | 6.47451726 | 2 | 2 |
| ENSG00000274958 | | -1.625845911 | 0.002500823 | -14.17391528 | 3.68E-19 | -15.79976119 | 8.797441463 | 9.461450882 | 8.65477198 | 7.481152682 | 7.35080119 | 2 | 2 |
| C14ORF166 | C14ORF166 | -0.562485462 | 0.441343032 | -15.23946381 | 3.39E-21 | -15.80194927 | 10.59268348 | 9.318399851 | 10.06083273 | 9.511620236 | 9.52849248 | 2 | 2 |
| FAM103A1 | | -1.696909959 | 0.066566843 | -14.10532119 | 1.77E-16 | -15.80223114 | 9.810482605 | 8.169643011 | 8.236946975 | 7.341208691 | 7.21862736 | 2 | 2 |
| ENSG00000146276 | GABRR1 | -6.462613674 | 0.241959192 | -9.340880047 | 0.074484213 | -15.80349372 | 5.866944937 | 2 | 2 | 2 | 2.14022021 | 2 | 2 |
| ENSG00000162571 | TTLL10 | -6.462613674 | 0.241959192 | -9.340880047 | 0.074484213 | -15.80349372 | 5.866944937 | 2 | 2 | 2 | 2.14022021 | 2 | 2 |
| ENSG00000169129 | AFAP1L2 | -13.361369 | 2.37E-16 | -2.442542977 | 8.11E-05 | -15.80391198 | 8.928409675 | 9.558016677 | 10.17589429 | 2 | 2 | 7.15911073 | 7.299480881 |
| ENSG00000160808 | MYL3 | -11.58268263 | 3.79E-08 | -4.233803653 | 0.005275183 | -15.81648629 | 4.815609471 | 7.503546079 | 8.960138144 | 2 | 2 | 3.56816602 | 4.376876162 |
| ENSG00000115307 | AUP1 | -14.40277027 | 8.03E-17 | -1.424359649 | 0.132806015 | -15.82712992 | 11.57661845 | 10.01303782 | 9.703463497 | 2 | 2 | 9.252303334 | 9.26976224 |
| ENSG00000206454 | | -7.997322373 | 3.56E-09 | -7.84874044 | 5.65E-15 | -15.84606281 | 6.816637977 | 6.04685525 | 7.225742779 | 2 | 2.2680111 | 2.139668783 | 2.172608095 |
| ENSG00000236346 | | -1.662790496 | 0.164155144 | -14.18504695 | 7.42E-15 | -15.84783745 | 10.09970294 | 7.954278496 | 7.818184403 | 7.326437704 | 7.45518898 | 2 | 2 |
| ENSG00000235712 | | -0.953021674 | 0.490247308 | -14.90219956 | 1.69E-15 | -15.85522123 | 7.53297911 | 9.328390744 | 10.76969939 | 8.767939462 | 8.82635531 | 2 | 2 |
| ENSG00000164691 | TAGAP | -9.892004321 | 5.72E-06 | -5.968529531 | 0.00012298 | -15.86053385 | 5.095959458 | 7.431218269 | 8.608698767 | 2 | 2.14022021 | 3.27687201 | 2.172608095 |
| ENSG00000234243 | | -2.783660372 | 0.019862551 | -13.08143963 | 2.32E-12 | -15.8651 | 6.96351253 | 6.602424674 | 9.014156722 | 5.178261742 | 5.39133044 | 2 | 2 |
| FAM96A | | -1.570265086 | 0.001142768 | -14.30283782 | 7.99E-20 | -15.8731029 | 9.460782017 | 8.717821467 | 9.180874355 | 7.638916932 | 7.55566492 | 2 | 2 |
| ENSG00000234951 | | -4.652508551 | 0.04670103 | -11.23298878 | 1.93E-05 | -15.88549733 | 7.33131396 | 5.595927442 | 2 | 2.695456494 | 2.77779907 | 2 | 2 |
| ENSG00000166415 | WDR72 | -6.004910762 | 0.029514146 | -9.88555653 | 0.0005163 | -15.89046729 | 5.866944937 | 2 | 4.874587708 | 2.141737246 | 2.14022021 | 2 | 2 |
| ENSG00000143549 | TPM3 | -15.29392071 | 2.89E-22 | -0.597023037 | 0.262103779 | -15.89094374 | 11.74093347 | 11.4868636 | 11.46280979 | 2 | 2 | 10.53282379 | 11.3098158 |
| ENSG00000162959 | MEMO1 | -12.23512577 | 1.37E-13 | -3.662169031 | 2.81E-08 | -15.89729481 | 9.081016759 | 7.819099893 | 8.408111533 | 2 | 2 | 5.180212365 | 4.870334174 |
| ENSG00000002586 | CD99 | 0.5849006 | 0.464793444 | -16.48426059 | 4.80E-24 | -15.89935999 | 12.01383685 | 10.79296235 | 10.7904855 | 11.90585331 | 11.9059382 | 2 | 2 |
| ENSG00000185666 | SYN3 | -7.235163111 | 0.187020863 | -8.673210621 | 0.101181698 | -15.90837373 | 2 | 3.686303414 | 4.874587708 | 2 | 2 | 2 | 2 |
| ENSG00000189068 | VSTM1 | -7.235163111 | 0.187020863 | -8.673210621 | 0.101181698 | -15.90837373 | 2 | 3.686303414 | 4.874587708 | 2 | 2 | 2 | 2 |
| ENSG00000143727 | ACP1 | -14.55557338 | 5.90E-20 | -1.362682189 | 0.011729059 | -15.91825557 | 11.32135895 | 10.4709455 | 10.54349105 | 2 | 2 | 9.482666068 | 9.465740058 |
| METTL12 | | -3.642177803 | 0.003766598 | -12.27826916 | 1.51E-10 | -15.92044696 | 5.70932728 | 6.228547981 | 8.236946975 | 3.997744386 | 3.82856379 | 2 | 2 |
| PRR25 | | -5.52740652 | 0.164314918 | -10.40803436 | 0.008442457 | -15.93544088 | 2 | 2 | 6.882775709 | 2.270786764 | 2.2680111 | 2 | 2 |
| ENSG00000234798 | | -2.017437175 | 0.016673089 | -13.94252247 | 1.54E-16 | -15.95995964 | 8.138771685 | 8.124485672 | 9.589475645 | 6.83085985 | 6.79406287 | 2 | 2 |
| ENSG00000170788 | DYDC1 | -6.042379204 | 0.006393932 | -9.927905237 | 3.36E-05 | -15.97028444 | 4.467311984 | 4.11352228 | 5.772716771 | 2.270786764 | 2 | 2 | 2 |
| ATP5J2 | ATP5J2 | -0.825717793 | 0.27841917 | -15.14871994 | 1.52E-20 | -15.97443774 | 10.65885801 | 9.532315021 | 9.439621147 | 9.167628414 | 9.16894939 | 2 | 2 |
| ENSG00000146166 | LGSN | -7.969309234 | 0.140578614 | -8.010108785 | 0.130419453 | -15.97941802 | 2 | 2 | 5.930865648 | 2 | 2 | 2.048074405 | 2 |
| ENSG00000198842 | DUSP27 | -7.969309234 | 0.140578614 | -8.010108785 | 0.130419453 | -15.97941802 | 2 | 2 | 5.930865648 | 2 | 2 | 2.048074405 | 2 |
| ENSG00000180432 | CYP8B1 | -8.769443765 | 0.006205289 | -7.218947353 | 0.013945355 | -15.98839112 | 2 | 3.076665209 | 6.629488387 | 2 | 2 | 2.094598369 | 2.059860178 |
| ATP5I | ATP5I | -1.280151129 | 0.070638909 | -14.70946185 | 1.50E-19 | -15.98961298 | 9.975371636 | 8.655694736 | 9.725218844 | 8.266399315 | 8.29975519 | 2 | 2 |
| ENSG00000163833 | FBXO40 | -7.281254634 | 0.042202744 | -8.719251418 | 0.012260343 | -16.00050605 | 4.00722531 | 3.076665209 | 4.523675391 | 2 | 2 | 2 | 2 |
| ENSG00000128709 | HOXD9 | -2.762806071 | 0.001048931 | -13.24392154 | 7.53E-15 | -16.00672761 | 7.709880939 | 7.232742069 | 8.865353883 | 5.552894743 | 5.34860058 | 2 | 2 |
| ENSG00000168092 | PAFAH1B2 | -15.18846877 | 5.18E-21 | -0.818283932 | 0.219386965 | -16.0067527 | 10.79750658 | 11.38523987 | 11.97287769 | 2 | 2 | 10.45290995 | 10.81806159 |
| UFD1L | | -0.882750565 | 0.338872911 | -15.1263472 | 2.70E-19 | -16.00909777 | 10.03472846 | 8.717821467 | 10.58572549 | 8.985700128 | 9.18624204 | 2 | 2 |
| ENSG00000181143 | MUC16 | -5.562608742 | 0.114438327 | -10.45056163 | 0.003368852 | -16.01317037 | 6.009025997 | 2 | 5.930865648 | 2.498691189 | 2 | 2 | 2 |
| ENSG00000224940 | PRRT4 | -10.28988296 | 0.000161254 | -5.742289316 | 0.013722868 | -16.03217227 | 7.667647223 | 6.464272742 | 2 | 2 | 2 | 2.775316226 | 2.225841198 |
| ENSG00000162882 | HAAO | -10.37356544 | 8.68E-18 | -5.668331318 | 1.54E-23 | -16.04189676 | 8.696033109 | 8.820493869 | 9.61300511 | 2.270786764 | 2 | 3.993419637 | 3.753786374 |
| LOC728485 | | -6.581760395 | 0.010454649 | -9.460934638 | 0.000119309 | -16.04269503 | 5.095959458 | 4.710564154 | 3.369317972 | 2 | 2.14022021 | 2 | 2 |
| ENSG00000125848 | FLRT3 | -6.409446965 | 5.95E-09 | -9.655944293 | 1.90E-16 | -16.06539126 | 6.366709671 | 9.687856435 | 9.014156722 | 3.349643765 | 3.15372745 | 2.267001569 | 2.117335275 |
| ENSG00000124260 | MAGEA10 | -6.597258651 | 0.013722176 | -9.476346418 | 0.000212929 | -16.07360507 | 4.815609471 | 3.076665209 | 5.156626401 | 2 | 2.14022021 | 2 | 2 |
| ENSG00000158089 | GALNT14 | -9.533616796 | 2.45E-06 | -6.543819863 | 7.73E-07 | -16.07743666 | 6.009025997 | 8.487557258 | 5.39244166 | 2 | 2.14022021 | 2.560105256 | 2.421067459 |
| ENSG00000274296 | | -2.285710391 | 0.113324339 | -13.79601156 | 2.55E-12 | -16.08172195 | 6.009025997 | 8.124485672 | 9.746650996 | 6.476767609 | 6.35006802 | 2 | 2 |
| ENSG00000145194 | ECE2 | -3.645858797 | 8.78E-08 | -12.44324343 | 1.20E-14 | -16.08910222 | 9.015175615 | 7.73146271 | 9.083172337 | 5.194621444 | 5.2591458 | 2.048074405 | 2 |
| ENSG00000184344 | GDF3 | -7.327341183 | 0.180987334 | -8.765389844 | 0.097152724 | -16.09273103 | 5.095959458 | 2 | 3.369317972 | 2 | 2 | 2 | 2 |
| ENSG00000164488 | DACT2 | -8.908890787 | 0.009085099 | -7.208443499 | 0.027721408 | -16.11733429 | 2 | 2 | 9.227351203 | 2.270786764 | 2 | 2.62493832 | 2.172608095 |
| ENSG00000235439 | | -0.254488501 | 0.764007023 | -15.86288542 | 2.39E-22 | -16.11737392 | 11.38138789 | 10.29361762 | 10.06936278 | 10.42990227 | 10.4653998 | 2 | 2 |
| ENSG00000126262 | FFAR2 | -8.121838862 | 0.131866271 | -7.999631488 | 0.129694026 | -16.12147035 | 2 | 2 | 6.073378139 | 2 | 2 | 2 | 2.059860178 |
| ENSG00000274580 | | -4.773665395 | 0.000112178 | -11.35523938 | 2.12E-09 | -16.12890477 | 6.366709671 | 4.710564154 | 6.958202216 | 2.784612485 | 2.68924543 | 2 | 2 |
| ENSG00000160654 | CD3G | -7.349329021 | 0.179452048 | -8.787377956 | 0.096127185 | -16.13670698 | 4.467311984 | 2 | 4.523675391 | 2 | 2 | 2 | 2 |
| ENSG00000143921 | ABCG8 | -9.445200745 | 0.018048848 | -6.692021648 | 0.075265997 | -16.13722239 | 2 | 2 | 7.342949392 | 2 | 2 | 2.225793514 | 2.117335275 |
| ENSG00000056291 | NPFFR2 | -5.635746983 | 0.000472147 | -10.51895583 | 3.06E-07 | -16.15470281 | 4.00722531 | 5.838937808 | 5.930865648 | 2.3892346 | 2.14022021 | 2 | 2 |
| ENSG00000118946 | PCDH17 | -4.055994607 | 0.085411097 | -12.10697349 | 3.81E-06 | -16.1629681 | 2 | 6.464272742 | 8.175061096 | 3.566290607 | 3.45167086 | 2 | 2 |
| APITD1 | APITD1 | -7.364994283 | 0.178432005 | -8.803043414 | 0.095461226 | -16.1680377 | 3.327711234 | 5.131603625 | 2 | 2 | 2 | 2 | 2 |
| TMEM261 | | -1.62692042 | 0.018869516 | -14.55829568 | 3.91E-19 | -16.1852161 | 10.04301211 | 8.993716073 | 8.865353883 | 7.717440541 | 7.86451142 | 2 | 2 |
| ENSG00000106410 | NOBOX | -7.381755626 | 0.177300996 | -8.819804966 | 0.094749097 | -16.20156059 | 2 | 2 | 5.39244166 | 2 | 2 | 2 | 2 |
| ENSG00000142319 | SLC6A3 | -7.381755626 | 0.177300996 | -8.819804966 | 0.094749097 | -16.20156059 | 2 | 2 | 5.39244166 | 2 | 2 | 2 | 2 |
| ENSG00000150244 | TRIM48 | -7.381755626 | 0.177300996 | -8.819804966 | 0.094749097 | -16.20156059 | 2 | 2 | 5.39244166 | 2 | 2 | 2 | 2 |
| ENSG00000185594 | SPATA8 | -7.381755626 | 0.177300996 | -8.819804966 | 0.094749097 | -16.20156059 | 2 | 2 | 5.39244166 | 2 | 2 | 2 | 2 |
| ENSG00000254709 | IGLL5 | -7.381755626 | 0.177300996 | -8.819804966 | 0.094749097 | -16.20156059 | 2 | 2 | 5.39244166 | 2 | 2 | 2 | 2 |
| PTPN20B | | -7.381755626 | 0.177300996 | -8.819804966 | 0.094749097 | -16.20156059 | 2 | 2 | 5.39244166 | 2 | 2 | 2 | 2 |
| ENSG00000165672 | PRDX3 | -14.74150712 | 5.04E-21 | -1.466016373 | 0.000902647 | -16.20752349 | 11.04852648 | 10.62073637 | 11.30562889 | 2 | 2 | 9.445288547 | 9.659568662 |
| ENSG00000178605 | GTPBP6 | -2.11681895 | 0.026099339 | -14.12113407 | 3.76E-16 | -16.23795302 | 8.042902253 | 8.255926468 | 9.868959282 | 6.820355514 | 6.95464102 | 2 | 2 |
| FAM213B | | -1.611264306 | 0.087402267 | -14.62711622 | 1.42E-17 | -16.23838053 | 10.32020829 | 8.414402626 | 8.99637432 | 7.797276258 | 7.95036812 | 2 | 2 |
| ENSG00000005108 | THSD7A | -6.179493001 | 0.07325688 | -10.06669174 | 0.003506974 | -16.24618474 | 2 | 5.946638857 | 5.156626401 | 2.270786764 | 2 | 2 | 2 |
| ENSG00000256463 | SALL3 | -5.688112566 | 0.101987045 | -10.57605157 | 0.002709551 | -16.26416414 | 6.653098118 | 5.131603625 | 2 | 2.498691189 | 2 | 2 | 2 |
| C11ORF30 | C11ORF30 | -7.41314878 | 0.175217153 | -8.851198513 | 0.09357979 | -16.26434729 | 2 | 4.11352228 | 4.874587708 | 2 | 2 | 2 | 2 |
| ENSG00000179636 | TPPP2 | -7.423107256 | 0.174626306 | -8.861157113 | 0.093248514 | -16.28426437 | 4.00722531 | 4.93638933 | 2 | 2 | 2 | 2 | 2 |
| GPR98 | | -7.423107256 | 0.174626306 | -8.861157113 | 0.093248514 | -16.28426437 | 4.00722531 | 4.93638933 | 2 | 2 | 2 | 2 | 2 |
| ENSG00000198075 | SULT1C4 | -7.629800564 | 9.09E-15 | -8.665768256 | 1.84E-19 | -16.29556882 | 10.56993315 | 9.506147172 | 7.691296561 | 3.162721474 | 2.86123014 | 2.656294034 | 2.509591799 |
| C9ORF9 | C9ORF9 | -7.429082904 | 0.174256431 | -8.867132836 | 0.093025295 | -16.29621574 | 5.095959458 | 3.686303414 | 2 | 2 | 2 | 2 | 2 |
| ENSG00000114124 | GRK7 | -5.046139498 | 0.002229055 | -11.25007781 | 1.65E-07 | -16.2962173 | 3.327711234 | 6.728495673 | 6.629488387 | 2.695456494 | 2.49394783 | 2 | 2 |
| ENSG00000126756 | UXT | -12.96046027 | 1.37E-15 | -3.348244472 | 8.59E-09 | -16.30870475 | 9.613202746 | 8.505280662 | 9.399592459 | 2 | 2 | 5.916072173 | 6.041166521 |
| ENSG00000135624 | CCT7 | -15.46893557 | 1.91E-21 | -0.846953049 | 0.231007098 | -16.31588862 | 12.37867048 | 11.28620006 | 11.2690689 | 2 | 2 | 10.76797168 | 11.01768743 |

| | | | | | | | | | | | | | |
|---|---|---|---|---|---|---|---|---|---|---|---|---|---|
| ENSG00000124749 | COL21A1 | -12.681129 | 2.65E-13 | -3.643063962 | 2.28E-05 | -16.32419296 | 7.667647223 | 9.506147172 | 9.149036566 | 2 | 2 | 5.056402493 | 5.759151667 |
| ENSG00000275903 | | -0.795244619 | 0.277753086 | -15.53415616 | 9.06E-22 | -16.32940078 | 11.00678643 | 10.01303782 | 9.798878489 | 9.504256136 | 9.65516634 | 2 | 2 |
| ENSG00000263290 | | -0.718687458 | 0.335781482 | -15.61266708 | 6.31E-22 | -16.33135453 | 9.657036304 | 10.35355093 | 11.02781271 | 9.749588203 | 9.72225407 | 2 | 2 |
| ENSG00000126233 | SLURP1 | -9.387649848 | 0.075817354 | -6.953521096 | 0.17318175 | -16.34117094 | 2 | 2 | 7.285536013 | 2 | 2 | 2.267001569 | 2 |
| ATP5SL | ATP5SL | -1.437817064 | 0.315574141 | -14.90701547 | 9.10E-15 | -16.34483253 | 10.73241055 | 9.549500232 | 7.029880454 | 8.30834735 | 8.33666436 | 2 | 2 |
| ENSG00000156096 | UGT2B4 | -6.738021241 | 0.011560674 | -9.621220923 | 0.000165086 | -16.35924216 | 4.815609471 | 3.076665209 | 5.39244166 | 2.141737246 | 2 | 2 | 2 |
| ENSG00000096171 | | -1.333707208 | 0.014307407 | -15.04103633 | 1.81E-21 | -16.37474354 | 10.34051281 | 9.497318007 | 9.670202709 | 8.447952961 | 8.6601233 | 2 | 2 |
| ENSG00000167770 | OTUB1 | -14.8844889 | 7.57E-20 | -1.492980728 | 0.027841943 | -16.37746963 | 10.4253394 | 11.02260251 | 11.73625717 | 2 | 2 | 9.487849724 | 9.834971282 |
| ENSG00000162624 | LHX8 | -6.253610945 | 0.243970533 | -10.12684244 | 0.050493994 | -16.38045338 | 2 | 6.602424674 | 2 | 2 | 2.2680111 | 2 | 2 |
| ANKRD32 | ANKRD32 | -7.473157107 | 0.171238625 | -8.911207589 | 0.091164266 | -16.3843647 | 2 | 5.457120953 | 2 | 2 | 2 | 2 | 2 |
| LOH12CR1 | | -7.473157107 | 0.171238625 | -8.911207589 | 0.091164266 | -16.3843647 | 2 | 5.457120953 | 2 | 2 | 2 | 2 | 2 |
| ENSG00000105618 | PRPF31 | -1.248581675 | 0.022867725 | -15.14984203 | 9.26E-22 | -16.39842371 | 9.87686338 | 9.514922632 | 10.43545992 | 8.695066731 | 8.80280692 | 2 | 2 |
| C1ORF106 | C1ORF106 | -5.109175678 | 1.90E-08 | -11.31266132 | 3.10E-11 | -16.42183699 | 6.468621203 | 6.228547981 | 5.930865648 | 2.695456494 | 2.49394783 | 2 | 2 |
| KIAA0141 | | -1.306821379 | 0.529869493 | -15.14877311 | 1.87E-11 | -16.45559449 | 11.22635546 | 9.28800521 | 4.874587708 | 8.684969711 | 8.69749358 | 2 | 2 |
| AIM1L | AIM1L | -5.291990678 | 0.168363493 | -11.17259994 | 0.003581429 | -16.46459062 | 7.624139768 | 2 | 2 | 2.498691189 | 2.49394783 | 2 | 2 |
| ENSG00000179709 | NLRP8 | -7.517831367 | 0.0310894 | -8.955828044 | 0.008318112 | -16.47365941 | 4.00722531 | 3.076665209 | 4.874587708 | 2 | 2 | 2 | 2 |
| ENSG00000134193 | REG4 | -6.797722995 | 0.085935609 | -9.682926943 | 0.009999334 | -16.48064994 | 3.327711234 | 6.04685525 | 2 | 2.141737246 | 2 | 2 | 2 |
| ENSG00000022355 | GABRA1 | -7.527218594 | 0.167701797 | -8.965269751 | 0.088907874 | -16.49248834 | 3.327711234 | 5.303523747 | 2 | 2 | 2 | 2 | 2 |
| ENSG00000184258 | CDR1 | -7.527218594 | 0.167701797 | -8.965269751 | 0.088907874 | -16.49248834 | 3.327711234 | 5.303523747 | 2 | 2 | 2 | 2 | 2 |
| ENSG00000100012 | SEC14L3 | -8.230785059 | 0.021848633 | -8.265731947 | 0.019975613 | -16.49651701 | 4.815609471 | 2 | 5.59507254 | 2 | 2 | 2.048074405 | 2 |
| ENSG00000203760 | CENPW | -11.95205247 | 5.41E-11 | -4.554166538 | 8.16E-06 | -16.50621901 | 9.219014269 | 7.572420273 | 6.958202216 | 2 | 2 | 3.930690651 | 4.188602707 |
| ENSG00000166405 | RIC3 | -7.745917373 | 1.25E-07 | -8.768964267 | 1.89E-10 | -16.51488164 | 5.330594134 | 6.228547981 | 8.206335842 | 2.270786764 | 2.14022021 | 2.094598369 | 2.117335275 |
| ENSG00000168081 | PNOC | -7.539321547 | 0.167050868 | -8.977372856 | 0.088390629 | -16.5166944 | 5.532353208 | 2 | 2 | 2 | 2 | 2 | 2 |
| ENSG00000236032 | OR5H14 | -7.539321547 | 0.167050868 | -8.977372856 | 0.088390629 | -16.5166944 | 5.532353208 | 2 | 2 | 2 | 2 | 2 | 2 |
| ENSG00000075043 | KCNQ2 | -6.033029888 | 0.090520336 | -10.5057349 | 0.003407034 | -16.53876479 | 2 | 6.728495673 | 4.523675391 | 2.3892346 | 2 | 2 | 2 |
| ATP5G3 | ATP5G3 | -0.776632518 | 0.355101172 | -15.76488637 | 1.37E-21 | -16.54151888 | 11.33828573 | 9.863030423 | 10.16797309 | 9.853054547 | 9.80623425 | 2 | 2 |
| ENSG00000182168 | UNC5C | -11.6802185 | 3.29E-11 | -4.863558103 | 4.55E-08 | -16.5437766 | 9.112843761 | 9.497318007 | 9.565556058 | 2.141737246 | 2 | 5.413723492 | 3.539239711 |
| ENSG00000124562 | SNRPC | -14.28585624 | 1.44E-19 | -2.259160623 | 1.17E-06 | -16.54501686 | 10.6695979 | 10.06773758 | 10.84614969 | 2 | 2 | 8.222994258 | 8.407389648 |
| ENSG00000224455 | | -1.66820197 | 0.00311339 | -14.88615665 | 7.94E-21 | -16.55435862 | 10.22876205 | 9.506147172 | 9.272377397 | 8.029321023 | 8.1187584 | 2 | 2 |
| ENSG00000197712 | FAM114A1 | -16.13812489 | 2.90E-24 | -0.416353402 | 0.49597846 | -16.55447829 | 11.95579124 | 12.24340894 | 12.87940139 | 2 | 2 | 11.85402479 | 12.12597882 |
| ENSG00000171236 | LRG1 | -6.848977939 | 0.008746159 | -9.728101464 | 0.000101151 | -16.5770794 | 4.00722531 | 3.686303414 | 5.772716771 | 2 | 2.14022021 | 2 | 2 |
| ENSG00000168952 | STXBP6 | -11.61753967 | 4.89E-21 | -4.960185646 | 1.50E-15 | -16.57772532 | 10.04301211 | 10.11467741 | 10.74860941 | 2.141737246 | 2.14022021 | 4.841348851 | 5.96101835 |
| ENSG00000275572 | GRIFIN | -9.015680226 | 0.010605213 | -7.563369916 | 0.021855645 | -16.57905014 | 5.866944937 | 2 | 6.073378139 | 2 | 2 | 2.139668783 | 2 |
| HN1 | | -1.613749404 | 0.483380008 | -14.96988625 | 5.72E-10 | -16.58363565 | 11.00254523 | 9.308339287 | 3.369317972 | 8.188900477 | 8.23280093 | 2 | 2 |
| ENSG00000215131 | C16orf90 | -8.799705346 | 0.033496988 | -7.788680822 | 0.049023059 | -16.58838617 | 2 | 2 | 6.718950348 | 2 | 2 | 2.048074405 | 2.059860178 |
| ENSG00000152795 | HNRNPDL | -15.34615832 | 5.54E-22 | -1.245552506 | 0.025960298 | -16.59171083 | 11.99474663 | 11.38523987 | 11.39656351 | 2 | 2 | 9.976966829 | 10.69176237 |
| ENSG00000132698 | RAB25 | -6.359425026 | 0.104994345 | -10.23319171 | 0.008232756 | -16.59261673 | 3.327711234 | 2 | 6.629488387 | 2 | 2.2680111 | 2 | 2 |
| ENSG00000186439 | TRDN | -7.577340319 | 0.164464069 | -9.015392102 | 0.086857455 | -16.59273242 | 4.467311984 | 2 | 4.874587708 | 2 | 2 | 2 | 2 |
| ENSG00000116857 | TMEM9 | -14.97784655 | 4.62E-21 | -1.617667864 | 0.002386617 | -16.59551442 | 11.26935334 | 10.67263725 | 11.65641302 | 2 | 2 | 9.592613586 | 9.687114391 |
| ENSG00000273707 | | -7.388721916 | 1.68E-23 | -9.217825479 | 2.45E-35 | -16.60654739 | 10.57565445 | 10.54992047 | 9.452720984 | 3.515101962 | 3.50452879 | 2.858551861 | 2.277179783 |
| ENSG00000213588 | ZBTB9 | -2.588545495 | 2.35E-08 | -14.03464276 | 3.83E-19 | -16.62318826 | 9.081016759 | 8.469613413 | 9.031722608 | 6.252910529 | 6.44777477 | 2 | 2 |
| ENSG00000171227 | TMEM37 | -11.82200887 | 5.52E-11 | -4.805041237 | 9.42E-07 | -16.62705011 | 7.750913381 | 7.979858146 | 8.511888463 | 2 | 2 | 4.331190017 | 2.885414003 |
| ENSG00000111144 | LTA4H | -14.46504117 | 1.19E-15 | -2.16261684 | 0.047129776 | -16.62765801 | 11.78851328 | 9.792607274 | 9.478569125 | 2 | 2 | 8.646918883 | 8.532164007 |
| ENSG00000183150 | GPR19 | -8.81223355 | 0.000157526 | -7.818526884 | 0.000187696 | -16.63076043 | 8.797441463 | 6.844429755 | 2 | 2 | 2.2680111 | 2.3070652 | 2.225841198 |
| ENSG00000128322 | IGLL1 | -5.882018696 | 0.132750246 | -10.76264904 | 0.005903997 | -16.64466773 | 2 | 2 | 7.225742779 | 2.270786764 | 2.2680111 | 2 | 2 |
| ENSG00000144290 | SLC4A10 | -7.605258208 | 0.162882455 | -9.04331034 | 0.085697496 | -16.64856855 | 2 | 2 | 5.59507254 | 2 | 2 | 2 | 2 |
| ENSG00000183034 | OTOP2 | -7.605258208 | 0.162882455 | -9.04331034 | 0.085697496 | -16.64856855 | 2 | 2 | 5.59507254 | 2 | 2 | 2 | 2 |
| ENSG00000185966 | LCE3E | -7.605258208 | 0.162882455 | -9.04331034 | 0.085697496 | -16.64856855 | 2 | 2 | 5.59507254 | 2 | 2 | 2 | 2 |
| ENSG00000276684 | | -7.605258208 | 0.162882455 | -9.04331034 | 0.085697496 | -16.64856855 | 2 | 2 | 5.59507254 | 2 | 2 | 2 | 2 |
| GPR133 | | -7.605258208 | 0.162882455 | -9.04331034 | 0.085697496 | -16.64856855 | 2 | 2 | 5.59507254 | 2 | 2 | 2 | 2 |
| PPAP2C | | -7.605258208 | 0.162882455 | -9.04331034 | 0.085697496 | -16.64856855 | 2 | 2 | 5.59507254 | 2 | 2 | 2 | 2 |
| ENSG00000234920 | | -3.949931237 | 0.007453224 | -12.71413449 | 5.12E-10 | -16.66406573 | 7.904282243 | 4.442710129 | 8.296287027 | 4.139692846 | 3.86943969 | 2 | 2 |
| ENSG00000164588 | HCN1 | -6.893584994 | 0.079982596 | -9.778779368 | 0.008998799 | -16.67236436 | 2 | 6.140560021 | 3.369317972 | 2.141737246 | 2 | 2 | 2 |
| ENSG00000103415 | HMOX2 | -1.8545673 | 0.012970946 | -14.83049012 | 2.41E-19 | -16.68505741 | 9.304185368 | 8.94259736 | 10.37481416 | 7.826439377 | 7.84670947 | 2 | 2 |
| ENSG00000118526 | TCF21 | -1.950013457 | 1.06E-06 | -14.74765076 | 1.89E-21 | -16.69766422 | 9.790941832 | 9.267380431 | 9.670202709 | 7.602596956 | 7.71833702 | 2 | 2 |
| GPR56 | | -7.631360786 | 0.161320596 | -9.069413244 | 0.08467269 | -16.70077403 | 2 | 4.11352228 | 5.156626401 | 2 | 2 | 2 | 2 |
| ENSG00000161905 | ALOX15 | -7.627171162 | 0.007409183 | -9.097276235 | 0.000960705 | -16.7244474 | 6.563806279 | 5.131603625 | 2 | 2 | 2.14022021 | 2.048074405 | 2 |
| ENSG00000113327 | GABRG2 | -8.434349714 | 0.115724282 | -8.312146243 | 0.113902532 | -16.74649596 | 6.366709671 | 2 | 2 | 2 | 2 | 2 | 2.059860178 |
| BRE | BRE | -2.216251242 | 5.37E-11 | -14.53151436 | 3.43E-21 | -16.7477656 | 9.544849385 | 9.267380431 | 9.301632639 | 7.19843704 | 7.18258783 | 2 | 2 |
| ENSG00000100385 | IL2RB | -6.859826352 | 2.73E-14 | -9.903645837 | 1.68E-21 | -16.76347219 | 8.075571721 | 8.487557258 | 9.049077186 | 3.023133209 | 2.68924543 | 2 | 2.277179783 |
| GBAS | | -1.847023831 | 2.80E-05 | -14.92005945 | 1.11E-21 | -16.76708328 | 10.1076228 | 9.443177293 | 9.658943166 | 7.959237999 | 7.90439517 | 2 | 2 |
| ATPIF1 | ATPIF1 | -1.203799463 | 0.018926992 | -15.56976991 | 3.46E-23 | -16.77356937 | 10.71696191 | 9.876711741 | 10.51254821 | 9.221512599 | 9.20032903 | 2 | 2 |
| ENSG00000077009 | NMRK2 | -9.973275412 | 0.057053589 | -6.804453028 | 0.180853 | -16.77772844 | 2 | 2 | 7.858115942 | 2 | 2 | 2.42098416 | 2 |
| ENSG00000141101 | NOB1 | -13.51643642 | 2.21E-17 | -3.264256948 | 5.41E-11 | -16.78069337 | 9.848787621 | 9.256956408 | 10.15199903 | 2 | 2 | 6.674237832 | 6.493217192 |
| C1ORF233 | C1ORF233 | -7.672979507 | 0.158698358 | -9.111032485 | 0.083004924 | -16.78401199 | 3.327711234 | 5.457120953 | 2 | 2 | 2 | 2 | 2 |
| ENSG00000233801 | | -1.435015151 | 2.58E-07 | -15.34914699 | 7.41E-24 | -16.78416214 | 10.15425007 | 10.160138 | 10.2528023 | 8.784246083 | 8.74082114 | 2 | 2 |
| ENSG00000189233 | NUGGC | -6.16552169 | 0.000727989 | -10.62943368 | 9.74E-07 | -16.79495537 | 5.330594134 | 3.686303414 | 6.534109959 | 2.141737246 | 2.2680111 | 2 | 2 |
| ENSG00000070886 | EPHA8 | -7.610567337 | 0.023911778 | -9.184523918 | 0.005293456 | -16.79509125 | 2 | 2 | 8.511888463 | 2.141737246 | 2.2680111 | 2.139668783 | 2 |
| ENSG00000214736 | TOMM6 | -1.644410552 | 0.003319963 | -15.15299243 | 1.42E-21 | -16.79740298 | 10.4253394 | 9.415325457 | 9.972637969 | 8.279879181 | 8.43865202 | 2 | 2 |
| ENSG00000122140 | MRPS2 | -14.40789173 | 1.43E-15 | -2.392467926 | 0.026069096 | -16.80035966 | 11.6462383 | 9.00621767 | 10.2528023 | 2 | 2 | 8.369127483 | 8.242012195 |
| C7ORF49 | C7ORF49 | -2.454658723 | 5.19E-05 | -14.34611829 | 5.23E-19 | -16.80077702 | 9.667789716 | 8.505280662 | 9.180874355 | 6.716553866 | 6.84106088 | 2 | 2 |
| CPSF3L | CPSF3L | -1.572216064 | 0.118300026 | -15.23400382 | 1.76E-18 | -16.80621989 | 11.01523161 | 9.148351514 | 9.227351203 | 8.486665614 | 8.53866332 | 2 | 2 |
| ENSG00000267855 | NDUFA7 | -11.42870315 | 9.09E-10 | -5.377529853 | 7.78E-07 | -16.806233 | 8.696033109 | 6.951736442 | 6.629488387 | 2 | 2 | 3.429858444 | 2.851942824 |
| ENSG00000196844 | PATE2 | -8.843041459 | 0.011524016 | -7.965875166 | 0.016562656 | -16.80891662 | 6.138360429 | 2 | 5.39244166 | 2 | 2 | 2.094598369 | 2 |
| ATP5D | ATP5D | -1.681069747 | 0.000402654 | -15.13587886 | 4.39E-22 | -16.8169486 | 10.31337612 | 9.981728319 | 9.528916236 | 8.241026137 | 8.37265277 | 2 | 2 |
| ENSG00000163956 | LRPAP1 | -15.63427014 | 1.45E-21 | -1.184926811 | 0.101824211 | -16.81919695 | 12.58137207 | 11.23134822 | 11.55376643 | 2 | 2 | 10.70681251 | 10.74487048 |
| ENSG00000230230 | | -0.863823127 | 0.183362892 | -15.95724924 | 1.68E-23 | -16.82107236 | 10.92846914 | 10.04973379 | 11.18521898 | 9.888054803 | 9.97960581 | 2 | 2 |
| ENSG00000089693 | MLF2 | -15.96100015 | 3.54E-23 | -0.861647162 | 0.185198772 | -16.82264732 | 12.40461712 | 11.47342019 | 12.59919771 | 2 | 2 | 11.20126917 | 11.52958355 |
| ENSG00000196591 | HDAC2 | -14.82179851 | 4.73E-22 | -2.002333599 | 6.83E-13 | -16.82413211 | 11.22635546 | 11.01334259 | 11.04521411 | 2 | 2 | 9.152412913 | 9.052154518 |
| NARFL | | -2.233834171 | 0.004148594 | -14.61390901 | 1.85E-18 | -16.84774319 | 9.922475255 | 8.297207175 | 9.69246155 | 7.190301983 | 7.31425651 | 2 | 2 |
| ENSG00000236222 | | -1.489081126 | 0.019227846 | -15.372549 | 1.13E-21 | -16.86163012 | 10.31337612 | 9.433952963 | 10.64399316 | 8.744518279 | 8.72001862 | 2 | 2 |
| LOC100996693 | | -5.403487965 | 0.16629284 | -11.45836736 | 0.003212725 | -16.86185533 | 7.904282243 | 2 | 2 | 2.784612485 | 2.2680111 | 2 | 2 |
| ENSG00000165795 | NDRG2 | -10.21026669 | 9.86E-54 | -6.659964216 | 6.66E-45 | -16.87023091 | 11.26581873 | 10.68435379 | 11.64504491 | 2.498691189 | 2.68924543 | 4.940196634 | 4.664017718 |
| LOC100130451 | | -6.998921973 | 0.001490774 | -9.881128031 | 3.32E-06 | -16.88005 | 5.330594134 | 4.442710129 | 4.874587708 | 2.141737246 | 2 | 2 | 2 |
| ENSG00000137161 | CNPY3 | -14.68622548 | 3.68E-20 | -2.201657697 | 3.78E-05 | -16.88788318 | 11.28340583 | 10.30378101 | 11.12511995 | 2 | 2 | 8.697250351 | 8.841275974 |
| ENSG00000228128 | | -1.839444256 | 0.196392556 | -15.04956737 | 7.47E-15 | -16.88901162 | 10.98545464 | 9.377333006 | 7.285536013 | 8.024753265 | 8.10812366 | 2 | 2 |
| MFSD4 | | -7.730973335 | 0.034783852 | -9.168975946 | 0.010097543 | -16.89994928 | 3.327711234 | 3.076665209 | 5.39244166 | 2 | 2 | 2 | 2 |
| CCBL1 | CCBL1 | -7.737316468 | 0.154010606 | -9.175370057 | 0.080179745 | -16.91268653 | 2 | 5.303523747 | 4.058998049 | 2 | 2 | 2 | 2 |
| TMEM194B | | -7.737316468 | 0.154010606 | -9.175370057 | 0.080179745 | -16.91268653 | 2 | 5.303523747 | 4.058998049 | 2 | 2 | 2 | 2 |
| ENSG00000064655 | EYA2 | -8.374401991 | 7.32E-25 | -8.5408363 | 2.16E-43 | -16.91523829 | 8.564112515 | 8.487557258 | 8.206335842 | 2.270786764 | 2.38539787 | 2.225793514 | 2.374681325 |
| ENSG00000111249 | CUX2 | -8.889340187 | 0.09564296 | -8.029919726 | 0.119677852 | -16.91925991 | 2 | 2 | 6.80318718 | 2 | 2 | 2.094598369 | 2 |
| ENSG00000159173 | TNNI1 | -9.438921799 | 7.20E-19 | -7.482617053 | 6.70E-22 | -16.92153885 | 8.737455672 | 9.125613817 | 9.528916236 | 2.3892346 | 2.14022021 | 2.42098416 | 3.183290849 |
| TCEB2 | | -1.009665968 | 0.140759233 | -15.91656627 | 4.98E-23 | -16.92623223 | 11.33828573 | 10.10308486 | 10.54960077 | 9.712435568 | 9.78435771 | 2 | 2 |
| ENSG00000159339 | PADI4 | -7.74496085 | 0.154010606 | -9.183014726 | 0.080201969 | -16.92797558 | 2 | 3.076665209 | 5.59507254 | 2 | 2 | 2 | 2 |
| ENSG00000140543 | DET1 | -11.87534931 | 7.26E-13 | -5.055073715 | 1.61E-14 | -16.93042303 | 8.631580145 | 7.954278496 | 7.777116083 | 2 | 2 | 3.336159116 | 3.908316285 |
| ENSG00000162998 | FRZB | -10.01457819 | 7.14E-115 | -6.918083699 | 8.28E-61 | -16.93266189 | 14.90935901 | 14.1054385 | 14.49069572 | 4.927308402 | 4.56059747 | 7.865372957 | 7.395535532 |
| ENSG00000133019 | CHRM3 | -5.862196443 | 0.001816155 | -11.0710254 | 1.14E-06 | -16.93322184 | 6.563806279 | 4.93638933 | 6.073378139 | 2.600425781 | 2 | 2 | 2 |
| ENSG00000023191 | RNH1 | -0.457354688 | 0.580917707 | -16.48912026 | 6.10E-24 | -16.94647495 | 12.02642475 | 10.8599287 | 10.71094342 | 10.81563567 | 10.9225423 | 2 | 2 |
| C19ORF83 | C19ORF83 | -7.757641634 | 0.151987999 | -9.195695109 | 0.078821897 | -16.95333674 | 5.095959458 | 2 | 4.523675391 | 2 | 2 | 2 | 2 |
| EFTUD1 | EFTUD1 | -7.760553034 | 0.15309083 | -9.198607105 | 0.079634469 | -16.95916014 | 2 | 5.72254323 | 2 | 2 | 2 | 2 | 2 |
| ENSG00000224774 | | -0.987915747 | 0.108262571 | -15.98011673 | 9.57E-24 | -16.96803247 | 11.33153883 | 10.27307349 | 10.64969256 | 9.861404292 | 9.80557618 | 2 | 2 |
| ENSG00000105583 | WDR83OS | -1.321650203 | 0.058965253 | -15.65578733 | 4.93E-22 | -16.97743753 | 11.08508709 | 9.733683991 | 10.35401885 | 9.138327453 | 9.21922971 | 2 | 2 |
| ENSG00000188487 | INSC | -5.89084461 | 0.138651767 | -11.09796558 | 0.005176626 | -16.98881019 | 2 | 2 | 7.552162999 | 2.498691189 | 2.14022021 | 2 | 2 |
| ENSG00000241186 | TDGF1 | -7.055786404 | 0.002404021 | -9.938294005 | 9.19E-06 | -16.99408041 | 4.815609471 | 5.595927442 | 4.058998049 | 2.141737246 | 2 | 2 | 2 |
| C14ORF2 | C14ORF2 | -1.84914422 | 0.00013089 | -15.1581486 | 4.64E-22 | -17.00729282 | 10.41260941 | 9.632483534 | 9.839340503 | 8.174538791 | 8.15434112 | 2 | 2 |
| LOC728392 | | -2.377717673 | 4.78E-05 | -14.65521679 | 6.04E-20 | -17.03293446 | 9.751047882 | 8.762721501 | 9.778213818 | 7.206526483 | 7.0991228 | 2 | 2 |
| ENSG00000145864 | GABRB2 | -8.572571233 | 0.000583912 | -8.462766456 | 5.03E-05 | -17.03533769 | 6.366709671 | 7.315436877 | 2 | 2.141737246 | 2 | 2.094598369 | 2.059860178 |

| | | | | | | | | | | | | | |
|---|---|---|---|---|---|---|---|---|---|---|---|---|---|
| ENSG00000122304 | PRM2 | -7.798747599 | 0.150705666 | -9.236802147 | 0.078206869 | -17.03554975 | 2 | 2 | 5.772716771 | 2 | 2 | 2 | 2 |
| ENSG00000166743 | ACSM1 | -7.798747599 | 0.150705666 | -9.236802147 | 0.078206869 | -17.03554975 | 2 | 2 | 5.772716771 | 2 | 2 | 2 | 2 |
| ENSG00000215547 | DEFB115 | -7.798747599 | 0.150705666 | -9.236802147 | 0.078206869 | -17.03554975 | 2 | 2 | 5.772716771 | 2 | 2 | 2 | 2 |
| ENSG00000236396 | SLC35G4 | -7.798747599 | 0.150705666 | -9.236802147 | 0.078206869 | -17.03554975 | 2 | 2 | 5.772716771 | 2 | 2 | 2 | 2 |
| SGOL2 | | -7.798747599 | 0.150705666 | -9.236802147 | 0.078206869 | -17.03554975 | 2 | 2 | 5.772716771 | 2 | 2 | 2 | 2 |
| ENSG00000162706 | CADM3 | -6.089848922 | 0.002287795 | -10.94601345 | 1.34E-07 | -17.03586237 | 2 | 8.004992147 | 10.15199903 | 3.615725078 | 3.2184422 | 2.048074405 | 2.117335275 |
| ENSG00000146282 | RARS2 | -14.54822994 | 1.35E-18 | -2.494566013 | 0.000351595 | -17.04279595 | 11.06489013 | 9.869887299 | 11.20857468 | 2 | 2 | 8.026277797 | 8.605550143 |
| FAM127A | | -0.874395563 | 0.391884695 | -16.20830731 | 3.43E-21 | -17.08270287 | 11.95579124 | 10.27823701 | 10.14394518 | 10.14060551 | 10.2069242 | 2 | 2 |
| ENSG00000145832 | SLC25A48 | -11.07440799 | 5.07E-05 | -6.017436359 | 0.010066815 | -17.09184434 | 8.444143041 | 6.78762695 | 5.39244166 | 2 | 2 | 3.172301926 | 2 |
| ENSG00000060709 | RIMBP2 | -9.054056141 | 0.002337322 | -8.045158834 | 0.003363005 | -17.09921498 | 2 | 4.11352228 | 6.80318718 | 2 | 2 | 2.048074405 | 2.059860178 |
| ENSG00000206262 | FOXL2NB | -8.92561242 | 5.27E-05 | -8.174748797 | 6.67E-05 | -17.10036122 | 2 | 6.04685525 | 9.714382178 | 2.141737246 | 2.2680111 | 2.384001718 | 2.277179783 |
| ENSG00000124783 | SSR1 | -15.89300937 | 1.22E-22 | -1.223278046 | 0.06883389 | -17.11628741 | 11.36496218 | 12.22881472 | 12.63555306 | 2 | 2 | 10.73886083 | 11.12688673 |
| ENSG00000226225 | | -1.017423177 | 0.381381705 | -16.10985322 | 1.78E-20 | -17.1272764 | 13.46777257 | 11.36354124 | 11.15748401 | 11.36437338 | 11.4085069 | 2 | 2.059860178 |
| ENSG00000232280 | | -1.197872574 | 0.03884132 | -15.93071867 | 9.42E-24 | -17.12859125 | 11.19324461 | 10.15453315 | 10.78012988 | 9.527851557 | 9.62191107 | 2 | 2 |
| ENSG00000168000 | BSCL2 | -12.65354496 | 5.31E-14 | -4.483754142 | 7.30E-10 | -17.1372991 | 8.107517749 | 9.624396729 | 8.65477198 | 2 | 2 | 4.572179576 | 4.793076545 |
| ENSG00000182004 | SNRPE | -12.9739239 | 4.02E-13 | -4.184639094 | 1.54E-05 | -17.15856299 | 10.23600589 | 8.414402626 | 8.175061096 | 2 | 2 | 5.333556395 | 5.108633494 |
| ENSG00000171794 | UTF1 | -7.860617787 | 0.146925888 | -9.298673108 | 0.075921834 | -17.1592909 | 5.70932728 | 3.076665209 | 2 | 2 | 2 | 2 | 2 |
| ENSG00000147274 | RBMX | -15.19934388 | 4.24E-23 | -1.963685261 | 1.07E-11 | -17.16302914 | 11.40728614 | 11.39715513 | 11.61039303 | 2 | 2 | 9.424950709 | 9.603161713 |
| ENSG00000141744 | PNMT | -9.085062307 | 1.61E-05 | -8.080859038 | 5.27E-06 | -17.16592135 | 4.815609471 | 4.442710129 | 6.43197674 | 2 | 2 | 2.048074405 | 2.059860178 |
| ENSG00000276463 | | -1.622202424 | 0.000194849 | -15.54511274 | 1.60E-23 | -17.16731516 | 10.09970294 | 10.23108619 | 10.74328839 | 8.767939462 | 8.77483902 | 2 | 2 |
| ENSG00000042304 | C2orf83 | -7.872067466 | 0.144467139 | -9.310122079 | 0.074292129 | -17.18218954 | 2 | 3.686303414 | 5.59507254 | 2 | 2 | 2 | 2 |
| ENSG00000137726 | FXYD6 | -8.344744409 | 1.58E-08 | -8.838659931 | 1.11E-09 | -17.18340434 | 11.48534568 | 10.60026656 | 9.287079172 | 3.840289928 | 2 | 3.10571689 | 2.746601469 |
| ENSG00000105402 | NAPA | -14.22996591 | 7.54E-17 | -2.954528176 | 0.000335973 | -17.18449409 | 11.31110672 | 10.10308486 | 9.503962296 | 2 | 2 | 7.386475997 | 7.749171054 |
| ATP5G1 | ATP5G1 | -2.488222925 | 1.22E-07 | -14.70902513 | 7.30E-21 | -17.19724805 | 9.949165863 | 9.377333006 | 9.227351203 | 7.161462517 | 7.03319408 | 2 | 2 |
| ENSG00000141698 | NT5C3B | -14.67694323 | 1.86E-18 | -2.524898987 | 0.000861059 | -17.20184221 | 11.57661845 | 9.849218119 | 10.94679967 | 2 | 2 | 8.387928811 | 8.494021883 |
| GUCY1A3 | | -6.01003437 | 0.128270761 | -11.21715427 | 0.004521991 | -17.22718863 | 7.667647223 | 2 | 2 | 2.498691189 | 2.14022021 | 2 | 2 |
| ENSG00000271043 | MTRNR2L2 | 0.238977344 | 0.74145361 | -17.46703197 | 5.47E-28 | -17.22805463 | 12.87380672 | 11.99322771 | 11.80682103 | 12.52089365 | 12.5632079 | 2 | 2 |
| ENSG00000232289 | | -7.902832044 | 0.144498138 | -9.340887892 | 0.074484213 | -17.24371994 | 5.866944937 | 2 | 2 | 2 | 2 | 2 | 2 |
| LECT1 | | -7.902832044 | 0.144498138 | -9.340887892 | 0.074484213 | -17.24371994 | 5.866944937 | 2 | 2 | 2 | 2 | 2 | 2 |
| ENSG00000214160 | ALG3 | -14.61191879 | 8.36E-20 | -2.644671196 | 1.58E-06 | -17.25658998 | 11.25872341 | 10.19331737 | 11.01462253 | 2 | 2 | 8.170345342 | 8.343344553 |
| ENSG00000124193 | SRSF6 | -15.38899009 | 8.63E-22 | -1.871093664 | 0.001171672 | -17.26008376 | 11.88382844 | 11.33904258 | 11.71738516 | 2 | 2 | 9.225442227 | 10.2062826 |
| ENSG00000182768 | NGRN | -15.27296411 | 1.35E-21 | -1.997459028 | 0.000337313 | -17.27042314 | 12.01804503 | 11.48909204 | 10.93751085 | 2 | 2 | 9.635529507 | 9.472638619 |
| ENSG00000100095 | SEZ6L | -6.699388816 | 0.039974799 | -10.57359291 | 0.001323662 | -17.27298172 | 2 | 5.946638857 | 6.203069917 | 2 | 2.2680111 | 2 | 2 |
| ENSG00000273594 | | -2.529119985 | 3.21E-07 | -14.75197018 | 1.79E-51 | -17.28109016 | 13.01313897 | 11.99636434 | 12.61966945 | 10.09121697 | 10.0618152 | 2.094598369 | 2.059860178 |
| ENSG00000124227 | ANKRD60 | -7.93245812 | 0.129721932 | -9.370508463 | 0.064527454 | -17.30296658 | 5.095959458 | 2 | 4.874587708 | 2 | 2 | 2 | 2 |
| ENSG00000221988 | PPT2 | -2.417288798 | 0.012512432 | -14.89024573 | 1.34E-17 | -17.30753453 | 8.107517749 | 9.799806309 | 10.43545992 | 7.330144653 | 7.35800033 | 2 | 2 |
| ENSG00000131914 | LIN28A | -7.104126063 | 2.06E-06 | -10.20344437 | 4.21E-09 | -17.30757043 | 8.674867122 | 6.140560021 | 5.39244166 | 2.3892346 | 2.38539787 | 2.094598369 | 2 |
| ENSG00000277134 | | -7.947227309 | 0.131244357 | -9.385278892 | 0.065559433 | -17.3325062 | 4.467311984 | 2 | 5.39244166 | 2 | 2 | 2 | 2 |
| ENSG00000173207 | CKS1B | -12.79322566 | 5.77E-12 | -4.545281146 | 3.27E-05 | -17.33850681 | 10.16187685 | 7.73146271 | 8.042706615 | 2 | 2 | 4.861667652 | 4.634793542 |
| ENSG00000151079 | KCNA6 | -7.967017724 | 0.129094584 | -9.405069063 | 0.064319529 | -17.37208679 | 4.00722531 | 5.595927442 | 2 | 2 | 2 | 2 | 2 |
| ENSG00000275837 | | -1.866386477 | 0.000962972 | -15.51030394 | 1.89E-22 | -17.37669041 | 10.41898845 | 10.74900803 | 9.681375057 | 8.516254096 | 8.47387021 | 2 | 2 |
| EMR1 | EMR1 | -7.969326963 | 0.140578614 | -9.407383641 | 0.072203243 | -17.3767106 | 2 | 2 | 5.930865648 | 2 | 2 | 2 | 2 |
| ENSG00000153822 | KCNJ16 | -7.969326963 | 0.140578614 | -9.407383641 | 0.072203243 | -17.3767106 | 2 | 2 | 5.930865648 | 2 | 2 | 2 | 2 |
| ENSG00000213822 | CEACAM18 | -7.969326963 | 0.140578614 | -9.407383641 | 0.072203243 | -17.3767106 | 2 | 2 | 5.930865648 | 2 | 2 | 2 | 2 |
| PYCRL | | -3.120029141 | 2.78E-15 | -14.25824813 | 3.95E-20 | -17.37827727 | 8.816894989 | 9.181799937 | 9.272377397 | 6.104969671 | 6.01871653 | 2 | 2 |
| ENSG00000223639 | | -0.138514409 | 0.770602177 | -17.24275302 | 3.01E-29 | -17.38126743 | 12.36055756 | 11.8287681 | 11.99766953 | 11.89275296 | 11.987549 | 2 | 2 |
| ENSG00000168843 | FSTL5 | -7.305532758 | 0.043742941 | -10.19038463 | 0.003202408 | -17.49591739 | 6.468621203 | 4.11352228 | 2 | 2.141737246 | 2 | 2 | 2 |
| ENSG00000133393 | FOPNL | -12.2200681 | 6.56E-11 | -5.286121991 | 1.73E-06 | -17.50619009 | 6.563806279 | 8.191702463 | 9.465702942 | 2 | 2 | 3.753554863 | 3.698365557 |
| ATP5G2 | ATP5G2 | -1.72197148 | 0.026151336 | -15.7930845 | 1.33E-21 | -17.51505597 | 11.36496218 | 9.842262059 | 10.23774519 | 8.934263273 | 8.90218458 | 2 | 2 |
| C2ORF71 | C2ORF71 | -6.826805378 | 0.198978044 | -10.70004343 | 0.037552988 | -17.52684881 | 2 | 2 | 7.163363831 | 2 | 2.2680111 | 2 | 2 |
| ENSG00000175497 | DPP10 | -6.322226464 | 0.058859905 | -11.21013608 | 0.001085539 | -17.53236255 | 5.532353208 | 7.315436877 | 2 | 2.498691189 | 2 | 2 | 2 |
| ENSG00000163288 | GABRB1 | -9.291966307 | 0.001246051 | -8.283340963 | 0.001696635 | -17.57530727 | 2 | 4.710564154 | 6.958202216 | 2 | 2 | 2.048074405 | 2.059860178 |
| ENSG00000169093 | ASMTL | -2.304980599 | 0.094222712 | -15.27700658 | 2.21E-15 | -17.58198718 | 11.18951828 | 9.672251278 | 7.502634626 | 7.836899669 | 7.8286851 | 2 | 2 |
| ENSG00000010932 | FMO1 | -9.742292799 | 0.013319182 | -7.884124253 | 0.033594955 | -17.62641705 | 2 | 3.076665209 | 7.600047325 | 2 | 2 | 2.183373728 | 2 |
| ENSG00000177324 | BEND2 | -7.381598027 | 0.004179068 | -10.26071984 | 3.51E-05 | -17.64231787 | 5.70932728 | 3.076665209 | 5.772716771 | 2 | 2.14022021 | 2 | 2 |
| ENSG00000154080 | CHST9 | -8.332303133 | 0.001958103 | -9.318421693 | 0.000297382 | -17.65072483 | 7.975256505 | 7.274681896 | 2 | 2.270786764 | 2 | 2.139668783 | 2 |
| ENSG00000147872 | PLIN2 | -15.43628074 | 2.80E-22 | -2.223124468 | 1.81E-05 | -17.65940521 | 11.35501623 | 11.53726652 | 12.12587352 | 2 | 2 | 9.744401079 | 9.191641004 |
| ENSG00000101282 | RSPO4 | -9.300632614 | 1.68E-08 | -8.365630704 | 2.09E-08 | -17.66626332 | 8.716893051 | 7.901717006 | 9.829330889 | 2.3892346 | 2.14022021 | 2.831336763 | 2 |
| ENSG00000124205 | EDN3 | -8.121858367 | 0.131866271 | -9.559916949 | 0.06701491 | -17.68177532 | 2 | 2 | 6.073378139 | 2 | 2 | 2 | 2 |
| ENSG00000160882 | CYP11B1 | -8.121858367 | 0.131866271 | -9.559916949 | 0.06701491 | -17.68177532 | 2 | 2 | 6.073378139 | 2 | 2 | 2 | 2 |
| ENSG00000164325 | TMEM174 | -8.121858367 | 0.131866271 | -9.559916949 | 0.06701491 | -17.68177532 | 2 | 2 | 6.073378139 | 2 | 2 | 2 | 2 |
| ENSG00000277092 | | -8.121858367 | 0.131866271 | -9.559916949 | 0.06701491 | -17.68177532 | 2 | 2 | 6.073378139 | 2 | 2 | 2 | 2 |
| ENSG00000142609 | CFAP74 | -8.138846373 | 0.111151568 | -9.576895343 | 0.052747312 | -17.71574172 | 2 | 4.11352228 | 5.772716771 | 2 | 2 | 2 | 2 |
| ENSG00000172889 | EGFL7 | -14.76442613 | 4.00E-20 | -2.954558168 | 1.08E-07 | -17.71898429 | 11.54780176 | 10.42977533 | 10.92816184 | 2 | 2 | 8.13252251 | 8.075273781 |
| C5ORF45 | C5ORF45 | -8.148467611 | 0.127775345 | -9.586525136 | 0.064293553 | -17.73499275 | 2 | 5.946638857 | 3.369317972 | 2 | 2 | 2 | 2 |
| ENSG00000143995 | MEIS1 | -1.26716305 | 0.076688241 | -16.49095442 | 3.09E-24 | -17.75811747 | 10.44422623 | 11.32663546 | 11.87409387 | 10.06920484 | 10.0596091 | 2 | 2 |
| ENSG00000088756 | ARHGAP28 | -12.77631581 | 8.66E-14 | -4.993537873 | 3.75E-10 | -17.76985369 | 7.750913381 | 9.470501606 | 9.399592459 | 2 | 2 | 4.220168345 | 4.500832793 |
| QTRTD1 | | -8.168723471 | 0.005631624 | -9.60671294 | 0.000863752 | -17.77543641 | 4.00722531 | 5.457120953 | 4.058998049 | 2 | 2 | 2 | 2 |
| ENSG00000204568 | MRPS18B | -2.326238557 | 4.58E-05 | -15.44960516 | 3.81E-22 | -17.77584371 | 10.83615308 | 9.890264534 | 9.935792933 | 7.910550164 | 8.04930085 | 2 | 2 |
| ENSG00000215217 | C5orf49 | -6.952123102 | 0.001092061 | -10.83276602 | 3.48E-06 | -17.78488912 | 4.467311984 | 3.686303414 | 7.029880454 | 2.141737246 | 2.14022021 | 2 | 2 |
| ENSG00000144331 | ZNF385B | -9.766182069 | 0.00640016 | -8.034836688 | 0.016160038 | -17.80101876 | 2 | 4.93638933 | 7.451345288 | 2 | 2 | 2 | 2.172608095 |
| ENSG00000161640 | SIGLEC11 | -8.971969342 | 0.092116635 | -8.849772584 | 0.090235542 | -17.82174193 | 2 | 2 | 6.882775709 | 2 | 2 | 2 | 2.059860178 |
| ENSG00000125378 | BMP4 | -14.09457691 | 9.48E-13 | -3.727906178 | 0.005664904 | -17.82248308 | 11.27990552 | 8.356994601 | 10.18377222 | 2 | 2 | 5.130840083 | 7.429968874 |
| ENSG00000146910 | CNPY1 | -8.192941984 | 0.12809176 | -9.631001455 | 0.064751221 | -17.82394344 | 6.138360429 | 2 | 2 | 2 | 2 | 2 | 2 |
| ENSG00000184330 | S100A7A | -8.192941984 | 0.12809176 | -9.631001455 | 0.064751221 | -17.82394344 | 6.138360429 | 2 | 2 | 2 | 2 | 2 | 2 |
| ENSG00000187730 | GABRD | -8.192941984 | 0.12809176 | -9.631001455 | 0.064751221 | -17.82394344 | 6.138360429 | 2 | 2 | 2 | 2 | 2 | 2 |
| ENSG00000188937 | NYX | -8.192941984 | 0.12809176 | -9.631001455 | 0.064751221 | -17.82394344 | 6.138360429 | 2 | 2 | 2 | 2 | 2 | 2 |
| ENSG00000277236 | | -8.192941984 | 0.12809176 | -9.631001455 | 0.064751221 | -17.82394344 | 6.138360429 | 2 | 2 | 2 | 2 | 2 | 2 |
| SLC22A20 | | -8.192941984 | 0.12809176 | -9.631001455 | 0.064751221 | -17.82394344 | 6.138360429 | 2 | 2 | 2 | 2 | 2 | 2 |
| ENSG00000047662 | FAM184B | -9.934776283 | 2.17E-12 | -7.907467101 | 1.92E-16 | -17.84224338 | 7.624139768 | 7.701025996 | 9.670202709 | 2.141737246 | 2.14022021 | 2.457042231 | 2.551896972 |
| ENSG00000163468 | CCT3 | -16.07062617 | 8.14E-24 | -1.772230508 | 0.001574623 | -17.84285668 | 12.87148291 | 12.01967418 | 11.95227171 | 2 | 2 | 10.44549716 | 10.69456799 |
| GPR125 | | -8.205434979 | 0.127606207 | -9.643494605 | 0.064416773 | -17.84892958 | 2 | 6.140560021 | 2 | 2 | 2 | 2 | 2 |
| C14ORF142 | C14ORF142 | -8.221339509 | 0.103241254 | -9.659387515 | 0.048008107 | -17.88072702 | 5.866944937 | 4.11352228 | 2 | 2 | 2 | 2 | 2 |
| ENSG00000142541 | RPL13A | -16.49529105 | 2.79E-22 | -1.395580101 | 0.116849792 | -17.89087115 | 13.6041805 | 12.27086722 | 11.79401137 | 2 | 2 | 11.25506317 | 11.48592732 |
| LOC100507462 | | -8.230994686 | 0.010847435 | -9.668991 | 0.002182721 | -17.89998569 | 3.327711234 | 5.72254323 | 4.058998049 | 2 | 2 | 2 | 2 |
| ENSG00000104043 | ATP8B4 | -11.85440194 | 3.39E-05 | -6.049195835 | 0.016287777 | -17.90359777 | 7.667647223 | 9.328390744 | 2 | 2 | 2 | 3.534812207 | 2.277179783 |
| ENSG00000140931 | CMTM3 | -15.36755098 | 7.97E-22 | -2.53688214 | 3.65E-06 | -17.90443312 | 11.14017917 | 11.41367288 | 12.17016655 | 2 | 2 | 9.162002347 | 9.064992397 |
| ENSG00000275744 | | -8.244930478 | 0.119108512 | -9.68298744 | 0.058685985 | -17.92791792 | 2 | 6.04685525 | 3.369317972 | 2 | 2 | 2 | 2 |
| ENSG00000237710 | | -5.149438521 | 0.005434951 | -12.78294029 | 2.88E-08 | -17.93237882 | 8.855032352 | 7.002537973 | 3.369317972 | 3.023133209 | 3.45167086 | 2 | 2 |
| ENSG00000136750 | GAD2 | -7.523469505 | 0.165845146 | -10.41005015 | 0.043736674 | -17.93351966 | 2 | 2 | 6.882775709 | 2.141737246 | 2 | 2 | 2 |
| KIAA1244 | | -8.253818902 | 0.124056351 | -9.691878688 | 0.062226406 | -17.94569759 | 3.327711234 | 2 | 6.073378139 | 2 | 2 | 2 | 2 |
| FAM179A | | -7.534847985 | 0.024868257 | -10.41353201 | 0.001209325 | -17.94837999 | 6.366709671 | 5.303523747 | 2 | 2 | 2.14022021 | 2 | 2 |
| ENSG00000146360 | GPR6 | -8.259796357 | 0.124626428 | -9.697856662 | 0.062683816 | -17.95765302 | 2 | 2 | 6.203069917 | 2 | 2 | 2 | 2 |
| ENSG00000205755 | CRLF2 | -8.26526465 | 0.009557939 | -9.703260214 | 0.001840609 | -17.96852486 | 3.327711234 | 4.11352228 | 5.772716771 | 2 | 2 | 2 | 2 |
| ENSG00000263163 | | -3.239362691 | 4.58E-05 | -14.74874395 | 1.94E-18 | -17.98810664 | 9.384607364 | 8.540087354 | 10.31867969 | 6.314087501 | 6.507263 | 2 | 2 |
| ENSG00000274587 | | -8.295400529 | 0.006714202 | -9.733393268 | 0.001125929 | -18.0287938 | 4.00722531 | 3.686303414 | 5.772716771 | 2 | 2 | 2 | 2 |
| ATP5F1 | ATP5F1 | -1.954548631 | 0.001462494 | -16.07960129 | 1.24E-23 | -18.03414992 | 11.51244431 | 10.54992047 | 10.43545992 | 8.990393018 | 8.95298909 | 2 | 2 |
| ENSG00000184305 | CCSER1 | -8.298092644 | 0.122588012 | -9.736153428 | 0.061494261 | -18.03424607 | 2 | 6.228547981 | 2 | 2 | 2 | 2 | 2 |
| ENSG00000276072 | | -2.229871531 | 0.027972615 | -15.80613529 | 2.29E-19 | -18.03600682 | 11.40085503 | 8.916342327 | 10.66102425 | 8.354580458 | 8.49850989 | 2 | 2 |
| ENSG00000088882 | CPXM1 | -10.2111067 | 8.96E-22 | -7.842992346 | 5.56E-19 | -18.05409905 | 10.84565509 | 10.1878402 | 10.44204362 | 2.498691189 | 2.2680111 | 3.904814898 | 2.509591799 |
| ENSG00000169397 | RNASE3 | -8.312681274 | 0.104130612 | -9.750732983 | 0.04896152 | -18.06341426 | 2 | 3.686303414 | 6.073378139 | 2 | 2 | 2 | 2 |
| ENSG00000135333 | EPHA7 | -11.12246529 | 1.44E-33 | -6.942691188 | 6.96E-20 | -18.06515648 | 10.51143645 | 12.4880879 | 11.89103474 | 2.600425781 | 2.38539787 | 4.652122235 | 5.408276315 |
| ENSG00000132518 | GUCY2D | -8.318699563 | 0.12143043 | -9.756760604 | 0.060818509 | -18.07546017 | 6.257048221 | 2 | 2 | 2 | 2 | 2 | 2 |
| LOC100652758 | | -8.321137138 | 0.091680275 | -9.759182639 | 0.041031569 | -18.08031978 | 2 | 5.838937808 | 4.523675391 | 2 | 2 | 2 | 2 |
| PPAPDC2 | | -8.329712391 | 0.112243988 | -9.76776888 | 0.054367856 | -18.09748127 | 3.327711234 | 6.140560021 | 2 | 2 | 2 | 2 | 2 |
| ENSG00000275181 | | -8.344562215 | 0.093742087 | -9.782609529 | 0.042358404 | -18.12717174 | 6.009025997 | 4.11352228 | 2 | 2 | 2 | 2 | 2 |
| ENSG00000226710 | | -1.932851372 | 0.057446139 | -16.22501126 | 1.69E-20 | -18.15786264 | 12.03890377 | 10.02537381 | 10.19160738 | 9.150957946 | 9.12426093 | 2 | 2 |

| | | | | | | | | | | | | | |
|---|---|---|---|---|---|---|---|---|---|---|---|---|---|
| CECR1 | CECR1 | -5.324029255 | 0.017718702 | -12.85007497 | 6.41E-07 | -18.17410423 | 2 | 8.337339476 | 8.236946975 | 3.349643765 | 3.01487888 | 2 | 2 |
| ENSG00000167741 | GGT6 | -8.374311791 | 0.10022545 | -9.812363655 | 0.046751733 | -18.18667545 | 4.00722531 | 2 | 6.073378139 | 2 | 2 | 2 | 2 |
| LINS | | -8.385159847 | 0.1181587 | -9.823221717 | 0.058719207 | -18.20838156 | 2 | 6.311476754 | 2 | 2 | 2 | 2 | 2 |
| ENSG00000120332 | TNN | -8.385689433 | 0.118145526 | -9.82375131 | 0.058715617 | -18.20944074 | 2 | 2 | 6.322058544 | 2 | 2 | 2 | 2 |
| PAPL | | -8.390140478 | 0.008335313 | -9.828136695 | 0.00155585 | -18.21827717 | 3.327711234 | 4.11352228 | 5.930865648 | 2 | 2 | 2 | 2 |
| ENSG00000278270 | | -1.253060345 | 0.200330183 | -16.97953015 | 3.46E-23 | -18.23259049 | 12.68425352 | 11.3756363 | 10.62675862 | 10.56855616 | 10.5624671 | 2 | 2 |
| ENSG00000101958 | GLRA2 | -9.238109834 | 1.50E-06 | -9.013861636 | 1.06E-10 | -18.25197147 | 7.384433967 | 7.274681896 | 5.930865648 | 2 | 2.14022021 | 2 | 2.172608095 |
| ENSG00000099203 | TMED1 | -14.29154645 | 3.37E-19 | -3.964046112 | 5.84E-15 | -18.25559256 | 10.58702944 | 10.06773758 | 10.92816184 | 2 | 2 | 6.405645065 | 6.874895611 |
| ENSG00000168143 | FAM83B | -7.237038323 | 7.28E-07 | -11.01978578 | 2.14E-08 | -18.2568241 | 5.330594134 | 6.311476754 | 8.486635645 | 2.3892346 | 2.2680111 | 2.048074405 | 2 |
| ENSG00000184009 | ACTG1 | -19.05103912 | 2.61E-35 | 0.791244403 | 0.041930843 | -18.25979472 | 15.33514107 | 15.18080079 | 15.44520605 | 2 | 2 | 15.86257079 | 16.33078418 |
| ENSG00000167755 | KLK6 | -9.572428669 | 0.003278093 | -8.691468347 | 0.004556654 | -18.26389702 | 6.89194353 | 5.946638857 | 2 | 2 | 2 | 2.094598369 | 2 |
| ENSG00000129744 | ART1 | -9.709988626 | 0.063965155 | -8.557806192 | 0.091732358 | -18.26779482 | 2 | 2 | 7.600047325 | 2 | 2 | 2 | 2.117335275 |
| KAL1 | | -8.415080636 | 0.105302052 | -9.853136731 | 0.050247697 | -18.26821737 | 3.327711234 | 6.228547981 | 2 | 2 | 2 | 2 | 2 |
| BAI2 | BAI2 | -8.420383463 | 0.104995707 | -9.858439568 | 0.050033331 | -18.27882303 | 2 | 6.228547981 | 3.369317972 | 2 | 2 | 2 | 2 |
| ENSG00000185559 | DLK1 | -6.583251447 | 0.000172096 | -11.70696517 | 5.27E-09 | -18.29021661 | 9.448362284 | 7.572420273 | 9.099922433 | 3.79808662 | 2 | 2.094598369 | 2 |
| ENSG00000142089 | IFITM3 | -17.40523805 | 2.39E-28 | -0.898746835 | 0.096892849 | -18.30398489 | 14.04407854 | 13.16759288 | 13.69420491 | 2 | 2 | 12.60303588 | 12.9385609 |
| ENSG00000163218 | PGLYRP4 | -8.434370083 | 0.115724282 | -9.872432568 | 0.05735638 | -18.30680265 | 6.366709671 | 2 | 2 | 2 | 2 | 2 | 2 |
| C6ORF57 | C6ORF57 | -8.436102222 | 0.006165262 | -9.874096389 | 0.001037164 | -18.31019861 | 5.095959458 | 3.076665209 | 5.59507254 | 2 | 2 | 2 | 2 |
| ENSG00000188848 | BEND4 | -8.444513083 | 0.089224711 | -9.882561482 | 0.040003152 | -18.32707457 | 4.00722531 | 6.140560021 | 2 | 2 | 2 | 2 | 2 |
| CECR6 | CECR6 | -5.327305181 | 0.023233848 | -13.01301752 | 9.51E-07 | -18.3403227 | 8.493334308 | 2 | 8.408111533 | 3.023133209 | 3.50452879 | 2 | 2 |
| ENSG00000140600 | SH3GL3 | -7.042690983 | 2.06E-06 | -11.31015343 | 1.98E-08 | -18.35284441 | 4.467311984 | 7.393647515 | 8.608698767 | 2.3892346 | 2.49394783 | 2.048074405 | 2 |
| ENSG00000159182 | PRAC1 | -8.473511096 | 0.001264306 | -9.911495393 | 0.00011836 | -18.38500649 | 5.532353208 | 4.11352228 | 4.874587708 | 2 | 2 | 2 | 2 |
| ENSG00000162881 | OXER1 | -12.25726192 | 3.16E-11 | -6.138496622 | 6.23E-09 | -18.39575854 | 9.048471759 | 6.844429755 | 8.904018013 | 2 | 2 | 3.534812207 | 2.782576693 |
| HOTS | | -7.758578741 | 0.000116576 | -10.63830691 | 5.37E-08 | -18.39688565 | 6.138360429 | 5.303523747 | 5.156626401 | 2 | 2.14022021 | 2 | 2 |
| ENSG00000016082 | ISL1 | -7.774653134 | 0.022393868 | -10.6592626 | 0.001096094 | -18.43391573 | 6.816637977 | 4.93638933 | 2 | 2.141737246 | 2 | 2 | 2 |
| C19ORF26 | C19ORF26 | -8.501475305 | 0.112649338 | -9.939538628 | 0.055406174 | -18.44101393 | 2 | 2 | 6.43197674 | 2 | 2 | 2 | 2 |
| ENSG00000188162 | OTOG | -8.501475305 | 0.112649338 | -9.939538628 | 0.055406174 | -18.44101393 | 2 | 2 | 6.43197674 | 2 | 2 | 2 | 2 |
| ENSG00000204978 | ERICH4 | -8.501475305 | 0.112649338 | -9.939538628 | 0.055406174 | -18.44101393 | 2 | 2 | 6.43197674 | 2 | 2 | 2 | 2 |
| ENSG00000258817 | OR4C13 | -8.501475305 | 0.112649338 | -9.939538628 | 0.055406174 | -18.44101393 | 2 | 2 | 6.43197674 | 2 | 2 | 2 | 2 |
| ENSG00000196497 | IPO4 | -12.03994363 | 7.79E-06 | -6.408707502 | 0.005140501 | -18.44865113 | 9.533135543 | 7.790474954 | 2 | 2 | 2 | 2.858551861 | 3.042026731 |
| ENSG00000127578 | WFIKKN1 | -12.40420012 | 1.78E-11 | -6.079398387 | 8.05E-09 | -18.48359851 | 9.247966931 | 9.055168103 | 6.629488387 | 2 | 2 | 3.483289517 | 3.15612297 |
| ENSG00000185070 | FLRT2 | -15.18259703 | 5.71E-22 | -3.317472497 | 2.58E-13 | -18.50006953 | 11.32135895 | 11.62702028 | 11.40669646 | 2 | 2 | 8.487172107 | 7.73202455 |
| C3ORF58 | C3ORF58 | -2.509618487 | 0.008410067 | -15.9933109 | 3.14E-20 | -18.50292939 | 9.112843761 | 11.26294702 | 11.2616444 | 8.408172799 | 8.2618769 | 2 | 2 |
| ENSG00000276701 | | -1.160255066 | 0.131821657 | -17.35087238 | 2.66E-26 | -18.51112744 | 12.9059531 | 11.6229672 | 11.62782301 | 11.02823523 | 11.028985 | 2 | 2 |
| ENSG00000276644 | DACH1 | -7.725357221 | 0.002571181 | -10.78961447 | 0.00010984 | -18.51497169 | 7.33131396 | 7.901717006 | 2 | 2.3892346 | 2 | 2.048074405 | 2 |
| ENSG00000272670 | | -8.546020467 | 0.0871768 | -9.984071224 | 0.039075168 | -18.53009169 | 2 | 3.686303414 | 6.322058544 | 2 | 2 | 2 | 2 |
| ENSG00000106384 | MOGAT3 | -9.315000474 | 0.008841057 | -9.224725909 | 0.008583972 | -18.53972638 | 4.467311984 | 2 | 7.029880454 | 2 | 2 | 2 | 2.059860178 |
| ENSG00000175928 | LRRN1 | -8.881614095 | 1.77E-11 | -9.680715954 | 7.72E-14 | -18.56233005 | 10.16187685 | 10.83561703 | 11.63070764 | 3.663521627 | 2 | 2.68698273 | 2.70970617 |
| ENSG00000226397 | C12orf77 | -8.565138657 | 0.000652799 | -10.00312101 | 4.88E-05 | -18.56825967 | 4.815609471 | 4.442710129 | 5.59507254 | 2 | 2 | 2 | 2 |
| ENSG00000149403 | GRIK4 | -10.08980752 | 0.000629169 | -8.538460183 | 0.001289371 | -18.6282677 | 7.867441304 | 4.442710129 | 2 | 2 | 2 | 2.094598369 | 2.059860178 |
| ENSG00000054803 | CBLN4 | -8.608653571 | 0.107655331 | -10.04671823 | 0.052627273 | -18.6553718 | 2 | 2 | 6.534109959 | 2 | 2 | 2 | 2 |
| ENSG00000154478 | GPR26 | -8.608653571 | 0.107655331 | -10.04671823 | 0.052627273 | -18.6553718 | 2 | 2 | 6.534109959 | 2 | 2 | 2 | 2 |
| SELV | | -8.608653571 | 0.107655331 | -10.04671823 | 0.052627273 | -18.6553718 | 2 | 2 | 6.534109959 | 2 | 2 | 2 | 2 |
| ENSG00000128052 | KDR | -8.664097429 | 1.21E-31 | -10.00498786 | 2.82E-43 | -18.66908529 | 9.810482605 | 10.95649913 | 9.757248847 | 2.600425781 | 3.01487888 | 2.34604626 | 2.421067459 |
| ENSG00000133710 | SPINK5 | -8.618664287 | 0.107207965 | -10.05672907 | 0.052348674 | -18.67539336 | 2 | 6.535001748 | 2 | 2 | 2 | 2 | 2 |
| ENSG00000131482 | G6PC | -8.619999762 | 0.07497303 | -10.05804639 | 0.03241055 | -18.67804615 | 2 | 4.11352228 | 6.322058544 | 2 | 2 | 2 | 2 |
| ATP5O | ATP5O | -2.279999815 | 4.65E-06 | -16.4024063 | 1.74E-25 | -18.68240612 | 11.70222964 | 10.84955954 | 11.02342937 | 8.94033939 | 8.99746255 | 2 | 2 |
| C9ORF117 | C9ORF117 | -8.643961865 | 0.000647135 | -10.08194506 | 4.95E-05 | -18.72590693 | 4.467311984 | 4.710564154 | 5.772716771 | 2 | 2 | 2 | 2 |
| ENSG00000164708 | PGAM2 | -7.422670465 | 0.00443527 | -11.30332091 | 3.81E-05 | -18.72599138 | 6.138360429 | 2 | 7.225742779 | 2.141737246 | 2.14022021 | 2 | 2 |
| ENSG00000113763 | UNC5A | -10.09882918 | 0.001900808 | -8.643341407 | 0.003902794 | -18.74217058 | 2 | 6.899080544 | 7.098165388 | 2 | 2 | 2.139668783 | 2 |
| ENSG00000273533 | | -7.939166874 | 0.133069745 | -10.82568944 | 0.032017989 | -18.76485631 | 2 | 2 | 7.285536013 | 2.141737246 | 2 | 2 | 2 |
| ENSG00000250479 | CHCHD10 | -2.947458444 | 1.68E-15 | -15.83641362 | 1.02E-24 | -18.78387207 | 10.41260941 | 10.61666558 | 10.94679967 | 7.739909082 | 7.76240929 | 2 | 2 |
| ENSG00000075891 | PAX2 | -9.95832395 | 0.000164922 | -8.843014537 | 0.00018628 | -18.80133849 | 3.327711234 | 5.72254323 | 7.451345288 | 2 | 2 | 2 | 2.117335275 |
| C4ORF29 | C4ORF29 | -8.69482045 | 0.001804089 | -10.13280943 | 0.000204802 | -18.82762988 | 4.00722531 | 6.04685525 | 4.523675391 | 2 | 2 | 2 | 2 |
| LOC729159 | | -8.703769476 | 0.090170759 | -10.14182752 | 0.041615684 | -18.84559699 | 3.327711234 | 2 | 6.534109959 | 2 | 2 | 2 | 2 |
| ENSG00000237727 | | -8.705434906 | 0.002690991 | -10.14342662 | 0.00035542 | -18.84886153 | 4.00722531 | 4.11352228 | 6.203069917 | 2 | 2 | 2 | 2 |
| ENSG00000105143 | SLC1A6 | -8.708418403 | 0.103202649 | -10.14648431 | 0.05002317 | -18.85490271 | 2 | 2 | 6.629488387 | 2 | 2 | 2 | 2 |
| ENSG00000170703 | TTLL6 | -9.470697606 | 4.37E-05 | -9.429043929 | 7.80E-07 | -18.89974154 | 4.467311984 | 6.228547981 | 8.460932916 | 2 | 2.14022021 | 2.139668783 | 2 |
| ENSG00000152254 | G6PC2 | -8.734363603 | 0.101987045 | -10.17242983 | 0.049380564 | -18.90679344 | 6.653098118 | 2 | 2 | 2 | 2 | 2 | 2 |
| ENSG00000164129 | NPY5R | -8.741016351 | 0.05466499 | -10.17905483 | 0.021735943 | -18.92007118 | 2 | 5.457120953 | 5.930865648 | 2 | 2 | 2 | 2 |
| CCDC176 | CCDC176 | -8.772157664 | 0.000311423 | -10.21013939 | 1.93E-05 | -18.98229706 | 4.467311984 | 5.72254323 | 5.156626401 | 2 | 2 | 2 | 2 |
| ENSG00000181234 | TMEM132C | -7.36625402 | 5.98E-19 | -11.62275743 | 3.13E-13 | -18.98901145 | 7.53297911 | 8.191702463 | 7.934808822 | 2.600425781 | 2.2680111 | 2.048074405 | 2 |
| GLTSCR2 | | -2.504498456 | 1.09E-05 | -16.50054817 | 4.13E-25 | -19.00504662 | 11.89759773 | 11.01334259 | 10.88048583 | 8.80304041 | 8.88359764 | 2 | 2 |
| ENSG00000188505 | NCCRP1 | -7.283632171 | 0.003718966 | -11.75136774 | 1.69E-05 | -19.03499991 | 3.327711234 | 3.076665209 | 8.143093325 | 2.270786764 | 2.14022021 | 2 | 2 |
| ENSG00000188282 | RUFY4 | -8.801728882 | 0.099210255 | -10.23979595 | 0.047728814 | -19.04152484 | 2 | 2 | 6.718950348 | 2 | 2 | 2 | 2 |
| ENSG00000213171 | LINGO4 | -8.801728882 | 0.099210255 | -10.23979595 | 0.047728814 | -19.04152484 | 2 | 2 | 6.718950348 | 2 | 2 | 2 | 2 |
| ENSG00000183878 | UTY | -6.791298827 | 3.31E-05 | -12.25275281 | 7.72E-09 | -19.04405163 | 4.467311984 | 7.874700259 | 7.342949392 | 2.141737246 | 2.59489879 | 2 | 2 |
| GATSL3 | | -4.658700633 | 0.019496517 | -14.40946795 | 1.44E-09 | -19.06816858 | 3.327711234 | 8.834582582 | 10.41552659 | 4.783303513 | 4.83276841 | 2 | 2 |
| ENSG00000165655 | ZNF503 | -14.73866975 | 8.96E-22 | -4.391152281 | 5.51E-54 | -19.12982203 | 10.98115017 | 10.89059638 | 11.16147894 | 2 | 2 | 6.670403488 | 6.685996416 |
| ENSG00000101198 | NKAIN4 | -11.62425798 | 1.57E-09 | -7.537462853 | 5.42E-10 | -19.16172083 | 8.631580145 | 6.228547981 | 8.007629717 | 2 | 2 | 2.592885947 | 2.172608095 |
| ENSG00000186297 | GABRA5 | -7.23138764 | 0.007699833 | -11.93365997 | 6.33E-05 | -19.16504761 | 9.699577239 | 3.076665209 | 2 | 2.3892346 | 2.68924543 | 2.048074405 | 2 |
| ENSG00000173157 | ADAMTS20 | -8.864372317 | 0.062511756 | -10.30241952 | 0.026212112 | -19.16679183 | 4.00722531 | 6.602424674 | 2 | 2 | 2 | 2 | 2 |
| ENSG00000144406 | UNC80 | -8.144713579 | 0.117647128 | -11.03119422 | 0.026673727 | -19.1759078 | 7.485143553 | 2 | 2 | 2.141737246 | 2 | 2 | 2 |
| KIAA1467 | | -8.87000732 | 0.000414663 | -10.30799118 | 2.90E-05 | -19.1779985 | 4.815609471 | 6.04685525 | 4.523675391 | 2 | 2 | 2 | 2 |
| ENSG00000118194 | TNNT2 | -14.30115987 | 7.00E-19 | -4.884731975 | 9.19E-19 | -19.18589185 | 10.85509492 | 10.39216648 | 10.44204362 | 2 | 2 | 5.253805068 | 6.192573806 |
| ENSG00000125861 | GFRA4 | -8.889368877 | 0.09564296 | -10.32743704 | 0.045651889 | -19.21680592 | 2 | 2 | 6.80318718 | 2 | 2 | 2 | 2 |
| LRRC48 | | -8.902866544 | 0.045222505 | -10.34090354 | 0.017099976 | -19.24377008 | 2 | 5.946638857 | 5.772716771 | 2 | 2 | 2 | 2 |
| ENSG00000100721 | TCL1A | -8.904498377 | 0.094997183 | -10.34256673 | 0.045276644 | -19.24706511 | 6.816637977 | 2 | 2 | 2 | 2 | 2 | 2 |
| ENSG00000136943 | CTSV | -9.374215877 | 7.95E-27 | -9.898868902 | 2.00E-34 | -19.27308478 | 11.28340583 | 10.06773758 | 9.888372618 | 2.784612485 | 2.49394783 | 2.267001569 | 2.671842496 |
| ENSG00000182459 | TEX19 | -8.932490228 | 0.050694512 | -10.37053173 | 0.019964283 | -19.30302196 | 2 | 4.710564154 | 6.534109959 | 2 | 2 | 2 | 2 |
| ENSG00000169314 | C22orf15 | -10.16027753 | 0.010529822 | -9.149354046 | 0.016231405 | -19.30963158 | 2 | 2 | 8.042706615 | 2 | 2 | 2.048074405 | 2.059860178 |
| ENSG00000198597 | ZNF536 | -8.212548147 | 0.113761161 | -11.09902418 | 0.025430364 | -19.31157233 | 2 | 2 | 7.552162999 | 2.141737246 | 2 | 2 | 2 |
| ENSG00000075673 | ATP12A | -8.9394932 | 0.093501828 | -10.37756199 | 0.044473202 | -19.31705519 | 2 | 6.844429755 | 2 | 2 | 2 | 2 | 2 |
| ENSG00000137070 | IL11RA | -14.67981199 | 1.35E-18 | -4.6404215 | 5.22E-11 | -19.32023349 | 11.70484245 | 10.46642862 | 10.23774519 | 2 | 2 | 6.377699085 | 6.389139248 |
| ENSG00000168671 | UGT3A2 | -8.948300064 | 0.044420522 | -10.38633777 | 0.016763071 | -19.33463783 | 5.330594134 | 6.311476754 | 2 | 2 | 2 | 2 | 2 |
| ENSG00000184983 | NDUFA6 | -2.777619073 | 2.30E-12 | -16.56564636 | 9.02E-27 | -19.34326543 | 11.4974524 | 11.01334259 | 11.63070764 | 8.601543953 | 8.673169 | 2 | 2 |
| ENSG00000101812 | H2BFM | -8.982588424 | 0.091680275 | -10.42065775 | 0.043476835 | -19.40324618 | 6.89194353 | 2 | 2 | 2 | 2 | 2 | 2 |
| LOC81691 | | -6.203793729 | 0.007422027 | -13.25779317 | 5.72E-07 | -19.4615869 | 2 | 8.276714465 | 9.014156722 | 3.227757282 | 2.59489879 | 2 | 2 |
| ENSG00000152208 | GRID2 | -11.09886057 | 1.04E-07 | -8.372214725 | 3.26E-08 | -19.47107529 | 7.904282243 | 7.901717006 | 4.874587708 | 2 | 2 | 2.267001569 | 2.059860178 |
| ENSG00000106031 | HOXA13 | -9.730346164 | 0.002128162 | -9.763179462 | 0.001837738 | -19.49352563 | 2 | 7.051611315 | 6.073378139 | 2 | 2 | 2.048074405 | 2 |
| ENSG00000234954 | | -9.050132697 | 0.088904811 | -10.48820287 | 0.042020739 | -19.53833557 | 2 | 2 | 6.958202216 | 2 | 2 | 2 | 2 |
| ENSG00000170786 | SDR16C5 | -9.056668317 | 0.088683678 | -10.49473857 | 0.041878072 | -19.55140689 | 6.96351253 | 2 | 2 | 2 | 2 | 2 | 2 |
| ENSG00000267206 | LCN6 | -9.056668317 | 0.088683678 | -10.49473857 | 0.041878072 | -19.55140689 | 6.96351253 | 2 | 2 | 2 | 2 | 2 | 2 |
| PTCHD2 | | -9.056668317 | 0.088683678 | -10.49473857 | 0.041878072 | -19.55140689 | 6.96351253 | 2 | 2 | 2 | 2 | 2 | 2 |
| EPT1 | | -9.078644091 | 0.000823063 | -10.51663356 | 7.83E-05 | -19.59527766 | 4.00722531 | 6.464272742 | 4.874587708 | 2 | 2 | 2 | 2 |
| ENSG00000067048 | DDX3Y | -4.900680942 | 1.90E-06 | -14.71217998 | 3.32E-16 | -19.61286092 | 7.53297911 | 9.733683991 | 10.2528023 | 4.887609717 | 4.83276841 | 2 | 2 |
| ENSG00000133980 | VRTN | -9.092239162 | 0.005680076 | -10.53024321 | 0.001064228 | -19.62248237 | 4.00722531 | 3.076665209 | 6.80318718 | 2 | 2 | 2 | 2 |
| ENSG00000198223 | CSF2RA | -8.386226549 | 0.000151624 | -11.2656531 | 1.67E-07 | -19.65187965 | 5.095959458 | 5.131603625 | 7.225742779 | 2 | 2.14022021 | 2 | 2 |
| ENSG00000160994 | CCDC105 | -9.124260452 | 0.085933312 | -10.56233155 | 0.040405933 | -19.68659201 | 2 | 2 | 7.029880454 | 2 | 2 | 2 | 2 |
| ENSG00000168032 | ENTPD3 | -10.62803211 | 6.61E-71 | -9.100711599 | 8.11E-112 | -19.72874371 | 11.95139721 | 11.36354124 | 11.62782301 | 2.498691189 | 2.68924543 | 3.235948948 | 3.383972333 |
| ENSG00000108953 | YWHAE | -1.512576916 | 0.006849738 | -18.24215915 | 5.88E-31 | -19.75473606 | 13.59294047 | 12.61696644 | 12.8350691 | 11.582282 | 11.5509701 | 2 | 2 |
| ENSG00000184454 | NCMAP | -8.450696697 | 0.000100793 | -11.33016966 | 8.40E-08 | -19.78086636 | 6.366709671 | 4.442710129 | 6.958202216 | 2 | 2.14022021 | 2 | 2 |
| ENSG00000164626 | KCNK5 | -9.186886152 | 0.000101604 | -10.6248683 | 4.87E-06 | -19.81175445 | 4.467311984 | 5.595927442 | 6.203069917 | 2 | 2 | 2 | 2 |
| ENSG00000186038 | HTR3E | -9.194765205 | 0.083130968 | -10.63283719 | 0.038886992 | -19.82760239 | 2 | 2 | 7.098165388 | 2 | 2 | 2 | 2 |
| ENSG00000184005 | ST6GALNAC3 | -10.90409024 | 1.19E-22 | -8.980666095 | 2.21E-34 | -19.88475634 | 9.317904228 | 9.883504052 | 10.94679967 | 2.270786764 | 2.14022021 | 2.656294034 | 2.671842496 |
| ENSG00000198088 | NUP62CL | -10.40011126 | 0.005896613 | -9.528453184 | 0.008390025 | -19.92856445 | 8.257427002 | 3.076665209 | 2 | 2 | 2 | 2.094598369 | 2 |
| C14ORF105 | C14ORF105 | -8.528398828 | 1.82E-05 | -11.41025039 | 4.19E-09 | -19.93864921 | 5.532353208 | 5.838937808 | 7.098165388 | 2.141737246 | 2 | 2 | 2 |

| | | | | | | | | | | | | | |
|---|---|---|---|---|---|---|---|---|---|---|---|---|---|
| ENSG00000168539 | CHRM1 | -9.261983895 | 0.080450078 | -10.70005672 | 0.037552988 | -19.96204061 | 2 | 2 | 7.163363831 | 2 | 2 | 2 | 2 |
| C14ORF1 | C14ORF1 | -3.570695546 | 2.85E-06 | -16.39223199 | 9.82E-23 | -19.96292754 | 12.00750147 | 10.81799725 | 10.32581712 | 7.668500375 | 7.70145652 | 2 | 2 |
| ENSG00000186329 | TMEM212 | -7.250715194 | 4.15E-07 | -12.71246834 | 1.42E-10 | -19.96318353 | 6.96351253 | 6.04685525 | 8.585098385 | 2.141737246 | 2.59489879 | 2 | 2 |
| ENSG00000152207 | CYSLTR2 | -11.5240395 | 1.29E-07 | -8.451207358 | 1.98E-07 | -19.97524686 | 8.75772933 | 7.761270542 | 4.874587708 | 2 | 2 | 2.34604626 | 2.059860178 |
| FAM212A | | -6.673269267 | 0.00022899 | -13.35488725 | 5.29E-09 | -20.02815651 | 9.371511179 | 7.572420273 | 5.156626401 | 2.141737246 | 3.2184422 | 2 | 2 |
| FAM115C | | -9.301954479 | 4.29E-05 | -10.73993521 | 1.59E-06 | -20.04188969 | 5.095959458 | 5.131603625 | 6.43197674 | 2 | 2 | 2 | 2 |
| ENSG00000182378 | PLCXD1 | -11.46388367 | 1.36E-07 | -8.58973016 | 1.27E-07 | -20.05361383 | 8.042902253 | 6.228547981 | 8.296287027 | 2 | 2 | 2.34604626 | 2 |
| C2ORF44 | C2ORF44 | -9.327466107 | 0.042243725 | -10.76551327 | 0.016346433 | -20.09297938 | 4.00722531 | 7.099070191 | 2 | 2 | 2 | 2 | 2 |
| ENSG00000157005 | SST | -7.109056351 | 5.17E-09 | -12.98969183 | 3.08E-12 | -20.09874819 | 6.96351253 | 8.862353853 | 6.629488387 | 2.498691189 | 2.49394783 | 2 | 2 |
| PTPLB | | -9.334460481 | 0.00169763 | -10.77245784 | 0.000219646 | -20.10691832 | 3.327711234 | 4.93638933 | 6.882775709 | 2 | 2 | 2 | 2 |
| ENSG00000145721 | LIX1 | -7.821176757 | 2.01E-06 | -12.29149971 | 1.87E-09 | -20.11267646 | 7.750913381 | 5.595927442 | 7.398165132 | 2.3892346 | 2 | 2 | 2 |
| ENSG00000145506 | NKD2 | -8.123948456 | 0.015869208 | -11.997996 | 0.000442844 | -20.12194446 | 8.340423424 | 4.710564154 | 2 | 2 | 2.2680111 | 2 | 2 |
| HRSP12 | | -9.355173271 | 0.050767575 | -10.79322794 | 0.020897808 | -20.14840121 | 3.327711234 | 7.189546432 | 2 | 2 | 2 | 2 | 2 |
| PVRL1 | | -9.355173271 | 0.050767575 | -10.79322794 | 0.020897808 | -20.14840121 | 3.327711234 | 7.189546432 | 2 | 2 | 2 | 2 | 2 |
| ERO1LB | | -9.363840984 | 7.30E-05 | -10.80182385 | 3.32E-06 | -20.16566483 | 4.467311984 | 6.389896633 | 5.772716771 | 2 | 2 | 2 | 2 |
| ENSG00000121570 | DPPA4 | -7.156910384 | 7.12E-07 | -13.04229455 | 9.03E-11 | -20.19920493 | 6.009025997 | 7.73146271 | 8.784770914 | 2.784612485 | 2.14022021 | 2 | 2 |
| METTL20 | | -9.39007844 | 1.89E-05 | -10.82805787 | 5.45E-07 | -20.21813631 | 5.095959458 | 5.457120953 | 6.43197674 | 2 | 2 | 2 | 2 |
| ENSG00000130222 | GADD45G | -9.999981771 | 2.30E-23 | -10.22174775 | 9.32E-26 | -20.22172952 | 13.44919347 | 11.89581018 | 11.39656351 | 3.462030052 | 3.08597292 | 3.600766115 | 2.509591799 |
| ENSG00000124440 | HIF3A | -7.510953038 | 7.80E-28 | -12.79390823 | 2.11E-42 | -20.30486126 | 11.22271378 | 10.41109345 | 12.0110149 | 4.17310023 | 4.22485516 | 2.183373728 | 2.059860178 |
| ENSG00000135750 | KCNK1 | -5.315944106 | 1.19E-12 | -15.04280323 | 2.14E-19 | -20.35874734 | 10.39976609 | 8.732943944 | 10.05225195 | 4.907595604 | 4.65713986 | 2 | 2 |
| ENSG00000198844 | ARHGEF15 | -9.490018491 | 0.072150422 | -10.92809416 | 0.033293021 | -20.41811265 | 7.384433967 | 2 | 2 | 2 | 2 | 2 | 2 |
| ENSG00000146151 | HMGCLL1 | -9.496896028 | 0.022954373 | -10.93493087 | 0.007443312 | -20.4318269 | 2 | 6.78762695 | 5.930865648 | 2 | 2 | 2 | 2 |
| ENSG00000275565 | | -8.279266899 | 6.69E-10 | -12.16177346 | 1.09E-12 | -20.44104036 | 6.89194353 | 6.844429755 | 7.342949392 | 2.270786764 | 2 | 2 | 2 |
| ENSG00000128713 | HOXD11 | -6.732475154 | 1.02E-23 | -13.78465159 | 1.13E-17 | -20.51712675 | 8.716893051 | 8.234834549 | 8.884815469 | 3.162721474 | 2.68924543 | 2 | 2 |
| ENSG00000196090 | PTPRT | -9.542041039 | 0.070313891 | -10.98011736 | 0.03241055 | -20.52215839 | 2 | 7.431218269 | 2 | 2 | 2 | 2 | 2 |
| ENSG00000145423 | SFRP2 | -8.573171855 | 6.69E-14 | -11.96216848 | 1.49E-16 | -20.53534033 | 10.09173937 | 8.623598428 | 8.677268317 | 2.784612485 | 2.14022021 | 2 | 2.117335275 |
| DHFRL1 | DHFRL1 | -9.55309056 | 0.025513953 | -10.99112953 | 0.008628998 | -20.54422009 | 2 | 5.131603625 | 7.163363831 | 2 | 2 | 2 | 2 |
| LRRC16B | | -9.557847315 | 0.069768927 | -10.99592383 | 0.032176413 | -20.55377115 | 2 | 2 | 7.451345288 | 2 | 2 | 2 | 2 |
| ENSG00000278488 | | -5.844026893 | 4.36E-13 | -14.72396844 | 3.58E-18 | -20.56799533 | 8.928409675 | 8.834582582 | 10.38167985 | 3.997744386 | 4.1930041 | 2 | 2 |
| ENSG00000171435 | KSR2 | -8.353432037 | 0.007376645 | -12.23409188 | 0.000130778 | -20.58752391 | 2 | 3.076665209 | 8.65477198 | 2.141737246 | 2.14022021 | 2 | 2 |
| ENSG00000130957 | FBP2 | -9.588034805 | 6.54E-05 | -11.02601957 | 3.10E-06 | -20.61405438 | 6.89194353 | 4.710564154 | 5.39244166 | 2 | 2 | 2 | 2 |
| ENSG00000219200 | RNASEK | -14.67949541 | 4.05E-16 | -5.942768304 | 1.84E-09 | -20.62226371 | 11.92922406 | 10.42046463 | 9.426401271 | 2 | 2 | 4.881704255 | 5.425444337 |
| ENSG00000174498 | IGDCC3 | -11.07468027 | 5.27E-15 | -9.547787613 | 8.96E-17 | -20.62246789 | 8.836089686 | 10.84260513 | 10.74328839 | 2.3892346 | 2 | 2.183373728 | 2.817676617 |
| PTPLA | | -9.603536806 | 9.39E-07 | -11.04151228 | 1.12E-08 | -20.64504908 | 6.009025997 | 6.228547981 | 5.59507254 | 2 | 2 | 2 | 2 |
| ENSG00000154856 | APCDD1 | -11.75875419 | 4.23E-21 | -8.893133048 | 1.26E-22 | -20.65188724 | 11.27287932 | 10.95649913 | 10.94216274 | 2.3892346 | 2 | 3.534812207 | 2.421067459 |
| ENSG00000169100 | SLC25A6 | -2.588920481 | 0.001748776 | -18.13253537 | 3.77E-27 | -20.72145585 | 13.67117821 | 12.90510119 | 11.7010103 | 10.37039102 | 10.3958393 | 2 | 2 |
| VPRBP | | -9.664670498 | 0.03944862 | -11.10272574 | 0.015480979 | -20.76739623 | 3.327711234 | 7.503546079 | 2 | 2 | 2 | 2 | 2 |
| ENSG00000276342 | | -8.977076536 | 0.006356279 | -11.85576123 | 0.00018896 | -20.83283776 | 8.075571721 | 5.595927442 | 2 | 2 | 2.14022021 | 2 | 2 |
| ENSG00000166863 | TAC3 | -10.44461438 | 2.94E-07 | -10.41378441 | 1.69E-07 | -20.85839879 | 7.709880939 | 6.140560021 | 5.59507254 | 2 | 2 | 2 | 2.059860178 |
| ENSG00000215864 | NBPF7 | -5.402593213 | 5.24E-16 | -15.55071555 | 2.08E-21 | -20.95330876 | 11.06489013 | 9.814097497 | 9.945092785 | 5.273747807 | 5.0259382 | 2 | 2 |
| INADL | | -9.774894812 | 7.10E-06 | -11.21287539 | 1.75E-07 | -20.9877702 | 5.866944937 | 5.131603625 | 6.882775709 | 2 | 2 | 2 | 2 |
| ENSG00000080709 | KCNN2 | -7.566794122 | 3.43E-09 | -13.44297111 | 1.61E-12 | -21.00976523 | 7.667647223 | 7.274681896 | 9.196533591 | 2.141737246 | 2.77779907 | 2 | 2 |
| ENSG00000105278 | ZFR2 | -9.792265109 | 0.015142983 | -11.23029801 | 0.004418378 | -21.02256312 | 2 | 6.602424674 | 6.80318718 | 2 | 2 | 2 | 2 |
| ENSG00000066468 | FGFR2 | -15.11410145 | 3.35E-18 | -5.913157217 | 5.61E-12 | -21.02725867 | 9.98400217 | 12.0991443 | 11.36918911 | 2 | 2 | 5.22272385 | 5.896817464 |
| C2ORF47 | C2ORF47 | -9.797962023 | 0.000195623 | -11.23595332 | 1.42E-05 | -21.03391534 | 4.00722531 | 5.595927442 | 7.225742779 | 2 | 2 | 2 | 2 |
| ENSG00000136944 | LMX1B | -9.082077356 | 0.001385303 | -11.96097036 | 1.40E-05 | -21.04304771 | 3.327711234 | 4.442710129 | 8.296287027 | 2 | 2.14022021 | 2 | 2 |
| ENSG00000126337 | KRT36 | -9.80324004 | 0.061920576 | -11.24131962 | 0.028164286 | -21.04455966 | 2 | 2 | 7.691296561 | 2 | 2 | 2 | 2 |
| ATP5B | ATP5B | -2.014757177 | 2.41E-07 | -19.03154174 | 2.63E-35 | -21.04629892 | 14.19387916 | 13.61548743 | 13.7248998 | 11.81304079 | 11.8929864 | 2 | 2 |
| C19ORF40 | C19ORF40 | -9.852327958 | 2.44E-06 | -11.29030699 | 4.43E-08 | -21.14263495 | 5.532353208 | 6.899080544 | 5.772716771 | 2 | 2 | 2 | 2 |
| ENSG00000170421 | KRT8 | -16.19683494 | 1.03E-19 | -4.947337218 | 3.05E-07 | -21.14417216 | 13.45696386 | 11.86353764 | 11.01019887 | 2 | 2 | 7.462723726 | 7.63942137 |
| KIAA2018 | | -9.871723216 | 1.12E-06 | -11.30970101 | 1.63E-08 | -21.18142422 | 5.330594134 | 6.602424674 | 6.43197674 | 2 | 2 | 2 | 2 |
| ENSG00000235302 | | -7.751109155 | 9.62E-16 | -13.43728743 | 2.45E-15 | -21.18839659 | 8.653385977 | 7.393647515 | 8.536706852 | 2.270786764 | 2.59489879 | 2 | 2 |
| ENSG00000118263 | KLF7 | -23.35027877 | 2.51E-15 | 2.157192766 | 0.484174515 | -21.19308601 | 3.327711234 | 2 | 9.25752426 | 2 | 2 | 9.916945014 | 9.75743869 |
| ENSG00000183317 | EPHA10 | -9.888380906 | 0.000112903 | -11.3263709 | 7.07E-06 | -21.21475181 | 4.815609471 | 4.93638933 | 7.398165132 | 2 | 2 | 2 | 2 |
| SGOL1 | | -9.89201281 | 1.97E-06 | -11.32999176 | 3.41E-08 | -21.22200457 | 6.96351253 | 5.595927442 | 5.772716771 | 2 | 2 | 2 | 2 |
| ENSG00000167664 | TMIGD2 | -11.12146884 | 0.000262746 | -10.11213177 | 0.000347179 | -21.23360061 | 2 | 4.442710129 | 8.941672936 | 2 | 2 | 2.048074405 | 2.059860178 |
| LOC113230 | | -9.905334214 | 0.059027778 | -11.34341507 | 0.026596694 | -21.24874928 | 7.790810999 | 2 | 2 | 2 | 2 | 2 | 2 |
| ENSG00000175206 | NPPA | -10.65413026 | 7.79E-10 | -10.64124784 | 2.78E-10 | -21.29537809 | 7.27616306 | 6.535001748 | 7.029880454 | 2 | 2 | 2 | 2.059860178 |
| KIAA1804 | | -9.929978997 | 0.024923147 | -11.36802777 | 0.008728946 | -21.29800677 | 4.00722531 | 7.73146271 | 2 | 2 | 2 | 2 | 2 |
| ENSG00000254772 | EEF1G | -2.306818845 | 0.003922264 | -18.9938981 | 3.17E-30 | -21.30071695 | 14.602198 | 13.48647061 | 12.83881593 | 11.49890168 | 11.5492022 | 2 | 2 |
| SLMO1 | | -9.944885292 | 0.05791763 | -11.38296664 | 0.026022069 | -21.32785193 | 7.82963488 | 2 | 2 | 2 | 2 | 2 | 2 |
| ENSG00000144278 | GALNT13 | -11.10661577 | 0.000640329 | -10.2268953 | 0.000840516 | -21.33351107 | 8.75772933 | 6.228547981 | 2 | 2 | 2 | 2.094598369 | 2 |
| ENSG00000125872 | LRRN4 | -10.4883222 | 7.32E-13 | -10.96944557 | 9.26E-19 | -21.45776777 | 7.975256505 | 8.655694736 | 10.14394518 | 2.270786764 | 2 | 2.139668783 | 2.059860178 |
| ENSG00000142449 | FBN3 | -12.33285749 | 9.70E-09 | -9.137317734 | 7.15E-09 | -21.47017522 | 9.87686338 | 7.638155693 | 5.39244166 | 2 | 2 | 2.183373728 | 2.277179783 |
| LPHN3 | | -10.03800639 | 0.011003457 | -11.47603889 | 0.002994622 | -21.51404528 | 7.218819781 | 6.602424674 | 2 | 2 | 2 | 2 | 2 |
| ENSG00000177519 | RPRM | -10.04771166 | 0.031076669 | -11.48577065 | 0.011727475 | -21.53348231 | 7.904282243 | 3.076665209 | 2 | 2 | 2 | 2 | 2 |
| ENSG00000074660 | SCARF1 | -10.05899509 | 0.030852431 | -11.49705419 | 0.011637524 | -21.55604927 | 7.904282243 | 2 | 3.369317972 | 2 | 2 | 2 | 2 |
| CCDC109B | CCDC109B | -10.06469241 | 0.021182597 | -11.50274038 | 0.007138053 | -21.56743279 | 7.867441304 | 4.11352228 | 2 | 2 | 2 | 2 | 2 |
| ENSG00000159251 | ACTC1 | -9.462665752 | 2.67E-06 | -12.10853513 | 1.55E-09 | -21.57120088 | 13.54270883 | 9.386925327 | 3.369317972 | 3.289987342 | 3.33976443 | 2.384001718 | 2.225841198 |
| ENSG00000188176 | SMTNL2 | -12.77260158 | 8.55E-13 | -8.810446796 | 9.20E-19 | -21.58304837 | 9.810482605 | 8.981105197 | 7.600047325 | 2 | 2 | 2.457042231 | 2.277179783 |
| MFI2 | | -10.0872416 | 0.010149381 | -11.52527373 | 0.002710501 | -21.61251533 | 6.816637977 | 7.145017454 | 2 | 2 | 2 | 2 | 2 |
| ACN9 | ACN9 | -10.10372782 | 0.011567328 | -11.54176283 | 0.00322719 | -21.64549065 | 7.579279349 | 6.04685525 | 2 | 2 | 2 | 2 | 2 |
| KIAA1919 | | -10.11223336 | 7.36E-07 | -11.55021229 | 1.03E-08 | -21.66244565 | 5.330594134 | 6.78762695 | 6.80318718 | 2 | 2 | 2 | 2 |
| LPPR3 | | -10.11507615 | 0.012786939 | -11.55311346 | 0.003681791 | -21.66818961 | 2 | 5.72254323 | 7.691296561 | 2 | 2 | 2 | 2 |
| ATP5A1 | ATP5A1 | -2.775581279 | 1.42E-08 | -18.90010363 | 1.18E-33 | -21.6756849 | 14.19108654 | 13.28254015 | 13.58169095 | 10.99300838 | 10.9316182 | 2 | 2 |
| CD97 | CD97 | -10.12185831 | 6.71E-07 | -11.55983717 | 9.19E-09 | -21.68169548 | 5.532353208 | 7.099070191 | 6.322058544 | 2 | 2 | 2 | 2 |
| ENSG00000095713 | CRTAC1 | -12.17590633 | 9.79E-12 | -9.52413193 | 3.69E-18 | -21.70003826 | 7.790810999 | 9.114109157 | 8.143093325 | 2 | 2 | 2.048074405 | 2.277179783 |
| ENSG00000185960 | SHOX | -10.1356927 | 4.99E-05 | -11.57368233 | 2.59E-06 | -21.70937503 | 5.866944937 | 4.442710129 | 7.551262999 | 2 | 2 | 2 | 2 |
| ENSG00000182256 | GABRG3 | -10.14414478 | 0.000618475 | -11.58214688 | 6.95E-05 | -21.72629166 | 3.327711234 | 5.303523747 | 7.777116083 | 2 | 2 | 2 | 2 |
| ENSG00000106038 | EVX1 | -10.18664241 | 0.000186661 | -11.62463825 | 1.49E-05 | -21.81128067 | 7.27616306 | 6.78762695 | 3.369317972 | 2 | 2 | 2 | 2 |
| SUV420H2 | | -10.19189991 | 3.71E-06 | -11.6298827 | 8.80E-08 | -21.82178261 | 6.89194353 | 7.002537973 | 4.874587708 | 2 | 2 | 2 | 2 |
| TMEM194A | | -10.19804134 | 0.008734471 | -11.63607326 | 0.002258289 | -21.8341146 | 7.096806373 | 7.099070191 | 2 | 2 | 2 | 2 | 2 |
| ENSG00000170689 | HOXB9 | -6.825409487 | 1.13E-08 | -15.02683286 | 2.39E-15 | -21.85224235 | 8.493334308 | 8.376385542 | 11.04521411 | 3.462030052 | 3.74316441 | 2 | 2 |
| ENSG00000112175 | BMP5 | -8.202125285 | 7.34E-06 | -13.67432478 | 1.75E-09 | -21.87645007 | 8.797441463 | 9.225224669 | 5.930865648 | 2.695456494 | 2 | 2 | 2 |
| ICT1 | | -10.23203685 | 0.008465185 | -11.67006903 | 0.00218059 | -21.90210588 | 7.33131396 | 6.899080544 | 2 | 2 | 2 | 2 | 2 |
| SELO | | -10.24974277 | 0.000161536 | -11.68773868 | 1.25E-05 | -21.93748145 | 7.218819781 | 7.002537973 | 3.369317972 | 2 | 2 | 2 | 2 |
| ENSG00000114374 | USP9Y | -8.151443702 | 1.37E-06 | -13.8347113 | 2.11E-10 | -21.986155 | 5.095959458 | 8.848535039 | 9.528916236 | 2.141737246 | 2.68924543 | 2 | 2 |
| C19ORF52 | C19ORF52 | -10.27840444 | 0.011601873 | -11.71644378 | 0.003311956 | -21.99484822 | 7.940205782 | 5.457120953 | 2 | 2 | 2 | 2 | 2 |
| ENSG00000169016 | E2F6 | -21.97088518 | 1.18E-10 | -0.074209531 | 0.985856566 | -22.04509472 | 2 | 7.790474954 | 2 | 2 | 2 | 6.216130818 | 6.165581817 |
| C10ORF2 | C10ORF2 | -10.34184862 | 1.89E-06 | -11.77983125 | 3.89E-08 | -22.12167987 | 7.435667355 | 6.602424674 | 5.156626401 | 2 | 2 | 2 | 2 |
| ENSG00000134716 | CYP2J2 | -10.34240695 | 0.024500134 | -11.78046739 | 0.008899899 | -22.12287434 | 8.19931886 | 3.076665209 | 2 | 2 | 2 | 2 | 2 |
| C1ORF168 | C1ORF168 | -10.36218567 | 1.43E-07 | -11.80016389 | 1.38E-09 | -22.16234955 | 6.257048221 | 6.04685525 | 7.398165132 | 2 | 2 | 2 | 2 |
| ENSG00000144227 | NXPH2 | -11.11538112 | 2.08E-05 | -11.05528182 | 1.63E-05 | -22.17066294 | 8.716893051 | 6.389896633 | 3.369317972 | 2 | 2 | 2 | 2.059860178 |
| CCDC23 | CCDC23 | -10.36976593 | 5.19E-07 | -11.80774629 | 7.36E-09 | -22.17751223 | 7.53297911 | 6.04685525 | 5.930865648 | 2 | 2 | 2 | 2 |
| N6AMT2 | | -10.37469909 | 1.11E-05 | -11.81268636 | 3.88E-07 | -22.18738545 | 7.27616306 | 4.442710129 | 7.098165388 | 2 | 2 | 2 | 2 |
| ATP5S | ATP5S | -10.39698909 | 2.03E-06 | -11.83497235 | 4.36E-08 | -22.23196145 | 5.095959458 | 7.467835385 | 6.718950348 | 2 | 2 | 2 | 2 |
| C5ORF28 | C5ORF28 | -10.41419386 | 2.54E-08 | -11.85217013 | 1.48E-10 | -22.26636399 | 7.27616306 | 6.535001748 | 6.203069917 | 2 | 2 | 2 | 2 |
| C9ORF41 | C9ORF41 | -10.4159536 | 4.88E-05 | -11.85394626 | 2.73E-06 | -22.26989986 | 4.00722531 | 6.535001748 | 7.734844356 | 2 | 2 | 2 | 2 |
| C9ORF91 | C9ORF91 | -10.45338291 | 0.000248027 | -11.89138377 | 2.29E-05 | -22.34476667 | 4.467311984 | 8.147241016 | 4.523675391 | 2 | 2 | 2 | 2 |
| ENSG00000275575 | | -7.106440501 | 2.61E-34 | -15.34477642 | 6.41E-22 | -22.45121692 | 10.37373127 | 9.532315021 | 10.48092713 | 3.663521627 | 3.60476711 | 2 | 2 |
| LYRM5 | | -10.50826638 | 1.36E-06 | -11.94624975 | 2.73E-08 | -22.45451613 | 7.579279349 | 6.844429755 | 5.156626401 | 2 | 2 | 2 | 2 |
| HMHA1 | | -10.55260862 | 1.06E-07 | -11.9905878 | 1.02E-09 | -22.54319642 | 6.366709671 | 6.140560021 | 7.646393253 | 2 | 2 | 2 | 2 |
| C7ORF60 | C7ORF60 | -10.55508139 | 0.005701427 | -11.99311422 | 0.00135936 | -22.54819561 | 2 | 7.274681896 | 7.600047325 | 2 | 2 | 2 | 2 |
| ENSG00000128714 | HOXD13 | -11.27849624 | 1.52E-05 | -11.30643314 | 1.13E-05 | -22.58492938 | 3.327711234 | 6.535001748 | 8.884815469 | 2 | 2 | 2.048074405 | 2 |
| OBFC1 | | -10.59428216 | 0.005267422 | -12.03231464 | 0.001236986 | -22.6265968 | 7.667647223 | 7.274681896 | 2 | 2 | 2 | 2 | 2 |
| CTAGE5 | CTAGE5 | -10.60078827 | 6.79E-08 | -12.03876716 | 5.89E-10 | -22.63955544 | 7.384433967 | 7.145017454 | 5.772716771 | 2 | 2 | 2 | 2 |
| ENSG00000256618 | MTRNR2L1 | -6.44598472 | 8.57E-36 | -16.30286171 | 5.86E-25 | -22.74884643 | 11.55070946 | 11.07694755 | 10.65536953 | 4.71689672 | 5.06165168 | 2 | 2 |

| | | | | | | | | | | | | |
|---|---|---|---|---|---|---|---|---|---|---|---|---|
| ENSG00000132906 | CASP9 | -22.72598336 | 4.37E-11 | -0.029813504 | 0.994412286 | -22.75579686 | 2 | 8.623598428 | 2 | 2 | 2 | 6.894026089 | 7.174740504 |
| PRKRIR | | -10.72335329 | 3.87E-05 | -12.16134859 | 2.20E-06 | -22.88470188 | 4.00722531 | 8.124485672 | 6.629488387 | 2 | 2 | 2 | 2 |
| ENSG00000174482 | LINGO2 | -9.216528345 | 2.15E-09 | -13.68657009 | 3.35E-12 | -22.90309843 | 8.836089686 | 6.78762695 | 9.099922433 | 2.3892346 | 2 | 2 | 2 |
| ENSG00000122756 | CNTFR | -11.20963142 | 7.15E-61 | -11.78815469 | 6.54E-97 | -22.99778611 | 11.78357814 | 11.65110249 | 11.76804571 | 2.600425781 | 2.2680111 | 2.42098416 | 2.172608095 |
| PPAP2A | | -10.78418322 | 0.005010769 | -12.22221916 | 0.001191046 | -23.00640238 | 2 | 6.728495673 | 8.236946975 | 2 | 2 | 2 | 2 |
| ENSG00000165246 | NLGN4Y | -10.79198587 | 2.26E-08 | -12.22996462 | 1.53E-10 | -23.02195048 | 7.096806373 | 7.701025996 | 6.073378139 | 2 | 2 | 2 | 2 |
| ENSG00000182196 | ARL6IP4 | -14.73665746 | 2.96E-19 | -8.295111068 | 2.00E-37 | -23.03176853 | 11.50646623 | 10.13189344 | 11.08361639 | 2 | 2 | 3.27687201 | 3.518055793 |
| ENSG00000105647 | PIK3R2 | -13.06116196 | 2.57E-11 | -9.980563693 | 4.14E-14 | -23.04172565 | 7.384433967 | 9.28800521 | 10.17589429 | 2 | 2 | 2.34604626 | 2.059860178 |
| ENSG00000081138 | CDH7 | -9.586585977 | 1.47E-06 | -13.46723453 | 1.77E-09 | -23.05382051 | 4.00722531 | 8.451443573 | 9.212024683 | 2.141737246 | 2.14022021 | 2 | 2 |
| ENSG00000153446 | C16orf89 | -22.42724748 | 2.64E-10 | -0.63413922 | 0.884155802 | -23.0613867 | 8.257427002 | 2 | 2 | 2 | 2 | 5.136410075 | 6.678855578 |
| ENSG00000164532 | TBX20 | -10.81441184 | 5.76E-10 | -12.25238673 | 1.35E-12 | -23.06679856 | 6.96351253 | 6.844429755 | 7.502634626 | 2 | 2 | 2 | 2 |
| ENSG00000085552 | IGSF9 | -10.79639359 | 9.02E-14 | -12.2924866 | 2.26E-17 | -23.08888019 | 10.34721797 | 9.125613817 | 8.434763955 | 2.270786764 | 2 | 2.094598369 | 2 |
| FAM178A | | -10.83909113 | 2.47E-06 | -12.27707889 | 6.59E-08 | -23.11617003 | 5.532353208 | 8.255926468 | 6.203069917 | 2 | 2 | 2 | 2 |
| CCDC101 | CCDC101 | -10.85710358 | 1.66E-06 | -12.29509054 | 4.00E-08 | -23.15219412 | 7.53297911 | 7.761270542 | 4.874587708 | 2 | 2 | 2 | 2 |
| ENSG00000166183 | ASPG | -12.89355673 | 9.93E-09 | -10.34194567 | 3.50E-09 | -23.23550239 | 10.52923584 | 7.761270542 | 5.39244166 | 2 | 2 | 2.183373728 | 2.117335275 |
| ENSG00000137252 | HCRTR2 | -10.92623444 | 0.003642609 | -12.36426857 | 0.000810633 | -23.29050301 | 2 | 7.431218269 | 8.110401118 | 2 | 2 | 2 | 2 |
| ENSG00000146013 | GFRA3 | -12.8238569 | 9.09E-15 | -10.46701675 | 1.43E-43 | -23.29087365 | 11.20436621 | 10.1878402 | 9.897982195 | 2.141737246 | 2 | 2.34604626 | 2.326754077 |
| ENSG00000179796 | LRRC3B | -10.90806492 | 7.62E-05 | -12.38346645 | 4.42E-06 | -23.29153136 | 9.473095744 | 8.916342327 | 2 | 2.141737246 | 2 | 2.048074405 | 2 |
| ENSG00000145808 | ADAMTS19 | -10.93012051 | 3.20E-06 | -12.36810983 | 9.51E-08 | -23.29823034 | 4.815609471 | 8.169643011 | 7.098165388 | 2 | 2 | 2 | 2 |
| ENSG00000164318 | EGFLAM | -15.34160129 | 6.05E-23 | -7.980884905 | 2.61E-95 | -23.3224862 | 11.38465071 | 11.86009815 | 11.56285656 | 2 | 2 | 3.904814898 | 4.161535617 |
| ENSG00000229637 | PRAC2 | -10.94890951 | 3.20E-07 | -12.38689372 | 5.00E-09 | -23.33580324 | 8.138771685 | 5.595927442 | 7.029880454 | 2 | 2 | 2 | 2 |
| FAM213A | | -7.941258762 | 2.55E-12 | -15.41222956 | 9.52E-17 | -23.35348832 | 11.00254523 | 9.267380431 | 9.954333071 | 3.663521627 | 2.38539787 | 2 | 2 |
| ENSG00000080224 | EPHA6 | -9.461683329 | 4.21E-10 | -13.92583328 | 8.33E-13 | -23.38751661 | 9.886101849 | 7.928237109 | 6.958202216 | 2.141737246 | 2.2680111 | 2 | 2 |
| VWA9 | | -10.99797717 | 1.29E-10 | -12.43595186 | 2.19E-13 | -23.43392903 | 7.624139768 | 7.145017454 | 7.098165388 | 2 | 2 | 2 | 2 |
| FAM73A | | -10.99906925 | 1.66E-08 | -12.43704907 | 1.14E-10 | -23.43611832 | 6.138360429 | 7.954278496 | 7.285536013 | 2 | 2 | 2 | 2 |
| C7ORF55 | C7ORF55 | -11.01194639 | 5.26E-06 | -12.44993793 | 1.84E-07 | -23.46188432 | 8.367060731 | 4.710564154 | 6.958202216 | 2 | 2 | 2 | 2 |
| TMEM180 | | -11.03674447 | 9.44E-10 | -12.47472121 | 2.86E-12 | -23.51146568 | 7.435667355 | 6.602424674 | 7.777116083 | 2 | 2 | 2 | 2 |
| ADRBK2 | ADRBK2 | -11.06441262 | 6.73E-08 | -12.50239501 | 7.11E-10 | -23.56680763 | 5.70932728 | 7.73146271 | 7.858115942 | 2 | 2 | 2 | 2 |
| MEF2BNB | | -11.0781609 | 4.33E-07 | -12.51614683 | 7.72E-09 | -23.59430773 | 8.444143041 | 5.946638857 | 6.534109959 | 2 | 2 | 2 | 2 |
| KIAA1377 | | -11.11049611 | 4.06E-06 | -12.54848789 | 1.36E-07 | -23.658984 | 4.467311984 | 7.701025996 | 8.143093325 | 2 | 2 | 2 | 2 |
| ENSG00000164093 | PITX2 | -8.152110933 | 1.87E-98 | -15.70118616 | 2.35E-63 | -23.8532971 | 13.83696579 | 13.25202848 | 13.47257427 | 5.514313187 | 5.52526112 | 2.139668783 | 2 |
| DDX26B | DDX26B | -11.2120349 | 1.28E-09 | -12.6500131 | 4.72E-12 | -23.862048 | 7.975256505 | 6.535001748 | 7.691296561 | 2 | 2 | 2 | 2 |
| KIAA0226 | | -11.26659221 | 3.62E-07 | -12.70457943 | 6.59E-09 | -23.97117163 | 5.70932728 | 7.002537973 | 8.608698767 | 2 | 2 | 2 | 2 |
| LPPR2 | | -11.31343199 | 4.87E-09 | -12.75141247 | 2.80E-11 | -24.06484446 | 6.257048221 | 7.954278496 | 8.042706615 | 2 | 2 | 2 | 2 |
| SEPP1 | | -11.31882617 | 3.59E-07 | -12.75681384 | 6.63E-09 | -24.07564002 | 8.737455672 | 6.602424674 | 6.073378139 | 2 | 2 | 2 | 2 |
| KIAA1598 | | -11.34516821 | 3.80E-05 | -12.78317105 | 2.53E-06 | -24.12833927 | 3.327711234 | 8.487557258 | 7.858115942 | 2 | 2 | 2 | 2 |
| HIATL1 | | -11.37926237 | 0.002308655 | -12.81729962 | 0.000477501 | -24.19656199 | 2 | 8.747909548 | 7.502634626 | 2 | 2 | 2 | 2 |
| ENSG00000110975 | SYT10 | -11.40301999 | 9.15E-09 | -12.84100196 | 6.48E-11 | -24.24402195 | 8.367060731 | 7.790474954 | 6.203069917 | 2 | 2 | 2 | 2 |
| PVRL3 | | -11.4044578 | 0.000145518 | -12.84246894 | 1.43E-05 | -24.24692674 | 3.327711234 | 5.946638857 | 9.116480286 | 2 | 2 | 2 | 2 |
| ENSG00000172987 | HPSE2 | -11.42759218 | 4.68E-11 | -12.88016586 | 2.28E-14 | -24.30775805 | 9.497412313 | 8.004992147 | 9.516493218 | 2.141737246 | 2 | 2.048074405 | 2 |
| LARGE | | -11.43627458 | 3.00E-05 | -12.87427749 | 1.91E-06 | -24.31055208 | 8.540903468 | 8.004992147 | 3.369317972 | 2 | 2 | 2 | 2 |
| C20ORF195 | C20ORF195 | -11.45160941 | 1.81E-05 | -12.88961027 | 9.97E-07 | -24.34121968 | 9.112843761 | 4.710564154 | 6.073378139 | 2 | 2 | 2 | 2 |
| ATHL1 | ATHL1 | -11.45443056 | 3.11E-11 | -12.89240675 | 4.53E-14 | -24.34683731 | 7.159102577 | 7.874700259 | 8.076950877 | 2 | 2 | 2 | 2 |
| SMEK1 | | -11.46466067 | 7.62E-09 | -12.90264284 | 5.30E-11 | -24.36730351 | 8.367060731 | 7.954278496 | 6.203069917 | 2 | 2 | 2 | 2 |
| FAM69B | | -9.955398621 | 3.10E-08 | -14.42276443 | 2.95E-11 | -24.37816305 | 5.330594134 | 8.929529568 | 10.36791565 | 2.270786764 | 2.14022021 | 2 | 2 |
| ERBB2IP | | -11.48510152 | 5.96E-07 | -12.9230919 | 1.34E-08 | -24.40819342 | 5.095959458 | 8.623598428 | 7.858115942 | 2 | 2 | 2 | 2 |
| NUPL1 | | -11.50857165 | 1.66E-08 | -12.94655529 | 1.44E-10 | -24.45512694 | 6.138360429 | 8.57407404 | 7.777116083 | 2 | 2 | 2 | 2 |
| ADCK4 | ADCK4 | -11.52193284 | 2.44E-12 | -12.95990751 | 1.80E-15 | -24.48184035 | 7.667647223 | 7.847167909 | 7.934808822 | 2 | 2 | 2 | 2 |
| CRAMP1L | CRAMP1L | -11.56030103 | 3.71E-05 | -12.99830657 | 2.59E-06 | -24.5586076 | 3.327711234 | 7.60566235 | 8.941672936 | 2 | 2 | 2 | 2 |
| ENSG00000130700 | GATA5 | -11.56116494 | 0.001626527 | -12.9992012 | 0.000312441 | -24.56036614 | 8.107517749 | 8.702538795 | 2 | 2 | 2 | 2 | 2 |
| DFNB31 | DFNB31 | -11.57616462 | 0.000208741 | -13.01418093 | 2.33E-05 | -24.59034555 | 5.70932728 | 3.076665209 | 9.330306409 | 2 | 2 | 2 | 2 |
| ENSG00000121871 | SLITRK3 | -11.5769108 | 7.55E-05 | -13.01492039 | 6.39E-06 | -24.59183119 | 3.327711234 | 6.666836837 | 9.212024683 | 2 | 2 | 2 | 2 |
| C17ORF59 | C17ORF59 | -11.58680195 | 9.84E-07 | -13.0247946 | 2.63E-08 | -24.61159655 | 5.330594134 | 7.189546432 | 9.031722608 | 2 | 2 | 2 | 2 |
| ADRBK1 | ADRBK1 | -11.62696502 | 4.95E-07 | -13.0649563 | 1.11E-08 | -24.69192132 | 5.095959458 | 8.747909548 | 8.042706615 | 2 | 2 | 2 | 2 |
| ADCK3 | ADCK3 | -11.63098806 | 1.03E-11 | -13.06896439 | 1.19E-14 | -24.69995244 | 8.313285031 | 7.928237109 | 7.398165132 | 2 | 2 | 2 | 2 |
| B3GALTL | B3GALTL | -11.64185093 | 2.53E-10 | -13.07983025 | 7.43E-13 | -24.72168117 | 6.89194353 | 7.979858146 | 8.511888463 | 2 | 2 | 2 | 2 |
| C2ORF43 | C2ORF43 | -11.66691361 | 1.14E-08 | -13.10489792 | 9.63E-11 | -24.77181152 | 8.444143041 | 8.376385542 | 6.073378139 | 2 | 2 | 2 | 2 |
| ENSG00000198692 | EIF1AY | -11.67150753 | 1.86E-09 | -13.10948933 | 9.56E-12 | -24.78099686 | 8.797441463 | 7.60566235 | 6.80318718 | 2 | 2 | 2 | 2 |
| ENSG00000240303 | ACAD11 | -12.89575782 | 2.22E-09 | -11.97712155 | 1.49E-10 | -24.87287936 | 6.009025997 | 9.479495904 | 9.907528187 | 2 | 2 | 2.094598369 | 2 |
| SETD8 | | -11.74724666 | 5.38E-10 | -13.1852275 | 2.04E-12 | -24.93247416 | 8.564112515 | 6.728495673 | 8.266922072 | 2 | 2 | 2 | 2 |
| STRA13 | | -11.779506 | 2.41E-12 | -13.21748209 | 2.08E-15 | -24.99698809 | 8.418903792 | 8.053983481 | 7.646393253 | 2 | 2 | 2 | 2 |
| C4ORF27 | C4ORF27 | -11.78415718 | 5.81E-09 | -13.22214138 | 4.35E-11 | -25.00629857 | 8.910412251 | 7.928237109 | 6.43197674 | 2 | 2 | 2 | 2 |
| ENSG00000136574 | GATA4 | -10.58429638 | 2.29E-15 | -14.4649279 | 1.81E-17 | -25.04922427 | 9.233563228 | 9.923599762 | 8.35328246 | 2.141737246 | 2.14022021 | 2 | 2 |
| ENSG00000184613 | NELL2 | -12.58666153 | 1.19E-12 | -12.46330428 | 1.70E-23 | -25.0499658 | 11.12465208 | 10.09139839 | 8.904018013 | 2.141737246 | 2 | 2.094598369 | 2.059860178 |
| ENSG00000163462 | TRIM46 | -23.11070279 | 1.39E-10 | -1.989224302 | 0.625290659 | -25.09992709 | 2 | 2 | 9.330306409 | 2 | 2 | 5.574182174 | 6.081436739 |
| ENSG00000260596 | DUX4 | -11.84406353 | 0.003332994 | -13.28211757 | 0.000813199 | -25.1261811 | 5.866944937 | 2 | 9.61300511 | 2 | 2 | 2 | 2 |
| TMEM35 | | -11.84569743 | 0.001257708 | -13.28373634 | 0.000235626 | -25.12943376 | 9.159305284 | 8.077869079 | 2 | 2 | 2 | 2 | 2 |
| UTP11L | | -11.90026946 | 7.45E-10 | -13.33825176 | 3.30E-12 | -25.23852122 | 8.928409675 | 8.124485672 | 6.80318718 | 2 | 2 | 2 | 2 |
| FAM195A | | -11.92591258 | 1.12E-11 | -13.36389074 | 1.59E-14 | -25.28980332 | 8.816894989 | 7.638155693 | 7.934808822 | 2 | 2 | 2 | 2 |
| LRRC16A | | -11.93524968 | 5.61E-11 | -13.3732294 | 1.29E-13 | -25.30847908 | 7.159102577 | 8.762721501 | 8.325066185 | 2 | 2 | 2 | 2 |
| CCDC132 | CCDC132 | -11.95388142 | 1.73E-11 | -13.39186013 | 2.82E-14 | -25.34574155 | 7.579279349 | 7.954278496 | 8.884815469 | 2 | 2 | 2 | 2 |
| C12ORF5 | C12ORF5 | -11.96022769 | 5.16E-10 | -13.39820996 | 2.14E-12 | -25.35843765 | 7.750913381 | 7.189546432 | 9.116480286 | 2 | 2 | 2 | 2 |
| ENSG00000241404 | EGFL8 | -12.00186859 | 6.50E-09 | -13.43985467 | 5.47E-11 | -25.44172326 | 7.867441304 | 6.602424674 | 9.25752426 | 2 | 2 | 2 | 2 |
| PCNXL4 | | -12.00302832 | 1.47E-05 | -13.4410352 | 8.73E-07 | -25.44406353 | 3.327711234 | 8.862353853 | 8.865353883 | 2 | 2 | 2 | 2 |
| ENSG00000164078 | MST1R | -22.34145118 | 5.92E-10 | -3.104974516 | 0.427095076 | -25.4464257 | 8.107517749 | 2 | 2 | 2 | 2 | 3.316666208 | 4.254128754 |
| C9ORF156 | C9ORF156 | -12.02592521 | 9.53E-07 | -13.46392229 | 2.89E-08 | -25.48984749 | 7.940205782 | 4.93638933 | 9.413059136 | 2 | 2 | 2 | 2 |
| ENSG00000169218 | RSPO1 | -12.03352364 | 0.001055988 | -13.47156424 | 0.000193765 | -25.50508788 | 9.397585735 | 8.147241016 | 2 | 2 | 2 | 2 | 2 |
| PCNXL2 | | -12.0539157 | 5.33E-10 | -13.49189868 | 2.33E-12 | -25.54581438 | 6.96351253 | 8.147241016 | 9.149036566 | 2 | 2 | 2 | 2 |
| SMEK2 | | -12.1076193 | 1.55E-11 | -13.54559888 | 2.68E-14 | -25.65321817 | 7.53297911 | 9.030900487 | 8.236946975 | 2 | 2 | 2 | 2 |
| SUV420H1 | | -12.16616999 | 2.98E-06 | -13.60417242 | 1.26E-07 | -25.77034241 | 4.00722531 | 9.090820894 | 8.941672936 | 2 | 2 | 2 | 2 |
| SEPN1 | | -12.23714484 | 3.43E-10 | -13.6751286 | 1.48E-12 | -25.91227344 | 8.107517749 | 9.405920729 | 7.225742779 | 2 | 2 | 2 | 2 |
| KIAA0020 | | -12.24295021 | 9.92E-12 | -13.68093022 | 1.61E-14 | -25.92388042 | 8.75772933 | 8.968383114 | 7.451345288 | 2 | 2 | 2 | 2 |
| C9ORF114 | C9ORF114 | -12.28082953 | 4.91E-11 | -13.71881139 | 1.29E-13 | -25.99964093 | 7.435667355 | 8.337339476 | 9.330306409 | 2 | 2 | 2 | 2 |
| WAPAL | | -12.28784399 | 2.76E-14 | -13.72581989 | 9.82E-18 | -26.01366388 | 8.367060731 | 8.639735838 | 8.699419242 | 2 | 2 | 2 | 2 |
| WIBG | | -12.31551821 | 8.50E-09 | -13.7535073 | 8.63E-11 | -26.06902551 | 6.257048221 | 8.607278467 | 9.452720984 | 2 | 2 | 2 | 2 |
| ENSG00000165556 | CDX2 | -12.34805451 | 1.07E-11 | -13.78603527 | 1.88E-14 | -26.13408978 | 7.579279349 | 9.267380431 | 8.585098385 | 2 | 2 | 2 | 2 |
| LOC389831 | | -12.39309353 | 3.83E-14 | -13.83107019 | 1.57E-17 | -26.22416372 | 8.998234871 | 8.395519306 | 8.585098385 | 2 | 2 | 2 | 2 |
| CCBL2 | CCBL2 | -12.4071666 | 4.17E-13 | -13.84514501 | 3.31E-16 | -26.25231162 | 9.143983763 | 8.004992147 | 8.721235202 | 2 | 2 | 2 | 2 |
| PCNX | | -12.42414437 | 4.57E-12 | -13.86212483 | 6.72E-15 | -26.2862692 | 9.384607364 | 8.505280662 | 7.818184403 | 2 | 2 | 2 | 2 |
| RNMTL1 | | -12.43683761 | 2.33E-11 | -13.87481973 | 5.35E-14 | -26.31165734 | 9.358295023 | 7.431218269 | 8.784770914 | 2 | 2 | 2 | 2 |
| GCN1L1 | | -12.46167973 | 1.09E-11 | -13.89966124 | 2.05E-14 | -26.36134097 | 7.940205782 | 9.532315021 | 8.236946975 | 2 | 2 | 2 | 2 |
| C21ORF59 | C21ORF59 | -12.46248833 | 1.21E-13 | -13.90046616 | 7.17E-17 | -26.36295448 | 9.233563228 | 8.451443573 | 8.408111533 | 2 | 2 | 2 | 2 |
| ENSG00000204569 | PPP1R10 | -12.46264544 | 8.61E-07 | -13.90064699 | 2.89E-08 | -26.36329243 | 4.467311984 | 9.067150531 | 9.516493218 | 2 | 2 | 2 | 2 |
| MLLT4 | | -12.47706858 | 8.15E-13 | -13.91504792 | 8.01E-16 | -26.3921165 | 8.107517749 | 9.367676479 | 8.511888463 | 2 | 2 | 2 | 2 |
| KIAA1279 | | -12.48706548 | 0.000641946 | -13.92510939 | 0.000111099 | -26.41217488 | 9.829762239 | 8.639735838 | 2 | 2 | 2 | 2 | 2 |
| FAM101B | | -12.50433697 | 1.51E-14 | -13.94231365 | 5.20E-18 | -26.44665062 | 8.493334308 | 8.777382925 | 9.049077186 | 2 | 2 | 2 | 2 |
| SELK | | -12.51040507 | 4.08E-08 | -13.94839897 | 6.65E-10 | -26.45880404 | 9.895281535 | 8.297207175 | 6.073378139 | 2 | 2 | 2 | 2 |
| ENTHD2 | | -12.51247594 | 3.69E-08 | -13.95046963 | 5.87E-10 | -26.46294557 | 6.138360429 | 8.255926468 | 9.907528187 | 2 | 2 | 2 | 2 |
| ENSG00000164128 | NPY1R | -12.8006308 | 5.02E-25 | -13.6697157 | 8.12E-38 | -26.4703465 | 12.08174942 | 11.1798568 | 11.06240811 | 2 | 2.2680111 | 2.094598369 | 2.059860178 |
| KIAA0195 | | -12.5472415 | 4.62E-12 | -13.98522275 | 7.30E-15 | -26.53246425 | 8.836089686 | 7.73146271 | 9.452720984 | 2 | 2 | 2 | 2 |
| C6ORF211 | C6ORF211 | -12.55523127 | 1.18E-13 | -13.9932096 | 7.24E-17 | -26.54844087 | 8.586954094 | 8.395519306 | 9.358421373 | 2 | 2 | 2 | 2 |
| CSRP2BP | CSRP2BP | -12.5570431 | 8.59E-11 | -13.9950275 | 3.00E-13 | -26.55207059 | 9.317904228 | 7.145017454 | 9.227351203 | 2 | 2 | 2 | 2 |
| C3ORF17 | C3ORF17 | -12.56589718 | 5.03E-13 | -14.00387667 | 4.60E-16 | -26.56977385 | 9.128497779 | 7.954278496 | 9.180874355 | 2 | 2 | 2 | 2 |
| ENSG00000187772 | LIN28B | -12.57306389 | 7.25E-07 | -14.01106611 | 2.41E-08 | -26.58413 | 4.467311984 | 9.479495904 | 9.358421373 | 2 | 2 | 2 | 2 |
| C9ORF89 | C9ORF89 | -12.58934739 | 6.33E-10 | -14.0273346 | 3.67E-12 | -26.61668199 | 8.963744667 | 6.844429755 | 9.636156974 | 2 | 2 | 2 | 2 |
| PCNXL3 | | -12.59219994 | 1.18E-13 | -14.03017848 | 7.44E-17 | -26.62237842 | 8.16936288 | 9.225224669 | 9.031722608 | 2 | 2 | 2 | 2 |
| MYEOV2 | | -12.61247676 | 8.12E-13 | -14.05045691 | 8.60E-16 | -26.66293367 | 9.410448395 | 7.954278496 | 8.978369997 | 2 | 2 | 2 | 2 |

| | | | | | | | | | | | | | |
|---|---|---|---|---|---|---|---|---|---|---|---|---|---|
| FAM92A1 | | -12.61706053 | 4.13E-12 | -14.05504212 | 6.56E-15 | -26.67210266 | 9.667789716 | 8.487557258 | 8.042706615 | 2 | 2 | 2 | 2 |
| WBP5 | | -12.62173261 | 5.81E-10 | -14.05971995 | 3.35E-12 | -26.68145256 | 9.848787621 | 8.57407404 | 7.098165388 | 2 | 2 | 2 | 2 |
| ENSG00000274488 | | -12.63758372 | 4.36E-11 | -14.0755679 | 1.32E-13 | -26.71315162 | 9.848787621 | 8.147241016 | 7.97167868 | 2 | 2 | 2 | 2 |
| DAK | DAK | -12.64447516 | 1.83E-13 | -14.08245433 | 1.34E-16 | -26.72692948 | 9.509418609 | 8.376385542 | 8.65477198 | 2 | 2 | 2 | 2 |
| C11ORF31 | C11ORF31 | -12.6562961 | 5.30E-12 | -14.09427818 | 9.12E-15 | -26.75057429 | 9.730679447 | 8.029695761 | 8.511888463 | 2 | 2 | 2 | 2 |
| C17ORF89 | C17ORF89 | -12.66907006 | 6.64E-13 | -14.10705039 | 6.91E-16 | -26.77612046 | 9.590776211 | 8.147241016 | 8.763901704 | 2 | 2 | 2 | 2 |
| SLMO2 | | -12.67136847 | 1.49E-13 | -14.10934764 | 1.05E-16 | -26.78071611 | 9.533135543 | 8.414402626 | 8.677268317 | 2 | 2 | 2 | 2 |
| AGPAT6 | AGPAT6 | -12.67698392 | 4.23E-11 | -14.11496835 | 1.29E-13 | -26.79195227 | 7.384433967 | 9.00621767 | 9.670202709 | 2 | 2 | 2 | 2 |
| PRUNE | | -12.69577154 | 3.57E-11 | -14.1337559 | 1.05E-13 | -26.82952744 | 7.82963488 | 8.414402626 | 9.888372618 | 2 | 2 | 2 | 2 |
| CARKD | CARKD | -12.69910746 | 2.08E-14 | -14.13708541 | 8.94E-18 | -26.83619287 | 9.262228249 | 8.433041974 | 9.132850258 | 2 | 2 | 2 | 2 |
| C11ORF73 | C11ORF73 | -12.71562004 | 1.49E-10 | -14.15360626 | 6.41E-13 | -26.8692263 | 10.00110988 | 7.761270542 | 8.206335842 | 2 | 2 | 2 | 2 |
| FAM96B | | -12.80151132 | 3.03E-15 | -14.23948862 | 8.44E-19 | -27.04099995 | 9.304185368 | 8.732943944 | 9.165043284 | 2 | 2 | 2 | 2 |
| TOMM70A | | -12.86486457 | 4.06E-08 | -14.30286186 | 7.37E-10 | -27.16772643 | 5.70932728 | 9.13702746 | 10.08627303 | 2 | 2 | 2 | 2 |
| ZMYM6NB | | -12.90697099 | 2.49E-14 | -14.34495023 | 1.26E-17 | -27.25192122 | 9.740899611 | 8.834582582 | 8.784770914 | 2 | 2 | 2 | 2 |
| ENSG00000149295 | DRD2 | -12.20157892 | 3.03E-13 | -15.08271317 | 8.06E-20 | -27.28429209 | 9.262228249 | 10.42512749 | 9.849281147 | 2.141737246 | 2 | 2 | 2 |
| ERO1L | | -12.9689958 | 2.20E-15 | -14.40697381 | 6.28E-19 | -27.37596961 | 8.816894989 | 9.540933215 | 9.301632639 | 2 | 2 | 2 | 2 |
| ENSG00000012817 | KDM5D | -12.97736636 | 2.48E-07 | -14.41536974 | 7.15E-09 | -27.3927361 | 4.815609471 | 9.549500232 | 10.05225195 | 2 | 2 | 2 | 2 |
| HIAT1 | | -13.01514472 | 1.32E-14 | -14.45312413 | 6.02E-18 | -27.46826885 | 9.829762239 | 8.791896851 | 9.06622548 | 2 | 2 | 2 | 2 |
| C17ORF85 | C17ORF85 | -13.04637344 | 1.67E-14 | -14.48435319 | 8.30E-18 | -27.53072663 | 9.128497779 | 8.747909548 | 9.878698603 | 2 | 2 | 2 | 2 |
| FTSJ2 | | -13.07568141 | 5.57E-11 | -14.51366908 | 2.22E-13 | -27.58935049 | 8.855032352 | 7.761270542 | 10.3329194 | 2 | 2 | 2 | 2 |
| LOC100129361 | | -13.08790059 | 3.48E-16 | -14.52587815 | 6.89E-20 | -27.61377875 | 9.371511179 | 9.114109157 | 9.589475645 | 2 | 2 | 2 | 2 |
| NDNL2 | | -13.09118813 | 4.71E-12 | -14.52917301 | 9.74E-15 | -27.62036114 | 8.340423424 | 8.717821467 | 10.28245311 | 2 | 2 | 2 | 2 |
| SRPR | | -13.10012501 | 1.14E-12 | -14.53810854 | 1.69E-15 | -27.63823355 | 10.22876205 | 8.687092496 | 8.608698767 | 2 | 2 | 2 | 2 |
| C10ORF32 | C10ORF32 | -13.12969921 | 3.76E-12 | -14.56768412 | 7.52E-15 | -27.69738333 | 10.1770105 | 9.308339287 | 7.97167868 | 2 | 2 | 2 | 2 |
| C1ORF86 | C1ORF86 | -13.14572891 | 1.48E-16 | -14.58370631 | 2.46E-20 | -27.72943522 | 9.544849385 | 9.488434476 | 9.227351203 | 2 | 2 | 2 | 2 |
| C17ORF70 | C17ORF70 | -13.14600796 | 1.83E-14 | -14.58398836 | 9.82E-18 | -27.72999631 | 8.737455672 | 9.267380431 | 9.999667074 | 2 | 2 | 2 | 2 |
| SELT | | -13.15445004 | 1.83E-14 | -14.59243049 | 9.85E-18 | -27.74688053 | 9.579430841 | 9.856140801 | 8.585098385 | 2 | 2 | 2 | 2 |
| HDHD1 | | -13.2311047 | 1.41E-11 | -14.6690918 | 4.12E-14 | -27.9001965 | 10.51739401 | 8.469613413 | 8.536706852 | 2 | 2 | 2 | 2 |
| LPHN1 | | -13.23840667 | 1.50E-15 | -14.67638592 | 4.66E-19 | -27.91479259 | 9.966689162 | 9.04308532 | 9.399592459 | 2 | 2 | 2 | 2 |
| CIRH1A | CIRH1A | -13.2490706 | 1.18E-14 | -14.68705128 | 5.98E-18 | -27.93612188 | 10.15425007 | 9.102512015 | 9.049077186 | 2 | 2 | 2 | 2 |
| ENSG00000263160 | | -13.26202203 | 6.83E-16 | -14.70000092 | 1.79E-19 | -27.96202295 | 9.048471759 | 9.583268459 | 9.878698603 | 2 | 2 | 2 | 2 |
| PPAPDC1B | | -13.3725813 | 2.76E-15 | -14.81056169 | 1.07E-18 | -28.18314299 | 9.064836028 | 9.461450882 | 10.19940021 | 2 | 2 | 2 | 2 |
| ENSG00000170290 | SLN | -14.15780965 | 6.07E-15 | -14.17029198 | 1.32E-15 | -28.32810163 | 11.32815351 | 9.856140801 | 9.358421373 | 2 | 2 | 2.048074405 | 2 |
| NRD1 | | -13.49671739 | 3.34E-15 | -14.93469863 | 1.45E-18 | -28.43141602 | 9.410448395 | 10.39692156 | 9.227351203 | 2 | 2 | 2 | 2 |
| KIAA1715 | | -13.50087648 | 1.36E-17 | -14.93885444 | 1.62E-21 | -28.43973092 | 9.678463567 | 9.726146444 | 9.926432745 | 2 | 2 | 2 | 2 |
| LEPREL2 | | -13.50112092 | 2.15E-12 | -14.93910783 | 4.53E-15 | -28.44022875 | 10.3868074 | 10.02537381 | 8.007629717 | 2 | 2 | 2 | 2 |
| GPR124 | | -13.52664051 | 3.52E-14 | -14.96462371 | 2.71E-17 | -28.49126422 | 8.797441463 | 9.632483534 | 10.49998282 | 2 | 2 | 2 | 2 |
| FAM115A | | -13.52867526 | 2.70E-14 | -14.96665826 | 1.96E-17 | -28.49533352 | 8.816894989 | 10.47993704 | 9.658943166 | 2 | 2 | 2 | 2 |
| ENSG00000233436 | BTBD18 | -22.97025283 | 1.88E-10 | -5.564174424 | 0.132010837 | -28.53442725 | 2 | 2 | 8.699419242 | 2 | 2 | 2.858551861 | 2.70970617 |
| MESDC2 | | -13.66881826 | 1.62E-15 | -15.1068 | 6.58E-19 | -28.77561826 | 10.57565445 | 9.632483534 | 9.330306409 | 2 | 2 | 2 | 2 |
| ENSG00000198759 | EGFL6 | -14.068026 | 3.62E-32 | -14.89456749 | 4.33E-50 | -28.96259348 | 13.04854016 | 12.46336633 | 12.78026686 | 2 | 2.2680111 | 2 | 2.172608095 |
| LPHN2 | | -13.77751417 | 4.44E-16 | -15.21549565 | 1.42E-19 | -28.99300982 | 9.509418609 | 10.64091981 | 9.757248847 | 2 | 2 | 2 | 2 |
| C16ORF13 | C16ORF13 | -13.77991618 | 5.39E-17 | -15.21789636 | 1.03E-20 | -28.99781253 | 10.51739401 | 9.763446433 | 9.757248847 | 2 | 2 | 2 | 2 |
| FAM195B | | -13.82531768 | 1.07E-17 | -15.2632972 | 1.47E-21 | -29.08861488 | 10.40620204 | 10.09139839 | 9.735974718 | 2 | 2 | 2 | 2 |
| ENSG00000271447 | MMP28 | -13.83000723 | 4.21E-15 | -15.26799062 | 2.31E-18 | -29.09799785 | 10.80239439 | 9.856140801 | 9.227351203 | 2 | 2 | 2 | 2 |
| PPP2R4 | | -13.89373379 | 2.10E-14 | -15.33171888 | 1.74E-17 | -29.22545267 | 10.72212986 | 8.820493869 | 10.36791565 | 2 | 2 | 2 | 2 |
| SEPW1 | | -13.89742018 | 1.78E-18 | -15.33539913 | 1.73E-22 | -29.23281932 | 10.3868074 | 10.03760521 | 10.07784268 | 2 | 2 | 2 | 2 |
| NHP2L1 | | -13.9046174 | 4.67E-17 | -15.34259818 | 9.51E-21 | -29.24721558 | 10.48127392 | 9.540933215 | 10.36098398 | 2 | 2 | 2 | 2 |
| APOA1BP | APOA1BP | -13.91850358 | 1.97E-18 | -15.35648269 | 1.99E-22 | -29.27498627 | 10.43795805 | 10.04973379 | 10.06936278 | 2 | 2 | 2 | 2 |
| ENSG00000132872 | SYT4 | -14.04861846 | 4.28E-09 | -15.48662231 | 6.73E-11 | -29.53524077 | 7.096806373 | 9.170736405 | 11.61622645 | 2 | 2 | 2 | 2 |
| WBSCR16 | | -14.07458129 | 1.24E-13 | -15.51256924 | 1.75E-16 | -29.58715053 | 11.26227543 | 9.018611864 | 9.888372618 | 2 | 2 | 2 | 2 |
| VIMP | | -14.09160741 | 8.82E-18 | -15.52958824 | 1.38E-21 | -29.62119565 | 10.54098126 | 9.792607274 | 10.63827116 | 2 | 2 | 2 | 2 |
| ENSG00000168830 | HTR1E | -14.18532242 | 1.60E-16 | -15.62330559 | 5.20E-20 | -29.80862801 | 10.95065381 | 9.540933215 | 10.56174313 | 2 | 2 | 2 | 2 |
| ENSG00000145824 | CXCL14 | -13.66376229 | 5.48E-51 | -16.16000093 | 7.52E-67 | -29.82376321 | 17.21220849 | 16.16443197 | 16.42045395 | 4.205751501 | 2.49394783 | 2.457042231 | 2.421067459 |
| FAM134C | | -14.32956124 | 1.01E-12 | -15.76755336 | 2.56E-15 | -30.0971146 | 8.777722034 | 10.10308486 | 11.61622645 | 2 | 2 | 2 | 2 |
| ENSG00000113196 | HAND1 | -15.06218693 | 1.50E-19 | -15.06926965 | 1.35E-20 | -30.13145658 | 11.93145677 | 10.82153847 | 11.00131064 | 2 | 2 | 2.048074405 | 2 |
| ENSG00000115361 | ACADL | -22.41199991 | 1.55E-09 | -7.733986089 | 0.040056024 | -30.145986 | 8.107517749 | 2 | 2 | 2 | 2 | 2.225793514 | 2.059860178 |
| RQCD1 | | -14.3910702 | 8.55E-20 | -15.82905019 | 5.79E-24 | -30.2201204 | 10.71177538 | 10.43440824 | 10.83118029 | 2 | 2 | 2 | 2 |
| ATP5L | ATP5L | -14.42959212 | 2.00E-17 | -15.86757532 | 4.56E-21 | -30.29716744 | 11.26581873 | 9.969011754 | 10.59756881 | 2 | 2 | 2 | 2 |
| GAREML | | -14.47164832 | 1.41E-18 | -15.90963018 | 1.88E-22 | -30.38127849 | 10.27872131 | 10.60026656 | 11.20857468 | 2 | 2 | 2 | 2 |
| ENSG00000188778 | ADRB3 | -22.27526278 | 1.00E-09 | -8.379109809 | 0.024265115 | -30.65437259 | 2 | 7.901717006 | 2 | 2 | 2 | 2.048074405 | 2.117335275 |
| ITFG3 | | -14.65138363 | 1.92E-16 | -16.08936974 | 8.29E-20 | -30.74075337 | 11.64895441 | 9.883504052 | 10.72180585 | 2 | 2 | 2 | 2 |
| ENSG00000165794 | SLC39A2 | -15.38605883 | 1.02E-22 | -15.3888278 | 4.29E-24 | -30.77488663 | 11.79588449 | 11.42535654 | 11.73625717 | 2 | 2 | 2.048074405 | 2 |
| ENSG00000159184 | HOXB13 | -14.71121418 | 2.71E-16 | -16.14920093 | 1.31E-19 | -30.8604151 | 11.81293891 | 10.3138733 | 10.2528023 | 2 | 2 | 2 | 2 |
| ENSG00000278229 | | -14.83882319 | 1.78E-16 | -16.27681041 | 8.33E-20 | -31.1156336 | 11.94919517 | 10.3387994 | 10.45512158 | 2 | 2 | 2 | 2 |
| PTPLAD1 | | -14.87283435 | 3.75E-20 | -16.31081643 | 2.85E-24 | -31.18365078 | 10.72727936 | 11.1062775 | 11.50743625 | 2 | 2 | 2 | 2 |
| SELM | | -14.99066729 | 2.47E-20 | -16.42864978 | 1.85E-24 | -31.41931708 | 11.4761985 | 10.73780697 | 11.46927014 | 2 | 2 | 2 | 2 |
| ENSG00000147655 | RSPO2 | -15.10581586 | 4.80E-22 | -16.54379706 | 1.55E-26 | -31.64961292 | 11.32815351 | 11.52862633 | 11.27276688 | 2 | 2 | 2 | 2 |
| ENSG00000187664 | HAPLN4 | -5.780628647 | 0.12640974 | -25.99482555 | 3.96E-13 | -31.7755542 | 8.797441463 | 2 | 2 | 2.868578001 | 2.59489879 | 2 | 2 |
| NGFRAP1 | | -15.3240478 | 6.60E-21 | -16.7620314 | 4.57E-25 | -32.0860792 | 12.04303957 | 11.13786633 | 11.46927014 | 2 | 2 | 2 | 2 |
| PVRL2 | | -15.44666811 | 1.34E-20 | -16.88465275 | 1.17E-24 | -32.33132085 | 12.27171276 | 11.31913965 | 11.36226358 | 2 | 2 | 2 | 2 |
| ENSG00000122585 | NPY | -13.7342139 | 7.39E-47 | -18.93995223 | 1.10E-30 | -32.67416613 | 13.82920345 | 13.39257657 | 14.03304396 | 2.600425781 | 2 | 2 | 2 |
| PPAP2B | | -15.84848928 | 1.70E-19 | -17.28647777 | 3.16E-23 | -33.13496705 | 11.20805444 | 12.84808039 | 11.82959564 | 2 | 2 | 2 | 2 |
| ENSG00000164946 | FREM1 | -18.62808917 | 8.61E-28 | -14.8769823 | 2.00E-63 | -33.50507148 | 13.93647517 | 14.74888626 | 15.5687961 | 2 | 2 | 2.183373728 | 2.466008509 |
| ENSG00000227739 | | -17.37665269 | 4.61E-25 | -18.81464269 | 1.34E-29 | -36.19129538 | 14.23241862 | 13.47302352 | 12.95729976 | 2 | 2 | 2 | 2 |
| GNB2L1 | | -17.4635956 | 7.36E-26 | -18.9015852 | 1.72E-30 | -36.3651808 | 14.29488152 | 13.51244688 | 13.16457284 | 2 | 2 | 2 | 2 |
| ENSG00000228854 | | -23.21968654 | 1.10E-06 | -27.64734231 | 3.91E-09 | -50.86702885 | 2 | 8.004992147 | 2 | 2 | 2 | 2 | 2 |
| ENSG00000149516 | MS4A3 | -22.69860806 | 1.98E-06 | -28.87441783 | 7.05E-10 | -51.57302588 | 8.107517749 | 2 | 2 | 2 | 2 | 2 | 2 |
| FAM73B | | -23.39256905 | 8.93E-07 | -28.56845822 | 1.08E-09 | -51.96102726 | 2 | 8.469613413 | 2 | 2 | 2 | 2 | 2 |
| ENSG00000181786 | ACTL9 | -22.90328409 | 1.57E-06 | -30.68964286 | 4.87E-11 | -53.59292696 | 8.138771685 | 2 | 2 | 2 | 2 | 2 | 2 |
| ENSG00000015413 | DPEP1 | -22.9392221 | 1.50E-06 | -33.05144846 | 1.13E-12 | -55.99067056 | 2 | 2 | 8.35328246 | 2 | 2 | 2 | 2 |
