## Supplementary Table 3 for "Identification of Cancer-Associated Fibroblasts in Glioblastoma and Defining Their Pro-tumoral Effects"

**SUPPLEMENTARY TABLE 3: CAF proliferation in response to various codnitions.**

**Sample Key**
A. Neurosphere media
B. Neurosphere media + anti-PDGF
C. Stem Cell CM
D. Stem Cell CM + anti-PDGF
E. Neurosphere media + anti-TGF-B
F. Stem Cell CM + anti-TGF-B
G. Neurosphere media + Both
H. Stem Cell CM + Both

**Cell Index at: 0:00:00**

| | 1 | 2 | 3 | 4 | 5 | 6 | 7 | 8 | 9 | 10 | 11 | 12 |
|---|---|---|---|---|---|---|---|---|---|---|---|---|
| A | | | | | | | | 0 | 0 | 0 | 0 | 0 |
| B | | | | | | | | 0 | 0 | 0 | 0 | 0 |
| C | | | | | | | | 0 | 0 | 0 | 0 | 0 |
| D | | | | | | | | 0 | 0 | 0 | 0 | 0 |
| E | | | | | | | | 0 | 0 | 0 | 0 | 0 |
| F | | | | | | | | 0 | 0 | 0 | 0 | 0 |
| G | | | | | | | | 0 | 0 | 0 | 0 | 0 |
| H | | | | | | | | 0 | 0 | 0 | 0 | 0 |

**Cell Index at: 0:00:21**

| | 1 | 2 | 3 | 4 | 5 | 6 | 7 | 8 | 9 | 10 | 11 | 12 |
|---|---|---|---|---|---|---|---|---|---|---|---|---|
| A | | | | | | | | 0.0002 | 0.0004 | -0.0015 | 0.0002 | -0.0053 |
| B | | | | | | | | -0.0008 | -0.0021 | -0.0047 | -0.0018 | -0.0007 |
| C | | | | | | | | 0.0068 | -0.0019 | -0.004 | 0.0002 | -0.0021 |
| D | | | | | | | | 0.0034 | 0.0065 | -0.0045 | -0.0015 | 0.0034 |
| E | | | | | | | | 0.0105 | -0.005 | -0.007 | -0.0049 | -0.0067 |
| F | | | | | | | | 0.0037 | -0.0026 | -0.0051 | 0.0058 | -0.0003 |
| G | | | | | | | | -0.0052 | 0.0036 | -0.0004 | -0.0006 | -0.0025 |
| H | | | | | | | | -0.0011 | -0.0023 | 0.0003 | 0.001 | 0.0013 |

**Cell Index at: 2:00:19**

| | 1 | 2 | 3 | 4 | 5 | 6 | 7 | 8 | 9 | 10 | 11 | 12 |
|---|---|---|---|---|---|---|---|---|---|---|---|---|
| A | | | | | | | | 0.0354 | 0.0381 | 0.0541 | 0.0476 | 0.0409 |
| B | | | | | | | | 0.021 | 0.0168 | 0.0256 | 0.0273 | 0.0427 |
| C | | | | | | | | 0.0082 | 0.0255 | 0.0357 | 0.0312 | 0.0352 |
| D | | | | | | | | 0.0192 | 0.0213 | 0.0136 | 0.0122 | 0.0274 |
| E | | | | | | | | 0.0018 | -0.0117 | -0.0149 | -0.0026 | -0.0132 |
| F | | | | | | | | 0.034 | 0.0321 | 0.0315 | 0.0302 | 0.0367 |
| G | | | | | | | | 0.0175 | 0.0284 | 0.0154 | 0.0209 | 0.0188 |
| H | | | | | | | | 0.054 | 0.0498 | 0.0592 | 0.0523 | 0.0384 |

**Cell Index at: 2:15:18**

| | 1 | 2 | 3 | 4 | 5 | 6 | 7 | 8 | 9 | 10 | 11 | 12 |
|---|---|---|---|---|---|---|---|---|---|---|---|---|
| A | | | | | | | | -0.0056 | -0.0034 | 0.0116 | 0.0022 | 0.0057 |
| B | | | | | | | | -0.0068 | -0.0022 | 0.0161 | 0.0131 | 0.0157 |
| C | | | | | | | | 0.0445 | 0.0435 | 0.059 | 0.0555 | 0.0489 |
| D | | | | | | | | 0.0482 | 0.0491 | 0.0496 | 0.0549 | 0.0534 |
| E | | | | | | | | 0.0146 | 0.0011 | 0.0089 | 0.009 | 0.008 |
| F | | | | | | | | 0.046 | 0.0501 | 0.0516 | 0.0551 | 0.0598 |
| G | | | | | | | | 0.0061 | 0.0139 | 0.0098 | 0.0152 | 0.0099 |
| H | | | | | | | | 0.045 | 0.0367 | 0.0527 | 0.0454 | 0.0335 |

**Cell Index at: 2:30:18**

| | 1 | 2 | 3 | 4 | 5 | 6 | 7 | 8 | 9 | 10 | 11 | 12 |
|---|---|---|---|---|---|---|---|---|---|---|---|---|
| A | | | | | | | | 0.0014 | -0.0012 | 0.0147 | 0.0104 | 0.0117 |
| B | | | | | | | | 0.0057 | 0.005 | 0.021 | 0.0186 | 0.0206 |
| C | | | | | | | | 0.0551 | 0.0592 | 0.0666 | 0.0762 | 0.0633 |
| D | | | | | | | | 0.0775 | 0.0838 | 0.0741 | 0.067 | 0.0754 |
| E | | | | | | | | 0.0338 | 0.0199 | 0.0195 | 0.0074 | 0.0255 |
| F | | | | | | | | 0.0668 | 0.066 | 0.0777 | 0.0681 | 0.0803 |
| G | | | | | | | | 0.0163 | 0.0198 | 0.0142 | 0.0295 | 0.0138 |
| H | | | | | | | | 0.0526 | 0.0487 | 0.0661 | 0.0618 | 0.0451 |

**Cell Index at: 2:45:18**

| | 1 | 2 | 3 | 4 | 5 | 6 | 7 | 8 | 9 | 10 | 11 | 12 |
|---|---|---|---|---|---|---|---|---|---|---|---|---|
| A | | | | | | | | 0.0024 | 0.0064 | 0.0235 | 0.018 | 0.0171 |
| B | | | | | | | | 0.0088 | 0.0158 | 0.0267 | 0.0278 | 0.0306 |
| C | | | | | | | | 0.0652 | 0.0799 | 0.0914 | 0.0945 | 0.0746 |
| D | | | | | | | | 0.0953 | 0.0875 | 0.0878 | 0.0823 | 0.0876 |
| E | | | | | | | | 0.0515 | 0.0297 | 0.0396 | 0.0228 | 0.036 |
| F | | | | | | | | 0.0795 | 0.0863 | 0.0934 | 0.0891 | 0.0871 |
| G | | | | | | | | 0.0249 | 0.0346 | 0.0224 | 0.0381 | 0.0234 |
| H | | | | | | | | 0.0606 | 0.0567 | 0.077 | 0.0671 | 0.0545 |

**Cell Index at: 3:00:18**

| | 1 | 2 | 3 | 4 | 5 | 6 | 7 | 8 | 9 | 10 | 11 | 12 |
|---|---|---|---|---|---|---|---|---|---|---|---|---|
| A | | | | | | | | 0.009 | 0.0142 | 0.0146 | 0.0251 | 0.023 |
| B | | | | | | | | 0.0158 | 0.0312 | 0.0406 | 0.0417 | 0.04 |
| C | | | | | | | | 0.0836 | 0.0812 | 0.0998 | 0.1008 | 0.0919 |
| D | | | | | | | | 0.1096 | 0.1074 | 0.1012 | 0.1089 | 0.1007 |
| E | | | | | | | | 0.0638 | 0.0435 | 0.0493 | 0.0356 | 0.0423 |
| F | | | | | | | | 0.0954 | 0.0974 | 0.101 | 0.0959 | 0.0998 |
| G | | | | | | | | 0.0362 | 0.0408 | 0.0332 | 0.0486 | 0.0319 |
| H | | | | | | | | 0.0698 | 0.0626 | 0.083 | 0.0725 | 0.0617 |

**Cell Index at: 3:15:19**

| | 1 | 2 | 3 | 4 | 5 | 6 | 7 | 8 | 9 | 10 | 11 | 12 |
|---|---|---|---|---|---|---|---|---|---|---|---|---|
| A | | | | | | | | 0.0147 | 0.0193 | 0.0142 | 0.0369 | 0.0324 |
| B | | | | | | | | 0.0229 | 0.0372 | 0.0439 | 0.0512 | 0.0412 |
| C | | | | | | | | 0.1043 | 0.0974 | 0.1092 | 0.1055 | 0.0934 |
| D | | | | | | | | 0.1159 | 0.117 | 0.1121 | 0.1127 | 0.1097 |
| E | | | | | | | | 0.0752 | 0.0535 | 0.0557 | 0.0432 | 0.0509 |
| F | | | | | | | | 0.0994 | 0.1058 | 0.1058 | 0.1041 | 0.1065 |
| G | | | | | | | | 0.0398 | 0.0475 | 0.0366 | 0.0546 | 0.0354 |
| H | | | | | | | | 0.0781 | 0.0687 | 0.0922 | 0.0792 | 0.0731 |

**Cell Index at: 3:30:19**

| | 1 | 2 | 3 | 4 | 5 | 6 | 7 | 8 | 9 | 10 | 11 | 12 |
|---|---|---|---|---|---|---|---|---|---|---|---|---|
| A | | | | | | | | 0.0296 | 0.0255 | 0.0189 | 0.0507 | 0.0398 |
| B | | | | | | | | 0.0251 | 0.0486 | 0.0488 | 0.0588 | 0.0526 |
| C | | | | | | | | 0.1009 | 0.1017 | 0.1124 | 0.1112 | 0.0969 |
| D | | | | | | | | 0.1185 | 0.1291 | 0.1102 | 0.1213 | 0.1053 |
| E | | | | | | | | 0.0824 | 0.0678 | 0.0629 | 0.0491 | 0.0536 |
| F | | | | | | | | 0.1101 | 0.1078 | 0.1142 | 0.1047 | 0.1153 |
| G | | | | | | | | 0.0464 | 0.0576 | 0.0477 | 0.0642 | 0.0478 |
| H | | | | | | | | 0.0851 | 0.0731 | 0.0975 | 0.083 | 0.0801 |

**Cell Index at: 3:45:19**

| | 1 | 2 | 3 | 4 | 5 | 6 | 7 | 8 | 9 | 10 | 11 | 12 |
|---|---|---|---|---|---|---|---|---|---|---|---|---|
| A | | | | | | | | 0.037 | 0.0304 | 0.0254 | 0.0628 | 0.0356 |
| B | | | | | | | | 0.0323 | 0.0568 | 0.0535 | 0.0641 | 0.0605 |
| C | | | | | | | | 0.1089 | 0.1095 | 0.124 | 0.1255 | 0.1 |
| D | | | | | | | | 0.1257 | 0.1328 | 0.121 | 0.1206 | 0.1178 |
| E | | | | | | | | 0.0832 | 0.0682 | 0.0766 | 0.0595 | 0.0633 |
| F | | | | | | | | 0.1042 | 0.1256 | 0.1163 | 0.1107 | 0.1188 |
| G | | | | | | | | 0.0567 | 0.0643 | 0.0564 | 0.0707 | 0.0531 |
| H | | | | | | | | 0.0881 | 0.0778 | 0.1022 | 0.0901 | 0.0851 |

**Cell Index at: 4:00:20**

| | 1 | 2 | 3 | 4 | 5 | 6 | 7 | 8 | 9 | 10 | 11 | 12 |
|---|---|---|---|---|---|---|---|---|---|---|---|---|
| A | | | | | | | | 0.0415 | 0.0343 | 0.0322 | 0.0664 | 0.04 |
| B | | | | | | | | 0.0361 | 0.0619 | 0.0571 | 0.0675 | 0.0683 |
| C | | | | | | | | 0.1148 | 0.1123 | 0.1281 | 0.1291 | 0.103 |
| D | | | | | | | | 0.133 | 0.1414 | 0.1333 | 0.1184 | 0.1172 |
| E | | | | | | | | 0.089 | 0.0761 | 0.0698 | 0.062 | 0.0678 |
| F | | | | | | | | 0.1127 | 0.1295 | 0.1254 | 0.1215 | 0.1249 |
| G | | | | | | | | 0.0591 | 0.0652 | 0.0569 | 0.0811 | 0.0592 |
| H | | | | | | | | 0.0869 | 0.0841 | 0.1048 | 0.0914 | 0.0887 |

**Cell Index at: 4:15:20**

| | 1 | 2 | 3 | 4 | 5 | 6 | 7 | 8 | 9 | 10 | 11 | 12 |
|---|---|---|---|---|---|---|---|---|---|---|---|---|
| A | | | | | | | | 0.0456 | 0.0418 | 0.0384 | 0.0733 | 0.0433 |
| B | | | | | | | | 0.0428 | 0.0656 | 0.0622 | 0.0747 | 0.0796 |
| C | | | | | | | | 0.1227 | 0.115 | 0.132 | 0.1386 | 0.1086 |
| D | | | | | | | | 0.1384 | 0.1478 | 0.1416 | 0.1302 | 0.1235 |
| E | | | | | | | | 0.0984 | 0.0823 | 0.0858 | 0.0667 | 0.0718 |
| F | | | | | | | | 0.108 | 0.1281 | 0.1288 | 0.122 | 0.1346 |
| G | | | | | | | | 0.0654 | 0.07 | 0.0671 | 0.0805 | 0.065 |

| | | | | | | | | | | | | |
|---|---|---|---|---|---|---|---|---|---|---|---|---|
| H | | | | | | | | 0.0897 | 0.0879 | 0.1106 | 0.0951 | 0.0905 |

**Cell Index at: 4:30:21**

| | 1 | 2 | 3 | 4 | 5 | 6 | 7 | 8 | 9 | 10 | 11 | 12 |
|---|---|---|---|---|---|---|---|---|---|---|---|---|
| A | | | | | | | | 0.0483 | 0.045 | 0.0399 | 0.0746 | 0.0517 |
| B | | | | | | | | 0.0435 | 0.0643 | 0.0689 | 0.0813 | 0.0817 |
| C | | | | | | | | 0.1234 | 0.1212 | 0.1359 | 0.1419 | 0.112 |
| D | | | | | | | | 0.1411 | 0.1527 | 0.1399 | 0.1305 | 0.133 |
| E | | | | | | | | 0.0978 | 0.0808 | 0.0909 | 0.0804 | 0.0681 |
| F | | | | | | | | 0.12 | 0.1386 | 0.1344 | 0.1217 | 0.1384 |
| G | | | | | | | | 0.0675 | 0.0766 | 0.0711 | 0.0877 | 0.0658 |
| H | | | | | | | | 0.0947 | 0.0934 | 0.1144 | 0.1002 | 0.092 |

**Cell Index at: 4:45:20**

| | 1 | 2 | 3 | 4 | 5 | 6 | 7 | 8 | 9 | 10 | 11 | 12 |
|---|---|---|---|---|---|---|---|---|---|---|---|---|
| A | | | | | | | | 0.0534 | 0.0476 | 0.0436 | 0.0777 | 0.0574 |
| B | | | | | | | | 0.0505 | 0.0792 | 0.0733 | 0.0824 | 0.0869 |
| C | | | | | | | | 0.1319 | 0.1198 | 0.1392 | 0.1452 | 0.1218 |
| D | | | | | | | | 0.1419 | 0.1536 | 0.1421 | 0.1304 | 0.1375 |
| E | | | | | | | | 0.1058 | 0.0889 | 0.0964 | 0.0815 | 0.0731 |
| F | | | | | | | | 0.1269 | 0.1404 | 0.1381 | 0.1282 | 0.1444 |
| G | | | | | | | | 0.0689 | 0.0769 | 0.0753 | 0.0924 | 0.0697 |
| H | | | | | | | | 0.098 | 0.0916 | 0.1226 | 0.1072 | 0.095 |

**Cell Index at: 5:00:21**

| | 1 | 2 | 3 | 4 | 5 | 6 | 7 | 8 | 9 | 10 | 11 | 12 |
|---|---|---|---|---|---|---|---|---|---|---|---|---|
| A | | | | | | | | 0.0551 | 0.0538 | 0.0447 | 0.083 | 0.065 |
| B | | | | | | | | 0.0527 | 0.0806 | 0.0747 | 0.0909 | 0.0933 |
| C | | | | | | | | 0.1365 | 0.1227 | 0.1502 | 0.153 | 0.1289 |
| D | | | | | | | | 0.1514 | 0.1595 | 0.1468 | 0.1314 | 0.139 |
| E | | | | | | | | 0.1121 | 0.0947 | 0.1045 | 0.0881 | 0.081 |
| F | | | | | | | | 0.1298 | 0.1457 | 0.1423 | 0.1298 | 0.148 |
| G | | | | | | | | 0.0714 | 0.0841 | 0.0782 | 0.0987 | 0.0784 |
| H | | | | | | | | 0.1041 | 0.0931 | 0.1251 | 0.1064 | 0.0974 |

**Cell Index at: 5:15:21**

| | 1 | 2 | 3 | 4 | 5 | 6 | 7 | 8 | 9 | 10 | 11 | 12 |
|---|---|---|---|---|---|---|---|---|---|---|---|---|
| A | | | | | | | | 0.053 | 0.0564 | 0.0481 | 0.0906 | 0.0709 |
| B | | | | | | | | 0.0587 | 0.085 | 0.0839 | 0.0998 | 0.0941 |
| C | | | | | | | | 0.1348 | 0.1277 | 0.1492 | 0.1616 | 0.1377 |
| D | | | | | | | | 0.1646 | 0.1621 | 0.1473 | 0.141 | 0.1395 |
| E | | | | | | | | 0.1203 | 0.1063 | 0.102 | 0.0878 | 0.0826 |
| F | | | | | | | | 0.1297 | 0.1459 | 0.1505 | 0.1385 | 0.1498 |
| G | | | | | | | | 0.0755 | 0.0901 | 0.0818 | 0.1036 | 0.0766 |
| H | | | | | | | | 0.1079 | 0.1016 | 0.1274 | 0.1066 | 0.0966 |

**Cell Index at: 5:30:21**

| | 1 | 2 | 3 | 4 | 5 | 6 | 7 | 8 | 9 | 10 | 11 | 12 |
|---|---|---|---|---|---|---|---|---|---|---|---|---|
| A | | | | | | | | 0.0511 | 0.0599 | 0.0567 | 0.0871 | 0.0737 |
| B | | | | | | | | 0.0604 | 0.0834 | 0.0839 | 0.0993 | 0.1006 |
| C | | | | | | | | 0.1439 | 0.1247 | 0.1574 | 0.1646 | 0.1388 |
| D | | | | | | | | 0.1492 | 0.1668 | 0.1525 | 0.145 | 0.1455 |
| E | | | | | | | | 0.1162 | 0.1111 | 0.1177 | 0.0858 | 0.0894 |
| F | | | | | | | | 0.1272 | 0.1543 | 0.1509 | 0.1417 | 0.1549 |
| G | | | | | | | | 0.0751 | 0.0995 | 0.0836 | 0.1018 | 0.0854 |
| H | | | | | | | | 0.1104 | 0.1008 | 0.1332 | 0.1146 | 0.0974 |

**Cell Index at: 5:45:22**

| | 1 | 2 | 3 | 4 | 5 | 6 | 7 | 8 | 9 | 10 | 11 | 12 |
|---|---|---|---|---|---|---|---|---|---|---|---|---|
| A | | | | | | | | 0.0606 | 0.0616 | 0.0589 | 0.0946 | 0.0783 |
| B | | | | | | | | 0.0631 | 0.0858 | 0.0842 | 0.0978 | 0.1015 |
| C | | | | | | | | 0.1413 | 0.1257 | 0.1551 | 0.1636 | 0.1415 |
| D | | | | | | | | 0.1552 | 0.1717 | 0.1571 | 0.1408 | 0.1495 |
| E | | | | | | | | 0.1163 | 0.1065 | 0.11 | 0.098 | 0.0885 |
| F | | | | | | | | 0.1317 | 0.1518 | 0.1591 | 0.1419 | 0.1587 |
| G | | | | | | | | 0.0796 | 0.0962 | 0.0862 | 0.1028 | 0.083 |
| H | | | | | | | | 0.1179 | 0.1034 | 0.1355 | 0.1166 | 0.1021 |

**Cell Index at: 6:00:22**

| | 1 | 2 | 3 | 4 | 5 | 6 | 7 | 8 | 9 | 10 | 11 | 12 |
|---|---|---|---|---|---|---|---|---|---|---|---|---|
| A | | | | | | | | 0.0586 | 0.0677 | 0.0627 | 0.0972 | 0.0844 |
| B | | | | | | | | 0.069 | 0.0918 | 0.0898 | 0.1068 | 0.1057 |
| C | | | | | | | | 0.1457 | 0.1263 | 0.1598 | 0.1724 | 0.1387 |
| D | | | | | | | | 0.1531 | 0.1774 | 0.1552 | 0.1542 | 0.1614 |
| E | | | | | | | | 0.1233 | 0.1215 | 0.1202 | 0.1046 | 0.0912 |
| F | | | | | | | | 0.141 | 0.1632 | 0.1613 | 0.1523 | 0.1577 |
| G | | | | | | | | 0.0845 | 0.1065 | 0.0907 | 0.1073 | 0.0896 |
| H | | | | | | | | 0.1178 | 0.1053 | 0.1371 | 0.117 | 0.104 |

**Cell Index at: 6:15:22**

| | 1 | 2 | 3 | 4 | 5 | 6 | 7 | 8 | 9 | 10 | 11 | 12 |
|---|---|---|---|---|---|---|---|---|---|---|---|---|
| A | | | | | | | | 0.066 | 0.0677 | 0.0654 | 0.1032 | 0.0884 |
| B | | | | | | | | 0.0731 | 0.0935 | 0.0951 | 0.1084 | 0.1055 |
| C | | | | | | | | 0.1456 | 0.1396 | 0.1537 | 0.1685 | 0.1515 |
| D | | | | | | | | 0.1624 | 0.1714 | 0.1549 | 0.1572 | 0.1678 |
| E | | | | | | | | 0.1172 | 0.1076 | 0.1255 | 0.1059 | 0.0964 |
| F | | | | | | | | 0.1367 | 0.1642 | 0.1613 | 0.154 | 0.1618 |
| G | | | | | | | | 0.08 | 0.1053 | 0.0934 | 0.1116 | 0.0875 |
| H | | | | | | | | 0.1166 | 0.1016 | 0.1384 | 0.1229 | 0.1042 |

**Cell Index at: 6:30:21**

| | 1 | 2 | 3 | 4 | 5 | 6 | 7 | 8 | 9 | 10 | 11 | 12 |
|---|---|---|---|---|---|---|---|---|---|---|---|---|
| A | | | | | | | | 0.0667 | 0.0706 | 0.0674 | 0.1021 | 0.0947 |
| B | | | | | | | | 0.0787 | 0.0894 | 0.0948 | 0.1146 | 0.1121 |
| C | | | | | | | | 0.1547 | 0.1372 | 0.1683 | 0.1738 | 0.1475 |
| D | | | | | | | | 0.1629 | 0.1759 | 0.1602 | 0.159 | 0.1604 |
| E | | | | | | | | 0.1282 | 0.1218 | 0.1233 | 0.1133 | 0.0934 |
| F | | | | | | | | 0.1519 | 0.16 | 0.1642 | 0.1576 | 0.1687 |
| G | | | | | | | | 0.082 | 0.1026 | 0.0988 | 0.1137 | 0.0894 |
| H | | | | | | | | 0.1235 | 0.1129 | 0.1399 | 0.1232 | 0.1074 |

**Cell Index at: 6:45:22**

| | 1 | 2 | 3 | 4 | 5 | 6 | 7 | 8 | 9 | 10 | 11 | 12 |
|---|---|---|---|---|---|---|---|---|---|---|---|---|
| A | | | | | | | | 0.0695 | 0.0733 | 0.0671 | 0.0969 | 0.0963 |
| B | | | | | | | | 0.0836 | 0.0923 | 0.0973 | 0.1163 | 0.1147 |
| C | | | | | | | | 0.1612 | 0.143 | 0.1719 | 0.1776 | 0.1582 |
| D | | | | | | | | 0.1624 | 0.1795 | 0.1583 | 0.1675 | 0.1583 |
| E | | | | | | | | 0.1219 | 0.1257 | 0.1209 | 0.1209 | 0.0909 |
| F | | | | | | | | 0.1463 | 0.1672 | 0.1692 | 0.1581 | 0.1675 |
| G | | | | | | | | 0.0869 | 0.1056 | 0.0987 | 0.1162 | 0.091 |
| H | | | | | | | | 0.1203 | 0.1099 | 0.1454 | 0.1305 | 0.1065 |

**Cell Index at: 7:00:21**

| | 1 | 2 | 3 | 4 | 5 | 6 | 7 | 8 | 9 | 10 | 11 | 12 |
|---|---|---|---|---|---|---|---|---|---|---|---|---|
| A | | | | | | | | 0.0692 | 0.0775 | 0.0727 | 0.0969 | 0.1027 |
| B | | | | | | | | 0.0906 | 0.094 | 0.1013 | 0.1186 | 0.1158 |
| C | | | | | | | | 0.1604 | 0.1395 | 0.1775 | 0.1719 | 0.1592 |
| D | | | | | | | | 0.1624 | 0.1786 | 0.1559 | 0.1664 | 0.1705 |
| E | | | | | | | | 0.1217 | 0.1307 | 0.1396 | 0.12 | 0.1005 |
| F | | | | | | | | 0.1502 | 0.169 | 0.1767 | 0.1602 | 0.172 |
| G | | | | | | | | 0.088 | 0.1069 | 0.0994 | 0.1178 | 0.0929 |
| H | | | | | | | | 0.1265 | 0.1131 | 0.1481 | 0.1292 | 0.1098 |

**Cell Index at: 7:15:21**

| | 1 | 2 | 3 | 4 | 5 | 6 | 7 | 8 | 9 | 10 | 11 | 12 |
|---|---|---|---|---|---|---|---|---|---|---|---|---|
| A | | | | | | | | 0.0731 | 0.0816 | 0.0747 | 0.1028 | 0.1045 |
| B | | | | | | | | 0.0907 | 0.1 | 0.0948 | 0.118 | 0.1202 |
| C | | | | | | | | 0.1607 | 0.1392 | 0.1743 | 0.1794 | 0.1579 |
| D | | | | | | | | 0.1686 | 0.1841 | 0.1688 | 0.159 | 0.1654 |
| E | | | | | | | | 0.1322 | 0.1499 | 0.1337 | 0.1223 | 0.1076 |
| F | | | | | | | | 0.1417 | 0.1733 | 0.1733 | 0.1615 | 0.178 |

| | | | | | | | | | | | | |
|---|---|---|---|---|---|---|---|---|---|---|---|---|
| G | | | | | | | | 0.0892 | 0.1102 | 0.1038 | 0.1156 | 0.0955 |
| H | | | | | | | | 0.1273 | 0.1156 | 0.1496 | 0.1302 | 0.1103 |

**Cell Index at: 7:30:22**

| | 1 | 2 | 3 | 4 | 5 | 6 | 7 | 8 | 9 | 10 | 11 | 12 |
|---|---|---|---|---|---|---|---|---|---|---|---|---|
| A | | | | | | | | 0.0765 | 0.0818 | 0.0749 | 0.1006 | 0.1068 |
| B | | | | | | | | 0.0947 | 0.102 | 0.1014 | 0.1227 | 0.1282 |
| C | | | | | | | | 0.1606 | 0.1458 | 0.1732 | 0.184 | 0.1605 |
| D | | | | | | | | 0.1668 | 0.196 | 0.1686 | 0.1655 | 0.1749 |
| E | | | | | | | | 0.1323 | 0.1342 | 0.1373 | 0.1268 | 0.1118 |
| F | | | | | | | | 0.1458 | 0.1717 | 0.177 | 0.1647 | 0.1776 |
| G | | | | | | | | 0.0849 | 0.1079 | 0.1097 | 0.1236 | 0.0962 |
| H | | | | | | | | 0.13 | 0.1148 | 0.1535 | 0.1324 | 0.1102 |

**Cell Index at: 7:45:23**

| | 1 | 2 | 3 | 4 | 5 | 6 | 7 | 8 | 9 | 10 | 11 | 12 |
|---|---|---|---|---|---|---|---|---|---|---|---|---|
| A | | | | | | | | 0.0742 | 0.0795 | 0.0769 | 0.0983 | 0.1097 |
| B | | | | | | | | 0.0971 | 0.109 | 0.105 | 0.1244 | 0.1217 |
| C | | | | | | | | 0.1568 | 0.1419 | 0.1783 | 0.1855 | 0.1686 |
| D | | | | | | | | 0.1696 | 0.1802 | 0.1729 | 0.1627 | 0.1796 |
| E | | | | | | | | 0.1354 | 0.137 | 0.1388 | 0.128 | 0.1104 |
| F | | | | | | | | 0.1569 | 0.1733 | 0.1821 | 0.1703 | 0.1789 |
| G | | | | | | | | 0.0915 | 0.1125 | 0.1069 | 0.1254 | 0.1006 |
| H | | | | | | | | 0.1314 | 0.117 | 0.1532 | 0.1327 | 0.1123 |

**Cell Index at: 8:00:24**

| | 1 | 2 | 3 | 4 | 5 | 6 | 7 | 8 | 9 | 10 | 11 | 12 |
|---|---|---|---|---|---|---|---|---|---|---|---|---|
| A | | | | | | | | 0.0802 | 0.0866 | 0.0817 | 0.1026 | 0.1061 |
| B | | | | | | | | 0.1014 | 0.1039 | 0.1064 | 0.1269 | 0.1217 |
| C | | | | | | | | 0.1558 | 0.1488 | 0.1761 | 0.186 | 0.1654 |
| D | | | | | | | | 0.1715 | 0.1845 | 0.1751 | 0.1617 | 0.1782 |
| E | | | | | | | | 0.1401 | 0.1323 | 0.1455 | 0.1359 | 0.1159 |
| F | | | | | | | | 0.1508 | 0.1777 | 0.1813 | 0.1745 | 0.1771 |
| G | | | | | | | | 0.0942 | 0.1135 | 0.1143 | 0.1302 | 0.1008 |
| H | | | | | | | | 0.1333 | 0.117 | 0.1584 | 0.1323 | 0.1087 |

**Cell Index at: 8:15:24**

| | 1 | 2 | 3 | 4 | 5 | 6 | 7 | 8 | 9 | 10 | 11 | 12 |
|---|---|---|---|---|---|---|---|---|---|---|---|---|
| A | | | | | | | | 0.0798 | 0.0865 | 0.0828 | 0.104 | 0.1099 |
| B | | | | | | | | 0.0989 | 0.1091 | 0.1049 | 0.1287 | 0.1183 |
| C | | | | | | | | 0.1623 | 0.1533 | 0.1762 | 0.1853 | 0.1632 |
| D | | | | | | | | 0.1725 | 0.1846 | 0.1747 | 0.163 | 0.1811 |
| E | | | | | | | | 0.1411 | 0.144 | 0.1487 | 0.1374 | 0.1275 |
| F | | | | | | | | 0.1562 | 0.1803 | 0.1768 | 0.176 | 0.1771 |
| G | | | | | | | | 0.096 | 0.1136 | 0.1145 | 0.1288 | 0.1019 |
| H | | | | | | | | 0.1341 | 0.1164 | 0.1583 | 0.1384 | 0.1101 |

**Cell Index at: 8:30:25**

| | 1 | 2 | 3 | 4 | 5 | 6 | 7 | 8 | 9 | 10 | 11 | 12 |
|---|---|---|---|---|---|---|---|---|---|---|---|---|
| A | | | | | | | | 0.0794 | 0.084 | 0.08 | 0.1077 | 0.1142 |
| B | | | | | | | | 0.1095 | 0.1099 | 0.1114 | 0.1332 | 0.126 |
| C | | | | | | | | 0.1655 | 0.1527 | 0.1817 | 0.1787 | 0.1592 |
| D | | | | | | | | 0.1724 | 0.1857 | 0.1775 | 0.1753 | 0.1823 |
| E | | | | | | | | 0.1546 | 0.154 | 0.1493 | 0.14 | 0.1246 |
| F | | | | | | | | 0.1595 | 0.1733 | 0.1865 | 0.1752 | 0.1774 |
| G | | | | | | | | 0.0944 | 0.1165 | 0.1163 | 0.1298 | 0.1011 |
| H | | | | | | | | 0.1369 | 0.1193 | 0.1596 | 0.1392 | 0.1136 |

**Cell Index at: 8:45:24**

| | 1 | 2 | 3 | 4 | 5 | 6 | 7 | 8 | 9 | 10 | 11 | 12 |
|---|---|---|---|---|---|---|---|---|---|---|---|---|
| A | | | | | | | | 0.0836 | 0.086 | 0.0778 | 0.1079 | 0.1122 |
| B | | | | | | | | 0.1107 | 0.1154 | 0.1113 | 0.1306 | 0.1225 |
| C | | | | | | | | 0.1685 | 0.1502 | 0.1847 | 0.1811 | 0.1631 |
| D | | | | | | | | 0.1786 | 0.1892 | 0.1827 | 0.1681 | 0.1877 |
| E | | | | | | | | 0.156 | 0.1524 | 0.1437 | 0.1418 | 0.118 |
| F | | | | | | | | 0.1544 | 0.1818 | 0.1839 | 0.1732 | 0.1826 |
| G | | | | | | | | 0.0979 | 0.1138 | 0.1183 | 0.132 | 0.0978 |
| H | | | | | | | | 0.1391 | 0.1207 | 0.1586 | 0.1402 | 0.1153 |

**Cell Index at: 9:00:25**

| | 1 | 2 | 3 | 4 | 5 | 6 | 7 | 8 | 9 | 10 | 11 | 12 |
|---|---|---|---|---|---|---|---|---|---|---|---|---|
| A | | | | | | | | 0.081 | 0.0885 | 0.0802 | 0.111 | 0.1134 |
| B | | | | | | | | 0.1112 | 0.1035 | 0.1126 | 0.1353 | 0.1269 |
| C | | | | | | | | 0.1708 | 0.1534 | 0.1836 | 0.1841 | 0.1707 |
| D | | | | | | | | 0.1829 | 0.1908 | 0.1887 | 0.1696 | 0.1852 |
| E | | | | | | | | 0.1472 | 0.1492 | 0.1547 | 0.1444 | 0.1299 |
| F | | | | | | | | 0.1573 | 0.178 | 0.1833 | 0.1705 | 0.1847 |
| G | | | | | | | | 0.0939 | 0.1207 | 0.1168 | 0.1318 | 0.1018 |
| H | | | | | | | | 0.1388 | 0.1232 | 0.1681 | 0.1392 | 0.1154 |

**Cell Index at: 9:15:25**

| | 1 | 2 | 3 | 4 | 5 | 6 | 7 | 8 | 9 | 10 | 11 | 12 |
|---|---|---|---|---|---|---|---|---|---|---|---|---|
| A | | | | | | | | 0.0883 | 0.0887 | 0.0804 | 0.1089 | 0.1106 |
| B | | | | | | | | 0.1145 | 0.1107 | 0.1168 | 0.1344 | 0.1244 |
| C | | | | | | | | 0.1629 | 0.1519 | 0.1811 | 0.1867 | 0.1725 |
| D | | | | | | | | 0.1749 | 0.1882 | 0.182 | 0.1711 | 0.1841 |
| E | | | | | | | | 0.151 | 0.1512 | 0.1497 | 0.1505 | 0.1314 |
| F | | | | | | | | 0.1604 | 0.1789 | 0.1894 | 0.173 | 0.1777 |
| G | | | | | | | | 0.0896 | 0.1225 | 0.1193 | 0.1363 | 0.1081 |
| H | | | | | | | | 0.1399 | 0.1259 | 0.1704 | 0.1427 | 0.1189 |

**Cell Index at: 9:30:24**

| | 1 | 2 | 3 | 4 | 5 | 6 | 7 | 8 | 9 | 10 | 11 | 12 |
|---|---|---|---|---|---|---|---|---|---|---|---|---|
| A | | | | | | | | 0.0844 | 0.0849 | 0.0826 | 0.105 | 0.1147 |
| B | | | | | | | | 0.1134 | 0.1056 | 0.1138 | 0.1332 | 0.1227 |
| C | | | | | | | | 0.1676 | 0.1436 | 0.19 | 0.1905 | 0.1697 |
| D | | | | | | | | 0.1832 | 0.1931 | 0.1869 | 0.1802 | 0.1855 |
| E | | | | | | | | 0.1471 | 0.155 | 0.1597 | 0.1453 | 0.1353 |
| F | | | | | | | | 0.1725 | 0.1831 | 0.1898 | 0.1768 | 0.181 |
| G | | | | | | | | 0.0891 | 0.1091 | 0.1206 | 0.1384 | 0.1053 |
| H | | | | | | | | 0.1411 | 0.1247 | 0.171 | 0.1473 | 0.1158 |

**Cell Index at: 9:45:24**

| | 1 | 2 | 3 | 4 | 5 | 6 | 7 | 8 | 9 | 10 | 11 | 12 |
|---|---|---|---|---|---|---|---|---|---|---|---|---|
| A | | | | | | | | 0.0835 | 0.0866 | 0.0829 | 0.1094 | 0.1148 |
| B | | | | | | | | 0.1121 | 0.1045 | 0.115 | 0.1383 | 0.125 |
| C | | | | | | | | 0.17 | 0.1473 | 0.1887 | 0.1971 | 0.1741 |
| D | | | | | | | | 0.1846 | 0.1807 | 0.1882 | 0.1797 | 0.1776 |
| E | | | | | | | | 0.1591 | 0.1571 | 0.1481 | 0.1474 | 0.1386 |
| F | | | | | | | | 0.1708 | 0.1835 | 0.196 | 0.1754 | 0.1773 |
| G | | | | | | | | 0.0912 | 0.1068 | 0.1302 | 0.143 | 0.1028 |
| H | | | | | | | | 0.1424 | 0.1278 | 0.1709 | 0.1486 | 0.1175 |

**Cell Index at: 10:00:24**

| | 1 | 2 | 3 | 4 | 5 | 6 | 7 | 8 | 9 | 10 | 11 | 12 |
|---|---|---|---|---|---|---|---|---|---|---|---|---|
| A | | | | | | | | 0.0877 | 0.088 | 0.0817 | 0.1101 | 0.1106 |
| B | | | | | | | | 0.1145 | 0.1076 | 0.113 | 0.1399 | 0.1272 |
| C | | | | | | | | 0.1753 | 0.1541 | 0.1919 | 0.1885 | 0.1761 |
| D | | | | | | | | 0.1863 | 0.1913 | 0.1821 | 0.1839 | 0.1866 |
| E | | | | | | | | 0.1492 | 0.1558 | 0.163 | 0.1509 | 0.1298 |
| F | | | | | | | | 0.1714 | 0.1895 | 0.191 | 0.1756 | 0.1803 |
| G | | | | | | | | 0.0902 | 0.1084 | 0.1267 | 0.1381 | 0.1119 |
| H | | | | | | | | 0.1464 | 0.125 | 0.1732 | 0.1511 | 0.117 |

**Cell Index at: 10:15:25**

| | 1 | 2 | 3 | 4 | 5 | 6 | 7 | 8 | 9 | 10 | 11 | 12 |
|---|---|---|---|---|---|---|---|---|---|---|---|---|
| A | | | | | | | | 0.0856 | 0.0934 | 0.0838 | 0.109 | 0.1138 |
| B | | | | | | | | 0.1116 | 0.1098 | 0.1135 | 0.1352 | 0.1276 |
| C | | | | | | | | 0.1667 | 0.1566 | 0.2008 | 0.1944 | 0.1663 |
| D | | | | | | | | 0.1788 | 0.1954 | 0.1878 | 0.1779 | 0.1822 |
| E | | | | | | | | 0.1507 | 0.1624 | 0.1515 | 0.1596 | 0.1325 |

| | | | | | | | | | | | | |
|---|---|---|---|---|---|---|---|---|---|---|---|---|
| F | | | | | | | | 0.1763 | 0.1839 | 0.2056 | 0.1779 | 0.188 |
| G | | | | | | | | 0.0946 | 0.1124 | 0.1253 | 0.1409 | 0.1064 |
| H | | | | | | | | 0.1443 | 0.1333 | 0.1729 | 0.1516 | 0.118 |

**Cell Index at: 10:30:26**

| | 1 | 2 | 3 | 4 | 5 | 6 | 7 | 8 | 9 | 10 | 11 | 12 |
|---|---|---|---|---|---|---|---|---|---|---|---|---|
| A | | | | | | | | 0.0821 | 0.091 | 0.0887 | 0.1107 | 0.1161 |
| B | | | | | | | | 0.1159 | 0.1074 | 0.11 | 0.1354 | 0.1309 |
| C | | | | | | | | 0.1735 | 0.1497 | 0.1941 | 0.1945 | 0.1716 |
| D | | | | | | | | 0.175 | 0.1881 | 0.193 | 0.1878 | 0.1832 |
| E | | | | | | | | 0.1507 | 0.1624 | 0.1543 | 0.155 | 0.1361 |
| F | | | | | | | | 0.1687 | 0.1869 | 0.1988 | 0.1843 | 0.1819 |
| G | | | | | | | | 0.0978 | 0.1119 | 0.1218 | 0.1442 | 0.1088 |
| H | | | | | | | | 0.1473 | 0.1307 | 0.1789 | 0.1545 | 0.1196 |

**Cell Index at: 10:45:27**

| | 1 | 2 | 3 | 4 | 5 | 6 | 7 | 8 | 9 | 10 | 11 | 12 |
|---|---|---|---|---|---|---|---|---|---|---|---|---|
| A | | | | | | | | 0.0799 | 0.0918 | 0.0938 | 0.1149 | 0.1201 |
| B | | | | | | | | 0.1127 | 0.112 | 0.1105 | 0.1338 | 0.1312 |
| C | | | | | | | | 0.1691 | 0.1591 | 0.1973 | 0.2005 | 0.172 |
| D | | | | | | | | 0.1757 | 0.1909 | 0.1955 | 0.1875 | 0.185 |
| E | | | | | | | | 0.1424 | 0.1511 | 0.1474 | 0.154 | 0.1371 |
| F | | | | | | | | 0.1681 | 0.1907 | 0.2051 | 0.1834 | 0.1778 |
| G | | | | | | | | 0.096 | 0.1094 | 0.1185 | 0.1462 | 0.105 |
| H | | | | | | | | 0.1486 | 0.1298 | 0.1796 | 0.1527 | 0.1156 |

**Cell Index at: 11:00:26**

| | 1 | 2 | 3 | 4 | 5 | 6 | 7 | 8 | 9 | 10 | 11 | 12 |
|---|---|---|---|---|---|---|---|---|---|---|---|---|
| A | | | | | | | | 0.0799 | 0.0935 | 0.0925 | 0.1069 | 0.122 |
| B | | | | | | | | 0.1167 | 0.1071 | 0.0984 | 0.1338 | 0.1326 |
| C | | | | | | | | 0.1703 | 0.1584 | 0.1932 | 0.2062 | 0.1771 |
| D | | | | | | | | 0.1659 | 0.1863 | 0.1962 | 0.1964 | 0.1892 |
| E | | | | | | | | 0.1541 | 0.1517 | 0.1673 | 0.1452 | 0.1377 |
| F | | | | | | | | 0.1672 | 0.1975 | 0.1893 | 0.1832 | 0.1817 |
| G | | | | | | | | 0.0917 | 0.1134 | 0.1223 | 0.1421 | 0.1059 |
| H | | | | | | | | 0.149 | 0.1327 | 0.1789 | 0.1561 | 0.1197 |

**Cell Index at: 11:15:25**

| | 1 | 2 | 3 | 4 | 5 | 6 | 7 | 8 | 9 | 10 | 11 | 12 |
|---|---|---|---|---|---|---|---|---|---|---|---|---|
| A | | | | | | | | 0.0862 | 0.0943 | 0.0926 | 0.1038 | 0.1185 |
| B | | | | | | | | 0.1156 | 0.105 | 0.1032 | 0.1336 | 0.1322 |
| C | | | | | | | | 0.1695 | 0.1573 | 0.1968 | 0.1988 | 0.176 |
| D | | | | | | | | 0.1869 | 0.1989 | 0.2016 | 0.1898 | 0.1905 |
| E | | | | | | | | 0.1509 | 0.1544 | 0.1622 | 0.1516 | 0.1415 |
| F | | | | | | | | 0.1727 | 0.1989 | 0.1848 | 0.1803 | 0.1792 |
| G | | | | | | | | 0.0983 | 0.1176 | 0.1229 | 0.1412 | 0.1022 |
| H | | | | | | | | 0.1521 | 0.1298 | 0.1826 | 0.1532 | 0.1155 |

**Cell Index at: 11:30:25**

| | 1 | 2 | 3 | 4 | 5 | 6 | 7 | 8 | 9 | 10 | 11 | 12 |
|---|---|---|---|---|---|---|---|---|---|---|---|---|
| A | | | | | | | | 0.0815 | 0.0874 | 0.0969 | 0.1047 | 0.1193 |
| B | | | | | | | | 0.122 | 0.1072 | 0.1063 | 0.1401 | 0.1274 |
| C | | | | | | | | 0.1737 | 0.1636 | 0.2 | 0.2011 | 0.1786 |
| D | | | | | | | | 0.1944 | 0.1954 | 0.2122 | 0.19 | 0.19 |
| E | | | | | | | | 0.155 | 0.1586 | 0.1635 | 0.1583 | 0.134 |
| F | | | | | | | | 0.1689 | 0.1909 | 0.1965 | 0.1845 | 0.1861 |
| G | | | | | | | | 0.0938 | 0.1156 | 0.1232 | 0.1439 | 0.1049 |
| H | | | | | | | | 0.1511 | 0.1278 | 0.1849 | 0.16 | 0.1171 |

**Cell Index at: 11:45:25**

| | 1 | 2 | 3 | 4 | 5 | 6 | 7 | 8 | 9 | 10 | 11 | 12 |
|---|---|---|---|---|---|---|---|---|---|---|---|---|
| A | | | | | | | | 0.0847 | 0.0906 | 0.0956 | 0.1042 | 0.1227 |
| B | | | | | | | | 0.1221 | 0.1063 | 0.1026 | 0.1405 | 0.1327 |
| C | | | | | | | | 0.1734 | 0.1694 | 0.2138 | 0.2099 | 0.178 |
| D | | | | | | | | 0.1821 | 0.1892 | 0.2126 | 0.1823 | 0.1929 |
| E | | | | | | | | 0.1558 | 0.1456 | 0.161 | 0.165 | 0.1333 |
| F | | | | | | | | 0.1741 | 0.1863 | 0.1977 | 0.1861 | 0.1952 |
| G | | | | | | | | 0.0938 | 0.1155 | 0.1164 | 0.1464 | 0.1105 |
| H | | | | | | | | 0.1509 | 0.1301 | 0.1845 | 0.1594 | 0.1231 |

**Cell Index at: 12:00:25**

| | 1 | 2 | 3 | 4 | 5 | 6 | 7 | 8 | 9 | 10 | 11 | 12 |
|---|---|---|---|---|---|---|---|---|---|---|---|---|
| A | | | | | | | | 0.0848 | 0.0868 | 0.0985 | 0.1042 | 0.1196 |
| B | | | | | | | | 0.1198 | 0.1066 | 0.1035 | 0.135 | 0.1335 |
| C | | | | | | | | 0.1713 | 0.1629 | 0.207 | 0.2151 | 0.1788 |
| D | | | | | | | | 0.1885 | 0.2063 | 0.2066 | 0.1922 | 0.1942 |
| E | | | | | | | | 0.1416 | 0.1535 | 0.1575 | 0.1592 | 0.1303 |
| F | | | | | | | | 0.1756 | 0.1874 | 0.1997 | 0.1891 | 0.1953 |
| G | | | | | | | | 0.095 | 0.1209 | 0.1222 | 0.1418 | 0.103 |
| H | | | | | | | | 0.1529 | 0.1316 | 0.1896 | 0.1595 | 0.121 |

**Cell Index at: 12:15:25**

| | 1 | 2 | 3 | 4 | 5 | 6 | 7 | 8 | 9 | 10 | 11 | 12 |
|---|---|---|---|---|---|---|---|---|---|---|---|---|
| A | | | | | | | | 0.0824 | 0.0876 | 0.098 | 0.1055 | 0.1194 |
| B | | | | | | | | 0.1207 | 0.1039 | 0.1033 | 0.1425 | 0.136 |
| C | | | | | | | | 0.1742 | 0.1702 | 0.2047 | 0.2127 | 0.1849 |
| D | | | | | | | | 0.1853 | 0.2077 | 0.1996 | 0.1914 | 0.1932 |
| E | | | | | | | | 0.1613 | 0.1588 | 0.1674 | 0.1648 | 0.1185 |
| F | | | | | | | | 0.1706 | 0.2051 | 0.2032 | 0.1921 | 0.1946 |
| G | | | | | | | | 0.0944 | 0.1209 | 0.1177 | 0.1474 | 0.1021 |
| H | | | | | | | | 0.1543 | 0.1312 | 0.1879 | 0.1621 | 0.1205 |

**Cell Index at: 12:30:26**

| | 1 | 2 | 3 | 4 | 5 | 6 | 7 | 8 | 9 | 10 | 11 | 12 |
|---|---|---|---|---|---|---|---|---|---|---|---|---|
| A | | | | | | | | 0.0853 | 0.0859 | 0.0969 | 0.1055 | 0.1165 |
| B | | | | | | | | 0.1244 | 0.1047 | 0.1001 | 0.1372 | 0.1321 |
| C | | | | | | | | 0.1734 | 0.1712 | 0.2108 | 0.206 | 0.1833 |
| D | | | | | | | | 0.1814 | 0.2103 | 0.2009 | 0.1941 | 0.191 |
| E | | | | | | | | 0.1516 | 0.1532 | 0.171 | 0.1547 | 0.1208 |
| F | | | | | | | | 0.174 | 0.1885 | 0.1923 | 0.1902 | 0.1959 |
| G | | | | | | | | 0.093 | 0.1207 | 0.1144 | 0.1447 | 0.1033 |
| H | | | | | | | | 0.1554 | 0.1305 | 0.1884 | 0.1621 | 0.1243 |

**Cell Index at: 12:45:26**

| | 1 | 2 | 3 | 4 | 5 | 6 | 7 | 8 | 9 | 10 | 11 | 12 |
|---|---|---|---|---|---|---|---|---|---|---|---|---|
| A | | | | | | | | 0.0781 | 0.0881 | 0.0976 | 0.1057 | 0.1202 |
| B | | | | | | | | 0.1239 | 0.1019 | 0.0926 | 0.1408 | 0.1345 |
| C | | | | | | | | 0.175 | 0.1714 | 0.2133 | 0.2155 | 0.1865 |
| D | | | | | | | | 0.1845 | 0.2074 | 0.2058 | 0.1933 | 0.19 |
| E | | | | | | | | 0.1552 | 0.1491 | 0.1628 | 0.1648 | 0.1227 |
| F | | | | | | | | 0.172 | 0.1901 | 0.2008 | 0.1927 | 0.196 |
| G | | | | | | | | 0.0952 | 0.113 | 0.1154 | 0.1461 | 0.0955 |
| H | | | | | | | | 0.1555 | 0.1374 | 0.1901 | 0.1627 | 0.1241 |

**Cell Index at: 13:00:26**

| | 1 | 2 | 3 | 4 | 5 | 6 | 7 | 8 | 9 | 10 | 11 | 12 |
|---|---|---|---|---|---|---|---|---|---|---|---|---|
| A | | | | | | | | 0.0787 | 0.0876 | 0.0932 | 0.1033 | 0.1263 |
| B | | | | | | | | 0.1214 | 0.0998 | 0.0948 | 0.1338 | 0.1301 |
| C | | | | | | | | 0.1686 | 0.1721 | 0.2091 | 0.2127 | 0.1886 |
| D | | | | | | | | 0.1829 | 0.212 | 0.2129 | 0.2005 | 0.1968 |
| E | | | | | | | | 0.1455 | 0.1472 | 0.1698 | 0.1615 | 0.1134 |
| F | | | | | | | | 0.1728 | 0.1862 | 0.1985 | 0.1893 | 0.1902 |
| G | | | | | | | | 0.1 | 0.1175 | 0.1129 | 0.1444 | 0.0978 |
| H | | | | | | | | 0.1583 | 0.1341 | 0.1919 | 0.16 | 0.1233 |

**Cell Index at: 13:15:25**

| | 1 | 2 | 3 | 4 | 5 | 6 | 7 | 8 | 9 | 10 | 11 | 12 |
|---|---|---|---|---|---|---|---|---|---|---|---|---|
| A | | | | | | | | 0.0747 | 0.0816 | 0.0957 | 0.0999 | 0.1217 |
| B | | | | | | | | 0.1225 | 0.1041 | 0.093 | 0.1394 | 0.1357 |
| C | | | | | | | | 0.1831 | 0.1741 | 0.2128 | 0.2192 | 0.1871 |
| D | | | | | | | | 0.1853 | 0.2035 | 0.1991 | 0.1947 | 0.2015 |

| | | | | | | | | | | | | |
|---|---|---|---|---|---|---|---|---|---|---|---|---|
| E | | | | | | | | 0.1494 | 0.1541 | 0.1605 | 0.1597 | 0.113 |
| F | | | | | | | | 0.185 | 0.1895 | 0.2035 | 0.1945 | 0.1973 |
| G | | | | | | | | 0.1025 | 0.118 | 0.1148 | 0.1437 | 0.0956 |
| H | | | | | | | | 0.1572 | 0.1385 | 0.1933 | 0.1633 | 0.1159 |

**Cell Index at: 13:30:25**

| | 1 | 2 | 3 | 4 | 5 | 6 | 7 | 8 | 9 | 10 | 11 | 12 |
|---|---|---|---|---|---|---|---|---|---|---|---|---|
| A | | | | | | | | 0.0731 | 0.0849 | 0.0977 | 0.0962 | 0.125 |
| B | | | | | | | | 0.1154 | 0.1095 | 0.0883 | 0.139 | 0.1355 |
| C | | | | | | | | 0.1695 | 0.1741 | 0.2142 | 0.2212 | 0.19 |
| D | | | | | | | | 0.1914 | 0.2077 | 0.2074 | 0.202 | 0.1974 |
| E | | | | | | | | 0.1467 | 0.1598 | 0.1658 | 0.1612 | 0.1178 |
| F | | | | | | | | 0.1797 | 0.1919 | 0.2079 | 0.1946 | 0.2052 |
| G | | | | | | | | 0.1043 | 0.1174 | 0.1155 | 0.1422 | 0.1005 |
| H | | | | | | | | 0.1561 | 0.1376 | 0.1922 | 0.1619 | 0.1166 |

**Cell Index at: 13:45:25**

| | 1 | 2 | 3 | 4 | 5 | 6 | 7 | 8 | 9 | 10 | 11 | 12 |
|---|---|---|---|---|---|---|---|---|---|---|---|---|
| A | | | | | | | | 0.0729 | 0.0805 | 0.096 | 0.0952 | 0.1181 |
| B | | | | | | | | 0.1177 | 0.1115 | 0.0793 | 0.1394 | 0.1322 |
| C | | | | | | | | 0.1774 | 0.1715 | 0.2142 | 0.2232 | 0.1847 |
| D | | | | | | | | 0.1938 | 0.2019 | 0.2121 | 0.201 | 0.1972 |
| E | | | | | | | | 0.1554 | 0.157 | 0.1726 | 0.1599 | 0.1193 |
| F | | | | | | | | 0.187 | 0.1958 | 0.2056 | 0.2025 | 0.2016 |
| G | | | | | | | | 0.104 | 0.1176 | 0.117 | 0.1409 | 0.0952 |
| H | | | | | | | | 0.1557 | 0.1377 | 0.192 | 0.1659 | 0.1186 |

**Cell Index at: 14:00:25**

| | 1 | 2 | 3 | 4 | 5 | 6 | 7 | 8 | 9 | 10 | 11 | 12 |
|---|---|---|---|---|---|---|---|---|---|---|---|---|
| A | | | | | | | | 0.0729 | 0.0819 | 0.0927 | 0.0979 | 0.1193 |
| B | | | | | | | | 0.1187 | 0.1108 | 0.0843 | 0.14 | 0.1335 |
| C | | | | | | | | 0.1718 | 0.1759 | 0.2135 | 0.2243 | 0.1882 |
| D | | | | | | | | 0.1959 | 0.2106 | 0.2148 | 0.2082 | 0.1953 |
| E | | | | | | | | 0.1491 | 0.152 | 0.1702 | 0.1627 | 0.1194 |
| F | | | | | | | | 0.1804 | 0.1866 | 0.1987 | 0.1913 | 0.2002 |
| G | | | | | | | | 0.1016 | 0.115 | 0.1146 | 0.1425 | 0.0989 |
| H | | | | | | | | 0.1541 | 0.1357 | 0.1959 | 0.1609 | 0.1184 |

**Cell Index at: 14:15:26**

| | 1 | 2 | 3 | 4 | 5 | 6 | 7 | 8 | 9 | 10 | 11 | 12 |
|---|---|---|---|---|---|---|---|---|---|---|---|---|
| A | | | | | | | | 0.071 | 0.0835 | 0.0963 | 0.0949 | 0.1221 |
| B | | | | | | | | 0.119 | 0.1095 | 0.0866 | 0.1372 | 0.1259 |
| C | | | | | | | | 0.1794 | 0.18 | 0.2106 | 0.2209 | 0.1859 |
| D | | | | | | | | 0.1913 | 0.2103 | 0.2115 | 0.2056 | 0.2012 |
| E | | | | | | | | 0.1379 | 0.1419 | 0.169 | 0.1617 | 0.1169 |
| F | | | | | | | | 0.1886 | 0.1922 | 0.204 | 0.1986 | 0.1999 |
| G | | | | | | | | 0.1026 | 0.1191 | 0.1196 | 0.1436 | 0.0995 |
| H | | | | | | | | 0.1581 | 0.1416 | 0.1971 | 0.1639 | 0.1149 |

**Cell Index at: 14:30:26**

| | 1 | 2 | 3 | 4 | 5 | 6 | 7 | 8 | 9 | 10 | 11 | 12 |
|---|---|---|---|---|---|---|---|---|---|---|---|---|
| A | | | | | | | | 0.0701 | 0.086 | 0.0913 | 0.0932 | 0.1185 |
| B | | | | | | | | 0.1136 | 0.1096 | 0.0891 | 0.1394 | 0.1334 |
| C | | | | | | | | 0.1775 | 0.1712 | 0.216 | 0.2279 | 0.1859 |
| D | | | | | | | | 0.1925 | 0.2094 | 0.2153 | 0.2037 | 0.1991 |
| E | | | | | | | | 0.1351 | 0.1495 | 0.1667 | 0.1572 | 0.1227 |
| F | | | | | | | | 0.1842 | 0.1943 | 0.2099 | 0.1917 | 0.2033 |
| G | | | | | | | | 0.0995 | 0.1135 | 0.119 | 0.1403 | 0.0988 |
| H | | | | | | | | 0.1572 | 0.139 | 0.1954 | 0.1658 | 0.1061 |

**Cell Index at: 14:45:27**

| | 1 | 2 | 3 | 4 | 5 | 6 | 7 | 8 | 9 | 10 | 11 | 12 |
|---|---|---|---|---|---|---|---|---|---|---|---|---|
| A | | | | | | | | 0.0677 | 0.0814 | 0.0962 | 0.0817 | 0.1195 |
| B | | | | | | | | 0.1107 | 0.1057 | 0.086 | 0.1307 | 0.1313 |
| C | | | | | | | | 0.1894 | 0.184 | 0.2157 | 0.2171 | 0.1912 |
| D | | | | | | | | 0.1875 | 0.2015 | 0.214 | 0.2018 | 0.197 |
| E | | | | | | | | 0.1473 | 0.1544 | 0.1696 | 0.1574 | 0.116 |
| F | | | | | | | | 0.1843 | 0.1946 | 0.2163 | 0.2005 | 0.2106 |
| G | | | | | | | | 0.095 | 0.1139 | 0.1174 | 0.1418 | 0.0994 |
| H | | | | | | | | 0.1565 | 0.1422 | 0.1999 | 0.1637 | 0.107 |

**Cell Index at: 15:00:27**

| | 1 | 2 | 3 | 4 | 5 | 6 | 7 | 8 | 9 | 10 | 11 | 12 |
|---|---|---|---|---|---|---|---|---|---|---|---|---|
| A | | | | | | | | 0.0638 | 0.0836 | 0.0972 | 0.0817 | 0.1177 |
| B | | | | | | | | 0.1159 | 0.1009 | 0.0913 | 0.1242 | 0.1313 |
| C | | | | | | | | 0.1806 | 0.1788 | 0.2213 | 0.2218 | 0.1887 |
| D | | | | | | | | 0.1854 | 0.2154 | 0.2111 | 0.2031 | 0.2075 |
| E | | | | | | | | 0.1393 | 0.1544 | 0.1694 | 0.1561 | 0.1152 |
| F | | | | | | | | 0.1841 | 0.1947 | 0.2123 | 0.1976 | 0.2109 |
| G | | | | | | | | 0.1006 | 0.1147 | 0.119 | 0.1458 | 0.1044 |
| H | | | | | | | | 0.1571 | 0.1366 | 0.1964 | 0.1643 | 0.1085 |

**Cell Index at: 15:15:27**

| | 1 | 2 | 3 | 4 | 5 | 6 | 7 | 8 | 9 | 10 | 11 | 12 |
|---|---|---|---|---|---|---|---|---|---|---|---|---|
| A | | | | | | | | 0.0622 | 0.083 | 0.095 | 0.0814 | 0.1188 |
| B | | | | | | | | 0.1115 | 0.1 | 0.0904 | 0.1279 | 0.1302 |
| C | | | | | | | | 0.1929 | 0.1787 | 0.2183 | 0.22 | 0.1908 |
| D | | | | | | | | 0.1877 | 0.2085 | 0.2149 | 0.1986 | 0.2053 |
| E | | | | | | | | 0.1446 | 0.1574 | 0.1759 | 0.1549 | 0.1079 |
| F | | | | | | | | 0.1838 | 0.1886 | 0.2128 | 0.2059 | 0.205 |
| G | | | | | | | | 0.0949 | 0.1185 | 0.1178 | 0.1368 | 0.1073 |
| H | | | | | | | | 0.1569 | 0.1412 | 0.2006 | 0.1618 | 0.1099 |

**Cell Index at: 15:30:26**

| | 1 | 2 | 3 | 4 | 5 | 6 | 7 | 8 | 9 | 10 | 11 | 12 |
|---|---|---|---|---|---|---|---|---|---|---|---|---|
| A | | | | | | | | 0.0628 | 0.0864 | 0.0944 | 0.0803 | 0.12 |
| B | | | | | | | | 0.114 | 0.1021 | 0.0862 | 0.1249 | 0.1259 |
| C | | | | | | | | 0.1801 | 0.1741 | 0.2172 | 0.2202 | 0.188 |
| D | | | | | | | | 0.1981 | 0.2087 | 0.2245 | 0.2106 | 0.204 |
| E | | | | | | | | 0.1456 | 0.1559 | 0.1705 | 0.1582 | 0.1186 |
| F | | | | | | | | 0.189 | 0.1915 | 0.2157 | 0.2048 | 0.2106 |
| G | | | | | | | | 0.0973 | 0.1144 | 0.1227 | 0.1407 | 0.1015 |
| H | | | | | | | | 0.1611 | 0.1363 | 0.2007 | 0.1611 | 0.1081 |

**Cell Index at: 15:45:26**

| | 1 | 2 | 3 | 4 | 5 | 6 | 7 | 8 | 9 | 10 | 11 | 12 |
|---|---|---|---|---|---|---|---|---|---|---|---|---|
| A | | | | | | | | 0.0666 | 0.0885 | 0.0961 | 0.0805 | 0.1155 |
| B | | | | | | | | 0.1162 | 0.1045 | 0.0909 | 0.1253 | 0.1308 |
| C | | | | | | | | 0.1815 | 0.1767 | 0.2156 | 0.2221 | 0.1924 |
| D | | | | | | | | 0.1945 | 0.2039 | 0.22 | 0.1995 | 0.2026 |
| E | | | | | | | | 0.1312 | 0.1472 | 0.1669 | 0.1555 | 0.1143 |
| F | | | | | | | | 0.1889 | 0.1909 | 0.2112 | 0.2006 | 0.2063 |
| G | | | | | | | | 0.0953 | 0.1158 | 0.1198 | 0.1385 | 0.1012 |
| H | | | | | | | | 0.1633 | 0.1368 | 0.1995 | 0.1493 | 0.1113 |

**Cell Index at: 16:00:27**

| | 1 | 2 | 3 | 4 | 5 | 6 | 7 | 8 | 9 | 10 | 11 | 12 |
|---|---|---|---|---|---|---|---|---|---|---|---|---|
| A | | | | | | | | 0.0598 | 0.0839 | 0.0926 | 0.0793 | 0.1178 |
| B | | | | | | | | 0.1077 | 0.1001 | 0.0949 | 0.1228 | 0.1336 |
| C | | | | | | | | 0.1825 | 0.1827 | 0.222 | 0.2228 | 0.1904 |
| D | | | | | | | | 0.1889 | 0.2115 | 0.2223 | 0.206 | 0.2144 |
| E | | | | | | | | 0.1409 | 0.1424 | 0.1751 | 0.1538 | 0.1208 |
| F | | | | | | | | 0.1977 | 0.1905 | 0.2088 | 0.2042 | 0.2096 |
| G | | | | | | | | 0.0919 | 0.1152 | 0.1132 | 0.1412 | 0.0998 |
| H | | | | | | | | 0.1582 | 0.1405 | 0.2017 | 0.1528 | 0.1095 |

**Cell Index at: 16:15:28**

| | 1 | 2 | 3 | 4 | 5 | 6 | 7 | 8 | 9 | 10 | 11 | 12 |
|---|---|---|---|---|---|---|---|---|---|---|---|---|
| A | | | | | | | | 0.0626 | 0.082 | 0.096 | 0.0786 | 0.1206 |
| B | | | | | | | | 0.1055 | 0.1004 | 0.091 | 0.1193 | 0.1347 |
| C | | | | | | | | 0.1831 | 0.1861 | 0.2201 | 0.2265 | 0.1905 |

| | | | | | | | | | | | | |
|---|---|---|---|---|---|---|---|---|---|---|---|---|
| D | | | | | | | | 0.1868 | 0.2065 | 0.2123 | 0.2149 | 0.2093 |
| E | | | | | | | | 0.141 | 0.1481 | 0.1822 | 0.1486 | 0.1241 |
| F | | | | | | | | 0.1918 | 0.1949 | 0.2176 | 0.2025 | 0.2148 |
| G | | | | | | | | 0.0916 | 0.1174 | 0.1186 | 0.1247 | 0.0996 |
| H | | | | | | | | 0.1624 | 0.1395 | 0.1985 | 0.1499 | 0.1118 |

**Cell Index at: 16:30:29**

| | 1 | 2 | 3 | 4 | 5 | 6 | 7 | 8 | 9 | 10 | 11 | 12 |
|---|---|---|---|---|---|---|---|---|---|---|---|---|
| A | | | | | | | | 0.0607 | 0.082 | 0.0933 | 0.0776 | 0.1212 |
| B | | | | | | | | 0.1051 | 0.1028 | 0.0864 | 0.1191 | 0.1353 |
| C | | | | | | | | 0.1869 | 0.1797 | 0.2205 | 0.2254 | 0.1959 |
| D | | | | | | | | 0.1944 | 0.2079 | 0.2184 | 0.2084 | 0.213 |
| E | | | | | | | | 0.1369 | 0.1457 | 0.1689 | 0.1544 | 0.1097 |
| F | | | | | | | | 0.1886 | 0.1958 | 0.2205 | 0.2039 | 0.2066 |
| G | | | | | | | | 0.0911 | 0.1128 | 0.1187 | 0.1286 | 0.0979 |
| H | | | | | | | | 0.1587 | 0.1412 | 0.2024 | 0.1515 | 0.1114 |

**Cell Index at: 16:45:20**

| | 1 | 2 | 3 | 4 | 5 | 6 | 7 | 8 | 9 | 10 | 11 | 12 |
|---|---|---|---|---|---|---|---|---|---|---|---|---|
| A | | | | | | | | 0.0659 | 0.0771 | 0.0963 | 0.0752 | 0.1223 |
| B | | | | | | | | 0.1055 | 0.1035 | 0.0877 | 0.1194 | 0.1402 |
| C | | | | | | | | 0.19 | 0.183 | 0.2178 | 0.2276 | 0.1938 |
| D | | | | | | | | 0.1937 | 0.2007 | 0.2207 | 0.21 | 0.2113 |
| E | | | | | | | | 0.1427 | 0.1487 | 0.1699 | 0.1482 | 0.1185 |
| F | | | | | | | | 0.1941 | 0.1923 | 0.2146 | 0.203 | 0.2102 |
| G | | | | | | | | 0.0892 | 0.1177 | 0.1197 | 0.1251 | 0.1014 |
| H | | | | | | | | 0.1606 | 0.1427 | 0.202 | 0.1533 | 0.1118 |

**Cell Index at: 17:00:18**

| | 1 | 2 | 3 | 4 | 5 | 6 | 7 | 8 | 9 | 10 | 11 | 12 |
|---|---|---|---|---|---|---|---|---|---|---|---|---|
| A | | | | | | | | 0.064 | 0.0845 | 0.0961 | 0.0766 | 0.1186 |
| B | | | | | | | | 0.1018 | 0.105 | 0.0899 | 0.1174 | 0.1262 |
| C | | | | | | | | 0.1791 | 0.1752 | 0.22 | 0.2339 | 0.1968 |
| D | | | | | | | | 0.2047 | 0.2038 | 0.2209 | 0.2041 | 0.212 |
| E | | | | | | | | 0.1541 | 0.1472 | 0.1721 | 0.1511 | 0.1195 |
| F | | | | | | | | 0.2047 | 0.1954 | 0.217 | 0.2001 | 0.2094 |
| G | | | | | | | | 0.0917 | 0.1107 | 0.1162 | 0.1244 | 0.1051 |
| H | | | | | | | | 0.1609 | 0.1384 | 0.1983 | 0.1512 | 0.1129 |

**Cell Index at: 17:15:16**

| | 1 | 2 | 3 | 4 | 5 | 6 | 7 | 8 | 9 | 10 | 11 | 12 |
|---|---|---|---|---|---|---|---|---|---|---|---|---|
| A | | | | | | | | 0.0611 | 0.0791 | 0.0933 | 0.0731 | 0.1233 |
| B | | | | | | | | 0.1054 | 0.1023 | 0.0933 | 0.1223 | 0.1265 |
| C | | | | | | | | 0.1762 | 0.1821 | 0.2208 | 0.229 | 0.1893 |
| D | | | | | | | | 0.1905 | 0.2006 | 0.215 | 0.2132 | 0.2091 |
| E | | | | | | | | 0.1382 | 0.1418 | 0.1671 | 0.1489 | 0.1162 |
| F | | | | | | | | 0.1953 | 0.1889 | 0.2106 | 0.203 | 0.2149 |
| G | | | | | | | | 0.0903 | 0.1134 | 0.1105 | 0.1297 | 0.1048 |
| H | | | | | | | | 0.159 | 0.1401 | 0.2018 | 0.1501 | 0.1125 |

**Cell Index at: 17:30:16**

| | 1 | 2 | 3 | 4 | 5 | 6 | 7 | 8 | 9 | 10 | 11 | 12 |
|---|---|---|---|---|---|---|---|---|---|---|---|---|
| A | | | | | | | | 0.0628 | 0.079 | 0.0893 | 0.0723 | 0.1184 |
| B | | | | | | | | 0.103 | 0.1073 | 0.0874 | 0.1185 | 0.1287 |
| C | | | | | | | | 0.1875 | 0.1855 | 0.2238 | 0.229 | 0.1985 |
| D | | | | | | | | 0.1945 | 0.1981 | 0.2246 | 0.2099 | 0.2072 |
| E | | | | | | | | 0.1459 | 0.1429 | 0.1746 | 0.1449 | 0.1109 |
| F | | | | | | | | 0.1978 | 0.1922 | 0.2192 | 0.2061 | 0.2145 |
| G | | | | | | | | 0.0903 | 0.1078 | 0.1182 | 0.1289 | 0.1024 |
| H | | | | | | | | 0.159 | 0.1414 | 0.1975 | 0.1513 | 0.1153 |

**Cell Index at: 17:45:17**

| | 1 | 2 | 3 | 4 | 5 | 6 | 7 | 8 | 9 | 10 | 11 | 12 |
|---|---|---|---|---|---|---|---|---|---|---|---|---|
| A | | | | | | | | 0.0622 | 0.0748 | 0.0791 | 0.0756 | 0.1169 |
| B | | | | | | | | 0.1084 | 0.1036 | 0.0889 | 0.1199 | 0.1288 |
| C | | | | | | | | 0.1815 | 0.1824 | 0.2258 | 0.2304 | 0.1956 |
| D | | | | | | | | 0.1964 | 0.1976 | 0.218 | 0.2093 | 0.2079 |
| E | | | | | | | | 0.1469 | 0.1384 | 0.1696 | 0.1522 | 0.1141 |
| F | | | | | | | | 0.1894 | 0.1923 | 0.216 | 0.2076 | 0.2091 |
| G | | | | | | | | 0.087 | 0.1109 | 0.114 | 0.1302 | 0.0964 |
| H | | | | | | | | 0.1562 | 0.1376 | 0.2012 | 0.15 | 0.114 |

**Cell Index at: 18:00:17**

| | 1 | 2 | 3 | 4 | 5 | 6 | 7 | 8 | 9 | 10 | 11 | 12 |
|---|---|---|---|---|---|---|---|---|---|---|---|---|
| A | | | | | | | | 0.0614 | 0.0762 | 0.0807 | 0.0745 | 0.1152 |
| B | | | | | | | | 0.1054 | 0.1027 | 0.0897 | 0.1182 | 0.1266 |
| C | | | | | | | | 0.1756 | 0.1849 | 0.2108 | 0.2243 | 0.1897 |
| D | | | | | | | | 0.1886 | 0.2038 | 0.2164 | 0.2103 | 0.2148 |
| E | | | | | | | | 0.1468 | 0.1387 | 0.1771 | 0.1463 | 0.1118 |
| F | | | | | | | | 0.1909 | 0.1916 | 0.2066 | 0.2073 | 0.2145 |
| G | | | | | | | | 0.0853 | 0.1078 | 0.1179 | 0.1269 | 0.0992 |
| H | | | | | | | | 0.1632 | 0.1407 | 0.2014 | 0.1485 | 0.1167 |

**Cell Index at: 18:15:18**

| | 1 | 2 | 3 | 4 | 5 | 6 | 7 | 8 | 9 | 10 | 11 | 12 |
|---|---|---|---|---|---|---|---|---|---|---|---|---|
| A | | | | | | | | 0.0617 | 0.0794 | 0.0782 | 0.079 | 0.1161 |
| B | | | | | | | | 0.105 | 0.0934 | 0.0915 | 0.1245 | 0.1272 |
| C | | | | | | | | 0.1917 | 0.1856 | 0.2241 | 0.2298 | 0.1942 |
| D | | | | | | | | 0.1957 | 0.205 | 0.2186 | 0.2132 | 0.2098 |
| E | | | | | | | | 0.1406 | 0.1381 | 0.1774 | 0.1461 | 0.1142 |
| F | | | | | | | | 0.1939 | 0.1923 | 0.2125 | 0.2047 | 0.2135 |
| G | | | | | | | | 0.0895 | 0.1049 | 0.1142 | 0.1293 | 0.0983 |
| H | | | | | | | | 0.1594 | 0.1395 | 0.2 | 0.1497 | 0.1163 |

**Cell Index at: 18:30:19**

| | 1 | 2 | 3 | 4 | 5 | 6 | 7 | 8 | 9 | 10 | 11 | 12 |
|---|---|---|---|---|---|---|---|---|---|---|---|---|
| A | | | | | | | | 0.0677 | 0.0757 | 0.0835 | 0.0721 | 0.11 |
| B | | | | | | | | 0.1056 | 0.0948 | 0.094 | 0.12 | 0.1265 |
| C | | | | | | | | 0.1869 | 0.1802 | 0.2228 | 0.2232 | 0.1936 |
| D | | | | | | | | 0.1936 | 0.218 | 0.2191 | 0.2171 | 0.215 |
| E | | | | | | | | 0.1446 | 0.1387 | 0.1788 | 0.141 | 0.112 |
| F | | | | | | | | 0.19 | 0.2 | 0.2121 | 0.2027 | 0.2109 |
| G | | | | | | | | 0.0843 | 0.1057 | 0.1167 | 0.1233 | 0.0972 |
| H | | | | | | | | 0.1598 | 0.1406 | 0.1973 | 0.1483 | 0.1128 |

**Cell Index at: 18:45:20**

| | 1 | 2 | 3 | 4 | 5 | 6 | 7 | 8 | 9 | 10 | 11 | 12 |
|---|---|---|---|---|---|---|---|---|---|---|---|---|
| A | | | | | | | | 0.062 | 0.0724 | 0.0852 | 0.0718 | 0.1095 |
| B | | | | | | | | 0.1056 | 0.0917 | 0.0909 | 0.1198 | 0.1272 |
| C | | | | | | | | 0.1843 | 0.1839 | 0.2202 | 0.2282 | 0.1946 |
| D | | | | | | | | 0.1825 | 0.2054 | 0.2209 | 0.206 | 0.212 |
| E | | | | | | | | 0.1472 | 0.1363 | 0.185 | 0.135 | 0.1047 |
| F | | | | | | | | 0.199 | 0.1944 | 0.2154 | 0.2062 | 0.2102 |
| G | | | | | | | | 0.0922 | 0.1116 | 0.1122 | 0.1312 | 0.1005 |
| H | | | | | | | | 0.1597 | 0.1393 | 0.2002 | 0.1478 | 0.1153 |

**Cell Index at: 19:00:20**

| | 1 | 2 | 3 | 4 | 5 | 6 | 7 | 8 | 9 | 10 | 11 | 12 |
|---|---|---|---|---|---|---|---|---|---|---|---|---|
| A | | | | | | | | 0.0577 | 0.0747 | 0.0814 | 0.0709 | 0.1097 |
| B | | | | | | | | 0.0999 | 0.0932 | 0.0881 | 0.1203 | 0.1185 |
| C | | | | | | | | 0.1811 | 0.1808 | 0.2221 | 0.2243 | 0.1909 |
| D | | | | | | | | 0.181 | 0.2058 | 0.2258 | 0.2093 | 0.2151 |
| E | | | | | | | | 0.1367 | 0.1338 | 0.1748 | 0.1484 | 0.1104 |
| F | | | | | | | | 0.1967 | 0.1876 | 0.2158 | 0.2086 | 0.2174 |
| G | | | | | | | | 0.0863 | 0.1031 | 0.1134 | 0.1266 | 0.0938 |
| H | | | | | | | | 0.1586 | 0.1409 | 0.2006 | 0.1527 | 0.1145 |

**Cell Index at: 19:15:20**

| | 1 | 2 | 3 | 4 | 5 | 6 | 7 | 8 | 9 | 10 | 11 | 12 |
|---|---|---|---|---|---|---|---|---|---|---|---|---|
| A | | | | | | | | 0.0591 | 0.0713 | 0.0779 | 0.0705 | 0.1137 |
| B | | | | | | | | 0.102 | 0.0994 | 0.0885 | 0.1171 | 0.1244 |

| | | | | | | | | | | | | |
|---|---|---|---|---|---|---|---|---|---|---|---|---|
| C | | | | | | | | 0.1872 | 0.1914 | 0.2293 | 0.2299 | 0.1877 |
| D | | | | | | | | 0.1795 | 0.2044 | 0.2219 | 0.2137 | 0.2197 |
| E | | | | | | | | 0.143 | 0.1405 | 0.1703 | 0.1481 | 0.1086 |
| F | | | | | | | | 0.1969 | 0.1924 | 0.2158 | 0.2071 | 0.2195 |
| G | | | | | | | | 0.0902 | 0.1048 | 0.1156 | 0.1267 | 0.0962 |
| H | | | | | | | | 0.1608 | 0.1413 | 0.2009 | 0.1499 | 0.1165 |

**Cell Index at: 19:30:20**

| | 1 | 2 | 3 | 4 | 5 | 6 | 7 | 8 | 9 | 10 | 11 | 12 |
|---|---|---|---|---|---|---|---|---|---|---|---|---|
| A | | | | | | | | 0.0561 | 0.0709 | 0.0806 | 0.0646 | 0.1134 |
| B | | | | | | | | 0.1003 | 0.0952 | 0.0873 | 0.1183 | 0.1264 |
| C | | | | | | | | 0.1896 | 0.1873 | 0.2254 | 0.2282 | 0.1958 |
| D | | | | | | | | 0.1956 | 0.2041 | 0.2139 | 0.203 | 0.2175 |
| E | | | | | | | | 0.1482 | 0.1258 | 0.168 | 0.1473 | 0.1169 |
| F | | | | | | | | 0.1921 | 0.1899 | 0.2147 | 0.2045 | 0.2164 |
| G | | | | | | | | 0.086 | 0.1008 | 0.1162 | 0.1199 | 0.095 |
| H | | | | | | | | 0.1642 | 0.1397 | 0.1983 | 0.1491 | 0.1169 |

**Cell Index at: 19:45:20**

| | 1 | 2 | 3 | 4 | 5 | 6 | 7 | 8 | 9 | 10 | 11 | 12 |
|---|---|---|---|---|---|---|---|---|---|---|---|---|
| A | | | | | | | | 0.0555 | 0.0752 | 0.0784 | 0.0629 | 0.1119 |
| B | | | | | | | | 0.0985 | 0.0927 | 0.09 | 0.113 | 0.1287 |
| C | | | | | | | | 0.1863 | 0.1903 | 0.2179 | 0.2322 | 0.1926 |
| D | | | | | | | | 0.1811 | 0.2091 | 0.2133 | 0.2067 | 0.2148 |
| E | | | | | | | | 0.1475 | 0.1296 | 0.1722 | 0.1395 | 0.1103 |
| F | | | | | | | | 0.1996 | 0.1936 | 0.2122 | 0.211 | 0.2163 |
| G | | | | | | | | 0.0883 | 0.1022 | 0.1113 | 0.1239 | 0.0908 |
| H | | | | | | | | 0.163 | 0.1423 | 0.1974 | 0.1494 | 0.1185 |

**Cell Index at: 20:00:21**

| | 1 | 2 | 3 | 4 | 5 | 6 | 7 | 8 | 9 | 10 | 11 | 12 |
|---|---|---|---|---|---|---|---|---|---|---|---|---|
| A | | | | | | | | 0.0555 | 0.0707 | 0.0865 | 0.0655 | 0.113 |
| B | | | | | | | | 0.1027 | 0.0884 | 0.0918 | 0.1176 | 0.1257 |
| C | | | | | | | | 0.1848 | 0.1879 | 0.2211 | 0.2337 | 0.1997 |
| D | | | | | | | | 0.191 | 0.2102 | 0.214 | 0.2103 | 0.2129 |
| E | | | | | | | | 0.1444 | 0.1291 | 0.1721 | 0.1439 | 0.1129 |
| F | | | | | | | | 0.1943 | 0.1908 | 0.2189 | 0.2077 | 0.2137 |
| G | | | | | | | | 0.0874 | 0.0984 | 0.1147 | 0.1256 | 0.096 |
| H | | | | | | | | 0.1656 | 0.1399 | 0.1994 | 0.1524 | 0.1186 |

**Cell Index at: 20:15:21**

| | 1 | 2 | 3 | 4 | 5 | 6 | 7 | 8 | 9 | 10 | 11 | 12 |
|---|---|---|---|---|---|---|---|---|---|---|---|---|
| A | | | | | | | | 0.0547 | 0.0733 | 0.0871 | 0.0668 | 0.1165 |
| B | | | | | | | | 0.1008 | 0.0945 | 0.0901 | 0.1151 | 0.1243 |
| C | | | | | | | | 0.1989 | 0.1871 | 0.2238 | 0.2319 | 0.1931 |
| D | | | | | | | | 0.1782 | 0.2109 | 0.2113 | 0.2086 | 0.2143 |
| E | | | | | | | | 0.1446 | 0.1358 | 0.1692 | 0.1421 | 0.116 |
| F | | | | | | | | 0.1983 | 0.1969 | 0.2178 | 0.1891 | 0.2162 |
| G | | | | | | | | 0.0869 | 0.0976 | 0.1125 | 0.1225 | 0.0949 |
| H | | | | | | | | 0.1624 | 0.1454 | 0.192 | 0.1515 | 0.1179 |

**Cell Index at: 20:30:22**

| | 1 | 2 | 3 | 4 | 5 | 6 | 7 | 8 | 9 | 10 | 11 | 12 |
|---|---|---|---|---|---|---|---|---|---|---|---|---|
| A | | | | | | | | 0.0544 | 0.0698 | 0.0809 | 0.0656 | 0.1103 |
| B | | | | | | | | 0.0956 | 0.0915 | 0.0891 | 0.115 | 0.1292 |
| C | | | | | | | | 0.1791 | 0.1878 | 0.2205 | 0.2329 | 0.1953 |
| D | | | | | | | | 0.1861 | 0.2097 | 0.2138 | 0.2058 | 0.2107 |
| E | | | | | | | | 0.1495 | 0.1284 | 0.176 | 0.136 | 0.1066 |
| F | | | | | | | | 0.1951 | 0.1967 | 0.2229 | 0.1927 | 0.2138 |
| G | | | | | | | | 0.0857 | 0.0959 | 0.0988 | 0.1262 | 0.0985 |
| H | | | | | | | | 0.167 | 0.141 | 0.1895 | 0.1547 | 0.1155 |

**Cell Index at: 20:45:22**

| | 1 | 2 | 3 | 4 | 5 | 6 | 7 | 8 | 9 | 10 | 11 | 12 |
|---|---|---|---|---|---|---|---|---|---|---|---|---|
| A | | | | | | | | 0.0525 | 0.069 | 0.0857 | 0.0684 | 0.11 |
| B | | | | | | | | 0.1042 | 0.0855 | 0.0857 | 0.1147 | 0.1271 |
| C | | | | | | | | 0.1773 | 0.1901 | 0.2273 | 0.2364 | 0.1983 |
| D | | | | | | | | 0.1853 | 0.214 | 0.2103 | 0.2136 | 0.2105 |
| E | | | | | | | | 0.1419 | 0.1291 | 0.1735 | 0.1476 | 0.1096 |
| F | | | | | | | | 0.1933 | 0.1957 | 0.2192 | 0.1921 | 0.2186 |
| G | | | | | | | | 0.0863 | 0.1027 | 0.0983 | 0.1226 | 0.0919 |
| H | | | | | | | | 0.1677 | 0.1407 | 0.1903 | 0.1505 | 0.1165 |

**Cell Index at: 21:00:22**

| | 1 | 2 | 3 | 4 | 5 | 6 | 7 | 8 | 9 | 10 | 11 | 12 |
|---|---|---|---|---|---|---|---|---|---|---|---|---|
| A | | | | | | | | 0.0504 | 0.0695 | 0.082 | 0.0652 | 0.1097 |
| B | | | | | | | | 0.0967 | 0.0915 | 0.0808 | 0.1088 | 0.1286 |
| C | | | | | | | | 0.1857 | 0.1891 | 0.228 | 0.2369 | 0.1975 |
| D | | | | | | | | 0.1868 | 0.2114 | 0.22 | 0.2091 | 0.2188 |
| E | | | | | | | | 0.1419 | 0.1358 | 0.1614 | 0.1507 | 0.1111 |
| F | | | | | | | | 0.1984 | 0.196 | 0.221 | 0.1957 | 0.2156 |
| G | | | | | | | | 0.0917 | 0.1017 | 0.1008 | 0.1251 | 0.0972 |
| H | | | | | | | | 0.1679 | 0.1469 | 0.1925 | 0.1542 | 0.1136 |

**Cell Index at: 21:15:23**

| | 1 | 2 | 3 | 4 | 5 | 6 | 7 | 8 | 9 | 10 | 11 | 12 |
|---|---|---|---|---|---|---|---|---|---|---|---|---|
| A | | | | | | | | 0.0541 | 0.0649 | 0.0798 | 0.0662 | 0.1095 |
| B | | | | | | | | 0.0971 | 0.0893 | 0.0813 | 0.1089 | 0.1218 |
| C | | | | | | | | 0.1855 | 0.1858 | 0.2228 | 0.2411 | 0.1938 |
| D | | | | | | | | 0.1776 | 0.2077 | 0.2113 | 0.212 | 0.2167 |
| E | | | | | | | | 0.1406 | 0.1249 | 0.1678 | 0.1374 | 0.1152 |
| F | | | | | | | | 0.2007 | 0.1944 | 0.2209 | 0.1951 | 0.2204 |
| G | | | | | | | | 0.0887 | 0.1017 | 0.1005 | 0.1248 | 0.0918 |
| H | | | | | | | | 0.1682 | 0.145 | 0.1929 | 0.1532 | 0.1152 |

**Cell Index at: 21:30:23**

| | 1 | 2 | 3 | 4 | 5 | 6 | 7 | 8 | 9 | 10 | 11 | 12 |
|---|---|---|---|---|---|---|---|---|---|---|---|---|
| A | | | | | | | | 0.0498 | 0.0628 | 0.0707 | 0.0675 | 0.1046 |
| B | | | | | | | | 0.0929 | 0.0894 | 0.0829 | 0.1032 | 0.1216 |
| C | | | | | | | | 0.1921 | 0.1822 | 0.2304 | 0.2313 | 0.1925 |
| D | | | | | | | | 0.1773 | 0.216 | 0.2116 | 0.2095 | 0.2106 |
| E | | | | | | | | 0.1362 | 0.1397 | 0.1663 | 0.1408 | 0.1124 |
| F | | | | | | | | 0.2004 | 0.1933 | 0.2234 | 0.1971 | 0.217 |
| G | | | | | | | | 0.0877 | 0.0998 | 0.0981 | 0.1247 | 0.089 |
| H | | | | | | | | 0.1651 | 0.1457 | 0.1974 | 0.1538 | 0.1152 |

**Cell Index at: 21:45:24**

| | 1 | 2 | 3 | 4 | 5 | 6 | 7 | 8 | 9 | 10 | 11 | 12 |
|---|---|---|---|---|---|---|---|---|---|---|---|---|
| A | | | | | | | | 0.05 | 0.0643 | 0.073 | 0.0689 | 0.1043 |
| B | | | | | | | | 0.0963 | 0.0918 | 0.087 | 0.109 | 0.1193 |
| C | | | | | | | | 0.188 | 0.1858 | 0.2296 | 0.233 | 0.1959 |
| D | | | | | | | | 0.1869 | 0.2111 | 0.211 | 0.2114 | 0.2149 |
| E | | | | | | | | 0.1439 | 0.1313 | 0.1661 | 0.1484 | 0.1027 |
| F | | | | | | | | 0.197 | 0.192 | 0.2177 | 0.1888 | 0.2198 |
| G | | | | | | | | 0.0854 | 0.1008 | 0.0989 | 0.1203 | 0.0923 |
| H | | | | | | | | 0.169 | 0.1443 | 0.1995 | 0.1562 | 0.1127 |

**Cell Index at: 22:00:24**

| | 1 | 2 | 3 | 4 | 5 | 6 | 7 | 8 | 9 | 10 | 11 | 12 |
|---|---|---|---|---|---|---|---|---|---|---|---|---|
| A | | | | | | | | 0.0471 | 0.0652 | 0.0709 | 0.0637 | 0.1061 |
| B | | | | | | | | 0.0912 | 0.0861 | 0.0796 | 0.1081 | 0.1235 |
| C | | | | | | | | 0.1836 | 0.1853 | 0.2312 | 0.2318 | 0.1975 |
| D | | | | | | | | 0.184 | 0.2142 | 0.2113 | 0.2101 | 0.2166 |
| E | | | | | | | | 0.1395 | 0.1313 | 0.1653 | 0.1421 | 0.1134 |
| F | | | | | | | | 0.1981 | 0.1951 | 0.2172 | 0.1963 | 0.2224 |
| G | | | | | | | | 0.0879 | 0.1005 | 0.0965 | 0.1211 | 0.0883 |
| H | | | | | | | | 0.1669 | 0.1426 | 0.1934 | 0.1554 | 0.1167 |

**Cell Index at: 22:15:25**

| | 1 | 2 | 3 | 4 | 5 | 6 | 7 | 8 | 9 | 10 | 11 | 12 |
|---|---|---|---|---|---|---|---|---|---|---|---|---|
| A | | | | | | | | 0.0481 | 0.0628 | 0.0723 | 0.0629 | 0.1021 |

| | | | | | | | | | | | | |
|---|---|---|---|---|---|---|---|---|---|---|---|---|
| B | | | | | | | | 0.0935 | 0.085 | 0.0802 | 0.1106 | 0.1251 |
| C | | | | | | | | 0.1899 | 0.1844 | 0.2321 | 0.2361 | 0.1944 |
| D | | | | | | | | 0.1898 | 0.2095 | 0.218 | 0.2098 | 0.2175 |
| E | | | | | | | | 0.1399 | 0.1365 | 0.1621 | 0.1369 | 0.1156 |
| F | | | | | | | | 0.2042 | 0.1893 | 0.2145 | 0.1951 | 0.2218 |
| G | | | | | | | | 0.0878 | 0.0968 | 0.0968 | 0.1209 | 0.0866 |
| H | | | | | | | | 0.1663 | 0.139 | 0.198 | 0.1529 | 0.1155 |

**Cell Index at: 22:30:26**

| | 1 | 2 | 3 | 4 | 5 | 6 | 7 | 8 | 9 | 10 | 11 | 12 |
|---|---|---|---|---|---|---|---|---|---|---|---|---|
| A | | | | | | | | 0.0467 | 0.0618 | 0.0734 | 0.0578 | 0.1042 |
| B | | | | | | | | 0.0938 | 0.0837 | 0.0768 | 0.1001 | 0.1168 |
| C | | | | | | | | 0.1905 | 0.1829 | 0.2305 | 0.2418 | 0.1881 |
| D | | | | | | | | 0.1838 | 0.2252 | 0.2161 | 0.2145 | 0.2134 |
| E | | | | | | | | 0.1293 | 0.1334 | 0.1729 | 0.1405 | 0.1166 |
| F | | | | | | | | 0.2006 | 0.1947 | 0.2168 | 0.195 | 0.2162 |
| G | | | | | | | | 0.0832 | 0.0967 | 0.0957 | 0.12 | 0.0843 |
| H | | | | | | | | 0.1646 | 0.1379 | 0.1933 | 0.1505 | 0.1137 |

**Cell Index at: 22:45:27**

| | 1 | 2 | 3 | 4 | 5 | 6 | 7 | 8 | 9 | 10 | 11 | 12 |
|---|---|---|---|---|---|---|---|---|---|---|---|---|
| A | | | | | | | | 0.0451 | 0.0606 | 0.0748 | 0.0583 | 0.1011 |
| B | | | | | | | | 0.0962 | 0.0785 | 0.0805 | 0.1043 | 0.1188 |
| C | | | | | | | | 0.1874 | 0.1861 | 0.2277 | 0.2429 | 0.1977 |
| D | | | | | | | | 0.1854 | 0.2159 | 0.2163 | 0.2098 | 0.212 |
| E | | | | | | | | 0.1253 | 0.128 | 0.1658 | 0.1325 | 0.1121 |
| F | | | | | | | | 0.1998 | 0.1938 | 0.2148 | 0.1941 | 0.2175 |
| G | | | | | | | | 0.0835 | 0.0976 | 0.0952 | 0.1187 | 0.0846 |
| H | | | | | | | | 0.1705 | 0.1396 | 0.1938 | 0.1555 | 0.1164 |

**Cell Index at: 23:00:27**

| | 1 | 2 | 3 | 4 | 5 | 6 | 7 | 8 | 9 | 10 | 11 | 12 |
|---|---|---|---|---|---|---|---|---|---|---|---|---|
| A | | | | | | | | 0.0486 | 0.0595 | 0.0737 | 0.0592 | 0.1023 |
| B | | | | | | | | 0.0941 | 0.0847 | 0.0793 | 0.1061 | 0.1154 |
| C | | | | | | | | 0.1894 | 0.1875 | 0.2256 | 0.234 | 0.1967 |
| D | | | | | | | | 0.1887 | 0.2198 | 0.2158 | 0.2045 | 0.2109 |
| E | | | | | | | | 0.1331 | 0.1281 | 0.1664 | 0.1265 | 0.1096 |
| F | | | | | | | | 0.1995 | 0.202 | 0.2218 | 0.1993 | 0.2118 |
| G | | | | | | | | 0.084 | 0.0973 | 0.0951 | 0.1151 | 0.086 |
| H | | | | | | | | 0.1662 | 0.138 | 0.1943 | 0.1535 | 0.1138 |

**Cell Index at: 23:15:27**

| | 1 | 2 | 3 | 4 | 5 | 6 | 7 | 8 | 9 | 10 | 11 | 12 |
|---|---|---|---|---|---|---|---|---|---|---|---|---|
| A | | | | | | | | 0.0471 | 0.058 | 0.0725 | 0.0588 | 0.1004 |
| B | | | | | | | | 0.0888 | 0.086 | 0.083 | 0.106 | 0.1197 |
| C | | | | | | | | 0.1937 | 0.1866 | 0.2314 | 0.2436 | 0.1973 |
| D | | | | | | | | 0.182 | 0.2063 | 0.2138 | 0.2088 | 0.2112 |
| E | | | | | | | | 0.146 | 0.1304 | 0.166 | 0.1329 | 0.1088 |
| F | | | | | | | | 0.1993 | 0.1966 | 0.2198 | 0.1934 | 0.2142 |
| G | | | | | | | | 0.0849 | 0.0968 | 0.097 | 0.1162 | 0.0856 |
| H | | | | | | | | 0.168 | 0.1291 | 0.1927 | 0.1492 | 0.1103 |

**Cell Index at: 23:30:28**

| | 1 | 2 | 3 | 4 | 5 | 6 | 7 | 8 | 9 | 10 | 11 | 12 |
|---|---|---|---|---|---|---|---|---|---|---|---|---|
| A | | | | | | | | 0.0468 | 0.0527 | 0.0747 | 0.058 | 0.0988 |
| B | | | | | | | | 0.0945 | 0.0836 | 0.0767 | 0.1082 | 0.1164 |
| C | | | | | | | | 0.1893 | 0.1877 | 0.2297 | 0.2452 | 0.1943 |
| D | | | | | | | | 0.1834 | 0.2146 | 0.2103 | 0.2191 | 0.2158 |
| E | | | | | | | | 0.1378 | 0.1311 | 0.1592 | 0.1386 | 0.1061 |
| F | | | | | | | | 0.2014 | 0.192 | 0.2197 | 0.1894 | 0.218 |
| G | | | | | | | | 0.0828 | 0.0944 | 0.0939 | 0.1136 | 0.0837 |
| H | | | | | | | | 0.1642 | 0.1345 | 0.1911 | 0.151 | 0.1141 |

**Cell Index at: 23:45:28**

| | 1 | 2 | 3 | 4 | 5 | 6 | 7 | 8 | 9 | 10 | 11 | 12 |
|---|---|---|---|---|---|---|---|---|---|---|---|---|
| A | | | | | | | | 0.0482 | 0.0542 | 0.0719 | 0.0567 | 0.0923 |
| B | | | | | | | | 0.0885 | 0.0857 | 0.0787 | 0.1027 | 0.1193 |
| C | | | | | | | | 0.1935 | 0.1864 | 0.2355 | 0.2428 | 0.1931 |
| D | | | | | | | | 0.1849 | 0.2104 | 0.2171 | 0.214 | 0.2101 |
| E | | | | | | | | 0.1318 | 0.1189 | 0.1629 | 0.137 | 0.1042 |
| F | | | | | | | | 0.2097 | 0.1954 | 0.2212 | 0.1946 | 0.2149 |
| G | | | | | | | | 0.0791 | 0.0927 | 0.0924 | 0.1156 | 0.0818 |
| H | | | | | | | | 0.1667 | 0.1339 | 0.1942 | 0.1513 | 0.1137 |

**Cell Index at: 24:00:29**

| | 1 | 2 | 3 | 4 | 5 | 6 | 7 | 8 | 9 | 10 | 11 | 12 |
|---|---|---|---|---|---|---|---|---|---|---|---|---|
| A | | | | | | | | 0.0464 | 0.0515 | 0.0743 | 0.0528 | 0.0953 |
| B | | | | | | | | 0.0942 | 0.0813 | 0.0787 | 0.1042 | 0.1139 |
| C | | | | | | | | 0.1942 | 0.1857 | 0.2289 | 0.2477 | 0.1976 |
| D | | | | | | | | 0.1815 | 0.2159 | 0.2095 | 0.2135 | 0.2046 |
| E | | | | | | | | 0.1361 | 0.1239 | 0.1637 | 0.1388 | 0.1131 |
| F | | | | | | | | 0.1986 | 0.1955 | 0.216 | 0.1933 | 0.2128 |
| G | | | | | | | | 0.0756 | 0.0918 | 0.0966 | 0.1138 | 0.0788 |
| H | | | | | | | | 0.1687 | 0.1352 | 0.1916 | 0.1512 | 0.1146 |

| | | | | | | | | | | | | | |
|---|---|---|---|---|---|---|---|---|---|---|---|---|---|
| A vs B | 0.030214 | B vs C | 0.000101 | C vs D | 0.676655 | D vs E | 0.00024 | E vs F | 0.000355 | F vs G | 2.68E-06 | G vs H | 0.006549 |
| A vs C | 1.43E-05 | B vs D | 2.07E-06 | C vs E | 0.001088 | D vs F | 0.826246 | E vs G | 0.004231 | F vs H | 0.01535 | | |
| A vs D | 4.13E-06 | B vs E | 0.006095 | C vs F | 0.574245 | D vs G | 1.91E-06 | E vs H | 0.313948 | | | | |
| A vs E | 0.000469 | B vs F | 2.61E-06 | C vs G | 8.22E-05 | D vs H | 0.012582 | | | | | | |
| A vs F | 1.08E-05 | B vs G | 0.751259 | C vs H | 0.011078 | | | | | | | | |
| A vs G | 0.046671 | B vs H | 0.008362 | | | | | | | | | | |
| A vs H | 0.000894 | | | | | | | | | | | | |

**Cell Index at: 24:15:28**

| | 1 | 2 | 3 | 4 | 5 | 6 | 7 | 8 | 9 | 10 | 11 | 12 |
|---|---|---|---|---|---|---|---|---|---|---|---|---|
| A | | | | | | | | 0.0449 | 0.0542 | 0.0723 | 0.0556 | 0.0925 |
| B | | | | | | | | 0.0887 | 0.0831 | 0.0814 | 0.1071 | 0.1099 |
| C | | | | | | | | 0.1853 | 0.184 | 0.2309 | 0.2494 | 0.1963 |
| D | | | | | | | | 0.1858 | 0.208 | 0.2201 | 0.2136 | 0.213 |
| E | | | | | | | | 0.1344 | 0.1201 | 0.1527 | 0.1326 | 0.1058 |
| F | | | | | | | | 0.1946 | 0.1987 | 0.2212 | 0.1933 | 0.2163 |
| G | | | | | | | | 0.0783 | 0.0922 | 0.0908 | 0.1138 | 0.0849 |
| H | | | | | | | | 0.1634 | 0.1349 | 0.1908 | 0.1524 | 0.1145 |

| *Count* | *Sum* | *Average* | *Variance* |
|---|---|---|---|
| 5 | 0.3203 | 0.06406 | 0.000419 |
| 5 | 0.4723 | 0.09446 | 0.000224 |
| 5 | 1.0541 | 0.21082 | 0.000692 |
| 5 | 1.025 | 0.205 | 0.000191 |
| 5 | 0.6756 | 0.13512 | 0.000361 |
| 5 | 1.0162 | 0.20324 | 0.000109 |
| 5 | 0.4566 | 0.09132 | 0.000234 |
| 5 | 0.7613 | 0.15226 | 0.000882 |

**Cell Index at: 24:30:29**

| | 1 | 2 | 3 | 4 | 5 | 6 | 7 | 8 | 9 | 10 | 11 | 12 |
|---|---|---|---|---|---|---|---|---|---|---|---|---|
| A | | | | | | | | 0.0473 | 0.0524 | 0.0736 | 0.0567 | 0.0917 |
| B | | | | | | | | 0.0873 | 0.0798 | 0.0803 | 0.1082 | 0.1085 |
| C | | | | | | | | 0.1891 | 0.1923 | 0.225 | 0.2449 | 0.1957 |
| D | | | | | | | | 0.1783 | 0.2138 | 0.2195 | 0.2161 | 0.2161 |
| E | | | | | | | | 0.1288 | 0.1274 | 0.1534 | 0.1391 | 0.1065 |
| F | | | | | | | | 0.1968 | 0.1931 | 0.2157 | 0.1955 | 0.2141 |
| G | | | | | | | | 0.0751 | 0.093 | 0.0904 | 0.1114 | 0.077 |
| H | | | | | | | | 0.1633 | 0.1363 | 0.1923 | 0.1532 | 0.1152 |

| *SS* | *df* | *MS* | *F* | *P-value* | *F crit* |
|---|---|---|---|---|---|
| 0.117299 | 7 | 0.016757 | 43.08214 | 1.62E-14 | 2.312741 |
| 0.012447 | 32 | 0.000389 | | | |
| 0.129746 | 39 | | | | |

**Cell Index at: 24:45:30**

| | 1 | 2 | 3 | 4 | 5 | 6 | 7 | 8 | 9 | 10 | 11 | 12 |
|---|---|---|---|---|---|---|---|---|---|---|---|---|
| A | | | | | | | | 0.0448 | 0.0543 | 0.0736 | 0.0552 | 0.0896 |
| B | | | | | | | | 0.0839 | 0.0767 | 0.0827 | 0.1064 | 0.1059 |
| C | | | | | | | | 0.1896 | 0.1918 | 0.2316 | 0.2402 | 0.1942 |
| D | | | | | | | | 0.1774 | 0.2144 | 0.2242 | 0.2091 | 0.2176 |
| E | | | | | | | | 0.1349 | 0.1231 | 0.1471 | 0.1331 | 0.1108 |
| F | | | | | | | | 0.2067 | 0.188 | 0.219 | 0.1897 | 0.2174 |
| G | | | | | | | | 0.0701 | 0.0948 | 0.0936 | 0.1129 | 0.0775 |
| H | | | | | | | | 0.1642 | 0.1277 | 0.1901 | 0.151 | 0.1123 |

**Cell Index at: 25:00:30**

| | 1 | 2 | 3 | 4 | 5 | 6 | 7 | 8 | 9 | 10 | 11 | 12 |
|---|---|---|---|---|---|---|---|---|---|---|---|---|
| A | | | | | | | | 0.0412 | 0.0535 | 0.0697 | 0.0538 | 0.0921 |
| B | | | | | | | | 0.0889 | 0.0775 | 0.0821 | 0.1017 | 0.1141 |
| C | | | | | | | | 0.1853 | 0.1905 | 0.2312 | 0.2392 | 0.1945 |
| D | | | | | | | | 0.1765 | 0.2156 | 0.2168 | 0.2137 | 0.2156 |
| E | | | | | | | | 0.1377 | 0.1274 | 0.1442 | 0.1277 | 0.1087 |
| F | | | | | | | | 0.1999 | 0.1879 | 0.2166 | 0.1962 | 0.2087 |
| G | | | | | | | | 0.0737 | 0.0933 | 0.0918 | 0.1135 | 0.0777 |
| H | | | | | | | | 0.1653 | 0.123 | 0.1925 | 0.1562 | 0.1124 |

**Cell Index at: 25:15:31**

| | 1 | 2 | 3 | 4 | 5 | 6 | 7 | 8 | 9 | 10 | 11 | 12 |
|---|---|---|---|---|---|---|---|---|---|---|---|---|
| A | | | | | | | | 0.0439 | 0.0565 | 0.0678 | 0.0515 | 0.0889 |
| B | | | | | | | | 0.0876 | 0.0774 | 0.0779 | 0.1004 | 0.1081 |
| C | | | | | | | | 0.1887 | 0.1892 | 0.2309 | 0.2464 | 0.1989 |
| D | | | | | | | | 0.1855 | 0.2097 | 0.2192 | 0.2096 | 0.2146 |
| E | | | | | | | | 0.1327 | 0.1338 | 0.1435 | 0.1353 | 0.1068 |
| F | | | | | | | | 0.2083 | 0.1924 | 0.2213 | 0.1955 | 0.2146 |
| G | | | | | | | | 0.07 | 0.0941 | 0.0867 | 0.1007 | 0.0714 |
| H | | | | | | | | 0.1653 | 0.1228 | 0.1903 | 0.1553 | 0.1088 |

**Cell Index at: 25:30:31**

| | 1 | 2 | 3 | 4 | 5 | 6 | 7 | 8 | 9 | 10 | 11 | 12 |
|---|---|---|---|---|---|---|---|---|---|---|---|---|
| A | | | | | | | | 0.0414 | 0.0568 | 0.0673 | 0.0556 | 0.0899 |
| B | | | | | | | | 0.0881 | 0.0771 | 0.0841 | 0.101 | 0.1085 |
| C | | | | | | | | 0.1867 | 0.1841 | 0.2388 | 0.2439 | 0.1944 |
| D | | | | | | | | 0.1808 | 0.2164 | 0.2142 | 0.2184 | 0.216 |
| E | | | | | | | | 0.1376 | 0.1276 | 0.1474 | 0.1339 | 0.1086 |
| F | | | | | | | | 0.1985 | 0.1907 | 0.2188 | 0.1952 | 0.2117 |
| G | | | | | | | | 0.0688 | 0.0882 | 0.0886 | 0.1029 | 0.0782 |
| H | | | | | | | | 0.164 | 0.1259 | 0.1944 | 0.1544 | 0.1073 |

**Cell Index at: 25:45:32**

| | 1 | 2 | 3 | 4 | 5 | 6 | 7 | 8 | 9 | 10 | 11 | 12 |
|---|---|---|---|---|---|---|---|---|---|---|---|---|
| A | | | | | | | | 0.0434 | 0.0562 | 0.0665 | 0.0537 | 0.085 |
| B | | | | | | | | 0.0905 | 0.0765 | 0.0864 | 0.0961 | 0.1082 |
| C | | | | | | | | 0.1881 | 0.1881 | 0.2352 | 0.2386 | 0.1916 |
| D | | | | | | | | 0.1813 | 0.2163 | 0.222 | 0.2143 | 0.2114 |
| E | | | | | | | | 0.132 | 0.1325 | 0.1453 | 0.1325 | 0.1032 |
| F | | | | | | | | 0.1945 | 0.1982 | 0.2208 | 0.1977 | 0.2152 |
| G | | | | | | | | 0.0705 | 0.094 | 0.0878 | 0.0959 | 0.0731 |
| H | | | | | | | | 0.1625 | 0.1218 | 0.1893 | 0.1553 | 0.1077 |

**Cell Index at: 26:00:31**

| | 1 | 2 | 3 | 4 | 5 | 6 | 7 | 8 | 9 | 10 | 11 | 12 |
|---|---|---|---|---|---|---|---|---|---|---|---|---|
| A | | | | | | | | 0.0399 | 0.0595 | 0.0687 | 0.0528 | 0.0845 |
| B | | | | | | | | 0.0898 | 0.0806 | 0.0808 | 0.1021 | 0.1114 |
| C | | | | | | | | 0.1895 | 0.1868 | 0.2317 | 0.2442 | 0.1944 |
| D | | | | | | | | 0.191 | 0.2149 | 0.2118 | 0.2152 | 0.2195 |
| E | | | | | | | | 0.1312 | 0.1277 | 0.1453 | 0.1332 | 0.1037 |
| F | | | | | | | | 0.2035 | 0.1961 | 0.2242 | 0.1962 | 0.2109 |
| G | | | | | | | | 0.0713 | 0.0938 | 0.0908 | 0.0976 | 0.0743 |
| H | | | | | | | | 0.1648 | 0.1228 | 0.1951 | 0.149 | 0.1061 |

**Cell Index at: 26:15:32**

| | 1 | 2 | 3 | 4 | 5 | 6 | 7 | 8 | 9 | 10 | 11 | 12 |
|---|---|---|---|---|---|---|---|---|---|---|---|---|
| A | | | | | | | | 0.0377 | 0.0579 | 0.0695 | 0.0531 | 0.0838 |
| B | | | | | | | | 0.0881 | 0.0795 | 0.0791 | 0.0971 | 0.1101 |
| C | | | | | | | | 0.1933 | 0.1884 | 0.2295 | 0.2397 | 0.1974 |
| D | | | | | | | | 0.177 | 0.2169 | 0.2168 | 0.2169 | 0.2213 |
| E | | | | | | | | 0.1311 | 0.1349 | 0.1431 | 0.1269 | 0.1107 |
| F | | | | | | | | 0.2067 | 0.2023 | 0.2147 | 0.1941 | 0.2137 |
| G | | | | | | | | 0.0761 | 0.0898 | 0.0888 | 0.1006 | 0.0759 |
| H | | | | | | | | 0.1618 | 0.1178 | 0.1938 | 0.1488 | 0.1061 |

**Cell Index at: 26:30:32**

| | 1 | 2 | 3 | 4 | 5 | 6 | 7 | 8 | 9 | 10 | 11 | 12 |
|---|---|---|---|---|---|---|---|---|---|---|---|---|
| A | | | | | | | | 0.0336 | 0.0574 | 0.0662 | 0.0555 | 0.0816 |
| B | | | | | | | | 0.0901 | 0.0833 | 0.0797 | 0.0953 | 0.1021 |
| C | | | | | | | | 0.1921 | 0.1904 | 0.2299 | 0.2356 | 0.1929 |
| D | | | | | | | | 0.1742 | 0.2121 | 0.2195 | 0.2116 | 0.211 |
| E | | | | | | | | 0.134 | 0.1314 | 0.1446 | 0.1312 | 0.0995 |
| F | | | | | | | | 0.2019 | 0.1907 | 0.2232 | 0.1974 | 0.2224 |
| G | | | | | | | | 0.075 | 0.0921 | 0.0853 | 0.0968 | 0.0696 |
| H | | | | | | | | 0.1647 | 0.1226 | 0.1922 | 0.1517 | 0.1026 |

**Cell Index at: 26:45:33**

| | 1 | 2 | 3 | 4 | 5 | 6 | 7 | 8 | 9 | 10 | 11 | 12 |
|---|---|---|---|---|---|---|---|---|---|---|---|---|
| A | | | | | | | | 0.0302 | 0.0556 | 0.0647 | 0.0559 | 0.0832 |
| B | | | | | | | | 0.0881 | 0.0807 | 0.0797 | 0.0949 | 0.1056 |
| C | | | | | | | | 0.1952 | 0.1882 | 0.2288 | 0.2379 | 0.1914 |
| D | | | | | | | | 0.1783 | 0.216 | 0.217 | 0.2168 | 0.2125 |
| E | | | | | | | | 0.1329 | 0.1159 | 0.1359 | 0.1319 | 0.1054 |
| F | | | | | | | | 0.1998 | 0.1867 | 0.2188 | 0.1944 | 0.2139 |
| G | | | | | | | | 0.0707 | 0.0861 | 0.0871 | 0.1008 | 0.0742 |
| H | | | | | | | | 0.1603 | 0.1179 | 0.1883 | 0.1516 | 0.1061 |

**Cell Index at: 27:00:34**

| | 1 | 2 | 3 | 4 | 5 | 6 | 7 | 8 | 9 | 10 | 11 | 12 |
|---|---|---|---|---|---|---|---|---|---|---|---|---|
| A | | | | | | | | 0.0315 | 0.0543 | 0.0662 | 0.0523 | 0.0837 |
| B | | | | | | | | 0.092 | 0.0765 | 0.0823 | 0.0908 | 0.1006 |
| C | | | | | | | | 0.1967 | 0.1874 | 0.232 | 0.2404 | 0.197 |
| D | | | | | | | | 0.1759 | 0.2101 | 0.2299 | 0.2243 | 0.209 |
| E | | | | | | | | 0.1324 | 0.1261 | 0.1347 | 0.135 | 0.0988 |
| F | | | | | | | | 0.1992 | 0.1908 | 0.2285 | 0.1952 | 0.2202 |
| G | | | | | | | | 0.0661 | 0.0902 | 0.0884 | 0.1031 | 0.0715 |
| H | | | | | | | | 0.1624 | 0.1181 | 0.1912 | 0.1525 | 0.1035 |

**Cell Index at: 27:15:34**

| | 1 | 2 | 3 | 4 | 5 | 6 | 7 | 8 | 9 | 10 | 11 | 12 |
|---|---|---|---|---|---|---|---|---|---|---|---|---|
| A | | | | | | | | 0.0304 | 0.0501 | 0.0629 | 0.054 | 0.0802 |
| B | | | | | | | | 0.0865 | 0.0794 | 0.079 | 0.0904 | 0.1006 |
| C | | | | | | | | 0.1895 | 0.1913 | 0.2293 | 0.2393 | 0.1901 |
| D | | | | | | | | 0.1825 | 0.2132 | 0.2194 | 0.2158 | 0.2146 |
| E | | | | | | | | 0.1236 | 0.1234 | 0.1399 | 0.1254 | 0.104 |
| F | | | | | | | | 0.1913 | 0.1902 | 0.2244 | 0.1921 | 0.2216 |
| G | | | | | | | | 0.0713 | 0.0886 | 0.0855 | 0.1026 | 0.0698 |
| H | | | | | | | | 0.1627 | 0.1146 | 0.1918 | 0.1551 | 0.1051 |

**Cell Index at: 27:30:35**

| | 1 | 2 | 3 | 4 | 5 | 6 | 7 | 8 | 9 | 10 | 11 | 12 |
|---|---|---|---|---|---|---|---|---|---|---|---|---|
| A | | | | | | | | 0.0262 | 0.0511 | 0.0606 | 0.0526 | 0.0775 |
| B | | | | | | | | 0.0853 | 0.0766 | 0.0784 | 0.0908 | 0.0977 |
| C | | | | | | | | 0.1935 | 0.1915 | 0.231 | 0.2424 | 0.193 |
| D | | | | | | | | 0.1758 | 0.2182 | 0.2195 | 0.2206 | 0.2147 |
| E | | | | | | | | 0.1193 | 0.1167 | 0.136 | 0.1291 | 0.1018 |
| F | | | | | | | | 0.1975 | 0.187 | 0.226 | 0.1973 | 0.2158 |
| G | | | | | | | | 0.0698 | 0.0862 | 0.0869 | 0.1004 | 0.0716 |
| H | | | | | | | | 0.1631 | 0.1191 | 0.1918 | 0.1517 | 0.1055 |

**Cell Index at: 27:45:34**

| | 1 | 2 | 3 | 4 | 5 | 6 | 7 | 8 | 9 | 10 | 11 | 12 |
|---|---|---|---|---|---|---|---|---|---|---|---|---|
| A | | | | | | | | 0.0287 | 0.0461 | 0.0579 | 0.0512 | 0.0803 |
| B | | | | | | | | 0.0756 | 0.0743 | 0.0802 | 0.0901 | 0.1014 |
| C | | | | | | | | 0.1985 | 0.1888 | 0.2242 | 0.2425 | 0.1937 |
| D | | | | | | | | 0.1775 | 0.2182 | 0.2228 | 0.2209 | 0.2007 |
| E | | | | | | | | 0.1257 | 0.1175 | 0.1413 | 0.1271 | 0.1064 |
| F | | | | | | | | 0.1995 | 0.1904 | 0.2219 | 0.1963 | 0.2137 |
| G | | | | | | | | 0.0758 | 0.0875 | 0.0888 | 0.102 | 0.0701 |
| H | | | | | | | | 0.1605 | 0.1179 | 0.1919 | 0.1538 | 0.1015 |

**Cell Index at: 28:00:35**

| | 1 | 2 | 3 | 4 | 5 | 6 | 7 | 8 | 9 | 10 | 11 | 12 |
|---|---|---|---|---|---|---|---|---|---|---|---|---|
| A | | | | | | | | 0.0274 | 0.0486 | 0.0606 | 0.048 | 0.0769 |
| B | | | | | | | | 0.0744 | 0.0771 | 0.0809 | 0.0889 | 0.1031 |
| C | | | | | | | | 0.2041 | 0.1884 | 0.2234 | 0.246 | 0.1929 |
| D | | | | | | | | 0.1738 | 0.2226 | 0.221 | 0.2133 | 0.2071 |
| E | | | | | | | | 0.1213 | 0.1127 | 0.1443 | 0.1267 | 0.1026 |
| F | | | | | | | | 0.2 | 0.1935 | 0.2231 | 0.1924 | 0.217 |
| G | | | | | | | | 0.0682 | 0.0863 | 0.0898 | 0.1061 | 0.0701 |
| H | | | | | | | | 0.1643 | 0.1187 | 0.1931 | 0.1563 | 0.1034 |

**Cell Index at: 28:15:35**

| | 1 | 2 | 3 | 4 | 5 | 6 | 7 | 8 | 9 | 10 | 11 | 12 |
|---|---|---|---|---|---|---|---|---|---|---|---|---|
| A | | | | | | | | 0.0313 | 0.0464 | 0.0618 | 0.0519 | 0.0804 |
| B | | | | | | | | 0.0796 | 0.0757 | 0.0804 | 0.0861 | 0.1031 |
| C | | | | | | | | 0.1953 | 0.1903 | 0.2214 | 0.243 | 0.1952 |
| D | | | | | | | | 0.179 | 0.2091 | 0.2239 | 0.2153 | 0.2088 |
| E | | | | | | | | 0.1288 | 0.1158 | 0.1358 | 0.1215 | 0.0898 |
| F | | | | | | | | 0.1957 | 0.1891 | 0.2275 | 0.1935 | 0.2128 |
| G | | | | | | | | 0.0636 | 0.0844 | 0.0858 | 0.1036 | 0.0676 |
| H | | | | | | | | 0.1639 | 0.1209 | 0.1906 | 0.1528 | 0.1027 |

**Cell Index at: 28:30:36**

| | 1 | 2 | 3 | 4 | 5 | 6 | 7 | 8 | 9 | 10 | 11 | 12 |
|---|---|---|---|---|---|---|---|---|---|---|---|---|
| A | | | | | | | | 0.031 | 0.0411 | 0.0588 | 0.0483 | 0.0783 |
| B | | | | | | | | 0.0795 | 0.0707 | 0.079 | 0.0844 | 0.1046 |
| C | | | | | | | | 0.1978 | 0.1912 | 0.222 | 0.2443 | 0.1981 |
| D | | | | | | | | 0.179 | 0.2026 | 0.227 | 0.2143 | 0.2094 |
| E | | | | | | | | 0.1189 | 0.126 | 0.1377 | 0.1217 | 0.0993 |
| F | | | | | | | | 0.1991 | 0.1916 | 0.2275 | 0.1923 | 0.2147 |
| G | | | | | | | | 0.071 | 0.0825 | 0.0864 | 0.1032 | 0.0711 |
| H | | | | | | | | 0.1617 | 0.1182 | 0.1917 | 0.1513 | 0.1018 |

**Cell Index at: 28:45:36**

| | 1 | 2 | 3 | 4 | 5 | 6 | 7 | 8 | 9 | 10 | 11 | 12 |
|---|---|---|---|---|---|---|---|---|---|---|---|---|
| A | | | | | | | | 0.0256 | 0.0438 | 0.0546 | 0.0498 | 0.0782 |
| B | | | | | | | | 0.0805 | 0.0716 | 0.0792 | 0.088 | 0.1035 |
| C | | | | | | | | 0.1926 | 0.1986 | 0.2217 | 0.2475 | 0.1991 |
| D | | | | | | | | 0.1768 | 0.2186 | 0.2124 | 0.2162 | 0.2071 |
| E | | | | | | | | 0.1178 | 0.1213 | 0.134 | 0.1308 | 0.0963 |
| F | | | | | | | | 0.1976 | 0.1899 | 0.2233 | 0.1935 | 0.2123 |
| G | | | | | | | | 0.0664 | 0.0822 | 0.0818 | 0.1028 | 0.0671 |
| H | | | | | | | | 0.1631 | 0.1179 | 0.1887 | 0.154 | 0.0987 |

**Cell Index at: 29:00:36**

| | 1 | 2 | 3 | 4 | 5 | 6 | 7 | 8 | 9 | 10 | 11 | 12 |
|---|---|---|---|---|---|---|---|---|---|---|---|---|
| A | | | | | | | | 0.0265 | 0.0454 | 0.0582 | 0.0461 | 0.0743 |
| B | | | | | | | | 0.0843 | 0.0761 | 0.079 | 0.0842 | 0.1035 |
| C | | | | | | | | 0.1981 | 0.1945 | 0.2131 | 0.245 | 0.1965 |
| D | | | | | | | | 0.1853 | 0.2142 | 0.2199 | 0.2226 | 0.2043 |
| E | | | | | | | | 0.1097 | 0.1195 | 0.1272 | 0.1217 | 0.091 |
| F | | | | | | | | 0.1938 | 0.189 | 0.2266 | 0.1908 | 0.2145 |
| G | | | | | | | | 0.0644 | 0.0835 | 0.0805 | 0.1018 | 0.0716 |
| H | | | | | | | | 0.164 | 0.1196 | 0.1935 | 0.1534 | 0.1024 |

**Cell Index at: 29:15:36**

| | 1 | 2 | 3 | 4 | 5 | 6 | 7 | 8 | 9 | 10 | 11 | 12 |
|---|---|---|---|---|---|---|---|---|---|---|---|---|
| A | | | | | | | | 0.0278 | 0.0451 | 0.055 | 0.0473 | 0.0704 |
| B | | | | | | | | 0.0767 | 0.0697 | 0.0739 | 0.0835 | 0.1039 |
| C | | | | | | | | 0.1873 | 0.1883 | 0.2112 | 0.2491 | 0.1969 |
| D | | | | | | | | 0.1828 | 0.2162 | 0.2232 | 0.2163 | 0.2081 |
| E | | | | | | | | 0.1167 | 0.1241 | 0.1355 | 0.1305 | 0.0947 |
| F | | | | | | | | 0.1993 | 0.1899 | 0.2315 | 0.1928 | 0.2134 |
| G | | | | | | | | 0.0636 | 0.0831 | 0.0826 | 0.0962 | 0.0706 |
| H | | | | | | | | 0.1657 | 0.1183 | 0.1899 | 0.1535 | 0.0995 |

**Cell Index at: 29:30:36**

| | 1 | 2 | 3 | 4 | 5 | 6 | 7 | 8 | 9 | 10 | 11 | 12 |
|---|---|---|---|---|---|---|---|---|---|---|---|---|
| A | | | | | | | | 0.0283 | 0.042 | 0.0557 | 0.0454 | 0.0727 |
| B | | | | | | | | 0.0768 | 0.0738 | 0.0754 | 0.0859 | 0.1038 |
| C | | | | | | | | 0.1932 | 0.1897 | 0.2111 | 0.2365 | 0.1881 |
| D | | | | | | | | 0.1852 | 0.2177 | 0.221 | 0.2147 | 0.1939 |
| E | | | | | | | | 0.114 | 0.1192 | 0.1375 | 0.1259 | 0.0898 |
| F | | | | | | | | 0.1957 | 0.1885 | 0.234 | 0.1928 | 0.2167 |
| G | | | | | | | | 0.0665 | 0.0845 | 0.0851 | 0.1044 | 0.071 |
| H | | | | | | | | 0.162 | 0.1189 | 0.1921 | 0.1557 | 0.1007 |

**Cell Index at: 29:45:37**

| | 1 | 2 | 3 | 4 | 5 | 6 | 7 | 8 | 9 | 10 | 11 | 12 |
|---|---|---|---|---|---|---|---|---|---|---|---|---|
| A | | | | | | | | 0.0256 | 0.0408 | 0.0538 | 0.0458 | 0.0713 |
| B | | | | | | | | 0.0782 | 0.0704 | 0.0779 | 0.0895 | 0.0997 |
| C | | | | | | | | 0.1902 | 0.1947 | 0.2078 | 0.244 | 0.1974 |
| D | | | | | | | | 0.1859 | 0.2208 | 0.2326 | 0.2146 | 0.2068 |
| E | | | | | | | | 0.1235 | 0.1188 | 0.136 | 0.1217 | 0.0894 |
| F | | | | | | | | 0.2065 | 0.1866 | 0.2227 | 0.1955 | 0.2121 |
| G | | | | | | | | 0.0628 | 0.0846 | 0.0814 | 0.1 | 0.0692 |
| H | | | | | | | | 0.1633 | 0.115 | 0.1889 | 0.1497 | 0.103 |

**Cell Index at: 30:00:36**

| | 1 | 2 | 3 | 4 | 5 | 6 | 7 | 8 | 9 | 10 | 11 | 12 |
|---|---|---|---|---|---|---|---|---|---|---|---|---|
| A | | | | | | | | 0.022 | 0.0402 | 0.0538 | 0.0435 | 0.0692 |
| B | | | | | | | | 0.0813 | 0.0682 | 0.0785 | 0.0816 | 0.0977 |
| C | | | | | | | | 0.1948 | 0.1961 | 0.2163 | 0.2442 | 0.1956 |
| D | | | | | | | | 0.1827 | 0.2252 | 0.2218 | 0.2122 | 0.2116 |
| E | | | | | | | | 0.1195 | 0.1217 | 0.1417 | 0.1186 | 0.099 |
| F | | | | | | | | 0.1993 | 0.1878 | 0.2301 | 0.1855 | 0.2181 |
| G | | | | | | | | 0.0595 | 0.0868 | 0.0762 | 0.0976 | 0.0737 |
| H | | | | | | | | 0.1638 | 0.1171 | 0.1922 | 0.1539 | 0.1047 |

**Cell Index at: 30:15:37**

| | 1 | 2 | 3 | 4 | 5 | 6 | 7 | 8 | 9 | 10 | 11 | 12 |
|---|---|---|---|---|---|---|---|---|---|---|---|---|
| A | | | | | | | | 0.0221 | 0.0413 | 0.0567 | 0.0461 | 0.0704 |
| B | | | | | | | | 0.0756 | 0.0686 | 0.0776 | 0.0825 | 0.0977 |
| C | | | | | | | | 0.1973 | 0.1985 | 0.2089 | 0.2524 | 0.201 |
| D | | | | | | | | 0.1799 | 0.2163 | 0.2235 | 0.2168 | 0.2024 |
| E | | | | | | | | 0.1173 | 0.1215 | 0.128 | 0.118 | 0.0897 |
| F | | | | | | | | 0.1954 | 0.1904 | 0.231 | 0.1959 | 0.218 |
| G | | | | | | | | 0.0644 | 0.0816 | 0.0825 | 0.0943 | 0.0695 |
| H | | | | | | | | 0.1629 | 0.1167 | 0.1878 | 0.1495 | 0.1071 |

**Cell Index at: 30:30:36**

| | 1 | 2 | 3 | 4 | 5 | 6 | 7 | 8 | 9 | 10 | 11 | 12 |
|---|---|---|---|---|---|---|---|---|---|---|---|---|
| A | | | | | | | | 0.017 | 0.0404 | 0.0554 | 0.0482 | 0.0716 |
| B | | | | | | | | 0.0743 | 0.071 | 0.0736 | 0.0811 | 0.1005 |
| C | | | | | | | | 0.1917 | 0.1943 | 0.2133 | 0.2481 | 0.1991 |
| D | | | | | | | | 0.1722 | 0.2184 | 0.2155 | 0.2139 | 0.1976 |
| E | | | | | | | | 0.1225 | 0.1162 | 0.139 | 0.1168 | 0.0976 |
| F | | | | | | | | 0.198 | 0.1843 | 0.2248 | 0.1922 | 0.2114 |
| G | | | | | | | | 0.0625 | 0.0833 | 0.082 | 0.0961 | 0.0699 |
| H | | | | | | | | 0.1608 | 0.1148 | 0.1771 | 0.1526 | 0.1048 |

**Cell Index at: 30:45:36**

| | 1 | 2 | 3 | 4 | 5 | 6 | 7 | 8 | 9 | 10 | 11 | 12 |
|---|---|---|---|---|---|---|---|---|---|---|---|---|
| A | | | | | | | | 0.0156 | 0.0374 | 0.0507 | 0.0437 | 0.0695 |
| B | | | | | | | | 0.0693 | 0.0669 | 0.0727 | 0.0826 | 0.0954 |
| C | | | | | | | | 0.197 | 0.195 | 0.2101 | 0.2476 | 0.1965 |
| D | | | | | | | | 0.1736 | 0.2267 | 0.2231 | 0.2095 | 0.2095 |
| E | | | | | | | | 0.115 | 0.1112 | 0.1384 | 0.1259 | 0.0887 |
| F | | | | | | | | 0.1866 | 0.1894 | 0.2231 | 0.1949 | 0.2114 |
| G | | | | | | | | 0.0589 | 0.0851 | 0.0796 | 0.1002 | 0.0656 |
| H | | | | | | | | 0.1645 | 0.1121 | 0.1783 | 0.1528 | 0.1029 |

**Cell Index at: 31:00:37**

| | 1 | 2 | 3 | 4 | 5 | 6 | 7 | 8 | 9 | 10 | 11 | 12 |
|---|---|---|---|---|---|---|---|---|---|---|---|---|
| A | | | | | | | | 0.0182 | 0.0353 | 0.0556 | 0.0434 | 0.0625 |
| B | | | | | | | | 0.0793 | 0.0674 | 0.0769 | 0.0788 | 0.0917 |
| C | | | | | | | | 0.189 | 0.1915 | 0.2149 | 0.2509 | 0.193 |
| D | | | | | | | | 0.1757 | 0.22 | 0.2326 | 0.2122 | 0.1904 |
| E | | | | | | | | 0.1177 | 0.1184 | 0.1384 | 0.1224 | 0.0949 |
| F | | | | | | | | 0.1921 | 0.1908 | 0.2335 | 0.1988 | 0.2078 |
| G | | | | | | | | 0.0573 | 0.0833 | 0.0802 | 0.0973 | 0.063 |
| H | | | | | | | | 0.1631 | 0.1163 | 0.1757 | 0.1557 | 0.0997 |

**Cell Index at: 31:15:37**

| | 1 | 2 | 3 | 4 | 5 | 6 | 7 | 8 | 9 | 10 | 11 | 12 |
|---|---|---|---|---|---|---|---|---|---|---|---|---|
| A | | | | | | | | 0.0178 | 0.0396 | 0.0511 | 0.043 | 0.0598 |
| B | | | | | | | | 0.07 | 0.063 | 0.0779 | 0.0763 | 0.0879 |
| C | | | | | | | | 0.1959 | 0.1893 | 0.2122 | 0.2485 | 0.193 |
| D | | | | | | | | 0.1783 | 0.2123 | 0.2276 | 0.2076 | 0.2012 |
| E | | | | | | | | 0.115 | 0.1078 | 0.1324 | 0.1217 | 0.096 |
| F | | | | | | | | 0.1954 | 0.1897 | 0.2271 | 0.2003 | 0.207 |
| G | | | | | | | | 0.0589 | 0.0796 | 0.0834 | 0.0909 | 0.0631 |
| H | | | | | | | | 0.1575 | 0.1143 | 0.1761 | 0.1528 | 0.0968 |

**Cell Index at: 31:30:38**

| | 1 | 2 | 3 | 4 | 5 | 6 | 7 | 8 | 9 | 10 | 11 | 12 |
|---|---|---|---|---|---|---|---|---|---|---|---|---|
| A | | | | | | | | 0.0159 | 0.0378 | 0.0464 | 0.0426 | 0.0631 |
| B | | | | | | | | 0.0721 | 0.0652 | 0.0729 | 0.0792 | 0.0898 |
| C | | | | | | | | 0.1903 | 0.1923 | 0.2137 | 0.249 | 0.1956 |
| D | | | | | | | | 0.1755 | 0.2133 | 0.2223 | 0.2158 | 0.2031 |
| E | | | | | | | | 0.1112 | 0.1131 | 0.1337 | 0.1228 | 0.0911 |
| F | | | | | | | | 0.1971 | 0.1873 | 0.2311 | 0.2012 | 0.2122 |
| G | | | | | | | | 0.0581 | 0.0817 | 0.0808 | 0.0924 | 0.0574 |
| H | | | | | | | | 0.1589 | 0.1143 | 0.1756 | 0.1541 | 0.098 |

**Cell Index at: 31:45:38**

| | 1 | 2 | 3 | 4 | 5 | 6 | 7 | 8 | 9 | 10 | 11 | 12 |
|---|---|---|---|---|---|---|---|---|---|---|---|---|
| A | | | | | | | | 0.0172 | 0.0374 | 0.0469 | 0.0407 | 0.0618 |
| B | | | | | | | | 0.073 | 0.0583 | 0.0711 | 0.0783 | 0.0841 |
| C | | | | | | | | 0.1951 | 0.1977 | 0.2173 | 0.2536 | 0.1961 |
| D | | | | | | | | 0.1749 | 0.2217 | 0.2279 | 0.2097 | 0.2104 |
| E | | | | | | | | 0.1116 | 0.1208 | 0.1352 | 0.1291 | 0.0953 |
| F | | | | | | | | 0.1962 | 0.1973 | 0.2225 | 0.1985 | 0.2054 |
| G | | | | | | | | 0.055 | 0.0771 | 0.081 | 0.0942 | 0.0604 |
| H | | | | | | | | 0.161 | 0.1162 | 0.1746 | 0.1526 | 0.0982 |

**Cell Index at: 32:00:39**

| | 1 | 2 | 3 | 4 | 5 | 6 | 7 | 8 | 9 | 10 | 11 | 12 |
|---|---|---|---|---|---|---|---|---|---|---|---|---|
| A | | | | | | | | 0.0138 | 0.0377 | 0.0485 | 0.0392 | 0.0582 |
| B | | | | | | | | 0.068 | 0.0657 | 0.071 | 0.0758 | 0.0907 |
| C | | | | | | | | 0.1929 | 0.1935 | 0.2111 | 0.2491 | 0.1981 |
| D | | | | | | | | 0.1743 | 0.2207 | 0.2232 | 0.2171 | 0.2016 |
| E | | | | | | | | 0.1099 | 0.111 | 0.1383 | 0.1226 | 0.092 |
| F | | | | | | | | 0.1996 | 0.1883 | 0.2269 | 0.196 | 0.2097 |
| G | | | | | | | | 0.0526 | 0.0798 | 0.0796 | 0.0956 | 0.0594 |
| H | | | | | | | | 0.1577 | 0.1199 | 0.1746 | 0.1526 | 0.0981 |

**Cell Index at: 32:15:38**

| | 1 | 2 | 3 | 4 | 5 | 6 | 7 | 8 | 9 | 10 | 11 | 12 |
|---|---|---|---|---|---|---|---|---|---|---|---|---|
| A | | | | | | | | 0.0137 | 0.0322 | 0.0463 | 0.0407 | 0.0619 |
| B | | | | | | | | 0.0735 | 0.0647 | 0.0689 | 0.0786 | 0.0874 |
| C | | | | | | | | 0.1972 | 0.1936 | 0.2135 | 0.2583 | 0.1947 |
| D | | | | | | | | 0.1735 | 0.2137 | 0.2316 | 0.2173 | 0.2068 |
| E | | | | | | | | 0.106 | 0.1142 | 0.1334 | 0.1217 | 0.0867 |
| F | | | | | | | | 0.193 | 0.187 | 0.2271 | 0.1973 | 0.2093 |
| G | | | | | | | | 0.0535 | 0.076 | 0.081 | 0.0904 | 0.0584 |
| H | | | | | | | | 0.1616 | 0.1153 | 0.1787 | 0.1535 | 0.0989 |

**Cell Index at: 32:30:39**

| | 1 | 2 | 3 | 4 | 5 | 6 | 7 | 8 | 9 | 10 | 11 | 12 |
|---|---|---|---|---|---|---|---|---|---|---|---|---|
| A | | | | | | | | 0.0116 | 0.033 | 0.0499 | 0.0375 | 0.0613 |
| B | | | | | | | | 0.0704 | 0.0683 | 0.0668 | 0.0772 | 0.0842 |
| C | | | | | | | | 0.1907 | 0.1993 | 0.2182 | 0.2498 | 0.1919 |
| D | | | | | | | | 0.1751 | 0.2169 | 0.224 | 0.2186 | 0.2131 |
| E | | | | | | | | 0.1146 | 0.1134 | 0.1268 | 0.1163 | 0.0951 |
| F | | | | | | | | 0.1962 | 0.1911 | 0.2289 | 0.1988 | 0.2086 |
| G | | | | | | | | 0.0493 | 0.0772 | 0.0814 | 0.0895 | 0.063 |
| H | | | | | | | | 0.1627 | 0.1151 | 0.1779 | 0.1525 | 0.1005 |

**Cell Index at: 32:45:38**

| | 1 | 2 | 3 | 4 | 5 | 6 | 7 | 8 | 9 | 10 | 11 | 12 |
|---|---|---|---|---|---|---|---|---|---|---|---|---|
| A | | | | | | | | 0.0174 | 0.0325 | 0.0473 | 0.0384 | 0.0561 |
| B | | | | | | | | 0.0661 | 0.0642 | 0.0692 | 0.079 | 0.0863 |
| C | | | | | | | | 0.1985 | 0.2008 | 0.2208 | 0.2501 | 0.1981 |
| D | | | | | | | | 0.1782 | 0.2139 | 0.2267 | 0.2181 | 0.1933 |
| E | | | | | | | | 0.1154 | 0.1197 | 0.1295 | 0.123 | 0.0845 |
| F | | | | | | | | 0.2007 | 0.1879 | 0.2248 | 0.2018 | 0.2027 |
| G | | | | | | | | 0.0529 | 0.0796 | 0.0739 | 0.0925 | 0.0647 |
| H | | | | | | | | 0.1597 | 0.1158 | 0.179 | 0.1528 | 0.0988 |

**Cell Index at: 33:00:38**

| | 1 | 2 | 3 | 4 | 5 | 6 | 7 | 8 | 9 | 10 | 11 | 12 |
|---|---|---|---|---|---|---|---|---|---|---|---|---|
| A | | | | | | | | 0.015 | 0.0324 | 0.0452 | 0.0384 | 0.0576 |
| B | | | | | | | | 0.062 | 0.067 | 0.0661 | 0.0795 | 0.0811 |
| C | | | | | | | | 0.1945 | 0.202 | 0.2143 | 0.2534 | 0.1942 |
| D | | | | | | | | 0.1811 | 0.2159 | 0.231 | 0.2138 | 0.2017 |
| E | | | | | | | | 0.1078 | 0.1157 | 0.1288 | 0.1142 | 0.0828 |
| F | | | | | | | | 0.1933 | 0.1914 | 0.2237 | 0.1975 | 0.2075 |
| G | | | | | | | | 0.0505 | 0.0714 | 0.0788 | 0.0938 | 0.059 |
| H | | | | | | | | 0.1593 | 0.1159 | 0.1787 | 0.1559 | 0.0983 |

**Cell Index at: 33:15:39**

| | 1 | 2 | 3 | 4 | 5 | 6 | 7 | 8 | 9 | 10 | 11 | 12 |
|---|---|---|---|---|---|---|---|---|---|---|---|---|
| A | | | | | | | | 0.0164 | 0.0353 | 0.0443 | 0.0368 | 0.059 |
| B | | | | | | | | 0.0588 | 0.0642 | 0.0675 | 0.0758 | 0.081 |
| C | | | | | | | | 0.1949 | 0.1938 | 0.2187 | 0.2464 | 0.1989 |
| D | | | | | | | | 0.1756 | 0.2236 | 0.2325 | 0.2081 | 0.2038 |
| E | | | | | | | | 0.1023 | 0.1127 | 0.1302 | 0.1215 | 0.0761 |
| F | | | | | | | | 0.1919 | 0.1933 | 0.2265 | 0.1963 | 0.2042 |
| G | | | | | | | | 0.0525 | 0.0707 | 0.0738 | 0.0944 | 0.0545 |
| H | | | | | | | | 0.1576 | 0.1126 | 0.1782 | 0.1535 | 0.1003 |

**Cell Index at: 33:30:39**

| | 1 | 2 | 3 | 4 | 5 | 6 | 7 | 8 | 9 | 10 | 11 | 12 |
|---|---|---|---|---|---|---|---|---|---|---|---|---|
| A | | | | | | | | 0.0153 | 0.0321 | 0.0442 | 0.0358 | 0.0565 |
| B | | | | | | | | 0.0661 | 0.0644 | 0.0608 | 0.0777 | 0.0819 |
| C | | | | | | | | 0.1995 | 0.1949 | 0.2149 | 0.2518 | 0.2005 |
| D | | | | | | | | 0.1763 | 0.215 | 0.2341 | 0.2125 | 0.2082 |
| E | | | | | | | | 0.0966 | 0.1112 | 0.1253 | 0.1146 | 0.0718 |
| F | | | | | | | | 0.1942 | 0.1871 | 0.2326 | 0.204 | 0.2019 |
| G | | | | | | | | 0.0489 | 0.072 | 0.0765 | 0.0908 | 0.0561 |
| H | | | | | | | | 0.1599 | 0.1144 | 0.1771 | 0.1522 | 0.1002 |

**Cell Index at: 33:45:40**

| | 1 | 2 | 3 | 4 | 5 | 6 | 7 | 8 | 9 | 10 | 11 | 12 |
|---|---|---|---|---|---|---|---|---|---|---|---|---|
| A | | | | | | | | 0.0113 | 0.0308 | 0.0389 | 0.0361 | 0.0567 |
| B | | | | | | | | 0.0667 | 0.0641 | 0.0675 | 0.0714 | 0.0843 |
| C | | | | | | | | 0.2 | 0.1987 | 0.2129 | 0.2474 | 0.1919 |
| D | | | | | | | | 0.1816 | 0.2216 | 0.2306 | 0.2189 | 0.2057 |
| E | | | | | | | | 0.1003 | 0.1087 | 0.1275 | 0.1158 | 0.0828 |
| F | | | | | | | | 0.1983 | 0.1909 | 0.2307 | 0.1998 | 0.2116 |
| G | | | | | | | | 0.0503 | 0.0707 | 0.0698 | 0.09 | 0.0581 |
| H | | | | | | | | 0.159 | 0.1174 | 0.1789 | 0.1548 | 0.1008 |

**Cell Index at: 34:00:39**

| | 1 | 2 | 3 | 4 | 5 | 6 | 7 | 8 | 9 | 10 | 11 | 12 |
|---|---|---|---|---|---|---|---|---|---|---|---|---|
| A | | | | | | | | 0.0112 | 0.0332 | 0.0403 | 0.0355 | 0.0557 |
| B | | | | | | | | 0.0625 | 0.063 | 0.0627 | 0.0705 | 0.082 |
| C | | | | | | | | 0.1994 | 0.2012 | 0.2181 | 0.2489 | 0.1923 |
| D | | | | | | | | 0.1927 | 0.2179 | 0.226 | 0.2171 | 0.2104 |
| E | | | | | | | | 0.1068 | 0.1088 | 0.1242 | 0.1086 | 0.08 |
| F | | | | | | | | 0.1963 | 0.1926 | 0.2267 | 0.1974 | 0.2085 |
| G | | | | | | | | 0.0494 | 0.0708 | 0.0694 | 0.0901 | 0.0543 |

| | | | | | | | | | | | | |
|---|---|---|---|---|---|---|---|---|---|---|---|---|
| H | | | | | | | | 0.1564 | 0.1159 | 0.1809 | 0.1501 | 0.0999 |

**Cell Index at: 34:15:40**

| | 1 | 2 | 3 | 4 | 5 | 6 | 7 | 8 | 9 | 10 | 11 | 12 |
|---|---|---|---|---|---|---|---|---|---|---|---|---|
| A | | | | | | | | 0.0119 | 0.0329 | 0.0406 | 0.036 | 0.0528 |
| B | | | | | | | | 0.0626 | 0.0629 | 0.0637 | 0.071 | 0.0803 |
| C | | | | | | | | 0.1919 | 0.1999 | 0.2197 | 0.2562 | 0.1896 |
| D | | | | | | | | 0.1898 | 0.2194 | 0.2408 | 0.2134 | 0.2061 |
| E | | | | | | | | 0.1023 | 0.1123 | 0.1193 | 0.1158 | 0.0844 |
| F | | | | | | | | 0.1981 | 0.1911 | 0.2306 | 0.2033 | 0.2032 |
| G | | | | | | | | 0.051 | 0.0704 | 0.0717 | 0.0906 | 0.0531 |
| H | | | | | | | | 0.1545 | 0.1139 | 0.1807 | 0.1525 | 0.097 |

**Cell Index at: 34:30:40**

| | 1 | 2 | 3 | 4 | 5 | 6 | 7 | 8 | 9 | 10 | 11 | 12 |
|---|---|---|---|---|---|---|---|---|---|---|---|---|
| A | | | | | | | | 0.0098 | 0.0344 | 0.0361 | 0.0359 | 0.0533 |
| B | | | | | | | | 0.0643 | 0.0596 | 0.0671 | 0.0681 | 0.0785 |
| C | | | | | | | | 0.1971 | 0.2032 | 0.2158 | 0.2585 | 0.1948 |
| D | | | | | | | | 0.1793 | 0.2193 | 0.2317 | 0.2108 | 0.2104 |
| E | | | | | | | | 0.094 | 0.1114 | 0.1302 | 0.1129 | 0.0871 |
| F | | | | | | | | 0.1996 | 0.1896 | 0.2373 | 0.1963 | 0.2092 |
| G | | | | | | | | 0.0484 | 0.0713 | 0.0747 | 0.0894 | 0.0555 |
| H | | | | | | | | 0.1609 | 0.1133 | 0.1791 | 0.1495 | 0.0965 |

**Cell Index at: 34:45:40**

| | 1 | 2 | 3 | 4 | 5 | 6 | 7 | 8 | 9 | 10 | 11 | 12 |
|---|---|---|---|---|---|---|---|---|---|---|---|---|
| A | | | | | | | | 0.0097 | 0.0291 | 0.0347 | 0.0347 | 0.0523 |
| B | | | | | | | | 0.0604 | 0.0606 | 0.0646 | 0.0699 | 0.0809 |
| C | | | | | | | | 0.1923 | 0.2 | 0.219 | 0.2561 | 0.1909 |
| D | | | | | | | | 0.1832 | 0.2129 | 0.2316 | 0.2134 | 0.2137 |
| E | | | | | | | | 0.1016 | 0.11 | 0.123 | 0.1102 | 0.0772 |
| F | | | | | | | | 0.1986 | 0.1962 | 0.2336 | 0.2012 | 0.2093 |
| G | | | | | | | | 0.0476 | 0.069 | 0.0729 | 0.0863 | 0.0529 |
| H | | | | | | | | 0.1595 | 0.1109 | 0.1813 | 0.1495 | 0.0978 |

**Cell Index at: 35:00:41**

| | 1 | 2 | 3 | 4 | 5 | 6 | 7 | 8 | 9 | 10 | 11 | 12 |
|---|---|---|---|---|---|---|---|---|---|---|---|---|
| A | | | | | | | | 0.007 | 0.0294 | 0.0369 | 0.0371 | 0.0505 |
| B | | | | | | | | 0.0617 | 0.0619 | 0.0652 | 0.0645 | 0.0773 |
| C | | | | | | | | 0.1817 | 0.1942 | 0.2177 | 0.2517 | 0.193 |
| D | | | | | | | | 0.1794 | 0.2155 | 0.2289 | 0.2094 | 0.2171 |
| E | | | | | | | | 0.0968 | 0.1173 | 0.1271 | 0.116 | 0.0849 |
| F | | | | | | | | 0.1983 | 0.1919 | 0.23 | 0.1993 | 0.2044 |
| G | | | | | | | | 0.0534 | 0.069 | 0.0676 | 0.0847 | 0.0501 |
| H | | | | | | | | 0.157 | 0.1108 | 0.1787 | 0.1474 | 0.0954 |

**Cell Index at: 35:15:41**

| | 1 | 2 | 3 | 4 | 5 | 6 | 7 | 8 | 9 | 10 | 11 | 12 |
|---|---|---|---|---|---|---|---|---|---|---|---|---|
| A | | | | | | | | 0.0071 | 0.032 | 0.0367 | 0.0373 | 0.0555 |
| B | | | | | | | | 0.0619 | 0.0574 | 0.0627 | 0.0682 | 0.0775 |
| C | | | | | | | | 0.19 | 0.1926 | 0.2244 | 0.2498 | 0.1928 |
| D | | | | | | | | 0.1804 | 0.2159 | 0.2357 | 0.2178 | 0.2057 |
| E | | | | | | | | 0.091 | 0.1185 | 0.1288 | 0.1131 | 0.0751 |
| F | | | | | | | | 0.2034 | 0.1942 | 0.2321 | 0.1962 | 0.207 |
| G | | | | | | | | 0.0496 | 0.0648 | 0.0679 | 0.0849 | 0.0528 |
| H | | | | | | | | 0.1559 | 0.1132 | 0.1798 | 0.1545 | 0.0987 |

**Cell Index at: 35:30:42**

| | 1 | 2 | 3 | 4 | 5 | 6 | 7 | 8 | 9 | 10 | 11 | 12 |
|---|---|---|---|---|---|---|---|---|---|---|---|---|
| A | | | | | | | | 0.0049 | 0.0295 | 0.0343 | 0.0332 | 0.0525 |
| B | | | | | | | | 0.0656 | 0.0541 | 0.0648 | 0.0669 | 0.0822 |
| C | | | | | | | | 0.1898 | 0.1932 | 0.2221 | 0.2525 | 0.1899 |
| D | | | | | | | | 0.1798 | 0.2221 | 0.2354 | 0.2167 | 0.2046 |
| E | | | | | | | | 0.1086 | 0.1145 | 0.1275 | 0.1102 | 0.08 |
| F | | | | | | | | 0.201 | 0.1914 | 0.2338 | 0.1985 | 0.2023 |
| G | | | | | | | | 0.0508 | 0.0725 | 0.0665 | 0.0864 | 0.0512 |
| H | | | | | | | | 0.1618 | 0.1094 | 0.1804 | 0.152 | 0.1001 |

**Cell Index at: 35:45:42**

| | 1 | 2 | 3 | 4 | 5 | 6 | 7 | 8 | 9 | 10 | 11 | 12 |
|---|---|---|---|---|---|---|---|---|---|---|---|---|
| A | | | | | | | | 0.0068 | 0.0278 | 0.0424 | 0.033 | 0.0496 |
| B | | | | | | | | 0.063 | 0.0524 | 0.0584 | 0.0675 | 0.0786 |
| C | | | | | | | | 0.1916 | 0.1891 | 0.2177 | 0.2505 | 0.1912 |
| D | | | | | | | | 0.179 | 0.225 | 0.2259 | 0.2172 | 0.2107 |
| E | | | | | | | | 0.1022 | 0.12 | 0.1266 | 0.1108 | 0.082 |
| F | | | | | | | | 0.2017 | 0.1925 | 0.2372 | 0.2057 | 0.2104 |
| G | | | | | | | | 0.0472 | 0.0671 | 0.0686 | 0.0777 | 0.0522 |
| H | | | | | | | | 0.1562 | 0.1013 | 0.1817 | 0.1516 | 0.101 |

**Cell Index at: 36:00:43**

| | 1 | 2 | 3 | 4 | 5 | 6 | 7 | 8 | 9 | 10 | 11 | 12 |
|---|---|---|---|---|---|---|---|---|---|---|---|---|
| A | | | | | | | | 0.0046 | 0.0305 | 0.0385 | 0.0328 | 0.0484 |
| B | | | | | | | | 0.0576 | 0.0559 | 0.0561 | 0.0661 | 0.0811 |
| C | | | | | | | | 0.1924 | 0.195 | 0.2183 | 0.2577 | 0.1935 |
| D | | | | | | | | 0.1865 | 0.2233 | 0.2312 | 0.2181 | 0.2058 |
| E | | | | | | | | 0.1032 | 0.1141 | 0.1244 | 0.1083 | 0.0808 |
| F | | | | | | | | 0.2056 | 0.1998 | 0.2327 | 0.202 | 0.2103 |
| G | | | | | | | | 0.0411 | 0.0677 | 0.0675 | 0.0773 | 0.0499 |
| H | | | | | | | | 0.1582 | 0.1031 | 0.1827 | 0.1539 | 0.102 |

**Cell Index at: 36:15:42**

| | 1 | 2 | 3 | 4 | 5 | 6 | 7 | 8 | 9 | 10 | 11 | 12 |
|---|---|---|---|---|---|---|---|---|---|---|---|---|
| A | | | | | | | | 0.0059 | 0.0303 | 0.0406 | 0.0316 | 0.0507 |
| B | | | | | | | | 0.0575 | 0.056 | 0.0548 | 0.0622 | 0.0799 |
| C | | | | | | | | 0.1931 | 0.1902 | 0.2248 | 0.2582 | 0.1973 |
| D | | | | | | | | 0.1829 | 0.221 | 0.2304 | 0.2145 | 0.208 |
| E | | | | | | | | 0.1016 | 0.1225 | 0.1217 | 0.1035 | 0.0769 |
| F | | | | | | | | 0.207 | 0.1963 | 0.2265 | 0.2044 | 0.2165 |
| G | | | | | | | | 0.0507 | 0.0648 | 0.067 | 0.077 | 0.0487 |
| H | | | | | | | | 0.1548 | 0.102 | 0.1824 | 0.1539 | 0.1014 |

**Cell Index at: 36:30:42**

| | 1 | 2 | 3 | 4 | 5 | 6 | 7 | 8 | 9 | 10 | 11 | 12 |
|---|---|---|---|---|---|---|---|---|---|---|---|---|
| A | | | | | | | | 0.0053 | 0.0267 | 0.041 | 0.0322 | 0.0486 |
| B | | | | | | | | 0.0601 | 0.0541 | 0.0551 | 0.0667 | 0.0823 |
| C | | | | | | | | 0.1937 | 0.1851 | 0.2223 | 0.2599 | 0.1922 |
| D | | | | | | | | 0.1835 | 0.2284 | 0.2248 | 0.2229 | 0.2054 |
| E | | | | | | | | 0.1139 | 0.1123 | 0.1217 | 0.1034 | 0.0829 |
| F | | | | | | | | 0.2067 | 0.1949 | 0.2344 | 0.2058 | 0.2163 |
| G | | | | | | | | 0.0406 | 0.0634 | 0.0656 | 0.074 | 0.0539 |
| H | | | | | | | | 0.1568 | 0.1033 | 0.1826 | 0.1552 | 0.1011 |

**Cell Index at: 36:45:42**

| | 1 | 2 | 3 | 4 | 5 | 6 | 7 | 8 | 9 | 10 | 11 | 12 |
|---|---|---|---|---|---|---|---|---|---|---|---|---|
| A | | | | | | | | 0.0053 | 0.0264 | 0.036 | 0.0313 | 0.0466 |
| B | | | | | | | | 0.0539 | 0.0544 | 0.0521 | 0.068 | 0.0821 |
| C | | | | | | | | 0.1871 | 0.1983 | 0.2201 | 0.2542 | 0.1924 |
| D | | | | | | | | 0.1869 | 0.2323 | 0.2344 | 0.215 | 0.2108 |
| E | | | | | | | | 0.1029 | 0.1097 | 0.1207 | 0.1027 | 0.0803 |
| F | | | | | | | | 0.2069 | 0.2005 | 0.2326 | 0.2024 | 0.2076 |
| G | | | | | | | | 0.0506 | 0.0668 | 0.0682 | 0.0764 | 0.0477 |
| H | | | | | | | | 0.1515 | 0.103 | 0.1812 | 0.1499 | 0.0977 |

**Cell Index at: 37:00:42**

| | 1 | 2 | 3 | 4 | 5 | 6 | 7 | 8 | 9 | 10 | 11 | 12 |
|---|---|---|---|---|---|---|---|---|---|---|---|---|
| A | | | | | | | | 0.0049 | 0.0261 | 0.0389 | 0.0262 | 0.0476 |
| B | | | | | | | | 0.0592 | 0.0533 | 0.0578 | 0.0658 | 0.0832 |
| C | | | | | | | | 0.1898 | 0.1935 | 0.2242 | 0.2564 | 0.1954 |
| D | | | | | | | | 0.1844 | 0.2283 | 0.2348 | 0.2221 | 0.206 |
| E | | | | | | | | 0.1043 | 0.1033 | 0.1209 | 0.1114 | 0.0773 |
| F | | | | | | | | 0.2049 | 0.1956 | 0.2319 | 0.2036 | 0.2161 |

| | | | | | | | | | | | |
|---|---|---|---|---|---|---|---|---|---|---|---|
| G | | | | | | | 0.0411 | 0.0671 | 0.0703 | 0.0695 | 0.0481 |
| H | | | | | | | 0.1555 | 0.1012 | 0.1839 | 0.1494 | 0.1012 |

**Cell Index at: 37:15:43**

| | 1 | 2 | 3 | 4 | 5 | 6 | 7 | 8 | 9 | 10 | 11 | 12 |
|---|---|---|---|---|---|---|---|---|---|---|---|---|
| A | | | | | | | | 0.0092 | 0.0257 | 0.0396 | 0.0293 | 0.0465 |
| B | | | | | | | | 0.0561 | 0.0529 | 0.0575 | 0.0621 | 0.081 |
| C | | | | | | | | 0.1908 | 0.1918 | 0.2191 | 0.2546 | 0.1946 |
| D | | | | | | | | 0.1825 | 0.2323 | 0.2291 | 0.2221 | 0.2019 |
| E | | | | | | | | 0.1013 | 0.1116 | 0.1194 | 0.1124 | 0.074 |
| F | | | | | | | | 0.2084 | 0.1922 | 0.2322 | 0.2029 | 0.2133 |
| G | | | | | | | | 0.0444 | 0.0635 | 0.0656 | 0.0714 | 0.0491 |
| H | | | | | | | | 0.1562 | 0.1026 | 0.1802 | 0.1521 | 0.1018 |

**Cell Index at: 37:30:43**

| | 1 | 2 | 3 | 4 | 5 | 6 | 7 | 8 | 9 | 10 | 11 | 12 |
|---|---|---|---|---|---|---|---|---|---|---|---|---|
| A | | | | | | | | 0.0064 | 0.0285 | 0.0416 | 0.0313 | 0.0461 |
| B | | | | | | | | 0.0553 | 0.055 | 0.0555 | 0.0617 | 0.0781 |
| C | | | | | | | | 0.1877 | 0.1946 | 0.2329 | 0.256 | 0.1876 |
| D | | | | | | | | 0.1791 | 0.2274 | 0.2276 | 0.2226 | 0.2056 |
| E | | | | | | | | 0.1007 | 0.1131 | 0.1122 | 0.1061 | 0.077 |
| F | | | | | | | | 0.2064 | 0.202 | 0.2327 | 0.2 | 0.2113 |
| G | | | | | | | | 0.0397 | 0.0613 | 0.063 | 0.0705 | 0.0521 |
| H | | | | | | | | 0.1589 | 0.1038 | 0.1784 | 0.1527 | 0.0991 |

**Cell Index at: 37:45:44**

| | 1 | 2 | 3 | 4 | 5 | 6 | 7 | 8 | 9 | 10 | 11 | 12 |
|---|---|---|---|---|---|---|---|---|---|---|---|---|
| A | | | | | | | | 0.0062 | 0.0279 | 0.0368 | 0.0284 | 0.0483 |
| B | | | | | | | | 0.0519 | 0.05 | 0.0544 | 0.065 | 0.0818 |
| C | | | | | | | | 0.1905 | 0.1876 | 0.2265 | 0.2584 | 0.1841 |
| D | | | | | | | | 0.185 | 0.2256 | 0.2341 | 0.225 | 0.2129 |
| E | | | | | | | | 0.099 | 0.1137 | 0.1126 | 0.1003 | 0.0712 |
| F | | | | | | | | 0.2083 | 0.1974 | 0.2297 | 0.208 | 0.2132 |
| G | | | | | | | | 0.0412 | 0.0645 | 0.0675 | 0.0733 | 0.049 |
| H | | | | | | | | 0.1575 | 0.1059 | 0.1825 | 0.1497 | 0.1031 |

**Cell Index at: 38:00:44**

| | 1 | 2 | 3 | 4 | 5 | 6 | 7 | 8 | 9 | 10 | 11 | 12 |
|---|---|---|---|---|---|---|---|---|---|---|---|---|
| A | | | | | | | | 0.0043 | 0.0266 | 0.0388 | 0.0308 | 0.0479 |
| B | | | | | | | | 0.0542 | 0.0529 | 0.0583 | 0.0646 | 0.0772 |
| C | | | | | | | | 0.1853 | 0.1863 | 0.2224 | 0.2606 | 0.1893 |
| D | | | | | | | | 0.184 | 0.2245 | 0.2305 | 0.2234 | 0.2062 |
| E | | | | | | | | 0.1101 | 0.1134 | 0.1161 | 0.1017 | 0.0684 |
| F | | | | | | | | 0.2057 | 0.198 | 0.2333 | 0.2009 | 0.2132 |
| G | | | | | | | | 0.0381 | 0.0642 | 0.0659 | 0.072 | 0.0481 |
| H | | | | | | | | 0.1594 | 0.1065 | 0.1864 | 0.1522 | 0.1004 |

**Cell Index at: 38:15:45**

| | 1 | 2 | 3 | 4 | 5 | 6 | 7 | 8 | 9 | 10 | 11 | 12 |
|---|---|---|---|---|---|---|---|---|---|---|---|---|
| A | | | | | | | | 0.0035 | 0.0259 | 0.0407 | 0.0265 | 0.0453 |
| B | | | | | | | | 0.0518 | 0.0497 | 0.0548 | 0.065 | 0.075 |
| C | | | | | | | | 0.1835 | 0.1914 | 0.2258 | 0.2592 | 0.1904 |
| D | | | | | | | | 0.1782 | 0.2182 | 0.2265 | 0.2298 | 0.2141 |
| E | | | | | | | | 0.1001 | 0.1115 | 0.1125 | 0.111 | 0.0751 |
| F | | | | | | | | 0.2163 | 0.1983 | 0.2265 | 0.2023 | 0.2108 |
| G | | | | | | | | 0.0413 | 0.0642 | 0.0672 | 0.0707 | 0.0469 |
| H | | | | | | | | 0.1592 | 0.1066 | 0.1809 | 0.1543 | 0.0985 |

**Cell Index at: 38:30:44**

| | 1 | 2 | 3 | 4 | 5 | 6 | 7 | 8 | 9 | 10 | 11 | 12 |
|---|---|---|---|---|---|---|---|---|---|---|---|---|
| A | | | | | | | | 0.002 | 0.0257 | 0.0356 | 0.0277 | 0.0451 |
| B | | | | | | | | 0.0503 | 0.052 | 0.0507 | 0.0653 | 0.0771 |
| C | | | | | | | | 0.1862 | 0.1868 | 0.2228 | 0.2581 | 0.1843 |
| D | | | | | | | | 0.1815 | 0.2212 | 0.2246 | 0.2234 | 0.2085 |
| E | | | | | | | | 0.1069 | 0.113 | 0.1097 | 0.104 | 0.077 |
| F | | | | | | | | 0.2085 | 0.1964 | 0.2391 | 0.1949 | 0.2097 |
| G | | | | | | | | 0.0434 | 0.0604 | 0.0693 | 0.0676 | 0.0493 |
| H | | | | | | | | 0.1595 | 0.1117 | 0.1805 | 0.1536 | 0.0991 |

**Cell Index at: 38:45:44**

| | 1 | 2 | 3 | 4 | 5 | 6 | 7 | 8 | 9 | 10 | 11 | 12 |
|---|---|---|---|---|---|---|---|---|---|---|---|---|
| A | | | | | | | | -0.0009 | 0.0212 | 0.0395 | 0.0264 | 0.0442 |
| B | | | | | | | | 0.0486 | 0.0505 | 0.0558 | 0.0651 | 0.0775 |
| C | | | | | | | | 0.187 | 0.1892 | 0.2216 | 0.2657 | 0.1865 |
| D | | | | | | | | 0.1821 | 0.2186 | 0.2331 | 0.2334 | 0.2094 |
| E | | | | | | | | 0.1001 | 0.1044 | 0.1101 | 0.1077 | 0.0721 |
| F | | | | | | | | 0.2159 | 0.2011 | 0.236 | 0.2041 | 0.2061 |
| G | | | | | | | | 0.0398 | 0.0617 | 0.063 | 0.0706 | 0.0515 |
| H | | | | | | | | 0.1541 | 0.1095 | 0.1829 | 0.1543 | 0.1011 |

**Cell Index at: 39:00:45**

| | 1 | 2 | 3 | 4 | 5 | 6 | 7 | 8 | 9 | 10 | 11 | 12 |
|---|---|---|---|---|---|---|---|---|---|---|---|---|
| A | | | | | | | | 0.001 | 0.0232 | 0.0369 | 0.0268 | 0.0486 |
| B | | | | | | | | 0.0493 | 0.0447 | 0.0546 | 0.0692 | 0.0763 |
| C | | | | | | | | 0.1814 | 0.1915 | 0.2199 | 0.2573 | 0.1907 |
| D | | | | | | | | 0.1861 | 0.2166 | 0.2312 | 0.2291 | 0.2095 |
| E | | | | | | | | 0.1053 | 0.109 | 0.1035 | 0.1106 | 0.0775 |
| F | | | | | | | | 0.2126 | 0.1968 | 0.2334 | 0.2066 | 0.2095 |
| G | | | | | | | | 0.0357 | 0.0637 | 0.0617 | 0.0737 | 0.05 |
| H | | | | | | | | 0.1618 | 0.1094 | 0.185 | 0.153 | 0.0979 |

**Cell Index at: 39:15:45**

| | 1 | 2 | 3 | 4 | 5 | 6 | 7 | 8 | 9 | 10 | 11 | 12 |
|---|---|---|---|---|---|---|---|---|---|---|---|---|
| A | | | | | | | | -0.0019 | 0.0236 | 0.0363 | 0.0221 | 0.0458 |
| B | | | | | | | | 0.0501 | 0.0452 | 0.0556 | 0.0646 | 0.0785 |
| C | | | | | | | | 0.1791 | 0.1847 | 0.227 | 0.2654 | 0.192 |
| D | | | | | | | | 0.1858 | 0.2241 | 0.2332 | 0.2266 | 0.2099 |
| E | | | | | | | | 0.1011 | 0.1031 | 0.1066 | 0.1089 | 0.0757 |
| F | | | | | | | | 0.2068 | 0.1986 | 0.2393 | 0.202 | 0.2145 |
| G | | | | | | | | 0.0398 | 0.0584 | 0.0572 | 0.071 | 0.0473 |
| H | | | | | | | | 0.1588 | 0.1082 | 0.1824 | 0.1532 | 0.0973 |

**Cell Index at: 39:30:46**

| | 1 | 2 | 3 | 4 | 5 | 6 | 7 | 8 | 9 | 10 | 11 | 12 |
|---|---|---|---|---|---|---|---|---|---|---|---|---|
| A | | | | | | | | -0.0023 | 0.0201 | 0.0384 | 0.0219 | 0.0451 |
| B | | | | | | | | 0.0466 | 0.0467 | 0.0579 | 0.0648 | 0.0775 |
| C | | | | | | | | 0.1846 | 0.1844 | 0.225 | 0.2615 | 0.1875 |
| D | | | | | | | | 0.1859 | 0.2252 | 0.2294 | 0.2273 | 0.2067 |
| E | | | | | | | | 0.1002 | 0.11 | 0.1144 | 0.1106 | 0.0801 |
| F | | | | | | | | 0.2063 | 0.1982 | 0.2419 | 0.2039 | 0.2121 |
| G | | | | | | | | 0.0386 | 0.0584 | 0.0659 | 0.0743 | 0.0469 |
| H | | | | | | | | 0.1617 | 0.1079 | 0.1884 | 0.1549 | 0.0961 |

**Cell Index at: 39:45:47**

| | 1 | 2 | 3 | 4 | 5 | 6 | 7 | 8 | 9 | 10 | 11 | 12 |
|---|---|---|---|---|---|---|---|---|---|---|---|---|
| A | | | | | | | | 0.0004 | 0.0189 | 0.0369 | 0.0233 | 0.045 |
| B | | | | | | | | 0.0496 | 0.0444 | 0.055 | 0.0708 | 0.084 |
| C | | | | | | | | 0.182 | 0.1881 | 0.2281 | 0.2606 | 0.1896 |
| D | | | | | | | | 0.1871 | 0.2305 | 0.23 | 0.2285 | 0.2089 |
| E | | | | | | | | 0.0955 | 0.1039 | 0.112 | 0.111 | 0.0772 |
| F | | | | | | | | 0.2049 | 0.2043 | 0.2337 | 0.203 | 0.2064 |
| G | | | | | | | | 0.0381 | 0.0611 | 0.0618 | 0.0693 | 0.0489 |
| H | | | | | | | | 0.1586 | 0.1077 | 0.185 | 0.1548 | 0.1008 |

**Cell Index at: 40:00:48**

| | 1 | 2 | 3 | 4 | 5 | 6 | 7 | 8 | 9 | 10 | 11 | 12 |
|---|---|---|---|---|---|---|---|---|---|---|---|---|
| A | | | | | | | | -0.0002 | 0.0194 | 0.0395 | 0.0214 | 0.0415 |
| B | | | | | | | | 0.0516 | 0.0441 | 0.0519 | 0.066 | 0.0798 |
| C | | | | | | | | 0.182 | 0.1761 | 0.2264 | 0.2604 | 0.1943 |
| D | | | | | | | | 0.1889 | 0.2301 | 0.2332 | 0.2317 | 0.2056 |
| E | | | | | | | | 0.1049 | 0.106 | 0.1037 | 0.1085 | 0.0776 |

| | | | | | | | | | | | |
|---|---|---|---|---|---|---|---|---|---|---|---|
| F | | | | | | | 0.2107 | 0.2 | 0.2367 | 0.2012 | 0.2099 |
| G | | | | | | | 0.0373 | 0.0591 | 0.0713 | 0.074 | 0.0496 |
| H | | | | | | | 0.1601 | 0.109 | 0.1869 | 0.1569 | 0.099 |

**Cell Index at: 40:15:48**

| | 1 | 2 | 3 | 4 | 5 | 6 | 7 | 8 | 9 | 10 | 11 | 12 |
|---|---|---|---|---|---|---|---|---|---|---|---|---|
| A | | | | | | | | 0.0003 | 0.0243 | 0.0381 | 0.0209 | 0.0401 |
| B | | | | | | | | 0.0493 | 0.0492 | 0.0484 | 0.0678 | 0.0824 |
| C | | | | | | | | 0.1778 | 0.1829 | 0.2254 | 0.2606 | 0.191 |
| D | | | | | | | | 0.1882 | 0.2271 | 0.2364 | 0.229 | 0.2078 |
| E | | | | | | | | 0.1017 | 0.1034 | 0.0994 | 0.1055 | 0.0768 |
| F | | | | | | | | 0.2122 | 0.2006 | 0.235 | 0.2019 | 0.2122 |
| G | | | | | | | | 0.0388 | 0.0624 | 0.062 | 0.0702 | 0.0481 |
| H | | | | | | | | 0.1608 | 0.1113 | 0.1846 | 0.1567 | 0.1028 |

**Cell Index at: 40:30:48**

| | 1 | 2 | 3 | 4 | 5 | 6 | 7 | 8 | 9 | 10 | 11 | 12 |
|---|---|---|---|---|---|---|---|---|---|---|---|---|
| A | | | | | | | | -0.0003 | 0.024 | 0.0395 | 0.0199 | 0.0404 |
| B | | | | | | | | 0.0484 | 0.0446 | 0.0494 | 0.0689 | 0.0782 |
| C | | | | | | | | 0.1845 | 0.189 | 0.2283 | 0.2647 | 0.1952 |
| D | | | | | | | | 0.1916 | 0.2297 | 0.2335 | 0.2308 | 0.2042 |
| E | | | | | | | | 0.1023 | 0.1028 | 0.1062 | 0.117 | 0.0836 |
| F | | | | | | | | 0.2105 | 0.205 | 0.2316 | 0.2041 | 0.2114 |
| G | | | | | | | | 0.0384 | 0.0615 | 0.0665 | 0.0689 | 0.0472 |
| H | | | | | | | | 0.1663 | 0.1115 | 0.1845 | 0.1595 | 0.0999 |

**Cell Index at: 40:45:49**

| | 1 | 2 | 3 | 4 | 5 | 6 | 7 | 8 | 9 | 10 | 11 | 12 |
|---|---|---|---|---|---|---|---|---|---|---|---|---|
| A | | | | | | | | -0.0006 | 0.0233 | 0.0413 | 0.0186 | 0.0396 |
| B | | | | | | | | 0.0467 | 0.0475 | 0.0541 | 0.0695 | 0.0762 |
| C | | | | | | | | 0.1829 | 0.1897 | 0.2276 | 0.263 | 0.1974 |
| D | | | | | | | | 0.1862 | 0.2259 | 0.2409 | 0.2354 | 0.2054 |
| E | | | | | | | | 0.0952 | 0.0955 | 0.1027 | 0.1062 | 0.0753 |
| F | | | | | | | | 0.2095 | 0.1994 | 0.2341 | 0.2024 | 0.2143 |
| G | | | | | | | | 0.0382 | 0.0624 | 0.0595 | 0.0685 | 0.0485 |
| H | | | | | | | | 0.1654 | 0.1111 | 0.1847 | 0.1484 | 0.1003 |

**Cell Index at: 41:00:48**

| | 1 | 2 | 3 | 4 | 5 | 6 | 7 | 8 | 9 | 10 | 11 | 12 |
|---|---|---|---|---|---|---|---|---|---|---|---|---|
| A | | | | | | | | -0.0031 | 0.0244 | 0.0358 | 0.0199 | 0.0377 |
| B | | | | | | | | 0.0479 | 0.0461 | 0.0546 | 0.0664 | 0.0781 |
| C | | | | | | | | 0.1873 | 0.1893 | 0.231 | 0.2673 | 0.1909 |
| D | | | | | | | | 0.1835 | 0.2255 | 0.2386 | 0.2326 | 0.2119 |
| E | | | | | | | | 0.0938 | 0.1027 | 0.107 | 0.1091 | 0.081 |
| F | | | | | | | | 0.2145 | 0.1966 | 0.2372 | 0.2009 | 0.2139 |
| G | | | | | | | | 0.0392 | 0.0608 | 0.0644 | 0.0692 | 0.0442 |
| H | | | | | | | | 0.1637 | 0.112 | 0.184 | 0.1503 | 0.1034 |

**Cell Index at: 41:15:47**

| | 1 | 2 | 3 | 4 | 5 | 6 | 7 | 8 | 9 | 10 | 11 | 12 |
|---|---|---|---|---|---|---|---|---|---|---|---|---|
| A | | | | | | | | -0.0031 | 0.0243 | 0.0376 | 0.0169 | 0.036 |
| B | | | | | | | | 0.0489 | 0.0488 | 0.0517 | 0.0693 | 0.0771 |
| C | | | | | | | | 0.1837 | 0.1867 | 0.2291 | 0.2676 | 0.1887 |
| D | | | | | | | | 0.1926 | 0.232 | 0.2268 | 0.2291 | 0.2106 |
| E | | | | | | | | 0.0973 | 0.1064 | 0.1094 | 0.1102 | 0.0783 |
| F | | | | | | | | 0.2064 | 0.2044 | 0.2374 | 0.2126 | 0.2168 |
| G | | | | | | | | 0.0391 | 0.0608 | 0.0676 | 0.066 | 0.0434 |
| H | | | | | | | | 0.1653 | 0.1129 | 0.1846 | 0.1468 | 0.1018 |

**Cell Index at: 41:30:47**

| | 1 | 2 | 3 | 4 | 5 | 6 | 7 | 8 | 9 | 10 | 11 | 12 |
|---|---|---|---|---|---|---|---|---|---|---|---|---|
| A | | | | | | | | -0.0006 | 0.0246 | 0.0367 | 0.0195 | 0.0357 |
| B | | | | | | | | 0.0438 | 0.0477 | 0.0492 | 0.0665 | 0.081 |
| C | | | | | | | | 0.1799 | 0.1866 | 0.2236 | 0.2648 | 0.1973 |
| D | | | | | | | | 0.1838 | 0.2302 | 0.2274 | 0.2294 | 0.2087 |
| E | | | | | | | | 0.0974 | 0.1068 | 0.1061 | 0.1076 | 0.0796 |
| F | | | | | | | | 0.2069 | 0.1995 | 0.2325 | 0.2048 | 0.2134 |
| G | | | | | | | | 0.0359 | 0.059 | 0.0597 | 0.0632 | 0.0429 |
| H | | | | | | | | 0.1644 | 0.1134 | 0.1852 | 0.1484 | 0.1043 |

**Cell Index at: 41:45:48**

| | 1 | 2 | 3 | 4 | 5 | 6 | 7 | 8 | 9 | 10 | 11 | 12 |
|---|---|---|---|---|---|---|---|---|---|---|---|---|
| A | | | | | | | | -0.0046 | 0.0244 | 0.037 | 0.0207 | 0.0339 |
| B | | | | | | | | 0.0466 | 0.0455 | 0.0494 | 0.0654 | 0.0792 |
| C | | | | | | | | 0.1772 | 0.1856 | 0.2261 | 0.2636 | 0.2017 |
| D | | | | | | | | 0.1822 | 0.2284 | 0.2304 | 0.227 | 0.2088 |
| E | | | | | | | | 0.0924 | 0.0941 | 0.106 | 0.1062 | 0.0715 |
| F | | | | | | | | 0.2092 | 0.2028 | 0.2318 | 0.2073 | 0.221 |
| G | | | | | | | | 0.0374 | 0.0598 | 0.0626 | 0.0624 | 0.0461 |
| H | | | | | | | | 0.1649 | 0.1117 | 0.1834 | 0.148 | 0.1019 |

**Cell Index at: 42:00:48**

| | 1 | 2 | 3 | 4 | 5 | 6 | 7 | 8 | 9 | 10 | 11 | 12 |
|---|---|---|---|---|---|---|---|---|---|---|---|---|
| A | | | | | | | | -0.0046 | 0.0217 | 0.0367 | 0.0237 | 0.0366 |
| B | | | | | | | | 0.0478 | 0.0454 | 0.0507 | 0.0672 | 0.0736 |
| C | | | | | | | | 0.1865 | 0.1803 | 0.227 | 0.2654 | 0.2021 |
| D | | | | | | | | 0.1934 | 0.2278 | 0.2275 | 0.2269 | 0.2087 |
| E | | | | | | | | 0.1001 | 0.0977 | 0.1105 | 0.1077 | 0.0804 |
| F | | | | | | | | 0.2105 | 0.2005 | 0.2312 | 0.2038 | 0.218 |
| G | | | | | | | | 0.0357 | 0.0612 | 0.0628 | 0.0627 | 0.0468 |
| H | | | | | | | | 0.166 | 0.1134 | 0.187 | 0.1504 | 0.1038 |

**Cell Index at: 42:15:48**

| | 1 | 2 | 3 | 4 | 5 | 6 | 7 | 8 | 9 | 10 | 11 | 12 |
|---|---|---|---|---|---|---|---|---|---|---|---|---|
| A | | | | | | | | -0.0043 | 0.0246 | 0.0399 | 0.0219 | 0.037 |
| B | | | | | | | | 0.0435 | 0.0475 | 0.049 | 0.0643 | 0.0796 |
| C | | | | | | | | 0.1866 | 0.1817 | 0.2299 | 0.267 | 0.203 |
| D | | | | | | | | 0.1897 | 0.2338 | 0.2306 | 0.2292 | 0.212 |
| E | | | | | | | | 0.0968 | 0.1 | 0.1082 | 0.1074 | 0.0822 |
| F | | | | | | | | 0.2093 | 0.1949 | 0.2334 | 0.2066 | 0.2192 |
| G | | | | | | | | 0.0343 | 0.0573 | 0.0586 | 0.0629 | 0.0456 |
| H | | | | | | | | 0.1656 | 0.1137 | 0.1863 | 0.1481 | 0.1024 |

**Cell Index at: 42:30:47**

| | 1 | 2 | 3 | 4 | 5 | 6 | 7 | 8 | 9 | 10 | 11 | 12 |
|---|---|---|---|---|---|---|---|---|---|---|---|---|
| A | | | | | | | | -0.006 | 0.0257 | 0.0353 | 0.0199 | 0.04 |
| B | | | | | | | | 0.0452 | 0.0413 | 0.0484 | 0.0696 | 0.078 |
| C | | | | | | | | 0.183 | 0.1845 | 0.2268 | 0.2654 | 0.2011 |
| D | | | | | | | | 0.1859 | 0.2367 | 0.237 | 0.2329 | 0.208 |
| E | | | | | | | | 0.1027 | 0.0947 | 0.1046 | 0.1107 | 0.0806 |
| F | | | | | | | | 0.2055 | 0.2037 | 0.2316 | 0.202 | 0.2209 |
| G | | | | | | | | 0.0394 | 0.0617 | 0.0587 | 0.0677 | 0.0471 |
| H | | | | | | | | 0.1653 | 0.115 | 0.1893 | 0.1488 | 0.1 |

**Cell Index at: 42:45:47**

| | 1 | 2 | 3 | 4 | 5 | 6 | 7 | 8 | 9 | 10 | 11 | 12 |
|---|---|---|---|---|---|---|---|---|---|---|---|---|
| A | | | | | | | | -0.0041 | 0.0228 | 0.0356 | 0.0155 | 0.038 |
| B | | | | | | | | 0.0427 | 0.046 | 0.0489 | 0.0656 | 0.0773 |
| C | | | | | | | | 0.1816 | 0.1804 | 0.2287 | 0.2662 | 0.1988 |
| D | | | | | | | | 0.1865 | 0.236 | 0.2346 | 0.228 | 0.2101 |
| E | | | | | | | | 0.0951 | 0.0989 | 0.1134 | 0.1099 | 0.0768 |
| F | | | | | | | | 0.2022 | 0.2004 | 0.2292 | 0.1974 | 0.2173 |
| G | | | | | | | | 0.0381 | 0.0613 | 0.0582 | 0.0655 | 0.043 |
| H | | | | | | | | 0.1624 | 0.1155 | 0.1883 | 0.1518 | 0.0988 |

**Cell Index at: 43:00:48**

| | 1 | 2 | 3 | 4 | 5 | 6 | 7 | 8 | 9 | 10 | 11 | 12 |
|---|---|---|---|---|---|---|---|---|---|---|---|---|
| A | | | | | | | | -0.0051 | 0.0261 | 0.0358 | 0.0203 | 0.0405 |
| B | | | | | | | | 0.0455 | 0.0499 | 0.0498 | 0.0684 | 0.0779 |
| C | | | | | | | | 0.1889 | 0.1874 | 0.2275 | 0.2671 | 0.2001 |
| D | | | | | | | | 0.1828 | 0.2325 | 0.2374 | 0.2331 | 0.2097 |

| | 1 | 2 | 3 | 4 | 5 | 6 | 7 | 8 | 9 | 10 | 11 | 12 |
|---|---|---|---|---|---|---|---|---|---|---|---|---|
| E | | | | | | | | 0.1008 | 0.0965 | 0.1153 | 0.1096 | 0.0807 |
| F | | | | | | | | 0.2097 | 0.2043 | 0.2342 | 0.202 | 0.2186 |
| G | | | | | | | | 0.0346 | 0.058 | 0.0547 | 0.0698 | 0.049 |
| H | | | | | | | | 0.1633 | 0.1144 | 0.1861 | 0.1532 | 0.1008 |

**Cell Index at: 43:15:48**

| | 1 | 2 | 3 | 4 | 5 | 6 | 7 | 8 | 9 | 10 | 11 | 12 |
|---|---|---|---|---|---|---|---|---|---|---|---|---|
| A | | | | | | | | -0.0053 | 0.0257 | 0.0345 | 0.0184 | 0.0389 |
| B | | | | | | | | 0.0454 | 0.0516 | 0.0514 | 0.0664 | 0.0834 |
| C | | | | | | | | 0.1887 | 0.1894 | 0.2255 | 0.2645 | 0.2007 |
| D | | | | | | | | 0.1839 | 0.2231 | 0.2339 | 0.2276 | 0.2124 |
| E | | | | | | | | 0.0991 | 0.0985 | 0.1179 | 0.1095 | 0.0834 |
| F | | | | | | | | 0.2125 | 0.1982 | 0.2322 | 0.2014 | 0.221 |
| G | | | | | | | | 0.0395 | 0.0589 | 0.0609 | 0.0685 | 0.0466 |
| H | | | | | | | | 0.1642 | 0.113 | 0.1833 | 0.1517 | 0.1 |

**Cell Index at: 43:30:49**

| | 1 | 2 | 3 | 4 | 5 | 6 | 7 | 8 | 9 | 10 | 11 | 12 |
|---|---|---|---|---|---|---|---|---|---|---|---|---|
| A | | | | | | | | -0.0049 | 0.023 | 0.034 | 0.0217 | 0.0399 |
| B | | | | | | | | 0.0453 | 0.0522 | 0.0446 | 0.0657 | 0.0792 |
| C | | | | | | | | 0.1899 | 0.178 | 0.2315 | 0.2614 | 0.206 |
| D | | | | | | | | 0.1797 | 0.228 | 0.2407 | 0.2382 | 0.2033 |
| E | | | | | | | | 0.0957 | 0.0947 | 0.1171 | 0.1016 | 0.0744 |
| F | | | | | | | | 0.2164 | 0.1944 | 0.2348 | 0.2012 | 0.2204 |
| G | | | | | | | | 0.0432 | 0.0603 | 0.0594 | 0.0638 | 0.0463 |
| H | | | | | | | | 0.1637 | 0.1169 | 0.1848 | 0.1505 | 0.0989 |

**Cell Index at: 43:45:49**

| | 1 | 2 | 3 | 4 | 5 | 6 | 7 | 8 | 9 | 10 | 11 | 12 |
|---|---|---|---|---|---|---|---|---|---|---|---|---|
| A | | | | | | | | -0.0056 | 0.0288 | 0.0302 | 0.0236 | 0.0376 |
| B | | | | | | | | 0.0496 | 0.044 | 0.0551 | 0.0642 | 0.0787 |
| C | | | | | | | | 0.1885 | 0.1873 | 0.2301 | 0.2668 | 0.2034 |
| D | | | | | | | | 0.1715 | 0.2292 | 0.2286 | 0.2362 | 0.2021 |
| E | | | | | | | | 0.0982 | 0.0964 | 0.1157 | 0.1099 | 0.086 |
| F | | | | | | | | 0.2113 | 0.1996 | 0.2334 | 0.2066 | 0.218 |
| G | | | | | | | | 0.0395 | 0.0607 | 0.0597 | 0.0676 | 0.0458 |
| H | | | | | | | | 0.1604 | 0.1173 | 0.1868 | 0.1487 | 0.0999 |

**Cell Index at: 44:00:50**

| | 1 | 2 | 3 | 4 | 5 | 6 | 7 | 8 | 9 | 10 | 11 | 12 |
|---|---|---|---|---|---|---|---|---|---|---|---|---|
| A | | | | | | | | -0.0059 | 0.0254 | 0.0345 | 0.0216 | 0.0385 |
| B | | | | | | | | 0.0447 | 0.0479 | 0.052 | 0.068 | 0.0837 |
| C | | | | | | | | 0.19 | 0.18 | 0.2279 | 0.2708 | 0.2069 |
| D | | | | | | | | 0.1818 | 0.2343 | 0.2365 | 0.2255 | 0.2088 |
| E | | | | | | | | 0.0969 | 0.0981 | 0.1174 | 0.1132 | 0.0817 |
| F | | | | | | | | 0.2195 | 0.1995 | 0.2328 | 0.2055 | 0.2184 |
| G | | | | | | | | 0.0389 | 0.0623 | 0.0612 | 0.0687 | 0.0477 |
| H | | | | | | | | 0.1605 | 0.1146 | 0.1872 | 0.1473 | 0.0993 |

**Cell Index at: 44:15:50**

| | 1 | 2 | 3 | 4 | 5 | 6 | 7 | 8 | 9 | 10 | 11 | 12 |
|---|---|---|---|---|---|---|---|---|---|---|---|---|
| A | | | | | | | | -0.0031 | 0.0268 | 0.0363 | 0.0178 | 0.0393 |
| B | | | | | | | | 0.0449 | 0.0445 | 0.0506 | 0.0703 | 0.0835 |
| C | | | | | | | | 0.1871 | 0.1789 | 0.2302 | 0.2725 | 0.2008 |
| D | | | | | | | | 0.1879 | 0.2307 | 0.2276 | 0.228 | 0.2048 |
| E | | | | | | | | 0.0997 | 0.0976 | 0.1125 | 0.11 | 0.0777 |
| F | | | | | | | | 0.2167 | 0.1996 | 0.234 | 0.2007 | 0.2182 |
| G | | | | | | | | 0.0407 | 0.0586 | 0.0623 | 0.0654 | 0.0451 |
| H | | | | | | | | 0.165 | 0.1179 | 0.1886 | 0.1478 | 0.0982 |

**Cell Index at: 44:30:50**

| | 1 | 2 | 3 | 4 | 5 | 6 | 7 | 8 | 9 | 10 | 11 | 12 |
|---|---|---|---|---|---|---|---|---|---|---|---|---|
| A | | | | | | | | -0.0035 | 0.0271 | 0.0347 | 0.0219 | 0.0397 |
| B | | | | | | | | 0.0464 | 0.045 | 0.0519 | 0.0677 | 0.0787 |
| C | | | | | | | | 0.1912 | 0.1826 | 0.2328 | 0.2685 | 0.2029 |
| D | | | | | | | | 0.1838 | 0.23 | 0.232 | 0.2292 | 0.2069 |
| E | | | | | | | | 0.0994 | 0.0977 | 0.1116 | 0.1173 | 0.078 |
| F | | | | | | | | 0.219 | 0.2071 | 0.2345 | 0.2045 | 0.2169 |
| G | | | | | | | | 0.0377 | 0.0585 | 0.0627 | 0.0693 | 0.0549 |
| H | | | | | | | | 0.1643 | 0.1181 | 0.1883 | 0.1465 | 0.1004 |

**Cell Index at: 44:45:51**

| | 1 | 2 | 3 | 4 | 5 | 6 | 7 | 8 | 9 | 10 | 11 | 12 |
|---|---|---|---|---|---|---|---|---|---|---|---|---|
| A | | | | | | | | -0.0032 | 0.0276 | 0.0415 | 0.0209 | 0.0409 |
| B | | | | | | | | 0.0423 | 0.0472 | 0.0477 | 0.0641 | 0.0808 |
| C | | | | | | | | 0.1953 | 0.1859 | 0.2346 | 0.2646 | 0.2047 |
| D | | | | | | | | 0.1869 | 0.2312 | 0.2304 | 0.2299 | 0.2068 |
| E | | | | | | | | 0.0947 | 0.103 | 0.1208 | 0.1219 | 0.0772 |
| F | | | | | | | | 0.221 | 0.1946 | 0.2269 | 0.2071 | 0.2144 |
| G | | | | | | | | 0.0389 | 0.0618 | 0.0601 | 0.0683 | 0.0494 |
| H | | | | | | | | 0.1608 | 0.1231 | 0.1865 | 0.1477 | 0.0993 |

**Cell Index at: 45:00:51**

| | 1 | 2 | 3 | 4 | 5 | 6 | 7 | 8 | 9 | 10 | 11 | 12 |
|---|---|---|---|---|---|---|---|---|---|---|---|---|
| A | | | | | | | | -0.0038 | 0.0258 | 0.0377 | 0.0186 | 0.0389 |
| B | | | | | | | | 0.0436 | 0.0457 | 0.0478 | 0.0644 | 0.0803 |
| C | | | | | | | | 0.194 | 0.1802 | 0.2302 | 0.2686 | 0.2032 |
| D | | | | | | | | 0.179 | 0.232 | 0.2288 | 0.2257 | 0.2093 |
| E | | | | | | | | 0.1084 | 0.0993 | 0.1174 | 0.1129 | 0.073 |
| F | | | | | | | | 0.2172 | 0.2015 | 0.2366 | 0.2063 | 0.2198 |
| G | | | | | | | | 0.0401 | 0.0542 | 0.0643 | 0.0697 | 0.0512 |
| H | | | | | | | | 0.1611 | 0.1218 | 0.1871 | 0.1479 | 0.0988 |

**Cell Index at: 45:15:52**

| | 1 | 2 | 3 | 4 | 5 | 6 | 7 | 8 | 9 | 10 | 11 | 12 |
|---|---|---|---|---|---|---|---|---|---|---|---|---|
| A | | | | | | | | -0.0047 | 0.0252 | 0.0414 | 0.0181 | 0.0363 |
| B | | | | | | | | 0.0411 | 0.0445 | 0.0485 | 0.0646 | 0.0798 |
| C | | | | | | | | 0.1945 | 0.1881 | 0.2339 | 0.2665 | 0.1988 |
| D | | | | | | | | 0.1779 | 0.2365 | 0.2265 | 0.2267 | 0.2112 |
| E | | | | | | | | 0.1067 | 0.0931 | 0.1152 | 0.115 | 0.0776 |
| F | | | | | | | | 0.2219 | 0.2029 | 0.2382 | 0.206 | 0.2185 |
| G | | | | | | | | 0.0345 | 0.0603 | 0.062 | 0.0749 | 0.0502 |
| H | | | | | | | | 0.164 | 0.1187 | 0.1859 | 0.1472 | 0.0974 |

**Cell Index at: 45:30:53**

| | 1 | 2 | 3 | 4 | 5 | 6 | 7 | 8 | 9 | 10 | 11 | 12 |
|---|---|---|---|---|---|---|---|---|---|---|---|---|
| A | | | | | | | | -0.0052 | 0.0233 | 0.0382 | 0.0207 | 0.0361 |
| B | | | | | | | | 0.0395 | 0.0483 | 0.0489 | 0.0674 | 0.0817 |
| C | | | | | | | | 0.1885 | 0.1892 | 0.2316 | 0.2662 | 0.2025 |
| D | | | | | | | | 0.1856 | 0.2412 | 0.2295 | 0.2278 | 0.2142 |
| E | | | | | | | | 0.1009 | 0.0979 | 0.1062 | 0.1125 | 0.0777 |
| F | | | | | | | | 0.2187 | 0.1994 | 0.2416 | 0.2042 | 0.217 |
| G | | | | | | | | 0.0405 | 0.0625 | 0.0635 | 0.0687 | 0.0487 |
| H | | | | | | | | 0.1646 | 0.1216 | 0.1885 | 0.1488 | 0.0972 |

**Cell Index at: 45:45:54**

| | 1 | 2 | 3 | 4 | 5 | 6 | 7 | 8 | 9 | 10 | 11 | 12 |
|---|---|---|---|---|---|---|---|---|---|---|---|---|
| A | | | | | | | | -0.0061 | 0.0249 | 0.0396 | 0.0178 | 0.0412 |
| B | | | | | | | | 0.0398 | 0.0439 | 0.047 | 0.0649 | 0.0767 |
| C | | | | | | | | 0.1922 | 0.1825 | 0.229 | 0.264 | 0.2015 |
| D | | | | | | | | 0.1802 | 0.2389 | 0.2299 | 0.2188 | 0.2127 |
| E | | | | | | | | 0.1005 | 0.0936 | 0.1153 | 0.1155 | 0.0757 |
| F | | | | | | | | 0.2179 | 0.2044 | 0.2383 | 0.206 | 0.2168 |
| G | | | | | | | | 0.0382 | 0.0633 | 0.0663 | 0.0746 | 0.0511 |
| H | | | | | | | | 0.1644 | 0.1215 | 0.1869 | 0.1455 | 0.0982 |

**Cell Index at: 46:00:54**

| | 1 | 2 | 3 | 4 | 5 | 6 | 7 | 8 | 9 | 10 | 11 | 12 |
|---|---|---|---|---|---|---|---|---|---|---|---|---|
| A | | | | | | | | -0.0091 | 0.0233 | 0.042 | 0.0073 | 0.0363 |
| B | | | | | | | | 0.0393 | 0.0422 | 0.0486 | 0.0667 | 0.0813 |
| C | | | | | | | | 0.189 | 0.1822 | 0.2261 | 0.266 | 0.1967 |

| | 1 | 2 | 3 | 4 | 5 | 6 | 7 | 8 | 9 | 10 | 11 | 12 |
|---|---|---|---|---|---|---|---|---|---|---|---|---|
| D | | | | | | | | 0.1834 | 0.2394 | 0.2311 | 0.2244 | 0.2103 |
| E | | | | | | | | 0.102 | 0.0963 | 0.1165 | 0.1103 | 0.0829 |
| F | | | | | | | | 0.2242 | 0.1986 | 0.2417 | 0.2001 | 0.213 |
| G | | | | | | | | 0.0359 | 0.0651 | 0.0685 | 0.0701 | 0.0512 |
| H | | | | | | | | 0.1624 | 0.1212 | 0.1885 | 0.1458 | 0.093 |

**Cell Index at: 46:15:51**

| | 1 | 2 | 3 | 4 | 5 | 6 | 7 | 8 | 9 | 10 | 11 | 12 |
|---|---|---|---|---|---|---|---|---|---|---|---|---|
| A | | | | | | | | -0.0073 | 0.0231 | 0.0396 | 0.0087 | 0.0388 |
| B | | | | | | | | 0.0372 | 0.0432 | 0.048 | 0.0649 | 0.0784 |
| C | | | | | | | | 0.1968 | 0.1855 | 0.226 | 0.2621 | 0.2017 |
| D | | | | | | | | 0.1912 | 0.2369 | 0.2309 | 0.228 | 0.2109 |
| E | | | | | | | | 0.1055 | 0.0966 | 0.1127 | 0.1114 | 0.0822 |
| F | | | | | | | | 0.2191 | 0.1991 | 0.2432 | 0.2002 | 0.2159 |
| G | | | | | | | | 0.0338 | 0.0617 | 0.0606 | 0.0714 | 0.0524 |
| H | | | | | | | | 0.1656 | 0.122 | 0.1894 | 0.1444 | 0.0942 |

**Cell Index at: 46:30:51**

| | 1 | 2 | 3 | 4 | 5 | 6 | 7 | 8 | 9 | 10 | 11 | 12 |
|---|---|---|---|---|---|---|---|---|---|---|---|---|
| A | | | | | | | | -0.0077 | 0.0219 | 0.0453 | 0.0097 | 0.0386 |
| B | | | | | | | | 0.0385 | 0.045 | 0.0476 | 0.0683 | 0.0791 |
| C | | | | | | | | 0.1933 | 0.1916 | 0.232 | 0.2698 | 0.1996 |
| D | | | | | | | | 0.1836 | 0.2322 | 0.2209 | 0.2291 | 0.204 |
| E | | | | | | | | 0.0976 | 0.0963 | 0.1122 | 0.1075 | 0.0779 |
| F | | | | | | | | 0.2102 | 0.2031 | 0.2411 | 0.2013 | 0.2117 |
| G | | | | | | | | 0.0368 | 0.0579 | 0.065 | 0.0754 | 0.0488 |
| H | | | | | | | | 0.1653 | 0.1215 | 0.1879 | 0.1449 | 0.0971 |

**Cell Index at: 46:45:52**

| | 1 | 2 | 3 | 4 | 5 | 6 | 7 | 8 | 9 | 10 | 11 | 12 |
|---|---|---|---|---|---|---|---|---|---|---|---|---|
| A | | | | | | | | -0.0075 | 0.0217 | 0.0429 | 0.0122 | 0.0394 |
| B | | | | | | | | 0.0375 | 0.0426 | 0.0471 | 0.0682 | 0.0826 |
| C | | | | | | | | 0.192 | 0.1867 | 0.2356 | 0.2601 | 0.2003 |
| D | | | | | | | | 0.189 | 0.2356 | 0.2282 | 0.2277 | 0.2066 |
| E | | | | | | | | 0.1048 | 0.0969 | 0.1093 | 0.1091 | 0.0856 |
| F | | | | | | | | 0.2113 | 0.2026 | 0.2419 | 0.2025 | 0.2089 |
| G | | | | | | | | 0.0337 | 0.0653 | 0.0642 | 0.0733 | 0.0477 |
| H | | | | | | | | 0.1658 | 0.1209 | 0.1886 | 0.1438 | 0.0968 |

**Cell Index at: 47:00:53**

| | 1 | 2 | 3 | 4 | 5 | 6 | 7 | 8 | 9 | 10 | 11 | 12 |
|---|---|---|---|---|---|---|---|---|---|---|---|---|
| A | | | | | | | | -0.0072 | 0.0148 | 0.0419 | 0.0092 | 0.0353 |
| B | | | | | | | | 0.0375 | 0.0409 | 0.0477 | 0.068 | 0.0788 |
| C | | | | | | | | 0.1957 | 0.1907 | 0.2309 | 0.2665 | 0.1952 |
| D | | | | | | | | 0.187 | 0.2397 | 0.2261 | 0.2221 | 0.2094 |
| E | | | | | | | | 0.0996 | 0.1014 | 0.113 | 0.1072 | 0.0717 |
| F | | | | | | | | 0.2136 | 0.2011 | 0.243 | 0.1989 | 0.2121 |
| G | | | | | | | | 0.0358 | 0.0641 | 0.07 | 0.0753 | 0.0489 |
| H | | | | | | | | 0.1673 | 0.1196 | 0.1888 | 0.1435 | 0.0974 |

**Cell Index at: 47:15:53**

| | 1 | 2 | 3 | 4 | 5 | 6 | 7 | 8 | 9 | 10 | 11 | 12 |
|---|---|---|---|---|---|---|---|---|---|---|---|---|
| A | | | | | | | | -0.006 | 0.0173 | 0.0425 | 0.0097 | 0.0381 |
| B | | | | | | | | 0.0406 | 0.0438 | 0.0449 | 0.068 | 0.0815 |
| C | | | | | | | | 0.1914 | 0.185 | 0.2353 | 0.264 | 0.1944 |
| D | | | | | | | | 0.1892 | 0.241 | 0.2344 | 0.2298 | 0.2086 |
| E | | | | | | | | 0.0986 | 0.0954 | 0.1059 | 0.1099 | 0.0774 |
| F | | | | | | | | 0.2197 | 0.2009 | 0.24 | 0.1992 | 0.2119 |
| G | | | | | | | | 0.0376 | 0.06 | 0.062 | 0.0764 | 0.0513 |
| H | | | | | | | | 0.1656 | 0.1227 | 0.1889 | 0.1466 | 0.0978 |

**Cell Index at: 47:30:53**

| | 1 | 2 | 3 | 4 | 5 | 6 | 7 | 8 | 9 | 10 | 11 | 12 |
|---|---|---|---|---|---|---|---|---|---|---|---|---|
| A | | | | | | | | -0.0085 | 0.0173 | 0.0406 | 0.0087 | 0.0364 |
| B | | | | | | | | 0.0408 | 0.0461 | 0.0436 | 0.07 | 0.0802 |
| C | | | | | | | | 0.1902 | 0.1825 | 0.2321 | 0.2617 | 0.2019 |
| D | | | | | | | | 0.1926 | 0.2299 | 0.2259 | 0.2308 | 0.2056 |
| E | | | | | | | | 0.1016 | 0.0882 | 0.1133 | 0.1097 | 0.0727 |
| F | | | | | | | | 0.2215 | 0.199 | 0.2405 | 0.2008 | 0.212 |
| G | | | | | | | | 0.0362 | 0.0585 | 0.065 | 0.0736 | 0.0464 |
| H | | | | | | | | 0.1651 | 0.1233 | 0.1822 | 0.1423 | 0.1021 |

**Cell Index at: 47:45:53**

| | 1 | 2 | 3 | 4 | 5 | 6 | 7 | 8 | 9 | 10 | 11 | 12 |
|---|---|---|---|---|---|---|---|---|---|---|---|---|
| A | | | | | | | | -0.0037 | 0.0184 | 0.043 | 0.0095 | 0.0362 |
| B | | | | | | | | 0.0361 | 0.0471 | 0.0452 | 0.0696 | 0.0803 |
| C | | | | | | | | 0.1938 | 0.1829 | 0.2352 | 0.264 | 0.1976 |
| D | | | | | | | | 0.1938 | 0.2392 | 0.2265 | 0.2325 | 0.213 |
| E | | | | | | | | 0.0979 | 0.0869 | 0.1167 | 0.1132 | 0.0832 |
| F | | | | | | | | 0.2176 | 0.2037 | 0.2364 | 0.2016 | 0.2133 |
| G | | | | | | | | 0.0388 | 0.0671 | 0.0617 | 0.0747 | 0.0445 |
| H | | | | | | | | 0.1677 | 0.1231 | 0.1812 | 0.1469 | 0.1013 |

**Cell Index at: 48:00:53**

| | 1 | 2 | 3 | 4 | 5 | 6 | 7 | 8 | 9 | 10 | 11 | 12 |
|---|---|---|---|---|---|---|---|---|---|---|---|---|
| A | | | | | | | | -0.0066 | 0.0185 | 0.0418 | 0.0074 | 0.0352 |
| B | | | | | | | | 0.0397 | 0.0478 | 0.0475 | 0.0676 | 0.0793 |
| C | | | | | | | | 0.2023 | 0.1823 | 0.2397 | 0.2657 | 0.2007 |
| D | | | | | | | | 0.186 | 0.2404 | 0.2249 | 0.2268 | 0.2142 |
| E | | | | | | | | 0.1071 | 0.0883 | 0.1125 | 0.1127 | 0.0806 |
| F | | | | | | | | 0.2205 | 0.2016 | 0.2429 | 0.1975 | 0.2131 |
| G | | | | | | | | 0.0399 | 0.0685 | 0.0659 | 0.0727 | 0.0458 |
| H | | | | | | | | 0.167 | 0.125 | 0.1838 | 0.1444 | 0.1006 |

| | | | | | | | | | | | | | |
|---|---|---|---|---|---|---|---|---|---|---|---|---|---|
| A vs B | 0.012755 | B vs C | 8.86E-05 | C vs D | 0.986073 | D vs E | 1.16E-05 | E vs F | 5.46E-06 | F vs G | 5.55E-07 | G vs H | 0.002364 |
| A vs C | 1.66E-05 | B vs D | 1.09E-06 | C vs E | 0.000562 | D vs F | 0.790801 | E vs G | 0.002104 | F vs H | 0.005191 | | |
| A vs D | 2.78E-07 | B vs E | 0.002262 | C vs F | 0.865714 | D vs G | 1.43E-06 | E vs H | 0.037767 | | | | |
| A vs E | 0.000122 | B vs F | 5.28E-07 | C vs G | 0.000121 | D vs H | 0.00408 | | | | | | |
| A vs F | 2.17E-07 | B vs G | 0.830674 | C vs.H | 0.008059 | | | | | | | | |
| A vs G | 0.008442 | B vs H | 0.001917 | | | | | | | | | | |
| A vs H | 0.000232 | | | | | | | | | | | | |

**Cell Index at: 48:15:53**

| | 1 | 2 | 3 | 4 | 5 | 6 | 7 | 8 | 9 | 10 | 11 | 12 |
|---|---|---|---|---|---|---|---|---|---|---|---|---|
| A | | | | | | | | -0.0062 | 0.019 | 0.0408 | 0.0087 | 0.0343 |
| B | | | | | | | | 0.0414 | 0.0438 | 0.0456 | 0.0646 | 0.0811 |
| C | | | | | | | | 0.1915 | 0.1841 | 0.2333 | 0.2658 | 0.2015 |
| D | | | | | | | | 0.1924 | 0.2378 | 0.2313 | 0.236 | 0.2117 |
| E | | | | | | | | 0.1006 | 0.0852 | 0.113 | 0.1108 | 0.0821 |
| F | | | | | | | | 0.2241 | 0.1986 | 0.2472 | 0.2096 | 0.2211 |
| G | | | | | | | | 0.0371 | 0.0651 | 0.0655 | 0.0738 | 0.0458 |
| H | | | | | | | | 0.1693 | 0.1196 | 0.1852 | 0.1434 | 0.103 |

| *Count* | *Sum* | *Average* | *Variance* |
|---|---|---|---|
| 5 | 0.0963 | 0.01926 | 0.000393 |
| 5 | 0.2819 | 0.05638 | 0.00027 |
| 5 | 1.0907 | 0.21814 | 0.001142 |
| 5 | 1.0923 | 0.21846 | 0.000416 |
| 5 | 0.5012 | 0.10024 | 0.00022 |
| 5 | 1.0756 | 0.21512 | 0.000325 |
| 5 | 0.2928 | 0.05856 | 0.000216 |
| 5 | 0.7208 | 0.14416 | 0.001089 |

**Cell Index at: 48:30:53**

| | 1 | 2 | 3 | 4 | 5 | 6 | 7 | 8 | 9 | 10 | 11 | 12 |
|---|---|---|---|---|---|---|---|---|---|---|---|---|
| A | | | | | | | | -0.0048 | 0.0218 | 0.0417 | 0.0097 | 0.0329 |
| B | | | | | | | | 0.0445 | 0.0459 | 0.0432 | 0.0687 | 0.0827 |
| C | | | | | | | | 0.1937 | 0.1842 | 0.2339 | 0.2647 | 0.1968 |
| D | | | | | | | | 0.1935 | 0.2452 | 0.221 | 0.2299 | 0.2073 |
| E | | | | | | | | 0.0937 | 0.0908 | 0.1178 | 0.1098 | 0.084 |
| F | | | | | | | | 0.2174 | 0.1961 | 0.2391 | 0.2016 | 0.2193 |
| G | | | | | | | | 0.041 | 0.0666 | 0.0595 | 0.0744 | 0.0458 |
| H | | | | | | | | 0.1659 | 0.1216 | 0.1839 | 0.1442 | 0.1014 |

| *SS* | *df* | *MS* | *F* | *P-value* | *F crit* |
|---|---|---|---|---|---|
| 0.233503 | 7 | 0.033358 | 65.54862 | 3.62E-17 | 2.312741 |
| 0.016285 | 32 | 0.000509 | | | |
| 0.249788 | 39 | | | | |

**Cell Index at: 48:45:54**

| | 1 | 2 | 3 | 4 | 5 | 6 | 7 | 8 | 9 | 10 | 11 | 12 |
|---|---|---|---|---|---|---|---|---|---|---|---|---|
| A | | | | | | | | -0.0066 | 0.018 | 0.0416 | 0.0091 | 0.0346 |
| B | | | | | | | | 0.0431 | 0.0438 | 0.0468 | 0.0672 | 0.0773 |
| C | | | | | | | | 0.1906 | 0.188 | 0.2403 | 0.2691 | 0.1992 |
| D | | | | | | | | 0.1875 | 0.2374 | 0.2168 | 0.2331 | 0.214 |
| E | | | | | | | | 0.101 | 0.09 | 0.1129 | 0.1066 | 0.0791 |
| F | | | | | | | | 0.2199 | 0.1981 | 0.2308 | 0.2016 | 0.2211 |
| G | | | | | | | | 0.0396 | 0.0628 | 0.0643 | 0.0726 | 0.0504 |
| H | | | | | | | | 0.1705 | 0.1232 | 0.1824 | 0.1474 | 0.1035 |

**Cell Index at: 49:00:55**

| | 1 | 2 | 3 | 4 | 5 | 6 | 7 | 8 | 9 | 10 | 11 | 12 |
|---|---|---|---|---|---|---|---|---|---|---|---|---|
| A | | | | | | | | -0.0056 | 0.0215 | 0.0405 | 0.0095 | 0.0343 |

| | | | | | | | | | | | | |
|---|---|---|---|---|---|---|---|---|---|---|---|---|
| B | | | | | | | | 0.0426 | 0.0413 | 0.0436 | 0.0696 | 0.0811 |
| C | | | | | | | | 0.1864 | 0.1853 | 0.2397 | 0.2644 | 0.1926 |
| D | | | | | | | | 0.1859 | 0.2371 | 0.2223 | 0.2333 | 0.2114 |
| E | | | | | | | | 0.0973 | 0.0869 | 0.115 | 0.1071 | 0.0809 |
| F | | | | | | | | 0.2183 | 0.2058 | 0.2246 | 0.1979 | 0.2207 |
| G | | | | | | | | 0.0418 | 0.0666 | 0.0644 | 0.0738 | 0.0461 |
| H | | | | | | | | 0.1722 | 0.122 | 0.1829 | 0.1431 | 0.1019 |

**Cell Index at: 49:15:56**

| | 1 | 2 | 3 | 4 | 5 | 6 | 7 | 8 | 9 | 10 | 11 | 12 |
|---|---|---|---|---|---|---|---|---|---|---|---|---|
| A | | | | | | | | -0.0074 | 0.0189 | 0.0404 | 0.0079 | 0.0342 |
| B | | | | | | | | 0.0411 | 0.0458 | 0.0451 | 0.0677 | 0.0755 |
| C | | | | | | | | 0.185 | 0.1829 | 0.2403 | 0.2655 | 0.1954 |
| D | | | | | | | | 0.1883 | 0.2431 | 0.2107 | 0.2256 | 0.2183 |
| E | | | | | | | | 0.0971 | 0.0963 | 0.1196 | 0.1077 | 0.0756 |
| F | | | | | | | | 0.2194 | 0.1982 | 0.2333 | 0.2005 | 0.2163 |
| G | | | | | | | | 0.0382 | 0.065 | 0.0604 | 0.0688 | 0.0473 |
| H | | | | | | | | 0.1669 | 0.1212 | 0.1831 | 0.1469 | 0.1018 |

**Cell Index at: 49:30:56**

| | 1 | 2 | 3 | 4 | 5 | 6 | 7 | 8 | 9 | 10 | 11 | 12 |
|---|---|---|---|---|---|---|---|---|---|---|---|---|
| A | | | | | | | | -0.0067 | 0.0191 | 0.039 | 0.0065 | 0.0353 |
| B | | | | | | | | 0.0417 | 0.0422 | 0.0485 | 0.0732 | 0.0711 |
| C | | | | | | | | 0.1866 | 0.1833 | 0.2384 | 0.262 | 0.198 |
| D | | | | | | | | 0.1925 | 0.2379 | 0.2202 | 0.228 | 0.2133 |
| E | | | | | | | | 0.1055 | 0.0888 | 0.117 | 0.1121 | 0.0844 |
| F | | | | | | | | 0.2165 | 0.2049 | 0.2287 | 0.2015 | 0.2216 |
| G | | | | | | | | 0.044 | 0.0651 | 0.0595 | 0.0693 | 0.0449 |
| H | | | | | | | | 0.17 | 0.1236 | 0.1829 | 0.1501 | 0.102 |

**Cell Index at: 49:45:57**

| | 1 | 2 | 3 | 4 | 5 | 6 | 7 | 8 | 9 | 10 | 11 | 12 |
|---|---|---|---|---|---|---|---|---|---|---|---|---|
| A | | | | | | | | -0.009 | 0.0185 | 0.0403 | 0.0081 | 0.0352 |
| B | | | | | | | | 0.0381 | 0.0347 | 0.0471 | 0.0696 | 0.0798 |
| C | | | | | | | | 0.1881 | 0.1848 | 0.2361 | 0.2616 | 0.1982 |
| D | | | | | | | | 0.187 | 0.2422 | 0.2134 | 0.2264 | 0.2069 |
| E | | | | | | | | 0.0941 | 0.0923 | 0.1173 | 0.1071 | 0.0821 |
| F | | | | | | | | 0.2171 | 0.2053 | 0.2291 | 0.2002 | 0.2158 |
| G | | | | | | | | 0.0446 | 0.0661 | 0.0618 | 0.0639 | 0.0456 |
| H | | | | | | | | 0.1688 | 0.1258 | 0.1816 | 0.1481 | 0.1022 |

**Cell Index at: 50:00:57**

| | 1 | 2 | 3 | 4 | 5 | 6 | 7 | 8 | 9 | 10 | 11 | 12 |
|---|---|---|---|---|---|---|---|---|---|---|---|---|
| A | | | | | | | | -0.001 | 0.0184 | 0.041 | 0.008 | 0.0315 |
| B | | | | | | | | 0.0454 | 0.0433 | 0.0475 | 0.069 | 0.0742 |
| C | | | | | | | | 0.1871 | 0.1858 | 0.2407 | 0.2627 | 0.1952 |
| D | | | | | | | | 0.1873 | 0.2434 | 0.2226 | 0.2328 | 0.2097 |
| E | | | | | | | | 0.1028 | 0.0883 | 0.1188 | 0.1018 | 0.0799 |
| F | | | | | | | | 0.2178 | 0.2016 | 0.2351 | 0.1991 | 0.2143 |
| G | | | | | | | | 0.0459 | 0.0602 | 0.0602 | 0.0664 | 0.0465 |
| H | | | | | | | | 0.1688 | 0.1185 | 0.1818 | 0.1498 | 0.1031 |

**Cell Index at: 50:15:57**

| | 1 | 2 | 3 | 4 | 5 | 6 | 7 | 8 | 9 | 10 | 11 | 12 |
|---|---|---|---|---|---|---|---|---|---|---|---|---|
| A | | | | | | | | -0.0032 | 0.018 | 0.0361 | 0.0057 | 0.0356 |
| B | | | | | | | | 0.0417 | 0.0409 | 0.0518 | 0.0689 | 0.0778 |
| C | | | | | | | | 0.1826 | 0.1879 | 0.2405 | 0.2593 | 0.2 |
| D | | | | | | | | 0.1865 | 0.2429 | 0.2211 | 0.2322 | 0.2117 |
| E | | | | | | | | 0.095 | 0.0914 | 0.118 | 0.1078 | 0.0741 |
| F | | | | | | | | 0.214 | 0.2022 | 0.2328 | 0.2085 | 0.2113 |
| G | | | | | | | | 0.0454 | 0.0628 | 0.0627 | 0.0649 | 0.0437 |
| H | | | | | | | | 0.1701 | 0.1182 | 0.1807 | 0.1494 | 0.1026 |

**Cell Index at: 50:30:58**

| | 1 | 2 | 3 | 4 | 5 | 6 | 7 | 8 | 9 | 10 | 11 | 12 |
|---|---|---|---|---|---|---|---|---|---|---|---|---|
| A | | | | | | | | -0.0034 | 0.017 | 0.0394 | 0.0097 | 0.0364 |
| B | | | | | | | | 0.0445 | 0.0419 | 0.0548 | 0.0694 | 0.0738 |
| C | | | | | | | | 0.1815 | 0.177 | 0.2326 | 0.2626 | 0.2046 |
| D | | | | | | | | 0.1844 | 0.2362 | 0.2172 | 0.2342 | 0.2119 |
| E | | | | | | | | 0.1017 | 0.0947 | 0.1189 | 0.1026 | 0.0868 |
| F | | | | | | | | 0.2154 | 0.2027 | 0.227 | 0.2003 | 0.2119 |
| G | | | | | | | | 0.0396 | 0.0627 | 0.0621 | 0.0651 | 0.0427 |
| H | | | | | | | | 0.1726 | 0.1219 | 0.182 | 0.1474 | 0.1071 |

**Cell Index at: 50:45:58**

| | 1 | 2 | 3 | 4 | 5 | 6 | 7 | 8 | 9 | 10 | 11 | 12 |
|---|---|---|---|---|---|---|---|---|---|---|---|---|
| A | | | | | | | | -0.0031 | 0.0204 | 0.0367 | 0.0054 | 0.0346 |
| B | | | | | | | | 0.0369 | 0.0427 | 0.0538 | 0.0706 | 0.0731 |
| C | | | | | | | | 0.1744 | 0.1759 | 0.2427 | 0.2626 | 0.2067 |
| D | | | | | | | | 0.1839 | 0.2402 | 0.2256 | 0.2298 | 0.2057 |
| E | | | | | | | | 0.1005 | 0.0953 | 0.1175 | 0.1027 | 0.0769 |
| F | | | | | | | | 0.2199 | 0.2046 | 0.234 | 0.2028 | 0.2052 |
| G | | | | | | | | 0.0412 | 0.0645 | 0.0624 | 0.0655 | 0.0435 |
| H | | | | | | | | 0.1706 | 0.1206 | 0.1824 | 0.1482 | 0.1038 |

**Cell Index at: 51:00:59**

| | 1 | 2 | 3 | 4 | 5 | 6 | 7 | 8 | 9 | 10 | 11 | 12 |
|---|---|---|---|---|---|---|---|---|---|---|---|---|
| A | | | | | | | | -0.0033 | 0.0186 | 0.0358 | 0.0106 | 0.0337 |
| B | | | | | | | | 0.0407 | 0.0434 | 0.0537 | 0.0675 | 0.0694 |
| C | | | | | | | | 0.1807 | 0.18 | 0.2405 | 0.2601 | 0.203 |
| D | | | | | | | | 0.1866 | 0.2387 | 0.2241 | 0.2268 | 0.2101 |
| E | | | | | | | | 0.0988 | 0.0863 | 0.1157 | 0.1031 | 0.0819 |
| F | | | | | | | | 0.2096 | 0.2011 | 0.2338 | 0.2128 | 0.2053 |
| G | | | | | | | | 0.042 | 0.0641 | 0.0644 | 0.0632 | 0.0455 |
| H | | | | | | | | 0.168 | 0.1225 | 0.1808 | 0.1473 | 0.1046 |

**Cell Index at: 51:16:00**

| | 1 | 2 | 3 | 4 | 5 | 6 | 7 | 8 | 9 | 10 | 11 | 12 |
|---|---|---|---|---|---|---|---|---|---|---|---|---|
| A | | | | | | | | -0.0003 | 0.0167 | 0.0364 | 0.01 | 0.0345 |
| B | | | | | | | | 0.0404 | 0.0405 | 0.0492 | 0.0643 | 0.0727 |
| C | | | | | | | | 0.1777 | 0.1811 | 0.2465 | 0.2659 | 0.2055 |
| D | | | | | | | | 0.1866 | 0.236 | 0.2165 | 0.2237 | 0.2161 |
| E | | | | | | | | 0.1006 | 0.0958 | 0.1172 | 0.1008 | 0.0826 |
| F | | | | | | | | 0.2169 | 0.2103 | 0.2363 | 0.2089 | 0.2042 |
| G | | | | | | | | 0.0412 | 0.0647 | 0.07 | 0.0623 | 0.0446 |
| H | | | | | | | | 0.1686 | 0.1229 | 0.1797 | 0.1474 | 0.1063 |

**Cell Index at: 51:31:01**

| | 1 | 2 | 3 | 4 | 5 | 6 | 7 | 8 | 9 | 10 | 11 | 12 |
|---|---|---|---|---|---|---|---|---|---|---|---|---|
| A | | | | | | | | -0.0033 | 0.0201 | 0.0369 | 0.0094 | 0.0355 |
| B | | | | | | | | 0.0389 | 0.0387 | 0.0481 | 0.0694 | 0.0685 |
| C | | | | | | | | 0.1815 | 0.1809 | 0.2444 | 0.2657 | 0.2003 |
| D | | | | | | | | 0.1972 | 0.2301 | 0.2207 | 0.2261 | 0.2186 |
| E | | | | | | | | 0.0996 | 0.0903 | 0.1091 | 0.1082 | 0.0819 |
| F | | | | | | | | 0.2165 | 0.2029 | 0.2302 | 0.2103 | 0.2035 |
| G | | | | | | | | 0.0417 | 0.0615 | 0.0667 | 0.0638 | 0.0468 |
| H | | | | | | | | 0.1746 | 0.1205 | 0.1768 | 0.1489 | 0.1068 |

**Cell Index at: 51:46:00**

| | 1 | 2 | 3 | 4 | 5 | 6 | 7 | 8 | 9 | 10 | 11 | 12 |
|---|---|---|---|---|---|---|---|---|---|---|---|---|
| A | | | | | | | | -0.0005 | 0.0184 | 0.0397 | 0.0049 | 0.0334 |
| B | | | | | | | | 0.0392 | 0.039 | 0.0458 | 0.0674 | 0.0677 |
| C | | | | | | | | 0.1804 | 0.1734 | 0.2451 | 0.2641 | 0.2039 |
| D | | | | | | | | 0.1881 | 0.2266 | 0.2195 | 0.2329 | 0.2121 |
| E | | | | | | | | 0.1016 | 0.0914 | 0.1134 | 0.1097 | 0.0808 |
| F | | | | | | | | 0.2138 | 0.2077 | 0.2342 | 0.2104 | 0.2005 |
| G | | | | | | | | 0.038 | 0.0637 | 0.0686 | 0.061 | 0.0437 |
| H | | | | | | | | 0.1725 | 0.1234 | 0.1809 | 0.1505 | 0.1095 |

**Cell Index at: 52:01:01**

| | 1 | 2 | 3 | 4 | 5 | 6 | 7 | 8 | 9 | 10 | 11 | 12 |
|---|---|---|---|---|---|---|---|---|---|---|---|---|

| | | | | | | | | | | | | |
|---|---|---|---|---|---|---|---|---|---|---|---|---|
| A | | | | | | | | -0.0014 | 0.0175 | 0.0417 | 0.0094 | 0.0334 |
| B | | | | | | | | 0.0399 | 0.0358 | 0.043 | 0.0641 | 0.0663 |
| C | | | | | | | | 0.1745 | 0.1727 | 0.246 | 0.2659 | 0.2035 |
| D | | | | | | | | 0.1867 | 0.2304 | 0.2216 | 0.2253 | 0.2115 |
| E | | | | | | | | 0.1064 | 0.0892 | 0.116 | 0.1049 | 0.0745 |
| F | | | | | | | | 0.2151 | 0.2085 | 0.2371 | 0.2159 | 0.2059 |
| G | | | | | | | | 0.0412 | 0.0685 | 0.0676 | 0.0647 | 0.0389 |
| H | | | | | | | | 0.1741 | 0.1233 | 0.1815 | 0.1474 | 0.1102 |

**Cell Index at: 52:16:01**

| | 1 | 2 | 3 | 4 | 5 | 6 | 7 | 8 | 9 | 10 | 11 | 12 |
|---|---|---|---|---|---|---|---|---|---|---|---|---|
| A | | | | | | | | -0.0064 | 0.015 | 0.0397 | 0.009 | 0.0336 |
| B | | | | | | | | 0.0383 | 0.0348 | 0.0496 | 0.0652 | 0.0735 |
| C | | | | | | | | 0.1744 | 0.1756 | 0.2525 | 0.2701 | 0.2039 |
| D | | | | | | | | 0.1813 | 0.2324 | 0.2226 | 0.2355 | 0.2148 |
| E | | | | | | | | 0.1045 | 0.0879 | 0.1094 | 0.111 | 0.0817 |
| F | | | | | | | | 0.2166 | 0.2064 | 0.2403 | 0.2177 | 0.2053 |
| G | | | | | | | | 0.0433 | 0.0638 | 0.0669 | 0.0689 | 0.0473 |
| H | | | | | | | | 0.1665 | 0.1254 | 0.1813 | 0.1517 | 0.1092 |

**Cell Index at: 52:31:01**

| | 1 | 2 | 3 | 4 | 5 | 6 | 7 | 8 | 9 | 10 | 11 | 12 |
|---|---|---|---|---|---|---|---|---|---|---|---|---|
| A | | | | | | | | -0.0043 | 0.0132 | 0.0362 | 0.0086 | 0.0344 |
| B | | | | | | | | 0.0372 | 0.0389 | 0.0457 | 0.0661 | 0.0734 |
| C | | | | | | | | 0.1788 | 0.1784 | 0.2536 | 0.2676 | 0.2009 |
| D | | | | | | | | 0.1935 | 0.2344 | 0.2234 | 0.2338 | 0.2172 |
| E | | | | | | | | 0.1014 | 0.0847 | 0.1141 | 0.1062 | 0.0805 |
| F | | | | | | | | 0.2177 | 0.2051 | 0.2395 | 0.2134 | 0.2122 |
| G | | | | | | | | 0.0372 | 0.0652 | 0.0718 | 0.0673 | 0.0449 |
| H | | | | | | | | 0.171 | 0.1248 | 0.1822 | 0.1519 | 0.1096 |

**Cell Index at: 52:46:02**

| | 1 | 2 | 3 | 4 | 5 | 6 | 7 | 8 | 9 | 10 | 11 | 12 |
|---|---|---|---|---|---|---|---|---|---|---|---|---|
| A | | | | | | | | -0.0033 | 0.0143 | 0.0386 | 0.0114 | 0.0375 |
| B | | | | | | | | 0.0405 | 0.0359 | 0.0447 | 0.0658 | 0.0708 |
| C | | | | | | | | 0.1783 | 0.1733 | 0.248 | 0.2686 | 0.2052 |
| D | | | | | | | | 0.1869 | 0.2288 | 0.2215 | 0.232 | 0.2195 |
| E | | | | | | | | 0.095 | 0.0882 | 0.1152 | 0.1074 | 0.0823 |
| F | | | | | | | | 0.216 | 0.2036 | 0.2398 | 0.2212 | 0.2079 |
| G | | | | | | | | 0.0403 | 0.0666 | 0.0662 | 0.0677 | 0.0391 |
| H | | | | | | | | 0.1712 | 0.1216 | 0.1817 | 0.1501 | 0.1102 |

**Cell Index at: 53:01:02**

| | 1 | 2 | 3 | 4 | 5 | 6 | 7 | 8 | 9 | 10 | 11 | 12 |
|---|---|---|---|---|---|---|---|---|---|---|---|---|
| A | | | | | | | | -0.0053 | 0.0143 | 0.0392 | 0.0093 | 0.0413 |
| B | | | | | | | | 0.0416 | 0.036 | 0.0419 | 0.0654 | 0.0685 |
| C | | | | | | | | 0.1829 | 0.1705 | 0.2497 | 0.2694 | 0.2039 |
| D | | | | | | | | 0.1923 | 0.2289 | 0.2248 | 0.2293 | 0.2205 |
| E | | | | | | | | 0.1028 | 0.0918 | 0.1102 | 0.1087 | 0.0799 |
| F | | | | | | | | 0.2185 | 0.2083 | 0.2374 | 0.2262 | 0.211 |
| G | | | | | | | | 0.0355 | 0.0635 | 0.0657 | 0.0659 | 0.047 |
| H | | | | | | | | 0.1708 | 0.1228 | 0.1847 | 0.1477 | 0.1077 |

**Cell Index at: 53:16:03**

| | 1 | 2 | 3 | 4 | 5 | 6 | 7 | 8 | 9 | 10 | 11 | 12 |
|---|---|---|---|---|---|---|---|---|---|---|---|---|
| A | | | | | | | | -0.0044 | 0.0153 | 0.0395 | 0.009 | 0.0387 |
| B | | | | | | | | 0.0363 | 0.0365 | 0.0471 | 0.0641 | 0.0666 |
| C | | | | | | | | 0.1733 | 0.179 | 0.2507 | 0.2667 | 0.2078 |
| D | | | | | | | | 0.1868 | 0.2294 | 0.2261 | 0.2368 | 0.2167 |
| E | | | | | | | | 0.0981 | 0.0931 | 0.1075 | 0.1109 | 0.0805 |
| F | | | | | | | | 0.2211 | 0.2088 | 0.2419 | 0.2269 | 0.2084 |
| G | | | | | | | | 0.0423 | 0.0633 | 0.0645 | 0.0676 | 0.0454 |
| H | | | | | | | | 0.1711 | 0.1234 | 0.1806 | 0.1511 | 0.1096 |

**Cell Index at: 53:31:04**

| | 1 | 2 | 3 | 4 | 5 | 6 | 7 | 8 | 9 | 10 | 11 | 12 |
|---|---|---|---|---|---|---|---|---|---|---|---|---|
| A | | | | | | | | -0.0057 | 0.0158 | 0.0386 | 0.0102 | 0.0365 |
| B | | | | | | | | 0.0407 | 0.0352 | 0.0468 | 0.0672 | 0.0689 |
| C | | | | | | | | 0.1749 | 0.1777 | 0.2515 | 0.2669 | 0.2098 |
| D | | | | | | | | 0.1884 | 0.2324 | 0.2301 | 0.2414 | 0.2157 |
| E | | | | | | | | 0.0999 | 0.0872 | 0.117 | 0.1113 | 0.0769 |
| F | | | | | | | | 0.2232 | 0.2047 | 0.2463 | 0.2209 | 0.2111 |
| G | | | | | | | | 0.0429 | 0.058 | 0.0613 | 0.0682 | 0.0448 |
| H | | | | | | | | 0.1727 | 0.1212 | 0.1815 | 0.1501 | 0.1088 |

**Cell Index at: 53:46:05**

| | 1 | 2 | 3 | 4 | 5 | 6 | 7 | 8 | 9 | 10 | 11 | 12 |
|---|---|---|---|---|---|---|---|---|---|---|---|---|
| A | | | | | | | | -0.0062 | 0.016 | 0.0389 | 0.0105 | 0.0394 |
| B | | | | | | | | 0.0381 | 0.0297 | 0.046 | 0.0669 | 0.0697 |
| C | | | | | | | | 0.1791 | 0.1796 | 0.2527 | 0.2683 | 0.2136 |
| D | | | | | | | | 0.1905 | 0.2302 | 0.2343 | 0.2337 | 0.2142 |
| E | | | | | | | | 0.1052 | 0.0961 | 0.1129 | 0.1084 | 0.0783 |
| F | | | | | | | | 0.2245 | 0.207 | 0.2453 | 0.219 | 0.2134 |
| G | | | | | | | | 0.0408 | 0.0593 | 0.0647 | 0.0636 | 0.0472 |
| H | | | | | | | | 0.1713 | 0.1253 | 0.1825 | 0.1507 | 0.1094 |

**Cell Index at: 54:01:05**

| | 1 | 2 | 3 | 4 | 5 | 6 | 7 | 8 | 9 | 10 | 11 | 12 |
|---|---|---|---|---|---|---|---|---|---|---|---|---|
| A | | | | | | | | -0.0061 | 0.0146 | 0.037 | 0.0107 | 0.0403 |
| B | | | | | | | | 0.0347 | 0.0372 | 0.0524 | 0.063 | 0.068 |
| C | | | | | | | | 0.1793 | 0.1853 | 0.2562 | 0.2602 | 0.2122 |
| D | | | | | | | | 0.1875 | 0.2356 | 0.2262 | 0.2391 | 0.2119 |
| E | | | | | | | | 0.0981 | 0.09 | 0.1056 | 0.1126 | 0.0651 |
| F | | | | | | | | 0.2219 | 0.2135 | 0.2495 | 0.2245 | 0.212 |
| G | | | | | | | | 0.0388 | 0.0602 | 0.065 | 0.0664 | 0.0442 |
| H | | | | | | | | 0.1737 | 0.1255 | 0.184 | 0.1488 | 0.1108 |

**Cell Index at: 54:16:05**

| | 1 | 2 | 3 | 4 | 5 | 6 | 7 | 8 | 9 | 10 | 11 | 12 |
|---|---|---|---|---|---|---|---|---|---|---|---|---|
| A | | | | | | | | -0.0045 | 0.0166 | 0.0408 | 0.0114 | 0.0399 |
| B | | | | | | | | 0.0342 | 0.0345 | 0.0488 | 0.0653 | 0.0662 |
| C | | | | | | | | 0.1738 | 0.1811 | 0.2588 | 0.271 | 0.2136 |
| D | | | | | | | | 0.1828 | 0.2375 | 0.2282 | 0.2392 | 0.2178 |
| E | | | | | | | | 0.0978 | 0.0941 | 0.1076 | 0.1114 | 0.0748 |
| F | | | | | | | | 0.2262 | 0.2157 | 0.2505 | 0.2249 | 0.215 |
| G | | | | | | | | 0.0425 | 0.0615 | 0.0634 | 0.0607 | 0.0445 |
| H | | | | | | | | 0.1739 | 0.1246 | 0.1847 | 0.1508 | 0.1136 |

**Cell Index at: 54:31:05**

| | 1 | 2 | 3 | 4 | 5 | 6 | 7 | 8 | 9 | 10 | 11 | 12 |
|---|---|---|---|---|---|---|---|---|---|---|---|---|
| A | | | | | | | | -0.0036 | 0.0166 | 0.0354 | 0.0128 | 0.0419 |
| B | | | | | | | | 0.0405 | 0.037 | 0.0468 | 0.0654 | 0.0637 |
| C | | | | | | | | 0.1786 | 0.1776 | 0.2604 | 0.2674 | 0.2115 |
| D | | | | | | | | 0.1877 | 0.2407 | 0.2311 | 0.2373 | 0.2225 |
| E | | | | | | | | 0.0969 | 0.0896 | 0.1119 | 0.1124 | 0.0789 |
| F | | | | | | | | 0.2265 | 0.2172 | 0.249 | 0.2235 | 0.2193 |
| G | | | | | | | | 0.04 | 0.0585 | 0.0608 | 0.062 | 0.0449 |
| H | | | | | | | | 0.1715 | 0.1225 | 0.1857 | 0.1503 | 0.1136 |

**Cell Index at: 54:46:05**

| | 1 | 2 | 3 | 4 | 5 | 6 | 7 | 8 | 9 | 10 | 11 | 12 |
|---|---|---|---|---|---|---|---|---|---|---|---|---|
| A | | | | | | | | -0.0066 | 0.015 | 0.0375 | 0.0123 | 0.0386 |
| B | | | | | | | | 0.0409 | 0.0337 | 0.0504 | 0.071 | 0.065 |
| C | | | | | | | | 0.1798 | 0.1863 | 0.2619 | 0.2717 | 0.2096 |
| D | | | | | | | | 0.1903 | 0.2384 | 0.2299 | 0.242 | 0.2181 |
| E | | | | | | | | 0.1059 | 0.0891 | 0.1097 | 0.12 | 0.0787 |
| F | | | | | | | | 0.2277 | 0.2186 | 0.2446 | 0.227 | 0.2152 |
| G | | | | | | | | 0.0428 | 0.0549 | 0.0629 | 0.0642 | 0.0458 |
| H | | | | | | | | 0.1693 | 0.1206 | 0.1849 | 0.1529 | 0.1135 |

**Cell Index at: 55:01:05**

| | 1 | 2 | 3 | 4 | 5 | 6 | 7 | 8 | 9 | 10 | 11 | 12 |
|---|---|---|---|---|---|---|---|---|---|---|---|---|
| A | | | | | | | | -0.0044 | 0.0143 | 0.0367 | 0.0126 | 0.0386 |
| B | | | | | | | | 0.0352 | 0.0291 | 0.0501 | 0.0641 | 0.068 |
| C | | | | | | | | 0.1777 | 0.1852 | 0.2602 | 0.2728 | 0.2106 |
| D | | | | | | | | 0.1941 | 0.2342 | 0.2273 | 0.2359 | 0.2248 |
| E | | | | | | | | 0.1006 | 0.094 | 0.1114 | 0.1125 | 0.0764 |
| F | | | | | | | | 0.2271 | 0.2118 | 0.246 | 0.2233 | 0.221 |
| G | | | | | | | | 0.0395 | 0.056 | 0.0641 | 0.0622 | 0.0423 |
| H | | | | | | | | 0.1716 | 0.1222 | 0.1832 | 0.1537 | 0.1144 |

**Cell Index at: 55:16:06**

| | 1 | 2 | 3 | 4 | 5 | 6 | 7 | 8 | 9 | 10 | 11 | 12 |
|---|---|---|---|---|---|---|---|---|---|---|---|---|
| A | | | | | | | | -0.0012 | 0.0144 | 0.0373 | 0.0107 | 0.0376 |
| B | | | | | | | | 0.0372 | 0.0307 | 0.0462 | 0.0669 | 0.0685 |
| C | | | | | | | | 0.1732 | 0.1874 | 0.2606 | 0.2688 | 0.2106 |
| D | | | | | | | | 0.1865 | 0.2357 | 0.2307 | 0.238 | 0.222 |
| E | | | | | | | | 0.1014 | 0.0959 | 0.1159 | 0.1141 | 0.0783 |
| F | | | | | | | | 0.2283 | 0.2187 | 0.2422 | 0.2235 | 0.2249 |
| G | | | | | | | | 0.0416 | 0.0537 | 0.0591 | 0.0616 | 0.0434 |
| H | | | | | | | | 0.1711 | 0.1213 | 0.1845 | 0.1497 | 0.1145 |

**Cell Index at: 55:31:07**

| | 1 | 2 | 3 | 4 | 5 | 6 | 7 | 8 | 9 | 10 | 11 | 12 |
|---|---|---|---|---|---|---|---|---|---|---|---|---|
| A | | | | | | | | -0.0057 | 0.0131 | 0.0354 | 0.0103 | 0.0377 |
| B | | | | | | | | 0.0405 | 0.0287 | 0.0487 | 0.0684 | 0.0636 |
| C | | | | | | | | 0.1748 | 0.1836 | 0.2638 | 0.2757 | 0.2131 |
| D | | | | | | | | 0.1863 | 0.239 | 0.2255 | 0.2389 | 0.2255 |
| E | | | | | | | | 0.0981 | 0.0975 | 0.1154 | 0.1088 | 0.0738 |
| F | | | | | | | | 0.223 | 0.2175 | 0.2528 | 0.2317 | 0.2227 |
| G | | | | | | | | 0.0425 | 0.0546 | 0.0632 | 0.0566 | 0.0408 |
| H | | | | | | | | 0.1756 | 0.1222 | 0.1848 | 0.1521 | 0.114 |

**Cell Index at: 55:46:07**

| | 1 | 2 | 3 | 4 | 5 | 6 | 7 | 8 | 9 | 10 | 11 | 12 |
|---|---|---|---|---|---|---|---|---|---|---|---|---|
| A | | | | | | | | -0.0041 | 0.0169 | 0.0369 | 0.0121 | 0.0376 |
| B | | | | | | | | 0.0381 | 0.0329 | 0.0487 | 0.0679 | 0.0632 |
| C | | | | | | | | 0.1737 | 0.1876 | 0.2684 | 0.2749 | 0.2135 |
| D | | | | | | | | 0.1942 | 0.2269 | 0.2268 | 0.2435 | 0.2211 |
| E | | | | | | | | 0.1007 | 0.0895 | 0.1074 | 0.1167 | 0.0776 |
| F | | | | | | | | 0.2251 | 0.2171 | 0.2487 | 0.2302 | 0.2242 |
| G | | | | | | | | 0.0392 | 0.0562 | 0.0621 | 0.0592 | 0.0416 |
| H | | | | | | | | 0.1756 | 0.1164 | 0.187 | 0.1531 | 0.1127 |

**Cell Index at: 56:01:07**

| | 1 | 2 | 3 | 4 | 5 | 6 | 7 | 8 | 9 | 10 | 11 | 12 |
|---|---|---|---|---|---|---|---|---|---|---|---|---|
| A | | | | | | | | -0.0033 | 0.0154 | 0.0374 | 0.0073 | 0.0386 |
| B | | | | | | | | 0.0345 | 0.0302 | 0.0504 | 0.0681 | 0.063 |
| C | | | | | | | | 0.175 | 0.186 | 0.2599 | 0.2778 | 0.2124 |
| D | | | | | | | | 0.1897 | 0.2335 | 0.2289 | 0.2454 | 0.2321 |
| E | | | | | | | | 0.1035 | 0.0933 | 0.1098 | 0.1095 | 0.0769 |
| F | | | | | | | | 0.228 | 0.2179 | 0.2528 | 0.2294 | 0.2276 |
| G | | | | | | | | 0.0404 | 0.0565 | 0.0614 | 0.0629 | 0.0457 |
| H | | | | | | | | 0.1733 | 0.1218 | 0.1801 | 0.1508 | 0.1162 |

**Cell Index at: 56:16:06**

| | 1 | 2 | 3 | 4 | 5 | 6 | 7 | 8 | 9 | 10 | 11 | 12 |
|---|---|---|---|---|---|---|---|---|---|---|---|---|
| A | | | | | | | | -0.0044 | 0.0161 | 0.0358 | 0.0102 | 0.0383 |
| B | | | | | | | | 0.034 | 0.0353 | 0.0507 | 0.0655 | 0.0648 |
| C | | | | | | | | 0.175 | 0.1887 | 0.2685 | 0.2783 | 0.2095 |
| D | | | | | | | | 0.1887 | 0.2356 | 0.2281 | 0.2443 | 0.2299 |
| E | | | | | | | | 0.0959 | 0.0889 | 0.1118 | 0.1124 | 0.0827 |
| F | | | | | | | | 0.2299 | 0.2188 | 0.2581 | 0.2277 | 0.2211 |
| G | | | | | | | | 0.0416 | 0.0544 | 0.0623 | 0.0565 | 0.0427 |
| H | | | | | | | | 0.1725 | 0.1216 | 0.1791 | 0.1556 | 0.1159 |

**Cell Index at: 56:31:07**

| | 1 | 2 | 3 | 4 | 5 | 6 | 7 | 8 | 9 | 10 | 11 | 12 |
|---|---|---|---|---|---|---|---|---|---|---|---|---|
| A | | | | | | | | -0.0018 | 0.0119 | 0.0372 | 0.0085 | 0.0388 |
| B | | | | | | | | 0.0361 | 0.037 | 0.045 | 0.0688 | 0.0669 |
| C | | | | | | | | 0.1764 | 0.1901 | 0.2668 | 0.2672 | 0.2094 |
| D | | | | | | | | 0.1896 | 0.2376 | 0.2313 | 0.2469 | 0.2267 |
| E | | | | | | | | 0.0977 | 0.0948 | 0.1121 | 0.1158 | 0.0784 |
| F | | | | | | | | 0.2286 | 0.2245 | 0.2585 | 0.2305 | 0.2307 |
| G | | | | | | | | 0.0441 | 0.0511 | 0.0629 | 0.0581 | 0.045 |
| H | | | | | | | | 0.1725 | 0.1193 | 0.1759 | 0.1533 | 0.1189 |

**Cell Index at: 56:46:08**

| | 1 | 2 | 3 | 4 | 5 | 6 | 7 | 8 | 9 | 10 | 11 | 12 |
|---|---|---|---|---|---|---|---|---|---|---|---|---|
| A | | | | | | | | -0.0027 | 0.016 | 0.0364 | 0.0091 | 0.0386 |
| B | | | | | | | | 0.0346 | 0.0341 | 0.0399 | 0.0691 | 0.0693 |
| C | | | | | | | | 0.1791 | 0.1852 | 0.2715 | 0.2725 | 0.2118 |
| D | | | | | | | | 0.184 | 0.2461 | 0.2305 | 0.2482 | 0.2201 |
| E | | | | | | | | 0.0968 | 0.0925 | 0.1091 | 0.1116 | 0.0765 |
| F | | | | | | | | 0.232 | 0.2229 | 0.2545 | 0.2256 | 0.227 |
| G | | | | | | | | 0.0441 | 0.0544 | 0.0589 | 0.0617 | 0.0415 |
| H | | | | | | | | 0.1736 | 0.1214 | 0.1742 | 0.1537 | 0.1169 |

**Cell Index at: 57:01:09**

| | 1 | 2 | 3 | 4 | 5 | 6 | 7 | 8 | 9 | 10 | 11 | 12 |
|---|---|---|---|---|---|---|---|---|---|---|---|---|
| A | | | | | | | | -0.0099 | 0.0147 | 0.0373 | 0.0066 | 0.035 |
| B | | | | | | | | 0.0371 | 0.0339 | 0.0439 | 0.0682 | 0.0673 |
| C | | | | | | | | 0.1771 | 0.1859 | 0.2731 | 0.2701 | 0.2134 |
| D | | | | | | | | 0.186 | 0.2473 | 0.2364 | 0.2429 | 0.2199 |
| E | | | | | | | | 0.0923 | 0.0913 | 0.105 | 0.1141 | 0.0784 |
| F | | | | | | | | 0.2332 | 0.2181 | 0.2624 | 0.2253 | 0.2225 |
| G | | | | | | | | 0.0395 | 0.0558 | 0.058 | 0.0605 | 0.0427 |
| H | | | | | | | | 0.1755 | 0.1208 | 0.1763 | 0.1591 | 0.1152 |

**Cell Index at: 57:16:10**

| | 1 | 2 | 3 | 4 | 5 | 6 | 7 | 8 | 9 | 10 | 11 | 12 |
|---|---|---|---|---|---|---|---|---|---|---|---|---|
| A | | | | | | | | -0.004 | 0.0121 | 0.0346 | 0.0067 | 0.0393 |
| B | | | | | | | | 0.0376 | 0.0348 | 0.0402 | 0.0713 | 0.063 |
| C | | | | | | | | 0.1818 | 0.1787 | 0.268 | 0.2686 | 0.2106 |
| D | | | | | | | | 0.1827 | 0.2431 | 0.2329 | 0.2477 | 0.227 |
| E | | | | | | | | 0.0962 | 0.0969 | 0.1064 | 0.1091 | 0.0826 |
| F | | | | | | | | 0.2294 | 0.2157 | 0.2572 | 0.2288 | 0.2248 |
| G | | | | | | | | 0.0445 | 0.0519 | 0.0549 | 0.0569 | 0.0442 |
| H | | | | | | | | 0.1787 | 0.1189 | 0.1795 | 0.16 | 0.1178 |

**Cell Index at: 57:31:10**

| | 1 | 2 | 3 | 4 | 5 | 6 | 7 | 8 | 9 | 10 | 11 | 12 |
|---|---|---|---|---|---|---|---|---|---|---|---|---|
| A | | | | | | | | -0.0032 | 0.0137 | 0.0392 | 0.0081 | 0.0364 |
| B | | | | | | | | 0.0361 | 0.0325 | 0.0446 | 0.0658 | 0.0599 |
| C | | | | | | | | 0.1867 | 0.1843 | 0.2702 | 0.272 | 0.2158 |
| D | | | | | | | | 0.186 | 0.248 | 0.2295 | 0.2483 | 0.2165 |
| E | | | | | | | | 0.096 | 0.0912 | 0.1124 | 0.1148 | 0.0792 |
| F | | | | | | | | 0.2335 | 0.2205 | 0.2551 | 0.2242 | 0.2277 |
| G | | | | | | | | 0.0383 | 0.0522 | 0.0564 | 0.0577 | 0.0414 |
| H | | | | | | | | 0.1778 | 0.1218 | 0.1799 | 0.1582 | 0.1173 |

**Cell Index at: 57:46:10**

| | 1 | 2 | 3 | 4 | 5 | 6 | 7 | 8 | 9 | 10 | 11 | 12 |
|---|---|---|---|---|---|---|---|---|---|---|---|---|
| A | | | | | | | | -0.0024 | 0.0135 | 0.0374 | 0.0043 | 0.0387 |
| B | | | | | | | | 0.0381 | 0.0345 | 0.0415 | 0.0655 | 0.0603 |
| C | | | | | | | | 0.1793 | 0.181 | 0.2675 | 0.2774 | 0.2154 |
| D | | | | | | | | 0.1885 | 0.2405 | 0.2322 | 0.2522 | 0.2208 |
| E | | | | | | | | 0.0958 | 0.0925 | 0.1109 | 0.1189 | 0.0849 |
| F | | | | | | | | 0.2247 | 0.2243 | 0.2564 | 0.235 | 0.226 |
| G | | | | | | | | 0.0412 | 0.0516 | 0.0562 | 0.0612 | 0.04 |
| H | | | | | | | | 0.1767 | 0.1193 | 0.1822 | 0.1645 | 0.1176 |

**Cell Index at: 58:01:10**

| | 1 | 2 | 3 | 4 | 5 | 6 | 7 | 8 | 9 | 10 | 11 | 12 |
|---|---|---|---|---|---|---|---|---|---|---|---|---|
| A | | | | | | | | -0.0049 | 0.0133 | 0.0376 | 0.0037 | 0.0399 |
| B | | | | | | | | 0.041 | 0.0311 | 0.0427 | 0.0711 | 0.0601 |
| C | | | | | | | | 0.1827 | 0.1895 | 0.2707 | 0.2715 | 0.2167 |
| D | | | | | | | | 0.1828 | 0.2486 | 0.2302 | 0.2576 | 0.2219 |
| E | | | | | | | | 0.0909 | 0.0925 | 0.1126 | 0.1149 | 0.0838 |
| F | | | | | | | | 0.2269 | 0.2217 | 0.2583 | 0.2312 | 0.2262 |
| G | | | | | | | | 0.0436 | 0.0479 | 0.0589 | 0.0571 | 0.0409 |
| H | | | | | | | | 0.176 | 0.1231 | 0.1816 | 0.1657 | 0.1197 |

**Cell Index at: 58:16:09**

| | 1 | 2 | 3 | 4 | 5 | 6 | 7 | 8 | 9 | 10 | 11 | 12 |
|---|---|---|---|---|---|---|---|---|---|---|---|---|
| A | | | | | | | | -0.0042 | 0.0128 | 0.0354 | 0.005 | 0.0399 |
| B | | | | | | | | 0.036 | 0.0286 | 0.0451 | 0.0684 | 0.0637 |
| C | | | | | | | | 0.1851 | 0.1873 | 0.2751 | 0.2717 | 0.2213 |
| D | | | | | | | | 0.1831 | 0.249 | 0.2394 | 0.2517 | 0.2352 |
| E | | | | | | | | 0.0931 | 0.0883 | 0.1115 | 0.1158 | 0.0779 |
| F | | | | | | | | 0.2364 | 0.2224 | 0.254 | 0.2317 | 0.2319 |
| G | | | | | | | | 0.0428 | 0.0511 | 0.0576 | 0.0599 | 0.0407 |
| H | | | | | | | | 0.1732 | 0.1216 | 0.1854 | 0.1666 | 0.1193 |

**Cell Index at: 58:31:09**

| | 1 | 2 | 3 | 4 | 5 | 6 | 7 | 8 | 9 | 10 | 11 | 12 |
|---|---|---|---|---|---|---|---|---|---|---|---|---|
| A | | | | | | | | -0.0027 | 0.0119 | 0.0351 | 0.0081 | 0.0418 |
| B | | | | | | | | 0.0358 | 0.0335 | 0.0415 | 0.0688 | 0.0641 |
| C | | | | | | | | 0.1812 | 0.1917 | 0.2722 | 0.2762 | 0.2234 |
| D | | | | | | | | 0.1982 | 0.2459 | 0.2336 | 0.2485 | 0.2295 |
| E | | | | | | | | 0.0967 | 0.0873 | 0.1192 | 0.1155 | 0.077 |
| F | | | | | | | | 0.2364 | 0.2273 | 0.2547 | 0.2294 | 0.2308 |
| G | | | | | | | | 0.0426 | 0.0502 | 0.056 | 0.0599 | 0.0413 |
| H | | | | | | | | 0.1761 | 0.1208 | 0.1845 | 0.1637 | 0.1187 |

**Cell Index at: 58:46:09**

| | 1 | 2 | 3 | 4 | 5 | 6 | 7 | 8 | 9 | 10 | 11 | 12 |
|---|---|---|---|---|---|---|---|---|---|---|---|---|
| A | | | | | | | | -0.0057 | 0.0125 | 0.0318 | 0.0059 | 0.0389 |
| B | | | | | | | | 0.0312 | 0.0295 | 0.0431 | 0.0691 | 0.0654 |
| C | | | | | | | | 0.1835 | 0.1932 | 0.2792 | 0.2762 | 0.2221 |
| D | | | | | | | | 0.1972 | 0.2497 | 0.2331 | 0.2551 | 0.229 |
| E | | | | | | | | 0.0966 | 0.0894 | 0.1191 | 0.1053 | 0.0779 |
| F | | | | | | | | 0.2293 | 0.2253 | 0.2639 | 0.2269 | 0.23 |
| G | | | | | | | | 0.0415 | 0.0555 | 0.0548 | 0.0617 | 0.039 |
| H | | | | | | | | 0.1743 | 0.1199 | 0.1844 | 0.1641 | 0.1142 |

**Cell Index at: 59:01:09**

| | 1 | 2 | 3 | 4 | 5 | 6 | 7 | 8 | 9 | 10 | 11 | 12 |
|---|---|---|---|---|---|---|---|---|---|---|---|---|
| A | | | | | | | | -0.0041 | 0.0129 | 0.0324 | 0.006 | 0.0405 |
| B | | | | | | | | 0.0353 | 0.0318 | 0.0431 | 0.0613 | 0.0677 |
| C | | | | | | | | 0.1804 | 0.1889 | 0.2736 | 0.2767 | 0.2184 |
| D | | | | | | | | 0.1947 | 0.2438 | 0.2392 | 0.2534 | 0.2317 |
| E | | | | | | | | 0.0916 | 0.0814 | 0.1088 | 0.1057 | 0.0734 |
| F | | | | | | | | 0.2332 | 0.2225 | 0.2587 | 0.2285 | 0.231 |
| G | | | | | | | | 0.0421 | 0.0547 | 0.0557 | 0.0609 | 0.0393 |
| H | | | | | | | | 0.181 | 0.12 | 0.1825 | 0.1644 | 0.1162 |

**Cell Index at: 59:16:09**

| | 1 | 2 | 3 | 4 | 5 | 6 | 7 | 8 | 9 | 10 | 11 | 12 |
|---|---|---|---|---|---|---|---|---|---|---|---|---|
| A | | | | | | | | -0.0014 | 0.0099 | 0.0316 | 0.0057 | 0.0411 |
| B | | | | | | | | 0.0357 | 0.0346 | 0.039 | 0.0673 | 0.0619 |
| C | | | | | | | | 0.1861 | 0.1924 | 0.2788 | 0.279 | 0.2251 |
| D | | | | | | | | 0.1918 | 0.2427 | 0.2327 | 0.2401 | 0.2295 |
| E | | | | | | | | 0.1011 | 0.0899 | 0.1184 | 0.114 | 0.078 |
| F | | | | | | | | 0.2394 | 0.2236 | 0.2566 | 0.2271 | 0.234 |
| G | | | | | | | | 0.0407 | 0.0531 | 0.053 | 0.0613 | 0.0375 |
| H | | | | | | | | 0.1807 | 0.1192 | 0.185 | 0.1666 | 0.1179 |

**Cell Index at: 59:31:10**

| | 1 | 2 | 3 | 4 | 5 | 6 | 7 | 8 | 9 | 10 | 11 | 12 |
|---|---|---|---|---|---|---|---|---|---|---|---|---|
| A | | | | | | | | -0.0031 | 0.0082 | 0.0298 | 0.0054 | 0.0402 |
| B | | | | | | | | 0.037 | 0.0276 | 0.0418 | 0.0681 | 0.066 |
| C | | | | | | | | 0.185 | 0.185 | 0.2791 | 0.2778 | 0.2205 |
| D | | | | | | | | 0.1843 | 0.2508 | 0.2379 | 0.2523 | 0.2355 |
| E | | | | | | | | 0.098 | 0.0904 | 0.1151 | 0.108 | 0.0783 |
| F | | | | | | | | 0.2463 | 0.2311 | 0.2619 | 0.2353 | 0.2314 |
| G | | | | | | | | 0.0465 | 0.0519 | 0.0592 | 0.0609 | 0.0358 |
| H | | | | | | | | 0.1746 | 0.1175 | 0.1847 | 0.1676 | 0.116 |

**Cell Index at: 59:46:10**

| | 1 | 2 | 3 | 4 | 5 | 6 | 7 | 8 | 9 | 10 | 11 | 12 |
|---|---|---|---|---|---|---|---|---|---|---|---|---|
| A | | | | | | | | -0.0035 | 0.0038 | 0.0275 | 0.0021 | 0.0386 |
| B | | | | | | | | 0.0312 | 0.0333 | 0.0429 | 0.065 | 0.0655 |
| C | | | | | | | | 0.1819 | 0.1902 | 0.2797 | 0.2807 | 0.2205 |
| D | | | | | | | | 0.1904 | 0.2452 | 0.2381 | 0.2458 | 0.2339 |
| E | | | | | | | | 0.0965 | 0.0878 | 0.1118 | 0.1069 | 0.0873 |
| F | | | | | | | | 0.2439 | 0.2232 | 0.2618 | 0.2291 | 0.2323 |
| G | | | | | | | | 0.0405 | 0.053 | 0.054 | 0.0586 | 0.0337 |
| H | | | | | | | | 0.1758 | 0.119 | 0.1828 | 0.1669 | 0.1186 |

**Cell Index at: 60:01:10**

| | 1 | 2 | 3 | 4 | 5 | 6 | 7 | 8 | 9 | 10 | 11 | 12 |
|---|---|---|---|---|---|---|---|---|---|---|---|---|
| A | | | | | | | | -0.0005 | 0.0055 | 0.0261 | 0.0033 | 0.0387 |
| B | | | | | | | | 0.038 | 0.0267 | 0.0415 | 0.0647 | 0.0635 |
| C | | | | | | | | 0.1874 | 0.1813 | 0.2727 | 0.2895 | 0.2185 |
| D | | | | | | | | 0.2074 | 0.2463 | 0.2347 | 0.2409 | 0.2323 |
| E | | | | | | | | 0.0895 | 0.0993 | 0.1184 | 0.1127 | 0.0763 |
| F | | | | | | | | 0.2397 | 0.2238 | 0.2597 | 0.2318 | 0.2316 |
| G | | | | | | | | 0.0371 | 0.0512 | 0.0545 | 0.0599 | 0.0348 |
| H | | | | | | | | 0.1776 | 0.1199 | 0.1804 | 0.17 | 0.118 |

**Cell Index at: 60:16:10**

| | 1 | 2 | 3 | 4 | 5 | 6 | 7 | 8 | 9 | 10 | 11 | 12 |
|---|---|---|---|---|---|---|---|---|---|---|---|---|
| A | | | | | | | | -0.0021 | -0.0004 | 0.0245 | 0.0041 | 0.0354 |
| B | | | | | | | | 0.0336 | 0.0303 | 0.0434 | 0.069 | 0.0645 |
| C | | | | | | | | 0.1871 | 0.1871 | 0.2826 | 0.2839 | 0.2277 |
| D | | | | | | | | 0.204 | 0.257 | 0.2315 | 0.244 | 0.2301 |
| E | | | | | | | | 0.1014 | 0.0935 | 0.1117 | 0.1092 | 0.0833 |
| F | | | | | | | | 0.2383 | 0.2235 | 0.2661 | 0.2339 | 0.232 |
| G | | | | | | | | 0.0453 | 0.0504 | 0.0568 | 0.0586 | 0.0391 |
| H | | | | | | | | 0.1763 | 0.1174 | 0.1826 | 0.1683 | 0.1213 |

**Cell Index at: 60:31:10**

| | 1 | 2 | 3 | 4 | 5 | 6 | 7 | 8 | 9 | 10 | 11 | 12 |
|---|---|---|---|---|---|---|---|---|---|---|---|---|
| A | | | | | | | | 0 | 0.0027 | 0.0293 | 0.0044 | 0.0371 |
| B | | | | | | | | 0.0311 | 0.032 | 0.0434 | 0.069 | 0.0652 |
| C | | | | | | | | 0.1833 | 0.1882 | 0.2812 | 0.2861 | 0.2228 |
| D | | | | | | | | 0.1977 | 0.2483 | 0.2356 | 0.2516 | 0.2411 |
| E | | | | | | | | 0.0993 | 0.0824 | 0.1143 | 0.109 | 0.0798 |
| F | | | | | | | | 0.2442 | 0.2256 | 0.2647 | 0.2354 | 0.2312 |
| G | | | | | | | | 0.0405 | 0.0479 | 0.0506 | 0.0602 | 0.0406 |
| H | | | | | | | | 0.1822 | 0.1177 | 0.1825 | 0.168 | 0.1207 |

**Cell Index at: 60:46:11**

| | 1 | 2 | 3 | 4 | 5 | 6 | 7 | 8 | 9 | 10 | 11 | 12 |
|---|---|---|---|---|---|---|---|---|---|---|---|---|
| A | | | | | | | | 0.002 | -0.0005 | 0.0285 | 0.0027 | 0.037 |
| B | | | | | | | | 0.0328 | 0.0327 | 0.0443 | 0.064 | 0.0624 |
| C | | | | | | | | 0.1829 | 0.1866 | 0.2775 | 0.2835 | 0.2277 |
| D | | | | | | | | 0.1971 | 0.2533 | 0.2383 | 0.258 | 0.2374 |
| E | | | | | | | | 0.1043 | 0.087 | 0.1091 | 0.1139 | 0.09 |
| F | | | | | | | | 0.2458 | 0.2268 | 0.2671 | 0.2366 | 0.2318 |
| G | | | | | | | | 0.0412 | 0.0483 | 0.056 | 0.0612 | 0.0409 |
| H | | | | | | | | 0.1785 | 0.1203 | 0.185 | 0.1721 | 0.1229 |

**Cell Index at: 61:01:11**

| | 1 | 2 | 3 | 4 | 5 | 6 | 7 | 8 | 9 | 10 | 11 | 12 |
|---|---|---|---|---|---|---|---|---|---|---|---|---|
| A | | | | | | | | -0.0008 | 0.0002 | 0.0278 | 0.0038 | 0.0366 |
| B | | | | | | | | 0.0333 | 0.0336 | 0.0425 | 0.0637 | 0.0661 |
| C | | | | | | | | 0.1824 | 0.1839 | 0.2819 | 0.2849 | 0.2278 |
| D | | | | | | | | 0.2037 | 0.2471 | 0.2343 | 0.2571 | 0.2378 |
| E | | | | | | | | 0.1031 | 0.0972 | 0.1138 | 0.1134 | 0.0794 |
| F | | | | | | | | 0.2449 | 0.2246 | 0.2689 | 0.2403 | 0.2295 |
| G | | | | | | | | 0.0394 | 0.0469 | 0.0513 | 0.0598 | 0.0416 |
| H | | | | | | | | 0.1772 | 0.1217 | 0.1826 | 0.1741 | 0.125 |

**Cell Index at: 61:16:12**

| | 1 | 2 | 3 | 4 | 5 | 6 | 7 | 8 | 9 | 10 | 11 | 12 |
|---|---|---|---|---|---|---|---|---|---|---|---|---|
| A | | | | | | | | 0.0017 | 0.0018 | 0.0318 | -0.0002 | 0.0394 |
| B | | | | | | | | 0.0352 | 0.0282 | 0.0387 | 0.0645 | 0.0632 |
| C | | | | | | | | 0.1884 | 0.186 | 0.2785 | 0.2794 | 0.2242 |
| D | | | | | | | | 0.2078 | 0.2442 | 0.2412 | 0.2532 | 0.2356 |
| E | | | | | | | | 0.1019 | 0.0931 | 0.1145 | 0.1167 | 0.0896 |
| F | | | | | | | | 0.2459 | 0.2268 | 0.2683 | 0.2359 | 0.224 |
| G | | | | | | | | 0.0397 | 0.0479 | 0.0542 | 0.0628 | 0.0399 |
| H | | | | | | | | 0.18 | 0.1225 | 0.1828 | 0.1735 | 0.1254 |

**Cell Index at: 61:31:13**

| | 1 | 2 | 3 | 4 | 5 | 6 | 7 | 8 | 9 | 10 | 11 | 12 |
|---|---|---|---|---|---|---|---|---|---|---|---|---|
| A | | | | | | | | -0.001 | 0.0024 | 0.0311 | 0.0012 | 0.0366 |
| B | | | | | | | | 0.0367 | 0.0348 | 0.0387 | 0.0649 | 0.0643 |
| C | | | | | | | | 0.1871 | 0.1866 | 0.2797 | 0.2859 | 0.224 |
| D | | | | | | | | 0.1967 | 0.24 | 0.2377 | 0.2588 | 0.2455 |
| E | | | | | | | | 0.0923 | 0.0914 | 0.1166 | 0.1083 | 0.0822 |
| F | | | | | | | | 0.2449 | 0.2332 | 0.2619 | 0.2398 | 0.2287 |
| G | | | | | | | | 0.0402 | 0.0488 | 0.0525 | 0.0611 | 0.0433 |
| H | | | | | | | | 0.1832 | 0.1238 | 0.1818 | 0.17 | 0.1296 |

**Cell Index at: 61:46:14**

| | 1 | 2 | 3 | 4 | 5 | 6 | 7 | 8 | 9 | 10 | 11 | 12 |
|---|---|---|---|---|---|---|---|---|---|---|---|---|
| A | | | | | | | | 0.0001 | -0.002 | 0.0321 | 0.0009 | 0.0351 |
| B | | | | | | | | 0.0391 | 0.0323 | 0.0391 | 0.0601 | 0.0628 |
| C | | | | | | | | 0.1905 | 0.1848 | 0.2786 | 0.2888 | 0.2253 |
| D | | | | | | | | 0.1981 | 0.2441 | 0.2398 | 0.2527 | 0.2467 |
| E | | | | | | | | 0.0979 | 0.099 | 0.1119 | 0.1141 | 0.0902 |
| F | | | | | | | | 0.2436 | 0.2235 | 0.2625 | 0.2433 | 0.2316 |
| G | | | | | | | | 0.0406 | 0.0453 | 0.0559 | 0.0615 | 0.0451 |
| H | | | | | | | | 0.1807 | 0.125 | 0.1879 | 0.1689 | 0.1274 |

**Cell Index at: 62:01:14**

| | 1 | 2 | 3 | 4 | 5 | 6 | 7 | 8 | 9 | 10 | 11 | 12 |
|---|---|---|---|---|---|---|---|---|---|---|---|---|
| A | | | | | | | | -0.0001 | 0.0017 | 0.0299 | 0.0041 | 0.035 |
| B | | | | | | | | 0.0342 | 0.0317 | 0.0403 | 0.0638 | 0.0672 |
| C | | | | | | | | 0.1902 | 0.1854 | 0.2817 | 0.2857 | 0.2309 |
| D | | | | | | | | 0.2024 | 0.2538 | 0.2398 | 0.2613 | 0.2361 |
| E | | | | | | | | 0.0944 | 0.0929 | 0.113 | 0.1112 | 0.078 |
| F | | | | | | | | 0.2534 | 0.2224 | 0.2668 | 0.2424 | 0.2276 |
| G | | | | | | | | 0.04 | 0.047 | 0.0573 | 0.0619 | 0.0446 |
| H | | | | | | | | 0.1814 | 0.1233 | 0.1899 | 0.1711 | 0.1299 |

**Cell Index at: 62:16:14**

| | 1 | 2 | 3 | 4 | 5 | 6 | 7 | 8 | 9 | 10 | 11 | 12 |
|---|---|---|---|---|---|---|---|---|---|---|---|---|
| A | | | | | | | | 0.0018 | 0.0027 | 0.0314 | 0.0045 | 0.0329 |
| B | | | | | | | | 0.0361 | 0.0327 | 0.0346 | 0.0662 | 0.0647 |
| C | | | | | | | | 0.1959 | 0.1873 | 0.2792 | 0.2926 | 0.2211 |
| D | | | | | | | | 0.2057 | 0.2481 | 0.2427 | 0.2594 | 0.2406 |
| E | | | | | | | | 0.1008 | 0.0908 | 0.1136 | 0.1072 | 0.0826 |
| F | | | | | | | | 0.2533 | 0.2176 | 0.265 | 0.2421 | 0.232 |
| G | | | | | | | | 0.0429 | 0.0523 | 0.0549 | 0.0615 | 0.0466 |
| H | | | | | | | | 0.1822 | 0.126 | 0.1911 | 0.1721 | 0.1284 |

**Cell Index at: 62:31:14**

| | 1 | 2 | 3 | 4 | 5 | 6 | 7 | 8 | 9 | 10 | 11 | 12 |
|---|---|---|---|---|---|---|---|---|---|---|---|---|
| A | | | | | | | | -0.0003 | 0.0038 | 0.0295 | 0.0023 | 0.0265 |
| B | | | | | | | | 0.0351 | 0.0337 | 0.0426 | 0.0666 | 0.0602 |
| C | | | | | | | | 0.1932 | 0.189 | 0.283 | 0.2928 | 0.2274 |
| D | | | | | | | | 0.2032 | 0.253 | 0.246 | 0.2571 | 0.2394 |
| E | | | | | | | | 0.0962 | 0.0918 | 0.1171 | 0.107 | 0.0824 |
| F | | | | | | | | 0.2447 | 0.2244 | 0.2693 | 0.2469 | 0.225 |
| G | | | | | | | | 0.0371 | 0.0454 | 0.0546 | 0.0597 | 0.0475 |
| H | | | | | | | | 0.1788 | 0.1266 | 0.189 | 0.1732 | 0.1286 |

**Cell Index at: 62:46:14**

| | 1 | 2 | 3 | 4 | 5 | 6 | 7 | 8 | 9 | 10 | 11 | 12 |
|---|---|---|---|---|---|---|---|---|---|---|---|---|
| A | | | | | | | | -0.0006 | 0.0042 | 0.0283 | 0.0038 | 0.0236 |
| B | | | | | | | | 0.0349 | 0.03 | 0.0415 | 0.0683 | 0.0614 |
| C | | | | | | | | 0.1935 | 0.1918 | 0.2861 | 0.2911 | 0.2274 |
| D | | | | | | | | 0.1979 | 0.2531 | 0.2341 | 0.2611 | 0.2449 |
| E | | | | | | | | 0.0987 | 0.089 | 0.1188 | 0.1092 | 0.0793 |
| F | | | | | | | | 0.2507 | 0.2261 | 0.2672 | 0.2456 | 0.2317 |
| G | | | | | | | | 0.035 | 0.0447 | 0.0564 | 0.0617 | 0.0465 |
| H | | | | | | | | 0.1852 | 0.1298 | 0.1871 | 0.1754 | 0.13 |

**Cell Index at: 63:01:14**

| | 1 | 2 | 3 | 4 | 5 | 6 | 7 | 8 | 9 | 10 | 11 | 12 |
|---|---|---|---|---|---|---|---|---|---|---|---|---|
| A | | | | | | | | 0.0008 | 0.0044 | 0.0334 | 0.0044 | 0.0262 |
| B | | | | | | | | 0.0383 | 0.0304 | 0.0379 | 0.0686 | 0.0618 |
| C | | | | | | | | 0.1988 | 0.1905 | 0.2821 | 0.2925 | 0.2296 |
| D | | | | | | | | 0.198 | 0.2484 | 0.2471 | 0.2585 | 0.2387 |
| E | | | | | | | | 0.0935 | 0.0918 | 0.1108 | 0.1158 | 0.0771 |
| F | | | | | | | | 0.2524 | 0.2182 | 0.2665 | 0.2498 | 0.2273 |
| G | | | | | | | | 0.0356 | 0.0465 | 0.0552 | 0.0592 | 0.0477 |
| H | | | | | | | | 0.1863 | 0.128 | 0.1913 | 0.1762 | 0.1301 |

**Cell Index at: 63:16:15**

| | 1 | 2 | 3 | 4 | 5 | 6 | 7 | 8 | 9 | 10 | 11 | 12 |
|---|---|---|---|---|---|---|---|---|---|---|---|---|
| A | | | | | | | | 0.004 | 0.0069 | 0.0311 | 0.0032 | 0.0272 |
| B | | | | | | | | 0.0286 | 0.0307 | 0.0372 | 0.0712 | 0.0629 |
| C | | | | | | | | 0.1995 | 0.189 | 0.289 | 0.2902 | 0.2324 |
| D | | | | | | | | 0.1912 | 0.2469 | 0.2467 | 0.2608 | 0.2424 |
| E | | | | | | | | 0.0976 | 0.0869 | 0.1088 | 0.1201 | 0.0766 |
| F | | | | | | | | 0.2518 | 0.2226 | 0.2694 | 0.251 | 0.2268 |
| G | | | | | | | | 0.0297 | 0.0449 | 0.0519 | 0.0587 | 0.0443 |
| H | | | | | | | | 0.1865 | 0.1328 | 0.1932 | 0.1725 | 0.1324 |

**Cell Index at: 63:31:16**

| | 1 | 2 | 3 | 4 | 5 | 6 | 7 | 8 | 9 | 10 | 11 | 12 |
|---|---|---|---|---|---|---|---|---|---|---|---|---|
| A | | | | | | | | 0.0018 | 0.0016 | 0.0337 | 0.0027 | 0.0269 |
| B | | | | | | | | 0.0302 | 0.0338 | 0.037 | 0.072 | 0.0619 |
| C | | | | | | | | 0.1999 | 0.1925 | 0.2863 | 0.2972 | 0.226 |
| D | | | | | | | | 0.1965 | 0.2477 | 0.24 | 0.2598 | 0.2413 |
| E | | | | | | | | 0.1005 | 0.0886 | 0.1148 | 0.1119 | 0.0773 |
| F | | | | | | | | 0.248 | 0.2232 | 0.2726 | 0.2453 | 0.2266 |
| G | | | | | | | | 0.0319 | 0.0472 | 0.0515 | 0.0575 | 0.047 |
| H | | | | | | | | 0.1878 | 0.13 | 0.1915 | 0.1777 | 0.1315 |

**Cell Index at: 63:46:16**

| | 1 | 2 | 3 | 4 | 5 | 6 | 7 | 8 | 9 | 10 | 11 | 12 |
|---|---|---|---|---|---|---|---|---|---|---|---|---|
| A | | | | | | | | 0.0055 | 0.0019 | 0.0345 | 0.0031 | 0.0296 |
| B | | | | | | | | 0.0345 | 0.0291 | 0.0378 | 0.0713 | 0.0634 |
| C | | | | | | | | 0.1955 | 0.1962 | 0.2863 | 0.2991 | 0.2275 |
| D | | | | | | | | 0.1995 | 0.2587 | 0.249 | 0.2716 | 0.2417 |
| E | | | | | | | | 0.0965 | 0.0875 | 0.1111 | 0.1044 | 0.0782 |
| F | | | | | | | | 0.2542 | 0.224 | 0.2682 | 0.2454 | 0.2301 |
| G | | | | | | | | 0.035 | 0.0458 | 0.0537 | 0.0598 | 0.0434 |

| | 1 | 2 | 3 | 4 | 5 | 6 | 7 | 8 | 9 | 10 | 11 | 12 |
|---|---|---|---|---|---|---|---|---|---|---|---|---|
| H | | | | | | | | 0.1853 | 0.1292 | 0.1924 | 0.1803 | 0.1334 |

**Cell Index at: 64:01:16**

| | 1 | 2 | 3 | 4 | 5 | 6 | 7 | 8 | 9 | 10 | 11 | 12 |
|---|---|---|---|---|---|---|---|---|---|---|---|---|
| A | | | | | | | | 0.0024 | 0.0019 | 0.0347 | 0.0037 | 0.0309 |
| B | | | | | | | | 0.0317 | 0.0293 | 0.0401 | 0.068 | 0.0641 |
| C | | | | | | | | 0.1934 | 0.1932 | 0.2887 | 0.2939 | 0.2291 |
| D | | | | | | | | 0.2032 | 0.2456 | 0.2467 | 0.2612 | 0.2461 |
| E | | | | | | | | 0.1006 | 0.0928 | 0.1137 | 0.1096 | 0.0815 |
| F | | | | | | | | 0.2557 | 0.2206 | 0.2642 | 0.247 | 0.2314 |
| G | | | | | | | | 0.0324 | 0.0433 | 0.0537 | 0.0525 | 0.0474 |
| H | | | | | | | | 0.1873 | 0.1296 | 0.1948 | 0.1811 | 0.1334 |

**Cell Index at: 64:16:16**

| | 1 | 2 | 3 | 4 | 5 | 6 | 7 | 8 | 9 | 10 | 11 | 12 |
|---|---|---|---|---|---|---|---|---|---|---|---|---|
| A | | | | | | | | 0.0038 | -0.0003 | 0.0336 | 0.0024 | 0.0294 |
| B | | | | | | | | 0.0285 | 0.0309 | 0.0389 | 0.0667 | 0.0625 |
| C | | | | | | | | 0.1981 | 0.1925 | 0.2871 | 0.2943 | 0.2286 |
| D | | | | | | | | 0.1963 | 0.2484 | 0.2383 | 0.2687 | 0.2408 |
| E | | | | | | | | 0.1003 | 0.0924 | 0.1124 | 0.1101 | 0.0783 |
| F | | | | | | | | 0.2492 | 0.2157 | 0.2665 | 0.2472 | 0.2329 |
| G | | | | | | | | 0.0355 | 0.0465 | 0.0508 | 0.0505 | 0.045 |
| H | | | | | | | | 0.1895 | 0.1302 | 0.1955 | 0.1808 | 0.1359 |

**Cell Index at: 64:31:16**

| | 1 | 2 | 3 | 4 | 5 | 6 | 7 | 8 | 9 | 10 | 11 | 12 |
|---|---|---|---|---|---|---|---|---|---|---|---|---|
| A | | | | | | | | 0.0032 | -0.0018 | 0.0334 | 0.0038 | 0.0309 |
| B | | | | | | | | 0.0312 | 0.0258 | 0.0394 | 0.0688 | 0.0631 |
| C | | | | | | | | 0.1951 | 0.1922 | 0.2908 | 0.293 | 0.2237 |
| D | | | | | | | | 0.1988 | 0.2579 | 0.2529 | 0.2681 | 0.2432 |
| E | | | | | | | | 0.0918 | 0.0836 | 0.1096 | 0.1029 | 0.0788 |
| F | | | | | | | | 0.2483 | 0.2107 | 0.2732 | 0.252 | 0.2331 |
| G | | | | | | | | 0.0337 | 0.0492 | 0.0512 | 0.0525 | 0.0446 |
| H | | | | | | | | 0.1899 | 0.1287 | 0.1961 | 0.1838 | 0.1362 |

**Cell Index at: 64:46:16**

| | 1 | 2 | 3 | 4 | 5 | 6 | 7 | 8 | 9 | 10 | 11 | 12 |
|---|---|---|---|---|---|---|---|---|---|---|---|---|
| A | | | | | | | | 0.0072 | -0.0024 | 0.033 | 0.0035 | 0.0296 |
| B | | | | | | | | 0.0321 | 0.0281 | 0.0386 | 0.0658 | 0.0646 |
| C | | | | | | | | 0.1925 | 0.1967 | 0.2994 | 0.2943 | 0.2257 |
| D | | | | | | | | 0.1952 | 0.2542 | 0.2513 | 0.261 | 0.2417 |
| E | | | | | | | | 0.0942 | 0.087 | 0.1199 | 0.1095 | 0.0742 |
| F | | | | | | | | 0.2511 | 0.2222 | 0.2694 | 0.2488 | 0.2383 |
| G | | | | | | | | 0.033 | 0.0486 | 0.0555 | 0.0549 | 0.042 |
| H | | | | | | | | 0.192 | 0.1246 | 0.1947 | 0.1844 | 0.1366 |

**Cell Index at: 65:01:16**

| | 1 | 2 | 3 | 4 | 5 | 6 | 7 | 8 | 9 | 10 | 11 | 12 |
|---|---|---|---|---|---|---|---|---|---|---|---|---|
| A | | | | | | | | 0.0064 | -0.0011 | 0.0323 | 0.0058 | 0.0256 |
| B | | | | | | | | 0.0308 | 0.0252 | 0.0389 | 0.068 | 0.0602 |
| C | | | | | | | | 0.1953 | 0.197 | 0.2912 | 0.2972 | 0.2245 |
| D | | | | | | | | 0.1957 | 0.2579 | 0.2594 | 0.2673 | 0.2458 |
| E | | | | | | | | 0.0874 | 0.0931 | 0.1136 | 0.1046 | 0.08 |
| F | | | | | | | | 0.2563 | 0.2215 | 0.2702 | 0.2494 | 0.2399 |
| G | | | | | | | | 0.037 | 0.0454 | 0.0571 | 0.0567 | 0.0466 |
| H | | | | | | | | 0.1928 | 0.1272 | 0.1987 | 0.1836 | 0.139 |

**Cell Index at: 65:16:17**

| | 1 | 2 | 3 | 4 | 5 | 6 | 7 | 8 | 9 | 10 | 11 | 12 |
|---|---|---|---|---|---|---|---|---|---|---|---|---|
| A | | | | | | | | 0.0076 | -0.0005 | 0.0342 | 0.0047 | 0.023 |
| B | | | | | | | | 0.0327 | 0.0275 | 0.0356 | 0.0703 | 0.0604 |
| C | | | | | | | | 0.1964 | 0.1949 | 0.2919 | 0.2954 | 0.227 |
| D | | | | | | | | 0.1946 | 0.2597 | 0.2485 | 0.2646 | 0.2354 |
| E | | | | | | | | 0.092 | 0.0929 | 0.1149 | 0.114 | 0.0732 |
| F | | | | | | | | 0.2446 | 0.2256 | 0.2678 | 0.2481 | 0.2355 |
| G | | | | | | | | 0.0345 | 0.0473 | 0.0567 | 0.0554 | 0.0454 |
| H | | | | | | | | 0.1942 | 0.1267 | 0.1995 | 0.1856 | 0.1366 |

**Cell Index at: 65:31:18**

| | 1 | 2 | 3 | 4 | 5 | 6 | 7 | 8 | 9 | 10 | 11 | 12 |
|---|---|---|---|---|---|---|---|---|---|---|---|---|
| A | | | | | | | | 0.008 | -0.0011 | 0.0344 | 0.0071 | 0.0261 |
| B | | | | | | | | 0.0337 | 0.0286 | 0.0425 | 0.0667 | 0.0647 |
| C | | | | | | | | 0.1915 | 0.1972 | 0.2908 | 0.2943 | 0.2281 |
| D | | | | | | | | 0.1919 | 0.2607 | 0.2545 | 0.269 | 0.2454 |
| E | | | | | | | | 0.0953 | 0.0839 | 0.1084 | 0.1149 | 0.0755 |
| F | | | | | | | | 0.2447 | 0.2228 | 0.266 | 0.2549 | 0.2372 |
| G | | | | | | | | 0.0355 | 0.0481 | 0.0571 | 0.0572 | 0.045 |
| H | | | | | | | | 0.1942 | 0.1233 | 0.1977 | 0.1871 | 0.141 |

**Cell Index at: 65:46:17**

| | 1 | 2 | 3 | 4 | 5 | 6 | 7 | 8 | 9 | 10 | 11 | 12 |
|---|---|---|---|---|---|---|---|---|---|---|---|---|
| A | | | | | | | | 0.0066 | -0.0021 | 0.0344 | 0.0044 | 0.0277 |
| B | | | | | | | | 0.0338 | 0.0268 | 0.041 | 0.0653 | 0.0616 |
| C | | | | | | | | 0.2003 | 0.194 | 0.2895 | 0.2994 | 0.2268 |
| D | | | | | | | | 0.1949 | 0.2702 | 0.2478 | 0.2711 | 0.241 |
| E | | | | | | | | 0.0972 | 0.091 | 0.0987 | 0.1188 | 0.0785 |
| F | | | | | | | | 0.2507 | 0.2234 | 0.2707 | 0.256 | 0.2402 |
| G | | | | | | | | 0.0389 | 0.0449 | 0.0585 | 0.0548 | 0.0442 |
| H | | | | | | | | 0.1937 | 0.1219 | 0.199 | 0.1874 | 0.1391 |

**Cell Index at: 66:01:17**

| | 1 | 2 | 3 | 4 | 5 | 6 | 7 | 8 | 9 | 10 | 11 | 12 |
|---|---|---|---|---|---|---|---|---|---|---|---|---|
| A | | | | | | | | 0.008 | -0.001 | 0.0344 | 0.0101 | 0.027 |
| B | | | | | | | | 0.037 | 0.027 | 0.0379 | 0.067 | 0.059 |
| C | | | | | | | | 0.2029 | 0.1967 | 0.2929 | 0.2927 | 0.2293 |
| D | | | | | | | | 0.2034 | 0.2562 | 0.2582 | 0.2724 | 0.2455 |
| E | | | | | | | | 0.1005 | 0.0881 | 0.1031 | 0.1195 | 0.0766 |
| F | | | | | | | | 0.2536 | 0.2185 | 0.2685 | 0.2528 | 0.2433 |
| G | | | | | | | | 0.04 | 0.0497 | 0.0589 | 0.0525 | 0.0454 |
| H | | | | | | | | 0.1961 | 0.122 | 0.2002 | 0.1889 | 0.1428 |

**Cell Index at: 66:16:17**

| | 1 | 2 | 3 | 4 | 5 | 6 | 7 | 8 | 9 | 10 | 11 | 12 |
|---|---|---|---|---|---|---|---|---|---|---|---|---|
| A | | | | | | | | 0.0075 | -0.0037 | 0.0322 | 0.0039 | 0.0309 |
| B | | | | | | | | 0.0353 | 0.0219 | 0.0368 | 0.0648 | 0.062 |
| C | | | | | | | | 0.2009 | 0.1952 | 0.2941 | 0.2915 | 0.2292 |
| D | | | | | | | | 0.1989 | 0.2697 | 0.2625 | 0.2681 | 0.2439 |
| E | | | | | | | | 0.0931 | 0.0824 | 0.1046 | 0.1144 | 0.071 |
| F | | | | | | | | 0.2488 | 0.2283 | 0.2694 | 0.2628 | 0.2488 |
| G | | | | | | | | 0.0423 | 0.0453 | 0.0569 | 0.055 | 0.0451 |
| H | | | | | | | | 0.1963 | 0.124 | 0.2 | 0.1866 | 0.1401 |

**Cell Index at: 66:31:16**

| | 1 | 2 | 3 | 4 | 5 | 6 | 7 | 8 | 9 | 10 | 11 | 12 |
|---|---|---|---|---|---|---|---|---|---|---|---|---|
| A | | | | | | | | 0.0043 | -0.0007 | 0.0336 | 0.0039 | 0.0297 |
| B | | | | | | | | 0.0388 | 0.0258 | 0.0376 | 0.0698 | 0.0623 |
| C | | | | | | | | 0.2054 | 0.1948 | 0.2906 | 0.2984 | 0.2234 |
| D | | | | | | | | 0.1991 | 0.2575 | 0.2641 | 0.2645 | 0.2511 |
| E | | | | | | | | 0.0867 | 0.0884 | 0.1046 | 0.1197 | 0.0781 |
| F | | | | | | | | 0.2537 | 0.2297 | 0.2658 | 0.2581 | 0.2434 |
| G | | | | | | | | 0.0359 | 0.0488 | 0.0574 | 0.051 | 0.0385 |
| H | | | | | | | | 0.1981 | 0.1241 | 0.2005 | 0.1911 | 0.1448 |

**Cell Index at: 66:46:17**

| | 1 | 2 | 3 | 4 | 5 | 6 | 7 | 8 | 9 | 10 | 11 | 12 |
|---|---|---|---|---|---|---|---|---|---|---|---|---|
| A | | | | | | | | 0.0066 | -0.0038 | 0.0335 | 0.0025 | 0.0332 |
| B | | | | | | | | 0.038 | 0.0279 | 0.0405 | 0.0665 | 0.0603 |
| C | | | | | | | | 0.2067 | 0.1984 | 0.2938 | 0.2983 | 0.2278 |
| D | | | | | | | | 0.204 | 0.2686 | 0.2607 | 0.2653 | 0.244 |
| E | | | | | | | | 0.0995 | 0.0953 | 0.1043 | 0.1135 | 0.0731 |
| F | | | | | | | | 0.2476 | 0.2283 | 0.2658 | 0.2541 | 0.2421 |

| | | | | | | | | | | | | |
|---|---|---|---|---|---|---|---|---|---|---|---|---|
| G | | | | | | | | 0.0363 | 0.0444 | 0.0592 | 0.056 | 0.0394 |
| H | | | | | | | | 0.1977 | 0.1292 | 0.2042 | 0.1894 | 0.1437 |

**Cell Index at: 67:01:18**

| | 1 | 2 | 3 | 4 | 5 | 6 | 7 | 8 | 9 | 10 | 11 | 12 |
|---|---|---|---|---|---|---|---|---|---|---|---|---|
| A | | | | | | | | 0.0055 | -0.0026 | 0.0325 | 0.0055 | 0.0274 |
| B | | | | | | | | 0.0366 | 0.0248 | 0.0402 | 0.0682 | 0.0629 |
| C | | | | | | | | 0.201 | 0.2005 | 0.2944 | 0.2979 | 0.2261 |
| D | | | | | | | | 0.2004 | 0.2695 | 0.2646 | 0.2656 | 0.2472 |
| E | | | | | | | | 0.094 | 0.0893 | 0.1093 | 0.1214 | 0.0757 |
| F | | | | | | | | 0.2508 | 0.2287 | 0.2661 | 0.2543 | 0.2431 |
| G | | | | | | | | 0.0352 | 0.0501 | 0.0579 | 0.0551 | 0.0339 |
| H | | | | | | | | 0.1982 | 0.1266 | 0.2025 | 0.1919 | 0.1414 |

**Cell Index at: 67:16:19**

| | 1 | 2 | 3 | 4 | 5 | 6 | 7 | 8 | 9 | 10 | 11 | 12 |
|---|---|---|---|---|---|---|---|---|---|---|---|---|
| A | | | | | | | | 0.0115 | -0.0011 | 0.0323 | 0.0046 | 0.0303 |
| B | | | | | | | | 0.0398 | 0.03 | 0.0427 | 0.065 | 0.0614 |
| C | | | | | | | | 0.2001 | 0.1972 | 0.2955 | 0.2927 | 0.2273 |
| D | | | | | | | | 0.2057 | 0.2631 | 0.2655 | 0.2659 | 0.2439 |
| E | | | | | | | | 0.098 | 0.0917 | 0.1144 | 0.117 | 0.0775 |
| F | | | | | | | | 0.2582 | 0.2275 | 0.2683 | 0.2521 | 0.2428 |
| G | | | | | | | | 0.032 | 0.0447 | 0.0624 | 0.0525 | 0.0339 |
| H | | | | | | | | 0.2025 | 0.1301 | 0.2047 | 0.1939 | 0.1428 |

**Cell Index at: 67:31:17**

| | 1 | 2 | 3 | 4 | 5 | 6 | 7 | 8 | 9 | 10 | 11 | 12 |
|---|---|---|---|---|---|---|---|---|---|---|---|---|
| A | | | | | | | | 0.004 | -0.0014 | 0.0328 | 0.0054 | 0.0262 |
| B | | | | | | | | 0.0392 | 0.0277 | 0.0401 | 0.0661 | 0.0616 |
| C | | | | | | | | 0.2014 | 0.2021 | 0.2941 | 0.2984 | 0.2217 |
| D | | | | | | | | 0.2038 | 0.2643 | 0.2704 | 0.2676 | 0.25 |
| E | | | | | | | | 0.0977 | 0.0919 | 0.1098 | 0.115 | 0.0798 |
| F | | | | | | | | 0.2535 | 0.2316 | 0.2663 | 0.2583 | 0.2418 |
| G | | | | | | | | 0.0395 | 0.0496 | 0.0618 | 0.0535 | 0.0399 |
| H | | | | | | | | 0.2047 | 0.1298 | 0.2011 | 0.1895 | 0.1438 |

**Cell Index at: 67:46:17**

| | 1 | 2 | 3 | 4 | 5 | 6 | 7 | 8 | 9 | 10 | 11 | 12 |
|---|---|---|---|---|---|---|---|---|---|---|---|---|
| A | | | | | | | | 0.0052 | -0.0018 | 0.0323 | 0.0048 | 0.0306 |
| B | | | | | | | | 0.0419 | 0.0261 | 0.0429 | 0.0653 | 0.0672 |
| C | | | | | | | | 0.1966 | 0.1965 | 0.2941 | 0.2951 | 0.228 |
| D | | | | | | | | 0.2044 | 0.2673 | 0.2683 | 0.2673 | 0.2545 |
| E | | | | | | | | 0.0965 | 0.0905 | 0.1128 | 0.118 | 0.0767 |
| F | | | | | | | | 0.2512 | 0.2335 | 0.2685 | 0.2508 | 0.246 |
| G | | | | | | | | 0.0343 | 0.0544 | 0.0628 | 0.0551 | 0.0375 |
| H | | | | | | | | 0.2073 | 0.126 | 0.2062 | 0.1895 | 0.1445 |

**Cell Index at: 68:01:18**

| | 1 | 2 | 3 | 4 | 5 | 6 | 7 | 8 | 9 | 10 | 11 | 12 |
|---|---|---|---|---|---|---|---|---|---|---|---|---|
| A | | | | | | | | 0.0045 | -0.0012 | 0.0345 | 0.0052 | 0.0273 |
| B | | | | | | | | 0.0372 | 0.0256 | 0.0379 | 0.0684 | 0.0608 |
| C | | | | | | | | 0.2056 | 0.1971 | 0.297 | 0.3004 | 0.2209 |
| D | | | | | | | | 0.1959 | 0.2695 | 0.2641 | 0.2671 | 0.2473 |
| E | | | | | | | | 0.0983 | 0.0948 | 0.112 | 0.1177 | 0.0773 |
| F | | | | | | | | 0.2455 | 0.2346 | 0.2666 | 0.258 | 0.2404 |
| G | | | | | | | | 0.0333 | 0.0491 | 0.0605 | 0.0505 | 0.0362 |
| H | | | | | | | | 0.2069 | 0.1254 | 0.2043 | 0.1895 | 0.147 |

**Cell Index at: 68:16:19**

| | 1 | 2 | 3 | 4 | 5 | 6 | 7 | 8 | 9 | 10 | 11 | 12 |
|---|---|---|---|---|---|---|---|---|---|---|---|---|
| A | | | | | | | | 0.0049 | -0.0018 | 0.036 | 0.0071 | 0.029 |
| B | | | | | | | | 0.0372 | 0.028 | 0.0421 | 0.0666 | 0.0638 |
| C | | | | | | | | 0.2 | 0.2008 | 0.2982 | 0.2965 | 0.2233 |
| D | | | | | | | | 0.1945 | 0.2679 | 0.2626 | 0.275 | 0.2471 |
| E | | | | | | | | 0.094 | 0.0908 | 0.1045 | 0.1111 | 0.0752 |
| F | | | | | | | | 0.2528 | 0.2359 | 0.2744 | 0.2586 | 0.2458 |
| G | | | | | | | | 0.0387 | 0.0514 | 0.0619 | 0.0519 | 0.0344 |
| H | | | | | | | | 0.2101 | 0.1283 | 0.2014 | 0.1935 | 0.1484 |

**Cell Index at: 68:31:20**

| | 1 | 2 | 3 | 4 | 5 | 6 | 7 | 8 | 9 | 10 | 11 | 12 |
|---|---|---|---|---|---|---|---|---|---|---|---|---|
| A | | | | | | | | 0.0055 | -0.0001 | 0.0336 | 0.0047 | 0.0287 |
| B | | | | | | | | 0.0396 | 0.0259 | 0.0399 | 0.0639 | 0.065 |
| C | | | | | | | | 0.2043 | 0.2007 | 0.2954 | 0.2979 | 0.2251 |
| D | | | | | | | | 0.2021 | 0.2713 | 0.2674 | 0.2746 | 0.2491 |
| E | | | | | | | | 0.1009 | 0.0885 | 0.1087 | 0.1143 | 0.0777 |
| F | | | | | | | | 0.2558 | 0.2372 | 0.2717 | 0.2576 | 0.2453 |
| G | | | | | | | | 0.0383 | 0.0488 | 0.063 | 0.0522 | 0.0373 |
| H | | | | | | | | 0.2083 | 0.1282 | 0.195 | 0.1925 | 0.1483 |

**Cell Index at: 68:46:21**

| | 1 | 2 | 3 | 4 | 5 | 6 | 7 | 8 | 9 | 10 | 11 | 12 |
|---|---|---|---|---|---|---|---|---|---|---|---|---|
| A | | | | | | | | 0.0079 | 0.0003 | 0.0331 | 0.0044 | 0.0266 |
| B | | | | | | | | 0.0403 | 0.0241 | 0.0396 | 0.0638 | 0.0634 |
| C | | | | | | | | 0.2063 | 0.1991 | 0.2969 | 0.2955 | 0.2208 |
| D | | | | | | | | 0.199 | 0.2633 | 0.2652 | 0.2737 | 0.245 |
| E | | | | | | | | 0.1071 | 0.0841 | 0.1148 | 0.1087 | 0.0773 |
| F | | | | | | | | 0.2421 | 0.239 | 0.2687 | 0.2628 | 0.2446 |
| G | | | | | | | | 0.0391 | 0.0499 | 0.063 | 0.0537 | 0.0348 |
| H | | | | | | | | 0.209 | 0.1255 | 0.1923 | 0.1955 | 0.1459 |

**Cell Index at: 69:01:22**

| | 1 | 2 | 3 | 4 | 5 | 6 | 7 | 8 | 9 | 10 | 11 | 12 |
|---|---|---|---|---|---|---|---|---|---|---|---|---|
| A | | | | | | | | 0.0063 | -0.0033 | 0.0338 | 0.0073 | 0.028 |
| B | | | | | | | | 0.0416 | 0.0273 | 0.0408 | 0.0645 | 0.066 |
| C | | | | | | | | 0.2025 | 0.2011 | 0.304 | 0.3007 | 0.226 |
| D | | | | | | | | 0.1969 | 0.2736 | 0.2725 | 0.2746 | 0.254 |
| E | | | | | | | | 0.1036 | 0.0839 | 0.1113 | 0.1076 | 0.0773 |
| F | | | | | | | | 0.2463 | 0.2367 | 0.2679 | 0.258 | 0.2478 |
| G | | | | | | | | 0.0364 | 0.0459 | 0.0631 | 0.0527 | 0.0341 |
| H | | | | | | | | 0.2115 | 0.127 | 0.1977 | 0.1924 | 0.1492 |

**Cell Index at: 69:16:23**

| | 1 | 2 | 3 | 4 | 5 | 6 | 7 | 8 | 9 | 10 | 11 | 12 |
|---|---|---|---|---|---|---|---|---|---|---|---|---|
| A | | | | | | | | 0.0031 | -0.0028 | 0.0332 | 0.0047 | 0.0268 |
| B | | | | | | | | 0.0393 | 0.0254 | 0.038 | 0.067 | 0.0597 |
| C | | | | | | | | 0.2078 | 0.1981 | 0.3021 | 0.3078 | 0.2282 |
| D | | | | | | | | 0.1954 | 0.2683 | 0.2711 | 0.2723 | 0.2458 |
| E | | | | | | | | 0.0961 | 0.0886 | 0.1141 | 0.1195 | 0.0744 |
| F | | | | | | | | 0.2494 | 0.2391 | 0.2656 | 0.2618 | 0.2519 |
| G | | | | | | | | 0.0354 | 0.0447 | 0.0616 | 0.0561 | 0.0351 |
| H | | | | | | | | 0.2082 | 0.1278 | 0.1998 | 0.1929 | 0.1513 |

**Cell Index at: 69:31:24**

| | 1 | 2 | 3 | 4 | 5 | 6 | 7 | 8 | 9 | 10 | 11 | 12 |
|---|---|---|---|---|---|---|---|---|---|---|---|---|
| A | | | | | | | | 0.0045 | -0.0047 | 0.0321 | 0.0048 | 0.0289 |
| B | | | | | | | | 0.0425 | 0.0268 | 0.0422 | 0.0673 | 0.0612 |
| C | | | | | | | | 0.2081 | 0.2014 | 0.3065 | 0.305 | 0.2244 |
| D | | | | | | | | 0.1999 | 0.2779 | 0.2691 | 0.273 | 0.2532 |
| E | | | | | | | | 0.1063 | 0.0896 | 0.1056 | 0.1182 | 0.0717 |
| F | | | | | | | | 0.2483 | 0.2325 | 0.2766 | 0.2603 | 0.2479 |
| G | | | | | | | | 0.0318 | 0.045 | 0.0619 | 0.053 | 0.0335 |
| H | | | | | | | | 0.2111 | 0.1284 | 0.1953 | 0.1937 | 0.1538 |

**Cell Index at: 69:46:23**

| | 1 | 2 | 3 | 4 | 5 | 6 | 7 | 8 | 9 | 10 | 11 | 12 |
|---|---|---|---|---|---|---|---|---|---|---|---|---|
| A | | | | | | | | 0.0024 | -0.0034 | 0.0332 | 0.0085 | 0.0246 |
| B | | | | | | | | 0.0394 | 0.0206 | 0.041 | 0.0658 | 0.0603 |
| C | | | | | | | | 0.2072 | 0.2072 | 0.3056 | 0.3095 | 0.2254 |
| D | | | | | | | | 0.1978 | 0.2731 | 0.2685 | 0.2694 | 0.2518 |
| E | | | | | | | | 0.1087 | 0.0824 | 0.1116 | 0.1203 | 0.0752 |

| | | | | | | | | | | | | |
|---|---|---|---|---|---|---|---|---|---|---|---|---|
| F | | | | | | | | 0.2473 | 0.2319 | 0.2791 | 0.265 | 0.2461 |
| G | | | | | | | | 0.033 | 0.0459 | 0.0608 | 0.0559 | 0.0309 |
| H | | | | | | | | 0.2113 | 0.1288 | 0.2003 | 0.1943 | 0.1516 |

**Cell Index at: 70:01:24**

| | 1 | 2 | 3 | 4 | 5 | 6 | 7 | 8 | 9 | 10 | 11 | 12 |
|---|---|---|---|---|---|---|---|---|---|---|---|---|
| A | | | | | | | | 0.0026 | -0.0037 | 0.0341 | 0.0039 | 0.0272 |
| B | | | | | | | | 0.0393 | 0.0264 | 0.044 | 0.0688 | 0.0595 |
| C | | | | | | | | 0.2051 | 0.2044 | 0.3012 | 0.3051 | 0.2294 |
| D | | | | | | | | 0.1932 | 0.2686 | 0.2727 | 0.2807 | 0.2552 |
| E | | | | | | | | 0.1002 | 0.0801 | 0.1152 | 0.1112 | 0.0753 |
| F | | | | | | | | 0.2507 | 0.2357 | 0.2761 | 0.2593 | 0.2489 |
| G | | | | | | | | 0.0326 | 0.0434 | 0.0587 | 0.0554 | 0.0309 |
| H | | | | | | | | 0.2138 | 0.1291 | 0.1967 | 0.1942 | 0.1507 |

**Cell Index at: 70:16:24**

| | 1 | 2 | 3 | 4 | 5 | 6 | 7 | 8 | 9 | 10 | 11 | 12 |
|---|---|---|---|---|---|---|---|---|---|---|---|---|
| A | | | | | | | | 0.0038 | -0.0055 | 0.0301 | 0.0076 | 0.0259 |
| B | | | | | | | | 0.0431 | 0.0278 | 0.0398 | 0.0724 | 0.0605 |
| C | | | | | | | | 0.2092 | 0.2028 | 0.3117 | 0.3026 | 0.2306 |
| D | | | | | | | | 0.1997 | 0.2734 | 0.2769 | 0.2764 | 0.2478 |
| E | | | | | | | | 0.0991 | 0.0839 | 0.1133 | 0.1167 | 0.0772 |
| F | | | | | | | | 0.2561 | 0.2326 | 0.2801 | 0.2655 | 0.2512 |
| G | | | | | | | | 0.033 | 0.0432 | 0.0585 | 0.0533 | 0.0318 |
| H | | | | | | | | 0.2125 | 0.1276 | 0.1958 | 0.1966 | 0.1511 |

**Cell Index at: 70:31:25**

| | 1 | 2 | 3 | 4 | 5 | 6 | 7 | 8 | 9 | 10 | 11 | 12 |
|---|---|---|---|---|---|---|---|---|---|---|---|---|
| A | | | | | | | | 0.0037 | -0.0028 | 0.0338 | 0.0062 | 0.0292 |
| B | | | | | | | | 0.0434 | 0.0293 | 0.0418 | 0.0707 | 0.0584 |
| C | | | | | | | | 0.211 | 0.2078 | 0.3077 | 0.3062 | 0.2248 |
| D | | | | | | | | 0.1978 | 0.2823 | 0.2772 | 0.2746 | 0.2469 |
| E | | | | | | | | 0.1072 | 0.0938 | 0.1118 | 0.1097 | 0.0779 |
| F | | | | | | | | 0.2546 | 0.2325 | 0.2786 | 0.2694 | 0.2552 |
| G | | | | | | | | 0.0253 | 0.0441 | 0.0591 | 0.0563 | 0.0304 |
| H | | | | | | | | 0.2128 | 0.1257 | 0.196 | 0.1979 | 0.1521 |

**Cell Index at: 70:46:25**

| | 1 | 2 | 3 | 4 | 5 | 6 | 7 | 8 | 9 | 10 | 11 | 12 |
|---|---|---|---|---|---|---|---|---|---|---|---|---|
| A | | | | | | | | -0.0028 | -0.0007 | 0.031 | 0.005 | 0.0231 |
| B | | | | | | | | 0.0433 | 0.0215 | 0.0441 | 0.0706 | 0.0608 |
| C | | | | | | | | 0.2094 | 0.1999 | 0.3042 | 0.3101 | 0.2254 |
| D | | | | | | | | 0.1958 | 0.277 | 0.2775 | 0.283 | 0.2507 |
| E | | | | | | | | 0.1091 | 0.0868 | 0.1065 | 0.1164 | 0.073 |
| F | | | | | | | | 0.253 | 0.2381 | 0.2797 | 0.2666 | 0.247 |
| G | | | | | | | | 0.0275 | 0.0434 | 0.0623 | 0.054 | 0.0314 |
| H | | | | | | | | 0.2118 | 0.1269 | 0.1945 | 0.1976 | 0.1514 |

**Cell Index at: 71:01:26**

| | 1 | 2 | 3 | 4 | 5 | 6 | 7 | 8 | 9 | 10 | 11 | 12 |
|---|---|---|---|---|---|---|---|---|---|---|---|---|
| A | | | | | | | | 0.0026 | -0.0047 | 0.0332 | 0.005 | 0.0275 |
| B | | | | | | | | 0.0415 | 0.0272 | 0.0409 | 0.0697 | 0.0655 |
| C | | | | | | | | 0.2136 | 0.2071 | 0.3057 | 0.308 | 0.2257 |
| D | | | | | | | | 0.1954 | 0.2723 | 0.2792 | 0.2798 | 0.2528 |
| E | | | | | | | | 0.1119 | 0.0844 | 0.1154 | 0.11 | 0.0732 |
| F | | | | | | | | 0.2522 | 0.2379 | 0.2834 | 0.2694 | 0.2526 |
| G | | | | | | | | 0.0286 | 0.0477 | 0.0589 | 0.0555 | 0.0308 |
| H | | | | | | | | 0.2092 | 0.129 | 0.1952 | 0.1982 | 0.1525 |

**Cell Index at: 71:16:27**

| | 1 | 2 | 3 | 4 | 5 | 6 | 7 | 8 | 9 | 10 | 11 | 12 |
|---|---|---|---|---|---|---|---|---|---|---|---|---|
| A | | | | | | | | 0.0013 | -0.0023 | 0.0311 | 0.0051 | 0.0266 |
| B | | | | | | | | 0.0442 | 0.0263 | 0.0402 | 0.0718 | 0.0573 |
| C | | | | | | | | 0.2056 | 0.2046 | 0.3059 | 0.3067 | 0.223 |
| D | | | | | | | | 0.2029 | 0.2729 | 0.2845 | 0.2768 | 0.2481 |
| E | | | | | | | | 0.1099 | 0.0855 | 0.1227 | 0.1082 | 0.0746 |
| F | | | | | | | | 0.2593 | 0.2323 | 0.2796 | 0.2675 | 0.2558 |
| G | | | | | | | | 0.0271 | 0.0419 | 0.0583 | 0.0537 | 0.0338 |
| H | | | | | | | | 0.2118 | 0.1311 | 0.1962 | 0.1987 | 0.1534 |

**Cell Index at: 71:31:28**

| | 1 | 2 | 3 | 4 | 5 | 6 | 7 | 8 | 9 | 10 | 11 | 12 |
|---|---|---|---|---|---|---|---|---|---|---|---|---|
| A | | | | | | | | 0.0033 | -0.0045 | 0.0344 | 0.0052 | 0.0235 |
| B | | | | | | | | 0.0478 | 0.0257 | 0.0439 | 0.0762 | 0.058 |
| C | | | | | | | | 0.2134 | 0.2085 | 0.3043 | 0.305 | 0.2307 |
| D | | | | | | | | 0.1919 | 0.2772 | 0.2741 | 0.287 | 0.2522 |
| E | | | | | | | | 0.1099 | 0.0921 | 0.1179 | 0.109 | 0.0814 |
| F | | | | | | | | 0.2514 | 0.2316 | 0.2786 | 0.267 | 0.2548 |
| G | | | | | | | | 0.0323 | 0.0376 | 0.0549 | 0.0509 | 0.0315 |
| H | | | | | | | | 0.2119 | 0.1296 | 0.1923 | 0.1979 | 0.1508 |

**Cell Index at: 71:46:29**

| | 1 | 2 | 3 | 4 | 5 | 6 | 7 | 8 | 9 | 10 | 11 | 12 |
|---|---|---|---|---|---|---|---|---|---|---|---|---|
| A | | | | | | | | -0.0023 | -0.0028 | 0.0306 | 0.0056 | 0.0277 |
| B | | | | | | | | 0.0519 | 0.0251 | 0.0462 | 0.0764 | 0.0557 |
| C | | | | | | | | 0.2135 | 0.1991 | 0.2927 | 0.3069 | 0.2274 |
| D | | | | | | | | 0.196 | 0.2823 | 0.2848 | 0.2814 | 0.2524 |
| E | | | | | | | | 0.1082 | 0.0876 | 0.1268 | 0.1129 | 0.0762 |
| F | | | | | | | | 0.2537 | 0.2246 | 0.2804 | 0.265 | 0.2549 |
| G | | | | | | | | 0.0325 | 0.0419 | 0.0604 | 0.0474 | 0.0338 |
| H | | | | | | | | 0.2097 | 0.1337 | 0.1988 | 0.198 | 0.1534 |

**Cell Index at: 72:01:29**

| | 1 | 2 | 3 | 4 | 5 | 6 | 7 | 8 | 9 | 10 | 11 | 12 |
|---|---|---|---|---|---|---|---|---|---|---|---|---|
| A | | | | | | | | 0.0012 | -0.0055 | 0.0327 | 0.0066 | 0.0261 |
| B | | | | | | | | 0.0477 | 0.028 | 0.0433 | 0.0754 | 0.0556 |
| C | | | | | | | | 0.2119 | 0.208 | 0.3049 | 0.3067 | 0.2272 |
| D | | | | | | | | 0.1928 | 0.2821 | 0.2827 | 0.2841 | 0.249 |
| E | | | | | | | | 0.1158 | 0.0797 | 0.1161 | 0.1097 | 0.0774 |
| F | | | | | | | | 0.2541 | 0.2281 | 0.2761 | 0.2624 | 0.2568 |
| G | | | | | | | | 0.0343 | 0.0313 | 0.0624 | 0.0528 | 0.0325 |
| H | | | | | | | | 0.2096 | 0.1325 | 0.198 | 0.1985 | 0.1597 |

| | | | | | | | | | | | | | |
|---|---|---|---|---|---|---|---|---|---|---|---|---|---|
| A vs B | 0.007764 | B vs C | 0.000377 | C vs D | 0.827768 | D vs E | 0.000221 | E vs F | 1.1E-06 | F vs G | 4.55E-08 | G vs H | 0.000217 |
| A vs C | 0.000183 | B vs D | 6.49E-05 | C vs E | 0.001242 | D vs F | 0.895914 | E vs G | 0.000991 | F vs H | 0.00347 | | |
| A vs D | 3.12E-05 | B vs E | 0.002874 | C vs F | 0.87988 | D vs G | 8.56E-05 | E vs H | 0.002568 | | | | |
| A vs E | 6.9E-05 | B vs F | 7.15E-08 | C vs G | 0.000411 | D vs H | 0.009333 | | | | | | |
| A vs F | 1.61E-08 | B vs G | 0.485343 | C vs H | 0.030785 | | | | | | | | |
| A vs G | 0.014106 | B vs H | 0.0002 | | | | | | | | | | |
| A vs H | 5.32E-05 | | | | | | | | | | | | |

**Cell Index at: 72:16:30**

| | 1 | 2 | 3 | 4 | 5 | 6 | 7 | 8 | 9 | 10 | 11 | 12 |
|---|---|---|---|---|---|---|---|---|---|---|---|---|
| A | | | | | | | | 0.0008 | -0.0014 | 0.0306 | 0.0042 | 0.0245 |
| B | | | | | | | | 0.0478 | 0.0138 | 0.0458 | 0.0754 | 0.0572 |
| C | | | | | | | | 0.2022 | 0.2089 | 0.2953 | 0.3101 | 0.2305 |
| D | | | | | | | | 0.2056 | 0.2755 | 0.2826 | 0.2785 | 0.2548 |
| E | | | | | | | | 0.1004 | 0.0833 | 0.1155 | 0.1133 | 0.0807 |
| F | | | | | | | | 0.2553 | 0.2357 | 0.2796 | 0.2666 | 0.2597 |
| G | | | | | | | | 0.0331 | 0.0373 | 0.0598 | 0.0525 | 0.031 |
| H | | | | | | | | 0.2119 | 0.1324 | 0.2022 | 0.2031 | 0.155 |

| *Count* | *Sum* | *Average* | *Variance* |
|---|---|---|---|
| 5 | 0.0611 | 0.01222 | 0.00027 |
| 5 | 0.25 | 0.05 | 0.000303 |
| 5 | 1.2587 | 0.25174 | 0.002487 |
| 5 | 1.2907 | 0.25814 | 0.001551 |
| 5 | 0.4987 | 0.09974 | 0.000381 |
| 5 | 1.2775 | 0.2555 | 0.000307 |
| 5 | 0.2133 | 0.04266 | 0.000199 |
| 5 | 0.8983 | 0.17966 | 0.001053 |

**Cell Index at: 72:31:30**

| | 1 | 2 | 3 | 4 | 5 | 6 | 7 | 8 | 9 | 10 | 11 | 12 |
|---|---|---|---|---|---|---|---|---|---|---|---|---|
| A | | | | | | | | -0.0005 | -0.0043 | 0.0351 | 0.003 | 0.0273 |
| B | | | | | | | | 0.0481 | 0.0202 | 0.0422 | 0.0748 | 0.06 |
| C | | | | | | | | 0.2077 | 0.2063 | 0.2963 | 0.3142 | 0.2293 |
| D | | | | | | | | 0.1982 | 0.2749 | 0.2814 | 0.2669 | 0.2547 |
| E | | | | | | | | 0.1048 | 0.0865 | 0.1084 | 0.1117 | 0.0816 |
| F | | | | | | | | 0.2587 | 0.2375 | 0.2765 | 0.2665 | 0.2574 |
| G | | | | | | | | 0.0343 | 0.037 | 0.0646 | 0.0539 | 0.0343 |
| H | | | | | | | | 0.2128 | 0.1373 | 0.2043 | 0.2012 | 0.1574 |

| *SS* | *df* | *MS* | *F* | *P-value* | *F crit* |
|---|---|---|---|---|---|
| 0.383849 | 7 | 0.054836 | 66.9754 | 2.63E-17 | 2.312741 |
| 0.0262 | 32 | 0.000819 | | | |
| 0.410049 | 39 | | | | |

**Cell Index at: 72:46:31**

| | 1 | 2 | 3 | 4 | 5 | 6 | 7 | 8 | 9 | 10 | 11 | 12 |
|---|---|---|---|---|---|---|---|---|---|---|---|---|
| A | | | | | | | | -0.0029 | -0.0047 | 0.0321 | 0.0034 | 0.0243 |
| B | | | | | | | | 0.0441 | 0.0152 | 0.0402 | 0.0763 | 0.0586 |
| C | | | | | | | | 0.2111 | 0.2045 | 0.3025 | 0.3166 | 0.2286 |

| | | | | | | | | | | | | |
|---|---|---|---|---|---|---|---|---|---|---|---|---|
| D | | | | | | | | 0.1953 | 0.2775 | 0.2782 | 0.285 | 0.2516 |
| E | | | | | | | | 0.1039 | 0.0908 | 0.1176 | 0.1099 | 0.0796 |
| F | | | | | | | | 0.256 | 0.2382 | 0.2747 | 0.2726 | 0.2554 |
| G | | | | | | | | 0.0369 | 0.0409 | 0.0648 | 0.0514 | 0.0368 |
| H | | | | | | | | 0.2129 | 0.1349 | 0.2053 | 0.2045 | 0.1549 |

**Cell Index at: 73:01:32**

| | 1 | 2 | 3 | 4 | 5 | 6 | 7 | 8 | 9 | 10 | 11 | 12 |
|---|---|---|---|---|---|---|---|---|---|---|---|---|
| A | | | | | | | | -0.0032 | -0.0011 | 0.0327 | 0.0023 | 0.0242 |
| B | | | | | | | | 0.0487 | 0.02 | 0.0398 | 0.0761 | 0.0596 |
| C | | | | | | | | 0.2128 | 0.2111 | 0.3005 | 0.3155 | 0.2261 |
| D | | | | | | | | 0.199 | 0.2742 | 0.2854 | 0.2774 | 0.2608 |
| E | | | | | | | | 0.1055 | 0.0887 | 0.1125 | 0.1033 | 0.0837 |
| F | | | | | | | | 0.2641 | 0.2451 | 0.2767 | 0.2703 | 0.2542 |
| G | | | | | | | | 0.037 | 0.0359 | 0.0632 | 0.0509 | 0.0347 |
| H | | | | | | | | 0.2143 | 0.138 | 0.2052 | 0.2036 | 0.1575 |

**Cell Index at: 73:16:33**

| | 1 | 2 | 3 | 4 | 5 | 6 | 7 | 8 | 9 | 10 | 11 | 12 |
|---|---|---|---|---|---|---|---|---|---|---|---|---|
| A | | | | | | | | 0.0033 | -0.0031 | 0.0365 | 0.0073 | 0.0221 |
| B | | | | | | | | 0.0473 | 0.0165 | 0.0375 | 0.0752 | 0.0576 |
| C | | | | | | | | 0.2035 | 0.2043 | 0.3025 | 0.3169 | 0.2251 |
| D | | | | | | | | 0.1995 | 0.279 | 0.2841 | 0.2911 | 0.2535 |
| E | | | | | | | | 0.112 | 0.0925 | 0.1141 | 0.1079 | 0.0858 |
| F | | | | | | | | 0.2618 | 0.2462 | 0.2774 | 0.2696 | 0.2544 |
| G | | | | | | | | 0.036 | 0.0383 | 0.0655 | 0.049 | 0.0378 |
| H | | | | | | | | 0.2135 | 0.1353 | 0.2026 | 0.1997 | 0.1547 |

**Cell Index at: 73:31:33**

| | 1 | 2 | 3 | 4 | 5 | 6 | 7 | 8 | 9 | 10 | 11 | 12 |
|---|---|---|---|---|---|---|---|---|---|---|---|---|
| A | | | | | | | | 0.0028 | -0.0027 | 0.0333 | 0.0044 | 0.0221 |
| B | | | | | | | | 0.0463 | 0.0233 | 0.0395 | 0.0757 | 0.0585 |
| C | | | | | | | | 0.2097 | 0.2073 | 0.3011 | 0.32 | 0.2318 |
| D | | | | | | | | 0.2033 | 0.2759 | 0.2887 | 0.2801 | 0.2585 |
| E | | | | | | | | 0.1125 | 0.0814 | 0.1146 | 0.1104 | 0.076 |
| F | | | | | | | | 0.2586 | 0.2487 | 0.2812 | 0.2704 | 0.256 |
| G | | | | | | | | 0.0348 | 0.0383 | 0.0613 | 0.0531 | 0.0345 |
| H | | | | | | | | 0.2133 | 0.1369 | 0.2046 | 0.2042 | 0.1577 |

**Cell Index at: 73:46:34**

| | 1 | 2 | 3 | 4 | 5 | 6 | 7 | 8 | 9 | 10 | 11 | 12 |
|---|---|---|---|---|---|---|---|---|---|---|---|---|
| A | | | | | | | | 0.0027 | -0.0034 | 0.0319 | 0.0023 | 0.0226 |
| B | | | | | | | | 0.0456 | 0.0207 | 0.0403 | 0.0789 | 0.054 |
| C | | | | | | | | 0.2079 | 0.2035 | 0.3056 | 0.3205 | 0.2291 |
| D | | | | | | | | 0.1992 | 0.2779 | 0.2778 | 0.2843 | 0.2607 |
| E | | | | | | | | 0.1026 | 0.0879 | 0.1133 | 0.1058 | 0.0809 |
| F | | | | | | | | 0.2585 | 0.2428 | 0.2786 | 0.2695 | 0.2573 |
| G | | | | | | | | 0.0342 | 0.0371 | 0.0643 | 0.0502 | 0.0354 |
| H | | | | | | | | 0.2168 | 0.1349 | 0.2026 | 0.2034 | 0.1555 |

**Cell Index at: 74:01:35**

| | 1 | 2 | 3 | 4 | 5 | 6 | 7 | 8 | 9 | 10 | 11 | 12 |
|---|---|---|---|---|---|---|---|---|---|---|---|---|
| A | | | | | | | | 0.0049 | -0.0011 | 0.0349 | 0.0066 | 0.0233 |
| B | | | | | | | | 0.0484 | 0.0206 | 0.0391 | 0.0774 | 0.0556 |
| C | | | | | | | | 0.214 | 0.2081 | 0.3073 | 0.3202 | 0.2318 |
| D | | | | | | | | 0.1995 | 0.2819 | 0.2835 | 0.2892 | 0.2525 |
| E | | | | | | | | 0.1122 | 0.0861 | 0.1238 | 0.1119 | 0.0783 |
| F | | | | | | | | 0.2602 | 0.245 | 0.2804 | 0.2688 | 0.2654 |
| G | | | | | | | | 0.0309 | 0.037 | 0.0624 | 0.0492 | 0.0361 |
| H | | | | | | | | 0.2147 | 0.1344 | 0.2091 | 0.2025 | 0.1575 |

**Cell Index at: 74:16:35**

| | 1 | 2 | 3 | 4 | 5 | 6 | 7 | 8 | 9 | 10 | 11 | 12 |
|---|---|---|---|---|---|---|---|---|---|---|---|---|
| A | | | | | | | | 0.0079 | -0.0062 | 0.0299 | 0.0079 | 0.022 |
| B | | | | | | | | 0.0487 | 0.0203 | 0.0412 | 0.0748 | 0.052 |
| C | | | | | | | | 0.2111 | 0.207 | 0.3072 | 0.3254 | 0.2301 |
| D | | | | | | | | 0.1968 | 0.2839 | 0.2808 | 0.291 | 0.2559 |
| E | | | | | | | | 0.1072 | 0.0819 | 0.1198 | 0.1139 | 0.0798 |
| F | | | | | | | | 0.2606 | 0.2445 | 0.2768 | 0.2689 | 0.2638 |
| G | | | | | | | | 0.0305 | 0.0371 | 0.0653 | 0.0533 | 0.0345 |
| H | | | | | | | | 0.2178 | 0.1324 | 0.2106 | 0.2017 | 0.159 |

**Cell Index at: 74:31:35**

| | 1 | 2 | 3 | 4 | 5 | 6 | 7 | 8 | 9 | 10 | 11 | 12 |
|---|---|---|---|---|---|---|---|---|---|---|---|---|
| A | | | | | | | | 0.0081 | -0.006 | 0.0334 | 0.0031 | 0.0253 |
| B | | | | | | | | 0.0457 | 0.0249 | 0.0402 | 0.0777 | 0.0553 |
| C | | | | | | | | 0.2138 | 0.2025 | 0.3004 | 0.3177 | 0.2298 |
| D | | | | | | | | 0.1999 | 0.2757 | 0.286 | 0.2938 | 0.2475 |
| E | | | | | | | | 0.106 | 0.091 | 0.1163 | 0.1123 | 0.0803 |
| F | | | | | | | | 0.2639 | 0.2444 | 0.2862 | 0.269 | 0.2653 |
| G | | | | | | | | 0.0293 | 0.0381 | 0.0687 | 0.049 | 0.0352 |
| H | | | | | | | | 0.221 | 0.1359 | 0.2129 | 0.2051 | 0.1617 |

**Cell Index at: 74:46:35**

| | 1 | 2 | 3 | 4 | 5 | 6 | 7 | 8 | 9 | 10 | 11 | 12 |
|---|---|---|---|---|---|---|---|---|---|---|---|---|
| A | | | | | | | | 0.0052 | -0.0057 | 0.0321 | 0.0055 | 0.0238 |
| B | | | | | | | | 0.0457 | 0.0189 | 0.0363 | 0.0737 | 0.0549 |
| C | | | | | | | | 0.2153 | 0.2078 | 0.3106 | 0.3227 | 0.2357 |
| D | | | | | | | | 0.1925 | 0.2682 | 0.2856 | 0.2881 | 0.254 |
| E | | | | | | | | 0.1127 | 0.0916 | 0.1179 | 0.112 | 0.0842 |
| F | | | | | | | | 0.2648 | 0.2428 | 0.2811 | 0.2731 | 0.2574 |
| G | | | | | | | | 0.0288 | 0.0354 | 0.0659 | 0.0501 | 0.0347 |
| H | | | | | | | | 0.2225 | 0.1318 | 0.2116 | 0.2041 | 0.1583 |

**Cell Index at: 75:01:35**

| | 1 | 2 | 3 | 4 | 5 | 6 | 7 | 8 | 9 | 10 | 11 | 12 |
|---|---|---|---|---|---|---|---|---|---|---|---|---|
| A | | | | | | | | 0.0074 | -0.0029 | 0.0356 | 0.0063 | 0.0253 |
| B | | | | | | | | 0.0489 | 0.0147 | 0.0361 | 0.0737 | 0.0554 |
| C | | | | | | | | 0.2105 | 0.2115 | 0.3063 | 0.3197 | 0.2348 |
| D | | | | | | | | 0.1918 | 0.2741 | 0.2852 | 0.2984 | 0.2507 |
| E | | | | | | | | 0.1057 | 0.0898 | 0.1163 | 0.1126 | 0.0801 |
| F | | | | | | | | 0.2722 | 0.2397 | 0.2904 | 0.271 | 0.2666 |
| G | | | | | | | | 0.0196 | 0.034 | 0.0634 | 0.049 | 0.035 |
| H | | | | | | | | 0.2209 | 0.1339 | 0.2164 | 0.208 | 0.16 |

**Cell Index at: 75:16:36**

| | 1 | 2 | 3 | 4 | 5 | 6 | 7 | 8 | 9 | 10 | 11 | 12 |
|---|---|---|---|---|---|---|---|---|---|---|---|---|
| A | | | | | | | | 0.0074 | -0.0041 | 0.0327 | 0.0046 | 0.0211 |
| B | | | | | | | | 0.0484 | 0.0159 | 0.0342 | 0.0772 | 0.0542 |
| C | | | | | | | | 0.215 | 0.2074 | 0.3022 | 0.3218 | 0.232 |
| D | | | | | | | | 0.2034 | 0.275 | 0.2885 | 0.2954 | 0.2591 |
| E | | | | | | | | 0.1114 | 0.1 | 0.1133 | 0.1103 | 0.0775 |
| F | | | | | | | | 0.2646 | 0.2419 | 0.2887 | 0.2699 | 0.267 |
| G | | | | | | | | 0.0203 | 0.0351 | 0.0628 | 0.0504 | 0.0338 |
| H | | | | | | | | 0.2284 | 0.1317 | 0.2181 | 0.2051 | 0.1579 |

**Cell Index at: 75:31:37**

| | 1 | 2 | 3 | 4 | 5 | 6 | 7 | 8 | 9 | 10 | 11 | 12 |
|---|---|---|---|---|---|---|---|---|---|---|---|---|
| A | | | | | | | | 0.0025 | -0.0064 | 0.0353 | 0.0075 | 0.0185 |
| B | | | | | | | | 0.0441 | 0.0214 | 0.0352 | 0.0765 | 0.059 |
| C | | | | | | | | 0.2141 | 0.2054 | 0.3062 | 0.3208 | 0.2349 |
| D | | | | | | | | 0.1965 | 0.279 | 0.2785 | 0.2934 | 0.2613 |
| E | | | | | | | | 0.1143 | 0.0938 | 0.115 | 0.1137 | 0.0757 |
| F | | | | | | | | 0.2653 | 0.2413 | 0.2861 | 0.2741 | 0.2649 |
| G | | | | | | | | 0.0194 | 0.0352 | 0.0653 | 0.0496 | 0.0345 |
| H | | | | | | | | 0.224 | 0.134 | 0.2171 | 0.2055 | 0.1566 |

**Cell Index at: 75:46:38**

| | 1 | 2 | 3 | 4 | 5 | 6 | 7 | 8 | 9 | 10 | 11 | 12 |
|---|---|---|---|---|---|---|---|---|---|---|---|---|
| A | | | | | | | | 0.0032 | -0.0031 | 0.0331 | 0.0061 | 0.0187 |
| B | | | | | | | | 0.0436 | 0.0263 | 0.0349 | 0.0745 | 0.0583 |

| | | | | | | | | | | | | |
|---|---|---|---|---|---|---|---|---|---|---|---|---|
| C | | | | | | | | 0.2154 | 0.2038 | 0.3047 | 0.3189 | 0.2375 |
| D | | | | | | | | 0.1996 | 0.2738 | 0.2828 | 0.299 | 0.2552 |
| E | | | | | | | | 0.1123 | 0.095 | 0.1215 | 0.1063 | 0.0748 |
| F | | | | | | | | 0.267 | 0.2474 | 0.287 | 0.2711 | 0.268 |
| G | | | | | | | | 0.023 | 0.0382 | 0.062 | 0.0482 | 0.0342 |
| H | | | | | | | | 0.2256 | 0.1363 | 0.2207 | 0.2056 | 0.1579 |

**Cell Index at: 76:01:38**

| | 1 | 2 | 3 | 4 | 5 | 6 | 7 | 8 | 9 | 10 | 11 | 12 |
|---|---|---|---|---|---|---|---|---|---|---|---|---|
| A | | | | | | | | 0.0039 | -0.0042 | 0.033 | 0.0062 | 0.019 |
| B | | | | | | | | 0.0435 | 0.0242 | 0.0363 | 0.0726 | 0.0608 |
| C | | | | | | | | 0.2054 | 0.2053 | 0.3067 | 0.3143 | 0.2366 |
| D | | | | | | | | 0.1946 | 0.2796 | 0.2895 | 0.3013 | 0.2577 |
| E | | | | | | | | 0.1132 | 0.0894 | 0.1064 | 0.1186 | 0.0763 |
| F | | | | | | | | 0.2697 | 0.2468 | 0.2889 | 0.2757 | 0.2633 |
| G | | | | | | | | 0.0199 | 0.0368 | 0.0614 | 0.0511 | 0.028 |
| H | | | | | | | | 0.2275 | 0.1355 | 0.2192 | 0.2052 | 0.1583 |

**Cell Index at: 76:16:39**

| | 1 | 2 | 3 | 4 | 5 | 6 | 7 | 8 | 9 | 10 | 11 | 12 |
|---|---|---|---|---|---|---|---|---|---|---|---|---|
| A | | | | | | | | 0.0045 | -0.0053 | 0.0316 | 0.0051 | 0.0176 |
| B | | | | | | | | 0.0421 | 0.023 | 0.0342 | 0.0716 | 0.0539 |
| C | | | | | | | | 0.2111 | 0.2045 | 0.3025 | 0.3204 | 0.2364 |
| D | | | | | | | | 0.2036 | 0.2753 | 0.2914 | 0.303 | 0.2537 |
| E | | | | | | | | 0.1125 | 0.0911 | 0.1159 | 0.1086 | 0.0699 |
| F | | | | | | | | 0.2636 | 0.2384 | 0.2891 | 0.2794 | 0.2632 |
| G | | | | | | | | 0.0184 | 0.0375 | 0.0667 | 0.045 | 0.0268 |
| H | | | | | | | | 0.2258 | 0.136 | 0.222 | 0.2095 | 0.1621 |

**Cell Index at: 76:31:39**

| | 1 | 2 | 3 | 4 | 5 | 6 | 7 | 8 | 9 | 10 | 11 | 12 |
|---|---|---|---|---|---|---|---|---|---|---|---|---|
| A | | | | | | | | 0.0022 | -0.0043 | 0.0279 | 0.0068 | 0.0158 |
| B | | | | | | | | 0.0423 | 0.0252 | 0.0327 | 0.0776 | 0.0545 |
| C | | | | | | | | 0.202 | 0.2041 | 0.3053 | 0.3265 | 0.2366 |
| D | | | | | | | | 0.2008 | 0.2795 | 0.2928 | 0.3031 | 0.2604 |
| E | | | | | | | | 0.1025 | 0.0892 | 0.1114 | 0.1123 | 0.0696 |
| F | | | | | | | | 0.267 | 0.2345 | 0.2877 | 0.277 | 0.2673 |
| G | | | | | | | | 0.0216 | 0.0346 | 0.0627 | 0.0441 | 0.0262 |
| H | | | | | | | | 0.2294 | 0.1373 | 0.22 | 0.2087 | 0.1645 |

**Cell Index at: 76:46:40**

| | 1 | 2 | 3 | 4 | 5 | 6 | 7 | 8 | 9 | 10 | 11 | 12 |
|---|---|---|---|---|---|---|---|---|---|---|---|---|
| A | | | | | | | | 0.0017 | -0.0036 | 0.0305 | 0.0056 | 0.0152 |
| B | | | | | | | | 0.0452 | 0.0211 | 0.034 | 0.0744 | 0.0533 |
| C | | | | | | | | 0.2085 | 0.2014 | 0.3051 | 0.3235 | 0.2375 |
| D | | | | | | | | 0.196 | 0.2763 | 0.2842 | 0.2999 | 0.2556 |
| E | | | | | | | | 0.1096 | 0.0832 | 0.1134 | 0.1176 | 0.0701 |
| F | | | | | | | | 0.2727 | 0.2394 | 0.2916 | 0.2813 | 0.2661 |
| G | | | | | | | | 0.0185 | 0.0394 | 0.0609 | 0.0465 | 0.0275 |
| H | | | | | | | | 0.2306 | 0.1373 | 0.2221 | 0.2104 | 0.1634 |

**Cell Index at: 77:01:40**

| | 1 | 2 | 3 | 4 | 5 | 6 | 7 | 8 | 9 | 10 | 11 | 12 |
|---|---|---|---|---|---|---|---|---|---|---|---|---|
| A | | | | | | | | 0.0016 | -0.0049 | 0.0333 | 0.0081 | 0.0081 |
| B | | | | | | | | 0.0431 | 0.0241 | 0.0318 | 0.074 | 0.0525 |
| C | | | | | | | | 0.2093 | 0.1999 | 0.3127 | 0.3253 | 0.2427 |
| D | | | | | | | | 0.1951 | 0.2758 | 0.29 | 0.2979 | 0.2645 |
| E | | | | | | | | 0.1001 | 0.0921 | 0.1147 | 0.1074 | 0.0699 |
| F | | | | | | | | 0.2735 | 0.2431 | 0.2897 | 0.2815 | 0.2656 |
| G | | | | | | | | 0.0191 | 0.0394 | 0.0601 | 0.0488 | 0.0277 |
| H | | | | | | | | 0.2317 | 0.1351 | 0.2231 | 0.2069 | 0.1656 |

**Cell Index at: 77:16:41**

| | 1 | 2 | 3 | 4 | 5 | 6 | 7 | 8 | 9 | 10 | 11 | 12 |
|---|---|---|---|---|---|---|---|---|---|---|---|---|
| A | | | | | | | | -0.0017 | -0.0036 | 0.0313 | 0.008 | 0.0094 |
| B | | | | | | | | 0.0457 | 0.0235 | 0.0316 | 0.0775 | 0.0506 |
| C | | | | | | | | 0.213 | 0.2037 | 0.3078 | 0.3236 | 0.247 |
| D | | | | | | | | 0.2048 | 0.2816 | 0.2827 | 0.2999 | 0.2589 |
| E | | | | | | | | 0.1068 | 0.0927 | 0.1088 | 0.1091 | 0.0669 |
| F | | | | | | | | 0.2733 | 0.241 | 0.2843 | 0.2844 | 0.2686 |
| G | | | | | | | | 0.0191 | 0.0403 | 0.0629 | 0.0494 | 0.0291 |
| H | | | | | | | | 0.2304 | 0.1382 | 0.2236 | 0.2139 | 0.1624 |

**Cell Index at: 77:31:41**

| | 1 | 2 | 3 | 4 | 5 | 6 | 7 | 8 | 9 | 10 | 11 | 12 |
|---|---|---|---|---|---|---|---|---|---|---|---|---|
| A | | | | | | | | 0.0003 | -0.0046 | 0.0305 | 0.0092 | 0.01 |
| B | | | | | | | | 0.0445 | 0.0253 | 0.0329 | 0.0782 | 0.0539 |
| C | | | | | | | | 0.2126 | 0.2033 | 0.307 | 0.3242 | 0.2418 |
| D | | | | | | | | 0.1975 | 0.2792 | 0.2896 | 0.3002 | 0.2555 |
| E | | | | | | | | 0.1067 | 0.0893 | 0.108 | 0.1108 | 0.0655 |
| F | | | | | | | | 0.2686 | 0.2422 | 0.2939 | 0.2781 | 0.2724 |
| G | | | | | | | | 0.018 | 0.0396 | 0.0613 | 0.0479 | 0.0261 |
| H | | | | | | | | 0.2287 | 0.1383 | 0.2204 | 0.2133 | 0.164 |

**Cell Index at: 77:46:42**

| | 1 | 2 | 3 | 4 | 5 | 6 | 7 | 8 | 9 | 10 | 11 | 12 |
|---|---|---|---|---|---|---|---|---|---|---|---|---|
| A | | | | | | | | -0.0006 | -0.0055 | 0.0323 | 0.0062 | 0.0104 |
| B | | | | | | | | 0.0468 | 0.0217 | 0.031 | 0.072 | 0.0542 |
| C | | | | | | | | 0.2186 | 0.2067 | 0.3089 | 0.3288 | 0.2425 |
| D | | | | | | | | 0.1958 | 0.2744 | 0.2932 | 0.2897 | 0.2569 |
| E | | | | | | | | 0.1027 | 0.0905 | 0.1142 | 0.1139 | 0.0659 |
| F | | | | | | | | 0.2811 | 0.2379 | 0.2931 | 0.2856 | 0.2699 |
| G | | | | | | | | 0.0209 | 0.0362 | 0.0609 | 0.0478 | 0.0283 |
| H | | | | | | | | 0.2329 | 0.1355 | 0.2174 | 0.2096 | 0.1658 |

**Cell Index at: 78:01:43**

| | 1 | 2 | 3 | 4 | 5 | 6 | 7 | 8 | 9 | 10 | 11 | 12 |
|---|---|---|---|---|---|---|---|---|---|---|---|---|
| A | | | | | | | | -0.002 | -0.0066 | 0.0332 | 0.0047 | 0.0149 |
| B | | | | | | | | 0.0455 | 0.0276 | 0.0324 | 0.0755 | 0.0515 |
| C | | | | | | | | 0.2085 | 0.2036 | 0.3114 | 0.3274 | 0.2474 |
| D | | | | | | | | 0.198 | 0.2875 | 0.2963 | 0.299 | 0.2541 |
| E | | | | | | | | 0.1005 | 0.0874 | 0.1129 | 0.1132 | 0.0719 |
| F | | | | | | | | 0.2685 | 0.2374 | 0.2965 | 0.2851 | 0.2722 |
| G | | | | | | | | 0.0256 | 0.0404 | 0.056 | 0.0488 | 0.0237 |
| H | | | | | | | | 0.2335 | 0.1326 | 0.2189 | 0.2119 | 0.1689 |

**Cell Index at: 78:16:43**

| | 1 | 2 | 3 | 4 | 5 | 6 | 7 | 8 | 9 | 10 | 11 | 12 |
|---|---|---|---|---|---|---|---|---|---|---|---|---|
| A | | | | | | | | -0.0029 | -0.0066 | 0.0321 | 0.0061 | 0.0115 |
| B | | | | | | | | 0.0467 | 0.0231 | 0.028 | 0.0755 | 0.0491 |
| C | | | | | | | | 0.2085 | 0.2052 | 0.3154 | 0.3295 | 0.2449 |
| D | | | | | | | | 0.195 | 0.2818 | 0.2957 | 0.2945 | 0.2605 |
| E | | | | | | | | 0.1072 | 0.0877 | 0.1149 | 0.1089 | 0.0682 |
| F | | | | | | | | 0.2734 | 0.2472 | 0.3044 | 0.2801 | 0.2792 |
| G | | | | | | | | 0.0207 | 0.0423 | 0.0618 | 0.0468 | 0.0302 |
| H | | | | | | | | 0.2343 | 0.1333 | 0.2189 | 0.2129 | 0.1662 |

**Cell Index at: 78:31:44**

| | 1 | 2 | 3 | 4 | 5 | 6 | 7 | 8 | 9 | 10 | 11 | 12 |
|---|---|---|---|---|---|---|---|---|---|---|---|---|
| A | | | | | | | | -0.0013 | -0.0035 | 0.0352 | 0.0066 | 0.013 |
| B | | | | | | | | 0.0489 | 0.0256 | 0.034 | 0.0717 | 0.0488 |
| C | | | | | | | | 0.2163 | 0.1982 | 0.3145 | 0.3304 | 0.2446 |
| D | | | | | | | | 0.192 | 0.2784 | 0.2892 | 0.2908 | 0.2532 |
| E | | | | | | | | 0.1 | 0.0922 | 0.1117 | 0.1162 | 0.0638 |
| F | | | | | | | | 0.2694 | 0.2392 | 0.2999 | 0.2858 | 0.276 |
| G | | | | | | | | 0.0231 | 0.0397 | 0.0636 | 0.0478 | 0.0293 |
| H | | | | | | | | 0.236 | 0.1344 | 0.2225 | 0.2115 | 0.1676 |

**Cell Index at: 78:46:44**

| | 1 | 2 | 3 | 4 | 5 | 6 | 7 | 8 | 9 | 10 | 11 | 12 |
|---|---|---|---|---|---|---|---|---|---|---|---|---|
| A | | | | | | | | -0.0036 | -0.0036 | 0.035 | 0.0076 | 0.0124 |

| | | | | | | | | | | | | |
|---|---|---|---|---|---|---|---|---|---|---|---|---|
| B | | | | | | | | 0.0444 | 0.0265 | 0.0362 | 0.0726 | 0.0513 |
| C | | | | | | | | 0.2096 | 0.2043 | 0.3179 | 0.3338 | 0.2421 |
| D | | | | | | | | 0.1979 | 0.2818 | 0.3004 | 0.3 | 0.2538 |
| E | | | | | | | | 0.1055 | 0.0856 | 0.1116 | 0.1134 | 0.071 |
| F | | | | | | | | 0.268 | 0.2451 | 0.302 | 0.2975 | 0.2756 |
| G | | | | | | | | 0.0235 | 0.0439 | 0.059 | 0.0505 | 0.0281 |
| H | | | | | | | | 0.2366 | 0.1336 | 0.2198 | 0.2134 | 0.17 |

**Cell Index at: 79:01:45**

| | 1 | 2 | 3 | 4 | 5 | 6 | 7 | 8 | 9 | 10 | 11 | 12 |
|---|---|---|---|---|---|---|---|---|---|---|---|---|
| A | | | | | | | | -0.0035 | -0.007 | 0.0319 | 0.0122 | 0.0119 |
| B | | | | | | | | 0.0423 | 0.0216 | 0.0317 | 0.0709 | 0.0563 |
| C | | | | | | | | 0.2128 | 0.2069 | 0.3134 | 0.3348 | 0.2476 |
| D | | | | | | | | 0.1929 | 0.2832 | 0.3081 | 0.2918 | 0.2576 |
| E | | | | | | | | 0.1079 | 0.0862 | 0.1196 | 0.1151 | 0.0676 |
| F | | | | | | | | 0.2703 | 0.2456 | 0.3038 | 0.2956 | 0.2775 |
| G | | | | | | | | 0.0181 | 0.0431 | 0.0617 | 0.0509 | 0.0267 |
| H | | | | | | | | 0.2399 | 0.1347 | 0.2237 | 0.214 | 0.1686 |

**Cell Index at: 79:16:45**

| | 1 | 2 | 3 | 4 | 5 | 6 | 7 | 8 | 9 | 10 | 11 | 12 |
|---|---|---|---|---|---|---|---|---|---|---|---|---|
| A | | | | | | | | -0.0064 | -0.0037 | 0.035 | 0.0092 | 0.0149 |
| B | | | | | | | | 0.0474 | 0.0231 | 0.0279 | 0.0722 | 0.0489 |
| C | | | | | | | | 0.2092 | 0.2053 | 0.3144 | 0.3362 | 0.2467 |
| D | | | | | | | | 0.1995 | 0.2841 | 0.2982 | 0.3019 | 0.2563 |
| E | | | | | | | | 0.0998 | 0.09 | 0.1182 | 0.1131 | 0.0655 |
| F | | | | | | | | 0.2743 | 0.2399 | 0.3037 | 0.2904 | 0.2785 |
| G | | | | | | | | 0.0202 | 0.0375 | 0.0604 | 0.0491 | 0.0311 |
| H | | | | | | | | 0.2366 | 0.1381 | 0.2236 | 0.2147 | 0.169 |

**Cell Index at: 79:31:46**

| | 1 | 2 | 3 | 4 | 5 | 6 | 7 | 8 | 9 | 10 | 11 | 12 |
|---|---|---|---|---|---|---|---|---|---|---|---|---|
| A | | | | | | | | -0.0069 | -0.0033 | 0.0353 | 0.0076 | 0.0142 |
| B | | | | | | | | 0.0457 | 0.0265 | 0.0303 | 0.0678 | 0.0495 |
| C | | | | | | | | 0.2079 | 0.2082 | 0.3124 | 0.3331 | 0.2463 |
| D | | | | | | | | 0.2017 | 0.2812 | 0.2936 | 0.3021 | 0.2522 |
| E | | | | | | | | 0.1066 | 0.0937 | 0.1181 | 0.1124 | 0.0664 |
| F | | | | | | | | 0.2782 | 0.2506 | 0.3019 | 0.2926 | 0.2745 |
| G | | | | | | | | 0.0224 | 0.038 | 0.0598 | 0.0469 | 0.031 |
| H | | | | | | | | 0.2378 | 0.1359 | 0.2229 | 0.2123 | 0.1723 |

**Cell Index at: 79:46:47**

| | 1 | 2 | 3 | 4 | 5 | 6 | 7 | 8 | 9 | 10 | 11 | 12 |
|---|---|---|---|---|---|---|---|---|---|---|---|---|
| A | | | | | | | | -0.0058 | -0.003 | 0.0311 | 0.009 | 0.0136 |
| B | | | | | | | | 0.0444 | 0.0238 | 0.0289 | 0.0689 | 0.0495 |
| C | | | | | | | | 0.2137 | 0.1999 | 0.3166 | 0.3347 | 0.2495 |
| D | | | | | | | | 0.1989 | 0.2877 | 0.301 | 0.3054 | 0.2504 |
| E | | | | | | | | 0.1032 | 0.0859 | 0.1124 | 0.1133 | 0.0696 |
| F | | | | | | | | 0.2817 | 0.2543 | 0.3102 | 0.2959 | 0.2753 |
| G | | | | | | | | 0.0229 | 0.0391 | 0.0612 | 0.0464 | 0.032 |
| H | | | | | | | | 0.2379 | 0.1405 | 0.225 | 0.2098 | 0.1698 |

**Cell Index at: 80:01:47**

| | 1 | 2 | 3 | 4 | 5 | 6 | 7 | 8 | 9 | 10 | 11 | 12 |
|---|---|---|---|---|---|---|---|---|---|---|---|---|
| A | | | | | | | | -0.0006 | -0.0023 | 0.0314 | 0.0093 | 0.0139 |
| B | | | | | | | | 0.0481 | 0.0276 | 0.027 | 0.07 | 0.0472 |
| C | | | | | | | | 0.2136 | 0.1987 | 0.3209 | 0.3382 | 0.2477 |
| D | | | | | | | | 0.1991 | 0.284 | 0.3002 | 0.2981 | 0.263 |
| E | | | | | | | | 0.1028 | 0.0939 | 0.118 | 0.1085 | 0.0707 |
| F | | | | | | | | 0.2796 | 0.2522 | 0.3096 | 0.2928 | 0.278 |
| G | | | | | | | | 0.0201 | 0.038 | 0.0607 | 0.0495 | 0.0308 |
| H | | | | | | | | 0.2368 | 0.1362 | 0.2279 | 0.213 | 0.1717 |

**Cell Index at: 80:16:48**

| | 1 | 2 | 3 | 4 | 5 | 6 | 7 | 8 | 9 | 10 | 11 | 12 |
|---|---|---|---|---|---|---|---|---|---|---|---|---|
| A | | | | | | | | -0.0037 | -0.0016 | 0.032 | 0.0093 | 0.0107 |
| B | | | | | | | | 0.0465 | 0.0284 | 0.0311 | 0.0683 | 0.0478 |
| C | | | | | | | | 0.2064 | 0.2032 | 0.3127 | 0.3332 | 0.2509 |
| D | | | | | | | | 0.1953 | 0.2868 | 0.297 | 0.3129 | 0.2595 |
| E | | | | | | | | 0.1009 | 0.0975 | 0.1128 | 0.1019 | 0.0693 |
| F | | | | | | | | 0.2752 | 0.2532 | 0.3111 | 0.2965 | 0.2797 |
| G | | | | | | | | 0.0237 | 0.0373 | 0.0609 | 0.0478 | 0.0293 |
| H | | | | | | | | 0.2394 | 0.1406 | 0.2274 | 0.2109 | 0.1724 |

**Cell Index at: 80:31:48**

| | 1 | 2 | 3 | 4 | 5 | 6 | 7 | 8 | 9 | 10 | 11 | 12 |
|---|---|---|---|---|---|---|---|---|---|---|---|---|
| A | | | | | | | | -0.0005 | -0.0029 | 0.0308 | 0.0087 | 0.013 |
| B | | | | | | | | 0.0426 | 0.0291 | 0.0252 | 0.0696 | 0.0504 |
| C | | | | | | | | 0.2163 | 0.1994 | 0.3178 | 0.3375 | 0.2514 |
| D | | | | | | | | 0.1928 | 0.2913 | 0.2968 | 0.3051 | 0.25 |
| E | | | | | | | | 0.0981 | 0.0864 | 0.1131 | 0.1088 | 0.0747 |
| F | | | | | | | | 0.2812 | 0.2569 | 0.3144 | 0.2949 | 0.2816 |
| G | | | | | | | | 0.0199 | 0.0411 | 0.057 | 0.0466 | 0.0291 |
| H | | | | | | | | 0.2388 | 0.1395 | 0.2264 | 0.2147 | 0.1732 |

**Cell Index at: 80:46:49**

| | 1 | 2 | 3 | 4 | 5 | 6 | 7 | 8 | 9 | 10 | 11 | 12 |
|---|---|---|---|---|---|---|---|---|---|---|---|---|
| A | | | | | | | | 0.0008 | -0.0048 | 0.032 | 0.0089 | 0.0146 |
| B | | | | | | | | 0.0437 | 0.0245 | 0.0303 | 0.0683 | 0.0487 |
| C | | | | | | | | 0.2161 | 0.1981 | 0.321 | 0.3447 | 0.2491 |
| D | | | | | | | | 0.1964 | 0.2861 | 0.299 | 0.3068 | 0.2574 |
| E | | | | | | | | 0.0987 | 0.085 | 0.1127 | 0.1044 | 0.0666 |
| F | | | | | | | | 0.2824 | 0.254 | 0.3104 | 0.294 | 0.2774 |
| G | | | | | | | | 0.0182 | 0.0387 | 0.0582 | 0.0417 | 0.0279 |
| H | | | | | | | | 0.237 | 0.142 | 0.2273 | 0.217 | 0.1726 |

**Cell Index at: 81:01:50**

| | 1 | 2 | 3 | 4 | 5 | 6 | 7 | 8 | 9 | 10 | 11 | 12 |
|---|---|---|---|---|---|---|---|---|---|---|---|---|
| A | | | | | | | | -0.0017 | -0.0041 | 0.0336 | 0.009 | 0.0098 |
| B | | | | | | | | 0.0355 | 0.022 | 0.0311 | 0.0682 | 0.0514 |
| C | | | | | | | | 0.2126 | 0.2045 | 0.3194 | 0.3377 | 0.2549 |
| D | | | | | | | | 0.1977 | 0.2874 | 0.3052 | 0.3077 | 0.262 |
| E | | | | | | | | 0.0907 | 0.0899 | 0.1154 | 0.1074 | 0.0674 |
| F | | | | | | | | 0.2794 | 0.2565 | 0.3101 | 0.2918 | 0.2752 |
| G | | | | | | | | 0.0189 | 0.0395 | 0.0613 | 0.0466 | 0.0317 |
| H | | | | | | | | 0.2391 | 0.1442 | 0.229 | 0.216 | 0.1708 |

**Cell Index at: 81:16:50**

| | 1 | 2 | 3 | 4 | 5 | 6 | 7 | 8 | 9 | 10 | 11 | 12 |
|---|---|---|---|---|---|---|---|---|---|---|---|---|
| A | | | | | | | | 0.0036 | -0.0066 | 0.033 | 0.0102 | 0.0142 |
| B | | | | | | | | 0.0422 | 0.0194 | 0.0299 | 0.0632 | 0.0455 |
| C | | | | | | | | 0.2122 | 0.2057 | 0.3222 | 0.3351 | 0.2559 |
| D | | | | | | | | 0.199 | 0.2875 | 0.3032 | 0.3077 | 0.2629 |
| E | | | | | | | | 0.0976 | 0.0923 | 0.1167 | 0.1094 | 0.0701 |
| F | | | | | | | | 0.2773 | 0.2483 | 0.3214 | 0.2992 | 0.2742 |
| G | | | | | | | | 0.019 | 0.0359 | 0.0643 | 0.0476 | 0.0328 |
| H | | | | | | | | 0.2362 | 0.1405 | 0.2264 | 0.2194 | 0.1724 |

**Cell Index at: 81:31:51**

| | 1 | 2 | 3 | 4 | 5 | 6 | 7 | 8 | 9 | 10 | 11 | 12 |
|---|---|---|---|---|---|---|---|---|---|---|---|---|
| A | | | | | | | | 0.0027 | -0.0024 | 0.032 | 0.0117 | 0.0121 |
| B | | | | | | | | 0.0412 | 0.0217 | 0.0268 | 0.0613 | 0.0497 |
| C | | | | | | | | 0.2179 | 0.2006 | 0.3223 | 0.332 | 0.2483 |
| D | | | | | | | | 0.2046 | 0.2892 | 0.3025 | 0.3074 | 0.2566 |
| E | | | | | | | | 0.0988 | 0.0956 | 0.1099 | 0.106 | 0.0702 |
| F | | | | | | | | 0.2835 | 0.2547 | 0.3168 | 0.2999 | 0.277 |
| G | | | | | | | | 0.0191 | 0.038 | 0.0626 | 0.0465 | 0.0316 |
| H | | | | | | | | 0.239 | 0.1418 | 0.2295 | 0.2176 | 0.1726 |

**Cell Index at: 81:46:52**

| | 1 | 2 | 3 | 4 | 5 | 6 | 7 | 8 | 9 | 10 | 11 | 12 |
|---|---|---|---|---|---|---|---|---|---|---|---|---|

| | | | | | | | | | | | | |
|---|---|---|---|---|---|---|---|---|---|---|---|---|
| A | | | | | | | | -0.0006 | -0.0029 | 0.0314 | 0.0077 | 0.0159 |
| B | | | | | | | | 0.0412 | 0.0231 | 0.0284 | 0.0647 | 0.049 |
| C | | | | | | | | 0.2203 | 0.2047 | 0.3271 | 0.3353 | 0.2481 |
| D | | | | | | | | 0.2014 | 0.2877 | 0.3087 | 0.3167 | 0.2617 |
| E | | | | | | | | 0.1003 | 0.0931 | 0.1127 | 0.1072 | 0.0697 |
| F | | | | | | | | 0.284 | 0.2559 | 0.3177 | 0.2999 | 0.2755 |
| G | | | | | | | | 0.0205 | 0.036 | 0.0627 | 0.0459 | 0.035 |
| H | | | | | | | | 0.2381 | 0.1438 | 0.2284 | 0.2165 | 0.1744 |

**Cell Index at: 82:01:52**

| | 1 | 2 | 3 | 4 | 5 | 6 | 7 | 8 | 9 | 10 | 11 | 12 |
|---|---|---|---|---|---|---|---|---|---|---|---|---|
| A | | | | | | | | 0.0019 | -0.0047 | 0.0342 | 0.0084 | 0.0133 |
| B | | | | | | | | 0.0446 | 0.0163 | 0.0322 | 0.0667 | 0.0513 |
| C | | | | | | | | 0.2243 | 0.203 | 0.3273 | 0.3383 | 0.2555 |
| D | | | | | | | | 0.199 | 0.2863 | 0.3063 | 0.3175 | 0.264 |
| E | | | | | | | | 0.1033 | 0.0897 | 0.1124 | 0.1093 | 0.0661 |
| F | | | | | | | | 0.2755 | 0.256 | 0.3228 | 0.2996 | 0.2802 |
| G | | | | | | | | 0.0222 | 0.0412 | 0.0657 | 0.0473 | 0.0316 |
| H | | | | | | | | 0.2403 | 0.1464 | 0.2285 | 0.2132 | 0.1736 |

**Cell Index at: 82:16:52**

| | 1 | 2 | 3 | 4 | 5 | 6 | 7 | 8 | 9 | 10 | 11 | 12 |
|---|---|---|---|---|---|---|---|---|---|---|---|---|
| A | | | | | | | | 0.0024 | -0.0042 | 0.0321 | 0.0083 | 0.0137 |
| B | | | | | | | | 0.0438 | 0.0203 | 0.026 | 0.0633 | 0.0532 |
| C | | | | | | | | 0.2166 | 0.2089 | 0.3238 | 0.346 | 0.2554 |
| D | | | | | | | | 0.1975 | 0.2913 | 0.3085 | 0.3193 | 0.2672 |
| E | | | | | | | | 0.102 | 0.0931 | 0.1112 | 0.1105 | 0.0741 |
| F | | | | | | | | 0.2864 | 0.2576 | 0.3079 | 0.3 | 0.2787 |
| G | | | | | | | | 0.0191 | 0.0376 | 0.0637 | 0.0475 | 0.0305 |
| H | | | | | | | | 0.239 | 0.1442 | 0.2303 | 0.2175 | 0.1759 |

**Cell Index at: 82:31:52**

| | 1 | 2 | 3 | 4 | 5 | 6 | 7 | 8 | 9 | 10 | 11 | 12 |
|---|---|---|---|---|---|---|---|---|---|---|---|---|
| A | | | | | | | | 0.0024 | -0.0019 | 0.0377 | 0.0082 | 0.0137 |
| B | | | | | | | | 0.0377 | 0.0162 | 0.0245 | 0.0616 | 0.0497 |
| C | | | | | | | | 0.2204 | 0.2076 | 0.3348 | 0.3436 | 0.2602 |
| D | | | | | | | | 0.2021 | 0.2971 | 0.3125 | 0.3186 | 0.273 |
| E | | | | | | | | 0.1023 | 0.0957 | 0.1073 | 0.1104 | 0.0632 |
| F | | | | | | | | 0.2854 | 0.2546 | 0.3199 | 0.2964 | 0.2793 |
| G | | | | | | | | 0.0216 | 0.0408 | 0.0597 | 0.0449 | 0.0272 |
| H | | | | | | | | 0.2393 | 0.1449 | 0.231 | 0.2175 | 0.1763 |

**Cell Index at: 82:46:53**

| | 1 | 2 | 3 | 4 | 5 | 6 | 7 | 8 | 9 | 10 | 11 | 12 |
|---|---|---|---|---|---|---|---|---|---|---|---|---|
| A | | | | | | | | 0.005 | -0.0032 | 0.0377 | 0.0122 | 0.0144 |
| B | | | | | | | | 0.0371 | 0.0155 | 0.0288 | 0.0606 | 0.0512 |
| C | | | | | | | | 0.2221 | 0.2001 | 0.3356 | 0.3413 | 0.2588 |
| D | | | | | | | | 0.2033 | 0.2865 | 0.3095 | 0.3138 | 0.2654 |
| E | | | | | | | | 0.1133 | 0.0991 | 0.1057 | 0.1119 | 0.0661 |
| F | | | | | | | | 0.2855 | 0.2587 | 0.3162 | 0.3005 | 0.2787 |
| G | | | | | | | | 0.0221 | 0.0399 | 0.0671 | 0.0444 | 0.0255 |
| H | | | | | | | | 0.2424 | 0.142 | 0.2284 | 0.2177 | 0.1725 |

**Cell Index at: 83:01:53**

| | 1 | 2 | 3 | 4 | 5 | 6 | 7 | 8 | 9 | 10 | 11 | 12 |
|---|---|---|---|---|---|---|---|---|---|---|---|---|
| A | | | | | | | | 0.007 | -0.0061 | 0.0372 | 0.0081 | 0.0108 |
| B | | | | | | | | 0.0374 | 0.0153 | 0.0282 | 0.0614 | 0.05 |
| C | | | | | | | | 0.2234 | 0.2086 | 0.3379 | 0.349 | 0.2605 |
| D | | | | | | | | 0.1969 | 0.2893 | 0.3088 | 0.3096 | 0.2709 |
| E | | | | | | | | 0.0979 | 0.0913 | 0.1131 | 0.1133 | 0.0688 |
| F | | | | | | | | 0.2844 | 0.2562 | 0.3159 | 0.3036 | 0.282 |
| G | | | | | | | | 0.0224 | 0.0428 | 0.0609 | 0.0466 | 0.0292 |
| H | | | | | | | | 0.2451 | 0.1423 | 0.2318 | 0.2143 | 0.1739 |

**Cell Index at: 83:16:53**

| | 1 | 2 | 3 | 4 | 5 | 6 | 7 | 8 | 9 | 10 | 11 | 12 |
|---|---|---|---|---|---|---|---|---|---|---|---|---|
| A | | | | | | | | 0.0052 | -0.0068 | 0.0394 | 0.0123 | 0.0126 |
| B | | | | | | | | 0.0346 | 0.018 | 0.0307 | 0.0617 | 0.0505 |
| C | | | | | | | | 0.2256 | 0.2092 | 0.338 | 0.3514 | 0.2568 |
| D | | | | | | | | 0.201 | 0.2927 | 0.3145 | 0.3166 | 0.2707 |
| E | | | | | | | | 0.1035 | 0.0915 | 0.1122 | 0.1152 | 0.0652 |
| F | | | | | | | | 0.2867 | 0.2607 | 0.3157 | 0.3108 | 0.2763 |
| G | | | | | | | | 0.0224 | 0.0402 | 0.0602 | 0.0455 | 0.0233 |
| H | | | | | | | | 0.2457 | 0.1462 | 0.2258 | 0.2159 | 0.1755 |

**Cell Index at: 83:31:53**

| | 1 | 2 | 3 | 4 | 5 | 6 | 7 | 8 | 9 | 10 | 11 | 12 |
|---|---|---|---|---|---|---|---|---|---|---|---|---|
| A | | | | | | | | 0.0039 | -0.0094 | 0.0381 | 0.0109 | 0.0113 |
| B | | | | | | | | 0.0352 | 0.0194 | 0.031 | 0.0604 | 0.0557 |
| C | | | | | | | | 0.2321 | 0.2027 | 0.3437 | 0.3552 | 0.2595 |
| D | | | | | | | | 0.2082 | 0.292 | 0.3159 | 0.3153 | 0.2667 |
| E | | | | | | | | 0.1066 | 0.0885 | 0.113 | 0.1124 | 0.0671 |
| F | | | | | | | | 0.2858 | 0.2599 | 0.3324 | 0.3061 | 0.2746 |
| G | | | | | | | | 0.0188 | 0.0422 | 0.0565 | 0.0456 | 0.0281 |
| H | | | | | | | | 0.2448 | 0.151 | 0.2315 | 0.2138 | 0.1742 |

**Cell Index at: 83:46:54**

| | 1 | 2 | 3 | 4 | 5 | 6 | 7 | 8 | 9 | 10 | 11 | 12 |
|---|---|---|---|---|---|---|---|---|---|---|---|---|
| A | | | | | | | | 0.0075 | -0.0076 | 0.0404 | 0.0124 | 0.0116 |
| B | | | | | | | | 0.0332 | 0.0202 | 0.0304 | 0.0605 | 0.0524 |
| C | | | | | | | | 0.227 | 0.2071 | 0.3389 | 0.3495 | 0.261 |
| D | | | | | | | | 0.2079 | 0.2915 | 0.3123 | 0.3216 | 0.2754 |
| E | | | | | | | | 0.1079 | 0.0893 | 0.1111 | 0.1117 | 0.0643 |
| F | | | | | | | | 0.2876 | 0.2639 | 0.3261 | 0.3084 | 0.2788 |
| G | | | | | | | | 0.023 | 0.0368 | 0.0567 | 0.045 | 0.0228 |
| H | | | | | | | | 0.2394 | 0.1498 | 0.2296 | 0.216 | 0.1742 |

**Cell Index at: 84:01:54**

| | 1 | 2 | 3 | 4 | 5 | 6 | 7 | 8 | 9 | 10 | 11 | 12 |
|---|---|---|---|---|---|---|---|---|---|---|---|---|
| A | | | | | | | | 0.0048 | -0.0098 | 0.0388 | 0.0114 | 0.0094 |
| B | | | | | | | | 0.0355 | 0.019 | 0.0328 | 0.0604 | 0.0536 |
| C | | | | | | | | 0.2271 | 0.2087 | 0.339 | 0.3508 | 0.265 |
| D | | | | | | | | 0.2195 | 0.2973 | 0.3133 | 0.3132 | 0.2691 |
| E | | | | | | | | 0.1065 | 0.0843 | 0.1003 | 0.1208 | 0.0641 |
| F | | | | | | | | 0.289 | 0.2604 | 0.3316 | 0.3105 | 0.2722 |
| G | | | | | | | | 0.0178 | 0.0431 | 0.0542 | 0.0485 | 0.0244 |
| H | | | | | | | | 0.2437 | 0.148 | 0.2317 | 0.2179 | 0.1732 |

**Cell Index at: 84:16:54**

| | 1 | 2 | 3 | 4 | 5 | 6 | 7 | 8 | 9 | 10 | 11 | 12 |
|---|---|---|---|---|---|---|---|---|---|---|---|---|
| A | | | | | | | | 0.009 | -0.0077 | 0.0376 | 0.0132 | 0.0093 |
| B | | | | | | | | 0.0415 | 0.0184 | 0.0304 | 0.0616 | 0.0605 |
| C | | | | | | | | 0.232 | 0.2113 | 0.3385 | 0.3548 | 0.2647 |
| D | | | | | | | | 0.2166 | 0.2891 | 0.3153 | 0.3139 | 0.2732 |
| E | | | | | | | | 0.095 | 0.0869 | 0.1045 | 0.1143 | 0.064 |
| F | | | | | | | | 0.285 | 0.2639 | 0.3247 | 0.3097 | 0.2759 |
| G | | | | | | | | 0.0146 | 0.0409 | 0.0522 | 0.0465 | 0.0273 |
| H | | | | | | | | 0.2438 | 0.1493 | 0.2288 | 0.218 | 0.1743 |

**Cell Index at: 84:31:55**

| | 1 | 2 | 3 | 4 | 5 | 6 | 7 | 8 | 9 | 10 | 11 | 12 |
|---|---|---|---|---|---|---|---|---|---|---|---|---|
| A | | | | | | | | 0.0073 | -0.0077 | 0.0412 | 0.0106 | 0.0091 |
| B | | | | | | | | 0.0423 | 0.0132 | 0.0329 | 0.0567 | 0.0614 |
| C | | | | | | | | 0.2294 | 0.2101 | 0.3448 | 0.351 | 0.2623 |
| D | | | | | | | | 0.2125 | 0.2875 | 0.3207 | 0.3216 | 0.2782 |
| E | | | | | | | | 0.1023 | 0.0932 | 0.1157 | 0.1081 | 0.067 |
| F | | | | | | | | 0.2863 | 0.2626 | 0.3323 | 0.3044 | 0.2789 |
| G | | | | | | | | 0.0206 | 0.0378 | 0.0485 | 0.0461 | 0.0252 |
| H | | | | | | | | 0.247 | 0.1527 | 0.2328 | 0.2181 | 0.1768 |

**Cell Index at: 84:46:56**

| | 1 | 2 | 3 | 4 | 5 | 6 | 7 | 8 | 9 | 10 | 11 | 12 |
|---|---|---|---|---|---|---|---|---|---|---|---|---|
| A | | | | | | | | 0.0041 | -0.0075 | 0.0427 | 0.0128 | 0.0105 |
| B | | | | | | | | 0.0377 | 0.0194 | 0.0314 | 0.0581 | 0.0623 |
| C | | | | | | | | 0.2315 | 0.2141 | 0.3402 | 0.3489 | 0.2604 |
| D | | | | | | | | 0.2125 | 0.2828 | 0.3135 | 0.3141 | 0.279 |
| E | | | | | | | | 0.1034 | 0.0915 | 0.1198 | 0.1175 | 0.0684 |
| F | | | | | | | | 0.2904 | 0.2599 | 0.333 | 0.3096 | 0.2765 |
| G | | | | | | | | 0.016 | 0.044 | 0.0468 | 0.0498 | 0.0287 |
| H | | | | | | | | 0.2494 | 0.1493 | 0.2328 | 0.213 | 0.1795 |

**Cell Index at: 85:01:56**

| | 1 | 2 | 3 | 4 | 5 | 6 | 7 | 8 | 9 | 10 | 11 | 12 |
|---|---|---|---|---|---|---|---|---|---|---|---|---|
| A | | | | | | | | 0.009 | -0.0094 | 0.0405 | 0.0127 | 0.0113 |
| B | | | | | | | | 0.0394 | 0.0205 | 0.0318 | 0.0586 | 0.0594 |
| C | | | | | | | | 0.2303 | 0.2099 | 0.3389 | 0.3551 | 0.2638 |
| D | | | | | | | | 0.2159 | 0.2906 | 0.3144 | 0.3172 | 0.2726 |
| E | | | | | | | | 0.1059 | 0.0816 | 0.1087 | 0.1122 | 0.0702 |
| F | | | | | | | | 0.2889 | 0.2624 | 0.3317 | 0.3177 | 0.2778 |
| G | | | | | | | | 0.02 | 0.0414 | 0.0483 | 0.0414 | 0.0231 |
| H | | | | | | | | 0.2515 | 0.1539 | 0.2332 | 0.215 | 0.1797 |

**Cell Index at: 85:16:57**

| | 1 | 2 | 3 | 4 | 5 | 6 | 7 | 8 | 9 | 10 | 11 | 12 |
|---|---|---|---|---|---|---|---|---|---|---|---|---|
| A | | | | | | | | 0.0068 | -0.0094 | 0.0414 | 0.0102 | 0.0099 |
| B | | | | | | | | 0.0403 | 0.0176 | 0.035 | 0.0625 | 0.0595 |
| C | | | | | | | | 0.2275 | 0.2146 | 0.3433 | 0.3514 | 0.2632 |
| D | | | | | | | | 0.214 | 0.288 | 0.3098 | 0.3143 | 0.2809 |
| E | | | | | | | | 0.1017 | 0.087 | 0.1046 | 0.0996 | 0.0653 |
| F | | | | | | | | 0.287 | 0.2652 | 0.331 | 0.3088 | 0.2743 |
| G | | | | | | | | 0.0212 | 0.0404 | 0.0538 | 0.0444 | 0.0258 |
| H | | | | | | | | 0.2489 | 0.1568 | 0.2315 | 0.2141 | 0.1781 |

**Cell Index at: 85:31:58**

| | 1 | 2 | 3 | 4 | 5 | 6 | 7 | 8 | 9 | 10 | 11 | 12 |
|---|---|---|---|---|---|---|---|---|---|---|---|---|
| A | | | | | | | | 0.0056 | -0.008 | 0.0416 | 0.0097 | 0.0093 |
| B | | | | | | | | 0.0405 | 0.0206 | 0.0333 | 0.0651 | 0.0592 |
| C | | | | | | | | 0.2232 | 0.2105 | 0.3443 | 0.3495 | 0.2684 |
| D | | | | | | | | 0.2201 | 0.2811 | 0.3087 | 0.3178 | 0.2704 |
| E | | | | | | | | 0.1035 | 0.0902 | 0.1093 | 0.1061 | 0.0727 |
| F | | | | | | | | 0.287 | 0.2622 | 0.3279 | 0.314 | 0.2723 |
| G | | | | | | | | 0.0222 | 0.0411 | 0.0527 | 0.0503 | 0.0299 |
| H | | | | | | | | 0.2546 | 0.1584 | 0.2345 | 0.2134 | 0.1799 |

**Cell Index at: 85:46:59**

| | 1 | 2 | 3 | 4 | 5 | 6 | 7 | 8 | 9 | 10 | 11 | 12 |
|---|---|---|---|---|---|---|---|---|---|---|---|---|
| A | | | | | | | | 0.0072 | -0.0079 | 0.0419 | 0.0111 | 0.013 |
| B | | | | | | | | 0.0387 | 0.0225 | 0.0311 | 0.0617 | 0.0583 |
| C | | | | | | | | 0.2264 | 0.2125 | 0.3385 | 0.3491 | 0.2673 |
| D | | | | | | | | 0.2201 | 0.2882 | 0.3118 | 0.3196 | 0.2733 |
| E | | | | | | | | 0.0985 | 0.0941 | 0.1115 | 0.1083 | 0.0696 |
| F | | | | | | | | 0.2882 | 0.264 | 0.3333 | 0.3153 | 0.2804 |
| G | | | | | | | | 0.0204 | 0.0405 | 0.0532 | 0.0523 | 0.0295 |
| H | | | | | | | | 0.255 | 0.1605 | 0.2384 | 0.2114 | 0.1801 |

**Cell Index at: 86:02:00**

| | 1 | 2 | 3 | 4 | 5 | 6 | 7 | 8 | 9 | 10 | 11 | 12 |
|---|---|---|---|---|---|---|---|---|---|---|---|---|
| A | | | | | | | | 0.008 | -0.0039 | 0.0384 | 0.007 | 0.0093 |
| B | | | | | | | | 0.0386 | 0.0237 | 0.0307 | 0.0639 | 0.0587 |
| C | | | | | | | | 0.2269 | 0.2145 | 0.3412 | 0.3546 | 0.2655 |
| D | | | | | | | | 0.2183 | 0.2865 | 0.3174 | 0.3174 | 0.2798 |
| E | | | | | | | | 0.1011 | 0.0947 | 0.1089 | 0.1037 | 0.0693 |
| F | | | | | | | | 0.2894 | 0.2633 | 0.3227 | 0.3114 | 0.2769 |
| G | | | | | | | | 0.0203 | 0.0374 | 0.0495 | 0.0476 | 0.0192 |
| H | | | | | | | | 0.2521 | 0.1594 | 0.2373 | 0.2138 | 0.1796 |

**Cell Index at: 86:17:00**

| | 1 | 2 | 3 | 4 | 5 | 6 | 7 | 8 | 9 | 10 | 11 | 12 |
|---|---|---|---|---|---|---|---|---|---|---|---|---|
| A | | | | | | | | 0.0048 | -0.0062 | 0.0368 | 0.0096 | 0.0099 |
| B | | | | | | | | 0.0371 | 0.0215 | 0.0333 | 0.0613 | 0.0609 |
| C | | | | | | | | 0.2201 | 0.216 | 0.3455 | 0.3528 | 0.2643 |
| D | | | | | | | | 0.2164 | 0.2864 | 0.3199 | 0.3176 | 0.2791 |
| E | | | | | | | | 0.1044 | 0.0956 | 0.1066 | 0.1073 | 0.0698 |
| F | | | | | | | | 0.2966 | 0.2642 | 0.326 | 0.3173 | 0.2789 |
| G | | | | | | | | 0.0208 | 0.0397 | 0.0529 | 0.0442 | 0.0307 |
| H | | | | | | | | 0.2531 | 0.161 | 0.2396 | 0.2185 | 0.1798 |

**Cell Index at: 86:32:01**

| | 1 | 2 | 3 | 4 | 5 | 6 | 7 | 8 | 9 | 10 | 11 | 12 |
|---|---|---|---|---|---|---|---|---|---|---|---|---|
| A | | | | | | | | 0.0034 | -0.0075 | 0.0394 | 0.0094 | 0.0074 |
| B | | | | | | | | 0.0423 | 0.0238 | 0.0316 | 0.0619 | 0.0604 |
| C | | | | | | | | 0.2243 | 0.2165 | 0.3507 | 0.3514 | 0.2662 |
| D | | | | | | | | 0.2171 | 0.2894 | 0.3194 | 0.3272 | 0.2747 |
| E | | | | | | | | 0.1009 | 0.0954 | 0.1074 | 0.1031 | 0.0768 |
| F | | | | | | | | 0.2925 | 0.2648 | 0.3241 | 0.315 | 0.2841 |
| G | | | | | | | | 0.019 | 0.039 | 0.0566 | 0.048 | 0.0274 |
| H | | | | | | | | 0.253 | 0.1603 | 0.2391 | 0.2205 | 0.1782 |

**Cell Index at: 86:47:01**

| | 1 | 2 | 3 | 4 | 5 | 6 | 7 | 8 | 9 | 10 | 11 | 12 |
|---|---|---|---|---|---|---|---|---|---|---|---|---|
| A | | | | | | | | 0.0044 | -0.005 | 0.0407 | 0.011 | 0.0123 |
| B | | | | | | | | 0.0416 | 0.0216 | 0.0315 | 0.0602 | 0.059 |
| C | | | | | | | | 0.22 | 0.2168 | 0.3529 | 0.3475 | 0.2668 |
| D | | | | | | | | 0.2148 | 0.2799 | 0.3184 | 0.3188 | 0.2764 |
| E | | | | | | | | 0.098 | 0.0854 | 0.1083 | 0.1027 | 0.0633 |
| F | | | | | | | | 0.3016 | 0.2632 | 0.3262 | 0.3191 | 0.2812 |
| G | | | | | | | | 0.0217 | 0.0403 | 0.053 | 0.0482 | 0.0324 |
| H | | | | | | | | 0.2558 | 0.1606 | 0.2395 | 0.2196 | 0.1828 |

**Cell Index at: 87:02:02**

| | 1 | 2 | 3 | 4 | 5 | 6 | 7 | 8 | 9 | 10 | 11 | 12 |
|---|---|---|---|---|---|---|---|---|---|---|---|---|
| A | | | | | | | | 0.0012 | -0.005 | 0.041 | 0.0107 | 0.0064 |
| B | | | | | | | | 0.0433 | 0.0238 | 0.0325 | 0.0584 | 0.0618 |
| C | | | | | | | | 0.2218 | 0.2188 | 0.3534 | 0.3632 | 0.2675 |
| D | | | | | | | | 0.2227 | 0.2906 | 0.324 | 0.3223 | 0.2758 |
| E | | | | | | | | 0.0949 | 0.0936 | 0.1118 | 0.1011 | 0.062 |
| F | | | | | | | | 0.2948 | 0.2618 | 0.3241 | 0.3215 | 0.2796 |
| G | | | | | | | | 0.0194 | 0.0402 | 0.0499 | 0.0505 | 0.0302 |
| H | | | | | | | | 0.2547 | 0.162 | 0.2366 | 0.2195 | 0.1806 |

**Cell Index at: 87:17:02**

| | 1 | 2 | 3 | 4 | 5 | 6 | 7 | 8 | 9 | 10 | 11 | 12 |
|---|---|---|---|---|---|---|---|---|---|---|---|---|
| A | | | | | | | | 0.0002 | -0.005 | 0.0418 | 0.012 | 0.009 |
| B | | | | | | | | 0.0428 | 0.0224 | 0.0328 | 0.0605 | 0.0595 |
| C | | | | | | | | 0.2275 | 0.2214 | 0.3503 | 0.3613 | 0.2695 |
| D | | | | | | | | 0.217 | 0.2902 | 0.3203 | 0.3191 | 0.2853 |
| E | | | | | | | | 0.0988 | 0.0842 | 0.1113 | 0.1013 | 0.0637 |
| F | | | | | | | | 0.2987 | 0.2611 | 0.3316 | 0.3269 | 0.283 |
| G | | | | | | | | 0.0237 | 0.0371 | 0.053 | 0.049 | 0.0269 |
| H | | | | | | | | 0.2517 | 0.1635 | 0.2399 | 0.2217 | 0.1831 |

**Cell Index at: 87:32:03**

| | 1 | 2 | 3 | 4 | 5 | 6 | 7 | 8 | 9 | 10 | 11 | 12 |
|---|---|---|---|---|---|---|---|---|---|---|---|---|
| A | | | | | | | | 0.0046 | -0.0033 | 0.0385 | 0.0112 | 0.0113 |
| B | | | | | | | | 0.0395 | 0.0242 | 0.0344 | 0.062 | 0.0604 |
| C | | | | | | | | 0.2289 | 0.2185 | 0.353 | 0.3659 | 0.272 |
| D | | | | | | | | 0.2204 | 0.295 | 0.3213 | 0.3253 | 0.2889 |
| E | | | | | | | | 0.0918 | 0.0882 | 0.1152 | 0.0963 | 0.066 |
| F | | | | | | | | 0.2908 | 0.2651 | 0.325 | 0.3222 | 0.2865 |
| G | | | | | | | | 0.0224 | 0.0429 | 0.0529 | 0.0485 | 0.0299 |
| H | | | | | | | | 0.2488 | 0.166 | 0.2401 | 0.2224 | 0.183 |

**Cell Index at: 87:47:03**

| | 1 | 2 | 3 | 4 | 5 | 6 | 7 | 8 | 9 | 10 | 11 | 12 |
|---|---|---|---|---|---|---|---|---|---|---|---|---|
| A | | | | | | | | 0.005 | -0.0049 | 0.038 | 0.0126 | 0.0145 |
| B | | | | | | | | 0.0427 | 0.0229 | 0.034 | 0.0616 | 0.0628 |
| C | | | | | | | | 0.2274 | 0.2172 | 0.3606 | 0.3624 | 0.277 |
| D | | | | | | | | 0.2139 | 0.2946 | 0.3283 | 0.3207 | 0.2787 |
| E | | | | | | | | 0.099 | 0.0862 | 0.1098 | 0.1034 | 0.0674 |
| F | | | | | | | | 0.2947 | 0.2736 | 0.3313 | 0.3239 | 0.2856 |
| G | | | | | | | | 0.0222 | 0.0427 | 0.0539 | 0.0482 | 0.0307 |
| H | | | | | | | | 0.2486 | 0.1638 | 0.2411 | 0.2261 | 0.1797 |

**Cell Index at: 88:02:04**

| | 1 | 2 | 3 | 4 | 5 | 6 | 7 | 8 | 9 | 10 | 11 | 12 |
|---|---|---|---|---|---|---|---|---|---|---|---|---|
| A | | | | | | | | 0.0057 | -0.0011 | 0.0393 | 0.0107 | 0.0146 |
| B | | | | | | | | 0.0413 | 0.0234 | 0.0376 | 0.0686 | 0.0575 |
| C | | | | | | | | 0.2308 | 0.223 | 0.3587 | 0.3667 | 0.2701 |
| D | | | | | | | | 0.2138 | 0.3006 | 0.3239 | 0.3201 | 0.2841 |
| E | | | | | | | | 0.104 | 0.0922 | 0.1136 | 0.0987 | 0.062 |
| F | | | | | | | | 0.2976 | 0.27 | 0.3315 | 0.3283 | 0.2876 |
| G | | | | | | | | 0.0246 | 0.047 | 0.0538 | 0.0504 | 0.0302 |
| H | | | | | | | | 0.2535 | 0.1648 | 0.2422 | 0.2264 | 0.1767 |

**Cell Index at: 88:17:05**

| | 1 | 2 | 3 | 4 | 5 | 6 | 7 | 8 | 9 | 10 | 11 | 12 |
|---|---|---|---|---|---|---|---|---|---|---|---|---|
| A | | | | | | | | 0.0036 | -0.0017 | 0.0415 | 0.0109 | 0.0136 |
| B | | | | | | | | 0.0401 | 0.025 | 0.0344 | 0.0721 | 0.0592 |
| C | | | | | | | | 0.2315 | 0.2181 | 0.3542 | 0.3717 | 0.2819 |
| D | | | | | | | | 0.2191 | 0.2937 | 0.3306 | 0.3204 | 0.2845 |
| E | | | | | | | | 0.1026 | 0.0882 | 0.1228 | 0.1054 | 0.0632 |
| F | | | | | | | | 0.2982 | 0.2722 | 0.3247 | 0.3265 | 0.2854 |
| G | | | | | | | | 0.0221 | 0.0406 | 0.0558 | 0.0502 | 0.0259 |
| H | | | | | | | | 0.2534 | 0.1614 | 0.246 | 0.2243 | 0.1806 |

**Cell Index at: 88:32:05**

| | 1 | 2 | 3 | 4 | 5 | 6 | 7 | 8 | 9 | 10 | 11 | 12 |
|---|---|---|---|---|---|---|---|---|---|---|---|---|
| A | | | | | | | | 0.0063 | -0.0037 | 0.0385 | 0.0103 | 0.014 |
| B | | | | | | | | 0.0414 | 0.0288 | 0.0353 | 0.0718 | 0.06 |
| C | | | | | | | | 0.2305 | 0.2185 | 0.3548 | 0.3674 | 0.2783 |
| D | | | | | | | | 0.2189 | 0.2979 | 0.3243 | 0.3275 | 0.2872 |
| E | | | | | | | | 0.0977 | 0.0961 | 0.1158 | 0.1062 | 0.0678 |
| F | | | | | | | | 0.3016 | 0.2773 | 0.3255 | 0.3281 | 0.2894 |
| G | | | | | | | | 0.0272 | 0.0456 | 0.0522 | 0.0537 | 0.0309 |
| H | | | | | | | | 0.2537 | 0.1635 | 0.2497 | 0.2249 | 0.1791 |

**Cell Index at: 88:47:05**

| | 1 | 2 | 3 | 4 | 5 | 6 | 7 | 8 | 9 | 10 | 11 | 12 |
|---|---|---|---|---|---|---|---|---|---|---|---|---|
| A | | | | | | | | 0.0064 | -0.0023 | 0.0387 | 0.0098 | 0.0173 |
| B | | | | | | | | 0.0451 | 0.0258 | 0.0357 | 0.0643 | 0.0609 |
| C | | | | | | | | 0.2333 | 0.2197 | 0.3578 | 0.3695 | 0.278 |
| D | | | | | | | | 0.2195 | 0.3115 | 0.3274 | 0.3296 | 0.2833 |
| E | | | | | | | | 0.098 | 0.0899 | 0.1182 | 0.1113 | 0.0666 |
| F | | | | | | | | 0.3038 | 0.2758 | 0.3301 | 0.3235 | 0.2852 |
| G | | | | | | | | 0.025 | 0.0461 | 0.0542 | 0.0515 | 0.0302 |
| H | | | | | | | | 0.257 | 0.1643 | 0.2464 | 0.226 | 0.1802 |

**Cell Index at: 89:02:05**

| | 1 | 2 | 3 | 4 | 5 | 6 | 7 | 8 | 9 | 10 | 11 | 12 |
|---|---|---|---|---|---|---|---|---|---|---|---|---|
| A | | | | | | | | 0.0072 | -0.0058 | 0.0409 | 0.0081 | 0.0155 |
| B | | | | | | | | 0.0378 | 0.023 | 0.0387 | 0.0701 | 0.063 |
| C | | | | | | | | 0.2331 | 0.2226 | 0.3576 | 0.3703 | 0.2815 |
| D | | | | | | | | 0.2241 | 0.2988 | 0.3248 | 0.3283 | 0.2841 |
| E | | | | | | | | 0.0966 | 0.093 | 0.1183 | 0.1107 | 0.0682 |
| F | | | | | | | | 0.2992 | 0.2782 | 0.3319 | 0.3275 | 0.2875 |
| G | | | | | | | | 0.0215 | 0.0435 | 0.059 | 0.0508 | 0.0318 |
| H | | | | | | | | 0.2578 | 0.1622 | 0.2476 | 0.2266 | 0.1821 |

**Cell Index at: 89:17:06**

| | 1 | 2 | 3 | 4 | 5 | 6 | 7 | 8 | 9 | 10 | 11 | 12 |
|---|---|---|---|---|---|---|---|---|---|---|---|---|
| A | | | | | | | | 0.0052 | -0.0013 | 0.0417 | 0.0077 | 0.0152 |
| B | | | | | | | | 0.04 | 0.0209 | 0.0394 | 0.0674 | 0.0613 |
| C | | | | | | | | 0.23 | 0.2206 | 0.3621 | 0.3703 | 0.2842 |
| D | | | | | | | | 0.22 | 0.3006 | 0.3235 | 0.329 | 0.2837 |
| E | | | | | | | | 0.0991 | 0.0867 | 0.1206 | 0.1071 | 0.0683 |
| F | | | | | | | | 0.3001 | 0.2798 | 0.3341 | 0.3245 | 0.2949 |
| G | | | | | | | | 0.0216 | 0.0428 | 0.0519 | 0.0511 | 0.0347 |
| H | | | | | | | | 0.2623 | 0.1652 | 0.2482 | 0.2265 | 0.1831 |

**Cell Index at: 89:32:06**

| | 1 | 2 | 3 | 4 | 5 | 6 | 7 | 8 | 9 | 10 | 11 | 12 |
|---|---|---|---|---|---|---|---|---|---|---|---|---|
| A | | | | | | | | 0.0098 | -0.0041 | 0.0387 | 0.011 | 0.0167 |
| B | | | | | | | | 0.0363 | 0.0248 | 0.0417 | 0.0669 | 0.0624 |
| C | | | | | | | | 0.2308 | 0.2187 | 0.3595 | 0.3702 | 0.283 |
| D | | | | | | | | 0.2233 | 0.2993 | 0.325 | 0.3311 | 0.2881 |
| E | | | | | | | | 0.097 | 0.0874 | 0.1181 | 0.1133 | 0.0656 |
| F | | | | | | | | 0.3043 | 0.2755 | 0.327 | 0.3275 | 0.2892 |
| G | | | | | | | | 0.0245 | 0.0405 | 0.057 | 0.0526 | 0.0278 |
| H | | | | | | | | 0.2622 | 0.1655 | 0.2468 | 0.2265 | 0.1822 |

**Cell Index at: 89:47:06**

| | 1 | 2 | 3 | 4 | 5 | 6 | 7 | 8 | 9 | 10 | 11 | 12 |
|---|---|---|---|---|---|---|---|---|---|---|---|---|
| A | | | | | | | | 0.0102 | -0.0048 | 0.0336 | 0.0091 | 0.0177 |
| B | | | | | | | | 0.0336 | 0.0282 | 0.0414 | 0.0737 | 0.0602 |
| C | | | | | | | | 0.2335 | 0.2223 | 0.3616 | 0.376 | 0.2779 |
| D | | | | | | | | 0.2176 | 0.3074 | 0.3347 | 0.3309 | 0.2912 |
| E | | | | | | | | 0.1014 | 0.0897 | 0.1191 | 0.109 | 0.0643 |
| F | | | | | | | | 0.3078 | 0.2753 | 0.3329 | 0.3302 | 0.2894 |
| G | | | | | | | | 0.022 | 0.0456 | 0.0569 | 0.0519 | 0.0295 |
| H | | | | | | | | 0.2625 | 0.1677 | 0.2485 | 0.2262 | 0.1883 |

**Cell Index at: 90:02:07**

| | 1 | 2 | 3 | 4 | 5 | 6 | 7 | 8 | 9 | 10 | 11 | 12 |
|---|---|---|---|---|---|---|---|---|---|---|---|---|
| A | | | | | | | | 0.0077 | -0.0043 | 0.0366 | 0.0076 | 0.0174 |
| B | | | | | | | | 0.0354 | 0.0261 | 0.0403 | 0.0688 | 0.0595 |
| C | | | | | | | | 0.2338 | 0.2215 | 0.3618 | 0.3712 | 0.2845 |
| D | | | | | | | | 0.222 | 0.3026 | 0.3314 | 0.3313 | 0.2887 |
| E | | | | | | | | 0.096 | 0.087 | 0.1172 | 0.1126 | 0.0707 |
| F | | | | | | | | 0.3083 | 0.283 | 0.3351 | 0.3297 | 0.2951 |
| G | | | | | | | | 0.0229 | 0.0456 | 0.0573 | 0.0516 | 0.033 |
| H | | | | | | | | 0.2651 | 0.1671 | 0.2508 | 0.2269 | 0.1851 |

**Cell Index at: 90:17:05**

| | 1 | 2 | 3 | 4 | 5 | 6 | 7 | 8 | 9 | 10 | 11 | 12 |
|---|---|---|---|---|---|---|---|---|---|---|---|---|
| A | | | | | | | | 0.0098 | -0.0054 | 0.038 | 0.0116 | 0.0169 |
| B | | | | | | | | 0.0374 | 0.0296 | 0.0404 | 0.073 | 0.0619 |
| C | | | | | | | | 0.2411 | 0.2256 | 0.3577 | 0.3654 | 0.2882 |
| D | | | | | | | | 0.218 | 0.3069 | 0.3359 | 0.3272 | 0.2912 |
| E | | | | | | | | 0.0976 | 0.0852 | 0.1232 | 0.1175 | 0.0685 |
| F | | | | | | | | 0.3025 | 0.2788 | 0.337 | 0.3299 | 0.297 |
| G | | | | | | | | 0.0242 | 0.0427 | 0.0583 | 0.053 | 0.0334 |
| H | | | | | | | | 0.2635 | 0.1691 | 0.2497 | 0.2292 | 0.1876 |

**Cell Index at: 90:32:06**

| | 1 | 2 | 3 | 4 | 5 | 6 | 7 | 8 | 9 | 10 | 11 | 12 |
|---|---|---|---|---|---|---|---|---|---|---|---|---|
| A | | | | | | | | 0.0113 | -0.0032 | 0.0393 | 0.0051 | 0.0168 |
| B | | | | | | | | 0.0358 | 0.0285 | 0.0461 | 0.0715 | 0.0631 |
| C | | | | | | | | 0.2423 | 0.2251 | 0.3632 | 0.3706 | 0.2887 |
| D | | | | | | | | 0.2193 | 0.307 | 0.3295 | 0.3393 | 0.2898 |
| E | | | | | | | | 0.1005 | 0.0956 | 0.1205 | 0.1148 | 0.0728 |
| F | | | | | | | | 0.3086 | 0.2841 | 0.3309 | 0.333 | 0.2904 |
| G | | | | | | | | 0.0237 | 0.0441 | 0.0558 | 0.0504 | 0.0316 |
| H | | | | | | | | 0.2705 | 0.1683 | 0.2479 | 0.2292 | 0.1893 |

**Cell Index at: 90:47:07**

| | 1 | 2 | 3 | 4 | 5 | 6 | 7 | 8 | 9 | 10 | 11 | 12 |
|---|---|---|---|---|---|---|---|---|---|---|---|---|
| A | | | | | | | | 0.0147 | -0.0012 | 0.0408 | 0.0069 | 0.0185 |
| B | | | | | | | | 0.0374 | 0.0314 | 0.0444 | 0.0678 | 0.0574 |
| C | | | | | | | | 0.2432 | 0.2285 | 0.3715 | 0.3737 | 0.2869 |
| D | | | | | | | | 0.2142 | 0.2973 | 0.328 | 0.3419 | 0.2881 |
| E | | | | | | | | 0.1014 | 0.089 | 0.1245 | 0.1176 | 0.068 |
| F | | | | | | | | 0.3049 | 0.2842 | 0.3337 | 0.3326 | 0.2945 |
| G | | | | | | | | 0.0243 | 0.0425 | 0.0578 | 0.0473 | 0.0329 |
| H | | | | | | | | 0.271 | 0.1687 | 0.2505 | 0.2307 | 0.1891 |

**Cell Index at: 91:02:07**

| | 1 | 2 | 3 | 4 | 5 | 6 | 7 | 8 | 9 | 10 | 11 | 12 |
|---|---|---|---|---|---|---|---|---|---|---|---|---|
| A | | | | | | | | 0.0131 | -0.0006 | 0.0383 | 0.0093 | 0.0207 |
| B | | | | | | | | 0.037 | 0.0298 | 0.0444 | 0.0668 | 0.0569 |
| C | | | | | | | | 0.2373 | 0.2262 | 0.3682 | 0.3738 | 0.2884 |
| D | | | | | | | | 0.2159 | 0.303 | 0.3303 | 0.3354 | 0.2869 |
| E | | | | | | | | 0.0935 | 0.0897 | 0.1233 | 0.1141 | 0.0677 |
| F | | | | | | | | 0.3078 | 0.2882 | 0.3318 | 0.3418 | 0.2963 |
| G | | | | | | | | 0.0234 | 0.0488 | 0.0569 | 0.0448 | 0.0309 |
| H | | | | | | | | 0.2722 | 0.1673 | 0.2506 | 0.2339 | 0.1904 |

**Cell Index at: 91:17:07**

| | 1 | 2 | 3 | 4 | 5 | 6 | 7 | 8 | 9 | 10 | 11 | 12 |
|---|---|---|---|---|---|---|---|---|---|---|---|---|
| A | | | | | | | | 0.0151 | -0.0006 | 0.0408 | 0.0093 | 0.0208 |
| B | | | | | | | | 0.0407 | 0.0288 | 0.0421 | 0.0685 | 0.0585 |
| C | | | | | | | | 0.234 | 0.2348 | 0.3679 | 0.3724 | 0.2916 |
| D | | | | | | | | 0.2248 | 0.3133 | 0.334 | 0.3333 | 0.2885 |
| E | | | | | | | | 0.1033 | 0.104 | 0.12 | 0.1103 | 0.075 |
| F | | | | | | | | 0.3107 | 0.2829 | 0.3413 | 0.3407 | 0.2923 |
| G | | | | | | | | 0.0269 | 0.0449 | 0.0608 | 0.0502 | 0.0276 |
| H | | | | | | | | 0.2721 | 0.1676 | 0.2529 | 0.2321 | 0.1877 |

**Cell Index at: 91:32:08**

| | 1 | 2 | 3 | 4 | 5 | 6 | 7 | 8 | 9 | 10 | 11 | 12 |
|---|---|---|---|---|---|---|---|---|---|---|---|---|
| A | | | | | | | | 0.0129 | 0.0002 | 0.0394 | 0.0087 | 0.0219 |
| B | | | | | | | | 0.0399 | 0.0334 | 0.0444 | 0.069 | 0.0581 |
| C | | | | | | | | 0.2306 | 0.232 | 0.3621 | 0.3733 | 0.296 |
| D | | | | | | | | 0.2238 | 0.3102 | 0.3331 | 0.3293 | 0.2924 |
| E | | | | | | | | 0.0919 | 0.0926 | 0.125 | 0.113 | 0.0661 |
| F | | | | | | | | 0.3079 | 0.2855 | 0.343 | 0.3378 | 0.2924 |
| G | | | | | | | | 0.0277 | 0.0467 | 0.0563 | 0.0498 | 0.03 |
| H | | | | | | | | 0.2712 | 0.1683 | 0.252 | 0.2352 | 0.1882 |

**Cell Index at: 91:47:09**

| | 1 | 2 | 3 | 4 | 5 | 6 | 7 | 8 | 9 | 10 | 11 | 12 |
|---|---|---|---|---|---|---|---|---|---|---|---|---|
| A | | | | | | | | 0.0107 | -0.0021 | 0.043 | 0.0109 | 0.0201 |
| B | | | | | | | | 0.0379 | 0.0331 | 0.0449 | 0.0681 | 0.0521 |
| C | | | | | | | | 0.2333 | 0.2353 | 0.3679 | 0.3775 | 0.2961 |
| D | | | | | | | | 0.2271 | 0.3072 | 0.3417 | 0.3372 | 0.2952 |
| E | | | | | | | | 0.1022 | 0.0923 | 0.1252 | 0.1133 | 0.0697 |
| F | | | | | | | | 0.3134 | 0.2869 | 0.3372 | 0.339 | 0.2982 |
| G | | | | | | | | 0.0297 | 0.0478 | 0.0557 | 0.0508 | 0.0253 |
| H | | | | | | | | 0.2738 | 0.1722 | 0.2527 | 0.2356 | 0.191 |

**Cell Index at: 92:02:10**

| | 1 | 2 | 3 | 4 | 5 | 6 | 7 | 8 | 9 | 10 | 11 | 12 |
|---|---|---|---|---|---|---|---|---|---|---|---|---|
| A | | | | | | | | 0.0131 | -0.0004 | 0.0415 | 0.0081 | 0.0209 |
| B | | | | | | | | 0.0433 | 0.0347 | 0.0447 | 0.069 | 0.0586 |
| C | | | | | | | | 0.2344 | 0.2373 | 0.3594 | 0.3773 | 0.2997 |
| D | | | | | | | | 0.2244 | 0.3006 | 0.3359 | 0.3355 | 0.2883 |
| E | | | | | | | | 0.1063 | 0.0919 | 0.1279 | 0.1222 | 0.0732 |
| F | | | | | | | | 0.3101 | 0.2912 | 0.3384 | 0.3413 | 0.2947 |
| G | | | | | | | | 0.0252 | 0.0446 | 0.0581 | 0.0503 | 0.0282 |
| H | | | | | | | | 0.2713 | 0.1686 | 0.2552 | 0.2363 | 0.1872 |

**Cell Index at: 92:17:11**

| | 1 | 2 | 3 | 4 | 5 | 6 | 7 | 8 | 9 | 10 | 11 | 12 |
|---|---|---|---|---|---|---|---|---|---|---|---|---|
| A | | | | | | | | 0.0162 | 0.0015 | 0.0425 | 0.0072 | 0.0221 |
| B | | | | | | | | 0.0394 | 0.0359 | 0.048 | 0.0742 | 0.0557 |
| C | | | | | | | | 0.2345 | 0.235 | 0.3625 | 0.3774 | 0.2985 |
| D | | | | | | | | 0.2207 | 0.3133 | 0.3283 | 0.3333 | 0.2987 |
| E | | | | | | | | 0.1047 | 0.0906 | 0.1258 | 0.1176 | 0.0718 |
| F | | | | | | | | 0.316 | 0.288 | 0.3393 | 0.3443 | 0.2943 |
| G | | | | | | | | 0.0298 | 0.048 | 0.0592 | 0.0492 | 0.0337 |
| H | | | | | | | | 0.2732 | 0.1737 | 0.2531 | 0.2346 | 0.1886 |

**Cell Index at: 92:32:11**

| | 1 | 2 | 3 | 4 | 5 | 6 | 7 | 8 | 9 | 10 | 11 | 12 |
|---|---|---|---|---|---|---|---|---|---|---|---|---|
| A | | | | | | | | 0.0145 | 0.0012 | 0.0426 | 0.0068 | 0.0199 |
| B | | | | | | | | 0.0467 | 0.0361 | 0.0456 | 0.0704 | 0.0575 |
| C | | | | | | | | 0.2323 | 0.2363 | 0.3574 | 0.3765 | 0.3006 |
| D | | | | | | | | 0.2257 | 0.3135 | 0.3294 | 0.338 | 0.2948 |
| E | | | | | | | | 0.104 | 0.0897 | 0.1259 | 0.12 | 0.0749 |
| F | | | | | | | | 0.3197 | 0.2877 | 0.3372 | 0.3461 | 0.3011 |
| G | | | | | | | | 0.0294 | 0.0498 | 0.0557 | 0.0494 | 0.0324 |
| H | | | | | | | | 0.2758 | 0.1719 | 0.2544 | 0.2372 | 0.1917 |

**Cell Index at: 92:47:12**

| | 1 | 2 | 3 | 4 | 5 | 6 | 7 | 8 | 9 | 10 | 11 | 12 |
|---|---|---|---|---|---|---|---|---|---|---|---|---|
| A | | | | | | | | 0.0169 | 0.0034 | 0.0416 | 0.0056 | 0.0221 |
| B | | | | | | | | 0.0477 | 0.0319 | 0.0439 | 0.0714 | 0.0549 |
| C | | | | | | | | 0.2375 | 0.2386 | 0.3638 | 0.3798 | 0.2968 |
| D | | | | | | | | 0.2325 | 0.3114 | 0.3251 | 0.3388 | 0.2905 |
| E | | | | | | | | 0.1 | 0.0909 | 0.1219 | 0.125 | 0.0702 |
| F | | | | | | | | 0.3241 | 0.2939 | 0.3463 | 0.3494 | 0.3059 |
| G | | | | | | | | 0.0306 | 0.0483 | 0.058 | 0.0481 | 0.0325 |
| H | | | | | | | | 0.2781 | 0.1736 | 0.2551 | 0.2356 | 0.1937 |

**Cell Index at: 93:02:12**

| | 1 | 2 | 3 | 4 | 5 | 6 | 7 | 8 | 9 | 10 | 11 | 12 |
|---|---|---|---|---|---|---|---|---|---|---|---|---|
| A | | | | | | | | 0.0152 | 0.0024 | 0.0432 | 0.0052 | 0.0221 |
| B | | | | | | | | 0.0422 | 0.0344 | 0.0467 | 0.0712 | 0.0544 |
| C | | | | | | | | 0.2377 | 0.2383 | 0.356 | 0.3792 | 0.3064 |
| D | | | | | | | | 0.236 | 0.315 | 0.3345 | 0.3365 | 0.3006 |
| E | | | | | | | | 0.1019 | 0.0925 | 0.1262 | 0.1217 | 0.0711 |
| F | | | | | | | | 0.3186 | 0.2984 | 0.3437 | 0.3453 | 0.305 |
| G | | | | | | | | 0.0277 | 0.049 | 0.0605 | 0.0544 | 0.0326 |
| H | | | | | | | | 0.2762 | 0.1724 | 0.2546 | 0.2359 | 0.1925 |

**Cell Index at: 93:17:13**

| | 1 | 2 | 3 | 4 | 5 | 6 | 7 | 8 | 9 | 10 | 11 | 12 |
|---|---|---|---|---|---|---|---|---|---|---|---|---|
| A | | | | | | | | 0.0143 | 0.0052 | 0.044 | 0.0059 | 0.0192 |
| B | | | | | | | | 0.0436 | 0.0334 | 0.045 | 0.0696 | 0.0556 |
| C | | | | | | | | 0.2505 | 0.2342 | 0.3616 | 0.3792 | 0.3059 |
| D | | | | | | | | 0.2311 | 0.3102 | 0.3252 | 0.3325 | 0.3017 |
| E | | | | | | | | 0.103 | 0.0993 | 0.1274 | 0.1227 | 0.0665 |
| F | | | | | | | | 0.326 | 0.2952 | 0.3464 | 0.3472 | 0.3024 |
| G | | | | | | | | 0.0315 | 0.0503 | 0.0606 | 0.0495 | 0.0324 |
| H | | | | | | | | 0.2779 | 0.1753 | 0.2579 | 0.235 | 0.1936 |

**Cell Index at: 93:32:13**

| | 1 | 2 | 3 | 4 | 5 | 6 | 7 | 8 | 9 | 10 | 11 | 12 |
|---|---|---|---|---|---|---|---|---|---|---|---|---|
| A | | | | | | | | 0.0138 | 0.0079 | 0.0407 | 0.0065 | 0.0213 |
| B | | | | | | | | 0.0426 | 0.033 | 0.0486 | 0.0753 | 0.0516 |
| C | | | | | | | | 0.2451 | 0.2355 | 0.3612 | 0.3809 | 0.311 |
| D | | | | | | | | 0.2312 | 0.317 | 0.3351 | 0.3411 | 0.2957 |
| E | | | | | | | | 0.0998 | 0.0951 | 0.1305 | 0.1208 | 0.0681 |
| F | | | | | | | | 0.329 | 0.2997 | 0.3458 | 0.3392 | 0.3052 |
| G | | | | | | | | 0.0319 | 0.0534 | 0.0578 | 0.0538 | 0.0332 |

| | 1 | 2 | 3 | 4 | 5 | 6 | 7 | 8 | 9 | 10 | 11 | 12 |
|---|---|---|---|---|---|---|---|---|---|---|---|---|
| H | | | | | | | | 0.2788 | 0.1713 | 0.2599 | 0.2362 | 0.1929 |

**Cell Index at: 93:47:14**

| | 1 | 2 | 3 | 4 | 5 | 6 | 7 | 8 | 9 | 10 | 11 | 12 |
|---|---|---|---|---|---|---|---|---|---|---|---|---|
| A | | | | | | | | 0.0134 | 0.0051 | 0.0437 | 0.0073 | 0.0225 |
| B | | | | | | | | 0.0479 | 0.0329 | 0.0476 | 0.0736 | 0.055 |
| C | | | | | | | | 0.2449 | 0.2336 | 0.3563 | 0.378 | 0.3047 |
| D | | | | | | | | 0.234 | 0.3127 | 0.3326 | 0.3353 | 0.3054 |
| E | | | | | | | | 0.0964 | 0.0977 | 0.1286 | 0.1196 | 0.069 |
| F | | | | | | | | 0.3247 | 0.297 | 0.3492 | 0.35 | 0.3104 |
| G | | | | | | | | 0.0317 | 0.0547 | 0.0582 | 0.0533 | 0.0339 |
| H | | | | | | | | 0.2791 | 0.1757 | 0.2601 | 0.2377 | 0.1944 |

**Cell Index at: 94:02:15**

| | 1 | 2 | 3 | 4 | 5 | 6 | 7 | 8 | 9 | 10 | 11 | 12 |
|---|---|---|---|---|---|---|---|---|---|---|---|---|
| A | | | | | | | | 0.0167 | 0.0065 | 0.0446 | 0.007 | 0.0237 |
| B | | | | | | | | 0.0419 | 0.0312 | 0.0467 | 0.0783 | 0.0554 |
| C | | | | | | | | 0.2485 | 0.2415 | 0.3551 | 0.3777 | 0.307 |
| D | | | | | | | | 0.237 | 0.3101 | 0.3357 | 0.3337 | 0.3035 |
| E | | | | | | | | 0.0951 | 0.0929 | 0.1243 | 0.1246 | 0.0712 |
| F | | | | | | | | 0.3211 | 0.2983 | 0.3466 | 0.3481 | 0.3116 |
| G | | | | | | | | 0.0298 | 0.0546 | 0.0548 | 0.052 | 0.0338 |
| H | | | | | | | | 0.2807 | 0.1785 | 0.259 | 0.2387 | 0.1961 |

**Cell Index at: 94:17:15**

| | 1 | 2 | 3 | 4 | 5 | 6 | 7 | 8 | 9 | 10 | 11 | 12 |
|---|---|---|---|---|---|---|---|---|---|---|---|---|
| A | | | | | | | | 0.0154 | 0.005 | 0.0426 | 0.0066 | 0.0247 |
| B | | | | | | | | 0.046 | 0.033 | 0.0455 | 0.0775 | 0.0562 |
| C | | | | | | | | 0.2481 | 0.2388 | 0.3563 | 0.3874 | 0.3047 |
| D | | | | | | | | 0.2312 | 0.306 | 0.3421 | 0.34 | 0.302 |
| E | | | | | | | | 0.101 | 0.097 | 0.1276 | 0.121 | 0.0779 |
| F | | | | | | | | 0.3282 | 0.2982 | 0.3456 | 0.3523 | 0.3057 |
| G | | | | | | | | 0.0293 | 0.0527 | 0.0573 | 0.0578 | 0.0344 |
| H | | | | | | | | 0.2812 | 0.1771 | 0.2598 | 0.239 | 0.1977 |

**Cell Index at: 94:32:16**

| | 1 | 2 | 3 | 4 | 5 | 6 | 7 | 8 | 9 | 10 | 11 | 12 |
|---|---|---|---|---|---|---|---|---|---|---|---|---|
| A | | | | | | | | 0.017 | 0.0037 | 0.0395 | 0.0078 | 0.026 |
| B | | | | | | | | 0.0445 | 0.0313 | 0.0458 | 0.0751 | 0.0576 |
| C | | | | | | | | 0.2445 | 0.2399 | 0.3591 | 0.3837 | 0.3109 |
| D | | | | | | | | 0.2395 | 0.3067 | 0.3429 | 0.333 | 0.302 |
| E | | | | | | | | 0.0938 | 0.0933 | 0.1254 | 0.1167 | 0.0768 |
| F | | | | | | | | 0.3273 | 0.2974 | 0.3466 | 0.3519 | 0.3047 |
| G | | | | | | | | 0.0315 | 0.054 | 0.0582 | 0.056 | 0.0352 |
| H | | | | | | | | 0.2823 | 0.1818 | 0.2605 | 0.2382 | 0.1997 |

**Cell Index at: 94:47:16**

| | 1 | 2 | 3 | 4 | 5 | 6 | 7 | 8 | 9 | 10 | 11 | 12 |
|---|---|---|---|---|---|---|---|---|---|---|---|---|
| A | | | | | | | | 0.0125 | 0.0029 | 0.0427 | 0.0053 | 0.0238 |
| B | | | | | | | | 0.0405 | 0.029 | 0.0474 | 0.0745 | 0.0573 |
| C | | | | | | | | 0.2466 | 0.2387 | 0.3601 | 0.3865 | 0.3077 |
| D | | | | | | | | 0.2401 | 0.3144 | 0.3445 | 0.3338 | 0.2989 |
| E | | | | | | | | 0.0953 | 0.0993 | 0.129 | 0.118 | 0.0761 |
| F | | | | | | | | 0.3279 | 0.2968 | 0.3542 | 0.3567 | 0.3062 |
| G | | | | | | | | 0.0379 | 0.0551 | 0.0596 | 0.0532 | 0.0397 |
| H | | | | | | | | 0.2829 | 0.1822 | 0.2636 | 0.2408 | 0.2017 |

**Cell Index at: 95:02:16**

| | 1 | 2 | 3 | 4 | 5 | 6 | 7 | 8 | 9 | 10 | 11 | 12 |
|---|---|---|---|---|---|---|---|---|---|---|---|---|
| A | | | | | | | | 0.0161 | 0.0039 | 0.0412 | 0.0078 | 0.0247 |
| B | | | | | | | | 0.0462 | 0.0316 | 0.0465 | 0.0757 | 0.0546 |
| C | | | | | | | | 0.2421 | 0.2421 | 0.3643 | 0.3839 | 0.3084 |
| D | | | | | | | | 0.2437 | 0.3146 | 0.3436 | 0.3333 | 0.3065 |
| E | | | | | | | | 0.0962 | 0.1028 | 0.1292 | 0.1236 | 0.0731 |
| F | | | | | | | | 0.3267 | 0.3002 | 0.3534 | 0.352 | 0.3023 |
| G | | | | | | | | 0.0338 | 0.053 | 0.0621 | 0.0588 | 0.0351 |
| H | | | | | | | | 0.2832 | 0.1787 | 0.2622 | 0.2418 | 0.2036 |

**Cell Index at: 95:17:17**

| | 1 | 2 | 3 | 4 | 5 | 6 | 7 | 8 | 9 | 10 | 11 | 12 |
|---|---|---|---|---|---|---|---|---|---|---|---|---|
| A | | | | | | | | 0.0185 | 0.0026 | 0.0425 | 0.006 | 0.0242 |
| B | | | | | | | | 0.0425 | 0.0298 | 0.0454 | 0.076 | 0.0551 |
| C | | | | | | | | 0.2461 | 0.2432 | 0.36 | 0.3934 | 0.3106 |
| D | | | | | | | | 0.2401 | 0.3099 | 0.3394 | 0.337 | 0.3082 |
| E | | | | | | | | 0.0959 | 0.0911 | 0.1271 | 0.1174 | 0.0781 |
| F | | | | | | | | 0.3333 | 0.2974 | 0.3565 | 0.3535 | 0.3041 |
| G | | | | | | | | 0.0304 | 0.0541 | 0.0603 | 0.0548 | 0.034 |
| H | | | | | | | | 0.2824 | 0.1833 | 0.2654 | 0.2388 | 0.2007 |

**Cell Index at: 95:32:18**

| | 1 | 2 | 3 | 4 | 5 | 6 | 7 | 8 | 9 | 10 | 11 | 12 |
|---|---|---|---|---|---|---|---|---|---|---|---|---|
| A | | | | | | | | 0.019 | 0.001 | 0.04 | 0.0072 | 0.0279 |
| B | | | | | | | | 0.0513 | 0.034 | 0.0492 | 0.0732 | 0.0525 |
| C | | | | | | | | 0.2471 | 0.2432 | 0.3636 | 0.3827 | 0.3088 |
| D | | | | | | | | 0.247 | 0.3077 | 0.3387 | 0.3406 | 0.3061 |
| E | | | | | | | | 0.106 | 0.0943 | 0.1309 | 0.118 | 0.0687 |
| F | | | | | | | | 0.3352 | 0.2959 | 0.3542 | 0.3534 | 0.3018 |
| G | | | | | | | | 0.0314 | 0.0549 | 0.0624 | 0.0628 | 0.037 |
| H | | | | | | | | 0.2875 | 0.1814 | 0.2687 | 0.2416 | 0.2032 |

**Cell Index at: 95:47:19**

| | 1 | 2 | 3 | 4 | 5 | 6 | 7 | 8 | 9 | 10 | 11 | 12 |
|---|---|---|---|---|---|---|---|---|---|---|---|---|
| A | | | | | | | | 0.0159 | 0.0002 | 0.0417 | 0.0089 | 0.0223 |
| B | | | | | | | | 0.0442 | 0.0342 | 0.0487 | 0.0747 | 0.0531 |
| C | | | | | | | | 0.2427 | 0.2493 | 0.368 | 0.3879 | 0.3103 |
| D | | | | | | | | 0.2477 | 0.3179 | 0.3465 | 0.3338 | 0.3079 |
| E | | | | | | | | 0.0941 | 0.0936 | 0.1372 | 0.1224 | 0.0691 |
| F | | | | | | | | 0.3388 | 0.2991 | 0.3492 | 0.3558 | 0.2953 |
| G | | | | | | | | 0.0323 | 0.0559 | 0.0573 | 0.0598 | 0.0378 |
| H | | | | | | | | 0.2845 | 0.1825 | 0.2663 | 0.2398 | 0.2033 |

**Cell Index at: 96:02:20**

| | 1 | 2 | 3 | 4 | 5 | 6 | 7 | 8 | 9 | 10 | 11 | 12 |
|---|---|---|---|---|---|---|---|---|---|---|---|---|
| A | | | | | | | | 0.0195 | 0.0008 | 0.041 | 0.0082 | 0.0244 |
| B | | | | | | | | 0.0456 | 0.0304 | 0.0455 | 0.073 | 0.0533 |
| C | | | | | | | | 0.2475 | 0.2423 | 0.368 | 0.3903 | 0.3124 |
| D | | | | | | | | 0.2479 | 0.3211 | 0.3482 | 0.3395 | 0.3086 |
| E | | | | | | | | 0.1039 | 0.0957 | 0.1283 | 0.1191 | 0.0687 |
| F | | | | | | | | 0.3329 | 0.3025 | 0.3583 | 0.3531 | 0.3019 |
| G | | | | | | | | 0.0378 | 0.0516 | 0.06 | 0.0547 | 0.0388 |
| H | | | | | | | | 0.2853 | 0.1834 | 0.2665 | 0.2383 | 0.2045 |

| | | | | | | | | | | | | | |
|---|---|---|---|---|---|---|---|---|---|---|---|---|---|
| A vs B | 0.013838 | B vs C | 0.000681 | C vs D | 0.978968 | D vs E | 3.2E-05 | E vs F | 6.92E-07 | F vs G | 3.26E-06 | G vs H | 0.000384 |
| A vs C | 0.000427 | B vs D | 2.63E-05 | C vs E | 0.001331 | D vs F | 0.460955 | E vs G | 0.003735 | F vs H | 0.004287 | | |
| A vs D | 1.5E-05 | B vs E | 0.003525 | C vs F | 0.610003 | D vs G | 6.04E-05 | E vs H | 0.000745 | | | | |
| A vs E | 0.000257 | B vs F | 4.9E-07 | C vs G | 0.000824 | D vs H | 0.017314 | | | | | | |
| A vs F | 2.54E-07 | B vs G | 0.908562 | C vs H | 0.070701 | | | | | | | | |
| A vs G | 0.008901 | B vs H | 0.000232 | | | | | | | | | | |
| A vs H | 0.000111 | | | | | | | | | | | | |

**Cell Index at: 96:17:21**

| | 1 | 2 | 3 | 4 | 5 | 6 | 7 | 8 | 9 | 10 | 11 | 12 |
|---|---|---|---|---|---|---|---|---|---|---|---|---|
| A | | | | | | | | 0.0181 | 0.0003 | 0.0414 | 0.0067 | 0.0223 |
| B | | | | | | | | 0.0442 | 0.0325 | 0.0447 | 0.0772 | 0.0525 |
| C | | | | | | | | 0.2536 | 0.2407 | 0.3676 | 0.3976 | 0.3152 |
| D | | | | | | | | 0.2458 | 0.3169 | 0.3404 | 0.3393 | 0.31 |
| E | | | | | | | | 0.1074 | 0.0974 | 0.1306 | 0.1217 | 0.0715 |
| F | | | | | | | | 0.338 | 0.2962 | 0.3572 | 0.3541 | 0.3134 |
| G | | | | | | | | 0.0348 | 0.0565 | 0.0609 | 0.0595 | 0.0377 |
| H | | | | | | | | 0.2863 | 0.1832 | 0.2688 | 0.2391 | 0.2041 |

| *Count* | *Sum* | *Average* | *Variance* |
|---|---|---|---|
| 5 | 0.0939 | 0.01878 | 0.00024 |
| 5 | 0.2478 | 0.04956 | 0.000241 |
| 5 | 1.5605 | 0.3121 | 0.004571 |
| 5 | 1.5653 | 0.31306 | 0.001566 |
| 5 | 0.5157 | 0.10314 | 0.000532 |
| 5 | 1.6487 | 0.32974 | 0.000722 |
| 5 | 0.2429 | 0.04858 | 9.72E-05 |
| 5 | 1.178 | 0.2356 | 0.001781 |

**Cell Index at: 96:32:21**

| | 1 | 2 | 3 | 4 | 5 | 6 | 7 | 8 | 9 | 10 | 11 | 12 |
|---|---|---|---|---|---|---|---|---|---|---|---|---|
| A | | | | | | | | 0.023 | 0.003 | 0.0383 | 0.0075 | 0.0238 |
| B | | | | | | | | 0.0387 | 0.0336 | 0.0506 | 0.0759 | 0.0498 |
| C | | | | | | | | 0.2511 | 0.2426 | 0.3691 | 0.39 | 0.3147 |
| D | | | | | | | | 0.2471 | 0.3226 | 0.3415 | 0.33 | 0.3079 |
| E | | | | | | | | 0.0981 | 0.0994 | 0.1269 | 0.1316 | 0.0681 |

| *SS* | *df* | *MS* | *F* | *P-value* | *F crit* |
|---|---|---|---|---|---|
| 0.633728 | 7 | 0.090533 | 74.2743 | 5.65E-18 | 2.312741 |
| 0.039005 | 32 | 0.001219 | | | |

| | | | | | | | | | | | | | | |
|---|---|---|---|---|---|---|---|---|---|---|---|---|---|---|
| F | | | | | | | | 0.3369 | 0.3047 | 0.3541 | 0.3638 | 0.3065 | | |
| G | | | | | | | | 0.0341 | 0.0564 | 0.0591 | 0.0604 | 0.0365 | 0.672733 | 39 |
| H | | | | | | | | 0.2822 | 0.1815 | 0.2691 | 0.2409 | 0.2042 | | |

**Cell Index at: 96:47:21**

| | 1 | 2 | 3 | 4 | 5 | 6 | 7 | 8 | 9 | 10 | 11 | 12 |
|---|---|---|---|---|---|---|---|---|---|---|---|---|
| A | | | | | | | | 0.023 | 0.0024 | 0.0405 | 0.0087 | 0.0247 |
| B | | | | | | | | 0.0457 | 0.0317 | 0.0497 | 0.0765 | 0.0549 |
| C | | | | | | | | 0.2518 | 0.2421 | 0.3715 | 0.3947 | 0.3126 |
| D | | | | | | | | 0.2417 | 0.3263 | 0.3413 | 0.3344 | 0.3105 |
| E | | | | | | | | 0.1067 | 0.0912 | 0.1297 | 0.131 | 0.0657 |
| F | | | | | | | | 0.342 | 0.3108 | 0.3509 | 0.3597 | 0.3062 |
| G | | | | | | | | 0.0361 | 0.0527 | 0.0655 | 0.0591 | 0.034 |
| H | | | | | | | | 0.2831 | 0.1828 | 0.2726 | 0.2433 | 0.2056 |

**Cell Index at: 97:02:21**

| | 1 | 2 | 3 | 4 | 5 | 6 | 7 | 8 | 9 | 10 | 11 | 12 |
|---|---|---|---|---|---|---|---|---|---|---|---|---|
| A | | | | | | | | 0.0243 | 0.0015 | 0.0408 | 0.0067 | 0.0252 |
| B | | | | | | | | 0.0483 | 0.0298 | 0.0558 | 0.0781 | 0.0497 |
| C | | | | | | | | 0.2538 | 0.2404 | 0.3712 | 0.3948 | 0.3137 |
| D | | | | | | | | 0.2435 | 0.3219 | 0.3499 | 0.3346 | 0.3032 |
| E | | | | | | | | 0.1063 | 0.0911 | 0.1364 | 0.1211 | 0.0708 |
| F | | | | | | | | 0.3439 | 0.3058 | 0.3541 | 0.3612 | 0.304 |
| G | | | | | | | | 0.0381 | 0.0517 | 0.0641 | 0.0574 | 0.0355 |
| H | | | | | | | | 0.2838 | 0.1864 | 0.272 | 0.2437 | 0.2064 |

**Cell Index at: 97:17:22**

| | 1 | 2 | 3 | 4 | 5 | 6 | 7 | 8 | 9 | 10 | 11 | 12 |
|---|---|---|---|---|---|---|---|---|---|---|---|---|
| A | | | | | | | | 0.0223 | -0.0006 | 0.0429 | 0.0087 | 0.0256 |
| B | | | | | | | | 0.0472 | 0.0351 | 0.0536 | 0.0768 | 0.0553 |
| C | | | | | | | | 0.2583 | 0.2436 | 0.3711 | 0.3977 | 0.3173 |
| D | | | | | | | | 0.2519 | 0.3191 | 0.3443 | 0.3359 | 0.3098 |
| E | | | | | | | | 0.1051 | 0.097 | 0.1401 | 0.1171 | 0.0706 |
| F | | | | | | | | 0.3429 | 0.3122 | 0.3479 | 0.3554 | 0.3076 |
| G | | | | | | | | 0.0401 | 0.0555 | 0.0608 | 0.0574 | 0.0364 |
| H | | | | | | | | 0.2871 | 0.1808 | 0.2728 | 0.2442 | 0.2066 |

**Cell Index at: 97:32:22**

| | 1 | 2 | 3 | 4 | 5 | 6 | 7 | 8 | 9 | 10 | 11 | 12 |
|---|---|---|---|---|---|---|---|---|---|---|---|---|
| A | | | | | | | | 0.0246 | 0.0021 | 0.0459 | 0.0079 | 0.0252 |
| B | | | | | | | | 0.045 | 0.0321 | 0.0553 | 0.0803 | 0.0528 |
| C | | | | | | | | 0.2538 | 0.2386 | 0.3679 | 0.3876 | 0.3065 |
| D | | | | | | | | 0.2531 | 0.3201 | 0.3479 | 0.3334 | 0.3063 |
| E | | | | | | | | 0.1005 | 0.0966 | 0.1292 | 0.1169 | 0.0714 |
| F | | | | | | | | 0.3441 | 0.3116 | 0.3558 | 0.3595 | 0.3119 |
| G | | | | | | | | 0.0402 | 0.0539 | 0.0618 | 0.0584 | 0.0368 |
| H | | | | | | | | 0.2873 | 0.1821 | 0.2709 | 0.2417 | 0.2035 |

**Cell Index at: 97:47:23**

| | 1 | 2 | 3 | 4 | 5 | 6 | 7 | 8 | 9 | 10 | 11 | 12 |
|---|---|---|---|---|---|---|---|---|---|---|---|---|
| A | | | | | | | | 0.0232 | 0.0003 | 0.0451 | 0.0094 | 0.0268 |
| B | | | | | | | | 0.0452 | 0.0308 | 0.0585 | 0.0754 | 0.0545 |
| C | | | | | | | | 0.2546 | 0.2402 | 0.3703 | 0.3893 | 0.3119 |
| D | | | | | | | | 0.25 | 0.3209 | 0.3481 | 0.3354 | 0.315 |
| E | | | | | | | | 0.1139 | 0.1007 | 0.1339 | 0.1204 | 0.0652 |
| F | | | | | | | | 0.344 | 0.3098 | 0.3556 | 0.3511 | 0.3067 |
| G | | | | | | | | 0.0373 | 0.0543 | 0.0658 | 0.0548 | 0.0388 |
| H | | | | | | | | 0.2847 | 0.1861 | 0.2732 | 0.2428 | 0.2104 |

**Cell Index at: 98:02:24**

| | 1 | 2 | 3 | 4 | 5 | 6 | 7 | 8 | 9 | 10 | 11 | 12 |
|---|---|---|---|---|---|---|---|---|---|---|---|---|
| A | | | | | | | | 0.0221 | 0.0004 | 0.047 | 0.0073 | 0.0247 |
| B | | | | | | | | 0.0451 | 0.0323 | 0.0598 | 0.0727 | 0.0546 |
| C | | | | | | | | 0.2556 | 0.246 | 0.3728 | 0.3905 | 0.3097 |
| D | | | | | | | | 0.257 | 0.319 | 0.3517 | 0.3338 | 0.3169 |
| E | | | | | | | | 0.1047 | 0.0995 | 0.1387 | 0.1206 | 0.0726 |
| F | | | | | | | | 0.349 | 0.3118 | 0.347 | 0.3572 | 0.3167 |
| G | | | | | | | | 0.0413 | 0.0542 | 0.0635 | 0.0614 | 0.041 |
| H | | | | | | | | 0.2834 | 0.185 | 0.2764 | 0.2427 | 0.209 |

**Cell Index at: 98:17:25**

| | 1 | 2 | 3 | 4 | 5 | 6 | 7 | 8 | 9 | 10 | 11 | 12 |
|---|---|---|---|---|---|---|---|---|---|---|---|---|
| A | | | | | | | | 0.0235 | 0.0004 | 0.0491 | 0.0081 | 0.0239 |
| B | | | | | | | | 0.0434 | 0.0328 | 0.0552 | 0.0761 | 0.0522 |
| C | | | | | | | | 0.2534 | 0.2463 | 0.3697 | 0.3928 | 0.3139 |
| D | | | | | | | | 0.2507 | 0.323 | 0.3443 | 0.329 | 0.3057 |
| E | | | | | | | | 0.0962 | 0.1019 | 0.1303 | 0.1143 | 0.0707 |
| F | | | | | | | | 0.3501 | 0.3111 | 0.3508 | 0.361 | 0.3116 |
| G | | | | | | | | 0.0447 | 0.0529 | 0.0644 | 0.0561 | 0.0374 |
| H | | | | | | | | 0.2858 | 0.1864 | 0.2745 | 0.2445 | 0.2103 |

**Cell Index at: 98:32:26**

| | 1 | 2 | 3 | 4 | 5 | 6 | 7 | 8 | 9 | 10 | 11 | 12 |
|---|---|---|---|---|---|---|---|---|---|---|---|---|
| A | | | | | | | | 0.0229 | -0.0008 | 0.0523 | 0.0085 | 0.0247 |
| B | | | | | | | | 0.0411 | 0.0317 | 0.0573 | 0.0737 | 0.054 |
| C | | | | | | | | 0.2577 | 0.2394 | 0.3725 | 0.3928 | 0.3161 |
| D | | | | | | | | 0.2527 | 0.3201 | 0.3412 | 0.3401 | 0.3092 |
| E | | | | | | | | 0.1147 | 0.0998 | 0.1325 | 0.1249 | 0.0716 |
| F | | | | | | | | 0.3456 | 0.3206 | 0.355 | 0.3585 | 0.3082 |
| G | | | | | | | | 0.0383 | 0.0524 | 0.0626 | 0.0584 | 0.0375 |
| H | | | | | | | | 0.2834 | 0.1873 | 0.2776 | 0.2465 | 0.2138 |

**Cell Index at: 98:47:26**

| | 1 | 2 | 3 | 4 | 5 | 6 | 7 | 8 | 9 | 10 | 11 | 12 |
|---|---|---|---|---|---|---|---|---|---|---|---|---|
| A | | | | | | | | 0.021 | -0.0004 | 0.0471 | 0.0079 | 0.0251 |
| B | | | | | | | | 0.0485 | 0.0287 | 0.0545 | 0.0724 | 0.0523 |
| C | | | | | | | | 0.2565 | 0.2449 | 0.3703 | 0.3945 | 0.3155 |
| D | | | | | | | | 0.252 | 0.3231 | 0.3472 | 0.3382 | 0.3083 |
| E | | | | | | | | 0.1025 | 0.0959 | 0.1375 | 0.1229 | 0.0712 |
| F | | | | | | | | 0.3528 | 0.3202 | 0.3617 | 0.3603 | 0.3035 |
| G | | | | | | | | 0.042 | 0.0525 | 0.0614 | 0.0592 | 0.037 |
| H | | | | | | | | 0.2848 | 0.1862 | 0.2755 | 0.2466 | 0.2116 |

**Cell Index at: 99:02:27**

| | 1 | 2 | 3 | 4 | 5 | 6 | 7 | 8 | 9 | 10 | 11 | 12 |
|---|---|---|---|---|---|---|---|---|---|---|---|---|
| A | | | | | | | | 0.0262 | 0.0034 | 0.0473 | 0.0118 | 0.0269 |
| B | | | | | | | | 0.0491 | 0.0293 | 0.058 | 0.0799 | 0.0544 |
| C | | | | | | | | 0.2595 | 0.2416 | 0.3685 | 0.3998 | 0.315 |
| D | | | | | | | | 0.2497 | 0.3265 | 0.3456 | 0.3369 | 0.312 |
| E | | | | | | | | 0.1123 | 0.1004 | 0.1423 | 0.1196 | 0.0729 |
| F | | | | | | | | 0.347 | 0.3173 | 0.3567 | 0.3629 | 0.3083 |
| G | | | | | | | | 0.0408 | 0.0546 | 0.0645 | 0.0586 | 0.0382 |
| H | | | | | | | | 0.2879 | 0.186 | 0.2747 | 0.2488 | 0.2157 |

**Cell Index at: 99:17:28**

| | 1 | 2 | 3 | 4 | 5 | 6 | 7 | 8 | 9 | 10 | 11 | 12 |
|---|---|---|---|---|---|---|---|---|---|---|---|---|
| A | | | | | | | | 0.0259 | 0.0035 | 0.0495 | 0.0115 | 0.0246 |
| B | | | | | | | | 0.0493 | 0.0323 | 0.0585 | 0.0771 | 0.0521 |
| C | | | | | | | | 0.2606 | 0.2451 | 0.3783 | 0.3961 | 0.3116 |
| D | | | | | | | | 0.2565 | 0.3215 | 0.3442 | 0.3451 | 0.3065 |
| E | | | | | | | | 0.1054 | 0.1067 | 0.1381 | 0.1192 | 0.0779 |
| F | | | | | | | | 0.3509 | 0.3122 | 0.3534 | 0.3674 | 0.3042 |
| G | | | | | | | | 0.0453 | 0.0575 | 0.0618 | 0.0579 | 0.0367 |
| H | | | | | | | | 0.2891 | 0.1843 | 0.2757 | 0.2486 | 0.2141 |

**Cell Index at: 99:32:28**

| | 1 | 2 | 3 | 4 | 5 | 6 | 7 | 8 | 9 | 10 | 11 | 12 |
|---|---|---|---|---|---|---|---|---|---|---|---|---|
| A | | | | | | | | 0.0241 | 0.0046 | 0.0453 | 0.0065 | 0.0249 |
| B | | | | | | | | 0.0495 | 0.0307 | 0.0545 | 0.0773 | 0.0543 |
| C | | | | | | | | 0.2612 | 0.2438 | 0.3821 | 0.3995 | 0.3169 |
| D | | | | | | | | 0.2549 | 0.32 | 0.3469 | 0.3353 | 0.3056 |

| | | | | | | | | | | | | |
|---|---|---|---|---|---|---|---|---|---|---|---|---|
| E | | | | | | | | 0.1052 | 0.0996 | 0.1391 | 0.117 | 0.0722 |
| F | | | | | | | | 0.3479 | 0.3174 | 0.3622 | 0.3655 | 0.3105 |
| G | | | | | | | | 0.0399 | 0.0573 | 0.0635 | 0.0566 | 0.0367 |
| H | | | | | | | | 0.2889 | 0.1853 | 0.2797 | 0.2501 | 0.2148 |

**Cell Index at: 99:47:29**

| | 1 | 2 | 3 | 4 | 5 | 6 | 7 | 8 | 9 | 10 | 11 | 12 |
|---|---|---|---|---|---|---|---|---|---|---|---|---|
| A | | | | | | | | 0.0257 | 0.0046 | 0.0492 | 0.0094 | 0.0274 |
| B | | | | | | | | 0.0484 | 0.0337 | 0.0586 | 0.0789 | 0.0481 |
| C | | | | | | | | 0.2571 | 0.2476 | 0.3758 | 0.3998 | 0.319 |
| D | | | | | | | | 0.2562 | 0.3241 | 0.347 | 0.3457 | 0.3073 |
| E | | | | | | | | 0.1021 | 0.0996 | 0.1379 | 0.1153 | 0.0725 |
| F | | | | | | | | 0.3547 | 0.3149 | 0.3655 | 0.3647 | 0.3096 |
| G | | | | | | | | 0.0444 | 0.0573 | 0.0703 | 0.056 | 0.0363 |
| H | | | | | | | | 0.284 | 0.1863 | 0.2793 | 0.2486 | 0.2169 |

**Cell Index at: 100:02:29**

| | 1 | 2 | 3 | 4 | 5 | 6 | 7 | 8 | 9 | 10 | 11 | 12 |
|---|---|---|---|---|---|---|---|---|---|---|---|---|
| A | | | | | | | | 0.0257 | 0.0074 | 0.047 | 0.0092 | 0.0247 |
| B | | | | | | | | 0.0512 | 0.0277 | 0.0593 | 0.0806 | 0.0528 |
| C | | | | | | | | 0.259 | 0.2507 | 0.3808 | 0.3982 | 0.317 |
| D | | | | | | | | 0.2621 | 0.3287 | 0.3522 | 0.3465 | 0.3099 |
| E | | | | | | | | 0.1099 | 0.1037 | 0.1367 | 0.1153 | 0.0709 |
| F | | | | | | | | 0.3453 | 0.3221 | 0.3645 | 0.3704 | 0.3071 |
| G | | | | | | | | 0.0431 | 0.0568 | 0.0685 | 0.0565 | 0.0363 |
| H | | | | | | | | 0.283 | 0.1851 | 0.2778 | 0.2481 | 0.218 |

**Cell Index at: 100:17:29**

| | 1 | 2 | 3 | 4 | 5 | 6 | 7 | 8 | 9 | 10 | 11 | 12 |
|---|---|---|---|---|---|---|---|---|---|---|---|---|
| A | | | | | | | | 0.0275 | 0.0052 | 0.0447 | 0.0101 | 0.0247 |
| B | | | | | | | | 0.0493 | 0.0303 | 0.0563 | 0.0813 | 0.0507 |
| C | | | | | | | | 0.2557 | 0.2445 | 0.3777 | 0.4021 | 0.3184 |
| D | | | | | | | | 0.2619 | 0.3273 | 0.3541 | 0.3483 | 0.3087 |
| E | | | | | | | | 0.1064 | 0.0984 | 0.1317 | 0.1211 | 0.068 |
| F | | | | | | | | 0.3512 | 0.3211 | 0.3646 | 0.3686 | 0.3128 |
| G | | | | | | | | 0.0464 | 0.0513 | 0.0636 | 0.0576 | 0.0343 |
| H | | | | | | | | 0.2875 | 0.1882 | 0.279 | 0.244 | 0.2161 |

**Cell Index at: 100:32:27**

| | 1 | 2 | 3 | 4 | 5 | 6 | 7 | 8 | 9 | 10 | 11 | 12 |
|---|---|---|---|---|---|---|---|---|---|---|---|---|
| A | | | | | | | | 0.0256 | 0.0092 | 0.048 | 0.0078 | 0.0255 |
| B | | | | | | | | 0.0501 | 0.031 | 0.0571 | 0.0831 | 0.0451 |
| C | | | | | | | | 0.2639 | 0.2471 | 0.3784 | 0.3975 | 0.3243 |
| D | | | | | | | | 0.2565 | 0.3238 | 0.3514 | 0.3464 | 0.3102 |
| E | | | | | | | | 0.1062 | 0.1013 | 0.133 | 0.1187 | 0.073 |
| F | | | | | | | | 0.3533 | 0.3214 | 0.3594 | 0.3717 | 0.3103 |
| G | | | | | | | | 0.0438 | 0.0508 | 0.0639 | 0.0487 | 0.0382 |
| H | | | | | | | | 0.2874 | 0.1886 | 0.2814 | 0.2457 | 0.2154 |

**Cell Index at: 100:47:28**

| | 1 | 2 | 3 | 4 | 5 | 6 | 7 | 8 | 9 | 10 | 11 | 12 |
|---|---|---|---|---|---|---|---|---|---|---|---|---|
| A | | | | | | | | 0.0255 | 0.0107 | 0.0466 | 0.0094 | 0.025 |
| B | | | | | | | | 0.0478 | 0.0357 | 0.0588 | 0.0793 | 0.0515 |
| C | | | | | | | | 0.2565 | 0.2587 | 0.3846 | 0.3993 | 0.3206 |
| D | | | | | | | | 0.2625 | 0.3275 | 0.3545 | 0.3515 | 0.3105 |
| E | | | | | | | | 0.1124 | 0.098 | 0.1394 | 0.1186 | 0.0718 |
| F | | | | | | | | 0.355 | 0.3263 | 0.3637 | 0.3657 | 0.3148 |
| G | | | | | | | | 0.0422 | 0.05 | 0.0617 | 0.0551 | 0.0391 |
| H | | | | | | | | 0.287 | 0.1908 | 0.2797 | 0.2493 | 0.2212 |

**Cell Index at: 101:02:28**

| | 1 | 2 | 3 | 4 | 5 | 6 | 7 | 8 | 9 | 10 | 11 | 12 |
|---|---|---|---|---|---|---|---|---|---|---|---|---|
| A | | | | | | | | 0.0271 | 0.013 | 0.0488 | 0.0099 | 0.0254 |
| B | | | | | | | | 0.0474 | 0.0348 | 0.0548 | 0.0802 | 0.0466 |
| C | | | | | | | | 0.2605 | 0.2536 | 0.3825 | 0.3961 | 0.3195 |
| D | | | | | | | | 0.2664 | 0.3393 | 0.3588 | 0.3552 | 0.3102 |
| E | | | | | | | | 0.1104 | 0.1012 | 0.1352 | 0.1197 | 0.0673 |
| F | | | | | | | | 0.3615 | 0.3182 | 0.3628 | 0.364 | 0.3187 |
| G | | | | | | | | 0.0451 | 0.0472 | 0.064 | 0.0505 | 0.0355 |
| H | | | | | | | | 0.2863 | 0.1906 | 0.2824 | 0.2519 | 0.2216 |

**Cell Index at: 101:17:29**

| | 1 | 2 | 3 | 4 | 5 | 6 | 7 | 8 | 9 | 10 | 11 | 12 |
|---|---|---|---|---|---|---|---|---|---|---|---|---|
| A | | | | | | | | 0.0294 | 0.0101 | 0.0494 | 0.0125 | 0.0258 |
| B | | | | | | | | 0.0497 | 0.0335 | 0.0554 | 0.0809 | 0.048 |
| C | | | | | | | | 0.2583 | 0.2535 | 0.3728 | 0.3954 | 0.3188 |
| D | | | | | | | | 0.2583 | 0.3242 | 0.3521 | 0.3504 | 0.3095 |
| E | | | | | | | | 0.1145 | 0.089 | 0.1349 | 0.117 | 0.0742 |
| F | | | | | | | | 0.3587 | 0.3256 | 0.3675 | 0.3722 | 0.3126 |
| G | | | | | | | | 0.0436 | 0.0478 | 0.062 | 0.0537 | 0.0357 |
| H | | | | | | | | 0.2858 | 0.193 | 0.28 | 0.2493 | 0.2203 |

**Cell Index at: 101:32:29**

| | 1 | 2 | 3 | 4 | 5 | 6 | 7 | 8 | 9 | 10 | 11 | 12 |
|---|---|---|---|---|---|---|---|---|---|---|---|---|
| A | | | | | | | | 0.0299 | 0.0108 | 0.0532 | 0.0123 | 0.0277 |
| B | | | | | | | | 0.049 | 0.0351 | 0.0587 | 0.0802 | 0.0481 |
| C | | | | | | | | 0.2573 | 0.2537 | 0.3795 | 0.3988 | 0.3187 |
| D | | | | | | | | 0.2635 | 0.329 | 0.35 | 0.3487 | 0.3113 |
| E | | | | | | | | 0.1184 | 0.0985 | 0.1351 | 0.1255 | 0.073 |
| F | | | | | | | | 0.3593 | 0.3181 | 0.3587 | 0.3729 | 0.3131 |
| G | | | | | | | | 0.0439 | 0.0488 | 0.0595 | 0.0486 | 0.0396 |
| H | | | | | | | | 0.2872 | 0.196 | 0.2831 | 0.2463 | 0.2244 |

**Cell Index at: 101:47:30**

| | 1 | 2 | 3 | 4 | 5 | 6 | 7 | 8 | 9 | 10 | 11 | 12 |
|---|---|---|---|---|---|---|---|---|---|---|---|---|
| A | | | | | | | | 0.0315 | 0.0094 | 0.0489 | 0.0115 | 0.0295 |
| B | | | | | | | | 0.0499 | 0.0341 | 0.0554 | 0.0772 | 0.0478 |
| C | | | | | | | | 0.2626 | 0.257 | 0.3786 | 0.4078 | 0.3203 |
| D | | | | | | | | 0.266 | 0.3273 | 0.3486 | 0.3535 | 0.3159 |
| E | | | | | | | | 0.1152 | 0.0976 | 0.1414 | 0.1239 | 0.0682 |
| F | | | | | | | | 0.3612 | 0.3178 | 0.3607 | 0.3693 | 0.3172 |
| G | | | | | | | | 0.0419 | 0.0521 | 0.0596 | 0.0461 | 0.0385 |
| H | | | | | | | | 0.2902 | 0.1925 | 0.2817 | 0.25 | 0.2236 |

**Cell Index at: 102:02:31**

| | 1 | 2 | 3 | 4 | 5 | 6 | 7 | 8 | 9 | 10 | 11 | 12 |
|---|---|---|---|---|---|---|---|---|---|---|---|---|
| A | | | | | | | | 0.0312 | 0.0088 | 0.0532 | 0.0124 | 0.0299 |
| B | | | | | | | | 0.0466 | 0.0406 | 0.057 | 0.0776 | 0.0505 |
| C | | | | | | | | 0.2635 | 0.2584 | 0.3821 | 0.4016 | 0.323 |
| D | | | | | | | | 0.2692 | 0.3312 | 0.3562 | 0.3602 | 0.3149 |
| E | | | | | | | | 0.1123 | 0.0958 | 0.1423 | 0.1191 | 0.076 |
| F | | | | | | | | 0.3613 | 0.3275 | 0.3639 | 0.3749 | 0.3171 |
| G | | | | | | | | 0.042 | 0.0504 | 0.0591 | 0.0531 | 0.0386 |
| H | | | | | | | | 0.2888 | 0.1949 | 0.2849 | 0.2499 | 0.2247 |

**Cell Index at: 102:17:31**

| | 1 | 2 | 3 | 4 | 5 | 6 | 7 | 8 | 9 | 10 | 11 | 12 |
|---|---|---|---|---|---|---|---|---|---|---|---|---|
| A | | | | | | | | 0.0318 | 0.0136 | 0.0539 | 0.0146 | 0.0297 |
| B | | | | | | | | 0.0438 | 0.0348 | 0.0579 | 0.071 | 0.048 |
| C | | | | | | | | 0.2626 | 0.2546 | 0.3745 | 0.4077 | 0.327 |
| D | | | | | | | | 0.2713 | 0.3266 | 0.3572 | 0.3533 | 0.3077 |
| E | | | | | | | | 0.113 | 0.1009 | 0.1427 | 0.1205 | 0.074 |
| F | | | | | | | | 0.3658 | 0.3228 | 0.3647 | 0.3741 | 0.3185 |
| G | | | | | | | | 0.0491 | 0.0526 | 0.0592 | 0.0499 | 0.0403 |
| H | | | | | | | | 0.2893 | 0.1892 | 0.2851 | 0.2515 | 0.2279 |

**Cell Index at: 102:32:31**

| | 1 | 2 | 3 | 4 | 5 | 6 | 7 | 8 | 9 | 10 | 11 | 12 |
|---|---|---|---|---|---|---|---|---|---|---|---|---|
| A | | | | | | | | 0.0349 | 0.0138 | 0.0536 | 0.0154 | 0.0285 |
| B | | | | | | | | 0.047 | 0.0388 | 0.057 | 0.0741 | 0.0492 |
| C | | | | | | | | 0.2585 | 0.2631 | 0.3779 | 0.4053 | 0.3182 |

| | | | | | | | | | | | | |
|---|---|---|---|---|---|---|---|---|---|---|---|---|
| D | | | | | | | | 0.2672 | 0.3275 | 0.3566 | 0.363 | 0.3085 |
| E | | | | | | | | 0.1073 | 0.0973 | 0.1346 | 0.1202 | 0.0791 |
| F | | | | | | | | 0.3699 | 0.3193 | 0.3613 | 0.3772 | 0.3141 |
| G | | | | | | | | 0.0435 | 0.0532 | 0.0609 | 0.0508 | 0.0394 |
| H | | | | | | | | 0.2888 | 0.1936 | 0.285 | 0.2523 | 0.2298 |

**Cell Index at: 102:47:31**

| | 1 | 2 | 3 | 4 | 5 | 6 | 7 | 8 | 9 | 10 | 11 | 12 |
|---|---|---|---|---|---|---|---|---|---|---|---|---|
| A | | | | | | | | 0.0333 | 0.0132 | 0.0558 | 0.0153 | 0.0278 |
| B | | | | | | | | 0.0501 | 0.0383 | 0.0542 | 0.0753 | 0.0476 |
| C | | | | | | | | 0.2634 | 0.2617 | 0.377 | 0.4058 | 0.3214 |
| D | | | | | | | | 0.2636 | 0.331 | 0.3522 | 0.3547 | 0.3144 |
| E | | | | | | | | 0.1137 | 0.1009 | 0.1372 | 0.1211 | 0.0729 |
| F | | | | | | | | 0.3674 | 0.3205 | 0.3635 | 0.3797 | 0.3093 |
| G | | | | | | | | 0.0412 | 0.0539 | 0.0621 | 0.0516 | 0.0398 |
| H | | | | | | | | 0.2877 | 0.1947 | 0.2843 | 0.2534 | 0.229 |

**Cell Index at: 103:02:31**

| | 1 | 2 | 3 | 4 | 5 | 6 | 7 | 8 | 9 | 10 | 11 | 12 |
|---|---|---|---|---|---|---|---|---|---|---|---|---|
| A | | | | | | | | 0.0369 | 0.0143 | 0.0559 | 0.0171 | 0.0275 |
| B | | | | | | | | 0.0461 | 0.0384 | 0.0568 | 0.0787 | 0.048 |
| C | | | | | | | | 0.2588 | 0.2546 | 0.3789 | 0.4179 | 0.3197 |
| D | | | | | | | | 0.2683 | 0.3328 | 0.3548 | 0.3552 | 0.3213 |
| E | | | | | | | | 0.1193 | 0.0983 | 0.1351 | 0.1194 | 0.0712 |
| F | | | | | | | | 0.367 | 0.3153 | 0.3635 | 0.3757 | 0.3187 |
| G | | | | | | | | 0.0437 | 0.0553 | 0.0638 | 0.0519 | 0.042 |
| H | | | | | | | | 0.2896 | 0.1953 | 0.2846 | 0.2537 | 0.2314 |

**Cell Index at: 103:17:31**

| | 1 | 2 | 3 | 4 | 5 | 6 | 7 | 8 | 9 | 10 | 11 | 12 |
|---|---|---|---|---|---|---|---|---|---|---|---|---|
| A | | | | | | | | 0.035 | 0.0159 | 0.059 | 0.0158 | 0.0257 |
| B | | | | | | | | 0.049 | 0.0382 | 0.0571 | 0.0745 | 0.0492 |
| C | | | | | | | | 0.2621 | 0.2587 | 0.3771 | 0.4127 | 0.3182 |
| D | | | | | | | | 0.2712 | 0.3297 | 0.3562 | 0.3653 | 0.3279 |
| E | | | | | | | | 0.1111 | 0.098 | 0.1354 | 0.115 | 0.0738 |
| F | | | | | | | | 0.3607 | 0.3239 | 0.3647 | 0.3764 | 0.3177 |
| G | | | | | | | | 0.044 | 0.0525 | 0.0644 | 0.0534 | 0.0416 |
| H | | | | | | | | 0.2904 | 0.1922 | 0.2822 | 0.2558 | 0.234 |

**Cell Index at: 103:32:32**

| | 1 | 2 | 3 | 4 | 5 | 6 | 7 | 8 | 9 | 10 | 11 | 12 |
|---|---|---|---|---|---|---|---|---|---|---|---|---|
| A | | | | | | | | 0.0374 | 0.0153 | 0.0572 | 0.0161 | 0.0268 |
| B | | | | | | | | 0.0472 | 0.0407 | 0.0512 | 0.0766 | 0.05 |
| C | | | | | | | | 0.2607 | 0.2615 | 0.3841 | 0.4078 | 0.3208 |
| D | | | | | | | | 0.2631 | 0.3339 | 0.3567 | 0.3533 | 0.3248 |
| E | | | | | | | | 0.1167 | 0.0944 | 0.1358 | 0.1236 | 0.0732 |
| F | | | | | | | | 0.3688 | 0.3282 | 0.3725 | 0.3776 | 0.3139 |
| G | | | | | | | | 0.0391 | 0.0579 | 0.0619 | 0.0536 | 0.0449 |
| H | | | | | | | | 0.2926 | 0.1941 | 0.2851 | 0.2539 | 0.232 |

**Cell Index at: 103:47:32**

| | 1 | 2 | 3 | 4 | 5 | 6 | 7 | 8 | 9 | 10 | 11 | 12 |
|---|---|---|---|---|---|---|---|---|---|---|---|---|
| A | | | | | | | | 0.0414 | 0.0189 | 0.0596 | 0.0148 | 0.0257 |
| B | | | | | | | | 0.0458 | 0.0413 | 0.0573 | 0.0787 | 0.0484 |
| C | | | | | | | | 0.2608 | 0.2631 | 0.3844 | 0.4103 | 0.3226 |
| D | | | | | | | | 0.2685 | 0.3347 | 0.3572 | 0.3538 | 0.3221 |
| E | | | | | | | | 0.1157 | 0.1011 | 0.1395 | 0.1195 | 0.0751 |
| F | | | | | | | | 0.3657 | 0.3209 | 0.3714 | 0.3828 | 0.3197 |
| G | | | | | | | | 0.0457 | 0.0595 | 0.0595 | 0.0513 | 0.0453 |
| H | | | | | | | | 0.2934 | 0.1954 | 0.2865 | 0.2561 | 0.2351 |

**Cell Index at: 104:02:33**

| | 1 | 2 | 3 | 4 | 5 | 6 | 7 | 8 | 9 | 10 | 11 | 12 |
|---|---|---|---|---|---|---|---|---|---|---|---|---|
| A | | | | | | | | 0.0417 | 0.0189 | 0.0561 | 0.013 | 0.0279 |
| B | | | | | | | | 0.0408 | 0.0423 | 0.0512 | 0.0763 | 0.0513 |
| C | | | | | | | | 0.2615 | 0.2621 | 0.3877 | 0.4031 | 0.3298 |
| D | | | | | | | | 0.2651 | 0.3361 | 0.3673 | 0.3629 | 0.319 |
| E | | | | | | | | 0.1121 | 0.0968 | 0.1345 | 0.1206 | 0.0719 |
| F | | | | | | | | 0.3672 | 0.3266 | 0.3753 | 0.376 | 0.3137 |
| G | | | | | | | | 0.0388 | 0.0588 | 0.0641 | 0.0551 | 0.0449 |
| H | | | | | | | | 0.2944 | 0.1977 | 0.2831 | 0.2588 | 0.239 |

**Cell Index at: 104:17:33**

| | 1 | 2 | 3 | 4 | 5 | 6 | 7 | 8 | 9 | 10 | 11 | 12 |
|---|---|---|---|---|---|---|---|---|---|---|---|---|
| A | | | | | | | | 0.0405 | 0.0192 | 0.054 | 0.0159 | 0.032 |
| B | | | | | | | | 0.0452 | 0.0424 | 0.0594 | 0.0784 | 0.0493 |
| C | | | | | | | | 0.2619 | 0.2614 | 0.3827 | 0.4068 | 0.3258 |
| D | | | | | | | | 0.2667 | 0.3369 | 0.3642 | 0.3588 | 0.3243 |
| E | | | | | | | | 0.112 | 0.095 | 0.1446 | 0.1223 | 0.0715 |
| F | | | | | | | | 0.3665 | 0.3297 | 0.3778 | 0.3809 | 0.3157 |
| G | | | | | | | | 0.0391 | 0.0561 | 0.0615 | 0.0495 | 0.0464 |
| H | | | | | | | | 0.2963 | 0.1984 | 0.2849 | 0.2559 | 0.2402 |

**Cell Index at: 104:32:33**

| | 1 | 2 | 3 | 4 | 5 | 6 | 7 | 8 | 9 | 10 | 11 | 12 |
|---|---|---|---|---|---|---|---|---|---|---|---|---|
| A | | | | | | | | 0.0405 | 0.0203 | 0.0565 | 0.016 | 0.0292 |
| B | | | | | | | | 0.0418 | 0.0429 | 0.057 | 0.0785 | 0.0495 |
| C | | | | | | | | 0.2619 | 0.2611 | 0.3851 | 0.4073 | 0.3223 |
| D | | | | | | | | 0.2626 | 0.3299 | 0.3533 | 0.3547 | 0.3179 |
| E | | | | | | | | 0.1123 | 0.0943 | 0.1359 | 0.1272 | 0.0733 |
| F | | | | | | | | 0.3696 | 0.3311 | 0.3724 | 0.3822 | 0.3143 |
| G | | | | | | | | 0.0429 | 0.0558 | 0.0613 | 0.0517 | 0.0497 |
| H | | | | | | | | 0.3005 | 0.198 | 0.2826 | 0.2587 | 0.2387 |

**Cell Index at: 104:47:33**

| | 1 | 2 | 3 | 4 | 5 | 6 | 7 | 8 | 9 | 10 | 11 | 12 |
|---|---|---|---|---|---|---|---|---|---|---|---|---|
| A | | | | | | | | 0.0395 | 0.02 | 0.0576 | 0.0171 | 0.0302 |
| B | | | | | | | | 0.0396 | 0.0433 | 0.0593 | 0.0795 | 0.0487 |
| C | | | | | | | | 0.261 | 0.2671 | 0.3898 | 0.4046 | 0.321 |
| D | | | | | | | | 0.2555 | 0.3286 | 0.357 | 0.3665 | 0.3115 |
| E | | | | | | | | 0.1203 | 0.0977 | 0.1445 | 0.1295 | 0.0701 |
| F | | | | | | | | 0.3763 | 0.3346 | 0.3714 | 0.3916 | 0.3192 |
| G | | | | | | | | 0.0457 | 0.0571 | 0.0605 | 0.0493 | 0.0482 |
| H | | | | | | | | 0.3003 | 0.1968 | 0.2811 | 0.2624 | 0.2404 |

**Cell Index at: 105:02:33**

| | 1 | 2 | 3 | 4 | 5 | 6 | 7 | 8 | 9 | 10 | 11 | 12 |
|---|---|---|---|---|---|---|---|---|---|---|---|---|
| A | | | | | | | | 0.0397 | 0.0206 | 0.0547 | 0.0165 | 0.0324 |
| B | | | | | | | | 0.0425 | 0.0414 | 0.0619 | 0.0775 | 0.048 |
| C | | | | | | | | 0.2604 | 0.2575 | 0.3914 | 0.4074 | 0.3269 |
| D | | | | | | | | 0.2642 | 0.3267 | 0.3612 | 0.3637 | 0.3151 |
| E | | | | | | | | 0.1216 | 0.1055 | 0.1363 | 0.1223 | 0.0738 |
| F | | | | | | | | 0.367 | 0.3299 | 0.3792 | 0.3882 | 0.3181 |
| G | | | | | | | | 0.0401 | 0.0589 | 0.0586 | 0.0492 | 0.0473 |
| H | | | | | | | | 0.3016 | 0.1984 | 0.2839 | 0.2621 | 0.2387 |

**Cell Index at: 105:17:34**

| | 1 | 2 | 3 | 4 | 5 | 6 | 7 | 8 | 9 | 10 | 11 | 12 |
|---|---|---|---|---|---|---|---|---|---|---|---|---|
| A | | | | | | | | 0.0396 | 0.0222 | 0.0524 | 0.018 | 0.0322 |
| B | | | | | | | | 0.0427 | 0.0403 | 0.0609 | 0.0743 | 0.0456 |
| C | | | | | | | | 0.2611 | 0.2657 | 0.3868 | 0.4085 | 0.3218 |
| D | | | | | | | | 0.2704 | 0.3343 | 0.3624 | 0.3664 | 0.3219 |
| E | | | | | | | | 0.1101 | 0.0994 | 0.1421 | 0.1191 | 0.0681 |
| F | | | | | | | | 0.3644 | 0.3271 | 0.3755 | 0.3892 | 0.3185 |
| G | | | | | | | | 0.0427 | 0.0576 | 0.0644 | 0.0494 | 0.0469 |
| H | | | | | | | | 0.2973 | 0.1998 | 0.2854 | 0.2653 | 0.2345 |

**Cell Index at: 105:32:34**

| | 1 | 2 | 3 | 4 | 5 | 6 | 7 | 8 | 9 | 10 | 11 | 12 |
|---|---|---|---|---|---|---|---|---|---|---|---|---|
| A | | | | | | | | 0.0395 | 0.026 | 0.0497 | 0.0176 | 0.031 |
| B | | | | | | | | 0.0407 | 0.0458 | 0.0601 | 0.0745 | 0.0489 |

| | | | | | | | | | | | | |
|---|---|---|---|---|---|---|---|---|---|---|---|---|
| C | | | | | | | | 0.2593 | 0.2615 | 0.3881 | 0.404 | 0.3248 |
| D | | | | | | | | 0.2602 | 0.3234 | 0.3543 | 0.3621 | 0.3176 |
| E | | | | | | | | 0.1086 | 0.106 | 0.1448 | 0.1218 | 0.0756 |
| F | | | | | | | | 0.3719 | 0.3325 | 0.3794 | 0.3886 | 0.3236 |
| G | | | | | | | | 0.0414 | 0.0562 | 0.0656 | 0.0513 | 0.0449 |
| H | | | | | | | | 0.2997 | 0.1969 | 0.2853 | 0.2646 | 0.2383 |

**Cell Index at: 105:47:35**

| | 1 | 2 | 3 | 4 | 5 | 6 | 7 | 8 | 9 | 10 | 11 | 12 |
|---|---|---|---|---|---|---|---|---|---|---|---|---|
| A | | | | | | | | 0.0359 | 0.0251 | 0.0504 | 0.0174 | 0.0329 |
| B | | | | | | | | 0.0401 | 0.044 | 0.0618 | 0.0738 | 0.0495 |
| C | | | | | | | | 0.2591 | 0.2648 | 0.3917 | 0.4096 | 0.3286 |
| D | | | | | | | | 0.2588 | 0.321 | 0.3588 | 0.358 | 0.3229 |
| E | | | | | | | | 0.1176 | 0.1079 | 0.1451 | 0.1221 | 0.0651 |
| F | | | | | | | | 0.3763 | 0.3323 | 0.3785 | 0.3894 | 0.3213 |
| G | | | | | | | | 0.0412 | 0.058 | 0.0632 | 0.0525 | 0.0441 |
| H | | | | | | | | 0.2985 | 0.1978 | 0.2854 | 0.2663 | 0.2385 |

**Cell Index at: 106:02:35**

| | 1 | 2 | 3 | 4 | 5 | 6 | 7 | 8 | 9 | 10 | 11 | 12 |
|---|---|---|---|---|---|---|---|---|---|---|---|---|
| A | | | | | | | | 0.0368 | 0.0249 | 0.0516 | 0.0219 | 0.0306 |
| B | | | | | | | | 0.0448 | 0.0433 | 0.067 | 0.0744 | 0.0501 |
| C | | | | | | | | 0.2648 | 0.2704 | 0.4071 | 0.4117 | 0.3239 |
| D | | | | | | | | 0.2627 | 0.3299 | 0.3641 | 0.3588 | 0.3231 |
| E | | | | | | | | 0.1119 | 0.1037 | 0.1443 | 0.1249 | 0.0697 |
| F | | | | | | | | 0.3728 | 0.334 | 0.3823 | 0.3913 | 0.3189 |
| G | | | | | | | | 0.0435 | 0.0636 | 0.0611 | 0.0523 | 0.0494 |
| H | | | | | | | | 0.3008 | 0.2003 | 0.284 | 0.2645 | 0.2398 |

**Cell Index at: 106:17:35**

| | 1 | 2 | 3 | 4 | 5 | 6 | 7 | 8 | 9 | 10 | 11 | 12 |
|---|---|---|---|---|---|---|---|---|---|---|---|---|
| A | | | | | | | | 0.0372 | 0.0255 | 0.0519 | 0.0229 | 0.0294 |
| B | | | | | | | | 0.0417 | 0.0441 | 0.0643 | 0.0764 | 0.0502 |
| C | | | | | | | | 0.2602 | 0.2702 | 0.4002 | 0.4144 | 0.323 |
| D | | | | | | | | 0.2634 | 0.325 | 0.3636 | 0.3672 | 0.3214 |
| E | | | | | | | | 0.1184 | 0.1064 | 0.1417 | 0.1228 | 0.0747 |
| F | | | | | | | | 0.3718 | 0.3348 | 0.3798 | 0.3949 | 0.3263 |
| G | | | | | | | | 0.0435 | 0.0519 | 0.0638 | 0.0524 | 0.0482 |
| H | | | | | | | | 0.3078 | 0.2005 | 0.2834 | 0.2642 | 0.2379 |

**Cell Index at: 106:32:36**

| | 1 | 2 | 3 | 4 | 5 | 6 | 7 | 8 | 9 | 10 | 11 | 12 |
|---|---|---|---|---|---|---|---|---|---|---|---|---|
| A | | | | | | | | 0.0326 | 0.0251 | 0.0487 | 0.02 | 0.0325 |
| B | | | | | | | | 0.0389 | 0.0463 | 0.0646 | 0.0774 | 0.0473 |
| C | | | | | | | | 0.2602 | 0.2691 | 0.4031 | 0.4131 | 0.3268 |
| D | | | | | | | | 0.265 | 0.3174 | 0.3615 | 0.3705 | 0.3264 |
| E | | | | | | | | 0.1185 | 0.1076 | 0.1438 | 0.1252 | 0.076 |
| F | | | | | | | | 0.375 | 0.3364 | 0.3759 | 0.3936 | 0.3185 |
| G | | | | | | | | 0.0476 | 0.0594 | 0.0646 | 0.05 | 0.0492 |
| H | | | | | | | | 0.3068 | 0.2002 | 0.2832 | 0.2673 | 0.2425 |

**Cell Index at: 106:47:37**

| | 1 | 2 | 3 | 4 | 5 | 6 | 7 | 8 | 9 | 10 | 11 | 12 |
|---|---|---|---|---|---|---|---|---|---|---|---|---|
| A | | | | | | | | 0.0334 | 0.0287 | 0.0501 | 0.0185 | 0.0316 |
| B | | | | | | | | 0.0368 | 0.0508 | 0.0654 | 0.0823 | 0.0506 |
| C | | | | | | | | 0.2621 | 0.2631 | 0.3999 | 0.4167 | 0.3225 |
| D | | | | | | | | 0.2635 | 0.3173 | 0.3725 | 0.3734 | 0.3237 |
| E | | | | | | | | 0.1146 | 0.1057 | 0.1376 | 0.1264 | 0.0746 |
| F | | | | | | | | 0.3736 | 0.3385 | 0.3794 | 0.398 | 0.3241 |
| G | | | | | | | | 0.0435 | 0.0533 | 0.0609 | 0.0554 | 0.0504 |
| H | | | | | | | | 0.3071 | 0.2024 | 0.2842 | 0.2674 | 0.2418 |

**Cell Index at: 107:02:37**

| | 1 | 2 | 3 | 4 | 5 | 6 | 7 | 8 | 9 | 10 | 11 | 12 |
|---|---|---|---|---|---|---|---|---|---|---|---|---|
| A | | | | | | | | 0.033 | 0.0283 | 0.0482 | 0.0179 | 0.0333 |
| B | | | | | | | | 0.0424 | 0.0466 | 0.0645 | 0.0731 | 0.0519 |
| C | | | | | | | | 0.2668 | 0.2664 | 0.4026 | 0.4195 | 0.3285 |
| D | | | | | | | | 0.2726 | 0.3201 | 0.3635 | 0.3683 | 0.3246 |
| E | | | | | | | | 0.127 | 0.104 | 0.141 | 0.1269 | 0.073 |
| F | | | | | | | | 0.383 | 0.3376 | 0.3888 | 0.395 | 0.3262 |
| G | | | | | | | | 0.0465 | 0.0582 | 0.0582 | 0.0559 | 0.0488 |
| H | | | | | | | | 0.305 | 0.2058 | 0.2865 | 0.2699 | 0.2417 |

**Cell Index at: 107:17:37**

| | 1 | 2 | 3 | 4 | 5 | 6 | 7 | 8 | 9 | 10 | 11 | 12 |
|---|---|---|---|---|---|---|---|---|---|---|---|---|
| A | | | | | | | | 0.0335 | 0.0279 | 0.0505 | 0.0198 | 0.0329 |
| B | | | | | | | | 0.0362 | 0.0453 | 0.0637 | 0.0775 | 0.0501 |
| C | | | | | | | | 0.2627 | 0.2708 | 0.3981 | 0.4246 | 0.3309 |
| D | | | | | | | | 0.2699 | 0.3302 | 0.3658 | 0.3695 | 0.3278 |
| E | | | | | | | | 0.127 | 0.1068 | 0.1439 | 0.1238 | 0.073 |
| F | | | | | | | | 0.3864 | 0.3328 | 0.3818 | 0.396 | 0.3229 |
| G | | | | | | | | 0.0466 | 0.0573 | 0.0618 | 0.0536 | 0.0486 |
| H | | | | | | | | 0.3046 | 0.2064 | 0.2878 | 0.2721 | 0.2437 |

**Cell Index at: 107:32:37**

| | 1 | 2 | 3 | 4 | 5 | 6 | 7 | 8 | 9 | 10 | 11 | 12 |
|---|---|---|---|---|---|---|---|---|---|---|---|---|
| A | | | | | | | | 0.032 | 0.0242 | 0.0505 | 0.0181 | 0.0318 |
| B | | | | | | | | 0.0386 | 0.0474 | 0.0604 | 0.0751 | 0.052 |
| C | | | | | | | | 0.2687 | 0.2683 | 0.4001 | 0.4211 | 0.3278 |
| D | | | | | | | | 0.2688 | 0.3273 | 0.363 | 0.3747 | 0.3291 |
| E | | | | | | | | 0.125 | 0.0957 | 0.1421 | 0.1256 | 0.0713 |
| F | | | | | | | | 0.3866 | 0.3423 | 0.3841 | 0.4011 | 0.331 |
| G | | | | | | | | 0.0505 | 0.0586 | 0.0682 | 0.0524 | 0.052 |
| H | | | | | | | | 0.3043 | 0.2061 | 0.2907 | 0.2675 | 0.2412 |

**Cell Index at: 107:47:37**

| | 1 | 2 | 3 | 4 | 5 | 6 | 7 | 8 | 9 | 10 | 11 | 12 |
|---|---|---|---|---|---|---|---|---|---|---|---|---|
| A | | | | | | | | 0.0287 | 0.0233 | 0.0529 | 0.0215 | 0.035 |
| B | | | | | | | | 0.04 | 0.046 | 0.0644 | 0.0795 | 0.0524 |
| C | | | | | | | | 0.2707 | 0.2738 | 0.4008 | 0.4251 | 0.3263 |
| D | | | | | | | | 0.2787 | 0.3291 | 0.3701 | 0.3749 | 0.3287 |
| E | | | | | | | | 0.1328 | 0.102 | 0.145 | 0.1243 | 0.0762 |
| F | | | | | | | | 0.3847 | 0.3492 | 0.3849 | 0.4027 | 0.3277 |
| G | | | | | | | | 0.0492 | 0.0556 | 0.0617 | 0.0575 | 0.0515 |
| H | | | | | | | | 0.3061 | 0.209 | 0.2943 | 0.2715 | 0.2441 |

**Cell Index at: 108:02:38**

| | 1 | 2 | 3 | 4 | 5 | 6 | 7 | 8 | 9 | 10 | 11 | 12 |
|---|---|---|---|---|---|---|---|---|---|---|---|---|
| A | | | | | | | | 0.0334 | 0.0225 | 0.0488 | 0.0223 | 0.036 |
| B | | | | | | | | 0.0411 | 0.0471 | 0.0658 | 0.0797 | 0.05 |
| C | | | | | | | | 0.2721 | 0.2672 | 0.4024 | 0.4293 | 0.3304 |
| D | | | | | | | | 0.2774 | 0.3251 | 0.3685 | 0.3772 | 0.3251 |
| E | | | | | | | | 0.124 | 0.1008 | 0.1378 | 0.1318 | 0.0771 |
| F | | | | | | | | 0.3892 | 0.3455 | 0.3827 | 0.4001 | 0.3323 |
| G | | | | | | | | 0.0454 | 0.0633 | 0.0664 | 0.0544 | 0.0533 |
| H | | | | | | | | 0.307 | 0.2084 | 0.2935 | 0.2683 | 0.2488 |

**Cell Index at: 108:17:39**

| | 1 | 2 | 3 | 4 | 5 | 6 | 7 | 8 | 9 | 10 | 11 | 12 |
|---|---|---|---|---|---|---|---|---|---|---|---|---|
| A | | | | | | | | 0.0309 | 0.0196 | 0.0519 | 0.0227 | 0.0342 |
| B | | | | | | | | 0.0405 | 0.0418 | 0.0609 | 0.0803 | 0.0523 |
| C | | | | | | | | 0.2706 | 0.2744 | 0.4052 | 0.424 | 0.3246 |
| D | | | | | | | | 0.2738 | 0.3314 | 0.3641 | 0.3835 | 0.3233 |
| E | | | | | | | | 0.126 | 0.101 | 0.1414 | 0.1249 | 0.0702 |
| F | | | | | | | | 0.3899 | 0.3475 | 0.3829 | 0.4022 | 0.3317 |
| G | | | | | | | | 0.0443 | 0.0566 | 0.0636 | 0.0562 | 0.0513 |
| H | | | | | | | | 0.3083 | 0.2128 | 0.2947 | 0.2706 | 0.2474 |

**Cell Index at: 108:32:39**

| | 1 | 2 | 3 | 4 | 5 | 6 | 7 | 8 | 9 | 10 | 11 | 12 |
|---|---|---|---|---|---|---|---|---|---|---|---|---|
| A | | | | | | | | 0.0294 | 0.0253 | 0.0503 | 0.0207 | 0.0391 |

| | | | | | | | | | | | | |
|---|---|---|---|---|---|---|---|---|---|---|---|---|
| B | | | | | | | | 0.0347 | 0.0456 | 0.0554 | 0.0799 | 0.0565 |
| C | | | | | | | | 0.2765 | 0.2755 | 0.4082 | 0.4229 | 0.3261 |
| D | | | | | | | | 0.2811 | 0.3367 | 0.3701 | 0.3791 | 0.3257 |
| E | | | | | | | | 0.1234 | 0.1058 | 0.1417 | 0.1333 | 0.0738 |
| F | | | | | | | | 0.3876 | 0.3495 | 0.3886 | 0.4059 | 0.3321 |
| G | | | | | | | | 0.0452 | 0.0588 | 0.0617 | 0.0553 | 0.05 |
| H | | | | | | | | 0.3102 | 0.2112 | 0.2971 | 0.2726 | 0.2485 |

**Cell Index at: 108:47:40**

| | 1 | 2 | 3 | 4 | 5 | 6 | 7 | 8 | 9 | 10 | 11 | 12 |
|---|---|---|---|---|---|---|---|---|---|---|---|---|
| A | | | | | | | | 0.0303 | 0.0225 | 0.0515 | 0.0183 | 0.0373 |
| B | | | | | | | | 0.035 | 0.0438 | 0.0557 | 0.08 | 0.0546 |
| C | | | | | | | | 0.2684 | 0.2759 | 0.4063 | 0.4284 | 0.3262 |
| D | | | | | | | | 0.2806 | 0.3352 | 0.3699 | 0.379 | 0.3257 |
| E | | | | | | | | 0.1201 | 0.0985 | 0.1384 | 0.1366 | 0.0742 |
| F | | | | | | | | 0.3909 | 0.3528 | 0.3876 | 0.4045 | 0.3278 |
| G | | | | | | | | 0.0452 | 0.0564 | 0.0648 | 0.0577 | 0.0483 |
| H | | | | | | | | 0.3115 | 0.213 | 0.2937 | 0.2763 | 0.2489 |

**Cell Index at: 109:02:40**

| | 1 | 2 | 3 | 4 | 5 | 6 | 7 | 8 | 9 | 10 | 11 | 12 |
|---|---|---|---|---|---|---|---|---|---|---|---|---|
| A | | | | | | | | 0.0314 | 0.0222 | 0.0556 | 0.0206 | 0.0391 |
| B | | | | | | | | 0.0372 | 0.0469 | 0.0536 | 0.0782 | 0.0572 |
| C | | | | | | | | 0.2778 | 0.2705 | 0.4105 | 0.4276 | 0.3234 |
| D | | | | | | | | 0.2829 | 0.3398 | 0.3801 | 0.3787 | 0.3287 |
| E | | | | | | | | 0.132 | 0.1009 | 0.1424 | 0.1332 | 0.081 |
| F | | | | | | | | 0.3878 | 0.3603 | 0.3944 | 0.402 | 0.3266 |
| G | | | | | | | | 0.0469 | 0.0602 | 0.0662 | 0.053 | 0.0431 |
| H | | | | | | | | 0.3104 | 0.2092 | 0.293 | 0.2721 | 0.2512 |

**Cell Index at: 109:17:41**

| | 1 | 2 | 3 | 4 | 5 | 6 | 7 | 8 | 9 | 10 | 11 | 12 |
|---|---|---|---|---|---|---|---|---|---|---|---|---|
| A | | | | | | | | 0.0283 | 0.0266 | 0.0569 | 0.0231 | 0.0368 |
| B | | | | | | | | 0.0379 | 0.0444 | 0.0565 | 0.0819 | 0.0551 |
| C | | | | | | | | 0.2709 | 0.2706 | 0.4093 | 0.4268 | 0.3286 |
| D | | | | | | | | 0.2885 | 0.341 | 0.3776 | 0.379 | 0.3334 |
| E | | | | | | | | 0.1229 | 0.0971 | 0.1371 | 0.1325 | 0.0754 |
| F | | | | | | | | 0.3845 | 0.3545 | 0.3894 | 0.4036 | 0.3261 |
| G | | | | | | | | 0.0417 | 0.0579 | 0.067 | 0.0586 | 0.0457 |
| H | | | | | | | | 0.3096 | 0.2101 | 0.2937 | 0.2761 | 0.2488 |

**Cell Index at: 109:32:41**

| | 1 | 2 | 3 | 4 | 5 | 6 | 7 | 8 | 9 | 10 | 11 | 12 |
|---|---|---|---|---|---|---|---|---|---|---|---|---|
| A | | | | | | | | 0.03 | 0.0295 | 0.0518 | 0.0234 | 0.0389 |
| B | | | | | | | | 0.034 | 0.0482 | 0.0551 | 0.0807 | 0.0592 |
| C | | | | | | | | 0.2779 | 0.2697 | 0.4132 | 0.427 | 0.3298 |
| D | | | | | | | | 0.2852 | 0.3349 | 0.3761 | 0.3773 | 0.333 |
| E | | | | | | | | 0.1253 | 0.1008 | 0.1364 | 0.1305 | 0.0748 |
| F | | | | | | | | 0.3894 | 0.3577 | 0.3917 | 0.4063 | 0.3268 |
| G | | | | | | | | 0.0486 | 0.0559 | 0.0671 | 0.0593 | 0.0491 |
| H | | | | | | | | 0.312 | 0.2118 | 0.2978 | 0.2763 | 0.2505 |

**Cell Index at: 109:47:41**

| | 1 | 2 | 3 | 4 | 5 | 6 | 7 | 8 | 9 | 10 | 11 | 12 |
|---|---|---|---|---|---|---|---|---|---|---|---|---|
| A | | | | | | | | 0.0326 | 0.0279 | 0.0528 | 0.0175 | 0.0405 |
| B | | | | | | | | 0.0333 | 0.0493 | 0.0534 | 0.0817 | 0.0536 |
| C | | | | | | | | 0.2805 | 0.2708 | 0.4053 | 0.4318 | 0.3302 |
| D | | | | | | | | 0.2827 | 0.3283 | 0.3786 | 0.3891 | 0.3273 |
| E | | | | | | | | 0.123 | 0.1034 | 0.1373 | 0.1324 | 0.0778 |
| F | | | | | | | | 0.384 | 0.3508 | 0.3902 | 0.401 | 0.3257 |
| G | | | | | | | | 0.0465 | 0.0621 | 0.0706 | 0.056 | 0.046 |
| H | | | | | | | | 0.3135 | 0.2091 | 0.2962 | 0.2777 | 0.2514 |

**Cell Index at: 110:02:41**

| | 1 | 2 | 3 | 4 | 5 | 6 | 7 | 8 | 9 | 10 | 11 | 12 |
|---|---|---|---|---|---|---|---|---|---|---|---|---|
| A | | | | | | | | 0.0305 | 0.0257 | 0.0544 | 0.017 | 0.0381 |
| B | | | | | | | | 0.0339 | 0.0453 | 0.0549 | 0.0819 | 0.0561 |
| C | | | | | | | | 0.2847 | 0.2655 | 0.4077 | 0.4343 | 0.3327 |
| D | | | | | | | | 0.282 | 0.332 | 0.3828 | 0.3793 | 0.3294 |
| E | | | | | | | | 0.129 | 0.1091 | 0.1339 | 0.1354 | 0.0764 |
| F | | | | | | | | 0.3897 | 0.3572 | 0.3945 | 0.4014 | 0.3278 |
| G | | | | | | | | 0.0447 | 0.0651 | 0.0681 | 0.053 | 0.0522 |
| H | | | | | | | | 0.3108 | 0.2099 | 0.2963 | 0.2747 | 0.2531 |

**Cell Index at: 110:17:40**

| | 1 | 2 | 3 | 4 | 5 | 6 | 7 | 8 | 9 | 10 | 11 | 12 |
|---|---|---|---|---|---|---|---|---|---|---|---|---|
| A | | | | | | | | 0.0315 | 0.0276 | 0.0561 | 0.0195 | 0.0393 |
| B | | | | | | | | 0.0373 | 0.0485 | 0.0559 | 0.083 | 0.0592 |
| C | | | | | | | | 0.2791 | 0.2715 | 0.4077 | 0.43 | 0.3312 |
| D | | | | | | | | 0.2907 | 0.3277 | 0.3824 | 0.3843 | 0.3296 |
| E | | | | | | | | 0.1296 | 0.1053 | 0.1398 | 0.129 | 0.0728 |
| F | | | | | | | | 0.3965 | 0.3512 | 0.3876 | 0.4094 | 0.328 |
| G | | | | | | | | 0.0475 | 0.0636 | 0.0698 | 0.0548 | 0.0488 |
| H | | | | | | | | 0.3049 | 0.2144 | 0.2972 | 0.2715 | 0.2536 |

**Cell Index at: 110:32:40**

| | 1 | 2 | 3 | 4 | 5 | 6 | 7 | 8 | 9 | 10 | 11 | 12 |
|---|---|---|---|---|---|---|---|---|---|---|---|---|
| A | | | | | | | | 0.028 | 0.0291 | 0.0566 | 0.0187 | 0.0402 |
| B | | | | | | | | 0.0354 | 0.0448 | 0.0536 | 0.0805 | 0.0569 |
| C | | | | | | | | 0.2774 | 0.2711 | 0.4163 | 0.4271 | 0.3355 |
| D | | | | | | | | 0.2925 | 0.3377 | 0.3815 | 0.3916 | 0.3365 |
| E | | | | | | | | 0.1199 | 0.1036 | 0.1302 | 0.134 | 0.0789 |
| F | | | | | | | | 0.3909 | 0.3607 | 0.3919 | 0.4117 | 0.3274 |
| G | | | | | | | | 0.0401 | 0.0636 | 0.067 | 0.0583 | 0.0482 |
| H | | | | | | | | 0.3041 | 0.2185 | 0.299 | 0.2718 | 0.2583 |

**Cell Index at: 110:47:40**

| | 1 | 2 | 3 | 4 | 5 | 6 | 7 | 8 | 9 | 10 | 11 | 12 |
|---|---|---|---|---|---|---|---|---|---|---|---|---|
| A | | | | | | | | 0.0278 | 0.0302 | 0.0578 | 0.0184 | 0.0384 |
| B | | | | | | | | 0.0334 | 0.0483 | 0.0588 | 0.0847 | 0.056 |
| C | | | | | | | | 0.2827 | 0.2772 | 0.4113 | 0.428 | 0.3355 |
| D | | | | | | | | 0.292 | 0.339 | 0.3825 | 0.3853 | 0.3396 |
| E | | | | | | | | 0.1273 | 0.1039 | 0.1441 | 0.1337 | 0.0794 |
| F | | | | | | | | 0.3936 | 0.3561 | 0.3874 | 0.413 | 0.3283 |
| G | | | | | | | | 0.0491 | 0.0673 | 0.0695 | 0.0578 | 0.0516 |
| H | | | | | | | | 0.3062 | 0.2139 | 0.2968 | 0.2729 | 0.258 |

**Cell Index at: 111:02:41**

| | 1 | 2 | 3 | 4 | 5 | 6 | 7 | 8 | 9 | 10 | 11 | 12 |
|---|---|---|---|---|---|---|---|---|---|---|---|---|
| A | | | | | | | | 0.0297 | 0.0324 | 0.0573 | 0.0183 | 0.0398 |
| B | | | | | | | | 0.032 | 0.0465 | 0.062 | 0.0807 | 0.057 |
| C | | | | | | | | 0.2757 | 0.273 | 0.4112 | 0.4319 | 0.3381 |
| D | | | | | | | | 0.2953 | 0.3346 | 0.3762 | 0.3907 | 0.3332 |
| E | | | | | | | | 0.1243 | 0.1063 | 0.1387 | 0.1294 | 0.0771 |
| F | | | | | | | | 0.4002 | 0.3559 | 0.3906 | 0.4077 | 0.3338 |
| G | | | | | | | | 0.0455 | 0.0656 | 0.0695 | 0.0553 | 0.0508 |
| H | | | | | | | | 0.3032 | 0.2169 | 0.2984 | 0.2715 | 0.2584 |

**Cell Index at: 111:17:42**

| | 1 | 2 | 3 | 4 | 5 | 6 | 7 | 8 | 9 | 10 | 11 | 12 |
|---|---|---|---|---|---|---|---|---|---|---|---|---|
| A | | | | | | | | 0.0329 | 0.0321 | 0.0572 | 0.0203 | 0.0402 |
| B | | | | | | | | 0.0339 | 0.0451 | 0.0552 | 0.0802 | 0.0567 |
| C | | | | | | | | 0.2818 | 0.2716 | 0.4021 | 0.4352 | 0.3374 |
| D | | | | | | | | 0.296 | 0.3375 | 0.3877 | 0.388 | 0.3416 |
| E | | | | | | | | 0.1288 | 0.1074 | 0.1374 | 0.1326 | 0.079 |
| F | | | | | | | | 0.4027 | 0.3549 | 0.3941 | 0.4079 | 0.3317 |
| G | | | | | | | | 0.0461 | 0.0664 | 0.0685 | 0.0608 | 0.0495 |
| H | | | | | | | | 0.3088 | 0.2173 | 0.3026 | 0.272 | 0.2596 |

**Cell Index at: 111:32:43**

| | 1 | 2 | 3 | 4 | 5 | 6 | 7 | 8 | 9 | 10 | 11 | 12 |
|---|---|---|---|---|---|---|---|---|---|---|---|---|

| | | | | | | | | | | | | |
|---|---|---|---|---|---|---|---|---|---|---|---|---|
| A | | | | | | | | 0.0296 | 0.0346 | 0.0606 | 0.0221 | 0.0385 |
| B | | | | | | | | 0.0356 | 0.0464 | 0.0563 | 0.0837 | 0.0537 |
| C | | | | | | | | 0.2852 | 0.2816 | 0.4119 | 0.43 | 0.3329 |
| D | | | | | | | | 0.2938 | 0.3436 | 0.3809 | 0.3979 | 0.337 |
| E | | | | | | | | 0.1306 | 0.1057 | 0.1342 | 0.1308 | 0.0733 |
| F | | | | | | | | 0.3995 | 0.3629 | 0.3948 | 0.4133 | 0.3332 |
| G | | | | | | | | 0.0523 | 0.0653 | 0.0726 | 0.0591 | 0.052 |
| H | | | | | | | | 0.3071 | 0.2158 | 0.3006 | 0.2771 | 0.2621 |

**Cell Index at: 111:47:44**

| | 1 | 2 | 3 | 4 | 5 | 6 | 7 | 8 | 9 | 10 | 11 | 12 |
|---|---|---|---|---|---|---|---|---|---|---|---|---|
| A | | | | | | | | 0.0317 | 0.0346 | 0.0583 | 0.0214 | 0.0381 |
| B | | | | | | | | 0.0362 | 0.0457 | 0.0573 | 0.0828 | 0.0567 |
| C | | | | | | | | 0.2881 | 0.2715 | 0.4065 | 0.438 | 0.3362 |
| D | | | | | | | | 0.2883 | 0.3308 | 0.3876 | 0.3863 | 0.3342 |
| E | | | | | | | | 0.1326 | 0.1081 | 0.1454 | 0.1403 | 0.0766 |
| F | | | | | | | | 0.4057 | 0.3574 | 0.3997 | 0.4121 | 0.3374 |
| G | | | | | | | | 0.0467 | 0.0649 | 0.0688 | 0.0585 | 0.0494 |
| H | | | | | | | | 0.3109 | 0.2197 | 0.2993 | 0.2716 | 0.2617 |

**Cell Index at: 112:02:45**

| | 1 | 2 | 3 | 4 | 5 | 6 | 7 | 8 | 9 | 10 | 11 | 12 |
|---|---|---|---|---|---|---|---|---|---|---|---|---|
| A | | | | | | | | 0.0279 | 0.0305 | 0.0596 | 0.021 | 0.0398 |
| B | | | | | | | | 0.0371 | 0.0453 | 0.0583 | 0.0842 | 0.0587 |
| C | | | | | | | | 0.2829 | 0.2703 | 0.4092 | 0.4424 | 0.3375 |
| D | | | | | | | | 0.2958 | 0.3396 | 0.3868 | 0.3894 | 0.3342 |
| E | | | | | | | | 0.1369 | 0.1026 | 0.1374 | 0.1388 | 0.0763 |
| F | | | | | | | | 0.4066 | 0.3674 | 0.4002 | 0.4194 | 0.333 |
| G | | | | | | | | 0.045 | 0.0657 | 0.0678 | 0.0584 | 0.0507 |
| H | | | | | | | | 0.3172 | 0.2171 | 0.2988 | 0.2759 | 0.2615 |

**Cell Index at: 112:17:46**

| | 1 | 2 | 3 | 4 | 5 | 6 | 7 | 8 | 9 | 10 | 11 | 12 |
|---|---|---|---|---|---|---|---|---|---|---|---|---|
| A | | | | | | | | 0.0319 | 0.0312 | 0.0561 | 0.0214 | 0.0397 |
| B | | | | | | | | 0.0342 | 0.0424 | 0.0567 | 0.0862 | 0.0572 |
| C | | | | | | | | 0.2894 | 0.2747 | 0.4091 | 0.4408 | 0.3378 |
| D | | | | | | | | 0.2873 | 0.3366 | 0.3836 | 0.3958 | 0.3399 |
| E | | | | | | | | 0.1354 | 0.1058 | 0.1356 | 0.1296 | 0.0756 |
| F | | | | | | | | 0.4077 | 0.3738 | 0.397 | 0.4152 | 0.3294 |
| G | | | | | | | | 0.0462 | 0.0635 | 0.0661 | 0.0565 | 0.0521 |
| H | | | | | | | | 0.314 | 0.2188 | 0.2991 | 0.2752 | 0.262 |

**Cell Index at: 112:32:46**

| | 1 | 2 | 3 | 4 | 5 | 6 | 7 | 8 | 9 | 10 | 11 | 12 |
|---|---|---|---|---|---|---|---|---|---|---|---|---|
| A | | | | | | | | 0.0329 | 0.0316 | 0.0589 | 0.0213 | 0.0438 |
| B | | | | | | | | 0.0334 | 0.0423 | 0.0583 | 0.0851 | 0.0596 |
| C | | | | | | | | 0.2907 | 0.2765 | 0.4164 | 0.4478 | 0.3355 |
| D | | | | | | | | 0.2949 | 0.3334 | 0.3882 | 0.3948 | 0.3428 |
| E | | | | | | | | 0.1247 | 0.1084 | 0.1366 | 0.1353 | 0.0815 |
| F | | | | | | | | 0.4045 | 0.3667 | 0.403 | 0.412 | 0.3319 |
| G | | | | | | | | 0.0448 | 0.0629 | 0.0697 | 0.0608 | 0.0583 |
| H | | | | | | | | 0.3222 | 0.2172 | 0.3002 | 0.2773 | 0.2607 |

**Cell Index at: 112:47:47**

| | 1 | 2 | 3 | 4 | 5 | 6 | 7 | 8 | 9 | 10 | 11 | 12 |
|---|---|---|---|---|---|---|---|---|---|---|---|---|
| A | | | | | | | | 0.0333 | 0.0319 | 0.0577 | 0.0235 | 0.0415 |
| B | | | | | | | | 0.0351 | 0.0452 | 0.0578 | 0.0843 | 0.0605 |
| C | | | | | | | | 0.2882 | 0.2702 | 0.4126 | 0.4447 | 0.3393 |
| D | | | | | | | | 0.2902 | 0.3318 | 0.3887 | 0.3926 | 0.3397 |
| E | | | | | | | | 0.13 | 0.1049 | 0.1448 | 0.1327 | 0.079 |
| F | | | | | | | | 0.4012 | 0.371 | 0.4017 | 0.4146 | 0.3292 |
| G | | | | | | | | 0.0465 | 0.0648 | 0.07 | 0.0622 | 0.0539 |
| H | | | | | | | | 0.3188 | 0.2174 | 0.3025 | 0.274 | 0.2613 |

**Cell Index at: 113:02:48**

| | 1 | 2 | 3 | 4 | 5 | 6 | 7 | 8 | 9 | 10 | 11 | 12 |
|---|---|---|---|---|---|---|---|---|---|---|---|---|
| A | | | | | | | | 0.0332 | 0.03 | 0.0592 | 0.0236 | 0.0438 |
| B | | | | | | | | 0.0401 | 0.0453 | 0.0582 | 0.0871 | 0.0584 |
| C | | | | | | | | 0.2898 | 0.271 | 0.4094 | 0.4472 | 0.3383 |
| D | | | | | | | | 0.2898 | 0.3365 | 0.3899 | 0.3846 | 0.3435 |
| E | | | | | | | | 0.1268 | 0.1028 | 0.1453 | 0.1407 | 0.0846 |
| F | | | | | | | | 0.4086 | 0.3702 | 0.4034 | 0.4203 | 0.3346 |
| G | | | | | | | | 0.0422 | 0.0647 | 0.0666 | 0.0623 | 0.0584 |
| H | | | | | | | | 0.3186 | 0.218 | 0.3025 | 0.2763 | 0.2642 |

**Cell Index at: 113:17:49**

| | 1 | 2 | 3 | 4 | 5 | 6 | 7 | 8 | 9 | 10 | 11 | 12 |
|---|---|---|---|---|---|---|---|---|---|---|---|---|
| A | | | | | | | | 0.0342 | 0.0302 | 0.0565 | 0.0243 | 0.0431 |
| B | | | | | | | | 0.0394 | 0.047 | 0.0584 | 0.0904 | 0.0611 |
| C | | | | | | | | 0.2899 | 0.2691 | 0.4124 | 0.4437 | 0.3323 |
| D | | | | | | | | 0.2993 | 0.3431 | 0.3867 | 0.3881 | 0.3375 |
| E | | | | | | | | 0.1362 | 0.1043 | 0.1367 | 0.1356 | 0.084 |
| F | | | | | | | | 0.4077 | 0.367 | 0.3985 | 0.4144 | 0.3405 |
| G | | | | | | | | 0.0451 | 0.064 | 0.07 | 0.0515 | 0.0526 |
| H | | | | | | | | 0.3231 | 0.2161 | 0.3063 | 0.2782 | 0.2629 |

**Cell Index at: 113:32:49**

| | 1 | 2 | 3 | 4 | 5 | 6 | 7 | 8 | 9 | 10 | 11 | 12 |
|---|---|---|---|---|---|---|---|---|---|---|---|---|
| A | | | | | | | | 0.0367 | 0.0317 | 0.0603 | 0.0264 | 0.0421 |
| B | | | | | | | | 0.0397 | 0.0441 | 0.0602 | 0.086 | 0.0591 |
| C | | | | | | | | 0.2915 | 0.2783 | 0.4131 | 0.4468 | 0.3331 |
| D | | | | | | | | 0.2982 | 0.3466 | 0.3931 | 0.3903 | 0.3373 |
| E | | | | | | | | 0.1323 | 0.097 | 0.1385 | 0.1391 | 0.0785 |
| F | | | | | | | | 0.4123 | 0.3673 | 0.4029 | 0.416 | 0.3429 |
| G | | | | | | | | 0.0438 | 0.0645 | 0.0679 | 0.0562 | 0.0547 |
| H | | | | | | | | 0.3232 | 0.2156 | 0.306 | 0.2762 | 0.2629 |

**Cell Index at: 113:47:50**

| | 1 | 2 | 3 | 4 | 5 | 6 | 7 | 8 | 9 | 10 | 11 | 12 |
|---|---|---|---|---|---|---|---|---|---|---|---|---|
| A | | | | | | | | 0.0372 | 0.0334 | 0.0571 | 0.0253 | 0.0409 |
| B | | | | | | | | 0.0434 | 0.0489 | 0.0574 | 0.0902 | 0.0629 |
| C | | | | | | | | 0.2966 | 0.2813 | 0.4099 | 0.45 | 0.3395 |
| D | | | | | | | | 0.2938 | 0.3388 | 0.3864 | 0.3932 | 0.3398 |
| E | | | | | | | | 0.1314 | 0.1063 | 0.1403 | 0.1348 | 0.0792 |
| F | | | | | | | | 0.4119 | 0.3701 | 0.4034 | 0.4122 | 0.3411 |
| G | | | | | | | | 0.0457 | 0.0636 | 0.067 | 0.0527 | 0.0567 |
| H | | | | | | | | 0.3204 | 0.2154 | 0.3065 | 0.2761 | 0.2644 |

**Cell Index at: 114:02:50**

| | 1 | 2 | 3 | 4 | 5 | 6 | 7 | 8 | 9 | 10 | 11 | 12 |
|---|---|---|---|---|---|---|---|---|---|---|---|---|
| A | | | | | | | | 0.0353 | 0.0337 | 0.0595 | 0.0262 | 0.0437 |
| B | | | | | | | | 0.0418 | 0.0453 | 0.0605 | 0.0902 | 0.0621 |
| C | | | | | | | | 0.2947 | 0.279 | 0.4085 | 0.4518 | 0.3388 |
| D | | | | | | | | 0.2981 | 0.3472 | 0.3825 | 0.3935 | 0.3453 |
| E | | | | | | | | 0.1319 | 0.1054 | 0.1445 | 0.1412 | 0.0812 |
| F | | | | | | | | 0.4118 | 0.3763 | 0.4037 | 0.4172 | 0.3407 |
| G | | | | | | | | 0.047 | 0.063 | 0.0691 | 0.0542 | 0.0551 |
| H | | | | | | | | 0.32 | 0.2152 | 0.3124 | 0.2795 | 0.266 |

**Cell Index at: 114:17:51**

| | 1 | 2 | 3 | 4 | 5 | 6 | 7 | 8 | 9 | 10 | 11 | 12 |
|---|---|---|---|---|---|---|---|---|---|---|---|---|
| A | | | | | | | | 0.0375 | 0.034 | 0.0575 | 0.0253 | 0.0429 |
| B | | | | | | | | 0.0409 | 0.0454 | 0.0577 | 0.0907 | 0.0599 |
| C | | | | | | | | 0.2986 | 0.2865 | 0.4113 | 0.447 | 0.3335 |
| D | | | | | | | | 0.2942 | 0.345 | 0.3953 | 0.3974 | 0.3392 |
| E | | | | | | | | 0.1337 | 0.1054 | 0.1449 | 0.1354 | 0.0785 |
| F | | | | | | | | 0.4106 | 0.3725 | 0.4013 | 0.4162 | 0.3458 |
| G | | | | | | | | 0.0459 | 0.0657 | 0.072 | 0.0526 | 0.0588 |
| H | | | | | | | | 0.3167 | 0.2129 | 0.3114 | 0.2779 | 0.2637 |

**Cell Index at: 114:32:51**

| | 1 | 2 | 3 | 4 | 5 | 6 | 7 | 8 | 9 | 10 | 11 | 12 |
|---|---|---|---|---|---|---|---|---|---|---|---|---|
| A | | | | | | | | 0.0383 | 0.0307 | 0.0592 | 0.0273 | 0.0429 |
| B | | | | | | | | 0.0406 | 0.0456 | 0.0555 | 0.0892 | 0.0595 |
| C | | | | | | | | 0.2911 | 0.2793 | 0.4087 | 0.4454 | 0.3395 |
| D | | | | | | | | 0.3003 | 0.3469 | 0.3931 | 0.398 | 0.348 |
| E | | | | | | | | 0.13 | 0.1058 | 0.1461 | 0.1417 | 0.0806 |
| F | | | | | | | | 0.4128 | 0.374 | 0.4023 | 0.4142 | 0.3391 |
| G | | | | | | | | 0.042 | 0.0683 | 0.0689 | 0.0579 | 0.0529 |
| H | | | | | | | | 0.3209 | 0.2163 | 0.3136 | 0.2806 | 0.264 |

**Cell Index at: 114:47:51**

| | 1 | 2 | 3 | 4 | 5 | 6 | 7 | 8 | 9 | 10 | 11 | 12 |
|---|---|---|---|---|---|---|---|---|---|---|---|---|
| A | | | | | | | | 0.0323 | 0.0346 | 0.0599 | 0.0242 | 0.0423 |
| B | | | | | | | | 0.0409 | 0.0445 | 0.0552 | 0.0882 | 0.0662 |
| C | | | | | | | | 0.2983 | 0.2811 | 0.4137 | 0.4545 | 0.3346 |
| D | | | | | | | | 0.3027 | 0.3449 | 0.3853 | 0.3928 | 0.3485 |
| E | | | | | | | | 0.1333 | 0.0985 | 0.1497 | 0.1411 | 0.0796 |
| F | | | | | | | | 0.4095 | 0.3738 | 0.3969 | 0.4175 | 0.3351 |
| G | | | | | | | | 0.0442 | 0.0675 | 0.0669 | 0.0557 | 0.0574 |
| H | | | | | | | | 0.3195 | 0.2144 | 0.3151 | 0.2784 | 0.2671 |

**Cell Index at: 115:02:51**

| | 1 | 2 | 3 | 4 | 5 | 6 | 7 | 8 | 9 | 10 | 11 | 12 |
|---|---|---|---|---|---|---|---|---|---|---|---|---|
| A | | | | | | | | 0.0339 | 0.0348 | 0.0588 | 0.0238 | 0.0434 |
| B | | | | | | | | 0.0396 | 0.0448 | 0.0545 | 0.09 | 0.0627 |
| C | | | | | | | | 0.2969 | 0.2832 | 0.4135 | 0.4616 | 0.3359 |
| D | | | | | | | | 0.2957 | 0.3402 | 0.3925 | 0.3948 | 0.3441 |
| E | | | | | | | | 0.1304 | 0.1048 | 0.1484 | 0.1385 | 0.0783 |
| F | | | | | | | | 0.4118 | 0.3742 | 0.4005 | 0.4186 | 0.336 |
| G | | | | | | | | 0.0454 | 0.068 | 0.0673 | 0.0574 | 0.0539 |
| H | | | | | | | | 0.3217 | 0.2161 | 0.3143 | 0.2789 | 0.2631 |

**Cell Index at: 115:17:51**

| | 1 | 2 | 3 | 4 | 5 | 6 | 7 | 8 | 9 | 10 | 11 | 12 |
|---|---|---|---|---|---|---|---|---|---|---|---|---|
| A | | | | | | | | 0.0342 | 0.0345 | 0.0625 | 0.0229 | 0.0424 |
| B | | | | | | | | 0.0427 | 0.0428 | 0.0594 | 0.0915 | 0.0656 |
| C | | | | | | | | 0.2917 | 0.2776 | 0.4087 | 0.4598 | 0.3427 |
| D | | | | | | | | 0.2891 | 0.3433 | 0.3872 | 0.3918 | 0.3431 |
| E | | | | | | | | 0.1399 | 0.104 | 0.1402 | 0.1308 | 0.0864 |
| F | | | | | | | | 0.4183 | 0.3755 | 0.4031 | 0.4223 | 0.3371 |
| G | | | | | | | | 0.047 | 0.0682 | 0.0711 | 0.056 | 0.0589 |
| H | | | | | | | | 0.3183 | 0.215 | 0.3162 | 0.2772 | 0.2695 |

**Cell Index at: 115:32:51**

| | 1 | 2 | 3 | 4 | 5 | 6 | 7 | 8 | 9 | 10 | 11 | 12 |
|---|---|---|---|---|---|---|---|---|---|---|---|---|
| A | | | | | | | | 0.0338 | 0.0347 | 0.0617 | 0.0262 | 0.0436 |
| B | | | | | | | | 0.0447 | 0.0447 | 0.0568 | 0.0937 | 0.0668 |
| C | | | | | | | | 0.2949 | 0.283 | 0.4167 | 0.4583 | 0.3414 |
| D | | | | | | | | 0.3004 | 0.3432 | 0.396 | 0.3978 | 0.3457 |
| E | | | | | | | | 0.138 | 0.1028 | 0.1448 | 0.1372 | 0.0861 |
| F | | | | | | | | 0.4142 | 0.3695 | 0.4071 | 0.4215 | 0.3408 |
| G | | | | | | | | 0.0453 | 0.0651 | 0.0686 | 0.0575 | 0.0584 |
| H | | | | | | | | 0.3174 | 0.2185 | 0.3161 | 0.2822 | 0.2661 |

**Cell Index at: 115:47:51**

| | 1 | 2 | 3 | 4 | 5 | 6 | 7 | 8 | 9 | 10 | 11 | 12 |
|---|---|---|---|---|---|---|---|---|---|---|---|---|
| A | | | | | | | | 0.0361 | 0.0351 | 0.0629 | 0.0205 | 0.0434 |
| B | | | | | | | | 0.0465 | 0.0461 | 0.0574 | 0.0896 | 0.0676 |
| C | | | | | | | | 0.2943 | 0.292 | 0.4164 | 0.4571 | 0.3384 |
| D | | | | | | | | 0.2901 | 0.3448 | 0.3988 | 0.4019 | 0.3448 |
| E | | | | | | | | 0.1361 | 0.1067 | 0.1478 | 0.1305 | 0.084 |
| F | | | | | | | | 0.4131 | 0.3713 | 0.4021 | 0.4215 | 0.3411 |
| G | | | | | | | | 0.0504 | 0.0657 | 0.0717 | 0.0604 | 0.0542 |
| H | | | | | | | | 0.3191 | 0.2186 | 0.3171 | 0.275 | 0.2662 |

**Cell Index at: 116:02:52**

| | 1 | 2 | 3 | 4 | 5 | 6 | 7 | 8 | 9 | 10 | 11 | 12 |
|---|---|---|---|---|---|---|---|---|---|---|---|---|
| A | | | | | | | | 0.0349 | 0.0385 | 0.0591 | 0.0225 | 0.0405 |
| B | | | | | | | | 0.0449 | 0.0499 | 0.0619 | 0.0929 | 0.0663 |
| C | | | | | | | | 0.2978 | 0.2873 | 0.4204 | 0.4615 | 0.3416 |
| D | | | | | | | | 0.3003 | 0.3404 | 0.3954 | 0.404 | 0.3514 |
| E | | | | | | | | 0.1408 | 0.1055 | 0.1442 | 0.1318 | 0.0844 |
| F | | | | | | | | 0.4097 | 0.3777 | 0.4094 | 0.4213 | 0.3409 |
| G | | | | | | | | 0.0471 | 0.0634 | 0.0667 | 0.054 | 0.0583 |
| H | | | | | | | | 0.319 | 0.2171 | 0.3127 | 0.2762 | 0.2679 |

**Cell Index at: 116:17:52**

| | 1 | 2 | 3 | 4 | 5 | 6 | 7 | 8 | 9 | 10 | 11 | 12 |
|---|---|---|---|---|---|---|---|---|---|---|---|---|
| A | | | | | | | | 0.0357 | 0.0387 | 0.0609 | 0.0234 | 0.0426 |
| B | | | | | | | | 0.0477 | 0.044 | 0.062 | 0.0872 | 0.0688 |
| C | | | | | | | | 0.2962 | 0.2901 | 0.4168 | 0.4655 | 0.3403 |
| D | | | | | | | | 0.3008 | 0.3461 | 0.3973 | 0.4094 | 0.3478 |
| E | | | | | | | | 0.1355 | 0.1098 | 0.1487 | 0.1334 | 0.0813 |
| F | | | | | | | | 0.4207 | 0.3706 | 0.4035 | 0.423 | 0.3441 |
| G | | | | | | | | 0.0472 | 0.0703 | 0.0707 | 0.0618 | 0.0563 |
| H | | | | | | | | 0.3193 | 0.2149 | 0.3114 | 0.2764 | 0.2692 |

**Cell Index at: 116:32:52**

| | 1 | 2 | 3 | 4 | 5 | 6 | 7 | 8 | 9 | 10 | 11 | 12 |
|---|---|---|---|---|---|---|---|---|---|---|---|---|
| A | | | | | | | | 0.0359 | 0.0352 | 0.0619 | 0.0207 | 0.0453 |
| B | | | | | | | | 0.0479 | 0.0431 | 0.0617 | 0.0924 | 0.0681 |
| C | | | | | | | | 0.2999 | 0.2824 | 0.4151 | 0.4614 | 0.3403 |
| D | | | | | | | | 0.2973 | 0.3515 | 0.3963 | 0.4066 | 0.3525 |
| E | | | | | | | | 0.1319 | 0.1096 | 0.1421 | 0.1347 | 0.0795 |
| F | | | | | | | | 0.4149 | 0.3758 | 0.4034 | 0.4232 | 0.3454 |
| G | | | | | | | | 0.0494 | 0.065 | 0.0676 | 0.0569 | 0.0557 |
| H | | | | | | | | 0.3231 | 0.2154 | 0.3173 | 0.2755 | 0.267 |

**Cell Index at: 116:47:53**

| | 1 | 2 | 3 | 4 | 5 | 6 | 7 | 8 | 9 | 10 | 11 | 12 |
|---|---|---|---|---|---|---|---|---|---|---|---|---|
| A | | | | | | | | 0.0353 | 0.0336 | 0.0582 | 0.0229 | 0.0404 |
| B | | | | | | | | 0.0464 | 0.0378 | 0.0631 | 0.0913 | 0.067 |
| C | | | | | | | | 0.3052 | 0.2889 | 0.4185 | 0.4621 | 0.3378 |
| D | | | | | | | | 0.299 | 0.3432 | 0.4032 | 0.4037 | 0.353 |
| E | | | | | | | | 0.1406 | 0.1074 | 0.1459 | 0.1397 | 0.0832 |
| F | | | | | | | | 0.4243 | 0.3749 | 0.4012 | 0.4282 | 0.3428 |
| G | | | | | | | | 0.048 | 0.0675 | 0.0697 | 0.0574 | 0.0538 |
| H | | | | | | | | 0.3232 | 0.2156 | 0.3145 | 0.2771 | 0.2691 |

**Cell Index at: 117:02:54**

| | 1 | 2 | 3 | 4 | 5 | 6 | 7 | 8 | 9 | 10 | 11 | 12 |
|---|---|---|---|---|---|---|---|---|---|---|---|---|
| A | | | | | | | | 0.0405 | 0.0331 | 0.0582 | 0.0207 | 0.0442 |
| B | | | | | | | | 0.0485 | 0.0405 | 0.0634 | 0.0916 | 0.0671 |
| C | | | | | | | | 0.3034 | 0.2864 | 0.4165 | 0.458 | 0.3413 |
| D | | | | | | | | 0.2995 | 0.3435 | 0.3954 | 0.4075 | 0.3533 |
| E | | | | | | | | 0.1314 | 0.1086 | 0.1455 | 0.1403 | 0.0818 |
| F | | | | | | | | 0.4204 | 0.3777 | 0.4071 | 0.424 | 0.3513 |
| G | | | | | | | | 0.0469 | 0.0677 | 0.0707 | 0.0572 | 0.0543 |
| H | | | | | | | | 0.3241 | 0.2139 | 0.3169 | 0.2803 | 0.2681 |

**Cell Index at: 117:17:55**

| | 1 | 2 | 3 | 4 | 5 | 6 | 7 | 8 | 9 | 10 | 11 | 12 |
|---|---|---|---|---|---|---|---|---|---|---|---|---|
| A | | | | | | | | 0.0398 | 0.0326 | 0.0631 | 0.0208 | 0.0431 |
| B | | | | | | | | 0.0483 | 0.0422 | 0.061 | 0.0891 | 0.0687 |
| C | | | | | | | | 0.3059 | 0.2905 | 0.4169 | 0.4617 | 0.3419 |
| D | | | | | | | | 0.3009 | 0.3491 | 0.3982 | 0.4012 | 0.3473 |
| E | | | | | | | | 0.1321 | 0.1109 | 0.1563 | 0.1375 | 0.0809 |
| F | | | | | | | | 0.4239 | 0.3825 | 0.4037 | 0.4248 | 0.346 |
| G | | | | | | | | 0.0473 | 0.0684 | 0.0707 | 0.0576 | 0.0554 |
| H | | | | | | | | 0.3255 | 0.2162 | 0.3186 | 0.2798 | 0.2704 |

**Cell Index at: 117:32:55**

| | 1 | 2 | 3 | 4 | 5 | 6 | 7 | 8 | 9 | 10 | 11 | 12 |
|---|---|---|---|---|---|---|---|---|---|---|---|---|
| A | | | | | | | | 0.0392 | 0.0314 | 0.0603 | 0.0203 | 0.0411 |
| B | | | | | | | | 0.0451 | 0.0442 | 0.0637 | 0.0921 | 0.0693 |
| C | | | | | | | | 0.3056 | 0.2904 | 0.4181 | 0.4651 | 0.3414 |
| D | | | | | | | | 0.2965 | 0.3504 | 0.4037 | 0.4035 | 0.3544 |
| E | | | | | | | | 0.1291 | 0.1101 | 0.1418 | 0.1387 | 0.081 |
| F | | | | | | | | 0.4257 | 0.3817 | 0.4104 | 0.4294 | 0.3526 |
| G | | | | | | | | 0.0504 | 0.0665 | 0.0697 | 0.0562 | 0.0576 |
| H | | | | | | | | 0.3267 | 0.2188 | 0.3185 | 0.2817 | 0.2692 |

**Cell Index at: 117:47:55**

| | 1 | 2 | 3 | 4 | 5 | 6 | 7 | 8 | 9 | 10 | 11 | 12 |
|---|---|---|---|---|---|---|---|---|---|---|---|---|
| A | | | | | | | | 0.0442 | 0.0324 | 0.0608 | 0.0222 | 0.044 |
| B | | | | | | | | 0.0459 | 0.0422 | 0.0651 | 0.0932 | 0.0685 |
| C | | | | | | | | 0.3106 | 0.2898 | 0.4225 | 0.4606 | 0.3427 |
| D | | | | | | | | 0.292 | 0.3568 | 0.3964 | 0.4036 | 0.3503 |
| E | | | | | | | | 0.1361 | 0.1112 | 0.1514 | 0.1352 | 0.0785 |
| F | | | | | | | | 0.4182 | 0.3798 | 0.406 | 0.4281 | 0.3502 |
| G | | | | | | | | 0.0466 | 0.0674 | 0.072 | 0.0597 | 0.0605 |
| H | | | | | | | | 0.3282 | 0.2203 | 0.3186 | 0.28 | 0.2694 |

**Cell Index at: 118:02:56**

| | 1 | 2 | 3 | 4 | 5 | 6 | 7 | 8 | 9 | 10 | 11 | 12 |
|---|---|---|---|---|---|---|---|---|---|---|---|---|
| A | | | | | | | | 0.0446 | 0.0333 | 0.0577 | 0.0234 | 0.045 |
| B | | | | | | | | 0.0422 | 0.0401 | 0.0606 | 0.09 | 0.0665 |
| C | | | | | | | | 0.3074 | 0.2915 | 0.4292 | 0.4653 | 0.3453 |
| D | | | | | | | | 0.3001 | 0.3467 | 0.4079 | 0.4056 | 0.3511 |
| E | | | | | | | | 0.1395 | 0.1076 | 0.15 | 0.1366 | 0.0805 |
| F | | | | | | | | 0.4225 | 0.3781 | 0.4076 | 0.4292 | 0.3502 |
| G | | | | | | | | 0.0504 | 0.0715 | 0.0738 | 0.0556 | 0.0582 |
| H | | | | | | | | 0.3264 | 0.222 | 0.3245 | 0.2802 | 0.2716 |

**Cell Index at: 118:17:57**

| | 1 | 2 | 3 | 4 | 5 | 6 | 7 | 8 | 9 | 10 | 11 | 12 |
|---|---|---|---|---|---|---|---|---|---|---|---|---|
| A | | | | | | | | 0.0439 | 0.032 | 0.06 | 0.0229 | 0.0452 |
| B | | | | | | | | 0.0472 | 0.0428 | 0.0629 | 0.0945 | 0.0696 |
| C | | | | | | | | 0.3055 | 0.2874 | 0.4354 | 0.4684 | 0.3447 |
| D | | | | | | | | 0.2975 | 0.3516 | 0.4013 | 0.4074 | 0.3496 |
| E | | | | | | | | 0.1386 | 0.1169 | 0.157 | 0.1383 | 0.079 |
| F | | | | | | | | 0.4253 | 0.3774 | 0.4094 | 0.4276 | 0.352 |
| G | | | | | | | | 0.0525 | 0.067 | 0.0749 | 0.0569 | 0.0589 |
| H | | | | | | | | 0.3294 | 0.2227 | 0.3242 | 0.2811 | 0.2708 |

**Cell Index at: 118:32:58**

| | 1 | 2 | 3 | 4 | 5 | 6 | 7 | 8 | 9 | 10 | 11 | 12 |
|---|---|---|---|---|---|---|---|---|---|---|---|---|
| A | | | | | | | | 0.0477 | 0.0379 | 0.0566 | 0.0236 | 0.0445 |
| B | | | | | | | | 0.0456 | 0.0405 | 0.0644 | 0.0906 | 0.0724 |
| C | | | | | | | | 0.3149 | 0.2903 | 0.4296 | 0.4686 | 0.3465 |
| D | | | | | | | | 0.2912 | 0.3478 | 0.3969 | 0.4119 | 0.3591 |
| E | | | | | | | | 0.1345 | 0.1087 | 0.1509 | 0.1314 | 0.0808 |
| F | | | | | | | | 0.4298 | 0.382 | 0.4123 | 0.4301 | 0.3527 |
| G | | | | | | | | 0.0515 | 0.0697 | 0.0746 | 0.0558 | 0.0524 |
| H | | | | | | | | 0.3284 | 0.2263 | 0.3239 | 0.2799 | 0.2733 |

**Cell Index at: 118:47:58**

| | 1 | 2 | 3 | 4 | 5 | 6 | 7 | 8 | 9 | 10 | 11 | 12 |
|---|---|---|---|---|---|---|---|---|---|---|---|---|
| A | | | | | | | | 0.0489 | 0.0371 | 0.0566 | 0.0242 | 0.0434 |
| B | | | | | | | | 0.0451 | 0.0394 | 0.0629 | 0.0882 | 0.0694 |
| C | | | | | | | | 0.3078 | 0.2872 | 0.4343 | 0.4664 | 0.3472 |
| D | | | | | | | | 0.2994 | 0.3457 | 0.4041 | 0.4065 | 0.3566 |
| E | | | | | | | | 0.1386 | 0.1157 | 0.1524 | 0.1368 | 0.0832 |
| F | | | | | | | | 0.4323 | 0.3839 | 0.4099 | 0.4303 | 0.3503 |
| G | | | | | | | | 0.0504 | 0.0658 | 0.0719 | 0.057 | 0.0553 |
| H | | | | | | | | 0.3285 | 0.2294 | 0.3231 | 0.278 | 0.2726 |

**Cell Index at: 119:02:58**

| | 1 | 2 | 3 | 4 | 5 | 6 | 7 | 8 | 9 | 10 | 11 | 12 |
|---|---|---|---|---|---|---|---|---|---|---|---|---|
| A | | | | | | | | 0.0458 | 0.0373 | 0.0566 | 0.0243 | 0.0418 |
| B | | | | | | | | 0.0393 | 0.0417 | 0.062 | 0.088 | 0.0697 |
| C | | | | | | | | 0.3141 | 0.2932 | 0.4295 | 0.4629 | 0.3439 |
| D | | | | | | | | 0.3105 | 0.3501 | 0.4032 | 0.4212 | 0.3573 |
| E | | | | | | | | 0.1358 | 0.118 | 0.1431 | 0.1376 | 0.0839 |
| F | | | | | | | | 0.4293 | 0.3777 | 0.4053 | 0.4333 | 0.349 |
| G | | | | | | | | 0.0487 | 0.0739 | 0.073 | 0.0554 | 0.0605 |
| H | | | | | | | | 0.3345 | 0.2262 | 0.3251 | 0.283 | 0.2724 |

**Cell Index at: 119:17:59**

| | 1 | 2 | 3 | 4 | 5 | 6 | 7 | 8 | 9 | 10 | 11 | 12 |
|---|---|---|---|---|---|---|---|---|---|---|---|---|
| A | | | | | | | | 0.0456 | 0.0367 | 0.0595 | 0.0287 | 0.0413 |
| B | | | | | | | | 0.0419 | 0.0423 | 0.0602 | 0.0912 | 0.0723 |
| C | | | | | | | | 0.3101 | 0.2908 | 0.4304 | 0.4611 | 0.3539 |
| D | | | | | | | | 0.2977 | 0.354 | 0.4118 | 0.4149 | 0.3563 |
| E | | | | | | | | 0.1415 | 0.1073 | 0.1507 | 0.1439 | 0.0811 |
| F | | | | | | | | 0.4258 | 0.384 | 0.4078 | 0.4274 | 0.3532 |
| G | | | | | | | | 0.0429 | 0.0686 | 0.0741 | 0.0554 | 0.0608 |
| H | | | | | | | | 0.3313 | 0.2229 | 0.3259 | 0.2799 | 0.273 |

**Cell Index at: 119:32:59**

| | 1 | 2 | 3 | 4 | 5 | 6 | 7 | 8 | 9 | 10 | 11 | 12 |
|---|---|---|---|---|---|---|---|---|---|---|---|---|
| A | | | | | | | | 0.0503 | 0.0357 | 0.0582 | 0.0279 | 0.04 |
| B | | | | | | | | 0.0445 | 0.0425 | 0.0633 | 0.0926 | 0.0754 |
| C | | | | | | | | 0.3106 | 0.2954 | 0.431 | 0.4645 | 0.3471 |
| D | | | | | | | | 0.302 | 0.3432 | 0.4094 | 0.4144 | 0.3563 |
| E | | | | | | | | 0.1436 | 0.1101 | 0.1527 | 0.1332 | 0.0793 |
| F | | | | | | | | 0.4286 | 0.381 | 0.4059 | 0.4397 | 0.3501 |
| G | | | | | | | | 0.0526 | 0.0719 | 0.0756 | 0.0509 | 0.0611 |
| H | | | | | | | | 0.3371 | 0.2229 | 0.3264 | 0.2811 | 0.274 |

**Cell Index at: 119:48:00**

| | 1 | 2 | 3 | 4 | 5 | 6 | 7 | 8 | 9 | 10 | 11 | 12 |
|---|---|---|---|---|---|---|---|---|---|---|---|---|
| A | | | | | | | | 0.0467 | 0.0367 | 0.0536 | 0.0273 | 0.0438 |
| B | | | | | | | | 0.0456 | 0.0425 | 0.0669 | 0.0891 | 0.066 |
| C | | | | | | | | 0.309 | 0.297 | 0.4316 | 0.4729 | 0.3537 |
| D | | | | | | | | 0.3081 | 0.3484 | 0.4082 | 0.4211 | 0.3679 |
| E | | | | | | | | 0.1447 | 0.1179 | 0.1529 | 0.1349 | 0.0764 |
| F | | | | | | | | 0.4331 | 0.3833 | 0.4149 | 0.4304 | 0.3566 |
| G | | | | | | | | 0.0448 | 0.0735 | 0.0721 | 0.0547 | 0.0622 |
| H | | | | | | | | 0.3363 | 0.2234 | 0.3296 | 0.286 | 0.2713 |

**Cell Index at: 120:03:01**

| | 1 | 2 | 3 | 4 | 5 | 6 | 7 | 8 | 9 | 10 | 11 | 12 |
|---|---|---|---|---|---|---|---|---|---|---|---|---|
| A | | | | | | | | 0.0449 | 0.037 | 0.056 | 0.0284 | 0.0374 |
| B | | | | | | | | 0.044 | 0.0401 | 0.0672 | 0.09 | 0.0707 |
| C | | | | | | | | 0.3165 | 0.2946 | 0.4377 | 0.4645 | 0.3562 |
| D | | | | | | | | 0.3037 | 0.3474 | 0.4074 | 0.414 | 0.3641 |
| E | | | | | | | | 0.1419 | 0.1128 | 0.1534 | 0.1464 | 0.0786 |
| F | | | | | | | | 0.4299 | 0.382 | 0.4159 | 0.4312 | 0.3525 |
| G | | | | | | | | 0.0439 | 0.0725 | 0.0722 | 0.0536 | 0.0664 |
| H | | | | | | | | 0.3333 | 0.2276 | 0.3266 | 0.2853 | 0.2737 |

| | | | | | | | | | | | | | |
|---|---|---|---|---|---|---|---|---|---|---|---|---|---|
| A vs B | 0.08066 | B vs C | 0.000425 | C vs D | 0.871053 | D vs E | 2.31E-05 | E vs F | 1.03E-06 | F vs G | 4.14E-06 | G vs H | 0.000139 |
| A vs C | 0.000481 | B vs D | 1.68E-05 | C vs E | 0.00077 | D vs F | 0.208677 | E vs G | 0.006546 | F vs H | 0.001997 | | |
| A vs D | 4.86E-05 | B vs E | 0.006277 | C vs F | 0.469646 | D vs G | 5.05E-05 | E vs H | 0.000201 | | | | |
| A vs E | 0.002192 | B vs F | 5.48E-07 | C vs G | 0.000584 | D vs H | 0.023605 | | | | | | |
| A vs F | 5.31E-06 | B vs G | 0.951531 | C vs H | 0.067283 | | | | | | | | |
| A vs G | 0.02121 | B vs H | 5.41E-05 | | | | | | | | | | |
| A vs H | 0.000119 | | | | | | | | | | | | |

**Cell Index at: 120:18:01**

| | 1 | 2 | 3 | 4 | 5 | 6 | 7 | 8 | 9 | 10 | 11 | 12 |
|---|---|---|---|---|---|---|---|---|---|---|---|---|
| A | | | | | | | | 0.0449 | 0.0363 | 0.0561 | 0.0278 | 0.0434 |
| B | | | | | | | | 0.0459 | 0.0427 | 0.0663 | 0.0907 | 0.0655 |
| C | | | | | | | | 0.3103 | 0.2932 | 0.4396 | 0.4638 | 0.3548 |
| D | | | | | | | | 0.2953 | 0.3521 | 0.4094 | 0.4155 | 0.3677 |
| E | | | | | | | | 0.1393 | 0.1188 | 0.1494 | 0.1367 | 0.0762 |
| F | | | | | | | | 0.4313 | 0.3809 | 0.4135 | 0.4315 | 0.3524 |
| G | | | | | | | | 0.0448 | 0.0731 | 0.0748 | 0.0523 | 0.0654 |

| *Count* | *Sum* | *Average* | *Variance* |
|---|---|---|---|
| 5 | 0.2037 | 0.04074 | 0.000107 |
| 5 | 0.312 | 0.0624 | 0.000422 |
| 5 | 1.8695 | 0.3739 | 0.005544 |
| 5 | 1.8366 | 0.36732 | 0.00206 |
| 5 | 0.6331 | 0.12662 | 0.00096 |

| | 1 | 2 | 3 | 4 | 5 | 6 | 7 | 8 | 9 | 10 | 11 | 12 |
|---|---|---|---|---|---|---|---|---|---|---|---|---|
| H | | | | | | | | 0.3389 | 0.2292 | 0.3289 | 0.2872 | 0.2711 |

| | | | |
|---|---|---|---|
| 5 | 2.0115 | 0.4023 | 0.001169 |
| 5 | 0.3086 | 0.06172 | 0.000158 |
| 5 | 1.4465 | 0.2893 | 0.001848 |

| *SS* | *df* | *MS* | *F* | *P-value* | *F crit* |
|---|---|---|---|---|---|
| 0.870046 | 7 | 0.124292 | 81.05397 | 1.53E-18 | 2.312741 |
| 0.04907 | 32 | 0.001533 | | | |
| 0.919116 | 39 | | | | |

**Cell Index at: 120:33:01**

| | 1 | 2 | 3 | 4 | 5 | 6 | 7 | 8 | 9 | 10 | 11 | 12 |
|---|---|---|---|---|---|---|---|---|---|---|---|---|
| A | | | | | | | | 0.0461 | 0.0382 | 0.061 | 0.0305 | 0.0456 |
| B | | | | | | | | 0.0445 | 0.0402 | 0.0675 | 0.0947 | 0.0673 |
| C | | | | | | | | 0.3114 | 0.2978 | 0.4348 | 0.4705 | 0.3488 |
| D | | | | | | | | 0.3086 | 0.3467 | 0.414 | 0.4131 | 0.3626 |
| E | | | | | | | | 0.1405 | 0.1141 | 0.1541 | 0.1417 | 0.0858 |
| F | | | | | | | | 0.4386 | 0.3839 | 0.4149 | 0.4311 | 0.3613 |
| G | | | | | | | | 0.0476 | 0.0775 | 0.0751 | 0.052 | 0.0662 |
| H | | | | | | | | 0.3382 | 0.2316 | 0.3302 | 0.2885 | 0.2717 |

**Cell Index at: 120:48:02**

| | 1 | 2 | 3 | 4 | 5 | 6 | 7 | 8 | 9 | 10 | 11 | 12 |
|---|---|---|---|---|---|---|---|---|---|---|---|---|
| A | | | | | | | | 0.0496 | 0.0335 | 0.0628 | 0.0285 | 0.0475 |
| B | | | | | | | | 0.0457 | 0.0454 | 0.0651 | 0.0891 | 0.0683 |
| C | | | | | | | | 0.3154 | 0.2985 | 0.4342 | 0.4677 | 0.3507 |
| D | | | | | | | | 0.3053 | 0.3547 | 0.4116 | 0.4204 | 0.3687 |
| E | | | | | | | | 0.146 | 0.112 | 0.1583 | 0.1435 | 0.0783 |
| F | | | | | | | | 0.4386 | 0.3907 | 0.4155 | 0.4325 | 0.3572 |
| G | | | | | | | | 0.0495 | 0.0733 | 0.0777 | 0.0508 | 0.0633 |
| H | | | | | | | | 0.344 | 0.2303 | 0.3286 | 0.291 | 0.2716 |

**Cell Index at: 121:03:02**

| | 1 | 2 | 3 | 4 | 5 | 6 | 7 | 8 | 9 | 10 | 11 | 12 |
|---|---|---|---|---|---|---|---|---|---|---|---|---|
| A | | | | | | | | 0.0481 | 0.0361 | 0.063 | 0.0296 | 0.0492 |
| B | | | | | | | | 0.0427 | 0.046 | 0.0695 | 0.0919 | 0.0678 |
| C | | | | | | | | 0.3115 | 0.296 | 0.4347 | 0.4644 | 0.354 |
| D | | | | | | | | 0.3088 | 0.3585 | 0.4186 | 0.4151 | 0.3688 |
| E | | | | | | | | 0.1373 | 0.1231 | 0.1579 | 0.1451 | 0.0777 |
| F | | | | | | | | 0.4444 | 0.3853 | 0.4173 | 0.4351 | 0.3641 |
| G | | | | | | | | 0.0461 | 0.0775 | 0.0777 | 0.0526 | 0.0661 |
| H | | | | | | | | 0.3392 | 0.2294 | 0.336 | 0.2905 | 0.2755 |

**Cell Index at: 121:18:02**

| | 1 | 2 | 3 | 4 | 5 | 6 | 7 | 8 | 9 | 10 | 11 | 12 |
|---|---|---|---|---|---|---|---|---|---|---|---|---|
| A | | | | | | | | 0.0501 | 0.0337 | 0.0602 | 0.0301 | 0.0485 |
| B | | | | | | | | 0.0422 | 0.0463 | 0.0632 | 0.089 | 0.0674 |
| C | | | | | | | | 0.3216 | 0.3033 | 0.44 | 0.4719 | 0.357 |
| D | | | | | | | | 0.3061 | 0.3567 | 0.4157 | 0.423 | 0.366 |
| E | | | | | | | | 0.1329 | 0.1138 | 0.1583 | 0.1411 | 0.0814 |
| F | | | | | | | | 0.4324 | 0.395 | 0.4184 | 0.4358 | 0.361 |
| G | | | | | | | | 0.0483 | 0.0679 | 0.0813 | 0.0501 | 0.0666 |
| H | | | | | | | | 0.3393 | 0.2277 | 0.336 | 0.2912 | 0.2733 |

**Cell Index at: 121:33:03**

| | 1 | 2 | 3 | 4 | 5 | 6 | 7 | 8 | 9 | 10 | 11 | 12 |
|---|---|---|---|---|---|---|---|---|---|---|---|---|
| A | | | | | | | | 0.0509 | 0.0372 | 0.0625 | 0.031 | 0.0489 |
| B | | | | | | | | 0.0492 | 0.0461 | 0.0667 | 0.0912 | 0.0702 |
| C | | | | | | | | 0.3214 | 0.308 | 0.4346 | 0.4715 | 0.3633 |
| D | | | | | | | | 0.3048 | 0.3663 | 0.4146 | 0.4202 | 0.3723 |
| E | | | | | | | | 0.1404 | 0.1131 | 0.149 | 0.1352 | 0.0869 |
| F | | | | | | | | 0.4415 | 0.3903 | 0.4136 | 0.4273 | 0.3523 |
| G | | | | | | | | 0.0451 | 0.0747 | 0.0749 | 0.05 | 0.0671 |
| H | | | | | | | | 0.3393 | 0.2304 | 0.3388 | 0.2923 | 0.2765 |

**Cell Index at: 121:48:04**

| | 1 | 2 | 3 | 4 | 5 | 6 | 7 | 8 | 9 | 10 | 11 | 12 |
|---|---|---|---|---|---|---|---|---|---|---|---|---|
| A | | | | | | | | 0.0536 | 0.0394 | 0.0636 | 0.0336 | 0.049 |
| B | | | | | | | | 0.0443 | 0.0466 | 0.0667 | 0.0903 | 0.0718 |
| C | | | | | | | | 0.329 | 0.3074 | 0.4381 | 0.4789 | 0.36 |
| D | | | | | | | | 0.3024 | 0.3585 | 0.4184 | 0.4273 | 0.372 |
| E | | | | | | | | 0.1376 | 0.114 | 0.1602 | 0.1433 | 0.083 |
| F | | | | | | | | 0.4328 | 0.3918 | 0.4086 | 0.4355 | 0.3573 |
| G | | | | | | | | 0.0455 | 0.0732 | 0.0692 | 0.0535 | 0.0659 |
| H | | | | | | | | 0.3343 | 0.2296 | 0.3429 | 0.2882 | 0.2803 |

**Cell Index at: 122:03:05**

| | 1 | 2 | 3 | 4 | 5 | 6 | 7 | 8 | 9 | 10 | 11 | 12 |
|---|---|---|---|---|---|---|---|---|---|---|---|---|
| A | | | | | | | | 0.0527 | 0.0422 | 0.0628 | 0.0343 | 0.0496 |
| B | | | | | | | | 0.0499 | 0.0453 | 0.0663 | 0.0896 | 0.0702 |
| C | | | | | | | | 0.3294 | 0.305 | 0.4411 | 0.4762 | 0.3689 |
| D | | | | | | | | 0.3035 | 0.3578 | 0.4144 | 0.4196 | 0.3732 |
| E | | | | | | | | 0.142 | 0.1203 | 0.1596 | 0.1374 | 0.0779 |
| F | | | | | | | | 0.4377 | 0.3904 | 0.4099 | 0.4344 | 0.3592 |
| G | | | | | | | | 0.0479 | 0.0754 | 0.0725 | 0.0519 | 0.0675 |
| H | | | | | | | | 0.3416 | 0.2336 | 0.3412 | 0.2885 | 0.2777 |

**Cell Index at: 122:18:06**

| | 1 | 2 | 3 | 4 | 5 | 6 | 7 | 8 | 9 | 10 | 11 | 12 |
|---|---|---|---|---|---|---|---|---|---|---|---|---|
| A | | | | | | | | 0.0543 | 0.045 | 0.0667 | 0.0315 | 0.0484 |
| B | | | | | | | | 0.0506 | 0.0467 | 0.0673 | 0.0904 | 0.0699 |
| C | | | | | | | | 0.3298 | 0.3067 | 0.4456 | 0.481 | 0.3687 |
| D | | | | | | | | 0.3059 | 0.3529 | 0.4201 | 0.4238 | 0.3747 |
| E | | | | | | | | 0.1333 | 0.1119 | 0.1522 | 0.1476 | 0.0816 |
| F | | | | | | | | 0.4387 | 0.3862 | 0.4103 | 0.4374 | 0.3579 |
| G | | | | | | | | 0.0467 | 0.0748 | 0.0745 | 0.0523 | 0.0628 |
| H | | | | | | | | 0.3353 | 0.2313 | 0.3413 | 0.2926 | 0.2797 |

**Cell Index at: 122:33:06**

| | 1 | 2 | 3 | 4 | 5 | 6 | 7 | 8 | 9 | 10 | 11 | 12 |
|---|---|---|---|---|---|---|---|---|---|---|---|---|
| A | | | | | | | | 0.0541 | 0.0489 | 0.0693 | 0.0322 | 0.0501 |
| B | | | | | | | | 0.0516 | 0.049 | 0.0643 | 0.09 | 0.0742 |
| C | | | | | | | | 0.3273 | 0.3058 | 0.4468 | 0.4803 | 0.3647 |
| D | | | | | | | | 0.3029 | 0.3664 | 0.4217 | 0.4294 | 0.3795 |
| E | | | | | | | | 0.1385 | 0.1168 | 0.1531 | 0.1451 | 0.081 |
| F | | | | | | | | 0.4384 | 0.3939 | 0.4127 | 0.4354 | 0.355 |
| G | | | | | | | | 0.0477 | 0.0731 | 0.0756 | 0.0588 | 0.0658 |
| H | | | | | | | | 0.3401 | 0.2327 | 0.3416 | 0.291 | 0.28 |

**Cell Index at: 122:48:06**

| | 1 | 2 | 3 | 4 | 5 | 6 | 7 | 8 | 9 | 10 | 11 | 12 |
|---|---|---|---|---|---|---|---|---|---|---|---|---|
| A | | | | | | | | 0.0573 | 0.0474 | 0.0709 | 0.0305 | 0.0476 |
| B | | | | | | | | 0.0493 | 0.0468 | 0.0677 | 0.0948 | 0.0684 |
| C | | | | | | | | 0.3228 | 0.3051 | 0.4472 | 0.4742 | 0.3642 |
| D | | | | | | | | 0.3076 | 0.3696 | 0.421 | 0.4281 | 0.3796 |
| E | | | | | | | | 0.1395 | 0.1195 | 0.1589 | 0.1378 | 0.0867 |
| F | | | | | | | | 0.4413 | 0.3882 | 0.4126 | 0.4337 | 0.358 |
| G | | | | | | | | 0.0465 | 0.0753 | 0.0752 | 0.0574 | 0.0677 |
| H | | | | | | | | 0.3397 | 0.2319 | 0.343 | 0.2938 | 0.2805 |

**Cell Index at: 123:03:07**

| | 1 | 2 | 3 | 4 | 5 | 6 | 7 | 8 | 9 | 10 | 11 | 12 |
|---|---|---|---|---|---|---|---|---|---|---|---|---|
| A | | | | | | | | 0.0531 | 0.0488 | 0.0687 | 0.0293 | 0.0483 |
| B | | | | | | | | 0.0482 | 0.0482 | 0.0676 | 0.0937 | 0.0728 |
| C | | | | | | | | 0.329 | 0.3066 | 0.446 | 0.487 | 0.3656 |
| D | | | | | | | | 0.3091 | 0.3624 | 0.4169 | 0.43 | 0.3798 |
| E | | | | | | | | 0.1477 | 0.114 | 0.1494 | 0.1428 | 0.0824 |
| F | | | | | | | | 0.4483 | 0.3834 | 0.4176 | 0.4382 | 0.3643 |
| G | | | | | | | | 0.0451 | 0.0788 | 0.0727 | 0.0533 | 0.0671 |
| H | | | | | | | | 0.3427 | 0.2319 | 0.3462 | 0.2931 | 0.2807 |

**Cell Index at: 123:18:07**

| | 1 | 2 | 3 | 4 | 5 | 6 | 7 | 8 | 9 | 10 | 11 | 12 |
|---|---|---|---|---|---|---|---|---|---|---|---|---|
| A | | | | | | | | 0.0597 | 0.0464 | 0.0708 | 0.0315 | 0.05 |
| B | | | | | | | | 0.0485 | 0.0475 | 0.0646 | 0.0955 | 0.0761 |
| C | | | | | | | | 0.3229 | 0.3018 | 0.4434 | 0.4867 | 0.3689 |
| D | | | | | | | | 0.3134 | 0.3598 | 0.412 | 0.4338 | 0.3824 |
| E | | | | | | | | 0.1411 | 0.1157 | 0.1459 | 0.1406 | 0.0765 |

| | | | | | | | | | | | | |
|---|---|---|---|---|---|---|---|---|---|---|---|---|
| F | | | | | | | | 0.4437 | 0.3908 | 0.4163 | 0.4399 | 0.3624 |
| G | | | | | | | | 0.0482 | 0.0753 | 0.0774 | 0.0529 | 0.0662 |
| H | | | | | | | | 0.3477 | 0.2338 | 0.3443 | 0.2903 | 0.2817 |

**Cell Index at: 123:33:08**

| | 1 | 2 | 3 | 4 | 5 | 6 | 7 | 8 | 9 | 10 | 11 | 12 |
|---|---|---|---|---|---|---|---|---|---|---|---|---|
| A | | | | | | | | 0.0601 | 0.0507 | 0.0709 | 0.0305 | 0.0519 |
| B | | | | | | | | 0.0486 | 0.0457 | 0.0661 | 0.097 | 0.0759 |
| C | | | | | | | | 0.3265 | 0.3057 | 0.4385 | 0.4827 | 0.3675 |
| D | | | | | | | | 0.3116 | 0.3605 | 0.419 | 0.4336 | 0.3752 |
| E | | | | | | | | 0.1351 | 0.1191 | 0.1503 | 0.1396 | 0.0838 |
| F | | | | | | | | 0.4468 | 0.3814 | 0.4247 | 0.4413 | 0.3646 |
| G | | | | | | | | 0.0548 | 0.0777 | 0.0787 | 0.052 | 0.0677 |
| H | | | | | | | | 0.3496 | 0.2409 | 0.3478 | 0.2888 | 0.281 |

**Cell Index at: 123:48:08**

| | 1 | 2 | 3 | 4 | 5 | 6 | 7 | 8 | 9 | 10 | 11 | 12 |
|---|---|---|---|---|---|---|---|---|---|---|---|---|
| A | | | | | | | | 0.0583 | 0.0497 | 0.0712 | 0.0301 | 0.0528 |
| B | | | | | | | | 0.0464 | 0.0511 | 0.0668 | 0.0922 | 0.0787 |
| C | | | | | | | | 0.3313 | 0.2987 | 0.4451 | 0.4751 | 0.3703 |
| D | | | | | | | | 0.3061 | 0.369 | 0.4191 | 0.4323 | 0.3736 |
| E | | | | | | | | 0.1347 | 0.1211 | 0.151 | 0.1443 | 0.0839 |
| F | | | | | | | | 0.4371 | 0.3887 | 0.4243 | 0.4444 | 0.364 |
| G | | | | | | | | 0.0495 | 0.0802 | 0.0758 | 0.0549 | 0.0701 |
| H | | | | | | | | 0.3482 | 0.2374 | 0.3488 | 0.2913 | 0.2802 |

**Cell Index at: 124:03:09**

| | 1 | 2 | 3 | 4 | 5 | 6 | 7 | 8 | 9 | 10 | 11 | 12 |
|---|---|---|---|---|---|---|---|---|---|---|---|---|
| A | | | | | | | | 0.0623 | 0.0472 | 0.0725 | 0.0309 | 0.0522 |
| B | | | | | | | | 0.0457 | 0.0474 | 0.0686 | 0.0945 | 0.0805 |
| C | | | | | | | | 0.3316 | 0.305 | 0.4482 | 0.4938 | 0.3678 |
| D | | | | | | | | 0.3138 | 0.3663 | 0.4296 | 0.4281 | 0.3756 |
| E | | | | | | | | 0.1403 | 0.1066 | 0.1431 | 0.1415 | 0.0845 |
| F | | | | | | | | 0.4443 | 0.3903 | 0.4201 | 0.4481 | 0.3697 |
| G | | | | | | | | 0.052 | 0.0789 | 0.0763 | 0.0538 | 0.0727 |
| H | | | | | | | | 0.3511 | 0.2398 | 0.3482 | 0.2935 | 0.2818 |

**Cell Index at: 124:18:09**

| | 1 | 2 | 3 | 4 | 5 | 6 | 7 | 8 | 9 | 10 | 11 | 12 |
|---|---|---|---|---|---|---|---|---|---|---|---|---|
| A | | | | | | | | 0.0635 | 0.0503 | 0.0749 | 0.0333 | 0.0532 |
| B | | | | | | | | 0.0489 | 0.0511 | 0.0674 | 0.0922 | 0.0744 |
| C | | | | | | | | 0.3345 | 0.302 | 0.449 | 0.4898 | 0.3738 |
| D | | | | | | | | 0.3116 | 0.3672 | 0.4266 | 0.4258 | 0.378 |
| E | | | | | | | | 0.1413 | 0.1199 | 0.1516 | 0.1433 | 0.0805 |
| F | | | | | | | | 0.4425 | 0.3828 | 0.4183 | 0.4485 | 0.3688 |
| G | | | | | | | | 0.0485 | 0.0786 | 0.0764 | 0.0569 | 0.0702 |
| H | | | | | | | | 0.3483 | 0.2393 | 0.3513 | 0.2912 | 0.2848 |

**Cell Index at: 124:33:08**

| | 1 | 2 | 3 | 4 | 5 | 6 | 7 | 8 | 9 | 10 | 11 | 12 |
|---|---|---|---|---|---|---|---|---|---|---|---|---|
| A | | | | | | | | 0.0664 | 0.0475 | 0.075 | 0.0314 | 0.0549 |
| B | | | | | | | | 0.0504 | 0.0535 | 0.0658 | 0.09 | 0.0774 |
| C | | | | | | | | 0.332 | 0.2909 | 0.4513 | 0.4873 | 0.3749 |
| D | | | | | | | | 0.3125 | 0.3669 | 0.4303 | 0.4297 | 0.3793 |
| E | | | | | | | | 0.1363 | 0.1188 | 0.1459 | 0.1329 | 0.0783 |
| F | | | | | | | | 0.4481 | 0.3909 | 0.4237 | 0.4466 | 0.3679 |
| G | | | | | | | | 0.0529 | 0.077 | 0.0732 | 0.0599 | 0.0714 |
| H | | | | | | | | 0.351 | 0.2394 | 0.3477 | 0.2907 | 0.285 |

**Cell Index at: 124:48:08**

| | 1 | 2 | 3 | 4 | 5 | 6 | 7 | 8 | 9 | 10 | 11 | 12 |
|---|---|---|---|---|---|---|---|---|---|---|---|---|
| A | | | | | | | | 0.0696 | 0.0476 | 0.0749 | 0.0292 | 0.055 |
| B | | | | | | | | 0.047 | 0.053 | 0.0699 | 0.0909 | 0.0779 |
| C | | | | | | | | 0.3324 | 0.2966 | 0.4489 | 0.4864 | 0.3729 |
| D | | | | | | | | 0.3141 | 0.3715 | 0.4233 | 0.421 | 0.3817 |
| E | | | | | | | | 0.1375 | 0.1201 | 0.1547 | 0.1315 | 0.0824 |
| F | | | | | | | | 0.4446 | 0.3867 | 0.4301 | 0.4458 | 0.3698 |
| G | | | | | | | | 0.0509 | 0.0741 | 0.0756 | 0.0636 | 0.0768 |
| H | | | | | | | | 0.3503 | 0.2419 | 0.3487 | 0.2924 | 0.2853 |

**Cell Index at: 125:03:08**

| | 1 | 2 | 3 | 4 | 5 | 6 | 7 | 8 | 9 | 10 | 11 | 12 |
|---|---|---|---|---|---|---|---|---|---|---|---|---|
| A | | | | | | | | 0.0712 | 0.0477 | 0.0789 | 0.0295 | 0.056 |
| B | | | | | | | | 0.0466 | 0.0553 | 0.0697 | 0.089 | 0.0807 |
| C | | | | | | | | 0.3354 | 0.3008 | 0.455 | 0.4903 | 0.3779 |
| D | | | | | | | | 0.3159 | 0.3766 | 0.4252 | 0.4241 | 0.3787 |
| E | | | | | | | | 0.1415 | 0.1255 | 0.1513 | 0.1337 | 0.0852 |
| F | | | | | | | | 0.4491 | 0.3905 | 0.4318 | 0.4547 | 0.3713 |
| G | | | | | | | | 0.05 | 0.0809 | 0.0769 | 0.0603 | 0.0732 |
| H | | | | | | | | 0.3512 | 0.242 | 0.3515 | 0.2905 | 0.2852 |

**Cell Index at: 125:18:09**

| | 1 | 2 | 3 | 4 | 5 | 6 | 7 | 8 | 9 | 10 | 11 | 12 |
|---|---|---|---|---|---|---|---|---|---|---|---|---|
| A | | | | | | | | 0.0731 | 0.0494 | 0.0795 | 0.0273 | 0.0549 |
| B | | | | | | | | 0.0477 | 0.0567 | 0.069 | 0.0947 | 0.0798 |
| C | | | | | | | | 0.3429 | 0.2991 | 0.4647 | 0.491 | 0.375 |
| D | | | | | | | | 0.3198 | 0.3764 | 0.4352 | 0.4302 | 0.377 |
| E | | | | | | | | 0.1356 | 0.1238 | 0.1539 | 0.1369 | 0.0839 |
| F | | | | | | | | 0.4481 | 0.3918 | 0.4338 | 0.4487 | 0.3714 |
| G | | | | | | | | 0.0483 | 0.0767 | 0.0765 | 0.0589 | 0.0758 |
| H | | | | | | | | 0.3533 | 0.2438 | 0.3525 | 0.294 | 0.2875 |

**Cell Index at: 125:33:10**

| | 1 | 2 | 3 | 4 | 5 | 6 | 7 | 8 | 9 | 10 | 11 | 12 |
|---|---|---|---|---|---|---|---|---|---|---|---|---|
| A | | | | | | | | 0.0719 | 0.0511 | 0.0782 | 0.028 | 0.058 |
| B | | | | | | | | 0.051 | 0.0517 | 0.0702 | 0.0965 | 0.0776 |
| C | | | | | | | | 0.3324 | 0.2926 | 0.4593 | 0.4895 | 0.373 |
| D | | | | | | | | 0.3088 | 0.3769 | 0.4305 | 0.4242 | 0.3731 |
| E | | | | | | | | 0.1411 | 0.1259 | 0.15 | 0.1334 | 0.0826 |
| F | | | | | | | | 0.4513 | 0.4028 | 0.4333 | 0.4419 | 0.3715 |
| G | | | | | | | | 0.0531 | 0.0766 | 0.0791 | 0.0551 | 0.072 |
| H | | | | | | | | 0.3541 | 0.2397 | 0.353 | 0.2923 | 0.29 |

**Cell Index at: 125:48:11**

| | 1 | 2 | 3 | 4 | 5 | 6 | 7 | 8 | 9 | 10 | 11 | 12 |
|---|---|---|---|---|---|---|---|---|---|---|---|---|
| A | | | | | | | | 0.0726 | 0.05 | 0.0784 | 0.0276 | 0.0581 |
| B | | | | | | | | 0.0519 | 0.0506 | 0.0628 | 0.0934 | 0.0796 |
| C | | | | | | | | 0.3328 | 0.296 | 0.4551 | 0.4888 | 0.3768 |
| D | | | | | | | | 0.3228 | 0.3699 | 0.4364 | 0.4293 | 0.3837 |
| E | | | | | | | | 0.1415 | 0.1251 | 0.1533 | 0.1279 | 0.0863 |
| F | | | | | | | | 0.447 | 0.4003 | 0.4336 | 0.4431 | 0.3716 |
| G | | | | | | | | 0.0522 | 0.0834 | 0.074 | 0.0597 | 0.0744 |
| H | | | | | | | | 0.3551 | 0.2423 | 0.3542 | 0.2914 | 0.2956 |

**Cell Index at: 126:03:12**

| | 1 | 2 | 3 | 4 | 5 | 6 | 7 | 8 | 9 | 10 | 11 | 12 |
|---|---|---|---|---|---|---|---|---|---|---|---|---|
| A | | | | | | | | 0.0725 | 0.0481 | 0.076 | 0.0297 | 0.0564 |
| B | | | | | | | | 0.0496 | 0.0508 | 0.0668 | 0.0923 | 0.0827 |
| C | | | | | | | | 0.3368 | 0.296 | 0.4587 | 0.4919 | 0.3812 |
| D | | | | | | | | 0.3192 | 0.3757 | 0.4403 | 0.427 | 0.3818 |
| E | | | | | | | | 0.1386 | 0.1312 | 0.1559 | 0.1428 | 0.0782 |
| F | | | | | | | | 0.447 | 0.3999 | 0.4282 | 0.4505 | 0.3702 |
| G | | | | | | | | 0.0529 | 0.082 | 0.0759 | 0.0611 | 0.0747 |
| H | | | | | | | | 0.3543 | 0.2412 | 0.3524 | 0.2942 | 0.2947 |

**Cell Index at: 126:18:13**

| | 1 | 2 | 3 | 4 | 5 | 6 | 7 | 8 | 9 | 10 | 11 | 12 |
|---|---|---|---|---|---|---|---|---|---|---|---|---|
| A | | | | | | | | 0.0741 | 0.0527 | 0.076 | 0.0311 | 0.0597 |
| B | | | | | | | | 0.0581 | 0.0514 | 0.0676 | 0.0931 | 0.0849 |
| C | | | | | | | | 0.3443 | 0.2981 | 0.4604 | 0.4909 | 0.3868 |
| D | | | | | | | | 0.3249 | 0.3745 | 0.4345 | 0.4188 | 0.3775 |

| | | | | | | | | | | | | |
|---|---|---|---|---|---|---|---|---|---|---|---|---|
| E | | | | | | | | 0.1461 | 0.1244 | 0.1499 | 0.1419 | 0.0803 |
| F | | | | | | | | 0.4568 | 0.3996 | 0.4362 | 0.45 | 0.3727 |
| G | | | | | | | | 0.0539 | 0.0822 | 0.0778 | 0.0594 | 0.0737 |
| H | | | | | | | | 0.3574 | 0.2394 | 0.3549 | 0.2938 | 0.2939 |

**Cell Index at: 126:33:14**

| | 1 | 2 | 3 | 4 | 5 | 6 | 7 | 8 | 9 | 10 | 11 | 12 |
|---|---|---|---|---|---|---|---|---|---|---|---|---|
| A | | | | | | | | 0.0727 | 0.0531 | 0.0758 | 0.0321 | 0.0582 |
| B | | | | | | | | 0.0522 | 0.0531 | 0.0658 | 0.0918 | 0.081 |
| C | | | | | | | | 0.3368 | 0.2944 | 0.4607 | 0.4896 | 0.3751 |
| D | | | | | | | | 0.3244 | 0.3619 | 0.4365 | 0.4299 | 0.385 |
| E | | | | | | | | 0.1381 | 0.1252 | 0.1596 | 0.1473 | 0.0814 |
| F | | | | | | | | 0.4454 | 0.3985 | 0.4335 | 0.4459 | 0.3775 |
| G | | | | | | | | 0.0547 | 0.0834 | 0.0763 | 0.0631 | 0.0759 |
| H | | | | | | | | 0.3574 | 0.2429 | 0.3592 | 0.2937 | 0.2972 |

**Cell Index at: 126:48:14**

| | 1 | 2 | 3 | 4 | 5 | 6 | 7 | 8 | 9 | 10 | 11 | 12 |
|---|---|---|---|---|---|---|---|---|---|---|---|---|
| A | | | | | | | | 0.0711 | 0.0504 | 0.081 | 0.033 | 0.0582 |
| B | | | | | | | | 0.0536 | 0.0543 | 0.0684 | 0.0909 | 0.0801 |
| C | | | | | | | | 0.3374 | 0.3025 | 0.459 | 0.4897 | 0.3842 |
| D | | | | | | | | 0.3346 | 0.3786 | 0.4395 | 0.4305 | 0.3837 |
| E | | | | | | | | 0.144 | 0.1234 | 0.1523 | 0.1393 | 0.0788 |
| F | | | | | | | | 0.4531 | 0.3972 | 0.4311 | 0.4459 | 0.3807 |
| G | | | | | | | | 0.0566 | 0.0874 | 0.0749 | 0.0631 | 0.0753 |
| H | | | | | | | | 0.3593 | 0.2418 | 0.3591 | 0.2947 | 0.2992 |

**Cell Index at: 127:03:15**

| | 1 | 2 | 3 | 4 | 5 | 6 | 7 | 8 | 9 | 10 | 11 | 12 |
|---|---|---|---|---|---|---|---|---|---|---|---|---|
| A | | | | | | | | 0.0727 | 0.053 | 0.0843 | 0.0337 | 0.0563 |
| B | | | | | | | | 0.0535 | 0.0537 | 0.067 | 0.0867 | 0.0837 |
| C | | | | | | | | 0.3402 | 0.2985 | 0.4635 | 0.4901 | 0.3845 |
| D | | | | | | | | 0.3357 | 0.3737 | 0.4381 | 0.4343 | 0.3847 |
| E | | | | | | | | 0.1437 | 0.116 | 0.1558 | 0.1417 | 0.0892 |
| F | | | | | | | | 0.4499 | 0.3987 | 0.4335 | 0.444 | 0.3796 |
| G | | | | | | | | 0.0566 | 0.0888 | 0.0738 | 0.0665 | 0.0734 |
| H | | | | | | | | 0.3626 | 0.2462 | 0.3611 | 0.2943 | 0.3006 |

**Cell Index at: 127:18:15**

| | 1 | 2 | 3 | 4 | 5 | 6 | 7 | 8 | 9 | 10 | 11 | 12 |
|---|---|---|---|---|---|---|---|---|---|---|---|---|
| A | | | | | | | | 0.0726 | 0.0537 | 0.0825 | 0.0332 | 0.0531 |
| B | | | | | | | | 0.0514 | 0.0512 | 0.0667 | 0.0957 | 0.0803 |
| C | | | | | | | | 0.3372 | 0.2965 | 0.4617 | 0.4883 | 0.389 |
| D | | | | | | | | 0.3254 | 0.3783 | 0.4302 | 0.4278 | 0.3779 |
| E | | | | | | | | 0.1362 | 0.1195 | 0.158 | 0.1406 | 0.0833 |
| F | | | | | | | | 0.452 | 0.3988 | 0.4279 | 0.4436 | 0.3785 |
| G | | | | | | | | 0.0542 | 0.0882 | 0.078 | 0.0635 | 0.0744 |
| H | | | | | | | | 0.357 | 0.246 | 0.363 | 0.2964 | 0.3017 |

**Cell Index at: 127:33:16**

| | 1 | 2 | 3 | 4 | 5 | 6 | 7 | 8 | 9 | 10 | 11 | 12 |
|---|---|---|---|---|---|---|---|---|---|---|---|---|
| A | | | | | | | | 0.0762 | 0.053 | 0.0853 | 0.0372 | 0.0531 |
| B | | | | | | | | 0.0506 | 0.0563 | 0.068 | 0.0938 | 0.087 |
| C | | | | | | | | 0.3437 | 0.3048 | 0.4609 | 0.4896 | 0.3843 |
| D | | | | | | | | 0.327 | 0.378 | 0.4369 | 0.4285 | 0.3834 |
| E | | | | | | | | 0.1478 | 0.1122 | 0.1542 | 0.1351 | 0.0858 |
| F | | | | | | | | 0.4557 | 0.4024 | 0.4283 | 0.4425 | 0.3833 |
| G | | | | | | | | 0.0531 | 0.0862 | 0.0777 | 0.0612 | 0.0738 |
| H | | | | | | | | 0.3564 | 0.2459 | 0.3648 | 0.2975 | 0.2995 |

**Cell Index at: 127:48:16**

| | 1 | 2 | 3 | 4 | 5 | 6 | 7 | 8 | 9 | 10 | 11 | 12 |
|---|---|---|---|---|---|---|---|---|---|---|---|---|
| A | | | | | | | | 0.0742 | 0.0518 | 0.0852 | 0.0308 | 0.0548 |
| B | | | | | | | | 0.052 | 0.0559 | 0.0665 | 0.0926 | 0.0833 |
| C | | | | | | | | 0.3403 | 0.2974 | 0.4581 | 0.487 | 0.3799 |
| D | | | | | | | | 0.3234 | 0.3756 | 0.4318 | 0.4339 | 0.3769 |
| E | | | | | | | | 0.1516 | 0.105 | 0.1599 | 0.1316 | 0.0877 |
| F | | | | | | | | 0.4483 | 0.4024 | 0.4354 | 0.4447 | 0.381 |
| G | | | | | | | | 0.0547 | 0.0881 | 0.0832 | 0.0648 | 0.0774 |
| H | | | | | | | | 0.3603 | 0.2472 | 0.3639 | 0.2994 | 0.2994 |

**Cell Index at: 128:03:17**

| | 1 | 2 | 3 | 4 | 5 | 6 | 7 | 8 | 9 | 10 | 11 | 12 |
|---|---|---|---|---|---|---|---|---|---|---|---|---|
| A | | | | | | | | 0.0732 | 0.0507 | 0.0837 | 0.0332 | 0.0539 |
| B | | | | | | | | 0.0532 | 0.0587 | 0.0682 | 0.0963 | 0.0786 |
| C | | | | | | | | 0.3433 | 0.2986 | 0.4641 | 0.4854 | 0.3852 |
| D | | | | | | | | 0.332 | 0.3841 | 0.4382 | 0.4313 | 0.3735 |
| E | | | | | | | | 0.1401 | 0.1024 | 0.163 | 0.1411 | 0.0834 |
| F | | | | | | | | 0.4494 | 0.4049 | 0.436 | 0.4438 | 0.3898 |
| G | | | | | | | | 0.0553 | 0.0843 | 0.0809 | 0.0679 | 0.0723 |
| H | | | | | | | | 0.3605 | 0.2454 | 0.3632 | 0.3 | 0.2976 |

**Cell Index at: 128:18:17**

| | 1 | 2 | 3 | 4 | 5 | 6 | 7 | 8 | 9 | 10 | 11 | 12 |
|---|---|---|---|---|---|---|---|---|---|---|---|---|
| A | | | | | | | | 0.0726 | 0.0511 | 0.0836 | 0.0354 | 0.0566 |
| B | | | | | | | | 0.0504 | 0.0573 | 0.0697 | 0.094 | 0.0801 |
| C | | | | | | | | 0.3382 | 0.2982 | 0.4642 | 0.4908 | 0.3851 |
| D | | | | | | | | 0.326 | 0.3788 | 0.4303 | 0.429 | 0.3888 |
| E | | | | | | | | 0.1441 | 0.1123 | 0.1613 | 0.1406 | 0.0865 |
| F | | | | | | | | 0.4535 | 0.403 | 0.4327 | 0.4499 | 0.3783 |
| G | | | | | | | | 0.052 | 0.0818 | 0.082 | 0.0678 | 0.0726 |
| H | | | | | | | | 0.363 | 0.2463 | 0.3612 | 0.2988 | 0.3002 |

**Cell Index at: 128:33:17**

| | 1 | 2 | 3 | 4 | 5 | 6 | 7 | 8 | 9 | 10 | 11 | 12 |
|---|---|---|---|---|---|---|---|---|---|---|---|---|
| A | | | | | | | | 0.0746 | 0.0526 | 0.0812 | 0.0344 | 0.0569 |
| B | | | | | | | | 0.0525 | 0.0575 | 0.0739 | 0.0938 | 0.0783 |
| C | | | | | | | | 0.3531 | 0.2999 | 0.4693 | 0.4938 | 0.3936 |
| D | | | | | | | | 0.3235 | 0.3818 | 0.4332 | 0.4229 | 0.3852 |
| E | | | | | | | | 0.1438 | 0.1107 | 0.1645 | 0.1413 | 0.0883 |
| F | | | | | | | | 0.4549 | 0.403 | 0.4329 | 0.4542 | 0.3836 |
| G | | | | | | | | 0.054 | 0.0889 | 0.0856 | 0.0716 | 0.0791 |
| H | | | | | | | | 0.3612 | 0.2473 | 0.3589 | 0.3018 | 0.2992 |

**Cell Index at: 128:48:18**

| | 1 | 2 | 3 | 4 | 5 | 6 | 7 | 8 | 9 | 10 | 11 | 12 |
|---|---|---|---|---|---|---|---|---|---|---|---|---|
| A | | | | | | | | 0.0749 | 0.0523 | 0.0836 | 0.0349 | 0.0555 |
| B | | | | | | | | 0.0494 | 0.0579 | 0.0744 | 0.0948 | 0.08 |
| C | | | | | | | | 0.3533 | 0.2997 | 0.4642 | 0.4945 | 0.3846 |
| D | | | | | | | | 0.3169 | 0.3687 | 0.4388 | 0.4247 | 0.3838 |
| E | | | | | | | | 0.1438 | 0.1118 | 0.1632 | 0.1419 | 0.0846 |
| F | | | | | | | | 0.4576 | 0.4098 | 0.4357 | 0.4465 | 0.3842 |
| G | | | | | | | | 0.0548 | 0.0873 | 0.0822 | 0.0691 | 0.0802 |
| H | | | | | | | | 0.3624 | 0.2496 | 0.3658 | 0.3035 | 0.2998 |

**Cell Index at: 129:03:19**

| | 1 | 2 | 3 | 4 | 5 | 6 | 7 | 8 | 9 | 10 | 11 | 12 |
|---|---|---|---|---|---|---|---|---|---|---|---|---|
| A | | | | | | | | 0.0744 | 0.0519 | 0.0863 | 0.0381 | 0.0584 |
| B | | | | | | | | 0.0471 | 0.0558 | 0.0709 | 0.0982 | 0.0844 |
| C | | | | | | | | 0.3542 | 0.2971 | 0.4647 | 0.4878 | 0.3877 |
| D | | | | | | | | 0.3318 | 0.3768 | 0.443 | 0.4296 | 0.3936 |
| E | | | | | | | | 0.1432 | 0.1167 | 0.1683 | 0.1423 | 0.089 |
| F | | | | | | | | 0.4533 | 0.396 | 0.4344 | 0.4609 | 0.3917 |
| G | | | | | | | | 0.0492 | 0.0915 | 0.0892 | 0.0697 | 0.0791 |
| H | | | | | | | | 0.3629 | 0.2481 | 0.3599 | 0.296 | 0.3043 |

**Cell Index at: 129:18:20**

| | 1 | 2 | 3 | 4 | 5 | 6 | 7 | 8 | 9 | 10 | 11 | 12 |
|---|---|---|---|---|---|---|---|---|---|---|---|---|
| A | | | | | | | | 0.0722 | 0.0494 | 0.0855 | 0.0301 | 0.0575 |
| B | | | | | | | | 0.0492 | 0.0542 | 0.0698 | 0.096 | 0.0819 |
| C | | | | | | | | 0.3505 | 0.3135 | 0.4727 | 0.4801 | 0.3915 |

| | | | | | | | | | | | | |
|---|---|---|---|---|---|---|---|---|---|---|---|---|
| D | | | | | | | | 0.3224 | 0.3812 | 0.4385 | 0.4258 | 0.392 |
| E | | | | | | | | 0.1481 | 0.1121 | 0.1608 | 0.1365 | 0.0829 |
| F | | | | | | | | 0.4521 | 0.4024 | 0.4309 | 0.4513 | 0.3829 |
| G | | | | | | | | 0.0544 | 0.0891 | 0.0844 | 0.072 | 0.0807 |
| H | | | | | | | | 0.3607 | 0.2469 | 0.3662 | 0.3016 | 0.3017 |

**Cell Index at: 129:33:20**

| | 1 | 2 | 3 | 4 | 5 | 6 | 7 | 8 | 9 | 10 | 11 | 12 |
|---|---|---|---|---|---|---|---|---|---|---|---|---|
| A | | | | | | | | 0.0737 | 0.0552 | 0.0831 | 0.0393 | 0.0601 |
| B | | | | | | | | 0.0524 | 0.0579 | 0.0706 | 0.0957 | 0.0859 |
| C | | | | | | | | 0.3616 | 0.2995 | 0.4666 | 0.4961 | 0.3885 |
| D | | | | | | | | 0.3244 | 0.378 | 0.4397 | 0.4337 | 0.3807 |
| E | | | | | | | | 0.1497 | 0.1107 | 0.1664 | 0.1385 | 0.0826 |
| F | | | | | | | | 0.4559 | 0.4079 | 0.4417 | 0.4556 | 0.389 |
| G | | | | | | | | 0.0533 | 0.0866 | 0.0854 | 0.0701 | 0.0798 |
| H | | | | | | | | 0.3651 | 0.2515 | 0.3659 | 0.3022 | 0.3068 |

**Cell Index at: 129:48:21**

| | 1 | 2 | 3 | 4 | 5 | 6 | 7 | 8 | 9 | 10 | 11 | 12 |
|---|---|---|---|---|---|---|---|---|---|---|---|---|
| A | | | | | | | | 0.0721 | 0.0533 | 0.0818 | 0.0344 | 0.0615 |
| B | | | | | | | | 0.0485 | 0.0592 | 0.0748 | 0.0958 | 0.0813 |
| C | | | | | | | | 0.3547 | 0.3093 | 0.475 | 0.499 | 0.3865 |
| D | | | | | | | | 0.331 | 0.3842 | 0.4394 | 0.4222 | 0.3823 |
| E | | | | | | | | 0.137 | 0.1086 | 0.1657 | 0.135 | 0.0879 |
| F | | | | | | | | 0.4538 | 0.413 | 0.4432 | 0.4572 | 0.3969 |
| G | | | | | | | | 0.0531 | 0.0892 | 0.0848 | 0.0689 | 0.0796 |
| H | | | | | | | | 0.3659 | 0.2491 | 0.3682 | 0.2997 | 0.3029 |

**Cell Index at: 130:03:21**

| | 1 | 2 | 3 | 4 | 5 | 6 | 7 | 8 | 9 | 10 | 11 | 12 |
|---|---|---|---|---|---|---|---|---|---|---|---|---|
| A | | | | | | | | 0.0739 | 0.0538 | 0.0846 | 0.0352 | 0.0614 |
| B | | | | | | | | 0.053 | 0.058 | 0.0754 | 0.0964 | 0.0842 |
| C | | | | | | | | 0.3531 | 0.3059 | 0.4786 | 0.509 | 0.3932 |
| D | | | | | | | | 0.3311 | 0.3897 | 0.4452 | 0.4308 | 0.3867 |
| E | | | | | | | | 0.138 | 0.1132 | 0.1656 | 0.1369 | 0.0816 |
| F | | | | | | | | 0.4563 | 0.4092 | 0.4393 | 0.4578 | 0.3857 |
| G | | | | | | | | 0.0535 | 0.0925 | 0.0858 | 0.0672 | 0.0802 |
| H | | | | | | | | 0.367 | 0.2532 | 0.3686 | 0.3032 | 0.3106 |

**Cell Index at: 130:18:22**

| | 1 | 2 | 3 | 4 | 5 | 6 | 7 | 8 | 9 | 10 | 11 | 12 |
|---|---|---|---|---|---|---|---|---|---|---|---|---|
| A | | | | | | | | 0.0759 | 0.0558 | 0.0886 | 0.0368 | 0.0663 |
| B | | | | | | | | 0.0542 | 0.0575 | 0.0764 | 0.0998 | 0.0835 |
| C | | | | | | | | 0.3551 | 0.303 | 0.4736 | 0.5063 | 0.3974 |
| D | | | | | | | | 0.3345 | 0.3835 | 0.4388 | 0.4324 | 0.3854 |
| E | | | | | | | | 0.137 | 0.1146 | 0.1609 | 0.1369 | 0.0876 |
| F | | | | | | | | 0.4566 | 0.4081 | 0.4405 | 0.4563 | 0.3879 |
| G | | | | | | | | 0.0548 | 0.0914 | 0.087 | 0.0678 | 0.0776 |
| H | | | | | | | | 0.371 | 0.2528 | 0.3669 | 0.3006 | 0.3096 |

**Cell Index at: 130:33:22**

| | 1 | 2 | 3 | 4 | 5 | 6 | 7 | 8 | 9 | 10 | 11 | 12 |
|---|---|---|---|---|---|---|---|---|---|---|---|---|
| A | | | | | | | | 0.0764 | 0.0573 | 0.0894 | 0.035 | 0.0634 |
| B | | | | | | | | 0.0508 | 0.0558 | 0.0766 | 0.0994 | 0.0873 |
| C | | | | | | | | 0.3511 | 0.3046 | 0.4713 | 0.506 | 0.3978 |
| D | | | | | | | | 0.3368 | 0.3855 | 0.4431 | 0.4368 | 0.3867 |
| E | | | | | | | | 0.1444 | 0.1088 | 0.1693 | 0.1351 | 0.0901 |
| F | | | | | | | | 0.4578 | 0.4088 | 0.4419 | 0.4589 | 0.3857 |
| G | | | | | | | | 0.0576 | 0.0918 | 0.0843 | 0.0665 | 0.08 |
| H | | | | | | | | 0.3695 | 0.2523 | 0.3676 | 0.3022 | 0.3105 |

**Cell Index at: 130:48:22**

| | 1 | 2 | 3 | 4 | 5 | 6 | 7 | 8 | 9 | 10 | 11 | 12 |
|---|---|---|---|---|---|---|---|---|---|---|---|---|
| A | | | | | | | | 0.0765 | 0.0563 | 0.088 | 0.0357 | 0.0621 |
| B | | | | | | | | 0.0562 | 0.0568 | 0.0752 | 0.0984 | 0.0825 |
| C | | | | | | | | 0.357 | 0.3051 | 0.4749 | 0.5039 | 0.3903 |
| D | | | | | | | | 0.3395 | 0.3818 | 0.4425 | 0.4302 | 0.3928 |
| E | | | | | | | | 0.1401 | 0.1088 | 0.1652 | 0.137 | 0.0853 |
| F | | | | | | | | 0.4548 | 0.4137 | 0.4399 | 0.4643 | 0.3888 |
| G | | | | | | | | 0.0508 | 0.0876 | 0.0844 | 0.0676 | 0.0767 |
| H | | | | | | | | 0.3685 | 0.2541 | 0.3672 | 0.3041 | 0.3145 |

**Cell Index at: 131:03:23**

| | 1 | 2 | 3 | 4 | 5 | 6 | 7 | 8 | 9 | 10 | 11 | 12 |
|---|---|---|---|---|---|---|---|---|---|---|---|---|
| A | | | | | | | | 0.08 | 0.0608 | 0.0853 | 0.0372 | 0.0618 |
| B | | | | | | | | 0.0543 | 0.0605 | 0.0762 | 0.0988 | 0.0841 |
| C | | | | | | | | 0.3589 | 0.301 | 0.4673 | 0.5058 | 0.399 |
| D | | | | | | | | 0.3371 | 0.3891 | 0.4437 | 0.4359 | 0.3902 |
| E | | | | | | | | 0.1448 | 0.108 | 0.1681 | 0.137 | 0.0836 |
| F | | | | | | | | 0.4543 | 0.419 | 0.4397 | 0.4616 | 0.3944 |
| G | | | | | | | | 0.0562 | 0.0872 | 0.0869 | 0.0684 | 0.0804 |
| H | | | | | | | | 0.3668 | 0.2591 | 0.3679 | 0.3064 | 0.3146 |

**Cell Index at: 131:18:23**

| | 1 | 2 | 3 | 4 | 5 | 6 | 7 | 8 | 9 | 10 | 11 | 12 |
|---|---|---|---|---|---|---|---|---|---|---|---|---|
| A | | | | | | | | 0.0776 | 0.0607 | 0.088 | 0.0403 | 0.0616 |
| B | | | | | | | | 0.0561 | 0.056 | 0.0734 | 0.0995 | 0.0876 |
| C | | | | | | | | 0.3624 | 0.3033 | 0.4741 | 0.5068 | 0.4022 |
| D | | | | | | | | 0.3299 | 0.3925 | 0.4449 | 0.4408 | 0.3922 |
| E | | | | | | | | 0.1365 | 0.1093 | 0.1698 | 0.1393 | 0.0805 |
| F | | | | | | | | 0.4557 | 0.4252 | 0.4483 | 0.4664 | 0.3993 |
| G | | | | | | | | 0.0535 | 0.0895 | 0.0909 | 0.0729 | 0.0804 |
| H | | | | | | | | 0.3705 | 0.2581 | 0.3654 | 0.3059 | 0.3132 |

**Cell Index at: 131:33:23**

| | 1 | 2 | 3 | 4 | 5 | 6 | 7 | 8 | 9 | 10 | 11 | 12 |
|---|---|---|---|---|---|---|---|---|---|---|---|---|
| A | | | | | | | | 0.0726 | 0.062 | 0.0861 | 0.0385 | 0.0636 |
| B | | | | | | | | 0.0602 | 0.0646 | 0.0742 | 0.0987 | 0.0828 |
| C | | | | | | | | 0.3607 | 0.3067 | 0.4818 | 0.5111 | 0.3981 |
| D | | | | | | | | 0.3344 | 0.3865 | 0.4459 | 0.4311 | 0.3851 |
| E | | | | | | | | 0.1392 | 0.1132 | 0.1624 | 0.1498 | 0.0865 |
| F | | | | | | | | 0.4582 | 0.4223 | 0.4469 | 0.4696 | 0.3961 |
| G | | | | | | | | 0.0514 | 0.0917 | 0.0818 | 0.066 | 0.0809 |
| H | | | | | | | | 0.3708 | 0.2612 | 0.3629 | 0.3076 | 0.317 |

**Cell Index at: 131:48:23**

| | 1 | 2 | 3 | 4 | 5 | 6 | 7 | 8 | 9 | 10 | 11 | 12 |
|---|---|---|---|---|---|---|---|---|---|---|---|---|
| A | | | | | | | | 0.078 | 0.0615 | 0.0845 | 0.0393 | 0.0594 |
| B | | | | | | | | 0.0594 | 0.06 | 0.08 | 0.1018 | 0.0852 |
| C | | | | | | | | 0.3545 | 0.305 | 0.4847 | 0.509 | 0.3968 |
| D | | | | | | | | 0.3381 | 0.3886 | 0.4392 | 0.4389 | 0.3841 |
| E | | | | | | | | 0.1422 | 0.1112 | 0.1606 | 0.1437 | 0.0857 |
| F | | | | | | | | 0.4554 | 0.4176 | 0.4523 | 0.4762 | 0.4007 |
| G | | | | | | | | 0.0541 | 0.0951 | 0.0843 | 0.0687 | 0.0833 |
| H | | | | | | | | 0.3735 | 0.264 | 0.3622 | 0.3094 | 0.3189 |

**Cell Index at: 132:03:24**

| | 1 | 2 | 3 | 4 | 5 | 6 | 7 | 8 | 9 | 10 | 11 | 12 |
|---|---|---|---|---|---|---|---|---|---|---|---|---|
| A | | | | | | | | 0.0747 | 0.0651 | 0.0853 | 0.0378 | 0.0623 |
| B | | | | | | | | 0.0583 | 0.0635 | 0.0782 | 0.0965 | 0.0849 |
| C | | | | | | | | 0.3527 | 0.3055 | 0.484 | 0.5158 | 0.4057 |
| D | | | | | | | | 0.3371 | 0.3956 | 0.4391 | 0.4387 | 0.3942 |
| E | | | | | | | | 0.1491 | 0.1203 | 0.1718 | 0.1451 | 0.0796 |
| F | | | | | | | | 0.4571 | 0.4154 | 0.454 | 0.4702 | 0.4012 |
| G | | | | | | | | 0.053 | 0.0889 | 0.0808 | 0.068 | 0.0838 |
| H | | | | | | | | 0.3729 | 0.2651 | 0.3635 | 0.3086 | 0.3203 |

**Cell Index at: 132:18:24**

| | 1 | 2 | 3 | 4 | 5 | 6 | 7 | 8 | 9 | 10 | 11 | 12 |
|---|---|---|---|---|---|---|---|---|---|---|---|---|
| A | | | | | | | | 0.078 | 0.065 | 0.089 | 0.0356 | 0.0625 |
| B | | | | | | | | 0.0601 | 0.0634 | 0.0793 | 0.0981 | 0.0849 |

| | | | | | | | | | | | | |
|---|---|---|---|---|---|---|---|---|---|---|---|---|
| C | | | | | | | | 0.358 | 0.308 | 0.4896 | 0.511 | 0.4004 |
| D | | | | | | | | 0.3308 | 0.3805 | 0.4439 | 0.4395 | 0.4042 |
| E | | | | | | | | 0.153 | 0.1213 | 0.1666 | 0.1432 | 0.0895 |
| F | | | | | | | | 0.4591 | 0.4216 | 0.4547 | 0.4757 | 0.3989 |
| G | | | | | | | | 0.0485 | 0.0874 | 0.0833 | 0.0694 | 0.0827 |
| H | | | | | | | | 0.3708 | 0.2631 | 0.3626 | 0.3128 | 0.3202 |

**Cell Index at: 132:33:24**

| | 1 | 2 | 3 | 4 | 5 | 6 | 7 | 8 | 9 | 10 | 11 | 12 |
|---|---|---|---|---|---|---|---|---|---|---|---|---|
| A | | | | | | | | 0.0739 | 0.0644 | 0.09 | 0.034 | 0.0657 |
| B | | | | | | | | 0.0648 | 0.0605 | 0.077 | 0.1004 | 0.0868 |
| C | | | | | | | | 0.3625 | 0.3078 | 0.4936 | 0.5168 | 0.4069 |
| D | | | | | | | | 0.3429 | 0.3818 | 0.4501 | 0.4472 | 0.395 |
| E | | | | | | | | 0.1398 | 0.1167 | 0.1689 | 0.1518 | 0.0864 |
| F | | | | | | | | 0.4641 | 0.4288 | 0.4478 | 0.4729 | 0.4032 |
| G | | | | | | | | 0.0543 | 0.0963 | 0.0816 | 0.0719 | 0.0781 |
| H | | | | | | | | 0.3747 | 0.2636 | 0.3615 | 0.3125 | 0.324 |

**Cell Index at: 132:48:24**

| | 1 | 2 | 3 | 4 | 5 | 6 | 7 | 8 | 9 | 10 | 11 | 12 |
|---|---|---|---|---|---|---|---|---|---|---|---|---|
| A | | | | | | | | 0.0751 | 0.0652 | 0.0876 | 0.0394 | 0.0662 |
| B | | | | | | | | 0.0639 | 0.057 | 0.0739 | 0.103 | 0.0855 |
| C | | | | | | | | 0.3607 | 0.3079 | 0.4922 | 0.5039 | 0.4017 |
| D | | | | | | | | 0.3451 | 0.3849 | 0.4475 | 0.4479 | 0.3905 |
| E | | | | | | | | 0.1491 | 0.1117 | 0.1696 | 0.1457 | 0.0775 |
| F | | | | | | | | 0.4624 | 0.4239 | 0.4515 | 0.4788 | 0.4023 |
| G | | | | | | | | 0.053 | 0.0942 | 0.0819 | 0.068 | 0.0814 |
| H | | | | | | | | 0.372 | 0.2657 | 0.3616 | 0.315 | 0.3244 |

**Cell Index at: 133:03:25**

| | 1 | 2 | 3 | 4 | 5 | 6 | 7 | 8 | 9 | 10 | 11 | 12 |
|---|---|---|---|---|---|---|---|---|---|---|---|---|
| A | | | | | | | | 0.0772 | 0.064 | 0.0844 | 0.0409 | 0.0657 |
| B | | | | | | | | 0.0608 | 0.0605 | 0.0793 | 0.0985 | 0.089 |
| C | | | | | | | | 0.3575 | 0.3116 | 0.4887 | 0.5103 | 0.4031 |
| D | | | | | | | | 0.3404 | 0.3853 | 0.4492 | 0.4455 | 0.392 |
| E | | | | | | | | 0.1463 | 0.1102 | 0.172 | 0.1512 | 0.0873 |
| F | | | | | | | | 0.46 | 0.4226 | 0.4535 | 0.4791 | 0.4019 |
| G | | | | | | | | 0.0543 | 0.095 | 0.0876 | 0.0693 | 0.0812 |
| H | | | | | | | | 0.3751 | 0.2673 | 0.3597 | 0.3123 | 0.3248 |

**Cell Index at: 133:18:25**

| | 1 | 2 | 3 | 4 | 5 | 6 | 7 | 8 | 9 | 10 | 11 | 12 |
|---|---|---|---|---|---|---|---|---|---|---|---|---|
| A | | | | | | | | 0.0742 | 0.0645 | 0.0872 | 0.0404 | 0.0703 |
| B | | | | | | | | 0.0613 | 0.0597 | 0.0713 | 0.1023 | 0.0866 |
| C | | | | | | | | 0.3672 | 0.3182 | 0.4969 | 0.5144 | 0.4075 |
| D | | | | | | | | 0.3498 | 0.3861 | 0.442 | 0.4566 | 0.3985 |
| E | | | | | | | | 0.1535 | 0.112 | 0.1749 | 0.1474 | 0.0846 |
| F | | | | | | | | 0.4673 | 0.4262 | 0.4565 | 0.4825 | 0.4036 |
| G | | | | | | | | 0.0542 | 0.0913 | 0.0842 | 0.0675 | 0.078 |
| H | | | | | | | | 0.3754 | 0.2653 | 0.358 | 0.3088 | 0.3269 |

**Cell Index at: 133:33:26**

| | 1 | 2 | 3 | 4 | 5 | 6 | 7 | 8 | 9 | 10 | 11 | 12 |
|---|---|---|---|---|---|---|---|---|---|---|---|---|
| A | | | | | | | | 0.0742 | 0.0672 | 0.0893 | 0.0403 | 0.068 |
| B | | | | | | | | 0.061 | 0.0559 | 0.0752 | 0.0966 | 0.0885 |
| C | | | | | | | | 0.3639 | 0.3141 | 0.4943 | 0.5212 | 0.4036 |
| D | | | | | | | | 0.341 | 0.3869 | 0.4402 | 0.4378 | 0.3985 |
| E | | | | | | | | 0.1451 | 0.1155 | 0.1739 | 0.1494 | 0.0782 |
| F | | | | | | | | 0.4578 | 0.4301 | 0.4579 | 0.4883 | 0.4013 |
| G | | | | | | | | 0.0566 | 0.0929 | 0.0848 | 0.0667 | 0.082 |
| H | | | | | | | | 0.3761 | 0.2682 | 0.3588 | 0.3097 | 0.3284 |

**Cell Index at: 133:48:27**

| | 1 | 2 | 3 | 4 | 5 | 6 | 7 | 8 | 9 | 10 | 11 | 12 |
|---|---|---|---|---|---|---|---|---|---|---|---|---|
| A | | | | | | | | 0.0738 | 0.0659 | 0.0882 | 0.0401 | 0.0703 |
| B | | | | | | | | 0.0654 | 0.0613 | 0.0739 | 0.0995 | 0.0925 |
| C | | | | | | | | 0.3709 | 0.3093 | 0.491 | 0.5159 | 0.4068 |
| D | | | | | | | | 0.3504 | 0.3826 | 0.4488 | 0.4534 | 0.4058 |
| E | | | | | | | | 0.154 | 0.1215 | 0.1742 | 0.145 | 0.0887 |
| F | | | | | | | | 0.4678 | 0.4273 | 0.4514 | 0.4866 | 0.4011 |
| G | | | | | | | | 0.054 | 0.0968 | 0.0879 | 0.0661 | 0.0825 |
| H | | | | | | | | 0.3763 | 0.2695 | 0.3602 | 0.3101 | 0.3294 |

**Cell Index at: 134:03:27**

| | 1 | 2 | 3 | 4 | 5 | 6 | 7 | 8 | 9 | 10 | 11 | 12 |
|---|---|---|---|---|---|---|---|---|---|---|---|---|
| A | | | | | | | | 0.0794 | 0.0647 | 0.0925 | 0.0409 | 0.0729 |
| B | | | | | | | | 0.0644 | 0.0599 | 0.08 | 0.0994 | 0.0878 |
| C | | | | | | | | 0.3667 | 0.3141 | 0.4978 | 0.5142 | 0.4081 |
| D | | | | | | | | 0.346 | 0.3915 | 0.4474 | 0.4476 | 0.4061 |
| E | | | | | | | | 0.143 | 0.1127 | 0.1735 | 0.1403 | 0.0781 |
| F | | | | | | | | 0.4681 | 0.4367 | 0.4619 | 0.485 | 0.4008 |
| G | | | | | | | | 0.054 | 0.0962 | 0.0885 | 0.0645 | 0.0809 |
| H | | | | | | | | 0.3805 | 0.2678 | 0.3617 | 0.3131 | 0.3267 |

**Cell Index at: 134:18:27**

| | 1 | 2 | 3 | 4 | 5 | 6 | 7 | 8 | 9 | 10 | 11 | 12 |
|---|---|---|---|---|---|---|---|---|---|---|---|---|
| A | | | | | | | | 0.0825 | 0.063 | 0.0947 | 0.0432 | 0.071 |
| B | | | | | | | | 0.0647 | 0.059 | 0.0799 | 0.1033 | 0.0863 |
| C | | | | | | | | 0.3653 | 0.3153 | 0.5024 | 0.519 | 0.4107 |
| D | | | | | | | | 0.3555 | 0.3908 | 0.46 | 0.4504 | 0.4059 |
| E | | | | | | | | 0.1528 | 0.1159 | 0.1741 | 0.15 | 0.0797 |
| F | | | | | | | | 0.4688 | 0.4265 | 0.4628 | 0.4887 | 0.3941 |
| G | | | | | | | | 0.0496 | 0.0968 | 0.0889 | 0.0635 | 0.0828 |
| H | | | | | | | | 0.3786 | 0.2739 | 0.3648 | 0.3125 | 0.3263 |

**Cell Index at: 134:33:27**

| | 1 | 2 | 3 | 4 | 5 | 6 | 7 | 8 | 9 | 10 | 11 | 12 |
|---|---|---|---|---|---|---|---|---|---|---|---|---|
| A | | | | | | | | 0.0787 | 0.0619 | 0.0915 | 0.0374 | 0.0746 |
| B | | | | | | | | 0.0623 | 0.0645 | 0.0781 | 0.0999 | 0.0872 |
| C | | | | | | | | 0.3623 | 0.3174 | 0.503 | 0.5153 | 0.4126 |
| D | | | | | | | | 0.3535 | 0.3871 | 0.4598 | 0.4537 | 0.4082 |
| E | | | | | | | | 0.1585 | 0.1088 | 0.1767 | 0.1476 | 0.0815 |
| F | | | | | | | | 0.4728 | 0.4366 | 0.4625 | 0.4843 | 0.3986 |
| G | | | | | | | | 0.0539 | 0.096 | 0.0898 | 0.0629 | 0.0843 |
| H | | | | | | | | 0.3754 | 0.2727 | 0.3669 | 0.3133 | 0.3255 |

**Cell Index at: 134:48:27**

| | 1 | 2 | 3 | 4 | 5 | 6 | 7 | 8 | 9 | 10 | 11 | 12 |
|---|---|---|---|---|---|---|---|---|---|---|---|---|
| A | | | | | | | | 0.0809 | 0.0658 | 0.0949 | 0.0416 | 0.0738 |
| B | | | | | | | | 0.0601 | 0.0649 | 0.0763 | 0.1039 | 0.0856 |
| C | | | | | | | | 0.3805 | 0.309 | 0.5043 | 0.5205 | 0.4123 |
| D | | | | | | | | 0.3556 | 0.3828 | 0.4502 | 0.4481 | 0.4093 |
| E | | | | | | | | 0.1497 | 0.1075 | 0.1731 | 0.149 | 0.0906 |
| F | | | | | | | | 0.4667 | 0.4333 | 0.4597 | 0.4855 | 0.4034 |
| G | | | | | | | | 0.0547 | 0.0953 | 0.0926 | 0.0662 | 0.0845 |
| H | | | | | | | | 0.3737 | 0.2716 | 0.3686 | 0.3127 | 0.3254 |

**Cell Index at: 135:03:28**

| | 1 | 2 | 3 | 4 | 5 | 6 | 7 | 8 | 9 | 10 | 11 | 12 |
|---|---|---|---|---|---|---|---|---|---|---|---|---|
| A | | | | | | | | 0.084 | 0.0657 | 0.0913 | 0.0362 | 0.0725 |
| B | | | | | | | | 0.0656 | 0.0631 | 0.0762 | 0.1022 | 0.0909 |
| C | | | | | | | | 0.3706 | 0.3157 | 0.5021 | 0.5125 | 0.4059 |
| D | | | | | | | | 0.3549 | 0.3914 | 0.458 | 0.4485 | 0.4116 |
| E | | | | | | | | 0.1527 | 0.117 | 0.1649 | 0.1534 | 0.0897 |
| F | | | | | | | | 0.4672 | 0.4351 | 0.4598 | 0.4938 | 0.4015 |
| G | | | | | | | | 0.0552 | 0.0978 | 0.0893 | 0.0642 | 0.0853 |
| H | | | | | | | | 0.378 | 0.2728 | 0.3716 | 0.3143 | 0.3225 |

**Cell Index at: 135:18:28**

| | 1 | 2 | 3 | 4 | 5 | 6 | 7 | 8 | 9 | 10 | 11 | 12 |
|---|---|---|---|---|---|---|---|---|---|---|---|---|
| A | | | | | | | | 0.082 | 0.069 | 0.0955 | 0.0369 | 0.0722 |

| | | | | | | | | | | | | |
|---|---|---|---|---|---|---|---|---|---|---|---|---|
| B | | | | | | | | 0.0629 | 0.0636 | 0.082 | 0.1003 | 0.0934 |
| C | | | | | | | | 0.3811 | 0.3211 | 0.4959 | 0.5175 | 0.4079 |
| D | | | | | | | | 0.3572 | 0.3966 | 0.4563 | 0.4486 | 0.4053 |
| E | | | | | | | | 0.152 | 0.1188 | 0.1688 | 0.15 | 0.0861 |
| F | | | | | | | | 0.4746 | 0.434 | 0.464 | 0.4926 | 0.3977 |
| G | | | | | | | | 0.0537 | 0.0972 | 0.0905 | 0.0717 | 0.0813 |
| H | | | | | | | | 0.3782 | 0.269 | 0.3745 | 0.3104 | 0.3235 |

**Cell Index at: 135:33:29**

| | 1 | 2 | 3 | 4 | 5 | 6 | 7 | 8 | 9 | 10 | 11 | 12 |
|---|---|---|---|---|---|---|---|---|---|---|---|---|
| A | | | | | | | | 0.079 | 0.0664 | 0.0986 | 0.0366 | 0.0733 |
| B | | | | | | | | 0.0635 | 0.0636 | 0.0775 | 0.1035 | 0.0895 |
| C | | | | | | | | 0.3743 | 0.32 | 0.4989 | 0.5211 | 0.4192 |
| D | | | | | | | | 0.3495 | 0.3929 | 0.4605 | 0.4416 | 0.4171 |
| E | | | | | | | | 0.1481 | 0.1169 | 0.164 | 0.1502 | 0.0799 |
| F | | | | | | | | 0.4679 | 0.435 | 0.4662 | 0.4939 | 0.4022 |
| G | | | | | | | | 0.0613 | 0.0956 | 0.0889 | 0.068 | 0.0814 |
| H | | | | | | | | 0.3868 | 0.2714 | 0.3702 | 0.3163 | 0.3293 |

**Cell Index at: 135:48:29**

| | 1 | 2 | 3 | 4 | 5 | 6 | 7 | 8 | 9 | 10 | 11 | 12 |
|---|---|---|---|---|---|---|---|---|---|---|---|---|
| A | | | | | | | | 0.0769 | 0.0712 | 0.0961 | 0.0395 | 0.0749 |
| B | | | | | | | | 0.0682 | 0.0658 | 0.0791 | 0.0986 | 0.0863 |
| C | | | | | | | | 0.3768 | 0.3113 | 0.4966 | 0.523 | 0.4132 |
| D | | | | | | | | 0.361 | 0.3907 | 0.4635 | 0.4425 | 0.4045 |
| E | | | | | | | | 0.1489 | 0.11 | 0.1726 | 0.1463 | 0.0782 |
| F | | | | | | | | 0.4746 | 0.4357 | 0.4748 | 0.4972 | 0.4017 |
| G | | | | | | | | 0.06 | 0.0977 | 0.0906 | 0.0669 | 0.084 |
| H | | | | | | | | 0.3857 | 0.2747 | 0.3717 | 0.3172 | 0.3279 |

**Cell Index at: 136:03:30**

| | 1 | 2 | 3 | 4 | 5 | 6 | 7 | 8 | 9 | 10 | 11 | 12 |
|---|---|---|---|---|---|---|---|---|---|---|---|---|
| A | | | | | | | | 0.0785 | 0.0726 | 0.0945 | 0.0424 | 0.0762 |
| B | | | | | | | | 0.0699 | 0.0667 | 0.0771 | 0.1004 | 0.0949 |
| C | | | | | | | | 0.3821 | 0.3197 | 0.4975 | 0.5253 | 0.4273 |
| D | | | | | | | | 0.3568 | 0.3996 | 0.4619 | 0.4439 | 0.4113 |
| E | | | | | | | | 0.1603 | 0.1099 | 0.1691 | 0.1481 | 0.0824 |
| F | | | | | | | | 0.472 | 0.4388 | 0.4681 | 0.4951 | 0.4077 |
| G | | | | | | | | 0.0646 | 0.0971 | 0.0927 | 0.0687 | 0.0778 |
| H | | | | | | | | 0.3844 | 0.2723 | 0.372 | 0.3156 | 0.3333 |

**Cell Index at: 136:18:30**

| | 1 | 2 | 3 | 4 | 5 | 6 | 7 | 8 | 9 | 10 | 11 | 12 |
|---|---|---|---|---|---|---|---|---|---|---|---|---|
| A | | | | | | | | 0.0826 | 0.0732 | 0.0969 | 0.0405 | 0.0764 |
| B | | | | | | | | 0.0678 | 0.0646 | 0.0803 | 0.0991 | 0.086 |
| C | | | | | | | | 0.3705 | 0.3211 | 0.4901 | 0.5276 | 0.4246 |
| D | | | | | | | | 0.3661 | 0.3972 | 0.4586 | 0.4412 | 0.4067 |
| E | | | | | | | | 0.1479 | 0.1176 | 0.1697 | 0.1525 | 0.0859 |
| F | | | | | | | | 0.4698 | 0.4385 | 0.4739 | 0.4987 | 0.4044 |
| G | | | | | | | | 0.0577 | 0.0979 | 0.0949 | 0.0703 | 0.0779 |
| H | | | | | | | | 0.3851 | 0.2712 | 0.371 | 0.3194 | 0.3314 |

**Cell Index at: 136:33:29**

| | 1 | 2 | 3 | 4 | 5 | 6 | 7 | 8 | 9 | 10 | 11 | 12 |
|---|---|---|---|---|---|---|---|---|---|---|---|---|
| A | | | | | | | | 0.0816 | 0.0757 | 0.0953 | 0.042 | 0.077 |
| B | | | | | | | | 0.0682 | 0.069 | 0.084 | 0.1042 | 0.0878 |
| C | | | | | | | | 0.3778 | 0.3227 | 0.496 | 0.53 | 0.4224 |
| D | | | | | | | | 0.3575 | 0.4 | 0.4704 | 0.4413 | 0.401 |
| E | | | | | | | | 0.1626 | 0.1127 | 0.1763 | 0.1482 | 0.0802 |
| F | | | | | | | | 0.4827 | 0.4398 | 0.4708 | 0.4911 | 0.4048 |
| G | | | | | | | | 0.0625 | 0.0996 | 0.0894 | 0.0632 | 0.0781 |
| H | | | | | | | | 0.3858 | 0.2652 | 0.3728 | 0.3185 | 0.3282 |

**Cell Index at: 136:48:30**

| | 1 | 2 | 3 | 4 | 5 | 6 | 7 | 8 | 9 | 10 | 11 | 12 |
|---|---|---|---|---|---|---|---|---|---|---|---|---|
| A | | | | | | | | 0.08 | 0.0789 | 0.095 | 0.0414 | 0.0729 |
| B | | | | | | | | 0.0692 | 0.0639 | 0.0837 | 0.1045 | 0.086 |
| C | | | | | | | | 0.3712 | 0.3184 | 0.4978 | 0.518 | 0.4197 |
| D | | | | | | | | 0.3531 | 0.4006 | 0.4614 | 0.4497 | 0.4082 |
| E | | | | | | | | 0.1554 | 0.1172 | 0.1684 | 0.1511 | 0.0863 |
| F | | | | | | | | 0.4745 | 0.435 | 0.4666 | 0.4939 | 0.4093 |
| G | | | | | | | | 0.0563 | 0.1012 | 0.0938 | 0.0681 | 0.0792 |
| H | | | | | | | | 0.3842 | 0.273 | 0.3735 | 0.3185 | 0.3302 |

**Cell Index at: 137:03:31**

| | 1 | 2 | 3 | 4 | 5 | 6 | 7 | 8 | 9 | 10 | 11 | 12 |
|---|---|---|---|---|---|---|---|---|---|---|---|---|
| A | | | | | | | | 0.0826 | 0.0776 | 0.0955 | 0.0427 | 0.0765 |
| B | | | | | | | | 0.0727 | 0.0638 | 0.0843 | 0.1029 | 0.0897 |
| C | | | | | | | | 0.373 | 0.321 | 0.4938 | 0.5223 | 0.4237 |
| D | | | | | | | | 0.3565 | 0.3929 | 0.4705 | 0.4491 | 0.4089 |
| E | | | | | | | | 0.1573 | 0.1189 | 0.1716 | 0.1578 | 0.0852 |
| F | | | | | | | | 0.4758 | 0.4375 | 0.4771 | 0.4942 | 0.4109 |
| G | | | | | | | | 0.0572 | 0.1013 | 0.0906 | 0.0707 | 0.0837 |
| H | | | | | | | | 0.3877 | 0.2723 | 0.3757 | 0.3212 | 0.328 |

**Cell Index at: 137:18:32**

| | 1 | 2 | 3 | 4 | 5 | 6 | 7 | 8 | 9 | 10 | 11 | 12 |
|---|---|---|---|---|---|---|---|---|---|---|---|---|
| A | | | | | | | | 0.0798 | 0.0785 | 0.0992 | 0.0433 | 0.0802 |
| B | | | | | | | | 0.072 | 0.0665 | 0.0855 | 0.1036 | 0.0906 |
| C | | | | | | | | 0.374 | 0.3175 | 0.5072 | 0.5237 | 0.4211 |
| D | | | | | | | | 0.3629 | 0.3998 | 0.4753 | 0.4594 | 0.4123 |
| E | | | | | | | | 0.1557 | 0.1202 | 0.1735 | 0.15 | 0.0815 |
| F | | | | | | | | 0.4775 | 0.4334 | 0.4745 | 0.4875 | 0.417 |
| G | | | | | | | | 0.0586 | 0.1021 | 0.0885 | 0.0707 | 0.083 |
| H | | | | | | | | 0.3858 | 0.273 | 0.3768 | 0.3195 | 0.3311 |

**Cell Index at: 137:33:32**

| | 1 | 2 | 3 | 4 | 5 | 6 | 7 | 8 | 9 | 10 | 11 | 12 |
|---|---|---|---|---|---|---|---|---|---|---|---|---|
| A | | | | | | | | 0.0804 | 0.0801 | 0.1008 | 0.0415 | 0.0784 |
| B | | | | | | | | 0.0683 | 0.0659 | 0.0803 | 0.0998 | 0.0897 |
| C | | | | | | | | 0.3786 | 0.3234 | 0.5049 | 0.5301 | 0.4203 |
| D | | | | | | | | 0.3543 | 0.3991 | 0.4721 | 0.4511 | 0.4167 |
| E | | | | | | | | 0.1579 | 0.1156 | 0.174 | 0.1503 | 0.084 |
| F | | | | | | | | 0.4714 | 0.4394 | 0.4761 | 0.4956 | 0.415 |
| G | | | | | | | | 0.0589 | 0.1065 | 0.0864 | 0.0723 | 0.0818 |
| H | | | | | | | | 0.3848 | 0.2735 | 0.3762 | 0.3217 | 0.3318 |

**Cell Index at: 137:48:33**

| | 1 | 2 | 3 | 4 | 5 | 6 | 7 | 8 | 9 | 10 | 11 | 12 |
|---|---|---|---|---|---|---|---|---|---|---|---|---|
| A | | | | | | | | 0.0833 | 0.0752 | 0.103 | 0.0399 | 0.0793 |
| B | | | | | | | | 0.0753 | 0.0674 | 0.0816 | 0.0991 | 0.0863 |
| C | | | | | | | | 0.3758 | 0.3248 | 0.5063 | 0.5269 | 0.4177 |
| D | | | | | | | | 0.36 | 0.3968 | 0.4791 | 0.4607 | 0.418 |
| E | | | | | | | | 0.1588 | 0.1194 | 0.1727 | 0.1577 | 0.08 |
| F | | | | | | | | 0.4814 | 0.4412 | 0.4809 | 0.498 | 0.4183 |
| G | | | | | | | | 0.056 | 0.1059 | 0.0895 | 0.0745 | 0.0821 |
| H | | | | | | | | 0.3869 | 0.2771 | 0.3801 | 0.3177 | 0.3354 |

**Cell Index at: 138:03:33**

| | 1 | 2 | 3 | 4 | 5 | 6 | 7 | 8 | 9 | 10 | 11 | 12 |
|---|---|---|---|---|---|---|---|---|---|---|---|---|
| A | | | | | | | | 0.0823 | 0.0764 | 0.0977 | 0.0405 | 0.0788 |
| B | | | | | | | | 0.0735 | 0.0713 | 0.0817 | 0.102 | 0.0866 |
| C | | | | | | | | 0.3779 | 0.3253 | 0.4914 | 0.5257 | 0.4216 |
| D | | | | | | | | 0.3675 | 0.399 | 0.4758 | 0.4498 | 0.4212 |
| E | | | | | | | | 0.153 | 0.1175 | 0.1653 | 0.1595 | 0.076 |
| F | | | | | | | | 0.4766 | 0.4413 | 0.4801 | 0.493 | 0.416 |
| G | | | | | | | | 0.0589 | 0.1062 | 0.0885 | 0.0693 | 0.0843 |
| H | | | | | | | | 0.3865 | 0.2767 | 0.3777 | 0.3213 | 0.3321 |

**Cell Index at: 138:18:34**

| | 1 | 2 | 3 | 4 | 5 | 6 | 7 | 8 | 9 | 10 | 11 | 12 |
|---|---|---|---|---|---|---|---|---|---|---|---|---|

| | | | | | | | | | | | | |
|---|---|---|---|---|---|---|---|---|---|---|---|---|
| A | | | | | | | | 0.0819 | 0.0824 | 0.0985 | 0.0377 | 0.0823 |
| B | | | | | | | | 0.0708 | 0.0679 | 0.085 | 0.1034 | 0.0881 |
| C | | | | | | | | 0.3747 | 0.3212 | 0.5007 | 0.5331 | 0.4227 |
| D | | | | | | | | 0.3726 | 0.4059 | 0.4748 | 0.4602 | 0.4141 |
| E | | | | | | | | 0.1524 | 0.1142 | 0.1651 | 0.1551 | 0.0778 |
| F | | | | | | | | 0.4832 | 0.4435 | 0.4764 | 0.4993 | 0.4132 |
| G | | | | | | | | 0.0541 | 0.1093 | 0.0875 | 0.0715 | 0.0824 |
| H | | | | | | | | 0.3884 | 0.2775 | 0.3777 | 0.3258 | 0.3364 |

**Cell Index at: 138:33:35**

| | 1 | 2 | 3 | 4 | 5 | 6 | 7 | 8 | 9 | 10 | 11 | 12 |
|---|---|---|---|---|---|---|---|---|---|---|---|---|
| A | | | | | | | | 0.0847 | 0.078 | 0.1022 | 0.0398 | 0.086 |
| B | | | | | | | | 0.0736 | 0.0705 | 0.0864 | 0.1011 | 0.0955 |
| C | | | | | | | | 0.3799 | 0.3296 | 0.5015 | 0.5268 | 0.4218 |
| D | | | | | | | | 0.3587 | 0.4101 | 0.4769 | 0.4525 | 0.4231 |
| E | | | | | | | | 0.1544 | 0.1218 | 0.1753 | 0.1654 | 0.0848 |
| F | | | | | | | | 0.4831 | 0.439 | 0.4842 | 0.4983 | 0.4108 |
| G | | | | | | | | 0.0591 | 0.1094 | 0.0892 | 0.0689 | 0.0811 |
| H | | | | | | | | 0.3868 | 0.2776 | 0.3812 | 0.3241 | 0.3365 |

**Cell Index at: 138:48:35**

| | 1 | 2 | 3 | 4 | 5 | 6 | 7 | 8 | 9 | 10 | 11 | 12 |
|---|---|---|---|---|---|---|---|---|---|---|---|---|
| A | | | | | | | | 0.0859 | 0.078 | 0.0992 | 0.0402 | 0.0849 |
| B | | | | | | | | 0.076 | 0.071 | 0.087 | 0.1 | 0.0919 |
| C | | | | | | | | 0.3781 | 0.3248 | 0.4969 | 0.5299 | 0.4291 |
| D | | | | | | | | 0.3749 | 0.4065 | 0.4745 | 0.4591 | 0.43 |
| E | | | | | | | | 0.16 | 0.1184 | 0.1723 | 0.1584 | 0.0862 |
| F | | | | | | | | 0.4817 | 0.44 | 0.4809 | 0.4923 | 0.4222 |
| G | | | | | | | | 0.0592 | 0.1106 | 0.0851 | 0.0715 | 0.0835 |
| H | | | | | | | | 0.3882 | 0.2816 | 0.3833 | 0.322 | 0.3358 |

**Cell Index at: 139:03:36**

| | 1 | 2 | 3 | 4 | 5 | 6 | 7 | 8 | 9 | 10 | 11 | 12 |
|---|---|---|---|---|---|---|---|---|---|---|---|---|
| A | | | | | | | | 0.0899 | 0.0787 | 0.1006 | 0.0406 | 0.0872 |
| B | | | | | | | | 0.0717 | 0.0661 | 0.0838 | 0.1037 | 0.0907 |
| C | | | | | | | | 0.373 | 0.3232 | 0.5021 | 0.5308 | 0.4245 |
| D | | | | | | | | 0.3768 | 0.4094 | 0.4797 | 0.4583 | 0.4279 |
| E | | | | | | | | 0.1549 | 0.1277 | 0.1659 | 0.1573 | 0.0922 |
| F | | | | | | | | 0.4807 | 0.441 | 0.4804 | 0.5023 | 0.4211 |
| G | | | | | | | | 0.0559 | 0.1059 | 0.0832 | 0.078 | 0.0845 |
| H | | | | | | | | 0.3894 | 0.2816 | 0.3855 | 0.3277 | 0.3358 |
