## Supplementary Figures and Supplementary Tables 2, 4, and 5 for "Identification of Cancer-Associated Fibroblasts in Glioblastoma and Defining Their Pro-tumoral Effects"

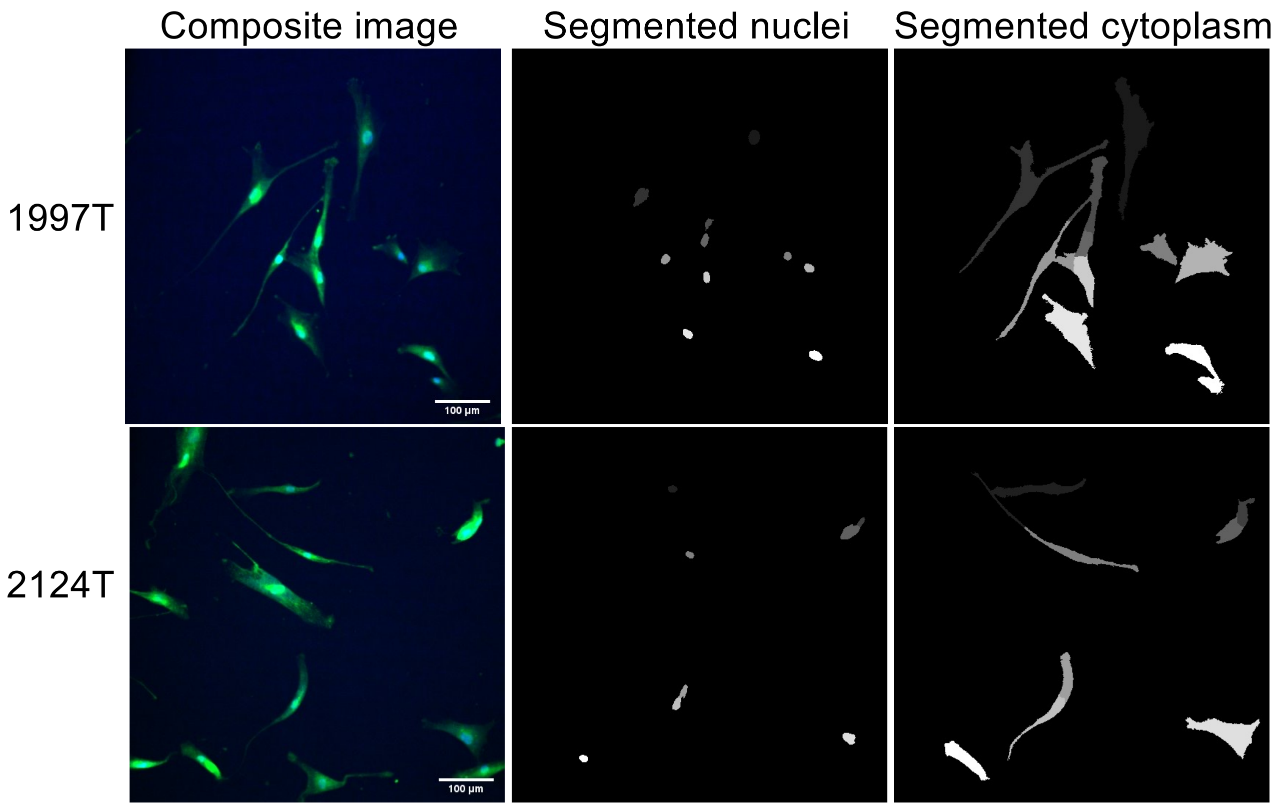


**Supplementary Figure 1. Representative segmented images from breast cancer CAFs. Related to Figure 1A.** Breast cancer CAF cells 1997T and 2124T were imaged (left) using CellTracker (green) and DAPI (blue). Segmentation was performed using ImageJ to split images into blue (nuclei, middle) and green (cytoplasm) channels. Segmented images were analyzed for morphology and results were used to generate a classifier, as described in the Methods section. 10x magnification; scale bar 100 μm.

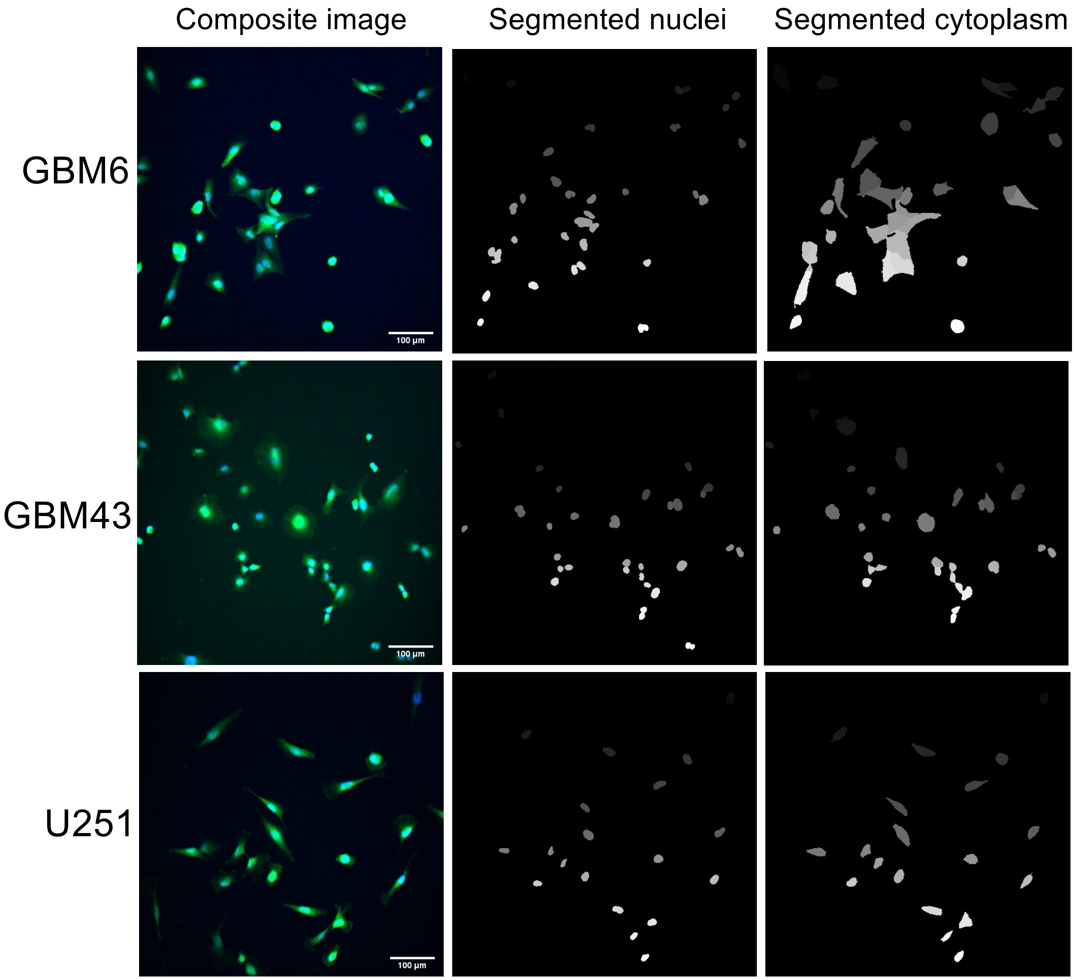


**Supplementary Figure 2. Representative segmented images from GBM cells. Related to Figure 1A.** GBM cells GBM6, GBM43, and U251 were imaged (left) using CellTracker (green) and DAPI (blue). Segmentation was performed using ImageJ to split images into blue (nuclei, middle) and green (cytoplasm, right) channels. Segmented images were analyzed for morphology and results were used to generate a classifier, as described in the Methods section. 10x magnification; scale bar 100 μm.

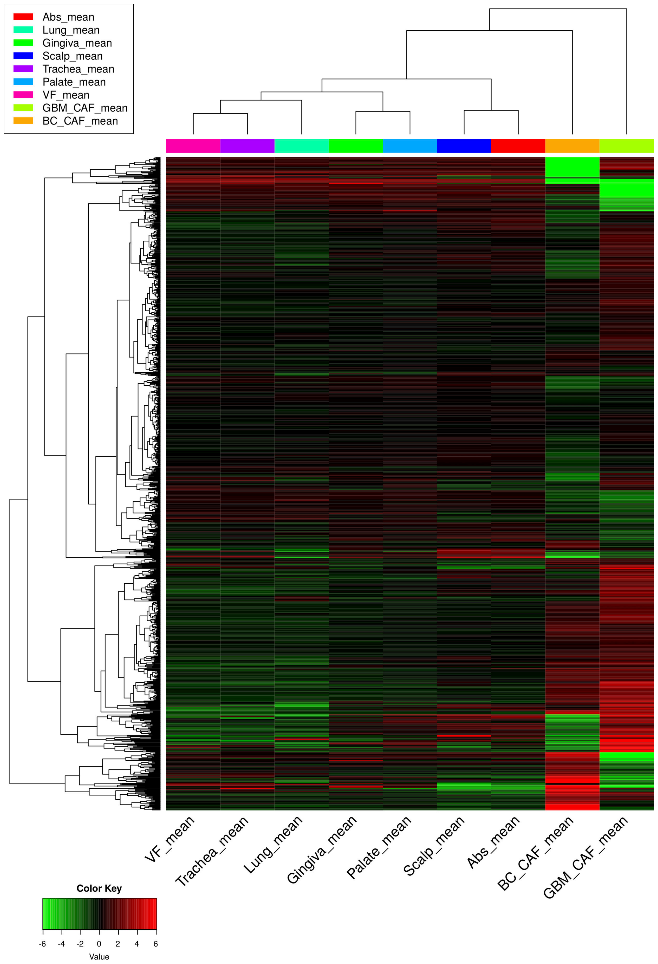


**Supplementary Figure 3. Transcriptomic profiling reveals that cells that emerge from serial trypsinization of primary GBMs resemble CAFs from other cancers. Related to Figure 1B.** RNA-seq was performed on cells isolated from GBM after serial trypsinization (n=1 case shown here from the two cases shown in **Fig. 1C**) which were then compared to archived RNA-seq results form breast cancer CAFs (n=1 case shown here from the two cases shown in **Fig. 1C**) and normal fibroblasts from seven different tissues (scalp dermis, soft palate, upper gingiva, vocal fold, trachea, lung, and abdomen dermis). Cells isolated from GBM after serial trypsinization more closely resembled breast cancer CAFs than normal fibroblasts. Of the 7 normal fibroblasts shown here, the GBM cells and breast cancer CAFs most closely resembled fibroblasts from abdomen dermis and scalp.

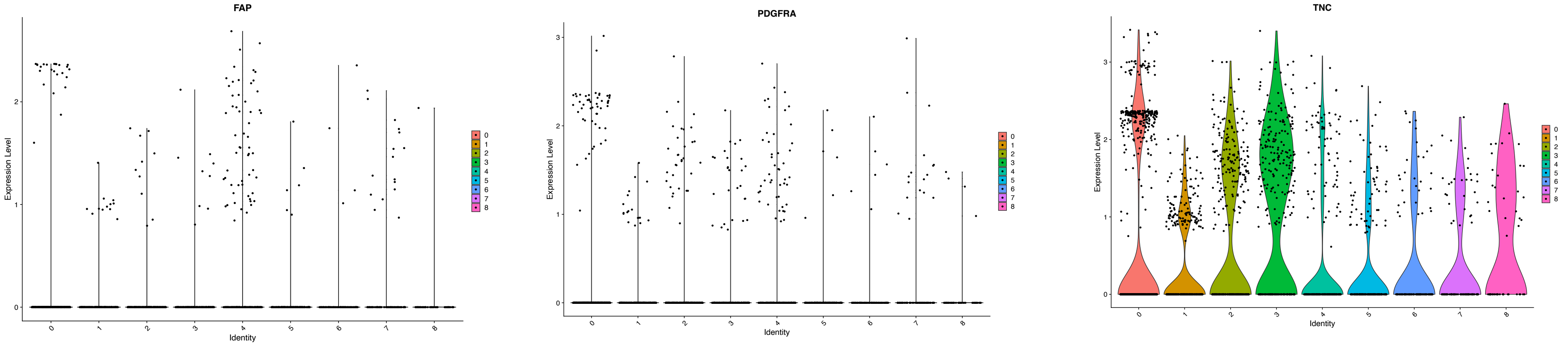


**Supplementary Figure 4. Single-cell analysis of cells emerging from serial trypsinization of patient GBM samples. Related to Figure 1C.** Single-cell RNA-seq was performed on 7,276 cells cells isolated from a single GBM after serial trypsinization. Shown are violin plots revealing expression of CAF markers FAP, PDGFA, and TNC in each of the 9 clusters identified in these cells.

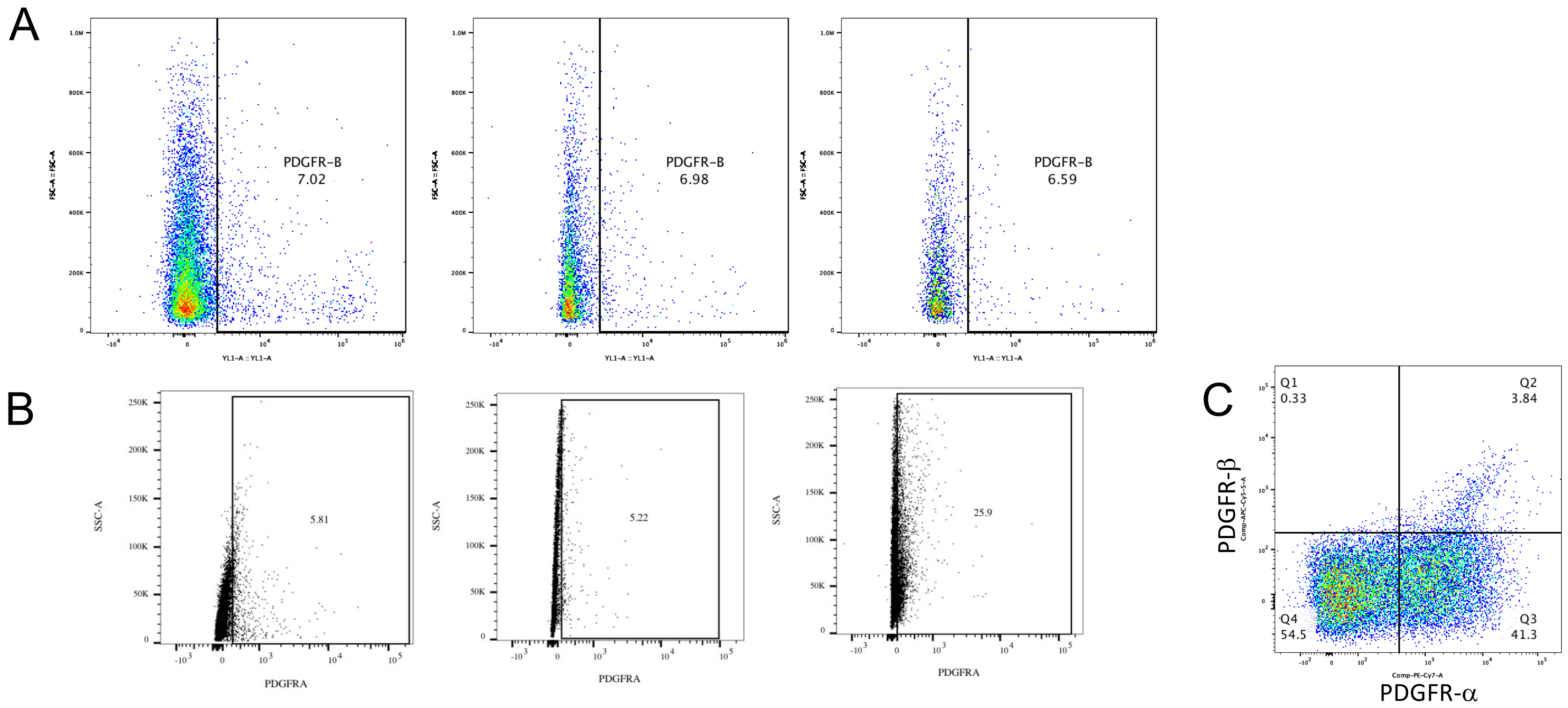


**Supplementary Figure 5. Flow cytometry of cells reveals no ubiquitous CAF markers. Related to Figure 1B.** (**A**) Flow cytometry of cells isolated from GBM after serial trypsinization reveals expression of CAF marker PDGFR-β in 6.59-7.02% of cells (n=3 cases). (**B**) Flow cytometry of cells isolated from GBM after serial trypsinization reveals expression of CAF marker PDGFR-α in 5.22-25.9% of cells (n=3 cases). (**C**) Costaining of a case of cells isolated from GBM after serial trypsinization reveals low levels of PDGFR-β alongside high levels of PDGFR-α in this case but with most of the PDGFR-β positivity occurring in PDGFR-α positive cells

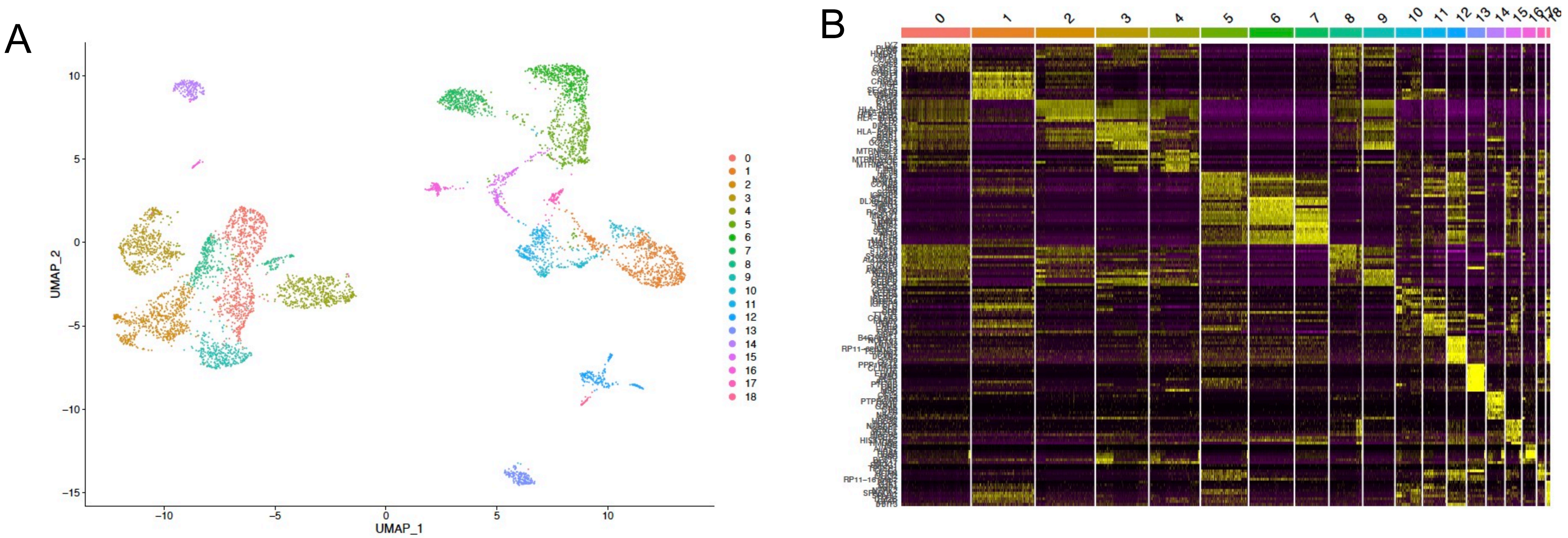


**Supplementary Figure 6. Single-cell RNA-seq reveals clusters of cells in GBM. Related to Figure 1D.** We analyzed profiles from 12074 single cells taken from 8 patient GBMs, revealing (**A**) 18 different clusters identified by a t-SNE plot and (**B**) transcriptomic differences between these 18 clusters as shown by a heatmap.

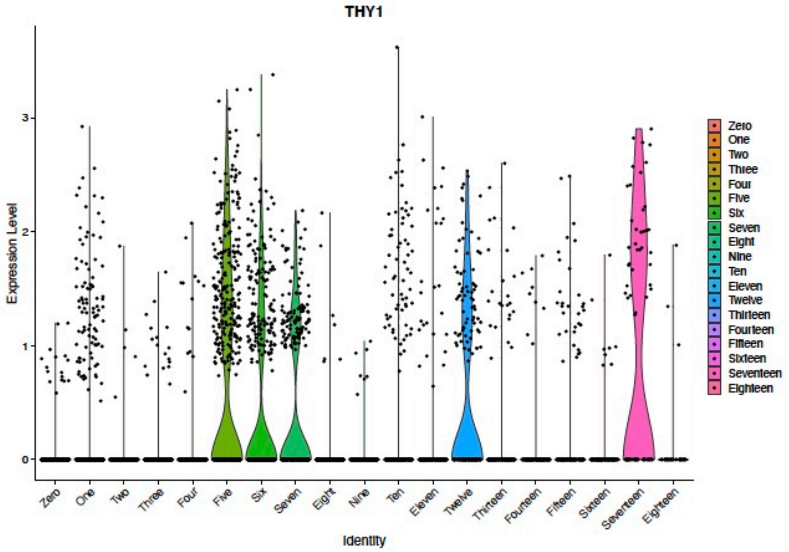


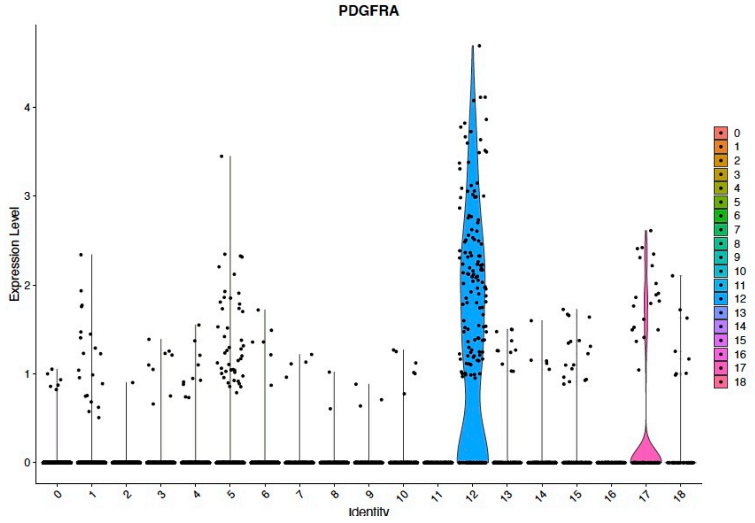


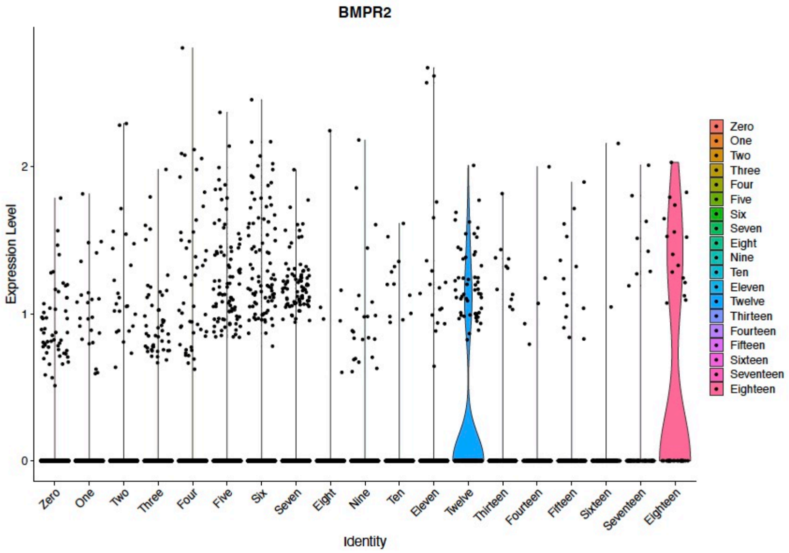


**Supplementary Figure 7. Expression of mesenchymal stem cell (MSC) markers in 18 clusters identified by single-cell RNA-seq of GBM. Related to Figure 1D.** Shown are violin plots revealing expression of MSC markers THY1, BMPR2, and PDGFRA in cells from 18 different clusters identified in patient GBMs. All 3 markers were expressed robustly in cluster 12. Smaller numbers of cells expressed these MSC markers in cluster 1, which exhibited robust pericyte marker expression and in cluster 13, which exhibited robust CAF marker expression.

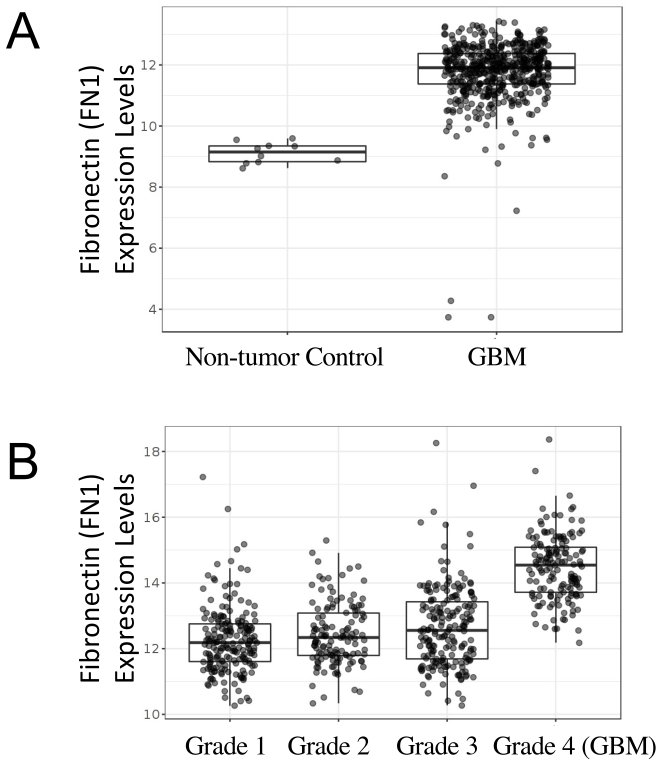


**Supplementary Figure 8. Transcriptomic data from GlioVis data set reveal more fibronectin expression in GBM compared to low-grade glioma or normal brain. Related to Figure 1D.** Population-based bioinformatic data was obtained from GlioVis (gliovis.bioinfo.cnis.es), and analyzed for fibronectin (*FN1*) expression, revealing that (**A**) GBM had significantly higher expression of *FN1* than non-tumor samples (P<0.001) and (**B**) GBM also had much higher expression of *FN1* than low grade gliomas (P<0.001).

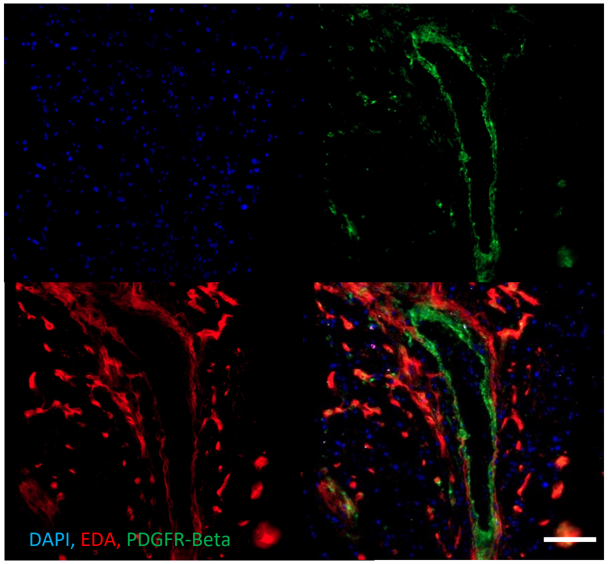


**Supplementary Figure 9. Co-localization of EDA with PDGFR-β staining in GBM. Related to Figure 1G.** Shown is another example of co-localization of EDA fibronectin (red) with PDFGR-β (green) staining, with DAPI nuclear staining in blue. 100x magnification, scale bar 30 μm.

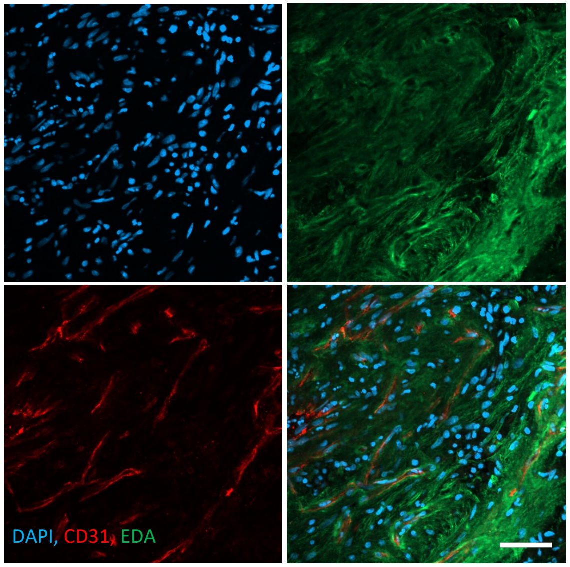


**Supplementary Figure 10. EDA staining is distinct from CD31 staining. Related to Figure 1G.** Shown is staining of EDA fibronectin (green) and CD31 endothelial cells (red) staining, with DAPI nuclear staining in blue. EDA deposition did not overlap with areas of CD31 staining. 100x magnification, scale bar 30 μm.

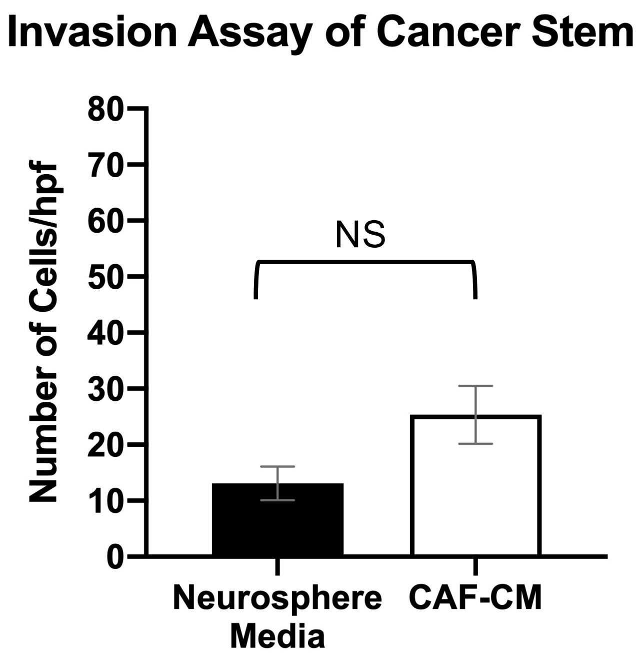


**Supplementary Figure 11. Lack of chemotactic attraction of GSCs towards CAFs. Related to Figure 2.** Neurospheres derived from GBM6 cells migrated equally towards control media or CAF CM (P=0.1=NS), indicating lack of chemotactic attraction.

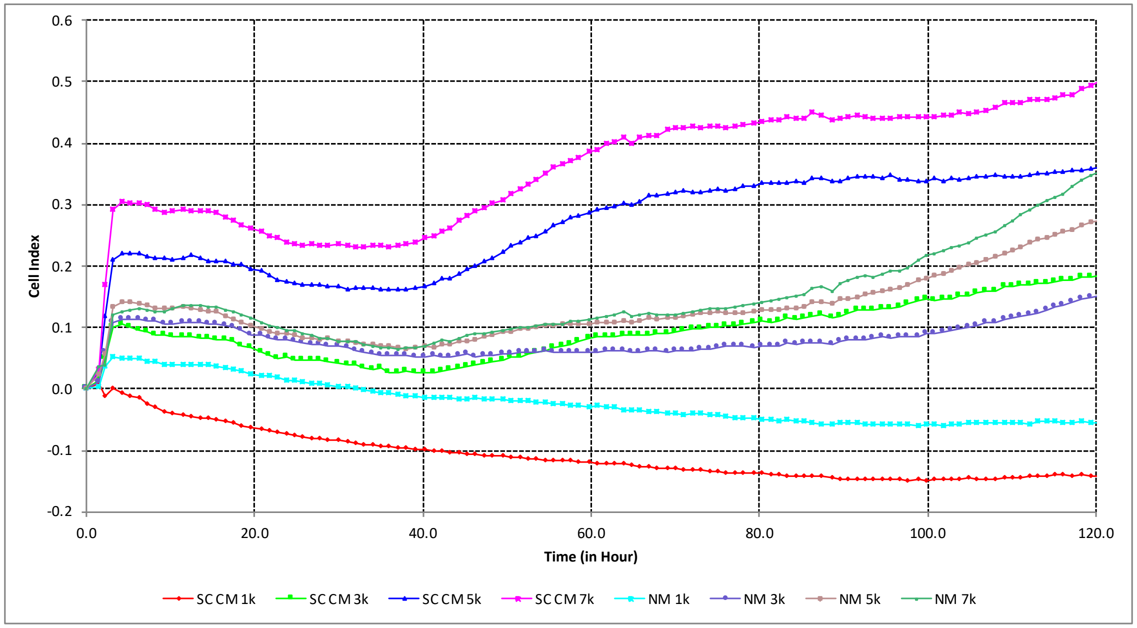


**Supplementary Figure 12. Optimization of number of cultured GBM CAFs for proliferation assay. Related to Figures 3B and 3E.** Shown is the continuous growth of CAFs plated at varying densities (1000, 3000, 5000, or 7000 cells per well in 96 well plates) in neuropshere media (NM) or stem cell conditioned media (SC CM). Proliferation was continuously assessed using the xCELLigence RTCA MP instrument to measure impedance as a surrogate for cell count over 120 hours. Based on these results, 1000 cells per well was chosen for experiments in **Figs. 3B and 3E**.

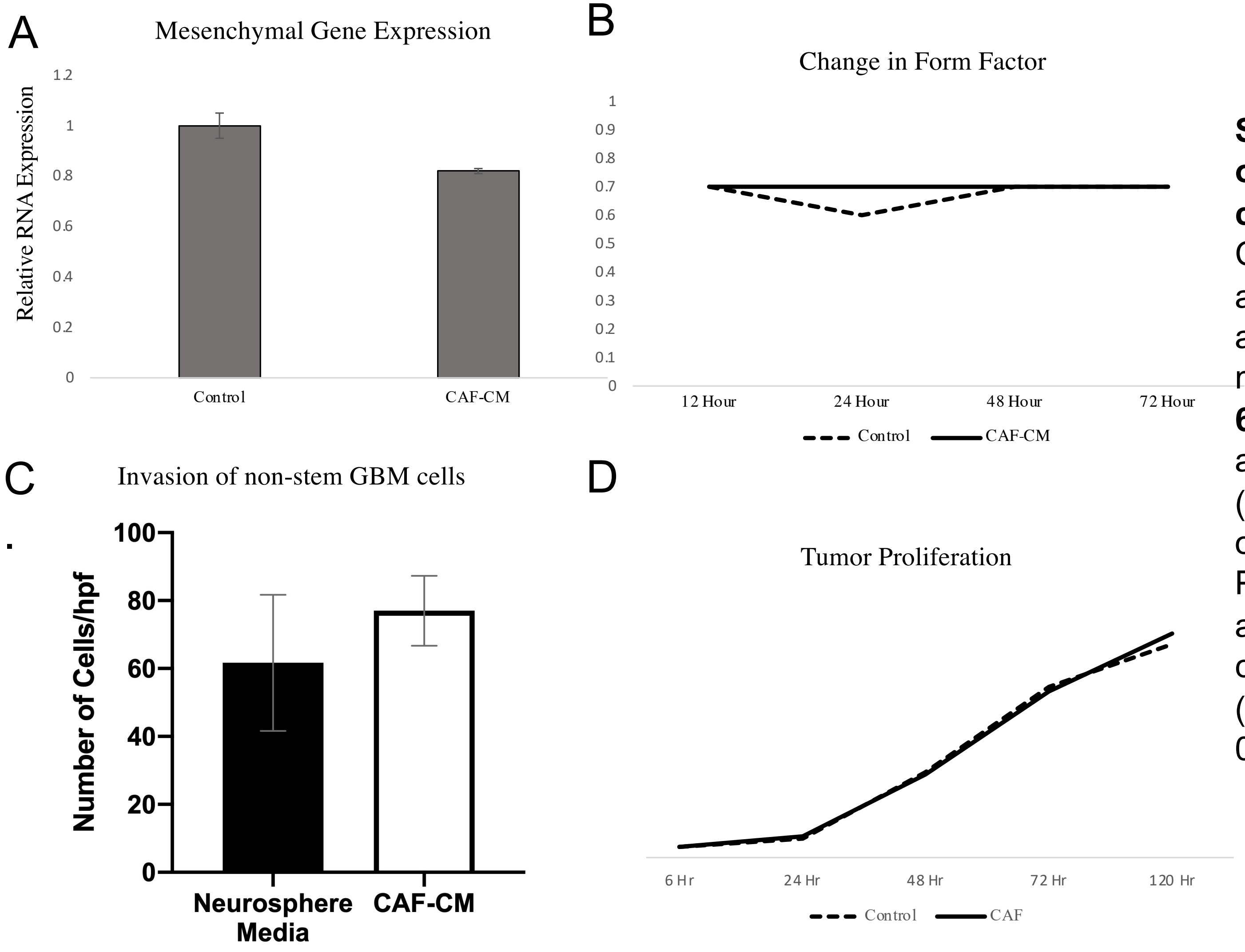


**Supplementary Figure 13. Effects of CAF CM on non-stem GBM cells. Related to Figure 2.** CAF CM did not alter properties of adherent GBM cells, including (**A**) aggregate expression of five mesenchymal genes (**Supp. Table 6**) as assessed by qPCR in adherent DBTRG-05MG cells (n=3/group; P=0.6); (**B**) morphology of adherent GBM6 cells (n=3/group; P=0.06-0.8); (**C**) invasion of adherent GBM6 cells in matrigel chambers (n=3/group; P=0.5); and (**D**) proliferation (n=3/group; P=0.3-0.9) of GBM6 cells.

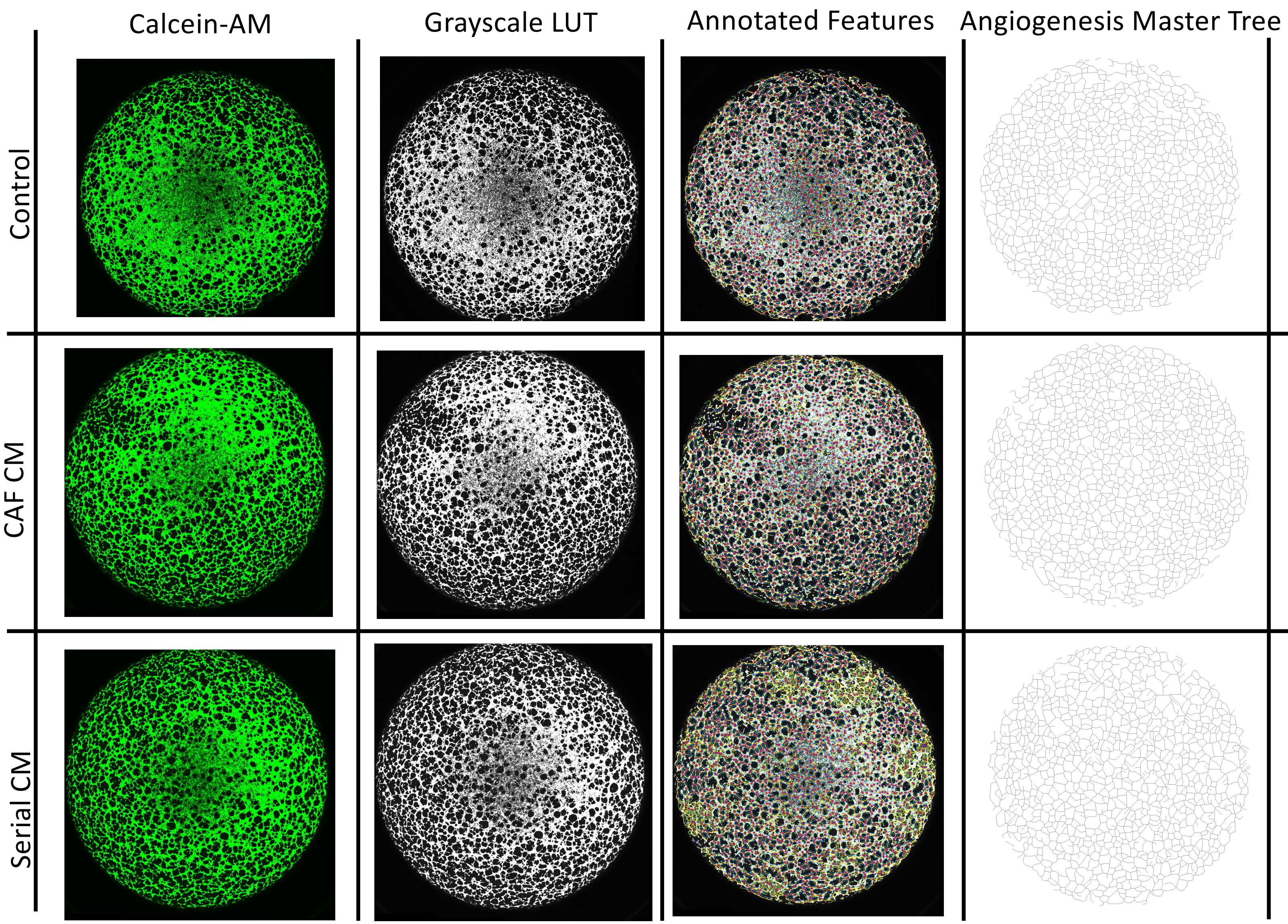


**Supplementary Figure 14. Representative images of HUVEC cells in various conditions at 4 hours. Related to Figure 4A.** Shown are pictures of cultured HUVEC cells labeled with Calcein-AM (left, green); the grayscale LUT (Look Up Tables) colorless image (second column); image with annotated features like nodes and master junctions (third column); and stripped down image of angiogenesis master tree representing quantified metrics without annotations.

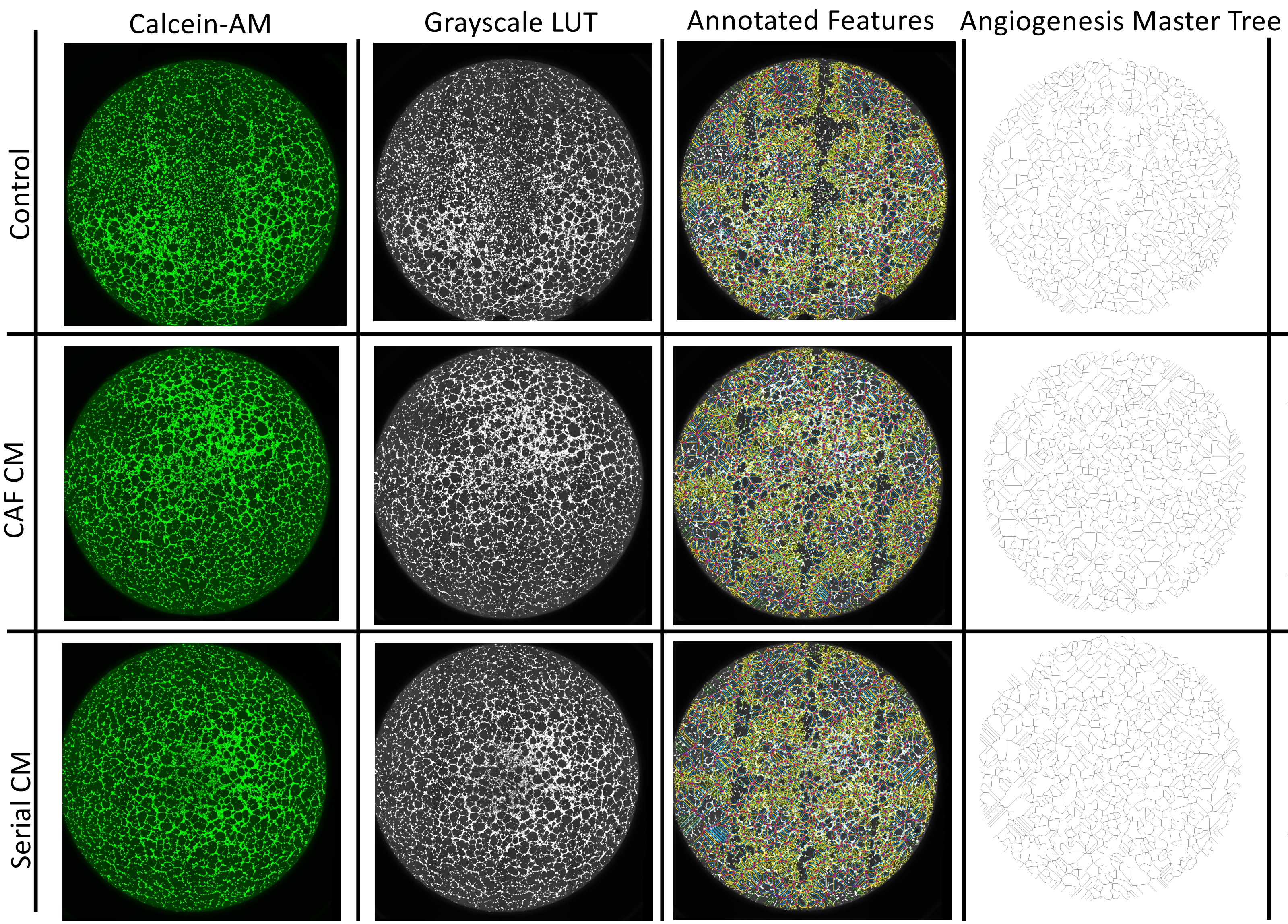


**Supplementary Figure 15. Representative images of HUVEC cells in various conditions at 8 hours. Related to Figure 4A.** Shown are pictures of cultured HUVEC cells labeled with Calcein-AM (left, green); the grayscale LUT (Look Up Tables) colorless image (second column); image with annotated features like nodes and master junctions (third column); and stripped down image of angiogenesis master tree representing quantified metrics without annotations.

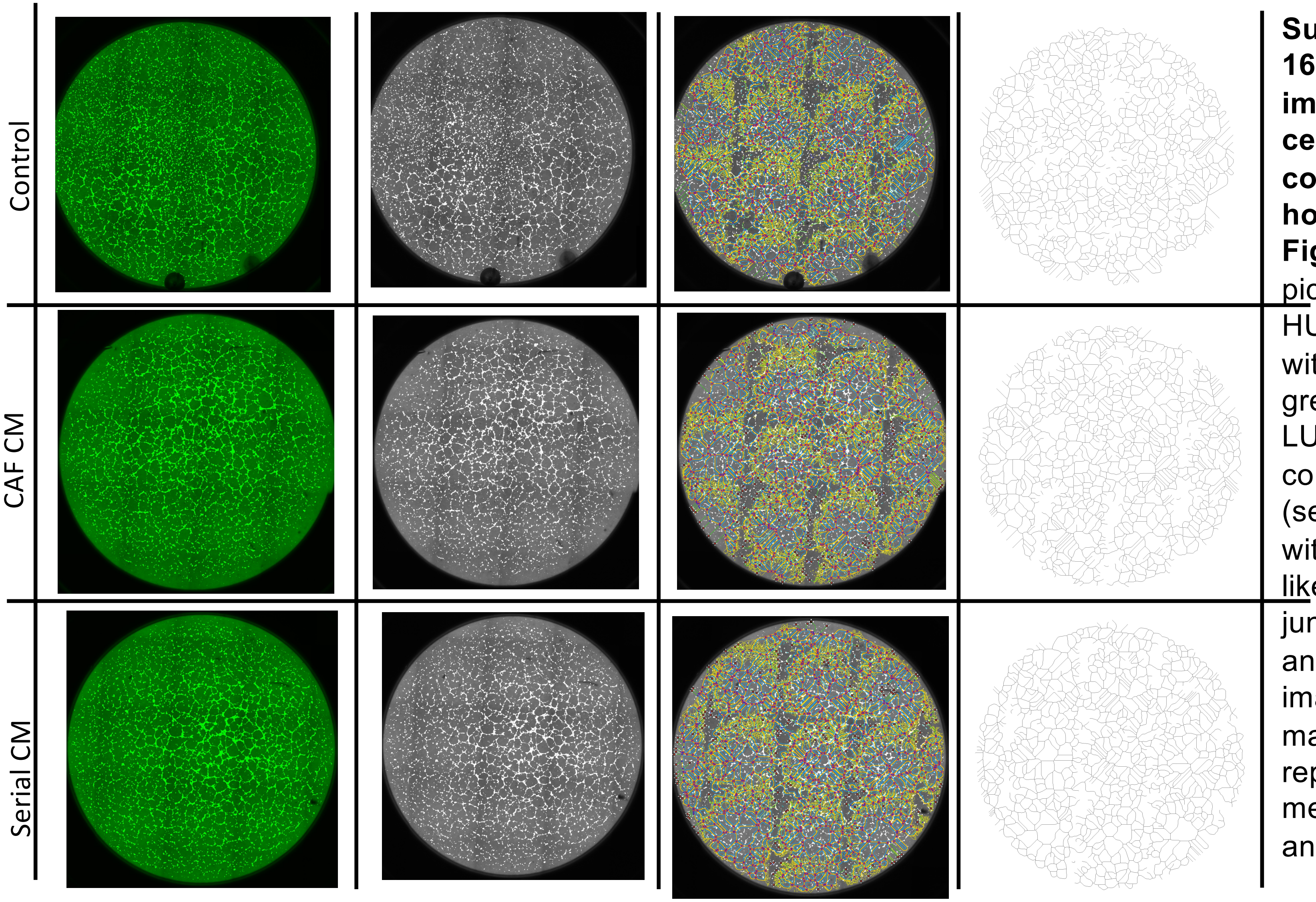


**Supplementary Figure 16. Representative images of HUVEC cells in various conditions at 16 hours. Related to Figure 4A.** Shown are pictures of cultured HUVEC cells labeled with Calcein-AM (left, green); the grayscale LUT (Look Up Tables) colorless image (second column); image with annotated features like nodes and master junctions (third column); and stripped down image of angiogenesis master tree representing quantified metrics without annotations.

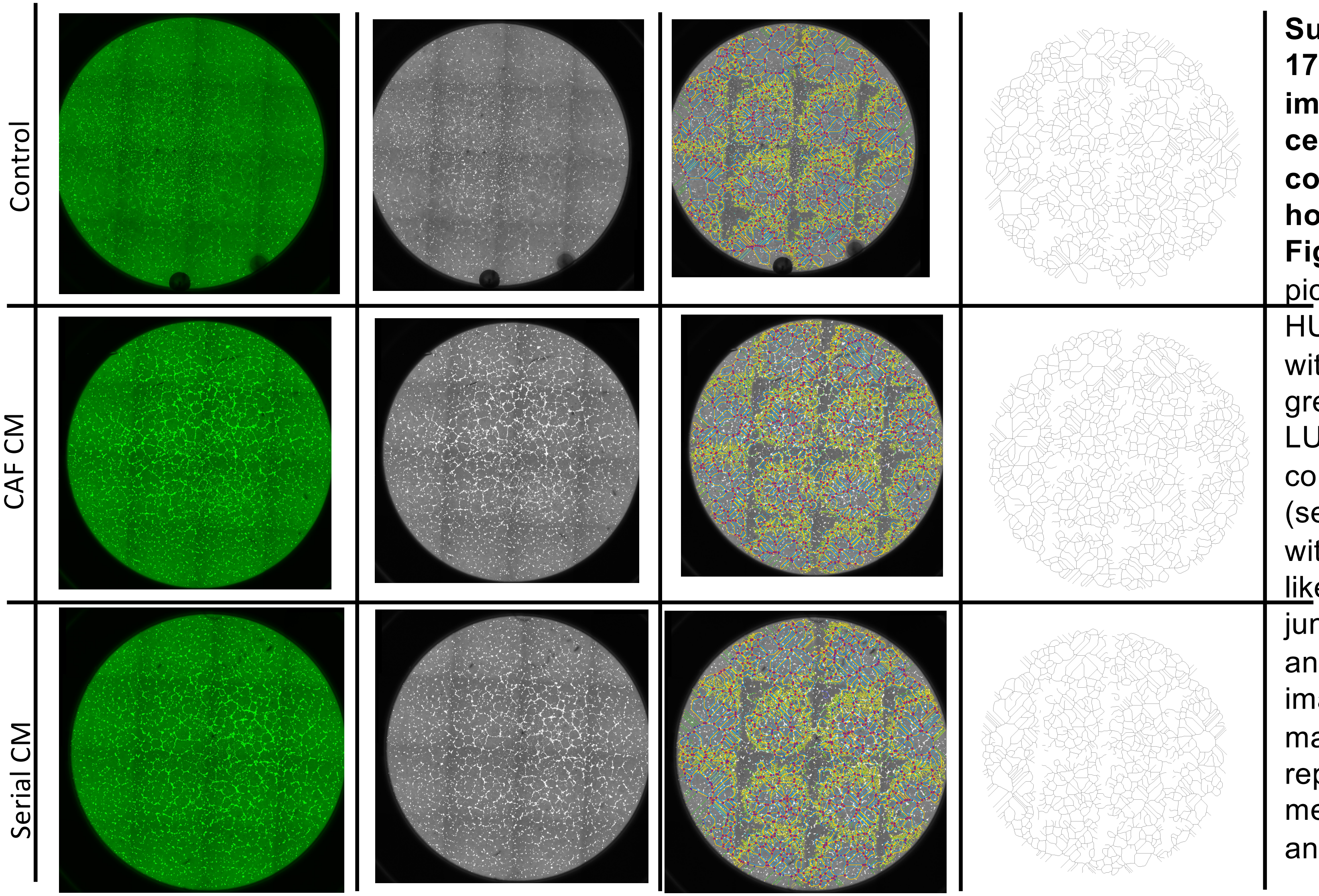


**Supplementary Figure 17. Representative images of HUVEC cells in various conditions at 24 hours. Related to Figure 4A.** Shown are pictures of cultured HUVEC cells labeled with Calcein-AM (left, green); the grayscale LUT (Look Up Tables) colorless image (second column); image with annotated features like nodes and master junctions (third column); and stripped down image of angiogenesis master tree representing quantified metrics without annotations.

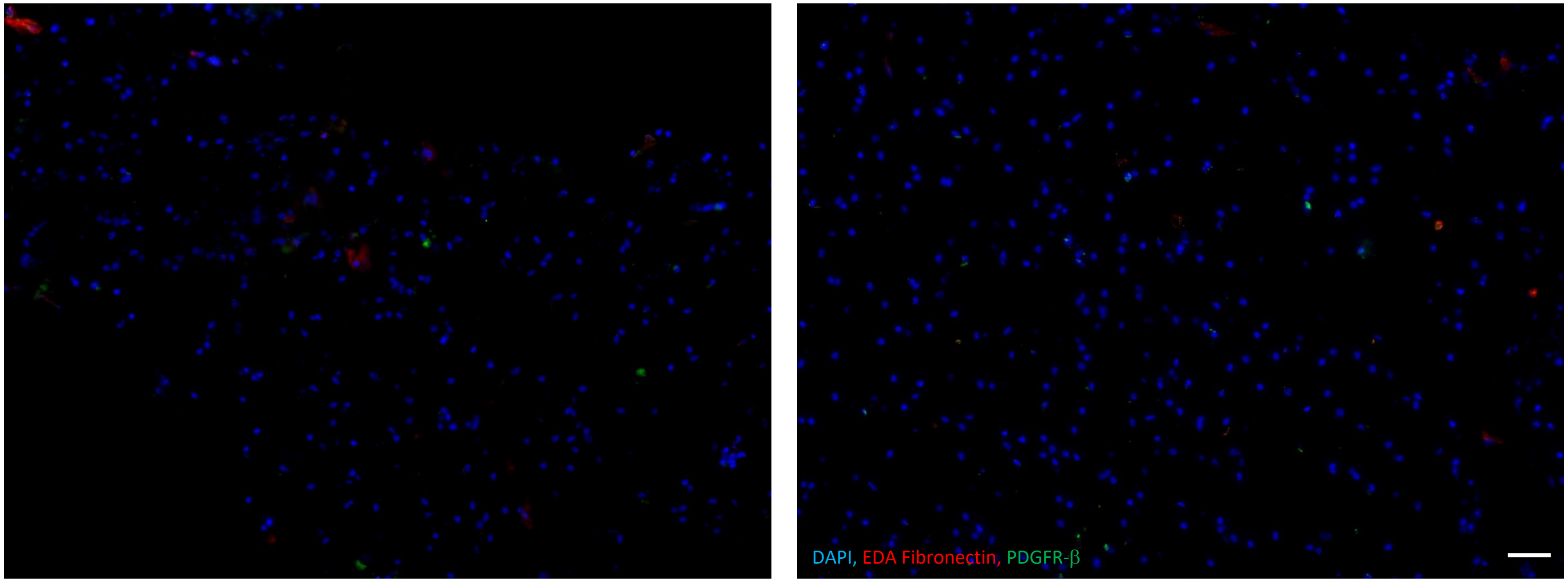


**Supplementary Figure 18. SVZ from tumor-free brain lacks PDGFR-β and EDA immunopositivity. Related to Figure 5E.** Shown is the SVZ from tumor-free brain tissue resected during epilepsy surgery revealing no PDGFR-β or EDA immunopositivity. 100x magnification, scale bar 30 μm.

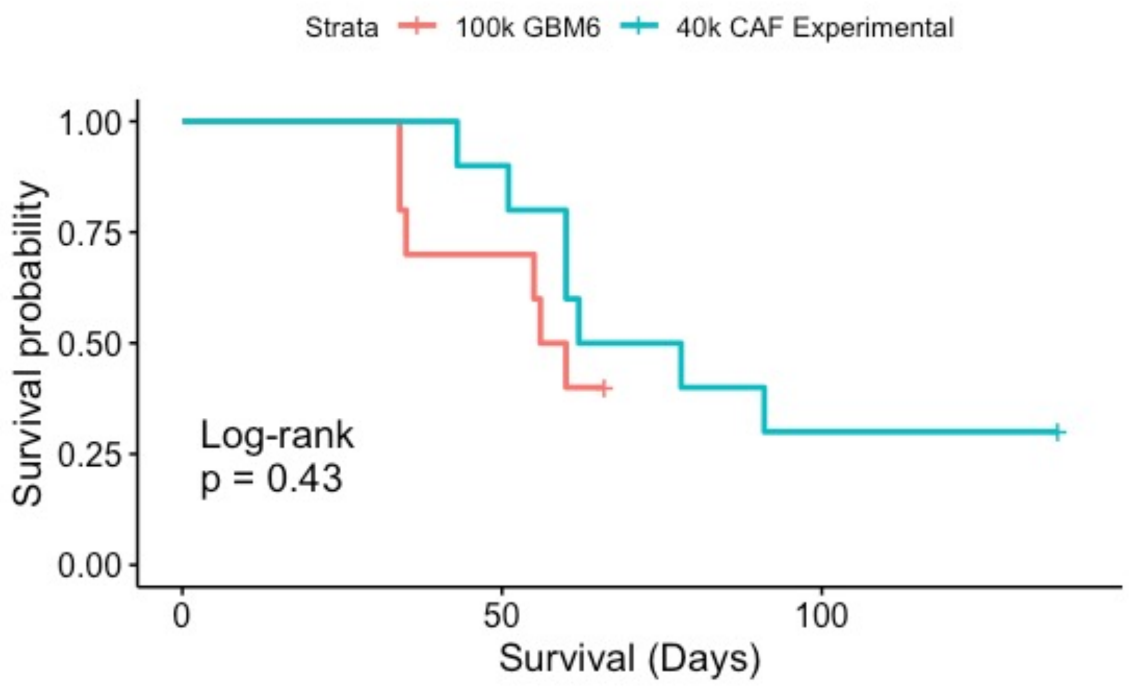


**Supplementary Figure 19. Addition of CAFs to GBM6 neurospheres accelerates tumor growth. Related to Figure 6A.** Shown are Kaplan-Meier curves revealing that the threshold for intracranial tumor formation in over half of mice with GBM6 neurospheres dropped from 100,000 GBM6 cells in the absence of CAFs (red curve) to 35,000 GBM6 cells when combined with 5,000 CAFs, creating comparable survival curves (P=0.4)

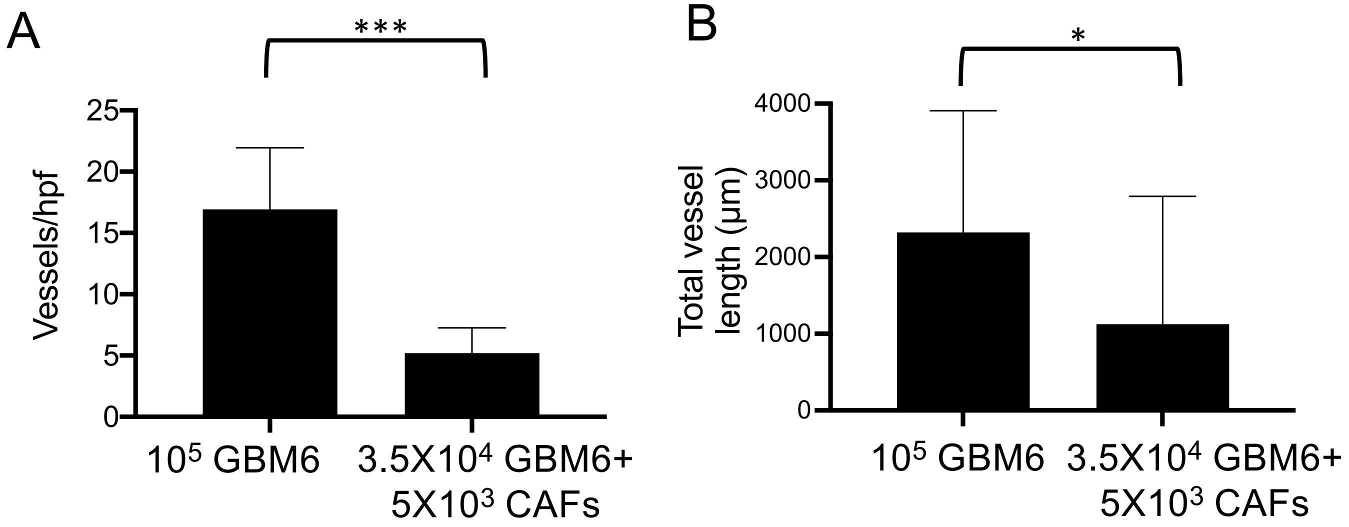


**Supplementary Figure 20. Addition of CAFs to GBM6 neurospheres alters tumor vessels *in vivo*. Related to Figures 6E-F.** In addition to the increased vessel diameter described in **Figs. 6E-F**, addition of CAFs to GBM6 neurospheres *in vivo* decreased vessel density ($P<0.001$), as shown here. The net effect of the former was greater than the latter, leading to increased total vessel surface area, as shown in **Figs. 6E-F**.

SUPPLEMENTARY TABLE 1 HAS BEEN UPLOADED SEPARATELY

**Supplementary Table 2: GSC-CAF receptor-ligand interactions documented by RNA-seq results**

| Receptor (expressed) in GBM_CAF | Ligand (expressed) in GSC | Read count of receptor in GBM_CAF1 | Read count of receptor in GBM_CAF2 | FPKM of ligand in GSC |
|---|---|---|---|---|
| NOTCH1 | DLK1 | 11112 | 6415 | 0 |
| NOTCH2 | DLK1 | 60689 | 51780 | 0 |
| NOTCH3 | DLK1 | 107836 | 69406 | 0 |
| NOTCH4 | DLK1 | 0 | 0 | 0 |
| NOTCH1 | JAG1 | 11112 | 6415 | 0 |
| NOTCH2 | JAG1 | 60689 | 51780 | 0 |
| NOTCH3 | JAG1 | 107836 | 69406 | 0 |
| NOTCH4 | JAG1 | 0 | 0 | 0 |
| NOTCH1 | JAG2 | 11112 | 6415 | 0.36549 |
| NOTCH2 | JAG2 | 60689 | 51780 | 0.36549 |
| NOTCH3 | JAG2 | 107836 | 69406 | 0.36549 |
| NOTCH4 | JAG2 | 0 | 0 | 0.36549 |
| NOTCH1 | DLL1 | 11112 | 6415 | 0.98157 |
| NOTCH2 | DLL1 | 60689 | 51780 | 0.98157 |
| NOTCH3 | DLL1 | 107836 | 69406 | 0.98157 |
| NOTCH4 | DLL1 | 0 | 0 | 0.98157 |
| NOTCH1 | DLL3 | 11112 | 6415 | 3.42671 |
| NOTCH2 | DLL3 | 60689 | 51780 | 3.42671 |
| NOTCH3 | DLL3 | 107836 | 69406 | 3.42671 |
| NOTCH4 | DLL3 | 0 | 0 | 3.42671 |
| NOTCH1 | DLL4 | 11112 | 6415 | 0.0703881 |
| NOTCH2 | DLL4 | 60689 | 51780 | 0.0703881 |
| NOTCH3 | DLL4 | 107836 | 69406 | 0.0703881 |
| NOTCH4 | DLL4 | 0 | 0 | 0.0703881 |
| EPHA1 | EFNA1 | 67 | 50 | 4.41594 |
| EPHA2 | EFNA1 | 10202 | 7810 | 4.41594 |
| EPHA3 | EFNA1 | 65 | 48 | 4.41594 |
| EPHA4 | EFNA1 | 42 | 32 | 4.41594 |
| EPHA5 | EFNA1 | 437 | 798 | 4.41594 |
| EPHA7 | EFNA1 | 156 | 227 | 4.41594 |
| EPHA8 | EFNA1 | 3 | 0 | 4.41594 |
| EPHA1 | EFNA2 | 67 | 50 | 1.30643 |
| EPHA2 | EFNA2 | 10202 | 7810 | 1.30643 |
| EPHA3 | EFNA2 | 65 | 48 | 1.30643 |
| EPHA4 | EFNA2 | 42 | 32 | 1.30643 |
| EPHA5 | EFNA2 | 437 | 798 | 1.30643 |
| EPHA7 | EFNA2 | 156 | 227 | 1.30643 |
| EPHA8 | EFNA2 | 3 | 0 | 1.30643 |
| EPHA1 | EFNA3 | 67 | 50 | 0 |

| | | | | |
|---|---|---|---|---|
| EPHA2 | EFNA3 | 10202 | 7810 | 0 |
| EPHA3 | EFNA3 | 65 | 48 | 0 |
| EPHA4 | EFNA3 | 42 | 32 | 0 |
| EPHA5 | EFNA3 | 437 | 798 | 0 |
| EPHA7 | EFNA3 | 156 | 227 | 0 |
| EPHA8 | EFNA3 | 3 | 0 | 0 |
| EPHA1 | EFNA4 | 67 | 50 | 4.61158 |
| EPHA2 | EFNA4 | 10202 | 7810 | 4.61158 |
| EPHA3 | EFNA4 | 65 | 48 | 4.61158 |
| EPHA4 | EFNA4 | 42 | 32 | 4.61158 |
| EPHA5 | EFNA4 | 437 | 798 | 4.61158 |
| EPHA7 | EFNA4 | 156 | 227 | 4.61158 |
| EPHA8 | EFNA4 | 3 | 0 | 4.61158 |
| EPHA1 | EFNA5 | 67 | 50 | 1.79451 |
| EPHA2 | EFNA5 | 10202 | 7810 | 1.79451 |
| EPHA3 | EFNA5 | 65 | 48 | 1.79451 |
| EPHA4 | EFNA5 | 42 | 32 | 1.79451 |
| EPHA5 | EFNA5 | 437 | 798 | 1.79451 |
| EPHA7 | EFNA5 | 156 | 227 | 1.79451 |
| EPHA8 | EFNA5 | 3 | 0 | 1.79451 |
| EPHB1 | EFNB1 | 67 | 50 | 12.6425 |
| EPHB2 | EFNB1 | 427 | 290 | 12.6425 |
| EPHB3 | EFNB1 | 117 | 27 | 12.6425 |
| EPHB4 | EFNB1 | 4383 | 3194 | 12.6425 |
| EPHB6 | EFNB1 | 28 | 39 | 12.6425 |
| EPHA4 | EFNB1 | 42 | 32 | 12.6425 |
| EPHB1 | EFNB2 | 67 | 50 | 12.2504 |
| EPHB2 | EFNB2 | 427 | 290 | 12.2504 |
| EPHB3 | EFNB2 | 117 | 27 | 12.2504 |
| EPHB4 | EFNB2 | 4383 | 3194 | 12.2504 |
| EPHB6 | EFNB2 | 28 | 39 | 12.2504 |
| EPHA4 | EFNB2 | 42 | 32 | 12.2504 |
| EPHB1 | EFNB3 | 67 | 50 | 5.66867 |
| EPHB2 | EFNB3 | 427 | 290 | 5.66867 |
| EPHB3 | EFNB3 | 117 | 27 | 5.66867 |
| EPHB4 | EFNB3 | 4383 | 3194 | 5.66867 |
| EPHB6 | EFNB3 | 28 | 39 | 5.66867 |
| EPHA4 | EFNB3 | 42 | 32 | 5.66867 |
| TNFRSF1A | TNF | 10094 | 10585 | 0.127431 |
| TNFRSF1B | TNF | 875 | 566 | 0.127431 |
| LTBR | TNF | 12006 | 10262 | 0.127431 |
| TNFRSF1A | LTA | 10094 | 10585 | 0.103095 |
| TNFRSF1B | LTA | 875 | 566 | 0.103095 |
| TNFRSF14 | LTA | 0 | 0 | 0.103095 |
| LTBR | LTA | 12006 | 10262 | 0.103095 |

| | | | | |
|---|---|---|---|---|
| TNFRSF8 | TNFSF8 | 0 | 0 | 0 |
| LTBR | LTB | 12006 | 10262 | 0.0581009 |
| TNFRSF10A | TNFSF10 | 350 | 271 | 0.0386981 |
| TNFRSF10B | TNFSF10 | 26190 | 21025 | 0.0386981 |
| TNFRSF10C | TNFSF10 | 1333 | 376 | 0.0386981 |
| TNFRSF10D | TNFSF10 | 16221 | 15578 | 0.0386981 |
| TNFRSF11B | TNFSF10 | 20976 | 41717 | 0.0386981 |
| TNFRSF11A | TNFSF11 | 27 | 20 | 0 |
| TNFRSF11B | TNFSF11 | 20976 | 41717 | 0 |
| TNFRSF9 | TNFSF9 | 53 | 15 | 0 |
| TNFRSF4 | TNFSF4 | 3 | 4 | 0.485994 |
| TNFRSF14 | TNFSF14 | 0 | 0 | 0.00968154 |
| TNFRSF6B | TNFSF14 | 0 | 0 | 0.00968154 |
| LTBR | TNFSF14 | 12006 | 10262 | 0.00968154 |
| TNFRSF18 | TNFSF18 | 15 | 7 | 0.134271 |
| IL2RA | IL2 | 32 | 12 | 0 |
| IL2RB | IL2 | 0 | 5 | 0 |
| IL2RG | IL2 | 20 | 8 | 0 |
| IL7R | IL7 | 172 | 132 | 0 |
| IL2RG | IL7 | 20 | 8 | 0 |
| IL9R | IL9 | 0 | 0 | 0 |
| IL2RG | IL9 | 20 | 8 | 0 |
| IL15RA | IL15 | 585 | 687 | 0.2739 |
| IL2RB | IL15 | 0 | 5 | 0.2739 |
| IL2RG | IL15 | 20 | 8 | 0.2739 |
| IL4R | IL4 | 10108 | 7306 | 1.33311 |
| IL2RG | IL4 | 20 | 8 | 1.33311 |
| IL13RA1 | IL4 | 13594 | 9479 | 1.33311 |
| IL13RA1 | IL13 | 13594 | 9479 | 0.0610946 |
| IL13RA2 | IL13 | 0 | 0 | 0.0610946 |
| IL4R | IL13 | 10108 | 7306 | 0.0610946 |
| IL2RG | IL13 | 20 | 8 | 0.0610946 |
| IL3RA | IL3 | 0 | 0 | 0.0278091 |
| CSF2RB | IL3 | 63 | 15 | 0.0278091 |
| IL5RA | IL5 | 1 | 0 | 0 |
| CSF2RB | IL5 | 63 | 15 | 0 |
| CSF2RA | CSF2 | 0 | 0 | 0 |
| CSF2RB | CSF2 | 63 | 15 | 0 |
| IFNGR1 | IFNG | 4144 | 2701 | 0.0533601 |
| IFNGR2 | IFNG | 142 | 123 | 0.0533601 |
| IL10RA | IL10 | 312 | 122 | 0.0175415 |
| IL10RB | IL10 | 61 | 49 | 0.0175415 |
| CD4 | IL16 | 20074 | 16693 | 0.791199 |
| IL12RB1 | IL12A | 91 | 39 | 1.29506 |
| IL12RB2 | IL12A | 5 | 4 | 1.29506 |

| | | | | |
|---|---|---|---|---|
| IL12RB1 | IL12B | 91 | 39 | 0 |
| IL12RB2 | IL12B | 5 | 4 | 0 |
| IFNAR1 | IFNA4 | 5037 | 3785 | 0 |
| IFNAR2 | IFNA4 | 223 | 248 | 0 |
| IFNAR1 | IFNA1 | 5037 | 3785 | 0 |
| IFNAR2 | IFNA1 | 223 | 248 | 0 |
| IFNAR1 | IFNA5 | 5037 | 3785 | 0 |
| IFNAR2 | IFNA5 | 223 | 248 | 0 |
| IFNAR1 | IFNA16 | 5037 | 3785 | 0 |
| IFNAR2 | IFNA16 | 223 | 248 | 0 |
| IFNAR1 | IFNA8 | 5037 | 3785 | 0 |
| IFNAR2 | IFNA8 | 223 | 248 | 0 |
| IFNAR1 | IFNA14 | 5037 | 3785 | 0 |
| IFNAR2 | IFNA14 | 223 | 248 | 0 |
| IFNAR1 | IFNA21 | 5037 | 3785 | 0 |
| IFNAR2 | IFNA21 | 223 | 248 | 0 |
| IFNAR1 | IFNA2 | 5037 | 3785 | 0 |
| IFNAR2 | IFNA2 | 223 | 248 | 0 |
| IFNAR1 | IFNA13 | 5037 | 3785 | 0 |
| IFNAR2 | IFNA13 | 223 | 248 | 0 |
| IFNAR1 | IFNA6 | 5037 | 3785 | 0 |
| IFNAR2 | IFNA6 | 223 | 248 | 0 |
| IFNAR1 | IFNA7 | 5037 | 3785 | 0 |
| IFNAR2 | IFNA7 | 223 | 248 | 0 |
| IFNAR1 | IFNA10 | 5037 | 3785 | 0 |
| IFNAR2 | IFNA10 | 223 | 248 | 0 |
| IFNAR1 | IFNA17 | 5037 | 3785 | 0 |
| IFNAR2 | IFNA17 | 223 | 248 | 0 |
| IFNAR1 | IFNW1 | 5037 | 3785 | 0 |
| IFNAR2 | IFNW1 | 223 | 248 | 0 |
| IFNAR1 | IFNB1 | 5037 | 3785 | 0 |
| IFNAR2 | IFNB1 | 223 | 248 | 0 |
| IL1R1 | IL1A | 3467 | 2856 | 0.0803254 |
| IL1R2 | IL1A | 6 | 0 | 0.0803254 |
| IL1R1 | IL1B | 3467 | 2856 | 0 |
| IL1R2 | IL1B | 6 | 0 | 0 |
| IL1R1 | IL1RN | 3467 | 2856 | 0 |
| IL1R2 | IL1RN | 6 | 0 | 0 |
| IL18R1 | IL18 | 16 | 2 | 0.265656 |
| IL18RAP | IL18 | 0 | 0 | 0.265656 |
| IGF1R | IGF1 | 23223 | 13852 | 0.445648 |
| IGF2R | IGF2 | 26729 | 24413 | 0.0390533 |
| INSR | INS | 2662 | 1842 | 0 |
| MET | HGF | 12653 | 12224 | 0.10775 |
| MST1R | MST1 | 44 | 89 | 0 |

| | | | | |
|---|---|---|---|---|
| FGFR1 | FGF1 | 10191 | 10818 | 2.83887 |
| FGFR2 | FGF1 | 246 | 328 | 2.83887 |
| FGFR3 | FGF1 | 1169 | 1988 | 2.83887 |
| FGFR4 | FGF1 | 190 | 188 | 2.83887 |
| FGFR1 | FGF2 | 10191 | 10818 | 13.6923 |
| FGFR2 | FGF2 | 246 | 328 | 13.6923 |
| FGFR3 | FGF2 | 1169 | 1988 | 13.6923 |
| FGFR4 | FGF2 | 190 | 188 | 13.6923 |
| FGFR1 | FGF3 | 10191 | 10818 | 0 |
| FGFR2 | FGF3 | 246 | 328 | 0 |
| FGFR1 | FGF4 | 10191 | 10818 | 0 |
| FGFR2 | FGF4 | 246 | 328 | 0 |
| FGFR3 | FGF4 | 1169 | 1988 | 0 |
| FGFR4 | FGF4 | 190 | 188 | 0 |
| FGFR1 | FGF5 | 10191 | 10818 | 0.425066 |
| FGFR2 | FGF5 | 246 | 328 | 0.425066 |
| FGFR3 | FGF5 | 1169 | 1988 | 0.425066 |
| FGFR1 | FGF6 | 10191 | 10818 | 0 |
| FGFR2 | FGF6 | 246 | 328 | 0 |
| FGFR4 | FGF6 | 190 | 188 | 0 |
| FGFR2 | FGF7 | 246 | 328 | 0.818809 |
| FGFR2 | FGF8 | 246 | 328 | 0 |
| FGFR3 | FGF8 | 1169 | 1988 | 0 |
| FGFR4 | FGF8 | 190 | 188 | 0 |
| FGFR1 | FGF9 | 10191 | 10818 | 0.0839945 |
| FGFR2 | FGF9 | 246 | 328 | 0.0839945 |
| FGFR3 | FGF9 | 1169 | 1988 | 0.0839945 |
| FGFR4 | FGF9 | 190 | 188 | 0.0839945 |
| FGFR1 | FGF10 | 10191 | 10818 | 0.042519 |
| FGFR2 | FGF10 | 246 | 328 | 0.042519 |
| FGFR3 | FGF10 | 1169 | 1988 | 0.042519 |
| FGFR4 | FGF10 | 190 | 188 | 0.042519 |
| FGFR1 | FGF11 | 10191 | 10818 | 3.2058 |
| FGFR2 | FGF11 | 246 | 328 | 3.2058 |
| FGFR3 | FGF11 | 1169 | 1988 | 3.2058 |
| FGFR4 | FGF11 | 190 | 188 | 3.2058 |
| FGFR1 | FGF12 | 10191 | 10818 | 5.89345 |
| FGFR2 | FGF12 | 246 | 328 | 5.89345 |
| FGFR3 | FGF12 | 1169 | 1988 | 5.89345 |
| FGFR4 | FGF12 | 190 | 188 | 5.89345 |
| FGFR1 | FGF13 | 10191 | 10818 | 0.297014 |
| FGFR2 | FGF13 | 246 | 328 | 0.297014 |
| FGFR3 | FGF13 | 1169 | 1988 | 0.297014 |
| FGFR4 | FGF13 | 190 | 188 | 0.297014 |
| FGFR1 | FGF14 | 10191 | 10818 | 10.0795 |

| | | | | |
|---|---|---|---|---|
| FGFR2 | FGF14 | 246 | 328 | 10.0795 |
| FGFR3 | FGF14 | 1169 | 1988 | 10.0795 |
| FGFR4 | FGF14 | 190 | 188 | 10.0795 |
| FGFR1 | FGF16 | 10191 | 10818 | 0 |
| FGFR2 | FGF16 | 246 | 328 | 0 |
| FGFR3 | FGF16 | 1169 | 1988 | 0 |
| FGFR4 | FGF16 | 190 | 188 | 0 |
| FGFR1 | FGF17 | 10191 | 10818 | 0.185556 |
| FGFR2 | FGF17 | 246 | 328 | 0.185556 |
| FGFR3 | FGF17 | 1169 | 1988 | 0.185556 |
| FGFR4 | FGF17 | 190 | 188 | 0.185556 |
| FGFR1 | FGF18 | 10191 | 10818 | 0.0453882 |
| FGFR2 | FGF18 | 246 | 328 | 0.0453882 |
| FGFR3 | FGF18 | 1169 | 1988 | 0.0453882 |
| FGFR4 | FGF18 | 190 | 188 | 0.0453882 |
| FGFR1 | FGF19 | 10191 | 10818 | 0 |
| FGFR2 | FGF19 | 246 | 328 | 0 |
| FGFR3 | FGF19 | 1169 | 1988 | 0 |
| FGFR4 | FGF19 | 190 | 188 | 0 |
| NTRK2 | BDNF | 12799 | 10507 | 2.43497 |
| NGFR | BDNF | 4 | 57 | 2.43497 |
| NTRK1 | NTF3 | 13 | 0 | 0 |
| NTRK2 | NTF3 | 12799 | 10507 | 0 |
| NTRK3 | NTF3 | 299 | 373 | 0 |
| NGFR | NTF3 | 4 | 57 | 0 |
| MPL | THPO | 7 | 9 | 0.0703603 |
| EPOR | EPO | 695 | 323 | 0 |
| TEK | ANGPT1 | 480 | 665 | 1.3593 |
| TEK | ANGPT2 | 480 | 665 | 0 |
| TEK | ANGPTL1 | 480 | 665 | 9.01043 |
| PDGFRA | PDGFA | 8662 | 8471 | 37.4467 |
| PDGFRA | PDGFB | 8662 | 8471 | 1.69045 |
| PDGFRB | PDGFB | 98286 | 53923 | 1.69045 |
| FLT1 | VEGFB | 104 | 39 | 21.0968 |
| NRP1 | VEGFB | 3090 | 1334 | 21.0968 |
| KDR | VEGFC | 8 | 8 | 0.0268396 |
| FLT4 | VEGFC | 6 | 4 | 0.0268396 |
| KDR | FIGF | 8 | 8 | 0.0286633 |
| FLT4 | FIGF | 6 | 4 | 0.0286633 |
| FLT1 | PGF | 104 | 39 | 2.00862 |
| NRP1 | PGF | 3090 | 1334 | 2.00862 |
| EGFR | EGF | 10281 | 11271 | 0.21968 |
| EGFR | TGFA | 10281 | 11271 | 3.27761 |
| EGFR | AREG | 10281 | 11271 | 0 |
| EGFR | BTC | 10281 | 11271 | 0.413846 |

| | | | | |
|---|---|---|---|---|
| ERBB3 | BTC | 6354 | 7391 | 0.413846 |
| ERBB4 | BTC | 8 | 4 | 0.413846 |
| ERBB4 | NRG1 | 8 | 4 | 2.43271 |
| ERBB3 | NRG1 | 6354 | 7391 | 2.43271 |
| ERBB2 | NRG1 | 7283 | 6644 | 2.43271 |
| EGFR | EREG | 10281 | 11271 | 0.0906336 |
| ERBB4 | EREG | 8 | 4 | 0.0906336 |
| IL11RA | IL11 | 584 | 471 | 0.966172 |
| IL6ST | IL11 | 18587 | 14035 | 0.966172 |
| CSF3R | CSF3 | 121 | 28 | 0 |
| LEPR | LEP | 2266 | 1870 | 0.0146018 |
| IL6R | IL6 | 988 | 1032 | 0.867347 |
| IL6ST | IL6 | 18587 | 14035 | 0.867347 |
| LIFR | LIF | 1176 | 931 | 2.53368 |
| IL6ST | LIF | 18587 | 14035 | 2.53368 |
| IL6ST | CTF1 | 18587 | 14035 | 0.896088 |
| LIFR | CTF1 | 1176 | 931 | 0.896088 |
| CNTFR | CNTF | 10 | 3 | 0 |
| IL6ST | CNTF | 18587 | 14035 | 0 |
| LIFR | CNTF | 1176 | 931 | 0 |
| OSMR | OSM | 20823 | 16891 | 0 |
| IL6ST | OSM | 18587 | 14035 | 0 |
| LIFR | OSM | 1176 | 931 | 0 |
| FLT3 | FLT3LG | 0 | 0 | 0 |
| CSF1R | CSF1 | 399 | 209 | 67.2675 |
| PTPRZ1 | MDK | 29 | 4 | 128.173 |
| PTPRB | MDK | 468 | 173 | 128.173 |
| PTPRZ1 | PTN | 29 | 4 | 251.53 |
| PTPRB | PTN | 468 | 173 | 251.53 |
| TGFBR1 | TGFB1 | 8114 | 6998 | 0 |
| TGFBR2 | TGFB1 | 7819 | 7775 | 0 |
| TGFBR3 | TGFB1 | 857 | 1254 | 0 |
| TGFBR1 | TGFB2 | 8114 | 6998 | 10.2392 |
| TGFBR2 | TGFB2 | 7819 | 7775 | 10.2392 |
| TGFBR3 | TGFB2 | 857 | 1254 | 10.2392 |
| TGFBR1 | TGFB3 | 8114 | 6998 | 4.49341 |
| TGFBR2 | TGFB3 | 7819 | 7775 | 4.49341 |
| TGFBR3 | TGFB3 | 857 | 1254 | 4.49341 |
| BMPR1A | BMP2 | 2617 | 1958 | 0.865568 |
| BMPR1B | BMP2 | 9 | 8 | 0.865568 |
| BMPR2 | BMP2 | 18786 | 14274 | 0.865568 |
| ACVR1 | BMP2 | 2883 | 2045 | 0.865568 |
| ACVR2B | BMP2 | 791 | 445 | 0.865568 |
| BMPR1A | BMP3 | 2617 | 1958 | 0.011317 |
| BMPR1B | BMP3 | 9 | 8 | 0.011317 |

| | | | | |
|---|---|---|---|---|
| BMPR2 | BMP3 | 18786 | 14274 | 0.011317 |
| ACVR1 | BMP3 | 2883 | 2045 | 0.011317 |
| ACVR2B | BMP3 | 791 | 445 | 0.011317 |
| BMPR1A | BMP4 | 2617 | 1958 | 3.56783 |
| BMPR1B | BMP4 | 9 | 8 | 3.56783 |
| BMPR2 | BMP4 | 18786 | 14274 | 3.56783 |
| ACVR1 | BMP4 | 2883 | 2045 | 3.56783 |
| ACVR2B | BMP4 | 791 | 445 | 3.56783 |
| BMPR1A | BMP5 | 2617 | 1958 | 0.00729969 |
| BMPR1B | BMP5 | 9 | 8 | 0.00729969 |
| BMPR2 | BMP5 | 18786 | 14274 | 0.00729969 |
| ACVR1 | BMP5 | 2883 | 2045 | 0.00729969 |
| ACVR2B | BMP5 | 791 | 445 | 0.00729969 |
| BMPR1A | BMP6 | 2617 | 1958 | 0 |
| BMPR1B | BMP6 | 9 | 8 | 0 |
| BMPR2 | BMP6 | 18786 | 14274 | 0 |
| ACVR1 | BMP6 | 2883 | 2045 | 0 |
| ACVR2B | BMP6 | 791 | 445 | 0 |
| BMPR1A | BMP7 | 2617 | 1958 | 16.4788 |
| BMPR1B | BMP7 | 9 | 8 | 16.4788 |
| BMPR2 | BMP7 | 18786 | 14274 | 16.4788 |
| ACVR1 | BMP7 | 2883 | 2045 | 16.4788 |
| ACVR2B | BMP7 | 791 | 445 | 16.4788 |
| BMPR1A | BMP10 | 2617 | 1958 | 0.033059 |
| BMPR1B | BMP10 | 9 | 8 | 0.033059 |
| BMPR2 | BMP10 | 18786 | 14274 | 0.033059 |
| ACVR1 | BMP10 | 2883 | 2045 | 0.033059 |
| ACVR2B | BMP10 | 791 | 445 | 0.033059 |
| BMPR1A | BMP15 | 2617 | 1958 | 0 |
| BMPR1B | BMP15 | 9 | 8 | 0 |
| BMPR2 | BMP15 | 18786 | 14274 | 0 |
| ACVR1 | BMP15 | 2883 | 2045 | 0 |
| ACVR2B | BMP15 | 791 | 445 | 0 |
| ACVR1 | INHBA | 2883 | 2045 | 0.0842814 |
| ACVR1B | INHBA | 2020 | 1906 | 0.0842814 |
| ACVR2B | INHBA | 791 | 445 | 0.0842814 |
| ACVR1 | INHBB | 2883 | 2045 | 0.744529 |
| ACVR1B | INHBB | 2020 | 1906 | 0.744529 |
| ACVR2B | INHBB | 791 | 445 | 0.744529 |
| ACVR1 | INHBC | 2883 | 2045 | 0.0573731 |
| ACVR1B | INHBC | 2020 | 1906 | 0.0573731 |
| ACVR2B | INHBC | 791 | 445 | 0.0573731 |
| ACVR1 | INHA | 2883 | 2045 | 0.186876 |
| ACVR1B | INHA | 2020 | 1906 | 0.186876 |
| ACVR2B | INHA | 791 | 445 | 0.186876 |

| | | | | |
|---|---|---|---|---|
| AMHR2 | AMH | 0 | 0 | 1.69292 |
| CD44 | COL1A1 | 47344 | 43823 | 36.441 |
| CD44 | COL1A2 | 47344 | 43823 | 149.164 |
| CD44 | COL3A1 | 47344 | 43823 | 3.12119 |
| CD44 | COL4A1 | 47344 | 43823 | 27.2355 |
| CD44 | COL4A2 | 47344 | 43823 | 35.8977 |
| CD44 | COL4A5 | 47344 | 43823 | 10.447 |
| CD44 | COL5A2 | 47344 | 43823 | 65.1758 |
| CD44 | COL5A3 | 47344 | 43823 | 70.4069 |
| CD44 | COL6A1 | 47344 | 43823 | 68.8776 |
| CD44 | COL6A2 | 47344 | 43823 | 29.2504 |
| CD44 | FN1 | 47344 | 43823 | 38.5671 |
| CD44 | IGFBP3 | 47344 | 43823 | 123.117 |
| CD44 | LAMA2 | 47344 | 43823 | 0.127887 |
| CD44 | LAMA4 | 47344 | 43823 | 39.3491 |
| CD44 | LAMA5 | 47344 | 43823 | 10.9562 |
| CD44 | LAMB1 | 47344 | 43823 | 92.5695 |
| CD44 | LAMB2 | 47344 | 43823 | 24.1174 |
| CD44 | LAMC1 | 47344 | 43823 | 57.7532 |
| CD44 | LAMC3 | 47344 | 43823 | 0.275787 |
| CD44 | SPP1 | 47344 | 43823 | 11.767 |
| CD44 | VCAN | 47344 | 43823 | 49.9514 |

SUPPLEMENTARY TABLE 3 HAS BEEN UPLOADED SEPARATELY

**Supplementary Table 4: Primers used in this manuscript for qPCR**

| Name | Species | Experimental Role | Forward | Reverse |
|---|---|---|---|---|
| FN | Human | Measure total fibronectin | CCACCCCCATAAGGCATAGG | GTAGGGGTCAAAGCACGAGTCATC |
| EDA-FN | Human | Measure EDA fibronectin | CCCAAGCTTAACATTGATCGCCCTAAAGGA | CCCGGTACCTGTGGACT GGGTTCCAATCAGG |
| Arg1 | Human | M2 macrophage gene expression | CAGAAGAATGGAAGAGTCAG | CAGATATGCAGGGAGTCACC |
| iNOS | Human | M1 macrophage gene expression | TGCATGGACCAGTATAAGGCAAGC | GCTTCTGGTCGATGTCATGAGCAA |
| MMP9 | Human | M2 macrophage gene expression | GATGCGTGGAGAGTCGAAAT | CACCAAACTGGATGACGATG |
| TGFB1 | Human | M2 macrophage gene expression | CCCAGCATCTGCAAAGCTC | GTCAATGTACAGCTGCCGCA |
| CXCL10 | Human | M1 macrophage gene expression | AGAACGGTGCGCTGCAC | CCTATGGCCCTGGGTCTA |
| Il1b | Human | M1 macrophage gene expression | CCACAGACCTTCCAGGAGAATG | GTGCAGTTCAGTGATCGTACAGG |
| GAPDH | Human | Housekeeping gene | CATGACAACTTTGGTATCGTGG | CCTGCTTCACCACCTTCTTG |
| ACTB | Human | Housekeeping gene | GAG CAC AGA GCC TCG CCT TT | ACA TGC CGG AGC CGT TGT C |
| NANOG | Human | Assess GSC gene expression | AGT CCC AAA GGC AAA CAA CCC ACT TC | TGC TGG AGG CTG AGG TAT TTC TGT CTC |
| Oct | Human | Assess GSC gene expression | GAC AGG GGG AGG GGA GGA GCT AGG | CTT CCC TCC AAC CAG TTG CCC CAA AC |
| SOX2 | Human | Assess GSC gene expression | GGG AAA TGG GAG GGG TGC AAA AGA GG | TTG CGT GAG TGT GGA TGG GAT TGG TG |
| LIF | Human | Assess mesenchymal gene expression | TGA ACC AGA TCA GGA GCC AA | AAG GTA CAC GAC TAT GCG GT |
| CHI3L1 | Human | Assess mesenchymal gene expression | CAG CAG CTA TGA CAT TGC CA | ATG CCC ATC ACC AGC TTA CT |
| COL4A2 | Human | Assess mesenchymal gene expression | TGT GGG CAT GAA AGG TCT CT | AAA ATC CAG CCT CGC CTT TG |
| FOSL2 | Human | Assess mesenchymal gene expression | AAG ACC TGG CGT GAT CAA GA | GCT CAG CAA TCT CCT TCT GC |
| TIMP1 | Human | Assess mesenchymal gene expression | TAC TTC CAC AGG TCC CAC AA | GCA GGG GAT GGA TAA ACA GG |
| SPOCD1 | Human | Assess mesenchymal gene expression | ATG GAG TGA AGC TTG TGT GC | TGG AAA ACC TGG CAC CCA |

**Supplementary Table 5: Antibodies used in this manuscript**

| Antigen | Species Source | Vendor |
|---|---|---|
| Mouse/Human CD11b | Rat | BioLegend |
| Mouse MHC Class II | Rat | eBioscience |
| Mouse CD206 | Rat | BioLegend |
| Human CD140a/PDGFR-$\alpha$ | Mouse | Invitrogen |
| Human CD140b/PDGFR-$\beta$ | Mouse | BioLegend |
| Human $\alpha$-SMA | Mouse | R&D Systems |
| Human PDGF | Rabbit | R&D Systems |
| Human TGF-$\beta$ | Mouse | R&D Systems |
| Human TLR4 | Goat | R&D Systems |
| Human Osteopontin | Goat | R&D Systems |
| Human HGF | Mouse | R&D Systems |
| Human and Mouse Fibronectin | Rabbit | Abcam |
| Human Nestin | Mouse | Abcam |
| Human and Mouse EDA | Mouse | Abcam |
| Human and Mouse CD31 | Rabbit | Abcam |
| Human PDGFR-B | Mouse | Santa Cruz Biotech |
| Human PDGFR-$\alpha$ | Rabbit | Abcam |
